# Direct Male Development in Chromosomally ZZ Zebrafish

**DOI:** 10.1101/2023.12.27.573483

**Authors:** Catherine A. Wilson, Peter Batzel, John H. Postlethwait

## Abstract

The genetics of sex determination varies across taxa, sometimes even within a species. Major domesticated strains of zebrafish (*Danio rerio*), including AB and TU, lack a strong genetic sex determining locus, but strains more recently derived from nature, like Nadia (NA), possess a ZZ male/ZW female chromosomal sex-determination system. AB strain fish pass through a juvenile ovary stage, forming oocytes that survive in fish that become females but die in fish that become males. To understand mechanisms of gonad development in NA zebrafish, we studied histology and single cell transcriptomics in developing ZZ and ZW fish. ZW fish developed oocytes by 22 days post-fertilization (dpf) but ZZ fish directly formed testes, avoiding a juvenile ovary phase. Gonads of some ZW and WW fish, however, developed oocytes that died as the gonad became a testis, mimicking AB fish, suggesting that the gynogenetically derived AB strain is chromosomally WW. Single-cell RNA-seq of 19dpf gonads showed similar cell types in ZZ and ZW fish, including germ cells, precursors of gonadal support cells, steroidogenic cells, interstitial/stromal cells, and immune cells, consistent with a bipotential juvenile gonad. In contrast, scRNA-seq of 30dpf gonads revealed that cells in ZZ gonads had transcriptomes characteristic of testicular Sertoli, Leydig, and germ cells while ZW gonads had granulosa cells, theca cells, and developing oocytes. Hematopoietic and vascular cells were similar in both sex genotypes. These results show that juvenile NA zebrafish initially develop a bipotential gonad; that a factor on the NA W chromosome or fewer than two Z chromosomes is essential to initiate oocyte development; and without the W factor or with two Z doses, NA gonads develop directly into testes without passing through the juvenile ovary stage. Sex determination in AB and TU strains mimics NA ZW and WW zebrafish, suggesting loss of the Z chromosome during domestication. Genetic analysis of the NA strain will facilitate our understanding of the evolution of sex determination mechanisms.

## Introduction

Vertebrates exhibit a wide variety of sex determination mechanisms, including genetic and environmental control factors (Nagahama et al., 2021). Among fishes with genetic sex determination, some species have a single major genetic sex determinant, but others use polygenic sex determination. Unlike mammals and birds, which have evolutionarily rather stable genetic sex determinants (Ioannidis et al., 2021, Terao et al., 2022), different fish lineages can have different major sex determination genes, sometimes even in the same species or genus (Matsuda and Sakaizumi, 2016, Pan et al., 2021, Song et al., 2021). Despite the popularity of zebrafish as a model organism (Bradford et al., 2017) and substantial knowledge about its gonadogenesis (Liew and Orban, 2014, Kossack and Draper, 2019, Aharon and Marlow, 2021), our understanding of the genetic regulation of zebrafish sex determination is insufficient.

Zebrafish gonads have been described as passing through a juvenile ovary stage, with all juveniles developing perinucleolar oocytes by about 20 days post-fertilization (dpf) (Takahashi, 1977, Uchida et al., 2002). In some individuals, these oocytes persist and the fish develops as a female, but in other individuals, oocytes die, the gonad transitions to a testis, and the fish becomes a male (Wang et al., 2007). Previous studies in zebrafish strains domesticated for mutagenesis research (Walker-Durchanek, 1980, Streisinger et al., 1981, Mullins et al., 1994) showed that germ cells, specifically oocytes, are essential for both the establishment and maintenance of female sex because zebrafish that lack germ cells become males (Slanchev et al., 2005), as do mutants that lose oocytes during meiosis (Rodriguez-Mari et al., 2010, Shive et al., 2010, Saito et al., 2011, Rodriguez-Mari et al., 2011, Beer and Draper, 2013, Dranow et al., 2013, Dranow et al., 2016, Ramanagoudr-Bhojappa et al., 2018, Takemoto et al., 2020, Blokhina et al., 2021, Islam et al., 2021). While zebrafish are gonochoristic, mutations that cause progressive oocyte loss in adults but retain some germ cells result in individuals that can first become females and later convert to a male phenotype (Dranow et al., 2013).

Mechanisms that cause some juvenile zebrafish to maintain oocytes and become females are not yet fully known, but oocyte survival and sex ratio are strongly influenced by both genetic and environmental factors. In general, stressful environments masculinize a clutch: an increased fraction of males occurs after hypoxia, high rearing density, high temperature, low nutrition, gamma rays, inbreeding, and exposure to exogenous cortisol (Walker and Streisinger, 1983, Shang et al., 2006, Lawrence et al., 2008, Abozaid et al., 2011, Ribas et al., 2017, Santos et al., 2017, Delomas and Dabrowski, 2018, Valdivieso et al., 2022). Sex phenotype has been mapped to different genetic locations in different zebrafish strains, and sex ratios are consistent in repeated matings of the same individual fish, consistent with polygenic sex determination in laboratory lines (Bradley et al., 2011, Anderson et al., 2012, Howe et al., 2013, Liew and Orban, 2014, Luzio et al., 2015).

Unlike laboratory strains, zebrafish lines not long adapted to laboratory life at the time of investigation, including the Nadia (NA) strain, have a genetic ZZ/ZW sex determination system, with the major Sex-Associated Region on chromosome 4 (*sar4*) located near the telomere of the right arm of chromosome-4 (chr4) (as displayed in Ensembl: (ensembl.org/Danio_rerio/Location/Chromosome?r=4)) (Tong et al., 2010, Anderson et al., 2012, Wilson et al., 2014). Individual NA strain zebrafish homozygous for Z alleles always develop as males and most ZW individuals become females, but some ZW individuals develop as males for all four strains investigated (Anderson et al., 2012, Wilson et al., 2014). In addition, WW fish in these less domesticated strains tend to become females, although some develop as males. The finding that ZZ fish do not become females suggests that the W allele is necessary but insufficient for female development, although the hypothesis that Z dosage regulates sex (two Z doses, always male; one or zero Z chromosomes, usually female) is not formally ruled out. Because ZW fish occasionally develop as males, either masculinizing environmental factors or segregating background genetic modifiers can affect zebrafish sex determination. These results raise the question: Do ZZ zebrafish pass through the juvenile ovary stage like laboratory strains, or alternatively, do they develop directly into males? Under the hypothesis that the W carries a factor that is necessary, but insufficient, for ovarian development, ZZ fish, lacking the hypothesized W-linked ovary factor, should not form juvenile ovaries with meiotic oocytes.

To test this hypothesis, we investigated gonad development in chromosomally ZW and ZZ zebrafish from the NA strain. At 19 days post-fertilization (dpf), gonads in ZW and ZZ fish were not distinguishable histologically. By about 22dpf, all NA ZW fish had begun to develop oocytes, like all fish of the domesticated AB and TU strains. Developing oocytes, however, died in some juvenile ZW fish as their gonads became testes. In contrast, gonads in ZZ fish did not pass through a juvenile ovary phase but developed directly into testes, as predicted if a W-linked factor is required for oocyte development or that two Z doses block female development. Single-cell RNA-seq (scRNA-seq) of 19dpf juvenile gonads showed that neither somatic cell clusters nor germ cell clusters were strongly differentiated by sex genotype, consistent with a bipotential gonad in both ZW and ZZ fish at this stage. Single cell transcriptomics at 30dpf, however, revealed that cells in ZW gonads had transcriptomes characteristic of theca and granulosa cells, maturing Stage IA and early Stage IB oocytes. ZZ gonads at 30dpf, however, showed developing Sertoli and Leydig cells, and some ZZ-specific interstitial cell types. Sex-specific hematopoietic and vascular cells were not detected. These results show that 1) juvenile NA zebrafish initially develop a bipotential gonad; 2) a factor on the W chromosome, or fewer than two Z chromosomes, is essential to trigger oocyte development; and 3) ZZ zebrafish gonads develop directly into testes. These results are as expected from the hypothesis that the W chromosome contains a factor that is necessary but insufficient for female development or the hypothesis that the Z chromosome contains a dosage-sensitive factor that inhibits female development, with fully penetrant inhibition of ovary development in ZZ fish. Under either hypothesis, standard laboratory strains produce both males and females, as do both ZW and WW NA zebrafish, suggesting loss of the Z chromosome leading to WW populations during domestication.

## RESULTS AND DISCUSSION

### Sex chromosomes and zebrafish sex development

To confirm the relationship of sex development and sex chromosomes, we crossed NA strain individuals genotyped for sex chromosomes. We evaluated 366 fish that were the offspring of eight different crosses of ZW females x ZZ males (five single pair crosses and three crosses with two males and two females). At sexual maturity, primers specific for Z- and W-chromosomes (Wilson et al., 2014) identified genotypic sex, while secondary sexual characteristics distinguished phenotypic sex (for females, a rounded abdomen, and pale ventral and anal fins, and for males, a streamlined body with yellow ventral and anal fins and sex tubercles). Offspring included 184 ZZ individuals (50.3%) and 182 ZW fish (49.7%), a 1:1 genotypic sex ratio (p=0.92, Chi-square test). This result shows first, that in meiosis, Z and W chromosomes segregate as homologous chromosomes, and second, that ZZ and ZW genotypes are equally likely to survive. All 184 ZZ individuals were phenotypic males. Of 182 ZW fish, 128 individuals (70.3%) were phenotypic females and 54 fish (29.7%) were phenotypic males, called neomales. Families varied widely in the percent of neomales, from 6.7% to 66.7% (Fig. 1A). This result would be explained either if sex determination in ZW NA strain zebrafish can be influenced by genetic factors in addition to *sar4* or if environmental factors increase the likelihood of male development in NA as in laboratory strains (Liew and Orban, 2014, Valdivieso et al., 2022).

**Figure 1.**
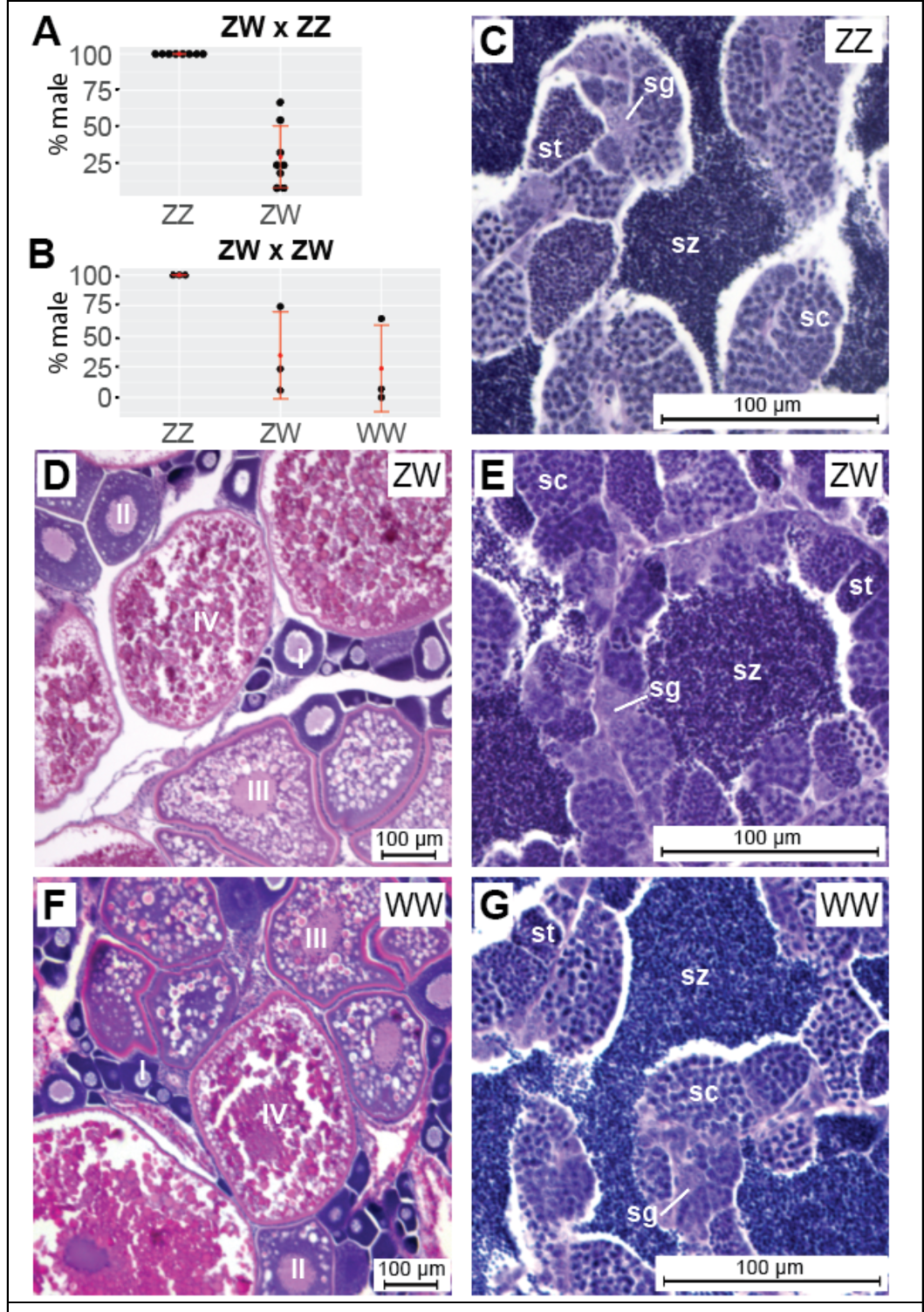
Sex ratios and gonad histology in adult ZW and ZZ zebrafish at 5 months post fertilization (mpf). A. Sex phenotypes in progeny of ZW females crossed to ZZ males. Within each sex genotype, each point represents a different cross. No ZZ individuals became adult females. B. Sex phenotypes in progeny of ZW females crossed to ZW neomales. No ZZ individuals became adult females. C-G. Cross sections of 5mpf adult gonads. C. ZZ adult testis. D. ZW ovary. E. ZW neomale testis. F. WW ovary. G. WW neomale testis. Ovary histology for ZW and WW fish showed no clear difference and testis histology of ZZ, ZW, and WW fish was the same. Abbreviations: o, oocyte; pc, pycnotic cell; sc, spermatocyte; sg, spermatogonium; sp, spermatid; sz, spermatozoa.

To study sex determination in chromosomally WW fish, we made three single pair matings between ZW phenotypic females and ZW neomales, resulting in a total of 158 progeny (Fig. 1B). Offspring included 43 (27.2%) ZZ individuals, 74 (46.9%) ZW fish, and 41 (25.9%) WW fish, approximating a 1:2:1 genotypic ratio (p=0.71, Chi square). All ZZ individuals were phenotypic males (43/43, 100%), while 30 of the 74 (40.5%) ZW fish and 10 of the 41 (25.9%) WW fish developed as neomales. The frequency of sex reversal was again variable among crosses, ranging between 5.9% and 74.2% (average 34.4%) for ZW and between 0% and 64.3% (average 23.7%) for WW (Fig. 1B). These results show that 1) homozygous W fish have normal viability, and 2) no Z-specific genes are essential to allow a fish to develop testes.

### Gonad histology in ZW, ZZ, and WW adults

Having shown that some ZW and WW fish become neomales, we wondered if adult neomale gonads differ from those in ZZ males. Histological sections showed that at 5mpf (months post fertilization), all adult NA ZZ fish examined (n=4) possessed testes with germ cells organized into cysts containing mitotic spermatogonia, meiotic spermatocytes, and post-meiotic spermatids, with mature spermatozoa present in the testis lumen (Fig. 1C). Abdominal sections of adult 5mpf NA ZW fish (n=4) with female secondary sexual characteristics possessed ovaries with oocytes in various stages of development (Selman et al., 1993), including meiotic perinucleolar oocytes (Stage I), larger oocytes containing cortical alveoli (Stage II), vitellogenic oocytes accumulating yolk (Stage III), and maturing oocytes in the final stages of oogenesis (Stage IV) (Fig. 1D). In contrast, in 5mpf genetic ZW neomales (n=4), testes had morphologies not detectably different from those of ZZ males; notably, 5mpf ZW adult neomales lacked any detectable oocytes (Fig.1E). Gonads in the phenotypically female WW fish (n=4) were also not detectably different from ovaries in ZW fish (Fig.1F), and gonads in phenotypically neomale WW fish (n=4) were no different from testes in ZZ fish (Fig. 1G). We conclude that NA strain neomales, whether ZW or WW, produce histologically normal testes with all stages of spermatogenesis like ZZ males, and by 5mpf, had no detectable oocytes.

### Gonad development in ZZ and ZW juveniles

All chromosomally ZZ NA fish had normal testes as adults, but because even phenotypically male adult laboratory zebrafish pass through a juvenile ovary stage (Takahashi, 1977, Uchida et al., 2002, Wang et al., 2007), we wondered whether ZZ zebrafish do too. The hypothesis that a factor on the W chromosome or that fewer than two Z chromosomes is necessary for ovary development predicts that ZZ fish will not pass through the juvenile ovary stage. To test this prediction, we analyzed differences in gonad development in ZZ and ZW NA zebrafish over developmental time in histological sections from two separate NA families at 10, 14, 19, 22, 26, and 30dpf.

Results showed that gonads in ZZ and ZW zebrafish were morphologically indistinguishable in cross-sections of 10, 14, and 19dpf larvae. In five ZW and four ZZ fish examined at 10dpf, bilateral gonads had clusters of germ cells surrounded by somatic cells (Fig. 2A, B), as for laboratory strain zebrafish (Takahashi, 1977, Uchida et al., 2002, Rodriguez-Mari et al., 2005, Wang et al., 2007, Dranow et al., 2013, Dranow et al., 2016). Gonads in both ZW and ZZ fish increased in size and in germ cell number at 14dpf and 19dpf, but gonads in ZW and ZZ fish remained morphologically indistinguishable (Fig. 2C-F), again as in laboratory strains. We conclude that the morphology of ZW and ZZ gonads appeared to be bipotential up to 19dpf.

**Figure 2.**
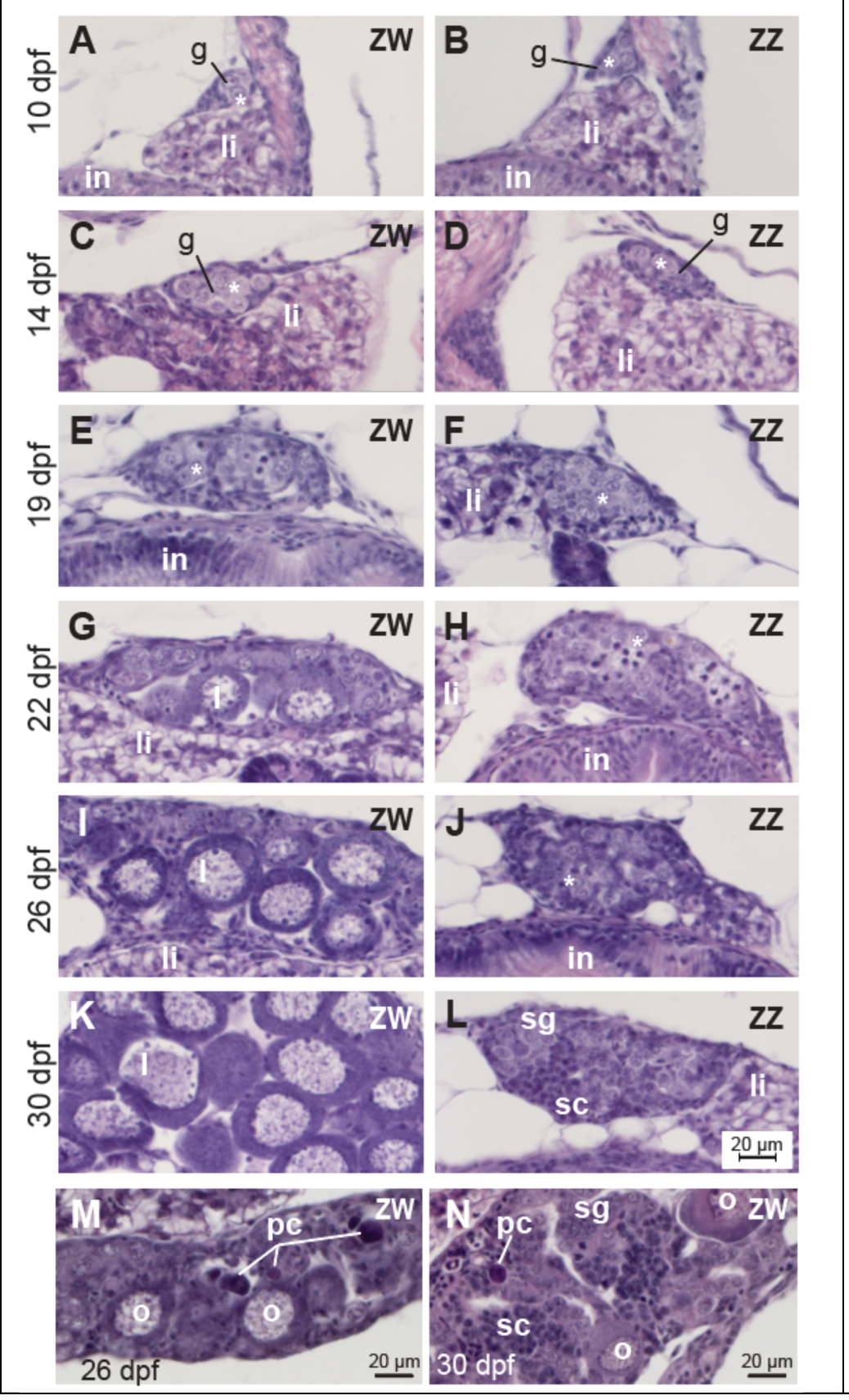
Developmental trajectory of gonads from ZW and ZZ fish. A, C, E, G, I, K, M, N: ZW gonads. B, D, F, H, J, L: ZZ gonads. A, B: 10dpf. C, D. 14dpf. E, F: 19dpf. G, H: 22dpf. I, J: 26dpf. K, L. 30dpf. At 19dpf and earlier, ZW and ZZ gonads were histologically similar, but at 22dpf and later, oocytes were developing in ZW, but not ZZ gonads. At 26dpf, some ZZ gonads showed clusters of spermatogonia, and at 30dpf, most ZZ gonads were still immature testes that were organized into cysts containing spermatogonia and spermatocytes. No ZZ gonad studied showed any oocytes at any developmental stage, but instead, showed direct development into testes. M. At 26dpf, some ZW gonads were mostly ovary with a few pycnotic cells. N. At 30dpf, some ZW gonads were mostly testes with a few pycnotic cells. Abbreviations: *, gonocytes; g, gonad; I and II, Stage I and Stage II oocytes; in, intestine; li, liver; o, oocyte; pc, pycnotic cell; sc, spermatocyte; sg, spermatogonium.

Between 19 and 22dpf, gonad development in ZW and ZZ sex genotypes began to differ. At 22dpf, gonads in nine of ten ZW individuals examined contained perinucleolar oocytes (Fig. 2G), while gonads in the tenth fish examined retained the indifferent morphology of younger fish. In contrast, all ten ZZ individuals at 22dpf still possessed indifferent gonads, failing to advance in developmental stage (Fig. 2H). By 26dpf, gonads of all 11 ZW fish studied had perinucleolar oocytes (Fig. 2I), but four of these 11 had gonads that also contained pyknotic cells representing dying oocytes (Fig. 2M). Similar pyknotic cells appear in juvenile gonads transforming to adult testis in laboratory strain zebrafish and in gonads of mutants experiencing female-to-male sex reversal (Takahashi, 1977, Uchida et al., 2002, Maack and Segner, 2003, Rodriguez-Mari et al., 2010, Rodriguez-Mari et al., 2011, Dranow et al., 2016, Blokhina et al., 2021, Xie et al., 2021). Of nine 26dpf ZZ fish, three contained largely undifferentiated gonads, and the other six had clusters of spermatogonia (Fig. 2J), including one with spermatocytes. At 30dpf, 10 of 12 ZW fish examined had a well-developed ovary full of perinucleolar oocytes (Fig. 2K). At 30dpf, oocytes had increased in size, but had not yet entered the cortical alveolar stage (Stage II, (Selman et al., 1993)). Two of these 12 ZW 30dpf fish, however, had gonads that were nearly all testis, including spermatogonia and spermatocytes, but also pyknotic oocyte-like cells and a few remaining oocytes (Fig. 2N). We interpret these two fish as ZW individuals that had gone through the juvenile ovary stage, but by 30dpf, were transitioning to neomales.

The developmental trajectory of ZZ males differed from that of ZW neomales. At 30 dpf, seven of 11 ZZ males investigated still had immature testes that were organized into cysts containing spermatogonia and spermatocytes (Fig. 2L); one ZZ individual still had largely undifferentiated gonads; three ZZ fish had maturing testes containing spermatogonia and spermatocytes; and one had spermatids. All three of these maturing male fish came from Family 1, which was fed paramecia instead of rotifers during development. Perinucleolar oocytes were never observed in any ZZ individual analyzed at any stage. We conclude that gonads in ZZ zebrafish of the NA strain: 1) are slower in developing sex-specific morphologies than ZW fish; 2) do not form immature oocytes; 3) do not pass through a juvenile hermaphrodite stage; and 4) develop directly into testes. This developmental trajectory differs from that of domesticated zebrafish, which all appear to pass through a juvenile hermaphrodite stage. Further, these findings are consistent with the hypothesis that a factor on the W chromosome, or alternatively, fewer than two Z chromosomes, is necessary for oocytes to develop in NA zebrafish gonads.

### 19dpf gonad gene expression patterns were similar in ZW and ZZ gonads

The histology investigations (Fig. 1, 2) showed that at 19dpf, the morphology of ZW and ZZ gonads could not be distinguished, but at 30dpf, gonads in both ZW and ZZ fish showed signs of differentiation. To learn whether gonad transcriptomes of ZW and ZZ fish had become different at 19dpf even though gonad morphologies had not, we performed scRNA-seq on 19dpf and 30dpf gonads dissected from ZW and ZZ animals. From resulting data, we removed red blood cells and contaminating non-gonad cells, and re-clustered remaining cells. We analyzed each time point separately and then merged all four datasets (two ages by two sex genotypes). Analysis of 19dpf gonads resulted in 783 ZW cells and 1270 ZZ cells grouped into 28 clusters (Fig. 3A, B, Supplementary Table S1).

**Figure 3.**
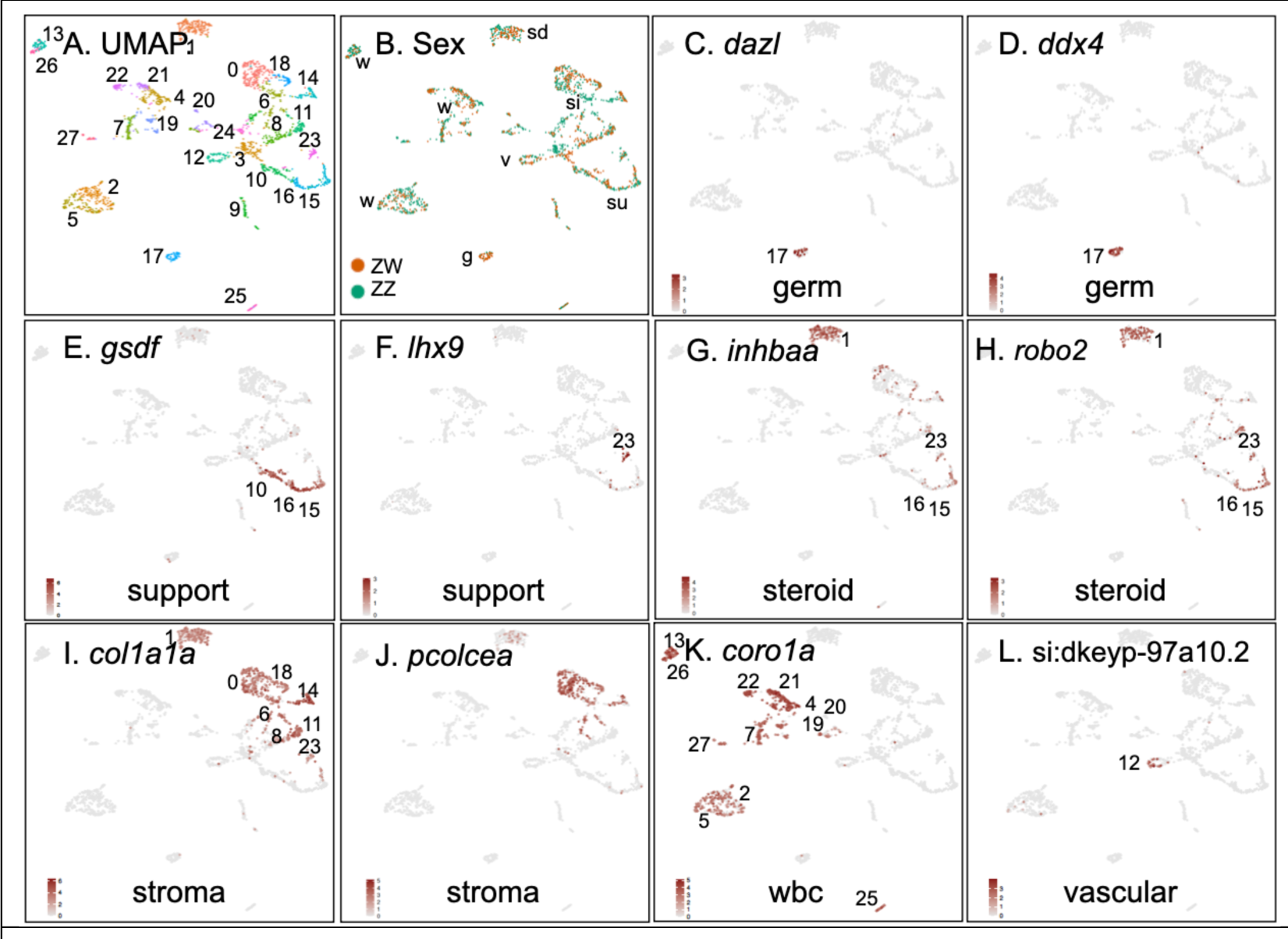
Cell-type marker genes for 19dpf ZW and ZZ gonads. A. UMAP plot for 19dpf gonads combining ZW and ZZ cells. B. Cells originating from ZW (red) or ZZ (blue) gonads, showing all clusters with mixtures of both sex genotypes. C, D. Germ cells marked by *dazl* and *ddx4* (*vasa*) were in cluster 19c17. E, F. Support cells (granulosa and Sertoli) or their precursors marked by *gsdf*, and *lhx9*. G, H. Steriogenic cells (theca, Leydig) or their precursors labeled with *inhbaa* and *robo2* expression. I, J. Stroma/interstitial cells marked by *col1a1a* and *pcolcea*. K. White blood cells identified by *coro1a* expression. L. Vasculature labeled with *si:dkeyp-97a10.2*. Abbreviations: g, germ cells; si, stroma/interstitial; sd, steroidogenic; su, support; v, vascular; w, white blood cells.

<u>Cell-type marker genes s</u> help characterize cluster identities (Qiu et al., 2021) (Supplementary Table S2). *Germ cells* are marked by the RNA-binding protein Dazl, which is essential before meiosis for germ cell cystogenesis and germline stem cell specification (Fu et al., 2015, Bertho et al., 2021). Expression of *dazl* identified 19dpf cluster 17 (19c17) as germ cells (Fig. 3C).

Additional markers germ cell markers included *ddx4* (Fig. 3D) and *dnd1*, as well as novel markers including *si:ch73-167f10.1* (Supplementary Table S2). *Support cells*, including granulosa cells in ovaries and Sertoli cells in testes, express the fish-specific TGFb-family protein Gsdf (gonadal soma derived factor) (Sawatari et al., 2007, Gautier et al., 2011, Rondeau et al., 2013, Yan et al., 2017, Hsu and Chung, 2021). At 19dpf, three clusters (19c10, c15, and c16) expressed *gsdf* (Fig. 3E), thus identifying support cell precursors. Cluster 19c23 strongly expressed *lhx9* (Fig. 3F), which is expressed in support cell precursors and is downregulated during differentiation into granulosa and Sertoli cells (Mazaud et al., 2002, Liu et al., 2022).

*Steroidogenic cells*, including theca cells in ovaries and Leydig cells in testes, arise from *Nr5a1*- expressing precursor cells (Stevant et al., 2019, Yan et al., 2020). Cluster 19c16 expressed *nr5a1b* as did 19c1, c15, and c23. Nearly all cells in Cluster 19c1 expressed *inhbaa* and *robo2* (Fig. 3G, H), which are markers of fetal Leydig cells in mouse (Archambeault and Yao, 2010, McClelland et al., 2015) and are necessary for steroidogenesis and folliculogenesis (Namwanje and Brown, 2016, Lu et al., 2020, Zhao et al., 2022), as well as *osr2* (Dickinson et al., 2008, McClelland et al., 2015, Martinot and Boerboom, 2021). *Interstitial and stromal cells* express the collagen gene *col1a1a* (Liu et al., 2022). Several clusters (19c0, c1, c6, c11, c14, and c18) expressed *col1a1a* at 19dpf and *pcolcea* (procollagen C-endopeptidase enhancer) (Fig.3I, J) defining them as stromal/interstitial cells. *Leukocytes* express the actin filament binding protein Coro1a (Song et al., 2004, Xavier et al., 2008). Gonads at 19dpf displayed a substantial variety of white blood cell types (Clusters 19c2, c4, c5, c7, c13, c19, c20, c21, c22, c26, c27) (Fig. 3K). *Vasculature* specific for the testis develop in XY mice coincident with the expression of *Sry* (Brennan et al., 2002) and appear early in zebrafish gonads (Kossack et al., 2023). In NA zebrafish, 19c12 expressed specifically the vascular marker *si:dkeyp-97a10.2* (Gomez et al., 2009) (Fig. 3L) and the lymphatic marker *lyve1b*, identifying gonadal vascular cells (Jackson et al., 2001, Prevo et al., 2001, Chen et al., 2013).

All 19dpf clusters contained both ZW and ZZ cells (Fig. 3B), which agrees with the histology findings, and suggests that ZW and ZZ gonads were morphologically and transcriptionally similar at 19dpf (Fig. 2E, F). Differential expression analysis of 19dpf clusters showed no significantly differentially expressed genes between ZZ and ZW genotypes in nealy all clusters; the most DE genes, mostly mitochondrial genes, appeared in 19c1, suggesting a slight difference in stress levels (Ilicic et al., 2016) in that cluster for ZW and ZZ samples (Supplementary Table S3), confirming strong similarity of ZZ and ZW cell types at 19dpf.

### 30dpf gonad gene expression patterns in ZW and ZZ gonads were distinct

The histology studies had shown that ZW and ZZ gonads were morphologically different by 30dpf (Fig. 2), so we wanted to identify gene expression patterns in 30dpf individuals with different chromosomal sexes by single cell transcriptomics. The 30dpf gonad sample gave 7787 ZW cells and 2843 ZZ cells in 48 clusters (Fig. 4A, Supplementary Table S4, 5). In contrast to 19dpf gonads, 30dpf gonads had several clusters specific for either ZW or ZZ cells (Fig. 4B).

**Figure 4.**
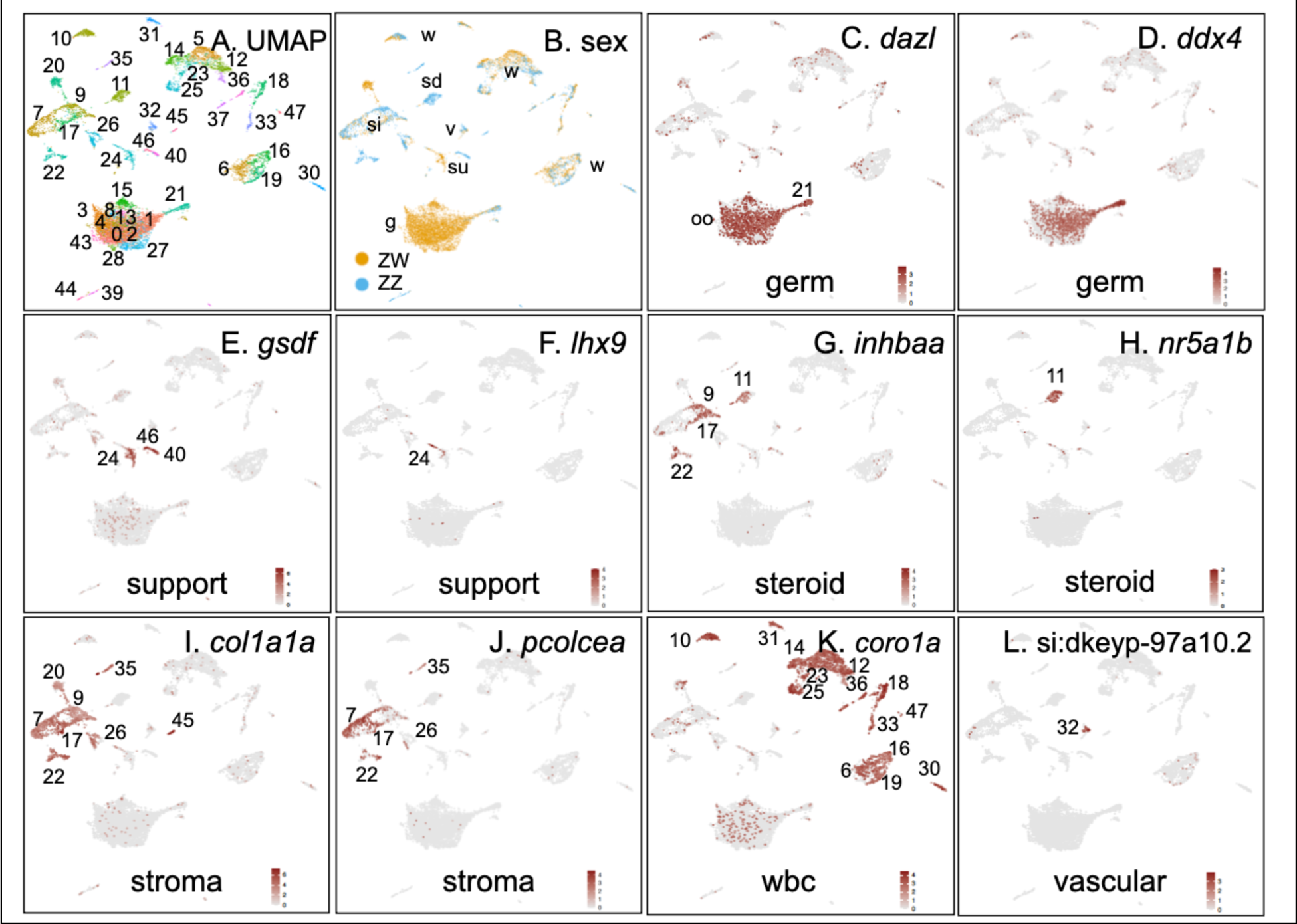
Cell-type marker genes for 30dpf ZW and ZZ gonads. A. UMAP plot for 30dpf gonads combining ZW and ZZ cells. B. Cells originating from ZW (gold) or ZZ (blue) gonads, showing several clusters with sex-specific genotypes. C, D. Germ cells marked by *dazl* and *ddx4* (*vasa*) expression. E, F. Support cells (granulosa and Sertoli) or their precursors marked by *gsdf* and *lhx9* expression. G, H. Steriogenic cells (theca, Leydig) or their precursors labeled with *inhbaa* and *nr5a1b* expression. I, J. Stroma/interstitial cells marked by *col1a1a* and *pcolcea* expresseion. K. White blood cells identified by *coro1a* expression. L. Vasculature labeled with *si:dkeyp-97a10.2* expression. Abbreviations: g, germ cells; oo, oocytes; si, stroma/interstitial; sd, steroidogenic; su, support; v, vascular; w, white blood cells.

*Germ cells* were clearly labeled by *dazl* and *ddx4* in a large group of ZW-specific clusters (30c0- c4, c8, c13, c15, c27, c28, c43) and in c21 (Fig. 4C, D), showing substantial proliferation of germ cells in ZW but much less in ZZ gonads. *Support cells* labeled by *gsdf* and *lhx9* identified mainly 30c24, c40, and c46. *Steroidogenic cells* marked by *inhbaa* and *nr5a1b* identified 30c11 for both markers, and 30c9, c17, and c22 specific for *inhbaa* (Fig. 4G, H). *Interstitial/stroma cells* expressed *col1a1a* in a variety of clusters (30c7, c9, c17, c20, c22, c26, c35, c45) and *pcolcea* in most of these (Fig. 4I, J). *Leukocytes* marked by *coro1a* included 14 clusters (Fig. 4K).

*Vascular cell* types denoted by *si:dkeyp-97a10.2* expression (Gomez et al., 2009) occupied 30c32. We conclude that by 30dpf, ZW and ZZ gonads had diverged considerably in terms of the cell types they contained, consistent with the histology results (Fig. 2).

### Combined datasets for age and sex genotype

Integrating all four samples (two ages, each with two sex genotypes) gave 40 clusters (Fig. 5A, Supplementary Table S6) with defined marker genes (Supplementary Table S7). Some clusters contained significant proportions of cells from both 19dpf and 30dpf, but many sorted out by age (Fig. 5B), suggesting cell type maturation. And, as expected from the 30dpf-only analysis (Fig. 5C), some clusters in the combined analysis were mostly ZW cells and others mainly ZZ cells, while many were still mixed. The analysis below provides a more detailed understanding of major cell types (Fig. 5D.

**Figure 5.**
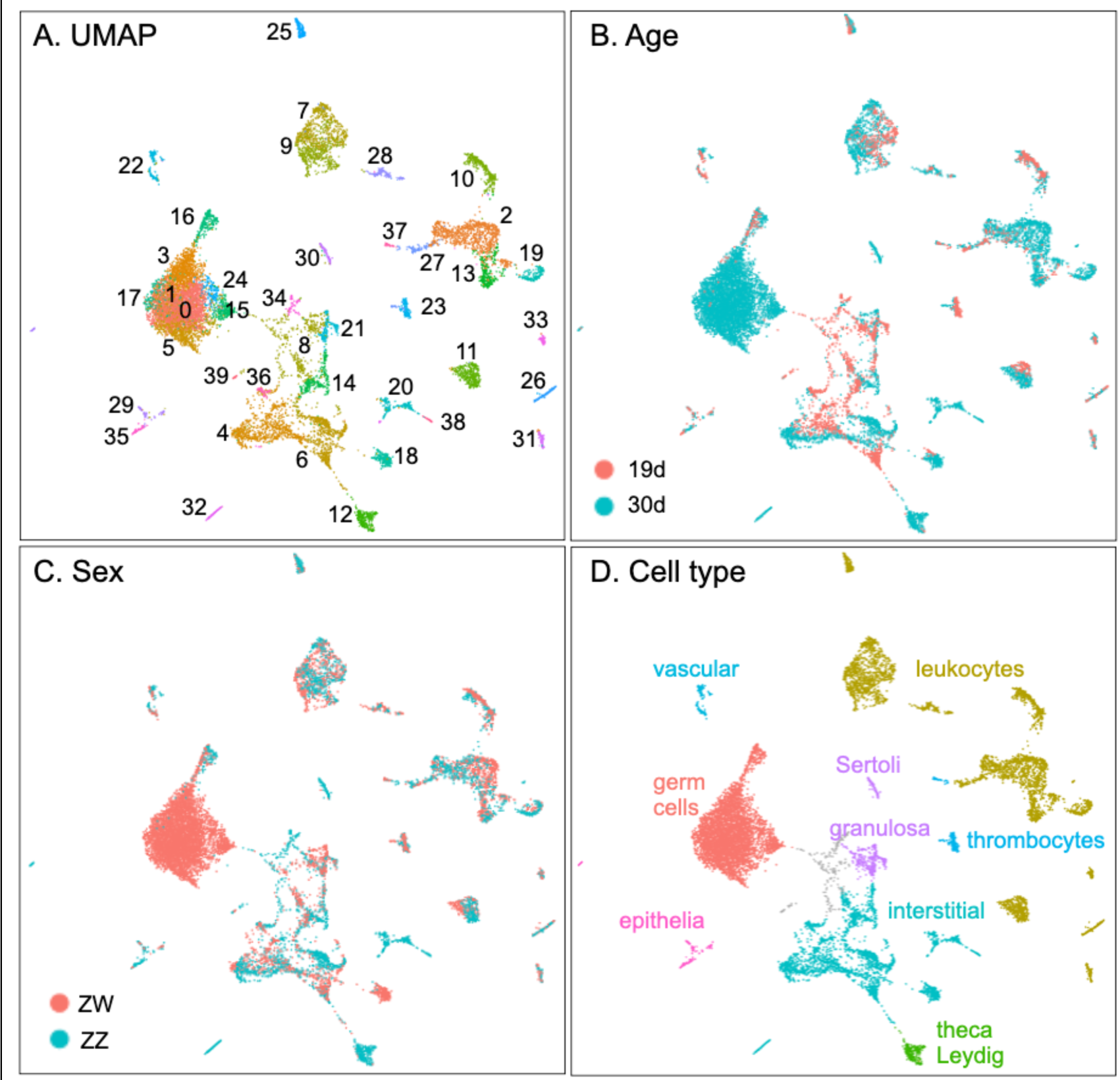
The combined dataset with two ages, each with two sex genotypes. A. UMAP displaying clusters. B. Samples displayed by age (19dpf, red; 30dpf, blue). C. Samples displayed by sex genotype (ZW, red; ZZ, blue). D. Inferred cel types.

### Germ cells

<u>Germ cell marker genes</u>, including *dazl*, *ddx4,* and *dnd1* (Olsen et al., 1997, Yoon et al., 1997, Maegawa et al., 1999, Howley and Ho, 2000, Tan et al., 2002, Ciruna et al., 2002, Weidinger et al., 2003, Krovel and Olsen, 2004, Slanchev et al., 2005, Houwing et al., 2008, Saito et al., 2011, Hartung et al., 2014, Hong et al., 2016, Bertho et al., 2021) were displayed in the merged 19dpf+30dpf (“1930”) data as expressed in several clusters (1930c0, c1, c3, c5, c15, c16, c17, c24) (Fig. 6D-J). Cluster 1930c16 contained 19dpf cells mixed with 30dpf cells, but the large group of *dnd1-*expressing clusters contained almost exclusively 30dpf ZW cells (Fig. 4B). We conclude that the large, several-cluster group of 30dpf ZW cells are developing oocytes and cluster 1930c16 cells are less mature germ cells that are generally similar in ZW and ZZ gonads.

**Figure 6.**
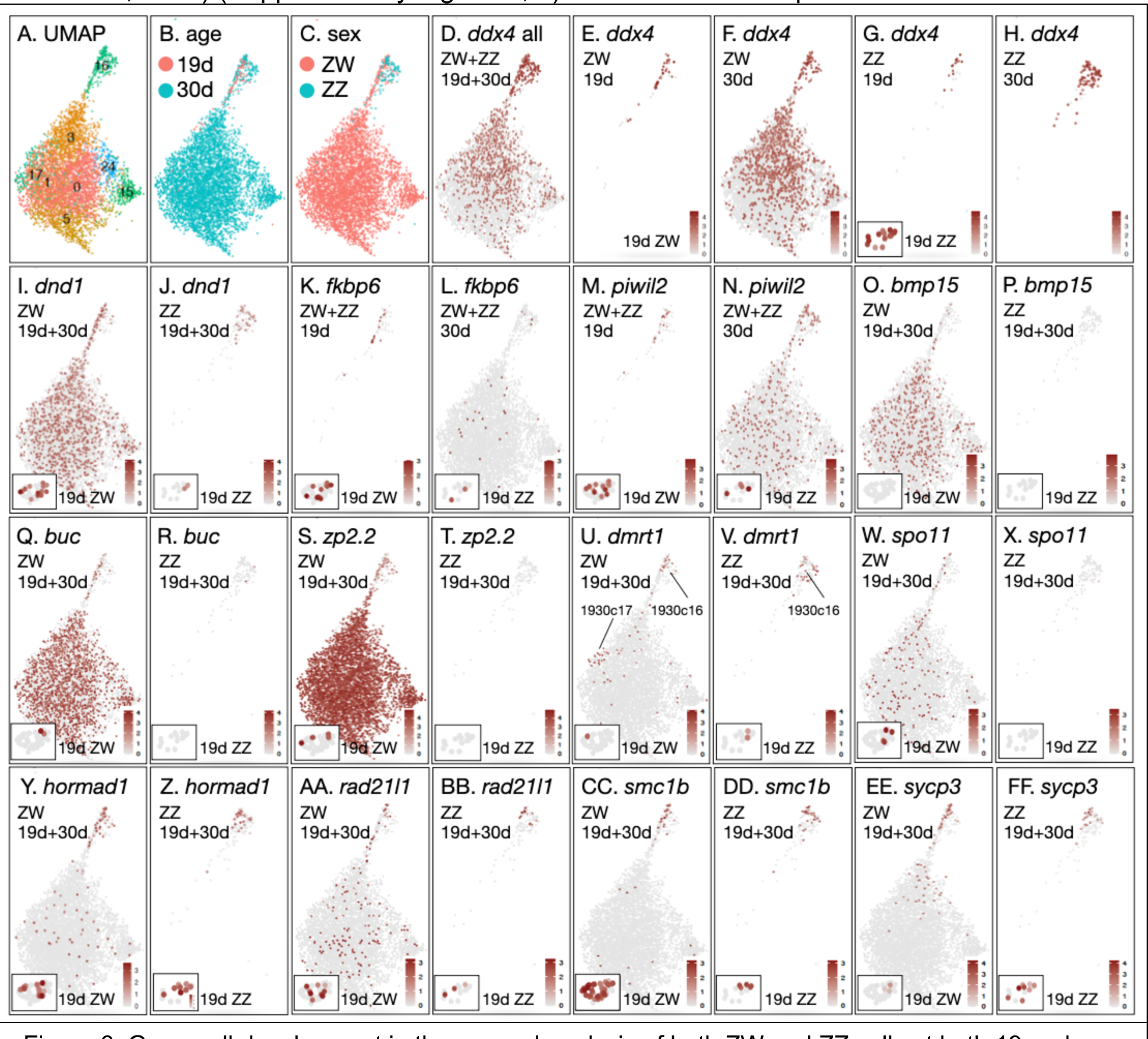
Germ cell development in the merged analysis of both ZW and ZZ cells at both 19 and 30dpf. A. UMAP isolating germ cell clusters. B. Germ cells marked by age (red, 19dpf; blue, 30dpf). C. Germ cells marked by sex genotype (red, ZW; blue, ZZ). D. *ddx4*(*vasa*) including all four conditions. E. *ddx4* showing only ZW cells at 19dpf. Insert in lower left of the panel shows *ddx4* expression in the ZW germ cell cluster 19c17 in the analysis of 19dpf cells only (see Fig. 3). F. *ddx4,* only ZW cells at 30dpf. G. *ddx4,* only ZZ cells at 19dpf. Insert in lower left of the panel shows *ddx4* expression in the ZZ germ cell cluster 19c17 from the analysis of 19dpf cells only. H. *ddx4,* only ZZ cells at 30dpf. I. *dnd1* ZW cells only at both 19dpf and 30dpf. J. *dnd1* ZZ cells only at both 19dpf and 30dpf. K. *fkbp6,* both ZW and ZZ cells only at 19dpf. L. *fkbp6,* both ZW and ZZ cells only at 30dpf. M. *piwil2,* ZW and ZZ cells at 19dpf. N. *piwil2,* ZW and ZZ cells at 30dpf. O. *bmp15,* ZW cells at both 19dpf and 30dpf. P. *bmp15,* ZZ cells at both 19dpf and 30dpf. Q. *buc,* ZW cells at both 19dpf and 30dpf. R. *buc,* ZZ cells at both 19dpf and 30dpf. S. *zp2.2,* ZW cells at both 19dpf and 30dpf. T *zp2.2,* ZZ cells at both 19dpf and 30dpf. U. *dmrt1,* ZW cells at both 19dpf and 30dpf. V. *dmrt1,* ZZ cells at both 19dpf and 30dpf. W. *spo11,* ZW cells at both 19dpf and 30dpf. X. *spo11,* ZZ cells at both 19dpf and 30dpf. Y. *hormad1,* ZW cells at both 19dpf and 30dpf. Z. *hormad1,* ZZ cells at both 19dpf and 30dpf. AA. r*ad21l1,* ZW cells at both 19dpf and 30dpf. BB. r*ad21l1,* ZZ cells at both 19dpf and 30dpf. CC. *smc1b,* ZW cells at both 19dpf and 30dpf. DD. s*mc1b,* ZZ cells at both 19dpf and 30dpf. EE. s*ycp3,* ZW cells at both 19dpf and 30dpf. FF. *sycp3* ZZ, cells at both 19dpf and 30dpf.

The Vasa-encoding gene *ddx4* was strongly expressed in young germ cells of 1930c16 and less strongly in oocytes (Fig. 6D), reflecting the observation that *ddx4* expression is higher in early stages of gametogenesis (Wang et al., 2007). Figures 6E-H show each of the four conditions separately in the combined 19+30dpf analysis. Results for germ cells (19c17 in the 19dpf-only analysis) are shown in the box at the lower left of the 19dpf panels in Figure 6. The distribution of *ddx4-*expressing ZZ cells at 19dpf and 30dpf (Fig. 6G, H) suggest that the transcriptomes of ZZ germ cells changed little between 19dpf and 30dpf, reflecting the delayed differentiation observed in ZZ gonads in our histological studies. This result also shows that ZZ germ cells did not develop an oocyte-like transcriptome, again consistent with the histology.

Expression of *dnd1* supports these conclusions (Fig. 6I, J,).

<u>Oocyte stages</u> recovered are limited by the scRNA-seq protocol, which passes cells through a 40µm filter. According to recent staging criteria (Bogoch et al., 2022), we would recover only oocytes from the earliest two stages, the symmetry-breaking stage (8–20 μm Selman Stage IA, oogonia and meiotic oocytes from leptotene to pachytene), and early Selman Stage IB, nuclear cleft, 15–50μm, oocytes from pachytene to mid-diplotene (Selman et al., 1993, Bogoch et al., 2022)). Comparison of gene expression patterns with other transcriptomic analyses of zebrafish oocytes (Wong et al., 2018, Zhu et al., 2018, Can et al., 2020, Bogoch et al., 2022) are difficult because at least some of the oocyte sizes we examined were not analyzed or were mixed with larger oocytes, and in other studies, oocyte sizes were listed only in relative terms, so comparison wasn’t possible (Cabrera-Quio et al., 2021).

<u>Cluster 1930c16</u> appears to consist of bipotential germ cells. Few genes were expressed at a higher level in 1930c16 at 19dpf than in 30dpf germ cells, but *fkbp6* was an exception (Fig. 6K, L, samples separated by age, not by sex genotype). Men homozygous for pathogenic variants of *FKBP6* show arrested spermatogenesis at the round-spermatid stage and have abnormal piRNA biogenesis like mouse *Fkbp6* mutants (Xiol et al., 2012, Wyrwoll et al., 2022). piRNAs interact with Piwi proteins, including Piwil2 in zebrafish, which aids in transposon defense (Houwing et al., 2008). Like *fkbp6, piwil2* transcripts accumulated in 1930c16 germ cells at 19dpf and 30dpf (Fig. 6M, N). The expression of *fkbp6* and *piwil2* in 19dpf germ cells shows that these cells represent an early step in the pathway of gametogenesis.

<u>Oocyte-specific genes</u> in NA gonads tended to label strongly the *dnd1*-expressing germ-cell clusters except for 1930c16. Accumulation of *bmp15* and *gdf9* transcripts was detected only in 30dpf ZW germ cells, but not in 19dpf ZW germ cells or ZZ germ cells (Fig. 6O, P, Supplementary Fig. S1A, B). The loss of either Bmp15 or Gdf9 function blocks ovarian follicle development in zebrafish and mouse (Dong et al., 1996, Su et al., 2004, Dranow et al., 2016,

Zhai et al., 2022, Chen et al., 2022). The 30dpf ZW cells also accumulated transcripts encoding the Balbiani body protein Bucky ball (*buc*), an early marker of oocyte asymmetry (Marlow and Mullins, 2008, Jamieson-Lucy et al., 2022), but few ZZ cells did, and only at a low level (Fig. 6Q, R). The low or undetected expression of *bmp15* and *gdf9* expression in 1930c16 suggests that these cells were at an earlier stage of differentiation than the large group of germ cell clusters (1930c0, c1, c3, c5, c15, c17, c24). Cells in the large group of 30dpf ZW germ cell clusters also expressed genes encoding zona pellucida egg coat proteins like Zp2.2 (Fig. 6S, T). At 19dpf, the germ cell cluster 19c17 had at least one ZW germ cell expressing a zona pellucida gene (*zp2.1, zp2.2, zp2.3, zp3, zp3.2, zp3a.1, zp3a.2, zp3b, zp3d.1, zp3d.2, zp3f.1*(*si:ch211- 14a17.7*), *zpcx*) but no *ZZ* germ cells expressed any *zp* gene at 19dpf; although this result was not statistically significant for any single gene (Supplemental Table 3), taken as a group, this finding would be expected if ZW germ cells were already beginning to differentiate as oocytes even before differences were apparent histologically. Other oocyte-specific genes like *zar1* (Miao et al., 2017) (Supplementary Fig. S1K, L) followed the same pattern.

Zebrafish orthologs of presumed fish vitellogenin receptors *vldlr* and *lrp13* (Hiramatsu N et al., 2013, Reading et al., 2014, Mushirobira et al., 2015, Morini et al., 2020) were expressed only in 30dpf ZW oocytes (Supplementary Fig. S1M-P). Vldlr may be associated with formation of yolk oil droplets and Lrp13 with vitellogenin uptake (Hiramatsu N et al., 2013). Other suggested vitellogenin receptor candidates, like *lrp1ab, lrp2a, lrp5* and *lrp6,* were not expressed in our dataset (Zhai et al., 2022).

<u>Dmrt1</u> is essential for normal testis differentiation in mammals and is expressed in both germ cells and Sertoli cells (Raymond et al., 1999, Raymond et al., 2000, Kim et al., 2007, Matson et al., 2010, Herpin et al., 2010, Kopp, 2012). In addition, variants of *dmrt1* are major sex determining genes in some amphibia, perhaps some snakes, in birds, and some fish (Nanda et al., 2002, Kobayashi et al., 2004, Smith et al., 2009, Janes et al., 2014, Cui et al., 2017, Mustapha et al., 2018, Ioannidis et al., 2021, Ogita et al., 2020). In NA gonads at 19dpf, *dmrt1* was expressed at low levels in both ZW and ZZ germ cells in 1930c16 (Fig. 6U, V). At 30dpf, *dmrt1* expression continued in 1930c16 germ cells in both ZW and ZZ genotypes, and in addition, appeared in 30dpf ZW germ cells in 1930c17 (Fig. 6U, V). Because *in situ* hybridization in adult TU laboratory strain zebrafish showed that *dmrt1* is expressed primarily in Stage IB oocytes (Webster et al., 2017), we conclude that that in our NA fish, the *dmrt1-* expressing ZW germ cells in 1930c17 represent stage 1B oocytes.

These results are consistent with the conclusion that 1) cluster 1930c16 represents largely undifferentiated germ cells that our histology studies (Fig. 2) showed were present in 19dpf ZW and ZZ gonads; 2) that some of the 19dpf ZW germ cells in 1930c16 appear to have had already started expressing weakly some oocyte genes; 3) that ZZ germ cells in 1930c16 remained largely undifferentiated at both 19dpf and 30dpf; 4) that most of the large group of 30dpf ZW-specific germ cell clusters represent Stage IA oocytes expressing strongly oocyte-specific genes; and 5) that 1930c17 represents early *dmrt1-*expressing Stage IB oocytes.

<u>Meiosis gene functions</u> are important for sex determination, as shown by mutations in meiosis genes that produce mostly neomale offspring (Rodriguez-Mari et al., 2010, Shive et al., 2010, Rodriguez-Mari et al., 2011, Saito et al., 2011, Ramanagoudr-Bhojappa et al., 2018, Takemoto et al., 2020, Blokhina et al., 2021, Islam et al., 2021). Meiotic recombination begins with the introduction of double-strand DNA breaks catalyzed by Spo11, Hormad1, and CCDC36 (IHO1) (Keeney, 2008, Shin et al., 2013, Stanzione et al., 2016). In zebrafish, *spo11* is necessary for homolog synapsis, for sperm production, and for preventing females from having abnormal offspring (Blokhina et al., 2019). In our samples, *spo11* expression appeared in ZW germ cells, but not in any ZZ cells, at both 19dpf and 30dpf (Fig. 6W, X), and *hormad1* was expressed in 1930c16 germ cells at both ages and both sex genotypes (Fig. 6Y, Z). Repair of Spo11-induced double strand breaks occurs by the DNA recombinases Dmc1 and Rad51 (Hunter, 2015) and Fanconi Anemia genes, including *fancl* and *brca2* (Grompe and D’Andrea, 2001). This process involves sister chromatid cohesion, achieved by cohesin components, including *rad21l1* in zebrafish, which is required for oogenesis but not spermatogenesis (Blokhina et al., 2021) and in our samples, was expressed at both ages in both sexes, in 1930c16 (Fig. 6AA, BB). Other meiosis genes, including *smc1b* (Fig. 6CC, DD, Supplementary Fig. S1C-F), which is essential for homolog pairing and synapsis (Islam et al., 2021), *rec8a*, and *rec8b,* and genes encoding synaptonemal complex proteins Sycp1, Sycp2, and Sycp3 (Fig. 6EE, FF, Supplementary Fig. S1G,H, Supp Table 7) are stronger markers for young germ cells in cluster 1930c16 than for oocytes in clusters 1930c0, c1, c3, c5, c15, c17, c24. The pattern for meiosis genes in general was opposite from the pattern for oocyte-specific genes, like *bmp15, buc,* and *zp2.2,* which increased from 1930c16 to the large group of oocyte clusters. How zebrafish regulate the switch from transcribing meiosis genes to maturing oocyte genes is unknown. The expression of meiosis genes in both ZW and ZZ germ cells (except for *spo11*) deepens the mystery of the mechanism in zebrafish that causes many meiotic gene mutants to tend to develop as males (*fancl, brca2, rad21l1, smc1b, sycp1, sycp2* (Rodriguez-Mari et al., 2010, Shive et al., 2010, Rodriguez-Mari et al., 2011, Saito et al., 2011, Ramanagoudr-Bhojappa et al., 2018, Takemoto et al., 2020, Blokhina et al., 2021, Islam et al., 2021), with *mlh1* and *spo11* being the exceptions (Feitsma et al., 2007, Leal et al., 2008, Blokhina et al., 2019). A hypothesis is that zebrafish females lack a synapsis checkpoint (Imai et al., 2021).

<u>Histone</u> H2ax phosphorylation results in foci at the sites of DNA breaks in meiosis (Hamer et al., 2003, Blokhina et al., 2021, Islam et al., 2021, Imai et al., 2021). Two H2ax histones had a reciprocal pattern of expression in NA gonads: *h2ax1*(*h2afx1*) was expressed strongly in nearly all somatic cells in all four samples and in both ZW and ZZ germ cells in 1930c16 but in few oocytes as defined by *buc* expression (Supplementary Fig. S1I, J). Reciprocally, *h2ax*(*h2afx*) was expressed strongly in 30dpf ZW oocytes but not in somatic cells and weakly in ZZ germ cells (Supplementary Fig. S1K, L). These results suggest that during oogenesis, histone H2ax replaces the somatic and early germ cell histone H2ax1 in zebrafish oogenesis.

#### Support cells

Support cells -- granulosa cells in the ovary and Sertoli cells in the testis -- control germ cell proliferation, differentiation, and maturation. Support cells in fish express *gsdf* in both ovaries and testes and *gsdf* is required for oocyte maturation (Sawatari et al., 2007, Gautier et al., 2011, Imai et al., 2015, Zhang et al., 2016, Yan et al., 2017, Jiang et al., 2022). Furthermore, a variant of *gsdf* is the major male sex determining gene in Luzon medaka and sablefish (Myosho et al., 2012, Herpin et al., 2021). In the combined analysis, *gsdf* was expressed strongly in just three cell clusters: 1930c30, c21, and c8, identifying support cells or their precursors (Fig. 7A, B). In 19dpf gonads, *gsdf* was expressed in 1930c8 and 1930c21 in both ZW and ZZ cells (Fig. 7A-C, E) suggesting that these are support cell precursors and, that ZW and ZZ pre-support cells at 19dpf are transcriptionally similar (Fig. 7C, E).

**Figure 7.**
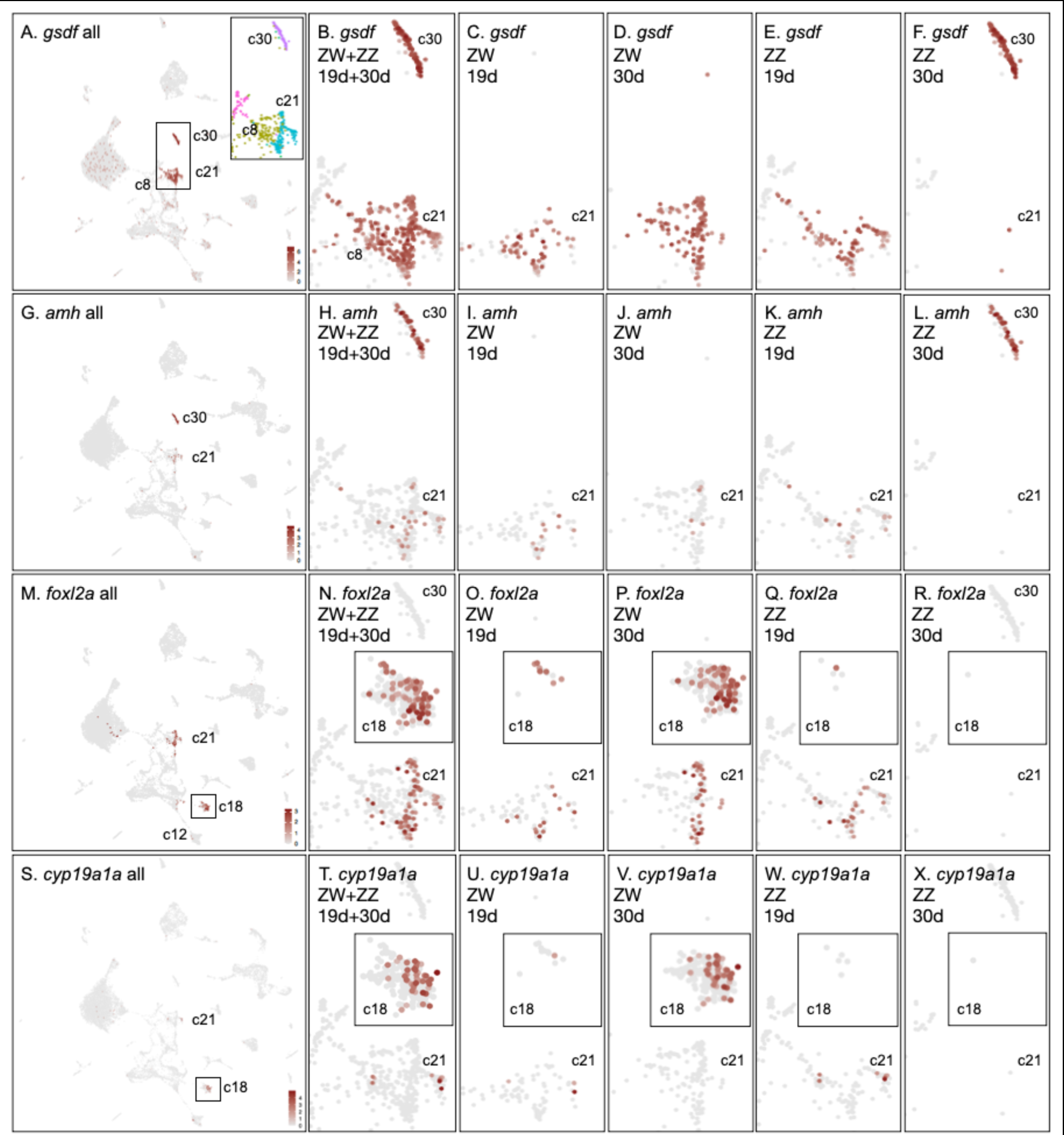
Support cell gene expression merging both time points and both sex genotypes. A. *gsdf* expression across the whole combined dataset. The rectangle marks *gsdf*-expressing support cells and their precursors, expanded to show cluster assignments in color. B. Enlargement of rectangle from part A, showing *gsdf* expression in clusters 1930c30, c21, and c8 combining all four conditions. C. *gsdf* expression in 19dpf ZW gonads. D. *gsdf* expression in 30dpf ZW cells. E. *gsdf* expression in 19dpf ZZ cells. F. *gsdf* expression in 30dpf ZZ cells. G-L. *amh* expression in samples as described for panels B-F. M-R. *foxl2a* expression. Box in M around 1930c18 is expanded as inserts in panels N-R. S-X. *cyp19a1a* expression. Boxed insert in panel S around 1930c18 appears as inserts in panels T-X.

At 30dpf, *gsdf* expression differed substantially in ZW and ZZ gonads. Cluster 1930c30 contained almost exclusively 30dpf ZZ cells (Fig. 7F) representing Sertoli cells. In contrast, ZW cells at 30dpf occupied 1930c21 and 1930c8 (Fig. 7D), suggesting that these clusters contain granulosa cells in addition to support cell precursors. Both *inha,* which is necessary for final oocyte maturation and ovulation (Lu et al., 2020), and *igf3*, which is essential for zebrafish ovarian follicles to develop beyond the primary growth stage (Xie et al., 2021), had virtually the same expression pattern as *gsdf* (Supplemental Fig. S2A-F). These results suggest: 1) *gsdf-* expressing 30dpf ZW cells in 1930c21 represent granulosa cells; 2) *gsdf-*expressing 30dpf-ZZ cells in 1930c30 represent Sertoli cells; 3) some *gsdf*-expressing cells of both sex genotypes at 19dpf in 1930c8 and 1930c21 likely represent bipotential precursors of support cells and at 30dpf, the ZW cells represent granulosa cell precursors; 4) substantial differentiation of sex-specific cell types occurs between 19dpf and 30dpf.

<u>Sertoli cells</u> in fish strongly express *amh, dmrt1*, and *sox9a* (Chiang et al., 2001a, Kobayashi et al., 2004, Rodriguez-Mari et al., 2005, Yan et al., 2005, Guo et al., 2005, Webster et al., 2017, Lin et al., 2017). AntiMüllerian Hormone (Amh), secreted by Sertoli cells in mammalian testes, destroys the developing female reproductive tract, and variants of *amh* or its receptor gene *amhr2* are male sex determinants in several species of fish (Morinaga et al., 2007, Kamiya et al., 2012, Hattori et al., 2012, Li et al., 2015, Ieda et al., 2018, Pan et al., 2021). At 19dpf, a few pre-support cells of both ZW and ZZ genotypes weakly expressed *amh* (Fig. 7G-I, K) and *dmrt1* (Supplemental Fig. S2G-I, K. Recall, *dmrt1* was also expressed in germ cells, Fig. 6U, V). At 30dpf, *amh* and *dmrt1* were still weakly expressed in just a few ZW cells but were strongly expressed in ZZ cells in 1930c30 (Fig. 7J, L and Supplemental Fig. S2J, L). Mammalian Sertoli cells express S*ox9,* which maintains testis development (Morais da Silva et al., 1996, Qin and Bishop, 2005, Barrionuevo et al., 2006). In our samples, *sox9a* was expressed strongly in ZZ Sertoli cells (1930c30) and was detected in some 1930c21 cells in ZW gonads and 19dpf ZZ gonads (Supplemental Fig. S2M-R). These results support the assignment of the 30dpf ZZ- specific cluster 1930c30 as Sertoli cells and 19dpf ZZ cells in 1930c21 as support cell precursors.

<u>Ovarian follicle cells</u> in mammals are multi-layered, with inner granulosa cells that form gap junctions with the oocyte that transport substances, and an outer granulosa cells that produce estradiol (Gilchrist et al., 2008). In teleosts, however, ovarian follicle cells form a single-cell epithelium over the oocyte, but it is unclear whether this single layer provides both the transport function through gap junctions and the endocrine function or if that single cell layer consists of two cell types, one that transports substances and one that provides estradiol (Devlin RH and Y., 2002). The mammalian granulosa gene *Foxl2* has two duplicates in zebrafish that arose in the teleost genome duplication (*foxl2a, foxl2b*) plus a third paralog, lost in mammals, that arose in a genome duplication event before the divergence of teleosts and mammals (*foxl2l,* alias *zgc:194189* or *foxl3*) (Crespo et al., 2013, Caulier et al., 2015, Bertho et al., 2016, Webster et al., 2017, Yang et al., 2017, Dai et al., 2021, Liu et al., 2022). In our analysis combining both ages and both sex genotypes, *foxl2a* was expressed in two clusters, in the support cell precursors in 1930c21 similar to the pattern of *gsdf* and *amh,* and additionally, in 1930c18 (Fig. 7M). At 19dpf in 1930c21, expression of *foxl2a* was similar to that of *gsdf*, but weaker, and appeared in both ZW and ZZ cells (Fig. 7O, Q), consistent with these cells being bipotential support cell precursors. At 30dpf, *foxl2a* was also expressed in ZW 1930c21, again consistent with these cells representing granulosa cells. A second major *foxl2a* expression domain appeared in cluster 1930c18 (Fig. 7M, box; N-R, insert). Cluster 1930c18 contained few 19dpf cells and only one 30dpf ZZ cell (Fig. 7N-R, insert), showing that it was a cell type that began to develop before 19d in both ZW and ZZ cells, but by 30dpf, was ZW-specific.

Expression of *foxl2b* was like that of *foxl2a* (Supplemental Fig. S2S-X). Expression of *foxl2l*, which is missing from mammals, showed scant expression limited to Stage IA and small Stage IB oocytes; studies show that oocytes at a more mature stage express *foxl2l* strongly (Kikuchi et al., 2020, Liu et al., 2022).

The *foxl2a-* and *foxl2b-*expressing cells in 1930c21 may be performing the transport function of mammalian granulosa cells because the gap junction connexin gene *gja11*(*cx34.5*) was specifically expressed in 1930c21 in a pattern like that of *gsdf* and *foxl2a* at 19dpf in presumed support cell precursors and at 30dpf in granulosa cells in ZW gonads and in Sertoli cells in ZZ gonads (Supplementary Fig. S2Y-DD). *In situ* hybridization had previously detected *gja11* expression in follicle cells surrounding Stage II oocytes (cortical alveolus stage, 140–340 μm (Bogoch et al., 2022), which were not present in our samples), but not in Stage IB oocytes (Liu et al., 2022); our data detected *gja11* expression at earlier stages presumably due to increased sensitivity. The oocyte partner of Gja11 could be encoded by *gjc4a.1*(*cx44.2*) or *gjc4b*(*cx43.4*), which were expressed strongly and rather specifically in 30dpf oocytes (Santos et al., 2007) (Supplementary Fig. S2KK, LL).

The estrogen-producing function of mammalian granulosa cells might be mediated by a cell type different from the 1930c21 cells that may perform the transport function. Gonadal aromatase, encoded by *cyp19a1a*, converts testosterone to estradiol or androstenedione to estrone (Chiang et al., 2001b, Chiang et al., 2001c, Tenugu et al., 2021). At 19dpf, a few *cyp19a1a*-expressing cells appeared with 1930c21 support cell precursors in both ZW and ZZ genotypes (Fig. 7S, U, W,), while at 30dpf, *cyp19a1a* was expressed in ZW cells in 1930c18 but not in ZZ cells (Fig. 7V, X). The strong expression of *gsdf, foxl2a,* and *gja11* in 1930c21 and *foxl2a* and *cyp19a1a* in 1930c18 suggests that the transport function and estrogen-production function of mammalian granulosa cells may be carried out by two different follicle cell types in zebrafish 30dpf ZW ovaries.

We conclude that expression patterns of *gsdf, foxl2a* and *foxl2b,* and *cyp19a1a* suggest that: 1) at 19dpf, support cell precursors were similar in ZW and ZZ gonads; 2) that 30dpf ZW but not ZZ gonads maintained these common support cell precursors; 3) that granulosa cells appeared in the UMAP near support cell precursors but Sertoli cells were distant; and 4) that the aromatase function of 30dpf ovarian follicle cells was provided by a ZW cell type different from the strongly *gsdf*-expressing 30dpf ZW follicle cell type.

#### Steroidogenic cells

Sex steroids are important for zebrafish gonad development (Menuet et al., 2004, de Waal et al., 2009, Caulier et al., 2015, Luzio et al., 2016, Lu et al., 2017, Zhai et al., 2018), but we know little about how steroidogenic cell development in ZW vs. ZZ zebrafish.

<u>Nr5a1</u> and Star initiate gonadal steroidogenesis (Fig. 8A, after (Tenugu et al., 2021)). Nr5a1 (Steroidogenic Factor-1, SF1) is a major regulator of genes that encode gonadal steroidogenesis enzymes in theca and Leydig cells and it is one of the earliest markers of gonadal differentiation, turning on in human gonads before *SRY* (Morohashi et al., 1995, Ikeda et al., 1996, Hanley et al., 1999). Zebrafish has two co-orthologs of *Nr5a1* (*nr5a1a* and *nr5a1b*) (von Hofsten et al., 2001, Kuo et al., 2005) from the teleost genome duplication (Taylor et al., 2003, Jaillon et al., 2004, Postlethwait et al., 1999, Amores et al., 1998). Loss-of-function mutations in *nr5a1a* disrupt development strongly in the interrenal and mutations in *nr5a1b* cause more severe phenotypes in the gonad (Yan et al., 2020). At 19dpf, *nr5a1b* was expressed in a few support precursor cells (1930c21) and a few 1930c6 cells, but at 30dpf, *nr5a1b* was strongly expressed in 1930c12, but only weakly in granulosa or Sertoli cells (Fig. 8B). We conclude that in 1930c12, ZW cells are theca cells and the ZZ cells are Leydig cells. Leydig cells in 1930c12 also specifically expressed *insl3*, *skor2*(*zgc:153395*), and *pcdh8* (Supplementary Table S7).

**Figure 8.**
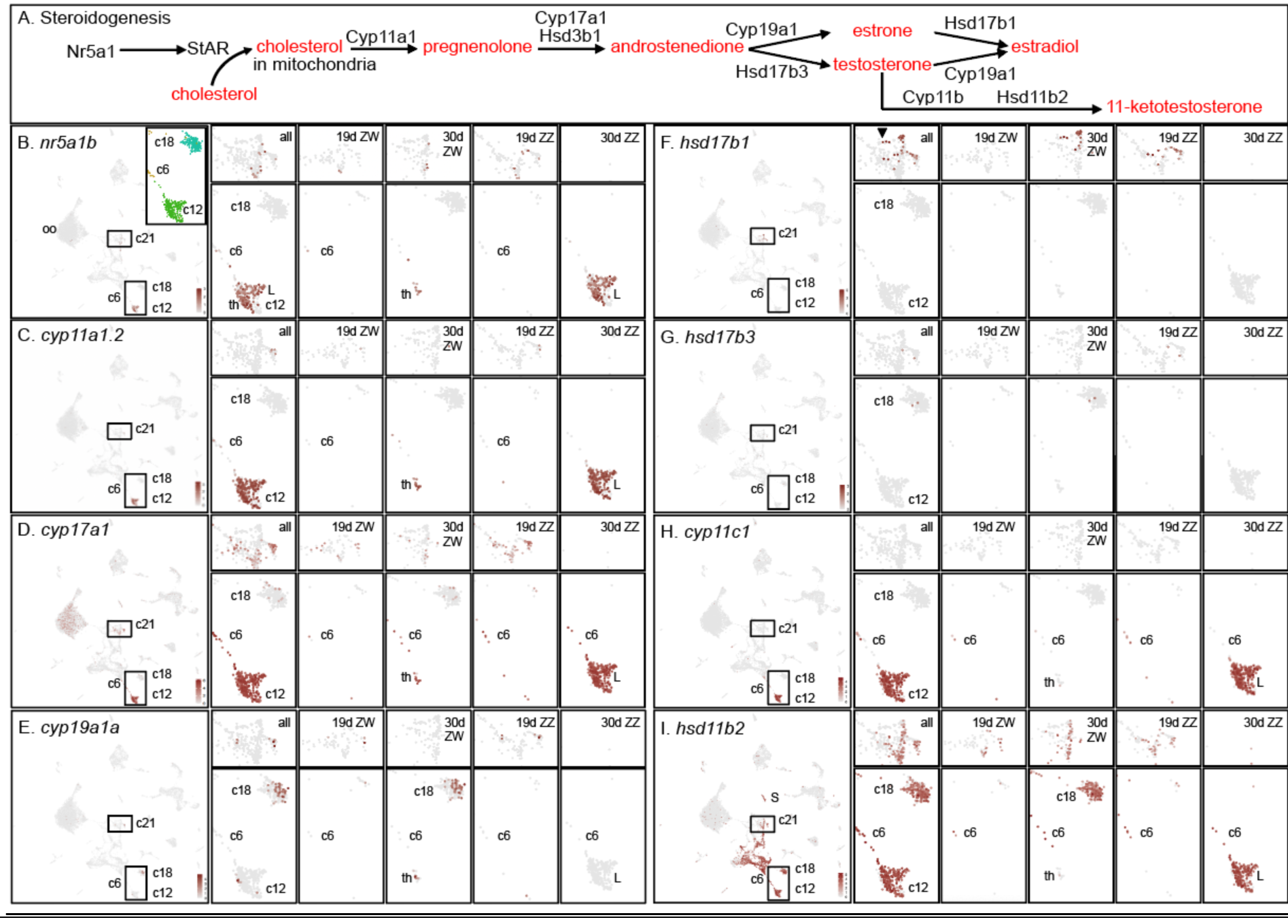
Steroidogenesis. A. An abbreviated pathway of steroidogenesis (after (Tenugu et al., 2021)). B. *nr5a1b* expression. The left panel shows expression in the entire dataset. The insert shows cluster demarcation for the most relevant steroidogenic enzymes. Subsequent panels show, at the top 1930c21, the support cell precursor and granulosa cell cluster and at the bottom the clusters corresponding to the insert in the left panel. Displayed from left to right are the enlargements of all four conditions: 19dpf ZW gonads, 30dpf ZW gonads, 19dpf ZZ gonads, and 30dpf ZZ gonads. Cluster numbers are indicated. C. cyp11a1.2 displayed as explained for *nr5a1b*. D. *cyp17a1*. E. *cyp19a1a*. F. *hsd17b1*. G. *hsd17b3*. H. *cyp11c1*. I. *hsd11b2*. Abbreviations: L, Leydig cells; th, theca cells.

<u>StAR</u> (steroidogenic acute regulatory protein) gene transcription is turned on by Nr5a1 (Sugawara et al., 1997). In an initial step (Fig. 8A), StAR brings cholesterol into the inner mitochondrial membrane (Miller and Auchus, 2011). Zebrafish adult gonads express *star* (Bauer et al., 2000, Ings and Van Der Kraak, 2006, de Waal et al., 2009), and although NA zebrafish at 19dpf had few *star-*expressing cells, at 30dpf, *star* was expressed strongly in theca and Leydig cells (1930c12) and in a few oocytes (Supplemental Fig. S3B).

<u>Cyp11a1</u> in mammals converts cholesterol to pregnenolone (Lai et al., 1998). Zebrafish has tandem duplicates of human *CYP11A1* (*cyp11a1.1* and *cyp11a1.2*, previously called *cyp11a1* and *cyp11a2*, respectively) (Hu et al., 2001, Goldstone et al., 2010, Parajes et al., 2013); originating in a duplication event that occurred after the divergence of zebrafish and carp lineages (gene tree ENSGT00940000158575). Disruption of *cyp11a1.2* results in androgen loss, and homozygous mutants develop into feminized males with disorganized testes and few spermatozoa (Parajes et al., 2013, Li et al., 2020a, Wang et al., 2022). At 19dpf, *cyp11a1.2* showed scant expression, but at 30dpf, it was widely expressed in ZW theca and ZZ Leydig cells and in 1930c6 (Fig. 8C). In contrast, expression of the tandem duplicate *cyp11a1.1* was not detected at 19dpf (Supplemental Fig. S3C) and at 30dpf, appeared in many ZW oocytes, as expected from the its known maternal expression, but not in ZZ germ cells (Hsu et al., 2002, Parajes et al., 2013, Wang et al., 2022). The *cyp11a1.1* gene was expressed weakly in ZZ Leydig cells but in no theca cells, and weakly in a few 1930c18 cells (Supplemental Fig. S2C). We conclude that the expression of *cyp11a1.1* and *cyp11a1.2* diverged after the tandem duplication event and that *cyp11a1.2* maintained the ancestral pattern in theca and Leydig cells but *cyp11a1.1* evolved stronger expression in the oocyte germ cell lineage, which doesn’t appear to be an ancestral expression domain because medaka, which lacks the tandem duplication, does not express *cyp11a1* in oocytes (Nakamoto et al., 2010). The regulatory mechanism causing different expression patterns for these two tightly linked tandemly duplicated genes is unknown.

<u>Cyp17a1 and Hsd3b</u> act on the product of Cyp11a1, pregnenolone, and convert it to androstenedione (Fig. 8A). Knockout of *cyp17a1* in AB zebrafish leads to loss of juvenile ovaries and an all-male population with reduced male secondary sex characteristics (Zhai et al., 2018). At 19dpf, *cyp17a1* expression was detected in many support cell precursors and in 1930c6, which may be theca/Leydig cell precursors (Fig. 8D). At 30dpf, *cyp17a1* was expressed weakly in granulosa and Sertoli cells, but at a high level in ZW theca and ZZ Leydig cells, and in many oocytes (Fig. 8D). Expression of *hsd3b1* was like that of *cyp17a1,* but at lower levels and in fewer cells (Supplemental Fig. S2D). Expression of *hsd3b2* (Lin et al., 2015) appeared almost exclusively in oocytes, like *cyp11a1.1* (Supplemental Fig. S2E).

<u>Cyp19a1 and Hsd17b</u> enzymes carry out the next step, converting androstenediol to estradiol (Miller and Auchus, 2011). In principle, these reactions can occur in either of two ways (Fig. 8A):1) Cyp19a1 can form estrone from which Hsd17b1 forms estradiol (E2); alternatively, 2) Hsd17b3 can form testosterone, which Cyp19a1 converts to estradiol (Fig. 8A). Cyp19a1 is important for zebrafish sex development because *cyp19a1a* mutants develop mostly as males (Dranow et al., 2016, Lau et al., 2016, Yin et al., 2017, Wu et al., 2020, Romano et al., 2020). In 19dpf NA zebrafish, *cyp19a1a* expression was found in a few support cell precursors (1930c21) in both ZW and ZZ gonads (Fig. 8E). At 30dpf, *cyp19a1a* expression was not detected in any ZZ cells, expected from its role in estrogen production. Transcript for *cyp19a1a* was present in some ZW theca cells (Fig. 8E, 1930c12), but the focus of *cyp19a1a* expression was in ZW 1930c18 cells (Fig. 8E), which also strongly expressed *foxl2a* and *foxl2b* but did not strongly express *gsdf* (Fig. 7A, M-R, Fig. S2S-X). Cluster 1930c18 ZW cells at both 19dpf and 30dpf also expressed specifically the transcription factor *arxa* (Supplemental Fig. S3F), whose mammalian ortholog *Arx* is expressed in, and is necessary for, the maintenance of fetal Leydig progenitors in mouse (Miyabayashi et al., 2013) and the formation of normal genitalia and sex steroid levels in humans (Gupta et al., 2019, Basa et al., 2021). The *cyp19a1a* cluster 1930c18 also expressed strongly and specifically the G protein-coupled receptors *s1pr3a, gpr17*, the junctional adhesion molecule *jam2b*, and the signaling gene *wnt11* (*wnt11r* (Postlethwait et al., 2019)). Expression of steroidogenic enzymes other than *cyp19a1a* in 1930c18 was either not detected or detected weakly, except for *hsd11b2*, which was broadly expressed in nearly all stromal/interstitial cell clusters marked by the collagen gene *col1a1a* (compare Fig. 8I to Fig. 9A). These results are consistent with zebrafish having two types of ovarian follicle cells, one strongly and specifically expressing *cyp19a1a, arxa,* foxl2a, *s1pr3a*, *gpr17,* and *wnt11* (1930c18), and the other expressing strongly *gsdf,* foxl2a, *lamb2l, inha,* and *igf3* (1930c21).

**Figure 9.**
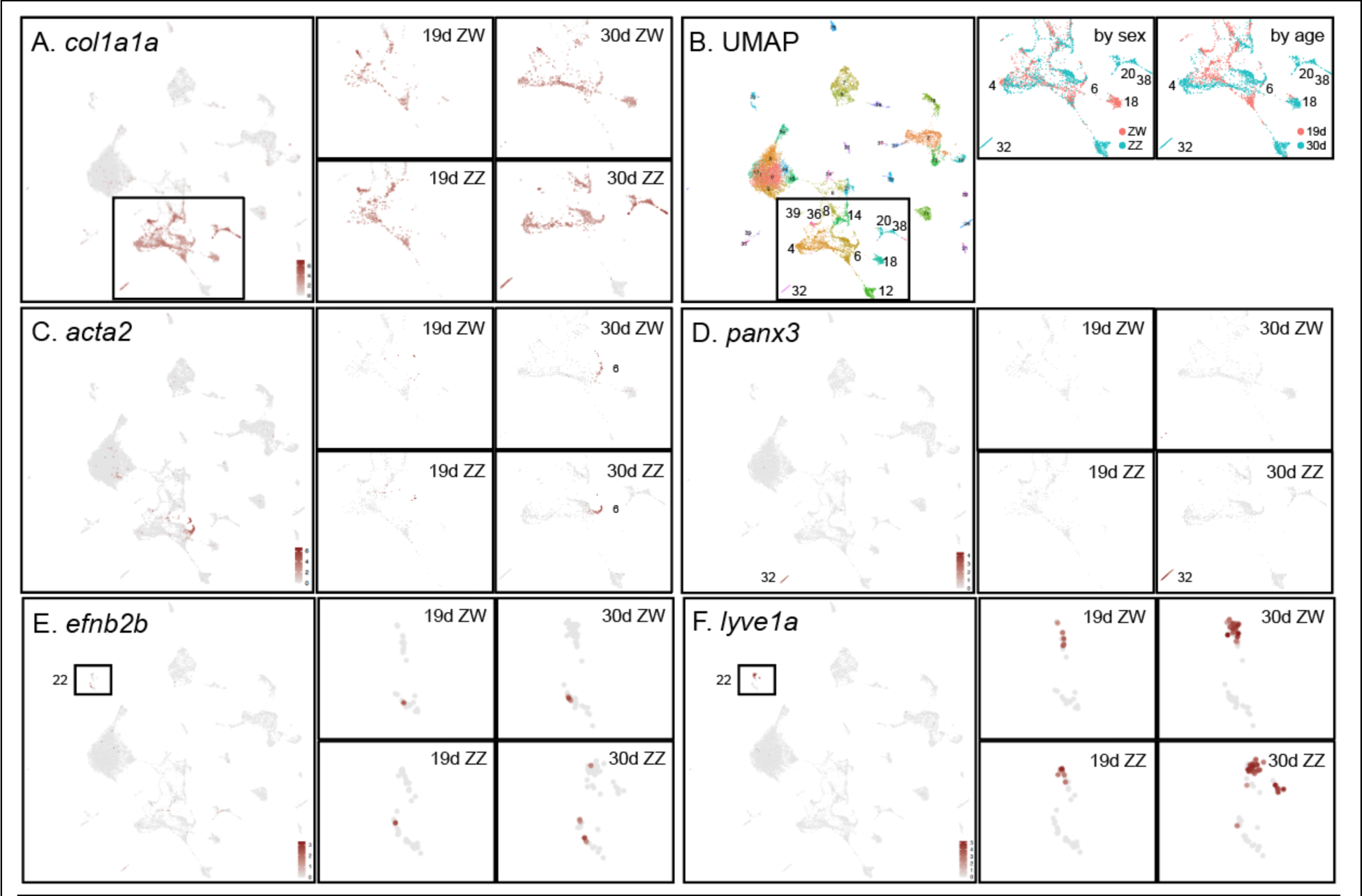
Stromal/interstitial clusters. A. Expression of *col1a1a* defines stromal/interstitial cells (Liu et al., 2022). The left large panel combines cells of all four conditions for the complete dataset and the four smaller panels to the right show an enlargement of the *col1a1a*-expressing domain for each condition independently. At 19dpf, ZW and ZZ expression patterns are similar but at 30dpf, some clusters are sex-genotype specific. B. The UMAP is in the left large panel; the middle panel shows the *col1a1a-*expressing region for samples separated by sex genotype (red, ZW; blue, ZZ); and the right panel shows samples separated by age (red, 19dpf; blue, 30dpf). C. *acta2* expression, with panels as described for panel A. D. *panx3* expression in the 30dpf ZZ-specific cluster 1930c32. E. *efn2b* expression, indicating likely arterial endothelial cells in 1930c22 boxed in the large panel and expanded in the four condition-specific panels to the right. F. *lyve1a* expression, a marker of lymphatics, labeling 1930c22.

Hsd17b, the second estrogen-producing enzyme activity (Fig. 8A), is the master sex determinant in *Seriola* fishes (Purcell et al., 2018, Koyama et al., 2019). Hsd17b activity could be provided in principle by either Hsd17b1 or Hsd17b3 (Mindnich et al., 2004, Mindnich et al., 2005). At 19dpf, *hsd17b1* expression appeared in ZZ pre-support cells but not in ZW cells in our samples (Fig. 8F). By 30dpf, however, *hsd17b1* expression was detected in *gsdf*-expressing ZW granulosa cells in 1930c21 (Fig. 8F) as in adult AB strain female zebrafish (Liu et al., 2022). Expression of *hsd17b1,* was not detected in the *cyp19a1a-*expressing 1930c18 ZW cells or in any 30dpf ZZ cells.

In the alternative pathway to estradiol (Fig. 8A) and in contrast to *hsd17b1, hsd17b3* expression was low at both 19dpf and 30dpf (Fig. 8G). We conclude that, as in adult AB strain zebrafish (Liu et al., 2022), *hsd17b3* appears to play a smaller role than *hsd17b1* in estrogen synthesis in juvenile zebrafish.

Our results suggest that three different transcriptomic clusters are important for estrogen production in 30dpf ZW zebrafish (Supplemental Fig. S3J-L): 1) theca cells (ZW cells in 1930c12) producing androstenedione using Cyp17a1 (Fig. 8D); 2) follicle cells in 1930c18 forming estrone catalyzed by Cyp19a1a (Fig. 8E); and 3) a different type of follicle cell in 1930c21 generating estradiol by Hsd17b1 (Fig. 6F). It is possible that the 1930c18 *cyp19a1a-* expressing cell type and the *hsd17b1-*expressing cell type are simply different states or stages of the same individual cells. Also, it could be that Hsd17b3 might become more important in older animals or in more mature ovarian follicles.

<u>Androgens</u> in fish gonads include testosterone and the principal teleost androgen 11-keto- testosterone (11KT) (Kusakabe et al., 2003, de Waal et al., 2008, Nelson et al., 2013, Tokarz et al., 2015, Tenugu et al., 2021). 11KT is produced by the enzymes Hsd17b3, Cyp11c, and Hsd11b2 (Fig. 8A). In 19dpf NA zebrafish, *cyp11c1* was expressed only in a few 1930c6 cells in both ZW and ZZ gonads (Fig. 8H), consistent with bi-potential gonads. At 30dpf, however, *cyp11c* was expressed strongly and specifically in ZZ Leydig cells but not in ZW theca cells (Fig. 8H). Within 1930c12, ZW cells, but not ZZ cells, expressed the “estrogen genes” *cyp19a1a* (Fig. 7S, Fig. 8E) and *foxl2a* (Fig. 7M); reciprocally, ZZ cells, but not ZW cells, expressed the “androgen gene” *cyp11c1* (Fig. 8H), confirming cell assignments as theca vs. Leydig cells.

Hsd11b2 can follow Cyp11c to produce 11KT (Fig. 8A). The *hsd11b2* gene was expressed more broadly than any of the other steroidogenic enzymes discussed here (Fig. 8I). At 19dpf, both ZW and ZZ gonads expressed *hsd11b2* both in pre-support cells and in 1930c6 cells (Fig. 8I), as well as in several stroma/interstitial cell clusters (see Fig. 9). At 30dpf, *hsd11b2* was expressed in ZW gonads in granulosa cells and *cyp19a1a-*expressing 1930c18 cells, and in ZZ gonads in Sertoli cells and Leydig cells, as well as in stromal/interstitial cells (Fig. 8I). *In vitro* enzyme assays show that zebrafish Hsd17b3 converts androstenedione to testosterone (Mindnich et al., 2005), but we did not specifically detect *hsd17b3* expression in ZZ NA gonads (Fig. 8G). Hsd17b1 is unlikely to perform this function because, although *hsd17b1* was expressed in pre-support cells in 19dpf ZZ animals, expression was not detected at 30dpf in ZZ gonads (Fig. 8F). This result suggests that at 30dpf, neither *hsd17b1* nor *hsd17b3* is likely to make a significant contribution to 11KT formation in ZZ zebrafish, although they might at later developmental stages. A different Hsd17b enzyme might Hsd17b3 activity in zebrafish testes or our samples may represent a stage too early for production of testosterone or 11KT.

<u>Steroid receptor</u> genes were expressed in several steroidogenic cell types. The estrogen receptor gene *esr1* was expressed in only a few cells in NA gonads at either 19 or 30dpf, consistent with a weak effect of *esr1* mutants on ovary development (Lu et al., 2017). At 19dpf in both ZW and ZZ gonads, *esr2a* was expressed in pre-support cells and some interstitial cells and at 30dpf, in ZW granulosa cells, in *cyp19a1a-*expressing 1930c18 cells, in a few oocytes, and in some ZZ Leydig cells, as well as in some interstitial/stromal cells in at both ages and sex genotypes (Supplemental Fig. S3G). The *esr2b* gene was expressed like *esr2a* in 30dpf ZW gonads, but also in 30dpf ZZ Sertoli cells (Supplemental Fig. S3H). In addition, unique among the steroid-related gene set examined, *esr2b* was expressed strongly in19dpf ZW cells in 1930c18 (the *cyp19a1a* cluster) (Fig. 8F). The androgen receptor gene *ar* was expressed with virtually the same pattern as *esr2b* (Supplemental Fig. S3I). Expression of *ar* in ZZ cells is in accordance with the expectation that zebrafish *ar* mutants would mostly develop as females, and *ar* expression in ZW cells could help explain why even mutant females have reduced fertility (Crowder et al., 2018, Yu et al., 2018).

#### Interstitial and stromal cells

<u>Interstitial and stromal cells</u> provide structural support, erect barriers, transmit forces, and supply signals, thus offering functions distinct from other gonadal cell types (Kinnear et al., 2020). Many interstitial and stromal cell types construct substantial intercellular matrix and are marked by expression of the collagen gene *col1a1a* (Liu et al., 2022) (Fig. 9A). Sex- and age-specific clusters were prominent among *col1a1a*-expressing interstitial/stromal cells (Fig. 9B). One conspicuous cluster was the 30dpf ZW-restricted *cyp19a1a-*expressing cluster 1930c18 discussed above, which has strong *col1a1a* expression (Fig. 9A) and low *gsdf* expression (Fig. 7A), further distinguishing 1930c18 from *hsd17b1*-expressing granulosa cells in 1930c21 (Fig. 8F). In contrast, clusters 1930c20 and 1930c38 were 30dpf ZZ-specific (Fig. 9B). These ZZ- specific clusters expressed many extracellular matrix components including the small leucine-rich proteoglycan gene *lumican* (Supplementary Fig. S4A), *nog2, ccn2b*(*ctgfb*), *ccn4b*(*wisp1b*), and *ogna* (Supplementary Table S7). Cluster 1930c32 had far more ZZ cells than ZW cells at 30dpf and no cells at 19dpf, and specifically expressed the channel-forming *pannexin* gene *panx3* (Fig. 9D), whose mammalian ortholog is a marker for the epididymis (Turmel et al., 2011), suggesting a role in zebrafish male gonadal ducts.

<u>Contractile cells</u> in 1930c6 specifically expressed the smooth muscle genes *acta2* (Fig. 9C), *csrp1b, cnn1b, tpm2,* and *mylkb*. This strong expression domain appeared in 30dpf gonads and a smaller domain in 19dpf gonads, but about equally in ZW and ZZ cells. In human testes, *ACTA2, CSRP1* and other contractile protein genes are expressed in peritubular myoid cells, a layer of flattened contractile cells around seminiferous tubules that help transport spermatozoa and testicular fluid (Maekawa et al., 1996). Peritubular myoid cells also inhabit the interstitial region of fish testes (Schulz and Nóbrega, 2011). ZW cells also expressed this set of smooth muscle cell genes, as do cells in human ovaries (Fan et al., 2019).

<u>Vascular endothelial cells</u> line arterials, venous vessels, and lymphatics (Wolf et al., 2019, Gurung et al., 2022) and enter zebrafish gonads early in gonadogenesis (Kossack et al., 2023). For arterials, Vegf-receptor-2 (*kdr*, *kdrl*) activates Notch1 signaling (*notch1a, notch1b, notchl, dll4*) to provide Ephrin-B2 (*efnb2a, efnb2b*) at the cell surface (Villa et al., 2001, Masumura et al., 2009, Scheppke et al., 2012, Nakayama et al., 2013). Both *efnb2a* and *efnb2b* (Fig. 7E), Vegf-receptor, and Notch signaling genes (e.g., *kdr,* Supplemental Fig. S4B) were expressed in 1930c22 at both ages and in both sex genotypes, identifying arterial endothelium. For venous endothelium, *smarca4a* acts via CouptfII (*nr2f2*) to place Ephb4 (*ephb4a* (Supplemental Fig.

S4C) and *ephb4b*) on the cell surface (Pereira et al., 2000, Wang et al., 2002, You et al., 2005, Muto et al., 2011, Davis et al., 2013, Model et al., 2014, Cui et al., 2015). The venous gene set was also expressed in 1930c22, suggesting that gonadal vein endothelial cells also appeared in this cluster. For lymphatics, *sox18* turns on Prox1 (*prox1a, prox1b*) to provide Vegfr3 (*flt4*) and thereby lymphatic identity supported by enriched expression of *lyve1a* (Fig. 7F) and *lyve1b* (Kaipainen et al., 1995, Prevo et al., 2001, Jackson et al., 2001, Francois et al., 2008, Lee et al., 2009, Aranguren et al., 2013). We conclude that NA gonads at both 19 and 30dpf had all three types of endothelia without age- or sex-specific endothelial cell types.

#### Gonad Immune Cells

Several types of white blood cells provide gonads with developmental and homeostatic functions. In mammalian testes, special lymphocytes are essential for normal testis function (Mossadegh-Keller and Sieweke, 2018, Garcia-Alonso et al., 2022). In the mammalian ovary, cytokine-mediated inflammation helps regulate ovulation (Field et al., 2014, Fan et al., 2019) and in laboratory zebrafish, activation of a specific macrophage population propels ovarian failure and masculinization (Bravo et al., 2023). To study leukocytes in developing NA gonads, we identified cells expressing the pan-leukocyte genes *coro1a,* (Fig. 10A), *ptprc* (*cd45*), and *lcp1* (*l-plastin*) (Zakrzewska et al., 2010, Torraca et al., 2014) (Table S7), all of which marked the same clusters and thus define the leukocytes in our dataset (Fig. 10A).

**Figure 10.**
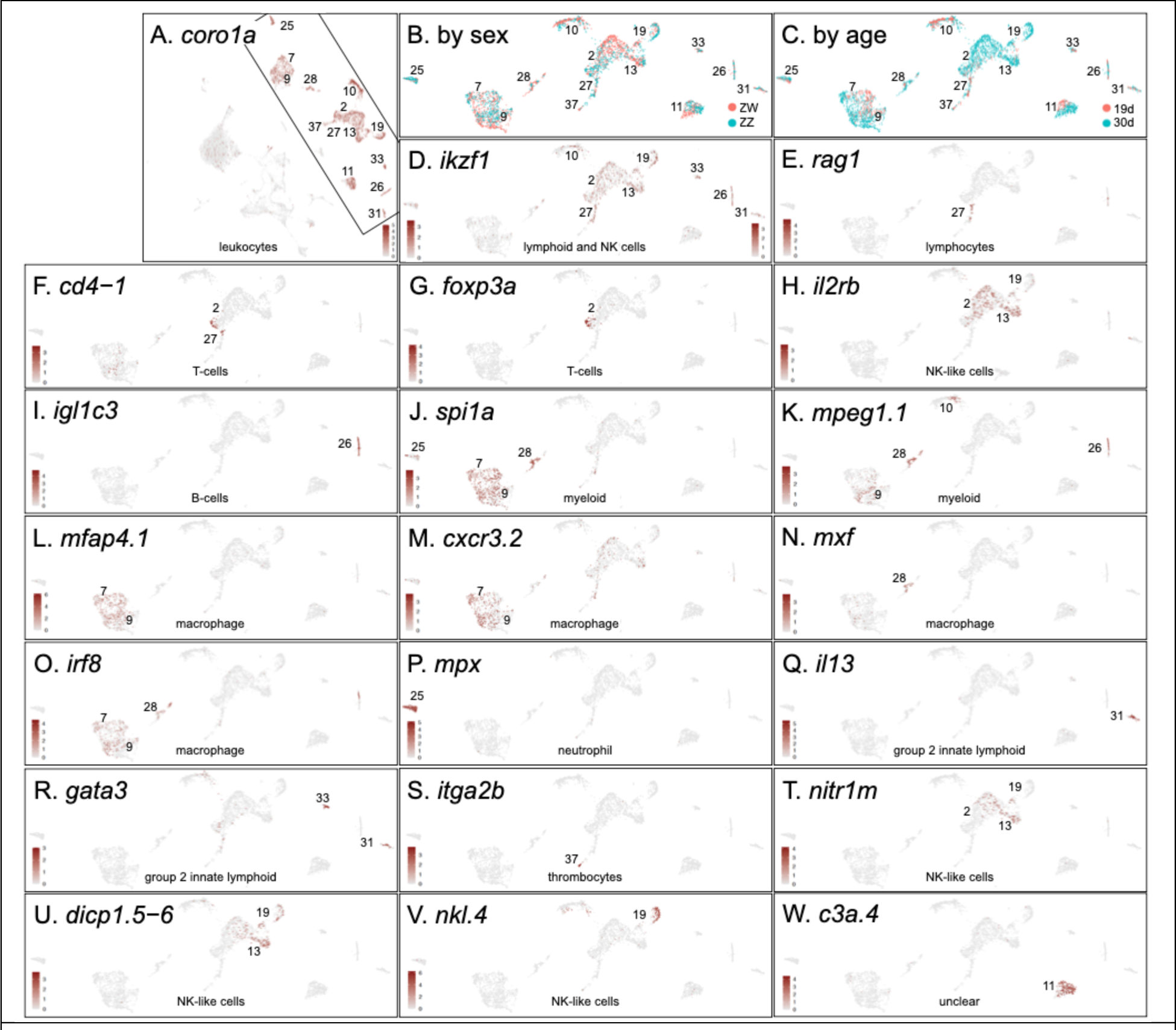
Immune cells. A. *coro1a* expression identifying leukocytes displayed for all four conditions analyzed together. The remainder of the panels show the boxed region rotated counterclockwise. B. Cells labeled according to sex genotype (red, ZW; blue, ZZ). C. Cells labeled according to age (red, 19dpf; blue, 30dpf). D. *ikzf1*. E. *rag1*. F. *cd4-1*. G. *foxp3a*. H. *il2rb*. I. *igl1c3*. J. *spi1a*. K. *mpeg1.1.* L. *mfap4.1*. M. *cxcf3.2.* N. *mxf.* O. *irf8.* P. *mpx*. Q. *il13.* R. *gata3*. S. *itga2b.* T. *nitr1m*. U. *dicp1.5-6*. V. *nkl.4*. W. *c3a.4*.

None of the leukocyte clusters were exclusively ZW or ZZ (Supplementary Table S6). Leukocyte cell types, however, partially sorted out by age, with some clusters being almost exclusively 30dpf (1930c2, c13, c19). These results suggest that at these stages, NA gonads may not have major sex-specific leukocyte populations and that between 19dpf and 30dpf, some new leukocyte populations appear in the gonads.

<u>Lymphocytes</u> and active immunity develop at about three weeks post-fertilization; the period our samples cover (Lam et al., 2004). Lymphocytes and NK cell development in mouse and zebrafish require Ikaros (*ikzf1*), which is expressed early in the lymphocyte lineage (Winandy et al., 1995, Wang et al., 1996, Willett et al., 2001, Schorpp et al., 2006). Both sex genotypes and both ages expressed *ikzf1* (Fig. 10D). In lymphocytes, Rag1 and Rag2 help rearrange immunoglobulin genes and T-cell receptor genes; in NA gonads, *rag1* and *rag2* were expressed in 1930c27 (Fig. 10E) even at 19dpf in both sex genotypes, identifying these cells as lymphoid. A few 1930c27 and adjacent 30dpf cells in 1930c2 expressed T-cell marker genes including *cd4-1* (Fig. 10F)*, cd8a, cd8b, zap70,* and *cd2*(*si:ch211-132g1*) (Blackburn et al., 2014, Moore et al., 2016b, Carmona et al., 2017, Shao et al., 2018). In NA gonads, the regulatory T-cell (Treg) gene *foxp3a* (Hui et al., 2017) was expressed in both ZW and ZZ gonads at 30dpf only (Fig. 10G), which contrasts to the testis-restricted expression of *foxp3a* in adult laboratory strain zebrafish (Li et al., 2020b)); this early *foxp3* expression in genetically female cells could explain why gonad function deteriorates in both ovary and testis in *foxp3a* mutants (Li et al., 2020b). Mixed with T-cells in 1930c2 and extending into 1930c13 were 30dpf cells expressing markers for some natural killer (NK)-like cells including *il2rb* (Carmona et al., 2017) (Fig. 10H), suggesting that in NA zebrafish, as in mammals, gene expression in NK-like cells is similar to that in some T-cells (Bezman et al., 2012).

<u>B-lymphocytes</u> also appeared in both 19dpf and 30dpf gonads of both sexes in 1930c26 marked by both light and heavy immunoglobulin genes including *igl1c3* (Fig. 10I), *igic1s1*, *igl3v5*, and *ighv1-4*, as well as the B-cell-specific transcription factor gene *pax5* (Cobaleda et al., 2007, Moore et al., 2016b). Cluster 1930c26 also expressed *cxcr4a,* encoding the receptor for the lymphocyte chemoattractant Cxcl12 (Bleul et al., 1996).

<u>Myeloid cells</u> include neutrophils, macrophages, monocytes, mast cells, eosinophils, and basophils (Wattrus and Zon, 2021). Early in myeloid development, zebrafish hematopoietic cells express *spi1a* and *spi1b* and these genes are required for macrophage specification (Bukrinsky et al., 2009, Roh-Johnson et al., 2017). NA gonads expressed *spi1a* in the 30dpf-specific cluster 1930c25 and the mixed-age clusters 1930c7, c9, and c28 (Fig.10J). Clusters 1930c7+9 expressed macrophage-specific genes like *mpeg1.1* and *mfap4.1*(*mfap4*, *zgc:77076*) (Fig. 10K, L) and *marco*. Macrophages in cluster 1930c7+9 were present in both sex genotypes.

Macrophage migration during infection and injury in zebrafish depend on Cxcr3, but of the three *cxcr3* paralogs in zebrafish, only *cxcr3.2* is required (Sommer et al., 2020); for the gonad, this may be because *cxcr3.2* is the only one of the three paralogs expressed strongly in most 1930c7+9 macrophages (Fig. 10M). Functionally different populations of macrophages have been identified in mammalian testes (Garcia-Alonso et al., 2022, DeFalco et al., 2015), but NA gonads did not appear to have macrophage sub-types expressing the relevant orthologs of the mammalian markers. Zebrafish gonads, however, do have multiple macrophage subtypes because 1930c28 represents a primarily 30dpf ZW population of *spi1^+^ mpeg1.1^+^* cells that uniquely expressed *mxf* (Fig. 10N) and other *myxovirus (influenza) resistance* paralogs, as well as the inflammation marker *tnfa.* Cells expressing the macrophage differentiation regulator *irf8* are present in 1930c7, c9, and c28 in both NA sex genotypes already by 19dpf (Fig. 10O) and so macrophages are likely in place in NA to help remodel ovarian follicles as in laboratory strain zebrafish (Bravo et al., 2023).

<u>Neutrophil</u>-specific markers, including *mpx* (Fig. 10P), *lyz,* and *mmp13a* were expressed strongly in 1930c25 in all four age/sex-genotype conditions. Other genes used as neutrophil markers, including *nccrp1, cebpa, cpa5, cma1, and prss1* (Tang et al., 2017, Rougeot et al., 2019), were expressed not only in 1930c25, but also strongly in several other clusters, decreasing utility as neutrophil-specific markers in NA gonads.

<u>Group 2 innate lymphoid cells</u> (ILC2s, nuocytes, Th2 cells) in mammals help provide innate immunity against helminth infection and play a role in allergic airway hyperreactivity (Neill et al., 2010, Moro et al., 2010, Barlow et al., 2012, Klein Wolterink et al., 2013). Key ILC2 genes include *IL4, IL5* (zebrafish ortholog: *csf2rb*), and *IL13* (Fig. 10Q) and in zebrafish, *il4* and *il13* are essential to suppress type-1 immune responses (Bottiglione et al., 2020) and were expressed specifically in 1930c31. Gata3 is essential for normal expression of *IL3* and *CSF2R* (but not *IL4*) in ILC2 cells (Zhu et al., 2004), and *gata3* was expressed specifically in 1930c31 and c33 (Fig. 10R). Genes strongly expressed in 1930c31 and c33 included *il11b*, *csf3a, gata3* and many genes annotated as ‘serine-type endopeptidase activity’ (including *si:dkey-78l4.2* and a dozen of its tandem duplicates), which are also strongly expressed in the 5dpf larval thymus (Farnsworth et al., 2019). We hypothesize that the ILC2-like cells in 1930c31 and c33 in zebrafish gonads may be acting to help regulate the extent of inflammation associated with developmental changes as gonads mature.

<u>Thrombocytes</u> appeared in 1930c37, which contained cells of both ages and sex genotypes, according to marker genes *itga2b(cd41)* (Fig. 10S*), mpl, gp1bb, zfpm1* and *coagulation factor V* (*f5)* (Lin et al., 2005, Khandekar et al., 2012, Tang et al., 2017).

<u>Natural killer (NK)-like cells</u> may be a variant type of cytotoxic innate lymphoid cell. Zebrafish NK-like cells express NK markers like *il2rb* (Carmona et al., 2017) (Fig. 10H). The 1930c2, c13, and c19 cells also expressed novel immune-type receptors (NITRs, e.g., *nitr1m*, Fig. 10T) that have properties like those of mammalian natural killer receptors (NKRs) (Yoder et al., 2010).

The *nitr* gene expression detected in the zebrafish ovary but not as maternal transcripts in oocytes (Yoder et al., 2010) could have been from the NK-like cells described here. Diverse immunoglobulin domain-containing protein (DICP) genes have been suggested to be associated with NK-like cells (Haire et al., 2012, Rodriguez-Nunez et al., 2016, Carmona et al., 2017) and the expression of *dicp* genes like *dicp1.5-6* (Fig. 10U) in 1930c2, c13, and c19 supports this notion. The same cell set expresses NK-lysin genes: *nkl.4* expression, which is up-regulated after viral infection (Pereiro et al., 2015) and increases ten-fold in *rag2-*deficient zebrafish (Moore et al., 2016a), was expressed specifically in NA gonads in 1930c19 (Fig. 10V).

<u>Cluster 1930c11</u> cells expressed the pan-leucocyte markers *coro1a, ptprc* (*cd45*), and *lcp1*, supporting twhite blood cell identity (Fig. 10A), but their specific cell type assignment is unclear. 1930c11 cells expressed many genes expressed almost exclusively in the thymus of 5dpf zebrafish (Farnsworth et al., 2019), including *c3a.4* (Fig. 10W), *si:ch73-160i9.2, mfap4.9(ENSDARG00000095746), si:dkey-203a12.7,* and *tuno4,2.* Cluster 1930c11 also expressed a number of *spink2* paralogs (*spink2.5, spink2.10, spink2.11*) that are expressed only in the zebrafish thymus at 5dpf (Farnsworth et al., 2019), while human *SPINK2* is proposed as a marker of hematopoietic stem cells (Calvanese et al., 2022). Nearly all 1930c11 genes were expressed in all four age/genotype conditions. We conclude that 1930c11 may represent an early-stage blood cell in gonads.

## CONCLUSIONS

Laboratory strains of zebrafish pass through a juvenile ovary phase in which all individuals initially form a gonad with oocytes that die in fish that become males but survive in individuals that become females (Takahashi, 1977, Uchida et al., 2002, Maack and Segner, 2003, Wang et al., 2007, Rodriguez-Mari et al., 2010) and have what appears to be a polygenic sex determination mechanism (Bradley et al., 2011, Anderson et al., 2012, Howe et al., 2013, Wilson et al., 2014, Luzio et al., 2015). In contrast, in NA strain zebrafish, which have a ZW female / ZZ male chromosomal sex determination mechanism (Anderson et al., 2012, Wilson et al., 2014), ZZ individuals do not develop juvenile ovaries but instead directly develop testes (Fig. 2). Although all NA strain ZW individuals initially form gonads with oocytes, some ZW fish lose these juvenile oocytes as their gonads become a testis, mimicking laboratory strains. Thus, the distinctive feature of the NA strain is the direct development of testis in ZZ fish by-passing the juvenile ovary phase.

Because gonads in some individuals carrying the W allele initially develop oocytes that undergo apoptosis like gonads in AB and TU strains, and because oocytes do not develop in fish that lack the W chromosome, we suspect that these laboratory strains, which were cleaned of background lethal mutations by either gynogenesis or repeated inbreeding (Walker- Durchanek, 1980, Streisinger et al., 1981, Chakrabarti et al., 1983, Mullins et al., 1994), are likely chromosomally WW.

The direct testis development in ZZ NA fish raises the question of whether they transit a transcriptomic state shared with developing gonads in ZW genetic females or alternatively, if gonads in the two sex genotypes are transcriptionally different even at 19dpf when they are morphologically indistinguishable. Single-cell RNA-seq showed that at 19dpf, all clusters consisted of an intermixture of ZW and ZZ cells, but by 30dpf, sex-genotype-specific clusters had developed. The intermingling of ZW and ZZ cells in 19dpf clusters shows that at this stage, ZW and ZZ gonads in NA fish were not only morphologically similar, but also had not differentiated sex-specific cell types according to gene expression patterns in our experiments.

Despite the general bipotential nature of 19dpf NA ZW and ZZ gonads, germ cells in 19dpf ZW gonads were already expressing low levels of oocyte-specific genes, for example, eggshell genes, that ZZ germ cells did not express. This result suggests that ovary-specific functions had already begun in ZW, but not ZZ, fish by 19dpf, but whether those differences are due to primary differences in germ cells or subtle differences in support cells, steroidogenic cells, or stromal/interstitial cells is not yet known. The findings that 30dfp ZZ germ cells clustered with 19dpf ZZ germ cells, and that ZZ germ cells did not express spermatogenesis-specific genes, showed that ZZ germ cells generally postponed development until after 30dpf, in contrast to ZW germ cells, many of which showed oocyte-specific expression by 30dpf.

Like germ cells, ZW and ZZ support cells had very similar transcriptomes at 19dpf, but by 30dpf, ZW and ZZ support cells had differentiated into granulosa cells and Sertoli cells, respectively. The differentiated nature of the 30dpf ZZ Sertoli cells vs. the relatively undifferentiated state of the 30dpf ZZ germ cells suggests that in ZZ fish, support cell differentiation preceded germ cell differentiation.

Analysis of steroidogenic cells showed that genes encoding enzymes that catalyze the last two steps in estrogen biosynthesis are expressed in three different cell clusters: theca cells in 1930c12 expressing *cyp17a1* can produce androstenedione, which follicle cells in 1930c18 expressing *cyp19a1a* can convert to estrone, which granulosa cells in 1930c21 expressing *hsd17b1* can convert to estradiol. Whether the *cyp19a1a*-expressing cells and the *hsd17b1-* expressing cells are different states of the same cells, or alternatively, are totally different cells is unclear. The failure to find significant expression of the testosterone-synthesizing enzyme gene *hsd17b3* in 30dpf NA gonads, mimicking findings with 40dpf AB ovaries (Liu et al., 2022), raises the question of how or whether gonads at these stages produce testosterone.

Histology supported the notion that the W chromosome in the four zebrafish strains that had a ZW / ZZ *sar-4* chromosomal sex determination mechanism (Wilson et al., 2014) has a locus necessary, but not sufficient, for gonads to initiate oocyte development. An alternative hypothesis is that the Z chromosome has a locus that is required in two doses to prevent oocyte development. The location of the genetic sex determinant at the telomere and the enormous number of repetitive elements in the region led to a poor quality genome assembly and the region lacks any of the ‘usual suspects’ for fish sex determination genes (e.g., Sox family, Dmrt1 variants, TGF-beta pathways) (Herpin and Schartl, 2015, Bertho et al., 2016). High quality PacBio genomic sequences from ZZ and WW individuals and detailed genetic analyses promise to lead to the molecular genetic identification of this elusive element.

## MATERIALS AND METHODS

### Fish Husbandry

The NA wild-type strain (NA, ZFIN ID: ZDB-GENO-030115-2), originating from the Nadia region of India, has been raised at the University of Oregon zebrafish facility since 1999, and has been selected for maintenance of the Z chromosome since about 2012. For developmental histology, juveniles of one NA family were raised in embryo medium E2 (Westerfield, 1993) and fed paramecia from 4-10 days post-fertilization (dpf) and other families were raised in E5 and fed rotifers from 4-10dpf (Best et al., 2010), but results from both were comparable and were combined. After 10dpf, fish were moved to a circulating fish water system (Westerfield, 1993) and fed Zeigler Adult Zebrafish Diet. Procedures were approved by the University of Oregon IACUC protocol #18-13.

### Genotyping

Animals were genotyped for Z and W chromosomes with PCR primers designed from RAD-tag sequences that contained a sex-linked marker from the published NA dataset (Fig. S1), using either the original published primer set (Wilson et al., 2014) or an improved primer set (F: CCGCGTTTATATCCTGGTAA and R: GTTGACCCAACTGGACTCTG), which amplified a 119-nucleotide (nt) fragment from NA genomic DNA. PCR conditions were: 6m 94°C; 45 cycles of 25s 94°C, 25s 61°C, 30s 72°C; followed by 10m at 72°C. The amplicon was digested with the restriction enzyme CviQI according to manufacturer’s instructions (New England Biolabs). The enzyme digested the 119nt amplicon from the Z allele to produce fragments of 43 and 76nt, but left the W allele uncut.

### Gonad Histology

Fish were euthanized in ice water and the caudal fin was removed for genotyping. Fish trunks from the gills to the urogenital pore were fixed in Bouin’s fixative (Sigma) for at least 48 hours, then washed repeatedly with 70% ethanol. Fixed tissues were embedded in paraffin, sectioned at 7 microns, and stained with hematoxylin and eosin.

### Single-cell RNA-Seq

Zebrafish were fin-clipped, genotyped for genetic sex, allowed to recover on the water system for a few days, and at 19 and 30 dpf, they were euthanized in ice water and gonads were dissected in chilled phosphate buffered saline (PBS) as described (Wang et al., 2017). Gonads from five fish per genotype were pooled in a well of a 24-well tissue culture plate and dissociated using a protocol adapted from (Covassin et al., 2006). PBS was replaced with 1.2mL of a solution containing 0.05% trypsin and 0.02% EDTA. Samples were incubated for 10 minutes at 28.5°C, pipetting up and down every two minutes with a 1000µl pipette tip to manually break up tissues. Adding 200µl of a solution containing 30% Fetal Bovine Serum and 6mM CaCl2 stopped digestion. Cells were transferred to a 1.5 mL microfuge tube and spun for 5 min at 2000 RPM (∼300 RCF) at 4°C. The supernatant was removed, and cells were resuspended in a solution containing 1% fetal bovine serum, 0.8 mM CaCl2, and 1x antibiotic/antimycotic (Sigma #A5955) in L-15 medium. Cells were spun a second time for 5 min at 2000 RPM at 4°C. The supernatant was removed, and cells were resuspended in the suspension solution, then filtered through a 40-micron strainer (Biologix #15-1040) to generate a suspension of single-cells. After microscopic visualization to check for dissociation, cells were counted with a Bio-RAD TC20 Automated Cell Counter and given to the University of Oregon Genomics & Cell Characterization Core Facility for library construction and sequencing.

Libraries were prepared with Chromium v2 chemistry (10x Genomics) targeting 10,000 cells. according to manufacturer’s instructions. Libraries were sequenced on an Illumina NextSeq 500.

Sequences were aligned to the zebrafish genome GRCZ11 using the Lawson zebrafish transcriptome annotation (Lawson et al., 2020) with Cell Ranger v. 6.1.1 (Zheng et al., 2017). Ambient RNA was removed with SoupX v 1.6.2 (Young and Behjati, 2020). For 19dpf individuals, ambient RNA derived from pancreas and erythrocytes, so we used the genes “hbaa1”, “cpa5”, “cpa1”, “cel.2”, “cel.1”, “amy2a”, “hbba1”, “ela2l”, “ela2”, “prss1”, “hbae5”, and “prss59.2” to estimate the contamination fraction. In 30dpf individuals, ambient RNA was derived from both erythrocytes and fragile oocytes, so we used the following genes to estimate the contamination fraction: hbaa1”, “hbaa2”, “hbba2”, “hbba1”, “hbae5”, “ddx4”, “dnd1”, “sycp1”, “sycp3”, and “dmc1”. Data were analyzed with Seurat v. 4.2.0 (Butler et al., 2018). Non-gonad clusters, including contaminating pancreatic cells from dissection errors and erythrocytes, were excluded and gonad cells were reclustered. Differential expression analysis was performed on 19dpf clusters comparing ZZ vs ZW genotypes using the Seurat “FindMarkers” function with the parameters min.pct=0.05, logfc.threshold=0.1, test.use=”MAST” (Finak et al., 2015).

## Author Contributions

CAW: Conceptualization, data curation, formal analysis, investigation, software, writing original draft. PB: Conceptualization, data curation, formal analysis, investigation, software, review and editing. JHP: Conceptualization, formal analysis, funding acquisition, investigation, project administration, supervision, writing original draft.

## Data Availability Statement

The datasets generated for this study can be found in the Sequence Read Archive (www.ncbi.nlm.nih.gov/sra) under the accession number PRJNA1055160.

## Supporting information

SuppTable S1 19dpf Cell Counts per Cluster

SuppTable S2 19dpf_AllMarkers_minpct10

SuppTable S3 19dpf ZZ vs ZW Differential Expression

SuppTable S4 30dpf Cell Counts per Cluster

SuppTable S5 30dpf_AllMarkers_minpct10

SuppTable S6 Merged 19dpf 30dpf Cell Counts per Cluster

SuppTable S7 Merged19and30dpf_AllMarkers_minpct10

SuppMethods Scripts Wilson

## Acknowledgements

We thank Ruth Bremiller and Poh Kheng Loi for help with histology, and the University of Oregon’s Genomics and Cell Characterization Core Facility (GC3F) for library preparation and sequencing. This work benefited from access to the University of Oregon high performance computing cluster, Talapas.

## Scope Statement

Our manuscript “Direct Male Development in Chromosomally ZZ Zebrafish” by Wilson, Bazel, and Postlethwait is submitted to *Frontiers in Cell and Developmental Biology* for the special volume entitled: *Proceedings of the 9th International Symposium on the Biology of Vertebrate Sex Determination 2023*. Our work uses histology and single-cell transcriptomics to probe the mechanisms of gonadal development in a zebrafish strain that maintains the chromosomal sex determination system found in the wild rather than in domesticated laboratory strains that lost the natural mechanism. Results show that gonads in chromosomally ZZ individuals develop directly into testes and avoid the juvenile ovary state found in gonads of all individuals in domesticated strains. Our scRNA-seq experiments revealed that gene expression in gonads at the histologically indifferent stage are highly similar in ZZ (all male) and ZW (mostly female) individuals. Eleven days later, however, when testes and ovaries are distinct morphologically, many transcriptional cell types are ZZ- or ZW-specific. This is the first work to identify genes that distinguish gonadal cell types at these early developmental states and the first to study gonad development in a zebrafish strain with the wild genetic sex determination system. This work is in the journal scope because it investigates “molecular and cellular reproduction” and “morphogenesis and patterning”.

## Supplementary Materials

### Supplementary Figures

**Supplementary Figure S1.**
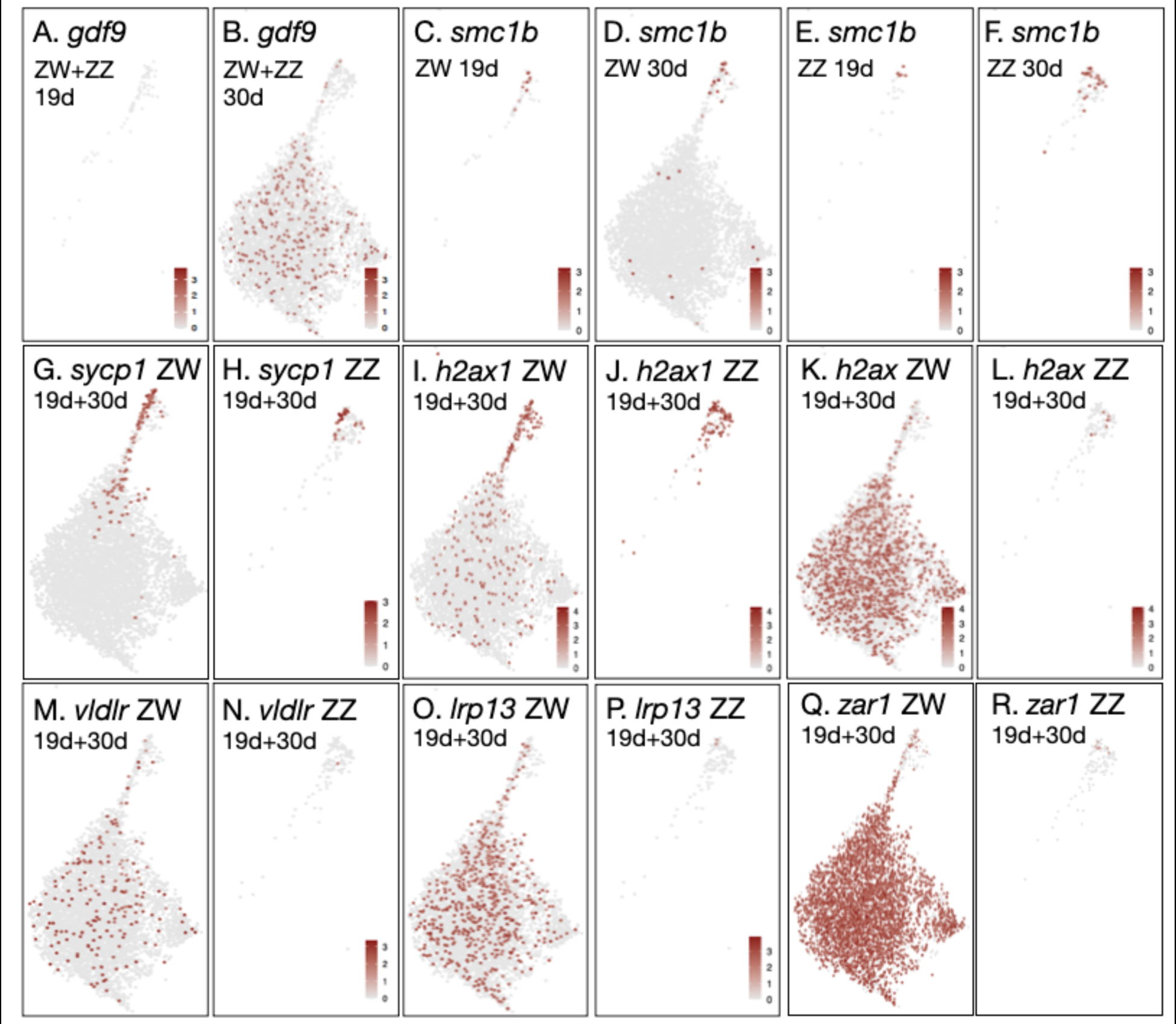
Germ cell gene expression in the merged analysis of both ZW and ZZ cells at both 19dpf and 30dpf. A. *gdf9,* ZW and ZZ cells at 19dpf only. B. *gdf9, ZW* and ZZ cells at 30dpf only. C. *smc1b,* ZW cells at 19dpf only. D. *smc1b,* ZW cells at 30dpf only. E *smc1b,* ZZ cells at 19dpf only. F. *smc1b,* ZZ cells at 30dpf only. G. *sycp1*, ZW cells at 19dpf and 30dpf. H. *sycp1*, ZZ cells at 19dpf and 30dpf. I. *h2ax1*, ZW cells at 19dpf and 30dpf. J. *h2ax1*, ZZ cells at 19dpf and 30dpf. K. *h2ax*, ZW cells at 19dpf and 30dpf. L. *h2ax*, ZZ cells at 19dpf and 30dpf. M. *vldlr*, ZW cells at 19dpf and 30dpf. N. *vldlr*, ZZ cells at 19dpf and 30dpf. O. *lrp13*, ZW cells at 19dpf and 30dpf. P. *lrp13*, ZZ cells at 19dpf and 30dpf. Q. *zar1*, ZW cells at 19dpf and 30dpf. R. *zar1*, ZZ cells at 19dpf and 30dpf.

**Supplementary Figure S2.**
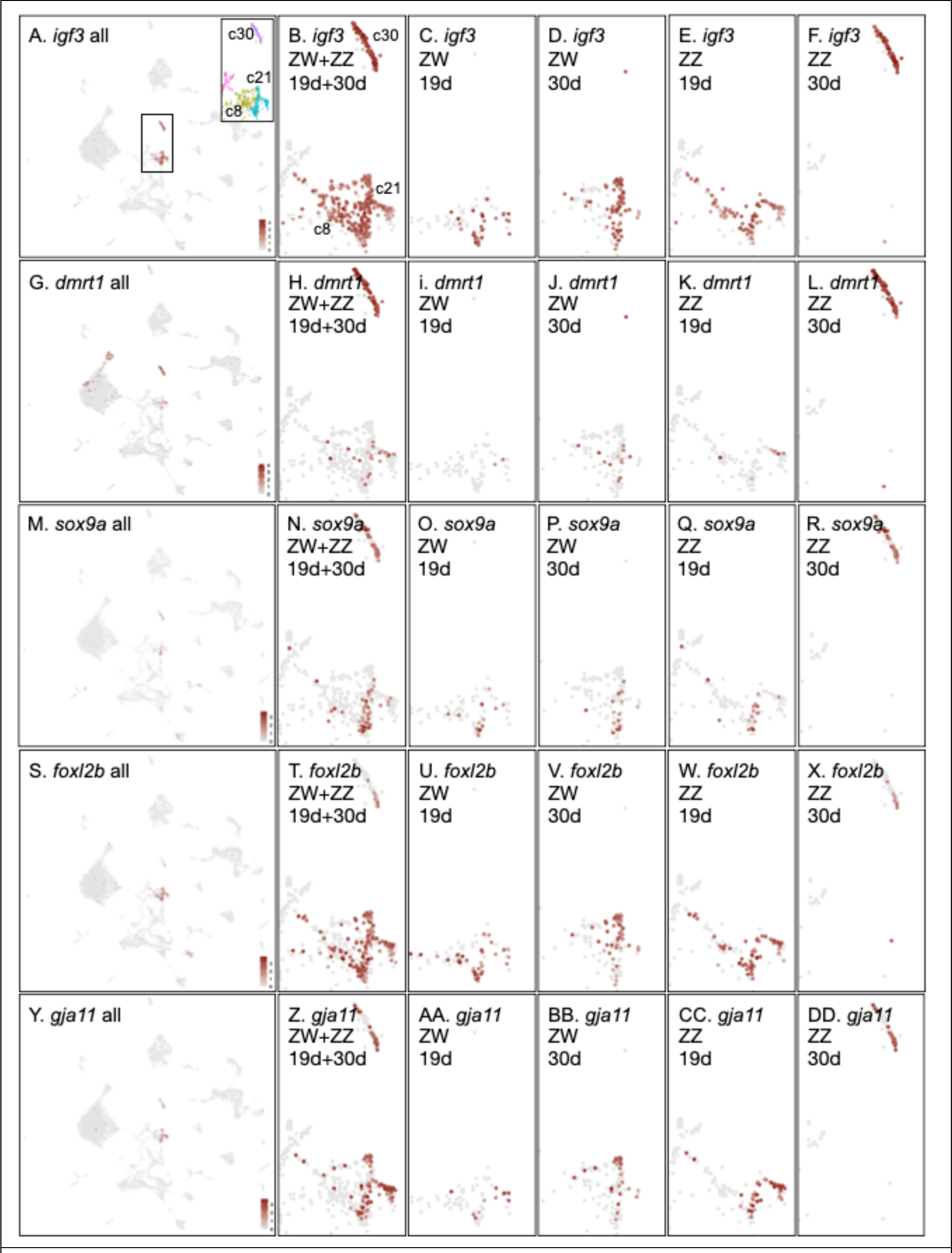
Support cell gene expression in the merged analysis of both ZW and ZZ cells at both 19dpf and 30dpf. A. *igf3* expression across the whole combined dataset. The rectangle marks *igf3*-expressing support cells and their precursors, expanded to show cluster assignments in color. B. Enlargement of rectangle from part A, showing *igf3* expression in clusters 1930c30, c21, and c8 combining all four conditions. C. *igf3* expression in 19dpf ZW gonads. D. *igf3* expression in 30dpf ZW cells. E. *igf3* expression in 19dpf ZZ cells. F. *igf3* expression in 30dpf ZZ cells. G-L. *dmrt1* expression in samples as described for panels A-F. M-R. *sox9a* expression. S-X, *foxl2b* expression. YDD. *gja11* expression. EE-JJ. *Lhx9* expression.

**Supplemental Figure S3.**
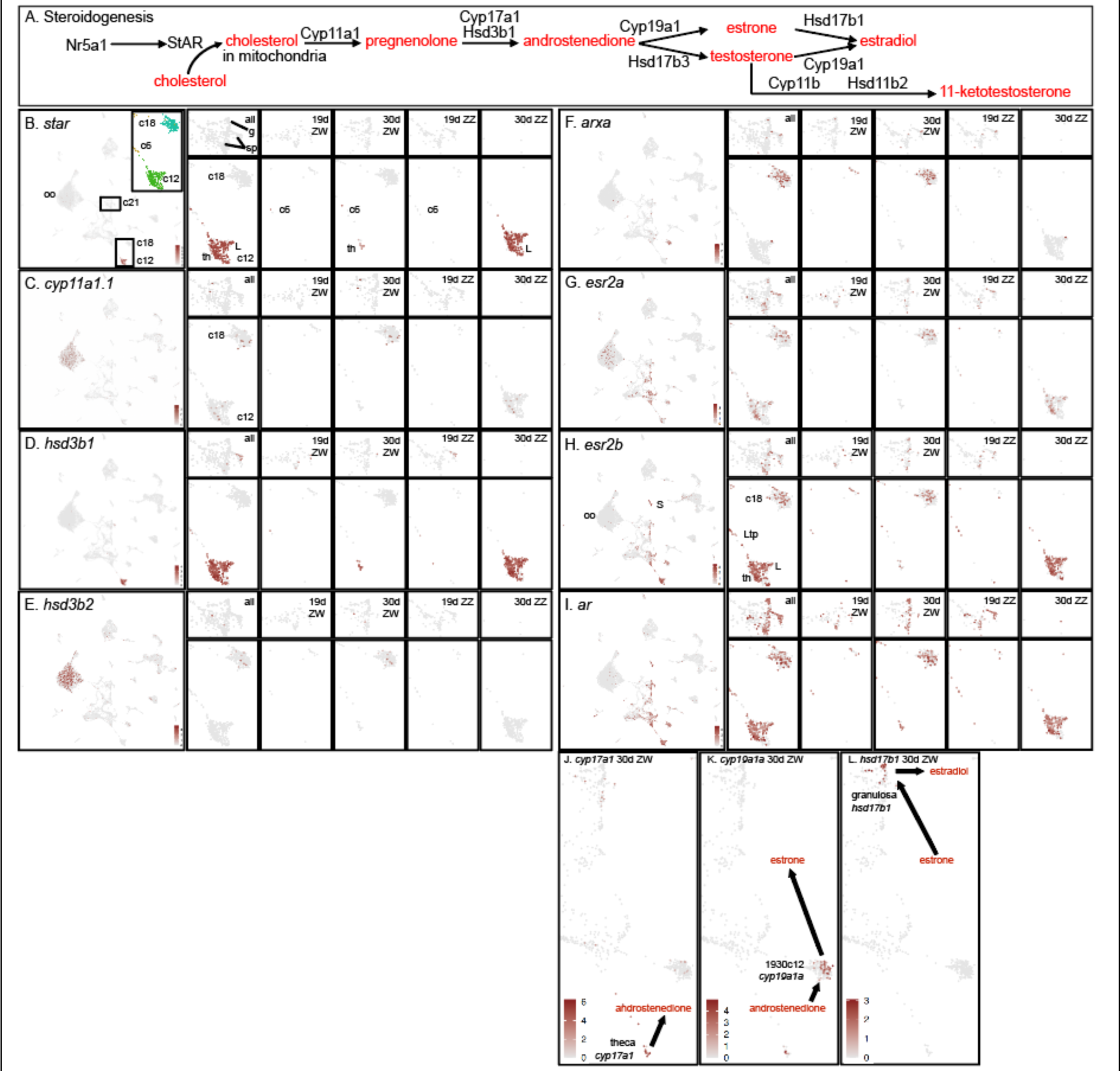
Steroidogenesis. A. An abbreviated pathway of steroidogenesis (after (Tenugu et al., 2021)). B. *star* expression. The left panel shows expression in the entire dataset. The insert shows cluster demarcation for the most relevant steroidogenic enzymes. Subsequent panels show, at the top 1930c21, the support cell precursor and granulosa cell cluster and at the bottom the clusters corresponding to the insert in the left-hand panel. Displayed from left to right are the enlargements of all four conditions, 19dpf ZW gonads, 30dpf ZW gonads, 19dpf ZZ gonads, and 30dpf ZZ gonads. Cluster numbers are indicated. L, Leydig cells; th, theca cells. C. *cyp11a1.1* displayed as explained for *star*. D. *hsd3b1*. E. *hsd3b2*. F. *arxa*. G. *esr2a*. H. *esr2b*. I. *ar*, androgen receptor. J-L. Three different cell types appear to be important for estrogen production in 30dpf ZW zebrafish: 1) theca cells in 1930c12 producing androstenedione using Cyp17a1; 2) follicle cells in 1930c18 forming estrone catalyzed by Cyp19a1a; and 3) a different type of follicle cell in 1930c21 generating estradiol by Hsd17b1.

**Supplementary Figure S4.**
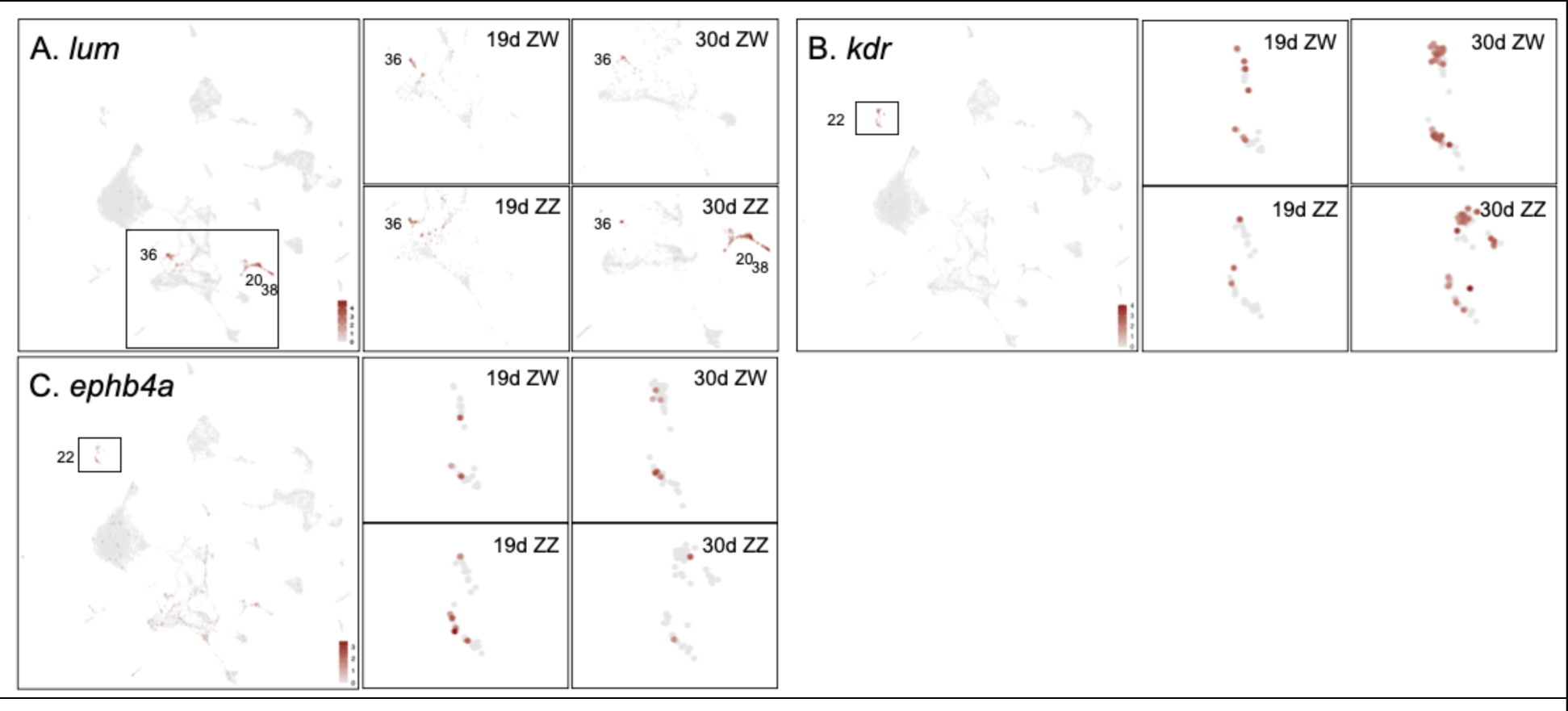
Stromal/interstitial/vascular clusters. A. *lum* expression mainly in the 30d ZZ-specific clusters 1930c20 and 1930c38. The left large panel combines cells of all four conditions and the four smaller panels show each condition independently. B. *ephb4a* expression, a venous endothelium gene, with the four smaller panels expanding cluster 1930c22. C. *kdr* expression with the four smaller panels expanding cluster 1930c22.

### Supplementary Tables

Supplementary Table S1. Cell counts per genotype per cluster for the 19dpf single-cell dataset

Supplementary Table S2. Gene expression markers for 19dpf clusters. We required genes to be expressed in a minimum of 10% of cells and a minimum log fold change of 0.25.

Supplementary Table S3. Differential expression analysis comparing ZZ cells to ZW cells for each cluster at 19dpf.

Supplementary Table S4. Cell counts per genotype per cluster for the 30dpf single-cell dataset

Supplementary Table S5. Gene expression markers for 30dpf clusters. We required genes to be expressed in a minimum of 10% of cells and a minimum log fold change of 0.25.

Supplementary Table S6. Cell counts per age per genotype per cluster for the combined 19dpf and 30dpf single-cell dataset

Supplementary Table S7. Gene expression markers for combined 19dpf and 30dpf clusters. We required genes to be expressed in a minimum of 10% of cells and a minimum log fold change of 0.25.

### Supplementary Methods

Supplementary Method 1. R scripts utilizing Seurat to analyze 19dpf single-cell data

Supplementary Method 2. R scripts utilizing Seurat to analyze 30dpf data

Supplementary Method 3. R scripts utilizing Seurat to analyze combined single-cell data

Supplementary Method 4. R scripts utilizing the SoupX package to remove ambient RNA.

