## Supplementary material for "Direct Male Development in Chromosomally ZZ Zebrafish": SuppTable S1 19dpf Cell Counts per Cluster

| Cluster | ZW cell number | ZZ cell number |
|---|---|---|
| 0 | 83 | 110 |
| 1 | 82 | 100 |
| 2 | 40 | 115 |
| 3 | 49 | 75 |
| 4 | 46 | 62 |
| 5 | 31 | 63 |
| 6 | 17 | 77 |
| 7 | 26 | 63 |
| 8 | 45 | 38 |
| 9 | 20 | 58 |
| 10 | 34 | 40 |
| 11 | 28 | 36 |
| 12 | 27 | 35 |
| 13 | 10 | 51 |
| 14 | 20 | 40 |
| 15 | 23 | 36 |
| 16 | 22 | 35 |
| 17 | 41 | 14 |
| 18 | 26 | 22 |
| 19 | 17 | 31 |
| 20 | 15 | 32 |
| 21 | 23 | 17 |
| 22 | 20 | 20 |
| 23 | 8 | 31 |
| 24 | 16 | 21 |
| 25 | 4 | 30 |
| 26 | 7 | 18 |
| 27 | 3 | 21 |
