## Supplementary material for "Direct Male Development in Chromosomally ZZ Zebrafish": SuppTable S2 19dpf_AllMarkers_minpct10

| | p_val | avg_log2FC | pct.1 | pct.2 | p_val_adj | cluster | gene |
|---|---|---|---|---|---|---|---|
| pcolcea | 1.3411541 | 1.519891235 | 0.798 | 0.095 | 2.70872898990793e-1 | 0 | pcolcea |
| scara3 | 1.1269219 | 0.925036739 | 0.42 | 0.02 | 2.27604426415497e-1 | 0 | scara3 |
| fn1a | 4.4906068 | 1.313103251 | 0.741 | 0.129 | 9.06967868478952e-1 | 0 | fn1a |
| igfbp5b | 4.2310578 | 1.470161863 | 0.725 | 0.124 | 8.54546744386896e-1 | 0 | igfbp5b |
| thbs3a | 3.5139065 | 0.820373987 | 0.326 | 0.007 | 7.09703714333451e-1 | 0 | thbs3a |
| slit3 | 1.3288986 | 0.999700448 | 0.492 | 0.044 | 2.68397664108791e-1 | 0 | slit3 |
| ccl25b | 4.9446429 | 1.482672395 | 0.72 | 0.136 | 9.98669539247061e-9 | 0 | ccl25b |
| plpp3 | 1.6856596 | 1.180835909 | 0.591 | 0.082 | 3.4045268930024e-93 | 0 | plpp3 |
| mfap5 | 1.1286465 | 1.156129411 | 0.668 | 0.116 | 2.27952734796551e-8 | 0 | mfap5 |
| crhbp | 5.1979432 | 0.708466969 | 0.275 | 0.006 | 1.04982859294666e-8 | 0 | crhbp |
| pmp22a | 4.1178283 | 1.192830999 | 0.876 | 0.27 | 8.31677800918026e-8 | 0 | pmp22a |
| spon2b | 3.7505252 | 0.76075324 | 0.254 | 0.005 | 7.57493575400409e-8 | 0 | spon2b |
| si:dkey-261h17.1 | 3.7941872 | 1.286889992 | 0.767 | 0.204 | 7.66312004857237e-8 | 0 | si:dkey-261h17.1 |
| FP102783.1 | 7.8978037 | 0.681313361 | 0.28 | 0.009 | 1.59511943166877e-8 | 0 | FP102783.1 |
| ptx3a | 3.1586302 | 1.062519916 | 0.409 | 0.037 | 6.37948548703345e-8 | 0 | ptx3a |
| mustn1a | 3.6203477 | 0.614234648 | 0.264 | 0.007 | 7.31201636315639e-8 | 0 | mustn1a |
| BX640522.2 | 4.4308027 | 1.0309582 | 0.539 | 0.081 | 8.94889229344171e-7 | 0 | BX640522.2 |
| osr1 | 1.8378590 | 1.02165878 | 0.539 | 0.095 | 3.71192384128675e-6 | 0 | osr1 |
| CR383676.1 | 1.3254026 | 0.72290987 | 0.933 | 0.637 | 2.67691581200878e-6 | 0 | CR383676.1 |
| col1a2 | 2.1057551 | 1.092457897 | 0.886 | 0.281 | 4.25299368715375e-6 | 0 | col1a2 |
| emid1 | 3.4439274 | 0.858482631 | 0.383 | 0.04 | 6.95570033643417e-6 | 0 | emid1 |
| emilin2a | 2.1519436 | 0.565091706 | 0.212 | 0.005 | 4.34628064195114e-6 | 0 | emilin2a |
| rbp4 | 1.1017985 | 1.1059081 | 0.72 | 0.174 | 2.2253025032111e-65 | 0 | rbp4 |
| si:ch211-227m13.1 | 7.4049152 | 0.519026463 | 0.233 | 0.009 | 1.49557073524688e-6 | 0 | si:ch211-227m13.1 |
| fstb | 1.7767422 | 0.700329727 | 0.306 | 0.024 | 3.58848625985326e-6 | 0 | fstb |
| neu3.5 | 3.6927189 | 0.561851206 | 0.202 | 0.005 | 7.45818451470439e-6 | 0 | neu3.5 |
| vasna | 8.1842254 | 0.864549635 | 0.451 | 0.069 | 1.65296800644449e-6 | 0 | vasna |
| klf2a | 1.7934762 | 1.045118019 | 0.601 | 0.141 | 3.62228404187252e-6 | 0 | klf2a |
| plpp1a | 4.4968845 | 1.02928473 | 0.487 | 0.09 | 9.08235763407949e-6 | 0 | plpp1a |
| zcchc24 | 4.1961094 | 0.743741426 | 0.368 | 0.045 | 8.47488234735914e-5 | 0 | zcchc24 |
| endouc | 8.8585007 | 0.910487824 | 0.492 | 0.088 | 1.78915139228872e-5 | 0 | endouc |
| tgfbr3 | 4.5922625 | 0.779746733 | 0.347 | 0.041 | 9.27499260365423e-5 | 0 | tgfbr3 |
| krt94 | 2.5852420 | 1.007352397 | 0.829 | 0.32 | 5.22141342065951e-5 | 0 | krt94 |
| col1a1b | 5.9329272 | 1.021814819 | 0.762 | 0.256 | 1.19827332326028e-5 | 0 | col1a1b |

| | | | | | | | |
|---|---|---|---|---|---|---|---|
| mgp | 7.5406976 | 0.622271932 | 0.238 | 0.014 | 1.52299470363411e-5 | 0 | mgp |
| htra1a | 1.0821246 | 0.698513198 | 0.285 | 0.024 | 2.18556707769117e-5 | 0 | htra1a |
| fstl1b | 4.4067332 | 0.973232567 | 0.72 | 0.238 | 8.90027906852657e-5 | 0 | fstl1b |
| col1a1a | 6.4306316 | 0.980108378 | 0.803 | 0.267 | 1.29879466822307e-5 | 0 | col1a1a |
| fgl2b | 6.4456512 | 0.405240235 | 0.161 | 0.003 | 1.30182817599192e-5 | 0 | fgl2b |
| si:ch73-173h19.3 | 3.7051058 | 0.446211434 | 0.171 | 0.005 | 7.48320219088335e-5 | 0 | si:ch73-173h19.3 |
| id2b | 2.8323719 | 0.799113244 | 0.358 | 0.05 | 5.72054157574644e-5 | 0 | id2b |
| sost | 5.2738051 | 0.727929886 | 0.342 | 0.045 | 1.0651504357219e-48 | 0 | sost |
| mfap2 | 1.2716651 | 0.931273945 | 0.668 | 0.211 | 2.56838201527394e-4 | 0 | mfap2 |
| icn | 3.7258607 | 1.024956231 | 0.72 | 0.216 | 7.5251209446539e-48 | 0 | icn |
| eda | 1.2252102 | 0.444072475 | 0.187 | 0.009 | 2.47455720946378e-4 | 0 | eda |
| colec12 | 1.7584537 | 0.800116143 | 0.425 | 0.083 | 3.55154898726431e-4 | 0 | colec12 |
| dcn | 2.0257007 | 0.94533375 | 0.71 | 0.24 | 4.09130772517637e-4 | 0 | dcn |
| CABZ01092746.1 | 1.2656820 | 0.901344598 | 0.601 | 0.173 | 2.55629807402115e-4 | 0 | CABZ01092746.1 |
| nova2 | 6.3099537 | 0.681362386 | 0.321 | 0.043 | 1.27442135779804e-4 | 0 | nova2 |
| fbln5 | 7.7316912 | 0.769452017 | 0.404 | 0.076 | 1.56156968630009e-4 | 0 | fbln5 |
| nfixb | 3.4371219 | 0.84050745 | 0.544 | 0.142 | 6.9419552259498e-44 | 0 | nfixb |
| AL732488.1 | 1.0126218 | 0.608183776 | 0.264 | 0.029 | 2.04519231179868e-4 | 0 | AL732488.1 |
| XLOC-016094 | 1.0213534 | 0.763407564 | 0.409 | 0.081 | 2.06282746578532e-4 | 0 | XLOC-016094 |
| adgrg6 | 4.0685817 | 0.378352385 | 0.135 | 0.003 | 8.21731452774024e-4 | 0 | adgrg6 |
| crispld2 | 5.2348928 | 0.511573796 | 0.202 | 0.014 | 1.0572913119866e-41 | 0 | crispld2 |
| LOC100537342 | 1.2690765 | 0.532449271 | 0.223 | 0.019 | 2.56315391204675e-4 | 0 | LOC100537342 |
| adrb3a | 4.9457161 | 0.32351855 | 0.13 | 0.003 | 9.98886295488211e-4 | 0 | adrb3a |
| CR855311.1 | 1.1577777 | 0.924694422 | 0.399 | 0.081 | 2.3383636826989e-40 | 0 | CR855311.1 |
| abi3bpa | 1.8750975 | 0.292556963 | 0.109 | 0.001 | 3.7871345847232e-40 | 0 | abi3bpa |
| BX663503.3 | 2.9752746 | 0.613519803 | 0.28 | 0.036 | 6.00916224541718e-4 | 0 | BX663503.3 |
| wnt7bb | 5.0447288 | 0.456876412 | 0.166 | 0.009 | 1.01888388670899e-3 | 0 | wnt7bb |
| igf2b | 3.2297233 | 0.849238802 | 0.492 | 0.135 | 6.52307230287957e-3 | 0 | igf2b |
| tsku | 3.2667256 | 0.639644357 | 0.337 | 0.057 | 6.59780588189085e-3 | 0 | tsku |
| efnb1 | 5.5963630 | 0.686609437 | 0.383 | 0.077 | 1.13029743921668e-3 | 0 | efnb1 |
| emp2 | 2.1863720 | 0.728612006 | 0.368 | 0.073 | 4.41581556295278e-3 | 0 | emp2 |
| dpysl3 | 2.3950801 | 0.48330489 | 0.212 | 0.019 | 4.83734337874495e-3 | 0 | dpysl3 |
| flnb | 2.4273380 | 0.577891892 | 0.259 | 0.032 | 4.90249471754176e-3 | 0 | flnb |
| fgfr2 | 9.6264864 | 0.598335287 | 0.264 | 0.035 | 1.94426147305228e-3 | 0 | fgfr2 |
| ddr2l | 3.9560214 | 0.501088366 | 0.228 | 0.024 | 7.98997659548361e-3 | 0 | ddr2l |

| | | | | | | | |
|---|---|---|---|---|---|---|---|
| lrp1aa | 6.2856613 | 0.753794725 | 0.415 | 0.099 | 1.26951501707082e-3 | 0 | lrp1aa |
| rspo3 | 1.3461697 | 0.488230683 | 0.218 | 0.022 | 2.71885896980957e-3 | 0 | rspo3 |
| nfia | 3.5356983 | 0.783415606 | 0.534 | 0.171 | 7.14105005328398e-3 | 0 | nfia |
| znf385a | 6.8278959 | 0.670196637 | 0.363 | 0.073 | 1.37903014056761e-3 | 0 | znf385a |
| nid1a | 8.9791662 | 0.534257262 | 0.254 | 0.033 | 1.81352220541177e-3 | 0 | nid1a |
| adamts15b | 1.3790930 | 0.510566293 | 0.212 | 0.021 | 2.78535420178679e-3 | 0 | adamts15b |
| col6a3 | 1.0737127 | 0.76969061 | 0.523 | 0.164 | 2.16857756457002e-3 | 0 | col6a3 |
| cdh10a | 3.8024469 | 0.369632809 | 0.15 | 0.009 | 7.67980215898135e-3 | 0 | cdh10a |
| si:ch211-105c13.3 | 5.6562746 | 0.657131132 | 0.383 | 0.085 | 1.14239779725858e-3 | 0 | si:ch211-105c13.3 |
| crabp1b | 1.5238306 | 0.288606018 | 0.109 | 0.003 | 3.07768068689367e-3 | 0 | crabp1b |
| rbp5 | 2.9685630 | 0.817479238 | 0.513 | 0.17 | 5.99560675218846e-3 | 0 | rbp5 |
| hsd11b2 | 6.9382900 | 0.818058675 | 0.674 | 0.258 | 1.40132645147875e-3 | 0 | hsd11b2 |
| sema3bl | 9.8310903 | 0.516580577 | 0.233 | 0.031 | 1.98558532261665e-3 | 0 | sema3bl |
| tgfbr2b | 1.3748891 | 0.490471886 | 0.212 | 0.025 | 2.776863617898e-31 | 0 | tgfbr2b |
| cdh6 | 2.6736648 | 0.596966956 | 0.326 | 0.063 | 5.40000082958935e-3 | 0 | cdh6 |
| col5a1 | 4.2421149 | 0.783290759 | 0.58 | 0.214 | 8.56779957572823e-3 | 0 | col5a1 |
| LOC108180269 | 1.0153091 | 0.613217519 | 0.306 | 0.06 | 2.05061979793387e-3 | 0 | LOC108180269 |
| zgc:195173 | 1.1908155 | 0.772314721 | 0.554 | 0.209 | 2.40509024394112e-3 | 0 | zgc:195173 |
| rgs5a | 2.0888073 | 0.641024728 | 0.347 | 0.073 | 4.21876428619632e-3 | 0 | rgs5a |
| loxl1 | 2.1844849 | 0.581076121 | 0.295 | 0.054 | 4.41200433401649e-3 | 0 | loxl1 |
| ugt5g1 | 2.2540077 | 0.355076069 | 0.15 | 0.011 | 4.5524194975615e-30 | 0 | ugt5g1 |
| clu | 7.2571041 | 0.578264694 | 0.264 | 0.044 | 1.46571731539936e-2 | 0 | clu |
| add3a | 2.0539266 | 0.753984628 | 0.523 | 0.195 | 4.14831562278498e-2 | 0 | add3a |
| rem1 | 4.1522356 | 0.382792842 | 0.161 | 0.014 | 8.38627036222979e-2 | 0 | rem1 |
| prrx1a | 6.9409374 | 0.35656624 | 0.166 | 0.015 | 1.40186114250368e-2 | 0 | prrx1a |
| mdka | 7.3514543 | 0.759493347 | 0.56 | 0.194 | 1.48477324352278e-2 | 0 | mdka |
| kirrel1a | 1.5168468 | 0.547213965 | 0.275 | 0.05 | 3.06357552152085e-2 | 0 | kirrel1a |
| fbn2b | 1.9023348 | 0.746351058 | 0.461 | 0.154 | 3.84214570500477e-2 | 0 | fbn2b |
| fbln2 | 1.2610470 | 0.442552861 | 0.212 | 0.029 | 2.54693679130774e-2 | 0 | fbln2 |
| s100a10b | 1.5432373 | 0.724422181 | 0.829 | 0.425 | 3.11687645226414e-2 | 0 | s100a10b |
| gpr78a | 7.6999629 | 0.339819713 | 0.145 | 0.012 | 1.55516151791322e-2 | 0 | gpr78a |
| akap12b | 1.2382383 | 0.680636752 | 0.43 | 0.128 | 2.50087009092822e-2 | 0 | akap12b |
| lrrc17 | 2.3000626 | 0.702373122 | 0.461 | 0.157 | 4.64543648663903e-2 | 0 | lrrc17 |
| hapln3 | 4.9099663 | 0.376786901 | 0.15 | 0.013 | 9.91665910262319e-2 | 0 | hapln3 |
| CU929237.1 | 6.4232342 | 0.509763967 | 0.238 | 0.04 | 1.2973006148658e-25 | 0 | CU929237.1 |

| | | | | | | | |
|---|---|---|---|---|---|---|---|
| si:dkey-7j14.5 | 7.8320466 | 0.308199371 | 0.135 | 0.01 | 1.58183846945435e-2 | 0 | si:dkey-7j14.5 |
| aadac | 1.5159904 | 0.262331159 | 0.109 | 0.005 | 3.06184594410057e-2 | 0 | aadac |
| fpr1 | 1.8913848 | 0.441081874 | 0.166 | 0.018 | 3.82002990820721e-2 | 0 | fpr1 |
| col6a2 | 4.2907151 | 0.613878704 | 0.378 | 0.105 | 8.66595743470665e-2 | 0 | col6a2 |
| CR847566.1 | 5.8160995 | 0.625977265 | 0.389 | 0.111 | 1.17467761724784e-2 | 0 | CR847566.1 |
| adcy6b | 5.8940838 | 0.266275958 | 0.104 | 0.005 | 1.19042812127376e-2 | 0 | adcy6b |
| slc6a4a | 7.6198058 | 0.340152257 | 0.15 | 0.014 | 1.5389721924851e-24 | 0 | slc6a4a |
| cyp1c2 | 2.7117399 | 0.457426375 | 0.187 | 0.025 | 5.47690120006223e-2 | 0 | cyp1c2 |
| marcksl1a | 3.8249912 | 0.652089553 | 0.782 | 0.491 | 7.7253348781128e-24 | 0 | marcksl1a |
| tmem100a | 6.0632924 | 0.444407164 | 0.218 | 0.036 | 1.2246031747716e-23 | 0 | tmem100a |
| tnfsf10l | 7.2061431 | 0.584367995 | 0.337 | 0.089 | 1.45542472471323e-2 | 0 | tnfsf10l |
| lrp1ab | 1.3250524 | 0.623553557 | 0.358 | 0.104 | 2.67620841231836e-2 | 0 | lrp1ab |
| asip2b | 1.7457623 | 0.5757923 | 0.264 | 0.053 | 3.52591621420041e-2 | 0 | asip2b |
| zgc:153704 | 4.2855599 | 0.479102769 | 0.197 | 0.029 | 8.65554552035225e-2 | 0 | zgc:153704 |
| wscd2 | 1.3266913 | 0.35862759 | 0.161 | 0.019 | 2.67951841972099e-2 | 0 | wscd2 |
| ablim1b | 2.9425854 | 0.543825621 | 0.269 | 0.06 | 5.94313985252734e-2 | 0 | ablim1b |
| sparc | 6.3040159 | 0.649282964 | 0.736 | 0.373 | 1.2732221096994e-21 | 0 | sparc |
| cebpd | 6.6066977 | 0.689644358 | 0.565 | 0.255 | 1.33435475307531e-2 | 0 | cebpd |
| abhd15a | 7.8199658 | 0.521902643 | 0.254 | 0.054 | 1.57939850187834e-2 | 0 | abhd15a |
| abca1a | 3.8484650 | 0.536631847 | 0.285 | 0.07 | 7.77274493536838e-2 | 0 | abca1a |
| cebpa | 1.2517035 | 0.644043669 | 0.43 | 0.152 | 2.52806557859676e-2 | 0 | cebpa |
| cxcl8b.1 | 2.3193274 | 0.692174164 | 0.337 | 0.094 | 4.68434573283114e-2 | 0 | cxcl8b.1 |
| slit2 | 2.6325837 | 0.347044729 | 0.15 | 0.019 | 5.31702938887338e-2 | 0 | slit2 |
| adamts15a | 4.2631706 | 0.548600586 | 0.326 | 0.093 | 8.61032585379807e-2 | 0 | adamts15a |
| kank2 | 6.7340849 | 0.480421258 | 0.238 | 0.051 | 1.36008312915396e-1 | 0 | kank2 |
| ptges | 1.0183590 | 0.397053579 | 0.161 | 0.022 | 2.05677975195873e-1 | 0 | ptges |
| col5a2a | 1.6172590 | 0.611039315 | 0.43 | 0.161 | 3.26637811827032e-1 | 0 | col5a2a |
| cdc42ep1b | 1.6736063 | 0.289207711 | 0.135 | 0.015 | 3.38018265700288e-1 | 0 | cdc42ep1b |
| si:ch1073-459j12.1 | 8.3741053 | 0.611011558 | 0.44 | 0.163 | 1.69131804999784e-1 | 0 | si:ch1073-459j12.1 |
| anxa1a | 3.2756826 | 0.627883073 | 0.575 | 0.253 | 6.61589621474836e-1 | 0 | anxa1a |
| kita | 4.1119167 | 0.336122321 | 0.135 | 0.016 | 8.30483820369744e-1 | 0 | kita |
| fbxo34 | 4.4953793 | 0.412937534 | 0.187 | 0.034 | 9.07931759433726e-1 | 0 | fbxo34 |
| c6 | 5.3528939 | 0.681038633 | 0.461 | 0.194 | 1.08112398871719e-1 | 0 | c6 |
| dhrs3b | 8.7177401 | 0.446937092 | 0.238 | 0.055 | 1.76072196900561e-1 | 0 | dhrs3b |
| cxcl18b | 8.7713860 | 0.617158576 | 0.358 | 0.117 | 1.77155684578322e-1 | 0 | cxcl18b |

| | | | | | | | |
|---|---|---|---|---|---|---|---|
| mfge8b | 9.1762601 | 0.460654253 | 0.181 | 0.033 | 1.85332926566356e-1 | 0 | mfge8b |
| cyp2n13 | 9.8373057 | 0.471434649 | 0.223 | 0.049 | 1.98684064036227e-1 | 0 | cyp2n13 |
| cyp2ad3 | 1.0682389 | 0.559269706 | 0.321 | 0.099 | 2.15752225055266e-1 | 0 | cyp2ad3 |
| NC-002333.4 | 1.7622386 | 0.555493783 | 0.876 | 0.561 | 3.55919342986394e-1 | 0 | NC-002333.4 |
| lxn | 1.9544016 | 0.642925363 | 0.508 | 0.247 | 3.94730497348067e-1 | 0 | lxn |
| agtr2 | 3.4380450 | 0.37528713 | 0.181 | 0.033 | 6.94381966655751e-1 | 0 | agtr2 |
| htra1b | 4.1764323 | 0.359142657 | 0.145 | 0.021 | 8.43514045293392e-1 | 0 | htra1b |
| id1 | 4.8958879 | 0.666702974 | 0.549 | 0.251 | 9.88822491846988e-1 | 0 | id1 |
| cyp2p7 | 9.9573793 | 0.406556171 | 0.212 | 0.046 | 2.01109191714738e-1 | 0 | cyp2p7 |
| tril | 1.1759121 | 0.405613337 | 0.197 | 0.04 | 2.37498985596738e-1 | 0 | tril |
| CABZ01039096.1 | 1.9657030 | 0.341535927 | 0.155 | 0.025 | 3.97013049289578e-1 | 0 | CABZ01039096.1 |
| RAMP1 | 3.5658918 | 0.305859153 | 0.124 | 0.015 | 7.20203171248593e-1 | 0 | RAMP1 |
| enpep | 4.1594667 | 0.293446142 | 0.13 | 0.017 | 8.40087507730752e-1 | 0 | enpep |
| ccdc80 | 6.2425309 | 0.439926446 | 0.233 | 0.058 | 1.26080397904159e-1 | 0 | ccdc80 |
| si:ch211-204a13.2 | 6.5498990 | 0.350607945 | 0.171 | 0.031 | 1.32288311440007e-1 | 0 | si:ch211-204a13.2 |
| ctsla | 7.5701235 | 0.567179335 | 0.565 | 0.288 | 1.52893786292011e-1 | 0 | ctsla |
| mmp14a | 1.0752637 | 0.441360295 | 0.285 | 0.085 | 2.17171018263926e-1 | 0 | mmp14a |
| col6a1 | 1.2440893 | 0.526661685 | 0.358 | 0.127 | 2.51268732685968e-1 | 0 | col6a1 |
| ramp2 | 1.3140127 | 0.515125849 | 0.207 | 0.047 | 2.65391160545237e-1 | 0 | ramp2 |
| cxcl12a | 1.4810198 | 0.520289507 | 0.28 | 0.08 | 2.99121586584924e-1 | 0 | cxcl12a |
| si:ch211-251b21.1 | 1.6624944 | 0.349951692 | 0.145 | 0.022 | 3.35773997320423e-1 | 0 | si:ch211-251b21.1 |
| im:7151449 | 2.2735941 | 0.381749507 | 0.155 | 0.027 | 4.59197808002947e-1 | 0 | im:7151449 |
| efemp2a | 4.7774911 | 0.325655111 | 0.166 | 0.03 | 9.6490988033991e-15 | 0 | efemp2a |
| c1qtnf2 | 5.6712122 | 0.341745702 | 0.176 | 0.035 | 1.14541473473631e-1 | 0 | c1qtnf2 |
| angptl1a | 6.9996467 | 0.456573051 | 0.238 | 0.061 | 1.41371865200589e-1 | 0 | angptl1a |
| cyp1c1 | 7.1886886 | 0.370784101 | 0.15 | 0.026 | 1.45189944206176e-1 | 0 | cyp1c1 |
| iqsec1b | 9.1414632 | 0.36655804 | 0.166 | 0.031 | 1.84630133895399e-1 | 0 | iqsec1b |
| TCIM (1 of many) | 1.6380199 | 0.328820029 | 0.124 | 0.017 | 3.30830883310541e-1 | 0 | TCIM (1 of many) |
| fgfr1a | 1.8968355 | 0.304463981 | 0.135 | 0.021 | 3.83103879132022e-1 | 0 | fgfr1a |
| AL954191.1 | 2.5483747 | 0.398045254 | 0.14 | 0.023 | 5.14695248647876e-1 | 0 | AL954191.1 |
| sostdc1a | 2.6080260 | 0.583228608 | 0.264 | 0.078 | 5.26743020822177e-1 | 0 | sostdc1a |
| loxa | 5.0058411 | 0.380877351 | 0.192 | 0.043 | 1.01102974033539e-1 | 0 | loxa |
| vim | 8.3036884 | 0.470988029 | 0.285 | 0.09 | 1.67709594795587e-1 | 0 | vim |
| cd82b | 9.2469215 | 0.415106228 | 0.254 | 0.074 | 1.86760073584381e-1 | 0 | cd82b |
| fzd7b | 9.8980269 | 0.48714173 | 0.28 | 0.091 | 1.99910450556818e-1 | 0 | fzd7b |

| | | | | | | | |
|---|---|---|---|---|---|---|---|
| dhrs3a | 1.8543649 | 0.43167948 | 0.238 | 0.067 | 3.74526085885071e-1 | 0 | dhrs3a |
| tmem176 | 6.1887283 | 0.46251341 | 0.358 | 0.138 | 1.24993747401514e-1 | 0 | tmem176 |
| tgfb1a | 6.4933449 | 0.318022123 | 0.166 | 0.035 | 1.3114608893793e-12 | 0 | tgfb1a |
| ppap2d | 6.9122020 | 0.326431279 | 0.15 | 0.029 | 1.3960574446661e-12 | 0 | ppap2d |
| si:ch211-198c19.3 | 7.5783638 | 0.590899052 | 0.404 | 0.166 | 1.53060215174607e-1 | 0 | si:ch211-198c19.3 |
| kitlga | 9.5521487 | 0.292720435 | 0.135 | 0.023 | 1.92924747550482e-1 | 0 | kitlga |
| lhfpl6 | 1.1745093 | 0.324212326 | 0.155 | 0.031 | 2.37215658757117e-1 | 0 | lhfpl6 |
| tent5c | 1.4749817 | 0.297766211 | 0.135 | 0.023 | 2.97902064771951e-1 | 0 | tent5c |
| antxr2a | 1.4907353 | 0.532317552 | 0.482 | 0.245 | 3.01083822960143e-1 | 0 | antxr2a |
| fgfr4 | 2.1084390 | 0.301679058 | 0.135 | 0.023 | 4.2584142884221e-12 | 0 | fgfr4 |
| fkbp1ab | 2.9639556 | 0.348294539 | 0.161 | 0.033 | 5.98630124269246e-1 | 0 | fkbp1ab |
| si:dkey-26m3.3 | 3.2945383 | 0.268241144 | 0.145 | 0.028 | 6.65397916552006e-1 | 0 | si:dkey-26m3.3 |
| si:ch73-46j18.5 | 5.1412340 | 0.502891898 | 0.674 | 0.373 | 1.03837504251471e-1 | 0 | si:ch73-46j18.5 |
| pdgfra | 6.1684188 | 0.475619063 | 0.316 | 0.118 | 1.24583554981533e-1 | 0 | pdgfra |
| LO018188.1 | 6.5777084 | 0.430370833 | 0.254 | 0.079 | 1.3284997836506e-11 | 0 | LO018188.1 |
| bag2 | 1.2180049 | 0.299110335 | 0.135 | 0.025 | 2.46000468375978e-1 | 0 | bag2 |
| c7a | 1.5638855 | 0.363129652 | 0.171 | 0.04 | 3.15857965926639e-1 | 0 | c7a |
| sc:d0202 | 1.9243070 | 0.347953978 | 0.15 | 0.031 | 3.88652293308039e-1 | 0 | sc:d0202 |
| slc7a1 | 2.0832781 | 0.286134551 | 0.14 | 0.027 | 4.20759683942079e-1 | 0 | slc7a1 |
| tead1b | 2.1203437 | 0.390460246 | 0.192 | 0.051 | 4.2824583445187e-11 | 0 | tead1b |
| LOC103909284 | 2.2008543 | 0.470217832 | 0.326 | 0.128 | 4.44506560759306e-1 | 0 | LOC103909284 |
| klf6a | 3.3633232 | 0.52158149 | 0.466 | 0.243 | 6.7929039049941e-11 | 0 | klf6a |
| ca5a | 3.6830476 | 0.269899213 | 0.135 | 0.026 | 7.43865137032778e-1 | 0 | ca5a |
| meis1a | 4.2610427 | 0.320306765 | 0.171 | 0.04 | 8.6060280979516e-11 | 0 | meis1a |
| gpc6a | 6.7488611 | 0.264495144 | 0.124 | 0.022 | 1.36306749215687e-1 | 0 | gpc6a |
| klf3 | 7.5404886 | 0.344682804 | 0.187 | 0.049 | 1.52295250047898e-1 | 0 | klf3 |
| dock9b | 7.9178628 | 0.275266499 | 0.104 | 0.015 | 1.59917075822635e-1 | 0 | dock9b |
| eva1ba | 8.4893773 | 0.375231306 | 0.223 | 0.067 | 1.71459954048812e-1 | 0 | eva1ba |
| gpc1b | 1.4574431 | 0.335289421 | 0.155 | 0.035 | 2.94359786869701e-1 | 0 | gpc1b |
| dpt | 1.6597264 | 0.303922683 | 0.119 | 0.021 | 3.35214957635653e-1 | 0 | dpt |
| egfra | 1.9294202 | 0.337860303 | 0.171 | 0.043 | 3.89685010990732e-1 | 0 | egfra |
| itga5 | 2.8141764 | 0.307857743 | 0.145 | 0.032 | 5.68379224901297e-1 | 0 | itga5 |
| si:ch1073-291c23.2 | 5.6371380 | 0.42817515 | 0.311 | 0.124 | 1.13853276917259e-0 | 0 | si:ch1073-291c23.2 |
| bicc1b | 5.7638511 | 0.32300339 | 0.15 | 0.035 | 1.16412502047178e-0 | 0 | bicc1b |
| col4a1 | 1.2392315 | 0.445240173 | 0.311 | 0.123 | 2.50287593673329e-0 | 0 | col4a1 |

| | | | | | | | |
|---|---|---|---|---|---|---|---|
| hoxb5b | 1.6872582 | 0.25064127 | 0.104 | 0.017 | 3.40775556275627e-0 | 0 | hoxb5b |
| june | 1.8304632 | 0.251618777 | 0.109 | 0.019 | 3.69698661902552e-0 | 0 | june |
| cd81a | 2.9618398 | 0.457346432 | 0.679 | 0.422 | 5.98202802418126e-0 | 0 | cd81a |
| tns1b | 3.2507124 | 0.306253233 | 0.155 | 0.038 | 6.56546388527292e-0 | 0 | tns1b |
| si:ch211-133n4.4 | 4.3369153 | 0.430225436 | 0.295 | 0.121 | 8.75926785948668e-0 | 0 | si:ch211-133n4.4 |
| s100v2 | 6.5342859 | 0.446157486 | 0.456 | 0.234 | 1.3197297390079e-08 | 0 | s100v2 |
| hic1 | 7.0193250 | 0.447179371 | 0.295 | 0.123 | 1.41769307997073e-0 | 0 | hic1 |
| cdon | 7.1771238 | 0.446797722 | 0.311 | 0.133 | 1.44956369592858e-0 | 0 | cdon |
| mmp14b | 7.3195827 | 0.442231297 | 0.269 | 0.105 | 1.47833613686986e-0 | 0 | mmp14b |
| rbpjb | 9.3448760 | 0.381241725 | 0.223 | 0.077 | 1.8873846186627e-08 | 0 | rbpjb |
| lamb1a | 1.1502635 | 0.425881605 | 0.301 | 0.127 | 2.32318726319011e-0 | 0 | lamb1a |
| cd28l | 1.5328360 | 0.307564906 | 0.161 | 0.043 | 3.09586895882334e-0 | 0 | cd28l |
| capn2a | 3.0591961 | 0.336317079 | 0.181 | 0.054 | 6.17865842421237e-0 | 0 | capn2a |
| pdgfrb | 3.1106477 | 0.338972056 | 0.181 | 0.054 | 6.28257527495327e-0 | 0 | pdgfrb |
| boc | 3.2790252 | 0.412214273 | 0.342 | 0.154 | 6.6226473345228e-08 | 0 | boc |
| si:ch211-39i22.1 | 3.3773935 | 0.509706537 | 0.352 | 0.17 | 6.82132167735808e-0 | 0 | si:ch211-39i22.1 |
| foxp4 | 3.6187457 | 0.455782071 | 0.363 | 0.177 | 7.30878087475906e-0 | 0 | foxp4 |
| zeb1b | 4.1945207 | 0.327845219 | 0.155 | 0.042 | 8.47167355208541e-0 | 0 | zeb1b |
| asic1b | 4.2466165 | 0.274376413 | 0.13 | 0.03 | 8.57689143681776e-0 | 0 | asic1b |
| lpar1 | 5.0909074 | 0.365109917 | 0.249 | 0.092 | 1.02821058000368e-0 | 0 | lpar1 |
| tgm1l1 | 5.1227291 | 0.297994715 | 0.145 | 0.037 | 1.0346376087541e-07 | 0 | tgm1l1 |
| CABZ01072614.1 | 5.6088570 | 0.467272857 | 0.342 | 0.162 | 1.13282085425893e-0 | 0 | CABZ01072614.1 |
| rpz5 | 1.0838716 | 0.416107495 | 0.461 | 0.24 | 2.18909549788825e-0 | 0 | rpz5 |
| mt-atp6 | 1.1749575 | 0.465755876 | 0.648 | 0.405 | 2.3730618166977e-07 | 0 | mt-atp6 |
| ank3b | 1.5190291 | 0.270377342 | 0.14 | 0.036 | 3.0679832157588e-07 | 0 | ank3b |
| bmp1a | 1.8430002 | 0.25640102 | 0.13 | 0.031 | 3.72230757288326e-0 | 0 | bmp1a |
| mt-nd2 | 1.9948559 | 0.421205103 | 0.596 | 0.356 | 4.02901055652551e-0 | 0 | mt-nd2 |
| rnd3b | 2.3723825 | 0.349990762 | 0.233 | 0.087 | 4.79150102834315e-0 | 0 | rnd3b |
| itgb1a | 2.8447500 | 0.334570048 | 0.207 | 0.072 | 5.74554166605473e-0 | 0 | itgb1a |
| atoh8 | 3.5394165 | 0.311330037 | 0.171 | 0.052 | 7.14855966725138e-0 | 0 | atoh8 |
| pitx2 | 5.3283751 | 0.310172031 | 0.135 | 0.035 | 1.07617193311658e-0 | 0 | pitx2 |
| sdc2 | 6.0356391 | 0.3685134 | 0.269 | 0.114 | 1.21901802950587e-0 | 0 | sdc2 |
| ghra | 8.1431644 | 0.315306874 | 0.155 | 0.046 | 1.64467491838536e-0 | 0 | ghra |
| junba | 8.6756922 | 0.444020084 | 0.58 | 0.391 | 1.75222955894158e-0 | 0 | junba |
| dedd1 | 1.0158667 | 0.458846327 | 0.368 | 0.201 | 2.05174604348166e-0 | 0 | dedd1 |

| | | | | | | | |
|---|---|---|---|---|---|---|---|
| mt-cyb | 1.0364724 | 0.433392146 | 0.606 | 0.359 | 2.09336331114906e-0 | 0 | mt-cyb |
| rbms3 | 1.1689471 | 0.285664851 | 0.15 | 0.044 | 2.36092263588614e-0 | 0 | rbms3 |
| si:dkey-238c7.16 | 1.3783621 | 0.343150099 | 0.259 | 0.106 | 2.78387794720989e-0 | 0 | si:dkey-238c7.16 |
| mt-co2 | 1.4982295 | 0.449821902 | 0.689 | 0.442 | 3.02597422387382e-0 | 0 | mt-co2 |
| stom | 1.5365766 | 0.260818499 | 0.124 | 0.031 | 3.10342395592225e-0 | 0 | stom |
| COX3 | 1.5460031 | 0.387007795 | 0.808 | 0.579 | 3.12246250367799e-0 | 0 | COX3 |
| arrdc3a | 2.0259875 | 0.285442591 | 0.181 | 0.06 | 4.09188705726563e-0 | 0 | arrdc3a |
| znf704 | 2.4292807 | 0.294007999 | 0.15 | 0.045 | 4.90641826347648e-0 | 0 | znf704 |
| lama4 | 2.6327038 | 0.36300775 | 0.228 | 0.091 | 5.3172718741016e-06 | 0 | lama4 |
| zfp36l1b | 2.9151401 | 0.412592437 | 0.363 | 0.188 | 5.88770858504577e-0 | 0 | zfp36l1b |
| rnd3a | 3.1896822 | 0.306938438 | 0.161 | 0.051 | 6.44220126710082e-0 | 0 | rnd3a |
| fxyd6l | 3.8335917 | 0.377284068 | 0.295 | 0.137 | 7.74270521358555e-0 | 0 | fxyd6l |
| eif4ebp3l | 3.9498778 | 0.42021706 | 0.363 | 0.189 | 7.97756833762113e-0 | 0 | eif4ebp3l |
| sik2b | 3.9674490 | 0.349368616 | 0.181 | 0.064 | 8.0130567680971e-06 | 0 | sik2b |
| ahnak | 4.4520598 | 0.392592012 | 0.363 | 0.189 | 8.99182525021865e-0 | 0 | ahnak |
| pabpc1b | 5.6021851 | 0.382107263 | 0.259 | 0.114 | 1.13147333725397e-0 | 0 | pabpc1b |
| jun | 7.1795240 | 0.412671735 | 0.591 | 0.369 | 1.45004847632032e-0 | 0 | jun |
| cadm4 | 1.1668890 | 0.260804379 | 0.15 | 0.047 | 2.35676573673173e-0 | 0 | cadm4 |
| tenm3 | 1.2061926 | 0.402811514 | 0.285 | 0.136 | 2.43614727855567e-0 | 0 | tenm3 |
| rgcc | 1.2385074 | 0.311130037 | 0.192 | 0.069 | 2.50141344666751e-0 | 0 | rgcc |
| fzd2 | 1.3467914 | 0.358744776 | 0.244 | 0.108 | 2.72011464935964e-0 | 0 | fzd2 |
| anxa2a | 1.8568730 | 0.405405622 | 0.352 | 0.175 | 3.7503264385456e-05 | 0 | anxa2a |
| col6a4a | 2.3682129 | 0.35703325 | 0.326 | 0.158 | 4.78307960366425e-0 | 0 | col6a4a |
| plekho2 | 2.4915249 | 0.347912763 | 0.212 | 0.086 | 5.03213285551649e-0 | 0 | plekho2 |
| slc38a5b | 2.6948488 | 0.414677417 | 0.347 | 0.195 | 5.44278624615497e-0 | 0 | slc38a5b |
| hoxc8a | 4.9912247 | 0.277636815 | 0.171 | 0.061 | 0.000100808 | 0 | hoxc8a |
| tnfaip6 | 6.8365982 | 0.266105243 | 0.119 | 0.033 | 0.000138079 | 0 | tnfaip6 |
| c1qtnf5 | 7.0500034 | 0.294198879 | 0.197 | 0.075 | 0.000142389 | 0 | c1qtnf5 |
| meis2b | 7.5959435 | 0.319630233 | 0.212 | 0.088 | 0.000153415 | 0 | meis2b |
| hspb1 | 8.3782955 | 0.408655249 | 0.321 | 0.169 | 0.000169216 | 0 | hspb1 |
| glipr2l | 9.3280546 | 0.28129709 | 0.15 | 0.051 | 0.000188399 | 0 | glipr2l |
| si:ch211-106h4.12 | 1.0787402 | 0.34028203 | 0.155 | 0.053 | 0.000217873 | 0 | si:ch211-106h4.12 |
| nfil3-6 | 1.6060868 | 0.347480337 | 0.275 | 0.133 | 0.000324381 | 0 | nfil3-6 |
| grb10a | 1.9479893 | 0.262471344 | 0.161 | 0.057 | 0.000393435 | 0 | grb10a |
| pbx1a | 2.0610623 | 0.359071113 | 0.233 | 0.107 | 0.000416273 | 0 | pbx1a |

| | | | | | | | |
|---|---|---|---|---|---|---|---|
| rbm5 | 2.3704111 | 0.328290989 | 0.218 | 0.096 | 0.000478752 | 0 | rbm5 |
| meis1b | 2.3907994 | 0.350287912 | 0.29 | 0.148 | 0.00048287 | 0 | meis1b |
| ddit3 | 2.6821875 | 0.407158018 | 0.337 | 0.193 | 0.000541721 | 0 | ddit3 |
| proza | 3.0135683 | 0.2574839 | 0.155 | 0.056 | 0.00060865 | 0 | proza |
| mt-nd3 | 3.0873816 | 0.330124729 | 0.446 | 0.259 | 0.000623558 | 0 | mt-nd3 |
| tmem176l.1 | 3.2136431 | 0.358619008 | 0.487 | 0.314 | 0.00064906 | 0 | tmem176l.1 |
| dlc1 | 3.5132979 | 0.251929099 | 0.119 | 0.036 | 0.000709581 | 0 | dlc1 |
| olfml3b | 3.5161941 | 0.318049457 | 0.228 | 0.103 | 0.000710166 | 0 | olfml3b |
| twist1a | 3.6690420 | 0.25501355 | 0.119 | 0.036 | 0.000741036 | 0 | twist1a |
| grb10b | 4.4295162 | 0.263541371 | 0.15 | 0.054 | 0.000894629 | 0 | grb10b |
| cast | 4.7180488 | 0.278926734 | 0.202 | 0.085 | 0.000952904 | 0 | cast |
| plod2 | 6.1217655 | 0.260069081 | 0.15 | 0.054 | 0.001236413 | 0 | plod2 |
| mt-nd1 | 6.6816521 | 0.358167244 | 0.528 | 0.342 | 0.001349493 | 0 | mt-nd1 |
| purab | 9.7091514 | 0.327640729 | 0.223 | 0.105 | 0.001960957 | 0 | purab |
| rarga | 1.0524536 | 0.328852951 | 0.187 | 0.08 | 0.002125641 | 0 | rarga |
| ip6k2b | 1.0586211 | 0.368038857 | 0.29 | 0.159 | 0.002138097 | 0 | ip6k2b |
| ccdc3b | 1.8853012 | 0.265750544 | 0.155 | 0.06 | 0.003807743 | 0 | ccdc3b |
| si:ch211-212k18.8 | 2.4663424 | 0.268921619 | 0.171 | 0.071 | 0.004981272 | 0 | si:ch211-212k18.8 |
| si:dkey-7j14.6 | 2.7247603 | 0.341278049 | 0.461 | 0.313 | 0.005503198 | 0 | si:dkey-7j14.6 |
| spry4 | 3.9415030 | 0.257342452 | 0.15 | 0.058 | 0.007960654 | 0 | spry4 |
| chd3 | 4.3278554 | 0.338031301 | 0.373 | 0.233 | 0.00874097 | 0 | chd3 |
| pmp22b | 4.3791397 | 0.25883849 | 0.14 | 0.052 | 0.008844549 | 0 | pmp22b |
| yap1 | 4.5168576 | 0.28389925 | 0.197 | 0.09 | 0.009122697 | 0 | yap1 |
| gng7 | 4.5869523 | 0.285815179 | 0.212 | 0.101 | 0.009264268 | 0 | gng7 |
| sptan1 | 4.6655151 | 0.281143394 | 0.197 | 0.09 | 0.009422941 | 0 | sptan1 |
| mt-nd4 | 4.9350839 | 0.323239671 | 0.606 | 0.426 | 0.009967389 | 0 | mt-nd4 |
| aoc2 | 5.5197216 | 0.337208188 | 0.249 | 0.126 | 0.011148182 | 0 | aoc2 |
| si:ch211-212k18.9 | 5.8011095 | 0.298033969 | 0.223 | 0.111 | 0.011716501 | 0 | si:ch211-212k18.9 |
| zfp36l1a | 7.9172303 | 0.347142384 | 0.44 | 0.303 | 0.01599043 | 0 | zfp36l1a |
| zgc:153372 | 8.0942372 | 0.269915887 | 0.135 | 0.051 | 0.016347931 | 0 | zgc:153372 |
| ackr3b | 1.0736351 | 0.265578813 | 0.166 | 0.07 | 0.021684208 | 0 | ackr3b |
| nrp1a | 1.4297302 | 0.29281628 | 0.218 | 0.106 | 0.028876262 | 0 | nrp1a |
| flna | 1.4804162 | 0.318736147 | 0.285 | 0.161 | 0.029899968 | 0 | flna |
| tuba8l3 | 1.5169952 | 0.273930834 | 0.347 | 0.196 | 0.030638754 | 0 | tuba8l3 |
| tspan4b | 1.5845421 | 0.267898451 | 0.207 | 0.1 | 0.032002998 | 0 | tspan4b |

| | | | | | | | |
|---|---|---|---|---|---|---|---|
| myl6 | 2.9589990 | 0.323918679 | 0.326 | 0.196 | 0.059762903 | 0 | myl6 |
| ckap4 | 3.6202747 | 0.301258021 | 0.254 | 0.142 | 0.073118689 | 0 | ckap4 |
| BX284696.1 | 4.0716392 | 0.308833927 | 0.259 | 0.151 | 0.082234898 | 0 | BX284696.1 |
| socs3a | 4.5310024 | 0.327131166 | 0.56 | 0.422 | 0.091512657 | 0 | socs3a |
| cald1a | 4.5347655 | 0.333528029 | 0.352 | 0.231 | 0.091588659 | 0 | cald1a |
| gstt1b | 5.1569374 | 0.285867621 | 0.295 | 0.175 | 0.104154666 | 0 | gstt1b |
| wu:fb16f03 | 6.1565968 | 0.264955319 | 0.202 | 0.103 | 0.124344787 | 0 | wu:fb16f03 |
| b2ml | 7.3617323 | 0.276722353 | 0.549 | 0.441 | 0.148684908 | 0 | b2ml |
| zgc:158463 | 7.4577453 | 0.2783923 | 0.218 | 0.112 | 0.150624083 | 0 | zgc:158463 |
| klf13 | 7.5994108 | 0.278458895 | 0.187 | 0.093 | 0.153485302 | 0 | klf13 |
| cnn3a | 8.2111507 | 0.26330735 | 0.238 | 0.13 | 0.165840612 | 0 | cnn3a |
| mgst1.1 | 9.4758410 | 0.266733337 | 0.228 | 0.125 | 0.191383562 | 0 | mgst1.1 |
| vat1 | 1.2577347 | 0.305923716 | 0.342 | 0.218 | 0.25402468 | 0 | vat1 |
| cyp3c1 | 2.1781059 | 0.251450407 | 0.171 | 0.085 | 0.439912067 | 0 | cyp3c1 |
| rbfox2 | 2.3045596 | 0.276838732 | 0.15 | 0.071 | 0.465451903 | 0 | rbfox2 |
| fosab | 2.3146149 | 0.310144182 | 0.56 | 0.456 | 0.467482789 | 0 | fosab |
| cldn5a | 2.5963615 | 0.286496545 | 0.223 | 0.121 | 0.524387138 | 0 | cldn5a |
| mt-co1 | 2.6189647 | 0.255878045 | 0.725 | 0.584 | 0.528952315 | 0 | mt-co1 |
| mmp16a | 3.6796211 | 0.268663018 | 0.218 | 0.119 | 0.743173092 | 0 | mmp16a |
| serinc5 | 4.2693980 | 0.256699971 | 0.202 | 0.111 | 0.862290327 | 0 | serinc5 |
| slc43a2a | 4.7545163 | 0.286117791 | 0.187 | 0.1 | 0.960269665 | 0 | slc43a2a |
| jpt1b | 5.4413401 | 0.261742487 | 0.451 | 0.327 | 1 | 0 | jpt1b |
| socs3b | 5.4430348 | 0.276645518 | 0.238 | 0.141 | 1 | 0 | socs3b |
| hdlbpa | 5.6454104 | 0.291003034 | 0.373 | 0.262 | 1 | 0 | hdlbpa |
| aldh9a1a.1 | 0.00011 | 0.288157831 | 0.43 | 0.335 | 1 | 0 | aldh9a1a.1 |
| celf2 | 0.000272 | 0.267315014 | 0.44 | 0.354 | 1 | 0 | celf2 |
| dnmt3ab | 0.000313 | 0.263659392 | 0.249 | 0.164 | 1 | 0 | dnmt3ab |
| ncor2 | 0.00032 | 0.252203038 | 0.197 | 0.119 | 1 | 0 | ncor2 |
| mafba | 0.000466 | 0.25921917 | 0.228 | 0.147 | 1 | 0 | mafba |
| adam10a | 0.000542 | 0.263161992 | 0.249 | 0.166 | 1 | 0 | adam10a |
| tspan14 | 0.000551 | 0.266338965 | 0.254 | 0.174 | 1 | 0 | tspan14 |
| sdc4 | 0.000588 | 0.250135389 | 0.347 | 0.261 | 1 | 0 | sdc4 |
| arl6ip5b | 0.001044 | 0.254375728 | 0.212 | 0.139 | 1 | 0 | arl6ip5b |
| gadd45ba | 0.001058 | 0.261413923 | 0.337 | 0.248 | 1 | 0 | gadd45ba |
| inhbaa | 3.3585063 | 1.430637634 | 0.83 | 0.056 | 6.78317522072511e-2 | 1 | inhbaa |

| | | | | | | | |
|---|---|---|---|---|---|---|---|
| nid2a | 2.3612289 | 1.320235484 | 0.813 | 0.057 | 4.76897412617855e-1 | 1 | nid2a |
| osr2 | 1.2473880 | 1.111420382 | 0.654 | 0.036 | 2.51934960004007e-1 | 1 | osr2 |
| rbp41 | 4.3431019 | 1.575396299 | 0.962 | 0.154 | 8.77176306774264e-1 | 1 | rbp4 |
| pdgfra1 | 3.2310767 | 1.056263697 | 0.742 | 0.078 | 6.52580563206426e-1 | 1 | pdgfra |
| nr2f1a | 1.5041147 | 1.09351316 | 0.747 | 0.099 | 3.03786046102e-114 | 1 | nr2f1a |
| lrrtm1 | 4.3230201 | 0.740022526 | 0.335 | 0.006 | 8.73120388277408e-1 | 1 | lrrtm1 |
| ndnf | 4.7860417 | 0.860424859 | 0.489 | 0.032 | 9.66636848045142e-1 | 1 | ndnf |
| robo2 | 2.3579399 | 0.884621781 | 0.571 | 0.052 | 4.76233140609926e-1 | 1 | robo2 |
| col18a1a | 1.3041505 | 0.935679227 | 0.555 | 0.05 | 2.63399282063747e-1 | 1 | col18a1a |
| hsd11b21 | 1.4825768 | 1.318064983 | 0.929 | 0.236 | 2.99436054442255e-9 | 1 | hsd11b2 |
| nid1b | 1.5303678 | 0.999205569 | 0.676 | 0.095 | 3.09088395423044e-9 | 1 | nid1b |
| c61 | 2.3465436 | 1.107286287 | 0.863 | 0.157 | 4.73931416606838e-9 | 1 | c6 |
| cd248a | 4.4171231 | 0.910110721 | 0.626 | 0.078 | 8.92126355879748e-9 | 1 | cd248a |
| dio2 | 5.3497505 | 0.643956981 | 0.374 | 0.019 | 1.08048911241685e-9 | 1 | dio2 |
| col6a4a1 | 4.3983042 | 1.018085797 | 0.725 | 0.121 | 8.88325504256788e-8 | 1 | col6a4a |
| nme2b.2 | 1.5302803 | 0.77347676 | 0.418 | 0.037 | 3.09070727632495e-7 | 1 | nme2b.2 |
| dkk3b | 3.9480565 | 0.85577076 | 0.703 | 0.119 | 7.97388983495131e-7 | 1 | dkk3b |
| caln2 | 1.2548872 | 0.479182237 | 0.258 | 0.007 | 2.534495834691e-77 | 1 | caln2 |
| tpbga | 1.1494537 | 0.450644143 | 0.253 | 0.007 | 2.32155171850283e-7 | 1 | tpbga |
| nlgn4xa | 3.5825065 | 0.431430088 | 0.22 | 0.004 | 7.23558841440659e-7 | 1 | nlgn4xa |
| mxra8a | 1.0210814 | 0.864260634 | 0.615 | 0.107 | 2.06227813370348e-7 | 1 | mxra8a |
| mdka1 | 7.6228361 | 1.021921445 | 0.78 | 0.174 | 1.5395842214253e-71 | 1 | mdka |
| cldn5a1 | 4.1453589 | 0.838140016 | 0.577 | 0.087 | 8.37238145297254e-7 | 1 | cldn5a |
| inka1a | 5.4848046 | 0.693014322 | 0.445 | 0.049 | 1.1077659882089e-70 | 1 | inka1a |
| postnb | 3.3334475 | 0.90181285 | 0.747 | 0.14 | 6.73256397005499e-6 | 1 | postnb |
| si:ch1073-291c23.2 | 3.6536857 | 0.79305704 | 0.593 | 0.098 | 7.3793490579687e-69 | 1 | si:ch1073-291c23.2 |
| cspg5a | 1.5988889 | 0.43670883 | 0.236 | 0.007 | 3.22927594650374e-6 | 1 | cspg5a |
| cd82a | 9.4832956 | 0.768295735 | 0.599 | 0.1 | 1.91534122076505e-6 | 1 | cd82a |
| fn1b | 1.0623023 | 0.903192825 | 0.692 | 0.135 | 2.14553211910828e-6 | 1 | fn1b |
| col27a1b | 2.7592546 | 0.669644588 | 0.407 | 0.044 | 5.57286651670038e-6 | 1 | col27a1b |
| atp1a3b | 4.1986523 | 0.577405648 | 0.346 | 0.029 | 8.48001810897025e-6 | 1 | atp1a3b |
| si:ch211-197h24.9 | 7.5997111 | 0.848625086 | 0.615 | 0.124 | 1.53491366275122e-6 | 1 | si:ch211-197h24.9 |
| vegfaa | 3.2238664 | 0.497813893 | 0.286 | 0.017 | 6.51124303655631e-6 | 1 | vegfaa |
| nrxn1a | 6.2002348 | 0.43067598 | 0.209 | 0.006 | 1.25226143715468e-6 | 1 | nrxn1a |
| slco2b1 | 2.0740487 | 0.68070702 | 0.418 | 0.051 | 4.1889562579522e-61 | 1 | slco2b1 |

| | | | | | | | |
|---|---|---|---|---|---|---|---|
| esr2b | 9.4873242 | 0.533083424 | 0.352 | 0.033 | 1.91615487623586e-6 | 1 | esr2b |
| CABZ01092746.11 | 3.7721136 | 0.876107031 | 0.725 | 0.164 | 7.61853793022403e-5 | 1 | CABZ01092746.1 |
| pcdh18b | 1.1384282 | 0.608439727 | 0.385 | 0.045 | 2.29928354724364e-5 | 1 | pcdh18b |
| zgc:1951731 | 1.4101001 | 0.84468973 | 0.758 | 0.191 | 2.84797920900969e-5 | 1 | zgc:195173 |
| lrrc171 | 1.1820248 | 0.772556673 | 0.659 | 0.14 | 2.38733561329275e-5 | 1 | lrrc17 |
| nrp1a1 | 1.3651150 | 0.695842221 | 0.495 | 0.08 | 2.75712278492859e-5 | 1 | nrp1a |
| pmp22a1 | 6.1128074 | 0.930571975 | 0.907 | 0.27 | 1.23460372123221e-5 | 1 | pmp22a |
| BX569787.1 | 1.0274320 | 0.673736366 | 0.467 | 0.074 | 2.07510459000999e-5 | 1 | BX569787.1 |
| zgc:158343 | 5.6973458 | 0.877318325 | 0.654 | 0.164 | 1.15069294982815e-5 | 1 | zgc:158343 |
| cd81a1 | 3.2985684 | 0.896691676 | 0.934 | 0.399 | 6.66211874551281e-5 | 1 | cd81a |
| si:dkey-9i23.16 | 6.3272404 | 0.626460629 | 0.451 | 0.071 | 1.2779127518729e-52 | 1 | si:dkey-9i23.16 |
| spon1b | 1.8566658 | 0.624446391 | 0.423 | 0.061 | 3.74990804527156e-5 | 1 | spon1b |
| tnfsf10 | 5.9556129 | 0.724056839 | 0.516 | 0.096 | 1.20285514628877e-5 | 1 | tnfsf10 |
| gstm.1 | 1.9103250 | 0.841436163 | 0.89 | 0.291 | 3.85828347556699e-5 | 1 | gstm.1 |
| ar | 3.9637963 | 0.633473262 | 0.467 | 0.079 | 8.00567950310947e-5 | 1 | ar |
| luzp2 | 1.0501998 | 0.417292421 | 0.231 | 0.014 | 2.12108860759043e-5 | 1 | luzp2 |
| grhl1 | 5.9774926 | 0.49345526 | 0.302 | 0.03 | 1.2072741893633e-49 | 1 | grhl1 |
| shisal1b | 3.6886569 | 0.309594452 | 0.176 | 0.006 | 7.44998050852833e-4 | 1 | shisal1b |
| col5a11 | 7.4773810 | 0.782158105 | 0.764 | 0.198 | 1.51020665826339e-4 | 1 | col5a1 |
| dcn1 | 8.0234589 | 0.872661564 | 0.83 | 0.231 | 1.620498001259e-48 | 1 | dcn |
| col1a1a1 | 3.5566957 | 0.84421683 | 0.912 | 0.26 | 7.18345848652234e-4 | 1 | col1a1a |
| si:ch211-106j24.1 | 1.5181950 | 0.431267906 | 0.236 | 0.017 | 3.06629853948967e-4 | 1 | si:ch211-106j24.1 |
| cxcl18b1 | 1.7555524 | 0.878500208 | 0.505 | 0.105 | 3.54568919909167e-4 | 1 | cxcl18b |
| gas1a | 2.8836282 | 0.700745511 | 0.516 | 0.106 | 5.82406392067286e-4 | 1 | gas1a |
| tmem1761 | 4.3746187 | 0.672080301 | 0.56 | 0.12 | 8.83541743623185e-4 | 1 | tmem176 |
| ptprz1b | 2.9822376 | 0.358280516 | 0.192 | 0.01 | 6.02322541288153e-4 | 1 | ptprz1b |
| nr2f2 | 2.0184497 | 0.619923986 | 0.418 | 0.07 | 4.07666303683597e-4 | 1 | nr2f2 |
| prkag2a | 2.1152225 | 0.351351895 | 0.187 | 0.009 | 4.2721150053876e-45 | 1 | prkag2a |
| ccbe1 | 4.5538287 | 0.486203567 | 0.269 | 0.026 | 9.19736782785113e-4 | 1 | ccbe1 |
| si:dkey-7j14.61 | 5.6148260 | 0.78997709 | 0.802 | 0.281 | 1.13402641131978e-4 | 1 | si:dkey-7j14.6 |
| cyp3c11 | 5.5322131 | 0.540756128 | 0.396 | 0.063 | 1.11734109881133e-4 | 1 | cyp3c1 |
| aebp1 | 1.2305410 | 0.565563515 | 0.418 | 0.071 | 2.48532375336642e-4 | 1 | aebp1 |
| myct1a | 3.6038014 | 0.329631595 | 0.225 | 0.017 | 7.27859782447108e-4 | 1 | myct1a |
| tmem119b | 5.3136909 | 0.504245152 | 0.269 | 0.029 | 1.07320615556727e-4 | 1 | tmem119b |
| sema4c | 5.6003711 | 0.349407205 | 0.198 | 0.013 | 1.13110695354879e-4 | 1 | sema4c |

| | | | | | | | |
|---|---|---|---|---|---|---|---|
| tmem176l.11 | 8.7143462 | 0.773026031 | 0.797 | 0.285 | 1.76003651734153e-4 | 1 | tmem176l.1 |
| fosab1 | 2.0068778 | 0.851962895 | 0.885 | 0.425 | 4.05329116961188e-4 | 1 | fosab |
| ece1 | 3.7176755 | 0.549670801 | 0.368 | 0.059 | 7.50858923042076e-4 | 1 | ece1 |
| adrb2a | 3.7285418 | 0.42341595 | 0.242 | 0.023 | 7.53053603713386e-4 | 1 | adrb2a |
| snai2 | 1.0224182 | 0.415064066 | 0.275 | 0.031 | 2.06497804938431e-4 | 1 | snai2 |
| pbx1b | 1.8018522 | 0.474067777 | 0.324 | 0.045 | 3.6392009366368e-40 | 1 | pbx1b |
| ptprga | 3.1535361 | 0.585250644 | 0.423 | 0.08 | 6.3691969992765e-40 | 1 | ptprga |
| srgap1b | 1.4119199 | 0.29933832 | 0.165 | 0.008 | 2.85165482378448e-3 | 1 | srgap1b |
| junba1 | 1.4593676 | 0.810338187 | 0.857 | 0.365 | 2.94748479242662e-3 | 1 | junba |
| si:ch211-195o20.7 | 2.4511396 | 0.428615301 | 0.291 | 0.036 | 4.95056669368541e-3 | 1 | si:ch211-195o20.7 |
| qkia | 4.3372666 | 0.745189177 | 0.808 | 0.321 | 8.75997745793763e-3 | 1 | qkia |
| col1a21 | 5.7618376 | 0.818941592 | 0.929 | 0.281 | 1.16371835855787e-3 | 1 | col1a2 |
| zgc:136930 | 1.0654796 | 0.282602373 | 0.115 | 0.002 | 2.15194920479268e-3 | 1 | zgc:136930 |
| loxl2b | 2.4835538 | 0.492273452 | 0.324 | 0.048 | 5.0160336390972e-38 | 1 | loxl2b |
| si:ch211-286o17.1 | 2.8697457 | 0.580361608 | 0.429 | 0.085 | 5.7960254679799e-38 | 1 | si:ch211-286o17.1 |
| scube1 | 5.1721209 | 0.362520223 | 0.214 | 0.018 | 1.04461326349804e-3 | 1 | scube1 |
| ntn1a | 7.8377975 | 0.309444001 | 0.176 | 0.011 | 1.58299996237298e-3 | 1 | ntn1a |
| pbx1a1 | 1.0470532 | 0.52385716 | 0.44 | 0.088 | 2.11473336224836e-3 | 1 | pbx1a |
| cpn1 | 4.4965175 | 0.679635282 | 0.588 | 0.161 | 9.08161648306004e-3 | 1 | cpn1 |
| gstm.3 | 5.8799556 | 0.774005946 | 0.819 | 0.241 | 1.18757464092274e-3 | 1 | gstm.3 |
| hic11 | 2.0408427 | 0.581824304 | 0.473 | 0.107 | 4.12189003394783e-3 | 1 | hic1 |
| cplx2l | 2.1347115 | 0.336648828 | 0.209 | 0.018 | 4.31147700246698e-3 | 1 | cplx2l |
| itga9 | 3.8787275 | 0.516938269 | 0.33 | 0.053 | 7.83386612628306e-3 | 1 | itga9 |
| cux1a | 4.6559517 | 0.556636639 | 0.401 | 0.084 | 9.40362565758325e-3 | 1 | cux1a |
| jun1 | 6.8734577 | 0.785329532 | 0.819 | 0.348 | 1.38823227142945e-3 | 1 | jun |
| mmp16a1 | 9.3507911 | 0.605699079 | 0.445 | 0.098 | 1.88857928147532e-3 | 1 | mmp16a |
| vstm4a | 1.9093192 | 0.35631271 | 0.231 | 0.025 | 3.85625207156155e-3 | 1 | vstm4a |
| c4 | 2.6802032 | 0.342269128 | 0.181 | 0.014 | 5.41320645929543e-3 | 1 | c4 |
| ch25h | 2.8593931 | 0.388785542 | 0.198 | 0.017 | 5.7751162652537e-34 | 1 | ch25h |
| si:ch211-212k18.9: | 3.4968436 | 0.523591347 | 0.429 | 0.092 | 7.06257506297467e-3 | 1 | si:ch211-212k18.9 |
| tshz3b | 5.9612714 | 0.429801856 | 0.291 | 0.043 | 1.20399800027073e-3 | 1 | tshz3b |
| tmem88b | 1.5723939 | 0.558494571 | 0.478 | 0.111 | 3.17576399675438e-3 | 1 | tmem88b |
| sptlc2a | 1.6108778 | 0.473013479 | 0.335 | 0.06 | 3.25349008151215e-3 | 1 | sptlc2a |
| cpox | 1.7218550 | 0.450598942 | 0.33 | 0.058 | 3.47763058059216e-3 | 1 | cpox |
| ier2a | 1.8232153 | 0.767748652 | 0.758 | 0.331 | 3.68234798816429e-3 | 1 | ier2a |

| | | | | | | | |
|---|---|---|---|---|---|---|---|
| mfap21 | 1.8573385 | 0.61968291 | 0.714 | 0.209 | 3.75126661765208e-3 | 1 | mfap2 |
| socs3a1 | 2.6285203 | 0.764662689 | 0.841 | 0.396 | 5.30882245317483e-3 | 1 | socs3a |
| ms4a17a.11 | 9.0619427 | 0.512595275 | 0.429 | 0.098 | 1.83024058257025e-3 | 1 | ms4a17a.11 |
| crispld1a | 1.3027090 | 0.25948002 | 0.148 | 0.008 | 2.63108155335844e-3 | 1 | crispld1a |
| nfic | 2.6565550 | 0.528006116 | 0.423 | 0.095 | 5.36544425409451e-3 | 1 | nfic |
| cyp27c1 | 3.4568366 | 0.433381419 | 0.291 | 0.045 | 6.98177297505434e-3 | 1 | cyp27c1 |
| ckba | 5.4173649 | 0.515617781 | 0.418 | 0.095 | 1.09414520184103e-3 | 1 | ckba |
| si:dkey-188i13.10 | 1.5613752 | 0.587653703 | 0.511 | 0.14 | 3.15350965097529e-3 | 1 | si:dkey-188i13.10 |
| nog1 | 4.1044938 | 0.361749215 | 0.192 | 0.018 | 8.28984623188611e-3 | 1 | nog1 |
| pgrmc1 | 4.6647809 | 0.609354139 | 0.599 | 0.191 | 9.42145813463389e-3 | 1 | pgrmc1 |
| snai1a | 9.3948993 | 0.37878596 | 0.275 | 0.041 | 1.89748782885037e-3 | 1 | snai1a |
| sh3gl1b | 2.6049559 | 0.398732651 | 0.291 | 0.048 | 5.26122944223769e-3 | 1 | sh3gl1b |
| CU459094.2 | 2.8999356 | 0.327443538 | 0.214 | 0.024 | 5.85699999728025e-3 | 1 | CU459094.2 |
| tmem47 | 3.0787521 | 0.383549898 | 0.236 | 0.031 | 6.21815571579093e-3 | 1 | tmem47 |
| fam198a | 7.1975823 | 0.369752438 | 0.253 | 0.036 | 1.45369570574094e-2 | 1 | fam198a |
| col12a1b | 1.4382377 | 0.434871919 | 0.286 | 0.046 | 2.90480875835649e-2 | 1 | col12a1b |
| nck1a | 2.1411018 | 0.369172072 | 0.291 | 0.05 | 4.32438342055234e-2 | 1 | nck1a |
| mtss1la | 3.6236170 | 0.329281163 | 0.192 | 0.021 | 7.31861935892744e-2 | 1 | mtss1la |
| spry2 | 4.9795717 | 0.450414229 | 0.407 | 0.096 | 1.00572411637208e-2 | 1 | spry2 |
| si:ch73-46j18.51 | 5.5743328 | 0.688854267 | 0.863 | 0.357 | 1.12584800194771e-2 | 1 | si:ch73-46j18.5 |
| col1a1b1 | 6.8714488 | 0.590571132 | 0.786 | 0.257 | 1.38782651919545e-2 | 1 | col1a1b |
| XLOC-000431 | 1.0956600 | 0.631112644 | 0.665 | 0.234 | 2.21290452608721e-2 | 1 | XLOC-000431 |
| lama5 | 1.2246864 | 0.444949632 | 0.269 | 0.045 | 2.47349920207995e-2 | 1 | lama5 |
| zgc:92744 | 1.7440379 | 0.662817253 | 0.56 | 0.178 | 3.52243352897952e-2 | 1 | zgc:92744 |
| rhoua | 4.5681367 | 0.395436549 | 0.275 | 0.047 | 9.22626589178543e-2 | 1 | rhoua |
| myl9b | 6.0747744 | 0.543988088 | 0.571 | 0.173 | 1.22692218728722e-2 | 1 | myl9b |
| chd31 | 6.8709680 | 0.592346879 | 0.604 | 0.211 | 1.3877294092297e-26 | 1 | chd3 |
| cdon1 | 1.1411824 | 0.493642659 | 0.456 | 0.12 | 2.30484614261313e-2 | 1 | cdon |
| ms4a17a.8 | 1.2073419 | 0.337090196 | 0.225 | 0.032 | 2.43846853670491e-2 | 1 | ms4a17a.8 |
| cald1a1 | 2.5928367 | 0.558760394 | 0.626 | 0.205 | 5.23675240078562e-2 | 1 | cald1a |
| loxl2a | 5.2198391 | 0.455731462 | 0.297 | 0.057 | 1.05425090805391e-2 | 1 | loxl2a |
| fstl1b1 | 3.1594243 | 0.557554649 | 0.72 | 0.241 | 6.38108929282059e-2 | 1 | fstl1b |
| enc1 | 3.3676431 | 0.431016557 | 0.308 | 0.063 | 6.80162889715363e-2 | 1 | enc1 |
| rgs7bpb | 3.9768135 | 0.305786649 | 0.209 | 0.029 | 8.03197037559097e-2 | 1 | rgs7bpb |
| eva1ba1 | 4.7983638 | 0.431024346 | 0.302 | 0.06 | 9.69125546158775e-2 | 1 | eva1ba |

| | | | | | | | |
|---|---|---|---|---|---|---|---|
| kif26ba | 5.2584643 | 0.283760741 | 0.154 | 0.014 | 1.06205203979881e-2 | 1 | kif26ba |
| irf1b | 6.2027892 | 0.255204925 | 0.148 | 0.013 | 1.25277734736316e-2 | 1 | irf1b |
| ctbp2a | 8.5511757 | 0.337833723 | 0.269 | 0.049 | 1.72708097162089e-2 | 1 | ctbp2a |
| si:ch211-132p1.3 | 9.7999307 | 0.505073443 | 0.434 | 0.123 | 1.97929201383564e-2 | 1 | si:ch211-132p1.3 |
| tenm31 | 1.0735620 | 0.496250254 | 0.445 | 0.122 | 2.16827322131801e-2 | 1 | tenm3 |
| cldn11a | 1.9611579 | 0.557506608 | 0.544 | 0.166 | 3.96095068625815e-2 | 1 | cldn11a |
| raraa | 2.0977481 | 0.387137719 | 0.286 | 0.055 | 4.23682198156169e-2 | 1 | raraa |
| tbx2a | 3.9388221 | 0.251867224 | 0.154 | 0.015 | 7.95523909462968e-2 | 1 | tbx2a |
| tagln | 4.3702503 | 0.558005035 | 0.423 | 0.113 | 8.82659472038807e-2 | 1 | tagln |
| h2afy2 | 6.7456179 | 0.562268711 | 0.593 | 0.228 | 1.36241246274944e-2 | 1 | h2afy2 |
| tnfrsf19 | 1.0642521 | 0.390710172 | 0.231 | 0.038 | 2.14947010055993e-2 | 1 | tnfrsf19 |
| sypl1 | 2.2274817 | 0.262788547 | 0.176 | 0.021 | 4.49884493484805e-2 | 1 | sypl1 |
| rrbp1b | 3.1792370 | 0.33719565 | 0.225 | 0.037 | 6.42110506185011e-2 | 1 | rrbp1b |
| si:ch1073-459j12.1 | 3.7626568 | 0.547227796 | 0.511 | 0.158 | 7.59943795454832e-2 | 1 | si:ch1073-459j12.1 |
| pkig | 8.0414636 | 0.421117058 | 0.396 | 0.105 | 1.62413442296902e-2 | 1 | pkig |
| mgst1.11 | 9.3338159 | 0.458837223 | 0.401 | 0.109 | 1.88515081358445e-2 | 1 | mgst1.1 |
| cpxm1a | 1.2906242 | 0.310783947 | 0.214 | 0.033 | 2.60667380179777e-2 | 1 | cpxm1a |
| socs3b1 | 1.3581160 | 0.513647434 | 0.423 | 0.124 | 2.7429870435909e-22 | 1 | socs3b |
| sult1st1 | 2.0002370 | 0.360929313 | 0.302 | 0.066 | 4.03987879643304e-2 | 1 | sult1st1 |
| marcksl1b | 2.0632269 | 0.577076059 | 0.665 | 0.248 | 4.16709951497748e-2 | 1 | marcksl1b |
| b2m | 3.5576409 | 0.583941469 | 0.731 | 0.323 | 7.1853675227823e-22 | 1 | b2m |
| mxra8b | 4.5604450 | 0.342917584 | 0.253 | 0.048 | 9.2107307943426e-22 | 1 | mxra8b |
| XLOC-005605 | 4.7208674 | 0.401969059 | 0.33 | 0.076 | 9.53473598813941e-2 | 1 | XLOC-005605 |
| ptenb | 5.6679813 | 0.451364304 | 0.385 | 0.106 | 1.14476218724547e-2 | 1 | ptenb |
| c1qtnf6a | 6.2170193 | 0.36533429 | 0.258 | 0.05 | 1.25565140455287e-2 | 1 | c1qtnf6a |
| esr2a | 6.7858217 | 0.290728217 | 0.192 | 0.027 | 1.37053242372746e-2 | 1 | esr2a |
| snx17 | 7.3307495 | 0.431245631 | 0.346 | 0.086 | 1.48059148356079e-2 | 1 | snx17 |
| col5a2a1 | 7.3353377 | 0.454746108 | 0.511 | 0.154 | 1.48151815915373e-2 | 1 | col5a2a |
| zgc:113531 | 1.2597608 | 0.410814653 | 0.258 | 0.051 | 2.54433890465463e-2 | 1 | zgc:113531 |
| cox4i2 | 1.8945317 | 0.324765617 | 0.242 | 0.044 | 3.82638579782135e-2 | 1 | cox4i2 |
| tob1b | 1.9545181 | 0.450897667 | 0.429 | 0.127 | 3.94754039989807e-2 | 1 | tob1b |
| aldh2.2 | 1.9708517 | 0.438952672 | 0.385 | 0.108 | 3.98052935468129e-2 | 1 | aldh2.2 |
| snap23.2 | 3.3025550 | 0.272524844 | 0.154 | 0.017 | 6.6701704816638e-21 | 1 | snap23.2 |
| ngdn | 7.8268417 | 0.398325701 | 0.374 | 0.105 | 1.58078722505843e-2 | 1 | ngdn |
| celf21 | 1.7863963 | 0.562940605 | 0.725 | 0.327 | 3.60798471143729e-2 | 1 | celf2 |

| | | | | | | | |
|---|---|---|---|---|---|---|---|
| cebpd1 | 2.1502939 | 0.544577899 | 0.654 | 0.248 | 4.34294860120952e-2 | 1 | cebpd |
| selenbp1 | 2.4007185 | 0.464424681 | 0.407 | 0.122 | 4.84873128528382e-2 | 1 | selenbp1 |
| fzd21 | 3.1371534 | 0.395028557 | 0.363 | 0.097 | 6.33610881463397e-2 | 1 | fzd2 |
| colec121 | 3.4619556 | 0.39100293 | 0.357 | 0.091 | 6.99211174324777e-2 | 1 | colec12 |
| osr11 | 3.6076477 | 0.462445069 | 0.396 | 0.111 | 7.28636615664812e-2 | 1 | osr1 |
| tcf12 | 4.0599989 | 0.432911137 | 0.368 | 0.103 | 8.19997985186114e-2 | 1 | tcf12 |
| nme3 | 4.0955033 | 0.424067671 | 0.445 | 0.138 | 8.27168819537163e-2 | 1 | nme3 |
| fzd7a | 1.1968105 | 0.362994538 | 0.225 | 0.043 | 2.41719826511561e-1 | 1 | fzd7a |
| fbn2b1 | 1.2972882 | 0.488402541 | 0.473 | 0.155 | 2.62013299608776e-1 | 1 | fbn2b |
| grb10a1 | 1.4430109 | 0.332932516 | 0.247 | 0.05 | 2.91444914423352e-1 | 1 | grb10a |
| smo | 2.2799593 | 0.343583182 | 0.242 | 0.049 | 4.60483388249188e-1 | 1 | smo |
| acss1 | 2.3894960 | 0.278714732 | 0.192 | 0.031 | 4.82606513785838e-1 | 1 | acss1 |
| ywhaz | 3.0962137 | 0.538134681 | 0.775 | 0.422 | 6.25342288287721e-1 | 1 | ywhaz |
| cnbpb | 3.2556987 | 0.496556258 | 0.67 | 0.297 | 6.57553478492383e-1 | 1 | cnbpb |
| CABZ01068366.1 | 3.2589283 | 0.379872786 | 0.335 | 0.087 | 6.58205751891392e-1 | 1 | CABZ01068366.1 |
| ghrb | 4.2866349 | 0.299676212 | 0.214 | 0.039 | 8.65771659754451e-1 | 1 | ghrb |
| hmgb1b | 4.6595239 | 0.503783314 | 0.665 | 0.273 | 9.41084054165314e-1 | 1 | hmgb1b |
| SCARF2 | 5.1417086 | 0.308808854 | 0.203 | 0.035 | 1.03847090192273e-1 | 1 | SCARF2 |
| cyp2ad31 | 5.5625380 | 0.454451406 | 0.352 | 0.097 | 1.12346580279937e-1 | 1 | cyp2ad3 |
| wls | 6.5245004 | 0.45842064 | 0.467 | 0.159 | 1.3177533619119e-18 | 1 | wls |
| c1qtnf51 | 6.9342834 | 0.439734262 | 0.286 | 0.067 | 1.40051722653044e-1 | 1 | c1qtnf5 |
| wfdc1 | 7.6601316 | 0.278095588 | 0.154 | 0.02 | 1.5471167829098e-18 | 1 | wfdc1 |
| serinc2 | 7.9600976 | 0.444660086 | 0.5 | 0.174 | 1.6077009134596e-18 | 1 | serinc2 |
| ets1 | 1.2225755 | 0.316363462 | 0.22 | 0.042 | 2.46923583765521e-1 | 1 | ets1 |
| boc1 | 1.3401721 | 0.466926214 | 0.451 | 0.145 | 2.70674577343633e-1 | 1 | boc |
| mmp14a1 | 1.3527691 | 0.37830181 | 0.319 | 0.082 | 2.73218779897688e-1 | 1 | mmp14a |
| si:dkey-261h17.11 | 1.4105005 | 0.481113992 | 0.61 | 0.222 | 2.84878797858568e-1 | 1 | si:dkey-261h17.1 |
| khdrbs1b | 1.6665159 | 0.428921464 | 0.401 | 0.128 | 3.36586226561634e-1 | 1 | khdrbs1b |
| ponzr1 | 1.6723129 | 0.542975482 | 0.791 | 0.36 | 3.37757037103017e-1 | 1 | ponzr1 |
| fkbp7 | 1.8695917 | 0.387598354 | 0.374 | 0.107 | 3.77601443788664e-1 | 1 | fkbp7 |
| olfml3b1 | 2.8539244 | 0.403173636 | 0.341 | 0.093 | 5.76407124830506e-1 | 1 | olfml3b |
| stxbp6 | 2.9408210 | 0.294267506 | 0.242 | 0.05 | 5.9395762343507e-18 | 1 | stxbp6 |
| mrc2 | 3.2100263 | 0.281584486 | 0.22 | 0.042 | 6.48329019320426e-1 | 1 | mrc2 |
| vdac1 | 3.5142847 | 0.411610401 | 0.467 | 0.159 | 7.09780097227109e-1 | 1 | vdac1 |
| col6a31 | 5.1228836 | 0.458235014 | 0.5 | 0.169 | 1.03466881040078e-1 | 1 | col6a3 |

| | | | | | | | |
|---|---|---|---|---|---|---|---|
| pde4ba | 5.7298305 | 0.465461622 | 0.429 | 0.142 | 1.1572538750841e-17 | 1 | pde4ba |
| tmem100a1 | 7.7440456 | 0.27892541 | 0.209 | 0.038 | 1.56406490149215e-1 | 1 | tmem100a |
| robo1 | 8.7539093 | 0.30576473 | 0.209 | 0.039 | 1.76802706914071e-1 | 1 | robo1 |
| jpt1b1 | 9.9714835 | 0.51138769 | 0.703 | 0.304 | 2.01394053494147e-1 | 1 | jpt1b |
| xirp2a | 1.2796513 | 0.355771094 | 0.275 | 0.066 | 2.584511899147e-17 | 1 | xirp2a |
| hoxc9a | 1.3272087 | 0.264890136 | 0.176 | 0.028 | 2.68056358539626e-1 | 1 | hoxc9a |
| lamb1a1 | 1.5509728 | 0.423474913 | 0.39 | 0.119 | 3.13249988603475e-1 | 1 | lamb1a |
| dhrs13l1 | 1.8846652 | 0.330005482 | 0.264 | 0.062 | 3.80645835231067e-1 | 1 | dhrs13l1 |
| jcada | 2.7099369 | 0.30673142 | 0.209 | 0.04 | 5.47325967063515e-1 | 1 | jcada |
| cxcl12a1 | 3.6440946 | 0.37711379 | 0.308 | 0.078 | 7.35997801549292e-1 | 1 | cxcl12a |
| eif4g3a | 5.4424275 | 0.255842865 | 0.143 | 0.019 | 1.09920709193084e-1 | 1 | eif4g3a |
| ifi45 | 6.5173106 | 0.513874055 | 0.489 | 0.176 | 1.31630122697351e-1 | 1 | ifi45 |
| tsc22d1 | 7.0233946 | 0.298477942 | 0.22 | 0.045 | 1.41851502714275e-1 | 1 | tsc22d1 |
| cnn3b | 8.3477831 | 0.277243755 | 0.198 | 0.037 | 1.68600175547061e-1 | 1 | cnn3b |
| CR383676.11 | 8.9795640 | 0.600217979 | 1 | 0.632 | 1.81360254668666e-1 | 1 | CR383676.1 |
| tmeff1b | 1.0277577 | 0.367043256 | 0.319 | 0.087 | 2.0757622617037e-16 | 1 | tmeff1b |
| nell2a | 1.4396997 | 0.284038617 | 0.17 | 0.027 | 2.90776152296928e-1 | 1 | nell2a |
| entpd1 | 1.5669297 | 0.384415942 | 0.401 | 0.127 | 3.1647280264319e-16 | 1 | entpd1 |
| yap11 | 2.3858639 | 0.339446082 | 0.302 | 0.081 | 4.81872948242252e-1 | 1 | yap1 |
| spred2b | 3.0844401 | 0.421046396 | 0.456 | 0.16 | 6.22964367063989e-1 | 1 | spred2b |
| antxr1c | 3.4664180 | 0.274427618 | 0.209 | 0.042 | 7.00112454976392e-1 | 1 | antxr1c |
| si:dkey-102m7.3 | 3.8403946 | 0.348006278 | 0.357 | 0.109 | 7.75644511254401e-1 | 1 | si:dkey-102m7.3 |
| igf2b1 | 6.6295518 | 0.48943734 | 0.418 | 0.144 | 1.33897059703297e-1 | 1 | igf2b |
| hmgb3a | 7.0649184 | 0.478561008 | 0.577 | 0.234 | 1.42690158336427e-1 | 1 | hmgb3a |
| mgst3b | 1.5095977 | 0.467243246 | 0.61 | 0.277 | 3.04893464629466e-1 | 1 | mgst3b |
| b2ml1 | 1.9821367 | 0.478000836 | 0.797 | 0.418 | 4.00332162756753e-1 | 1 | b2ml |
| lama41 | 1.9921859 | 0.385116068 | 0.302 | 0.085 | 4.02361790401909e-1 | 1 | lama4 |
| map2k6 | 2.5922484 | 0.392479064 | 0.39 | 0.132 | 5.23556412370875e-1 | 1 | map2k6 |
| nop58 | 4.1087672 | 0.405680043 | 0.505 | 0.197 | 8.29847720629165e-1 | 1 | nop58 |
| cdh2 | 4.7837394 | 0.358739819 | 0.357 | 0.109 | 9.66171861207506e-1 | 1 | cdh2 |
| aldh9a1a.11 | 5.4857619 | 0.451275755 | 0.676 | 0.312 | 1.10795933284915e-1 | 1 | aldh9a1a.1 |
| FO744854.1 | 6.4655338 | 0.254112802 | 0.132 | 0.018 | 1.30584387988476e-1 | 1 | FO744854.1 |
| lhfpl2a | 6.8636810 | 0.351911952 | 0.275 | 0.074 | 1.38625765849652e-1 | 1 | lhfpl2a |
| lgals17 | 7.2019524 | 0.363276362 | 0.242 | 0.059 | 1.45457834098707e-1 | 1 | lgals17 |
| slc38a5b1 | 9.4515805 | 0.385306944 | 0.484 | 0.183 | 1.90893572618066e-1 | 1 | slc38a5b |

| | | | | | | | |
|---|---|---|---|---|---|---|---|
| mvp | 2.3775634 | 0.355243453 | 0.297 | 0.088 | 4.8019648034789e-14 | 1 | mvp |
| zgc:174863 | 2.8324032 | 0.375944372 | 0.407 | 0.147 | 5.72060479637811e-1 | 1 | zgc:174863 |
| mycb | 3.3704937 | 0.392511982 | 0.401 | 0.142 | 6.80738623179754e-1 | 1 | mycb |
| ms4a17a.9 | 3.4035694 | 0.346646247 | 0.374 | 0.122 | 6.8741892208505e-14 | 1 | ms4a17a.9 |
| ctdsp2 | 3.4796085 | 0.367713658 | 0.423 | 0.154 | 7.02776532731045e-1 | 1 | ctdsp2 |
| ctdspla | 3.7422477 | 0.287629989 | 0.225 | 0.053 | 7.55821770317734e-1 | 1 | ctdspla |
| adh5 | 5.0510533 | 0.432250954 | 0.687 | 0.335 | 1.02016124258546e-1 | 1 | adh5 |
| h3f3c | 5.8492473 | 0.480806388 | 0.83 | 0.505 | 1.18137249645312e-1 | 1 | h3f3c |
| cnn3a1 | 6.6760135 | 0.332734352 | 0.368 | 0.118 | 1.34835444750257e-1 | 1 | cnn3a |
| si:ch211-39i22.11 | 7.5954554 | 0.376378103 | 0.451 | 0.162 | 1.53405414465493e-1 | 1 | si:ch211-39i22.1 |
| agrn | 9.1254187 | 0.342786435 | 0.319 | 0.097 | 1.8430608232392e-13 | 1 | agrn |
| BX322530.1 | 1.1639348 | 0.291551549 | 0.247 | 0.063 | 2.35079931685171e-1 | 1 | BX322530.1 |
| f10 | 1.4256537 | 0.35008574 | 0.33 | 0.103 | 2.87939280395943e-1 | 1 | f10 |
| si:dkey-239h2.3 | 1.5510306 | 0.35763381 | 0.396 | 0.143 | 3.13261667217095e-1 | 1 | si:dkey-239h2.3 |
| XLOC-0160941 | 4.7349005 | 0.318814116 | 0.308 | 0.092 | 9.56307855258099e-1 | 1 | XLOC-016094 |
| si:ch211-222l21.1 | 5.0419438 | 0.485845325 | 0.874 | 0.504 | 1.01832140834523e-1 | 1 | si:ch211-222l21.1 |
| dhrs3b1 | 5.5517291 | 0.28422655 | 0.231 | 0.057 | 1.12128273711397e-1 | 1 | dhrs3b |
| rbfox21 | 5.8930971 | 0.276114946 | 0.242 | 0.063 | 1.19022882861561e-1 | 1 | rbfox2 |
| tgif1 | 7.0362643 | 0.352875071 | 0.379 | 0.135 | 1.42111431259793e-1 | 1 | tgif1 |
| col6a11 | 7.4682331 | 0.339118552 | 0.379 | 0.126 | 1.50835904345652e-1 | 1 | col6a1 |
| tmem98 | 8.3778621 | 0.250233624 | 0.176 | 0.036 | 1.69207682257766e-1 | 1 | tmem98 |
| cyb5r1 | 8.6894077 | 0.335360001 | 0.264 | 0.077 | 1.75499967442777e-1 | 1 | cyb5r1 |
| ifi46 | 9.4075095 | 0.401330202 | 0.445 | 0.172 | 1.90003469975581e-1 | 1 | ifi46 |
| add3a1 | 9.9005571 | 0.391631694 | 0.495 | 0.199 | 1.99961553739934e-1 | 1 | add3a |
| grpel2 | 1.1169217 | 0.301492039 | 0.346 | 0.117 | 2.25584683731779e-1 | 1 | grpel2 |
| meis2b1 | 1.6090910 | 0.327168957 | 0.28 | 0.082 | 3.24988121919552e-1 | 1 | meis2b |
| lpar11 | 1.7994643 | 0.346860957 | 0.291 | 0.089 | 3.63437807090251e-1 | 1 | lpar1 |
| wu:fi37e09 | 1.8199523 | 0.372688216 | 0.44 | 0.174 | 3.67575773476391e-1 | 1 | wu:fi37e09 |
| si:ch211-212k18.8 | 1.9500389 | 0.287736836 | 0.242 | 0.064 | 3.93849374230982e-1 | 1 | si:ch211-212k18.8 |
| tmem50a | 2.0475815 | 0.361243116 | 0.451 | 0.178 | 4.13550043532726e-1 | 1 | tmem50a |
| tomm20a | 2.2127564 | 0.383566629 | 0.478 | 0.2 | 4.46910423256805e-1 | 1 | tomm20a |
| rab13 | 2.3763678 | 0.273954327 | 0.242 | 0.065 | 4.79955023079985e-1 | 1 | rab13 |
| sept10 | 2.4609383 | 0.278079894 | 0.247 | 0.068 | 4.9703572383481e-12 | 1 | sept10 |
| ppib | 2.7502883 | 0.427654417 | 0.923 | 0.621 | 5.55475747576268e-1 | 1 | ppib |
| nop56 | 3.7658500 | 0.407402442 | 0.56 | 0.26 | 7.60588736301154e-1 | 1 | nop56 |

| | | | | | | | |
|---|---|---|---|---|---|---|---|
| si:ch211-133n4.41 | 4.9119708 | 0.342677368 | 0.346 | 0.117 | 9.92070761935418e-1 | 1 | si:ch211-133n4.4 |
| ube2e2 | 5.0443763 | 0.268365009 | 0.247 | 0.069 | 1.01881269783674e-1 | 1 | ube2e2 |
| cbx1a | 5.1105344 | 0.346170503 | 0.357 | 0.128 | 1.03217465001435e-1 | 1 | cbx1a |
| foxo3b | 5.3400348 | 0.343532193 | 0.269 | 0.082 | 1.07852683691474e-1 | 1 | foxo3b |
| si:ch73-281n10.2 | 6.6864897 | 0.44101106 | 0.78 | 0.406 | 1.35047034344012e-1 | 1 | si:ch73-281n10.2 |
| ier5l | 7.6767777 | 0.288309651 | 0.203 | 0.05 | 1.55047881192066e-1 | 1 | ier5l |
| ephb3 | 8.1522146 | 0.329823912 | 0.297 | 0.094 | 1.64650279530776e-1 | 1 | ephb3 |
| parp12b | 1.7551643 | 0.310402952 | 0.236 | 0.067 | 3.54490550751873e-1 | 1 | parp12b |
| atf4b | 1.8685869 | 0.422570576 | 0.643 | 0.339 | 3.77398496640099e-1 | 1 | atf4b |
| col4a11 | 2.0682719 | 0.346476072 | 0.352 | 0.121 | 4.17728895437256e-1 | 1 | col4a1 |
| lrp1aa1 | 2.6581512 | 0.300331801 | 0.33 | 0.109 | 5.36866816444838e-1 | 1 | lrp1aa |
| isg15 | 2.8280870 | 0.265163474 | 0.176 | 0.039 | 5.71188741999366e-1 | 1 | isg15 |
| gmfb | 2.8752280 | 0.335475396 | 0.401 | 0.155 | 5.80709813697847e-1 | 1 | gmfb |
| aldh2.1 | 3.4680120 | 0.260312057 | 0.22 | 0.06 | 7.00434399309535e-1 | 1 | aldh2.1 |
| dedd11 | 3.5130175 | 0.390049787 | 0.456 | 0.194 | 7.09524148457938e-1 | 1 | dedd1 |
| pfn2l | 4.0393235 | 0.377278937 | 0.549 | 0.24 | 8.15822186177031e-1 | 1 | pfn2l |
| nt5dc2 | 4.3544501 | 0.267522994 | 0.253 | 0.075 | 8.79468302910133e-1 | 1 | nt5dc2 |
| marcksl1a1 | 5.4068131 | 0.44332218 | 0.846 | 0.486 | 1.09201404742936e-1 | 1 | marcksl1a |
| snx6 | 5.4485752 | 0.312438291 | 0.269 | 0.086 | 1.10044873787095e-1 | 1 | snx6 |
| XLOC-013964 | 6.4146311 | 0.284505892 | 0.39 | 0.143 | 1.29556306040572e-1 | 1 | XLOC-013964 |
| lipf | 7.1693720 | 0.276875169 | 0.236 | 0.068 | 1.44799807888865e-1 | 1 | lipf |
| sulf2b | 8.5636031 | 0.278208323 | 0.198 | 0.05 | 1.72959092692391e-1 | 1 | sulf2b |
| tuba8l31 | 8.6119620 | 0.333833186 | 0.473 | 0.185 | 1.73935797119713e-1 | 1 | tuba8l3 |
| ciarta | 9.7135331 | 0.337324804 | 0.423 | 0.169 | 1.96184229414076e-1 | 1 | ciarta |
| hmgn6 | 1.1966045 | 0.42412768 | 0.846 | 0.55 | 2.41678225959052e-1 | 1 | hmgn6 |
| mmp2 | 1.3782372 | 0.345974428 | 0.489 | 0.196 | 2.78362571448988e-1 | 1 | mmp2 |
| cbx7a | 1.4063110 | 0.294259241 | 0.181 | 0.044 | 2.84032651626651e-1 | 1 | cbx7a |
| etv4 | 1.6300641 | 0.258801258 | 0.198 | 0.05 | 3.29224061056685e-1 | 1 | etv4 |
| fam168a | 1.8131281 | 0.320982633 | 0.368 | 0.138 | 3.66197495359767e-1 | 1 | fam168a |
| ctsla1 | 2.1712896 | 0.34145119 | 0.615 | 0.285 | 4.3853537558335e-10 | 1 | ctsla |
| myclb | 2.3729626 | 0.284814983 | 0.253 | 0.077 | 4.79267273475762e-1 | 1 | myclb |
| tpm4a | 3.3383123 | 0.364291165 | 0.522 | 0.225 | 6.74238955010115e-1 | 1 | tpm4a |
| lgals9l3 | 3.4807912 | 0.331870332 | 0.418 | 0.169 | 7.03015409907411e-1 | 1 | lgals9l3 |
| si:dkey-6n21.12 | 4.0087941 | 0.280814468 | 0.231 | 0.067 | 8.09656148040324e-1 | 1 | si:dkey-6n21.12 |
| nenf | 5.3510841 | 0.259981705 | 0.269 | 0.087 | 1.08075847010687e-0 | 1 | nenf |

| | | | | | | | |
|---|---|---|---|---|---|---|---|
| si:ch211-222k6.3 | 5.3519439 | 0.260295685 | 0.203 | 0.055 | 1.08093212867631e-0 | 1 | si:ch211-222k6.3 |
| hmgn7 | 5.8243338 | 0.409519436 | 0.797 | 0.51 | 1.17634069847463e-0 | 1 | hmgn7 |
| si:ch211-286b5.5 | 7.1508644 | 0.327834144 | 0.478 | 0.206 | 1.44426009630342e-0 | 1 | si:ch211-286b5.5 |
| serinc51 | 8.0271894 | 0.338228691 | 0.291 | 0.103 | 1.6212514483654e-09 | 1 | serinc5 |
| cab39l | 1.3572239 | 0.273347423 | 0.247 | 0.08 | 2.74118515292813e-0 | 1 | cab39l |
| cpne3 | 1.4805048 | 0.27663379 | 0.22 | 0.065 | 2.99017570476638e-0 | 1 | cpne3 |
| ctnnb2 | 1.5822748 | 0.304724396 | 0.302 | 0.109 | 3.19572060959375e-0 | 1 | ctnnb2 |
| sesn1 | 1.6904528 | 0.396968336 | 0.445 | 0.199 | 3.41420759862365e-0 | 1 | sesn1 |
| sparc1 | 1.9588073 | 0.453288933 | 0.83 | 0.366 | 3.9562031706542e-09 | 1 | sparc |
| rad21a | 2.1403959 | 0.360723774 | 0.489 | 0.232 | 4.32295768945952e-0 | 1 | rad21a |
| hspe1 | 2.2491005 | 0.376469973 | 0.566 | 0.271 | 4.54250828688096e-0 | 1 | hspe1 |
| hist2h2l | 2.3538834 | 0.370943731 | 0.44 | 0.193 | 4.75413845540868e-0 | 1 | hist2h2l |
| rnf13 | 3.1984998 | 0.259067935 | 0.209 | 0.06 | 6.46001010775654e-0 | 1 | rnf13 |
| id11 | 3.3369587 | 0.389590286 | 0.549 | 0.253 | 6.73965561263242e-0 | 1 | id1 |
| calm2a | 4.0722024 | 0.348847439 | 0.5 | 0.233 | 8.22462722281557e-0 | 1 | calm2a |
| sox11a | 4.7400256 | 0.268390423 | 0.214 | 0.063 | 9.57342976719962e-0 | 1 | sox11a |
| hnrnpa0b | 4.9449607 | 0.392901793 | 0.813 | 0.49 | 9.98733723597498e-0 | 1 | hnrnpa0b |
| nfyba | 9.1007520 | 0.316549916 | 0.462 | 0.211 | 1.8380788998677e-08 | 1 | nfyba |
| ptena | 9.4570403 | 0.293893717 | 0.313 | 0.12 | 1.91003844587749e-0 | 1 | ptena |
| emd | 1.0117759 | 0.278835856 | 0.352 | 0.138 | 2.04348384153639e-0 | 1 | emd |
| smarce1 | 1.0726013 | 0.310045762 | 0.489 | 0.228 | 2.16633285767977e-0 | 1 | smarce1 |
| zgc:112437 | 1.0740322 | 0.287460727 | 0.192 | 0.054 | 2.16922292259164e-0 | 1 | zgc:112437 |
| antxr2a1 | 1.1727296 | 0.367818643 | 0.511 | 0.243 | 2.36856201083407e-0 | 1 | antxr2a |
| tnfrsf1a | 1.1803256 | 0.268401421 | 0.247 | 0.082 | 2.38390366312981e-0 | 1 | tnfrsf1a |
| flot1b | 1.3650309 | 0.285651337 | 0.418 | 0.18 | 2.75695293618352e-0 | 1 | flot1b |
| cxxc5a | 1.3873794 | 0.252060328 | 0.236 | 0.076 | 2.8020902459985e-08 | 1 | cxxc5a |
| cygb1 | 1.6005260 | 0.288439745 | 0.269 | 0.091 | 3.23258252954743e-0 | 1 | cygb1 |
| col6a21 | 1.8416682 | 0.279613151 | 0.313 | 0.113 | 3.71961732396913e-0 | 1 | col6a2 |
| si:dkey-11f4.14 | 2.0985070 | 0.345582258 | 0.418 | 0.186 | 4.23835463850412e-0 | 1 | si:dkey-11f4.14 |
| aldh6a1 | 2.8578978 | 0.261970245 | 0.308 | 0.121 | 5.77209633814844e-0 | 1 | aldh6a1 |
| grinab | 2.9610833 | 0.304830667 | 0.385 | 0.163 | 5.98049999563513e-0 | 1 | grinab |
| csnk1a1 | 2.9664032 | 0.32438267 | 0.516 | 0.251 | 5.991244585577e-08 | 1 | csnk1a1 |
| FO082781.1 | 3.0618109 | 0.270391744 | 0.253 | 0.086 | 6.18393947505432e-0 | 1 | FO082781.1 |
| crim1 | 4.2883298 | 0.277317425 | 0.198 | 0.059 | 8.66113974766208e-0 | 1 | crim1 |
| aip | 4.4522725 | 0.285713469 | 0.335 | 0.136 | 8.99225482914681e-0 | 1 | aip |

| | | | | | | | |
|---|---|---|---|---|---|---|---|
| appb | 5.5759318 | 0.253296253 | 0.231 | 0.075 | 1.12617095902085e-0 | 1 | appb |
| vdrb | 6.0836016 | 0.290226851 | 0.253 | 0.089 | 1.22870501682246e-0 | 1 | vdrb |
| bsg | 7.1176700 | 0.360057226 | 0.665 | 0.366 | 1.43755582103652e-0 | 1 | bsg |
| prdx2 | 7.9798581 | 0.351631889 | 0.791 | 0.465 | 1.61169194845908e-0 | 1 | prdx2 |
| nfia1 | 8.1876309 | 0.318744347 | 0.423 | 0.184 | 1.65365582247661e-0 | 1 | nfia |
| snu13b | 9.3602815 | 0.288857558 | 0.555 | 0.272 | 1.890496059174e-07 | 1 | snu13b |
| NC-002333.41 | 9.8261660 | 0.431474607 | 0.912 | 0.559 | 1.98459076031369e-0 | 1 | NC-002333.4 |
| pdia3 | 9.9363205 | 0.375930774 | 0.775 | 0.462 | 2.006838670389e-07 | 1 | pdia3 |
| pabpn1 | 1.0015657 | 0.352204457 | 0.527 | 0.276 | 2.02286238717922e-0 | 1 | pabpn1 |
| pik3r3a | 1.5046788 | 0.263465146 | 0.33 | 0.131 | 3.03899996757233e-0 | 1 | pik3r3a |
| foxp41 | 1.5611677 | 0.328437154 | 0.396 | 0.174 | 3.1530904293507e-07 | 1 | foxp4 |
| sept2 | 2.0904144 | 0.322855972 | 0.533 | 0.265 | 4.22201013457309e-0 | 1 | sept2 |
| zfp36l1a1 | 2.0971001 | 0.348722345 | 0.571 | 0.291 | 4.23551321348586e-0 | 1 | zfp36l1a |
| flot2a | 2.1288570 | 0.276828509 | 0.335 | 0.138 | 4.29965258636668e-0 | 1 | flot2a |
| cbx5 | 2.6540711 | 0.299405424 | 0.44 | 0.205 | 5.36042756463255e-0 | 1 | cbx5 |
| mafba1 | 4.4161853 | 0.27093835 | 0.335 | 0.137 | 8.91936952027686e-0 | 1 | mafba |
| tax1bp3 | 5.3140717 | 0.266465898 | 0.363 | 0.155 | 1.0732830713747e-06 | 1 | tax1bp3 |
| il6st | 5.4461836 | 0.277220587 | 0.214 | 0.073 | 1.09996570820436e-0 | 1 | il6st |
| si:ch73-335l21.4 | 5.6763355 | 0.367341824 | 0.269 | 0.105 | 1.14644949411821e-0 | 1 | si:ch73-335l21.4 |
| lman2 | 6.8096611 | 0.277759139 | 0.39 | 0.178 | 1.37534726974231e-0 | 1 | lman2 |
| lxn1 | 6.8843671 | 0.353392895 | 0.495 | 0.25 | 1.39043563821288e-0 | 1 | lxn |
| rnf213a | 7.5921203 | 0.288676563 | 0.302 | 0.123 | 1.53338055104371e-0 | 1 | rnf213a |
| adamts15a1 | 8.1504119 | 0.259341492 | 0.269 | 0.1 | 1.64613870548928e-0 | 1 | adamts15a |
| nucks1b | 8.8744194 | 0.289329714 | 0.341 | 0.149 | 1.79236650411765e-0 | 1 | nucks1b |
| tegt | 9.1745287 | 0.323362594 | 0.621 | 0.343 | 1.85297957851353e-0 | 1 | tegt |
| jdp2b | 9.7148140 | 0.316453076 | 0.324 | 0.138 | 1.96210098501368e-0 | 1 | jdp2b |
| laptm4a | 1.2093893 | 0.273251218 | 0.39 | 0.178 | 2.44260366952549e-0 | 1 | laptm4a |
| atp2b2 | 1.3819763 | 0.27996139 | 0.324 | 0.137 | 2.79117756228786e-0 | 1 | atp2b2 |
| midn | 1.4967824 | 0.288063223 | 0.368 | 0.166 | 3.02305157596775e-0 | 1 | midn |
| rpl27a | 1.6166338 | 0.325621629 | 0.918 | 0.669 | 3.26511529514958e-0 | 1 | rpl27a |
| eif4a1b | 1.6658042 | 0.348452839 | 0.736 | 0.469 | 3.36442477679758e-0 | 1 | eif4a1b |
| etf1b | 1.9546139 | 0.28685075 | 0.401 | 0.188 | 3.94773385631772e-0 | 1 | etf1b |
| ftr82 | 2.0050739 | 0.274001199 | 0.253 | 0.095 | 4.04964790660586e-0 | 1 | ftr82 |
| egr1 | 2.0793626 | 0.3190267 | 0.341 | 0.152 | 4.19968869711383e-0 | 1 | egr1 |
| LOC103910438 | 2.3670098 | 0.273883683 | 0.374 | 0.171 | 4.78064980706938e-0 | 1 | LOC103910438 |

| | | | | | | | |
|---|---|---|---|---|---|---|---|
| anxa11b | 2.3954923 | 0.286516275 | 0.566 | 0.291 | 4.83817598605101e-0 | 1 | anxa11b |
| sumo2b | 2.5840809 | 0.333621384 | 0.736 | 0.438 | 5.21906823804809e-0 | 1 | sumo2b |
| khsrp | 3.0244676 | 0.25534634 | 0.363 | 0.165 | 6.10851737667979e-0 | 1 | khsrp |
| hmgb1a | 3.0329543 | 0.337365283 | 0.692 | 0.408 | 6.12565780756559e-0 | 1 | hmgb1a |
| fzd7b1 | 3.1013166 | 0.255823375 | 0.253 | 0.095 | 6.26372932579825e-0 | 1 | fzd7b |
| dcaf7 | 3.2635300 | 0.25762009 | 0.286 | 0.118 | 6.59135158409716e-0 | 1 | dcaf7 |
| hnrnpaba | 3.5878229 | 0.342968819 | 0.808 | 0.506 | 7.24632604478667e-0 | 1 | hnrnpaba |
| CABZ01072614.11 | 3.9436839 | 0.321314199 | 0.352 | 0.162 | 7.96505848276401e-0 | 1 | CABZ01072614.1 |
| rsrp1 | 4.0184416 | 0.258522842 | 0.302 | 0.125 | 8.11604655282958e-0 | 1 | rsrp1 |
| srsf5a | 4.0470242 | 0.309438369 | 0.654 | 0.368 | 8.17377490572191e-0 | 1 | srsf5a |
| pnn | 5.5276852 | 0.254384756 | 0.319 | 0.139 | 1.1164265954606e-05 | 1 | pnn |
| pik3r2 | 5.9311137 | 0.251929477 | 0.247 | 0.094 | 1.1979070486726e-05 | 1 | pik3r2 |
| cct8 | 8.9529749 | 0.289720329 | 0.593 | 0.322 | 1.80823234967729e-0 | 1 | cct8 |
| bccip | 9.3576796 | 0.257840141 | 0.445 | 0.225 | 1.88997054960094e-0 | 1 | bccip |
| tomm5 | 1.0390152 | 0.284736007 | 0.555 | 0.303 | 2.09849913977895e-0 | 1 | tomm5 |
| dnmt3ab1 | 1.0643294 | 0.286182034 | 0.341 | 0.156 | 2.14962613268081e-0 | 1 | dnmt3ab |
| ncl | 1.2524409 | 0.310165808 | 0.681 | 0.397 | 2.52955495549508e-0 | 1 | ncl |
| eif3g | 1.2661148 | 0.30102735 | 0.637 | 0.379 | 2.55717214831826e-0 | 1 | eif3g |
| hp1bp3 | 1.3093159 | 0.268766318 | 0.456 | 0.228 | 2.64442540776834e-0 | 1 | hp1bp3 |
| si:ch211-288g17.3 | 1.5691945 | 0.298149502 | 0.527 | 0.276 | 3.16930229740697e-0 | 1 | si:ch211-288g17.3 |
| si:ch73-376l24.2 | 1.6897221 | 0.286055181 | 0.335 | 0.154 | 3.4127319174547e-05 | 1 | si:ch73-376l24.2 |
| sf3b5 | 2.1849064 | 0.252794698 | 0.379 | 0.183 | 4.41285560785342e-0 | 1 | sf3b5 |
| ywhaqb | 2.5614154 | 0.302217834 | 0.489 | 0.264 | 5.17329081225784e-0 | 1 | ywhaqb |
| nucks1a | 2.5819286 | 0.282551103 | 0.429 | 0.218 | 5.21472138521395e-0 | 1 | nucks1a |
| acin1a | 3.0274124 | 0.308423084 | 0.511 | 0.282 | 6.11446496891304e-0 | 1 | acin1a |
| pin1 | 3.1688855 | 0.255966863 | 0.412 | 0.206 | 6.40019814295256e-0 | 1 | pin1 |
| efnb11 | 3.4679868 | 0.261930506 | 0.236 | 0.092 | 7.00429306147719e-0 | 1 | efnb1 |
| serbp1a | 3.5214984 | 0.315752497 | 0.885 | 0.637 | 7.11237041974691e-0 | 1 | serbp1a |
| irf2bp2b | 4.3891620 | 0.263851543 | 0.374 | 0.179 | 8.86479068244529e-0 | 1 | irf2bp2b |
| ube2e3 | 4.5150195 | 0.287309352 | 0.462 | 0.242 | 9.11898493351168e-0 | 1 | ube2e3 |
| ddx27 | 4.7351375 | 0.255838548 | 0.275 | 0.121 | 9.56355734258847e-0 | 1 | ddx27 |
| rtn4a | 5.0929023 | 0.26278873 | 0.451 | 0.235 | 0.000102861 | 1 | rtn4a |
| npm1a | 5.3293644 | 0.263983005 | 0.533 | 0.292 | 0.000107637 | 1 | npm1a |
| krt941 | 5.4965366 | 0.289317163 | 0.676 | 0.338 | 0.000111014 | 1 | krt94 |
| si:ch73-376l24.3 | 6.4095054 | 0.251086161 | 0.341 | 0.159 | 0.000129453 | 1 | si:ch73-376l24.3 |

| | | | | | | | |
|---|---|---|---|---|---|---|---|
| rab2a | 7.0982026 | 0.256036547 | 0.407 | 0.205 | 0.000143362 | 1 | rab2a |
| ywhae1 | 7.4564497 | 0.259647309 | 0.654 | 0.358 | 0.000150598 | 1 | ywhae1 |
| tnrc6c1 | 9.0871073 | 0.297218858 | 0.429 | 0.228 | 0.000183532 | 1 | tnrc6c1 |
| hnrnpa1b | 9.8330680 | 0.257959281 | 0.44 | 0.232 | 0.000198598 | 1 | hnrnpa1b |
| gtpbp4 | 1.1845606 | 0.259229161 | 0.538 | 0.298 | 0.000239246 | 1 | gtpbp4 |
| paip2b | 1.4962230 | 0.265254546 | 0.571 | 0.326 | 0.000302192 | 1 | paip2b |
| itm2ba | 1.5075376 | 0.293580333 | 0.659 | 0.394 | 0.000304477 | 1 | itm2ba |
| flna1 | 1.6775915 | 0.265381411 | 0.33 | 0.157 | 0.000338823 | 1 | flna |
| gadd45ba1 | 1.7098595 | 0.306875837 | 0.44 | 0.238 | 0.00034534 | 1 | gadd45ba |
| skp1 | 1.9089900 | 0.274531461 | 0.588 | 0.343 | 0.000385559 | 1 | skp1 |
| si:ch211-196l7.4 | 2.0239036 | 0.285044741 | 0.654 | 0.395 | 0.000408768 | 1 | si:ch211-196l7.4 |
| nop53 | 2.3886007 | 0.272915216 | 0.582 | 0.337 | 0.000482426 | 1 | nop53 |
| rnf10 | 2.8431656 | 0.261624642 | 0.385 | 0.202 | 0.000574234 | 1 | rnf10 |
| eef1db | 2.8491004 | 0.258349378 | 0.599 | 0.343 | 0.000575433 | 1 | eef1db |
| tsc22d3 | 4.5168083 | 0.255634625 | 0.44 | 0.236 | 0.00091226 | 1 | tsc22d3 |
| syncrip | 4.7971374 | 0.28320559 | 0.626 | 0.375 | 0.000968878 | 1 | syncrip |
| selenow1 | 5.7575435 | 0.300333251 | 0.868 | 0.575 | 0.001162851 | 1 | selenow1 |
| fkbp1aa | 6.0350485 | 0.29845842 | 0.83 | 0.56 | 0.001218899 | 1 | fkbp1aa |
| nsa2 | 6.2352349 | 0.290408177 | 0.742 | 0.506 | 0.00125933 | 1 | nsa2 |
| seta | 8.0786215 | 0.255557517 | 0.582 | 0.339 | 0.001631639 | 1 | seta |
| cct2 | 8.3692468 | 0.257363038 | 0.632 | 0.381 | 0.001690337 | 1 | cct2 |
| ubc | 1.7857814 | 0.272910331 | 0.863 | 0.639 | 0.003606743 | 1 | ubc |
| snrpb | 1.8265132 | 0.263958513 | 0.654 | 0.405 | 0.003689009 | 1 | snrpb |
| zgc:162730 | 1.9380977 | 0.327862159 | 0.709 | 0.476 | 0.003914376 | 1 | zgc:162730 |
| rsl24d1 | 1.9897405 | 0.293171726 | 0.786 | 0.558 | 0.004018679 | 1 | rsl24d1 |
| gnb1b | 7.4377765 | 0.253842317 | 0.533 | 0.326 | 0.015022077 | 1 | gnb1b |
| eif5a | 8.8482622 | 0.27106935 | 0.791 | 0.586 | 0.017870835 | 1 | eif5a |
| rpl22l1 | 7.2653564 | 0.258108004 | 0.852 | 0.603 | 0.146738405 | 1 | rpl22l1 |
| COX31 | 0.000287 | 0.255616173 | 0.841 | 0.578 | 1 | 1 | COX3 |
| grn1 | 2.3007906 | 1.890091402 | 0.845 | 0.048 | 4.64690696101113e-2 | 2 | grn1 |
| si:dkey-5n18.1 | 7.2917731 | 1.498080164 | 0.677 | 0.051 | 1.4727194318383e-13 | 2 | si:dkey-5n18.1 |
| c1qb | 8.6545974 | 1.416957526 | 0.626 | 0.05 | 1.74796905064593e-1 | 2 | c1qb |
| c1qc | 1.0046920 | 1.336405698 | 0.6 | 0.052 | 2.02917645510371e-1 | 2 | c1qc |
| zgc:153118 | 1.1115949 | 1.336996171 | 0.548 | 0.045 | 2.24508822164913e-1 | 2 | zgc:153118 |
| NPC2 (1 of many) | 5.7857802 | 1.24176252 | 0.523 | 0.041 | 1.16855403781385e-1 | 2 | NPC2 (1 of many) |

| | | | | | | | |
|---|---|---|---|---|---|---|---|
| cd74a | 2.4648976 | 1.610478533 | 0.845 | 0.165 | 4.97835388230717e-1 | 2 | cd74a |
| ccl34b.8 | 8.1475789 | 1.212709899 | 0.471 | 0.036 | 1.64556651826245e-9 | 2 | ccl34b.8 |
| marco | 8.3939803 | 1.142101663 | 0.497 | 0.042 | 1.6953322094276e-90 | 2 | marco |
| si:dkey-7i4.24 | 2.6518115 | 1.254782087 | 0.406 | 0.025 | 5.35586388505299e-8 | 2 | si:dkey-7i4.24 |
| cd74b | 9.3850971 | 1.429804479 | 0.845 | 0.207 | 1.89550806504274e-8 | 2 | cd74b |
| si:busm1-266f07.2 | 2.0896254 | 1.338003322 | 0.703 | 0.121 | 4.22041642078859e-8 | 2 | si:busm1-266f07.2 |
| lgals2a | 2.4897753 | 1.354912493 | 0.774 | 0.168 | 5.02859931888153e-8 | 2 | lgals2a |
| lygl1 | 9.3648261 | 1.187540532 | 0.6 | 0.083 | 1.89141393447439e-8 | 2 | lygl1 |
| ctss2.2 | 8.7484092 | 1.080864721 | 0.503 | 0.069 | 1.76691621781029e-6 | 2 | ctss2.2 |
| LOC564660 | 7.4032838 | 1.014027089 | 0.335 | 0.027 | 1.49524123566364e-6 | 2 | LOC564660 |
| c1qa | 7.7741970 | 0.988091403 | 0.452 | 0.057 | 1.57015458283104e-6 | 2 | c1qa |
| ms4a17a.10 | 5.8304324 | 0.9047993 | 0.394 | 0.043 | 1.17757244873582e-5 | 2 | ms4a17a.10 |
| jac3 | 1.3656455 | 0.962501651 | 0.323 | 0.026 | 2.75819427192643e-5 | 2 | jac3 |
| CU929150.1 | 7.6754898 | 0.807718609 | 0.29 | 0.022 | 1.55021868346523e-5 | 2 | CU929150.1 |
| zgc:103700 | 1.0291387 | 1.076994258 | 0.523 | 0.1 | 2.07855151590265e-5 | 2 | zgc:103700 |
| lgals9l1 | 2.1052852 | 0.870303777 | 0.323 | 0.033 | 4.25204466300883e-5 | 2 | lgals9l1 |
| tmsb4x | 2.6340715 | 0.393687248 | 1 | 0.937 | 5.32003433041774e-5 | 2 | tmsb4x |
| spi1a | 2.4000555 | 0.80075383 | 0.329 | 0.036 | 4.84739210604305e-4 | 2 | spi1a |
| fcer1gl | 1.2140995 | 1.082719116 | 0.8 | 0.219 | 2.45211684012138e-4 | 2 | fcer1gl |
| lgals3bpb | 1.8433565 | 0.886386679 | 0.323 | 0.035 | 3.72302726881306e-4 | 2 | lgals3bpb |
| ccl39.1 | 4.2693985 | 0.77310539 | 0.232 | 0.015 | 8.62290426219274e-4 | 2 | ccl39.1 |
| actb2 | 4.0757399 | 0.434680533 | 1 | 0.922 | 8.23177207763371e-4 | 2 | actb2 |
| pfn1 | 2.1819613 | 1.049591421 | 0.987 | 0.379 | 4.4069072763975e-38 | 2 | pfn1 |
| txn | 3.8608769 | 0.897465037 | 0.387 | 0.07 | 7.79781311987363e-3 | 2 | txn |
| slc3a2a | 5.8352346 | 0.727702875 | 0.297 | 0.039 | 1.17854233904286e-3 | 2 | slc3a2a |
| ccl39.3 | 1.8086300 | 0.596926907 | 0.187 | 0.012 | 3.65289010234959e-3 | 2 | ccl39.3 |
| mhc2b | 2.3220438 | 0.900720498 | 0.394 | 0.073 | 4.68983187981862e-3 | 2 | mhc2b |
| mafbb | 3.1358201 | 0.661667309 | 0.265 | 0.031 | 6.33341599420216e-3 | 2 | mafbb |
| BX649485.2 | 2.3921583 | 0.820748433 | 0.297 | 0.04 | 4.83144219137292e-3 | 2 | BX649485.2 |
| grna | 8.1909941 | 0.749229255 | 0.329 | 0.053 | 1.65433508852807e-3 | 2 | grna |
| zgc:64065 | 1.8122780 | 0.689525995 | 0.271 | 0.035 | 3.66025801798119e-3 | 2 | zgc:64065 |
| CU499330.1 | 1.3163745 | 0.802592437 | 0.368 | 0.066 | 2.65868169861897e-3 | 2 | CU499330.1 |
| ctsba | 2.9378941 | 1.017639661 | 0.587 | 0.223 | 5.93366486473028e-3 | 2 | ctsba |
| mhc2dab | 3.4861106 | 0.803183131 | 0.355 | 0.068 | 7.04089769104682e-3 | 2 | mhc2dab |
| f13a1b | 6.5041620 | 0.605832717 | 0.219 | 0.024 | 1.31364561580568e-3 | 2 | f13a1b |

| | | | | | | | |
|---|---|---|---|---|---|---|---|
| gstt1a | 5.8262364 | 0.640751929 | 0.271 | 0.041 | 1.17672496881668e-2 | 2 | gstt1a |
| havcr1 | 7.5770693 | 0.617455139 | 0.265 | 0.038 | 1.53034068847999e-2 | 2 | havcr1 |
| gng10 | 1.6719808 | 0.748658427 | 0.361 | 0.077 | 3.37689974287764e-2 | 2 | gng10 |
| krt18b | 2.1654602 | 0.888439234 | 0.658 | 0.298 | 4.37358014831834e-2 | 2 | krt18b |
| ifi30 | 2.7051662 | 0.918473093 | 0.639 | 0.286 | 5.46362425976477e-2 | 2 | ifi30 |
| gpx1a | 8.3637685 | 0.844924695 | 0.587 | 0.212 | 1.68923032731482e-2 | 2 | gpx1a |
| mfap4 | 1.1614106 | 0.690787674 | 0.277 | 0.044 | 2.34570104192408e-2 | 2 | mfap4 |
| cndp2 | 1.3364040 | 0.801856551 | 0.387 | 0.099 | 2.69913520829096e-2 | 2 | cndp2 |
| cotl1 | 2.1901687 | 0.811138633 | 0.826 | 0.384 | 4.42348389133683e-2 | 2 | cotl1 |
| si:ch211-147m6.1 | 1.0829345 | 0.547519859 | 0.206 | 0.024 | 2.18720291362601e-2 | 2 | si:ch211-147m6.1 |
| si:ch211-194m7.3 | 6.7340838 | 0.518273211 | 0.187 | 0.02 | 1.36008291498857e-2 | 2 | si:ch211-194m7.3 |
| si:ch211-283g2.3 | 1.6405435 | 0.56497321 | 0.187 | 0.021 | 3.31340574046705e-2 | 2 | si:ch211-283g2.3 |
| ccl35.1 | 2.5811253 | 0.81625528 | 0.342 | 0.076 | 5.21309879146247e-2 | 2 | ccl35.1 |
| zgc:174904 | 6.4094846 | 0.703791665 | 0.284 | 0.055 | 1.29452362403646e-2 | 2 | zgc:174904 |
| ckbb | 6.9681396 | 0.757366116 | 0.529 | 0.168 | 1.40735517015306e-2 | 2 | ckbb |
| ccr12b.2 | 7.8429798 | 0.520941792 | 0.2 | 0.026 | 1.58404664570748e-2 | 2 | ccr12b.2 |
| si:ch211-165i18.2 | 2.0483263 | 0.614134601 | 0.258 | 0.044 | 4.13700477702944e-2 | 2 | si:ch211-165i18.2 |
| psap | 7.7398457 | 0.903016869 | 0.561 | 0.259 | 1.56321664642891e-2 | 2 | psap |
| ccl34b.1 | 9.5093019 | 0.496947029 | 0.161 | 0.017 | 1.92059372163035e-2 | 2 | ccl34b.1 |
| jac9 | 1.3405707 | 0.492642067 | 0.168 | 0.019 | 2.70755077944792e-2 | 2 | jac9 |
| LOC100536946 | 2.2493329 | 0.571076022 | 0.219 | 0.034 | 4.54297775264193e-2 | 2 | LOC100536946 |
| arpc1b | 3.1649157 | 0.74820954 | 0.755 | 0.335 | 6.39218025046915e-2 | 2 | arpc1b |
| slc40a1 | 5.0677643 | 0.63164316 | 0.258 | 0.051 | 1.02353636361783e-2 | 2 | slc40a1 |
| me1 | 1.8948290 | 0.36127416 | 0.135 | 0.013 | 3.82698631777041e-2 | 2 | me1 |
| rnaset2l | 4.6758746 | 0.633748248 | 0.297 | 0.069 | 9.44386410223814e-2 | 2 | rnaset2l |
| actb1 | 6.5019044 | 0.497260546 | 0.916 | 0.73 | 1.31318965161584e-1 | 2 | actb1 |
| itgae.1 | 1.4569256 | 0.566405756 | 0.239 | 0.045 | 2.94255270811375e-1 | 2 | itgae.1 |
| atp6v1g1 | 1.7207726 | 0.778465729 | 0.452 | 0.177 | 3.47544451822e-19 | 2 | atp6v1g1 |
| LOC101885706 | 2.9161997 | 0.420585403 | 0.148 | 0.017 | 5.88984867315288e-1 | 2 | LOC101885706 |
| ccl39.2 | 3.7878993 | 0.304125977 | 0.11 | 0.008 | 7.6504202303749e-19 | 2 | ccl39.2 |
| snx10b | 1.2137449 | 0.339081561 | 0.123 | 0.011 | 2.45140072596819e-1 | 2 | snx10b |
| si:zfos-2330d3.3 | 2.0543670 | 0.429924476 | 0.148 | 0.017 | 4.14920516503293e-1 | 2 | si:zfos-2330d3.3 |
| cmklr1 | 7.0965539 | 0.487238622 | 0.194 | 0.032 | 1.43329099266468e-1 | 2 | cmklr1 |
| ehd4 | 9.3253707 | 0.523022489 | 0.226 | 0.045 | 1.88344512545999e-1 | 2 | ehd4 |
| ccl34a.4 | 1.7652998 | 0.474912909 | 0.187 | 0.03 | 3.56537608890448e-1 | 2 | ccl34a.4 |

| | | | | | | | |
|---|---|---|---|---|---|---|---|
| BX323596.2 | 3.4085884 | 0.437073075 | 0.129 | 0.015 | 6.88432607588016e-1 | 2 | BX323596.2 |
| ctsd | 5.0332476 | 0.794222077 | 0.426 | 0.173 | 1.01656502166399e-1 | 2 | ctsd |
| si:ch211-1o7.3 | 9.6598391 | 0.376343991 | 0.142 | 0.019 | 1.95099771102665e-1 | 2 | si:ch211-1o7.3 |
| si:ch211-198c19.3: | 4.1219663 | 0.756391993 | 0.432 | 0.169 | 8.3251353960639e-15 | 2 | si:ch211-198c19.3 |
| tspan36 | 4.1483654 | 0.716122484 | 0.348 | 0.121 | 8.37845363500661e-1 | 2 | tspan36 |
| hspa8 | 1.2282012 | 0.28357506 | 0.89 | 0.839 | 2.48059806804745e-1 | 2 | hspa8 |
| fabp11a | 8.3385681 | 0.801112096 | 0.316 | 0.097 | 1.68414061617091e-1 | 2 | fabp11a |
| cd9b | 1.0855377 | 0.584104516 | 0.174 | 0.032 | 2.19246054860291e-1 | 2 | cd9b |
| si:dkey-27i16.2 | 1.2134436 | 0.571276139 | 0.29 | 0.087 | 2.45079213959839e-1 | 2 | si:dkey-27i16.2 |
| cebpb | 3.9265545 | 0.687412407 | 0.548 | 0.28 | 7.93046215880576e-1 | 2 | cebpb |
| si:busm1-194e12.1 | 4.0786896 | 0.541761004 | 0.232 | 0.059 | 8.23772945407541e-1 | 2 | si:busm1-194e12.11 |
| gnpda1 | 7.9880282 | 0.463400218 | 0.206 | 0.048 | 1.61334206104166e-1 | 2 | gnpda1 |
| map3k14a | 1.5856699 | 0.539580087 | 0.2 | 0.047 | 3.20257749825039e-1 | 2 | map3k14a |
| ms4a17a.6 | 1.9226063 | 0.51129199 | 0.226 | 0.058 | 3.88308803097579e-1 | 2 | ms4a17a.6 |
| mpeg1.1 | 1.9972121 | 0.349663332 | 0.135 | 0.021 | 4.03376940433055e-1 | 2 | mpeg1.1 |
| ctsk | 2.4255254 | 0.648524174 | 0.335 | 0.126 | 4.89883377423876e-1 | 2 | ctsk |
| laptm5 | 2.5965020 | 0.591272239 | 0.471 | 0.2 | 5.2441551159755e-12 | 2 | laptm5 |
| CR318588.1 | 4.0157868 | 0.660139544 | 0.561 | 0.248 | 8.11068477171112e-1 | 2 | CR318588.1 |
| epdl1 | 1.6761497 | 0.363284092 | 0.116 | 0.016 | 3.38531963533749e-1 | 2 | epdl1 |
| cx32.2 | 1.8972399 | 0.382645408 | 0.168 | 0.034 | 3.83185544521937e-1 | 2 | cx32.2 |
| si:dkey-56f14.7 | 2.4769062 | 0.295861047 | 0.11 | 0.015 | 5.00260750243182e-1 | 2 | si:dkey-56f14.7 |
| irf8 | 4.8241962 | 0.444334546 | 0.206 | 0.053 | 9.74342920524825e-1 | 2 | irf8 |
| lta4h | 5.7314749 | 0.511014522 | 0.271 | 0.085 | 1.15758599977246e-1 | 2 | lta4h |
| npc2 | 8.3478270 | 0.704071529 | 0.465 | 0.26 | 1.68601063295533e-1 | 2 | npc2 |
| LOC110437731 | 9.0537840 | 0.498316569 | 0.29 | 0.086 | 1.82859275784872e-1 | 2 | LOC110437731 |
| LO018493.1 | 9.9105748 | 0.399003277 | 0.155 | 0.031 | 2.0016388123047e-10 | 2 | LO018493.1 |
| mrc1b | 1.1672423 | 0.447317532 | 0.206 | 0.051 | 2.35747929214988e-1 | 2 | mrc1b |
| apoeb | 4.9778098 | 0.513609486 | 0.155 | 0.032 | 1.00536825388045e-0 | 2 | apoeb |
| mcl1a | 5.6399746 | 0.479164516 | 0.742 | 0.534 | 1.13910567144501e-0 | 2 | mcl1a |
| cd63 | 7.3132747 | 0.467647293 | 0.703 | 0.562 | 1.47706209383607e-0 | 2 | cd63 |
| hmox1a | 2.0226807 | 0.429243942 | 0.129 | 0.024 | 4.08520827575787e-0 | 2 | hmox1a |
| clic2 | 3.1330424 | 0.524242927 | 0.258 | 0.089 | 6.32780576304019e-0 | 2 | clic2 |
| gsdmea | 8.3718235 | 0.289603891 | 0.116 | 0.02 | 1.69085721020358e-0 | 2 | gsdmea |
| rac2 | 1.1173878 | 0.537795311 | 0.484 | 0.249 | 2.2567881685034e-08 | 2 | rac2 |
| mapre1a | 1.1873994 | 0.546524134 | 0.394 | 0.188 | 2.39819065632779e-0 | 2 | mapre1a |

| | | | | | | | |
|---|---|---|---|---|---|---|---|
| scpep1 | 1.8757245 | 0.528938083 | 0.29 | 0.116 | 3.78840079619441e-0 | 2 | scpep1 |
| si:ch73-158p21.3 | 1.9840030 | 0.299827572 | 0.123 | 0.023 | 4.00709097461575e-0 | 2 | si:ch73-158p21.3 |
| atp6v1e1b | 2.3140590 | 0.523012716 | 0.303 | 0.124 | 4.67370500046322e-0 | 2 | atp6v1e1b |
| zgc:152830 | 3.6340424 | 0.461175786 | 0.219 | 0.071 | 7.339675468311e-08 | 2 | zgc:152830 |
| si:cabz01074946.1 | 5.9064313 | 0.458465055 | 0.206 | 0.064 | 1.19292193679104e-0 | 2 | si:cabz01074946.1 |
| sftpbb | 6.7682271 | 0.522893122 | 0.245 | 0.086 | 1.36697883308744e-0 | 2 | sftpbb |
| atp6v0ca | 9.3106506 | 0.585805993 | 0.413 | 0.226 | 1.88047211542239e-0 | 2 | atp6v0ca |
| cst14b.1 | 9.4889654 | 0.542282458 | 0.271 | 0.107 | 1.91648634197094e-0 | 2 | cst14b.1 |
| wasb | 1.7858752 | 0.51853666 | 0.335 | 0.149 | 3.60693225080634e-0 | 2 | wasb |
| cfl1 | 2.2881301 | 0.300464765 | 0.884 | 0.76 | 4.62133641981538e-0 | 2 | cfl1 |
| tuba8l2 | 4.2156077 | 0.506214667 | 0.252 | 0.097 | 8.51426295751021e-0 | 2 | tuba8l2 |
| coro1a | 5.4078434 | 0.459360983 | 0.555 | 0.265 | 1.09222213555805e-0 | 2 | coro1a |
| cap1 | 5.9917630 | 0.519279561 | 0.51 | 0.32 | 1.21015638003307e-0 | 2 | cap1 |
| arl6ip1 | 6.1289379 | 0.559474414 | 0.452 | 0.279 | 1.23786159871998e-0 | 2 | arl6ip1 |
| dpp7 | 1.5785174 | 0.503798308 | 0.265 | 0.112 | 3.18813178167693e-0 | 2 | dpp7 |
| sec61g | 1.9545665 | 0.310906967 | 0.813 | 0.722 | 3.94763798157512e-0 | 2 | sec61g |
| mdh1aa | 2.2825626 | 0.494900723 | 0.516 | 0.349 | 4.610091770703e-06 | 2 | mdh1aa |
| tpp1 | 9.8898811 | 0.453444079 | 0.226 | 0.088 | 1.99745929116618e-0 | 2 | tpp1 |
| gabarapb | 1.0255134 | 0.44801957 | 0.561 | 0.447 | 2.07122955845232e-0 | 2 | gabarapb |
| si:ch211-212k18.7 | 1.3162344 | 0.491802044 | 0.342 | 0.178 | 2.65839868988199e-0 | 2 | si:ch211-212k18.7 |
| zgc:1627301 | 1.3787175 | 0.42754746 | 0.671 | 0.482 | 2.78459577309222e-0 | 2 | zgc:162730 |
| alas1 | 1.5884672 | 0.503711653 | 0.271 | 0.122 | 3.20822731579444e-0 | 2 | alas1 |
| naga | 2.2796949 | 0.410002177 | 0.206 | 0.076 | 4.60429993982553e-0 | 2 | naga |
| fgl2a | 3.8572543 | 0.442125094 | 0.277 | 0.121 | 7.79049666956783e-0 | 2 | fgl2a |
| glrx | 4.0798819 | 0.431862489 | 0.548 | 0.421 | 8.24013750800474e-0 | 2 | glrx |
| fth1a | 5.6181957 | 0.479420565 | 0.581 | 0.446 | 0.000113471 | 2 | fth1a |
| cfp | 6.8707780 | 0.380527882 | 0.213 | 0.078 | 0.000138769 | 2 | cfp |
| enpp1 | 1.0151440 | 0.282965558 | 0.123 | 0.032 | 0.000205029 | 2 | enpp1 |
| smox | 1.2964935 | 0.473755536 | 0.226 | 0.097 | 0.000261853 | 2 | smox |
| pepd | 4.9472234 | 0.40919805 | 0.226 | 0.1 | 0.000999191 | 2 | pepd |
| si:dkey-262k9.4 | 6.2107231 | 0.382574057 | 0.213 | 0.089 | 0.00125438 | 2 | si:dkey-262k9.4 |
| grapa | 7.1666727 | 0.255857088 | 0.103 | 0.026 | 0.001447453 | 2 | grapa |
| capgb | 9.5824405 | 0.377909073 | 0.265 | 0.12 | 0.001935366 | 2 | capgb |
| zgc:64051 | 1.0831834 | 0.426757548 | 0.323 | 0.169 | 0.002187706 | 2 | zgc:64051 |
| spi1b | 1.0958460 | 0.37185604 | 0.265 | 0.12 | 0.00221328 | 2 | spi1b |

| | | | | | | | |
|---|---|---|---|---|---|---|---|
| ctsz | 1.3738706 | 0.438195854 | 0.239 | 0.112 | 0.002774806 | 2 | ctsz |
| pycard | 1.7798087 | 0.389177738 | 0.277 | 0.139 | 0.00359468 | 2 | pycard |
| BX901920.1 | 1.9776480 | 0.35693746 | 0.187 | 0.074 | 0.003994256 | 2 | BX901920.1 |
| XLOC-028370 | 3.5304608 | 0.369682912 | 0.148 | 0.053 | 0.007130472 | 2 | XLOC-028370 |
| samsn1a | 3.7467866 | 0.304578805 | 0.142 | 0.048 | 0.007567385 | 2 | samsn1a |
| ctsh | 4.5305432 | 0.436938789 | 0.252 | 0.128 | 0.009150338 | 2 | ctsh |
| nccrp1 | 6.0951724 | 0.436696651 | 0.355 | 0.215 | 0.01231042 | 2 | nccrp1 |
| CR392341.1 | 6.6169795 | 0.317778483 | 0.155 | 0.058 | 0.013364314 | 2 | CR392341.1 |
| ctsa | 7.8879151 | 0.442295746 | 0.245 | 0.127 | 0.015931222 | 2 | ctsa |
| si:ch211-102c2.4 | 1.1858749 | 0.345189068 | 0.329 | 0.168 | 0.023951117 | 2 | si:ch211-102c2.4 |
| rap1b | 1.6994931 | 0.38431028 | 0.445 | 0.322 | 0.034324662 | 2 | rap1b |
| fuca2 | 2.0564575 | 0.36884812 | 0.194 | 0.089 | 0.041534272 | 2 | fuca2 |
| gm2a | 2.0910080 | 0.404980362 | 0.245 | 0.127 | 0.042232089 | 2 | gm2a |
| CR855311.11 | 2.4372458 | 0.303164671 | 0.226 | 0.101 | 0.049225054 | 2 | CR855311.1 |
| cxcr3.2 | 3.6544628 | 0.262001977 | 0.116 | 0.039 | 0.073809185 | 2 | cxcr3.2 |
| atp6v1ba | 6.2002338 | 0.351218969 | 0.174 | 0.079 | 0.125226124 | 2 | atp6v1ba |
| kctd12.2 | 6.4104485 | 0.400020048 | 0.329 | 0.206 | 0.129471829 | 2 | kctd12.2 |
| slc11a2 | 6.9989805 | 0.294227049 | 0.142 | 0.056 | 0.14135841 | 2 | slc11a2 |
| si:dkey-53k12.2 | 7.6567070 | 0.266778283 | 0.116 | 0.041 | 0.154642513 | 2 | si:dkey-53k12.2 |
| asah1b | 1.2504924 | 0.337713338 | 0.206 | 0.105 | 0.252561954 | 2 | asah1b |
| arpc3 | 1.3230766 | 0.334083724 | 0.555 | 0.479 | 0.267221791 | 2 | arpc3 |
| dnase2 | 1.4945786 | 0.252470785 | 0.103 | 0.035 | 0.301860043 | 2 | dnase2 |
| rgs2 | 1.6051395 | 0.253816724 | 0.11 | 0.038 | 0.324190036 | 2 | rgs2 |
| arpc2 | 2.3586539 | 0.300941811 | 0.548 | 0.485 | 0.476377343 | 2 | arpc2 |
| cebpa1 | 3.1061736 | 0.375358061 | 0.284 | 0.169 | 0.627353889 | 2 | cebpa |
| lipf1 | 3.7346067 | 0.338695515 | 0.161 | 0.076 | 0.75427852 | 2 | lipf |
| XLOC-025423 | 4.1535634 | 0.359894062 | 0.31 | 0.187 | 0.838895203 | 2 | XLOC-025423 |
| arpc5b | 4.2982115 | 0.351165452 | 0.348 | 0.226 | 0.868109788 | 2 | arpc5b |
| si:dkey-69o16.5 | 6.6383703 | 0.272437575 | 0.142 | 0.064 | 1 | 2 | si:dkey-69o16.5 |
| selenop | 6.9236904 | 0.26525008 | 0.129 | 0.055 | 1 | 2 | selenop |
| mbd2 | 7.9021073 | 0.25153235 | 0.135 | 0.059 | 1 | 2 | mbd2 |
| ecm1b | 0.000114 | 0.458796165 | 0.226 | 0.131 | 1 | 2 | ecm1b |
| atp6v1d | 0.000128 | 0.261218624 | 0.116 | 0.048 | 1 | 2 | atp6v1d |
| rab8b | 0.000146 | 0.324066827 | 0.2 | 0.113 | 1 | 2 | rab8b |
| grap2b | 0.000152 | 0.308362087 | 0.303 | 0.189 | 1 | 2 | grap2b |

| | | | | | | | | |
|---|---|---|---|---|---|---|---|---|
| si:ch211-15b10.6 | 0.000153 | 0.326206705 | 0.187 | 0.103 | | 1 | 2 | si:ch211-15b10.6 |
| tmsb1 | 0.000164 | 0.255101686 | 0.323 | 0.188 | | 1 | 2 | tmsb1 |
| ccl39.7 | 0.000292 | 0.417041308 | 0.148 | 0.075 | | 1 | 2 | ccl39.7 |
| vsir | 0.000325 | 0.282701963 | 0.194 | 0.107 | | 1 | 2 | vsir |
| zgc:110239 | 0.000329 | 0.311918386 | 0.206 | 0.122 | | 1 | 2 | zgc:110239 |
| lamtor5 | 0.000358 | 0.330331441 | 0.213 | 0.128 | | 1 | 2 | lamtor5 |
| sat1a.2 | 0.000367 | 0.323562427 | 0.406 | 0.335 | | 1 | 2 | sat1a.2 |
| pkma | 0.000424 | 0.348040466 | 0.277 | 0.195 | | 1 | 2 | pkma |
| XLOC-023492 | 0.000454 | 0.262002942 | 0.168 | 0.089 | | 1 | 2 | XLOC-023492 |
| arpc4l | 0.00059 | 0.306934417 | 0.419 | 0.349 | | 1 | 2 | arpc4l |
| atp5l | 0.000691 | 0.250914989 | 0.561 | 0.526 | | 1 | 2 | atp5l |
| sdcbp2 | 0.001015 | 0.298165753 | 0.4 | 0.342 | | 1 | 2 | sdcbp2 |
| derl2 | 0.001404 | 0.336574269 | 0.194 | 0.122 | | 1 | 2 | derl2 |
| rnaseka | 0.001744 | 0.308756873 | 0.2 | 0.128 | | 1 | 2 | rnaseka |
| lgmn | 0.00273 | 0.285878125 | 0.2 | 0.128 | | 1 | 2 | lgmn |
| cyba | 0.003452 | 0.268613921 | 0.2 | 0.131 | | 1 | 2 | cyba |
| capza1a | 0.003737 | 0.268963883 | 0.387 | 0.344 | | 1 | 2 | capza1a |
| litaf | 0.004516 | 0.278122275 | 0.271 | 0.206 | | 1 | 2 | litaf |
| mcl1b | 0.004879 | 0.271142949 | 0.374 | 0.316 | | 1 | 2 | mcl1b |
| gabarapa | 0.006696 | 0.309544465 | 0.31 | 0.258 | | 1 | 2 | gabarapa |
| lgals9l31 | 0.006914 | 0.294235444 | 0.245 | 0.186 | | 1 | 2 | lgals9l3 |
| bloc1s6 | 0.007447 | 0.27673157 | 0.194 | 0.134 | | 1 | 2 | bloc1s6 |
| ssr3 | 0.008049 | 0.259211957 | 0.439 | 0.386 | | 1 | 2 | ssr3 |
| bhmt | 1.5495760 | 0.426646169 | 0.177 | 0.017 | 3.12967878884949e-2 | | 3 | bhmt |
| rplp0 | 6.1811549 | 0.333893225 | 0.992 | 0.882 | 1.24840785951414e-2 | | 3 | rplp0 |
| rps12 | 1.2451495 | 0.305410461 | 0.984 | 0.886 | 2.5148285223329e-21 | | 3 | rps12 |
| rps7 | 1.9043791 | 0.263689833 | 0.976 | 0.908 | 3.84627448828244e-2 | | 3 | rps7 |
| rps2 | 5.5083168 | 0.254152293 | 0.984 | 0.918 | 1.11251475089239e-1 | | 3 | rps2 |
| si:dkey-14d8.6 | 2.4190545 | 0.513568876 | 0.194 | 0.025 | 4.88576438057898e-1 | | 3 | si:dkey-14d8.6 |
| ppdpfa | 7.3479643 | 0.540478008 | 0.21 | 0.03 | 1.48406835009789e-1 | | 3 | ppdpfa |
| rpl35 | 5.4923575 | 0.260187722 | 0.984 | 0.901 | 1.10929144881251e-1 | | 3 | rpl35 |
| cel.2 | 3.0097348 | 0.26085603 | 0.105 | 0.012 | 6.07876140945145e-1 | | 3 | cel.2 |
| rpl7 | 4.4775981 | 0.253028857 | 0.944 | 0.866 | 9.04340506297141e-1 | | 3 | rpl7 |
| aldob | 2.0821617 | 0.295665081 | 0.121 | 0.019 | 4.20534202487883e-0 | | 3 | aldob |
| cpa4 | 5.3562756 | 0.285446144 | 0.113 | 0.017 | 1.08180699655901e-0 | | 3 | cpa4 |

| | | | | | | | |
|---|---|---|---|---|---|---|---|
| eif4ebp3 | 1.5040641 | 0.299528088 | 0.105 | 0.017 | 3.03775833664605e-0 | 3 | eif4ebp3 |
| gamt | 2.7828682 | 0.33563612 | 0.137 | 0.03 | 5.62055890980338e-0 | 3 | gamt |
| cpb1 | 3.1976336 | 0.264430012 | 0.121 | 0.023 | 6.45826072850646e-0 | 3 | cpb1 |
| rpl22l11 | 5.7049042 | 0.346980969 | 0.653 | 0.623 | 0.000115222 | 3 | rpl22l1 |
| prdx21 | 0.001463 | 0.29559286 | 0.484 | 0.494 | 1 | 3 | prdx2 |
| gstm.31 | 0.002382 | 0.372504011 | 0.387 | 0.286 | 1 | 3 | gstm.3 |
| zgc:927441 | 0.003662 | 0.303672603 | 0.298 | 0.206 | 1 | 3 | zgc:92744 |
| fth1a1 | 0.004504 | 0.289678904 | 0.476 | 0.455 | 1 | 3 | fth1a |
| ccl33.3 | 3.8111374 | 2.376691915 | 0.889 | 0.054 | 7.69735421125931e-1 | 4 | ccl33.3 |
| pfn11 | 2.5567072 | 1.347984635 | 1 | 0.393 | 5.16378165975664e-7 | 4 | pfn1 |
| vaspa | 2.5045451 | 1.242612267 | 0.62 | 0.096 | 5.05842975361017e-6 | 4 | vaspa |
| coro1a1 | 3.0907639 | 1.277937684 | 0.88 | 0.254 | 6.24241598081446e-5 | 4 | coro1a |
| arpc1b1 | 7.7453537 | 1.253398382 | 0.926 | 0.335 | 1.56432910135416e-5 | 4 | arpc1b |
| fcer1gl1 | 4.9258054 | 1.275220631 | 0.88 | 0.229 | 9.9486493284444e-52 | 4 | fcer1gl |
| sftpba | 6.8425827 | 1.00795561 | 0.454 | 0.055 | 1.38199642942189e-4 | 4 | sftpba |
| actb21 | 4.2860821 | 0.552872024 | 1 | 0.924 | 8.65660005024387e-4 | 4 | actb2 |
| si:ch211-214p16.1 | 9.5490727 | 1.249158379 | 0.769 | 0.166 | 1.92862622416455e-4 | 4 | si:ch211-214p16.1 |
| actb11 | 3.8766995 | 0.790223315 | 1 | 0.73 | 7.82977009102995e-4 | 4 | actb1 |
| cotl11 | 5.9399922 | 1.127460201 | 0.898 | 0.39 | 1.19970023016347e-4 | 4 | cotl1 |
| tmsb4x1 | 6.1219895 | 0.43189027 | 1 | 0.938 | 1.23645821964075e-4 | 4 | tmsb4x |
| si:ch211-220f21.3 | 7.4412310 | 0.724755872 | 0.269 | 0.02 | 1.50290544332123e-4 | 4 | si:ch211-220f21.3 |
| ckbb1 | 4.2509455 | 1.170275228 | 0.639 | 0.171 | 8.58563473366679e-3 | 4 | ckbb |
| arpc5b1 | 4.0005481 | 1.114551988 | 0.667 | 0.212 | 8.07990719449708e-3 | 4 | arpc5b |
| arpc31 | 4.2412626 | 0.901748703 | 0.806 | 0.467 | 8.56607819591803e-2 | 4 | arpc3 |
| wipf1a | 2.9789204 | 0.825224587 | 0.38 | 0.071 | 6.01652560185698e-2 | 4 | wipf1a |
| cfl11 | 1.4247761 | 0.554044286 | 0.917 | 0.761 | 2.87762042507334e-2 | 4 | cfl1 |
| ptpn6 | 1.0567816 | 0.90512693 | 0.472 | 0.131 | 2.13438193251457e-2 | 4 | ptpn6 |
| rac21 | 3.3816522 | 0.858038102 | 0.639 | 0.246 | 6.82992303537412e-2 | 4 | rac2 |
| arpc4 | 6.1113877 | 0.91080879 | 0.648 | 0.303 | 1.23431698923245e-2 | 4 | arpc4 |
| grap2b1 | 9.9768001 | 0.866959666 | 0.546 | 0.178 | 2.01501432922915e-2 | 4 | grap2b |
| LOC100536213 | 1.4770705 | 0.884579459 | 0.472 | 0.135 | 2.98323947690162e-2 | 4 | LOC100536213 |
| eomesa | 1.5196405 | 0.609934187 | 0.25 | 0.035 | 3.0692180995777e-21 | 4 | eomesa |
| rplp2 | 2.3453740 | 0.282790367 | 0.991 | 0.926 | 4.73695191902391e-2 | 4 | rplp2 |
| fam49ba | 1.4159853 | 0.862320036 | 0.417 | 0.111 | 2.85986557761818e-2 | 4 | fam49ba |
| arpc21 | 3.9261528 | 0.790966188 | 0.741 | 0.476 | 7.9296508611725e-20 | 4 | arpc2 |

| | | | | | | | |
|---|---|---|---|---|---|---|---|
| myl12.1 | 1.1018293 | 0.614874964 | 0.796 | 0.625 | 2.22536481355697e-1 | 4 | myl12.1 |
| cd74a1 | 3.6389390 | 0.772355468 | 0.611 | 0.194 | 7.34956529351797e-1 | 4 | cd74a |
| kctd12.21 | 4.3447149 | 0.827615486 | 0.519 | 0.199 | 8.77502082785503e-1 | 4 | kctd12.2 |
| wasa | 4.6458885 | 0.748475937 | 0.389 | 0.107 | 9.38330111609813e-1 | 4 | wasa |
| spi1b1 | 1.8231240 | 0.784735706 | 0.398 | 0.116 | 3.682163690743e-16 | 4 | spi1b |
| hcls1 | 4.2836985 | 0.715200881 | 0.37 | 0.101 | 8.65178595016724e-1 | 4 | hcls1 |
| ptmaa | 5.1915436 | 0.556556096 | 0.833 | 0.65 | 1.04853607130036e-1 | 4 | ptmaa |
| fcer1g | 5.3068803 | 0.583526168 | 0.241 | 0.043 | 1.07183061880131e-1 | 4 | fcer1g |
| mef2b | 2.7100206 | 0.464440256 | 0.176 | 0.023 | 5.4734287818322e-15 | 4 | mef2b |
| LOC101882805 | 1.1024828 | 0.322159452 | 0.102 | 0.007 | 2.2266846856112e-14 | 4 | LOC101882805 |
| c6ast3 | 2.0280944 | 0.301647828 | 0.148 | 0.018 | 4.09614228733627e-1 | 4 | c6ast3 |
| abracl | 4.2910501 | 0.775396594 | 0.509 | 0.231 | 8.66663402925482e-1 | 4 | abracl |
| LOC110438870 | 4.9613873 | 0.277643475 | 0.102 | 0.008 | 1.00205139466052e-1 | 4 | LOC110438870 |
| si:dkey-222f2.7 | 9.7729928 | 0.513607407 | 0.204 | 0.037 | 1.97385136319258e-1 | 4 | si:dkey-222f2.7 |
| CR753876.1 | 1.2668409 | 0.67316432 | 0.324 | 0.093 | 2.55863868489906e-1 | 4 | CR753876.1 |
| si:ch211-102c2.41 | 9.0770137 | 0.698784129 | 0.435 | 0.166 | 1.83328446329473e-1 | 4 | si:ch211-102c2.4 |
| rps17 | 5.6165085 | 0.259476181 | 0.963 | 0.877 | 1.13436624149344e-0 | 4 | rps17 |
| abi3a | 7.4165735 | 0.533482603 | 0.241 | 0.061 | 1.49792536558785e-0 | 4 | abi3a |
| dicp1.1 | 8.4898490 | 0.4082395 | 0.139 | 0.021 | 1.71469481290489e-0 | 4 | dicp1.1 |
| si:dkey-172o19.2 | 1.0401280 | 0.314363163 | 0.111 | 0.013 | 2.10074665834497e-0 | 4 | si:dkey-172o19.2 |
| lcp2b | 4.9394561 | 0.371958863 | 0.148 | 0.025 | 9.9762195097763e-09 | 4 | lcp2b |
| arpc4l1 | 5.2948712 | 0.651372707 | 0.565 | 0.343 | 1.06940513842366e-0 | 4 | arpc4l |
| LOC108190565 | 6.2242563 | 0.337255384 | 0.12 | 0.017 | 1.25711306279396e-0 | 4 | LOC108190565 |
| h1fx | 6.4110646 | 0.790420613 | 0.472 | 0.229 | 1.29484272467776e-0 | 4 | h1fx |
| ccl35.11 | 2.1097150 | 0.592920409 | 0.287 | 0.085 | 4.26099147267439e-0 | 4 | ccl35.1 |
| csk | 2.0158116 | 0.590451768 | 0.306 | 0.112 | 4.0713348392221e-07 | 4 | csk |
| cd74b1 | 2.2773577 | 0.588478245 | 0.546 | 0.239 | 4.59957951188776e-0 | 4 | cd74b |
| zgc:153867 | 4.0569680 | 0.383897258 | 0.722 | 0.638 | 8.19385836036819e-0 | 4 | zgc:153867 |
| si:dkey-33i11.9 | 1.7898889 | 0.416687055 | 0.167 | 0.039 | 3.6150387414601e-06 | 4 | si:dkey-33i11.9 |
| CR381673.1 | 3.6772478 | 0.482363186 | 0.194 | 0.053 | 7.42693754140921e-0 | 4 | CR381673.1 |
| skap2 | 4.2765756 | 0.569706479 | 0.287 | 0.109 | 8.63739983844465e-0 | 4 | skap2 |
| ephx2 | 7.4371824 | 0.440459328 | 0.185 | 0.05 | 1.50208774827676e-0 | 4 | ephx2 |
| hmgn2 | 1.4614047 | 0.545829463 | 0.741 | 0.63 | 2.95159916540462e-0 | 4 | hmgn2 |
| cap11 | 2.8179065 | 0.598232419 | 0.509 | 0.325 | 5.69132585024406e-0 | 4 | cap1 |
| arhgdig | 3.7226756 | 0.569926231 | 0.528 | 0.362 | 7.51868794355014e-0 | 4 | arhgdig |

| | | | | | | | |
|---|---|---|---|---|---|---|---|
| yrk | 1.3655096 | 0.457474721 | 0.241 | 0.086 | 0.000275792 | 4 | yrk |
| si:ch73-264p11.1 | 3.9816364 | 0.354038263 | 0.13 | 0.031 | 0.000804171 | 4 | si:ch73-264p11.1 |
| mcl1a1 | 6.8051767 | 0.431617804 | 0.667 | 0.543 | 0.001374442 | 4 | mcl1a |
| mapre1a1 | 7.8470308 | 0.556398057 | 0.361 | 0.195 | 0.001584865 | 4 | mapre1a |
| CU467110.1 | 2.3389883 | 0.367166885 | 0.13 | 0.034 | 0.004724055 | 4 | CU467110.1 |
| si:ch73-380l3.4 | 2.6500902 | 0.306868642 | 0.12 | 0.03 | 0.005352387 | 4 | si:ch73-380l3.4 |
| zgc:640511 | 3.5020296 | 0.484069869 | 0.343 | 0.172 | 0.007073049 | 4 | zgc:64051 |
| rhogb | 3.5644892 | 0.501046812 | 0.324 | 0.166 | 0.007199199 | 4 | rhogb |
| ikzf2 | 4.3385150 | 0.350910495 | 0.167 | 0.054 | 0.008762499 | 4 | ikzf2 |
| AL954142.1 | 4.3392274 | 0.314209943 | 0.12 | 0.031 | 0.008763938 | 4 | AL954142.1 |
| rgs18 | 6.9399778 | 0.329209348 | 0.12 | 0.032 | 0.014016673 | 4 | rgs18 |
| itga1 | 1.6871518 | 0.384947049 | 0.167 | 0.057 | 0.034075407 | 4 | itga1 |
| washc2c | 7.7423296 | 0.375229944 | 0.157 | 0.057 | 0.156371832 | 4 | washc2c |
| hmgb2a | 1.1707176 | 0.411808067 | 0.63 | 0.585 | 0.236449837 | 4 | hmgb2a |
| rnaset2l1 | 1.6502652 | 0.351192194 | 0.194 | 0.08 | 0.333304069 | 4 | rnaset2l |
| calm1b | 3.8406324 | 0.416805262 | 0.269 | 0.144 | 0.775692529 | 4 | calm1b |
| stk17b | 6.3006639 | 0.443989366 | 0.25 | 0.136 | 1 | 4 | stk17b |
| sept6 | 6.7473829 | 0.435119535 | 0.306 | 0.189 | 1 | 4 | sept6 |
| si:zfos-741a10.3 | 0.000106 | 0.295817657 | 0.139 | 0.053 | 1 | 4 | si:zfos-741a10.3 |
| laptm51 | 0.000127 | 0.37698408 | 0.352 | 0.213 | 1 | 4 | laptm5 |
| lcp1 | 0.000151 | 0.367870793 | 0.157 | 0.068 | 1 | 4 | lcp1 |
| ppp1r18 | 0.000152 | 0.302816342 | 0.157 | 0.067 | 1 | 4 | ppp1r18 |
| ipcef1 | 0.000179 | 0.258398914 | 0.102 | 0.034 | 1 | 4 | ipcef1 |
| sh3bgrl | 0.00026 | 0.407587991 | 0.315 | 0.213 | 1 | 4 | sh3bgrl |
| fybb | 0.000326 | 0.281161958 | 0.139 | 0.058 | 1 | 4 | fybb |
| wdr1 | 0.000543 | 0.418215778 | 0.333 | 0.237 | 1 | 4 | wdr1 |
| capza1b | 0.000803 | 0.370256883 | 0.426 | 0.365 | 1 | 4 | capza1b |
| cox8b | 0.001206 | 0.394929664 | 0.222 | 0.133 | 1 | 4 | cox8b |
| capzb | 0.001279 | 0.38101365 | 0.426 | 0.389 | 1 | 4 | capzb |
| si:ch211-214p16.2 | 0.002071 | 0.284373437 | 0.176 | 0.091 | 1 | 4 | si:ch211-214p16.2 |
| myo1f | 0.002275 | 0.263235159 | 0.13 | 0.061 | 1 | 4 | myo1f |
| idh2 | 0.004109 | 0.391928082 | 0.417 | 0.386 | 1 | 4 | idh2 |
| LOC103909115 | 0.004681 | 0.342026887 | 0.278 | 0.2 | 1 | 4 | LOC103909115 |
| syk | 0.004889 | 0.276784512 | 0.148 | 0.079 | 1 | 4 | syk |
| dtnbp1a | 0.006896 | 0.322688527 | 0.167 | 0.099 | 1 | 4 | dtnbp1a |

| | | | | | | | |
|---|---|---|---|---|---|---|---|
| zgc:1627302 | 0.00713 | 0.25601094 | 0.546 | 0.493 | 1 | 4 | zgc:162730 |
| lasp1 | 0.007748 | 0.323073656 | 0.389 | 0.361 | 1 | 4 | lasp1 |
| mrc1b1 | 1.5790629 | 1.727864962 | 0.915 | 0.022 | 3.18923351605073e-2 | 5 | mrc1b |
| si:ch211-165i18.21 | 2.0360730 | 1.668006392 | 0.883 | 0.021 | 4.11225666383325e-2 | 5 | si:ch211-165i18.2 |
| itgae.11 | 9.5784779 | 1.477148064 | 0.862 | 0.021 | 1.93456518805086e-2 | 5 | itgae.1 |
| mfap41 | 1.0645372 | 1.663210273 | 0.872 | 0.023 | 2.15004598248695e-2 | 5 | mfap4 |
| CU499330.11 | 7.9119931 | 1.827589228 | 0.979 | 0.046 | 1.59798526534437e-2 | 5 | CU499330.1 |
| c1qb1 | 4.1743676 | 1.878156956 | 0.989 | 0.05 | 8.43097028280381e-2 | 5 | c1qb |
| havcr11 | 1.8489913 | 1.309791034 | 0.755 | 0.021 | 3.73440785106626e-1 | 5 | havcr1 |
| c1qa1 | 8.8561921 | 1.616132982 | 0.936 | 0.046 | 1.78868511943827e-1 | 5 | c1qa |
| cx32.21 | 3.4897421 | 1.115318722 | 0.66 | 0.015 | 7.04823213904198e-1 | 5 | cx32.2 |
| si:dkey-5n18.11 | 8.5334206 | 1.892260404 | 0.979 | 0.056 | 1.72349496323279e-1 | 5 | si:dkey-5n18.1 |
| c1qc1 | 3.8502749 | 1.683143908 | 0.957 | 0.052 | 7.77640027541167e-1 | 5 | c1qc |
| marco1 | 1.2381731 | 1.555944846 | 0.862 | 0.039 | 2.50073823601731e-1 | 5 | marco |
| ms4a17a.101 | 1.6598636 | 1.416355313 | 0.798 | 0.034 | 3.35242668080429e-1 | 5 | ms4a17a.10 |
| ccl34a.41 | 1.0746111 | 1.197418072 | 0.606 | 0.015 | 2.17039207213773e-1 | 5 | ccl34a.4 |
| ctss2.21 | 1.0998564 | 1.576083837 | 0.936 | 0.062 | 2.22137999550531e-1 | 5 | ctss2.2 |
| cfp1 | 9.9245617 | 1.398668428 | 0.883 | 0.051 | 2.00446373906411e-1 | 5 | cfp |
| BX649485.21 | 2.0063071 | 1.53125991 | 0.713 | 0.028 | 4.05213851561392e-1 | 5 | BX649485.2 |
| LOC1005369461 | 8.2045826 | 1.355871847 | 0.638 | 0.02 | 1.65707955145406e-1 | 5 | LOC100536946 |
| zgc:1531181 | 5.1456355 | 1.640227659 | 0.83 | 0.047 | 1.03926401190673e-1 | 5 | zgc:153118 |
| NPC2 (1 of many)1 | 1.3088526 | 1.388107648 | 0.809 | 0.042 | 2.64348961959772e-1 | 5 | NPC2 (1 of many) |
| lgals3bpb1 | 3.4379494 | 1.339979244 | 0.681 | 0.027 | 6.94362650512441e-1 | 5 | lgals3bpb |
| LO018493.11 | 2.7096097 | 0.905202082 | 0.574 | 0.015 | 5.47259882199517e-1 | 5 | LO018493.1 |
| havcr2 | 1.9202987 | 0.679013827 | 0.457 | 0.006 | 3.87842740030152e-1 | 5 | havcr2 |
| cmklr11 | 1.9158549 | 1.038601704 | 0.585 | 0.018 | 3.86945215739731e-1 | 5 | cmklr1 |
| ms4a17a.61 | 7.5719823 | 1.143889945 | 0.734 | 0.039 | 1.52931328438464e-1 | 5 | ms4a17a.6 |
| ccl34b.81 | 1.4016333 | 1.449135228 | 0.723 | 0.037 | 2.83087890888055e-1 | 5 | ccl34b.8 |
| grna1 | 2.5994979 | 1.1267684 | 0.755 | 0.041 | 5.25020597025387e-1 | 5 | grna |
| lygl11 | 1.3638639 | 1.633738849 | 0.926 | 0.083 | 2.7545959614536e-13 | 5 | lygl1 |
| zgc:640651 | 1.0914811 | 1.015629992 | 0.628 | 0.025 | 2.20446454380007e-1 | 5 | zgc:64065 |
| slc7a7 | 7.5851914 | 0.658232782 | 0.394 | 0.005 | 1.53198111946417e-1 | 5 | slc7a7 |
| spi1a1 | 9.5599139 | 0.991408184 | 0.649 | 0.03 | 1.93081581430285e-1 | 5 | spi1a |
| mafbb1 | 1.7632333 | 0.977506768 | 0.585 | 0.023 | 3.56120243897261e-1 | 5 | mafbb |
| ccr12b.21 | 2.8133395 | 0.883911017 | 0.511 | 0.017 | 5.68210191644027e-1 | 5 | ccr12b.2 |

| | | | | | | | |
|---|---|---|---|---|---|---|---|
| lect2l | 5.1533112 | 1.173970338 | 0.638 | 0.032 | 1.04081427629834e-1 | 5 | lect2l |
| LOC1104377311 | 2.5965115 | 1.594037804 | 0.819 | 0.067 | 5.24417429009478e-1 | 5 | LOC110437731 |
| slc3a2a1 | 4.6443030 | 0.953090255 | 0.628 | 0.031 | 9.3800988368426e-12 | 5 | slc3a2a |
| grn11 | 3.7312421 | 1.571814166 | 0.851 | 0.073 | 7.53598972963759e-1 | 5 | grn1 |
| f13a1b1 | 2.0185423 | 0.746999716 | 0.5 | 0.017 | 4.07684997742418e-1 | 5 | f13a1b |
| si:ch73-343l4.2 | 2.1853873 | 0.560891867 | 0.351 | 0.005 | 4.41382689398312e-1 | 5 | si:ch73-343l4.2 |
| si:ch211-147m6.11 | 2.5771421 | 0.926892116 | 0.489 | 0.017 | 5.20505403834444e-1 | 5 | si:ch211-147m6.1 |
| cxcr3.21 | 9.2362123 | 0.763367298 | 0.532 | 0.022 | 1.86543780363021e-1 | 5 | cxcr3.2 |
| zgc:1749041 | 4.8017092 | 1.161831486 | 0.67 | 0.044 | 9.69801217064957e-1 | 5 | zgc:174904 |
| zgc:1528301 | 8.3593167 | 1.016719605 | 0.713 | 0.053 | 1.68833120926829e-1 | 5 | zgc:152830 |
| smox1 | 5.5499439 | 1.117350348 | 0.798 | 0.074 | 1.12092218014512e-1 | 5 | smox |
| si:ch211-39i2.2 | 3.9757884 | 0.744327194 | 0.479 | 0.019 | 8.02989997795512e-1 | 5 | si:ch211-39i2.2 |
| BX649485.1 | 9.2875377 | 0.742661375 | 0.447 | 0.015 | 1.87580399699434e-1 | 5 | BX649485.1 |
| jac31 | 8.1649077 | 1.141337895 | 0.521 | 0.026 | 1.64906641604108e-1 | 5 | jac3 |
| tgfbi | 6.7399698 | 0.907475014 | 0.617 | 0.041 | 1.3612717148073e-98 | 5 | tgfbi |
| FO907110.1 | 2.2850367 | 0.546424689 | 0.287 | 0.003 | 4.6150886297111e-98 | 5 | FO907110.1 |
| samsn1a1 | 1.7551851 | 0.799611874 | 0.553 | 0.032 | 3.54494748998885e-9 | 5 | samsn1a |
| tcnbb | 3.3096637 | 0.789146506 | 0.436 | 0.017 | 6.68452779645327e-9 | 5 | tcnbb |
| sftpbb1 | 6.4280385 | 1.102769622 | 0.745 | 0.067 | 1.29827094117019e-9 | 5 | sftpbb |
| mhc2dab1 | 2.1207692 | 1.119371446 | 0.702 | 0.061 | 4.28331756649884e-9 | 5 | mhc2dab |
| csf2rb | 3.5694874 | 0.701186315 | 0.457 | 0.02 | 7.20929388299268e-9 | 5 | csf2rb |
| si:ch211-283g2.31 | 7.2018277 | 0.766877545 | 0.415 | 0.015 | 1.45455315761103e-9 | 5 | si:ch211-283g2.3 |
| zgc:136870 | 2.0959224 | 0.551373721 | 0.351 | 0.01 | 4.23313449847802e-9 | 5 | zgc:136870 |
| si:ch211-194m7.31 | 2.5347870 | 0.750716224 | 0.404 | 0.015 | 5.11950943302606e-8 | 5 | si:ch211-194m7.3 |
| naga1 | 2.5574526 | 0.946795069 | 0.66 | 0.059 | 5.16528704616018e-8 | 5 | naga |
| jac91 | 4.9839819 | 0.755818119 | 0.383 | 0.014 | 1.00661484316603e-8 | 5 | jac9 |
| gng101 | 1.1655858 | 0.949385284 | 0.713 | 0.069 | 2.35413382941714e-8 | 5 | gng10 |
| LOC5646601 | 4.0459658 | 1.191255967 | 0.489 | 0.029 | 8.17163728941438e-8 | 5 | LOC564660 |
| si:busm1-194e12.1 | 7.8503619 | 0.958532273 | 0.596 | 0.047 | 1.58553759908367e-8 | 5 | si:busm1-194e12.11 |
| zgc:1037001 | 8.6222730 | 1.287863477 | 0.809 | 0.099 | 1.74144047969631e-8 | 5 | zgc:103700 |
| tpp11 | 1.0103094 | 1.017024325 | 0.691 | 0.07 | 2.04052198793069e-8 | 5 | tpp1 |
| hdac9b | 3.7822045 | 0.603387833 | 0.447 | 0.023 | 7.638918503224e-83 | 5 | hdac9b |
| AL928650.1 | 9.0444160 | 0.526590693 | 0.277 | 0.005 | 1.82670070347481e-8 | 5 | AL928650.1 |
| si:dkey-102g19.3 | 1.5146431 | 0.810423026 | 0.521 | 0.036 | 3.05912470673622e-7 | 5 | si:dkey-102g19.3 |
| vsir1 | 2.9369405 | 1.050456039 | 0.734 | 0.084 | 5.93173886353493e-7 | 5 | vsir |

| | | | | | | | |
|---|---|---|---|---|---|---|---|
| ehd41 | 1.8555450 | 0.720808931 | 0.521 | 0.036 | 3.74764433736845e-7 | 5 | ehd4 |
| si:zfos-2330d3.31 | 1.3832058 | 0.804705357 | 0.34 | 0.012 | 2.79366077969873e-7 | 5 | si:zfos-2330d3.3 |
| il4r.1 | 3.4551330 | 0.881000936 | 0.564 | 0.047 | 6.9783322005598e-77 | 5 | il4r.1 |
| CR855311.12 | 9.5326093 | 1.101968754 | 0.723 | 0.081 | 1.92530111736125e-7 | 5 | CR855311.1 |
| CABZ01030278.1 | 5.2872834 | 0.465419417 | 0.234 | 0.003 | 1.067872642954e-75 | 5 | CABZ01030278.1 |
| clic21 | 1.2651513 | 0.955812005 | 0.681 | 0.074 | 2.55522623676588e-7 | 5 | clic2 |
| parvg | 1.4734571 | 0.91336101 | 0.723 | 0.082 | 2.97594131865883e-7 | 5 | parvg |
| entpd2a.1 | 5.4515951 | 0.468921936 | 0.266 | 0.006 | 1.10105866288242e-7 | 5 | entpd2a.1 |
| abca1b | 6.1842055 | 0.564781694 | 0.351 | 0.014 | 1.24902399024086e-7 | 5 | abca1b |
| cd22 | 1.1341640 | 0.374433895 | 0.223 | 0.003 | 2.29067104451407e-7 | 5 | cd22 |
| txn1 | 1.2974348 | 0.977900959 | 0.66 | 0.067 | 2.62042923849093e-7 | 5 | txn |
| fgl2a1 | 4.7110967 | 1.17723647 | 0.766 | 0.103 | 9.51500218259496e-7 | 5 | fgl2a |
| atp1a3a | 6.4960786 | 0.54240545 | 0.383 | 0.019 | 1.31201301460598e-7 | 5 | atp1a3a |
| si:busm1-266f07.2: | 7.6751804 | 1.412665922 | 0.84 | 0.132 | 1.55015618860872e-7 | 5 | si:busm1-266f07.2 |
| moxd1 | 6.4429537 | 0.526819578 | 0.277 | 0.007 | 1.3012833741216e-72 | 5 | moxd1 |
| m17 | 7.1934414 | 0.51112272 | 0.277 | 0.007 | 1.45285936154589e-7 | 5 | m17 |
| CABZ01001434.1 | 6.5797309 | 0.64198395 | 0.34 | 0.014 | 1.32890825462629e-7 | 5 | CABZ01001434.1 |
| lgals2a1 | 1.5876699 | 1.386771071 | 0.957 | 0.178 | 3.20661691415533e-7 | 5 | lgals2a |
| si:cabz01074946.1 | 5.4671400 | 0.816649509 | 0.564 | 0.052 | 1.10419828036524e-6 | 5 | si:cabz01074946.1 |
| calhm6 | 1.8606594 | 0.546887408 | 0.372 | 0.019 | 3.75797384724529e-6 | 5 | calhm6 |
| mhc2b1 | 8.2820550 | 1.085966014 | 0.638 | 0.072 | 1.6727266675893e-68 | 5 | mhc2b |
| fosl2 | 2.6766060 | 0.847134699 | 0.585 | 0.06 | 5.4059412724221e-68 | 5 | fosl2 |
| tcima | 3.3809897 | 0.904059312 | 0.628 | 0.068 | 6.82858504878699e-6 | 5 | tcima |
| scpep11 | 4.9661773 | 1.007653949 | 0.723 | 0.101 | 1.00301884600429e-6 | 5 | scpep1 |
| rnaset2l2 | 1.3795928 | 0.861512654 | 0.606 | 0.062 | 2.78636369946281e-6 | 5 | rnaset2l |
| dnase21 | 2.3670573 | 0.576175198 | 0.394 | 0.023 | 4.78074567590573e-6 | 5 | dnase2 |
| si:ch211-264f5.2 | 4.2158095 | 0.448579651 | 0.255 | 0.007 | 8.51467045584201e-6 | 5 | si:ch211-264f5.2 |
| si:dkey-7i4.241 | 2.4371658 | 1.070109482 | 0.457 | 0.034 | 4.92234386168254e-6 | 5 | si:dkey-7i4.24 |
| gsdmea1 | 6.8359038 | 0.505746041 | 0.319 | 0.014 | 1.38064749234836e-6 | 5 | gsdmea |
| ccl34b.11 | 1.1064985 | 0.577550517 | 0.319 | 0.014 | 2.23479516420693e-6 | 5 | ccl34b.1 |
| plekho1b | 3.2415859 | 0.67124133 | 0.468 | 0.037 | 6.54703122151741e-6 | 5 | plekho1b |
| tspan361 | 3.4549103 | 1.092414665 | 0.723 | 0.11 | 6.97788248948489e-6 | 5 | tspan36 |
| stoml3b | 3.7759528 | 0.39851118 | 0.213 | 0.004 | 7.62629190297452e-6 | 5 | stoml3b |
| dhrs9 | 1.5264780 | 0.680216513 | 0.447 | 0.034 | 3.08302761807339e-6 | 5 | dhrs9 |
| tspan37 | 2.7291385 | 0.615203096 | 0.468 | 0.037 | 5.51204104821576e-6 | 5 | tspan37 |

| | | | | | | | |
|---|---|---|---|---|---|---|---|
| ctsba1 | 6.1441573 | 1.242922823 | 0.936 | 0.217 | 1.2409354584362e-61 | 5 | ctsba |
| si:ch211-74m13.3 | 8.4655720 | 0.887692253 | 0.521 | 0.054 | 1.70979159609476e-6 | 5 | si:ch211-74m13.3 |
| gpx1a1 | 3.5701272 | 1.211183378 | 0.926 | 0.208 | 7.2105859654135e-60 | 5 | gpx1a |
| jac2 | 4.0965903 | 0.457282084 | 0.245 | 0.007 | 8.27388347091618e-6 | 5 | jac2 |
| cd74b2 | 8.2666989 | 1.399245203 | 0.947 | 0.222 | 1.66962519341399e-5 | 5 | cd74b |
| CR392341.11 | 9.9085912 | 0.748959013 | 0.489 | 0.045 | 2.00123818454953e-5 | 5 | CR392341.1 |
| cybb | 1.6736706 | 0.434684619 | 0.298 | 0.013 | 3.38031263565517e-5 | 5 | cybb |
| lta4h1 | 3.9134019 | 0.811067769 | 0.617 | 0.075 | 7.90389795539706e-5 | 5 | lta4h |
| ms4a17a.91 | 4.7263134 | 0.935657028 | 0.734 | 0.116 | 9.54573534968881e-5 | 5 | ms4a17a.9 |
| si:ch1073-392o20.[illegible] | 9.3235729 | 0.712013761 | 0.521 | 0.054 | 1.88308203574824e-5 | 5 | si:ch1073-392o20.2 |
| jac8 | 2.0818001 | 0.571585245 | 0.245 | 0.008 | 4.20461184728532e-5 | 5 | jac8 |
| si:busm1-71b9.3 | 2.7907213 | 0.29140539 | 0.16 | 0.001 | 5.6364199437596e-58 | 5 | si:busm1-71b9.3 |
| BX323596.21 | 6.7445262 | 0.605908116 | 0.277 | 0.011 | 1.36219196539582e-5 | 5 | BX323596.2 |
| acsl4b | 1.2760551 | 0.57207954 | 0.394 | 0.028 | 2.57724854650638e-5 | 5 | acsl4b |
| hmha1b | 1.4729779 | 0.693037922 | 0.532 | 0.057 | 2.97497360526846e-5 | 5 | hmha1b |
| plxnc1 | 1.5516569 | 0.679051835 | 0.489 | 0.047 | 3.13388160605123e-5 | 5 | plxnc1 |
| psap1 | 6.8617549 | 1.234991542 | 0.936 | 0.251 | 1.38586864376092e-5 | 5 | psap |
| plin2 | 9.8120321 | 0.507592688 | 0.266 | 0.011 | 1.98173612961328e-5 | 5 | plin2 |
| glb1l | 2.1364846 | 0.58423728 | 0.426 | 0.035 | 4.31505807229084e-5 | 5 | glb1l |
| ctsd1 | 3.6699778 | 1.086468378 | 0.819 | 0.162 | 7.41225417739857e-5 | 5 | ctsd |
| si:dkey-262k9.41 | 5.1904606 | 0.698704049 | 0.606 | 0.074 | 1.04831733043032e-5 | 5 | si:dkey-262k9.4 |
| ets2 | 5.2684841 | 0.688177445 | 0.511 | 0.054 | 1.06407573851853e-5 | 5 | ets2 |
| ctsc | 1.4710545 | 0.737444935 | 0.457 | 0.043 | 2.97108891608894e-5 | 5 | ctsc |
| trpv1 | 1.6426166 | 0.378816087 | 0.255 | 0.01 | 3.31759280788625e-5 | 5 | trpv1 |
| gm2a1 | 2.7520293 | 0.880558643 | 0.713 | 0.108 | 5.55827358591206e-5 | 5 | gm2a |
| laptm52 | 4.1569324 | 1.11912469 | 0.915 | 0.187 | 8.39575653343389e-5 | 5 | laptm5 |
| slc16a9a | 5.9685046 | 0.360563072 | 0.202 | 0.005 | 1.20545888651925e-5 | 5 | slc16a9a |
| cndp21 | 2.2569442 | 0.86701516 | 0.649 | 0.095 | 4.55835038525761e-5 | 5 | cndp2 |
| cxcr4b | 2.3713790 | 1.083920829 | 0.777 | 0.132 | 4.78947428549855e-5 | 5 | cxcr4b |
| si:ch211-283g2.2 | 3.4987250 | 0.36425308 | 0.223 | 0.007 | 7.06637503708713e-5 | 5 | si:ch211-283g2.2 |
| si:ch73-158p21.31 | 4.5633050 | 0.595147384 | 0.309 | 0.017 | 9.21650730306706e-5 | 5 | si:ch73-158p21.3 |
| fcer1gl2 | 8.4083215 | 1.323033522 | 0.979 | 0.229 | 1.69822870395668e-5 | 5 | fcer1gl |
| slc6a6b | 9.1347075 | 0.653449225 | 0.5 | 0.052 | 1.84493687925227e-5 | 5 | slc6a6b |
| gnpda11 | 1.8389930 | 0.618115728 | 0.447 | 0.041 | 3.71421433100662e-5 | 5 | gnpda1 |
| ctsa1 | 2.2043949 | 0.871252836 | 0.681 | 0.11 | 4.45221647019195e-5 | 5 | ctsa |

| | | | | | | | |
|---|---|---|---|---|---|---|---|
| LOC1018857061 | 2.7852054 | 0.491606057 | 0.287 | 0.014 | 5.62527940985037e-5 | 5 | LOC101885706 |
| lyn | 5.0172487 | 0.56259958 | 0.415 | 0.035 | 1.01333372266229e-5 | 5 | lyn |
| cebpb1 | 2.9630273 | 1.25969277 | 0.936 | 0.27 | 5.98442627673114e-5 | 5 | cebpb |
| dpp71 | 2.1618454 | 0.835291076 | 0.638 | 0.098 | 4.36627919590014e-5 | 5 | dpp7 |
| si:ch211-286b5.9 | 3.3976764 | 0.277709003 | 0.181 | 0.004 | 6.86228703735619e-5 | 5 | si:ch211-286b5.9 |
| FO704661.1 | 7.4161346 | 0.341839697 | 0.202 | 0.006 | 1.49783671876231e-5 | 5 | FO704661.1 |
| adgrg3 | 1.3451757 | 0.390279811 | 0.213 | 0.007 | 2.71685143168502e-5 | 5 | adgrg3 |
| LOC100538255 | 1.5334182 | 0.39006758 | 0.223 | 0.008 | 3.09704487141055e-5 | 5 | LOC100538255 |
| ctsh1 | 9.3632882 | 0.871480027 | 0.67 | 0.112 | 1.89110333423242e-4 | 5 | ctsh |
| ptgis | 3.0798127 | 0.432262524 | 0.245 | 0.011 | 6.22029780240322e-4 | 5 | ptgis |
| si:ch211-102c2.42 | 5.2372732 | 0.93822934 | 0.83 | 0.149 | 1.0577720848559e-47 | 5 | si:ch211-102c2.4 |
| selp | 6.6510631 | 0.359481327 | 0.223 | 0.009 | 1.343315225998e-47 | 5 | selp |
| cd74a2 | 8.1414937 | 1.360746091 | 0.809 | 0.187 | 1.64433749108359e-4 | 5 | cd74a |
| si:ch73-248e21.5 | 1.3024890 | 0.521456988 | 0.394 | 0.035 | 2.63063720207389e-4 | 5 | si:ch73-248e21.5 |
| si:ch211-212k18.7: | 2.0091328 | 0.926661945 | 0.787 | 0.162 | 4.05784560037811e-4 | 5 | si:ch211-212k18.7 |
| si:dkey-56f14.71 | 2.5800720 | 0.370880116 | 0.245 | 0.011 | 5.21097147995161e-4 | 5 | si:dkey-56f14.7 |
| csf3r | 2.7372567 | 0.303820912 | 0.17 | 0.004 | 5.52843744435223e-4 | 5 | csf3r |
| ctsz1 | 7.2976481 | 0.923402395 | 0.606 | 0.098 | 1.47390599279305e-4 | 5 | ctsz |
| ly6m5 | 7.6810253 | 0.430787086 | 0.277 | 0.016 | 1.55133669543539e-4 | 5 | ly6m5 |
| nrros | 1.3350029 | 0.577255767 | 0.404 | 0.039 | 2.6963053663288e-46 | 5 | nrros |
| fuca21 | 1.6330512 | 0.749074817 | 0.543 | 0.075 | 3.29827363545705e-4 | 5 | fuca2 |
| zgc:640512 | 5.7412689 | 0.935946123 | 0.787 | 0.152 | 1.15956409968923e-4 | 5 | zgc:64051 |
| irf81 | 2.0385150 | 0.565452346 | 0.436 | 0.046 | 4.11718881057605e-4 | 5 | irf8 |
| itgae.2 | 2.0779790 | 0.532509217 | 0.351 | 0.029 | 4.19689434672097e-4 | 5 | itgae.2 |
| CT027744.2 | 3.4557219 | 0.520878707 | 0.404 | 0.039 | 6.97952167163401e-4 | 5 | CT027744.2 |
| enpp11 | 1.1613158 | 0.47487108 | 0.33 | 0.025 | 2.34550961981933e-4 | 5 | enpp1 |
| pmaip1 | 1.2354973 | 0.7528895 | 0.489 | 0.065 | 2.4953340978971e-44 | 5 | pmaip1 |
| ckbb2 | 3.4467748 | 0.910835047 | 0.84 | 0.165 | 6.96145111280418e-4 | 5 | ckbb |
| slc40a11 | 4.1229172 | 0.665140594 | 0.436 | 0.049 | 8.32705587360276e-4 | 5 | slc40a1 |
| il10ra | 7.5962427 | 0.568828225 | 0.383 | 0.037 | 1.53421314463001e-4 | 5 | il10ra |
| lgals9l11 | 8.0471603 | 0.620217334 | 0.394 | 0.038 | 1.6252849765349e-43 | 5 | lgals9l1 |
| si:zfos-1069f5.1 | 8.0799973 | 0.276636898 | 0.149 | 0.003 | 1.6319170618812e-43 | 5 | si:zfos-1069f5.1 |
| cdaa | 8.4905972 | 0.436362368 | 0.287 | 0.019 | 1.71484593579688e-4 | 5 | cdaa |
| slc35f6 | 3.1374003 | 0.661008687 | 0.436 | 0.052 | 6.33660744696649e-4 | 5 | slc35f6 |
| XLOC-005448 | 5.0365893 | 0.542476635 | 0.277 | 0.018 | 1.0172399467859e-42 | 5 | XLOC-005448 |

| | | | | | | | |
|---|---|---|---|---|---|---|---|
| mapk11 | 1.3252692 | 0.361666 | 0.255 | 0.015 | 2.6766463892775e-42 | 5 | mapk11 |
| atp6v1e1b1 | 1.5575064 | 0.787984991 | 0.638 | 0.114 | 3.1456957941721e-42 | 5 | atp6v1e1b |
| zbtb16a | 2.2895755 | 0.363548785 | 0.223 | 0.011 | 4.6242557228561e-42 | 5 | zbtb16a |
| aplp2 | 2.4447605 | 0.711924778 | 0.564 | 0.087 | 4.93768284486097e-4 | 5 | aplp2 |
| gnsb | 3.0012239 | 0.490860182 | 0.351 | 0.032 | 6.06157203184766e-4 | 5 | gnsb |
| grapa1 | 4.0128456 | 0.417415033 | 0.287 | 0.02 | 8.10474442276905e-4 | 5 | grapa |
| mfng | 4.6451181 | 0.554694192 | 0.426 | 0.047 | 9.38174513209378e-4 | 5 | mfng |
| xpnpep2 | 9.6300018 | 0.287948099 | 0.202 | 0.008 | 1.94497147848142e-4 | 5 | xpnpep2 |
| gpr183a | 3.4763644 | 0.573360465 | 0.436 | 0.051 | 7.02121335818347e-4 | 5 | gpr183a |
| itgb2 | 4.0643320 | 0.855703926 | 0.734 | 0.142 | 8.20873145163055e-4 | 5 | itgb2 |
| galk1 | 4.9818261 | 0.359192966 | 0.234 | 0.013 | 1.00617942606171e-4 | 5 | galk1 |
| zgc:63831 | 9.9756014 | 0.675753259 | 0.585 | 0.094 | 2.01477223330218e-4 | 5 | zgc:63831 |
| nccrp11 | 1.6195073 | 0.910333101 | 0.809 | 0.198 | 3.27091889706577e-4 | 5 | nccrp1 |
| si:zfos-741a10.31 | 4.0207534 | 0.545011804 | 0.394 | 0.042 | 8.12071569219766e-4 | 5 | si:zfos-741a10.3 |
| mrc1a | 4.7953071 | 0.377334989 | 0.202 | 0.009 | 9.68508191305162e-4 | 5 | mrc1a |
| prcp | 6.6854929 | 0.359278086 | 0.255 | 0.016 | 1.35026900710905e-3 | 5 | prcp |
| tcirg1b | 9.4530725 | 0.437896624 | 0.319 | 0.028 | 1.90923706380128e-3 | 5 | tcirg1b |
| atp6v1ba1 | 9.7085291 | 0.622507836 | 0.479 | 0.067 | 1.96083164064422e-3 | 5 | atp6v1ba |
| lpar5a | 5.2533491 | 0.55923407 | 0.415 | 0.049 | 1.06101892395956e-3 | 5 | lpar5a |
| sc:d0284 | 1.4804188 | 0.352171965 | 0.245 | 0.015 | 2.99000187744358e-3 | 5 | sc:d0284 |
| si:ch211-9d9.8 | 1.6256197 | 0.289304473 | 0.213 | 0.011 | 3.28326420451373e-3 | 5 | si:ch211-9d9.8 |
| me2 | 2.7118832 | 0.439957701 | 0.287 | 0.023 | 5.47719067588526e-3 | 5 | me2 |
| si:ch211-193e13.5 | 6.9140690 | 0.320231711 | 0.234 | 0.014 | 1.39643453218715e-3 | 5 | si:ch211-193e13.5 |
| atp6v1ab | 1.0169646 | 0.38733622 | 0.266 | 0.019 | 2.05396351034553e-3 | 5 | atp6v1ab |
| arl6ip11 | 1.0581044 | 0.924831905 | 0.872 | 0.264 | 2.13705351601611e-3 | 5 | arl6ip1 |
| tuba8l21 | 1.0659581 | 0.615917706 | 0.543 | 0.088 | 2.15291575259712e-3 | 5 | tuba8l2 |
| CABZ01020840.1 | 1.1631603 | 0.270335362 | 0.181 | 0.007 | 2.3492350364476e-37 | 5 | CABZ01020840.1 |
| asah1b1 | 1.3552902 | 0.71612366 | 0.543 | 0.092 | 2.73727969649175e-3 | 5 | asah1b |
| atp6v0ca1 | 1.4437312 | 0.946424174 | 0.787 | 0.214 | 2.91590398177073e-3 | 5 | atp6v0ca |
| LOC101886073 | 2.4191312 | 0.290116714 | 0.213 | 0.011 | 4.88591940714531e-3 | 5 | LOC101886073 |
| si:ch211-1o7.31 | 2.5191239 | 0.325272134 | 0.255 | 0.017 | 5.08787465619827e-3 | 5 | si:ch211-1o7.3 |
| si:ch211-243a20.4 | 3.5538622 | 0.427752233 | 0.277 | 0.021 | 7.17773566153867e-3 | 5 | si:ch211-243a20.4 |
| zmp:0000000936 | 3.9904053 | 0.254392102 | 0.17 | 0.006 | 8.05942164951922e-3 | 5 | zmp:0000000936 |
| si:dkey-148a17.6 | 4.0865811 | 0.262473042 | 0.17 | 0.006 | 8.2536678851314e-37 | 5 | si:dkey-148a17.6 |
| cd631 | 9.6511234 | 0.915715037 | 0.979 | 0.553 | 1.9492374098006e-36 | 5 | cd63 |

| | | | | | | | |
|---|---|---|---|---|---|---|---|
| hlx1 | 2.3818823 | 0.508525205 | 0.351 | 0.037 | 4.81068769383515e-3 | 5 | hlx1 |
| krt18b1 | 3.5024356 | 0.986730525 | 0.904 | 0.297 | 7.0738692253239e-36 | 5 | krt18b |
| cyba1 | 4.1407233 | 0.710262452 | 0.606 | 0.114 | 8.36301903614123e-3 | 5 | cyba |
| crema | 4.1938429 | 0.691758256 | 0.66 | 0.127 | 8.47030463108396e-3 | 5 | crema |
| ncf2 | 4.5064998 | 0.382368651 | 0.266 | 0.02 | 9.10177778396849e-3 | 5 | ncf2 |
| sb:cb81 | 5.9026026 | 0.439999462 | 0.309 | 0.028 | 1.19214864842063e-3 | 5 | sb:cb81 |
| hmox1a1 | 1.7973422 | 0.489397106 | 0.266 | 0.021 | 3.63009219293055e-3 | 5 | hmox1a |
| snx10b1 | 4.4095402 | 0.314085276 | 0.202 | 0.011 | 8.90594840086423e-3 | 5 | snx10b |
| ltb4r | 5.0094811 | 0.388863336 | 0.245 | 0.017 | 1.01176490215294e-3 | 5 | ltb4r |
| mapre1a2 | 7.4222216 | 0.758185756 | 0.766 | 0.177 | 1.49906611254129e-3 | 5 | mapre1a |
| rgn | 9.9918836 | 0.259736848 | 0.234 | 0.016 | 2.01806074632197e-3 | 5 | rgn |
| CU929150.11 | 1.1519641 | 0.619464144 | 0.309 | 0.029 | 2.32662200597999e-3 | 5 | CU929150.1 |
| tmsb11 | 4.1496304 | 0.815315713 | 0.766 | 0.171 | 8.38100858282567e-3 | 5 | tmsb1 |
| LOC100148254 | 6.0283997 | 0.430918716 | 0.277 | 0.024 | 1.21755589090827e-3 | 5 | LOC100148254 |
| nfil3-61 | 8.1023646 | 0.717915517 | 0.617 | 0.124 | 1.6364345879861e-33 | 5 | nfil3-6 |
| rnaseka1 | 1.5535625 | 0.678947956 | 0.574 | 0.113 | 3.13773018691635e-3 | 5 | rnaseka |
| CABZ01073834.1 | 1.6177106 | 0.485323566 | 0.383 | 0.047 | 3.26729024612439e-3 | 5 | CABZ01073834.1 |
| stx11a | 3.5006446 | 0.362660993 | 0.255 | 0.02 | 7.07025204455151e-3 | 5 | stx11a |
| BX901920.11 | 3.5830083 | 0.594333255 | 0.447 | 0.065 | 7.23660194848383e-3 | 5 | BX901920.1 |
| atp6v1g11 | 3.6457945 | 0.769096588 | 0.713 | 0.173 | 7.36341132556352e-3 | 5 | atp6v1g1 |
| ctsl.1 | 4.3545680 | 0.280062492 | 0.149 | 0.005 | 8.79492101457936e-3 | 5 | ctsl.1 |
| zgc:153317 | 4.4798102 | 0.395134529 | 0.277 | 0.025 | 9.04787277016594e-3 | 5 | zgc:153317 |
| srgn | 5.5682082 | 0.884623417 | 0.851 | 0.222 | 1.12461101086922e-3 | 5 | srgn |
| slc39a1 | 6.6013308 | 0.403117676 | 0.234 | 0.017 | 1.33327079158608e-3 | 5 | slc39a1 |
| dusp1 | 7.6216435 | 0.68999857 | 0.511 | 0.089 | 1.53934334777745e-3 | 5 | dusp1 |
| thy1 | 7.9130623 | 0.551371356 | 0.351 | 0.04 | 1.59820119477591e-3 | 5 | thy1 |
| ccl39.11 | 1.1224606 | 0.602535404 | 0.255 | 0.021 | 2.26703374450923e-3 | 5 | ccl39.1 |
| arhgef5 | 1.1231816 | 0.463796657 | 0.34 | 0.038 | 2.26848996120184e-3 | 5 | arhgef5 |
| snx27a | 1.5914905 | 0.43698674 | 0.319 | 0.034 | 3.21433337880169e-3 | 5 | snx27a |
| atp6v1h | 2.4864966 | 0.499035619 | 0.404 | 0.056 | 5.02197735782462e-3 | 5 | atp6v1h |
| tec | 3.0094419 | 0.428083443 | 0.33 | 0.037 | 6.07816995500436e-3 | 5 | tec |
| CU638870.1 | 3.1108872 | 0.381170376 | 0.266 | 0.023 | 6.28305891897859e-3 | 5 | CU638870.1 |
| LOC797949 | 3.4647845 | 0.254479495 | 0.16 | 0.007 | 6.9978253354967e-32 | 5 | LOC797949 |
| zgc:101715 | 4.3833564 | 0.368560921 | 0.277 | 0.025 | 8.85306496156625e-3 | 5 | zgc:101715 |
| g6pd | 5.9560823 | 0.350529181 | 0.277 | 0.026 | 1.20294996028064e-3 | 5 | g6pd |

| | | | | | | | |
|---|---|---|---|---|---|---|---|
| fermt3b | 6.8309021 | 0.381556032 | 0.277 | 0.025 | 1.37963730319682e-3 | 5 | fermt3b |
| rab44 | 1.4385034 | 0.523036505 | 0.394 | 0.053 | 2.90534544424e-31 | 5 | rab44 |
| klf2b | 1.6787189 | 0.302724839 | 0.202 | 0.013 | 3.39050875751885e-3 | 5 | klf2b |
| illr3 | 3.4648873 | 0.269812366 | 0.117 | 0.003 | 6.99803301355406e-3 | 5 | illr3 |
| tent5ab | 4.1422280 | 0.297648776 | 0.181 | 0.01 | 8.36605795842918e-3 | 5 | tent5ab |
| si:ch73-237c6.1 | 4.2326093 | 0.525508493 | 0.394 | 0.055 | 8.54860119288284e-3 | 5 | si:ch73-237c6.1 |
| ctss2.1 | 4.8915595 | 0.612787528 | 0.415 | 0.061 | 9.87948275667863e-3 | 5 | ctss2.1 |
| lamp1a | 5.3685177 | 0.643871982 | 0.479 | 0.084 | 1.08427953241721e-3 | 5 | lamp1a |
| gadd45bb | 8.8401807 | 0.525750977 | 0.372 | 0.049 | 1.78545131224489e-3 | 5 | gadd45bb |
| XLOC-0283701 | 2.1602137 | 0.653198226 | 0.351 | 0.046 | 4.362983805586e-30 | 5 | XLOC-028370 |
| ripk3 | 2.2712984 | 0.257025568 | 0.17 | 0.009 | 4.58734156596256e-3 | 5 | ripk3 |
| zgc:123107 | 3.0792454 | 0.352850717 | 0.191 | 0.012 | 6.21915206758407e-3 | 5 | zgc:123107 |
| aatka | 4.8395517 | 0.281238419 | 0.149 | 0.006 | 9.77444257994191e-3 | 5 | aatka |
| bach1b | 7.0525986 | 0.364895848 | 0.266 | 0.025 | 1.42441334456702e-2 | 5 | bach1b |
| lifrb | 2.3463074 | 0.295098063 | 0.16 | 0.008 | 4.73883714748159e-2 | 5 | lifrb |
| pdk2b | 2.4283104 | 0.705938432 | 0.553 | 0.117 | 4.90445866854676e-2 | 5 | pdk2b |
| wasb1 | 4.9218044 | 0.62618533 | 0.66 | 0.139 | 9.94056844692381e-2 | 5 | wasb |
| vmp1 | 7.5063129 | 0.491447341 | 0.34 | 0.044 | 1.51605002204761e-2 | 5 | vmp1 |
| pycard1 | 7.9303817 | 0.644768604 | 0.606 | 0.127 | 1.60169920213258e-2 | 5 | pycard |
| cd83 | 9.3526729 | 0.555020799 | 0.319 | 0.039 | 1.88895936095108e-2 | 5 | cd83 |
| alas11 | 1.3717369 | 0.602801144 | 0.553 | 0.113 | 2.77049712491304e-2 | 5 | alas1 |
| cebpa2 | 1.9286065 | 0.765410758 | 0.649 | 0.155 | 3.89520663706867e-2 | 5 | cebpa |
| mpp1 | 3.8491688 | 0.374758331 | 0.287 | 0.031 | 7.77416625768368e-2 | 5 | mpp1 |
| pkma1 | 4.2231131 | 0.683247861 | 0.691 | 0.178 | 8.52942171217248e-2 | 5 | pkma |
| si:ch73-248e21.7 | 6.9494727 | 0.519617751 | 0.404 | 0.063 | 1.40358501095311e-2 | 5 | si:ch73-248e21.7 |
| ifi301 | 7.0479144 | 0.961574857 | 0.809 | 0.289 | 1.42346727272672e-2 | 5 | ifi30 |
| mgrn1b | 1.0241250 | 0.343808629 | 0.234 | 0.021 | 2.06842538239402e-2 | 5 | mgrn1b |
| zmp:0000001082 | 1.1675797 | 0.402953349 | 0.319 | 0.039 | 2.35816087862956e-2 | 5 | zmp:0000001082 |
| ccl38.1 | 2.3526958 | 0.284078846 | 0.234 | 0.02 | 4.75173971566233e-2 | 5 | ccl38.1 |
| si:dkey-53k12.21 | 2.8304832 | 0.375168276 | 0.298 | 0.034 | 5.71672701919199e-2 | 5 | si:dkey-53k12.2 |
| rab8b1 | 3.6357100 | 0.555065203 | 0.511 | 0.101 | 7.34304352239369e-2 | 5 | rab8b |
| pepd1 | 5.3840013 | 0.543296148 | 0.479 | 0.091 | 1.08740675585746e-2 | 5 | pepd |
| si:ch211-152p11.4 | 6.7338703 | 0.262892383 | 0.16 | 0.009 | 1.36003978759848e-2 | 5 | si:ch211-152p11.4 |
| entpd11 | 8.5719149 | 0.67758588 | 0.574 | 0.131 | 1.73126966513718e-2 | 5 | entpd1 |
| ndrg1a | 1.1356135 | 0.415986862 | 0.362 | 0.053 | 2.29359876901026e-2 | 5 | ndrg1a |

| | | | | | | | |
|---|---|---|---|---|---|---|---|
| tram1 | 1.7086264 | 0.58041714 | 0.543 | 0.118 | 3.45091291890683e-2 | 5 | tram1 |
| si:ch211-204c21.1 | 2.1244566 | 0.547601319 | 0.426 | 0.073 | 4.29076500632457e-2 | 5 | si:ch211-204c21.1 |
| lamtor51 | 2.4758434 | 0.570924515 | 0.543 | 0.115 | 5.00046092720042e-2 | 5 | lamtor5 |
| lrrc15 | 3.0760885 | 0.356025157 | 0.255 | 0.026 | 6.21277609593985e-2 | 5 | lrrc15 |
| wee2 | 3.7393250 | 0.310862328 | 0.245 | 0.024 | 7.55231485217655e-2 | 5 | wee2 |
| abcg2a | 7.2003361 | 0.287362792 | 0.149 | 0.008 | 1.45425189046248e-2 | 5 | abcg2a |
| atf3 | 8.7013823 | 0.433733362 | 0.266 | 0.029 | 1.75741820080202e-2 | 5 | atf3 |
| jak1 | 1.2284858 | 0.556810366 | 0.521 | 0.109 | 2.48117290215255e-2 | 5 | jak1 |
| ptprc | 1.2418864 | 0.553138032 | 0.521 | 0.104 | 2.50823802799031e-2 | 5 | ptprc |
| nr4a1 | 1.5433654 | 0.543082772 | 0.543 | 0.11 | 3.11713514625222e-2 | 5 | nr4a1 |
| pnp5a | 2.5821201 | 0.460918918 | 0.362 | 0.056 | 5.21510807044245e-2 | 5 | pnp5a |
| glipr1a | 2.6749733 | 0.52827244 | 0.447 | 0.08 | 5.40264376142585e-2 | 5 | glipr1a |
| cxcl8b.11 | 4.6874930 | 0.96663491 | 0.468 | 0.1 | 9.46732968920781e-2 | 5 | cxcl8b.1 |
| b3gnt3.4 | 4.8484202 | 0.372359824 | 0.266 | 0.03 | 9.79235440553412e-2 | 5 | b3gnt3.4 |
| ccl39.31 | 6.7002883 | 0.47484782 | 0.202 | 0.017 | 1.3532572317807e-24 | 5 | ccl39.3 |
| dnase1l4.1 | 8.2586064 | 0.251335468 | 0.16 | 0.01 | 1.66799074496022e-2 | 5 | dnase1l4.1 |
| ccdc88b | 9.3183649 | 0.315569048 | 0.255 | 0.028 | 1.88203016117908e-2 | 5 | ccdc88b |
| si:ch211-209f23.6 | 9.8853885 | 0.255452278 | 0.16 | 0.01 | 1.99655192926461e-2 | 5 | si:ch211-209f23.6 |
| zgc:1627303 | 1.3221828 | 0.831987416 | 0.936 | 0.475 | 2.67041265252335e-2 | 5 | zgc:162730 |
| glmp | 1.4899619 | 0.435886611 | 0.362 | 0.057 | 3.00927620004856e-2 | 5 | glmp |
| rac22 | 1.5619277 | 0.794740796 | 0.84 | 0.239 | 3.15462547591571e-2 | 5 | rac2 |
| sdc41 | 1.6965190 | 0.82059743 | 0.734 | 0.247 | 3.42645960898402e-2 | 5 | sdc4 |
| CU499336.2 | 2.1683793 | 0.251228824 | 0.181 | 0.013 | 4.37947581803992e-2 | 5 | CU499336.2 |
| CABZ01076275.1 | 2.5991215 | 0.335877532 | 0.191 | 0.015 | 5.24944574516351e-2 | 5 | CABZ01076275.1 |
| zgc:1102391 | 2.6561823 | 0.569635123 | 0.511 | 0.111 | 5.36469139432314e-2 | 5 | zgc:110239 |
| dennd4a | 3.6021027 | 0.399359551 | 0.287 | 0.036 | 7.27516700393671e-2 | 5 | dennd4a |
| paox1 | 5.3523134 | 0.351234414 | 0.223 | 0.022 | 1.08100674604998e-2 | 5 | paox1 |
| xbp1 | 1.0352730 | 0.784516685 | 0.926 | 0.416 | 2.0909408823092e-23 | 5 | xbp1 |
| si:cabz01102082.1 | 1.2841430 | 0.283316224 | 0.17 | 0.012 | 2.59358380480709e-2 | 5 | si:cabz01102082.1 |
| timp2b | 1.3169292 | 0.422626797 | 0.277 | 0.035 | 2.65980207248012e-2 | 5 | timp2b |
| CABZ01078737.1 | 1.3473624 | 0.252789744 | 0.149 | 0.009 | 2.72126784362685e-2 | 5 | CABZ01078737.1 |
| npc21 | 1.8541776 | 0.799523561 | 0.745 | 0.253 | 3.74488258452835e-2 | 5 | npc2 |
| mbd21 | 2.3676738 | 0.368995484 | 0.34 | 0.052 | 4.78199095480893e-2 | 5 | mbd2 |
| cotl12 | 2.8289310 | 0.88713506 | 0.979 | 0.39 | 5.71359198911779e-2 | 5 | cotl1 |
| XLOC-042595 | 3.7991883 | 0.446315379 | 0.479 | 0.092 | 7.67322070916946e-2 | 5 | XLOC-042595 |

| | | | | | | | |
|---|---|---|---|---|---|---|---|
| bcl10 | 4.5073097 | 0.29256879 | 0.191 | 0.016 | 9.10341348266735e-2 | 5 | bcl10 |
| stat3 | 5.3928457 | 0.419025343 | 0.394 | 0.068 | 1.08919306400275e-2 | 5 | stat3 |
| spi1b2 | 5.8555548 | 0.544495595 | 0.532 | 0.112 | 1.18264640476906e-2 | 5 | spi1b |
| npl | 6.2660975 | 0.364696097 | 0.266 | 0.033 | 1.26556371491571e-2 | 5 | npl |
| ssh2a | 7.2417230 | 0.283854941 | 0.202 | 0.018 | 1.46261079473688e-2 | 5 | ssh2a |
| cd9b1 | 8.1713118 | 0.494936018 | 0.266 | 0.032 | 1.65035985938598e-2 | 5 | cd9b |
| atp6ap2 | 8.6346967 | 0.599632381 | 0.511 | 0.122 | 1.74394969476501e-2 | 5 | atp6ap2 |
| derl21 | 9.9303463 | 0.510868251 | 0.5 | 0.11 | 2.00563204252564e-2 | 5 | derl2 |
| mid1ip1l | 1.0991822 | 0.414138881 | 0.394 | 0.07 | 2.22001839787645e-2 | 5 | mid1ip1l |
| canx | 1.2765014 | 0.582985178 | 0.617 | 0.161 | 2.57814996291624e-2 | 5 | canx |
| rgs21 | 1.7914489 | 0.405722226 | 0.266 | 0.033 | 3.61818943174867e-2 | 5 | rgs2 |
| atp6v1d1 | 2.2107297 | 0.363213815 | 0.298 | 0.042 | 4.46501084973384e-2 | 5 | atp6v1d |
| pim1 | 2.7806487 | 0.597023185 | 0.585 | 0.149 | 5.61607631763243e-2 | 5 | pim1 |
| caspa | 3.6687811 | 0.297199497 | 0.234 | 0.026 | 7.40983737569311e-2 | 5 | caspa |
| si:zfos-2326c3.2 | 3.9311800 | 0.313071024 | 0.202 | 0.019 | 7.93980443893676e-2 | 5 | si:zfos-2326c3.2 |
| CR936442.1 | 4.2426145 | 0.489724721 | 0.415 | 0.077 | 8.56880853585642e-2 | 5 | CR936442.1 |
| cd99 | 5.0032505 | 0.397320997 | 0.298 | 0.042 | 1.01050651939809e-2 | 5 | cd99 |
| si:dkey-27i16.21 | 5.4511906 | 0.623899765 | 0.426 | 0.087 | 1.10097698174477e-2 | 5 | si:dkey-27i16.2 |
| FERMT3 (1 of many) | 8.2957797 | 0.408155872 | 0.34 | 0.055 | 1.67549863666801e-2 | 5 | FERMT3 (1 of many) |
| mfsd1 | 8.7537260 | 0.334879948 | 0.234 | 0.026 | 1.76799005402342e-2 | 5 | mfsd1 |
| slc15a4 | 1.7293704 | 0.31889404 | 0.223 | 0.024 | 3.49280959204262e-2 | 5 | slc15a4 |
| gnao1a | 2.0176453 | 0.318649373 | 0.234 | 0.027 | 4.07503839801818e-2 | 5 | gnao1a |
| stk17b1 | 2.2006106 | 0.516104681 | 0.532 | 0.124 | 4.44457323900871e-2 | 5 | stk17b |
| lrrc59 | 2.3014841 | 0.425306468 | 0.426 | 0.086 | 4.64830751873025e-2 | 5 | lrrc59 |
| arhgap4b | 3.1951858 | 0.300835875 | 0.223 | 0.025 | 6.4533168950249e-21 | 5 | arhgap4b |
| tmsb4x2 | 3.6481938 | 0.328296091 | 1 | 0.939 | 7.36825714975837e-2 | 5 | tmsb4x |
| prss16 | 3.6758254 | 0.388034555 | 0.277 | 0.038 | 7.4240646428834e-21 | 5 | prss16 |
| bri3 | 3.7892731 | 0.374395791 | 0.298 | 0.044 | 7.65319504520988e-2 | 5 | bri3 |
| prdx5 | 6.5503876 | 0.695475645 | 0.745 | 0.275 | 1.32298180250454e-2 | 5 | prdx5 |
| FP102158.1 | 1.2546673 | 0.257769418 | 0.191 | 0.018 | 2.53405172314267e-2 | 5 | FP102158.1 |
| si:dkey-222f2.71 | 1.3138269 | 0.360940083 | 0.266 | 0.035 | 2.6535362658338e-20 | 5 | si:dkey-222f2.7 |
| cyfip1 | 1.7081889 | 0.443944993 | 0.404 | 0.079 | 3.45002917609806e-2 | 5 | cyfip1 |
| dnase1l1 | 1.7755779 | 0.514748079 | 0.404 | 0.082 | 3.58613478997382e-2 | 5 | dnase1l1 |
| suclg1 | 2.8963308 | 0.453798697 | 0.617 | 0.172 | 5.8497194091335e-20 | 5 | suclg1 |
| atp6v1aa | 2.9460498 | 0.333500288 | 0.287 | 0.042 | 5.95013681924446e-2 | 5 | atp6v1aa |

| | | | | | | | |
|---|---|---|---|---|---|---|---|
| XLOC-0234921 | 3.1917111 | 0.497618678 | 0.404 | 0.08 | 6.44629896859928e-2 | 5 | XLOC-023492 |
| rhogb1 | 4.0566419 | 0.548503025 | 0.596 | 0.154 | 8.19319965761713e-2 | 5 | rhogb |
| si:dkey-192g7.3 | 4.9164013 | 0.30105138 | 0.213 | 0.023 | 9.92965589279312e-2 | 5 | si:dkey-192g7.3 |
| adcy7 | 6.3513594 | 0.336237516 | 0.234 | 0.029 | 1.28278405970814e-1 | 5 | adcy7 |
| rnf144b | 6.6401675 | 0.326160507 | 0.223 | 0.026 | 1.34111464613902e-1 | 5 | rnf144b |
| zgc:77486 | 8.3968769 | 0.357592721 | 0.287 | 0.043 | 1.69591724032407e-1 | 5 | zgc:77486 |
| ptpn61 | 8.4953534 | 0.460897111 | 0.553 | 0.13 | 1.71580653944684e-1 | 5 | ptpn6 |
| cst14b.11 | 1.0994339 | 0.526966171 | 0.457 | 0.103 | 2.22052680833712e-1 | 5 | cst14b.1 |
| glipr2 | 1.3312248 | 0.302027472 | 0.213 | 0.024 | 2.68867475183489e-1 | 5 | glipr2 |
| git2b | 1.3449170 | 0.299075179 | 0.223 | 0.026 | 2.71632902405159e-1 | 5 | git2b |
| atp6v1c1a | 1.8255638 | 0.296272161 | 0.255 | 0.034 | 3.68709124494713e-1 | 5 | atp6v1c1a |
| tagln2 | 1.8995576 | 0.687000858 | 0.862 | 0.319 | 3.83653648671463e-1 | 5 | tagln2 |
| mir21-1 | 2.0702451 | 0.513999744 | 0.447 | 0.1 | 4.18127420583289e-1 | 5 | mir21-1 |
| si:ch211-15b10.61 | 3.1435100 | 0.451293699 | 0.436 | 0.094 | 6.3489471682059e-19 | 5 | si:ch211-15b10.6 |
| ctsk1 | 4.2409537 | 0.655183191 | 0.489 | 0.125 | 8.56545430375362e-1 | 5 | ctsk |
| slc43a2a1 | 4.5414872 | 0.527722916 | 0.426 | 0.093 | 9.17244176285413e-1 | 5 | slc43a2a |
| rhbdf1a | 4.6318237 | 0.324647507 | 0.277 | 0.041 | 9.35489440888839e-1 | 5 | rhbdf1a |
| rab32a | 5.0250340 | 0.311295178 | 0.223 | 0.027 | 1.01490612804472e-1 | 5 | rab32a |
| csrnp1b | 5.8598659 | 0.459902166 | 0.426 | 0.092 | 1.1835171310422e-18 | 5 | csrnp1b |
| adipor2 | 1.0368123 | 0.411955011 | 0.351 | 0.066 | 2.09404997344925e-1 | 5 | adipor2 |
| si:dkey-11f4.141 | 1.0517435 | 0.699017074 | 0.596 | 0.188 | 2.12420652855175e-1 | 5 | si:dkey-11f4.14 |
| egr11 | 1.1128421 | 0.57578166 | 0.553 | 0.151 | 2.24760719540324e-1 | 5 | egr1 |
| osbp | 1.3115453 | 0.310259526 | 0.245 | 0.033 | 2.6489281202558e-18 | 5 | osbp |
| XLOC-0004311 | 2.1165039 | 0.593257718 | 0.777 | 0.248 | 4.27470310047988e-1 | 5 | XLOC-000431 |
| tmem189 | 2.2526457 | 0.382576514 | 0.319 | 0.057 | 4.54966862404702e-1 | 5 | tmem189 |
| atp2b21 | 2.7961942 | 0.517513919 | 0.521 | 0.136 | 5.6474735762655e-18 | 5 | atp2b2 |
| arl5c | 2.9116520 | 0.431438301 | 0.394 | 0.082 | 5.88066359418964e-1 | 5 | arl5c |
| wipf1b | 3.7822337 | 0.439289431 | 0.34 | 0.066 | 7.63897741028745e-1 | 5 | wipf1b |
| pgd | 4.4356812 | 0.475802702 | 0.436 | 0.103 | 8.95874540899501e-1 | 5 | pgd |
| si:dkey-69o16.51 | 1.0259815 | 0.356880772 | 0.319 | 0.058 | 2.07217484371562e-1 | 5 | si:dkey-69o16.5 |
| grap2b2 | 1.1437430 | 0.50945414 | 0.649 | 0.176 | 2.31001790311187e-1 | 5 | grap2b |
| atp6v0b | 1.1823709 | 0.472843644 | 0.436 | 0.103 | 2.38803458880312e-1 | 5 | atp6v0b |
| ppargc1b | 1.3794736 | 0.283689866 | 0.202 | 0.024 | 2.7861228451895e-17 | 5 | ppargc1b |
| cycsb | 1.4462149 | 0.589979848 | 0.723 | 0.253 | 2.9209203108449e-17 | 5 | cycsb |
| pfn12 | 1.4775996 | 0.963564278 | 1 | 0.397 | 2.98430800515864e-1 | 5 | pfn1 |

| | | | | | | | |
|---|---|---|---|---|---|---|---|
| tmem86a | 1.9136007 | 0.250083148 | 0.138 | 0.011 | 3.86489944177504e-1 | 5 | tmem86a |
| nfe2l2a | 2.1864787 | 0.391513776 | 0.33 | 0.062 | 4.41603103283776e-1 | 5 | nfe2l2a |
| atp6ap1b | 2.6316511 | 0.330656532 | 0.234 | 0.033 | 5.31514583046877e-1 | 5 | atp6ap1b |
| DHDH | 2.7582539 | 0.383463674 | 0.436 | 0.102 | 5.57084556149553e-1 | 5 | DHDH |
| lnpep | 3.3181834 | 0.302835249 | 0.255 | 0.038 | 6.7017351198495e-17 | 5 | lnpep |
| slc30a1a | 5.0493430 | 0.291480164 | 0.223 | 0.03 | 1.01981581160144e-1 | 5 | slc30a1a |
| si:dkey-25e12.3 | 5.9556140 | 0.34563335 | 0.223 | 0.03 | 1.2028553676091e-16 | 5 | si:dkey-25e12.3 |
| nfkbiaa | 6.0984421 | 0.609521216 | 0.532 | 0.151 | 1.2317023589137e-16 | 5 | nfkbiaa |
| slc20a1b | 6.4164298 | 0.399334578 | 0.309 | 0.056 | 1.29592633315966e-1 | 5 | slc20a1b |
| alox5ap | 7.2093864 | 0.283658571 | 0.223 | 0.03 | 1.45607977825057e-1 | 5 | alox5ap |
| actb22 | 8.5665825 | 0.362536832 | 1 | 0.924 | 1.73019267029749e-1 | 5 | actb2 |
| XLOC-005449 | 9.2532417 | 0.428730954 | 0.34 | 0.066 | 1.86887724597082e-1 | 5 | XLOC-005449 |
| efhd2 | 1.0554763 | 0.365873381 | 0.309 | 0.056 | 2.13174560983407e-1 | 5 | efhd2 |
| cyp4f3 | 1.1320745 | 0.258622385 | 0.191 | 0.022 | 2.28645094726916e-1 | 5 | cyp4f3 |
| elovl1b | 1.2966489 | 0.504937593 | 0.543 | 0.151 | 2.61884197854352e-1 | 5 | elovl1b |
| bhlhe40 | 1.3223732 | 0.609575283 | 0.617 | 0.199 | 2.67079725094012e-1 | 5 | bhlhe40 |
| lima1a | 1.4070960 | 0.375823941 | 0.255 | 0.04 | 2.84191191559739e-1 | 5 | lima1a |
| abcc3 | 1.5533107 | 0.274751264 | 0.213 | 0.028 | 3.13722167199503e-1 | 5 | abcc3 |
| LO018340.1 | 1.6428234 | 0.318098598 | 0.255 | 0.039 | 3.31801051805589e-1 | 5 | LO018340.1 |
| vma21 | 1.7646164 | 0.438141714 | 0.489 | 0.128 | 3.56399582153193e-1 | 5 | vma21 |
| slc7a8a | 2.0508713 | 0.285689182 | 0.223 | 0.031 | 4.1421447980724e-16 | 5 | slc7a8a |
| foxo1a | 2.1547821 | 0.351275936 | 0.255 | 0.04 | 4.35201356992598e-1 | 5 | foxo1a |
| rp2 | 2.6347184 | 0.278003165 | 0.245 | 0.036 | 5.32134088047578e-1 | 5 | rp2 |
| txnipa | 3.1088950 | 0.6948307 | 0.777 | 0.355 | 6.27903531972053e-1 | 5 | txnipa |
| dusp5 | 3.3274605 | 0.359646436 | 0.255 | 0.04 | 6.72047200504629e-1 | 5 | dusp5 |
| trib3 | 3.5764352 | 0.483856564 | 0.447 | 0.114 | 7.22332633091047e-1 | 5 | trib3 |
| hcls11 | 3.6719802 | 0.415652028 | 0.436 | 0.1 | 7.41629851420357e-1 | 5 | hcls1 |
| myo1f1 | 5.1619088 | 0.351930792 | 0.298 | 0.054 | 1.04255073649575e-1 | 5 | myo1f |
| coro1a2 | 5.6384978 | 0.649861024 | 0.84 | 0.261 | 1.13880741360045e-1 | 5 | coro1a |
| cap12 | 7.2302201 | 0.635345328 | 0.798 | 0.313 | 1.46028757022383e-1 | 5 | cap1 |
| abracl1 | 7.9524614 | 0.520537818 | 0.691 | 0.225 | 1.60615864135406e-1 | 5 | abracl |
| COX7A2 (1 of many) | 8.2853462 | 0.527410304 | 0.787 | 0.295 | 1.67339138259903e-1 | 5 | COX7A2 (1 of many) |
| rab20 | 8.8823693 | 0.253509425 | 0.213 | 0.029 | 1.79397212797693e-1 | 5 | rab20 |
| CABZ01080702.1 | 8.8947830 | 0.537141426 | 0.553 | 0.162 | 1.79647932636951e-1 | 5 | CABZ01080702.1 |
| tnfrsf1a1 | 9.6976213 | 0.426554022 | 0.372 | 0.083 | 1.95862858216565e-1 | 5 | tnfrsf1a |

| | | | | | | | |
|---|---|---|---|---|---|---|---|
| myh9a | 9.9267259 | 0.30615368 | 0.223 | 0.032 | 2.00490083227695e-1 | 5 | myh9a |
| dram2b | 1.0126198 | 0.362956942 | 0.287 | 0.052 | 2.04518831157979e-1 | 5 | dram2b |
| sgpl1 | 1.0674846 | 0.275622243 | 0.17 | 0.019 | 2.15599878379446e-1 | 5 | sgpl1 |
| pcyox1 | 1.0801158 | 0.475648846 | 0.404 | 0.099 | 2.18151004992857e-1 | 5 | pcyox1 |
| porb | 1.5759593 | 0.30400258 | 0.245 | 0.038 | 3.18296517843491e-1 | 5 | porb |
| swap70b | 1.9683713 | 0.254153365 | 0.213 | 0.029 | 3.9755195847339e-15 | 5 | swap70b |
| m6pr | 2.0377405 | 0.401403985 | 0.319 | 0.064 | 4.11562450340046e-1 | 5 | m6pr |
| dse | 2.2019752 | 0.280617568 | 0.213 | 0.029 | 4.44732938728673e-1 | 5 | dse |
| sh2b3 | 2.4240077 | 0.400545987 | 0.298 | 0.057 | 4.89576837386804e-1 | 5 | sh2b3 |
| galt | 2.5994100 | 0.263646902 | 0.202 | 0.027 | 5.25002855882372e-1 | 5 | galt |
| plek | 2.6994340 | 0.331477577 | 0.266 | 0.045 | 5.45204698575807e-1 | 5 | plek |
| arpc1b2 | 2.7081279 | 0.725619115 | 0.936 | 0.339 | 5.46960611393925e-1 | 5 | arpc1b |
| ccl35.12 | 2.7888266 | 0.588731418 | 0.372 | 0.083 | 5.63259310423378e-1 | 5 | ccl35.1 |
| crip1 | 2.9291292 | 0.650593349 | 0.904 | 0.358 | 5.91596241576532e-1 | 5 | crip1 |
| sdr16c5b | 2.9694835 | 0.393588698 | 0.351 | 0.077 | 5.99746588298508e-1 | 5 | sdr16c5b |
| mboat2a | 3.5719056 | 0.257260911 | 0.202 | 0.027 | 7.2141778500259e-15 | 5 | mboat2a |
| cox17 | 7.9405614 | 0.450067713 | 0.574 | 0.173 | 1.6037551888368e-14 | 5 | cox17 |
| lmcd1 | 8.7520894 | 0.263775535 | 0.255 | 0.042 | 1.76765951579646e-1 | 5 | lmcd1 |
| ssr2 | 1.0070382 | 0.61189425 | 0.851 | 0.403 | 2.03391511506046e-1 | 5 | ssr2 |
| ifi461 | 1.2092317 | 0.42308367 | 0.606 | 0.176 | 2.442285396262e-14 | 5 | ifi46 |
| LOC100151196 | 1.3729841 | 0.275715466 | 0.223 | 0.034 | 2.7730161419876e-14 | 5 | LOC100151196 |
| mcl1a2 | 1.4074093 | 0.634437113 | 0.947 | 0.53 | 2.84254461909035e-1 | 5 | mcl1a |
| atp6v0d1 | 1.5612560 | 0.316363967 | 0.287 | 0.054 | 3.15326883484681e-1 | 5 | atp6v0d1 |
| ktn1 | 1.7100372 | 0.387861809 | 0.34 | 0.074 | 3.45376231902042e-1 | 5 | ktn1 |
| ostf1 | 1.7792192 | 0.314751479 | 0.277 | 0.05 | 3.59348911111356e-1 | 5 | ostf1 |
| wdr11 | 1.8606646 | 0.502418683 | 0.66 | 0.222 | 3.75798448988708e-1 | 5 | wdr1 |
| tfe3a | 2.0248645 | 0.314330834 | 0.255 | 0.044 | 4.08961884053245e-1 | 5 | tfe3a |
| cpox1 | 2.1166755 | 0.326633396 | 0.33 | 0.07 | 4.27504961627654e-1 | 5 | cpox |
| itm2bb | 2.1219931 | 0.586162762 | 0.872 | 0.337 | 4.28578955528135e-1 | 5 | itm2bb |
| RASA2 | 2.2978372 | 0.333416438 | 0.266 | 0.047 | 4.64094184720658e-1 | 5 | RASA2 |
| grinab1 | 2.7879028 | 0.455886137 | 0.553 | 0.165 | 5.63072732046267e-1 | 5 | grinab |
| echdc2 | 6.2205336 | 0.320876352 | 0.287 | 0.057 | 1.25636118364236e-1 | 5 | echdc2 |
| unc93b1 | 6.5918591 | 0.263321958 | 0.181 | 0.023 | 1.33135779359832e-1 | 5 | unc93b1 |
| zfr2 | 8.5262218 | 0.321504585 | 0.277 | 0.053 | 1.72204103111882e-1 | 5 | zfr2 |
| tnfb | 8.6115368 | 0.395624506 | 0.202 | 0.029 | 1.73927208825887e-1 | 5 | tnfb |

| | | | | | | | |
|---|---|---|---|---|---|---|---|
| bzw1b | 9.8140187 | 0.584047845 | 0.681 | 0.272 | 1.98213737059554e-1 | 5 | bzw1b |
| rgs19 | 1.0269301 | 0.297204528 | 0.234 | 0.039 | 2.07409081624328e-1 | 5 | rgs19 |
| klf131 | 1.0561664 | 0.39918455 | 0.372 | 0.088 | 2.13313929442824e-1 | 5 | klf13 |
| wasf2 | 1.2280134 | 0.358284144 | 0.404 | 0.102 | 2.48021880243568e-1 | 5 | wasf2 |
| slc11a21 | 1.2364351 | 0.257536942 | 0.277 | 0.053 | 2.49722811863905e-1 | 5 | slc11a2 |
| ube2l3a | 1.5467786 | 0.439715766 | 0.521 | 0.161 | 3.12402878751322e-1 | 5 | ube2l3a |
| SBNO2 | 1.6788114 | 0.277250292 | 0.213 | 0.033 | 3.39069548132202e-1 | 5 | SBNO2 |
| lgmn1 | 1.9436459 | 0.519579568 | 0.426 | 0.12 | 3.92558182363024e-1 | 5 | lgmn |
| si:ch1073-429i10.3 | 1.9872884 | 0.536109711 | 0.809 | 0.334 | 4.01372643901177e-1 | 5 | si:ch1073-429i10.3.1 |
| nol4lb | 2.0128090 | 0.276553304 | 0.223 | 0.036 | 4.06527037907819e-1 | 5 | nol4lb |
| syngr2a | 2.8719902 | 0.290725914 | 0.287 | 0.058 | 5.80055862620748e-1 | 5 | syngr2a |
| atp1b1a | 3.1558796 | 0.499971781 | 0.745 | 0.278 | 6.37393005259295e-1 | 5 | atp1b1a |
| ppt1 | 3.5548640 | 0.539883824 | 0.702 | 0.272 | 7.17975897168625e-1 | 5 | ppt1 |
| commd7 | 4.0111224 | 0.332390854 | 0.426 | 0.111 | 8.10126408859666e-1 | 5 | commd7 |
| sat1a.21 | 4.3619521 | 0.560523876 | 0.777 | 0.319 | 8.80983477768863e-1 | 5 | sat1a.2 |
| ssr31 | 4.5733124 | 0.634251201 | 0.777 | 0.371 | 9.2367191567439e-13 | 5 | ssr3 |
| fkbp2 | 4.9420014 | 0.425254811 | 0.596 | 0.199 | 9.98136036239574e-1 | 5 | fkbp2 |
| atp2a2b | 5.5434794 | 0.335048028 | 0.266 | 0.052 | 1.11961655297818e-1 | 5 | atp2a2b |
| abca1a1 | 5.7000246 | 0.381738131 | 0.34 | 0.078 | 1.15123397115707e-1 | 5 | abca1a |
| arpc5b2 | 6.1343419 | 0.513154557 | 0.638 | 0.216 | 1.23895304672327e-1 | 5 | arpc5b |
| ubl3b | 6.8395599 | 0.274552993 | 0.245 | 0.044 | 1.38138591851936e-1 | 5 | ubl3b |
| zgc:162698 | 7.4429531 | 0.362283643 | 0.309 | 0.068 | 1.50325325753019e-1 | 5 | zgc:162698 |
| il13ra1 | 7.4927293 | 0.261000369 | 0.181 | 0.025 | 1.51330655151972e-1 | 5 | il13ra1 |
| plp2 | 7.5918946 | 0.336236408 | 0.298 | 0.063 | 1.53333495633304e-1 | 5 | plp2 |
| tmed5 | 1.1243464 | 0.385500215 | 0.298 | 0.066 | 2.27084244711867e-1 | 5 | tmed5 |
| LOC1039091151 | 1.1864540 | 0.461082952 | 0.574 | 0.186 | 2.39628115933816e-1 | 5 | LOC103909115 |
| fam49bb | 1.6061896 | 0.299847019 | 0.287 | 0.059 | 3.24402116698623e-1 | 5 | fam49bb |
| map3k14a1 | 1.6450367 | 0.343929581 | 0.255 | 0.049 | 3.32248065644494e-1 | 5 | map3k14a |
| BX908750.1 | 2.1532856 | 0.259135584 | 0.181 | 0.025 | 4.34899098998189e-1 | 5 | BX908750.1 |
| lgals9l32 | 2.2715792 | 0.490105392 | 0.532 | 0.174 | 4.58790851185552e-1 | 5 | lgals9l3 |
| ecm1b1 | 2.4637885 | 0.685179729 | 0.415 | 0.125 | 4.97611368294357e-1 | 5 | ecm1b |
| ccl39.21 | 3.0181384 | 0.306592357 | 0.117 | 0.011 | 6.09573419673569e-1 | 5 | ccl39.2 |
| rassf2a | 3.3497244 | 0.262968473 | 0.234 | 0.041 | 6.76543845369593e-1 | 5 | rassf2a |
| tpi1b | 3.6292706 | 0.472446825 | 0.628 | 0.232 | 7.33003798058513e-1 | 5 | tpi1b |
| abi2b | 3.7430570 | 0.315545207 | 0.287 | 0.061 | 7.55985222820465e-1 | 5 | abi2b |

| | | | | | | | |
|---|---|---|---|---|---|---|---|
| aoc21 | 3.9617277 | 0.365140196 | 0.447 | 0.123 | 8.00150148971142e-1 | 5 | aoc2 |
| mob1a | 4.3849732 | 0.455479399 | 0.596 | 0.205 | 8.85633037330531e-1 | 5 | mob1a |
| si:dkey-112e7.2 | 5.0684446 | 0.498188737 | 0.67 | 0.258 | 1.02367376812144e-1 | 5 | si:dkey-112e7.2 |
| grb2b | 6.1569517 | 0.372699321 | 0.362 | 0.092 | 1.2435195501219e-11 | 5 | grb2b |
| zc3h12b | 6.7408858 | 0.25666097 | 0.17 | 0.024 | 1.36145672368977e-1 | 5 | zc3h12b |
| capgb1 | 6.8106721 | 0.387515388 | 0.436 | 0.116 | 1.37555146116974e-1 | 5 | capgb |
| wbp2nl | 7.2035813 | 0.348553494 | 0.404 | 0.109 | 1.45490732905623e-1 | 5 | wbp2nl |
| bloc1s61 | 8.2846736 | 0.323235561 | 0.436 | 0.124 | 1.67325553026227e-1 | 5 | bloc1s6 |
| eno3 | 9.3070702 | 0.505937487 | 0.702 | 0.28 | 1.87974897903675e-1 | 5 | eno3 |
| si:ch1073-325m22. | 1.2394519 | 0.499766936 | 0.766 | 0.345 | 2.50332101610526e-1 | 5 | si:ch1073-325m22.2 |
| blcap | 1.2637685 | 0.359345286 | 0.468 | 0.138 | 2.55243336733269e-1 | 5 | blcap |
| cbr1l | 1.3136187 | 0.356608267 | 0.479 | 0.143 | 2.65311584581421e-1 | 5 | cbr1l |
| ipo7 | 1.3568018 | 0.371647579 | 0.468 | 0.142 | 2.74033269330499e-1 | 5 | ipo7 |
| gmppb | 1.3848023 | 0.344846533 | 0.351 | 0.09 | 2.79688525544278e-1 | 5 | gmppb |
| selplg | 1.3992578 | 0.309223257 | 0.213 | 0.037 | 2.82608107882054e-1 | 5 | selplg |
| arpc4l2 | 1.4723272 | 0.546157325 | 0.755 | 0.335 | 2.97365943812151e-1 | 5 | arpc4l |
| serp1 | 2.2240374 | 0.435809418 | 0.479 | 0.157 | 4.4918883648296e-11 | 5 | serp1 |
| si:ch211-198c19.3 | 2.3182252 | 0.572331145 | 0.521 | 0.173 | 4.68211949709647e-1 | 5 | si:ch211-198c19.3 |
| msna | 2.6340668 | 0.571643816 | 0.819 | 0.388 | 5.32002479103252e-1 | 5 | msna |
| zgc:1583431 | 3.0608825 | 0.520796199 | 0.543 | 0.191 | 6.18206455637527e-1 | 5 | zgc:158343 |
| gnai2a | 3.6952434 | 0.455407277 | 0.511 | 0.174 | 7.46328326760918e-1 | 5 | gnai2a |
| arhgdig1 | 5.3464655 | 0.555674667 | 0.819 | 0.349 | 1.07982563786324e-1 | 5 | arhgdig |
| litaf1 | 6.1865072 | 0.420960289 | 0.564 | 0.194 | 1.24948886531523e-1 | 5 | litaf |
| skap21 | 6.7996017 | 0.331183308 | 0.394 | 0.106 | 1.37331556951002e-1 | 5 | skap2 |
| mef2cb | 6.8713475 | 0.280617829 | 0.234 | 0.045 | 1.38780607284274e-1 | 5 | mef2cb |
| rap1b1 | 8.0990764 | 0.506523155 | 0.745 | 0.311 | 1.63577046965069e-1 | 5 | rap1b |
| lipf2 | 8.3866586 | 0.407899185 | 0.298 | 0.072 | 1.69385345476559e-1 | 5 | lipf |
| junba2 | 1.0667910 | 0.599795856 | 0.84 | 0.388 | 2.15459791501581e-1 | 5 | junba |
| si:ch211-66e2.3 | 1.4292384 | 0.367611694 | 0.319 | 0.077 | 2.88663285900709e-1 | 5 | si:ch211-66e2.3 |
| antxr2a2 | 2.2174259 | 0.574726223 | 0.606 | 0.251 | 4.47853515575394e-1 | 5 | antxr2a |
| XLOC-0254231 | 2.9732833 | 0.417534129 | 0.543 | 0.179 | 6.00514042849399e-1 | 5 | XLOC-025423 |
| hexb | 5.1818603 | 0.385017644 | 0.309 | 0.08 | 1.0465803287629e-09 | 5 | hexb |
| MYADM | 5.2228003 | 0.290610305 | 0.266 | 0.059 | 1.05484898615661e-0 | 5 | MYADM |
| cpvl | 6.9139502 | 0.303681397 | 0.277 | 0.065 | 1.39641053694978e-0 | 5 | cpvl |
| arrdc2 | 9.4374180 | 0.28431122 | 0.245 | 0.053 | 1.90607531998769e-0 | 5 | arrdc2 |

| | | | | | | | |
|---|---|---|---|---|---|---|---|
| tradd | 1.1173768 | 0.266043678 | 0.255 | 0.057 | 2.25676604250949e-0 | 5 | tradd |
| tmed2 | 1.2214208 | 0.414157845 | 0.723 | 0.296 | 2.46690376199149e-0 | 5 | tmed2 |
| park7 | 1.3867015 | 0.258163258 | 0.319 | 0.082 | 2.80072112695821e-0 | 5 | park7 |
| itga4 | 1.8120276 | 0.304938049 | 0.255 | 0.058 | 3.65975223820287e-0 | 5 | itga4 |
| srgap2 | 1.8458302 | 0.283590379 | 0.223 | 0.046 | 3.72802339532658e-0 | 5 | srgap2 |
| gstt1a1 | 1.9147789 | 0.305700785 | 0.234 | 0.049 | 3.86727899682783e-0 | 5 | gstt1a |
| rin3 | 2.0230353 | 0.276547393 | 0.255 | 0.057 | 4.08592441527888e-0 | 5 | rin3 |
| degs1 | 2.0671268 | 0.349530348 | 0.383 | 0.114 | 4.17497600064695e-0 | 5 | degs1 |
| triqk | 2.1330720 | 0.301895307 | 0.298 | 0.076 | 4.30816552507056e-0 | 5 | triqk |
| serpinb1l3 | 2.5122955 | 0.267195171 | 0.372 | 0.105 | 5.07408333993582e-0 | 5 | serpinb1l3 |
| clptm1l | 3.0122113 | 0.25565535 | 0.234 | 0.05 | 6.08376321793960e-0 | 5 | clptm1l |
| uqcrq | 3.5586876 | 0.477383445 | 0.745 | 0.359 | 7.18748139070814e-0 | 5 | uqcrq |
| sec61b | 3.7806120 | 0.503827334 | 0.926 | 0.555 | 7.63570223014786e-0 | 5 | sec61b |
| elmo1 | 5.2034608 | 0.27785297 | 0.266 | 0.063 | 1.05094298051311e-0 | 5 | elmo1 |
| rpn1 | 5.6691588 | 0.417701941 | 0.574 | 0.224 | 1.14500001807954e-0 | 5 | rpn1 |
| ier2b | 5.9667437 | 0.487004385 | 0.723 | 0.312 | 1.20510323691897e-0 | 5 | ier2b |
| ubl7b | 6.7868673 | 0.299907749 | 0.298 | 0.078 | 1.37074359871857e-0 | 5 | ubl7b |
| ddit31 | 9.6022089 | 0.459440997 | 0.511 | 0.192 | 1.93935813444766e-0 | 5 | ddit3 |
| sec61g1 | 1.1506579 | 0.425963735 | 0.936 | 0.719 | 2.32398393211482e-0 | 5 | sec61g |
| actr2a | 1.1949974 | 0.356722533 | 0.457 | 0.153 | 2.41353626957605e-0 | 5 | actr2a |
| npepl1 | 1.2133323 | 0.338209988 | 0.351 | 0.103 | 2.45056741270951e-0 | 5 | npepl1 |
| hyou1 | 1.2341216 | 0.31213259 | 0.287 | 0.075 | 2.4925554976666e-08 | 5 | hyou1 |
| hsp90b1 | 1.2452271 | 0.394249293 | 0.468 | 0.165 | 2.51498530217526e-0 | 5 | hsp90b1 |
| mbnl1 | 1.3901047 | 0.353920064 | 0.5 | 0.175 | 2.80759453939273e-0 | 5 | mbnl1 |
| clta | 2.3810794 | 0.336211502 | 0.532 | 0.198 | 4.80906621123249e-0 | 5 | clta |
| irf2 | 2.7965147 | 0.291017728 | 0.34 | 0.099 | 5.64812086718876e-0 | 5 | irf2 |
| ost4 | 2.9718322 | 0.383544522 | 0.681 | 0.284 | 6.00220968944431e-0 | 5 | ost4 |
| mt-co11 | 3.0534848 | 0.541661863 | 0.904 | 0.582 | 6.16712344334463e-0 | 5 | mt-co1 |
| cox7b | 3.2349960 | 0.480655194 | 0.872 | 0.492 | 6.53372157937346e-0 | 5 | cox7b |
| glrx1 | 3.3419054 | 0.468133151 | 0.819 | 0.412 | 6.74964648585672e-0 | 5 | glrx |
| nfkbie | 3.3688324 | 0.337402807 | 0.362 | 0.11 | 6.80403094371699e-0 | 5 | nfkbie |
| atp5f1d | 3.6941214 | 0.402231064 | 0.745 | 0.353 | 7.46101715644174e-0 | 5 | atp5f1d |
| zgc:123258 | 4.0387905 | 0.269049271 | 0.298 | 0.079 | 8.15714523993552e-0 | 5 | zgc:123258 |
| pgam1a | 4.3001780 | 0.381888527 | 0.426 | 0.148 | 8.68506970123883e-0 | 5 | pgam1a |
| arpc22 | 4.4415452 | 0.477617165 | 0.862 | 0.472 | 8.97058900313177e-0 | 5 | arpc2 |

| | | | | | | | |
|---|---|---|---|---|---|---|---|
| atp5if1b | 4.9981072 | 0.330046703 | 0.649 | 0.261 | 1.00946772686827e-0 | 5 | atp5if1b |
| tln1 | 5.5498598 | 0.399629222 | 0.553 | 0.213 | 1.12090519206677e-0 | 5 | tln1 |
| rab5c | 5.7045219 | 0.330253183 | 0.5 | 0.178 | 1.15214229968847e-0 | 5 | rab5c |
| vps29 | 5.8450279 | 0.313041423 | 0.5 | 0.181 | 1.18052028854733e-0 | 5 | vps29 |
| gapdhs | 6.6164046 | 0.506737137 | 0.83 | 0.464 | 1.33631525664998e-0 | 5 | gapdhs |
| anxa5b | 8.8075201 | 0.361313553 | 0.457 | 0.16 | 1.778854838975e-07 | 5 | anxa5b |
| hspa5 | 9.1195298 | 0.506394682 | 0.511 | 0.212 | 1.84187144621268e-0 | 5 | hspa5 |
| COX5B | 9.9143907 | 0.459813296 | 0.84 | 0.467 | 2.00240950742948e-0 | 5 | COX5B |
| gdi2 | 1.1253919 | 0.407636075 | 0.66 | 0.291 | 2.27295414535292e-0 | 5 | gdi2 |
| ugp2b | 1.2482952 | 0.272957489 | 0.255 | 0.065 | 2.52118197490104e-0 | 5 | ugp2b |
| aldoaa | 1.2563998 | 0.426128357 | 0.638 | 0.276 | 2.53755075899662e-0 | 5 | aldoaa |
| cox7a2l | 1.5194288 | 0.331172045 | 0.489 | 0.18 | 3.06879048989758e-0 | 5 | cox7a2l |
| cdc42l | 1.5322609 | 0.441710895 | 0.713 | 0.333 | 3.09470734675931e-0 | 5 | cdc42l |
| dusp2 | 1.5796790 | 0.323581042 | 0.319 | 0.092 | 3.19047771272813e-0 | 5 | dusp2 |
| LOC1005362131 | 1.6110321 | 0.318475416 | 0.436 | 0.139 | 3.25380169086231e-0 | 5 | LOC100536213 |
| uqcr10 | 2.0022104 | 0.375680205 | 0.702 | 0.311 | 4.04386438044235e-0 | 5 | uqcr10 |
| XLOC-010899 | 2.2383776 | 0.361446645 | 0.447 | 0.158 | 4.52085126328236e-0 | 5 | XLOC-010899 |
| smim15 | 2.3686998 | 0.369895552 | 0.511 | 0.201 | 4.78406306305128e-0 | 5 | smim15 |
| ldhba | 3.2608974 | 0.396563442 | 0.713 | 0.325 | 6.58603461004089e-0 | 5 | ldhba |
| ccr9a | 4.3666444 | 0.391312665 | 0.309 | 0.091 | 8.81931188337506e-0 | 5 | ccr9a |
| gnai1 | 4.5942295 | 0.289602444 | 0.362 | 0.116 | 9.27896532783664e-0 | 5 | gnai1 |
| calr3a | 5.5344306 | 0.366880095 | 0.564 | 0.225 | 1.1177889674327e-06 | 5 | calr3a |
| rnf213a1 | 5.8003945 | 0.383473314 | 0.372 | 0.127 | 1.17150568833466e-0 | 5 | rnf213a |
| actb12 | 5.9113403 | 0.438382815 | 0.989 | 0.732 | 1.19391340268386e-0 | 5 | actb1 |
| atf4a | 6.5806094 | 0.353214777 | 0.521 | 0.204 | 1.32908569523065e-0 | 5 | atf4a |
| cox6a1 | 6.7854285 | 0.439482063 | 0.872 | 0.482 | 1.37045301028946e-0 | 5 | cox6a1 |
| pak2a | 7.1230695 | 0.25979745 | 0.33 | 0.098 | 1.43864636261449e-0 | 5 | pak2a |
| tmem176l.3a | 7.4761441 | 0.295234689 | 0.298 | 0.088 | 1.50995682517175e-0 | 5 | tmem176l.3a |
| mbnl3 | 7.6487755 | 0.287731385 | 0.309 | 0.092 | 1.54482320648216e-0 | 5 | mbnl3 |
| sdhb | 8.3937083 | 0.284182318 | 0.404 | 0.139 | 1.69527728086854e-0 | 5 | sdhb |
| rab1ab | 8.4129543 | 0.362582386 | 0.532 | 0.216 | 1.69916438899862e-0 | 5 | rab1ab |
| si:ch73-234b20.5 | 9.0231599 | 0.271857181 | 0.383 | 0.127 | 1.82240762283007e-0 | 5 | si:ch73-234b20.5 |
| snx1a | 9.4851971 | 0.319330196 | 0.436 | 0.159 | 1.91572527434859e-0 | 5 | snx1a |
| slc44a2 | 1.1865684 | 0.257736454 | 0.266 | 0.073 | 2.39651238721859e-0 | 5 | slc44a2 |
| arf2a | 1.3003081 | 0.349933524 | 0.415 | 0.152 | 2.62623246790493e-0 | 5 | arf2a |

| | | | | | | | |
|---|---|---|---|---|---|---|---|
| chmp2bb | 1.5293429 | 0.252066653 | 0.277 | 0.079 | 3.08881403770982e-0 | 5 | chmp2bb |
| si:dkey-180p18.9 | 2.4982645 | 0.309629024 | 0.415 | 0.151 | 5.04574488254031e-0 | 5 | si:dkey-180p18.9 |
| manf | 2.6962380 | 0.315485701 | 0.34 | 0.114 | 5.44559198842815e-0 | 5 | manf |
| oaz1b | 2.9227379 | 0.432925712 | 0.894 | 0.511 | 5.90305393279823e-0 | 5 | oaz1b |
| tagapb | 3.0032179 | 0.264307194 | 0.277 | 0.079 | 6.06559923406165e-0 | 5 | tagapb |
| dnajc1 | 3.1057343 | 0.28478078 | 0.372 | 0.128 | 6.27265174583508e-0 | 5 | dnajc1 |
| cltca | 3.4285863 | 0.268428857 | 0.351 | 0.114 | 6.9247158413744e-06 | 5 | cltca |
| sec61a1 | 3.4693811 | 0.29436664 | 0.521 | 0.216 | 7.00710910486676e-0 | 5 | sec61a1 |
| elf1 | 3.4749606 | 0.281080675 | 0.319 | 0.1 | 7.01837795618448e-0 | 5 | elf1 |
| lxn2 | 4.8833132 | 0.422764187 | 0.585 | 0.257 | 9.86282778041377e-0 | 5 | lxn |
| cfl12 | 5.0370559 | 0.3693299 | 0.989 | 0.759 | 1.01733418353114e-0 | 5 | cfl1 |
| higd2a | 6.1100040 | 0.267927112 | 0.351 | 0.118 | 1.23403752367053e-0 | 5 | higd2a |
| dad1 | 6.2830793 | 0.42418774 | 0.777 | 0.407 | 1.26899352988524e-0 | 5 | dad1 |
| pfkfb4b | 6.3626814 | 0.321868252 | 0.521 | 0.21 | 1.28507078040103e-0 | 5 | pfkfb4b |
| abcc5 | 6.6936004 | 0.259172915 | 0.298 | 0.091 | 1.35190648487412e-0 | 5 | abcc5 |
| slc30a7 | 7.0877919 | 0.263724051 | 0.277 | 0.084 | 1.43152134018417e-0 | 5 | slc30a7 |
| cox5aa | 7.0912435 | 0.400152235 | 0.755 | 0.398 | 1.4322184649769e-05 | 5 | cox5aa |
| prdx6 | 7.4119996 | 0.355275034 | 0.489 | 0.206 | 1.49700156575395e-0 | 5 | prdx6 |
| g3bp1 | 7.5589260 | 0.313210684 | 0.33 | 0.112 | 1.52667630230182e-0 | 5 | g3bp1 |
| arhgdia | 7.7759415 | 0.384669742 | 0.766 | 0.36 | 1.57050691202615e-0 | 5 | arhgdia |
| p4hb | 8.3865550 | 0.346545134 | 0.5 | 0.209 | 1.69383252819056e-0 | 5 | p4hb |
| mrpl33 | 8.5545657 | 0.277304427 | 0.394 | 0.145 | 1.72776563624094e-0 | 5 | mrpl33 |
| zgc:153713 | 1.0396585 | 0.360177152 | 0.585 | 0.264 | 2.09979827905017e-0 | 5 | zgc:153713 |
| rgs5b | 1.0622003 | 0.301081289 | 0.213 | 0.055 | 2.14532600232902e-0 | 5 | rgs5b |
| dbi | 1.1315497 | 0.399403251 | 0.755 | 0.391 | 2.28539096991986e-0 | 5 | dbi |
| gpia | 1.2957765 | 0.303151895 | 0.362 | 0.131 | 2.61707989585338e-0 | 5 | gpia |
| snd1 | 1.2987478 | 0.304901879 | 0.436 | 0.176 | 2.62308097558951e-0 | 5 | snd1 |
| sptssa | 1.4047485 | 0.288462193 | 0.34 | 0.116 | 2.83717064691791e-0 | 5 | sptssa |
| scamp2 | 1.4492651 | 0.268404476 | 0.298 | 0.095 | 2.92708079222798e-0 | 5 | scamp2 |
| si:dkeyp-27c8.2 | 1.7766243 | 0.258954321 | 0.213 | 0.057 | 3.58824811521979e-0 | 5 | si:dkeyp-27c8.2 |
| tspan141 | 1.7914816 | 0.284489796 | 0.447 | 0.169 | 3.61825541539221e-0 | 5 | tspan14 |
| arpc41 | 2.0015579 | 0.355045747 | 0.66 | 0.305 | 4.04254662128695e-0 | 5 | arpc4 |
| tmed7 | 2.2302186 | 0.368252503 | 0.638 | 0.296 | 4.50437266173936e-0 | 5 | tmed7 |
| midn1 | 2.6282746 | 0.324293514 | 0.436 | 0.172 | 5.30832629452575e-0 | 5 | midn |
| eno1a | 3.9025418 | 0.308742365 | 0.447 | 0.18 | 7.88196379268221e-0 | 5 | eno1a |

| | | | | | | | |
|---|---|---|---|---|---|---|---|
| f11r.1 | 3.9052039 | 0.280107661 | 0.298 | 0.097 | 7.88734049725833e-0 | 5 | f11r.1 |
| rhoab | 3.9350124 | 0.37610596 | 0.628 | 0.303 | 7.9475447373465e-05 | 5 | rhoab |
| eif4ebp1 | 3.9636791 | 0.376705284 | 0.66 | 0.332 | 8.0054427482749e-05 | 5 | eif4ebp1 |
| ddx271 | 3.9739229 | 0.271215337 | 0.351 | 0.124 | 8.02613226963044e-0 | 5 | ddx27 |
| ube2v2 | 4.0199522 | 0.369026641 | 0.702 | 0.353 | 8.11909753826757e-0 | 5 | ube2v2 |
| eif3c | 5.0632122 | 0.316342278 | 0.617 | 0.279 | 0.000102262 | 5 | eif3c |
| mtdha | 5.1884100 | 0.29426067 | 0.468 | 0.192 | 0.00010479 | 5 | mtdha |
| capza1b1 | 6.1605037 | 0.366338511 | 0.713 | 0.352 | 0.000124424 | 5 | capza1b |
| clic1 | 8.2972828 | 0.33332331 | 0.489 | 0.207 | 0.00016758 | 5 | clic1 |
| atp5mf | 8.9080902 | 0.268109168 | 0.638 | 0.296 | 0.000179917 | 5 | atp5mf |
| sec11a | 9.6431015 | 0.334626671 | 0.564 | 0.262 | 0.000194762 | 5 | sec11a |
| gabarapa1 | 1.3331366 | 0.290540364 | 0.564 | 0.247 | 0.000269254 | 5 | gabarapa |
| flna2 | 1.3883718 | 0.321769166 | 0.404 | 0.161 | 0.000280409 | 5 | flna |
| atp1a1a.1 | 1.4668360 | 0.254486728 | 0.351 | 0.129 | 0.000296257 | 5 | atp1a1a.1 |
| atp5l1 | 1.5440406 | 0.382830227 | 0.851 | 0.514 | 0.00031185 | 5 | atp5l |
| cox6b2 | 1.6859476 | 0.370146642 | 0.936 | 0.655 | 0.000340511 | 5 | cox6b2 |
| cox7c | 1.7694280 | 0.389176715 | 0.851 | 0.572 | 0.000357371 | 5 | cox7c |
| pdcd10a | 1.7901158 | 0.26114684 | 0.266 | 0.087 | 0.00036155 | 5 | pdcd10a |
| celf22 | 1.8722457 | 0.401198437 | 0.67 | 0.347 | 0.000378137 | 5 | celf2 |
| ponzr11 | 1.8900153 | 0.364834981 | 0.787 | 0.379 | 0.000381726 | 5 | ponzr1 |
| ccl39.71 | 1.9621204 | 0.364644949 | 0.234 | 0.073 | 0.000396289 | 5 | ccl39.7 |
| atp5po | 2.0199593 | 0.283558503 | 0.649 | 0.307 | 0.000407971 | 5 | atp5po |
| ist1 | 2.1092852 | 0.252832665 | 0.277 | 0.092 | 0.000426012 | 5 | ist1 |
| mdh1aa1 | 2.5595414 | 0.356013325 | 0.66 | 0.347 | 0.000516951 | 5 | mdh1aa |
| reep5 | 2.5856818 | 0.32096097 | 0.553 | 0.256 | 0.00052223 | 5 | reep5 |
| krtcap2 | 2.6385505 | 0.357287778 | 0.596 | 0.296 | 0.000532908 | 5 | krtcap2 |
| ostc | 2.8089914 | 0.327185998 | 0.532 | 0.243 | 0.000567332 | 5 | ostc |
| fosab2 | 2.8698538 | 0.437759504 | 0.83 | 0.448 | 0.000579624 | 5 | fosab |
| fam120c | 3.3209178 | 0.283787805 | 0.362 | 0.141 | 0.000670726 | 5 | fam120c |
| higd1a | 4.0066076 | 0.324392963 | 0.564 | 0.265 | 0.000809215 | 5 | higd1a |
| nfkbiab | 4.0825064 | 0.344971188 | 0.606 | 0.291 | 0.000824544 | 5 | nfkbiab |
| cnp | 4.2557295 | 0.318827206 | 0.489 | 0.223 | 0.00085953 | 5 | cnp |
| arpc32 | 4.5642589 | 0.382911888 | 0.83 | 0.468 | 0.000921843 | 5 | arpc3 |
| mcl1b1 | 4.8174425 | 0.332561075 | 0.638 | 0.306 | 0.000972979 | 5 | mcl1b |
| cdk2ap2 | 4.9128603 | 0.288821462 | 0.319 | 0.121 | 0.00099225 | 5 | cdk2ap2 |

| | | | | | | | |
|---|---|---|---|---|---|---|---|
| uqcrh | 5.1778965 | 0.284258143 | 0.628 | 0.313 | 0.00104578 | 5 | uqcrh |
| magt1 | 5.5342285 | 0.28949827 | 0.457 | 0.201 | 0.001117748 | 5 | magt1 |
| capza1a1 | 7.1616074 | 0.332697393 | 0.67 | 0.332 | 0.00144643 | 5 | capza1a |
| btg1 | 8.7400886 | 0.390408514 | 0.777 | 0.455 | 0.001765236 | 5 | btg1 |
| nop10 | 9.0579745 | 0.30569609 | 0.511 | 0.239 | 0.001829439 | 5 | nop10 |
| arf5 | 9.0689847 | 0.295039966 | 0.553 | 0.262 | 0.001831663 | 5 | arf5 |
| cst14a.2 | 9.2602124 | 0.31391049 | 0.617 | 0.298 | 0.001870285 | 5 | cst14a.2 |
| hspa81 | 9.4510385 | 0.252540099 | 0.968 | 0.837 | 0.001908826 | 5 | hspa8 |
| max | 1.0214636 | 0.252166178 | 0.447 | 0.192 | 0.00206305 | 5 | max |
| anxa11b1 | 1.1089714 | 0.28932649 | 0.638 | 0.3 | 0.00223979 | 5 | anxa11b |
| metrnla | 1.2090938 | 0.250845651 | 0.298 | 0.107 | 0.002442007 | 5 | metrnla |
| zgc:162944 | 1.2458901 | 0.331638166 | 0.734 | 0.389 | 0.002516324 | 5 | zgc:162944 |
| odc1 | 1.5500508 | 0.263837018 | 0.309 | 0.116 | 0.003130638 | 5 | odc1 |
| cox4i1 | 1.5725783 | 0.364019762 | 0.894 | 0.566 | 0.003176137 | 5 | cox4i1 |
| sdcbp21 | 1.9108028 | 0.332392085 | 0.649 | 0.332 | 0.003859249 | 5 | sdcbp2 |
| cox6c | 2.1293450 | 0.301856707 | 0.745 | 0.405 | 0.004300638 | 5 | cox6c |
| socs3b2 | 2.3488211 | 0.323633162 | 0.34 | 0.141 | 0.004743914 | 5 | socs3b |
| GCA | 3.0507077 | 0.297018431 | 0.606 | 0.3 | 0.006161514 | 5 | GCA |
| si:ch211-235e9.6 | 3.0689696 | 0.253386575 | 0.457 | 0.208 | 0.006198398 | 5 | si:ch211-235e9.6 |
| rsl1d1 | 3.6331993 | 0.262322132 | 0.5 | 0.233 | 0.007337973 | 5 | rsl1d1 |
| spcs2 | 3.6629593 | 0.277044924 | 0.521 | 0.245 | 0.007398079 | 5 | spcs2 |
| atp5f1c | 4.4799842 | 0.260008668 | 0.66 | 0.346 | 0.009048224 | 5 | atp5f1c |
| bsg1 | 7.0099131 | 0.321299919 | 0.702 | 0.377 | 0.014157922 | 5 | bsg |
| jdp2b1 | 8.4140800 | 0.304867489 | 0.34 | 0.145 | 0.016993917 | 5 | jdp2b |
| rab5ab | 8.5290932 | 0.251580339 | 0.33 | 0.137 | 0.01722621 | 5 | rab5ab |
| atp5md | 8.5668046 | 0.281845125 | 0.638 | 0.344 | 0.017302375 | 5 | atp5md |
| zfp36l1a2 | 1.2584132 | 0.324868557 | 0.585 | 0.303 | 0.025416172 | 5 | zfp36l1a |
| btg2 | 1.7220772 | 0.286103668 | 0.511 | 0.253 | 0.034780794 | 5 | btg2 |
| ier2a1 | 2.0283686 | 0.322871167 | 0.66 | 0.355 | 0.040966963 | 5 | ier2a |
| ndufb3 | 2.1635015 | 0.252365975 | 0.617 | 0.328 | 0.043696242 | 5 | ndufb3 |
| COX32 | 2.2455678 | 0.417176233 | 0.809 | 0.591 | 0.045353735 | 5 | COX3 |
| vat11 | 2.7676415 | 0.257306132 | 0.468 | 0.218 | 0.055898056 | 5 | vat1 |
| pdia31 | 3.0262887 | 0.337362643 | 0.777 | 0.476 | 0.061121954 | 5 | pdia3 |
| calr3b | 3.5215490 | 0.274114046 | 0.383 | 0.178 | 0.071124727 | 5 | calr3b |
| pdia6 | 3.7706660 | 0.25524112 | 0.362 | 0.166 | 0.076156141 | 5 | pdia6 |

| | | | | | | | |
|---|---|---|---|---|---|---|---|
| rer1 | 3.7960272 | 0.256321268 | 0.372 | 0.171 | 0.076668362 | 5 | rer1 |
| socs3a2 | 5.1580227 | 0.297252751 | 0.777 | 0.419 | 0.104176586 | 5 | socs3a |
| atp5pf | 8.8438680 | 0.302140426 | 0.681 | 0.423 | 0.178619603 | 5 | atp5pf |
| zgc:56493 | 1.2718608 | 0.304157038 | 0.957 | 0.711 | 0.256877737 | 5 | zgc:56493 |
| tmem258 | 1.6645897 | 0.27971632 | 0.713 | 0.417 | 0.336197202 | 5 | tmem258 |
| ssr4 | 1.7203652 | 0.262769503 | 0.777 | 0.474 | 0.347462172 | 5 | ssr4 |
| itm2ba1 | 2.2918199 | 0.255242545 | 0.734 | 0.403 | 0.462878871 | 5 | itm2ba |
| fthl27 | 2.6014767 | 0.328610849 | 0.872 | 0.621 | 0.525420264 | 5 | fthl27 |
| ucp2 | 2.8800090 | 0.282596902 | 0.713 | 0.416 | 0.581675421 | 5 | ucp2 |
| atp5mc3b | 3.0138504 | 0.299806554 | 0.84 | 0.628 | 0.608707368 | 5 | atp5mc3b |
| icn1 | 3.9444891 | 0.429722711 | 0.447 | 0.254 | 0.796668475 | 5 | icn |
| ppdpfb | 7.3822374 | 0.302981514 | 0.894 | 0.611 | 1 | 5 | ppdpfb |
| selenof | 0.000143 | 0.263037669 | 0.681 | 0.417 | 1 | 5 | selenof |
| rpl27a1 | 0.000607 | 0.25333563 | 0.936 | 0.679 | 1 | 5 | rpl27a |
| zgc:1537041 | 4.9856815 | 0.895996723 | 0.319 | 0.031 | 1.00695810496235e-3 | 6 | zgc:153704 |
| cfd | 3.3374736 | 1.192925493 | 0.468 | 0.076 | 6.74069556266792e-3 | 6 | cfd |
| tagln1 | 4.0686037 | 1.029990283 | 0.596 | 0.118 | 8.2173588942544e-36 | 6 | tagln |
| cxcl12a2 | 1.0795409 | 0.90013524 | 0.489 | 0.08 | 2.18034881072678e-3 | 6 | cxcl12a |
| ccl25b1 | 1.7551182 | 1.158394503 | 0.681 | 0.167 | 3.54481241483976e-3 | 6 | ccl25b |
| acta2 | 1.9353427 | 0.563884459 | 0.181 | 0.009 | 3.90881175687037e-3 | 6 | acta2 |
| col1a22 | 1.6197592 | 1.017481842 | 0.904 | 0.311 | 3.27142782114124e-3 | 6 | col1a2 |
| tcf21 | 2.2451041 | 0.323059694 | 0.202 | 0.013 | 4.53443689547168e-3 | 6 | tcf21 |
| epha4b | 3.0295129 | 0.48703592 | 0.223 | 0.017 | 6.11870736087947e-3 | 6 | epha4b |
| angptl1a1 | 5.1885510 | 0.743292529 | 0.404 | 0.062 | 1.04793165642988e-2 | 6 | angptl1a |
| rgs5a1 | 1.1130219 | 0.815102886 | 0.457 | 0.082 | 2.24797043226957e-2 | 6 | rgs5a |
| pcolcea1 | 3.8965187 | 0.886474313 | 0.617 | 0.139 | 7.86979900777204e-2 | 6 | pcolcea |
| asip2b1 | 7.7090231 | 0.806172876 | 0.383 | 0.058 | 1.55699141534629e-2 | 6 | asip2b |
| col1a1b2 | 5.0901311 | 0.976579753 | 0.819 | 0.279 | 1.02805378247503e-2 | 6 | col1a1b |
| postnb1 | 1.2526388 | 0.980412467 | 0.638 | 0.172 | 2.52995466945663e-2 | 6 | postnb |
| col5a12 | 1.3710182 | 0.861508393 | 0.745 | 0.224 | 2.76904547546354e-2 | 6 | col5a1 |
| col1a1a2 | 1.5402920 | 0.942487941 | 0.83 | 0.292 | 3.11092794802228e-2 | 6 | col1a1a |
| col6a12 | 6.5092347 | 0.764775413 | 0.543 | 0.13 | 1.31467014152923e-2 | 6 | col6a1 |
| tspan18a | 7.5825787 | 0.400896222 | 0.223 | 0.022 | 1.53145343225365e-2 | 6 | tspan18a |
| cyp1c21 | 1.1331641 | 0.525174178 | 0.255 | 0.03 | 2.28865161940172e-2 | 6 | cyp1c2 |
| dcn2 | 1.7719485 | 0.872754105 | 0.777 | 0.26 | 3.57880441901697e-2 | 6 | dcn |

| | | | | | | | |
|---|---|---|---|---|---|---|---|
| rspo1 | 6.1126259 | 0.536564521 | 0.223 | 0.023 | 1.23456705640464e-2 | 6 | rspo1 |
| dpt1 | 7.6945767 | 0.621069563 | 0.213 | 0.021 | 1.55407366984534e-2 | 6 | dpt |
| sparc2 | 7.7911090 | 0.832649198 | 0.872 | 0.384 | 1.57357028515573e-2 | 6 | sparc |
| hand2 | 9.5816986 | 0.309254386 | 0.138 | 0.008 | 1.93521567315075e-2 | 6 | hand2 |
| si:ch1073-459j12.1 | 2.8657617 | 0.808936591 | 0.585 | 0.17 | 5.78797906673034e-2 | 6 | si:ch1073-459j12.1 |
| rbp51 | 5.1141923 | 0.806625554 | 0.596 | 0.183 | 1.03291342389424e-1 | 6 | rbp5 |
| sfrp1a | 8.5286436 | 0.64510116 | 0.351 | 0.066 | 1.72253014803407e-1 | 6 | sfrp1a |
| mdka2 | 9.5160744 | 0.813308305 | 0.649 | 0.208 | 1.92196155964848e-1 | 6 | mdka |
| atoh81 | 2.3196864 | 0.493391169 | 0.309 | 0.052 | 4.68507068829595e-1 | 6 | atoh8 |
| fhl3b | 4.8827233 | 0.398349458 | 0.16 | 0.014 | 9.86163637670377e-1 | 6 | fhl3b |
| rap1gap | 6.8374663 | 0.330105387 | 0.149 | 0.012 | 1.38096308796334e-1 | 6 | rap1gap |
| birc6-as2 | 8.1785559 | 0.260182287 | 0.106 | 0.005 | 1.65182295141379e-1 | 6 | birc6-as2 |
| six1b | 1.4757810 | 0.272254188 | 0.117 | 0.007 | 2.98063494037833e-1 | 6 | six1b |
| si:ch73-46j18.52 | 1.8555319 | 0.733962318 | 0.84 | 0.38 | 3.74761790480496e-1 | 6 | si:ch73-46j18.5 |
| mfap22 | 2.0260091 | 0.761355784 | 0.66 | 0.234 | 4.0919307277954e-17 | 6 | mfap2 |
| ephb31 | 5.1769776 | 0.580984826 | 0.404 | 0.097 | 1.04559416890399e-1 | 6 | ephb3 |
| mdkb | 2.9018566 | 0.517090007 | 0.213 | 0.029 | 5.86087983812749e-1 | 6 | mdkb |
| hoxd9a | 1.8818428 | 0.252203024 | 0.138 | 0.012 | 3.80075808880666e-1 | 6 | hoxd9a |
| tpm1 | 1.9394756 | 0.738998946 | 0.543 | 0.188 | 3.91715897607244e-1 | 6 | tpm1 |
| angpt2b | 5.5101595 | 0.305026295 | 0.138 | 0.013 | 1.11288693404488e-1 | 6 | angpt2b |
| bmper | 7.9553709 | 0.553236305 | 0.362 | 0.085 | 1.60674627849184e-1 | 6 | bmper |
| olfml3a | 9.4264604 | 0.399852689 | 0.245 | 0.041 | 1.90386220870455e-1 | 6 | olfml3a |
| myl9b1 | 1.0616007 | 0.645129002 | 0.564 | 0.191 | 2.14411494120053e-1 | 6 | myl9b |
| prss35 | 1.1151951 | 0.544411167 | 0.309 | 0.064 | 2.2523595717389e-14 | 6 | prss35 |
| c1qtnf52 | 3.5707905 | 0.587894224 | 0.33 | 0.075 | 7.21192558644014e-1 | 6 | c1qtnf5 |
| smoc1 | 9.5552556 | 0.255342777 | 0.106 | 0.008 | 1.92987498705494e-1 | 6 | smoc1 |
| epha3 | 1.3575061 | 0.462119471 | 0.255 | 0.048 | 2.74175511246688e-1 | 6 | epha3 |
| LO018188.11 | 1.8000466 | 0.506900673 | 0.351 | 0.083 | 3.63555414032287e-1 | 6 | LO018188.1 |
| rai14 | 1.9865474 | 0.277498753 | 0.181 | 0.024 | 4.01222994230075e-1 | 6 | rai14 |
| wt1b | 2.3052240 | 0.54711142 | 0.287 | 0.062 | 4.65586100833107e-1 | 6 | wt1b |
| c62 | 2.7795833 | 0.680993821 | 0.574 | 0.202 | 5.61392456512668e-1 | 6 | c6 |
| nfixb1 | 3.6454937 | 0.620336365 | 0.511 | 0.164 | 7.36280375931429e-1 | 6 | nfixb |
| tenm32 | 9.2138086 | 0.561108404 | 0.447 | 0.136 | 1.86091294042212e-1 | 6 | tenm3 |
| cxcl12b | 1.2473313 | 0.451275031 | 0.138 | 0.015 | 2.51923503992808e-1 | 6 | cxcl12b |
| si:dkey-9i23.161 | 1.5428310 | 0.449948881 | 0.362 | 0.092 | 3.11605591785592e-1 | 6 | si:dkey-9i23.16 |

| | | | | | | | |
|---|---|---|---|---|---|---|---|
| ednraa | 1.6508397 | 0.419037453 | 0.255 | 0.05 | 3.33420110829013e-1 | 6 | ednraa |
| fstl1b2 | 2.2322977 | 0.653455349 | 0.67 | 0.265 | 4.50857171318114e-1 | 6 | fstl1b |
| jpt1b2 | 2.4564545 | 0.61478864 | 0.713 | 0.321 | 4.96130130520511e-1 | 6 | jpt1b |
| foxp42 | 3.0558110 | 0.610749942 | 0.511 | 0.179 | 6.17182153119392e-1 | 6 | foxp4 |
| zgc:1135311 | 6.3796561 | 0.431500871 | 0.277 | 0.059 | 1.28849916135037e-1 | 6 | zgc:113531 |
| col12a1b1 | 8.9262119 | 0.530370091 | 0.266 | 0.058 | 1.8028270337703e-11 | 6 | col12a1b |
| cald1a2 | 1.4197608 | 0.607051849 | 0.585 | 0.226 | 2.86749097379219e-1 | 6 | cald1a |
| pabpc1b1 | 2.6132772 | 0.506787896 | 0.394 | 0.115 | 5.27803596276051e-1 | 6 | pabpc1b |
| MDFI | 5.3579796 | 0.356081062 | 0.223 | 0.042 | 1.08215114024375e-1 | 6 | MDFI |
| mmp16a2 | 9.4203070 | 0.596640844 | 0.383 | 0.116 | 1.90261940911886e-1 | 6 | mmp16a |
| fxyd6l1 | 9.5471449 | 0.526328972 | 0.426 | 0.139 | 1.92823686828228e-1 | 6 | fxyd6l |
| BX322618.1 | 1.1999971 | 0.649601745 | 0.266 | 0.063 | 2.42363431917786e-1 | 6 | BX322618.1 |
| cxcl8b.3 | 3.6812648 | 0.274657141 | 0.128 | 0.015 | 7.43505071725514e-1 | 6 | cxcl8b.3 |
| gata5 | 3.7135696 | 0.309067476 | 0.16 | 0.024 | 7.50029660070809e-1 | 6 | gata5 |
| tmem471 | 4.0144512 | 0.348288861 | 0.213 | 0.041 | 8.10798714540301e-1 | 6 | tmem47 |
| bhlhe41 | 4.4099655 | 0.317695727 | 0.213 | 0.041 | 8.90680737816098e-1 | 6 | bhlhe41 |
| snai1a1 | 4.4846161 | 0.365265752 | 0.245 | 0.053 | 9.05757920596686e-1 | 6 | snai1a |
| MMP23B | 4.6238993 | 0.319922207 | 0.191 | 0.034 | 9.3388895402546e-10 | 6 | MMP23B |
| col6a32 | 6.8486479 | 0.572208448 | 0.5 | 0.183 | 1.38322142738058e-0 | 6 | col6a3 |
| pdgfrb1 | 6.8793401 | 0.395468088 | 0.255 | 0.057 | 1.38942033878376e-0 | 6 | pdgfrb |
| col6a22 | 6.9524167 | 0.52689147 | 0.383 | 0.119 | 1.40417960200033e-0 | 6 | col6a2 |
| sh3bgrl2 | 8.5776541 | 0.273649176 | 0.149 | 0.021 | 1.73242880507726e-0 | 6 | sh3bgrl2 |
| si:ch211-212k18.9: | 1.3954696 | 0.407690386 | 0.372 | 0.11 | 2.81843002221104e-0 | 6 | si:ch211-212k18.9 |
| CABZ01092746.12 | 1.9864313 | 0.585776639 | 0.521 | 0.198 | 4.01199530632721e-0 | 6 | CABZ01092746.1 |
| hsd11b22 | 2.0109674 | 0.607467564 | 0.67 | 0.279 | 4.06155101147159e-0 | 6 | hsd11b2 |
| fn1b1 | 2.4445852 | 0.549046321 | 0.479 | 0.17 | 4.93732881418798e-0 | 6 | fn1b |
| s1pr5a | 2.5498712 | 0.314252717 | 0.181 | 0.032 | 5.14997500494178e-0 | 6 | s1pr5a |
| meis1b1 | 2.6273572 | 0.532333847 | 0.426 | 0.148 | 5.3064734058574e-09 | 6 | meis1b |
| cd81a2 | 2.6544165 | 0.597971808 | 0.809 | 0.429 | 5.36112502662061e-0 | 6 | cd81a |
| myl61 | 3.0015568 | 0.558941421 | 0.511 | 0.194 | 6.06224435285194e-0 | 6 | myl6 |
| tpm4b | 3.5229809 | 0.349980694 | 0.213 | 0.043 | 7.11536471552287e-0 | 6 | tpm4b |
| mex3b | 4.0643650 | 0.379254854 | 0.266 | 0.064 | 8.20879814372619e-0 | 6 | mex3b |
| col12a1a | 4.8880013 | 0.475705056 | 0.223 | 0.049 | 9.87229640265694e-0 | 6 | col12a1a |
| jun2 | 6.5713302 | 0.601329536 | 0.734 | 0.373 | 1.32721157516492e-0 | 6 | jun |
| sdc21 | 8.1431689 | 0.429035363 | 0.372 | 0.117 | 1.64467583510129e-0 | 6 | sdc2 |

| | | | | | | | |
|---|---|---|---|---|---|---|---|
| tmem176l.12 | 9.3788270 | 0.54918787 | 0.67 | 0.314 | 1.8942416972719e-08 | 6 | tmem176l.1 |
| cxcl8b.12 | 1.1364478 | 0.670985445 | 0.34 | 0.106 | 2.29528363562948e-0 | 6 | cxcl8b.1 |
| meis1a1 | 1.5180080 | 0.327557463 | 0.213 | 0.045 | 3.06592084008044e-0 | 6 | meis1a |
| krt8 | 1.7915785 | 0.571161399 | 0.851 | 0.512 | 3.61845126468619e-0 | 6 | krt8 |
| cnn3a2 | 2.3641374 | 0.411525099 | 0.394 | 0.128 | 4.77484832126145e-0 | 6 | cnn3a |
| tns1b1 | 3.3458890 | 0.311067294 | 0.202 | 0.042 | 6.75769205448288e-0 | 6 | tns1b |
| aoc22 | 3.4638765 | 0.666850336 | 0.362 | 0.127 | 6.9959913933359e-08 | 6 | aoc2 |
| tmem119b1 | 6.5023958 | 0.307038825 | 0.202 | 0.043 | 1.31328889054457e-0 | 6 | tmem119b |
| vasna1 | 8.4210353 | 0.423788744 | 0.319 | 0.094 | 1.70079651110966e-0 | 6 | vasna |
| sema3fa | 1.2973196 | 0.278988113 | 0.16 | 0.029 | 2.62019644476322e-0 | 6 | sema3fa |
| mfap51 | 1.3436147 | 0.541831745 | 0.426 | 0.155 | 2.71369864600217e-0 | 6 | mfap5 |
| nid2a1 | 1.4045478 | 0.440855123 | 0.351 | 0.112 | 2.83676533691489e-0 | 6 | nid2a |
| plpp31 | 1.5584448 | 0.392669012 | 0.372 | 0.118 | 3.14759114006903e-0 | 6 | plpp3 |
| sostdc1a1 | 1.8149266 | 0.583104267 | 0.287 | 0.086 | 3.66560744403382e-0 | 6 | sostdc1a |
| cox4i21 | 1.8670859 | 0.344833907 | 0.223 | 0.054 | 3.7709534076695e-07 | 6 | cox4i2 |
| sox4a | 1.9633013 | 0.450080773 | 0.415 | 0.15 | 3.9652797795919e-07 | 6 | sox4a |
| col5a2a2 | 2.5347428 | 0.474783557 | 0.457 | 0.173 | 5.11942017888563e-0 | 6 | col5a2a |
| CABZ01072614.12 | 2.6289911 | 0.488835808 | 0.436 | 0.167 | 5.30977338530537e-0 | 6 | CABZ01072614.1 |
| h3f3c1 | 2.9168760 | 0.490492274 | 0.819 | 0.52 | 5.89121452049375e-0 | 6 | h3f3c |
| rbfox1 | 6.2277223 | 0.311800928 | 0.202 | 0.046 | 1.25781307420241e-0 | 6 | rbfox1 |
| pbx1a2 | 6.2565069 | 0.430544375 | 0.33 | 0.109 | 1.26362669883277e-0 | 6 | pbx1a |
| serpinh1b | 7.0802267 | 0.353380582 | 0.245 | 0.065 | 1.42999340463417e-0 | 6 | serpinh1b |
| rbpms2b | 9.5194311 | 0.35689191 | 0.202 | 0.048 | 1.92263950717408e-0 | 6 | rbpms2b |
| mxra8b1 | 1.3488226 | 0.336757945 | 0.223 | 0.058 | 2.72421701746757e-0 | 6 | mxra8b |
| myl9a | 1.4525517 | 0.483749212 | 0.33 | 0.11 | 2.93371873488267e-0 | 6 | myl9a |
| fhl2a | 1.5796768 | 0.266955928 | 0.138 | 0.024 | 3.19047327774849e-0 | 6 | fhl2a |
| wnt5b | 1.6112963 | 0.259568091 | 0.16 | 0.031 | 3.25433518912188e-0 | 6 | wnt5b |
| emilin1a | 1.6567261 | 0.29881191 | 0.213 | 0.053 | 3.34608988793881e-0 | 6 | emilin1a |
| edil3a | 1.9465878 | 0.277230062 | 0.149 | 0.028 | 3.93152338370832e-0 | 6 | edil3a |
| meis2b2 | 2.3581141 | 0.406635647 | 0.287 | 0.09 | 4.76268314500365e-0 | 6 | meis2b |
| dnmt3ab2 | 2.7003137 | 0.445579374 | 0.404 | 0.161 | 5.4538237025231e-06 | 6 | dnmt3ab |
| oip5-as1 | 2.8257835 | 0.305729211 | 0.202 | 0.049 | 5.70723499098216e-0 | 6 | oip5-as1 |
| zfp36l2 | 2.9614929 | 0.314618441 | 0.223 | 0.058 | 5.98132738600201e-0 | 6 | zfp36l2 |
| s100a10b1 | 2.9649863 | 0.49332488 | 0.851 | 0.444 | 5.98838299695566e-0 | 6 | s100a10b |
| stau2 | 4.5226761 | 0.435299363 | 0.383 | 0.149 | 9.13444903506894e-0 | 6 | stau2 |

| | | | | | | | |
|---|---|---|---|---|---|---|---|
| angptl4 | 5.5092332 | 0.340052595 | 0.277 | 0.084 | 1.11269984325787e-0 | 6 | angptl4 |
| vim1 | 5.7668291 | 0.380934855 | 0.309 | 0.098 | 1.16472647562361e-0 | 6 | vim |
| loxa1 | 6.1544868 | 0.33576191 | 0.202 | 0.05 | 1.24302170280132e-0 | 6 | loxa |
| s100v21 | 8.3824123 | 0.43971357 | 0.543 | 0.241 | 1.69299582901628e-0 | 6 | s100v2 |
| svep1 | 9.7308845 | 0.296110747 | 0.128 | 0.023 | 1.9653467525281e-05 | 6 | svep1 |
| atf5b | 9.7457105 | 0.29065513 | 0.234 | 0.065 | 1.96834115015084e-0 | 6 | atf5b |
| frzb | 9.9174179 | 0.251274075 | 0.128 | 0.022 | 2.00302090552609e-0 | 6 | frzb |
| igfbp5b1 | 1.0682462 | 0.498797113 | 0.426 | 0.169 | 2.1575369583176e-05 | 6 | igfbp5b |
| tnfsf10l1 | 1.2612854 | 0.458146788 | 0.298 | 0.103 | 2.54741813742578e-0 | 6 | tnfsf10l |
| si:dkey-40c11.2 | 1.3599683 | 0.271085605 | 0.149 | 0.031 | 2.74672812802606e-0 | 6 | si:dkey-40c11.2 |
| pdia32 | 1.4972567 | 0.446695185 | 0.798 | 0.475 | 3.02400946012524e-0 | 6 | pdia3 |
| tnfrsf191 | 2.5286891 | 0.299957679 | 0.191 | 0.048 | 5.10719354615985e-0 | 6 | tnfrsf19 |
| net1 | 2.8062646 | 0.31876411 | 0.255 | 0.078 | 5.66781264300267e-0 | 6 | net1 |
| si:ch1073-291c23.2 | 3.0610880 | 0.399292529 | 0.351 | 0.131 | 6.18247951560037e-0 | 6 | si:ch1073-291c23.2 |
| col16a1 | 4.8160053 | 0.354427115 | 0.245 | 0.076 | 9.72688592598695e-0 | 6 | col16a1 |
| si:ch211-133n4.42 | 5.9047329 | 0.394413117 | 0.34 | 0.128 | 0.000119258 | 6 | si:ch211-133n4.4 |
| gsta.1 | 6.1016203 | 0.368967133 | 0.277 | 0.094 | 0.000123234 | 6 | gsta.1 |
| ppib1 | 6.8979080 | 0.401542995 | 0.894 | 0.636 | 0.000139317 | 6 | ppib |
| fkbp9 | 7.1174626 | 0.374515556 | 0.266 | 0.088 | 0.000143751 | 6 | fkbp9 |
| sgce | 7.3615763 | 0.268511979 | 0.202 | 0.055 | 0.000148682 | 6 | sgce |
| chd32 | 8.2405895 | 0.447018151 | 0.489 | 0.234 | 0.000166435 | 6 | chd3 |
| vdac3 | 8.2911428 | 0.425238727 | 0.5 | 0.246 | 0.000167456 | 6 | vdac3 |
| id3 | 9.7082064 | 0.38894298 | 0.362 | 0.14 | 0.000196077 | 6 | id3 |
| f101 | 9.9891932 | 0.399193171 | 0.309 | 0.114 | 0.000201752 | 6 | f10 |
| si:ch211-39i22.12 | 1.2479713 | 0.41287427 | 0.415 | 0.176 | 0.000252053 | 6 | si:ch211-39i22.1 |
| cd248a1 | 1.2912476 | 0.368788583 | 0.319 | 0.117 | 0.000260793 | 6 | cd248a |
| adh8a | 1.3296771 | 0.514910875 | 0.213 | 0.064 | 0.000268555 | 6 | adh8a |
| pcdh18b1 | 1.6447653 | 0.339272756 | 0.223 | 0.068 | 0.000332193 | 6 | pcdh18b |
| cdh21 | 1.8146939 | 0.414620574 | 0.319 | 0.122 | 0.000366514 | 6 | cdh2 |
| adh51 | 1.8767776 | 0.429510417 | 0.606 | 0.355 | 0.000379053 | 6 | adh5 |
| si:ch211-212k18.82 | 1.8788617 | 0.29907236 | 0.234 | 0.073 | 0.000379474 | 6 | si:ch211-212k18.8 |
| crtap | 2.2017380 | 0.319560051 | 0.255 | 0.084 | 0.000444685 | 6 | crtap |
| nell2a1 | 2.7031545 | 0.322001486 | 0.149 | 0.035 | 0.000545956 | 6 | nell2a |
| pgrmc11 | 2.7930305 | 0.358880557 | 0.479 | 0.215 | 0.000564108 | 6 | pgrmc1 |
| gas1a1 | 2.9338044 | 0.363916928 | 0.34 | 0.132 | 0.00059254 | 6 | gas1a |

| | | | | | | | |
|---|---|---|---|---|---|---|---|
| fkbp71 | 3.6162372 | 0.320316446 | 0.319 | 0.122 | 0.000730371 | 6 | fkbp7 |
| osr12 | 3.7848226 | 0.320743211 | 0.34 | 0.126 | 0.000764421 | 6 | osr1 |
| hmgb3a1 | 4.5490978 | 0.414234918 | 0.521 | 0.252 | 0.000918781 | 6 | hmgb3a |
| adamts15a2 | 5.1600374 | 0.374561033 | 0.287 | 0.107 | 0.001042173 | 6 | adamts15a |
| cadm2a | 5.5813853 | 0.309728842 | 0.16 | 0.041 | 0.001127272 | 6 | cadm2a |
| CR383676.12 | 5.7462720 | 0.392660337 | 0.851 | 0.656 | 0.001160575 | 6 | CR383676.1 |
| timp2a | 6.0050046 | 0.373464447 | 0.287 | 0.107 | 0.001212831 | 6 | timp2a |
| cyp2p71 | 6.2023069 | 0.310789329 | 0.191 | 0.056 | 0.00125268 | 6 | cyp2p7 |
| tmem1762 | 6.6277439 | 0.404860606 | 0.351 | 0.149 | 0.001338605 | 6 | tmem176 |
| si:dkey-56m19.5 | 6.9507872 | 0.380460961 | 0.266 | 0.097 | 0.00140385 | 6 | si:dkey-56m19.5 |
| si:dkey-238c7.161 | 8.0352607 | 0.351184139 | 0.298 | 0.112 | 0.001622882 | 6 | si:dkey-238c7.16 |
| clu1 | 1.3353331 | 0.397442851 | 0.191 | 0.058 | 0.002696972 | 6 | clu |
| krt942 | 1.4688021 | 0.45857597 | 0.638 | 0.355 | 0.00296654 | 6 | krt94 |
| ftr821 | 1.6774540 | 0.302552368 | 0.277 | 0.101 | 0.003387954 | 6 | ftr82 |
| zgc:110339 | 1.8752392 | 0.286265113 | 0.362 | 0.152 | 0.003787421 | 6 | zgc:110339 |
| rbms31 | 1.9127824 | 0.277200399 | 0.17 | 0.048 | 0.003863247 | 6 | rbms3 |
| LOC1039092841 | 1.9573502 | 0.383650099 | 0.33 | 0.137 | 0.00395326 | 6 | LOC103909284 |
| fzd4 | 2.1009247 | 0.264469654 | 0.191 | 0.057 | 0.004243238 | 6 | fzd4 |
| calua | 2.2115849 | 0.291390286 | 0.213 | 0.07 | 0.004466738 | 6 | calua |
| gng71 | 2.5936200 | 0.301818915 | 0.277 | 0.104 | 0.005238334 | 6 | gng7 |
| kdm6bb | 2.6907881 | 0.33032716 | 0.277 | 0.107 | 0.005434585 | 6 | kdm6bb |
| rbp42 | 2.7691173 | 0.413124537 | 0.468 | 0.214 | 0.005592786 | 6 | rbp4 |
| fblim1 | 3.0665585 | 0.2790402 | 0.255 | 0.092 | 0.006193528 | 6 | fblim1 |
| nanos1 | 3.2394117 | 0.30087237 | 0.255 | 0.091 | 0.00654264 | 6 | nanos1 |
| pmp22a2 | 3.4950301 | 0.410756344 | 0.617 | 0.312 | 0.007058912 | 6 | pmp22a |
| sept101 | 3.6031876 | 0.290185622 | 0.223 | 0.077 | 0.007277358 | 6 | sept10 |
| col6a4a2 | 4.7024703 | 0.386105735 | 0.372 | 0.164 | 0.009497579 | 6 | col6a4a |
| gtf2a1l | 6.0966332 | 0.268225541 | 0.128 | 0.031 | 0.01231337 | 6 | gtf2a1l |
| XLOC-0254232 | 6.2709606 | 0.605576552 | 0.372 | 0.187 | 0.012665459 | 6 | XLOC-025423 |
| krt18a.1 | 6.4233284 | 0.371989053 | 0.543 | 0.273 | 0.012973196 | 6 | krt18a.1 |
| lamb1a2 | 7.9032562 | 0.335446323 | 0.319 | 0.135 | 0.015962207 | 6 | lamb1a |
| krt15 | 8.1280864 | 0.37270367 | 0.372 | 0.159 | 0.016416296 | 6 | krt15 |
| tm2d3 | 1.0753303 | 0.262369133 | 0.255 | 0.099 | 0.021718447 | 6 | tm2d3 |
| CR847566.11 | 1.1153340 | 0.32981924 | 0.309 | 0.128 | 0.022526402 | 6 | CR847566.1 |
| add3a2 | 1.2850938 | 0.371866625 | 0.426 | 0.216 | 0.02595504 | 6 | add3a |

| | | | | | | | |
|---|---|---|---|---|---|---|---|
| XLOC-041269 | 1.4623912 | 0.264100768 | 0.181 | 0.057 | 0.029535917 | 6 | XLOC-041269 |
| tspan7 | 1.6727281 | 0.270824975 | 0.255 | 0.098 | 0.033784091 | 6 | tspan7 |
| olfml3b2 | 1.7199049 | 0.328959366 | 0.266 | 0.108 | 0.034736921 | 6 | olfml3b |
| fn1a1 | 1.9198813 | 0.343669986 | 0.383 | 0.176 | 0.038775843 | 6 | fn1a |
| ptp4a2a | 2.0432469 | 0.270321289 | 0.266 | 0.109 | 0.041267459 | 6 | ptp4a2a |
| selenbp11 | 2.3502239 | 0.30244014 | 0.319 | 0.138 | 0.047467473 | 6 | selenbp1 |
| FO082781.11 | 2.3777298 | 0.266771185 | 0.245 | 0.094 | 0.048023009 | 6 | FO082781.1 |
| svild | 2.4252594 | 0.332123581 | 0.245 | 0.097 | 0.048982965 | 6 | svild |
| cygb11 | 2.4354930 | 0.321490428 | 0.255 | 0.099 | 0.049189653 | 6 | cygb1 |
| atrx | 2.4624458 | 0.339432231 | 0.457 | 0.238 | 0.049734018 | 6 | atrx |
| rnd3b1 | 2.5598995 | 0.281003271 | 0.245 | 0.093 | 0.051702292 | 6 | rnd3b |
| tuba8l32 | 2.9423381 | 0.298651912 | 0.426 | 0.2 | 0.059426403 | 6 | tuba8l3 |
| si:ch211-286o17.1 | 3.0553436 | 0.287053623 | 0.266 | 0.108 | 0.061708776 | 6 | si:ch211-286o17.1 |
| loxl2b1 | 3.1989895 | 0.29139336 | 0.191 | 0.067 | 0.064609993 | 6 | loxl2b |
| efnb12 | 3.8706255 | 0.332268015 | 0.245 | 0.098 | 0.078175025 | 6 | efnb1 |
| pleca | 3.9808298 | 0.287200359 | 0.213 | 0.078 | 0.080400821 | 6 | pleca |
| BX569787.11 | 4.0314522 | 0.276156432 | 0.255 | 0.102 | 0.081423241 | 6 | BX569787.1 |
| zfp36l1b1 | 4.5365699 | 0.356466646 | 0.394 | 0.195 | 0.091625104 | 6 | zfp36l1b |
| ciarta1 | 4.6079320 | 0.326039716 | 0.372 | 0.183 | 0.093066404 | 6 | ciarta |
| BX640522.21 | 5.0146813 | 0.420179335 | 0.277 | 0.116 | 0.101281519 | 6 | BX640522.2 |
| BX322530.11 | 5.1451389 | 0.260319008 | 0.202 | 0.074 | 0.103916371 | 6 | BX322530.1 |
| nme31 | 5.1498433 | 0.332903271 | 0.33 | 0.157 | 0.104011385 | 6 | nme3 |
| hdlbpa1 | 5.5466231 | 0.329953238 | 0.479 | 0.262 | 0.112025148 | 6 | hdlbpa |
| mmp21 | 5.8869784 | 0.34595388 | 0.415 | 0.213 | 0.118899303 | 6 | mmp2 |
| fzd22 | 7.7652428 | 0.284628385 | 0.266 | 0.114 | 0.156834611 | 6 | fzd2 |
| CABZ01045617.1 | 8.5647720 | 0.29350305 | 0.277 | 0.12 | 0.172982701 | 6 | CABZ01045617.1 |
| selenow11 | 8.6829722 | 0.321216658 | 0.83 | 0.59 | 0.175369991 | 6 | selenow1 |
| si:ch73-281n10.21 | 8.7488146 | 0.318897785 | 0.723 | 0.425 | 0.176699809 | 6 | si:ch73-281n10.2 |
| ier5l1 | 9.4913627 | 0.250246292 | 0.17 | 0.058 | 0.191697054 | 6 | ier5l |
| si:ch211-191i18.2 | 1.0092472 | 0.25872863 | 0.117 | 0.032 | 0.203837671 | 6 | si:ch211-191i18.2 |
| timm23a | 1.2128554 | 0.286294961 | 0.351 | 0.174 | 0.244960413 | 6 | timm23a |
| colec122 | 1.4229588 | 0.282547074 | 0.255 | 0.108 | 0.28739499 | 6 | colec12 |
| gna11b | 1.4286199 | 0.281081939 | 0.202 | 0.078 | 0.288538375 | 6 | gna11b |
| ms4a17a.92 | 1.4741952 | 0.28971747 | 0.298 | 0.136 | 0.29774322 | 6 | ms4a17a.9 |
| mgst3b1 | 1.5643556 | 0.319965623 | 0.5 | 0.297 | 0.315952912 | 6 | mgst3b |

| | | | | | | | |
|---|---|---|---|---|---|---|---|
| aldh9a1a.12 | 1.7028491 | 0.347585225 | 0.543 | 0.335 | 0.343924442 | 6 | aldh9a1a.1 |
| dkk3b1 | 1.8744050 | 0.293340952 | 0.34 | 0.163 | 0.378573583 | 6 | dkk3b |
| dedd12 | 1.8756722 | 0.327610773 | 0.394 | 0.209 | 0.378829527 | 6 | dedd1 |
| si:dkey-7j14.62 | 1.9094128 | 0.328061789 | 0.543 | 0.316 | 0.385644106 | 6 | si:dkey-7j14.6 |
| ip6k2a | 1.9819450 | 0.293651748 | 0.255 | 0.114 | 0.400293439 | 6 | ip6k2a |
| hmgn61 | 2.0522291 | 0.324777097 | 0.777 | 0.566 | 0.414488729 | 6 | hmgn6 |
| boc2 | 2.0581692 | 0.300576267 | 0.34 | 0.164 | 0.415688441 | 6 | boc |
| pfdn1 | 2.1069669 | 0.295609585 | 0.404 | 0.222 | 0.425544112 | 6 | pfdn1 |
| mknk2b | 2.2358126 | 0.278016694 | 0.415 | 0.216 | 0.45156708 | 6 | mknk2b |
| fbn2b2 | 2.5199229 | 0.302775437 | 0.351 | 0.175 | 0.508948828 | 6 | fbn2b |
| cyp2ad32 | 2.6436692 | 0.314983094 | 0.255 | 0.113 | 0.533941875 | 6 | cyp2ad3 |
| XLOC-041870 | 2.6635591 | 0.335624425 | 0.138 | 0.045 | 0.537959033 | 6 | XLOC-041870 |
| ctsla2 | 2.7208356 | 0.269280291 | 0.543 | 0.303 | 0.549527181 | 6 | ctsla |
| dbn1 | 2.7500886 | 0.276776019 | 0.266 | 0.12 | 0.555435397 | 6 | dbn1 |
| wls1 | 3.2366151 | 0.311847129 | 0.34 | 0.179 | 0.653699171 | 6 | wls |
| sost1 | 3.2700715 | 0.294244199 | 0.181 | 0.067 | 0.660456344 | 6 | sost |
| arl6ip5b1 | 3.2797384 | 0.256319456 | 0.298 | 0.139 | 0.662408769 | 6 | arl6ip5b |
| CABZ01080702.11 | 4.4240727 | 0.33222276 | 0.33 | 0.173 | 0.893529982 | 6 | CABZ01080702.1 |
| slc38a5b2 | 4.4276438 | 0.273223521 | 0.383 | 0.201 | 0.894251223 | 6 | slc38a5b |
| sept7a | 4.7576276 | 0.281522075 | 0.245 | 0.111 | 0.960898047 | 6 | sept7a |
| aes | 4.8938254 | 0.250098533 | 0.181 | 0.071 | 0.98840592 | 6 | aes |
| si:dkey-28b4.7 | 5.7982308 | 0.260843113 | 0.372 | 0.191 | 1 | 6 | si:dkey-28b4.7 |
| pfn2l1 | 7.2577357 | 0.291285234 | 0.457 | 0.259 | 1 | 6 | pfn2l |
| flna3 | 7.4654843 | 0.325140204 | 0.319 | 0.165 | 1 | 6 | flna |
| cd82a1 | 7.7475367 | 0.251032389 | 0.287 | 0.137 | 1 | 6 | cd82a |
| h2afy21 | 7.9849072 | 0.279257538 | 0.436 | 0.252 | 1 | 6 | h2afy2 |
| XLOC-0054491 | 8.4776629 | 0.316943553 | 0.181 | 0.074 | 1 | 6 | XLOC-005449 |
| hmgb1b1 | 9.5342390 | 0.284104542 | 0.511 | 0.297 | 1 | 6 | hmgb1b |
| mycb1 | 0.000102 | 0.280842049 | 0.309 | 0.158 | 1 | 6 | mycb |
| col4a12 | 0.000115 | 0.307828799 | 0.277 | 0.134 | 1 | 6 | col4a1 |
| arhgef1b | 0.000128 | 0.26506518 | 0.255 | 0.122 | 1 | 6 | arhgef1b |
| ahnak1 | 0.000129 | 0.252451109 | 0.372 | 0.197 | 1 | 6 | ahnak |
| fosb | 0.000149 | 0.324294331 | 0.234 | 0.111 | 1 | 6 | fosb |
| zgc:1584631 | 0.00015 | 0.28936379 | 0.245 | 0.116 | 1 | 6 | zgc:158463 |
| slc3a2b | 0.000155 | 0.268766878 | 0.319 | 0.169 | 1 | 6 | slc3a2b |

| | | | | | | | |
|---|---|---|---|---|---|---|---|
| tnrc6c11 | 0.000156 | 0.296381456 | 0.404 | 0.238 | 1 | 6 | tnrc6c1 |
| id12 | 0.000189 | 0.367177906 | 0.447 | 0.271 | 1 | 6 | id1 |
| pbx4 | 0.000216 | 0.263185721 | 0.223 | 0.107 | 1 | 6 | pbx4 |
| mcl1b2 | 0.000218 | 0.261192385 | 0.521 | 0.311 | 1 | 6 | mcl1b |
| BX901920.12 | 0.000256 | 0.31139202 | 0.181 | 0.078 | 1 | 6 | BX901920.1 |
| tpm4a1 | 0.000348 | 0.276807202 | 0.415 | 0.243 | 1 | 6 | tpm4a |
| irf2bp2a | 0.000386 | 0.264786725 | 0.415 | 0.245 | 1 | 6 | irf2bp2a |
| anxa1a1 | 0.000443 | 0.279535359 | 0.468 | 0.274 | 1 | 6 | anxa1a |
| fam133b | 0.000484 | 0.274249731 | 0.351 | 0.209 | 1 | 6 | fam133b |
| fosab3 | 0.000645 | 0.320656426 | 0.617 | 0.458 | 1 | 6 | fosab |
| anxa5b1 | 0.000804 | 0.262356483 | 0.298 | 0.167 | 1 | 6 | anxa5b |
| NC-002333.42 | 0.001073 | 0.291106543 | 0.777 | 0.581 | 1 | 6 | NC-002333.4 |
| ier2a2 | 0.001335 | 0.293068438 | 0.511 | 0.362 | 1 | 6 | ier2a |
| FP236356.1 | 2.7111307 | 1.220215424 | 0.528 | 0.012 | 5.47567078468366e-1 | 7 | FP236356.1 |
| XLOC-0425951 | 2.4278203 | 1.901972474 | 0.899 | 0.074 | 4.90346881476151e-1 | 7 | XLOC-042595 |
| LOC103910871 | 1.2169701 | 1.664480499 | 0.876 | 0.069 | 2.45791465010569e-1 | 7 | LOC103910871 |
| XLOC-042596 | 1.0457393 | 0.867947557 | 0.472 | 0.009 | 2.11207983137749e-1 | 7 | XLOC-042596 |
| CR936442.11 | 3.2633806 | 1.53758689 | 0.787 | 0.061 | 6.59104984651786e-1 | 7 | CR936442.1 |
| si:dkey-183p4.10 | 2.3102342 | 0.687648726 | 0.382 | 0.006 | 4.66598004249026e-1 | 7 | si:dkey-183p4.10 |
| ccl38.6 | 1.0431279 | 1.297317466 | 0.404 | 0.008 | 2.10680547473398e-1 | 7 | ccl38.6 |
| LOC110439931 | 2.0127870 | 0.543374213 | 0.292 | 0.002 | 4.06522602636916e-1 | 7 | LOC110439931 |
| zbtb32 | 1.1725705 | 0.630773802 | 0.315 | 0.003 | 2.36824080491143e-1 | 7 | zbtb32 |
| si:rp71-81e14.2 | 1.5300640 | 0.537187513 | 0.281 | 0.002 | 3.09027026545692e-1 | 7 | si:rp71-81e14.2 |
| si:ch211-132g1.4 | 2.7993211 | 0.605480443 | 0.281 | 0.002 | 5.65378899206053e-1 | 7 | si:ch211-132g1.4 |
| zap70 | 2.3560861 | 0.563522193 | 0.27 | 0.002 | 4.75858710592636e-1 | 7 | zap70 |
| si:ch73-223p23.2 | 1.0044725 | 0.447246975 | 0.247 | 0.001 | 2.02873322230405e-1 | 7 | si:ch73-223p23.2 |
| ccl36.1 | 3.1083705 | 1.00174439 | 0.292 | 0.003 | 6.27797602770115e-1 | 7 | ccl36.1 |
| si:dkey-183i3.6 | 1.1133731 | 0.603784517 | 0.27 | 0.002 | 2.248679843586e-97 | 7 | si:dkey-183i3.6 |
| itk | 2.1123319 | 0.493065195 | 0.27 | 0.002 | 4.26627674941425e-9 | 7 | itk |
| XLOC-0139641 | 1.2912115 | 1.828905318 | 0.899 | 0.131 | 2.60786000376661e-9 | 7 | XLOC-013964 |
| si:dkey-260g12.1 | 1.8236719 | 0.62748298 | 0.348 | 0.008 | 3.68327018296423e-9 | 7 | si:dkey-260g12.1 |
| bcl11ba | 1.2836879 | 0.658054296 | 0.292 | 0.004 | 2.59266453971168e-9 | 7 | bcl11ba |
| XLOC-0108991 | 6.6766868 | 1.776168272 | 0.899 | 0.139 | 1.34849044226699e-9 | 7 | XLOC-010899 |
| ccl38a.5 | 6.8956414 | 0.897217221 | 0.258 | 0.002 | 1.39271269397752e-9 | 7 | ccl38a.5 |
| sla2 | 8.2801062 | 0.737783463 | 0.36 | 0.01 | 1.6723330552603e-92 | 7 | sla2 |

| | | | | | | | |
|---|---|---|---|---|---|---|---|
| LOC101885675 | 2.4694805 | 0.514351697 | 0.27 | 0.003 | 4.98760990207024e-9 | 7 | LOC101885675 |
| BX005105.1 | 5.6078160 | 0.38709462 | 0.236 | 0.002 | 1.13261061271341e-8 | 7 | BX005105.1 |
| dusp21 | 1.3795650 | 1.414431164 | 0.708 | 0.075 | 2.78630756815775e-8 | 7 | dusp2 |
| si:ch211-196f2.6 | 3.1999051 | 0.453605811 | 0.225 | 0.001 | 6.46284839041986e-8 | 7 | si:ch211-196f2.6 |
| tagapa | 1.0840025 | 1.068259117 | 0.573 | 0.044 | 2.18936001874994e-8 | 7 | tagapa |
| tcf7 | 6.0517907 | 0.606462052 | 0.292 | 0.007 | 1.22228018539452e-7 | 7 | tcf7 |
| XLOC-043043 | 3.3636404 | 1.230141591 | 0.506 | 0.035 | 6.79354469928316e-7 | 7 | XLOC-043043 |
| FP016106.1 | 5.2463563 | 0.380737033 | 0.191 | 0.001 | 1.05960659393145e-7 | 7 | FP016106.1 |
| ncaldb | 1.0085073 | 0.860477769 | 0.494 | 0.035 | 2.03688232793275e-7 | 7 | ncaldb |
| XLOC-0004312 | 7.6022139 | 1.790741506 | 0.966 | 0.241 | 1.53541915851254e-7 | 7 | XLOC-000431 |
| gzmk | 1.3443365 | 0.428660965 | 0.213 | 0.002 | 2.71515659398864e-7 | 7 | gzmk |
| cd40lg | 2.4971479 | 0.369655251 | 0.202 | 0.002 | 5.0434897993824e-72 | 7 | cd40lg |
| lbh | 1.0027647 | 0.859994397 | 0.551 | 0.048 | 2.02528395569105e-7 | 7 | lbh |
| nkl.2 | 1.7685232 | 0.750551172 | 0.225 | 0.003 | 3.57188640209567e-7 | 7 | nkl.2 |
| si:ch211-214p16.1 | 7.3697198 | 1.54773658 | 0.921 | 0.165 | 1.48846232609522e-7 | 7 | si:ch211-214p16.1 |
| si:ch211-167j9.5 | 3.3716376 | 0.389362398 | 0.213 | 0.003 | 6.80969647529129e-7 | 7 | si:ch211-167j9.5 |
| XLOC-039757 | 5.7814961 | 0.684663031 | 0.348 | 0.015 | 1.16768877539254e-6 | 7 | XLOC-039757 |
| cxcr3.1 | 3.1608143 | 0.410887825 | 0.225 | 0.004 | 6.38389673417439e-6 | 7 | cxcr3.1 |
| si:dkey-109a10.2 | 2.4014012 | 0.545700949 | 0.247 | 0.006 | 4.85011004282166e-6 | 7 | si:dkey-109a10.2 |
| si:ch1073-220m6.1 | 9.6888005 | 0.503239677 | 0.247 | 0.006 | 1.95684704015938e-6 | 7 | si:ch1073-220m6.1 |
| abo | 2.6949171 | 0.441941582 | 0.225 | 0.005 | 5.44292424637926e-6 | 7 | abo |
| ccr2 | 3.8880568 | 0.790738093 | 0.416 | 0.028 | 7.85270835589745e-6 | 7 | ccr2 |
| CR786582.2 | 5.0004085 | 0.403720966 | 0.225 | 0.005 | 1.00993250852074e-6 | 7 | CR786582.2 |
| lck | 7.0181727 | 0.37962722 | 0.18 | 0.002 | 1.41746034559972e-6 | 7 | lck |
| si:dkeyp-85e10.3 | 7.8964466 | 0.432641212 | 0.236 | 0.006 | 1.59484533041996e-6 | 7 | si:dkeyp-85e10.3 |
| si:ch73-285p12.4 | 2.0200726 | 0.377558672 | 0.202 | 0.003 | 4.079940821934e-62 | 7 | si:ch73-285p12.4 |
| ccl38a.4 | 6.7020681 | 0.513433967 | 0.169 | 0.001 | 1.3536167107607e-61 | 7 | ccl38a.4 |
| CR848749.2 | 1.4464459 | 0.463661187 | 0.191 | 0.003 | 2.92138688568878e-6 | 7 | CR848749.2 |
| si:ch211-119e14.1 | 2.4911080 | 0.792262465 | 0.472 | 0.041 | 5.03129085477945e-5 | 7 | si:ch211-119e14.1 |
| si:ch211-93g21.1 | 1.2279911 | 0.518478023 | 0.281 | 0.011 | 2.48017368223847e-5 | 7 | si:ch211-93g21.1 |
| zgc:136605 | 1.1817427 | 0.445070212 | 0.202 | 0.004 | 2.3867658545931e-56 | 7 | zgc:136605 |
| irf4b | 1.2869079 | 0.478470445 | 0.236 | 0.007 | 2.59916795458762e-5 | 7 | irf4b |
| tnfsf14 | 1.8347087 | 0.250172933 | 0.135 | 0 | 3.70556122577347e-5 | 7 | tnfsf14 |
| ccl32b.3 | 2.4265197 | 0.443051842 | 0.213 | 0.005 | 4.90084197424133e-5 | 7 | ccl32b.3 |
| cytip | 2.4506023 | 0.400945809 | 0.225 | 0.006 | 4.94948150092393e-5 | 7 | cytip |

| | | | | | | | |
|---|---|---|---|---|---|---|---|
| hpdb | 5.2343618 | 0.45061657 | 0.18 | 0.003 | 1.05718405276414e-5 | 7 | hpdb |
| grap2a | 1.0186479 | 0.654400446 | 0.438 | 0.041 | 2.0573631860903e-51 | 7 | grap2a |
| bcl2l10 | 4.9599405 | 1.031540632 | 0.697 | 0.12 | 1.0017591843145e-50 | 7 | bcl2l10 |
| si:ch211-132g1.1 | 2.6786965 | 0.349613841 | 0.18 | 0.004 | 5.41016334210182e-5 | 7 | si:ch211-132g1.1 |
| ikzf1 | 3.3059394 | 0.584955334 | 0.36 | 0.027 | 6.67700600415113e-4 | 7 | ikzf1 |
| arl5c1 | 3.0431482 | 0.897714643 | 0.551 | 0.076 | 6.14624656114248e-4 | 7 | arl5c |
| skap1 | 1.1216497 | 0.400376694 | 0.213 | 0.008 | 2.265395993978e-46 | 7 | skap1 |
| rorc | 1.2242071 | 0.356419739 | 0.112 | 0 | 2.47253119754039e-4 | 7 | rorc |
| XLOC-014079 | 9.7121166 | 0.64062067 | 0.292 | 0.018 | 1.96155620446018e-4 | 7 | XLOC-014079 |
| im:7142702 | 2.9038614 | 0.355082145 | 0.146 | 0.002 | 5.86492888636294e-4 | 7 | im:7142702 |
| rag1 | 3.4476103 | 0.508652143 | 0.146 | 0.002 | 6.96313867753555e-4 | 7 | rag1 |
| faslg | 1.5801054 | 0.364495366 | 0.213 | 0.008 | 3.19133904559978e-4 | 7 | faslg |
| arpp21 | 8.1978131 | 0.389615311 | 0.135 | 0.002 | 1.65571231423484e-4 | 7 | arpp21 |
| themis | 1.0735734 | 0.251276443 | 0.135 | 0.002 | 2.16829639282855e-4 | 7 | themis |
| smpd3 | 1.8677102 | 0.332197715 | 0.169 | 0.004 | 3.77221429466952e-4 | 7 | smpd3 |
| satb1b | 1.5028055 | 0.27794065 | 0.124 | 0.001 | 3.03521630558311e-4 | 7 | satb1b |
| id31 | 2.9136664 | 1.132698506 | 0.64 | 0.128 | 5.88473221253976e-4 | 7 | id3 |
| il7r | 2.4144729 | 0.471084669 | 0.236 | 0.013 | 4.87651096668967e-4 | 7 | il7r |
| scinlb | 5.3753281 | 0.78615003 | 0.539 | 0.081 | 1.08565501962091e-3 | 7 | scinlb |
| ccr9b | 9.4228947 | 0.376260681 | 0.146 | 0.003 | 1.90314205500938e-3 | 7 | ccr9b |
| gfi1ab | 3.6274103 | 0.489185758 | 0.27 | 0.019 | 7.32628062297522e-3 | 7 | gfi1ab |
| il21r.1 | 4.6331741 | 0.403347392 | 0.236 | 0.013 | 9.35762185561483e-3 | 7 | il21r.1 |
| CABZ01084575.1 | 9.5190515 | 0.51655442 | 0.27 | 0.019 | 1.92256284346196e-3 | 7 | CABZ01084575.1 |
| si:dkey-171c9.3 | 1.2513502 | 0.433215262 | 0.258 | 0.017 | 2.52735212954886e-3 | 7 | si:dkey-171c9.3 |
| tnfb1 | 3.2395627 | 0.677378623 | 0.303 | 0.025 | 6.54294491544658e-3 | 7 | tnfb |
| rgs13 | 4.3900398 | 0.783777511 | 0.539 | 0.085 | 8.86656354701242e-3 | 7 | rgs13 |
| runx3 | 7.2873510 | 0.514139506 | 0.247 | 0.016 | 1.47182630112469e-3 | 7 | runx3 |
| LOC101882375 | 7.7792123 | 0.311118963 | 0.135 | 0.003 | 1.5711675262339e-37 | 7 | LOC101882375 |
| sh2d1ab | 1.0171547 | 0.280526583 | 0.135 | 0.003 | 2.05434745482671e-3 | 7 | sh2d1ab |
| si:ch211-25d12.7 | 2.9229587 | 0.285086093 | 0.157 | 0.005 | 5.90349974955588e-3 | 7 | si:ch211-25d12.7 |
| wasb2 | 7.1879829 | 0.949117623 | 0.663 | 0.141 | 1.45175692052595e-3 | 7 | wasb |
| cyth4a | 9.6488989 | 0.347751414 | 0.213 | 0.011 | 1.94878811726824e-3 | 7 | cyth4a |
| nr4a11 | 1.9577649 | 0.867805034 | 0.596 | 0.108 | 3.95409787827394e-3 | 7 | nr4a1 |
| cd247l | 7.4473737 | 0.321202148 | 0.191 | 0.009 | 1.50414607187286e-3 | 7 | cd247l |
| LOC1005362132 | 7.6580440 | 0.91200773 | 0.652 | 0.13 | 1.54669515591108e-3 | 7 | LOC100536213 |

| | | | | | | | |
|---|---|---|---|---|---|---|---|
| nuggc.3 | 1.0575505 | 0.267121388 | 0.169 | 0.006 | 2.13593488893795e-3 | 7 | nuggc.3 |
| si:dkey-21e2.12 | 3.0086245 | 0.429851556 | 0.135 | 0.003 | 6.07651895547443e-3 | 7 | si:dkey-21e2.12 |
| coro1a3 | 4.4748444 | 1.038466485 | 0.933 | 0.258 | 9.03784330833451e-3 | 7 | coro1a |
| tbc1d10c | 1.4039949 | 0.320022093 | 0.169 | 0.007 | 2.8356486211936e-34 | 7 | tbc1d10c |
| arhgef3l | 1.6782065 | 0.424097217 | 0.258 | 0.02 | 3.38947368504193e-3 | 7 | arhgef3l |
| CABZ01079663.1 | 5.4890533 | 0.413282773 | 0.27 | 0.022 | 1.10862410637991e-3 | 7 | CABZ01079663.1 |
| si:dkey-27i16.22 | 6.2285464 | 0.725219772 | 0.506 | 0.084 | 1.25797953426569e-3 | 7 | si:dkey-27i16.2 |
| tagapb1 | 3.1561152 | 0.658448828 | 0.461 | 0.071 | 6.37440588039551e-3 | 7 | tagapb |
| wdr19 | 4.0970783 | 0.480331556 | 0.27 | 0.023 | 8.27486911818358e-3 | 7 | wdr19 |
| si:ch211-277c7.6 | 2.4616876 | 0.601735632 | 0.258 | 0.022 | 4.97187054463286e-3 | 7 | si:ch211-277c7.6 |
| BX120005.1 | 2.4846589 | 0.430655448 | 0.281 | 0.025 | 5.01826561374428e-3 | 7 | BX120005.1 |
| CABZ01054965.1 | 9.8801631 | 0.340585647 | 0.157 | 0.007 | 1.99549654484688e-3 | 7 | CABZ01054965.1 |
| CR392341.12 | 1.4985982 | 0.607380752 | 0.382 | 0.051 | 3.0267188742628e-30 | 7 | CR392341.1 |
| prkacba | 2.9985751 | 0.356230309 | 0.236 | 0.018 | 6.05622214956923e-3 | 7 | prkacba |
| cabz01093075.1 | 9.5598823 | 0.311560715 | 0.146 | 0.006 | 1.93080943320831e-2 | 7 | cabz01093075.1 |
| map4k6 | 3.3755843 | 0.262204364 | 0.157 | 0.007 | 6.81766768072655e-2 | 7 | map4k6 |
| si:ch211-207n23.2 | 5.9582386 | 0.495657408 | 0.213 | 0.015 | 1.2033854566491e-28 | 7 | si:ch211-207n23.2 |
| rasgrp3 | 6.1768869 | 0.284497887 | 0.18 | 0.01 | 1.24754586586767e-2 | 7 | rasgrp3 |
| FQ323156.1 | 3.3791873 | 0.614367787 | 0.393 | 0.061 | 6.82494462573465e-2 | 7 | FQ323156.1 |
| XLOC-026836 | 3.4752835 | 0.70400503 | 0.438 | 0.071 | 7.01903012083244e-2 | 7 | XLOC-026836 |
| rac23 | 1.1262814 | 0.888510407 | 0.831 | 0.241 | 2.27475056428973e-2 | 7 | rac2 |
| zgc:152863 | 2.2440084 | 0.342913602 | 0.191 | 0.013 | 4.53222378565674e-2 | 7 | zgc:152863 |
| ccl34b.4 | 5.5979717 | 0.502338026 | 0.169 | 0.01 | 1.13062234983747e-2 | 7 | ccl34b.4 |
| p2ry10 | 1.9173617 | 0.341627899 | 0.225 | 0.019 | 3.87249556867005e-2 | 7 | p2ry10 |
| mbnl11 | 2.6707703 | 0.744889527 | 0.652 | 0.169 | 5.39415489670397e-2 | 7 | mbnl1 |
| samsn1b | 9.7036963 | 0.350066146 | 0.191 | 0.014 | 1.95985554655858e-2 | 7 | samsn1b |
| XLOC-019150 | 5.6705452 | 0.322516982 | 0.169 | 0.011 | 1.14528002691287e-2 | 7 | XLOC-019150 |
| ptprc1 | 6.6220522 | 0.626225135 | 0.517 | 0.105 | 1.33745589811105e-2 | 7 | ptprc |
| arhgdig2 | 1.3341098 | 0.871169487 | 0.854 | 0.349 | 2.69450164893519e-2 | 7 | arhgdig |
| si:ch211-105j21.9 | 1.3476111 | 0.437374708 | 0.27 | 0.03 | 2.72177021694408e-2 | 7 | si:ch211-105j21.9 |
| traf1 | 1.3534731 | 0.255619177 | 0.157 | 0.009 | 2.73360966809274e-2 | 7 | traf1 |
| dub | 2.0493403 | 0.564091114 | 0.427 | 0.077 | 4.13905275903487e-2 | 7 | dub |
| lcp2a | 8.7619307 | 0.446024259 | 0.292 | 0.038 | 1.76964715877009e-2 | 7 | lcp2a |
| agap2 | 2.1074516 | 0.301351492 | 0.202 | 0.017 | 4.25642002667109e-2 | 7 | agap2 |
| scgn | 2.2506354 | 0.341942934 | 0.202 | 0.017 | 4.54560840880972e-2 | 7 | scgn |

| | | | | | | | |
|---|---|---|---|---|---|---|---|
| si:ch73-226l13.2 | 2.6489840 | 0.290541315 | 0.135 | 0.007 | 5.35015309881897e-2 | 7 | si:ch73-226l13.2 |
| capgb2 | 3.9613220 | 0.75618786 | 0.506 | 0.114 | 8.00068223517767e-2 | 7 | capgb |
| dok2 | 6.8060313 | 0.455021074 | 0.303 | 0.041 | 1.37461414441819e-2 | 7 | dok2 |
| si:dkey-185m8.2 | 8.6100963 | 0.420486681 | 0.281 | 0.036 | 1.73898115099998e-2 | 7 | si:dkey-185m8.2 |
| stk17al | 1.0803429 | 0.48746985 | 0.315 | 0.045 | 2.18196874986481e-2 | 7 | stk17al |
| unc119b | 1.1452136 | 0.278728197 | 0.157 | 0.01 | 2.31298808329475e-2 | 7 | unc119b |
| pgam1a1 | 1.3169600 | 0.700509829 | 0.551 | 0.143 | 2.65986415726058e-2 | 7 | pgam1a |
| cd44b | 1.9233200 | 0.351473728 | 0.191 | 0.016 | 3.88452946455584e-2 | 7 | cd44b |
| h1fx1 | 5.7032992 | 0.905338301 | 0.697 | 0.222 | 1.15189535326277e-2 | 7 | h1fx |
| egr3 | 6.6838430 | 0.442673688 | 0.236 | 0.026 | 1.34993577384521e-2 | 7 | egr3 |
| CU019662.1 | 1.1885761 | 0.353964828 | 0.213 | 0.021 | 2.40056720076498e-2 | 7 | CU019662.1 |
| zgc:152774 | 1.2441558 | 0.324789771 | 0.213 | 0.021 | 2.51282162653795e-2 | 7 | zgc:152774 |
| si:ch211-209l18.2 | 2.7357875 | 0.280136191 | 0.124 | 0.006 | 5.52547008074462e-2 | 7 | si:ch211-209l18.2 |
| cd4-1 | 3.0372605 | 0.338759266 | 0.124 | 0.006 | 6.13435522141792e-2 | 7 | cd4-1 |
| CABZ01117603.1 | 8.5150108 | 0.257446461 | 0.157 | 0.011 | 1.71977673496062e-2 | 7 | CABZ01117603.1 |
| batf3 | 1.0073161 | 0.27281433 | 0.169 | 0.013 | 2.03447639365643e-2 | 7 | batf3 |
| egr2b | 1.1788053 | 0.404393825 | 0.146 | 0.01 | 2.38083309808611e-2 | 7 | egr2b |
| FERMT3 (1 of many) | 1.5385705 | 0.455180438 | 0.337 | 0.055 | 3.10745096804573e-2 | 7 | FERMT3 (1 of many) |
| rel | 1.9955527 | 0.355385993 | 0.213 | 0.023 | 4.03041793942667e-2 | 7 | rel |
| il34 | 2.4807482 | 0.425724469 | 0.225 | 0.025 | 5.01036723682906e-2 | 7 | il34 |
| si:dkey-53k12.22 | 2.7958273 | 0.398395744 | 0.27 | 0.036 | 5.64673257836071e-2 | 7 | si:dkey-53k12.2 |
| gnai2b | 1.6469862 | 0.399317971 | 0.247 | 0.032 | 3.32641822420754e-1 | 7 | gnai2b |
| sept9b | 2.7353256 | 0.425045419 | 0.292 | 0.045 | 5.52453729969245e-1 | 7 | sept9b |
| zgc:174945 | 3.9018203 | 0.253725104 | 0.157 | 0.012 | 7.88050651985169e-1 | 7 | zgc:174945 |
| pfn13 | 5.4060207 | 0.965681921 | 0.989 | 0.399 | 1.09185400245733e-1 | 7 | pfn1 |
| wasa1 | 6.7708612 | 0.54407389 | 0.472 | 0.106 | 1.36751085656255e-1 | 7 | wasa |
| si:cabz01074946.1 | 9.2904517 | 0.469065195 | 0.348 | 0.062 | 1.87639254082343e-1 | 7 | si:cabz01074946.1 |
| srgn1 | 1.3389936 | 0.738552261 | 0.719 | 0.229 | 2.70436544045938e-1 | 7 | srgn |
| histh1l | 1.5734215 | 0.749666006 | 0.528 | 0.147 | 3.17783957545587e-1 | 7 | histh1l |
| rplp21 | 2.0244053 | 0.280158654 | 1 | 0.926 | 4.08869141151934e-1 | 7 | rplp2 |
| ccr9a1 | 2.2109681 | 0.556084957 | 0.416 | 0.087 | 4.46549234894694e-1 | 7 | ccr9a |
| si:dkey-103e21.5 | 2.2601671 | 0.259065455 | 0.169 | 0.015 | 4.5648596799819e-18 | 7 | si:dkey-103e21.5 |
| fli1b | 2.6207422 | 0.381442744 | 0.236 | 0.031 | 5.29311320342387e-1 | 7 | fli1b |
| rps171 | 3.5758486 | 0.345068612 | 0.989 | 0.877 | 7.22214151839142e-1 | 7 | rps17 |
| si:ch211-67e16.3 | 3.6969578 | 0.337199133 | 0.169 | 0.015 | 7.466745824349e-18 | 7 | si:ch211-67e16.3 |

| | | | | | | | |
|---|---|---|---|---|---|---|---|
| arpc1b3 | 1.1497874 | 0.826408827 | 0.854 | 0.344 | 2.32222578870821e-1 | 7 | arpc1b |
| ptk2ba | 1.3983750 | 0.34856371 | 0.191 | 0.021 | 2.82429801654138e-1 | 7 | ptk2ba |
| XLOC-043760 | 4.1018457 | 0.366470805 | 0.169 | 0.016 | 8.28449776474747e-1 | 7 | XLOC-043760 |
| calm1b1 | 6.3198348 | 0.555337015 | 0.517 | 0.135 | 1.2764170529258e-16 | 7 | calm1b |
| il10ra1 | 9.5789764 | 0.4134545 | 0.27 | 0.043 | 1.934665872731e-16 | 7 | il10ra |
| pim2 | 1.7980627 | 0.444084918 | 0.337 | 0.065 | 3.63154737947618e-1 | 7 | pim2 |
| lpar5b | 1.9179812 | 0.268908674 | 0.169 | 0.017 | 3.87374669407227e-1 | 7 | lpar5b |
| gadd45ga | 3.0602216 | 0.480924042 | 0.27 | 0.045 | 6.18072958500591e-1 | 7 | gadd45ga |
| rps6ka1 | 7.7584582 | 0.439178823 | 0.27 | 0.045 | 1.5669758180503e-15 | 7 | rps6ka1 |
| gpr183a1 | 1.7873371 | 0.491376214 | 0.303 | 0.058 | 3.60988492693146e-1 | 7 | gpr183a |
| hcst | 3.7180228 | 0.255363563 | 0.157 | 0.016 | 7.50929067662833e-1 | 7 | hcst |
| ca2 | 3.7665513 | 0.431405303 | 0.225 | 0.033 | 7.60730376927448e-1 | 7 | ca2 |
| plk3 | 4.0052388 | 0.424248829 | 0.281 | 0.05 | 8.08938085940124e-1 | 7 | plk3 |
| LOC1039091152 | 4.1415619 | 0.560964863 | 0.596 | 0.186 | 8.36471261593697e-1 | 7 | LOC103909115 |
| spint1a | 4.4472525 | 0.37337095 | 0.202 | 0.027 | 8.98211607564436e-1 | 7 | spint1a |
| zgc:640513 | 8.1483173 | 0.624564139 | 0.551 | 0.164 | 1.6457156514485e-14 | 7 | zgc:64051 |
| vsir2 | 1.1778827 | 0.499902604 | 0.416 | 0.1 | 2.37896983999295e-1 | 7 | vsir |
| unc13d | 1.2815648 | 0.341673807 | 0.213 | 0.03 | 2.58837646507836e-1 | 7 | unc13d |
| zgc:194312 | 1.5202897 | 0.283368136 | 0.191 | 0.024 | 3.07052921417912e-1 | 7 | zgc:194312 |
| laptm53 | 1.7230663 | 0.659801182 | 0.629 | 0.202 | 3.48007708640681e-1 | 7 | laptm5 |
| sept12 | 2.1239206 | 0.446355271 | 0.348 | 0.077 | 4.28968244997448e-1 | 7 | sept12 |
| tec1 | 2.3790537 | 0.373984823 | 0.247 | 0.041 | 4.80497487781948e-1 | 7 | tec |
| rps27.1 | 2.4819509 | 0.287289695 | 0.989 | 0.894 | 5.01279628149722e-1 | 7 | rps27.1 |
| rsl24d11 | 3.3671038 | 0.593599222 | 0.921 | 0.563 | 6.8005396864087e-14 | 7 | rsl24d1 |
| LOC1001511961 | 3.5480999 | 0.314199953 | 0.225 | 0.034 | 7.16609747195338e-1 | 7 | LOC100151196 |
| grap2b3 | 3.8525589 | 0.563623266 | 0.596 | 0.179 | 7.78101331843529e-1 | 7 | grap2b |
| si:dkey-1h24.6 | 4.1571758 | 0.271025671 | 0.112 | 0.008 | 8.39624807172686e-1 | 7 | si:dkey-1h24.6 |
| btg11 | 4.5582742 | 0.677105877 | 0.843 | 0.452 | 9.20634655268963e-1 | 7 | btg1 |
| si:dkeyp-68b7.12 | 4.6319666 | 0.418488686 | 0.258 | 0.045 | 9.35518310237313e-1 | 7 | si:dkeyp-68b7.12 |
| si:ch211-241f5.3 | 5.8009537 | 0.358659439 | 0.213 | 0.031 | 1.17161862866884e-1 | 7 | si:ch211-241f5.3 |
| aldocb | 5.9343375 | 0.417291403 | 0.281 | 0.053 | 1.19855815209364e-1 | 7 | aldocb |
| AL954142.11 | 8.0811489 | 0.29564077 | 0.202 | 0.028 | 1.63214964386908e-1 | 7 | AL954142.1 |
| ikzf21 | 1.2592057 | 0.450390297 | 0.27 | 0.05 | 2.54321795300914e-1 | 7 | ikzf2 |
| elf11 | 1.7123768 | 0.547774051 | 0.382 | 0.098 | 3.45848750545156e-1 | 7 | elf1 |
| samd11 | 2.2820727 | 0.277985792 | 0.146 | 0.015 | 4.60910229835157e-1 | 7 | samd11 |

| | | | | | | | |
|---|---|---|---|---|---|---|---|
| znrf1 | 2.7069443 | 0.255648017 | 0.112 | 0.009 | 5.46721545626157e-1 | 7 | znrf1 |
| uck2a | 4.5159700 | 0.443038476 | 0.315 | 0.069 | 9.12090471810492e-1 | 7 | uck2a |
| clic11 | 4.9148061 | 0.551279496 | 0.596 | 0.203 | 9.92643399683861e-1 | 7 | clic1 |
| CU207241.1 | 5.9580132 | 0.281133381 | 0.169 | 0.021 | 1.20333993875736e-1 | 7 | CU207241.1 |
| dtnba | 6.1345125 | 0.29605872 | 0.213 | 0.033 | 1.23898750536981e-1 | 7 | dtnba |
| si:rp71-68n21.9 | 6.2416887 | 0.385186967 | 0.281 | 0.055 | 1.26063388124828e-1 | 7 | si:rp71-68n21.9 |
| RASSF5 | 1.1266338 | 0.457826143 | 0.315 | 0.07 | 2.27546229269373e-1 | 7 | RASSF5 |
| CR381673.11 | 1.2567564 | 0.411938493 | 0.27 | 0.051 | 2.53827107430084e-1 | 7 | CR381673.1 |
| si:dkey-78l4.4 | 1.3097340 | 0.362699995 | 0.146 | 0.016 | 2.64526979710025e-1 | 7 | si:dkey-78l4.4 |
| b3gnt5a | 2.7309648 | 0.346390743 | 0.258 | 0.049 | 5.51572976403651e-1 | 7 | b3gnt5a |
| si:ch73-208g10.1 | 2.8340479 | 0.317107746 | 0.191 | 0.028 | 5.72392671635711e-1 | 7 | si:ch73-208g10.1 |
| sept61 | 4.4850727 | 0.552757802 | 0.528 | 0.18 | 9.05850145965889e-1 | 7 | sept6 |
| pik3r1 | 4.5218447 | 0.481861547 | 0.371 | 0.096 | 9.13276979817729e-1 | 7 | pik3r1 |
| XLOC-033137 | 5.0004648 | 0.310453729 | 0.18 | 0.025 | 1.00994389012109e-1 | 7 | XLOC-033137 |
| ppifb | 9.7260530 | 0.51557189 | 0.472 | 0.157 | 1.96437093995178e-1 | 7 | ppifb |
| nfkbiaa1 | 1.0873755 | 0.616962146 | 0.472 | 0.155 | 2.19617231558745e-1 | 7 | nfkbiaa |
| imp4 | 1.8607475 | 0.293939508 | 0.281 | 0.058 | 3.75815174642153e-1 | 7 | imp4 |
| dtnbp1a1 | 2.8175060 | 0.470052643 | 0.348 | 0.091 | 5.69051705043265e-1 | 7 | dtnbp1a |
| rhof | 2.9400990 | 0.293275557 | 0.202 | 0.033 | 5.93811795466155e-1 | 7 | rhof |
| si:dkey-262k9.42 | 3.0738329 | 0.400253078 | 0.348 | 0.087 | 6.20822048724709e-1 | 7 | si:dkey-262k9.4 |
| fam49ba1 | 3.1097452 | 0.415893751 | 0.416 | 0.114 | 6.28075254308713e-1 | 7 | fam49ba |
| XLOC-0254233 | 4.3572599 | 0.776012959 | 0.494 | 0.182 | 8.80035783783029e-1 | 7 | XLOC-025423 |
| lpar5a1 | 5.4203193 | 0.350170015 | 0.27 | 0.056 | 1.09474189032538e-1 | 7 | lpar5a |
| sat1a.22 | 8.4616408 | 0.585719376 | 0.719 | 0.323 | 1.70899760532485e-1 | 7 | sat1a.2 |
| cxcr4b1 | 1.0577727 | 0.57347246 | 0.472 | 0.147 | 2.13638369147567e-1 | 7 | cxcr4b |
| gsdmeb | 1.7259844 | 0.269833311 | 0.157 | 0.022 | 3.48597073469951e-1 | 7 | gsdmeb |
| socs1a | 1.7570258 | 0.369365108 | 0.225 | 0.043 | 3.54866515191007e-1 | 7 | socs1a |
| XLOC-011085 | 1.8015941 | 0.346327141 | 0.202 | 0.035 | 3.63867980002858e-1 | 7 | XLOC-011085 |
| msna1 | 2.5054895 | 0.600168394 | 0.73 | 0.393 | 5.06033722770407e-1 | 7 | msna |
| orai1a | 2.5375144 | 0.252765779 | 0.135 | 0.016 | 5.12501802779414e-1 | 7 | orai1a |
| ptpn62 | 4.3508455 | 0.511061828 | 0.438 | 0.136 | 8.78740279142533e-1 | 7 | ptpn6 |
| wipf1a1 | 4.9625976 | 0.419098026 | 0.315 | 0.077 | 1.0022958420707e-09 | 7 | wipf1a |
| ddx49 | 7.3654712 | 0.309582301 | 0.169 | 0.026 | 1.48760423071442e-0 | 7 | ddx49 |
| serbp1a1 | 1.1716349 | 0.450198689 | 0.91 | 0.648 | 2.36635109286124e-0 | 7 | serbp1a |
| mfng1 | 1.4621185 | 0.29498161 | 0.258 | 0.056 | 2.95304092530838e-0 | 7 | mfng |

| | | | | | | | |
|---|---|---|---|---|---|---|---|
| RASA21 | 1.8233001 | 0.333809666 | 0.236 | 0.049 | 3.68251932135843e-0 | 7 | RASA2 |
| si:ch211-226m16.2 | 1.8884751 | 0.358817589 | 0.281 | 0.068 | 3.81415329206787e-0 | 7 | si:ch211-226m16.2 |
| litaf2 | 2.8355621 | 0.472340844 | 0.528 | 0.196 | 5.72698485871915e-0 | 7 | litaf |
| XLOC-0054492 | 3.5893361 | 0.42949094 | 0.281 | 0.07 | 7.24938218237276e-0 | 7 | XLOC-005449 |
| XLOC-032002 | 4.5025961 | 0.277801236 | 0.18 | 0.03 | 9.09389349273814e-0 | 7 | XLOC-032002 |
| wipf1b1 | 5.4404442 | 0.336176977 | 0.281 | 0.069 | 1.09880651968545e-0 | 7 | wipf1b |
| hmha1b1 | 5.7110853 | 0.345591227 | 0.281 | 0.069 | 1.15346791452856e-0 | 7 | hmha1b |
| csk1 | 7.6724618 | 0.423583919 | 0.371 | 0.111 | 1.54960711121245e-0 | 7 | csk |
| XLOC-0234922 | 7.7198985 | 0.41761436 | 0.315 | 0.085 | 1.55918790139588e-0 | 7 | XLOC-023492 |
| slc25a5 | 9.5465621 | 0.25742519 | 0.966 | 0.896 | 1.92811915329089e-0 | 7 | slc25a5 |
| ppp1r181 | 1.0628533 | 0.260921236 | 0.27 | 0.062 | 2.14664501191887e-0 | 7 | ppp1r18 |
| ccdc50 | 1.8242395 | 0.309200227 | 0.146 | 0.022 | 3.68441651907402e-0 | 7 | ccdc50 |
| mcl1a3 | 1.9147665 | 0.544237877 | 0.865 | 0.535 | 3.86725404995257e-0 | 7 | mcl1a |
| cox171 | 2.1410626 | 0.457371967 | 0.483 | 0.178 | 4.32430430628537e-0 | 7 | cox17 |
| fam49a | 2.3604342 | 0.319378358 | 0.258 | 0.062 | 4.76736915019726e-0 | 7 | fam49a |
| arpc5a | 4.1024129 | 0.469467057 | 0.483 | 0.18 | 8.28564353047972e-0 | 7 | arpc5a |
| syk1 | 4.1250863 | 0.407553892 | 0.281 | 0.074 | 8.33143693565859e-0 | 7 | syk |
| prkacab | 4.1913902 | 0.264027746 | 0.236 | 0.053 | 8.46535096814263e-0 | 7 | prkacab |
| rps23 | 9.9499496 | 0.263851325 | 1 | 0.878 | 2.00959132925811e-0 | 7 | rps23 |
| hmga1a | 9.9993454 | 0.560412389 | 0.753 | 0.426 | 2.01956779452861e-0 | 7 | hmga1a |
| si:ch73-264p11.11 | 1.8100135 | 0.261046121 | 0.169 | 0.03 | 3.65568433714934e-0 | 7 | si:ch73-264p11.1 |
| hcls12 | 2.1337159 | 0.387799613 | 0.348 | 0.105 | 4.30946601474394e-0 | 7 | hcls1 |
| fmnl1a | 2.7804264 | 0.293329922 | 0.18 | 0.036 | 5.61562732773731e-0 | 7 | fmnl1a |
| sik1 | 4.4101162 | 0.356338523 | 0.292 | 0.084 | 8.90711183178712e-0 | 7 | sik1 |
| ascc1 | 5.3080717 | 0.320942397 | 0.202 | 0.045 | 1.07207125285695e-0 | 7 | ascc1 |
| abracl2 | 5.4669727 | 0.466497137 | 0.551 | 0.232 | 1.10416447680965e-0 | 7 | abracl |
| sesn11 | 6.7299805 | 0.482931146 | 0.506 | 0.208 | 1.35925416493881e-0 | 7 | sesn1 |
| cap13 | 8.8367604 | 0.502516623 | 0.663 | 0.32 | 1.78476049997511e-0 | 7 | cap1 |
| mbpb | 1.0277088 | 0.324217547 | 0.292 | 0.085 | 2.07566360817386e-0 | 7 | mbpb |
| ubash3ba | 1.0835349 | 0.263062537 | 0.124 | 0.019 | 2.18841557540707e-0 | 7 | ubash3ba |
| iqgap1 | 1.0892231 | 0.307728713 | 0.281 | 0.08 | 2.19990403764818e-0 | 7 | iqgap1 |
| cox6b1 | 1.1050513 | 0.320707254 | 0.382 | 0.126 | 2.23187224203479e-0 | 7 | cox6b1 |
| ier2b1 | 1.5149048 | 0.473089348 | 0.674 | 0.315 | 3.05965323449536e-0 | 7 | ier2b |
| plcg2 | 1.5370389 | 0.310898723 | 0.191 | 0.042 | 3.10435765546191e-0 | 7 | plcg2 |
| dusp11 | 1.7479537 | 0.409566045 | 0.315 | 0.099 | 3.53034219879555e-0 | 7 | dusp1 |

| | | | | | | | |
|---|---|---|---|---|---|---|---|
| arrdc3b | 2.0748640 | 0.28170536 | 0.169 | 0.034 | 4.19060294972196e-0 | 7 | arrdc3b |
| mef2b1 | 2.2083381 | 0.318382475 | 0.146 | 0.026 | 4.46018051067987e-0 | 7 | mef2b |
| glipr1a1 | 2.5220455 | 0.287252793 | 0.303 | 0.088 | 5.09377540099567e-0 | 7 | glipr1a |
| crlf3 | 2.7656327 | 0.333753466 | 0.315 | 0.1 | 5.58574853433937e-0 | 7 | crlf3 |
| cotl13 | 2.9882281 | 0.544620225 | 0.798 | 0.399 | 6.03532440107812e-0 | 7 | cotl1 |
| nfkbie1 | 3.2109128 | 0.361179152 | 0.337 | 0.112 | 6.48508077354775e-0 | 7 | nfkbie |
| vaspa1 | 3.8874800 | 0.322425262 | 0.36 | 0.113 | 7.85154347110872e-0 | 7 | vaspa |
| ctss2.11 | 4.3011267 | 0.2674006 | 0.258 | 0.069 | 8.68698565996511e-0 | 7 | ctss2.1 |
| tnip1 | 4.7709471 | 0.370595112 | 0.247 | 0.07 | 9.63588189525617e-0 | 7 | tnip1 |
| h2afx | 5.5316603 | 0.453948302 | 0.191 | 0.046 | 1.11722944625998e-0 | 7 | h2afx |
| ccni | 5.5954453 | 0.47101382 | 0.719 | 0.406 | 1.13011209129446e-0 | 7 | ccni |
| XLOC-0283702 | 5.6183508 | 0.3359693 | 0.213 | 0.053 | 1.13473832243608e-0 | 7 | XLOC-028370 |
| si:dkeyp-19e1.3 | 8.4771192 | 0.299041728 | 0.18 | 0.041 | 1.71212376749132e-0 | 7 | si:dkeyp-19e1.3 |
| cdc42l1 | 9.7080447 | 0.446693424 | 0.652 | 0.337 | 1.96073378905149e-0 | 7 | cdc42l |
| arhgdia1 | 1.0291781 | 0.455546633 | 0.697 | 0.364 | 2.07863111400008e-0 | 7 | arhgdia |
| mob1a1 | 1.3331421 | 0.362693677 | 0.506 | 0.21 | 2.69254711145961e-0 | 7 | mob1a |
| abi3a1 | 1.5006127 | 0.263422253 | 0.236 | 0.062 | 3.03078764134221e-0 | 7 | abi3a |
| si:dkey-118j18.2 | 1.6302595 | 0.271626468 | 0.247 | 0.07 | 3.29263528245117e-0 | 7 | si:dkey-118j18.2 |
| hmgn21 | 1.7940516 | 0.531151658 | 0.899 | 0.624 | 3.62344621576247e-0 | 7 | hmgn2 |
| ezrb | 3.5114193 | 0.258208878 | 0.225 | 0.061 | 7.09201366637541e-0 | 7 | ezrb |
| dock8 | 4.4004645 | 0.26455522 | 0.157 | 0.034 | 8.88761832212626e-0 | 7 | dock8 |
| stim1a | 4.9114366 | 0.28590899 | 0.18 | 0.044 | 9.91962857783034e-0 | 7 | stim1a |
| atp2b1a | 5.4625726 | 0.378927825 | 0.213 | 0.059 | 0.000110328 | 7 | atp2b1a |
| mibp2 | 5.8493406 | 0.412132515 | 0.899 | 0.622 | 0.000118139 | 7 | mibp2 |
| XLOC-0418701 | 7.4265579 | 0.284873682 | 0.18 | 0.043 | 0.000149994 | 7 | XLOC-041870 |
| BX901920.13 | 1.0646900 | 0.346568793 | 0.247 | 0.075 | 0.000215035 | 7 | BX901920.1 |
| nrros1 | 1.1714087 | 0.262516857 | 0.191 | 0.049 | 0.000236589 | 7 | nrros |
| midn2 | 1.2799209 | 0.400309964 | 0.416 | 0.174 | 0.000258506 | 7 | midn |
| slc25a55a | 1.4159210 | 0.334433571 | 0.315 | 0.114 | 0.000285974 | 7 | slc25a55a |
| eif4ebp2 | 1.4912556 | 0.385497454 | 0.584 | 0.285 | 0.000301189 | 7 | eif4ebp2 |
| twf1a | 1.5274708 | 0.367597482 | 0.449 | 0.195 | 0.000308503 | 7 | twf1a |
| itga41 | 1.5301283 | 0.284110299 | 0.213 | 0.06 | 0.00030904 | 7 | itga4 |
| cd831 | 2.6777261 | 0.306443922 | 0.18 | 0.046 | 0.00054082 | 7 | cd83 |
| arpc33 | 2.6887212 | 0.43255219 | 0.764 | 0.472 | 0.000543041 | 7 | arpc3 |
| wu:fb55g09 | 3.2230206 | 0.474825391 | 0.562 | 0.305 | 0.000650953 | 7 | wu:fb55g09 |

| | | | | | | | |
|---|---|---|---|---|---|---|---|
| sept21 | 3.8577155 | 0.386560898 | 0.562 | 0.276 | 0.000779143 | 7 | sept2 |
| si:dkey-112e7.21 | 4.6028249 | 0.411423154 | 0.517 | 0.265 | 0.000929633 | 7 | si:dkey-112e7.2 |
| crip11 | 5.0113526 | 0.43262922 | 0.697 | 0.368 | 0.001012143 | 7 | crip1 |
| si:ch211-132p1.31 | 5.5761121 | 0.320630313 | 0.36 | 0.141 | 0.001126207 | 7 | si:ch211-132p1.3 |
| itgae.21 | 6.0967560 | 0.295440676 | 0.157 | 0.039 | 0.001231362 | 7 | itgae.2 |
| stxbp2 | 7.0783918 | 0.282203706 | 0.213 | 0.065 | 0.001429623 | 7 | stxbp2 |
| sept7b | 7.3715224 | 0.316432015 | 0.371 | 0.152 | 0.001488826 | 7 | sept7b |
| gnai2a1 | 7.6427636 | 0.333003482 | 0.416 | 0.179 | 0.001543609 | 7 | gnai2a |
| eif3c1 | 7.8817871 | 0.376266593 | 0.562 | 0.282 | 0.001591885 | 7 | eif3c |
| hnrnpa1b1 | 8.3789140 | 0.336147441 | 0.506 | 0.239 | 0.001692289 | 7 | hnrnpa1b |
| wdr12 | 1.0612299 | 0.319214458 | 0.506 | 0.23 | 0.002143366 | 7 | wdr1 |
| si:dkey-112a7.4 | 1.0932420 | 0.251187683 | 0.258 | 0.087 | 0.002208021 | 7 | si:dkey-112a7.4 |
| pnrc2 | 1.1662474 | 0.337605849 | 0.944 | 0.72 | 0.00235547 | 7 | pnrc2 |
| ucp21 | 1.2521832 | 0.387095932 | 0.753 | 0.415 | 0.002529035 | 7 | ucp2 |
| ap1s1 | 1.3581696 | 0.298206057 | 0.281 | 0.103 | 0.002743095 | 7 | ap1s1 |
| nutf2l | 1.3699806 | 0.312801047 | 0.393 | 0.168 | 0.00276695 | 7 | nutf2l |
| fosl21 | 1.3821182 | 0.300476174 | 0.236 | 0.077 | 0.002791464 | 7 | fosl2 |
| kmt5aa | 1.6840890 | 0.275244474 | 0.371 | 0.153 | 0.003401355 | 7 | kmt5aa |
| rnf44 | 1.8348232 | 0.3061929 | 0.337 | 0.132 | 0.003705792 | 7 | rnf44 |
| eno1a1 | 2.0163670 | 0.350239051 | 0.404 | 0.183 | 0.004072457 | 7 | eno1a |
| BX322618.11 | 2.0471185 | 0.317556305 | 0.213 | 0.066 | 0.004134565 | 7 | BX322618.1 |
| tpi1b1 | 2.1808341 | 0.329776708 | 0.494 | 0.239 | 0.004404631 | 7 | tpi1b |
| hnrnph1l | 2.2445808 | 0.251225693 | 0.326 | 0.125 | 0.00453338 | 7 | hnrnph1l |
| mob3a | 2.2881650 | 0.320695314 | 0.416 | 0.183 | 0.004621407 | 7 | mob3a |
| ran | 2.4676818 | 0.342441049 | 0.888 | 0.657 | 0.004983977 | 7 | ran |
| eif3ba | 2.5945684 | 0.351790497 | 0.708 | 0.386 | 0.00524025 | 7 | eif3ba |
| si:ch211-196l7.41 | 2.8184752 | 0.371122337 | 0.697 | 0.406 | 0.005692474 | 7 | si:ch211-196l7.4 |
| bptf | 2.9196885 | 0.262551395 | 0.202 | 0.063 | 0.005896895 | 7 | bptf |
| sh3kbp1 | 3.7405121 | 0.313141983 | 0.281 | 0.106 | 0.007554712 | 7 | sh3kbp1 |
| atp5if1b1 | 5.7501212 | 0.384570299 | 0.506 | 0.269 | 0.01161352 | 7 | atp5if1b |
| myca | 6.7303129 | 0.289028976 | 0.191 | 0.06 | 0.013593213 | 7 | myca |
| kctd5a | 7.9237033 | 0.34252891 | 0.483 | 0.242 | 0.016003504 | 7 | kctd5a |
| CR753876.11 | 8.9840229 | 0.294835447 | 0.27 | 0.098 | 0.018145031 | 7 | CR753876.1 |
| actb13 | 1.0995407 | 0.331573756 | 0.899 | 0.737 | 0.022207424 | 7 | actb1 |
| arpc42 | 1.1298089 | 0.349791154 | 0.573 | 0.309 | 0.022818751 | 7 | arpc4 |

| | | | | | | | |
|---|---|---|---|---|---|---|---|
| actr2a1 | 1.1308525 | 0.321378993 | 0.36 | 0.158 | 0.022839829 | 7 | actr2a |
| capza1b2 | 1.2255361 | 0.338393493 | 0.64 | 0.356 | 0.024752153 | 7 | capza1b |
| elmo11 | 1.3976079 | 0.251743427 | 0.202 | 0.066 | 0.028227488 | 7 | elmo1 |
| eif2s1b | 1.9424853 | 0.330760217 | 0.528 | 0.293 | 0.039232376 | 7 | eif2s1b |
| ddx61 | 1.9969003 | 0.331921921 | 0.607 | 0.336 | 0.040331396 | 7 | ddx61 |
| nfkbiab1 | 2.0333059 | 0.353261633 | 0.539 | 0.295 | 0.041066681 | 7 | nfkbiab |
| anp32a | 2.1386152 | 0.292492651 | 0.573 | 0.3 | 0.043193613 | 7 | anp32a |
| tefa | 2.3853789 | 0.268520303 | 0.337 | 0.145 | 0.048177499 | 7 | tefa |
| eif3f | 2.3929109 | 0.32614293 | 0.876 | 0.638 | 0.048329623 | 7 | eif3f |
| gng5 | 2.7601627 | 0.352661426 | 0.629 | 0.377 | 0.055747007 | 7 | gng5 |
| ppp1caa | 2.9798925 | 0.292616753 | 0.427 | 0.209 | 0.060184891 | 7 | ppp1caa |
| ddx21 | 3.2872616 | 0.287095162 | 0.461 | 0.228 | 0.066392823 | 7 | ddx21 |
| srsf2a | 3.4942437 | 0.313617413 | 0.506 | 0.268 | 0.070573241 | 7 | srsf2a |
| arf6a | 3.8600462 | 0.300041663 | 0.348 | 0.159 | 0.077961355 | 7 | arf6a |
| cox5aa1 | 4.0455617 | 0.331204408 | 0.663 | 0.403 | 0.081708212 | 7 | cox5aa |
| si:ch211-222l21.11 | 4.2971476 | 0.382467851 | 0.798 | 0.524 | 0.086789491 | 7 | si:ch211-222l21.1 |
| ldhba1 | 4.6656011 | 0.305379186 | 0.596 | 0.331 | 0.094231146 | 7 | ldhba |
| arf6b | 5.3839456 | 0.26464895 | 0.371 | 0.172 | 0.10873955 | 7 | arf6b |
| zfp36l1a3 | 6.0055569 | 0.366072077 | 0.539 | 0.306 | 0.121294235 | 7 | zfp36l1a |
| ponzr12 | 6.7665344 | 0.345747351 | 0.663 | 0.386 | 0.136663696 | 7 | ponzr1 |
| LOC1039104381 | 7.7732732 | 0.26765575 | 0.382 | 0.18 | 0.1569968 | 7 | LOC103910438 |
| nap1l1 | 8.9604834 | 0.373071458 | 0.584 | 0.354 | 0.180974885 | 7 | nap1l1 |
| ywhaqa | 9.6232596 | 0.280758082 | 0.483 | 0.248 | 0.194360976 | 7 | ywhaqa |
| capzb1 | 1.2416432 | 0.320684125 | 0.64 | 0.379 | 0.250774678 | 7 | capzb |
| nsa21 | 1.3588425 | 0.332447722 | 0.753 | 0.516 | 0.27444542 | 7 | nsa2 |
| nhp2 | 1.5706174 | 0.257191131 | 0.461 | 0.237 | 0.317217611 | 7 | nhp2 |
| zgc:1629441 | 1.9061016 | 0.298806898 | 0.663 | 0.393 | 0.384975344 | 7 | zgc:162944 |
| ilf2 | 2.1460056 | 0.277956792 | 0.461 | 0.244 | 0.433428753 | 7 | ilf2 |
| ddx39aa | 2.8633408 | 0.250858864 | 0.393 | 0.199 | 0.578308952 | 7 | ddx39aa |
| eif3s10 | 3.0774107 | 0.281421651 | 0.461 | 0.255 | 0.621544643 | 7 | eif3s10 |
| cox7b1 | 3.5399955 | 0.284286381 | 0.764 | 0.498 | 0.714972903 | 7 | cox7b |
| btf3 | 4.0956871 | 0.257748011 | 0.899 | 0.723 | 0.827205926 | 7 | btf3 |
| brd2a | 4.6887178 | 0.253784843 | 0.303 | 0.143 | 0.946980348 | 7 | brd2a |
| ube2na | 4.7612389 | 0.26474888 | 0.427 | 0.229 | 0.961627431 | 7 | ube2na |
| GCA1 | 4.8049827 | 0.281013799 | 0.528 | 0.304 | 0.97046237 | 7 | GCA |

| | | | | | | | |
|---|---|---|---|---|---|---|---|
| eif3ea | 4.9474566 | 0.299188397 | 0.685 | 0.444 | 0.999237818 | 7 | eif3ea |
| mych | 4.9656029 | 0.2579683 | 0.337 | 0.166 | 1 | 7 | mych |
| slc25a3b | 5.0327688 | 0.301424374 | 0.831 | 0.566 | 1 | 7 | slc25a3b |
| cct81 | 5.1013412 | 0.284851577 | 0.562 | 0.336 | 1 | 7 | cct8 |
| eif3d | 5.3103736 | 0.278175282 | 0.663 | 0.414 | 1 | 7 | eif3d |
| nop531 | 5.5031180 | 0.290976212 | 0.573 | 0.349 | 1 | 7 | nop53 |
| si:ch211-102c2.43 | 5.9620245 | 0.356478276 | 0.348 | 0.173 | 1 | 7 | si:ch211-102c2.4 |
| aldoaa1 | 7.0838740 | 0.280980003 | 0.483 | 0.284 | 1 | 7 | aldoaa |
| eef2b | 7.6710805 | 0.256273806 | 0.933 | 0.773 | 1 | 7 | eef2b |
| arpc23 | 8.0193322 | 0.288362031 | 0.73 | 0.479 | 1 | 7 | arpc2 |
| irf2bp2a1 | 8.0440307 | 0.292956723 | 0.438 | 0.245 | 1 | 7 | irf2bp2a |
| ywhabl | 8.6422683 | 0.288062804 | 0.674 | 0.435 | 1 | 7 | ywhabl |
| si:dkey-42i9.4 | 9.4742532 | 0.355645561 | 0.36 | 0.195 | 1 | 7 | si:dkey-42i9.4 |
| hmgb2a1 | 9.5354367 | 0.367452811 | 0.73 | 0.581 | 1 | 7 | hmgb2a |
| glrx2 | 9.5434999 | 0.29619597 | 0.64 | 0.421 | 1 | 7 | glrx |
| egr12 | 0.000106 | 0.315658252 | 0.315 | 0.162 | 1 | 7 | egr1 |
| cebpb2 | 0.000116 | 0.390956284 | 0.472 | 0.292 | 1 | 7 | cebpb |
| spcs3 | 0.000119 | 0.275084557 | 0.427 | 0.241 | 1 | 7 | spcs3 |
| arpc4l3 | 0.000135 | 0.251342139 | 0.584 | 0.344 | 1 | 7 | arpc4l |
| myl12.11 | 0.000135 | 0.288051342 | 0.82 | 0.626 | 1 | 7 | myl12.1 |
| zgc:1627304 | 0.000161 | 0.325359909 | 0.697 | 0.487 | 1 | 7 | zgc:162730 |
| eif3g1 | 0.000346 | 0.252610284 | 0.607 | 0.392 | 1 | 7 | eif3g |
| lasp11 | 0.000371 | 0.253323101 | 0.562 | 0.354 | 1 | 7 | lasp1 |
| ptmaa1 | 0.000391 | 0.279888299 | 0.843 | 0.651 | 1 | 7 | ptmaa |
| eef1da | 0.000462 | 0.262579295 | 0.809 | 0.609 | 1 | 7 | eef1da |
| gtpbp41 | 0.000491 | 0.270327001 | 0.483 | 0.311 | 1 | 7 | gtpbp4 |
| trib31 | 0.000559 | 0.271527945 | 0.247 | 0.123 | 1 | 7 | trib3 |
| krt151 | 1.4813757 | 1.693782173 | 0.867 | 0.14 | 2.99193455991876e-7 | 8 | krt15 |
| krt4 | 1.9212518 | 1.504513106 | 0.687 | 0.081 | 3.88035231492333e-7 | 8 | krt4 |
| gstm.32 | 6.3054651 | 1.557258374 | 0.964 | 0.264 | 1.27351479624227e-5 | 8 | gstm.3 |
| anxa1a2 | 1.4893990 | 1.334229111 | 0.855 | 0.259 | 3.00813935788586e-4 | 8 | anxa1a |
| s100a10b2 | 3.6130401 | 1.235846164 | 0.952 | 0.442 | 7.2972572290169e-41 | 8 | s100a10b |
| si:ch211-213d14.2 | 1.6809255 | 0.722264627 | 0.265 | 0.017 | 3.39496535787175e-3 | 8 | si:ch211-213d14.2 |
| krt81 | 5.5271200 | 1.050897596 | 0.928 | 0.51 | 1.11631243624258e-3 | 8 | krt8 |
| s100a10a | 2.5439705 | 0.668400697 | 0.253 | 0.019 | 5.13805740793556e-3 | 8 | s100a10a |

| | | | | | | | |
|---|---|---|---|---|---|---|---|
| krt91 | 1.4982159 | 1.074887568 | 0.47 | 0.083 | 3.02594680653698e-2 | 8 | krt91 |
| icn2 | 1.8072452 | 1.098663464 | 0.759 | 0.242 | 3.6500931578749e-24 | 8 | icn |
| cygb12 | 2.2176355 | 0.907919555 | 0.446 | 0.092 | 4.47895851451112e-2 | 8 | cygb1 |
| cavin1b | 5.5545565 | 0.816216973 | 0.373 | 0.063 | 1.12185379202734e-2 | 8 | cavin1b |
| anxa2a1 | 6.0443081 | 0.988055824 | 0.614 | 0.174 | 1.22076891456718e-2 | 8 | anxa2a |
| tmsb12 | 7.1100538 | 1.042537799 | 0.602 | 0.181 | 1.43601758480651e-2 | 8 | tmsb1 |
| ucmaa | 3.7580932 | 0.627912158 | 0.241 | 0.029 | 7.59022102599999e-2 | 8 | ucmaa |
| cldnc | 6.9257789 | 0.759240748 | 0.361 | 0.064 | 1.39879957282891e-1 | 8 | cldnc |
| krt18a.11 | 4.2159551 | 1.016480286 | 0.663 | 0.269 | 8.51496456730948e-1 | 8 | krt18a.1 |
| ppdpfa1 | 2.0033348 | 0.487014511 | 0.229 | 0.033 | 4.04613535878504e-1 | 8 | ppdpfa |
| postnb2 | 6.3779955 | 0.882583557 | 0.53 | 0.179 | 1.28816375741084e-1 | 8 | postnb |
| krt943 | 4.3765033 | 0.830053459 | 0.711 | 0.353 | 8.83922379120076e-1 | 8 | krt94 |
| si:ch73-46j18.53 | 5.5897178 | 0.676328311 | 0.711 | 0.388 | 1.12895531943897e-1 | 8 | si:ch73-46j18.5 |
| hspb11 | 1.8366812 | 0.695433937 | 0.458 | 0.171 | 3.70954511875894e-0 | 8 | hspb1 |
| gstm.11 | 3.5565654 | 0.707118503 | 0.614 | 0.332 | 7.18319523277038e-0 | 8 | gstm.1 |
| aldh1a2 | 1.3770842 | 0.669279348 | 0.506 | 0.195 | 2.78129715559115e-0 | 8 | aldh1a2 |
| podxl | 8.2156104 | 0.600175042 | 0.337 | 0.108 | 1.65930685029879e-0 | 8 | podxl |
| ybx1 | 1.5560941 | 0.257061165 | 0.928 | 0.859 | 3.14284344887047e-0 | 8 | ybx1 |
| eif4ebp3l1 | 5.0862911 | 0.66889063 | 0.41 | 0.196 | 0.000102728 | 8 | eif4ebp3l |
| si:ch211-105c13.31 | 1.4691205 | 0.527261322 | 0.289 | 0.105 | 0.000296718 | 8 | si:ch211-105c13.3 |
| zgc:110182 | 2.1509522 | 0.472052546 | 0.229 | 0.078 | 0.004344278 | 8 | zgc:110182 |
| fgf24 | 4.2734549 | 0.364432264 | 0.133 | 0.031 | 0.008631097 | 8 | fgf24 |
| adh8a1 | 6.5645843 | 0.448488945 | 0.205 | 0.065 | 0.013258491 | 8 | adh8a |
| si:dkey-16p21.8 | 8.4119994 | 0.333951295 | 0.133 | 0.032 | 0.016989715 | 8 | si:dkey-16p21.8 |
| selenow12 | 4.5769782 | 0.361257061 | 0.663 | 0.598 | 0.092441229 | 8 | selenow1 |
| fth1a2 | 5.6260125 | 0.421570112 | 0.614 | 0.45 | 0.113628576 | 8 | fth1a |
| cavin2b | 6.6263779 | 0.502188354 | 0.301 | 0.145 | 0.133832956 | 8 | cavin2b |
| nupr1b | 6.7021626 | 0.266028871 | 0.12 | 0.03 | 0.135363578 | 8 | nupr1b |
| rnd3b2 | 1.7568430 | 0.390299166 | 0.229 | 0.095 | 0.354829593 | 8 | rnd3b |
| si:ch211-106h4.12 | 2.8781027 | 0.365062533 | 0.169 | 0.058 | 0.581290417 | 8 | si:ch211-106h4.12 |
| s100v22 | 4.3102924 | 0.47025636 | 0.398 | 0.249 | 0.870549768 | 8 | s100v2 |
| id13 | 5.8239250 | 0.458818428 | 0.446 | 0.272 | 1 | 8 | id1 |
| sparc3 | 6.8776737 | 0.387565645 | 0.59 | 0.399 | 1 | 8 | sparc |
| capns1a | 7.7823036 | 0.461579551 | 0.386 | 0.249 | 1 | 8 | capns1a |
| jam2b | 0.000103 | 0.31256175 | 0.12 | 0.038 | 1 | 8 | jam2b |

| | | | | | | | |
|---|---|---|---|---|---|---|---|
| si:ch211-198c19.3: | 0.000206 | 0.412386813 | 0.337 | 0.182 | 1 | 8 | si:ch211-198c19.3 |
| hdlbpa2 | 0.000207 | 0.456608058 | 0.386 | 0.267 | 1 | 8 | hdlbpa |
| tagln3 | 0.000214 | 0.441100502 | 0.265 | 0.135 | 1 | 8 | tagln |
| hmgn71 | 0.000531 | 0.295938618 | 0.59 | 0.532 | 1 | 8 | hmgn7 |
| wt1b1 | 0.00091 | 0.331392984 | 0.157 | 0.068 | 1 | 8 | wt1b |
| cebpd2 | 0.000925 | 0.375321214 | 0.41 | 0.279 | 1 | 8 | cebpd |
| gchfr | 0.001282 | 0.27439904 | 0.12 | 0.046 | 1 | 8 | gchfr |
| akt2l | 0.001619 | 0.255679133 | 0.108 | 0.04 | 1 | 8 | akt2l |
| myl9a1 | 0.001943 | 0.332677298 | 0.217 | 0.116 | 1 | 8 | myl9a |
| fn1b2 | 0.002288 | 0.407722938 | 0.289 | 0.18 | 1 | 8 | fn1b |
| lgals3b | 0.002373 | 0.408661958 | 0.301 | 0.207 | 1 | 8 | lgals3b |
| aldh9a1a.13 | 0.004237 | 0.382078632 | 0.41 | 0.342 | 1 | 8 | aldh9a1a.1 |
| hsbp1a | 0.004365 | 0.31621019 | 0.157 | 0.078 | 1 | 8 | hsbp1a |
| dbi1 | 0.004776 | 0.329901222 | 0.482 | 0.405 | 1 | 8 | dbi |
| cav1 | 0.004917 | 0.311678321 | 0.157 | 0.078 | 1 | 8 | cav1 |
| ahcy | 0.005523 | 0.270396117 | 0.205 | 0.112 | 1 | 8 | ahcy |
| si:ch73-281n10.22 | 0.006935 | 0.3286886 | 0.494 | 0.436 | 1 | 8 | si:ch73-281n10.2 |
| cldn5a2 | 0.007866 | 0.291495547 | 0.217 | 0.127 | 1 | 8 | cldn5a |
| col1a23 | 0.008653 | 0.286459821 | 0.482 | 0.331 | 1 | 8 | col1a2 |
| cldnb | 2.6260503 | 1.209954235 | 0.41 | 0.003 | 5.30383384619122e-1 | 9 | cldnb |
| tnks1bp1 | 6.5361562 | 0.804859629 | 0.359 | 0.001 | 1.32010747707206e-1 | 9 | tnks1bp1 |
| epcam | 6.7259371 | 1.327678561 | 0.526 | 0.013 | 1.35843752406425e-1 | 9 | epcam |
| wu:fb18f06 | 1.8926373 | 1.284634304 | 0.41 | 0.007 | 3.82255970330632e-1 | 9 | wu:fb18f06 |
| krt5 | 1.7531680 | 1.236015134 | 0.295 | 0.002 | 3.54087346726954e-1 | 9 | krt5 |
| si:dkey-193p11.2 | 4.2152814 | 1.173659648 | 0.282 | 0.002 | 8.51360401758265e-1 | 9 | si:dkey-193p11.2 |
| zgc:193505 | 3.3368994 | 0.940275369 | 0.269 | 0.001 | 6.73953576653636e-1 | 9 | zgc:193505 |
| tmem176l.4 | 6.1365474 | 0.770615408 | 0.308 | 0.004 | 1.23939849436723e-1 | 9 | tmem176l.4 |
| krt17 | 1.4221224 | 1.111924195 | 0.256 | 0.001 | 2.87226079879986e-9 | 9 | krt17 |
| icn21 | 9.9330052 | 1.263898389 | 0.282 | 0.003 | 2.00616907147494e-9 | 9 | icn2 |
| si:dkey-247k7.2 | 6.7240620 | 0.990918728 | 0.256 | 0.002 | 1.35805881785802e-9 | 9 | si:dkey-247k7.2 |
| spint2 | 6.2537852 | 0.998847815 | 0.423 | 0.015 | 1.26307701350177e-9 | 9 | spint2 |
| anxa1c | 6.3641152 | 0.767493018 | 0.244 | 0.001 | 1.28536035046416e-9 | 9 | anxa1c |
| selenow2b | 2.4456388 | 0.410955975 | 0.218 | 0 | 4.9394567928339e-93 | 9 | selenow2b |
| znf185 | 5.3501005 | 0.5769216 | 0.244 | 0.002 | 1.08055981030616e-8 | 9 | znf185 |
| si:dkey-87o1.2 | 1.2188417 | 0.693435227 | 0.295 | 0.005 | 2.46169468046255e-8 | 9 | si:dkey-87o1.2 |

| | | | | | | | |
|---|---|---|---|---|---|---|---|
| si:dkeyp-86h10.3 | 2.3637701 | 0.596057801 | 0.231 | 0.001 | 4.77410656257017e-8 | 9 | si:dkeyp-86h10.3 |
| soul2 | 2.6553138 | 0.628288474 | 0.231 | 0.001 | 5.36293730797481e-8 | 9 | soul2 |
| cldne | 1.1085131 | 0.630383165 | 0.205 | 0 | 2.23886401804331e-8 | 9 | cldne |
| evpla | 1.1085131 | 0.480164338 | 0.205 | 0 | 2.23886401804331e-8 | 9 | evpla |
| hrc | 7.7883907 | 0.605928165 | 0.244 | 0.002 | 1.57302127719408e-8 | 9 | hrc |
| ppl | 1.4063980 | 0.605125538 | 0.231 | 0.002 | 2.8405022416428e-83 | 9 | ppl |
| zgc:111983 | 4.9715522 | 0.828812432 | 0.192 | 0 | 1.00410441422622e-8 | 9 | zgc:111983 |
| abcb5 | 4.9715522 | 0.524871642 | 0.192 | 0 | 1.00410441422622e-8 | 9 | abcb5 |
| pkp3a | 3.4104466 | 0.575692519 | 0.269 | 0.004 | 6.88807909609956e-8 | 9 | pkp3a |
| rhcgl1 | 1.7722007 | 0.637150629 | 0.192 | 0.001 | 3.57931383213744e-7 | 9 | rhcgl1 |
| cldnh | 2.2068188 | 0.548525751 | 0.179 | 0 | 4.45711205057064e-7 | 9 | cldnh |
| anxa3a | 2.2068188 | 0.349381646 | 0.179 | 0 | 4.45711205057064e-7 | 9 | anxa3a |
| tmem45b | 2.5483873 | 0.333681537 | 0.192 | 0.001 | 5.14697795409724e-7 | 9 | tmem45b |
| zgc:153284 | 7.7966188 | 0.764829482 | 0.269 | 0.005 | 1.5746831168668e-75 | 9 | zgc:153284 |
| anxa1b | 9.7943355 | 0.67194986 | 0.205 | 0.002 | 1.97816195049892e-7 | 9 | anxa1b |
| oclna | 6.0452710 | 0.462454117 | 0.231 | 0.003 | 1.22096338594845e-7 | 9 | oclna |
| eps8l1a | 7.5378722 | 0.340942272 | 0.179 | 0.001 | 1.5224240616117e-70 | 9 | eps8l1a |
| elf3 | 8.5033859 | 0.390463739 | 0.179 | 0.001 | 1.71742887016953e-7 | 9 | elf3 |
| zgc:174935 | 9.6986515 | 0.385934124 | 0.167 | 0 | 1.95883664382819e-7 | 9 | zgc:174935 |
| cldnf | 9.6986515 | 0.358945847 | 0.167 | 0 | 1.95883664382819e-7 | 9 | cldnf |
| mid1ip1a | 7.0497149 | 0.916851069 | 0.282 | 0.008 | 1.42383092773354e-6 | 9 | mid1ip1a |
| tmem54a | 3.5390472 | 0.31888024 | 0.205 | 0.002 | 7.14781373406247e-6 | 9 | tmem54a |
| krt1-19d | 3.9491542 | 0.810153835 | 0.192 | 0.002 | 7.97610679088718e-6 | 9 | krt1-19d |
| si:ch73-52f15.5 | 5.4854796 | 0.461033358 | 0.179 | 0.001 | 1.10790232610205e-6 | 9 | si:ch73-52f15.5 |
| ccl20b | 1.1299690 | 0.562431953 | 0.231 | 0.004 | 2.28219844750569e-6 | 9 | ccl20b |
| cldn7b | 1.1501492 | 0.391875489 | 0.231 | 0.004 | 2.32295642812454e-6 | 9 | cldn7b |
| elovl7a | 2.3424311 | 0.532473698 | 0.256 | 0.006 | 4.73100814018797e-6 | 9 | elovl7a |
| CABZ01027551.1 | 3.1493752 | 0.442566098 | 0.167 | 0.001 | 6.36079318202905e-6 | 9 | CABZ01027551.1 |
| si:ch211-125o16.4 | 4.2218834 | 0.443391828 | 0.154 | 0 | 8.52693806955553e-6 | 9 | si:ch211-125o16.4 |
| si:dkey-248g15.3 | 4.3004808 | 0.546767576 | 0.167 | 0.001 | 8.68568117249449e-6 | 9 | si:dkey-248g15.3 |
| LOC101882729 | 4.3004808 | 0.403114454 | 0.167 | 0.001 | 8.68568117249449e-6 | 9 | LOC101882729 |
| dhrs13a.2 | 2.9590887 | 0.650874384 | 0.192 | 0.002 | 5.97647155429168e-6 | 9 | dhrs13a.2 |
| scel | 1.8858941 | 0.546223099 | 0.167 | 0.001 | 3.80894037963854e-6 | 9 | scel |
| si:dkey-7c18.24 | 8.5825650 | 0.95615691 | 0.192 | 0.003 | 1.7334206561836e-59 | 9 | si:dkey-7c18.24 |
| si:ch211-157c3.4 | 1.4197399 | 0.4713639 | 0.154 | 0.001 | 2.86744872670748e-5 | 9 | si:ch211-157c3.4 |

| | | | | | | | |
|---|---|---|---|---|---|---|---|
| si:ch211-217k17.9 | 1.4889918 | 0.4999787 | 0.154 | 0.001 | 3.00731686811392e-5 | 9 | si:ch211-217k17.9 |
| oclnb | 1.6377554 | 0.302519527 | 0.154 | 0.001 | 3.30777473048428e-5 | 9 | oclnb |
| CR926130.1 | 1.8212813 | 0.338034889 | 0.141 | 0 | 3.67844191734114e-5 | 9 | CR926130.1 |
| si:ch73-347e22.8 | 1.8212813 | 0.276681123 | 0.141 | 0 | 3.67844191734114e-5 | 9 | si:ch73-347e22.8 |
| gpa33a | 1.8212813 | 0.261061731 | 0.141 | 0 | 3.67844191734114e-5 | 9 | gpa33a |
| dlx3b | 1.8212813 | 0.257951628 | 0.141 | 0 | 3.67844191734114e-5 | 9 | dlx3b |
| tlcd1 | 1.8212813 | 0.250974832 | 0.141 | 0 | 3.67844191734114e-5 | 9 | tlcd1 |
| lmo7a | 1.9255203 | 0.411310391 | 0.192 | 0.003 | 3.88897338345934e-5 | 9 | lmo7a |
| si:ch211-25g7.5 | 4.5262893 | 0.380060806 | 0.167 | 0.002 | 9.14174660630903e-5 | 9 | si:ch211-25g7.5 |
| zgc:154054 | 8.8601375 | 0.333717498 | 0.154 | 0.001 | 1.78948198164439e-5 | 9 | zgc:154054 |
| erbb3a | 8.8601375 | 0.262280539 | 0.154 | 0.001 | 1.78948198164439e-5 | 9 | erbb3a |
| LOC100333235 | 9.0576755 | 0.320779061 | 0.154 | 0.001 | 1.82937872867884e-5 | 9 | LOC100333235 |
| hmgcra | 9.4660098 | 0.327784674 | 0.154 | 0.001 | 1.91185001237149e-5 | 9 | hmgcra |
| krt97 | 5.3029031 | 0.435556491 | 0.141 | 0.001 | 1.07102734517095e-5 | 9 | krt97 |
| si:ch73-204p21.2 | 5.3029031 | 0.320481319 | 0.141 | 0.001 | 1.07102734517095e-5 | 9 | si:ch73-204p21.2 |
| stx11b.1 | 6.1109627 | 0.351142651 | 0.141 | 0.001 | 1.23423113995837e-5 | 9 | stx11b.1 |
| gdpd3a | 6.4066966 | 0.299877848 | 0.141 | 0.001 | 1.2939605318059e-53 | 9 | gdpd3a |
| si:dkey-221j11.3 | 7.7913394 | 0.33527959 | 0.128 | 0 | 1.57361682758258e-5 | 9 | si:dkey-221j11.3 |
| CR854839.1 | 7.7913394 | 0.303110504 | 0.128 | 0 | 1.57361682758258e-5 | 9 | CR854839.1 |
| cldn23a | 7.7913394 | 0.299894838 | 0.128 | 0 | 1.57361682758258e-5 | 9 | cldn23a |
| st14b | 7.7913394 | 0.273054694 | 0.128 | 0 | 1.57361682758258e-5 | 9 | st14b |
| bicdl2 | 2.4211816 | 0.301519457 | 0.167 | 0.002 | 4.89006053662143e-5 | 9 | bicdl2 |
| si:ch211-202f5.3 | 3.0033224 | 0.358255807 | 0.167 | 0.002 | 6.06581035816066e-5 | 9 | si:ch211-202f5.3 |
| cd9b2 | 1.1923394 | 0.882694369 | 0.397 | 0.029 | 2.40816795832191e-5 | 9 | cd9b |
| stard14 | 2.3323224 | 0.543642483 | 0.218 | 0.006 | 4.71059171495877e-5 | 9 | stard14 |
| lye | 8.5904373 | 0.48275753 | 0.154 | 0.002 | 1.73501063625552e-5 | 9 | lye |
| cyldb | 1.5580665 | 0.305082593 | 0.154 | 0.002 | 3.1468269724816e-50 | 9 | cyldb |
| agr1 | 1.2387177 | 0.500285684 | 0.179 | 0.004 | 2.50183829085936e-4 | 9 | agr1 |
| AL929237.1 | 2.0773012 | 0.435310348 | 0.128 | 0.001 | 4.1955254000996e-48 | 9 | AL929237.1 |
| ANXA1 (1 of many) | 3.3081782 | 0.365302228 | 0.115 | 0 | 6.68152764491391e-4 | 9 | ANXA1 (1 of many) |
| hepacam2 | 3.3081782 | 0.304337046 | 0.115 | 0 | 6.68152764491391e-4 | 9 | hepacam2 |
| XLOC-041945 | 3.3081782 | 0.29245849 | 0.115 | 0 | 6.68152764491391e-4 | 9 | XLOC-041945 |
| ponzr2 | 3.3081782 | 0.280932396 | 0.115 | 0 | 6.68152764491391e-4 | 9 | ponzr2 |
| krt96 | 3.3081782 | 0.271333386 | 0.115 | 0 | 6.68152764491391e-4 | 9 | krt96 |
| zgc:110333 | 3.3081782 | 0.266554671 | 0.115 | 0 | 6.68152764491391e-4 | 9 | zgc:110333 |

| | | | | | | | |
|---|---|---|---|---|---|---|---|
| CABZ01048956.1 | 3.3081782 | 0.259277478 | 0.115 | 0 | 6.68152764491391e-4 | 9 | CABZ01048956.1 |
| zgc:100868 | 3.3081782 | 0.258677614 | 0.115 | 0 | 6.68152764491391e-4 | 9 | zgc:100868 |
| prrg2 | 6.5602376 | 0.457149102 | 0.282 | 0.014 | 1.32497120130005e-4 | 9 | prrg2 |
| tmem30b | 7.5388466 | 0.301393589 | 0.128 | 0.001 | 1.52262086252103e-4 | 9 | tmem30b |
| rhbg | 8.2131440 | 0.383752867 | 0.128 | 0.001 | 1.65880870155752e-4 | 9 | rhbg |
| hunk | 8.8923183 | 0.289849256 | 0.154 | 0.003 | 1.79598152733052e-4 | 9 | hunk |
| arrdc1a | 3.1469783 | 0.303762043 | 0.167 | 0.004 | 6.35595214315301e-4 | 9 | arrdc1a |
| mir375-2 | 8.1017144 | 0.302839559 | 0.115 | 0.001 | 1.63630327144758e-4 | 9 | mir375-2 |
| LOC100331080 | 8.6853877 | 0.317961846 | 0.115 | 0.001 | 1.75418775495678e-4 | 9 | LOC100331080 |
| krtt1c19e | 8.8890988 | 0.496194271 | 0.115 | 0.001 | 1.79533130272779e-4 | 9 | krtt1c19e |
| pof1b | 1.2117270 | 0.512373629 | 0.244 | 0.012 | 2.44732518309077e-4 | 9 | pof1b |
| zgc:92313 | 1.2095267 | 0.510230281 | 0.282 | 0.017 | 2.44288112075514e-4 | 9 | zgc:92313 |
| rassf7b | 1.1500052 | 0.358982151 | 0.154 | 0.004 | 2.32266555196337e-3 | 9 | rassf7b |
| myod1 | 2.1284946 | 0.285178104 | 0.115 | 0.001 | 4.29892069657111e-3 | 9 | myod1 |
| si:ch211-69b7.6 | 2.2198593 | 0.323428679 | 0.115 | 0.001 | 4.48344997591511e-3 | 9 | si:ch211-69b7.6 |
| lgals2b | 2.2669940 | 0.591914772 | 0.115 | 0.001 | 4.57864797981175e-3 | 9 | lgals2b |
| pou2f3 | 2.4144548 | 0.275712341 | 0.115 | 0.001 | 4.87647442125443e-3 | 9 | pou2f3 |
| si:ch211-79k12.1 | 4.6860810 | 0.454266224 | 0.218 | 0.01 | 9.46447794576027e-3 | 9 | si:ch211-79k12.1 |
| zgc:113279 | 3.0551006 | 0.272951835 | 0.103 | 0.001 | 6.17038685277046e-3 | 9 | zgc:113279 |
| cx30.3 | 3.1258475 | 0.286878282 | 0.103 | 0.001 | 6.31327437265822e-3 | 9 | cx30.3 |
| col18a1b | 1.2132180 | 0.281105096 | 0.128 | 0.002 | 2.45033648396508e-3 | 9 | col18a1b |
| s100v1 | 1.4826142 | 0.260935328 | 0.128 | 0.002 | 2.99443608998575e-3 | 9 | s100v1 |
| rassf10a | 5.3128300 | 0.28832034 | 0.141 | 0.003 | 1.07303228171087e-3 | 9 | rassf10a |
| FP017215.1 | 9.1816228 | 0.259270703 | 0.179 | 0.007 | 1.85441237586394e-3 | 9 | FP017215.1 |
| frmd4ba | 9.2472127 | 0.361477345 | 0.218 | 0.011 | 1.86765954918784e-3 | 9 | frmd4ba |
| cdh1 | 3.0573104 | 0.658822823 | 0.346 | 0.034 | 6.1748499788551e-35 | 9 | cdh1 |
| spint1a1 | 2.5410698 | 0.524855054 | 0.295 | 0.024 | 5.13219873602493e-3 | 9 | spint1a |
| b3gnt7l | 6.7692077 | 0.289417019 | 0.128 | 0.003 | 1.36717688746401e-3 | 9 | b3gnt7l |
| si:dkey-38p12.3 | 2.9900273 | 0.282750797 | 0.167 | 0.006 | 6.03895826605523e-3 | 9 | si:dkey-38p12.3 |
| st14a | 3.5440566 | 0.257900804 | 0.167 | 0.006 | 7.15793131094708e-3 | 9 | st14a |
| spaca4l | 3.6962222 | 0.486539934 | 0.179 | 0.008 | 7.46526015086387e-3 | 9 | spaca4l |
| si:dkey-96g2.1 | 5.2842135 | 0.509436473 | 0.103 | 0.001 | 1.06725261327889e-3 | 9 | si:dkey-96g2.1 |
| myf5 | 6.3565073 | 0.28221129 | 0.103 | 0.001 | 1.28382379590468e-3 | 9 | myf5 |
| si:ch1073-345a8.1 | 2.1114397 | 0.308932558 | 0.154 | 0.005 | 4.26447477555389e-3 | 9 | si:ch1073-345a8.1 |
| tnk1 | 9.7055280 | 0.250912045 | 0.167 | 0.007 | 1.9602254954747e-31 | 9 | tnk1 |

| | | | | | | | |
|---|---|---|---|---|---|---|---|
| capn9 | 1.6366566 | 0.273083805 | 0.115 | 0.002 | 3.30555548856773e-3 | 9 | capn9 |
| sh3yl1 | 2.0121482 | 0.291512207 | 0.128 | 0.003 | 4.06393590749853e-3 | 9 | sh3yl1 |
| chst6 | 2.3586787 | 0.279276023 | 0.128 | 0.003 | 4.76382355162248e-3 | 9 | chst6 |
| tm4sf4 | 1.5549820 | 0.430059251 | 0.103 | 0.002 | 3.14059719856497e-2 | 9 | tm4sf4 |
| zgc:153665 | 1.7386401 | 0.264335407 | 0.103 | 0.002 | 3.51153144843513e-2 | 9 | zgc:153665 |
| s100w | 2.5705106 | 0.559627157 | 0.244 | 0.021 | 5.1916603135021e-27 | 9 | s100w |
| si:dkeyp-1h4.6 | 3.2976102 | 0.353261485 | 0.128 | 0.004 | 6.66018350472713e-2 | 9 | si:dkeyp-1h4.6 |
| cfl1l | 6.7005091 | 0.988047379 | 0.385 | 0.058 | 1.35330183338322e-2 | 9 | cfl1l |
| rnf128a | 1.0520923 | 0.404232463 | 0.244 | 0.021 | 2.12491091520593e-2 | 9 | rnf128a |
| f2rl1.2 | 1.5371588 | 0.25130631 | 0.103 | 0.002 | 3.10459978708741e-2 | 9 | f2rl1.2 |
| elovl1a | 3.9879024 | 0.380881919 | 0.179 | 0.011 | 8.05436662355591e-2 | 9 | elovl1a |
| ankrd22 | 2.5332392 | 0.332319487 | 0.218 | 0.017 | 5.11638341345863e-2 | 9 | ankrd22 |
| bik | 1.8181345 | 0.328449871 | 0.192 | 0.014 | 3.67208626178624e-2 | 9 | bik |
| cmah | 1.8676990 | 0.267999694 | 0.141 | 0.007 | 3.7721918195212e-22 | 9 | cmah |
| xkr8.2 | 5.1453977 | 0.293902524 | 0.103 | 0.003 | 1.03921598910699e-2 | 9 | xkr8.2 |
| perp | 5.4742564 | 0.698669867 | 0.436 | 0.085 | 1.10563558262663e-2 | 9 | perp |
| arfgap3 | 8.7775041 | 0.312745151 | 0.179 | 0.013 | 1.77279251064021e-2 | 9 | arfgap3 |
| zgc:92242 | 1.1672263 | 0.587323138 | 0.282 | 0.036 | 2.35744705307902e-2 | 9 | zgc:92242 |
| aldob1 | 1.6249777 | 0.572351937 | 0.205 | 0.018 | 3.28196748357246e-2 | 9 | aldob |
| si:busm1-57f23.1 | 2.2078950 | 0.437950034 | 0.218 | 0.021 | 4.45928560972516e-2 | 9 | si:busm1-57f23.1 |
| ap1s3b | 2.6029371 | 0.352965873 | 0.192 | 0.016 | 5.25715226131368e-2 | 9 | ap1s3b |
| CABZ01079427.1 | 4.4607952 | 0.363661845 | 0.231 | 0.024 | 9.00946824297382e-2 | 9 | CABZ01079427.1 |
| nqo1 | 2.1559367 | 0.407203873 | 0.179 | 0.015 | 4.35434539817466e-1 | 9 | nqo1 |
| vwde | 4.4029871 | 0.326697202 | 0.141 | 0.009 | 8.89271311871924e-1 | 9 | vwde |
| six1b1 | 1.0234964 | 0.310702016 | 0.128 | 0.007 | 2.06715587521627e-1 | 9 | six1b |
| cbx3b | 1.1975116 | 0.384668798 | 0.192 | 0.018 | 2.41861419969405e-1 | 9 | cbx3b |
| casz1 | 1.7319687 | 0.307098394 | 0.141 | 0.009 | 3.49805718970348e-1 | 9 | casz1 |
| jupa | 1.9432769 | 0.58353227 | 0.423 | 0.087 | 3.92483640237147e-1 | 9 | jupa |
| mgll | 2.2074031 | 0.257668269 | 0.141 | 0.009 | 4.45829221092386e-1 | 9 | mgll |
| gipc2 | 2.3973472 | 0.338918142 | 0.192 | 0.018 | 4.84192229149789e-1 | 9 | gipc2 |
| notch3 | 2.0666912 | 0.449044949 | 0.244 | 0.031 | 4.17409631311145e-1 | 9 | notch3 |
| zgc:92380 | 2.5761485 | 0.453931316 | 0.154 | 0.012 | 5.20304731599382e-1 | 9 | zgc:92380 |
| si:ch211-207n23.2 | 5.8907336 | 0.705200023 | 0.179 | 0.018 | 1.18975148427226e-1 | 9 | si:ch211-207n23.2 |
| mgst2 | 2.0705865 | 0.251379263 | 0.154 | 0.013 | 4.18196357976349e-1 | 9 | mgst2 |
| dsg2.1 | 2.3298998 | 0.285693329 | 0.154 | 0.013 | 4.70569879991713e-1 | 9 | dsg2.1 |

| | | | | | | | |
|---|---|---|---|---|---|---|---|
| phlda2 | 4.4563171 | 0.597654369 | 0.321 | 0.058 | 9.00042373183857e-1 | 9 | phlda2 |
| ptprfb | 6.7387805 | 0.313651003 | 0.218 | 0.027 | 1.36103151043662e-1 | 9 | ptprfb |
| tdh | 1.9584952 | 0.570733373 | 0.333 | 0.066 | 3.95557292687526e-1 | 9 | tdh |
| f11r.11 | 2.5548393 | 0.586098466 | 0.397 | 0.095 | 5.16000899243089e-1 | 9 | f11r.1 |
| CABZ01092156.1 | 3.0518632 | 0.279706305 | 0.205 | 0.027 | 6.16384820259041e-1 | 9 | CABZ01092156.1 |
| arhgef51 | 8.9284747 | 0.426590364 | 0.256 | 0.044 | 1.80328405430021e-1 | 9 | arhgef5 |
| ezrb1 | 2.5922042 | 0.412061256 | 0.295 | 0.059 | 5.23547493556114e-1 | 9 | ezrb |
| xiap | 3.1474360 | 0.40602819 | 0.269 | 0.05 | 6.35687650211048e-1 | 9 | xiap |
| si:dkey-85k7.7 | 5.2726623 | 0.26784658 | 0.141 | 0.014 | 1.06491960951024e-1 | 9 | si:dkey-85k7.7 |
| sec23b | 1.1906550 | 0.454763405 | 0.205 | 0.032 | 2.40476602445743e-1 | 9 | sec23b |
| fuca1.1 | 1.7510284 | 0.443603018 | 0.269 | 0.056 | 3.53655214897876e-1 | 9 | fuca1.1 |
| rasgef1ba | 1.8921308 | 0.306899594 | 0.218 | 0.037 | 3.82153671365202e-1 | 9 | rasgef1ba |
| fabp3 | 2.3128556 | 0.617356272 | 0.5 | 0.169 | 4.67127450662936e-1 | 9 | fabp3 |
| dspa | 2.9758781 | 0.336702703 | 0.231 | 0.042 | 6.01038112102453e-1 | 9 | dspa |
| fxyd1 | 5.4226058 | 0.57112511 | 0.359 | 0.098 | 1.09520369803879e-0 | 9 | fxyd1 |
| si:dkey-16p21.81 | 6.0618947 | 0.299044174 | 0.192 | 0.03 | 1.22432088133911e-0 | 9 | si:dkey-16p21.8 |
| krt911 | 1.2107343 | 1.138613183 | 0.321 | 0.09 | 2.44532009774498e-0 | 9 | krt91 |
| hsd11b1la | 1.4533029 | 0.271164254 | 0.167 | 0.024 | 2.93523603336285e-0 | 9 | hsd11b1la |
| icn3 | 2.4079903 | 1.064373668 | 0.526 | 0.253 | 4.86341802956931e-0 | 9 | icn |
| rgs22 | 3.2135938 | 0.5200254 | 0.205 | 0.037 | 6.49049557049583e-0 | 9 | rgs2 |
| tead3b | 3.4918746 | 0.25634958 | 0.179 | 0.028 | 7.05253921392532e-0 | 9 | tead3b |
| si:ch211-137i24.10 | 5.3383936 | 0.366890023 | 0.115 | 0.012 | 1.07819535664528e-0 | 9 | si:ch211-137i24.10 |
| pam | 6.5742431 | 0.350886828 | 0.192 | 0.033 | 1.32779987969602e-0 | 9 | pam |
| tmsb | 7.7792826 | 0.523758807 | 0.244 | 0.052 | 1.57118172462426e-0 | 9 | tmsb |
| si:ch211-251p5.5 | 1.2365459 | 0.30164835 | 0.167 | 0.026 | 2.49745177615623e-0 | 9 | si:ch211-251p5.5 |
| mpc1 | 1.8107681 | 0.374908987 | 0.282 | 0.068 | 3.65720848193345e-0 | 9 | mpc1 |
| net11 | 2.7161955 | 0.391686928 | 0.308 | 0.078 | 5.48590022354909e-0 | 9 | net1 |
| ank3b1 | 8.0850800 | 0.276052392 | 0.205 | 0.039 | 1.63294362685773e-0 | 9 | ank3b |
| mid1ip1b | 1.5852994 | 0.385523961 | 0.256 | 0.061 | 3.20182936429953e-0 | 9 | mid1ip1b |
| ak1 | 4.8186409 | 0.453680988 | 0.423 | 0.146 | 9.73220912169576e-0 | 9 | ak1 |
| dsc2l | 5.2651405 | 0.277392379 | 0.179 | 0.033 | 1.06340043424008e-0 | 9 | dsc2l |
| wu:fi04e12 | 7.4353494 | 0.323803484 | 0.154 | 0.026 | 1.5017175347099e-06 | 9 | wu:fi04e12 |
| krt41 | 7.4866409 | 0.942624345 | 0.308 | 0.097 | 1.51207686863998e-0 | 9 | krt4 |
| tmbim4 | 1.7016418 | 0.3218761 | 0.244 | 0.06 | 3.43680597923529e-0 | 9 | tmbim4 |
| cxadr | 2.2093597 | 0.275537859 | 0.167 | 0.031 | 4.46224389976694e-0 | 9 | cxadr |

| | | | | | | | |
|---|---|---|---|---|---|---|---|
| slc20a1b1 | 2.3476155 | 0.338067607 | 0.244 | 0.061 | 4.74147922723074e-0 | 9 | slc20a1b |
| plecb | 2.5696496 | 0.388334106 | 0.244 | 0.062 | 5.18992148699537e-0 | 9 | plecb |
| cavin2a | 2.8193317 | 0.289882004 | 0.167 | 0.031 | 5.6942042873643e-06 | 9 | cavin2a |
| tp53inp1 | 3.2987146 | 0.348418875 | 0.256 | 0.067 | 6.66241393177446e-0 | 9 | tp53inp1 |
| tax1bp1b | 4.2052204 | 0.386621285 | 0.321 | 0.1 | 8.49328371565739e-0 | 9 | tax1bp1b |
| mat2aa | 9.8891046 | 0.343101526 | 0.308 | 0.093 | 1.99730246785184e-0 | 9 | mat2aa |
| si:dkey-204f11.64 | 1.0121327 | 0.30939066 | 0.205 | 0.048 | 2.04420448893873e-0 | 9 | si:dkey-204f11.64 |
| vps28 | 1.1129660 | 0.321011403 | 0.359 | 0.118 | 2.24785756710468e-0 | 9 | vps28 |
| btbd10b | 2.8348408 | 0.405282828 | 0.308 | 0.1 | 5.72552797976095e-0 | 9 | btbd10b |
| rpz | 3.3386114 | 0.294053972 | 0.179 | 0.039 | 6.7429934814503e-05 | 9 | rpz |
| s100v23 | 4.6270836 | 0.593991772 | 0.5 | 0.245 | 9.34532087718258e-0 | 9 | s100v2 |
| rnasekb | 4.8406108 | 0.322974029 | 0.192 | 0.046 | 9.77658169957262e-0 | 9 | rnasekb |
| sox11a1 | 5.0119860 | 0.428720982 | 0.244 | 0.07 | 0.000101227 | 9 | sox11a |
| si:ch211-166a6.5 | 5.8159300 | 0.330305801 | 0.231 | 0.062 | 0.000117464 | 9 | si:ch211-166a6.5 |
| galnt7 | 5.8191586 | 0.342356123 | 0.231 | 0.063 | 0.00011753 | 9 | galnt7 |
| pleca1 | 6.5721312 | 0.298359513 | 0.269 | 0.077 | 0.000132737 | 9 | pleca |
| tcnbb1 | 7.5699857 | 0.371568148 | 0.154 | 0.031 | 0.000152891 | 9 | tcnbb |
| tuba8l22 | 1.0566453 | 0.356181787 | 0.308 | 0.101 | 0.000213411 | 9 | tuba8l2 |
| CT573383.1 | 1.2276642 | 0.450737835 | 0.115 | 0.019 | 0.000247951 | 9 | CT573383.1 |
| exosc1 | 1.3224965 | 0.258873783 | 0.218 | 0.058 | 0.000267105 | 9 | exosc1 |
| b3gnt2l | 1.5144386 | 0.252991828 | 0.128 | 0.023 | 0.000305871 | 9 | b3gnt2l |
| cadm2a1 | 1.5935990 | 0.283048256 | 0.179 | 0.041 | 0.000321859 | 9 | cadm2a |
| ucp22 | 1.9359684 | 0.516188734 | 0.692 | 0.419 | 0.000391008 | 9 | ucp2 |
| nectin1b | 4.1882882 | 0.276492521 | 0.218 | 0.06 | 0.000845909 | 9 | nectin1b |
| tspan9a | 4.2266203 | 0.304174374 | 0.141 | 0.029 | 0.000853651 | 9 | tspan9a |
| reep51 | 5.3843475 | 0.428058418 | 0.526 | 0.26 | 0.001087477 | 9 | reep5 |
| hopx | 6.6193451 | 0.433563406 | 0.218 | 0.065 | 0.001336909 | 9 | hopx |
| gabarapl2 | 1.0816085 | 0.348996648 | 0.308 | 0.112 | 0.002184525 | 9 | gabarapl2 |
| ccnd1 | 1.1521071 | 0.36374153 | 0.282 | 0.099 | 0.002326911 | 9 | ccnd1 |
| smarcd1 | 1.2097872 | 0.26324691 | 0.256 | 0.082 | 0.002443407 | 9 | smarcd1 |
| reep3b | 1.4066815 | 0.287228303 | 0.269 | 0.089 | 0.002841075 | 9 | reep3b |
| sqstm1 | 1.4939707 | 0.335814038 | 0.244 | 0.078 | 0.003017373 | 9 | sqstm1 |
| cav2 | 1.9785386 | 0.257155528 | 0.154 | 0.036 | 0.003996054 | 9 | cav2 |
| cox7a2l1 | 2.4593367 | 0.34442284 | 0.423 | 0.185 | 0.004967122 | 9 | cox7a2l |
| chmp4bb | 2.4928668 | 0.311757597 | 0.282 | 0.098 | 0.005034843 | 9 | chmp4bb |

| | | | | | | | |
|---|---|---|---|---|---|---|---|
| mgst1.2 | 2.6629368 | 0.43143756 | 0.397 | 0.176 | 0.005378334 | 9 | mgst1.2 |
| chd4b | 2.7180581 | 0.351788169 | 0.346 | 0.137 | 0.005489662 | 9 | chd4b |
| mpc2 | 4.9354978 | 0.287258499 | 0.308 | 0.116 | 0.009968225 | 9 | mpc2 |
| bcl2l13 | 5.5929937 | 0.276482105 | 0.167 | 0.044 | 0.011296169 | 9 | bcl2l13 |
| chmp2bb1 | 5.9632894 | 0.300404833 | 0.244 | 0.082 | 0.012044056 | 9 | chmp2bb |
| psmd13 | 6.1402187 | 0.30534947 | 0.359 | 0.147 | 0.0124014 | 9 | psmd13 |
| cnpy1 | 6.5222054 | 0.39738994 | 0.282 | 0.105 | 0.013172898 | 9 | cnpy1 |
| mgst3b2 | 1.0889562 | 0.377121616 | 0.564 | 0.296 | 0.021993649 | 9 | mgst3b |
| ctsf | 1.1284741 | 0.289888099 | 0.218 | 0.07 | 0.022791792 | 9 | ctsf |
| yipf4 | 1.1475932 | 0.281073675 | 0.321 | 0.126 | 0.02317794 | 9 | yipf4 |
| tuba1a | 1.1528463 | 0.270994944 | 0.141 | 0.034 | 0.023284037 | 9 | tuba1a |
| tubb2b | 1.2960993 | 0.411880472 | 0.577 | 0.316 | 0.026177318 | 9 | tubb2b |
| vdac2 | 1.3980990 | 0.401046387 | 0.705 | 0.44 | 0.028237407 | 9 | vdac2 |
| tmem165 | 1.7101881 | 0.28793607 | 0.244 | 0.086 | 0.034540669 | 9 | tmem165 |
| pfn2 | 1.7812638 | 0.356585301 | 0.603 | 0.313 | 0.035976186 | 9 | pfn2 |
| calr | 1.7820645 | 0.373536064 | 0.333 | 0.145 | 0.035992358 | 9 | calr |
| tmed71 | 1.8451222 | 0.362588217 | 0.564 | 0.302 | 0.037265934 | 9 | tmed7 |
| arpc1a | 2.0078365 | 0.288702638 | 0.359 | 0.149 | 0.040552275 | 9 | arpc1a |
| eml2 | 2.1083470 | 0.277986499 | 0.128 | 0.031 | 0.042582286 | 9 | eml2 |
| tmem59 | 2.2313124 | 0.378276016 | 0.423 | 0.206 | 0.045065817 | 9 | tmem59 |
| crema1 | 2.5423436 | 0.398948185 | 0.333 | 0.144 | 0.051347714 | 9 | crema |
| luc7l | 3.1637256 | 0.270067039 | 0.205 | 0.067 | 0.063897767 | 9 | luc7l |
| kras | 3.4227770 | 0.250895422 | 0.192 | 0.06 | 0.069129828 | 9 | kras |
| tjp2b | 3.5491821 | 0.253142105 | 0.231 | 0.078 | 0.071682832 | 9 | tjp2b |
| cnbpa | 3.5646186 | 0.292270625 | 0.5 | 0.248 | 0.071994604 | 9 | cnbpa |
| lyrm4 | 3.5757876 | 0.262918224 | 0.295 | 0.116 | 0.072220182 | 9 | lyrm4 |
| lsm12b | 3.5919043 | 0.326800032 | 0.513 | 0.267 | 0.072545692 | 9 | lsm12b |
| cd991 | 3.9205677 | 0.293247835 | 0.167 | 0.05 | 0.079183707 | 9 | cd99 |
| tegt1 | 3.9374759 | 0.371522699 | 0.628 | 0.357 | 0.079525201 | 9 | tegt |
| si:dkey-23i12.5 | 4.6342180 | 0.30962908 | 0.333 | 0.145 | 0.093597302 | 9 | si:dkey-23i12.5 |
| arhgef1b1 | 5.5594471 | 0.316275337 | 0.295 | 0.121 | 0.112284154 | 9 | arhgef1b |
| CABZ01072614.13 | 5.5988338 | 0.271306269 | 0.385 | 0.171 | 0.113079648 | 9 | CABZ01072614.1 |
| csnk1a11 | 5.6123979 | 0.305444177 | 0.526 | 0.265 | 0.113353602 | 9 | csnk1a1 |
| CU929070.1 | 5.8151121 | 0.316933412 | 0.231 | 0.085 | 0.11744782 | 9 | CU929070.1 |
| ccl19a.1 | 7.0064076 | 0.330663624 | 0.115 | 0.027 | 0.141508415 | 9 | ccl19a.1 |

| | | | | | | | |
|---|---|---|---|---|---|---|---|
| slc3a2b1 | 7.2742643 | 0.305360528 | 0.372 | 0.168 | 0.146918316 | 9 | slc3a2b |
| arf2b | 7.5973495 | 0.306808105 | 0.423 | 0.204 | 0.153443669 | 9 | arf2b |
| stk35 | 8.3775507 | 0.286502465 | 0.244 | 0.093 | 0.169201392 | 9 | stk35 |
| ddt | 8.6172855 | 0.265000972 | 0.321 | 0.136 | 0.174043317 | 9 | ddt |
| ppp2cb | 8.7890825 | 0.288510959 | 0.474 | 0.236 | 0.177513101 | 9 | ppp2cb |
| s100u | 8.8474101 | 0.266152872 | 0.192 | 0.064 | 0.178691142 | 9 | s100u |
| sf3a2 | 8.9920621 | 0.302962837 | 0.359 | 0.165 | 0.181612679 | 9 | sf3a2 |
| si:dkey-28b4.71 | 9.8281870 | 0.294459009 | 0.41 | 0.191 | 0.198499894 | 9 | si:dkey-28b4.7 |
| vkorc1l1 | 1.0016677 | 0.253627208 | 0.192 | 0.064 | 0.202306827 | 9 | vkorc1l1 |
| ndufs5 | 1.1071456 | 0.3085389 | 0.449 | 0.232 | 0.223610203 | 9 | ndufs5 |
| abcf1 | 1.3560965 | 0.334627212 | 0.385 | 0.187 | 0.273890828 | 9 | abcf1 |
| si:ch211-286b5.51 | 1.3679354 | 0.330209705 | 0.436 | 0.222 | 0.276281929 | 9 | si:ch211-286b5.5 |
| fbp1a | 1.4378225 | 0.279042129 | 0.179 | 0.06 | 0.290397026 | 9 | fbp1a |
| ndufb9 | 1.6294425 | 0.322494054 | 0.564 | 0.32 | 0.329098516 | 9 | ndufb9 |
| hyou11 | 1.7064068 | 0.254800665 | 0.218 | 0.079 | 0.344643001 | 9 | hyou1 |
| ppifb1 | 1.8066782 | 0.307404093 | 0.346 | 0.163 | 0.364894798 | 9 | ppifb |
| nupr1a | 1.8550170 | 0.272357324 | 0.359 | 0.167 | 0.374657787 | 9 | nupr1a |
| srsf5a1 | 1.9483988 | 0.320062363 | 0.667 | 0.383 | 0.393518106 | 9 | srsf5a |
| cst14b.12 | 1.9698559 | 0.335991247 | 0.269 | 0.113 | 0.397851802 | 9 | cst14b.1 |
| si:ch73-281n10.23 | 2.0250871 | 0.34219792 | 0.718 | 0.428 | 0.409006853 | 9 | si:ch73-281n10.2 |
| nucb2b | 2.1157690 | 0.269564767 | 0.141 | 0.041 | 0.427321871 | 9 | nucb2b |
| anp32a1 | 2.1515137 | 0.334987281 | 0.526 | 0.303 | 0.434541229 | 9 | anp32a |
| matr3l1.1 | 2.6108808 | 0.261403587 | 0.256 | 0.105 | 0.527319606 | 9 | matr3l1.1 |
| ywhah | 2.6161574 | 0.297071237 | 0.295 | 0.129 | 0.528385315 | 9 | ywhah |
| cntfr | 2.7117749 | 0.250826937 | 0.141 | 0.041 | 0.547697183 | 9 | cntfr |
| surf4l | 3.1704338 | 0.261542064 | 0.205 | 0.076 | 0.640332533 | 9 | surf4l |
| prelid3b | 3.3070709 | 0.319392583 | 0.359 | 0.175 | 0.667929121 | 9 | prelid3b |
| smdt1b | 3.3321920 | 0.321160703 | 0.449 | 0.245 | 0.673002829 | 9 | smdt1b |
| atp5pf1 | 3.3640698 | 0.337842853 | 0.667 | 0.426 | 0.67944118 | 9 | atp5pf |
| mcl1b3 | 3.6374716 | 0.429853598 | 0.5 | 0.314 | 0.734660149 | 9 | mcl1b |
| gpm6ab | 3.7432541 | 0.257627915 | 0.103 | 0.025 | 0.756025035 | 9 | gpm6ab |
| atp1b1a1 | 3.8737664 | 0.300668075 | 0.526 | 0.29 | 0.782384615 | 9 | atp1b1a |
| ndufa3 | 4.0613393 | 0.291868445 | 0.449 | 0.24 | 0.820268712 | 9 | ndufa3 |
| tefa1 | 4.7789122 | 0.258354046 | 0.321 | 0.147 | 0.9651969 | 9 | tefa |
| hmga1a1 | 4.8766194 | 0.369072563 | 0.641 | 0.432 | 0.984930835 | 9 | hmga1a |

| | | | | | | | |
|---|---|---|---|---|---|---|---|
| si:ch1073-429i10.3 | 4.9868143 | 0.322230957 | 0.577 | 0.347 | 1 | 9 | si:ch1073-429i10.3.1 |
| zgc:1748631 | 5.9759609 | 0.30613703 | 0.333 | 0.164 | 1 | 9 | zgc:174863 |
| dedd13 | 6.0549730 | 0.289865729 | 0.41 | 0.209 | 1 | 9 | dedd1 |
| atp5f1b | 6.0967022 | 0.325664478 | 0.782 | 0.526 | 1 | 9 | atp5f1b |
| si:dkey-42i9.41 | 6.2050189 | 0.279780671 | 0.385 | 0.195 | 1 | 9 | si:dkey-42i9.4 |
| ppp3ccb | 6.2828297 | 0.272498115 | 0.295 | 0.133 | 1 | 9 | ppp3ccb |
| chmp1b | 6.3734101 | 0.265782055 | 0.372 | 0.184 | 1 | 9 | chmp1b |
| NC-002333.43 | 6.4508429 | 0.363412386 | 0.846 | 0.58 | 1 | 9 | NC-002333.4 |
| flot2a1 | 6.9148172 | 0.253548509 | 0.321 | 0.149 | 1 | 9 | flot2a |
| erp44 | 7.2468733 | 0.253896697 | 0.308 | 0.144 | 1 | 9 | erp44 |
| midn3 | 7.3828711 | 0.34682085 | 0.346 | 0.178 | 1 | 9 | midn |
| ppa1b | 7.6731133 | 0.250480941 | 0.244 | 0.104 | 1 | 9 | ppa1b |
| fam32a | 7.6806122 | 0.297155761 | 0.538 | 0.313 | 1 | 9 | fam32a |
| ndufa6 | 7.9383744 | 0.256880579 | 0.538 | 0.309 | 1 | 9 | ndufa6 |
| sgk1 | 8.9368146 | 0.252535508 | 0.192 | 0.073 | 1 | 9 | sgk1 |
| rbm51 | 9.0629904 | 0.250268236 | 0.244 | 0.102 | 1 | 9 | rbm5 |
| atp5if1b2 | 9.8170283 | 0.271034055 | 0.487 | 0.271 | 1 | 9 | atp5if1b |
| ndufa5 | 0.000106 | 0.253146001 | 0.436 | 0.236 | 1 | 9 | ndufa5 |
| ndufb2 | 0.00011 | 0.279786523 | 0.436 | 0.238 | 1 | 9 | ndufb2 |
| selenot2 | 0.000111 | 0.288179169 | 0.244 | 0.11 | 1 | 9 | selenot2 |
| vps4b | 0.000113 | 0.271472545 | 0.295 | 0.139 | 1 | 9 | vps4b |
| klf2a1 | 0.000114 | 0.495041522 | 0.321 | 0.179 | 1 | 9 | klf2a |
| calr3a1 | 0.000115 | 0.336714159 | 0.423 | 0.233 | 1 | 9 | calr3a |
| gpatch8 | 0.000117 | 0.269830041 | 0.244 | 0.106 | 1 | 9 | gpatch8 |
| tob1b1 | 0.000121 | 0.297489318 | 0.308 | 0.148 | 1 | 9 | tob1b |
| csnk1da | 0.000125 | 0.265207968 | 0.244 | 0.106 | 1 | 9 | csnk1da |
| ndufb5 | 0.000131 | 0.298874079 | 0.385 | 0.21 | 1 | 9 | ndufb5 |
| fam162a | 0.000139 | 0.269898967 | 0.244 | 0.109 | 1 | 9 | fam162a |
| si:ch211-39i22.13 | 0.000146 | 0.298832795 | 0.359 | 0.18 | 1 | 9 | si:ch211-39i22.1 |
| hmgn3 | 0.00015 | 0.25890346 | 0.333 | 0.164 | 1 | 9 | hmgn3 |
| btg21 | 0.000159 | 0.289572378 | 0.462 | 0.257 | 1 | 9 | btg2 |
| ldhba2 | 0.000181 | 0.302439917 | 0.551 | 0.335 | 1 | 9 | ldhba |
| cdk2ap21 | 0.000182 | 0.266610773 | 0.269 | 0.125 | 1 | 9 | cdk2ap2 |
| psma1 | 0.000215 | 0.273551284 | 0.641 | 0.391 | 1 | 9 | psma1 |
| gnb1b1 | 0.000237 | 0.260093927 | 0.577 | 0.335 | 1 | 9 | gnb1b |

| | | | | | | | |
|---|---|---|---|---|---|---|---|
| NDUFB1 | 0.000275 | 0.256697708 | 0.474 | 0.275 | 1 | 9 | NDUFB1 |
| cox7b2 | 0.00028 | 0.310267758 | 0.731 | 0.501 | 1 | 9 | cox7b |
| ralbp1 | 0.000295 | 0.255991596 | 0.244 | 0.112 | 1 | 9 | ralbp1 |
| jun3 | 0.000311 | 0.336545126 | 0.603 | 0.381 | 1 | 9 | jun |
| atp5mf1 | 0.000331 | 0.25008967 | 0.513 | 0.304 | 1 | 9 | atp5mf |
| nop532 | 0.000355 | 0.250392568 | 0.59 | 0.349 | 1 | 9 | nop53 |
| spry21 | 0.000367 | 0.264429214 | 0.256 | 0.118 | 1 | 9 | spry2 |
| capns1a1 | 0.000461 | 0.364673828 | 0.397 | 0.248 | 1 | 9 | capns1a |
| chmp2a | 0.000462 | 0.254097547 | 0.346 | 0.188 | 1 | 9 | chmp2a |
| gnb1a | 0.000465 | 0.322366536 | 0.423 | 0.263 | 1 | 9 | gnb1a |
| marcksl1b1 | 0.000469 | 0.365182559 | 0.449 | 0.279 | 1 | 9 | marcksl1b |
| ahnak2 | 0.000491 | 0.294396399 | 0.359 | 0.199 | 1 | 9 | ahnak |
| sdf2l1 | 0.000495 | 0.300565517 | 0.192 | 0.084 | 1 | 9 | sdf2l1 |
| psmb4 | 0.000529 | 0.275949461 | 0.462 | 0.283 | 1 | 9 | psmb4 |
| calm2b | 0.000529 | 0.261631572 | 0.705 | 0.442 | 1 | 9 | calm2b |
| anxa4 | 0.000586 | 0.466429498 | 0.385 | 0.236 | 1 | 9 | anxa4 |
| zgc:91910 | 0.000656 | 0.255551806 | 0.487 | 0.297 | 1 | 9 | zgc:91910 |
| ier2b2 | 0.000689 | 0.335623913 | 0.487 | 0.324 | 1 | 9 | ier2b |
| slc38a5b3 | 0.000698 | 0.279593318 | 0.359 | 0.203 | 1 | 9 | slc38a5b |
| ywhabl1 | 0.000729 | 0.277502247 | 0.628 | 0.438 | 1 | 9 | ywhabl |
| calm1b2 | 0.000797 | 0.304664537 | 0.282 | 0.146 | 1 | 9 | calm1b |
| fth1a3 | 0.001018 | 0.26646707 | 0.667 | 0.448 | 1 | 9 | fth1a |
| COX5B1 | 0.001051 | 0.262144061 | 0.667 | 0.477 | 1 | 9 | COX5B |
| cebpd3 | 0.00128 | 0.307552315 | 0.449 | 0.278 | 1 | 9 | cebpd |
| b2ml2 | 0.001539 | 0.270579658 | 0.641 | 0.443 | 1 | 9 | b2ml |
| pycard2 | 0.001628 | 0.28744766 | 0.269 | 0.144 | 1 | 9 | pycard |
| atf4b1 | 0.001708 | 0.256073379 | 0.564 | 0.358 | 1 | 9 | atf4b |
| si:dkey-56m19.51 | 0.001817 | 0.298344394 | 0.205 | 0.101 | 1 | 9 | si:dkey-56m19.5 |
| gadd45ba2 | 0.001915 | 0.254570016 | 0.41 | 0.25 | 1 | 9 | gadd45ba |
| ier5 | 0.001918 | 0.284767314 | 0.346 | 0.209 | 1 | 9 | ier5 |
| tmsb13 | 0.002291 | 0.475452921 | 0.308 | 0.194 | 1 | 9 | tmsb1 |
| hspa51 | 0.002388 | 0.256936392 | 0.372 | 0.22 | 1 | 9 | hspa5 |
| ckap41 | 0.002845 | 0.252571066 | 0.269 | 0.148 | 1 | 9 | ckap4 |
| h2afx1 | 0.003061 | 0.251067415 | 0.564 | 0.378 | 1 | 9 | h2afx1 |
| selenow13 | 0.003674 | 0.27856266 | 0.769 | 0.594 | 1 | 9 | selenow1 |

| | | | | | | | |
|---|---|---|---|---|---|---|---|
| qkia1 | 0.003904 | 0.298144245 | 0.5 | 0.359 | 1 | 9 | qkia |
| agr2 | 0.005143 | 0.3026328 | 0.103 | 0.039 | 1 | 9 | agr2 |
| gsdf | 2.0394468 | 1.588939885 | 0.662 | 0.066 | 4.11907074679548e-7 | 10 | gsdf |
| hspe11 | 5.6064716 | 1.393185969 | 0.905 | 0.274 | 1.13233908325559e-4 | 10 | hspe1 |
| aldh1a21 | 2.5421583 | 1.298887518 | 0.743 | 0.188 | 5.13439729875942e-3 | 10 | aldh1a2 |
| cahz | 1.2677654 | 0.534106054 | 0.216 | 0.011 | 2.56050596213984e-3 | 10 | cahz |
| CR318588.11 | 2.2458493 | 1.307676799 | 0.878 | 0.249 | 4.53594194064736e-3 | 10 | CR318588.1 |
| zgc:927442 | 4.4866817 | 1.14110261 | 0.757 | 0.191 | 9.06175117760671e-3 | 10 | zgc:92744 |
| CR318588.3 | 8.1976035 | 0.880305899 | 0.419 | 0.072 | 1.65566998866839e-2 | 10 | CR318588.3 |
| ponzr13 | 4.7104304 | 1.010960887 | 0.811 | 0.382 | 9.51365647814515e-2 | 10 | ponzr1 |
| ppdpfa2 | 1.4410086 | 0.693246862 | 0.27 | 0.032 | 2.91040514433933e-2 | 10 | ppdpfa |
| jam2a | 3.3021993 | 1.017297834 | 0.473 | 0.112 | 6.66945198481114e-1 | 10 | jam2a |
| igf3 | 1.0896393 | 0.744861217 | 0.297 | 0.043 | 2.20074468444811e-1 | 10 | igf3 |
| foxl2b | 2.3906018 | 0.706504092 | 0.311 | 0.048 | 4.82829854825013e-1 | 10 | foxl2b |
| gstm.33 | 2.1510352 | 1.03804664 | 0.743 | 0.275 | 4.34444588954988e-1 | 10 | gstm.3 |
| cyp17a1 | 1.8468500 | 0.61592294 | 0.243 | 0.035 | 3.73008299811257e-1 | 10 | cyp17a1 |
| rpz51 | 2.7114872 | 0.918614936 | 0.595 | 0.248 | 5.4763908047824e-13 | 10 | rpz5 |
| ecrg4a | 2.3663875 | 0.68711144 | 0.311 | 0.062 | 4.77939289659281e-1 | 10 | ecrg4a |
| ifi451 | 5.6883676 | 0.924179501 | 0.527 | 0.192 | 1.1488796223665e-11 | 10 | ifi45 |
| itm2bb1 | 8.0560598 | 0.853050128 | 0.676 | 0.35 | 1.62708241081369e-1 | 10 | itm2bb |
| ccdc187 | 9.8846639 | 0.723960294 | 0.311 | 0.071 | 1.99640556866068e-1 | 10 | ccdc187 |
| fkbp1aa1 | 1.6975708 | 0.587356231 | 0.77 | 0.577 | 3.42858383064301e-1 | 10 | fkbp1aa |
| asb11 | 3.3366467 | 0.484759828 | 0.162 | 0.023 | 6.73902549900818e-0 | 10 | asb11 |
| anxa13l | 9.3120346 | 0.452299346 | 0.149 | 0.02 | 1.88075163153362e-0 | 10 | anxa13l |
| fez1 | 1.3136120 | 0.471508196 | 0.176 | 0.03 | 2.65310226088222e-0 | 10 | fez1 |
| glulb | 8.0984880 | 0.742058757 | 0.284 | 0.082 | 1.63565162596681e-0 | 10 | glulb |
| XLOC-0056051 | 1.7900270 | 0.615857205 | 0.297 | 0.09 | 3.61531753548447e-0 | 10 | XLOC-005605 |
| bcam | 3.2483933 | 0.620754176 | 0.297 | 0.096 | 6.56078000004399e-0 | 10 | bcam |
| bhmt1 | 4.6687066 | 0.321413203 | 0.135 | 0.023 | 9.42938672403669e-0 | 10 | bhmt |
| fthl271 | 6.4676683 | 0.439427875 | 0.743 | 0.628 | 0.000130627 | 10 | fthl27 |
| nr0b1 | 6.8014082 | 0.490853214 | 0.176 | 0.04 | 0.000137368 | 10 | nr0b1 |
| ifi462 | 8.1932830 | 0.674896955 | 0.405 | 0.188 | 0.00016548 | 10 | ifi46 |
| cldn11a1 | 8.7646880 | 0.638253437 | 0.432 | 0.191 | 0.00017702 | 10 | cldn11a |
| man2c1 | 1.4692475 | 0.416990125 | 0.149 | 0.03 | 0.000296744 | 10 | man2c1 |
| mpzl2b | 1.5630838 | 0.494807015 | 0.216 | 0.059 | 0.000315696 | 10 | mpzl2b |

| | | | | | | | |
|---|---|---|---|---|---|---|---|
| cx43 | 1.9495803 | 0.546392546 | 0.257 | 0.078 | 0.000393757 | 10 | cx43 |
| tuba8l33 | 3.1068901 | 0.616340173 | 0.419 | 0.202 | 0.000627499 | 10 | tuba8l3 |
| si:ch73-1a9.3 | 4.1800000 | 0.465320546 | 0.689 | 0.589 | 0.000844235 | 10 | si:ch73-1a9.3 |
| gamt1 | 7.9512646 | 0.365649283 | 0.149 | 0.032 | 0.001605917 | 10 | gamt |
| phlda1 | 8.4584969 | 0.283438172 | 0.108 | 0.018 | 0.001708363 | 10 | phlda1 |
| rasd1 | 9.5186048 | 0.285068153 | 0.108 | 0.018 | 0.001922473 | 10 | rasd1 |
| odc11 | 2.8348814 | 0.59191578 | 0.297 | 0.118 | 0.00572561 | 10 | odc1 |
| ttll12 | 3.5574111 | 0.393446565 | 0.122 | 0.024 | 0.007184903 | 10 | ttll12 |
| rpia | 5.5078552 | 0.548643264 | 0.243 | 0.087 | 0.011124215 | 10 | rpia |
| LOC103910903 | 5.6972443 | 0.337772597 | 0.162 | 0.04 | 0.011506724 | 10 | LOC103910903 |
| si:dkey-188i13.11 | 8.1160948 | 0.643391077 | 0.365 | 0.182 | 0.016392077 | 10 | si:dkey-188i13.11 |
| fkbp1b | 8.6659309 | 0.389680532 | 0.135 | 0.032 | 0.017502581 | 10 | fkbp1b |
| si:dkey-26c10.5 | 2.0659199 | 0.327077485 | 0.108 | 0.022 | 0.041725385 | 10 | si:dkey-26c10.5 |
| h3f3d | 4.7217985 | 0.290258037 | 0.703 | 0.676 | 0.095366166 | 10 | h3f3d |
| ap2m1a | 7.9207416 | 0.588112366 | 0.446 | 0.304 | 0.15997522 | 10 | ap2m1a |
| hsd3b7 | 1.0992291 | 0.425423085 | 0.203 | 0.072 | 0.22201131 | 10 | hsd3b7 |
| cdh4 | 1.1133438 | 0.317337581 | 0.122 | 0.03 | 0.224862049 | 10 | cdh4 |
| nupr1b1 | 1.3361800 | 0.30828711 | 0.122 | 0.03 | 0.269868279 | 10 | nupr1b |
| emx2 | 1.6014792 | 0.457875393 | 0.176 | 0.06 | 0.323450755 | 10 | emx2 |
| si:dkey-6n6.1 | 1.7731563 | 0.351625417 | 0.108 | 0.025 | 0.358124384 | 10 | si:dkey-6n6.1 |
| atp1b3b | 1.8207336 | 0.422306849 | 0.176 | 0.06 | 0.367733573 | 10 | atp1b3b |
| pfn21 | 2.0926929 | 0.525896935 | 0.459 | 0.319 | 0.422661193 | 10 | pfn2 |
| cpb11 | 3.0504064 | 0.279792211 | 0.108 | 0.026 | 0.6160906 | 10 | cpb1 |
| gpx1b | 4.0656691 | 0.369087063 | 0.122 | 0.033 | 0.821143199 | 10 | gpx1b |
| XLOC-009815 | 4.2691006 | 0.34137963 | 0.176 | 0.062 | 0.86223026 | 10 | XLOC-009815 |
| gstm.12 | 4.4659745 | 0.477019126 | 0.486 | 0.338 | 0.901992883 | 10 | gstm.1 |
| tmsb2 | 6.5086153 | 0.381640017 | 0.162 | 0.055 | 1 | 10 | tmsb |
| si:ch211-117c9.5 | 7.2174977 | 0.32140988 | 0.122 | 0.035 | 1 | 10 | si:ch211-117c9.5 |
| alcamb | 0.000121 | 0.34576655 | 0.149 | 0.051 | 1 | 10 | alcamb |
| dpp3 | 0.000136 | 0.461122126 | 0.203 | 0.087 | 1 | 10 | dpp3 |
| prdx22 | 0.000197 | 0.377864906 | 0.581 | 0.49 | 1 | 10 | prdx2 |
| bambia | 0.000229 | 0.348150469 | 0.162 | 0.06 | 1 | 10 | bambia |
| oaz1a | 0.000242 | 0.321543436 | 0.622 | 0.597 | 1 | 10 | oaz1a |
| alcama | 0.00043 | 0.352872595 | 0.149 | 0.056 | 1 | 10 | alcama |
| isg151 | 0.000484 | 0.290176452 | 0.135 | 0.048 | 1 | 10 | isg15 |

| | | | | | | | |
|---|---|---|---|---|---|---|---|
| si:ch211-222l21.12 | 0.000505 | 0.343837406 | 0.608 | 0.533 | 1 | 10 | si:ch211-222l21.1 |
| CABZ01044746.1 | 0.000629 | 0.259499018 | 0.108 | 0.034 | 1 | 10 | CABZ01044746.1 |
| eif4ebp11 | 0.000717 | 0.444207428 | 0.432 | 0.344 | 1 | 10 | eif4ebp1 |
| gstt1b1 | 0.000972 | 0.436397038 | 0.297 | 0.182 | 1 | 10 | gstt1b |
| pi15a | 0.001449 | 0.286628858 | 0.216 | 0.1 | 1 | 10 | pi15a |
| glrx5 | 0.001514 | 0.391145944 | 0.176 | 0.082 | 1 | 10 | glrx5 |
| calm3b | 0.001533 | 0.372443217 | 0.486 | 0.41 | 1 | 10 | calm3b |
| si:ch73-281n10.24 | 0.001551 | 0.398198027 | 0.5 | 0.436 | 1 | 10 | si:ch73-281n10.2 |
| slc25a3b1 | 0.001796 | 0.258507332 | 0.568 | 0.578 | 1 | 10 | slc25a3b |
| pnp5a1 | 0.002963 | 0.330836092 | 0.149 | 0.066 | 1 | 10 | pnp5a |
| hmgb3a2 | 0.003679 | 0.34945211 | 0.365 | 0.26 | 1 | 10 | hmgb3a |
| krt18a.12 | 0.00484 | 0.359468295 | 0.405 | 0.28 | 1 | 10 | krt18a.1 |
| rdh10a | 0.005273 | 0.349968879 | 0.176 | 0.089 | 1 | 10 | rdh10a |
| naxe | 0.005677 | 0.270634061 | 0.108 | 0.044 | 1 | 10 | naxe |
| ywhae11 | 0.005835 | 0.366927072 | 0.432 | 0.382 | 1 | 10 | ywhae1 |
| mmp22 | 0.007563 | 0.357989985 | 0.311 | 0.218 | 1 | 10 | mmp2 |
| sumo3a | 0.00922 | 0.339834273 | 0.446 | 0.412 | 1 | 10 | sumo3a |
| phc2a | 0.00929 | 0.318736463 | 0.176 | 0.094 | 1 | 10 | phc2a |
| edem1 | 0.009311 | 0.284781876 | 0.122 | 0.056 | 1 | 10 | edem1 |
| nr2f1a1 | 0.009698 | 0.345318727 | 0.243 | 0.153 | 1 | 10 | nr2f1a |
| marcksl1b2 | 0.009945 | 0.368784929 | 0.378 | 0.281 | 1 | 10 | marcksl1b |
| slc29a1b | 3.3057389 | 1.087335155 | 0.594 | 0.011 | 6.67660088591426e-1 | 11 | slc29a1b |
| cavin1b1 | 3.3515227 | 1.633179792 | 0.906 | 0.049 | 6.76907053549492e-1 | 11 | cavin1b |
| s100a10a1 | 3.9659305 | 1.103067724 | 0.531 | 0.012 | 8.00998998325203e-1 | 11 | s100a10a |
| jam2b1 | 1.8869944 | 0.988103069 | 0.625 | 0.022 | 3.81116263868424e-1 | 11 | jam2b |
| podxl1 | 1.9159086 | 1.67216239 | 0.953 | 0.091 | 3.86956068893013e-1 | 11 | podxl |
| si:ch211-156j16.1 | 1.2500280 | 1.145954381 | 0.75 | 0.047 | 2.52468173774943e-1 | 11 | si:ch211-156j16.1 |
| cav11 | 3.1830965 | 1.256685627 | 0.797 | 0.058 | 6.42890005079435e-9 | 11 | cav1 |
| cldnc1 | 7.2838133 | 1.523268494 | 0.734 | 0.055 | 1.47111178805771e-9 | 11 | cldnc |
| si:ch211-106h4.12 | 8.5503615 | 1.251687095 | 0.656 | 0.043 | 1.72691652593461e-8 | 11 | si:ch211-106h4.12 |
| lurap1 | 2.5433977 | 0.6965919 | 0.391 | 0.011 | 5.13690038716722e-8 | 11 | lurap1 |
| frzb1 | 7.7044039 | 1.012193177 | 0.422 | 0.014 | 1.55605847358352e-8 | 11 | frzb |
| eppk1 | 4.6819605 | 1.063310305 | 0.625 | 0.041 | 9.45615577730356e-8 | 11 | eppk1 |
| pcdh10b | 1.0394142 | 0.597047562 | 0.328 | 0.007 | 2.09930501835932e-8 | 11 | pcdh10b |
| krt42 | 3.7149174 | 1.460283687 | 0.844 | 0.082 | 7.50301875151894e-8 | 11 | krt4 |

| | | | | | | | |
|---|---|---|---|---|---|---|---|
| krt152 | 1.4925858 | 1.786277094 | 0.984 | 0.143 | 3.01457554231636e-7 | 11 | krt15 |
| serpine3 | 3.4238096 | 0.712699793 | 0.375 | 0.012 | 6.91506836839434e-7 | 11 | serpine3 |
| sfrp1a1 | 8.7827973 | 1.087934972 | 0.703 | 0.059 | 1.77386158805088e-7 | 11 | sfrp1a |
| postnb3 | 9.7070902 | 1.735471761 | 1 | 0.168 | 1.9605410272846e-73 | 11 | postnb |
| gcga | 1.6639700 | 0.710932785 | 0.453 | 0.021 | 3.36072036424892e-7 | 11 | gcga |
| bnc2 | 9.5598517 | 1.094689793 | 0.688 | 0.068 | 1.93080326596126e-6 | 11 | bnc2 |
| slc43a3b | 1.1825077 | 0.847000315 | 0.547 | 0.038 | 2.38831087062648e-6 | 11 | slc43a3b |
| frya | 9.8920224 | 0.30972737 | 0.188 | 0.001 | 1.99789177984357e-6 | 11 | frya |
| im:71514491 | 4.0667523 | 0.626922955 | 0.453 | 0.025 | 8.21361972241984e-6 | 11 | im:7151449 |
| igfbp2a | 1.8972164 | 0.377888186 | 0.25 | 0.005 | 3.83180805625936e-6 | 11 | igfbp2a |
| cyp2ad2 | 1.2161296 | 0.630640981 | 0.328 | 0.012 | 2.45621702723595e-5 | 11 | cyp2ad2 |
| paplna | 1.2791753 | 0.82071655 | 0.516 | 0.037 | 2.5835504491232e-59 | 11 | paplna |
| BNC1 | 1.9545915 | 0.454890726 | 0.297 | 0.009 | 3.94768845618313e-5 | 11 | BNC1 |
| si:ch211-63p21.1 | 2.7475222 | 0.58093804 | 0.312 | 0.011 | 5.54917065991087e-5 | 11 | si:ch211-63p21.1 |
| ehd2b | 2.7660933 | 0.840095826 | 0.516 | 0.039 | 5.58667879157267e-5 | 11 | ehd2b |
| tmem981 | 2.1781329 | 0.719303957 | 0.484 | 0.035 | 4.39917503885824e-5 | 11 | tmem98 |
| cldn5a3 | 3.0028351 | 1.228920551 | 0.766 | 0.11 | 6.06482617229053e-5 | 11 | cldn5a |
| cavin2b1 | 2.0212312 | 1.195511037 | 0.828 | 0.13 | 4.08228070662648e-5 | 11 | cavin2b |
| hspb12 | 2.7775417 | 1.258595353 | 0.891 | 0.16 | 5.60980098309202e-5 | 11 | hspb1 |
| pcsk9 | 3.6000045 | 0.345943969 | 0.172 | 0.002 | 7.27092914916781e-5 | 11 | pcsk9 |
| tgm2b | 2.5348322 | 0.359111788 | 0.219 | 0.005 | 5.11960069742637e-4 | 11 | tgm2b |
| pleca2 | 2.1686957 | 1.020456577 | 0.594 | 0.068 | 4.38011489620554e-4 | 11 | pleca |
| myrf | 3.1847842 | 0.349783788 | 0.188 | 0.003 | 6.43230870066117e-4 | 11 | myrf |
| rhpn2 | 2.0271283 | 0.667738067 | 0.391 | 0.026 | 4.0941911975705e-47 | 11 | rhpn2 |
| fsta | 6.8333554 | 0.511240249 | 0.281 | 0.012 | 1.3801327970352e-46 | 11 | fsta |
| fn1b3 | 1.1590826 | 1.287710612 | 0.859 | 0.163 | 2.34099919960403e-4 | 11 | fn1b |
| krt912 | 3.6755481 | 1.268290583 | 0.641 | 0.082 | 7.42350463732316e-4 | 11 | krt91 |
| cygb2 | 9.4243077 | 0.321314745 | 0.156 | 0.002 | 1.90342743785877e-4 | 11 | cygb2 |
| cxadr1 | 1.5292415 | 0.589192485 | 0.375 | 0.025 | 3.08860918677785e-4 | 11 | cxadr |
| zgc:1124371 | 2.1143677 | 0.862210376 | 0.516 | 0.052 | 4.2703886425519e-45 | 11 | zgc:112437 |
| si:ch211-66e2.31 | 6.2933010 | 1.012994009 | 0.594 | 0.072 | 1.27105800366342e-4 | 11 | si:ch211-66e2.3 |
| efna1b | 9.8305882 | 0.598545152 | 0.344 | 0.021 | 1.98548390119708e-4 | 11 | efna1b |
| gstm.34 | 5.4386597 | 1.505648941 | 0.984 | 0.27 | 1.09844611136422e-4 | 11 | gstm.3 |
| col12a1a1 | 3.5183657 | 0.816822969 | 0.469 | 0.044 | 7.10604328592823e-4 | 11 | col12a1a |
| itga6b | 7.8302301 | 0.382518321 | 0.188 | 0.004 | 1.58147159099715e-4 | 11 | itga6b |

| | | | | | | | |
|---|---|---|---|---|---|---|---|
| sema3fa1 | 1.9429088 | 0.58287896 | 0.359 | 0.024 | 3.92409294310489e-4 | 11 | sema3fa |
| epha4l | 2.7632961 | 0.536652312 | 0.328 | 0.02 | 5.58102919168651e-4 | 11 | epha4l |
| flnb1 | 5.4876750 | 0.680306526 | 0.453 | 0.04 | 1.10834573821671e-4 | 11 | flnb |
| zgc:1101821 | 2.4903344 | 0.771616523 | 0.578 | 0.069 | 5.02972847078017e-4 | 11 | zgc:110182 |
| endouc1 | 3.3106445 | 1.169937769 | 0.688 | 0.107 | 6.68650876225422e-4 | 11 | endouc |
| wnt5b1 | 1.3816350 | 0.637500688 | 0.359 | 0.027 | 2.79048826047283e-3 | 11 | wnt5b |
| anxa1a3 | 4.4917586 | 1.317225879 | 0.938 | 0.262 | 9.07200491226434e-3 | 11 | anxa1a |
| CU984579.1 | 2.4600739 | 0.476208532 | 0.297 | 0.018 | 4.96861131689985e-3 | 11 | CU984579.1 |
| aldh1a22 | 2.5028511 | 1.15803818 | 0.875 | 0.186 | 5.05500847245323e-3 | 11 | aldh1a2 |
| pkp3b | 2.7021489 | 0.748485479 | 0.547 | 0.069 | 5.45753021776908e-3 | 11 | pkp3b |
| cygb13 | 2.8317367 | 1.022130894 | 0.609 | 0.091 | 5.71925871795345e-3 | 11 | cygb1 |
| plod21 | 6.4783160 | 0.656742056 | 0.469 | 0.05 | 1.30842548641098e-3 | 11 | plod2 |
| col1a1b3 | 7.1205773 | 1.243701816 | 0.984 | 0.282 | 1.43814300744061e-3 | 11 | col1a1b |
| ucmaa1 | 1.7126503 | 0.752663734 | 0.344 | 0.027 | 3.45903998348602e-3 | 11 | ucmaa |
| myo1cb | 1.7380322 | 0.604220994 | 0.391 | 0.036 | 3.51030363989451e-3 | 11 | myo1cb |
| sox6 | 1.7889472 | 0.640621206 | 0.406 | 0.04 | 3.61313666044228e-3 | 11 | sox6 |
| aplp1 | 2.2392502 | 0.463830622 | 0.281 | 0.017 | 4.52261379578504e-3 | 11 | aplp1 |
| bmper1 | 2.7174301 | 0.850197111 | 0.578 | 0.082 | 5.48839358365291e-3 | 11 | bmper |
| slc43a1b | 4.0059677 | 0.396067449 | 0.203 | 0.008 | 8.0908531334228e-35 | 11 | slc43a1b |
| si:ch211-105c13.3: | 9.3329767 | 0.826486049 | 0.641 | 0.096 | 1.88498131204214e-3 | 11 | si:ch211-105c13.3 |
| s100a10b3 | 1.7891001 | 1.244242418 | 0.984 | 0.446 | 3.61344556352962e-3 | 11 | s100a10b |
| dkk3b2 | 2.5330451 | 1.005501852 | 0.75 | 0.152 | 5.11599136815783e-3 | 11 | dkk3b |
| cgnl1 | 6.6849307 | 0.647444084 | 0.438 | 0.048 | 1.35015547100067e-3 | 11 | cgnl1 |
| CABZ01040998.1 | 2.0022028 | 0.280106215 | 0.141 | 0.003 | 4.04384910394595e-3 | 11 | CABZ01040998.1 |
| prss351 | 2.0827595 | 0.720639201 | 0.5 | 0.062 | 4.20654938269745e-3 | 11 | prss35 |
| kcnq5a | 3.5551112 | 0.415605884 | 0.219 | 0.01 | 7.18025817886413e-3 | 11 | kcnq5a |
| FP016056.1 | 4.2157572 | 0.429960227 | 0.219 | 0.01 | 8.514564935573e-33 | 11 | FP016056.1 |
| ugt1b5 | 6.8812346 | 0.683015721 | 0.531 | 0.072 | 1.3898029556103e-32 | 11 | ugt1b5 |
| col6a4a3 | 3.1983594 | 1.076396467 | 0.734 | 0.156 | 6.45972652311767e-3 | 11 | col6a4a |
| col1a24 | 3.6819327 | 1.191686036 | 0.984 | 0.317 | 7.43639963662203e-3 | 11 | col1a2 |
| dchs1b | 5.4666714 | 0.396681355 | 0.234 | 0.013 | 1.10410362733924e-3 | 11 | dchs1b |
| col1a1a3 | 2.2904273 | 1.168850256 | 0.969 | 0.296 | 4.62597606409552e-3 | 11 | col1a1a |
| emilin1a1 | 3.1825780 | 0.65639073 | 0.422 | 0.048 | 6.42785281640785e-3 | 11 | emilin1a |
| cxcl8b.31 | 5.4601497 | 0.48313307 | 0.234 | 0.013 | 1.10278643693637e-3 | 11 | cxcl8b.3 |
| anxa2a2 | 9.5620371 | 1.117823792 | 0.766 | 0.174 | 1.93124463929973e-3 | 11 | anxa2a |

| | | | | | | | |
|---|---|---|---|---|---|---|---|
| c7a1 | 1.2940397 | 0.623681722 | 0.391 | 0.041 | 2.61357204861349e-3 | 11 | c7a |
| cdh22 | 8.7968935 | 0.858907294 | 0.625 | 0.115 | 1.77670860007898e-2 | 11 | cdh2 |
| prss23 | 5.2353335 | 0.580864734 | 0.359 | 0.037 | 1.05738030898328e-2 | 11 | prss23 |
| lrtm2a | 6.0354966 | 0.291294922 | 0.188 | 0.008 | 1.21898926512507e-2 | 11 | lrtm2a |
| krt82 | 7.5397058 | 1.053468152 | 0.969 | 0.513 | 1.52279439767217e-2 | 11 | krt8 |
| si:ch211-213d14.2 | 1.5171584 | 0.507174822 | 0.266 | 0.019 | 3.06420497893475e-2 | 11 | si:ch211-213d14.2 |
| col4a13 | 3.1549248 | 0.917775528 | 0.641 | 0.125 | 6.3720016478788e-28 | 11 | col4a1 |
| clic4 | 4.9806004 | 0.724770195 | 0.562 | 0.095 | 1.00593188204076e-2 | 11 | clic4 |
| ildr2 | 6.6271903 | 0.40031943 | 0.266 | 0.02 | 1.3384936312941e-27 | 11 | ildr2 |
| ptprk | 9.4780710 | 0.554594583 | 0.359 | 0.038 | 1.91428600755124e-2 | 11 | ptprk |
| si:ch211-250c4.4 | 1.3409654 | 0.475069553 | 0.312 | 0.028 | 2.7083478426785e-27 | 11 | si:ch211-250c4.4 |
| meis1b2 | 1.6116922 | 0.852349025 | 0.688 | 0.144 | 3.2551348262704e-27 | 11 | meis1b |
| brinp2 | 8.0716920 | 0.330526173 | 0.188 | 0.009 | 1.63023965006316e-2 | 11 | brinp2 |
| kank1a | 8.0716920 | 0.316081543 | 0.188 | 0.009 | 1.63023965006316e-2 | 11 | kank1a |
| epb41l4a | 9.0785823 | 0.353655459 | 0.203 | 0.011 | 1.83360128103344e-2 | 11 | epb41l4a |
| si:ch211-170d8.8 | 9.2888394 | 0.292728407 | 0.156 | 0.006 | 1.87606691179153e-2 | 11 | si:ch211-170d8.8 |
| npr2 | 1.2161802 | 0.296745144 | 0.188 | 0.009 | 2.45631934009867e-2 | 11 | npr2 |
| fgf241 | 2.0590637 | 0.521403524 | 0.297 | 0.027 | 4.15869111151023e-2 | 11 | fgf24 |
| adh8a2 | 7.1727972 | 0.67059138 | 0.438 | 0.059 | 1.44868986623815e-2 | 11 | adh8a |
| LOC100334706 | 7.2946843 | 0.31186777 | 0.219 | 0.014 | 1.47330739081491e-2 | 11 | LOC100334706 |
| mob2a | 8.9038570 | 0.591214817 | 0.344 | 0.038 | 1.79831200034297e-2 | 11 | mob2a |
| sostdc1a2 | 1.2457585 | 0.797833862 | 0.516 | 0.082 | 2.51605855734517e-2 | 11 | sostdc1a |
| tmem88b1 | 1.5203620 | 0.811744737 | 0.641 | 0.127 | 3.07067523969418e-2 | 11 | tmem88b |
| sorbs1 | 1.5951683 | 0.496729544 | 0.328 | 0.034 | 3.22176160716086e-2 | 11 | sorbs1 |
| cgna | 1.8059311 | 0.255074551 | 0.125 | 0.003 | 3.64743905916917e-2 | 11 | cgna |
| si:ch1073-459j12.1 | 2.0199382 | 0.875777048 | 0.75 | 0.171 | 4.0796692993331e-25 | 11 | si:ch1073-459j12.1 |
| fxyd11 | 2.4300193 | 0.634317536 | 0.547 | 0.094 | 4.9079100093881e-25 | 11 | fxyd1 |
| mmp23 | 3.6117038 | 0.89271733 | 0.797 | 0.203 | 7.29455828337254e-2 | 11 | mmp2 |
| pdlim3b | 5.1163495 | 0.365121433 | 0.203 | 0.012 | 1.03334912643856e-2 | 11 | pdlim3b |
| col6a33 | 1.0937580 | 0.913926685 | 0.734 | 0.181 | 2.20906312026969e-2 | 11 | col6a3 |
| ghra1 | 2.3860309 | 0.522904041 | 0.375 | 0.046 | 4.81906664788923e-2 | 11 | ghra |
| epb41l5 | 2.4049214 | 0.559654523 | 0.391 | 0.051 | 4.8572199388525e-24 | 11 | epb41l5 |
| krt18a.13 | 2.8325186 | 1.093609416 | 0.844 | 0.267 | 5.72083781674815e-2 | 11 | krt18a.1 |
| wt1a | 3.2462316 | 0.695455209 | 0.484 | 0.079 | 6.55641413264402e-2 | 11 | wt1a |
| rdh10a1 | 4.2588390 | 0.620285351 | 0.5 | 0.079 | 8.60157727806014e-2 | 11 | rdh10a |

| | | | | | | | |
|---|---|---|---|---|---|---|---|
| entpd12 | 5.2337423 | 0.821297015 | 0.625 | 0.136 | 1.05705894493214e-2 | 11 | entpd1 |
| ptgs1 | 6.2339490 | 0.3641127 | 0.25 | 0.02 | 1.25907068104443e-2 | 11 | ptgs1 |
| si:ch73-46j18.54 | 9.3871237 | 0.952857992 | 0.922 | 0.385 | 1.89591738996645e-2 | 11 | si:ch73-46j18.5 |
| itgb4 | 1.6520307 | 0.43647978 | 0.219 | 0.016 | 3.33660645230043e-2 | 11 | itgb4 |
| ahnak3 | 1.6795106 | 0.853789577 | 0.734 | 0.189 | 3.39210768250052e-2 | 11 | ahnak |
| cast1 | 8.3847574 | 0.677272009 | 0.484 | 0.084 | 1.69346947079202e-2 | 11 | cast |
| tagln4 | 9.8843839 | 0.852253804 | 0.609 | 0.125 | 1.99634902014195e-2 | 11 | tagln |
| cdon2 | 9.9921467 | 0.765680495 | 0.625 | 0.134 | 2.01811387286916e-2 | 11 | cdon |
| col5a2a3 | 1.7227670 | 0.824381957 | 0.703 | 0.169 | 3.47947259170847e-2 | 11 | col5a2a |
| cyp2ad33 | 2.9521453 | 0.739171607 | 0.547 | 0.106 | 5.96244791536377e-2 | 11 | cyp2ad3 |
| col5a13 | 4.5370581 | 0.900700105 | 0.812 | 0.23 | 9.16349632735121e-2 | 11 | col5a1 |
| ptprsa | 7.9466892 | 0.478307434 | 0.328 | 0.039 | 1.60499282743792e-2 | 11 | ptprsa |
| col21a1 | 1.0553848 | 0.449143829 | 0.281 | 0.029 | 2.13156079006069e-2 | 11 | col21a1 |
| si:ch211-198c19.3 | 1.3388661 | 0.941040748 | 0.703 | 0.172 | 2.70410792322888e-2 | 11 | si:ch211-198c19.3 |
| ehd1b | 5.4509999 | 0.544912703 | 0.422 | 0.066 | 1.10093845885072e-2 | 11 | ehd1b |
| oaz2b | 5.5336600 | 0.426707283 | 0.281 | 0.03 | 1.11763332450366e-2 | 11 | oaz2b |
| sccpdha | 6.7825946 | 0.302792308 | 0.141 | 0.006 | 1.36988065001434e-2 | 11 | sccpdha |
| cebpd4 | 9.7364350 | 0.891830524 | 0.844 | 0.266 | 1.96646779272771e-2 | 11 | cebpd |
| boc3 | 1.6812664 | 0.770237918 | 0.656 | 0.156 | 3.39565378160254e-2 | 11 | boc |
| gata6 | 2.1235321 | 0.479076606 | 0.375 | 0.053 | 4.28889790501551e-2 | 11 | gata6 |
| sparc4 | 2.7482363 | 0.925535562 | 0.953 | 0.389 | 5.55061285556061e-2 | 11 | sparc |
| nt5e | 4.1841998 | 0.328851923 | 0.172 | 0.011 | 8.45082847807766e-2 | 11 | nt5e |
| icn4 | 5.6914727 | 1.081188507 | 0.797 | 0.246 | 1.14950675806738e-1 | 11 | icn |
| si:ch211-137i24.10 | 5.8756843 | 0.336928948 | 0.172 | 0.011 | 1.18671196117862e-1 | 11 | si:ch211-137i24.10 |
| c63 | 5.8926056 | 0.827349548 | 0.766 | 0.201 | 1.19012955975804e-1 | 11 | c6 |
| arvcfb | 7.8497961 | 0.331169173 | 0.188 | 0.013 | 1.5854233378662e-19 | 11 | arvcfb |
| rtn3 | 8.2445743 | 0.604973922 | 0.438 | 0.076 | 1.66515668914711e-1 | 11 | rtn3 |
| tnfsf10l2 | 9.6740914 | 0.712918749 | 0.5 | 0.1 | 1.95387625032202e-1 | 11 | tnfsf10l |
| nrp1a2 | 1.1557591 | 0.696271893 | 0.516 | 0.104 | 2.33428678677271e-1 | 11 | nrp1a |
| scarb2a | 1.3083821 | 0.541537499 | 0.359 | 0.052 | 2.64253948838818e-1 | 11 | scarb2a |
| ftr822 | 1.7881164 | 0.628233055 | 0.5 | 0.097 | 3.61145887896914e-1 | 11 | ftr82 |
| id14 | 1.8993302 | 0.949579049 | 0.828 | 0.261 | 3.8360772414666e-19 | 11 | id1 |
| krt944 | 1.8995249 | 0.97871972 | 0.891 | 0.351 | 3.83647058826943e-1 | 11 | krt94 |
| usp43b | 1.9624318 | 0.354075754 | 0.203 | 0.016 | 3.9635237066037e-19 | 11 | usp43b |
| si:dkey-16l2.20 | 3.0035184 | 0.44661981 | 0.297 | 0.036 | 6.06620617425481e-1 | 11 | si:dkey-16l2.20 |

| | | | | | | | |
|---|---|---|---|---|---|---|---|
| fat2 | 3.7415662 | 0.328886204 | 0.172 | 0.011 | 7.5568414218629e-19 | 11 | fat2 |
| myl9a2 | 4.1033633 | 0.629641806 | 0.531 | 0.107 | 8.28756298230786e-1 | 11 | myl9a |
| cav21 | 4.4677195 | 0.449430398 | 0.281 | 0.032 | 9.0234531524789e-19 | 11 | cav2 |
| cyp27c11 | 4.9519974 | 0.520315224 | 0.375 | 0.057 | 1.00015493485961e-1 | 11 | cyp27c1 |
| cd151 | 9.9233752 | 0.430218708 | 0.312 | 0.04 | 2.0042240962371e-18 | 11 | cd151 |
| itgb3b | 1.2057842 | 0.379588056 | 0.234 | 0.023 | 2.43532241476482e-1 | 11 | itgb3b |
| rab3db | 1.4663170 | 0.375811232 | 0.234 | 0.023 | 2.96152045851705e-1 | 11 | rab3db |
| tjp1a | 2.3049962 | 0.537081372 | 0.438 | 0.078 | 4.65540091131158e-1 | 11 | tjp1a |
| klf2a2 | 2.8043850 | 0.802926252 | 0.656 | 0.169 | 5.66401647959854e-1 | 11 | klf2a |
| mfap52 | 3.1214051 | 0.772647784 | 0.641 | 0.152 | 6.30430195452902e-1 | 11 | mfap5 |
| fstl1b3 | 6.2998811 | 0.853930557 | 0.828 | 0.266 | 1.272386991881e-17 | 11 | fstl1b |
| cfd1 | 7.0069668 | 0.912205257 | 0.438 | 0.083 | 1.41519709415919e-1 | 11 | cfd |
| loxl2a1 | 9.4068974 | 0.508909524 | 0.406 | 0.068 | 1.89991107776458e-1 | 11 | loxl2a |
| DST | 1.2510902 | 0.333066673 | 0.203 | 0.018 | 2.52682690156754e-1 | 11 | DST |
| tmsb14 | 1.6600196 | 0.972944488 | 0.641 | 0.184 | 3.35274163010472e-1 | 11 | tmsb1 |
| tlr5b | 2.2276215 | 0.401482294 | 0.297 | 0.039 | 4.49912726544871e-1 | 11 | tlr5b |
| wt1b2 | 2.5141513 | 0.573714808 | 0.375 | 0.062 | 5.07783142556233e-1 | 11 | wt1b |
| slco2b11 | 2.5642807 | 0.594763171 | 0.406 | 0.073 | 5.17907788742911e-1 | 11 | slco2b1 |
| atp10a | 3.9653458 | 0.542108651 | 0.391 | 0.068 | 8.00880905617825e-1 | 11 | atp10a |
| map2k1 | 3.9984551 | 0.339190498 | 0.25 | 0.028 | 8.0756798751914e-17 | 11 | map2k1 |
| sema3d | 4.9356139 | 0.422452266 | 0.266 | 0.032 | 9.96845957992721e-1 | 11 | sema3d |
| rap1gap1 | 4.9737051 | 0.310387888 | 0.172 | 0.013 | 1.00453923210139e-1 | 11 | rap1gap |
| angptl1a2 | 5.0192497 | 0.643381968 | 0.391 | 0.067 | 1.01373787416342e-1 | 11 | angptl1a |
| tjp2b1 | 5.6394419 | 0.525445947 | 0.406 | 0.074 | 1.13899809572886e-1 | 11 | tjp2b |
| ppp1r3ca | 6.3003809 | 0.280334639 | 0.141 | 0.008 | 1.27248794402969e-1 | 11 | ppp1r3ca |
| hoxc10a | 9.5568257 | 0.260701695 | 0.141 | 0.008 | 1.93019210242877e-1 | 11 | hoxc10a |
| nat16 | 1.1083844 | 0.254180507 | 0.125 | 0.006 | 2.23860404202607e-1 | 11 | nat16 |
| myof | 1.3696221 | 0.301918331 | 0.188 | 0.016 | 2.76622587275769e-1 | 11 | myof |
| olfml3b3 | 1.6599978 | 0.539369744 | 0.5 | 0.102 | 3.35269756397431e-1 | 11 | olfml3b |
| ppfibp1a | 1.7133654 | 0.425397311 | 0.312 | 0.045 | 3.46048411199634e-1 | 11 | ppfibp1a |
| sult1st6 | 1.9281444 | 0.456889785 | 0.25 | 0.03 | 3.89427331660521e-1 | 11 | sult1st6 |
| BX640522.22 | 1.9712650 | 0.784628494 | 0.5 | 0.111 | 3.98136409020789e-1 | 11 | BX640522.2 |
| mical2a | 9.5294017 | 0.296596556 | 0.156 | 0.011 | 1.92465327081766e-1 | 11 | mical2a |
| mmp11b | 1.0711523 | 0.280447318 | 0.156 | 0.011 | 2.1634063668712e-15 | 11 | mmp11b |
| bmp2b | 2.3600941 | 0.332139979 | 0.188 | 0.017 | 4.76668212979046e-1 | 11 | bmp2b |

| | | | | | | | |
|---|---|---|---|---|---|---|---|
| rbpms2b1 | 3.0975342 | 0.375913156 | 0.312 | 0.047 | 6.25608986032055e-1 | 11 | rbpms2b |
| hdlbpa3 | 3.7889690 | 0.754975066 | 0.734 | 0.257 | 7.65258087127795e-1 | 11 | hdlbpa |
| tspan4a | 3.8207515 | 0.544467887 | 0.344 | 0.06 | 7.71677194004211e-1 | 11 | tspan4a |
| tnfsf101 | 4.0419044 | 0.635279801 | 0.516 | 0.121 | 8.1634344594936e-15 | 11 | tnfsf10 |
| fstb1 | 4.1038406 | 0.432460712 | 0.297 | 0.042 | 8.28852698677773e-1 | 11 | fstb |
| ntd5 | 4.6867279 | 0.294052727 | 0.188 | 0.017 | 9.46578451767304e-1 | 11 | ntd5 |
| edil3a1 | 5.1589464 | 0.33428654 | 0.234 | 0.027 | 1.0419524195059e-14 | 11 | edil3a |
| fam171a2a | 6.0211943 | 0.423920354 | 0.297 | 0.044 | 1.21610062099719e-1 | 11 | fam171a2a |
| reep3a | 8.6601800 | 0.405869096 | 0.297 | 0.044 | 1.74909657198577e-1 | 11 | reep3a |
| si:ch73-375g18.1 | 1.1320210 | 0.343944843 | 0.234 | 0.028 | 2.28634287536376e-1 | 11 | si:ch73-375g18.1 |
| c1qtnf21 | 2.0376793 | 0.394291193 | 0.281 | 0.04 | 4.11550091841228e-1 | 11 | c1qtnf2 |
| hsbp1a1 | 2.3556178 | 0.535533657 | 0.375 | 0.072 | 4.75764133793745e-1 | 11 | hsbp1a |
| hsd11b23 | 2.7790188 | 0.82316312 | 0.828 | 0.28 | 5.61278434129832e-1 | 11 | hsd11b2 |
| slmapb | 2.9179859 | 0.351520068 | 0.234 | 0.029 | 5.89345631890645e-1 | 11 | slmapb |
| notch31 | 3.3603692 | 0.358708743 | 0.25 | 0.032 | 6.78693775369023e-1 | 11 | notch3 |
| ackr3b1 | 3.8737707 | 0.499563558 | 0.375 | 0.07 | 7.8238548028025e-14 | 11 | ackr3b |
| LOC101884451 | 7.8585182 | 0.625244079 | 0.484 | 0.117 | 1.58718493190432e-1 | 11 | LOC101884451 |
| itgav | 8.7475436 | 0.546137094 | 0.406 | 0.086 | 1.7667413883693e-13 | 11 | itgav |
| col6a13 | 9.1206696 | 0.647283007 | 0.531 | 0.136 | 1.84210165391059e-1 | 11 | col6a1 |
| ctsf1 | 1.0941264 | 0.417218212 | 0.359 | 0.067 | 2.20980723773929e-1 | 11 | ctsf |
| col6a23 | 1.3083048 | 0.577275771 | 0.5 | 0.119 | 2.64238332469265e-1 | 11 | col6a2 |
| gpr155a | 1.3792640 | 0.327018335 | 0.188 | 0.019 | 2.78569952271363e-1 | 11 | gpr155a |
| myh11a | 1.6425810 | 0.544343059 | 0.438 | 0.096 | 3.31752104135261e-1 | 11 | myh11a |
| dcn3 | 2.4923507 | 0.779353674 | 0.812 | 0.267 | 5.03380071733743e-1 | 11 | dcn |
| f102 | 3.1848345 | 0.579609628 | 0.469 | 0.111 | 6.43241031254413e-1 | 11 | f10 |
| myl62 | 4.1503439 | 0.652753173 | 0.656 | 0.194 | 8.38244968261889e-1 | 11 | myl6 |
| cyp2n131 | 4.5932627 | 0.426461495 | 0.328 | 0.057 | 9.27701269380135e-1 | 11 | cyp2n13 |
| fibinb | 4.9716904 | 0.499675948 | 0.328 | 0.059 | 1.00413232366434e-1 | 11 | fibinb |
| fxyd6l2 | 8.4525930 | 0.624766482 | 0.516 | 0.14 | 1.7071702224125e-12 | 11 | fxyd6l |
| znf385a1 | 8.7341507 | 0.482884237 | 0.422 | 0.09 | 1.76403641849383e-1 | 11 | znf385a |
| xirp2a1 | 9.3454502 | 0.466937585 | 0.375 | 0.075 | 1.88750058244892e-1 | 11 | xirp2a |
| tmem1763 | 9.7057526 | 0.585012577 | 0.547 | 0.146 | 1.96027086225027e-1 | 11 | tmem176 |
| cd81a3 | 1.0395987 | 0.74985466 | 0.906 | 0.431 | 2.09967751033738e-1 | 11 | cd81a |
| selenbp12 | 1.2881445 | 0.648112677 | 0.5 | 0.135 | 2.6016655967339e-12 | 11 | selenbp1 |
| gstm.13 | 1.7934724 | 0.712409367 | 0.828 | 0.328 | 3.6222762452628e-12 | 11 | gstm.1 |

| | | | | | | | |
|---|---|---|---|---|---|---|---|
| tenm33 | 1.9154871 | 0.623899539 | 0.516 | 0.138 | 3.86870938559261e-1 | 11 | tenm3 |
| phactr4a | 2.3571305 | 0.396905018 | 0.312 | 0.055 | 4.76069661514472e-1 | 11 | phactr4a |
| si:dkey-56m19.52 | 2.4073714 | 0.519596792 | 0.422 | 0.095 | 4.86216810783682e-1 | 11 | si:dkey-56m19.5 |
| zgc:1135312 | 2.6349572 | 0.465948784 | 0.328 | 0.061 | 5.32182310409306e-1 | 11 | zgc:113531 |
| lama42 | 2.6398255 | 0.488396809 | 0.422 | 0.094 | 5.33165566348321e-1 | 11 | lama4 |
| CR847566.12 | 5.2524365 | 0.626590311 | 0.484 | 0.125 | 1.06083461805983e-1 | 11 | CR847566.1 |
| LOC101882334 | 5.8369017 | 0.304789435 | 0.156 | 0.015 | 1.17887904702558e-1 | 11 | LOC101882334 |
| LOC101886030 | 6.1989138 | 0.292589501 | 0.188 | 0.021 | 1.25199462381714e-1 | 11 | LOC101886030 |
| zgc:154058 | 7.2056185 | 0.329389527 | 0.203 | 0.025 | 1.45531877764358e-1 | 11 | zgc:154058 |
| cald1a3 | 7.3010253 | 0.65814162 | 0.688 | 0.228 | 1.47458809639335e-1 | 11 | cald1a |
| CABZ01080702.12 | 1.0068954 | 0.661284049 | 0.547 | 0.168 | 2.0336268045618e-11 | 11 | CABZ01080702.1 |
| si:ch211-1a19.3 | 1.5289976 | 0.535404415 | 0.359 | 0.076 | 3.08811654774733e-1 | 11 | si:ch211-1a19.3 |
| rsu1 | 1.5725840 | 0.468392381 | 0.391 | 0.086 | 3.17614807804951e-1 | 11 | rsu1 |
| CU459094.21 | 2.0646033 | 0.38336975 | 0.234 | 0.035 | 4.16987931206828e-1 | 11 | CU459094.2 |
| zfp36l1b2 | 2.2276280 | 0.630970692 | 0.609 | 0.192 | 4.49914045695359e-1 | 11 | zfp36l1b |
| palm1b | 2.3077669 | 0.395192636 | 0.312 | 0.058 | 4.66099682682062e-1 | 11 | palm1b |
| vim2 | 2.4991712 | 0.525460845 | 0.422 | 0.098 | 5.0475762638599e-11 | 11 | vim |
| rai141 | 2.7375046 | 0.25231398 | 0.203 | 0.026 | 5.52893815667159e-1 | 11 | rai14 |
| rnd3b3 | 2.8064056 | 0.449117626 | 0.406 | 0.091 | 5.66809747924473e-1 | 11 | rnd3b |
| aoc23 | 3.1766526 | 0.717987875 | 0.469 | 0.127 | 6.41588536066852e-1 | 11 | aoc2 |
| si:ch73-138n13.1 | 3.4079904 | 0.450595971 | 0.344 | 0.071 | 6.88311837509830e-1 | 11 | si:ch73-138n13.1 |
| ptpn21 | 3.9382882 | 0.373242864 | 0.281 | 0.049 | 7.9541606925438e-11 | 11 | ptpn21 |
| aldh2.21 | 3.9779239 | 0.479540633 | 0.469 | 0.122 | 8.03421309145188e-1 | 11 | aldh2.2 |
| nanos11 | 4.1612832 | 0.556816789 | 0.391 | 0.089 | 8.40454372655674e-1 | 11 | nanos1 |
| map7d1a | 5.1471000 | 0.260758401 | 0.156 | 0.016 | 1.03955980451582e-1 | 11 | map7d1a |
| CABZ01072614.14 | 6.3432859 | 0.590966803 | 0.547 | 0.167 | 1.2811534607891e-10 | 11 | CABZ01072614.1 |
| s100v24 | 7.4950965 | 0.643369783 | 0.688 | 0.241 | 1.51378465653847e-1 | 11 | s100v2 |
| dbpb | 1.0137996 | 0.406490824 | 0.297 | 0.056 | 2.04757106753488e-1 | 11 | dbpb |
| eif4ebp3l2 | 1.0585351 | 0.684330679 | 0.578 | 0.193 | 2.13792351566466e-1 | 11 | eif4ebp3l |
| si:dkey-9i23.162 | 1.3925037 | 0.441796266 | 0.406 | 0.095 | 2.81243974614482e-1 | 11 | si:dkey-9i23.16 |
| itga51 | 1.7562669 | 0.341513638 | 0.234 | 0.036 | 3.54713244436077e-1 | 11 | itga5 |
| cxcl12b1 | 1.7775827 | 0.292160118 | 0.156 | 0.016 | 3.590183786668e-10 | 11 | cxcl12b |
| LOC103910015 | 1.8327823 | 0.362958142 | 0.25 | 0.042 | 3.70167057161486e-1 | 11 | LOC103910015 |
| nfixb2 | 2.6501798 | 0.599409971 | 0.562 | 0.168 | 5.35256832106511e-1 | 11 | nfixb |
| tspan18b | 2.9446064 | 0.470538803 | 0.312 | 0.065 | 5.9472217186335e-10 | 11 | tspan18b |

| | | | | | | | |
|---|---|---|---|---|---|---|---|
| col27a1b1 | 2.9762350 | 0.43572911 | 0.328 | 0.068 | 6.01110202979448e-1 | 11 | col27a1b |
| ccdc801 | 3.0093956 | 0.401692393 | 0.328 | 0.066 | 6.07807641276272e-1 | 11 | ccdc80 |
| quo | 6.6331270 | 0.3744466 | 0.172 | 0.021 | 1.3396926726011e-09 | 11 | quo |
| itga91 | 9.4028353 | 0.416268383 | 0.328 | 0.07 | 1.89909066469588e-0 | 11 | itga9 |
| pmp22a3 | 1.0916509 | 0.665635443 | 0.844 | 0.309 | 2.20480737681194e-0 | 11 | pmp22a |
| mink1 | 1.2304628 | 0.274397272 | 0.172 | 0.021 | 2.48516580831267e-0 | 11 | mink1 |
| epha31 | 1.3199478 | 0.507408487 | 0.266 | 0.051 | 2.66589860909551e-0 | 11 | epha3 |
| fbln51 | 2.6058998 | 0.530123386 | 0.391 | 0.098 | 5.26313601578337e-0 | 11 | fbln5 |
| hspb8 | 2.9859735 | 0.271945912 | 0.156 | 0.018 | 6.03077087799333e-0 | 11 | hspb8 |
| fap | 3.3284557 | 0.307956611 | 0.172 | 0.022 | 6.72248209181606e-0 | 11 | fap |
| ppp1r13l | 3.3284557 | 0.293377556 | 0.172 | 0.022 | 6.72248209181606e-0 | 11 | ppp1r13l |
| aida | 3.9190603 | 0.300307499 | 0.25 | 0.045 | 7.91532615493469e-0 | 11 | aida |
| si:ch211-166a6.51 | 4.2508638 | 0.389586624 | 0.297 | 0.061 | 8.58546969527407e-0 | 11 | si:ch211-166a6.5 |
| cxcl18b2 | 4.7792445 | 0.543613631 | 0.469 | 0.129 | 9.6526402067531e-09 | 11 | cxcl18b |
| fbn2b3 | 6.3427104 | 0.493352146 | 0.562 | 0.171 | 1.2810372241244e-08 | 11 | fbn2b |
| hoxb3a | 8.7004818 | 0.285193633 | 0.188 | 0.027 | 1.75723631819273e-0 | 11 | hoxb3a |
| piezo2a.2 | 9.9776545 | 0.270359258 | 0.188 | 0.027 | 2.01518688857691e-0 | 11 | piezo2a.2 |
| gpc6a1 | 1.1096605 | 0.267341664 | 0.188 | 0.027 | 2.24118144526819e-0 | 11 | gpc6a |
| zgc:1533721 | 1.2258770 | 0.386080726 | 0.266 | 0.052 | 2.47590392261347e-0 | 11 | zgc:153372 |
| si:dkey-28b4.72 | 1.3148316 | 0.530526638 | 0.562 | 0.188 | 2.65556542771235e-0 | 11 | si:dkey-28b4.7 |
| fgl2a2 | 1.3875231 | 0.45110573 | 0.453 | 0.123 | 2.80238054502525e-0 | 11 | fgl2a |
| wnt4a | 1.4961359 | 0.280535337 | 0.172 | 0.023 | 3.02174582394636e-0 | 11 | wnt4a |
| mdkb1 | 2.0574841 | 0.395645235 | 0.203 | 0.032 | 4.15550063949134e-0 | 11 | mdkb |
| flna4 | 2.2867060 | 0.48306508 | 0.516 | 0.161 | 4.61846020101512e-0 | 11 | flna |
| slc38a5b4 | 2.5133151 | 0.50216327 | 0.578 | 0.198 | 5.07614261543381e-0 | 11 | slc38a5b |
| cpne31 | 2.6926092 | 0.398071765 | 0.312 | 0.071 | 5.43826283429527e-0 | 11 | cpne3 |
| jupa1 | 2.9387899 | 0.451000183 | 0.359 | 0.091 | 5.9354741134146e-08 | 11 | jupa |
| llgl1 | 3.0069331 | 0.272777898 | 0.156 | 0.02 | 6.07310281917213e-0 | 11 | llgl1 |
| csrp1a | 3.0938912 | 0.450257884 | 0.484 | 0.147 | 6.24873219995452e-0 | 11 | csrp1a |
| selenow14 | 3.7138709 | 0.559570362 | 0.922 | 0.591 | 7.5009051451503e-08 | 11 | selenow1 |
| slc12a7b | 3.8690539 | 0.299341188 | 0.203 | 0.033 | 7.81432829390869e-0 | 11 | slc12a7b |
| si:ch211-132b12.7 | 4.3536896 | 0.375601207 | 0.281 | 0.06 | 8.79314692237845e-0 | 11 | si:ch211-132b12.7 |
| hoxc3a | 4.4442331 | 0.380918933 | 0.297 | 0.065 | 8.9760176533211e-08 | 11 | hoxc3a |
| tril1 | 4.5800989 | 0.372011398 | 0.25 | 0.048 | 9.25042590164393e-0 | 11 | tril |
| irf2bpl | 4.9585243 | 0.355013204 | 0.266 | 0.055 | 1.00147315460313e-0 | 11 | irf2bpl |

| | | | | | | | |
|---|---|---|---|---|---|---|---|
| pik3r21 | 5.3304969 | 0.441274744 | 0.375 | 0.099 | 1.07660047025447e-0 | 11 | pik3r2 |
| dbn11 | 5.7678443 | 0.50611892 | 0.406 | 0.118 | 1.16493152411009e-0 | 11 | dbn1 |
| lgals3b1 | 6.2955315 | 0.558858397 | 0.547 | 0.2 | 1.27150850938254e-0 | 11 | lgals3b |
| elovl1b1 | 6.7646947 | 0.533400789 | 0.484 | 0.159 | 1.36626540468927e-0 | 11 | elovl1b |
| utrn | 7.9288797 | 0.347076882 | 0.219 | 0.039 | 1.60139583442327e-0 | 11 | utrn |
| LOC1039092842 | 8.9298163 | 0.504392219 | 0.453 | 0.136 | 1.80355500466179e-0 | 11 | LOC103909284 |
| pmp22b1 | 1.0838511 | 0.300624218 | 0.266 | 0.054 | 2.18905415631229e-0 | 11 | pmp22b |
| sesn12 | 1.2233542 | 0.651433422 | 0.547 | 0.21 | 2.47080867718889e-0 | 11 | sesn1 |
| f3b | 1.2667589 | 0.462053213 | 0.344 | 0.089 | 2.55847300182586e-0 | 11 | f3b |
| LO018188.12 | 1.3575977 | 0.488443977 | 0.344 | 0.088 | 2.74194019763889e-0 | 11 | LO018188.1 |
| gsta.11 | 1.4194154 | 0.406702933 | 0.359 | 0.094 | 2.86679340496399e-0 | 11 | gsta.1 |
| gna12a | 1.4844517 | 0.326354921 | 0.25 | 0.05 | 2.99814724796199e-0 | 11 | gna12a |
| aqp8a.1 | 1.6942821 | 0.306395098 | 0.109 | 0.01 | 3.42194160683278e-0 | 11 | aqp8a.1 |
| gata51 | 1.7198509 | 0.264247318 | 0.172 | 0.025 | 3.47358304538082e-0 | 11 | gata5 |
| meis3 | 1.8622310 | 0.357335583 | 0.203 | 0.035 | 3.76114801630592e-0 | 11 | meis3 |
| tpm4a2 | 1.9830370 | 0.538158799 | 0.641 | 0.238 | 4.00513991300945e-0 | 11 | tpm4a |
| dag1 | 2.1047335 | 0.40655381 | 0.328 | 0.082 | 4.25093030705998e-0 | 11 | dag1 |
| zgc:1584632 | 2.1556037 | 0.482271775 | 0.391 | 0.113 | 4.35367289216505e-0 | 11 | zgc:158463 |
| rspo31 | 2.2145964 | 0.325517895 | 0.203 | 0.035 | 4.47282037594607e-0 | 11 | rspo3 |
| fzd7b2 | 2.3689913 | 0.414464617 | 0.375 | 0.1 | 4.78465188493373e-0 | 11 | fzd7b |
| jun4 | 2.4962678 | 0.631945073 | 0.828 | 0.376 | 5.0417121639864e-07 | 11 | jun |
| aldh9a1a.14 | 4.4726935 | 0.56476196 | 0.734 | 0.332 | 9.03349918041712e-0 | 11 | aldh9a1a.1 |
| her9 | 5.0844911 | 0.358011252 | 0.234 | 0.046 | 1.0269146864002e-06 | 11 | her9 |
| map1lc3a | 5.9255984 | 0.437341531 | 0.391 | 0.114 | 1.1967931257453e-06 | 11 | map1lc3a |
| pabpc1b2 | 6.6562381 | 0.420594396 | 0.406 | 0.119 | 1.34436042494435e-0 | 11 | pabpc1b |
| esr2a1 | 7.1502756 | 0.299593082 | 0.203 | 0.037 | 1.44414117952701e-0 | 11 | esr2a |
| sypl2a | 7.3699865 | 0.370110407 | 0.359 | 0.097 | 1.48851618247475e-0 | 11 | sypl2a |
| BX322530.12 | 8.1935213 | 0.389524871 | 0.297 | 0.073 | 1.65484550141018e-0 | 11 | BX322530.1 |
| fzd7a1 | 8.4844715 | 0.327558236 | 0.25 | 0.053 | 1.71360871046117e-0 | 11 | fzd7a |
| tspan71 | 9.8481907 | 0.390532062 | 0.359 | 0.098 | 1.98903907927602e-0 | 11 | tspan7 |
| dhrs13l11 | 1.0195827 | 0.375602197 | 0.297 | 0.073 | 2.05925133210666e-0 | 11 | dhrs13l1 |
| mxra8b2 | 1.1170657 | 0.316643027 | 0.266 | 0.059 | 2.25613764918115e-0 | 11 | mxra8b |
| dcbld2 | 2.0001789 | 0.28560591 | 0.219 | 0.043 | 4.03976146639057e-0 | 11 | dcbld2 |
| dspa1 | 2.2430649 | 0.303474512 | 0.219 | 0.043 | 4.53031824688009e-0 | 11 | dspa |
| si:ch73-22o12.1 | 2.3845509 | 0.324677638 | 0.25 | 0.055 | 4.81607751926509e-0 | 11 | si:ch73-22o12.1 |

| | | | | | | | |
|---|---|---|---|---|---|---|---|
| slc7a11 | 3.0123027 | 0.250403456 | 0.188 | 0.033 | 6.08394779268579e-0 | 11 | slc7a1 |
| tns1b2 | 3.1338382 | 0.318857965 | 0.219 | 0.044 | 6.3294130735449e-06 | 11 | tns1b |
| gpm6ab1 | 3.2715836 | 0.278122908 | 0.156 | 0.024 | 6.60761740320779e-0 | 11 | gpm6ab |
| stau21 | 3.2799693 | 0.446755114 | 0.453 | 0.15 | 6.62455411097089e-0 | 11 | stau2 |
| daam1b | 3.3353633 | 0.283991677 | 0.188 | 0.034 | 6.73643341755835e-0 | 11 | daam1b |
| phyh | 3.6592731 | 0.29571801 | 0.219 | 0.045 | 7.39063401617571e-0 | 11 | phyh |
| cyp2p6 | 3.8633396 | 0.31328519 | 0.188 | 0.034 | 7.80278715044242e-0 | 11 | cyp2p6 |
| tmtc1 | 3.9904039 | 0.272146127 | 0.172 | 0.029 | 8.05941887942439e-0 | 11 | tmtc1 |
| coro1ca | 4.4014920 | 0.337369046 | 0.297 | 0.075 | 8.88969351222715e-0 | 11 | coro1ca |
| cry2 | 4.5506761 | 0.320442728 | 0.281 | 0.069 | 9.19100053604465e-0 | 11 | cry2 |
| sort1a | 5.0151876 | 0.260683836 | 0.156 | 0.024 | 1.01291745970037e-0 | 11 | sort1a |
| dsc2l1 | 5.1962841 | 0.267574253 | 0.188 | 0.034 | 1.04949351721795e-0 | 11 | dsc2l |
| CU459056.1 | 5.3962601 | 0.309202292 | 0.297 | 0.074 | 1.08988266564248e-0 | 11 | CU459056.1 |
| capns1a2 | 5.6555992 | 0.487108255 | 0.594 | 0.243 | 1.14226137797393e-0 | 11 | capns1a |
| plecb1 | 5.8142065 | 0.315744595 | 0.266 | 0.062 | 1.17429528965185e-0 | 11 | plecb |
| b2ml3 | 5.9075764 | 0.526062561 | 0.844 | 0.438 | 1.19315321810538e-0 | 11 | b2ml |
| cd151l | 6.3139955 | 0.313036131 | 0.328 | 0.087 | 1.27523768880357e-0 | 11 | cd151l |
| arl4d | 8.7599221 | 0.335052278 | 0.281 | 0.07 | 1.76924147308934e-0 | 11 | arl4d |
| si:dkey-238c7.162 | 1.6315994 | 0.379471475 | 0.375 | 0.112 | 3.29534136400047e-0 | 11 | si:dkey-238c7.16 |
| tmem176l.13 | 1.7298845 | 0.523652429 | 0.719 | 0.317 | 3.49384780623347e-0 | 11 | tmem176l.1 |
| nck1b | 1.7664842 | 0.367072512 | 0.312 | 0.088 | 3.56776814114245e-0 | 11 | nck1b |
| flii | 1.8629458 | 0.251964508 | 0.203 | 0.041 | 3.76259170997189e-0 | 11 | flii |
| hbegfa | 2.0017017 | 0.300606895 | 0.25 | 0.058 | 4.04283699131244e-0 | 11 | hbegfa |
| hoxc9a1 | 2.0066888 | 0.282760395 | 0.188 | 0.036 | 4.05290943938702e-0 | 11 | hoxc9a |
| nfia2 | 2.1400662 | 0.44971 | 0.531 | 0.195 | 4.32229188980792e-0 | 11 | nfia |
| asip2b2 | 2.2108961 | 0.46815295 | 0.266 | 0.067 | 4.46534705504941e-0 | 11 | asip2b |
| plk31 | 2.3210614 | 0.367938185 | 0.234 | 0.054 | 4.6878477951844e-05 | 11 | plk3 |
| mrtfab | 2.3507033 | 0.301324213 | 0.219 | 0.047 | 4.74771550037244e-0 | 11 | mrtfab |
| mgst1.12 | 2.4459557 | 0.423482625 | 0.391 | 0.126 | 4.94009676442668e-0 | 11 | mgst1.1 |
| BX569787.12 | 2.7508615 | 0.37858072 | 0.344 | 0.101 | 5.55591504860863e-0 | 11 | BX569787.1 |
| ptges1 | 2.8422952 | 0.260360167 | 0.172 | 0.031 | 5.74058364742286e-0 | 11 | ptges |
| si:ch211-39i22.14 | 2.8680066 | 0.483235461 | 0.484 | 0.178 | 5.79251311240467e-0 | 11 | si:ch211-39i22.1 |
| mknk2b1 | 2.8926640 | 0.410059548 | 0.562 | 0.214 | 5.84231357428878e-0 | 11 | mknk2b |
| glud1a | 3.0238620 | 0.4288866 | 0.406 | 0.139 | 6.10729419623191e-0 | 11 | glud1a |
| stxbp61 | 3.5724188 | 0.325632278 | 0.25 | 0.061 | 7.21521442745015e-0 | 11 | stxbp6 |

| | | | | | | | |
|---|---|---|---|---|---|---|---|
| gpc4 | 5.3959412 | 0.264993031 | 0.219 | 0.048 | 0.000108982 | 11 | gpc4 |
| gli3 | 6.1334637 | 0.285911629 | 0.234 | 0.055 | 0.000123878 | 11 | gli3 |
| fat1a | 7.6648666 | 0.295651107 | 0.203 | 0.044 | 0.000154807 | 11 | fat1a |
| nkd1 | 7.9234754 | 0.274466073 | 0.203 | 0.044 | 0.00016003 | 11 | nkd1 |
| mt-nd41 | 8.0466910 | 0.520166659 | 0.812 | 0.431 | 0.000162519 | 11 | mt-nd4 |
| fzd41 | 8.1061285 | 0.375068337 | 0.234 | 0.058 | 0.000163719 | 11 | fzd4 |
| cdr2l | 8.7080060 | 0.250465585 | 0.172 | 0.033 | 0.000175876 | 11 | cdr2l |
| chd33 | 9.1636036 | 0.473526658 | 0.562 | 0.236 | 0.000185077 | 11 | chd3 |
| calm2b1 | 9.3115365 | 0.495218355 | 0.828 | 0.44 | 0.000188065 | 11 | calm2b |
| gulp1a | 9.3687762 | 0.308303623 | 0.25 | 0.062 | 0.000189221 | 11 | gulp1a |
| gchfr1 | 9.8536829 | 0.295350724 | 0.203 | 0.044 | 0.000199015 | 11 | gchfr |
| fam3c | 1.0131016 | 0.311442383 | 0.234 | 0.058 | 0.000204616 | 11 | fam3c |
| acin1a1 | 1.0505378 | 0.485066639 | 0.625 | 0.292 | 0.000212177 | 11 | acin1a |
| wu:fb16f031 | 1.0701658 | 0.309512265 | 0.344 | 0.104 | 0.000216141 | 11 | wu:fb16f03 |
| XLOC-0412691 | 1.2024051 | 0.351750677 | 0.234 | 0.057 | 0.00024285 | 11 | XLOC-041269 |
| vat12 | 1.2265218 | 0.497273965 | 0.531 | 0.22 | 0.000247721 | 11 | vat1 |
| zgc:1951732 | 1.4192882 | 0.443895701 | 0.578 | 0.23 | 0.000286654 | 11 | zgc:195173 |
| CR383676.13 | 1.4599370 | 0.467628436 | 0.891 | 0.657 | 0.000294863 | 11 | CR383676.1 |
| cyp2p72 | 1.5630841 | 0.27264608 | 0.234 | 0.056 | 0.000315696 | 11 | cyp2p7 |
| adam10a1 | 1.5770723 | 0.394053806 | 0.453 | 0.165 | 0.000318521 | 11 | adam10a |
| stat1a | 1.7264978 | 0.378124301 | 0.406 | 0.143 | 0.000348701 | 11 | stat1a |
| kdm6bb1 | 1.7796793 | 0.338966828 | 0.344 | 0.107 | 0.000359442 | 11 | kdm6bb |
| snx171 | 1.9158887 | 0.377801247 | 0.328 | 0.101 | 0.000386952 | 11 | snx17 |
| rbms2a | 1.9200716 | 0.255172742 | 0.219 | 0.051 | 0.000387797 | 11 | rbms2a |
| CU539058.1 | 1.9508676 | 0.265788355 | 0.156 | 0.029 | 0.000394017 | 11 | CU539058.1 |
| gas1a2 | 2.1450968 | 0.427008081 | 0.391 | 0.134 | 0.000433245 | 11 | gas1a |
| pdlim1 | 2.2524510 | 0.303865176 | 0.312 | 0.093 | 0.000454928 | 11 | pdlim1 |
| pfn22 | 2.4665139 | 0.453364609 | 0.672 | 0.313 | 0.000498162 | 11 | pfn2 |
| dedd14 | 2.9509006 | 0.456582922 | 0.5 | 0.208 | 0.000595993 | 11 | dedd1 |
| lxn3 | 3.4317778 | 0.477106018 | 0.594 | 0.261 | 0.000693116 | 11 | lxn |
| loxa2 | 3.5871170 | 0.279444105 | 0.219 | 0.052 | 0.00072449 | 11 | loxa |
| gadd45ba3 | 3.6356825 | 0.437544917 | 0.578 | 0.246 | 0.000734299 | 11 | gadd45ba |
| aebp11 | 3.7264349 | 0.341259081 | 0.312 | 0.095 | 0.000752628 | 11 | aebp1 |
| mphosph8 | 3.8349897 | 0.398410237 | 0.578 | 0.246 | 0.000774553 | 11 | mphosph8 |
| cpn11 | 4.0190682 | 0.391753568 | 0.5 | 0.189 | 0.000811731 | 11 | cpn1 |

| | | | | | | | |
|---|---|---|---|---|---|---|---|
| si:ch1073-291c23.2 | 4.1278565 | 0.375064953 | 0.391 | 0.133 | 0.000833703 | 11 | si:ch1073-291c23.2 |
| mir199-3 | 6.3933631 | 0.306838195 | 0.188 | 0.042 | 0.001291268 | 11 | mir199-3 |
| mt-nd11 | 6.3969123 | 0.554090351 | 0.672 | 0.35 | 0.001291984 | 11 | mt-nd1 |
| serinc21 | 6.8669284 | 0.406003136 | 0.484 | 0.194 | 0.001386914 | 11 | serinc2 |
| aldh2.11 | 7.0506623 | 0.265090463 | 0.25 | 0.068 | 0.001424022 | 11 | aldh2.1 |
| cd248a2 | 7.5784022 | 0.264243289 | 0.375 | 0.118 | 0.00153061 | 11 | cd248a |
| pbx41 | 9.6547631 | 0.309473032 | 0.328 | 0.105 | 0.001949973 | 11 | pbx4 |
| capns1b | 1.0756294 | 0.440575259 | 0.547 | 0.249 | 0.002172449 | 11 | capns1b |
| cstf2 | 1.0765372 | 0.285491951 | 0.312 | 0.099 | 0.002174282 | 11 | cstf2 |
| ckap42 | 1.1527830 | 0.339286574 | 0.406 | 0.144 | 0.002328276 | 11 | ckap4 |
| srpx | 1.2732301 | 0.250278617 | 0.141 | 0.026 | 0.002571543 | 11 | srpx |
| wu:fb55g091 | 1.3311686 | 0.451645126 | 0.625 | 0.306 | 0.002688561 | 11 | wu:fb55g09 |
| si:ch211-286o17.1 | 1.4892639 | 0.345953896 | 0.328 | 0.108 | 0.003007866 | 11 | si:ch211-286o17.1 |
| kmt2bb | 1.5438278 | 0.399085319 | 0.391 | 0.148 | 0.003118069 | 11 | kmt2bb |
| si:ch211-133n4.43 | 1.5645118 | 0.347647199 | 0.375 | 0.13 | 0.003159845 | 11 | si:ch211-133n4.4 |
| arpc1a1 | 1.6453952 | 0.341768736 | 0.406 | 0.149 | 0.003323205 | 11 | arpc1a |
| cxcl8b.13 | 1.6564201 | 0.266759013 | 0.344 | 0.109 | 0.003345472 | 11 | cxcl8b.1 |
| meis2b3 | 1.7134453 | 0.344616789 | 0.297 | 0.093 | 0.003460645 | 11 | meis2b |
| fkbp72 | 1.8113010 | 0.325931778 | 0.359 | 0.123 | 0.003658285 | 11 | fkbp7 |
| ak11 | 1.9225084 | 0.286198629 | 0.406 | 0.149 | 0.00388289 | 11 | ak1 |
| si:ch211-212k18.9 | 2.0358449 | 0.315486218 | 0.344 | 0.114 | 0.004111796 | 11 | si:ch211-212k18.9 |
| fzd23 | 2.1266097 | 0.300519469 | 0.344 | 0.113 | 0.004295114 | 11 | fzd2 |
| c1qtnf6a1 | 2.1330503 | 0.257084857 | 0.234 | 0.063 | 0.004308122 | 11 | c1qtnf6a |
| ncam1a | 2.1714258 | 0.275770206 | 0.156 | 0.033 | 0.004385629 | 11 | ncam1a |
| sox4a1 | 2.1895073 | 0.327878385 | 0.422 | 0.154 | 0.004422148 | 11 | sox4a |
| ctnnb1 | 2.2083452 | 0.392398593 | 0.5 | 0.209 | 0.004460195 | 11 | ctnnb1 |
| si:ch211-214j24.10 | 2.2470698 | 0.328263784 | 0.422 | 0.162 | 0.004538407 | 11 | si:ch211-214j24.10 |
| s100u1 | 2.4853728 | 0.265748913 | 0.234 | 0.064 | 0.005019707 | 11 | s100u |
| pdk2b1 | 2.5465567 | 0.358007373 | 0.359 | 0.13 | 0.005143281 | 11 | pdk2b |
| CABZ01045617.11 | 2.6599360 | 0.302068936 | 0.344 | 0.12 | 0.005372273 | 11 | CABZ01045617.1 |
| ehd3 | 2.7850422 | 0.333502146 | 0.141 | 0.028 | 0.00562495 | 11 | ehd3 |
| lrrc58b | 2.8261809 | 0.253109847 | 0.156 | 0.033 | 0.005708038 | 11 | lrrc58b |
| syncrip1 | 3.1566829 | 0.423970682 | 0.734 | 0.386 | 0.006375553 | 11 | syncrip |
| qkia2 | 3.3825557 | 0.416887975 | 0.719 | 0.353 | 0.006831748 | 11 | qkia |
| ppp1cb | 4.2221981 | 0.315942314 | 0.281 | 0.091 | 0.008527574 | 11 | ppp1cb |

| | | | | | | | |
|---|---|---|---|---|---|---|---|
| LO018148.1 | 4.2537866 | 0.255145828 | 0.156 | 0.034 | 0.008591373 | 11 | LO018148.1 |
| csrnp1b1 | 4.3164747 | 0.251987185 | 0.312 | 0.1 | 0.008717984 | 11 | csrnp1b |
| crim11 | 4.7400343 | 0.283603095 | 0.234 | 0.066 | 0.009573447 | 11 | crim1 |
| lamb1a3 | 5.3574379 | 0.330288289 | 0.375 | 0.136 | 0.010820417 | 11 | lamb1a |
| tsr1 | 5.4224515 | 0.254912611 | 0.188 | 0.047 | 0.010951725 | 11 | tsr1 |
| tagln21 | 5.9196480 | 0.448265103 | 0.656 | 0.333 | 0.011955913 | 11 | tagln2 |
| si:ch211-196l7.42 | 6.0941020 | 0.440105155 | 0.719 | 0.408 | 0.012308258 | 11 | si:ch211-196l7.4 |
| ctnnb21 | 6.2878119 | 0.272353083 | 0.344 | 0.119 | 0.012699494 | 11 | ctnnb2 |
| ctsk2 | 6.5302432 | 0.289640358 | 0.375 | 0.134 | 0.013189132 | 11 | ctsk |
| colec123 | 6.7244510 | 0.269147324 | 0.328 | 0.108 | 0.013581374 | 11 | colec12 |
| si:dkey-51e6.1 | 6.7824166 | 0.275315154 | 0.359 | 0.129 | 0.013698447 | 11 | si:dkey-51e6.1 |
| arl6ip5b2 | 6.8762405 | 0.30894548 | 0.375 | 0.139 | 0.013887943 | 11 | arl6ip5b |
| rhoua1 | 7.1468258 | 0.326113674 | 0.219 | 0.062 | 0.014434444 | 11 | rhoua |
| rnf131 | 7.3904178 | 0.27875975 | 0.234 | 0.068 | 0.014926427 | 11 | rnf13 |
| cirbpb | 7.9735036 | 0.310446735 | 0.984 | 0.809 | 0.016104085 | 11 | cirbpb |
| ndfip1l | 8.9349943 | 0.378378737 | 0.328 | 0.121 | 0.018046008 | 11 | ndfip1l |
| ctnna1 | 9.1519754 | 0.251135317 | 0.234 | 0.068 | 0.018484245 | 11 | ctnna1 |
| myl9b2 | 9.1746144 | 0.359900119 | 0.484 | 0.199 | 0.018529969 | 11 | myl9b |
| tpm11 | 1.0003881 | 0.400550469 | 0.453 | 0.196 | 0.020204839 | 11 | tpm1 |
| adamts15a3 | 1.0140501 | 0.338891665 | 0.312 | 0.108 | 0.020480771 | 11 | adamts15a |
| zyx | 1.0708720 | 0.281122798 | 0.344 | 0.124 | 0.021628403 | 11 | zyx |
| foxp1b | 1.0940948 | 0.289536163 | 0.203 | 0.056 | 0.022097434 | 11 | foxp1b |
| usp36 | 1.2699618 | 0.319898024 | 0.328 | 0.12 | 0.02564942 | 11 | usp36 |
| zmiz1a | 1.2909875 | 0.274151649 | 0.203 | 0.056 | 0.026074076 | 11 | zmiz1a |
| cebpa3 | 1.3907004 | 0.356554587 | 0.422 | 0.17 | 0.028087978 | 11 | cebpa |
| f11r.12 | 1.4912046 | 0.286451689 | 0.297 | 0.1 | 0.030117861 | 11 | f11r.1 |
| pabpn11 | 1.5548155 | 0.375132543 | 0.594 | 0.289 | 0.03140261 | 11 | pabpn1 |
| pfn2l2 | 1.6777189 | 0.347229741 | 0.578 | 0.258 | 0.03388489 | 11 | pfn2l |
| notch2 | 1.7284569 | 0.326519583 | 0.375 | 0.147 | 0.034909645 | 11 | notch2 |
| akap12b1 | 1.7382683 | 0.339604894 | 0.391 | 0.149 | 0.035107806 | 11 | akap12b |
| plscr3b | 1.7497128 | 0.31600756 | 0.344 | 0.129 | 0.03533895 | 11 | plscr3b |
| rmdn1 | 1.8621675 | 0.274603141 | 0.312 | 0.111 | 0.037610199 | 11 | rmdn1 |
| timp2a1 | 1.8835682 | 0.294283123 | 0.312 | 0.109 | 0.038042427 | 11 | timp2a |
| foxp43 | 2.0549285 | 0.3824306 | 0.438 | 0.186 | 0.041503391 | 11 | foxp4 |
| wls2 | 2.1326859 | 0.349293581 | 0.422 | 0.179 | 0.043073859 | 11 | wls |

| | | | | | | | |
|---|---|---|---|---|---|---|---|
| vdac11 | 2.1581883 | 0.282546238 | 0.438 | 0.178 | 0.043588929 | 11 | vdac1 |
| fblim11 | 2.5150951 | 0.267744385 | 0.281 | 0.094 | 0.050797376 | 11 | fblim1 |
| znf609a | 2.8071929 | 0.251056253 | 0.172 | 0.044 | 0.056696876 | 11 | znf609a |
| dynll1 | 3.0034104 | 0.419595355 | 0.703 | 0.392 | 0.06065988 | 11 | dynll1 |
| si:ch211-140m22.7 | 3.0584333 | 0.253792231 | 0.234 | 0.074 | 0.061771179 | 11 | si:ch211-140m22.7 |
| akt1s1 | 3.0889529 | 0.257489783 | 0.234 | 0.073 | 0.062387584 | 11 | akt1s1 |
| sgta | 3.3131328 | 0.258327194 | 0.297 | 0.106 | 0.066915343 | 11 | sgta |
| mycbp | 3.3917542 | 0.343116238 | 0.484 | 0.223 | 0.06850326 | 11 | mycbp |
| flot1b1 | 3.4424385 | 0.288077384 | 0.453 | 0.193 | 0.069526932 | 11 | flot1b |
| ncor21 | 3.6564559 | 0.255592143 | 0.328 | 0.119 | 0.073849441 | 11 | ncor2 |
| fam168a1 | 3.7033484 | 0.314074463 | 0.375 | 0.152 | 0.074796528 | 11 | fam168a |
| map2k61 | 3.7889544 | 0.293164179 | 0.375 | 0.148 | 0.076525513 | 11 | map2k6 |
| socs3a3 | 3.8195572 | 0.44016695 | 0.766 | 0.424 | 0.077143598 | 11 | socs3a |
| smg7 | 3.9346459 | 0.256115971 | 0.188 | 0.052 | 0.079468044 | 11 | smg7 |
| lrp1ab1 | 4.3697229 | 0.291726634 | 0.328 | 0.121 | 0.088255295 | 11 | lrp1ab |
| myh10 | 4.4113954 | 0.264546987 | 0.234 | 0.073 | 0.089096955 | 11 | myh10 |
| robo21 | 4.8795261 | 0.313337659 | 0.266 | 0.092 | 0.098551789 | 11 | robo2 |
| ctsla3 | 5.1484271 | 0.289416388 | 0.641 | 0.303 | 0.103982782 | 11 | ctsla |
| usp9 | 5.5558052 | 0.289080804 | 0.25 | 0.084 | 0.112210598 | 11 | usp9 |
| XLOC-000946 | 5.5845508 | 0.287435094 | 0.219 | 0.07 | 0.112791174 | 11 | XLOC-000946 |
| si:dkey-7j14.63 | 5.7654459 | 0.404316641 | 0.609 | 0.317 | 0.116444712 | 11 | si:dkey-7j14.6 |
| tspan9b | 5.8037907 | 0.268067737 | 0.25 | 0.082 | 0.117219162 | 11 | tspan9b |
| gstt1b2 | 5.8268629 | 0.369889636 | 0.406 | 0.179 | 0.117685151 | 11 | gstt1b |
| NC-002333.44 | 5.9483032 | 0.436186734 | 0.812 | 0.583 | 0.120137881 | 11 | NC-002333.4 |
| mfap23 | 6.0270438 | 0.325846673 | 0.562 | 0.244 | 0.121728206 | 11 | mfap2 |
| rerea | 6.8902099 | 0.251387273 | 0.219 | 0.068 | 0.13916157 | 11 | rerea |
| elof1 | 6.9095292 | 0.314873413 | 0.438 | 0.197 | 0.139551762 | 11 | elof1 |
| csnk1g2b | 8.1927409 | 0.257460956 | 0.219 | 0.07 | 0.165468788 | 11 | csnk1g2b |
| klf11a | 8.3124788 | 0.253044258 | 0.203 | 0.062 | 0.167887135 | 11 | klf11a |
| abcc51 | 8.8327271 | 0.302642117 | 0.266 | 0.095 | 0.17839459 | 11 | abcc5 |
| fosb1 | 8.9387660 | 0.320003856 | 0.297 | 0.11 | 0.180536258 | 11 | fosb |
| mt-nd21 | 9.3406086 | 0.419624829 | 0.656 | 0.37 | 0.188652272 | 11 | mt-nd2 |
| bhlhe401 | 9.3680175 | 0.365571156 | 0.453 | 0.21 | 0.18920585 | 11 | bhlhe40 |
| si:ch211-288g17.3[illegible] | 9.5591666 | 0.360761993 | 0.578 | 0.289 | 0.193066489 | 11 | si:ch211-288g17.3 |
| klf6a1 | 9.9196652 | 0.338547489 | 0.547 | 0.255 | 0.20034748 | 11 | klf6a |

| | | | | | | | |
|---|---|---|---|---|---|---|---|
| lamp2 | 1.0040393 | 0.295460305 | 0.391 | 0.165 | 0.202785832 | 11 | lamp2 |
| anxa5b2 | 1.0382366 | 0.312785637 | 0.391 | 0.166 | 0.209692648 | 11 | anxa5b |
| mgst1.21 | 1.0633875 | 0.286673829 | 0.406 | 0.177 | 0.214772375 | 11 | mgst1.2 |
| mych1 | 1.1049543 | 0.286799689 | 0.391 | 0.166 | 0.223167623 | 11 | mych |
| ywhaqa1 | 1.1484452 | 0.327612963 | 0.516 | 0.25 | 0.231951496 | 11 | ywhaqa |
| pfkfb4b1 | 1.2788080 | 0.342156275 | 0.453 | 0.217 | 0.258280854 | 11 | pfkfb4b |
| cux1a1 | 1.3216324 | 0.311991182 | 0.281 | 0.106 | 0.266930111 | 11 | cux1a |
| eva1ba2 | 1.3225728 | 0.256560826 | 0.234 | 0.077 | 0.26712004 | 11 | eva1ba |
| bsg2 | 1.3252068 | 0.39043323 | 0.656 | 0.384 | 0.267652037 | 11 | bsg |
| celf23 | 1.3359946 | 0.378153201 | 0.656 | 0.352 | 0.269830832 | 11 | celf2 |
| jpt1b3 | 1.3421397 | 0.365499896 | 0.656 | 0.329 | 0.271071964 | 11 | jpt1b |
| mgst3b3 | 1.3929959 | 0.299758015 | 0.609 | 0.297 | 0.281343385 | 11 | mgst3b |
| smdt1b1 | 1.4315429 | 0.320258827 | 0.5 | 0.245 | 0.289128727 | 11 | smdt1b |
| igf2b2 | 1.4675380 | 0.305698433 | 0.391 | 0.161 | 0.29639866 | 11 | igf2b |
| rbm39a | 1.4745464 | 0.310478718 | 0.453 | 0.208 | 0.297814152 | 11 | rbm39a |
| tgfb3 | 1.5532917 | 0.255776988 | 0.25 | 0.086 | 0.313718341 | 11 | tgfb3 |
| nfil3-62 | 1.6330818 | 0.29575406 | 0.344 | 0.14 | 0.329833531 | 11 | nfil3-6 |
| nucb2a | 1.7138047 | 0.293792568 | 0.312 | 0.124 | 0.346137149 | 11 | nucb2a |
| ctgfa | 2.0244586 | 0.266457915 | 0.203 | 0.064 | 0.40887992 | 11 | ctgfa |
| si:ch211-222l21.13 | 2.3475738 | 0.395842092 | 0.859 | 0.526 | 0.474139482 | 11 | si:ch211-222l21.1 |
| fkbp5 | 2.3753826 | 0.274724978 | 0.359 | 0.151 | 0.479756042 | 11 | fkbp5 |
| zfp36l1a4 | 2.6266662 | 0.403431171 | 0.547 | 0.308 | 0.530507783 | 11 | zfp36l1a |
| fam120c1 | 2.7118660 | 0.29075826 | 0.344 | 0.145 | 0.547715582 | 11 | fam120c |
| macf1a | 2.7963743 | 0.263574459 | 0.234 | 0.082 | 0.564783723 | 11 | macf1a |
| rnf7 | 3.7030540 | 0.262444595 | 0.344 | 0.146 | 0.747905831 | 11 | rnf7 |
| pfdn11 | 3.8899197 | 0.25866623 | 0.469 | 0.222 | 0.785647095 | 11 | pfdn1 |
| csnk1db | 4.3823969 | 0.280550712 | 0.312 | 0.132 | 0.885112702 | 11 | csnk1db |
| ndufb91 | 4.6166445 | 0.270012053 | 0.609 | 0.32 | 0.932423695 | 11 | ndufb9 |
| foxo3b1 | 4.9162399 | 0.259587114 | 0.25 | 0.094 | 0.992932992 | 11 | foxo3b |
| uqcr101 | 5.5282180 | 0.283295541 | 0.609 | 0.32 | 1 | 11 | uqcr10 |
| mycb2 | 5.5910858 | 0.289571626 | 0.359 | 0.159 | 1 | 11 | mycb |
| tob1a | 5.7436352 | 0.266363545 | 0.438 | 0.204 | 1 | 11 | tob1a |
| tmem50a1 | 5.7957332 | 0.26916294 | 0.422 | 0.195 | 1 | 11 | tmem50a |
| pbx1a3 | 6.1713063 | 0.284957639 | 0.281 | 0.113 | 1 | 11 | pbx1a |
| si:ch211-213a13.1 | 6.5048058 | 0.25996334 | 0.344 | 0.149 | 1 | 11 | si:ch211-213a13.1 |

| | | | | | | | |
|---|---|---|---|---|---|---|---|
| fosab4 | 6.5091571 | 0.426919309 | 0.734 | 0.457 | 1 | 11 | fosab |
| dap1b | 6.6377883 | 0.332000268 | 0.484 | 0.252 | 1 | 11 | dap1b |
| h1f0 | 8.0561447 | 0.256991544 | 0.297 | 0.119 | 1 | 11 | h1f0 |
| eif4a2 | 8.3291030 | 0.252462401 | 0.281 | 0.116 | 1 | 11 | eif4a2 |
| ctdsp21 | 0.000113 | 0.314771669 | 0.359 | 0.172 | 1 | 11 | ctdsp2 |
| hmgb1a1 | 0.000116 | 0.293432112 | 0.734 | 0.423 | 1 | 11 | hmgb1a |
| col12a1b2 | 0.00012 | 0.253426016 | 0.188 | 0.063 | 1 | 11 | col12a1b |
| vdrb1 | 0.000125 | 0.258543686 | 0.25 | 0.099 | 1 | 11 | vdrb |
| mdh1aa2 | 0.000129 | 0.270520156 | 0.641 | 0.352 | 1 | 11 | mdh1aa |
| ogt.1 | 0.000134 | 0.269097 | 0.234 | 0.092 | 1 | 11 | ogt.1 |
| add1 | 0.000138 | 0.285587904 | 0.266 | 0.111 | 1 | 11 | add1 |
| krt18b2 | 0.000138 | 0.297516918 | 0.609 | 0.316 | 1 | 11 | krt18b |
| adh52 | 0.000143 | 0.302033721 | 0.625 | 0.358 | 1 | 11 | adh5 |
| junba3 | 0.000165 | 0.398389952 | 0.656 | 0.4 | 1 | 11 | junba |
| si:ch73-376l24.31 | 0.00017 | 0.266957192 | 0.359 | 0.169 | 1 | 11 | si:ch73-376l24.3 |
| irf2bp2b1 | 0.000214 | 0.269582644 | 0.391 | 0.19 | 1 | 11 | irf2bp2b |
| mmp16a3 | 0.000219 | 0.328850489 | 0.281 | 0.123 | 1 | 11 | mmp16a |
| mdka3 | 0.00022 | 0.287959724 | 0.453 | 0.22 | 1 | 11 | mdka |
| mcl1b4 | 0.000227 | 0.286639546 | 0.578 | 0.312 | 1 | 11 | mcl1b |
| id2a | 0.00024 | 0.311967202 | 0.344 | 0.167 | 1 | 11 | id2a |
| LOC1039104382 | 0.000246 | 0.309162395 | 0.359 | 0.183 | 1 | 11 | LOC103910438 |
| mt-atp61 | 0.000252 | 0.421799367 | 0.641 | 0.42 | 1 | 11 | mt-atp6 |
| itm2ba2 | 0.000253 | 0.326417969 | 0.656 | 0.41 | 1 | 11 | itm2ba |
| atox1 | 0.000253 | 0.264369731 | 0.375 | 0.188 | 1 | 11 | atox1 |
| cnbpb1 | 0.000263 | 0.287533682 | 0.578 | 0.322 | 1 | 11 | cnbpb |
| add3a3 | 0.00029 | 0.266198516 | 0.438 | 0.218 | 1 | 11 | add3a |
| zgc:1538671 | 0.000342 | 0.279329688 | 0.797 | 0.638 | 1 | 11 | zgc:153867 |
| rhoaa | 0.000369 | 0.291620324 | 0.516 | 0.296 | 1 | 11 | rhoaa |
| ccl25b2 | 0.000379 | 0.362642546 | 0.375 | 0.185 | 1 | 11 | ccl25b |
| spred2b1 | 0.000385 | 0.253315209 | 0.375 | 0.18 | 1 | 11 | spred2b |
| h2afy22 | 0.000396 | 0.258668927 | 0.484 | 0.253 | 1 | 11 | h2afy2 |
| mt-cyb1 | 0.000415 | 0.45133771 | 0.547 | 0.377 | 1 | 11 | mt-cyb |
| hmgn72 | 0.000415 | 0.317162191 | 0.766 | 0.527 | 1 | 11 | hmgn7 |
| tpm3 | 0.0005 | 0.294752976 | 0.734 | 0.454 | 1 | 11 | tpm3 |
| dynlt3 | 0.000522 | 0.250797425 | 0.422 | 0.22 | 1 | 11 | dynlt3 |

| | | | | | | | |
|---|---|---|---|---|---|---|---|
| mt-nd5 | 0.000546 | 0.307744699 | 0.344 | 0.181 | 1 | 11 | mt-nd5 |
| crip12 | 0.000559 | 0.346947208 | 0.594 | 0.376 | 1 | 11 | crip1 |
| btg22 | 0.000573 | 0.310912189 | 0.453 | 0.259 | 1 | 11 | btg2 |
| CABZ01092746.13 | 0.000614 | 0.262498261 | 0.422 | 0.206 | 1 | 11 | CABZ01092746.1 |
| hist2h2l1 | 0.00064 | 0.250939939 | 0.406 | 0.209 | 1 | 11 | hist2h2l |
| ndufa4l | 0.000683 | 0.298198813 | 0.844 | 0.569 | 1 | 11 | ndufa4l |
| sem1 | 0.00072 | 0.279549722 | 0.625 | 0.389 | 1 | 11 | sem1 |
| tefa2 | 0.000997 | 0.253937605 | 0.297 | 0.149 | 1 | 11 | tefa |
| kctd5a1 | 0.001178 | 0.252803051 | 0.438 | 0.246 | 1 | 11 | kctd5a |
| ywhaqb1 | 0.001528 | 0.278322257 | 0.469 | 0.278 | 1 | 11 | ywhaqb |
| marcksl1a2 | 0.002334 | 0.282193795 | 0.781 | 0.509 | 1 | 11 | marcksl1a |
| msna2 | 0.002435 | 0.252089786 | 0.656 | 0.4 | 1 | 11 | msna |
| COX33 | 0.004018 | 0.316019263 | 0.766 | 0.596 | 1 | 11 | COX3 |
| ppib2 | 0.004816 | 0.28137616 | 0.875 | 0.64 | 1 | 11 | ppib |
| ppdpfb1 | 0.007365 | 0.25008674 | 0.875 | 0.616 | 1 | 11 | ppdpfb |
| h3f3c2 | 0.00792 | 0.264928727 | 0.797 | 0.525 | 1 | 11 | h3f3c |
| cldn5b | 3.9110598 | 1.161947147 | 0.435 | 0.003 | 7.89916762962428e-1 | 12 | cldn5b |
| she | 2.3623895 | 0.984130247 | 0.403 | 0.003 | 4.77131820169276e-1 | 12 | she |
| pecam1 | 1.6579451 | 0.656982713 | 0.226 | 0 | 3.34855176276956e-9 | 12 | pecam1 |
| lyve1b | 2.1130476 | 0.700762292 | 0.226 | 0 | 4.2677223243625e-90 | 12 | lyve1b |
| si:ch211-145b13.6 | 1.2301159 | 0.906919955 | 0.355 | 0.007 | 2.48446515277483e-8 | 12 | si:ch211-145b13.6 |
| si:dkeyp-97a10.2 | 5.2408146 | 0.782377807 | 0.306 | 0.004 | 1.05848734142715e-8 | 12 | si:dkeyp-97a10.2 |
| cdh5 | 2.4891728 | 0.681909948 | 0.21 | 0 | 5.02738243228736e-8 | 12 | cdh5 |
| si:ch211-152c2.3 | 3.2840835 | 0.794597902 | 0.242 | 0.002 | 6.63286361494239e-8 | 12 | si:ch211-152c2.3 |
| erg | 1.3744318 | 0.902160478 | 0.323 | 0.007 | 2.77593992654749e-7 | 12 | erg |
| exoc3l2a | 2.3805608 | 0.695504307 | 0.21 | 0.001 | 4.80801875200602e-7 | 12 | exoc3l2a |
| cd81b | 4.6504143 | 0.640764411 | 0.21 | 0.001 | 9.39244180438254e-7 | 12 | cd81b |
| rasip1 | 2.3507737 | 0.532701645 | 0.194 | 0.001 | 4.74785781866992e-7 | 12 | rasip1 |
| LOC100535225 | 3.6140780 | 0.558186945 | 0.177 | 0 | 7.29935351545241e-6 | 12 | LOC100535225 |
| bcl6b | 3.6140780 | 0.454082196 | 0.177 | 0 | 7.29935351545241e-6 | 12 | bcl6b |
| si:ch211-33e4.2 | 6.5156774 | 0.651884515 | 0.242 | 0.003 | 1.31597137944476e-6 | 12 | si:ch211-33e4.2 |
| igfbp7 | 1.2001992 | 1.362582276 | 0.581 | 0.049 | 2.4240425131324e-63 | 12 | igfbp7 |
| sele | 2.5049497 | 0.492066785 | 0.177 | 0.001 | 5.05924693926604e-6 | 12 | sele |
| kdr | 4.2933689 | 0.480990411 | 0.161 | 0 | 8.67131734853897e-6 | 12 | kdr |
| si:dkey-126g1.9 | 8.0838519 | 0.7253067 | 0.274 | 0.007 | 1.63269558322464e-6 | 12 | si:dkey-126g1.9 |

| | | | | | | | |
|---|---|---|---|---|---|---|---|
| plvapb | 2.3505983 | 1.079361246 | 0.355 | 0.016 | 4.74750347512368e-5 | 12 | plvapb |
| stab1 | 8.5667036 | 0.515051243 | 0.194 | 0.002 | 1.7302171402587e-56 | 12 | stab1 |
| lyve1a | 2.3424119 | 0.581074695 | 0.161 | 0.001 | 4.73096945691973e-5 | 12 | lyve1a |
| notchl | 1.3629878 | 0.36403791 | 0.129 | 0 | 2.7528266352834e-54 | 12 | notchl |
| dab2 | 5.1871599 | 0.449011264 | 0.177 | 0.002 | 1.04765069762008e-5 | 12 | dab2 |
| si:ch73-334d15.2 | 1.7468375 | 0.622672022 | 0.258 | 0.009 | 3.52808772740614e-4 | 12 | si:ch73-334d15.2 |
| ldb2a | 2.7186129 | 0.596593017 | 0.21 | 0.005 | 5.49078259222763e-4 | 12 | ldb2a |
| etv2 | 7.0762179 | 0.500453653 | 0.177 | 0.003 | 1.42918374426373e-4 | 12 | etv2 |
| plvapa | 5.6850287 | 0.484795194 | 0.145 | 0.001 | 1.14820525567011e-4 | 12 | plvapa |
| clec14a | 5.2458658 | 0.603051181 | 0.21 | 0.006 | 1.0595075222895e-43 | 12 | clec14a |
| flt4 | 6.9662860 | 0.659561359 | 0.226 | 0.008 | 1.40698079198838e-4 | 12 | flt4 |
| rbp2a | 8.2042983 | 0.901228072 | 0.226 | 0.008 | 1.65702213627659e-4 | 12 | rbp2a |
| gpr182 | 6.3106590 | 0.326657829 | 0.113 | 0 | 1.27456380194759e-4 | 12 | gpr182 |
| dysf | 6.3106590 | 0.317507853 | 0.113 | 0 | 1.27456380194759e-4 | 12 | dysf |
| si:ch211-14k19.8 | 7.4800779 | 0.30891119 | 0.113 | 0 | 1.51075134424458e-4 | 12 | si:ch211-14k19.8 |
| plekhg5a | 3.9449337 | 0.470541245 | 0.177 | 0.004 | 7.96758266855979e-3 | 12 | plekhg5a |
| sox7 | 3.1640419 | 0.483275256 | 0.194 | 0.006 | 6.3904155206033e-38 | 12 | sox7 |
| akap12a | 7.7771304 | 0.302192891 | 0.113 | 0.001 | 1.57074703090866e-3 | 12 | akap12a |
| fabp11a1 | 8.7232821 | 1.480036344 | 0.613 | 0.098 | 1.76184129644661e-3 | 12 | fabp11a |
| arhgef9b | 9.9965025 | 0.296881362 | 0.113 | 0.001 | 2.01899362902022e-3 | 12 | arhgef9b |
| plk2b | 1.0830516 | 0.465502094 | 0.161 | 0.004 | 2.18743946975232e-3 | 12 | plk2b |
| tie1 | 6.7420893 | 0.492667146 | 0.177 | 0.005 | 1.36169978685823e-3 | 12 | tie1 |
| slc9a3r1b | 1.2798002 | 0.332289627 | 0.129 | 0.002 | 2.58481249972674e-3 | 12 | slc9a3r1b |
| ccm2l | 1.3358275 | 0.322467681 | 0.129 | 0.002 | 2.69797097324813e-3 | 12 | ccm2l |
| ecscr | 4.9180201 | 0.465451639 | 0.194 | 0.008 | 9.93292522900132e-3 | 12 | ecscr |
| fkbp1ab1 | 4.6354009 | 0.870431429 | 0.355 | 0.035 | 9.36211933788288e-3 | 12 | fkbp1ab |
| myct1a1 | 6.6010098 | 0.826898587 | 0.306 | 0.027 | 1.33320596890917e-2 | 12 | myct1a |
| aqp1a.1 | 2.1665496 | 0.4010167 | 0.113 | 0.002 | 4.3757803420951e-28 | 12 | aqp1a.1 |
| ccl38.11 | 2.3108944 | 0.878576935 | 0.274 | 0.022 | 4.66731360620703e-2 | 12 | ccl38.1 |
| tm4sf18 | 1.1415615 | 0.43198031 | 0.161 | 0.006 | 2.30561183704915e-2 | 12 | tm4sf18 |
| aqp8a.11 | 5.0035660 | 0.620320137 | 0.177 | 0.008 | 1.01057024125788e-2 | 12 | aqp8a.1 |
| kdrl | 1.3979276 | 0.377077843 | 0.113 | 0.002 | 2.82339454343431e-2 | 12 | kdrl |
| ramp21 | 6.6399828 | 0.946630115 | 0.387 | 0.052 | 1.3410773368003e-24 | 12 | ramp2 |
| wasf3b | 2.4002600 | 0.586867987 | 0.21 | 0.014 | 4.84780530914951e-2 | 12 | wasf3b |
| fgd5a | 3.4998027 | 0.562635245 | 0.21 | 0.014 | 7.06855171383817e-2 | 12 | fgd5a |

| | | | | | | | |
|---|---|---|---|---|---|---|---|
| egfl7 | 1.4488000 | 0.554344911 | 0.21 | 0.014 | 2.92614151671045e-2 | 12 | egfl7 |
| mrc1a1 | 1.4892102 | 0.565838719 | 0.194 | 0.012 | 3.00775798238014e-2 | 12 | mrc1a |
| sox18 | 6.0129104 | 0.440437113 | 0.161 | 0.008 | 1.21442752085373e-2 | 12 | sox18 |
| parvab | 7.6702062 | 0.587201574 | 0.21 | 0.015 | 1.54915155649882e-2 | 12 | parvab |
| tmem88a | 1.7500763 | 0.609134634 | 0.258 | 0.024 | 3.53462917897849e-2 | 12 | tmem88a |
| pcdh12 | 2.3302175 | 0.304132695 | 0.129 | 0.005 | 4.70634046675708e-2 | 12 | pcdh12 |
| si:ch211-248e11.2 | 2.3632390 | 0.423129111 | 0.129 | 0.005 | 4.77303399391065e-2 | 12 | si:ch211-248e11.2 |
| postna | 1.2289375 | 0.418519948 | 0.161 | 0.009 | 2.48208519267801e-2 | 12 | postna |
| calcrlb | 5.4789032 | 0.363998547 | 0.129 | 0.005 | 1.10657409394815e-1 | 12 | calcrlb |
| fam174b | 9.7418304 | 0.507069309 | 0.21 | 0.017 | 1.9675575009707e-19 | 12 | fam174b |
| ushbp1 | 1.2412623 | 0.361967306 | 0.145 | 0.008 | 2.5069775661379e-18 | 12 | ushbp1 |
| cpn12 | 3.8628748 | 1.025539862 | 0.597 | 0.186 | 7.80184830176514e-1 | 12 | cpn1 |
| tmem108 | 1.3936722 | 0.446270156 | 0.161 | 0.012 | 2.81479985706834e-1 | 12 | tmem108 |
| reep1 | 1.4379170 | 0.335734917 | 0.129 | 0.007 | 2.90416097015344e-1 | 12 | reep1 |
| lfng | 1.6431444 | 0.4564892 | 0.177 | 0.015 | 3.31865877538121e-1 | 12 | lfng |
| rab11bb | 7.6316094 | 0.63155436 | 0.242 | 0.033 | 1.54135615656736e-1 | 12 | rab11bb |
| si:ch211-156j16.11 | 3.6629917 | 0.765165037 | 0.323 | 0.061 | 7.39814438466049e-1 | 12 | si:ch211-156j16.1 |
| afap1l1b | 2.1555648 | 0.470703885 | 0.177 | 0.019 | 4.35359442361533e-1 | 12 | afap1l1b |
| rgcc1 | 2.6751435 | 0.88262748 | 0.339 | 0.072 | 5.40298733486282e-1 | 12 | rgcc |
| ednraa1 | 2.9068853 | 0.698886224 | 0.29 | 0.052 | 5.87103624465341e-1 | 12 | ednraa |
| rbp7b | 8.0802993 | 0.545868604 | 0.145 | 0.013 | 1.63197806205181e-1 | 12 | rbp7b |
| npdc1a | 1.1719081 | 0.323466454 | 0.113 | 0.007 | 2.36690283156697e-1 | 12 | npdc1a |
| mcamb | 3.1597291 | 0.342781987 | 0.129 | 0.01 | 6.38170498713036e-1 | 12 | mcamb |
| hapln31 | 1.1046032 | 0.557560487 | 0.177 | 0.021 | 2.23096727428388e-1 | 12 | hapln3 |
| krt18a.14 | 1.6793652 | 0.872322293 | 0.661 | 0.273 | 3.39181394487788e-1 | 12 | krt18a.1 |
| vwa1 | 4.2951248 | 0.490264468 | 0.161 | 0.018 | 8.67486357701857e-1 | 12 | vwa1 |
| lmo4a | 5.0298084 | 0.438209972 | 0.194 | 0.026 | 1.01587041991755e-0 | 12 | lmo4a |
| vat13 | 5.3161243 | 0.828902812 | 0.565 | 0.219 | 1.07369762532752e-0 | 12 | vat1 |
| si:ch211-66e2.32 | 1.2383019 | 0.679423977 | 0.339 | 0.08 | 2.50099842793256e-0 | 12 | si:ch211-66e2.3 |
| fkbp1aa2 | 1.1479472 | 0.60479577 | 0.806 | 0.577 | 2.31850908335324e-0 | 12 | fkbp1aa |
| marcksl1b3 | 2.1600552 | 0.898079286 | 0.581 | 0.276 | 4.3626635315239e-08 | 12 | marcksl1b |
| lmo2 | 1.8135197 | 0.409917532 | 0.161 | 0.023 | 3.66276584574321e-0 | 12 | lmo2 |
| rhocb | 1.9540870 | 0.759446556 | 0.468 | 0.178 | 3.94666958667095e-0 | 12 | rhocb |
| fli1a | 1.0810184 | 0.340020171 | 0.145 | 0.02 | 2.18333299549769e-0 | 12 | fli1a |
| carhsp1 | 2.1306038 | 0.636670587 | 0.323 | 0.095 | 4.30318063192079e-0 | 12 | carhsp1 |

| | | | | | | | |
|---|---|---|---|---|---|---|---|
| crip2 | 2.8050874 | 0.286208266 | 0.129 | 0.016 | 5.66543518580257e-0 | 12 | crip2 |
| palm2 | 4.5527484 | 0.411234616 | 0.145 | 0.021 | 9.19518598490919e-0 | 12 | palm2 |
| krt83 | 4.6864508 | 0.626359213 | 0.79 | 0.519 | 9.46522484486851e-0 | 12 | krt8 |
| mob2b | 1.2103265 | 0.389487266 | 0.145 | 0.022 | 2.4444964963088e-05 | 12 | mob2b |
| cxcl12a3 | 1.3553525 | 0.775105161 | 0.306 | 0.092 | 2.7374056270298e-05 | 12 | cxcl12a |
| mycn | 4.0584250 | 0.370325735 | 0.129 | 0.019 | 8.1968010751037e-05 | 12 | mycn |
| cdh61 | 1.1721927 | 0.627576877 | 0.274 | 0.082 | 0.000236748 | 12 | cdh6 |
| thy11 | 1.6925033 | 0.563914016 | 0.21 | 0.05 | 0.000341835 | 12 | thy1 |
| si:ch211-284f22.3 | 2.3423731 | 0.401657008 | 0.177 | 0.037 | 0.000473089 | 12 | si:ch211-284f22.3 |
| tspan4b1 | 2.8766257 | 0.55131098 | 0.306 | 0.104 | 0.000580992 | 12 | tspan4b |
| zfp36l21 | 3.3195142 | 0.559896919 | 0.226 | 0.061 | 0.000670442 | 12 | zfp36l2 |
| tagln22 | 3.9405138 | 0.640495703 | 0.565 | 0.336 | 0.000795866 | 12 | tagln2 |
| oaz2a | 8.4788033 | 0.376742399 | 0.113 | 0.017 | 0.001712464 | 12 | oaz2a |
| il13ra11 | 1.3775890 | 0.344761684 | 0.145 | 0.028 | 0.002782317 | 12 | il13ra1 |
| kank3 | 1.6846383 | 0.276674843 | 0.113 | 0.018 | 0.003402464 | 12 | kank3 |
| ponzr14 | 1.9027908 | 0.585815433 | 0.629 | 0.391 | 0.003843067 | 12 | ponzr1 |
| slc22a31 | 1.9380179 | 0.347230949 | 0.129 | 0.023 | 0.003914215 | 12 | slc22a31 |
| rhoca | 3.8205777 | 0.554795352 | 0.306 | 0.114 | 0.007716421 | 12 | rhoca |
| vamp5 | 4.0950551 | 0.355299705 | 0.161 | 0.036 | 0.008270783 | 12 | vamp5 |
| limch1b | 4.1730625 | 0.276730301 | 0.145 | 0.03 | 0.008428334 | 12 | limch1b |
| cirbpb1 | 4.4587311 | 0.315529033 | 0.935 | 0.811 | 0.009005299 | 12 | cirbpb |
| rps6ka3b | 4.5092205 | 0.268650409 | 0.113 | 0.019 | 0.009107273 | 12 | rps6ka3b |
| ckba1 | 5.5509034 | 0.568433226 | 0.306 | 0.118 | 0.01121116 | 12 | ckba |
| tuba8l34 | 5.9114544 | 0.616136078 | 0.419 | 0.204 | 0.011939364 | 12 | tuba8l3 |
| arl4aa | 6.3333066 | 0.274640948 | 0.113 | 0.019 | 0.012791379 | 12 | arl4aa |
| glula | 1.0753993 | 0.512127542 | 0.242 | 0.079 | 0.02171984 | 12 | glula |
| ehd42 | 1.3613234 | 0.461250988 | 0.194 | 0.054 | 0.027494649 | 12 | ehd4 |
| tcima1 | 1.5294373 | 0.556198604 | 0.258 | 0.088 | 0.030890047 | 12 | tcima |
| tcf4 | 2.6665611 | 0.319884003 | 0.145 | 0.034 | 0.053856536 | 12 | tcf4 |
| myl9a3 | 2.7248095 | 0.575908609 | 0.29 | 0.115 | 0.055032979 | 12 | myl9a |
| mmp14b1 | 3.0099151 | 0.513904597 | 0.29 | 0.115 | 0.060791256 | 12 | mmp14b |
| yrk1 | 3.2103572 | 0.444557421 | 0.258 | 0.089 | 0.064839585 | 12 | yrk |
| fscn1a | 3.4168841 | 0.365738677 | 0.145 | 0.034 | 0.069010808 | 12 | fscn1a |
| lgmn2 | 3.5354952 | 0.558911517 | 0.306 | 0.128 | 0.071406398 | 12 | lgmn |
| nova21 | 3.6132237 | 0.428809391 | 0.21 | 0.064 | 0.072976281 | 12 | nova2 |

| | | | | | | | |
|---|---|---|---|---|---|---|---|
| oaz2b1 | 3.8643733 | 0.321938791 | 0.145 | 0.034 | 0.078048749 | 12 | oaz2b |
| rasgef1bb | 4.8819483 | 0.301526928 | 0.161 | 0.041 | 0.098600712 | 12 | rasgef1bb |
| piezo2a.21 | 5.9972959 | 0.310765468 | 0.129 | 0.029 | 0.121127386 | 12 | piezo2a.2 |
| calm2a1 | 6.0082648 | 0.593183209 | 0.435 | 0.251 | 0.121348926 | 12 | calm2a |
| ephb4a | 6.8758612 | 0.30047367 | 0.145 | 0.035 | 0.13887177 | 12 | ephb4a |
| znf423 | 1.2001735 | 0.319870998 | 0.129 | 0.03 | 0.242399058 | 12 | znf423 |
| marcksl1a3 | 1.5336021 | 0.469958678 | 0.645 | 0.514 | 0.309741633 | 12 | marcksl1a |
| loxl11 | 1.6202204 | 0.460433879 | 0.21 | 0.072 | 0.327235932 | 12 | loxl1 |
| grb10a2 | 2.1875037 | 0.409208055 | 0.194 | 0.063 | 0.441810122 | 12 | grb10a |
| si:ch211-1a19.31 | 2.3406689 | 0.393483401 | 0.226 | 0.08 | 0.472744913 | 12 | si:ch211-1a19.3 |
| pfn23 | 2.4851310 | 0.549595182 | 0.484 | 0.319 | 0.501921918 | 12 | pfn2 |
| hspg2 | 3.0151931 | 0.398024884 | 0.161 | 0.049 | 0.608978555 | 12 | hspg2 |
| txn2 | 3.0773927 | 0.422071378 | 0.242 | 0.089 | 0.621541005 | 12 | txn |
| si:ch211-204c21.1 | 3.4786599 | 0.416349724 | 0.226 | 0.085 | 0.702584946 | 12 | si:ch211-204c21.1 |
| arpc1a2 | 3.6772759 | 0.45807114 | 0.323 | 0.152 | 0.742699419 | 12 | arpc1a |
| exoc6 | 4.4803145 | 0.323561502 | 0.161 | 0.048 | 0.90488913 | 12 | exoc6 |
| tpm4a3 | 6.5086255 | 0.481575476 | 0.419 | 0.246 | 1 | 12 | tpm4a |
| si:ch211-71n6.4 | 6.7062968 | 0.341190696 | 0.145 | 0.042 | 1 | 12 | si:ch211-71n6.4 |
| LOC1018844511 | 6.9025332 | 0.459180993 | 0.274 | 0.124 | 1 | 12 | LOC101884451 |
| myo1ca | 7.4555253 | 0.415128279 | 0.161 | 0.052 | 1 | 12 | myo1ca |
| fzd42 | 8.0595492 | 0.359898464 | 0.177 | 0.06 | 1 | 12 | fzd4 |
| hmgn22 | 8.5842556 | 0.44472086 | 0.742 | 0.632 | 1 | 12 | hmgn2 |
| myo1cb1 | 0.000101 | 0.332894211 | 0.145 | 0.044 | 1 | 12 | myo1cb |
| cd276 | 0.000125 | 0.365513614 | 0.194 | 0.071 | 1 | 12 | cd276 |
| tuba8l4 | 0.000128 | 0.464921216 | 0.581 | 0.453 | 1 | 12 | tuba8l4 |
| ehd31 | 0.000132 | 0.307769986 | 0.113 | 0.029 | 1 | 12 | ehd3 |
| macf1a1 | 0.000163 | 0.393229457 | 0.21 | 0.083 | 1 | 12 | macf1a |
| tpte | 0.000183 | 0.342995238 | 0.145 | 0.046 | 1 | 12 | tpte |
| gas6 | 0.000303 | 0.300691457 | 0.177 | 0.065 | 1 | 12 | gas6 |
| me3 | 0.000355 | 0.327240945 | 0.129 | 0.04 | 1 | 12 | me3 |
| cav12 | 0.000359 | 0.429037846 | 0.194 | 0.078 | 1 | 12 | cav1 |
| CU929150.12 | 0.000386 | 0.359114127 | 0.129 | 0.039 | 1 | 12 | CU929150.1 |
| ivns1abpa | 0.000396 | 0.298385879 | 0.113 | 0.032 | 1 | 12 | ivns1abpa |
| selenop1 | 0.000402 | 0.345298018 | 0.161 | 0.057 | 1 | 12 | selenop |
| ctsla4 | 0.00043 | 0.551385969 | 0.435 | 0.31 | 1 | 12 | ctsla |

| | | | | | | | | |
|---|---|---|---|---|---|---|---|---|
| prkd2 | 0.000514 | 0.261817739 | 0.129 | 0.041 | | 1 | 12 | prkd2 |
| cyfip11 | 0.000566 | 0.345337636 | 0.21 | 0.09 | | 1 | 12 | cyfip1 |
| pros1 | 0.000664 | 0.286495518 | 0.145 | 0.05 | | 1 | 12 | pros1 |
| acsbg2 | 0.000768 | 0.255071693 | 0.113 | 0.034 | | 1 | 12 | acsbg2 |
| hmgb2a2 | 0.000803 | 0.382679918 | 0.661 | 0.585 | | 1 | 12 | hmgb2a |
| erlin1 | 0.000968 | 0.27339664 | 0.145 | 0.052 | | 1 | 12 | erlin1 |
| rgl1 | 0.001225 | 0.318708236 | 0.145 | 0.054 | | 1 | 12 | rgl1 |
| svild1 | 0.001586 | 0.394215144 | 0.21 | 0.1 | | 1 | 12 | svild |
| ms4a17a.81 | 0.001594 | 0.305639788 | 0.129 | 0.046 | | 1 | 12 | ms4a17a.8 |
| gtf2h5 | 0.001742 | 0.359076597 | 0.226 | 0.109 | | 1 | 12 | gtf2h5 |
| fli1b1 | 0.001857 | 0.257863831 | 0.113 | 0.037 | | 1 | 12 | fli1b |
| cavin2b2 | 0.001934 | 0.374695617 | 0.274 | 0.148 | | 1 | 12 | cavin2b |
| pora | 0.002294 | 0.28124193 | 0.145 | 0.058 | | 1 | 12 | pora |
| CT030188.1 | 0.002354 | 0.321697251 | 0.177 | 0.077 | | 1 | 12 | CT030188.1 |
| galnt1 | 0.002409 | 0.256772537 | 0.129 | 0.047 | | 1 | 12 | galnt1 |
| cgnl11 | 0.002513 | 0.279817768 | 0.145 | 0.057 | | 1 | 12 | cgnl1 |
| dut | 0.002844 | 0.337967431 | 0.21 | 0.099 | | 1 | 12 | dut |
| fam171a2a1 | 0.002928 | 0.270400706 | 0.129 | 0.049 | | 1 | 12 | fam171a2a |
| tmem88b2 | 0.003056 | 0.359990178 | 0.258 | 0.14 | | 1 | 12 | tmem88b |
| hnrnpa0b1 | 0.003584 | 0.301781726 | 0.581 | 0.517 | | 1 | 12 | hnrnpa0b |
| snx9b | 0.00444 | 0.300463269 | 0.161 | 0.072 | | 1 | 12 | snx9b |
| gstr | 0.00511 | 0.388453606 | 0.29 | 0.178 | | 1 | 12 | gstr |
| pfn2l3 | 0.005177 | 0.385147857 | 0.371 | 0.264 | | 1 | 12 | pfn2l |
| ms4a17a.111 | 0.006608 | 0.339213279 | 0.226 | 0.124 | | 1 | 12 | ms4a17a.11 |
| myl63 | 0.006764 | 0.344801546 | 0.323 | 0.205 | | 1 | 12 | myl6 |
| pdlim11 | 0.007079 | 0.27667125 | 0.194 | 0.096 | | 1 | 12 | pdlim1 |
| clic22 | 0.007267 | 0.335365048 | 0.194 | 0.098 | | 1 | 12 | clic2 |
| krt945 | 0.007306 | 0.402043002 | 0.468 | 0.364 | | 1 | 12 | krt94 |
| cavin1b2 | 0.007757 | 0.280263322 | 0.161 | 0.073 | | 1 | 12 | cavin1b |
| phlda21 | 0.008542 | 0.299790534 | 0.145 | 0.065 | | 1 | 12 | phlda2 |
| cdc42 | 0.008597 | 0.354476375 | 0.435 | 0.362 | | 1 | 12 | cdc42 |
| sept7b1 | 0.008821 | 0.377637701 | 0.258 | 0.158 | | 1 | 12 | sept7b |
| si:dkey-203a12.2 | 0 | 2.813516733 | 1 | 0.013 | | 0 | 13 | si:dkey-203a12.2 |
| si:dkeyp-75b4.10 | 0 | 2.80364611 | 1 | 0.012 | | 0 | 13 | si:dkeyp-75b4.10 |
| si:ch211-14a17.10 | 0 | 2.618035529 | 1 | 0.014 | | 0 | 13 | si:ch211-14a17.10 |

| | | | | | | | |
|---|---|---|---|---|---|---|---|
| spink2.3 | 0 | 2.401896022 | 1 | 0.014 | 0 | 13 | spink2.3 |
| itln1 | 0 | 2.310512651 | 1 | 0.014 | 0 | 13 | itln1 |
| si:ch211-194m7.8 | 0 | 2.279633668 | 1 | 0.009 | 0 | 13 | si:ch211-194m7.8 |
| LOC110437886 | 0 | 2.058184012 | 0.984 | 0.008 | 0 | 13 | LOC110437886 |
| clca1 | 0 | 2.037263823 | 0.984 | 0.006 | 0 | 13 | clca1 |
| LOC110438970 | 0 | 2.032877047 | 0.984 | 0.009 | 0 | 13 | LOC110438970 |
| CABZ01112049.1 | 0 | 2.026490309 | 0.967 | 0.005 | 0 | 13 | CABZ01112049.1 |
| cfi | 0 | 1.856743392 | 0.984 | 0.006 | 0 | 13 | cfi |
| c3a.5 | 0 | 1.792036133 | 0.934 | 0.01 | 0 | 13 | c3a.5 |
| cfhl2 | 0 | 1.462999979 | 0.82 | 0.004 | 0 | 13 | cfhl2 |
| c3a.4 | 0 | 1.401672068 | 0.787 | 0.003 | 0 | 13 | c3a.4 |
| si:ch1073-110a20.3 | 7.2840943 | 2.184942975 | 0.967 | 0.012 | 1.47116853042513e-3 | 13 | si:ch1073-110a20.3 |
| si:dkey-203a12.5 | 1.3519474 | 1.992147768 | 0.934 | 0.011 | 2.73052824789975e-3 | 13 | si:dkey-203a12.5 |
| si:dkey-77g12.4 | 7.6003513 | 2.216864265 | 0.967 | 0.014 | 1.53504295941653e-2 | 13 | si:dkey-77g12.4 |
| si:zfos-2330d3.1 | 1.5351072 | 2.038067502 | 0.967 | 0.014 | 3.10045602823225e-2 | 13 | si:zfos-2330d3.1 |
| ccl34b.9 | 1.7140447 | 2.168618472 | 0.984 | 0.016 | 3.46185614140744e-2 | 13 | ccl34b.9 |
| si:ch1073-376c22.1 | 6.7449533 | 1.472569266 | 0.803 | 0.006 | 1.3622782229585e-28 | 13 | si:ch1073-376c22.1 |
| LOC108179162 | 2.1536678 | 1.461487889 | 0.77 | 0.005 | 4.34976293978031e-2 | 13 | LOC108179162 |
| si:dkey-203a12.7 | 2.0729651 | 2.218011787 | 0.918 | 0.013 | 4.18676761602464e-2 | 13 | si:dkey-203a12.7 |
| spink2.2 | 5.7154066 | 2.749449237 | 1 | 0.019 | 1.15434068447527e-2 | 13 | spink2.2 |
| agr21 | 2.6403811 | 2.037572149 | 0.934 | 0.014 | 5.33277782006744e-2 | 13 | agr2 |
| zmp:0000001323 | 1.2005489 | 2.412608053 | 0.967 | 0.018 | 2.42474867966951e-2 | 13 | zmp:0000001323 |
| MFAP4 (1 of many) | 8.9020767 | 2.492213717 | 1 | 0.022 | 1.79795244160672e-2 | 13 | MFAP4 (1 of many) |
| FO203432.1 | 6.6226935 | 1.843340157 | 0.869 | 0.014 | 1.33758541800047e-2 | 13 | FO203432.1 |
| spink2.4 | 5.3282005 | 2.370357695 | 0.869 | 0.014 | 1.07613667194817e-2 | 13 | spink2.4 |
| si:ch73-359m17.2 | 1.6181033 | 2.438507656 | 1 | 0.025 | 3.26808338216991e-2 | 13 | si:ch73-359m17.2 |
| BX322787.1 | 1.1677813 | 2.450328101 | 0.984 | 0.027 | 2.35856803376641e-2 | 13 | BX322787.1 |
| spink2.5 | 5.8162843 | 2.01603461 | 0.77 | 0.012 | 1.17471494655155e-2 | 13 | spink2.5 |
| LOC110438722 | 6.0789420 | 1.015574267 | 0.557 | 0.002 | 1.22776391703584e-2 | 13 | LOC110438722 |
| LOC103909452 | 1.2242102 | 1.098209594 | 0.656 | 0.008 | 2.47253753852814e-1 | 13 | LOC103909452 |
| LOC101883382 | 1.1332354 | 1.059705883 | 0.574 | 0.006 | 2.28879573289586e-1 | 13 | LOC101883382 |
| nfe2 | 1.3036125 | 1.07342836 | 0.607 | 0.009 | 2.63290632885601e-1 | 13 | nfe2 |
| cfh | 1.0540628 | 1.090510905 | 0.59 | 0.008 | 2.12889064536228e-1 | 13 | cfh |
| cst14b.2 | 1.6093031 | 1.667921953 | 0.656 | 0.014 | 3.25030955874164e-1 | 13 | cst14b.2 |
| si:dkey-203a12.4 | 1.6142603 | 1.397927697 | 0.475 | 0.004 | 3.26032169696806e-1 | 13 | si:dkey-203a12.4 |

| | | | | | | | |
|---|---|---|---|---|---|---|---|
| timp4.2 | 9.6199316 | 1.080002021 | 0.557 | 0.008 | 1.94293760101466e-1 | 13 | timp4.2 |
| spink2.1 | 8.8249967 | 2.180312935 | 0.967 | 0.055 | 1.78238458521096e-1 | 13 | spink2.1 |
| LOC110437920 | 1.7547045 | 0.611947191 | 0.328 | 0 | 3.54397670966522e-1 | 13 | LOC110437920 |
| si:ch211-212d10.1 | 2.0354942 | 1.237503674 | 0.607 | 0.016 | 4.11108778512281e-1 | 13 | si:ch211-212d10.1 |
| cfbl | 7.6000218 | 1.960490502 | 0.984 | 0.073 | 1.53497640504996e-1 | 13 | cfbl |
| viml | 5.7657423 | 0.802661823 | 0.426 | 0.005 | 1.16450699200047e-1 | 13 | viml |
| LOC1104377312 | 1.1616492 | 1.753596153 | 0.967 | 0.075 | 2.34618301893288e-1 | 13 | LOC110437731 |
| si:dkey-203a12.9 | 4.2384968 | 1.091517022 | 0.344 | 0.003 | 8.56049217785535e-1 | 13 | si:dkey-203a12.9 |
| BX908782.3 | 4.2142704 | 1.147848547 | 0.639 | 0.027 | 8.51156201800952e-1 | 13 | BX908782.3 |
| adgre13 | 8.2728211 | 0.554983405 | 0.344 | 0.004 | 1.67086168917527e-1 | 13 | adgre13 |
| cfp2 | 9.7326126 | 1.405007909 | 0.869 | 0.065 | 1.96569577034921e-1 | 13 | cfp |
| CABZ01072950.1 | 1.8749745 | 0.489372811 | 0.279 | 0.001 | 3.78688604349483e-1 | 13 | CABZ01072950.1 |
| si:ch211-194m7.5 | 4.4148330 | 0.500773348 | 0.262 | 0.001 | 8.91663827789821e-1 | 13 | si:ch211-194m7.5 |
| si:dkey-27h10.2 | 2.4852900 | 0.89593664 | 0.475 | 0.015 | 5.01954033380743e-9 | 13 | si:dkey-27h10.2 |
| si:dkey-102g19.31 | 1.8699965 | 1.123545002 | 0.672 | 0.039 | 3.77683198954738e-9 | 13 | si:dkey-102g19.3 |
| si:ch211-214p16.2 | 4.1967747 | 1.495894569 | 0.852 | 0.072 | 8.47622591230642e-9 | 13 | si:ch211-214p16.2 |
| adam8a | 1.1920591 | 0.945504686 | 0.59 | 0.029 | 2.40760182255479e-8 | 13 | adam8a |
| si:ch1073-67j19.1 | 2.4307035 | 0.794972645 | 0.246 | 0.001 | 4.90929200878445e-8 | 13 | si:ch1073-67j19.1 |
| cdaa1 | 1.3283098 | 0.716430704 | 0.475 | 0.018 | 2.68278740779658e-8 | 13 | cdaa |
| CABZ01058647.1 | 1.8730969 | 1.047268668 | 0.279 | 0.003 | 3.78309400959069e-8 | 13 | CABZ01058647.1 |
| sult5a1 | 5.9057955 | 0.682919573 | 0.41 | 0.012 | 1.19279353142013e-8 | 13 | sult5a1 |
| ca6 | 9.7233875 | 0.787505697 | 0.393 | 0.011 | 1.96383258388379e-8 | 13 | ca6 |
| osmr | 1.9760321 | 0.527789615 | 0.262 | 0.003 | 3.99099217744192e-8 | 13 | osmr |
| cxcr2 | 1.7736070 | 0.392693887 | 0.23 | 0.002 | 3.58215425734213e-7 | 13 | cxcr2 |
| si:dkeyp-66d1.7 | 3.2656328 | 0.749847976 | 0.426 | 0.018 | 6.59559872551318e-7 | 13 | si:dkeyp-66d1.7 |
| gata2a | 3.0893421 | 0.856856284 | 0.541 | 0.033 | 6.2395443820372e-71 | 13 | gata2a |
| si:ch73-160i9.2 | 2.6616879 | 0.321572239 | 0.18 | 0 | 5.37581113766e-70 | 13 | si:ch73-160i9.2 |
| BX936393.1 | 4.7545752 | 0.317526107 | 0.164 | 0 | 9.6028156076456e-70 | 13 | BX936393.1 |
| BX511132.1 | 5.1792195 | 0.445057385 | 0.246 | 0.003 | 1.04604698144931e-6 | 13 | BX511132.1 |
| si:ch211-210b2.4 | 1.8608825 | 0.476104017 | 0.213 | 0.002 | 3.75842439770183e-6 | 13 | si:ch211-210b2.4 |
| si:ch211-276a23.5 | 4.3626700 | 0.67030415 | 0.393 | 0.016 | 8.81128469794617e-6 | 13 | si:ch211-276a23.5 |
| cfhl1 | 3.8130857 | 0.337645537 | 0.197 | 0.001 | 7.70128930013657e-6 | 13 | cfhl1 |
| si:ch211-270n8.1 | 4.0187077 | 0.386830901 | 0.197 | 0.001 | 8.11658410056938e-6 | 13 | si:ch211-270n8.1 |
| CU075735.1 | 1.6829913 | 0.420671939 | 0.262 | 0.005 | 3.39913762797537e-6 | 13 | CU075735.1 |
| cd59 | 4.0622498 | 0.449007772 | 0.262 | 0.006 | 8.20452605926226e-6 | 13 | cd59 |

| | | | | | | | |
|---|---|---|---|---|---|---|---|
| si:ch211-214p16.1: | 1.7153812 | 1.640775975 | 1 | 0.173 | 3.46455555037409e-5 | 13 | si:ch211-214p16.1 |
| pigrl2.3 | 4.5888885 | 0.360365277 | 0.213 | 0.003 | 9.26817819864296e-5 | 13 | pigrl2.3 |
| lrrc8da | 1.3950418 | 0.671279695 | 0.41 | 0.024 | 2.81756595445238e-5 | 13 | lrrc8da |
| h6pd | 4.4669356 | 0.423650023 | 0.246 | 0.007 | 9.02186999308911e-5 | 13 | h6pd |
| mctp2b | 1.3520928 | 0.422959397 | 0.213 | 0.005 | 2.73082184126491e-4 | 13 | mctp2b |
| itgb3a | 6.1223437 | 0.373409156 | 0.197 | 0.004 | 1.23652976570811e-4 | 13 | itgb3a |
| zgc:77112 | 1.8874597 | 0.301026704 | 0.18 | 0.003 | 3.8121025549759e-47 | 13 | zgc:77112 |
| tspan2b | 2.7897469 | 0.376260955 | 0.213 | 0.005 | 5.63445200997147e-4 | 13 | tspan2b |
| myh11a1 | 6.9511681 | 1.034086794 | 0.672 | 0.089 | 1.40392742562885e-4 | 13 | myh11a |
| XLOC-019062 | 9.1228805 | 0.33787202 | 0.18 | 0.004 | 1.84254818138573e-4 | 13 | XLOC-019062 |
| si:ch73-237c6.11 | 9.2295083 | 0.81802469 | 0.525 | 0.057 | 1.86408380637123e-4 | 13 | si:ch73-237c6.1 |
| spns3 | 4.9977074 | 0.575805295 | 0.328 | 0.02 | 1.00938697198889e-4 | 13 | spns3 |
| atp1a3a1 | 1.7325996 | 0.526321307 | 0.361 | 0.025 | 3.49933152798204e-4 | 13 | atp1a3a |
| zgc:172053 | 1.9670278 | 0.38399047 | 0.213 | 0.007 | 3.97280609748676e-4 | 13 | zgc:172053 |
| vwa11 | 1.2482949 | 0.380298046 | 0.164 | 0.003 | 2.52118121099984e-3 | 13 | vwa11 |
| limd2 | 1.2532171 | 0.782749072 | 0.492 | 0.053 | 2.53112261870038e-3 | 13 | limd2 |
| illr4 | 1.7106169 | 0.389776129 | 0.213 | 0.007 | 3.45493297860416e-3 | 13 | illr4 |
| myofl | 2.3615131 | 0.527544145 | 0.361 | 0.027 | 4.76954816166403e-3 | 13 | myofl |
| mir223 | 2.5902760 | 0.739544224 | 0.459 | 0.045 | 5.23158055974942e-3 | 13 | mir223 |
| calm1b3 | 9.0885420 | 1.035543295 | 0.754 | 0.133 | 1.83561284072297e-3 | 13 | calm1b |
| CT030188.11 | 2.2068615 | 0.813995167 | 0.541 | 0.066 | 4.45719832230802e-3 | 13 | CT030188.1 |
| si:ch211-74f19.2 | 3.0304239 | 0.597283253 | 0.328 | 0.023 | 6.12054729211169e-3 | 13 | si:ch211-74f19.2 |
| ptprr | 4.1534634 | 0.424601576 | 0.246 | 0.012 | 8.38875009179339e-3 | 13 | ptprr |
| cd7al | 1.4332704 | 0.55706762 | 0.377 | 0.032 | 2.89477632508978e-3 | 13 | cd7al |
| itgb21 | 1.0744589 | 0.99257503 | 0.803 | 0.15 | 2.17008471088005e-3 | 13 | itgb2 |
| CR848841.1 | 5.7531770 | 0.379911553 | 0.213 | 0.009 | 1.16196916847401e-3 | 13 | CR848841.1 |
| si:ch1073-443f11.2 | 5.7830336 | 0.660870361 | 0.492 | 0.058 | 1.16799930540962e-3 | 13 | si:ch1073-443f11.2 |
| gpr183b | 7.4088728 | 0.309487892 | 0.164 | 0.004 | 1.49637004137358e-3 | 13 | gpr183b |
| qsox1 | 1.1729013 | 0.620026536 | 0.41 | 0.041 | 2.36890894418837e-3 | 13 | qsox1 |
| slc44a4 | 7.0159866 | 0.26306235 | 0.164 | 0.005 | 1.41701881702966e-3 | 13 | slc44a4 |
| rgcc2 | 2.0118940 | 0.691699126 | 0.508 | 0.067 | 4.06342237887809e-3 | 13 | rgcc |
| pygb | 2.3286261 | 0.643970045 | 0.393 | 0.042 | 4.70312632284196e-3 | 13 | pygb |
| myh9a1 | 1.0550495 | 0.509257479 | 0.344 | 0.031 | 2.13088365966004e-2 | 13 | myh9a |
| il16 | 2.8464990 | 0.611881503 | 0.377 | 0.039 | 5.74907415417549e-2 | 13 | il16 |
| glipr1a2 | 3.9531646 | 0.822309103 | 0.541 | 0.083 | 7.98420671207531e-2 | 13 | glipr1a |

| | | | | | | | |
|---|---|---|---|---|---|---|---|
| capgb3 | 6.0619006 | 0.939146775 | 0.639 | 0.115 | 1.22432207206703e-2 | 13 | capgb |
| p4hb1 | 1.8593105 | 1.01834261 | 0.803 | 0.204 | 3.755249612902e-28 | 13 | p4hb |
| atp2a3 | 2.7512135 | 0.591158405 | 0.393 | 0.044 | 5.55662609793356e-2 | 13 | atp2a3 |
| prkcbb | 1.3141492 | 0.428160798 | 0.311 | 0.027 | 2.65418718005492e-2 | 13 | prkcbb |
| cyp4f31 | 3.6390953 | 0.522211475 | 0.279 | 0.022 | 7.34988086691417e-2 | 13 | cyp4f3 |
| hacd3 | 3.8720488 | 0.554962387 | 0.377 | 0.041 | 7.82037714222907e-2 | 13 | hacd3 |
| si:dkey-261l7.2 | 4.1205241 | 0.282968756 | 0.131 | 0.003 | 8.32222260399185e-2 | 13 | si:dkey-261l7.2 |
| gfi1ab1 | 4.8438731 | 0.381064209 | 0.279 | 0.022 | 9.78317061587851e-2 | 13 | gfi1ab |
| ikzf22 | 1.1458892 | 0.571565579 | 0.41 | 0.049 | 2.31435257215301e-2 | 13 | ikzf2 |
| itpr1b | 1.6037382 | 0.345091736 | 0.18 | 0.008 | 3.23907007113605e-2 | 13 | itpr1b |
| srgn2 | 1.4312851 | 1.023336971 | 0.852 | 0.232 | 2.89076663034166e-2 | 13 | srgn |
| scinlb1 | 3.6787070 | 0.669886181 | 0.541 | 0.087 | 7.42988454249855e-2 | 13 | scinlb |
| alox5a | 3.7829402 | 0.544243112 | 0.295 | 0.027 | 7.64040442111399e-2 | 13 | alox5a |
| clstn1 | 5.5487265 | 0.465109689 | 0.279 | 0.025 | 1.12067629703267e-2 | 13 | clstn1 |
| cdkn3 | 3.3501962 | 0.324815338 | 0.213 | 0.014 | 6.76639133659639e-2 | 13 | cdkn3 |
| selplg1 | 5.5642821 | 0.486915998 | 0.328 | 0.036 | 1.12381807136873e-2 | 13 | selplg |
| ccr9a2 | 7.3082534 | 0.679806024 | 0.525 | 0.088 | 1.47604794085984e-2 | 13 | ccr9a |
| cox6b11 | 1.2448677 | 0.732113862 | 0.607 | 0.123 | 2.51425941996738e-2 | 13 | cox6b1 |
| def6c | 2.7242029 | 0.251532802 | 0.213 | 0.015 | 5.50207270087153e-2 | 13 | def6c |
| samsn1b1 | 5.6954291 | 0.392884609 | 0.213 | 0.015 | 1.15030582255856e-2 | 13 | samsn1b |
| coro1a4 | 6.1035018 | 1.061677609 | 0.852 | 0.27 | 1.23272427642558e-2 | 13 | coro1a |
| si:dkey-21o19.2 | 7.7563450 | 0.320671304 | 0.197 | 0.013 | 1.56654900506163e-2 | 13 | si:dkey-21o19.2 |
| LOC1001482541 | 1.7493158 | 0.450332034 | 0.279 | 0.028 | 3.53309312772656e-2 | 13 | LOC100148254 |
| si:ch73-208g10.11 | 1.9061136 | 0.412967109 | 0.279 | 0.028 | 3.84977769463795e-2 | 13 | si:ch73-208g10.1 |
| ssr32 | 2.2906955 | 1.006544463 | 0.869 | 0.375 | 4.62651770410799e-2 | 13 | ssr3 |
| galnt12 | 4.3791594 | 0.411907358 | 0.246 | 0.022 | 8.84458840113206e-2 | 13 | galnt12 |
| si:dkey-42l23.2 | 2.0624948 | 0.260223798 | 0.18 | 0.011 | 4.16562084782734e-2 | 13 | si:dkey-42l23.2 |
| pla2g4aa | 3.9298901 | 0.405086141 | 0.23 | 0.02 | 7.93719917282013e-2 | 13 | pla2g4aa |
| unc13d1 | 2.5314335 | 0.443400701 | 0.279 | 0.031 | 5.11273627423897e-1 | 13 | unc13d |
| slc6a6b1 | 4.7442988 | 0.643671264 | 0.393 | 0.063 | 9.58206036427254e-1 | 13 | slc6a6b |
| syngr2a1 | 7.2063411 | 0.56265204 | 0.377 | 0.059 | 1.45546471228181e-1 | 13 | syngr2a |
| alox5ap1 | 1.4025904 | 0.406435881 | 0.279 | 0.031 | 2.83281184763107e-1 | 13 | alox5ap |
| pdia4 | 2.8773756 | 0.789219476 | 0.443 | 0.085 | 5.81143559699019e-1 | 13 | pdia4 |
| si:cabz01036022.1 | 6.0113374 | 0.319657774 | 0.148 | 0.008 | 1.21410983198377e-1 | 13 | si:cabz01036022.1 |
| peak3 | 7.5957821 | 0.256145203 | 0.131 | 0.006 | 1.53412012765823e-1 | 13 | peak3 |

| | | | | | | | |
|---|---|---|---|---|---|---|---|
| grap2b4 | 8.2632936 | 0.882031172 | 0.672 | 0.183 | 1.66893741397965e-1 | 13 | grap2b |
| nucb2b1 | 1.4714396 | 0.426114842 | 0.295 | 0.037 | 2.97186656898482e-1 | 13 | nucb2b |
| bach1b1 | 1.5343599 | 0.360302032 | 0.262 | 0.029 | 3.09894679607159e-1 | 13 | bach1b |
| rogdi | 1.5569526 | 0.32734965 | 0.23 | 0.022 | 3.14457718070662e-1 | 13 | rogdi |
| cyba2 | 2.2524199 | 0.721935251 | 0.541 | 0.124 | 4.54921256125159e-1 | 13 | cyba |
| rasal3 | 2.7874167 | 0.382637515 | 0.23 | 0.023 | 5.62974565185005e-1 | 13 | rasal3 |
| si:dkey-126g1.7 | 4.5772442 | 0.287026741 | 0.148 | 0.009 | 9.24466020455051e-1 | 13 | si:dkey-126g1.7 |
| si:dkey-24p1.7 | 1.0515581 | 0.256390998 | 0.164 | 0.011 | 2.12383205493932e-1 | 13 | si:dkey-24p1.7 |
| ehd1a | 1.0739304 | 0.36291688 | 0.279 | 0.035 | 2.16901735902921e-1 | 13 | ehd1a |
| fmnl1a1 | 1.1324195 | 0.411491683 | 0.279 | 0.035 | 2.28714768476794e-1 | 13 | fmnl1a |
| nfkbiaa2 | 1.4013475 | 0.850563332 | 0.59 | 0.155 | 2.83030173826874e-1 | 13 | nfkbiaa |
| vaspa2 | 2.9189564 | 0.567572283 | 0.541 | 0.111 | 5.89541642169517e-1 | 13 | vaspa |
| CT573383.11 | 3.1440550 | 0.319209735 | 0.197 | 0.017 | 6.35004788617531e-1 | 13 | CT573383.1 |
| atf6 | 3.8500504 | 0.588811504 | 0.443 | 0.088 | 7.775946835074e-16 | 13 | atf6 |
| lpcat2 | 4.5574528 | 0.355629371 | 0.246 | 0.028 | 9.20468749282662e-1 | 13 | lpcat2 |
| sec24d | 4.5809000 | 0.335116005 | 0.213 | 0.021 | 9.25204385289475e-1 | 13 | sec24d |
| zmp:00000010821 | 6.0493456 | 0.428501276 | 0.311 | 0.044 | 1.2217863399025e-15 | 13 | zmp:0000001082 |
| camk1b | 1.5548006 | 0.312142282 | 0.213 | 0.021 | 3.1402308564094e-15 | 13 | camk1b |
| gsr | 1.8538605 | 0.51679925 | 0.311 | 0.048 | 3.74424208643205e-1 | 13 | gsr |
| ltb4r1 | 2.7402377 | 0.400085399 | 0.213 | 0.022 | 5.53445821697407e-1 | 13 | ltb4r |
| xbp11 | 2.8482503 | 0.80042603 | 0.902 | 0.425 | 5.7526112689582e-15 | 13 | xbp1 |
| rac24 | 3.4667985 | 0.81450006 | 0.82 | 0.249 | 7.00189297514531e-1 | 13 | rac2 |
| fybb1 | 5.0208268 | 0.375023353 | 0.344 | 0.054 | 1.01405639854208e-1 | 13 | fybb |
| btk | 6.4695356 | 0.349480494 | 0.295 | 0.041 | 1.30665211088005e-1 | 13 | btk |
| dhrs13a.1 | 1.1477429 | 0.408311849 | 0.23 | 0.027 | 2.31809648009503e-1 | 13 | dhrs13a.1 |
| eno1a2 | 1.5831000 | 0.740546132 | 0.607 | 0.18 | 3.19738719419257e-1 | 13 | eno1a |
| myo1f2 | 1.6393464 | 0.410380095 | 0.344 | 0.056 | 3.31098803573987e-1 | 13 | myo1f |
| glipr21 | 1.7628580 | 0.354475772 | 0.23 | 0.026 | 3.56044433818015e-1 | 13 | glipr2 |
| dnajc3a | 2.4426292 | 0.421031443 | 0.246 | 0.031 | 4.9333783475817e-14 | 13 | dnajc3a |
| nr4a12 | 3.3199139 | 0.647080524 | 0.508 | 0.118 | 6.70523016561533e-1 | 13 | nr4a1 |
| syk2 | 3.3242870 | 0.493075382 | 0.393 | 0.074 | 6.71406260863909e-1 | 13 | syk |
| LO017700.1 | 4.3389656 | 0.25251391 | 0.131 | 0.008 | 8.76340883255226e-1 | 13 | LO017700.1 |
| si:ch211-136m16.8 | 8.2317820 | 0.310638847 | 0.246 | 0.031 | 1.66257301161755e-1 | 13 | si:ch211-136m16.8 |
| aldocb1 | 1.1023626 | 0.467239735 | 0.328 | 0.055 | 2.22644182780812e-1 | 13 | aldocb |
| si:zfos-741a10.32 | 1.7723568 | 0.473684675 | 0.311 | 0.05 | 3.57962913218112e-1 | 13 | si:zfos-741a10.3 |

| | | | | | | | |
|---|---|---|---|---|---|---|---|
| ctss1 | 2.4317291 | 0.253973292 | 0.197 | 0.02 | 4.91136332164729e-1 | 13 | ctss1 |
| aspm | 2.6595059 | 0.25354409 | 0.148 | 0.011 | 5.37140410805311e-1 | 13 | aspm |
| fcer1gl3 | 2.8739307 | 0.805940456 | 0.885 | 0.244 | 5.8044778540508e-13 | 13 | fcer1gl |
| rgs14a | 3.4432595 | 0.280343638 | 0.213 | 0.024 | 6.95435127424525e-1 | 13 | rgs14a |
| moxd11 | 9.0360242 | 0.280104376 | 0.164 | 0.015 | 1.82500580928292e-1 | 13 | moxd1 |
| camkk1a | 1.2068247 | 0.25868929 | 0.115 | 0.007 | 2.43742399097629e-1 | 13 | camkk1a |
| cnpy11 | 1.3241496 | 0.629305438 | 0.426 | 0.102 | 2.67438494771476e-1 | 13 | cnpy1 |
| slc2a3b | 4.1485717 | 0.262248248 | 0.18 | 0.019 | 8.37887045763401e-1 | 13 | slc2a3b |
| gyg1a | 4.8169112 | 0.503719878 | 0.377 | 0.079 | 9.72871566726553e-1 | 13 | gyg1a |
| rasgrp4 | 4.9173052 | 0.260743162 | 0.18 | 0.019 | 9.931481471346e-12 | 13 | rasgrp4 |
| si:dkey-185m8.21 | 6.1040884 | 0.381766696 | 0.262 | 0.04 | 1.2328427467017e-11 | 13 | si:dkey-185m8.2 |
| XLOC-006268 | 8.9586232 | 0.286196608 | 0.148 | 0.013 | 1.80937312940221e-1 | 13 | XLOC-006268 |
| pfn14 | 2.6269393 | 0.947316044 | 1 | 0.407 | 5.30562933056113e-1 | 13 | pfn1 |
| tmsb15 | 3.6133746 | 0.698965545 | 0.623 | 0.185 | 7.29793285983095e-1 | 13 | tmsb1 |
| cebpb3 | 3.8106821 | 0.739052654 | 0.787 | 0.285 | 7.69643464175893e-1 | 13 | cebpb |
| hsp90b11 | 4.8847085 | 0.725680109 | 0.541 | 0.168 | 9.86564585478964e-1 | 13 | hsp90b1 |
| CU019662.11 | 5.5916739 | 0.292517791 | 0.197 | 0.024 | 1.12935039252289e-1 | 13 | CU019662.1 |
| hspa52 | 6.0263387 | 0.875401776 | 0.59 | 0.215 | 1.21713962968019e-1 | 13 | hspa5 |
| cbfa2t3 | 8.6912704 | 0.46642851 | 0.262 | 0.044 | 1.75537588775642e-1 | 13 | cbfa2t3 |
| actr2a2 | 9.2995251 | 0.577012148 | 0.541 | 0.155 | 1.87822509388819e-1 | 13 | actr2a |
| lrrfip1a | 1.0079027 | 0.385439288 | 0.361 | 0.074 | 2.03566117649392e-1 | 13 | lrrfip1a |
| si:ch73-248e21.71 | 1.4126102 | 0.440977096 | 0.344 | 0.07 | 2.85304884140609e-1 | 13 | si:ch73-248e21.7 |
| ostf11 | 1.7688593 | 0.368259691 | 0.295 | 0.053 | 3.57256518295724e-1 | 13 | ostf1 |
| dock81 | 1.7879402 | 0.307023959 | 0.23 | 0.034 | 3.6111028957576e-10 | 13 | dock8 |
| b4galt4 | 2.6568044 | 0.256872996 | 0.131 | 0.011 | 5.36594796102267e-1 | 13 | b4galt4 |
| pycard3 | 2.9864834 | 0.612615703 | 0.492 | 0.139 | 6.03180071728392e-1 | 13 | pycard |
| syvn1 | 4.6209151 | 0.337117277 | 0.197 | 0.026 | 9.33286227687834e-1 | 13 | syvn1 |
| hk1 | 4.6209151 | 0.318776156 | 0.197 | 0.026 | 9.33286227687834e-1 | 13 | hk1 |
| cotl14 | 4.6507459 | 0.789653563 | 0.902 | 0.402 | 9.39311159971385e-1 | 13 | cotl1 |
| mbnl12 | 8.4859498 | 0.589301848 | 0.557 | 0.178 | 1.71390729817155e-0 | 13 | mbnl1 |
| ptpn63 | 9.0170236 | 0.518444476 | 0.508 | 0.138 | 1.82116826816363e-0 | 13 | ptpn6 |
| camk2g2 | 1.0013185 | 0.370617848 | 0.262 | 0.046 | 2.02236313819404e-0 | 13 | camk2g2 |
| far1 | 1.0065590 | 0.406804336 | 0.246 | 0.041 | 2.0329472748293e-09 | 13 | far1 |
| mylk5 | 1.2444418 | 0.260930188 | 0.164 | 0.019 | 2.51339922530707e-0 | 13 | mylk5 |
| rab441 | 1.5399632 | 0.361004628 | 0.311 | 0.061 | 3.11026386827033e-0 | 13 | rab44 |

| | | | | | | | |
|---|---|---|---|---|---|---|---|
| plxnc11 | 2.5129542 | 0.473691953 | 0.295 | 0.06 | 5.0754136591047e-09 | 13 | plxnc1 |
| p2ry101 | 2.7075413 | 0.293542103 | 0.18 | 0.023 | 5.46842116877243e-0 | 13 | p2ry10 |
| yrk2 | 3.3552435 | 0.393580823 | 0.377 | 0.086 | 6.77658530854842e-0 | 13 | yrk |
| syne2b | 3.7308810 | 0.277521072 | 0.148 | 0.016 | 7.53526052124645e-0 | 13 | syne2b |
| prkcda | 6.7654854 | 0.314006569 | 0.213 | 0.033 | 1.36642510056381e-0 | 13 | prkcda |
| tram11 | 7.9442266 | 0.522246485 | 0.443 | 0.128 | 1.60449545555926e-0 | 13 | tram1 |
| LOC103911430 | 8.3716086 | 0.332670191 | 0.23 | 0.038 | 1.69081378948896e-0 | 13 | LOC103911430 |
| arhgap4b1 | 1.0716228 | 0.287171243 | 0.197 | 0.029 | 2.16435667385907e-0 | 13 | arhgap4b |
| arhgap20b | 1.0963075 | 0.295096669 | 0.164 | 0.02 | 2.21421240508823e-0 | 13 | arhgap20b |
| si:dkeyp-68b7.121 | 1.2843705 | 0.312663827 | 0.262 | 0.048 | 2.59404329856584e-0 | 13 | si:dkeyp-68b7.12 |
| lta4h2 | 1.5867121 | 0.41735061 | 0.377 | 0.091 | 3.20468246497636e-0 | 13 | lta4h |
| itgae.22 | 1.7855005 | 0.289707594 | 0.23 | 0.038 | 3.60617544764893e-0 | 13 | itgae.2 |
| sec61a1l | 4.8894828 | 0.314288334 | 0.295 | 0.063 | 9.87528857758289e-0 | 13 | sec61a1l |
| MYADM1 | 5.1248756 | 0.345077851 | 0.295 | 0.062 | 1.03507114011656e-0 | 13 | MYADM |
| acap1 | 5.4979713 | 0.316135516 | 0.197 | 0.031 | 1.11042527487733e-0 | 13 | acap1 |
| sgk11 | 8.2264660 | 0.383730784 | 0.311 | 0.071 | 1.66149934870073e-0 | 13 | sgk1 |
| si:ch211-226m16.2 | 9.2279825 | 0.338703638 | 0.311 | 0.07 | 1.86377562923037e-0 | 13 | si:ch211-226m16.2 |
| spi1b3 | 1.2153325 | 0.463734079 | 0.443 | 0.122 | 2.4546070942877e-07 | 13 | spi1b |
| cxcr4b2 | 1.3052084 | 0.520756224 | 0.508 | 0.151 | 2.63612943578826e-0 | 13 | cxcr4b |
| slc9a3r1a | 1.3916818 | 0.357949971 | 0.279 | 0.059 | 2.81077976864486e-0 | 13 | slc9a3r1a |
| arhgdig3 | 1.5262990 | 0.638009668 | 0.787 | 0.358 | 3.08266618186654e-0 | 13 | arhgdig |
| grasp | 1.9269013 | 0.411365762 | 0.262 | 0.054 | 3.89176256908525e-0 | 13 | grasp |
| hephl1a | 1.9630722 | 0.275568076 | 0.18 | 0.027 | 3.96481712372084e-0 | 13 | hephl1a |
| h1fx2 | 2.0846615 | 0.580707867 | 0.639 | 0.23 | 4.21039096831465e-0 | 13 | h1fx |
| sec11a1 | 2.3266577 | 0.600856879 | 0.623 | 0.265 | 4.69915065502721e-0 | 13 | sec11a |
| rgs191 | 2.9212690 | 0.271536545 | 0.23 | 0.042 | 5.90008715130898e-0 | 13 | rgs19 |
| cd632 | 2.9525172 | 0.570264643 | 0.885 | 0.563 | 5.96319902248734e-0 | 13 | cd63 |
| wdr13 | 3.6646694 | 0.589986288 | 0.59 | 0.231 | 7.40153291771282e-0 | 13 | wdr1 |
| sec22ba | 4.4301286 | 0.279869486 | 0.246 | 0.049 | 8.9475308816421e-07 | 13 | sec22ba |
| wasa2 | 4.5547552 | 0.426841416 | 0.41 | 0.113 | 9.1992390859178e-07 | 13 | wasa |
| pip4k2aa | 5.5684009 | 0.260216813 | 0.18 | 0.028 | 1.12464993921486e-0 | 13 | pip4k2aa |
| zgc:638311 | 6.8237165 | 0.385030462 | 0.393 | 0.108 | 1.37818602589155e-0 | 13 | zgc:63831 |
| ddost | 6.8518335 | 0.493431934 | 0.459 | 0.152 | 1.3838648266289e-06 | 13 | ddost |
| CABZ01076758.1 | 7.4029926 | 0.252366538 | 0.164 | 0.024 | 1.49518243389423e-0 | 13 | CABZ01076758.1 |
| rhogb2 | 7.4376335 | 0.521483832 | 0.492 | 0.164 | 1.50217885240371e-0 | 13 | rhogb |

| | | | | | | | |
|---|---|---|---|---|---|---|---|
| mgst1.22 | 1.1368494 | 0.467724001 | 0.508 | 0.174 | 2.29609490213883e-0 | 13 | mgst1.2 |
| ank3b2 | 1.3743261 | 0.351089276 | 0.213 | 0.04 | 2.77572658632902e-0 | 13 | ank3b |
| tpmt.1 | 1.5963569 | 0.273756226 | 0.213 | 0.04 | 3.22416220475826e-0 | 13 | tpmt.1 |
| erlec1 | 2.0423023 | 0.317703973 | 0.295 | 0.071 | 4.12483810913871e-0 | 13 | erlec1 |
| CABZ01073834.11 | 2.1704689 | 0.29489112 | 0.262 | 0.057 | 4.38369616888289e-0 | 13 | CABZ01073834.1 |
| prdx4 | 2.3802798 | 0.54011855 | 0.557 | 0.231 | 4.80745123863351e-0 | 13 | prdx4 |
| fli1b2 | 2.8963662 | 0.251768637 | 0.197 | 0.035 | 5.84979095328046e-0 | 13 | fli1b |
| arpc1b4 | 2.9743886 | 0.65389162 | 0.852 | 0.351 | 6.00737278209162e-0 | 13 | arpc1b |
| gmds | 4.6265517 | 0.316505049 | 0.246 | 0.054 | 9.34424660484514e-0 | 13 | gmds |
| abi3a2 | 5.2271233 | 0.286244778 | 0.279 | 0.064 | 1.05572210662744e-0 | 13 | abi3a |
| actn1 | 5.8170201 | 0.31537153 | 0.279 | 0.066 | 1.17486356101097e-0 | 13 | actn1 |
| hck | 6.0961345 | 0.262000738 | 0.197 | 0.036 | 1.23123629548974e-0 | 13 | hck |
| furina | 6.7619566 | 0.372489764 | 0.311 | 0.082 | 1.36571238608779e-0 | 13 | furina |
| elmo12 | 7.5821998 | 0.310463246 | 0.279 | 0.066 | 1.53137691223117e-0 | 13 | elmo1 |
| snd11 | 7.8153320 | 0.440963184 | 0.492 | 0.178 | 1.57846260689219e-0 | 13 | snd1 |
| laptm54 | 8.1605362 | 0.529878893 | 0.59 | 0.209 | 1.64818350630425e-0 | 13 | laptm5 |
| dnajb9b | 9.1355427 | 0.354584981 | 0.148 | 0.022 | 1.84510556754278e-0 | 13 | dnajb9b |
| gmip | 1.1113366 | 0.264425367 | 0.131 | 0.018 | 2.24456673226718e-0 | 13 | gmip |
| spcs21 | 1.1711198 | 0.51822349 | 0.59 | 0.248 | 2.36531067526143e-0 | 13 | spcs2 |
| ssr21 | 1.2312743 | 0.537114767 | 0.77 | 0.413 | 2.48680486116316e-0 | 13 | ssr2 |
| cds2 | 1.5385753 | 0.297243277 | 0.197 | 0.038 | 3.10746055976312e-0 | 13 | cds2 |
| pkma2 | 2.0380432 | 0.479630737 | 0.508 | 0.192 | 4.11623593399789e-0 | 13 | pkma |
| si:dkey-118j18.21 | 3.1332029 | 0.341341026 | 0.279 | 0.071 | 6.32812999436147e-0 | 13 | si:dkey-118j18.2 |
| manf1 | 3.2049903 | 0.486025025 | 0.361 | 0.117 | 6.47311901564819e-0 | 13 | manf |
| wasb3 | 3.4321559 | 0.459357411 | 0.459 | 0.154 | 6.9319254294664e-05 | 13 | wasb |
| msna3 | 3.7355458 | 0.539976885 | 0.77 | 0.396 | 7.5446819964289e-05 | 13 | msna |
| hmha1b2 | 4.3169101 | 0.347343403 | 0.279 | 0.072 | 8.71886347037661e-0 | 13 | hmha1b |
| dnajb11 | 6.0667393 | 0.349797588 | 0.262 | 0.067 | 0.00012253 | 13 | dnajb11 |
| sec61g2 | 6.4580971 | 0.409774608 | 0.869 | 0.725 | 0.000130434 | 13 | sec61g |
| sdf2l11 | 7.4131316 | 0.359315948 | 0.295 | 0.082 | 0.000149723 | 13 | sdf2l1 |
| si:dkeyp-27c8.21 | 7.6456792 | 0.256291412 | 0.246 | 0.058 | 0.00015442 | 13 | si:dkeyp-27c8.2 |
| pdia33 | 1.0254607 | 0.518444995 | 0.787 | 0.48 | 0.000207112 | 13 | pdia3 |
| gng2 | 1.0792894 | 0.258755711 | 0.295 | 0.077 | 0.000217984 | 13 | gng2 |
| mrtfab1 | 1.2314312 | 0.313994673 | 0.213 | 0.048 | 0.000248712 | 13 | mrtfab |
| ost41 | 1.3476747 | 0.517667682 | 0.607 | 0.293 | 0.00027219 | 13 | ost4 |

| | | | | | | | |
|---|---|---|---|---|---|---|---|
| tes | 1.4925746 | 0.333476273 | 0.279 | 0.075 | 0.000301455 | 13 | tes |
| ldha | 1.6422457 | 0.287792926 | 0.23 | 0.054 | 0.000331684 | 13 | ldha |
| inpp5d | 1.6487818 | 0.256116464 | 0.197 | 0.041 | 0.000333004 | 13 | inpp5d |
| ssr41 | 1.7171910 | 0.538408749 | 0.738 | 0.48 | 0.000346821 | 13 | ssr4 |
| cd81a4 | 1.9132352 | 0.557712029 | 0.803 | 0.435 | 0.000386416 | 13 | cd81a |
| cyth1a | 2.1567781 | 0.279702748 | 0.246 | 0.062 | 0.000435604 | 13 | cyth1a |
| selenos | 2.2234277 | 0.32378631 | 0.246 | 0.063 | 0.000449066 | 13 | selenos |
| nccrp12 | 2.5526747 | 0.405978383 | 0.541 | 0.216 | 0.000515564 | 13 | nccrp1 |
| sh3kbp11 | 2.6547948 | 0.324491309 | 0.344 | 0.107 | 0.000536189 | 13 | sh3kbp1 |
| herpud1 | 3.6966570 | 0.291558869 | 0.246 | 0.064 | 0.000746614 | 13 | herpud1 |
| arpc5a1 | 4.2931819 | 0.428120615 | 0.475 | 0.184 | 0.000867094 | 13 | arpc5a |
| zgc:640514 | 4.7553071 | 0.413239142 | 0.475 | 0.172 | 0.000960429 | 13 | zgc:64051 |
| lrrc591 | 4.9497016 | 0.320427717 | 0.311 | 0.095 | 0.000999691 | 13 | lrrc59 |
| dnajc11 | 5.1493217 | 0.389679998 | 0.377 | 0.132 | 0.001040009 | 13 | dnajc1 |
| ptprc2 | 5.5175002 | 0.355456406 | 0.361 | 0.116 | 0.00111437 | 13 | ptprc |
| agap21 | 6.4707696 | 0.264354434 | 0.131 | 0.022 | 0.001306901 | 13 | agap2 |
| calr3a2 | 9.1109439 | 0.438971558 | 0.541 | 0.231 | 0.001840137 | 13 | calr3a |
| srp54 | 1.8907417 | 0.273328407 | 0.279 | 0.082 | 0.003818731 | 13 | srp54 |
| si:dkey-73n10.1 | 1.9432267 | 0.271112558 | 0.164 | 0.035 | 0.003924735 | 13 | si:dkey-73n10.1 |
| rab11fip1b | 1.9659240 | 0.260410616 | 0.262 | 0.074 | 0.003970577 | 13 | rab11fip1b |
| actb14 | 1.9795196 | 0.421817858 | 0.984 | 0.737 | 0.003998036 | 13 | actb1 |
| stk17b2 | 2.0959465 | 0.387846392 | 0.377 | 0.135 | 0.004233183 | 13 | stk17b |
| LOC1039091153 | 2.3190755 | 0.428503944 | 0.475 | 0.196 | 0.004683837 | 13 | LOC103909115 |
| hmgn31 | 2.5662275 | 0.285769818 | 0.443 | 0.162 | 0.00518301 | 13 | hmgn3 |
| si:dkey-56i24.1 | 2.5787122 | 0.266591922 | 0.131 | 0.024 | 0.005208225 | 13 | si:dkey-56i24.1 |
| hyou12 | 2.7362942 | 0.364758341 | 0.262 | 0.079 | 0.005526494 | 13 | hyou1 |
| zgc:172106 | 2.7438056 | 0.310908614 | 0.213 | 0.055 | 0.005541664 | 13 | zgc:172106 |
| hlx11 | 2.7520939 | 0.259649305 | 0.197 | 0.047 | 0.005558404 | 13 | hlx1 |
| si:ch1073-392o20.[illegible] | 3.7121045 | 0.29318213 | 0.246 | 0.07 | 0.007497338 | 13 | si:ch1073-392o20.2 |
| si:ch211-102c2.44 | 3.9815937 | 0.329864271 | 0.475 | 0.171 | 0.008041625 | 13 | si:ch211-102c2.4 |
| clint1a | 5.5620142 | 0.280708624 | 0.279 | 0.086 | 0.0112336 | 13 | clint1a |
| arrb2b | 6.8791929 | 0.366833058 | 0.344 | 0.125 | 0.013893906 | 13 | arrb2b |
| cebpa4 | 7.8554039 | 0.333524728 | 0.443 | 0.169 | 0.015865559 | 13 | cebpa |
| ssr1 | 9.0014283 | 0.46336065 | 0.541 | 0.272 | 0.018180185 | 13 | ssr1 |
| samsn1a2 | 1.2600779 | 0.275437181 | 0.197 | 0.051 | 0.025449795 | 13 | samsn1a |

| | | | | | | | |
|---|---|---|---|---|---|---|---|
| calr3b1 | 1.8657867 | 0.535241828 | 0.393 | 0.181 | 0.037683295 | 13 | calr3b |
| cap14 | 2.0359892 | 0.451774144 | 0.623 | 0.326 | 0.041120875 | 13 | cap1 |
| cdipt | 2.0984856 | 0.279502004 | 0.279 | 0.091 | 0.042383115 | 13 | cdipt |
| rab5aa | 2.6293863 | 0.314626794 | 0.328 | 0.119 | 0.053105715 | 13 | rab5aa |
| tmed10 | 2.8396676 | 0.389212115 | 0.492 | 0.239 | 0.057352767 | 13 | tmed10 |
| cox8a | 2.8776200 | 0.420646575 | 0.705 | 0.47 | 0.058119291 | 13 | cox8a |
| nnt | 3.2527733 | 0.258981321 | 0.213 | 0.063 | 0.065696264 | 13 | nnt |
| pgd1 | 3.3821110 | 0.304921137 | 0.311 | 0.112 | 0.068308496 | 13 | pgd |
| si:dkey-262k9.43 | 3.4748718 | 0.27316418 | 0.279 | 0.092 | 0.070181986 | 13 | si:dkey-262k9.4 |
| si:ch73-234b20.51 | 4.1584557 | 0.303590517 | 0.344 | 0.133 | 0.08398833 | 13 | si:ch73-234b20.5 |
| parp8 | 4.2010328 | 0.298287094 | 0.164 | 0.042 | 0.08484826 | 13 | parp8 |
| zbtb20 | 5.4857349 | 0.395953019 | 0.361 | 0.148 | 0.110795388 | 13 | zbtb20 |
| hcls13 | 6.8282512 | 0.25779294 | 0.311 | 0.109 | 0.13791019 | 13 | hcls1 |
| rpn11 | 9.4291065 | 0.382045981 | 0.475 | 0.233 | 0.190439665 | 13 | rpn1 |
| arpc34 | 1.2574521 | 0.384969323 | 0.803 | 0.475 | 0.253967617 | 13 | arpc3 |
| arhgdia2 | 1.3151427 | 0.399704382 | 0.656 | 0.37 | 0.265619389 | 13 | arhgdia |
| krtcap21 | 1.5438908 | 0.40544304 | 0.541 | 0.303 | 0.311819637 | 13 | krtcap2 |
| rca2.1 | 1.8090103 | 0.290549202 | 0.246 | 0.087 | 0.365365816 | 13 | rca2.1 |
| ppib3 | 1.8234676 | 0.386620045 | 0.918 | 0.639 | 0.368285765 | 13 | ppib |
| rnf130 | 1.8437951 | 0.297533564 | 0.246 | 0.086 | 0.372391297 | 13 | rnf130 |
| crip13 | 1.8764138 | 0.414089333 | 0.705 | 0.373 | 0.378979307 | 13 | crip1 |
| arpc43 | 2.4686635 | 0.354392367 | 0.59 | 0.312 | 0.498595979 | 13 | arpc4 |
| hm13 | 3.1017370 | 0.309479386 | 0.311 | 0.127 | 0.626457834 | 13 | hm13 |
| dbi2 | 3.1232443 | 0.397326128 | 0.656 | 0.4 | 0.630801667 | 13 | dbi |
| clic12 | 3.8976377 | 0.299182247 | 0.459 | 0.213 | 0.787205891 | 13 | clic1 |
| pdia61 | 4.5318653 | 0.385644149 | 0.361 | 0.169 | 0.915300852 | 13 | pdia6 |
| rrbp1a | 4.9078959 | 0.252879592 | 0.262 | 0.1 | 0.991247754 | 13 | rrbp1a |
| selenof1 | 5.9655151 | 0.360254431 | 0.689 | 0.421 | 1 | 13 | selenof |
| arf1 | 6.7515238 | 0.316550073 | 0.311 | 0.133 | 1 | 13 | arf1 |
| itgav1 | 7.0203676 | 0.258216277 | 0.246 | 0.091 | 1 | 13 | itgav |
| tmed9 | 7.4245860 | 0.294901642 | 0.361 | 0.162 | 1 | 13 | tmed9 |
| gapdhs1 | 7.7762061 | 0.370938353 | 0.721 | 0.473 | 1 | 13 | gapdhs |
| capzb2 | 0.000103 | 0.351820912 | 0.639 | 0.383 | 1 | 13 | capzb |
| hmgb2a3 | 0.000108 | 0.373682694 | 0.82 | 0.581 | 1 | 13 | hmgb2a |
| mt-co12 | 0.000169 | 0.377728696 | 0.836 | 0.59 | 1 | 13 | mt-co1 |

| | | | | | | | |
|---|---|---|---|---|---|---|---|
| twf1a1 | 0.000187 | 0.317365038 | 0.393 | 0.2 | 1 | 13 | twf1a |
| RASA22 | 0.000242 | 0.262588584 | 0.164 | 0.054 | 1 | 13 | RASA2 |
| ostc1 | 0.000327 | 0.29965964 | 0.459 | 0.25 | 1 | 13 | ostc |
| cfl13 | 0.000409 | 0.25202646 | 0.902 | 0.766 | 1 | 13 | cfl1 |
| dad11 | 0.000417 | 0.333971811 | 0.656 | 0.417 | 1 | 13 | dad1 |
| spcs1 | 0.00053 | 0.310805033 | 0.557 | 0.354 | 1 | 13 | spcs1 |
| tln11 | 0.000827 | 0.28687185 | 0.41 | 0.223 | 1 | 13 | tln1 |
| lxn4 | 0.001141 | 0.280649678 | 0.475 | 0.265 | 1 | 13 | lxn |
| atf4a1 | 0.001236 | 0.2622228 | 0.393 | 0.213 | 1 | 13 | atf4a |
| tmem2581 | 0.002551 | 0.266100582 | 0.623 | 0.424 | 1 | 13 | tmem258 |
| si:ch73-91k6.2 | 0.00261 | 0.267771032 | 0.197 | 0.088 | 1 | 13 | si:ch73-91k6.2 |
| mt-atp62 | 0.004122 | 0.364546966 | 0.574 | 0.423 | 1 | 13 | mt-atp6 |
| lasp12 | 0.004802 | 0.256080905 | 0.541 | 0.357 | 1 | 13 | lasp1 |
| si:ch211-243a20.3 | 1.6470021 | 1.54218109 | 0.717 | 0.007 | 3.32645015996768e-2 | 14 | si:ch211-243a20.3 |
| wfdc2 | 4.1921718 | 1.443239637 | 0.683 | 0.012 | 8.46692954502127e-1 | 14 | wfdc2 |
| tnmd | 2.6224141 | 0.998233782 | 0.367 | 0.001 | 5.29648983490153e-1 | 14 | tnmd |
| lum | 8.1384602 | 1.234546625 | 0.617 | 0.02 | 1.64372481994351e-1 | 14 | lum |
| sfrp2 | 8.6017661 | 0.944371618 | 0.383 | 0.005 | 1.73729871498026e-1 | 14 | sfrp2 |
| si:dkey-12l12.1 | 5.1976370 | 0.921885938 | 0.417 | 0.009 | 1.04976675320412e-1 | 14 | si:dkey-12l12.1 |
| pcolce2b | 5.6499892 | 1.162346256 | 0.6 | 0.03 | 1.14112832008173e-9 | 14 | pcolce2b |
| sfrp1b | 1.2355351 | 1.029528554 | 0.45 | 0.014 | 2.49541024845238e-9 | 14 | sfrp1b |
| cyp26b1 | 6.1438275 | 0.98987073 | 0.367 | 0.007 | 1.24086884814731e-9 | 14 | cyp26b1 |
| crabp2b | 1.7219896 | 0.939105723 | 0.483 | 0.02 | 3.47790244866727e-8 | 14 | crabp2b |
| twist1a1 | 1.1260693 | 1.086848057 | 0.55 | 0.028 | 2.27432222560078e-8 | 14 | twist1a |
| col11a1b | 3.3726974 | 1.026369019 | 0.483 | 0.024 | 6.81183699135772e-7 | 14 | col11a1b |
| asip1 | 1.5211378 | 0.960797479 | 0.383 | 0.013 | 3.07224213786493e-7 | 14 | asip1 |
| pdgfrl | 2.2342883 | 1.17265439 | 0.633 | 0.047 | 4.51259227717758e-7 | 14 | pdgfrl |
| tnc | 5.0338167 | 0.635797136 | 0.217 | 0.002 | 1.01667997547814e-6 | 14 | tnc |
| ism1 | 7.1868413 | 0.748101093 | 0.3 | 0.008 | 1.45152635663992e-6 | 14 | ism1 |
| pbx3b | 1.6528962 | 1.218694996 | 0.633 | 0.055 | 3.33835457269536e-6 | 14 | pbx3b |
| thbs2b | 9.5455505 | 0.674254682 | 0.317 | 0.01 | 1.92791484259209e-6 | 14 | thbs2b |
| efnb2a | 1.0550984 | 1.012496697 | 0.483 | 0.03 | 2.1309822392805e-62 | 14 | efnb2a |
| thbs4b | 1.6171294 | 0.569507709 | 0.267 | 0.006 | 3.26611637252436e-6 | 14 | thbs4b |
| cyr61 | 6.2353672 | 1.145781005 | 0.617 | 0.055 | 1.25935713217277e-6 | 14 | cyr61 |
| apcdd1l | 6.9312001 | 0.671204068 | 0.317 | 0.01 | 1.39989449756228e-6 | 14 | apcdd1l |

| | | | | | | | |
|---|---|---|---|---|---|---|---|
| CR855375.3 | 5.3216987 | 0.402948427 | 0.15 | 0 | 1.07482349200381e-5 | 14 | CR855375.3 |
| tmem26b | 5.3216987 | 0.396138928 | 0.15 | 0 | 1.07482349200381e-5 | 14 | tmem26b |
| itga11a | 4.5236056 | 0.435005074 | 0.183 | 0.002 | 9.13632628227386e-5 | 14 | itga11a |
| cntfr1 | 4.5801461 | 0.956251355 | 0.467 | 0.032 | 9.2505211116739e-55 | 14 | cntfr |
| crabp1a | 1.4406167 | 0.755061918 | 0.35 | 0.016 | 2.90961374442453e-5 | 14 | crabp1a |
| CU929237.11 | 4.0658814 | 0.977108176 | 0.517 | 0.045 | 8.21186075227573e-5 | 14 | CU929237.1 |
| col2a1b | 7.2058935 | 0.829965647 | 0.367 | 0.021 | 1.45537432498105e-4 | 14 | col2a1b |
| palm1a | 3.3543961 | 0.719988392 | 0.383 | 0.024 | 6.7748738786395e-48 | 14 | palm1a |
| wnt11r | 1.9634335 | 0.797636236 | 0.317 | 0.015 | 3.96554680690494e-4 | 14 | wnt11r |
| trabd2b | 1.7379243 | 0.306521528 | 0.167 | 0.002 | 3.51008590548536e-4 | 14 | trabd2b |
| mkxa | 2.3998086 | 0.357575361 | 0.117 | 0 | 4.84689350901363e-4 | 14 | mkxa |
| cpe | 6.0992134 | 0.480262106 | 0.2 | 0.005 | 1.23185814450135e-4 | 14 | cpe |
| cd248b | 9.2221383 | 0.518974785 | 0.2 | 0.005 | 1.86259528309557e-4 | 14 | cd248b |
| col1a1a4 | 2.5775768 | 1.422375398 | 1 | 0.296 | 5.20593197605945e-4 | 14 | col1a1a |
| lhfpl61 | 3.3475460 | 0.773731564 | 0.4 | 0.032 | 6.76103874087787e-4 | 14 | lhfpl6 |
| adh8a3 | 7.6394180 | 1.211360182 | 0.517 | 0.057 | 1.54293325746638e-4 | 14 | adh8a |
| col1a25 | 1.3841557 | 1.409581628 | 1 | 0.318 | 2.79557937523788e-4 | 14 | col1a2 |
| vwa12 | 1.7132635 | 0.684106337 | 0.283 | 0.015 | 3.46027841800975e-3 | 14 | vwa1 |
| col5a2a4 | 2.8073376 | 1.186670315 | 0.8 | 0.167 | 5.66997983245663e-3 | 14 | col5a2a |
| zfhx4 | 4.0950748 | 0.720423896 | 0.35 | 0.026 | 8.27082276479335e-3 | 14 | zfhx4 |
| alx4a | 1.7597176 | 0.505235514 | 0.233 | 0.01 | 3.55410169406191e-3 | 14 | alx4a |
| timp2a2 | 3.6570122 | 1.021183743 | 0.633 | 0.1 | 7.38606772495637e-3 | 14 | timp2a |
| ebf3a | 7.3698530 | 0.528604381 | 0.233 | 0.01 | 1.48848921620249e-3 | 14 | ebf3a |
| mfap24 | 3.8922561 | 1.241998327 | 0.883 | 0.235 | 7.86118975324414e-3 | 14 | mfap2 |
| cdh11 | 9.1387147 | 0.880404041 | 0.4 | 0.039 | 1.84574622267212e-3 | 14 | cdh11 |
| FP102018.1 | 9.2946864 | 0.748349437 | 0.35 | 0.029 | 1.87724781984359e-3 | 14 | FP102018.1 |
| efna2a | 1.2942365 | 0.438279407 | 0.233 | 0.011 | 2.61396965568094e-3 | 14 | efna2a |
| bcl11aa | 7.0044332 | 0.52253464 | 0.233 | 0.011 | 1.4146853862259e-33 | 14 | bcl11aa |
| gulp1a1 | 1.0813596 | 0.856196108 | 0.467 | 0.056 | 2.18402210145214e-3 | 14 | gulp1a |
| hhipl2 | 1.6068574 | 0.354434404 | 0.15 | 0.003 | 3.24536996664368e-3 | 14 | hhipl2 |
| LOC1005373421 | 2.7040132 | 0.671694169 | 0.35 | 0.029 | 5.46129548832023e-3 | 14 | LOC100537342 |
| cav22 | 2.3340240 | 0.761334187 | 0.35 | 0.031 | 4.71402842939372e-3 | 14 | cav2 |
| aqp3a | 6.8328046 | 0.999673557 | 0.317 | 0.025 | 1.38002155446403e-3 | 14 | aqp3a |
| adgra2 | 7.8333728 | 0.417554305 | 0.2 | 0.008 | 1.58210630449307e-3 | 14 | adgra2 |
| cspg4 | 5.2567383 | 0.448227206 | 0.167 | 0.005 | 1.06170344366558e-3 | 14 | cspg4 |

| | | | | | | | |
|---|---|---|---|---|---|---|---|
| endouc2 | 1.4037035 | 1.178433819 | 0.617 | 0.111 | 2.83506003009052e-3 | 14 | endouc |
| cacna1g | 1.6348440 | 0.481852403 | 0.217 | 0.011 | 3.30189461975442e-3 | 14 | cacna1g |
| aoc24 | 2.5980920 | 1.086594002 | 0.65 | 0.122 | 5.24736660388631e-3 | 14 | aoc2 |
| XLOC-042917 | 3.4285421 | 0.587573744 | 0.267 | 0.018 | 6.92462651917719e-3 | 14 | XLOC-042917 |
| si:dkey-282h22.5 | 3.5238206 | 0.30597486 | 0.1 | 0.001 | 7.1170605608631e-30 | 14 | si:dkey-282h22.5 |
| sfrp1a2 | 4.8183915 | 0.968298715 | 0.483 | 0.067 | 9.73170541290848e-3 | 14 | sfrp1a |
| plpp32 | 7.9051445 | 1.081638065 | 0.633 | 0.114 | 1.5966020449434e-29 | 14 | plpp3 |
| fstl1b4 | 1.7935444 | 1.15093103 | 0.9 | 0.265 | 3.62242175731774e-2 | 14 | fstl1b |
| efcc1 | 2.0004543 | 0.440217435 | 0.2 | 0.009 | 4.04031759289374e-2 | 14 | efcc1 |
| col1a1b4 | 2.9654074 | 1.256797431 | 0.9 | 0.286 | 5.98923345664398e-2 | 14 | col1a1b |
| col5a14 | 7.6769107 | 1.090930993 | 0.867 | 0.229 | 1.55050567129754e-2 | 14 | col5a1 |
| col12a1b3 | 1.1015339 | 0.965383725 | 0.433 | 0.056 | 2.22476811270522e-2 | 14 | col12a1b |
| mmp24 | 1.8570714 | 1.075015783 | 0.783 | 0.205 | 3.75072722706718e-2 | 14 | mmp2 |
| vwde1 | 2.7826958 | 0.634598309 | 0.183 | 0.008 | 5.6202107911066e-27 | 14 | vwde |
| cdon3 | 3.3663118 | 1.019919813 | 0.633 | 0.135 | 6.79893997685326e-2 | 14 | cdon |
| anos1b | 3.8330585 | 0.386869649 | 0.183 | 0.008 | 7.74162837866681e-2 | 14 | anos1b |
| hsp70.1 | 2.8303960 | 0.279950893 | 0.1 | 0.001 | 5.71655099371424e-2 | 14 | hsp70.1 |
| zgc:162612 | 1.3858996 | 0.301839834 | 0.133 | 0.004 | 2.79910142947771e-2 | 14 | zgc:162612 |
| LO018188.13 | 3.4465462 | 0.92151725 | 0.5 | 0.083 | 6.96098946503654e-2 | 14 | LO018188.1 |
| bmp3 | 4.0471860 | 0.497313401 | 0.217 | 0.014 | 8.1741017432176e-25 | 14 | bmp3 |
| col16a11 | 6.9148500 | 0.787134083 | 0.467 | 0.072 | 1.39659226628588e-2 | 14 | col16a1 |
| enpp2 | 7.8839558 | 0.350749029 | 0.167 | 0.007 | 1.59232255334418e-2 | 14 | enpp2 |
| serpinf1 | 1.4940357 | 0.682699771 | 0.383 | 0.049 | 3.0175039096139e-24 | 14 | serpinf1 |
| ctgfa1 | 4.2777249 | 0.8451932 | 0.417 | 0.058 | 8.63972109415292e-2 | 14 | ctgfa |
| tpm4b1 | 1.6466774 | 0.690761628 | 0.35 | 0.042 | 3.32579450704857e-2 | 14 | tpm4b |
| metrnla1 | 2.0186397 | 0.90859388 | 0.533 | 0.103 | 4.07704669372212e-2 | 14 | metrnla |
| abi3bpb | 6.3813064 | 0.665250318 | 0.333 | 0.038 | 1.28883247053706e-2 | 14 | abi3bpb |
| tmeff2a | 7.4063895 | 0.488142666 | 0.2 | 0.013 | 1.49586849914135e-2 | 14 | tmeff2a |
| dcn4 | 1.2187743 | 1.037337569 | 0.867 | 0.266 | 2.46155864768187e-2 | 14 | dcn |
| boc4 | 2.6587495 | 0.930445475 | 0.65 | 0.157 | 5.36987638591636e-2 | 14 | boc |
| srpx2 | 6.1703821 | 0.396377972 | 0.183 | 0.011 | 1.24623207703764e-2 | 14 | srpx2 |
| efna5b | 9.1837753 | 0.559688918 | 0.217 | 0.016 | 1.8548471083294e-21 | 14 | efna5b |
| glis2a | 5.4988604 | 0.496963184 | 0.267 | 0.026 | 1.110604847991e-20 | 14 | glis2a |
| vipr1a | 6.5618216 | 0.45794873 | 0.217 | 0.017 | 1.32529111972457e-2 | 14 | vipr1a |
| tfpia | 6.8914597 | 0.484881374 | 0.233 | 0.02 | 1.39186811999537e-2 | 14 | tfpia |

| | | | | | | | |
|---|---|---|---|---|---|---|---|
| marcksl1b4 | 7.9140070 | 1.037983185 | 0.8 | 0.27 | 1.59839201371447e-2 | 14 | marcksl1b |
| lepr | 9.4274763 | 0.341066575 | 0.133 | 0.005 | 1.90406739568003e-2 | 14 | lepr |
| sparc5 | 1.7329597 | 0.979093341 | 0.933 | 0.391 | 3.50005882981418e-2 | 14 | sparc |
| CABZ01085139.1 | 1.9141447 | 0.348543321 | 0.15 | 0.007 | 3.86599811721433e-2 | 14 | CABZ01085139.1 |
| chrna5 | 2.3690617 | 0.382979312 | 0.117 | 0.004 | 4.78479403217191e-2 | 14 | chrna5 |
| mfap53 | 3.5178659 | 1.01125456 | 0.617 | 0.154 | 7.10503380154194e-2 | 14 | mfap5 |
| fibinb1 | 3.9361986 | 0.702385152 | 0.383 | 0.058 | 7.94994034730106e-2 | 14 | fibinb |
| si:ch211-251b21.1 | 4.6633183 | 0.546584162 | 0.267 | 0.027 | 9.4185040624788e-20 | 14 | si:ch211-251b21.1 |
| CABZ01074363.1 | 5.7118427 | 0.522506619 | 0.217 | 0.018 | 1.15362088682884e-1 | 14 | CABZ01074363.1 |
| si:dkey-261h17.12 | 1.3226813 | 1.083464567 | 0.75 | 0.241 | 2.67141953512561e-1 | 14 | si:dkey-261h17.1 |
| ecrg4a1 | 1.3302313 | 0.857981708 | 0.4 | 0.062 | 2.68666835474124e-1 | 14 | ecrg4a |
| lrp1aa2 | 1.5853857 | 0.829279835 | 0.533 | 0.116 | 3.20200369227334e-1 | 14 | lrp1aa |
| si:ch211-39i22.15 | 1.6809678 | 1.006093673 | 0.633 | 0.174 | 3.39505078203093e-1 | 14 | si:ch211-39i22.1 |
| pmp22a4 | 1.9157054 | 1.010857627 | 0.85 | 0.31 | 3.86915027021898e-1 | 14 | pmp22a |
| olfml3b4 | 2.6067291 | 0.806377645 | 0.5 | 0.103 | 5.26481090085163e-1 | 14 | olfml3b |
| sema3d1 | 3.0280713 | 0.535776342 | 0.283 | 0.032 | 6.11579578740677e-1 | 14 | sema3d |
| tuft1a | 4.9601264 | 0.393513277 | 0.167 | 0.01 | 1.00179674712959e-1 | 14 | tuft1a |
| col6a34 | 1.4628895 | 0.91202476 | 0.667 | 0.184 | 2.95459803907883e-1 | 14 | col6a3 |
| lpl | 1.5804312 | 0.451006283 | 0.25 | 0.025 | 3.19199696784466e-1 | 14 | lpl |
| emilin1b | 3.8656481 | 0.451440687 | 0.217 | 0.019 | 7.80744957037179e-1 | 14 | emilin1b |
| si:dkey-238c7.163 | 7.4702319 | 0.813746591 | 0.5 | 0.109 | 1.50876275380577e-1 | 14 | si:dkey-238c7.16 |
| fam69c | 1.9388346 | 0.586128195 | 0.3 | 0.04 | 3.9158643162438e-17 | 14 | fam69c |
| paplna1 | 2.3662292 | 0.627004565 | 0.317 | 0.044 | 4.77907313890747e-1 | 14 | paplna |
| col6a14 | 3.4057316 | 0.854126483 | 0.55 | 0.137 | 6.87855626861222e-1 | 14 | col6a1 |
| aspn | 1.0999439 | 0.274984593 | 0.1 | 0.003 | 2.2215567509532e-16 | 14 | aspn |
| rbfox11 | 1.1740183 | 0.587495493 | 0.317 | 0.046 | 2.37116495992868e-1 | 14 | rbfox1 |
| p4ha2 | 3.3420238 | 0.466216897 | 0.217 | 0.022 | 6.74988564552825e-1 | 14 | p4ha2 |
| ogna | 4.5193483 | 0.374135118 | 0.117 | 0.005 | 9.12772795447629e-1 | 14 | ogna |
| susd5 | 4.7536188 | 0.312417913 | 0.117 | 0.005 | 9.60088398551134e-1 | 14 | susd5 |
| igf1 | 4.7536188 | 0.295893426 | 0.117 | 0.005 | 9.60088398551134e-1 | 14 | igf1 |
| col21a11 | 8.8735715 | 0.593324525 | 0.25 | 0.03 | 1.79219524790207e-1 | 14 | col21a1 |
| qsox11 | 1.0692366 | 0.599476686 | 0.3 | 0.044 | 2.15953725122878e-1 | 14 | qsox1 |
| si:ch1073-459j12.1 | 1.0696993 | 0.878637636 | 0.617 | 0.176 | 2.16047183134777e-1 | 14 | si:ch1073-459j12.1 |
| cemip | 1.4763552 | 0.507899911 | 0.183 | 0.016 | 2.98179473190032e-1 | 14 | cemip |
| pigr | 2.7077248 | 0.5111146 | 0.267 | 0.035 | 5.46879185427836e-1 | 14 | pigr |

| | | | | | | | |
|---|---|---|---|---|---|---|---|
| col12a1a2 | 3.2003284 | 0.681852697 | 0.317 | 0.05 | 6.46370343745768e-1 | 14 | col12a1a |
| wnt16 | 4.6857581 | 0.326116746 | 0.15 | 0.01 | 9.46382568424365e-1 | 14 | wnt16 |
| col6a24 | 4.7232083 | 0.761361923 | 0.5 | 0.12 | 9.5394639341275e-15 | 14 | col6a2 |
| thbs2a | 6.6349242 | 0.407323435 | 0.183 | 0.016 | 1.34005564597617e-1 | 14 | thbs2a |
| nuak1a | 1.1769927 | 0.325545188 | 0.183 | 0.016 | 2.37717235093769e-1 | 14 | nuak1a |
| emx21 | 2.1878373 | 0.576874579 | 0.333 | 0.056 | 4.41877510800049e-1 | 14 | emx2 |
| loxl12 | 5.3017496 | 0.626561105 | 0.367 | 0.068 | 1.07079437981012e-1 | 14 | loxl1 |
| s100v25 | 7.2259517 | 0.833680705 | 0.683 | 0.242 | 1.45942546699191e-1 | 14 | s100v2 |
| coch | 7.5791023 | 0.48420434 | 0.25 | 0.032 | 1.53075130852159e-1 | 14 | coch |
| tpm4a4 | 8.2353686 | 0.860613641 | 0.683 | 0.238 | 1.66329741015268e-1 | 14 | tpm4a |
| meis2b4 | 9.8142959 | 0.671280512 | 0.417 | 0.09 | 1.98219334572858e-1 | 14 | meis2b |
| auts2a | 1.0914803 | 0.4235361 | 0.2 | 0.021 | 2.2044629140204e-13 | 14 | auts2a |
| ecm1b2 | 1.1503083 | 0.687679625 | 0.517 | 0.127 | 2.32327773510026e-1 | 14 | ecm1b |
| chad | 1.3374957 | 0.530910149 | 0.217 | 0.026 | 2.70134026091303e-1 | 14 | chad |
| sox181 | 1.8118010 | 0.271005134 | 0.133 | 0.009 | 3.65929453277618e-1 | 14 | sox18 |
| fbln52 | 2.3513529 | 0.701287399 | 0.433 | 0.097 | 4.74902754882917e-1 | 14 | fbln5 |
| fbln21 | 3.6446819 | 0.555663088 | 0.267 | 0.039 | 7.36116411411846e-1 | 14 | fbln2 |
| loxa3 | 8.6030917 | 0.559658743 | 0.3 | 0.05 | 1.73756644865826e-1 | 14 | loxa |
| ntn1b | 9.1073977 | 0.369714626 | 0.15 | 0.012 | 1.83942111947182e-1 | 14 | ntn1b |
| ctsla5 | 9.6718976 | 0.76900665 | 0.767 | 0.3 | 1.95343317792261e-1 | 14 | ctsla |
| ucmaa2 | 1.2063349 | 0.640421566 | 0.233 | 0.031 | 2.43643472789697e-1 | 14 | ucmaa |
| tgfb2 | 1.2517185 | 0.312306353 | 0.15 | 0.012 | 2.5280959767064e-12 | 14 | tgfb2 |
| calhm2 | 1.5783174 | 0.425371563 | 0.233 | 0.031 | 3.18772783734965e-1 | 14 | calhm2 |
| cpn13 | 1.8666559 | 0.804516308 | 0.583 | 0.187 | 3.77008497794819e-1 | 14 | cpn1 |
| palld | 2.8828467 | 0.4217986 | 0.233 | 0.032 | 5.82248548053741e-1 | 14 | palld |
| krt913 | 5.2000337 | 0.631511943 | 0.417 | 0.089 | 1.05025082077919e-1 | 14 | krt91 |
| zfhx3 | 8.8979618 | 0.606355659 | 0.333 | 0.067 | 1.79712135000995e-1 | 14 | zfhx3 |
| mmp14a2 | 9.8679826 | 0.646378889 | 0.4 | 0.094 | 1.99303646559133e-1 | 14 | mmp14a |
| hmcn1 | 1.0443059 | 0.279845417 | 0.133 | 0.01 | 2.10918476531096e-1 | 14 | hmcn1 |
| pcdh11 | 1.1196784 | 0.355443881 | 0.15 | 0.013 | 2.26141462975683e-1 | 14 | pcdh11 |
| pcdh7b | 2.8754807 | 0.392582408 | 0.183 | 0.021 | 5.80760838288513e-1 | 14 | pcdh7b |
| pcsk6 | 3.4113504 | 0.33937847 | 0.15 | 0.014 | 6.88990443651719e-1 | 14 | pcsk6 |
| stom1 | 5.8320025 | 0.422741411 | 0.233 | 0.034 | 1.17788955751436e-1 | 14 | stom |
| ahnak4 | 7.0562387 | 0.780655154 | 0.567 | 0.195 | 1.42514853309117e-1 | 14 | ahnak |
| ccl19a.11 | 7.8469916 | 0.528181779 | 0.2 | 0.025 | 1.58485690542358e-1 | 14 | ccl19a.1 |

| | | | | | | | |
|---|---|---|---|---|---|---|---|
| cd81a5 | 8.1366357 | 0.743670226 | 0.85 | 0.434 | 1.64335633185524e-1 | 14 | cd81a |
| zgc:1537042 | 8.5392864 | 0.516391784 | 0.25 | 0.038 | 1.72467968285763e-1 | 14 | zgc:153704 |
| CR383676.14 | 1.0958765 | 0.616152633 | 0.983 | 0.655 | 2.21334194966004e-1 | 14 | CR383676.1 |
| ckba2 | 1.1184907 | 0.651165515 | 0.433 | 0.114 | 2.25901572205163e-1 | 14 | ckba |
| grm4 | 1.5421763 | 0.415009153 | 0.2 | 0.027 | 3.11473353382183e-1 | 14 | grm4 |
| AL954322.2 | 1.5652995 | 0.27263278 | 0.133 | 0.011 | 3.16143554712218e-1 | 14 | AL954322.2 |
| twsg1a | 1.8297063 | 0.373187773 | 0.183 | 0.022 | 3.69545787507814e-1 | 14 | twsg1a |
| rnd3b4 | 2.0476409 | 0.61976452 | 0.383 | 0.092 | 4.13562034818851e-1 | 14 | rnd3b |
| tgfbr31 | 2.5610105 | 0.52199981 | 0.317 | 0.062 | 5.17247297985315e-1 | 14 | tgfbr3 |
| twist1b | 3.4779145 | 0.283205736 | 0.15 | 0.015 | 7.02434395254975e-1 | 14 | twist1b |
| ehd2b1 | 4.0462206 | 0.50572475 | 0.267 | 0.047 | 8.17215182581674e-1 | 14 | ehd2b |
| cd1511 | 4.3514682 | 0.50782688 | 0.25 | 0.043 | 8.78866045885596e-1 | 14 | cd151 |
| BX088707.3 | 4.4003654 | 0.328019152 | 0.133 | 0.012 | 8.88741816833274e-1 | 14 | BX088707.3 |
| col8a2 | 6.0068410 | 0.269244334 | 0.1 | 0.006 | 1.21320168744826e-0 | 14 | col8a2 |
| si:ch211-105c13.3 | 7.7433875 | 0.683749227 | 0.4 | 0.104 | 1.5639319780093e-09 | 14 | si:ch211-105c13.3 |
| sox11a2 | 1.1974947 | 0.600769211 | 0.317 | 0.07 | 2.4185801523092e-09 | 14 | sox11a |
| fbn2b4 | 1.2258931 | 0.667090549 | 0.533 | 0.172 | 2.47593634372625e-0 | 14 | fbn2b |
| cnpy4 | 1.2627057 | 0.411470356 | 0.267 | 0.049 | 2.5502867601957e-09 | 14 | cnpy4 |
| htra1a1 | 1.2813767 | 0.558723415 | 0.25 | 0.043 | 2.58799654113198e-0 | 14 | htra1a |
| mir214a | 1.3370640 | 0.373893796 | 0.167 | 0.02 | 2.70046828069748e-0 | 14 | mir214a |
| plxdc2 | 1.5277964 | 0.356493739 | 0.183 | 0.024 | 3.08569039284683e-0 | 14 | plxdc2 |
| fgf18a | 1.7269785 | 0.250753278 | 0.117 | 0.009 | 3.48797858453139e-0 | 14 | fgf18a |
| agmo | 1.7446646 | 0.36131794 | 0.167 | 0.02 | 3.52369926433666e-0 | 14 | agmo |
| LOC101885478 | 2.9008934 | 0.264109408 | 0.1 | 0.007 | 5.8589345901695e-09 | 14 | LOC101885478 |
| cavin2b3 | 3.9131527 | 0.70241917 | 0.467 | 0.142 | 7.90339468207579e-0 | 14 | cavin2b |
| pitx21 | 4.2242381 | 0.449385007 | 0.233 | 0.039 | 8.53169382196268e-0 | 14 | pitx2 |
| hmbox1a | 4.6332125 | 0.525827819 | 0.333 | 0.077 | 9.35769928769602e-0 | 14 | hmbox1a |
| si:ch211-195m9.3 | 5.8793872 | 0.399347443 | 0.167 | 0.021 | 1.18745984132312e-0 | 14 | si:ch211-195m9.3 |
| adamts15a4 | 5.8929683 | 0.580391527 | 0.4 | 0.106 | 1.19020281584798e-0 | 14 | adamts15a |
| rasl12 | 8.5844955 | 0.334939005 | 0.15 | 0.017 | 1.73381056113681e-0 | 14 | rasl12 |
| fzd7a2 | 9.2944415 | 0.496208179 | 0.267 | 0.053 | 1.87719835058555e-0 | 14 | fzd7a |
| sdc22 | 1.0240535 | 0.625511861 | 0.417 | 0.12 | 2.06828093326237e-0 | 14 | sdc2 |
| actn11 | 1.1948722 | 0.504572799 | 0.3 | 0.065 | 2.41328346008839e-0 | 14 | actn1 |
| ppib4 | 1.4547895 | 0.550575906 | 0.917 | 0.64 | 2.93823842572102e-0 | 14 | ppib |
| mmp14b2 | 1.4901575 | 0.611819713 | 0.4 | 0.112 | 3.00967115260095e-0 | 14 | mmp14b |

| | | | | | | | |
|---|---|---|---|---|---|---|---|
| hspa12a | 1.8005635 | 0.315854671 | 0.15 | 0.017 | 3.63659821714428e-0 | 14 | hspa12a |
| ppfibp1b | 2.0146329 | 0.305266568 | 0.167 | 0.021 | 4.06895413629786e-0 | 14 | ppfibp1b |
| slit31 | 2.2820965 | 0.560162303 | 0.333 | 0.078 | 4.60915037240496e-0 | 14 | slit3 |
| akap12b2 | 2.3127190 | 0.684859308 | 0.467 | 0.147 | 4.67099870095824e-0 | 14 | akap12b |
| prss352 | 2.3537752 | 0.64940759 | 0.3 | 0.069 | 4.75391993210422e-0 | 14 | prss35 |
| meox1 | 2.7695137 | 0.352067899 | 0.15 | 0.018 | 5.59358694311408e-0 | 14 | meox1 |
| jag1b | 2.7721069 | 0.497407201 | 0.233 | 0.043 | 5.59882449722998e-0 | 14 | jag1b |
| fkbp91 | 2.9684216 | 0.547779448 | 0.35 | 0.089 | 5.99532113003403e-0 | 14 | fkbp9 |
| BX640522.23 | 3.0058905 | 0.664137497 | 0.417 | 0.115 | 6.07099718557568e-0 | 14 | BX640522.2 |
| efna1b1 | 3.0375519 | 0.356629755 | 0.183 | 0.026 | 6.13494376715204e-0 | 14 | efna1b |
| ppfibp1a1 | 3.3849197 | 0.400090787 | 0.25 | 0.047 | 6.83652240850583e-0 | 14 | ppfibp1a |
| ptprk1 | 3.7948789 | 0.46522462 | 0.233 | 0.043 | 7.66451708406966e-0 | 14 | ptprk |
| cfd2 | 5.2507041 | 0.614293715 | 0.35 | 0.086 | 1.06048471750451e-0 | 14 | cfd |
| wnt5b2 | 7.2863443 | 0.418566807 | 0.2 | 0.032 | 1.4716229782174e-07 | 14 | wnt5b |
| tril2 | 9.5224188 | 0.443521103 | 0.25 | 0.049 | 1.92324292762635e-0 | 14 | tril |
| si:ch211-250c4.41 | 9.5459462 | 0.370221201 | 0.2 | 0.032 | 1.9279947585149e-07 | 14 | si:ch211-250c4.4 |
| mt-nd22 | 9.5625945 | 0.669960705 | 0.783 | 0.366 | 1.93135722154932e-0 | 14 | mt-nd2 |
| fpr11 | 1.4443349 | 0.346116377 | 0.183 | 0.027 | 2.91712334662535e-0 | 14 | fpr1 |
| si:ch73-46j18.55 | 1.8219132 | 0.653481872 | 0.783 | 0.39 | 3.67971809346653e-0 | 14 | si:ch73-46j18.5 |
| marcksl1a4 | 2.0778839 | 0.648318956 | 0.817 | 0.509 | 4.19670219086244e-0 | 14 | marcksl1a |
| svep11 | 2.4168139 | 0.31550276 | 0.167 | 0.023 | 4.88123918724226e-0 | 14 | svep1 |
| si:dkey-16l2.201 | 2.5318435 | 0.383323069 | 0.217 | 0.039 | 5.11356449820418e-0 | 14 | si:dkey-16l2.20 |
| atrn | 3.8389627 | 0.296171068 | 0.15 | 0.02 | 7.75355314388947e-0 | 14 | atrn |
| LOC1018823341 | 5.5497690 | 0.304885655 | 0.133 | 0.016 | 1.12088686245588e-0 | 14 | LOC101882334 |
| vat14 | 6.0978168 | 0.628408495 | 0.567 | 0.219 | 1.23157606117426e-0 | 14 | vat1 |
| cercam | 6.5238604 | 0.329101203 | 0.2 | 0.035 | 1.31762409264201e-0 | 14 | cercam |
| myocd | 7.2122592 | 0.269784819 | 0.117 | 0.012 | 1.45666000656105e-0 | 14 | myocd |
| myl9a4 | 7.3234232 | 0.580197485 | 0.383 | 0.112 | 1.47911179566509e-0 | 14 | myl9a |
| gas1a3 | 7.4071298 | 0.574586817 | 0.417 | 0.134 | 1.49601800996826e-0 | 14 | gas1a |
| lsp1 | 7.4099868 | 0.422944406 | 0.217 | 0.041 | 1.49659504253443e-0 | 14 | lsp1 |
| LOC108190717 | 8.4691045 | 0.260131072 | 0.133 | 0.016 | 1.71050505446078e-0 | 14 | LOC108190717 |
| tnfsf12 | 1.0281003 | 0.39326949 | 0.233 | 0.047 | 2.07645436459215e-0 | 14 | tnfsf12 |
| p3h1 | 1.0542976 | 0.323092907 | 0.167 | 0.025 | 2.12936487082281e-0 | 14 | p3h1 |
| rbp52 | 1.0806636 | 0.570558411 | 0.533 | 0.192 | 2.18261638059545e-0 | 14 | rbp5 |
| cav13 | 1.7124789 | 0.520271858 | 0.3 | 0.074 | 3.45869364061677e-0 | 14 | cav1 |

| | | | | | | | |
|---|---|---|---|---|---|---|---|
| mgst1.23 | 2.3927976 | 0.572147272 | 0.483 | 0.175 | 4.83273334319439e-0 | 14 | mgst1.2 |
| cd28l1 | 2.6252401 | 0.392084012 | 0.233 | 0.049 | 5.30219753295225e-0 | 14 | cd28l |
| pawr | 3.1828936 | 0.328699779 | 0.183 | 0.031 | 6.42849025035179e-0 | 14 | pawr |
| nfixb3 | 3.5427469 | 0.593559925 | 0.483 | 0.171 | 7.15528602356841e-0 | 14 | nfixb |
| amotl2b | 3.9878796 | 0.270010789 | 0.133 | 0.017 | 8.05432061560506e-0 | 14 | amotl2b |
| hsd11b24 | 4.3024797 | 0.705721747 | 0.65 | 0.286 | 8.68971840057944e-0 | 14 | hsd11b2 |
| pabpc1b3 | 4.4178893 | 0.531198002 | 0.383 | 0.12 | 8.92281104511818e-0 | 14 | pabpc1b |
| meis1b3 | 4.4772382 | 0.588551422 | 0.433 | 0.153 | 9.0426780170453e-06 | 14 | meis1b |
| CR847566.13 | 4.9117662 | 0.573117134 | 0.4 | 0.129 | 9.92029429060583e-0 | 14 | CR847566.1 |
| tmem88a1 | 5.0474492 | 0.322229041 | 0.167 | 0.027 | 1.01943333407926e-0 | 14 | tmem88a |
| mgp1 | 5.3668382 | 0.297731091 | 0.183 | 0.031 | 1.08394031225444e-0 | 14 | mgp |
| hoxa9b | 5.6720879 | 0.31937595 | 0.15 | 0.022 | 1.14559160107383e-0 | 14 | hoxa9b |
| mt-nd31 | 5.7329696 | 0.572110099 | 0.633 | 0.266 | 1.15788788579872e-0 | 14 | mt-nd3 |
| itgb1a1 | 5.9628669 | 0.451293667 | 0.3 | 0.078 | 1.20432023343408e-0 | 14 | itgb1a |
| sc:d02021 | 6.2910538 | 0.405986178 | 0.2 | 0.038 | 1.27060414076953e-0 | 14 | sc:d0202 |
| dbn12 | 6.7919447 | 0.591528491 | 0.367 | 0.12 | 1.37176907454482e-0 | 14 | dbn1 |
| mt-nd42 | 8.1565759 | 0.580984795 | 0.817 | 0.431 | 1.64738363817628e-0 | 14 | mt-nd4 |
| ctbp1 | 1.1889700 | 0.532252025 | 0.45 | 0.166 | 2.40136286141078e-0 | 14 | ctbp1 |
| pdcd6 | 1.2331485 | 0.487482188 | 0.383 | 0.124 | 2.49059019976037e-0 | 14 | pdcd6 |
| plpp2b | 1.2523307 | 0.331488774 | 0.15 | 0.023 | 2.52933248302878e-0 | 14 | plpp2b |
| SCARF21 | 1.2936754 | 0.351619226 | 0.217 | 0.045 | 2.61283639073048e-0 | 14 | SCARF2 |
| gpc6a2 | 1.3534078 | 0.299957488 | 0.167 | 0.028 | 2.73347787941554e-0 | 14 | gpc6a |
| zfp36l22 | 1.6705728 | 0.468231467 | 0.25 | 0.06 | 3.37405594799978e-0 | 14 | zfp36l2 |
| hmgb3a3 | 1.9863826 | 0.601328775 | 0.567 | 0.255 | 4.01189704482219e-0 | 14 | hmgb3a |
| pdia5 | 2.2638218 | 0.442282127 | 0.3 | 0.083 | 4.57224105452233e-0 | 14 | pdia5 |
| pdgfrb2 | 2.4386455 | 0.44934892 | 0.25 | 0.061 | 4.92533233594287e-0 | 14 | pdgfrb |
| sostdc1a3 | 2.7274089 | 0.579483328 | 0.317 | 0.088 | 5.50854782124106e-0 | 14 | sostdc1a |
| XLOC-040108 | 2.8711084 | 0.282143644 | 0.1 | 0.011 | 5.7987777287051e-05 | 14 | XLOC-040108 |
| chd34 | 3.3503655 | 0.546423734 | 0.55 | 0.237 | 6.76673333572415e-0 | 14 | chd3 |
| sgcb | 3.8562988 | 0.295187436 | 0.15 | 0.024 | 7.78856686825288e-0 | 14 | sgcb |
| emp21 | 5.3669100 | 0.504190762 | 0.317 | 0.094 | 0.000108395 | 14 | emp2 |
| ckap43 | 6.7349172 | 0.519047074 | 0.4 | 0.145 | 0.000136025 | 14 | ckap4 |
| fhod3b | 7.1776816 | 0.278592809 | 0.133 | 0.02 | 0.000144968 | 14 | fhod3b |
| loxl3b | 8.0296752 | 0.299078406 | 0.133 | 0.02 | 0.000162175 | 14 | loxl3b |
| glipr2l1 | 8.3546542 | 0.347122087 | 0.233 | 0.055 | 0.000168739 | 14 | glipr2l |

| | | | | | | | |
|---|---|---|---|---|---|---|---|
| hic12 | 9.1851691 | 0.484172397 | 0.383 | 0.132 | 0.000185513 | 14 | hic1 |
| jun5 | 9.8029982 | 0.621382491 | 0.7 | 0.38 | 0.000197991 | 14 | jun |
| filip1l | 1.1315311 | 0.360960243 | 0.2 | 0.043 | 0.000228535 | 14 | filip1l |
| mt-cyb2 | 1.2582421 | 0.65687768 | 0.7 | 0.373 | 0.000254127 | 14 | mt-cyb |
| fat1a1 | 1.2900018 | 0.430296428 | 0.2 | 0.045 | 0.000260542 | 14 | fat1a |
| plod1a | 1.3549522 | 0.356391771 | 0.183 | 0.038 | 0.00027366 | 14 | plod1a |
| lrrc172 | 1.4814138 | 0.577220118 | 0.45 | 0.178 | 0.000299201 | 14 | lrrc17 |
| cd248a3 | 1.5846909 | 0.542585975 | 0.35 | 0.12 | 0.00032006 | 14 | cd248a |
| kank21 | 1.6455517 | 0.367401848 | 0.25 | 0.063 | 0.000332352 | 14 | kank2 |
| LO018340.11 | 1.7222062 | 0.384462716 | 0.2 | 0.044 | 0.000347834 | 14 | LO018340.1 |
| gstr1 | 1.9054044 | 0.439442581 | 0.45 | 0.173 | 0.000384835 | 14 | gstr |
| lrp5 | 1.9202628 | 0.257007623 | 0.133 | 0.021 | 0.000387835 | 14 | lrp5 |
| arl6ip5b3 | 1.9527736 | 0.508740261 | 0.383 | 0.139 | 0.000394402 | 14 | arl6ip5b |
| marcksa | 1.9729262 | 0.375206452 | 0.217 | 0.051 | 0.000398472 | 14 | marcksa |
| kdelr2a | 2.0103552 | 0.451794945 | 0.317 | 0.1 | 0.000406031 | 14 | kdelr2a |
| pcolcea2 | 2.3984636 | 0.544122996 | 0.433 | 0.152 | 0.000484418 | 14 | pcolcea |
| plod3 | 2.5742991 | 0.445829077 | 0.233 | 0.061 | 0.000519931 | 14 | plod3 |
| srpx1 | 2.6151102 | 0.286072761 | 0.15 | 0.026 | 0.000528174 | 14 | srpx |
| COX34 | 2.6807033 | 0.543649916 | 0.933 | 0.591 | 0.000541422 | 14 | COX3 |
| myl64 | 2.7115357 | 0.55517773 | 0.483 | 0.2 | 0.000547649 | 14 | myl6 |
| cd99l2 | 3.4737254 | 0.420850411 | 0.283 | 0.084 | 0.000701588 | 14 | cd99l2 |
| efemp2a1 | 3.6699806 | 0.342142826 | 0.183 | 0.039 | 0.000741226 | 14 | efemp2a |
| mgst1.13 | 3.8994144 | 0.446572141 | 0.367 | 0.128 | 0.000787565 | 14 | mgst1.1 |
| fkbp73 | 4.2385932 | 0.490242821 | 0.35 | 0.124 | 0.000856069 | 14 | fkbp7 |
| tmem119b2 | 4.3335080 | 0.351866258 | 0.2 | 0.046 | 0.000875239 | 14 | tmem119b |
| flna5 | 4.3786238 | 0.564180744 | 0.417 | 0.165 | 0.000884351 | 14 | flna |
| mt-co21 | 4.8702134 | 0.620042424 | 0.767 | 0.456 | 0.000983637 | 14 | mt-co2 |
| XLOC-004556 | 5.0346416 | 0.292409713 | 0.167 | 0.033 | 0.001016847 | 14 | XLOC-004556 |
| crhbp1 | 6.2482979 | 0.40633925 | 0.15 | 0.028 | 0.001261969 | 14 | crhbp |
| egfl71 | 6.4875658 | 0.250158623 | 0.117 | 0.017 | 0.001310294 | 14 | egfl7 |
| mt-nd12 | 6.5036488 | 0.602740789 | 0.65 | 0.351 | 0.001313542 | 14 | mt-nd1 |
| slc7a12 | 7.0014549 | 0.344381012 | 0.167 | 0.034 | 0.001414084 | 14 | slc7a1 |
| klf6a2 | 7.3104980 | 0.558050028 | 0.533 | 0.256 | 0.001476501 | 14 | klf6a |
| lrp1ab2 | 8.3010833 | 0.460232541 | 0.35 | 0.121 | 0.00167657 | 14 | lrp1ab |
| RAMP11 | 8.5447267 | 0.272036771 | 0.133 | 0.022 | 0.001725778 | 14 | RAMP1 |

| | | | | | | | |
|---|---|---|---|---|---|---|---|
| cald1a4 | 9.1587240 | 0.544563843 | 0.517 | 0.234 | 0.001849788 | 14 | cald1a |
| tspan4b2 | 9.2130604 | 0.437487613 | 0.317 | 0.104 | 0.001860762 | 14 | tspan4b |
| scara31 | 9.2313771 | 0.365102437 | 0.217 | 0.053 | 0.001864461 | 14 | scara3 |
| CABZ01092746.14 | 9.5147084 | 0.519086745 | 0.5 | 0.205 | 0.001921686 | 14 | CABZ01092746.1 |
| fhl1b | 9.9619937 | 0.275752534 | 0.167 | 0.034 | 0.002012024 | 14 | fhl1b |
| castor2 | 1.0756327 | 0.302244313 | 0.167 | 0.035 | 0.002172455 | 14 | castor2 |
| gng72 | 1.0832051 | 0.417582253 | 0.317 | 0.105 | 0.002187749 | 14 | gng7 |
| si:ch211-71n6.41 | 1.2574113 | 0.315129814 | 0.183 | 0.041 | 0.002539594 | 14 | si:ch211-71n6.4 |
| BX663503.31 | 1.3312637 | 0.337506044 | 0.217 | 0.054 | 0.002688753 | 14 | BX663503.3 |
| pleca3 | 1.5025206 | 0.401803948 | 0.267 | 0.079 | 0.003034641 | 14 | pleca |
| mt-atp63 | 1.6075284 | 0.613068324 | 0.717 | 0.419 | 0.003246725 | 14 | mt-atp6 |
| lpar12 | 1.6507761 | 0.476691743 | 0.3 | 0.101 | 0.003334073 | 14 | lpar1 |
| myl9b3 | 1.8778168 | 0.510269174 | 0.467 | 0.2 | 0.003792627 | 14 | myl9b |
| tspan72 | 1.9543958 | 0.414763498 | 0.3 | 0.1 | 0.003947293 | 14 | tspan7 |
| nfia3 | 2.0547745 | 0.486242894 | 0.467 | 0.197 | 0.004150028 | 14 | nfia |
| adamts15b1 | 2.0888881 | 0.34352032 | 0.167 | 0.035 | 0.004218927 | 14 | adamts15b |
| si:ch211-133n4.44 | 3.8056393 | 0.468315772 | 0.35 | 0.131 | 0.00768625 | 14 | si:ch211-133n4.4 |
| hoxc8a1 | 4.0330859 | 0.374537192 | 0.233 | 0.067 | 0.008145624 | 14 | hoxc8a |
| rarga1 | 4.5367763 | 0.411522321 | 0.267 | 0.084 | 0.009162927 | 14 | rarga |
| krt43 | 4.7412822 | 0.316353156 | 0.317 | 0.099 | 0.009575968 | 14 | krt4 |
| hsd3b71 | 4.8488316 | 0.339031676 | 0.25 | 0.071 | 0.009793185 | 14 | hsd3b7 |
| dedd15 | 5.3732941 | 0.510289058 | 0.45 | 0.21 | 0.010852442 | 14 | dedd1 |
| tmeff1b1 | 6.5196857 | 0.381813875 | 0.3 | 0.102 | 0.013167809 | 14 | tmeff1b |
| fermt2 | 6.5250380 | 0.335941138 | 0.183 | 0.045 | 0.013178619 | 14 | fermt2 |
| fkbp10b | 6.7278181 | 0.28920955 | 0.15 | 0.032 | 0.013588174 | 14 | fkbp10b |
| dusp8a | 6.9547555 | 0.302390566 | 0.133 | 0.026 | 0.01404652 | 14 | dusp8a |
| serpinh1b1 | 7.6431029 | 0.376586547 | 0.233 | 0.068 | 0.015436775 | 14 | serpinh1b |
| h2afx11 | 8.3665256 | 0.504432951 | 0.667 | 0.377 | 0.016897872 | 14 | h2afx1 |
| krt18b3 | 8.9715161 | 0.475726428 | 0.633 | 0.316 | 0.018119771 | 14 | krt18b |
| kras1 | 9.4367767 | 0.315321619 | 0.217 | 0.061 | 0.019059458 | 14 | kras |
| ephb32 | 9.5240357 | 0.420895708 | 0.3 | 0.106 | 0.019235695 | 14 | ephb3 |
| stk17al1 | 1.0483915 | 0.299452615 | 0.2 | 0.052 | 0.021174364 | 14 | stk17al |
| znf385a2 | 1.1249974 | 0.400245618 | 0.283 | 0.095 | 0.022721575 | 14 | znf385a |
| calua1 | 1.1770368 | 0.387405331 | 0.233 | 0.071 | 0.023772613 | 14 | calua |
| si:ch211-106j24.11 | 1.2863735 | 0.2796211 | 0.15 | 0.033 | 0.025980886 | 14 | si:ch211-106j24.1 |

| | | | | | | | |
|---|---|---|---|---|---|---|---|
| znf1035 | 1.3563610 | 0.330722694 | 0.2 | 0.055 | 0.027394425 | 14 | znf1035 |
| adamts2 | 1.3810941 | 0.307186778 | 0.167 | 0.04 | 0.027893958 | 14 | adamts2 |
| NC-002333.45 | 1.7431842 | 0.484099814 | 0.9 | 0.581 | 0.035207093 | 14 | NC-002333.4 |
| hspg21 | 1.7593491 | 0.338452672 | 0.183 | 0.048 | 0.035533574 | 14 | hspg2 |
| leng8 | 1.7832950 | 0.281101495 | 0.167 | 0.04 | 0.03601721 | 14 | leng8 |
| ddit32 | 1.8179845 | 0.487294532 | 0.433 | 0.2 | 0.036717833 | 14 | ddit3 |
| XLOC-0320021 | 1.8820233 | 0.308016765 | 0.15 | 0.033 | 0.038011226 | 14 | XLOC-032002 |
| si:dkey-16l2.16 | 1.9031790 | 0.292491658 | 0.2 | 0.055 | 0.038438508 | 14 | si:dkey-16l2.16 |
| zyx1 | 1.9032541 | 0.441397352 | 0.317 | 0.125 | 0.038440023 | 14 | zyx |
| mt-co13 | 2.0133788 | 0.464665393 | 0.883 | 0.588 | 0.040664212 | 14 | mt-co1 |
| flot2a2 | 2.2207513 | 0.428753381 | 0.35 | 0.149 | 0.044852515 | 14 | flot2a |
| utrn1 | 2.5813380 | 0.276974176 | 0.167 | 0.041 | 0.052135284 | 14 | utrn |
| jpt1b4 | 2.6417353 | 0.471311691 | 0.6 | 0.331 | 0.053355128 | 14 | jpt1b |
| gli31 | 2.8645263 | 0.305852355 | 0.2 | 0.056 | 0.057854839 | 14 | gli3 |
| net12 | 3.0094853 | 0.356538037 | 0.25 | 0.081 | 0.060782575 | 14 | net1 |
| acvr1l | 3.0136000 | 0.290354227 | 0.15 | 0.035 | 0.06086568 | 14 | acvr1l |
| LOC1039092843 | 3.0339479 | 0.440069671 | 0.35 | 0.14 | 0.061276647 | 14 | LOC103909284 |
| cnn3a3 | 3.1692518 | 0.425390774 | 0.333 | 0.135 | 0.06400938 | 14 | cnn3a |
| map4k4 | 3.3101368 | 0.335769841 | 0.2 | 0.058 | 0.066854834 | 14 | map4k4 |
| meis1a2 | 3.5235059 | 0.306938868 | 0.183 | 0.049 | 0.071164249 | 14 | meis1a |
| tbx2b | 3.5520983 | 0.303816395 | 0.167 | 0.042 | 0.07174173 | 14 | tbx2b |
| palm21 | 3.6832868 | 0.25440574 | 0.117 | 0.022 | 0.074391344 | 14 | palm2 |
| ephb4a1 | 4.7070722 | 0.261971376 | 0.15 | 0.035 | 0.095068739 | 14 | ephb4a |
| parvaa | 4.8336641 | 0.313379362 | 0.217 | 0.066 | 0.097625515 | 14 | parvaa |
| crispld21 | 4.9899766 | 0.273612888 | 0.133 | 0.029 | 0.100782558 | 14 | crispld2 |
| ackr3b2 | 5.4344490 | 0.371137773 | 0.233 | 0.074 | 0.109759568 | 14 | ackr3b |
| slc43a3b1 | 5.8958300 | 0.296654414 | 0.183 | 0.05 | 0.119078079 | 14 | slc43a3b |
| mir199-31 | 6.0386826 | 0.318881757 | 0.167 | 0.043 | 0.121963274 | 14 | mir199-3 |
| prss231 | 6.3000506 | 0.277008375 | 0.167 | 0.043 | 0.127242123 | 14 | prss23 |
| ppap2d1 | 6.4103007 | 0.3022147 | 0.15 | 0.037 | 0.129468844 | 14 | ppap2d |
| klf7b | 6.9204804 | 0.309873013 | 0.2 | 0.059 | 0.139772943 | 14 | klf7b |
| tead1b1 | 7.3332844 | 0.334313794 | 0.2 | 0.06 | 0.148110345 | 14 | tead1b |
| myo1ca1 | 7.4151206 | 0.30078322 | 0.183 | 0.051 | 0.149763192 | 14 | myo1ca |
| lgals2a2 | 7.8095750 | 0.564945363 | 0.433 | 0.207 | 0.157729988 | 14 | lgals2a |
| reck | 8.9030087 | 0.286575077 | 0.167 | 0.045 | 0.179814068 | 14 | reck |

| | | | | | | | |
|---|---|---|---|---|---|---|---|
| tsku1 | 9.3299915 | 0.386815649 | 0.233 | 0.078 | 0.18843784 | 14 | tsku |
| edil3a2 | 9.7252421 | 0.317071409 | 0.133 | 0.031 | 0.196420716 | 14 | edil3a |
| tpst1 | 1.1029795 | 0.326312239 | 0.217 | 0.071 | 0.222768783 | 14 | tpst1 |
| angptl1a3 | 1.2662304 | 0.487798315 | 0.217 | 0.073 | 0.255740567 | 14 | angptl1a |
| lims1 | 1.2833653 | 0.318827231 | 0.2 | 0.062 | 0.25920129 | 14 | lims1 |
| wu:fi46h04 | 1.5826892 | 0.284057459 | 0.167 | 0.047 | 0.319655739 | 14 | wu:fi46h04 |
| mylka | 1.6210210 | 0.287792127 | 0.15 | 0.039 | 0.327397626 | 14 | mylka |
| tuba1b | 1.8665566 | 0.286083999 | 0.233 | 0.078 | 0.376988447 | 14 | tuba1b |
| tmem1764 | 1.9272684 | 0.391257831 | 0.35 | 0.153 | 0.389250402 | 14 | tmem176 |
| her6 | 2.0083599 | 0.307006713 | 0.15 | 0.039 | 0.405628459 | 14 | her6 |
| calr1 | 2.3074634 | 0.37530736 | 0.333 | 0.146 | 0.466038384 | 14 | calr |
| gstt1b3 | 2.6942603 | 0.38026912 | 0.383 | 0.18 | 0.544159772 | 14 | gstt1b |
| golt1bb | 2.7147231 | 0.296660915 | 0.217 | 0.074 | 0.548292636 | 14 | golt1bb |
| colec124 | 2.7633161 | 0.344802481 | 0.283 | 0.11 | 0.558106968 | 14 | colec12 |
| ptbp1b | 2.8883181 | 0.266073351 | 0.2 | 0.065 | 0.583353624 | 14 | ptbp1b |
| rcan1a | 2.8894714 | 0.267048864 | 0.183 | 0.056 | 0.583586551 | 14 | rcan1a |
| FAT4 | 3.6737193 | 0.276588434 | 0.15 | 0.041 | 0.741981103 | 14 | FAT4 |
| ddx6 | 3.7033599 | 0.298992916 | 0.25 | 0.093 | 0.747967611 | 14 | ddx6 |
| myh101 | 3.9223478 | 0.297779204 | 0.217 | 0.074 | 0.792196586 | 14 | myh10 |
| tgfb31 | 3.9652470 | 0.348568445 | 0.233 | 0.086 | 0.800860942 | 14 | tgfb3 |
| igfbp5b2 | 4.3610686 | 0.432221425 | 0.383 | 0.174 | 0.880805038 | 14 | igfbp5b |
| lrpap1 | 4.4257006 | 0.319089039 | 0.267 | 0.105 | 0.893858768 | 14 | lrpap1 |
| kirrel1a1 | 5.3144647 | 0.322922674 | 0.2 | 0.067 | 1 | 14 | kirrel1a |
| col15a1a | 5.3166686 | 0.263053591 | 0.15 | 0.042 | 1 | 14 | col15a1a |
| cldn11a2 | 5.4784943 | 0.458998162 | 0.4 | 0.194 | 1 | 14 | cldn11a |
| sdc42 | 5.5204357 | 0.398465404 | 0.483 | 0.263 | 1 | 14 | sdc4 |
| adh8b | 5.5440069 | 0.308316961 | 0.25 | 0.097 | 1 | 14 | adh8b |
| tmem176l.14 | 5.6135740 | 0.414784688 | 0.567 | 0.323 | 1 | 14 | tmem176l.1 |
| ier2b3 | 6.1289274 | 0.474365709 | 0.517 | 0.325 | 1 | 14 | ier2b |
| daam1b1 | 6.6001363 | 0.258554072 | 0.133 | 0.036 | 1 | 14 | daam1b |
| per3 | 7.6527288 | 0.294765501 | 0.2 | 0.07 | 1 | 14 | per3 |
| hbegfa1 | 7.6910081 | 0.331029714 | 0.183 | 0.061 | 1 | 14 | hbegfa |
| tubb5 | 7.7544343 | 0.326527603 | 0.217 | 0.078 | 1 | 14 | tubb5 |
| acadm | 8.3595343 | 0.369767806 | 0.333 | 0.16 | 1 | 14 | acadm |
| dhrs3a1 | 8.9707929 | 0.343754615 | 0.217 | 0.079 | 1 | 14 | dhrs3a |

| | | | | | | | |
|---|---|---|---|---|---|---|---|
| bsg3 | 9.1481276 | 0.403966071 | 0.6 | 0.386 | 1 | 14 | bsg |
| enpp12 | 9.5148034 | 0.27528627 | 0.133 | 0.036 | 1 | 14 | enpp1 |
| s1pr2 | 9.6287060 | 0.305487383 | 0.183 | 0.062 | 1 | 14 | s1pr2 |
| notch32 | 9.8535731 | 0.267250752 | 0.133 | 0.036 | 1 | 14 | notch3 |
| aebp12 | 9.9021634 | 0.287559738 | 0.25 | 0.097 | 1 | 14 | aebp1 |
| ctnnb22 | 9.9999892 | 0.325618277 | 0.283 | 0.121 | 1 | 14 | ctnnb2 |
| vasnb | 0.000109 | 0.268812587 | 0.183 | 0.061 | 1 | 14 | vasnb |
| fxyd6l3 | 0.00011 | 0.388138367 | 0.317 | 0.147 | 1 | 14 | fxyd6l |
| vasna2 | 0.000111 | 0.375021211 | 0.25 | 0.1 | 1 | 14 | vasna |
| ssr11 | 0.000118 | 0.364526636 | 0.483 | 0.274 | 1 | 14 | ssr1 |
| tenm34 | 0.00012 | 0.363188578 | 0.317 | 0.145 | 1 | 14 | tenm3 |
| mex3b1 | 0.000122 | 0.281898294 | 0.2 | 0.07 | 1 | 14 | mex3b |
| LOC1018844512 | 0.000125 | 0.364662773 | 0.283 | 0.124 | 1 | 14 | LOC101884451 |
| fosb2 | 0.000126 | 0.418029791 | 0.267 | 0.112 | 1 | 14 | fosb |
| chd6 | 0.00013 | 0.32798361 | 0.233 | 0.094 | 1 | 14 | chd6 |
| sept7a1 | 0.000131 | 0.318268916 | 0.267 | 0.113 | 1 | 14 | sept7a |
| XLOC-0268361 | 0.000135 | 0.461310522 | 0.217 | 0.083 | 1 | 14 | XLOC-026836 |
| mrc21 | 0.000141 | 0.2648568 | 0.167 | 0.054 | 1 | 14 | mrc2 |
| epha32 | 0.000143 | 0.308474448 | 0.167 | 0.054 | 1 | 14 | epha3 |
| tnrc6c12 | 0.000145 | 0.38645472 | 0.433 | 0.24 | 1 | 14 | tnrc6c1 |
| dnmt3ab3 | 0.000148 | 0.395773486 | 0.333 | 0.167 | 1 | 14 | dnmt3ab |
| s100a10b4 | 0.000153 | 0.409951451 | 0.717 | 0.455 | 1 | 14 | s100a10b |
| si:dkey-7j14.64 | 0.000179 | 0.365811983 | 0.55 | 0.32 | 1 | 14 | si:dkey-7j14.6 |
| brd3b | 0.000186 | 0.329872172 | 0.267 | 0.115 | 1 | 14 | brd3b |
| fzd7b3 | 0.000199 | 0.347168524 | 0.25 | 0.105 | 1 | 14 | fzd7b |
| selenom | 0.000211 | 0.306719286 | 0.233 | 0.096 | 1 | 14 | selenom |
| ywhah1 | 0.000231 | 0.339107898 | 0.283 | 0.131 | 1 | 14 | ywhah |
| BX322530.13 | 0.000247 | 0.306389638 | 0.2 | 0.076 | 1 | 14 | BX322530.1 |
| creb3l3l | 0.000256 | 0.289061548 | 0.233 | 0.095 | 1 | 14 | creb3l3l |
| hdlbpa4 | 0.00026 | 0.393349816 | 0.45 | 0.267 | 1 | 14 | hdlbpa |
| loxl2a2 | 0.000268 | 0.295045823 | 0.2 | 0.074 | 1 | 14 | loxl2a |
| si:ch211-286o17.1 | 0.000269 | 0.385838758 | 0.25 | 0.111 | 1 | 14 | si:ch211-286o17.1 |
| hif1al | 0.000291 | 0.285129287 | 0.233 | 0.096 | 1 | 14 | hif1al |
| surf4l1 | 0.000301 | 0.322189992 | 0.2 | 0.077 | 1 | 14 | surf4l |
| zgc:1135313 | 0.000323 | 0.284655532 | 0.183 | 0.066 | 1 | 14 | zgc:113531 |

| | | | | | | | |
|---|---|---|---|---|---|---|---|
| crtap1 | 0.000333 | 0.331089693 | 0.217 | 0.088 | 1 | 14 | crtap |
| CR753876.12 | 0.000346 | 0.273321763 | 0.25 | 0.101 | 1 | 14 | CR753876.1 |
| mt-nd51 | 0.000353 | 0.343153875 | 0.35 | 0.181 | 1 | 14 | mt-nd5 |
| fn1a2 | 0.000359 | 0.413802567 | 0.35 | 0.181 | 1 | 14 | fn1a |
| sept8a | 0.000361 | 0.258642731 | 0.217 | 0.086 | 1 | 14 | sept8a |
| pard3bb | 0.000385 | 0.299729759 | 0.183 | 0.069 | 1 | 14 | pard3bb |
| agrn1 | 0.00041 | 0.356268634 | 0.25 | 0.112 | 1 | 14 | agrn |
| zgc:919101 | 0.000422 | 0.342699233 | 0.483 | 0.299 | 1 | 14 | zgc:91910 |
| postnb4 | 0.000424 | 0.433265913 | 0.367 | 0.188 | 1 | 14 | postnb |
| antxr2a3 | 0.000431 | 0.362077833 | 0.45 | 0.261 | 1 | 14 | antxr2a |
| pho | 0.000461 | 0.319209985 | 0.267 | 0.123 | 1 | 14 | pho |
| hmgb1a2 | 0.000466 | 0.353754083 | 0.633 | 0.427 | 1 | 14 | hmgb1a |
| FP101882.1 | 0.000468 | 0.283902475 | 0.15 | 0.05 | 1 | 14 | FP101882.1 |
| grb10b1 | 0.000518 | 0.261365508 | 0.167 | 0.06 | 1 | 14 | grb10b |
| sdha | 0.000524 | 0.281457838 | 0.233 | 0.1 | 1 | 14 | sdha |
| zgc:1584633 | 0.00053 | 0.255003971 | 0.267 | 0.118 | 1 | 14 | zgc:158463 |
| mgst3a | 0.000546 | 0.396328892 | 0.283 | 0.144 | 1 | 14 | mgst3a |
| angptl41 | 0.000576 | 0.288638536 | 0.217 | 0.089 | 1 | 14 | angptl4 |
| gsta.12 | 0.000591 | 0.260794132 | 0.233 | 0.098 | 1 | 14 | gsta.1 |
| dhrs3b2 | 0.000592 | 0.277200894 | 0.183 | 0.069 | 1 | 14 | dhrs3b |
| gna11b1 | 0.000598 | 0.277891492 | 0.2 | 0.08 | 1 | 14 | gna11b |
| rhoab1 | 0.00063 | 0.369130776 | 0.483 | 0.312 | 1 | 14 | rhoab |
| ndfip1 | 0.000648 | 0.346243839 | 0.383 | 0.223 | 1 | 14 | ndfip1 |
| col6a4a4 | 0.000662 | 0.366199964 | 0.333 | 0.169 | 1 | 14 | col6a4a |
| nabp1a | 0.000684 | 0.282525075 | 0.2 | 0.082 | 1 | 14 | nabp1a |
| sept102 | 0.000695 | 0.251772577 | 0.2 | 0.08 | 1 | 14 | sept10 |
| serinc1 | 0.000704 | 0.289339383 | 0.25 | 0.113 | 1 | 14 | serinc1 |
| pfn2l4 | 0.000799 | 0.338947025 | 0.45 | 0.262 | 1 | 14 | pfn2l |
| slc38a5b5 | 0.000866 | 0.336403936 | 0.367 | 0.205 | 1 | 14 | slc38a5b |
| hmga2 | 0.000899 | 0.259123525 | 0.15 | 0.053 | 1 | 14 | hmga2 |
| zbtb37 | 0.000904 | 0.261804629 | 0.133 | 0.045 | 1 | 14 | zbtb37 |
| asph | 0.000981 | 0.277614766 | 0.2 | 0.083 | 1 | 14 | asph |
| ctsk3 | 0.000987 | 0.288839113 | 0.283 | 0.137 | 1 | 14 | ctsk |
| selenof2 | 0.001137 | 0.32461555 | 0.617 | 0.423 | 1 | 14 | selenof |
| gadd45ba4 | 0.00114 | 0.352790498 | 0.417 | 0.251 | 1 | 14 | gadd45ba |

| | | | | | | | |
|---|---|---|---|---|---|---|---|
| rbm4.1 | 0.001155 | 0.27156351 | 0.217 | 0.095 | 1 | 14 | rbm4.1 |
| krt18a.15 | 0.001221 | 0.2980007 | 0.5 | 0.279 | 1 | 14 | krt18a.1 |
| cygb14 | 0.001243 | 0.297850548 | 0.233 | 0.103 | 1 | 14 | cygb1 |
| ssr22 | 0.001269 | 0.296898223 | 0.633 | 0.417 | 1 | 14 | ssr2 |
| bnc21 | 0.0014 | 0.262552268 | 0.2 | 0.084 | 1 | 14 | bnc2 |
| rab2a1 | 0.00143 | 0.33106559 | 0.367 | 0.218 | 1 | 14 | rab2a |
| add11 | 0.001474 | 0.331605243 | 0.233 | 0.113 | 1 | 14 | add1 |
| socs3b3 | 0.001485 | 0.356095975 | 0.283 | 0.146 | 1 | 14 | socs3b |
| si:ch211-212k18.9 | 0.001565 | 0.26069076 | 0.25 | 0.118 | 1 | 14 | si:ch211-212k18.9 |
| plecb2 | 0.001578 | 0.253186352 | 0.167 | 0.066 | 1 | 14 | plecb |
| puraa | 0.001579 | 0.26044254 | 0.217 | 0.098 | 1 | 14 | puraa |
| cxcl12a4 | 0.00189 | 0.285009046 | 0.217 | 0.095 | 1 | 14 | cxcl12a |
| ctnnb11 | 0.001996 | 0.315514361 | 0.367 | 0.214 | 1 | 14 | ctnnb1 |
| ip6k2b1 | 0.002007 | 0.365619112 | 0.3 | 0.167 | 1 | 14 | ip6k2b |
| zgc:77086 | 0.002156 | 0.25195481 | 0.25 | 0.122 | 1 | 14 | zgc:77086 |
| spred2b2 | 0.002163 | 0.290138968 | 0.333 | 0.181 | 1 | 14 | spred2b |
| mcl1b5 | 0.002508 | 0.316982822 | 0.5 | 0.315 | 1 | 14 | mcl1b |
| u2surp | 0.002514 | 0.310467079 | 0.283 | 0.153 | 1 | 14 | u2surp |
| tegt2 | 0.002624 | 0.289514641 | 0.533 | 0.362 | 1 | 14 | tegt |
| anxa1a4 | 0.002671 | 0.3265713 | 0.467 | 0.277 | 1 | 14 | anxa1a |
| ndufc2 | 0.002683 | 0.319411061 | 0.4 | 0.25 | 1 | 14 | ndufc2 |
| srsf9 | 0.002714 | 0.280094338 | 0.317 | 0.177 | 1 | 14 | srsf9 |
| pdgfra2 | 0.002806 | 0.264380354 | 0.267 | 0.133 | 1 | 14 | pdgfra |
| fthl272 | 0.002901 | 0.286679274 | 0.817 | 0.627 | 1 | 14 | fthl27 |
| rbp43 | 0.002949 | 0.277426036 | 0.417 | 0.219 | 1 | 14 | rbp4 |
| si:ch73-376l24.32 | 0.002965 | 0.312383834 | 0.3 | 0.171 | 1 | 14 | si:ch73-376l24.3 |
| higd1a1 | 0.003101 | 0.297971254 | 0.433 | 0.274 | 1 | 14 | higd1a |
| sost2 | 0.003113 | 0.313078235 | 0.167 | 0.07 | 1 | 14 | sost |
| ndnf1 | 0.003133 | 0.254570735 | 0.167 | 0.07 | 1 | 14 | ndnf |
| rap1b2 | 0.003192 | 0.304545157 | 0.483 | 0.326 | 1 | 14 | rap1b |
| lamp21 | 0.003221 | 0.286566239 | 0.3 | 0.168 | 1 | 14 | lamp2 |
| wls3 | 0.00336 | 0.305663637 | 0.317 | 0.182 | 1 | 14 | wls |
| fzd24 | 0.003685 | 0.254871665 | 0.233 | 0.117 | 1 | 14 | fzd2 |
| tspan142 | 0.003837 | 0.268210582 | 0.317 | 0.177 | 1 | 14 | tspan14 |
| ccl25b3 | 0.003856 | 0.380408733 | 0.333 | 0.186 | 1 | 14 | ccl25b |

| | | | | | | | | |
|---|---|---|---|---|---|---|---|---|
| CABZ01080702.13 | 0.004154 | 0.269815065 | 0.317 | 0.176 | | 1 | 14 | CABZ01080702.1 |
| tagln5 | 0.004294 | 0.427529067 | 0.25 | 0.137 | | 1 | 14 | tagln |
| mcfd2 | 0.004343 | 0.254945308 | 0.183 | 0.084 | | 1 | 14 | mcfd2 |
| tubb4b | 0.004417 | 0.250744831 | 0.817 | 0.669 | | 1 | 14 | tubb4b |
| add3a4 | 0.00465 | 0.296894168 | 0.367 | 0.221 | | 1 | 14 | add3a |
| appb1 | 0.004831 | 0.286454033 | 0.183 | 0.085 | | 1 | 14 | appb |
| fam107b | 0.005187 | 0.341318037 | 0.283 | 0.165 | | 1 | 14 | fam107b |
| alyref | 0.005266 | 0.255659276 | 0.25 | 0.132 | | 1 | 14 | alyref |
| dynll11 | 0.005515 | 0.292068003 | 0.567 | 0.397 | | 1 | 14 | dynll1 |
| si:ch73-335l21.41 | 0.005613 | 0.274135397 | 0.233 | 0.116 | | 1 | 14 | si:ch73-335l21.4 |
| srrm1 | 0.005633 | 0.282334348 | 0.25 | 0.139 | | 1 | 14 | srrm1 |
| syncrip2 | 0.005645 | 0.278586821 | 0.55 | 0.392 | | 1 | 14 | syncrip |
| acin1a2 | 0.005802 | 0.307270921 | 0.433 | 0.298 | | 1 | 14 | acin1a |
| anxa5b3 | 0.005813 | 0.270485319 | 0.3 | 0.169 | | 1 | 14 | anxa5b |
| si:dkey-239h2.31 | 0.005858 | 0.273151665 | 0.283 | 0.161 | | 1 | 14 | si:dkey-239h2.3 |
| ier51 | 0.006234 | 0.302491341 | 0.333 | 0.211 | | 1 | 14 | ier5 |
| kmt2bb1 | 0.006656 | 0.285651283 | 0.267 | 0.152 | | 1 | 14 | kmt2bb |
| h3f3c3 | 0.006699 | 0.271803951 | 0.683 | 0.529 | | 1 | 14 | h3f3c |
| mycb3 | 0.006723 | 0.260841822 | 0.283 | 0.161 | | 1 | 14 | mycb |
| calm2b2 | 0.006767 | 0.263350226 | 0.633 | 0.446 | | 1 | 14 | calm2b |
| si:ch211-213a13.1: | 0.006832 | 0.266761466 | 0.267 | 0.151 | | 1 | 14 | si:ch211-213a13.1 |
| tcf121 | 0.007557 | 0.261284599 | 0.233 | 0.123 | | 1 | 14 | tcf12 |
| hist2h2l2 | 0.008001 | 0.267835869 | 0.35 | 0.211 | | 1 | 14 | hist2h2l |
| nucks1a1 | 0.008041 | 0.271798231 | 0.367 | 0.232 | | 1 | 14 | nucks1a |
| sec11a2 | 0.00822 | 0.258647478 | 0.417 | 0.272 | | 1 | 14 | sec11a |
| sox4a2 | 0.008237 | 0.259421246 | 0.283 | 0.158 | | 1 | 14 | sox4a |
| notch21 | 0.008242 | 0.269101022 | 0.267 | 0.151 | | 1 | 14 | notch2 |
| rsrp11 | 0.008558 | 0.256109388 | 0.25 | 0.138 | | 1 | 14 | rsrp1 |
| sar1b | 0.009464 | 0.250114539 | 0.25 | 0.141 | | 1 | 14 | sar1b |
| col4a14 | 0.009473 | 0.280219117 | 0.25 | 0.138 | | 1 | 14 | col4a1 |
| cx34.5 | 5.7620390 | 1.186445132 | 0.559 | 0.013 | 1.16375902098054e-1 | | 15 | cx34.5 |
| gpr17 | 9.0890418 | 0.569082982 | 0.322 | 0.001 | 1.83571378064603e-1 | | 15 | gpr17 |
| sema7a | 3.4554842 | 1.110734296 | 0.593 | 0.02 | 6.97904145275001e-1 | | 15 | sema7a |
| ecrg4a2 | 3.8057049 | 1.589395906 | 0.814 | 0.05 | 7.68638223079042e-1 | | 15 | ecrg4a |
| rasd11 | 5.0228905 | 0.870719198 | 0.441 | 0.009 | 1.01447320729872e-1 | | 15 | rasd1 |

| | | | | | | | |
|---|---|---|---|---|---|---|---|
| ackr4b | 8.9377082 | 1.024664383 | 0.542 | 0.025 | 1.80514893946612e-8 | 15 | ackr4b |
| pik3ip1 | 1.9755259 | 0.986837316 | 0.559 | 0.031 | 3.98996969853893e-7 | 15 | pik3ip1 |
| man2c11 | 9.7274187 | 0.94714286 | 0.475 | 0.021 | 1.9646467628373e-76 | 15 | man2c1 |
| inha | 3.6368241 | 0.612016594 | 0.305 | 0.007 | 7.34529373789289e-6 | 15 | inha |
| ghrb1 | 4.2218951 | 0.842986056 | 0.576 | 0.039 | 8.52696163216841e-6 | 15 | ghrb |
| cyp17a11 | 4.3221646 | 1.098466488 | 0.508 | 0.029 | 8.72947602965045e-6 | 15 | cyp17a1 |
| XLOC-0056052 | 5.2671120 | 1.180323296 | 0.763 | 0.078 | 1.0637986245773e-65 | 15 | XLOC-005605 |
| dag11 | 1.8456314 | 1.140537958 | 0.712 | 0.071 | 3.72762179904424e-6 | 15 | dag1 |
| pi15a1 | 2.6856477 | 1.339241507 | 0.78 | 0.084 | 5.4242027880128e-64 | 15 | pi15a |
| igf31 | 8.9164196 | 1.166934595 | 0.542 | 0.037 | 1.80084926955055e-6 | 15 | igf3 |
| COLEC10 | 9.4071383 | 0.504180307 | 0.254 | 0.005 | 1.89995973363667e-6 | 15 | COLEC10 |
| asb111 | 2.4400977 | 0.724020132 | 0.39 | 0.017 | 4.9282654242013e-61 | 15 | asb11 |
| met | 7.2916607 | 0.620472782 | 0.339 | 0.012 | 1.47269672484626e-6 | 15 | met |
| tmtops2b | 2.5077842 | 0.32592178 | 0.169 | 0.001 | 5.06497190148648e-5 | 15 | tmtops2b |
| ecm1b3 | 8.1717865 | 1.369193302 | 0.847 | 0.118 | 1.65045572882126e-5 | 15 | ecm1b |
| anxa13l1 | 1.7315728 | 0.595468532 | 0.356 | 0.015 | 3.49725772393298e-5 | 15 | anxa13l |
| podxl2 | 1.0457158 | 1.198286394 | 0.814 | 0.097 | 2.11203236859983e-5 | 15 | podxl |
| slc7a3a | 1.1681645 | 0.987821517 | 0.61 | 0.059 | 2.35934185364739e-5 | 15 | slc7a3a |
| apoeb1 | 4.0697915 | 0.742988384 | 0.458 | 0.029 | 8.21975802225011e-5 | 15 | apoeb |
| gsdf1 | 1.6071879 | 1.176295405 | 0.678 | 0.07 | 3.2460374969924e-53 | 15 | gsdf |
| rn7sk | 2.8721921 | 1.219777425 | 0.712 | 0.089 | 5.80096653278408e-5 | 15 | rn7sk |
| cldn11a3 | 4.0297797 | 1.383181884 | 0.966 | 0.177 | 8.13894606151217e-5 | 15 | cldn11a |
| LOC1039109031 | 1.2756095 | 1.037015151 | 0.458 | 0.032 | 2.57634860344114e-5 | 15 | LOC103910903 |
| robo11 | 2.4834973 | 0.755157245 | 0.508 | 0.041 | 5.01591953276414e-5 | 15 | robo1 |
| jam2a1 | 6.4728172 | 1.088699647 | 0.797 | 0.106 | 1.30731489766485e-5 | 15 | jam2a |
| ptprga1 | 1.0955496 | 1.068180847 | 0.712 | 0.092 | 2.21268156854142e-4 | 15 | ptprga |
| cyp19a1a | 1.5457612 | 0.491636566 | 0.186 | 0.003 | 3.121973961959e-49 | 15 | cyp19a1a |
| CABZ01068366.11 | 1.8479277 | 1.054166621 | 0.695 | 0.091 | 3.73225966678623e-4 | 15 | CABZ01068366.1 |
| tmem88b3 | 4.2049818 | 1.257548747 | 0.78 | 0.125 | 8.49280177840937e-4 | 15 | tmem88b |
| CABZ01068367.1 | 1.3057019 | 0.79715287 | 0.458 | 0.037 | 2.63712627551684e-4 | 15 | CABZ01068367.1 |
| crim12 | 1.3796606 | 0.835890329 | 0.559 | 0.057 | 2.78650052223697e-4 | 15 | crim1 |
| dpp31 | 1.5272029 | 0.872437902 | 0.627 | 0.075 | 3.08449178133436e-4 | 15 | dpp3 |
| islr2 | 8.8750617 | 0.629508798 | 0.322 | 0.017 | 1.79249622548077e-4 | 15 | islr2 |
| hsd3b1 | 2.2309696 | 0.323128441 | 0.153 | 0.002 | 4.50588931570273e-4 | 15 | hsd3b1 |
| drd2a | 2.6908442 | 0.31757023 | 0.153 | 0.002 | 5.43469811201517e-4 | 15 | drd2a |

| | | | | | | | |
|---|---|---|---|---|---|---|---|
| foxl2b1 | 1.8550473 | 0.812856544 | 0.492 | 0.045 | 3.74663922213506e-4 | 15 | foxl2b |
| musk | 3.5403624 | 0.419263239 | 0.22 | 0.007 | 7.1504701259563e-42 | 15 | musk |
| rxfp3.2a | 2.7905403 | 0.2739655 | 0.136 | 0.001 | 5.63605431213531e-4 | 15 | rxfp3.2a |
| greb1 | 3.7351407 | 0.300643503 | 0.169 | 0.003 | 7.54386387203463e-4 | 15 | greb1 |
| CABZ01068358.1 | 2.3037901 | 0.914119334 | 0.576 | 0.073 | 4.65296490989947e-3 | 15 | CABZ01068358.1 |
| aldh1a23 | 4.4359956 | 1.20328952 | 0.915 | 0.187 | 8.9593804184447e-39 | 15 | aldh1a2 |
| cx431 | 4.5819728 | 1.004939372 | 0.576 | 0.07 | 9.25421056103274e-3 | 15 | cx43 |
| bcam1 | 9.7552448 | 1.019702974 | 0.644 | 0.087 | 1.97026681082573e-3 | 15 | bcam |
| XLOC-0098151 | 1.6918141 | 0.845490629 | 0.492 | 0.053 | 3.41695706949143e-3 | 15 | XLOC-009815 |
| tnfaip61 | 2.7031872 | 0.701058111 | 0.39 | 0.031 | 5.45962731271726e-3 | 15 | tnfaip6 |
| itm2bb2 | 4.0308884 | 1.335666719 | 0.983 | 0.343 | 8.14118531301568e-3 | 15 | itm2bb |
| glulb1 | 5.8580096 | 0.943717187 | 0.576 | 0.075 | 1.18314221560752e-3 | 15 | glulb |
| dusp14 | 1.1330253 | 0.565229325 | 0.305 | 0.018 | 2.28837130292719e-3 | 15 | dusp14 |
| nr2f1a2 | 1.1332717 | 1.087484955 | 0.763 | 0.138 | 2.28886891657341e-3 | 15 | nr2f1a |
| oprd1a | 1.0217526 | 0.257045135 | 0.153 | 0.003 | 2.06363374558051e-3 | 15 | oprd1a |
| phc2a1 | 1.3025225 | 0.980004748 | 0.576 | 0.083 | 2.63070484257972e-3 | 15 | phc2a |
| slc38a4 | 4.0866526 | 0.307830507 | 0.203 | 0.007 | 8.25381229275896e-3 | 15 | slc38a4 |
| foxl2a | 4.6325430 | 0.476892863 | 0.288 | 0.017 | 9.35634720912228e-3 | 15 | foxl2a |
| XLOC-018328 | 5.2870455 | 0.439228957 | 0.186 | 0.006 | 1.06782459728345e-3 | 15 | XLOC-018328 |
| tox3 | 8.2650788 | 0.334228322 | 0.186 | 0.006 | 1.66929798227406e-3 | 15 | tox3 |
| trim47 | 1.0386585 | 0.484505474 | 0.305 | 0.02 | 2.09777865839126e-3 | 15 | trim47 |
| atp1a3b1 | 1.6834677 | 0.733499118 | 0.441 | 0.046 | 3.40009972185938e-3 | 15 | atp1a3b |
| appa | 1.9506640 | 0.968163314 | 0.712 | 0.131 | 3.93975616744102e-3 | 15 | appa |
| hspe12 | 2.9516891 | 1.238552922 | 0.932 | 0.278 | 5.96152654344892e-3 | 15 | hspe1 |
| r3hdml | 3.9136615 | 0.256545196 | 0.169 | 0.005 | 7.90442215355495e-3 | 15 | r3hdml |
| arrdc3a1 | 8.7224271 | 0.717753648 | 0.492 | 0.059 | 1.76166861479359e-3 | 15 | arrdc3a |
| si:dkey-42i9.7 | 1.1619718 | 0.713665149 | 0.407 | 0.042 | 2.34683458838171e-3 | 15 | si:dkey-42i9.7 |
| il15 | 2.0449203 | 0.788687748 | 0.508 | 0.068 | 4.13012571696666e-3 | 15 | il15 |
| si:ch211-225b11.1 | 2.6868177 | 0.652131241 | 0.424 | 0.047 | 5.42656579793412e-3 | 15 | si:ch211-225b11.1 |
| htr1aa | 5.6629278 | 0.254899515 | 0.119 | 0.002 | 1.14374154195617e-2 | 15 | htr1aa |
| mpzl2b1 | 1.0020309 | 0.813868516 | 0.441 | 0.053 | 2.02380197795496e-2 | 15 | mpzl2b |
| hsd17b1 | 1.9908243 | 0.267783765 | 0.153 | 0.004 | 4.02086793149353e-2 | 15 | hsd17b1 |
| atp1b3b1 | 2.2322463 | 0.716661011 | 0.441 | 0.053 | 4.5084680038882e-29 | 15 | atp1b3b |
| CR318588.31 | 2.7486623 | 0.718739479 | 0.525 | 0.071 | 5.55147325497334e-2 | 15 | CR318588.3 |
| XLOC-042229 | 3.9521562 | 0.48448124 | 0.271 | 0.019 | 7.98217004137781e-2 | 15 | XLOC-042229 |

| | | | | | | | |
|---|---|---|---|---|---|---|---|
| nid1b1 | 6.3486269 | 0.935923499 | 0.678 | 0.13 | 1.28223219060635e-2 | 15 | nid1b |
| lamb1a4 | 1.8538160 | 0.873710639 | 0.678 | 0.128 | 3.74415234258034e-2 | 15 | lamb1a |
| ar1 | 2.0555708 | 0.766376403 | 0.593 | 0.099 | 4.15163643593718e-2 | 15 | ar |
| wu:fc75a09 | 3.3819267 | 0.450579213 | 0.271 | 0.019 | 6.83047740891421e-2 | 15 | wu:fc75a09 |
| fam102ab | 1.7410161 | 0.612580197 | 0.356 | 0.036 | 3.51633040917321e-2 | 15 | fam102ab |
| hsd3b72 | 2.7157096 | 0.773561816 | 0.475 | 0.065 | 5.48491873619924e-2 | 15 | hsd3b7 |
| bambia1 | 4.3273838 | 0.808378435 | 0.424 | 0.054 | 8.74001711477342e-2 | 15 | bambia |
| mxra8a1 | 5.4045785 | 0.824830218 | 0.695 | 0.136 | 1.0915627372646e-26 | 15 | mxra8a |
| gata61 | 6.0268150 | 0.640519182 | 0.424 | 0.052 | 1.21723584293687e-2 | 15 | gata6 |
| znf1104 | 7.1694938 | 0.477566485 | 0.254 | 0.018 | 1.44802266316873e-2 | 15 | znf1104 |
| slc4a4a | 3.9318680 | 0.38928949 | 0.22 | 0.013 | 7.94119384962326e-2 | 15 | slc4a4a |
| cox7a1 | 5.0145109 | 0.518597946 | 0.407 | 0.05 | 1.01278077809038e-2 | 15 | cox7a1 |
| pgrmc12 | 5.9261451 | 0.970633619 | 0.797 | 0.21 | 1.19690353207636e-2 | 15 | pgrmc1 |
| ifi463 | 1.5791410 | 0.919567659 | 0.763 | 0.179 | 3.18939125911447e-2 | 15 | ifi46 |
| CR318588.12 | 2.0315569 | 1.230053072 | 0.898 | 0.253 | 4.10313548915158e-2 | 15 | CR318588.1 |
| pdgfaa | 8.2655117 | 0.594550285 | 0.407 | 0.052 | 1.66938541203997e-2 | 15 | pdgfaa |
| zgc:927443 | 1.1829250 | 1.064840281 | 0.763 | 0.195 | 2.38915376734779e-2 | 15 | zgc:92744 |
| dmrt1 | 1.3174149 | 0.272005823 | 0.119 | 0.003 | 2.66078296514318e-2 | 15 | dmrt1 |
| cdh12 | 2.8514370 | 0.590657562 | 0.339 | 0.037 | 5.75904747502703e-2 | 15 | cdh1 |
| ctsla6 | 2.5589311 | 0.920361866 | 0.932 | 0.296 | 5.16827331312058e-2 | 15 | ctsla |
| si:ch73-61d6.3 | 7.2963967 | 0.451183838 | 0.305 | 0.031 | 1.47365324429827e-2 | 15 | si:ch73-61d6.3 |
| XLOC-034863 | 1.0700164 | 0.260015144 | 0.169 | 0.008 | 2.16111229929569e-2 | 15 | XLOC-034863 |
| plxna4 | 1.1936201 | 0.514692486 | 0.271 | 0.025 | 2.41075455338595e-2 | 15 | plxna4 |
| tnfrsfa | 1.5960063 | 0.516079965 | 0.356 | 0.043 | 3.22345393103861e-2 | 15 | tnfrsfa |
| grip1 | 2.7271396 | 0.332521591 | 0.186 | 0.011 | 5.50800392233405e-2 | 15 | grip1 |
| akap12b3 | 3.3626105 | 0.910854051 | 0.644 | 0.142 | 6.79146451274085e-2 | 15 | akap12b |
| phactr4a1 | 3.7759698 | 0.601766934 | 0.39 | 0.053 | 7.62632623373967e-2 | 15 | phactr4a |
| myclb1 | 5.1103112 | 0.746977344 | 0.475 | 0.081 | 1.03212955367593e-2 | 15 | myclb |
| msi2a | 9.4142067 | 0.541575498 | 0.373 | 0.049 | 1.9013873380263e-21 | 15 | msi2a |
| rftn2 | 1.9560965 | 0.296450929 | 0.186 | 0.011 | 3.95072811948503e-2 | 15 | rftn2 |
| si:dkey-22o12.2 | 3.8298735 | 0.484042839 | 0.254 | 0.023 | 7.73519570382395e-2 | 15 | si:dkey-22o12.2 |
| XLOC-015291 | 5.1636557 | 0.417790989 | 0.271 | 0.026 | 1.04290354749073e-2 | 15 | XLOC-015291 |
| krt18a.16 | 5.5891870 | 1.065880551 | 0.847 | 0.268 | 1.1288481069699e-20 | 15 | krt18a.1 |
| parp12a | 7.1494010 | 0.540388687 | 0.322 | 0.038 | 1.44396452570068e-2 | 15 | parp12a |
| col4a5 | 9.1583461 | 0.422598832 | 0.254 | 0.023 | 1.8497111808014e-20 | 15 | col4a5 |

| | | | | | | | |
|---|---|---|---|---|---|---|---|
| hspd1 | 9.5092430 | 0.776099888 | 0.661 | 0.158 | 1.92058182026159e-2 | 15 | hspd1 |
| pde4ba1 | 1.0253673 | 0.834773858 | 0.644 | 0.153 | 2.07093445722327e-2 | 15 | pde4ba |
| alcama1 | 1.1189940 | 0.592160068 | 0.373 | 0.05 | 2.26003236808632e-2 | 15 | alcama |
| hs3st3b1b | 1.6249982 | 0.375542554 | 0.22 | 0.017 | 3.28200895387503e-2 | 15 | hs3st3b1b |
| glula1 | 2.1151381 | 0.724118725 | 0.441 | 0.073 | 4.27194452921023e-2 | 15 | glula |
| ldlrap1a | 2.2897908 | 0.414916648 | 0.288 | 0.03 | 4.62469055090278e-2 | 15 | ldlrap1a |
| fitm2 | 4.1326133 | 0.321536197 | 0.203 | 0.015 | 8.34663919276856e-2 | 15 | fitm2 |
| CR751602.2 | 1.0478520 | 0.413614961 | 0.237 | 0.021 | 2.11634684665194e-1 | 15 | CR751602.2 |
| epb41l51 | 1.6707040 | 0.500825037 | 0.373 | 0.052 | 3.37432088594923e-1 | 15 | epb41l5 |
| zfpm1 | 1.7492215 | 0.414130035 | 0.271 | 0.028 | 3.53290277113604e-1 | 15 | zfpm1 |
| tcf122 | 1.9144077 | 0.70737764 | 0.542 | 0.114 | 3.86652935427149e-1 | 15 | tcf12 |
| gdpd4a | 4.7051621 | 0.327079984 | 0.186 | 0.013 | 9.50301601601342e-1 | 15 | gdpd4a |
| si:dkey-283j8.1 | 8.6672922 | 0.317962594 | 0.153 | 0.008 | 1.75053301890355e-1 | 15 | si:dkey-283j8.1 |
| fdxr | 8.9839681 | 0.4987731 | 0.288 | 0.033 | 1.81449203862133e-1 | 15 | fdxr |
| atp1b1a2 | 1.0646440 | 0.844051444 | 0.847 | 0.283 | 2.15026162764823e-1 | 15 | atp1b1a |
| aff2 | 1.2606302 | 0.336053909 | 0.186 | 0.013 | 2.5460948866773e-18 | 15 | aff2 |
| lgals171 | 1.4656706 | 0.629927763 | 0.407 | 0.065 | 2.96021495607493e-1 | 15 | lgals17 |
| col4a6 | 1.5179090 | 0.514941021 | 0.288 | 0.034 | 3.06572084024013e-1 | 15 | col4a6 |
| cpn14 | 1.7254630 | 0.82849801 | 0.695 | 0.184 | 3.48491767498597e-1 | 15 | cpn1 |
| zeb2b | 2.2386813 | 0.31046699 | 0.169 | 0.011 | 4.52146477481185e-1 | 15 | zeb2b |
| ifi452 | 3.4112613 | 0.839109258 | 0.729 | 0.188 | 6.88972461112274e-1 | 15 | ifi45 |
| si:dkey-188i13.9 | 4.2323844 | 0.441931428 | 0.305 | 0.037 | 8.54814682137978e-1 | 15 | si:dkey-188i13.9 |
| si:dkey-7j14.65 | 4.2713498 | 0.832119739 | 0.898 | 0.31 | 8.62684537543958e-1 | 15 | si:dkey-7j14.6 |
| si:ch211-136a13.1 | 4.3565403 | 0.44089087 | 0.203 | 0.017 | 8.79890464305079e-1 | 15 | si:ch211-136a13.1 |
| tcf41 | 4.8434212 | 0.431730392 | 0.271 | 0.03 | 9.78225791284349e-1 | 15 | tcf4 |
| tp53inp11 | 1.5594677 | 0.639142861 | 0.39 | 0.065 | 3.14965697198615e-1 | 15 | tp53inp1 |
| si:dkey-187e18.1 | 3.0284378 | 0.597555542 | 0.441 | 0.082 | 6.11653588931764e-1 | 15 | si:dkey-187e18.1 |
| mfge8b1 | 4.5556138 | 0.431942376 | 0.305 | 0.039 | 9.20097323780997e-1 | 15 | mfge8b |
| hpn | 4.6280073 | 0.347370869 | 0.169 | 0.012 | 9.34718649891081e-1 | 15 | hpn |
| tmem176l.15 | 6.4495517 | 0.835300784 | 0.864 | 0.314 | 1.30261597334634e-1 | 15 | tmem176l.1 |
| zmp:0000000711 | 6.9684964 | 0.251759012 | 0.153 | 0.009 | 1.40742722332554e-1 | 15 | zmp:0000000711 |
| elavl4 | 7.9622493 | 0.319562313 | 0.169 | 0.012 | 1.60813551106395e-1 | 15 | elavl4 |
| f103 | 8.1960969 | 0.639277841 | 0.525 | 0.111 | 1.65536569124405e-1 | 15 | f10 |
| si:dkey-73n10.11 | 1.0991465 | 0.393040861 | 0.271 | 0.032 | 2.21994624026684e-1 | 15 | si:dkey-73n10.1 |
| zgc:92140 | 3.0311772 | 0.50451889 | 0.356 | 0.056 | 6.12206869495738e-1 | 15 | zgc:92140 |

| | | | | | | | |
|---|---|---|---|---|---|---|---|
| rhoq | 3.4707444 | 0.377541334 | 0.237 | 0.025 | 7.00986265724598e-1 | 15 | rhoq |
| CABZ01021532.1 | 3.9107977 | 0.439061342 | 0.22 | 0.022 | 7.89863820622994e-1 | 15 | CABZ01021532.1 |
| lama51 | 4.1199984 | 0.546212392 | 0.356 | 0.057 | 8.32116085319251e-1 | 15 | lama5 |
| chtopa | 4.6711106 | 0.680021639 | 0.559 | 0.136 | 9.43424210625203e-1 | 15 | chtopa |
| ap2m1a1 | 6.7899364 | 0.798298872 | 0.814 | 0.295 | 1.37136346394747e-1 | 15 | ap2m1a |
| cebpd5 | 7.8419151 | 0.800619773 | 0.847 | 0.267 | 1.58383160534294e-1 | 15 | cebpd |
| rpia1 | 7.9124834 | 0.670555074 | 0.424 | 0.083 | 1.59808429001536e-1 | 15 | rpia |
| mov10b.1 | 8.1982072 | 0.370159714 | 0.237 | 0.026 | 1.65579191637432e-1 | 15 | mov10b.1 |
| scn4bb | 9.3832043 | 0.353002423 | 0.203 | 0.019 | 1.89512579207709e-1 | 15 | scn4bb |
| odc12 | 9.5009572 | 0.708924194 | 0.508 | 0.113 | 1.91890833526211e-1 | 15 | odc1 |
| gstm.35 | 9.6862673 | 0.951648793 | 0.864 | 0.275 | 1.95633542548753e-1 | 15 | gstm.3 |
| mmp25 | 1.2811351 | 0.766966278 | 0.712 | 0.207 | 2.58750867850362e-1 | 15 | mmp2 |
| pde7a | 1.3536479 | 0.563420441 | 0.441 | 0.088 | 2.7339627948984e-15 | 15 | pde7a |
| enc11 | 1.9386919 | 0.585027875 | 0.407 | 0.075 | 3.91557604955172e-1 | 15 | enc1 |
| svild2 | 2.3219481 | 0.636896255 | 0.458 | 0.093 | 4.6896387761548e-15 | 15 | svild |
| angptl42 | 2.9411657 | 0.638472845 | 0.424 | 0.083 | 5.94027238031334e-1 | 15 | angptl4 |
| LOC553316 | 4.0961580 | 0.56034334 | 0.356 | 0.061 | 8.27301044068413e-1 | 15 | LOC553316 |
| si:dkey-21a6.5 | 4.5575345 | 0.292728127 | 0.203 | 0.019 | 9.20485245282027e-1 | 15 | si:dkey-21a6.5 |
| si:ch73-86n18.1 | 4.6210906 | 0.280145856 | 0.169 | 0.013 | 9.3332168317053e-15 | 15 | si:ch73-86n18.1 |
| ccdc1871 | 4.8976800 | 0.568474671 | 0.39 | 0.07 | 9.89184432217643e-1 | 15 | ccdc187 |
| jam3b | 6.3373491 | 0.492516441 | 0.356 | 0.059 | 1.27995440506069e-1 | 15 | jam3b |
| glrx51 | 8.4170909 | 0.478532971 | 0.407 | 0.076 | 1.69999985656706e-1 | 15 | glrx5 |
| sox4a3 | 1.0464903 | 0.78805032 | 0.559 | 0.15 | 2.11359657287387e-1 | 15 | sox4a |
| rhoj | 1.1225884 | 0.35046837 | 0.203 | 0.02 | 2.26729191742151e-1 | 15 | rhoj |
| hspg22 | 1.2722611 | 0.444867864 | 0.305 | 0.045 | 2.56958574881762e-1 | 15 | hspg2 |
| ptprfb1 | 1.4655185 | 0.377642367 | 0.237 | 0.028 | 2.95990785306137e-1 | 15 | ptprfb |
| sema3b | 1.6199285 | 0.356436085 | 0.203 | 0.02 | 3.27176962296161e-1 | 15 | sema3b |
| fam234a | 1.8359908 | 0.38768124 | 0.237 | 0.028 | 3.70815073929671e-1 | 15 | fam234a |
| uxs1 | 1.9025275 | 0.433558316 | 0.288 | 0.041 | 3.84253499133732e-1 | 15 | uxs1 |
| vasnb1 | 2.0814077 | 0.52463324 | 0.339 | 0.057 | 4.20381914464824e-1 | 15 | vasnb |
| zgc:100829 | 2.3272003 | 0.546022917 | 0.39 | 0.073 | 4.70024658521247e-1 | 15 | zgc:100829 |
| mtus1b | 2.9045153 | 0.388062345 | 0.237 | 0.028 | 5.86624968704163e-1 | 15 | mtus1b |
| si:ch211-117c9.51 | 2.9721945 | 0.392008056 | 0.254 | 0.032 | 6.00294136377377e-1 | 15 | si:ch211-117c9.5 |
| alpl | 5.2817760 | 0.2787118 | 0.153 | 0.011 | 1.06676030741121e-1 | 15 | alpl |
| robo22 | 6.4290314 | 0.603712025 | 0.424 | 0.088 | 1.29847147695193e-1 | 15 | robo2 |

| | | | | | | | |
|---|---|---|---|---|---|---|---|
| tmsb3 | 1.2738357 | 0.487751827 | 0.322 | 0.051 | 2.57276596422548e-1 | 15 | tmsb |
| hbp1 | 1.3167373 | 0.500095697 | 0.356 | 0.065 | 2.65941441286106e-1 | 15 | hbp1 |
| ece2b | 1.4053994 | 0.467146138 | 0.322 | 0.052 | 2.83848517671621e-1 | 15 | ece2b |
| chd7 | 2.2544439 | 0.405459591 | 0.288 | 0.043 | 4.55330048060484e-1 | 15 | chd7 |
| c64 | 2.2609805 | 0.788745905 | 0.678 | 0.205 | 4.56650233946332e-1 | 15 | c6 |
| nav3 | 3.0482018 | 0.426533152 | 0.254 | 0.034 | 6.15645318448488e-1 | 15 | nav3 |
| antxr1c1 | 3.0790363 | 0.484653761 | 0.305 | 0.049 | 6.21872977408443e-1 | 15 | antxr1c |
| irs1 | 3.7153668 | 0.396691734 | 0.288 | 0.043 | 7.50392645963586e-1 | 15 | irs1 |
| zgc:1951733 | 6.9916677 | 0.72663164 | 0.712 | 0.227 | 1.41210713800688e-1 | 15 | zgc:195173 |
| junba4 | 1.2306964 | 0.795352616 | 0.898 | 0.394 | 2.4856376565821e-12 | 15 | junba |
| socs3a4 | 1.7005642 | 0.79471427 | 0.898 | 0.421 | 3.43462953597003e-1 | 15 | socs3a |
| si:ch211-199g17.2 | 3.2093622 | 0.531529654 | 0.39 | 0.081 | 6.48194899526555e-1 | 15 | si:ch211-199g17.2 |
| rnf128a1 | 4.1388645 | 0.334260248 | 0.203 | 0.024 | 8.35926483054326e-1 | 15 | rnf128a |
| si:dkey-188i13.101 | 4.2347336 | 0.617775259 | 0.576 | 0.161 | 8.55289158049302e-1 | 15 | si:dkey-188i13.10 |
| cd1512 | 5.3623035 | 0.469070566 | 0.271 | 0.042 | 1.08302443964443e-1 | 15 | cd151 |
| si:zfos-464b6.2 | 6.0549418 | 0.371459982 | 0.305 | 0.052 | 1.22291661142238e-1 | 15 | si:zfos-464b6.2 |
| mknk2b2 | 6.5168342 | 0.733360452 | 0.627 | 0.213 | 1.3162050162119e-11 | 15 | mknk2b |
| itga3b | 8.5935849 | 0.250127051 | 0.153 | 0.013 | 1.7356463461111e-11 | 15 | itga3b |
| chka | 9.6602193 | 0.388584882 | 0.288 | 0.047 | 1.95107449735158e-1 | 15 | chka |
| rnf213b | 1.0630521 | 0.39946464 | 0.186 | 0.021 | 2.14704648972271e-1 | 15 | rnf213b |
| alcamb1 | 1.0702081 | 0.471607937 | 0.288 | 0.047 | 2.16149935550007e-1 | 15 | alcamb |
| cldn12 | 1.1125003 | 0.417888244 | 0.288 | 0.048 | 2.2469169133811e-11 | 15 | cldn12 |
| prkar2aa | 1.2108103 | 0.568071139 | 0.525 | 0.143 | 2.44547361428854e-1 | 15 | prkar2aa |
| hvcn1 | 1.7025266 | 0.425652109 | 0.305 | 0.054 | 3.4385931183914e-11 | 15 | hvcn1 |
| wt1a1 | 1.7480828 | 0.522541137 | 0.39 | 0.082 | 3.53060298558726e-1 | 15 | wt1a |
| cdh23 | 1.7482366 | 0.586281344 | 0.475 | 0.121 | 3.53091348072453e-1 | 15 | cdh2 |
| slc43a2a2 | 1.8120969 | 0.603075856 | 0.424 | 0.099 | 3.65989219886793e-1 | 15 | slc43a2a |
| nxn | 2.4289149 | 0.339918993 | 0.22 | 0.029 | 4.90567952116122e-1 | 15 | nxn |
| LOC110438024 | 2.8577519 | 0.308905538 | 0.169 | 0.017 | 5.77180156985667e-1 | 15 | LOC110438024 |
| hist2h2l3 | 2.9133726 | 0.719171632 | 0.61 | 0.203 | 5.88413877144046e-1 | 15 | hist2h2l |
| zgc:1101822 | 3.0964714 | 0.482605717 | 0.373 | 0.076 | 6.25394340542782e-1 | 15 | zgc:110182 |
| b2m1 | 3.8866337 | 0.752040581 | 0.797 | 0.346 | 7.8498342233871e-11 | 15 | b2m |
| rpz52 | 3.9230947 | 0.749553958 | 0.695 | 0.248 | 7.92347445954686e-1 | 15 | rpz5 |
| irf2bp2b2 | 6.1715114 | 0.620318447 | 0.593 | 0.185 | 1.24646017244764e-1 | 15 | irf2bp2b |
| mbnl31 | 6.2271389 | 0.490067103 | 0.407 | 0.093 | 1.25769526099599e-1 | 15 | mbnl3 |

| | | | | | | | |
|---|---|---|---|---|---|---|---|
| sema5bb | 6.4520866 | 0.289965203 | 0.153 | 0.014 | 1.30312794172804e-1 | 15 | sema5bb |
| steap4 | 6.6991840 | 0.285272148 | 0.153 | 0.014 | 1.35303419751822e-1 | 15 | steap4 |
| wu:fc27b11 | 7.2777135 | 0.360777626 | 0.237 | 0.034 | 1.46987980905846e-1 | 15 | wu:fc27b11 |
| ace | 7.4847569 | 0.396377693 | 0.237 | 0.035 | 1.51169635248099e-1 | 15 | ace |
| plcl1 | 8.4383725 | 0.325886097 | 0.169 | 0.018 | 1.70429810212071e-1 | 15 | plcl1 |
| cox6a2 | 1.1399201 | 0.374353058 | 0.254 | 0.04 | 2.30229680379034e-1 | 15 | cox6a2 |
| lsr | 1.3657921 | 0.327625222 | 0.254 | 0.039 | 2.75849044563752e-1 | 15 | lsr |
| stat1a1 | 1.6265501 | 0.52159171 | 0.508 | 0.14 | 3.2851434100719e-10 | 15 | stat1a |
| myo10l3 | 1.7014310 | 0.305628188 | 0.186 | 0.022 | 3.43638021294472e-1 | 15 | myo10l3 |
| crybg1a | 1.7961908 | 0.294439258 | 0.203 | 0.026 | 3.62776666217762e-1 | 15 | crybg1a |
| si:ch211-264f5.6 | 1.8127832 | 0.294285775 | 0.153 | 0.015 | 3.66127841023824e-1 | 15 | si:ch211-264f5.6 |
| atp1a1a.11 | 1.8496442 | 0.573397595 | 0.475 | 0.13 | 3.73572643359096e-1 | 15 | atp1a1a.1 |
| sema3ab | 2.4677176 | 0.429249287 | 0.186 | 0.023 | 4.98404941120602e-1 | 15 | sema3ab |
| lamc1 | 2.6567879 | 0.504179181 | 0.339 | 0.069 | 5.36591467250835e-1 | 15 | lamc1 |
| kremen1 | 2.9296164 | 0.349206503 | 0.169 | 0.019 | 5.91694627419002e-1 | 15 | kremen1 |
| cd82a2 | 5.1760298 | 0.62013518 | 0.475 | 0.134 | 1.045402744945e-09 | 15 | cd82a |
| slc16a10 | 5.2054628 | 0.36863461 | 0.271 | 0.048 | 1.05134733397758e-0 | 15 | slc16a10 |
| zfp36l1b3 | 5.3039966 | 0.611715867 | 0.61 | 0.193 | 1.07124821195177e-0 | 15 | zfp36l1b |
| aebp2 | 5.5124254 | 0.400093303 | 0.305 | 0.059 | 1.11334456945539e-0 | 15 | aebp2 |
| nos1apa | 6.3611658 | 0.39795624 | 0.322 | 0.064 | 1.28476467503429e-0 | 15 | nos1apa |
| atp10a1 | 6.4513726 | 0.43562071 | 0.339 | 0.07 | 1.30298373555449e-0 | 15 | atp10a |
| sppl3 | 7.3898904 | 0.398244671 | 0.305 | 0.059 | 1.49253617797865e-0 | 15 | sppl3 |
| mafba2 | 7.6169394 | 0.610047395 | 0.492 | 0.145 | 1.53839326617249e-0 | 15 | mafba |
| gstm.14 | 7.6654128 | 0.68778739 | 0.814 | 0.33 | 1.5481834361058e-09 | 15 | gstm.1 |
| wbp1la | 7.8054049 | 0.331090541 | 0.203 | 0.028 | 1.57645763178018e-0 | 15 | wbp1la |
| COX35 | 9.0014938 | 0.70158203 | 0.966 | 0.59 | 1.81803171218073e-0 | 15 | COX3 |
| slc29a1a | 9.2839816 | 0.400366755 | 0.254 | 0.043 | 1.87508577252702e-0 | 15 | slc29a1a |
| sigmar1 | 9.7802791 | 0.342559377 | 0.288 | 0.053 | 1.97532297178993e-0 | 15 | sigmar1 |
| zgc:66433 | 1.3755133 | 0.407000577 | 0.271 | 0.049 | 2.7781242821584e-09 | 15 | zgc:66433 |
| sox11b | 1.7265862 | 0.279964038 | 0.169 | 0.02 | 3.48718617310498e-0 | 15 | sox11b |
| ywhaz1 | 1.9850382 | 0.650629633 | 0.864 | 0.441 | 4.00918177516798e-0 | 15 | ywhaz |
| arf2b1 | 1.9978537 | 0.594880043 | 0.593 | 0.201 | 4.03506512956483e-0 | 15 | arf2b |
| stat2 | 2.1873828 | 0.415967035 | 0.322 | 0.066 | 4.41785714746006e-0 | 15 | stat2 |
| sort1a1 | 2.4104588 | 0.275314771 | 0.186 | 0.024 | 4.86840375689176e-0 | 15 | sort1a |
| si:dkey-220k22.1 | 2.7811956 | 0.259884094 | 0.169 | 0.02 | 5.61718093063773e-0 | 15 | si:dkey-220k22.1 |

| | | | | | | | |
|---|---|---|---|---|---|---|---|
| pfkfb4b2 | 2.8188823 | 0.704934048 | 0.576 | 0.214 | 5.69329670055956e-0 | 15 | pfkfb4b |
| gdpd5b | 2.8966960 | 0.314990047 | 0.203 | 0.029 | 5.85045696222063e-0 | 15 | gdpd5b |
| slc1a3a | 3.0717323 | 0.384671041 | 0.22 | 0.034 | 6.2039778826707e-09 | 15 | slc1a3a |
| cmtm6 | 3.4281890 | 0.473510553 | 0.356 | 0.085 | 6.92391347283678e-0 | 15 | cmtm6 |
| NC-002333.46 | 5.0667161 | 0.69596759 | 1 | 0.578 | 1.02332466362922e-0 | 15 | NC-002333.4 |
| nr0b11 | 6.2377990 | 0.341511867 | 0.237 | 0.039 | 1.25984828040268e-0 | 15 | nr0b1 |
| hdac4 | 6.8333086 | 0.373769692 | 0.237 | 0.04 | 1.38012335323734e-0 | 15 | hdac4 |
| pcbp2 | 6.9164886 | 0.519618528 | 0.542 | 0.175 | 1.39692320601311e-0 | 15 | pcbp2 |
| FP016056.11 | 6.9760907 | 0.253093289 | 0.136 | 0.013 | 1.40896105820447e-0 | 15 | FP016056.1 |
| sema6bb | 7.8307216 | 0.257229226 | 0.136 | 0.013 | 1.58157084531865e-0 | 15 | sema6bb |
| ifi302 | 1.0204522 | 0.641424655 | 0.746 | 0.3 | 2.06100750713692e-0 | 15 | ifi30 |
| aldh2.22 | 1.0488095 | 0.50139189 | 0.441 | 0.124 | 2.11828069595705e-0 | 15 | aldh2.2 |
| thbs3b | 1.1320012 | 0.250028751 | 0.119 | 0.01 | 2.28630289256259e-0 | 15 | thbs3b |
| ppp1r14bb | 1.1727850 | 0.362384963 | 0.322 | 0.069 | 2.36867406031403e-0 | 15 | ppp1r14bb |
| acat1 | 1.2257905 | 0.36467095 | 0.254 | 0.046 | 2.47572925068278e-0 | 15 | acat1 |
| hmgb3a4 | 1.2441438 | 0.621017592 | 0.678 | 0.252 | 2.51279741585264e-0 | 15 | hmgb3a |
| si:dkey-17m8.1 | 1.2670049 | 0.400185866 | 0.305 | 0.064 | 2.55896981961938e-0 | 15 | si:dkey-17m8.1 |
| iscub | 1.9806877 | 0.467112591 | 0.475 | 0.138 | 4.00039514011944e-0 | 15 | iscub |
| lgals9l33 | 2.2542813 | 0.518071251 | 0.559 | 0.18 | 4.55297209278965e-0 | 15 | lgals9l3 |
| txnipa1 | 2.3404920 | 0.650684548 | 0.797 | 0.361 | 4.72709179260678e-0 | 15 | txnipa |
| agrn2 | 2.5842347 | 0.50992236 | 0.407 | 0.108 | 5.21937897270743e-0 | 15 | agrn |
| hmgb1b2 | 2.8070153 | 0.579868938 | 0.763 | 0.294 | 5.66932897273905e-0 | 15 | hmgb1b |
| dnmt3bb.1 | 2.8917640 | 0.392296875 | 0.254 | 0.049 | 5.84049577928253e-0 | 15 | dnmt3bb.1 |
| golph3 | 2.9345577 | 0.305011646 | 0.254 | 0.047 | 5.92692625705744e-0 | 15 | golph3 |
| nectin1b1 | 3.0508195 | 0.425123288 | 0.288 | 0.059 | 6.16174023718927e-0 | 15 | nectin1b |
| col18a1a1 | 3.9836496 | 0.538122748 | 0.356 | 0.087 | 8.04577713944148e-0 | 15 | col18a1a |
| tusc3 | 4.0905461 | 0.310576425 | 0.203 | 0.032 | 8.26167600855204e-0 | 15 | tusc3 |
| ponzr15 | 4.6230982 | 0.659387815 | 0.847 | 0.385 | 9.33727149786513e-0 | 15 | ponzr1 |
| sptbn2 | 4.8101338 | 0.42000658 | 0.288 | 0.062 | 9.71502739364297e-0 | 15 | sptbn2 |
| pdia62 | 5.5596274 | 0.523306115 | 0.508 | 0.165 | 1.12287795298864e-0 | 15 | pdia6 |
| pdxka | 6.7489994 | 0.279598862 | 0.203 | 0.032 | 1.36309541075274e-0 | 15 | pdxka |
| tuba8l35 | 8.6594959 | 0.640564472 | 0.559 | 0.2 | 1.74895838770984e-0 | 15 | tuba8l3 |
| chd2 | 8.8265818 | 0.314082517 | 0.237 | 0.043 | 1.78270473173835e-0 | 15 | chd2 |
| adamts21 | 1.1871637 | 0.368059722 | 0.22 | 0.039 | 2.39771461188788e-0 | 15 | adamts2 |
| cpxm1a1 | 1.2073460 | 0.358259953 | 0.237 | 0.044 | 2.4384769066016e-07 | 15 | cpxm1a |

| | | | | | | | |
|---|---|---|---|---|---|---|---|
| CR383676.15 | 1.2534711 | 0.604825908 | 1 | 0.655 | 2.53163573767618e-0 | 15 | CR383676.1 |
| cd99l21 | 1.5723194 | 0.439468811 | 0.339 | 0.083 | 3.17561368021812e-0 | 15 | cd99l2 |
| btg23 | 1.8715253 | 0.62647849 | 0.627 | 0.254 | 3.77991979492109e-0 | 15 | btg2 |
| dlc11 | 1.8772468 | 0.359599759 | 0.22 | 0.039 | 3.79147555016392e-0 | 15 | dlc1 |
| dynll2a | 2.2047275 | 0.275553411 | 0.186 | 0.028 | 4.45288826097769e-0 | 15 | dynll2a |
| sparc6 | 2.2072999 | 0.67862253 | 0.949 | 0.391 | 4.45808364719475e-0 | 15 | sparc |
| inhbaa1 | 2.2718321 | 0.496690695 | 0.424 | 0.115 | 4.58841931395223e-0 | 15 | inhbaa |
| sumo3a1 | 2.3975978 | 0.586211093 | 0.831 | 0.401 | 4.84242839898838e-0 | 15 | sumo3a |
| golga4 | 3.2721657 | 0.276963534 | 0.169 | 0.024 | 6.6087932192767e-07 | 15 | golga4 |
| FP236812.2 | 3.6485869 | 0.277327862 | 0.169 | 0.024 | 7.36905096647826e-0 | 15 | FP236812.2 |
| junbb | 3.7384946 | 0.387618268 | 0.271 | 0.059 | 7.55063772764628e-0 | 15 | junbb |
| h3f3d1 | 3.9599670 | 0.481558738 | 0.949 | 0.669 | 7.99794536532791e-0 | 15 | h3f3d |
| pik3r22 | 3.9936116 | 0.454903014 | 0.373 | 0.1 | 8.06589737237879e-0 | 15 | pik3r2 |
| znf292a | 4.4425858 | 0.327145155 | 0.237 | 0.046 | 8.9726905562401e-07 | 15 | znf292a |
| cyyr1 | 4.9549183 | 0.299709981 | 0.203 | 0.035 | 1.00074486378857e-0 | 15 | cyyr1 |
| oga | 5.0019802 | 0.328726404 | 0.22 | 0.04 | 1.0102499410889e-06 | 15 | oga |
| bmp4 | 5.2282556 | 0.306632934 | 0.169 | 0.025 | 1.05595079498012e-0 | 15 | bmp4 |
| nktr | 5.2462221 | 0.337893005 | 0.271 | 0.058 | 1.0595794785764e-06 | 15 | nktr |
| selenof3 | 5.8037494 | 0.573077894 | 0.831 | 0.417 | 1.17218327389163e-0 | 15 | selenof |
| si:dkey-42i9.42 | 6.1057374 | 0.52546066 | 0.542 | 0.192 | 1.23317579388641e-0 | 15 | si:dkey-42i9.4 |
| reep3b1 | 6.3532388 | 0.420708267 | 0.339 | 0.089 | 1.28316364147709e-0 | 15 | reep3b |
| arl4d1 | 6.4921703 | 0.343730782 | 0.305 | 0.069 | 1.311223647169e-06 | 15 | arl4d |
| ptp4a2b | 8.6286482 | 0.487105112 | 0.458 | 0.144 | 1.74272808719713e-0 | 15 | ptp4a2b |
| tegt3 | 8.7576693 | 0.581812184 | 0.763 | 0.356 | 1.76878647327485e-0 | 15 | tegt |
| mt-co14 | 8.9727767 | 0.613882886 | 0.983 | 0.586 | 1.81223172444728e-0 | 15 | mt-co1 |
| h2afx12 | 9.5647578 | 0.622976238 | 0.78 | 0.374 | 1.93179413959326e-0 | 15 | h2afx1 |
| ing4 | 1.0301914 | 0.344340506 | 0.322 | 0.078 | 2.08067766931839e-0 | 15 | ing4 |
| ivns1abpa1 | 1.1673685 | 0.269024844 | 0.186 | 0.03 | 2.35773417595704e-0 | 15 | ivns1abpa |
| tspan3a | 1.2348483 | 0.411611505 | 0.39 | 0.111 | 2.49402316015537e-0 | 15 | tspan3a |
| sh3gl3b | 1.3801027 | 0.257019755 | 0.169 | 0.026 | 2.78739357568416e-0 | 15 | sh3gl3b |
| eppk11 | 1.3857300 | 0.392001953 | 0.254 | 0.054 | 2.79875894431024e-0 | 15 | eppk1 |
| tsc22d11 | 1.5050356 | 0.395239725 | 0.254 | 0.055 | 3.03972056575526e-0 | 15 | tsc22d1 |
| fosab5 | 1.7653843 | 0.649302092 | 0.864 | 0.454 | 3.56554686717991e-0 | 15 | fosab |
| h1f01 | 1.7834894 | 0.464796958 | 0.407 | 0.116 | 3.60211364497632e-0 | 15 | h1f0 |
| desi1a | 2.3210737 | 0.268676106 | 0.169 | 0.026 | 4.68787258863971e-0 | 15 | desi1a |

| | | | | | | | |
|---|---|---|---|---|---|---|---|
| smurf2 | 2.6622350 | 0.33313847 | 0.186 | 0.032 | 5.37691616617735e-0 | 15 | smurf2 |
| wu:fj29h11 | 2.9306675 | 0.297878847 | 0.169 | 0.027 | 5.91906916326453e-0 | 15 | wu:fj29h11 |
| ralgps2 | 3.0516646 | 0.289180293 | 0.186 | 0.032 | 6.1634470040834e-06 | 15 | ralgps2 |
| si:ch73-376l24.33 | 3.4891085 | 0.505509978 | 0.475 | 0.166 | 7.0469524996619e-06 | 15 | si:ch73-376l24.3 |
| gclm | 3.5290497 | 0.300162213 | 0.237 | 0.049 | 7.12762177429073e-0 | 15 | gclm |
| igflr1 | 3.5838651 | 0.398525603 | 0.288 | 0.069 | 7.23833236720751e-0 | 15 | igflr1 |
| zgc:195175 | 3.6999138 | 0.331424569 | 0.22 | 0.044 | 7.47271596279455e-0 | 15 | zgc:195175 |
| tmem50a2 | 3.7379055 | 0.489947261 | 0.525 | 0.192 | 7.54944779404065e-0 | 15 | tmem50a |
| c4b | 3.7915090 | 0.272347939 | 0.136 | 0.017 | 7.65771089462491e-0 | 15 | c4b |
| tm2d1 | 3.8961845 | 0.303918232 | 0.186 | 0.032 | 7.86912387996761e-0 | 15 | tm2d1 |
| ch25hl3 | 4.0270239 | 0.351679567 | 0.271 | 0.062 | 8.1333801844161e-06 | 15 | ch25hl3 |
| znf609b | 4.1439514 | 0.35077642 | 0.237 | 0.05 | 8.36953868210794e-0 | 15 | znf609b |
| si:ch211-197h24.9 | 4.5303760 | 0.522899424 | 0.458 | 0.158 | 9.15000058284834e-0 | 15 | si:ch211-197h24.9 |
| si:ch73-22o12.11 | 4.5556593 | 0.327261928 | 0.254 | 0.056 | 9.20106519262595e-0 | 15 | si:ch73-22o12.1 |
| fez11 | 4.6669025 | 0.278463481 | 0.186 | 0.031 | 9.42574301920539e-0 | 15 | fez1 |
| lpar13 | 4.7102787 | 0.451501126 | 0.356 | 0.099 | 9.51334992273312e-0 | 15 | lpar1 |
| fgfrl1a | 4.7600250 | 0.36468913 | 0.237 | 0.05 | 9.61382260016611e-0 | 15 | fgfrl1a |
| gas61 | 4.9045397 | 0.357138639 | 0.271 | 0.063 | 9.90569901242763e-0 | 15 | gas6 |
| fam20b | 5.7862666 | 0.296737732 | 0.169 | 0.028 | 1.16865228400259e-0 | 15 | fam20b |
| vapb | 7.3836551 | 0.386306106 | 0.339 | 0.094 | 1.49127682123858e-0 | 15 | vapb |
| zc3h11a | 8.0555837 | 0.339710318 | 0.254 | 0.058 | 1.62698624401338e-0 | 15 | zc3h11a |
| chd4a | 9.1827857 | 0.476612063 | 0.508 | 0.188 | 1.85464724749129e-0 | 15 | chd4a |
| eif4ebp12 | 1.0194722 | 0.51941846 | 0.712 | 0.336 | 2.05902820050276e-0 | 15 | eif4ebp1 |
| plod22 | 1.1147984 | 0.358473716 | 0.254 | 0.058 | 2.25155836129584e-0 | 15 | plod2 |
| ndrg2 | 1.2010036 | 0.341938239 | 0.356 | 0.099 | 2.42566715669418e-0 | 15 | ndrg2 |
| schip1 | 1.2633548 | 0.258479715 | 0.186 | 0.033 | 2.55159769018858e-0 | 15 | schip1 |
| csgalnact1a | 1.2828462 | 0.305054529 | 0.203 | 0.039 | 2.59096466879904e-0 | 15 | csgalnact1a |
| snx61 | 1.4195018 | 0.371822892 | 0.339 | 0.095 | 2.86696791289724e-0 | 15 | snx6 |
| cd164 | 1.4786639 | 0.418771098 | 0.339 | 0.098 | 2.98645767845055e-0 | 15 | cd164 |
| mt-nd43 | 1.6971883 | 0.587255027 | 0.814 | 0.432 | 3.4278112540805e-05 | 15 | mt-nd4 |
| midn4 | 1.6984985 | 0.500251081 | 0.475 | 0.176 | 3.43045743821794e-0 | 15 | midn |
| atrx1 | 1.8872236 | 0.518193397 | 0.576 | 0.238 | 3.81162556006871e-0 | 15 | atrx |
| pi4kb | 1.9677403 | 0.326798816 | 0.203 | 0.041 | 3.97424524336787e-0 | 15 | pi4kb |
| bmpr2a | 2.0045599 | 0.283589507 | 0.203 | 0.039 | 4.04860976302078e-0 | 15 | bmpr2a |
| adipor21 | 2.5131701 | 0.331361961 | 0.288 | 0.073 | 5.0758497277302e-05 | 15 | adipor2 |

| | | | | | | | |
|---|---|---|---|---|---|---|---|
| eef1db1 | 2.5314994 | 0.511177575 | 0.746 | 0.354 | 5.11286950781381e-0 | 15 | eef1db |
| wu:fb55g092 | 2.5934650 | 0.521397195 | 0.678 | 0.305 | 5.23802128384789e-0 | 15 | wu:fb55g09 |
| zgc:86598 | 2.6606154 | 0.411949458 | 0.424 | 0.142 | 5.3736450569649e-05 | 15 | zgc:86598 |
| msi2b | 2.7122331 | 0.304816886 | 0.237 | 0.053 | 5.47789719815525e-0 | 15 | msi2b |
| mmp14a3 | 2.7968905 | 0.38834272 | 0.339 | 0.096 | 5.64887978375419e-0 | 15 | mmp14a |
| grinab2 | 3.0296770 | 0.465728175 | 0.475 | 0.174 | 6.11903872482722e-0 | 15 | grinab |
| nacc2 | 3.1367906 | 0.273994846 | 0.203 | 0.041 | 6.33537611542358e-0 | 15 | nacc2 |
| foxn3 | 3.1620253 | 0.262079522 | 0.22 | 0.046 | 6.38634267931667e-0 | 15 | foxn3 |
| nr2f21 | 3.1910734 | 0.355186024 | 0.339 | 0.093 | 6.44501108537784e-0 | 15 | nr2f2 |
| ucp23 | 3.2218736 | 0.538595537 | 0.814 | 0.418 | 6.50721821779461e-0 | 15 | ucp2 |
| tnrc6c13 | 3.2592743 | 0.468615303 | 0.593 | 0.235 | 6.58275649315413e-0 | 15 | tnrc6c1 |
| dad12 | 3.2679290 | 0.511822286 | 0.814 | 0.412 | 6.60023627919565e-0 | 15 | dad1 |
| spcs22 | 3.4265807 | 0.480578572 | 0.593 | 0.248 | 6.92066520549387e-0 | 15 | spcs2 |
| nme32 | 3.5671797 | 0.420581865 | 0.458 | 0.156 | 7.20463296067426e-0 | 15 | nme3 |
| adgrl2a | 3.6062128 | 0.315036587 | 0.203 | 0.041 | 7.28346818865471e-0 | 15 | adgrl2a |
| ttll121 | 3.7767416 | 0.303878236 | 0.153 | 0.024 | 7.62788511650332e-0 | 15 | ttll12 |
| papolg | 3.8334955 | 0.250113091 | 0.186 | 0.035 | 7.74251087253407e-0 | 15 | papolg |
| serinc52 | 3.9739203 | 0.519613575 | 0.356 | 0.112 | 8.02612699523792e-0 | 15 | serinc5 |
| dynll2b | 4.0281949 | 0.37732518 | 0.39 | 0.122 | 8.13574537214482e-0 | 15 | dynll2b |
| pdk2a | 4.4249709 | 0.303605612 | 0.186 | 0.036 | 8.93711387330531e-0 | 15 | pdk2a |
| map1lc3a1 | 4.6045423 | 0.387458609 | 0.373 | 0.115 | 9.29979422643121e-0 | 15 | map1lc3a |
| fn1b4 | 4.9278952 | 0.65870415 | 0.458 | 0.176 | 9.95286994909523e-0 | 15 | fn1b |
| emc10 | 5.4457051 | 0.401492929 | 0.373 | 0.12 | 0.000109987 | 15 | emc10 |
| mphosph81 | 5.9151546 | 0.467151951 | 0.593 | 0.247 | 0.000119468 | 15 | mphosph8 |
| tlk1b | 6.0812599 | 0.264808979 | 0.203 | 0.042 | 0.000122823 | 15 | tlk1b |
| hnrnpl2 | 6.6673981 | 0.393179637 | 0.441 | 0.153 | 0.000134661 | 15 | hnrnpl2 |
| arglu1a | 6.6844979 | 0.42012124 | 0.356 | 0.11 | 0.000135007 | 15 | arglu1a |
| kmt2cb | 7.5564139 | 0.404417712 | 0.322 | 0.094 | 0.000152617 | 15 | kmt2cb |
| adgrl3.1 | 7.6481748 | 0.36530389 | 0.237 | 0.055 | 0.00015447 | 15 | adgrl3.1 |
| clec11a | 8.7847355 | 0.255284491 | 0.169 | 0.03 | 0.000177425 | 15 | clec11a |
| ldb1a | 8.8042831 | 0.284418036 | 0.203 | 0.043 | 0.00017782 | 15 | ldb1a |
| tjp2b2 | 9.0105663 | 0.375675772 | 0.288 | 0.078 | 0.000181986 | 15 | tjp2b |
| matr3l1.11 | 1.0333769 | 0.376590377 | 0.339 | 0.104 | 0.000208711 | 15 | matr3l1.1 |
| cox8b1 | 1.1382494 | 0.270629387 | 0.407 | 0.13 | 0.000229892 | 15 | cox8b |
| wwtr1 | 1.1614884 | 0.256953701 | 0.22 | 0.048 | 0.000234586 | 15 | wwtr1 |

| | | | | | | | |
|---|---|---|---|---|---|---|---|
| pcyox11 | 1.1794744 | 0.411255397 | 0.339 | 0.106 | 0.000238218 | 15 | pcyox1 |
| jcada1 | 1.2387369 | 0.345457294 | 0.22 | 0.05 | 0.000250188 | 15 | jcada |
| ndufa4l1 | 1.2434098 | 0.473150574 | 0.898 | 0.568 | 0.000251131 | 15 | ndufa4l |
| si:dkey-177p2.6 | 1.2467209 | 0.389874009 | 0.373 | 0.118 | 0.0002518 | 15 | si:dkey-177p2.6 |
| adam10a2 | 1.2617569 | 0.43182768 | 0.458 | 0.165 | 0.000254837 | 15 | adam10a |
| pgrmc2 | 1.3291598 | 0.331593818 | 0.271 | 0.071 | 0.00026845 | 15 | pgrmc2 |
| ogdha | 1.4311556 | 0.369439121 | 0.339 | 0.102 | 0.000289051 | 15 | ogdha |
| ube2d4 | 1.5033893 | 0.301405161 | 0.22 | 0.05 | 0.00030364 | 15 | ube2d4 |
| si:dkey-56m19.53 | 1.5762458 | 0.348945657 | 0.339 | 0.098 | 0.000318354 | 15 | si:dkey-56m19.5 |
| rbm4.11 | 1.6209728 | 0.321650654 | 0.322 | 0.092 | 0.000327388 | 15 | rbm4.1 |
| rbm39a1 | 1.6408915 | 0.461246784 | 0.508 | 0.207 | 0.000331411 | 15 | rbm39a |
| slc38a2 | 1.6865978 | 0.279993626 | 0.203 | 0.044 | 0.000340642 | 15 | slc38a2 |
| tmem134 | 1.8109121 | 0.260202238 | 0.254 | 0.062 | 0.00036575 | 15 | tmem134 |
| selenbp13 | 1.8338899 | 0.431666007 | 0.407 | 0.139 | 0.000370391 | 15 | selenbp1 |
| lman21 | 1.8771081 | 0.400854918 | 0.492 | 0.188 | 0.00037912 | 15 | lman2 |
| anxa2a3 | 1.9398235 | 0.664115065 | 0.458 | 0.184 | 0.000391786 | 15 | anxa2a |
| tram2 | 2.0812688 | 0.324440714 | 0.271 | 0.071 | 0.000420354 | 15 | tram2 |
| zgc:56095 | 2.1000071 | 0.289592034 | 0.288 | 0.079 | 0.000424138 | 15 | zgc:56095 |
| wu:fb16f032 | 2.2534461 | 0.376505828 | 0.339 | 0.105 | 0.000455129 | 15 | wu:fb16f03 |
| jun6 | 2.2941215 | 0.605877329 | 0.695 | 0.381 | 0.000463344 | 15 | jun |
| ndufs4 | 2.3264196 | 0.408285709 | 0.441 | 0.167 | 0.000469867 | 15 | ndufs4 |
| cald1a5 | 2.4111695 | 0.452817242 | 0.593 | 0.232 | 0.000486984 | 15 | cald1a |
| tle2b | 2.4800832 | 0.261151611 | 0.169 | 0.032 | 0.000500902 | 15 | tle2b |
| egfra1 | 2.4966359 | 0.273527848 | 0.22 | 0.05 | 0.000504246 | 15 | egfra |
| uqcrc1 | 2.6231212 | 0.354946563 | 0.492 | 0.182 | 0.000529792 | 15 | uqcrc1 |
| rnf213a2 | 2.6900153 | 0.394827176 | 0.39 | 0.131 | 0.000543302 | 15 | rnf213a |
| foxo1a1 | 3.1035371 | 0.349657553 | 0.203 | 0.045 | 0.000626821 | 15 | foxo1a |
| ubb | 3.3665730 | 0.533762125 | 0.678 | 0.371 | 0.000679947 | 15 | ubb |
| gchfr2 | 3.4185766 | 0.311808684 | 0.203 | 0.045 | 0.00069045 | 15 | gchfr |
| krt946 | 3.5029977 | 0.484240833 | 0.847 | 0.353 | 0.0007075 | 15 | krt94 |
| si:dkeyp-69e1.8 | 3.5253632 | 0.278363454 | 0.186 | 0.039 | 0.000712018 | 15 | si:dkeyp-69e1.8 |
| ier2a3 | 3.7104293 | 0.553566708 | 0.712 | 0.358 | 0.000749395 | 15 | ier2a |
| s1pr21 | 3.8092348 | 0.344034905 | 0.237 | 0.06 | 0.000769351 | 15 | s1pr2 |
| zgc:162509 | 3.9650368 | 0.274320604 | 0.22 | 0.052 | 0.000800818 | 15 | zgc:162509 |
| mt-co22 | 4.0193679 | 0.626081264 | 0.847 | 0.454 | 0.000811792 | 15 | mt-co2 |

| | | | | | | | |
|---|---|---|---|---|---|---|---|
| tomm20a1 | 4.7892980 | 0.432154298 | 0.525 | 0.216 | 0.000967295 | 15 | tomm20a |
| arfrp1 | 4.8441976 | 0.26083815 | 0.22 | 0.053 | 0.000978383 | 15 | arfrp1 |
| edem11 | 4.9092991 | 0.302628103 | 0.22 | 0.053 | 0.000991531 | 15 | edem1 |
| si:dkey-188i13.111 | 5.1476545 | 0.470005107 | 0.458 | 0.181 | 0.001039672 | 15 | si:dkey-188i13.11 |
| eci2 | 5.4148008 | 0.261424529 | 0.22 | 0.053 | 0.001093627 | 15 | eci2 |
| atf4b2 | 5.6421325 | 0.473467167 | 0.712 | 0.355 | 0.001139542 | 15 | atf4b |
| nfic1 | 5.9352502 | 0.384762243 | 0.356 | 0.117 | 0.001198742 | 15 | nfic |
| tomm6 | 6.0420316 | 0.386694199 | 0.542 | 0.222 | 0.001220309 | 15 | tomm6 |
| kdm2ba | 6.5377313 | 0.289766489 | 0.237 | 0.06 | 0.001320426 | 15 | kdm2ba |
| si:dkey-11f4.142 | 6.7242014 | 0.427565221 | 0.508 | 0.198 | 0.001358087 | 15 | si:dkey-11f4.14 |
| rfxap | 6.7835116 | 0.291873869 | 0.288 | 0.083 | 0.001370066 | 15 | rfxap |
| srsf10b | 7.3210475 | 0.307565893 | 0.322 | 0.099 | 0.001478632 | 15 | srsf10b |
| cdc34b | 7.3926594 | 0.258570212 | 0.237 | 0.06 | 0.001493095 | 15 | cdc34b |
| tnrc6c2 | 8.1665943 | 0.26082301 | 0.186 | 0.04 | 0.001649407 | 15 | tnrc6c2 |
| meis1b4 | 8.6376246 | 0.419930543 | 0.424 | 0.153 | 0.001744541 | 15 | meis1b |
| FO082781.12 | 8.8190812 | 0.392823702 | 0.305 | 0.095 | 0.00178119 | 15 | FO082781.1 |
| ganab | 8.9071567 | 0.260774046 | 0.169 | 0.035 | 0.001798978 | 15 | ganab |
| ddx3xa | 9.8554802 | 0.326185601 | 0.288 | 0.084 | 0.001990511 | 15 | ddx3xa |
| nt5c2b | 1.0136432 | 0.305702148 | 0.22 | 0.055 | 0.002047255 | 15 | nt5c2b |
| rbbp6 | 1.0428667 | 0.411379262 | 0.373 | 0.132 | 0.002106278 | 15 | rbbp6 |
| si:ch211-191j22.3 | 1.0475871 | 0.360164928 | 0.458 | 0.175 | 0.002115812 | 15 | si:ch211-191j22.3 |
| snx172 | 1.0480442 | 0.398032633 | 0.322 | 0.102 | 0.002116735 | 15 | snx17 |
| adamts15a5 | 1.0568594 | 0.376426929 | 0.339 | 0.108 | 0.002134539 | 15 | adamts15a |
| rnf24 | 1.1302463 | 0.254336622 | 0.186 | 0.041 | 0.002282759 | 15 | rnf24 |
| pin4 | 1.1598082 | 0.338723818 | 0.373 | 0.13 | 0.002342465 | 15 | pin4 |
| nrp2a | 1.1699154 | 0.285421487 | 0.169 | 0.035 | 0.002362878 | 15 | nrp2a |
| gorasp2 | 1.2030981 | 0.298754072 | 0.288 | 0.085 | 0.002429897 | 15 | gorasp2 |
| neo1a | 1.2269729 | 0.260149028 | 0.22 | 0.054 | 0.002478117 | 15 | neo1a |
| zgc:113208 | 1.2768683 | 0.345064523 | 0.254 | 0.071 | 0.002578891 | 15 | zgc:113208 |
| txndc9 | 1.3426765 | 0.397460923 | 0.458 | 0.184 | 0.002711804 | 15 | txndc9 |
| wls4 | 1.3551153 | 0.416323964 | 0.458 | 0.178 | 0.002736926 | 15 | wls |
| sumf2 | 1.3659001 | 0.251925507 | 0.153 | 0.029 | 0.002758709 | 15 | sumf2 |
| si:ch211-286b5.52 | 1.3722250 | 0.420745467 | 0.525 | 0.221 | 0.002771483 | 15 | si:ch211-286b5.5 |
| ube2e31 | 1.5648086 | 0.447512534 | 0.559 | 0.253 | 0.003160444 | 15 | ube2e3 |
| klf11b | 1.6018523 | 0.333982275 | 0.305 | 0.093 | 0.003235261 | 15 | klf11b |

| | | | | | | | |
|---|---|---|---|---|---|---|---|
| pfn2l5 | 1.6892178 | 0.410050549 | 0.61 | 0.258 | 0.003411713 | 15 | pfn2l |
| cx43.4 | 1.6943091 | 0.314319791 | 0.22 | 0.056 | 0.003421996 | 15 | cx43.4 |
| LEPROTL1 | 1.7085178 | 0.373663855 | 0.356 | 0.122 | 0.003450693 | 15 | LEPROTL1 |
| hnrnpabb | 1.7682113 | 0.474876072 | 0.746 | 0.401 | 0.003571256 | 15 | hnrnpabb |
| nucks1a2 | 1.8024064 | 0.404496373 | 0.542 | 0.227 | 0.00364032 | 15 | nucks1a |
| sdc43 | 1.8352381 | 0.449776431 | 0.593 | 0.26 | 0.00370663 | 15 | sdc4 |
| fbxl6 | 1.8705183 | 0.360694683 | 0.356 | 0.123 | 0.003777886 | 15 | fbxl6 |
| si:ch73-1a9.31 | 1.8731007 | 0.485358874 | 0.966 | 0.582 | 0.003783102 | 15 | si:ch73-1a9.3 |
| kmt2bb2 | 1.8742623 | 0.356406062 | 0.407 | 0.148 | 0.003785448 | 15 | kmt2bb |
| sod2 | 1.9618735 | 0.331842609 | 0.39 | 0.142 | 0.003962396 | 15 | sod2 |
| tmed91 | 1.9659687 | 0.350642731 | 0.424 | 0.16 | 0.003970667 | 15 | tmed9 |
| fthl273 | 2.0351395 | 0.439375381 | 0.915 | 0.624 | 0.004110371 | 15 | fthl27 |
| zgc:919102 | 2.1009541 | 0.443163748 | 0.61 | 0.295 | 0.004243297 | 15 | zgc:91910 |
| ndufc21 | 2.1763720 | 0.397673022 | 0.559 | 0.245 | 0.004395619 | 15 | ndufc2 |
| oaz1a1 | 2.2509144 | 0.424460201 | 0.864 | 0.59 | 0.004546172 | 15 | oaz1a |
| top2b | 2.2597634 | 0.352145698 | 0.237 | 0.067 | 0.004564044 | 15 | top2b |
| gstt1b4 | 2.3385279 | 0.382640785 | 0.458 | 0.178 | 0.004723125 | 15 | gstt1b |
| si:ch73-335l21.42 | 2.3544271 | 0.427326843 | 0.339 | 0.113 | 0.004755236 | 15 | si:ch73-335l21.4 |
| ndfip11 | 2.3877147 | 0.414096989 | 0.508 | 0.219 | 0.004822468 | 15 | ndfip1 |
| grhl11 | 2.4113317 | 0.315669753 | 0.203 | 0.05 | 0.004870167 | 15 | grhl1 |
| erbb2 | 2.5591778 | 0.256921538 | 0.203 | 0.049 | 0.005168772 | 15 | erbb2 |
| oaz1b1 | 2.5857215 | 0.448141902 | 0.847 | 0.519 | 0.005222382 | 15 | oaz1b |
| cux1a2 | 2.6360638 | 0.350869748 | 0.322 | 0.106 | 0.005324058 | 15 | cux1a |
| ctnnb12 | 2.6750033 | 0.408511188 | 0.508 | 0.21 | 0.005402704 | 15 | ctnnb1 |
| cd992 | 2.7851831 | 0.270251671 | 0.203 | 0.05 | 0.005625234 | 15 | cd99 |
| tgif11 | 2.8540015 | 0.418282003 | 0.39 | 0.15 | 0.005764227 | 15 | tgif1 |
| itga92 | 3.0767379 | 0.338008565 | 0.254 | 0.072 | 0.006214088 | 15 | itga9 |
| si:ch211-166g5.4 | 3.0768677 | 0.268182407 | 0.169 | 0.037 | 0.00621435 | 15 | si:ch211-166g5.4 |
| prkcsh | 3.1053124 | 0.320702395 | 0.322 | 0.105 | 0.006271799 | 15 | prkcsh |
| psmd3 | 3.1132791 | 0.394266561 | 0.39 | 0.15 | 0.00628789 | 15 | psmd3 |
| osr21 | 3.1161602 | 0.327005203 | 0.288 | 0.084 | 0.006293709 | 15 | osr2 |
| fundc1 | 3.1931122 | 0.281340408 | 0.288 | 0.088 | 0.006449129 | 15 | fundc1 |
| top1l | 3.3896195 | 0.396792392 | 0.441 | 0.175 | 0.006846015 | 15 | top1l |
| sumo1 | 3.4272148 | 0.380849231 | 0.441 | 0.179 | 0.006921946 | 15 | sumo1 |
| BX548028.1 | 3.4690613 | 0.303437979 | 0.22 | 0.058 | 0.007006463 | 15 | BX548028.1 |

| | | | | | | | |
|---|---|---|---|---|---|---|---|
| erbin | 3.5798607 | 0.26878786 | 0.237 | 0.065 | 0.007230245 | 15 | erbin |
| ndufs7 | 3.6072212 | 0.352922484 | 0.441 | 0.176 | 0.007285505 | 15 | ndufs7 |
| uqcrc2b | 3.6468219 | 0.310331648 | 0.322 | 0.107 | 0.007365486 | 15 | uqcrc2b |
| arhgap5 | 3.8872610 | 0.267802791 | 0.22 | 0.058 | 0.007851101 | 15 | arhgap5 |
| nabp1a1 | 4.1027467 | 0.275720767 | 0.271 | 0.08 | 0.008286318 | 15 | nabp1a |
| prdx3 | 4.2521498 | 0.315930575 | 0.458 | 0.181 | 0.008588067 | 15 | prdx3 |
| tnrc6a | 4.7917944 | 0.319730752 | 0.322 | 0.108 | 0.009677987 | 15 | tnrc6a |
| pkig1 | 4.7977115 | 0.333177115 | 0.356 | 0.124 | 0.009689938 | 15 | pkig |
| dap1b1 | 5.0074039 | 0.411791476 | 0.559 | 0.251 | 0.010113454 | 15 | dap1b |
| coa3a | 5.1342092 | 0.314617946 | 0.288 | 0.093 | 0.010369563 | 15 | coa3a |
| tspan9b1 | 5.1405422 | 0.315422939 | 0.271 | 0.082 | 0.010382353 | 15 | tspan9b |
| atp6v0b1 | 5.1751281 | 0.349287864 | 0.322 | 0.112 | 0.010452206 | 15 | atp6v0b |
| nudcd2 | 5.1926519 | 0.272564688 | 0.288 | 0.09 | 0.010487599 | 15 | nudcd2 |
| b2ml4 | 5.5329107 | 0.442653811 | 0.814 | 0.44 | 0.01117482 | 15 | b2ml |
| BX005309.1 | 5.5562918 | 0.310588522 | 0.22 | 0.059 | 0.011222043 | 15 | BX005309.1 |
| BX569787.13 | 5.8086523 | 0.403104123 | 0.305 | 0.103 | 0.011731735 | 15 | BX569787.1 |
| rras | 6.2183145 | 0.320088618 | 0.339 | 0.117 | 0.01255913 | 15 | rras |
| hmgb2b | 6.3213223 | 0.442228506 | 0.831 | 0.434 | 0.012767175 | 15 | hmgb2b |
| chmp5b | 6.5622833 | 0.383587059 | 0.424 | 0.172 | 0.013253844 | 15 | chmp5b |
| serinc11 | 6.9657853 | 0.358062499 | 0.322 | 0.111 | 0.014068797 | 15 | serinc1 |
| chmp2a1 | 7.5222420 | 0.349792402 | 0.458 | 0.186 | 0.015192672 | 15 | chmp2a |
| ncalda | 7.5630663 | 0.290918737 | 0.237 | 0.068 | 0.015275125 | 15 | ncalda |
| hnrpkl | 7.5797836 | 0.363619325 | 0.373 | 0.14 | 0.015308889 | 15 | hnrpkl |
| si:dkey-19a16.7 | 7.6148344 | 0.306799793 | 0.22 | 0.061 | 0.015379681 | 15 | si:dkey-19a16.7 |
| ormdl1 | 7.8818997 | 0.280582163 | 0.22 | 0.061 | 0.015919073 | 15 | ormdl1 |
| ostc2 | 7.9418590 | 0.441276818 | 0.525 | 0.248 | 0.016040173 | 15 | ostc |
| ctnna11 | 8.0393354 | 0.289975738 | 0.237 | 0.068 | 0.016237046 | 15 | ctnna1 |
| ier52 | 8.1210921 | 0.405629978 | 0.475 | 0.207 | 0.01640217 | 15 | ier5 |
| brd3b1 | 8.2049297 | 0.352465472 | 0.322 | 0.114 | 0.016571497 | 15 | brd3b |
| hnrnpc | 8.2663764 | 0.324415781 | 0.305 | 0.102 | 0.016695601 | 15 | hnrnpc |
| psenen | 8.3323803 | 0.378622676 | 0.373 | 0.146 | 0.016828909 | 15 | psenen |
| kdm1a | 8.4980500 | 0.307900877 | 0.339 | 0.119 | 0.017163512 | 15 | kdm1a |
| eif4g2a | 8.5575058 | 0.317591391 | 0.288 | 0.093 | 0.017283595 | 15 | eif4g2a |
| clk4a | 8.9501305 | 0.305612402 | 0.237 | 0.068 | 0.018076579 | 15 | clk4a |
| tubb51 | 8.9929745 | 0.360861679 | 0.254 | 0.077 | 0.018163111 | 15 | tubb5 |

| | | | | | | | |
|---|---|---|---|---|---|---|---|
| si:dkey-28b4.73 | 9.1214066 | 0.358217939 | 0.475 | 0.191 | 0.018422505 | 15 | si:dkey-28b4.7 |
| slc3a2b2 | 9.1899161 | 0.38529218 | 0.424 | 0.169 | 0.018560874 | 15 | slc3a2b |
| dpm2 | 9.4585537 | 0.326139194 | 0.271 | 0.086 | 0.019103441 | 15 | dpm2 |
| fundc2 | 9.4646015 | 0.341753221 | 0.356 | 0.132 | 0.019115656 | 15 | fundc2 |
| bnc22 | 9.7266912 | 0.304260399 | 0.271 | 0.082 | 0.019644998 | 15 | bnc2 |
| tmem106ba | 9.8099145 | 0.257662645 | 0.22 | 0.06 | 0.019813084 | 15 | tmem106ba |
| sufu | 1.0044976 | 0.263739386 | 0.22 | 0.06 | 0.020287839 | 15 | sufu |
| ddx24 | 1.0254521 | 0.307766089 | 0.271 | 0.086 | 0.020711057 | 15 | ddx24 |
| zgc:152791 | 1.1074626 | 0.280552034 | 0.237 | 0.067 | 0.022367423 | 15 | zgc:152791 |
| selenok | 1.1086239 | 0.387049366 | 0.458 | 0.2 | 0.022390878 | 15 | selenok |
| golga7 | 1.1153398 | 0.388680101 | 0.424 | 0.177 | 0.022526519 | 15 | golga7 |
| ap1s2 | 1.1772566 | 0.34814452 | 0.407 | 0.16 | 0.023777053 | 15 | ap1s2 |
| si:ch211-204c21.1: | 1.1882290 | 0.297560417 | 0.271 | 0.084 | 0.023998663 | 15 | si:ch211-204c21.1 |
| flot1b2 | 1.1958791 | 0.355463048 | 0.458 | 0.193 | 0.024153171 | 15 | flot1b |
| hnrnpa0b2 | 1.2004029 | 0.435366827 | 0.881 | 0.508 | 0.024244538 | 15 | hnrnpa0b |
| esd | 1.2163718 | 0.308626456 | 0.441 | 0.174 | 0.024567062 | 15 | esd |
| bud31 | 1.2246075 | 0.322191157 | 0.542 | 0.238 | 0.0247334 | 15 | bud31 |
| hmgb1a3 | 1.3081216 | 0.46481312 | 0.729 | 0.424 | 0.026420133 | 15 | hmgb1a |
| fkbp51 | 1.3792078 | 0.342574586 | 0.39 | 0.151 | 0.02785586 | 15 | fkbp5 |
| txndc15 | 1.4162371 | 0.326011971 | 0.305 | 0.106 | 0.028603743 | 15 | txndc15 |
| brd2a1 | 1.4475063 | 0.348293252 | 0.373 | 0.143 | 0.029235285 | 15 | brd2a |
| si:ch211-212k18.7: | 1.5138190 | 0.38236382 | 0.441 | 0.183 | 0.030574603 | 15 | si:ch211-212k18.7 |
| nol4lb1 | 1.5442025 | 0.268655409 | 0.169 | 0.041 | 0.031188259 | 15 | nol4lb |
| itga42 | 1.5498045 | 0.290038624 | 0.22 | 0.062 | 0.031301403 | 15 | itga4 |
| eif4ebp3l3 | 1.5956585 | 0.349695306 | 0.475 | 0.197 | 0.032227516 | 15 | eif4ebp3l |
| ppp2cb1 | 1.5959682 | 0.362382742 | 0.525 | 0.237 | 0.032233772 | 15 | ppp2cb |
| aebp13 | 1.6205870 | 0.331577225 | 0.288 | 0.096 | 0.032730997 | 15 | aebp1 |
| bzw1b1 | 1.6290158 | 0.397580736 | 0.576 | 0.282 | 0.032901234 | 15 | bzw1b |
| trib32 | 1.6676521 | 0.32818135 | 0.339 | 0.123 | 0.033681571 | 15 | trib3 |
| si:ch211-15b10.62 | 1.8095908 | 0.302122027 | 0.305 | 0.104 | 0.036548306 | 15 | si:ch211-15b10.6 |
| crebzf | 1.8841364 | 0.378558909 | 0.373 | 0.147 | 0.038053905 | 15 | crebzf |
| cd151l1 | 1.9543824 | 0.352180721 | 0.271 | 0.089 | 0.039472662 | 15 | cd151l |
| sap30l | 2.0205048 | 0.338304624 | 0.492 | 0.218 | 0.040808137 | 15 | sap30l |
| fkbp21 | 2.1250429 | 0.358721344 | 0.475 | 0.21 | 0.042919492 | 15 | fkbp2 |
| pgd2 | 2.3276949 | 0.268501264 | 0.322 | 0.112 | 0.047012455 | 15 | pgd |

| | | | | | | | |
|---|---|---|---|---|---|---|---|
| lrp1aa3 | 2.3608362 | 0.348127468 | 0.339 | 0.122 | 0.047681809 | 15 | lrp1aa |
| tsc22d31 | 2.3651912 | 0.40012896 | 0.525 | 0.246 | 0.047769769 | 15 | tsc22d3 |
| tma7 | 2.3816447 | 0.410080664 | 0.814 | 0.467 | 0.04810208 | 15 | tma7 |
| eci1 | 2.4658638 | 0.272475226 | 0.271 | 0.089 | 0.049803053 | 15 | eci1 |
| CABZ01026681.1 | 2.4666731 | 0.301250328 | 0.203 | 0.058 | 0.049819398 | 15 | CABZ01026681.1 |
| gnb1b2 | 2.4737105 | 0.405345694 | 0.644 | 0.335 | 0.049961531 | 15 | gnb1b |
| hp1bp31 | 2.4742269 | 0.385023282 | 0.508 | 0.241 | 0.049971961 | 15 | hp1bp3 |
| psme1 | 2.4922622 | 0.363196283 | 0.492 | 0.226 | 0.050336221 | 15 | psme1 |
| cxxc5a1 | 2.6047490 | 0.384038134 | 0.254 | 0.085 | 0.052608116 | 15 | cxxc5a |
| col27a1b2 | 2.6412724 | 0.304334156 | 0.237 | 0.071 | 0.053345781 | 15 | col27a1b |
| chac1 | 2.6792206 | 0.337090503 | 0.593 | 0.283 | 0.054112219 | 15 | chac1 |
| fzd7b4 | 2.6809370 | 0.300018826 | 0.305 | 0.103 | 0.054146885 | 15 | fzd7b |
| pnp5a2 | 2.6858866 | 0.333463081 | 0.22 | 0.065 | 0.054246853 | 15 | pnp5a |
| wdr46 | 2.7056804 | 0.266806735 | 0.22 | 0.065 | 0.054646628 | 15 | wdr46 |
| tmed72 | 2.8276650 | 0.392889313 | 0.593 | 0.304 | 0.057110351 | 15 | tmed7 |
| fzd25 | 2.8647514 | 0.310451031 | 0.322 | 0.115 | 0.057859386 | 15 | fzd2 |
| txnl4a | 2.8818270 | 0.254221026 | 0.288 | 0.097 | 0.05820426 | 15 | txnl4a |
| atf7ip | 2.9801486 | 0.306591315 | 0.305 | 0.108 | 0.060190062 | 15 | atf7ip |
| pbx42 | 3.0009368 | 0.303429249 | 0.305 | 0.106 | 0.060609921 | 15 | pbx4 |
| ctsf2 | 3.0254913 | 0.256373381 | 0.237 | 0.071 | 0.061105848 | 15 | ctsf |
| si:ch73-376l24.21 | 3.0995255 | 0.325848377 | 0.407 | 0.163 | 0.062601118 | 15 | si:ch73-376l24.2 |
| ier3ip1 | 3.1105749 | 0.352708792 | 0.492 | 0.222 | 0.062824283 | 15 | ier3ip1 |
| taf13 | 3.1521740 | 0.354680258 | 0.576 | 0.272 | 0.06366446 | 15 | taf13 |
| rsf1b.1 | 3.1712373 | 0.320850216 | 0.288 | 0.101 | 0.064049481 | 15 | rsf1b.1 |
| selenot21 | 3.2133277 | 0.301828534 | 0.305 | 0.109 | 0.064899581 | 15 | selenot2 |
| ugt1b51 | 3.2493510 | 0.302322127 | 0.254 | 0.081 | 0.065627144 | 15 | ugt1b5 |
| fmr1 | 3.3776062 | 0.259262008 | 0.237 | 0.072 | 0.068217513 | 15 | fmr1 |
| mmgt1 | 3.5932072 | 0.271260778 | 0.39 | 0.151 | 0.072572007 | 15 | mmgt1 |
| vma211 | 3.6217847 | 0.319292327 | 0.356 | 0.138 | 0.073149186 | 15 | vma21 |
| smarce11 | 3.7542315 | 0.37473269 | 0.508 | 0.243 | 0.075824215 | 15 | smarce1 |
| ddx611 | 3.8031510 | 0.382118053 | 0.661 | 0.338 | 0.076812242 | 15 | ddx61 |
| zgc:770861 | 3.8223057 | 0.369591268 | 0.322 | 0.12 | 0.077199109 | 15 | zgc:77086 |
| ppib5 | 4.0312706 | 0.412464902 | 0.966 | 0.638 | 0.081419574 | 15 | ppib |
| foxp44 | 4.2692738 | 0.379302986 | 0.441 | 0.187 | 0.086226524 | 15 | foxp4 |
| uck2a1 | 4.3471052 | 0.305378704 | 0.237 | 0.075 | 0.087798485 | 15 | uck2a |

| | | | | | | | |
|---|---|---|---|---|---|---|---|
| si:ch73-46j18.56 | 4.3516254 | 0.478753218 | 0.797 | 0.39 | 0.08788978 | 15 | si:ch73-46j18.5 |
| akt1s11 | 4.3910709 | 0.284035757 | 0.237 | 0.073 | 0.08868646 | 15 | akt1s1 |
| CABZ01080702.14 | 4.5703555 | 0.351702531 | 0.424 | 0.173 | 0.09230747 | 15 | CABZ01080702.1 |
| wsb1 | 4.6109185 | 0.25620017 | 0.254 | 0.081 | 0.093126723 | 15 | wsb1 |
| lsm4 | 4.6641422 | 0.326137948 | 0.559 | 0.258 | 0.094201681 | 15 | lsm4 |
| med28 | 4.7006195 | 0.3054643 | 0.288 | 0.102 | 0.094938412 | 15 | med28 |
| tob1b2 | 4.7054878 | 0.33519446 | 0.373 | 0.147 | 0.095036738 | 15 | tob1b |
| chd61 | 5.2619625 | 0.302364166 | 0.271 | 0.093 | 0.106275859 | 15 | chd6 |
| zgc:114200 | 5.3115422 | 0.25392131 | 0.186 | 0.05 | 0.107277218 | 15 | zgc:114200 |
| rab11fip1b1 | 5.4523576 | 0.295495249 | 0.237 | 0.075 | 0.110121267 | 15 | rab11fip1b |
| ddx5 | 5.6118947 | 0.399366445 | 0.661 | 0.35 | 0.113343437 | 15 | ddx5 |
| aldh7a1 | 5.6502466 | 0.29388711 | 0.288 | 0.103 | 0.114118032 | 15 | aldh7a1 |
| arhgef1b2 | 5.6702413 | 0.358021921 | 0.322 | 0.122 | 0.114521865 | 15 | arhgef1b |
| gkap1 | 5.7202248 | 0.305350475 | 0.322 | 0.121 | 0.115531381 | 15 | gkap1 |
| cdc14b | 5.7559104 | 0.306545552 | 0.237 | 0.075 | 0.116252124 | 15 | cdc14b |
| tia1 | 5.9127741 | 0.308973445 | 0.271 | 0.094 | 0.119420299 | 15 | tia1 |
| rbmx | 6.2586484 | 0.328676189 | 0.373 | 0.152 | 0.126405922 | 15 | rbmx |
| casc4 | 6.4306729 | 0.281335153 | 0.169 | 0.044 | 0.129880301 | 15 | casc4 |
| ssbp3b | 6.5310206 | 0.28312697 | 0.237 | 0.076 | 0.131907024 | 15 | ssbp3b |
| krt84 | 6.6207132 | 0.452117013 | 0.864 | 0.517 | 0.133718546 | 15 | krt8 |
| shc1 | 6.6514726 | 0.277828233 | 0.169 | 0.044 | 0.134339792 | 15 | shc1 |
| tor1 | 6.7623989 | 0.288718368 | 0.271 | 0.093 | 0.136580172 | 15 | tor1 |
| baz2ba | 6.8721330 | 0.286143629 | 0.271 | 0.093 | 0.138796471 | 15 | baz2ba |
| rsrp12 | 6.9223227 | 0.374604748 | 0.339 | 0.135 | 0.139810152 | 15 | rsrp1 |
| lama43 | 7.0080523 | 0.276200534 | 0.288 | 0.098 | 0.141541634 | 15 | lama4 |
| timp2a3 | 7.0833258 | 0.312220933 | 0.305 | 0.11 | 0.143061932 | 15 | timp2a |
| snrpd3l | 7.6925340 | 0.364199734 | 0.373 | 0.161 | 0.15536611 | 15 | snrpd3l |
| tmed101 | 7.7368767 | 0.348567285 | 0.508 | 0.239 | 0.156261699 | 15 | tmed10 |
| si:ch211-194k22.8 | 8.0730608 | 0.277608893 | 0.237 | 0.076 | 0.16305161 | 15 | si:ch211-194k22.8 |
| h2afva | 8.1891108 | 0.357032598 | 0.492 | 0.23 | 0.165395471 | 15 | h2afva |
| ube2q2 | 8.8301773 | 0.30084104 | 0.254 | 0.086 | 0.178343091 | 15 | ube2q2 |
| bcap31 | 9.0466196 | 0.295480954 | 0.271 | 0.096 | 0.182714577 | 15 | bcap31 |
| dido1 | 9.0982411 | 0.354832396 | 0.373 | 0.156 | 0.183757176 | 15 | dido1 |
| sik2b1 | 9.5191140 | 0.353053566 | 0.22 | 0.07 | 0.192257547 | 15 | sik2b |
| LOC1018844513 | 9.5258266 | 0.323917496 | 0.322 | 0.123 | 0.192393122 | 15 | LOC101884451 |

| | | | | | | | |
|---|---|---|---|---|---|---|---|
| parp12b1 | 9.8024996 | 0.276233143 | 0.237 | 0.077 | 0.197981086 | 15 | parp12b |
| aplp21 | 9.8060253 | 0.308411523 | 0.288 | 0.103 | 0.198052294 | 15 | aplp2 |
| mvp1 | 9.8707884 | 0.257747377 | 0.288 | 0.101 | 0.199360313 | 15 | mvp |
| wdr83os | 1.0049694 | 0.326055684 | 0.407 | 0.18 | 0.202973686 | 15 | wdr83os |
| ftr823 | 1.0208168 | 0.323374494 | 0.288 | 0.104 | 0.206174381 | 15 | ftr82 |
| psme2 | 1.0705297 | 0.316749033 | 0.441 | 0.192 | 0.216214889 | 15 | psme2 |
| polr2i | 1.0715737 | 0.261405061 | 0.254 | 0.087 | 0.216425745 | 15 | polr2i |
| elocb | 1.0757653 | 0.348220448 | 0.576 | 0.288 | 0.217272329 | 15 | elocb |
| psmb7 | 1.0805049 | 0.352689274 | 0.593 | 0.302 | 0.21822958 | 15 | psmb7 |
| nudt21 | 1.1070768 | 0.257951453 | 0.339 | 0.13 | 0.223596308 | 15 | nudt21 |
| psma6a | 1.1138726 | 0.299541573 | 0.373 | 0.155 | 0.224968857 | 15 | psma6a |
| ndnf2 | 1.1164443 | 0.293548452 | 0.22 | 0.068 | 0.225488257 | 15 | ndnf |
| atp5f1b1 | 1.1211673 | 0.380722326 | 0.847 | 0.527 | 0.226442178 | 15 | atp5f1b |
| rca2.11 | 1.1486513 | 0.271262899 | 0.254 | 0.087 | 0.231993107 | 15 | rca2.1 |
| asah1b2 | 1.1510094 | 0.339077453 | 0.288 | 0.108 | 0.232469382 | 15 | asah1b |
| selenot1a | 1.1832915 | 0.287400166 | 0.288 | 0.108 | 0.238989385 | 15 | selenot1a |
| anp32b | 1.2231468 | 0.331094437 | 0.492 | 0.235 | 0.247038979 | 15 | anp32b |
| arhgef4 | 1.2283265 | 0.273680833 | 0.169 | 0.046 | 0.248085108 | 15 | arhgef4 |
| anxa13 | 1.2792704 | 0.267999817 | 0.237 | 0.078 | 0.25837425 | 15 | anxa13 |
| trappc5 | 1.2912244 | 0.273739467 | 0.254 | 0.089 | 0.260788608 | 15 | trappc5 |
| hnrnph1l1 | 1.3264789 | 0.302547778 | 0.322 | 0.129 | 0.267908957 | 15 | hnrnph1l |
| ginm1 | 1.3352770 | 0.317426796 | 0.305 | 0.118 | 0.269685902 | 15 | ginm1 |
| aif1l | 1.3535366 | 0.267056425 | 0.254 | 0.084 | 0.273373799 | 15 | aif1l |
| sec11a3 | 1.4998865 | 0.375414772 | 0.525 | 0.268 | 0.302932088 | 15 | sec11a |
| txndc5 | 1.5823413 | 0.274316323 | 0.254 | 0.087 | 0.319585482 | 15 | txndc5 |
| flot2a3 | 1.5936605 | 0.318549376 | 0.356 | 0.149 | 0.321871614 | 15 | flot2a |
| ankrd11 | 1.6207710 | 0.313610304 | 0.356 | 0.147 | 0.327347122 | 15 | ankrd11 |
| ndufa51 | 1.6287983 | 0.314193567 | 0.492 | 0.236 | 0.328968403 | 15 | ndufa5 |
| glrx3 | 1.7084651 | 0.263420079 | 0.288 | 0.108 | 0.345058708 | 15 | glrx3 |
| csnk1a12 | 1.7106902 | 0.361683597 | 0.525 | 0.267 | 0.345508113 | 15 | csnk1a1 |
| krtcap22 | 1.7359735 | 0.396503083 | 0.559 | 0.302 | 0.350614578 | 15 | krtcap2 |
| mesd | 1.7375542 | 0.262213821 | 0.254 | 0.089 | 0.350933839 | 15 | mesd |
| csnk1da1 | 1.8714589 | 0.275516967 | 0.288 | 0.106 | 0.377978564 | 15 | csnk1da |
| mt-nd23 | 1.9023854 | 0.420132127 | 0.678 | 0.37 | 0.384224799 | 15 | mt-nd2 |
| sec62 | 1.9249701 | 0.280428891 | 0.288 | 0.109 | 0.388786217 | 15 | sec62 |

| | | | | | | | |
|---|---|---|---|---|---|---|---|
| tspan31 | 1.9388308 | 0.296943559 | 0.203 | 0.064 | 0.391585663 | 15 | tspan31 |
| si:ch211-74m13.31 | 2.0061094 | 0.289729249 | 0.22 | 0.071 | 0.405173926 | 15 | si:ch211-74m13.3 |
| znfx1 | 2.1679030 | 0.2652004 | 0.169 | 0.047 | 0.43785138 | 15 | znfx1 |
| rcan1a1 | 2.2329282 | 0.252494403 | 0.186 | 0.056 | 0.450984526 | 15 | rcan1a |
| hif1ab | 2.2645315 | 0.343479958 | 0.39 | 0.174 | 0.45736744 | 15 | hif1ab |
| ptges3b | 2.2741320 | 0.313401637 | 0.475 | 0.225 | 0.459306455 | 15 | ptges3b |
| kmt5aa1 | 2.2834581 | 0.26528218 | 0.373 | 0.156 | 0.461190041 | 15 | kmt5aa |
| mt-nd13 | 2.4365547 | 0.408029796 | 0.644 | 0.351 | 0.492110968 | 15 | mt-nd1 |
| dlst | 2.4540568 | 0.305681117 | 0.373 | 0.162 | 0.495645862 | 15 | dlst |
| eif4a1a | 2.5288280 | 0.328089295 | 0.644 | 0.329 | 0.510747394 | 15 | eif4a1a |
| rmdn11 | 2.5668450 | 0.315440014 | 0.288 | 0.113 | 0.518425695 | 15 | rmdn1 |
| psmg1 | 2.5674541 | 0.296405083 | 0.254 | 0.093 | 0.518548719 | 15 | psmg1 |
| itm2ba3 | 2.6331179 | 0.399689814 | 0.678 | 0.41 | 0.531810826 | 15 | itm2ba |
| psmb1 | 2.7103051 | 0.328065585 | 0.644 | 0.336 | 0.54740034 | 15 | psmb1 |
| zgc:111986 | 2.7430278 | 0.329010947 | 0.475 | 0.228 | 0.554009343 | 15 | zgc:111986 |
| paip2b1 | 2.7736743 | 0.333176078 | 0.644 | 0.338 | 0.560199001 | 15 | paip2b |
| canx1 | 2.8687636 | 0.342190833 | 0.39 | 0.175 | 0.579404194 | 15 | canx |
| mir21-11 | 2.9054376 | 0.3202158 | 0.288 | 0.111 | 0.586811247 | 15 | mir21-1 |
| sdhc | 2.9260547 | 0.324081401 | 0.373 | 0.168 | 0.590975282 | 15 | sdhc |
| dcaf71 | 3.0347186 | 0.265612611 | 0.322 | 0.128 | 0.61292213 | 15 | dcaf7 |
| copg2 | 3.2159670 | 0.268518763 | 0.254 | 0.092 | 0.64952886 | 15 | copg2 |
| si:ch211-196l7.43 | 3.2930173 | 0.373149879 | 0.695 | 0.41 | 0.665090707 | 15 | si:ch211-196l7.4 |
| sf3b51 | 3.3979681 | 0.371545332 | 0.407 | 0.195 | 0.68628763 | 15 | sf3b5 |
| acadm1 | 3.5705837 | 0.26517932 | 0.373 | 0.159 | 0.721150794 | 15 | acadm |
| prkab1b | 3.6529144 | 0.330320699 | 0.424 | 0.206 | 0.737779122 | 15 | prkab1b |
| ndufa11 | 3.6651445 | 0.279088009 | 0.441 | 0.205 | 0.740249254 | 15 | ndufa11 |
| tardbp | 3.7573254 | 0.27961762 | 0.288 | 0.114 | 0.758867012 | 15 | tardbp |
| zgc:1748632 | 3.8212740 | 0.313714978 | 0.373 | 0.164 | 0.77178271 | 15 | zgc:174863 |
| esr2b1 | 3.8286995 | 0.253084037 | 0.186 | 0.058 | 0.773282442 | 15 | esr2b |
| pigx | 3.8840345 | 0.290455765 | 0.186 | 0.059 | 0.784458468 | 15 | pigx |
| ndufs3 | 3.8981124 | 0.263022721 | 0.356 | 0.152 | 0.787301776 | 15 | ndufs3 |
| polr2k | 3.9950328 | 0.26540356 | 0.288 | 0.112 | 0.806876793 | 15 | polr2k |
| nop101 | 4.1740062 | 0.329496951 | 0.492 | 0.244 | 0.843024037 | 15 | nop10 |
| ctdsp22 | 4.5523984 | 0.278758239 | 0.39 | 0.171 | 0.919447923 | 15 | ctdsp2 |
| scarb2a1 | 4.5885688 | 0.265821562 | 0.186 | 0.058 | 0.926753256 | 15 | scarb2a |

| | | | | | | | |
|---|---|---|---|---|---|---|---|
| fam120c2 | 4.7255844 | 0.296208943 | 0.339 | 0.146 | 0.954426282 | 15 | fam120c |
| tob1a1 | 4.7973172 | 0.292922569 | 0.441 | 0.204 | 0.968914168 | 15 | tob1a |
| slc38a5b6 | 4.8699728 | 0.293645237 | 0.441 | 0.202 | 0.983588424 | 15 | slc38a5b |
| srsf1b | 5.0990416 | 0.262076726 | 0.237 | 0.085 | 1 | 15 | srsf1b |
| edf1 | 5.1611015 | 0.306694035 | 0.678 | 0.363 | 1 | 15 | edf1 |
| pbdc1 | 5.1839144 | 0.304371337 | 0.424 | 0.206 | 1 | 15 | pbdc1 |
| emc6 | 5.2417724 | 0.270883736 | 0.305 | 0.124 | 1 | 15 | emc6 |
| polr2l | 5.3583012 | 0.280460101 | 0.475 | 0.226 | 1 | 15 | polr2l |
| lamp22 | 5.4496207 | 0.290577633 | 0.373 | 0.166 | 1 | 15 | lamp2 |
| ptbp3 | 5.5142010 | 0.293252484 | 0.237 | 0.087 | 1 | 15 | ptbp3 |
| rpn12 | 5.5288726 | 0.314194982 | 0.475 | 0.233 | 1 | 15 | rpn1 |
| ISCU (1 of many) | 5.5510852 | 0.253774562 | 0.39 | 0.171 | 1 | 15 | ISCU (1 of many) |
| ssr12 | 5.6707513 | 0.304994434 | 0.525 | 0.273 | 1 | 15 | ssr1 |
| sar1b1 | 5.9355134 | 0.311816779 | 0.322 | 0.139 | 1 | 15 | sar1b |
| vdac31 | 5.9701064 | 0.292356951 | 0.508 | 0.25 | 1 | 15 | vdac3 |
| tnrc6b | 6.5527992 | 0.257869502 | 0.237 | 0.085 | 1 | 15 | tnrc6b |
| psmd1 | 6.7523175 | 0.279348953 | 0.356 | 0.156 | 1 | 15 | psmd1 |
| PYURF | 7.0173517 | 0.257248848 | 0.22 | 0.078 | 1 | 15 | PYURF |
| adh53 | 7.1993447 | 0.33310052 | 0.644 | 0.358 | 1 | 15 | adh5 |
| CABZ01072614.15 | 7.3279593 | 0.271720834 | 0.39 | 0.173 | 1 | 15 | CABZ01072614.1 |
| fkbp1aa3 | 7.9062152 | 0.375094175 | 0.881 | 0.575 | 1 | 15 | fkbp1aa |
| khdrbs1a | 7.9417944 | 0.354645701 | 0.932 | 0.59 | 1 | 15 | khdrbs1a |
| cox8a1 | 8.6115773 | 0.346836187 | 0.78 | 0.468 | 1 | 15 | cox8a |
| calm1a | 8.6488905 | 0.31349411 | 0.458 | 0.226 | 1 | 15 | calm1a |
| acin1a3 | 9.0023603 | 0.315976291 | 0.559 | 0.294 | 1 | 15 | acin1a |
| cemip2 | 9.1728217 | 0.258047362 | 0.136 | 0.037 | 1 | 15 | cemip2 |
| eif5b | 9.2495819 | 0.339812511 | 0.407 | 0.203 | 1 | 15 | eif5b |
| stat31 | 9.3536201 | 0.259408501 | 0.22 | 0.078 | 1 | 15 | stat3 |
| gatad2b | 9.3798818 | 0.292935437 | 0.288 | 0.118 | 1 | 15 | gatad2b |
| tm2d31 | 0.00011 | 0.264763614 | 0.254 | 0.102 | 1 | 15 | tm2d3 |
| trappc1 | 0.000111 | 0.267018769 | 0.424 | 0.202 | 1 | 15 | trappc1 |
| hmgb3b | 0.000127 | 0.308691028 | 0.322 | 0.145 | 1 | 15 | hmgb3b |
| atp5mc1 | 0.00013 | 0.269455467 | 0.627 | 0.333 | 1 | 15 | atp5mc1 |
| LOC1039104383 | 0.00014 | 0.332486296 | 0.373 | 0.183 | 1 | 15 | LOC103910438 |
| si:ch73-281n10.25 | 0.000147 | 0.343624349 | 0.746 | 0.43 | 1 | 15 | si:ch73-281n10.2 |

| | | | | | | | |
|---|---|---|---|---|---|---|---|
| tnfsf102 | 0.000148 | 0.296388207 | 0.305 | 0.128 | 1 | 15 | tnfsf10 |
| golt1bb1 | 0.000153 | 0.274258442 | 0.203 | 0.074 | 1 | 15 | golt1bb |
| jam2b2 | 0.000155 | 0.282284241 | 0.136 | 0.038 | 1 | 15 | jam2b |
| asph1 | 0.000158 | 0.278625374 | 0.22 | 0.083 | 1 | 15 | asph |
| nhp21 | 0.000167 | 0.277601699 | 0.475 | 0.24 | 1 | 15 | nhp2 |
| psmc1a | 0.000168 | 0.255646112 | 0.339 | 0.153 | 1 | 15 | psmc1a |
| gpx4b | 0.000175 | 0.293047872 | 0.542 | 0.297 | 1 | 15 | gpx4b |
| tram12 | 0.000176 | 0.272023709 | 0.305 | 0.132 | 1 | 15 | tram1 |
| rcc2 | 0.00019 | 0.287363462 | 0.356 | 0.167 | 1 | 15 | rcc2 |
| cpne1 | 0.000191 | 0.282192471 | 0.39 | 0.189 | 1 | 15 | cpne1 |
| rer11 | 0.000198 | 0.3267374 | 0.356 | 0.175 | 1 | 15 | rer1 |
| rtn4a1 | 0.000208 | 0.297981984 | 0.458 | 0.248 | 1 | 15 | rtn4a |
| rbm4.3 | 0.000214 | 0.307360763 | 0.593 | 0.333 | 1 | 15 | rbm4.3 |
| smarcd11 | 0.000224 | 0.257719532 | 0.22 | 0.084 | 1 | 15 | smarcd1 |
| srsf5a2 | 0.000229 | 0.324965154 | 0.644 | 0.386 | 1 | 15 | srsf5a |
| rab10 | 0.000239 | 0.277546322 | 0.593 | 0.317 | 1 | 15 | rab10 |
| eif3ja | 0.000247 | 0.260484643 | 0.458 | 0.232 | 1 | 15 | eif3ja |
| srsf3b | 0.000269 | 0.336132045 | 0.525 | 0.305 | 1 | 15 | srsf3b |
| psmb6 | 0.000274 | 0.284911195 | 0.525 | 0.291 | 1 | 15 | psmb6 |
| rab2a2 | 0.000296 | 0.329971509 | 0.407 | 0.217 | 1 | 15 | rab2a |
| mtdha1 | 0.000301 | 0.254790427 | 0.407 | 0.199 | 1 | 15 | mtdha |
| tmed21 | 0.000301 | 0.299555704 | 0.559 | 0.309 | 1 | 15 | tmed2 |
| lsm6 | 0.000314 | 0.274377005 | 0.407 | 0.205 | 1 | 15 | lsm6 |
| pdha1a | 0.000317 | 0.256975312 | 0.288 | 0.128 | 1 | 15 | pdha1a |
| echs1 | 0.000326 | 0.266761964 | 0.305 | 0.139 | 1 | 15 | echs1 |
| nsfl1c | 0.000326 | 0.273857868 | 0.305 | 0.141 | 1 | 15 | nsfl1c |
| gmfb1 | 0.000335 | 0.274924384 | 0.356 | 0.172 | 1 | 15 | gmfb |
| surf4l2 | 0.000355 | 0.275237351 | 0.203 | 0.077 | 1 | 15 | surf4l |
| si:ch211-51e12.7 | 0.000382 | 0.25220652 | 0.39 | 0.191 | 1 | 15 | si:ch211-51e12.7 |
| ndufb10 | 0.000389 | 0.255455002 | 0.441 | 0.229 | 1 | 15 | ndufb10 |
| laptm4a1 | 0.000389 | 0.252220862 | 0.39 | 0.191 | 1 | 15 | laptm4a |
| sfpq | 0.000392 | 0.267903763 | 0.288 | 0.129 | 1 | 15 | sfpq |
| thoc2 | 0.000393 | 0.272178502 | 0.271 | 0.12 | 1 | 15 | thoc2 |
| spcs31 | 0.000398 | 0.274453819 | 0.458 | 0.243 | 1 | 15 | spcs3 |
| psip1b | 0.0004 | 0.266704041 | 0.271 | 0.119 | 1 | 15 | psip1b |

| | | | | | | | |
|---|---|---|---|---|---|---|---|
| xbp12 | 0.000429 | 0.263591306 | 0.746 | 0.43 | 1 | 15 | xbp1 |
| ndufb92 | 0.000431 | 0.28382143 | 0.559 | 0.322 | 1 | 15 | ndufb9 |
| ola1 | 0.000434 | 0.315271147 | 0.542 | 0.327 | 1 | 15 | ola1 |
| slco2b12 | 0.000441 | 0.309226775 | 0.203 | 0.079 | 1 | 15 | slco2b1 |
| si:ch211-222l21.14 | 0.000446 | 0.351858337 | 0.831 | 0.528 | 1 | 15 | si:ch211-222l21.1 |
| pfn24 | 0.000479 | 0.282688616 | 0.576 | 0.317 | 1 | 15 | pfn2 |
| si:ch211-132p1.32 | 0.000521 | 0.293514862 | 0.305 | 0.145 | 1 | 15 | si:ch211-132p1.3 |
| uqcr102 | 0.000525 | 0.305088362 | 0.542 | 0.323 | 1 | 15 | uqcr10 |
| anapc16 | 0.000531 | 0.254664965 | 0.339 | 0.165 | 1 | 15 | anapc16 |
| socs3b4 | 0.000535 | 0.346170693 | 0.305 | 0.145 | 1 | 15 | socs3b |
| mycb4 | 0.000539 | 0.329085001 | 0.322 | 0.16 | 1 | 15 | mycb |
| zgc:1629442 | 0.000548 | 0.284635953 | 0.678 | 0.397 | 1 | 15 | zgc:162944 |
| sap18 | 0.000588 | 0.25058802 | 0.441 | 0.238 | 1 | 15 | sap18 |
| hnrnpa1a | 0.000613 | 0.287198571 | 0.39 | 0.206 | 1 | 15 | hnrnpa1a |
| smim13 | 0.000623 | 0.256913016 | 0.22 | 0.092 | 1 | 15 | smim13 |
| serinc22 | 0.000626 | 0.265320219 | 0.39 | 0.198 | 1 | 15 | serinc2 |
| mif | 0.000709 | 0.279774078 | 0.525 | 0.313 | 1 | 15 | mif |
| rnf101 | 0.00072 | 0.265682712 | 0.407 | 0.212 | 1 | 15 | rnf10 |
| sdcbp22 | 0.000805 | 0.261668859 | 0.61 | 0.338 | 1 | 15 | sdcbp2 |
| skp11 | 0.000836 | 0.262507878 | 0.61 | 0.357 | 1 | 15 | skp1 |
| atp5f1c1 | 0.00095 | 0.268367898 | 0.576 | 0.354 | 1 | 15 | atp5f1c |
| hint1 | 0.000993 | 0.281223667 | 0.508 | 0.303 | 1 | 15 | hint1 |
| sat1a.23 | 0.000998 | 0.281789499 | 0.576 | 0.333 | 1 | 15 | sat1a.2 |
| pin11 | 0.001005 | 0.265660921 | 0.407 | 0.219 | 1 | 15 | pin1 |
| chd35 | 0.001007 | 0.254654221 | 0.458 | 0.24 | 1 | 15 | chd3 |
| rad21a1 | 0.001195 | 0.289845134 | 0.441 | 0.249 | 1 | 15 | rad21a |
| h2afy23 | 0.001259 | 0.259704474 | 0.458 | 0.255 | 1 | 15 | h2afy2 |
| ywhae12 | 0.001369 | 0.265676753 | 0.627 | 0.377 | 1 | 15 | ywhae1 |
| hspa53 | 0.001736 | 0.262571816 | 0.407 | 0.22 | 1 | 15 | hspa5 |
| qkia3 | 0.001745 | 0.278772975 | 0.61 | 0.357 | 1 | 15 | qkia |
| atp5fa1 | 0.00199 | 0.254883199 | 0.729 | 0.459 | 1 | 15 | atp5fa1 |
| ndufb4 | 0.002002 | 0.25080984 | 0.508 | 0.315 | 1 | 15 | ndufb4 |
| hypk | 0.002181 | 0.254707278 | 0.475 | 0.283 | 1 | 15 | hypk |
| snrpb1 | 0.00225 | 0.268766264 | 0.644 | 0.42 | 1 | 15 | snrpb |
| cnbpb2 | 0.002365 | 0.270571329 | 0.525 | 0.324 | 1 | 15 | cnbpb |

| | | | | | | | |
|---|---|---|---|---|---|---|---|
| ddit33 | 0.003146 | 0.292143579 | 0.356 | 0.202 | 1 | 15 | ddit3 |
| pim11 | 0.003804 | 0.271219123 | 0.305 | 0.165 | 1 | 15 | pim1 |
| prdx23 | 0.004503 | 0.271529853 | 0.695 | 0.487 | 1 | 15 | prdx2 |
| ubc1 | 0.004894 | 0.258337816 | 0.864 | 0.653 | 1 | 15 | ubc |
| calr3a3 | 0.004962 | 0.272339314 | 0.39 | 0.236 | 1 | 15 | calr3a |
| mgst3b4 | 0.005706 | 0.253454213 | 0.475 | 0.301 | 1 | 15 | mgst3b |
| notch22 | 0.006887 | 0.259747151 | 0.271 | 0.151 | 1 | 15 | notch2 |
| si:dkey-6n6.11 | 5.6284660 | 1.443139582 | 0.702 | 0.009 | 1.13678129421761e-2 | 16 | si:dkey-6n6.1 |
| LOC1039109032 | 8.8346095 | 1.658694504 | 0.825 | 0.022 | 1.78432609442825e-1 | 16 | LOC103910903 |
| fgfr3 | 3.2794584 | 1.400526986 | 0.807 | 0.021 | 6.62352225435192e-1 | 16 | fgfr3 |
| igf32 | 1.2336189 | 1.601902984 | 0.86 | 0.029 | 2.49154013487025e-1 | 16 | igf3 |
| alcama2 | 2.2177779 | 1.45828974 | 0.842 | 0.037 | 4.47924605391585e-1 | 16 | alcama |
| LOC1104380241 | 2.8841788 | 0.926682674 | 0.509 | 0.008 | 5.82517601952401e-1 | 16 | LOC110438024 |
| cdh41 | 1.3479482 | 1.075481331 | 0.632 | 0.017 | 2.72245113147912e-1 | 16 | cdh4 |
| foxl2b2 | 3.5501743 | 1.371862225 | 0.825 | 0.036 | 7.17028718462096e-1 | 16 | foxl2b |
| atp1b3b2 | 3.9796611 | 1.325518949 | 0.825 | 0.042 | 8.03772171348682e-1 | 16 | atp1b3b |
| phlda11 | 7.2427967 | 0.836301229 | 0.474 | 0.008 | 1.46282766619878e-1 | 16 | phlda1 |
| znf536 | 2.9192961 | 0.795307811 | 0.421 | 0.006 | 5.89610246439451e-1 | 16 | znf536 |
| fez12 | 4.7476058 | 1.082728637 | 0.596 | 0.02 | 9.58873953214605e-1 | 16 | fez1 |
| pcdh1b | 9.9627471 | 1.013835061 | 0.614 | 0.022 | 2.01217604065718e-1 | 16 | pcdh1b |
| bcam2 | 5.0671875 | 1.729385122 | 0.965 | 0.079 | 1.02341986033964e-1 | 16 | bcam |
| gsdf2 | 8.8375578 | 1.861773227 | 0.895 | 0.064 | 1.78492156425212e-1 | 16 | gsdf |
| islr21 | 2.7201107 | 0.816937872 | 0.491 | 0.013 | 5.49380761463656e-1 | 16 | islr2 |
| mpzl2b2 | 3.6262686 | 1.256166913 | 0.754 | 0.045 | 7.32397471952446e-1 | 16 | mpzl2b |
| si:ch211-117c9.52 | 1.3026912 | 0.937018649 | 0.579 | 0.023 | 2.63104553189256e-9 | 16 | si:ch211-117c9.5 |
| cx432 | 9.7856978 | 1.613546074 | 0.842 | 0.063 | 1.97641739602707e-9 | 16 | cx43 |
| alcamb2 | 1.0193425 | 1.065678978 | 0.684 | 0.036 | 2.05876618530834e-9 | 16 | alcamb |
| pi15a2 | 2.2033764 | 1.659783867 | 0.912 | 0.081 | 4.45015938265906e-9 | 16 | pi15a |
| si:dkey-26c10.51 | 2.3724984 | 0.87417013 | 0.456 | 0.013 | 4.79173503212114e-9 | 16 | si:dkey-26c10.5 |
| jam2a2 | 4.8727266 | 1.822172424 | 0.965 | 0.102 | 9.84144605426821e-9 | 16 | jam2a |
| fgfr41 | 2.4326285 | 0.915705213 | 0.526 | 0.02 | 4.91317980589322e-9 | 16 | fgfr4 |
| cadm2a2 | 1.6332753 | 1.01218891 | 0.614 | 0.03 | 3.29872628522373e-9 | 16 | cadm2a |
| XLOC-0422291 | 4.3352130 | 0.710918891 | 0.456 | 0.014 | 8.75582971255815e-9 | 16 | XLOC-042229 |
| spock1 | 3.0756163 | 0.61849027 | 0.333 | 0.005 | 6.21182235185589e-8 | 16 | spock1 |
| slc7a3a1 | 8.6286316 | 1.139708892 | 0.754 | 0.056 | 1.74272473407889e-8 | 16 | slc7a3a |

| | | | | | | | |
|---|---|---|---|---|---|---|---|
| metrn | 1.4563446 | 0.851345008 | 0.509 | 0.022 | 2.9413793206896e-82 | 16 | metrn |
| gpx1b1 | 1.3090956 | 1.148085362 | 0.509 | 0.023 | 2.64398057991261e-8 | 16 | gpx1b |
| corin | 3.2657042 | 0.784728058 | 0.421 | 0.014 | 6.59574282557449e-8 | 16 | corin |
| bambia2 | 4.7028796 | 1.121484411 | 0.684 | 0.047 | 9.49840600600151e-8 | 16 | bambia |
| hsd3b73 | 1.3351385 | 1.266562602 | 0.737 | 0.058 | 2.69657931337281e-7 | 16 | hsd3b7 |
| lamb2l | 2.0045201 | 0.511944796 | 0.281 | 0.004 | 4.0485293524287e-78 | 16 | lamb2l |
| ndp | 8.2623317 | 0.566353937 | 0.333 | 0.008 | 1.66874313705803e-7 | 16 | ndp |
| arhgef18a | 6.2534869 | 0.643082844 | 0.386 | 0.013 | 1.26301675142844e-7 | 16 | arhgef18a |
| itga3b1 | 1.9127070 | 0.558504058 | 0.333 | 0.008 | 3.86309449859106e-7 | 16 | itga3b |
| tgfb2l | 1.5747384 | 0.605465599 | 0.368 | 0.011 | 3.1804993312335e-71 | 16 | tgfb2l |
| hpn1 | 1.0044443 | 0.512945724 | 0.316 | 0.008 | 2.0286761928835e-68 | 16 | hpn |
| si:ch211-225b11.1 | 5.1591227 | 0.963013606 | 0.596 | 0.043 | 1.04198802766947e-6 | 16 | si:ch211-225b11.1 |
| CABZ01021532.11 | 7.4755476 | 0.8231354 | 0.404 | 0.017 | 1.50983635616671e-6 | 16 | CABZ01021532.1 |
| ece2b1 | 4.0466286 | 1.003219443 | 0.596 | 0.045 | 8.17297590114076e-6 | 16 | ece2b |
| atp11a | 5.3676557 | 0.545654984 | 0.298 | 0.007 | 1.08410543965878e-6 | 16 | atp11a |
| tent5ba | 6.6785070 | 0.590284005 | 0.368 | 0.013 | 1.34885805896873e-6 | 16 | tent5ba |
| smoc2 | 6.5866252 | 0.675325877 | 0.298 | 0.008 | 1.33030070508374e-6 | 16 | smoc2 |
| si:ch211-276k2.1 | 6.6532506 | 0.543137439 | 0.263 | 0.005 | 1.34375702582445e-6 | 16 | si:ch211-276k2.1 |
| fzd10 | 2.2723693 | 0.412040579 | 0.263 | 0.005 | 4.58950428724977e-6 | 16 | fzd10 |
| scn4bb1 | 2.7807335 | 0.64713226 | 0.368 | 0.014 | 5.61624748025425e-6 | 16 | scn4bb |
| cx34.51 | 4.5863843 | 0.831926562 | 0.404 | 0.018 | 9.26312048781132e-6 | 16 | cx34.5 |
| cldn11a4 | 1.3967352 | 1.541979578 | 1 | 0.177 | 2.82098616421479e-6 | 16 | cldn11a |
| nectin1b2 | 3.6870640 | 0.897131878 | 0.614 | 0.05 | 7.44676318139693e-6 | 16 | nectin1b |
| col4a51 | 2.9830983 | 0.640490217 | 0.404 | 0.019 | 6.02496371351287e-5 | 16 | col4a5 |
| rdh10a2 | 5.4590873 | 1.311298598 | 0.702 | 0.075 | 1.10257186825425e-5 | 16 | rdh10a |
| col4a61 | 2.6637791 | 0.743396441 | 0.474 | 0.029 | 5.3800347450745e-58 | 16 | col4a6 |
| CABZ01049847.1 | 4.5089870 | 0.565063429 | 0.298 | 0.009 | 9.10680108731287e-5 | 16 | CABZ01049847.1 |
| inhbb | 1.1623854 | 0.491602428 | 0.246 | 0.005 | 2.34766984724749e-5 | 16 | inhbb |
| f104 | 1.8705001 | 1.134679339 | 0.807 | 0.103 | 3.77784920558806e-5 | 16 | f10 |
| cemip21 | 4.5711164 | 0.715698958 | 0.456 | 0.028 | 9.23228391640104e-5 | 16 | cemip2 |
| bmp6 | 4.9621041 | 0.669187094 | 0.368 | 0.017 | 1.0021961846041e-54 | 16 | bmp6 |
| nav31 | 7.9188552 | 0.753801922 | 0.456 | 0.029 | 1.59937119222723e-5 | 16 | nav3 |
| slc2a1b | 3.0590134 | 0.566447608 | 0.316 | 0.012 | 6.17828942568181e-5 | 16 | slc2a1b |
| foxl2a1 | 1.7592334 | 0.613203383 | 0.351 | 0.016 | 3.55312370087628e-5 | 16 | foxl2a |
| nat8l | 2.6548465 | 0.631288527 | 0.351 | 0.016 | 5.36199360546596e-5 | 16 | nat8l |

| | | | | | | | |
|---|---|---|---|---|---|---|---|
| LOC5533161 | 2.6597989 | 0.877867247 | 0.596 | 0.054 | 5.37199591610823e-5 | 16 | LOC553316 |
| si:dkey-188i13.91 | 4.5601916 | 0.81844445 | 0.474 | 0.033 | 9.21021908448775e-5 | 16 | si:dkey-188i13.9 |
| nr0b12 | 9.2641545 | 0.749561687 | 0.474 | 0.033 | 1.87108129329406e-5 | 16 | nr0b1 |
| lsr1 | 3.8436453 | 0.724498165 | 0.474 | 0.033 | 7.76301056290408e-5 | 16 | lsr |
| sox9a | 4.1241287 | 0.544755421 | 0.316 | 0.012 | 8.32950284531215e-5 | 16 | sox9a |
| plxna41 | 4.5758468 | 0.694930653 | 0.386 | 0.022 | 9.24183795159323e-5 | 16 | plxna4 |
| CR318588.32 | 1.0455598 | 1.006416175 | 0.649 | 0.068 | 2.11171731209493e-4 | 16 | CR318588.3 |
| bmpr2a1 | 1.6430929 | 0.763430325 | 0.456 | 0.032 | 3.31855478355273e-4 | 16 | bmpr2a |
| hkdc1 | 2.3946987 | 0.438985966 | 0.246 | 0.007 | 4.83657316345717e-4 | 16 | hkdc1 |
| ldlrap1a1 | 3.0781504 | 0.624601263 | 0.421 | 0.027 | 6.21694054983774e-4 | 16 | ldlrap1a |
| gdpd4a1 | 1.3178855 | 0.574806043 | 0.281 | 0.01 | 2.66173343270062e-4 | 16 | gdpd4a |
| fam184b | 8.7633093 | 0.380302328 | 0.228 | 0.006 | 1.76992559091956e-4 | 16 | fam184b |
| agrn3 | 1.0907474 | 1.057211453 | 0.737 | 0.099 | 2.20298262020138e-4 | 16 | agrn |
| rpz53 | 1.0989370 | 1.450846573 | 0.982 | 0.24 | 2.2195231591423e-47 | 16 | rpz5 |
| prr13 | 1.6841740 | 0.726667828 | 0.421 | 0.029 | 3.40152637233036e-4 | 16 | prr13 |
| etv41 | 1.7822386 | 0.810220284 | 0.544 | 0.05 | 3.59958741086769e-4 | 16 | etv4 |
| mbnl32 | 3.4480293 | 0.970471913 | 0.684 | 0.085 | 6.96398491204441e-4 | 16 | mbnl3 |
| appa1 | 5.7417532 | 1.168199297 | 0.807 | 0.129 | 1.15966191200041e-4 | 16 | appa |
| XLOC-0056053 | 3.0611715 | 1.013387492 | 0.667 | 0.082 | 6.18264825692273e-4 | 16 | XLOC-005605 |
| slc38a21 | 6.8447928 | 0.705917577 | 0.456 | 0.037 | 1.38244281532224e-4 | 16 | slc38a2 |
| CABZ01068366.12 | 1.2857466 | 0.915547126 | 0.702 | 0.092 | 2.5968225591479e-44 | 16 | CABZ01068366.1 |
| efhd1 | 2.8795960 | 0.452411692 | 0.263 | 0.01 | 5.81592010070961e-4 | 16 | efhd1 |
| XLOC-002129 | 3.1758092 | 0.477893245 | 0.246 | 0.008 | 6.41418192017496e-4 | 16 | XLOC-002129 |
| pcdh10a | 7.8535503 | 0.400219345 | 0.246 | 0.008 | 1.58618157149349e-4 | 16 | pcdh10a |
| aldh1a24 | 1.6388480 | 1.397600042 | 0.912 | 0.187 | 3.30998147012378e-4 | 16 | aldh1a2 |
| ecrg4a3 | 2.1281505 | 1.072202729 | 0.561 | 0.058 | 4.29822570092871e-4 | 16 | ecrg4a |
| prkar2aa1 | 7.6379669 | 1.028941183 | 0.789 | 0.136 | 1.54264017674686e-4 | 16 | prkar2aa |
| fgd5a1 | 8.2893077 | 0.449736318 | 0.281 | 0.012 | 1.67419149291195e-4 | 16 | fgd5a |
| gata62 | 1.6136157 | 0.821227846 | 0.509 | 0.05 | 3.25901980023526e-4 | 16 | gata6 |
| emilin1a2 | 4.3306296 | 0.812194989 | 0.491 | 0.048 | 8.74657266854862e-4 | 16 | emilin1a |
| asb112 | 9.0786748 | 0.549160428 | 0.333 | 0.019 | 1.83361996561315e-4 | 16 | asb11 |
| amh | 3.1264533 | 0.354566699 | 0.193 | 0.005 | 6.31449786149179e-4 | 16 | amh |
| si:ch211-207d6.2 | 2.2678746 | 0.951242585 | 0.509 | 0.053 | 4.58042651254805e-3 | 16 | si:ch211-207d6.2 |
| slc38a3a | 5.5076214 | 0.284450523 | 0.175 | 0.004 | 1.1123743099404e-38 | 16 | slc38a3a |
| ifi464 | 3.2475289 | 1.200392918 | 0.842 | 0.177 | 6.55903431195555e-3 | 16 | ifi46 |

| | | | | | | | |
|---|---|---|---|---|---|---|---|
| chrdl2 | 3.9047114 | 0.361202287 | 0.211 | 0.007 | 7.88634565900692e-3 | 16 | chrdl2 |
| zgc:92275 | 5.1085370 | 0.265753588 | 0.14 | 0.002 | 1.03177122341527e-3 | 16 | zgc:92275 |
| wu:fc27b111 | 6.2204339 | 0.597796361 | 0.386 | 0.03 | 1.25634104492115e-3 | 16 | wu:fc27b11 |
| si:dkey-42i9.71 | 2.8994689 | 0.836118213 | 0.439 | 0.041 | 5.85605737764689e-3 | 16 | si:dkey-42i9.7 |
| ccdc1872 | 8.8066621 | 0.897963723 | 0.544 | 0.066 | 1.778681563363e-36 | 16 | ccdc187 |
| itm2bb3 | 1.6241691 | 1.356348311 | 0.982 | 0.344 | 3.28033444865319e-3 | 16 | itm2bb |
| lgals172 | 4.2247504 | 0.898772292 | 0.526 | 0.062 | 8.53272854339441e-3 | 16 | lgals17 |
| myh102 | 1.6419087 | 0.74910807 | 0.544 | 0.065 | 3.31616312315978e-3 | 16 | myh10 |
| XLOC-0098152 | 1.7241565 | 0.750826516 | 0.491 | 0.054 | 3.48227907314101e-3 | 16 | XLOC-009815 |
| nr5a1b | 1.8521602 | 0.280772068 | 0.158 | 0.003 | 3.74080799810772e-3 | 16 | nr5a1b |
| chka1 | 9.3372422 | 0.623853162 | 0.439 | 0.043 | 1.88584282529688e-3 | 16 | chka |
| hey2 | 1.2106335 | 0.289278233 | 0.14 | 0.002 | 2.44511657808321e-3 | 16 | hey2 |
| trim35-14 | 1.2839996 | 0.3434603 | 0.175 | 0.005 | 2.59329405508529e-3 | 16 | trim35-14 |
| ifi453 | 2.2262823 | 1.306490414 | 0.825 | 0.186 | 4.49642247878631e-3 | 16 | ifi45 |
| si:dkey-178e17.3 | 3.5088641 | 0.439218912 | 0.281 | 0.017 | 7.08685282831462e-3 | 16 | si:dkey-178e17.3 |
| emx22 | 3.6047681 | 0.729442858 | 0.474 | 0.053 | 7.28055020534743e-3 | 16 | emx2 |
| gig2e | 3.7688156 | 0.493743832 | 0.298 | 0.02 | 7.61187695750377e-3 | 16 | gig2e |
| si:ch211-198m17.1 | 4.2662897 | 0.305473332 | 0.158 | 0.004 | 8.61662536393116e-3 | 16 | si:ch211-198m17.1 |
| nos1apa1 | 4.6448351 | 0.750527232 | 0.491 | 0.059 | 9.38117353462579e-3 | 16 | nos1apa |
| col14a1a | 6.0479944 | 0.303084471 | 0.158 | 0.004 | 1.22151344591618e-3 | 16 | col14a1a |
| CABZ01084501.3 | 8.1593289 | 0.41712923 | 0.193 | 0.007 | 1.64793966123259e-3 | 16 | CABZ01084501.3 |
| edem12 | 2.8592894 | 0.650582305 | 0.439 | 0.047 | 5.77490688259094e-3 | 16 | edem1 |
| si:ch73-61d6.31 | 2.0569521 | 0.539705205 | 0.351 | 0.03 | 4.15442634361296e-3 | 16 | si:ch73-61d6.3 |
| msi2a1 | 2.5306459 | 0.619515361 | 0.439 | 0.047 | 5.11114567018047e-3 | 16 | msi2a |
| trim35-33 | 6.1765427 | 0.287920769 | 0.158 | 0.004 | 1.24747634110583e-3 | 16 | trim35-33 |
| zgc:1527911 | 7.2622352 | 0.721333832 | 0.491 | 0.06 | 1.46675365755374e-3 | 16 | zgc:152791 |
| adamtsl5 | 8.0078829 | 0.294049929 | 0.158 | 0.004 | 1.61735211435557e-3 | 16 | adamtsl5 |
| isg152 | 1.5406498 | 0.692235116 | 0.404 | 0.041 | 3.11165040584453e-3 | 16 | isg15 |
| CR318588.13 | 4.2759405 | 1.344238019 | 0.93 | 0.253 | 8.63611717796807e-3 | 16 | CR318588.1 |
| vasnb2 | 5.8335599 | 0.680124507 | 0.456 | 0.054 | 1.17820409460152e-2 | 16 | vasnb |
| zgc:158482 | 7.5903112 | 0.306658105 | 0.211 | 0.01 | 1.5330151533357e-29 | 16 | zgc:158482 |
| si:dkey-44g23.5 | 1.5514890 | 0.44465118 | 0.298 | 0.022 | 3.13354243921909e-2 | 16 | si:dkey-44g23.5 |
| ponzr16 | 1.9351722 | 1.148703021 | 0.965 | 0.382 | 3.90846731735429e-2 | 16 | ponzr1 |
| si:ch211-199g17.2 | 2.1075726 | 0.689770062 | 0.544 | 0.077 | 4.25666444512143e-2 | 16 | si:ch211-199g17.2 |
| si:dkey-188i13.102 | 2.8172823 | 0.931149193 | 0.754 | 0.156 | 5.69006513158688e-2 | 16 | si:dkey-188i13.10 |

| | | | | | | | |
|---|---|---|---|---|---|---|---|
| tmtc2a | 6.6649530 | 0.293756038 | 0.158 | 0.005 | 1.34612057595829e-2 | 16 | tmtc2a |
| hspe13 | 1.0350758 | 1.12845321 | 0.947 | 0.279 | 2.0905426157006e-28 | 16 | hspe1 |
| thbs3b1 | 1.5281985 | 0.361637873 | 0.193 | 0.008 | 3.08650261884495e-2 | 16 | thbs3b |
| sema3ab1 | 2.3261344 | 0.561531338 | 0.281 | 0.02 | 4.69809365213664e-2 | 16 | sema3ab |
| mpped2a | 2.4618601 | 0.484374835 | 0.333 | 0.029 | 4.97221884571545e-2 | 16 | mpped2a |
| lamc11 | 2.8354150 | 0.728072029 | 0.491 | 0.065 | 5.72668777829905e-2 | 16 | lamc1 |
| errfi1a | 3.1258985 | 0.480752794 | 0.263 | 0.018 | 6.31337724196239e-2 | 16 | errfi1a |
| aplp22 | 1.4887998 | 0.792681289 | 0.579 | 0.095 | 3.0069290255868e-27 | 16 | aplp2 |
| arrdc3a2 | 1.7493030 | 0.808465414 | 0.456 | 0.06 | 3.53306745105459e-2 | 16 | arrdc3a |
| aif1l1 | 2.5997974 | 0.71370366 | 0.526 | 0.077 | 5.25081082605653e-2 | 16 | aif1l |
| adgrl3.11 | 8.8342833 | 0.624964761 | 0.421 | 0.05 | 1.78426021197203e-2 | 16 | adgrl3.1 |
| adam10a3 | 1.3808393 | 0.908540621 | 0.719 | 0.158 | 2.78888132756631e-2 | 16 | adam10a |
| decr1 | 1.8914434 | 0.637829868 | 0.509 | 0.076 | 3.82014842285362e-2 | 16 | decr1 |
| arf2b2 | 1.9584205 | 0.931516036 | 0.789 | 0.196 | 3.95542199460933e-2 | 16 | arf2b |
| sat1a.1 | 2.2094976 | 0.412322713 | 0.193 | 0.009 | 4.46252231645128e-2 | 16 | sat1a.1 |
| sv2bb | 3.2408352 | 0.366593965 | 0.228 | 0.014 | 6.54551495283017e-2 | 16 | sv2bb |
| dusp4 | 4.9077211 | 0.463623199 | 0.246 | 0.017 | 9.91212449590948e-2 | 16 | dusp4 |
| XLOC-0152911 | 7.2637990 | 0.513904233 | 0.298 | 0.026 | 1.46706950271402e-2 | 16 | XLOC-015291 |
| ace1 | 7.3656393 | 0.533419129 | 0.333 | 0.032 | 1.48763817153768e-2 | 16 | ace |
| phactr4a2 | 7.8974313 | 0.642134522 | 0.421 | 0.053 | 1.5950442186225e-25 | 16 | phactr4a |
| LOC110437926 | 1.2489839 | 0.252529942 | 0.123 | 0.003 | 2.52257281954148e-2 | 16 | LOC110437926 |
| XLOC-004335 | 1.7294507 | 0.319530413 | 0.158 | 0.006 | 3.49297164479104e-2 | 16 | XLOC-004335 |
| ptprfb2 | 2.1118523 | 0.482993982 | 0.298 | 0.026 | 4.26530816122745e-2 | 16 | ptprfb |
| jam3b1 | 2.2516187 | 0.603831903 | 0.439 | 0.057 | 4.54759443500238e-2 | 16 | jam3b |
| b2m2 | 2.6512567 | 1.071294103 | 0.93 | 0.343 | 5.35474330400828e-2 | 16 | b2m |
| igsf3 | 4.9086669 | 0.365990448 | 0.211 | 0.012 | 9.91403469967811e-2 | 16 | igsf3 |
| plxna2 | 5.9905726 | 0.417034281 | 0.246 | 0.017 | 1.20991595105153e-2 | 16 | plxna2 |
| znfx11 | 8.9926667 | 0.588329297 | 0.368 | 0.042 | 1.81624889857625e-2 | 16 | znfx1 |
| tgfb32 | 3.9221788 | 0.65631059 | 0.509 | 0.079 | 7.92162471293525e-2 | 16 | tgfb3 |
| lama52 | 5.8677296 | 0.623353079 | 0.421 | 0.055 | 1.18510536242516e-2 | 16 | lama5 |
| pik3c2a | 6.5226894 | 0.460351469 | 0.316 | 0.031 | 1.31738758647087e-2 | 16 | pik3c2a |
| large2 | 9.1951743 | 0.270849739 | 0.175 | 0.008 | 1.85714936721568e-2 | 16 | large2 |
| robo12 | 1.8541439 | 0.644788964 | 0.368 | 0.045 | 3.74481453943683e-2 | 16 | robo1 |
| mxa | 2.6509622 | 0.37411969 | 0.211 | 0.013 | 5.35414847723413e-2 | 16 | mxa |
| inha1 | 2.6595960 | 0.413798821 | 0.193 | 0.011 | 5.37158612279385e-2 | 16 | inha |

| | | | | | | | |
|---|---|---|---|---|---|---|---|
| CABZ01068367.11 | 1.1662678 | 0.54660069 | 0.351 | 0.04 | 2.35551121377843e-2 | 16 | CABZ01068367.1 |
| ndrg3a | 1.4521109 | 0.462478994 | 0.281 | 0.026 | 2.93282840143169e-2 | 16 | ndrg3a |
| cyp17a12 | 1.7850994 | 0.435403229 | 0.333 | 0.035 | 3.60536530478685e-2 | 16 | cyp17a1 |
| phc2a2 | 1.7943580 | 0.701402594 | 0.509 | 0.085 | 3.62406493380901e-2 | 16 | phc2a |
| slc4a4a1 | 1.9408713 | 0.363721342 | 0.211 | 0.014 | 3.91997796370722e-2 | 16 | slc4a4a |
| ache | 3.2652191 | 0.276190731 | 0.193 | 0.011 | 6.59476306002493e-2 | 16 | ache |
| zgc:921401 | 3.9739297 | 0.580560421 | 0.404 | 0.055 | 8.02614595643842e-2 | 16 | zgc:92140 |
| fgfrl1a1 | 4.2103672 | 0.648396723 | 0.368 | 0.047 | 8.50367865158452e-2 | 16 | fgfrl1a |
| hspd11 | 4.9976299 | 0.771751396 | 0.684 | 0.158 | 1.00937132429574e-2 | 16 | hspd1 |
| cdh13 | 5.9832013 | 0.522032766 | 0.333 | 0.037 | 1.20842718465347e-2 | 16 | cdh1 |
| adamts9 | 8.2053109 | 0.325427584 | 0.175 | 0.009 | 1.65722666175124e-2 | 16 | adamts9 |
| notch1b | 8.9485032 | 0.361507429 | 0.211 | 0.014 | 1.80732920717701e-2 | 16 | notch1b |
| tbx18 | 9.3840768 | 0.373816755 | 0.158 | 0.007 | 1.89530199804831e-2 | 16 | tbx18 |
| XLOC-040516 | 1.0732470 | 0.313536025 | 0.175 | 0.009 | 2.16763710332937e-2 | 16 | XLOC-040516 |
| nanos12 | 1.1557800 | 0.759627063 | 0.509 | 0.087 | 2.33432898665884e-2 | 16 | nanos1 |
| hmcn2 | 1.3232284 | 0.331873022 | 0.193 | 0.012 | 2.67252449136926e-2 | 16 | hmcn2 |
| adgrg1 | 1.3660142 | 0.344155963 | 0.193 | 0.012 | 2.75893889920805e-2 | 16 | adgrg1 |
| tspan9b2 | 2.2118054 | 0.631532836 | 0.474 | 0.076 | 4.46718352895985e-2 | 16 | tspan9b |
| odc13 | 3.8050708 | 0.835974927 | 0.561 | 0.112 | 7.68510161825956e-2 | 16 | odc1 |
| CU984600.2 | 4.5906830 | 0.392081995 | 0.263 | 0.024 | 9.271802461184e-21 | 16 | CU984600.2 |
| zgc:175088 | 9.6324272 | 0.691724471 | 0.561 | 0.111 | 1.94546132517375e-2 | 16 | zgc:175088 |
| dynll2b1 | 9.8749894 | 0.698828105 | 0.579 | 0.117 | 1.99445162498212e-2 | 16 | dynll2b |
| slc16a101 | 1.5817660 | 0.558560578 | 0.351 | 0.046 | 3.19469295077693e-2 | 16 | slc16a10 |
| zgc:560951 | 1.7533695 | 0.534939439 | 0.456 | 0.075 | 3.5412804906271e-20 | 16 | zgc:56095 |
| slit21 | 1.8855503 | 0.422957055 | 0.263 | 0.024 | 3.80824605849331e-2 | 16 | slit2 |
| podxl3 | 2.4283089 | 0.816837015 | 0.561 | 0.105 | 4.90445550808695e-2 | 16 | podxl |
| ttll122 | 2.7959985 | 0.571190443 | 0.246 | 0.022 | 5.64707827236772e-2 | 16 | ttll12 |
| CABZ01068358.11 | 2.8863080 | 0.699855529 | 0.456 | 0.077 | 5.82947645841458e-2 | 16 | CABZ01068358.1 |
| dnmt3bb.11 | 2.9391782 | 0.541215183 | 0.351 | 0.046 | 5.93625838762748e-2 | 16 | dnmt3bb.1 |
| irge4 | 4.8924608 | 0.281526577 | 0.175 | 0.01 | 9.88130314575003e-2 | 16 | irge4 |
| ehd32 | 5.0663904 | 0.459722327 | 0.263 | 0.025 | 1.02325887022719e-1 | 16 | ehd3 |
| zgc:927444 | 5.7229835 | 0.922491998 | 0.754 | 0.196 | 1.1558709926196e-19 | 16 | zgc:92744 |
| sulf1 | 6.1290078 | 0.370613999 | 0.228 | 0.018 | 1.23787572446893e-1 | 16 | sulf1 |
| CABZ01086029.1 | 8.7514813 | 0.440752926 | 0.298 | 0.033 | 1.76753668916141e-1 | 16 | CABZ01086029.1 |
| znf11041 | 9.5352319 | 0.445037547 | 0.228 | 0.019 | 1.92583080148879e-1 | 16 | znf1104 |

| | | | | | | | |
|---|---|---|---|---|---|---|---|
| zgc:123010 | 1.1248373 | 0.509072124 | 0.298 | 0.033 | 2.27183408061871e-1 | 16 | zgc:123010 |
| gstm.36 | 1.4297613 | 1.153034033 | 0.86 | 0.276 | 2.88768899846091e-1 | 16 | gstm.3 |
| steap41 | 1.4766818 | 0.397026589 | 0.193 | 0.013 | 2.98245442429882e-1 | 16 | steap4 |
| tle2b1 | 1.6232790 | 0.421473894 | 0.281 | 0.029 | 3.27853673946433e-1 | 16 | tle2b |
| rsad2 | 1.8086694 | 0.293002376 | 0.158 | 0.008 | 3.652969730144e-19 | 16 | rsad2 |
| map6a | 2.3324103 | 0.258447405 | 0.14 | 0.006 | 4.7107691142975e-19 | 16 | map6a |
| mxra8a2 | 2.7930747 | 0.780325459 | 0.614 | 0.139 | 5.64117308576656e-1 | 16 | mxra8a |
| slc44a5b | 2.8480730 | 0.410012758 | 0.263 | 0.026 | 5.75225312598607e-1 | 16 | slc44a5b |
| parp12a1 | 9.0291645 | 0.481239377 | 0.316 | 0.039 | 1.82362036845206e-1 | 16 | parp12a |
| si:dkey-238i5.2 | 1.3828718 | 0.315281504 | 0.193 | 0.014 | 2.79298629536218e-1 | 16 | si:dkey-238i5.2 |
| pde7a1 | 1.6565286 | 0.61208519 | 0.474 | 0.087 | 3.3456908680734e-18 | 16 | pde7a |
| cxxc5a2 | 1.7898976 | 0.592682261 | 0.456 | 0.079 | 3.61505624830897e-1 | 16 | cxxc5a |
| metrnla2 | 2.1779714 | 0.675510798 | 0.526 | 0.104 | 4.39884902133705e-1 | 16 | metrnla |
| svild3 | 3.9824654 | 0.668135768 | 0.491 | 0.093 | 8.04338556527518e-1 | 16 | svild |
| tnfrsfa1 | 4.6249090 | 0.454821034 | 0.333 | 0.044 | 9.34092879169061e-1 | 16 | tnfrsfa |
| mmp26 | 6.4269323 | 0.782443222 | 0.772 | 0.206 | 1.29804752103728e-1 | 16 | mmp2 |
| tmem50a3 | 7.1648126 | 0.744873494 | 0.684 | 0.188 | 1.44707721669452e-1 | 16 | tmem50a |
| tmed102 | 9.1103046 | 0.736837982 | 0.772 | 0.232 | 1.84000822554e-17 | 16 | tmed10 |
| tmed92 | 1.2975366 | 0.681231512 | 0.632 | 0.155 | 2.62063477870775e-1 | 16 | tmed9 |
| uxs11 | 1.5648211 | 0.444597581 | 0.316 | 0.041 | 3.16046926394613e-1 | 16 | uxs1 |
| slc29a1a1 | 2.4459643 | 0.491259132 | 0.316 | 0.041 | 4.94011415744743e-1 | 16 | slc29a1a |
| wu:fj29h111 | 2.5658123 | 0.355989985 | 0.246 | 0.025 | 5.18217121664333e-1 | 16 | wu:fj29h11 |
| dpp32 | 6.3500452 | 0.635437233 | 0.439 | 0.081 | 1.2825186391122e-16 | 16 | dpp3 |
| spry22 | 7.0029486 | 0.640354569 | 0.526 | 0.112 | 1.41438553543401e-1 | 16 | spry2 |
| mgll1 | 7.3214813 | 0.269189921 | 0.158 | 0.01 | 1.47871958613643e-1 | 16 | mgll |
| hbegfb | 8.3625651 | 0.325617297 | 0.193 | 0.015 | 1.68898728163057e-1 | 16 | hbegfb |
| atp6v0cb | 8.7606204 | 0.396185935 | 0.298 | 0.037 | 1.76938251574902e-1 | 16 | atp6v0cb |
| dag12 | 1.4612671 | 0.579717883 | 0.439 | 0.08 | 2.95132122577321e-1 | 16 | dag1 |
| pgrmc13 | 1.6528896 | 0.811915414 | 0.702 | 0.214 | 3.33834131169506e-1 | 16 | pgrmc1 |
| tmem88b4 | 1.6902666 | 0.710918045 | 0.579 | 0.131 | 3.41383155125897e-1 | 16 | tmem88b |
| fstl3 | 1.8239790 | 0.470370636 | 0.228 | 0.023 | 3.68389038746273e-1 | 16 | fstl3 |
| rpia2 | 2.2824733 | 0.648500779 | 0.439 | 0.083 | 4.6099114289265e-16 | 16 | rpia |
| man1b1b | 2.2931224 | 0.459942011 | 0.333 | 0.048 | 4.6314194479612e-16 | 16 | man1b1b |
| rnf1301 | 2.5928704 | 0.556258763 | 0.439 | 0.081 | 5.2368204760931e-16 | 16 | rnf130 |
| fkbp1aa4 | 2.6577752 | 0.749508714 | 0.965 | 0.573 | 5.36790862621872e-1 | 16 | fkbp1aa |

| | | | | | | | |
|---|---|---|---|---|---|---|---|
| tbck | 4.0117347 | 0.269037005 | 0.158 | 0.01 | 8.10250070812685e-1 | 16 | tbck |
| atp1b1a3 | 4.0875326 | 0.787407263 | 0.825 | 0.284 | 8.25558964627649e-1 | 16 | atp1b1a |
| mafa | 1.9134710 | 0.272163794 | 0.158 | 0.011 | 3.86463750896395e-1 | 16 | mafa |
| ar2 | 1.9945321 | 0.60035026 | 0.491 | 0.102 | 4.02835662120863e-1 | 16 | ar |
| ap2m1a2 | 2.1453604 | 0.798986337 | 0.807 | 0.295 | 4.33298456388754e-1 | 16 | ap2m1a |
| cldnc2 | 3.0813457 | 0.503787632 | 0.404 | 0.066 | 6.22339406426184e-1 | 16 | cldnc |
| rhoj1 | 3.1141062 | 0.330137068 | 0.211 | 0.02 | 6.28956041988507e-1 | 16 | rhoj |
| sparc7 | 3.1489720 | 0.878020057 | 0.982 | 0.39 | 6.35997875718894e-1 | 16 | sparc |
| spred2b3 | 4.2623125 | 0.698755896 | 0.649 | 0.173 | 8.60859270984009e-1 | 16 | spred2b |
| pfn25 | 4.4589261 | 0.805024047 | 0.842 | 0.309 | 9.00569322195942e-1 | 16 | pfn2 |
| sigmar11 | 6.7139899 | 0.441748255 | 0.333 | 0.052 | 1.35602454890843e-1 | 16 | sigmar1 |
| selenof4 | 7.3505516 | 0.782933555 | 0.895 | 0.415 | 1.48459091726402e-1 | 16 | selenof |
| sdc44 | 8.8679286 | 0.741551894 | 0.789 | 0.254 | 1.79105555471232e-1 | 16 | sdc4 |
| ppfibp1a2 | 9.1211785 | 0.425101904 | 0.316 | 0.046 | 1.84220443045734e-1 | 16 | ppfibp1a |
| tjp1a1 | 9.4035019 | 0.538795613 | 0.421 | 0.079 | 1.89922528308021e-1 | 16 | tjp1a |
| arl4d2 | 9.5019765 | 0.525655045 | 0.386 | 0.067 | 1.91911420004277e-1 | 16 | arl4d |
| mtm1 | 1.2620818 | 0.427142263 | 0.281 | 0.038 | 2.54902680271134e-1 | 16 | mtm1 |
| ccdc125 | 1.5116501 | 0.282946488 | 0.175 | 0.014 | 3.0530797305569e-14 | 16 | ccdc125 |
| si:ch211-214p13.9 | 1.6428047 | 0.402056645 | 0.246 | 0.029 | 3.31797281345971e-1 | 16 | si:ch211-214p13.9 |
| rilpl2 | 1.6713295 | 0.391616993 | 0.263 | 0.033 | 3.37558419693417e-1 | 16 | rilpl2 |
| tegt4 | 1.7637822 | 0.777608414 | 0.842 | 0.354 | 3.5623110416408e-14 | 16 | tegt |
| wu:fc75a091 | 1.8429820 | 0.377145294 | 0.211 | 0.021 | 3.72227083293011e-1 | 16 | wu:fc75a09 |
| ccl19a.12 | 1.8641098 | 0.532804288 | 0.228 | 0.025 | 3.76494267553897e-1 | 16 | ccl19a.1 |
| tnika | 2.3102574 | 0.308718676 | 0.193 | 0.018 | 4.66602687227344e-1 | 16 | tnika |
| myo10l31 | 2.6305158 | 0.358068807 | 0.211 | 0.022 | 5.31285285119663e-1 | 16 | myo10l3 |
| zgc:114181 | 3.9396515 | 0.471364767 | 0.281 | 0.039 | 7.95691428098369e-1 | 16 | zgc:114181 |
| asph2 | 5.2996776 | 0.532203401 | 0.404 | 0.078 | 1.07037589300481e-1 | 16 | asph |
| gdi1 | 8.1931165 | 0.409490956 | 0.263 | 0.035 | 1.65476375552523e-1 | 16 | gdi1 |
| adgrl2a1 | 8.8650781 | 0.414722233 | 0.281 | 0.039 | 1.79047983756804e-1 | 16 | adgrl2a |
| abhd15a1 | 1.3527627 | 0.529508934 | 0.368 | 0.064 | 2.73217492742701e-1 | 16 | abhd15a |
| afap1l1a | 1.4296710 | 0.297897332 | 0.158 | 0.012 | 2.88750671886727e-1 | 16 | afap1l1a |
| arhgap51 | 1.4414328 | 0.427855195 | 0.333 | 0.055 | 2.91126200044952e-1 | 16 | arhgap5 |
| zgc:1008291 | 1.6401753 | 0.569304247 | 0.386 | 0.074 | 3.31266216548674e-1 | 16 | zgc:100829 |
| gas62 | 1.6548607 | 0.484340262 | 0.351 | 0.06 | 3.34232216022734e-1 | 16 | gas6 |
| zgc:1625091 | 1.8017814 | 0.411891271 | 0.316 | 0.049 | 3.63905791783188e-1 | 16 | zgc:162509 |

| | | | | | | | |
|---|---|---|---|---|---|---|---|
| afap1l1b1 | 1.8523537 | 0.317528958 | 0.193 | 0.019 | 3.74119892340108e-1 | 16 | afap1l1b |
| zgc:171686 | 2.4865172 | 0.280417094 | 0.158 | 0.012 | 5.02201880620611e-1 | 16 | zgc:171686 |
| hmgb3a5 | 2.8204850 | 0.760408002 | 0.737 | 0.251 | 5.69653369350543e-1 | 16 | hmgb3a |
| emc101 | 3.0481157 | 0.553698683 | 0.491 | 0.117 | 6.15627941178943e-1 | 16 | emc10 |
| abl2 | 3.2525349 | 0.288488234 | 0.175 | 0.016 | 6.56914491253791e-1 | 16 | abl2 |
| sdf2l12 | 3.4931681 | 0.458848931 | 0.404 | 0.079 | 7.0551517964195e-13 | 16 | sdf2l1 |
| hist2h2l4 | 3.9951228 | 0.710980717 | 0.667 | 0.202 | 8.06894966705737e-1 | 16 | hist2h2l |
| amd1 | 4.6802368 | 0.442072687 | 0.281 | 0.041 | 9.45267432319115e-1 | 16 | amd1 |
| si:dkey-188i13.112 | 5.0559644 | 0.678650579 | 0.614 | 0.176 | 1.02115313409569e-1 | 16 | si:dkey-188i13.11 |
| prnprs3 | 5.2569072 | 0.298301073 | 0.158 | 0.013 | 1.06173755985095e-1 | 16 | prnprs3 |
| ociad2 | 5.9240913 | 0.272048344 | 0.158 | 0.013 | 1.19648872316538e-1 | 16 | ociad2 |
| cnih1 | 6.9148377 | 0.525742282 | 0.421 | 0.089 | 1.39658978742919e-1 | 16 | cnih1 |
| lhfpl2a1 | 7.1125105 | 0.558024295 | 0.404 | 0.083 | 1.43651376534048e-1 | 16 | lhfpl2a |
| cd99l22 | 9.2242907 | 0.51708838 | 0.404 | 0.081 | 1.86303000098445e-1 | 16 | cd99l2 |
| dad13 | 9.9230581 | 0.71817748 | 0.895 | 0.411 | 2.00416006171103e-1 | 16 | dad1 |
| dynll2a1 | 1.0138377 | 0.325618605 | 0.228 | 0.027 | 2.04764801069847e-1 | 16 | dynll2a |
| ifi35 | 1.0770082 | 0.42999523 | 0.298 | 0.048 | 2.17523364509409e-1 | 16 | ifi35 |
| st5 | 1.2021120 | 0.435239468 | 0.263 | 0.037 | 2.42790563766132e-1 | 16 | st5 |
| mphosph82 | 1.2720412 | 0.703313371 | 0.702 | 0.244 | 2.5691417318101e-12 | 16 | mphosph8 |
| csrnp2 | 1.3986912 | 0.328614773 | 0.211 | 0.024 | 2.82493675539484e-1 | 16 | csrnp2 |
| si:dkey-73n10.12 | 1.4559789 | 0.418464591 | 0.246 | 0.033 | 2.9406407523232e-12 | 16 | si:dkey-73n10.1 |
| pdia63 | 1.5261140 | 0.633520319 | 0.579 | 0.163 | 3.0822925504457e-12 | 16 | pdia6 |
| nrp2a1 | 1.8334680 | 0.436313024 | 0.246 | 0.033 | 3.70305546307507e-1 | 16 | nrp2a |
| zfp36l1b4 | 1.9053858 | 0.686651735 | 0.649 | 0.192 | 3.84830782502477e-1 | 16 | zfp36l1b |
| top1l1 | 2.1376126 | 0.640263872 | 0.596 | 0.171 | 4.31733627249908e-1 | 16 | top1l |
| serpinh1a | 2.4168996 | 0.308565366 | 0.193 | 0.02 | 4.88141221591785e-1 | 16 | serpinh1a |
| cox7a11 | 2.5897801 | 0.353114028 | 0.316 | 0.053 | 5.23057904447768e-1 | 16 | cox7a1 |
| lhfpl2b | 2.8306190 | 0.421923673 | 0.298 | 0.048 | 5.71700125236039e-1 | 16 | lhfpl2b |
| ctsla7 | 2.8421254 | 0.667752816 | 0.842 | 0.299 | 5.74024084656847e-1 | 16 | ctsla |
| txndc12 | 2.9462926 | 0.529807117 | 0.404 | 0.085 | 5.95062735688932e-1 | 16 | txndc12 |
| sik11 | 3.9899094 | 0.527235231 | 0.404 | 0.084 | 8.05842010328169e-1 | 16 | sik1 |
| rnf11b | 4.0114714 | 0.319772532 | 0.211 | 0.025 | 8.101968847014e-12 | 16 | rnf11b |
| sema3b1 | 4.4045889 | 0.343171267 | 0.193 | 0.021 | 8.89594822270804e-1 | 16 | sema3b |
| calr3a4 | 4.5134228 | 0.695400477 | 0.684 | 0.228 | 9.11576012086837e-1 | 16 | calr3a |
| si:ch211-106e7.2 | 4.9745201 | 0.313583951 | 0.193 | 0.021 | 1.00470382893875e-1 | 16 | si:ch211-106e7.2 |

| | | | | | | | |
|---|---|---|---|---|---|---|---|
| FAT41 | 5.4872811 | 0.381363573 | 0.263 | 0.038 | 1.10826617535649e-1 | 16 | FAT4 |
| tmem176l.16 | 5.7315862 | 0.735465266 | 0.825 | 0.316 | 1.1576084801431e-11 | 16 | tmem176l.1 |
| zgc:664331 | 5.8540362 | 0.394751859 | 0.298 | 0.048 | 1.18233969853156e-1 | 16 | zgc:66433 |
| si:ch211-236l14.4 | 6.0362826 | 0.287496563 | 0.158 | 0.014 | 1.21914801570655e-1 | 16 | si:ch211-236l14.4 |
| gpx4b1 | 6.5176788 | 0.667320103 | 0.772 | 0.291 | 1.31637559235608e-1 | 16 | gpx4b |
| ackr4b1 | 6.8585354 | 0.424519509 | 0.246 | 0.034 | 1.38521841329814e-1 | 16 | ackr4b |
| pip4p1a | 7.2731545 | 0.255653927 | 0.14 | 0.011 | 1.46895902443944e-1 | 16 | pip4p1a |
| zgc:1101823 | 7.3625978 | 0.481428791 | 0.386 | 0.076 | 1.48702387858025e-1 | 16 | zgc:110182 |
| tmem30ab | 8.1105336 | 0.506340672 | 0.404 | 0.087 | 1.63808447371853e-1 | 16 | tmem30ab |
| fkbp1b1 | 1.1989208 | 0.350175139 | 0.228 | 0.03 | 2.42146050187999e-1 | 16 | fkbp1b |
| hspa54 | 1.3120805 | 0.682576067 | 0.667 | 0.213 | 2.65000906120726e-1 | 16 | hspa5 |
| slc20a2 | 1.9978415 | 0.375667307 | 0.246 | 0.035 | 4.03504066360591e-1 | 16 | slc20a2 |
| lamp23 | 2.0288289 | 0.519496467 | 0.579 | 0.16 | 4.09762587298731e-1 | 16 | lamp2 |
| crybg1a1 | 2.3154115 | 0.342870383 | 0.211 | 0.026 | 4.67643668604597e-1 | 16 | crybg1a |
| phlda22 | 2.3740214 | 0.427627229 | 0.333 | 0.06 | 4.79481110456755e-1 | 16 | phlda2 |
| pgr | 2.9358361 | 0.343481422 | 0.193 | 0.022 | 5.92950820275102e-1 | 16 | pgr |
| h2afx13 | 2.9719683 | 0.747461692 | 0.842 | 0.372 | 6.00248446204051e-1 | 16 | h2afx1 |
| fkbp22 | 3.0312499 | 0.6165653 | 0.632 | 0.206 | 6.12221543517417e-1 | 16 | fkbp2 |
| si:dkey-7j14.66 | 3.2720917 | 0.708164498 | 0.807 | 0.313 | 6.60864379311361e-1 | 16 | si:dkey-7j14.6 |
| plcl11 | 3.5181093 | 0.286006268 | 0.175 | 0.018 | 7.1055255478382e-11 | 16 | plcl1 |
| XLOC-035202 | 3.7440590 | 0.310997134 | 0.158 | 0.015 | 7.56187606684484e-1 | 16 | XLOC-035202 |
| rhoq1 | 3.9334005 | 0.333108561 | 0.211 | 0.026 | 7.94428901327298e-1 | 16 | rhoq |
| smurf21 | 4.1718732 | 0.360368802 | 0.228 | 0.031 | 8.42593233572655e-1 | 16 | smurf2 |
| farp2 | 4.2732473 | 0.264805404 | 0.175 | 0.018 | 8.63067759918209e-1 | 16 | farp2 |
| ca8 | 4.8743633 | 0.266836762 | 0.175 | 0.018 | 9.84475157814555e-1 | 16 | ca8 |
| btr09 | 5.3789122 | 0.273280183 | 0.193 | 0.022 | 1.08637890781194e-1 | 16 | btr09 |
| anxa131 | 6.3910084 | 0.437460527 | 0.368 | 0.074 | 1.29079197215769e-1 | 16 | anxa13 |
| tuba8l36 | 6.7623443 | 0.725070516 | 0.614 | 0.199 | 1.36579068524582e-1 | 16 | tuba8l3 |
| zgc:923131 | 7.8121034 | 0.280715758 | 0.193 | 0.022 | 1.57781053622018e-1 | 16 | zgc:92313 |
| marcksl1b5 | 7.9532798 | 0.822094317 | 0.702 | 0.273 | 1.60632393947581e-1 | 16 | marcksl1b |
| srpra | 7.9685783 | 0.482462199 | 0.351 | 0.072 | 1.60941376836129e-1 | 16 | srpra |
| prdx41 | 8.8552316 | 0.544293157 | 0.684 | 0.229 | 1.78849113963014e-1 | 16 | prdx4 |
| f11r.13 | 9.4059938 | 0.517534402 | 0.421 | 0.098 | 1.89972858180518e-1 | 16 | f11r.1 |
| hp1bp32 | 1.0633891 | 0.608169355 | 0.684 | 0.236 | 2.14772702240409e-1 | 16 | hp1bp3 |
| hs3st3b1b1 | 1.1684942 | 0.265151847 | 0.175 | 0.019 | 2.36000790002856e-1 | 16 | hs3st3b1b |

| | | | | | | | |
|---|---|---|---|---|---|---|---|
| sema7a1 | 1.2633021 | 0.380700258 | 0.228 | 0.031 | 2.55149134011888e-1 | 16 | sema7a |
| fas | 1.2750479 | 0.341397088 | 0.228 | 0.032 | 2.57521436608381e-1 | 16 | fas |
| tpcn2 | 1.3105565 | 0.25768579 | 0.175 | 0.019 | 2.6469310351885e-10 | 16 | tpcn2 |
| nsfl1c1 | 1.4945984 | 0.459306291 | 0.509 | 0.135 | 3.01864046910845e-1 | 16 | nsfl1c |
| wu:fa19b12 | 1.6007926 | 0.28893411 | 0.193 | 0.023 | 3.23312091877254e-1 | 16 | wu:fa19b12 |
| ywhaz2 | 1.6290535 | 0.668592477 | 0.93 | 0.44 | 3.29019938325797e-1 | 16 | ywhaz |
| smad5 | 1.8607039 | 0.419491163 | 0.281 | 0.049 | 3.75806381869842e-1 | 16 | smad5 |
| ndrg21 | 1.9072334 | 0.4659151 | 0.421 | 0.097 | 3.85203940730866e-1 | 16 | ndrg2 |
| inka1a1 | 2.0424116 | 0.449262724 | 0.368 | 0.076 | 4.12505888304856e-1 | 16 | inka1a |
| fam107b1 | 2.0953715 | 0.679785742 | 0.526 | 0.159 | 4.23202195218419e-1 | 16 | fam107b |
| serpinf11 | 2.2981823 | 0.387859679 | 0.298 | 0.052 | 4.64163898941188e-1 | 16 | serpinf1 |
| atf5b1 | 2.4224318 | 0.428804515 | 0.333 | 0.065 | 4.89258560742889e-1 | 16 | atf5b |
| chd4b1 | 2.5206001 | 0.562702454 | 0.491 | 0.135 | 5.09085608366613e-1 | 16 | chd4b |
| wbp1la1 | 2.5485192 | 0.334390744 | 0.211 | 0.028 | 5.14724424244298e-1 | 16 | wbp1la |
| ifi303 | 2.6537792 | 0.718377108 | 0.772 | 0.299 | 5.3598378616313e-10 | 16 | ifi30 |
| reep3b2 | 2.9372154 | 0.47950739 | 0.386 | 0.088 | 5.93229413871368e-1 | 16 | reep3b |
| mydgf | 2.9385007 | 0.443651658 | 0.386 | 0.086 | 5.9348899232723e-10 | 16 | mydgf |
| nars | 3.6911902 | 0.498302111 | 0.491 | 0.133 | 7.45509686013653e-1 | 16 | nars |
| appb2 | 5.2817400 | 0.476546905 | 0.368 | 0.08 | 1.06675304543852e-0 | 16 | appb |
| prkd21 | 5.3129799 | 0.342415466 | 0.246 | 0.038 | 1.07306256874612e-0 | 16 | prkd2 |
| rn7sk1 | 6.5694605 | 0.660140136 | 0.404 | 0.099 | 1.32683393790574e-0 | 16 | rn7sk |
| cd151l2 | 6.7267410 | 0.432816008 | 0.386 | 0.086 | 1.35859989087856e-0 | 16 | cd151l |
| ttpal | 6.8863964 | 0.31634586 | 0.228 | 0.033 | 1.39084548523084e-0 | 16 | ttpal |
| pam1 | 7.0525544 | 0.384381127 | 0.228 | 0.034 | 1.42440443130452e-0 | 16 | pam |
| chd71 | 7.6905014 | 0.384926372 | 0.263 | 0.044 | 1.55325057624296e-0 | 16 | chd7 |
| phgdh | 7.9858459 | 0.258903938 | 0.158 | 0.016 | 1.61290131165497e-0 | 16 | phgdh |
| ralaa | 8.1381505 | 0.271005021 | 0.193 | 0.024 | 1.643662274445e-09 | 16 | ralaa |
| auh | 8.5533226 | 0.356079862 | 0.228 | 0.034 | 1.72751457671678e-0 | 16 | auh |
| elavl41 | 9.8295625 | 0.25898914 | 0.14 | 0.013 | 1.98527675232997e-0 | 16 | elavl4 |
| jak2a | 1.0499842 | 0.291348115 | 0.193 | 0.025 | 2.12065325002106e-0 | 16 | jak2a |
| tbca | 1.0659118 | 0.590800719 | 0.754 | 0.289 | 2.15282217648846e-0 | 16 | tbca |
| lzic | 1.4286200 | 0.338727839 | 0.246 | 0.04 | 2.88538396133324e-0 | 16 | lzic |
| si:ch73-42p12.2 | 1.4386603 | 0.534314226 | 0.544 | 0.168 | 2.90566237680734e-0 | 16 | si:ch73-42p12.2 |
| pcyox12 | 1.6480360 | 0.458226082 | 0.421 | 0.104 | 3.32853849506554e-0 | 16 | pcyox1 |
| lgals9l34 | 1.7571375 | 0.614014287 | 0.561 | 0.18 | 3.54889069785695e-0 | 16 | lgals9l3 |

| | | | | | | | |
|---|---|---|---|---|---|---|---|
| esd1 | 1.8179150 | 0.514634 | 0.561 | 0.171 | 3.67164304361873e-C | 16 | esd |
| gpcpd1 | 1.8272234 | 0.288876585 | 0.175 | 0.021 | 3.69044328706765e-C | 16 | gpcpd1 |
| btbd6b | 2.1797801 | 0.291445734 | 0.193 | 0.025 | 4.40250192800349e-C | 16 | btbd6b |
| pde4ba2 | 2.4179068 | 0.598038806 | 0.526 | 0.157 | 4.88344639716371e-C | 16 | pde4ba |
| si:ch73-1a9.32 | 2.4530324 | 0.649535873 | 0.982 | 0.582 | 4.95438972088697e-C | 16 | si:ch73-1a9.3 |
| gstt1b5 | 2.4552840 | 0.526287495 | 0.579 | 0.175 | 4.9589371895798e-09 | 16 | gstt1b |
| fdxr1 | 2.6566631 | 0.395746052 | 0.228 | 0.035 | 5.36566252049676e-C | 16 | fdxr |
| psmd8 | 2.6968606 | 0.500110385 | 0.509 | 0.146 | 5.4468494459916e-09 | 16 | psmd8 |
| chrac1 | 2.9727181 | 0.433110387 | 0.386 | 0.091 | 6.00399877008196e-C | 16 | chrac1 |
| lamb1a5 | 3.7585254 | 0.587060125 | 0.474 | 0.134 | 7.59109390461118e-C | 16 | lamb1a |
| sdcbp23 | 3.8665293 | 0.622953699 | 0.807 | 0.333 | 7.80922925006441e-C | 16 | sdcbp2 |
| LOC566587 | 4.0496313 | 0.257178218 | 0.175 | 0.021 | 8.17904048227914e-C | 16 | LOC566587 |
| epsti1 | 4.1333463 | 0.258487116 | 0.158 | 0.017 | 8.34811963821265e-C | 16 | epsti1 |
| ppib6 | 4.4295945 | 0.571366955 | 0.947 | 0.639 | 8.94645209802692e-C | 16 | ppib |
| rnf213b1 | 4.9744404 | 0.273576806 | 0.175 | 0.021 | 1.00468773315939e-C | 16 | rnf213b |
| nhsl1a | 5.2322846 | 0.252083271 | 0.14 | 0.014 | 1.05676453320443e-C | 16 | nhsl1a |
| hspg23 | 5.4943262 | 0.367980574 | 0.263 | 0.046 | 1.10968906591198e-C | 16 | hspg2 |
| blmh | 5.5141244 | 0.378024691 | 0.368 | 0.086 | 1.11368770750677e-C | 16 | blmh |
| tmem30aa | 5.5398293 | 0.510845235 | 0.421 | 0.11 | 1.11887933026929e-C | 16 | tmem30aa |
| nab1b | 6.2801385 | 0.320097849 | 0.175 | 0.022 | 1.26839958107748e-C | 16 | nab1b |
| tmem33 | 6.5645959 | 0.442456545 | 0.333 | 0.074 | 1.32585143992285e-C | 16 | tmem33 |
| txndc151 | 7.0822611 | 0.458795182 | 0.404 | 0.104 | 1.43040429264009e-C | 16 | txndc15 |
| tm2d32 | 7.2307104 | 0.4840459 | 0.386 | 0.098 | 1.46038659337109e-C | 16 | tm2d3 |
| ndfip12 | 7.6728583 | 0.608762342 | 0.596 | 0.217 | 1.54968720539597e-C | 16 | ndfip1 |
| tp53 | 9.6554592 | 0.551645399 | 0.544 | 0.175 | 1.95011311028137e-C | 16 | tp53 |
| eef1e1 | 9.8955858 | 0.280493501 | 0.211 | 0.031 | 1.99861148331694e-C | 16 | eef1e1 |
| rer12 | 1.0586983 | 0.511577343 | 0.544 | 0.17 | 2.13825307687463e-C | 16 | rer1 |
| fam13b | 1.0856428 | 0.38909595 | 0.281 | 0.053 | 2.19267284523104e-C | 16 | fam13b |
| emc61 | 1.1140270 | 0.483197253 | 0.439 | 0.12 | 2.2500004996773e-08 | 16 | emc6 |
| bace2 | 1.1287384 | 0.294617971 | 0.211 | 0.032 | 2.27971303412841e-C | 16 | bace2 |
| si:ch211-114c12.2 | 1.2453716 | 0.399820522 | 0.263 | 0.049 | 2.51527714145938e-C | 16 | si:ch211-114c12.2 |
| zgc:1142001 | 1.2705051 | 0.399607236 | 0.263 | 0.048 | 2.56603925471982e-C | 16 | zgc:114200 |
| ccdc6b | 1.3608968 | 0.256090466 | 0.193 | 0.027 | 2.74860345506567e-C | 16 | ccdc6b |
| bcl7ba | 1.4125564 | 0.288461761 | 0.211 | 0.032 | 2.85294020492371e-C | 16 | bcl7ba |
| irf2bp2b3 | 1.4917496 | 0.584855478 | 0.561 | 0.186 | 3.01288676683284e-C | 16 | irf2bp2b |

| | | | | | | | |
|---|---|---|---|---|---|---|---|
| chtopa1 | 1.5221741 | 0.518255855 | 0.474 | 0.139 | 3.0743351041402e-08 | 16 | chtopa |
| swap70b1 | 2.0520891 | 0.379538929 | 0.211 | 0.033 | 4.1446045508953e-08 | 16 | swap70b |
| rbx1 | 2.2203531 | 0.575190841 | 0.86 | 0.375 | 4.48444724366133e-0 | 16 | rbx1 |
| derl3 | 2.5893914 | 0.25305881 | 0.158 | 0.019 | 5.22979390757896e-0 | 16 | derl3 |
| glulb2 | 2.7148154 | 0.487663746 | 0.351 | 0.082 | 5.48311269907268e-0 | 16 | glulb |
| gstm.15 | 2.7231672 | 0.644841008 | 0.807 | 0.331 | 5.49998096153854e-0 | 16 | gstm.1 |
| sumo3a2 | 2.7706124 | 0.610650974 | 0.86 | 0.401 | 5.59580604379628e-0 | 16 | sumo3a |
| zgc:92606 | 2.8976117 | 0.436905635 | 0.386 | 0.096 | 5.85230651861837e-0 | 16 | zgc:92606 |
| tnfrsf9a | 2.9574834 | 0.34665727 | 0.193 | 0.027 | 5.97322940653082e-0 | 16 | tnfrsf9a |
| dhps | 3.0437174 | 0.323697497 | 0.228 | 0.038 | 6.14739613038174e-0 | 16 | dhps |
| itm2ba4 | 3.1192411 | 0.614435169 | 0.86 | 0.405 | 6.29993134475168e-0 | 16 | itm2ba |
| crema2 | 3.5954947 | 0.510338143 | 0.491 | 0.141 | 7.26182083450667e-0 | 16 | crema |
| fmr11 | 4.5646533 | 0.394681173 | 0.316 | 0.07 | 9.2192303740345e-08 | 16 | fmr1 |
| mrpl38 | 4.8253598 | 0.317177898 | 0.246 | 0.045 | 9.74577919937829e-0 | 16 | mrpl38 |
| CABZ01088149.1 | 5.0255702 | 0.287570632 | 0.228 | 0.038 | 1.01501441386585e-0 | 16 | CABZ01088149.1 |
| ucp24 | 5.1189798 | 0.599138864 | 0.93 | 0.415 | 1.03388035525841e-0 | 16 | ucp2 |
| si:ch211-286b5.53 | 5.3438084 | 0.563741121 | 0.614 | 0.219 | 1.07928898941515e-0 | 16 | si:ch211-286b5.5 |
| glod5 | 5.8106709 | 0.361774699 | 0.281 | 0.057 | 1.17358121113163e-0 | 16 | glod5 |
| h1f02 | 6.0600967 | 0.575084599 | 0.421 | 0.116 | 1.22395773896393e-0 | 16 | h1f0 |
| sde2 | 6.0950552 | 0.322831278 | 0.246 | 0.045 | 1.23101831437902e-0 | 16 | sde2 |
| si:ch211-1a19.32 | 6.1122887 | 0.527020552 | 0.333 | 0.077 | 1.23449896392295e-0 | 16 | si:ch211-1a19.3 |
| si:ch211-264f5.61 | 6.4226980 | 0.273539805 | 0.14 | 0.015 | 1.29719233424806e-0 | 16 | si:ch211-264f5.6 |
| zgc:1584634 | 6.4254031 | 0.448059745 | 0.421 | 0.114 | 1.29773867972495e-0 | 16 | zgc:158463 |
| CABZ01072614.16 | 7.0717961 | 0.441970172 | 0.544 | 0.169 | 1.4282906642979e-07 | 16 | CABZ01072614.1 |
| rnf213a3 | 7.2322722 | 0.533439836 | 0.439 | 0.13 | 1.46070201835944e-0 | 16 | rnf213a |
| ext2 | 7.8600882 | 0.363035717 | 0.263 | 0.051 | 1.58750201423753e-0 | 16 | ext2 |
| wnt4a1 | 8.9733552 | 0.276232481 | 0.175 | 0.024 | 1.81234855478328e-0 | 16 | wnt4a |
| pag1 | 9.0676290 | 0.289764521 | 0.211 | 0.033 | 1.83138904812271e-0 | 16 | pag1 |
| limch1b1 | 9.2582473 | 0.310328576 | 0.193 | 0.029 | 1.86988820778072e-0 | 16 | limch1b |
| psme21 | 9.9347491 | 0.575822398 | 0.544 | 0.189 | 2.00652128024887e-0 | 16 | psme2 |
| stat1b | 1.0318563 | 0.292888173 | 0.158 | 0.02 | 2.08404030207103e-0 | 16 | stat1b |
| tob1a2 | 1.4102635 | 0.585927618 | 0.561 | 0.201 | 2.84830938915692e-0 | 16 | tob1a |
| robo23 | 1.4213889 | 0.424803073 | 0.368 | 0.09 | 2.87077929356534e-0 | 16 | robo2 |
| arid1b | 1.4345765 | 0.255186587 | 0.175 | 0.024 | 2.89741423532091e-0 | 16 | arid1b |
| txnipa2 | 1.4389264 | 0.64421782 | 0.772 | 0.362 | 2.90619973832253e-0 | 16 | txnipa |

| | | | | | | | |
|---|---|---|---|---|---|---|---|
| btf3l4 | 1.4449439 | 0.491487752 | 0.561 | 0.195 | 2.91835319946164e-0 | 16 | btf3l4 |
| rpe | 1.6123637 | 0.385280479 | 0.333 | 0.079 | 3.25649105863903e-0 | 16 | rpe |
| rras1 | 1.6960886 | 0.420129754 | 0.421 | 0.115 | 3.42559014665992e-0 | 16 | rras |
| nme33 | 1.7786363 | 0.519079952 | 0.491 | 0.156 | 3.59231177467228e-0 | 16 | nme3 |
| ewsr1b | 1.7798599 | 0.437359084 | 0.544 | 0.176 | 3.59478322645301e-0 | 16 | ewsr1b |
| fn1a3 | 1.9481328 | 0.460076712 | 0.579 | 0.175 | 3.93464387058414e-0 | 16 | fn1a |
| prdx31 | 1.9930164 | 0.49300963 | 0.526 | 0.179 | 4.02529525305146e-0 | 16 | prdx3 |
| hsd11b1la1 | 2.0209999 | 0.277080797 | 0.175 | 0.025 | 4.08181363562585e-0 | 16 | hsd11b1la |
| dynlt31 | 2.2068081 | 0.521371486 | 0.596 | 0.216 | 4.45709048116136e-0 | 16 | dynlt3 |
| si:ch211-218c6.8 | 2.3020846 | 0.405776456 | 0.386 | 0.101 | 4.64952027746893e-0 | 16 | si:ch211-218c6.8 |
| purab1 | 2.3490744 | 0.423905305 | 0.404 | 0.108 | 4.74442560670868e-0 | 16 | purab |
| sdf2 | 2.4093480 | 0.333490306 | 0.228 | 0.041 | 4.86616022980196e-0 | 16 | sdf2 |
| ilf21 | 2.4581717 | 0.550310633 | 0.632 | 0.242 | 4.96476953075754e-0 | 16 | ilf2 |
| ddx39b | 2.5466334 | 0.278282011 | 0.211 | 0.035 | 5.14343555026571e-0 | 16 | ddx39b |
| sf3b52 | 2.5849222 | 0.520107006 | 0.544 | 0.191 | 5.22076748146265e-0 | 16 | sf3b5 |
| msi2b1 | 2.7236918 | 0.327269428 | 0.263 | 0.052 | 5.50104044817706e-0 | 16 | msi2b |
| med4 | 2.7905432 | 0.33517199 | 0.228 | 0.041 | 5.63606026591732e-0 | 16 | med4 |
| calub | 2.9414832 | 0.317620311 | 0.211 | 0.035 | 5.94091368275728e-0 | 16 | calub |
| CR376783.1 | 3.0159175 | 0.281499354 | 0.175 | 0.025 | 6.09124874079936e-0 | 16 | CR376783.1 |
| fthl274 | 3.3326401 | 0.554607312 | 0.947 | 0.624 | 6.73093335228927e-0 | 16 | fthl27 |
| flot1b3 | 3.3572726 | 0.515331956 | 0.544 | 0.191 | 6.78068347981373e-0 | 16 | flot1b |
| golga71 | 3.5872378 | 0.47812143 | 0.526 | 0.174 | 7.24514425427671e-0 | 16 | golga7 |
| tmed73 | 3.6416684 | 0.564984874 | 0.702 | 0.301 | 7.35507783185194e-0 | 16 | tmed7 |
| ywhaqb2 | 3.9694577 | 0.573577366 | 0.667 | 0.273 | 8.01711386030302e-0 | 16 | ywhaqb |
| fundc21 | 4.0690643 | 0.443964053 | 0.439 | 0.13 | 8.21828921336029e-0 | 16 | fundc2 |
| psme11 | 4.1632294 | 0.524163563 | 0.596 | 0.223 | 8.40847442385728e-0 | 16 | psme1 |
| tgif12 | 4.6357256 | 0.478159031 | 0.474 | 0.148 | 9.36277511988003e-0 | 16 | tgif1 |
| arf4a | 4.7541567 | 0.292658877 | 0.193 | 0.031 | 9.60197029854023e-0 | 16 | arf4a |
| si:dkey-42i9.6 | 4.8467123 | 0.413949225 | 0.298 | 0.068 | 9.78890498713103e-0 | 16 | si:dkey-42i9.6 |
| b4galt6 | 4.8483678 | 0.382038195 | 0.316 | 0.073 | 9.79224856922561e-0 | 16 | b4galt6 |
| nid1b2 | 4.9957267 | 0.455435059 | 0.474 | 0.136 | 1.00898692919297e-0 | 16 | nid1b |
| mif1 | 5.0314323 | 0.533822664 | 0.719 | 0.308 | 1.01619839328223e-0 | 16 | mif |
| tomm61 | 5.2354632 | 0.499044546 | 0.596 | 0.221 | 1.05740650303345e-0 | 16 | tomm6 |
| atp1a1a.12 | 5.3140315 | 0.469915193 | 0.439 | 0.131 | 1.07327495400236e-0 | 16 | atp1a1a.1 |
| cfap36 | 5.7234842 | 0.307214844 | 0.193 | 0.031 | 1.15597211005049e-0 | 16 | cfap36 |

| | | | | | | | |
|---|---|---|---|---|---|---|---|
| ewsr1a | 5.8494353 | 0.416626603 | 0.351 | 0.09 | 1.18141045444626e-C | 16 | ewsr1a |
| gnb1b3 | 5.9438860 | 0.575570748 | 0.719 | 0.333 | 1.20048666763177e-C | 16 | gnb1b |
| id2b1 | 6.1031766 | 0.418759546 | 0.316 | 0.072 | 1.23265859088545e-C | 16 | id2b |
| ndufc22 | 6.4119258 | 0.514103511 | 0.632 | 0.243 | 1.29501667363258e-C | 16 | ndufc2 |
| ckap44 | 6.5675226 | 0.464775984 | 0.474 | 0.143 | 1.32644255572742e-C | 16 | ckap4 |
| mgst3b5 | 6.5958440 | 0.569837273 | 0.684 | 0.295 | 1.33216263092523e-C | 16 | mgst3b |
| si:ch211-195o20.7 | 7.1817172 | 0.319925733 | 0.263 | 0.053 | 1.45049142660245e-C | 16 | si:ch211-195o20.7 |
| ccnd11 | 7.3142304 | 0.443966736 | 0.368 | 0.098 | 1.47725512641711e-C | 16 | ccnd1 |
| tmem14ca | 7.5329400 | 0.385802267 | 0.404 | 0.116 | 1.52142791101064e-C | 16 | tmem14ca |
| nucb2a1 | 7.6324385 | 0.460883117 | 0.421 | 0.122 | 1.54152361371463e-C | 16 | nucb2a |
| si:dkey-22o12.21 | 7.7603316 | 0.257247434 | 0.175 | 0.026 | 1.56735418734793e-C | 16 | si:dkey-22o12.2 |
| canx2 | 8.2138459 | 0.500616111 | 0.509 | 0.172 | 1.65895046250534e-C | 16 | canx |
| hmg20b | 8.4803577 | 0.310658368 | 0.263 | 0.055 | 1.71277785845269e-C | 16 | hmg20b |
| rbmx1 | 8.5351888 | 0.45670823 | 0.474 | 0.15 | 1.72385209326205e-C | 16 | rbmx |
| nedd8l | 8.7608719 | 0.37815205 | 0.351 | 0.091 | 1.76943329800031e-C | 16 | nedd8l |
| igflr11 | 9.7208674 | 0.418546303 | 0.298 | 0.069 | 1.96332359047399e-C | 16 | igflr1 |
| manf2 | 9.8518144 | 0.44248186 | 0.404 | 0.116 | 1.98977095517617e-C | 16 | manf |
| ostc3 | 1.0403297 | 0.573156931 | 0.614 | 0.246 | 2.10115406624679e-C | 16 | ostc |
| timm44 | 1.0517311 | 0.288908372 | 0.193 | 0.032 | 2.12418136876666e-C | 16 | timm44 |
| sptbn21 | 1.0656367 | 0.352447019 | 0.281 | 0.062 | 2.15226661973199e-C | 16 | sptbn2 |
| ndufb6 | 1.0989278 | 0.429451112 | 0.544 | 0.19 | 2.21950459420565e-C | 16 | ndufb6 |
| pomp | 1.1477961 | 0.551885394 | 0.86 | 0.409 | 2.31820388380353e-C | 16 | pomp |
| actr1 | 1.1770881 | 0.350851276 | 0.298 | 0.068 | 2.37736496067669e-C | 16 | actr1 |
| psmc5 | 1.1801351 | 0.473744861 | 0.526 | 0.184 | 2.38351903577424e-C | 16 | psmc5 |
| rps6ka3a | 1.2371154 | 0.298229729 | 0.246 | 0.048 | 2.49860216863765e-C | 16 | rps6ka3a |
| si:dkey-187e18.11 | 1.3025211 | 0.427063623 | 0.333 | 0.085 | 2.63070201238817e-C | 16 | si:dkey-187e18.1 |
| grinab3 | 1.3063692 | 0.496948115 | 0.509 | 0.173 | 2.63847396711221e-C | 16 | grinab |
| trip12 | 1.3238806 | 0.282816342 | 0.228 | 0.043 | 2.67384184517407e-C | 16 | trip12 |
| krtcap23 | 1.3842922 | 0.541708504 | 0.702 | 0.298 | 2.79585502407901e-C | 16 | krtcap2 |
| cpxm1a2 | 1.4448506 | 0.376982458 | 0.228 | 0.044 | 2.91816485293898e-C | 16 | cpxm1a |
| snrpd1 | 1.4772633 | 0.492935721 | 0.702 | 0.27 | 2.98362871125512e-C | 16 | snrpd1 |
| hnrnpl21 | 1.5310434 | 0.442225547 | 0.474 | 0.152 | 3.09224854360154e-C | 16 | hnrnpl2 |
| psmb10 | 1.5362636 | 0.363684777 | 0.316 | 0.077 | 3.10279163415828e-C | 16 | psmb10 |
| sec11a4 | 1.5718922 | 0.533088223 | 0.649 | 0.265 | 3.17475076191038e-C | 16 | sec11a |
| ppil1 | 1.5768729 | 0.30828247 | 0.246 | 0.049 | 3.18481026619323e-C | 16 | ppil1 |

| | | | | | | | |
|---|---|---|---|---|---|---|---|
| si:ch211-212k18.7: | 1.6025281 | 0.515947841 | 0.526 | 0.18 | 3.23662604137413e-0 | 16 | si:ch211-212k18.7 |
| wls5 | 1.6320251 | 0.477917385 | 0.526 | 0.176 | 3.29620122671406e-0 | 16 | wls |
| npc22 | 1.6591824 | 0.451920893 | 0.702 | 0.263 | 3.35105081285666e-0 | 16 | npc2 |
| pigx1 | 1.7832920 | 0.346470888 | 0.263 | 0.057 | 3.6017149549264e-06 | 16 | pigx |
| dspa2 | 1.9554885 | 0.286053457 | 0.228 | 0.044 | 3.94950022745161e-0 | 16 | dspa |
| fam168a2 | 2.1661690 | 0.44060993 | 0.474 | 0.15 | 4.37501173072526e-0 | 16 | fam168a |
| ndufs2 | 2.2185773 | 0.42493724 | 0.456 | 0.145 | 4.48086058239712e-0 | 16 | ndufs2 |
| tmed22 | 2.2769071 | 0.561826218 | 0.667 | 0.306 | 4.5986693896401e-06 | 16 | tmed2 |
| wt1a2 | 2.3537230 | 0.418367712 | 0.333 | 0.084 | 4.75381436697798e-0 | 16 | wt1a |
| lman1 | 2.4124275 | 0.328511358 | 0.263 | 0.057 | 4.87237990534638e-0 | 16 | lman1 |
| znf609b1 | 2.4382014 | 0.337950378 | 0.246 | 0.05 | 4.92443539503788e-0 | 16 | znf609b |
| rpn13 | 2.4572942 | 0.503389056 | 0.596 | 0.23 | 4.96299728189452e-0 | 16 | rpn1 |
| tnfrsf192 | 2.4876534 | 0.345379163 | 0.246 | 0.05 | 5.02431359699575e-0 | 16 | tnfrsf19 |
| ecm1b4 | 2.5252263 | 0.463871464 | 0.439 | 0.13 | 5.10019963764612e-0 | 16 | ecm1b |
| ndufa7 | 2.5952210 | 0.43724107 | 0.439 | 0.138 | 5.24156804580532e-0 | 16 | ndufa7 |
| csnk1g2a | 3.0024573 | 0.339266387 | 0.263 | 0.058 | 6.06406313309134e-0 | 16 | csnk1g2a |
| srp541 | 3.2950414 | 0.403329745 | 0.316 | 0.081 | 6.65499525810888e-0 | 16 | srp54 |
| rab101 | 3.3702751 | 0.513006356 | 0.737 | 0.313 | 6.8069446773067e-06 | 16 | rab10 |
| oaz1b2 | 3.4759840 | 0.529500112 | 0.93 | 0.517 | 7.02044504467192e-0 | 16 | oaz1b |
| psmb11 | 3.5522647 | 0.524150931 | 0.754 | 0.333 | 7.174509044579e-06 | 16 | psmb1 |
| golt1bb2 | 3.5874240 | 0.3504208 | 0.298 | 0.072 | 7.24552040857114e-0 | 16 | golt1bb |
| pmpcb | 3.8011631 | 0.328517184 | 0.263 | 0.058 | 7.67720912861266e-0 | 16 | pmpcb |
| oaz1a2 | 3.8058859 | 0.525987612 | 0.947 | 0.588 | 7.68674775962066e-0 | 16 | oaz1a |
| sap181 | 3.8721134 | 0.475191915 | 0.596 | 0.234 | 7.82050751735856e-0 | 16 | sap18 |
| CABZ01044746.11 | 4.0166551 | 0.266132843 | 0.193 | 0.033 | 8.11243831183514e-0 | 16 | CABZ01044746.1 |
| usf1l | 4.0951634 | 0.424335912 | 0.368 | 0.106 | 8.27100168469487e-0 | 16 | usf1l |
| atrx2 | 4.3905106 | 0.52417508 | 0.596 | 0.238 | 8.86751438736475e-0 | 16 | atrx |
| rap2c | 4.6901809 | 0.471151668 | 0.404 | 0.125 | 9.47275842851684e-0 | 16 | rap2c |
| atp5f1c2 | 4.8313627 | 0.483791819 | 0.789 | 0.348 | 9.75790343697895e-0 | 16 | atp5f1c |
| crlf31 | 5.1854610 | 0.363658158 | 0.368 | 0.102 | 1.04730756054388e-0 | 16 | crlf3 |
| lman22 | 5.1876288 | 0.498160671 | 0.509 | 0.188 | 1.04774539466548e-0 | 16 | lman2 |
| adamts22 | 6.0509806 | 0.286098179 | 0.211 | 0.039 | 1.22211656994334e-0 | 16 | adamts2 |
| LOC108190056 | 6.0888566 | 0.251971198 | 0.175 | 0.028 | 1.22976637476106e-0 | 16 | LOC108190056 |
| enc12 | 6.3350804 | 0.351453079 | 0.316 | 0.078 | 1.27949618954649e-0 | 16 | enc1 |
| fhl1b1 | 6.5507208 | 0.286499381 | 0.193 | 0.033 | 1.32304908691591e-0 | 16 | fhl1b |

| | | | | | | | |
|---|---|---|---|---|---|---|---|
| pgam5 | 6.6024433 | 0.349757094 | 0.351 | 0.094 | 1.33349547642085e-0 | 16 | pgam5 |
| si:dkey-17m8.11 | 6.7797809 | 0.334319485 | 0.281 | 0.065 | 1.36931234933146e-0 | 16 | si:dkey-17m8.1 |
| tapbp.2 | 6.7944265 | 0.287744427 | 0.158 | 0.024 | 1.37227032699545e-0 | 16 | tapbp.2 |
| mff | 6.9353405 | 0.29911199 | 0.298 | 0.071 | 1.40073072432274e-0 | 16 | mff |
| psmb3 | 7.7246783 | 0.477300509 | 0.632 | 0.251 | 1.5601532826093e-05 | 16 | psmb3 |
| marcksb | 8.1038647 | 0.398263794 | 0.333 | 0.089 | 1.63673755420122e-0 | 16 | marcksb |
| eif2b1 | 8.3606066 | 0.297911312 | 0.246 | 0.053 | 1.68859172784019e-0 | 16 | eif2b1 |
| sumf21 | 8.4461645 | 0.257394908 | 0.175 | 0.029 | 1.70587185175973e-0 | 16 | sumf2 |
| mrgbp | 8.5738520 | 0.294627881 | 0.263 | 0.059 | 1.7316609078229e-05 | 16 | mrgbp |
| dpm3 | 8.8216208 | 0.274405956 | 0.228 | 0.046 | 1.7817027559626e-05 | 16 | dpm3 |
| calr3b2 | 9.1112976 | 0.480865929 | 0.509 | 0.178 | 1.84020878812358e-0 | 16 | calr3b |
| zbtb46 | 1.0309598 | 0.288994803 | 0.211 | 0.04 | 2.08222964315369e-0 | 16 | zbtb46 |
| zgc:1748633 | 1.0328288 | 0.49609945 | 0.456 | 0.162 | 2.08600434758232e-0 | 16 | zgc:174863 |
| selenot22 | 1.0442293 | 0.397162908 | 0.368 | 0.108 | 2.10903000275822e-0 | 16 | selenot2 |
| arhgap35a | 1.0842971 | 0.257663597 | 0.193 | 0.034 | 2.18995493306472e-0 | 16 | arhgap35a |
| stat21 | 1.1018355 | 0.38432437 | 0.281 | 0.067 | 2.22537727685329e-0 | 16 | stat2 |
| slc39a7 | 1.1050344 | 0.376302579 | 0.333 | 0.091 | 2.23183812311019e-0 | 16 | slc39a7 |
| il151 | 1.1434133 | 0.386361549 | 0.298 | 0.075 | 2.30935190424019e-0 | 16 | il15 |
| ptges3b1 | 1.1526196 | 0.467625628 | 0.579 | 0.223 | 2.32794590924588e-0 | 16 | ptges3b |
| dhrs3a2 | 1.2745448 | 0.467809583 | 0.298 | 0.077 | 2.5741982583905e-05 | 16 | dhrs3a |
| psma3 | 1.2778312 | 0.513827136 | 0.649 | 0.28 | 2.5808358435163e-05 | 16 | psma3 |
| zranb2 | 1.2849832 | 0.359353823 | 0.351 | 0.097 | 2.59528076505837e-0 | 16 | zranb2 |
| mmgt11 | 1.2873093 | 0.467278304 | 0.439 | 0.15 | 2.59997874317112e-0 | 16 | mmgt1 |
| yif1a | 1.3750533 | 0.322585787 | 0.263 | 0.06 | 2.77719525351171e-0 | 16 | yif1a |
| ctnna12 | 1.3852832 | 0.327871174 | 0.281 | 0.067 | 2.79785666701784e-0 | 16 | ctnna1 |
| rbb4l | 1.3982794 | 0.365988831 | 0.368 | 0.107 | 2.82410494409984e-0 | 16 | rbb4l |
| nfybb | 1.6065974 | 0.331883819 | 0.246 | 0.054 | 3.24484488750238e-0 | 16 | nfybb |
| tmem256 | 1.6943876 | 0.38507112 | 0.316 | 0.085 | 3.42215466447062e-0 | 16 | tmem256 |
| CABZ01113812.1 | 1.7364626 | 0.286386884 | 0.193 | 0.036 | 3.50713363123483e-0 | 16 | CABZ01113812.1 |
| cmpk | 1.8190179 | 0.393185528 | 0.439 | 0.142 | 3.67387052623041e-0 | 16 | cmpk |
| anapc11 | 1.8588345 | 0.373228372 | 0.333 | 0.093 | 3.75428808741918e-0 | 16 | anapc11 |
| prr12a | 1.9899363 | 0.259474146 | 0.175 | 0.03 | 4.01907435686378e-0 | 16 | prr12a |
| tcima2 | 2.0149305 | 0.303061388 | 0.333 | 0.086 | 4.06955523604973e-0 | 16 | tcima |
| slc38a5b7 | 2.0167642 | 0.411690494 | 0.561 | 0.199 | 4.07325868131259e-0 | 16 | slc38a5b |
| psmc6 | 2.0941053 | 0.414183957 | 0.439 | 0.145 | 4.22946459744254e-0 | 16 | psmc6 |

| | | | | | | | |
|---|---|---|---|---|---|---|---|
| nacc21 | 2.1029084 | 0.252957039 | 0.211 | 0.041 | 4.24724419058603e-0 | 16 | nacc2 |
| nmt1a | 2.1334034 | 0.334374573 | 0.333 | 0.091 | 4.30883485448715e-0 | 16 | nmt1a |
| si:ch211-197h24.9: | 2.3411334 | 0.495506338 | 0.456 | 0.159 | 4.72838728460935e-0 | 16 | si:ch211-197h24.9 |
| spcs32 | 2.3589930 | 0.495682693 | 0.579 | 0.24 | 4.76445826174385e-0 | 16 | spcs3 |
| lsm61 | 2.4447761 | 0.381100485 | 0.561 | 0.201 | 4.93771432689062e-0 | 16 | lsm6 |
| cops4 | 2.4554184 | 0.377690904 | 0.351 | 0.102 | 4.9592085893175e-05 | 16 | cops4 |
| sar1b2 | 2.4816911 | 0.407845984 | 0.421 | 0.136 | 5.01227161613574e-0 | 16 | sar1b |
| ralbp11 | 2.6018726 | 0.384463883 | 0.368 | 0.11 | 5.25500217283515e-0 | 16 | ralbp1 |
| ciapin1 | 2.6731272 | 0.293528753 | 0.281 | 0.068 | 5.39891514951199e-0 | 16 | ciapin1 |
| NC-002333.47 | 2.6848669 | 0.585695064 | 0.965 | 0.58 | 5.42262580442018e-0 | 16 | NC-002333.4 |
| ch25hl31 | 2.7417594 | 0.359958944 | 0.263 | 0.062 | 5.53753156750473e-0 | 16 | ch25hl3 |
| psmd131 | 2.9889431 | 0.410238511 | 0.439 | 0.147 | 6.03676847044408e-0 | 16 | psmd13 |
| rnaseh1 | 3.0314577 | 0.329737828 | 0.246 | 0.057 | 6.12263524279302e-0 | 16 | rnaseh1 |
| selenos1 | 3.3814365 | 0.33782 | 0.263 | 0.062 | 6.82948750006974e-0 | 16 | selenos |
| ppp1cab | 3.4512634 | 0.337870256 | 0.386 | 0.116 | 6.97051688307076e-0 | 16 | ppp1cab |
| psmb71 | 3.5565355 | 0.466115031 | 0.702 | 0.299 | 7.18313475068797e-0 | 16 | psmb7 |
| LEPROTL11 | 3.5932391 | 0.425735843 | 0.386 | 0.121 | 7.25726517646642e-0 | 16 | LEPROTL1 |
| LOC1018844514 | 3.6155010 | 0.460728165 | 0.386 | 0.121 | 7.30222748294188e-0 | 16 | LOC101884451 |
| gpia1 | 3.7934824 | 0.358496236 | 0.421 | 0.134 | 7.66169659091977e-0 | 16 | gpia |
| cdc37 | 3.9476721 | 0.333631395 | 0.263 | 0.063 | 7.9731135041009e-05 | 16 | cdc37 |
| mef2cb1 | 4.0434334 | 0.323040224 | 0.228 | 0.049 | 8.1665225195357e-05 | 16 | mef2cb |
| wu:fu71h07 | 4.1027600 | 0.314715043 | 0.228 | 0.05 | 8.28634448050275e-0 | 16 | wu:fu71h07 |
| bcap311 | 4.1501223 | 0.369788519 | 0.333 | 0.095 | 8.38200217110842e-0 | 16 | bcap31 |
| zgc:193742 | 4.1982752 | 0.261085638 | 0.211 | 0.043 | 8.47925646028382e-0 | 16 | zgc:193742 |
| ywhae2 | 4.3329317 | 0.291469337 | 0.228 | 0.049 | 8.75122229368961e-0 | 16 | ywhae2 |
| ptgr1 | 4.4955534 | 0.42413648 | 0.368 | 0.115 | 9.07966931105147e-0 | 16 | ptgr1 |
| hnrnpa0b3 | 4.5233601 | 0.519935973 | 0.877 | 0.509 | 9.13583050019313e-0 | 16 | hnrnpa0b |
| gmfb2 | 4.5706729 | 0.451722827 | 0.474 | 0.169 | 9.23138811627709e-0 | 16 | gmfb |
| hmgb1a4 | 4.7543647 | 0.538319146 | 0.825 | 0.421 | 9.60239046338336e-0 | 16 | hmgb1a |
| chd4a1 | 4.7903841 | 0.44671717 | 0.509 | 0.188 | 9.67513893246815e-0 | 16 | chd4a |
| eif1axa | 4.9958966 | 0.348777266 | 0.404 | 0.125 | 0.000100902 | 16 | eif1axa |
| ccdc85b | 5.0738441 | 0.328951539 | 0.263 | 0.063 | 0.000102476 | 16 | ccdc85b |
| ap1s21 | 5.1698244 | 0.42054752 | 0.456 | 0.159 | 0.000104415 | 16 | ap1s2 |
| ier3ip11 | 5.3122626 | 0.495039282 | 0.544 | 0.221 | 0.000107292 | 16 | ier3ip1 |
| rbm4.12 | 5.4284599 | 0.341714936 | 0.333 | 0.092 | 0.000109639 | 16 | rbm4.1 |

| | | | | | | | |
|---|---|---|---|---|---|---|---|
| tbl2 | 5.5577994 | 0.252689572 | 0.175 | 0.031 | 0.000112251 | 16 | tbl2 |
| zgc:1629443 | 5.6229314 | 0.493281081 | 0.807 | 0.393 | 0.000113566 | 16 | zgc:162944 |
| axin1 | 5.9372817 | 0.262775128 | 0.193 | 0.037 | 0.000119915 | 16 | axin1 |
| ddah2 | 6.0716738 | 0.363033365 | 0.281 | 0.072 | 0.00012263 | 16 | ddah2 |
| spcs11 | 6.1343408 | 0.539017572 | 0.684 | 0.351 | 0.000123895 | 16 | spcs1 |
| sec22ba1 | 6.1468951 | 0.25834996 | 0.228 | 0.05 | 0.000124149 | 16 | sec22ba |
| kmt2cb1 | 6.6623173 | 0.342569102 | 0.333 | 0.094 | 0.000134559 | 16 | kmt2cb |
| chac11 | 7.0251205 | 0.462614941 | 0.649 | 0.282 | 0.000141886 | 16 | chac1 |
| ube2d2l | 7.0591325 | 0.35708896 | 0.263 | 0.065 | 0.000142573 | 16 | ube2d2l |
| si:ch73-281n10.26 | 7.3200894 | 0.566996406 | 0.842 | 0.427 | 0.000147844 | 16 | si:ch73-281n10.2 |
| bckdk | 7.4226751 | 0.365285426 | 0.316 | 0.088 | 0.000149916 | 16 | bckdk |
| sod21 | 7.5420929 | 0.35966754 | 0.421 | 0.142 | 0.000152328 | 16 | sod2 |
| tnfaip62 | 7.5499796 | 0.285919126 | 0.193 | 0.037 | 0.000152487 | 16 | tnfaip6 |
| mtx2 | 7.7233369 | 0.292234306 | 0.228 | 0.051 | 0.000155988 | 16 | mtx2 |
| stat1a2 | 7.8457256 | 0.431343333 | 0.421 | 0.143 | 0.00015846 | 16 | stat1a |
| rrp36 | 8.0461193 | 0.268618268 | 0.246 | 0.057 | 0.000162507 | 16 | rrp36 |
| hsp90b12 | 8.1401748 | 0.445420126 | 0.474 | 0.171 | 0.000164407 | 16 | hsp90b1 |
| sdhdb | 8.1470987 | 0.359165604 | 0.386 | 0.121 | 0.000164547 | 16 | sdhdb |
| crebzf1 | 8.2083838 | 0.383315566 | 0.439 | 0.146 | 0.000165785 | 16 | crebzf |
| atp10a2 | 8.7136630 | 0.351534238 | 0.281 | 0.072 | 0.00017599 | 16 | atp10a |
| cbx3a | 8.9839180 | 0.432279929 | 0.667 | 0.275 | 0.000181448 | 16 | cbx3a |
| rtn4a2 | 9.0492747 | 0.454555122 | 0.596 | 0.244 | 0.000182768 | 16 | rtn4a |
| rab11fip1b2 | 9.0805414 | 0.393657614 | 0.281 | 0.074 | 0.0001834 | 16 | rab11fip1b |
| ndufab1b | 9.2785563 | 0.378227367 | 0.526 | 0.2 | 0.000187399 | 16 | ndufab1b |
| pkp3b1 | 9.3881341 | 0.323546489 | 0.298 | 0.078 | 0.000189612 | 16 | pkp3b |
| cox5ab | 9.4269837 | 0.408531841 | 0.684 | 0.298 | 0.000190397 | 16 | cox5ab |
| lama44 | 9.5711458 | 0.433354111 | 0.333 | 0.097 | 0.000193308 | 16 | lama4 |
| emd1 | 1.0384851 | 0.4021404 | 0.439 | 0.149 | 0.000209743 | 16 | emd |
| ddx51 | 1.0750204 | 0.514841828 | 0.737 | 0.349 | 0.000217122 | 16 | ddx5 |
| pnp5a3 | 1.0879972 | 0.285438857 | 0.263 | 0.064 | 0.000219743 | 16 | pnp5a |
| rrbp1a1 | 1.0920789 | 0.378453224 | 0.333 | 0.098 | 0.000220567 | 16 | rrbp1a |
| eif4h | 1.1573893 | 0.329511146 | 0.404 | 0.13 | 0.000233758 | 16 | eif4h |
| psmg11 | 1.1587433 | 0.375483711 | 0.316 | 0.091 | 0.000234031 | 16 | psmg1 |
| ndufb41 | 1.1640017 | 0.419200465 | 0.702 | 0.309 | 0.000235093 | 16 | ndufb4 |
| eny2 | 1.1851441 | 0.426403745 | 0.439 | 0.157 | 0.000239364 | 16 | eny2 |

| | | | | | | | |
|---|---|---|---|---|---|---|---|
| vapal | 1.1890574 | 0.31918253 | 0.263 | 0.065 | 0.000240154 | 16 | vapal |
| anp32b1 | 1.3104936 | 0.455042371 | 0.561 | 0.233 | 0.00026468 | 16 | anp32b |
| psmd4b | 1.3303424 | 0.372591755 | 0.386 | 0.124 | 0.000268689 | 16 | psmd4b |
| rnf102 | 1.3512800 | 0.418334164 | 0.544 | 0.209 | 0.000272918 | 16 | rnf10 |
| sec24a | 1.3994808 | 0.269679504 | 0.246 | 0.058 | 0.000282653 | 16 | sec24a |
| tm9sf4 | 1.4768877 | 0.382567098 | 0.281 | 0.077 | 0.000298287 | 16 | tm9sf4 |
| gng73 | 1.4933014 | 0.378724119 | 0.351 | 0.105 | 0.000301602 | 16 | gng7 |
| med281 | 1.4974591 | 0.390909977 | 0.333 | 0.101 | 0.000302442 | 16 | med28 |
| hnrnpl | 1.5536513 | 0.304069482 | 0.281 | 0.073 | 0.000313791 | 16 | hnrnpl |
| smad1 | 1.5653090 | 0.254807829 | 0.175 | 0.032 | 0.000316145 | 16 | smad1 |
| apex1 | 1.6766669 | 0.335525068 | 0.298 | 0.082 | 0.000338636 | 16 | apex1 |
| CABZ01110881.1 | 1.7683187 | 0.324002291 | 0.246 | 0.06 | 0.000357147 | 16 | CABZ01110881.1 |
| mcfd21 | 1.7935149 | 0.309649725 | 0.298 | 0.081 | 0.000362236 | 16 | mcfd2 |
| tor11 | 1.8790697 | 0.364667445 | 0.316 | 0.092 | 0.000379516 | 16 | tor1 |
| lsm41 | 1.9008614 | 0.429458277 | 0.614 | 0.256 | 0.000383917 | 16 | lsm4 |
| zgc:92818 | 2.0208101 | 0.322973244 | 0.263 | 0.067 | 0.000408143 | 16 | zgc:92818 |
| psma11 | 2.1358350 | 0.48000047 | 0.772 | 0.39 | 0.000431375 | 16 | psma1 |
| CABZ01112656.1 | 2.1541987 | 0.318531369 | 0.263 | 0.068 | 0.000435084 | 16 | CABZ01112656.1 |
| sod1 | 2.1739883 | 0.469792751 | 0.807 | 0.416 | 0.00043908 | 16 | sod1 |
| zgc:1232581 | 2.1816557 | 0.341559149 | 0.298 | 0.083 | 0.000440629 | 16 | zgc:123258 |
| etnk1 | 2.2215862 | 0.321452119 | 0.228 | 0.053 | 0.000448694 | 16 | etnk1 |
| si:zfos-464b6.21 | 2.3799807 | 0.317190753 | 0.228 | 0.054 | 0.000480685 | 16 | si:zfos-464b6.2 |
| coil | 2.3942097 | 0.271221174 | 0.211 | 0.046 | 0.000483559 | 16 | coil |
| eif4g2a1 | 2.6540015 | 0.34268015 | 0.316 | 0.092 | 0.000536029 | 16 | eif4g2a |
| si:ch211-132p1.33 | 2.8907174 | 0.465235398 | 0.404 | 0.143 | 0.000583838 | 16 | si:ch211-132p1.3 |
| zgc:770862 | 2.9191033 | 0.393744256 | 0.368 | 0.119 | 0.000589571 | 16 | zgc:77086 |
| chd62 | 3.0574911 | 0.352541479 | 0.316 | 0.092 | 0.000617521 | 16 | chd6 |
| eif4ebp13 | 3.0717126 | 0.450149664 | 0.719 | 0.336 | 0.000620394 | 16 | eif4ebp1 |
| si:dkey-172h23.2 | 3.1814745 | 0.315935215 | 0.228 | 0.054 | 0.000642562 | 16 | si:dkey-172h23.2 |
| MRPS12 | 3.2046775 | 0.324487906 | 0.386 | 0.129 | 0.000647249 | 16 | MRPS12 |
| csgalnact1a1 | 3.2735753 | 0.265842281 | 0.193 | 0.04 | 0.000661164 | 16 | csgalnact1a |
| tpm4a5 | 3.5195948 | 0.507295921 | 0.579 | 0.241 | 0.000710853 | 16 | tpm4a |
| sox4a4 | 3.5332538 | 0.444899494 | 0.439 | 0.154 | 0.000713611 | 16 | sox4a |
| ctnnb13 | 3.6350203 | 0.448181091 | 0.526 | 0.21 | 0.000734165 | 16 | ctnnb1 |
| epha33 | 3.8434395 | 0.287170556 | 0.228 | 0.053 | 0.000776259 | 16 | epha3 |

| | | | | | | | |
|---|---|---|---|---|---|---|---|
| ssr42 | 3.9885378 | 0.47891868 | 0.842 | 0.478 | 0.000805565 | 16 | ssr4 |
| dpy30 | 4.0923844 | 0.362052108 | 0.561 | 0.223 | 0.000826539 | 16 | dpy30 |
| rbm8a | 4.1625064 | 0.431920141 | 0.772 | 0.366 | 0.000840701 | 16 | rbm8a |
| tvp23b | 4.3029044 | 0.270917794 | 0.193 | 0.041 | 0.000869058 | 16 | tvp23b |
| ndufs51 | 4.3338126 | 0.395134726 | 0.561 | 0.232 | 0.0008753 | 16 | ndufs5 |
| atp5meb | 4.5344717 | 0.428982098 | 0.825 | 0.419 | 0.000915827 | 16 | atp5meb |
| med10 | 4.6183119 | 0.412573407 | 0.456 | 0.175 | 0.00093276 | 16 | med10 |
| zfand3 | 4.6358457 | 0.317845102 | 0.368 | 0.117 | 0.000936302 | 16 | zfand3 |
| eif4e1c | 4.7409173 | 0.336196515 | 0.316 | 0.094 | 0.000957523 | 16 | eif4e1c |
| mtpn | 4.7804322 | 0.456047996 | 0.579 | 0.257 | 0.000965504 | 16 | mtpn |
| dpm21 | 4.9513107 | 0.315857175 | 0.298 | 0.085 | 0.001000016 | 16 | dpm2 |
| eif4g1a | 5.1277708 | 0.416169395 | 0.404 | 0.143 | 0.001035656 | 16 | eif4g1a |
| kpna4 | 5.2343904 | 0.261758971 | 0.246 | 0.061 | 0.00105719 | 16 | kpna4 |
| rcan1a2 | 5.2812566 | 0.277584849 | 0.228 | 0.055 | 0.001066655 | 16 | rcan1a |
| prdx61 | 5.3161655 | 0.432859399 | 0.509 | 0.21 | 0.001073706 | 16 | prdx6 |
| lrrc8aa | 5.4107947 | 0.291798609 | 0.263 | 0.069 | 0.001092818 | 16 | lrrc8aa |
| selenok1 | 5.4714082 | 0.396683327 | 0.509 | 0.199 | 0.00110506 | 16 | selenok |
| slka | 5.6464422 | 0.313215957 | 0.228 | 0.055 | 0.001140412 | 16 | slka |
| rnf121 | 5.9452435 | 0.296024835 | 0.193 | 0.043 | 0.001200761 | 16 | rnf121 |
| pfn2l6 | 6.0271184 | 0.452451801 | 0.614 | 0.258 | 0.001217297 | 16 | pfn2l |
| pin12 | 6.3868181 | 0.377484394 | 0.544 | 0.215 | 0.001289946 | 16 | pin1 |
| dnajc8 | 6.5677263 | 0.327940021 | 0.333 | 0.104 | 0.001326484 | 16 | dnajc8 |
| reep52 | 6.7526190 | 0.473856233 | 0.579 | 0.261 | 0.001363826 | 16 | reep5 |
| mtdha2 | 6.9712117 | 0.41980897 | 0.491 | 0.197 | 0.001407976 | 16 | mtdha |
| naa38 | 7.0170818 | 0.30930859 | 0.246 | 0.063 | 0.00141724 | 16 | naa38 |
| psma4 | 7.0441860 | 0.441188067 | 0.632 | 0.284 | 0.001422714 | 16 | psma4 |
| srsf11 | 7.1375761 | 0.380624118 | 0.474 | 0.178 | 0.001441576 | 16 | srsf11 |
| glrx52 | 7.2510938 | 0.332715039 | 0.281 | 0.08 | 0.001464503 | 16 | glrx5 |
| cdv3 | 7.5162428 | 0.391948284 | 0.509 | 0.205 | 0.001518056 | 16 | cdv3 |
| matr3l1.12 | 7.5196124 | 0.336799552 | 0.333 | 0.104 | 0.001518736 | 16 | matr3l1.1 |
| rpz2 | 7.8669409 | 0.307381672 | 0.298 | 0.087 | 0.001588886 | 16 | rpz2 |
| mafba3 | 8.1463934 | 0.38853384 | 0.421 | 0.147 | 0.001645327 | 16 | mafba |
| ube2v21 | 8.2252571 | 0.445678366 | 0.737 | 0.358 | 0.001661255 | 16 | ube2v2 |
| si:ch211-271b14.1 | 8.3694634 | 0.264675289 | 0.211 | 0.049 | 0.001690381 | 16 | si:ch211-271b14.1 |
| tjp2b3 | 8.6244887 | 0.312618771 | 0.281 | 0.078 | 0.001741888 | 16 | tjp2b |

| | | | | | | | |
|---|---|---|---|---|---|---|---|
| golgb1 | 8.7982862 | 0.332888282 | 0.298 | 0.088 | 0.00177699 | 16 | golgb1 |
| calm2b3 | 9.3302661 | 0.500258148 | 0.825 | 0.441 | 0.001884434 | 16 | calm2b |
| hmgb1b3 | 9.4767088 | 0.458281049 | 0.667 | 0.297 | 0.001914011 | 16 | hmgb1b |
| ctbp2a1 | 9.4888483 | 0.294829622 | 0.246 | 0.063 | 0.001916463 | 16 | ctbp2a |
| frg1 | 9.5609830 | 0.280133819 | 0.246 | 0.063 | 0.001931032 | 16 | frg1 |
| coq10b | 1.0354420 | 0.269335813 | 0.211 | 0.05 | 0.002091282 | 16 | coq10b |
| iscub1 | 1.0384430 | 0.340635099 | 0.404 | 0.14 | 0.002097343 | 16 | iscub |
| cgnl12 | 1.0473727 | 0.264978768 | 0.228 | 0.055 | 0.002115379 | 16 | cgnl1 |
| ube2v1 | 1.0903358 | 0.45235077 | 0.649 | 0.314 | 0.002202151 | 16 | ube2v1 |
| stk35l | 1.0956701 | 0.303164746 | 0.246 | 0.064 | 0.002212925 | 16 | stk35l |
| adam19a | 1.1211717 | 0.264736224 | 0.211 | 0.049 | 0.002264431 | 16 | adam19a |
| prrc2a | 1.1263029 | 0.279267197 | 0.228 | 0.056 | 0.002274794 | 16 | prrc2a |
| tmem39a | 1.1315321 | 0.285171523 | 0.263 | 0.072 | 0.002285356 | 16 | tmem39a |
| bcat2 | 1.1612091 | 0.269137255 | 0.175 | 0.036 | 0.002345294 | 16 | bcat2 |
| mgst3a1 | 1.1989437 | 0.353885319 | 0.404 | 0.141 | 0.002421507 | 16 | mgst3a |
| yipf3 | 1.2614341 | 0.264728047 | 0.228 | 0.057 | 0.002547719 | 16 | yipf3 |
| xaf1 | 1.2669695 | 0.324230781 | 0.246 | 0.065 | 0.002558898 | 16 | xaf1 |
| trappc51 | 1.2674396 | 0.293291751 | 0.298 | 0.088 | 0.002559848 | 16 | trappc5 |
| nudcd21 | 1.2871976 | 0.320159737 | 0.298 | 0.09 | 0.002599753 | 16 | nudcd2 |
| ndufa31 | 1.3194248 | 0.409722875 | 0.544 | 0.239 | 0.002664842 | 16 | ndufa3 |
| cltcb | 1.3264363 | 0.259655538 | 0.211 | 0.05 | 0.002679004 | 16 | cltcb |
| CABZ01034698.2 | 1.3390936 | 0.29863066 | 0.246 | 0.065 | 0.002704567 | 16 | CABZ01034698.2 |
| CZQB01141835.1 | 1.4207845 | 0.316468151 | 0.298 | 0.09 | 0.002869558 | 16 | CZQB01141835.1 |
| ankrd111 | 1.4534207 | 0.388251944 | 0.404 | 0.146 | 0.002935474 | 16 | ankrd11 |
| os9 | 1.4629296 | 0.318680847 | 0.281 | 0.081 | 0.002954679 | 16 | os9 |
| ppt11 | 1.4701138 | 0.424576026 | 0.632 | 0.282 | 0.002969189 | 16 | ppt1 |
| erlin11 | 1.4886446 | 0.290396303 | 0.211 | 0.051 | 0.003006616 | 16 | erlin1 |
| atp5mc11 | 1.5317767 | 0.439504274 | 0.702 | 0.331 | 0.003093729 | 16 | atp5mc1 |
| tnrc6a1 | 1.5529564 | 0.336365736 | 0.333 | 0.108 | 0.003136506 | 16 | tnrc6a |
| ssr13 | 1.5923391 | 0.407556415 | 0.596 | 0.271 | 0.003216047 | 16 | ssr1 |
| ube2e32 | 1.6285279 | 0.465445357 | 0.561 | 0.253 | 0.003289138 | 16 | ube2e3 |
| hint11 | 1.6316518 | 0.390993988 | 0.667 | 0.298 | 0.003295447 | 16 | hint1 |
| cnpy41 | 1.6529614 | 0.278833204 | 0.211 | 0.051 | 0.003338486 | 16 | cnpy4 |
| ctsd2 | 1.6598870 | 0.311249491 | 0.491 | 0.183 | 0.003352474 | 16 | ctsd |
| rasl11a | 1.6944133 | 0.258326659 | 0.175 | 0.037 | 0.003422207 | 16 | rasl11a |

| | | | | | | | |
|---|---|---|---|---|---|---|---|
| atf4b3 | 1.7289891 | 0.463690483 | 0.719 | 0.355 | 0.003492039 | 16 | atf4b |
| si:dkeyp-75h12.5 | 1.8059261 | 0.296115877 | 0.263 | 0.073 | 0.003647429 | 16 | si:dkeyp-75h12.5 |
| zgc:193541 | 1.8078168 | 0.438614389 | 0.86 | 0.467 | 0.003651248 | 16 | zgc:193541 |
| eif5a2 | 1.8505799 | 0.427110413 | 0.877 | 0.455 | 0.003737616 | 16 | eif5a2 |
| ap2b1 | 1.8659248 | 0.342923184 | 0.263 | 0.074 | 0.003768608 | 16 | ap2b1 |
| edf11 | 1.8695305 | 0.43733921 | 0.719 | 0.362 | 0.003775891 | 16 | edf1 |
| arl6ip12 | 1.8869268 | 0.412231847 | 0.632 | 0.282 | 0.003811026 | 16 | arl6ip1 |
| ppp2r5d | 1.9425962 | 0.272837602 | 0.228 | 0.057 | 0.003923462 | 16 | ppp2r5d |
| uqcrc11 | 2.0043109 | 0.412526849 | 0.456 | 0.183 | 0.004048107 | 16 | uqcrc1 |
| ddt1 | 2.0186996 | 0.327143509 | 0.386 | 0.136 | 0.004077168 | 16 | ddt |
| rps6kb1b | 2.1047574 | 0.324107377 | 0.333 | 0.109 | 0.004250979 | 16 | rps6kb1b |
| ptenb1 | 2.1279632 | 0.339817843 | 0.368 | 0.124 | 0.004297847 | 16 | ptenb |
| ddx23 | 2.2021142 | 0.327285482 | 0.281 | 0.084 | 0.00444761 | 16 | ddx23 |
| h2afva1 | 2.2830847 | 0.393471893 | 0.544 | 0.229 | 0.004611146 | 16 | h2afva |
| atp5mea | 2.3098396 | 0.37391856 | 0.456 | 0.18 | 0.004665183 | 16 | atp5mea |
| thtpa | 2.3304911 | 0.268214516 | 0.228 | 0.058 | 0.004706893 | 16 | thtpa |
| imp3 | 2.4296129 | 0.275033255 | 0.263 | 0.074 | 0.004907089 | 16 | imp3 |
| vat15 | 2.4418902 | 0.413273472 | 0.544 | 0.221 | 0.004931886 | 16 | vat1 |
| lcorl | 2.5174752 | 0.279925784 | 0.211 | 0.052 | 0.005084545 | 16 | lcorl |
| lsm5 | 2.5400238 | 0.376473969 | 0.544 | 0.232 | 0.005130086 | 16 | lsm5 |
| nufip2 | 2.5555930 | 0.273654899 | 0.211 | 0.052 | 0.005161531 | 16 | nufip2 |
| ik | 2.6004052 | 0.324920313 | 0.281 | 0.085 | 0.005252038 | 16 | ik |
| hnrnpk | 2.6328488 | 0.372015746 | 0.491 | 0.197 | 0.005317565 | 16 | hnrnpk |
| dock7 | 2.7625766 | 0.261027712 | 0.175 | 0.038 | 0.005579576 | 16 | dock7 |
| ddost1 | 2.7694159 | 0.403815704 | 0.404 | 0.154 | 0.005593389 | 16 | ddost |
| si:rp71-45k5.4 | 2.8704010 | 0.379587871 | 0.526 | 0.221 | 0.005797349 | 16 | si:rp71-45k5.4 |
| selenot1b | 2.9461433 | 0.321439667 | 0.351 | 0.12 | 0.005950326 | 16 | selenot1b |
| rtraf | 2.9818579 | 0.325019424 | 0.281 | 0.085 | 0.006022459 | 16 | rtraf |
| bsg4 | 3.0063734 | 0.444980689 | 0.772 | 0.381 | 0.006071972 | 16 | bsg |
| ppp2cb2 | 3.0819341 | 0.434703102 | 0.526 | 0.237 | 0.006224582 | 16 | ppp2cb |
| zgc:919103 | 3.1180809 | 0.411368627 | 0.632 | 0.295 | 0.006297588 | 16 | zgc:91910 |
| zgc:55733 | 3.1264333 | 0.265154011 | 0.211 | 0.052 | 0.006314457 | 16 | zgc:55733 |
| parp12b2 | 3.1459736 | 0.335730951 | 0.263 | 0.077 | 0.006353923 | 16 | parp12b |
| lypla2 | 3.2575167 | 0.299420463 | 0.263 | 0.075 | 0.006579207 | 16 | lypla2 |
| ndufa4l2 | 3.6832861 | 0.442906209 | 0.912 | 0.568 | 0.007439133 | 16 | ndufa4l |

| | | | | | | | |
|---|---|---|---|---|---|---|---|
| nudt211 | 3.8623682 | 0.327561972 | 0.368 | 0.129 | 0.007800825 | 16 | nudt21 |
| lsm7 | 3.9386270 | 0.374995431 | 0.544 | 0.227 | 0.007954845 | 16 | lsm7 |
| arl14ep | 4.0229496 | 0.286835124 | 0.263 | 0.076 | 0.008125151 | 16 | arl14ep |
| gabpa | 4.1162559 | 0.259216887 | 0.246 | 0.067 | 0.008313602 | 16 | gabpa |
| si:ch211-222l21.15 | 4.1679834 | 0.497526531 | 0.912 | 0.526 | 0.008418076 | 16 | si:ch211-222l21.1 |
| chd9 | 4.4684364 | 0.309564743 | 0.246 | 0.069 | 0.009024901 | 16 | chd9 |
| midn5 | 4.5429282 | 0.442703782 | 0.439 | 0.177 | 0.009175352 | 16 | midn |
| ndufa8 | 4.5672439 | 0.33042218 | 0.509 | 0.212 | 0.009224463 | 16 | ndufa8 |
| serinc12 | 4.6434913 | 0.336596726 | 0.333 | 0.111 | 0.00937846 | 16 | serinc1 |
| cox6a11 | 4.6758730 | 0.437565211 | 0.86 | 0.49 | 0.009443861 | 16 | cox6a1 |
| hsbp1a2 | 4.6918628 | 0.297301308 | 0.263 | 0.076 | 0.009476155 | 16 | hsbp1a |
| mfap1 | 5.1526673 | 0.281188869 | 0.263 | 0.076 | 0.010406842 | 16 | mfap1 |
| ctsa2 | 5.3134335 | 0.327003477 | 0.368 | 0.129 | 0.010731542 | 16 | ctsa |
| uqcr103 | 5.4520019 | 0.420211838 | 0.649 | 0.32 | 0.011011408 | 16 | uqcr10 |
| rhoca1 | 5.7536264 | 0.365016921 | 0.333 | 0.114 | 0.011620599 | 16 | rhoca |
| atp5mf2 | 5.7935932 | 0.389389282 | 0.632 | 0.302 | 0.01170132 | 16 | atp5mf |
| manbal | 5.9501788 | 0.266467562 | 0.211 | 0.054 | 0.012017576 | 16 | manbal |
| ndufa61 | 5.9928200 | 0.315565267 | 0.667 | 0.308 | 0.012103699 | 16 | ndufa6 |
| pkig2 | 6.0798264 | 0.373670585 | 0.351 | 0.124 | 0.012279425 | 16 | pkig |
| ndufs71 | 6.4670262 | 0.29268822 | 0.456 | 0.176 | 0.013061453 | 16 | ndufs7 |
| adssl | 6.4924709 | 0.326287711 | 0.386 | 0.138 | 0.013112843 | 16 | adssl |
| arpin | 6.5195232 | 0.252410223 | 0.263 | 0.076 | 0.013167481 | 16 | arpin |
| tars | 6.5397501 | 0.251513648 | 0.281 | 0.084 | 0.013208333 | 16 | tars |
| selenot1a1 | 6.7196811 | 0.327977918 | 0.316 | 0.107 | 0.01357174 | 16 | selenot1a |
| tspan3a1 | 6.7525180 | 0.319242131 | 0.333 | 0.113 | 0.013638061 | 16 | tspan3a |
| ndufb8 | 6.7679621 | 0.424607434 | 0.649 | 0.334 | 0.013669253 | 16 | ndufb8 |
| psmc4 | 6.8338230 | 0.361586685 | 0.421 | 0.164 | 0.013802272 | 16 | psmc4 |
| smarca4a | 6.9296480 | 0.321095179 | 0.298 | 0.098 | 0.01399581 | 16 | smarca4a |
| capns1b1 | 6.9567836 | 0.399945758 | 0.561 | 0.25 | 0.014050616 | 16 | capns1b |
| cux1a3 | 7.1732326 | 0.348255656 | 0.316 | 0.106 | 0.014487778 | 16 | cux1a |
| etfa | 7.1756594 | 0.295416918 | 0.404 | 0.151 | 0.014492679 | 16 | etfa |
| s100v26 | 7.2010415 | 0.373197929 | 0.596 | 0.245 | 0.014543944 | 16 | s100v2 |
| ebag9 | 7.2308750 | 0.290160901 | 0.316 | 0.105 | 0.014604198 | 16 | ebag9 |
| mesd1 | 7.5698499 | 0.304815436 | 0.281 | 0.088 | 0.015288826 | 16 | mesd |
| akirin1 | 7.6813328 | 0.367504315 | 0.333 | 0.12 | 0.015513988 | 16 | akirin1 |

| | | | | | | | |
|---|---|---|---|---|---|---|---|
| perp1 | 7.8206558 | 0.282096811 | 0.298 | 0.093 | 0.015795379 | 16 | perp |
| h3f3d2 | 7.8630725 | 0.402728898 | 0.965 | 0.669 | 0.015881048 | 16 | h3f3d |
| uqcrfs1 | 7.9167570 | 0.330081146 | 0.544 | 0.234 | 0.015989474 | 16 | uqcrfs1 |
| arpc1a3 | 8.1230497 | 0.342847451 | 0.404 | 0.15 | 0.016406123 | 16 | arpc1a |
| abcf11 | 8.1401782 | 0.38308353 | 0.456 | 0.187 | 0.016440718 | 16 | abcf1 |
| purba | 8.6924235 | 0.31927739 | 0.298 | 0.098 | 0.017556088 | 16 | purba |
| sympk | 8.7705923 | 0.258087154 | 0.175 | 0.041 | 0.017713965 | 16 | sympk |
| rab5c1 | 8.7965060 | 0.361214732 | 0.456 | 0.185 | 0.017766303 | 16 | rab5c |
| ap2a1 | 9.1566167 | 0.298807584 | 0.246 | 0.072 | 0.018493619 | 16 | ap2a1 |
| ost42 | 9.3623811 | 0.378514809 | 0.632 | 0.293 | 0.018909201 | 16 | ost4 |
| STMP1 | 9.3770116 | 0.344276546 | 0.526 | 0.23 | 0.01893875 | 16 | STMP1 |
| ubb1 | 9.6018821 | 0.43107537 | 0.719 | 0.37 | 0.019392921 | 16 | ubb |
| rhoua2 | 9.6358738 | 0.301614066 | 0.228 | 0.062 | 0.019461574 | 16 | rhoua |
| csnk1a13 | 9.6483077 | 0.419850348 | 0.561 | 0.266 | 0.019486687 | 16 | csnk1a1 |
| srp19 | 9.8424028 | 0.365527343 | 0.351 | 0.128 | 0.019878701 | 16 | srp19 |
| BX284696.11 | 1.0105632 | 0.348059292 | 0.404 | 0.154 | 0.020410347 | 16 | BX284696.1 |
| pgd3 | 1.0757443 | 0.283821428 | 0.333 | 0.112 | 0.021726809 | 16 | pgd |
| mapre1b | 1.0774962 | 0.292417135 | 0.333 | 0.115 | 0.021762192 | 16 | mapre1b |
| dido11 | 1.0825464 | 0.357503407 | 0.404 | 0.156 | 0.021864191 | 16 | dido1 |
| yipf5 | 1.0893467 | 0.292698739 | 0.263 | 0.08 | 0.022001536 | 16 | yipf5 |
| hyou13 | 1.0984230 | 0.2843666 | 0.263 | 0.079 | 0.02218485 | 16 | hyou1 |
| ap2s1 | 1.1107136 | 0.296727598 | 0.351 | 0.125 | 0.022433083 | 16 | ap2s1 |
| lmf2b | 1.1249209 | 0.27863433 | 0.228 | 0.064 | 0.022720028 | 16 | lmf2b |
| ctbp11 | 1.1314121 | 0.378502048 | 0.421 | 0.167 | 0.02285113 | 16 | ctbp1 |
| trappc6b | 1.1439572 | 0.269473177 | 0.193 | 0.049 | 0.023104504 | 16 | trappc6b |
| si:ch211-51e12.71 | 1.1484028 | 0.377566663 | 0.456 | 0.189 | 0.023194293 | 16 | si:ch211-51e12.7 |
| atp5f1d1 | 1.2224483 | 0.344363048 | 0.719 | 0.36 | 0.024689789 | 16 | atp5f1d |
| ptbp1a | 1.2420092 | 0.348367424 | 0.404 | 0.158 | 0.02508486 | 16 | ptbp1a |
| trappc11 | 1.2591376 | 0.372688555 | 0.474 | 0.201 | 0.025430803 | 16 | trappc1 |
| gcsha | 1.2669132 | 0.25208285 | 0.228 | 0.063 | 0.025587847 | 16 | gcsha |
| mt-nd52 | 1.2748261 | 0.3278995 | 0.439 | 0.178 | 0.025747663 | 16 | mt-nd5 |
| atl3 | 1.2989599 | 0.264086165 | 0.211 | 0.056 | 0.026235095 | 16 | atl3 |
| tmem214 | 1.3102191 | 0.266032624 | 0.228 | 0.064 | 0.026462496 | 16 | tmem214 |
| glula2 | 1.3124606 | 0.298507903 | 0.263 | 0.079 | 0.026507767 | 16 | glula |
| rab6a | 1.3562429 | 0.26406264 | 0.246 | 0.071 | 0.02739204 | 16 | rab6a |

| | | | | | | | |
|---|---|---|---|---|---|---|---|
| zc4h2 | 1.3659334 | 0.321394426 | 0.298 | 0.101 | 0.027587758 | 16 | zc4h2 |
| tmem45a | 1.6242145 | 0.264995766 | 0.175 | 0.042 | 0.032804262 | 16 | tmem45a |
| calm1a1 | 1.6290214 | 0.346548983 | 0.526 | 0.225 | 0.032901347 | 16 | calm1a |
| wdr83os1 | 1.6485891 | 0.342244024 | 0.439 | 0.179 | 0.033296555 | 16 | wdr83os |
| zbtb201 | 1.6574371 | 0.359159322 | 0.386 | 0.148 | 0.033475258 | 16 | zbtb20 |
| atp5pd | 1.7187354 | 0.360132753 | 0.737 | 0.371 | 0.034713299 | 16 | atp5pd |
| hsdl2 | 1.7565280 | 0.285798373 | 0.298 | 0.1 | 0.035476597 | 16 | hsdl2 |
| tmem591 | 1.7649360 | 0.389280478 | 0.474 | 0.207 | 0.035646414 | 16 | tmem59 |
| gtf2h51 | 1.7903196 | 0.270865369 | 0.316 | 0.107 | 0.036159086 | 16 | gtf2h5 |
| hnrnpabb1 | 1.8116812 | 0.444952498 | 0.754 | 0.401 | 0.036590525 | 16 | hnrnpabb |
| ptp4a2a1 | 1.8201435 | 0.301443078 | 0.316 | 0.11 | 0.036761439 | 16 | ptp4a2a |
| praf2 | 1.8821142 | 0.267064178 | 0.211 | 0.058 | 0.038013061 | 16 | praf2 |
| mrps11 | 1.9007011 | 0.263807109 | 0.298 | 0.099 | 0.03838846 | 16 | mrps11 |
| puf60a | 1.9502501 | 0.287411891 | 0.351 | 0.124 | 0.039389202 | 16 | puf60a |
| irf9 | 1.9716748 | 0.270752592 | 0.175 | 0.042 | 0.039821918 | 16 | irf9 |
| inip | 1.9906207 | 0.266039538 | 0.263 | 0.082 | 0.040204568 | 16 | inip |
| hdac1 | 2.1142524 | 0.388277844 | 0.579 | 0.273 | 0.042701556 | 16 | hdac1 |
| calm3b1 | 2.1263526 | 0.427693615 | 0.737 | 0.404 | 0.042945945 | 16 | calm3b |
| krt18a.17 | 2.1532215 | 0.39444954 | 0.614 | 0.276 | 0.043488616 | 16 | krt18a.1 |
| rsf1b.11 | 2.1573771 | 0.307010714 | 0.298 | 0.101 | 0.043572547 | 16 | rsf1b.1 |
| ddx39ab | 2.2375309 | 0.296143573 | 0.456 | 0.186 | 0.045191412 | 16 | ddx39ab |
| tma71 | 2.2415673 | 0.410238648 | 0.86 | 0.466 | 0.045272935 | 16 | tma7 |
| pfdn12 | 2.3069373 | 0.328652468 | 0.509 | 0.222 | 0.046593213 | 16 | pfdn1 |
| chmp6b | 2.3291424 | 0.324205389 | 0.281 | 0.095 | 0.04704169 | 16 | chmp6b |
| fubp1 | 2.3374289 | 0.329362296 | 0.368 | 0.139 | 0.047209052 | 16 | fubp1 |
| fam133b1 | 2.3635881 | 0.313907465 | 0.491 | 0.208 | 0.047737389 | 16 | fam133b |
| uri1 | 2.4346162 | 0.308383353 | 0.246 | 0.076 | 0.049171944 | 16 | uri1 |
| mrpl30 | 2.5195915 | 0.295775904 | 0.351 | 0.131 | 0.050888191 | 16 | mrpl30 |
| ddx612 | 2.6314600 | 0.403879569 | 0.667 | 0.338 | 0.0531476 | 16 | ddx61 |
| tnrc6c14 | 2.6443255 | 0.378096415 | 0.526 | 0.237 | 0.053407442 | 16 | tnrc6c1 |
| wu:fi37e091 | 2.7349546 | 0.339249792 | 0.456 | 0.19 | 0.055237879 | 16 | wu:fi37e09 |
| laptm4a2 | 2.7390179 | 0.326002213 | 0.456 | 0.189 | 0.055319945 | 16 | laptm4a |
| eif4a1a1 | 2.7486178 | 0.379030691 | 0.667 | 0.329 | 0.055513834 | 16 | eif4a1a |
| ube2d2 | 2.9937318 | 0.392271088 | 0.544 | 0.264 | 0.060464402 | 16 | ube2d2 |
| sf3a21 | 3.0035890 | 0.355965782 | 0.404 | 0.166 | 0.060663488 | 16 | sf3a2 |

| | | | | | | | |
|---|---|---|---|---|---|---|---|
| sypl2a1 | 3.0372059 | 0.281734886 | 0.298 | 0.099 | 0.061342448 | 16 | sypl2a |
| atp6ap21 | 3.0470617 | 0.333437969 | 0.351 | 0.134 | 0.061541507 | 16 | atp6ap2 |
| dda1 | 3.0961924 | 0.258047006 | 0.246 | 0.074 | 0.0625338 | 16 | dda1 |
| si:ch73-138n13.11 | 3.1034165 | 0.281117166 | 0.246 | 0.074 | 0.062679703 | 16 | si:ch73-138n13.1 |
| chchd2 | 3.1693582 | 0.299232414 | 0.316 | 0.111 | 0.064011528 | 16 | chchd2 |
| gabarapb1 | 3.3171299 | 0.411094631 | 0.789 | 0.446 | 0.066996073 | 16 | gabarapb |
| yeats4 | 3.4218827 | 0.298760829 | 0.281 | 0.095 | 0.069111765 | 16 | yeats4 |
| si:dkey-88p24.11 | 3.4562137 | 0.265265017 | 0.263 | 0.084 | 0.06980515 | 16 | si:dkey-88p24.11 |
| cmtm61 | 3.5456353 | 0.316014196 | 0.263 | 0.088 | 0.071611197 | 16 | cmtm6 |
| rab14 | 3.5571570 | 0.294913983 | 0.333 | 0.121 | 0.071843901 | 16 | rab14 |
| hbp11 | 3.5777823 | 0.312761114 | 0.228 | 0.068 | 0.072260471 | 16 | hbp1 |
| ndufs6 | 3.6232468 | 0.334138592 | 0.456 | 0.198 | 0.073178716 | 16 | ndufs6 |
| si:ch211-85n16.3 | 3.6560261 | 0.257499301 | 0.211 | 0.06 | 0.073840761 | 16 | si:ch211-85n16.3 |
| srsf5a3 | 3.7207887 | 0.398690922 | 0.719 | 0.384 | 0.075148771 | 16 | srsf5a |
| si:dkeyp-27c8.22 | 3.7857747 | 0.27696528 | 0.211 | 0.059 | 0.076461293 | 16 | si:dkeyp-27c8.2 |
| tle2a | 3.7946517 | 0.255598257 | 0.158 | 0.037 | 0.07664058 | 16 | tle2a |
| gstk1 | 3.9817640 | 0.314846821 | 0.316 | 0.115 | 0.080419689 | 16 | gstk1 |
| dctn3 | 4.1459893 | 0.33426601 | 0.404 | 0.166 | 0.083736548 | 16 | dctn3 |
| eif1axb | 4.1988168 | 0.390314037 | 0.632 | 0.336 | 0.084803504 | 16 | eif1axb |
| txndc17 | 4.3654499 | 0.274360709 | 0.246 | 0.077 | 0.088168994 | 16 | txndc17 |
| txnl4a1 | 4.4734132 | 0.303541778 | 0.281 | 0.097 | 0.090349528 | 16 | txnl4a |
| erp441 | 4.6057364 | 0.320048935 | 0.368 | 0.144 | 0.093022059 | 16 | erp44 |
| hmga1a2 | 4.6933158 | 0.399730704 | 0.807 | 0.429 | 0.094790899 | 16 | hmga1a |
| zgc:1951751 | 4.7179837 | 0.258743727 | 0.175 | 0.045 | 0.095289118 | 16 | zgc:195175 |
| zgc:100918 | 4.7358864 | 0.271722875 | 0.246 | 0.077 | 0.095650698 | 16 | zgc:100918 |
| psmd14 | 4.7539435 | 0.343618245 | 0.386 | 0.159 | 0.096015397 | 16 | psmd14 |
| rsrc2 | 4.8769111 | 0.259025295 | 0.246 | 0.077 | 0.098498975 | 16 | rsrc2 |
| si:ch211-213a13.1[illegible] | 5.1947409 | 0.359069745 | 0.368 | 0.149 | 0.104918183 | 16 | si:ch211-213a13.1 |
| cdh24 | 5.1957792 | 0.347371199 | 0.333 | 0.125 | 0.104939153 | 16 | cdh2 |
| sf3b6 | 5.2179898 | 0.359220577 | 0.632 | 0.31 | 0.105387741 | 16 | sf3b6 |
| gstr2 | 5.2397083 | 0.293060456 | 0.421 | 0.175 | 0.105826389 | 16 | gstr |
| tm2d2 | 5.3475583 | 0.256907785 | 0.211 | 0.061 | 0.108004637 | 16 | tm2d2 |
| gstm.2 | 5.5087486 | 0.275292312 | 0.316 | 0.113 | 0.111260196 | 16 | gstm.2 |
| rca2.12 | 5.6112708 | 0.282056588 | 0.263 | 0.087 | 0.113330838 | 16 | rca2.1 |
| prkcsh1 | 5.6439189 | 0.302747476 | 0.298 | 0.106 | 0.113990231 | 16 | prkcsh |

| | | | | | | | |
|---|---|---|---|---|---|---|---|
| mob4 | 5.6903062 | 0.306454263 | 0.316 | 0.117 | 0.114927115 | 16 | mob4 |
| ppp2r1ba | 5.7132388 | 0.310531476 | 0.351 | 0.137 | 0.115390286 | 16 | ppp2r1ba |
| tpp12 | 5.7881951 | 0.285024892 | 0.281 | 0.093 | 0.116904177 | 16 | tpp1 |
| si:dkey-46a10.3 | 5.7973305 | 0.250718161 | 0.228 | 0.069 | 0.117088686 | 16 | si:dkey-46a10.3 |
| khdrbs1a1 | 5.8657737 | 0.405411913 | 0.93 | 0.59 | 0.118471032 | 16 | khdrbs1a |
| cnpy12 | 5.8825642 | 0.294800849 | 0.298 | 0.107 | 0.118810151 | 16 | cnpy1 |
| ssuh2rs1 | 5.8999189 | 0.267002303 | 0.246 | 0.078 | 0.119160663 | 16 | ssuh2rs1 |
| snx12 | 5.9660012 | 0.271369917 | 0.333 | 0.12 | 0.120495327 | 16 | snx12 |
| znf593 | 6.0520668 | 0.281194453 | 0.316 | 0.116 | 0.122233594 | 16 | znf593 |
| mpc21 | 6.4971536 | 0.302490362 | 0.316 | 0.118 | 0.131223012 | 16 | mpc2 |
| lrpap11 | 6.5903685 | 0.278525104 | 0.298 | 0.105 | 0.133105673 | 16 | lrpap1 |
| ndufb21 | 7.0137505 | 0.344685739 | 0.509 | 0.238 | 0.141656719 | 16 | ndufb2 |
| kmt2bb3 | 7.0400216 | 0.343329253 | 0.368 | 0.149 | 0.142187317 | 16 | kmt2bb |
| yap12 | 7.1041299 | 0.289397608 | 0.281 | 0.095 | 0.143482113 | 16 | yap1 |
| psmc2 | 7.5889839 | 0.280839978 | 0.386 | 0.153 | 0.153274709 | 16 | psmc2 |
| mrps36 | 7.8195244 | 0.304128859 | 0.368 | 0.151 | 0.157930936 | 16 | mrps36 |
| aurkaip1 | 7.9198093 | 0.276476098 | 0.228 | 0.072 | 0.159956389 | 16 | aurkaip1 |
| si:dkey-6n21.121 | 8.1290797 | 0.252917862 | 0.246 | 0.077 | 0.164183023 | 16 | si:dkey-6n21.12 |
| nucks1a3 | 8.2601213 | 0.388928028 | 0.491 | 0.229 | 0.16682967 | 16 | nucks1a |
| paip2b2 | 8.3715799 | 0.379167499 | 0.649 | 0.339 | 0.1690808 | 16 | paip2b |
| sept7b2 | 8.3972319 | 0.285925261 | 0.386 | 0.155 | 0.169598894 | 16 | sept7b |
| eif5b1 | 8.7949588 | 0.350617336 | 0.456 | 0.202 | 0.177631783 | 16 | eif5b |
| psmg3 | 9.0744591 | 0.302069602 | 0.351 | 0.139 | 0.183276851 | 16 | psmg3 |
| eif5 | 9.0795708 | 0.369726548 | 0.526 | 0.26 | 0.183380093 | 16 | eif5 |
| calr2 | 9.6333340 | 0.303564129 | 0.368 | 0.146 | 0.194564447 | 16 | calr |
| jagn1b | 9.6497295 | 0.291850791 | 0.316 | 0.118 | 0.194895587 | 16 | jagn1b |
| mrps16 | 9.7327890 | 0.262806838 | 0.246 | 0.081 | 0.19657314 | 16 | mrps16 |
| snx62 | 9.7928119 | 0.259941876 | 0.281 | 0.097 | 0.197785424 | 16 | snx6 |
| ap1m1 | 9.8687062 | 0.302226196 | 0.298 | 0.109 | 0.19931826 | 16 | ap1m1 |
| echs11 | 9.9000158 | 0.292086675 | 0.351 | 0.138 | 0.199950621 | 16 | echs1 |
| ndufs8a | 9.9299352 | 0.26097964 | 0.298 | 0.108 | 0.200554902 | 16 | ndufs8a |
| dynll12 | 9.9557869 | 0.383222166 | 0.754 | 0.392 | 0.20107703 | 16 | dynll1 |
| elocb1 | 9.9789511 | 0.375438458 | 0.561 | 0.289 | 0.201544876 | 16 | elocb |
| mrpl36 | 1.0123379 | 0.317715177 | 0.421 | 0.183 | 0.204461892 | 16 | mrpl36 |
| nutf2 | 1.0157470 | 0.27484474 | 0.298 | 0.108 | 0.205150423 | 16 | nutf2 |

| | | | | | | | |
|---|---|---|---|---|---|---|---|
| smim151 | 1.0577822 | 0.29918876 | 0.474 | 0.207 | 0.213640288 | 16 | smim15 |
| snd12 | 1.0610260 | 0.278231947 | 0.421 | 0.181 | 0.214295422 | 16 | snd1 |
| p4hb2 | 1.1168820 | 0.337879043 | 0.474 | 0.215 | 0.225576665 | 16 | p4hb |
| snrnp70 | 1.1594989 | 0.262285577 | 0.298 | 0.107 | 0.234184008 | 16 | snrnp70 |
| cdc421 | 1.1821551 | 0.353149384 | 0.684 | 0.355 | 0.238759882 | 16 | cdc42 |
| higd1a2 | 1.1853039 | 0.369310413 | 0.561 | 0.271 | 0.239395847 | 16 | higd1a |
| cyfip12 | 1.2071294 | 0.271656216 | 0.263 | 0.089 | 0.243803928 | 16 | cyfip1 |
| helz | 1.2409756 | 0.259496286 | 0.228 | 0.072 | 0.250639844 | 16 | helz |
| CR383676.16 | 1.3034552 | 0.530755528 | 1 | 0.655 | 0.263258848 | 16 | CR383676.1 |
| cebpd6 | 1.3052973 | 0.41207091 | 0.579 | 0.276 | 0.263630901 | 16 | cebpd |
| emc7 | 1.3210565 | 0.267434109 | 0.281 | 0.101 | 0.266813794 | 16 | emc7 |
| rbm4.31 | 1.4091137 | 0.384634339 | 0.632 | 0.332 | 0.284598703 | 16 | rbm4.3 |
| ilf3b | 1.4159245 | 0.293406197 | 0.333 | 0.128 | 0.285974275 | 16 | ilf3b |
| snrpd3l1 | 1.4171620 | 0.294003535 | 0.386 | 0.161 | 0.286224225 | 16 | snrpd3l |
| ing41 | 1.4177036 | 0.254145556 | 0.246 | 0.081 | 0.286333609 | 16 | ing4 |
| snrpe | 1.5055884 | 0.316843243 | 0.561 | 0.271 | 0.3040837 | 16 | snrpe |
| psma5 | 1.5437574 | 0.333726362 | 0.544 | 0.268 | 0.311792691 | 16 | psma5 |
| sltm | 1.5617148 | 0.307111179 | 0.316 | 0.121 | 0.315419548 | 16 | sltm |
| fam162a1 | 1.5720979 | 0.255224971 | 0.298 | 0.109 | 0.317516626 | 16 | fam162a |
| spry41 | 1.5939874 | 0.255594138 | 0.211 | 0.063 | 0.321937653 | 16 | spry4 |
| eef1db2 | 1.6146397 | 0.356440677 | 0.684 | 0.356 | 0.3261088 | 16 | eef1db |
| sem11 | 1.6291984 | 0.361216372 | 0.719 | 0.387 | 0.329049208 | 16 | sem1 |
| ktn11 | 1.6899998 | 0.265260326 | 0.246 | 0.082 | 0.341329269 | 16 | ktn1 |
| hif1ab1 | 1.7319963 | 0.327657591 | 0.404 | 0.174 | 0.34981131 | 16 | hif1ab |
| ndufab1a | 1.7767370 | 0.343401571 | 0.404 | 0.183 | 0.358847574 | 16 | ndufab1a |
| atp6v1e1a | 1.7913401 | 0.262727755 | 0.281 | 0.1 | 0.36179697 | 16 | atp6v1e1a |
| ube2l3b | 1.8085642 | 0.331944983 | 0.474 | 0.223 | 0.36527573 | 16 | ube2l3b |
| hacd2 | 1.8224682 | 0.258607686 | 0.211 | 0.066 | 0.36808392 | 16 | hacd2 |
| rab2a3 | 1.8469173 | 0.31056123 | 0.474 | 0.215 | 0.373021895 | 16 | rab2a |
| il6st1 | 1.8650114 | 0.253316375 | 0.246 | 0.081 | 0.376676362 | 16 | il6st |
| psenen1 | 1.9591467 | 0.324926968 | 0.351 | 0.147 | 0.395688876 | 16 | psenen |
| serinc23 | 2.0861615 | 0.370145643 | 0.421 | 0.197 | 0.42134204 | 16 | serinc2 |
| slc25a3b2 | 2.1524275 | 0.380165684 | 0.965 | 0.567 | 0.4347258 | 16 | slc25a3b |
| h3f3c4 | 2.1868203 | 0.416214592 | 0.877 | 0.524 | 0.441672103 | 16 | h3f3c |
| ptena1 | 2.2144247 | 0.281588224 | 0.333 | 0.131 | 0.44724737 | 16 | ptena |

| | | | | | | | |
|---|---|---|---|---|---|---|---|
| creb3l3l1 | 2.2507567 | 0.298197668 | 0.263 | 0.095 | 0.454585335 | 16 | creb3l3l |
| aldh9a1a.15 | 2.2736732 | 0.44648186 | 0.579 | 0.338 | 0.459213779 | 16 | aldh9a1a.1 |
| glud1a1 | 2.3139273 | 0.277502507 | 0.351 | 0.142 | 0.467343911 | 16 | glud1a |
| vangl2 | 2.3237232 | 0.252459338 | 0.158 | 0.042 | 0.469322379 | 16 | vangl2 |
| si:ch211-191j22.31 | 2.3767197 | 0.312584589 | 0.404 | 0.177 | 0.480026094 | 16 | si:ch211-191j22.3 |
| rmdn12 | 2.4583591 | 0.254053008 | 0.298 | 0.113 | 0.496514789 | 16 | rmdn1 |
| zc3h15 | 2.4822951 | 0.257957012 | 0.263 | 0.095 | 0.501349155 | 16 | zc3h15 |
| sap30l1 | 2.4983715 | 0.293047251 | 0.474 | 0.219 | 0.504596093 | 16 | sap30l |
| eif1b | 2.5078249 | 0.376232157 | 0.807 | 0.501 | 0.506505404 | 16 | eif1b |
| hm131 | 2.5386799 | 0.32002008 | 0.316 | 0.127 | 0.512737184 | 16 | hm13 |
| zgc:77650 | 2.5583253 | 0.25638936 | 0.281 | 0.102 | 0.516704965 | 16 | zgc:77650 |
| tax1bp31 | 2.5984548 | 0.324005026 | 0.386 | 0.167 | 0.524809927 | 16 | tax1bp3 |
| sdhb1 | 2.6294594 | 0.2891245 | 0.351 | 0.145 | 0.531071916 | 16 | sdhb |
| hmgb3b1 | 2.6921407 | 0.283559003 | 0.351 | 0.145 | 0.543731661 | 16 | hmgb3b |
| commd3 | 2.7014905 | 0.260104812 | 0.316 | 0.123 | 0.545620037 | 16 | commd3 |
| gorasp21 | 2.8890081 | 0.272871525 | 0.246 | 0.086 | 0.583492978 | 16 | gorasp2 |
| pdgfaa1 | 2.9062733 | 0.276627892 | 0.193 | 0.059 | 0.586980019 | 16 | pdgfaa |
| ogdha1 | 3.0593611 | 0.290590951 | 0.281 | 0.104 | 0.61789918 | 16 | ogdha |
| ndufa12 | 3.0625041 | 0.312130503 | 0.421 | 0.194 | 0.618533969 | 16 | ndufa12 |
| smdt1b2 | 3.1142077 | 0.313607237 | 0.509 | 0.246 | 0.628976533 | 16 | smdt1b |
| puraa1 | 3.1992979 | 0.297713641 | 0.263 | 0.097 | 0.646162207 | 16 | puraa |
| ndufa1 | 3.2164672 | 0.277879436 | 0.596 | 0.307 | 0.6496299 | 16 | ndufa1 |
| hnrnph1l2 | 3.2390632 | 0.299897589 | 0.316 | 0.129 | 0.654193597 | 16 | hnrnph1l |
| etfb | 3.2716862 | 0.270213148 | 0.368 | 0.157 | 0.660782468 | 16 | etfb |
| glrx31 | 3.4235747 | 0.296982053 | 0.281 | 0.108 | 0.691459399 | 16 | glrx3 |
| psmb5 | 3.5319838 | 0.286343376 | 0.333 | 0.135 | 0.713354787 | 16 | psmb5 |
| cpne11 | 3.5514106 | 0.30263578 | 0.421 | 0.188 | 0.717278402 | 16 | cpne1 |
| cox8a2 | 3.7730310 | 0.362045384 | 0.825 | 0.467 | 0.762039088 | 16 | cox8a |
| qkia4 | 3.8751953 | 0.355008267 | 0.684 | 0.355 | 0.782673206 | 16 | qkia |
| syap1 | 3.9388807 | 0.265383565 | 0.281 | 0.105 | 0.795535745 | 16 | syap1 |
| ube2e1 | 4.0287892 | 0.299246067 | 0.439 | 0.202 | 0.813694567 | 16 | ube2e1 |
| arf2a1 | 4.1095708 | 0.299146973 | 0.368 | 0.158 | 0.830010032 | 16 | arf2a |
| si:ch211-235e9.61 | 4.2063565 | 0.283534639 | 0.456 | 0.212 | 0.849557823 | 16 | si:ch211-235e9.6 |
| ndufb7 | 4.2087685 | 0.265384173 | 0.614 | 0.304 | 0.850044979 | 16 | ndufb7 |
| mat2aa1 | 4.3300601 | 0.26557215 | 0.263 | 0.096 | 0.874542251 | 16 | mat2aa |

| | | | | | | | |
|---|---|---|---|---|---|---|---|
| tomm20a2 | 4.4058476 | 0.327760809 | 0.456 | 0.218 | 0.889849055 | 16 | tomm20a |
| pum1 | 4.4520690 | 0.25627084 | 0.211 | 0.068 | 0.899184379 | 16 | pum1 |
| csnk1da2 | 4.5089942 | 0.280085544 | 0.281 | 0.107 | 0.910681576 | 16 | csnk1da |
| ndufb31 | 4.5225485 | 0.337705983 | 0.614 | 0.334 | 0.913419137 | 16 | ndufb3 |
| COX36 | 4.6459470 | 0.444040851 | 0.895 | 0.592 | 0.938341936 | 16 | COX3 |
| atp5mc3a | 4.7881679 | 0.34843049 | 0.614 | 0.343 | 0.967066276 | 16 | atp5mc3a |
| dnaja2a | 4.7954919 | 0.311140558 | 0.544 | 0.272 | 0.968545508 | 16 | dnaja2a |
| si:ch73-37h15.2 | 4.8546244 | 0.278249973 | 0.211 | 0.07 | 0.980488501 | 16 | si:ch73-37h15.2 |
| ppm1g | 4.8619263 | 0.266185813 | 0.316 | 0.126 | 0.981963257 | 16 | ppm1g |
| rbm19 | 4.9850587 | 0.255246607 | 0.246 | 0.089 | 1 | 16 | rbm19 |
| jpt2 | 5.0321959 | 0.302589607 | 0.491 | 0.238 | 1 | 16 | jpt2 |
| gatad2b1 | 5.0636679 | 0.29034524 | 0.298 | 0.118 | 1 | 16 | gatad2b |
| grpel21 | 5.0762087 | 0.303029556 | 0.316 | 0.132 | 1 | 16 | grpel2 |
| rbms2a1 | 5.1036091 | 0.273166684 | 0.175 | 0.053 | 1 | 16 | rbms2a |
| hspa4b | 5.1841923 | 0.253369774 | 0.298 | 0.118 | 1 | 16 | hspa4b |
| rtf1 | 5.2360737 | 0.299937643 | 0.298 | 0.121 | 1 | 16 | rtf1 |
| sdha1 | 5.3300034 | 0.273051136 | 0.263 | 0.099 | 1 | 16 | sdha |
| chchd6b | 5.3535866 | 0.270457038 | 0.263 | 0.1 | 1 | 16 | chchd6b |
| rab5if | 5.4463722 | 0.271241556 | 0.298 | 0.121 | 1 | 16 | rab5if |
| ndufa9a | 5.5628555 | 0.259039542 | 0.316 | 0.129 | 1 | 16 | ndufa9a |
| atf7ip1 | 5.7109657 | 0.275857352 | 0.281 | 0.109 | 1 | 16 | atf7ip |
| tm9sf2 | 6.0149300 | 0.260577805 | 0.281 | 0.108 | 1 | 16 | tm9sf2 |
| atox11 | 6.1751747 | 0.305766806 | 0.404 | 0.188 | 1 | 16 | atox1 |
| rab11a | 6.1801906 | 0.274004599 | 0.404 | 0.179 | 1 | 16 | rab11a |
| ssr23 | 6.2404456 | 0.334713614 | 0.719 | 0.415 | 1 | 16 | ssr2 |
| CU694197.1 | 6.2924798 | 0.282185491 | 0.456 | 0.216 | 1 | 16 | CU694197.1 |
| bud311 | 6.3001905 | 0.279732889 | 0.491 | 0.24 | 1 | 16 | bud31 |
| hnrnpaba1 | 6.3973278 | 0.37907049 | 0.86 | 0.524 | 1 | 16 | hnrnpaba |
| ywhabl2 | 6.5052772 | 0.360728581 | 0.719 | 0.438 | 1 | 16 | ywhabl |
| zgc:865981 | 6.6027289 | 0.316589008 | 0.333 | 0.145 | 1 | 16 | zgc:86598 |
| mt-nd24 | 6.6323508 | 0.375370666 | 0.667 | 0.37 | 1 | 16 | mt-nd2 |
| fkbp74 | 6.7383509 | 0.262636315 | 0.316 | 0.125 | 1 | 16 | fkbp7 |
| brd2a2 | 6.7972625 | 0.320728433 | 0.333 | 0.144 | 1 | 16 | brd2a |
| stau22 | 6.8335982 | 0.323752298 | 0.351 | 0.154 | 1 | 16 | stau2 |
| acadvl | 6.9439279 | 0.28063652 | 0.281 | 0.112 | 1 | 16 | acadvl |

| | | | | | | | |
|---|---|---|---|---|---|---|---|
| srsf3b1 | 7.0100810 | 0.337710113 | 0.561 | 0.304 | 1 | 16 | srsf3b |
| psmd31 | 7.2749819 | 0.266684355 | 0.351 | 0.151 | 1 | 16 | psmd3 |
| cpsf6 | 7.3719287 | 0.287825459 | 0.351 | 0.154 | 1 | 16 | cpsf6 |
| anxa5b4 | 7.4918841 | 0.279064347 | 0.386 | 0.167 | 1 | 16 | anxa5b |
| eif4a1b1 | 7.5371731 | 0.357134281 | 0.789 | 0.484 | 1 | 16 | eif4a1b |
| atp5pb | 7.5907550 | 0.338188937 | 0.789 | 0.487 | 1 | 16 | atp5pb |
| acin1a4 | 7.7797182 | 0.332066239 | 0.561 | 0.294 | 1 | 16 | acin1a |
| aimp2 | 7.9332930 | 0.276526757 | 0.439 | 0.208 | 1 | 16 | aimp2 |
| rab11ba | 8.0210256 | 0.284622773 | 0.368 | 0.166 | 1 | 16 | rab11ba |
| drap1 | 8.3564825 | 0.28884879 | 0.404 | 0.189 | 1 | 16 | drap1 |
| cox5aa2 | 8.3933103 | 0.294078682 | 0.737 | 0.405 | 1 | 16 | cox5aa |
| ube2ib | 8.4431946 | 0.297716849 | 0.474 | 0.234 | 1 | 16 | ube2ib |
| crk | 8.4624991 | 0.256025838 | 0.193 | 0.062 | 1 | 16 | crk |
| ndufa10 | 8.6673434 | 0.256481577 | 0.333 | 0.142 | 1 | 16 | ndufa10 |
| snu13b1 | 8.6888744 | 0.27592336 | 0.579 | 0.289 | 1 | 16 | snu13b |
| timm23a1 | 8.8478714 | 0.281932468 | 0.386 | 0.176 | 1 | 16 | timm23a |
| b2ml5 | 8.9927912 | 0.357308209 | 0.754 | 0.442 | 1 | 16 | b2ml |
| pfdn5 | 9.0038565 | 0.25259103 | 0.456 | 0.218 | 1 | 16 | pfdn5 |
| degs11 | 9.1265694 | 0.266792242 | 0.298 | 0.121 | 1 | 16 | degs1 |
| gabarapa2 | 9.3028687 | 0.299590164 | 0.509 | 0.255 | 1 | 16 | gabarapa |
| eif4ebp21 | 9.3813241 | 0.304109756 | 0.561 | 0.291 | 1 | 16 | eif4ebp2 |
| pdap1a | 9.4799459 | 0.254156366 | 0.298 | 0.122 | 1 | 16 | pdap1a |
| hmgn62 | 9.6295880 | 0.373243588 | 0.877 | 0.567 | 1 | 16 | hmgn6 |
| cox7a2l2 | 9.8237510 | 0.278703265 | 0.404 | 0.188 | 1 | 16 | cox7a2l |
| vdac12 | 0.0001 | 0.307790774 | 0.386 | 0.18 | 1 | 16 | vdac1 |
| dlst1 | 0.000105 | 0.25551995 | 0.368 | 0.162 | 1 | 16 | dlst |
| cdc42l2 | 0.000105 | 0.308462921 | 0.649 | 0.342 | 1 | 16 | cdc42l |
| taf15 | 0.000107 | 0.314150573 | 0.386 | 0.186 | 1 | 16 | taf15 |
| hnrnpc1 | 0.000108 | 0.270323186 | 0.263 | 0.104 | 1 | 16 | hnrnpc |
| tomm70a | 0.000113 | 0.257332151 | 0.281 | 0.114 | 1 | 16 | tomm70a |
| ywhae13 | 0.000114 | 0.336969953 | 0.649 | 0.376 | 1 | 16 | ywhae1 |
| si:ch73-376l24.22 | 0.000117 | 0.281911695 | 0.368 | 0.165 | 1 | 16 | si:ch73-376l24.2 |
| gtf2e2 | 0.000124 | 0.262815678 | 0.246 | 0.094 | 1 | 16 | gtf2e2 |
| tufm | 0.000128 | 0.254920982 | 0.281 | 0.114 | 1 | 16 | tufm |
| prdx24 | 0.000129 | 0.349099515 | 0.807 | 0.484 | 1 | 16 | prdx2 |

| | | | | | | | | |
|---|---|---|---|---|---|---|---|---|
| flot2a4 | 0.00013 | 0.297710668 | 0.333 | 0.15 | | 1 | 16 | flot2a |
| sat1a.24 | 0.000131 | 0.341649755 | 0.596 | 0.333 | | 1 | 16 | sat1a.2 |
| caprin1b | 0.000132 | 0.265229786 | 0.526 | 0.259 | | 1 | 16 | caprin1b |
| psma6a1 | 0.000134 | 0.25035679 | 0.351 | 0.156 | | 1 | 16 | psma6a |
| ywhah2 | 0.000135 | 0.337210206 | 0.298 | 0.131 | | 1 | 16 | ywhah |
| hnrnpa0a | 0.000135 | 0.279240272 | 0.333 | 0.144 | | 1 | 16 | hnrnpa0a |
| csnk2b | 0.000136 | 0.324547924 | 0.544 | 0.3 | | 1 | 16 | csnk2b |
| LOC1039104384 | 0.000148 | 0.291683723 | 0.386 | 0.183 | | 1 | 16 | LOC103910438 |
| arhgef1b3 | 0.000159 | 0.263701703 | 0.298 | 0.123 | | 1 | 16 | arhgef1b |
| irf21 | 0.000161 | 0.259548993 | 0.263 | 0.106 | | 1 | 16 | irf2 |
| cope | 0.000162 | 0.286756378 | 0.404 | 0.196 | | 1 | 16 | cope |
| vdac32 | 0.00017 | 0.252218182 | 0.509 | 0.25 | | 1 | 16 | vdac3 |
| rad21a2 | 0.000171 | 0.265639754 | 0.509 | 0.247 | | 1 | 16 | rad21a |
| sub1b | 0.000177 | 0.264921303 | 0.404 | 0.19 | | 1 | 16 | sub1b |
| map1lc3b | 0.000178 | 0.261874308 | 0.421 | 0.201 | | 1 | 16 | map1lc3b |
| ndufa111 | 0.000181 | 0.258930311 | 0.421 | 0.206 | | 1 | 16 | ndufa11 |
| tomm51 | 0.000181 | 0.27980831 | 0.596 | 0.317 | | 1 | 16 | tomm5 |
| snrpb2 | 0.000189 | 0.319432022 | 0.719 | 0.418 | | 1 | 16 | snrpb |
| pnn1 | 0.0002 | 0.25975519 | 0.333 | 0.15 | | 1 | 16 | pnn |
| spcs23 | 0.000203 | 0.322416175 | 0.474 | 0.252 | | 1 | 16 | spcs2 |
| pabpn12 | 0.000204 | 0.29944859 | 0.544 | 0.291 | | 1 | 16 | pabpn1 |
| lrrc592 | 0.000212 | 0.25082234 | 0.246 | 0.098 | | 1 | 16 | lrrc59 |
| sub1a | 0.000222 | 0.296930221 | 0.649 | 0.362 | | 1 | 16 | sub1a |
| cyc1 | 0.000227 | 0.25825244 | 0.456 | 0.23 | | 1 | 16 | cyc1 |
| pbx43 | 0.000232 | 0.266788104 | 0.263 | 0.108 | | 1 | 16 | pbx4 |
| vamp3 | 0.00024 | 0.285738386 | 0.439 | 0.221 | | 1 | 16 | vamp3 |
| mycbp1 | 0.000242 | 0.258913932 | 0.456 | 0.225 | | 1 | 16 | mycbp |
| si:ch1073-429i10.3 | 0.000245 | 0.27466221 | 0.649 | 0.348 | | 1 | 16 | si:ch1073-429i10.3.1 |
| rhoaa1 | 0.000247 | 0.308099178 | 0.544 | 0.296 | | 1 | 16 | rhoaa |
| copz1 | 0.000249 | 0.26419095 | 0.404 | 0.199 | | 1 | 16 | copz1 |
| ube2na1 | 0.000254 | 0.290359588 | 0.456 | 0.232 | | 1 | 16 | ube2na |
| psmb61 | 0.000274 | 0.289972051 | 0.526 | 0.292 | | 1 | 16 | psmb6 |
| cycsb1 | 0.000278 | 0.274651578 | 0.509 | 0.268 | | 1 | 16 | cycsb |
| gstp1 | 0.00029 | 0.290599325 | 0.526 | 0.278 | | 1 | 16 | gstp1 |
| u2surp1 | 0.000322 | 0.262230993 | 0.333 | 0.152 | | 1 | 16 | u2surp |

| | | | | | | | | |
|---|---|---|---|---|---|---|---|---|
| si:dkey-177p2.61 | 0.000332 | 0.258663894 | 0.281 | 0.12 | | 1 | 16 | si:dkey-177p2.6 |
| mrps6 | 0.000348 | 0.251271335 | 0.263 | 0.113 | | 1 | 16 | mrps6 |
| ube2ia | 0.000379 | 0.277745164 | 0.368 | 0.18 | | 1 | 16 | ube2ia |
| syf2 | 0.000387 | 0.26286734 | 0.404 | 0.201 | | 1 | 16 | syf2 |
| arf51 | 0.000388 | 0.31599216 | 0.474 | 0.269 | | 1 | 16 | arf5 |
| ptmaa2 | 0.000403 | 0.327887299 | 0.912 | 0.652 | | 1 | 16 | ptmaa |
| psmb41 | 0.000405 | 0.295200139 | 0.509 | 0.283 | | 1 | 16 | psmb4 |
| mt-co15 | 0.000406 | 0.3654811 | 0.807 | 0.591 | | 1 | 16 | mt-co1 |
| hnrnpub | 0.00041 | 0.257397317 | 0.298 | 0.134 | | 1 | 16 | hnrnpub |
| taf131 | 0.000416 | 0.266436487 | 0.509 | 0.275 | | 1 | 16 | taf13 |
| rac1a | 0.000427 | 0.263500625 | 0.263 | 0.112 | | 1 | 16 | rac1a |
| fam120c3 | 0.000434 | 0.286484222 | 0.316 | 0.147 | | 1 | 16 | fam120c |
| atp5f1b2 | 0.000437 | 0.302300441 | 0.807 | 0.528 | | 1 | 16 | atp5f1b |
| eif4a3 | 0.000453 | 0.253276889 | 0.421 | 0.216 | | 1 | 16 | eif4a3 |
| snrpg | 0.000455 | 0.297000563 | 0.491 | 0.268 | | 1 | 16 | snrpg |
| mt-nd44 | 0.000457 | 0.318817883 | 0.737 | 0.434 | | 1 | 16 | mt-nd4 |
| mcts1 | 0.000466 | 0.253410554 | 0.333 | 0.16 | | 1 | 16 | mcts1 |
| tra2a | 0.00048 | 0.337274026 | 0.404 | 0.222 | | 1 | 16 | tra2a |
| snrpd2 | 0.00048 | 0.276178502 | 0.491 | 0.264 | | 1 | 16 | snrpd2 |
| hmgb2b1 | 0.000541 | 0.330471424 | 0.702 | 0.438 | | 1 | 16 | hmgb2b |
| sf1 | 0.000545 | 0.251798548 | 0.281 | 0.126 | | 1 | 16 | sf1 |
| eif3ba1 | 0.000556 | 0.291133046 | 0.667 | 0.392 | | 1 | 16 | eif3ba |
| hnrnpa0l | 0.000557 | 0.317550141 | 0.965 | 0.701 | | 1 | 16 | hnrnpa0l |
| psma2 | 0.000594 | 0.257980574 | 0.439 | 0.231 | | 1 | 16 | psma2 |
| rad23b | 0.000628 | 0.281462959 | 0.474 | 0.26 | | 1 | 16 | rad23b |
| COX5B2 | 0.000642 | 0.269469934 | 0.772 | 0.476 | | 1 | 16 | COX5B |
| gng51 | 0.000691 | 0.298642288 | 0.632 | 0.381 | | 1 | 16 | gng5 |
| txndc91 | 0.000708 | 0.258005332 | 0.368 | 0.187 | | 1 | 16 | txndc9 |
| safb | 0.000716 | 0.251695555 | 0.211 | 0.085 | | 1 | 16 | safb |
| si:ch211-288g17.3[illegible] | 0.000728 | 0.337477752 | 0.509 | 0.292 | | 1 | 16 | si:ch211-288g17.3 |
| fkbp3 | 0.000739 | 0.276314458 | 0.456 | 0.252 | | 1 | 16 | fkbp3 |
| elavl1a | 0.000786 | 0.284075607 | 0.281 | 0.131 | | 1 | 16 | elavl1a |
| dynlrb1 | 0.000819 | 0.269524423 | 0.474 | 0.267 | | 1 | 16 | dynlrb1 |
| slirp | 0.000842 | 0.260075773 | 0.333 | 0.17 | | 1 | 16 | slirp |
| snx1a1 | 0.000851 | 0.262927066 | 0.333 | 0.167 | | 1 | 16 | snx1a |

| | | | | | | | | |
|---|---|---|---|---|---|---|---|---|
| hsbp1b | 0.000972 | 0.261553296 | 0.456 | 0.25 | | 1 | 16 | hsbp1b |
| st13 | 0.001024 | 0.251543502 | 0.298 | 0.146 | | 1 | 16 | st13 |
| prmt1 | 0.00106 | 0.266590584 | 0.421 | 0.229 | | 1 | 16 | prmt1 |
| wu:fb55g093 | 0.001071 | 0.277641197 | 0.544 | 0.309 | | 1 | 16 | wu:fb55g09 |
| mknk2b3 | 0.001089 | 0.311030942 | 0.404 | 0.22 | | 1 | 16 | mknk2b |
| smarce12 | 0.001094 | 0.264514006 | 0.439 | 0.245 | | 1 | 16 | smarce1 |
| ssr33 | 0.001133 | 0.287403271 | 0.614 | 0.383 | | 1 | 16 | ssr3 |
| atp5po1 | 0.001203 | 0.259994248 | 0.544 | 0.316 | | 1 | 16 | atp5po |
| brk1 | 0.001264 | 0.264392624 | 0.439 | 0.244 | | 1 | 16 | brk1 |
| hnrnpa1b2 | 0.001399 | 0.255589757 | 0.439 | 0.245 | | 1 | 16 | hnrnpa1b |
| ranbp1 | 0.001606 | 0.260386404 | 0.544 | 0.324 | | 1 | 16 | ranbp1 |
| klf6a3 | 0.001648 | 0.254174574 | 0.474 | 0.258 | | 1 | 16 | klf6a |
| si:dkey-42i9.43 | 0.001688 | 0.277511721 | 0.368 | 0.197 | | 1 | 16 | si:dkey-42i9.4 |
| xbp13 | 0.001864 | 0.252111564 | 0.702 | 0.432 | | 1 | 16 | xbp1 |
| si:ch73-46j18.57 | 0.002125 | 0.311562407 | 0.772 | 0.391 | | 1 | 16 | si:ch73-46j18.5 |
| atp5f1e | 0.00269 | 0.258232875 | 0.807 | 0.543 | | 1 | 16 | atp5f1e |
| cox7c1 | 0.002798 | 0.268861932 | 0.825 | 0.578 | | 1 | 16 | cox7c |
| pdia34 | 0.002946 | 0.281818797 | 0.754 | 0.482 | | 1 | 16 | pdia3 |
| psmd11 | 0.003187 | 0.262465588 | 0.298 | 0.158 | | 1 | 16 | psmd1 |
| mt-nd14 | 0.003283 | 0.279491772 | 0.579 | 0.353 | | 1 | 16 | mt-nd1 |
| eif5a1 | 0.008151 | 0.250977912 | 0.912 | 0.595 | | 1 | 16 | eif5a |
| ddx4 | 0 | 1.863883519 | 0.891 | 0.002 | | 0 | 17 | ddx4 |
| si:ch73-167f10.1 | 1.4098618 | 1.30105425 | 0.655 | 0.002 | 2.84749807844936e-2 | | 17 | si:ch73-167f10.1 |
| hsp90aa1.2 | 2.0792140 | 1.516778729 | 0.709 | 0.008 | 4.19938860079024e-2 | | 17 | hsp90aa1.2 |
| elavl2 | 7.6183421 | 0.988836115 | 0.455 | 0.001 | 1.53867656527929e-1 | | 17 | elavl2 |
| si:ch211-226l4.6 | 5.8680168 | 0.982250697 | 0.455 | 0.002 | 1.18516336102935e-1 | | 17 | si:ch211-226l4.6 |
| zgc:193690 | 2.0269815 | 0.909195674 | 0.418 | 0.001 | 4.09389462849226e-1 | | 17 | zgc:193690 |
| zgc:56699 | 1.4567450 | 1.031937762 | 0.473 | 0.004 | 2.94218802473399e-1 | | 17 | zgc:56699 |
| chtopb | 1.4977767 | 0.95009256 | 0.436 | 0.002 | 3.0250596832816e-15 | | 17 | chtopb |
| zgc:173742 | 3.4195824 | 0.862602044 | 0.364 | 0 | 6.90653069725214e-1 | | 17 | zgc:173742 |
| zgc:103482 | 3.7978645 | 0.83505541 | 0.364 | 0 | 7.67054703052171e-1 | | 17 | zgc:103482 |
| hormad1 | 3.0610783 | 0.837005968 | 0.4 | 0.002 | 6.18245993524144e-1 | | 17 | hormad1 |
| dazl | 3.9029809 | 0.924574345 | 0.345 | 0 | 7.88285054480906e-1 | | 17 | dazl |
| zgc:55461 | 3.6726572 | 1.257608126 | 0.564 | 0.011 | 7.41766585428483e-1 | | 17 | zgc:55461 |
| syce2 | 5.9672358 | 0.749635952 | 0.327 | 0 | 1.20520263040341e-1 | | 17 | syce2 |

| | | | | | | | |
|---|---|---|---|---|---|---|---|
| adad1 | 7.0996017 | 0.803613447 | 0.327 | 0 | 1.43390655904091e-1 | 17 | adad1 |
| pld6 | 8.6455175 | 0.888420628 | 0.382 | 0.002 | 1.74613518798419e-1 | 17 | pld6 |
| zgc:92287 | 1.3647927 | 0.916404843 | 0.418 | 0.004 | 2.75647201281872e-1 | 17 | zgc:92287 |
| org | 1.4211949 | 0.809633984 | 0.345 | 0.001 | 2.87038752297114e-1 | 17 | org |
| si:dkey-189g17.2 | 1.7267079 | 0.914168158 | 0.382 | 0.003 | 3.48743208594296e-1 | 17 | si:dkey-189g17.2 |
| sycp1 | 1.0875934 | 0.748353627 | 0.291 | 0.001 | 2.19661239891419e-1 | 17 | sycp1 |
| tdrd9 | 1.7857166 | 0.589244707 | 0.255 | 0 | 3.60661198046384e-1 | 17 | tdrd9 |
| npm2b | 7.6829664 | 0.850742688 | 0.309 | 0.002 | 1.55172873025293e-1 | 17 | npm2b |
| dnd1 | 8.3971942 | 0.813016582 | 0.309 | 0.002 | 1.69598132357124e-1 | 17 | dnd1 |
| piwil2 | 1.7556625 | 0.671050999 | 0.291 | 0.001 | 3.54591164420754e-1 | 17 | piwil2 |
| smc1b | 6.3798153 | 0.632580293 | 0.273 | 0.001 | 1.28853130978802e-1 | 17 | smc1b |
| LOC101883959 | 6.5893319 | 0.678482573 | 0.273 | 0.001 | 1.33084736540351e-1 | 17 | LOC101883959 |
| si:dkey-40g16.5 | 8.5324598 | 0.613198866 | 0.273 | 0.001 | 1.72330091416129e-1 | 17 | si:dkey-40g16.5 |
| CABZ01083945.1 | 1.1891565 | 0.671891354 | 0.309 | 0.002 | 2.40173947556358e-1 | 17 | CABZ01083945.1 |
| pabpn1l | 1.3315041 | 0.703971061 | 0.309 | 0.002 | 2.68923902712379e-1 | 17 | pabpn1l |
| pcp4l1 | 4.1313967 | 0.682794237 | 0.291 | 0.002 | 8.34418200519841e-1 | 17 | pcp4l1 |
| exd1 | 1.3614288 | 0.684075729 | 0.273 | 0.001 | 2.74967788117769e-9 | 17 | exd1 |
| eif4e1b | 7.7703058 | 0.601243404 | 0.255 | 0.001 | 1.56936868033853e-9 | 17 | eif4e1b |
| ccnd3 | 9.1187436 | 0.650206347 | 0.255 | 0.001 | 1.84171266261211e-9 | 17 | ccnd3 |
| zgc:161973 | 1.3385275 | 0.563549242 | 0.255 | 0.001 | 2.70342411074913e-9 | 17 | zgc:161973 |
| fkbp6 | 1.1976977 | 0.604604191 | 0.236 | 0 | 2.41899024506908e-9 | 17 | fkbp6 |
| rad21l1 | 2.6378078 | 0.541750669 | 0.218 | 0 | 5.32758059592913e-9 | 17 | rad21l1 |
| piwil1 | 7.3883846 | 0.648317813 | 0.236 | 0.001 | 1.49223205639844e-8 | 17 | piwil1 |
| si:ch211-176l24.4 | 2.2238105 | 0.707594116 | 0.273 | 0.002 | 4.49143026371617e-8 | 17 | si:ch211-176l24.4 |
| CR382383.2 | 1.3261602 | 0.536557465 | 0.218 | 0 | 2.67844585893685e-8 | 17 | CR382383.2 |
| sycp3 | 1.3717207 | 0.496106339 | 0.218 | 0 | 2.7704644189831e-86 | 17 | sycp3 |
| stag3 | 1.4675790 | 0.527144279 | 0.218 | 0 | 2.96406937408701e-8 | 17 | stag3 |
| asz1 | 4.9833952 | 0.579927515 | 0.218 | 0.001 | 1.00649632957151e-7 | 17 | asz1 |
| arl15a | 1.5730689 | 0.468090114 | 0.2 | 0 | 3.17712726173212e-7 | 17 | arl15a |
| rbm46 | 3.6782918 | 0.478253297 | 0.182 | 0 | 7.429046128109e-78 | 17 | rbm46 |
| s100a1 | 9.0583126 | 0.605805717 | 0.236 | 0.002 | 1.82950741493185e-7 | 17 | s100a1 |
| nme4 | 5.0138278 | 0.580409703 | 0.255 | 0.003 | 1.01264280164068e-7 | 17 | nme4 |
| hnrnpa3 | 4.1636202 | 0.458598095 | 0.2 | 0.001 | 8.40926384786357e-7 | 17 | hnrnpa3 |
| crp2 | 1.4242314 | 0.533441323 | 0.182 | 0 | 2.87652028707732e-7 | 17 | crp2 |
| rad9b | 4.2585260 | 0.53261528 | 0.164 | 0 | 8.60094501163513e-7 | 17 | rad9b |

| | | | | | | | |
|---|---|---|---|---|---|---|---|
| ccdc106b | 4.2585260 | 0.454491743 | 0.164 | 0 | 8.60094501163513e-7 | 17 | ccdc106b |
| gpat2 | 4.2585260 | 0.434277572 | 0.164 | 0 | 8.60094501163513e-7 | 17 | gpat2 |
| si:dkeyp-93d12.1 | 4.2585260 | 0.39981087 | 0.164 | 0 | 8.60094501163513e-7 | 17 | si:dkeyp-93d12.1 |
| si:ch211-14a17.7 | 1.4620615 | 0.46920094 | 0.164 | 0 | 2.95292571797994e-6 | 17 | si:ch211-14a17.7 |
| cdca9 | 1.5109466 | 0.407241078 | 0.164 | 0 | 3.05165896515355e-6 | 17 | cdca9 |
| spdya | 1.6675933 | 0.384827569 | 0.164 | 0 | 3.36803835602934e-6 | 17 | spdya |
| pabpc1l | 1.7809142 | 0.485797304 | 0.164 | 0 | 3.59691260559716e-6 | 17 | pabpc1l |
| zar1l | 4.8729081 | 0.384298373 | 0.145 | 0 | 9.84181256644164e-6 | 17 | zar1l |
| LOC110440179 | 4.8729081 | 0.365368545 | 0.145 | 0 | 9.84181256644164e-6 | 17 | LOC110440179 |
| stk31 | 4.8729081 | 0.354603207 | 0.145 | 0 | 9.84181256644164e-6 | 17 | stk31 |
| tdrd1 | 9.1426754 | 0.66650851 | 0.255 | 0.005 | 1.84654615157889e-6 | 17 | tdrd1 |
| odf3l2a | 6.3686606 | 0.522685281 | 0.218 | 0.003 | 1.28627840099116e-6 | 17 | odf3l2a |
| e2f5 | 3.3059765 | 0.687632623 | 0.255 | 0.005 | 6.67708075475926e-5 | 17 | e2f5 |
| spata6l | 2.2258744 | 0.442858242 | 0.2 | 0.002 | 4.49559860025264e-5 | 17 | spata6l |
| zglp1 | 1.5226735 | 0.364413873 | 0.145 | 0 | 3.07534370136188e-5 | 17 | zglp1 |
| moto | 1.6247887 | 0.387202079 | 0.145 | 0 | 3.28158587397267e-5 | 17 | moto |
| cpeb1b | 5.5201287 | 0.316279423 | 0.127 | 0 | 1.11490040958516e-5 | 17 | cpeb1b |
| adad2 | 5.5201287 | 0.304798444 | 0.127 | 0 | 1.11490040958516e-5 | 17 | adad2 |
| si:ch211-241j8.2 | 5.5201287 | 0.299323478 | 0.127 | 0 | 1.11490040958516e-5 | 17 | si:ch211-241j8.2 |
| tbpl2 | 5.5201287 | 0.29757944 | 0.127 | 0 | 1.11490040958516e-5 | 17 | tbpl2 |
| tdrkh | 4.4724301 | 0.615736478 | 0.255 | 0.007 | 9.03296712333568e-5 | 17 | tdrkh |
| si:ch211-173m16.2 | 1.1993821 | 0.431359482 | 0.145 | 0.001 | 2.42239221402957e-4 | 17 | si:ch211-173m16.2 |
| nfr | 1.4709415 | 0.332891757 | 0.145 | 0.001 | 2.97086063242945e-4 | 17 | nfr |
| esco2 | 3.7921640 | 0.85021179 | 0.327 | 0.015 | 7.65903365719816e-4 | 17 | esco2 |
| noxo1b | 7.0055540 | 0.440187932 | 0.164 | 0.002 | 1.41491174825791e-4 | 17 | noxo1b |
| BX571701.1 | 1.1876950 | 0.380539515 | 0.127 | 0 | 2.3987876643232e-46 | 17 | BX571701.1 |
| si:dkey-241l7.4 | 1.1876950 | 0.365063556 | 0.127 | 0 | 2.3987876643232e-46 | 17 | si:dkey-241l7.4 |
| nanog | 1.1876950 | 0.362852723 | 0.127 | 0 | 2.3987876643232e-46 | 17 | nanog |
| spata22 | 1.2262590 | 0.317937744 | 0.127 | 0 | 2.47667547874412e-4 | 17 | spata22 |
| CR933730.1 | 1.3933874 | 0.335603531 | 0.127 | 0 | 2.8142246217607e-46 | 17 | CR933730.1 |
| ndufs8b | 1.4852685 | 0.328552771 | 0.127 | 0 | 2.99979698334041e-4 | 17 | ndufs8b |
| akap1a | 6.2056202 | 0.390561191 | 0.109 | 0 | 1.25334911292777e-4 | 17 | akap1a |
| si:ch211-226h8.14 | 6.2056202 | 0.30510431 | 0.109 | 0 | 1.25334911292777e-4 | 17 | si:ch211-226h8.14 |
| CCNB1IP1 | 6.2056202 | 0.297812152 | 0.109 | 0 | 1.25334911292777e-4 | 17 | CCNB1IP1 |
| si:ch211-57b15.2 | 6.2056202 | 0.284886704 | 0.109 | 0 | 1.25334911292777e-4 | 17 | si:ch211-57b15.2 |

| | | | | | | | |
|---|---|---|---|---|---|---|---|
| MAJIN | 6.2056202 | 0.273107722 | 0.109 | 0 | 1.25334911292777e-4 | 17 | MAJIN |
| zgc:92591 | 6.2056202 | 0.251255801 | 0.109 | 0 | 1.25334911292777e-4 | 17 | zgc:92591 |
| kpna2 | 9.5068675 | 0.897220541 | 0.382 | 0.024 | 1.9201020333751e-44 | 17 | kpna2 |
| si:ch211-227n13.3 | 1.0196552 | 0.381376724 | 0.145 | 0.001 | 2.05939776504742e-4 | 17 | si:ch211-227n13.3 |
| si:dkey-92i17.2 | 5.0449224 | 0.526138005 | 0.218 | 0.006 | 1.01892299441764e-4 | 17 | si:dkey-92i17.2 |
| wasf3a | 6.3843015 | 0.34742137 | 0.127 | 0.001 | 1.28943738500872e-4 | 17 | wasf3a |
| ccdc9b | 6.3843015 | 0.290476577 | 0.127 | 0.001 | 1.28943738500872e-4 | 17 | ccdc9b |
| si:dkeyp-50f7.2 | 6.5677449 | 0.391761568 | 0.127 | 0.001 | 1.32648744981245e-4 | 17 | si:dkeyp-50f7.2 |
| stmn1b | 1.0929323 | 0.624334814 | 0.236 | 0.008 | 2.20739544941589e-4 | 17 | stmn1b |
| lmnl3 | 1.1896524 | 0.362258579 | 0.145 | 0.002 | 2.40274115217522e-3 | 17 | lmnl3 |
| alg13 | 9.7171576 | 0.347300744 | 0.109 | 0 | 1.96257432292674e-3 | 17 | alg13 |
| si:ch211-286b5.2 | 9.7171576 | 0.319098786 | 0.109 | 0 | 1.96257432292674e-3 | 17 | si:ch211-286b5.2 |
| arf3a | 1.0026031 | 0.304913854 | 0.109 | 0 | 2.02495748558181e-3 | 17 | arf3a |
| LOC110440049 | 1.0026031 | 0.278194316 | 0.109 | 0 | 2.02495748558181e-3 | 17 | LOC110440049 |
| si:ch211-10a23.2 | 1.0344669 | 0.325257371 | 0.109 | 0 | 2.08931285713856e-3 | 17 | si:ch211-10a23.2 |
| pm20d1.1 | 1.0344669 | 0.26168315 | 0.109 | 0 | 2.08931285713856e-3 | 17 | pm20d1.1 |
| LOC101886261 | 1.0673379 | 0.257761928 | 0.109 | 0 | 2.15570244525511e-3 | 17 | LOC101886261 |
| rnf17 | 1.1362291 | 0.271854147 | 0.109 | 0 | 2.29484205546468e-3 | 17 | rnf17 |
| nsg2 | 3.3389425 | 0.477061036 | 0.182 | 0.004 | 6.74366234305411e-3 | 17 | nsg2 |
| zgc:65894 | 4.8750781 | 0.423110487 | 0.164 | 0.003 | 9.84619540796367e-3 | 17 | zgc:65894 |
| ube2c | 1.1869652 | 0.87164492 | 0.4 | 0.034 | 2.39731374378727e-3 | 17 | ube2c |
| zar1 | 2.8181979 | 0.487582842 | 0.145 | 0.002 | 5.69191432828785e-3 | 17 | zar1 |
| dctpp1 | 9.6084801 | 0.778582748 | 0.327 | 0.024 | 1.9406247420483e-33 | 17 | dctpp1 |
| birc5a | 1.1825929 | 0.741950823 | 0.309 | 0.021 | 2.38848305559473e-3 | 17 | birc5a |
| hpf1 | 1.5824949 | 0.937422488 | 0.418 | 0.042 | 3.19616513112662e-3 | 17 | hpf1 |
| slbp2 | 2.4373626 | 0.329032838 | 0.109 | 0.001 | 4.9227413506151e-33 | 17 | slbp2 |
| alox5b.3 | 2.5741497 | 0.335173764 | 0.109 | 0.001 | 5.19901016628652e-3 | 17 | alox5b.3 |
| si:dkey-37o8.1 | 2.8710297 | 0.351327688 | 0.109 | 0.001 | 5.79861879169811e-3 | 17 | si:dkey-37o8.1 |
| sncb | 2.8710297 | 0.261761542 | 0.109 | 0.001 | 5.79861879169811e-3 | 17 | sncb |
| tmsb5 | 3.2019192 | 0.274731482 | 0.109 | 0.001 | 6.46691634596064e-3 | 17 | tmsb5 |
| zgc:162297 | 1.4294066 | 0.324079934 | 0.127 | 0.002 | 2.88697262841229e-3 | 17 | zgc:162297 |
| trip13 | 2.2050900 | 0.452695809 | 0.164 | 0.004 | 4.45362033617051e-3 | 17 | trip13 |
| slbp | 4.4071442 | 0.776387601 | 0.345 | 0.028 | 8.90110918076815e-3 | 17 | slbp |
| dut1 | 1.0772874 | 1.120714576 | 0.582 | 0.089 | 2.17579751523778e-3 | 17 | dut |
| chek1 | 2.1563714 | 0.531060663 | 0.218 | 0.01 | 4.35522347086565e-3 | 17 | chek1 |

| | | | | | | | |
|---|---|---|---|---|---|---|---|
| gra | 1.3247072 | 0.478650639 | 0.182 | 0.006 | 2.67551119538864e-3 | 17 | gra |
| LOC101883594 | 2.9055213 | 0.403270622 | 0.164 | 0.005 | 5.8682813760964e-30 | 17 | LOC101883594 |
| rad51 | 1.5858554 | 0.6442011 | 0.273 | 0.018 | 3.20295216630547e-2 | 17 | rad51 |
| si:dkey-6d5.1 | 1.8816002 | 0.346322191 | 0.127 | 0.002 | 3.80026807193458e-2 | 17 | si:dkey-6d5.1 |
| nsmce1 | 3.0657152 | 0.66282752 | 0.273 | 0.019 | 6.19182501846247e-2 | 17 | nsmce1 |
| hells | 1.3804715 | 1.01240751 | 0.527 | 0.082 | 2.78813842282541e-2 | 17 | hells |
| rrm2 | 5.5307354 | 1.003594731 | 0.473 | 0.063 | 1.11704264358762e-2 | 17 | rrm2 |
| terb1 | 8.5481082 | 0.317853275 | 0.127 | 0.003 | 1.72646141422286e-2 | 17 | terb1 |
| ddx39b1 | 4.7266155 | 0.783785806 | 0.327 | 0.032 | 9.54634547185938e-2 | 17 | ddx39b |
| stmn1a | 1.3946857 | 1.13198803 | 0.564 | 0.101 | 2.81684686885105e-2 | 17 | stmn1a |
| wdr76 | 4.6550636 | 0.404583575 | 0.164 | 0.007 | 9.40183208718664e-2 | 17 | wdr76 |
| zgc:153681 | 5.2870422 | 0.312894904 | 0.109 | 0.002 | 1.06782391853891e-2 | 17 | zgc:153681 |
| si:ch211-147d7.5 | 5.6085919 | 0.256983745 | 0.109 | 0.002 | 1.13276731021781e-2 | 17 | si:ch211-147d7.5 |
| eef1a1a | 5.7200263 | 0.318166634 | 0.109 | 0.002 | 1.1552737192373e-22 | 17 | eef1a1a |
| eif4eb | 6.1894135 | 0.767268324 | 0.345 | 0.04 | 1.25007585977755e-2 | 17 | eif4eb |
| cox5b2 | 6.9368093 | 0.415378262 | 0.145 | 0.005 | 1.40102738715504e-2 | 17 | cox5b2 |
| sub1b1 | 1.0689108 | 1.155298714 | 0.673 | 0.183 | 2.15887922528169e-2 | 17 | sub1b |
| lim2.2 | 1.1929529 | 0.397413498 | 0.127 | 0.004 | 2.40940704621164e-2 | 17 | lim2.2 |
| gcshb | 1.9452611 | 0.484625847 | 0.218 | 0.015 | 3.92884398991411e-2 | 17 | gcshb |
| si:ch211-212d10.2 | 4.4343067 | 0.596314004 | 0.218 | 0.015 | 8.95596929650715e-2 | 17 | si:ch211-212d10.2 |
| hddc3 | 8.7801385 | 0.70635851 | 0.327 | 0.038 | 1.77332457404023e-2 | 17 | hddc3 |
| mgmt | 1.7934916 | 0.815866737 | 0.4 | 0.058 | 3.62231503897725e-2 | 17 | mgmt |
| tomm20a3 | 2.8738683 | 1.128405953 | 0.691 | 0.212 | 5.80435200721315e-2 | 17 | tomm20a |
| morc3a | 5.7346974 | 0.634577962 | 0.236 | 0.019 | 1.15823683931923e-2 | 17 | morc3a |
| ube2t | 9.1996540 | 0.563739422 | 0.218 | 0.016 | 1.85805413129603e-2 | 17 | ube2t |
| dnajc18 | 1.6582029 | 0.523951037 | 0.218 | 0.016 | 3.34907254794325e-2 | 17 | dnajc18 |
| cdca8 | 3.4528998 | 0.583224369 | 0.236 | 0.02 | 6.97382180828234e-2 | 17 | cdca8 |
| hmgb2b2 | 6.0921951 | 1.073028895 | 0.891 | 0.433 | 1.23044065525396e-1 | 17 | hmgb2b |
| cdk1 | 8.4643453 | 0.689324583 | 0.291 | 0.031 | 1.70954383824386e-1 | 17 | cdk1 |
| lsm62 | 1.2540945 | 1.121569761 | 0.655 | 0.199 | 2.5328947335049e-19 | 17 | lsm6 |
| mindy3 | 2.4522792 | 0.369228901 | 0.127 | 0.005 | 4.95286839726806e-1 | 17 | mindy3 |
| tnnt2d | 2.5627266 | 0.333210658 | 0.127 | 0.005 | 5.1759390590788e-19 | 17 | tnnt2d |
| idh3g | 3.3342425 | 0.769250123 | 0.327 | 0.043 | 6.7341697448821e-19 | 17 | idh3g |
| nasp | 3.4044618 | 0.915825761 | 0.436 | 0.077 | 6.87599165398945e-1 | 17 | nasp |
| hmgb1b4 | 7.7406401 | 1.088238766 | 0.764 | 0.295 | 1.56337710104561e-1 | 17 | hmgb1b |

| | | | | | | | |
|---|---|---|---|---|---|---|---|
| mrnip | 9.4304080 | 0.263213881 | 0.109 | 0.003 | 1.90465950845533e-1 | 17 | mrnip |
| si:ch211-126c2.4 | 1.0588515 | 0.276406988 | 0.109 | 0.003 | 2.13856246823417e-1 | 17 | si:ch211-126c2.4 |
| mad2l2 | 1.1683974 | 0.643842855 | 0.236 | 0.022 | 2.35981239630681e-1 | 17 | mad2l2 |
| ppid | 2.7697131 | 0.975724047 | 0.491 | 0.106 | 5.59398967938547e-1 | 17 | ppid |
| nutf2l1 | 2.8178695 | 1.009652341 | 0.6 | 0.166 | 5.69125118179387e-1 | 17 | nutf2l |
| spata4 | 5.8525953 | 0.287082144 | 0.127 | 0.005 | 1.18204868346047e-1 | 17 | spata4 |
| snrpg1 | 9.7655189 | 1.089155213 | 0.709 | 0.263 | 1.97234186365131e-1 | 17 | snrpg |
| arl2bp | 1.4781818 | 0.450494753 | 0.2 | 0.016 | 2.98548379742425e-1 | 17 | arl2bp |
| nsmce2 | 2.7156537 | 0.662703401 | 0.273 | 0.032 | 5.48480587170002e-1 | 17 | nsmce2 |
| psmc3ip | 2.7677270 | 0.455556257 | 0.164 | 0.01 | 5.58997822252106e-1 | 17 | psmc3ip |
| dcun1d5 | 3.1057628 | 0.780581471 | 0.4 | 0.07 | 6.27270921996417e-1 | 17 | dcun1d5 |
| cdc6 | 3.5755983 | 0.51638375 | 0.2 | 0.016 | 7.22163596635298e-1 | 17 | cdc6 |
| orc2 | 5.1581294 | 0.294385869 | 0.109 | 0.004 | 1.04178739861792e-1 | 17 | orc2 |
| dynll2b2 | 5.2336250 | 0.978018076 | 0.509 | 0.119 | 1.05703524894881e-1 | 17 | dynll2b |
| spc24 | 6.1262852 | 0.458331683 | 0.182 | 0.013 | 1.23732582595516e-1 | 17 | spc24 |
| gins3 | 1.4208356 | 0.496503914 | 0.164 | 0.011 | 2.8696617314623e-16 | 17 | gins3 |
| phb2b | 2.3911165 | 0.686877549 | 0.345 | 0.054 | 4.8293380730985e-16 | 17 | phb2b |
| rbm8a1 | 2.5582198 | 1.074900968 | 0.764 | 0.367 | 5.16683662797462e-1 | 17 | rbm8a |
| aurkb | 2.5651117 | 0.564130199 | 0.236 | 0.024 | 5.18075623917398e-1 | 17 | aurkb |
| cse1l | 3.1382432 | 0.611384863 | 0.255 | 0.029 | 6.33830998595222e-1 | 17 | cse1l |
| gtf2b | 6.4984620 | 0.886560521 | 0.455 | 0.099 | 1.3124943811463e-15 | 17 | gtf2b |
| cks2 | 6.7159939 | 0.577619667 | 0.236 | 0.025 | 1.35642929840854e-1 | 17 | cks2 |
| fen1 | 8.7974870 | 0.789296966 | 0.382 | 0.068 | 1.77682845711355e-1 | 17 | fen1 |
| uchl1 | 9.0638203 | 0.66687412 | 0.291 | 0.039 | 1.83061979655733e-1 | 17 | uchl1 |
| si:ch73-95l15.5 | 1.0060314 | 0.385951635 | 0.127 | 0.006 | 2.03188179751952e-1 | 17 | si:ch73-95l15.5 |
| mad2l1 | 1.2287042 | 0.58480254 | 0.236 | 0.025 | 2.48161392471256e-1 | 17 | mad2l1 |
| snrpa1 | 1.4155424 | 0.92381534 | 0.473 | 0.112 | 2.85897118160942e-1 | 17 | snrpa1 |
| atf7ip2 | 3.6723881 | 0.395645477 | 0.164 | 0.012 | 7.41712239592922e-1 | 17 | atf7ip2 |
| sms | 3.6798964 | 0.700287649 | 0.327 | 0.05 | 7.43228677684757e-1 | 17 | sms |
| hirip3 | 4.6600045 | 0.833765878 | 0.4 | 0.079 | 9.41181109859185e-1 | 17 | hirip3 |
| larp7 | 6.9597630 | 0.64440711 | 0.291 | 0.041 | 1.40566334740482e-1 | 17 | larp7 |
| gtpbp1l | 9.9223291 | 0.395504676 | 0.145 | 0.009 | 2.00401282629175e-1 | 17 | gtpbp1l |
| brf2 | 1.1595500 | 0.355910133 | 0.145 | 0.009 | 2.34194315370116e-1 | 17 | brf2 |
| ssrp1a | 1.4215980 | 0.764753563 | 0.345 | 0.06 | 2.87120158238419e-1 | 17 | ssrp1a |
| dpy301 | 2.2342732 | 1.023793521 | 0.618 | 0.222 | 4.51256170604919e-1 | 17 | dpy30 |

| | | | | | | | |
|---|---|---|---|---|---|---|---|
| CABZ01080568.1 | 2.2812454 | 0.282178274 | 0.109 | 0.005 | 4.6074313514351e-14 | 17 | CABZ01080568.1 |
| ankrd10a | 2.3554485 | 0.510140307 | 0.218 | 0.023 | 4.75729944237338e-1 | 17 | ankrd10a |
| hibadha | 2.4384421 | 0.260669461 | 0.109 | 0.005 | 4.92492150958053e-1 | 17 | hibadha |
| chaf1a | 2.5672534 | 0.816172329 | 0.382 | 0.074 | 5.18508171596668e-1 | 17 | chaf1a |
| grhpra | 2.5718597 | 0.275426886 | 0.109 | 0.005 | 5.19438521867236e-1 | 17 | grhpra |
| parp3 | 2.8589986 | 0.703274293 | 0.309 | 0.049 | 5.77431964475798e-1 | 17 | parp3 |
| nuf2 | 6.1782157 | 0.344387312 | 0.145 | 0.01 | 1.24781423244362e-1 | 17 | nuf2 |
| pane1 | 6.7159998 | 0.408208507 | 0.164 | 0.013 | 1.35643048283871e-1 | 17 | pane1 |
| calm2b4 | 6.8695246 | 0.878789208 | 0.818 | 0.442 | 1.38743789153865e-1 | 17 | calm2b |
| lsm71 | 1.1951961 | 1.013993743 | 0.618 | 0.225 | 2.41393760367084e-1 | 17 | lsm7 |
| LOC110438491 | 1.8448807 | 0.352722036 | 0.164 | 0.013 | 3.72610561386901e-1 | 17 | LOC110438491 |
| LOC103908877 | 1.9101481 | 0.395553573 | 0.164 | 0.013 | 3.85792615169455e-1 | 17 | LOC103908877 |
| zgc:194246 | 2.0471848 | 0.644060706 | 0.291 | 0.044 | 4.13469921422616e-1 | 17 | zgc:194246 |
| prdx1 | 2.1074287 | 0.754679294 | 0.345 | 0.063 | 4.25637379996841e-1 | 17 | prdx1 |
| smfn | 2.4701776 | 0.762051434 | 0.364 | 0.071 | 4.98901786654592e-1 | 17 | smfn |
| srsf7b | 3.1243789 | 0.263414638 | 0.109 | 0.005 | 6.3103081519837e-13 | 17 | srsf7b |
| hddc2 | 6.1191746 | 0.63312294 | 0.273 | 0.04 | 1.23588970091359e-1 | 17 | hddc2 |
| ap1m2 | 1.0623244 | 0.335602366 | 0.145 | 0.011 | 2.14557667606605e-1 | 17 | ap1m2 |
| srsf2b | 1.7386040 | 0.503536478 | 0.218 | 0.026 | 3.51145857646132e-1 | 17 | srsf2b |
| cenpw | 1.9384357 | 0.583985749 | 0.255 | 0.036 | 3.91505862436392e-1 | 17 | cenpw |
| rtca | 2.0130537 | 0.72872736 | 0.364 | 0.074 | 4.06576459970239e-1 | 17 | rtca |
| trmt112 | 2.0275508 | 0.941962856 | 0.564 | 0.191 | 4.09504452533779e-1 | 17 | trmt112 |
| erh | 2.3046163 | 0.911583673 | 0.727 | 0.38 | 4.65463355895806e-1 | 17 | erh |
| mycbp2 | 2.7279199 | 0.949427857 | 0.6 | 0.221 | 5.50957986975994e-1 | 17 | mycbp |
| mrto4 | 3.0155050 | 0.710215014 | 0.364 | 0.075 | 6.09041549135623e-1 | 17 | mrto4 |
| phax | 3.1154331 | 0.635357391 | 0.273 | 0.042 | 6.2922403782052e-12 | 17 | phax |
| ttf2 | 3.2396581 | 0.283710566 | 0.109 | 0.006 | 6.54313755718997e-1 | 17 | ttf2 |
| si:dkey-148h10.5 | 3.2779970 | 0.274973003 | 0.109 | 0.006 | 6.62057073015279e-1 | 17 | si:dkey-148h10.5 |
| zgc:92594 | 3.2779970 | 0.260145479 | 0.109 | 0.006 | 6.62057073015279e-1 | 17 | zgc:92594 |
| atp5mc3b1 | 8.3615613 | 0.662337302 | 0.855 | 0.632 | 1.68878453667901e-1 | 17 | atp5mc3b |
| rabif | 8.9814987 | 0.728016847 | 0.364 | 0.077 | 1.81399330454607e-1 | 17 | rabif |
| sbds | 1.1242695 | 0.59096613 | 0.273 | 0.043 | 2.27068730788783e-1 | 17 | sbds |
| qdprb1 | 1.2255206 | 0.501948162 | 0.236 | 0.033 | 2.47518410146696e-1 | 17 | qdprb1 |
| cmss1 | 1.2535561 | 0.602600058 | 0.273 | 0.044 | 2.53180742738514e-1 | 17 | cmss1 |
| top1mt | 1.3086461 | 0.362870488 | 0.127 | 0.009 | 2.64307270707326e-1 | 17 | top1mt |

| | | | | | | | |
|---|---|---|---|---|---|---|---|
| pgam51 | 1.3173215 | 0.737650968 | 0.4 | 0.093 | 2.66059433345737e-1 | 17 | pgam5 |
| snrnp27 | 1.5717568 | 0.820935541 | 0.455 | 0.123 | 3.1744773876952e-11 | 17 | snrnp27 |
| timm23a2 | 2.3671597 | 0.851353781 | 0.527 | 0.173 | 4.78095247071538e-1 | 17 | timm23a |
| pus10 | 2.4309014 | 0.291637111 | 0.109 | 0.006 | 4.90969158095672e-1 | 17 | pus10 |
| tyms | 3.6557903 | 0.620367546 | 0.291 | 0.05 | 7.38359985049215e-1 | 17 | tyms |
| ubc2 | 5.2812772 | 0.628928197 | 0.836 | 0.654 | 1.06665956424207e-1 | 17 | ubc |
| setb | 6.4250555 | 0.844592974 | 0.727 | 0.387 | 1.29766846259902e-1 | 17 | setb |
| rpa2 | 7.1346307 | 0.757558566 | 0.4 | 0.095 | 1.44098137994703e-1 | 17 | rpa2 |
| lsm51 | 8.7108466 | 0.894132312 | 0.582 | 0.231 | 1.75932969301563e-1 | 17 | lsm5 |
| nap1l11 | 8.7746306 | 0.853698516 | 0.709 | 0.354 | 1.77221215290237e-1 | 17 | nap1l1 |
| lnpk | 1.0824842 | 0.401774619 | 0.145 | 0.013 | 2.18629339803939e-1 | 17 | lnpk |
| mob1bb | 1.4644464 | 0.6076241 | 0.255 | 0.041 | 2.95774256252295e-1 | 17 | mob1bb |
| cenpp | 1.4924487 | 0.292134735 | 0.109 | 0.007 | 3.01429876478882e-1 | 17 | cenpp |
| rcor2 | 1.5731369 | 0.268876517 | 0.109 | 0.007 | 3.17726477830079e-1 | 17 | rcor2 |
| dhx16 | 2.4260992 | 0.577006283 | 0.291 | 0.053 | 4.89999257536517e-1 | 17 | dhx16 |
| snrpd11 | 2.5863300 | 0.930537002 | 0.618 | 0.272 | 5.22361081890721e-1 | 17 | snrpd1 |
| zgc:110130 | 2.6496092 | 0.285886477 | 0.127 | 0.01 | 5.35141578360167e-1 | 17 | zgc:110130 |
| npm1a1 | 2.8625604 | 0.956043867 | 0.636 | 0.304 | 5.78151329849541e-1 | 17 | npm1a |
| h2afvb | 3.4745544 | 0.757994306 | 0.818 | 0.527 | 7.01755757584353e-1 | 17 | h2afvb |
| top2a | 3.6816267 | 0.430654208 | 0.2 | 0.025 | 7.43578150613837e-1 | 17 | top2a |
| unga | 3.9338396 | 0.502481774 | 0.2 | 0.026 | 7.94517596511109e-1 | 17 | unga |
| xrcc5 | 5.2223562 | 0.39302876 | 0.164 | 0.017 | 1.05475928327678e-0 | 17 | xrcc5 |
| si:dkey-30c15.10 | 7.6378205 | 0.33631583 | 0.109 | 0.007 | 1.54261061261764e-0 | 17 | si:dkey-30c15.10 |
| chchd3b | 8.0296292 | 0.331915054 | 0.109 | 0.007 | 1.62174422859909e-0 | 17 | chchd3b |
| zswim7 | 8.1103292 | 0.331443473 | 0.109 | 0.007 | 1.63804319696192e-0 | 17 | zswim7 |
| nsl1 | 1.0590771 | 0.320045776 | 0.127 | 0.01 | 2.13901802243087e-0 | 17 | nsl1 |
| alkbh4 | 1.2146268 | 0.354717687 | 0.145 | 0.014 | 2.45318184901535e-0 | 17 | alkbh4 |
| ruvbl1 | 1.5465166 | 0.679804987 | 0.327 | 0.073 | 3.12349972762477e-0 | 17 | ruvbl1 |
| rpf1 | 2.4613264 | 0.761878419 | 0.4 | 0.111 | 4.97114098184961e-0 | 17 | rpf1 |
| ppp1r14aa | 2.7105642 | 0.314095953 | 0.127 | 0.011 | 5.47452660666636e-0 | 17 | ppp1r14aa |
| alkbh3 | 3.0210955 | 0.436898849 | 0.182 | 0.023 | 6.10170662419457e-0 | 17 | alkbh3 |
| hint12 | 3.1785917 | 0.863089617 | 0.636 | 0.3 | 6.41980171518822e-0 | 17 | hint1 |
| sh3glb2b | 3.3310897 | 0.359377499 | 0.164 | 0.018 | 6.72780205497593e-0 | 17 | sh3glb2b |
| haus4 | 3.4699249 | 0.333460628 | 0.109 | 0.008 | 7.00820750114299e-0 | 17 | haus4 |
| cenps | 3.5593438 | 0.368821714 | 0.164 | 0.018 | 7.18880679523631e-0 | 17 | cenps |

| | | | | | | | |
|---|---|---|---|---|---|---|---|
| zgc:66472 | 4.0730546 | 0.42699649 | 0.182 | 0.023 | 8.22634846813986e-0 | 17 | zgc:66472 |
| ddx18 | 5.3679483 | 0.725919304 | 0.382 | 0.101 | 1.08416453065223e-0 | 17 | ddx18 |
| COX7A2 (1 of many) | 5.6817843 | 0.852031023 | 0.636 | 0.309 | 1.14754997787296e-0 | 17 | COX7A2 (1 of many) |
| mibp | 7.6721002 | 0.473672851 | 0.182 | 0.023 | 1.54953408777096e-0 | 17 | mibp |
| chtf18 | 8.7839201 | 0.320075201 | 0.127 | 0.011 | 1.77408834532367e-0 | 17 | chtf18 |
| zgc:103499 | 9.0708022 | 0.320443866 | 0.127 | 0.011 | 1.83202992701045e-0 | 17 | zgc:103499 |
| lsm3 | 9.3625363 | 0.81306895 | 0.473 | 0.158 | 1.89095146391718e-0 | 17 | lsm3 |
| snu13b2 | 1.0191059 | 0.822517058 | 0.618 | 0.288 | 2.05828827529452e-0 | 17 | snu13b |
| srrm2 | 1.0952254 | 0.757793576 | 0.382 | 0.105 | 2.21202690027656e-0 | 17 | srrm2 |
| auh1 | 1.2253525 | 0.506541835 | 0.218 | 0.035 | 2.47484456352989e-0 | 17 | auh |
| ndc1 | 1.2897743 | 0.594566015 | 0.236 | 0.041 | 2.60495729117735e-0 | 17 | ndc1 |
| si:ch211-222l21.16 | 1.3804774 | 0.697013626 | 0.873 | 0.527 | 2.78815021684564e-0 | 17 | si:ch211-222l21.1 |
| dnajc9 | 1.3975917 | 0.641044932 | 0.309 | 0.068 | 2.82271601766869e-0 | 17 | dnajc9 |
| snrpe1 | 1.4381211 | 0.863515426 | 0.6 | 0.27 | 2.90457337514648e-0 | 17 | snrpe |
| krcp | 1.6577618 | 0.251625786 | 0.109 | 0.008 | 3.34818164623478e-0 | 17 | krcp |
| lsm42 | 1.6739821 | 0.815529642 | 0.582 | 0.258 | 3.38094176534473e-0 | 17 | lsm4 |
| faub | 1.7477619 | 0.659769021 | 0.364 | 0.094 | 3.52995486245569e-0 | 17 | faub |
| zgc:162025 | 1.9580788 | 0.698134087 | 0.345 | 0.087 | 3.95473180850452e-0 | 17 | zgc:162025 |
| lsm12a | 2.3356927 | 0.590125532 | 0.291 | 0.062 | 4.71739862109943e-0 | 17 | lsm12a |
| mief2 | 2.3663349 | 0.287286667 | 0.127 | 0.012 | 4.77928665187945e-0 | 17 | mief2 |
| dguok | 2.8884528 | 0.375216714 | 0.145 | 0.016 | 5.83380815426588e-0 | 17 | dguok |
| psmd10 | 2.8953253 | 0.566864617 | 0.273 | 0.055 | 5.8476886273445e-08 | 17 | psmd10 |
| ralaa1 | 3.3214574 | 0.394368173 | 0.182 | 0.025 | 6.70834762867063e-0 | 17 | ralaa |
| ccdc130 | 3.5146478 | 0.404790553 | 0.164 | 0.02 | 7.09853417758975e-0 | 17 | ccdc130 |
| ppp4r2b | 3.5815419 | 0.483144773 | 0.2 | 0.03 | 7.23364027405115e-0 | 17 | ppp4r2b |
| rcc21 | 3.6017299 | 0.776086574 | 0.473 | 0.164 | 7.27441405469144e-0 | 17 | rcc2 |
| strap | 4.6144588 | 0.721371042 | 0.364 | 0.099 | 9.3198225593743e-08 | 17 | strap |
| nudc | 5.0301593 | 0.838520786 | 0.491 | 0.187 | 1.01594128441129e-0 | 17 | nudc |
| si:dkey-109l4.3 | 5.8163739 | 0.311754786 | 0.127 | 0.012 | 1.17473304891918e-0 | 17 | si:dkey-109l4.3 |
| zgc:101744 | 6.4121050 | 0.305582258 | 0.127 | 0.012 | 1.29505285815242e-0 | 17 | zgc:101744 |
| xrcc1 | 6.7172376 | 0.347989674 | 0.145 | 0.016 | 1.35668047920719e-0 | 17 | xrcc1 |
| minos1 | 7.0371811 | 0.775116759 | 0.473 | 0.168 | 1.42129947166713e-0 | 17 | minos1 |
| parp1 | 9.0458273 | 0.56841404 | 0.273 | 0.057 | 1.82698575593565e-0 | 17 | parp1 |
| ppa2 | 9.8525885 | 0.433297465 | 0.182 | 0.026 | 1.98992729997367e-0 | 17 | ppa2 |
| cct7 | 1.1094858 | 0.779560038 | 0.636 | 0.336 | 2.2408286299058e-07 | 17 | cct7 |

| | | | | | | | |
|---|---|---|---|---|---|---|---|
| gmnn | 1.1623404 | 0.480996769 | 0.182 | 0.026 | 2.3475790838425e-07 | 17 | gmnn |
| cpsf2 | 1.2609985 | 0.490239776 | 0.236 | 0.044 | 2.54683881803008e-0 | 17 | cpsf2 |
| hnrnpabb2 | 1.2741477 | 0.7524494 | 0.691 | 0.403 | 2.57339624886587e-0 | 17 | hnrnpabb |
| si:ch211-51e12.72 | 2.0812167 | 0.756077258 | 0.491 | 0.188 | 4.20343337587639e-0 | 17 | si:ch211-51e12.7 |
| hspa4a | 2.8019730 | 0.502703443 | 0.236 | 0.045 | 5.65914504076077e-0 | 17 | hspa4a |
| wrb | 2.8387243 | 0.580346885 | 0.236 | 0.046 | 5.73337155011561e-0 | 17 | wrb |
| dlgap5 | 2.9386540 | 0.372968893 | 0.164 | 0.022 | 5.93519966076858e-0 | 17 | dlgap5 |
| coil1 | 3.2344754 | 0.530786059 | 0.236 | 0.046 | 6.53267016549163e-0 | 17 | coil |
| cops5 | 3.2711310 | 0.581537528 | 0.291 | 0.068 | 6.60670345938871e-0 | 17 | cops5 |
| polr2l1 | 3.2731755 | 0.821021423 | 0.527 | 0.225 | 6.6108327061082e-07 | 17 | polr2l |
| lmo4a1 | 3.4627913 | 0.46894021 | 0.182 | 0.027 | 6.99379971508263e-0 | 17 | lmo4a |
| gtf2f2a | 3.5044709 | 0.737277364 | 0.4 | 0.128 | 7.07797999313964e-0 | 17 | gtf2f2a |
| chek2 | 3.7323169 | 0.319520119 | 0.127 | 0.013 | 7.53816045075801e-0 | 17 | chek2 |
| tubb2b1 | 3.7794548 | 0.812886961 | 0.636 | 0.317 | 7.63336488316199e-0 | 17 | tubb2b |
| pcgf6 | 4.3822178 | 0.357003856 | 0.109 | 0.01 | 8.85076539893757e-0 | 17 | pcgf6 |
| lsm1 | 4.9376739 | 0.622734903 | 0.309 | 0.076 | 9.97262014136495e-0 | 17 | lsm1 |
| cdc23 | 4.9778830 | 0.297554331 | 0.109 | 0.01 | 1.00538303639106e-0 | 17 | cdc23 |
| polr2f | 6.0040751 | 0.763106205 | 0.473 | 0.182 | 1.21264305849917e-0 | 17 | polr2f |
| atp5f1b3 | 6.1200988 | 0.668701484 | 0.745 | 0.53 | 1.23607635931393e-0 | 17 | atp5f1b |
| prpf31 | 6.2901744 | 0.52481749 | 0.218 | 0.04 | 1.27042653389944e-0 | 17 | prpf31 |
| aprt | 6.5121939 | 0.558002532 | 0.236 | 0.047 | 1.3152678111581e-06 | 17 | aprt |
| smarcad1a | 6.8362252 | 0.373758392 | 0.164 | 0.023 | 1.38071242317242e-0 | 17 | smarcad1a |
| rgp1 | 7.4896411 | 0.365919428 | 0.127 | 0.014 | 1.51268283202741e-0 | 17 | rgp1 |
| ndel1a | 7.8551103 | 0.314874895 | 0.127 | 0.014 | 1.58649664264373e-0 | 17 | ndel1a |
| si:dkey-185e18.7 | 8.9550665 | 0.370870134 | 0.145 | 0.018 | 1.80865479168511e-0 | 17 | si:dkey-185e18.7 |
| nelfcd | 9.9744400 | 0.319171283 | 0.145 | 0.018 | 2.01453765487127e-0 | 17 | nelfcd |
| mcm7 | 1.0480753 | 0.698888052 | 0.345 | 0.098 | 2.11679777335245e-0 | 17 | mcm7 |
| zgc:101851 | 1.1190023 | 0.547054582 | 0.236 | 0.048 | 2.26004897878564e-0 | 17 | zgc:101851 |
| ndufab1b1 | 1.1531487 | 0.781106538 | 0.491 | 0.201 | 2.32901443694396e-0 | 17 | ndufab1b |
| igbp1 | 1.2340836 | 0.337105036 | 0.182 | 0.028 | 2.49247872114692e-0 | 17 | igbp1 |
| eloca | 1.4588195 | 0.596219602 | 0.273 | 0.063 | 2.94637780690249e-0 | 17 | eloca |
| gcsha1 | 1.5287063 | 0.556231961 | 0.273 | 0.062 | 3.08752823330824e-0 | 17 | gcsha |
| snrpd3l2 | 1.6483160 | 0.742190999 | 0.436 | 0.159 | 3.32910401775586e-0 | 17 | snrpd3l |
| orc3 | 1.7932238 | 0.336585525 | 0.145 | 0.019 | 3.62177422773338e-0 | 17 | orc3 |
| orc4 | 2.1211407 | 0.417300701 | 0.182 | 0.029 | 4.28406787364538e-0 | 17 | orc4 |

| | | | | | | | |
|---|---|---|---|---|---|---|---|
| rpa3 | 2.1839829 | 0.67251077 | 0.364 | 0.109 | 4.41099044802278e-0 | 17 | rpa3 |
| ppa1b1 | 2.8992354 | 0.671447198 | 0.345 | 0.103 | 5.85558577822007e-0 | 17 | ppa1b |
| selenou1a | 2.9255529 | 0.533321073 | 0.236 | 0.05 | 5.90873926095608e-0 | 17 | selenou1a |
| ddx52 | 3.1214938 | 0.412709908 | 0.182 | 0.03 | 6.3044811213974e-06 | 17 | ddx52 |
| samd1a | 3.1465007 | 0.335616546 | 0.127 | 0.015 | 6.3549874889799e-06 | 17 | samd1a |
| h3f3a | 3.2581317 | 0.729367738 | 0.673 | 0.388 | 6.58044872665959e-0 | 17 | h3f3a |
| rnaseh2b | 3.6608882 | 0.479730267 | 0.218 | 0.043 | 7.39389609251605e-0 | 17 | rnaseh2b |
| si:ch211-210c8.6 | 3.6925322 | 0.325906102 | 0.127 | 0.015 | 7.45780744585082e-0 | 17 | si:ch211-210c8.6 |
| cenph | 3.7050090 | 0.282487594 | 0.109 | 0.011 | 7.483006747862e-06 | 17 | cenph |
| smim8 | 4.6603828 | 0.49791198 | 0.255 | 0.057 | 9.41257518141742e-0 | 17 | smim8 |
| nup50 | 6.9528257 | 0.592537147 | 0.327 | 0.093 | 1.40426221922734e-0 | 17 | nup50 |
| cxxc1b | 7.6361146 | 0.341066267 | 0.109 | 0.011 | 1.54226607891825e-0 | 17 | cxxc1b |
| sart1 | 9.2296788 | 0.732816228 | 0.364 | 0.119 | 1.86411823148011e-0 | 17 | sart1 |
| apooa | 9.2474161 | 0.442181499 | 0.2 | 0.037 | 1.86770063313429e-0 | 17 | apooa |
| snrpf | 1.0160887 | 0.709415096 | 0.636 | 0.388 | 2.05219451306428e-0 | 17 | snrpf |
| vrk1 | 1.2432441 | 0.555901536 | 0.255 | 0.06 | 2.51098022706822e-0 | 17 | vrk1 |
| ptgr11 | 1.2926559 | 0.584193098 | 0.364 | 0.115 | 2.61077732078674e-0 | 17 | ptgr1 |
| mcmbp | 1.4597050 | 0.354603006 | 0.182 | 0.032 | 2.94816633266494e-0 | 17 | mcmbp |
| nedd8 | 1.5486431 | 0.690629368 | 0.382 | 0.131 | 3.12779455388297e-0 | 17 | nedd8 |
| si:ch211-102c2.7 | 1.5633031 | 0.358758683 | 0.145 | 0.021 | 3.15740338873377e-0 | 17 | si:ch211-102c2.7 |
| smu1a | 1.6393248 | 0.575177572 | 0.273 | 0.069 | 3.31094444041581e-0 | 17 | smu1a |
| rpia3 | 1.7321556 | 0.629767673 | 0.309 | 0.087 | 3.4984347589046e-05 | 17 | rpia |
| zcchc2 | 1.7404671 | 0.256225448 | 0.127 | 0.016 | 3.51522155039116e-0 | 17 | zcchc2 |
| alkbh5 | 1.7595462 | 0.428664178 | 0.182 | 0.032 | 3.5537555641903e-05 | 17 | alkbh5 |
| si:ch211-127m7.2 | 1.8415848 | 0.306333191 | 0.109 | 0.012 | 3.71944886860336e-0 | 17 | si:ch211-127m7.2 |
| zgc:56106 | 1.9922397 | 0.583861909 | 0.273 | 0.07 | 4.02372656159138e-0 | 17 | zgc:56106 |
| purbb | 2.2668388 | 0.363965943 | 0.164 | 0.026 | 4.57833442638307e-0 | 17 | purbb |
| fam117ab | 2.5604526 | 0.317373471 | 0.127 | 0.016 | 5.17134612401626e-0 | 17 | fam117ab |
| cbx3a1 | 2.5799150 | 0.732395327 | 0.545 | 0.279 | 5.21065451784135e-0 | 17 | cbx3a |
| pcna | 4.0502843 | 0.638598873 | 0.473 | 0.171 | 8.18035929552975e-0 | 17 | pcna |
| aaas | 4.2940984 | 0.372297294 | 0.164 | 0.027 | 8.67279070190022e-0 | 17 | aaas |
| psma51 | 4.8343999 | 0.721298859 | 0.527 | 0.269 | 9.76403757272685e-0 | 17 | psma5 |
| zgc:101569 | 6.5506630 | 0.358260676 | 0.164 | 0.028 | 0.000132304 | 17 | zgc:101569 |
| eif4e1c1 | 6.5656111 | 0.654607056 | 0.309 | 0.094 | 0.000132606 | 17 | eif4e1c |
| pcmt | 7.1829477 | 0.501207201 | 0.236 | 0.056 | 0.000145074 | 17 | pcmt |

| | | | | | | | |
|---|---|---|---|---|---|---|---|
| rbx11 | 8.2004244 | 0.733632496 | 0.6 | 0.382 | 0.000165624 | 17 | rbx1 |
| zgc:136564 | 8.3293686 | 0.417668182 | 0.145 | 0.023 | 0.000168228 | 17 | zgc:136564 |
| polr2h | 8.9045538 | 0.533594515 | 0.255 | 0.065 | 0.000179845 | 17 | polr2h |
| calm3b2 | 8.9432465 | 0.67341661 | 0.636 | 0.407 | 0.000180627 | 17 | calm3b |
| atp5mc12 | 9.0162091 | 0.656123515 | 0.618 | 0.333 | 0.0001821 | 17 | atp5mc1 |
| imp31 | 1.0045453 | 0.543995101 | 0.273 | 0.074 | 0.000202888 | 17 | imp3 |
| ppih | 1.0121447 | 0.630357353 | 0.273 | 0.076 | 0.000204423 | 17 | ppih |
| sod22 | 1.0879129 | 0.68114517 | 0.382 | 0.143 | 0.000219726 | 17 | sod2 |
| edc4 | 1.0896266 | 0.349449936 | 0.145 | 0.023 | 0.000220072 | 17 | edc4 |
| si:dkey-87o1.21 | 1.0959254 | 0.269619571 | 0.109 | 0.013 | 0.000221344 | 17 | si:dkey-87o1.2 |
| gyg1a1 | 1.1625724 | 0.575111064 | 0.291 | 0.082 | 0.000234805 | 17 | gyg1a |
| si:dkey-6i22.5 | 1.2378129 | 0.400115167 | 0.145 | 0.023 | 0.000250001 | 17 | si:dkey-6i22.5 |
| gid8a | 1.2647118 | 0.357873346 | 0.164 | 0.029 | 0.000255434 | 17 | gid8a |
| txlna | 1.2924819 | 0.459287627 | 0.182 | 0.036 | 0.000261043 | 17 | txlna |
| seta1 | 1.3136526 | 0.693483651 | 0.582 | 0.355 | 0.000265318 | 17 | seta |
| btr04 | 1.4120650 | 0.358808238 | 0.109 | 0.013 | 0.000285195 | 17 | btr04 |
| psmd4a | 1.4160829 | 0.552925495 | 0.291 | 0.084 | 0.000286006 | 17 | psmd4a |
| zgc:114104 | 1.4472054 | 0.300182831 | 0.109 | 0.013 | 0.000292292 | 17 | zgc:114104 |
| sf3b61 | 1.4747687 | 0.675792116 | 0.564 | 0.312 | 0.000297859 | 17 | sf3b6 |
| gnl3 | 1.4921235 | 0.682973021 | 0.345 | 0.12 | 0.000301364 | 17 | gnl3 |
| ing2 | 1.6111992 | 0.477466624 | 0.2 | 0.043 | 0.000325414 | 17 | ing2 |
| senp3a | 1.6161332 | 0.26202405 | 0.109 | 0.013 | 0.00032641 | 17 | senp3a |
| cript | 1.6991780 | 0.415533057 | 0.2 | 0.043 | 0.000343183 | 17 | cript |
| rcbtb1 | 1.7181012 | 0.287593312 | 0.109 | 0.013 | 0.000347005 | 17 | rcbtb1 |
| dynll2a2 | 1.8003180 | 0.381568594 | 0.164 | 0.029 | 0.00036361 | 17 | dynll2a |
| rsl1d11 | 1.9241349 | 0.696407283 | 0.491 | 0.239 | 0.000388618 | 17 | rsl1d1 |
| epcam1 | 1.9714306 | 0.383077104 | 0.164 | 0.028 | 0.00039817 | 17 | epcam |
| mcm3 | 2.0941098 | 0.541740721 | 0.255 | 0.067 | 0.000422947 | 17 | mcm3 |
| polr3f | 2.3525550 | 0.378109994 | 0.182 | 0.036 | 0.000475146 | 17 | polr3f |
| tk2 | 2.5809624 | 0.381829622 | 0.182 | 0.036 | 0.000521277 | 17 | tk2 |
| wdr82 | 2.6845225 | 0.588735988 | 0.309 | 0.098 | 0.000542193 | 17 | wdr82 |
| ciapin11 | 2.7969321 | 0.551300566 | 0.255 | 0.069 | 0.000564896 | 17 | ciapin1 |
| utp23 | 2.9217631 | 0.510537997 | 0.255 | 0.068 | 0.000590109 | 17 | utp23 |
| fkbp8 | 3.1116487 | 0.290472335 | 0.127 | 0.019 | 0.00062846 | 17 | fkbp8 |
| snrpd21 | 3.1544055 | 0.680208198 | 0.509 | 0.263 | 0.000637095 | 17 | snrpd2 |

| | | | | | | | |
|---|---|---|---|---|---|---|---|
| tubb4b1 | 3.2239004 | 0.463146121 | 0.764 | 0.671 | 0.000651131 | 17 | tubb4b |
| ap3s2 | 3.2935675 | 0.424814405 | 0.2 | 0.044 | 0.000665202 | 17 | ap3s2 |
| atp5mea1 | 3.6574426 | 0.714891301 | 0.418 | 0.182 | 0.000738694 | 17 | atp5mea |
| wee21 | 3.7495444 | 0.371453224 | 0.164 | 0.03 | 0.000757295 | 17 | wee2 |
| h2afx14 | 4.6695859 | 0.633773585 | 0.618 | 0.379 | 0.000943116 | 17 | h2afx1 |
| brdt | 4.7152742 | 0.584193005 | 0.273 | 0.08 | 0.000952344 | 17 | brdt |
| psma8 | 4.9342325 | 0.646203921 | 0.564 | 0.324 | 0.000996567 | 17 | psma8 |
| drg2 | 5.2004293 | 0.555770671 | 0.309 | 0.1 | 0.001050331 | 17 | drg2 |
| tktb | 5.4345754 | 0.639546089 | 0.345 | 0.125 | 0.001097621 | 17 | tktb |
| bnip3la | 5.6276250 | 0.311185859 | 0.127 | 0.019 | 0.001136611 | 17 | bnip3la |
| dnph1 | 5.9143024 | 0.41359865 | 0.2 | 0.045 | 0.001194512 | 17 | dnph1 |
| ss18 | 6.1134141 | 0.564683848 | 0.291 | 0.091 | 0.001234726 | 17 | ss18 |
| skp12 | 6.3143994 | 0.624401955 | 0.6 | 0.358 | 0.001275319 | 17 | skp1 |
| mrpl47 | 6.6608946 | 0.553821144 | 0.273 | 0.081 | 0.001345301 | 17 | mrpl47 |
| pold1 | 6.7994457 | 0.35461184 | 0.145 | 0.025 | 0.001373284 | 17 | pold1 |
| baz1a | 8.2789775 | 0.486421719 | 0.273 | 0.08 | 0.001672105 | 17 | baz1a |
| hnrnph3 | 8.6032373 | 0.272456363 | 0.127 | 0.02 | 0.001737596 | 17 | hnrnph3 |
| them4 | 8.6899498 | 0.255514058 | 0.109 | 0.015 | 0.001755109 | 17 | them4 |
| prmt5 | 8.8269271 | 0.466125298 | 0.2 | 0.047 | 0.001782774 | 17 | prmt5 |
| pttg1 | 8.9816338 | 0.363896092 | 0.164 | 0.032 | 0.001814021 | 17 | pttg1 |
| ppie | 1.0268385 | 0.397934066 | 0.164 | 0.032 | 0.002073906 | 17 | ppie |
| cdc20 | 1.0406880 | 0.298379088 | 0.127 | 0.02 | 0.002101878 | 17 | cdc20 |
| hnrnpa0l1 | 1.1519772 | 0.358832307 | 0.782 | 0.706 | 0.002326648 | 17 | hnrnpa0l |
| xrcc6 | 1.1949361 | 0.419950426 | 0.2 | 0.047 | 0.002413413 | 17 | xrcc6 |
| uqcrc12 | 1.1978484 | 0.626773565 | 0.418 | 0.185 | 0.002419295 | 17 | uqcrc1 |
| bud312 | 1.2364947 | 0.658665184 | 0.473 | 0.241 | 0.002497348 | 17 | bud31 |
| hspd12 | 1.2547076 | 0.621473036 | 0.4 | 0.166 | 0.002534133 | 17 | hspd1 |
| pop5 | 1.4137428 | 0.436463383 | 0.218 | 0.055 | 0.002855337 | 17 | pop5 |
| pfdn6 | 1.4920303 | 0.620868112 | 0.382 | 0.156 | 0.003013454 | 17 | pfdn6 |
| zgc:865982 | 1.6226068 | 0.597772835 | 0.364 | 0.145 | 0.003277179 | 17 | zgc:86598 |
| hadhaa | 1.6715097 | 0.380601794 | 0.182 | 0.04 | 0.003375948 | 17 | hadhaa |
| rbb4l1 | 1.7407461 | 0.605339743 | 0.309 | 0.108 | 0.003515785 | 17 | rbb4l |
| lin37 | 1.8934417 | 0.442322099 | 0.182 | 0.041 | 0.003824184 | 17 | lin37 |
| parp2 | 1.9744901 | 0.397937135 | 0.182 | 0.04 | 0.003987878 | 17 | parp2 |
| hspa9 | 2.0304014 | 0.546729801 | 0.309 | 0.105 | 0.004100802 | 17 | hspa9 |

| | | | | | | | |
|---|---|---|---|---|---|---|---|
| cdk2 | 2.1463950 | 0.470411698 | 0.182 | 0.041 | 0.004335074 | 17 | cdk2 |
| rpp21 | 2.2839881 | 0.441754126 | 0.2 | 0.049 | 0.004612971 | 17 | rpp21 |
| tcp1 | 2.3783914 | 0.615126486 | 0.564 | 0.348 | 0.004803637 | 17 | tcp1 |
| hnrnph1 | 2.4195581 | 0.283408575 | 0.109 | 0.016 | 0.004886782 | 17 | hnrnph1 |
| tsn | 2.4318193 | 0.357063252 | 0.145 | 0.027 | 0.004911546 | 17 | tsn |
| dis3 | 2.4322951 | 0.286805592 | 0.109 | 0.016 | 0.004912507 | 17 | dis3 |
| nxt2 | 2.4575753 | 0.514482947 | 0.255 | 0.077 | 0.004963565 | 17 | nxt2 |
| wdr121 | 2.5528493 | 0.470811195 | 0.236 | 0.066 | 0.00515599 | 17 | wdr12 |
| mcm2 | 2.5790625 | 0.528185993 | 0.291 | 0.094 | 0.005208933 | 17 | mcm2 |
| rnf4 | 2.6542003 | 0.331273019 | 0.164 | 0.034 | 0.005360688 | 17 | rnf4 |
| ndufa81 | 2.7310728 | 0.66154783 | 0.436 | 0.214 | 0.005515948 | 17 | ndufa8 |
| rtf2 | 2.8167843 | 0.517734623 | 0.273 | 0.086 | 0.005689059 | 17 | rtf2 |
| supt3h | 3.1220424 | 0.500704283 | 0.236 | 0.068 | 0.006305589 | 17 | supt3h |
| mybbp1a | 3.2200856 | 0.397734759 | 0.182 | 0.042 | 0.006503607 | 17 | mybbp1a |
| abcf2a | 3.3340188 | 0.552247364 | 0.255 | 0.078 | 0.006733718 | 17 | abcf2a |
| cct5 | 3.3604870 | 0.597701775 | 0.6 | 0.411 | 0.006787176 | 17 | cct5 |
| dcaf13 | 3.4456694 | 0.436795437 | 0.182 | 0.042 | 0.006959219 | 17 | dcaf13 |
| mbd3b | 3.4912694 | 0.455673614 | 0.236 | 0.066 | 0.007051317 | 17 | mbd3b |
| selenoh | 3.5176713 | 0.612635891 | 0.382 | 0.159 | 0.007104641 | 17 | selenoh |
| prpsap1 | 3.5478109 | 0.402968336 | 0.2 | 0.049 | 0.007165514 | 17 | prpsap1 |
| nup93 | 3.6824451 | 0.430902375 | 0.182 | 0.042 | 0.007437434 | 17 | nup93 |
| usp1 | 4.1043790 | 0.495446129 | 0.218 | 0.059 | 0.008289614 | 17 | usp1 |
| magoh | 4.1583735 | 0.552565453 | 0.327 | 0.12 | 0.008398667 | 17 | magoh |
| ldlrap1a2 | 4.3965924 | 0.306994584 | 0.164 | 0.034 | 0.008879798 | 17 | ldlrap1a |
| rai142 | 4.6018815 | 0.400137939 | 0.145 | 0.028 | 0.00929442 | 17 | rai14 |
| mfap11 | 4.6046713 | 0.473889379 | 0.255 | 0.077 | 0.009300055 | 17 | mfap1 |
| ndufab1a1 | 4.9936122 | 0.637439983 | 0.4 | 0.184 | 0.010085599 | 17 | ndufab1a |
| timm13 | 5.1286034 | 0.541266318 | 0.273 | 0.09 | 0.01035824 | 17 | timm13 |
| si:dkey-46a10.31 | 5.1826518 | 0.526489 | 0.236 | 0.069 | 0.010467402 | 17 | si:dkey-46a10.3 |
| ilf22 | 5.2208108 | 0.638772595 | 0.473 | 0.247 | 0.010544472 | 17 | ilf2 |
| shmt1 | 5.5068586 | 0.457174974 | 0.182 | 0.044 | 0.011122202 | 17 | shmt1 |
| LOC103909972 | 5.7778813 | 0.565923367 | 0.327 | 0.124 | 0.011669587 | 17 | LOC103909972 |
| CABZ01044053.1 | 5.8603732 | 0.432199643 | 0.218 | 0.059 | 0.011836196 | 17 | CABZ01044053.1 |
| ndufv2 | 5.9119794 | 0.546199607 | 0.291 | 0.103 | 0.011940425 | 17 | ndufv2 |
| cirbpa | 6.1730436 | 0.359396136 | 0.8 | 0.703 | 0.012467696 | 17 | cirbpa |

| | | | | | | | |
|---|---|---|---|---|---|---|---|
| puf60a1 | 6.4554429 | 0.582304912 | 0.327 | 0.125 | 0.013038058 | 17 | puf60a |
| gtf2h3 | 6.4848123 | 0.312076482 | 0.145 | 0.029 | 0.013097375 | 17 | gtf2h3 |
| psmd132 | 6.6404431 | 0.574158865 | 0.364 | 0.149 | 0.013411703 | 17 | psmd13 |
| si:dkey-276j7.1 | 6.6679401 | 0.505426332 | 0.255 | 0.079 | 0.013467239 | 17 | si:dkey-276j7.1 |
| riok2 | 6.8322781 | 0.49777482 | 0.218 | 0.061 | 0.013799152 | 17 | riok2 |
| mtx21 | 7.0685108 | 0.413860447 | 0.2 | 0.052 | 0.014276271 | 17 | mtx2 |
| mrpl21 | 7.0956938 | 0.399341964 | 0.164 | 0.036 | 0.014331173 | 17 | mrpl21 |
| stip1 | 7.2040307 | 0.581968377 | 0.327 | 0.127 | 0.014549981 | 17 | stip1 |
| ufc1 | 7.5606689 | 0.515109505 | 0.291 | 0.1 | 0.015270283 | 17 | ufc1 |
| ankrd40 | 7.8190108 | 0.31271655 | 0.127 | 0.023 | 0.015792056 | 17 | ankrd40 |
| rangrf | 8.2518666 | 0.328683447 | 0.145 | 0.029 | 0.016666295 | 17 | rangrf |
| nudcd22 | 8.5292053 | 0.502377665 | 0.273 | 0.091 | 0.017226436 | 17 | nudcd2 |
| thop1 | 9.3655200 | 0.395162898 | 0.182 | 0.045 | 0.018915541 | 17 | thop1 |
| atp5md1 | 9.5161836 | 0.620929734 | 0.564 | 0.352 | 0.019219836 | 17 | atp5md |
| lin52 | 9.8987525 | 0.495942578 | 0.255 | 0.081 | 0.01999251 | 17 | lin52 |
| si:ch211-276i12.11 | 1.0197397 | 0.443080358 | 0.236 | 0.07 | 0.020595684 | 17 | si:ch211-276i12.11 |
| cdca7a | 1.0561838 | 0.491881247 | 0.2 | 0.053 | 0.021331744 | 17 | cdca7a |
| rps19bp1 | 1.1045987 | 0.512354317 | 0.255 | 0.082 | 0.022309581 | 17 | rps19bp1 |
| aldh7a11 | 1.1334258 | 0.509097176 | 0.291 | 0.104 | 0.022891802 | 17 | aldh7a1 |
| gstp11 | 1.1934535 | 0.523002229 | 0.545 | 0.278 | 0.024104182 | 17 | gstp1 |
| taf151 | 1.2300515 | 0.614417421 | 0.4 | 0.186 | 0.024843351 | 17 | taf15 |
| ppme1 | 1.3538582 | 0.478496136 | 0.236 | 0.072 | 0.027343875 | 17 | ppme1 |
| gpn1 | 1.5230635 | 0.37700362 | 0.145 | 0.031 | 0.030761315 | 17 | gpn1 |
| ndufb22 | 1.6233343 | 0.607921619 | 0.455 | 0.24 | 0.032786485 | 17 | ndufb2 |
| nol12 | 1.6379456 | 0.442355998 | 0.218 | 0.063 | 0.033081589 | 17 | nol12 |
| mcm4 | 1.7252646 | 0.433106061 | 0.236 | 0.072 | 0.03484517 | 17 | mcm4 |
| psmd7 | 1.9012471 | 0.605464861 | 0.382 | 0.177 | 0.038399489 | 17 | psmd7 |
| CABZ01092156.11 | 1.9215871 | 0.356328237 | 0.145 | 0.031 | 0.038810296 | 17 | CABZ01092156.1 |
| lyrm9 | 1.9625523 | 0.349889533 | 0.2 | 0.054 | 0.039637669 | 17 | lyrm9 |
| rtcb | 1.9779423 | 0.306142497 | 0.145 | 0.031 | 0.039948503 | 17 | rtcb |
| dele1 | 1.9943380 | 0.411759207 | 0.145 | 0.031 | 0.040279645 | 17 | dele1 |
| cct3 | 2.0009553 | 0.607754493 | 0.527 | 0.333 | 0.040413295 | 17 | cct3 |
| csnk2b1 | 2.0339619 | 0.590807275 | 0.509 | 0.302 | 0.041079929 | 17 | csnk2b |
| smarca1 | 2.1418538 | 0.33028767 | 0.109 | 0.018 | 0.043259023 | 17 | smarca1 |
| rbbp4 | 2.1768327 | 0.589553004 | 0.327 | 0.132 | 0.04396549 | 17 | rbbp4 |

| | | | | | | | |
|---|---|---|---|---|---|---|---|
| tfdp1b | 2.1792294 | 0.2941177 | 0.127 | 0.024 | 0.044013898 | 17 | tfdp1b |
| ptges3b2 | 2.2078713 | 0.643742821 | 0.436 | 0.227 | 0.044592377 | 17 | ptges3b |
| zgc:923132 | 2.2501956 | 0.287373904 | 0.127 | 0.024 | 0.045447201 | 17 | zgc:92313 |
| atp5fa11 | 2.4151202 | 0.525246163 | 0.636 | 0.462 | 0.048778184 | 17 | atp5fa1 |
| tuba8l | 2.5511120 | 0.564170088 | 0.545 | 0.312 | 0.05152481 | 17 | tuba8l |
| clns1a | 2.5754941 | 0.524674648 | 0.255 | 0.086 | 0.052017256 | 17 | clns1a |
| eif2s2 | 2.5997626 | 0.520998354 | 0.655 | 0.466 | 0.052507407 | 17 | eif2s2 |
| trmt6 | 2.7451880 | 0.284618784 | 0.127 | 0.025 | 0.055444563 | 17 | trmt6 |
| acaa1 | 2.7677135 | 0.29272261 | 0.145 | 0.031 | 0.055899511 | 17 | acaa1 |
| CABZ01005379.1 | 2.7873431 | 0.390635604 | 0.164 | 0.039 | 0.056295969 | 17 | CABZ01005379.1 |
| ttc9c | 3.0643982 | 0.293985142 | 0.127 | 0.025 | 0.061891651 | 17 | ttc9c |
| uhrf1 | 3.3127017 | 0.289990077 | 0.109 | 0.019 | 0.066906638 | 17 | uhrf1 |
| si:ch211-214j24.1( | 3.4404511 | 0.566768406 | 0.364 | 0.165 | 0.069486791 | 17 | si:ch211-214j24.10 |
| gstm.16 | 3.4637766 | 0.534815769 | 0.582 | 0.337 | 0.069957896 | 17 | gstm.1 |
| pdcd4b | 3.8160587 | 0.254943327 | 0.109 | 0.019 | 0.077072938 | 17 | pdcd4b |
| slmapb1 | 3.8252914 | 0.359885406 | 0.145 | 0.032 | 0.077259411 | 17 | slmapb |
| paip2b3 | 3.8401178 | 0.548861469 | 0.545 | 0.342 | 0.07755886 | 17 | paip2b |
| nub1 | 4.0418551 | 0.306538211 | 0.145 | 0.032 | 0.081633348 | 17 | nub1 |
| cox5aa3 | 4.1588024 | 0.60096205 | 0.582 | 0.41 | 0.083995333 | 17 | cox5aa |
| bmi1b | 4.1923889 | 0.292400254 | 0.127 | 0.025 | 0.08467368 | 17 | bmi1b |
| tsr2 | 4.2844718 | 0.50410348 | 0.255 | 0.088 | 0.086533478 | 17 | tsr2 |
| pdrg1 | 4.6864861 | 0.341622444 | 0.182 | 0.048 | 0.09465296 | 17 | pdrg1 |
| tchp | 4.7075926 | 0.261839773 | 0.109 | 0.019 | 0.095079249 | 17 | tchp |
| synrg | 5.0007991 | 0.257389141 | 0.109 | 0.019 | 0.10100114 | 17 | synrg |
| sap182 | 5.0254217 | 0.607202329 | 0.436 | 0.238 | 0.101498443 | 17 | sap18 |
| rnf19a | 5.3690005 | 0.33263961 | 0.164 | 0.04 | 0.108437703 | 17 | rnf19a |
| gnpda12 | 5.4766831 | 0.3603562 | 0.2 | 0.056 | 0.11061257 | 17 | gnpda1 |
| utp15 | 5.7101421 | 0.347613835 | 0.164 | 0.041 | 0.115327741 | 17 | utp15 |
| tuba8l41 | 5.8097667 | 0.541698774 | 0.636 | 0.452 | 0.11733986 | 17 | tuba8l4 |
| aldh3a2a | 6.1843447 | 0.263660047 | 0.109 | 0.02 | 0.124905212 | 17 | aldh3a2a |
| cetn3 | 6.2481014 | 0.346359905 | 0.145 | 0.033 | 0.126192906 | 17 | cetn3 |
| eif4a31 | 6.5399875 | 0.56903279 | 0.418 | 0.216 | 0.132088128 | 17 | eif4a3 |
| med282 | 6.5642166 | 0.477310336 | 0.273 | 0.103 | 0.132577484 | 17 | med28 |
| prim1 | 6.8141987 | 0.26824909 | 0.109 | 0.02 | 0.137626372 | 17 | prim1 |
| ss18l2 | 6.8348567 | 0.453338312 | 0.236 | 0.078 | 0.138043601 | 17 | ss18l2 |

| | | | | | | | |
|---|---|---|---|---|---|---|---|
| vrk3 | 7.1667270 | 0.251444864 | 0.109 | 0.02 | 0.144746385 | 17 | vrk3 |
| plrg1 | 7.5040545 | 0.426249765 | 0.2 | 0.059 | 0.15155939 | 17 | plrg1 |
| pgls | 8.5455020 | 0.279444485 | 0.109 | 0.02 | 0.172593505 | 17 | pgls |
| brk11 | 8.5485488 | 0.585832334 | 0.436 | 0.245 | 0.172655041 | 17 | brk1 |
| dynlrb11 | 8.5935794 | 0.598761194 | 0.455 | 0.268 | 0.173564525 | 17 | dynlrb1 |
| taf132 | 8.6501929 | 0.562433878 | 0.473 | 0.276 | 0.174707946 | 17 | taf13 |
| pus1 | 8.8179230 | 0.349573655 | 0.164 | 0.042 | 0.178095591 | 17 | pus1 |
| XLOC-044204 | 9.3081237 | 0.300860751 | 0.145 | 0.034 | 0.187996176 | 17 | XLOC-044204 |
| chchd10 | 9.4403368 | 0.506523717 | 0.182 | 0.052 | 0.190666484 | 17 | chchd10 |
| trappc3 | 9.6253059 | 0.383455106 | 0.2 | 0.059 | 0.194402304 | 17 | trappc3 |
| coq9 | 1.0127777 | 0.373695458 | 0.2 | 0.06 | 0.204550718 | 17 | coq9 |
| ttc5 | 1.0329255 | 0.301617072 | 0.127 | 0.027 | 0.208619967 | 17 | ttc5 |
| mrps9 | 1.0702368 | 0.454819864 | 0.182 | 0.052 | 0.21615574 | 17 | mrps9 |
| bbip1 | 1.0825515 | 0.493917486 | 0.255 | 0.092 | 0.218642937 | 17 | bbip1 |
| mapre1a3 | 1.0892067 | 0.499105677 | 0.418 | 0.198 | 0.219987087 | 17 | mapre1a |
| fkbp31 | 1.0942457 | 0.619773364 | 0.436 | 0.253 | 0.221004811 | 17 | fkbp3 |
| rpa1 | 1.1488820 | 0.469635266 | 0.255 | 0.092 | 0.232039707 | 17 | rpa1 |
| srsf3b2 | 1.2539582 | 0.566500816 | 0.491 | 0.307 | 0.253261956 | 17 | srsf3b |
| wdr83 | 1.2880204 | 0.256953707 | 0.109 | 0.021 | 0.260141485 | 17 | wdr83 |
| oaz1a3 | 1.3016572 | 0.403386731 | 0.709 | 0.595 | 0.262895718 | 17 | oaz1a |
| ppat | 1.3472157 | 0.294439114 | 0.127 | 0.028 | 0.272097165 | 17 | ppat |
| h2afva2 | 1.3491152 | 0.621530166 | 0.418 | 0.233 | 0.2724808 | 17 | h2afva |
| gtf2e21 | 1.4576720 | 0.465164617 | 0.255 | 0.094 | 0.294406029 | 17 | gtf2e2 |
| ahcy1 | 1.4822615 | 0.455350543 | 0.291 | 0.11 | 0.299372359 | 17 | ahcy |
| cd2bp2 | 1.4990046 | 0.392141439 | 0.2 | 0.061 | 0.302753979 | 17 | cd2bp2 |
| eftud2 | 1.5075168 | 0.324199633 | 0.127 | 0.028 | 0.304473172 | 17 | eftud2 |
| chrac11 | 1.5701099 | 0.488317113 | 0.255 | 0.095 | 0.317115107 | 17 | chrac1 |
| atp5pb1 | 1.6120061 | 0.501950618 | 0.618 | 0.492 | 0.325576887 | 17 | atp5pb |
| atp13a1 | 1.6546900 | 0.304872395 | 0.127 | 0.028 | 0.334197758 | 17 | atp13a1 |
| psmf1 | 1.7079155 | 0.450108581 | 0.2 | 0.063 | 0.344947709 | 17 | psmf1 |
| ndufs72 | 1.7658506 | 0.54212266 | 0.364 | 0.178 | 0.356648861 | 17 | ndufs7 |
| nolc1 | 2.0281638 | 0.484958734 | 0.236 | 0.085 | 0.409628255 | 17 | nolc1 |
| gstr3 | 2.0722794 | 0.525996976 | 0.364 | 0.176 | 0.418538271 | 17 | gstr |
| spoplb | 2.1091568 | 0.299635602 | 0.145 | 0.036 | 0.425986416 | 17 | spoplb |
| glud1b | 2.1225129 | 0.400673253 | 0.218 | 0.074 | 0.428683935 | 17 | glud1b |

| | | | | | | | |
|---|---|---|---|---|---|---|---|
| psmc1a1 | 2.1477029 | 0.5618081 | 0.327 | 0.154 | 0.433771573 | 17 | psmc1a |
| COX5B3 | 2.1505549 | 0.507292989 | 0.6 | 0.481 | 0.434347591 | 17 | COX5B |
| map7d3 | 2.2096714 | 0.28200085 | 0.109 | 0.022 | 0.44628734 | 17 | map7d3 |
| mrpl14 | 2.2492973 | 0.429676379 | 0.255 | 0.095 | 0.454290577 | 17 | mrpl14 |
| rcc1l | 2.2500805 | 0.264654566 | 0.127 | 0.029 | 0.454448766 | 17 | rcc1l |
| ufd1l | 2.2973944 | 0.478257999 | 0.255 | 0.096 | 0.46400475 | 17 | ufd1l |
| dnaja2b | 2.3063608 | 0.469904083 | 0.255 | 0.096 | 0.465815693 | 17 | dnaja2b |
| psmc41 | 2.3529613 | 0.5601034 | 0.345 | 0.166 | 0.475227605 | 17 | psmc4 |
| thap12a | 2.4720084 | 0.276911504 | 0.127 | 0.029 | 0.499271539 | 17 | thap12a |
| tomm34 | 2.5082397 | 0.316121904 | 0.127 | 0.029 | 0.506589181 | 17 | tomm34 |
| rexo4 | 2.5170686 | 0.455373942 | 0.218 | 0.074 | 0.508372358 | 17 | rexo4 |
| asf1ba | 2.5879962 | 0.268521005 | 0.145 | 0.036 | 0.522697604 | 17 | asf1ba |
| cct21 | 2.6306267 | 0.513768525 | 0.564 | 0.399 | 0.531307689 | 17 | cct2 |
| zgc:171480 | 2.6339649 | 0.445041941 | 0.236 | 0.086 | 0.531981891 | 17 | zgc:171480 |
| si:ch211-276i12.9 | 2.6896763 | 0.417980151 | 0.182 | 0.055 | 0.54323394 | 17 | si:ch211-276i12.9 |
| ccng1 | 2.7257543 | 0.596117672 | 0.545 | 0.408 | 0.550520599 | 17 | ccng1 |
| hat1 | 2.7992895 | 0.354293432 | 0.164 | 0.045 | 0.565372514 | 17 | hat1 |
| smc5 | 2.8470791 | 0.325690546 | 0.145 | 0.037 | 0.575024572 | 17 | smc5 |
| rfc4 | 2.8914229 | 0.41086711 | 0.2 | 0.064 | 0.583980695 | 17 | rfc4 |
| slc25a25a | 2.9403113 | 0.42702787 | 0.182 | 0.055 | 0.593854683 | 17 | slc25a25a |
| hmgb2a4 | 2.9772676 | 0.399891692 | 0.782 | 0.582 | 0.601318753 | 17 | hmgb2a |
| txn21 | 2.9969708 | 0.5036884 | 0.291 | 0.123 | 0.605298193 | 17 | txn2 |
| uqcc2 | 2.9988330 | 0.476891283 | 0.255 | 0.1 | 0.605674301 | 17 | uqcc2 |
| slc25a12 | 3.0359880 | 0.279632111 | 0.109 | 0.022 | 0.613178507 | 17 | slc25a12 |
| mcm6 | 3.2896586 | 0.405075502 | 0.236 | 0.085 | 0.664412349 | 17 | mcm6 |
| sae1 | 3.3673325 | 0.42213801 | 0.182 | 0.056 | 0.68010016 | 17 | sae1 |
| timm10 | 3.4142886 | 0.454986803 | 0.255 | 0.098 | 0.689583876 | 17 | timm10 |
| sigmar12 | 3.6044003 | 0.423413634 | 0.182 | 0.056 | 0.727980742 | 17 | sigmar1 |
| acot13 | 3.6683490 | 0.342287572 | 0.164 | 0.046 | 0.740896465 | 17 | acot13 |
| prdx32 | 3.8807770 | 0.51578716 | 0.364 | 0.184 | 0.783800538 | 17 | prdx3 |
| gins4 | 3.9140305 | 0.259470259 | 0.109 | 0.023 | 0.790516753 | 17 | gins4 |
| cstf21 | 4.4228730 | 0.467129402 | 0.255 | 0.102 | 0.893287675 | 17 | cstf2 |
| eif5a21 | 4.4418370 | 0.474315305 | 0.6 | 0.463 | 0.897117827 | 17 | eif5a2 |
| naa10 | 4.5512214 | 0.542465657 | 0.436 | 0.265 | 0.919210204 | 17 | naa10 |
| rcc1 | 4.5541446 | 0.337569258 | 0.145 | 0.039 | 0.919800595 | 17 | rcc1 |

| | | | | | | | |
|---|---|---|---|---|---|---|---|
| mre11a | 4.6527557 | 0.267683605 | 0.109 | 0.023 | 0.939717074 | 17 | mre11a |
| sub1a1 | 4.6899491 | 0.556708865 | 0.509 | 0.367 | 0.947229035 | 17 | sub1a |
| pole4 | 4.8603160 | 0.406581481 | 0.236 | 0.088 | 0.981638032 | 17 | pole4 |
| rfc5 | 4.9585197 | 0.432034358 | 0.2 | 0.067 | 1 | 17 | rfc5 |
| zmat5 | 5.2792053 | 0.304146446 | 0.127 | 0.031 | 1 | 17 | zmat5 |
| cct6a | 6.0736235 | 0.527151711 | 0.473 | 0.297 | 1 | 17 | cct6a |
| fgfr1op | 6.0958230 | 0.308020999 | 0.127 | 0.031 | 1 | 17 | fgfr1op |
| mcm5 | 6.4852011 | 0.394127669 | 0.218 | 0.078 | 1 | 17 | mcm5 |
| nop16 | 6.9384203 | 0.446018589 | 0.236 | 0.092 | 1 | 17 | nop16 |
| znhit1 | 7.3796658 | 0.361134577 | 0.164 | 0.049 | 1 | 17 | znhit1 |
| llph | 7.4573731 | 0.502163926 | 0.291 | 0.132 | 1 | 17 | llph |
| prpf6 | 7.5962537 | 0.333458999 | 0.164 | 0.049 | 1 | 17 | prpf6 |
| api5 | 7.6498906 | 0.421449452 | 0.236 | 0.092 | 1 | 17 | api5 |
| usf1 | 7.8129039 | 0.332540839 | 0.164 | 0.049 | 1 | 17 | usf1 |
| ran1 | 7.8504244 | 0.291486853 | 0.691 | 0.666 | 1 | 17 | ran |
| pacsin3 | 8.0724837 | 0.255038722 | 0.109 | 0.024 | 1 | 17 | pacsin3 |
| uba2 | 9.1622555 | 0.41320708 | 0.218 | 0.081 | 1 | 17 | uba2 |
| si:ch211-210c8.7 | 9.2120191 | 0.372266373 | 0.182 | 0.059 | 1 | 17 | si:ch211-210c8.7 |
| uchl3 | 9.4527155 | 0.393896609 | 0.182 | 0.059 | 1 | 17 | uchl3 |
| polr1d | 9.4952632 | 0.34171349 | 0.182 | 0.059 | 1 | 17 | polr1d |
| rae1 | 9.5475628 | 0.349951751 | 0.164 | 0.05 | 1 | 17 | rae1 |
| ccdc137 | 9.9290281 | 0.266123407 | 0.127 | 0.032 | 1 | 17 | ccdc137 |
| pde6d | 0.000101 | 0.252247258 | 0.109 | 0.025 | 1 | 17 | pde6d |
| psmd141 | 0.000103 | 0.49045918 | 0.327 | 0.161 | 1 | 17 | psmd14 |
| gid8b | 0.000115 | 0.355254906 | 0.182 | 0.06 | 1 | 17 | gid8b |
| sympk1 | 0.000115 | 0.353746299 | 0.145 | 0.042 | 1 | 17 | sympk |
| rnf8 | 0.000115 | 0.334914047 | 0.145 | 0.041 | 1 | 17 | rnf8 |
| ctnnbl1 | 0.000116 | 0.280220708 | 0.127 | 0.033 | 1 | 17 | ctnnbl1 |
| uba3 | 0.000116 | 0.309520789 | 0.145 | 0.041 | 1 | 17 | uba3 |
| rnf175 | 0.000116 | 0.250209851 | 0.109 | 0.025 | 1 | 17 | rnf175 |
| ube2l3a1 | 0.000121 | 0.464799913 | 0.345 | 0.173 | 1 | 17 | ube2l3a |
| ndufb51 | 0.000122 | 0.48673675 | 0.382 | 0.212 | 1 | 17 | ndufb5 |
| arl6ip4 | 0.000123 | 0.345490922 | 0.164 | 0.051 | 1 | 17 | arl6ip4 |
| zw10 | 0.000126 | 0.269120836 | 0.109 | 0.025 | 1 | 17 | zw10 |
| nsun2 | 0.00013 | 0.302226592 | 0.127 | 0.033 | 1 | 17 | nsun2 |

| | | | | | | | |
|---|---|---|---|---|---|---|---|
| hnrnpul1 | 0.000141 | 0.322583892 | 0.2 | 0.071 | 1 | 17 | hnrnpul1 |
| si:dkey-188i13.113 | 0.000142 | 0.534173484 | 0.345 | 0.184 | 1 | 17 | si:dkey-188i13.11 |
| nars1 | 0.000142 | 0.510998451 | 0.291 | 0.139 | 1 | 17 | nars |
| prpf4 | 0.000147 | 0.288995242 | 0.127 | 0.034 | 1 | 17 | prpf4 |
| eef1e11 | 0.000149 | 0.267367652 | 0.127 | 0.034 | 1 | 17 | eef1e1 |
| pcnp | 0.000149 | 0.32048749 | 0.164 | 0.051 | 1 | 17 | pcnp |
| nelfb | 0.000151 | 0.255983879 | 0.109 | 0.026 | 1 | 17 | nelfb |
| wu:fa19b121 | 0.000152 | 0.262490046 | 0.109 | 0.026 | 1 | 17 | wu:fa19b12 |
| asf1bb | 0.000152 | 0.378125573 | 0.2 | 0.072 | 1 | 17 | asf1bb |
| cops3 | 0.000158 | 0.436381511 | 0.2 | 0.073 | 1 | 17 | cops3 |
| thoc7 | 0.000159 | 0.365786254 | 0.2 | 0.072 | 1 | 17 | thoc7 |
| sf3b53 | 0.000163 | 0.480359226 | 0.364 | 0.196 | 1 | 17 | sf3b5 |
| itpa | 0.000164 | 0.325411237 | 0.164 | 0.052 | 1 | 17 | itpa |
| eif2b3 | 0.000179 | 0.278017725 | 0.127 | 0.034 | 1 | 17 | eif2b3 |
| cfl1l1 | 0.000183 | 0.265404577 | 0.2 | 0.067 | 1 | 17 | cfl1l |
| sod11 | 0.000184 | 0.491331389 | 0.527 | 0.424 | 1 | 17 | sod1 |
| p4ha1b | 0.000185 | 0.266339055 | 0.109 | 0.026 | 1 | 17 | p4ha1b |
| mrpl9 | 0.000188 | 0.26977302 | 0.127 | 0.034 | 1 | 17 | mrpl9 |
| gkap11 | 0.000192 | 0.416462629 | 0.273 | 0.123 | 1 | 17 | gkap1 |
| siva1 | 0.000192 | 0.333861768 | 0.164 | 0.053 | 1 | 17 | siva1 |
| hax1 | 0.0002 | 0.338363804 | 0.145 | 0.044 | 1 | 17 | hax1 |
| apex11 | 0.000203 | 0.397211518 | 0.218 | 0.085 | 1 | 17 | apex1 |
| ebna1bp2 | 0.000203 | 0.55025435 | 0.273 | 0.128 | 1 | 17 | ebna1bp2 |
| uba1 | 0.000207 | 0.470318872 | 0.327 | 0.17 | 1 | 17 | uba1 |
| atp5pd1 | 0.000212 | 0.494164245 | 0.509 | 0.377 | 1 | 17 | atp5pd |
| cox7a2a | 0.000216 | 0.541330379 | 0.455 | 0.305 | 1 | 17 | cox7a2a |
| slirp1 | 0.000218 | 0.496850611 | 0.327 | 0.17 | 1 | 17 | slirp |
| mdm2 | 0.000218 | 0.36921994 | 0.127 | 0.035 | 1 | 17 | mdm2 |
| dhfr | 0.000222 | 0.285099241 | 0.127 | 0.035 | 1 | 17 | dhfr |
| ttf1 | 0.000227 | 0.333185313 | 0.164 | 0.053 | 1 | 17 | ttf1 |
| mrpl57 | 0.000228 | 0.508271227 | 0.291 | 0.144 | 1 | 17 | mrpl57 |
| gapdhs2 | 0.00023 | 0.392309287 | 0.618 | 0.477 | 1 | 17 | gapdhs |
| tma72 | 0.000232 | 0.415261845 | 0.6 | 0.474 | 1 | 17 | tma7 |
| leng1 | 0.000241 | 0.321624078 | 0.145 | 0.044 | 1 | 17 | leng1 |
| usp5 | 0.000242 | 0.37757047 | 0.182 | 0.064 | 1 | 17 | usp5 |

| | | | | | | | |
|---|---|---|---|---|---|---|---|
| ndufa32 | 0.000243 | 0.516506242 | 0.4 | 0.244 | 1 | 17 | ndufa3 |
| lyar | 0.000253 | 0.436312526 | 0.2 | 0.077 | 1 | 17 | lyar |
| gatd3a | 0.000256 | 0.328748185 | 0.145 | 0.045 | 1 | 17 | gatd3a |
| dda11 | 0.000259 | 0.358726069 | 0.2 | 0.076 | 1 | 17 | dda1 |
| c1d | 0.000264 | 0.400016708 | 0.2 | 0.076 | 1 | 17 | c1d |
| cenpx | 0.000271 | 0.38854147 | 0.2 | 0.075 | 1 | 17 | cenpx |
| rabl3 | 0.000276 | 0.269253756 | 0.127 | 0.035 | 1 | 17 | rabl3 |
| emg1 | 0.000285 | 0.382400178 | 0.182 | 0.065 | 1 | 17 | emg1 |
| etfa1 | 0.000286 | 0.400209075 | 0.309 | 0.154 | 1 | 17 | etfa |
| polr2i1 | 0.000288 | 0.432867866 | 0.218 | 0.089 | 1 | 17 | polr2i |
| mrps25 | 0.00029 | 0.440997765 | 0.255 | 0.115 | 1 | 17 | mrps25 |
| rad23b1 | 0.000296 | 0.470279239 | 0.418 | 0.262 | 1 | 17 | rad23b |
| polr1c | 0.000297 | 0.269493673 | 0.127 | 0.036 | 1 | 17 | polr1c |
| timm8b | 0.000303 | 0.465474478 | 0.236 | 0.103 | 1 | 17 | timm8b |
| ubtf | 0.000308 | 0.430575715 | 0.236 | 0.1 | 1 | 17 | ubtf |
| sart3 | 0.000311 | 0.337858538 | 0.127 | 0.036 | 1 | 17 | sart3 |
| prmt11 | 0.000311 | 0.50090604 | 0.382 | 0.23 | 1 | 17 | prmt1 |
| utp11l | 0.000311 | 0.308976344 | 0.164 | 0.054 | 1 | 17 | utp11l |
| polr2k1 | 0.000311 | 0.396857718 | 0.255 | 0.113 | 1 | 17 | polr2k |
| tomm62 | 0.000318 | 0.492222277 | 0.382 | 0.227 | 1 | 17 | tomm6 |
| bida | 0.000318 | 0.346471433 | 0.182 | 0.065 | 1 | 17 | bida |
| psma6a2 | 0.000318 | 0.427895189 | 0.309 | 0.158 | 1 | 17 | psma6a |
| riox1 | 0.000327 | 0.25039936 | 0.127 | 0.036 | 1 | 17 | riox1 |
| gpia2 | 0.000328 | 0.411524522 | 0.291 | 0.138 | 1 | 17 | gpia |
| tprb | 0.000334 | 0.267776467 | 0.127 | 0.036 | 1 | 17 | tprb |
| fam168b | 0.000338 | 0.364984705 | 0.218 | 0.088 | 1 | 17 | fam168b |
| mtap | 0.000354 | 0.419998044 | 0.255 | 0.116 | 1 | 17 | mtap |
| mki67 | 0.000355 | 0.254556623 | 0.109 | 0.028 | 1 | 17 | mki67 |
| h3f3d3 | 0.000368 | 0.282288531 | 0.727 | 0.676 | 1 | 17 | h3f3d |
| psmb72 | 0.000377 | 0.477158352 | 0.455 | 0.307 | 1 | 17 | psmb7 |
| tcea2 | 0.000379 | 0.293347563 | 0.109 | 0.028 | 1 | 17 | tcea2 |
| ttc33 | 0.000392 | 0.297363789 | 0.127 | 0.037 | 1 | 17 | ttc33 |
| ddx47 | 0.000395 | 0.409921288 | 0.2 | 0.079 | 1 | 17 | ddx47 |
| cct82 | 0.000414 | 0.462408307 | 0.491 | 0.342 | 1 | 17 | cct8 |
| smn1 | 0.000416 | 0.33043469 | 0.164 | 0.056 | 1 | 17 | smn1 |

| | | | | | | | |
|---|---|---|---|---|---|---|---|
| uxt | 0.000419 | 0.271989711 | 0.127 | 0.037 | 1 | 17 | uxt |
| msh6 | 0.000421 | 0.250082284 | 0.109 | 0.028 | 1 | 17 | msh6 |
| krr1 | 0.000432 | 0.30999212 | 0.127 | 0.037 | 1 | 17 | krr1 |
| mrps23 | 0.000437 | 0.250200533 | 0.145 | 0.046 | 1 | 17 | mrps23 |
| g3bp2 | 0.000439 | 0.309966687 | 0.145 | 0.046 | 1 | 17 | g3bp2 |
| ube2ia1 | 0.00044 | 0.485576465 | 0.327 | 0.182 | 1 | 17 | ube2ia |
| med101 | 0.000448 | 0.45809277 | 0.327 | 0.178 | 1 | 17 | med10 |
| nudt212 | 0.00045 | 0.459635837 | 0.273 | 0.132 | 1 | 17 | nudt21 |
| med11 | 0.000454 | 0.395836088 | 0.218 | 0.091 | 1 | 17 | med11 |
| sf3a22 | 0.000454 | 0.502603612 | 0.309 | 0.168 | 1 | 17 | sf3a2 |
| vdac21 | 0.000462 | 0.404954556 | 0.582 | 0.447 | 1 | 17 | vdac2 |
| si:ch211-284b7.3 | 0.000473 | 0.395648011 | 0.2 | 0.08 | 1 | 17 | si:ch211-284b7.3 |
| ube2f | 0.000496 | 0.295201135 | 0.145 | 0.047 | 1 | 17 | ube2f |
| aifm1 | 0.000504 | 0.25194862 | 0.127 | 0.037 | 1 | 17 | aifm1 |
| mrps33 | 0.000508 | 0.369847976 | 0.218 | 0.091 | 1 | 17 | mrps33 |
| STMP11 | 0.000511 | 0.489739489 | 0.382 | 0.234 | 1 | 17 | STMP1 |
| cpsf3 | 0.000516 | 0.269672132 | 0.145 | 0.047 | 1 | 17 | cpsf3 |
| pole3 | 0.000548 | 0.361419924 | 0.182 | 0.069 | 1 | 17 | pole3 |
| rnf168 | 0.000551 | 0.269217655 | 0.109 | 0.029 | 1 | 17 | rnf168 |
| ubtfl | 0.000553 | 0.32206284 | 0.145 | 0.048 | 1 | 17 | ubtfl |
| ndufs61 | 0.000554 | 0.506281271 | 0.345 | 0.202 | 1 | 17 | ndufs6 |
| emc9 | 0.000586 | 0.296902654 | 0.145 | 0.048 | 1 | 17 | emc9 |
| mrpl17 | 0.000601 | 0.366454403 | 0.182 | 0.069 | 1 | 17 | mrpl17 |
| rrp9 | 0.000602 | 0.34452299 | 0.145 | 0.048 | 1 | 17 | rrp9 |
| vdac13 | 0.000604 | 0.457665154 | 0.327 | 0.182 | 1 | 17 | vdac1 |
| phf5a | 0.000606 | 0.438379138 | 0.327 | 0.182 | 1 | 17 | phf5a |
| ppp1r2 | 0.000619 | 0.30712989 | 0.127 | 0.039 | 1 | 17 | ppp1r2 |
| polr2c | 0.000622 | 0.316016796 | 0.164 | 0.058 | 1 | 17 | polr2c |
| atp5if1a | 0.000627 | 0.453959132 | 0.509 | 0.398 | 1 | 17 | atp5if1a |
| nudt1 | 0.00063 | 0.29026882 | 0.109 | 0.03 | 1 | 17 | nudt1 |
| prpf40a | 0.000633 | 0.406134527 | 0.182 | 0.07 | 1 | 17 | prpf40a |
| FQ976913.1 | 0.000647 | 0.361223857 | 0.164 | 0.059 | 1 | 17 | FQ976913.1 |
| med8 | 0.000649 | 0.33962558 | 0.145 | 0.048 | 1 | 17 | med8 |
| ube2g1a | 0.000652 | 0.35385594 | 0.2 | 0.08 | 1 | 17 | ube2g1a |
| gmppaa | 0.000683 | 0.263290191 | 0.109 | 0.03 | 1 | 17 | gmppaa |

| | | | | | | | |
|---|---|---|---|---|---|---|---|
| pa2g4a | 0.000684 | 0.407777432 | 0.291 | 0.147 | 1 | 17 | pa2g4a |
| rnaseh11 | 0.000689 | 0.327093407 | 0.164 | 0.059 | 1 | 17 | rnaseh1 |
| ndufa62 | 0.000708 | 0.466514299 | 0.455 | 0.314 | 1 | 17 | ndufa6 |
| ndufb71 | 0.000712 | 0.497175396 | 0.436 | 0.309 | 1 | 17 | ndufb7 |
| psmb51 | 0.000712 | 0.444487955 | 0.273 | 0.137 | 1 | 17 | psmb5 |
| wu:fi46h041 | 0.000745 | 0.251370174 | 0.145 | 0.048 | 1 | 17 | wu:fi46h04 |
| cox7b3 | 0.000772 | 0.383015814 | 0.6 | 0.507 | 1 | 17 | cox7b |
| ddx613 | 0.000778 | 0.442465056 | 0.473 | 0.344 | 1 | 17 | ddx61 |
| eaf2 | 0.000786 | 0.255340523 | 0.127 | 0.039 | 1 | 17 | eaf2 |
| xpot | 0.000815 | 0.303157015 | 0.145 | 0.049 | 1 | 17 | xpot |
| cnot9 | 0.000844 | 0.409584685 | 0.2 | 0.084 | 1 | 17 | cnot9 |
| mrps18c | 0.000857 | 0.34078143 | 0.2 | 0.083 | 1 | 17 | mrps18c |
| ercc1 | 0.000869 | 0.278908215 | 0.127 | 0.04 | 1 | 17 | ercc1 |
| nop581 | 0.000904 | 0.436727726 | 0.364 | 0.22 | 1 | 17 | nop58 |
| pfdn13 | 0.000942 | 0.453989262 | 0.364 | 0.226 | 1 | 17 | pfdn1 |
| pdap1a1 | 0.00095 | 0.418317209 | 0.255 | 0.123 | 1 | 17 | pdap1a |
| mrpl53 | 0.000967 | 0.354428702 | 0.182 | 0.072 | 1 | 17 | mrpl53 |
| idh3b | 0.000978 | 0.310076102 | 0.164 | 0.06 | 1 | 17 | idh3b |
| naa40 | 0.000988 | 0.350451139 | 0.182 | 0.072 | 1 | 17 | naa40 |
| rpl7l1 | 0.001031 | 0.424963139 | 0.255 | 0.124 | 1 | 17 | rpl7l1 |
| sf3a1 | 0.001048 | 0.318724212 | 0.164 | 0.061 | 1 | 17 | sf3a1 |
| mrpl331 | 0.001083 | 0.412589523 | 0.291 | 0.153 | 1 | 17 | mrpl33 |
| hp1bp33 | 0.0011 | 0.431147764 | 0.382 | 0.245 | 1 | 17 | hp1bp3 |
| mrpl44 | 0.001109 | 0.271186661 | 0.145 | 0.05 | 1 | 17 | mrpl44 |
| kat5b | 0.001165 | 0.305909012 | 0.145 | 0.051 | 1 | 17 | kat5b |
| dynll13 | 0.001174 | 0.405810516 | 0.527 | 0.398 | 1 | 17 | dynll1 |
| znhit3 | 0.001192 | 0.307527802 | 0.127 | 0.041 | 1 | 17 | znhit3 |
| tsen34 | 0.001206 | 0.307049292 | 0.145 | 0.052 | 1 | 17 | tsen34 |
| ndufb61 | 0.001209 | 0.485655522 | 0.327 | 0.196 | 1 | 17 | ndufb6 |
| csnk2a1 | 0.001222 | 0.434007538 | 0.345 | 0.209 | 1 | 17 | csnk2a1 |
| swt1 | 0.001227 | 0.251134029 | 0.127 | 0.04 | 1 | 17 | swt1 |
| maco1a | 0.001256 | 0.315521476 | 0.145 | 0.052 | 1 | 17 | maco1a |
| dhps1 | 0.001266 | 0.266930047 | 0.127 | 0.041 | 1 | 17 | dhps |
| ccdc58 | 0.001282 | 0.302291452 | 0.145 | 0.051 | 1 | 17 | ccdc58 |
| atp5meb1 | 0.001326 | 0.466034744 | 0.509 | 0.428 | 1 | 17 | atp5meb |

| | | | | | | | |
|---|---|---|---|---|---|---|---|
| anp32b2 | 0.001331 | 0.508440049 | 0.364 | 0.239 | 1 | 17 | anp32b |
| LOC110437782 | 0.001331 | 0.250883494 | 0.109 | 0.032 | 1 | 17 | LOC110437782 |
| ddx231 | 0.001367 | 0.335005116 | 0.2 | 0.086 | 1 | 17 | ddx23 |
| yars | 0.001384 | 0.25572076 | 0.127 | 0.042 | 1 | 17 | yars |
| vcp | 0.001385 | 0.453065415 | 0.327 | 0.191 | 1 | 17 | vcp |
| pfdn2 | 0.001411 | 0.426675546 | 0.273 | 0.146 | 1 | 17 | pfdn2 |
| zgc:56304 | 0.001415 | 0.412602272 | 0.309 | 0.173 | 1 | 17 | zgc:56304 |
| odc14 | 0.001428 | 0.368636527 | 0.255 | 0.121 | 1 | 17 | odc1 |
| ik1 | 0.001434 | 0.360781662 | 0.2 | 0.087 | 1 | 17 | ik |
| sumo3a3 | 0.00144 | 0.420688764 | 0.509 | 0.411 | 1 | 17 | sumo3a |
| mrps14 | 0.001478 | 0.422543674 | 0.236 | 0.115 | 1 | 17 | mrps14 |
| rtraf1 | 0.001508 | 0.352551861 | 0.2 | 0.087 | 1 | 17 | rtraf |
| flot1b4 | 0.001549 | 0.472494342 | 0.327 | 0.197 | 1 | 17 | flot1b |
| dnm1l | 0.001565 | 0.295240124 | 0.109 | 0.033 | 1 | 17 | dnm1l |
| ndufb101 | 0.001611 | 0.455875134 | 0.364 | 0.231 | 1 | 17 | ndufb10 |
| rbbp5 | 0.001614 | 0.311279311 | 0.145 | 0.053 | 1 | 17 | rbbp5 |
| rbm191 | 0.001691 | 0.445493356 | 0.2 | 0.09 | 1 | 17 | rbm19 |
| snrpb3 | 0.001696 | 0.404502258 | 0.509 | 0.424 | 1 | 17 | snrpb |
| mif2 | 0.001701 | 0.508190886 | 0.418 | 0.316 | 1 | 17 | mif |
| prim2 | 0.001803 | 0.270880378 | 0.109 | 0.033 | 1 | 17 | prim2 |
| polr2b | 0.001861 | 0.331100017 | 0.164 | 0.064 | 1 | 17 | polr2b |
| rpp14 | 0.001909 | 0.32515831 | 0.145 | 0.053 | 1 | 17 | rpp14 |
| hmgb3b2 | 0.001917 | 0.420286214 | 0.273 | 0.147 | 1 | 17 | hmgb3b |
| dck | 0.002003 | 0.265925049 | 0.127 | 0.043 | 1 | 17 | dck |
| pno1 | 0.002054 | 0.467911945 | 0.309 | 0.186 | 1 | 17 | pno1 |
| zgc:112496 | 0.002058 | 0.26796373 | 0.127 | 0.043 | 1 | 17 | zgc:112496 |
| rchy1 | 0.002099 | 0.303651676 | 0.145 | 0.054 | 1 | 17 | rchy1 |
| polr2d | 0.002108 | 0.43628799 | 0.327 | 0.198 | 1 | 17 | polr2d |
| si:ch211-156b7.4 | 0.002136 | 0.361234731 | 0.145 | 0.054 | 1 | 17 | si:ch211-156b7.4 |
| nfs1 | 0.002145 | 0.311033808 | 0.145 | 0.054 | 1 | 17 | nfs1 |
| nmt2 | 0.002234 | 0.265471021 | 0.109 | 0.034 | 1 | 17 | nmt2 |
| ptrh2 | 0.002423 | 0.281111047 | 0.127 | 0.044 | 1 | 17 | ptrh2 |
| stk35l1 | 0.002431 | 0.309770607 | 0.164 | 0.066 | 1 | 17 | stk35l |
| ube2d1b | 0.002441 | 0.395974533 | 0.345 | 0.211 | 1 | 17 | ube2d1b |
| fundc11 | 0.002485 | 0.368571073 | 0.2 | 0.091 | 1 | 17 | fundc1 |

| | | | | | | | |
|---|---|---|---|---|---|---|---|
| ncln | 0.002489 | 0.37272684 | 0.164 | 0.067 | 1 | 17 | ncln |
| cops7a | 0.002493 | 0.356992577 | 0.182 | 0.079 | 1 | 17 | cops7a |
| dnajc81 | 0.0025 | 0.429155681 | 0.218 | 0.107 | 1 | 17 | dnajc8 |
| zgc:158803 | 0.002563 | 0.370565022 | 0.182 | 0.079 | 1 | 17 | zgc:158803 |
| got2b | 0.002579 | 0.364265525 | 0.2 | 0.091 | 1 | 17 | got2b |
| atrxl | 0.002583 | 0.298688217 | 0.127 | 0.045 | 1 | 17 | atrxl |
| uqcrfs11 | 0.002589 | 0.437462219 | 0.364 | 0.239 | 1 | 17 | uqcrfs1 |
| atp5f1c3 | 0.002602 | 0.454531767 | 0.455 | 0.358 | 1 | 17 | atp5f1c |
| zc3h14 | 0.002642 | 0.311242247 | 0.127 | 0.045 | 1 | 17 | zc3h14 |
| cox6c1 | 0.002694 | 0.420456104 | 0.509 | 0.418 | 1 | 17 | cox6c |
| trappc12 | 0.002737 | 0.454805883 | 0.327 | 0.205 | 1 | 17 | trappc1 |
| mphosph83 | 0.002748 | 0.406527551 | 0.382 | 0.253 | 1 | 17 | mphosph8 |
| ube2v22 | 0.002759 | 0.412407976 | 0.473 | 0.366 | 1 | 17 | ube2v2 |
| txndc171 | 0.002767 | 0.320477101 | 0.182 | 0.079 | 1 | 17 | txndc17 |
| afmid | 0.002785 | 0.254989579 | 0.109 | 0.035 | 1 | 17 | afmid |
| khdrbs1a2 | 0.002828 | 0.264381124 | 0.618 | 0.599 | 1 | 17 | khdrbs1a |
| pqbp1 | 0.002835 | 0.268676887 | 0.127 | 0.045 | 1 | 17 | pqbp1 |
| cdk7 | 0.002873 | 0.256532147 | 0.127 | 0.045 | 1 | 17 | cdk7 |
| atp5mc3a1 | 0.002879 | 0.454512398 | 0.436 | 0.348 | 1 | 17 | atp5mc3a |
| rbm22 | 0.002886 | 0.394002548 | 0.2 | 0.094 | 1 | 17 | rbm22 |
| hspe14 | 0.00303 | 0.424412053 | 0.436 | 0.293 | 1 | 17 | hspe1 |
| tbca1 | 0.003034 | 0.439517253 | 0.418 | 0.299 | 1 | 17 | tbca |
| psmc51 | 0.003055 | 0.450663381 | 0.309 | 0.19 | 1 | 17 | psmc5 |
| cops41 | 0.003078 | 0.334279226 | 0.218 | 0.106 | 1 | 17 | cops4 |
| wu:fc33b09 | 0.00308 | 0.254180022 | 0.127 | 0.045 | 1 | 17 | wu:fc33b09 |
| ubap2b | 0.00318 | 0.335795145 | 0.2 | 0.092 | 1 | 17 | ubap2b |
| commd71 | 0.00329 | 0.3668346 | 0.236 | 0.122 | 1 | 17 | commd7 |
| eif1ad | 0.003316 | 0.33840392 | 0.273 | 0.147 | 1 | 17 | eif1ad |
| caprin1a | 0.003323 | 0.279379069 | 0.182 | 0.078 | 1 | 17 | caprin1a |
| eno31 | 0.003344 | 0.386790718 | 0.436 | 0.295 | 1 | 17 | eno3 |
| arpp19b | 0.003451 | 0.356155625 | 0.236 | 0.12 | 1 | 17 | arpp19b |
| med21 | 0.003502 | 0.319822041 | 0.164 | 0.068 | 1 | 17 | med21 |
| pgk1 | 0.003588 | 0.345815938 | 0.218 | 0.108 | 1 | 17 | pgk1 |
| ndufa2 | 0.003597 | 0.436427644 | 0.327 | 0.205 | 1 | 17 | ndufa2 |
| cox7c2 | 0.003644 | 0.294329189 | 0.618 | 0.583 | 1 | 17 | cox7c |

| | | | | | | | |
|---|---|---|---|---|---|---|---|
| mrps28 | 0.003646 | 0.302334939 | 0.145 | 0.057 | 1 | 17 | mrps28 |
| supt4h1 | 0.003704 | 0.349872006 | 0.236 | 0.122 | 1 | 17 | supt4h1 |
| ruvbl2 | 0.003799 | 0.25350551 | 0.109 | 0.036 | 1 | 17 | ruvbl2 |
| tmem131 | 0.003821 | 0.271372011 | 0.127 | 0.046 | 1 | 17 | tmem131 |
| glrx32 | 0.003887 | 0.403043161 | 0.218 | 0.11 | 1 | 17 | glrx3 |
| pitpnaa | 0.003915 | 0.292100659 | 0.164 | 0.068 | 1 | 17 | pitpnaa |
| bola1 | 0.003951 | 0.300580023 | 0.127 | 0.047 | 1 | 17 | bola1 |
| psmd81 | 0.004042 | 0.352228559 | 0.273 | 0.153 | 1 | 17 | psmd8 |
| hnrnpl22 | 0.004127 | 0.406600159 | 0.273 | 0.158 | 1 | 17 | hnrnpl2 |
| adrm1 | 0.004139 | 0.373721387 | 0.2 | 0.097 | 1 | 17 | adrm1 |
| zgc:63470 | 0.004458 | 0.316053586 | 0.127 | 0.048 | 1 | 17 | zgc:63470 |
| dcun1d1 | 0.004568 | 0.268522203 | 0.109 | 0.037 | 1 | 17 | dcun1d1 |
| cmc2 | 0.004684 | 0.265409636 | 0.127 | 0.048 | 1 | 17 | cmc2 |
| tuba8l23 | 0.004789 | 0.311223619 | 0.218 | 0.105 | 1 | 17 | tuba8l2 |
| taf11 | 0.005019 | 0.291428627 | 0.164 | 0.071 | 1 | 17 | taf11 |
| thap4 | 0.005034 | 0.256053429 | 0.109 | 0.038 | 1 | 17 | thap4 |
| u2af1 | 0.005036 | 0.39713438 | 0.345 | 0.232 | 1 | 17 | u2af1 |
| srsf4 | 0.005039 | 0.354195069 | 0.2 | 0.099 | 1 | 17 | srsf4 |
| fdx1 | 0.00504 | 0.321410746 | 0.2 | 0.096 | 1 | 17 | fdx1 |
| prpf39 | 0.005054 | 0.348285117 | 0.218 | 0.111 | 1 | 17 | prpf39 |
| uqcc3 | 0.00512 | 0.315105262 | 0.145 | 0.06 | 1 | 17 | uqcc3 |
| ppig | 0.005186 | 0.356625573 | 0.182 | 0.086 | 1 | 17 | ppig |
| yeats41 | 0.005204 | 0.3028981 | 0.2 | 0.098 | 1 | 17 | yeats4 |
| psmc3 | 0.005266 | 0.364669967 | 0.255 | 0.143 | 1 | 17 | psmc3 |
| rbmx2 | 0.005297 | 0.356096644 | 0.273 | 0.156 | 1 | 17 | rbmx |
| ube2d21 | 0.005549 | 0.382410265 | 0.382 | 0.268 | 1 | 17 | ube2d2 |
| sf3b4 | 0.005624 | 0.378872386 | 0.182 | 0.087 | 1 | 17 | sf3b4 |
| fbl | 0.005644 | 0.427644277 | 0.309 | 0.2 | 1 | 17 | fbl |
| nsfl1c2 | 0.00566 | 0.371807503 | 0.255 | 0.143 | 1 | 17 | nsfl1c |
| nip7 | 0.005845 | 0.344424134 | 0.218 | 0.114 | 1 | 17 | nip7 |
| u2af2b | 0.005989 | 0.371011224 | 0.291 | 0.177 | 1 | 17 | u2af2b |
| exosc5 | 0.006072 | 0.257473302 | 0.109 | 0.039 | 1 | 17 | exosc5 |
| ubxn1 | 0.006248 | 0.344571712 | 0.182 | 0.086 | 1 | 17 | ubxn1 |
| npepl11 | 0.006424 | 0.30059405 | 0.218 | 0.111 | 1 | 17 | npepl1 |
| hnrnpa0b4 | 0.006425 | 0.296719709 | 0.6 | 0.517 | 1 | 17 | hnrnpa0b |

| | | | | | | | | |
|---|---|---|---|---|---|---|---|---|
| ssu72 | 0.0065 | 0.338119436 | 0.2 | 0.099 | | 1 | 17 | ssu72 |
| gtf2a2 | 0.006554 | 0.340258055 | 0.182 | 0.087 | | 1 | 17 | gtf2a2 |
| cycsb2 | 0.006667 | 0.389700266 | 0.382 | 0.271 | | 1 | 17 | cycsb |
| srsf5a4 | 0.006729 | 0.37006607 | 0.473 | 0.391 | | 1 | 17 | srsf5a |
| lig1 | 0.006832 | 0.364429832 | 0.182 | 0.087 | | 1 | 17 | lig1 |
| AK6 | 0.006888 | 0.343743635 | 0.145 | 0.062 | | 1 | 17 | AK6 |
| tia1l | 0.007006 | 0.377229777 | 0.218 | 0.117 | | 1 | 17 | tia1l |
| ndufa9a1 | 0.00715 | 0.359899654 | 0.236 | 0.132 | | 1 | 17 | ndufa9a |
| si:dkey-180p18.91 | 0.00723 | 0.349741262 | 0.273 | 0.159 | | 1 | 17 | si:dkey-180p18.9 |
| ccdc25 | 0.007236 | 0.401329794 | 0.273 | 0.164 | | 1 | 17 | ccdc25 |
| ywhaqa2 | 0.007257 | 0.371073959 | 0.364 | 0.256 | | 1 | 17 | ywhaqa |
| exosc8 | 0.00726 | 0.309670163 | 0.127 | 0.051 | | 1 | 17 | exosc8 |
| myl12.2 | 0.00738 | 0.390382646 | 0.218 | 0.116 | | 1 | 17 | myl12.2 |
| pabpc1b4 | 0.007812 | 0.303419747 | 0.236 | 0.125 | | 1 | 17 | pabpc1b |
| elocb2 | 0.00808 | 0.41832744 | 0.382 | 0.294 | | 1 | 17 | elocb |
| ube2nb | 0.008129 | 0.347222065 | 0.255 | 0.147 | | 1 | 17 | ube2nb |
| stau1 | 0.008211 | 0.299148642 | 0.2 | 0.1 | | 1 | 17 | stau1 |
| grpel22 | 0.008418 | 0.405303604 | 0.236 | 0.135 | | 1 | 17 | grpel2 |
| ctsf3 | 0.008532 | 0.263493663 | 0.164 | 0.073 | | 1 | 17 | ctsf |
| gnb1a1 | 0.008547 | 0.326625363 | 0.382 | 0.265 | | 1 | 17 | gnb1a |
| cox4i11 | 0.008548 | 0.273440741 | 0.6 | 0.58 | | 1 | 17 | cox4i1 |
| ctps1a | 0.008596 | 0.251935802 | 0.145 | 0.062 | | 1 | 17 | ctps1a |
| gtf2f1 | 0.008707 | 0.340786322 | 0.2 | 0.104 | | 1 | 17 | gtf2f1 |
| mrpl24 | 0.00927 | 0.331947378 | 0.145 | 0.064 | | 1 | 17 | mrpl24 |
| pim3 | 0.009443 | 0.255927628 | 0.145 | 0.062 | | 1 | 17 | pim3 |
| hsdl21 | 0.009522 | 0.319326071 | 0.2 | 0.103 | | 1 | 17 | hsdl2 |
| mrpl18 | 0.009549 | 0.302539844 | 0.2 | 0.104 | | 1 | 17 | mrpl18 |
| ddx3xb | 0.009553 | 0.304199976 | 0.2 | 0.103 | | 1 | 17 | ddx3xb |
| psip1b1 | 0.009791 | 0.380990042 | 0.218 | 0.121 | | 1 | 17 | psip1b |
| psmc61 | 0.009959 | 0.329623796 | 0.255 | 0.15 | | 1 | 17 | psmc6 |
| coch1 | 8.4957593 | 1.677945023 | 0.771 | 0.021 | 1.71588851074802e-1 | | 18 | coch |
| cald1b | 8.2896278 | 1.411531099 | 0.729 | 0.019 | 1.67425612846295e-1 | | 18 | cald1b |
| pvalb9 | 3.2852763 | 0.895952062 | 0.438 | 0.004 | 6.63527268523644e-1 | | 18 | pvalb9 |
| rspo11 | 4.2764990 | 1.134878183 | 0.583 | 0.019 | 8.63724511247184e-1 | | 18 | rspo1 |
| rasl121 | 5.3563787 | 0.816170316 | 0.438 | 0.011 | 1.08182782244704e-8 | | 18 | rasl12 |

| | | | | | | | |
|---|---|---|---|---|---|---|---|
| sned1 | 1.3438490 | 0.847402396 | 0.5 | 0.016 | 2.71417199383316e-8 | 18 | sned1 |
| mylkb | 3.4511889 | 0.62892275 | 0.333 | 0.005 | 6.97036626267662e-8 | 18 | mylkb |
| spon1b1 | 2.7538472 | 1.280506773 | 0.875 | 0.074 | 5.56194536869604e-7 | 18 | spon1b |
| sost3 | 1.5531825 | 1.236950755 | 0.729 | 0.057 | 3.13696288290171e-6 | 18 | sost |
| cdh62 | 4.2314550 | 1.299714642 | 0.792 | 0.071 | 8.54626978897295e-6 | 18 | cdh6 |
| hoxb5b1 | 2.6792857 | 0.77046654 | 0.417 | 0.016 | 5.41135352365672e-6 | 18 | hoxb5b |
| prrx1b | 1.7338730 | 0.537430568 | 0.312 | 0.009 | 3.50190348928406e-5 | 18 | prrx1b |
| mex3b2 | 2.3814131 | 1.218996546 | 0.667 | 0.059 | 4.80974004827964e-5 | 18 | mex3b |
| rgs4 | 1.0718359 | 0.872021461 | 0.5 | 0.029 | 2.16478700168864e-5 | 18 | rgs4 |
| pitx22 | 2.1995187 | 0.941655415 | 0.521 | 0.033 | 4.44236797754445e-5 | 18 | pitx2 |
| fgf242 | 5.5992186 | 0.802015826 | 0.458 | 0.025 | 1.13087418840307e-5 | 18 | fgf24 |
| elnb | 5.8026079 | 0.474262077 | 0.208 | 0.003 | 1.17195273644583e-5 | 18 | elnb |
| htra1a2 | 5.1529802 | 0.884360645 | 0.542 | 0.037 | 1.04074742176353e-5 | 18 | htra1a |
| CABZ01039096.11 | 6.9318446 | 0.774742762 | 0.458 | 0.027 | 1.40002466548639e-5 | 18 | CABZ01039096.1 |
| tmeff2a1 | 7.5191718 | 0.531596529 | 0.312 | 0.011 | 1.51864713605075e-4 | 18 | tmeff2a |
| cdc42ep1b1 | 1.1430137 | 0.626366262 | 0.375 | 0.018 | 2.3085449449336e-48 | 18 | cdc42ep1b |
| barx1 | 1.5204921 | 0.653640489 | 0.312 | 0.012 | 3.07093793532788e-4 | 18 | barx1 |
| rgs5a2 | 2.5094858 | 1.167699661 | 0.729 | 0.084 | 5.06840856794203e-4 | 18 | rgs5a |
| mamdc2a | 8.0372682 | 0.266496195 | 0.167 | 0.002 | 1.62328707758014e-4 | 18 | mamdc2a |
| isl2b | 7.1861425 | 0.586817244 | 0.333 | 0.015 | 1.45138520889834e-4 | 18 | isl2b |
| phyhiplb | 2.5798759 | 0.436284211 | 0.25 | 0.007 | 5.21057547311071e-4 | 18 | phyhiplb |
| c1qtnf53 | 1.2799463 | 1.039423028 | 0.667 | 0.073 | 2.58510769276609e-4 | 18 | c1qtnf5 |
| nfixb4 | 3.4579574 | 1.327965318 | 0.917 | 0.162 | 6.98403674927856e-4 | 18 | nfixb |
| angpt2b1 | 5.4680402 | 0.534623003 | 0.292 | 0.012 | 1.10438007928007e-4 | 18 | angpt2b |
| nppc | 6.1917226 | 0.456739995 | 0.208 | 0.005 | 1.25054222853828e-4 | 18 | nppc |
| BX640522.24 | 1.9607968 | 1.13915345 | 0.792 | 0.108 | 3.96022139032194e-4 | 18 | BX640522.2 |
| CR847566.14 | 1.3246685 | 1.13545884 | 0.792 | 0.121 | 2.6754330883192e-38 | 18 | CR847566.1 |
| shisa2b | 2.8140832 | 0.421325599 | 0.229 | 0.007 | 5.68360403613981e-3 | 18 | shisa2b |
| igfbp5b3 | 3.2897522 | 1.30458207 | 0.917 | 0.163 | 6.64431259178029e-3 | 18 | igfbp5b |
| LOC1039092844 | 3.4784322 | 1.10495443 | 0.812 | 0.13 | 7.02538957058486e-3 | 18 | LOC103909284 |
| abi3bpb1 | 4.6753648 | 0.788643027 | 0.458 | 0.037 | 9.44283439575548e-3 | 18 | abi3bpb |
| rcan2 | 2.5216048 | 0.528330804 | 0.312 | 0.016 | 5.09288538467138e-3 | 18 | rcan2 |
| vim3 | 7.8733506 | 1.094142229 | 0.688 | 0.094 | 1.59018062325443e-3 | 18 | vim |
| ccdc3b1 | 1.0605461 | 0.82455332 | 0.542 | 0.057 | 2.14198508051225e-3 | 18 | ccdc3b |
| bicc1b1 | 1.3505158 | 0.712535975 | 0.438 | 0.036 | 2.72763686511917e-3 | 18 | bicc1b |

| | | | | | | | |
|---|---|---|---|---|---|---|---|
| nuak1a1 | 2.4574135 | 0.52098629 | 0.292 | 0.015 | 4.96323823176971e-3 | 18 | nuak1a |
| si:dkey-26m3.31 | 8.9058963 | 0.63307515 | 0.396 | 0.03 | 1.79872388037438e-3 | 18 | si:dkey-26m3.3 |
| ece11 | 4.9931900 | 0.874201901 | 0.583 | 0.074 | 1.0084745960335e-31 | 18 | ece1 |
| bgna | 7.9044124 | 0.533128935 | 0.271 | 0.014 | 1.59645418117167e-3 | 18 | bgna |
| igfbp2b | 1.4280435 | 0.602764814 | 0.354 | 0.026 | 2.88421951150441e-3 | 18 | igfbp2b |
| kcnq5b | 8.6344114 | 0.298020657 | 0.167 | 0.004 | 1.74389208592852e-3 | 18 | kcnq5b |
| fhl1a | 1.2318281 | 0.601319005 | 0.292 | 0.018 | 2.48792340566501e-2 | 18 | fhl1a |
| net13 | 1.7416000 | 0.843412944 | 0.562 | 0.075 | 3.51750954483998e-2 | 18 | net1 |
| si:ch211-207d6.21 | 2.9825052 | 0.731649242 | 0.5 | 0.055 | 6.02376582545469e-2 | 18 | si:ch211-207d6.2 |
| atoh82 | 3.5109348 | 0.793214347 | 0.479 | 0.053 | 7.09103511556103e-2 | 18 | atoh8 |
| mn1a | 1.1260363 | 0.410933128 | 0.229 | 0.011 | 2.27425562077236e-2 | 18 | mn1a |
| BX927333.1 | 1.4041908 | 0.253876349 | 0.167 | 0.005 | 2.83604434156454e-2 | 18 | BX927333.1 |
| cdh10a1 | 5.9120816 | 0.49416976 | 0.271 | 0.016 | 1.19406313454958e-2 | 18 | cdh10a |
| ctgfa2 | 9.5911661 | 1.069602702 | 0.479 | 0.058 | 1.93712783524751e-2 | 18 | ctgfa |
| pcolcea3 | 3.7435016 | 1.067124018 | 0.792 | 0.146 | 7.56075031394379e-2 | 18 | pcolcea |
| dcn5 | 5.7586086 | 1.178260157 | 0.979 | 0.267 | 1.16306619135494e-2 | 18 | dcn |
| cacna1g1 | 1.4758199 | 0.386114499 | 0.229 | 0.012 | 2.98071364395379e-2 | 18 | cacna1g |
| fibinb2 | 1.8420146 | 0.728345686 | 0.479 | 0.058 | 3.72031692621253e-2 | 18 | fibinb |
| loxa4 | 3.8387600 | 0.70160929 | 0.438 | 0.048 | 7.75314359345968e-2 | 18 | loxa |
| foxp45 | 4.2198872 | 1.065768583 | 0.792 | 0.18 | 8.5229063507783e-26 | 18 | foxp4 |
| mmrn2a | 7.2028857 | 0.431332676 | 0.25 | 0.015 | 1.45476683945392e-2 | 18 | mmrn2a |
| relt | 2.1806571 | 0.278196165 | 0.167 | 0.006 | 4.40427323510837e-2 | 18 | relt |
| adgrd1 | 2.5259441 | 0.327829484 | 0.167 | 0.006 | 5.10164946721955e-2 | 18 | adgrd1 |
| CU929237.12 | 9.6890023 | 0.676207715 | 0.438 | 0.049 | 1.9568877947594e-24 | 18 | CU929237.1 |
| si:ch73-335l21.43 | 1.7257723 | 0.986702223 | 0.625 | 0.107 | 3.48554251576096e-2 | 18 | si:ch73-335l21.4 |
| loxl13 | 2.6040473 | 0.757738189 | 0.5 | 0.066 | 5.25939437586991e-2 | 18 | loxl1 |
| id15 | 3.4842562 | 1.226758758 | 0.917 | 0.264 | 7.0371523404677e-24 | 18 | id1 |
| osr13 | 4.3484987 | 0.890998891 | 0.688 | 0.123 | 8.78266298446196e-2 | 18 | osr1 |
| ptx3a1 | 5.8674155 | 0.878355507 | 0.479 | 0.062 | 1.18504191542365e-2 | 18 | ptx3a |
| ryk | 8.1080962 | 0.496951602 | 0.312 | 0.027 | 1.63759220657716e-2 | 18 | ryk |
| fn1a4 | 1.0242179 | 1.013810128 | 0.792 | 0.171 | 2.06861303476718e-2 | 18 | fn1a |
| aga | 1.0846930 | 0.671455885 | 0.438 | 0.055 | 2.19075461544l3e-23 | 18 | aga |
| igflr12 | 1.2100557 | 0.754967198 | 0.479 | 0.066 | 2.44394959805923e-2 | 18 | igflr1 |
| sox12 | 2.2063498 | 0.430327867 | 0.229 | 0.014 | 4.45616481143552e-2 | 18 | sox12 |
| lum1 | 5.5168414 | 0.562059909 | 0.333 | 0.031 | 1.11423647322198e-2 | 18 | lum |

| | | | | | | | |
|---|---|---|---|---|---|---|---|
| mfap25 | 1.4274070 | 1.132204582 | 0.833 | 0.24 | 2.88293394142478e-2 | 18 | mfap2 |
| pdlim3a | 1.6414110 | 0.308594455 | 0.167 | 0.007 | 3.31515793456261e-2 | 18 | pdlim3a |
| hspa12b | 2.7177253 | 0.566377852 | 0.354 | 0.037 | 5.4889899095337e-22 | 18 | hspa12b |
| tnfsf103 | 2.8009094 | 0.871718453 | 0.646 | 0.121 | 5.65699682167685e-2 | 18 | tnfsf10 |
| sertad4 | 5.6857074 | 0.464165513 | 0.292 | 0.025 | 1.14834232555581e-2 | 18 | sertad4 |
| cx43.41 | 5.6875391 | 0.633679475 | 0.417 | 0.052 | 1.14871227943688e-2 | 18 | cx43.4 |
| mfap54 | 1.2020935 | 0.94540192 | 0.75 | 0.154 | 2.42786838410669e-2 | 18 | mfap5 |
| mmp16a4 | 3.2614519 | 0.864413635 | 0.625 | 0.116 | 6.58715460348674e-2 | 18 | mmp16a |
| rasl11b | 7.0698398 | 0.442728907 | 0.229 | 0.016 | 1.42789556332866e-2 | 18 | rasl11b |
| hmcn11 | 7.5734077 | 0.28092039 | 0.188 | 0.01 | 1.52960115860311e-2 | 18 | hmcn1 |
| mir214a1 | 8.3480253 | 0.426223687 | 0.25 | 0.019 | 1.68605068374356e-2 | 18 | mir214a |
| XLOC-0412692 | 2.6882963 | 0.652160031 | 0.417 | 0.054 | 5.42955216255201e-2 | 18 | XLOC-041269 |
| tnfsf10l3 | 4.2349404 | 0.703429556 | 0.583 | 0.101 | 8.55330924457197e-2 | 18 | tnfsf10l |
| tagln6 | 4.8371295 | 0.937219381 | 0.646 | 0.128 | 9.76955058827222e-2 | 18 | tagln |
| CABZ01092746.15 | 5.0028731 | 0.996811592 | 0.792 | 0.199 | 1.01043028074168e-1 | 18 | CABZ01092746.1 |
| wif1 | 5.9763564 | 0.29172621 | 0.146 | 0.006 | 1.20704470858509e-1 | 18 | wif1 |
| si:ch1073-459j12.1 | 6.1910975 | 0.906225019 | 0.771 | 0.175 | 1.2504159712523e-19 | 18 | si:ch1073-459j12.1 |
| tgfb33 | 6.8951093 | 0.728943933 | 0.5 | 0.081 | 1.39260522615355e-1 | 18 | tgfb3 |
| cald1a6 | 2.0192955 | 0.947455791 | 0.833 | 0.228 | 4.07837119485464e-1 | 18 | cald1a |
| BX663503.32 | 2.7055322 | 0.614054306 | 0.396 | 0.051 | 5.46436348984472e-1 | 18 | BX663503.3 |
| pcdh7b1 | 2.9843150 | 0.475128995 | 0.25 | 0.02 | 6.02742110295405e-1 | 18 | pcdh7b |
| foxc1b | 3.0425310 | 0.330584735 | 0.25 | 0.02 | 6.14499993588597e-1 | 18 | foxc1b |
| pim31 | 3.7666836 | 0.551682046 | 0.417 | 0.056 | 7.60757099380116e-1 | 18 | pim3 |
| tmsb21 | 5.7221644 | 0.320498121 | 0.208 | 0.014 | 1.15570556161308e-1 | 18 | tmsb2 |
| abhd15a2 | 5.7350606 | 0.725765466 | 0.438 | 0.064 | 1.15831019766538e-1 | 18 | abhd15a |
| col1a26 | 7.6727339 | 1.059119929 | 1 | 0.322 | 1.5496620770087e-18 | 18 | col1a2 |
| boc5 | 9.8535669 | 0.893148164 | 0.688 | 0.159 | 1.99012492295072e-1 | 18 | boc |
| rspo32 | 4.3167368 | 0.524934504 | 0.312 | 0.034 | 8.71851347046819e-1 | 18 | rspo3 |
| f3b1 | 5.3718303 | 0.718678511 | 0.5 | 0.087 | 1.08494857585842e-1 | 18 | f3b |
| sh3bgrl21 | 7.9788815 | 0.427760347 | 0.25 | 0.022 | 1.61149470061188e-1 | 18 | sh3bgrl2 |
| pbx1a4 | 8.3372220 | 0.732233109 | 0.562 | 0.108 | 1.6838687395293e-17 | 18 | pbx1a |
| LOC110439623 | 3.3713653 | 0.334173867 | 0.208 | 0.015 | 6.80914651950733e-1 | 18 | LOC110439623 |
| filip1l1 | 3.5721436 | 0.539702786 | 0.333 | 0.04 | 7.21465862415068e-1 | 18 | filip1l |
| fstl1b5 | 3.8642337 | 0.939868264 | 0.896 | 0.269 | 7.80459283938835e-1 | 18 | fstl1b |
| jag1b1 | 4.6620511 | 0.568491881 | 0.333 | 0.041 | 9.4159447338383e-17 | 18 | jag1b |

| | | | | | | | |
|---|---|---|---|---|---|---|---|
| fbn2b5 | 4.9893087 | 0.815260136 | 0.708 | 0.17 | 1.00769068001368e-1 | 18 | fbn2b |
| lrrc173 | 6.1422884 | 0.842451135 | 0.708 | 0.173 | 1.24055799767249e-1 | 18 | lrrc17 |
| rergla | 1.2058960 | 0.388516168 | 0.167 | 0.01 | 2.43554823504431e-1 | 18 | rergla |
| mxra8b3 | 1.3268649 | 0.555152691 | 0.396 | 0.058 | 2.67986923883164e-1 | 18 | mxra8b |
| dkk3b3 | 1.4117212 | 0.822781514 | 0.667 | 0.159 | 2.85125338731258e-1 | 18 | dkk3b |
| elovl6 | 2.4950825 | 0.299489106 | 0.188 | 0.013 | 5.03931827413243e-1 | 18 | elovl6 |
| LOC110438055 | 2.5759701 | 0.260152456 | 0.146 | 0.007 | 5.20268693107876e-1 | 18 | LOC110438055 |
| lgals2a3 | 4.2900848 | 0.904384417 | 0.792 | 0.2 | 8.66468431293831e-1 | 18 | lgals2a |
| fkbp92 | 4.7415741 | 0.686980219 | 0.479 | 0.087 | 9.57655731303047e-1 | 18 | fkbp9 |
| dhrs3b3 | 5.8684169 | 0.583492557 | 0.417 | 0.064 | 1.18524417023992e-1 | 18 | dhrs3b |
| pi15b | 8.0867261 | 0.377159416 | 0.167 | 0.01 | 1.63327608467203e-1 | 18 | pi15b |
| twist1a2 | 1.3308228 | 0.443940759 | 0.312 | 0.037 | 2.68786293368349e-1 | 18 | twist1a |
| gata52 | 1.3407991 | 0.422022037 | 0.25 | 0.025 | 2.70801207419768e-1 | 18 | gata5 |
| c1qtnf6a2 | 1.9336049 | 0.570792708 | 0.396 | 0.06 | 3.90530190761714e-1 | 18 | c1qtnf6a |
| hmga21 | 2.3337769 | 0.495192501 | 0.354 | 0.049 | 4.71352933863243e-1 | 18 | hmga2 |
| bmper2 | 2.4642199 | 0.697309929 | 0.479 | 0.088 | 4.97698512587511e-1 | 18 | bmper |
| sept9a | 3.9424295 | 0.536024028 | 0.396 | 0.062 | 7.96252493729954e-1 | 18 | sept9a |
| col1a1a5 | 4.4402187 | 0.920166895 | 0.958 | 0.302 | 8.96790987691359e-1 | 18 | col1a1a |
| col5a15 | 6.8794300 | 0.886310186 | 0.792 | 0.235 | 1.38943848435307e-1 | 18 | col5a1 |
| mmp15b | 2.1621754 | 0.497886415 | 0.292 | 0.036 | 4.36694569557406e-1 | 18 | mmp15b |
| efna2b | 3.7894142 | 0.342244996 | 0.188 | 0.015 | 7.65347993470423e-1 | 18 | efna2b |
| si:dkey-261h17.13 | 3.8107219 | 0.900523897 | 0.792 | 0.243 | 7.69651511255908e-1 | 18 | si:dkey-261h17.1 |
| pmp22a5 | 7.1963213 | 0.90031572 | 0.917 | 0.312 | 1.45344103090655e-1 | 18 | pmp22a |
| cacnb2a | 1.0778036 | 0.261267219 | 0.167 | 0.012 | 2.17684010555174e-1 | 18 | cacnb2a |
| dpysl31 | 1.8000125 | 0.401764665 | 0.271 | 0.032 | 3.63548539262167e-1 | 18 | dpysl3 |
| efemp2a2 | 2.2878435 | 0.427528585 | 0.292 | 0.037 | 4.62075770569626e-1 | 18 | efemp2a |
| bckdhbl | 2.6595717 | 0.395759685 | 0.292 | 0.038 | 5.37153703115701e-1 | 18 | bckdhbl |
| lmo4b | 3.3818323 | 0.265389484 | 0.208 | 0.019 | 6.83028674233862e-1 | 18 | lmo4b |
| znf503 | 4.3295804 | 0.512908703 | 0.312 | 0.044 | 8.74445371566833e-1 | 18 | znf503 |
| rbp53 | 8.6382192 | 0.746635912 | 0.688 | 0.19 | 1.74466114315965e-1 | 18 | rbp5 |
| col6a35 | 1.0111989 | 0.708548308 | 0.708 | 0.186 | 2.04231853060745e-1 | 18 | col6a3 |
| col15a1a1 | 2.4008650 | 0.412940535 | 0.292 | 0.039 | 4.8490271873133e-12 | 18 | col15a1a |
| tmem45a1 | 3.6355599 | 0.408457777 | 0.292 | 0.04 | 7.34274048470354e-1 | 18 | tmem45a |
| hoxb5a | 5.9043826 | 0.331410962 | 0.188 | 0.017 | 1.19250817334523e-1 | 18 | hoxb5a |
| glipr2l2 | 6.8572079 | 0.527139951 | 0.333 | 0.053 | 1.38495028508501e-1 | 18 | glipr2l |

| | | | | | | | |
|---|---|---|---|---|---|---|---|
| lrp1ab3 | 7.1767918 | 0.60751612 | 0.521 | 0.118 | 1.4494966463991e-11 | 18 | lrp1ab |
| rassf8a | 1.1661320 | 0.352679452 | 0.208 | 0.022 | 2.35523681636991e-1 | 18 | rassf8a |
| snai1a2 | 2.0655905 | 0.528167769 | 0.333 | 0.055 | 4.17187320066653e-1 | 18 | snai1a |
| myl65 | 2.5223926 | 0.755664296 | 0.667 | 0.197 | 5.09447647168552e-1 | 18 | myl6 |
| vasna3 | 2.6056699 | 0.610685964 | 0.458 | 0.096 | 5.26267150764549e-1 | 18 | vasna |
| hic13 | 3.5936617 | 0.590160309 | 0.542 | 0.129 | 7.25811870402713e-1 | 18 | hic1 |
| grb10b2 | 3.8532928 | 0.515882241 | 0.333 | 0.056 | 7.78249556647897e-1 | 18 | grb10b |
| tsku2 | 4.0016473 | 0.555111588 | 0.396 | 0.076 | 8.08212705281335e-1 | 18 | tsku |
| mylka1 | 4.1395100 | 0.363219026 | 0.271 | 0.037 | 8.36056845993624e-1 | 18 | mylka |
| wnt5b3 | 4.1557744 | 0.404173935 | 0.25 | 0.032 | 8.39341761897961e-1 | 18 | wnt5b |
| sparc8 | 4.3599332 | 0.841464432 | 0.938 | 0.394 | 8.80575715522311e-1 | 18 | sparc |
| cdh111 | 8.1337217 | 0.46882854 | 0.292 | 0.044 | 1.64276777438496e-1 | 18 | cdh11 |
| ccl25b4 | 9.8383274 | 0.820348293 | 0.646 | 0.18 | 1.98704698666297e-1 | 18 | ccl25b |
| tuba1a1 | 1.4551945 | 0.408728556 | 0.25 | 0.033 | 2.93905641876162e-1 | 18 | tuba1a |
| cnn3a4 | 1.6296781 | 0.625494036 | 0.521 | 0.131 | 3.29146104982643e-1 | 18 | cnn3a |
| hmgn32 | 1.7119294 | 0.697968904 | 0.562 | 0.161 | 3.45758386489923e-1 | 18 | hmgn3 |
| nuak1b | 1.8626728 | 0.29912186 | 0.188 | 0.019 | 3.76204038632156e-1 | 18 | nuak1b |
| col5a2a5 | 2.0723653 | 0.688609813 | 0.625 | 0.175 | 4.18555637834426e-1 | 18 | col5a2a |
| hoxc8a2 | 2.0968970 | 0.513944736 | 0.354 | 0.065 | 4.23510297142813e-1 | 18 | hoxc8a |
| FP102018.11 | 2.1000351 | 0.358387791 | 0.25 | 0.033 | 4.24144094851194e-1 | 18 | FP102018.1 |
| gas1a4 | 2.5306981 | 0.643449383 | 0.521 | 0.133 | 5.11125108918307e-1 | 18 | gas1a |
| mrc22 | 2.5913409 | 0.424720209 | 0.312 | 0.051 | 5.23373123426964e-1 | 18 | mrc2 |
| hoxc3a1 | 2.6242693 | 0.497716042 | 0.354 | 0.066 | 5.30023674995723e-1 | 18 | hoxc3a |
| serpinh1b2 | 2.8859700 | 0.582483292 | 0.354 | 0.066 | 5.82879371951728e-1 | 18 | serpinh1b |
| nfia4 | 3.0499390 | 0.686714321 | 0.646 | 0.194 | 6.15996195760736e-1 | 18 | nfia |
| tpm4b2 | 3.5790964 | 0.438018667 | 0.292 | 0.045 | 7.22870108622868e-1 | 18 | tpm4b |
| rab131 | 4.1806055 | 0.477903991 | 0.375 | 0.074 | 8.44356899494026e-1 | 18 | rab13 |
| ackr3b3 | 4.6362942 | 0.51577105 | 0.375 | 0.072 | 9.36392343438976e-1 | 18 | ackr3b |
| crabp2b1 | 4.8023278 | 0.326733259 | 0.229 | 0.029 | 9.69926162098118e-1 | 18 | crabp2b |
| emid11 | 4.9764410 | 0.58514599 | 0.354 | 0.066 | 1.00509180282854e-0 | 18 | emid1 |
| tspan4b3 | 6.0462482 | 0.503849288 | 0.458 | 0.102 | 1.22116076783855e-0 | 18 | tspan4b |
| mical3b | 6.0727869 | 0.333444367 | 0.208 | 0.025 | 1.22652077921764e-0 | 18 | mical3b |
| zgc:171566 | 6.9614111 | 0.271473447 | 0.167 | 0.016 | 1.40599620045671e-0 | 18 | zgc:171566 |
| fkbp1ab2 | 7.5632237 | 0.389838595 | 0.271 | 0.039 | 1.52754429431967e-0 | 18 | fkbp1ab |
| zgc:153675 | 8.7643731 | 0.470965234 | 0.333 | 0.062 | 1.77014044663956e-0 | 18 | zgc:153675 |

| | | | | | | | |
|---|---|---|---|---|---|---|---|
| kdm7ab | 9.1660133 | 0.398734393 | 0.25 | 0.036 | 1.85125970803277e-0 | 18 | kdm7ab |
| LO018188.14 | 1.0598251 | 0.542370101 | 0.417 | 0.088 | 2.14052887975512e-0 | 18 | LO018188.1 |
| tspan73 | 1.3543671 | 0.500990018 | 0.438 | 0.098 | 2.73541537463158e-0 | 18 | tspan7 |
| pora1 | 1.7158273 | 0.394160366 | 0.312 | 0.054 | 3.46545645495878e-0 | 18 | pora |
| zfhx31 | 1.8975719 | 0.462234538 | 0.354 | 0.068 | 3.83252604943362e-0 | 18 | zfhx3 |
| epha4l1 | 2.0593694 | 0.278611978 | 0.208 | 0.025 | 4.15930850767659e-0 | 18 | epha4l |
| fblim12 | 2.2580503 | 0.51086291 | 0.417 | 0.092 | 4.56058434536508e-0 | 18 | fblim1 |
| hmgb3a6 | 2.8179572 | 0.75397888 | 0.688 | 0.254 | 5.69142824267948e-0 | 18 | hmgb3a |
| zgc:92664 | 2.9847519 | 0.472800554 | 0.333 | 0.064 | 6.02830355876176e-0 | 18 | zgc:92664 |
| mmp11b1 | 3.1493137 | 0.290137106 | 0.146 | 0.013 | 6.36066902972911e-0 | 18 | mmp11b |
| ppib7 | 4.9219447 | 0.602094716 | 0.979 | 0.64 | 9.94085188180185e-0 | 18 | ppib |
| nid1a1 | 4.9425414 | 0.374582817 | 0.292 | 0.048 | 9.98245090485516e-0 | 18 | nid1a |
| myl9b4 | 7.5964177 | 0.671537748 | 0.625 | 0.198 | 1.5342485009879e-08 | 18 | myl9b |
| tenm35 | 8.0626893 | 0.606713282 | 0.521 | 0.141 | 1.62842136888878e-0 | 18 | tenm3 |
| rcn1 | 8.5750253 | 0.339699042 | 0.229 | 0.032 | 1.73189787775012e-0 | 18 | rcn1 |
| myl9a5 | 9.3208046 | 0.578963112 | 0.458 | 0.112 | 1.8825229235137e-08 | 18 | myl9a |
| fosab6 | 1.1304206 | 0.793207774 | 0.917 | 0.455 | 2.28311048722799e-0 | 18 | fosab |
| tead1b2 | 1.3253533 | 0.455613199 | 0.312 | 0.058 | 2.6768161804793e-08 | 18 | tead1b |
| cadm41 | 1.3916818 | 0.404088653 | 0.292 | 0.051 | 2.81077988735998e-0 | 18 | cadm4 |
| quo1 | 2.3033070 | 0.269912152 | 0.188 | 0.022 | 4.65198919402755e-0 | 18 | quo |
| fzd43 | 2.5112787 | 0.396328791 | 0.312 | 0.057 | 5.07202966666708e-0 | 18 | fzd4 |
| kif26ba1 | 2.8612559 | 0.294373025 | 0.188 | 0.023 | 5.77887861416724e-0 | 18 | kif26ba |
| si:ch211-204a13.2: | 3.2741929 | 0.398350661 | 0.25 | 0.039 | 6.61288751601379e-0 | 18 | si:ch211-204a13.2 |
| CR383676.17 | 3.6573288 | 0.630829161 | 1 | 0.656 | 7.38670698556344e-0 | 18 | CR383676.1 |
| epha4b1 | 4.0033664 | 0.282875801 | 0.188 | 0.023 | 8.08559925548833e-0 | 18 | epha4b |
| LOC1005373422 | 4.2870006 | 0.389836097 | 0.229 | 0.034 | 8.65845511948473e-0 | 18 | LOC100537342 |
| atrx3 | 4.7059928 | 0.617585351 | 0.667 | 0.238 | 9.50469373250864e-0 | 18 | atrx |
| crtap2 | 5.6369598 | 0.510405243 | 0.375 | 0.085 | 1.13849677412464e-0 | 18 | crtap |
| si:ch211-267e7.3 | 5.9519336 | 0.252137332 | 0.146 | 0.014 | 1.20211204483361e-0 | 18 | si:ch211-267e7.3 |
| xirp2a2 | 7.0609411 | 0.498257708 | 0.354 | 0.078 | 1.42609827865781e-0 | 18 | xirp2a |
| h3f3c5 | 7.8753381 | 0.650420376 | 0.833 | 0.527 | 1.59058204992915e-0 | 18 | h3f3c |
| h1f03 | 8.1794308 | 0.670761405 | 0.438 | 0.117 | 1.6519996579898e-07 | 18 | h1f0 |
| ube2e33 | 8.7996936 | 0.648754674 | 0.667 | 0.252 | 1.77727411838881e-0 | 18 | ube2e3 |
| ssr14 | 1.0108963 | 0.555968012 | 0.729 | 0.269 | 2.04170735013118e-0 | 18 | ssr1 |
| efna5b1 | 1.0655486 | 0.250300677 | 0.167 | 0.019 | 2.15208857309228e-0 | 18 | efna5b |

| | | | | | | | |
|---|---|---|---|---|---|---|---|
| jpt1b5 | 1.0945657 | 0.681576008 | 0.771 | 0.329 | 2.21069443633966e-0 | 18 | jpt1b |
| col1a1b5 | 1.0998464 | 0.717981386 | 0.792 | 0.292 | 2.22135996369209e-0 | 18 | col1a1b |
| fzd7b5 | 1.1084499 | 0.519065873 | 0.417 | 0.102 | 2.23873630056249e-0 | 18 | fzd7b |
| si:dkey-222b8.1 | 1.3541606 | 0.328363022 | 0.188 | 0.025 | 2.73499829326182e-0 | 18 | si:dkey-222b8.1 |
| marcksb1 | 1.6787852 | 0.501272506 | 0.375 | 0.089 | 3.39064248337838e-0 | 18 | marcksb |
| col6a4a5 | 1.9114316 | 0.599251777 | 0.562 | 0.164 | 3.86051851389374e-0 | 18 | col6a4a |
| pdgfc | 2.0618749 | 0.273421303 | 0.146 | 0.015 | 4.16436879614809e-0 | 18 | pdgfc |
| stau11 | 2.2050263 | 0.459825039 | 0.396 | 0.096 | 4.45349174089509e-0 | 18 | stau1 |
| igf2b3 | 2.7058736 | 0.600516292 | 0.542 | 0.159 | 5.46505305037578e-0 | 18 | igf2b |
| marcksl1a5 | 4.8369143 | 0.664056024 | 0.958 | 0.507 | 9.76911593813412e-0 | 18 | marcksl1a |
| fkbp10b1 | 5.0211846 | 0.346841478 | 0.208 | 0.031 | 1.01412866511499e-0 | 18 | fkbp10b |
| bcl9 | 5.0587890 | 0.32864712 | 0.188 | 0.026 | 1.02172362106763e-0 | 18 | bcl9 |
| serpine1 | 5.1176940 | 0.287570258 | 0.146 | 0.016 | 1.03362067628732e-0 | 18 | serpine1 |
| thbs2b1 | 5.2988046 | 0.306483292 | 0.146 | 0.016 | 1.0701995678701e-06 | 18 | thbs2b |
| srd5a3 | 5.5012392 | 0.256172241 | 0.188 | 0.026 | 1.11108528819148e-0 | 18 | srd5a3 |
| ckap45 | 5.6632637 | 0.588584097 | 0.479 | 0.145 | 1.1438093763403e-06 | 18 | ckap4 |
| s100v27 | 6.5716274 | 0.576629653 | 0.708 | 0.244 | 1.32727160007988e-0 | 18 | s100v2 |
| sat1b | 6.6321406 | 0.317179663 | 0.188 | 0.026 | 1.33949344082245e-0 | 18 | sat1b |
| si:ch211-286o17.1 | 8.2174020 | 0.509020456 | 0.417 | 0.108 | 1.6596687009417e-06 | 18 | si:ch211-286o17.1 |
| cyr611 | 1.3433589 | 0.429789226 | 0.312 | 0.066 | 2.71318201973878e-0 | 18 | cyr61 |
| cyp27c12 | 1.4943665 | 0.488473335 | 0.292 | 0.062 | 3.01817220120361e-0 | 18 | cyp27c1 |
| efnb13 | 1.5219494 | 0.456810093 | 0.396 | 0.098 | 3.0738813061116e-06 | 18 | efnb1 |
| nme34 | 2.1735369 | 0.52399322 | 0.5 | 0.157 | 4.38989255975892e-0 | 18 | nme3 |
| asip2b3 | 2.5656075 | 0.501403515 | 0.312 | 0.067 | 5.18175750808147e-0 | 18 | asip2b |
| smo1 | 2.6994917 | 0.398543351 | 0.292 | 0.061 | 5.45216340254086e-0 | 18 | smo |
| meis1b5 | 2.7139762 | 0.537987808 | 0.5 | 0.153 | 5.48141773358713e-0 | 18 | meis1b |
| id32 | 3.0265689 | 0.552808558 | 0.479 | 0.143 | 6.11276126539077e-0 | 18 | id3 |
| cldn11a5 | 3.0785551 | 0.613093932 | 0.583 | 0.191 | 6.21775778045791e-0 | 18 | cldn11a |
| lhfpl2a2 | 3.3762147 | 0.434692952 | 0.354 | 0.085 | 6.81894098609358e-0 | 18 | lhfpl2a |
| bambia3 | 3.6410641 | 0.358132689 | 0.292 | 0.059 | 7.35385727089401e-0 | 18 | bambia |
| pdgfrb3 | 4.2564485 | 0.395367521 | 0.292 | 0.061 | 8.59674913286618e-0 | 18 | pdgfrb |
| rhpn21 | 4.4259579 | 0.32745012 | 0.208 | 0.034 | 8.9391072571823e-06 | 18 | rhpn2 |
| palld1 | 4.6621157 | 0.297530576 | 0.208 | 0.034 | 9.41607520645549e-0 | 18 | palld |
| calua2 | 4.7138540 | 0.423619785 | 0.312 | 0.071 | 9.52057095267516e-0 | 18 | calua |
| flot2a5 | 4.8627002 | 0.483390031 | 0.479 | 0.148 | 9.82119574943605e-0 | 18 | flot2a |

| | | | | | | | |
|---|---|---|---|---|---|---|---|
| rasgef1bb1 | 5.2737459 | 0.354525234 | 0.229 | 0.04 | 1.06513846453548e-0 | 18 | rasgef1bb |
| zgc:162939 | 5.6420494 | 0.300906619 | 0.208 | 0.034 | 1.13952473696521e-0 | 18 | zgc:162939 |
| cd81a6 | 6.2178048 | 0.657748678 | 0.896 | 0.435 | 1.25581005532054e-0 | 18 | cd81a |
| selenon | 6.3112299 | 0.255664129 | 0.167 | 0.023 | 1.27467911410406e-0 | 18 | selenon |
| cd82a3 | 6.6408509 | 0.517671857 | 0.458 | 0.137 | 1.34125266322217e-0 | 18 | cd82a |
| ankrd46a | 7.1315503 | 0.31399712 | 0.208 | 0.035 | 1.4403592150366e-05 | 18 | ankrd46a |
| rnd3a1 | 7.4794010 | 0.411160129 | 0.271 | 0.056 | 1.51061463562737e-0 | 18 | rnd3a |
| meis2b5 | 7.8008431 | 0.394590453 | 0.375 | 0.093 | 1.57553629063565e-0 | 18 | meis2b |
| tpm4a6 | 9.8591671 | 0.591121987 | 0.646 | 0.241 | 1.99125598720659e-0 | 18 | tpm4a |
| adgrl3.12 | 1.0265357 | 0.358050846 | 0.271 | 0.055 | 2.07329418747962e-0 | 18 | adgrl3.1 |
| ptbp1b1 | 1.0542278 | 0.351465216 | 0.292 | 0.064 | 2.12922394126709e-0 | 18 | ptbp1b |
| dpysl2b | 1.0636845 | 0.321154336 | 0.229 | 0.041 | 2.14832378557411e-0 | 18 | dpysl2b |
| col6a15 | 1.1201162 | 0.480580041 | 0.479 | 0.141 | 2.26229887879055e-0 | 18 | col6a1 |
| pfn2l7 | 1.1934406 | 0.571489342 | 0.667 | 0.258 | 2.41039212031851e-0 | 18 | pfn2l |
| jun7 | 1.3177607 | 0.664482825 | 0.812 | 0.38 | 2.66148139233285e-0 | 18 | jun |
| fam168a3 | 1.4183807 | 0.481195476 | 0.479 | 0.151 | 2.86470356666968e-0 | 18 | fam168a |
| tmem100a2 | 1.4851399 | 0.313895892 | 0.25 | 0.048 | 2.99953715091717e-0 | 18 | tmem100a |
| praf21 | 1.5490483 | 0.338934172 | 0.271 | 0.057 | 3.12861293354923e-0 | 18 | praf2 |
| gulp1a2 | 1.6411190 | 0.343609501 | 0.292 | 0.063 | 3.31456808825116e-0 | 18 | gulp1a |
| hnrnpa0a1 | 1.6753869 | 0.493936305 | 0.458 | 0.142 | 3.38377892560788e-0 | 18 | hnrnpa0a |
| cldn11b | 1.7364323 | 0.275037784 | 0.188 | 0.03 | 3.50707232625628e-0 | 18 | cldn11b |
| tmeff1b2 | 1.9096934 | 0.456878157 | 0.375 | 0.101 | 3.85700787966838e-0 | 18 | tmeff1b |
| nova22 | 1.9193190 | 0.387062291 | 0.292 | 0.063 | 3.87644860013256e-0 | 18 | nova2 |
| rbfox12 | 2.1620739 | 0.323185902 | 0.25 | 0.049 | 4.36674069543812e-0 | 18 | rbfox1 |
| dvl2 | 2.2641199 | 0.32666397 | 0.25 | 0.05 | 4.57284298094954e-0 | 18 | dvl2 |
| si:dkeyp-113d7.1 | 2.4178626 | 0.366827171 | 0.375 | 0.097 | 4.8833572009349e-05 | 18 | si:dkeyp-113d7.1 |
| rem11 | 2.4445251 | 0.308683169 | 0.167 | 0.024 | 4.93720744843238e-0 | 18 | rem1 |
| antxr2a4 | 2.5594145 | 0.58362236 | 0.646 | 0.258 | 5.16924959327069e-0 | 18 | antxr2a |
| mbd3b1 | 2.5689211 | 0.359850315 | 0.292 | 0.066 | 5.18845001181859e-0 | 18 | mbd3b |
| parvaa1 | 2.6006068 | 0.359142283 | 0.292 | 0.065 | 5.25244566113241e-0 | 18 | parvaa |
| rerea1 | 2.6751638 | 0.399839178 | 0.292 | 0.067 | 5.40302850374073e-0 | 18 | rerea |
| hmgn73 | 2.8742629 | 0.57119776 | 0.896 | 0.526 | 5.80514887407703e-0 | 18 | hmgn7 |
| nr2f1a3 | 2.9331525 | 0.516935872 | 0.479 | 0.149 | 5.92408813406813e-0 | 18 | nr2f1a |
| kirrel1a2 | 3.1396919 | 0.387181387 | 0.292 | 0.066 | 6.34123576492795e-0 | 18 | kirrel1a |
| lpar2a | 3.2328373 | 0.266107093 | 0.229 | 0.042 | 6.52936155918593e-0 | 18 | lpar2a |

| | | | | | | | |
|---|---|---|---|---|---|---|---|
| gatad2b2 | 3.2507623 | 0.481267596 | 0.396 | 0.116 | 6.5655646641709e-05 | 18 | gatad2b |
| ecm2 | 3.3476573 | 0.257325296 | 0.146 | 0.019 | 6.76126346061013e-0 | 18 | ecm2 |
| tubb52 | 4.0603552 | 0.449097754 | 0.312 | 0.077 | 8.2006995570014e-05 | 18 | tubb5 |
| MMP23B1 | 4.6574707 | 0.293316027 | 0.208 | 0.037 | 9.40669367401776e-0 | 18 | MMP23B |
| fam198a1 | 4.7981427 | 0.291098839 | 0.25 | 0.05 | 9.69080883519933e-0 | 18 | fam198a |
| pbx3b1 | 4.9614887 | 0.397046296 | 0.292 | 0.067 | 0.000100207 | 18 | pbx3b |
| rbm39a2 | 5.0297319 | 0.501901129 | 0.562 | 0.207 | 0.000101585 | 18 | rbm39a |
| sav1 | 5.5213564 | 0.31510757 | 0.271 | 0.059 | 0.000111515 | 18 | sav1 |
| pdgfra3 | 5.5239743 | 0.449844364 | 0.438 | 0.129 | 0.000111568 | 18 | pdgfra |
| luzp1 | 7.5167517 | 0.281126591 | 0.188 | 0.032 | 0.000151816 | 18 | luzp1 |
| pcdh18b2 | 7.6827251 | 0.422207401 | 0.292 | 0.07 | 0.000155168 | 18 | pcdh18b |
| angptl43 | 8.4740876 | 0.442980467 | 0.333 | 0.087 | 0.000171151 | 18 | angptl4 |
| aebp14 | 9.0027970 | 0.414145377 | 0.354 | 0.096 | 0.000181829 | 18 | aebp1 |
| fxyd6l4 | 9.2927929 | 0.452706264 | 0.458 | 0.145 | 0.000187687 | 18 | fxyd6l |
| ldlrad3 | 1.0033084 | 0.285610148 | 0.188 | 0.032 | 0.000202638 | 18 | ldlrad3 |
| mgst1.14 | 1.0945181 | 0.449158524 | 0.417 | 0.128 | 0.00022106 | 18 | mgst1.1 |
| col6a25 | 1.1518997 | 0.459950425 | 0.417 | 0.124 | 0.000232649 | 18 | col6a2 |
| olfml3a1 | 1.2799013 | 0.32975449 | 0.229 | 0.046 | 0.000258502 | 18 | olfml3a |
| zcchc241 | 1.2846699 | 0.397601143 | 0.292 | 0.07 | 0.000259465 | 18 | zcchc24 |
| si:ch211-133n4.45 | 1.3328505 | 0.49255409 | 0.417 | 0.131 | 0.000269196 | 18 | si:ch211-133n4.4 |
| arid5b | 1.3848626 | 0.324907053 | 0.167 | 0.027 | 0.000279701 | 18 | arid5b |
| anxa41 | 1.6074822 | 0.492649533 | 0.583 | 0.234 | 0.000324663 | 18 | anxa4 |
| si:ch73-46j18.58 | 1.7311504 | 0.60914204 | 0.792 | 0.392 | 0.00034964 | 18 | si:ch73-46j18.5 |
| ctsla8 | 1.8164547 | 0.440546715 | 0.771 | 0.303 | 0.000366869 | 18 | ctsla |
| zgc:55943 | 2.2567194 | 0.284803836 | 0.25 | 0.055 | 0.00045579 | 18 | zgc:55943 |
| hp1bp34 | 2.3293752 | 0.509559163 | 0.583 | 0.24 | 0.000470464 | 18 | hp1bp3 |
| scara32 | 2.4212102 | 0.323221401 | 0.25 | 0.053 | 0.000489012 | 18 | scara3 |
| fzd26 | 2.4782201 | 0.456738173 | 0.375 | 0.115 | 0.000500526 | 18 | fzd2 |
| tmem119b3 | 2.5462650 | 0.272740599 | 0.229 | 0.046 | 0.000514269 | 18 | tmem119b |
| si:ch211-288g17.3: | 2.7928944 | 0.570046318 | 0.667 | 0.289 | 0.000564081 | 18 | si:ch211-288g17.3 |
| olfml3b5 | 2.8253744 | 0.424790585 | 0.375 | 0.109 | 0.000570641 | 18 | olfml3b |
| mgst3b6 | 3.1738379 | 0.523303781 | 0.667 | 0.298 | 0.00064102 | 18 | mgst3b |
| fkbp75 | 3.1843931 | 0.448204074 | 0.396 | 0.124 | 0.000643152 | 18 | fkbp7 |
| tnfsf121 | 3.2060149 | 0.318198716 | 0.229 | 0.048 | 0.000647519 | 18 | tnfsf12 |
| her91 | 3.2097902 | 0.360395425 | 0.229 | 0.048 | 0.000648281 | 18 | her9 |

| | | | | | | | |
|---|---|---|---|---|---|---|---|
| cdon4 | 3.3089407 | 0.486261885 | 0.438 | 0.143 | 0.000668307 | 18 | cdon |
| s100a10b5 | 3.3759641 | 0.617074619 | 0.896 | 0.453 | 0.000681843 | 18 | s100a10b |
| zgc:770863 | 3.5270334 | 0.389354236 | 0.396 | 0.119 | 0.000712355 | 18 | zgc:77086 |
| rab40b | 3.5958356 | 0.26124251 | 0.167 | 0.028 | 0.000726251 | 18 | rab40b |
| rgs12b | 3.7772220 | 0.250701132 | 0.125 | 0.016 | 0.000762886 | 18 | rgs12b |
| arhgap31 | 3.8629885 | 0.297609476 | 0.229 | 0.048 | 0.000780208 | 18 | arhgap31 |
| icn5 | 4.0484005 | 0.628366497 | 0.667 | 0.253 | 0.000817655 | 18 | icn |
| txndc51 | 5.5415789 | 0.433226952 | 0.312 | 0.087 | 0.001119233 | 18 | txndc5 |
| adam19a1 | 5.7980599 | 0.292140002 | 0.229 | 0.049 | 0.001171034 | 18 | adam19a |
| si:ch73-281n10.27 | 6.0668127 | 0.576333542 | 0.854 | 0.429 | 0.001225314 | 18 | si:ch73-281n10.2 |
| rrbp1b1 | 6.2397978 | 0.275443903 | 0.229 | 0.049 | 0.001260252 | 18 | rrbp1b |
| ctnnb14 | 6.8267224 | 0.484864194 | 0.542 | 0.211 | 0.001378793 | 18 | ctnnb1 |
| wfdc11 | 6.9341823 | 0.291383188 | 0.167 | 0.029 | 0.001400497 | 18 | wfdc1 |
| chd36 | 7.3018287 | 0.509156456 | 0.583 | 0.238 | 0.00147475 | 18 | chd3 |
| znf385a3 | 7.4339278 | 0.431885783 | 0.333 | 0.095 | 0.00150143 | 18 | znf385a |
| hmgb1b5 | 7.9663116 | 0.50068111 | 0.688 | 0.298 | 0.001608956 | 18 | hmgb1b |
| cep290 | 8.4804823 | 0.257784662 | 0.125 | 0.017 | 0.001712803 | 18 | cep290 |
| hmgn63 | 9.2791623 | 0.50573281 | 0.896 | 0.568 | 0.001874112 | 18 | hmgn6 |
| arl6ip5b4 | 9.6558011 | 0.433401639 | 0.417 | 0.14 | 0.001950182 | 18 | arl6ip5b |
| stau23 | 9.9826584 | 0.459135242 | 0.438 | 0.153 | 0.002016198 | 18 | stau2 |
| si:ch211-39i22.16 | 1.0122725 | 0.474572341 | 0.5 | 0.18 | 0.002044487 | 18 | si:ch211-39i22.1 |
| smarce13 | 1.0663677 | 0.46670401 | 0.583 | 0.243 | 0.002153743 | 18 | smarce1 |
| meaf6 | 1.1441093 | 0.395720787 | 0.417 | 0.141 | 0.002310758 | 18 | meaf6 |
| si:ch211-212k18.83 | 1.2720903 | 0.316724155 | 0.292 | 0.075 | 0.002569241 | 18 | si:ch211-212k18.8 |
| aip1 | 1.4314163 | 0.35797802 | 0.438 | 0.147 | 0.002891032 | 18 | aip |
| XLOC-035413 | 1.5058589 | 0.331231135 | 0.292 | 0.079 | 0.003041383 | 18 | XLOC-035413 |
| mmp14a4 | 1.7196885 | 0.388578198 | 0.333 | 0.098 | 0.003473255 | 18 | mmp14a |
| ephb33 | 1.8138496 | 0.381088273 | 0.354 | 0.106 | 0.003663432 | 18 | ephb3 |
| ash1l | 1.9117200 | 0.338506269 | 0.25 | 0.061 | 0.003861101 | 18 | ash1l |
| zranb21 | 1.9309523 | 0.351141454 | 0.333 | 0.098 | 0.003899944 | 18 | zranb2 |
| si:ch211-105c13.34 | 1.9519810 | 0.441631303 | 0.354 | 0.107 | 0.003942416 | 18 | si:ch211-105c13.3 |
| ss18l21 | 1.9905840 | 0.282345477 | 0.292 | 0.077 | 0.004020383 | 18 | ss18l2 |
| hmgb1a5 | 2.0023930 | 0.568982509 | 0.75 | 0.425 | 0.004044233 | 18 | hmgb1a |
| pdgfrl1 | 2.0625334 | 0.308092971 | 0.25 | 0.06 | 0.004165699 | 18 | pdgfrl |
| nucb2a2 | 2.1370501 | 0.443711902 | 0.375 | 0.124 | 0.0043162 | 18 | nucb2a |

| | | | | | | | |
|---|---|---|---|---|---|---|---|
| pdia64 | 2.1511220 | 0.433984924 | 0.458 | 0.168 | 0.004344621 | 18 | pdia6 |
| ybx11 | 2.5582802 | 0.281884187 | 0.979 | 0.859 | 0.005166959 | 18 | ybx1 |
| meis31 | 2.6234207 | 0.255469525 | 0.188 | 0.037 | 0.005298523 | 18 | meis3 |
| ndrg22 | 2.6424070 | 0.381948655 | 0.333 | 0.101 | 0.00533687 | 18 | ndrg2 |
| colec125 | 2.7517426 | 0.417088126 | 0.354 | 0.109 | 0.005557695 | 18 | colec12 |
| nkd11 | 3.1079424 | 0.261488152 | 0.208 | 0.045 | 0.006277111 | 18 | nkd1 |
| cd248a4 | 3.5128694 | 0.424103249 | 0.375 | 0.12 | 0.007094942 | 18 | cd248a |
| si:ch73-1a9.33 | 3.5612832 | 0.519809745 | 0.875 | 0.586 | 0.007192724 | 18 | si:ch73-1a9.3 |
| jund | 4.0865971 | 0.365806199 | 0.292 | 0.082 | 0.0082537 | 18 | jund |
| ube2ib1 | 4.1678048 | 0.418502461 | 0.562 | 0.232 | 0.008417715 | 18 | ube2ib |
| erlec11 | 4.4886805 | 0.295287931 | 0.271 | 0.073 | 0.009065788 | 18 | erlec1 |
| esd2 | 4.5568059 | 0.429725638 | 0.458 | 0.175 | 0.009203381 | 18 | esd |
| usp14 | 4.5865809 | 0.260547424 | 0.25 | 0.062 | 0.009263517 | 18 | usp14 |
| sar1b3 | 4.8369799 | 0.395026344 | 0.396 | 0.138 | 0.009769248 | 18 | sar1b |
| dbn13 | 4.8631310 | 0.363284193 | 0.375 | 0.121 | 0.009822066 | 18 | dbn1 |
| zc4h21 | 5.6271477 | 0.290167432 | 0.333 | 0.101 | 0.01136515 | 18 | zc4h2 |
| fosb3 | 5.9747055 | 0.634196399 | 0.333 | 0.111 | 0.012067113 | 18 | fosb |
| phf2 | 6.0050681 | 0.276970579 | 0.188 | 0.039 | 0.012128436 | 18 | phf2 |
| ptenb2 | 6.0346049 | 0.381102067 | 0.375 | 0.125 | 0.012188092 | 18 | ptenb |
| med111 | 6.4703184 | 0.257195236 | 0.312 | 0.089 | 0.013068102 | 18 | med11 |
| smarca2 | 6.5171422 | 0.309753546 | 0.271 | 0.074 | 0.013162672 | 18 | smarca2 |
| zgc:1951734 | 6.9144360 | 0.451178719 | 0.583 | 0.233 | 0.013965087 | 18 | zgc:195173 |
| rbm4.13 | 6.9984571 | 0.348089977 | 0.312 | 0.093 | 0.014134784 | 18 | rbm4.1 |
| homer3b | 7.6003941 | 0.300419949 | 0.229 | 0.057 | 0.015350516 | 18 | homer3b |
| spred2b4 | 7.6063292 | 0.459048497 | 0.458 | 0.179 | 0.015362503 | 18 | spred2b |
| meis1a3 | 7.6563485 | 0.367676241 | 0.208 | 0.049 | 0.015463527 | 18 | meis1a |
| pfn26 | 8.0030718 | 0.431102419 | 0.708 | 0.315 | 0.016163804 | 18 | pfn2 |
| ahnak5 | 8.1848388 | 0.452614939 | 0.5 | 0.198 | 0.016530919 | 18 | ahnak |
| mmp14b3 | 8.6751129 | 0.394554222 | 0.354 | 0.115 | 0.017521126 | 18 | mmp14b |
| BX569787.14 | 9.8235327 | 0.351706334 | 0.333 | 0.103 | 0.019840589 | 18 | BX569787.1 |
| ddit34 | 1.0479010 | 0.529386135 | 0.479 | 0.2 | 0.021164458 | 18 | ddit3 |
| prkab1b1 | 1.0770413 | 0.356342544 | 0.521 | 0.205 | 0.021753004 | 18 | prkab1b |
| slit32 | 1.0779207 | 0.312374993 | 0.292 | 0.08 | 0.021770766 | 18 | slit3 |
| ppdpfb2 | 1.0977823 | 0.432945568 | 0.854 | 0.618 | 0.02217191 | 18 | ppdpfb |
| ncor22 | 1.1864154 | 0.387066087 | 0.354 | 0.12 | 0.023962033 | 18 | ncor2 |

| | | | | | | | |
|---|---|---|---|---|---|---|---|
| tshz2 | 1.2100450 | 0.31675738 | 0.188 | 0.042 | 0.024439281 | 18 | tshz2 |
| frs2b | 1.2408465 | 0.256756096 | 0.208 | 0.048 | 0.025061378 | 18 | frs2b |
| fam168b1 | 1.2971113 | 0.334156961 | 0.292 | 0.087 | 0.026197759 | 18 | fam168b |
| ppfibp1a3 | 1.3001759 | 0.292702934 | 0.208 | 0.049 | 0.026259654 | 18 | ppfibp1a |
| tuba1b1 | 1.5379809 | 0.362380633 | 0.271 | 0.078 | 0.031062602 | 18 | tuba1b |
| cntfr2 | 1.5420244 | 0.314527952 | 0.188 | 0.041 | 0.031144269 | 18 | cntfr |
| zgc:110158 | 1.5987398 | 0.274275933 | 0.229 | 0.058 | 0.032289749 | 18 | zgc:110158 |
| bsg5 | 1.7267444 | 0.486458661 | 0.708 | 0.385 | 0.034875058 | 18 | bsg |
| cd82b1 | 1.8050282 | 0.340935642 | 0.292 | 0.086 | 0.036456155 | 18 | cd82b |
| junba5 | 1.8761235 | 0.529291835 | 0.771 | 0.4 | 0.037892066 | 18 | junba |
| krt947 | 1.9070414 | 0.52462442 | 0.75 | 0.358 | 0.038516516 | 18 | krt94 |
| ptprga2 | 1.9441509 | 0.331900801 | 0.333 | 0.105 | 0.039266016 | 18 | ptprga |
| akap12b4 | 2.1467972 | 0.426166193 | 0.417 | 0.15 | 0.043358864 | 18 | akap12b |
| rbms32 | 2.2147001 | 0.264383421 | 0.208 | 0.05 | 0.044730299 | 18 | rbms3 |
| zc3h18 | 2.2472361 | 0.255156605 | 0.188 | 0.043 | 0.045387428 | 18 | zc3h18 |
| ccnd12 | 2.4805438 | 0.361895168 | 0.312 | 0.101 | 0.050099544 | 18 | ccnd1 |
| cmtm3 | 2.5264302 | 0.330926792 | 0.25 | 0.072 | 0.051026312 | 18 | cmtm3 |
| etv42 | 2.5775584 | 0.296205145 | 0.229 | 0.059 | 0.052058948 | 18 | etv4 |
| lsp11 | 2.6315516 | 0.261141989 | 0.188 | 0.043 | 0.053149449 | 18 | lsp1 |
| myef2 | 2.6370589 | 0.2577562 | 0.25 | 0.069 | 0.053260679 | 18 | myef2 |
| pafah1b1b | 2.8110678 | 0.299870489 | 0.229 | 0.061 | 0.056775137 | 18 | pafah1b1b |
| abl1 | 2.8660551 | 0.304528032 | 0.229 | 0.061 | 0.057885715 | 18 | abl1 |
| pin13 | 3.0842720 | 0.363556881 | 0.521 | 0.217 | 0.062293043 | 18 | pin1 |
| cd2761 | 3.3729270 | 0.300302072 | 0.25 | 0.071 | 0.068123008 | 18 | cd276 |
| si:ch211-51e12.73 | 3.4440920 | 0.361594978 | 0.479 | 0.19 | 0.069560326 | 18 | si:ch211-51e12.7 |
| syf21 | 3.6748128 | 0.382946154 | 0.479 | 0.2 | 0.074220194 | 18 | syf2 |
| TPGS1 | 3.6810489 | 0.250253224 | 0.167 | 0.036 | 0.074346145 | 18 | TPGS1 |
| gdf10a | 3.8635817 | 0.272070524 | 0.104 | 0.016 | 0.07803276 | 18 | gdf10a |
| CABZ01080702.15 | 3.8891777 | 0.36990676 | 0.458 | 0.173 | 0.078549724 | 18 | CABZ01080702.1 |
| smarcc2 | 4.0549734 | 0.266367418 | 0.188 | 0.044 | 0.081898299 | 18 | smarcc2 |
| acin1a5 | 4.2982083 | 0.452367336 | 0.604 | 0.295 | 0.086810915 | 18 | acin1a |
| fthl275 | 4.3616500 | 0.426009783 | 0.896 | 0.626 | 0.088092245 | 18 | fthl27 |
| yif1a1 | 4.5946796 | 0.270663504 | 0.229 | 0.062 | 0.092798746 | 18 | yif1a |
| sult6b1 | 4.6372726 | 0.292268946 | 0.271 | 0.082 | 0.093658997 | 18 | sult6b1 |
| FP102052.2 | 5.1066792 | 0.254372497 | 0.188 | 0.044 | 0.103139601 | 18 | FP102052.2 |

| | | | | | | | |
|---|---|---|---|---|---|---|---|
| irf2bp2b4 | 5.3495805 | 0.416656034 | 0.458 | 0.19 | 0.108045479 | 18 | irf2bp2b |
| hsd11b25 | 5.4489773 | 0.51166 | 0.646 | 0.288 | 0.110052996 | 18 | hsd11b2 |
| si:ch211-196l7.44 | 5.6717300 | 0.42845422 | 0.771 | 0.41 | 0.114551932 | 18 | si:ch211-196l7.4 |
| zbtb202 | 5.6842172 | 0.373378931 | 0.396 | 0.149 | 0.114804135 | 18 | zbtb20 |
| pkp3b2 | 6.3127209 | 0.255472293 | 0.271 | 0.079 | 0.127498025 | 18 | pkp3b |
| mxra8a3 | 6.3909844 | 0.370263438 | 0.396 | 0.146 | 0.129078712 | 18 | mxra8a |
| bnc23 | 6.9026168 | 0.343936245 | 0.271 | 0.083 | 0.139412152 | 18 | bnc2 |
| si:ch1073-291c23.2 | 7.2050196 | 0.369488409 | 0.375 | 0.136 | 0.145519782 | 18 | si:ch1073-291c23.2 |
| manbal1 | 7.2197649 | 0.26560149 | 0.208 | 0.055 | 0.145817592 | 18 | manbal |
| serpina10a | 7.2582223 | 0.278816718 | 0.208 | 0.054 | 0.146594317 | 18 | serpina10a |
| dnase1l11 | 7.5354688 | 0.276953125 | 0.292 | 0.092 | 0.152193864 | 18 | dnase1l1 |
| klf2a3 | 7.5551718 | 0.377971284 | 0.458 | 0.178 | 0.152591806 | 18 | klf2a |
| neo1a1 | 7.5948186 | 0.279917921 | 0.208 | 0.055 | 0.153392553 | 18 | neo1a |
| CU104709.1 | 7.7590888 | 0.312115829 | 0.208 | 0.056 | 0.156710318 | 18 | CU104709.1 |
| syncrip3 | 7.9022476 | 0.439672873 | 0.729 | 0.389 | 0.159601695 | 18 | syncrip |
| ssr43 | 8.6186790 | 0.428296632 | 0.812 | 0.48 | 0.174071461 | 18 | ssr4 |
| pdcd61 | 8.6430475 | 0.319325115 | 0.354 | 0.126 | 0.174563632 | 18 | pdcd6 |
| fbxl61 | 8.6550874 | 0.295163739 | 0.354 | 0.124 | 0.174806802 | 18 | fbxl6 |
| ier53 | 8.7619051 | 0.395480107 | 0.479 | 0.208 | 0.176964198 | 18 | ier5 |
| selenom1 | 8.7733937 | 0.305325593 | 0.292 | 0.095 | 0.177196233 | 18 | selenom |
| hspa4b1 | 9.6029634 | 0.296760008 | 0.333 | 0.118 | 0.193951052 | 18 | hspa4b |
| tmem263 | 9.6325622 | 0.264660592 | 0.292 | 0.093 | 0.19454886 | 18 | tmem263 |
| chmp6b1 | 1.0529532 | 0.273517991 | 0.292 | 0.095 | 0.212664976 | 18 | chmp6b |
| calr3 | 1.0739306 | 0.382848664 | 0.375 | 0.147 | 0.216901779 | 18 | calr |
| krt18b4 | 1.0773992 | 0.482684273 | 0.625 | 0.318 | 0.217602324 | 18 | krt18b |
| mak16 | 1.1763875 | 0.337876591 | 0.292 | 0.101 | 0.237595001 | 18 | mak16 |
| sdc45 | 1.1779710 | 0.360145266 | 0.604 | 0.261 | 0.237914817 | 18 | sdc4 |
| dio21 | 1.2719127 | 0.257184429 | 0.188 | 0.047 | 0.256888224 | 18 | dio2 |
| si:ch211-204c21.1: | 1.2746516 | 0.294051383 | 0.271 | 0.085 | 0.257441391 | 18 | si:ch211-204c21.1 |
| si:ch211-194k22.8: | 1.2905607 | 0.321187977 | 0.25 | 0.077 | 0.260654563 | 18 | si:ch211-194k22.8 |
| tspan4a1 | 1.3249927 | 0.2641873 | 0.229 | 0.065 | 0.267608781 | 18 | tspan4a |
| kank22 | 1.3693199 | 0.271661382 | 0.229 | 0.065 | 0.276561553 | 18 | kank2 |
| fam171a2a2 | 1.4551486 | 0.258482432 | 0.188 | 0.048 | 0.29389638 | 18 | fam171a2a |
| nucks1a4 | 1.4726051 | 0.422366837 | 0.5 | 0.23 | 0.297422059 | 18 | nucks1a |
| pdlim12 | 1.5489399 | 0.256056643 | 0.292 | 0.095 | 0.312839395 | 18 | pdlim1 |

| | | | | | | | |
|---|---|---|---|---|---|---|---|
| coro1ca1 | 1.6987175 | 0.287325507 | 0.25 | 0.078 | 0.343089992 | 18 | coro1ca |
| pa2g4b | 1.7753742 | 0.353933644 | 0.396 | 0.163 | 0.358572345 | 18 | pa2g4b |
| irf2bpl1 | 1.8244493 | 0.283357616 | 0.208 | 0.058 | 0.368484041 | 18 | irf2bpl |
| si:ch211-246m6.4 | 1.9995731 | 0.311341468 | 0.375 | 0.143 | 0.403853788 | 18 | si:ch211-246m6.4 |
| ssuh2rs11 | 2.2651283 | 0.267237001 | 0.25 | 0.078 | 0.45748797 | 18 | ssuh2rs1 |
| u2af2b1 | 2.8095413 | 0.329840063 | 0.417 | 0.175 | 0.567443061 | 18 | u2af2b |
| ctbp12 | 2.9532819 | 0.310507992 | 0.417 | 0.168 | 0.596474364 | 18 | ctbp1 |
| aldh2.12 | 2.9828386 | 0.257887345 | 0.229 | 0.07 | 0.602443924 | 18 | aldh2.1 |
| pard3bb1 | 3.0675831 | 0.26547131 | 0.229 | 0.069 | 0.619559765 | 18 | pard3bb |
| h2afva3 | 3.0900126 | 0.359952683 | 0.5 | 0.231 | 0.624089854 | 18 | h2afva |
| col16a12 | 3.1245631 | 0.308752688 | 0.25 | 0.079 | 0.631068022 | 18 | col16a1 |
| si:dkey-28b4.74 | 3.2512909 | 0.337114155 | 0.458 | 0.193 | 0.656663232 | 18 | si:dkey-28b4.7 |
| psmb52 | 3.2590390 | 0.306339744 | 0.354 | 0.135 | 0.658228116 | 18 | psmb5 |
| kmt2bb4 | 3.3631713 | 0.322361021 | 0.375 | 0.15 | 0.679259708 | 18 | kmt2bb |
| ube2v11 | 3.4127027 | 0.363915887 | 0.625 | 0.316 | 0.689263566 | 18 | ube2v1 |
| plpp33 | 3.4706172 | 0.391740923 | 0.333 | 0.124 | 0.70096057 | 18 | plpp3 |
| pik3r3a1 | 3.5267407 | 0.281845327 | 0.375 | 0.143 | 0.712295822 | 18 | pik3r3a |
| pkig3 | 3.6231909 | 0.322537766 | 0.333 | 0.126 | 0.731775885 | 18 | pkig |
| ier5l2 | 3.6386012 | 0.260845299 | 0.208 | 0.06 | 0.734888301 | 18 | ier5l |
| cpsf61 | 3.8322838 | 0.344783431 | 0.375 | 0.154 | 0.774006361 | 18 | cpsf6 |
| nap1l12 | 3.9191330 | 0.386981455 | 0.667 | 0.356 | 0.791547299 | 18 | nap1l1 |
| qkia5 | 3.9856246 | 0.399257161 | 0.688 | 0.356 | 0.804976601 | 18 | qkia |
| tax1bp32 | 4.0342015 | 0.379294487 | 0.396 | 0.168 | 0.814787695 | 18 | tax1bp3 |
| cd151l3 | 4.0857095 | 0.273284655 | 0.271 | 0.09 | 0.825190768 | 18 | cd151l |
| eef1g | 4.1192745 | 0.251214219 | 0.958 | 0.838 | 0.83196989 | 18 | eef1g |
| dedd16 | 4.8825723 | 0.334791002 | 0.479 | 0.211 | 0.986133133 | 18 | dedd1 |
| spon2b1 | 5.1474051 | 0.296335363 | 0.125 | 0.026 | 1 | 18 | spon2b |
| sept22 | 5.3134603 | 0.416741747 | 0.542 | 0.282 | 1 | 18 | sept2 |
| hdlbpa5 | 5.3186660 | 0.326404574 | 0.562 | 0.265 | 1 | 18 | hdlbpa |
| sult1st11 | 5.3778342 | 0.274495217 | 0.25 | 0.083 | 1 | 18 | sult1st1 |
| socs3a5 | 5.4336726 | 0.43706484 | 0.75 | 0.427 | 1 | 18 | socs3a |
| ptmab | 5.6481357 | 0.320578521 | 0.938 | 0.761 | 1 | 18 | ptmab |
| taf133 | 5.6840075 | 0.369781666 | 0.542 | 0.275 | 1 | 18 | taf13 |
| zgc:1583432 | 5.7612456 | 0.373379958 | 0.458 | 0.201 | 1 | 18 | zgc:158343 |
| anxa2a4 | 5.9431593 | 0.395836324 | 0.438 | 0.186 | 1 | 18 | anxa2a |

| | | | | | | | | |
|---|---|---|---|---|---|---|---|---|
| zgc:77849 | 6.1229880 | 0.323866498 | 0.417 | 0.182 | | 1 | 18 | zgc:77849 |
| kdelr2b | 6.1304935 | 0.301352748 | 0.375 | 0.153 | | 1 | 18 | kdelr2b |
| zmat2 | 6.2256153 | 0.274855412 | 0.438 | 0.183 | | 1 | 18 | zmat2 |
| cdh25 | 6.3498039 | 0.303283817 | 0.333 | 0.126 | | 1 | 18 | cdh2 |
| col18a1a2 | 6.9180126 | 0.268204996 | 0.271 | 0.09 | | 1 | 18 | col18a1a |
| nop533 | 7.2831550 | 0.342796708 | 0.688 | 0.35 | | 1 | 18 | nop53 |
| ywhaqb3 | 8.0993969 | 0.355432571 | 0.562 | 0.277 | | 1 | 18 | ywhaqb |
| sgk12 | 8.2712664 | 0.263108306 | 0.229 | 0.074 | | 1 | 18 | sgk1 |
| rbbp41 | 8.5692045 | 0.307672224 | 0.333 | 0.132 | | 1 | 18 | rbbp4 |
| dynlt32 | 8.8091205 | 0.316524688 | 0.479 | 0.221 | | 1 | 18 | dynlt3 |
| yap13 | 9.4030856 | 0.274228899 | 0.271 | 0.096 | | 1 | 18 | yap1 |
| krtcap24 | 9.4476984 | 0.360329279 | 0.583 | 0.303 | | 1 | 18 | krtcap2 |
| cebpg | 9.4908904 | 0.274005389 | 0.25 | 0.087 | | 1 | 18 | cebpg |
| tbca2 | 9.5774963 | 0.35087808 | 0.562 | 0.296 | | 1 | 18 | tbca |
| psma31 | 9.6192480 | 0.346325735 | 0.562 | 0.284 | | 1 | 18 | psma3 |
| si:ch211-212k18.9! | 9.9548248 | 0.286702848 | 0.312 | 0.117 | | 1 | 18 | si:ch211-212k18.9 |
| arpp19b1 | 0.000102 | 0.259456239 | 0.312 | 0.119 | | 1 | 18 | arpp19b |
| btg3 | 0.000103 | 0.262816952 | 0.229 | 0.077 | | 1 | 18 | btg3 |
| rras2 | 0.000106 | 0.264731072 | 0.312 | 0.119 | | 1 | 18 | rras |
| gtf2a21 | 0.000106 | 0.251889168 | 0.25 | 0.085 | | 1 | 18 | gtf2a2 |
| hdac11 | 0.000107 | 0.351943417 | 0.542 | 0.275 | | 1 | 18 | hdac1 |
| arf2b3 | 0.000108 | 0.301545078 | 0.458 | 0.207 | | 1 | 18 | arf2b |
| fkbp32 | 0.000109 | 0.307398123 | 0.521 | 0.251 | | 1 | 18 | fkbp3 |
| h2afvb1 | 0.000114 | 0.393872391 | 0.812 | 0.528 | | 1 | 18 | h2afvb |
| rarga2 | 0.000114 | 0.266404303 | 0.25 | 0.086 | | 1 | 18 | rarga |
| vapb1 | 0.000116 | 0.250311867 | 0.271 | 0.097 | | 1 | 18 | vapb |
| foxo3b2 | 0.000118 | 0.250242078 | 0.271 | 0.094 | | 1 | 18 | foxo3b |
| mcl1b6 | 0.000119 | 0.379745445 | 0.604 | 0.314 | | 1 | 18 | mcl1b |
| smndc1 | 0.000123 | 0.294776016 | 0.396 | 0.174 | | 1 | 18 | smndc1 |
| id2b2 | 0.000127 | 0.290690304 | 0.229 | 0.075 | | 1 | 18 | id2b |
| tns1b3 | 0.000129 | 0.269197039 | 0.167 | 0.046 | | 1 | 18 | tns1b |
| nucks1b1 | 0.00013 | 0.295338723 | 0.375 | 0.16 | | 1 | 18 | nucks1b |
| ssbp3b1 | 0.00013 | 0.262697033 | 0.229 | 0.077 | | 1 | 18 | ssbp3b |
| pbx44 | 0.000131 | 0.252658557 | 0.292 | 0.108 | | 1 | 18 | pbx4 |
| gstt1b6 | 0.000135 | 0.290937064 | 0.417 | 0.181 | | 1 | 18 | gstt1b |

| | | | | | | | | |
|---|---|---|---|---|---|---|---|---|
| tcima3 | 0.00014 | 0.377338394 | 0.25 | 0.089 | | 1 | 18 | tcima |
| cirbpb2 | 0.000142 | 0.279533295 | 0.979 | 0.811 | | 1 | 18 | cirbpb |
| plpp1a1 | 0.000144 | 0.345061939 | 0.312 | 0.123 | | 1 | 18 | plpp1a |
| cdc14b1 | 0.000144 | 0.252647353 | 0.229 | 0.077 | | 1 | 18 | cdc14b |
| jdp2b2 | 0.000145 | 0.378301184 | 0.354 | 0.15 | | 1 | 18 | jdp2b |
| paip2b4 | 0.000147 | 0.348702945 | 0.625 | 0.341 | | 1 | 18 | paip2b |
| notch23 | 0.000157 | 0.310204731 | 0.354 | 0.15 | | 1 | 18 | notch2 |
| ywhabb | 0.000162 | 0.352283082 | 0.708 | 0.401 | | 1 | 18 | ywhabb |
| hspb13 | 0.000162 | 0.302111866 | 0.417 | 0.177 | | 1 | 18 | hspb1 |
| rad21a3 | 0.000166 | 0.349923892 | 0.5 | 0.248 | | 1 | 18 | rad21a |
| dynlrb12 | 0.000166 | 0.294637457 | 0.542 | 0.267 | | 1 | 18 | dynlrb1 |
| anxa5b5 | 0.000174 | 0.285880417 | 0.396 | 0.168 | | 1 | 18 | anxa5b |
| cbx51 | 0.000176 | 0.378483168 | 0.438 | 0.221 | | 1 | 18 | cbx5 |
| tra2a1 | 0.000188 | 0.328215628 | 0.458 | 0.222 | | 1 | 18 | tra2a |
| ptp4a1 | 0.000189 | 0.328743674 | 0.438 | 0.21 | | 1 | 18 | ptp4a1 |
| zyx2 | 0.000189 | 0.297933588 | 0.312 | 0.126 | | 1 | 18 | zyx |
| khdrbs1a3 | 0.000195 | 0.352706392 | 0.792 | 0.595 | | 1 | 18 | khdrbs1a |
| timp2a4 | 0.000217 | 0.274641702 | 0.292 | 0.111 | | 1 | 18 | timp2a |
| rpl27a2 | 0.000225 | 0.337680624 | 0.938 | 0.685 | | 1 | 18 | rpl27a |
| sntb2 | 0.000225 | 0.261263933 | 0.25 | 0.091 | | 1 | 18 | sntb2 |
| pabpc1b5 | 0.000238 | 0.263259544 | 0.312 | 0.123 | | 1 | 18 | pabpc1b |
| dad14 | 0.000238 | 0.377236393 | 0.688 | 0.418 | | 1 | 18 | dad1 |
| vdac33 | 0.00026 | 0.315778627 | 0.5 | 0.252 | | 1 | 18 | vdac3 |
| ddx39aa1 | 0.000263 | 0.264836517 | 0.438 | 0.202 | | 1 | 18 | ddx39aa |
| sf11 | 0.000267 | 0.257543178 | 0.312 | 0.126 | | 1 | 18 | sf1 |
| sec61b1 | 0.000301 | 0.355861237 | 0.812 | 0.566 | | 1 | 18 | sec61b |
| psip1b2 | 0.000326 | 0.3062049 | 0.292 | 0.119 | | 1 | 18 | psip1b |
| nr3c1 | 0.000327 | 0.280643186 | 0.229 | 0.082 | | 1 | 18 | nr3c1 |
| tmem167a | 0.000342 | 0.298568832 | 0.375 | 0.173 | | 1 | 18 | tmem167a |
| si:ch211-194b1.1 | 0.000345 | 0.276512035 | 0.146 | 0.041 | | 1 | 18 | si:ch211-194b1.1 |
| hbegfa2 | 0.000351 | 0.260563353 | 0.188 | 0.061 | | 1 | 18 | hbegfa |
| sesn13 | 0.000365 | 0.304457699 | 0.458 | 0.215 | | 1 | 18 | sesn1 |
| ccdc802 | 0.00037 | 0.260861307 | 0.208 | 0.071 | | 1 | 18 | ccdc80 |
| calm2a2 | 0.000396 | 0.307148897 | 0.5 | 0.251 | | 1 | 18 | calm2a |
| rab1ab1 | 0.000396 | 0.317444248 | 0.438 | 0.225 | | 1 | 18 | rab1ab |

| | | | | | | | |
|---|---|---|---|---|---|---|---|
| sdcbp24 | 0.0004 | 0.330135273 | 0.625 | 0.34 | 1 | 18 | sdcbp2 |
| fam133b2 | 0.0004 | 0.273767683 | 0.438 | 0.21 | 1 | 18 | fam133b |
| hmgb2b3 | 0.000403 | 0.414725452 | 0.688 | 0.439 | 1 | 18 | hmgb2b |
| flna6 | 0.000413 | 0.292868581 | 0.375 | 0.167 | 1 | 18 | flna |
| tomm63 | 0.000413 | 0.291877835 | 0.458 | 0.226 | 1 | 18 | tomm6 |
| nr2f22 | 0.000427 | 0.305488817 | 0.25 | 0.097 | 1 | 18 | nr2f2 |
| csnk1a14 | 0.000433 | 0.301535144 | 0.521 | 0.269 | 1 | 18 | csnk1a1 |
| h3f3a1 | 0.000434 | 0.363481183 | 0.646 | 0.39 | 1 | 18 | h3f3a |
| hspa55 | 0.000438 | 0.285972076 | 0.458 | 0.22 | 1 | 18 | hspa5 |
| rhoab2 | 0.00044 | 0.311357607 | 0.583 | 0.311 | 1 | 18 | rhoab |
| NC-002333.48 | 0.000462 | 0.414005954 | 0.875 | 0.583 | 1 | 18 | NC-002333.4 |
| ier2a4 | 0.000471 | 0.391533095 | 0.625 | 0.362 | 1 | 18 | ier2a |
| dnmt3ab4 | 0.000495 | 0.264523237 | 0.375 | 0.167 | 1 | 18 | dnmt3ab |
| bcas2 | 0.000499 | 0.253785755 | 0.396 | 0.185 | 1 | 18 | bcas2 |
| sltm1 | 0.000503 | 0.27902522 | 0.292 | 0.123 | 1 | 18 | sltm |
| snrpb4 | 0.000503 | 0.331778461 | 0.708 | 0.42 | 1 | 18 | snrpb |
| calr3b3 | 0.00051 | 0.351167285 | 0.375 | 0.183 | 1 | 18 | calr3b |
| psmb73 | 0.000511 | 0.27529703 | 0.583 | 0.304 | 1 | 18 | psmb7 |
| brd7 | 0.000515 | 0.261769734 | 0.25 | 0.099 | 1 | 18 | brd7 |
| pnn2 | 0.000526 | 0.284372806 | 0.333 | 0.151 | 1 | 18 | pnn |
| add3a5 | 0.000533 | 0.291834705 | 0.458 | 0.22 | 1 | 18 | add3a |
| tram21 | 0.000548 | 0.256480531 | 0.208 | 0.074 | 1 | 18 | tram2 |
| ndfip1l1 | 0.00058 | 0.296641389 | 0.292 | 0.123 | 1 | 18 | ndfip1l |
| arhgef1b4 | 0.000588 | 0.295018793 | 0.292 | 0.124 | 1 | 18 | arhgef1b |
| h2afy24 | 0.000624 | 0.325190511 | 0.479 | 0.255 | 1 | 18 | h2afy2 |
| nosip | 0.00063 | 0.254608634 | 0.188 | 0.065 | 1 | 18 | nosip |
| mt-nd25 | 0.000638 | 0.357384032 | 0.646 | 0.372 | 1 | 18 | mt-nd2 |
| fam32a1 | 0.000653 | 0.310869376 | 0.562 | 0.315 | 1 | 18 | fam32a |
| surf4l3 | 0.00076 | 0.28406411 | 0.208 | 0.077 | 1 | 18 | surf4l |
| ndufb93 | 0.000769 | 0.28484725 | 0.583 | 0.323 | 1 | 18 | ndufb9 |
| angptl1a4 | 0.000807 | 0.269236609 | 0.208 | 0.074 | 1 | 18 | angptl1a |
| mt-atp64 | 0.000819 | 0.388164507 | 0.729 | 0.42 | 1 | 18 | mt-atp6 |
| add12 | 0.000861 | 0.253475535 | 0.271 | 0.113 | 1 | 18 | add1 |
| anapc161 | 0.000907 | 0.271464176 | 0.354 | 0.166 | 1 | 18 | anapc16 |
| psma12 | 0.000969 | 0.315399529 | 0.646 | 0.395 | 1 | 18 | psma1 |

| | | | | | | | |
|---|---|---|---|---|---|---|---|
| ppp2cb3 | 0.000976 | 0.284160957 | 0.458 | 0.24 | 1 | 18 | ppp2cb |
| tagln23 | 0.001023 | 0.322483834 | 0.604 | 0.337 | 1 | 18 | tagln2 |
| calm3b3 | 0.001025 | 0.309368998 | 0.688 | 0.407 | 1 | 18 | calm3b |
| nfyba1 | 0.001039 | 0.28019406 | 0.438 | 0.228 | 1 | 18 | nfyba |
| gadd45ba5 | 0.001085 | 0.304527505 | 0.479 | 0.251 | 1 | 18 | gadd45ba |
| btg24 | 0.00112 | 0.293261131 | 0.5 | 0.259 | 1 | 18 | btg2 |
| crip14 | 0.001191 | 0.366030639 | 0.625 | 0.377 | 1 | 18 | crip1 |
| calr3a5 | 0.001249 | 0.323118056 | 0.438 | 0.236 | 1 | 18 | calr3a |
| serinc53 | 0.001278 | 0.261968741 | 0.271 | 0.115 | 1 | 18 | serinc5 |
| eif4a1b2 | 0.001326 | 0.334914798 | 0.708 | 0.487 | 1 | 18 | eif4a1b |
| actn12 | 0.001449 | 0.252760859 | 0.188 | 0.069 | 1 | 18 | actn1 |
| eif2s3 | 0.001499 | 0.31755755 | 0.688 | 0.434 | 1 | 18 | eif2s3 |
| sumo2b1 | 0.001537 | 0.32315349 | 0.688 | 0.459 | 1 | 18 | sumo2b |
| sumo3a4 | 0.001597 | 0.309915946 | 0.667 | 0.407 | 1 | 18 | sumo3a |
| mt-nd15 | 0.001641 | 0.381625058 | 0.583 | 0.354 | 1 | 18 | mt-nd1 |
| hsf2 | 0.001697 | 0.250940405 | 0.25 | 0.109 | 1 | 18 | hsf2 |
| zgc:1538672 | 0.001718 | 0.301888064 | 0.875 | 0.637 | 1 | 18 | zgc:153867 |
| hsp90b13 | 0.00172 | 0.279222735 | 0.354 | 0.175 | 1 | 18 | hsp90b1 |
| lman23 | 0.001741 | 0.266828656 | 0.375 | 0.192 | 1 | 18 | lman2 |
| anp32b3 | 0.001815 | 0.27493172 | 0.438 | 0.237 | 1 | 18 | anp32b |
| hnrnpaba2 | 0.001818 | 0.31432326 | 0.771 | 0.527 | 1 | 18 | hnrnpaba |
| tmem176l.17 | 0.001876 | 0.309927096 | 0.583 | 0.324 | 1 | 18 | tmem176l.1 |
| atf4b4 | 0.001896 | 0.317502429 | 0.583 | 0.36 | 1 | 18 | atf4b |
| ctdsp23 | 0.001929 | 0.25137528 | 0.354 | 0.173 | 1 | 18 | ctdsp2 |
| ubc3 | 0.001997 | 0.285402574 | 0.875 | 0.654 | 1 | 18 | ubc |
| klf6a4 | 0.002108 | 0.334267356 | 0.458 | 0.259 | 1 | 18 | klf6a |
| adh54 | 0.002125 | 0.281863156 | 0.604 | 0.36 | 1 | 18 | adh5 |
| sod12 | 0.002217 | 0.285670174 | 0.667 | 0.421 | 1 | 18 | sod1 |
| dap1b2 | 0.00227 | 0.283235354 | 0.458 | 0.255 | 1 | 18 | dap1b |
| chd4a2 | 0.002574 | 0.267588898 | 0.375 | 0.192 | 1 | 18 | chd4a |
| btf3l41 | 0.002773 | 0.256993682 | 0.375 | 0.201 | 1 | 18 | btf3l4 |
| rpl22l12 | 0.002781 | 0.310533432 | 0.854 | 0.619 | 1 | 18 | rpl22l1 |
| eif3ba2 | 0.002989 | 0.260108854 | 0.667 | 0.393 | 1 | 18 | eif3ba |
| srsf5a5 | 0.003025 | 0.269135638 | 0.646 | 0.387 | 1 | 18 | srsf5a |
| rhoaa2 | 0.003113 | 0.272713874 | 0.521 | 0.298 | 1 | 18 | rhoaa |

| | | | | | | | |
|---|---|---|---|---|---|---|---|
| ywhae14 | 0.003226 | 0.256753217 | 0.646 | 0.378 | 1 | 18 | ywhae1 |
| tpm12 | 0.003488 | 0.303641102 | 0.375 | 0.2 | 1 | 18 | tpm1 |
| si:dkey-42i9.44 | 0.003565 | 0.255633858 | 0.375 | 0.198 | 1 | 18 | si:dkey-42i9.4 |
| eif1b1 | 0.0036 | 0.292347907 | 0.729 | 0.504 | 1 | 18 | eif1b |
| mmp27 | 0.003934 | 0.264603965 | 0.417 | 0.217 | 1 | 18 | mmp2 |
| magt11 | 0.004319 | 0.26368698 | 0.375 | 0.208 | 1 | 18 | magt1 |
| mdka4 | 0.004319 | 0.25883084 | 0.438 | 0.223 | 1 | 18 | mdka |
| ostc4 | 0.004753 | 0.267898209 | 0.438 | 0.252 | 1 | 18 | ostc |
| tmem2582 | 0.005751 | 0.250408048 | 0.688 | 0.424 | 1 | 18 | tmem258 |
| phf5a1 | 0.006463 | 0.262128054 | 0.333 | 0.182 | 1 | 18 | phf5a |
| tpm31 | 0.007121 | 0.257251277 | 0.708 | 0.457 | 1 | 18 | tpm3 |
| il11b | 1.9394123 | 1.324756597 | 0.521 | 0.003 | 3.91703116499701e-1 | 19 | il11b |
| si:dkey-78l4.41 | 1.5315431 | 1.574927187 | 0.562 | 0.008 | 3.0932576876384e-15 | 19 | si:dkey-78l4.4 |
| si:dkey-78l4.6 | 1.4712448 | 1.579081571 | 0.375 | 0.001 | 2.9714731610474e-14 | 19 | si:dkey-78l4.6 |
| si:dkey-21e2.13 | 2.5600875 | 1.621837636 | 0.354 | 0 | 5.17060891701522e-1 | 19 | si:dkey-21e2.13 |
| il4 | 1.3447945 | 1.325839383 | 0.396 | 0.002 | 2.71608154488738e-1 | 19 | il4 |
| csf3a | 2.6996585 | 0.915556857 | 0.354 | 0.001 | 5.45250037093909e-1 | 19 | csf3a |
| si:dkey-78l4.2 | 1.5844386 | 1.484816004 | 0.438 | 0.004 | 3.20009065969878e-1 | 19 | si:dkey-78l4.2 |
| il13 | 4.0783688 | 1.165554335 | 0.333 | 0.001 | 8.23708146713762e-1 | 19 | il13 |
| si:dkey-21e2.16 | 1.0019911 | 1.477100976 | 0.292 | 0.001 | 2.02372152603959e-1 | 19 | si:dkey-21e2.16 |
| si:dkey-21e2.10 | 3.4005408 | 1.382119667 | 0.271 | 0 | 6.86807243193929e-1 | 19 | si:dkey-21e2.10 |
| si:dkey-21e2.14 | 9.6869668 | 0.793018823 | 0.25 | 0 | 1.9564766850827e-10 | 19 | si:dkey-21e2.14 |
| si:dkey-21e2.15 | 1.3238711 | 1.507906406 | 0.333 | 0.003 | 2.67382264546331e-1 | 19 | si:dkey-21e2.15 |
| CR847844.1 | 2.6662405 | 0.684315737 | 0.271 | 0.001 | 5.38500596873937e-1 | 19 | CR847844.1 |
| si:dkey-21e2.3 | 6.7967505 | 0.746154719 | 0.271 | 0.002 | 1.37273970673748e-8 | 19 | si:dkey-21e2.3 |
| XLOC-0430431 | 6.1832477 | 1.422289223 | 0.667 | 0.041 | 1.24883054834955e-7 | 19 | XLOC-043043 |
| si:dkey-21e2.8 | 2.7585160 | 1.110883411 | 0.208 | 0.001 | 5.57137494839747e-7 | 19 | si:dkey-21e2.8 |
| si:ch211-241f5.31 | 3.5653244 | 1.130405038 | 0.542 | 0.027 | 7.20088571339142e-7 | 19 | si:ch211-241f5.3 |
| faslg1 | 3.4255850 | 0.750413267 | 0.333 | 0.009 | 6.9186541043758e-63 | 19 | faslg |
| adgre14 | 9.8763760 | 0.454132049 | 0.208 | 0.002 | 1.99473167347962e-6 | 19 | adgre14 |
| LOC100150882 | 2.0524258 | 1.47236406 | 0.354 | 0.011 | 4.14528446977684e-6 | 19 | LOC100150882 |
| LOC103911745 | 2.1367672 | 0.397754109 | 0.146 | 0 | 4.31562880477128e-6 | 19 | LOC103911745 |
| si:dkey-9i23.4 | 1.0858550 | 1.230670321 | 0.292 | 0.007 | 2.19310139817848e-6 | 19 | si:dkey-9i23.4 |
| gata3 | 2.5486408 | 0.701821599 | 0.354 | 0.012 | 5.14748997531368e-5 | 19 | gata3 |
| CR847936.10 | 5.0068492 | 0.365544139 | 0.146 | 0 | 1.01123334271e-53 | 19 | CR847936.10 |

| | | | | | | | |
|---|---|---|---|---|---|---|---|
| XLOC-0140791 | 1.8070880 | 0.905848843 | 0.417 | 0.021 | 3.64977573875289e-5 | 19 | XLOC-014079 |
| cd247l1 | 1.6835784 | 0.55493014 | 0.292 | 0.01 | 3.40032347457725e-4 | 19 | cd247l |
| si:dkey-8e10.2 | 1.9168417 | 0.382911814 | 0.146 | 0.001 | 3.87144532040474e-4 | 19 | si:dkey-8e10.2 |
| ca21 | 9.8362461 | 1.016533864 | 0.458 | 0.031 | 1.98662662535531e-4 | 19 | ca2 |
| si:dkey-78l4.7 | 6.7730838 | 0.434493766 | 0.104 | 0 | 1.36795973959901e-4 | 19 | si:dkey-78l4.7 |
| BX571675.1 | 2.9843371 | 0.342799468 | 0.146 | 0.001 | 6.02746566874367e-4 | 19 | BX571675.1 |
| sox13 | 5.4391811 | 0.549745447 | 0.271 | 0.01 | 1.0985514074165e-41 | 19 | sox13 |
| si:ch211-212d10.1 | 1.6598469 | 1.29411948 | 0.375 | 0.025 | 3.35239290758927e-3 | 19 | si:ch211-212d10.1 |
| si:ch211-191c10.1 | 4.4636117 | 0.730870401 | 0.354 | 0.022 | 9.01515666546924e-3 | 19 | si:ch211-191c10.1 |
| LOC103911004 | 5.5945723 | 0.425224377 | 0.146 | 0.002 | 1.12993577326988e-3 | 19 | LOC103911004 |
| CABZ01084575.11 | 5.7987164 | 0.744584317 | 0.354 | 0.022 | 1.17116676815828e-3 | 19 | CABZ01084575.1 |
| XLOC-0397571 | 3.3126472 | 0.657567875 | 0.354 | 0.022 | 6.69055371929053e-3 | 19 | XLOC-039757 |
| tnfrsf9a1 | 1.0676802 | 0.863564483 | 0.354 | 0.024 | 2.15639385487015e-3 | 19 | tnfrsf9a |
| si:dkey-33i11.91 | 3.0218302 | 0.714490125 | 0.417 | 0.037 | 6.10319049090104e-3 | 19 | si:dkey-33i11.9 |
| FP236356.11 | 1.0953503 | 0.671658887 | 0.354 | 0.026 | 2.21227913019634e-3 | 19 | FP236356.1 |
| laptm55 | 8.5189262 | 1.382950244 | 0.812 | 0.206 | 1.72056753420007e-2 | 19 | laptm5 |
| ctss11 | 2.5128849 | 0.570951959 | 0.292 | 0.019 | 5.07527378434205e-2 | 19 | ctss1 |
| cldni | 3.0485894 | 0.418993005 | 0.188 | 0.007 | 6.15723611798622e-2 | 19 | cldni |
| ccl34b.41 | 3.9068195 | 0.696181181 | 0.229 | 0.011 | 7.89060342609788e-2 | 19 | ccl34b.4 |
| si:dkey-12f6.5 | 2.9974295 | 0.352472116 | 0.125 | 0.003 | 6.05390854914892e-2 | 19 | si:dkey-12f6.5 |
| acp5b | 1.7218851 | 0.344044878 | 0.167 | 0.007 | 3.47769140838064e-2 | 19 | acp5b |
| cfbl1 | 7.8087344 | 1.183084648 | 0.521 | 0.089 | 1.57713010388284e-2 | 19 | cfbl |
| sla21 | 1.5206568 | 0.516689107 | 0.25 | 0.019 | 3.07127069005277e-2 | 19 | sla2 |
| capgb4 | 3.8742661 | 0.924529623 | 0.583 | 0.12 | 7.82485535412336e-1 | 19 | capgb |
| batf31 | 1.2395047 | 0.446329527 | 0.208 | 0.015 | 2.50342772279746e-1 | 19 | batf3 |
| sult5a11 | 4.8581074 | 0.572566065 | 0.229 | 0.019 | 9.81191960786179e-1 | 19 | sult5a1 |
| BX322787.11 | 6.3186531 | 0.636755872 | 0.375 | 0.048 | 1.27617838606822e-1 | 19 | BX322787.1 |
| ogfrl1 | 1.7968634 | 0.344045521 | 0.146 | 0.007 | 3.62912506429373e-1 | 19 | ogfrl1 |
| AL954142.12 | 2.2078907 | 0.578761576 | 0.271 | 0.03 | 4.4592769960396e-15 | 19 | AL954142.1 |
| pfn15 | 2.2613520 | 1.073668556 | 1 | 0.411 | 4.56725273890659e-1 | 19 | pfn1 |
| FO203432.11 | 6.6985199 | 0.531248852 | 0.292 | 0.033 | 1.3529000730626e-14 | 19 | FO203432.1 |
| calm1b4 | 4.0114846 | 0.819331999 | 0.583 | 0.141 | 8.10199564512582e-1 | 19 | calm1b |
| il341 | 3.3741402 | 0.536266771 | 0.25 | 0.029 | 6.81475101583239e-1 | 19 | il34 |
| zgc:1366051 | 3.7263311 | 0.367558543 | 0.146 | 0.009 | 7.52607100854102e-1 | 19 | zgc:136605 |
| srgn3 | 4.1306206 | 0.921911827 | 0.75 | 0.238 | 8.34261450796665e-1 | 19 | srgn |

| | | | | | | | |
|---|---|---|---|---|---|---|---|
| cass4 | 1.1564624 | 0.27392195 | 0.125 | 0.007 | 2.33570720627704e-1 | 19 | cass4 |
| CU075735.11 | 1.5804977 | 0.377672872 | 0.146 | 0.01 | 3.19213121387646e-1 | 19 | CU075735.1 |
| tagapa1 | 3.3628262 | 0.722212846 | 0.354 | 0.06 | 6.79190021285041e-1 | 19 | tagapa |
| arpc1b5 | 4.3987112 | 0.95091047 | 0.854 | 0.354 | 8.88407704495301e-1 | 19 | arpc1b |
| dusp51 | 1.0675297 | 0.645489838 | 0.292 | 0.044 | 2.15608979646179e-1 | 19 | dusp5 |
| CABZ01073834.12 | 2.2671468 | 0.578614647 | 0.333 | 0.056 | 4.57895639468346e-1 | 19 | CABZ01073834.1 |
| si:ch211-277c7.61 | 2.8017018 | 0.566379789 | 0.229 | 0.027 | 5.65859725380703e-1 | 19 | si:ch211-277c7.6 |
| CU207241.11 | 3.1705764 | 0.457150558 | 0.208 | 0.023 | 6.40361318520765e-1 | 19 | CU207241.1 |
| rasal31 | 3.6275885 | 0.451794702 | 0.208 | 0.025 | 7.3266406119127e-10 | 19 | rasal3 |
| eomesa1 | 8.5130398 | 0.567095868 | 0.271 | 0.041 | 1.71937865449462e-0 | 19 | eomesa |
| hck1 | 9.0183302 | 0.444610046 | 0.25 | 0.036 | 1.82143216747892e-0 | 19 | hck |
| ndufa4 | 1.0844000 | 0.425840697 | 0.208 | 0.026 | 2.19016273918746e-0 | 19 | ndufa4 |
| ncaldb1 | 1.2298887 | 0.538725217 | 0.292 | 0.049 | 2.48400628713505e-0 | 19 | ncaldb |
| scgn1 | 3.6653171 | 0.338766315 | 0.188 | 0.021 | 7.40284112256317e-0 | 19 | scgn |
| bhlhe402 | 4.2477244 | 0.819231649 | 0.583 | 0.209 | 8.57912911628428e-0 | 19 | bhlhe40 |
| pik3cd | 6.6039151 | 0.288981884 | 0.167 | 0.017 | 1.33379274008584e-0 | 19 | pik3cd |
| elf12 | 9.2604053 | 0.660700097 | 0.417 | 0.103 | 1.87032406153642e-0 | 19 | elf1 |
| grap2b5 | 1.0924371 | 0.726912709 | 0.604 | 0.188 | 2.20639536496817e-0 | 19 | grap2b |
| coro1a5 | 1.4380996 | 0.806039547 | 0.771 | 0.275 | 2.90452993029978e-0 | 19 | coro1a |
| cox172 | 1.5348319 | 0.713473148 | 0.562 | 0.182 | 3.09990008189521e-0 | 19 | cox17 |
| sik12 | 3.7707805 | 0.563233238 | 0.375 | 0.086 | 7.61584553409686e-0 | 19 | sik1 |
| si:ch211-102c2.45 | 5.7586995 | 0.756276547 | 0.562 | 0.171 | 1.16308454161124e-0 | 19 | si:ch211-102c2.4 |
| ptprc3 | 7.1900914 | 0.645397573 | 0.438 | 0.115 | 1.45218277476117e-0 | 19 | ptprc |
| sh3bp2 | 7.9558485 | 0.324073249 | 0.146 | 0.015 | 1.60684274021376e-0 | 19 | sh3bp2 |
| pgam1a2 | 1.8348208 | 0.593135251 | 0.5 | 0.153 | 3.70578765036431e-0 | 19 | pgam1a |
| si:ch73-208g10.12 | 3.0922654 | 0.376900785 | 0.208 | 0.031 | 6.2454485844886e-07 | 19 | si:ch73-208g10.1 |
| CABZ01041494.1 | 3.7273929 | 0.273838455 | 0.104 | 0.008 | 7.52821560504818e-0 | 19 | CABZ01041494.1 |
| si:ch211-207n23.2 | 4.6276235 | 0.429563471 | 0.167 | 0.02 | 9.34641130196173e-0 | 19 | si:ch211-207n23.2 |
| wasb4 | 6.0930608 | 0.678537942 | 0.5 | 0.155 | 1.23061549636906e-0 | 19 | wasb |
| wasa3 | 9.8089384 | 0.639267246 | 0.417 | 0.115 | 1.98111128871791e-0 | 19 | wasa |
| ccr21 | 1.3006393 | 0.530600985 | 0.229 | 0.04 | 2.62690124395808e-0 | 19 | ccr2 |
| tnip11 | 3.0091934 | 0.502801606 | 0.312 | 0.072 | 6.07766802992093e-0 | 19 | tnip1 |
| cotl15 | 4.7593925 | 0.728693094 | 0.854 | 0.406 | 9.61254513555486e-0 | 19 | cotl1 |
| dnajb9b1 | 4.8189892 | 0.340436386 | 0.167 | 0.023 | 9.73291262785065e-0 | 19 | dnajb9b |
| zgc:640515 | 4.9341396 | 0.586984444 | 0.542 | 0.172 | 9.96548180036862e-0 | 19 | zgc:64051 |

| | | | | | | | |
|---|---|---|---|---|---|---|---|
| hhex | 6.7058429 | 0.304684076 | 0.125 | 0.013 | 1.35437909281346e-0 | 19 | hhex |
| far11 | 9.1847985 | 0.441258716 | 0.229 | 0.043 | 1.85505376770546e-0 | 19 | far1 |
| atp2a31 | 1.3448510 | 0.365873595 | 0.25 | 0.05 | 2.71619568779747e-0 | 19 | atp2a3 |
| MFAP4 (1 of many)1 | 1.4452660 | 0.343302684 | 0.25 | 0.046 | 2.91900384403821e-0 | 19 | MFAP4 (1 of many) |
| ptpn64 | 1.4763117 | 0.597820508 | 0.458 | 0.142 | 2.98170693104043e-0 | 19 | ptpn6 |
| btk1 | 3.7496681 | 0.400337924 | 0.229 | 0.044 | 7.57320482076739e-0 | 19 | btk |
| ldha1 | 4.8397506 | 0.465683415 | 0.25 | 0.055 | 9.7748443781389e-05 | 19 | ldha |
| id33 | 5.1761773 | 0.628117631 | 0.438 | 0.144 | 0.000104543 | 19 | id3 |
| actb15 | 6.4319764 | 0.488841091 | 0.958 | 0.739 | 0.000129907 | 19 | actb1 |
| mylk51 | 6.8832696 | 0.313552044 | 0.146 | 0.02 | 0.000139021 | 19 | mylk5 |
| CABZ01117603.11 | 6.8875083 | 0.25651474 | 0.125 | 0.015 | 0.000139107 | 19 | CABZ01117603.1 |
| spns31 | 7.1547569 | 0.344557237 | 0.167 | 0.026 | 0.000144505 | 19 | spns3 |
| rac25 | 1.0711473 | 0.660057448 | 0.625 | 0.258 | 0.00021634 | 19 | rac2 |
| slc2a3b1 | 1.1406126 | 0.318317578 | 0.146 | 0.021 | 0.00023037 | 19 | slc2a3b |
| fermt3b1 | 1.7173774 | 0.316533504 | 0.188 | 0.033 | 0.000346859 | 19 | fermt3b |
| crip15 | 2.2120168 | 0.648512698 | 0.708 | 0.375 | 0.000446761 | 19 | crip1 |
| itgb22 | 2.2863373 | 0.713926259 | 0.438 | 0.163 | 0.000461772 | 19 | itgb2 |
| tes1 | 2.6733455 | 0.475688202 | 0.292 | 0.076 | 0.000539936 | 19 | tes |
| wipf1a2 | 3.0711363 | 0.470028472 | 0.312 | 0.082 | 0.000620277 | 19 | wipf1a |
| cyb5r2 | 3.6035610 | 0.264427883 | 0.125 | 0.016 | 0.000727811 | 19 | cyb5r2 |
| ppp1r182 | 3.6405142 | 0.476958501 | 0.271 | 0.067 | 0.000735275 | 19 | ppp1r18 |
| def6a | 3.6626534 | 0.336058747 | 0.167 | 0.028 | 0.000739746 | 19 | def6a |
| si:dkey-53k12.23 | 3.8164942 | 0.410522649 | 0.208 | 0.042 | 0.000770817 | 19 | si:dkey-53k12.2 |
| alox5ap2 | 4.0041175 | 0.380459497 | 0.188 | 0.035 | 0.000808712 | 19 | alox5ap |
| aldocb2 | 4.8513130 | 0.394712966 | 0.25 | 0.059 | 0.00097982 | 19 | aldocb |
| fybb2 | 5.3400606 | 0.384074728 | 0.25 | 0.058 | 0.001078532 | 19 | fybb |
| rps6ka11 | 5.3450945 | 0.405655323 | 0.229 | 0.051 | 0.001079549 | 19 | rps6ka1 |
| rgs131 | 5.5439398 | 0.5683681 | 0.333 | 0.099 | 0.00111971 | 19 | rgs13 |
| lcp2b1 | 5.9349205 | 0.339262967 | 0.167 | 0.028 | 0.001198676 | 19 | lcp2b |
| si:dkeyp-68b7.122 | 6.1612769 | 0.367072669 | 0.229 | 0.05 | 0.001244393 | 19 | si:dkeyp-68b7.12 |
| zgc:101583 | 6.2635387 | 0.316907238 | 0.146 | 0.023 | 0.001265047 | 19 | zgc:101583 |
| LOC1081907171 | 6.3913293 | 0.285409952 | 0.125 | 0.017 | 0.001290857 | 19 | LOC108190717 |
| tmc8 | 1.0357509 | 0.260357166 | 0.167 | 0.029 | 0.002091906 | 19 | tmc8 |
| cox6b12 | 1.4083581 | 0.467476281 | 0.396 | 0.131 | 0.002844461 | 19 | cox6b1 |
| h1fx3 | 1.5103922 | 0.625238264 | 0.562 | 0.234 | 0.003050539 | 19 | h1fx |

| | | | | | | | |
|---|---|---|---|---|---|---|---|
| il161 | 1.5891497 | 0.380980872 | 0.208 | 0.045 | 0.003209606 | 19 | il16 |
| tagapb2 | 1.6124528 | 0.493896201 | 0.292 | 0.083 | 0.003256671 | 19 | tagapb |
| efhd21 | 1.6733840 | 0.429378874 | 0.25 | 0.063 | 0.003379734 | 19 | efhd2 |
| sla1 | 1.8029693 | 0.345497056 | 0.188 | 0.038 | 0.003641457 | 19 | sla1 |
| vaspa3 | 4.0119896 | 0.476036242 | 0.375 | 0.117 | 0.008103016 | 19 | vaspa |
| gapdhs3 | 4.0702013 | 0.537876206 | 0.771 | 0.474 | 0.008220586 | 19 | gapdhs |
| XLOC-0268362 | 4.0758972 | 0.475727278 | 0.292 | 0.082 | 0.00823209 | 19 | XLOC-026836 |
| xbp14 | 4.8328378 | 0.580103436 | 0.708 | 0.433 | 0.009760883 | 19 | xbp1 |
| ctss2.12 | 4.9021276 | 0.377791173 | 0.271 | 0.073 | 0.009900827 | 19 | ctss2.1 |
| cfl14 | 5.5051197 | 0.372625479 | 0.917 | 0.766 | 0.01111869 | 19 | cfl1 |
| si:dkey-27h10.21 | 8.1034579 | 0.279872923 | 0.146 | 0.026 | 0.016366554 | 19 | si:dkey-27h10.2 |
| hyou14 | 9.6986122 | 0.443688549 | 0.271 | 0.08 | 0.019588287 | 19 | hyou1 |
| scinlb2 | 1.0068458 | 0.420828706 | 0.312 | 0.095 | 0.020335266 | 19 | scinlb |
| fcer1gl4 | 1.0218091 | 0.682455876 | 0.562 | 0.256 | 0.020637479 | 19 | fcer1gl |
| selplg2 | 1.2058368 | 0.307306514 | 0.188 | 0.041 | 0.024354286 | 19 | selplg |
| arhgdig4 | 1.4847320 | 0.567424194 | 0.646 | 0.364 | 0.029987133 | 19 | arhgdig |
| arhgef3l1 | 1.6624434 | 0.319695521 | 0.146 | 0.028 | 0.033576371 | 19 | arhgef3l |
| limd21 | 2.0037213 | 0.43195109 | 0.229 | 0.062 | 0.04046916 | 19 | limd2 |
| hcls14 | 2.3071007 | 0.435898651 | 0.333 | 0.11 | 0.046596514 | 19 | hcls1 |
| si:dkey-237j10.2 | 2.3337465 | 0.289648349 | 0.125 | 0.021 | 0.04713468 | 19 | si:dkey-237j10.2 |
| glipr1a3 | 2.5034270 | 0.434674345 | 0.292 | 0.092 | 0.050561717 | 19 | glipr1a |
| CR392341.13 | 2.6435769 | 0.378132905 | 0.229 | 0.061 | 0.053392324 | 19 | CR392341.1 |
| pik3r11 | 2.7131621 | 0.419417682 | 0.312 | 0.103 | 0.054797735 | 19 | pik3r1 |
| cd633 | 2.7602692 | 0.472108905 | 0.833 | 0.566 | 0.055749158 | 19 | cd63 |
| tpi1b2 | 3.7991132 | 0.526561391 | 0.5 | 0.244 | 0.076730691 | 19 | tpi1b |
| dusp22 | 3.9444432 | 0.581529057 | 0.292 | 0.098 | 0.07966592 | 19 | dusp2 |
| derl22 | 3.9574110 | 0.476268379 | 0.333 | 0.122 | 0.079927831 | 19 | derl2 |
| si:ch211-226m16.2 | 5.4821385 | 0.298496177 | 0.25 | 0.073 | 0.110722753 | 19 | si:ch211-226m16.2 |
| hmha1b3 | 6.0611958 | 0.361613964 | 0.25 | 0.074 | 0.122417973 | 19 | hmha1b |
| abracl3 | 6.5685455 | 0.52661065 | 0.5 | 0.24 | 0.132664914 | 19 | abracl |
| stxbp21 | 7.1038185 | 0.399082258 | 0.229 | 0.068 | 0.143475824 | 19 | stxbp2 |
| ikzf11 | 8.2996708 | 0.314988966 | 0.167 | 0.038 | 0.167628453 | 19 | ikzf1 |
| RASSF51 | 8.3367452 | 0.386626605 | 0.25 | 0.077 | 0.168377244 | 19 | RASSF5 |
| arpc5a2 | 9.7662393 | 0.521064012 | 0.417 | 0.188 | 0.197248737 | 19 | arpc5a |
| reep53 | 1.0628184 | 0.488413927 | 0.521 | 0.264 | 0.214657441 | 19 | reep5 |

| | | | | | | | |
|---|---|---|---|---|---|---|---|
| pygb1 | 1.4345092 | 0.339031981 | 0.188 | 0.049 | 0.289727842 | 19 | pygb |
| sec61g3 | 1.5891659 | 0.371279594 | 0.875 | 0.726 | 0.320963849 | 19 | sec61g |
| atp5if1b3 | 1.6196255 | 0.496470729 | 0.521 | 0.273 | 0.327115765 | 19 | atp5if1b |
| yrk3 | 1.7638635 | 0.424650353 | 0.271 | 0.09 | 0.356247522 | 19 | yrk |
| fam49ba2 | 1.9003572 | 0.390637881 | 0.333 | 0.122 | 0.383815147 | 19 | fam49ba |
| nr4a13 | 1.9016382 | 0.441057063 | 0.333 | 0.124 | 0.384073871 | 19 | nr4a1 |
| skap22 | 2.1296487 | 0.404166863 | 0.312 | 0.114 | 0.430125165 | 19 | skap2 |
| LOC1001482542 | 2.2071126 | 0.265692262 | 0.146 | 0.033 | 0.445770551 | 19 | LOC100148254 |
| LOC1039114301 | 2.4831936 | 0.252422125 | 0.167 | 0.04 | 0.501530626 | 19 | LOC103911430 |
| syk3 | 2.5202598 | 0.326711507 | 0.25 | 0.079 | 0.509016884 | 19 | syk |
| si:zfos-2330d3.11 | 2.6888094 | 0.274575611 | 0.167 | 0.039 | 0.543058835 | 19 | si:zfos-2330d3.1 |
| BX120005.11 | 3.5240248 | 0.300237789 | 0.146 | 0.034 | 0.711747308 | 19 | BX120005.1 |
| pim21 | 3.8885029 | 0.383577689 | 0.229 | 0.074 | 0.785360934 | 19 | pim2 |
| inpp5d1 | 4.5046951 | 0.261266171 | 0.167 | 0.043 | 0.909813285 | 19 | inpp5d |
| ptger4b | 4.7589638 | 0.264617098 | 0.125 | 0.027 | 0.961167922 | 19 | ptger4b |
| cox6a21 | 5.1016183 | 0.272496477 | 0.167 | 0.043 | 1 | 19 | cox6a2 |
| si:ch211-66e2.33 | 6.7758182 | 0.388290828 | 0.25 | 0.084 | 1 | 19 | si:ch211-66e2.3 |
| sh2b31 | 7.5005564 | 0.288179726 | 0.208 | 0.065 | 1 | 19 | sh2b3 |
| wdr14 | 8.7446290 | 0.456714439 | 0.458 | 0.236 | 1 | 19 | wdr1 |
| lasp13 | 0.000101 | 0.439307695 | 0.583 | 0.357 | 1 | 19 | lasp1 |
| ms4a17a.62 | 0.000107 | 0.406099901 | 0.208 | 0.067 | 1 | 19 | ms4a17a.6 |
| rhogb3 | 0.000115 | 0.41800984 | 0.375 | 0.169 | 1 | 19 | rhogb |
| XLOC-018356 | 0.000124 | 0.312071572 | 0.146 | 0.037 | 1 | 19 | XLOC-018356 |
| csf2rb1 | 0.000137 | 0.262011858 | 0.146 | 0.038 | 1 | 19 | csf2rb |
| XLOC-0234923 | 0.000143 | 0.368858411 | 0.25 | 0.091 | 1 | 19 | XLOC-023492 |
| sat1a.25 | 0.000144 | 0.407536097 | 0.583 | 0.334 | 1 | 19 | sat1a.2 |
| ostf12 | 0.00017 | 0.272750695 | 0.188 | 0.057 | 1 | 19 | ostf1 |
| csk2 | 0.000173 | 0.368351958 | 0.292 | 0.118 | 1 | 19 | csk |
| BX005183.3 | 0.000182 | 0.301734664 | 0.188 | 0.058 | 1 | 19 | BX005183.3 |
| arl5c2 | 0.0002 | 0.332198664 | 0.25 | 0.093 | 1 | 19 | arl5c |
| lcp11 | 0.000207 | 0.322240542 | 0.208 | 0.07 | 1 | 19 | lcp1 |
| cisd2 | 0.000242 | 0.283449078 | 0.188 | 0.06 | 1 | 19 | cisd2 |
| ugcg | 0.000309 | 0.253494031 | 0.125 | 0.032 | 1 | 19 | ugcg |
| mapre1a4 | 0.000323 | 0.357392146 | 0.417 | 0.198 | 1 | 19 | mapre1a |
| bcl2l101 | 0.000356 | 0.417817799 | 0.312 | 0.141 | 1 | 19 | bcl2l10 |

| | | | | | | | |
|---|---|---|---|---|---|---|---|
| lpcat21 | 0.000371 | 0.251289032 | 0.125 | 0.032 | 1 | 19 | lpcat2 |
| XLOC-0425952 | 0.000405 | 0.338841527 | 0.271 | 0.106 | 1 | 19 | XLOC-042595 |
| arrb2b1 | 0.000408 | 0.37550111 | 0.292 | 0.128 | 1 | 19 | arrb2b |
| serpine2 | 0.000409 | 0.271768708 | 0.167 | 0.051 | 1 | 19 | serpine2 |
| cebpb4 | 0.000416 | 0.424520712 | 0.542 | 0.294 | 1 | 19 | cebpb |
| sept62 | 0.00044 | 0.417501819 | 0.375 | 0.191 | 1 | 19 | sept6 |
| egr31 | 0.000485 | 0.294576373 | 0.125 | 0.033 | 1 | 19 | egr3 |
| mcl1a4 | 0.000486 | 0.393653924 | 0.708 | 0.545 | 1 | 19 | mcl1a |
| rnaset2l3 | 0.000528 | 0.297949944 | 0.229 | 0.083 | 1 | 19 | rnaset2l |
| samsn1a3 | 0.000548 | 0.284586626 | 0.167 | 0.053 | 1 | 19 | samsn1a |
| gpr183a2 | 0.000565 | 0.330572754 | 0.188 | 0.066 | 1 | 19 | gpr183a |
| lpar5a2 | 0.000573 | 0.261923424 | 0.188 | 0.063 | 1 | 19 | lpar5a |
| plxnc12 | 0.000618 | 0.266681761 | 0.188 | 0.064 | 1 | 19 | plxnc1 |
| ccr9a3 | 0.000626 | 0.313985952 | 0.25 | 0.098 | 1 | 19 | ccr9a |
| bri3bp | 0.000664 | 0.281067576 | 0.167 | 0.054 | 1 | 19 | bri3bp |
| arpc24 | 0.000676 | 0.376920872 | 0.646 | 0.486 | 1 | 19 | arpc2 |
| hif1ab2 | 0.00068 | 0.368867796 | 0.354 | 0.176 | 1 | 19 | hif1ab |
| scamp21 | 0.000684 | 0.271843275 | 0.25 | 0.101 | 1 | 19 | scamp2 |
| GCA2 | 0.000763 | 0.423447093 | 0.479 | 0.31 | 1 | 19 | GCA |
| dub1 | 0.000805 | 0.297664608 | 0.229 | 0.088 | 1 | 19 | dub |
| nptna | 0.000846 | 0.279616487 | 0.167 | 0.055 | 1 | 19 | nptna |
| lbh1 | 0.000852 | 0.300146254 | 0.188 | 0.067 | 1 | 19 | lbh |
| actr2a3 | 0.000899 | 0.372449047 | 0.333 | 0.163 | 1 | 19 | actr2a |
| cyba3 | 0.00095 | 0.349625593 | 0.292 | 0.133 | 1 | 19 | cyba |
| mob3a1 | 0.000957 | 0.332303408 | 0.375 | 0.189 | 1 | 19 | mob3a |
| LOC1005362133 | 0.001001 | 0.322321434 | 0.333 | 0.149 | 1 | 19 | LOC100536213 |
| LOC1039091154 | 0.001012 | 0.39354953 | 0.375 | 0.2 | 1 | 19 | LOC103909115 |
| ptpn22 | 0.001026 | 0.283828377 | 0.104 | 0.027 | 1 | 19 | ptpn22 |
| fosl22 | 0.001088 | 0.332287512 | 0.208 | 0.081 | 1 | 19 | fosl2 |
| mlec | 0.001102 | 0.280624718 | 0.167 | 0.057 | 1 | 19 | mlec |
| CR855389.1 | 0.001305 | 0.271250768 | 0.125 | 0.037 | 1 | 19 | CR855389.1 |
| gna13b | 0.001441 | 0.25307302 | 0.167 | 0.058 | 1 | 19 | gna13b |
| myl12.12 | 0.001448 | 0.30527462 | 0.75 | 0.631 | 1 | 19 | myl12.1 |
| sept7b3 | 0.001517 | 0.36009174 | 0.312 | 0.157 | 1 | 19 | sept7b |
| si:dkey-262k9.2 | 0.001547 | 0.274062789 | 0.104 | 0.028 | 1 | 19 | si:dkey-262k9.2 |

| | | | | | | | |
|---|---|---|---|---|---|---|---|
| zgc:112356 | 0.001733 | 0.260765668 | 0.208 | 0.082 | 1 | 19 | zgc:112356 |
| pkma3 | 0.001747 | 0.310321593 | 0.375 | 0.197 | 1 | 19 | pkma |
| XLOC-0254234 | 0.001772 | 0.370919456 | 0.375 | 0.192 | 1 | 19 | XLOC-025423 |
| si:ch73-248e21.51 | 0.002123 | 0.279430493 | 0.146 | 0.049 | 1 | 19 | si:ch73-248e21.5 |
| mbnl13 | 0.002234 | 0.414042309 | 0.333 | 0.186 | 1 | 19 | mbnl1 |
| dok21 | 0.002256 | 0.30556722 | 0.146 | 0.05 | 1 | 19 | dok2 |
| rarga3 | 0.002289 | 0.298173378 | 0.208 | 0.087 | 1 | 19 | rarga |
| sept121 | 0.002488 | 0.283379484 | 0.208 | 0.086 | 1 | 19 | sept12 |
| furina1 | 0.002774 | 0.258128059 | 0.208 | 0.086 | 1 | 19 | furina |
| rnf1302 | 0.002862 | 0.273637825 | 0.208 | 0.088 | 1 | 19 | rnf130 |
| capzb3 | 0.002876 | 0.348056786 | 0.562 | 0.386 | 1 | 19 | capzb |
| park71 | 0.003023 | 0.313941176 | 0.208 | 0.09 | 1 | 19 | park7 |
| myd88 | 0.003114 | 0.259784636 | 0.208 | 0.088 | 1 | 19 | myd88 |
| pef1 | 0.003353 | 0.278792898 | 0.25 | 0.116 | 1 | 19 | pef1 |
| higd1a3 | 0.003594 | 0.326961165 | 0.458 | 0.274 | 1 | 19 | higd1a |
| cox7a2a1 | 0.003709 | 0.387610891 | 0.458 | 0.305 | 1 | 19 | cox7a2a |
| lypla21 | 0.003729 | 0.294590416 | 0.188 | 0.078 | 1 | 19 | lypla2 |
| arpc44 | 0.004195 | 0.288366086 | 0.521 | 0.316 | 1 | 19 | arpc4 |
| CR753876.13 | 0.004266 | 0.334578585 | 0.229 | 0.102 | 1 | 19 | CR753876.1 |
| arpc35 | 0.004328 | 0.345488999 | 0.625 | 0.481 | 1 | 19 | arpc3 |
| sh3bgrl1 | 0.004522 | 0.315086853 | 0.375 | 0.215 | 1 | 19 | sh3bgrl |
| med19a | 0.004988 | 0.283757663 | 0.229 | 0.106 | 1 | 19 | med19a |
| txn3 | 0.005334 | 0.298566631 | 0.208 | 0.091 | 1 | 19 | txn |
| cap15 | 0.005367 | 0.324324348 | 0.521 | 0.33 | 1 | 19 | cap1 |
| eno1a3 | 0.006774 | 0.305767542 | 0.333 | 0.189 | 1 | 19 | eno1a |
| nfkbiaa3 | 0.007 | 0.421967079 | 0.292 | 0.165 | 1 | 19 | nfkbiaa |
| edf12 | 0.007257 | 0.35351234 | 0.5 | 0.369 | 1 | 19 | edf1 |
| arpc4l4 | 0.007394 | 0.280342319 | 0.542 | 0.35 | 1 | 19 | arpc4l |
| tomm7 | 0.007817 | 0.31294932 | 0.396 | 0.251 | 1 | 19 | tomm7 |
| arhgdia3 | 0.008574 | 0.320234808 | 0.521 | 0.375 | 1 | 19 | arhgdia |
| gp1bb | 1.8425681 | 1.041944272 | 0.319 | 0 | 3.72143482698437e-1 | 20 | gp1bb |
| si:ch211-161c3.6 | 2.7573182 | 1.069394696 | 0.426 | 0.005 | 5.56895564828249e-1 | 20 | si:ch211-161c3.6 |
| itga2b | 2.3749601 | 0.676273094 | 0.213 | 0 | 4.79670694778779e-8 | 20 | itga2b |
| blvrb | 1.0240748 | 0.695228987 | 0.319 | 0.007 | 2.06832395039069e-6 | 20 | blvrb |
| spi2 | 8.3916422 | 0.303652497 | 0.149 | 0 | 1.69485999034605e-6 | 20 | spi2 |

| | | | | | | | |
|---|---|---|---|---|---|---|---|
| myb | 4.1642488 | 0.674189508 | 0.404 | 0.016 | 8.41053338547968e-6 | 20 | myb |
| blf | 1.3435911 | 0.439688495 | 0.17 | 0.001 | 2.7136509847847e-57 | 20 | blf |
| znfl1l | 6.4623578 | 0.31754029 | 0.213 | 0.005 | 1.30520240605945e-4 | 20 | znfl1l |
| sb:cb811 | 1.9514137 | 0.801373445 | 0.447 | 0.032 | 3.94127025163345e-4 | 20 | sb:cb81 |
| zgc:1716861 | 6.6289838 | 0.63104902 | 0.277 | 0.01 | 1.3388558651888e-41 | 20 | zgc:171686 |
| tnr | 1.1164374 | 0.396412212 | 0.128 | 0.001 | 2.25486872809377e-3 | 20 | tnr |
| plac8l1 | 6.1491693 | 0.502081646 | 0.149 | 0.002 | 1.24194773480889e-3 | 20 | plac8l1 |
| CABZ01066694.1 | 1.0266810 | 0.397095917 | 0.277 | 0.012 | 2.07358775739854e-3 | 20 | CABZ01066694.1 |
| si:dkey-217l24.1 | 1.6962878 | 0.474073342 | 0.255 | 0.01 | 3.42599262966354e-3 | 20 | si:dkey-217l24.1 |
| wnt2 | 9.7613284 | 0.33275576 | 0.106 | 0 | 1.97149551311706e-3 | 20 | wnt2 |
| apln | 1.3414713 | 0.590501305 | 0.17 | 0.003 | 2.70936959424354e-3 | 20 | apln |
| csf1rb | 1.3718334 | 0.350551301 | 0.234 | 0.009 | 2.7706919708381e-34 | 20 | csf1rb |
| parvb | 1.7023787 | 0.551164821 | 0.191 | 0.006 | 3.43829444727062e-3 | 20 | parvb |
| thbs1b | 3.6031933 | 0.770094457 | 0.213 | 0.008 | 7.27736960459657e-3 | 20 | thbs1b |
| fli1a1 | 3.9817613 | 0.576161305 | 0.298 | 0.017 | 8.04196339750711e-3 | 20 | fli1a |
| gcnt4a | 1.2160097 | 0.544655818 | 0.255 | 0.013 | 2.45597482269196e-2 | 20 | gcnt4a |
| hhex1 | 2.5406594 | 0.315067938 | 0.234 | 0.011 | 5.13136998967301e-2 | 20 | hhex |
| dub2 | 3.8307920 | 0.745711279 | 0.574 | 0.08 | 7.73705074524096e-2 | 20 | dub |
| nfe21 | 6.5134528 | 0.466183314 | 0.277 | 0.021 | 1.31552207287311e-2 | 20 | nfe2 |
| mir2231 | 2.4233392 | 0.716804972 | 0.404 | 0.049 | 4.89441825640748e-2 | 20 | mir223 |
| si:ch211-226m16.2 | 5.8014875 | 0.670641675 | 0.468 | 0.068 | 1.17172643411226e-1 | 20 | si:ch211-226m16.2 |
| si:ch211-214p16.2 | 5.9272247 | 0.854310932 | 0.532 | 0.085 | 1.19712157596487e-1 | 20 | si:ch211-214p16.2 |
| pmp22b2 | 8.2321608 | 0.712787814 | 0.404 | 0.052 | 1.66264951729441e-1 | 20 | pmp22b |
| lmo21 | 2.1302637 | 0.44446458 | 0.255 | 0.022 | 4.30249374697145e-1 | 20 | lmo2 |
| hbegfb1 | 9.1953793 | 0.672837922 | 0.213 | 0.016 | 1.85719076675416e-1 | 20 | hbegfb |
| plek1 | 1.5488861 | 0.782042689 | 0.362 | 0.048 | 3.12828533506399e-1 | 20 | plek |
| sept122 | 2.0321295 | 0.72974902 | 0.468 | 0.08 | 4.10429209008736e-1 | 20 | sept12 |
| p2ry13 | 2.9209735 | 0.363539872 | 0.106 | 0.003 | 5.89949035503333e-1 | 20 | p2ry13 |
| hacd31 | 4.2621504 | 0.803439431 | 0.319 | 0.045 | 8.60826536210454e-1 | 20 | hacd3 |
| cd7al1 | 7.5272797 | 0.265114243 | 0.298 | 0.036 | 1.52028468252269e-1 | 20 | cd7al |
| itgb3b1 | 1.0479022 | 0.719140062 | 0.234 | 0.025 | 2.1164480831934e-13 | 20 | itgb3b |
| cxcr4b3 | 1.8830344 | 0.887873585 | 0.617 | 0.15 | 3.8031645894594e-13 | 20 | cxcr4b |
| boka | 5.7608769 | 0.404118562 | 0.149 | 0.01 | 1.16352432148189e-1 | 20 | boka |
| zgc:171506 | 8.9031316 | 0.28773432 | 0.149 | 0.01 | 1.79816550248543e-1 | 20 | zgc:171506 |
| inka1b | 2.3050331 | 0.524327978 | 0.213 | 0.022 | 4.65547535789827e-1 | 20 | inka1b |

| | | | | | | | |
|---|---|---|---|---|---|---|---|
| DIPK2B | 3.6856060 | 0.29638994 | 0.106 | 0.005 | 7.44381857209594e-1 | 20 | DIPK2B |
| mef2aa | 4.9365806 | 0.728103048 | 0.447 | 0.093 | 9.97041184298455e-1 | 20 | mef2aa |
| ncf21 | 6.7278396 | 0.399803398 | 0.234 | 0.027 | 1.35882177891472e-1 | 20 | ncf2 |
| celf4 | 1.4101959 | 0.415675544 | 0.128 | 0.008 | 2.84817283212807e-1 | 20 | celf4 |
| rhag | 1.7854542 | 0.379832948 | 0.128 | 0.008 | 3.60608188005572e-1 | 20 | rhag |
| si:ch211-33e4.21 | 2.4697649 | 0.345010435 | 0.128 | 0.008 | 4.98818427121028e-1 | 20 | si:ch211-33e4.2 |
| ptpn11b | 3.0668815 | 0.380711567 | 0.17 | 0.015 | 6.19418071473766e-1 | 20 | ptpn11b |
| XLOC-014336 | 4.0780537 | 0.3011724 | 0.277 | 0.037 | 8.23644509377712e-1 | 20 | XLOC-014336 |
| prkacba1 | 4.1672289 | 0.398007829 | 0.213 | 0.023 | 8.41655235096967e-1 | 20 | prkacba |
| nup88 | 4.4041923 | 0.26953809 | 0.106 | 0.005 | 8.89514737979753e-1 | 20 | nup88 |
| ahcy2 | 5.9052351 | 0.685405277 | 0.468 | 0.107 | 1.19268033608295e-1 | 20 | ahcy |
| riox2 | 6.1447989 | 0.262126253 | 0.234 | 0.028 | 1.2410650463557e-10 | 20 | riox2 |
| myca1 | 1.0112226 | 0.51690657 | 0.34 | 0.059 | 2.04236638111621e-1 | 20 | myca |
| cbfa2t31 | 1.0166362 | 0.391604831 | 0.298 | 0.045 | 2.05330016976963e-1 | 20 | cbfa2t3 |
| susd1 | 1.3673172 | 0.358870303 | 0.149 | 0.012 | 2.7615706052986e-10 | 20 | susd1 |
| CU499330.12 | 2.1516506 | 0.483894923 | 0.426 | 0.08 | 4.34568872896851e-1 | 20 | CU499330.1 |
| dennd2c | 5.2911190 | 0.322098225 | 0.128 | 0.01 | 1.06864730907567e-0 | 20 | dennd2c |
| XLOC-0190621 | 1.4272188 | 0.267621566 | 0.106 | 0.007 | 2.88255399005527e-0 | 20 | XLOC-019062 |
| gbp | 1.6667145 | 0.622360609 | 0.617 | 0.198 | 3.36626337997355e-0 | 20 | gbp |
| si:ch211-214p16.1 | 1.7372326 | 0.76808309 | 0.66 | 0.186 | 3.50868877148818e-0 | 20 | si:ch211-214p16.1 |
| tp53rk | 3.2351542 | 0.253143872 | 0.277 | 0.045 | 6.53404100418159e-0 | 20 | tp53rk |
| rac3a | 3.4292911 | 0.462213865 | 0.149 | 0.015 | 6.92613928000317e-0 | 20 | rac3a |
| ppan | 4.4298384 | 0.376837579 | 0.277 | 0.047 | 8.94694462761163e-0 | 20 | ppan |
| cdkn2c | 4.9619800 | 0.304167591 | 0.106 | 0.007 | 1.00217111662453e-0 | 20 | cdkn2c |
| si:dkey-185m8.22 | 6.8162373 | 0.405220111 | 0.255 | 0.041 | 1.3766754487696e-07 | 20 | si:dkey-185m8.2 |
| mybbp1a1 | 7.3846157 | 0.258639779 | 0.255 | 0.04 | 1.49147084057482e-0 | 20 | mybbp1a |
| dnmt3bb.12 | 1.0315509 | 0.414312536 | 0.277 | 0.049 | 2.08342336923213e-0 | 20 | dnmt3bb.1 |
| si:ch211-222l21.17 | 1.3307850 | 0.714888461 | 0.894 | 0.528 | 2.68778662567948e-0 | 20 | si:ch211-222l21.1 |
| AL954142.13 | 1.4656820 | 0.48155754 | 0.213 | 0.031 | 2.96023799004155e-0 | 20 | AL954142.1 |
| dhx37 | 2.0628561 | 0.265066455 | 0.17 | 0.02 | 4.16635052985268e-0 | 20 | dhx37 |
| gnai2b1 | 4.7236735 | 0.284175065 | 0.234 | 0.037 | 9.54040349727615e-0 | 20 | gnai2b |
| limd22 | 6.1508881 | 0.508899054 | 0.298 | 0.06 | 1.24229486998446e-0 | 20 | limd2 |
| suv39h1b | 6.6994582 | 0.490494116 | 0.149 | 0.017 | 1.35308958222415e-0 | 20 | suv39h1b |
| spns2 | 6.7573716 | 0.324049353 | 0.106 | 0.008 | 1.3647863485908e-06 | 20 | spns2 |
| MCMDC2 | 8.9397052 | 0.295497677 | 0.362 | 0.08 | 1.80555226663284e-0 | 20 | MCMDC2 |

| | | | | | | | |
|---|---|---|---|---|---|---|---|
| kdm6bb2 | 1.0698742 | 0.540453956 | 0.404 | 0.108 | 2.16082504931077e-0 | 20 | kdm6bb |
| bmp16 | 2.2711292 | 0.488556472 | 0.149 | 0.018 | 4.58699970302942e-0 | 20 | bmp16 |
| cxcr3.22 | 2.2796019 | 0.385908771 | 0.234 | 0.04 | 4.60411199384444e-0 | 20 | cxcr3.2 |
| pim22 | 2.3956483 | 0.494154975 | 0.319 | 0.072 | 4.83849092423462e-0 | 20 | pim2 |
| soga1 | 2.4264049 | 0.462914786 | 0.234 | 0.041 | 4.9006100243541e-06 | 20 | soga1 |
| wasa4 | 2.5610624 | 0.385629462 | 0.447 | 0.114 | 5.17257777143365e-0 | 20 | wasa |
| ppifb2 | 4.3903325 | 0.488678047 | 0.511 | 0.162 | 8.86715465860592e-0 | 20 | ppifb |
| rap1gap2a | 7.6284700 | 0.420964534 | 0.128 | 0.014 | 1.54072209040921e-0 | 20 | rap1gap2a |
| nudt9 | 8.1846003 | 0.25671913 | 0.213 | 0.035 | 1.65304372749964e-0 | 20 | nudt9 |
| fabp31 | 8.8894961 | 0.616477635 | 0.511 | 0.174 | 1.79541154396573e-0 | 20 | fabp3 |
| ophn1 | 9.3480803 | 0.370071644 | 0.255 | 0.049 | 1.88803179524252e-0 | 20 | ophn1 |
| ptmaa3 | 1.2807686 | 0.590906787 | 0.957 | 0.652 | 2.58676853014202e-0 | 20 | ptmaa |
| adgrg11 | 1.9236864 | 0.341963956 | 0.128 | 0.014 | 3.88526949726216e-0 | 20 | adgrg1 |
| rabgap1l | 2.2570214 | 0.511676466 | 0.234 | 0.046 | 4.55850623211948e-0 | 20 | rabgap1l |
| tmem110l | 2.3422143 | 0.276484423 | 0.128 | 0.014 | 4.73057026961059e-0 | 20 | tmem110l |
| XLOC-0425953 | 2.8758974 | 0.547459075 | 0.383 | 0.103 | 5.80845004672205e-0 | 20 | XLOC-042595 |
| ppid1 | 3.4796181 | 0.318737216 | 0.404 | 0.11 | 7.02778486652492e-0 | 20 | ppid |
| tagapb3 | 3.9736227 | 0.33324815 | 0.34 | 0.082 | 8.02552577531823e-0 | 20 | tagapb |
| glcci1a | 4.1658117 | 0.266436383 | 0.128 | 0.015 | 8.41368998383118e-0 | 20 | glcci1a |
| aldoaa2 | 4.6368006 | 0.601770548 | 0.638 | 0.284 | 9.36494617756637e-0 | 20 | aldoaa |
| pfn16 | 6.0289675 | 0.756312683 | 0.979 | 0.411 | 0.000121767 | 20 | pfn1 |
| laptm56 | 6.0349642 | 0.454402989 | 0.638 | 0.211 | 0.000121888 | 20 | laptm5 |
| selenoh1 | 1.3347479 | 0.404844565 | 0.489 | 0.157 | 0.000269579 | 20 | selenoh |
| gatad2b3 | 1.4094578 | 0.397895134 | 0.404 | 0.116 | 0.000284668 | 20 | gatad2b |
| rsl24d12 | 1.7085926 | 0.510333307 | 0.83 | 0.572 | 0.000345084 | 20 | rsl24d1 |
| stmn1a1 | 2.1683855 | 0.424666118 | 0.383 | 0.107 | 0.000437949 | 20 | stmn1a |
| utp4 | 3.6272349 | 0.254982705 | 0.213 | 0.041 | 0.000732593 | 20 | utp4 |
| LOC1039091155 | 4.1568934 | 0.513105382 | 0.532 | 0.196 | 0.000839568 | 20 | LOC103909115 |
| vamp51 | 4.2540506 | 0.441514439 | 0.191 | 0.037 | 0.000859191 | 20 | vamp5 |
| hells1 | 4.3877763 | 0.264504764 | 0.34 | 0.088 | 0.000886199 | 20 | hells |
| fbp1a1 | 7.1023601 | 0.328424378 | 0.255 | 0.06 | 0.001434464 | 20 | fbp1a |
| si:dkey-112e7.22 | 1.1879265 | 0.543181182 | 0.596 | 0.269 | 0.002399255 | 20 | si:dkey-112e7.2 |
| mrpl42 | 1.2405521 | 0.289218872 | 0.362 | 0.103 | 0.002505543 | 20 | mrpl42 |
| rap1b3 | 1.4481753 | 0.699658339 | 0.66 | 0.323 | 0.00292488 | 20 | rap1b |
| FP074902.2 | 1.5805317 | 0.265785162 | 0.277 | 0.07 | 0.0031922 | 20 | FP074902.2 |

| | | | | | | | |
|---|---|---|---|---|---|---|---|
| parvg1 | 1.6685252 | 0.360742641 | 0.362 | 0.105 | 0.00336992 | 20 | parvg |
| mcm71 | 2.1137686 | 0.357014581 | 0.34 | 0.099 | 0.004269179 | 20 | mcm7 |
| si:dkey-23i12.51 | 2.2130770 | 0.370782093 | 0.426 | 0.146 | 0.004469752 | 20 | si:dkey-23i12.5 |
| ikzf12 | 2.4051740 | 0.250134433 | 0.191 | 0.038 | 0.00485773 | 20 | ikzf1 |
| myh11a2 | 2.9443152 | 0.607035789 | 0.319 | 0.101 | 0.005946633 | 20 | myh11a |
| rbm28 | 2.9792375 | 0.355611688 | 0.426 | 0.143 | 0.006017166 | 20 | rbm28 |
| eef1g1 | 3.1594800 | 0.262215897 | 0.894 | 0.839 | 0.006381202 | 20 | eef1g |
| cox6b13 | 3.3278721 | 0.363800152 | 0.404 | 0.131 | 0.006721303 | 20 | cox6b1 |
| lrrc8da1 | 3.3331904 | 0.393376792 | 0.17 | 0.033 | 0.006732045 | 20 | lrrc8da |
| adssl1 | 3.3415045 | 0.334789296 | 0.426 | 0.139 | 0.006748837 | 20 | adssl |
| ap1s11 | 3.5219659 | 0.451868428 | 0.34 | 0.105 | 0.007113315 | 20 | ap1s1 |
| nr4a14 | 3.5394374 | 0.45695967 | 0.383 | 0.123 | 0.007148602 | 20 | nr4a1 |
| nop582 | 4.2209956 | 0.445277662 | 0.532 | 0.217 | 0.008525145 | 20 | nop58 |
| psmg2 | 4.4022757 | 0.295255241 | 0.34 | 0.101 | 0.008891276 | 20 | psmg2 |
| dera | 4.5555487 | 0.262850259 | 0.149 | 0.026 | 0.009200842 | 20 | dera |
| fermt3b2 | 4.6569215 | 0.439662732 | 0.17 | 0.034 | 0.009405584 | 20 | fermt3b |
| tln12 | 4.6988912 | 0.636352059 | 0.511 | 0.222 | 0.009490351 | 20 | tln1 |
| id2a1 | 5.1392035 | 0.494927298 | 0.447 | 0.166 | 0.010379649 | 20 | id2a |
| nutf2l2 | 7.0308226 | 0.35658583 | 0.468 | 0.171 | 0.014200152 | 20 | nutf2l |
| ivns1abpb | 7.3988085 | 0.270799175 | 0.191 | 0.041 | 0.014943374 | 20 | ivns1abpb |
| pgam1a3 | 7.5185795 | 0.316362099 | 0.447 | 0.154 | 0.015185275 | 20 | pgam1a |
| dcps | 7.9299074 | 0.313137669 | 0.149 | 0.027 | 0.016016034 | 20 | dcps |
| npm1b | 8.0766715 | 0.297612427 | 0.234 | 0.057 | 0.016312454 | 20 | npm1b |
| gfi1ab2 | 9.4590258 | 0.289560871 | 0.149 | 0.027 | 0.019104395 | 20 | gfi1ab |
| tktb1 | 9.5578387 | 0.312065455 | 0.383 | 0.125 | 0.019303967 | 20 | tktb |
| serbp1a2 | 1.1371182 | 0.381210877 | 0.809 | 0.656 | 0.022966376 | 20 | serbp1a |
| pik3r12 | 1.2729508 | 0.276191032 | 0.34 | 0.103 | 0.025709789 | 20 | pik3r1 |
| hmgb3b3 | 1.3278312 | 0.258816062 | 0.426 | 0.144 | 0.026818208 | 20 | hmgb3b |
| arl6ip13 | 1.6774552 | 0.468056991 | 0.617 | 0.284 | 0.033879563 | 20 | arl6ip1 |
| cnbpa1 | 1.7151679 | 0.44720101 | 0.553 | 0.251 | 0.034641248 | 20 | cnbpa |
| ncaldb2 | 1.8251258 | 0.276251798 | 0.213 | 0.051 | 0.036862067 | 20 | ncaldb |
| arpc1b6 | 1.9767507 | 0.514242005 | 0.787 | 0.356 | 0.039924434 | 20 | arpc1b |
| gtpbp42 | 2.1417911 | 0.520679527 | 0.596 | 0.312 | 0.043257756 | 20 | gtpbp4 |
| ddx211 | 2.1816982 | 0.427432662 | 0.532 | 0.231 | 0.044063759 | 20 | ddx21 |
| si:dkey-239h2.32 | 2.2950095 | 0.377668566 | 0.426 | 0.159 | 0.046352308 | 20 | si:dkey-239h2.3 |

| | | | | | | | |
|---|---|---|---|---|---|---|---|
| creg1 | 2.5950559 | 0.322993742 | 0.191 | 0.044 | 0.052412345 | 20 | creg1 |
| capgb5 | 2.6905153 | 0.289492744 | 0.383 | 0.125 | 0.054340339 | 20 | capgb |
| atp2b1a1 | 2.7675953 | 0.335360268 | 0.234 | 0.062 | 0.055897124 | 20 | atp2b1a |
| plk32 | 2.9964023 | 0.495265192 | 0.213 | 0.056 | 0.060518339 | 20 | plk3 |
| naa401 | 3.6256070 | 0.26621875 | 0.255 | 0.071 | 0.073226386 | 20 | naa40 |
| thrap3b | 3.8532415 | 0.370594596 | 0.383 | 0.136 | 0.07782392 | 20 | thrap3b |
| zgc:640516 | 3.8824013 | 0.341208254 | 0.468 | 0.174 | 0.07841286 | 20 | zgc:64051 |
| prps1a | 4.0131382 | 0.273606474 | 0.362 | 0.12 | 0.081053352 | 20 | prps1a |
| wu:fb55g094 | 5.8536211 | 0.553445574 | 0.596 | 0.309 | 0.118225586 | 20 | wu:fb55g09 |
| mob2b1 | 5.9827300 | 0.386464491 | 0.128 | 0.024 | 0.120833198 | 20 | mob2b |
| si:ch211-284f22.31 | 5.9855161 | 0.331731052 | 0.17 | 0.038 | 0.12088947 | 20 | si:ch211-284f22.3 |
| tfpia1 | 6.8461153 | 0.306749567 | 0.128 | 0.024 | 0.138270992 | 20 | tfpia |
| myg1 | 7.0063370 | 0.277965375 | 0.319 | 0.106 | 0.14150699 | 20 | myg1 |
| derl23 | 7.1214561 | 0.645646539 | 0.319 | 0.123 | 0.143832049 | 20 | derl2 |
| pdcd4a | 9.0377638 | 0.254370004 | 0.234 | 0.067 | 0.182535716 | 20 | pdcd4a |
| bcl2l102 | 1.0076464 | 0.485425794 | 0.362 | 0.14 | 0.203514355 | 20 | bcl2l10 |
| twistnb | 1.0270206 | 0.2889137 | 0.277 | 0.087 | 0.207427367 | 20 | twistnb |
| carhsp11 | 1.0771108 | 0.310292196 | 0.298 | 0.097 | 0.217544081 | 20 | carhsp1 |
| gspt1l | 1.1495646 | 0.279408564 | 0.34 | 0.119 | 0.232177571 | 20 | gspt1l |
| rgs23 | 1.1533931 | 0.389492808 | 0.17 | 0.04 | 0.232950823 | 20 | rgs2 |
| mcl1a5 | 1.1581612 | 0.509159985 | 0.809 | 0.543 | 0.233913829 | 20 | mcl1a |
| hectd3 | 1.1981836 | 0.258891278 | 0.191 | 0.048 | 0.24199715 | 20 | hectd3 |
| si:ch73-234b20.52 | 1.2320091 | 0.510857212 | 0.34 | 0.134 | 0.248828882 | 20 | si:ch73-234b20.5 |
| cct31 | 1.6960205 | 0.455526482 | 0.596 | 0.333 | 0.342545277 | 20 | cct3 |
| lpp | 1.7666432 | 0.381192867 | 0.191 | 0.051 | 0.356808939 | 20 | lpp |
| ncf1 | 1.8744236 | 0.285821325 | 0.149 | 0.033 | 0.378577342 | 20 | ncf1 |
| XLOC-0268363 | 1.9281772 | 0.475443536 | 0.255 | 0.083 | 0.389433966 | 20 | XLOC-026836 |
| si:ch211-14a17.11 | 2.0102088 | 0.288128106 | 0.298 | 0.1 | 0.406001875 | 20 | si:ch211-14a17.11 |
| metap2b | 2.4256355 | 0.38270887 | 0.553 | 0.261 | 0.489905617 | 20 | metap2b |
| si:ch211-203d1.3 | 2.6273943 | 0.261258537 | 0.106 | 0.019 | 0.530654846 | 20 | si:ch211-203d1.3 |
| eif3ba3 | 2.9983766 | 0.435107652 | 0.66 | 0.394 | 0.605582131 | 20 | eif3ba |
| si:cabz01074946.1 | 3.0426000 | 0.323986673 | 0.234 | 0.071 | 0.614513924 | 20 | si:cabz01074946.1 |
| cdc34b1 | 3.1248862 | 0.288267811 | 0.213 | 0.062 | 0.631133281 | 20 | cdc34b |
| fhl2a1 | 3.1575136 | 0.446625191 | 0.128 | 0.027 | 0.637723022 | 20 | fhl2a |
| sept9b1 | 3.2032748 | 0.304437015 | 0.191 | 0.052 | 0.646965424 | 20 | sept9b |

| | | | | | | | |
|---|---|---|---|---|---|---|---|
| btg12 | 3.7046212 | 0.531482697 | 0.723 | 0.463 | 0.748222356 | 20 | btg1 |
| MZT1 | 3.9147007 | 0.315365966 | 0.234 | 0.073 | 0.790652119 | 20 | MZT1 |
| dusp6 | 4.6264324 | 0.379078126 | 0.277 | 0.098 | 0.934400553 | 20 | dusp6 |
| ptpn2b | 4.6556931 | 0.270497239 | 0.191 | 0.053 | 0.940310337 | 20 | ptpn2b |
| kmt5aa2 | 4.7847851 | 0.366154938 | 0.383 | 0.157 | 0.966383064 | 20 | kmt5aa |
| RASSF52 | 5.2620715 | 0.438553759 | 0.234 | 0.077 | 1 | 20 | RASSF5 |
| hmga1a3 | 5.7375764 | 0.475397736 | 0.681 | 0.434 | 1 | 20 | hmga1a |
| fgl2a3 | 5.8200981 | 0.754944885 | 0.298 | 0.129 | 1 | 20 | fgl2a |
| bloc1s5 | 6.0054355 | 0.256551731 | 0.128 | 0.028 | 1 | 20 | bloc1s5 |
| msi2a2 | 6.4204322 | 0.288225223 | 0.191 | 0.055 | 1 | 20 | msi2a |
| nfyba2 | 6.7723243 | 0.348860215 | 0.489 | 0.227 | 1 | 20 | nfyba |
| pinx1 | 7.1949952 | 0.281581355 | 0.447 | 0.194 | 1 | 20 | pinx1 |
| sept63 | 7.2036796 | 0.296675873 | 0.447 | 0.189 | 1 | 20 | sept6 |
| pde7a2 | 7.5843799 | 0.263456293 | 0.277 | 0.094 | 1 | 20 | pde7a |
| ddx272 | 7.8958867 | 0.271869482 | 0.34 | 0.13 | 1 | 20 | ddx27 |
| ranbp11 | 8.0445459 | 0.35941494 | 0.617 | 0.323 | 1 | 20 | ranbp1 |
| XLOC-0139642 | 8.1692348 | 0.371190511 | 0.383 | 0.159 | 1 | 20 | XLOC-013964 |
| mgst1.24 | 9.6654747 | 0.363836201 | 0.404 | 0.179 | 1 | 20 | mgst1.2 |
| tab1 | 0.000102 | 0.255486193 | 0.277 | 0.097 | 1 | 20 | tab1 |
| actn13 | 0.000109 | 0.351827247 | 0.213 | 0.069 | 1 | 20 | actn1 |
| ldhba3 | 0.000127 | 0.396292892 | 0.596 | 0.337 | 1 | 20 | ldhba |
| si:ch211-105j21.91 | 0.000129 | 0.268492041 | 0.149 | 0.038 | 1 | 20 | si:ch211-105j21.9 |
| b3gnt3.41 | 0.00013 | 0.284767264 | 0.149 | 0.038 | 1 | 20 | b3gnt3.4 |
| rsrc21 | 0.000133 | 0.262435227 | 0.234 | 0.078 | 1 | 20 | rsrc2 |
| nop561 | 0.000136 | 0.411040996 | 0.511 | 0.281 | 1 | 20 | nop56 |
| dkc1 | 0.000138 | 0.332046816 | 0.404 | 0.184 | 1 | 20 | dkc1 |
| grcc10 | 0.000159 | 0.251508299 | 0.298 | 0.111 | 1 | 20 | grcc10 |
| atp5if1b4 | 0.00016 | 0.358088641 | 0.532 | 0.273 | 1 | 20 | atp5if1b |
| sesn14 | 0.000167 | 0.32449924 | 0.468 | 0.215 | 1 | 20 | sesn1 |
| ddx39ab1 | 0.000177 | 0.264412957 | 0.426 | 0.188 | 1 | 20 | ddx39ab |
| CABZ01079663.11 | 0.000202 | 0.254523857 | 0.128 | 0.031 | 1 | 20 | CABZ01079663.1 |
| arhgap20b1 | 0.000203 | 0.286613335 | 0.106 | 0.023 | 1 | 20 | arhgap20b |
| eif3k | 0.000205 | 0.353423925 | 0.574 | 0.328 | 1 | 20 | eif3k |
| snrpb5 | 0.00026 | 0.383661174 | 0.66 | 0.421 | 1 | 20 | snrpb |
| si:ch211-214p13.9 | 0.000297 | 0.370259718 | 0.128 | 0.033 | 1 | 20 | si:ch211-214p13.9 |

| | | | | | | | |
|---|---|---|---|---|---|---|---|
| hmgn23 | 0.000307 | 0.398908829 | 0.894 | 0.63 | 1 | 20 | hmgn2 |
| si:dkey-51e6.11 | 0.00037 | 0.277806355 | 0.319 | 0.132 | 1 | 20 | si:dkey-51e6.1 |
| cct83 | 0.000417 | 0.350926831 | 0.574 | 0.34 | 1 | 20 | cct8 |
| clic13 | 0.000432 | 0.30031072 | 0.447 | 0.215 | 1 | 20 | clic1 |
| bmp61 | 0.000457 | 0.255738834 | 0.106 | 0.025 | 1 | 20 | bmp6 |
| atp5mc3b2 | 0.000478 | 0.323244357 | 0.723 | 0.636 | 1 | 20 | atp5mc3b |
| tefa3 | 0.000492 | 0.281147168 | 0.34 | 0.149 | 1 | 20 | tefa |
| vdac22 | 0.000494 | 0.369887344 | 0.66 | 0.445 | 1 | 20 | vdac2 |
| mrpl361 | 0.000509 | 0.252061151 | 0.404 | 0.185 | 1 | 20 | mrpl36 |
| sumo3b | 0.000548 | 0.259373073 | 0.191 | 0.065 | 1 | 20 | sumo3b |
| twf1a2 | 0.000619 | 0.295309342 | 0.426 | 0.201 | 1 | 20 | twf1a |
| prdx25 | 0.00064 | 0.370297497 | 0.723 | 0.488 | 1 | 20 | prdx2 |
| cct4 | 0.000648 | 0.356522454 | 0.574 | 0.356 | 1 | 20 | cct4 |
| eif2s31 | 0.000732 | 0.338534485 | 0.681 | 0.434 | 1 | 20 | eif2s3 |
| gapdhs4 | 0.000761 | 0.402810591 | 0.702 | 0.476 | 1 | 20 | gapdhs |
| zeb2a | 0.000799 | 0.342525062 | 0.17 | 0.057 | 1 | 20 | zeb2a |
| si:ch211-196l7.45 | 0.000801 | 0.38234078 | 0.66 | 0.412 | 1 | 20 | si:ch211-196l7.4 |
| mibp21 | 0.000869 | 0.314551214 | 0.745 | 0.631 | 1 | 20 | mibp2 |
| ccnd2a | 0.000994 | 0.394661584 | 0.319 | 0.147 | 1 | 20 | ccnd2a |
| siah1 | 0.001047 | 0.345743647 | 0.149 | 0.048 | 1 | 20 | siah1 |
| ncl1 | 0.0011 | 0.377102376 | 0.596 | 0.418 | 1 | 20 | ncl |
| rnf441 | 0.00119 | 0.331339716 | 0.298 | 0.138 | 1 | 20 | rnf44 |
| si:dkey-261h17.14 | 0.001194 | 0.302859236 | 0.489 | 0.251 | 1 | 20 | si:dkey-261h17.1 |
| pdcd10b | 0.001209 | 0.267107858 | 0.234 | 0.093 | 1 | 20 | pdcd10b |
| ucp25 | 0.001302 | 0.340718865 | 0.66 | 0.424 | 1 | 20 | ucp2 |
| eif3g2 | 0.001457 | 0.315101932 | 0.638 | 0.396 | 1 | 20 | eif3g |
| dido12 | 0.001489 | 0.260653288 | 0.34 | 0.158 | 1 | 20 | dido1 |
| svila | 0.001574 | 0.283236417 | 0.128 | 0.038 | 1 | 20 | svila |
| tagln24 | 0.00174 | 0.341979187 | 0.574 | 0.338 | 1 | 20 | tagln2 |
| rhof1 | 0.001746 | 0.269889206 | 0.128 | 0.038 | 1 | 20 | rhof |
| pfdn4 | 0.001764 | 0.307215683 | 0.489 | 0.284 | 1 | 20 | pfdn4 |
| capza1b3 | 0.001807 | 0.343785633 | 0.574 | 0.364 | 1 | 20 | capza1b |
| CR753876.14 | 0.001848 | 0.422289405 | 0.234 | 0.102 | 1 | 20 | CR753876.1 |
| eif3ha | 0.001977 | 0.342633524 | 0.532 | 0.354 | 1 | 20 | eif3ha |
| mdh2 | 0.001998 | 0.307682791 | 0.596 | 0.373 | 1 | 20 | mdh2 |

| | | | | | | | | |
|---|---|---|---|---|---|---|---|---|
| si:ch211-260e23.9 | 0.00202 | 0.356032131 | 0.213 | 0.088 | | 1 | 20 | si:ch211-260e23.9 |
| arpc25 | 0.002147 | 0.301835381 | 0.766 | 0.483 | | 1 | 20 | arpc2 |
| ctgfa3 | 0.002211 | 0.582172045 | 0.17 | 0.066 | | 1 | 20 | ctgfa |
| cct51 | 0.002392 | 0.308675111 | 0.617 | 0.411 | | 1 | 20 | cct5 |
| mtap1 | 0.002842 | 0.275926634 | 0.255 | 0.116 | | 1 | 20 | mtap |
| ndfip13 | 0.002859 | 0.273787671 | 0.426 | 0.223 | | 1 | 20 | ndfip1 |
| atp2a32 | 0.003054 | 0.279976527 | 0.149 | 0.052 | | 1 | 20 | atp2a3 |
| eif5a22 | 0.003144 | 0.322513105 | 0.638 | 0.463 | | 1 | 20 | eif5a2 |
| nhp22 | 0.003365 | 0.282004156 | 0.426 | 0.243 | | 1 | 20 | nhp2 |
| celf24 | 0.003379 | 0.290248819 | 0.596 | 0.356 | | 1 | 20 | celf2 |
| eif3d1 | 0.003571 | 0.310379198 | 0.596 | 0.42 | | 1 | 20 | eif3d |
| zgc:1625092 | 0.003838 | 0.299328241 | 0.149 | 0.054 | | 1 | 20 | zgc:162509 |
| dynlrb13 | 0.004105 | 0.270587879 | 0.468 | 0.268 | | 1 | 20 | dynlrb1 |
| rpl22l13 | 0.004118 | 0.263006972 | 0.702 | 0.623 | | 1 | 20 | rpl22l1 |
| eif3f1 | 0.00423 | 0.267575532 | 0.766 | 0.646 | | 1 | 20 | eif3f |
| nupr1a1 | 0.00429 | 0.326336526 | 0.319 | 0.171 | | 1 | 20 | nupr1a |
| mob1a2 | 0.004339 | 0.284334626 | 0.404 | 0.218 | | 1 | 20 | mob1a |
| rfx5 | 0.004348 | 0.259558745 | 0.149 | 0.054 | | 1 | 20 | rfx5 |
| ola11 | 0.004936 | 0.292816632 | 0.511 | 0.329 | | 1 | 20 | ola1 |
| cox5aa4 | 0.004977 | 0.274051536 | 0.638 | 0.409 | | 1 | 20 | cox5aa |
| cct71 | 0.005059 | 0.276355884 | 0.532 | 0.339 | | 1 | 20 | cct7 |
| alox5a1 | 0.00516 | 0.279049978 | 0.106 | 0.033 | | 1 | 20 | alox5a |
| eif3m | 0.006433 | 0.268075601 | 0.66 | 0.524 | | 1 | 20 | eif3m |
| ak2 | 0.006679 | 0.288547768 | 0.553 | 0.38 | | 1 | 20 | ak2 |
| nap1l13 | 0.006818 | 0.282281959 | 0.553 | 0.359 | | 1 | 20 | nap1l1 |
| usf1l1 | 0.007043 | 0.253644391 | 0.234 | 0.11 | | 1 | 20 | usf1l |
| fam53b | 0.007165 | 0.268942125 | 0.213 | 0.097 | | 1 | 20 | fam53b |
| sept23 | 0.008704 | 0.275871876 | 0.468 | 0.284 | | 1 | 20 | sept2 |
| myl12.13 | 0.009031 | 0.257157207 | 0.851 | 0.629 | | 1 | 20 | myl12.1 |
| tubb4b2 | 0.009631 | 0.254721939 | 0.83 | 0.67 | | 1 | 20 | tubb4b |
| prdx51 | 0.009668 | 0.25415579 | 0.468 | 0.292 | | 1 | 20 | prdx5 |
| btg25 | 0.009744 | 0.251823573 | 0.447 | 0.26 | | 1 | 20 | btg2 |
| ccl33.31 | 3.2380845 | 2.443729146 | 1 | 0.08 | 6.53995929763437e-8 | | 21 | ccl33.3 |
| si:ch211-220f21.31 | 3.4038536 | 1.123437136 | 0.525 | 0.023 | 6.87476321842307e-6 | | 21 | si:ch211-220f21.3 |
| lcp2b2 | 1.0806878 | 1.035196709 | 0.5 | 0.022 | 2.18266519918896e-6 | | 21 | lcp2b |

| | | | | | | | |
|---|---|---|---|---|---|---|---|
| sftpba1 | 1.8737400 | 1.428119416 | 0.775 | 0.062 | 3.78439276592287e-6 | 21 | sftpba |
| vaspa4 | 6.5848394 | 1.571205512 | 0.9 | 0.108 | 1.32994002597599e-5 | 21 | vaspa |
| LOC1081905651 | 1.2561837 | 0.734587601 | 0.35 | 0.016 | 2.53711433732627e-4 | 21 | LOC108190565 |
| eomesa2 | 1.7709129 | 1.042284111 | 0.5 | 0.037 | 3.57671295153614e-3 | 21 | eomesa |
| si:ch211-102c2.46 | 3.3218358 | 1.463801467 | 0.925 | 0.166 | 6.70911188294655e-3 | 21 | si:ch211-102c2.4 |
| ckbb3 | 2.8754436 | 1.464596015 | 0.925 | 0.181 | 5.80753353110036e-3 | 21 | ckbb |
| abi3a3 | 4.6716589 | 1.033114009 | 0.575 | 0.06 | 9.43534957286168e-3 | 21 | abi3a |
| fcer1g1 | 1.4197984 | 0.966444566 | 0.5 | 0.044 | 2.86756693717318e-3 | 21 | fcer1g |
| LOC1005362134 | 2.4379983 | 1.27188882 | 0.85 | 0.139 | 4.92402519017186e-3 | 21 | LOC100536213 |
| fcer1gl5 | 2.8393028 | 1.498343309 | 1 | 0.248 | 5.73453995853558e-3 | 21 | fcer1gl |
| mef2b2 | 3.6648288 | 0.765507468 | 0.375 | 0.025 | 7.40185491663498e-3 | 21 | mef2b |
| fam49ba3 | 1.1664218 | 1.254364417 | 0.725 | 0.116 | 2.35582213975364e-3 | 21 | fam49ba |
| dicp1.11 | 8.2044422 | 0.707372001 | 0.325 | 0.021 | 1.65705120784928e-2 | 21 | dicp1.1 |
| si:ch211-215e19.4 | 1.3395628 | 0.522439781 | 0.225 | 0.009 | 2.70551504926883e-2 | 21 | si:ch211-215e19.4 |
| hcls15 | 2.8856848 | 1.101054273 | 0.675 | 0.104 | 5.82821777703927e-2 | 21 | hcls1 |
| wipf1a3 | 6.2325539 | 1.068230572 | 0.575 | 0.078 | 1.25878892430833e-2 | 21 | wipf1a |
| CR381673.12 | 1.6786595 | 0.951316132 | 0.475 | 0.053 | 3.39038863063046e-2 | 21 | CR381673.1 |
| pfn17 | 1.8209359 | 1.301291574 | 1 | 0.413 | 3.67774428159301e-2 | 21 | pfn1 |
| coro1a6 | 2.7201227 | 1.307687739 | 0.925 | 0.274 | 5.49383196166301e-2 | 21 | coro1a |
| si:dkey-33i11.92 | 7.6487745 | 0.796165183 | 0.4 | 0.038 | 1.54482300031591e-2 | 21 | si:dkey-33i11.9 |
| ptpn65 | 8.9923115 | 1.183373824 | 0.725 | 0.138 | 1.8161771549163e-23 | 21 | ptpn6 |
| FO393424.2 | 3.6739114 | 0.421079221 | 0.175 | 0.006 | 7.42019903688002e-2 | 21 | FO393424.2 |
| arpc1b7 | 4.9220457 | 1.27796153 | 0.975 | 0.354 | 9.9410558704933e-23 | 21 | arpc1b |
| si:ch73-264p11.12 | 6.1285290 | 0.668008882 | 0.35 | 0.03 | 1.23777901234804e-2 | 21 | si:ch73-264p11.1 |
| washc2c1 | 7.2770461 | 0.859796422 | 0.45 | 0.055 | 1.46974501039376e-2 | 21 | washc2c |
| CR753876.15 | 1.1469344 | 1.050610414 | 0.6 | 0.095 | 2.31646342207847e-2 | 21 | CR753876.1 |
| itga11 | 2.0805507 | 0.905258617 | 0.45 | 0.056 | 4.20208830148286e-2 | 21 | itga1 |
| cd74a3 | 1.1755887 | 1.166084757 | 0.925 | 0.202 | 2.37433666775001e-2 | 21 | cd74a |
| lcp12 | 4.2810168 | 0.851442526 | 0.475 | 0.065 | 8.64636970898665e-2 | 21 | lcp1 |
| cd74b3 | 1.6012326 | 1.179977426 | 0.95 | 0.241 | 3.23400962133232e-1 | 21 | cd74b |
| si:ch211-188c18.1 | 3.9886447 | 0.286139874 | 0.1 | 0.002 | 8.05586578425504e-1 | 21 | si:ch211-188c18.1 |
| actb23 | 1.1092920 | 0.5753307 | 1 | 0.926 | 2.24043714090113e-1 | 21 | actb2 |
| cotl16 | 4.0551012 | 1.123773853 | 0.925 | 0.407 | 8.19008800870647e-1 | 21 | cotl1 |
| actb16 | 6.8288148 | 0.789123842 | 1 | 0.739 | 1.37921573803192e-1 | 21 | actb1 |
| si:ch73-380l3.41 | 7.0323340 | 0.656451977 | 0.3 | 0.029 | 1.42032051569585e-1 | 21 | si:ch73-380l3.4 |

| | | | | | | | |
|---|---|---|---|---|---|---|---|
| TSTA3 (1 of many) | 9.6681933 | 0.390176662 | 0.175 | 0.009 | 1.95268501535825e-1 | 21 | TSTA3 (1 of many) |
| tmsb4x3 | 1.0560189 | 0.426549915 | 1 | 0.941 | 2.13284137921545e-1 | 21 | tmsb4x |
| arpc5b3 | 1.2831308 | 1.142368527 | 0.75 | 0.225 | 2.59153941216643e-1 | 21 | arpc5b |
| itgb23 | 9.7688020 | 1.042568583 | 0.675 | 0.159 | 1.97300494692006e-1 | 21 | itgb2 |
| arpc26 | 5.1060190 | 0.982224229 | 0.925 | 0.481 | 1.03126266437937e-1 | 21 | arpc2 |
| ephx21 | 7.0954104 | 0.699997088 | 0.375 | 0.051 | 1.43306004264237e-1 | 21 | ephx2 |
| grap2b6 | 7.6228287 | 0.995530026 | 0.725 | 0.187 | 1.53958271967646e-1 | 21 | grap2b |
| CT573231.1 | 4.3118552 | 0.645635711 | 0.275 | 0.029 | 8.70865399524089e-1 | 21 | CT573231.1 |
| rac26 | 2.9714258 | 1.010546389 | 0.8 | 0.256 | 6.00138874462787e-1 | 21 | rac2 |
| si:ch211-214p16.1 | 4.1454570 | 1.09063765 | 0.75 | 0.186 | 8.37257967283082e-1 | 21 | si:ch211-214p16.1 |
| arpc4l5 | 7.2743897 | 0.98653527 | 0.825 | 0.345 | 1.46920850665605e-1 | 21 | arpc4l |
| spi1b4 | 2.2910644 | 0.899339246 | 0.55 | 0.123 | 4.62726291864055e-1 | 21 | spi1b |
| wasa5 | 4.6156126 | 0.834022918 | 0.525 | 0.114 | 9.32215286085659e-1 | 21 | wasa |
| LOC1018828051 | 7.5238822 | 0.373283112 | 0.15 | 0.009 | 1.51959850361353e-1 | 21 | LOC101882805 |
| ccl35.13 | 7.7461612 | 0.826867306 | 0.475 | 0.088 | 1.56449218994427e-1 | 21 | ccl35.1 |
| arpc45 | 2.1351405 | 0.994427258 | 0.775 | 0.312 | 4.31234334711058e-1 | 21 | arpc4 |
| si:dkey-12f6.51 | 2.2972763 | 0.261466033 | 0.1 | 0.004 | 4.63980909171798e-1 | 21 | si:dkey-12f6.5 |
| LOC108181220 | 2.3060985 | 0.364774911 | 0.15 | 0.01 | 4.6576271806483e-11 | 21 | LOC108181220 |
| arpc36 | 3.6695345 | 0.893898957 | 0.875 | 0.477 | 7.41135891271753e-1 | 21 | arpc3 |
| LOC100535743 | 5.0552372 | 0.467412022 | 0.2 | 0.018 | 1.02100627549508e-1 | 21 | LOC100535743 |
| si:ch211-214k5.3 | 5.9389089 | 0.543855957 | 0.25 | 0.029 | 1.19948144467375e-1 | 21 | si:ch211-214k5.3 |
| rnaset2l4 | 1.2068731 | 0.720335023 | 0.425 | 0.08 | 2.43752167421421e-1 | 21 | rnaset2l |
| ppp1r183 | 1.2338159 | 0.701936051 | 0.375 | 0.065 | 2.4919380293144e-10 | 21 | ppp1r18 |
| XLOC-0268364 | 2.0589763 | 0.742394778 | 0.425 | 0.08 | 4.1585145470718e-10 | 21 | XLOC-026836 |
| cfl15 | 7.3827021 | 0.579181911 | 0.975 | 0.765 | 1.49108435294927e-0 | 21 | cfl1 |
| si:dkey-172o19.21 | 9.9531879 | 0.392575374 | 0.175 | 0.015 | 2.0102453674623e-09 | 21 | si:dkey-172o19.2 |
| mapre1a5 | 1.1127144 | 0.815216152 | 0.65 | 0.195 | 2.2473493386025e-09 | 21 | mapre1a |
| ptmaa4 | 3.3927187 | 0.668978169 | 0.95 | 0.653 | 6.85227415991678e-0 | 21 | ptmaa |
| gm2a2 | 3.5566376 | 0.831403057 | 0.5 | 0.128 | 7.18334113202589e-0 | 21 | gm2a |
| CABZ01046427.2 | 3.8731895 | 0.358185297 | 0.15 | 0.012 | 7.82268097130528e-0 | 21 | CABZ01046427.2 |
| pxna | 4.0833991 | 0.706472999 | 0.4 | 0.084 | 8.24724129523436e-0 | 21 | pxna |
| si:ch211-214p16.2 | 5.9194561 | 0.783665406 | 0.425 | 0.088 | 1.19555255233628e-0 | 21 | si:ch211-214p16.2 |
| skap23 | 6.1075378 | 0.684044441 | 0.475 | 0.112 | 1.23353941347762e-0 | 21 | skap2 |
| sftpbb2 | 6.4803490 | 0.76277749 | 0.425 | 0.091 | 1.30883610017817e-0 | 21 | sftpbb |
| kctd12.22 | 9.9279068 | 0.860667342 | 0.625 | 0.207 | 2.00513935328019e-0 | 21 | kctd12.2 |

| | | | | | | | |
|---|---|---|---|---|---|---|---|
| BX682234.1 | 1.2780755 | 0.323413549 | 0.15 | 0.012 | 2.58132915500158e-0 | 21 | BX682234.1 |
| rgs132 | 3.8708017 | 0.769060309 | 0.425 | 0.098 | 7.81785837842821e-0 | 21 | rgs13 |
| zgc:1037002 | 4.9091953 | 0.769861853 | 0.5 | 0.124 | 9.91510184101081e-0 | 21 | zgc:103700 |
| tmc6a | 6.8537190 | 0.345409304 | 0.15 | 0.013 | 1.38424562788387e-0 | 21 | tmc6a |
| syk4 | 7.4401621 | 0.580826745 | 0.375 | 0.077 | 1.50268955816746e-0 | 21 | syk |
| myl12.14 | 7.6425446 | 0.663151472 | 0.875 | 0.629 | 1.54356473537379e-0 | 21 | myl12.1 |
| sla11 | 7.9074577 | 0.502565089 | 0.25 | 0.037 | 1.59706924973567e-0 | 21 | sla1 |
| gpr132b | 1.0106981 | 0.503368423 | 0.25 | 0.036 | 2.04130704326618e-0 | 21 | gpr132b |
| gpam | 1.4304234 | 0.277648007 | 0.1 | 0.006 | 2.88902627043922e-0 | 21 | gpam |
| zgc:152891 | 3.6903315 | 0.324813911 | 0.15 | 0.014 | 7.45336264604173e-0 | 21 | zgc:152891 |
| h1fx4 | 5.3443712 | 0.871119126 | 0.65 | 0.234 | 1.07940266053046e-0 | 21 | h1fx |
| mcl1a6 | 6.6661731 | 0.713661694 | 0.9 | 0.542 | 1.34636699370462e-0 | 21 | mcl1a |
| si:dkey-222f2.72 | 1.5487359 | 0.591761732 | 0.25 | 0.041 | 3.12798208658442e-0 | 21 | si:dkey-222f2.7 |
| CU972454.4 | 1.6445926 | 0.389316695 | 0.15 | 0.015 | 3.32158385883131e-0 | 21 | CU972454.4 |
| rplp22 | 2.4298295 | 0.250017956 | 0.975 | 0.928 | 4.90752673943722e-0 | 21 | rplp2 |
| mdkb2 | 2.4973167 | 0.499003342 | 0.225 | 0.034 | 5.04383061920274e-0 | 21 | mdkb |
| yrk4 | 2.9700796 | 0.653387951 | 0.375 | 0.089 | 5.99866990889035e-0 | 21 | yrk |
| SLA (1 of many) | 3.1026363 | 0.257426397 | 0.1 | 0.007 | 6.26639464956915e-0 | 21 | SLA (1 of many) |
| arhgdig5 | 3.5046795 | 0.79387529 | 0.725 | 0.363 | 7.07840129877236e-0 | 21 | arhgdig |
| CABZ01046432.1 | 4.8886215 | 0.358299201 | 0.15 | 0.016 | 9.87354899981694e-0 | 21 | CABZ01046432.1 |
| si:dkey-9i23.41 | 7.3056977 | 0.368982124 | 0.125 | 0.011 | 1.47553177182327e-0 | 21 | si:dkey-9i23.4 |
| ctss12 | 1.0335228 | 0.411160247 | 0.175 | 0.023 | 2.08740617141836e-0 | 21 | ctss1 |
| LOC101884267 | 1.7808473 | 0.480233056 | 0.15 | 0.017 | 3.59677740189949e-0 | 21 | LOC101884267 |
| grasp1 | 2.5847020 | 0.579854049 | 0.275 | 0.056 | 5.220322737218e-05 | 21 | grasp |
| si:ch73-27e22.7 | 3.2226392 | 0.322213003 | 0.125 | 0.012 | 6.50876446653247e-0 | 21 | si:ch73-27e22.7 |
| si:busm1-266f07.2 | 6.8449812 | 0.661895444 | 0.525 | 0.157 | 0.000138248 | 21 | si:busm1-266f07.2 |
| dok22 | 7.1742483 | 0.505542039 | 0.25 | 0.048 | 0.000144898 | 21 | dok2 |
| cox173 | 7.2170430 | 0.626377327 | 0.525 | 0.184 | 0.000145763 | 21 | cox17 |
| npsn | 7.2809659 | 0.337495196 | 0.2 | 0.03 | 0.000147054 | 21 | npsn |
| capzb4 | 7.7734409 | 0.703407408 | 0.7 | 0.384 | 0.000157 | 21 | capzb |
| laptm57 | 1.0420277 | 0.678830995 | 0.6 | 0.213 | 0.000210458 | 21 | laptm5 |
| cfbl2 | 1.5231543 | 0.552472942 | 0.375 | 0.094 | 0.000307631 | 21 | cfbl |
| capza1b4 | 1.5800128 | 0.719965471 | 0.675 | 0.362 | 0.000319115 | 21 | capza1b |
| cap16 | 1.8763611 | 0.728716268 | 0.675 | 0.328 | 0.000378969 | 21 | cap1 |
| zgc:640517 | 2.1039473 | 0.701584446 | 0.5 | 0.175 | 0.000424934 | 21 | zgc:64051 |

| | | | | | | | |
|---|---|---|---|---|---|---|---|
| cdkn1bb | 2.5523135 | 0.620721259 | 0.475 | 0.165 | 0.000515491 | 21 | cdkn1bb |
| parvg2 | 2.6996869 | 0.5676101 | 0.375 | 0.106 | 0.000545256 | 21 | parvg |
| rhogb4 | 2.9223630 | 0.690314617 | 0.475 | 0.168 | 0.00059023 | 21 | rhogb |
| wdr15 | 3.7856526 | 0.734423103 | 0.55 | 0.235 | 0.000764588 | 21 | wdr1 |
| si:ch211-198c19.1 | 4.5532947 | 0.276433104 | 0.125 | 0.014 | 0.000919629 | 21 | si:ch211-198c19.1 |
| rap1gap2a1 | 4.8419796 | 0.282861601 | 0.125 | 0.014 | 0.000977935 | 21 | rap1gap2a |
| LOC110438754 | 7.1894165 | 0.299147372 | 0.125 | 0.015 | 0.001452046 | 21 | LOC110438754 |
| XLOC-0183561 | 1.4672182 | 0.437493189 | 0.2 | 0.036 | 0.002963341 | 21 | XLOC-018356 |
| cnn2 | 2.3643974 | 0.63267296 | 0.6 | 0.282 | 0.004775374 | 21 | cnn2 |
| csk3 | 2.8170082 | 0.534099599 | 0.375 | 0.118 | 0.005689512 | 21 | csk |
| stk17b3 | 3.4511357 | 0.617438045 | 0.4 | 0.137 | 0.006970259 | 21 | stk17b |
| abracl4 | 4.1893123 | 0.702552687 | 0.525 | 0.24 | 0.008461154 | 21 | abracl |
| zgc:1627305 | 5.1984660 | 0.582382283 | 0.925 | 0.488 | 0.010499342 | 21 | zgc:162730 |
| tln13 | 7.1248094 | 0.659607257 | 0.5 | 0.223 | 0.014389978 | 21 | tln1 |
| idh21 | 8.4538437 | 0.651792418 | 0.65 | 0.382 | 0.017074228 | 21 | idh2 |
| mob1a3 | 1.0683418 | 0.593927786 | 0.5 | 0.217 | 0.021577301 | 21 | mob1a |
| AL954142.14 | 1.3603826 | 0.327609029 | 0.175 | 0.032 | 0.027475648 | 21 | AL954142.1 |
| mob3a2 | 1.4543234 | 0.607984188 | 0.45 | 0.188 | 0.029372971 | 21 | mob3a |
| ccr9a4 | 1.4581260 | 0.538843748 | 0.325 | 0.097 | 0.029449772 | 21 | ccr9a |
| zgc:1538673 | 1.5251353 | 0.426461611 | 0.775 | 0.64 | 0.030803158 | 21 | zgc:153867 |
| nptna1 | 2.8818664 | 0.403976825 | 0.225 | 0.054 | 0.058205056 | 21 | nptna |
| eno32 | 3.4503657 | 0.627290006 | 0.575 | 0.294 | 0.069687037 | 21 | eno3 |
| ipcef11 | 3.5286192 | 0.308366722 | 0.175 | 0.035 | 0.071267522 | 21 | ipcef1 |
| CR855389.11 | 4.1834459 | 0.353189557 | 0.175 | 0.036 | 0.084493057 | 21 | CR855389.1 |
| si:ch73-380l3.2 | 5.4578715 | 0.452341786 | 0.225 | 0.058 | 0.110232632 | 21 | si:ch73-380l3.2 |
| btk2 | 5.6716514 | 0.367472313 | 0.2 | 0.046 | 0.114550344 | 21 | btk |
| fhit | 7.0551307 | 0.397447243 | 0.225 | 0.058 | 0.142492476 | 21 | fhit |
| twf1a3 | 7.5713040 | 0.563320577 | 0.45 | 0.201 | 0.152917628 | 21 | twf1a |
| fybb3 | 7.6576031 | 0.45151473 | 0.225 | 0.059 | 0.15466061 | 21 | fybb |
| clic14 | 8.1584286 | 0.535942879 | 0.475 | 0.215 | 0.164775783 | 21 | clic1 |
| glipr1a4 | 9.2321919 | 0.422864067 | 0.3 | 0.093 | 0.186462582 | 21 | glipr1a |
| si:ch211-93g21.11 | 1.0268574 | 0.256931999 | 0.125 | 0.021 | 0.207394404 | 21 | si:ch211-93g21.1 |
| zdhhc23a | 1.5302809 | 0.255523693 | 0.1 | 0.014 | 0.309070839 | 21 | zdhhc23a |
| cdc42l3 | 2.1975477 | 0.528164961 | 0.6 | 0.345 | 0.443838728 | 21 | cdc42l |
| ppp2cab | 2.2749646 | 0.402082306 | 0.175 | 0.041 | 0.459474603 | 21 | ppp2cab |

| | | | | | | | |
|---|---|---|---|---|---|---|---|
| wdr191 | 2.4745708 | 0.390682831 | 0.15 | 0.031 | 0.499789065 | 21 | wdr19 |
| mhc2dab2 | 3.4822423 | 0.439762548 | 0.275 | 0.086 | 0.703308486 | 21 | mhc2dab |
| RASSF53 | 3.5281116 | 0.425515474 | 0.25 | 0.077 | 0.712572702 | 21 | RASSF5 |
| mibp22 | 4.2989354 | 0.420862945 | 0.8 | 0.63 | 0.868255988 | 21 | mibp2 |
| ptprc4 | 4.4723308 | 0.49747364 | 0.325 | 0.119 | 0.903276663 | 21 | ptprc |
| stk17al2 | 4.7057775 | 0.358635089 | 0.2 | 0.054 | 0.95042589 | 21 | stk17al |
| lgals9l12 | 6.1324924 | 0.273283235 | 0.2 | 0.052 | 1 | 21 | lgals9l1 |
| BX120005.12 | 6.5144864 | 0.411808134 | 0.15 | 0.034 | 1 | 21 | BX120005.1 |
| pcmtd1 | 7.6553377 | 0.384457997 | 0.2 | 0.057 | 1 | 21 | pcmtd1 |
| cox6a22 | 7.7121789 | 0.280061956 | 0.175 | 0.044 | 1 | 21 | cox6a2 |
| CU207241.12 | 9.8571128 | 0.255043944 | 0.125 | 0.025 | 1 | 21 | CU207241.1 |
| srgn4 | 0.0001 | 0.500798554 | 0.525 | 0.245 | 1 | 21 | srgn |
| dtnbp1a2 | 0.000104 | 0.461473319 | 0.275 | 0.099 | 1 | 21 | dtnbp1a |
| XLOC-0320022 | 0.000105 | 0.330039291 | 0.15 | 0.034 | 1 | 21 | XLOC-032002 |
| XLOC-0254235 | 0.000117 | 0.537092277 | 0.425 | 0.191 | 1 | 21 | XLOC-025423 |
| lasp14 | 0.000127 | 0.480791508 | 0.575 | 0.358 | 1 | 21 | lasp1 |
| ikzf23 | 0.000143 | 0.319271603 | 0.2 | 0.057 | 1 | 21 | ikzf2 |
| XLOC-0418702 | 0.000153 | 0.373124768 | 0.175 | 0.047 | 1 | 21 | XLOC-041870 |
| pnrc21 | 0.000164 | 0.293977406 | 0.85 | 0.727 | 1 | 21 | pnrc2 |
| cryl1 | 0.000184 | 0.323790743 | 0.175 | 0.048 | 1 | 21 | cryl1 |
| vamp8 | 0.000217 | 0.367629382 | 0.25 | 0.088 | 1 | 21 | vamp8 |
| sh3bgrl4 | 0.000244 | 0.479664996 | 0.425 | 0.214 | 1 | 21 | sh3bgrl |
| tradd1 | 0.000278 | 0.347077426 | 0.2 | 0.063 | 1 | 21 | tradd |
| ctss2.13 | 0.0003 | 0.395107334 | 0.225 | 0.074 | 1 | 21 | ctss2.1 |
| LOC1039091156 | 0.000327 | 0.490910133 | 0.4 | 0.2 | 1 | 21 | LOC103909115 |
| si:ch211-66e2.34 | 0.000328 | 0.326045563 | 0.25 | 0.085 | 1 | 21 | si:ch211-66e2.3 |
| FERMT3 (1 of many) | 0.000406 | 0.384223412 | 0.2 | 0.065 | 1 | 21 | FERMT3 (1 of many) |
| actr2a4 | 0.000427 | 0.467914262 | 0.35 | 0.163 | 1 | 21 | actr2a |
| lrrfip1a1 | 0.00049 | 0.409038577 | 0.225 | 0.08 | 1 | 21 | lrrfip1a |
| sh3bgrl3 | 0.000502 | 0.48483651 | 0.4 | 0.213 | 1 | 21 | sh3bgrl3 |
| rps6ka12 | 0.000612 | 0.312458847 | 0.175 | 0.053 | 1 | 21 | rps6ka1 |
| si:ch211-277c7.62 | 0.000711 | 0.274942293 | 0.125 | 0.03 | 1 | 21 | si:ch211-277c7.6 |
| itm2bb4 | 0.000785 | 0.410559044 | 0.625 | 0.356 | 1 | 21 | itm2bb |
| arhgdia4 | 0.000826 | 0.455153098 | 0.55 | 0.375 | 1 | 21 | arhgdia |
| cnpy3 | 0.000949 | 0.363715289 | 0.225 | 0.085 | 1 | 21 | cnpy3 |

| | | | | | | | |
|---|---|---|---|---|---|---|---|
| ccdc28a | 0.001073 | 0.396370204 | 0.25 | 0.103 | 1 | 21 | ccdc28a |
| atp6v1e1b2 | 0.001206 | 0.390517901 | 0.3 | 0.134 | 1 | 21 | atp6v1e1b |
| crlf32 | 0.001207 | 0.428931095 | 0.25 | 0.106 | 1 | 21 | crlf3 |
| ak12 | 0.001266 | 0.378662143 | 0.325 | 0.153 | 1 | 21 | ak1 |
| si:busm1-194e12.1 | 0.001554 | 0.296639711 | 0.2 | 0.069 | 1 | 21 | si:busm1-194e12.11 |
| alox5a2 | 0.001559 | 0.259323415 | 0.125 | 0.033 | 1 | 21 | alox5a |
| ucp26 | 0.002623 | 0.41385008 | 0.6 | 0.426 | 1 | 21 | ucp2 |
| ccni1 | 0.002707 | 0.371687108 | 0.575 | 0.416 | 1 | 21 | ccni |
| myo1f3 | 0.002841 | 0.324051848 | 0.175 | 0.062 | 1 | 21 | myo1f |
| gapdhs5 | 0.002866 | 0.397158824 | 0.625 | 0.478 | 1 | 21 | gapdhs |
| fam210b | 0.003196 | 0.303612991 | 0.15 | 0.05 | 1 | 21 | fam210b |
| si:rp71-68n21.91 | 0.003215 | 0.28141725 | 0.175 | 0.063 | 1 | 21 | si:rp71-68n21.9 |
| limd23 | 0.003733 | 0.295755858 | 0.175 | 0.063 | 1 | 21 | limd2 |
| tpm32 | 0.003912 | 0.342284604 | 0.625 | 0.46 | 1 | 21 | tpm3 |
| CR318588.14 | 0.004078 | 0.369581246 | 0.5 | 0.267 | 1 | 21 | CR318588.1 |
| eif4ebp14 | 0.004228 | 0.357778703 | 0.525 | 0.343 | 1 | 21 | eif4ebp1 |
| mcu | 0.004393 | 0.306977318 | 0.125 | 0.039 | 1 | 21 | mcu |
| atp5md2 | 0.004577 | 0.387290243 | 0.525 | 0.354 | 1 | 21 | atp5md |
| mhc2b2 | 0.004615 | 0.384125588 | 0.225 | 0.095 | 1 | 21 | mhc2b |
| rasal32 | 0.00533 | 0.265367324 | 0.1 | 0.028 | 1 | 21 | rasal3 |
| pkma4 | 0.006682 | 0.341385745 | 0.35 | 0.198 | 1 | 21 | pkma |
| eif4ebp22 | 0.006828 | 0.354773472 | 0.45 | 0.295 | 1 | 21 | eif4ebp2 |
| ypel5 | 0.007064 | 0.286027105 | 0.2 | 0.086 | 1 | 21 | ypel5 |
| cnp1 | 0.007274 | 0.38952052 | 0.375 | 0.232 | 1 | 21 | cnp |
| BX005340.1 | 0.007337 | 0.287347047 | 0.175 | 0.07 | 1 | 21 | BX005340.1 |
| si:ch1073-443f11.2 | 0.007382 | 0.276644524 | 0.175 | 0.069 | 1 | 21 | si:ch1073-443f11.2 |
| sat1a.26 | 0.008049 | 0.367212538 | 0.475 | 0.337 | 1 | 21 | sat1a.2 |
| si:zfos-741a10.33 | 0.009864 | 0.264075146 | 0.15 | 0.056 | 1 | 21 | si:zfos-741a10.3 |
| XLOC-0183562 | 1.5132332 | 1.851541534 | 0.875 | 0.023 | 3.05627721840742e-1 | 22 | XLOC-018356 |
| gpr132b1 | 7.2852818 | 1.55957537 | 0.85 | 0.025 | 1.47140836829078e-1 | 22 | gpr132b |
| BX120005.13 | 5.8751725 | 1.339258061 | 0.7 | 0.023 | 1.18660859800796e-1 | 22 | BX120005.1 |
| CABZ01046427.21 | 5.9140051 | 0.846963227 | 0.425 | 0.006 | 1.19445162222134e-1 | 22 | CABZ01046427.2 |
| LOC1005357431 | 9.0944429 | 1.037604393 | 0.5 | 0.012 | 1.83680464937645e-9 | 22 | LOC100535743 |
| si:ch73-380l3.42 | 4.3032823 | 1.203305205 | 0.625 | 0.023 | 8.69133928453416e-9 | 22 | si:ch73-380l3.4 |
| ccl33.32 | 6.6045445 | 2.473434869 | 1 | 0.08 | 1.33391985346101e-8 | 22 | ccl33.3 |

| | | | | | | | |
|---|---|---|---|---|---|---|---|
| CT573231.11 | 3.0050769 | 1.039427299 | 0.575 | 0.023 | 6.06935391062412e-7 | 22 | CT573231.1 |
| CU855821.1 | 1.4079146 | 0.781537248 | 0.35 | 0.006 | 2.84356528546809e-7 | 22 | CU855821.1 |
| LOC1018842671 | 2.1994907 | 0.889913141 | 0.425 | 0.012 | 4.44231151615771e-7 | 22 | LOC101884267 |
| eomesa3 | 6.1677848 | 1.188814444 | 0.65 | 0.034 | 1.24570750282026e-6 | 22 | eomesa |
| sftpba2 | 3.4876385 | 1.482068517 | 0.825 | 0.061 | 7.0439834889336e-69 | 22 | sftpba |
| LOC1081905652 | 8.6307464 | 0.844136247 | 0.425 | 0.014 | 1.74315186810531e-6 | 22 | LOC108190565 |
| XLOC-0268365 | 5.4385856 | 1.529725435 | 0.825 | 0.072 | 1.09843115164e-62 | 22 | XLOC-026836 |
| si:ch211-215e19.41 | 3.2538802 | 0.6801274 | 0.325 | 0.007 | 6.57186191949861e-6 | 22 | si:ch211-215e19.4 |
| si:ch211-277c7.63 | 8.1868430 | 1.138394589 | 0.5 | 0.023 | 1.65349669469308e-6 | 22 | si:ch211-277c7.6 |
| ccl35.14 | 1.5433539 | 1.546395024 | 0.85 | 0.081 | 3.11711202491871e-5 | 22 | ccl35.1 |
| pou2f2a | 2.4332907 | 0.719931995 | 0.35 | 0.01 | 4.91451728006183e-5 | 22 | pou2f2a |
| CR753876.16 | 6.4680298 | 1.466153501 | 0.875 | 0.09 | 1.30634799584461e-5 | 22 | CR753876.1 |
| fcer1g2 | 2.2451874 | 1.156513163 | 0.625 | 0.042 | 4.53460505049851e-5 | 22 | fcer1g |
| LOC1104388701 | 1.4395485 | 0.655693889 | 0.3 | 0.007 | 2.90745624024459e-5 | 22 | LOC110438870 |
| CU972454.41 | 5.6536227 | 0.7872556 | 0.35 | 0.011 | 1.1418621924875e-53 | 22 | CU972454.4 |
| CABZ01046425.1 | 2.0800467 | 0.488514348 | 0.2 | 0.002 | 4.20107033601379e-5 | 22 | CABZ01046425.1 |
| scdb | 8.5554047 | 0.644990441 | 0.3 | 0.008 | 1.72793510685509e-5 | 22 | scdb |
| si:cabz01036006.1 | 6.3701069 | 0.711727468 | 0.325 | 0.011 | 1.28657051048127e-4 | 22 | si:cabz01036006.1 |
| si:ch73-27e22.71 | 1.5480538 | 0.684799054 | 0.3 | 0.009 | 3.12660429606587e-4 | 22 | si:ch73-27e22.7 |
| nr4a15 | 9.7158361 | 1.436908343 | 0.875 | 0.115 | 1.96230743129314e-4 | 22 | nr4a1 |
| si:ch211-220f21.32 | 5.0810526 | 0.977532317 | 0.45 | 0.025 | 1.02622020130641e-4 | 22 | si:ch211-220f21.3 |
| si:ch211-214k5.31 | 6.1451501 | 0.796195563 | 0.45 | 0.025 | 1.24113597742327e-4 | 22 | si:ch211-214k5.3 |
| dicp1.12 | 9.7590614 | 0.823622355 | 0.4 | 0.02 | 1.97103764173402e-4 | 22 | dicp1.1 |
| si:dkey-172o19.22 | 7.8724433 | 0.645168738 | 0.325 | 0.012 | 1.58999739080157e-4 | 22 | si:dkey-172o19.2 |
| tagapb4 | 9.5840753 | 1.196251619 | 0.7 | 0.076 | 1.93569569541017e-4 | 22 | tagapb |
| si:ch211-102c2.47 | 3.3509250 | 1.541706669 | 0.95 | 0.165 | 6.76786339776693e-4 | 22 | si:ch211-102c2.4 |
| vaspa5 | 9.7566886 | 1.368320378 | 0.825 | 0.11 | 1.97055839801692e-4 | 22 | vaspa |
| ptprc5 | 7.6609367 | 1.304174801 | 0.8 | 0.11 | 1.5472793982793e-39 | 22 | ptprc |
| gpr137ba | 7.8931490 | 0.581375662 | 0.275 | 0.01 | 1.59417931011609e-3 | 22 | gpr137ba |
| itgb24 | 1.1983349 | 1.436011241 | 0.9 | 0.155 | 2.42027716141468e-3 | 22 | itgb2 |
| mybpha | 7.3454289 | 0.284186644 | 0.125 | 0.001 | 1.48355627749749e-3 | 22 | mybpha |
| themis2 | 3.5623861 | 0.537020496 | 0.225 | 0.007 | 7.19495122996203e-3 | 22 | themis2 |
| si:dkey-262k9.21 | 4.2258884 | 0.68402715 | 0.375 | 0.023 | 8.53502683965644e-3 | 22 | si:dkey-262k9.2 |
| LOC1005362135 | 4.9009161 | 1.355770052 | 0.85 | 0.139 | 9.89838044404762e-3 | 22 | LOC100536213 |
| hck2 | 2.0951198 | 0.803982126 | 0.425 | 0.033 | 4.23151355098709e-3 | 22 | hck |

| | | | | | | | |
|---|---|---|---|---|---|---|---|
| tagapa2 | 2.5221483 | 1.093272264 | 0.55 | 0.058 | 5.09398293923973e-3 | 22 | tagapa |
| BX005183.31 | 4.7349421 | 0.945321369 | 0.525 | 0.052 | 9.5631625736345e-32 | 22 | BX005183.3 |
| XLOC-0320023 | 7.0748599 | 0.906943877 | 0.4 | 0.029 | 1.42890945884865e-3 | 22 | XLOC-032002 |
| si:dkeyp-104b3.21 | 1.1485847 | 0.305248752 | 0.125 | 0.001 | 2.31979656553352e-3 | 22 | si:dkeyp-104b3.21 |
| mir2232 | 4.0741610 | 0.962478745 | 0.5 | 0.048 | 8.22858306725544e-3 | 22 | mir223 |
| BX682234.11 | 8.4979732 | 0.469781487 | 0.25 | 0.01 | 1.71633564966918e-3 | 22 | BX682234.1 |
| wipf1a4 | 1.2802924 | 1.036370438 | 0.625 | 0.077 | 2.58580661101424e-3 | 22 | wipf1a |
| oxct1a | 2.4215699 | 0.415760179 | 0.175 | 0.005 | 4.89084487871311e-2 | 22 | oxct1a |
| ckbb4 | 2.5639817 | 1.299675887 | 0.9 | 0.181 | 5.17847392086374e-2 | 22 | ckbb |
| fgd4a | 3.6535693 | 0.556609492 | 0.25 | 0.012 | 7.37911408615206e-2 | 22 | fgd4a |
| gm2a3 | 3.6691380 | 1.223892461 | 0.725 | 0.124 | 7.41055802863383e-2 | 22 | gm2a |
| CR391910.2 | 4.8393609 | 0.408393531 | 0.2 | 0.007 | 9.77405733934594e-2 | 22 | CR391910.2 |
| ephx22 | 1.3062358 | 0.891458074 | 0.475 | 0.049 | 2.63820446136061e-2 | 22 | ephx2 |
| ctss2.14 | 2.8422084 | 0.987149841 | 0.55 | 0.068 | 5.74040840829318e-2 | 22 | ctss2.1 |
| LOC101886459 | 4.4300571 | 0.286630081 | 0.15 | 0.003 | 8.9473864621779e-27 | 22 | LOC101886459 |
| si:ch73-264p11.13 | 6.2870871 | 0.684162802 | 0.375 | 0.029 | 1.26980299459291e-2 | 22 | si:ch73-264p11.1 |
| rgs133 | 4.8184999 | 1.164192849 | 0.625 | 0.094 | 9.73192425682393e-2 | 22 | rgs13 |
| fcer1gl6 | 4.8367055 | 1.381386738 | 0.95 | 0.249 | 9.76869422740354e-2 | 22 | fcer1gl |
| ptpn66 | 7.5292558 | 1.18829275 | 0.75 | 0.137 | 1.5206837997636e-25 | 22 | ptpn6 |
| CABZ01111959.1 | 1.2435958 | 0.676394121 | 0.3 | 0.02 | 2.51169045643695e-2 | 22 | CABZ01111959.1 |
| arl11 | 4.0275880 | 0.389016658 | 0.15 | 0.004 | 8.13451957990088e-2 | 22 | arl11 |
| pstpip1b | 5.1943663 | 0.323232396 | 0.15 | 0.004 | 1.04910616270405e-2 | 22 | pstpip1b |
| inpp5f | 5.3055445 | 0.364495421 | 0.15 | 0.004 | 1.07156083682905e-2 | 22 | inpp5f |
| mpeg1.11 | 9.6401689 | 0.607860121 | 0.325 | 0.024 | 1.94702493063158e-2 | 22 | mpeg1.1 |
| cd74b4 | 2.3457242 | 1.289591588 | 1 | 0.24 | 4.73765929793276e-2 | 22 | cd74b |
| spi1b5 | 3.0902173 | 1.100718036 | 0.7 | 0.12 | 6.24131199143247e-2 | 22 | spi1b |
| cnpy31 | 3.8787685 | 0.957772422 | 0.55 | 0.078 | 7.83394884585517e-2 | 22 | cnpy3 |
| cd74a4 | 5.7467216 | 1.26720166 | 0.95 | 0.201 | 1.16066537639743e-2 | 22 | cd74a |
| LOC1018828052 | 7.9733063 | 0.392198944 | 0.2 | 0.008 | 1.61036868193529e-2 | 22 | LOC101882805 |
| sftpbb3 | 1.7510715 | 0.980940361 | 0.6 | 0.088 | 3.5366391572605e-23 | 22 | sftpbb |
| nptna2 | 1.8931347 | 0.770274869 | 0.45 | 0.05 | 3.8235643514016e-23 | 22 | nptna |
| si:dkey-33i11.93 | 2.0441863 | 0.744932402 | 0.4 | 0.038 | 4.1286432304274e-23 | 22 | si:dkey-33i11.9 |
| b3gnt2l1 | 4.0766193 | 0.534776559 | 0.3 | 0.022 | 8.23354800463151e-2 | 22 | b3gnt2l |
| si:ch211-195b13.1 | 7.5474306 | 0.564582499 | 0.275 | 0.018 | 1.52435456333178e-2 | 22 | si:ch211-195b13.1 |
| zdhhc23a1 | 1.1659857 | 0.491981368 | 0.225 | 0.012 | 2.35494137654189e-2 | 22 | zdhhc23a |

| | | | | | | | |
|---|---|---|---|---|---|---|---|
| coro1a7 | 1.3623479 | 1.236567051 | 0.975 | 0.273 | 2.75153415961702e-2 | 22 | coro1a |
| si:ch211-119e14.1: | 2.0297905 | 0.812000831 | 0.45 | 0.052 | 4.09956802789038e-2 | 22 | si:ch211-119e14.1 |
| rab442 | 6.3172926 | 0.89441636 | 0.475 | 0.06 | 1.27590359684022e-2 | 22 | rab44 |
| si:ch211-214p16.2 | 1.2366008 | 1.034213996 | 0.575 | 0.086 | 2.49756278566368e-2 | 22 | si:ch211-214p16.2 |
| arpc1b8 | 1.2420683 | 1.228278614 | 1 | 0.353 | 2.50860539194954e-2 | 22 | arpc1b |
| kctd12.23 | 1.3859756 | 1.127383307 | 0.825 | 0.204 | 2.79925493984381e-2 | 22 | kctd12.2 |
| XLOC-0418703 | 2.2350042 | 0.764842501 | 0.4 | 0.042 | 4.51403810260924e-2 | 22 | XLOC-041870 |
| washc2c2 | 2.4235845 | 0.801567855 | 0.45 | 0.055 | 4.8949137443928e-21 | 22 | washc2c |
| ptgs2b | 3.8757348 | 0.485971037 | 0.2 | 0.01 | 7.82782172117382e-2 | 22 | ptgs2b |
| zgc:1528911 | 4.6665872 | 0.485280445 | 0.225 | 0.013 | 9.42510616829227e-2 | 22 | zgc:152891 |
| itga12 | 6.7936876 | 0.837570822 | 0.45 | 0.056 | 1.3721210995614e-20 | 22 | itga1 |
| dok23 | 5.2039606 | 0.757697218 | 0.4 | 0.045 | 1.05104393481742e-1 | 22 | dok2 |
| lcp13 | 1.1842622 | 0.802646417 | 0.475 | 0.065 | 2.39185447590269e-1 | 22 | lcp1 |
| lrrfip1a2 | 2.0004423 | 0.847753511 | 0.5 | 0.074 | 4.04029350140698e-1 | 22 | lrrfip1a |
| zgc:1037003 | 2.4111421 | 0.940505025 | 0.675 | 0.12 | 4.8697838825109e-19 | 22 | zgc:103700 |
| pfn18 | 3.8146542 | 1.192315223 | 1 | 0.413 | 7.7044570920522e-19 | 22 | pfn1 |
| si:dkey-9i23.5 | 4.1917559 | 0.271651643 | 0.1 | 0.002 | 8.4660894110566e-19 | 22 | si:dkey-9i23.5 |
| il7r1 | 4.5211343 | 0.495200574 | 0.25 | 0.018 | 9.13133498975215e-1 | 22 | il7r |
| XLOC-0331371 | 4.6531823 | 0.643829377 | 0.3 | 0.027 | 9.39803232766881e-1 | 22 | XLOC-033137 |
| zgc:640518 | 7.0326369 | 1.112266034 | 0.725 | 0.17 | 1.42038168619915e-1 | 22 | zgc:64051 |
| CU207241.13 | 1.5308277 | 0.515656197 | 0.275 | 0.022 | 3.09181274463826e-1 | 22 | CU207241.1 |
| ptprea | 2.0997464 | 0.707603934 | 0.35 | 0.038 | 4.24085785440198e-1 | 22 | ptprea |
| hps5 | 2.7766363 | 0.42937714 | 0.225 | 0.016 | 5.60797237789855e-1 | 22 | hps5 |
| itga43 | 3.7760147 | 0.745670352 | 0.425 | 0.059 | 7.62641700169895e-1 | 22 | itga4 |
| il10ra2 | 4.0357901 | 0.726688724 | 0.375 | 0.046 | 8.1510854527239e-17 | 22 | il10ra |
| stk17b4 | 4.2424266 | 0.962737415 | 0.625 | 0.133 | 8.568429067972e-17 | 22 | stk17b |
| pstpip1a | 4.9916027 | 0.387191693 | 0.2 | 0.012 | 1.00815400759734e-1 | 22 | pstpip1a |
| si:ch211-214p16.1! | 5.7420057 | 1.198247243 | 0.8 | 0.185 | 1.15971290808818e-1 | 22 | si:ch211-214p16.1 |
| CMKLR1 (1 of many) | 8.3729298 | 0.378186537 | 0.175 | 0.009 | 1.69108063685251e-1 | 22 | CMKLR1 (1 of many) |
| si:busm1-266f07.2: | 1.1212713 | 1.02753858 | 0.725 | 0.153 | 2.26463165354689e-1 | 22 | si:busm1-266f07.2 |
| egr32 | 2.2949057 | 0.591006914 | 0.3 | 0.03 | 4.63502104670069e-1 | 22 | egr3 |
| fam49ba4 | 3.1039721 | 0.881146187 | 0.6 | 0.118 | 6.26909245656192e-1 | 22 | fam49ba |
| FO393424.21 | 5.1960585 | 0.347692911 | 0.15 | 0.007 | 1.04944794689664e-1 | 22 | FO393424.2 |
| cfbl3 | 6.0105078 | 0.755913942 | 0.55 | 0.09 | 1.21394227680877e-1 | 22 | cfbl |
| cotl17 | 6.1574849 | 1.05273048 | 0.975 | 0.406 | 1.24362724385695e-1 | 22 | cotl1 |

| | | | | | | | |
|---|---|---|---|---|---|---|---|
| lcp2b3 | 7.5194177 | 0.55426807 | 0.275 | 0.027 | 1.51869680839024e-1 | 22 | lcp2b |
| mef2b3 | 1.6683447 | 0.465321217 | 0.275 | 0.027 | 3.36955592345479e-1 | 22 | mef2b |
| CABZ01109481.1 | 3.8216648 | 0.554698065 | 0.275 | 0.029 | 7.71861655180606e-1 | 22 | CABZ01109481.1 |
| XLOC-0437601 | 4.2696596 | 0.47480191 | 0.225 | 0.019 | 8.62343163659791e-1 | 22 | XLOC-043760 |
| zgc:1749451 | 5.2799679 | 0.397242138 | 0.2 | 0.015 | 1.06639513320144e-1 | 22 | zgc:174945 |
| ccr9a5 | 6.6046998 | 0.886726327 | 0.5 | 0.093 | 1.33395123313982e-1 | 22 | ccr9a |
| mcl1a7 | 7.2684352 | 0.888490866 | 0.975 | 0.541 | 1.46800585977782e-1 | 22 | mcl1a |
| arl5c3 | 9.3116987 | 0.841292818 | 0.475 | 0.089 | 1.88068378976727e-1 | 22 | arl5c |
| CABZ01046432.11 | 1.6022469 | 0.390977818 | 0.2 | 0.015 | 3.23605809427845e-1 | 22 | CABZ01046432.1 |
| nocta | 2.2156876 | 0.550616259 | 0.25 | 0.025 | 4.47502438454953e-1 | 22 | nocta |
| CR381673.13 | 3.1973229 | 0.681763279 | 0.375 | 0.055 | 6.45763318485781e-1 | 22 | CR381673.1 |
| yrk5 | 3.6141724 | 0.787902898 | 0.475 | 0.087 | 7.29954407772518e-1 | 22 | yrk |
| plekho21 | 6.1714088 | 0.763022252 | 0.475 | 0.09 | 1.2464394498274e-12 | 22 | plekho2 |
| aldoab | 7.6092843 | 0.360687277 | 0.2 | 0.016 | 1.53684716984974e-1 | 22 | aldoab |
| laptm58 | 9.8694184 | 0.952486783 | 0.75 | 0.21 | 1.99332645049753e-1 | 22 | laptm5 |
| abi3a4 | 1.2096063 | 0.661928032 | 0.4 | 0.063 | 2.44304184992214e-1 | 22 | abi3a |
| myo1f4 | 1.3031433 | 0.682100732 | 0.375 | 0.059 | 2.63195857062763e-1 | 22 | myo1f |
| runx31 | 4.8462748 | 0.508414588 | 0.225 | 0.022 | 9.78802133396331e-1 | 22 | runx3 |
| srgn5 | 6.1673939 | 0.915151498 | 0.8 | 0.239 | 1.24562856577519e-1 | 22 | srgn |
| arhgap4a | 1.0310854 | 0.372432652 | 0.175 | 0.013 | 2.08248319976358e-1 | 22 | arhgap4a |
| cfl16 | 1.3057191 | 0.60153419 | 0.975 | 0.765 | 2.63716098318121e-1 | 22 | cfl1 |
| rap1gap2a2 | 1.8595071 | 0.343300332 | 0.175 | 0.013 | 3.75564666195777e-1 | 22 | rap1gap2a |
| actb24 | 1.9086395 | 0.45509314 | 1 | 0.926 | 3.85487937287576e-1 | 22 | actb2 |
| msna4 | 2.1574714 | 0.854250732 | 0.9 | 0.398 | 4.35744499792222e-1 | 22 | msna |
| wasa6 | 6.3335642 | 0.862626708 | 0.5 | 0.115 | 1.27918996226031e-1 | 22 | wasa |
| grasp2 | 7.5503622 | 0.564148207 | 0.35 | 0.055 | 1.52494666188312e-1 | 22 | grasp |
| csk4 | 8.4106133 | 0.803256636 | 0.5 | 0.115 | 1.69869158203618e-1 | 22 | csk |
| si:ch211-199g17.2[illegible] | 9.0577690 | 0.700390728 | 0.425 | 0.084 | 1.82939760509944e-1 | 22 | si:ch211-199g17.2 |
| grap2b7 | 1.2710234 | 0.911924526 | 0.65 | 0.188 | 2.5670861242862e-10 | 22 | grap2b |
| cxcr4b4 | 1.7832725 | 0.893241691 | 0.6 | 0.152 | 3.60167551508354e-1 | 22 | cxcr4b |
| cdkn1bb1 | 3.6038147 | 0.844332851 | 0.575 | 0.163 | 7.27862462522783e-1 | 22 | cdkn1bb |
| arpc5b4 | 4.2093016 | 0.880289726 | 0.7 | 0.226 | 8.50152663952055e-1 | 22 | arpc5b |
| fmnl1a2 | 6.6590111 | 0.453082728 | 0.275 | 0.037 | 1.34492048566384e-0 | 22 | fmnl1a |
| arpc46 | 1.0525255 | 0.910200844 | 0.75 | 0.312 | 2.12578587642902e-0 | 22 | arpc4 |
| OSBPL8 | 1.2527366 | 0.299737239 | 0.15 | 0.011 | 2.53015214326143e-0 | 22 | OSBPL8 |

| | | | | | | | |
|---|---|---|---|---|---|---|---|
| cblb | 1.3736369 | 0.462977057 | 0.2 | 0.021 | 2.77433451955476e-0 | 22 | cblb |
| XLOC-0430432 | 1.5494158 | 0.623574027 | 0.325 | 0.05 | 3.12935521809939e-0 | 22 | XLOC-043043 |
| efhd22 | 2.0306296 | 0.577116052 | 0.35 | 0.061 | 4.10126277116454e-0 | 22 | efhd2 |
| pdpk1a | 2.0757350 | 0.289357455 | 0.125 | 0.008 | 4.19236200244241e-0 | 22 | pdpk1a |
| hcls16 | 2.2191947 | 0.74167696 | 0.475 | 0.108 | 4.48210772672845e-0 | 22 | hcls1 |
| zgc:1627306 | 2.3476151 | 0.823159029 | 0.925 | 0.488 | 4.74147823078423e-0 | 22 | zgc:162730 |
| syk5 | 2.4441348 | 0.578107457 | 0.4 | 0.077 | 4.93641920891536e-0 | 22 | syk |
| tmsb4x4 | 2.4976552 | 0.34452542 | 1 | 0.941 | 5.04451422038882e-0 | 22 | tmsb4x |
| rgs181 | 2.9931481 | 0.473655974 | 0.25 | 0.032 | 6.0452613685925e-09 | 22 | rgs18 |
| unc93b11 | 4.0886720 | 0.406518182 | 0.225 | 0.027 | 8.25789090890569e-0 | 22 | unc93b1 |
| XLOC-0140792 | 4.0889613 | 0.418397433 | 0.225 | 0.026 | 8.25847529440898e-0 | 22 | XLOC-014079 |
| actb17 | 4.6724829 | 0.627256225 | 1 | 0.739 | 9.43701391141377e-0 | 22 | actb1 |
| si:ch73-380l3.21 | 5.9585890 | 0.603347986 | 0.325 | 0.056 | 1.20345622636201e-0 | 22 | si:ch73-380l3.2 |
| ipcef12 | 6.2604921 | 0.485899314 | 0.25 | 0.033 | 1.26443159559713e-0 | 22 | ipcef1 |
| arpc27 | 6.2861385 | 0.795753 | 0.85 | 0.482 | 1.26961140455821e-0 | 22 | arpc2 |
| RASSF54 | 7.5563119 | 0.688598262 | 0.375 | 0.075 | 1.52614832603933e-0 | 22 | RASSF5 |
| si:dkey-206d17.12 | 2.1652437 | 0.548878514 | 0.275 | 0.043 | 4.37314288191136e-0 | 22 | si:dkey-206d17.12 |
| skap24 | 2.3266142 | 0.613525159 | 0.475 | 0.112 | 4.69906284310778e-0 | 22 | skap2 |
| si:dkey-222f2.73 | 2.3389570 | 0.507187608 | 0.275 | 0.041 | 4.72399159592591e-0 | 22 | si:dkey-222f2.7 |
| arhgap22 | 2.3668476 | 0.263506448 | 0.1 | 0.005 | 4.78032223350987e-0 | 22 | arhgap22 |
| hyal3 | 2.7668409 | 0.362124443 | 0.15 | 0.013 | 5.5881886349989e-08 | 22 | hyal3 |
| vipas39 | 3.0511418 | 0.307557145 | 0.15 | 0.013 | 6.16239121611184e-0 | 22 | vipas39 |
| rac27 | 3.0568702 | 0.838227649 | 0.725 | 0.257 | 6.17396092008215e-0 | 22 | rac2 |
| si:ch211-243a20.4 | 3.3512134 | 0.432914289 | 0.225 | 0.029 | 6.76844585550168e-0 | 22 | si:ch211-243a20.4 |
| sema4ab | 3.6536985 | 0.359607339 | 0.225 | 0.029 | 7.37937504026826e-0 | 22 | sema4ab |
| si:rp71-68n21.92 | 4.2845397 | 0.579305597 | 0.325 | 0.06 | 8.65348490745537e-0 | 22 | si:rp71-68n21.9 |
| itm2bb5 | 4.4323145 | 0.760365149 | 0.875 | 0.352 | 8.95194567715435e-0 | 22 | itm2bb |
| bpifcl | 4.6590495 | 0.414105754 | 0.2 | 0.023 | 9.40988240266896e-0 | 22 | bpifcl |
| dusp12 | 5.7672625 | 0.734762897 | 0.425 | 0.102 | 1.16481402247064e-0 | 22 | dusp1 |
| tmc6a1 | 6.3971493 | 0.345185557 | 0.15 | 0.013 | 1.29203225592319e-0 | 22 | tmc6a |
| vsir3 | 6.4501155 | 0.673920777 | 0.45 | 0.107 | 1.30272984025494e-0 | 22 | vsir |
| mfng2 | 6.8612235 | 0.543884916 | 0.325 | 0.059 | 1.38576131378667e-0 | 22 | mfng |
| arhgap33 | 7.7307508 | 0.251211593 | 0.15 | 0.013 | 1.56137974165575e-0 | 22 | arhgap33 |
| si:ch211-105j21.92 | 7.9138939 | 0.466643007 | 0.25 | 0.036 | 1.59836916008911e-0 | 22 | si:ch211-105j21.9 |
| mapre1a6 | 7.9950406 | 0.730007474 | 0.625 | 0.195 | 1.61475835646035e-0 | 22 | mapre1a |

| | | | | | | | |
|---|---|---|---|---|---|---|---|
| plek2 | 9.3204993 | 0.456643364 | 0.3 | 0.05 | 1.88246125506165e-0 | 22 | plek |
| myh11a3 | 1.1447152 | 0.677748208 | 0.425 | 0.1 | 2.31198143999703e-0 | 22 | myh11a |
| dapp1 | 1.2992298 | 0.257009182 | 0.125 | 0.009 | 2.62405451320788e-0 | 22 | dapp1 |
| si:zfos-741a10.34 | 1.5121963 | 0.59781619 | 0.3 | 0.053 | 3.05418301051273e-0 | 22 | si:zfos-741a10.3 |
| sla12 | 1.5577625 | 0.420210135 | 0.25 | 0.037 | 3.14621294247341e-0 | 22 | sla1 |
| rhogb5 | 2.4111318 | 0.731594858 | 0.55 | 0.167 | 4.86976292334144e-0 | 22 | rhogb |
| crema3 | 3.5545660 | 0.725072841 | 0.5 | 0.144 | 7.17915697841441e-0 | 22 | crema |
| ikzf24 | 5.5528417 | 0.540529192 | 0.3 | 0.055 | 1.12150744493215e-0 | 22 | ikzf2 |
| adgre5b.3 | 5.8144168 | 0.406229404 | 0.2 | 0.026 | 1.17433776303698e-0 | 22 | adgre5b.3 |
| tln14 | 6.1020084 | 0.770902359 | 0.6 | 0.221 | 1.23242264564315e-0 | 22 | tln1 |
| si:ch211-197k17.3 | 6.2437364 | 0.471156172 | 0.225 | 0.033 | 1.26104745776548e-0 | 22 | si:ch211-197k17.3 |
| si:ch211-153b23.7 | 8.2920352 | 0.434574708 | 0.2 | 0.026 | 1.67474235201954e-0 | 22 | si:ch211-153b23.7 |
| rnaset2l5 | 9.4871122 | 0.586245832 | 0.375 | 0.081 | 1.91611206459289e-0 | 22 | rnaset2l |
| TSTA3 (1 of many)1 | 1.0328808 | 0.263951832 | 0.125 | 0.01 | 2.08610949630618e-0 | 22 | TSTA3 (1 of many) |
| selplg3 | 1.4252870 | 0.457230489 | 0.25 | 0.041 | 2.87865226849027e-0 | 22 | selplg |
| h1fx5 | 1.5509875 | 0.842693656 | 0.65 | 0.234 | 3.13252952988793e-0 | 22 | h1fx |
| fam43a | 1.5850051 | 0.378405223 | 0.175 | 0.021 | 3.20123499830438e-0 | 22 | fam43a |
| tnfb2 | 2.4593413 | 0.459810421 | 0.225 | 0.033 | 4.96713168529565e-0 | 22 | tnfb |
| si:dkey-32n7.4 | 2.6704500 | 0.302930884 | 0.1 | 0.007 | 5.39350800912062e-0 | 22 | si:dkey-32n7.4 |
| mrtfaa | 2.7815588 | 0.353835607 | 0.15 | 0.016 | 5.61791432936055e-0 | 22 | mrtfaa |
| pnrc22 | 2.8082702 | 0.473363291 | 0.95 | 0.725 | 5.67186335651477e-0 | 22 | pnrc2 |
| st8sia4 | 2.8175871 | 0.271846865 | 0.1 | 0.007 | 5.69068075707712e-0 | 22 | st8sia4 |
| arhgdig6 | 4.4058625 | 0.749769466 | 0.775 | 0.362 | 8.89852057305457e-0 | 22 | arhgdig |
| st3gal5 | 4.7082704 | 0.354076703 | 0.2 | 0.028 | 9.50929373138568e-0 | 22 | st3gal5 |
| pparg | 5.8353948 | 0.315958721 | 0.175 | 0.022 | 1.17857470313173e-0 | 22 | pparg |
| pik3ap1 | 6.2903151 | 0.286044815 | 0.15 | 0.016 | 1.27045494268158e-0 | 22 | pik3ap1 |
| agap22 | 6.3203685 | 0.3513733 | 0.175 | 0.022 | 1.27652483798171e-0 | 22 | agap2 |
| arpc37 | 7.9392821 | 0.707571438 | 0.825 | 0.478 | 1.6034968071887e-05 | 22 | arpc3 |
| dusp22b | 8.6911484 | 0.400510188 | 0.2 | 0.029 | 1.75535124291734e-0 | 22 | dusp22b |
| CT027744.21 | 1.0487870 | 0.416580715 | 0.275 | 0.052 | 2.11823527010232e-0 | 22 | CT027744.2 |
| rsrp13 | 1.1134804 | 0.663399323 | 0.45 | 0.135 | 2.24889636589008e-0 | 22 | rsrp1 |
| arpc5a3 | 1.1591567 | 0.646519617 | 0.55 | 0.186 | 2.34114879392135e-0 | 22 | arpc5a |
| LOC1081907172 | 1.4319616 | 0.295387005 | 0.15 | 0.017 | 2.89213296751321e-0 | 22 | LOC108190717 |
| mcu1 | 2.0632746 | 0.371483404 | 0.225 | 0.037 | 4.16719577652716e-0 | 22 | mcu |
| mtmr12 | 2.0688689 | 0.310569207 | 0.15 | 0.017 | 4.17849453940616e-0 | 22 | mtmr12 |

| | | | | | | | |
|---|---|---|---|---|---|---|---|
| LOC1018860731 | 2.9173765 | 0.348845295 | 0.15 | 0.018 | 5.89222546736798e-0 | 22 | LOC101886073 |
| myl12.15 | 3.2965946 | 0.563432433 | 0.9 | 0.629 | 6.65813222891824e-0 | 22 | myl12.1 |
| cers1 | 3.5421516 | 0.385351116 | 0.175 | 0.024 | 7.15408365273824e-0 | 22 | cers1 |
| ampd3b | 3.8552559 | 0.294766985 | 0.15 | 0.018 | 7.78646050060337e-0 | 22 | ampd3b |
| capzb5 | 4.0802934 | 0.666154486 | 0.75 | 0.383 | 8.2409686947575e-05 | 22 | capzb |
| wdr192 | 4.1301547 | 0.366067832 | 0.2 | 0.03 | 8.34167362485609e-0 | 22 | wdr19 |
| tox2 | 5.9424807 | 0.3438184 | 0.175 | 0.025 | 0.00012002 | 22 | tox2 |
| lipia | 6.0924457 | 0.267237791 | 0.125 | 0.013 | 0.000123049 | 22 | lipia |
| npsn1 | 6.1184013 | 0.417864003 | 0.2 | 0.03 | 0.000123573 | 22 | npsn |
| si:ch211-203d1.31 | 7.2512935 | 0.288445325 | 0.15 | 0.018 | 0.000146454 | 22 | si:ch211-203d1.3 |
| fam210b1 | 1.0001568 | 0.340406199 | 0.25 | 0.048 | 0.000202002 | 22 | fam210b |
| fam57ba | 1.2181303 | 0.27357417 | 0.125 | 0.013 | 0.000246026 | 22 | fam57ba |
| fam49al | 1.2363540 | 0.295783192 | 0.15 | 0.019 | 0.000249706 | 22 | fam49al |
| zfp36l1a5 | 1.3886777 | 0.730161126 | 0.65 | 0.309 | 0.000280471 | 22 | zfp36l1a |
| mhc2dab3 | 1.4635813 | 0.498506984 | 0.35 | 0.085 | 0.0002956 | 22 | mhc2dab |
| ptpn221 | 1.6801786 | 0.293211908 | 0.175 | 0.026 | 0.000339346 | 22 | ptpn22 |
| fybb4 | 1.7638618 | 0.429032227 | 0.275 | 0.058 | 0.000356247 | 22 | fybb |
| ccr22 | 2.1439685 | 0.44135152 | 0.225 | 0.041 | 0.000433017 | 22 | ccr2 |
| malt3 | 2.1909137 | 0.311579153 | 0.15 | 0.02 | 0.000442499 | 22 | malt3 |
| si:busm1-194e12.1 | 3.2146602 | 0.429380835 | 0.3 | 0.067 | 0.000649265 | 22 | si:busm1-194e12.11 |
| LOC1039091157 | 3.5506143 | 0.643681762 | 0.525 | 0.198 | 0.000717118 | 22 | LOC103909115 |
| si:dkey-234h16.7 | 3.7932908 | 0.340204733 | 0.175 | 0.027 | 0.000766131 | 22 | si:dkey-234h16.7 |
| glipr1a5 | 3.9524513 | 0.505220542 | 0.35 | 0.092 | 0.000798277 | 22 | glipr1a |
| sh2d3ca | 4.3847334 | 0.281597493 | 0.15 | 0.02 | 0.000885585 | 22 | sh2d3ca |
| si:ch73-334d15.21 | 4.5845277 | 0.296674143 | 0.125 | 0.014 | 0.000925937 | 22 | si:ch73-334d15.2 |
| ubash3ba1 | 5.7376891 | 0.36036849 | 0.15 | 0.021 | 0.001158841 | 22 | ubash3ba |
| mdkb3 | 7.2041336 | 0.301085461 | 0.2 | 0.034 | 0.001455019 | 22 | mdkb |
| btg13 | 1.0160098 | 0.601572145 | 0.8 | 0.463 | 0.002052035 | 22 | btg1 |
| heca | 1.0244101 | 0.259019193 | 0.125 | 0.015 | 0.002069001 | 22 | heca |
| efna2a1 | 1.1658619 | 0.259677881 | 0.125 | 0.015 | 0.002354691 | 22 | efna2a |
| LOC110440216 | 1.1672899 | 0.255005435 | 0.1 | 0.01 | 0.002357576 | 22 | LOC110440216 |
| arrb2b2 | 1.1729002 | 0.534969705 | 0.4 | 0.126 | 0.002368907 | 22 | arrb2b |
| si:dkey-192g7.31 | 1.2491350 | 0.353973829 | 0.175 | 0.029 | 0.002522878 | 22 | si:dkey-192g7.3 |
| osgn1 | 1.4481050 | 0.37216131 | 0.2 | 0.037 | 0.002924738 | 22 | osgn1 |
| dtnbp1a3 | 1.4785441 | 0.444710971 | 0.35 | 0.097 | 0.002986216 | 22 | dtnbp1a |

| | | | | | | | |
|---|---|---|---|---|---|---|---|
| mbnl14 | 1.5057813 | 0.647559268 | 0.475 | 0.184 | 0.003041227 | 22 | mbnl1 |
| kdm7ab1 | 1.5470293 | 0.369124598 | 0.2 | 0.037 | 0.003124535 | 22 | kdm7ab |
| nlrc3 | 1.6979905 | 0.331952105 | 0.125 | 0.016 | 0.003429432 | 22 | nlrc3 |
| rab11fip4b | 2.1168624 | 0.346977048 | 0.175 | 0.03 | 0.004275427 | 22 | rab11fip4b |
| arhgef6 | 2.3460560 | 0.306132821 | 0.2 | 0.037 | 0.004738329 | 22 | arhgef6 |
| mych2 | 2.3873989 | 0.615432653 | 0.45 | 0.168 | 0.00482183 | 22 | mych |
| ncalda1 | 2.4734320 | 0.449121262 | 0.275 | 0.069 | 0.004995591 | 22 | ncalda |
| sat1a.27 | 3.0426997 | 0.643231647 | 0.65 | 0.334 | 0.006145341 | 22 | sat1a.2 |
| mob3a3 | 3.9210128 | 0.57274608 | 0.475 | 0.187 | 0.00791927 | 22 | mob3a |
| ptp4a2b1 | 3.9630968 | 0.565445351 | 0.425 | 0.148 | 0.008004267 | 22 | ptp4a2b |
| sox5 | 4.5819362 | 0.276609545 | 0.125 | 0.017 | 0.009254137 | 22 | sox5 |
| pygb2 | 4.5954136 | 0.400589468 | 0.225 | 0.049 | 0.009281357 | 22 | pygb |
| LOC1039114302 | 4.9937181 | 0.43854032 | 0.2 | 0.04 | 0.010085813 | 22 | LOC103911430 |
| FP101882.11 | 5.2307989 | 0.445645785 | 0.225 | 0.05 | 0.010564645 | 22 | FP101882.1 |
| XLOC-0254236 | 5.6600432 | 0.753244575 | 0.475 | 0.19 | 0.011431589 | 22 | XLOC-025423 |
| si:dkey-262k9.44 | 5.7772752 | 0.486083727 | 0.325 | 0.093 | 0.011668363 | 22 | si:dkey-262k9.4 |
| pak2a1 | 6.4238845 | 0.450910953 | 0.35 | 0.104 | 0.01297432 | 22 | pak2a |
| cnp2 | 7.1185702 | 0.575545184 | 0.525 | 0.229 | 0.014377376 | 22 | cnp |
| cap17 | 7.7334972 | 0.6236318 | 0.65 | 0.328 | 0.015619344 | 22 | cap1 |
| capza1b5 | 7.7411138 | 0.608933656 | 0.675 | 0.362 | 0.015634728 | 22 | capza1b |
| ucp27 | 9.5235157 | 0.570751142 | 0.75 | 0.423 | 0.019234645 | 22 | ucp2 |
| sh3bgrl5 | 1.0549785 | 0.583218342 | 0.5 | 0.213 | 0.021307402 | 22 | sh3bgrl |
| rps6ka13 | 1.0958422 | 0.433457779 | 0.225 | 0.052 | 0.022132726 | 22 | rps6ka1 |
| plxnc13 | 1.7100744 | 0.421455469 | 0.25 | 0.063 | 0.034538373 | 22 | plxnc1 |
| si:dkey-185m8.23 | 1.7435322 | 0.389056324 | 0.2 | 0.043 | 0.035214122 | 22 | si:dkey-185m8.2 |
| ptk2ba1 | 1.7456738 | 0.33163446 | 0.15 | 0.026 | 0.035257374 | 22 | ptk2ba |
| AL954142.15 | 1.7916755 | 0.27127739 | 0.175 | 0.032 | 0.036186471 | 22 | AL954142.1 |
| mob1a4 | 1.7976618 | 0.563415652 | 0.5 | 0.217 | 0.036307376 | 22 | mob1a |
| myd881 | 1.9628906 | 0.4273692 | 0.3 | 0.087 | 0.039644503 | 22 | myd88 |
| si:ch211-136m16.8 | 2.0048486 | 0.391040134 | 0.175 | 0.034 | 0.040491928 | 22 | si:ch211-136m16.8 |
| fermt3b3 | 2.0365566 | 0.331456236 | 0.175 | 0.034 | 0.041132334 | 22 | fermt3b |
| pxna1 | 2.4419540 | 0.395315601 | 0.3 | 0.086 | 0.049320145 | 22 | pxna |
| ccdc88b1 | 2.9111647 | 0.375702519 | 0.175 | 0.035 | 0.058796794 | 22 | ccdc88b |
| cdc42l4 | 2.9642794 | 0.54298493 | 0.65 | 0.344 | 0.059869552 | 22 | cdc42l |
| crfb4 | 3.3056020 | 0.412359101 | 0.25 | 0.066 | 0.066763244 | 22 | crfb4 |

| | | | | | | | |
|---|---|---|---|---|---|---|---|
| fam49bb1 | 3.8291386 | 0.380389001 | 0.25 | 0.066 | 0.077337114 | 22 | fam49bb |
| dock82 | 5.9083445 | 0.349485418 | 0.175 | 0.037 | 0.119330835 | 22 | dock8 |
| cyba4 | 6.6228359 | 0.460478386 | 0.375 | 0.132 | 0.133761417 | 22 | cyba |
| ppp1r184 | 6.7969114 | 0.422761353 | 0.25 | 0.068 | 0.137277221 | 22 | ppp1r18 |
| tescb | 6.8840686 | 0.281290346 | 0.125 | 0.02 | 0.139037534 | 22 | tescb |
| actn4 | 6.9195731 | 0.461602063 | 0.35 | 0.122 | 0.139754619 | 22 | actn4 |
| apbb1ip | 7.4797838 | 0.34654476 | 0.15 | 0.029 | 0.151069194 | 22 | apbb1ip |
| il21r.11 | 8.6820176 | 0.312226997 | 0.125 | 0.021 | 0.175350711 | 22 | il21r.1 |
| tyk2 | 1.1861151 | 0.283323693 | 0.125 | 0.021 | 0.239559677 | 22 | tyk2 |
| XLOC-0108992 | 1.1871657 | 0.494230755 | 0.425 | 0.166 | 0.239771874 | 22 | XLOC-010899 |
| LOC101886015 | 1.5391892 | 0.261019779 | 0.15 | 0.029 | 0.31087006 | 22 | LOC101886015 |
| gadd45bb1 | 1.5423497 | 0.425230394 | 0.225 | 0.061 | 0.311508388 | 22 | gadd45bb |
| si:dkey-52l18.4 | 1.6980506 | 0.332334842 | 0.2 | 0.05 | 0.34295528 | 22 | si:dkey-52l18.4 |
| clic15 | 1.7377365 | 0.505624098 | 0.475 | 0.215 | 0.350970657 | 22 | clic1 |
| glipr22 | 1.7803329 | 0.280487546 | 0.15 | 0.03 | 0.359573844 | 22 | glipr2 |
| abracl5 | 1.8308405 | 0.564379218 | 0.5 | 0.241 | 0.369774862 | 22 | abracl |
| sik13 | 1.8452719 | 0.475071268 | 0.275 | 0.089 | 0.372689585 | 22 | sik1 |
| cox6b14 | 1.8478902 | 0.371566195 | 0.375 | 0.133 | 0.37321839 | 22 | cox6b1 |
| actr2a5 | 1.9624275 | 0.521398096 | 0.4 | 0.162 | 0.396351489 | 22 | actr2a |
| wdr16 | 2.3810846 | 0.513456494 | 0.5 | 0.236 | 0.480907661 | 22 | wdr1 |
| mrtfab2 | 2.3919522 | 0.299412675 | 0.2 | 0.05 | 0.483102586 | 22 | mrtfab |
| CABZ01079663.12 | 2.3922223 | 0.260396501 | 0.15 | 0.03 | 0.483157142 | 22 | CABZ01079663.1 |
| csrnp1b2 | 2.8024651 | 0.431414362 | 0.3 | 0.103 | 0.566013889 | 22 | csrnp1b |
| gpr155a1 | 2.8973552 | 0.259389426 | 0.125 | 0.023 | 0.58517884 | 22 | gpr155a |
| atp2a33 | 3.3149567 | 0.315467509 | 0.2 | 0.052 | 0.669521809 | 22 | atp2a3 |
| scinlb3 | 3.6527322 | 0.352329841 | 0.3 | 0.096 | 0.737742331 | 22 | scinlb |
| ppp2cab1 | 3.9338838 | 0.252874602 | 0.175 | 0.041 | 0.794526527 | 22 | ppp2cab |
| si:ch211-212k18.74 | 4.0420636 | 0.484362655 | 0.425 | 0.185 | 0.816375588 | 22 | si:ch211-212k18.7 |
| plekho1b1 | 4.6388928 | 0.375909014 | 0.2 | 0.054 | 0.936917194 | 22 | plekho1b |
| lasp15 | 4.8544916 | 0.465786341 | 0.625 | 0.357 | 0.980461674 | 22 | lasp1 |
| sept64 | 4.9770333 | 0.480045641 | 0.425 | 0.191 | 1 | 22 | sept6 |
| per1b | 5.6943492 | 0.474358211 | 0.275 | 0.097 | 1 | 22 | per1b |
| cox8b2 | 5.9498160 | 0.384067968 | 0.35 | 0.133 | 1 | 22 | cox8b |
| FP085398.1 | 6.0967011 | 0.274229044 | 0.125 | 0.024 | 1 | 22 | FP085398.1 |
| arhgdia5 | 6.1200890 | 0.472828972 | 0.65 | 0.373 | 1 | 22 | arhgdia |

| | | | | | | | |
|---|---|---|---|---|---|---|---|
| cmip | 6.4289856 | 0.344284215 | 0.225 | 0.067 | 1 | 22 | cmip |
| XLOC-0234924 | 7.7436627 | 0.363177951 | 0.275 | 0.091 | 1 | 22 | XLOC-023492 |
| mast2 | 9.0885695 | 0.269342005 | 0.2 | 0.055 | 1 | 22 | mast2 |
| si:dkeyp-19e1.31 | 0.000103 | 0.252102823 | 0.175 | 0.044 | 1 | 22 | si:dkeyp-19e1.3 |
| BX901920.14 | 0.000104 | 0.35099746 | 0.25 | 0.079 | 1 | 22 | BX901920.1 |
| ldha2 | 0.000104 | 0.319462057 | 0.2 | 0.057 | 1 | 22 | ldha |
| pcmtd11 | 0.000105 | 0.329675757 | 0.2 | 0.057 | 1 | 22 | pcmtd1 |
| slc6a6b2 | 0.000114 | 0.388157771 | 0.225 | 0.069 | 1 | 22 | slc6a6b |
| sema7a2 | 0.000123 | 0.260795719 | 0.15 | 0.034 | 1 | 22 | sema7a |
| si:ch1073-443f11.2 | 0.000123 | 0.300792904 | 0.225 | 0.068 | 1 | 22 | si:ch1073-443f11.2 |
| tmx4 | 0.000133 | 0.381613918 | 0.175 | 0.048 | 1 | 22 | tmx4 |
| gna13b1 | 0.000134 | 0.322782892 | 0.2 | 0.058 | 1 | 22 | gna13b |
| plk33 | 0.000151 | 0.296708975 | 0.2 | 0.057 | 1 | 22 | plk3 |
| ptmaa5 | 0.00016 | 0.389318365 | 0.825 | 0.656 | 1 | 22 | ptmaa |
| irs11 | 0.000162 | 0.337215653 | 0.175 | 0.048 | 1 | 22 | irs1 |
| si:ch73-269m14.2 | 0.000188 | 0.416473126 | 0.25 | 0.09 | 1 | 22 | si:ch73-269m14.2 |
| n4bp1 | 0.000193 | 0.261264691 | 0.125 | 0.027 | 1 | 22 | n4bp1 |
| arid4a | 0.000289 | 0.351631834 | 0.225 | 0.076 | 1 | 22 | arid4a |
| chac12 | 0.000292 | 0.445604797 | 0.525 | 0.287 | 1 | 22 | chac1 |
| pkn1a | 0.000307 | 0.266511233 | 0.125 | 0.028 | 1 | 22 | pkn1a |
| csf2rb2 | 0.000312 | 0.26410632 | 0.15 | 0.038 | 1 | 22 | csf2rb |
| cd832 | 0.000315 | 0.317313089 | 0.175 | 0.049 | 1 | 22 | cd83 |
| XLOC-043861 | 0.000315 | 0.26442421 | 0.175 | 0.049 | 1 | 22 | XLOC-043861 |
| uckl1b | 0.000333 | 0.26921226 | 0.125 | 0.029 | 1 | 22 | uckl1b |
| limd24 | 0.000379 | 0.349847231 | 0.2 | 0.063 | 1 | 22 | limd2 |
| commd31 | 0.000386 | 0.383166397 | 0.3 | 0.125 | 1 | 22 | commd3 |
| arf6a1 | 0.00049 | 0.421080723 | 0.35 | 0.164 | 1 | 22 | arf6a |
| crlf33 | 0.000491 | 0.310229264 | 0.275 | 0.106 | 1 | 22 | crlf3 |
| mylipb | 0.000496 | 0.260704987 | 0.175 | 0.051 | 1 | 22 | mylipb |
| arhgap15 | 0.000557 | 0.330685208 | 0.15 | 0.041 | 1 | 22 | arhgap15 |
| tmem56a | 0.000606 | 0.262142008 | 0.125 | 0.03 | 1 | 22 | tmem56a |
| unm-hu7910 | 0.000622 | 0.260018793 | 0.175 | 0.052 | 1 | 22 | unm-hu7910 |
| herpud11 | 0.000657 | 0.319401101 | 0.2 | 0.066 | 1 | 22 | herpud1 |
| parvg3 | 0.00068 | 0.34689569 | 0.275 | 0.108 | 1 | 22 | parvg |
| idh22 | 0.000785 | 0.435166305 | 0.575 | 0.384 | 1 | 22 | idh2 |

| | | | | | | | |
|---|---|---|---|---|---|---|---|
| srgap3 | 0.000928 | 0.318022038 | 0.175 | 0.056 | 1 | 22 | srgap3 |
| ankrd112 | 0.000951 | 0.382773411 | 0.325 | 0.15 | 1 | 22 | ankrd11 |
| dnajb1b | 0.000989 | 0.290862862 | 0.175 | 0.056 | 1 | 22 | dnajb1b |
| tpi1b3 | 0.001111 | 0.38707769 | 0.45 | 0.246 | 1 | 22 | tpi1b |
| jdp2b3 | 0.00114 | 0.410135004 | 0.325 | 0.151 | 1 | 22 | jdp2b |
| nr1d2a | 0.001249 | 0.417789535 | 0.3 | 0.138 | 1 | 22 | nr1d2a |
| XLOC-011107 | 0.001352 | 0.288761392 | 0.125 | 0.033 | 1 | 22 | XLOC-011107 |
| alox5a3 | 0.00138 | 0.297082254 | 0.125 | 0.033 | 1 | 22 | alox5a |
| bin2b | 0.001464 | 0.284281794 | 0.175 | 0.058 | 1 | 22 | bin2b |
| cox174 | 0.001625 | 0.3706805 | 0.375 | 0.187 | 1 | 22 | cox17 |
| notch24 | 0.001671 | 0.344416232 | 0.325 | 0.151 | 1 | 22 | notch2 |
| pycard4 | 0.001694 | 0.339162633 | 0.325 | 0.146 | 1 | 22 | pycard |
| capza1a2 | 0.001731 | 0.381050981 | 0.55 | 0.344 | 1 | 22 | capza1a |
| slc25a33 | 0.001735 | 0.263062889 | 0.175 | 0.059 | 1 | 22 | slc25a33 |
| twf1a4 | 0.00186 | 0.411380422 | 0.375 | 0.203 | 1 | 22 | twf1a |
| sept123 | 0.002053 | 0.277891623 | 0.225 | 0.086 | 1 | 22 | sept12 |
| btk3 | 0.002278 | 0.263408189 | 0.15 | 0.047 | 1 | 22 | btk |
| cib1 | 0.002311 | 0.269909585 | 0.25 | 0.105 | 1 | 22 | cib1 |
| prelid3b1 | 0.00268 | 0.340773109 | 0.35 | 0.179 | 1 | 22 | prelid3b |
| adcy71 | 0.002787 | 0.27928116 | 0.125 | 0.036 | 1 | 22 | adcy7 |
| eno33 | 0.002957 | 0.371814725 | 0.5 | 0.295 | 1 | 22 | eno3 |
| selenot1a2 | 0.00322 | 0.292018863 | 0.25 | 0.11 | 1 | 22 | selenot1a |
| atp6v1g12 | 0.003403 | 0.321501118 | 0.375 | 0.194 | 1 | 22 | atp6v1g1 |
| elf13 | 0.003423 | 0.30232959 | 0.25 | 0.107 | 1 | 22 | elf1 |
| kdm2ba1 | 0.003458 | 0.269906206 | 0.175 | 0.062 | 1 | 22 | kdm2ba |
| si:ch73-248e21.72 | 0.003723 | 0.250388269 | 0.2 | 0.076 | 1 | 22 | si:ch73-248e21.7 |
| sh3kbp12 | 0.003961 | 0.302882102 | 0.25 | 0.111 | 1 | 22 | sh3kbp1 |
| ppifb3 | 0.00424 | 0.330784054 | 0.325 | 0.167 | 1 | 22 | ppifb |
| rhoab3 | 0.004737 | 0.320195824 | 0.525 | 0.313 | 1 | 22 | rhoab |
| FERMT3 (1 of many) | 0.004896 | 0.277634909 | 0.175 | 0.065 | 1 | 22 | FERMT3 (1 of many) |
| ier2b4 | 0.004995 | 0.363636818 | 0.525 | 0.326 | 1 | 22 | ier2b |
| klf132 | 0.005658 | 0.299354004 | 0.225 | 0.099 | 1 | 22 | klf13 |
| zgc:638312 | 0.005748 | 0.282712885 | 0.25 | 0.114 | 1 | 22 | zgc:63831 |
| wasb5 | 0.005909 | 0.317914305 | 0.325 | 0.16 | 1 | 22 | wasb |
| gapdhs6 | 0.005958 | 0.335375718 | 0.7 | 0.476 | 1 | 22 | gapdhs |

| | | | | | | | |
|---|---|---|---|---|---|---|---|
| mt-co16 | 0.005981 | 0.34130489 | 0.825 | 0.592 | 1 | 22 | mt-co1 |
| cebpb5 | 0.006131 | 0.396099548 | 0.5 | 0.296 | 1 | 22 | cebpb |
| CR383676.18 | 0.006283 | 0.463203214 | 0.975 | 0.658 | 1 | 22 | CR383676.1 |
| calm1b5 | 0.006736 | 0.319433673 | 0.3 | 0.148 | 1 | 22 | calm1b |
| mhc2b3 | 0.007573 | 0.277193163 | 0.225 | 0.095 | 1 | 22 | mhc2b |
| rnf1303 | 0.00833 | 0.307234133 | 0.2 | 0.088 | 1 | 22 | rnf130 |
| prkar2aa2 | 0.008713 | 0.263204199 | 0.3 | 0.151 | 1 | 22 | prkar2aa |
| znf10351 | 0.009785 | 0.276571889 | 0.15 | 0.058 | 1 | 22 | znf1035 |
| lhx9 | 3.2079186 | 1.126229374 | 0.615 | 0.006 | 6.4790332553262e-17 | 23 | lhx9 |
| nog11 | 3.1646107 | 1.190081798 | 0.692 | 0.021 | 6.39156431979473e-1 | 23 | nog1 |
| frem2b | 1.8746572 | 0.825228355 | 0.462 | 0.011 | 3.78624531048567e-8 | 23 | frem2b |
| lrrc51 | 1.5412229 | 0.354298206 | 0.205 | 0 | 3.11280806345299e-7 | 23 | lrrc51 |
| bmp62 | 6.1090798 | 0.830402009 | 0.462 | 0.018 | 1.23385086099961e-6 | 23 | bmp6 |
| nanos13 | 1.6542584 | 1.498985348 | 0.821 | 0.085 | 3.34110571675505e-5 | 23 | nanos1 |
| cldnc3 | 2.9969444 | 1.393141544 | 0.744 | 0.063 | 6.05292876187992e-5 | 23 | cldnc |
| wdr93 | 9.8003313 | 0.323887087 | 0.205 | 0.002 | 1.97937291669554e-5 | 23 | wdr93 |
| zgc:1101824 | 9.9653884 | 1.159198144 | 0.769 | 0.071 | 2.01270951154403e-5 | 23 | zgc:110182 |
| usp43b1 | 5.1406591 | 0.557863571 | 0.359 | 0.016 | 1.0382589382819e-42 | 23 | usp43b |
| slc4a4a2 | 8.3319435 | 0.527081748 | 0.333 | 0.013 | 1.68280263694565e-4 | 23 | slc4a4a |
| lrriq1 | 1.2762694 | 0.379381081 | 0.154 | 0.001 | 2.5776814187621e-42 | 23 | lrriq1 |
| pax2a | 3.7820136 | 0.40699649 | 0.231 | 0.005 | 7.63853294614221e-4 | 23 | pax2a |
| neurl1ab | 5.2827589 | 0.353160224 | 0.205 | 0.004 | 1.06695881526145e-4 | 23 | neurl1ab |
| cdh63 | 2.1353919 | 0.986011346 | 0.718 | 0.075 | 4.31285119499673e-3 | 23 | cdh6 |
| BX005003.1 | 6.4727951 | 0.282475819 | 0.128 | 0.001 | 1.30731044301569e-3 | 23 | BX005003.1 |
| lingo2a | 7.0200982 | 0.293147917 | 0.154 | 0.002 | 1.41784924422332e-3 | 23 | lingo2a |
| ralyl | 7.4641166 | 0.357875901 | 0.154 | 0.002 | 1.50752763964679e-3 | 23 | ralyl |
| si:ch211-207d6.22 | 7.8485005 | 1.142585641 | 0.59 | 0.056 | 1.58516166125788e-3 | 23 | si:ch211-207d6.2 |
| lamb2 | 3.1128266 | 0.435250685 | 0.231 | 0.007 | 6.28697589527506e-3 | 23 | lamb2 |
| snx10a | 1.0803413 | 0.378640182 | 0.179 | 0.004 | 2.18196548421625e-3 | 23 | snx10a |
| ackr3b4 | 5.1577010 | 0.978174872 | 0.615 | 0.069 | 1.04170087659855e-3 | 23 | ackr3b |
| cfl1l2 | 2.4371206 | 0.843238308 | 0.564 | 0.061 | 4.92225254492267e-2 | 23 | cfl1l |
| tmem238a | 3.6234336 | 0.551872752 | 0.385 | 0.028 | 7.31824885765019e-2 | 23 | tmem238a |
| fryb | 6.1661534 | 0.301136876 | 0.179 | 0.005 | 1.24537801872352e-2 | 23 | fryb |
| zgc:153311 | 1.5704955 | 0.309395211 | 0.128 | 0.002 | 3.17192994406551e-2 | 23 | zgc:153311 |
| bcam3 | 1.9131615 | 1.148627487 | 0.667 | 0.092 | 3.86401247218588e-2 | 23 | bcam |

| | | | | | | | |
|---|---|---|---|---|---|---|---|
| net14 | 1.2266734 | 0.805177237 | 0.615 | 0.076 | 2.47751246065117e-2 | 23 | net1 |
| gpx1b2 | 5.9561073 | 0.588874912 | 0.385 | 0.029 | 1.20295500347641e-2 | 23 | gpx1b |
| efnb3a | 1.2152834 | 0.251099868 | 0.154 | 0.004 | 2.454508055144e-25 | 23 | efnb3a |
| hoxb8a | 1.2770009 | 0.449234318 | 0.308 | 0.02 | 2.57915877954154e-2 | 23 | hoxb8a |
| emx23 | 2.8081847 | 0.663008941 | 0.513 | 0.056 | 5.67169068348938e-2 | 23 | emx2 |
| hsbp1a3 | 1.0255705 | 0.768411364 | 0.564 | 0.072 | 2.07134493760577e-2 | 23 | hsbp1a |
| arvcfb1 | 2.8429346 | 0.460166098 | 0.256 | 0.014 | 5.74187513260358e-2 | 23 | arvcfb |
| wu:fc27b112 | 2.9063931 | 0.541554917 | 0.385 | 0.033 | 5.87004230024181e-2 | 23 | wu:fc27b11 |
| rbfox3a | 5.4041509 | 0.322951339 | 0.179 | 0.006 | 1.09147635869443e-2 | 23 | rbfox3a |
| tnika1 | 6.8901173 | 0.43315362 | 0.282 | 0.018 | 1.39159701096615e-2 | 23 | tnika |
| si:ch211-156j16.12 | 1.2870912 | 0.739373991 | 0.513 | 0.06 | 2.59953818209064e-2 | 23 | si:ch211-156j16.1 |
| f11r.14 | 7.7752043 | 0.776000991 | 0.641 | 0.096 | 1.57035801682797e-2 | 23 | f11r.1 |
| lama53 | 9.5930427 | 0.690210937 | 0.487 | 0.057 | 1.93750684864009e-2 | 23 | lama5 |
| si:dkey-178e17.31 | 1.6078111 | 0.558802334 | 0.282 | 0.019 | 3.24729617864383e-2 | 23 | si:dkey-178e17.3 |
| tmeff1b3 | 2.2118949 | 0.93539621 | 0.615 | 0.098 | 4.46736430147675e-2 | 23 | tmeff1b |
| rgrb | 5.5009175 | 0.350837478 | 0.359 | 0.031 | 1.11102032602433e-2 | 23 | rgrb |
| fat1a2 | 7.9790903 | 0.610601606 | 0.41 | 0.042 | 1.61153688400657e-2 | 23 | fat1a |
| esr2b2 | 8.7556016 | 0.647133491 | 0.462 | 0.054 | 1.76836886308056e-2 | 23 | esr2b |
| fosb4 | 1.8103880 | 0.971482509 | 0.641 | 0.106 | 3.65644065355599e-2 | 23 | fosb |
| ifi454 | 1.6313199 | 1.209505394 | 0.821 | 0.192 | 3.29477681135599e-2 | 23 | ifi45 |
| enkur | 4.6567067 | 0.314744159 | 0.179 | 0.008 | 9.4051505778724e-20 | 23 | enkur |
| jam2b3 | 6.3643298 | 0.605386974 | 0.359 | 0.035 | 1.28540370009926e-1 | 23 | jam2b |
| mmp15b1 | 1.5117076 | 0.577599037 | 0.359 | 0.036 | 3.05319595465349e-1 | 23 | mmp15b |
| podxl4 | 9.4568487 | 0.894136547 | 0.641 | 0.108 | 1.90999973333351e-18 | 23 | podxl |
| fstl31 | 6.8412845 | 0.483780224 | 0.282 | 0.024 | 1.38173424780347e-1 | 23 | fstl3 |
| pcdh1b1 | 8.9076561 | 0.473428532 | 0.333 | 0.032 | 1.79907931474084e-1 | 23 | pcdh1b |
| id2a2 | 1.0861541 | 1.000507561 | 0.718 | 0.162 | 2.19370546101709e-1 | 23 | id2a |
| trdn | 1.1643821 | 0.2714967 | 0.154 | 0.006 | 2.351702678068e-17 | 23 | trdn |
| MDFI1 | 1.1705421 | 0.585681892 | 0.385 | 0.044 | 2.36414406115141e-1 | 23 | MDFI |
| LOC5533162 | 1.2133465 | 0.591323239 | 0.462 | 0.061 | 2.45059604308884e-1 | 23 | LOC553316 |
| rdh10a3 | 2.2363608 | 0.733624834 | 0.538 | 0.084 | 4.5167779720629e-17 | 23 | rdh10a |
| rpz54 | 3.0671505 | 1.070236014 | 0.872 | 0.249 | 6.19472405198113e-1 | 23 | rpz5 |
| cdh42 | 4.2200542 | 0.470702869 | 0.308 | 0.029 | 8.52324358624899e-1 | 23 | cdh4 |
| CABZ01086029.11 | 4.7329433 | 0.484379486 | 0.333 | 0.034 | 9.55912575327682e-1 | 23 | CABZ01086029.1 |
| tbx2b1 | 4.7387343 | 0.569744483 | 0.359 | 0.04 | 9.57082180266945e-1 | 23 | tbx2b |

| | | | | | | | |
|---|---|---|---|---|---|---|---|
| sox61 | 5.6254725 | 0.507431813 | 0.385 | 0.045 | 1.13617669648523e-1 | 23 | sox6 |
| meis32 | 1.5443615 | 0.525756282 | 0.333 | 0.035 | 3.11914706937165e-1 | 23 | meis3 |
| cxxc5a3 | 2.1864482 | 0.649311402 | 0.513 | 0.082 | 4.41596948219846e-1 | 23 | cxxc5a |
| wt1a3 | 2.8463778 | 0.722711937 | 0.513 | 0.083 | 5.74882936579775e-1 | 23 | wt1a |
| grhl12 | 3.4260018 | 0.567117653 | 0.385 | 0.048 | 6.91949590356144e-1 | 23 | grhl1 |
| emilin1a3 | 4.3590475 | 0.543050278 | 0.41 | 0.053 | 8.803968356453e-16 | 23 | emilin1a |
| tmsb4 | 4.9190275 | 0.560070994 | 0.41 | 0.052 | 9.93495993584775e-1 | 23 | tmsb |
| pi15a3 | 5.1264912 | 0.825073303 | 0.564 | 0.095 | 1.0353974390562e-15 | 23 | pi15a |
| angptl44 | 1.2308010 | 0.707825163 | 0.513 | 0.085 | 2.48584885178739e-1 | 23 | angptl4 |
| epb41l4a1 | 1.5548828 | 0.33317429 | 0.205 | 0.014 | 3.14039699056249e-1 | 23 | epb41l4a |
| pdgfaa2 | 1.7182047 | 0.579990377 | 0.41 | 0.056 | 3.47025810270051e-1 | 23 | pdgfaa |
| rbpms2b2 | 1.8791127 | 0.531385002 | 0.385 | 0.049 | 3.79524395907881e-1 | 23 | rbpms2b |
| jdp2b4 | 2.0171222 | 0.824590992 | 0.667 | 0.144 | 4.07398176876694e-1 | 23 | jdp2b |
| ncam1a1 | 2.0940601 | 0.443898727 | 0.308 | 0.031 | 4.22937330950562e-1 | 23 | ncam1a |
| glod51 | 2.3746047 | 0.53748679 | 0.41 | 0.056 | 4.79598930488718e-1 | 23 | glod5 |
| si:dkey-28b4.75 | 3.1092437 | 0.865586983 | 0.744 | 0.189 | 6.27973953010878e-1 | 23 | si:dkey-28b4.7 |
| ftr824 | 3.3736397 | 0.673445862 | 0.564 | 0.1 | 6.81374030093726e-1 | 23 | ftr82 |
| apex12 | 3.7469528 | 0.64014846 | 0.487 | 0.081 | 7.56772063804206e-1 | 23 | apex1 |
| jun8 | 3.7854878 | 1.021780035 | 0.974 | 0.378 | 7.64554979166695e-1 | 23 | jun |
| mdka5 | 6.6995125 | 1.043213026 | 0.795 | 0.217 | 1.35310054341864e-1 | 23 | mdka |
| si:ch73-335l21.44 | 8.2761910 | 0.91494165 | 0.564 | 0.111 | 1.67154231469435e-1 | 23 | si:ch73-335l21.4 |
| f3b2 | 1.1466931 | 0.67727914 | 0.513 | 0.088 | 2.31597610162914e-1 | 23 | f3b |
| tead3a | 1.8860925 | 0.517473754 | 0.282 | 0.029 | 3.80934118882633e-1 | 23 | tead3a |
| zgc:1141811 | 2.5814856 | 0.537995775 | 0.333 | 0.04 | 5.21382661701291e-1 | 23 | zgc:114181 |
| XLOC-0152912 | 2.6389407 | 0.414542707 | 0.282 | 0.029 | 5.32986869064722e-1 | 23 | XLOC-015291 |
| ube2e34 | 4.4361563 | 0.831885945 | 0.846 | 0.25 | 8.95970501943439e-1 | 23 | ube2e3 |
| cdh26 | 4.9986078 | 0.659702677 | 0.615 | 0.122 | 1.00956883061714e-1 | 23 | cdh2 |
| agrn4 | 1.0428283 | 0.665789323 | 0.564 | 0.108 | 2.10620038185623e-1 | 23 | agrn |
| ifi465 | 1.1479795 | 0.876774328 | 0.718 | 0.186 | 2.31857425021167e-1 | 23 | ifi46 |
| ndrg1a1 | 1.1957644 | 0.469823713 | 0.41 | 0.06 | 2.41508540842525e-1 | 23 | ndrg1a |
| AL954322.21 | 1.7963966 | 0.300444713 | 0.179 | 0.012 | 3.6281823630201e-13 | 23 | AL954322.2 |
| prss353 | 2.0566685 | 0.640263183 | 0.436 | 0.068 | 4.15385340065621e-1 | 23 | prss35 |
| pleca4 | 2.4138391 | 0.640754011 | 0.462 | 0.077 | 4.87523098935939e-1 | 23 | pleca |
| junba6 | 2.4767550 | 0.960443088 | 0.974 | 0.398 | 5.00230218930792e-1 | 23 | junba |
| hoxc9a2 | 2.9753619 | 0.437698158 | 0.308 | 0.036 | 6.00933860520459e-1 | 23 | hoxc9a |

| | | | | | | | |
|---|---|---|---|---|---|---|---|
| fn1b5 | 3.6592687 | 0.856700104 | 0.718 | 0.174 | 7.39062503209516e-1 | 23 | fn1b |
| gadd45ba6 | 4.1177282 | 0.873504598 | 0.821 | 0.245 | 8.31657575132788e-1 | 23 | gadd45ba |
| h1f04 | 5.6320275 | 0.77000048 | 0.564 | 0.116 | 1.13750059460498e-1 | 23 | h1f0 |
| esr2a2 | 7.8843279 | 0.466637157 | 0.308 | 0.037 | 1.59239771176878e-1 | 23 | esr2a |
| bnc24 | 1.0534171 | 0.650351346 | 0.462 | 0.08 | 2.12758671448086e-1 | 23 | bnc2 |
| CR318588.15 | 1.0551837 | 1.136479174 | 0.821 | 0.261 | 2.13115468887126e-1 | 23 | CR318588.1 |
| pcbd1 | 1.1695172 | 0.268473395 | 0.333 | 0.042 | 2.3620739166516e-12 | 23 | pcbd1 |
| fosab7 | 1.2212216 | 0.957158526 | 1 | 0.455 | 2.46650138134551e-1 | 23 | fosab |
| si:ch211-108c6.2 | 1.7112240 | 0.324827468 | 0.205 | 0.017 | 3.45615929226003e-1 | 23 | si:ch211-108c6.2 |
| BX957317.2 | 1.9756570 | 0.2768502 | 0.179 | 0.013 | 3.9902345136752e-12 | 23 | BX957317.2 |
| tspan9b3 | 2.0393055 | 0.565352359 | 0.462 | 0.08 | 4.11878550883761e-1 | 23 | tspan9b |
| zbtb203 | 2.4372390 | 0.673296984 | 0.641 | 0.145 | 4.92249170447065e-1 | 23 | zbtb20 |
| chd4a3 | 3.9011976 | 0.712615046 | 0.718 | 0.187 | 7.87924883338479e-1 | 23 | chd4a |
| krt18a.18 | 4.6261386 | 0.965937679 | 0.846 | 0.274 | 9.34341228789142e-1 | 23 | krt18a.1 |
| cavin1b3 | 4.7141153 | 0.494089394 | 0.436 | 0.068 | 9.52109884980407e-1 | 23 | cavin1b |
| hoxb6a | 5.5019187 | 0.295625695 | 0.205 | 0.017 | 1.11122253663101e-1 | 23 | hoxb6a |
| itsn2a | 8.3849182 | 0.263320194 | 0.179 | 0.013 | 1.6935019466647e-11 | 23 | itsn2a |
| hoxb5a1 | 9.0590558 | 0.356317346 | 0.205 | 0.018 | 1.82965750250543e-1 | 23 | hoxb5a |
| si:ch211-286b5.54 | 9.7611144 | 0.747872612 | 0.769 | 0.22 | 1.97145227968454e-1 | 23 | si:ch211-286b5.5 |
| socs3a6 | 9.8304042 | 0.929000878 | 0.974 | 0.425 | 1.98544675014665e-1 | 23 | socs3a |
| oaz2b2 | 1.1041988 | 0.389398482 | 0.282 | 0.033 | 2.23015043618777e-1 | 23 | oaz2b |
| epha34 | 2.3148265 | 0.434351085 | 0.359 | 0.052 | 4.67525526733265e-1 | 23 | epha3 |
| hmcn21 | 2.5169621 | 0.285905412 | 0.179 | 0.014 | 5.08350849672121e-1 | 23 | hmcn2 |
| pik3r3b | 2.6587126 | 0.565290962 | 0.333 | 0.048 | 5.36980184345608e-1 | 23 | pik3r3b |
| lhfpl2a3 | 4.1794164 | 0.539176343 | 0.462 | 0.085 | 8.44116745341653e-1 | 23 | lhfpl2a |
| gig2e1 | 5.0185284 | 0.356201258 | 0.231 | 0.024 | 1.01359220104637e-1 | 23 | gig2e |
| NPAS3 | 5.8152442 | 0.262756587 | 0.154 | 0.01 | 1.17450488168091e-1 | 23 | NPAS3 |
| ptges3b3 | 6.1812949 | 0.682701125 | 0.769 | 0.222 | 1.24843614351202e-1 | 23 | ptges3b |
| mphosph84 | 6.5430588 | 0.816057255 | 0.769 | 0.247 | 1.32150158815392e-1 | 23 | mphosph8 |
| ptk7a | 7.0811316 | 0.426692132 | 0.308 | 0.041 | 1.43017616267807e-1 | 23 | ptk7a |
| cd151l4 | 7.1142362 | 0.540726732 | 0.462 | 0.087 | 1.43686228727172e-1 | 23 | cd151l |
| alcama3 | 8.0746091 | 0.451061707 | 0.359 | 0.053 | 1.63082881289269e-1 | 23 | alcama |
| ugt1b52 | 1.3640553 | 0.514217255 | 0.436 | 0.079 | 2.75498260781268e-1 | 23 | ugt1b5 |
| mto1 | 1.7602629 | 0.281652187 | 0.154 | 0.011 | 3.55520316247697e-1 | 23 | mto1 |
| si:ch211-222l21.18 | 2.1592682 | 0.811991876 | 1 | 0.527 | 4.36107414186456e-1 | 23 | si:ch211-222l21.1 |

| | | | | | | | |
|---|---|---|---|---|---|---|---|
| cyp26a1 | 2.4118286 | 0.258144529 | 0.154 | 0.011 | 4.87117038484706e-1 | 23 | cyp26a1 |
| tcta | 3.1395165 | 0.440239758 | 0.359 | 0.057 | 6.34088152627061e-1 | 23 | tcta |
| fgf243 | 3.1875526 | 0.501383935 | 0.256 | 0.031 | 6.43790003561908e-1 | 23 | fgf24 |
| ptprsa1 | 4.0144259 | 0.413316957 | 0.308 | 0.043 | 8.10793612337901e-1 | 23 | ptprsa |
| pdlim3b1 | 4.3279735 | 0.315754963 | 0.179 | 0.015 | 8.74120821904284e-1 | 23 | pdlim3b |
| tmem982 | 5.0353231 | 0.39989158 | 0.308 | 0.044 | 1.01698422579756e-0 | 23 | tmem98 |
| zgc:1527912 | 6.5656822 | 0.536185942 | 0.385 | 0.066 | 1.32607083442781e-0 | 23 | zgc:152791 |
| si:dkey-250d21.1 | 6.8663601 | 0.543073365 | 0.538 | 0.124 | 1.38679875187918e-0 | 23 | si:dkey-250d21.1 |
| tmem237b | 7.3275044 | 0.285286841 | 0.205 | 0.02 | 1.47993607464448e-0 | 23 | tmem237b |
| klf11b1 | 8.7347586 | 0.547216671 | 0.462 | 0.092 | 1.76415921079036e-0 | 23 | klf11b |
| si:dkey-85k7.12 | 1.2486328 | 0.258153972 | 0.179 | 0.016 | 2.52186385536128e-0 | 23 | si:dkey-85k7.12 |
| tm2d21 | 1.6462262 | 0.408009258 | 0.359 | 0.059 | 3.32488325496355e-0 | 23 | tm2d2 |
| b2m3 | 1.7881561 | 0.834568365 | 0.846 | 0.35 | 3.61153897400334e-0 | 23 | b2m |
| ccnd13 | 2.5659355 | 0.588974706 | 0.462 | 0.099 | 5.18242006863055e-0 | 23 | ccnd1 |
| wls6 | 2.6796672 | 0.665792547 | 0.641 | 0.177 | 5.41212391545763e-0 | 23 | wls |
| cux1a4 | 2.7073853 | 0.560855629 | 0.487 | 0.105 | 5.46810629193739e-0 | 23 | cux1a |
| tjp2b4 | 2.7101189 | 0.48571035 | 0.41 | 0.078 | 5.47362725499725e-0 | 23 | tjp2b |
| ier2b5 | 2.7270661 | 0.778641359 | 0.872 | 0.32 | 5.50785547743776e-0 | 23 | ier2b |
| ngdn1 | 2.9962488 | 0.571522486 | 0.513 | 0.121 | 6.05152378382938e-0 | 23 | ngdn |
| her92 | 3.2967269 | 0.564659433 | 0.308 | 0.047 | 6.65839936403136e-0 | 23 | her9 |
| ftr14 | 3.5194745 | 0.266506619 | 0.205 | 0.021 | 7.10828266884676e-0 | 23 | ftr14 |
| sv2bb1 | 4.1705352 | 0.298703198 | 0.179 | 0.017 | 8.42322999891211e-0 | 23 | sv2bb |
| arl4d3 | 4.9954782 | 0.50231628 | 0.385 | 0.07 | 1.00893673864663e-08 | 23 | arl4d |
| itgav2 | 5.4124605 | 0.540371112 | 0.436 | 0.089 | 1.09315465850812e-0 | 23 | itgav |
| ehmt1b | 6.8904030 | 0.498852873 | 0.385 | 0.072 | 1.39165469822379e-0 | 23 | ehmt1b |
| ctnnb23 | 7.8113956 | 0.541236003 | 0.513 | 0.118 | 1.57766756981146e-0 | 23 | ctnnb2 |
| marcksb2 | 8.4849008 | 0.514967788 | 0.436 | 0.089 | 1.71369542527301e-0 | 23 | marcksb |
| hsd11b26 | 9.2566474 | 0.832824092 | 0.872 | 0.286 | 1.86956508778764e-0 | 23 | hsd11b2 |
| NAV1 (1 of many) | 1.0284294 | 0.445959719 | 0.308 | 0.049 | 2.07711888254331e-0 | 23 | NAV1 (1 of many) |
| tpm4a7 | 1.0322385 | 0.720539356 | 0.769 | 0.241 | 2.08481213660226e-0 | 23 | tpm4a |
| nucks1a5 | 1.0698291 | 0.711507818 | 0.718 | 0.227 | 2.1607339394517e-08 | 23 | nucks1a |
| ucmaa3 | 1.2002211 | 0.353650986 | 0.256 | 0.033 | 2.42408660074203e-0 | 23 | ucmaa |
| tspan3a2 | 1.3905591 | 0.50283263 | 0.487 | 0.112 | 2.80851240918191e-0 | 23 | tspan3a |
| hoxb3a1 | 1.5300913 | 0.320460808 | 0.231 | 0.028 | 3.09032540749451e-0 | 23 | hoxb3a |
| sumf1 | 1.5834578 | 0.31670872 | 0.256 | 0.034 | 3.19810973847594e-0 | 23 | sumf1 |

| | | | | | | | |
|---|---|---|---|---|---|---|---|
| si:dkey-42i9.72 | 1.6220600 | 0.378373614 | 0.308 | 0.047 | 3.27607473475192e-0 | 23 | si:dkey-42i9.7 |
| lsr2 | 2.2777653 | 0.334927246 | 0.282 | 0.041 | 4.60040265354341e-0 | 23 | lsr |
| dis3l2 | 2.5239753 | 0.288307795 | 0.231 | 0.029 | 5.09767297997814e-0 | 23 | dis3l2 |
| thtpa1 | 2.5768706 | 0.436953846 | 0.333 | 0.057 | 5.2045055508228e-08 | 23 | thtpa |
| epcam2 | 2.5790683 | 0.35390193 | 0.231 | 0.028 | 5.20894436317065e-0 | 23 | epcam |
| cd99l23 | 3.6461304 | 0.503746142 | 0.41 | 0.084 | 7.36408962480727e-0 | 23 | cd99l2 |
| nfia5 | 3.9284965 | 0.630547639 | 0.692 | 0.196 | 7.9343845259083e-08 | 23 | nfia |
| tgif13 | 4.3696794 | 0.60177784 | 0.564 | 0.149 | 8.8254415068142e-08 | 23 | tgif1 |
| ptmaa6 | 6.3578526 | 0.675313203 | 1 | 0.653 | 1.28409549135826e-0 | 23 | ptmaa |
| sox4a5 | 6.7887673 | 0.676066415 | 0.564 | 0.154 | 1.37112734501255e-0 | 23 | sox4a |
| zgc:923133 | 6.8348542 | 0.3011861 | 0.205 | 0.024 | 1.38043551234453e-0 | 23 | zgc:92313 |
| si:ch211-269e2.1 | 8.4688539 | 0.264953685 | 0.154 | 0.014 | 1.71045443833007e-0 | 23 | si:ch211-269e2.1 |
| nkd2a | 9.1070583 | 0.333602965 | 0.179 | 0.019 | 1.83935257321756e-0 | 23 | nkd2a |
| egr13 | 1.0483438 | 0.603984597 | 0.59 | 0.161 | 2.11734003137444e-0 | 23 | egr1 |
| si:dkey-188i13.114 | 1.6371297 | 0.61218425 | 0.615 | 0.18 | 3.30651086348474e-0 | 23 | si:dkey-188i13.11 |
| gcga1 | 1.7224054 | 0.362865987 | 0.231 | 0.031 | 3.47874223347033e-0 | 23 | gcga |
| hspb14 | 1.7452412 | 0.56429693 | 0.641 | 0.174 | 3.52486383758487e-0 | 23 | hspb1 |
| med13a | 2.0032790 | 0.275516538 | 0.179 | 0.019 | 4.04602271880973e-0 | 23 | med13a |
| psma6l | 2.0204294 | 0.414078358 | 0.359 | 0.07 | 4.08066141778451e-0 | 23 | psma6l |
| pard3bb2 | 2.0452108 | 0.385238763 | 0.359 | 0.067 | 4.13071234787326e-0 | 23 | pard3bb |
| vasnb3 | 2.0514449 | 0.408513584 | 0.333 | 0.059 | 4.14330343569516e-0 | 23 | vasnb |
| psmd142 | 2.0596303 | 0.543649854 | 0.564 | 0.158 | 4.15983544107766e-0 | 23 | psmd14 |
| spint1a2 | 2.4899065 | 0.259327106 | 0.231 | 0.03 | 5.02886434635108e-0 | 23 | spint1a |
| XLOC-0098153 | 3.3492857 | 0.416364771 | 0.333 | 0.06 | 6.7645525182683e-07 | 23 | XLOC-009815 |
| cyp27c13 | 3.8769847 | 0.442971647 | 0.333 | 0.062 | 7.83034606845706e-0 | 23 | cyp27c1 |
| si:ch211-276i12.11 | 4.3008079 | 0.355239848 | 0.359 | 0.069 | 8.68634182148674e-0 | 23 | si:ch211-276i12.11 |
| zgc:1533722 | 4.3950997 | 0.401410558 | 0.308 | 0.054 | 8.87678299221869e-0 | 23 | zgc:153372 |
| cldn11a6 | 4.6348664 | 0.699544009 | 0.641 | 0.191 | 9.36103979328644e-0 | 23 | cldn11a |
| zgc:770864 | 4.7267987 | 0.485908094 | 0.487 | 0.119 | 9.54671548098947e-0 | 23 | zgc:77086 |
| hmgb2b4 | 5.5392013 | 0.743153514 | 0.949 | 0.435 | 1.1187524886072e-06 | 23 | hmgb2b |
| rsad21 | 5.8166176 | 0.284267095 | 0.128 | 0.01 | 1.17478226273436e-0 | 23 | rsad2 |
| fosl1a | 5.8334696 | 0.438096395 | 0.282 | 0.048 | 1.17818585652843e-0 | 23 | fosl1a |
| si:ch211-1a19.33 | 6.4831245 | 0.451648438 | 0.385 | 0.079 | 1.30939666008064e-0 | 23 | si:ch211-1a19.3 |
| btbd6b1 | 6.7736026 | 0.36838775 | 0.205 | 0.027 | 1.36806451837583e-0 | 23 | btbd6b |
| hmgn64 | 6.8064178 | 0.637433889 | 0.949 | 0.569 | 1.37469221513755e-0 | 23 | hmgn6 |

| | | | | | | | |
|---|---|---|---|---|---|---|---|
| ctgfa4 | 7.2490504 | 0.493065527 | 0.333 | 0.063 | 1.46409071720705e-0 | 23 | ctgfa |
| tle2b2 | 7.2732755 | 0.334710798 | 0.231 | 0.032 | 1.46898346352749e-0 | 23 | tle2b |
| ywhah3 | 7.4997706 | 0.571914904 | 0.487 | 0.129 | 1.51472867156029e-0 | 23 | ywhah |
| btg26 | 8.3347999 | 0.647276612 | 0.744 | 0.256 | 1.68337954500661e-0 | 23 | btg2 |
| fam133b3 | 8.9558812 | 0.449975204 | 0.692 | 0.206 | 1.80881932819272e-0 | 23 | fam133b |
| ier2a5 | 9.2600615 | 0.755884231 | 0.846 | 0.359 | 1.87025462519203e-0 | 23 | ier2a |
| tia11 | 1.4336643 | 0.449736386 | 0.41 | 0.093 | 2.89557180537224e-0 | 23 | tia1 |
| CU984600.21 | 1.4878703 | 0.318610879 | 0.205 | 0.027 | 3.00505180706535e-0 | 23 | CU984600.2 |
| ppfibp1a4 | 1.4887048 | 0.409475391 | 0.282 | 0.049 | 3.0067372052568e-06 | 23 | ppfibp1a |
| CABZ01087560.1 | 1.5323884 | 0.433521796 | 0.462 | 0.113 | 3.09496497456103e-0 | 23 | CABZ01087560.1 |
| lzts2a | 2.3605589 | 0.28446764 | 0.205 | 0.028 | 4.76762094412494e-0 | 23 | lzts2a |
| coro1ca2 | 2.5495310 | 0.48039624 | 0.359 | 0.077 | 5.14928790611903e-0 | 23 | coro1ca |
| prdx11 | 2.7192670 | 0.381488473 | 0.333 | 0.065 | 5.49210359947576e-0 | 23 | prdx1 |
| nucks1b2 | 2.7759416 | 0.472773816 | 0.564 | 0.158 | 5.60656930947579e-0 | 23 | nucks1b |
| hmgb1b6 | 2.9140953 | 0.621301639 | 0.821 | 0.297 | 5.88559828512799e-0 | 23 | hmgb1b |
| lnx2b | 3.2306218 | 0.282947984 | 0.231 | 0.034 | 6.5248870248745e-06 | 23 | lnx2b |
| fam168a4 | 3.6474397 | 0.505740627 | 0.538 | 0.151 | 7.36673413618599e-0 | 23 | fam168a |
| zdhhc13 | 3.7412597 | 0.345249931 | 0.256 | 0.042 | 7.55622232892576e-0 | 23 | zdhhc13 |
| dachc | 4.5026728 | 0.322119795 | 0.179 | 0.022 | 9.09404830802358e-0 | 23 | dachc |
| XLOC-0412693 | 4.6371350 | 0.417081268 | 0.308 | 0.058 | 9.36562167528665e-0 | 23 | XLOC-041269 |
| cldn121 | 4.6763103 | 0.358508216 | 0.282 | 0.051 | 9.44474393395244e-0 | 23 | cldn12 |
| itpa1 | 4.8042073 | 0.327273875 | 0.282 | 0.05 | 9.70305755375371e-0 | 23 | itpa |
| si:ch73-61d6.32 | 4.8869159 | 0.31324703 | 0.231 | 0.035 | 9.87010419814642e-0 | 23 | si:ch73-61d6.3 |
| si:ch211-199g17.2: | 5.2718918 | 0.396496593 | 0.385 | 0.085 | 1.06476398736852e-0 | 23 | si:ch211-199g17.2 |
| ccdc90b | 5.3179658 | 0.356228494 | 0.282 | 0.051 | 1.07406955541214e-0 | 23 | ccdc90b |
| mov10b.11 | 5.4746819 | 0.288357982 | 0.205 | 0.029 | 1.10572151101513e-0 | 23 | mov10b.1 |
| dnajc7 | 6.2574806 | 0.291989367 | 0.256 | 0.043 | 1.2638233663131e-05 | 23 | dnajc7 |
| phactr4b | 6.8709058 | 0.289942678 | 0.179 | 0.023 | 1.38771685603484e-0 | 23 | phactr4b |
| cnpy42 | 7.3427637 | 0.330573652 | 0.282 | 0.051 | 1.48301799273453e-0 | 23 | cnpy4 |
| si:dkey-6n21.122 | 7.9082921 | 0.391758322 | 0.359 | 0.076 | 1.59723776230535e-0 | 23 | si:dkey-6n21.12 |
| dpyda.1 | 8.4388840 | 0.260331631 | 0.179 | 0.023 | 1.70440141108361e-0 | 23 | dpyda.1 |
| afdna | 8.6695975 | 0.323538486 | 0.256 | 0.043 | 1.75099861613719e-0 | 23 | afdna |
| hist2h2l5 | 9.2855943 | 0.625955399 | 0.615 | 0.207 | 1.87541149904244e-0 | 23 | hist2h2l |
| hmgn74 | 9.5187139 | 0.638058135 | 0.949 | 0.527 | 1.92249466062252e-0 | 23 | hmgn7 |
| dhrs3a3 | 9.5741063 | 0.445367059 | 0.359 | 0.078 | 1.93368225895057e-0 | 23 | dhrs3a |

| | | | | | | | |
|---|---|---|---|---|---|---|---|
| sema3b2 | 1.0226693 | 0.263050187 | 0.179 | 0.023 | 2.06548524617577e-0 | 23 | sema3b |
| psmc52 | 1.0797555 | 0.481516188 | 0.59 | 0.186 | 2.18078231426961e-0 | 23 | psmc5 |
| hnrnpabb3 | 1.0983630 | 0.623034219 | 0.949 | 0.4 | 2.21836394695636e-0 | 23 | hnrnpabb |
| cntrl | 1.1442758 | 0.29670463 | 0.154 | 0.017 | 2.31109397755254e-0 | 23 | cntrl |
| ptprz1b1 | 1.1859566 | 0.252252586 | 0.179 | 0.023 | 2.39527669164394e-0 | 23 | ptprz1b |
| nkd12 | 1.1870807 | 0.383846664 | 0.256 | 0.045 | 2.39754709003384e-0 | 23 | nkd1 |
| dchs1b1 | 1.1932645 | 0.285783574 | 0.154 | 0.017 | 2.41003638091502e-0 | 23 | dchs1b |
| si:ch211-154o6.3 | 1.2100439 | 0.265814415 | 0.154 | 0.017 | 2.44392576763363e-0 | 23 | si:ch211-154o6.3 |
| foxo4 | 1.2141783 | 0.356340436 | 0.256 | 0.045 | 2.45227595513023e-0 | 23 | foxo4 |
| pias4a | 1.2300771 | 0.302366266 | 0.256 | 0.044 | 2.48438685153095e-0 | 23 | pias4a |
| LOC1039092845 | 1.2533456 | 0.496178043 | 0.513 | 0.139 | 2.53138216714146e-0 | 23 | LOC103909284 |
| chd4b2 | 1.3034096 | 0.495660943 | 0.487 | 0.139 | 2.63249652319624e-0 | 23 | chd4b |
| BX957297.2 | 1.3922396 | 0.281012974 | 0.179 | 0.023 | 2.8119064377827e-05 | 23 | BX957297.2 |
| tle2a1 | 1.5267272 | 0.325520218 | 0.231 | 0.037 | 3.08353098269641e-0 | 23 | tle2a |
| spop | 1.6721455 | 0.320352909 | 0.256 | 0.044 | 3.37723226994533e-0 | 23 | spop |
| rbbp61 | 2.0766961 | 0.426046216 | 0.487 | 0.132 | 4.19430329951985e-0 | 23 | rbbp6 |
| ponzr17 | 2.4783303 | 0.677661787 | 0.872 | 0.389 | 5.00548376852973e-0 | 23 | ponzr1 |
| s100v28 | 2.4975480 | 0.619894125 | 0.692 | 0.247 | 5.04429781689035e-0 | 23 | s100v2 |
| smarcd12 | 2.5244569 | 0.395928936 | 0.359 | 0.083 | 5.09864560898296e-0 | 23 | smarcd1 |
| nme35 | 2.6341508 | 0.489130231 | 0.538 | 0.158 | 5.32019451063688e-0 | 23 | nme3 |
| si:ch73-376l24.23 | 2.7280910 | 0.430147905 | 0.564 | 0.163 | 5.50992557768033e-0 | 23 | si:ch73-376l24.2 |
| psmd133 | 2.8518311 | 0.529845576 | 0.487 | 0.148 | 5.7598433832038e-05 | 23 | psmd13 |
| fcho2 | 3.2173290 | 0.265241789 | 0.231 | 0.037 | 6.49803951830017e-0 | 23 | fcho2 |
| ar3 | 3.4171256 | 0.512689821 | 0.41 | 0.107 | 6.90156873698912e-0 | 23 | ar |
| CABZ01092156.12 | 3.5851211 | 0.267019843 | 0.205 | 0.03 | 7.24086923523816e-0 | 23 | CABZ01092156.1 |
| cfap45 | 3.6700591 | 0.36442257 | 0.205 | 0.031 | 7.41241856169781e-0 | 23 | cfap45 |
| tsc22d12 | 3.7261102 | 0.419458833 | 0.282 | 0.056 | 7.52562487343206e-0 | 23 | tsc22d1 |
| znf5031 | 3.8322910 | 0.364706498 | 0.256 | 0.047 | 7.74007823775314e-0 | 23 | znf503 |
| seh1l | 3.9111799 | 0.41403238 | 0.333 | 0.075 | 7.89941006556378e-0 | 23 | seh1l |
| CT573476.1 | 4.4421335 | 0.286316024 | 0.205 | 0.031 | 8.97177707910091e-0 | 23 | CT573476.1 |
| nfic2 | 4.9951712 | 0.504723026 | 0.436 | 0.118 | 0.000100887 | 23 | nfic |
| irs12 | 5.3356943 | 0.302878789 | 0.256 | 0.046 | 0.000107765 | 23 | irs1 |
| prkd22 | 5.8048264 | 0.356516802 | 0.231 | 0.04 | 0.00011724 | 23 | prkd2 |
| ddah21 | 6.0505228 | 0.353874869 | 0.333 | 0.073 | 0.000122202 | 23 | ddah2 |
| tango2 | 6.2231171 | 0.351013826 | 0.256 | 0.048 | 0.000125688 | 23 | tango2 |

| | | | | | | | |
|---|---|---|---|---|---|---|---|
| si:ch73-281n10.28 | 6.6410035 | 0.657438398 | 0.897 | 0.43 | 0.000134128 | 23 | si:ch73-281n10.2 |
| si:dkey-188i13.103 | 6.6687770 | 0.521526644 | 0.538 | 0.166 | 0.000134689 | 23 | si:dkey-188i13.10 |
| pbx45 | 6.9531177 | 0.426927766 | 0.41 | 0.106 | 0.000140432 | 23 | pbx4 |
| sumo3a5 | 7.1824531 | 0.619346807 | 0.846 | 0.405 | 0.000145064 | 23 | sumo3a |
| fthl276 | 7.4572160 | 0.568332671 | 0.974 | 0.626 | 0.000150613 | 23 | fthl27 |
| mycb5 | 7.5738742 | 0.535386546 | 0.513 | 0.158 | 0.00015297 | 23 | mycb |
| pak4 | 7.7466392 | 0.255640424 | 0.179 | 0.025 | 0.000156459 | 23 | pak4 |
| naa30 | 7.8871884 | 0.252943721 | 0.205 | 0.032 | 0.000159298 | 23 | naa30 |
| ptmab1 | 8.0729976 | 0.471531978 | 0.974 | 0.761 | 0.00016305 | 23 | ptmab |
| elof11 | 8.4103423 | 0.474086383 | 0.59 | 0.197 | 0.000169864 | 23 | elof1 |
| wnt4a2 | 8.4261897 | 0.29886775 | 0.179 | 0.025 | 0.000170184 | 23 | wnt4a |
| rgs5b1 | 9.0868547 | 0.461671855 | 0.282 | 0.058 | 0.000183527 | 23 | rgs5b |
| cox8b3 | 9.9409503 | 0.409941562 | 0.462 | 0.131 | 0.000200777 | 23 | cox8b |
| atrx4 | 1.0010923 | 0.56702777 | 0.641 | 0.24 | 0.000202191 | 23 | atrx |
| st51 | 1.0012101 | 0.327512243 | 0.231 | 0.04 | 0.000202214 | 23 | st5 |
| psmc62 | 1.0935025 | 0.448181077 | 0.487 | 0.146 | 0.000220855 | 23 | psmc6 |
| magi1b | 1.2776742 | 0.254198004 | 0.205 | 0.032 | 0.000258052 | 23 | magi1b |
| stat22 | 1.3266764 | 0.441930226 | 0.308 | 0.069 | 0.000267949 | 23 | stat2 |
| otub1b | 1.3749874 | 0.346330458 | 0.308 | 0.067 | 0.000277706 | 23 | otub1b |
| ttc27 | 1.3948559 | 0.292733793 | 0.179 | 0.026 | 0.000281719 | 23 | ttc27 |
| si:ch73-46j18.59 | 1.3968853 | 0.677382788 | 0.923 | 0.391 | 0.000282129 | 23 | si:ch73-46j18.5 |
| BX284638.1 | 1.3979221 | 0.259154776 | 0.179 | 0.026 | 0.000282338 | 23 | BX284638.1 |
| sart11 | 1.4597021 | 0.390819418 | 0.436 | 0.12 | 0.000294816 | 23 | sart1 |
| ubb2 | 1.4652602 | 0.560037404 | 0.872 | 0.371 | 0.000295939 | 23 | ubb |
| sub1a2 | 1.5463371 | 0.564090925 | 0.821 | 0.362 | 0.000312314 | 23 | sub1a |
| golga72 | 1.6575946 | 0.49824915 | 0.538 | 0.177 | 0.000334784 | 23 | golga7 |
| gpc41 | 1.7142531 | 0.343964635 | 0.256 | 0.05 | 0.000346228 | 23 | gpc4 |
| erbb21 | 1.7142531 | 0.325350432 | 0.256 | 0.05 | 0.000346228 | 23 | erbb2 |
| fkbp1aa5 | 1.7705893 | 0.574054889 | 0.949 | 0.577 | 0.000357606 | 23 | fkbp1aa |
| psmc1b | 1.7818520 | 0.340651718 | 0.333 | 0.077 | 0.000359881 | 23 | psmc1b |
| si:ch211-22i13.2 | 1.8834474 | 0.388622443 | 0.385 | 0.1 | 0.0003804 | 23 | si:ch211-22i13.2 |
| wwtr11 | 1.9478215 | 0.318588638 | 0.256 | 0.049 | 0.000393402 | 23 | wwtr1 |
| tnfrsfa2 | 1.9807402 | 0.284644252 | 0.256 | 0.048 | 0.00040005 | 23 | tnfrsfa |
| mycn1 | 2.0106676 | 0.253231942 | 0.154 | 0.02 | 0.000406095 | 23 | mycn |
| aldh1a25 | 2.1231545 | 0.4438299 | 0.641 | 0.199 | 0.000428814 | 23 | aldh1a2 |

| | | | | | | | |
|---|---|---|---|---|---|---|---|
| parp12b3 | 2.1274296 | 0.339398082 | 0.333 | 0.077 | 0.000429677 | 23 | parp12b |
| zgc:174906 | 2.1374925 | 0.368457294 | 0.256 | 0.051 | 0.000431709 | 23 | zgc:174906 |
| tspan18b1 | 2.2087785 | 0.366036382 | 0.308 | 0.068 | 0.000446107 | 23 | tspan18b |
| CR376783.11 | 2.2300663 | 0.254832466 | 0.179 | 0.026 | 0.000450406 | 23 | CR376783.1 |
| psmd4b1 | 2.4693845 | 0.424901133 | 0.436 | 0.126 | 0.000498742 | 23 | psmd4b |
| AK61 | 2.4728419 | 0.348246744 | 0.282 | 0.06 | 0.00049944 | 23 | AK6 |
| rmdn13 | 2.5436839 | 0.38147782 | 0.41 | 0.112 | 0.000513748 | 23 | rmdn1 |
| schip11 | 2.5656407 | 0.341186064 | 0.205 | 0.034 | 0.000518182 | 23 | schip1 |
| tardbpl | 2.7542345 | 0.45950355 | 0.487 | 0.15 | 0.000556273 | 23 | tardbpl |
| dnajc91 | 2.7930028 | 0.385133233 | 0.308 | 0.07 | 0.000564103 | 23 | dnajc9 |
| sf3b2 | 3.2034875 | 0.414656697 | 0.436 | 0.126 | 0.000647008 | 23 | sf3b2 |
| ehd2b2 | 3.2928467 | 0.320497118 | 0.256 | 0.05 | 0.000665056 | 23 | ehd2b |
| hmgb3a7 | 3.4634197 | 0.618011131 | 0.641 | 0.257 | 0.000699507 | 23 | hmgb3a |
| si:ch1073-157b13.1 | 3.5263870 | 0.263947995 | 0.205 | 0.034 | 0.000712224 | 23 | si:ch1073-157b13.1 |
| gna11b2 | 3.5648033 | 0.363254175 | 0.333 | 0.079 | 0.000719983 | 23 | gna11b |
| mir21-12 | 3.8706684 | 0.389816361 | 0.41 | 0.11 | 0.000781759 | 23 | mir21-1 |
| anks1b | 4.0085351 | 0.274378196 | 0.179 | 0.028 | 0.000809604 | 23 | anks1b |
| hdac12 | 4.3072541 | 0.550285763 | 0.667 | 0.274 | 0.000869936 | 23 | hdac1 |
| nsd3 | 4.4264536 | 0.27765676 | 0.205 | 0.034 | 0.000894011 | 23 | nsd3 |
| ctnnd1 | 4.5774831 | 0.289916786 | 0.231 | 0.043 | 0.000924514 | 23 | ctnnd1 |
| si:dkey-42i9.45 | 4.6271665 | 0.496122251 | 0.564 | 0.195 | 0.000934549 | 23 | si:dkey-42i9.4 |
| hmga22 | 4.7478891 | 0.362596238 | 0.256 | 0.052 | 0.000958931 | 23 | hmga2 |
| zfand5b | 5.1977880 | 0.403832644 | 0.41 | 0.116 | 0.001049797 | 23 | zfand5b |
| plod23 | 5.3082713 | 0.286436095 | 0.282 | 0.059 | 0.001072112 | 23 | plod2 |
| tshz3b1 | 5.4305959 | 0.339843264 | 0.282 | 0.06 | 0.001096817 | 23 | tshz3b |
| chtopa2 | 5.4511517 | 0.453652712 | 0.462 | 0.143 | 0.001100969 | 23 | chtopa |
| emd2 | 5.5257932 | 0.427189149 | 0.487 | 0.15 | 0.001116044 | 23 | emd |
| cfap20 | 5.7234221 | 0.298870993 | 0.282 | 0.061 | 0.00115596 | 23 | cfap20 |
| nfil3 | 6.0250629 | 0.296468335 | 0.205 | 0.035 | 0.001216882 | 23 | nfil3 |
| h2afy25 | 6.1825233 | 0.473473943 | 0.692 | 0.252 | 0.001248684 | 23 | h2afy2 |
| serinc24 | 6.1842776 | 0.491601133 | 0.564 | 0.196 | 0.001249039 | 23 | serinc2 |
| hnrnpl1 | 6.2324046 | 0.406115174 | 0.308 | 0.075 | 0.001258759 | 23 | hnrnpl |
| tp531 | 6.3284668 | 0.523485893 | 0.513 | 0.179 | 0.00127816 | 23 | tp53 |
| stat1b1 | 6.8952655 | 0.276931162 | 0.154 | 0.021 | 0.001392637 | 23 | stat1b |
| srsf3b3 | 7.0303707 | 0.530852535 | 0.744 | 0.303 | 0.001419924 | 23 | srsf3b |

| | | | | | | | |
|---|---|---|---|---|---|---|---|
| phc2a3 | 7.1757499 | 0.380064167 | 0.359 | 0.092 | 0.001449286 | 23 | phc2a |
| si:ch211-251p5.51 | 7.2540037 | 0.269653346 | 0.179 | 0.028 | 0.001465091 | 23 | si:ch211-251p5.5 |
| clic41 | 7.4567934 | 0.421244814 | 0.385 | 0.104 | 0.001506049 | 23 | clic4 |
| rn7sk2 | 7.6434847 | 0.381896436 | 0.385 | 0.102 | 0.001543755 | 23 | rn7sk |
| ywhaqa3 | 8.4911927 | 0.524325337 | 0.641 | 0.251 | 0.001714966 | 23 | ywhaqa |
| mrpl20 | 8.5359117 | 0.369529593 | 0.41 | 0.118 | 0.001723998 | 23 | mrpl20 |
| psma6a3 | 1.0028437 | 0.400179317 | 0.487 | 0.155 | 0.002025444 | 23 | psma6a |
| atp1b1a4 | 1.0491507 | 0.588008867 | 0.667 | 0.292 | 0.00211897 | 23 | atp1b1a |
| mycbp3 | 1.0638363 | 0.457987524 | 0.615 | 0.224 | 0.00214863 | 23 | mycbp |
| h2afvb2 | 1.1170414 | 0.571819581 | 0.974 | 0.526 | 0.002256089 | 23 | h2afvb |
| smarce14 | 1.1532135 | 0.50925072 | 0.615 | 0.244 | 0.002329145 | 23 | smarce1 |
| snrpe2 | 1.1771472 | 0.457401072 | 0.692 | 0.271 | 0.002377484 | 23 | snrpe |
| si:ch211-288g17.34 | 1.3271955 | 0.623698509 | 0.667 | 0.291 | 0.002680537 | 23 | si:ch211-288g17.3 |
| snrpg2 | 1.3931069 | 0.472788844 | 0.692 | 0.266 | 0.002813658 | 23 | snrpg |
| ulk2 | 1.4042775 | 0.29430152 | 0.333 | 0.082 | 0.002836219 | 23 | ulk2 |
| kmt5aa3 | 1.4089119 | 0.390166175 | 0.487 | 0.156 | 0.002845579 | 23 | kmt5aa |
| zgc:103692 | 1.4224430 | 0.256283936 | 0.205 | 0.037 | 0.002872908 | 23 | zgc:103692 |
| elavl1b | 1.4556548 | 0.253725154 | 0.179 | 0.029 | 0.002939986 | 23 | elavl1b |
| tjp1a2 | 1.6331791 | 0.36232075 | 0.333 | 0.084 | 0.003298532 | 23 | tjp1a |
| stxbp62 | 1.6507347 | 0.292295014 | 0.282 | 0.063 | 0.003333989 | 23 | stxbp6 |
| myh103 | 1.7796087 | 0.347248951 | 0.308 | 0.074 | 0.003594276 | 23 | myh10 |
| psat1 | 1.7895506 | 0.251943328 | 0.179 | 0.029 | 0.003614355 | 23 | psat1 |
| vgll4b | 1.8918453 | 0.257456464 | 0.205 | 0.038 | 0.00382096 | 23 | vgll4b |
| hnrnpa0a2 | 1.9013904 | 0.40096742 | 0.462 | 0.143 | 0.003840238 | 23 | hnrnpa0a |
| wscd21 | 2.0440049 | 0.275211408 | 0.179 | 0.029 | 0.004128277 | 23 | wscd2 |
| traf4a | 2.0562088 | 0.254535515 | 0.128 | 0.016 | 0.004152925 | 23 | traf4a |
| psmb12 | 2.0744903 | 0.52515285 | 0.744 | 0.337 | 0.004189848 | 23 | psmb1 |
| fbxl62 | 2.1123447 | 0.421947373 | 0.41 | 0.124 | 0.004266303 | 23 | fbxl6 |
| sipa1l2 | 2.1766708 | 0.261147862 | 0.205 | 0.037 | 0.004396222 | 23 | sipa1l2 |
| si:ch1073-429i10.3 | 2.1791201 | 0.471741085 | 0.821 | 0.347 | 0.004401169 | 23 | si:ch1073-429i10.3.1 |
| rad21a4 | 2.2576137 | 0.482729121 | 0.641 | 0.247 | 0.004559702 | 23 | rad21a |
| dbn14 | 2.3141253 | 0.405654476 | 0.41 | 0.121 | 0.004673839 | 23 | dbn1 |
| tax1bp33 | 2.6683695 | 0.4033641 | 0.513 | 0.167 | 0.005389306 | 23 | tax1bp3 |
| rab28 | 2.6695433 | 0.253239773 | 0.179 | 0.03 | 0.005391677 | 23 | rab28 |
| txndc172 | 2.7775916 | 0.325650366 | 0.308 | 0.077 | 0.005609902 | 23 | txndc17 |

| | | | | | | | |
|---|---|---|---|---|---|---|---|
| CU459056.11 | 2.7858132 | 0.35854887 | 0.308 | 0.077 | 0.005626507 | 23 | CU459056.1 |
| znfx12 | 2.8500243 | 0.31586378 | 0.231 | 0.047 | 0.005756194 | 23 | znfx1 |
| mrpl52 | 2.8546227 | 0.265704561 | 0.333 | 0.085 | 0.005765482 | 23 | mrpl52 |
| actn14 | 2.9509178 | 0.393682366 | 0.282 | 0.068 | 0.005959969 | 23 | actn1 |
| si:ch211-197h24.9 | 3.0687694 | 0.429408191 | 0.487 | 0.161 | 0.006197994 | 23 | si:ch211-197h24.9 |
| nos1apa2 | 3.3014980 | 0.339587825 | 0.282 | 0.067 | 0.006668036 | 23 | nos1apa |
| ywhaqb4 | 3.3344906 | 0.485823874 | 0.692 | 0.276 | 0.006734671 | 23 | ywhaqb |
| serbp1b | 3.3671115 | 0.355209309 | 0.41 | 0.121 | 0.006800555 | 23 | serbp1b |
| brd3a | 3.5491019 | 0.371752392 | 0.308 | 0.079 | 0.007168121 | 23 | brd3a |
| usp7 | 3.5654390 | 0.295929797 | 0.282 | 0.067 | 0.007201117 | 23 | usp7 |
| pfn2l8 | 3.5765871 | 0.501604798 | 0.667 | 0.26 | 0.007223633 | 23 | pfn2l |
| rrbp1a2 | 3.5866835 | 0.370873996 | 0.359 | 0.1 | 0.007244025 | 23 | rrbp1a |
| isg153 | 3.7809251 | 0.327085675 | 0.231 | 0.048 | 0.007636335 | 23 | isg15 |
| rps27l | 3.7929825 | 0.485603902 | 0.692 | 0.301 | 0.007660687 | 23 | rps27l |
| calcoco1a | 3.8114041 | 0.279331651 | 0.256 | 0.057 | 0.007697893 | 23 | calcoco1a |
| prdx26 | 4.0843044 | 0.551378636 | 0.872 | 0.486 | 0.00824907 | 23 | prdx2 |
| tardbp1 | 4.2192197 | 0.37096805 | 0.385 | 0.114 | 0.008521558 | 23 | tardbp |
| qkia6 | 4.3199041 | 0.570514578 | 0.744 | 0.357 | 0.00872491 | 23 | qkia |
| esd3 | 4.6841238 | 0.459614023 | 0.487 | 0.175 | 0.009460525 | 23 | esd |
| sumo11 | 4.8258184 | 0.40816097 | 0.513 | 0.18 | 0.009746706 | 23 | sumo1 |
| top1l2 | 5.2682440 | 0.426266824 | 0.513 | 0.176 | 0.010640272 | 23 | top1l |
| cfdp1 | 5.4421345 | 0.371545303 | 0.282 | 0.071 | 0.010991479 | 23 | cfdp1 |
| rhogd | 5.4890333 | 0.258729953 | 0.256 | 0.057 | 0.011086201 | 23 | rhogd |
| gstm.37 | 5.5948195 | 0.596226416 | 0.718 | 0.284 | 0.011299857 | 23 | gstm.3 |
| baz1a1 | 5.6726619 | 0.343988794 | 0.308 | 0.081 | 0.011457075 | 23 | baz1a |
| zgc:194981 | 5.7302410 | 0.295107919 | 0.282 | 0.068 | 0.011573368 | 23 | zgc:194981 |
| chd37 | 5.7612248 | 0.454771783 | 0.641 | 0.238 | 0.011635946 | 23 | chd3 |
| ada2a | 5.9686301 | 0.254114708 | 0.179 | 0.032 | 0.012054842 | 23 | ada2a |
| cald1a7 | 6.5949822 | 0.456689721 | 0.641 | 0.234 | 0.013319886 | 23 | cald1a |
| cavin2b4 | 6.7292661 | 0.404885352 | 0.462 | 0.145 | 0.013591099 | 23 | cavin2b |
| rbm39a3 | 6.8310398 | 0.433250937 | 0.564 | 0.209 | 0.013796651 | 23 | rbm39a |
| pomp1 | 7.4327492 | 0.503776341 | 0.846 | 0.413 | 0.015011924 | 23 | pomp |
| pabpn13 | 7.4619652 | 0.489077724 | 0.667 | 0.291 | 0.015070931 | 23 | pabpn1 |
| nop583 | 7.5147207 | 0.460006451 | 0.564 | 0.218 | 0.015177482 | 23 | nop58 |
| ncoa5 | 7.5371303 | 0.278328568 | 0.282 | 0.069 | 0.015222742 | 23 | ncoa5 |

| | | | | | | | |
|---|---|---|---|---|---|---|---|
| dpy302 | 7.9867833 | 0.391797235 | 0.59 | 0.226 | 0.016130906 | 23 | dpy30 |
| lgals173 | 8.0179821 | 0.416318877 | 0.282 | 0.071 | 0.016193918 | 23 | lgals17 |
| foxo3b3 | 8.0962747 | 0.401169985 | 0.333 | 0.094 | 0.016352046 | 23 | foxo3b |
| si:ch211-194b1.11 | 8.3604835 | 0.285481596 | 0.205 | 0.041 | 0.016885669 | 23 | si:ch211-194b1.1 |
| psma41 | 8.6499268 | 0.446808789 | 0.692 | 0.286 | 0.017470257 | 23 | psma4 |
| mcm72 | 8.9998628 | 0.308620979 | 0.359 | 0.1 | 0.018177023 | 23 | mcm7 |
| atp2b22 | 9.1601651 | 0.437793835 | 0.436 | 0.148 | 0.018500786 | 23 | atp2b2 |
| myl12.21 | 9.1672607 | 0.299905276 | 0.385 | 0.114 | 0.018515117 | 23 | myl12.2 |
| sparc9 | 1.0166210 | 0.584801807 | 0.923 | 0.397 | 0.020532696 | 23 | sparc |
| nckap1 | 1.0402285 | 0.298948905 | 0.256 | 0.06 | 0.021009495 | 23 | nckap1 |
| etf1b1 | 1.1117220 | 0.380099503 | 0.564 | 0.2 | 0.02245345 | 23 | etf1b |
| tomm22 | 1.1228734 | 0.302861225 | 0.41 | 0.126 | 0.022678675 | 23 | tomm22 |
| psmc1a2 | 1.1320238 | 0.367972805 | 0.462 | 0.153 | 0.022863487 | 23 | psmc1a |
| efnb14 | 1.1357735 | 0.330969394 | 0.359 | 0.1 | 0.022939219 | 23 | efnb1 |
| srrm11 | 1.1679416 | 0.430159748 | 0.41 | 0.137 | 0.023588917 | 23 | srrm1 |
| wdr45 | 1.1901090 | 0.302867268 | 0.256 | 0.061 | 0.024036632 | 23 | wdr45 |
| si:ch73-1a9.34 | 1.2100901 | 0.523572066 | 0.923 | 0.586 | 0.02444019 | 23 | si:ch73-1a9.3 |
| chka2 | 1.2114156 | 0.268224786 | 0.231 | 0.05 | 0.024466962 | 23 | chka |
| calr4 | 1.2605107 | 0.410496634 | 0.436 | 0.146 | 0.025458535 | 23 | calr |
| ilf3b1 | 1.2969442 | 0.355582518 | 0.41 | 0.129 | 0.026194382 | 23 | ilf3b |
| CABZ01021532.12 | 1.3414986 | 0.253573556 | 0.154 | 0.025 | 0.027094249 | 23 | CABZ01021532.1 |
| mgst3a2 | 1.3558270 | 0.357570602 | 0.436 | 0.143 | 0.027383639 | 23 | mgst3a |
| gstm.17 | 1.4609221 | 0.509304281 | 0.769 | 0.336 | 0.029506245 | 23 | gstm.1 |
| psip1b3 | 1.5683722 | 0.366930263 | 0.385 | 0.118 | 0.031676414 | 23 | psip1b |
| usp51 | 1.7396321 | 0.326109409 | 0.256 | 0.063 | 0.035135351 | 23 | usp5 |
| ash1l1 | 1.7626145 | 0.295739123 | 0.256 | 0.061 | 0.035599527 | 23 | ash1l |
| hnrnpa0b5 | 1.7844903 | 0.521464416 | 0.923 | 0.511 | 0.036041351 | 23 | hnrnpa0b |
| khdrbs1a4 | 1.8167772 | 0.492848784 | 0.949 | 0.593 | 0.036693449 | 23 | khdrbs1a |
| rbm4.32 | 1.8238949 | 0.483677276 | 0.718 | 0.333 | 0.036837207 | 23 | rbm4.3 |
| atp1a1a.13 | 1.9129695 | 0.380241517 | 0.41 | 0.134 | 0.038636246 | 23 | atp1a1a.1 |
| chd63 | 1.9148690 | 0.332088312 | 0.333 | 0.094 | 0.03867461 | 23 | chd6 |
| meis1b6 | 1.9261169 | 0.41043537 | 0.462 | 0.155 | 0.038901784 | 23 | meis1b |
| edf13 | 1.9350798 | 0.448029878 | 0.795 | 0.364 | 0.039082808 | 23 | edf1 |
| hsp90ab1 | 1.9545780 | 0.255123525 | 1 | 0.914 | 0.039476612 | 23 | hsp90ab1 |
| rnf442 | 1.9881914 | 0.412656689 | 0.41 | 0.136 | 0.040155503 | 23 | rnf44 |

| | | | | | | | |
|---|---|---|---|---|---|---|---|
| psmd82 | 2.0157094 | 0.394524727 | 0.436 | 0.151 | 0.040711285 | 23 | psmd8 |
| ddx614 | 2.0367426 | 0.486635738 | 0.718 | 0.34 | 0.041136091 | 23 | ddx61 |
| krt85 | 2.1932593 | 0.534428698 | 0.897 | 0.52 | 0.04429726 | 23 | krt8 |
| bckdhb | 2.2393260 | 0.285412602 | 0.256 | 0.064 | 0.045227669 | 23 | bckdhb |
| pet100 | 2.3832736 | 0.303553802 | 0.308 | 0.085 | 0.048134978 | 23 | pet100 |
| ube2ia2 | 2.4052857 | 0.405128274 | 0.487 | 0.18 | 0.048579557 | 23 | ube2ia |
| tnrc6c15 | 2.4824818 | 0.445813517 | 0.59 | 0.239 | 0.050138687 | 23 | tnrc6c1 |
| ext21 | 2.5648454 | 0.291082547 | 0.231 | 0.053 | 0.051802183 | 23 | ext2 |
| si:ch211-51e12.74 | 2.5880193 | 0.390996381 | 0.513 | 0.19 | 0.052270227 | 23 | si:ch211-51e12.7 |
| txnipa3 | 2.6127286 | 0.45364478 | 0.821 | 0.365 | 0.052769281 | 23 | txnipa |
| psma32 | 2.6427582 | 0.430408531 | 0.667 | 0.283 | 0.053375789 | 23 | psma3 |
| srgap31 | 2.6547282 | 0.366573259 | 0.231 | 0.055 | 0.053617546 | 23 | srgap3 |
| march6 | 2.6935091 | 0.286885669 | 0.205 | 0.044 | 0.054400804 | 23 | march6 |
| psmb31 | 2.7030015 | 0.420485327 | 0.615 | 0.255 | 0.054592521 | 23 | psmb3 |
| pde4ba3 | 2.8116103 | 0.44423812 | 0.462 | 0.161 | 0.056786095 | 23 | pde4ba |
| lsm43 | 2.8466625 | 0.43626215 | 0.615 | 0.259 | 0.057494043 | 23 | lsm4 |
| ppig1 | 2.8565975 | 0.271443141 | 0.308 | 0.084 | 0.057694701 | 23 | ppig |
| snrpd3l3 | 2.9043384 | 0.339485521 | 0.462 | 0.161 | 0.058658923 | 23 | snrpd3l |
| eif1axa1 | 3.0962115 | 0.384402851 | 0.385 | 0.128 | 0.062534185 | 23 | eif1axa |
| snrpd12 | 3.3201321 | 0.448830168 | 0.641 | 0.275 | 0.067056709 | 23 | snrpd1 |
| h2afx15 | 3.5812804 | 0.509255699 | 0.769 | 0.378 | 0.072331122 | 23 | h2afx1 |
| brd9 | 4.0460032 | 0.272808343 | 0.282 | 0.074 | 0.081717128 | 23 | brd9 |
| psmb74 | 4.1429890 | 0.419205869 | 0.692 | 0.303 | 0.08367595 | 23 | psmb7 |
| ptbp1b2 | 4.2831685 | 0.294537905 | 0.256 | 0.065 | 0.086507155 | 23 | ptbp1b |
| elovl1b2 | 4.3276419 | 0.381547502 | 0.462 | 0.163 | 0.087405384 | 23 | elovl1b |
| nt5c2b1 | 4.5406923 | 0.331122826 | 0.231 | 0.056 | 0.091708363 | 23 | nt5c2b |
| prps1a1 | 4.6534317 | 0.277789289 | 0.385 | 0.121 | 0.09398536 | 23 | prps1a |
| serbp1a3 | 4.6638535 | 0.414743101 | 0.949 | 0.654 | 0.094195849 | 23 | serbp1a |
| fam84a | 4.7237957 | 0.26370427 | 0.128 | 0.02 | 0.095406503 | 23 | fam84a |
| hnrpkl1 | 4.8163609 | 0.413902492 | 0.41 | 0.142 | 0.097276041 | 23 | hnrpkl |
| inka1a2 | 4.8212406 | 0.410239742 | 0.282 | 0.08 | 0.097374597 | 23 | inka1a |
| mrps111 | 5.0372236 | 0.276545876 | 0.333 | 0.1 | 0.101736806 | 23 | mrps11 |
| tubb53 | 5.1783868 | 0.366115374 | 0.282 | 0.079 | 0.104587879 | 23 | tubb5 |
| phf5a2 | 5.2536994 | 0.373641205 | 0.487 | 0.18 | 0.106108969 | 23 | phf5a |
| sae11 | 5.4489596 | 0.289275803 | 0.231 | 0.056 | 0.110052638 | 23 | sae1 |

| | | | | | | | |
|---|---|---|---|---|---|---|---|
| xaf11 | 5.5417467 | 0.307753249 | 0.256 | 0.066 | 0.11192666 | 23 | xaf1 |
| zmynd11 | 5.5982485 | 0.255452744 | 0.231 | 0.056 | 0.113067826 | 23 | zmynd11 |
| hmga1a4 | 5.7105852 | 0.536576526 | 0.769 | 0.433 | 0.115336691 | 23 | hmga1a |
| polr2eb | 5.7768825 | 0.3049101 | 0.41 | 0.139 | 0.116675696 | 23 | polr2eb |
| sf3b62 | 5.8233729 | 0.396571306 | 0.718 | 0.311 | 0.117614663 | 23 | sf3b6 |
| yy1a | 5.8846065 | 0.37669443 | 0.333 | 0.107 | 0.118851398 | 23 | yy1a |
| chmp1b1 | 5.9675614 | 0.374487288 | 0.487 | 0.186 | 0.120526839 | 23 | chmp1b |
| meaf61 | 5.9910206 | 0.33313096 | 0.41 | 0.143 | 0.121000644 | 23 | meaf6 |
| cdc26 | 6.0512915 | 0.355622844 | 0.41 | 0.144 | 0.122217935 | 23 | cdc26 |
| ssuh2rs12 | 6.1323421 | 0.298306596 | 0.282 | 0.079 | 0.123854915 | 23 | ssuh2rs1 |
| caprin1a1 | 6.1404112 | 0.300052682 | 0.282 | 0.077 | 0.124017886 | 23 | caprin1a |
| rnf103 | 6.6298734 | 0.454624331 | 0.513 | 0.212 | 0.133903555 | 23 | rnf10 |
| mkrn1 | 6.7776412 | 0.344872627 | 0.282 | 0.08 | 0.13688802 | 23 | mkrn1 |
| psme12 | 7.0458002 | 0.384824296 | 0.564 | 0.227 | 0.142304027 | 23 | psme1 |
| psmd71 | 7.3144676 | 0.3960015 | 0.462 | 0.177 | 0.147730302 | 23 | psmd7 |
| med112 | 7.5240875 | 0.294098849 | 0.308 | 0.09 | 0.151963996 | 23 | med11 |
| pbx1b1 | 7.8039607 | 0.268058594 | 0.256 | 0.066 | 0.157616596 | 23 | pbx1b |
| faf2 | 7.9026484 | 0.278756566 | 0.256 | 0.069 | 0.159609792 | 23 | faf2 |
| psma13 | 7.9340149 | 0.461538589 | 0.744 | 0.394 | 0.1602433 | 23 | psma1 |
| ctnnb15 | 7.9784018 | 0.466141908 | 0.513 | 0.213 | 0.161139782 | 23 | ctnnb1 |
| mvp2 | 8.0181977 | 0.312354202 | 0.333 | 0.102 | 0.161943539 | 23 | mvp |
| ubtfl1 | 8.4955431 | 0.254723519 | 0.205 | 0.047 | 0.171584485 | 23 | ubtfl |
| znf865 | 8.5097243 | 0.261831273 | 0.231 | 0.057 | 0.171870903 | 23 | znf865 |
| zc4h22 | 8.5648308 | 0.266496213 | 0.333 | 0.102 | 0.172983888 | 23 | zc4h2 |
| dido13 | 8.6440577 | 0.375109153 | 0.436 | 0.157 | 0.174584035 | 23 | dido1 |
| spred2b5 | 8.8082685 | 0.394812843 | 0.487 | 0.18 | 0.177900599 | 23 | spred2b |
| serinc54 | 9.1020085 | 0.339339493 | 0.359 | 0.114 | 0.183833267 | 23 | serinc5 |
| hnrnpc2 | 9.4418486 | 0.291709745 | 0.333 | 0.104 | 0.190697017 | 23 | hnrnpc |
| mxra8a4 | 9.5552848 | 0.345797746 | 0.436 | 0.146 | 0.192988089 | 23 | mxra8a |
| si:ch211-39i22.17 | 9.6547195 | 0.408819696 | 0.487 | 0.181 | 0.19499637 | 23 | si:ch211-39i22.1 |
| lrpap12 | 1.0155885 | 0.340324147 | 0.333 | 0.106 | 0.205118427 | 23 | lrpap1 |
| si:ch73-22o12.12 | 1.0583719 | 0.282745246 | 0.231 | 0.058 | 0.213759385 | 23 | si:ch73-22o12.1 |
| hnrnph1l3 | 1.0618476 | 0.309628561 | 0.385 | 0.129 | 0.21446136 | 23 | hnrnph1l |
| taf152 | 1.1176328 | 0.344892698 | 0.487 | 0.186 | 0.225728301 | 23 | taf15 |
| igf2bp2b | 1.1350152 | 0.255148274 | 0.256 | 0.068 | 0.229239036 | 23 | igf2bp2b |

| | | | | | | | |
|---|---|---|---|---|---|---|---|
| slc25a55a1 | 1.1814580 | 0.320031404 | 0.359 | 0.118 | 0.238619078 | 23 | slc25a55a |
| ehmt1a | 1.1851767 | 0.25970097 | 0.179 | 0.039 | 0.239370151 | 23 | ehmt1a |
| kctd12.24 | 1.2420742 | 0.390300816 | 0.538 | 0.209 | 0.250861729 | 23 | kctd12.2 |
| tcf3b | 1.2575041 | 0.260854975 | 0.231 | 0.058 | 0.253978118 | 23 | tcf3b |
| smim131 | 1.2579194 | 0.283083027 | 0.308 | 0.091 | 0.254061984 | 23 | smim13 |
| huwe1 | 1.2744971 | 0.299582732 | 0.231 | 0.06 | 0.25741019 | 23 | huwe1 |
| prdx62 | 1.3224937 | 0.303236976 | 0.538 | 0.212 | 0.267104053 | 23 | prdx6 |
| smarcb1b | 1.3515081 | 0.271905908 | 0.282 | 0.081 | 0.272964092 | 23 | smarcb1b |
| polr2d1 | 1.4828004 | 0.385327568 | 0.487 | 0.196 | 0.299481215 | 23 | polr2d |
| wt1b3 | 1.5662290 | 0.263063723 | 0.256 | 0.068 | 0.316331274 | 23 | wt1b |
| zgc:919104 | 1.6026227 | 0.379763005 | 0.667 | 0.297 | 0.323681719 | 23 | zgc:91910 |
| rpl22l14 | 1.6099827 | 0.456054653 | 0.974 | 0.618 | 0.325168221 | 23 | rpl22l1 |
| pde7a3 | 1.6604842 | 0.29294947 | 0.308 | 0.094 | 0.335368004 | 23 | pde7a |
| csnk1g2a1 | 1.6949570 | 0.261442539 | 0.231 | 0.06 | 0.342330466 | 23 | csnk1g2a |
| rbbp42 | 1.7407193 | 0.341640616 | 0.385 | 0.132 | 0.351573082 | 23 | rbbp4 |
| si:ch211-196l7.46 | 1.8366440 | 0.400070975 | 0.846 | 0.41 | 0.370946993 | 23 | si:ch211-196l7.4 |
| flot2a6 | 1.9130016 | 0.356891376 | 0.41 | 0.15 | 0.386368952 | 23 | flot2a |
| ndufa33 | 1.9288045 | 0.339361208 | 0.564 | 0.242 | 0.389560665 | 23 | ndufa3 |
| cct22 | 1.9620436 | 0.4213155 | 0.795 | 0.396 | 0.396273953 | 23 | cct2 |
| snrpa | 2.0013923 | 0.301298903 | 0.385 | 0.135 | 0.404221207 | 23 | snrpa |
| gata63 | 2.1224038 | 0.266983571 | 0.231 | 0.059 | 0.428661897 | 23 | gata6 |
| aurkaip11 | 2.1457279 | 0.260791444 | 0.256 | 0.073 | 0.433372665 | 23 | aurkaip1 |
| stat1a3 | 2.1678868 | 0.413796591 | 0.385 | 0.146 | 0.437848112 | 23 | stat1a |
| ctsf4 | 2.1760035 | 0.287822867 | 0.256 | 0.072 | 0.43948744 | 23 | ctsf |
| psmb42 | 2.1896632 | 0.394220225 | 0.615 | 0.283 | 0.442246278 | 23 | psmb4 |
| wdr61 | 2.1947948 | 0.257236941 | 0.256 | 0.072 | 0.443282707 | 23 | wdr61 |
| snrpd22 | 2.2104063 | 0.395382572 | 0.59 | 0.264 | 0.446435779 | 23 | snrpd2 |
| jund1 | 2.2495961 | 0.294901894 | 0.282 | 0.083 | 0.454350933 | 23 | jund |
| ppid2 | 2.2773309 | 0.303121259 | 0.333 | 0.112 | 0.45995253 | 23 | ppid |
| pfn27 | 2.2818214 | 0.447932578 | 0.667 | 0.317 | 0.460859469 | 23 | pfn2 |
| hnrnpk1 | 2.4032288 | 0.390525558 | 0.487 | 0.2 | 0.48538013 | 23 | hnrnpk |
| dhrs13a.3 | 2.4453074 | 0.277869046 | 0.205 | 0.051 | 0.493878742 | 23 | dhrs13a.3 |
| tcp11 | 2.4503570 | 0.350788684 | 0.769 | 0.346 | 0.494898606 | 23 | tcp1 |
| acadl | 2.4603818 | 0.259347578 | 0.231 | 0.061 | 0.496923319 | 23 | acadl |
| psme22 | 2.5814393 | 0.360705982 | 0.487 | 0.194 | 0.521373309 | 23 | psme2 |

| | | | | | | | |
|---|---|---|---|---|---|---|---|
| asah1b3 | 2.7364265 | 0.297182088 | 0.333 | 0.109 | 0.552676079 | 23 | asah1b |
| rtn31 | 2.7578757 | 0.262421467 | 0.282 | 0.084 | 0.557008156 | 23 | rtn3 |
| cmtm31 | 2.7621911 | 0.251274568 | 0.256 | 0.073 | 0.557879757 | 23 | cmtm3 |
| gemin8 | 2.8709028 | 0.262858266 | 0.282 | 0.086 | 0.579836245 | 23 | gemin8 |
| pfdn21 | 2.9523688 | 0.3515953 | 0.385 | 0.144 | 0.596289946 | 23 | pfdn2 |
| kmt2cb2 | 3.0222584 | 0.296028782 | 0.308 | 0.097 | 0.610405534 | 23 | kmt2cb |
| elocb3 | 3.0299465 | 0.33770449 | 0.667 | 0.289 | 0.611958303 | 23 | elocb |
| cpne32 | 3.0324790 | 0.310015056 | 0.256 | 0.075 | 0.61246979 | 23 | cpne3 |
| si:ch211-195m9.31 | 3.0419100 | 0.271152948 | 0.128 | 0.023 | 0.614374569 | 23 | si:ch211-195m9.3 |
| hnrnpl23 | 3.0568019 | 0.343615816 | 0.41 | 0.156 | 0.617382296 | 23 | hnrnpl2 |
| nars2 | 3.0610737 | 0.306934226 | 0.385 | 0.139 | 0.618245061 | 23 | nars |
| seta2 | 3.0978105 | 0.41876557 | 0.718 | 0.354 | 0.625664798 | 23 | seta |
| h3f3d4 | 3.1080824 | 0.414865051 | 0.974 | 0.671 | 0.62773942 | 23 | h3f3d |
| add13 | 3.2626900 | 0.324546352 | 0.333 | 0.112 | 0.658965515 | 23 | add1 |
| kpna41 | 3.2710222 | 0.274937772 | 0.231 | 0.063 | 0.660648362 | 23 | kpna4 |
| si:ch211-235e9.62 | 3.2778981 | 0.349970623 | 0.513 | 0.213 | 0.662037096 | 23 | si:ch211-235e9.6 |
| hdgfl2 | 3.2886078 | 0.293184058 | 0.256 | 0.075 | 0.664200137 | 23 | hdgfl2 |
| oaz1a4 | 3.3027730 | 0.413497492 | 0.897 | 0.592 | 0.667061064 | 23 | oaz1a |
| atp6ap22 | 3.3102027 | 0.287629666 | 0.385 | 0.135 | 0.668561651 | 23 | atp6ap2 |
| tnip12 | 3.3943330 | 0.287190412 | 0.256 | 0.074 | 0.685553443 | 23 | tnip1 |
| brd3b2 | 3.4325818 | 0.359992115 | 0.333 | 0.115 | 0.69327855 | 23 | brd3b |
| psmb2 | 3.4658107 | 0.369261581 | 0.59 | 0.263 | 0.699989797 | 23 | psmb2 |
| ginm11 | 3.5391230 | 0.254277486 | 0.359 | 0.118 | 0.714796688 | 23 | ginm1 |
| ddx53 | 3.6204730 | 0.463231949 | 0.692 | 0.353 | 0.731226948 | 23 | ddx5 |
| rac1a1 | 3.6687126 | 0.310386747 | 0.333 | 0.112 | 0.740969892 | 23 | rac1a |
| eif51 | 3.6699711 | 0.350404064 | 0.59 | 0.261 | 0.741224064 | 23 | eif5 |
| ralab | 3.7714825 | 0.282220693 | 0.205 | 0.053 | 0.761726321 | 23 | ralab |
| polr2b1 | 3.7757975 | 0.284334471 | 0.231 | 0.064 | 0.762597827 | 23 | polr2b |
| snrpb6 | 3.9434581 | 0.401877645 | 0.821 | 0.419 | 0.796460252 | 23 | snrpb |
| ppp6c | 3.9713128 | 0.260445356 | 0.41 | 0.148 | 0.802086055 | 23 | ppp6c |
| ddx241 | 4.0240421 | 0.286452715 | 0.282 | 0.088 | 0.812735783 | 23 | ddx24 |
| brd71 | 4.0830526 | 0.27270584 | 0.308 | 0.099 | 0.824654152 | 23 | brd7 |
| zgc:1124372 | 4.1237476 | 0.292106445 | 0.231 | 0.063 | 0.832873321 | 23 | zgc:112437 |
| uba11 | 4.1304156 | 0.322961599 | 0.436 | 0.17 | 0.834220054 | 23 | uba1 |
| gatc | 4.1841531 | 0.33288733 | 0.282 | 0.09 | 0.84507341 | 23 | gatc |

| | | | | | | | |
|---|---|---|---|---|---|---|---|
| sptbn22 | 4.1909947 | 0.296067458 | 0.231 | 0.065 | 0.846455207 | 23 | sptbn2 |
| prelid1a | 4.2314861 | 0.273779269 | 0.333 | 0.113 | 0.854633251 | 23 | prelid1a |
| bcas21 | 4.2573879 | 0.320179673 | 0.462 | 0.184 | 0.859864636 | 23 | bcas2 |
| cox14 | 4.2921430 | 0.276144805 | 0.385 | 0.14 | 0.866884141 | 23 | cox14 |
| zgc:1620251 | 4.4035322 | 0.308778308 | 0.282 | 0.09 | 0.889381401 | 23 | zgc:162025 |
| sf3b54 | 4.4075079 | 0.377313621 | 0.462 | 0.196 | 0.890184375 | 23 | sf3b5 |
| akap12b5 | 4.5024628 | 0.384044404 | 0.41 | 0.151 | 0.909362423 | 23 | akap12b |
| tgfb34 | 4.5824405 | 0.295738409 | 0.282 | 0.087 | 0.925515527 | 23 | tgfb3 |
| trib33 | 4.9093911 | 0.289018941 | 0.359 | 0.124 | 0.991549732 | 23 | trib3 |
| atp10a3 | 4.9657879 | 0.262443932 | 0.256 | 0.075 | 1 | 23 | atp10a |
| snrnp271 | 4.9916454 | 0.288905363 | 0.359 | 0.127 | 1 | 23 | snrnp27 |
| btf3l42 | 5.0072402 | 0.30238686 | 0.487 | 0.2 | 1 | 23 | btf3l4 |
| lrp1ab4 | 5.0235834 | 0.306000362 | 0.359 | 0.123 | 1 | 23 | lrp1ab |
| zgc:1951735 | 5.0469602 | 0.339773649 | 0.59 | 0.234 | 1 | 23 | zgc:195173 |
| gnb1b4 | 5.1349547 | 0.409319709 | 0.692 | 0.337 | 1 | 23 | gnb1b |
| tpd52l2a | 5.3039606 | 0.276213662 | 0.231 | 0.065 | 1 | 23 | tpd52l2a |
| eif4g2b | 5.6055528 | 0.277121146 | 0.282 | 0.09 | 1 | 23 | eif4g2b |
| psmd32 | 5.7053853 | 0.268211829 | 0.41 | 0.152 | 1 | 23 | psmd3 |
| ndufs62 | 5.7395481 | 0.304383345 | 0.487 | 0.2 | 1 | 23 | ndufs6 |
| ppdpfb3 | 5.7564741 | 0.426186876 | 0.949 | 0.618 | 1 | 23 | ppdpfb |
| appa2 | 5.7935751 | 0.362228854 | 0.385 | 0.143 | 1 | 23 | appa |
| ap1s22 | 5.8555981 | 0.356993715 | 0.41 | 0.162 | 1 | 23 | ap1s2 |
| aip2 | 6.0867609 | 0.335056274 | 0.385 | 0.149 | 1 | 23 | aip |
| pfkfb4b3 | 6.1879985 | 0.354767424 | 0.513 | 0.219 | 1 | 23 | pfkfb4b |
| ndufa63 | 6.2656863 | 0.344878102 | 0.641 | 0.312 | 1 | 23 | ndufa6 |
| gatad2b4 | 6.4403115 | 0.328266774 | 0.333 | 0.119 | 1 | 23 | gatad2b |
| hm132 | 6.5375639 | 0.272353827 | 0.359 | 0.128 | 1 | 23 | hm13 |
| bptf1 | 6.6608551 | 0.260490719 | 0.231 | 0.066 | 1 | 23 | bptf |
| rab11a1 | 6.6979232 | 0.28773689 | 0.462 | 0.18 | 1 | 23 | rab11a |
| puf60a2 | 6.7473297 | 0.267685266 | 0.359 | 0.126 | 1 | 23 | puf60a |
| ppm1aa | 6.8596143 | 0.268834426 | 0.308 | 0.101 | 1 | 23 | ppm1aa |
| sec63 | 6.8886826 | 0.260870674 | 0.256 | 0.077 | 1 | 23 | sec63 |
| neo1a2 | 7.5192833 | 0.297298916 | 0.205 | 0.056 | 1 | 23 | neo1a |
| cebpd7 | 7.5904448 | 0.399680567 | 0.641 | 0.277 | 1 | 23 | cebpd |
| polr2f1 | 7.6912023 | 0.289992574 | 0.462 | 0.185 | 1 | 23 | polr2f |

| | | | | | | | | |
|---|---|---|---|---|---|---|---|---|
| si:ch211-105c13.3! | 7.8082116 | 0.267342138 | 0.333 | 0.108 | | 1 | 23 | si:ch211-105c13.3 |
| tegt5 | 7.9056272 | 0.410511222 | 0.692 | 0.361 | | 1 | 23 | tegt |
| tax1bp1b1 | 8.1371753 | 0.27613339 | 0.308 | 0.104 | | 1 | 23 | tax1bp1b |
| sap30l2 | 8.1411010 | 0.320853178 | 0.513 | 0.221 | | 1 | 23 | sap30l |
| llph1 | 8.2016814 | 0.284259803 | 0.359 | 0.132 | | 1 | 23 | llph |
| sfpq1 | 8.2950064 | 0.288099347 | 0.359 | 0.129 | | 1 | 23 | sfpq |
| churc1 | 8.3453795 | 0.255392574 | 0.359 | 0.128 | | 1 | 23 | churc1 |
| gulp1a3 | 8.3589505 | 0.251462661 | 0.231 | 0.065 | | 1 | 23 | gulp1a |
| elavl1a1 | 8.4136821 | 0.276967986 | 0.359 | 0.131 | | 1 | 23 | elavl1a |
| ube2ib2 | 9.0145705 | 0.328593785 | 0.538 | 0.234 | | 1 | 23 | ube2ib |
| sem12 | 9.4444437 | 0.372615085 | 0.769 | 0.389 | | 1 | 23 | sem1 |
| trmt1121 | 9.7071855 | 0.326024321 | 0.462 | 0.196 | | 1 | 23 | trmt112 |
| znf318 | 9.9689812 | 0.250152523 | 0.179 | 0.046 | | 1 | 23 | znf318 |
| banf1 | 9.9790391 | 0.324353167 | 0.487 | 0.207 | | 1 | 23 | banf1 |
| hmgb1a6 | 0.000102 | 0.399795809 | 0.795 | 0.426 | | 1 | 23 | hmgb1a |
| nmi | 0.000102 | 0.30501549 | 0.333 | 0.122 | | 1 | 23 | nmi |
| eif2s32 | 0.000103 | 0.399417624 | 0.769 | 0.433 | | 1 | 23 | eif2s3 |
| trappc13 | 0.000104 | 0.307568124 | 0.487 | 0.203 | | 1 | 23 | trappc1 |
| b2ml6 | 0.000109 | 0.408884959 | 0.846 | 0.443 | | 1 | 23 | b2ml |
| khsrp1 | 0.000111 | 0.306794173 | 0.436 | 0.178 | | 1 | 23 | khsrp |
| u2af11 | 0.000117 | 0.296456889 | 0.538 | 0.229 | | 1 | 23 | u2af1 |
| eif4g2a2 | 0.000121 | 0.279710111 | 0.282 | 0.095 | | 1 | 23 | eif4g2a |
| arf11 | 0.000125 | 0.298456301 | 0.359 | 0.134 | | 1 | 23 | arf1 |
| plk34 | 0.000126 | 0.278848388 | 0.205 | 0.057 | | 1 | 23 | plk3 |
| flna7 | 0.000126 | 0.252753916 | 0.436 | 0.167 | | 1 | 23 | flna |
| eif3m1 | 0.000128 | 0.407012742 | 0.846 | 0.521 | | 1 | 23 | eif3m |
| acin1a6 | 0.000131 | 0.363924571 | 0.615 | 0.296 | | 1 | 23 | acin1a |
| add3a6 | 0.000131 | 0.354909354 | 0.513 | 0.22 | | 1 | 23 | add3a |
| ptprga3 | 0.000134 | 0.305073638 | 0.308 | 0.106 | | 1 | 23 | ptprga |
| ube2v12 | 0.000136 | 0.363359278 | 0.641 | 0.317 | | 1 | 23 | ube2v1 |
| nedd81 | 0.00014 | 0.254978012 | 0.359 | 0.134 | | 1 | 23 | nedd8 |
| zgc:928181 | 0.00014 | 0.253890524 | 0.231 | 0.069 | | 1 | 23 | zgc:92818 |
| snrnp701 | 0.000143 | 0.289391855 | 0.308 | 0.108 | | 1 | 23 | snrnp70 |
| aplp23 | 0.000143 | 0.262342544 | 0.308 | 0.105 | | 1 | 23 | aplp2 |
| hspa91 | 0.000144 | 0.27883823 | 0.308 | 0.107 | | 1 | 23 | hspa9 |

| | | | | | | | |
|---|---|---|---|---|---|---|---|
| ak21 | 0.000149 | 0.256965527 | 0.795 | 0.376 | 1 | 23 | ak2 |
| ndufb23 | 0.00015 | 0.293979308 | 0.538 | 0.24 | 1 | 23 | ndufb2 |
| ndufb42 | 0.00015 | 0.324428764 | 0.641 | 0.314 | 1 | 23 | ndufb4 |
| nsfl1c3 | 0.00015 | 0.317524695 | 0.359 | 0.142 | 1 | 23 | nsfl1c |
| ddx39ab2 | 0.000151 | 0.259116683 | 0.462 | 0.188 | 1 | 23 | ddx39ab |
| cct41 | 0.000154 | 0.338133678 | 0.718 | 0.354 | 1 | 23 | cct4 |
| eif4ebp15 | 0.000156 | 0.33947825 | 0.692 | 0.34 | 1 | 23 | eif4ebp1 |
| selenom2 | 0.000158 | 0.276730167 | 0.282 | 0.096 | 1 | 23 | selenom |
| cirbpa1 | 0.000159 | 0.389368207 | 0.974 | 0.701 | 1 | 23 | cirbpa |
| CR318588.33 | 0.000159 | 0.301326938 | 0.256 | 0.081 | 1 | 23 | CR318588.3 |
| brdt1 | 0.000159 | 0.261312781 | 0.256 | 0.082 | 1 | 23 | brdt |
| vps281 | 0.000172 | 0.275293077 | 0.333 | 0.123 | 1 | 23 | vps28 |
| bzw1b2 | 0.000176 | 0.356962136 | 0.59 | 0.285 | 1 | 23 | bzw1b |
| foxp46 | 0.000176 | 0.331190565 | 0.462 | 0.189 | 1 | 23 | foxp4 |
| psmc21 | 0.000178 | 0.302798962 | 0.385 | 0.155 | 1 | 23 | psmc2 |
| tomm20b | 0.000182 | 0.321416209 | 0.641 | 0.299 | 1 | 23 | tomm20b |
| anxa1a5 | 0.000182 | 0.42427488 | 0.59 | 0.277 | 1 | 23 | anxa1a |
| cct32 | 0.000183 | 0.323974635 | 0.692 | 0.332 | 1 | 23 | cct3 |
| snrpf1 | 0.000185 | 0.354165068 | 0.744 | 0.388 | 1 | 23 | snrpf |
| stmn1a2 | 0.000193 | 0.336875709 | 0.308 | 0.109 | 1 | 23 | stmn1a |
| CABZ01115113.1 | 0.000193 | 0.283649664 | 0.359 | 0.14 | 1 | 23 | CABZ01115113.1 |
| pcyox13 | 0.000194 | 0.268294644 | 0.308 | 0.109 | 1 | 23 | pcyox1 |
| XLOC-0056054 | 0.000195 | 0.293595364 | 0.282 | 0.094 | 1 | 23 | XLOC-005605 |
| ncor23 | 0.000197 | 0.264292502 | 0.333 | 0.122 | 1 | 23 | ncor2 |
| rad23b2 | 0.000198 | 0.328838302 | 0.564 | 0.26 | 1 | 23 | rad23b |
| cd2762 | 0.000201 | 0.26702562 | 0.231 | 0.072 | 1 | 23 | cd276 |
| atp5if1a1 | 0.000202 | 0.347383322 | 0.769 | 0.394 | 1 | 23 | atp5if1a |
| fam107b2 | 0.000207 | 0.28227294 | 0.41 | 0.164 | 1 | 23 | fam107b |
| aimp21 | 0.000213 | 0.277249439 | 0.487 | 0.209 | 1 | 23 | aimp2 |
| anxa5b6 | 0.000213 | 0.328167862 | 0.41 | 0.169 | 1 | 23 | anxa5b |
| srsf2a1 | 0.000219 | 0.300348277 | 0.59 | 0.272 | 1 | 23 | srsf2a |
| txndc92 | 0.000229 | 0.297406357 | 0.436 | 0.187 | 1 | 23 | txndc9 |
| ddx273 | 0.000232 | 0.335599919 | 0.333 | 0.131 | 1 | 23 | ddx27 |
| vdac14 | 0.000234 | 0.289448456 | 0.436 | 0.181 | 1 | 23 | vdac1 |
| si:dkey-11f4.143 | 0.000246 | 0.275308567 | 0.487 | 0.201 | 1 | 23 | si:dkey-11f4.14 |

| | | | | | | | | |
|---|---|---|---|---|---|---|---|---|
| rps27.2 | 0.000247 | 0.321176906 | 0.974 | 0.765 | | 1 | 23 | rps27.2 |
| pik3r3a2 | 0.00025 | 0.324688026 | 0.359 | 0.144 | | 1 | 23 | pik3r3a |
| zgc:778491 | 0.000252 | 0.282568925 | 0.436 | 0.183 | | 1 | 23 | zgc:77849 |
| cct6a1 | 0.000252 | 0.324253733 | 0.615 | 0.295 | | 1 | 23 | cct6a |
| cxcl18b3 | 0.000252 | 0.331766466 | 0.359 | 0.136 | | 1 | 23 | cxcl18b |
| fzd44 | 0.000256 | 0.274183328 | 0.205 | 0.06 | | 1 | 23 | fzd4 |
| nhp23 | 0.000257 | 0.338761228 | 0.513 | 0.242 | | 1 | 23 | nhp2 |
| ap2m1a3 | 0.000263 | 0.30511836 | 0.641 | 0.303 | | 1 | 23 | ap2m1a |
| serinc13 | 0.000263 | 0.304376872 | 0.308 | 0.113 | | 1 | 23 | serinc1 |
| rnf213a4 | 0.000268 | 0.398646496 | 0.333 | 0.135 | | 1 | 23 | rnf213a |
| krt153 | 0.00027 | 0.384416909 | 0.41 | 0.164 | | 1 | 23 | krt15 |
| rhoaa3 | 0.000274 | 0.343810443 | 0.615 | 0.297 | | 1 | 23 | rhoaa |
| atf4b5 | 0.000281 | 0.392493731 | 0.667 | 0.36 | | 1 | 23 | atf4b |
| uba52 | 0.000287 | 0.261103774 | 1 | 0.845 | | 1 | 23 | uba52 |
| col27a1b3 | 0.000297 | 0.268095529 | 0.231 | 0.073 | | 1 | 23 | col27a1b |
| dkk3b4 | 0.000299 | 0.319381207 | 0.41 | 0.166 | | 1 | 23 | dkk3b |
| srsf7a | 0.000303 | 0.353437592 | 0.487 | 0.236 | | 1 | 23 | srsf7a |
| zgc:1119861 | 0.000306 | 0.349058534 | 0.487 | 0.23 | | 1 | 23 | zgc:111986 |
| ddost2 | 0.000307 | 0.262413687 | 0.385 | 0.156 | | 1 | 23 | ddost |
| psmc31 | 0.000323 | 0.258979519 | 0.359 | 0.142 | | 1 | 23 | psmc3 |
| fam120c4 | 0.000323 | 0.325778524 | 0.359 | 0.147 | | 1 | 23 | fam120c |
| sesn15 | 0.000331 | 0.402577445 | 0.462 | 0.216 | | 1 | 23 | sesn1 |
| selenof5 | 0.000334 | 0.341065882 | 0.821 | 0.421 | | 1 | 23 | selenof |
| rtn4a3 | 0.000348 | 0.33233897 | 0.513 | 0.249 | | 1 | 23 | rtn4a |
| psmb62 | 0.000348 | 0.317905185 | 0.59 | 0.292 | | 1 | 23 | psmb6 |
| krt948 | 0.000349 | 0.424438401 | 0.718 | 0.361 | | 1 | 23 | krt94 |
| dcaf72 | 0.000361 | 0.274593765 | 0.333 | 0.129 | | 1 | 23 | dcaf7 |
| anxa2a5 | 0.000362 | 0.395581379 | 0.436 | 0.187 | | 1 | 23 | anxa2a |
| dkc11 | 0.000362 | 0.269336674 | 0.436 | 0.184 | | 1 | 23 | dkc1 |
| nop534 | 0.000362 | 0.376930684 | 0.641 | 0.353 | | 1 | 23 | nop53 |
| med102 | 0.000366 | 0.291384455 | 0.41 | 0.178 | | 1 | 23 | med10 |
| mcts11 | 0.000366 | 0.262283807 | 0.385 | 0.161 | | 1 | 23 | mcts1 |
| tmpob | 0.000386 | 0.254380704 | 0.231 | 0.076 | | 1 | 23 | tmpob |
| cdkn1a | 0.000392 | 0.314902131 | 0.282 | 0.103 | | 1 | 23 | cdkn1a |
| myclb2 | 0.000396 | 0.296387888 | 0.256 | 0.089 | | 1 | 23 | myclb |

| | | | | | | | |
|---|---|---|---|---|---|---|---|
| dnaja2a1 | 0.000399 | 0.282424234 | 0.59 | 0.274 | 1 | 23 | dnaja2a |
| pik3r23 | 0.000399 | 0.319010367 | 0.282 | 0.104 | 1 | 23 | pik3r2 |
| vamp31 | 0.000401 | 0.281545782 | 0.487 | 0.222 | 1 | 23 | vamp3 |
| h2afva4 | 0.000404 | 0.341835192 | 0.487 | 0.233 | 1 | 23 | h2afva |
| csnk2b2 | 0.000405 | 0.306732845 | 0.615 | 0.301 | 1 | 23 | csnk2b |
| supt16h | 0.000409 | 0.29189531 | 0.231 | 0.077 | 1 | 23 | supt16h |
| cbx52 | 0.000412 | 0.308709712 | 0.487 | 0.221 | 1 | 23 | cbx5 |
| tomm52 | 0.000414 | 0.341410691 | 0.615 | 0.319 | 1 | 23 | tomm5 |
| calr3b4 | 0.000418 | 0.262311337 | 0.436 | 0.182 | 1 | 23 | calr3b |
| ubr7 | 0.000427 | 0.250561086 | 0.256 | 0.089 | 1 | 23 | ubr7 |
| prkab1b2 | 0.000431 | 0.281492203 | 0.462 | 0.208 | 1 | 23 | prkab1b |
| lrrc174 | 0.000434 | 0.378274207 | 0.41 | 0.181 | 1 | 23 | lrrc17 |
| mbd3a | 0.000434 | 0.25256675 | 0.282 | 0.103 | 1 | 23 | mbd3a |
| hspa56 | 0.000448 | 0.305625596 | 0.487 | 0.221 | 1 | 23 | hspa5 |
| ube2e11 | 0.000469 | 0.265163289 | 0.462 | 0.204 | 1 | 23 | ube2e1 |
| hif1ab3 | 0.000472 | 0.297066363 | 0.41 | 0.175 | 1 | 23 | hif1ab |
| si:ch211-212k18.7[illegible] | 0.000478 | 0.277299088 | 0.436 | 0.185 | 1 | 23 | si:ch211-212k18.7 |
| cct72 | 0.000488 | 0.300475054 | 0.692 | 0.337 | 1 | 23 | cct7 |
| si:rp71-45k5.41 | 0.000492 | 0.285368614 | 0.487 | 0.224 | 1 | 23 | si:rp71-45k5.4 |
| mif3 | 0.000502 | 0.305761229 | 0.615 | 0.314 | 1 | 23 | mif |
| cebpb6 | 0.000504 | 0.386975733 | 0.59 | 0.294 | 1 | 23 | cebpb |
| khdrbs1b1 | 0.000508 | 0.284380276 | 0.359 | 0.148 | 1 | 23 | khdrbs1b |
| mmp28 | 0.000524 | 0.328696706 | 0.487 | 0.217 | 1 | 23 | mmp2 |
| calm2b5 | 0.000543 | 0.345201031 | 0.821 | 0.445 | 1 | 23 | calm2b |
| hnrnpa1a1 | 0.000544 | 0.277987848 | 0.462 | 0.206 | 1 | 23 | hnrnpa1a |
| ctsla9 | 0.000553 | 0.268197532 | 0.641 | 0.308 | 1 | 23 | ctsla |
| rtca1 | 0.000558 | 0.277687994 | 0.231 | 0.079 | 1 | 23 | rtca |
| ankrd113 | 0.000569 | 0.289610879 | 0.359 | 0.149 | 1 | 23 | ankrd11 |
| rpl5b | 0.000593 | 0.311555774 | 0.974 | 0.759 | 1 | 23 | rpl5b |
| paip2b5 | 0.000602 | 0.315448044 | 0.667 | 0.341 | 1 | 23 | paip2b |
| ppp2cb4 | 0.000617 | 0.326273121 | 0.487 | 0.24 | 1 | 23 | ppp2cb |
| si:ch73-308l14.2 | 0.00062 | 0.275787521 | 0.231 | 0.081 | 1 | 23 | si:ch73-308l14.2 |
| caprin1b1 | 0.000622 | 0.260899612 | 0.564 | 0.261 | 1 | 23 | caprin1b |
| sf3a23 | 0.000625 | 0.285677652 | 0.385 | 0.168 | 1 | 23 | sf3a2 |
| sltm2 | 0.000669 | 0.295880531 | 0.308 | 0.123 | 1 | 23 | sltm |

| | | | | | | | |
|---|---|---|---|---|---|---|---|
| eif3ja1 | 0.000718 | 0.278082356 | 0.487 | 0.234 | 1 | 23 | eif3ja |
| sap183 | 0.000739 | 0.292016897 | 0.487 | 0.239 | 1 | 23 | sap18 |
| cnih11 | 0.00074 | 0.276299134 | 0.256 | 0.095 | 1 | 23 | cnih1 |
| smdt1b3 | 0.000762 | 0.28323812 | 0.513 | 0.248 | 1 | 23 | smdt1b |
| nudc1 | 0.000764 | 0.296864803 | 0.41 | 0.191 | 1 | 23 | nudc |
| hypk1 | 0.00077 | 0.261601114 | 0.59 | 0.283 | 1 | 23 | hypk |
| pdcd5 | 0.000776 | 0.260360959 | 0.359 | 0.153 | 1 | 23 | pdcd5 |
| anxa42 | 0.000776 | 0.250285636 | 0.513 | 0.237 | 1 | 23 | anxa4 |
| adh55 | 0.000782 | 0.370388542 | 0.615 | 0.361 | 1 | 23 | adh5 |
| taf9 | 0.000796 | 0.259526962 | 0.333 | 0.137 | 1 | 23 | taf9 |
| LEPROTL12 | 0.000797 | 0.295600879 | 0.308 | 0.125 | 1 | 23 | LEPROTL1 |
| max1 | 0.000833 | 0.261726059 | 0.436 | 0.199 | 1 | 23 | max |
| nop562 | 0.000866 | 0.317506173 | 0.564 | 0.281 | 1 | 23 | nop56 |
| h3f3c6 | 0.000883 | 0.388591833 | 0.821 | 0.528 | 1 | 23 | h3f3c |
| lsm72 | 0.000907 | 0.281488603 | 0.487 | 0.23 | 1 | 23 | lsm7 |
| pim12 | 0.000923 | 0.257052424 | 0.385 | 0.165 | 1 | 23 | pim1 |
| si:ch73-42p12.21 | 0.000931 | 0.286539559 | 0.385 | 0.174 | 1 | 23 | si:ch73-42p12.2 |
| rpl4 | 0.000941 | 0.260509287 | 1 | 0.823 | 1 | 23 | rpl4 |
| sod13 | 0.00096 | 0.328603641 | 0.744 | 0.421 | 1 | 23 | sod1 |
| fbl1 | 0.001049 | 0.263945912 | 0.436 | 0.198 | 1 | 23 | fbl |
| cpsf62 | 0.001054 | 0.260789438 | 0.359 | 0.155 | 1 | 23 | cpsf6 |
| cirbpb3 | 0.001067 | 0.271799719 | 0.949 | 0.812 | 1 | 23 | cirbpb |
| drap11 | 0.001075 | 0.256026321 | 0.41 | 0.191 | 1 | 23 | drap1 |
| itm2bb6 | 0.001121 | 0.325359964 | 0.667 | 0.356 | 1 | 23 | itm2bb |
| purab2 | 0.001134 | 0.279115322 | 0.282 | 0.113 | 1 | 23 | purab |
| map1lc3b1 | 0.00114 | 0.273462619 | 0.436 | 0.203 | 1 | 23 | map1lc3b |
| syncripl | 0.00126 | 0.26383798 | 0.282 | 0.112 | 1 | 23 | syncripl |
| ptenb3 | 0.001266 | 0.306553362 | 0.308 | 0.127 | 1 | 23 | ptenb |
| inhbaa2 | 0.00128 | 0.282105834 | 0.308 | 0.12 | 1 | 23 | inhbaa |
| wbp1 | 0.001317 | 0.256679188 | 0.154 | 0.046 | 1 | 23 | wbp1 |
| flot1b5 | 0.001378 | 0.297511678 | 0.41 | 0.197 | 1 | 23 | flot1b |
| eif3g3 | 0.001395 | 0.291255266 | 0.744 | 0.395 | 1 | 23 | eif3g |
| kctd5a2 | 0.00142 | 0.26814741 | 0.513 | 0.247 | 1 | 23 | kctd5a |
| abcf12 | 0.001445 | 0.2796754 | 0.41 | 0.191 | 1 | 23 | abcf1 |
| boc6 | 0.001465 | 0.289822882 | 0.385 | 0.168 | 1 | 23 | boc |

| | | | | | | | |
|---|---|---|---|---|---|---|---|
| ndufc23 | 0.001473 | 0.257466355 | 0.513 | 0.249 | 1 | 23 | ndufc2 |
| ube2v23 | 0.001491 | 0.28232614 | 0.692 | 0.362 | 1 | 23 | ube2v2 |
| psma81 | 0.001524 | 0.281282152 | 0.615 | 0.325 | 1 | 23 | psma8 |
| alyref1 | 0.001531 | 0.291341898 | 0.308 | 0.132 | 1 | 23 | alyref |
| si:dkey-26c10.52 | 0.001604 | 0.263329454 | 0.103 | 0.024 | 1 | 23 | si:dkey-26c10.5 |
| tm9sf3 | 0.001659 | 0.271975028 | 0.205 | 0.073 | 1 | 23 | tm9sf3 |
| cldn5a4 | 0.001693 | 0.2912988 | 0.308 | 0.127 | 1 | 23 | cldn5a |
| mcl1b7 | 0.001708 | 0.329404671 | 0.59 | 0.315 | 1 | 23 | mcl1b |
| nfkbiab2 | 0.001722 | 0.311979013 | 0.564 | 0.3 | 1 | 23 | nfkbiab |
| ranbp12 | 0.001766 | 0.309931465 | 0.59 | 0.325 | 1 | 23 | ranbp1 |
| pin14 | 0.001822 | 0.298307307 | 0.436 | 0.22 | 1 | 23 | pin1 |
| ndufa121 | 0.002017 | 0.251305011 | 0.41 | 0.196 | 1 | 23 | ndufa12 |
| psenen2 | 0.002064 | 0.250937443 | 0.333 | 0.149 | 1 | 23 | psenen |
| prdx52 | 0.002193 | 0.275450805 | 0.564 | 0.291 | 1 | 23 | prdx5 |
| NC-002333.49 | 0.002221 | 0.359898458 | 0.897 | 0.584 | 1 | 23 | NC-002333.4 |
| naa101 | 0.002235 | 0.260780398 | 0.513 | 0.265 | 1 | 23 | naa10 |
| rbm8a2 | 0.002393 | 0.259340471 | 0.692 | 0.371 | 1 | 23 | rbm8a |
| tmem50a4 | 0.002517 | 0.275469726 | 0.41 | 0.198 | 1 | 23 | tmem50a |
| ppib8 | 0.002691 | 0.355400814 | 0.974 | 0.641 | 1 | 23 | ppib |
| celf25 | 0.002741 | 0.300002709 | 0.667 | 0.356 | 1 | 23 | celf2 |
| eif3ba4 | 0.002767 | 0.306784971 | 0.667 | 0.395 | 1 | 23 | eif3ba |
| cnbpa2 | 0.002775 | 0.252446214 | 0.487 | 0.253 | 1 | 23 | cnbpa |
| hnrnpa1b3 | 0.002858 | 0.257199531 | 0.487 | 0.246 | 1 | 23 | hnrnpa1b |
| eif3c2 | 0.002878 | 0.279587542 | 0.538 | 0.289 | 1 | 23 | eif3c |
| bzw1a | 0.00288 | 0.267177119 | 0.359 | 0.167 | 1 | 23 | bzw1a |
| eif1b2 | 0.003067 | 0.317944356 | 0.846 | 0.503 | 1 | 23 | eif1b |
| mgst3b7 | 0.003176 | 0.277950841 | 0.564 | 0.301 | 1 | 23 | mgst3b |
| eif3i | 0.003443 | 0.296694842 | 0.667 | 0.389 | 1 | 23 | eif3i |
| pcbp21 | 0.003668 | 0.307648524 | 0.359 | 0.182 | 1 | 23 | pcbp2 |
| dad15 | 0.003682 | 0.287352306 | 0.718 | 0.418 | 1 | 23 | dad1 |
| eny21 | 0.003833 | 0.2795593 | 0.333 | 0.161 | 1 | 23 | eny2 |
| gmfb3 | 0.003935 | 0.253467078 | 0.359 | 0.173 | 1 | 23 | gmfb |
| arhgef1b5 | 0.004008 | 0.268463177 | 0.282 | 0.125 | 1 | 23 | arhgef1b |
| hnrnpa0l2 | 0.004047 | 0.276048782 | 0.897 | 0.704 | 1 | 23 | hnrnpa0l |
| tmem176l.18 | 0.004079 | 0.27050579 | 0.641 | 0.324 | 1 | 23 | tmem176l.1 |

| | | | | | | | |
|---|---|---|---|---|---|---|---|
| rab102 | 0.004184 | 0.292685998 | 0.564 | 0.32 | 1 | 23 | rab10 |
| hnrnpaba3 | 0.004479 | 0.319153445 | 0.795 | 0.528 | 1 | 23 | hnrnpaba |
| ncl2 | 0.004537 | 0.27951513 | 0.718 | 0.416 | 1 | 23 | ncl |
| si:dkey-7j14.67 | 0.004917 | 0.295690785 | 0.59 | 0.321 | 1 | 23 | si:dkey-7j14.6 |
| id16 | 0.004918 | 0.310103867 | 0.513 | 0.274 | 1 | 23 | id1 |
| ier54 | 0.005152 | 0.25897931 | 0.41 | 0.211 | 1 | 23 | ier5 |
| selenow15 | 0.005497 | 0.319190237 | 0.923 | 0.595 | 1 | 23 | selenow1 |
| jpt1b6 | 0.005725 | 0.27075071 | 0.641 | 0.333 | 1 | 23 | jpt1b |
| h3f3a2 | 0.006131 | 0.258441676 | 0.667 | 0.391 | 1 | 23 | h3f3a |
| atp5f1b4 | 0.006278 | 0.278250669 | 0.846 | 0.53 | 1 | 23 | atp5f1b |
| mcm21 | 0.006372 | 0.265253844 | 0.231 | 0.097 | 1 | 23 | mcm2 |
| snu13b3 | 0.0064 | 0.27181151 | 0.513 | 0.292 | 1 | 23 | snu13b |
| eif3ea1 | 0.006977 | 0.268079018 | 0.744 | 0.449 | 1 | 23 | eif3ea |
| gng52 | 0.007163 | 0.253409838 | 0.667 | 0.383 | 1 | 23 | gng5 |
| ccng11 | 0.00748 | 0.263484308 | 0.667 | 0.406 | 1 | 23 | ccng1 |
| ywhae15 | 0.007644 | 0.273933832 | 0.615 | 0.379 | 1 | 23 | ywhae1 |
| sumo2b2 | 0.00856 | 0.273409382 | 0.769 | 0.458 | 1 | 23 | sumo2b |
| slc25a3b3 | 0.009498 | 0.296807993 | 0.846 | 0.572 | 1 | 23 | slc25a3b |
| stmn1a3 | 3.5257168 | 1.431086582 | 0.811 | 0.1 | 7.1208903841866e-41 | 24 | stmn1a |
| rrm21 | 7.8615050 | 1.146685154 | 0.541 | 0.066 | 1.58778816769066e-2 | 24 | rrm2 |
| mad2l11 | 1.1073778 | 0.659720137 | 0.297 | 0.026 | 2.23657095761791e-1 | 24 | mad2l1 |
| dut2 | 1.4514430 | 1.028928557 | 0.541 | 0.094 | 2.93147958241419e-1 | 24 | dut |
| dsn1 | 4.6645602 | 0.347563903 | 0.108 | 0.003 | 9.42101230427483e-1 | 24 | dsn1 |
| hmgb2a5 | 2.7085792 | 1.029004608 | 0.973 | 0.581 | 5.4705174426004e-15 | 24 | hmgb2a |
| rpa21 | 1.9281154 | 0.978136067 | 0.514 | 0.095 | 3.89421479789074e-1 | 24 | rpa2 |
| pcna1 | 2.8119201 | 1.108632633 | 0.676 | 0.17 | 5.67923512271234e-1 | 24 | pcna |
| tert | 1.2157327 | 0.256219427 | 0.108 | 0.004 | 2.45541537227125e-1 | 24 | tert |
| si:ch211-156b7.41 | 6.3627002 | 0.740539222 | 0.351 | 0.052 | 1.28507456916649e-1 | 24 | si:ch211-156b7.4 |
| zgc:110540 | 3.2775355 | 0.582471425 | 0.243 | 0.026 | 6.61963853155656e-1 | 24 | zgc:110540 |
| hmga1a5 | 1.1197014 | 0.944352807 | 0.865 | 0.432 | 2.26146098110089e-1 | 24 | hmga1a |
| slbp1 | 1.5616857 | 0.545832425 | 0.27 | 0.032 | 3.15413678903529e-1 | 24 | slbp |
| tubb2b2 | 1.9733230 | 1.082630595 | 0.757 | 0.318 | 3.9855204757349e-1C | 24 | tubb2b |
| rplp2l | 5.9472070 | 0.280488862 | 1 | 0.94 | 1.20115740848636e-C | 24 | rplp2l |
| hmgn24 | 6.2676057 | 0.870424305 | 1 | 0.629 | 1.26586833499772e-C | 24 | hmgn2 |
| asf1ba1 | 6.6214756 | 0.571424867 | 0.27 | 0.034 | 1.33733943768337e-C | 24 | asf1ba |

| | | | | | | | |
|---|---|---|---|---|---|---|---|
| bub1bb | 2.0771670 | 0.275017232 | 0.108 | 0.005 | 4.19525419970672e-0 | 24 | bub1bb |
| mibp1 | 7.4188006 | 0.575583974 | 0.216 | 0.024 | 1.49837516441235e-0 | 24 | mibp |
| arhgef39 | 4.0294405 | 0.320503695 | 0.135 | 0.01 | 8.13826098517319e-0 | 24 | arhgef39 |
| tank | 4.3841023 | 0.297081165 | 0.162 | 0.014 | 8.85457154406708e-0 | 24 | tank |
| h3f3a3 | 1.9956085 | 0.912892509 | 0.757 | 0.389 | 4.03053063010921e-0 | 24 | h3f3a |
| banf11 | 3.8943429 | 0.857352559 | 0.595 | 0.205 | 7.86540439222516e-0 | 24 | banf1 |
| cdk11 | 4.1444522 | 0.504688727 | 0.243 | 0.034 | 8.37055010938997e-0 | 24 | cdk1 |
| dscc1 | 5.9149366 | 0.416888484 | 0.162 | 0.016 | 1.19463975078991e-0 | 24 | dscc1 |
| sumo3b1 | 6.8888473 | 0.662630482 | 0.324 | 0.063 | 1.39134050141806e-0 | 24 | sumo3b |
| hmgb2b5 | 7.5302044 | 0.859679021 | 0.811 | 0.438 | 1.52087539601571e-0 | 24 | hmgb2b |
| tpx2 | 8.6201465 | 0.357495931 | 0.162 | 0.016 | 1.74101100577632e-0 | 24 | tpx2 |
| si:ch73-281n10.29 | 2.2837686 | 0.814355443 | 0.811 | 0.432 | 4.61252761328128e-0 | 24 | si:ch73-281n10.2 |
| dck1 | 1.9086315 | 0.504941744 | 0.243 | 0.041 | 3.85486309020129e-0 | 24 | dck |
| gins2 | 3.6489488 | 0.497903916 | 0.216 | 0.034 | 7.36978202047e-05 | 24 | gins2 |
| LOC100330864 | 4.1074721 | 0.297011693 | 0.108 | 0.009 | 8.29586142852616e-0 | 24 | LOC100330864 |
| si:ch211-222l21.19 | 4.2700405 | 0.691914561 | 0.865 | 0.53 | 8.62420086685719e-0 | 24 | si:ch211-222l21.1 |
| rfc41 | 1.0871805 | 0.529913066 | 0.297 | 0.064 | 0.000219578 | 24 | rfc4 |
| dnph11 | 1.2108272 | 0.481268037 | 0.243 | 0.045 | 0.000244551 | 24 | dnph1 |
| anp32b4 | 1.2434884 | 0.778599197 | 0.568 | 0.236 | 0.000251147 | 24 | anp32b |
| tuba8l42 | 1.5820509 | 0.737125462 | 0.73 | 0.452 | 0.000319527 | 24 | tuba8l4 |
| esco21 | 2.1528384 | 0.337114174 | 0.162 | 0.021 | 0.000434809 | 24 | esco2 |
| cirbpa2 | 2.7369012 | 0.518858256 | 0.892 | 0.703 | 0.000552772 | 24 | cirbpa |
| gid8a1 | 3.5638435 | 0.372522686 | 0.189 | 0.029 | 0.000719789 | 24 | gid8a |
| spc25 | 5.2146660 | 0.258452006 | 0.135 | 0.016 | 0.001053206 | 24 | spc25 |
| orc41 | 7.4528250 | 0.318938232 | 0.189 | 0.03 | 0.001505247 | 24 | orc4 |
| vrk11 | 1.5273675 | 0.478762804 | 0.27 | 0.061 | 0.003084824 | 24 | vrk1 |
| tuba8l1 | 2.1793944 | 0.749631092 | 0.622 | 0.312 | 0.004401723 | 24 | tuba8l |
| atic | 2.6802216 | 0.440245197 | 0.189 | 0.033 | 0.005413244 | 24 | atic |
| ccna2 | 3.1290507 | 0.405360533 | 0.162 | 0.025 | 0.006319744 | 24 | ccna2 |
| siva11 | 3.2009488 | 0.39186731 | 0.243 | 0.052 | 0.006464956 | 24 | siva1 |
| snrpd13 | 8.9188474 | 0.670702211 | 0.568 | 0.276 | 0.018013396 | 24 | snrpd1 |
| smc4 | 1.4146247 | 0.297056444 | 0.135 | 0.02 | 0.028571177 | 24 | smc4 |
| dnajc92 | 3.5019699 | 0.408066335 | 0.27 | 0.071 | 0.070729287 | 24 | dnajc9 |
| fabp32 | 6.6431575 | 0.686427291 | 0.432 | 0.177 | 0.134171853 | 24 | fabp3 |
| cbx53 | 7.8951923 | 0.631867352 | 0.486 | 0.221 | 0.159459199 | 24 | cbx5 |

| | | | | | | | |
|---|---|---|---|---|---|---|---|
| ube2c1 | 9.7556590 | 0.385835034 | 0.189 | 0.041 | 0.197035046 | 24 | ube2c |
| ranbp13 | 9.9653854 | 0.632334117 | 0.568 | 0.325 | 0.201270891 | 24 | ranbp1 |
| h2afvb3 | 1.0594943 | 0.554933485 | 0.757 | 0.531 | 0.213986077 | 24 | h2afvb |
| LOC101885938 | 1.0843170 | 0.259657613 | 0.135 | 0.023 | 0.218999507 | 24 | LOC101885938 |
| dtymk | 1.1160448 | 0.316619038 | 0.162 | 0.032 | 0.225407577 | 24 | dtymk |
| rrm1 | 1.3379921 | 0.333220428 | 0.189 | 0.042 | 0.270234269 | 24 | rrm1 |
| calm2b6 | 1.4685062 | 0.61044832 | 0.676 | 0.448 | 0.296594206 | 24 | calm2b |
| cks1b | 1.6804883 | 0.418578828 | 0.189 | 0.044 | 0.339408229 | 24 | cks1b |
| rpa31 | 1.9672441 | 0.519386874 | 0.324 | 0.112 | 0.397324303 | 24 | rpa3 |
| dlgap51 | 1.9699330 | 0.262235513 | 0.135 | 0.024 | 0.397867381 | 24 | dlgap5 |
| swi5 | 1.9830103 | 0.430956966 | 0.162 | 0.034 | 0.400508597 | 24 | swi5 |
| hmgn75 | 2.0543509 | 0.512888 | 0.73 | 0.531 | 0.41491726 | 24 | hmgn7 |
| tmsb6 | 2.3918577 | 0.505697182 | 0.216 | 0.056 | 0.483083504 | 24 | tmsb |
| si:dkey-16p21.82 | 2.5027422 | 0.361596007 | 0.162 | 0.034 | 0.505478849 | 24 | si:dkey-16p21.8 |
| si:dkey-6i22.51 | 2.5661056 | 0.295351488 | 0.135 | 0.025 | 0.51827635 | 24 | si:dkey-6i22.5 |
| rfc2 | 2.9907034 | 0.372330533 | 0.162 | 0.035 | 0.604032371 | 24 | rfc2 |
| tubb4b3 | 3.2195519 | 0.433748724 | 0.865 | 0.67 | 0.650252903 | 24 | tubb4b |
| snrpg3 | 3.2871046 | 0.611405149 | 0.514 | 0.27 | 0.663896531 | 24 | snrpg |
| snrpc | 3.5390187 | 0.526166181 | 0.378 | 0.153 | 0.714775625 | 24 | snrpc |
| chtf8 | 3.7964595 | 0.303432151 | 0.189 | 0.045 | 0.766770926 | 24 | chtf8 |
| pola2 | 4.1210115 | 0.288589031 | 0.108 | 0.017 | 0.832320697 | 24 | pola2 |
| lig11 | 4.1845563 | 0.465500893 | 0.27 | 0.086 | 0.845154847 | 24 | lig1 |
| zgc:564931 | 5.9574442 | 0.347757286 | 0.865 | 0.72 | 1 | 24 | zgc:56493 |
| fth1a4 | 6.0551709 | 0.507628079 | 0.73 | 0.451 | 1 | 24 | fth1a |
| prdx27 | 8.2387750 | 0.485270648 | 0.676 | 0.49 | 1 | 24 | prdx2 |
| suv39h1b1 | 8.6157637 | 0.288292666 | 0.108 | 0.018 | 1 | 24 | suv39h1b |
| ppil3 | 8.7123743 | 0.302650471 | 0.162 | 0.037 | 1 | 24 | ppil3 |
| atp5pf2 | 0.000105 | 0.528485662 | 0.649 | 0.431 | 1 | 24 | atp5pf |
| rpl41 | 0.000133 | 0.256168722 | 0.946 | 0.824 | 1 | 24 | rpl4 |
| cks21 | 0.000134 | 0.351784276 | 0.135 | 0.028 | 1 | 24 | cks2 |
| arl14ep1 | 0.000141 | 0.376488594 | 0.243 | 0.078 | 1 | 24 | arl14ep |
| unga1 | 0.000147 | 0.294275208 | 0.135 | 0.028 | 1 | 24 | unga |
| cenpw1 | 0.000149 | 0.337877957 | 0.162 | 0.039 | 1 | 24 | cenpw |
| polr2h1 | 0.000178 | 0.448961831 | 0.216 | 0.067 | 1 | 24 | polr2h |
| rfc3 | 0.000196 | 0.372853714 | 0.189 | 0.054 | 1 | 24 | rfc3 |

| | | | | | | | |
|---|---|---|---|---|---|---|---|
| rbbp43 | 0.000202 | 0.511851116 | 0.324 | 0.134 | 1 | 24 | rbbp4 |
| rpl22l15 | 0.000232 | 0.370658551 | 0.73 | 0.623 | 1 | 24 | rpl22l1 |
| rpl71 | 0.000246 | 0.258219734 | 1 | 0.868 | 1 | 24 | rpl7 |
| cenps1 | 0.000276 | 0.282921483 | 0.108 | 0.021 | 1 | 24 | cenps |
| si:dkey-88p24.111 | 0.000292 | 0.443681946 | 0.243 | 0.086 | 1 | 24 | si:dkey-88p24.11 |
| hnrnpabb4 | 0.000305 | 0.490292777 | 0.595 | 0.407 | 1 | 24 | hnrnpabb |
| cdca7a1 | 0.000305 | 0.343054307 | 0.189 | 0.054 | 1 | 24 | cdca7a |
| kif22 | 0.000391 | 0.278619867 | 0.135 | 0.031 | 1 | 24 | kif22 |
| si:ch211-288g17.3! | 0.000514 | 0.506372745 | 0.514 | 0.294 | 1 | 24 | si:ch211-288g17.3 |
| armc1l | 0.000522 | 0.262034414 | 0.108 | 0.022 | 1 | 24 | armc1l |
| pttg11 | 0.000614 | 0.32753475 | 0.135 | 0.033 | 1 | 24 | pttg1 |
| mcm73 | 0.000632 | 0.363625569 | 0.27 | 0.102 | 1 | 24 | mcm7 |
| sumo3a6 | 0.000687 | 0.472162708 | 0.595 | 0.41 | 1 | 24 | sumo3a |
| fen11 | 0.000726 | 0.401341355 | 0.216 | 0.074 | 1 | 24 | fen1 |
| nup62l | 0.000741 | 0.392705393 | 0.243 | 0.089 | 1 | 24 | nup62l |
| hmgb1b7 | 0.000765 | 0.488830441 | 0.514 | 0.303 | 1 | 24 | hmgb1b |
| dynll14 | 0.000806 | 0.483574563 | 0.595 | 0.398 | 1 | 24 | dynll1 |
| lsm73 | 0.000947 | 0.444338819 | 0.432 | 0.232 | 1 | 24 | lsm7 |
| snrpf2 | 0.00112 | 0.458882216 | 0.568 | 0.391 | 1 | 24 | snrpf |
| rpa11 | 0.001189 | 0.364734138 | 0.243 | 0.094 | 1 | 24 | rpa1 |
| seta3 | 0.00128 | 0.481535824 | 0.514 | 0.358 | 1 | 24 | seta |
| zgc:194627 | 0.001282 | 0.307123821 | 0.108 | 0.025 | 1 | 24 | zgc:194627 |
| nup501 | 0.001378 | 0.393586025 | 0.243 | 0.097 | 1 | 24 | nup50 |
| rps27.21 | 0.001466 | 0.267797993 | 0.919 | 0.766 | 1 | 24 | rps27.2 |
| cdc25b | 0.001469 | 0.277655352 | 0.108 | 0.025 | 1 | 24 | cdc25b |
| sub1b2 | 0.001607 | 0.43962308 | 0.378 | 0.192 | 1 | 24 | sub1b |
| snrpe3 | 0.001708 | 0.483201317 | 0.459 | 0.276 | 1 | 24 | snrpe |
| dek | 0.001841 | 0.272861131 | 0.162 | 0.05 | 1 | 24 | dek |
| tubb6 | 0.002335 | 0.262609029 | 0.108 | 0.027 | 1 | 24 | tubb6 |
| wdr461 | 0.00257 | 0.301724475 | 0.189 | 0.067 | 1 | 24 | wdr46 |
| hmgn65 | 0.002636 | 0.343127897 | 0.757 | 0.572 | 1 | 24 | hmgn6 |
| sub1a3 | 0.003875 | 0.466421612 | 0.486 | 0.368 | 1 | 24 | sub1a |
| tyms1 | 0.004109 | 0.283066696 | 0.162 | 0.054 | 1 | 24 | tyms |
| tfam | 0.004294 | 0.296453209 | 0.216 | 0.087 | 1 | 24 | tfam |
| ran2 | 0.004741 | 0.259575499 | 0.73 | 0.666 | 1 | 24 | ran |

| | | | | | | | |
|---|---|---|---|---|---|---|---|
| atp5md3 | 0.005004 | 0.39564756 | 0.541 | 0.354 | 1 | 24 | atp5md |
| cbx3a2 | 0.0056 | 0.458842264 | 0.432 | 0.283 | 1 | 24 | cbx3a |
| luc7l3 | 0.005887 | 0.261565585 | 0.216 | 0.088 | 1 | 24 | luc7l3 |
| atp5mc13 | 0.006066 | 0.448678105 | 0.486 | 0.338 | 1 | 24 | atp5mc1 |
| mcm41 | 0.006316 | 0.285248269 | 0.189 | 0.075 | 1 | 24 | mcm4 |
| psmd101 | 0.007174 | 0.259529392 | 0.162 | 0.059 | 1 | 24 | psmd10 |
| ahcy3 | 0.008427 | 0.371830028 | 0.243 | 0.113 | 1 | 24 | ahcy |
| cenpx1 | 0.00864 | 0.282445045 | 0.189 | 0.076 | 1 | 24 | cenpx |
| eif2s21 | 0.008883 | 0.362855944 | 0.568 | 0.469 | 1 | 24 | eif2s2 |
| kpna21 | 0.009442 | 0.292320248 | 0.108 | 0.032 | 1 | 24 | kpna2 |
| mibp23 | 0.009531 | 0.274284252 | 0.757 | 0.631 | 1 | 24 | mibp2 |
| sinhcafl | 0.009901 | 0.332639868 | 0.216 | 0.099 | 1 | 24 | sinhcafl |
| lyz | 0 | 2.899976624 | 0.912 | 0 | 0 | 25 | lyz |
| mpx | 0 | 2.288244016 | 0.971 | 0 | 0 | 25 | mpx |
| ch25hl2 | 0 | 2.023974133 | 0.912 | 0.004 | 0 | 25 | ch25hl2 |
| mmp13a | 4.3338186 | 1.906629972 | 0.824 | 0.003 | 8.75301354447324e-2 | 25 | mmp13a |
| scpp8 | 6.4773939 | 1.711743857 | 0.794 | 0.005 | 1.30823924614206e-2 | 25 | scpp8 |
| npsn2 | 1.3033301 | 2.435671829 | 1 | 0.017 | 2.63233590487593e-2 | 25 | npsn |
| BX908782.2 | 2.1584370 | 2.162864716 | 0.941 | 0.017 | 4.35939535613178e-2 | 25 | BX908782.2 |
| si:ch1073-429i10.1 | 1.4776459 | 1.495411561 | 0.559 | 0.003 | 2.98440149919033e-1 | 25 | si:ch1073-429i10.1 |
| chia.6 | 8.7868525 | 0.854854207 | 0.412 | 0.002 | 1.7746806103347e-13 | 25 | chia.6 |
| BX908782.31 | 1.5462263 | 2.091809591 | 0.912 | 0.03 | 3.12291333396829e-1 | 25 | BX908782.3 |
| il6r | 1.8287355 | 1.337338388 | 0.588 | 0.01 | 3.69349721933539e-1 | 25 | il6r |
| lect2l1 | 3.8594227 | 2.596774098 | 1 | 0.044 | 7.79487603390976e-1 | 25 | lect2l |
| cnksr2b | 8.3856513 | 0.593402596 | 0.294 | 0 | 1.69364999765991e-1 | 25 | cnksr2b |
| si:ch211-284o19.8 | 3.1060380 | 0.831159318 | 0.353 | 0.002 | 6.27326496804527e-1 | 25 | si:ch211-284o19.8 |
| si:ch211-260e23.8 | 1.3063535 | 0.691949322 | 0.294 | 0.001 | 2.63844219830022e-1 | 25 | si:ch211-260e23.8 |
| si:dkey-19b23.14 | 5.2727869 | 0.707312016 | 0.294 | 0.001 | 1.06494477710417e-9 | 25 | si:dkey-19b23.14 |
| cd7al2 | 1.9848504 | 1.556030869 | 0.735 | 0.03 | 4.00880252285975e-9 | 25 | cd7al |
| ly6m51 | 1.5193980 | 1.281067612 | 0.588 | 0.018 | 3.06872832692308e-8 | 25 | ly6m5 |
| il1b | 2.0338986 | 1.121566706 | 0.412 | 0.007 | 4.10786502381415e-8 | 25 | il1b |
| si:ch211-286b5.8 | 1.8109621 | 0.867602082 | 0.353 | 0.004 | 3.6576001871304e-86 | 25 | si:ch211-286b5.8 |
| si:ch211-193e13.5 | 2.0429536 | 1.197850244 | 0.529 | 0.015 | 4.12615348731517e-8 | 25 | si:ch211-193e13.5 |
| f8 | 2.9256314 | 0.659888726 | 0.235 | 0.001 | 5.90889793245927e-8 | 25 | f8 |
| abcb9 | 1.9420029 | 0.87442188 | 0.324 | 0.004 | 3.92226343705505e-8 | 25 | abcb9 |

| | | | | | | | |
|---|---|---|---|---|---|---|---|
| si:ch211-67e16.31 | 3.7770860 | 1.144769313 | 0.5 | 0.014 | 7.62858077724634e-8 | 25 | si:ch211-67e16.3 |
| CR848841.11 | 2.3206537 | 1.07415914 | 0.412 | 0.008 | 4.68702442788296e-7 | 25 | CR848841.1 |
| thy12 | 3.4132222 | 1.658343992 | 0.765 | 0.043 | 6.8936850057544e-76 | 25 | thy1 |
| si:ch211-9d9.81 | 7.3387460 | 0.961951293 | 0.441 | 0.013 | 1.482206532966e-67 | 25 | si:ch211-9d9.8 |
| abcc13 | 6.5854443 | 0.389554019 | 0.176 | 0 | 1.33006220184977e-6 | 25 | abcc13 |
| CR354540.1 | 2.5373081 | 0.427944052 | 0.147 | 0 | 5.12460121985131e-6 | 25 | CR354540.1 |
| ncf11 | 4.4059441 | 1.095880918 | 0.559 | 0.026 | 8.89868530855099e-5 | 25 | ncf1 |
| si:ch211-9d9.1 | 1.8791699 | 0.573320167 | 0.206 | 0.002 | 3.7953595906541e-55 | 25 | si:ch211-9d9.1 |
| mmp9 | 2.2661321 | 0.768975304 | 0.206 | 0.002 | 4.57690702323604e-5 | 25 | mmp9 |
| si:ch211-276a23.5 | 2.9009328 | 1.031895418 | 0.471 | 0.02 | 5.85901414698298e-5 | 25 | si:ch211-276a23.5 |
| si:ch73-248e21.73 | 1.3132093 | 1.432152699 | 0.765 | 0.067 | 2.65228890021736e-5 | 25 | si:ch73-248e21.7 |
| sult2st1 | 7.4187703 | 1.275103155 | 0.588 | 0.037 | 1.4983690529309e-50 | 25 | sult2st1 |
| ca61 | 3.0387226 | 0.991744556 | 0.412 | 0.016 | 6.13730814652976e-5 | 25 | ca6 |
| si:dkey-163m14.6 | 3.4022721 | 0.325152689 | 0.118 | 0 | 6.87156896264258e-5 | 25 | si:dkey-163m14.6 |
| alox5ap3 | 2.1412183 | 1.105154449 | 0.529 | 0.03 | 4.32461865354815e-4 | 25 | alox5ap |
| sms1 | 2.8451704 | 1.24522569 | 0.618 | 0.048 | 5.74639082393305e-4 | 25 | sms |
| illr41 | 1.2703092 | 0.681398137 | 0.294 | 0.009 | 2.56564359935664e-4 | 25 | illr4 |
| si:dkey-42l23.4 | 2.1023607 | 0.527332116 | 0.206 | 0.003 | 4.24613796641139e-4 | 25 | si:dkey-42l23.4 |
| adam8a1 | 4.5831356 | 1.172843908 | 0.529 | 0.038 | 9.25655900151209e-4 | 25 | adam8a |
| hbxb | 1.2219578 | 0.325552835 | 0.176 | 0.002 | 2.467988199336e-39 | 25 | hbxb |
| dhrs91 | 1.6392274 | 1.10724761 | 0.559 | 0.045 | 3.31074775887543e-3 | 25 | dhrs9 |
| CT030188.12 | 1.6969624 | 1.33723777 | 0.676 | 0.07 | 3.42735504642236e-3 | 25 | CT030188.1 |
| lta4h3 | 2.1233522 | 1.564618409 | 0.735 | 0.089 | 4.28853458178766e-3 | 25 | lta4h |
| srgn6 | 9.1777840 | 1.878630133 | 1 | 0.238 | 1.85363704927585e-3 | 25 | srgn |
| cfl1l3 | 4.7381434 | 1.21704248 | 0.618 | 0.061 | 9.5696283845955e-34 | 25 | cfl1l |
| ltb4r2 | 4.7807121 | 0.885862594 | 0.382 | 0.022 | 9.65560442186677e-3 | 25 | ltb4r |
| cxcr4b5 | 4.2206239 | 1.56890883 | 0.882 | 0.149 | 8.52439426528674e-3 | 25 | cxcr4b |
| gsto1 | 7.7060236 | 0.392575449 | 0.176 | 0.003 | 1.55638559054998e-3 | 25 | gsto1 |
| prkcda1 | 2.0699228 | 0.943071897 | 0.441 | 0.031 | 4.18062308160464e-3 | 25 | prkcda |
| LOC101882843 | 3.0485996 | 0.726894021 | 0.294 | 0.013 | 6.1572566914217e-32 | 25 | LOC101882843 |
| si:ch211-264f5.21 | 2.8557754 | 0.734930369 | 0.294 | 0.013 | 5.76780971664421e-3 | 25 | si:ch211-264f5.2 |
| ryr3 | 1.1420874 | 0.420229983 | 0.147 | 0.002 | 2.30667397418053e-2 | 25 | ryr3 |
| aldh8a1 | 1.4863751 | 0.30957803 | 0.147 | 0.002 | 3.00203189548516e-2 | 25 | aldh8a1 |
| zgc:638313 | 2.0367439 | 1.23796786 | 0.735 | 0.106 | 4.11361171908651e-2 | 25 | zgc:63831 |
| lrrc151 | 3.5840482 | 0.920747469 | 0.412 | 0.03 | 7.23870230710058e-2 | 25 | lrrc15 |

| | | | | | | | |
|---|---|---|---|---|---|---|---|
| LOC101885065 | 8.0624124 | 0.655011925 | 0.324 | 0.018 | 1.62836543946199e-2 | 25 | LOC101885065 |
| hyal31 | 2.0238592 | 0.527651278 | 0.265 | 0.011 | 4.08758847697975e-2 | 25 | hyal3 |
| CT573383.12 | 2.0618791 | 0.572201241 | 0.324 | 0.018 | 4.16437726248634e-2 | 25 | CT573383.1 |
| HACD4 | 2.4043506 | 0.966147044 | 0.471 | 0.042 | 4.85606706407002e-2 | 25 | HACD4 |
| scinlb4 | 2.5953001 | 1.221590897 | 0.676 | 0.091 | 5.24172780170874e-2 | 25 | scinlb |
| ponzr6 | 4.5140611 | 0.414691645 | 0.206 | 0.006 | 9.1170493759386e-28 | 25 | ponzr6 |
| il342 | 1.5427701 | 0.834007855 | 0.382 | 0.028 | 3.11593283762819e-2 | 25 | il34 |
| vsir4 | 7.1625107 | 1.225580065 | 0.676 | 0.104 | 1.44661229357405e-2 | 25 | vsir |
| ctss13 | 8.7366457 | 0.641161174 | 0.324 | 0.021 | 1.7645403423975e-24 | 25 | ctss1 |
| asah1a | 3.0549311 | 0.604512915 | 0.265 | 0.014 | 6.17004444491663e-2 | 25 | asah1a |
| vwa111 | 4.4813314 | 0.463550338 | 0.176 | 0.005 | 9.05094509332092e-2 | 25 | vwa11 |
| si:dkey-222f2.74 | 9.0419613 | 0.913343489 | 0.412 | 0.039 | 1.82620493939083e-2 | 25 | si:dkey-222f2.7 |
| nccrp13 | 2.1671280 | 1.446814662 | 0.824 | 0.216 | 4.37694858447455e-2 | 25 | nccrp1 |
| CU499336.21 | 2.6810628 | 0.572027191 | 0.265 | 0.017 | 5.41494254067458e-2 | 25 | CU499336.2 |
| foxq1a | 6.4875501 | 0.326328731 | 0.147 | 0.004 | 1.31029050000578e-1 | 25 | foxq1a |
| si:ch73-343l4.8 | 7.4393397 | 0.466182599 | 0.176 | 0.007 | 1.50252344045225e-1 | 25 | si:ch73-343l4.8 |
| XLOC-039817 | 1.0200181 | 0.488190708 | 0.206 | 0.01 | 2.06013057770169e-1 | 25 | XLOC-039817 |
| cfbl4 | 1.9153859 | 0.985885123 | 0.618 | 0.091 | 3.86850497288463e-1 | 25 | cfbl |
| glipr1a6 | 1.3936107 | 1.093593544 | 0.559 | 0.089 | 2.81467553895505e-1 | 25 | glipr1a |
| timp2b1 | 7.7296116 | 0.806122859 | 0.382 | 0.04 | 1.56114966931054e-1 | 25 | timp2b |
| capgb6 | 9.9513047 | 1.137699483 | 0.647 | 0.122 | 2.00986502656149e-1 | 25 | capgb |
| ssuh2.3 | 1.0427571 | 0.263325272 | 0.118 | 0.003 | 2.10605662942614e-1 | 25 | ssuh2.3 |
| CABZ01001434.11 | 1.2263426 | 0.937277418 | 0.294 | 0.024 | 2.47684416488287e-1 | 25 | CABZ01001434.1 |
| fmo5 | 1.5565833 | 0.59140157 | 0.265 | 0.019 | 3.14383137074441e-1 | 25 | fmo5 |
| si:ch1073-443f11.2 | 4.9429534 | 0.862099583 | 0.471 | 0.064 | 9.98328298542886e-1 | 25 | si:ch1073-443f11.2 |
| tcnbb2 | 9.2620961 | 1.019222223 | 0.324 | 0.031 | 1.8706655508695e-16 | 25 | tcnbb |
| lpar5a3 | 3.5290792 | 0.861013896 | 0.441 | 0.059 | 7.12768145549522e-1 | 25 | lpar5a |
| cebpb7 | 4.1467342 | 1.235623288 | 0.912 | 0.29 | 8.37515920410845e-1 | 25 | cebpb |
| mir2233 | 9.6476696 | 0.777159876 | 0.412 | 0.051 | 1.94853984198145e-1 | 25 | mir223 |
| si:ch211-136m16.8 | 1.0491951 | 0.71394191 | 0.324 | 0.032 | 2.11905943675132e-1 | 25 | si:ch211-136m16.8 |
| mmp25b | 1.0558484 | 0.317958819 | 0.147 | 0.006 | 2.13249712125421e-1 | 25 | mmp25b |
| elovl5 | 1.3851410 | 0.264640296 | 0.147 | 0.006 | 2.79756935582663e-1 | 25 | elovl5 |
| bdh1 | 2.2874658 | 0.431069478 | 0.235 | 0.017 | 4.61999471092039e-1 | 25 | bdh1 |
| vamp81 | 2.7991945 | 0.997870212 | 0.5 | 0.084 | 5.65353329782002e-1 | 25 | vamp8 |
| spi1b6 | 2.8542580 | 1.080429081 | 0.618 | 0.123 | 5.76474507554125e-1 | 25 | spi1b |

| | | | | | | | |
|---|---|---|---|---|---|---|---|
| sc:d156 | 1.1696110 | 0.481721307 | 0.147 | 0.006 | 2.36226338640784e-1 | 25 | sc:d156 |
| BX649411.1 | 1.5856333 | 0.312879346 | 0.147 | 0.006 | 3.20250370139896e-1 | 25 | BX649411.1 |
| pgd4 | 2.4295557 | 0.983848728 | 0.559 | 0.111 | 4.90697366501034e-1 | 25 | pgd |
| fcer1gl7 | 3.6210057 | 1.174151478 | 0.912 | 0.252 | 7.31334533347323e-1 | 25 | fcer1gl |
| si:ch211-8c17.2 | 3.6245880 | 0.317935023 | 0.118 | 0.004 | 7.32058055351665e-1 | 25 | si:ch211-8c17.2 |
| agtrap | 4.7305932 | 0.338271365 | 0.176 | 0.01 | 9.55437908892135e-1 | 25 | agtrap |
| pxk | 3.4085436 | 0.512685502 | 0.235 | 0.019 | 6.88423557830275e-1 | 25 | pxk |
| triobpa | 6.2243583 | 0.54161355 | 0.235 | 0.02 | 1.25713364983335e-1 | 25 | triobpa |
| cyp4f32 | 2.5554936 | 0.552951475 | 0.265 | 0.026 | 5.16133045963739e-1 | 25 | cyp4f3 |
| ppp1r14aa1 | 4.8722058 | 0.279190758 | 0.176 | 0.011 | 9.84039408736408e-1 | 25 | ppp1r14aa |
| camk1db | 9.3297728 | 0.30894879 | 0.147 | 0.008 | 1.88433421721822e-1 | 25 | camk1db |
| CABZ01073834.13 | 1.4350516 | 0.78486099 | 0.382 | 0.057 | 2.89837373230564e-1 | 25 | CABZ01073834.1 |
| mapk111 | 2.2389612 | 0.573519278 | 0.235 | 0.022 | 4.52202995568138e-1 | 25 | mapk11 |
| itgb25 | 3.7240716 | 0.981578342 | 0.647 | 0.161 | 7.52150752915266e-1 | 25 | itgb2 |
| gsto2 | 6.2768005 | 1.038023798 | 0.471 | 0.097 | 1.26772540279734e-1 | 25 | gsto2 |
| mid1ip1l1 | 6.5054135 | 0.782986498 | 0.441 | 0.079 | 1.31389836818894e-1 | 25 | mid1ip1l |
| spag1a | 1.7851824 | 0.545833279 | 0.294 | 0.036 | 3.60553295762507e-1 | 25 | spag1a |
| gra1 | 2.2475117 | 0.351974711 | 0.147 | 0.009 | 4.53929956606612e-1 | 25 | gra |
| plxnc14 | 2.4363926 | 0.713407865 | 0.382 | 0.062 | 4.92078228002895e-1 | 25 | plxnc1 |
| actb25 | 7.0978834 | 0.475116417 | 1 | 0.926 | 1.43355951089332e-0 | 25 | actb2 |
| jakmip1 | 1.2892033 | 0.341760208 | 0.118 | 0.006 | 2.60380399259468e-0 | 25 | jakmip1 |
| si:ch73-248e21.52 | 1.3141399 | 0.709485487 | 0.324 | 0.047 | 2.65416854166148e-0 | 25 | si:ch73-248e21.5 |
| si:dkey-27h10.22 | 1.7848506 | 0.559239785 | 0.235 | 0.025 | 3.60486281232685e-0 | 25 | si:dkey-27h10.2 |
| CABZ01117603.12 | 7.2046571 | 0.382564855 | 0.176 | 0.015 | 1.45512460026689e-0 | 25 | CABZ01117603.1 |
| sema4aa | 7.5348908 | 0.34842402 | 0.118 | 0.006 | 1.52182190318566e-0 | 25 | sema4aa |
| nrros2 | 1.1791627 | 0.695146297 | 0.324 | 0.051 | 2.38155506162584e-0 | 25 | nrros |
| zgc:55262 | 2.2494814 | 0.603045094 | 0.294 | 0.043 | 4.54327776738413e-0 | 25 | zgc:55262 |
| hpgd | 3.4619795 | 0.353829422 | 0.206 | 0.021 | 6.99216003229181e-0 | 25 | hpgd |
| si:ch211-66e2.35 | 3.8290630 | 0.764269049 | 0.412 | 0.082 | 7.73355874012191e-0 | 25 | si:ch211-66e2.3 |
| s1pr4 | 4.1170372 | 0.304117102 | 0.118 | 0.007 | 8.31518008579799e-0 | 25 | s1pr4 |
| zgc:175264 | 5.7456135 | 0.531473749 | 0.265 | 0.036 | 1.16044156276671e-0 | 25 | zgc:175264 |
| ctss2.15 | 5.8369085 | 0.685922318 | 0.382 | 0.072 | 1.17888042698374e-0 | 25 | ctss2.1 |
| atf31 | 6.5305971 | 0.611200081 | 0.265 | 0.036 | 1.3189847058827e-07 | 25 | atf3 |
| CABZ01111959.11 | 6.9428940 | 0.535260379 | 0.206 | 0.022 | 1.40225631941337e-0 | 25 | CABZ01111959.1 |
| rnaset2 | 1.3796051 | 0.634326348 | 0.294 | 0.046 | 2.78638845464095e-0 | 25 | rnaset2 |

| | | | | | | | |
|---|---|---|---|---|---|---|---|
| myh9a2 | 1.5759176 | 0.515879845 | 0.265 | 0.037 | 3.18288088801016e-0 | 25 | myh9a |
| cebpa5 | 1.6616890 | 0.861971816 | 0.588 | 0.171 | 3.35611336589107e-0 | 25 | cebpa |
| mapre1a7 | 1.9919754 | 0.857670335 | 0.618 | 0.197 | 4.02319280263084e-0 | 25 | mapre1a |
| gpr183b1 | 2.0477707 | 0.264768594 | 0.118 | 0.007 | 4.13588266900945e-0 | 25 | gpr183b |
| cybb1 | 2.7075491 | 0.397169826 | 0.206 | 0.023 | 5.46843704594071e-0 | 25 | cybb |
| selenow16 | 3.2125065 | 0.730656236 | 0.853 | 0.597 | 6.4882995637494e-07 | 25 | selenow1 |
| zgc:114045 | 3.4474625 | 0.34427019 | 0.206 | 0.023 | 6.9628401820519e-07 | 25 | zgc:114045 |
| zmp:00000010822 | 4.4451802 | 0.644021204 | 0.294 | 0.048 | 8.9779304834118e-07 | 25 | zmp:0000001082 |
| ampd3b1 | 5.9416986 | 0.368175021 | 0.176 | 0.018 | 1.20004488024376e-0 | 25 | ampd3b |
| slc7a111 | 6.5318385 | 0.316795954 | 0.118 | 0.008 | 1.31923542228724e-0 | 25 | slc7a11 |
| laptm59 | 7.2577808 | 0.825512304 | 0.676 | 0.213 | 1.46585399704468e-0 | 25 | laptm5 |
| f5 | 7.4025316 | 0.349875328 | 0.118 | 0.008 | 1.49508932109513e-0 | 25 | f5 |
| gde1 | 8.3416091 | 0.259117227 | 0.176 | 0.018 | 1.68475479539803e-0 | 25 | gde1 |
| CR381673.14 | 9.6073159 | 0.563627209 | 0.324 | 0.056 | 1.94038960351424e-0 | 25 | CR381673.1 |
| LOC110438237 | 1.0227027 | 0.663112948 | 0.324 | 0.061 | 2.06555272845183e-0 | 25 | LOC110438237 |
| pfn19 | 1.3763994 | 0.973189248 | 1 | 0.415 | 2.77991398875783e-0 | 25 | pfn1 |
| slc9a3r1a1 | 2.1416798 | 0.606653672 | 0.324 | 0.061 | 4.32555080403626e-0 | 25 | slc9a3r1a |
| spns32 | 2.4041783 | 0.557589566 | 0.206 | 0.026 | 4.85571893117202e-0 | 25 | spns3 |
| rac28 | 2.8443091 | 0.836010107 | 0.706 | 0.259 | 5.74465110565565e-0 | 25 | rac2 |
| adam19b | 2.9468128 | 0.343244038 | 0.147 | 0.013 | 5.95167798003728e-0 | 25 | adam19b |
| wasb6 | 3.3641438 | 0.816062992 | 0.529 | 0.157 | 6.79456128373673e-0 | 25 | wasb |
| gsr1 | 3.4181767 | 0.51634642 | 0.294 | 0.051 | 6.90369162445399e-0 | 25 | gsr |
| amd11 | 6.8719078 | 0.583263033 | 0.265 | 0.044 | 1.38791922234191e-0 | 25 | amd1 |
| rgs14a1 | 7.2903821 | 0.451347684 | 0.206 | 0.027 | 1.47243848033606e-0 | 25 | rgs14a |
| limd25 | 7.6046014 | 0.526719973 | 0.324 | 0.061 | 1.53590134488872e-0 | 25 | limd2 |
| tnfaip8l2a | 8.7213383 | 0.422779278 | 0.235 | 0.034 | 1.76144871162215e-0 | 25 | tnfaip8l2a |
| dusp23 | 1.7074780 | 0.665456548 | 0.412 | 0.097 | 3.44859344308677e-0 | 25 | dusp2 |
| si:ch211-105j21.93 | 2.8412369 | 0.581868579 | 0.235 | 0.037 | 5.73844618960952e-0 | 25 | si:ch211-105j21.9 |
| COX37 | 3.9529369 | 0.700534313 | 0.941 | 0.595 | 7.98374677454074e-0 | 25 | COX3 |
| zgc:100846 | 3.9695131 | 0.337445693 | 0.176 | 0.022 | 8.01722571670325e-0 | 25 | zgc:100846 |
| adam28 | 3.9770283 | 0.375464586 | 0.118 | 0.01 | 8.03240422555237e-0 | 25 | adam28 |
| cd634 | 4.1417787 | 0.690443384 | 0.824 | 0.568 | 8.3651505283701e-05 | 25 | cd63 |
| gapdhs7 | 4.3522042 | 0.750854853 | 0.794 | 0.475 | 8.79014684164089e-0 | 25 | gapdhs |
| raraa1 | 4.3887291 | 0.714333747 | 0.324 | 0.072 | 8.8639161668582e-05 | 25 | raraa |
| CABZ01058046.1 | 4.5546120 | 0.297281303 | 0.147 | 0.015 | 9.19894999044593e-0 | 25 | CABZ01058046.1 |

| | | | | | | | |
|---|---|---|---|---|---|---|---|
| ptprc6 | 7.2993528 | 0.647442719 | 0.441 | 0.118 | 0.000147425 | 25 | ptprc |
| papss2b | 1.2487330 | 0.395249722 | 0.206 | 0.031 | 0.000252207 | 25 | papss2b |
| odc15 | 1.4522661 | 0.598012431 | 0.441 | 0.119 | 0.000293314 | 25 | odc1 |
| cotl18 | 1.4639032 | 0.782892935 | 0.824 | 0.41 | 0.000295665 | 25 | cotl1 |
| hbegfa3 | 1.7930590 | 0.559718778 | 0.294 | 0.06 | 0.000362144 | 25 | hbegfa |
| si:cabz01036022.1 | 2.1059688 | 0.382950247 | 0.118 | 0.011 | 0.000425343 | 25 | si:cabz01036022.1 |
| mrps15 | 2.1586904 | 0.52013545 | 0.294 | 0.062 | 0.000435991 | 25 | mrps15 |
| mt-co17 | 2.3117162 | 0.668380828 | 0.912 | 0.592 | 0.000466897 | 25 | mt-co1 |
| itgae.23 | 3.4950128 | 0.416692559 | 0.235 | 0.041 | 0.000705888 | 25 | itgae.2 |
| alox5a4 | 3.5244661 | 0.434298926 | 0.206 | 0.032 | 0.000711836 | 25 | alox5a |
| rab443 | 5.0633992 | 0.635866743 | 0.294 | 0.065 | 0.001022655 | 25 | rab44 |
| cybc1 | 5.0733585 | 0.463572375 | 0.265 | 0.051 | 0.001024666 | 25 | cybc1 |
| madd | 6.9265110 | 0.2814877 | 0.147 | 0.018 | 0.001398947 | 25 | madd |
| zgc:1528631 | 7.0176253 | 0.470185262 | 0.147 | 0.018 | 0.00141735 | 25 | zgc:152863 |
| mt-co23 | 7.4876249 | 0.76759681 | 0.824 | 0.459 | 0.001512276 | 25 | mt-co2 |
| ank3b3 | 7.8117984 | 0.413313128 | 0.235 | 0.042 | 0.001577749 | 25 | ank3b |
| BX664618.1 | 8.4577275 | 0.317690367 | 0.118 | 0.012 | 0.001708207 | 25 | BX664618.1 |
| zdhhc20a | 9.2699551 | 0.496941626 | 0.235 | 0.045 | 0.001872253 | 25 | zdhhc20a |
| si:dkey-102g19.32 | 1.1945721 | 0.560668024 | 0.265 | 0.054 | 0.002412677 | 25 | si:dkey-102g19.3 |
| si:zfos-741a10.35 | 1.2086449 | 0.514769183 | 0.265 | 0.054 | 0.0024411 | 25 | si:zfos-741a10.3 |
| samsn1b2 | 2.1749096 | 0.342737731 | 0.147 | 0.019 | 0.004392665 | 25 | samsn1b |
| klf2b1 | 2.3573811 | 0.305184979 | 0.147 | 0.019 | 0.004761203 | 25 | klf2b |
| si:ch211-226m16.2 | 3.0946912 | 0.677341726 | 0.294 | 0.074 | 0.006250348 | 25 | si:ch211-226m16.2 |
| il162 | 3.7687847 | 0.388963424 | 0.235 | 0.046 | 0.007611814 | 25 | il16 |
| zgc:640519 | 4.4672036 | 0.68058566 | 0.5 | 0.175 | 0.009022411 | 25 | zgc:64051 |
| f11r.15 | 4.6194175 | 0.634002565 | 0.353 | 0.102 | 0.009329838 | 25 | f11r.1 |
| ets21 | 5.5146136 | 0.48922314 | 0.294 | 0.071 | 0.011137865 | 25 | ets2 |
| arpc1b9 | 5.8992483 | 0.728185138 | 0.765 | 0.359 | 0.011914712 | 25 | arpc1b |
| cyba5 | 8.3668601 | 0.600809748 | 0.412 | 0.132 | 0.016898547 | 25 | cyba |
| ncf22 | 9.9406799 | 0.295619247 | 0.176 | 0.029 | 0.020077191 | 25 | ncf2 |
| si:ch211-102c2.48 | 1.0892954 | 0.662173855 | 0.5 | 0.175 | 0.022000501 | 25 | si:ch211-102c2.4 |
| mrtfab3 | 1.1083190 | 0.437386098 | 0.235 | 0.05 | 0.02238472 | 25 | mrtfab |
| stk17b5 | 1.6672007 | 0.627985108 | 0.412 | 0.138 | 0.033672454 | 25 | stk17b |
| cited4b | 2.3122595 | 0.517224137 | 0.294 | 0.077 | 0.046700706 | 25 | cited4b |
| erg1 | 3.0461328 | 0.254810448 | 0.118 | 0.015 | 0.061522745 | 25 | erg |

| | | | | | | | |
|---|---|---|---|---|---|---|---|
| id34 | 3.7566419 | 0.635543414 | 0.412 | 0.146 | 0.075872897 | 25 | id3 |
| lpcat22 | 3.7772928 | 0.332817665 | 0.176 | 0.032 | 0.076289984 | 25 | lpcat2 |
| srd5a2a | 4.1302036 | 0.432752225 | 0.265 | 0.067 | 0.083417723 | 25 | srd5a2a |
| cbx7a1 | 5.8757974 | 0.334840705 | 0.235 | 0.053 | 0.118673482 | 25 | cbx7a |
| cox6a23 | 6.4604240 | 0.348229642 | 0.206 | 0.044 | 0.130481185 | 25 | cox6a2 |
| atg16l1 | 6.4712257 | 0.410360059 | 0.147 | 0.025 | 0.130699347 | 25 | atg16l1 |
| gpia3 | 6.7299625 | 0.646264771 | 0.382 | 0.138 | 0.135925054 | 25 | gpia |
| gadd45aa | 6.8640829 | 0.357094356 | 0.118 | 0.016 | 0.138633884 | 25 | gadd45aa |
| lxn5 | 8.6093431 | 0.627952364 | 0.559 | 0.267 | 0.173882903 | 25 | lxn |
| si:dkey-185m8.24 | 8.6325135 | 0.315720918 | 0.206 | 0.044 | 0.174350877 | 25 | si:dkey-185m8.2 |
| rps6ka5 | 9.0029864 | 0.293272836 | 0.147 | 0.025 | 0.181833317 | 25 | rps6ka5 |
| rgs192 | 9.6007142 | 0.360879749 | 0.206 | 0.045 | 0.193905626 | 25 | rgs19 |
| sdr16c5b1 | 9.6194267 | 0.469215098 | 0.294 | 0.086 | 0.194283561 | 25 | sdr16c5b |
| LOC1018857062 | 1.1279452 | 0.273821812 | 0.147 | 0.025 | 0.227811109 | 25 | LOC101885706 |
| apmap | 1.1833035 | 0.327958719 | 0.206 | 0.045 | 0.238991813 | 25 | apmap |
| mt-cyb3 | 1.2108308 | 0.727411509 | 0.618 | 0.378 | 0.244551514 | 25 | mt-cyb |
| ccdc88b2 | 1.3850147 | 0.38444709 | 0.176 | 0.036 | 0.279731433 | 25 | ccdc88b |
| coro1a8 | 1.4622438 | 0.621669577 | 0.647 | 0.281 | 0.295329389 | 25 | coro1a |
| tktb2 | 1.6227990 | 0.584858175 | 0.353 | 0.127 | 0.327756716 | 25 | tktb |
| pxna2 | 1.8573788 | 0.466133734 | 0.294 | 0.087 | 0.375134798 | 25 | pxna |
| larp6a | 1.9632300 | 0.489192425 | 0.206 | 0.05 | 0.396513577 | 25 | larp6a |
| FAR2 | 2.1251578 | 0.253443183 | 0.147 | 0.026 | 0.429218123 | 25 | FAR2 |
| tpst11 | 2.2554109 | 0.339463086 | 0.265 | 0.072 | 0.455525344 | 25 | tpst1 |
| pgp | 3.4871534 | 0.386560363 | 0.235 | 0.063 | 0.704300379 | 25 | pgp |
| RASSF55 | 4.1113362 | 0.453300873 | 0.265 | 0.077 | 0.830366587 | 25 | RASSF5 |
| mt-nd45 | 4.2215957 | 0.602037429 | 0.706 | 0.438 | 0.852635702 | 25 | mt-nd4 |
| rhogb6 | 4.3405103 | 0.571617666 | 0.412 | 0.17 | 0.876652882 | 25 | rhogb |
| gpr183a3 | 4.6371869 | 0.489850801 | 0.235 | 0.066 | 0.936572648 | 25 | gpr183a |
| arhgef3l2 | 5.6426296 | 0.291091047 | 0.147 | 0.028 | 1 | 25 | arhgef3l |
| arhgdig7 | 5.6710742 | 0.555957499 | 0.647 | 0.366 | 1 | 25 | arhgdig |
| tprg1 | 5.9088034 | 0.314888289 | 0.147 | 0.029 | 1 | 25 | tprg1 |
| si:dkey-262k9.45 | 6.2269449 | 0.446713054 | 0.294 | 0.095 | 1 | 25 | si:dkey-262k9.4 |
| gmip1 | 6.6265958 | 0.280718293 | 0.118 | 0.02 | 1 | 25 | gmip |
| cnp3 | 6.6462053 | 0.602299376 | 0.471 | 0.231 | 1 | 25 | cnp |
| zgc:776501 | 7.9145705 | 0.580953644 | 0.294 | 0.103 | 1 | 25 | zgc:77650 |

| | | | | | | | |
|---|---|---|---|---|---|---|---|
| lrp10 | 7.9620169 | 0.372249916 | 0.235 | 0.066 | 1 | 25 | lrp10 |
| rps6ka14 | 7.9835349 | 0.355013508 | 0.206 | 0.052 | 1 | 25 | rps6ka1 |
| si:dkey-222f8.3 | 8.1632957 | 0.461957706 | 0.294 | 0.099 | 1 | 25 | si:dkey-222f8.3 |
| twf1a5 | 8.9200708 | 0.545534786 | 0.441 | 0.202 | 1 | 25 | twf1a |
| XLOC-0110851 | 9.0261780 | 0.284235711 | 0.176 | 0.04 | 1 | 25 | XLOC-011085 |
| CU570894.1 | 9.0479251 | 0.278774732 | 0.118 | 0.02 | 1 | 25 | CU570894.1 |
| zgc:1749042 | 9.5101003 | 0.563240631 | 0.235 | 0.07 | 1 | 25 | zgc:174904 |
| si:dkey-260g12.11 | 9.9340401 | 0.357170272 | 0.118 | 0.021 | 1 | 25 | si:dkey-260g12.1 |
| dhrs13a.11 | 9.9471131 | 0.359544059 | 0.147 | 0.031 | 1 | 25 | dhrs13a.1 |
| rab11fip4b1 | 0.000113 | 0.333403354 | 0.147 | 0.031 | 1 | 25 | rab11fip4b |
| BX005340.11 | 0.000125 | 0.387965388 | 0.235 | 0.07 | 1 | 25 | BX005340.1 |
| arpc5b5 | 0.00013 | 0.522520509 | 0.5 | 0.231 | 1 | 25 | arpc5b |
| pkn1b | 0.000147 | 0.264803643 | 0.147 | 0.031 | 1 | 25 | pkn1b |
| nme2b.21 | 0.000158 | 0.357948934 | 0.235 | 0.068 | 1 | 25 | nme2b.2 |
| pik3r5 | 0.00016 | 0.250005595 | 0.118 | 0.021 | 1 | 25 | pik3r5 |
| lcp14 | 0.000165 | 0.404950799 | 0.235 | 0.07 | 1 | 25 | lcp1 |
| selplg4 | 0.000174 | 0.3021926 | 0.176 | 0.043 | 1 | 25 | selplg |
| xbp15 | 0.000189 | 0.521005316 | 0.676 | 0.435 | 1 | 25 | xbp1 |
| tmsb16 | 0.000192 | 0.578146165 | 0.441 | 0.194 | 1 | 25 | tmsb1 |
| dub3 | 0.000249 | 0.432630999 | 0.265 | 0.089 | 1 | 25 | dub |
| actn41 | 0.000268 | 0.488680478 | 0.324 | 0.124 | 1 | 25 | actn4 |
| si:rp71-68n21.93 | 0.000415 | 0.408592655 | 0.206 | 0.063 | 1 | 25 | si:rp71-68n21.9 |
| tmem30ab1 | 0.000435 | 0.382151977 | 0.265 | 0.093 | 1 | 25 | tmem30ab |
| fermt3b4 | 0.000541 | 0.251197792 | 0.147 | 0.035 | 1 | 25 | fermt3b |
| clic16 | 0.000558 | 0.461458828 | 0.441 | 0.216 | 1 | 25 | clic1 |
| abracl6 | 0.000655 | 0.48618633 | 0.471 | 0.242 | 1 | 25 | abracl |
| ATP11B | 0.000661 | 0.35385073 | 0.176 | 0.05 | 1 | 25 | ATP11B |
| sh3kbp13 | 0.000668 | 0.377174631 | 0.294 | 0.111 | 1 | 25 | sh3kbp1 |
| farsa | 0.000683 | 0.335451174 | 0.235 | 0.078 | 1 | 25 | farsa |
| myo1f5 | 0.000704 | 0.289102879 | 0.206 | 0.062 | 1 | 25 | myo1f |
| syngr2a2 | 0.000802 | 0.380205372 | 0.206 | 0.066 | 1 | 25 | syngr2a |
| si:ch73-234b20.53 | 0.00082 | 0.426863958 | 0.324 | 0.136 | 1 | 25 | si:ch73-234b20.5 |
| ncf4 | 0.000926 | 0.257854249 | 0.118 | 0.025 | 1 | 25 | ncf4 |
| pisd | 0.001142 | 0.285866399 | 0.118 | 0.026 | 1 | 25 | pisd |
| si:dkey-69o16.52 | 0.001142 | 0.336868585 | 0.206 | 0.067 | 1 | 25 | si:dkey-69o16.5 |

| | | | | | | | |
|---|---|---|---|---|---|---|---|
| wbp2nl1 | 0.001212 | 0.416430009 | 0.294 | 0.12 | 1 | 25 | wbp2nl |
| erlin12 | 0.00123 | 0.341756349 | 0.176 | 0.053 | 1 | 25 | erlin1 |
| LOC1039091158 | 0.001274 | 0.440457904 | 0.412 | 0.2 | 1 | 25 | LOC103909115 |
| si:ch1073-340j19.1 | 0.001327 | 0.319347409 | 0.147 | 0.04 | 1 | 25 | si:ch1073-340j19.1 |
| CT027744.22 | 0.00151 | 0.333144323 | 0.176 | 0.054 | 1 | 25 | CT027744.2 |
| slc6a6b3 | 0.001569 | 0.374610652 | 0.206 | 0.07 | 1 | 25 | slc6a6b |
| lasp16 | 0.001574 | 0.467215469 | 0.529 | 0.36 | 1 | 25 | lasp1 |
| naga2 | 0.001701 | 0.323366569 | 0.235 | 0.083 | 1 | 25 | naga |
| tagapb5 | 0.001741 | 0.350955007 | 0.235 | 0.085 | 1 | 25 | tagapb |
| tmem592 | 0.001757 | 0.561485768 | 0.382 | 0.212 | 1 | 25 | tmem59 |
| plek3 | 0.001864 | 0.258660929 | 0.176 | 0.053 | 1 | 25 | plek |
| LOC1001511962 | 0.00191 | 0.254988651 | 0.147 | 0.041 | 1 | 25 | LOC100151196 |
| fam107b3 | 0.002071 | 0.418519531 | 0.353 | 0.166 | 1 | 25 | fam107b |
| mia3 | 0.00215 | 0.331668713 | 0.147 | 0.042 | 1 | 25 | mia3 |
| wdr17 | 0.002177 | 0.440118314 | 0.441 | 0.238 | 1 | 25 | wdr1 |
| mbnl15 | 0.002212 | 0.411157548 | 0.382 | 0.186 | 1 | 25 | mbnl1 |
| rca2.13 | 0.002233 | 0.349489629 | 0.235 | 0.089 | 1 | 25 | rca2.1 |
| itm2bb7 | 0.002358 | 0.414108066 | 0.588 | 0.358 | 1 | 25 | itm2bb |
| ugp2b1 | 0.00238 | 0.302968332 | 0.206 | 0.071 | 1 | 25 | ugp2b |
| actb18 | 0.002487 | 0.335204607 | 0.912 | 0.741 | 1 | 25 | actb1 |
| si:dkey-172h23.21 | 0.002528 | 0.271481961 | 0.176 | 0.056 | 1 | 25 | si:dkey-172h23.2 |
| zgc:1721061 | 0.002563 | 0.32491572 | 0.176 | 0.057 | 1 | 25 | zgc:172106 |
| pim23 | 0.0026 | 0.426609509 | 0.206 | 0.075 | 1 | 25 | pim2 |
| pak2a2 | 0.002633 | 0.330196529 | 0.265 | 0.106 | 1 | 25 | pak2a |
| acsl4b1 | 0.002685 | 0.327601064 | 0.147 | 0.043 | 1 | 25 | acsl4b |
| LO018102.1 | 0.003283 | 0.326786704 | 0.176 | 0.06 | 1 | 25 | LO018102.1 |
| CR855337.1 | 0.00341 | 0.289085175 | 0.118 | 0.03 | 1 | 25 | CR855337.1 |
| mt-atp65 | 0.003685 | 0.440841864 | 0.647 | 0.424 | 1 | 25 | mt-atp6 |
| tsc22d32 | 0.003715 | 0.476897027 | 0.441 | 0.25 | 1 | 25 | tsc22d3 |
| fybb5 | 0.004224 | 0.315752998 | 0.176 | 0.06 | 1 | 25 | fybb |
| plpp1a2 | 0.004263 | 0.300326637 | 0.294 | 0.125 | 1 | 25 | plpp1a |
| cul1b | 0.004384 | 0.289948581 | 0.118 | 0.032 | 1 | 25 | cul1b |
| skap25 | 0.004568 | 0.396685879 | 0.265 | 0.116 | 1 | 25 | skap2 |
| elovl1b3 | 0.004779 | 0.4243832 | 0.324 | 0.166 | 1 | 25 | elovl1b |
| tmem1311 | 0.005406 | 0.276268314 | 0.147 | 0.047 | 1 | 25 | tmem131 |

| | | | | | | | |
|---|---|---|---|---|---|---|---|
| mcl1b8 | 0.005408 | 0.460960839 | 0.471 | 0.318 | 1 | 25 | mcl1b |
| cap18 | 0.005749 | 0.442913854 | 0.5 | 0.332 | 1 | 25 | cap1 |
| lrrfip1a3 | 0.006151 | 0.330681327 | 0.206 | 0.08 | 1 | 25 | lrrfip1a |
| ctsd3 | 0.006349 | 0.443480123 | 0.353 | 0.189 | 1 | 25 | ctsd |
| cst14b.13 | 0.006561 | 0.336517752 | 0.265 | 0.117 | 1 | 25 | cst14b.1 |
| CR383676.19 | 0.00669 | 0.431003219 | 0.941 | 0.66 | 1 | 25 | CR383676.1 |
| psen1 | 0.007022 | 0.287704014 | 0.176 | 0.064 | 1 | 25 | psen1 |
| zgc:1748634 | 0.007123 | 0.396039503 | 0.324 | 0.168 | 1 | 25 | zgc:174863 |
| sult6b11 | 0.007653 | 0.29223359 | 0.206 | 0.084 | 1 | 25 | sult6b1 |
| eno34 | 0.00843 | 0.46322593 | 0.441 | 0.297 | 1 | 25 | eno3 |
| zgc:1017151 | 0.008656 | 0.28183653 | 0.118 | 0.035 | 1 | 25 | zgc:101715 |
| arhgef52 | 0.008838 | 0.340728232 | 0.147 | 0.05 | 1 | 25 | arhgef5 |
| slc40a12 | 0.00885 | 0.274497838 | 0.176 | 0.065 | 1 | 25 | slc40a1 |
| si:dkey-203a12.21 | 5.9963960 | 2.54491029 | 1 | 0.03 | 1.21109210202692e-1 | 26 | si:dkey-203a12.2 |
| si:dkeyp-75b4.101 | 5.9378724 | 2.457444979 | 0.96 | 0.03 | 1.19927209422902e-1 | 26 | si:dkeyp-75b4.10 |
| si:dkey-77g12.41 | 8.2888645 | 1.883593505 | 0.96 | 0.031 | 1.67410196378549e-1 | 26 | si:dkey-77g12.4 |
| spink2.21 | 1.9864008 | 2.417046429 | 1 | 0.037 | 4.01193378564063e-1 | 26 | spink2.2 |
| BX322787.12 | 6.2371903 | 2.287275747 | 0.96 | 0.044 | 1.25972532820629e-8 | 26 | BX322787.1 |
| MFAP4 (1 of many)2 | 1.0126173 | 2.148408604 | 0.92 | 0.04 | 2.04518320512075e-8 | 26 | MFAP4 (1 of many) |
| itln11 | 2.7919353 | 1.842211678 | 0.84 | 0.033 | 5.63887177850835e-8 | 26 | itln1 |
| spink2.41 | 5.0515861 | 2.013272925 | 0.8 | 0.03 | 1.02026885123782e-8 | 26 | spink2.4 |
| si:dkey-203a12.71 | 6.3827207 | 1.823067073 | 0.8 | 0.03 | 1.28911810689714e-8 | 26 | si:dkey-203a12.7 |
| si:ch73-359m17.21 | 8.3738724 | 2.19152558 | 0.92 | 0.043 | 1.69127101418351e-7 | 26 | si:ch73-359m17.2 |
| ccl34b.91 | 6.1027363 | 1.74156629 | 0.84 | 0.035 | 1.23256966311327e-7 | 26 | ccl34b.9 |
| si:ch211-194m7.81 | 5.4367662 | 1.690692191 | 0.76 | 0.03 | 1.09806367442239e-7 | 26 | si:ch211-194m7.8 |
| spink2.31 | 3.2130421 | 1.811175602 | 0.8 | 0.034 | 6.48938113223761e-7 | 26 | spink2.3 |
| zmp:00000013231 | 9.5276259 | 1.963421773 | 0.8 | 0.037 | 1.92429461516428e-6 | 26 | zmp:0000001323 |
| spink2.51 | 5.8861601 | 1.692202119 | 0.68 | 0.026 | 1.18882776807587e-6 | 26 | spink2.5 |
| si:ch211-14a17.101 | 8.9486280 | 1.925162006 | 0.76 | 0.035 | 1.80735441549363e-6 | 26 | si:ch211-14a17.10 |
| si:dkey-203a12.51 | 1.7317955 | 1.36803798 | 0.64 | 0.031 | 3.49770746503771e-5 | 26 | si:dkey-203a12.5 |
| si:ch1073-110a20.3 | 5.3890969 | 1.505444286 | 0.64 | 0.033 | 1.08843590239035e-4 | 26 | si:ch1073-110a20.3 |
| agr22 | 2.8976718 | 1.482882754 | 0.64 | 0.034 | 5.85242782046647e-4 | 26 | agr2 |
| spink2.11 | 6.4486961 | 1.681916491 | 0.84 | 0.073 | 1.30244317038541e-4 | 26 | spink2.1 |
| LOC1104378861 | 1.3474732 | 1.301782571 | 0.56 | 0.031 | 2.72149165947621e-3 | 26 | LOC110437886 |
| si:zfos-2330d3.12 | 5.7102359 | 1.31510238 | 0.6 | 0.036 | 1.15329636173458e-3 | 26 | si:zfos-2330d3.1 |

| | | | | | | | |
|---|---|---|---|---|---|---|---|
| cst14b.21 | 3.1400969 | 1.08361907 | 0.44 | 0.028 | 6.34205377118855e-2 | 26 | cst14b.2 |
| clca11 | 4.3309273 | 1.176318943 | 0.44 | 0.03 | 8.74717398495543e-2 | 26 | clca1 |
| si:ch1073-376c22.1 | 7.0364948 | 0.991463344 | 0.4 | 0.025 | 1.4211608649033e-24 | 26 | si:ch1073-376c22.1 |
| si:ch211-214p16.2 | 1.9142458 | 1.340411299 | 0.72 | 0.087 | 3.86620226168408e-2 | 26 | si:ch211-214p16.2 |
| pdia2 | 3.3302742 | 0.621902794 | 0.28 | 0.012 | 6.72615494494616e-2 | 26 | pdia2 |
| LOC1104377313 | 1.5530864 | 1.47615149 | 0.72 | 0.094 | 3.13676866708552e-2 | 26 | LOC110437731 |
| LOC1104389701 | 1.0169866 | 1.038851918 | 0.44 | 0.033 | 2.05400786825689e-2 | 26 | LOC110438970 |
| CABZ01112049.11 | 7.6331059 | 0.955635844 | 0.4 | 0.029 | 1.54165840068213e-1 | 26 | CABZ01112049.1 |
| si:dkey-203a12.91 | 3.8387075 | 0.983615607 | 0.24 | 0.011 | 7.75303761922013e-1 | 26 | si:dkey-203a12.9 |
| si:ch211-214p16.1 | 7.4199265 | 1.567188512 | 0.88 | 0.189 | 1.49860256117353e-1 | 26 | si:ch211-214p16.1 |
| LOC1081791621 | 9.4437016 | 0.852093021 | 0.32 | 0.024 | 1.90734441762847e-1 | 26 | LOC108179162 |
| si:dkey-203a12.41 | 2.8830358 | 0.824969634 | 0.24 | 0.015 | 5.82286746790489e-1 | 26 | si:dkey-203a12.4 |
| LOC100329433 | 5.0885255 | 0.351853086 | 0.12 | 0.003 | 1.02772950808872e-1 | 26 | LOC100329433 |
| cfbl5 | 1.0711418 | 1.210363605 | 0.56 | 0.094 | 2.16338514090112e-1 | 26 | cfbl |
| cfi1 | 3.8125025 | 0.808572523 | 0.32 | 0.031 | 7.7001113991096e-11 | 26 | cfi |
| si:dkey-102g19.33 | 1.0786363 | 0.975027356 | 0.4 | 0.054 | 2.17852192207404e-1 | 26 | si:dkey-102g19.3 |
| FO203432.12 | 2.5922799 | 0.845556119 | 0.32 | 0.036 | 5.23562782789458e-0 | 26 | FO203432.1 |
| sult5a12 | 5.5889255 | 0.735972977 | 0.24 | 0.021 | 1.12879529793309e-0 | 26 | sult5a1 |
| si:ch211-212d10.1 | 4.4329482 | 0.696656323 | 0.28 | 0.03 | 8.95322566083726e-0 | 26 | si:ch211-212d10.1 |
| LOC1104387221 | 1.0638284 | 0.654808731 | 0.2 | 0.017 | 2.1486143717564e-07 | 26 | LOC110438722 |
| BX908782.32 | 1.9454604 | 0.746618727 | 0.32 | 0.041 | 3.92924655386722e-0 | 26 | BX908782.3 |
| calm1b6 | 4.7754826 | 1.045562977 | 0.56 | 0.146 | 9.64504232108388e-0 | 26 | calm1b |
| CU075735.12 | 4.9230673 | 0.458306119 | 0.16 | 0.011 | 9.94311921424726e-0 | 26 | CU075735.1 |
| cfhl21 | 8.1699827 | 0.640490982 | 0.24 | 0.026 | 1.65009141025317e-0 | 26 | cfhl2 |
| coro1a9 | 6.8486193 | 0.985423848 | 0.8 | 0.281 | 1.38321564688824e-0 | 26 | coro1a |
| zgc:91860 | 3.7577337 | 0.414782817 | 0.12 | 0.008 | 7.58949481040257e-0 | 26 | zgc:91860 |
| ssr34 | 6.4795077 | 0.915158753 | 0.8 | 0.385 | 0.000130867 | 26 | ssr3 |
| capgb7 | 6.8320252 | 0.937977811 | 0.48 | 0.126 | 0.000137986 | 26 | capgb |
| c6ast31 | 7.8664737 | 0.540471739 | 0.2 | 0.022 | 0.000158879 | 26 | c6ast3 |
| c3a.41 | 2.3898412 | 0.588161309 | 0.2 | 0.024 | 0.000482676 | 26 | c3a.4 |
| pfn110 | 5.5753574 | 0.933905593 | 0.88 | 0.419 | 0.001126055 | 26 | pfn1 |
| mctp2b1 | 1.4783436 | 0.340331135 | 0.12 | 0.01 | 0.002985811 | 26 | mctp2b |
| rflnb | 2.7153259 | 0.379549104 | 0.12 | 0.01 | 0.005484144 | 26 | rflnb |
| acat2 | 9.0391624 | 0.374353784 | 0.12 | 0.011 | 0.018256396 | 26 | acat2 |
| tmsb17 | 9.1592144 | 0.837285327 | 0.56 | 0.194 | 0.018498865 | 26 | tmsb1 |

| | | | | | | | |
|---|---|---|---|---|---|---|---|
| prss59.1 | 1.0792480 | 0.486201299 | 0.12 | 0.011 | 0.021797574 | 26 | prss59.1 |
| lta4h4 | 1.7379777 | 0.813793797 | 0.36 | 0.096 | 0.035101936 | 26 | lta4h |
| pycard5 | 3.0855134 | 0.757783951 | 0.44 | 0.145 | 0.062318116 | 26 | pycard |
| cip2a | 4.2378010 | 0.383735716 | 0.12 | 0.013 | 0.085590868 | 26 | cip2a |
| timp4.21 | 7.5517721 | 0.432957029 | 0.16 | 0.023 | 0.152523142 | 26 | timp4.2 |
| c3a.51 | 1.1372817 | 0.5493787 | 0.2 | 0.035 | 0.229696802 | 26 | c3a.5 |
| actb19 | 1.5195052 | 0.480698378 | 0.88 | 0.742 | 0.30689447 | 26 | actb1 |
| CT030188.13 | 6.6112193 | 0.647290561 | 0.28 | 0.077 | 1 | 26 | CT030188.1 |
| si:dkeyp-66d1.71 | 8.0621953 | 0.468273187 | 0.16 | 0.028 | 1 | 26 | si:dkeyp-66d1.7 |
| ctrl | 8.3805595 | 0.518384987 | 0.12 | 0.017 | 1 | 26 | ctrl |
| arpc1b10 | 0.00014 | 0.656909312 | 0.72 | 0.362 | 1 | 26 | arpc1b |
| gyg1a2 | 0.000159 | 0.673244288 | 0.28 | 0.085 | 1 | 26 | gyg1a |
| ly75 | 0.000173 | 0.394318623 | 0.12 | 0.018 | 1 | 26 | ly75 |
| cotl19 | 0.000208 | 0.657792392 | 0.68 | 0.413 | 1 | 26 | cotl1 |
| nbas | 0.000224 | 0.349181808 | 0.12 | 0.019 | 1 | 26 | nbas |
| cfp3 | 0.000286 | 0.609749531 | 0.28 | 0.086 | 1 | 26 | cfp |
| si:ch73-208g10.13 | 0.000385 | 0.529372376 | 0.16 | 0.034 | 1 | 26 | si:ch73-208g10.1 |
| si:ch1073-443f11.2 | 0.000403 | 0.534022204 | 0.24 | 0.069 | 1 | 26 | si:ch1073-443f11.2 |
| rspry1 | 0.000414 | 0.454487429 | 0.16 | 0.034 | 1 | 26 | rspry1 |
| eno1a4 | 0.000685 | 0.695248083 | 0.4 | 0.19 | 1 | 26 | eno1a |
| unc13d2 | 0.000812 | 0.504385415 | 0.16 | 0.037 | 1 | 26 | unc13d |
| rac29 | 0.000829 | 0.58916517 | 0.52 | 0.263 | 1 | 26 | rac2 |
| pstpip2 | 0.000898 | 0.402814828 | 0.12 | 0.022 | 1 | 26 | pstpip2 |
| LOC1018833821 | 0.001041 | 0.333614389 | 0.12 | 0.022 | 1 | 26 | LOC101883382 |
| slc2a3b2 | 0.001058 | 0.392980347 | 0.12 | 0.022 | 1 | 26 | slc2a3b |
| srgn7 | 0.001355 | 0.605191697 | 0.48 | 0.247 | 1 | 26 | srgn |
| sort1b | 0.001448 | 0.409420821 | 0.12 | 0.023 | 1 | 26 | sort1b |
| serp11 | 0.001454 | 0.671240129 | 0.36 | 0.169 | 1 | 26 | serp1 |
| si:dkey-163m14.2 | 0.001636 | 0.319826742 | 0.16 | 0.038 | 1 | 26 | si:dkey-163m14.2 |
| cfl17 | 0.001788 | 0.291153546 | 0.8 | 0.769 | 1 | 26 | cfl1 |
| grap2b8 | 0.001859 | 0.62240378 | 0.4 | 0.195 | 1 | 26 | grap2b |
| ehd1a1 | 0.001864 | 0.442662416 | 0.16 | 0.041 | 1 | 26 | ehd1a |
| cfh1 | 0.002034 | 0.358274493 | 0.12 | 0.024 | 1 | 26 | cfh |
| crip16 | 0.002289 | 0.549698139 | 0.56 | 0.38 | 1 | 26 | crip1 |
| si:ch211-276a23.5: | 0.003204 | 0.344268069 | 0.12 | 0.026 | 1 | 26 | si:ch211-276a23.5 |

| | | | | | | | |
|---|---|---|---|---|---|---|---|
| rogdi1 | 0.00418 | 0.384641331 | 0.12 | 0.027 | 1 | 26 | rogdi |
| dub4 | 0.004678 | 0.484358342 | 0.24 | 0.09 | 1 | 26 | dub |
| adam8a2 | 0.004695 | 0.384217595 | 0.16 | 0.044 | 1 | 26 | adam8a |
| ppp1r185 | 0.006134 | 0.479256682 | 0.2 | 0.07 | 1 | 26 | ppp1r18 |
| cpb12 | 0.006203 | 0.285517969 | 0.12 | 0.028 | 1 | 26 | cpb1 |
| btk4 | 0.006584 | 0.387794883 | 0.16 | 0.047 | 1 | 26 | btk |
| cdaa2 | 0.00788 | 0.364051247 | 0.12 | 0.03 | 1 | 26 | cdaa |
| hacd32 | 0.009189 | 0.404695766 | 0.16 | 0.05 | 1 | 26 | hacd3 |
| CR855337.11 | 0.009503 | 0.318953243 | 0.12 | 0.031 | 1 | 26 | CR855337.1 |
| LOC101886766 | 0 | 1.977748278 | 0.958 | 0.003 | 0 | 27 | LOC101886766 |
| LOC103908723 | 0 | 1.970961701 | 0.958 | 0 | 0 | 27 | LOC103908723 |
| igl4v8 | 0 | 1.941431003 | 1 | 0.001 | 0 | 27 | igl4v8 |
| zgc:194275 | 0 | 1.936041546 | 0.958 | 0 | 0 | 27 | zgc:194275 |
| cd37 | 0 | 1.901943465 | 0.958 | 0.001 | 0 | 27 | cd37 |
| igl3v5 | 0 | 1.69235369 | 0.958 | 0 | 0 | 27 | igl3v5 |
| CU896602.1 | 0 | 1.597909661 | 0.792 | 0 | 0 | 27 | CU896602.1 |
| BX571825.5 | 0 | 1.493681185 | 0.75 | 0 | 0 | 27 | BX571825.5 |
| igl1c3 | 0 | 1.409398601 | 0.75 | 0 | 0 | 27 | igl1c3 |
| igic1s1 | 0 | 1.404518762 | 0.792 | 0 | 0 | 27 | igic1s1 |
| pax5 | 0 | 1.40000569 | 0.792 | 0 | 0 | 27 | pax5 |
| XLOC-026705 | 2.6637096 | 1.37368532 | 0.667 | 0 | 5.37989435851292e-2 | 27 | XLOC-026705 |
| CT573342.2 | 4.3754987 | 1.248186482 | 0.625 | 0.001 | 8.83719483249283e-2 | 27 | CT573342.2 |
| CU571382.4 | 1.4643275 | 0.817257737 | 0.417 | 0 | 2.95750229928306e-1 | 27 | CU571382.4 |
| LOC101882264 | 8.4713970 | 1.390373186 | 0.708 | 0.007 | 1.71096806121024e-1 | 27 | LOC101882264 |
| LOC101886979 | 2.0074023 | 0.959767179 | 0.417 | 0 | 4.05435055703285e-1 | 27 | LOC101886979 |
| BX571825.3 | 2.1662469 | 1.0280132 | 0.417 | 0 | 4.37516904372676e-1 | 27 | BX571825.3 |
| ighv1-4 | 1.7979005 | 1.153366159 | 0.5 | 0.002 | 3.63121974282534e-1 | 27 | ighv1-4 |
| LOC108179343 | 4.1254432 | 0.705514224 | 0.333 | 0 | 8.33215778472133e-1 | 27 | LOC108179343 |
| cxcr4a | 2.2741074 | 1.349502835 | 0.667 | 0.008 | 4.59301487214493e-1 | 27 | cxcr4a |
| capga | 6.5577412 | 0.800292735 | 0.333 | 0.001 | 1.32446699157741e-1 | 27 | capga |
| sp6 | 2.0773307 | 0.681476491 | 0.375 | 0.002 | 4.19558483767446e-1 | 27 | sp6 |
| FP016018.1 | 1.0166530 | 0.463941576 | 0.25 | 0 | 2.05333419930427e-1 | 27 | FP016018.1 |
| CR944667.2 | 4.8343600 | 0.381220374 | 0.208 | 0 | 9.76395697735372e-9 | 27 | CR944667.2 |
| si:dkey-24p1.1 | 2.9689774 | 0.553342249 | 0.25 | 0.001 | 5.9964437145589e-81 | 27 | si:dkey-24p1.1 |
| LOC110438240 | 3.3651953 | 0.503275178 | 0.25 | 0.001 | 6.79668502275038e-8 | 27 | LOC110438240 |

| | | | | | | | |
|---|---|---|---|---|---|---|---|
| cobll1a | 4.3588025 | 0.525234487 | 0.292 | 0.002 | 8.80347351365429e-8 | 27 | cobll1a |
| epb41a | 8.1985882 | 1.0943338 | 0.708 | 0.023 | 1.65586887289806e-7 | 27 | epb41a |
| ncf12 | 3.1378153 | 1.440345262 | 0.75 | 0.027 | 6.33744566457437e-7 | 27 | ncf1 |
| tspan2a | 5.0285916 | 0.916093622 | 0.542 | 0.013 | 1.01562466375858e-7 | 27 | tspan2a |
| CU207301.2 | 2.2507419 | 0.377449745 | 0.167 | 0 | 4.54582355893897e-7 | 27 | CU207301.2 |
| irf4b1 | 5.6628971 | 0.797520812 | 0.5 | 0.011 | 1.14373534682782e-7 | 27 | irf4b |
| cldng | 1.2898454 | 0.381267455 | 0.25 | 0.001 | 2.60510084055391e-7 | 27 | cldng |
| sh2d4a | 9.9004489 | 0.607912877 | 0.333 | 0.004 | 1.99959368433436e-6 | 27 | sh2d4a |
| CT573231.12 | 9.1090855 | 1.197428662 | 0.667 | 0.026 | 1.83976200951791e-6 | 27 | CT573231.1 |
| swap70a | 7.8804136 | 0.693733742 | 0.375 | 0.007 | 1.59160714738674e-6 | 27 | swap70a |
| pou2f2a1 | 1.6609868 | 0.782937679 | 0.458 | 0.012 | 3.35469515619935e-5 | 27 | pou2f2a |
| XLOC-016788 | 1.0380567 | 0.304751984 | 0.125 | 0 | 2.09656323481029e-5 | 27 | XLOC-016788 |
| si:ch73-380l3.22 | 1.1932190 | 1.262931156 | 0.792 | 0.052 | 2.40994444183945e-4 | 27 | si:ch73-380l3.2 |
| si:ch211-145b13.6 | 3.1993449 | 0.798651593 | 0.417 | 0.013 | 6.46171699873609e-4 | 27 | si:ch211-145b13.6 |
| ccr9a6 | 1.3706394 | 1.601064056 | 0.917 | 0.092 | 2.76828042780699e-4 | 27 | ccr9a |
| si:ch211-119e14.1: | 1.6905674 | 1.265215924 | 0.708 | 0.052 | 3.41443915406372e-3 | 27 | si:ch211-119e14.1 |
| rtn4rl2b | 1.8370951 | 0.320633004 | 0.125 | 0 | 3.71038101465923e-3 | 27 | rtn4rl2b |
| tlr20.2 | 2.2170187 | 0.261360997 | 0.125 | 0 | 4.47771280032381e-3 | 27 | tlr20.2 |
| blnk | 3.8920749 | 0.757230446 | 0.458 | 0.02 | 7.86082386073466e-3 | 27 | blnk |
| si:ch73-7i4.3 | 1.3676345 | 0.471856493 | 0.208 | 0.003 | 2.76221149635704e-3 | 27 | si:ch73-7i4.3 |
| mcph1 | 1.8660647 | 0.425876272 | 0.208 | 0.003 | 3.76889102575131e-3 | 27 | mcph1 |
| arhgap151 | 3.0432005 | 0.914540212 | 0.583 | 0.037 | 6.1463520715485e-35 | 27 | arhgap15 |
| snx18b | 2.2446110 | 0.311739543 | 0.167 | 0.002 | 4.53344098950621e-3 | 27 | snx18b |
| chrna1 | 2.5429118 | 0.384168462 | 0.167 | 0.002 | 5.1359191391514e-34 | 27 | chrna1 |
| zgc:136254 | 3.4011127 | 0.357814415 | 0.167 | 0.002 | 6.86922748239394e-3 | 27 | zgc:136254 |
| si:dkey-78k11.9 | 1.2328139 | 0.466855172 | 0.25 | 0.006 | 2.48991428814989e-3 | 27 | si:dkey-78k11.9 |
| snx20 | 9.0350902 | 0.505801495 | 0.333 | 0.012 | 1.82481718155984e-3 | 27 | snx20 |
| cxcr4b6 | 1.1949189 | 1.829467475 | 1 | 0.151 | 2.4133778658896e-32 | 27 | cxcr4b |
| XLOC-0425961 | 1.3001120 | 0.771224709 | 0.458 | 0.024 | 2.62583633276804e-3 | 27 | XLOC-042596 |
| XLOC-0425954 | 5.7958640 | 1.603391129 | 0.875 | 0.1 | 1.17059066891504e-3 | 27 | XLOC-042595 |
| rbbp8l | 2.6766284 | 0.610627912 | 0.292 | 0.009 | 5.40598647426855e-3 | 27 | rbbp8l |
| slc46a2 | 3.3413784 | 0.593947667 | 0.292 | 0.009 | 6.74858204640793e-3 | 27 | slc46a2 |
| ighd | 7.0040588 | 0.341779346 | 0.125 | 0.001 | 1.4146097621465e-30 | 27 | ighd |
| si:ch1073-220m6.1 | 1.0072204 | 0.572935241 | 0.333 | 0.013 | 2.03428314246403e-3 | 27 | si:ch1073-220m6.1 |
| si:dkeyp-117h8.2 | 4.8073861 | 0.382055053 | 0.167 | 0.002 | 9.70947779837006e-3 | 27 | si:dkeyp-117h8.2 |

| | | | | | | | |
|---|---|---|---|---|---|---|---|
| pacsin31 | 1.7949014 | 0.793684705 | 0.417 | 0.022 | 3.6251623868726e-29 | 27 | pacsin3 |
| XLOC-0139643 | 1.5958347 | 1.720465602 | 0.958 | 0.155 | 3.22310751158564e-2 | 27 | XLOC-013964 |
| XLOC-0183563 | 1.6867949 | 1.049280088 | 0.5 | 0.034 | 3.40681968682478e-2 | 27 | XLOC-018356 |
| znf296 | 1.8261824 | 0.558750619 | 0.292 | 0.011 | 3.68834065625778e-2 | 27 | znf296 |
| LOC110437938 | 1.0105003 | 0.317755467 | 0.167 | 0.003 | 2.0409075273177e-26 | 27 | LOC110437938 |
| mrap2a | 1.2328511 | 0.351871686 | 0.167 | 0.003 | 2.48998948122559e-2 | 27 | mrap2a |
| rgs134 | 1.8340424 | 1.240504489 | 0.792 | 0.096 | 3.70421561024221e-2 | 27 | rgs13 |
| XLOC-028371 | 2.1961314 | 0.592073444 | 0.292 | 0.012 | 4.43552669573932e-2 | 27 | XLOC-028371 |
| cytip1 | 1.9954303 | 0.536658272 | 0.292 | 0.012 | 4.03017068739845e-2 | 27 | cytip |
| srrm4 | 4.8498027 | 0.420610815 | 0.167 | 0.003 | 9.79514666086118e-2 | 27 | srrm4 |
| si:ch211-198c19.1 | 2.2206029 | 0.504684812 | 0.292 | 0.013 | 4.48495187890511e-2 | 27 | si:ch211-198c19.1 |
| si:dkey-183p4.101 | 3.2570729 | 0.652470579 | 0.333 | 0.018 | 6.57831023519547e-2 | 27 | si:dkey-183p4.10 |
| grap2b9 | 4.1769991 | 1.352438281 | 1 | 0.188 | 8.43628517871216e-2 | 27 | grap2b |
| LOC1039094521 | 6.0978599 | 0.808129215 | 0.375 | 0.023 | 1.2315847644851e-21 | 27 | LOC103909452 |
| foxm1 | 1.3721581 | 0.266793537 | 0.167 | 0.004 | 2.7713477476195e-21 | 27 | foxm1 |
| ikzf13 | 9.8110710 | 0.678274251 | 0.458 | 0.037 | 1.98154201100514e-2 | 27 | ikzf1 |
| sftpba3 | 1.7015454 | 1.125739508 | 0.625 | 0.069 | 3.43661134564486e-2 | 27 | sftpba |
| dtx4a | 2.3806544 | 0.485797109 | 0.292 | 0.015 | 4.8082078587628e-20 | 27 | dtx4a |
| XLOC-0108993 | 2.5497195 | 1.468442917 | 0.875 | 0.163 | 5.14966854575188e-1 | 27 | XLOC-010899 |
| rel1 | 2.5896948 | 0.700863122 | 0.375 | 0.027 | 5.23040672620366e-1 | 27 | rel |
| syk6 | 3.5344531 | 1.013240184 | 0.625 | 0.077 | 7.13853509737941e-1 | 27 | syk |
| tlr19 | 6.2999723 | 0.258615787 | 0.125 | 0.002 | 1.27240541812797e-1 | 27 | tlr19 |
| BX649442.1 | 7.1327671 | 0.265767542 | 0.125 | 0.002 | 1.44060498985077e-1 | 27 | BX649442.1 |
| traf5 | 9.1435302 | 0.252704728 | 0.167 | 0.005 | 1.84671880689123e-1 | 27 | traf5 |
| LOC1039108711 | 3.7934032 | 1.153303211 | 0.667 | 0.097 | 7.66153658073031e-1 | 27 | LOC103910871 |
| si:busm1-266f07.2 | 4.3015935 | 1.152190962 | 0.917 | 0.156 | 8.68792840418766e-1 | 27 | si:busm1-266f07.2 |
| bpifcl1 | 8.9638168 | 0.50097916 | 0.333 | 0.023 | 1.81042209553073e-1 | 27 | bpifcl |
| CR936442.12 | 9.2410337 | 1.143918289 | 0.625 | 0.086 | 1.86641157787695e-1 | 27 | CR936442.1 |
| csrnp1a | 1.1228177 | 0.546470768 | 0.375 | 0.03 | 2.26775494685602e-1 | 27 | csrnp1a |
| lmo22 | 2.2159513 | 0.431784457 | 0.333 | 0.023 | 4.47555684386061e-1 | 27 | lmo2 |
| ascc11 | 3.8633631 | 0.786154565 | 0.458 | 0.047 | 7.80283452401427e-1 | 27 | ascc1 |
| XLOC-0004313 | 1.4365859 | 1.492988986 | 0.958 | 0.264 | 2.90147270004799e-1 | 27 | XLOC-000431 |
| nr4a16 | 1.6006934 | 1.024107143 | 0.75 | 0.122 | 3.23292059189145e-1 | 27 | nr4a1 |
| zgc:195282 | 4.5491682 | 0.440708784 | 0.25 | 0.014 | 9.18795504843556e-1 | 27 | zgc:195282 |
| prkcha | 8.2449992 | 0.428565053 | 0.25 | 0.014 | 1.66524250660504e-1 | 27 | prkcha |

| | | | | | | | |
|---|---|---|---|---|---|---|---|
| mpeg1.12 | 8.9119136 | 0.602604502 | 0.333 | 0.026 | 1.79993919783219e-1 | 27 | mpeg1.1 |
| klhl6 | 1.1108174 | 0.309616865 | 0.208 | 0.01 | 2.24351790382201e-1 | 27 | klhl6 |
| illr1 | 5.2172399 | 0.37780146 | 0.167 | 0.006 | 1.05372595286379e-1 | 27 | illr1 |
| mhc2b4 | 9.2933191 | 0.909262765 | 0.625 | 0.091 | 1.87697166210416e-1 | 27 | mhc2b |
| ptp4a3 | 9.4450272 | 0.817380894 | 0.458 | 0.054 | 1.90761215993758e-1 | 27 | ptp4a3 |
| cited4b1 | 1.0013118 | 0.843537928 | 0.542 | 0.076 | 2.02234958076025e-1 | 27 | cited4b |
| smc2 | 1.5737278 | 0.403099297 | 0.292 | 0.021 | 3.17845809239175e-1 | 27 | smc2 |
| tagapa3 | 2.2326573 | 0.841388538 | 0.5 | 0.062 | 4.50929806147404e-1 | 27 | tagapa |
| herpud12 | 3.1510166 | 0.697298082 | 0.5 | 0.064 | 6.36410841485212e-1 | 27 | herpud1 |
| rhoh | 3.7068290 | 0.350942499 | 0.167 | 0.007 | 7.48668271936492e-1 | 27 | rhoh |
| si:dkey-92i15.4 | 3.9149913 | 0.323923965 | 0.167 | 0.007 | 7.90710793452266e-1 | 27 | si:dkey-92i15.4 |
| s1pr41 | 4.2105722 | 0.349436545 | 0.167 | 0.007 | 8.50409281428833e-1 | 27 | s1pr4 |
| cd74b5 | 1.8444723 | 1.197677666 | 1 | 0.246 | 3.72528084790625e-1 | 27 | cd74b |
| BX323082.1 | 4.2700218 | 0.265174166 | 0.125 | 0.004 | 8.62416309854486e-1 | 27 | BX323082.1 |
| ccr9b1 | 4.3448696 | 0.360543998 | 0.167 | 0.007 | 8.7753332784266e-12 | 27 | ccr9b |
| si:ch211-114l13.12 | 4.3670978 | 0.31843047 | 0.125 | 0.004 | 8.82022756947242e-1 | 27 | si:ch211-114l13.12 |
| CT027980.1 | 4.8860663 | 0.294234081 | 0.125 | 0.004 | 9.86838829265153e-1 | 27 | CT027980.1 |
| mbnl16 | 1.3109628 | 0.85797553 | 0.833 | 0.182 | 2.64775170473289e-1 | 27 | mbnl1 |
| egr33 | 1.4165649 | 0.579703966 | 0.333 | 0.032 | 2.86103617007252e-1 | 27 | egr3 |
| CU207241.14 | 1.8057190 | 0.422843569 | 0.292 | 0.024 | 3.64701076701968e-1 | 27 | CU207241.1 |
| tex30 | 2.3257087 | 0.294147151 | 0.167 | 0.008 | 4.69723405737918e-1 | 27 | tex30 |
| si:rp71-68n21.94 | 3.2412843 | 0.676508904 | 0.458 | 0.06 | 6.54642210047269e-1 | 27 | si:rp71-68n21.9 |
| si:dkeyp-68b7.123 | 3.4334938 | 0.643046591 | 0.417 | 0.05 | 6.93462746971804e-1 | 27 | si:dkeyp-68b7.12 |
| cd74a5 | 4.2159881 | 1.211524531 | 0.875 | 0.208 | 8.51503127580002e-1 | 27 | cd74a |
| ralgps21 | 5.5914186 | 0.489615955 | 0.333 | 0.033 | 1.12929882008235e-1 | 27 | ralgps2 |
| cmtm8a | 9.8862180 | 0.265736038 | 0.125 | 0.004 | 1.99671945150287e-1 | 27 | cmtm8a |
| pdpk1a1 | 1.3825519 | 0.29274525 | 0.167 | 0.008 | 2.79234026404229e-1 | 27 | pdpk1a |
| tnip2 | 1.4806322 | 0.336328071 | 0.208 | 0.013 | 2.99043298023702e-1 | 27 | tnip2 |
| cenpn | 1.6995654 | 0.259158365 | 0.208 | 0.013 | 3.43261227760511e-1 | 27 | cenpn |
| hmbsa | 3.0782674 | 0.359555356 | 0.208 | 0.014 | 6.21717676128084e-1 | 27 | hmbsa |
| mki671 | 3.4638331 | 0.521440327 | 0.292 | 0.027 | 6.99590390356589e-1 | 27 | mki67 |
| SIPA1 | 5.6049408 | 0.320338038 | 0.167 | 0.009 | 1.13202990822211e-0 | 27 | SIPA1 |
| LOC1003308641 | 6.5924452 | 0.301782158 | 0.167 | 0.009 | 1.33147615763883e-0 | 27 | LOC100330864 |
| parvg4 | 6.9394847 | 0.831420389 | 0.583 | 0.105 | 1.4015677437366e-09 | 27 | parvg |
| ncapg | 9.6150292 | 0.329531663 | 0.208 | 0.014 | 1.9419474504351e-09 | 27 | ncapg |

| | | | | | | | |
|---|---|---|---|---|---|---|---|
| nfkb2 | 1.2672052 | 0.482036983 | 0.333 | 0.036 | 2.55937453650571e-0 | 27 | nfkb2 |
| fli1b3 | 1.4104519 | 0.483051422 | 0.333 | 0.036 | 2.84868971971818e-0 | 27 | fli1b |
| dap1b3 | 1.6740889 | 0.972076752 | 0.833 | 0.253 | 3.3811573825182e-09 | 27 | dap1b |
| ncf23 | 1.7796283 | 0.487427166 | 0.292 | 0.028 | 3.59431540625118e-0 | 27 | ncf2 |
| trim44 | 3.0421584 | 0.450625467 | 0.25 | 0.021 | 6.14424750653376e-0 | 27 | trim44 |
| pane11 | 3.6976132 | 0.288996234 | 0.208 | 0.015 | 7.46806943455894e-0 | 27 | pane1 |
| zgc:165555.3 | 5.7699850 | 0.485173573 | 0.25 | 0.022 | 1.16536388093642e-0 | 27 | zgc:165555.3 |
| LOC1005362136 | 5.9575483 | 1.016210902 | 0.667 | 0.147 | 1.20324603542945e-0 | 27 | LOC100536213 |
| rac210 | 6.5592828 | 1.012339386 | 0.917 | 0.259 | 1.32477836187048e-0 | 27 | rac2 |
| dapp11 | 7.9745172 | 0.352520209 | 0.167 | 0.01 | 1.61061323889341e-0 | 27 | dapp1 |
| vav2 | 1.0027472 | 0.296946463 | 0.167 | 0.01 | 2.02524853894962e-0 | 27 | vav2 |
| spdl1 | 1.0163200 | 0.29456781 | 0.167 | 0.01 | 2.05266158653316e-0 | 27 | spdl1 |
| pacsin2 | 1.0254775 | 0.697576542 | 0.375 | 0.051 | 2.07115694164795e-0 | 27 | pacsin2 |
| tacc3 | 1.0869353 | 0.298972035 | 0.167 | 0.01 | 2.1952832949908e-08 | 27 | tacc3 |
| cd7al3 | 1.1077661 | 0.484796068 | 0.333 | 0.039 | 2.237355279796e-08 | 27 | cd7al |
| bcl11aa1 | 1.6825311 | 0.33776901 | 0.208 | 0.016 | 3.39820821281173e-0 | 27 | bcl11aa |
| dck2 | 1.7656511 | 0.713431586 | 0.333 | 0.041 | 3.56608567094015e-0 | 27 | dck |
| wu:fb55g095 | 1.8988430 | 0.979643966 | 0.917 | 0.309 | 3.83509338241805e-0 | 27 | wu:fb55g09 |
| sc:d02841 | 1.9661623 | 0.465173352 | 0.25 | 0.023 | 3.97105803321036e-0 | 27 | sc:d0284 |
| kif221 | 2.2527142 | 0.384285382 | 0.292 | 0.03 | 4.54980703060385e-0 | 27 | kif22 |
| xpc | 2.6597778 | 0.51411506 | 0.292 | 0.031 | 5.3719533882664e-08 | 27 | xpc |
| XLOC-0430433 | 2.7229277 | 0.966383574 | 0.375 | 0.052 | 5.4994972591303e-08 | 27 | XLOC-043043 |
| rab11fip2 | 3.0969754 | 0.262850006 | 0.167 | 0.01 | 6.2549612328534e-08 | 27 | rab11fip2 |
| anln | 3.7567047 | 0.266622437 | 0.167 | 0.01 | 7.58741656415321e-0 | 27 | anln |
| rassf2b | 4.7001806 | 0.261193292 | 0.208 | 0.016 | 9.49295488186501e-0 | 27 | rassf2b |
| h1fx6 | 6.4421979 | 1.113296331 | 0.792 | 0.236 | 1.30113071330197e-0 | 27 | h1fx |
| smchd1 | 6.5357833 | 0.309398361 | 0.25 | 0.023 | 1.32003216453479e-0 | 27 | smchd1 |
| mibp3 | 1.0162990 | 0.683342432 | 0.25 | 0.025 | 2.05261918772794e-0 | 27 | mibp |
| fam53b1 | 1.2580286 | 0.691770951 | 0.5 | 0.095 | 2.54084056001269e-0 | 27 | fam53b |
| tfdp2 | 1.4926383 | 0.635181851 | 0.375 | 0.055 | 3.01468160786692e-0 | 27 | tfdp2 |
| stx2a | 1.7353853 | 0.253844743 | 0.208 | 0.017 | 3.50495784012075e-0 | 27 | stx2a |
| kctd5a3 | 2.6416174 | 0.879531427 | 0.792 | 0.246 | 5.33527468289181e-0 | 27 | kctd5a |
| ccna21 | 2.6914438 | 0.421618164 | 0.25 | 0.025 | 5.43590913485888e-0 | 27 | ccna2 |
| tnfaip8l2a1 | 3.9416493 | 0.46249776 | 0.292 | 0.035 | 7.96094913887271e-0 | 27 | tnfaip8l2a |
| plk35 | 5.1303387 | 0.589600584 | 0.375 | 0.056 | 1.03617452092409e-0 | 27 | plk3 |

| | | | | | | | |
|---|---|---|---|---|---|---|---|
| nuf21 | 7.8739946 | 0.2677052 | 0.167 | 0.012 | 1.59031069536264e-0 | 27 | nuf2 |
| mier1a | 8.0448767 | 0.388019164 | 0.25 | 0.026 | 1.62482376250558e-0 | 27 | mier1a |
| nfkbiab3 | 1.4851177 | 0.891611055 | 0.833 | 0.299 | 2.99949222608844e-0 | 27 | nfkbiab |
| sftpbb4 | 1.8742662 | 0.626639784 | 0.5 | 0.093 | 3.78545562754993e-0 | 27 | sftpbb |
| cd833 | 2.3756666 | 0.712980476 | 0.333 | 0.048 | 4.7981338814599e-06 | 27 | cd83 |
| hdhd2 | 2.4812487 | 0.406109309 | 0.25 | 0.028 | 5.01137807691851e-0 | 27 | hdhd2 |
| top2a1 | 3.4155357 | 0.359684862 | 0.25 | 0.027 | 6.89835760574996e-0 | 27 | top2a |
| mad2l12 | 8.5936685 | 0.314460306 | 0.25 | 0.028 | 1.73566323148983e-0 | 27 | mad2l1 |
| si:dkey-42i9.73 | 8.8487104 | 0.455971805 | 0.333 | 0.049 | 1.78717405443248e-0 | 27 | si:dkey-42i9.7 |
| creb1b | 1.1523123 | 0.341189769 | 0.208 | 0.021 | 2.32732516139564e-0 | 27 | creb1b |
| arhgdig8 | 1.5453830 | 0.883682324 | 0.875 | 0.364 | 3.12121022790965e-0 | 27 | arhgdig |
| zgc:153409 | 1.6470318 | 0.358931289 | 0.25 | 0.03 | 3.32651028218785e-0 | 27 | zgc:153409 |
| armc1l1 | 1.7877658 | 0.356015995 | 0.208 | 0.021 | 3.61075078803661e-0 | 27 | armc1l |
| gpr55a | 1.9103487 | 0.285909718 | 0.125 | 0.008 | 3.85833136920721e-0 | 27 | gpr55a |
| fam43a1 | 2.4049440 | 0.362919058 | 0.208 | 0.021 | 4.85726553300426e-0 | 27 | fam43a |
| rgcc3 | 2.5604358 | 0.523428386 | 0.417 | 0.076 | 5.1713123347815e-05 | 27 | rgcc |
| SLC7A1 (1 of many) | 2.9262419 | 0.261953364 | 0.167 | 0.014 | 5.91013084678059e-0 | 27 | SLC7A1 (1 of many) |
| ccl35.15 | 3.1406933 | 0.783432235 | 0.458 | 0.092 | 6.34325841033967e-0 | 27 | ccl35.1 |
| tmc6a2 | 3.1912087 | 0.332688775 | 0.167 | 0.014 | 6.44528426731892e-0 | 27 | tmc6a |
| rrm11 | 3.3377159 | 0.445246114 | 0.292 | 0.042 | 6.74118488823458e-0 | 27 | rrm1 |
| sh3d21 | 3.5323277 | 0.338653916 | 0.208 | 0.022 | 7.13424236501745e-0 | 27 | sh3d21 |
| inpp5d2 | 4.8438516 | 0.524972009 | 0.292 | 0.043 | 9.78312716204087e-0 | 27 | inpp5d |
| cad | 5.0697107 | 0.43599519 | 0.25 | 0.032 | 0.000102393 | 27 | cad |
| ciita | 5.5662682 | 0.270294105 | 0.125 | 0.008 | 0.000112422 | 27 | ciita |
| id35 | 5.7952827 | 0.682249483 | 0.583 | 0.145 | 0.000117047 | 27 | id3 |
| si:ch211-214p16.1 | 5.8408684 | 1.044520059 | 0.667 | 0.192 | 0.000117968 | 27 | si:ch211-214p16.1 |
| dtymk1 | 7.0487629 | 0.31877803 | 0.25 | 0.032 | 0.000142364 | 27 | dtymk |
| rnf216 | 9.0678450 | 0.292703025 | 0.208 | 0.023 | 0.000183143 | 27 | rnf216 |
| cks1b1 | 1.0010487 | 0.44306069 | 0.292 | 0.043 | 0.000202182 | 27 | cks1b |
| sypl2b | 1.0544218 | 0.586680269 | 0.417 | 0.086 | 0.000212962 | 27 | sypl2b |
| gpx1a2 | 1.0629886 | 0.773712547 | 0.75 | 0.234 | 0.000214692 | 27 | gpx1a |
| celf26 | 1.3348442 | 0.773191865 | 0.875 | 0.356 | 0.000269598 | 27 | celf2 |
| ikzf25 | 1.6463190 | 0.510880808 | 0.333 | 0.057 | 0.000332507 | 27 | ikzf2 |
| wasa7 | 2.0024115 | 0.653798203 | 0.5 | 0.118 | 0.000404427 | 27 | wasa |
| dub5 | 2.1891793 | 0.600221125 | 0.417 | 0.088 | 0.000442149 | 27 | dub |

| | | | | | | | |
|---|---|---|---|---|---|---|---|
| lnpa | 2.9091233 | 0.429753787 | 0.25 | 0.035 | 0.000587556 | 27 | lnpa |
| spc251 | 2.9343947 | 0.309546449 | 0.167 | 0.016 | 0.00059266 | 27 | spc25 |
| plcg21 | 2.9778987 | 0.425577124 | 0.292 | 0.046 | 0.000601446 | 27 | plcg2 |
| brms1 | 2.9936868 | 0.311084232 | 0.25 | 0.034 | 0.000604635 | 27 | brms1 |
| pim24 | 3.5336860 | 0.527259227 | 0.375 | 0.074 | 0.000713699 | 27 | pim2 |
| nfkbiaa4 | 3.6129101 | 0.735984095 | 0.583 | 0.163 | 0.000729699 | 27 | nfkbiaa |
| btg27 | 4.2307416 | 0.715068977 | 0.75 | 0.259 | 0.000854483 | 27 | btg2 |
| zgc:1037004 | 4.3545361 | 0.542069778 | 0.542 | 0.126 | 0.000879486 | 27 | zgc:103700 |
| crip17 | 4.3577409 | 0.774896374 | 0.958 | 0.376 | 0.000880133 | 27 | crip1 |
| ptmaa7 | 4.4385248 | 0.667238049 | 1 | 0.655 | 0.000896449 | 27 | ptmaa |
| smox2 | 5.1629247 | 0.488891505 | 0.458 | 0.102 | 0.001042756 | 27 | smox |
| derl24 | 5.7842017 | 0.541913863 | 0.5 | 0.123 | 0.001168235 | 27 | derl2 |
| h2afx2 | 6.8645298 | 0.587052285 | 0.292 | 0.049 | 0.001386429 | 27 | h2afx |
| cdk12 | 7.3383529 | 0.396572575 | 0.25 | 0.036 | 0.001482127 | 27 | cdk1 |
| ran3 | 7.8151033 | 0.606701524 | 1 | 0.663 | 0.001578416 | 27 | ran |
| klf133 | 7.9580061 | 0.659957124 | 0.417 | 0.098 | 0.001607278 | 27 | klf13 |
| tsc22d33 | 8.1461558 | 0.731775605 | 0.708 | 0.248 | 0.001645279 | 27 | tsc22d3 |
| tpx21 | 8.6570831 | 0.335259684 | 0.167 | 0.017 | 0.001748471 | 27 | tpx2 |
| pycard6 | 9.5313327 | 0.645549614 | 0.542 | 0.144 | 0.001925043 | 27 | pycard |
| lyn1 | 9.8711277 | 0.464996049 | 0.292 | 0.05 | 0.001993672 | 27 | lyn |
| si:ch73-234b20.54 | 1.2404069 | 0.636051931 | 0.5 | 0.135 | 0.00250525 | 27 | si:ch73-234b20.5 |
| im:7138535 | 1.2712972 | 0.333589621 | 0.208 | 0.027 | 0.002567639 | 27 | im:7138535 |
| si:ch211-147a11.3 | 1.4204527 | 0.385330897 | 0.292 | 0.05 | 0.002868888 | 27 | si:ch211-147a11.3 |
| trappc6bl | 1.4566940 | 0.313758696 | 0.292 | 0.049 | 0.002942085 | 27 | trappc6bl |
| dusp24 | 1.5144103 | 0.755898227 | 0.417 | 0.099 | 0.003058655 | 27 | dusp2 |
| rasal33 | 1.6226855 | 0.302150096 | 0.208 | 0.027 | 0.003277338 | 27 | rasal3 |
| zgc:1105401 | 1.6562254 | 0.424472596 | 0.208 | 0.027 | 0.003345079 | 27 | zgc:110540 |
| si:ch211-105j21.94 | 1.7004554 | 0.411954459 | 0.25 | 0.038 | 0.00343441 | 27 | si:ch211-105j21.9 |
| histh1l1 | 1.8088979 | 0.732220232 | 0.542 | 0.159 | 0.003653431 | 27 | histh1l |
| XLOC-0140793 | 1.8882075 | 0.525678233 | 0.208 | 0.028 | 0.003813613 | 27 | XLOC-014079 |
| RASSF56 | 2.0965489 | 0.431324164 | 0.375 | 0.077 | 0.0042344 | 27 | RASSF5 |
| skap26 | 2.1041918 | 0.611804546 | 0.458 | 0.115 | 0.004249836 | 27 | skap2 |
| si:ch211-153b23.7 | 2.2561505 | 0.330109338 | 0.208 | 0.027 | 0.004556747 | 27 | si:ch211-153b23.7 |
| atp2a34 | 2.3825897 | 0.430388628 | 0.292 | 0.052 | 0.004812116 | 27 | atp2a3 |
| bcl7bb | 2.8646041 | 0.485967017 | 0.375 | 0.082 | 0.005785641 | 27 | bcl7bb |

| | | | | | | | |
|---|---|---|---|---|---|---|---|
| ube2w | 2.9114498 | 0.347647379 | 0.25 | 0.039 | 0.005880255 | 27 | ube2w |
| cyba6 | 3.0679063 | 0.561383228 | 0.5 | 0.132 | 0.00619625 | 27 | cyba |
| zgc:6405110 | 3.1764414 | 0.585535458 | 0.625 | 0.176 | 0.006415459 | 27 | zgc:64051 |
| mbd3a1 | 3.2069927 | 0.515548713 | 0.417 | 0.102 | 0.006477163 | 27 | mbd3a |
| nusap1 | 3.4203672 | 0.303511652 | 0.167 | 0.019 | 0.006908116 | 27 | nusap1 |
| kmt2a | 3.6957932 | 0.342519205 | 0.25 | 0.04 | 0.007464394 | 27 | kmt2a |
| cks22 | 3.7895209 | 0.344806769 | 0.208 | 0.028 | 0.007653695 | 27 | cks2 |
| pi4k2b | 4.4898492 | 0.275026176 | 0.208 | 0.029 | 0.009068149 | 27 | pi4k2b |
| ncaldb3 | 4.7763873 | 0.355143982 | 0.292 | 0.052 | 0.009646869 | 27 | ncaldb |
| si:ch211-161h7.4 | 4.8909927 | 0.258144393 | 0.167 | 0.019 | 0.009878338 | 27 | si:ch211-161h7.4 |
| calm1b7 | 4.8983643 | 0.521804506 | 0.542 | 0.146 | 0.009893226 | 27 | calm1b |
| dgkzb | 5.2035479 | 0.316814367 | 0.125 | 0.011 | 0.010509606 | 27 | dgkzb |
| mcl1a8 | 5.2524734 | 0.667241131 | 0.958 | 0.544 | 0.010608421 | 27 | mcl1a |
| pigv | 5.3997488 | 0.291777735 | 0.125 | 0.011 | 0.010905873 | 27 | pigv |
| dgkg | 5.6550458 | 0.266142826 | 0.125 | 0.011 | 0.011421496 | 27 | dgkg |
| hopx1 | 5.8613104 | 0.468732847 | 0.333 | 0.068 | 0.011838089 | 27 | hopx |
| zgc:1943121 | 6.1317886 | 0.306732488 | 0.208 | 0.029 | 0.012384374 | 27 | zgc:194312 |
| tagapb6 | 6.3433827 | 0.50826424 | 0.375 | 0.084 | 0.01281173 | 27 | tagapb |
| elmo13 | 6.4687365 | 0.448162591 | 0.333 | 0.069 | 0.013064907 | 27 | elmo1 |
| bcl2l103 | 6.7963016 | 0.598995733 | 0.5 | 0.14 | 0.01372649 | 27 | bcl2l10 |
| tmc81 | 6.8776745 | 0.40863392 | 0.208 | 0.03 | 0.013890839 | 27 | tmc8 |
| tram13 | 7.9415417 | 0.644971353 | 0.458 | 0.133 | 0.016039532 | 27 | tram1 |
| CR855337.12 | 8.1990300 | 0.345398654 | 0.208 | 0.03 | 0.016559581 | 27 | CR855337.1 |
| sesn16 | 8.2082815 | 0.699442129 | 0.625 | 0.216 | 0.016578266 | 27 | sesn1 |
| hcst1 | 8.3646664 | 0.306947666 | 0.167 | 0.02 | 0.016894117 | 27 | hcst |
| ccr23 | 8.4787508 | 0.491926994 | 0.25 | 0.042 | 0.017124533 | 27 | ccr2 |
| lipea | 8.8138974 | 0.295155375 | 0.125 | 0.012 | 0.017801429 | 27 | lipea |
| nuggc.31 | 1.0251957 | 0.251199054 | 0.125 | 0.012 | 0.020705879 | 27 | nuggc.3 |
| ube2c2 | 1.0308001 | 0.303489358 | 0.25 | 0.041 | 0.020819071 | 27 | ube2c |
| wipf1b2 | 1.0314068 | 0.579263635 | 0.333 | 0.075 | 0.020831324 | 27 | wipf1b |
| rps27.11 | 1.2631954 | 0.310454101 | 1 | 0.897 | 0.025512759 | 27 | rps27.1 |
| si:ch211-243a20.4: | 1.2919064 | 0.335105214 | 0.208 | 0.031 | 0.026092635 | 27 | si:ch211-243a20.4 |
| ppp4r2a | 1.5796254 | 0.31254957 | 0.208 | 0.031 | 0.031903695 | 27 | ppp4r2a |
| pik3r51 | 1.7143678 | 0.335187403 | 0.167 | 0.021 | 0.034625088 | 27 | pik3r5 |
| rsl24d13 | 1.7280455 | 0.603590242 | 0.958 | 0.574 | 0.034901335 | 27 | rsl24d1 |

| | | | | | | | |
|---|---|---|---|---|---|---|---|
| msna5 | 1.8409437 | 0.633384599 | 0.917 | 0.401 | 0.037181542 | 27 | msna |
| ccni2 | 1.8598373 | 0.641950452 | 0.875 | 0.414 | 0.037563134 | 27 | ccni |
| si:ch211-93g21.12 | 2.0894517 | 0.283280294 | 0.167 | 0.021 | 0.042200658 | 27 | si:ch211-93g21.1 |
| tmed74 | 2.3071865 | 0.592617275 | 0.75 | 0.307 | 0.046598246 | 27 | tmed7 |
| si:ch211-195b13.1 | 2.5192823 | 0.319911239 | 0.167 | 0.021 | 0.050881945 | 27 | si:ch211-195b13.1 |
| rabl6b | 2.6926009 | 0.31712486 | 0.208 | 0.033 | 0.054382462 | 27 | rabl6b |
| wdfy4 | 3.0824505 | 0.250833363 | 0.125 | 0.013 | 0.062256254 | 27 | wdfy4 |
| mef2d | 3.4165040 | 0.457146764 | 0.375 | 0.092 | 0.069003132 | 27 | mef2d |
| thrap3b1 | 3.4945975 | 0.560133626 | 0.458 | 0.138 | 0.070580386 | 27 | thrap3b |
| rpl10 | 3.7953475 | 0.285898539 | 1 | 0.9 | 0.076654634 | 27 | rpl10 |
| coro1a10 | 3.8261153 | 0.633326286 | 0.833 | 0.28 | 0.077276052 | 27 | coro1a |
| evla | 4.1054593 | 0.433035707 | 0.208 | 0.035 | 0.082917962 | 27 | evla |
| saraf | 4.2863409 | 0.380356436 | 0.25 | 0.047 | 0.086571228 | 27 | saraf |
| gm2a4 | 4.3077212 | 0.566470634 | 0.458 | 0.132 | 0.087003046 | 27 | gm2a |
| btk5 | 4.5231121 | 0.377480207 | 0.25 | 0.046 | 0.091353297 | 27 | btk |
| mych3 | 4.5925934 | 0.625181602 | 0.5 | 0.169 | 0.09275661 | 27 | mych |
| rpl72 | 5.1285711 | 0.338528488 | 1 | 0.869 | 0.103581752 | 27 | rpl7 |
| glo1 | 6.1006384 | 0.610088931 | 0.667 | 0.28 | 0.123214595 | 27 | glo1 |
| hnrnpul1l | 6.1667820 | 0.388048149 | 0.333 | 0.078 | 0.124550498 | 27 | hnrnpul1l |
| dusp13 | 7.0570548 | 0.658332195 | 0.375 | 0.105 | 0.142531337 | 27 | dusp1 |
| LOC1039091159 | 7.2984289 | 0.553839168 | 0.583 | 0.2 | 0.147406369 | 27 | LOC103909115 |
| rplp23 | 8.1168924 | 0.259891244 | 1 | 0.928 | 0.163936878 | 27 | rplp2 |
| polr3glb | 8.1420262 | 0.313727593 | 0.292 | 0.062 | 0.164444504 | 27 | polr3glb |
| ccnd2a1 | 8.9225883 | 0.597472243 | 0.458 | 0.147 | 0.180209517 | 27 | ccnd2a |
| syngr2a3 | 1.0086792 | 0.393174155 | 0.292 | 0.065 | 0.203722956 | 27 | syngr2a |
| dhfr1 | 1.0181737 | 0.263507868 | 0.208 | 0.035 | 0.205640557 | 27 | dhfr |
| rpl19 | 1.0821639 | 0.287036664 | 1 | 0.894 | 0.218564654 | 27 | rpl19 |
| si:dkeyp-85e10.31 | 1.1137300 | 0.256330345 | 0.125 | 0.014 | 0.224940065 | 27 | si:dkeyp-85e10.3 |
| cybb2 | 1.1184419 | 0.345502098 | 0.167 | 0.024 | 0.225891725 | 27 | cybb |
| srsf7a1 | 1.1270736 | 0.597407664 | 0.583 | 0.237 | 0.227635056 | 27 | srsf7a |
| imp41 | 1.1408778 | 0.396201914 | 0.292 | 0.065 | 0.230423094 | 27 | imp4 |
| ube2e12 | 1.1483003 | 0.485693413 | 0.583 | 0.204 | 0.231922225 | 27 | ube2e1 |
| dlgap52 | 1.1846075 | 0.255702561 | 0.167 | 0.024 | 0.239255182 | 27 | dlgap5 |
| arf6a2 | 1.2242372 | 0.502422708 | 0.5 | 0.163 | 0.24725919 | 27 | arf6a |
| rrm22 | 1.2595979 | 0.806727769 | 0.292 | 0.072 | 0.254400994 | 27 | rrm2 |

| | | | | | | | |
|---|---|---|---|---|---|---|---|
| cnpy32 | 1.3332908 | 0.457499297 | 0.333 | 0.084 | 0.269284744 | 27 | cnpy3 |
| rps3 | 1.4721357 | 0.271691915 | 0.958 | 0.883 | 0.297327254 | 27 | rps3 |
| si:ch211-239j9.1 | 1.5555526 | 0.260582354 | 0.125 | 0.015 | 0.314174978 | 27 | si:ch211-239j9.1 |
| tp53inp12 | 1.6366764 | 0.56703679 | 0.292 | 0.071 | 0.330559544 | 27 | tp53inp1 |
| nosip1 | 1.7400457 | 0.307427301 | 0.292 | 0.065 | 0.35143703 | 27 | nosip |
| seta4 | 1.8232457 | 0.635862032 | 0.708 | 0.357 | 0.368240951 | 27 | seta |
| sri | 1.8245944 | 0.533969243 | 0.583 | 0.224 | 0.368513344 | 27 | sri |
| prcp1 | 1.8652542 | 0.314234757 | 0.167 | 0.025 | 0.376725411 | 27 | prcp |
| rpl15 | 1.9551687 | 0.279010389 | 1 | 0.9 | 0.394885431 | 27 | rpl15 |
| atf4a2 | 1.9674316 | 0.511854396 | 0.583 | 0.214 | 0.39736218 | 27 | atf4a |
| si:busm1-194e12.1 | 1.9688543 | 0.528842155 | 0.292 | 0.069 | 0.397649522 | 27 | si:busm1-194e12.11 |
| zgc:640652 | 2.0058408 | 0.358310159 | 0.25 | 0.05 | 0.405119673 | 27 | zgc:64065 |
| rps231 | 2.1736434 | 0.297338401 | 1 | 0.882 | 0.439010764 | 27 | rps23 |
| luc7l1 | 2.2019906 | 0.412805995 | 0.292 | 0.07 | 0.444736055 | 27 | luc7l |
| mknk2b4 | 2.2904932 | 0.545323663 | 0.583 | 0.221 | 0.462610928 | 27 | mknk2b |
| CABZ01090890.1 | 2.3546484 | 0.414716387 | 0.333 | 0.086 | 0.47556835 | 27 | CABZ01090890.1 |
| znf7041 | 2.4420284 | 0.372266118 | 0.25 | 0.052 | 0.493216481 | 27 | znf704 |
| selenow2a | 2.4514879 | 0.311766219 | 0.25 | 0.052 | 0.495127015 | 27 | selenow2a |
| lbr | 2.4890981 | 0.556122514 | 0.292 | 0.073 | 0.502723155 | 27 | lbr |
| higd2a1 | 2.5414649 | 0.431616423 | 0.417 | 0.125 | 0.513299685 | 27 | higd2a |
| rhogb7 | 2.6368630 | 0.542250553 | 0.5 | 0.17 | 0.532567231 | 27 | rhogb |
| si:ch211-260e23.9: | 2.7813420 | 0.449120915 | 0.333 | 0.088 | 0.561747652 | 27 | si:ch211-260e23.9 |
| btg14 | 2.8475231 | 0.593588869 | 0.833 | 0.465 | 0.575114251 | 27 | btg1 |
| baxa | 3.1566972 | 0.449424105 | 0.292 | 0.072 | 0.63755815 | 27 | baxa |
| eif3c3 | 3.1710741 | 0.558679033 | 0.667 | 0.29 | 0.64046184 | 27 | eif3c |
| egr2a | 3.3275858 | 0.26079265 | 0.125 | 0.016 | 0.67207252 | 27 | egr2a |
| pik3r13 | 3.8783840 | 0.404645541 | 0.375 | 0.105 | 0.783317225 | 27 | pik3r1 |
| lsm8 | 4.3305493 | 0.381104665 | 0.417 | 0.129 | 0.874641044 | 27 | lsm8 |
| rpl18a | 4.4025330 | 0.256118713 | 1 | 0.903 | 0.889179591 | 27 | rpl18a |
| fmnl1a3 | 4.4065123 | 0.331379129 | 0.208 | 0.04 | 0.889983304 | 27 | fmnl1a |
| rps5 | 4.4322434 | 0.288238467 | 1 | 0.89 | 0.895180218 | 27 | rps5 |
| birc5a1 | 4.4503518 | 0.295398423 | 0.167 | 0.027 | 0.898837561 | 27 | birc5a |
| CABZ01005379.11 | 4.7183165 | 0.325854366 | 0.208 | 0.04 | 0.952958388 | 27 | CABZ01005379.1 |
| RASA23 | 4.7550383 | 0.372218463 | 0.25 | 0.055 | 0.960375097 | 27 | RASA2 |
| XLOC-0397572 | 4.8708005 | 0.398799765 | 0.167 | 0.028 | 0.983755582 | 27 | XLOC-039757 |

| | | | | | | | | |
|---|---|---|---|---|---|---|---|---|
| ago2 | 5.1971432 | 0.290983111 | 0.208 | 0.04 | | 1 | 27 | ago2 |
| ucp28 | 5.9387343 | 0.535439777 | 0.875 | 0.424 | | 1 | 27 | ucp2 |
| pde4ba4 | 6.0680541 | 0.549356184 | 0.458 | 0.163 | | 1 | 27 | pde4ba |
| nutf2l3 | 6.3800382 | 0.584145814 | 0.458 | 0.175 | | 1 | 27 | nutf2l |
| si:ch1073-429i10.3 | 6.4072490 | 0.577082837 | 0.708 | 0.352 | | 1 | 27 | si:ch1073-429i10.3.1 |
| cry1ab | 7.3293658 | 0.310571386 | 0.208 | 0.042 | | 1 | 27 | cry1ab |
| XLOC-0283703 | 7.5148199 | 0.442223242 | 0.25 | 0.058 | | 1 | 27 | XLOC-028370 |
| casc3 | 7.7534477 | 0.309115471 | 0.208 | 0.042 | | 1 | 27 | casc3 |
| calm3a | 7.9401311 | 0.412481074 | 0.292 | 0.078 | | 1 | 27 | calm3a |
| laptm510 | 8.0817415 | 0.474433293 | 0.625 | 0.216 | | 1 | 27 | laptm5 |
| kifc1 | 8.3331488 | 0.256187889 | 0.125 | 0.017 | | 1 | 27 | kifc1 |
| tcea1 | 8.3559924 | 0.423952176 | 0.375 | 0.115 | | 1 | 27 | tcea1 |
| pfn111 | 8.3737949 | 0.830779418 | 1 | 0.418 | | 1 | 27 | pfn1 |
| cd40 | 8.7145390 | 0.357843397 | 0.167 | 0.03 | | 1 | 27 | cd40 |
| tubb2b3 | 8.8583986 | 0.727208728 | 0.625 | 0.322 | | 1 | 27 | tubb2b |
| srsf111 | 9.0721170 | 0.437635841 | 0.5 | 0.182 | | 1 | 27 | srsf11 |
| ivns1abpb1 | 9.2078524 | 0.26270751 | 0.208 | 0.042 | | 1 | 27 | ivns1abpb |
| tpst1l | 9.3752566 | 0.315521 | 0.208 | 0.043 | | 1 | 27 | tpst1l |
| eif3f2 | 9.5655575 | 0.485868599 | 1 | 0.644 | | 1 | 27 | eif3f |
| rpl27 | 9.5892659 | 0.258003308 | 1 | 0.897 | | 1 | 27 | rpl27 |
| mtmr11 | 0.00011 | 0.250039494 | 0.125 | 0.018 | | 1 | 27 | mtmr11 |
| grasp3 | 0.000112 | 0.327247777 | 0.25 | 0.058 | | 1 | 27 | grasp |
| spi1b7 | 0.000113 | 0.391925417 | 0.417 | 0.128 | | 1 | 27 | spi1b |
| ccdc6b1 | 0.000114 | 0.294214033 | 0.167 | 0.03 | | 1 | 27 | ccdc6b |
| cdc27 | 0.000114 | 0.323736265 | 0.167 | 0.03 | | 1 | 27 | cdc27 |
| slc25a55a2 | 0.000117 | 0.44430526 | 0.375 | 0.12 | | 1 | 27 | slc25a55a |
| gpia4 | 0.000122 | 0.394174001 | 0.417 | 0.139 | | 1 | 27 | gpia |
| mbd4 | 0.000136 | 0.268553325 | 0.208 | 0.043 | | 1 | 27 | mbd4 |
| CABZ01088428.1 | 0.00014 | 0.347258715 | 0.167 | 0.031 | | 1 | 27 | CABZ01088428.1 |
| zgc:638314 | 0.000143 | 0.362731473 | 0.375 | 0.113 | | 1 | 27 | zgc:63831 |
| sinhcafl1 | 0.000145 | 0.339857947 | 0.333 | 0.098 | | 1 | 27 | sinhcafl |
| clic17 | 0.000152 | 0.482513615 | 0.542 | 0.216 | | 1 | 27 | clic1 |
| rap1aa | 0.00016 | 0.388564178 | 0.458 | 0.163 | | 1 | 27 | rap1aa |
| rpl10a | 0.000161 | 0.26732409 | 1 | 0.899 | | 1 | 27 | rpl10a |
| plekho22 | 0.000165 | 0.354198024 | 0.333 | 0.095 | | 1 | 27 | plekho2 |

| | | | | | | | |
|---|---|---|---|---|---|---|---|
| srsf1a | 0.000168 | 0.285700696 | 0.292 | 0.077 | 1 | 27 | srsf1a |
| rps4x | 0.000171 | 0.258456274 | 1 | 0.899 | 1 | 27 | rps4x |
| sbno1 | 0.000177 | 0.263120308 | 0.167 | 0.031 | 1 | 27 | sbno1 |
| setbp1 | 0.000177 | 0.252453905 | 0.167 | 0.031 | 1 | 27 | setbp1 |
| zgc:1123561 | 0.000183 | 0.41343648 | 0.292 | 0.082 | 1 | 27 | zgc:112356 |
| stk351 | 0.000187 | 0.320453154 | 0.333 | 0.096 | 1 | 27 | stk35 |
| pkn1b1 | 0.000191 | 0.281220718 | 0.167 | 0.031 | 1 | 27 | pkn1b |
| gdi21 | 0.000197 | 0.540863231 | 0.625 | 0.304 | 1 | 27 | gdi2 |
| ptprc7 | 0.000208 | 0.433635377 | 0.375 | 0.12 | 1 | 27 | ptprc |
| mob1a5 | 0.000222 | 0.456629197 | 0.542 | 0.219 | 1 | 27 | mob1a |
| slc15a41 | 0.000223 | 0.281648315 | 0.167 | 0.032 | 1 | 27 | slc15a4 |
| lcp2a1 | 0.000223 | 0.332582456 | 0.208 | 0.047 | 1 | 27 | lcp2a |
| rcc22 | 0.000232 | 0.41694869 | 0.458 | 0.169 | 1 | 27 | rcc2 |
| CABZ01109481.11 | 0.000241 | 0.291263457 | 0.167 | 0.032 | 1 | 27 | CABZ01109481.1 |
| cdkn1bb2 | 0.000249 | 0.425197809 | 0.458 | 0.167 | 1 | 27 | cdkn1bb |
| si:dkey-42i9.46 | 0.00025 | 0.461911059 | 0.5 | 0.199 | 1 | 27 | si:dkey-42i9.4 |
| naca | 0.000256 | 0.336296594 | 1 | 0.832 | 1 | 27 | naca |
| si:dkey-27i16.23 | 0.000257 | 0.390761018 | 0.333 | 0.1 | 1 | 27 | si:dkey-27i16.2 |
| hmgn25 | 0.000291 | 0.628116006 | 0.875 | 0.633 | 1 | 27 | hmgn2 |
| rad21a5 | 0.000294 | 0.523095198 | 0.542 | 0.251 | 1 | 27 | rad21a |
| uba21 | 0.0003 | 0.318573224 | 0.292 | 0.082 | 1 | 27 | uba2 |
| lpar5a4 | 0.000304 | 0.312805324 | 0.25 | 0.063 | 1 | 27 | lpar5a |
| thoc5 | 0.000324 | 0.311710421 | 0.25 | 0.065 | 1 | 27 | thoc5 |
| rpa22 | 0.000328 | 0.323229972 | 0.333 | 0.1 | 1 | 27 | rpa2 |
| araf | 0.000332 | 0.252886172 | 0.167 | 0.033 | 1 | 27 | araf |
| srrt | 0.00035 | 0.254857604 | 0.208 | 0.048 | 1 | 27 | srrt |
| cnot91 | 0.000356 | 0.314383028 | 0.292 | 0.085 | 1 | 27 | cnot9 |
| ap1s12 | 0.000358 | 0.385663257 | 0.333 | 0.108 | 1 | 27 | ap1s1 |
| rnaset21 | 0.000362 | 0.298326703 | 0.208 | 0.048 | 1 | 27 | rnaset2 |
| gpr183a4 | 0.000364 | 0.364551154 | 0.25 | 0.066 | 1 | 27 | gpr183a |
| jpt21 | 0.000367 | 0.394263243 | 0.583 | 0.241 | 1 | 27 | jpt2 |
| iqgap11 | 0.000371 | 0.362568255 | 0.292 | 0.086 | 1 | 27 | iqgap1 |
| zgpat | 0.0004 | 0.257735567 | 0.167 | 0.034 | 1 | 27 | zgpat |
| gnai2a2 | 0.000408 | 0.473413966 | 0.458 | 0.186 | 1 | 27 | gnai2a |
| fbxl3a | 0.000419 | 0.282064813 | 0.208 | 0.049 | 1 | 27 | fbxl3a |

| | | | | | | | |
|---|---|---|---|---|---|---|---|
| tefa4 | 0.00043 | 0.377791538 | 0.417 | 0.15 | 1 | 27 | tefa |
| ier55 | 0.000467 | 0.448689041 | 0.5 | 0.211 | 1 | 27 | ier5 |
| ppp3r1b | 0.000552 | 0.290858027 | 0.375 | 0.124 | 1 | 27 | ppp3r1b |
| dek1 | 0.000556 | 0.255916947 | 0.208 | 0.05 | 1 | 27 | dek |
| psmd9 | 0.000558 | 0.267120396 | 0.208 | 0.051 | 1 | 27 | psmd9 |
| rangap1a | 0.000571 | 0.305983892 | 0.25 | 0.069 | 1 | 27 | rangap1a |
| ddit35 | 0.00063 | 0.567162656 | 0.458 | 0.203 | 1 | 27 | ddit3 |
| rps172 | 0.000631 | 0.262792959 | 1 | 0.88 | 1 | 27 | rps17 |
| rbb4l2 | 0.000647 | 0.336943424 | 0.333 | 0.111 | 1 | 27 | rbb4l |
| BX120005.14 | 0.000671 | 0.269781754 | 0.167 | 0.035 | 1 | 27 | BX120005.1 |
| wipf1a5 | 0.000676 | 0.321590623 | 0.292 | 0.085 | 1 | 27 | wipf1a |
| fam49a1 | 0.000684 | 0.258077262 | 0.25 | 0.069 | 1 | 27 | fam49a |
| il10ra3 | 0.000713 | 0.25810727 | 0.208 | 0.051 | 1 | 27 | il10ra |
| gltpa | 0.00072 | 0.252948327 | 0.125 | 0.022 | 1 | 27 | gltpa |
| gadd45ga1 | 0.000801 | 0.278282858 | 0.208 | 0.053 | 1 | 27 | gadd45ga |
| ptbp31 | 0.000832 | 0.291957364 | 0.292 | 0.089 | 1 | 27 | ptbp3 |
| sept65 | 0.000833 | 0.44037237 | 0.458 | 0.192 | 1 | 27 | sept6 |
| mef2cb2 | 0.000845 | 0.257089295 | 0.208 | 0.052 | 1 | 27 | mef2cb |
| zgc:85858 | 0.000877 | 0.268945163 | 0.208 | 0.053 | 1 | 27 | zgc:85858 |
| mzt2b | 0.0009 | 0.277009864 | 0.208 | 0.054 | 1 | 27 | mzt2b |
| kmt2bb5 | 0.000922 | 0.339742014 | 0.417 | 0.152 | 1 | 27 | kmt2bb |
| nop535 | 0.000978 | 0.466993646 | 0.667 | 0.355 | 1 | 27 | nop53 |
| hmga1a6 | 0.001024 | 0.590950852 | 0.667 | 0.437 | 1 | 27 | hmga1a |
| stk17al3 | 0.001044 | 0.346524858 | 0.208 | 0.055 | 1 | 27 | stk17al |
| LOC1039104385 | 0.001095 | 0.400795939 | 0.458 | 0.185 | 1 | 27 | LOC103910438 |
| cox7a2a2 | 0.001103 | 0.398910366 | 0.625 | 0.305 | 1 | 27 | cox7a2a |
| eif3ea2 | 0.001104 | 0.448840254 | 0.75 | 0.451 | 1 | 27 | eif3ea |
| ahcy4 | 0.001113 | 0.31647892 | 0.333 | 0.113 | 1 | 27 | ahcy |
| rfc31 | 0.001126 | 0.260343504 | 0.208 | 0.054 | 1 | 27 | rfc3 |
| chmp2a2 | 0.001153 | 0.399087467 | 0.458 | 0.191 | 1 | 27 | chmp2a |
| vsir5 | 0.001183 | 0.324247997 | 0.333 | 0.111 | 1 | 27 | vsir |
| ier2a6 | 0.001189 | 0.484474419 | 0.708 | 0.364 | 1 | 27 | ier2a |
| tyms2 | 0.001294 | 0.278110534 | 0.208 | 0.054 | 1 | 27 | tyms |
| tmem106a | 0.001304 | 0.273551139 | 0.208 | 0.055 | 1 | 27 | tmem106a |
| bri3bp1 | 0.001322 | 0.268915475 | 0.208 | 0.055 | 1 | 27 | bri3bp |

| | | | | | | | |
|---|---|---|---|---|---|---|---|
| dut3 | 0.001337 | 0.501188797 | 0.292 | 0.1 | 1 | 27 | dut |
| rhof2 | 0.001444 | 0.267560953 | 0.167 | 0.039 | 1 | 27 | rhof |
| eif3ha1 | 0.001457 | 0.425727931 | 0.667 | 0.355 | 1 | 27 | eif3ha |
| yrk6 | 0.001505 | 0.28913525 | 0.292 | 0.092 | 1 | 27 | yrk |
| cenpx2 | 0.001547 | 0.330990529 | 0.25 | 0.076 | 1 | 27 | cenpx |
| ctss2.16 | 0.001553 | 0.331086696 | 0.25 | 0.075 | 1 | 27 | ctss2.1 |
| zfand5b1 | 0.001627 | 0.326435594 | 0.333 | 0.12 | 1 | 27 | zfand5b |
| ptmab2 | 0.001653 | 0.347878258 | 0.958 | 0.763 | 1 | 27 | ptmab |
| lsm52 | 0.001657 | 0.40710675 | 0.5 | 0.237 | 1 | 27 | lsm5 |
| sdhaf4 | 0.001669 | 0.251822823 | 0.208 | 0.057 | 1 | 27 | sdhaf4 |
| sat1a.28 | 0.001673 | 0.416870657 | 0.667 | 0.336 | 1 | 27 | sat1a.2 |
| slka1 | 0.001706 | 0.315058239 | 0.208 | 0.058 | 1 | 27 | slka |
| si:dkey-172h23.22 | 0.001718 | 0.272404578 | 0.208 | 0.057 | 1 | 27 | si:dkey-172h23.2 |
| nucks1b3 | 0.001781 | 0.303341946 | 0.417 | 0.162 | 1 | 27 | nucks1b |
| pde7a4 | 0.001823 | 0.279407826 | 0.292 | 0.096 | 1 | 27 | pde7a |
| snrpd23 | 0.001826 | 0.432640694 | 0.542 | 0.267 | 1 | 27 | snrpd2 |
| sh3bgrl31 | 0.001828 | 0.329026266 | 0.5 | 0.213 | 1 | 27 | sh3bgrl3 |
| eif3m2 | 0.001831 | 0.4284102 | 0.875 | 0.523 | 1 | 27 | eif3m |
| taf10 | 0.001834 | 0.307696194 | 0.25 | 0.078 | 1 | 27 | taf10 |
| tnip13 | 0.001844 | 0.280582663 | 0.25 | 0.075 | 1 | 27 | tnip1 |
| hmha1b4 | 0.001948 | 0.277290191 | 0.25 | 0.076 | 1 | 27 | hmha1b |
| ier2b6 | 0.001977 | 0.464146863 | 0.625 | 0.327 | 1 | 27 | ier2b |
| dnaja2a2 | 0.001983 | 0.382750105 | 0.583 | 0.276 | 1 | 27 | dnaja2a |
| marcksl1a6 | 0.00199 | 0.448645114 | 0.917 | 0.513 | 1 | 27 | marcksl1a |
| cirbpb4 | 0.00203 | 0.296443989 | 0.958 | 0.813 | 1 | 27 | cirbpb |
| stmn1a4 | 0.00213 | 0.620948303 | 0.292 | 0.111 | 1 | 27 | stmn1a |
| gng21 | 0.002181 | 0.369410079 | 0.25 | 0.082 | 1 | 27 | gng2 |
| magt12 | 0.002386 | 0.38905213 | 0.458 | 0.209 | 1 | 27 | magt1 |
| ddx615 | 0.002388 | 0.435863065 | 0.625 | 0.344 | 1 | 27 | ddx61 |
| foxp1b1 | 0.002425 | 0.259457347 | 0.208 | 0.059 | 1 | 27 | foxp1b |
| zgc:1627307 | 0.002433 | 0.491145935 | 0.75 | 0.493 | 1 | 27 | zgc:162730 |
| ppp1caa1 | 0.00258 | 0.409035397 | 0.458 | 0.215 | 1 | 27 | ppp1caa |
| pdcd51 | 0.002602 | 0.34306549 | 0.375 | 0.155 | 1 | 27 | pdcd5 |
| si:ch211-22i13.21 | 0.002609 | 0.326909468 | 0.292 | 0.103 | 1 | 27 | si:ch211-22i13.2 |
| parp21 | 0.002741 | 0.275218524 | 0.167 | 0.042 | 1 | 27 | parp2 |

| | | | | | | | | |
|---|---|---|---|---|---|---|---|---|
| grb2b1 | 0.002815 | 0.287143546 | 0.292 | 0.102 | | 1 | 27 | grb2b |
| slc44a21 | 0.002856 | 0.2741965 | 0.25 | 0.08 | | 1 | 27 | slc44a2 |
| trappc31 | 0.002942 | 0.295452415 | 0.208 | 0.061 | | 1 | 27 | trappc3 |
| eif3k1 | 0.002964 | 0.372149349 | 0.625 | 0.33 | | 1 | 27 | eif3k |
| smad3a | 0.003052 | 0.267008916 | 0.167 | 0.043 | | 1 | 27 | smad3a |
| nt5dc21 | 0.003226 | 0.408927202 | 0.25 | 0.089 | | 1 | 27 | nt5dc2 |
| arpc5a4 | 0.003251 | 0.292304463 | 0.458 | 0.19 | | 1 | 27 | arpc5a |
| ddx39ab3 | 0.003294 | 0.393773375 | 0.417 | 0.191 | | 1 | 27 | ddx39ab |
| brdt2 | 0.003341 | 0.310656608 | 0.25 | 0.083 | | 1 | 27 | brdt |
| u2af12 | 0.003495 | 0.334716168 | 0.5 | 0.232 | | 1 | 27 | u2af1 |
| ripor1.1 | 0.003518 | 0.253079781 | 0.125 | 0.027 | | 1 | 27 | ripor1.1 |
| cul4b | 0.003726 | 0.273241202 | 0.208 | 0.063 | | 1 | 27 | cul4b |
| med19a1 | 0.003983 | 0.288960237 | 0.292 | 0.107 | | 1 | 27 | med19a |
| eed | 0.004369 | 0.270159274 | 0.208 | 0.065 | | 1 | 27 | eed |
| capzb6 | 0.004435 | 0.373097457 | 0.708 | 0.387 | | 1 | 27 | capzb |
| cct73 | 0.004596 | 0.414336715 | 0.583 | 0.341 | | 1 | 27 | cct7 |
| tpt1 | 0.004624 | 0.255860141 | 1 | 0.879 | | 1 | 27 | tpt1 |
| si:dkey-185m8.25 | 0.004635 | 0.274425549 | 0.167 | 0.045 | | 1 | 27 | si:dkey-185m8.2 |
| sumo3b2 | 0.004654 | 0.329376082 | 0.208 | 0.066 | | 1 | 27 | sumo3b |
| ndufa82 | 0.004898 | 0.322795165 | 0.458 | 0.218 | | 1 | 27 | ndufa8 |
| gemin81 | 0.004929 | 0.296458383 | 0.25 | 0.088 | | 1 | 27 | gemin8 |
| eif2s1b1 | 0.00499 | 0.336271721 | 0.583 | 0.3 | | 1 | 27 | eif2s1b |
| acin1a7 | 0.005213 | 0.403416455 | 0.542 | 0.299 | | 1 | 27 | acin1a |
| tp532 | 0.005226 | 0.316519771 | 0.417 | 0.182 | | 1 | 27 | tp53 |
| snx5 | 0.005555 | 0.302882593 | 0.292 | 0.109 | | 1 | 27 | snx5 |
| ada | 0.005592 | 0.255555279 | 0.208 | 0.066 | | 1 | 27 | ada |
| aldoaa3 | 0.005607 | 0.336898503 | 0.583 | 0.289 | | 1 | 27 | aldoaa |
| ufm1 | 0.005771 | 0.298181644 | 0.292 | 0.113 | | 1 | 27 | ufm1 |
| galnt71 | 0.005981 | 0.27975896 | 0.208 | 0.067 | | 1 | 27 | galnt7 |
| taf91 | 0.006075 | 0.313251164 | 0.333 | 0.139 | | 1 | 27 | taf9 |
| lamtor4 | 0.006577 | 0.352658729 | 0.292 | 0.12 | | 1 | 27 | lamtor4 |
| atp5po2 | 0.006629 | 0.348826354 | 0.583 | 0.319 | | 1 | 27 | atp5po |
| eif3s6ip | 0.006918 | 0.300119209 | 0.5 | 0.243 | | 1 | 27 | eif3s6ip |
| pfdn41 | 0.007189 | 0.343095934 | 0.542 | 0.286 | | 1 | 27 | pfdn4 |
| zgc:778492 | 0.007195 | 0.295432435 | 0.417 | 0.185 | | 1 | 27 | zgc:77849 |

| | | | | | | | |
|---|---|---|---|---|---|---|---|
| psmb63 | 0.007406 | 0.347298487 | 0.542 | 0.295 | 1 | 27 | psmb6 |
| nfyba3 | 0.007529 | 0.360218869 | 0.458 | 0.23 | 1 | 27 | nfyba |
| sec11a5 | 0.007703 | 0.331865281 | 0.542 | 0.273 | 1 | 27 | sec11a |
| cbx1b | 0.007731 | 0.27769208 | 0.25 | 0.091 | 1 | 27 | cbx1b |
| ywhaba | 0.007933 | 0.366992256 | 0.667 | 0.392 | 1 | 27 | ywhaba |
| ppifb4 | 0.007986 | 0.285123787 | 0.375 | 0.168 | 1 | 27 | ppifb |
| cap19 | 0.00815 | 0.416868368 | 0.583 | 0.332 | 1 | 27 | cap1 |
| rpl5b1 | 0.008253 | 0.286652404 | 0.958 | 0.761 | 1 | 27 | rpl5b |
| psmg31 | 0.008576 | 0.272806363 | 0.333 | 0.143 | 1 | 27 | psmg3 |
| hnrnpa1b4 | 0.008724 | 0.382391524 | 0.458 | 0.248 | 1 | 27 | hnrnpa1b |
| puf60b | 0.008725 | 0.265125015 | 0.292 | 0.116 | 1 | 27 | puf60b |
| ncoa51 | 0.009312 | 0.271647324 | 0.208 | 0.071 | 1 | 27 | ncoa5 |
| wasb7 | 0.009394 | 0.312498628 | 0.375 | 0.16 | 1 | 27 | wasb |
| anxa11b2 | 0.009757 | 0.311064576 | 0.583 | 0.312 | 1 | 27 | anxa11b |
| nfkbie2 | 0.009834 | 0.283042875 | 0.292 | 0.12 | 1 | 27 | nfkbie |
