## Supplementary material for "Direct Male Development in Chromosomally ZZ Zebrafish": SuppTable S3 19dpf ZZ vs ZW Differential Expression

| gene | p_val | avg_log2FC | pct.1 (ZZ) | pct.2 (ZW) | p_val_adj | cluster |
|---|---|---|---|---|---|---|
| gpx1a | 1.881000 | -0.34879294 | 0 | 0.133 | 0.3799057 | 0 |
| gpx4a | 4.900109 | -0.25006359 | 0 | 0.12 | 0.9896752 | 0 |
| ptx3a | 0.00012 | -0.61609224 | 0.282 | 0.578 | 1 | 0 |
| ms4a17a.8 | 0.00013 | -0.2418373 | 0 | 0.108 | 1 | 0 |
| ssbp3b | 0.00015 | -0.33247283 | 0.009 | 0.157 | 1 | 0 |
| abhd15a | 0.00028 | -0.48589062 | 0.145 | 0.398 | 1 | 0 |
| igfbp5b | 0.00048 | -0.35674652 | 0.673 | 0.795 | 1 | 0 |
| ddr1 | 0.00052 | -0.35223849 | 0.018 | 0.169 | 1 | 0 |
| fth1a | 0.00079 | -0.48362649 | 0.218 | 0.434 | 1 | 0 |
| ube2j2 | 0.00084 | -0.1557213 | 0 | 0.084 | 1 | 0 |
| stub1 | 0.00084 | -0.17441134 | 0 | 0.084 | 1 | 0 |
| wnt5b | 0.00084 | -0.20095202 | 0 | 0.084 | 1 | 0 |
| id2b | 0.00094 | -0.48111273 | 0.282 | 0.458 | 1 | 0 |
| pdzrn3b | 0.00107 | 0.205079491 | 0.091 | 0 | 1 | 0 |
| uap1 | 0.00155 | -0.41020891 | 0.045 | 0.205 | 1 | 0 |
| ranbp1 | 0.00178 | 0.394256284 | 0.291 | 0.108 | 1 | 0 |
| c4b | 0.00185 | 0.373883584 | 0.136 | 0.012 | 1 | 0 |
| cops4 | 0.00203 | 0.177391102 | 0.082 | 0 | 1 | 0 |
| hat1 | 0.00203 | 0.164207085 | 0.082 | 0 | 1 | 0 |
| pagr1 | 0.00203 | 0.191255021 | 0.082 | 0 | 1 | 0 |
| vbp1 | 0.00203 | 0.162602869 | 0.082 | 0 | 1 | 0 |
| cfd | 0.00203 | 0.142846745 | 0.082 | 0 | 1 | 0 |
| fbxo25 | 0.00217 | -0.1836497 | 0 | 0.072 | 1 | 0 |
| lta4h | 0.00217 | -0.15492709 | 0 | 0.072 | 1 | 0 |
| clptm1l | 0.00217 | -0.15973533 | 0 | 0.072 | 1 | 0 |
| wu:fi36c02 | 0.00217 | -0.1659772 | 0 | 0.072 | 1 | 0 |
| elfn1a | 0.00217 | -0.20332041 | 0 | 0.072 | 1 | 0 |
| elnb | 0.00217 | -0.2222945 | 0 | 0.072 | 1 | 0 |
| h1fx | 0.00232 | -0.31982151 | 0.018 | 0.096 | 1 | 0 |
| si:ch211-198c19.3 | 0.00235 | 0.509327529 | 0.491 | 0.289 | 1 | 0 |
| snrpd1 | 0.00242 | 0.318196845 | 0.182 | 0.084 | 1 | 0 |
| igf2b | 0.00277 | 0.431023234 | 0.582 | 0.373 | 1 | 0 |
| hmgb1b | 0.00295 | 0.414742566 | 0.345 | 0.169 | 1 | 0 |
| rgs5b | 0.00302 | 0.129616758 | 0.082 | 0.012 | 1 | 0 |
| mt-atp6 | 0.00308 | -0.24508149 | 0.555 | 0.771 | 1 | 0 |
| col6a4a | 0.00316 | 0.455090419 | 0.418 | 0.205 | 1 | 0 |
| plpp1a | 0.00326 | -0.41557513 | 0.382 | 0.627 | 1 | 0 |
| sub1b | 0.00329 | 0.322292766 | 0.164 | 0.036 | 1 | 0 |
| im:6904045 | 0.00385 | 0.188281774 | 0.073 | 0 | 1 | 0 |
| si:dkey-110g7.8 | 0.00385 | 0.197530102 | 0.073 | 0 | 1 | 0 |
| ppih | 0.00385 | 0.125236236 | 0.073 | 0 | 1 | 0 |
| zmynd11 | 0.00385 | 0.197574977 | 0.073 | 0 | 1 | 0 |
| nlrc3l | 0.00385 | 0.204999615 | 0.073 | 0 | 1 | 0 |
| kdm5c | 0.00385 | 0.204668683 | 0.073 | 0 | 1 | 0 |
| chaf1a | 0.00385 | 0.175118388 | 0.073 | 0 | 1 | 0 |
| ZNF512B | 0.00385 | 0.210986096 | 0.073 | 0 | 1 | 0 |
| pfdn4 | 0.00463 | 0.33514552 | 0.255 | 0.096 | 1 | 0 |

| | | | | | | |
|---|---|---|---|---|---|---|
| hnrnpabb | 0.00473 | 0.370328594 | 0.436 | 0.277 | 1 | 0 |
| CABZ01039096.1 | 0.00488 | -0.36558914 | 0.082 | 0.253 | 1 | 0 |
| XLOC-032112 | 0.00559 | -0.12051001 | 0 | 0.06 | 1 | 0 |
| mgrn1b | 0.00559 | -0.17878024 | 0 | 0.06 | 1 | 0 |
| mettl6 | 0.00559 | -0.10636909 | 0 | 0.06 | 1 | 0 |
| nudt8 | 0.00559 | -0.13568831 | 0 | 0.06 | 1 | 0 |
| cd2ap | 0.00559 | -0.15595966 | 0 | 0.06 | 1 | 0 |
| si:dkey-52l18.4 | 0.00559 | -0.14826108 | 0 | 0.06 | 1 | 0 |
| CU633984.1 | 0.00559 | -0.16456936 | 0 | 0.06 | 1 | 0 |
| c3a.1 | 0.00559 | -0.17500629 | 0 | 0.06 | 1 | 0 |
| cds2 | 0.00559 | -0.15191093 | 0 | 0.06 | 1 | 0 |
| slco2a1 | 0.00559 | -0.1542939 | 0 | 0.06 | 1 | 0 |
| txn | 0.00559 | -0.18215329 | 0 | 0.06 | 1 | 0 |
| zmp:0000001082 | 0.00559 | -0.15639508 | 0 | 0.06 | 1 | 0 |
| rsl1d1 | 0.00566 | 0.35766164 | 0.282 | 0.133 | 1 | 0 |
| col1a1a | 0.00634 | 0.277272096 | 0.864 | 0.723 | 1 | 0 |
| EML4 | 0.00671 | 0.230853539 | 0.118 | 0.012 | 1 | 0 |
| mt-co2 | 0.00684 | -0.28687845 | 0.6 | 0.807 | 1 | 0 |
| XLOC-013964 | 0.00721 | 0.183422249 | 0.109 | 0.012 | 1 | 0 |
| pmpcb | 0.00723 | -0.21734674 | 0.009 | 0.096 | 1 | 0 |
| cplx2 | 0.00728 | 0.18904101 | 0.064 | 0 | 1 | 0 |
| si:ch211-106j24.1 | 0.00728 | 0.166686932 | 0.064 | 0 | 1 | 0 |
| dera | 0.00728 | 0.164030534 | 0.064 | 0 | 1 | 0 |
| asf1bb | 0.00728 | 0.158557389 | 0.064 | 0 | 1 | 0 |
| nagk | 0.00728 | 0.171822922 | 0.064 | 0 | 1 | 0 |
| utp15 | 0.00728 | 0.123463524 | 0.064 | 0 | 1 | 0 |
| c6 | 0.00734 | 0.299791671 | 0.5 | 0.41 | 1 | 0 |
| coq6 | 0.0074 | -0.15675329 | 0.009 | 0.096 | 1 | 0 |
| fap | 0.00809 | 0.269659838 | 0.118 | 0.012 | 1 | 0 |
| ednraa | 0.0088 | 0.362852055 | 0.191 | 0.048 | 1 | 0 |
| mdka | 0.00883 | 0.374173848 | 0.636 | 0.458 | 1 | 0 |
| thoc2 | 0.00905 | 0.322302888 | 0.182 | 0.048 | 1 | 0 |
| jund | 0.00911 | -0.38245662 | 0.091 | 0.241 | 1 | 0 |
| ugt5g1 | 0.00924 | -0.34990069 | 0.082 | 0.241 | 1 | 0 |
| ndufb9 | 0.0096 | 0.343863643 | 0.364 | 0.169 | 1 | 0 |
| tmem33 | 0.01024 | 0.234328481 | 0.127 | 0.024 | 1 | 0 |
| tubb2b | 0.01096 | 0.329444989 | 0.182 | 0.06 | 1 | 0 |
| mfge8b | 0.01098 | -0.39159989 | 0.109 | 0.277 | 1 | 0 |
| aoc2 | 0.01108 | 0.425318803 | 0.327 | 0.145 | 1 | 0 |
| znf219 | 0.0111 | -0.21317106 | 0.009 | 0.096 | 1 | 0 |
| cyp1c2 | 0.01158 | 0.414226249 | 0.255 | 0.096 | 1 | 0 |
| si:ch211-194b1.1 | 0.01186 | -0.2186982 | 0.018 | 0.12 | 1 | 0 |
| col1a2 | 0.0127 | 0.202465083 | 0.936 | 0.819 | 1 | 0 |
| ncbp2 | 0.01304 | -0.16708005 | 0.009 | 0.096 | 1 | 0 |
| arf6b | 0.01319 | -0.33578064 | 0.073 | 0.205 | 1 | 0 |
| cdc5l | 0.01329 | -0.19821362 | 0.009 | 0.096 | 1 | 0 |
| fam20b | 0.01348 | -0.20251886 | 0.009 | 0.096 | 1 | 0 |
| sec61a1 | 0.01355 | -0.19330077 | 0.245 | 0.253 | 1 | 0 |

| | | | | | | |
|---|---|---|---|---|---|---|
| emilin2a | 0.01361 | 0.414576242 | 0.282 | 0.12 | 1 | 0 |
| sra1 | 0.01382 | 0.117171146 | 0.055 | 0 | 1 | 0 |
| rxfp2a | 0.01382 | 0.157851134 | 0.055 | 0 | 1 | 0 |
| chp2 | 0.01382 | 0.115853052 | 0.055 | 0 | 1 | 0 |
| tfcp2 | 0.01382 | 0.132960785 | 0.055 | 0 | 1 | 0 |
| uchl3 | 0.01382 | 0.10147699 | 0.055 | 0 | 1 | 0 |
| dcaf13 | 0.01382 | 0.10836241 | 0.055 | 0 | 1 | 0 |
| si:dkeyp-115e12.7 | 0.01382 | 0.160180579 | 0.055 | 0 | 1 | 0 |
| rnd1b | 0.01382 | 0.14641805 | 0.055 | 0 | 1 | 0 |
| akt2l | 0.01382 | 0.117682468 | 0.055 | 0 | 1 | 0 |
| ptgdsb.2 | 0.01382 | 0.121607129 | 0.055 | 0 | 1 | 0 |
| taf6 | 0.01382 | 0.131639202 | 0.055 | 0 | 1 | 0 |
| virma | 0.01382 | 0.1642403 | 0.055 | 0 | 1 | 0 |
| mfap2 | 0.01395 | 0.279265882 | 0.755 | 0.554 | 1 | 0 |
| gstm.1 | 0.01397 | 0.329695677 | 0.291 | 0.12 | 1 | 0 |
| bzw1b | 0.01399 | 0.153525858 | 0.227 | 0.108 | 1 | 0 |
| npm1a | 0.01405 | 0.274950723 | 0.291 | 0.12 | 1 | 0 |
| aldh1a2 | 0.01439 | 0.296512738 | 0.155 | 0.036 | 1 | 0 |
| hmgb1a | 0.01459 | 0.302888664 | 0.464 | 0.361 | 1 | 0 |
| spcs1 | 0.0146 | 0.345932021 | 0.382 | 0.193 | 1 | 0 |
| cd63 | 0.01471 | 0.100979353 | 0.682 | 0.542 | 1 | 0 |
| adh8a | 0.01504 | 0.32966378 | 0.2 | 0.06 | 1 | 0 |
| has2 | 0.01545 | -0.24519378 | 0.036 | 0.145 | 1 | 0 |
| cebpb | 0.01559 | -0.33839221 | 0.064 | 0.193 | 1 | 0 |
| adh5 | 0.016 | 0.297414995 | 0.364 | 0.181 | 1 | 0 |
| emd | 0.01623 | 0.280451186 | 0.155 | 0.036 | 1 | 0 |
| map4k4 | 0.01687 | -0.2925959 | 0.045 | 0.169 | 1 | 0 |
| ifngr1l | 0.01715 | -0.29858178 | 0.045 | 0.157 | 1 | 0 |
| tcima | 0.01738 | -0.41106117 | 0.055 | 0.181 | 1 | 0 |
| dhrs3a | 0.01768 | -0.32272485 | 0.191 | 0.301 | 1 | 0 |
| wnt7bb | 0.01841 | 0.345557949 | 0.209 | 0.108 | 1 | 0 |
| calub | 0.01842 | -0.24278591 | 0.018 | 0.108 | 1 | 0 |
| hmgn7 | 0.01852 | 0.311875382 | 0.527 | 0.325 | 1 | 0 |
| coa6 | 0.01904 | 0.173494634 | 0.1 | 0.012 | 1 | 0 |
| ms4a17a.9 | 0.01921 | 0.259412415 | 0.191 | 0.06 | 1 | 0 |
| selenoh | 0.01964 | 0.228893731 | 0.136 | 0.048 | 1 | 0 |
| pcna | 0.01992 | 0.276420362 | 0.127 | 0.024 | 1 | 0 |
| canx | 0.02017 | -0.3054901 | 0.064 | 0.193 | 1 | 0 |
| june | 0.02044 | -0.29754414 | 0.055 | 0.181 | 1 | 0 |
| si:zfos-169g10.3 | 0.02071 | 0.243749871 | 0.209 | 0.072 | 1 | 0 |
| htra1a | 0.02083 | -0.33049402 | 0.236 | 0.349 | 1 | 0 |
| spry2 | 0.02141 | 0.248528488 | 0.236 | 0.096 | 1 | 0 |
| camk2g1 | 0.02188 | 0.239237458 | 0.127 | 0.024 | 1 | 0 |
| ewsr1a | 0.02214 | 0.246803688 | 0.127 | 0.024 | 1 | 0 |
| fkbp1ab | 0.0222 | -0.37509537 | 0.1 | 0.241 | 1 | 0 |
| ruvbl1 | 0.02234 | 0.197899325 | 0.127 | 0.024 | 1 | 0 |
| inpp5l | 0.02278 | -0.21828236 | 0.018 | 0.108 | 1 | 0 |
| atrnl1b | 0.02296 | -0.26040352 | 0.018 | 0.108 | 1 | 0 |

| | | | | | | |
|---|---|---|---|---|---|---|
| ndufv3 | 0.02303 | -0.23553097 | 0.027 | 0.12 | 1 | 0 |
| reep3b | 0.0233 | 0.243742078 | 0.118 | 0.024 | 1 | 0 |
| fabp3 | 0.02348 | -0.24586918 | 0.064 | 0.193 | 1 | 0 |
| ppp6r3 | 0.02351 | 0.230527531 | 0.1 | 0.012 | 1 | 0 |
| gnai1 | 0.02352 | -0.19745055 | 0.009 | 0.084 | 1 | 0 |
| sik1 | 0.02362 | -0.27276392 | 0.036 | 0.145 | 1 | 0 |
| lgals2a | 0.02381 | 0.402554355 | 0.364 | 0.193 | 1 | 0 |
| mt-cyb | 0.02386 | -0.22342921 | 0.527 | 0.711 | 1 | 0 |
| mphosph6 | 0.02465 | 0.238519966 | 0.145 | 0.036 | 1 | 0 |
| sod1 | 0.02517 | 0.125681005 | 0.345 | 0.205 | 1 | 0 |
| arl6ip1 | 0.02543 | -0.2586759 | 0.127 | 0.205 | 1 | 0 |
| si:ch211-67e16.11 | 0.02549 | -0.30351502 | 0.018 | 0.096 | 1 | 0 |
| kdelr3 | 0.02581 | -0.15492721 | 0.009 | 0.084 | 1 | 0 |
| wu:fj16a03 | 0.02654 | -0.2124566 | 0.009 | 0.084 | 1 | 0 |
| thbs3a | 0.02725 | 0.264502829 | 0.355 | 0.289 | 1 | 0 |
| XLOC-016094 | 0.02768 | 0.318339982 | 0.491 | 0.301 | 1 | 0 |
| mmp14b | 0.02775 | -0.36985968 | 0.2 | 0.361 | 1 | 0 |
| asip2b | 0.02775 | 0.361365895 | 0.336 | 0.169 | 1 | 0 |
| eva1ba | 0.02794 | 0.323899072 | 0.291 | 0.133 | 1 | 0 |
| adcyap1b | 0.0283 | -0.24481742 | 0.009 | 0.084 | 1 | 0 |
| RPL37A | 0.02875 | -0.1908448 | 0.009 | 0.084 | 1 | 0 |
| ca8 | 0.02892 | -0.17013945 | 0.009 | 0.084 | 1 | 0 |
| nppc | 0.02896 | -0.2407048 | 0.009 | 0.084 | 1 | 0 |
| cnih1 | 0.02897 | -0.15407953 | 0.009 | 0.084 | 1 | 0 |
| ier2b | 0.02897 | -0.36869828 | 0.282 | 0.434 | 1 | 0 |
| krt91 | 0.02956 | 0.253708283 | 0.145 | 0.084 | 1 | 0 |
| spon2b | 0.03068 | -0.39492771 | 0.182 | 0.349 | 1 | 0 |
| ndufa4l | 0.0314 | 0.230256674 | 0.491 | 0.422 | 1 | 0 |
| cmpk | 0.0322 | -0.22295657 | 0.027 | 0.12 | 1 | 0 |
| hells | 0.03282 | 0.239197976 | 0.082 | 0.012 | 1 | 0 |
| coa3a | 0.03323 | -0.10707429 | 0.018 | 0.096 | 1 | 0 |
| ptbp1b | 0.0335 | 0.232269983 | 0.118 | 0.024 | 1 | 0 |
| blmh | 0.03436 | 0.159707565 | 0.091 | 0.012 | 1 | 0 |
| nsa2 | 0.03475 | 0.210239893 | 0.464 | 0.301 | 1 | 0 |
| atp5mea | 0.03491 | -0.21110707 | 0.055 | 0.169 | 1 | 0 |
| cdc42ep2 | 0.03557 | -0.19048335 | 0.045 | 0.145 | 1 | 0 |
| adcy6b | 0.03561 | -0.28264668 | 0.055 | 0.169 | 1 | 0 |
| vasna | 0.03604 | -0.30028642 | 0.391 | 0.53 | 1 | 0 |
| laptm4a | 0.03677 | -0.30529167 | 0.091 | 0.217 | 1 | 0 |
| sec61b | 0.03752 | 0.232227002 | 0.627 | 0.446 | 1 | 0 |
| nupr1a | 0.03768 | 0.273449918 | 0.245 | 0.108 | 1 | 0 |
| rpl27a | 0.03788 | -0.25538156 | 0.518 | 0.614 | 1 | 0 |
| psmc3 | 0.03832 | 0.168504662 | 0.091 | 0.012 | 1 | 0 |
| ppa1b | 0.03842 | 0.122999603 | 0.091 | 0.012 | 1 | 0 |
| cyp2ad2 | 0.03878 | 0.280543062 | 0.091 | 0.012 | 1 | 0 |
| zcrb1 | 0.04059 | 0.187307305 | 0.082 | 0.012 | 1 | 0 |
| insra | 0.04122 | -0.1796249 | 0.009 | 0.072 | 1 | 0 |
| dub | 0.04287 | -0.29665483 | 0.055 | 0.157 | 1 | 0 |

| | | | | | | |
|---|---|---|---|---|---|---|
| rps8a | 0.04326 | 0.135926775 | 0.855 | 0.855 | 1 | 0 |
| adh8b | 0.04352 | -0.24585044 | 0.1 | 0.181 | 1 | 0 |
| tnfaip6 | 0.04402 | -0.22294612 | 0.073 | 0.181 | 1 | 0 |
| gadd45ab | 0.04405 | -0.15285022 | 0.009 | 0.072 | 1 | 0 |
| cavin2a | 0.04407 | -0.16674879 | 0.027 | 0.108 | 1 | 0 |
| mboat7 | 0.04416 | 0.123868928 | 0.073 | 0.012 | 1 | 0 |
| copz1 | 0.04462 | 0.204795167 | 0.155 | 0.048 | 1 | 0 |
| twistnb | 0.04479 | 0.212630829 | 0.109 | 0.024 | 1 | 0 |
| mfge8a | 0.04485 | 0.243356283 | 0.109 | 0.024 | 1 | 0 |
| col4a1 | 0.04506 | 0.332772222 | 0.382 | 0.217 | 1 | 0 |
| dock9b | 0.04517 | -0.18966339 | 0.064 | 0.157 | 1 | 0 |
| eif1axa | 0.0454 | 0.140378766 | 0.1 | 0.024 | 1 | 0 |
| abi3bpb | 0.04545 | -0.26486839 | 0.045 | 0.133 | 1 | 0 |
| rgcc | 0.04586 | -0.33289171 | 0.136 | 0.265 | 1 | 0 |
| rspo1 | 0.0467 | -0.19631648 | 0.018 | 0.096 | 1 | 0 |
| si:dkey-10o6.2 | 0.04724 | -0.21138819 | 0.018 | 0.096 | 1 | 0 |
| si:dkey-9i23.16 | 0.04757 | 0.256328235 | 0.227 | 0.096 | 1 | 0 |
| hspd1 | 0.04767 | 0.237827941 | 0.145 | 0.048 | 1 | 0 |
| snx16 | 0.04775 | -0.17612391 | 0.018 | 0.096 | 1 | 0 |
| cilp | 0.04796 | -0.22573421 | 0.018 | 0.096 | 1 | 0 |
| wdr21 | 0.04865 | -0.11829057 | 0.009 | 0.072 | 1 | 0 |
| cyp2ad3 | 0.04938 | 0.292924669 | 0.391 | 0.229 | 1 | 0 |
| itga5 | 0.04945 | -0.28549383 | 0.091 | 0.217 | 1 | 0 |
| pho | 0.04948 | 0.312634867 | 0.2 | 0.084 | 1 | 0 |
| ppiab | 0.05027 | 0.205737892 | 0.791 | 0.699 | 1 | 0 |
| ykt6 | 0.05089 | -0.18204515 | 0.009 | 0.072 | 1 | 0 |
| rgl1 | 0.05104 | 0.223287084 | 0.109 | 0.024 | 1 | 0 |
| hnrnpa0l | 0.05131 | 0.171281358 | 0.627 | 0.47 | 1 | 0 |
| cish | 0.05223 | -0.28588997 | 0.1 | 0.205 | 1 | 0 |
| LOC101883507 | 0.05257 | -0.14935316 | 0.009 | 0.072 | 1 | 0 |
| shq1 | 0.0526 | -0.17086283 | 0.009 | 0.072 | 1 | 0 |
| c1qtnf5 | 0.05263 | 0.26952928 | 0.255 | 0.12 | 1 | 0 |
| kcnd3 | 0.05276 | -0.20504947 | 0.009 | 0.072 | 1 | 0 |
| LO018340.1 | 0.05311 | -0.24228116 | 0.045 | 0.145 | 1 | 0 |
| hmgb3a | 0.05328 | 0.296967954 | 0.291 | 0.157 | 1 | 0 |
| cotl1 | 0.05386 | -0.19765088 | 0.045 | 0.145 | 1 | 0 |
| srprb | 0.05433 | 0.257863264 | 0.227 | 0.108 | 1 | 0 |
| ggnbp2 | 0.05459 | 0.154354939 | 0.082 | 0.012 | 1 | 0 |
| gxylt2 | 0.05502 | -0.16265893 | 0.009 | 0.072 | 1 | 0 |
| krt15 | 0.05513 | -0.26057392 | 0.164 | 0.229 | 1 | 0 |
| LOC108180204 | 0.0553 | -0.18469543 | 0.009 | 0.072 | 1 | 0 |
| slc25a20 | 0.05532 | 0.117633209 | 0.127 | 0.048 | 1 | 0 |
| tpm3 | 0.05542 | 0.112457871 | 0.418 | 0.301 | 1 | 0 |
| si:ch73-375g18.1 | 0.0562 | -0.16015789 | 0.009 | 0.072 | 1 | 0 |
| emc2 | 0.05657 | -0.24460687 | 0.064 | 0.169 | 1 | 0 |
| dpy30 | 0.05717 | 0.211717929 | 0.164 | 0.06 | 1 | 0 |
| ccdc187 | 0.0576 | -0.16273207 | 0.009 | 0.072 | 1 | 0 |
| sox4a | 0.05769 | 0.28613229 | 0.255 | 0.12 | 1 | 0 |

| | | | | | | |
|---|---|---|---|---|---|---|
| srrt | 0.05826 | -0.13390508 | 0.009 | 0.072 | 1 | 0 |
| lin52 | 0.05834 | -0.10054225 | 0.009 | 0.072 | 1 | 0 |
| ada2a | 0.05836 | -0.16251898 | 0.009 | 0.072 | 1 | 0 |
| rps24 | 0.05848 | 0.177673806 | 0.909 | 0.819 | 1 | 0 |
| ppib | 0.05909 | 0.227161302 | 0.773 | 0.651 | 1 | 0 |
| cxcl12a | 0.05917 | 0.321541518 | 0.345 | 0.193 | 1 | 0 |
| si:dkey-250d21.1 | 0.05918 | 0.216811645 | 0.127 | 0.036 | 1 | 0 |
| arhgef6 | 0.05981 | -0.16301263 | 0.009 | 0.072 | 1 | 0 |
| elp6 | 0.06005 | -0.13733946 | 0.009 | 0.072 | 1 | 0 |
| stag2a | 0.06012 | -0.15766412 | 0.009 | 0.072 | 1 | 0 |
| mmp2 | 0.06024 | 0.308031224 | 0.373 | 0.217 | 1 | 0 |
| eif2a | 0.0604 | -0.16308579 | 0.009 | 0.072 | 1 | 0 |
| amotl2b | 0.06042 | 0.167218962 | 0.082 | 0.012 | 1 | 0 |
| rbfox1 | 0.06043 | -0.17282837 | 0.009 | 0.072 | 1 | 0 |
| mfn1b | 0.06046 | -0.140972 | 0.009 | 0.072 | 1 | 0 |
| col4a2 | 0.06142 | 0.18117348 | 0.082 | 0.012 | 1 | 0 |
| zbtb20 | 0.0622 | -0.30061118 | 0.136 | 0.253 | 1 | 0 |
| vegfaa | 0.06225 | -0.2769219 | 0.064 | 0.169 | 1 | 0 |
| akap8l | 0.06244 | -0.25238695 | 0.064 | 0.169 | 1 | 0 |
| sh3gl1b | 0.06274 | -0.25011671 | 0.064 | 0.169 | 1 | 0 |
| sec62 | 0.0629 | 0.259299416 | 0.173 | 0.072 | 1 | 0 |
| slc6a4a | 0.06335 | 0.236383075 | 0.2 | 0.084 | 1 | 0 |
| gabarapl2 | 0.06405 | -0.24916886 | 0.064 | 0.169 | 1 | 0 |
| cdc42ep3 | 0.06416 | 0.163010905 | 0.082 | 0.012 | 1 | 0 |
| XLOC-016198 | 0.06425 | 0.190383066 | 0.082 | 0.012 | 1 | 0 |
| rpl23a | 0.0643 | 0.243836069 | 0.773 | 0.651 | 1 | 0 |
| c1s | 0.06438 | 0.165789585 | 0.082 | 0.012 | 1 | 0 |
| spon1b | 0.06462 | -0.24601674 | 0.109 | 0.181 | 1 | 0 |
| pgpep1 | 0.06471 | 0.19840409 | 0.127 | 0.036 | 1 | 0 |
| atf7ip | 0.06601 | 0.219429316 | 0.145 | 0.048 | 1 | 0 |
| si:ch211-213a13.1 | 0.06626 | 0.216866927 | 0.164 | 0.06 | 1 | 0 |
| btd | 0.06632 | 0.210394643 | 0.127 | 0.036 | 1 | 0 |
| ptmaa | 0.06634 | 0.317266868 | 0.364 | 0.217 | 1 | 0 |
| hmgb2a | 0.06651 | -0.28502914 | 0.345 | 0.458 | 1 | 0 |
| gnaia | 0.06745 | -0.17236245 | 0.027 | 0.108 | 1 | 0 |
| si:ch211-114c12.2 | 0.06765 | 0.203675324 | 0.1 | 0.024 | 1 | 0 |
| ccl25b | 0.06826 | 0.11264552 | 0.718 | 0.723 | 1 | 0 |
| actr2a | 0.06956 | 0.14813136 | 0.1 | 0.024 | 1 | 0 |
| hm13 | 0.07088 | 0.108511747 | 0.118 | 0.12 | 1 | 0 |
| cd82a | 0.07121 | 0.244060772 | 0.164 | 0.06 | 1 | 0 |
| chd4a | 0.0713 | 0.229621201 | 0.164 | 0.06 | 1 | 0 |
| BX284696.1 | 0.07202 | -0.29543022 | 0.2 | 0.337 | 1 | 0 |
| dctn2 | 0.0724 | 0.20172097 | 0.145 | 0.048 | 1 | 0 |
| slc25a3b | 0.07273 | 0.20820956 | 0.455 | 0.301 | 1 | 0 |
| ptmab | 0.07314 | 0.218201268 | 0.582 | 0.422 | 1 | 0 |
| calua | 0.07371 | -0.22098332 | 0.073 | 0.181 | 1 | 0 |
| ccdc97 | 0.07375 | 0.179728721 | 0.073 | 0.012 | 1 | 0 |
| ddt | 0.07416 | 0.223284399 | 0.182 | 0.072 | 1 | 0 |

| | | | | | | |
|---|---|---|---|---|---|---|
| khdrbs1b | 0.07456 | 0.245205038 | 0.182 | 0.072 | 1 | 0 |
| zgc:77650 | 0.07464 | -0.16966354 | 0.055 | 0.145 | 1 | 0 |
| nfia | 0.07467 | 0.101218392 | 0.582 | 0.47 | 1 | 0 |
| gm2a | 0.07508 | -0.12016015 | 0.018 | 0.084 | 1 | 0 |
| psenen | 0.07572 | 0.140112738 | 0.118 | 0.036 | 1 | 0 |
| crabp2a | 0.07574 | -0.22717669 | 0.036 | 0.12 | 1 | 0 |
| helz | 0.07576 | -0.17289647 | 0.036 | 0.12 | 1 | 0 |
| col6a1 | 0.07595 | 0.211421676 | 0.418 | 0.277 | 1 | 0 |
| rps27.1 | 0.07688 | 0.209181418 | 0.809 | 0.711 | 1 | 0 |
| rpl14 | 0.07725 | 0.104207721 | 0.809 | 0.807 | 1 | 0 |
| gstp1 | 0.07736 | -0.13557026 | 0.345 | 0.361 | 1 | 0 |
| pwwp2b | 0.0775 | 0.217895693 | 0.109 | 0.048 | 1 | 0 |
| commd1 | 0.07759 | 0.169595542 | 0.073 | 0.012 | 1 | 0 |
| rfc3 | 0.0776 | 0.145813527 | 0.064 | 0.012 | 1 | 0 |
| pwp2h | 0.07799 | 0.155309873 | 0.073 | 0.012 | 1 | 0 |
| cox7a2a | 0.07834 | 0.223375909 | 0.191 | 0.084 | 1 | 0 |
| rps16 | 0.07867 | 0.183363766 | 0.864 | 0.771 | 1 | 0 |
| aph1b | 0.07929 | 0.186777604 | 0.091 | 0.024 | 1 | 0 |
| rps19 | 0.08016 | 0.159443593 | 0.927 | 0.831 | 1 | 0 |
| timm17a | 0.08032 | 0.208654694 | 0.136 | 0.072 | 1 | 0 |
| ypel5 | 0.08034 | -0.20215596 | 0.036 | 0.12 | 1 | 0 |
| CABZ01072614.1 | 0.0807 | -0.19237754 | 0.282 | 0.422 | 1 | 0 |
| srsf1a | 0.08079 | 0.147043018 | 0.073 | 0.012 | 1 | 0 |
| cdkn1bb | 0.08094 | -0.26089553 | 0.073 | 0.169 | 1 | 0 |
| gramd4b | 0.08144 | -0.10629411 | 0.009 | 0.06 | 1 | 0 |
| aldh7a1 | 0.08162 | 0.201304926 | 0.136 | 0.048 | 1 | 0 |
| fbxo28 | 0.08175 | -0.17464121 | 0.036 | 0.12 | 1 | 0 |
| hcar1-4 | 0.08187 | -0.11388813 | 0.009 | 0.06 | 1 | 0 |
| chd4b | 0.08194 | 0.199401127 | 0.118 | 0.036 | 1 | 0 |
| rnf169 | 0.08206 | -0.20283603 | 0.036 | 0.12 | 1 | 0 |
| sparc | 0.08259 | 0.186188743 | 0.773 | 0.687 | 1 | 0 |
| junbb | 0.08276 | -0.22193507 | 0.036 | 0.12 | 1 | 0 |
| gpatch4 | 0.08313 | 0.199573593 | 0.109 | 0.036 | 1 | 0 |
| nat10 | 0.08314 | 0.168508067 | 0.073 | 0.012 | 1 | 0 |
| lrig2 | 0.08344 | 0.197678789 | 0.1 | 0.024 | 1 | 0 |
| tuba8l2 | 0.08427 | 0.166892924 | 0.1 | 0.024 | 1 | 0 |
| cox4i2 | 0.08448 | 0.210322974 | 0.145 | 0.06 | 1 | 0 |
| herpud2 | 0.08459 | 0.163258918 | 0.1 | 0.024 | 1 | 0 |
| xirp2a | 0.08459 | 0.180313693 | 0.173 | 0.072 | 1 | 0 |
| gnb1b | 0.08469 | -0.28074058 | 0.245 | 0.373 | 1 | 0 |
| ciao2b | 0.0848 | 0.144655131 | 0.073 | 0.012 | 1 | 0 |
| rem1 | 0.08502 | 0.27606406 | 0.209 | 0.096 | 1 | 0 |
| mxra8a | 0.08614 | 0.23977622 | 0.191 | 0.084 | 1 | 0 |
| mmgt1 | 0.08639 | 0.220884306 | 0.155 | 0.06 | 1 | 0 |
| capn2b | 0.08657 | -0.18657474 | 0.018 | 0.084 | 1 | 0 |
| kdm1a | 0.08754 | 0.180872547 | 0.118 | 0.036 | 1 | 0 |
| nme3 | 0.0883 | 0.26780336 | 0.236 | 0.12 | 1 | 0 |
| id3 | 0.08832 | 0.292188485 | 0.264 | 0.145 | 1 | 0 |

| | | | | | | |
|---|---|---|---|---|---|---|
| rrp36 | 0.08851 | 0.164586491 | 0.1 | 0.024 | 1 | 0 |
| si:dkey-266m15.6 | 0.08878 | 0.189309909 | 0.1 | 0.024 | 1 | 0 |
| wbp2nl | 0.08904 | -0.20088317 | 0.055 | 0.145 | 1 | 0 |
| kpnb1 | 0.08906 | 0.190841523 | 0.118 | 0.036 | 1 | 0 |
| hoxc3a | 0.09013 | 0.269586281 | 0.182 | 0.084 | 1 | 0 |
| si:dkey-17o15.2 | 0.09022 | -0.20860326 | 0.027 | 0.096 | 1 | 0 |
| parp16 | 0.09048 | -0.17919549 | 0.027 | 0.096 | 1 | 0 |
| zc3h11a | 0.09108 | 0.144813911 | 0.073 | 0.012 | 1 | 0 |
| si:ch73-25f10.6 | 0.09124 | 0.150182136 | 0.073 | 0.012 | 1 | 0 |
| arid1b | 0.09133 | 0.159179046 | 0.073 | 0.012 | 1 | 0 |
| efemp2a | 0.0914 | -0.26729546 | 0.118 | 0.229 | 1 | 0 |
| net1 | 0.09145 | -0.13778004 | 0.018 | 0.084 | 1 | 0 |
| ap2a1 | 0.09193 | -0.18342875 | 0.073 | 0.12 | 1 | 0 |
| prpf40a | 0.09223 | -0.12587769 | 0.018 | 0.084 | 1 | 0 |
| foxo6b | 0.0923 | -0.17290001 | 0.018 | 0.084 | 1 | 0 |
| angptl2a | 0.09259 | -0.15759476 | 0.018 | 0.084 | 1 | 0 |
| rpl36a | 0.09265 | 0.160207836 | 0.882 | 0.819 | 1 | 0 |
| tob1a | 0.09268 | -0.28482005 | 0.218 | 0.349 | 1 | 0 |
| cyr61 | 0.09297 | -0.2545133 | 0.109 | 0.181 | 1 | 0 |
| bgnb | 0.09302 | 0.217540234 | 0.109 | 0.036 | 1 | 0 |
| gpc1b | 0.09307 | 0.197388258 | 0.173 | 0.133 | 1 | 0 |
| ptpn1 | 0.09377 | 0.173952174 | 0.064 | 0.012 | 1 | 0 |
| leng8 | 0.09439 | -0.14801978 | 0.018 | 0.084 | 1 | 0 |
| plpp3 | 0.09448 | -0.25352183 | 0.527 | 0.675 | 1 | 0 |
| mir21-1 | 0.09473 | 0.166624273 | 0.173 | 0.084 | 1 | 0 |
| ddx61 | 0.09479 | 0.261751642 | 0.345 | 0.205 | 1 | 0 |
| usp6nl | 0.09494 | -0.15546329 | 0.018 | 0.084 | 1 | 0 |
| vdrb | 0.09509 | 0.222903628 | 0.173 | 0.072 | 1 | 0 |
| myo18ab | 0.09511 | -0.21573649 | 0.045 | 0.133 | 1 | 0 |
| ube2d1a | 0.09603 | 0.129825911 | 0.073 | 0.012 | 1 | 0 |
| si:ch211-251b21.1 | 0.09612 | 0.285582468 | 0.191 | 0.084 | 1 | 0 |
| mark3a | 0.09738 | 0.145804582 | 0.073 | 0.012 | 1 | 0 |
| mrpl12 | 0.09757 | 0.128823465 | 0.145 | 0.06 | 1 | 0 |
| sirt7 | 0.09819 | 0.16720698 | 0.109 | 0.036 | 1 | 0 |
| lamb1a | 0.09858 | 0.203518359 | 0.327 | 0.265 | 1 | 0 |
| dkc1 | 0.09943 | 0.200559091 | 0.2 | 0.145 | 1 | 0 |
| ak3 | 0.09959 | -0.13763807 | 0.009 | 0.06 | 1 | 0 |
| rsl24d1 | 0.0996 | 0.210347122 | 0.509 | 0.434 | 1 | 0 |
| usp24 | 0.09971 | -0.13083593 | 0.009 | 0.06 | 1 | 0 |
| cdc42ep5 | 0.09973 | 0.199691207 | 0.118 | 0.048 | 1 | 0 |
| rpl13 | 0.09973 | 0.145963842 | 0.945 | 0.88 | 1 | 0 |
| DDX17 | 0.09979 | -0.20150888 | 0.073 | 0.169 | 1 | 0 |
| bag2 | 0.10024 | -0.20786231 | 0.091 | 0.193 | 1 | 0 |
| capns1b | 0.10034 | 0.232607332 | 0.264 | 0.169 | 1 | 0 |
| LOC110438108 | 0.10096 | -0.13090031 | 0.009 | 0.06 | 1 | 0 |
| abce1 | 0.10096 | 0.16041694 | 0.091 | 0.024 | 1 | 0 |
| fryl | 0.10125 | -0.204543 | 0.027 | 0.096 | 1 | 0 |
| sdha | 0.10131 | -0.18044915 | 0.027 | 0.096 | 1 | 0 |

| | | | | | | |
|---|---|---|---|---|---|---|
| rpl24 | 0.10144 | 0.17784905 | 0.864 | 0.783 | 1 | 0 |
| cfl1 | 0.1017 | 0.236221959 | 0.6 | 0.446 | 1 | 0 |
| ptk7a | 0.10182 | 0.139294545 | 0.073 | 0.012 | 1 | 0 |
| rraga | 0.10207 | 0.162222323 | 0.064 | 0.012 | 1 | 0 |
| ddx46 | 0.1022 | 0.135298143 | 0.073 | 0.012 | 1 | 0 |
| csnk1g2b | 0.10241 | -0.22350563 | 0.045 | 0.12 | 1 | 0 |
| ybx1 | 0.10295 | 0.176611628 | 0.809 | 0.735 | 1 | 0 |
| zgc:92744 | 0.10297 | 0.186887944 | 0.091 | 0.048 | 1 | 0 |
| bod1 | 0.10321 | 0.163452516 | 0.082 | 0.024 | 1 | 0 |
| irf2bp1 | 0.10381 | 0.163221769 | 0.073 | 0.012 | 1 | 0 |
| hoxb5b | 0.104 | -0.22196068 | 0.064 | 0.157 | 1 | 0 |
| si:dkey-9i23.15 | 0.10451 | -0.19698745 | 0.027 | 0.096 | 1 | 0 |
| acap1 | 0.1049 | -0.16516007 | 0.009 | 0.06 | 1 | 0 |
| rnf7 | 0.10501 | 0.17748947 | 0.173 | 0.072 | 1 | 0 |
| rpz2 | 0.10514 | 0.171021904 | 0.091 | 0.036 | 1 | 0 |
| higd2a | 0.10547 | 0.195987079 | 0.136 | 0.048 | 1 | 0 |
| spns1 | 0.10558 | 0.13408992 | 0.091 | 0.024 | 1 | 0 |
| SBNO2 | 0.10592 | -0.15836845 | 0.009 | 0.06 | 1 | 0 |
| pdlim3a | 0.10596 | -0.12647513 | 0.009 | 0.06 | 1 | 0 |
| rbm39b | 0.10628 | 0.172600659 | 0.136 | 0.048 | 1 | 0 |
| ran | 0.10629 | 0.194178752 | 0.518 | 0.446 | 1 | 0 |
| CABZ01068358.1 | 0.10701 | -0.19183653 | 0.064 | 0.157 | 1 | 0 |
| mrps25 | 0.10743 | -0.16679486 | 0.091 | 0.12 | 1 | 0 |
| ccm2 | 0.10787 | -0.12463787 | 0.009 | 0.06 | 1 | 0 |
| CABZ01088428.1 | 0.10788 | -0.15206507 | 0.009 | 0.06 | 1 | 0 |
| krt4 | 0.10824 | -0.23456359 | 0.064 | 0.145 | 1 | 0 |
| ugt1b5 | 0.10866 | -0.24156868 | 0.073 | 0.169 | 1 | 0 |
| klf2a | 0.1091 | -0.23766357 | 0.545 | 0.675 | 1 | 0 |
| myl9b | 0.10927 | 0.257068609 | 0.355 | 0.217 | 1 | 0 |
| si:ch73-367f21.4 | 0.10946 | -0.17142471 | 0.027 | 0.096 | 1 | 0 |
| spen | 0.11001 | -0.17850149 | 0.027 | 0.096 | 1 | 0 |
| tet2 | 0.11007 | -0.15940882 | 0.009 | 0.06 | 1 | 0 |
| eef1akmt1 | 0.11041 | 0.163940722 | 0.064 | 0.012 | 1 | 0 |
| atf5b | 0.11054 | -0.20316688 | 0.073 | 0.169 | 1 | 0 |
| kif1b | 0.11062 | -0.12065904 | 0.009 | 0.06 | 1 | 0 |
| rac3b | 0.11127 | 0.100507737 | 0.064 | 0.012 | 1 | 0 |
| dcaf7 | 0.11145 | -0.20227752 | 0.064 | 0.157 | 1 | 0 |
| ddx3xa | 0.11153 | -0.20174457 | 0.064 | 0.157 | 1 | 0 |
| pcnp | 0.11154 | -0.11336502 | 0.027 | 0.084 | 1 | 0 |
| stk17b | 0.11168 | -0.1487319 | 0.018 | 0.06 | 1 | 0 |
| sri | 0.11234 | 0.221480977 | 0.136 | 0.06 | 1 | 0 |
| pole4 | 0.11296 | 0.132981247 | 0.064 | 0.012 | 1 | 0 |
| BX663503.3 | 0.11341 | -0.28722141 | 0.227 | 0.349 | 1 | 0 |
| ip6k2b | 0.11377 | 0.21643963 | 0.345 | 0.217 | 1 | 0 |
| psmb3 | 0.11391 | 0.22475413 | 0.218 | 0.108 | 1 | 0 |
| tmem176 | 0.11395 | 0.221139203 | 0.418 | 0.277 | 1 | 0 |
| marcksl1a | 0.11408 | 0.167159553 | 0.836 | 0.711 | 1 | 0 |
| glyr1 | 0.11486 | -0.14505908 | 0.009 | 0.06 | 1 | 0 |

| | | | | | | |
|---|---|---|---|---|---|---|
| ms4a17a.14 | 0.11537 | -0.12936041 | 0.009 | 0.06 | 1 | 0 |
| XLOC-012680 | 0.11555 | -0.11279454 | 0.009 | 0.06 | 1 | 0 |
| rhbdf1a | 0.11596 | -0.12012017 | 0.009 | 0.06 | 1 | 0 |
| si:ch211-212k18.7 | 0.11616 | -0.14672649 | 0.009 | 0.06 | 1 | 0 |
| sec11a | 0.11646 | 0.146830788 | 0.236 | 0.133 | 1 | 0 |
| txnl4a | 0.11664 | -0.17055747 | 0.027 | 0.096 | 1 | 0 |
| npr2 | 0.1167 | -0.14619312 | 0.009 | 0.06 | 1 | 0 |
| alkbh5 | 0.11697 | 0.138316034 | 0.064 | 0.012 | 1 | 0 |
| mical2a | 0.11712 | -0.15506954 | 0.009 | 0.06 | 1 | 0 |
| arhgef5 | 0.11749 | 0.172562417 | 0.091 | 0.024 | 1 | 0 |
| rab5c | 0.11757 | 0.204124979 | 0.127 | 0.048 | 1 | 0 |
| srgn | 0.11777 | -0.14778147 | 0.236 | 0.253 | 1 | 0 |
| psmd12 | 0.11808 | -0.10788474 | 0.009 | 0.06 | 1 | 0 |
| fpr1 | 0.11809 | 0.266434601 | 0.2 | 0.12 | 1 | 0 |
| unk | 0.11815 | -0.12611146 | 0.009 | 0.06 | 1 | 0 |
| pcyt2 | 0.1184 | -0.13430051 | 0.009 | 0.06 | 1 | 0 |
| tegt | 0.1185 | -0.25479785 | 0.255 | 0.373 | 1 | 0 |
| hspb8 | 0.11872 | -0.11518675 | 0.009 | 0.06 | 1 | 0 |
| cyth1a | 0.11884 | -0.16735514 | 0.036 | 0.108 | 1 | 0 |
| cops7a | 0.11896 | 0.171491975 | 0.109 | 0.048 | 1 | 0 |
| HACD4 | 0.11911 | 0.178510941 | 0.091 | 0.024 | 1 | 0 |
| cercam | 0.11922 | -0.17719959 | 0.027 | 0.096 | 1 | 0 |
| sema5bb | 0.1194 | -0.14530331 | 0.009 | 0.06 | 1 | 0 |
| syncrip | 0.11961 | -0.22690462 | 0.309 | 0.398 | 1 | 0 |
| tm9sf1 | 0.11996 | -0.11470127 | 0.009 | 0.06 | 1 | 0 |
| sfrp1a | 0.12008 | 0.131670575 | 0.118 | 0.048 | 1 | 0 |
| xrn1 | 0.12059 | -0.12175447 | 0.009 | 0.06 | 1 | 0 |
| zmynd19 | 0.12114 | -0.12572886 | 0.009 | 0.06 | 1 | 0 |
| tnrc6c1 | 0.12138 | 0.254684176 | 0.309 | 0.181 | 1 | 0 |
| tmem198a | 0.12164 | -0.11952234 | 0.009 | 0.06 | 1 | 0 |
| slc39a6 | 0.1219 | -0.1819502 | 0.027 | 0.096 | 1 | 0 |
| wwp1 | 0.12199 | -0.14866599 | 0.027 | 0.096 | 1 | 0 |
| p4hb | 0.122 | -0.19000147 | 0.227 | 0.277 | 1 | 0 |
| USP53 (1 of many) | 0.12225 | -0.11612944 | 0.009 | 0.06 | 1 | 0 |
| dnajc12 | 0.12228 | -0.11068228 | 0.009 | 0.06 | 1 | 0 |
| fam160a2 | 0.1223 | -0.13796232 | 0.009 | 0.06 | 1 | 0 |
| meis2b | 0.12265 | 0.237878196 | 0.264 | 0.145 | 1 | 0 |
| tgm1l1 | 0.12314 | 0.191895596 | 0.164 | 0.12 | 1 | 0 |
| ccdc3b | 0.12325 | -0.22492082 | 0.127 | 0.193 | 1 | 0 |
| atp5mf | 0.12344 | -0.20118508 | 0.1 | 0.205 | 1 | 0 |
| synm | 0.12344 | -0.15312086 | 0.073 | 0.157 | 1 | 0 |
| psmd4b | 0.12363 | 0.157060482 | 0.082 | 0.024 | 1 | 0 |
| mrps21 | 0.12393 | 0.16824163 | 0.082 | 0.036 | 1 | 0 |
| rdh14b | 0.12406 | 0.151409603 | 0.064 | 0.012 | 1 | 0 |
| ddx24 | 0.12436 | 0.173955788 | 0.1 | 0.06 | 1 | 0 |
| zgc:92818 | 0.1248 | 0.164337122 | 0.1 | 0.06 | 1 | 0 |
| ssr3 | 0.12552 | 0.216588864 | 0.336 | 0.205 | 1 | 0 |
| ptp4a2a | 0.12586 | 0.176437159 | 0.136 | 0.06 | 1 | 0 |

| | | | | | | |
|---|---|---|---|---|---|---|
| gstm.2 | 0.12627 | -0.16363643 | 0.018 | 0.072 | 1 | 0 |
| tgfb3 | 0.12647 | -0.15576103 | 0.073 | 0.157 | 1 | 0 |
| cmasa | 0.12654 | 0.156617818 | 0.082 | 0.024 | 1 | 0 |
| gas6 | 0.12765 | 0.249565813 | 0.127 | 0.048 | 1 | 0 |
| si:dkey-239h2.3 | 0.12778 | 0.183772283 | 0.227 | 0.12 | 1 | 0 |
| rpl34 | 0.12888 | 0.141527728 | 0.873 | 0.819 | 1 | 0 |
| eif4g3a | 0.12895 | 0.1554631 | 0.055 | 0.012 | 1 | 0 |
| wu:fb16f03 | 0.12963 | 0.15733079 | 0.245 | 0.145 | 1 | 0 |
| myh10 | 0.12971 | 0.222179953 | 0.109 | 0.036 | 1 | 0 |
| zgc:158343 | 0.12972 | -0.27827451 | 0.182 | 0.301 | 1 | 0 |
| ddx28 | 0.13095 | -0.11036801 | 0.018 | 0.072 | 1 | 0 |
| uqcc2 | 0.13098 | 0.187985994 | 0.136 | 0.06 | 1 | 0 |
| ssbp1 | 0.13136 | 0.149241174 | 0.073 | 0.024 | 1 | 0 |
| si:ch211-106h4.12 | 0.1316 | 0.284559455 | 0.2 | 0.096 | 1 | 0 |
| gstt1b | 0.13162 | 0.167667297 | 0.345 | 0.229 | 1 | 0 |
| rgs5a | 0.13193 | -0.18482565 | 0.327 | 0.373 | 1 | 0 |
| zgc:113208 | 0.13221 | 0.16023091 | 0.091 | 0.024 | 1 | 0 |
| chac1 | 0.13264 | -0.27674739 | 0.164 | 0.265 | 1 | 0 |
| serinc1 | 0.13341 | -0.14779437 | 0.136 | 0.229 | 1 | 0 |
| loxl1 | 0.13347 | -0.2548111 | 0.255 | 0.349 | 1 | 0 |
| hadh | 0.13436 | 0.150303842 | 0.145 | 0.06 | 1 | 0 |
| uprt | 0.13484 | 0.127421789 | 0.064 | 0.012 | 1 | 0 |
| mgst3b | 0.13492 | 0.255847542 | 0.309 | 0.205 | 1 | 0 |
| tex261 | 0.13513 | 0.162854447 | 0.082 | 0.024 | 1 | 0 |
| pdcd10a | 0.1358 | 0.115928209 | 0.064 | 0.012 | 1 | 0 |
| fam198a | 0.13634 | 0.182809668 | 0.127 | 0.048 | 1 | 0 |
| cab39l | 0.13651 | 0.125439499 | 0.109 | 0.036 | 1 | 0 |
| svbp | 0.13732 | 0.128126299 | 0.064 | 0.012 | 1 | 0 |
| gabarapb | 0.13741 | 0.219691086 | 0.4 | 0.265 | 1 | 0 |
| chka | 0.13793 | 0.151549394 | 0.064 | 0.012 | 1 | 0 |
| trmt112 | 0.1391 | 0.231916324 | 0.236 | 0.133 | 1 | 0 |
| nutf2l | 0.13928 | 0.107029722 | 0.064 | 0.012 | 1 | 0 |
| eef1g | 0.13943 | 0.132378336 | 0.818 | 0.699 | 1 | 0 |
| srsf10b | 0.14016 | 0.15666931 | 0.109 | 0.036 | 1 | 0 |
| tmem106ba | 0.14037 | -0.15855141 | 0.036 | 0.108 | 1 | 0 |
| metap2b | 0.14076 | 0.236881657 | 0.236 | 0.133 | 1 | 0 |
| ppp1r14bb | 0.14127 | 0.193280595 | 0.1 | 0.036 | 1 | 0 |
| barx1 | 0.14129 | -0.20012307 | 0.036 | 0.108 | 1 | 0 |
| sdad1 | 0.14144 | 0.161679052 | 0.064 | 0.012 | 1 | 0 |
| peli1a | 0.14179 | -0.10361973 | 0.036 | 0.096 | 1 | 0 |
| furinb | 0.14191 | -0.16253236 | 0.036 | 0.108 | 1 | 0 |
| eef1da | 0.14224 | 0.116282607 | 0.591 | 0.47 | 1 | 0 |
| ephb2b | 0.1424 | 0.19006855 | 0.064 | 0.012 | 1 | 0 |
| il13ra2 | 0.14271 | -0.19272204 | 0.036 | 0.108 | 1 | 0 |
| alas1 | 0.1428 | -0.15002256 | 0.036 | 0.108 | 1 | 0 |
| htatip2 | 0.14289 | -0.18389421 | 0.036 | 0.108 | 1 | 0 |
| ep300b | 0.14295 | -0.16637309 | 0.036 | 0.108 | 1 | 0 |
| rbx1 | 0.1432 | 0.221551262 | 0.336 | 0.253 | 1 | 0 |

| | | | | | | |
|---|---|---|---|---|---|---|
| txnrd3 | 0.14361 | 0.161402515 | 0.109 | 0.036 | 1 | 0 |
| rbp5 | 0.14383 | 0.21275741 | 0.545 | 0.47 | 1 | 0 |
| ngrn | 0.14473 | 0.135075288 | 0.064 | 0.012 | 1 | 0 |
| LOC110439623 | 0.14514 | 0.170447995 | 0.109 | 0.036 | 1 | 0 |
| bco2a | 0.14555 | -0.12335282 | 0.018 | 0.072 | 1 | 0 |
| si:dkeyp-23e4.3 | 0.14565 | -0.12269698 | 0.045 | 0.108 | 1 | 0 |
| sema3d | 0.14687 | -0.17210605 | 0.055 | 0.133 | 1 | 0 |
| chmp7 | 0.14716 | 0.12009996 | 0.064 | 0.012 | 1 | 0 |
| dapk3 | 0.14746 | -0.13844545 | 0.018 | 0.072 | 1 | 0 |
| hoxb3a | 0.14818 | 0.188574861 | 0.082 | 0.024 | 1 | 0 |
| rabac1 | 0.14839 | 0.164823469 | 0.091 | 0.036 | 1 | 0 |
| atg5 | 0.14874 | -0.13694982 | 0.018 | 0.072 | 1 | 0 |
| mid2 | 0.14908 | -0.18613365 | 0.036 | 0.096 | 1 | 0 |
| cdc42 | 0.14922 | -0.22053044 | 0.273 | 0.361 | 1 | 0 |
| tmx1 | 0.15021 | 0.104286999 | 0.064 | 0.012 | 1 | 0 |
| uri1 | 0.15051 | 0.124035155 | 0.064 | 0.012 | 1 | 0 |
| shrprbck1r | 0.15084 | 0.126296784 | 0.064 | 0.012 | 1 | 0 |
| praf2 | 0.15128 | -0.17598557 | 0.045 | 0.12 | 1 | 0 |
| s1pr2 | 0.15159 | -0.17799524 | 0.1 | 0.145 | 1 | 0 |
| ppfibp1b | 0.15159 | -0.16480561 | 0.018 | 0.072 | 1 | 0 |
| si:ch211-204a13.2 | 0.15166 | -0.23870479 | 0.127 | 0.229 | 1 | 0 |
| lmnb2 | 0.15177 | 0.171137678 | 0.127 | 0.048 | 1 | 0 |
| slit3 | 0.1519 | -0.23530185 | 0.436 | 0.566 | 1 | 0 |
| naa20 | 0.15197 | 0.172177923 | 0.127 | 0.048 | 1 | 0 |
| crabp1b | 0.15198 | 0.207138448 | 0.145 | 0.06 | 1 | 0 |
| hmga1b | 0.15232 | 0.132195013 | 0.064 | 0.012 | 1 | 0 |
| thoc7 | 0.15234 | 0.128892779 | 0.1 | 0.036 | 1 | 0 |
| ndufs2 | 0.15294 | 0.180625568 | 0.145 | 0.06 | 1 | 0 |
| sec61g | 0.15336 | 0.19435762 | 0.773 | 0.663 | 1 | 0 |
| nop10 | 0.15347 | 0.23916153 | 0.282 | 0.169 | 1 | 0 |
| kctd5a | 0.15352 | 0.241296508 | 0.218 | 0.12 | 1 | 0 |
| epha4b | 0.15355 | 0.179633756 | 0.1 | 0.036 | 1 | 0 |
| epb41l5 | 0.15401 | 0.143302986 | 0.064 | 0.012 | 1 | 0 |
| phf21aa | 0.15412 | -0.16474792 | 0.018 | 0.06 | 1 | 0 |
| rpl19 | 0.15492 | 0.197975324 | 0.773 | 0.651 | 1 | 0 |
| cnot1 | 0.15585 | -0.16590208 | 0.045 | 0.12 | 1 | 0 |
| sppl2 | 0.15647 | 0.124782422 | 0.064 | 0.012 | 1 | 0 |
| tardbpl | 0.15695 | -0.23588014 | 0.136 | 0.217 | 1 | 0 |
| abi3bpa | 0.15733 | -0.18199752 | 0.073 | 0.157 | 1 | 0 |
| nktr | 0.15766 | -0.17759792 | 0.045 | 0.12 | 1 | 0 |
| junba | 0.15802 | -0.1300221 | 0.573 | 0.59 | 1 | 0 |
| rpf1 | 0.15918 | -0.14679776 | 0.082 | 0.169 | 1 | 0 |
| si:rp71-79p20.2 | 0.15952 | 0.114415171 | 0.1 | 0.036 | 1 | 0 |
| cdr2a | 0.15996 | 0.160003953 | 0.055 | 0.012 | 1 | 0 |
| dnajb12a | 0.16019 | -0.14761926 | 0.018 | 0.06 | 1 | 0 |
| si:ch211-218c6.8 | 0.16036 | 0.125212382 | 0.1 | 0.036 | 1 | 0 |
| fcho2 | 0.16081 | 0.136368875 | 0.064 | 0.012 | 1 | 0 |
| nr2f1a | 0.16104 | -0.18842364 | 0.1 | 0.157 | 1 | 0 |

| | | | | | | |
|---|---|---|---|---|---|---|
| chtf8 | 0.16145 | 0.115326727 | 0.064 | 0.012 | 1 | 0 |
| lgals3b | 0.16153 | 0.222774538 | 0.255 | 0.145 | 1 | 0 |
| ddit3 | 0.16162 | -0.25628701 | 0.291 | 0.398 | 1 | 0 |
| si:ch211-213d14.2 | 0.1625 | 0.116787499 | 0.064 | 0.012 | 1 | 0 |
| slc6a6a | 0.1635 | 0.163157444 | 0.064 | 0.012 | 1 | 0 |
| ptena | 0.16374 | 0.197072164 | 0.209 | 0.108 | 1 | 0 |
| AL954191.1 | 0.16383 | 0.275887927 | 0.173 | 0.096 | 1 | 0 |
| ilk | 0.16436 | 0.134360176 | 0.118 | 0.048 | 1 | 0 |
| elf2b | 0.16453 | 0.12533167 | 0.064 | 0.012 | 1 | 0 |
| clic2 | 0.16453 | -0.19925658 | 0.082 | 0.157 | 1 | 0 |
| acvr2aa | 0.16468 | 0.14693311 | 0.064 | 0.012 | 1 | 0 |
| psmb7 | 0.16484 | 0.215231247 | 0.264 | 0.181 | 1 | 0 |
| plagx | 0.16512 | 0.146559304 | 0.064 | 0.012 | 1 | 0 |
| pin1 | 0.16526 | 0.193896758 | 0.209 | 0.108 | 1 | 0 |
| si:ch73-59f11.3 | 0.16527 | 0.152631885 | 0.082 | 0.024 | 1 | 0 |
| faf2 | 0.16545 | -0.10258294 | 0.018 | 0.072 | 1 | 0 |
| mrpl32 | 0.1656 | 0.125250097 | 0.118 | 0.048 | 1 | 0 |
| crabp1a | 0.166 | 0.10340075 | 0.136 | 0.072 | 1 | 0 |
| eif6 | 0.16608 | 0.180969819 | 0.182 | 0.096 | 1 | 0 |
| dnajc8 | 0.1661 | -0.15314973 | 0.027 | 0.084 | 1 | 0 |
| msra | 0.16615 | 0.115241449 | 0.082 | 0.024 | 1 | 0 |
| si:dkey-89b17.4 | 0.16658 | -0.1442899 | 0.027 | 0.084 | 1 | 0 |
| zgc:112982 | 0.16677 | -0.12690169 | 0.018 | 0.072 | 1 | 0 |
| tmc6b | 0.16683 | -0.14186985 | 0.018 | 0.072 | 1 | 0 |
| surf4l | 0.16731 | -0.17997302 | 0.055 | 0.133 | 1 | 0 |
| usp9 | 0.16776 | 0.195306671 | 0.145 | 0.072 | 1 | 0 |
| tomm20a | 0.1679 | 0.198256346 | 0.182 | 0.108 | 1 | 0 |
| arhgef1b | 0.16817 | 0.200759618 | 0.173 | 0.084 | 1 | 0 |
| si:dkey-16l2.20 | 0.16822 | -0.20147195 | 0.045 | 0.108 | 1 | 0 |
| myl12.1 | 0.16874 | 0.121004311 | 0.536 | 0.422 | 1 | 0 |
| immp2l | 0.16878 | -0.10882842 | 0.018 | 0.072 | 1 | 0 |
| tm9sf4 | 0.16879 | -0.11417174 | 0.045 | 0.108 | 1 | 0 |
| serbp1b | 0.16884 | 0.172762637 | 0.118 | 0.048 | 1 | 0 |
| sdr16c5b | 0.16923 | 0.156639803 | 0.082 | 0.024 | 1 | 0 |
| prrx1b | 0.16931 | -0.15153731 | 0.018 | 0.072 | 1 | 0 |
| ttyh2 | 0.16969 | -0.15984964 | 0.018 | 0.072 | 1 | 0 |
| mrps16 | 0.16987 | -0.10694021 | 0.018 | 0.072 | 1 | 0 |
| c7a | 0.17011 | -0.12674079 | 0.164 | 0.181 | 1 | 0 |
| rplp2l | 0.1702 | 0.113603225 | 0.9 | 0.867 | 1 | 0 |
| COX7A2 (1 of many) | 0.17036 | 0.167941694 | 0.264 | 0.157 | 1 | 0 |
| itpkcb | 0.17059 | -0.20525639 | 0.073 | 0.157 | 1 | 0 |
| lzts2a | 0.1708 | -0.12662057 | 0.018 | 0.072 | 1 | 0 |
| eif1ad | 0.17094 | 0.143935397 | 0.136 | 0.06 | 1 | 0 |
| fosl2 | 0.17094 | -0.10535272 | 0.018 | 0.072 | 1 | 0 |
| cndp2 | 0.17206 | 0.117369125 | 0.082 | 0.024 | 1 | 0 |
| fbxo45 | 0.17243 | -0.13732094 | 0.018 | 0.072 | 1 | 0 |
| l3hypdh | 0.17247 | -0.10591365 | 0.018 | 0.072 | 1 | 0 |
| ptprfa | 0.17263 | -0.12234625 | 0.018 | 0.072 | 1 | 0 |

| | | | | | | |
|---|---|---|---|---|---|---|
| cs | 0.17263 | -0.17294377 | 0.064 | 0.145 | 1 | 0 |
| tmem59l | 0.17299 | -0.10781025 | 0.018 | 0.072 | 1 | 0 |
| sppl3 | 0.17338 | -0.20359071 | 0.073 | 0.145 | 1 | 0 |
| elmo3 | 0.17358 | -0.14350741 | 0.018 | 0.072 | 1 | 0 |
| XLOC-000431 | 0.17401 | 0.220882305 | 0.236 | 0.133 | 1 | 0 |
| psd3l | 0.17447 | -0.13749972 | 0.018 | 0.072 | 1 | 0 |
| shisa9b | 0.17448 | -0.15114378 | 0.018 | 0.072 | 1 | 0 |
| gbp | 0.17487 | 0.23451123 | 0.273 | 0.169 | 1 | 0 |
| npb | 0.17599 | 0.144154177 | 0.055 | 0.012 | 1 | 0 |
| si:ch211-235e9.6 | 0.17671 | 0.174115201 | 0.191 | 0.096 | 1 | 0 |
| lsm5 | 0.17714 | -0.15865769 | 0.064 | 0.145 | 1 | 0 |
| tuba1b | 0.17734 | -0.1877525 | 0.064 | 0.145 | 1 | 0 |
| bms1 | 0.17743 | 0.121328073 | 0.1 | 0.036 | 1 | 0 |
| rbms2a | 0.17772 | 0.164288992 | 0.109 | 0.072 | 1 | 0 |
| pard3bb | 0.17814 | -0.22087991 | 0.091 | 0.181 | 1 | 0 |
| vcp | 0.17823 | 0.224846009 | 0.236 | 0.145 | 1 | 0 |
| slc10a3 | 0.17902 | -0.12020894 | 0.027 | 0.084 | 1 | 0 |
| eif4a2 | 0.17906 | 0.226509987 | 0.182 | 0.096 | 1 | 0 |
| arl5a | 0.18001 | 0.100502588 | 0.082 | 0.024 | 1 | 0 |
| atpv0e2 | 0.18009 | -0.11700688 | 0.036 | 0.096 | 1 | 0 |
| si:ch73-173h19.3 | 0.18072 | -0.22488284 | 0.127 | 0.229 | 1 | 0 |
| fen1 | 0.18164 | 0.134307996 | 0.082 | 0.024 | 1 | 0 |
| mmp14a | 0.1823 | 0.221909266 | 0.336 | 0.217 | 1 | 0 |
| foxp4 | 0.18234 | -0.11872315 | 0.355 | 0.373 | 1 | 0 |
| mt-co1 | 0.18287 | -0.17752931 | 0.7 | 0.759 | 1 | 0 |
| FP102052.2 | 0.18289 | -0.17971547 | 0.073 | 0.157 | 1 | 0 |
| zgc:114188 | 0.18313 | 0.144287805 | 0.891 | 0.807 | 1 | 0 |
| COX3 | 0.18324 | -0.13538387 | 0.764 | 0.867 | 1 | 0 |
| rac1b | 0.18354 | 0.154461121 | 0.091 | 0.036 | 1 | 0 |
| bnip3lb | 0.18384 | -0.18153853 | 0.091 | 0.181 | 1 | 0 |
| neu3.5 | 0.18428 | -0.25559345 | 0.164 | 0.253 | 1 | 0 |
| eif4bb | 0.18466 | -0.10648077 | 0.118 | 0.133 | 1 | 0 |
| cnpy1 | 0.18466 | -0.10640867 | 0.027 | 0.084 | 1 | 0 |
| svep1 | 0.18562 | 0.208994023 | 0.127 | 0.06 | 1 | 0 |
| ppap2d | 0.18605 | -0.21226057 | 0.109 | 0.205 | 1 | 0 |
| prdx4 | 0.18625 | -0.14256145 | 0.218 | 0.241 | 1 | 0 |
| ftsj3 | 0.18669 | 0.171133633 | 0.118 | 0.048 | 1 | 0 |
| cyp2p7 | 0.18696 | -0.20707473 | 0.182 | 0.253 | 1 | 0 |
| cirbpa | 0.18758 | 0.106577466 | 0.655 | 0.542 | 1 | 0 |
| vkorc1l1 | 0.18804 | 0.115332605 | 0.082 | 0.024 | 1 | 0 |
| npc2 | 0.18805 | 0.190930393 | 0.264 | 0.157 | 1 | 0 |
| ca5a | 0.18821 | -0.19199861 | 0.109 | 0.169 | 1 | 0 |
| wu:fu71h07 | 0.18832 | -0.12603161 | 0.055 | 0.12 | 1 | 0 |
| casc4 | 0.18909 | 0.148768589 | 0.082 | 0.024 | 1 | 0 |
| ddah2 | 0.18935 | -0.16683851 | 0.045 | 0.096 | 1 | 0 |
| bckdhb | 0.18961 | 0.148415151 | 0.091 | 0.036 | 1 | 0 |
| asap1b | 0.18966 | -0.13038642 | 0.027 | 0.084 | 1 | 0 |
| snrpf | 0.18984 | -0.13816135 | 0.2 | 0.229 | 1 | 0 |

| | | | | | | |
|---|---|---|---|---|---|---|
| rbm8a | 0.19001 | 0.199094861 | 0.245 | 0.145 | 1 | 0 |
| gata2a | 0.19083 | 0.108060041 | 0.127 | 0.06 | 1 | 0 |
| ccng2 | 0.19092 | -0.17679479 | 0.045 | 0.096 | 1 | 0 |
| zgc:171480 | 0.19127 | 0.120093151 | 0.082 | 0.024 | 1 | 0 |
| slc39a7 | 0.19178 | 0.14565304 | 0.082 | 0.024 | 1 | 0 |
| fam168a | 0.19198 | 0.228286517 | 0.264 | 0.157 | 1 | 0 |
| si:ch211-246m6.4 | 0.19199 | -0.21458077 | 0.118 | 0.205 | 1 | 0 |
| zgc:56304 | 0.1925 | 0.1783477 | 0.164 | 0.084 | 1 | 0 |
| FO082781.1 | 0.19262 | 0.214774725 | 0.182 | 0.108 | 1 | 0 |
| actb1 | 0.19319 | -0.22232376 | 0.464 | 0.566 | 1 | 0 |
| srpx2 | 0.19408 | 0.138673471 | 0.082 | 0.024 | 1 | 0 |
| cdv3 | 0.19447 | -0.20287047 | 0.164 | 0.241 | 1 | 0 |
| si:dkey-238c7.16 | 0.19497 | 0.140359888 | 0.3 | 0.205 | 1 | 0 |
| atp5mc3a | 0.19497 | -0.10141656 | 0.291 | 0.301 | 1 | 0 |
| palld | 0.19506 | -0.15200941 | 0.027 | 0.084 | 1 | 0 |
| ccnt1 | 0.19507 | 0.186877957 | 0.1 | 0.036 | 1 | 0 |
| myzap | 0.1952 | -0.19546846 | 0.055 | 0.12 | 1 | 0 |
| rpl35 | 0.19612 | 0.166018233 | 0.8 | 0.687 | 1 | 0 |
| usp22 | 0.19659 | 0.143743862 | 0.091 | 0.036 | 1 | 0 |
| nup58 | 0.19681 | 0.154052028 | 0.091 | 0.036 | 1 | 0 |
| slain2 | 0.19691 | -0.1238373 | 0.027 | 0.084 | 1 | 0 |
| lrp1aa | 0.19696 | 0.159708903 | 0.464 | 0.349 | 1 | 0 |
| rimkla | 0.19711 | 0.100098472 | 0.073 | 0.024 | 1 | 0 |
| gchfr | 0.1978 | 0.123105717 | 0.082 | 0.024 | 1 | 0 |
| ndufb3 | 0.19784 | 0.207744125 | 0.318 | 0.205 | 1 | 0 |
| klf15 | 0.19844 | 0.156609865 | 0.082 | 0.024 | 1 | 0 |
| srsf2b | 0.19879 | 0.125200907 | 0.055 | 0.012 | 1 | 0 |
| pa2g4a | 0.19883 | 0.170580153 | 0.118 | 0.048 | 1 | 0 |
| ptk2aa | 0.199 | -0.17301132 | 0.082 | 0.133 | 1 | 0 |
| mmp15a | 0.19964 | 0.179654377 | 0.109 | 0.048 | 1 | 0 |
| gps1 | 0.19987 | 0.126443028 | 0.082 | 0.024 | 1 | 0 |
| kdm6al | 0.19991 | 0.147320683 | 0.082 | 0.024 | 1 | 0 |
| rbp7b | 0.19994 | 0.190573048 | 0.109 | 0.048 | 1 | 0 |
| hspa12b | 0.19998 | -0.11760483 | 0.073 | 0.145 | 1 | 0 |
| naa38 | 0.20001 | -0.11181945 | 0.036 | 0.096 | 1 | 0 |
| fam133b | 0.20022 | 0.194701859 | 0.264 | 0.193 | 1 | 0 |
| patl1 | 0.20037 | 0.107894886 | 0.082 | 0.024 | 1 | 0 |
| CABZ01074363.1 | 0.2004 | 0.137475517 | 0.082 | 0.024 | 1 | 0 |
| ppdpfb | 0.20049 | 0.138303821 | 0.673 | 0.554 | 1 | 0 |
| arl8a | 0.2005 | 0.131412015 | 0.082 | 0.024 | 1 | 0 |
| cox7c | 0.20054 | 0.144456104 | 0.582 | 0.458 | 1 | 0 |
| mylka | 0.20075 | 0.130809033 | 0.082 | 0.024 | 1 | 0 |
| aida | 0.20095 | 0.146287318 | 0.1 | 0.036 | 1 | 0 |
| ptprk | 0.2015 | 0.195008961 | 0.118 | 0.048 | 1 | 0 |
| alg5 | 0.20214 | 0.157597006 | 0.155 | 0.072 | 1 | 0 |
| ctsc | 0.20227 | -0.13096801 | 0.027 | 0.084 | 1 | 0 |
| prss35 | 0.20334 | -0.22746926 | 0.091 | 0.169 | 1 | 0 |
| rps15 | 0.20386 | 0.138163447 | 0.864 | 0.807 | 1 | 0 |

| | | | | | | |
|---|---|---|---|---|---|---|
| eef1a1l1 | 0.2041 | -0.10696782 | 0.891 | 0.94 | 1 | 0 |
| fxyd6l | 0.20538 | 0.223744231 | 0.345 | 0.229 | 1 | 0 |
| mmrn2a | 0.20542 | -0.14073813 | 0.027 | 0.084 | 1 | 0 |
| lsm6 | 0.20566 | 0.157968186 | 0.118 | 0.048 | 1 | 0 |
| pleca | 0.20606 | 0.157651603 | 0.1 | 0.036 | 1 | 0 |
| dnajb6a | 0.20649 | 0.120628009 | 0.055 | 0.012 | 1 | 0 |
| fkbp1aa | 0.20754 | 0.199712072 | 0.464 | 0.337 | 1 | 0 |
| prps1b | 0.20776 | -0.12818458 | 0.027 | 0.084 | 1 | 0 |
| LOC108180269 | 0.20845 | -0.22286276 | 0.255 | 0.373 | 1 | 0 |
| gpr78a | 0.20907 | -0.11934142 | 0.136 | 0.157 | 1 | 0 |
| u2af2b | 0.21015 | 0.179465501 | 0.173 | 0.108 | 1 | 0 |
| ctcf | 0.21018 | -0.1334988 | 0.027 | 0.084 | 1 | 0 |
| si:ch211-284f22.3 | 0.2112 | 0.1504101 | 0.1 | 0.036 | 1 | 0 |
| ccdc59 | 0.21143 | 0.117364341 | 0.055 | 0.012 | 1 | 0 |
| zgc:171679 | 0.21259 | 0.120729457 | 0.055 | 0.012 | 1 | 0 |
| pus1 | 0.21264 | 0.142297851 | 0.073 | 0.024 | 1 | 0 |
| plxna1a | 0.21298 | 0.170277249 | 0.1 | 0.036 | 1 | 0 |
| ap3s1.1 | 0.21317 | 0.155171357 | 0.1 | 0.036 | 1 | 0 |
| wls | 0.21387 | 0.196746801 | 0.273 | 0.205 | 1 | 0 |
| bsdc1 | 0.21391 | 0.151150811 | 0.091 | 0.048 | 1 | 0 |
| snrpd3l | 0.21418 | 0.120779934 | 0.082 | 0.06 | 1 | 0 |
| snrpa | 0.21453 | 0.133393296 | 0.1 | 0.036 | 1 | 0 |
| cacna1c | 0.21462 | 0.114928937 | 0.055 | 0.012 | 1 | 0 |
| hnrnpa1b | 0.21507 | -0.1957454 | 0.155 | 0.217 | 1 | 0 |
| gng12a | 0.21551 | 0.154144331 | 0.091 | 0.036 | 1 | 0 |
| glula | 0.21555 | 0.164549152 | 0.118 | 0.048 | 1 | 0 |
| ramp2 | 0.21606 | -0.25052686 | 0.173 | 0.253 | 1 | 0 |
| dhrs3b | 0.21625 | 0.112523225 | 0.245 | 0.229 | 1 | 0 |
| dda1 | 0.21674 | 0.166257863 | 0.118 | 0.06 | 1 | 0 |
| waca | 0.2173 | -0.16264987 | 0.036 | 0.096 | 1 | 0 |
| gas1a | 0.2175 | 0.114103692 | 0.264 | 0.181 | 1 | 0 |
| mycbp | 0.21767 | 0.191040038 | 0.264 | 0.193 | 1 | 0 |
| rbm4.2 | 0.21774 | 0.188155985 | 0.145 | 0.072 | 1 | 0 |
| ifngr2 | 0.21787 | -0.12435624 | 0.036 | 0.096 | 1 | 0 |
| dctn4 | 0.21817 | 0.122089129 | 0.055 | 0.012 | 1 | 0 |
| flrt2 | 0.21842 | -0.1461686 | 0.036 | 0.096 | 1 | 0 |
| snrpe | 0.21919 | 0.169476283 | 0.127 | 0.072 | 1 | 0 |
| rab1ba | 0.21921 | -0.1321314 | 0.036 | 0.096 | 1 | 0 |
| FP102783.1 | 0.21932 | 0.206583287 | 0.327 | 0.217 | 1 | 0 |
| lamtor4 | 0.21973 | 0.169953903 | 0.109 | 0.048 | 1 | 0 |
| rpl23 | 0.22064 | 0.142455927 | 0.864 | 0.771 | 1 | 0 |
| anxa11b | 0.22104 | 0.153092087 | 0.255 | 0.217 | 1 | 0 |
| bloc1s6 | 0.22166 | -0.11839812 | 0.082 | 0.108 | 1 | 0 |
| zgc:77849 | 0.22244 | 0.124760292 | 0.218 | 0.193 | 1 | 0 |
| CU929418.1 | 0.22424 | 0.111032603 | 0.055 | 0.012 | 1 | 0 |
| pgam1a | 0.22461 | 0.142758575 | 0.109 | 0.048 | 1 | 0 |
| si:dkey-79d12.5 | 0.22527 | -0.12055367 | 0.018 | 0.06 | 1 | 0 |
| setdb1a | 0.22546 | 0.111900834 | 0.055 | 0.012 | 1 | 0 |

| | | | | | | |
|---|---|---|---|---|---|---|
| krt18a.1 | 0.22553 | 0.205982281 | 0.255 | 0.157 | 1 | 0 |
| pik3r2 | 0.22653 | -0.14937925 | 0.109 | 0.145 | 1 | 0 |
| CABZ01045617.1 | 0.22698 | 0.162935379 | 0.191 | 0.157 | 1 | 0 |
| hikeshi | 0.22772 | -0.10446977 | 0.036 | 0.096 | 1 | 0 |
| pbdc1 | 0.22816 | -0.1967264 | 0.155 | 0.241 | 1 | 0 |
| fhit | 0.22831 | 0.101094177 | 0.055 | 0.012 | 1 | 0 |
| cyc1 | 0.22837 | 0.164032548 | 0.182 | 0.096 | 1 | 0 |
| lsm12b | 0.22923 | 0.162982114 | 0.282 | 0.181 | 1 | 0 |
| ak2 | 0.22933 | 0.219842328 | 0.355 | 0.253 | 1 | 0 |
| fstl1a | 0.22941 | -0.13881571 | 0.018 | 0.06 | 1 | 0 |
| lig1 | 0.22949 | 0.124094177 | 0.055 | 0.012 | 1 | 0 |
| rbm4.1 | 0.23004 | 0.158486991 | 0.136 | 0.096 | 1 | 0 |
| trappc6b | 0.23016 | 0.112442695 | 0.055 | 0.012 | 1 | 0 |
| cct2 | 0.23023 | 0.205402099 | 0.318 | 0.229 | 1 | 0 |
| fbl | 0.23064 | 0.175993631 | 0.182 | 0.096 | 1 | 0 |
| tcp11l2 | 0.23065 | -0.12756767 | 0.036 | 0.096 | 1 | 0 |
| prdx5 | 0.23135 | 0.142995223 | 0.173 | 0.133 | 1 | 0 |
| ska2 | 0.23191 | 0.140146723 | 0.073 | 0.024 | 1 | 0 |
| smarcd2 | 0.23211 | -0.15027773 | 0.036 | 0.096 | 1 | 0 |
| fstb | 0.23241 | 0.225135469 | 0.355 | 0.241 | 1 | 0 |
| ralbp1 | 0.23261 | -0.14643653 | 0.073 | 0.145 | 1 | 0 |
| cry2 | 0.2329 | -0.14052231 | 0.036 | 0.096 | 1 | 0 |
| tmem131 | 0.23294 | -0.1025555 | 0.018 | 0.06 | 1 | 0 |
| ndufs8a | 0.23307 | -0.1246456 | 0.036 | 0.096 | 1 | 0 |
| gsta.1 | 0.2332 | 0.178945805 | 0.182 | 0.096 | 1 | 0 |
| eef1b2 | 0.23401 | 0.14664729 | 0.718 | 0.602 | 1 | 0 |
| ANKRD50 | 0.23435 | -0.13922639 | 0.036 | 0.096 | 1 | 0 |
| fosl1a | 0.23457 | -0.14044756 | 0.036 | 0.096 | 1 | 0 |
| CABZ01110881.1 | 0.23474 | 0.112419005 | 0.055 | 0.012 | 1 | 0 |
| ahnak | 0.23492 | 0.207895099 | 0.4 | 0.313 | 1 | 0 |
| ywhaz | 0.23518 | 0.170387936 | 0.309 | 0.253 | 1 | 0 |
| mier3a | 0.23536 | 0.117239925 | 0.055 | 0.012 | 1 | 0 |
| plxna4 | 0.23562 | 0.115266762 | 0.055 | 0.012 | 1 | 0 |
| ndufb2 | 0.2357 | 0.200827301 | 0.236 | 0.145 | 1 | 0 |
| ssuh2rs1 | 0.23604 | -0.11249292 | 0.045 | 0.108 | 1 | 0 |
| sypl2a | 0.23621 | 0.126604924 | 0.155 | 0.133 | 1 | 0 |
| MMP23B | 0.23667 | 0.158459497 | 0.118 | 0.06 | 1 | 0 |
| egr1 | 0.23747 | 0.120180501 | 0.191 | 0.12 | 1 | 0 |
| nudt17 | 0.23829 | -0.15694734 | 0.055 | 0.096 | 1 | 0 |
| pbx1a | 0.23836 | 0.21381967 | 0.264 | 0.193 | 1 | 0 |
| creg2 | 0.23884 | -0.17841505 | 0.091 | 0.169 | 1 | 0 |
| zgc:153713 | 0.24067 | 0.143830923 | 0.155 | 0.12 | 1 | 0 |
| tnfrsf19 | 0.24069 | 0.172199823 | 0.127 | 0.06 | 1 | 0 |
| atp6v1g1 | 0.24205 | -0.14621342 | 0.045 | 0.096 | 1 | 0 |
| tars | 0.24261 | 0.127093217 | 0.073 | 0.024 | 1 | 0 |
| birc2 | 0.24314 | -0.15919623 | 0.073 | 0.145 | 1 | 0 |
| elocb | 0.24329 | 0.185637534 | 0.282 | 0.181 | 1 | 0 |
| fbn2b | 0.24344 | 0.103816286 | 0.5 | 0.41 | 1 | 0 |

| | | | | | | |
|---|---|---|---|---|---|---|
| fam162a | 0.24348 | 0.106599125 | 0.064 | 0.024 | 1 | 0 |
| ttyh3a | 0.24354 | 0.125929565 | 0.055 | 0.012 | 1 | 0 |
| adamts15b | 0.24415 | -0.23383664 | 0.173 | 0.265 | 1 | 0 |
| wdr1 | 0.2444 | -0.11704494 | 0.045 | 0.108 | 1 | 0 |
| ptgr1 | 0.24453 | 0.134531578 | 0.109 | 0.048 | 1 | 0 |
| sumo2b | 0.24548 | 0.205594628 | 0.427 | 0.325 | 1 | 0 |
| pcyox1 | 0.24661 | -0.13861416 | 0.045 | 0.108 | 1 | 0 |
| cdc42l | 0.24789 | -0.15031782 | 0.245 | 0.289 | 1 | 0 |
| idh2 | 0.24821 | 0.115480152 | 0.3 | 0.205 | 1 | 0 |
| arrb2b | 0.24892 | -0.13671896 | 0.055 | 0.084 | 1 | 0 |
| aldoaa | 0.24918 | -0.15626 | 0.127 | 0.217 | 1 | 0 |
| si:dkey-17m8.1 | 0.24922 | -0.16377846 | 0.045 | 0.108 | 1 | 0 |
| stau1 | 0.24965 | 0.179737171 | 0.136 | 0.072 | 1 | 0 |
| lgals9l3 | 0.24991 | -0.1904476 | 0.118 | 0.205 | 1 | 0 |
| gtpbp4 | 0.24999 | -0.18909795 | 0.218 | 0.325 | 1 | 0 |
| kat5b | 0.25007 | 0.101124332 | 0.055 | 0.012 | 1 | 0 |
| hspb1 | 0.25009 | -0.22668874 | 0.273 | 0.386 | 1 | 0 |
| commd6 | 0.25061 | -0.11275078 | 0.018 | 0.06 | 1 | 0 |
| purab | 0.25078 | -0.17595505 | 0.2 | 0.253 | 1 | 0 |
| zgc:66160 | 0.25101 | -0.14344196 | 0.045 | 0.108 | 1 | 0 |
| si:ch211-204c21.1 | 0.25111 | 0.139997925 | 0.173 | 0.096 | 1 | 0 |
| dusp1 | 0.25127 | -0.13688572 | 0.045 | 0.108 | 1 | 0 |
| uqcrc1 | 0.25128 | 0.174354105 | 0.164 | 0.084 | 1 | 0 |
| col12a1b | 0.25154 | 0.146888119 | 0.055 | 0.012 | 1 | 0 |
| efnb2a | 0.25163 | -0.16788936 | 0.064 | 0.133 | 1 | 0 |
| midn | 0.25178 | -0.16333313 | 0.064 | 0.108 | 1 | 0 |
| hnrnpul1l | 0.25254 | 0.106086997 | 0.091 | 0.036 | 1 | 0 |
| btbd9 | 0.2529 | 0.113204109 | 0.055 | 0.012 | 1 | 0 |
| zgc:110782 | 0.25319 | 0.121434852 | 0.055 | 0.012 | 1 | 0 |
| SCARF2 | 0.25422 | 0.122549894 | 0.073 | 0.024 | 1 | 0 |
| pabpc1a | 0.25455 | 0.166654048 | 0.636 | 0.518 | 1 | 0 |
| kmt5aa | 0.25507 | -0.13835996 | 0.036 | 0.072 | 1 | 0 |
| sept9a | 0.25529 | -0.1551243 | 0.091 | 0.169 | 1 | 0 |
| dlat | 0.2553 | -0.12729127 | 0.018 | 0.06 | 1 | 0 |
| tspan18a | 0.25532 | 0.117447271 | 0.055 | 0.012 | 1 | 0 |
| rexo1 | 0.25538 | 0.102557191 | 0.055 | 0.012 | 1 | 0 |
| pbx1b | 0.25569 | 0.153087099 | 0.109 | 0.06 | 1 | 0 |
| vapb | 0.25591 | 0.138012957 | 0.127 | 0.06 | 1 | 0 |
| ryk | 0.25705 | -0.12440322 | 0.055 | 0.12 | 1 | 0 |
| gnai2a | 0.25727 | 0.204017876 | 0.155 | 0.084 | 1 | 0 |
| tpm4b | 0.25744 | 0.157287642 | 0.109 | 0.048 | 1 | 0 |
| atp5fa1 | 0.25758 | 0.182681123 | 0.364 | 0.253 | 1 | 0 |
| cdc37 | 0.25798 | -0.12640286 | 0.018 | 0.06 | 1 | 0 |
| si:ch211-250c4.4 | 0.25802 | 0.151628803 | 0.127 | 0.06 | 1 | 0 |
| phyh | 0.25807 | -0.13989665 | 0.018 | 0.06 | 1 | 0 |
| tmem198b | 0.25848 | -0.15100648 | 0.055 | 0.12 | 1 | 0 |
| tsku | 0.25851 | -0.21438824 | 0.291 | 0.398 | 1 | 0 |
| ppp1r37 | 0.25853 | 0.134539646 | 0.073 | 0.024 | 1 | 0 |

| | | | | | | |
|---|---|---|---|---|---|---|
| nucks1a | 0.25879 | 0.186024419 | 0.209 | 0.12 | 1 | 0 |
| zgc:92242 | 0.25886 | 0.127773012 | 0.073 | 0.024 | 1 | 0 |
| CR855311.1 | 0.2589 | -0.10273276 | 0.4 | 0.398 | 1 | 0 |
| chchd1 | 0.2597 | 0.117288611 | 0.064 | 0.024 | 1 | 0 |
| nkain1 | 0.26019 | 0.141331174 | 0.082 | 0.036 | 1 | 0 |
| gng5 | 0.2605 | 0.161317145 | 0.264 | 0.169 | 1 | 0 |
| fndc3ba | 0.26066 | -0.12705857 | 0.018 | 0.06 | 1 | 0 |
| tcf3b | 0.26073 | -0.14337668 | 0.055 | 0.12 | 1 | 0 |
| jcada | 0.26357 | -0.16090153 | 0.064 | 0.133 | 1 | 0 |
| npm3 | 0.26365 | 0.170465441 | 0.118 | 0.06 | 1 | 0 |
| pin4 | 0.26369 | 0.164317904 | 0.145 | 0.072 | 1 | 0 |
| ap1g1 | 0.2643 | 0.126701506 | 0.064 | 0.024 | 1 | 0 |
| zgc:92335 | 0.26465 | -0.15279409 | 0.036 | 0.084 | 1 | 0 |
| actr1 | 0.26514 | 0.160663954 | 0.091 | 0.036 | 1 | 0 |
| ppp3cca | 0.26538 | -0.10937971 | 0.018 | 0.06 | 1 | 0 |
| cnpy4 | 0.26825 | 0.108306034 | 0.073 | 0.024 | 1 | 0 |
| nat9 | 0.26835 | 0.126067369 | 0.064 | 0.024 | 1 | 0 |
| selenot1a | 0.27027 | -0.12316441 | 0.055 | 0.084 | 1 | 0 |
| cybrd1 | 0.27101 | -0.13268877 | 0.027 | 0.06 | 1 | 0 |
| ywhaqa | 0.2712 | -0.18837443 | 0.127 | 0.205 | 1 | 0 |
| atp5pb | 0.27163 | 0.19114597 | 0.4 | 0.289 | 1 | 0 |
| cbx5 | 0.27217 | 0.182743246 | 0.227 | 0.145 | 1 | 0 |
| pbx3b | 0.273 | -0.20311724 | 0.091 | 0.169 | 1 | 0 |
| si:dkey-188i13.10 | 0.27377 | 0.131833716 | 0.091 | 0.048 | 1 | 0 |
| mef2d | 0.27387 | -0.18942358 | 0.082 | 0.157 | 1 | 0 |
| tomm20b | 0.2744 | 0.178891667 | 0.236 | 0.145 | 1 | 0 |
| b2m | 0.27465 | 0.193630433 | 0.264 | 0.169 | 1 | 0 |
| eps15 | 0.27512 | -0.13447477 | 0.018 | 0.06 | 1 | 0 |
| wdr46 | 0.27516 | -0.11369157 | 0.036 | 0.084 | 1 | 0 |
| plod2 | 0.27565 | 0.197033002 | 0.182 | 0.108 | 1 | 0 |
| atxn1a | 0.27595 | 0.133198761 | 0.073 | 0.024 | 1 | 0 |
| senp2 | 0.27602 | 0.101305514 | 0.073 | 0.024 | 1 | 0 |
| rab14 | 0.27658 | 0.101814439 | 0.164 | 0.096 | 1 | 0 |
| arid4a | 0.2767 | 0.113474591 | 0.073 | 0.024 | 1 | 0 |
| vdac1 | 0.27738 | -0.16685399 | 0.091 | 0.157 | 1 | 0 |
| poldip2 | 0.27769 | 0.106974313 | 0.073 | 0.024 | 1 | 0 |
| tcf25 | 0.27795 | 0.165379391 | 0.109 | 0.06 | 1 | 0 |
| gars | 0.27798 | 0.104690209 | 0.091 | 0.036 | 1 | 0 |
| pttg1ipb | 0.27856 | 0.160985181 | 0.091 | 0.036 | 1 | 0 |
| si:ch211-191j22.3 | 0.2786 | 0.146090721 | 0.191 | 0.108 | 1 | 0 |
| psmg3 | 0.27895 | 0.130013535 | 0.127 | 0.06 | 1 | 0 |
| ube2ia | 0.281 | 0.17960989 | 0.191 | 0.108 | 1 | 0 |
| agpat3 | 0.28123 | 0.136560801 | 0.073 | 0.024 | 1 | 0 |
| nfyba | 0.28158 | 0.155167087 | 0.191 | 0.108 | 1 | 0 |
| rps7 | 0.2818 | 0.134516856 | 0.809 | 0.711 | 1 | 0 |
| myl9a | 0.28182 | 0.148862856 | 0.145 | 0.108 | 1 | 0 |
| gulp1a | 0.28194 | 0.169300205 | 0.191 | 0.108 | 1 | 0 |
| jpt1b | 0.28228 | -0.14476544 | 0.427 | 0.482 | 1 | 0 |

| | | | | | | |
|---|---|---|---|---|---|---|
| sssca1 | 0.28234 | 0.109608228 | 0.073 | 0.024 | 1 | 0 |
| zgc:77056 | 0.28241 | 0.118932443 | 0.091 | 0.036 | 1 | 0 |
| dynll1 | 0.28247 | -0.16872585 | 0.264 | 0.313 | 1 | 0 |
| bloc1s2 | 0.28394 | 0.106507615 | 0.073 | 0.036 | 1 | 0 |
| tmem17 | 0.28395 | -0.10072771 | 0.018 | 0.06 | 1 | 0 |
| cpsf6 | 0.28403 | -0.16955639 | 0.1 | 0.157 | 1 | 0 |
| yaf2 | 0.28409 | 0.14559338 | 0.127 | 0.06 | 1 | 0 |
| rsf1b.1 | 0.28538 | 0.151471212 | 0.1 | 0.048 | 1 | 0 |
| slc35e1 | 0.28559 | 0.122832995 | 0.064 | 0.024 | 1 | 0 |
| add3a | 0.28582 | -0.18565641 | 0.482 | 0.578 | 1 | 0 |
| atrx | 0.28637 | 0.192351473 | 0.273 | 0.193 | 1 | 0 |
| krtcap2 | 0.28637 | -0.15140704 | 0.227 | 0.325 | 1 | 0 |
| sem1 | 0.28795 | 0.182642866 | 0.345 | 0.241 | 1 | 0 |
| smad7 | 0.2881 | 0.12987428 | 0.073 | 0.024 | 1 | 0 |
| jagn1b | 0.28823 | 0.144068857 | 0.118 | 0.072 | 1 | 0 |
| fam208b | 0.28859 | -0.11308571 | 0.018 | 0.06 | 1 | 0 |
| tnfsf10l | 0.28971 | -0.20289724 | 0.291 | 0.398 | 1 | 0 |
| jupa | 0.29014 | 0.185255718 | 0.155 | 0.084 | 1 | 0 |
| pwp1 | 0.29017 | 0.132797551 | 0.091 | 0.036 | 1 | 0 |
| srpk1a | 0.29026 | -0.11530631 | 0.018 | 0.06 | 1 | 0 |
| zc4h2 | 0.29034 | 0.1288533 | 0.064 | 0.024 | 1 | 0 |
| xkr9 | 0.29043 | -0.10403529 | 0.055 | 0.108 | 1 | 0 |
| tbl1xr1b | 0.29064 | 0.129387836 | 0.073 | 0.024 | 1 | 0 |
| pdcd11 | 0.29082 | 0.121275807 | 0.073 | 0.024 | 1 | 0 |
| zfhx4 | 0.29177 | -0.11128139 | 0.018 | 0.06 | 1 | 0 |
| sox11a | 0.29187 | 0.116450634 | 0.073 | 0.024 | 1 | 0 |
| ctps1a | 0.29192 | 0.112960181 | 0.073 | 0.024 | 1 | 0 |
| hapln3 | 0.29245 | -0.17585786 | 0.127 | 0.181 | 1 | 0 |
| slc22a31 | 0.29261 | -0.1279576 | 0.018 | 0.06 | 1 | 0 |
| spred2b | 0.29265 | 0.193073614 | 0.291 | 0.217 | 1 | 0 |
| vim | 0.29291 | 0.212552758 | 0.327 | 0.229 | 1 | 0 |
| itgb1bp1 | 0.29333 | 0.140683364 | 0.064 | 0.024 | 1 | 0 |
| dicer1 | 0.29358 | -0.12265092 | 0.018 | 0.06 | 1 | 0 |
| zgc:55943 | 0.29409 | -0.1388267 | 0.045 | 0.096 | 1 | 0 |
| ppp2r1ba | 0.2947 | 0.129062736 | 0.109 | 0.048 | 1 | 0 |
| rgs3a | 0.29518 | -0.1522654 | 0.055 | 0.096 | 1 | 0 |
| krt94 | 0.29519 | -0.12088386 | 0.8 | 0.867 | 1 | 0 |
| mrps10 | 0.29522 | 0.131559097 | 0.091 | 0.036 | 1 | 0 |
| sema3fa | 0.29546 | 0.141789354 | 0.109 | 0.048 | 1 | 0 |
| arl4cb | 0.2959 | 0.133902588 | 0.091 | 0.036 | 1 | 0 |
| tcf7l1a | 0.29602 | 0.137419461 | 0.091 | 0.036 | 1 | 0 |
| gemin7 | 0.29686 | 0.134414121 | 0.091 | 0.048 | 1 | 0 |
| tomm7 | 0.29712 | 0.176700652 | 0.2 | 0.12 | 1 | 0 |
| abcc5 | 0.29725 | -0.16466245 | 0.109 | 0.157 | 1 | 0 |
| lnpep | 0.29756 | -0.12351368 | 0.018 | 0.06 | 1 | 0 |
| si:ch1073-170o4.1 | 0.29804 | -0.14889422 | 0.036 | 0.072 | 1 | 0 |
| atp2b2 | 0.29818 | -0.19666839 | 0.145 | 0.229 | 1 | 0 |
| pla2g15 | 0.29867 | -0.13761474 | 0.045 | 0.096 | 1 | 0 |

| | | | | | | |
|---|---|---|---|---|---|---|
| bhlhe40 | 0.29912 | 0.204332574 | 0.273 | 0.181 | 1 | 0 |
| ssr4 | 0.29964 | -0.11734142 | 0.355 | 0.373 | 1 | 0 |
| smg6 | 0.30003 | 0.127395681 | 0.091 | 0.036 | 1 | 0 |
| sephs2 | 0.30011 | -0.13094281 | 0.036 | 0.072 | 1 | 0 |
| si:ch211-262e15.1 | 0.30087 | 0.107034392 | 0.136 | 0.072 | 1 | 0 |
| fam49a | 0.3011 | -0.13391462 | 0.045 | 0.084 | 1 | 0 |
| ube2d2l | 0.3024 | 0.149788562 | 0.118 | 0.06 | 1 | 0 |
| rpa3 | 0.30258 | 0.127673076 | 0.073 | 0.036 | 1 | 0 |
| rab6ba | 0.30327 | -0.11501191 | 0.018 | 0.06 | 1 | 0 |
| snx7 | 0.30366 | -0.10096529 | 0.018 | 0.06 | 1 | 0 |
| ckap5 | 0.30394 | -0.11124533 | 0.018 | 0.06 | 1 | 0 |
| zgc:77086 | 0.30439 | -0.11495066 | 0.109 | 0.181 | 1 | 0 |
| si:ch211-288d18.1 | 0.30484 | -0.11762524 | 0.018 | 0.06 | 1 | 0 |
| gnpat | 0.30503 | -0.11405131 | 0.018 | 0.06 | 1 | 0 |
| cmtm6 | 0.30555 | 0.1295096 | 0.091 | 0.048 | 1 | 0 |
| top1l | 0.30578 | 0.143397641 | 0.2 | 0.12 | 1 | 0 |
| reep3a | 0.30711 | -0.11170854 | 0.064 | 0.12 | 1 | 0 |
| rasl11a | 0.30712 | -0.15877253 | 0.064 | 0.12 | 1 | 0 |
| ddx19 | 0.30746 | -0.13221917 | 0.027 | 0.072 | 1 | 0 |
| gfpt2 | 0.30794 | -0.10327229 | 0.027 | 0.072 | 1 | 0 |
| scara3 | 0.30803 | -0.17384441 | 0.373 | 0.482 | 1 | 0 |
| col5a1 | 0.30865 | -0.17660416 | 0.536 | 0.639 | 1 | 0 |
| dedd1 | 0.30866 | 0.205848626 | 0.409 | 0.313 | 1 | 0 |
| psmg2 | 0.30945 | -0.12922121 | 0.045 | 0.084 | 1 | 0 |
| ankrd12 | 0.30959 | 0.152927673 | 0.118 | 0.06 | 1 | 0 |
| tlr5b | 0.30971 | -0.11137383 | 0.091 | 0.157 | 1 | 0 |
| spag9a | 0.31035 | 0.111544346 | 0.118 | 0.06 | 1 | 0 |
| dusp6 | 0.31058 | -0.12740388 | 0.082 | 0.108 | 1 | 0 |
| nars | 0.31283 | 0.14372516 | 0.1 | 0.048 | 1 | 0 |
| drap1 | 0.31419 | 0.110318356 | 0.136 | 0.072 | 1 | 0 |
| gabarapa | 0.31423 | -0.10547527 | 0.2 | 0.217 | 1 | 0 |
| rpl10a | 0.31434 | 0.13981307 | 0.836 | 0.759 | 1 | 0 |
| osr2 | 0.31496 | 0.194698592 | 0.155 | 0.096 | 1 | 0 |
| ccbe1 | 0.31532 | 0.165781159 | 0.136 | 0.072 | 1 | 0 |
| si:rp71-45k5.4 | 0.31557 | 0.158425606 | 0.164 | 0.096 | 1 | 0 |
| ptp4a1 | 0.3159 | 0.140923005 | 0.2 | 0.12 | 1 | 0 |
| zdhhc8b | 0.31693 | 0.148551537 | 0.082 | 0.036 | 1 | 0 |
| wdr43 | 0.3175 | 0.165746288 | 0.155 | 0.084 | 1 | 0 |
| ret | 0.31882 | -0.15898292 | 0.055 | 0.108 | 1 | 0 |
| CU929259.1 | 0.32002 | 0.124411221 | 0.064 | 0.036 | 1 | 0 |
| fbln1 | 0.32049 | 0.152519176 | 0.091 | 0.048 | 1 | 0 |
| itsn1 | 0.32069 | 0.112299838 | 0.055 | 0.024 | 1 | 0 |
| cox6b2 | 0.32114 | 0.151450761 | 0.591 | 0.482 | 1 | 0 |
| ppm1la | 0.32148 | -0.12730859 | 0.036 | 0.084 | 1 | 0 |
| api5 | 0.32253 | 0.100645395 | 0.073 | 0.036 | 1 | 0 |
| creb3l2 | 0.32307 | 0.171325274 | 0.155 | 0.084 | 1 | 0 |
| twist3 | 0.32428 | -0.15462781 | 0.036 | 0.084 | 1 | 0 |
| smndc1 | 0.32429 | -0.17217953 | 0.145 | 0.229 | 1 | 0 |

| | | | | | | |
|---|---|---|---|---|---|---|
| mibp2 | 0.32436 | -0.18397498 | 0.382 | 0.458 | 1 | 0 |
| wscd2 | 0.32476 | 0.161344061 | 0.182 | 0.133 | 1 | 0 |
| gar1 | 0.32509 | 0.109043669 | 0.082 | 0.036 | 1 | 0 |
| rrp15 | 0.32536 | 0.129963671 | 0.082 | 0.036 | 1 | 0 |
| si:dkeyp-69b9.6 | 0.32599 | 0.149043122 | 0.082 | 0.036 | 1 | 0 |
| slc25a5 | 0.3263 | -0.10288535 | 0.8 | 0.88 | 1 | 0 |
| gtf2f2a | 0.32641 | 0.146343268 | 0.127 | 0.072 | 1 | 0 |
| tktb | 0.32661 | -0.10271556 | 0.045 | 0.06 | 1 | 0 |
| si:dkey-163m14.2 | 0.32677 | 0.120026894 | 0.118 | 0.06 | 1 | 0 |
| pno1 | 0.32682 | 0.148546748 | 0.164 | 0.108 | 1 | 0 |
| cemip | 0.32692 | 0.105844859 | 0.082 | 0.036 | 1 | 0 |
| kmt2ca | 0.32927 | -0.1007028 | 0.027 | 0.072 | 1 | 0 |
| ccdc25 | 0.32981 | 0.140389444 | 0.155 | 0.084 | 1 | 0 |
| nap1l1 | 0.33058 | -0.17172644 | 0.218 | 0.313 | 1 | 0 |
| eif1axb | 0.331 | -0.13326175 | 0.273 | 0.313 | 1 | 0 |
| fbxl7 | 0.33136 | 0.146355769 | 0.1 | 0.048 | 1 | 0 |
| dnajc1 | 0.33224 | 0.134611214 | 0.091 | 0.06 | 1 | 0 |
| LOC566587 | 0.33229 | -0.11827966 | 0.036 | 0.084 | 1 | 0 |
| anxa1a | 0.33269 | 0.176488558 | 0.618 | 0.518 | 1 | 0 |
| mboat2b | 0.33277 | 0.124602257 | 0.064 | 0.024 | 1 | 0 |
| parn | 0.33426 | 0.1157529 | 0.064 | 0.024 | 1 | 0 |
| dlg3 | 0.33475 | -0.12302565 | 0.027 | 0.072 | 1 | 0 |
| pfdn1 | 0.33599 | 0.165342693 | 0.255 | 0.169 | 1 | 0 |
| ext2 | 0.33612 | 0.140916312 | 0.1 | 0.048 | 1 | 0 |
| fblim1 | 0.33638 | 0.16949342 | 0.182 | 0.12 | 1 | 0 |
| LOC101886030 | 0.3372 | 0.133878159 | 0.073 | 0.036 | 1 | 0 |
| cdc42bpb | 0.33728 | -0.15029444 | 0.064 | 0.12 | 1 | 0 |
| cipcb | 0.33828 | -0.16079879 | 0.045 | 0.096 | 1 | 0 |
| si:ch73-46j18.5 | 0.33865 | 0.14452213 | 0.709 | 0.627 | 1 | 0 |
| sfxn5b | 0.3395 | -0.10644374 | 0.027 | 0.072 | 1 | 0 |
| brms1la | 0.34019 | 0.103364713 | 0.1 | 0.048 | 1 | 0 |
| taf13 | 0.34132 | 0.15513765 | 0.309 | 0.217 | 1 | 0 |
| ephb4a | 0.34268 | -0.12549797 | 0.045 | 0.072 | 1 | 0 |
| bxdc2 | 0.34303 | 0.159665607 | 0.118 | 0.06 | 1 | 0 |
| ppm1aa | 0.34366 | 0.158492413 | 0.136 | 0.072 | 1 | 0 |
| zgc:63587 | 0.34375 | -0.10427329 | 0.027 | 0.072 | 1 | 0 |
| ndfip1 | 0.34388 | 0.163992792 | 0.182 | 0.12 | 1 | 0 |
| slco2b1 | 0.34392 | -0.14997301 | 0.127 | 0.169 | 1 | 0 |
| slc12a4 | 0.34394 | -0.11803505 | 0.027 | 0.072 | 1 | 0 |
| tefb | 0.34414 | -0.12079915 | 0.027 | 0.072 | 1 | 0 |
| tp53 | 0.34416 | -0.13971481 | 0.055 | 0.108 | 1 | 0 |
| ak4 | 0.3444 | -0.10585187 | 0.027 | 0.072 | 1 | 0 |
| ankrd10b | 0.34468 | -0.1120019 | 0.027 | 0.072 | 1 | 0 |
| fam120c | 0.34469 | 0.158504912 | 0.164 | 0.096 | 1 | 0 |
| rpl35a | 0.34482 | 0.127846434 | 0.818 | 0.771 | 1 | 0 |
| pdcd5 | 0.34522 | 0.143055564 | 0.155 | 0.096 | 1 | 0 |
| sgce | 0.34629 | 0.126975862 | 0.091 | 0.048 | 1 | 0 |
| usf2 | 0.3474 | -0.13139321 | 0.027 | 0.06 | 1 | 0 |

| | | | | | | |
|---|---|---|---|---|---|---|
| csnk1db | 0.34925 | 0.154755085 | 0.164 | 0.096 | 1 | 0 |
| PRRC2B | 0.34954 | -0.1106193 | 0.055 | 0.108 | 1 | 0 |
| dio1 | 0.34966 | -0.13494549 | 0.036 | 0.084 | 1 | 0 |
| col6a2 | 0.34981 | -0.12137626 | 0.364 | 0.398 | 1 | 0 |
| xbp1 | 0.35018 | -0.14439882 | 0.391 | 0.494 | 1 | 0 |
| rnd3a | 0.3503 | -0.16769654 | 0.127 | 0.205 | 1 | 0 |
| nupr1b | 0.35077 | 0.127534465 | 0.127 | 0.084 | 1 | 0 |
| mgp | 0.3508 | -0.16699501 | 0.218 | 0.265 | 1 | 0 |
| tbl1xr1a | 0.35144 | -0.11764271 | 0.064 | 0.12 | 1 | 0 |
| adamts15a | 0.35164 | 0.19349439 | 0.364 | 0.277 | 1 | 0 |
| eda | 0.3518 | -0.16089742 | 0.164 | 0.217 | 1 | 0 |
| cnbpa | 0.35207 | 0.144569872 | 0.164 | 0.096 | 1 | 0 |
| rbbp6 | 0.35265 | 0.15624991 | 0.182 | 0.108 | 1 | 0 |
| si:ch211-222k6.3 | 0.35287 | 0.149964073 | 0.136 | 0.072 | 1 | 0 |
| aadac | 0.35324 | 0.147742544 | 0.136 | 0.072 | 1 | 0 |
| vps37b | 0.35328 | 0.11198281 | 0.055 | 0.024 | 1 | 0 |
| dars | 0.35458 | -0.10400775 | 0.045 | 0.096 | 1 | 0 |
| csad | 0.35474 | -0.10812788 | 0.027 | 0.06 | 1 | 0 |
| tspan4a | 0.35509 | 0.144098761 | 0.145 | 0.084 | 1 | 0 |
| coa5 | 0.35521 | -0.13777012 | 0.055 | 0.096 | 1 | 0 |
| slc20a2 | 0.35527 | 0.104443602 | 0.1 | 0.048 | 1 | 0 |
| ndufaf5 | 0.35534 | 0.116567953 | 0.082 | 0.036 | 1 | 0 |
| zgc:110239 | 0.35582 | -0.15295893 | 0.1 | 0.169 | 1 | 0 |
| rps27a | 0.35701 | 0.101466158 | 0.882 | 0.855 | 1 | 0 |
| LOC100536647 | 0.35794 | -0.14456131 | 0.064 | 0.108 | 1 | 0 |
| mrpl17 | 0.35803 | 0.116525088 | 0.082 | 0.036 | 1 | 0 |
| rab2a | 0.35807 | 0.162091061 | 0.218 | 0.157 | 1 | 0 |
| slit2 | 0.35856 | 0.168552465 | 0.182 | 0.108 | 1 | 0 |
| bicc1b | 0.35886 | -0.17792371 | 0.118 | 0.193 | 1 | 0 |
| tmub2 | 0.35959 | -0.10871819 | 0.036 | 0.084 | 1 | 0 |
| ucp2 | 0.35959 | -0.12687265 | 0.173 | 0.253 | 1 | 0 |
| mustn1a | 0.35981 | -0.19031128 | 0.227 | 0.313 | 1 | 0 |
| traf7 | 0.36021 | 0.141429313 | 0.082 | 0.036 | 1 | 0 |
| exoc6 | 0.36045 | 0.147746801 | 0.082 | 0.036 | 1 | 0 |
| polr2k | 0.36053 | 0.123258038 | 0.118 | 0.06 | 1 | 0 |
| mtbl | 0.3611 | -0.13064087 | 0.036 | 0.084 | 1 | 0 |
| nufip2 | 0.36166 | -0.14469105 | 0.055 | 0.108 | 1 | 0 |
| slc37a2 | 0.36216 | -0.11148096 | 0.036 | 0.084 | 1 | 0 |
| ptpra | 0.3633 | -0.10245552 | 0.045 | 0.096 | 1 | 0 |
| eif5a | 0.36342 | 0.16776608 | 0.518 | 0.434 | 1 | 0 |
| clic4 | 0.3635 | -0.1508034 | 0.073 | 0.133 | 1 | 0 |
| sertad2b | 0.36355 | -0.11487752 | 0.036 | 0.084 | 1 | 0 |
| selenot1b | 0.36396 | -0.13707868 | 0.091 | 0.133 | 1 | 0 |
| frg1 | 0.36441 | -0.11806567 | 0.036 | 0.084 | 1 | 0 |
| rbm39a | 0.36508 | -0.10974075 | 0.255 | 0.277 | 1 | 0 |
| sav1 | 0.36563 | -0.10540568 | 0.064 | 0.12 | 1 | 0 |
| psma1 | 0.3659 | -0.16279215 | 0.245 | 0.313 | 1 | 0 |
| cyb5r3 | 0.36603 | -0.13639553 | 0.064 | 0.12 | 1 | 0 |

| | | | | | | |
|---|---|---|---|---|---|---|
| tgfb2 | 0.3664 | -0.11616324 | 0.036 | 0.084 | 1 | 0 |
| mbtps2 | 0.36663 | -0.10631488 | 0.036 | 0.084 | 1 | 0 |
| plod3 | 0.36691 | -0.14760617 | 0.082 | 0.145 | 1 | 0 |
| slc4a10a | 0.36727 | -0.13316839 | 0.036 | 0.084 | 1 | 0 |
| rap1b | 0.36728 | -0.18576456 | 0.209 | 0.289 | 1 | 0 |
| letm1 | 0.36789 | -0.10289941 | 0.036 | 0.084 | 1 | 0 |
| os9 | 0.36792 | -0.10244956 | 0.036 | 0.084 | 1 | 0 |
| mid1ip1l | 0.36804 | -0.12733077 | 0.036 | 0.084 | 1 | 0 |
| phyhiplb | 0.3681 | -0.13597054 | 0.036 | 0.084 | 1 | 0 |
| sept6 | 0.3681 | -0.10441942 | 0.036 | 0.084 | 1 | 0 |
| arpc3 | 0.36818 | -0.14626844 | 0.155 | 0.205 | 1 | 0 |
| rras2 | 0.3683 | 0.132106229 | 0.118 | 0.084 | 1 | 0 |
| sh3pxd2b | 0.36841 | -0.15587762 | 0.109 | 0.181 | 1 | 0 |
| lamc1 | 0.36877 | -0.16962374 | 0.109 | 0.181 | 1 | 0 |
| seh1l | 0.36882 | 0.118936389 | 0.1 | 0.06 | 1 | 0 |
| LO018188.1 | 0.3691 | -0.14196784 | 0.218 | 0.301 | 1 | 0 |
| ripor3 | 0.36926 | 0.135214679 | 0.118 | 0.06 | 1 | 0 |
| ing4 | 0.36945 | 0.125128887 | 0.1 | 0.048 | 1 | 0 |
| commd3 | 0.37015 | 0.118860133 | 0.091 | 0.048 | 1 | 0 |
| sulf2b | 0.37109 | 0.159935878 | 0.109 | 0.06 | 1 | 0 |
| dlc1 | 0.37113 | -0.15037146 | 0.091 | 0.157 | 1 | 0 |
| c1qtnf2 | 0.3715 | 0.166669876 | 0.209 | 0.133 | 1 | 0 |
| XLOC-042917 | 0.37156 | 0.14548475 | 0.082 | 0.036 | 1 | 0 |
| rpl22 | 0.37161 | 0.139259908 | 0.764 | 0.699 | 1 | 0 |
| ncoa4 | 0.37184 | -0.14748787 | 0.073 | 0.12 | 1 | 0 |
| rasgef1bb | 0.37185 | -0.12306439 | 0.055 | 0.108 | 1 | 0 |
| eif3jb | 0.37213 | -0.13894114 | 0.155 | 0.193 | 1 | 0 |
| hsd20b2 | 0.37257 | 0.139151009 | 0.109 | 0.072 | 1 | 0 |
| tspan31 | 0.37401 | 0.113005251 | 0.118 | 0.06 | 1 | 0 |
| zgc:162944 | 0.37476 | -0.12295744 | 0.164 | 0.193 | 1 | 0 |
| cbx1a | 0.37527 | 0.121158231 | 0.118 | 0.06 | 1 | 0 |
| srsf6a | 0.37543 | -0.1412878 | 0.082 | 0.145 | 1 | 0 |
| prdx6 | 0.37618 | 0.133005564 | 0.236 | 0.157 | 1 | 0 |
| chchd6b | 0.37638 | -0.11034066 | 0.045 | 0.096 | 1 | 0 |
| ssu72 | 0.37787 | 0.101960439 | 0.073 | 0.048 | 1 | 0 |
| pcdh11 | 0.3781 | -0.14576381 | 0.045 | 0.096 | 1 | 0 |
| zfp36l1b | 0.37921 | 0.184596267 | 0.4 | 0.313 | 1 | 0 |
| nkap | 0.38007 | -0.12581626 | 0.055 | 0.108 | 1 | 0 |
| snw1 | 0.38013 | 0.119522838 | 0.109 | 0.06 | 1 | 0 |
| ckap4 | 0.38018 | -0.18151617 | 0.218 | 0.301 | 1 | 0 |
| prrc2c | 0.3802 | 0.104604752 | 0.064 | 0.024 | 1 | 0 |
| tent4b | 0.38024 | -0.12561979 | 0.045 | 0.096 | 1 | 0 |
| si:ch211-108d22.2 | 0.38088 | -0.14617076 | 0.064 | 0.12 | 1 | 0 |
| rpl8 | 0.381 | 0.113786069 | 0.764 | 0.675 | 1 | 0 |
| rpl31 | 0.38145 | 0.114728267 | 0.827 | 0.771 | 1 | 0 |
| ddx18 | 0.38158 | -0.10508195 | 0.045 | 0.096 | 1 | 0 |
| cyp2n13 | 0.3818 | 0.189499214 | 0.255 | 0.181 | 1 | 0 |
| fgfr2 | 0.3843 | -0.19142642 | 0.227 | 0.313 | 1 | 0 |

| | | | | | | |
|---|---|---|---|---|---|---|
| rbm34 | 0.38432 | 0.105707119 | 0.082 | 0.036 | 1 | 0 |
| ppp1cb | 0.38438 | -0.15052169 | 0.064 | 0.108 | 1 | 0 |
| nono | 0.38446 | -0.10924939 | 0.082 | 0.108 | 1 | 0 |
| LOC103910871 | 0.38458 | 0.118271897 | 0.082 | 0.036 | 1 | 0 |
| eno1a | 0.38459 | -0.10474187 | 0.109 | 0.133 | 1 | 0 |
| mrpl53 | 0.38462 | 0.115515614 | 0.082 | 0.036 | 1 | 0 |
| amotl2a | 0.38475 | -0.14567683 | 0.055 | 0.108 | 1 | 0 |
| rybpb | 0.38499 | 0.115351171 | 0.1 | 0.048 | 1 | 0 |
| snai1a | 0.38541 | -0.1405086 | 0.055 | 0.108 | 1 | 0 |
| h3f3d | 0.38576 | 0.132811827 | 0.664 | 0.566 | 1 | 0 |
| rasa1a | 0.38628 | 0.133694719 | 0.073 | 0.036 | 1 | 0 |
| si:ch211-133n4.4 | 0.38645 | 0.106409029 | 0.327 | 0.253 | 1 | 0 |
| foxo3b | 0.38699 | -0.10226294 | 0.127 | 0.145 | 1 | 0 |
| ino80c | 0.38787 | -0.13025205 | 0.064 | 0.108 | 1 | 0 |
| lonrf1l | 0.38797 | 0.113864731 | 0.064 | 0.024 | 1 | 0 |
| rps18 | 0.38798 | 0.118707643 | 0.836 | 0.783 | 1 | 0 |
| pitpnbl | 0.38806 | -0.11110002 | 0.055 | 0.108 | 1 | 0 |
| stxbp6 | 0.38824 | -0.1303476 | 0.045 | 0.084 | 1 | 0 |
| ebna1bp2 | 0.38851 | -0.10588859 | 0.055 | 0.108 | 1 | 0 |
| syncripl | 0.38884 | 0.13856216 | 0.164 | 0.096 | 1 | 0 |
| bcl2l10 | 0.389 | -0.12404519 | 0.064 | 0.12 | 1 | 0 |
| calhm2 | 0.38932 | 0.13316358 | 0.091 | 0.048 | 1 | 0 |
| rtn3 | 0.38966 | -0.14838978 | 0.073 | 0.133 | 1 | 0 |
| hmgb2b | 0.38997 | 0.10022417 | 0.436 | 0.349 | 1 | 0 |
| mgst1.1 | 0.39018 | 0.161955962 | 0.264 | 0.181 | 1 | 0 |
| evi5b | 0.39021 | -0.1057134 | 0.036 | 0.06 | 1 | 0 |
| ldlrad4a | 0.39024 | -0.14201251 | 0.073 | 0.133 | 1 | 0 |
| seta | 0.39061 | 0.149604372 | 0.291 | 0.205 | 1 | 0 |
| FP102018.1 | 0.39092 | -0.13502982 | 0.064 | 0.12 | 1 | 0 |
| gtf2f1 | 0.39099 | -0.14043325 | 0.064 | 0.12 | 1 | 0 |
| rps5 | 0.39147 | 0.14643464 | 0.736 | 0.663 | 1 | 0 |
| rae1 | 0.39462 | 0.10622082 | 0.064 | 0.024 | 1 | 0 |
| pbxip1b | 0.39488 | 0.104280328 | 0.055 | 0.024 | 1 | 0 |
| ctbp1 | 0.39499 | 0.146021042 | 0.273 | 0.229 | 1 | 0 |
| tmem165 | 0.39518 | 0.100261953 | 0.082 | 0.036 | 1 | 0 |
| lix1l | 0.39557 | 0.128216741 | 0.1 | 0.048 | 1 | 0 |
| hnrnpul1 | 0.39589 | 0.101399173 | 0.082 | 0.036 | 1 | 0 |
| rpl17 | 0.39722 | 0.102238172 | 0.818 | 0.783 | 1 | 0 |
| ldb2b | 0.39733 | -0.12200755 | 0.045 | 0.072 | 1 | 0 |
| ctdspl2b | 0.39786 | -0.12987238 | 0.036 | 0.072 | 1 | 0 |
| gabpa | 0.39798 | 0.116904142 | 0.073 | 0.036 | 1 | 0 |
| hsdl2 | 0.39827 | 0.108808343 | 0.073 | 0.036 | 1 | 0 |
| eif2b4 | 0.3987 | -0.11240533 | 0.036 | 0.072 | 1 | 0 |
| sumo3a | 0.39886 | 0.129036628 | 0.391 | 0.301 | 1 | 0 |
| sap30bp | 0.39949 | 0.111226461 | 0.082 | 0.036 | 1 | 0 |
| sypl2b | 0.39978 | -0.12146408 | 0.073 | 0.108 | 1 | 0 |
| metrnla | 0.39978 | -0.16582194 | 0.109 | 0.169 | 1 | 0 |
| anp32b | 0.39998 | -0.12629224 | 0.118 | 0.169 | 1 | 0 |

| | | | | | | |
|---|---|---|---|---|---|---|
| fkbp7 | 0.40025 | 0.165817466 | 0.236 | 0.169 | 1 | 0 |
| boc | 0.40075 | 0.161260706 | 0.382 | 0.289 | 1 | 0 |
| rbm14a | 0.40437 | 0.104761161 | 0.082 | 0.048 | 1 | 0 |
| rasgef1ba | 0.40498 | 0.127997525 | 0.073 | 0.036 | 1 | 0 |
| tnfsf12 | 0.40507 | 0.131651166 | 0.091 | 0.048 | 1 | 0 |
| rad23b | 0.40613 | -0.10504034 | 0.145 | 0.169 | 1 | 0 |
| fkbp9 | 0.40615 | -0.11444855 | 0.127 | 0.193 | 1 | 0 |
| baiap2a | 0.40652 | 0.10587916 | 0.082 | 0.036 | 1 | 0 |
| yy1b | 0.40653 | 0.10932592 | 0.082 | 0.036 | 1 | 0 |
| smdt1b | 0.40678 | 0.141316077 | 0.218 | 0.169 | 1 | 0 |
| taf10 | 0.4075 | 0.105576547 | 0.082 | 0.036 | 1 | 0 |
| si:ch211-196l7.4 | 0.40778 | -0.17032712 | 0.373 | 0.458 | 1 | 0 |
| amd1 | 0.40779 | 0.100458728 | 0.064 | 0.024 | 1 | 0 |
| fxyd1 | 0.40838 | -0.13294461 | 0.091 | 0.133 | 1 | 0 |
| lsm4 | 0.40841 | 0.145375982 | 0.245 | 0.193 | 1 | 0 |
| si:ch211-198a12.6 | 0.40944 | 0.125746796 | 0.082 | 0.036 | 1 | 0 |
| zranb2 | 0.40999 | 0.134707608 | 0.091 | 0.048 | 1 | 0 |
| mast2 | 0.4103 | 0.150960545 | 0.145 | 0.084 | 1 | 0 |
| srsf5a | 0.41031 | 0.165016831 | 0.355 | 0.265 | 1 | 0 |
| nipblb | 0.41183 | 0.104406952 | 0.064 | 0.024 | 1 | 0 |
| clu | 0.41388 | 0.188622621 | 0.3 | 0.217 | 1 | 0 |
| erbb2 | 0.41422 | 0.121456591 | 0.073 | 0.036 | 1 | 0 |
| NDUFB1 | 0.41443 | 0.131799512 | 0.245 | 0.169 | 1 | 0 |
| si:dkey-67c22.2 | 0.4145 | 0.116985988 | 0.155 | 0.12 | 1 | 0 |
| glipr2l | 0.41528 | 0.162360543 | 0.173 | 0.12 | 1 | 0 |
| ormdl2 | 0.41534 | -0.10775599 | 0.027 | 0.06 | 1 | 0 |
| hipk3b | 0.41551 | 0.133194032 | 0.082 | 0.048 | 1 | 0 |
| rtn4b | 0.41699 | 0.107962669 | 0.082 | 0.048 | 1 | 0 |
| cbx1b | 0.41828 | 0.132201039 | 0.127 | 0.072 | 1 | 0 |
| rnd3b | 0.41978 | -0.16983892 | 0.2 | 0.277 | 1 | 0 |
| scaf4b | 0.42015 | -0.10426931 | 0.027 | 0.06 | 1 | 0 |
| cnn3a | 0.42146 | 0.145378683 | 0.273 | 0.193 | 1 | 0 |
| col5a2a | 0.42182 | 0.137900142 | 0.455 | 0.398 | 1 | 0 |
| hic1 | 0.42192 | 0.178611203 | 0.327 | 0.253 | 1 | 0 |
| ogt.1 | 0.42213 | -0.15564841 | 0.1 | 0.157 | 1 | 0 |
| srsf2a | 0.42291 | -0.13434889 | 0.2 | 0.241 | 1 | 0 |
| nkain4 | 0.42346 | -0.11147337 | 0.027 | 0.06 | 1 | 0 |
| arpc4 | 0.42378 | -0.14144561 | 0.164 | 0.217 | 1 | 0 |
| ndfip2 | 0.4241 | -0.12023795 | 0.091 | 0.12 | 1 | 0 |
| tmem258 | 0.42544 | -0.13054471 | 0.318 | 0.361 | 1 | 0 |
| sestd1 | 0.42639 | 0.10152438 | 0.055 | 0.036 | 1 | 0 |
| rps19bp1 | 0.42655 | 0.107897817 | 0.091 | 0.048 | 1 | 0 |
| rab5if | 0.42751 | 0.121021085 | 0.127 | 0.072 | 1 | 0 |
| thbs2a | 0.42825 | 0.132481125 | 0.109 | 0.06 | 1 | 0 |
| vps29 | 0.42917 | 0.127246158 | 0.127 | 0.072 | 1 | 0 |
| ppp2r2d | 0.43219 | -0.12700048 | 0.064 | 0.108 | 1 | 0 |
| pim1 | 0.43254 | -0.15388557 | 0.164 | 0.217 | 1 | 0 |
| rps21 | 0.43298 | 0.115076428 | 0.827 | 0.759 | 1 | 0 |

| | | | | | | |
|---|---|---|---|---|---|---|
| tubb5 | 0.433 | 0.12948295 | 0.118 | 0.084 | 1 | 0 |
| mrps36 | 0.4335 | 0.112430375 | 0.127 | 0.072 | 1 | 0 |
| ppp3ccb | 0.43414 | -0.14498421 | 0.109 | 0.157 | 1 | 0 |
| eif3i | 0.43421 | 0.134181771 | 0.264 | 0.217 | 1 | 0 |
| mapk9 | 0.43467 | -0.11285594 | 0.027 | 0.06 | 1 | 0 |
| skia | 0.43935 | -0.10221579 | 0.1 | 0.157 | 1 | 0 |
| lhfpl2a | 0.43993 | -0.11813458 | 0.064 | 0.096 | 1 | 0 |
| slc30a5 | 0.44095 | -0.10173031 | 0.036 | 0.06 | 1 | 0 |
| dynlt3 | 0.44276 | -0.1441537 | 0.109 | 0.157 | 1 | 0 |
| notch3 | 0.44347 | 0.109836397 | 0.109 | 0.06 | 1 | 0 |
| cst14b.1 | 0.44369 | 0.13507281 | 0.173 | 0.108 | 1 | 0 |
| twist1b | 0.444 | 0.139375577 | 0.109 | 0.06 | 1 | 0 |
| metap1 | 0.4453 | 0.103197034 | 0.064 | 0.036 | 1 | 0 |
| flot2a | 0.4453 | 0.12749024 | 0.173 | 0.108 | 1 | 0 |
| rpl27 | 0.44649 | 0.122948605 | 0.791 | 0.723 | 1 | 0 |
| zgc:114045 | 0.44653 | -0.10617167 | 0.036 | 0.06 | 1 | 0 |
| olfml3a | 0.44692 | -0.144926 | 0.091 | 0.145 | 1 | 0 |
| dusp7 | 0.44758 | -0.12392229 | 0.045 | 0.084 | 1 | 0 |
| trib3 | 0.44822 | -0.14431034 | 0.091 | 0.145 | 1 | 0 |
| pigx | 0.44829 | 0.116917653 | 0.091 | 0.048 | 1 | 0 |
| dio2 | 0.44845 | 0.115043524 | 0.127 | 0.072 | 1 | 0 |
| neo1a | 0.44856 | 0.111084047 | 0.127 | 0.096 | 1 | 0 |
| rab41 | 0.44914 | -0.10173878 | 0.036 | 0.072 | 1 | 0 |
| hmgn2 | 0.44982 | -0.14969502 | 0.391 | 0.482 | 1 | 0 |
| zfand5b | 0.44983 | -0.13757606 | 0.1 | 0.157 | 1 | 0 |
| tet3 | 0.45093 | -0.11857402 | 0.036 | 0.072 | 1 | 0 |
| dgcr2 | 0.45118 | 0.119852879 | 0.082 | 0.048 | 1 | 0 |
| smarca2 | 0.45135 | 0.11109505 | 0.127 | 0.072 | 1 | 0 |
| hnrpkl | 0.45204 | -0.12781815 | 0.064 | 0.108 | 1 | 0 |
| ndufs4 | 0.453 | 0.100303359 | 0.127 | 0.108 | 1 | 0 |
| selenbp1 | 0.45428 | 0.142463414 | 0.2 | 0.145 | 1 | 0 |
| gigyf1a | 0.45442 | 0.111128944 | 0.055 | 0.024 | 1 | 0 |
| trappc1 | 0.45505 | 0.126324427 | 0.173 | 0.133 | 1 | 0 |
| ctsba | 0.45595 | -0.12862742 | 0.191 | 0.265 | 1 | 0 |
| dcn | 0.45638 | 0.119298611 | 0.736 | 0.675 | 1 | 0 |
| zfand5a | 0.45704 | -0.13405732 | 0.1 | 0.157 | 1 | 0 |
| yif1a | 0.45773 | 0.115419156 | 0.109 | 0.06 | 1 | 0 |
| akt1s1 | 0.45864 | -0.12594317 | 0.064 | 0.108 | 1 | 0 |
| lhfpl6 | 0.45872 | -0.1371023 | 0.127 | 0.193 | 1 | 0 |
| mkrn1 | 0.45921 | 0.114572511 | 0.109 | 0.06 | 1 | 0 |
| fnbp1l | 0.45982 | 0.123246049 | 0.109 | 0.06 | 1 | 0 |
| rhoua | 0.46138 | 0.119029861 | 0.109 | 0.06 | 1 | 0 |
| pfdn6 | 0.46297 | 0.106631956 | 0.127 | 0.096 | 1 | 0 |
| ppp4cb | 0.46343 | -0.11595353 | 0.073 | 0.108 | 1 | 0 |
| si:ch211-214j24.10 | 0.46426 | 0.140457514 | 0.155 | 0.096 | 1 | 0 |
| lrrn2 | 0.46483 | -0.13567366 | 0.045 | 0.084 | 1 | 0 |
| rnf44 | 0.46493 | -0.13798487 | 0.127 | 0.193 | 1 | 0 |
| ndufb5 | 0.4653 | -0.11008137 | 0.127 | 0.193 | 1 | 0 |

| | | | | | | |
|---|---|---|---|---|---|---|
| fam208ab | 0.46565 | 0.1012035 | 0.055 | 0.024 | 1 | 0 |
| hnrnpc | 0.46652 | -0.10351794 | 0.073 | 0.096 | 1 | 0 |
| cstf2 | 0.4668 | -0.10574942 | 0.036 | 0.072 | 1 | 0 |
| galns | 0.46756 | -0.10912876 | 0.027 | 0.06 | 1 | 0 |
| ash1l | 0.46757 | -0.12127068 | 0.064 | 0.108 | 1 | 0 |
| tmed9 | 0.46865 | 0.110915549 | 0.145 | 0.108 | 1 | 0 |
| tia1l | 0.46866 | -0.1105483 | 0.055 | 0.096 | 1 | 0 |
| txndc15 | 0.47033 | 0.128454559 | 0.127 | 0.084 | 1 | 0 |
| dhx8 | 0.47215 | -0.1069908 | 0.027 | 0.06 | 1 | 0 |
| ctdsp2 | 0.47262 | 0.125885176 | 0.255 | 0.217 | 1 | 0 |
| mycbp2 | 0.47326 | 0.118810169 | 0.073 | 0.036 | 1 | 0 |
| btf3l4 | 0.47414 | 0.134675816 | 0.155 | 0.096 | 1 | 0 |
| sdcbp2 | 0.47433 | -0.12340621 | 0.382 | 0.422 | 1 | 0 |
| arrdc3a | 0.47449 | -0.10040589 | 0.155 | 0.217 | 1 | 0 |
| si:dkey-33c12.4 | 0.47488 | 0.13077597 | 0.2 | 0.157 | 1 | 0 |
| calr3b | 0.47537 | -0.14103294 | 0.1 | 0.157 | 1 | 0 |
| rab8b | 0.47878 | 0.114223068 | 0.1 | 0.06 | 1 | 0 |
| fzd7b | 0.47917 | 0.107616427 | 0.309 | 0.241 | 1 | 0 |
| zfhx3 | 0.4801 | -0.10253497 | 0.091 | 0.145 | 1 | 0 |
| hp1bp3 | 0.4801 | 0.123041421 | 0.155 | 0.096 | 1 | 0 |
| rpl7a | 0.48123 | 0.100134483 | 0.845 | 0.795 | 1 | 0 |
| selenof | 0.48152 | 0.12311021 | 0.318 | 0.241 | 1 | 0 |
| ano10a | 0.4829 | 0.104886642 | 0.073 | 0.036 | 1 | 0 |
| osbpl10b | 0.48427 | -0.10104923 | 0.027 | 0.06 | 1 | 0 |
| gtf2a2 | 0.48513 | 0.109006535 | 0.091 | 0.06 | 1 | 0 |
| cep350 | 0.48528 | 0.101308889 | 0.055 | 0.024 | 1 | 0 |
| tent5c | 0.48552 | -0.14169483 | 0.109 | 0.169 | 1 | 0 |
| c4 | 0.48634 | -0.12718295 | 0.082 | 0.12 | 1 | 0 |
| emid1 | 0.4889 | -0.16128301 | 0.355 | 0.422 | 1 | 0 |
| prdx2 | 0.48953 | -0.14023409 | 0.236 | 0.313 | 1 | 0 |
| psmd1 | 0.49064 | -0.11669883 | 0.064 | 0.108 | 1 | 0 |
| pdgfra | 0.49065 | -0.11900831 | 0.3 | 0.337 | 1 | 0 |
| ndufs3 | 0.49107 | 0.104681446 | 0.136 | 0.108 | 1 | 0 |
| lmbr1l | 0.49231 | -0.11663212 | 0.055 | 0.096 | 1 | 0 |
| si:dkey-188i13.11 | 0.49406 | 0.136896026 | 0.182 | 0.133 | 1 | 0 |
| gnav1 | 0.49486 | -0.11098668 | 0.036 | 0.072 | 1 | 0 |
| zgc:110158 | 0.49597 | -0.13927627 | 0.073 | 0.12 | 1 | 0 |
| nell2a | 0.49948 | -0.13761516 | 0.073 | 0.108 | 1 | 0 |
| tob1b | 0.50016 | -0.10914443 | 0.155 | 0.181 | 1 | 0 |
| reck | 0.50026 | -0.11925612 | 0.1 | 0.157 | 1 | 0 |
| tuba8l3 | 0.5003 | 0.132172404 | 0.382 | 0.301 | 1 | 0 |
| tspan35 | 0.50097 | -0.10346666 | 0.064 | 0.108 | 1 | 0 |
| clint1a | 0.50105 | -0.12759803 | 0.1 | 0.157 | 1 | 0 |
| serf2 | 0.50126 | 0.134670615 | 0.564 | 0.482 | 1 | 0 |
| xrn2 | 0.50131 | 0.110589666 | 0.091 | 0.048 | 1 | 0 |
| XLOC-041269 | 0.5023 | -0.1331843 | 0.1 | 0.157 | 1 | 0 |
| supt6h | 0.50298 | 0.12431115 | 0.118 | 0.072 | 1 | 0 |
| ywhaba | 0.50298 | -0.13962654 | 0.227 | 0.277 | 1 | 0 |

| | | | | | | |
|---|---|---|---|---|---|---|
| m6pr | 0.50397 | -0.11404233 | 0.045 | 0.084 | 1 | 0 |
| ostc | 0.50402 | 0.138094569 | 0.209 | 0.157 | 1 | 0 |
| smarcc2 | 0.50675 | -0.10611796 | 0.036 | 0.072 | 1 | 0 |
| tmsb2 | 0.50713 | -0.10698024 | 0.027 | 0.06 | 1 | 0 |
| inip | 0.50797 | 0.104451759 | 0.091 | 0.048 | 1 | 0 |
| PAXBP1 | 0.50806 | 0.11330872 | 0.091 | 0.048 | 1 | 0 |
| polr2f | 0.50816 | 0.114669012 | 0.155 | 0.12 | 1 | 0 |
| si:ch211-160o17.4 | 0.50822 | 0.117348281 | 0.091 | 0.048 | 1 | 0 |
| XLOC-010899 | 0.50853 | 0.101170465 | 0.091 | 0.048 | 1 | 0 |
| ap2m1b | 0.50937 | -0.10170373 | 0.091 | 0.145 | 1 | 0 |
| slc29a1a | 0.50965 | 0.104766859 | 0.091 | 0.048 | 1 | 0 |
| fn1a | 0.50968 | -0.10377009 | 0.718 | 0.771 | 1 | 0 |
| lmf2b | 0.50981 | 0.109330054 | 0.082 | 0.048 | 1 | 0 |
| spcs3 | 0.51064 | 0.101098788 | 0.182 | 0.157 | 1 | 0 |
| pkp3b | 0.51073 | -0.11703335 | 0.091 | 0.145 | 1 | 0 |
| anp32e | 0.51136 | 0.111318819 | 0.091 | 0.048 | 1 | 0 |
| ylpm1 | 0.51325 | 0.1068451 | 0.091 | 0.048 | 1 | 0 |
| clec11a | 0.51367 | 0.104831252 | 0.091 | 0.048 | 1 | 0 |
| si:ch211-66k16.27 | 0.51536 | -0.11478617 | 0.118 | 0.157 | 1 | 0 |
| nr1d2a | 0.51577 | 0.113703459 | 0.218 | 0.157 | 1 | 0 |
| papola | 0.51645 | -0.10349073 | 0.036 | 0.072 | 1 | 0 |
| kremen1 | 0.51682 | 0.105441276 | 0.055 | 0.024 | 1 | 0 |
| tmem45a | 0.51682 | 0.106974535 | 0.1 | 0.06 | 1 | 0 |
| txndc9 | 0.5182 | 0.112404867 | 0.118 | 0.072 | 1 | 0 |
| il6st | 0.51825 | 0.109860585 | 0.145 | 0.096 | 1 | 0 |
| epha4l | 0.51873 | -0.12856188 | 0.073 | 0.12 | 1 | 0 |
| suco | 0.51889 | 0.105034867 | 0.073 | 0.036 | 1 | 0 |
| ktn1 | 0.51998 | -0.11887366 | 0.082 | 0.133 | 1 | 0 |
| si:ch211-15b10.6 | 0.52096 | -0.11231093 | 0.064 | 0.108 | 1 | 0 |
| acadm | 0.5216 | -0.12414865 | 0.155 | 0.193 | 1 | 0 |
| sult1st1 | 0.52203 | -0.10442318 | 0.145 | 0.205 | 1 | 0 |
| yy1a | 0.52215 | -0.11073916 | 0.045 | 0.084 | 1 | 0 |
| endouc | 0.5222 | -0.13971158 | 0.464 | 0.53 | 1 | 0 |
| fgl2b | 0.52434 | -0.10512399 | 0.136 | 0.193 | 1 | 0 |
| plecb | 0.52459 | 0.129328447 | 0.118 | 0.072 | 1 | 0 |
| ece2b | 0.5249 | -0.1181608 | 0.045 | 0.084 | 1 | 0 |
| rcan2 | 0.52584 | -0.12017778 | 0.045 | 0.084 | 1 | 0 |
| asah1b | 0.52823 | 0.116418292 | 0.1 | 0.06 | 1 | 0 |
| atp5f1b | 0.52844 | 0.142257819 | 0.409 | 0.337 | 1 | 0 |
| ctsd | 0.52869 | -0.11327254 | 0.1 | 0.133 | 1 | 0 |
| gmfb | 0.52949 | 0.131050938 | 0.2 | 0.145 | 1 | 0 |
| pcolce2b | 0.53269 | -0.11330405 | 0.073 | 0.12 | 1 | 0 |
| nedd4l | 0.53276 | 0.102906305 | 0.073 | 0.036 | 1 | 0 |
| gpr27 | 0.53295 | 0.103599764 | 0.073 | 0.036 | 1 | 0 |
| col15a1a | 0.53364 | 0.100990271 | 0.073 | 0.048 | 1 | 0 |
| sccpdhb | 0.53542 | -0.1190036 | 0.064 | 0.108 | 1 | 0 |
| rnf185 | 0.53559 | 0.104153274 | 0.118 | 0.072 | 1 | 0 |
| tpm4a | 0.53639 | 0.142660451 | 0.3 | 0.229 | 1 | 0 |

| | | | | | | |
|---|---|---|---|---|---|---|
| apcdd1l | 0.53683 | -0.10168902 | 0.055 | 0.096 | 1 | 0 |
| ctnnb1 | 0.5369 | 0.137284286 | 0.3 | 0.229 | 1 | 0 |
| tmem176l.1 | 0.53755 | -0.13452527 | 0.455 | 0.53 | 1 | 0 |
| kank2 | 0.53762 | -0.12820117 | 0.209 | 0.277 | 1 | 0 |
| rbm5 | 0.53952 | 0.119724703 | 0.245 | 0.181 | 1 | 0 |
| serinc5 | 0.53965 | 0.14302249 | 0.227 | 0.169 | 1 | 0 |
| cacybp | 0.53966 | -0.10386113 | 0.055 | 0.084 | 1 | 0 |
| zgc:77486 | 0.53977 | -0.10293258 | 0.045 | 0.084 | 1 | 0 |
| polr2l | 0.53982 | 0.106149843 | 0.164 | 0.108 | 1 | 0 |
| cct8 | 0.54015 | -0.13297112 | 0.2 | 0.265 | 1 | 0 |
| tln1 | 0.54089 | 0.113815678 | 0.1 | 0.06 | 1 | 0 |
| lrpap1 | 0.54093 | -0.1036877 | 0.055 | 0.096 | 1 | 0 |
| plat | 0.54246 | -0.10093714 | 0.055 | 0.096 | 1 | 0 |
| eif3d | 0.54264 | -0.13009228 | 0.264 | 0.313 | 1 | 0 |
| rpl7 | 0.54273 | 0.117445936 | 0.727 | 0.675 | 1 | 0 |
| rpz | 0.5447 | -0.10787798 | 0.055 | 0.096 | 1 | 0 |
| ar | 0.54496 | 0.115400018 | 0.145 | 0.096 | 1 | 0 |
| prrx1a | 0.54585 | 0.115643952 | 0.191 | 0.133 | 1 | 0 |
| pdcd6 | 0.54668 | -0.12332222 | 0.136 | 0.181 | 1 | 0 |
| lrrc8aa | 0.54704 | 0.109107225 | 0.109 | 0.084 | 1 | 0 |
| nova2 | 0.54825 | -0.11623882 | 0.291 | 0.361 | 1 | 0 |
| ptenb | 0.54852 | 0.127066846 | 0.164 | 0.12 | 1 | 0 |
| elof1 | 0.54907 | 0.112835186 | 0.191 | 0.133 | 1 | 0 |
| fam168b | 0.54974 | -0.11372881 | 0.118 | 0.157 | 1 | 0 |
| scpep1 | 0.55043 | 0.106374644 | 0.118 | 0.084 | 1 | 0 |
| csrp1a | 0.5514 | 0.111760197 | 0.136 | 0.096 | 1 | 0 |
| ndufa3 | 0.55195 | -0.12629665 | 0.2 | 0.265 | 1 | 0 |
| rnf10 | 0.55245 | 0.114584966 | 0.191 | 0.133 | 1 | 0 |
| kdm4b | 0.55275 | 0.116321909 | 0.118 | 0.072 | 1 | 0 |
| FAT4 | 0.55309 | -0.10768048 | 0.045 | 0.072 | 1 | 0 |
| aldh9a1a.1 | 0.55327 | 0.131998931 | 0.464 | 0.386 | 1 | 0 |
| tcp1 | 0.55351 | 0.126281348 | 0.245 | 0.205 | 1 | 0 |
| gstk1 | 0.55826 | -0.10359711 | 0.082 | 0.12 | 1 | 0 |
| psma8 | 0.55944 | 0.118332226 | 0.2 | 0.145 | 1 | 0 |
| isg20 | 0.56048 | 0.10093825 | 0.109 | 0.072 | 1 | 0 |
| kdelr2a | 0.56134 | -0.10419148 | 0.127 | 0.181 | 1 | 0 |
| cntfr | 0.56341 | -0.11781948 | 0.055 | 0.084 | 1 | 0 |
| eif3ha | 0.56368 | 0.113951427 | 0.227 | 0.181 | 1 | 0 |
| mafba | 0.56436 | -0.12766299 | 0.2 | 0.265 | 1 | 0 |
| cxcl8b.1 | 0.56475 | -0.1742489 | 0.309 | 0.373 | 1 | 0 |
| nop53 | 0.56693 | 0.1120154 | 0.309 | 0.241 | 1 | 0 |
| mmp16a | 0.5673 | -0.12150797 | 0.191 | 0.253 | 1 | 0 |
| CABZ01090890.1 | 0.56751 | -0.11957545 | 0.073 | 0.108 | 1 | 0 |
| myo1ca | 0.56844 | 0.106083198 | 0.136 | 0.108 | 1 | 0 |
| si:ch73-42p12.2 | 0.56995 | 0.105450165 | 0.1 | 0.06 | 1 | 0 |
| ehmt1b | 0.57355 | 0.117690397 | 0.109 | 0.072 | 1 | 0 |
| inhbaa | 0.57356 | 0.139964381 | 0.127 | 0.084 | 1 | 0 |
| arhgap5 | 0.57699 | 0.105095866 | 0.1 | 0.06 | 1 | 0 |

| | | | | | | |
|---|---|---|---|---|---|---|
| adgrl3.1 | 0.57953 | 0.111412488 | 0.082 | 0.048 | 1 | 0 |
| tm9sf3 | 0.58217 | 0.10182638 | 0.1 | 0.06 | 1 | 0 |
| erh | 0.58225 | 0.103987235 | 0.255 | 0.193 | 1 | 0 |
| psmc6 | 0.58305 | -0.11189679 | 0.109 | 0.145 | 1 | 0 |
| ywhabl | 0.58386 | 0.129245022 | 0.318 | 0.253 | 1 | 0 |
| psap | 0.58437 | 0.119357581 | 0.227 | 0.169 | 1 | 0 |
| pdia5 | 0.585 | -0.11027194 | 0.118 | 0.157 | 1 | 0 |
| hsp90b1 | 0.58561 | 0.117844546 | 0.182 | 0.133 | 1 | 0 |
| map1lc3b | 0.58777 | 0.124052356 | 0.255 | 0.193 | 1 | 0 |
| per3 | 0.58898 | -0.10700011 | 0.1 | 0.133 | 1 | 0 |
| sh3kbp1 | 0.58987 | 0.105391835 | 0.1 | 0.06 | 1 | 0 |
| emp2 | 0.59091 | 0.127867671 | 0.391 | 0.337 | 1 | 0 |
| tra2a | 0.59418 | 0.125801296 | 0.255 | 0.193 | 1 | 0 |
| gadd45ba | 0.59438 | -0.14110634 | 0.309 | 0.373 | 1 | 0 |
| spry4 | 0.59588 | 0.115457736 | 0.173 | 0.12 | 1 | 0 |
| celf2 | 0.5972 | -0.12649937 | 0.409 | 0.482 | 1 | 0 |
| f10 | 0.59817 | -0.11695972 | 0.109 | 0.157 | 1 | 0 |
| cavin2b | 0.6012 | -0.12699343 | 0.182 | 0.241 | 1 | 0 |
| si:ch211-222l21.1 | 0.60206 | 0.137948648 | 0.236 | 0.181 | 1 | 0 |
| baz2ba | 0.60239 | 0.112858484 | 0.145 | 0.108 | 1 | 0 |
| ap1m1 | 0.6025 | -0.11687841 | 0.109 | 0.157 | 1 | 0 |
| dnase1l1 | 0.60651 | 0.105890346 | 0.109 | 0.072 | 1 | 0 |
| CU929237.1 | 0.60733 | 0.138591639 | 0.264 | 0.205 | 1 | 0 |
| pik3r3a | 0.60822 | 0.110158867 | 0.127 | 0.084 | 1 | 0 |
| si:ch1073-291c23.2 | 0.60834 | 0.10756882 | 0.327 | 0.289 | 1 | 0 |
| angptl1a | 0.60984 | 0.112957436 | 0.264 | 0.205 | 1 | 0 |
| eif3ja | 0.61094 | 0.117008954 | 0.182 | 0.133 | 1 | 0 |
| hnrnpa0b | 0.61508 | 0.104814525 | 0.464 | 0.422 | 1 | 0 |
| btg2 | 0.62024 | -0.11086121 | 0.291 | 0.325 | 1 | 0 |
| sptan1 | 0.6212 | -0.11885724 | 0.173 | 0.229 | 1 | 0 |
| cox6a1 | 0.62966 | 0.115885097 | 0.345 | 0.289 | 1 | 0 |
| col18a1a | 0.63164 | 0.111619916 | 0.127 | 0.084 | 1 | 0 |
| rarga | 0.6322 | 0.129608603 | 0.209 | 0.157 | 1 | 0 |
| mcl1b | 0.63693 | 0.120725586 | 0.391 | 0.325 | 1 | 0 |
| si:ch73-335l21.4 | 0.63869 | -0.11808126 | 0.173 | 0.205 | 1 | 0 |
| rsrp1 | 0.64151 | -0.10584113 | 0.1 | 0.145 | 1 | 0 |
| dap1b | 0.64164 | 0.120441711 | 0.355 | 0.301 | 1 | 0 |
| kirrel1a | 0.6426 | -0.12005566 | 0.255 | 0.301 | 1 | 0 |
| il11ra | 0.64533 | -0.1087326 | 0.091 | 0.133 | 1 | 0 |
| eif3c | 0.64634 | -0.11269773 | 0.145 | 0.193 | 1 | 0 |
| slc6a2 | 0.64658 | -0.10796132 | 0.073 | 0.108 | 1 | 0 |
| crhbp | 0.64812 | 0.114966532 | 0.3 | 0.241 | 1 | 0 |
| mt-nd1 | 0.65009 | -0.1126368 | 0.5 | 0.566 | 1 | 0 |
| tagln | 0.65084 | 0.100947488 | 0.091 | 0.06 | 1 | 0 |
| cldn5a | 0.65642 | -0.11337324 | 0.209 | 0.241 | 1 | 0 |
| arf2b | 0.66756 | 0.102255746 | 0.191 | 0.145 | 1 | 0 |
| ube2v2 | 0.67039 | -0.11139199 | 0.209 | 0.253 | 1 | 0 |
| cpn1 | 0.67131 | 0.111616484 | 0.327 | 0.277 | 1 | 0 |

| | | | | | | |
|---|---|---|---|---|---|---|
| akap12b | 0.67268 | 0.117253043 | 0.455 | 0.398 | 1 | 0 |
| slc38a5b | 0.67405 | 0.100756008 | 0.364 | 0.325 | 1 | 0 |
| plscr3b | 0.68308 | -0.10937065 | 0.191 | 0.241 | 1 | 0 |
| srsf9 | 0.684 | 0.10150102 | 0.191 | 0.145 | 1 | 0 |
| snx12 | 0.68483 | 0.103695712 | 0.136 | 0.096 | 1 | 0 |
| capza1a | 0.68503 | -0.10831157 | 0.182 | 0.229 | 1 | 0 |
| atf4b | 0.68509 | -0.11216437 | 0.264 | 0.313 | 1 | 0 |
| cyp3c1 | 0.6883 | 0.108601712 | 0.191 | 0.145 | 1 | 0 |
| nop56 | 0.69213 | 0.108134602 | 0.255 | 0.205 | 1 | 0 |
| cst14a.2 | 0.69309 | 0.106159074 | 0.264 | 0.217 | 1 | 0 |
| rhoab | 0.69352 | -0.10890167 | 0.255 | 0.301 | 1 | 0 |
| cxcl18b | 0.71864 | 0.112227303 | 0.382 | 0.325 | 1 | 0 |
| ghra | 0.71928 | 0.105470895 | 0.173 | 0.133 | 1 | 0 |
| loxa | 0.72633 | -0.10382835 | 0.173 | 0.217 | 1 | 0 |
| socs3b | 0.72768 | -0.10788907 | 0.218 | 0.265 | 1 | 0 |
| pi15a | 0.73083 | 0.110482813 | 0.155 | 0.12 | 1 | 0 |
| h2afy2 | 0.73291 | 0.103090313 | 0.236 | 0.193 | 1 | 0 |
| FP101882.1 | 0.74522 | -0.10295329 | 0.1 | 0.133 | 1 | 0 |
| nrp1a | 0.75155 | 0.101864177 | 0.236 | 0.193 | 1 | 0 |
| myl6 | 0.75304 | 0.103042594 | 0.345 | 0.301 | 1 | 0 |
| ms4a17a.8 | 6.473712 | -0.79697607 | 0 | 0.5 | 1.30749581 | 1 |
| mt-cyb | 3.077123 | -1.00183051 | 0.15 | 0.78 | 6.21486665 | 1 |
| mt-co2 | 5.334717 | -1.08258982 | 0.26 | 0.89 | 1.07745297 | 1 |
| rpl27a | 1.101782 | -0.39436795 | 0.88 | 0.963 | 2.22527093 | 1 |
| mt-atp6 | 4.446166 | -0.84183519 | 0.27 | 0.793 | 8.97992259 | 1 |
| mt-co1 | 6.798324 | -0.69336471 | 0.54 | 0.951 | 1.37305766 | 1 |
| gpx1a | 2.238877 | -0.34329878 | 0.01 | 0.317 | 4.52186094 | 1 |
| COX3 | 4.829062 | -0.54119392 | 0.71 | 1 | 9.75325703 | 1 |
| mt-nd1 | 2.502836 | -0.73120137 | 0.3 | 0.78 | 5.05497950 | 1 |
| mt-nd3 | 2.900638 | -0.61578324 | 0.14 | 0.585 | 5.85841992 | 1 |
| CR855311.1 | 9.214428 | -0.54889033 | 0 | 0.268 | 1.86103804 | 1 |
| tspo | 1.184674 | -0.43742721 | 0.01 | 0.317 | 2.39268647 | 1 |
| fth1a | 3.979737 | -0.56727427 | 0.32 | 0.744 | 0.0008038 | 1 |
| XLOC-000431 | 4.726231 | -0.60687461 | 0.55 | 0.805 | 0.0095456 | 1 |
| mt-nd4 | 4.779948 | -0.5314163 | 0.49 | 0.854 | 0.0096541 | 1 |
| mt-nd2 | 4.206842 | -0.45465756 | 0.37 | 0.732 | 0.0849656 | 1 |
| zgc:92744 | 4.770126 | 0.617274306 | 0.69 | 0.402 | 0.0963423 | 1 |
| notum1b | 1.289119 | -0.31817339 | 0 | 0.146 | 0.2603634 | 1 |
| si:ch211-212k18.8 | 1.589459 | -0.42600037 | 0.11 | 0.402 | 0.3210232 | 1 |
| LOC101885938 | 3.217151 | -0.22172233 | 0 | 0.134 | 0.649768 | 1 |
| atp1a3b | 9.942387 | -0.48787845 | 0.23 | 0.488 | 1 | 1 |
| ghrb | 0.00015 | -0.4062687 | 0.1 | 0.354 | 1 | 1 |
| fabp3 | 0.00016 | -0.40711918 | 0.16 | 0.415 | 1 | 1 |
| BX908782.2 | 0.0002 | -0.21180779 | 0 | 0.11 | 1 | 1 |
| diabloa | 0.0002 | -0.17612145 | 0 | 0.11 | 1 | 1 |
| mtor | 0.0002 | -0.16344771 | 0 | 0.11 | 1 | 1 |
| si:dkey-16p21.8 | 0.00022 | -0.30300989 | 0.02 | 0.195 | 1 | 1 |
| ch25h | 0.00024 | -0.45844647 | 0.09 | 0.329 | 1 | 1 |

| | | | | | | |
|---|---|---|---|---|---|---|
| rpz2 | 0.00026 | -0.3084895 | 0.08 | 0.305 | 1 | 1 |
| si:dkey-188i13.10 | 0.00031 | 0.297403825 | 0.56 | 0.451 | 1 | 1 |
| actb1 | 0.00032 | -0.42226797 | 0.57 | 0.817 | 1 | 1 |
| rhoaa | 0.00042 | 0.196743051 | 0.45 | 0.415 | 1 | 1 |
| XLOC-013964 | 0.00046 | -0.48552769 | 0.29 | 0.512 | 1 | 1 |
| ppa2 | 0.00049 | -0.15201158 | 0 | 0.098 | 1 | 1 |
| si:ch73-193i22.1 | 0.00071 | 0.190455824 | 0.1 | 0 | 1 | 1 |
| prkacbb | 0.00073 | -0.10237754 | 0.01 | 0.098 | 1 | 1 |
| osr1 | 0.00084 | 0.414278034 | 0.47 | 0.305 | 1 | 1 |
| eif1ad | 0.00088 | 0.281706844 | 0.31 | 0.195 | 1 | 1 |
| zgc:174904 | 0.00089 | -0.17663303 | 0.01 | 0.146 | 1 | 1 |
| junbb | 0.00091 | -0.33951417 | 0.09 | 0.305 | 1 | 1 |
| slc25a55a | 0.00104 | -0.28306679 | 0.1 | 0.305 | 1 | 1 |
| erh | 0.00105 | 0.250253771 | 0.57 | 0.5 | 1 | 1 |
| scube1 | 0.00106 | -0.22803652 | 0.13 | 0.317 | 1 | 1 |
| sbno1 | 0.00119 | -0.15142598 | 0 | 0.085 | 1 | 1 |
| cib2 | 0.00119 | -0.12882258 | 0 | 0.085 | 1 | 1 |
| gatb | 0.00119 | -0.11381256 | 0 | 0.085 | 1 | 1 |
| fmnl2a | 0.00119 | -0.15995639 | 0 | 0.085 | 1 | 1 |
| si:ch73-236c18.8 | 0.00119 | -0.17522867 | 0 | 0.085 | 1 | 1 |
| timm23a | 0.00183 | -0.25873994 | 0.22 | 0.451 | 1 | 1 |
| smox | 0.00188 | -0.16854236 | 0.01 | 0.134 | 1 | 1 |
| zmiz2 | 0.00192 | -0.21308166 | 0.01 | 0.134 | 1 | 1 |
| ptbp1b | 0.00202 | -0.3116321 | 0.1 | 0.305 | 1 | 1 |
| ep300b | 0.00219 | -0.25494119 | 0.02 | 0.159 | 1 | 1 |
| adamts2 | 0.0023 | -0.18539339 | 0.05 | 0.195 | 1 | 1 |
| ube2d2 | 0.00231 | -0.3050847 | 0.21 | 0.451 | 1 | 1 |
| tssc4 | 0.00248 | 0.276138705 | 0.2 | 0.073 | 1 | 1 |
| zgc:136930 | 0.00248 | 0.384769719 | 0.18 | 0.037 | 1 | 1 |
| adam17b | 0.00262 | -0.19584281 | 0.01 | 0.122 | 1 | 1 |
| BX322530.1 | 0.00265 | -0.13230929 | 0.18 | 0.329 | 1 | 1 |
| dhrs3b | 0.00267 | 0.325091022 | 0.29 | 0.159 | 1 | 1 |
| wdr36 | 0.00269 | -0.16956918 | 0.04 | 0.171 | 1 | 1 |
| itga9 | 0.00271 | -0.27749563 | 0.23 | 0.451 | 1 | 1 |
| LOC571912 | 0.00276 | 0.136301872 | 0.08 | 0 | 1 | 1 |
| lrrtm1 | 0.00284 | 0.450120998 | 0.44 | 0.207 | 1 | 1 |
| ccdc130 | 0.00292 | -0.1240511 | 0 | 0.073 | 1 | 1 |
| ctdnep1b | 0.00292 | -0.12032849 | 0 | 0.073 | 1 | 1 |
| ftr63 | 0.00292 | -0.12851824 | 0 | 0.073 | 1 | 1 |
| dnajb14 | 0.00292 | -0.10474077 | 0 | 0.073 | 1 | 1 |
| CABZ01102240.1 | 0.00292 | -0.10586564 | 0 | 0.073 | 1 | 1 |
| cdkn1d | 0.00292 | -0.13546224 | 0 | 0.073 | 1 | 1 |
| e2f8 | 0.00292 | -0.11293579 | 0 | 0.073 | 1 | 1 |
| CT030188.1 | 0.00292 | -0.1072393 | 0 | 0.073 | 1 | 1 |
| si:dkey-3h3.3 | 0.00292 | -0.1177422 | 0 | 0.073 | 1 | 1 |
| cdh11 | 0.00305 | -0.33526863 | 0.08 | 0.256 | 1 | 1 |
| hmgb1a | 0.00342 | 0.291979296 | 0.74 | 0.634 | 1 | 1 |
| tspan18b | 0.00374 | -0.31002227 | 0.1 | 0.293 | 1 | 1 |

| | | | | | | |
|---|---|---|---|---|---|---|
| aldh2.1 | 0.00394 | 0.123758256 | 0.22 | 0.22 | 1 | 1 |
| glo1 | 0.00404 | 0.206001522 | 0.42 | 0.366 | 1 | 1 |
| ahr2 | 0.00404 | -0.18719155 | 0.02 | 0.134 | 1 | 1 |
| pgr | 0.00411 | -0.28492771 | 0.07 | 0.244 | 1 | 1 |
| tfpia | 0.00422 | -0.19228861 | 0.01 | 0.122 | 1 | 1 |
| fbxo9 | 0.00448 | -0.24402846 | 0.05 | 0.207 | 1 | 1 |
| hiat1b | 0.00459 | -0.11376286 | 0.01 | 0.098 | 1 | 1 |
| polr3glb | 0.00463 | 0.271106592 | 0.26 | 0.134 | 1 | 1 |
| h3f3c | 0.00468 | 0.19320333 | 0.86 | 0.793 | 1 | 1 |
| homer3b | 0.00479 | -0.14005838 | 0.1 | 0.244 | 1 | 1 |
| csrnp1b | 0.00481 | -0.17750963 | 0.08 | 0.22 | 1 | 1 |
| tmem56a | 0.00508 | -0.10680846 | 0.01 | 0.085 | 1 | 1 |
| ppp4cb | 0.00509 | -0.13242935 | 0.13 | 0.268 | 1 | 1 |
| dnajc5aa | 0.00516 | 0.21322834 | 0.13 | 0.012 | 1 | 1 |
| ccl25b | 0.00522 | -0.33975767 | 0.3 | 0.537 | 1 | 1 |
| sh3glb1a | 0.00545 | 0.14645403 | 0.07 | 0 | 1 | 1 |
| lias | 0.00545 | 0.144596166 | 0.07 | 0 | 1 | 1 |
| zgc:162945 | 0.00545 | 0.128458999 | 0.07 | 0 | 1 | 1 |
| ndst3 | 0.00545 | 0.146906633 | 0.07 | 0 | 1 | 1 |
| pnpla8 | 0.00545 | 0.148612908 | 0.07 | 0 | 1 | 1 |
| NC-002333.4 | 0.00557 | -0.21308141 | 0.86 | 0.976 | 1 | 1 |
| ptprga | 0.0056 | -0.21314187 | 0.33 | 0.537 | 1 | 1 |
| si:rp71-36a1.2 | 0.00577 | -0.12549154 | 0.03 | 0.122 | 1 | 1 |
| ndfip1l | 0.00578 | -0.29874153 | 0.17 | 0.378 | 1 | 1 |
| sat1a.2 | 0.00587 | 0.132447704 | 0.37 | 0.366 | 1 | 1 |
| csnk1db | 0.00588 | -0.16102312 | 0.17 | 0.329 | 1 | 1 |
| stxbp3 | 0.00595 | 0.225115757 | 0.15 | 0.037 | 1 | 1 |
| ppdpfb | 0.00628 | 0.147182936 | 0.83 | 0.829 | 1 | 1 |
| cxcl12a | 0.00636 | -0.40291286 | 0.22 | 0.415 | 1 | 1 |
| atp5mf | 0.0064 | -0.32133211 | 0.32 | 0.512 | 1 | 1 |
| mphosph8 | 0.00663 | 0.292963923 | 0.46 | 0.329 | 1 | 1 |
| otub1b | 0.00701 | -0.2024381 | 0.06 | 0.207 | 1 | 1 |
| git2a | 0.00717 | -0.11692071 | 0 | 0.061 | 1 | 1 |
| narf | 0.00717 | -0.11322376 | 0 | 0.061 | 1 | 1 |
| BX511061.1 | 0.00717 | -0.12408059 | 0 | 0.061 | 1 | 1 |
| uvrag | 0.00717 | -0.11548795 | 0 | 0.061 | 1 | 1 |
| adam9 | 0.00717 | -0.10524295 | 0 | 0.061 | 1 | 1 |
| kdm6a | 0.00717 | -0.10924307 | 0 | 0.061 | 1 | 1 |
| angel2 | 0.00717 | -0.11359866 | 0 | 0.061 | 1 | 1 |
| bmp8a | 0.00717 | -0.12319494 | 0 | 0.061 | 1 | 1 |
| cnn1b | 0.00717 | -0.11188696 | 0 | 0.061 | 1 | 1 |
| gnpda1 | 0.00717 | -0.10045748 | 0 | 0.061 | 1 | 1 |
| tmem42b | 0.00717 | -0.12137626 | 0 | 0.061 | 1 | 1 |
| txn | 0.00717 | -0.12074847 | 0 | 0.061 | 1 | 1 |
| cdkn1ba | 0.00717 | -0.13610574 | 0 | 0.061 | 1 | 1 |
| cyp27a1.4 | 0.00717 | -0.14475831 | 0 | 0.061 | 1 | 1 |
| senp3b | 0.00717 | -0.10621129 | 0 | 0.061 | 1 | 1 |
| csnk2a2a | 0.00717 | -0.10402798 | 0 | 0.061 | 1 | 1 |

| | | | | | | |
|---|---|---|---|---|---|---|
| odc1 | 0.00717 | -0.14804326 | 0 | 0.061 | 1 | 1 |
| si:ch211-183d5.2 | 0.00717 | -0.13010162 | 0 | 0.061 | 1 | 1 |
| eif4g1a | 0.00732 | -0.13304227 | 0.14 | 0.28 | 1 | 1 |
| inip | 0.00748 | -0.15937438 | 0.11 | 0.256 | 1 | 1 |
| BX957362.1 | 0.00749 | -0.21426039 | 0.01 | 0.11 | 1 | 1 |
| cmtm7 | 0.00755 | -0.23985864 | 0.05 | 0.195 | 1 | 1 |
| stard7 | 0.00765 | -0.13978866 | 0.02 | 0.122 | 1 | 1 |
| tmem98 | 0.00809 | -0.24859481 | 0.1 | 0.268 | 1 | 1 |
| tpm4a | 0.0081 | -0.29394822 | 0.46 | 0.598 | 1 | 1 |
| ZNF512B | 0.00829 | -0.12474732 | 0.02 | 0.11 | 1 | 1 |
| fhl2a | 0.00833 | -0.20206169 | 0.02 | 0.134 | 1 | 1 |
| id1 | 0.00837 | 0.276650317 | 0.65 | 0.427 | 1 | 1 |
| zgc:110239 | 0.00845 | -0.16130322 | 0.12 | 0.268 | 1 | 1 |
| fabp11a | 0.00847 | -0.23665043 | 0.02 | 0.134 | 1 | 1 |
| cdk2ap2 | 0.00847 | -0.18932173 | 0.05 | 0.183 | 1 | 1 |
| loxl2b | 0.00857 | -0.28687384 | 0.23 | 0.439 | 1 | 1 |
| ccnd2a | 0.00862 | -0.20407698 | 0.21 | 0.39 | 1 | 1 |
| XLOC-010899 | 0.00873 | -0.35173802 | 0.32 | 0.5 | 1 | 1 |
| tra2a | 0.00875 | -0.28133847 | 0.29 | 0.512 | 1 | 1 |
| adamts1 | 0.00896 | -0.15919898 | 0.01 | 0.11 | 1 | 1 |
| osr2 | 0.00897 | 0.235718706 | 0.69 | 0.61 | 1 | 1 |
| selenoh | 0.00905 | 0.301183122 | 0.37 | 0.244 | 1 | 1 |
| cnn2 | 0.00958 | -0.29496199 | 0.28 | 0.5 | 1 | 1 |
| pop5 | 0.00964 | -0.23239948 | 0.06 | 0.195 | 1 | 1 |
| XLOC-025423 | 0.01035 | -0.38109543 | 0.21 | 0.415 | 1 | 1 |
| rapsn | 0.01078 | 0.142801057 | 0.06 | 0 | 1 | 1 |
| plcg1 | 0.01078 | 0.126031594 | 0.06 | 0 | 1 | 1 |
| cln6a | 0.01078 | 0.103083119 | 0.06 | 0 | 1 | 1 |
| dhx9 | 0.01078 | 0.126236638 | 0.06 | 0 | 1 | 1 |
| stag2a | 0.01078 | 0.134244101 | 0.06 | 0 | 1 | 1 |
| appl2 | 0.01078 | 0.116248575 | 0.06 | 0 | 1 | 1 |
| tap1 | 0.011 | -0.14874492 | 0.01 | 0.098 | 1 | 1 |
| hsp90b1 | 0.01121 | -0.18405807 | 0.17 | 0.341 | 1 | 1 |
| luzp2 | 0.01131 | 0.263376549 | 0.28 | 0.171 | 1 | 1 |
| pum3 | 0.0115 | -0.25729918 | 0.07 | 0.22 | 1 | 1 |
| prps1a | 0.0118 | -0.11550723 | 0.14 | 0.268 | 1 | 1 |
| mbd3a | 0.01181 | 0.118598392 | 0.18 | 0.159 | 1 | 1 |
| XLOC-042595 | 0.01226 | -0.1108363 | 0.01 | 0.085 | 1 | 1 |
| sfrp1a | 0.01238 | -0.19392753 | 0.08 | 0.22 | 1 | 1 |
| gnai2a | 0.01334 | -0.15848374 | 0.16 | 0.317 | 1 | 1 |
| carhsp1 | 0.01341 | -0.22984899 | 0.12 | 0.293 | 1 | 1 |
| wdr5 | 0.01357 | -0.13487387 | 0.06 | 0.171 | 1 | 1 |
| nosip | 0.01358 | -0.12331643 | 0.04 | 0.146 | 1 | 1 |
| cyp3c1 | 0.01359 | -0.2035558 | 0.31 | 0.5 | 1 | 1 |
| yap1 | 0.01398 | 0.299500286 | 0.38 | 0.207 | 1 | 1 |
| wls | 0.01414 | 0.224123737 | 0.5 | 0.427 | 1 | 1 |
| sec23ip | 0.01523 | -0.11140382 | 0.01 | 0.085 | 1 | 1 |
| bhlhe41 | 0.01544 | -0.18204636 | 0.02 | 0.122 | 1 | 1 |

| | | | | | | |
|---|---|---|---|---|---|---|
| ube2ib | 0.01547 | 0.129338604 | 0.43 | 0.415 | 1 | 1 |
| rpz | 0.01562 | 0.207369604 | 0.14 | 0.061 | 1 | 1 |
| mrpl53 | 0.01579 | -0.11398421 | 0.07 | 0.183 | 1 | 1 |
| pik3r3b | 0.01597 | -0.20067679 | 0.06 | 0.195 | 1 | 1 |
| hsd11b2 | 0.01606 | 0.174950801 | 0.97 | 0.878 | 1 | 1 |
| col6a3 | 0.01611 | -0.10378915 | 0.43 | 0.585 | 1 | 1 |
| CABZ01066926.1 | 0.01612 | 0.147942773 | 0.11 | 0.012 | 1 | 1 |
| mrpl4 | 0.01618 | -0.1729983 | 0.08 | 0.22 | 1 | 1 |
| si:ch211-210g13.5 | 0.01621 | 0.177555347 | 0.11 | 0.012 | 1 | 1 |
| eif3s10 | 0.01654 | -0.27089249 | 0.24 | 0.439 | 1 | 1 |
| ptp4a2a | 0.01654 | 0.22059084 | 0.31 | 0.207 | 1 | 1 |
| smarcb1b | 0.01668 | 0.10858263 | 0.24 | 0.232 | 1 | 1 |
| zc3h13 | 0.01742 | -0.2547134 | 0.09 | 0.244 | 1 | 1 |
| CABZ01074307.1 | 0.01743 | -0.18461425 | 0.01 | 0.098 | 1 | 1 |
| LOC108191614 | 0.01754 | -0.27850041 | 0.09 | 0.232 | 1 | 1 |
| mcts1 | 0.01766 | -0.11549757 | 0.17 | 0.305 | 1 | 1 |
| ehmt1b | 0.01775 | 0.229881848 | 0.16 | 0.073 | 1 | 1 |
| cul1b | 0.01778 | -0.1106298 | 0.03 | 0.122 | 1 | 1 |
| bin2b | 0.0181 | -0.13421219 | 0.01 | 0.098 | 1 | 1 |
| cry4 | 0.01812 | -0.10644965 | 0.01 | 0.098 | 1 | 1 |
| dbn1 | 0.01826 | -0.19522691 | 0.12 | 0.268 | 1 | 1 |
| nfyba | 0.01829 | 0.118300584 | 0.46 | 0.463 | 1 | 1 |
| ndufa1 | 0.01834 | -0.27857628 | 0.38 | 0.537 | 1 | 1 |
| si:ch211-181d7.3 | 0.01843 | -0.14955401 | 0.01 | 0.098 | 1 | 1 |
| zgc:86598 | 0.01845 | 0.222306319 | 0.27 | 0.183 | 1 | 1 |
| usp18 | 0.0186 | -0.15170795 | 0.01 | 0.098 | 1 | 1 |
| uba2 | 0.01862 | 0.187373473 | 0.15 | 0.085 | 1 | 1 |
| fbl | 0.01864 | -0.2961008 | 0.28 | 0.476 | 1 | 1 |
| prss35 | 0.01872 | 0.208448922 | 0.1 | 0.012 | 1 | 1 |
| gm2a | 0.01877 | -0.15500357 | 0.01 | 0.098 | 1 | 1 |
| chmp4bb | 0.01892 | 0.149126057 | 0.16 | 0.122 | 1 | 1 |
| bmt2 | 0.01896 | -0.137564 | 0.01 | 0.098 | 1 | 1 |
| ptmab | 0.01904 | 0.15239551 | 0.98 | 0.89 | 1 | 1 |
| akirin2 | 0.01931 | -0.2118501 | 0.06 | 0.195 | 1 | 1 |
| nkap | 0.01934 | -0.15080663 | 0.17 | 0.317 | 1 | 1 |
| yeats4 | 0.0195 | -0.14326617 | 0.11 | 0.244 | 1 | 1 |
| myocd | 0.01952 | -0.18131676 | 0.04 | 0.159 | 1 | 1 |
| anxa5b | 0.01959 | 0.146649669 | 0.31 | 0.28 | 1 | 1 |
| psma2 | 0.01964 | 0.15869139 | 0.36 | 0.317 | 1 | 1 |
| qkia | 0.01982 | 0.135813507 | 0.81 | 0.805 | 1 | 1 |
| si:ch73-22o12.1 | 0.01987 | -0.16078646 | 0.08 | 0.207 | 1 | 1 |
| si:ch211-195h23.3 | 0.01987 | -0.22107275 | 0.05 | 0.171 | 1 | 1 |
| reep5 | 0.0199 | 0.171642091 | 0.4 | 0.366 | 1 | 1 |
| anapc15 | 0.02022 | -0.12947023 | 0.03 | 0.134 | 1 | 1 |
| rgs4 | 0.0203 | -0.21616894 | 0.05 | 0.171 | 1 | 1 |
| zgc:171497 | 0.02068 | -0.14739917 | 0.06 | 0.171 | 1 | 1 |
| XLOC-032256 | 0.02078 | -0.21706327 | 0.02 | 0.098 | 1 | 1 |
| erbin | 0.02094 | -0.13783758 | 0.02 | 0.11 | 1 | 1 |

| | | | | | | |
|---|---|---|---|---|---|---|
| ldb1a | 0.021 | -0.1692221 | 0.04 | 0.159 | 1 | 1 |
| cmss1 | 0.02109 | -0.17558441 | 0.04 | 0.159 | 1 | 1 |
| acin1a | 0.02117 | -0.11140091 | 0.44 | 0.598 | 1 | 1 |
| pcdh20 | 0.02128 | 0.22227161 | 0.09 | 0.012 | 1 | 1 |
| si:dkey-261h17.1 | 0.02131 | 0.193703214 | 0.63 | 0.585 | 1 | 1 |
| paf1 | 0.02138 | 0.107249589 | 0.05 | 0 | 1 | 1 |
| pbxip1a | 0.02138 | 0.111932112 | 0.05 | 0 | 1 | 1 |
| ptprda | 0.02138 | 0.1020911 | 0.05 | 0 | 1 | 1 |
| dnmbp | 0.02138 | 0.133860923 | 0.05 | 0 | 1 | 1 |
| metrnlb | 0.02138 | 0.114042705 | 0.05 | 0 | 1 | 1 |
| peak1 | 0.02138 | 0.116023721 | 0.05 | 0 | 1 | 1 |
| twist1a | 0.02138 | 0.103316138 | 0.05 | 0 | 1 | 1 |
| colec11 | 0.02138 | 0.105705124 | 0.05 | 0 | 1 | 1 |
| alkbh1 | 0.02138 | 0.11175469 | 0.05 | 0 | 1 | 1 |
| plxna2 | 0.02138 | 0.113576049 | 0.05 | 0 | 1 | 1 |
| cbx7a | 0.02195 | -0.30086708 | 0.11 | 0.268 | 1 | 1 |
| polr3f | 0.02213 | -0.13673988 | 0.05 | 0.159 | 1 | 1 |
| tagln | 0.02221 | -0.36274699 | 0.36 | 0.5 | 1 | 1 |
| chpt1 | 0.02268 | 0.141657509 | 0.1 | 0.012 | 1 | 1 |
| nop16 | 0.02304 | -0.21967481 | 0.08 | 0.207 | 1 | 1 |
| chmp1a | 0.02318 | -0.11058967 | 0.05 | 0.146 | 1 | 1 |
| gstm.2 | 0.02322 | 0.149851689 | 0.28 | 0.244 | 1 | 1 |
| mylipb | 0.0236 | 0.230653404 | 0.16 | 0.049 | 1 | 1 |
| tktb | 0.02362 | -0.24778645 | 0.12 | 0.28 | 1 | 1 |
| CABZ01032488.1 | 0.02362 | -0.14730367 | 0.03 | 0.134 | 1 | 1 |
| asic1b | 0.02374 | 0.275887673 | 0.21 | 0.073 | 1 | 1 |
| atic | 0.02374 | -0.11510627 | 0.03 | 0.122 | 1 | 1 |
| chchd3a | 0.02384 | 0.22239246 | 0.19 | 0.085 | 1 | 1 |
| bud31 | 0.02411 | 0.112228595 | 0.37 | 0.378 | 1 | 1 |
| pin1 | 0.02418 | 0.113972275 | 0.41 | 0.415 | 1 | 1 |
| spag1a | 0.02422 | -0.16499945 | 0.02 | 0.11 | 1 | 1 |
| ephb4a | 0.02426 | 0.211473852 | 0.13 | 0.061 | 1 | 1 |
| u2af1 | 0.02441 | -0.17201695 | 0.24 | 0.402 | 1 | 1 |
| prrc2c | 0.02463 | -0.17441102 | 0.03 | 0.134 | 1 | 1 |
| si:rp71-36a1.3 | 0.025 | -0.18423075 | 0.05 | 0.171 | 1 | 1 |
| egln1b | 0.025 | -0.15304352 | 0.07 | 0.195 | 1 | 1 |
| fzd2 | 0.02505 | -0.15731092 | 0.29 | 0.451 | 1 | 1 |
| nfia | 0.02516 | -0.1596902 | 0.35 | 0.512 | 1 | 1 |
| CABZ01092746.1 | 0.02529 | 0.240178627 | 0.78 | 0.659 | 1 | 1 |
| arf5 | 0.0257 | 0.164745459 | 0.44 | 0.39 | 1 | 1 |
| myca | 0.026 | -0.19183609 | 0.05 | 0.146 | 1 | 1 |
| arid1ab | 0.02607 | 0.175803277 | 0.09 | 0.012 | 1 | 1 |
| sarnp | 0.02612 | -0.26434187 | 0.18 | 0.354 | 1 | 1 |
| egfra | 0.02622 | -0.21368275 | 0.1 | 0.244 | 1 | 1 |
| dynll2b | 0.0264 | 0.136814551 | 0.23 | 0.22 | 1 | 1 |
| syf2 | 0.02644 | 0.271747442 | 0.45 | 0.268 | 1 | 1 |
| btg1 | 0.02669 | 0.25786305 | 0.7 | 0.524 | 1 | 1 |
| znf385a | 0.02669 | 0.235418008 | 0.18 | 0.085 | 1 | 1 |

| | | | | | | |
|---|---|---|---|---|---|---|
| psmb3 | 0.02685 | -0.20984956 | 0.3 | 0.488 | 1 | 1 |
| nol7 | 0.0269 | -0.15343508 | 0.03 | 0.134 | 1 | 1 |
| mafba | 0.02723 | -0.12548782 | 0.27 | 0.415 | 1 | 1 |
| tmem50a | 0.02731 | -0.18145499 | 0.37 | 0.549 | 1 | 1 |
| bod1 | 0.02761 | 0.205200844 | 0.14 | 0.049 | 1 | 1 |
| myo10l3 | 0.02776 | -0.14736153 | 0.02 | 0.11 | 1 | 1 |
| isg15 | 0.02777 | -0.30041317 | 0.11 | 0.256 | 1 | 1 |
| atp6ap1b | 0.02792 | -0.18202066 | 0.02 | 0.11 | 1 | 1 |
| angptl2b | 0.02805 | -0.12778888 | 0.05 | 0.146 | 1 | 1 |
| fam84a | 0.02805 | 0.224437156 | 0.15 | 0.037 | 1 | 1 |
| si:dkey-180p18.9 | 0.02806 | 0.188405979 | 0.3 | 0.22 | 1 | 1 |
| rcl1 | 0.02829 | -0.2001859 | 0.07 | 0.183 | 1 | 1 |
| cygb1 | 0.02834 | -0.28674089 | 0.19 | 0.366 | 1 | 1 |
| atp5mc3a | 0.0284 | 0.195833601 | 0.53 | 0.463 | 1 | 1 |
| hic1 | 0.02868 | 0.14143975 | 0.48 | 0.463 | 1 | 1 |
| rad21a | 0.02877 | -0.12117692 | 0.42 | 0.573 | 1 | 1 |
| crebzf | 0.02882 | -0.27811388 | 0.18 | 0.354 | 1 | 1 |
| nr0b1 | 0.0304 | -0.21818178 | 0.09 | 0.232 | 1 | 1 |
| zc3h7a | 0.03047 | -0.12764655 | 0.01 | 0.085 | 1 | 1 |
| si:dkey-100n23.5 | 0.03049 | -0.1224533 | 0.09 | 0.195 | 1 | 1 |
| ets1 | 0.03133 | -0.218865 | 0.15 | 0.305 | 1 | 1 |
| pbx1a | 0.03186 | 0.14764218 | 0.45 | 0.427 | 1 | 1 |
| itga1 | 0.032 | -0.12844968 | 0.02 | 0.11 | 1 | 1 |
| col4a1 | 0.0321 | -0.24948526 | 0.27 | 0.451 | 1 | 1 |
| man2b1 | 0.03226 | -0.181187 | 0.02 | 0.11 | 1 | 1 |
| stox1 | 0.03257 | -0.18913003 | 0.06 | 0.183 | 1 | 1 |
| drap1 | 0.03323 | -0.15643382 | 0.28 | 0.439 | 1 | 1 |
| si:dkey-225k4.1 | 0.03325 | -0.12761522 | 0.01 | 0.085 | 1 | 1 |
| srpx | 0.03352 | -0.17151524 | 0.07 | 0.183 | 1 | 1 |
| capns1b | 0.03373 | 0.159742043 | 0.46 | 0.427 | 1 | 1 |
| ldhba | 0.03446 | -0.24676191 | 0.29 | 0.439 | 1 | 1 |
| cdv3 | 0.03449 | -0.15786058 | 0.29 | 0.451 | 1 | 1 |
| reck | 0.03461 | -0.15432973 | 0.05 | 0.159 | 1 | 1 |
| fuca2 | 0.03466 | -0.15850175 | 0.05 | 0.159 | 1 | 1 |
| nup93 | 0.03476 | 0.184501921 | 0.13 | 0.037 | 1 | 1 |
| cfd | 0.03488 | 0.251828643 | 0.22 | 0.085 | 1 | 1 |
| si:dkey-110g7.8 | 0.03495 | 0.166660575 | 0.12 | 0.024 | 1 | 1 |
| usp43b | 0.03497 | -0.17532675 | 0.02 | 0.11 | 1 | 1 |
| ms4a17a.14 | 0.03503 | -0.20951344 | 0.04 | 0.146 | 1 | 1 |
| pex7 | 0.0355 | -0.14404136 | 0.02 | 0.11 | 1 | 1 |
| ptges | 0.03573 | -0.10138512 | 0.05 | 0.134 | 1 | 1 |
| sdhaf4 | 0.03578 | -0.1254105 | 0.02 | 0.11 | 1 | 1 |
| papolg | 0.0358 | -0.14870246 | 0.02 | 0.11 | 1 | 1 |
| ahcy | 0.03598 | -0.18075038 | 0.05 | 0.159 | 1 | 1 |
| si:dkey-12j5.1 | 0.03616 | -0.14167964 | 0.06 | 0.171 | 1 | 1 |
| si:dkey-29h14.10 | 0.03616 | -0.14145778 | 0.01 | 0.085 | 1 | 1 |
| actn1 | 0.03627 | -0.19593694 | 0.07 | 0.195 | 1 | 1 |
| pomp | 0.03628 | 0.110742809 | 0.55 | 0.561 | 1 | 1 |

| | | | | | | |
|---|---|---|---|---|---|---|
| tlr5b | 0.03668 | -0.14330643 | 0.04 | 0.134 | 1 | 1 |
| tnfrsf1a | 0.03688 | -0.19278656 | 0.18 | 0.329 | 1 | 1 |
| ss18l2 | 0.03691 | 0.161190008 | 0.17 | 0.134 | 1 | 1 |
| sestd1 | 0.03705 | -0.12927223 | 0.06 | 0.159 | 1 | 1 |
| snu13b | 0.03729 | -0.12573239 | 0.48 | 0.646 | 1 | 1 |
| khsrp | 0.03744 | -0.23934273 | 0.28 | 0.463 | 1 | 1 |
| palld | 0.03745 | -0.1291191 | 0.04 | 0.134 | 1 | 1 |
| BX957297.2 | 0.03761 | -0.16568965 | 0.01 | 0.085 | 1 | 1 |
| CU655842.1 | 0.03779 | -0.14366716 | 0.01 | 0.085 | 1 | 1 |
| rbms1b | 0.03785 | -0.19851021 | 0.07 | 0.195 | 1 | 1 |
| scrn3 | 0.03812 | -0.12599477 | 0.01 | 0.085 | 1 | 1 |
| LOC103909972 | 0.03814 | -0.25402991 | 0.18 | 0.341 | 1 | 1 |
| midn | 0.03818 | -0.20965236 | 0.29 | 0.463 | 1 | 1 |
| selenoo1 | 0.03827 | -0.11248385 | 0.01 | 0.085 | 1 | 1 |
| fbln2 | 0.03851 | 0.175505836 | 0.12 | 0.024 | 1 | 1 |
| lgals3b | 0.03868 | 0.135040329 | 0.31 | 0.293 | 1 | 1 |
| gab1 | 0.03878 | -0.1481849 | 0.01 | 0.085 | 1 | 1 |
| cyp27c1 | 0.03886 | 0.267725164 | 0.35 | 0.22 | 1 | 1 |
| tbca | 0.03894 | -0.24199456 | 0.35 | 0.537 | 1 | 1 |
| ttf1 | 0.03901 | -0.1932473 | 0.07 | 0.195 | 1 | 1 |
| fstl3 | 0.03924 | -0.20343611 | 0.04 | 0.146 | 1 | 1 |
| abcd3a | 0.03926 | -0.13443 | 0.02 | 0.098 | 1 | 1 |
| irf8 | 0.03951 | -0.28404615 | 0.13 | 0.232 | 1 | 1 |
| mrto4 | 0.03991 | -0.1807741 | 0.07 | 0.195 | 1 | 1 |
| sulf2b | 0.03992 | -0.25811823 | 0.13 | 0.28 | 1 | 1 |
| bnc2 | 0.03995 | -0.2337237 | 0.07 | 0.195 | 1 | 1 |
| mrpl46 | 0.04018 | -0.21426931 | 0.05 | 0.159 | 1 | 1 |
| si:ch211-132p1.3 | 0.04031 | -0.26356611 | 0.35 | 0.537 | 1 | 1 |
| tcf12 | 0.04036 | 0.229213863 | 0.41 | 0.317 | 1 | 1 |
| eif3i | 0.04044 | 0.231078289 | 0.6 | 0.476 | 1 | 1 |
| cadpsb | 0.04151 | 0.247485772 | 0.14 | 0.037 | 1 | 1 |
| mrps14 | 0.04154 | -0.16118023 | 0.09 | 0.22 | 1 | 1 |
| sltm | 0.04157 | -0.23365624 | 0.15 | 0.305 | 1 | 1 |
| znf598 | 0.04245 | 0.158147459 | 0.11 | 0.024 | 1 | 1 |
| mrpl54 | 0.04245 | 0.17443493 | 0.35 | 0.28 | 1 | 1 |
| si:ch211-213a13.1 | 0.0432 | -0.24422815 | 0.24 | 0.366 | 1 | 1 |
| plekha1a | 0.04359 | 0.234760147 | 0.17 | 0.061 | 1 | 1 |
| fbxl7 | 0.04371 | 0.193093922 | 0.15 | 0.049 | 1 | 1 |
| bida | 0.0438 | -0.13173497 | 0.04 | 0.134 | 1 | 1 |
| hsd17b10 | 0.04391 | -0.13616422 | 0.08 | 0.195 | 1 | 1 |
| gyg1a | 0.04403 | 0.107679393 | 0.12 | 0.11 | 1 | 1 |
| thoc7 | 0.04458 | -0.19830281 | 0.08 | 0.207 | 1 | 1 |
| tagln2 | 0.04491 | -0.17491524 | 0.28 | 0.439 | 1 | 1 |
| wt1a | 0.04501 | -0.28185515 | 0.14 | 0.28 | 1 | 1 |
| adh5 | 0.04502 | 0.134374611 | 0.69 | 0.683 | 1 | 1 |
| gpr108 | 0.04504 | -0.19406991 | 0.05 | 0.146 | 1 | 1 |
| tuba8l | 0.04525 | -0.27009023 | 0.36 | 0.488 | 1 | 1 |
| dnajc7 | 0.04528 | -0.1582665 | 0.03 | 0.11 | 1 | 1 |

| | | | | | | |
|---|---|---|---|---|---|---|
| brd8 | 0.04552 | -0.10330979 | 0.02 | 0.098 | 1 | 1 |
| hadh | 0.04558 | -0.16954761 | 0.1 | 0.232 | 1 | 1 |
| XLOC-041269 | 0.04565 | 0.266088515 | 0.19 | 0.085 | 1 | 1 |
| brdt | 0.04659 | 0.162665391 | 0.2 | 0.159 | 1 | 1 |
| eif3m | 0.04717 | 0.145810231 | 0.74 | 0.695 | 1 | 1 |
| si:ch211-74m13.3 | 0.04734 | -0.13394719 | 0.08 | 0.183 | 1 | 1 |
| pigt | 0.04741 | -0.1686007 | 0.02 | 0.098 | 1 | 1 |
| mau2 | 0.04755 | -0.1382216 | 0.03 | 0.122 | 1 | 1 |
| psmd3 | 0.04759 | -0.18690749 | 0.16 | 0.305 | 1 | 1 |
| si:ch211-150o23.3 | 0.04779 | -0.17476101 | 0.03 | 0.122 | 1 | 1 |
| mibp2 | 0.04795 | -0.15809509 | 0.75 | 0.89 | 1 | 1 |
| cped1 | 0.04833 | -0.12757219 | 0.06 | 0.159 | 1 | 1 |
| papola | 0.0485 | 0.170062876 | 0.16 | 0.11 | 1 | 1 |
| paip2b | 0.04883 | 0.15465143 | 0.59 | 0.549 | 1 | 1 |
| ncln | 0.049 | 0.212820319 | 0.19 | 0.073 | 1 | 1 |
| rpp25l | 0.04915 | -0.11938173 | 0.03 | 0.122 | 1 | 1 |
| wbp1 | 0.04926 | -0.1404132 | 0.03 | 0.122 | 1 | 1 |
| trir | 0.04973 | 0.108070894 | 0.15 | 0.134 | 1 | 1 |
| pno1 | 0.04994 | -0.15355158 | 0.27 | 0.427 | 1 | 1 |
| mrps9 | 0.05046 | -0.10216226 | 0.08 | 0.171 | 1 | 1 |
| hsp90ab1 | 0.05049 | 0.101576773 | 0.99 | 0.951 | 1 | 1 |
| zgc:162730 | 0.05057 | -0.22132505 | 0.67 | 0.756 | 1 | 1 |
| eif3ba | 0.05062 | 0.197581432 | 0.6 | 0.524 | 1 | 1 |
| thap11 | 0.05108 | -0.11531604 | 0.02 | 0.098 | 1 | 1 |
| chmp5b | 0.05131 | 0.205294547 | 0.29 | 0.195 | 1 | 1 |
| eif2s1a | 0.05155 | -0.12141237 | 0.18 | 0.305 | 1 | 1 |
| zgc:85789 | 0.05158 | -0.14032197 | 0.03 | 0.122 | 1 | 1 |
| sema4ba | 0.05183 | 0.113423202 | 0.14 | 0.061 | 1 | 1 |
| ywhaz | 0.05227 | 0.14249239 | 0.79 | 0.756 | 1 | 1 |
| snu13a | 0.05266 | -0.13935315 | 0.03 | 0.122 | 1 | 1 |
| boc | 0.05286 | -0.15682348 | 0.38 | 0.537 | 1 | 1 |
| p4hb | 0.05313 | -0.21660043 | 0.29 | 0.329 | 1 | 1 |
| oxsr1a | 0.05328 | -0.17149466 | 0.07 | 0.183 | 1 | 1 |
| chd4a | 0.0534 | -0.21149857 | 0.24 | 0.402 | 1 | 1 |
| krt94 | 0.05349 | 0.255496715 | 0.74 | 0.598 | 1 | 1 |
| bmpr2a | 0.05387 | 0.173664806 | 0.11 | 0.024 | 1 | 1 |
| pitrm1 | 0.05425 | -0.11522771 | 0.02 | 0.098 | 1 | 1 |
| med8 | 0.05443 | 0.164439411 | 0.14 | 0.073 | 1 | 1 |
| mmp16a | 0.05447 | -0.14582137 | 0.38 | 0.524 | 1 | 1 |
| rab18a | 0.05481 | -0.19308862 | 0.06 | 0.171 | 1 | 1 |
| chico | 0.05514 | 0.156856072 | 0.08 | 0.012 | 1 | 1 |
| serinc5 | 0.05532 | -0.22696375 | 0.22 | 0.378 | 1 | 1 |
| krt8 | 0.05538 | -0.21703708 | 0.68 | 0.829 | 1 | 1 |
| tbk1 | 0.05564 | -0.14102296 | 0.02 | 0.098 | 1 | 1 |
| si:ch73-347e22.4 | 0.05601 | 0.190251104 | 0.16 | 0.073 | 1 | 1 |
| pop4 | 0.0564 | -0.10433617 | 0.02 | 0.098 | 1 | 1 |
| sh3bp5b | 0.05643 | 0.169243798 | 0.13 | 0.037 | 1 | 1 |
| igfbp7 | 0.05656 | -0.25290974 | 0.12 | 0.256 | 1 | 1 |

| | | | | | | |
|---|---|---|---|---|---|---|
| zgc:77650 | 0.05667 | -0.17760806 | 0.04 | 0.134 | 1 | 1 |
| hspa9 | 0.0568 | -0.12879954 | 0.13 | 0.244 | 1 | 1 |
| si:dkey-23i12.7 | 0.05707 | -0.22879796 | 0.04 | 0.134 | 1 | 1 |
| hnrnpk | 0.0571 | 0.180075407 | 0.33 | 0.256 | 1 | 1 |
| rxfp3.3b | 0.05722 | -0.13212409 | 0.05 | 0.146 | 1 | 1 |
| fblim1 | 0.0574 | 0.159374042 | 0.25 | 0.207 | 1 | 1 |
| sema4c | 0.05759 | 0.220982806 | 0.24 | 0.146 | 1 | 1 |
| rasl11a | 0.05775 | -0.13154403 | 0.12 | 0.232 | 1 | 1 |
| rtn3 | 0.05791 | -0.13133168 | 0.02 | 0.098 | 1 | 1 |
| spag7 | 0.05795 | 0.166632694 | 0.13 | 0.073 | 1 | 1 |
| scamp4 | 0.058 | -0.16762803 | 0.08 | 0.195 | 1 | 1 |
| ndufs5 | 0.05821 | -0.23031331 | 0.23 | 0.378 | 1 | 1 |
| ppp1r10 | 0.05848 | -0.2297959 | 0.14 | 0.28 | 1 | 1 |
| adamts15a | 0.05873 | -0.22133057 | 0.2 | 0.354 | 1 | 1 |
| seta | 0.05873 | 0.104915709 | 0.58 | 0.585 | 1 | 1 |
| nfya | 0.05875 | 0.123252278 | 0.08 | 0.049 | 1 | 1 |
| tmem176l.1 | 0.05898 | -0.18865306 | 0.75 | 0.854 | 1 | 1 |
| faub | 0.0591 | 0.104516716 | 0.12 | 0.11 | 1 | 1 |
| sept12 | 0.05921 | -0.11681287 | 0.02 | 0.098 | 1 | 1 |
| impact | 0.06032 | -0.11277756 | 0.02 | 0.098 | 1 | 1 |
| dcaf7 | 0.06041 | 0.121693287 | 0.3 | 0.268 | 1 | 1 |
| nkain1 | 0.06042 | 0.154336521 | 0.2 | 0.146 | 1 | 1 |
| si:dkey-28b4.7 | 0.06052 | 0.198256214 | 0.39 | 0.317 | 1 | 1 |
| CABZ01072614.1 | 0.06063 | -0.18399108 | 0.28 | 0.439 | 1 | 1 |
| bola1 | 0.06076 | 0.125410038 | 0.11 | 0.024 | 1 | 1 |
| ggh | 0.061 | 0.125666607 | 0.12 | 0.073 | 1 | 1 |
| prpf19 | 0.06107 | -0.13538505 | 0.09 | 0.195 | 1 | 1 |
| kmt5aa | 0.06129 | 0.167981795 | 0.28 | 0.22 | 1 | 1 |
| tmem69 | 0.06218 | -0.11897565 | 0.02 | 0.098 | 1 | 1 |
| tcp11l2 | 0.0625 | 0.167011798 | 0.13 | 0.037 | 1 | 1 |
| zgc:152791 | 0.06282 | -0.2778084 | 0.13 | 0.256 | 1 | 1 |
| si:dkey-192l18.9 | 0.06284 | 0.104208277 | 0.08 | 0.012 | 1 | 1 |
| cherp | 0.06287 | 0.130630045 | 0.07 | 0.012 | 1 | 1 |
| cox4i2 | 0.06314 | 0.196114289 | 0.27 | 0.207 | 1 | 1 |
| kif5ba | 0.06317 | -0.10171555 | 0.07 | 0.159 | 1 | 1 |
| cul4b | 0.0637 | -0.13369555 | 0.07 | 0.171 | 1 | 1 |
| nat9 | 0.06376 | 0.134524534 | 0.19 | 0.146 | 1 | 1 |
| ube2d4 | 0.06419 | -0.18522402 | 0.08 | 0.195 | 1 | 1 |
| si:dkey-33c12.4 | 0.06425 | 0.178501216 | 0.28 | 0.22 | 1 | 1 |
| irf7 | 0.06436 | -0.14198433 | 0.02 | 0.073 | 1 | 1 |
| rab11fip1b | 0.06463 | -0.1898761 | 0.04 | 0.11 | 1 | 1 |
| kctd12.2 | 0.06476 | 0.102449008 | 0.26 | 0.268 | 1 | 1 |
| ino80e | 0.06481 | -0.13765606 | 0.05 | 0.146 | 1 | 1 |
| ctsd | 0.06492 | 0.177149416 | 0.17 | 0.061 | 1 | 1 |
| hmgn7 | 0.06497 | -0.10383929 | 0.74 | 0.866 | 1 | 1 |
| chtopa | 0.06507 | -0.21057812 | 0.16 | 0.305 | 1 | 1 |
| ispd | 0.06519 | 0.136223208 | 0.09 | 0.024 | 1 | 1 |
| si:rp71-36a1.5 | 0.0652 | -0.184435 | 0.03 | 0.11 | 1 | 1 |

| | | | | | | |
|---|---|---|---|---|---|---|
| gtf2a1l | 0.06532 | -0.15718942 | 0.04 | 0.134 | 1 | 1 |
| LOC101885444 | 0.06537 | -0.13594711 | 0.03 | 0.11 | 1 | 1 |
| polr2k | 0.06628 | 0.226672575 | 0.25 | 0.159 | 1 | 1 |
| brd2a | 0.06646 | -0.22833349 | 0.21 | 0.366 | 1 | 1 |
| ulk2 | 0.06649 | -0.19107504 | 0.1 | 0.171 | 1 | 1 |
| trim35-14 | 0.06656 | -0.13635986 | 0.01 | 0.073 | 1 | 1 |
| cxcl8b.1 | 0.06676 | -0.13614814 | 0.04 | 0.122 | 1 | 1 |
| rarga | 0.06695 | -0.16068315 | 0.04 | 0.134 | 1 | 1 |
| atp5mc1 | 0.06702 | -0.257319 | 0.3 | 0.451 | 1 | 1 |
| mrpl33 | 0.06704 | -0.1539628 | 0.17 | 0.305 | 1 | 1 |
| ago4 | 0.06722 | -0.1337375 | 0.02 | 0.098 | 1 | 1 |
| CABZ01073166.1 | 0.06726 | -0.1287766 | 0.02 | 0.098 | 1 | 1 |
| micall1a | 0.06727 | -0.12328837 | 0.02 | 0.098 | 1 | 1 |
| map4k4 | 0.0673 | -0.22315671 | 0.06 | 0.159 | 1 | 1 |
| jam3b | 0.0678 | -0.10855238 | 0.08 | 0.171 | 1 | 1 |
| cox5ab | 0.06813 | -0.21618933 | 0.29 | 0.451 | 1 | 1 |
| cct6a | 0.06823 | 0.119364288 | 0.47 | 0.463 | 1 | 1 |
| CT573476.1 | 0.06855 | -0.13972397 | 0.04 | 0.134 | 1 | 1 |
| arhgap42b | 0.0687 | -0.15693977 | 0.04 | 0.134 | 1 | 1 |
| BX569787.1 | 0.06876 | -0.24543008 | 0.39 | 0.561 | 1 | 1 |
| rrs1 | 0.06905 | 0.167718215 | 0.24 | 0.171 | 1 | 1 |
| nudt22 | 0.06918 | -0.11101808 | 0.01 | 0.073 | 1 | 1 |
| CU571074.1 | 0.06922 | 0.169821499 | 0.12 | 0.037 | 1 | 1 |
| CABZ01068358.1 | 0.07001 | -0.14860121 | 0.12 | 0.232 | 1 | 1 |
| cox6c | 0.07016 | -0.22896008 | 0.45 | 0.585 | 1 | 1 |
| gnb1b | 0.07047 | -0.20508156 | 0.49 | 0.585 | 1 | 1 |
| brd4 | 0.07053 | -0.13506077 | 0.06 | 0.159 | 1 | 1 |
| jagn1b | 0.07091 | -0.16398763 | 0.14 | 0.268 | 1 | 1 |
| sh3glb1b | 0.07107 | 0.159609369 | 0.15 | 0.049 | 1 | 1 |
| si:zfos-464b6.2 | 0.07116 | -0.15378177 | 0.04 | 0.11 | 1 | 1 |
| igf2b | 0.07125 | 0.203478415 | 0.44 | 0.39 | 1 | 1 |
| btg2 | 0.07126 | 0.26315032 | 0.47 | 0.305 | 1 | 1 |
| rbpjb | 0.07127 | -0.11634893 | 0.07 | 0.159 | 1 | 1 |
| clic4 | 0.07136 | -0.14089438 | 0.17 | 0.293 | 1 | 1 |
| tmem43 | 0.07155 | -0.11473629 | 0.03 | 0.11 | 1 | 1 |
| kpnb1 | 0.07166 | 0.196859743 | 0.2 | 0.085 | 1 | 1 |
| lama4 | 0.07172 | -0.10231678 | 0.25 | 0.366 | 1 | 1 |
| fkbp2 | 0.07214 | -0.18078003 | 0.27 | 0.427 | 1 | 1 |
| casp8ap2 | 0.07231 | -0.10670785 | 0.06 | 0.146 | 1 | 1 |
| st8sia6 | 0.07247 | -0.1185156 | 0.01 | 0.073 | 1 | 1 |
| si:dkey-10c21.1 | 0.07259 | -0.14213998 | 0.01 | 0.073 | 1 | 1 |
| abcc3 | 0.07267 | -0.11165351 | 0.02 | 0.085 | 1 | 1 |
| sumo2b | 0.07269 | 0.195194924 | 0.78 | 0.683 | 1 | 1 |
| usf1l | 0.07284 | -0.13206998 | 0.15 | 0.268 | 1 | 1 |
| ifi46 | 0.07289 | -0.25190843 | 0.37 | 0.537 | 1 | 1 |
| si:dkeyp-9d4.2 | 0.07293 | -0.14065294 | 0.03 | 0.11 | 1 | 1 |
| cnot9 | 0.07311 | 0.172018908 | 0.15 | 0.061 | 1 | 1 |
| lhfpl2b | 0.07315 | -0.17981008 | 0.1 | 0.22 | 1 | 1 |

| | | | | | | |
|---|---|---|---|---|---|---|
| XLOC-024248 | 0.0736 | -0.14014096 | 0.01 | 0.073 | 1 | 1 |
| si:dkey-23f9.4 | 0.0738 | -0.1147465 | 0.01 | 0.073 | 1 | 1 |
| dr1 | 0.07386 | -0.10435314 | 0.04 | 0.122 | 1 | 1 |
| hibadhb | 0.07399 | 0.201582298 | 0.19 | 0.098 | 1 | 1 |
| wbp2nl | 0.07402 | -0.22696762 | 0.09 | 0.207 | 1 | 1 |
| mir214a | 0.0741 | -0.11309142 | 0.01 | 0.073 | 1 | 1 |
| ube2g1b | 0.07449 | 0.205704251 | 0.28 | 0.183 | 1 | 1 |
| eif2b2 | 0.0745 | 0.183436392 | 0.19 | 0.098 | 1 | 1 |
| larsb | 0.07465 | -0.10189682 | 0.01 | 0.073 | 1 | 1 |
| si:ch211-245h14.1 | 0.0747 | -0.11900781 | 0.06 | 0.146 | 1 | 1 |
| CABZ01029938.1 | 0.07473 | -0.14826818 | 0.01 | 0.073 | 1 | 1 |
| nars | 0.07488 | 0.165803961 | 0.22 | 0.146 | 1 | 1 |
| afap1l2 | 0.0749 | -0.10368772 | 0.01 | 0.073 | 1 | 1 |
| tmem17 | 0.07506 | -0.15177631 | 0.06 | 0.159 | 1 | 1 |
| BX005392.2 | 0.07541 | -0.10846793 | 0.01 | 0.073 | 1 | 1 |
| itpkb | 0.07575 | -0.1145761 | 0.01 | 0.073 | 1 | 1 |
| rab1ab | 0.07578 | -0.23645209 | 0.25 | 0.402 | 1 | 1 |
| csnk2a2b | 0.0758 | -0.11691041 | 0.01 | 0.073 | 1 | 1 |
| tmem30ab | 0.07595 | -0.14104972 | 0.13 | 0.244 | 1 | 1 |
| fli1a | 0.076 | -0.13404954 | 0.01 | 0.073 | 1 | 1 |
| snai2 | 0.07617 | -0.18517369 | 0.21 | 0.354 | 1 | 1 |
| gigyf1a | 0.0762 | 0.123033932 | 0.11 | 0.085 | 1 | 1 |
| serinc2 | 0.07623 | -0.15294204 | 0.43 | 0.585 | 1 | 1 |
| kdelr2a | 0.07624 | -0.11077221 | 0.14 | 0.244 | 1 | 1 |
| atp5if1a | 0.07632 | -0.19497095 | 0.43 | 0.598 | 1 | 1 |
| parga | 0.07635 | -0.10213143 | 0.01 | 0.073 | 1 | 1 |
| usp48 | 0.07647 | -0.13269892 | 0.01 | 0.073 | 1 | 1 |
| bmi1a | 0.07649 | -0.12548444 | 0.01 | 0.073 | 1 | 1 |
| map1lc3a | 0.07651 | 0.200857039 | 0.3 | 0.207 | 1 | 1 |
| fam210ab | 0.07701 | -0.16707281 | 0.06 | 0.159 | 1 | 1 |
| mhc1zea | 0.07705 | -0.1092468 | 0.01 | 0.073 | 1 | 1 |
| tmem201 | 0.07716 | -0.11354848 | 0.01 | 0.073 | 1 | 1 |
| purba | 0.07731 | 0.151178955 | 0.2 | 0.146 | 1 | 1 |
| riox2 | 0.07753 | 0.155266516 | 0.1 | 0.024 | 1 | 1 |
| rnaseh2c | 0.07779 | -0.16458096 | 0.09 | 0.207 | 1 | 1 |
| pfdn5 | 0.07787 | 0.12060378 | 0.4 | 0.378 | 1 | 1 |
| eif1axa | 0.07817 | -0.17589839 | 0.18 | 0.317 | 1 | 1 |
| cox5aa | 0.07833 | -0.20349819 | 0.43 | 0.512 | 1 | 1 |
| zgc:158320 | 0.07855 | 0.206721134 | 0.17 | 0.073 | 1 | 1 |
| fthl27 | 0.07858 | 0.175327816 | 0.88 | 0.793 | 1 | 1 |
| morf4l1 | 0.07868 | 0.195921531 | 0.47 | 0.378 | 1 | 1 |
| lims1 | 0.07871 | -0.17309882 | 0.05 | 0.146 | 1 | 1 |
| nme2b.2 | 0.07886 | 0.22641512 | 0.49 | 0.329 | 1 | 1 |
| rab11ba | 0.079 | -0.15932971 | 0.13 | 0.256 | 1 | 1 |
| hnrnpm | 0.07902 | 0.19082746 | 0.19 | 0.122 | 1 | 1 |
| myl9a | 0.0791 | -0.21927707 | 0.09 | 0.207 | 1 | 1 |
| fam49a | 0.07913 | 0.112568889 | 0.08 | 0.012 | 1 | 1 |
| acvrl1 | 0.07914 | -0.20934045 | 0.09 | 0.207 | 1 | 1 |

| | | | | | | |
|---|---|---|---|---|---|---|
| ctdsp2 | 0.07916 | -0.13394923 | 0.36 | 0.5 | 1 | 1 |
| zdbf2 | 0.07982 | 0.113337255 | 0.08 | 0.012 | 1 | 1 |
| vdac1 | 0.08106 | 0.159547077 | 0.49 | 0.439 | 1 | 1 |
| XLOC-004556 | 0.08143 | -0.14140854 | 0.09 | 0.195 | 1 | 1 |
| gpr85 | 0.08161 | 0.184930444 | 0.11 | 0.037 | 1 | 1 |
| agap3 | 0.08162 | -0.14181762 | 0.05 | 0.146 | 1 | 1 |
| crip1 | 0.08163 | 0.220925515 | 0.2 | 0.085 | 1 | 1 |
| svbp | 0.08176 | 0.121092813 | 0.1 | 0.024 | 1 | 1 |
| steap4 | 0.08177 | -0.14227216 | 0.03 | 0.11 | 1 | 1 |
| dusp6 | 0.08214 | -0.15861762 | 0.03 | 0.11 | 1 | 1 |
| ctbp2a | 0.0824 | -0.10362011 | 0.22 | 0.329 | 1 | 1 |
| prmt1 | 0.08265 | -0.15312964 | 0.34 | 0.488 | 1 | 1 |
| robo1 | 0.08274 | -0.12580368 | 0.16 | 0.268 | 1 | 1 |
| utp25 | 0.083 | -0.17300403 | 0.03 | 0.098 | 1 | 1 |
| lsm12a | 0.08309 | 0.140306392 | 0.15 | 0.098 | 1 | 1 |
| c4b | 0.0841 | -0.10565524 | 0.01 | 0.061 | 1 | 1 |
| ost4 | 0.08414 | -0.23953858 | 0.29 | 0.415 | 1 | 1 |
| pes | 0.08437 | -0.16838477 | 0.14 | 0.268 | 1 | 1 |
| cdca4 | 0.08447 | 0.141159731 | 0.26 | 0.207 | 1 | 1 |
| faf1 | 0.08466 | 0.128020295 | 0.13 | 0.098 | 1 | 1 |
| psma6l | 0.08532 | -0.14093161 | 0.06 | 0.159 | 1 | 1 |
| lmnb1 | 0.08582 | -0.13791683 | 0.08 | 0.183 | 1 | 1 |
| nifk | 0.08611 | -0.116467 | 0.15 | 0.256 | 1 | 1 |
| si:dkey-89b17.4 | 0.08644 | 0.189318335 | 0.16 | 0.061 | 1 | 1 |
| cotl1 | 0.08655 | -0.20072478 | 0.13 | 0.256 | 1 | 1 |
| flna | 0.08684 | -0.24613292 | 0.26 | 0.415 | 1 | 1 |
| si:ch73-37h15.2 | 0.08699 | -0.1476496 | 0.07 | 0.171 | 1 | 1 |
| fbn2b | 0.087 | 0.21276068 | 0.51 | 0.427 | 1 | 1 |
| cdr2l | 0.0872 | -0.15449999 | 0.03 | 0.11 | 1 | 1 |
| asb12a | 0.08794 | -0.10971325 | 0.04 | 0.122 | 1 | 1 |
| aars | 0.08802 | -0.16583785 | 0.12 | 0.244 | 1 | 1 |
| alas1 | 0.08822 | 0.177824827 | 0.2 | 0.098 | 1 | 1 |
| nfkbiaa | 0.08854 | -0.20719599 | 0.17 | 0.232 | 1 | 1 |
| ndfip1 | 0.08889 | -0.1415253 | 0.28 | 0.415 | 1 | 1 |
| cdh2 | 0.08906 | -0.17779908 | 0.29 | 0.439 | 1 | 1 |
| etf1b | 0.08909 | -0.19601208 | 0.33 | 0.488 | 1 | 1 |
| pdcd10b | 0.08911 | 0.158388765 | 0.18 | 0.11 | 1 | 1 |
| zgc:92664 | 0.08956 | 0.143697158 | 0.2 | 0.159 | 1 | 1 |
| rab2a | 0.08979 | 0.139770624 | 0.42 | 0.39 | 1 | 1 |
| tm2d3 | 0.09 | 0.18983452 | 0.18 | 0.098 | 1 | 1 |
| rnf213b | 0.09036 | -0.14630502 | 0.03 | 0.11 | 1 | 1 |
| tmem11 | 0.0906 | 0.186836102 | 0.18 | 0.085 | 1 | 1 |
| g3bp1 | 0.09064 | -0.14647013 | 0.07 | 0.171 | 1 | 1 |
| sdr16c5a | 0.09067 | 0.118544101 | 0.07 | 0.012 | 1 | 1 |
| wt1b | 0.09068 | -0.19672213 | 0.06 | 0.159 | 1 | 1 |
| zdhhc20a | 0.09092 | 0.166434056 | 0.12 | 0.037 | 1 | 1 |
| vim | 0.09105 | 0.20625798 | 0.26 | 0.134 | 1 | 1 |
| cyp11c1 | 0.09109 | 0.198828601 | 0.14 | 0.073 | 1 | 1 |

| | | | | | | |
|---|---|---|---|---|---|---|
| ms4a17a.11 | 0.09138 | 0.216006698 | 0.48 | 0.366 | 1 | 1 |
| zgc:161969 | 0.09153 | 0.117257495 | 0.12 | 0.098 | 1 | 1 |
| pcyox1 | 0.0916 | -0.1473844 | 0.09 | 0.195 | 1 | 1 |
| gart | 0.09194 | -0.1049505 | 0.03 | 0.11 | 1 | 1 |
| sf3b1 | 0.09197 | 0.225627174 | 0.21 | 0.098 | 1 | 1 |
| lin7a | 0.09218 | -0.18745752 | 0.07 | 0.171 | 1 | 1 |
| itprip | 0.09268 | -0.13762012 | 0.03 | 0.11 | 1 | 1 |
| ube2l3a | 0.09291 | -0.17724134 | 0.16 | 0.293 | 1 | 1 |
| p3h2 | 0.09294 | -0.17016585 | 0.06 | 0.159 | 1 | 1 |
| gdi1 | 0.09304 | -0.12851616 | 0.03 | 0.11 | 1 | 1 |
| lysmd3 | 0.09307 | 0.17108996 | 0.12 | 0.049 | 1 | 1 |
| mfge8a | 0.09328 | 0.16835739 | 0.13 | 0.049 | 1 | 1 |
| ssr1 | 0.09334 | -0.18621063 | 0.4 | 0.561 | 1 | 1 |
| hnrnpl2 | 0.09338 | -0.14746339 | 0.21 | 0.341 | 1 | 1 |
| tubb2b | 0.09342 | -0.12970028 | 0.39 | 0.537 | 1 | 1 |
| stat3 | 0.09343 | 0.151873863 | 0.11 | 0.061 | 1 | 1 |
| srek1ip1 | 0.09353 | -0.14107162 | 0.03 | 0.11 | 1 | 1 |
| lamp2 | 0.09373 | -0.20304173 | 0.22 | 0.366 | 1 | 1 |
| ywhabl | 0.09415 | 0.152967383 | 0.55 | 0.5 | 1 | 1 |
| ctnna1 | 0.09425 | -0.11619431 | 0.1 | 0.195 | 1 | 1 |
| jpt2 | 0.09443 | -0.12321537 | 0.25 | 0.378 | 1 | 1 |
| tab1 | 0.09448 | 0.123553758 | 0.19 | 0.159 | 1 | 1 |
| spns1 | 0.09448 | -0.15000601 | 0.06 | 0.159 | 1 | 1 |
| tspan9b | 0.09449 | -0.20702293 | 0.13 | 0.256 | 1 | 1 |
| aldh9a1a.1 | 0.09475 | 0.144765089 | 0.69 | 0.659 | 1 | 1 |
| smarcc2 | 0.09499 | -0.15057999 | 0.06 | 0.159 | 1 | 1 |
| c4 | 0.09519 | -0.17416196 | 0.13 | 0.244 | 1 | 1 |
| si:ch73-237c6.1 | 0.09545 | -0.15615078 | 0.04 | 0.122 | 1 | 1 |
| sf3b2 | 0.09561 | 0.22325245 | 0.31 | 0.207 | 1 | 1 |
| arpc4l | 0.09606 | -0.19425405 | 0.33 | 0.488 | 1 | 1 |
| mtpn | 0.09654 | -0.13450447 | 0.37 | 0.512 | 1 | 1 |
| rbmx | 0.09733 | -0.21012727 | 0.21 | 0.354 | 1 | 1 |
| utp4 | 0.09757 | 0.138004564 | 0.1 | 0.037 | 1 | 1 |
| sox11a | 0.0976 | 0.139295634 | 0.23 | 0.195 | 1 | 1 |
| nsrp1 | 0.09795 | -0.14580093 | 0.03 | 0.085 | 1 | 1 |
| rpe | 0.09806 | 0.138032841 | 0.17 | 0.073 | 1 | 1 |
| caprin1b | 0.09821 | 0.183688938 | 0.44 | 0.341 | 1 | 1 |
| cdkn1bb | 0.09822 | 0.153215586 | 0.21 | 0.146 | 1 | 1 |
| BX663503.3 | 0.09823 | 0.139983623 | 0.07 | 0.037 | 1 | 1 |
| dldh | 0.09854 | 0.141210519 | 0.3 | 0.256 | 1 | 1 |
| zgc:112356 | 0.09872 | 0.105720995 | 0.16 | 0.146 | 1 | 1 |
| zdhhc4 | 0.09881 | 0.16824655 | 0.15 | 0.073 | 1 | 1 |
| cbx8a | 0.09935 | -0.1230612 | 0.04 | 0.122 | 1 | 1 |
| psmb7 | 0.0996 | 0.209569127 | 0.49 | 0.378 | 1 | 1 |
| maco1a | 0.09966 | -0.1141768 | 0.09 | 0.183 | 1 | 1 |
| arpc3 | 0.09968 | -0.20625604 | 0.4 | 0.549 | 1 | 1 |
| ndufa11 | 0.09973 | -0.17048665 | 0.25 | 0.39 | 1 | 1 |
| p3h4 | 0.0998 | -0.10008187 | 0.02 | 0.085 | 1 | 1 |

| | | | | | | |
|---|---|---|---|---|---|---|
| rpp21 | 0.1001 | -0.12800248 | 0.07 | 0.159 | 1 | 1 |
| smco4 | 0.10019 | 0.123324098 | 0.12 | 0.037 | 1 | 1 |
| ino80c | 0.10083 | 0.163247328 | 0.16 | 0.061 | 1 | 1 |
| vrk1 | 0.10104 | -0.1693718 | 0.07 | 0.159 | 1 | 1 |
| cald1a | 0.10108 | -0.20359174 | 0.56 | 0.707 | 1 | 1 |
| antxr2a | 0.10117 | 0.220313472 | 0.56 | 0.451 | 1 | 1 |
| cfl1 | 0.10127 | -0.16947984 | 0.71 | 0.841 | 1 | 1 |
| c1qbp | 0.10148 | -0.1485 | 0.19 | 0.317 | 1 | 1 |
| rnf13 | 0.10268 | 0.123551001 | 0.22 | 0.195 | 1 | 1 |
| hsdl2 | 0.10292 | -0.14729299 | 0.07 | 0.171 | 1 | 1 |
| calr | 0.10309 | -0.21617339 | 0.25 | 0.354 | 1 | 1 |
| sort1a | 0.10323 | -0.14755542 | 0.02 | 0.085 | 1 | 1 |
| ndfip2 | 0.10346 | 0.151864441 | 0.11 | 0.037 | 1 | 1 |
| rab40c | 0.10401 | -0.14108764 | 0.04 | 0.122 | 1 | 1 |
| rpz5 | 0.10402 | -0.19361201 | 0.38 | 0.537 | 1 | 1 |
| pias2 | 0.10454 | -0.12193542 | 0.04 | 0.122 | 1 | 1 |
| tmem59 | 0.10464 | -0.14602593 | 0.29 | 0.427 | 1 | 1 |
| si:ch73-380l3.2 | 0.10485 | -0.15343987 | 0.08 | 0.183 | 1 | 1 |
| CDK2AP1 | 0.10496 | 0.139493454 | 0.12 | 0.061 | 1 | 1 |
| tia1l | 0.10546 | -0.12013883 | 0.13 | 0.232 | 1 | 1 |
| zgc:66433 | 0.10556 | -0.11478894 | 0.09 | 0.183 | 1 | 1 |
| zgc:162509 | 0.10557 | -0.10446267 | 0.08 | 0.171 | 1 | 1 |
| epha3 | 0.10568 | 0.155210756 | 0.12 | 0.037 | 1 | 1 |
| prpf38b | 0.10604 | -0.14854915 | 0.07 | 0.171 | 1 | 1 |
| ctnnb1 | 0.10658 | -0.22428102 | 0.3 | 0.451 | 1 | 1 |
| grpel2 | 0.10702 | 0.118653216 | 0.36 | 0.329 | 1 | 1 |
| irf1b | 0.10711 | -0.20436861 | 0.1 | 0.207 | 1 | 1 |
| grhl1 | 0.10728 | -0.18338095 | 0.24 | 0.378 | 1 | 1 |
| tubb4b | 0.10757 | 0.171279787 | 0.85 | 0.793 | 1 | 1 |
| golt1bb | 0.10759 | -0.18629197 | 0.09 | 0.195 | 1 | 1 |
| si:dkey-11f4.14 | 0.1077 | 0.218845759 | 0.46 | 0.366 | 1 | 1 |
| marcksl1b | 0.10797 | -0.15478134 | 0.6 | 0.744 | 1 | 1 |
| cgnl1 | 0.10809 | -0.13163006 | 0.09 | 0.183 | 1 | 1 |
| capn2b | 0.10839 | -0.13582789 | 0.02 | 0.085 | 1 | 1 |
| psmc1a | 0.10885 | -0.14808912 | 0.19 | 0.317 | 1 | 1 |
| map6a | 0.10899 | -0.11793611 | 0.02 | 0.085 | 1 | 1 |
| si:rp71-36a1.1 | 0.10915 | 0.144129292 | 0.13 | 0.073 | 1 | 1 |
| cnot8 | 0.10918 | 0.163167661 | 0.12 | 0.049 | 1 | 1 |
| cmtr1 | 0.1092 | -0.11086968 | 0.02 | 0.085 | 1 | 1 |
| CABZ01110881.1 | 0.10934 | -0.12520168 | 0.07 | 0.159 | 1 | 1 |
| rbx1 | 0.10947 | 0.160422397 | 0.5 | 0.451 | 1 | 1 |
| lnx2b | 0.10952 | 0.183502715 | 0.17 | 0.085 | 1 | 1 |
| ywhaba | 0.10977 | -0.12810591 | 0.41 | 0.549 | 1 | 1 |
| gsto2 | 0.10983 | 0.176686657 | 0.21 | 0.098 | 1 | 1 |
| XLOC-043857 | 0.11034 | -0.16079841 | 0.08 | 0.183 | 1 | 1 |
| sypl2b | 0.11167 | -0.16986988 | 0.08 | 0.183 | 1 | 1 |
| dhfr | 0.1117 | -0.13479554 | 0.04 | 0.122 | 1 | 1 |
| si:ch73-86n18.1 | 0.1119 | -0.13941035 | 0.02 | 0.085 | 1 | 1 |

| | | | | | | |
|---|---|---|---|---|---|---|
| cyfip1 | 0.11193 | -0.19499459 | 0.08 | 0.183 | 1 | 1 |
| zgc:163098 | 0.11201 | -0.14816083 | 0.04 | 0.122 | 1 | 1 |
| myef2 | 0.1122 | -0.16707156 | 0.12 | 0.195 | 1 | 1 |
| eef1e1 | 0.11221 | -0.10887388 | 0.02 | 0.085 | 1 | 1 |
| bcl2l1 | 0.11266 | 0.15806523 | 0.14 | 0.085 | 1 | 1 |
| plk2a | 0.11272 | 0.111589931 | 0.1 | 0.037 | 1 | 1 |
| phactr2 | 0.11325 | -0.12011586 | 0.02 | 0.085 | 1 | 1 |
| cyp17a1 | 0.11325 | 0.29309151 | 0.1 | 0.037 | 1 | 1 |
| psat1 | 0.11333 | -0.10833078 | 0.02 | 0.085 | 1 | 1 |
| si:dkey-286j15.3 | 0.11343 | -0.10851918 | 0.07 | 0.159 | 1 | 1 |
| si:dkey-260j18.2 | 0.11386 | 0.112735053 | 0.07 | 0.012 | 1 | 1 |
| mob1ba | 0.11416 | -0.10948045 | 0.04 | 0.122 | 1 | 1 |
| ubqln4 | 0.11418 | -0.15351103 | 0.08 | 0.146 | 1 | 1 |
| nansa | 0.11421 | -0.15288968 | 0.08 | 0.183 | 1 | 1 |
| si:dkey-239h2.3 | 0.11425 | -0.20230713 | 0.35 | 0.451 | 1 | 1 |
| dhrs4 | 0.11437 | 0.142642157 | 0.19 | 0.085 | 1 | 1 |
| nsun2 | 0.11446 | -0.1351634 | 0.04 | 0.122 | 1 | 1 |
| tecrb | 0.11448 | 0.132878188 | 0.09 | 0.024 | 1 | 1 |
| pmp22a | 0.1145 | -0.10004259 | 0.9 | 0.915 | 1 | 1 |
| atp6ap2 | 0.11455 | 0.143823103 | 0.19 | 0.134 | 1 | 1 |
| wu:fi04e12 | 0.11468 | 0.138812442 | 0.12 | 0.049 | 1 | 1 |
| slbp | 0.11481 | 0.113484464 | 0.06 | 0.012 | 1 | 1 |
| smc1al | 0.11483 | 0.130486374 | 0.35 | 0.305 | 1 | 1 |
| mast2 | 0.11502 | -0.16184729 | 0.04 | 0.122 | 1 | 1 |
| thoc1 | 0.11544 | -0.12490754 | 0.04 | 0.122 | 1 | 1 |
| CABZ01058650.1 | 0.11569 | 0.138555647 | 0.19 | 0.134 | 1 | 1 |
| rasl11b | 0.11578 | -0.10298012 | 0.03 | 0.098 | 1 | 1 |
| ptpn9a | 0.11593 | 0.146556772 | 0.09 | 0.024 | 1 | 1 |
| slc7a10a | 0.1161 | -0.14343811 | 0.04 | 0.122 | 1 | 1 |
| pfdn4 | 0.11611 | -0.11803849 | 0.38 | 0.512 | 1 | 1 |
| rasa3 | 0.11625 | -0.10623429 | 0.01 | 0.061 | 1 | 1 |
| sugt1 | 0.11626 | -0.13773402 | 0.04 | 0.122 | 1 | 1 |
| ift80 | 0.11696 | 0.106476199 | 0.07 | 0.012 | 1 | 1 |
| ndufs2 | 0.11718 | -0.11795488 | 0.15 | 0.256 | 1 | 1 |
| si:ch211-22i13.2 | 0.11721 | 0.147063096 | 0.17 | 0.122 | 1 | 1 |
| wu:fb74b10 | 0.11727 | -0.15430534 | 0.05 | 0.134 | 1 | 1 |
| eif1b | 0.11738 | 0.17265373 | 0.78 | 0.683 | 1 | 1 |
| phyh | 0.1174 | 0.129772765 | 0.15 | 0.098 | 1 | 1 |
| atpaf2 | 0.11755 | 0.123462669 | 0.08 | 0.024 | 1 | 1 |
| si:ch211-196c10.15 | 0.11785 | -0.11598421 | 0.02 | 0.085 | 1 | 1 |
| ywhaqb | 0.11793 | -0.19401543 | 0.42 | 0.573 | 1 | 1 |
| si:dkeyp-118h3.6 | 0.11814 | 0.116087436 | 0.07 | 0.012 | 1 | 1 |
| mepce | 0.11836 | 0.125950384 | 0.08 | 0.024 | 1 | 1 |
| sdhaf2 | 0.11868 | -0.10827209 | 0.02 | 0.085 | 1 | 1 |
| pbxip1b | 0.11907 | -0.15439692 | 0.04 | 0.098 | 1 | 1 |
| zfp36l1a | 0.11921 | -0.12843186 | 0.51 | 0.646 | 1 | 1 |
| adhfe1 | 0.11924 | -0.11910542 | 0.02 | 0.085 | 1 | 1 |
| cwf19l2 | 0.11934 | -0.1108946 | 0.03 | 0.098 | 1 | 1 |

| | | | | | | |
|---|---|---|---|---|---|---|
| ms4a17a.9 | 0.11944 | 0.22241222 | 0.43 | 0.305 | 1 | 1 |
| kctd5a | 0.11952 | 0.183875346 | 0.37 | 0.293 | 1 | 1 |
| cstf2 | 0.11965 | 0.161120726 | 0.21 | 0.146 | 1 | 1 |
| socs1b | 0.11972 | -0.11652873 | 0.02 | 0.085 | 1 | 1 |
| mospd1 | 0.1203 | -0.10878186 | 0.02 | 0.085 | 1 | 1 |
| loxl2a | 0.12031 | -0.24755565 | 0.24 | 0.366 | 1 | 1 |
| parvab | 0.12032 | -0.10468242 | 0.02 | 0.085 | 1 | 1 |
| det1 | 0.12038 | -0.10506321 | 0.02 | 0.085 | 1 | 1 |
| rhoac | 0.12039 | 0.164435094 | 0.41 | 0.354 | 1 | 1 |
| kirrel3l | 0.12069 | 0.205948095 | 0.17 | 0.073 | 1 | 1 |
| si:ch211-136a13.1 | 0.12075 | -0.13222598 | 0.06 | 0.146 | 1 | 1 |
| mych | 0.12113 | -0.11877799 | 0.26 | 0.378 | 1 | 1 |
| tm9sf4 | 0.1214 | -0.1840684 | 0.11 | 0.207 | 1 | 1 |
| glipr2l | 0.12154 | -0.12161961 | 0.06 | 0.146 | 1 | 1 |
| si:ch211-132g1.3 | 0.12221 | -0.13395038 | 0.02 | 0.085 | 1 | 1 |
| pdlim1 | 0.1224 | 0.144208276 | 0.16 | 0.098 | 1 | 1 |
| tpbgb | 0.12242 | -0.14176834 | 0.02 | 0.085 | 1 | 1 |
| zgc:92818 | 0.12257 | 0.144644022 | 0.16 | 0.11 | 1 | 1 |
| idh2 | 0.12276 | 0.199975379 | 0.6 | 0.5 | 1 | 1 |
| atp5f1e | 0.12282 | 0.123689423 | 0.72 | 0.707 | 1 | 1 |
| rbfa | 0.12282 | -0.10510176 | 0.02 | 0.085 | 1 | 1 |
| arrb2a | 0.12285 | -0.13159369 | 0.02 | 0.085 | 1 | 1 |
| setdb1b | 0.12291 | -0.12484039 | 0.02 | 0.085 | 1 | 1 |
| scarb2a | 0.12293 | -0.11624219 | 0.01 | 0.061 | 1 | 1 |
| gna11b | 0.12295 | 0.180830696 | 0.15 | 0.061 | 1 | 1 |
| nucks1b | 0.12295 | -0.10329428 | 0.29 | 0.402 | 1 | 1 |
| impdh1b | 0.12334 | 0.149175946 | 0.14 | 0.061 | 1 | 1 |
| gmfb | 0.12353 | 0.160426082 | 0.43 | 0.366 | 1 | 1 |
| appb | 0.12363 | -0.13387939 | 0.18 | 0.293 | 1 | 1 |
| rbpms2b | 0.12377 | 0.127346929 | 0.09 | 0.049 | 1 | 1 |
| si:ch211-133n4.4 | 0.12384 | -0.13504281 | 0.29 | 0.415 | 1 | 1 |
| slco2b1 | 0.12397 | -0.22380511 | 0.35 | 0.5 | 1 | 1 |
| tmem178b | 0.12416 | 0.211283457 | 0.14 | 0.061 | 1 | 1 |
| gnai1 | 0.12424 | -0.10919381 | 0.09 | 0.183 | 1 | 1 |
| fzd7b | 0.12425 | -0.15024871 | 0.2 | 0.317 | 1 | 1 |
| hypk | 0.12444 | -0.19391396 | 0.41 | 0.561 | 1 | 1 |
| dpm2 | 0.12445 | 0.175237895 | 0.17 | 0.073 | 1 | 1 |
| rsrp1 | 0.12449 | -0.18756003 | 0.24 | 0.378 | 1 | 1 |
| lpar1 | 0.12471 | -0.23474773 | 0.23 | 0.366 | 1 | 1 |
| clu | 0.12513 | -0.17883257 | 0.1 | 0.207 | 1 | 1 |
| klf11a | 0.12541 | 0.160845124 | 0.13 | 0.049 | 1 | 1 |
| tubb5 | 0.12586 | -0.17271071 | 0.15 | 0.268 | 1 | 1 |
| CU459056.1 | 0.1259 | 0.196686713 | 0.22 | 0.122 | 1 | 1 |
| zgc:66160 | 0.12732 | -0.1217987 | 0.07 | 0.159 | 1 | 1 |
| CT573342.1 | 0.12738 | -0.11266928 | 0.01 | 0.061 | 1 | 1 |
| ap3s2 | 0.12792 | 0.111844713 | 0.09 | 0.024 | 1 | 1 |
| DDX17 | 0.12803 | -0.18337488 | 0.1 | 0.207 | 1 | 1 |
| rbck1 | 0.1281 | -0.13307447 | 0.05 | 0.134 | 1 | 1 |

| | | | | | | |
|---|---|---|---|---|---|---|
| adi1 | 0.12828 | -0.15426335 | 0.14 | 0.256 | 1 | 1 |
| znf362b | 0.12828 | 0.117204689 | 0.11 | 0.085 | 1 | 1 |
| cish | 0.12852 | 0.142742269 | 0.11 | 0.061 | 1 | 1 |
| skib | 0.12857 | -0.16490348 | 0.13 | 0.244 | 1 | 1 |
| eno3 | 0.12874 | -0.18147171 | 0.27 | 0.366 | 1 | 1 |
| snrnp200 | 0.12876 | -0.10239437 | 0.09 | 0.171 | 1 | 1 |
| noc3l | 0.12878 | 0.132814876 | 0.11 | 0.037 | 1 | 1 |
| lsm5 | 0.12917 | -0.14321978 | 0.27 | 0.402 | 1 | 1 |
| ddit3 | 0.12922 | 0.177064044 | 0.36 | 0.293 | 1 | 1 |
| plpp3 | 0.12969 | -0.18800783 | 0.08 | 0.171 | 1 | 1 |
| cux1a | 0.1299 | 0.1754861 | 0.43 | 0.366 | 1 | 1 |
| nckap1 | 0.1302 | -0.10224002 | 0.11 | 0.195 | 1 | 1 |
| kmt2a | 0.13042 | 0.164466497 | 0.13 | 0.061 | 1 | 1 |
| ccl39.7 | 0.13057 | -0.11344384 | 0.15 | 0.171 | 1 | 1 |
| cdk7 | 0.1306 | 0.121204186 | 0.1 | 0.037 | 1 | 1 |
| cpne3 | 0.13083 | -0.14099055 | 0.17 | 0.28 | 1 | 1 |
| ap1m1 | 0.13086 | 0.124037939 | 0.12 | 0.073 | 1 | 1 |
| pknox1.1 | 0.13086 | 0.146923061 | 0.11 | 0.037 | 1 | 1 |
| si:ch211-276c2.4 | 0.1309 | -0.12324892 | 0.03 | 0.098 | 1 | 1 |
| clns1a | 0.13114 | 0.129970763 | 0.21 | 0.11 | 1 | 1 |
| mapre1b | 0.13122 | 0.189050803 | 0.26 | 0.146 | 1 | 1 |
| ndufv1 | 0.13144 | 0.171911151 | 0.21 | 0.122 | 1 | 1 |
| ephb3 | 0.1315 | -0.16052415 | 0.24 | 0.366 | 1 | 1 |
| sptlc2a | 0.13224 | -0.21917505 | 0.28 | 0.402 | 1 | 1 |
| ago1 | 0.13229 | 0.126344569 | 0.09 | 0.024 | 1 | 1 |
| NDUFAF8 | 0.13268 | -0.10367166 | 0.01 | 0.061 | 1 | 1 |
| col27a1b | 0.13272 | 0.237419554 | 0.47 | 0.329 | 1 | 1 |
| znf330 | 0.13284 | 0.158043107 | 0.14 | 0.061 | 1 | 1 |
| ier5l | 0.13352 | -0.11531458 | 0.16 | 0.256 | 1 | 1 |
| pxna | 0.13376 | -0.12871996 | 0.05 | 0.134 | 1 | 1 |
| gnl2 | 0.13408 | 0.139545787 | 0.11 | 0.037 | 1 | 1 |
| pawr | 0.13423 | -0.16313023 | 0.03 | 0.098 | 1 | 1 |
| mtss1la | 0.13444 | -0.18494511 | 0.14 | 0.256 | 1 | 1 |
| tmpob | 0.13471 | -0.13902106 | 0.05 | 0.134 | 1 | 1 |
| cnp | 0.13489 | 0.175939152 | 0.3 | 0.22 | 1 | 1 |
| slc7a6os | 0.13499 | -0.14735394 | 0.05 | 0.134 | 1 | 1 |
| gnl3 | 0.13505 | -0.20763381 | 0.18 | 0.293 | 1 | 1 |
| si:ch211-286b5.5 | 0.13512 | 0.143019703 | 0.5 | 0.451 | 1 | 1 |
| si:ch211-212k18.7 | 0.1353 | -0.22890955 | 0.17 | 0.28 | 1 | 1 |
| rarab | 0.1354 | -0.11409936 | 0.01 | 0.061 | 1 | 1 |
| rab7a | 0.13568 | -0.10485697 | 0.12 | 0.207 | 1 | 1 |
| ppme1 | 0.1357 | 0.160680862 | 0.16 | 0.073 | 1 | 1 |
| baxa | 0.13617 | 0.155800923 | 0.13 | 0.049 | 1 | 1 |
| jund | 0.13658 | 0.185581928 | 0.18 | 0.085 | 1 | 1 |
| cpsf3 | 0.13667 | -0.1003754 | 0.03 | 0.098 | 1 | 1 |
| CU459094.2 | 0.13756 | -0.17954253 | 0.16 | 0.28 | 1 | 1 |
| ldb1b | 0.1381 | -0.1662637 | 0.05 | 0.11 | 1 | 1 |
| mir21-1 | 0.13815 | -0.20074971 | 0.13 | 0.244 | 1 | 1 |

| | | | | | | |
|---|---|---|---|---|---|---|
| aldh1a2 | 0.13817 | -0.17364392 | 0.29 | 0.427 | 1 | 1 |
| nenf | 0.13847 | -0.18764249 | 0.21 | 0.341 | 1 | 1 |
| amdhd2 | 0.1391 | 0.139221818 | 0.11 | 0.037 | 1 | 1 |
| ndufs6 | 0.13915 | -0.18271306 | 0.19 | 0.317 | 1 | 1 |
| si:dkey-27i16.2 | 0.13963 | 0.118400905 | 0.15 | 0.122 | 1 | 1 |
| atp5fa1 | 0.13987 | -0.18676098 | 0.58 | 0.683 | 1 | 1 |
| ubr5 | 0.14002 | 0.157294808 | 0.15 | 0.061 | 1 | 1 |
| ctdspl2a | 0.14037 | -0.16819971 | 0.13 | 0.244 | 1 | 1 |
| emilin1a | 0.14041 | -0.14791929 | 0.03 | 0.098 | 1 | 1 |
| c1qtnf5 | 0.14065 | 0.26082387 | 0.34 | 0.22 | 1 | 1 |
| stk17b | 0.14067 | 0.101584354 | 0.11 | 0.037 | 1 | 1 |
| ilk | 0.14095 | -0.18735147 | 0.2 | 0.329 | 1 | 1 |
| rabac1 | 0.14118 | 0.14241398 | 0.13 | 0.073 | 1 | 1 |
| copb2 | 0.14136 | -0.12425767 | 0.13 | 0.232 | 1 | 1 |
| bnip4 | 0.1419 | 0.1324735 | 0.09 | 0.024 | 1 | 1 |
| lipf | 0.14222 | -0.20305493 | 0.18 | 0.305 | 1 | 1 |
| acadm | 0.14237 | -0.18071748 | 0.19 | 0.317 | 1 | 1 |
| ankhb | 0.14241 | -0.10385162 | 0.01 | 0.061 | 1 | 1 |
| ndufs4 | 0.14243 | 0.151013334 | 0.28 | 0.22 | 1 | 1 |
| sdhb | 0.14258 | -0.11688114 | 0.14 | 0.244 | 1 | 1 |
| BX537342.1 | 0.14266 | -0.11657553 | 0.01 | 0.061 | 1 | 1 |
| lsp1 | 0.14294 | -0.12456097 | 0.09 | 0.183 | 1 | 1 |
| hist2h2l | 0.14331 | -0.18877912 | 0.41 | 0.476 | 1 | 1 |
| rcc2 | 0.14332 | -0.12671481 | 0.22 | 0.329 | 1 | 1 |
| cycsb | 0.14345 | -0.2016042 | 0.28 | 0.402 | 1 | 1 |
| tfcp2 | 0.144 | 0.139337146 | 0.11 | 0.037 | 1 | 1 |
| zbtb46 | 0.14414 | -0.15647602 | 0.03 | 0.098 | 1 | 1 |
| sdad1 | 0.14437 | 0.155067263 | 0.13 | 0.073 | 1 | 1 |
| pa2g4a | 0.1445 | -0.18475906 | 0.21 | 0.329 | 1 | 1 |
| ralba | 0.14453 | 0.167952545 | 0.15 | 0.061 | 1 | 1 |
| XLOC-014336 | 0.14465 | -0.10846912 | 0.04 | 0.11 | 1 | 1 |
| idh1 | 0.14485 | -0.10341905 | 0.02 | 0.073 | 1 | 1 |
| mrps30 | 0.14488 | -0.13538514 | 0.08 | 0.134 | 1 | 1 |
| LOC100334937 | 0.14503 | -0.17116866 | 0.04 | 0.11 | 1 | 1 |
| col18a1a | 0.14522 | -0.20556345 | 0.49 | 0.634 | 1 | 1 |
| palm3 | 0.1456 | 0.120410827 | 0.07 | 0.024 | 1 | 1 |
| ntn1b | 0.14566 | 0.131250346 | 0.09 | 0.024 | 1 | 1 |
| cdc26 | 0.1457 | 0.148517943 | 0.27 | 0.22 | 1 | 1 |
| eif4a1a | 0.1463 | -0.20259408 | 0.47 | 0.61 | 1 | 1 |
| siah1 | 0.14686 | -0.15114446 | 0.07 | 0.159 | 1 | 1 |
| LOC110439840 | 0.14702 | 0.108081709 | 0.09 | 0.024 | 1 | 1 |
| fam172a | 0.14771 | -0.12669735 | 0.09 | 0.183 | 1 | 1 |
| clpp | 0.14792 | 0.130024232 | 0.16 | 0.098 | 1 | 1 |
| zgc:113921 | 0.14793 | -0.10583854 | 0.01 | 0.061 | 1 | 1 |
| castor2 | 0.14862 | 0.146182061 | 0.1 | 0.037 | 1 | 1 |
| atp5po | 0.14872 | 0.107644878 | 0.43 | 0.415 | 1 | 1 |
| rnf126 | 0.14875 | -0.10165353 | 0.01 | 0.061 | 1 | 1 |
| inhbaa | 0.14879 | 0.101256608 | 0.83 | 0.829 | 1 | 1 |

| | | | | | | |
|---|---|---|---|---|---|---|
| zgc:162025 | 0.14925 | 0.114867707 | 0.23 | 0.134 | 1 | 1 |
| rab5if | 0.14938 | 0.180612275 | 0.25 | 0.146 | 1 | 1 |
| hddc3 | 0.14974 | 0.106180988 | 0.07 | 0.024 | 1 | 1 |
| pak1ip1 | 0.14974 | -0.12878395 | 0.06 | 0.146 | 1 | 1 |
| sdha | 0.14991 | 0.159865599 | 0.18 | 0.122 | 1 | 1 |
| c1galt1b | 0.15091 | -0.145834 | 0.06 | 0.146 | 1 | 1 |
| pnrc2 | 0.15112 | 0.121108105 | 0.93 | 0.89 | 1 | 1 |
| eci2 | 0.15114 | 0.109401131 | 0.11 | 0.037 | 1 | 1 |
| krt91 | 0.15124 | 0.132965916 | 0.06 | 0.037 | 1 | 1 |
| wwp2 | 0.15133 | 0.108136596 | 0.06 | 0.012 | 1 | 1 |
| fam129bb | 0.15135 | -0.11000161 | 0.02 | 0.073 | 1 | 1 |
| ttpal | 0.15156 | 0.107451898 | 0.08 | 0.024 | 1 | 1 |
| tfap4 | 0.15167 | 0.181077528 | 0.17 | 0.085 | 1 | 1 |
| si:ch1073-429i10.3.1 | 0.15176 | 0.149254302 | 0.4 | 0.354 | 1 | 1 |
| smarcad1a | 0.15208 | 0.115950976 | 0.09 | 0.024 | 1 | 1 |
| ifi35 | 0.15214 | -0.16280724 | 0.07 | 0.159 | 1 | 1 |
| sgk3 | 0.15217 | -0.11274367 | 0.03 | 0.098 | 1 | 1 |
| gnsa | 0.15223 | 0.1200281 | 0.07 | 0.024 | 1 | 1 |
| phldb2b | 0.15271 | 0.17689697 | 0.14 | 0.073 | 1 | 1 |
| hsd11b1la | 0.15286 | -0.10717099 | 0.03 | 0.098 | 1 | 1 |
| cops2 | 0.153 | -0.1313265 | 0.14 | 0.244 | 1 | 1 |
| trit1 | 0.15303 | 0.141465173 | 0.1 | 0.037 | 1 | 1 |
| nr3c1 | 0.15347 | -0.11205042 | 0.03 | 0.098 | 1 | 1 |
| spoplb | 0.15392 | 0.119812364 | 0.08 | 0.024 | 1 | 1 |
| polr1c | 0.15398 | 0.11632135 | 0.1 | 0.049 | 1 | 1 |
| eif2b4 | 0.154 | 0.12369242 | 0.09 | 0.024 | 1 | 1 |
| pgap2 | 0.15429 | 0.157431868 | 0.11 | 0.049 | 1 | 1 |
| nr2f2 | 0.15434 | -0.14071685 | 0.36 | 0.488 | 1 | 1 |
| tmem39a | 0.15435 | -0.10317622 | 0.07 | 0.146 | 1 | 1 |
| celf5a | 0.15438 | 0.147686094 | 0.13 | 0.049 | 1 | 1 |
| adrb2a | 0.15439 | 0.152712484 | 0.26 | 0.22 | 1 | 1 |
| si:ch211-194c3.5 | 0.15471 | -0.1450785 | 0.03 | 0.098 | 1 | 1 |
| u2af2b | 0.15485 | 0.181148948 | 0.36 | 0.268 | 1 | 1 |
| pitx2 | 0.15496 | 0.14128281 | 0.11 | 0.049 | 1 | 1 |
| rrbp1b | 0.15525 | -0.19610243 | 0.18 | 0.28 | 1 | 1 |
| pgam5 | 0.1553 | -0.13218439 | 0.07 | 0.159 | 1 | 1 |
| wacb | 0.1554 | 0.129550676 | 0.09 | 0.024 | 1 | 1 |
| LO017872.1 | 0.15542 | 0.136652559 | 0.09 | 0.024 | 1 | 1 |
| thoc3 | 0.15587 | -0.10993053 | 0.03 | 0.098 | 1 | 1 |
| mapk7 | 0.15655 | 0.109735181 | 0.08 | 0.024 | 1 | 1 |
| rasgef1ba | 0.1571 | -0.11754704 | 0.03 | 0.098 | 1 | 1 |
| efnb1 | 0.15741 | 0.223943848 | 0.29 | 0.171 | 1 | 1 |
| ccni | 0.15758 | 0.150691754 | 0.57 | 0.5 | 1 | 1 |
| rbms2b | 0.1577 | 0.12410275 | 0.11 | 0.037 | 1 | 1 |
| si:dkey-217f16.1 | 0.1586 | -0.12386623 | 0.03 | 0.098 | 1 | 1 |
| srgap2 | 0.15865 | 0.184800247 | 0.16 | 0.085 | 1 | 1 |
| snx5 | 0.15867 | 0.131307641 | 0.16 | 0.098 | 1 | 1 |
| si:ch211-212d10.2 | 0.15986 | 0.107468062 | 0.07 | 0.024 | 1 | 1 |

| | | | | | | |
|---|---|---|---|---|---|---|
| myg1 | 0.16002 | -0.11167608 | 0.12 | 0.22 | 1 | 1 |
| acadl | 0.16034 | -0.10439767 | 0.05 | 0.122 | 1 | 1 |
| ndst2b | 0.16052 | 0.118815475 | 0.06 | 0.012 | 1 | 1 |
| adam19a | 0.16119 | 0.167401191 | 0.16 | 0.085 | 1 | 1 |
| lpgat1 | 0.16173 | -0.13793432 | 0.02 | 0.061 | 1 | 1 |
| map1lc3b | 0.16199 | 0.107508402 | 0.34 | 0.317 | 1 | 1 |
| ddx3xa | 0.16244 | 0.112598181 | 0.11 | 0.073 | 1 | 1 |
| rhobtb4 | 0.16293 | -0.1325274 | 0.03 | 0.085 | 1 | 1 |
| gadd45ga | 0.16317 | -0.12121553 | 0.07 | 0.159 | 1 | 1 |
| ppp6c | 0.1633 | 0.168321661 | 0.25 | 0.171 | 1 | 1 |
| pkig | 0.16378 | 0.186246486 | 0.44 | 0.341 | 1 | 1 |
| CABZ01038709.1 | 0.16394 | 0.126428038 | 0.08 | 0.037 | 1 | 1 |
| wipf2a | 0.16408 | -0.12163428 | 0.02 | 0.073 | 1 | 1 |
| sh3bgrl3 | 0.16414 | 0.159099967 | 0.27 | 0.207 | 1 | 1 |
| igf2bp3 | 0.16432 | -0.1487545 | 0.07 | 0.159 | 1 | 1 |
| rasgef1bb | 0.16433 | 0.152957113 | 0.14 | 0.073 | 1 | 1 |
| abca1a | 0.1646 | -0.16614364 | 0.07 | 0.159 | 1 | 1 |
| zgc:112962 | 0.16468 | -0.11450905 | 0.02 | 0.073 | 1 | 1 |
| sinhcafl | 0.16538 | -0.12858423 | 0.14 | 0.244 | 1 | 1 |
| morc2 | 0.16545 | 0.144710688 | 0.11 | 0.049 | 1 | 1 |
| map2k6 | 0.16551 | -0.16374144 | 0.33 | 0.463 | 1 | 1 |
| dusp5 | 0.16729 | 0.102153585 | 0.06 | 0.012 | 1 | 1 |
| psmd12 | 0.16788 | -0.1397688 | 0.08 | 0.171 | 1 | 1 |
| psme4b | 0.16824 | 0.115276645 | 0.06 | 0.012 | 1 | 1 |
| atp5f1b | 0.16858 | -0.17920316 | 0.65 | 0.756 | 1 | 1 |
| psma3 | 0.16905 | -0.18661285 | 0.35 | 0.488 | 1 | 1 |
| prrc2a | 0.16929 | 0.129806395 | 0.14 | 0.061 | 1 | 1 |
| farsb | 0.16929 | 0.139328715 | 0.13 | 0.061 | 1 | 1 |
| parp9 | 0.16943 | -0.11953333 | 0.02 | 0.073 | 1 | 1 |
| mia3 | 0.16944 | 0.126996898 | 0.07 | 0.024 | 1 | 1 |
| mcrs1 | 0.17033 | -0.14993025 | 0.05 | 0.122 | 1 | 1 |
| tm2d2 | 0.17078 | -0.1038599 | 0.04 | 0.11 | 1 | 1 |
| raraa | 0.17083 | 0.203193934 | 0.33 | 0.232 | 1 | 1 |
| nfyc | 0.17098 | -0.16949203 | 0.09 | 0.171 | 1 | 1 |
| jdp2b | 0.17176 | 0.161901002 | 0.34 | 0.305 | 1 | 1 |
| arl1 | 0.1719 | -0.13666235 | 0.1 | 0.195 | 1 | 1 |
| rnasekb | 0.17193 | -0.13945869 | 0.1 | 0.195 | 1 | 1 |
| gab2 | 0.17202 | 0.116986127 | 0.08 | 0.037 | 1 | 1 |
| s1pr2 | 0.17229 | 0.125399209 | 0.21 | 0.171 | 1 | 1 |
| hmgcrb | 0.17299 | 0.145765981 | 0.09 | 0.037 | 1 | 1 |
| arnt2 | 0.17317 | 0.103434453 | 0.06 | 0.012 | 1 | 1 |
| ecscr | 0.17353 | -0.12084474 | 0.04 | 0.11 | 1 | 1 |
| prr12a | 0.17366 | 0.122733164 | 0.11 | 0.061 | 1 | 1 |
| si:dkey-172m14.1 | 0.17393 | -0.10599004 | 0.05 | 0.122 | 1 | 1 |
| selenot2 | 0.17441 | 0.182260636 | 0.27 | 0.159 | 1 | 1 |
| phkg2 | 0.17473 | 0.114907774 | 0.09 | 0.037 | 1 | 1 |
| eif2s2 | 0.17536 | -0.17066517 | 0.59 | 0.72 | 1 | 1 |
| phf11 | 0.17586 | -0.10003126 | 0.07 | 0.146 | 1 | 1 |

| | | | | | | |
|---|---|---|---|---|---|---|
| ap3s1.1 | 0.17691 | 0.109170521 | 0.13 | 0.098 | 1 | 1 |
| cpt1aa | 0.17706 | 0.108770253 | 0.06 | 0.012 | 1 | 1 |
| ctsh | 0.17707 | -0.14069387 | 0.11 | 0.207 | 1 | 1 |
| ddx46 | 0.17715 | -0.10425417 | 0.04 | 0.11 | 1 | 1 |
| atpaf1 | 0.17749 | -0.11711041 | 0.04 | 0.11 | 1 | 1 |
| usp25 | 0.17759 | 0.111346894 | 0.06 | 0.012 | 1 | 1 |
| pcdh18b | 0.17772 | 0.101432741 | 0.39 | 0.378 | 1 | 1 |
| cnn3b | 0.17795 | 0.100987093 | 0.21 | 0.183 | 1 | 1 |
| copg2 | 0.17797 | -0.16555582 | 0.09 | 0.183 | 1 | 1 |
| myct1a | 0.17799 | 0.184412029 | 0.26 | 0.183 | 1 | 1 |
| CU694197.1 | 0.17814 | 0.175933033 | 0.38 | 0.293 | 1 | 1 |
| egfl6 | 0.17821 | 0.147680757 | 0.14 | 0.073 | 1 | 1 |
| pabpc1b | 0.17829 | 0.17221873 | 0.25 | 0.171 | 1 | 1 |
| fbxo28 | 0.17901 | 0.160190935 | 0.19 | 0.11 | 1 | 1 |
| arfgap2 | 0.17903 | 0.157559513 | 0.13 | 0.061 | 1 | 1 |
| ralbb | 0.17993 | -0.13473104 | 0.09 | 0.183 | 1 | 1 |
| mtbl | 0.18029 | -0.10984615 | 0.05 | 0.085 | 1 | 1 |
| ptpn21 | 0.1804 | 0.161150041 | 0.11 | 0.049 | 1 | 1 |
| sec22bb | 0.18046 | 0.115809946 | 0.08 | 0.024 | 1 | 1 |
| fam107b | 0.18053 | 0.193798367 | 0.3 | 0.183 | 1 | 1 |
| dnm3a | 0.18067 | 0.12591534 | 0.1 | 0.037 | 1 | 1 |
| tmem189 | 0.18068 | 0.116475308 | 0.1 | 0.037 | 1 | 1 |
| cox4i1 | 0.18109 | 0.160947303 | 0.81 | 0.72 | 1 | 1 |
| RPL37A | 0.18125 | -0.15022372 | 0.09 | 0.183 | 1 | 1 |
| zc3h18 | 0.18129 | -0.11681029 | 0.04 | 0.11 | 1 | 1 |
| axin2 | 0.18131 | -0.16367226 | 0.09 | 0.183 | 1 | 1 |
| zgc:162780 | 0.18291 | 0.12718654 | 0.1 | 0.037 | 1 | 1 |
| arl8a | 0.18301 | 0.143607318 | 0.16 | 0.073 | 1 | 1 |
| si:ch1073-340j19.1 | 0.1833 | -0.12795401 | 0.04 | 0.11 | 1 | 1 |
| si:dkeyp-68b7.7 | 0.18359 | 0.106343558 | 0.08 | 0.037 | 1 | 1 |
| gmps | 0.18391 | -0.11123922 | 0.04 | 0.11 | 1 | 1 |
| si:ch73-386h18.1 | 0.18453 | 0.100500762 | 0.06 | 0.012 | 1 | 1 |
| atp5pb | 0.18547 | 0.170861589 | 0.72 | 0.622 | 1 | 1 |
| ogdha | 0.18584 | -0.1811607 | 0.13 | 0.22 | 1 | 1 |
| dhx15 | 0.18664 | -0.15814555 | 0.17 | 0.28 | 1 | 1 |
| tspan36 | 0.18701 | -0.18520588 | 0.26 | 0.354 | 1 | 1 |
| ppp1cab | 0.18742 | 0.131248085 | 0.2 | 0.146 | 1 | 1 |
| tnrc6a | 0.18768 | 0.162808668 | 0.19 | 0.134 | 1 | 1 |
| cox20 | 0.18797 | -0.11261514 | 0.05 | 0.122 | 1 | 1 |
| cox7a2l | 0.18814 | -0.10904495 | 0.18 | 0.28 | 1 | 1 |
| hadhb | 0.18815 | -0.15402193 | 0.11 | 0.207 | 1 | 1 |
| CR407547.1 | 0.18822 | 0.163243667 | 0.16 | 0.073 | 1 | 1 |
| setdb2 | 0.18852 | 0.102609522 | 0.08 | 0.037 | 1 | 1 |
| mrpl57 | 0.18883 | -0.13503365 | 0.16 | 0.268 | 1 | 1 |
| sf3a1 | 0.18893 | -0.11463384 | 0.05 | 0.122 | 1 | 1 |
| pold3 | 0.18905 | -0.104298 | 0.04 | 0.11 | 1 | 1 |
| lmnb2 | 0.18906 | -0.17208811 | 0.12 | 0.22 | 1 | 1 |
| si:dkeyp-75h12.5 | 0.18931 | 0.157053514 | 0.19 | 0.122 | 1 | 1 |

| | | | | | | |
|---|---|---|---|---|---|---|
| dph7 | 0.18939 | 0.103781448 | 0.06 | 0.012 | 1 | 1 |
| filip1l | 0.18973 | -0.11915323 | 0.04 | 0.11 | 1 | 1 |
| mgst1.2 | 0.18984 | -0.15769232 | 0.31 | 0.341 | 1 | 1 |
| xbp1 | 0.18986 | -0.18962435 | 0.57 | 0.646 | 1 | 1 |
| ddx3xb | 0.19001 | -0.10537041 | 0.09 | 0.171 | 1 | 1 |
| pdk4 | 0.19014 | -0.1221882 | 0.04 | 0.11 | 1 | 1 |
| srpk1a | 0.19014 | -0.11939256 | 0.04 | 0.11 | 1 | 1 |
| setx | 0.19018 | -0.12875917 | 0.04 | 0.11 | 1 | 1 |
| hmgb3b | 0.19101 | -0.1369112 | 0.16 | 0.268 | 1 | 1 |
| nop56 | 0.19117 | -0.16515494 | 0.5 | 0.634 | 1 | 1 |
| zfand3 | 0.1913 | -0.15218128 | 0.15 | 0.256 | 1 | 1 |
| si:dkey-12l12.1 | 0.19134 | 0.135499471 | 0.06 | 0.012 | 1 | 1 |
| cxxc5a | 0.19148 | -0.12780048 | 0.19 | 0.293 | 1 | 1 |
| alyref | 0.19165 | -0.18247755 | 0.14 | 0.232 | 1 | 1 |
| anxa4 | 0.19203 | -0.16293649 | 0.32 | 0.451 | 1 | 1 |
| slc43a2a | 0.1922 | -0.11364045 | 0.07 | 0.146 | 1 | 1 |
| eif3k | 0.19289 | -0.12806537 | 0.42 | 0.549 | 1 | 1 |
| chpf2 | 0.19491 | -0.1037425 | 0.02 | 0.073 | 1 | 1 |
| mrps11 | 0.19531 | 0.108077635 | 0.12 | 0.049 | 1 | 1 |
| sirt2 | 0.19537 | 0.104545719 | 0.12 | 0.049 | 1 | 1 |
| rnf44 | 0.19545 | 0.166831551 | 0.31 | 0.232 | 1 | 1 |
| cst14b.1 | 0.19551 | 0.115513693 | 0.09 | 0.037 | 1 | 1 |
| arl4d | 0.19567 | 0.115323563 | 0.08 | 0.024 | 1 | 1 |
| ism1 | 0.19584 | -0.11167648 | 0.02 | 0.073 | 1 | 1 |
| pfkfb4b | 0.19609 | 0.166291965 | 0.34 | 0.268 | 1 | 1 |
| gatad2b | 0.19623 | 0.193859738 | 0.24 | 0.146 | 1 | 1 |
| taf7 | 0.19627 | 0.126023323 | 0.23 | 0.183 | 1 | 1 |
| tmem203 | 0.19635 | 0.116739863 | 0.1 | 0.049 | 1 | 1 |
| ngrn | 0.19642 | 0.123498941 | 0.13 | 0.073 | 1 | 1 |
| afg3l2 | 0.19732 | -0.13211168 | 0.04 | 0.098 | 1 | 1 |
| si:ch211-66k16.27 | 0.19748 | 0.145869604 | 0.33 | 0.268 | 1 | 1 |
| mrpl20 | 0.19882 | -0.14698923 | 0.12 | 0.22 | 1 | 1 |
| col15a1a | 0.19887 | -0.1670838 | 0.12 | 0.22 | 1 | 1 |
| acadvl | 0.19906 | -0.14208406 | 0.12 | 0.22 | 1 | 1 |
| uchl3 | 0.1992 | -0.128657 | 0.05 | 0.122 | 1 | 1 |
| zgc:65873 | 0.19945 | -0.14896712 | 0.14 | 0.244 | 1 | 1 |
| snrpd1 | 0.19958 | -0.15512973 | 0.29 | 0.415 | 1 | 1 |
| btbd6a | 0.19979 | 0.121294258 | 0.07 | 0.024 | 1 | 1 |
| fam234a | 0.2001 | 0.115047836 | 0.08 | 0.037 | 1 | 1 |
| ubap2l | 0.20029 | 0.101094698 | 0.11 | 0.085 | 1 | 1 |
| igf2bp2a | 0.20049 | 0.159330252 | 0.14 | 0.061 | 1 | 1 |
| zgc:56304 | 0.20103 | -0.124203 | 0.2 | 0.305 | 1 | 1 |
| myl6 | 0.2013 | -0.13350637 | 0.3 | 0.415 | 1 | 1 |
| meis1b | 0.20131 | -0.14395904 | 0.23 | 0.341 | 1 | 1 |
| zfp36l1b | 0.20167 | -0.10027879 | 0.28 | 0.378 | 1 | 1 |
| CZQB01141835.1 | 0.20181 | -0.11352711 | 0.14 | 0.232 | 1 | 1 |
| matr3l1.1 | 0.20197 | -0.16573939 | 0.14 | 0.244 | 1 | 1 |
| nipblb | 0.20263 | 0.123005676 | 0.11 | 0.049 | 1 | 1 |

| | | | | | | |
|---|---|---|---|---|---|---|
| si:ch73-299h12.6 | 0.20265 | 0.104305599 | 0.08 | 0.024 | 1 | 1 |
| vps26c | 0.20271 | 0.128379719 | 0.12 | 0.073 | 1 | 1 |
| lama5 | 0.20333 | 0.215552082 | 0.32 | 0.207 | 1 | 1 |
| tcima | 0.20349 | 0.12703001 | 0.06 | 0.024 | 1 | 1 |
| klf7b | 0.20407 | 0.143226762 | 0.14 | 0.085 | 1 | 1 |
| vps26bl | 0.20434 | 0.102764922 | 0.06 | 0.012 | 1 | 1 |
| mbnl2 | 0.20486 | 0.118616068 | 0.07 | 0.024 | 1 | 1 |
| stx6 | 0.20512 | 0.108714834 | 0.08 | 0.024 | 1 | 1 |
| CABZ01113812.1 | 0.20574 | -0.11464573 | 0.03 | 0.085 | 1 | 1 |
| cflara | 0.20607 | -0.14714319 | 0.15 | 0.207 | 1 | 1 |
| ctps1a | 0.20608 | 0.144544024 | 0.15 | 0.073 | 1 | 1 |
| hbegfa | 0.2073 | 0.122406108 | 0.11 | 0.061 | 1 | 1 |
| nab1b | 0.20745 | -0.10682499 | 0.02 | 0.073 | 1 | 1 |
| sppl3 | 0.20746 | 0.154268331 | 0.15 | 0.073 | 1 | 1 |
| coa6 | 0.20809 | 0.127662448 | 0.17 | 0.122 | 1 | 1 |
| mbd3b | 0.20823 | 0.180569861 | 0.21 | 0.122 | 1 | 1 |
| phf10 | 0.20848 | -0.10886931 | 0.05 | 0.122 | 1 | 1 |
| pgpep1 | 0.20921 | -0.12637766 | 0.07 | 0.134 | 1 | 1 |
| oaz2a | 0.20956 | 0.100645842 | 0.1 | 0.073 | 1 | 1 |
| capns1a | 0.2096 | -0.14556014 | 0.36 | 0.488 | 1 | 1 |
| gna11a | 0.20993 | 0.128784117 | 0.08 | 0.024 | 1 | 1 |
| col6a1 | 0.20995 | -0.11476279 | 0.33 | 0.439 | 1 | 1 |
| magoh | 0.21014 | -0.11068869 | 0.07 | 0.146 | 1 | 1 |
| nudt8 | 0.21029 | 0.116765722 | 0.08 | 0.024 | 1 | 1 |
| dnajc8 | 0.21071 | -0.10204556 | 0.14 | 0.232 | 1 | 1 |
| mrps23 | 0.21107 | -0.11025382 | 0.08 | 0.159 | 1 | 1 |
| ano5b | 0.21163 | -0.11938335 | 0.05 | 0.122 | 1 | 1 |
| fam110b | 0.21171 | -0.12351254 | 0.05 | 0.122 | 1 | 1 |
| arfrp1 | 0.21186 | -0.1009193 | 0.11 | 0.195 | 1 | 1 |
| LOC797168 | 0.21192 | 0.14620683 | 0.06 | 0.012 | 1 | 1 |
| nup98 | 0.21196 | -0.1185463 | 0.03 | 0.085 | 1 | 1 |
| glud1a | 0.21259 | -0.13328636 | 0.21 | 0.317 | 1 | 1 |
| slc25a24 | 0.21274 | -0.10603066 | 0.05 | 0.122 | 1 | 1 |
| cldn11a | 0.21292 | 0.201947923 | 0.6 | 0.476 | 1 | 1 |
| CABZ01045617.1 | 0.21315 | 0.115260235 | 0.17 | 0.159 | 1 | 1 |
| mcu | 0.21322 | -0.11557625 | 0.05 | 0.122 | 1 | 1 |
| psmd1 | 0.21322 | -0.13763306 | 0.24 | 0.354 | 1 | 1 |
| actr2a | 0.21323 | -0.13303595 | 0.05 | 0.122 | 1 | 1 |
| gpc4 | 0.21532 | 0.162413336 | 0.17 | 0.085 | 1 | 1 |
| fundc2 | 0.21562 | -0.11130802 | 0.23 | 0.329 | 1 | 1 |
| tmem30aa | 0.21609 | 0.132034261 | 0.17 | 0.085 | 1 | 1 |
| eif2s3 | 0.21638 | 0.136333618 | 0.67 | 0.61 | 1 | 1 |
| cxcl18b | 0.21639 | -0.23495699 | 0.45 | 0.573 | 1 | 1 |
| zgc:110158 | 0.21645 | -0.10830727 | 0.08 | 0.159 | 1 | 1 |
| chdh | 0.21669 | -0.10286873 | 0.09 | 0.171 | 1 | 1 |
| snrpb | 0.21673 | -0.16037493 | 0.6 | 0.72 | 1 | 1 |
| tomm7 | 0.21751 | -0.10666501 | 0.27 | 0.378 | 1 | 1 |
| cd40 | 0.21829 | 0.133890536 | 0.1 | 0.049 | 1 | 1 |

| | | | | | | |
|---|---|---|---|---|---|---|
| pcmtl | 0.21858 | 0.119382871 | 0.08 | 0.037 | 1 | 1 |
| si:dkey-112a7.4 | 0.21859 | -0.10260645 | 0.1 | 0.183 | 1 | 1 |
| sec24b | 0.21878 | 0.123065895 | 0.09 | 0.037 | 1 | 1 |
| mob2a | 0.21908 | -0.11358964 | 0.06 | 0.134 | 1 | 1 |
| gbp1 | 0.21927 | -0.1452962 | 0.04 | 0.085 | 1 | 1 |
| snip1 | 0.22054 | 0.103860147 | 0.09 | 0.049 | 1 | 1 |
| pom121 | 0.22059 | -0.13544316 | 0.03 | 0.085 | 1 | 1 |
| tfg | 0.22112 | 0.109418824 | 0.22 | 0.183 | 1 | 1 |
| rab1ba | 0.22159 | -0.10742213 | 0.03 | 0.085 | 1 | 1 |
| nap1l4b | 0.22181 | 0.120253102 | 0.12 | 0.073 | 1 | 1 |
| si:ch211-262h13.3 | 0.22217 | -0.13366537 | 0.06 | 0.134 | 1 | 1 |
| atl2 | 0.22309 | 0.119516864 | 0.12 | 0.049 | 1 | 1 |
| zgc:158803 | 0.22339 | -0.11633579 | 0.11 | 0.195 | 1 | 1 |
| tmco1 | 0.22345 | -0.13788412 | 0.14 | 0.195 | 1 | 1 |
| btg3 | 0.22379 | -0.1151852 | 0.19 | 0.232 | 1 | 1 |
| ano6 | 0.22396 | 0.105896377 | 0.07 | 0.024 | 1 | 1 |
| bmp1a | 0.22426 | 0.100610146 | 0.08 | 0.024 | 1 | 1 |
| neo1b | 0.2243 | -0.10583954 | 0.04 | 0.098 | 1 | 1 |
| si:dkey-42i9.7 | 0.22484 | -0.11975612 | 0.07 | 0.085 | 1 | 1 |
| psmf1 | 0.2266 | -0.10332884 | 0.07 | 0.146 | 1 | 1 |
| kmt5b | 0.22685 | 0.113047648 | 0.1 | 0.049 | 1 | 1 |
| arpin | 0.22752 | -0.11254402 | 0.08 | 0.159 | 1 | 1 |
| pfn2l | 0.22764 | -0.16063723 | 0.51 | 0.598 | 1 | 1 |
| ddx18 | 0.22764 | 0.116895003 | 0.19 | 0.159 | 1 | 1 |
| pdcd4a | 0.22825 | 0.113251248 | 0.11 | 0.061 | 1 | 1 |
| pnpla7a | 0.22906 | 0.102566097 | 0.07 | 0.024 | 1 | 1 |
| zgc:174863 | 0.22923 | -0.16649469 | 0.35 | 0.476 | 1 | 1 |
| icn | 0.22944 | 0.191840955 | 0.07 | 0.024 | 1 | 1 |
| orc6 | 0.22945 | 0.111500367 | 0.1 | 0.037 | 1 | 1 |
| si:ch211-71n6.4 | 0.23011 | 0.121696634 | 0.1 | 0.061 | 1 | 1 |
| hmgcl | 0.23052 | -0.11239018 | 0.04 | 0.085 | 1 | 1 |
| tmem199 | 0.23068 | 0.109121129 | 0.08 | 0.024 | 1 | 1 |
| man1b1b | 0.23113 | 0.142804658 | 0.1 | 0.049 | 1 | 1 |
| gorasp2 | 0.23127 | 0.14359241 | 0.17 | 0.085 | 1 | 1 |
| slc22a18 | 0.23154 | -0.10922711 | 0.03 | 0.085 | 1 | 1 |
| PDCL3 | 0.23208 | -0.11254126 | 0.06 | 0.134 | 1 | 1 |
| sumf2 | 0.23222 | 0.103242667 | 0.08 | 0.024 | 1 | 1 |
| anapc16 | 0.23235 | 0.118450536 | 0.24 | 0.195 | 1 | 1 |
| si:ch211-197h24.9 | 0.23262 | -0.16109873 | 0.56 | 0.683 | 1 | 1 |
| cct3 | 0.23266 | -0.17099728 | 0.45 | 0.573 | 1 | 1 |
| igfbp5b | 0.23309 | 0.210640266 | 0.35 | 0.268 | 1 | 1 |
| anp32e | 0.23333 | 0.155310155 | 0.25 | 0.171 | 1 | 1 |
| XLOC-009815 | 0.23362 | -0.17710246 | 0.16 | 0.244 | 1 | 1 |
| bms1 | 0.23398 | 0.100979272 | 0.14 | 0.11 | 1 | 1 |
| nr1d2b | 0.23427 | 0.114779379 | 0.08 | 0.037 | 1 | 1 |
| eva1ba | 0.23433 | -0.17391594 | 0.25 | 0.366 | 1 | 1 |
| sema3d | 0.23483 | 0.140649233 | 0.1 | 0.037 | 1 | 1 |
| anapc10 | 0.2349 | 0.106101456 | 0.09 | 0.037 | 1 | 1 |

| | | | | | | |
|---|---|---|---|---|---|---|
| s100u | 0.23502 | -0.12145592 | 0.13 | 0.22 | 1 | 1 |
| flcn | 0.23512 | 0.106664669 | 0.08 | 0.024 | 1 | 1 |
| dek | 0.23532 | 0.115331036 | 0.13 | 0.098 | 1 | 1 |
| rbm14a | 0.23629 | 0.114506315 | 0.11 | 0.049 | 1 | 1 |
| tnfsf12 | 0.23646 | 0.124723116 | 0.15 | 0.11 | 1 | 1 |
| tshz3b | 0.23652 | 0.163143123 | 0.32 | 0.256 | 1 | 1 |
| senp2 | 0.23673 | -0.11307792 | 0.08 | 0.159 | 1 | 1 |
| itgav | 0.23692 | -0.13350994 | 0.07 | 0.146 | 1 | 1 |
| sept10 | 0.23729 | -0.13829884 | 0.2 | 0.305 | 1 | 1 |
| cry-dash | 0.23737 | 0.112555483 | 0.12 | 0.085 | 1 | 1 |
| dvl2 | 0.23744 | 0.128118738 | 0.15 | 0.073 | 1 | 1 |
| igf1rb | 0.23814 | 0.108117473 | 0.06 | 0.024 | 1 | 1 |
| maff | 0.23865 | -0.14211168 | 0.08 | 0.159 | 1 | 1 |
| mmp14b | 0.23907 | 0.178284048 | 0.29 | 0.207 | 1 | 1 |
| ncoa5 | 0.23932 | 0.102048885 | 0.12 | 0.073 | 1 | 1 |
| nabp1a | 0.23961 | -0.11667304 | 0.12 | 0.207 | 1 | 1 |
| ccdc47 | 0.23987 | 0.130418509 | 0.18 | 0.122 | 1 | 1 |
| eif3ha | 0.2401 | -0.13237758 | 0.51 | 0.634 | 1 | 1 |
| ddb2 | 0.24093 | 0.144666424 | 0.18 | 0.11 | 1 | 1 |
| tars | 0.2418 | -0.12081497 | 0.09 | 0.171 | 1 | 1 |
| nol11 | 0.24181 | 0.113177898 | 0.1 | 0.049 | 1 | 1 |
| tbx2a | 0.24203 | 0.121299272 | 0.17 | 0.134 | 1 | 1 |
| aaas | 0.24209 | 0.121732449 | 0.09 | 0.037 | 1 | 1 |
| pla2g15 | 0.24298 | -0.11562283 | 0.09 | 0.171 | 1 | 1 |
| arv1 | 0.24309 | 0.104248669 | 0.05 | 0.024 | 1 | 1 |
| lsm14ab | 0.24315 | 0.155631477 | 0.2 | 0.122 | 1 | 1 |
| h2afva | 0.24322 | -0.110788 | 0.33 | 0.439 | 1 | 1 |
| atp1b1a | 0.24376 | -0.15566452 | 0.28 | 0.341 | 1 | 1 |
| ranbp3b | 0.24376 | -0.11597305 | 0.13 | 0.22 | 1 | 1 |
| txn2 | 0.24457 | -0.12273204 | 0.17 | 0.268 | 1 | 1 |
| surf4l | 0.24472 | -0.14141902 | 0.08 | 0.159 | 1 | 1 |
| pds5b | 0.2453 | 0.114757791 | 0.11 | 0.061 | 1 | 1 |
| snw1 | 0.24563 | -0.13264544 | 0.18 | 0.28 | 1 | 1 |
| zbtb20 | 0.24609 | -0.18468341 | 0.19 | 0.293 | 1 | 1 |
| rars | 0.24654 | -0.1120626 | 0.07 | 0.146 | 1 | 1 |
| cep85l | 0.2468 | 0.101060743 | 0.08 | 0.037 | 1 | 1 |
| si:dkey-187e18.1 | 0.24769 | -0.14542424 | 0.13 | 0.195 | 1 | 1 |
| txnrd3 | 0.24817 | -0.14366758 | 0.11 | 0.195 | 1 | 1 |
| eif4a2 | 0.24905 | 0.151306224 | 0.3 | 0.22 | 1 | 1 |
| tgfb3 | 0.24984 | 0.140187675 | 0.21 | 0.122 | 1 | 1 |
| rnf111 | 0.25002 | 0.104243032 | 0.05 | 0.012 | 1 | 1 |
| ptgr1 | 0.25021 | 0.11247958 | 0.18 | 0.134 | 1 | 1 |
| cdc14b | 0.2506 | 0.143363756 | 0.13 | 0.073 | 1 | 1 |
| mrpl15 | 0.25134 | 0.101119033 | 0.1 | 0.061 | 1 | 1 |
| si:ch211-204a13.2 | 0.25235 | 0.101068422 | 0.05 | 0.012 | 1 | 1 |
| ablim1b | 0.25265 | -0.10120521 | 0.18 | 0.268 | 1 | 1 |
| serpinh1a | 0.25319 | -0.10289417 | 0.03 | 0.085 | 1 | 1 |
| psmg1 | 0.25331 | -0.15998406 | 0.14 | 0.232 | 1 | 1 |

| | | | | | | |
|---|---|---|---|---|---|---|
| hmgb2a | 0.25339 | -0.12508041 | 0.54 | 0.659 | 1 | 1 |
| gbp | 0.25364 | 0.183115101 | 0.42 | 0.317 | 1 | 1 |
| nolc1 | 0.2541 | -0.14226653 | 0.1 | 0.183 | 1 | 1 |
| rp9 | 0.25466 | 0.10600398 | 0.12 | 0.061 | 1 | 1 |
| tmtc1 | 0.25485 | 0.113707472 | 0.09 | 0.037 | 1 | 1 |
| brd9 | 0.25613 | -0.14213887 | 0.08 | 0.159 | 1 | 1 |
| psmb4 | 0.25613 | -0.15145353 | 0.32 | 0.439 | 1 | 1 |
| nsun5 | 0.25659 | 0.101907996 | 0.08 | 0.037 | 1 | 1 |
| tbl1xr1a | 0.25666 | 0.168436506 | 0.24 | 0.146 | 1 | 1 |
| ppm1aa | 0.25674 | 0.133103862 | 0.2 | 0.146 | 1 | 1 |
| rap1b | 0.25699 | 0.126543837 | 0.33 | 0.28 | 1 | 1 |
| ikbkg | 0.25749 | -0.11673316 | 0.05 | 0.11 | 1 | 1 |
| aldh7a1 | 0.25794 | -0.13718158 | 0.2 | 0.305 | 1 | 1 |
| nucks1a | 0.258 | -0.11388284 | 0.38 | 0.488 | 1 | 1 |
| surf6 | 0.25803 | 0.117648883 | 0.09 | 0.049 | 1 | 1 |
| LOC108180269 | 0.25856 | 0.150312025 | 0.15 | 0.073 | 1 | 1 |
| si:rp71-45k5.4 | 0.2586 | -0.16650684 | 0.29 | 0.402 | 1 | 1 |
| zgc:172139 | 0.25872 | 0.100000984 | 0.05 | 0.012 | 1 | 1 |
| nccrp1 | 0.25879 | 0.137113897 | 0.25 | 0.183 | 1 | 1 |
| tns3.2 | 0.25906 | 0.112268565 | 0.17 | 0.134 | 1 | 1 |
| phc2a | 0.25927 | 0.129480659 | 0.13 | 0.085 | 1 | 1 |
| sec61g | 0.25946 | 0.122891334 | 0.9 | 0.829 | 1 | 1 |
| eif2a | 0.26058 | 0.12134804 | 0.13 | 0.061 | 1 | 1 |
| si:ch73-269m14.2 | 0.26136 | -0.10342559 | 0.17 | 0.256 | 1 | 1 |
| hadhab | 0.2614 | -0.10091855 | 0.03 | 0.085 | 1 | 1 |
| lonrf1l | 0.2615 | -0.14025961 | 0.05 | 0.11 | 1 | 1 |
| yy1a | 0.26174 | 0.159372238 | 0.29 | 0.207 | 1 | 1 |
| evi5b | 0.26184 | -0.11299256 | 0.03 | 0.085 | 1 | 1 |
| entpd1 | 0.26198 | -0.13156705 | 0.35 | 0.463 | 1 | 1 |
| ube2f | 0.26227 | 0.111949077 | 0.11 | 0.061 | 1 | 1 |
| fbxl3a | 0.26236 | 0.133563257 | 0.1 | 0.049 | 1 | 1 |
| hs3st3b1b | 0.2629 | -0.11392233 | 0.03 | 0.085 | 1 | 1 |
| ralab | 0.26329 | 0.116734692 | 0.11 | 0.049 | 1 | 1 |
| eif4e3 | 0.26335 | 0.146728201 | 0.14 | 0.073 | 1 | 1 |
| selenot1a | 0.2637 | -0.11566381 | 0.09 | 0.171 | 1 | 1 |
| gabarapb | 0.26416 | 0.151546436 | 0.67 | 0.573 | 1 | 1 |
| dnajc19 | 0.26417 | 0.144547688 | 0.29 | 0.22 | 1 | 1 |
| denr | 0.26466 | 0.161472156 | 0.32 | 0.232 | 1 | 1 |
| ubiad1 | 0.26496 | -0.10524717 | 0.05 | 0.11 | 1 | 1 |
| itm2bb | 0.26562 | 0.180507029 | 0.49 | 0.378 | 1 | 1 |
| tardbp | 0.26592 | 0.138849054 | 0.22 | 0.146 | 1 | 1 |
| dnajc4 | 0.26594 | -0.13509942 | 0.09 | 0.171 | 1 | 1 |
| mdh1aa | 0.26595 | -0.15812435 | 0.37 | 0.488 | 1 | 1 |
| slc29a4 | 0.26608 | -0.14646853 | 0.08 | 0.146 | 1 | 1 |
| add3a | 0.26612 | 0.15900277 | 0.53 | 0.451 | 1 | 1 |
| ap2b1 | 0.2665 | -0.14872213 | 0.1 | 0.183 | 1 | 1 |
| arhgap35a | 0.26682 | -0.13644455 | 0.08 | 0.146 | 1 | 1 |
| abcc5 | 0.26731 | -0.14883596 | 0.14 | 0.232 | 1 | 1 |

| | | | | | | |
|---|---|---|---|---|---|---|
| fubp1 | 0.26779 | 0.142976867 | 0.31 | 0.207 | 1 | 1 |
| atp5md | 0.26856 | -0.12023274 | 0.37 | 0.488 | 1 | 1 |
| sipa1l2.1 | 0.26879 | 0.135037527 | 0.13 | 0.061 | 1 | 1 |
| lrrc17 | 0.26938 | 0.135904414 | 0.69 | 0.622 | 1 | 1 |
| arhgap5 | 0.26947 | -0.14004376 | 0.09 | 0.159 | 1 | 1 |
| dcn | 0.27162 | 0.125226121 | 0.86 | 0.793 | 1 | 1 |
| gli3 | 0.27181 | -0.15057136 | 0.1 | 0.183 | 1 | 1 |
| MDFI | 0.27227 | -0.16245596 | 0.13 | 0.22 | 1 | 1 |
| abhd17b | 0.27298 | -0.11598029 | 0.05 | 0.11 | 1 | 1 |
| sult1st1 | 0.27405 | -0.16683268 | 0.26 | 0.354 | 1 | 1 |
| cbr1 | 0.27417 | 0.103499661 | 0.07 | 0.037 | 1 | 1 |
| ociad1 | 0.27433 | 0.12166605 | 0.12 | 0.061 | 1 | 1 |
| ostc | 0.27447 | -0.16359911 | 0.36 | 0.463 | 1 | 1 |
| mrps24 | 0.27468 | -0.12742788 | 0.11 | 0.195 | 1 | 1 |
| prrc1 | 0.27495 | 0.141179142 | 0.12 | 0.061 | 1 | 1 |
| ephb2a | 0.27507 | -0.12871589 | 0.06 | 0.122 | 1 | 1 |
| serbp1a | 0.27508 | 0.101294446 | 0.91 | 0.854 | 1 | 1 |
| rhogc | 0.27521 | -0.14002863 | 0.12 | 0.195 | 1 | 1 |
| zeb1b | 0.27555 | -0.14764606 | 0.13 | 0.22 | 1 | 1 |
| tia1 | 0.27557 | -0.14023123 | 0.11 | 0.195 | 1 | 1 |
| aida | 0.27594 | 0.133150637 | 0.18 | 0.098 | 1 | 1 |
| anxa1a | 0.27668 | 0.127519619 | 0.47 | 0.451 | 1 | 1 |
| ndufa8 | 0.27823 | 0.112420421 | 0.33 | 0.293 | 1 | 1 |
| dnmt3bb.1 | 0.27936 | 0.112462444 | 0.11 | 0.049 | 1 | 1 |
| cyb5a | 0.27974 | -0.1317688 | 0.15 | 0.244 | 1 | 1 |
| CR847566.1 | 0.28011 | 0.143381418 | 0.23 | 0.183 | 1 | 1 |
| marcksb | 0.2805 | -0.1565702 | 0.13 | 0.22 | 1 | 1 |
| bcas3 | 0.28106 | -0.14808435 | 0.08 | 0.146 | 1 | 1 |
| mrpl13 | 0.28179 | -0.11487921 | 0.09 | 0.146 | 1 | 1 |
| znf706 | 0.28358 | 0.102385462 | 0.24 | 0.207 | 1 | 1 |
| gpatch1 | 0.28369 | 0.135065675 | 0.13 | 0.061 | 1 | 1 |
| tnrc18 | 0.28428 | -0.10122472 | 0.15 | 0.232 | 1 | 1 |
| prpf4bb | 0.28551 | 0.115852055 | 0.13 | 0.061 | 1 | 1 |
| dhrs13a.3 | 0.28569 | 0.107902418 | 0.16 | 0.134 | 1 | 1 |
| kdm5bb | 0.28582 | -0.12468634 | 0.05 | 0.11 | 1 | 1 |
| rasa1b | 0.2861 | -0.10406617 | 0.02 | 0.061 | 1 | 1 |
| CU929070.1 | 0.28659 | -0.1071971 | 0.21 | 0.305 | 1 | 1 |
| adamts10 | 0.28672 | 0.106585922 | 0.11 | 0.049 | 1 | 1 |
| prkag2a | 0.2868 | 0.141680928 | 0.21 | 0.159 | 1 | 1 |
| zmynd19 | 0.28708 | 0.100878494 | 0.08 | 0.037 | 1 | 1 |
| ccdc3b | 0.28827 | -0.11280908 | 0.17 | 0.256 | 1 | 1 |
| txndc9 | 0.2892 | 0.120490888 | 0.37 | 0.317 | 1 | 1 |
| rfxank | 0.28954 | -0.10855411 | 0.04 | 0.098 | 1 | 1 |
| rin3 | 0.29029 | 0.153071855 | 0.17 | 0.098 | 1 | 1 |
| srpra | 0.29172 | 0.108427643 | 0.16 | 0.122 | 1 | 1 |
| eef1db | 0.29269 | -0.10172954 | 0.55 | 0.659 | 1 | 1 |
| tmem134 | 0.29317 | 0.103397363 | 0.15 | 0.11 | 1 | 1 |
| rhoua | 0.29349 | 0.165779244 | 0.31 | 0.232 | 1 | 1 |

| | | | | | | |
|---|---|---|---|---|---|---|
| srsf1b | 0.29355 | 0.109024538 | 0.19 | 0.159 | 1 | 1 |
| ddx47 | 0.2942 | 0.144015785 | 0.19 | 0.122 | 1 | 1 |
| jupa | 0.29431 | 0.100408278 | 0.14 | 0.11 | 1 | 1 |
| mcrip1 | 0.29458 | 0.139065108 | 0.21 | 0.146 | 1 | 1 |
| NC-002333.17 | 0.29586 | -0.10775551 | 0.04 | 0.098 | 1 | 1 |
| pabpn1 | 0.29599 | -0.10149276 | 0.48 | 0.585 | 1 | 1 |
| pdhb | 0.29604 | 0.125475596 | 0.17 | 0.11 | 1 | 1 |
| hpgd | 0.29612 | -0.11028604 | 0.05 | 0.11 | 1 | 1 |
| adipor1a | 0.29614 | 0.125683843 | 0.12 | 0.061 | 1 | 1 |
| supt6h | 0.29718 | -0.11536452 | 0.1 | 0.146 | 1 | 1 |
| arhgef2 | 0.29766 | 0.118532228 | 0.09 | 0.037 | 1 | 1 |
| ppp1r12a | 0.29775 | -0.12249016 | 0.04 | 0.098 | 1 | 1 |
| prelid1a | 0.2978 | 0.127504624 | 0.15 | 0.085 | 1 | 1 |
| lhfpl2a | 0.29848 | -0.13976787 | 0.23 | 0.329 | 1 | 1 |
| si:dkey-6n21.12 | 0.29866 | 0.131191644 | 0.25 | 0.207 | 1 | 1 |
| col21a1 | 0.29897 | -0.10622018 | 0.09 | 0.159 | 1 | 1 |
| smc3 | 0.29975 | 0.107244524 | 0.12 | 0.073 | 1 | 1 |
| cope | 0.30115 | -0.14638405 | 0.26 | 0.341 | 1 | 1 |
| anp32b | 0.30254 | -0.10280356 | 0.41 | 0.439 | 1 | 1 |
| LOC103910871 | 0.30254 | -0.15920259 | 0.11 | 0.183 | 1 | 1 |
| ddt | 0.30326 | 0.156387037 | 0.31 | 0.22 | 1 | 1 |
| psap | 0.30417 | 0.132780845 | 0.28 | 0.232 | 1 | 1 |
| cipcb | 0.30442 | 0.101253739 | 0.09 | 0.061 | 1 | 1 |
| mark3b | 0.30477 | 0.104080579 | 0.09 | 0.049 | 1 | 1 |
| msi2a | 0.30566 | 0.140282613 | 0.16 | 0.098 | 1 | 1 |
| FQ323156.1 | 0.3059 | -0.14761842 | 0.11 | 0.183 | 1 | 1 |
| eif6 | 0.30601 | -0.12559287 | 0.23 | 0.329 | 1 | 1 |
| uqcc1 | 0.30624 | 0.100954719 | 0.09 | 0.049 | 1 | 1 |
| proser1 | 0.30692 | -0.10667303 | 0.03 | 0.073 | 1 | 1 |
| zgc:193541 | 0.30732 | -0.14002164 | 0.51 | 0.622 | 1 | 1 |
| pnisr | 0.30784 | 0.146329508 | 0.2 | 0.134 | 1 | 1 |
| sfxn5b | 0.30832 | -0.10188871 | 0.08 | 0.146 | 1 | 1 |
| esf1 | 0.30861 | -0.11091006 | 0.17 | 0.256 | 1 | 1 |
| ivd | 0.31025 | 0.133559923 | 0.21 | 0.134 | 1 | 1 |
| tuba8l3 | 0.31071 | 0.135493475 | 0.5 | 0.439 | 1 | 1 |
| itpkcb | 0.31097 | -0.11728724 | 0.05 | 0.11 | 1 | 1 |
| pitpnb | 0.31158 | 0.107941732 | 0.11 | 0.049 | 1 | 1 |
| meaf6 | 0.31208 | -0.14728251 | 0.21 | 0.305 | 1 | 1 |
| hoxc3a | 0.31337 | 0.107706433 | 0.22 | 0.183 | 1 | 1 |
| tgfb1a | 0.31475 | -0.11374716 | 0.05 | 0.11 | 1 | 1 |
| selenof | 0.31534 | -0.13312656 | 0.51 | 0.622 | 1 | 1 |
| sertad2a | 0.31654 | -0.11035842 | 0.05 | 0.11 | 1 | 1 |
| uqcrc2b | 0.318 | 0.105949092 | 0.15 | 0.11 | 1 | 1 |
| svild | 0.3184 | 0.121148066 | 0.24 | 0.195 | 1 | 1 |
| tsku | 0.31939 | -0.10512798 | 0.1 | 0.171 | 1 | 1 |
| ube2q2 | 0.31996 | -0.10064155 | 0.13 | 0.207 | 1 | 1 |
| mpc2 | 0.32007 | -0.10531943 | 0.15 | 0.232 | 1 | 1 |
| xpot | 0.32023 | -0.10547537 | 0.05 | 0.11 | 1 | 1 |

| | | | | | | |
|---|---|---|---|---|---|---|
| nfe2l3 | 0.32042 | -0.10831111 | 0.05 | 0.11 | 1 | 1 |
| si:ch1073-392o20.2 | 0.32052 | -0.11038621 | 0.05 | 0.11 | 1 | 1 |
| ANKRD50 | 0.32147 | 0.109851499 | 0.11 | 0.073 | 1 | 1 |
| kars | 0.32219 | -0.13754917 | 0.24 | 0.341 | 1 | 1 |
| nup107 | 0.3222 | -0.103094 | 0.05 | 0.11 | 1 | 1 |
| si:ch211-235e9.6 | 0.3247 | -0.12852227 | 0.23 | 0.329 | 1 | 1 |
| caln2 | 0.32601 | -0.16869735 | 0.22 | 0.305 | 1 | 1 |
| ctdsp1 | 0.32644 | 0.134660512 | 0.21 | 0.134 | 1 | 1 |
| selenom | 0.32654 | -0.10693146 | 0.16 | 0.244 | 1 | 1 |
| bmper | 0.32715 | 0.156997881 | 0.24 | 0.171 | 1 | 1 |
| rbm39b | 0.3273 | 0.122260499 | 0.17 | 0.122 | 1 | 1 |
| sumo1 | 0.32752 | -0.10429109 | 0.25 | 0.341 | 1 | 1 |
| cmc2 | 0.32788 | -0.11395607 | 0.06 | 0.122 | 1 | 1 |
| ndufab1b | 0.32824 | -0.13325065 | 0.23 | 0.329 | 1 | 1 |
| yipf4 | 0.32874 | -0.14165672 | 0.22 | 0.317 | 1 | 1 |
| sod2 | 0.32887 | -0.12848432 | 0.23 | 0.329 | 1 | 1 |
| vhl | 0.3293 | 0.101432004 | 0.08 | 0.049 | 1 | 1 |
| abhd4 | 0.32959 | 0.10002147 | 0.06 | 0.024 | 1 | 1 |
| spina | 0.32973 | -0.10154151 | 0.08 | 0.146 | 1 | 1 |
| RSBN1 | 0.33178 | 0.100415228 | 0.09 | 0.037 | 1 | 1 |
| rtraf | 0.33268 | 0.108335206 | 0.14 | 0.085 | 1 | 1 |
| txnl4a | 0.33289 | -0.13300648 | 0.13 | 0.207 | 1 | 1 |
| il6st | 0.33302 | 0.167802835 | 0.25 | 0.171 | 1 | 1 |
| gng5 | 0.33323 | 0.149785269 | 0.52 | 0.427 | 1 | 1 |
| calm3b | 0.33324 | -0.10201825 | 0.47 | 0.573 | 1 | 1 |
| apopt1 | 0.33476 | 0.103366079 | 0.08 | 0.037 | 1 | 1 |
| fxyd1 | 0.33884 | -0.10123829 | 0.06 | 0.122 | 1 | 1 |
| ddx51 | 0.34001 | -0.11873834 | 0.06 | 0.122 | 1 | 1 |
| b3glcta | 0.34028 | -0.11275875 | 0.06 | 0.122 | 1 | 1 |
| cbr1l | 0.34119 | 0.133195309 | 0.28 | 0.22 | 1 | 1 |
| h3f3a | 0.34147 | -0.13846202 | 0.54 | 0.646 | 1 | 1 |
| calr3a | 0.34153 | -0.16859053 | 0.31 | 0.39 | 1 | 1 |
| fkbp10b | 0.3417 | -0.11923286 | 0.06 | 0.122 | 1 | 1 |
| CABZ01077217.1 | 0.34191 | 0.12813283 | 0.12 | 0.061 | 1 | 1 |
| serpinh1b | 0.34273 | 0.130713551 | 0.14 | 0.098 | 1 | 1 |
| PDPR | 0.34273 | -0.11022329 | 0.04 | 0.085 | 1 | 1 |
| camk2g1 | 0.34399 | -0.12284085 | 0.07 | 0.134 | 1 | 1 |
| mesd | 0.34439 | -0.11257314 | 0.09 | 0.134 | 1 | 1 |
| EML4 | 0.34548 | 0.103776901 | 0.14 | 0.11 | 1 | 1 |
| wbp1lb | 0.34587 | 0.124108382 | 0.13 | 0.073 | 1 | 1 |
| rpl7l1 | 0.34652 | -0.14434417 | 0.15 | 0.232 | 1 | 1 |
| poldip2 | 0.34654 | 0.111052721 | 0.13 | 0.073 | 1 | 1 |
| med30 | 0.34675 | -0.107481 | 0.1 | 0.159 | 1 | 1 |
| dnajc1 | 0.34746 | -0.12033887 | 0.19 | 0.28 | 1 | 1 |
| pwp1 | 0.34751 | -0.10601754 | 0.13 | 0.207 | 1 | 1 |
| si:dkeyp-69b9.6 | 0.34796 | -0.11239615 | 0.07 | 0.134 | 1 | 1 |
| lamtor4 | 0.34982 | -0.1370148 | 0.14 | 0.207 | 1 | 1 |
| ankrd11 | 0.35056 | -0.15084918 | 0.16 | 0.244 | 1 | 1 |

| | | | | | | |
|---|---|---|---|---|---|---|
| ankrd52a | 0.35085 | -0.11372073 | 0.07 | 0.134 | 1 | 1 |
| timmdc1 | 0.35178 | -0.10055121 | 0.07 | 0.134 | 1 | 1 |
| ctcf | 0.35184 | 0.13142976 | 0.14 | 0.085 | 1 | 1 |
| si:dkey-7j14.6 | 0.35209 | -0.1072315 | 0.78 | 0.829 | 1 | 1 |
| eif4g3a | 0.3532 | -0.15052742 | 0.11 | 0.183 | 1 | 1 |
| ewsr1a | 0.35365 | -0.13408975 | 0.11 | 0.183 | 1 | 1 |
| ssr3 | 0.354 | -0.15524251 | 0.4 | 0.5 | 1 | 1 |
| lysmd2 | 0.35404 | -0.10943525 | 0.13 | 0.207 | 1 | 1 |
| smim13 | 0.35451 | -0.10008553 | 0.11 | 0.183 | 1 | 1 |
| ap2a1 | 0.35484 | -0.11027698 | 0.09 | 0.159 | 1 | 1 |
| klf8 | 0.35508 | -0.11230003 | 0.07 | 0.134 | 1 | 1 |
| pwp2h | 0.35528 | -0.10780434 | 0.1 | 0.171 | 1 | 1 |
| ubc | 0.35534 | 0.100182893 | 0.88 | 0.841 | 1 | 1 |
| uqcrh | 0.35596 | 0.142903122 | 0.56 | 0.488 | 1 | 1 |
| arl6ip1 | 0.35702 | 0.142861968 | 0.38 | 0.28 | 1 | 1 |
| mzt2b | 0.35757 | 0.100903435 | 0.13 | 0.073 | 1 | 1 |
| nrp1a | 0.35761 | 0.162023019 | 0.54 | 0.439 | 1 | 1 |
| tmed7 | 0.35842 | 0.121391301 | 0.45 | 0.39 | 1 | 1 |
| map7d1b | 0.35923 | 0.124810621 | 0.2 | 0.122 | 1 | 1 |
| ahnak | 0.36007 | -0.15993813 | 0.28 | 0.366 | 1 | 1 |
| zgc:114181 | 0.36071 | -0.10618087 | 0.08 | 0.146 | 1 | 1 |
| cst14a.2 | 0.36238 | 0.108390243 | 0.42 | 0.39 | 1 | 1 |
| fam32a | 0.36265 | 0.139899197 | 0.54 | 0.463 | 1 | 1 |
| foxo3b | 0.36309 | -0.1641482 | 0.23 | 0.317 | 1 | 1 |
| ccnl1a | 0.36365 | -0.10428447 | 0.09 | 0.159 | 1 | 1 |
| bzw1a | 0.36439 | -0.11313949 | 0.23 | 0.317 | 1 | 1 |
| wfdc1 | 0.36608 | 0.109454009 | 0.17 | 0.134 | 1 | 1 |
| zmp:0000000912 | 0.36624 | -0.10017887 | 0.03 | 0.073 | 1 | 1 |
| ncl | 0.36658 | -0.12713731 | 0.64 | 0.732 | 1 | 1 |
| thtpa | 0.36662 | -0.13251551 | 0.1 | 0.171 | 1 | 1 |
| rgs5a | 0.36731 | -0.14955212 | 0.15 | 0.232 | 1 | 1 |
| ndufb6 | 0.36795 | -0.12971485 | 0.24 | 0.317 | 1 | 1 |
| si:dkey-67c22.2 | 0.36815 | 0.146732873 | 0.3 | 0.22 | 1 | 1 |
| uqcc2 | 0.36952 | -0.10084146 | 0.11 | 0.183 | 1 | 1 |
| slc35a1 | 0.36971 | 0.106338511 | 0.1 | 0.049 | 1 | 1 |
| rsrc2 | 0.37055 | -0.10950676 | 0.12 | 0.195 | 1 | 1 |
| cnbpa | 0.3726 | -0.11174647 | 0.41 | 0.512 | 1 | 1 |
| ska2 | 0.37395 | 0.118791581 | 0.14 | 0.085 | 1 | 1 |
| psmc2 | 0.37484 | -0.10534368 | 0.23 | 0.317 | 1 | 1 |
| endouc | 0.375 | -0.12162454 | 0.1 | 0.171 | 1 | 1 |
| glrx | 0.3752 | 0.115266579 | 0.62 | 0.573 | 1 | 1 |
| smad5 | 0.37662 | -0.11655422 | 0.11 | 0.183 | 1 | 1 |
| itgb1a | 0.37735 | -0.13036525 | 0.13 | 0.207 | 1 | 1 |
| hectd1 | 0.37813 | 0.115965156 | 0.13 | 0.073 | 1 | 1 |
| cap1 | 0.38209 | -0.11918091 | 0.22 | 0.268 | 1 | 1 |
| arhgef1b | 0.3838 | 0.148253325 | 0.18 | 0.11 | 1 | 1 |
| minos1 | 0.38431 | -0.10349906 | 0.23 | 0.256 | 1 | 1 |
| tomm5 | 0.38465 | -0.13207113 | 0.51 | 0.61 | 1 | 1 |

| | | | | | | |
|---|---|---|---|---|---|---|
| kirrel1a | 0.38585 | 0.129928474 | 0.16 | 0.098 | 1 | 1 |
| stat1a | 0.38591 | -0.13279363 | 0.18 | 0.256 | 1 | 1 |
| col12a1b | 0.38686 | -0.16416051 | 0.25 | 0.329 | 1 | 1 |
| cavin2b | 0.38716 | 0.114518607 | 0.26 | 0.22 | 1 | 1 |
| ptk7a | 0.38819 | 0.10484634 | 0.11 | 0.061 | 1 | 1 |
| gnb1a | 0.38898 | 0.112252384 | 0.4 | 0.354 | 1 | 1 |
| fbp1a | 0.38902 | 0.110264396 | 0.15 | 0.098 | 1 | 1 |
| ddost | 0.38948 | -0.12139136 | 0.16 | 0.232 | 1 | 1 |
| ppifb | 0.39115 | 0.103036708 | 0.23 | 0.171 | 1 | 1 |
| rfxap | 0.39176 | 0.108280613 | 0.21 | 0.134 | 1 | 1 |
| meis1a | 0.392 | -0.11110725 | 0.03 | 0.073 | 1 | 1 |
| tlk1b | 0.39327 | 0.10942532 | 0.12 | 0.085 | 1 | 1 |
| lrp10 | 0.39329 | 0.112783203 | 0.15 | 0.085 | 1 | 1 |
| aldh2.2 | 0.39371 | -0.13740046 | 0.34 | 0.439 | 1 | 1 |
| csde1 | 0.39482 | -0.11392549 | 0.21 | 0.268 | 1 | 1 |
| cldn5a | 0.39503 | -0.149601 | 0.54 | 0.622 | 1 | 1 |
| klf11b | 0.39683 | 0.123931288 | 0.14 | 0.098 | 1 | 1 |
| mbtps1 | 0.39687 | 0.114832025 | 0.13 | 0.073 | 1 | 1 |
| rca2.1 | 0.39814 | 0.105893659 | 0.15 | 0.085 | 1 | 1 |
| si:dkey-188i13.9 | 0.40098 | -0.12458505 | 0.06 | 0.11 | 1 | 1 |
| f10 | 0.40145 | 0.151490113 | 0.37 | 0.28 | 1 | 1 |
| CABZ01068367.1 | 0.40163 | 0.12429641 | 0.2 | 0.134 | 1 | 1 |
| ywhaqa | 0.40231 | -0.13203329 | 0.31 | 0.378 | 1 | 1 |
| si:dkey-42i9.6 | 0.40269 | -0.10218147 | 0.1 | 0.134 | 1 | 1 |
| si:ch211-1a19.3 | 0.40282 | -0.15948004 | 0.16 | 0.22 | 1 | 1 |
| tnfrsf19 | 0.40346 | 0.124462615 | 0.25 | 0.207 | 1 | 1 |
| zc3h7bb | 0.40532 | -0.12643652 | 0.09 | 0.146 | 1 | 1 |
| taf13 | 0.40551 | 0.135047825 | 0.51 | 0.427 | 1 | 1 |
| park7 | 0.40707 | 0.104779728 | 0.18 | 0.11 | 1 | 1 |
| si:dkeyp-44a8.2 | 0.40767 | 0.102793373 | 0.12 | 0.073 | 1 | 1 |
| med29 | 0.41212 | 0.117396667 | 0.17 | 0.122 | 1 | 1 |
| arid4a | 0.41249 | 0.100897287 | 0.17 | 0.122 | 1 | 1 |
| socs1a | 0.41388 | -0.10581922 | 0.05 | 0.085 | 1 | 1 |
| si:dkey-112e7.2 | 0.41528 | 0.124815154 | 0.37 | 0.305 | 1 | 1 |
| atf4b | 0.41544 | 0.117521802 | 0.67 | 0.61 | 1 | 1 |
| oaz1b | 0.41736 | 0.12789022 | 0.7 | 0.622 | 1 | 1 |
| tead1b | 0.41764 | 0.109809635 | 0.1 | 0.061 | 1 | 1 |
| ctdspl2b | 0.41886 | -0.10764131 | 0.07 | 0.122 | 1 | 1 |
| etv4 | 0.41888 | -0.13484531 | 0.17 | 0.232 | 1 | 1 |
| trnau1apb | 0.4189 | -0.11130045 | 0.21 | 0.293 | 1 | 1 |
| cdkn1a | 0.42014 | 0.120350466 | 0.27 | 0.22 | 1 | 1 |
| acss1 | 0.42149 | 0.12455461 | 0.22 | 0.159 | 1 | 1 |
| ddx5 | 0.42207 | -0.1328723 | 0.5 | 0.573 | 1 | 1 |
| si:ch73-281n10.2 | 0.42269 | 0.120864897 | 0.81 | 0.744 | 1 | 1 |
| rtf1 | 0.42401 | 0.11217913 | 0.27 | 0.22 | 1 | 1 |
| postnb | 0.42594 | 0.103993287 | 0.76 | 0.732 | 1 | 1 |
| mfap5 | 0.42598 | 0.135665852 | 0.25 | 0.171 | 1 | 1 |
| adh8b | 0.42641 | -0.11878382 | 0.22 | 0.28 | 1 | 1 |

| | | | | | | |
|---|---|---|---|---|---|---|
| hoxc8a | 0.42686 | 0.104508951 | 0.2 | 0.159 | 1 | 1 |
| cacybp | 0.42805 | 0.11034483 | 0.16 | 0.098 | 1 | 1 |
| zmat2 | 0.43007 | -0.1078139 | 0.29 | 0.378 | 1 | 1 |
| rer1 | 0.43136 | 0.125286587 | 0.33 | 0.268 | 1 | 1 |
| arrb2b | 0.43138 | -0.1185204 | 0.14 | 0.195 | 1 | 1 |
| aldoaa | 0.43195 | 0.112825423 | 0.42 | 0.366 | 1 | 1 |
| atf3 | 0.4322 | -0.10864523 | 0.05 | 0.098 | 1 | 1 |
| cyba | 0.43285 | -0.10507128 | 0.09 | 0.146 | 1 | 1 |
| emx2 | 0.43333 | 0.120539687 | 0.12 | 0.073 | 1 | 1 |
| pagr1 | 0.43753 | -0.11826766 | 0.21 | 0.293 | 1 | 1 |
| arhgdia | 0.44006 | -0.12503562 | 0.3 | 0.39 | 1 | 1 |
| fzd7a | 0.44042 | -0.13503036 | 0.2 | 0.256 | 1 | 1 |
| si:ch211-15b10.6 | 0.44074 | 0.101460847 | 0.25 | 0.207 | 1 | 1 |
| exosc6 | 0.44181 | -0.10550154 | 0.1 | 0.159 | 1 | 1 |
| ccdc90b | 0.44182 | 0.117789866 | 0.14 | 0.085 | 1 | 1 |
| eif3g | 0.44468 | 0.10719386 | 0.66 | 0.61 | 1 | 1 |
| s100a10b | 0.44528 | 0.137952124 | 0.64 | 0.549 | 1 | 1 |
| parp12b | 0.44578 | -0.1273371 | 0.2 | 0.28 | 1 | 1 |
| wtap | 0.44657 | 0.117434748 | 0.18 | 0.122 | 1 | 1 |
| atp2b2 | 0.44729 | 0.122508347 | 0.35 | 0.293 | 1 | 1 |
| pfn2 | 0.44773 | -0.12317245 | 0.48 | 0.573 | 1 | 1 |
| sept7b | 0.44842 | -0.13843282 | 0.23 | 0.305 | 1 | 1 |
| ankrd12 | 0.45135 | 0.13071756 | 0.2 | 0.134 | 1 | 1 |
| anp32a | 0.45449 | -0.13067112 | 0.47 | 0.549 | 1 | 1 |
| ddx27 | 0.45455 | 0.120561997 | 0.3 | 0.244 | 1 | 1 |
| nop10 | 0.45466 | 0.131677563 | 0.46 | 0.378 | 1 | 1 |
| mcl1a | 0.45541 | -0.13495254 | 0.59 | 0.659 | 1 | 1 |
| leng8 | 0.45912 | -0.10355156 | 0.06 | 0.11 | 1 | 1 |
| golgb1 | 0.45948 | 0.116798776 | 0.13 | 0.085 | 1 | 1 |
| rab18b | 0.46143 | 0.111239555 | 0.14 | 0.098 | 1 | 1 |
| ldlrad4a | 0.46409 | 0.116684793 | 0.16 | 0.11 | 1 | 1 |
| cyp2n13 | 0.47259 | 0.127684124 | 0.18 | 0.122 | 1 | 1 |
| zgc:110339 | 0.47406 | 0.127084281 | 0.37 | 0.293 | 1 | 1 |
| net1 | 0.47418 | 0.101410122 | 0.09 | 0.061 | 1 | 1 |
| rbm4.3 | 0.47505 | -0.12362758 | 0.46 | 0.549 | 1 | 1 |
| mrps6 | 0.47536 | 0.101970641 | 0.23 | 0.159 | 1 | 1 |
| nog1 | 0.47736 | 0.143803299 | 0.22 | 0.159 | 1 | 1 |
| rps27l | 0.479 | 0.119641528 | 0.49 | 0.402 | 1 | 1 |
| zgc:113531 | 0.47932 | 0.151675299 | 0.29 | 0.22 | 1 | 1 |
| myh11a | 0.48017 | -0.10836654 | 0.15 | 0.22 | 1 | 1 |
| gpx4b | 0.48651 | -0.11544359 | 0.47 | 0.549 | 1 | 1 |
| pard3bb | 0.48857 | -0.10549258 | 0.12 | 0.183 | 1 | 1 |
| atf7ip | 0.48869 | 0.11879278 | 0.25 | 0.195 | 1 | 1 |
| rbms2a | 0.49069 | -0.1023431 | 0.08 | 0.134 | 1 | 1 |
| pdgfaa | 0.49108 | 0.111590143 | 0.21 | 0.159 | 1 | 1 |
| kif26ba | 0.49192 | 0.104717072 | 0.17 | 0.134 | 1 | 1 |
| scaf1 | 0.49335 | -0.10133206 | 0.11 | 0.171 | 1 | 1 |
| smdt1b | 0.49456 | 0.118371844 | 0.41 | 0.354 | 1 | 1 |

| | | | | | | |
|---|---|---|---|---|---|---|
| sema3bl | 0.49595 | 0.107299519 | 0.1 | 0.061 | 1 | 1 |
| CR318588.1 | 0.4975 | -0.11954123 | 0.36 | 0.366 | 1 | 1 |
| spty2d1 | 0.4976 | 0.102979242 | 0.11 | 0.073 | 1 | 1 |
| cox8b | 0.49823 | -0.12008052 | 0.3 | 0.378 | 1 | 1 |
| ches1 | 0.49887 | 0.107972396 | 0.11 | 0.061 | 1 | 1 |
| ndufa6 | 0.50023 | -0.10988341 | 0.39 | 0.476 | 1 | 1 |
| si:ch211-140m22.7 | 0.50072 | -0.10362791 | 0.14 | 0.195 | 1 | 1 |
| gas1a | 0.50329 | 0.118102248 | 0.54 | 0.488 | 1 | 1 |
| foxp4 | 0.50339 | 0.136544234 | 0.43 | 0.354 | 1 | 1 |
| kank2 | 0.50368 | 0.10879312 | 0.14 | 0.11 | 1 | 1 |
| lgals2a | 0.5058 | 0.129367628 | 0.09 | 0.061 | 1 | 1 |
| bahcc1b | 0.50828 | 0.112105501 | 0.16 | 0.11 | 1 | 1 |
| npc2 | 0.50893 | -0.10186284 | 0.32 | 0.402 | 1 | 1 |
| si:ch211-106h4.12 | 0.50964 | 0.112082221 | 0.11 | 0.085 | 1 | 1 |
| ifi45 | 0.50967 | -0.13612598 | 0.47 | 0.512 | 1 | 1 |
| wu:fb55g09 | 0.51106 | 0.119602381 | 0.53 | 0.463 | 1 | 1 |
| tax1bp1b | 0.51305 | 0.106899627 | 0.19 | 0.134 | 1 | 1 |
| tsc22d1 | 0.51353 | 0.12096591 | 0.25 | 0.183 | 1 | 1 |
| pitpnbl | 0.51359 | 0.102184023 | 0.2 | 0.146 | 1 | 1 |
| capza1b | 0.51393 | -0.11065146 | 0.32 | 0.402 | 1 | 1 |
| tpst1 | 0.51411 | 0.104501641 | 0.18 | 0.134 | 1 | 1 |
| hspa5 | 0.518 | -0.13601325 | 0.29 | 0.366 | 1 | 1 |
| serp1 | 0.51885 | -0.10886096 | 0.17 | 0.232 | 1 | 1 |
| zgc:158463 | 0.51886 | -0.11517159 | 0.23 | 0.305 | 1 | 1 |
| cltca | 0.51923 | 0.106263175 | 0.21 | 0.146 | 1 | 1 |
| jam2a | 0.51987 | 0.113722543 | 0.36 | 0.28 | 1 | 1 |
| grb10a | 0.52267 | 0.111844679 | 0.28 | 0.207 | 1 | 1 |
| txnipa | 0.5297 | -0.12455877 | 0.42 | 0.488 | 1 | 1 |
| psme2 | 0.53063 | -0.10286661 | 0.3 | 0.378 | 1 | 1 |
| fbxl6 | 0.53379 | -0.11320481 | 0.22 | 0.293 | 1 | 1 |
| mdh2 | 0.53547 | -0.10743883 | 0.53 | 0.61 | 1 | 1 |
| eif4ebp1 | 0.5392 | -0.10651583 | 0.48 | 0.561 | 1 | 1 |
| ar | 0.54182 | -0.10680115 | 0.43 | 0.512 | 1 | 1 |
| banf1 | 0.54226 | -0.12226206 | 0.29 | 0.366 | 1 | 1 |
| lgals17 | 0.54432 | -0.12582778 | 0.21 | 0.28 | 1 | 1 |
| emp2 | 0.54448 | 0.113477041 | 0.21 | 0.159 | 1 | 1 |
| nfixb | 0.55013 | 0.12615664 | 0.33 | 0.268 | 1 | 1 |
| serbp1b | 0.55665 | 0.100395403 | 0.23 | 0.183 | 1 | 1 |
| mtdha | 0.5573 | 0.117860575 | 0.4 | 0.329 | 1 | 1 |
| meis3 | 0.55736 | 0.109743572 | 0.12 | 0.073 | 1 | 1 |
| LOC103910438 | 0.55833 | -0.11584382 | 0.34 | 0.415 | 1 | 1 |
| snd1 | 0.56016 | 0.101192834 | 0.29 | 0.244 | 1 | 1 |
| tmem47 | 0.5606 | 0.116945762 | 0.26 | 0.207 | 1 | 1 |
| elocb | 0.5667 | 0.104808565 | 0.46 | 0.39 | 1 | 1 |
| hnrnpa1a | 0.56789 | -0.11664729 | 0.25 | 0.317 | 1 | 1 |
| myclb | 0.56899 | 0.113965048 | 0.28 | 0.22 | 1 | 1 |
| ugt1b5 | 0.57134 | -0.11179861 | 0.16 | 0.22 | 1 | 1 |
| mcm2 | 0.5741 | 0.121964194 | 0.2 | 0.146 | 1 | 1 |

| | | | | | | |
|---|---|---|---|---|---|---|
| srrm1 | 0.57708 | -0.10408974 | 0.17 | 0.232 | 1 | 1 |
| cspg5a | 0.57899 | -0.12010808 | 0.21 | 0.268 | 1 | 1 |
| tmem88b | 0.57932 | 0.116622516 | 0.51 | 0.439 | 1 | 1 |
| fxyd6l | 0.58009 | -0.10184037 | 0.27 | 0.341 | 1 | 1 |
| klf2a | 0.58141 | -0.12512616 | 0.3 | 0.354 | 1 | 1 |
| acta2 | 0.59821 | -0.10204782 | 0.03 | 0.061 | 1 | 1 |
| smo | 0.60466 | 0.107590264 | 0.27 | 0.207 | 1 | 1 |
| cyp2ad3 | 0.61463 | -0.11251858 | 0.32 | 0.39 | 1 | 1 |
| selenok | 0.61485 | 0.102339598 | 0.3 | 0.244 | 1 | 1 |
| krt18a.1 | 0.63525 | -0.11408272 | 0.47 | 0.537 | 1 | 1 |
| nfil3-6 | 0.64434 | -0.10180752 | 0.21 | 0.268 | 1 | 1 |
| ccbe1 | 0.65045 | -0.10267801 | 0.25 | 0.293 | 1 | 1 |
| apoc1 | 7.265025 | -0.54643431 | 0 | 0.15 | 1 | 2 |
| LOC100536946 | 0.00045 | -0.69068168 | 0.165 | 0.375 | 1 | 2 |
| pcf11 | 0.00154 | -0.33070495 | 0 | 0.1 | 1 | 2 |
| mycbp | 0.0016 | 0.381927271 | 0.148 | 0 | 1 | 2 |
| zgc:92744 | 0.0023 | 0.413790598 | 0.139 | 0 | 1 | 2 |
| commd1 | 0.00319 | -0.5656589 | 0.07 | 0.25 | 1 | 2 |
| cpb1 | 0.00472 | 0.372432387 | 0.122 | 0 | 1 | 2 |
| jac8 | 0.00475 | -0.69801723 | 0.043 | 0.225 | 1 | 2 |
| rpl27a | 0.00475 | -0.27494508 | 0.548 | 0.625 | 1 | 2 |
| XLOC-000431 | 0.005 | 0.317602813 | 0.235 | 0.075 | 1 | 2 |
| psme2 | 0.00626 | 0.422151548 | 0.235 | 0.05 | 1 | 2 |
| rpl18 | 0.0067 | 0.220065696 | 0.957 | 0.875 | 1 | 2 |
| CABZ01032488.1 | 0.00705 | -0.17728601 | 0 | 0.075 | 1 | 2 |
| slc44a1b | 0.00705 | -0.21900237 | 0 | 0.075 | 1 | 2 |
| CU694532.1 | 0.00705 | -0.17267915 | 0 | 0.075 | 1 | 2 |
| PYURF | 0.00705 | -0.27939693 | 0 | 0.075 | 1 | 2 |
| slc16a9a | 0.00705 | -0.20521556 | 0 | 0.075 | 1 | 2 |
| thoc3 | 0.00705 | -0.18233814 | 0 | 0.075 | 1 | 2 |
| zpr1 | 0.00705 | -0.24365258 | 0 | 0.075 | 1 | 2 |
| ctnnbl1 | 0.00705 | -0.23393081 | 0 | 0.075 | 1 | 2 |
| LOC101886073 | 0.00705 | -0.22426682 | 0 | 0.075 | 1 | 2 |
| cc2d1b | 0.00705 | -0.22361869 | 0 | 0.075 | 1 | 2 |
| eif4e2rs1 | 0.00705 | -0.22489308 | 0 | 0.075 | 1 | 2 |
| rbp2a | 0.00705 | -0.22713852 | 0 | 0.075 | 1 | 2 |
| si:dkey-126g1.7 | 0.00705 | -0.22119453 | 0 | 0.075 | 1 | 2 |
| nudt3a | 0.00705 | -0.24858064 | 0 | 0.075 | 1 | 2 |
| prpsap1 | 0.00705 | -0.1908137 | 0 | 0.075 | 1 | 2 |
| srcap | 0.00705 | -0.19682709 | 0 | 0.075 | 1 | 2 |
| tgm1l1 | 0.00705 | -0.21099606 | 0 | 0.075 | 1 | 2 |
| ctsla | 0.00735 | -0.49746503 | 0.07 | 0.2 | 1 | 2 |
| rps3a | 0.00762 | 0.21371992 | 0.957 | 0.9 | 1 | 2 |
| nifk | 0.00926 | -0.30787658 | 0.017 | 0.15 | 1 | 2 |
| march5l | 0.00963 | 0.294342352 | 0.104 | 0 | 1 | 2 |
| sept8a | 0.00963 | 0.31264194 | 0.104 | 0 | 1 | 2 |
| slc3a2b | 0.00963 | 0.282624985 | 0.104 | 0 | 1 | 2 |
| timm10 | 0.00963 | 0.301579692 | 0.104 | 0 | 1 | 2 |

| | | | | | | |
|---|---|---|---|---|---|---|
| gkap1 | 0.00963 | 0.270959903 | 0.104 | 0 | 1 | 2 |
| gstm.2 | 0.00963 | 0.283641178 | 0.104 | 0 | 1 | 2 |
| ppp1cab | 0.00963 | 0.265899059 | 0.104 | 0 | 1 | 2 |
| sod2 | 0.00963 | 0.28599438 | 0.104 | 0 | 1 | 2 |
| mt-atp6 | 0.00987 | -0.110923 | 0.243 | 0.375 | 1 | 2 |
| si:ch73-335l21.4 | 0.00995 | -0.31546865 | 0.009 | 0.125 | 1 | 2 |
| zgc:193541 | 0.01007 | -0.34110542 | 0.383 | 0.5 | 1 | 2 |
| ypel1 | 0.01111 | -0.39097321 | 0.009 | 0.125 | 1 | 2 |
| eef2b | 0.0129 | -0.13493073 | 0.765 | 0.75 | 1 | 2 |
| gamt | 0.01375 | 0.209900702 | 0.096 | 0 | 1 | 2 |
| sh3kbp1 | 0.01486 | -0.38015358 | 0.026 | 0.15 | 1 | 2 |
| nck1a | 0.01591 | -0.45377123 | 0.061 | 0.2 | 1 | 2 |
| coro1a | 0.01718 | -0.38725661 | 0.504 | 0.7 | 1 | 2 |
| rpn1 | 0.01864 | 0.433277258 | 0.174 | 0.025 | 1 | 2 |
| si:dkey-72l14.4 | 0.01962 | 0.232523141 | 0.087 | 0 | 1 | 2 |
| si:dkey-188i13.11 | 0.01962 | 0.21830885 | 0.087 | 0 | 1 | 2 |
| sowahd | 0.01962 | 0.23506634 | 0.087 | 0 | 1 | 2 |
| acp1 | 0.01962 | 0.22010589 | 0.087 | 0 | 1 | 2 |
| plekhb2 | 0.01962 | 0.240419713 | 0.087 | 0 | 1 | 2 |
| plekho2 | 0.01962 | 0.248917535 | 0.087 | 0 | 1 | 2 |
| tuba8l | 0.02208 | 0.433577125 | 0.174 | 0.025 | 1 | 2 |
| rab14 | 0.02298 | -0.36633764 | 0.035 | 0.175 | 1 | 2 |
| arpc2 | 0.02323 | 0.365965177 | 0.591 | 0.425 | 1 | 2 |
| COX5B | 0.02493 | -0.12471287 | 0.391 | 0.375 | 1 | 2 |
| ucp2 | 0.02596 | 0.511931752 | 0.383 | 0.175 | 1 | 2 |
| c6ast3 | 0.028 | 0.175851577 | 0.078 | 0 | 1 | 2 |
| rap2c | 0.028 | 0.213803639 | 0.078 | 0 | 1 | 2 |
| XLOC-005448 | 0.028 | 0.246564746 | 0.078 | 0 | 1 | 2 |
| cadm4 | 0.028 | 0.24703901 | 0.078 | 0 | 1 | 2 |
| ccl39.6 | 0.028 | 0.301872203 | 0.078 | 0 | 1 | 2 |
| cel.1 | 0.028 | 0.203856825 | 0.078 | 0 | 1 | 2 |
| cel.2 | 0.028 | 0.174132535 | 0.078 | 0 | 1 | 2 |
| cpa1 | 0.028 | 0.246750908 | 0.078 | 0 | 1 | 2 |
| endou | 0.028 | 0.165106495 | 0.078 | 0 | 1 | 2 |
| pdcd5 | 0.028 | 0.196138736 | 0.078 | 0 | 1 | 2 |
| rhebl1 | 0.028 | 0.177242585 | 0.078 | 0 | 1 | 2 |
| rtn3 | 0.028 | 0.210938032 | 0.078 | 0 | 1 | 2 |
| sco1 | 0.028 | 0.220735512 | 0.078 | 0 | 1 | 2 |
| uqcrc2a | 0.028 | 0.172667429 | 0.078 | 0 | 1 | 2 |
| vps26c | 0.028 | 0.234229899 | 0.078 | 0 | 1 | 2 |
| si:ch211-106a19.1 | 0.02827 | -0.36567899 | 0.026 | 0.15 | 1 | 2 |
| smim4 | 0.0284 | -0.33785583 | 0.026 | 0.15 | 1 | 2 |
| psmb2 | 0.02959 | 0.315783581 | 0.304 | 0.125 | 1 | 2 |
| CABZ01046432.1 | 0.02962 | -0.30995026 | 0.009 | 0.1 | 1 | 2 |
| ppp6r3 | 0.02973 | -0.31792856 | 0.017 | 0.125 | 1 | 2 |
| rfc5 | 0.03234 | -0.27153342 | 0.009 | 0.1 | 1 | 2 |
| rpl28 | 0.03238 | 0.204346443 | 0.939 | 0.85 | 1 | 2 |
| BX072576.2 | 0.03279 | -0.19427439 | 0 | 0.05 | 1 | 2 |

| | | | | | | |
|---|---|---|---|---|---|---|
| CU633745.1 | 0.03279 | -0.14442045 | 0 | 0.05 | 1 | 2 |
| LOC110437969 | 0.03279 | -0.20082857 | 0 | 0.05 | 1 | 2 |
| XLOC-041980 | 0.03279 | -0.15487771 | 0 | 0.05 | 1 | 2 |
| aga | 0.03279 | -0.15145937 | 0 | 0.05 | 1 | 2 |
| ak4 | 0.03279 | -0.18210851 | 0 | 0.05 | 1 | 2 |
| babam2 | 0.03279 | -0.14909347 | 0 | 0.05 | 1 | 2 |
| bbc3 | 0.03279 | -0.17222726 | 0 | 0.05 | 1 | 2 |
| brcc3 | 0.03279 | -0.15212885 | 0 | 0.05 | 1 | 2 |
| cd99l2 | 0.03279 | -0.16274431 | 0 | 0.05 | 1 | 2 |
| dctn1a | 0.03279 | -0.15553655 | 0 | 0.05 | 1 | 2 |
| dnajc9 | 0.03279 | -0.20549863 | 0 | 0.05 | 1 | 2 |
| dnase1l4.2 | 0.03279 | -0.18246714 | 0 | 0.05 | 1 | 2 |
| eif4e2 | 0.03279 | -0.19405655 | 0 | 0.05 | 1 | 2 |
| emc9 | 0.03279 | -0.1708121 | 0 | 0.05 | 1 | 2 |
| enc2 | 0.03279 | -0.13631181 | 0 | 0.05 | 1 | 2 |
| iqcc | 0.03279 | -0.17469393 | 0 | 0.05 | 1 | 2 |
| junbb | 0.03279 | -0.19798197 | 0 | 0.05 | 1 | 2 |
| kbtbd11 | 0.03279 | -0.20078031 | 0 | 0.05 | 1 | 2 |
| mtg2 | 0.03279 | -0.14257535 | 0 | 0.05 | 1 | 2 |
| mtmr14 | 0.03279 | -0.14850165 | 0 | 0.05 | 1 | 2 |
| nudcd2 | 0.03279 | -0.1510464 | 0 | 0.05 | 1 | 2 |
| oxsm | 0.03279 | -0.12554565 | 0 | 0.05 | 1 | 2 |
| rassf4a | 0.03279 | -0.18870765 | 0 | 0.05 | 1 | 2 |
| rbck1 | 0.03279 | -0.19686789 | 0 | 0.05 | 1 | 2 |
| sh3bp5la | 0.03279 | -0.17537731 | 0 | 0.05 | 1 | 2 |
| si:dkey-78l4.2 | 0.03279 | -0.10311516 | 0 | 0.05 | 1 | 2 |
| smn1 | 0.03279 | -0.15645865 | 0 | 0.05 | 1 | 2 |
| suv39h1b | 0.03279 | -0.18609619 | 0 | 0.05 | 1 | 2 |
| tmem131 | 0.03279 | -0.1380473 | 0 | 0.05 | 1 | 2 |
| trim35-1 | 0.03279 | -0.18595625 | 0 | 0.05 | 1 | 2 |
| vps41 | 0.03279 | -0.15836323 | 0 | 0.05 | 1 | 2 |
| zgc:110540 | 0.03279 | -0.14195592 | 0 | 0.05 | 1 | 2 |
| zgc:85858 | 0.03279 | -0.15675514 | 0 | 0.05 | 1 | 2 |
| zmp:0000000936 | 0.03279 | -0.18054209 | 0 | 0.05 | 1 | 2 |
| znf410 | 0.03279 | -0.15150859 | 0 | 0.05 | 1 | 2 |
| znf704 | 0.03279 | -0.14313876 | 0 | 0.05 | 1 | 2 |
| golt1bb | 0.03279 | -0.16156435 | 0 | 0.05 | 1 | 2 |
| stx10 | 0.03279 | -0.16757128 | 0 | 0.05 | 1 | 2 |
| CU104709.1 | 0.03279 | -0.15081315 | 0 | 0.05 | 1 | 2 |
| aldh3a2b | 0.03279 | -0.16406912 | 0 | 0.05 | 1 | 2 |
| cnot7 | 0.03279 | -0.11989165 | 0 | 0.05 | 1 | 2 |
| daxx | 0.03279 | -0.13797454 | 0 | 0.05 | 1 | 2 |
| ganab | 0.03279 | -0.17984445 | 0 | 0.05 | 1 | 2 |
| gins4 | 0.03279 | -0.16629386 | 0 | 0.05 | 1 | 2 |
| hic1l | 0.03279 | -0.1685497 | 0 | 0.05 | 1 | 2 |
| irge4 | 0.03279 | -0.1938591 | 0 | 0.05 | 1 | 2 |
| lmbrd1 | 0.03279 | -0.15931611 | 0 | 0.05 | 1 | 2 |
| med22 | 0.03279 | -0.15894977 | 0 | 0.05 | 1 | 2 |

| | | | | | | |
|---|---|---|---|---|---|---|
| med28 | 0.03279 | -0.15857935 | 0 | 0.05 | 1 | 2 |
| mpzl2b | 0.03279 | -0.17534631 | 0 | 0.05 | 1 | 2 |
| otud5a | 0.03279 | -0.2015093 | 0 | 0.05 | 1 | 2 |
| plecb | 0.03279 | -0.13436294 | 0 | 0.05 | 1 | 2 |
| reck | 0.03279 | -0.13592721 | 0 | 0.05 | 1 | 2 |
| si:ch1073-358c10.1 | 0.03279 | -0.18359994 | 0 | 0.05 | 1 | 2 |
| si:ch211-171h4.5 | 0.03279 | -0.14920455 | 0 | 0.05 | 1 | 2 |
| si:ch211-76m11.3 | 0.03279 | -0.17362183 | 0 | 0.05 | 1 | 2 |
| si:dkey-30g5.1 | 0.03279 | -0.19495444 | 0 | 0.05 | 1 | 2 |
| tmx4 | 0.03279 | -0.19109527 | 0 | 0.05 | 1 | 2 |
| ttc32 | 0.03279 | -0.13400259 | 0 | 0.05 | 1 | 2 |
| utp15 | 0.03279 | -0.14495314 | 0 | 0.05 | 1 | 2 |
| cpn1 | 0.03279 | -0.16038787 | 0 | 0.05 | 1 | 2 |
| tnika | 0.03279 | -0.18292423 | 0 | 0.05 | 1 | 2 |
| LOC101885014 | 0.03279 | -0.16625991 | 0 | 0.05 | 1 | 2 |
| ddx56 | 0.03279 | -0.17825368 | 0 | 0.05 | 1 | 2 |
| ncln | 0.03279 | -0.1653571 | 0 | 0.05 | 1 | 2 |
| rbm7 | 0.03279 | -0.15279723 | 0 | 0.05 | 1 | 2 |
| si:ch211-253b8.2 | 0.03279 | -0.1676187 | 0 | 0.05 | 1 | 2 |
| tnfrsfa | 0.03279 | -0.15113859 | 0 | 0.05 | 1 | 2 |
| wu:fb16f03 | 0.03279 | -0.16475503 | 0 | 0.05 | 1 | 2 |
| eif5a | 0.03296 | 0.358769544 | 0.557 | 0.325 | 1 | 2 |
| mrps27 | 0.03407 | -0.26118253 | 0.009 | 0.1 | 1 | 2 |
| NC-002333.4 | 0.03442 | 0.569798286 | 0.391 | 0.175 | 1 | 2 |
| ilf2 | 0.03453 | -0.15683294 | 0.174 | 0.175 | 1 | 2 |
| cebpb | 0.03463 | -0.38922315 | 0.496 | 0.7 | 1 | 2 |
| mcrip1 | 0.0348 | -0.31419428 | 0.017 | 0.125 | 1 | 2 |
| si:ch211-122f10.4 | 0.03513 | -0.322933 | 0.017 | 0.125 | 1 | 2 |
| mych | 0.03565 | -0.26604604 | 0.009 | 0.1 | 1 | 2 |
| zgc:162730 | 0.03642 | -0.31888625 | 0.617 | 0.825 | 1 | 2 |
| rgs19 | 0.03703 | -0.27207747 | 0.009 | 0.1 | 1 | 2 |
| si:ch211-15b10.6 | 0.03713 | 0.209521783 | 0.217 | 0.1 | 1 | 2 |
| scarb2c | 0.03714 | -0.28935383 | 0.009 | 0.1 | 1 | 2 |
| gstp1 | 0.0379 | -0.35843413 | 0.183 | 0.275 | 1 | 2 |
| gng10 | 0.03945 | -0.3338745 | 0.33 | 0.45 | 1 | 2 |
| ebna1bp2 | 0.03999 | 0.164876955 | 0.07 | 0 | 1 | 2 |
| rras | 0.03999 | 0.185264859 | 0.07 | 0 | 1 | 2 |
| CELA1 (1 of many) | 0.03999 | 0.325506074 | 0.07 | 0 | 1 | 2 |
| pla2g1b | 0.03999 | 0.160047814 | 0.07 | 0 | 1 | 2 |
| selenot1a | 0.03999 | 0.175121683 | 0.07 | 0 | 1 | 2 |
| ssuh2rs1 | 0.03999 | 0.218350747 | 0.07 | 0 | 1 | 2 |
| cops5 | 0.03999 | 0.157611106 | 0.07 | 0 | 1 | 2 |
| gyg1a | 0.03999 | 0.18713334 | 0.07 | 0 | 1 | 2 |
| atp6v0b | 0.03999 | 0.228565666 | 0.07 | 0 | 1 | 2 |
| dhrs9 | 0.03999 | 0.206270067 | 0.07 | 0 | 1 | 2 |
| si:dkey-222f8.3 | 0.03999 | 0.182352945 | 0.07 | 0 | 1 | 2 |
| zgc:175088 | 0.03999 | 0.204227131 | 0.07 | 0 | 1 | 2 |
| arl14ep | 0.03999 | 0.221024255 | 0.07 | 0 | 1 | 2 |

| | | | | | | |
|---|---|---|---|---|---|---|
| lrrc8aa | 0.03999 | 0.188275027 | 0.07 | 0 | 1 | 2 |
| si:dkey-31f5.8 | 0.03999 | 0.207235134 | 0.07 | 0 | 1 | 2 |
| tmem131l | 0.03999 | 0.184547423 | 0.07 | 0 | 1 | 2 |
| zgc:92590 | 0.03999 | 0.223251885 | 0.07 | 0 | 1 | 2 |
| kctd12.2 | 0.04186 | 0.303020314 | 0.374 | 0.2 | 1 | 2 |
| ccl34a.4 | 0.04254 | -0.45869486 | 0.139 | 0.325 | 1 | 2 |
| BX649485.2 | 0.04484 | -0.49906065 | 0.252 | 0.425 | 1 | 2 |
| mhc2dab | 0.04603 | 0.475364015 | 0.409 | 0.2 | 1 | 2 |
| jac9 | 0.04625 | -0.45238232 | 0.122 | 0.3 | 1 | 2 |
| romo1 | 0.0475 | 0.398139705 | 0.409 | 0.2 | 1 | 2 |
| ywhaba | 0.04914 | -0.17953979 | 0.322 | 0.35 | 1 | 2 |
| rps26l | 0.0508 | 0.163984352 | 0.974 | 0.875 | 1 | 2 |
| pgam1a | 0.05089 | 0.388345781 | 0.2 | 0.05 | 1 | 2 |
| dusp6 | 0.05126 | -0.2842452 | 0.035 | 0.15 | 1 | 2 |
| ppifb | 0.05626 | 0.364098198 | 0.191 | 0.05 | 1 | 2 |
| gabarapb | 0.05673 | 0.370274295 | 0.617 | 0.4 | 1 | 2 |
| alyref | 0.05719 | 0.164495488 | 0.061 | 0 | 1 | 2 |
| cpa5 | 0.05719 | 0.241651733 | 0.061 | 0 | 1 | 2 |
| ctrl | 0.05719 | 0.240111893 | 0.061 | 0 | 1 | 2 |
| gng2 | 0.05719 | 0.138254938 | 0.061 | 0 | 1 | 2 |
| isoc2 | 0.05719 | 0.176807827 | 0.061 | 0 | 1 | 2 |
| lman2 | 0.05719 | 0.164240024 | 0.061 | 0 | 1 | 2 |
| ncf1 | 0.05719 | 0.189848478 | 0.061 | 0 | 1 | 2 |
| psma6l | 0.05719 | 0.157655235 | 0.061 | 0 | 1 | 2 |
| rps6kb1b | 0.05719 | 0.16601061 | 0.061 | 0 | 1 | 2 |
| sla1 | 0.05719 | 0.162106367 | 0.061 | 0 | 1 | 2 |
| zfp36l1b | 0.05719 | 0.17699379 | 0.061 | 0 | 1 | 2 |
| cul4b | 0.05719 | 0.129953534 | 0.061 | 0 | 1 | 2 |
| eif4g2b | 0.05719 | 0.175040509 | 0.061 | 0 | 1 | 2 |
| ell | 0.05719 | 0.180563203 | 0.061 | 0 | 1 | 2 |
| lmnb1 | 0.05719 | 0.174464119 | 0.061 | 0 | 1 | 2 |
| spock3 | 0.05719 | 0.195006976 | 0.061 | 0 | 1 | 2 |
| tgif1 | 0.05719 | 0.168849843 | 0.061 | 0 | 1 | 2 |
| chaf1a | 0.05719 | 0.180330198 | 0.061 | 0 | 1 | 2 |
| erp27 | 0.05719 | 0.152928483 | 0.061 | 0 | 1 | 2 |
| lcorl | 0.05719 | 0.186050393 | 0.061 | 0 | 1 | 2 |
| mrpl28 | 0.05719 | 0.163541083 | 0.061 | 0 | 1 | 2 |
| mrps33 | 0.05719 | 0.178463389 | 0.061 | 0 | 1 | 2 |
| si:ch211-240l19.8 | 0.05719 | 0.24749814 | 0.061 | 0 | 1 | 2 |
| si:ch73-71c20.5 | 0.05719 | 0.162801805 | 0.061 | 0 | 1 | 2 |
| si:dkey-33i11.9 | 0.05719 | 0.2318935 | 0.061 | 0 | 1 | 2 |
| smim29 | 0.05719 | 0.152275629 | 0.061 | 0 | 1 | 2 |
| stoml3b | 0.05719 | 0.182793052 | 0.061 | 0 | 1 | 2 |
| tardbpl | 0.05719 | 0.176356142 | 0.061 | 0 | 1 | 2 |
| timm13 | 0.05719 | 0.188213382 | 0.061 | 0 | 1 | 2 |
| tnip1 | 0.05719 | 0.164723851 | 0.061 | 0 | 1 | 2 |
| uri1 | 0.05719 | 0.170559863 | 0.061 | 0 | 1 | 2 |
| zc4h2 | 0.05719 | 0.176933231 | 0.061 | 0 | 1 | 2 |

| | | | | | | |
|---|---|---|---|---|---|---|
| anp32e | 0.05719 | 0.166229683 | 0.061 | 0 | 1 | 2 |
| chchd3a | 0.05719 | 0.183991646 | 0.061 | 0 | 1 | 2 |
| cpa4 | 0.05719 | 0.141554294 | 0.061 | 0 | 1 | 2 |
| cwc25 | 0.05719 | 0.15765326 | 0.061 | 0 | 1 | 2 |
| ppid | 0.05719 | 0.170069749 | 0.061 | 0 | 1 | 2 |
| rnf25 | 0.05719 | 0.124701814 | 0.061 | 0 | 1 | 2 |
| slc38a5b | 0.05719 | 0.140084479 | 0.061 | 0 | 1 | 2 |
| dpm2 | 0.05719 | 0.164959296 | 0.061 | 0 | 1 | 2 |
| manbal | 0.05719 | 0.194475484 | 0.061 | 0 | 1 | 2 |
| si:ch211-195b11.3 | 0.05719 | 0.151584516 | 0.061 | 0 | 1 | 2 |
| zgc:85789 | 0.05719 | 0.136598091 | 0.061 | 0 | 1 | 2 |
| cybrd1 | 0.05885 | -0.34328287 | 0.035 | 0.15 | 1 | 2 |
| selenow1 | 0.05888 | -0.38863972 | 0.339 | 0.55 | 1 | 2 |
| anxa1a | 0.0593 | -0.32618395 | 0.078 | 0.225 | 1 | 2 |
| dynlrb1 | 0.05934 | 0.342946097 | 0.165 | 0.05 | 1 | 2 |
| slc7a7 | 0.06001 | -0.25599648 | 0.026 | 0.125 | 1 | 2 |
| ccdc124 | 0.06004 | -0.33339289 | 0.087 | 0.175 | 1 | 2 |
| mcl1a | 0.06045 | -0.17381247 | 0.73 | 0.775 | 1 | 2 |
| itm2bb | 0.06115 | 0.411887245 | 0.452 | 0.25 | 1 | 2 |
| atf3 | 0.06165 | -0.37733767 | 0.043 | 0.15 | 1 | 2 |
| col4a3bpb | 0.06202 | -0.29059102 | 0.017 | 0.1 | 1 | 2 |
| cox4i1 | 0.06234 | 0.124183249 | 0.574 | 0.425 | 1 | 2 |
| map1lc3b | 0.06288 | -0.36534724 | 0.096 | 0.2 | 1 | 2 |
| ddost | 0.0645 | 0.197023738 | 0.122 | 0.025 | 1 | 2 |
| higd1a | 0.06609 | -0.38336796 | 0.096 | 0.25 | 1 | 2 |
| cyc1 | 0.06691 | 0.294613407 | 0.139 | 0.025 | 1 | 2 |
| lgmn | 0.06731 | -0.17122764 | 0.2 | 0.2 | 1 | 2 |
| rab11ba | 0.06911 | 0.339782727 | 0.139 | 0.025 | 1 | 2 |
| eif4a1b | 0.07058 | -0.27729651 | 0.339 | 0.425 | 1 | 2 |
| taf13 | 0.07101 | 0.271368793 | 0.139 | 0.025 | 1 | 2 |
| cox8a | 0.07196 | 0.252008254 | 0.443 | 0.275 | 1 | 2 |
| jun | 0.072 | -0.28794742 | 0.026 | 0.125 | 1 | 2 |
| znf706 | 0.07261 | -0.36654284 | 0.087 | 0.225 | 1 | 2 |
| anapc16 | 0.07408 | 0.227697072 | 0.191 | 0.075 | 1 | 2 |
| erh | 0.07463 | 0.363965763 | 0.226 | 0.075 | 1 | 2 |
| eny2 | 0.07576 | 0.277628724 | 0.139 | 0.025 | 1 | 2 |
| zbtb2b | 0.07647 | -0.29637423 | 0.026 | 0.125 | 1 | 2 |
| gdi2 | 0.07674 | 0.384648304 | 0.261 | 0.1 | 1 | 2 |
| txnrd3 | 0.07814 | -0.29283155 | 0.026 | 0.125 | 1 | 2 |
| dusp2 | 0.07852 | -0.39846641 | 0.07 | 0.175 | 1 | 2 |
| nt5c2a | 0.07953 | -0.2354181 | 0.017 | 0.1 | 1 | 2 |
| XLOC-013964 | 0.08024 | 0.320594288 | 0.139 | 0.025 | 1 | 2 |
| srsf3b | 0.08105 | 0.292607437 | 0.139 | 0.025 | 1 | 2 |
| twf1a | 0.08136 | 0.325080472 | 0.183 | 0.05 | 1 | 2 |
| DDX17 | 0.08201 | 0.143143836 | 0.052 | 0 | 1 | 2 |
| abce1 | 0.08201 | 0.141595367 | 0.052 | 0 | 1 | 2 |
| amy2a | 0.08201 | 0.160457407 | 0.052 | 0 | 1 | 2 |
| apoda.2 | 0.08201 | 0.146410147 | 0.052 | 0 | 1 | 2 |

| | | | | | | |
|---|---|---|---|---|---|---|
| atp7a | 0.08201 | 0.156563507 | 0.052 | 0 | 1 | 2 |
| bloc1s1 | 0.08201 | 0.128076734 | 0.052 | 0 | 1 | 2 |
| fcf1 | 0.08201 | 0.142818749 | 0.052 | 0 | 1 | 2 |
| flvcr2b | 0.08201 | 0.145927407 | 0.052 | 0 | 1 | 2 |
| git2b | 0.08201 | 0.140859432 | 0.052 | 0 | 1 | 2 |
| hnrnpm | 0.08201 | 0.123154197 | 0.052 | 0 | 1 | 2 |
| hsd17b10 | 0.08201 | 0.119322525 | 0.052 | 0 | 1 | 2 |
| rab42b | 0.08201 | 0.137195568 | 0.052 | 0 | 1 | 2 |
| rtn4ip1 | 0.08201 | 0.131130228 | 0.052 | 0 | 1 | 2 |
| si:ch211-243a20.4 | 0.08201 | 0.151701413 | 0.052 | 0 | 1 | 2 |
| SMIM10 | 0.08201 | 0.145598673 | 0.052 | 0 | 1 | 2 |
| dap | 0.08201 | 0.15583919 | 0.052 | 0 | 1 | 2 |
| eif4g2a | 0.08201 | 0.160108551 | 0.052 | 0 | 1 | 2 |
| plp2 | 0.08201 | 0.146588031 | 0.052 | 0 | 1 | 2 |
| CABZ01020840.1 | 0.08201 | 0.160641994 | 0.052 | 0 | 1 | 2 |
| ackr3b | 0.08201 | 0.127009077 | 0.052 | 0 | 1 | 2 |
| arl8a | 0.08201 | 0.117841263 | 0.052 | 0 | 1 | 2 |
| asb8 | 0.08201 | 0.104377628 | 0.052 | 0 | 1 | 2 |
| cdk11b | 0.08201 | 0.159236678 | 0.052 | 0 | 1 | 2 |
| eif4ebp3 | 0.08201 | 0.107422852 | 0.052 | 0 | 1 | 2 |
| hp1bp3 | 0.08201 | 0.131128588 | 0.052 | 0 | 1 | 2 |
| nucb2a | 0.08201 | 0.149168992 | 0.052 | 0 | 1 | 2 |
| ophn1 | 0.08201 | 0.174214901 | 0.052 | 0 | 1 | 2 |
| rbmx | 0.08201 | 0.14552096 | 0.052 | 0 | 1 | 2 |
| si:ch211-22i13.2 | 0.08201 | 0.141130762 | 0.052 | 0 | 1 | 2 |
| si:dkey-27b3.2 | 0.08201 | 0.180979935 | 0.052 | 0 | 1 | 2 |
| slc25a28 | 0.08201 | 0.142577126 | 0.052 | 0 | 1 | 2 |
| slc9a3r1a | 0.08201 | 0.157903653 | 0.052 | 0 | 1 | 2 |
| txndc12 | 0.08201 | 0.167919143 | 0.052 | 0 | 1 | 2 |
| ube2c | 0.08201 | 0.133447948 | 0.052 | 0 | 1 | 2 |
| ap1b1 | 0.08201 | 0.141427738 | 0.052 | 0 | 1 | 2 |
| cd164 | 0.08201 | 0.109697193 | 0.052 | 0 | 1 | 2 |
| dnaja2b | 0.08201 | 0.134586464 | 0.052 | 0 | 1 | 2 |
| naxd | 0.08201 | 0.136719375 | 0.052 | 0 | 1 | 2 |
| CABZ01068251.1 | 0.08201 | 0.112356792 | 0.052 | 0 | 1 | 2 |
| cfdp1 | 0.08201 | 0.150518202 | 0.052 | 0 | 1 | 2 |
| cish | 0.08201 | 0.148939105 | 0.052 | 0 | 1 | 2 |
| coa1 | 0.08201 | 0.150563963 | 0.052 | 0 | 1 | 2 |
| hbp1 | 0.08201 | 0.124731818 | 0.052 | 0 | 1 | 2 |
| impa1 | 0.08201 | 0.170422117 | 0.052 | 0 | 1 | 2 |
| ivd | 0.08201 | 0.131070579 | 0.052 | 0 | 1 | 2 |
| mrpl52 | 0.08201 | 0.13194625 | 0.052 | 0 | 1 | 2 |
| psmb10 | 0.08201 | 0.150160513 | 0.052 | 0 | 1 | 2 |
| si:ch211-214p16.2 | 0.08201 | 0.180652034 | 0.052 | 0 | 1 | 2 |
| si:dkey-14d8.6 | 0.08201 | 0.169328872 | 0.052 | 0 | 1 | 2 |
| trappc4 | 0.08201 | 0.138997637 | 0.052 | 0 | 1 | 2 |
| trir | 0.08201 | 0.14398262 | 0.052 | 0 | 1 | 2 |
| u2surp | 0.08201 | 0.151084375 | 0.052 | 0 | 1 | 2 |

| | | | | | | |
|---|---|---|---|---|---|---|
| wipi2 | 0.08201 | 0.119697803 | 0.052 | 0 | 1 | 2 |
| yy1a | 0.08201 | 0.117352046 | 0.052 | 0 | 1 | 2 |
| zgc:112160 | 0.08201 | 0.198182508 | 0.052 | 0 | 1 | 2 |
| bsdc1 | 0.08201 | 0.148435504 | 0.052 | 0 | 1 | 2 |
| LOC103910136 | 0.08201 | 0.189625564 | 0.052 | 0 | 1 | 2 |
| gnaia | 0.08201 | 0.160618117 | 0.052 | 0 | 1 | 2 |
| golph3 | 0.08201 | 0.118195766 | 0.052 | 0 | 1 | 2 |
| pltp | 0.08201 | 0.138890761 | 0.052 | 0 | 1 | 2 |
| si:dkey-6e2.2 | 0.08201 | 0.140984755 | 0.052 | 0 | 1 | 2 |
| strap | 0.08201 | 0.127273545 | 0.052 | 0 | 1 | 2 |
| zgc:194627 | 0.08201 | 0.143454752 | 0.052 | 0 | 1 | 2 |
| hyou1 | 0.08201 | 0.156561458 | 0.052 | 0 | 1 | 2 |
| ppig | 0.08201 | 0.133553648 | 0.052 | 0 | 1 | 2 |
| prss59.1 | 0.08201 | 0.249285176 | 0.052 | 0 | 1 | 2 |
| aimp1 | 0.08223 | -0.30988322 | 0.078 | 0.175 | 1 | 2 |
| ppp4r3b | 0.08536 | -0.19271961 | 0.009 | 0.075 | 1 | 2 |
| cox6c | 0.08725 | 0.341552137 | 0.47 | 0.275 | 1 | 2 |
| hnrnpa0l | 0.08814 | 0.172761272 | 0.661 | 0.5 | 1 | 2 |
| crema | 0.0883 | -0.3637739 | 0.122 | 0.225 | 1 | 2 |
| mt-co1 | 0.08902 | 0.159170074 | 0.452 | 0.3 | 1 | 2 |
| klf13 | 0.08904 | -0.26869512 | 0.017 | 0.1 | 1 | 2 |
| txnipa | 0.08942 | -0.26903037 | 0.278 | 0.35 | 1 | 2 |
| txn | 0.09062 | 0.36702056 | 0.417 | 0.3 | 1 | 2 |
| arpc4l | 0.09177 | 0.36800549 | 0.47 | 0.275 | 1 | 2 |
| sirt7 | 0.09215 | -0.19738072 | 0.009 | 0.075 | 1 | 2 |
| XLOC-025423 | 0.09418 | 0.375871198 | 0.339 | 0.225 | 1 | 2 |
| sh3bp1 | 0.09447 | -0.23569026 | 0.017 | 0.1 | 1 | 2 |
| eif3i | 0.09619 | 0.22976992 | 0.313 | 0.175 | 1 | 2 |
| sesn1 | 0.09707 | 0.312628631 | 0.122 | 0.025 | 1 | 2 |
| si:ch211-132p1.3 | 0.09791 | 0.302727622 | 0.174 | 0.05 | 1 | 2 |
| hsbp1b | 0.09931 | 0.235633198 | 0.226 | 0.1 | 1 | 2 |
| ncor1 | 0.09952 | -0.22732256 | 0.009 | 0.075 | 1 | 2 |
| rnft1 | 0.10017 | -0.21987194 | 0.009 | 0.075 | 1 | 2 |
| dnttip2 | 0.10102 | -0.23869119 | 0.017 | 0.1 | 1 | 2 |
| pef1 | 0.10109 | -0.31008555 | 0.043 | 0.15 | 1 | 2 |
| ociad1 | 0.10122 | -0.21128775 | 0.009 | 0.075 | 1 | 2 |
| derl2 | 0.10154 | 0.3890382 | 0.226 | 0.1 | 1 | 2 |
| naa38 | 0.10282 | -0.24225959 | 0.017 | 0.1 | 1 | 2 |
| LOC110437731 | 0.10332 | -0.34629586 | 0.261 | 0.375 | 1 | 2 |
| hnrnpk | 0.10368 | 0.289184696 | 0.13 | 0.025 | 1 | 2 |
| st3gal5 | 0.10496 | -0.21294055 | 0.009 | 0.075 | 1 | 2 |
| sarnp | 0.10541 | -0.32549359 | 0.078 | 0.2 | 1 | 2 |
| mtrex | 0.10701 | -0.17564887 | 0.009 | 0.075 | 1 | 2 |
| rps9 | 0.10703 | 0.145944838 | 0.965 | 0.875 | 1 | 2 |
| mif | 0.10723 | 0.334044043 | 0.174 | 0.05 | 1 | 2 |
| ythdc1 | 0.1096 | -0.20321591 | 0.009 | 0.075 | 1 | 2 |
| fth1a | 0.11063 | -0.312278 | 0.548 | 0.675 | 1 | 2 |
| mboat2a | 0.11063 | -0.24299181 | 0.009 | 0.075 | 1 | 2 |

| | | | | | | |
|---|---|---|---|---|---|---|
| wasb | 0.11161 | 0.246613385 | 0.374 | 0.225 | 1 | 2 |
| abcf1 | 0.11307 | 0.264453614 | 0.122 | 0.025 | 1 | 2 |
| ntan1 | 0.11526 | -0.2278539 | 0.009 | 0.075 | 1 | 2 |
| rpl13 | 0.11554 | -0.15540763 | 0.878 | 0.95 | 1 | 2 |
| nek6 | 0.11651 | -0.1863204 | 0.009 | 0.075 | 1 | 2 |
| si:dkey-25e12.3 | 0.11696 | -0.22557571 | 0.07 | 0.175 | 1 | 2 |
| ttc4 | 0.11723 | -0.20745659 | 0.009 | 0.075 | 1 | 2 |
| alox5ap | 0.11732 | -0.22633012 | 0.009 | 0.075 | 1 | 2 |
| XLOC-001102 | 0.11772 | -0.22245149 | 0.009 | 0.075 | 1 | 2 |
| rtn4a | 0.1181 | 0.210592189 | 0.122 | 0.025 | 1 | 2 |
| rnf44 | 0.11872 | -0.2100084 | 0.009 | 0.075 | 1 | 2 |
| zgc:92161 | 0.11877 | -0.20717267 | 0.009 | 0.075 | 1 | 2 |
| ier5 | 0.11957 | 0.174264899 | 0.104 | 0.025 | 1 | 2 |
| slc25a22a | 0.11961 | -0.20055922 | 0.009 | 0.075 | 1 | 2 |
| ube2g1a | 0.11965 | -0.18055318 | 0.009 | 0.075 | 1 | 2 |
| hipk3a | 0.1197 | -0.23615663 | 0.009 | 0.075 | 1 | 2 |
| zgc:66160 | 0.11982 | -0.20442448 | 0.009 | 0.075 | 1 | 2 |
| CABZ01078737.1 | 0.12028 | -0.26941098 | 0.026 | 0.1 | 1 | 2 |
| tomm7 | 0.12036 | 0.327555773 | 0.243 | 0.1 | 1 | 2 |
| klf6a | 0.12202 | -0.34035023 | 0.104 | 0.225 | 1 | 2 |
| eif3d | 0.12262 | 0.311480633 | 0.339 | 0.175 | 1 | 2 |
| ctsba | 0.1266 | -0.27641064 | 0.557 | 0.675 | 1 | 2 |
| rpl27 | 0.12738 | 0.143031529 | 0.922 | 0.8 | 1 | 2 |
| bzw1b | 0.12837 | -0.36417288 | 0.235 | 0.4 | 1 | 2 |
| ndufa10 | 0.12876 | 0.240450817 | 0.122 | 0.025 | 1 | 2 |
| si:ch73-158p21.3 | 0.12933 | 0.211245645 | 0.148 | 0.05 | 1 | 2 |
| rhoac | 0.12959 | 0.31099783 | 0.27 | 0.125 | 1 | 2 |
| zgc:64051 | 0.13407 | 0.37290476 | 0.365 | 0.2 | 1 | 2 |
| sdhb | 0.13494 | 0.324629202 | 0.2 | 0.075 | 1 | 2 |
| gmppb | 0.13514 | 0.276102794 | 0.165 | 0.05 | 1 | 2 |
| ms4a17a.6 | 0.13556 | 0.298253591 | 0.261 | 0.125 | 1 | 2 |
| hmgb2b | 0.1364 | 0.290703486 | 0.304 | 0.15 | 1 | 2 |
| FO704821.1 | 0.13799 | -0.3349075 | 0.07 | 0.175 | 1 | 2 |
| qkia | 0.13927 | 0.193440794 | 0.113 | 0.025 | 1 | 2 |
| rpl10a | 0.14018 | 0.18382666 | 0.896 | 0.8 | 1 | 2 |
| atp6v0ca | 0.14084 | -0.147177 | 0.409 | 0.425 | 1 | 2 |
| slc25a33 | 0.14092 | -0.23239258 | 0.035 | 0.125 | 1 | 2 |
| rhoab | 0.14107 | -0.3496159 | 0.235 | 0.375 | 1 | 2 |
| emc1 | 0.14138 | -0.14954713 | 0.009 | 0.05 | 1 | 2 |
| hs1bp3 | 0.14138 | -0.14278525 | 0.009 | 0.05 | 1 | 2 |
| psip1a | 0.14138 | -0.13468999 | 0.009 | 0.05 | 1 | 2 |
| si:ch211-210g13.5 | 0.14138 | -0.13419455 | 0.009 | 0.05 | 1 | 2 |
| tpm1 | 0.14138 | -0.19222209 | 0.009 | 0.05 | 1 | 2 |
| ddx47 | 0.14138 | -0.13147746 | 0.009 | 0.05 | 1 | 2 |
| rpz2 | 0.14138 | -0.15160293 | 0.009 | 0.05 | 1 | 2 |
| snapc2 | 0.14138 | -0.18165862 | 0.009 | 0.05 | 1 | 2 |
| atat1 | 0.14138 | -0.14993946 | 0.009 | 0.05 | 1 | 2 |
| golga2 | 0.14138 | -0.13855392 | 0.009 | 0.05 | 1 | 2 |

| | | | | | | |
|---|---|---|---|---|---|---|
| gpd1l | 0.14138 | -0.15217522 | 0.009 | 0.05 | 1 | 2 |
| hnrnph3 | 0.14138 | -0.15495916 | 0.009 | 0.05 | 1 | 2 |
| senp2 | 0.14138 | -0.10278052 | 0.009 | 0.05 | 1 | 2 |
| si:dkey-42l23.2 | 0.14138 | -0.15989759 | 0.009 | 0.05 | 1 | 2 |
| smpd2b | 0.14138 | -0.15740916 | 0.009 | 0.05 | 1 | 2 |
| tfe3b | 0.14138 | -0.15921531 | 0.009 | 0.05 | 1 | 2 |
| tm2d1 | 0.14138 | -0.12935619 | 0.009 | 0.05 | 1 | 2 |
| xpot | 0.14138 | -0.13366824 | 0.009 | 0.05 | 1 | 2 |
| egr2a | 0.14138 | -0.14809652 | 0.009 | 0.05 | 1 | 2 |
| gnpda2 | 0.14138 | -0.13779148 | 0.009 | 0.05 | 1 | 2 |
| irak4 | 0.14138 | -0.14936067 | 0.009 | 0.05 | 1 | 2 |
| itga4 | 0.14138 | -0.15799698 | 0.009 | 0.05 | 1 | 2 |
| leng8 | 0.14138 | -0.11383307 | 0.009 | 0.05 | 1 | 2 |
| map2k4b | 0.14138 | -0.17107703 | 0.009 | 0.05 | 1 | 2 |
| scinla | 0.14138 | -0.15462864 | 0.009 | 0.05 | 1 | 2 |
| shq1 | 0.14138 | -0.12362973 | 0.009 | 0.05 | 1 | 2 |
| si:cabz01036006.1 | 0.14138 | -0.17312833 | 0.009 | 0.05 | 1 | 2 |
| tvp23b | 0.14138 | -0.17364295 | 0.009 | 0.05 | 1 | 2 |
| zgc:101016 | 0.14138 | -0.14417449 | 0.009 | 0.05 | 1 | 2 |
| zgc:174877 | 0.14138 | -0.14885998 | 0.009 | 0.05 | 1 | 2 |
| znhit3 | 0.14138 | -0.17604141 | 0.009 | 0.05 | 1 | 2 |
| CU984600.2 | 0.14138 | -0.16104209 | 0.009 | 0.05 | 1 | 2 |
| dcp2 | 0.14138 | -0.12671329 | 0.009 | 0.05 | 1 | 2 |
| pdk4 | 0.14138 | -0.1835043 | 0.009 | 0.05 | 1 | 2 |
| si:ch211-255f4.7 | 0.14138 | -0.10534168 | 0.009 | 0.05 | 1 | 2 |
| FO704661.1 | 0.14138 | -0.17115688 | 0.009 | 0.05 | 1 | 2 |
| mrpl15 | 0.14138 | -0.10742663 | 0.009 | 0.05 | 1 | 2 |
| npepps | 0.14138 | -0.17993958 | 0.009 | 0.05 | 1 | 2 |
| rprd1b | 0.14138 | -0.16256842 | 0.009 | 0.05 | 1 | 2 |
| si:ch211-125a15.1 | 0.14138 | -0.13944557 | 0.009 | 0.05 | 1 | 2 |
| unc13d | 0.14138 | -0.12604536 | 0.009 | 0.05 | 1 | 2 |
| ppp1r9ba | 0.14138 | -0.13809102 | 0.009 | 0.05 | 1 | 2 |
| serinc5 | 0.14138 | -0.13101996 | 0.009 | 0.05 | 1 | 2 |
| flot2b | 0.14138 | -0.15725467 | 0.009 | 0.05 | 1 | 2 |
| rsu1 | 0.14138 | -0.11348522 | 0.009 | 0.05 | 1 | 2 |
| aaas | 0.14138 | -0.10349495 | 0.009 | 0.05 | 1 | 2 |
| chpfa | 0.14138 | -0.15456139 | 0.009 | 0.05 | 1 | 2 |
| dpf2 | 0.14138 | -0.12167694 | 0.009 | 0.05 | 1 | 2 |
| jac6 | 0.14138 | -0.1173227 | 0.009 | 0.05 | 1 | 2 |
| nudt8 | 0.14138 | -0.13546234 | 0.009 | 0.05 | 1 | 2 |
| plxnb2a | 0.14138 | -0.15655587 | 0.009 | 0.05 | 1 | 2 |
| tbcd | 0.14138 | -0.11563478 | 0.009 | 0.05 | 1 | 2 |
| CR855865.1 | 0.14138 | -0.11531858 | 0.009 | 0.05 | 1 | 2 |
| LOC110437917 | 0.14138 | -0.10974447 | 0.009 | 0.05 | 1 | 2 |
| TPGS1 | 0.14138 | -0.12744113 | 0.009 | 0.05 | 1 | 2 |
| fnta | 0.14138 | -0.14489252 | 0.009 | 0.05 | 1 | 2 |
| glg1a | 0.14138 | -0.16731508 | 0.009 | 0.05 | 1 | 2 |
| rpp38 | 0.14138 | -0.10713106 | 0.009 | 0.05 | 1 | 2 |

| | | | | | | |
|---|---|---|---|---|---|---|
| si:ch73-41e3.7 | 0.14138 | -0.16474135 | 0.009 | 0.05 | 1 | 2 |
| slc35c2 | 0.14138 | -0.14397419 | 0.009 | 0.05 | 1 | 2 |
| trappc6bl | 0.14138 | -0.1096779 | 0.009 | 0.05 | 1 | 2 |
| agfg1a | 0.14138 | -0.14863106 | 0.009 | 0.05 | 1 | 2 |
| cep170aa | 0.14138 | -0.1523994 | 0.009 | 0.05 | 1 | 2 |
| crls1 | 0.14138 | -0.14517225 | 0.009 | 0.05 | 1 | 2 |
| exosc8 | 0.14138 | -0.11909979 | 0.009 | 0.05 | 1 | 2 |
| foxj3 | 0.14138 | -0.13201601 | 0.009 | 0.05 | 1 | 2 |
| frrs1b | 0.14138 | -0.15479113 | 0.009 | 0.05 | 1 | 2 |
| gpr65 | 0.14138 | -0.1372162 | 0.009 | 0.05 | 1 | 2 |
| klhdc4 | 0.14138 | -0.15919559 | 0.009 | 0.05 | 1 | 2 |
| ldha | 0.14138 | -0.11469028 | 0.009 | 0.05 | 1 | 2 |
| rab8a | 0.14138 | -0.14961687 | 0.009 | 0.05 | 1 | 2 |
| sbds | 0.14138 | -0.11628282 | 0.009 | 0.05 | 1 | 2 |
| si:ch211-264f5.2 | 0.14138 | -0.12411179 | 0.009 | 0.05 | 1 | 2 |
| si:ch211-276a23.5 | 0.14138 | -0.13501075 | 0.009 | 0.05 | 1 | 2 |
| suz12b | 0.14138 | -0.14167625 | 0.009 | 0.05 | 1 | 2 |
| tmem141 | 0.14138 | -0.16385355 | 0.009 | 0.05 | 1 | 2 |
| ubash3ba | 0.14138 | -0.13362087 | 0.009 | 0.05 | 1 | 2 |
| znf1010 | 0.14138 | -0.12552042 | 0.009 | 0.05 | 1 | 2 |
| vps26b | 0.14138 | -0.1090705 | 0.009 | 0.05 | 1 | 2 |
| btbd10a | 0.14138 | -0.11741927 | 0.009 | 0.05 | 1 | 2 |
| lpar5a | 0.14141 | 0.211888044 | 0.113 | 0.025 | 1 | 2 |
| ifi46 | 0.14144 | -0.3259959 | 0.148 | 0.25 | 1 | 2 |
| slc25a25a | 0.14175 | -0.26334924 | 0.035 | 0.125 | 1 | 2 |
| tp53 | 0.14285 | -0.259548 | 0.035 | 0.125 | 1 | 2 |
| abracl | 0.14311 | -0.25013032 | 0.235 | 0.3 | 1 | 2 |
| ecm1b | 0.14314 | 0.458643091 | 0.261 | 0.125 | 1 | 2 |
| lgals3bpb | 0.14386 | -0.36907526 | 0.278 | 0.45 | 1 | 2 |
| tmem214 | 0.14607 | -0.26721974 | 0.035 | 0.125 | 1 | 2 |
| cndp2 | 0.14681 | -0.21736508 | 0.348 | 0.5 | 1 | 2 |
| suclg1 | 0.14802 | -0.1391683 | 0.235 | 0.25 | 1 | 2 |
| bloc1s6 | 0.14908 | 0.152613171 | 0.217 | 0.125 | 1 | 2 |
| smarce1 | 0.15067 | -0.19952743 | 0.113 | 0.15 | 1 | 2 |
| rps3 | 0.15227 | 0.162164705 | 0.939 | 0.875 | 1 | 2 |
| rnf185 | 0.15259 | 0.247854187 | 0.113 | 0.025 | 1 | 2 |
| ap1m1 | 0.15522 | -0.29778246 | 0.07 | 0.15 | 1 | 2 |
| rab2a | 0.15525 | -0.24878468 | 0.113 | 0.175 | 1 | 2 |
| ndfip1l | 0.15559 | -0.27322128 | 0.052 | 0.125 | 1 | 2 |
| ifi30 | 0.15624 | 0.29167779 | 0.678 | 0.525 | 1 | 2 |
| rpl17 | 0.15674 | 0.184082812 | 0.878 | 0.775 | 1 | 2 |
| vps25 | 0.15747 | -0.27486966 | 0.026 | 0.1 | 1 | 2 |
| capns1a | 0.15951 | 0.3014888 | 0.157 | 0.05 | 1 | 2 |
| ctss2.1 | 0.16021 | -0.19852115 | 0.122 | 0.15 | 1 | 2 |
| atp5pb | 0.1608 | 0.291734339 | 0.443 | 0.275 | 1 | 2 |
| sf3a2 | 0.16115 | 0.238313211 | 0.157 | 0.05 | 1 | 2 |
| si:busm1-194e12.11 | 0.1617 | 0.350198486 | 0.261 | 0.15 | 1 | 2 |
| hmha1b | 0.16214 | 0.238286756 | 0.113 | 0.025 | 1 | 2 |

| | | | | | | |
|---|---|---|---|---|---|---|
| rpl14 | 0.16433 | 0.159762634 | 0.861 | 0.725 | 1 | 2 |
| si:ch211-39i2.2 | 0.16504 | -0.30234409 | 0.052 | 0.15 | 1 | 2 |
| rpl34 | 0.16537 | 0.137256141 | 0.965 | 0.9 | 1 | 2 |
| zgc:163080 | 0.16542 | 0.278462904 | 0.157 | 0.05 | 1 | 2 |
| sec61a1 | 0.16603 | 0.303840627 | 0.261 | 0.125 | 1 | 2 |
| eif2s2 | 0.16945 | -0.28419586 | 0.287 | 0.45 | 1 | 2 |
| bccip | 0.17049 | -0.30144448 | 0.087 | 0.2 | 1 | 2 |
| cxcr3.2 | 0.17154 | -0.14859 | 0.113 | 0.125 | 1 | 2 |
| uba52 | 0.17182 | -0.16745048 | 0.757 | 0.825 | 1 | 2 |
| spcs1 | 0.17312 | 0.334716989 | 0.261 | 0.125 | 1 | 2 |
| fkbp5 | 0.17356 | -0.28025264 | 0.052 | 0.15 | 1 | 2 |
| si:dkey-262k9.4 | 0.17432 | -0.21725876 | 0.2 | 0.25 | 1 | 2 |
| cope | 0.17435 | 0.225853865 | 0.113 | 0.025 | 1 | 2 |
| ccl39.7 | 0.17513 | 0.422896006 | 0.174 | 0.075 | 1 | 2 |
| ier2b | 0.17659 | -0.31907207 | 0.096 | 0.2 | 1 | 2 |
| CU694197.1 | 0.17785 | 0.297573522 | 0.191 | 0.075 | 1 | 2 |
| si:ch73-1a9.3 | 0.17812 | 0.322807763 | 0.383 | 0.225 | 1 | 2 |
| rhocb | 0.17853 | -0.29427891 | 0.087 | 0.175 | 1 | 2 |
| rpl10 | 0.18006 | 0.120758576 | 0.93 | 0.825 | 1 | 2 |
| si:ch211-283g2.3 | 0.18026 | 0.301215498 | 0.217 | 0.1 | 1 | 2 |
| pdcd6 | 0.18059 | -0.19138708 | 0.139 | 0.175 | 1 | 2 |
| clic2 | 0.18073 | 0.324682114 | 0.287 | 0.175 | 1 | 2 |
| ms4a17a.10 | 0.1811 | 0.185149873 | 0.426 | 0.3 | 1 | 2 |
| si:ch211-79k12.1 | 0.18326 | 0.232786405 | 0.087 | 0.025 | 1 | 2 |
| selenok | 0.18424 | -0.28482287 | 0.104 | 0.225 | 1 | 2 |
| cebpa | 0.18461 | -0.32682884 | 0.252 | 0.375 | 1 | 2 |
| gm2a | 0.18525 | 0.269809924 | 0.278 | 0.15 | 1 | 2 |
| wu:fi37e09 | 0.18604 | 0.114477587 | 0.087 | 0.025 | 1 | 2 |
| spcs3 | 0.18709 | -0.11195343 | 0.122 | 0.125 | 1 | 2 |
| mef2cb | 0.18801 | -0.2784356 | 0.061 | 0.125 | 1 | 2 |
| rps18 | 0.1881 | 0.132712399 | 0.957 | 0.9 | 1 | 2 |
| il4r.1 | 0.18822 | -0.30417836 | 0.096 | 0.2 | 1 | 2 |
| tmem263 | 0.18837 | -0.23194799 | 0.026 | 0.1 | 1 | 2 |
| gpd1c | 0.1892 | -0.23456567 | 0.078 | 0.125 | 1 | 2 |
| taf12 | 0.18961 | 0.200132218 | 0.104 | 0.025 | 1 | 2 |
| gtf2h5 | 0.19128 | -0.23275158 | 0.026 | 0.1 | 1 | 2 |
| glrx | 0.19163 | 0.267105824 | 0.591 | 0.425 | 1 | 2 |
| rbm4.3 | 0.192 | -0.30087072 | 0.113 | 0.225 | 1 | 2 |
| fam120c | 0.19357 | -0.21246734 | 0.035 | 0.075 | 1 | 2 |
| flna | 0.19372 | -0.22923632 | 0.026 | 0.075 | 1 | 2 |
| ube2ib | 0.19497 | -0.17418963 | 0.026 | 0.1 | 1 | 2 |
| manf | 0.19562 | -0.19957037 | 0.026 | 0.1 | 1 | 2 |
| gnsb | 0.19627 | -0.29687789 | 0.043 | 0.125 | 1 | 2 |
| fuca1.1 | 0.1965 | -0.20339971 | 0.026 | 0.1 | 1 | 2 |
| rad21a | 0.19695 | -0.22821938 | 0.096 | 0.15 | 1 | 2 |
| si:busm1-266f07.2 | 0.19753 | 0.207560718 | 0.722 | 0.65 | 1 | 2 |
| nfkbiaa | 0.19867 | -0.29619665 | 0.113 | 0.2 | 1 | 2 |
| dennd4a | 0.19873 | -0.23717471 | 0.026 | 0.1 | 1 | 2 |

| | | | | | | |
|---|---|---|---|---|---|---|
| zgpat | 0.19917 | -0.20823479 | 0.026 | 0.1 | 1 | 2 |
| lgals3a | 0.19919 | -0.20256254 | 0.096 | 0.2 | 1 | 2 |
| ankrd11 | 0.19938 | -0.19223952 | 0.026 | 0.1 | 1 | 2 |
| nucks1b | 0.20068 | -0.19346201 | 0.026 | 0.1 | 1 | 2 |
| rfc4 | 0.20079 | -0.19430983 | 0.026 | 0.1 | 1 | 2 |
| tcnbb | 0.20082 | -0.23245376 | 0.026 | 0.1 | 1 | 2 |
| si:ch211-199g17.2 | 0.20087 | -0.19919005 | 0.026 | 0.1 | 1 | 2 |
| csrnp1b | 0.20137 | -0.29756817 | 0.078 | 0.175 | 1 | 2 |
| wasf2 | 0.20158 | -0.26940691 | 0.078 | 0.175 | 1 | 2 |
| spns1 | 0.20177 | -0.22524235 | 0.026 | 0.075 | 1 | 2 |
| atf4b | 0.20221 | -0.13407965 | 0.113 | 0.125 | 1 | 2 |
| ccl35.1 | 0.20384 | 0.33871612 | 0.365 | 0.275 | 1 | 2 |
| adh5 | 0.20387 | 0.142723847 | 0.191 | 0.1 | 1 | 2 |
| cst14a.2 | 0.20431 | 0.326952886 | 0.313 | 0.175 | 1 | 2 |
| snrnp27 | 0.20436 | 0.193287655 | 0.104 | 0.025 | 1 | 2 |
| ndufs3 | 0.20535 | 0.162742132 | 0.104 | 0.025 | 1 | 2 |
| mcl1b | 0.20553 | -0.26399444 | 0.348 | 0.45 | 1 | 2 |
| rhof | 0.20553 | -0.13128467 | 0.017 | 0.075 | 1 | 2 |
| rbm19 | 0.2066 | -0.23876638 | 0.017 | 0.075 | 1 | 2 |
| eif2s3 | 0.2076 | -0.29166607 | 0.278 | 0.4 | 1 | 2 |
| eif4bb | 0.20782 | -0.25894731 | 0.104 | 0.175 | 1 | 2 |
| si:ch1073-325m22.2 | 0.20839 | 0.269105269 | 0.374 | 0.225 | 1 | 2 |
| mhc2b | 0.20904 | 0.222187598 | 0.409 | 0.35 | 1 | 2 |
| dars | 0.20954 | -0.30712529 | 0.113 | 0.225 | 1 | 2 |
| zgc:114188 | 0.2096 | 0.170531677 | 0.896 | 0.8 | 1 | 2 |
| ISCU (1 of many) | 0.20972 | 0.248362223 | 0.104 | 0.025 | 1 | 2 |
| LO018493.1 | 0.21043 | 0.279356023 | 0.183 | 0.075 | 1 | 2 |
| ncbp2 | 0.2131 | 0.202555754 | 0.104 | 0.025 | 1 | 2 |
| si:dkey-192g7.3 | 0.21474 | 0.216436838 | 0.104 | 0.025 | 1 | 2 |
| snx5 | 0.21539 | 0.272588985 | 0.183 | 0.075 | 1 | 2 |
| tmsb1 | 0.21696 | 0.29571022 | 0.348 | 0.25 | 1 | 2 |
| ubxn1 | 0.21804 | -0.14043284 | 0.017 | 0.075 | 1 | 2 |
| cyth1b | 0.21905 | -0.25249916 | 0.035 | 0.1 | 1 | 2 |
| mrps17 | 0.21935 | -0.19258047 | 0.017 | 0.075 | 1 | 2 |
| atp5f1c | 0.2195 | 0.300511015 | 0.278 | 0.15 | 1 | 2 |
| rpa3 | 0.21967 | -0.19723105 | 0.035 | 0.075 | 1 | 2 |
| myl12.1 | 0.22019 | 0.225136661 | 0.626 | 0.525 | 1 | 2 |
| sept6 | 0.22059 | 0.229213644 | 0.104 | 0.025 | 1 | 2 |
| ms4a17a.9 | 0.22082 | 0.309366932 | 0.217 | 0.1 | 1 | 2 |
| cpeb4b | 0.22108 | -0.21458496 | 0.017 | 0.075 | 1 | 2 |
| CR392341.1 | 0.2211 | 0.275126838 | 0.183 | 0.075 | 1 | 2 |
| ndufa3 | 0.22151 | 0.23951716 | 0.122 | 0.05 | 1 | 2 |
| icn | 0.22332 | 0.215678391 | 0.261 | 0.15 | 1 | 2 |
| eif1ad | 0.22335 | 0.127447136 | 0.087 | 0.025 | 1 | 2 |
| mdh2 | 0.22346 | 0.205731932 | 0.296 | 0.175 | 1 | 2 |
| srsf1b | 0.22455 | -0.16028878 | 0.017 | 0.075 | 1 | 2 |
| zgc:123107 | 0.22549 | 0.246755687 | 0.104 | 0.025 | 1 | 2 |
| zgc:158803 | 0.22731 | -0.2003493 | 0.026 | 0.075 | 1 | 2 |

| | | | | | | |
|---|---|---|---|---|---|---|
| rack1 | 0.22762 | 0.167471265 | 0.861 | 0.775 | 1 | 2 |
| zgc:103700 | 0.22958 | 0.24856436 | 0.548 | 0.45 | 1 | 2 |
| zgc:136439 | 0.22992 | -0.22576807 | 0.043 | 0.125 | 1 | 2 |
| pitpnaa | 0.23028 | -0.24114061 | 0.061 | 0.125 | 1 | 2 |
| aars | 0.23159 | -0.24414845 | 0.052 | 0.125 | 1 | 2 |
| atg12 | 0.23172 | -0.16514441 | 0.017 | 0.075 | 1 | 2 |
| fosl2 | 0.23203 | -0.27578782 | 0.061 | 0.15 | 1 | 2 |
| nop58 | 0.23228 | 0.230102759 | 0.096 | 0.025 | 1 | 2 |
| trappc5 | 0.23241 | 0.190170733 | 0.096 | 0.025 | 1 | 2 |
| txndc9 | 0.23377 | -0.26355377 | 0.078 | 0.175 | 1 | 2 |
| ndufb7 | 0.23442 | -0.26819834 | 0.183 | 0.275 | 1 | 2 |
| dctn5 | 0.235 | -0.16059119 | 0.017 | 0.075 | 1 | 2 |
| bloc1s2 | 0.23584 | 0.139122193 | 0.087 | 0.025 | 1 | 2 |
| prdx2 | 0.23671 | -0.26987954 | 0.304 | 0.4 | 1 | 2 |
| LOC101883999 | 0.23702 | -0.1762977 | 0.017 | 0.075 | 1 | 2 |
| ehd4 | 0.23729 | -0.28394236 | 0.191 | 0.325 | 1 | 2 |
| sh3glb1b | 0.23746 | -0.23594327 | 0.043 | 0.125 | 1 | 2 |
| si:ch211-66e2.3 | 0.23786 | -0.23576425 | 0.043 | 0.125 | 1 | 2 |
| pde4ba | 0.23888 | 0.105553559 | 0.078 | 0.025 | 1 | 2 |
| mrpl33 | 0.23898 | -0.28499484 | 0.113 | 0.225 | 1 | 2 |
| sgpl1 | 0.23915 | -0.19598891 | 0.035 | 0.075 | 1 | 2 |
| ctsa | 0.23919 | 0.294966612 | 0.278 | 0.15 | 1 | 2 |
| rin1a | 0.23976 | -0.21268698 | 0.017 | 0.075 | 1 | 2 |
| hprt1 | 0.24043 | -0.24337388 | 0.043 | 0.1 | 1 | 2 |
| klf2a | 0.24076 | -0.13756542 | 0.017 | 0.075 | 1 | 2 |
| slc3a2a | 0.24082 | -0.25764988 | 0.261 | 0.4 | 1 | 2 |
| snrpb | 0.24108 | 0.274182362 | 0.278 | 0.15 | 1 | 2 |
| lsm5 | 0.24152 | -0.25113495 | 0.104 | 0.175 | 1 | 2 |
| rasgrp4 | 0.24409 | -0.1887582 | 0.017 | 0.075 | 1 | 2 |
| atp5f1e | 0.2446 | 0.238538532 | 0.443 | 0.3 | 1 | 2 |
| rbm14a | 0.24646 | -0.19822864 | 0.017 | 0.075 | 1 | 2 |
| snrpd2 | 0.24682 | 0.216723126 | 0.096 | 0.025 | 1 | 2 |
| gnai2b | 0.24827 | -0.20266756 | 0.017 | 0.075 | 1 | 2 |
| uqcrfs1 | 0.24846 | -0.24719315 | 0.174 | 0.25 | 1 | 2 |
| arpc5b | 0.24852 | 0.233384709 | 0.383 | 0.25 | 1 | 2 |
| atp6v1g1 | 0.24955 | 0.211228651 | 0.487 | 0.35 | 1 | 2 |
| letm1 | 0.2498 | -0.19913777 | 0.017 | 0.075 | 1 | 2 |
| hsd17b14 | 0.24993 | -0.20736325 | 0.017 | 0.075 | 1 | 2 |
| cdc34a | 0.25016 | -0.20250083 | 0.017 | 0.075 | 1 | 2 |
| dusp5 | 0.25035 | -0.21949097 | 0.035 | 0.1 | 1 | 2 |
| rars | 0.25114 | -0.20358854 | 0.017 | 0.075 | 1 | 2 |
| rpl24 | 0.25249 | 0.127192026 | 0.939 | 0.875 | 1 | 2 |
| urm1 | 0.25302 | -0.12750161 | 0.017 | 0.075 | 1 | 2 |
| rsf1b.1 | 0.25302 | -0.20310592 | 0.017 | 0.075 | 1 | 2 |
| CR855389.1 | 0.25364 | -0.23454477 | 0.035 | 0.1 | 1 | 2 |
| atp8a1 | 0.25494 | -0.18344994 | 0.017 | 0.075 | 1 | 2 |
| chmp5b | 0.25631 | 0.236021965 | 0.139 | 0.05 | 1 | 2 |
| vps35 | 0.25828 | -0.22804489 | 0.035 | 0.1 | 1 | 2 |

| | | | | | | |
|---|---|---|---|---|---|---|
| ehd1b | 0.25892 | -0.14741844 | 0.017 | 0.075 | 1 | 2 |
| nars | 0.25953 | -0.24635841 | 0.061 | 0.125 | 1 | 2 |
| apbb1ip | 0.26029 | -0.16860402 | 0.017 | 0.075 | 1 | 2 |
| epdl1 | 0.26046 | 0.293788657 | 0.139 | 0.05 | 1 | 2 |
| vamp8 | 0.26063 | -0.23780592 | 0.052 | 0.125 | 1 | 2 |
| rps17 | 0.26113 | 0.121396918 | 0.896 | 0.85 | 1 | 2 |
| cdc42l | 0.26176 | 0.128422589 | 0.261 | 0.175 | 1 | 2 |
| trim25 | 0.26242 | -0.18278386 | 0.026 | 0.075 | 1 | 2 |
| polr2f | 0.26331 | 0.165087014 | 0.096 | 0.025 | 1 | 2 |
| prdx3 | 0.26341 | 0.222810978 | 0.139 | 0.05 | 1 | 2 |
| wdfy1 | 0.26357 | -0.17695365 | 0.017 | 0.075 | 1 | 2 |
| atp1b1a | 0.26373 | 0.101033063 | 0.313 | 0.225 | 1 | 2 |
| ndufa8 | 0.26376 | 0.225834053 | 0.139 | 0.05 | 1 | 2 |
| eif4a1a | 0.264 | -0.30142044 | 0.183 | 0.3 | 1 | 2 |
| mtmr8 | 0.26412 | -0.1514011 | 0.017 | 0.075 | 1 | 2 |
| selenot1b | 0.26487 | 0.186495131 | 0.096 | 0.025 | 1 | 2 |
| ift46 | 0.26557 | -0.16236177 | 0.017 | 0.075 | 1 | 2 |
| mlec | 0.26612 | 0.207548649 | 0.096 | 0.025 | 1 | 2 |
| acadm | 0.27049 | -0.13986634 | 0.087 | 0.1 | 1 | 2 |
| glo1 | 0.27099 | 0.264874771 | 0.209 | 0.1 | 1 | 2 |
| znf593 | 0.27102 | -0.23191085 | 0.096 | 0.175 | 1 | 2 |
| bcas2 | 0.27114 | 0.1791393 | 0.096 | 0.025 | 1 | 2 |
| mknk2a | 0.27118 | 0.219064465 | 0.096 | 0.025 | 1 | 2 |
| tnfrsf11b | 0.2719 | -0.25804372 | 0.035 | 0.1 | 1 | 2 |
| CABZ01055347.1 | 0.27343 | -0.19401971 | 0.026 | 0.075 | 1 | 2 |
| gosr2 | 0.2749 | -0.13810869 | 0.061 | 0.075 | 1 | 2 |
| top1l | 0.27493 | 0.168232174 | 0.087 | 0.025 | 1 | 2 |
| ufm1 | 0.27547 | 0.169792881 | 0.096 | 0.025 | 1 | 2 |
| u2af1 | 0.27555 | 0.247744292 | 0.183 | 0.1 | 1 | 2 |
| tob1b | 0.27557 | 0.170942651 | 0.096 | 0.025 | 1 | 2 |
| glmp | 0.276 | 0.106898739 | 0.139 | 0.075 | 1 | 2 |
| si:ch211-214p16.1 | 0.27676 | 0.263592078 | 0.096 | 0.025 | 1 | 2 |
| pgp | 0.27796 | 0.175695062 | 0.096 | 0.025 | 1 | 2 |
| psmd8 | 0.28019 | 0.153680993 | 0.087 | 0.025 | 1 | 2 |
| nt5dc2 | 0.28049 | 0.150429702 | 0.087 | 0.025 | 1 | 2 |
| FERMT3 (1 of many) | 0.28063 | 0.111072642 | 0.113 | 0.05 | 1 | 2 |
| tdh | 0.28107 | -0.1973466 | 0.052 | 0.125 | 1 | 2 |
| eef1da | 0.28191 | 0.234530176 | 0.617 | 0.475 | 1 | 2 |
| trappc6b | 0.28197 | -0.19383577 | 0.043 | 0.1 | 1 | 2 |
| bcl2l10 | 0.28284 | 0.20861504 | 0.096 | 0.025 | 1 | 2 |
| zgc:152830 | 0.28297 | -0.17796134 | 0.209 | 0.25 | 1 | 2 |
| psmc3 | 0.28403 | 0.15039215 | 0.087 | 0.025 | 1 | 2 |
| eif5a2 | 0.28426 | 0.199978147 | 0.374 | 0.25 | 1 | 2 |
| si:ch211-260e23.9 | 0.28546 | 0.195397002 | 0.087 | 0.025 | 1 | 2 |
| gnb1a | 0.28841 | -0.19329184 | 0.2 | 0.25 | 1 | 2 |
| ctsd | 0.29056 | -0.28041668 | 0.391 | 0.525 | 1 | 2 |
| cct8 | 0.29378 | 0.140667731 | 0.235 | 0.15 | 1 | 2 |
| tmcc3 | 0.29464 | -0.185754 | 0.026 | 0.075 | 1 | 2 |

| | | | | | | |
|---|---|---|---|---|---|---|
| snrpd1 | 0.29511 | -0.19748924 | 0.165 | 0.275 | 1 | 2 |
| naga | 0.2956 | 0.282818323 | 0.235 | 0.125 | 1 | 2 |
| fthl27 | 0.29623 | -0.14237306 | 0.652 | 0.675 | 1 | 2 |
| si:dkey-17m8.1 | 0.29713 | -0.19742438 | 0.026 | 0.075 | 1 | 2 |
| glrx5 | 0.2982 | -0.17288884 | 0.026 | 0.075 | 1 | 2 |
| mknk2b | 0.299 | 0.116464975 | 0.139 | 0.075 | 1 | 2 |
| psmd4a | 0.29901 | 0.148524085 | 0.087 | 0.025 | 1 | 2 |
| LOC100151196 | 0.29961 | 0.147523543 | 0.087 | 0.025 | 1 | 2 |
| mat2aa | 0.30006 | -0.21981055 | 0.035 | 0.1 | 1 | 2 |
| xpnpep2 | 0.30111 | -0.19563953 | 0.035 | 0.075 | 1 | 2 |
| si:ch211-102c2.4 | 0.30227 | -0.14428939 | 0.322 | 0.35 | 1 | 2 |
| prelid1a | 0.30277 | -0.20417533 | 0.035 | 0.1 | 1 | 2 |
| grna | 0.30351 | -0.1508214 | 0.322 | 0.35 | 1 | 2 |
| si:ch211-191j22.3 | 0.30368 | 0.196737992 | 0.13 | 0.05 | 1 | 2 |
| ppt1 | 0.30474 | -0.23167665 | 0.287 | 0.375 | 1 | 2 |
| cirbpa | 0.30481 | 0.243646593 | 0.574 | 0.45 | 1 | 2 |
| eif3c | 0.3058 | 0.195199915 | 0.191 | 0.1 | 1 | 2 |
| ttyh3a | 0.30581 | -0.15263965 | 0.017 | 0.05 | 1 | 2 |
| cnbpb | 0.30618 | 0.118367537 | 0.148 | 0.075 | 1 | 2 |
| psmb4 | 0.30663 | 0.255802669 | 0.235 | 0.125 | 1 | 2 |
| hmgn2 | 0.30788 | 0.299406141 | 0.452 | 0.325 | 1 | 2 |
| commd3 | 0.30788 | -0.10258217 | 0.148 | 0.15 | 1 | 2 |
| ap2m1a | 0.30875 | 0.220982549 | 0.261 | 0.15 | 1 | 2 |
| chac1 | 0.30892 | -0.20957846 | 0.174 | 0.225 | 1 | 2 |
| adam10a | 0.30922 | 0.181805711 | 0.087 | 0.025 | 1 | 2 |
| mpp1 | 0.30952 | 0.197050183 | 0.087 | 0.025 | 1 | 2 |
| jpt1b | 0.30957 | -0.22412835 | 0.035 | 0.1 | 1 | 2 |
| sgk3 | 0.30962 | -0.16323505 | 0.035 | 0.1 | 1 | 2 |
| rgs2 | 0.30988 | -0.27128176 | 0.087 | 0.175 | 1 | 2 |
| fam162a | 0.31091 | 0.182104844 | 0.087 | 0.025 | 1 | 2 |
| AL928650.1 | 0.31181 | 0.127970502 | 0.078 | 0.025 | 1 | 2 |
| grn1 | 0.31183 | 0.167293268 | 0.87 | 0.775 | 1 | 2 |
| dnajc19 | 0.31252 | 0.103938794 | 0.078 | 0.025 | 1 | 2 |
| jac2 | 0.3128 | -0.2770721 | 0.07 | 0.15 | 1 | 2 |
| morf4l1 | 0.31315 | -0.18615922 | 0.157 | 0.2 | 1 | 2 |
| nfkbiab | 0.31403 | 0.263703246 | 0.2 | 0.1 | 1 | 2 |
| btg1 | 0.31496 | 0.186060746 | 0.417 | 0.3 | 1 | 2 |
| csnk2a1 | 0.31503 | 0.18533436 | 0.122 | 0.075 | 1 | 2 |
| rps13 | 0.31507 | -0.11004606 | 0.904 | 0.925 | 1 | 2 |
| ndufa4l | 0.31507 | 0.195953737 | 0.452 | 0.325 | 1 | 2 |
| fbl | 0.31718 | 0.189234335 | 0.13 | 0.05 | 1 | 2 |
| trnau1apb | 0.31824 | -0.18233858 | 0.035 | 0.1 | 1 | 2 |
| il13ra1 | 0.31826 | -0.17503726 | 0.035 | 0.1 | 1 | 2 |
| tmem18 | 0.31917 | -0.2235162 | 0.052 | 0.125 | 1 | 2 |
| ponzr1 | 0.32006 | 0.15702812 | 0.322 | 0.225 | 1 | 2 |
| dap1b | 0.32006 | -0.23593455 | 0.078 | 0.15 | 1 | 2 |
| prkab1b | 0.32006 | 0.226090263 | 0.13 | 0.05 | 1 | 2 |
| pim2 | 0.32118 | 0.184572914 | 0.078 | 0.025 | 1 | 2 |

| | | | | | | |
|---|---|---|---|---|---|---|
| rgs5b | 0.32123 | -0.16551731 | 0.035 | 0.1 | 1 | 2 |
| ifi45 | 0.32127 | -0.18690333 | 0.035 | 0.1 | 1 | 2 |
| hypk | 0.32139 | 0.19844565 | 0.13 | 0.05 | 1 | 2 |
| ptpn2b | 0.32196 | -0.19892208 | 0.035 | 0.1 | 1 | 2 |
| mtpn | 0.32213 | -0.26655608 | 0.165 | 0.275 | 1 | 2 |
| h2afva | 0.32234 | -0.20828057 | 0.043 | 0.1 | 1 | 2 |
| ms4a17a.11 | 0.3231 | 0.139244128 | 0.087 | 0.025 | 1 | 2 |
| mrps14 | 0.32316 | -0.19063235 | 0.035 | 0.1 | 1 | 2 |
| surf2 | 0.32331 | -0.1709856 | 0.035 | 0.1 | 1 | 2 |
| cstf2 | 0.32404 | -0.1581617 | 0.035 | 0.1 | 1 | 2 |
| sgta | 0.32414 | -0.15608604 | 0.052 | 0.125 | 1 | 2 |
| nsa2 | 0.32591 | -0.24273884 | 0.383 | 0.5 | 1 | 2 |
| atp5mc3a | 0.32596 | 0.217678368 | 0.287 | 0.175 | 1 | 2 |
| eif5b | 0.32643 | 0.153240239 | 0.087 | 0.025 | 1 | 2 |
| elmo1 | 0.32806 | 0.150790235 | 0.087 | 0.025 | 1 | 2 |
| ywhabb | 0.3285 | -0.24237612 | 0.261 | 0.35 | 1 | 2 |
| tnfrsf1a | 0.32932 | -0.24139769 | 0.061 | 0.125 | 1 | 2 |
| sptssa | 0.33117 | 0.233922364 | 0.122 | 0.05 | 1 | 2 |
| srsf5a | 0.33257 | -0.25497566 | 0.165 | 0.275 | 1 | 2 |
| rhogb | 0.33336 | -0.25658722 | 0.174 | 0.275 | 1 | 2 |
| celf2 | 0.33387 | -0.23022285 | 0.139 | 0.2 | 1 | 2 |
| cox7a2l | 0.33528 | 0.158151408 | 0.157 | 0.075 | 1 | 2 |
| nr1d2a | 0.33595 | 0.152308028 | 0.087 | 0.025 | 1 | 2 |
| si:ch211-217k17.10 | 0.33602 | -0.28383276 | 0.07 | 0.125 | 1 | 2 |
| ndufa9a | 0.3387 | -0.24435965 | 0.078 | 0.15 | 1 | 2 |
| slc25a11 | 0.33925 | -0.19137344 | 0.078 | 0.125 | 1 | 2 |
| ccl39.2 | 0.33994 | -0.22431422 | 0.087 | 0.175 | 1 | 2 |
| gda | 0.34027 | -0.17889332 | 0.026 | 0.075 | 1 | 2 |
| si:dkey-27i16.2 | 0.34114 | -0.19093264 | 0.261 | 0.375 | 1 | 2 |
| LOC100538255 | 0.34139 | 0.195882463 | 0.087 | 0.025 | 1 | 2 |
| si:ch73-376l24.3 | 0.34265 | -0.23115897 | 0.078 | 0.15 | 1 | 2 |
| ncor2 | 0.34271 | -0.20163315 | 0.052 | 0.125 | 1 | 2 |
| tomm5 | 0.34293 | -0.10767955 | 0.191 | 0.2 | 1 | 2 |
| iscub | 0.34373 | 0.178037007 | 0.087 | 0.025 | 1 | 2 |
| dram2b | 0.34381 | -0.21404724 | 0.07 | 0.15 | 1 | 2 |
| pmaip1 | 0.3441 | -0.21791683 | 0.07 | 0.15 | 1 | 2 |
| bcl7bb | 0.34429 | -0.21848451 | 0.052 | 0.125 | 1 | 2 |
| map3k14a | 0.34573 | 0.287985193 | 0.226 | 0.125 | 1 | 2 |
| lsm8 | 0.34587 | 0.157234338 | 0.087 | 0.025 | 1 | 2 |
| rnaseka | 0.34762 | 0.229506914 | 0.226 | 0.125 | 1 | 2 |
| chchd2 | 0.3478 | 0.163843245 | 0.087 | 0.025 | 1 | 2 |
| elovl1b | 0.34783 | -0.24369968 | 0.139 | 0.225 | 1 | 2 |
| entpd1 | 0.34811 | 0.252210373 | 0.191 | 0.1 | 1 | 2 |
| tmed10 | 0.34814 | -0.24001204 | 0.096 | 0.175 | 1 | 2 |
| rpl7l1 | 0.34833 | -0.19130947 | 0.043 | 0.1 | 1 | 2 |
| mcts1 | 0.35025 | -0.19541777 | 0.061 | 0.125 | 1 | 2 |
| si:cabz01074946.1 | 0.3512 | 0.133977088 | 0.226 | 0.15 | 1 | 2 |
| wdr1 | 0.35202 | -0.2457832 | 0.209 | 0.3 | 1 | 2 |

| | | | | | | |
|---|---|---|---|---|---|---|
| zc3h14 | 0.35234 | -0.18811971 | 0.026 | 0.075 | 1 | 2 |
| wipf1b | 0.3527 | 0.185030717 | 0.122 | 0.05 | 1 | 2 |
| mrpl32 | 0.35276 | 0.163996153 | 0.087 | 0.025 | 1 | 2 |
| fam49ba | 0.35302 | 0.225043444 | 0.139 | 0.075 | 1 | 2 |
| fgl2a | 0.35432 | -0.14852041 | 0.27 | 0.3 | 1 | 2 |
| sept7b | 0.35499 | -0.19938059 | 0.052 | 0.1 | 1 | 2 |
| tra2a | 0.3553 | -0.13545424 | 0.13 | 0.15 | 1 | 2 |
| zgc:110591 | 0.35534 | 0.138857941 | 0.078 | 0.025 | 1 | 2 |
| psmg1 | 0.35605 | 0.172906192 | 0.087 | 0.025 | 1 | 2 |
| atp6v1ab | 0.35605 | 0.155895079 | 0.087 | 0.025 | 1 | 2 |
| irf2 | 0.35631 | 0.190960387 | 0.087 | 0.025 | 1 | 2 |
| gnai2a | 0.35776 | 0.113382882 | 0.139 | 0.075 | 1 | 2 |
| minos1 | 0.35818 | 0.169794589 | 0.157 | 0.075 | 1 | 2 |
| bcam | 0.36085 | -0.19818724 | 0.043 | 0.1 | 1 | 2 |
| snd1 | 0.36101 | -0.19351312 | 0.104 | 0.15 | 1 | 2 |
| sept12 | 0.36144 | 0.148087224 | 0.113 | 0.05 | 1 | 2 |
| lipf | 0.36392 | 0.189790924 | 0.183 | 0.1 | 1 | 2 |
| zgc:77849 | 0.36433 | -0.1124401 | 0.165 | 0.175 | 1 | 2 |
| ywhaqb | 0.3652 | -0.14750669 | 0.07 | 0.1 | 1 | 2 |
| akirin1 | 0.36538 | -0.16086049 | 0.026 | 0.075 | 1 | 2 |
| BX323596.1 | 0.36568 | 0.228857668 | 0.078 | 0.025 | 1 | 2 |
| si:rp71-45k5.4 | 0.3666 | 0.15204754 | 0.148 | 0.075 | 1 | 2 |
| gng5 | 0.36674 | 0.109608937 | 0.252 | 0.175 | 1 | 2 |
| rps19bp1 | 0.3669 | -0.17698913 | 0.026 | 0.075 | 1 | 2 |
| tm2d3 | 0.36881 | -0.11447411 | 0.026 | 0.075 | 1 | 2 |
| parvg | 0.36909 | -0.10668688 | 0.165 | 0.175 | 1 | 2 |
| XLOC-023492 | 0.37014 | 0.238081372 | 0.191 | 0.1 | 1 | 2 |
| rsrp1 | 0.37057 | -0.17294138 | 0.043 | 0.075 | 1 | 2 |
| cox7a2a | 0.37103 | 0.165356858 | 0.235 | 0.2 | 1 | 2 |
| atp6v1e1b | 0.37124 | 0.183375302 | 0.33 | 0.225 | 1 | 2 |
| ndufb6 | 0.37231 | 0.109640548 | 0.139 | 0.075 | 1 | 2 |
| ssr4 | 0.37402 | -0.16129698 | 0.374 | 0.425 | 1 | 2 |
| rps10 | 0.37546 | 0.11509768 | 0.913 | 0.85 | 1 | 2 |
| cdk5rap3 | 0.37665 | 0.113119743 | 0.078 | 0.025 | 1 | 2 |
| elocb | 0.37746 | -0.23958936 | 0.157 | 0.25 | 1 | 2 |
| sec61b | 0.37751 | 0.197789905 | 0.626 | 0.5 | 1 | 2 |
| zgc:56676 | 0.37881 | 0.222752064 | 0.191 | 0.1 | 1 | 2 |
| fundc2 | 0.37917 | 0.175274584 | 0.078 | 0.025 | 1 | 2 |
| chkb | 0.37932 | -0.15423914 | 0.017 | 0.05 | 1 | 2 |
| elf2b | 0.38207 | -0.15065097 | 0.017 | 0.05 | 1 | 2 |
| snx1a | 0.38244 | -0.15590228 | 0.139 | 0.175 | 1 | 2 |
| ngdn | 0.3869 | -0.13860226 | 0.026 | 0.075 | 1 | 2 |
| lgalsla | 0.38876 | -0.15317221 | 0.017 | 0.05 | 1 | 2 |
| snx20 | 0.38974 | -0.16565369 | 0.026 | 0.075 | 1 | 2 |
| naca | 0.38999 | 0.122973715 | 0.887 | 0.8 | 1 | 2 |
| capns1b | 0.39224 | -0.23174645 | 0.104 | 0.175 | 1 | 2 |
| srgap3 | 0.39271 | -0.19018342 | 0.026 | 0.075 | 1 | 2 |
| caprin1b | 0.3934 | 0.214709598 | 0.165 | 0.1 | 1 | 2 |

| | | | | | | |
|---|---|---|---|---|---|---|
| eif4h | 0.39528 | 0.128146777 | 0.113 | 0.05 | 1 | 2 |
| cuedc1b | 0.39602 | -0.16571197 | 0.026 | 0.075 | 1 | 2 |
| srsf9 | 0.39645 | -0.15787876 | 0.035 | 0.075 | 1 | 2 |
| sh3pxd2b | 0.39658 | -0.18059146 | 0.035 | 0.075 | 1 | 2 |
| fybb | 0.39664 | 0.135754087 | 0.078 | 0.025 | 1 | 2 |
| rpl7 | 0.39874 | 0.119899918 | 0.904 | 0.825 | 1 | 2 |
| hspa9 | 0.40108 | 0.171220702 | 0.07 | 0.025 | 1 | 2 |
| ifngr1l | 0.40117 | -0.14481415 | 0.017 | 0.05 | 1 | 2 |
| MED9 | 0.40148 | -0.14069712 | 0.026 | 0.05 | 1 | 2 |
| si:zfos-2326c3.2 | 0.40179 | -0.16442599 | 0.026 | 0.075 | 1 | 2 |
| atp6v1f | 0.40197 | 0.101587737 | 0.07 | 0.025 | 1 | 2 |
| dnaja2a | 0.40222 | 0.165572403 | 0.191 | 0.15 | 1 | 2 |
| smox | 0.40241 | -0.25459218 | 0.2 | 0.3 | 1 | 2 |
| dkc1 | 0.40335 | 0.108071436 | 0.104 | 0.05 | 1 | 2 |
| eif3s6ip | 0.40424 | -0.21986494 | 0.13 | 0.2 | 1 | 2 |
| tuba8l2 | 0.40454 | 0.225656856 | 0.278 | 0.175 | 1 | 2 |
| jdp2b | 0.40614 | -0.16444694 | 0.043 | 0.075 | 1 | 2 |
| sri | 0.40647 | -0.2115507 | 0.113 | 0.2 | 1 | 2 |
| setb | 0.40702 | 0.227920244 | 0.217 | 0.125 | 1 | 2 |
| glipr2 | 0.40756 | -0.16471518 | 0.026 | 0.075 | 1 | 2 |
| mgst1.2 | 0.40799 | 0.120842898 | 0.078 | 0.025 | 1 | 2 |
| tmem258 | 0.40828 | 0.131540147 | 0.313 | 0.225 | 1 | 2 |
| atp5if1a | 0.40862 | 0.151870094 | 0.348 | 0.25 | 1 | 2 |
| rnf7 | 0.40916 | 0.167117401 | 0.113 | 0.05 | 1 | 2 |
| vsir | 0.40919 | 0.238282129 | 0.217 | 0.125 | 1 | 2 |
| ncaldb | 0.41037 | -0.16220216 | 0.026 | 0.075 | 1 | 2 |
| scpep1 | 0.41151 | 0.143851791 | 0.313 | 0.225 | 1 | 2 |
| ndufb10 | 0.41153 | 0.103968133 | 0.139 | 0.075 | 1 | 2 |
| arpc4 | 0.41189 | 0.13157264 | 0.365 | 0.275 | 1 | 2 |
| socs3b | 0.41194 | -0.14127198 | 0.078 | 0.15 | 1 | 2 |
| gmppab | 0.41214 | -0.12116177 | 0.026 | 0.075 | 1 | 2 |
| acin1a | 0.41306 | -0.19672944 | 0.061 | 0.125 | 1 | 2 |
| smndc1 | 0.41354 | -0.14066846 | 0.061 | 0.125 | 1 | 2 |
| bzw1a | 0.41494 | -0.18074164 | 0.096 | 0.175 | 1 | 2 |
| dpm1 | 0.4161 | 0.136532156 | 0.078 | 0.025 | 1 | 2 |
| dctn3 | 0.41633 | 0.186428295 | 0.113 | 0.05 | 1 | 2 |
| tk2 | 0.41726 | -0.14393269 | 0.026 | 0.075 | 1 | 2 |
| hmox2b | 0.41773 | -0.13648611 | 0.026 | 0.075 | 1 | 2 |
| ppp6c | 0.41785 | 0.130777053 | 0.078 | 0.025 | 1 | 2 |
| eef1db | 0.41823 | -0.14545072 | 0.243 | 0.275 | 1 | 2 |
| psmd1 | 0.41896 | 0.154233075 | 0.078 | 0.025 | 1 | 2 |
| galk2 | 0.41906 | -0.13726886 | 0.026 | 0.075 | 1 | 2 |
| gabarapl2 | 0.41955 | 0.158950021 | 0.078 | 0.025 | 1 | 2 |
| u2af2a | 0.42012 | -0.14894999 | 0.017 | 0.05 | 1 | 2 |
| lsm6 | 0.42043 | 0.126150179 | 0.07 | 0.05 | 1 | 2 |
| ccdc12 | 0.42141 | -0.13281806 | 0.026 | 0.075 | 1 | 2 |
| si:dkeyp-97b10.3 | 0.42188 | 0.153994767 | 0.052 | 0.025 | 1 | 2 |
| naa10 | 0.42208 | -0.23185047 | 0.139 | 0.225 | 1 | 2 |

| | | | | | | |
|---|---|---|---|---|---|---|
| zgc:173915 | 0.42233 | -0.1880169 | 0.026 | 0.075 | 1 | 2 |
| ogdha | 0.42234 | 0.153284802 | 0.078 | 0.025 | 1 | 2 |
| chd4a | 0.42332 | 0.130469814 | 0.078 | 0.025 | 1 | 2 |
| ptpn6 | 0.42463 | 0.199706312 | 0.183 | 0.125 | 1 | 2 |
| cited4b | 0.42469 | -0.20915051 | 0.061 | 0.125 | 1 | 2 |
| CABZ01073834.1 | 0.42546 | -0.10144283 | 0.113 | 0.125 | 1 | 2 |
| atp5l | 0.42583 | 0.161500981 | 0.591 | 0.475 | 1 | 2 |
| jagn1b | 0.42606 | 0.155019296 | 0.07 | 0.025 | 1 | 2 |
| STMP1 | 0.42633 | 0.214338191 | 0.183 | 0.1 | 1 | 2 |
| FBXO48 | 0.4267 | -0.18903164 | 0.061 | 0.125 | 1 | 2 |
| psma1 | 0.42716 | -0.22258441 | 0.243 | 0.35 | 1 | 2 |
| kpnb1 | 0.42797 | -0.19763753 | 0.061 | 0.125 | 1 | 2 |
| mid1ip1l | 0.42916 | -0.2172413 | 0.104 | 0.175 | 1 | 2 |
| gpia | 0.42927 | 0.196051726 | 0.13 | 0.075 | 1 | 2 |
| copb2 | 0.42936 | -0.14641992 | 0.026 | 0.075 | 1 | 2 |
| dnajc1 | 0.42952 | 0.187207347 | 0.113 | 0.05 | 1 | 2 |
| si:ch211-193e13.5 | 0.4301 | -0.16371144 | 0.026 | 0.075 | 1 | 2 |
| rmdn1 | 0.43066 | 0.148642609 | 0.078 | 0.025 | 1 | 2 |
| ppp1cbl | 0.43115 | -0.17498086 | 0.026 | 0.075 | 1 | 2 |
| pcyox1 | 0.43195 | -0.20395592 | 0.061 | 0.125 | 1 | 2 |
| tegt | 0.43227 | -0.19006364 | 0.122 | 0.175 | 1 | 2 |
| porb | 0.43228 | -0.14246985 | 0.026 | 0.075 | 1 | 2 |
| ube2e3 | 0.43238 | 0.10559127 | 0.113 | 0.05 | 1 | 2 |
| BX901920.1 | 0.43252 | 0.236789013 | 0.209 | 0.125 | 1 | 2 |
| si:dkey-112a7.4 | 0.43266 | 0.151503101 | 0.078 | 0.025 | 1 | 2 |
| si:ch73-269m14.2 | 0.43357 | 0.147165107 | 0.078 | 0.025 | 1 | 2 |
| rit1 | 0.43479 | -0.14019386 | 0.026 | 0.075 | 1 | 2 |
| ciapin1 | 0.43482 | -0.17559787 | 0.052 | 0.1 | 1 | 2 |
| fosab | 0.435 | -0.22749479 | 0.209 | 0.275 | 1 | 2 |
| wasa | 0.435 | 0.156933312 | 0.078 | 0.025 | 1 | 2 |
| ttc1 | 0.43506 | -0.16545526 | 0.026 | 0.075 | 1 | 2 |
| utp3 | 0.43547 | -0.1824217 | 0.043 | 0.1 | 1 | 2 |
| ndufa2 | 0.43553 | -0.2168018 | 0.139 | 0.225 | 1 | 2 |
| mgst3b | 0.43575 | 0.133412295 | 0.113 | 0.05 | 1 | 2 |
| map7d1b | 0.43611 | 0.179277697 | 0.078 | 0.025 | 1 | 2 |
| rbm8a | 0.43619 | -0.2017034 | 0.157 | 0.25 | 1 | 2 |
| eif1b | 0.43655 | 0.184832611 | 0.383 | 0.275 | 1 | 2 |
| asna1 | 0.43666 | -0.12868799 | 0.026 | 0.075 | 1 | 2 |
| ptges3a | 0.43666 | -0.19932152 | 0.078 | 0.15 | 1 | 2 |
| ctps1a | 0.43694 | -0.12302823 | 0.026 | 0.075 | 1 | 2 |
| pdk2b | 0.43746 | -0.11428763 | 0.183 | 0.2 | 1 | 2 |
| atox1 | 0.43823 | 0.136077684 | 0.139 | 0.075 | 1 | 2 |
| atp6v1c1b | 0.43854 | 0.144892342 | 0.078 | 0.025 | 1 | 2 |
| ube2l3a | 0.43951 | -0.20197647 | 0.183 | 0.25 | 1 | 2 |
| paip2b | 0.43984 | -0.17383677 | 0.104 | 0.15 | 1 | 2 |
| mrps10 | 0.43999 | -0.11554271 | 0.087 | 0.1 | 1 | 2 |
| prss16 | 0.44004 | 0.134246618 | 0.078 | 0.025 | 1 | 2 |
| bin2b | 0.44019 | 0.136898653 | 0.078 | 0.025 | 1 | 2 |

| | | | | | | |
|---|---|---|---|---|---|---|
| spsb4b | 0.44308 | -0.19686935 | 0.043 | 0.1 | 1 | 2 |
| hnrnpa0b | 0.44318 | 0.202864831 | 0.322 | 0.25 | 1 | 2 |
| ddx23 | 0.44354 | -0.13433033 | 0.052 | 0.075 | 1 | 2 |
| snw1 | 0.44371 | -0.19074381 | 0.043 | 0.1 | 1 | 2 |
| bhlhe40 | 0.44379 | 0.229673259 | 0.243 | 0.15 | 1 | 2 |
| cct6a | 0.44397 | 0.200606044 | 0.183 | 0.1 | 1 | 2 |
| mdm2 | 0.44433 | -0.14557062 | 0.017 | 0.05 | 1 | 2 |
| nfyba | 0.44626 | -0.17302683 | 0.078 | 0.15 | 1 | 2 |
| nop53 | 0.4463 | -0.21189541 | 0.174 | 0.25 | 1 | 2 |
| idh1 | 0.44696 | -0.17566239 | 0.035 | 0.075 | 1 | 2 |
| dusp1 | 0.4474 | -0.13844892 | 0.078 | 0.1 | 1 | 2 |
| rnaset2l | 0.44799 | 0.237309312 | 0.322 | 0.225 | 1 | 2 |
| tbcb | 0.44819 | -0.1812728 | 0.078 | 0.15 | 1 | 2 |
| csnk2b | 0.4493 | -0.14463949 | 0.165 | 0.2 | 1 | 2 |
| samsn1a | 0.44956 | -0.22327149 | 0.122 | 0.2 | 1 | 2 |
| lamp1a | 0.45033 | -0.18173659 | 0.061 | 0.125 | 1 | 2 |
| elf1 | 0.45055 | -0.14955184 | 0.07 | 0.1 | 1 | 2 |
| idh3b | 0.45265 | -0.15662513 | 0.017 | 0.05 | 1 | 2 |
| h3f3a | 0.45266 | 0.15450459 | 0.174 | 0.1 | 1 | 2 |
| ube2l3b | 0.45268 | -0.16647568 | 0.113 | 0.15 | 1 | 2 |
| cxcl20 | 0.45289 | 0.125370686 | 0.07 | 0.025 | 1 | 2 |
| ctbp1 | 0.45296 | -0.17198986 | 0.061 | 0.125 | 1 | 2 |
| srsf10b | 0.45327 | -0.19900126 | 0.061 | 0.125 | 1 | 2 |
| mtbl | 0.45345 | 0.128111709 | 0.113 | 0.05 | 1 | 2 |
| larp4b | 0.45412 | -0.13392486 | 0.017 | 0.05 | 1 | 2 |
| sumo1 | 0.45436 | -0.16637423 | 0.061 | 0.125 | 1 | 2 |
| tfg | 0.45447 | -0.16703458 | 0.061 | 0.125 | 1 | 2 |
| commd8 | 0.45489 | -0.13917424 | 0.026 | 0.05 | 1 | 2 |
| hcls1 | 0.45511 | 0.19664405 | 0.165 | 0.1 | 1 | 2 |
| ppme1 | 0.45565 | -0.15532114 | 0.043 | 0.075 | 1 | 2 |
| selenol | 0.45646 | -0.14002516 | 0.017 | 0.05 | 1 | 2 |
| si:ch73-281n10.2 | 0.46 | 0.212517068 | 0.174 | 0.1 | 1 | 2 |
| tollip | 0.46048 | -0.19236604 | 0.07 | 0.125 | 1 | 2 |
| hif1ab | 0.46058 | 0.169369831 | 0.113 | 0.05 | 1 | 2 |
| aamp | 0.46126 | -0.15681195 | 0.043 | 0.075 | 1 | 2 |
| rxrga | 0.46142 | -0.17085885 | 0.052 | 0.1 | 1 | 2 |
| mt-nd1 | 0.46222 | -0.23639575 | 0.183 | 0.275 | 1 | 2 |
| snx9b | 0.46247 | 0.17293641 | 0.113 | 0.05 | 1 | 2 |
| ppp2cb | 0.46275 | 0.178863233 | 0.148 | 0.075 | 1 | 2 |
| BX323596.2 | 0.46291 | 0.239985665 | 0.148 | 0.075 | 1 | 2 |
| CU467110.1 | 0.46538 | 0.142059753 | 0.061 | 0.025 | 1 | 2 |
| zgc:101715 | 0.46654 | 0.170302808 | 0.113 | 0.05 | 1 | 2 |
| ndufc2 | 0.46755 | 0.15524321 | 0.113 | 0.05 | 1 | 2 |
| h2afvb | 0.46786 | 0.222211015 | 0.374 | 0.275 | 1 | 2 |
| vps29 | 0.46839 | 0.207852735 | 0.174 | 0.1 | 1 | 2 |
| msna | 0.46886 | 0.223682838 | 0.27 | 0.175 | 1 | 2 |
| zgc:152791 | 0.46942 | -0.1445415 | 0.017 | 0.05 | 1 | 2 |
| rrp7a | 0.47362 | -0.14892159 | 0.017 | 0.05 | 1 | 2 |

| | | | | | | |
|---|---|---|---|---|---|---|
| mycbp2 | 0.47451 | -0.13783029 | 0.017 | 0.05 | 1 | 2 |
| zgc:56304 | 0.47734 | -0.13324837 | 0.087 | 0.15 | 1 | 2 |
| bbip1 | 0.47737 | -0.16110311 | 0.052 | 0.1 | 1 | 2 |
| psap | 0.47754 | -0.182288 | 0.539 | 0.625 | 1 | 2 |
| jac3 | 0.47957 | -0.22074631 | 0.296 | 0.4 | 1 | 2 |
| xpr1b | 0.47997 | -0.16860101 | 0.052 | 0.1 | 1 | 2 |
| nudc | 0.48039 | -0.10506974 | 0.087 | 0.1 | 1 | 2 |
| si:dkey-51e6.1 | 0.48085 | -0.1726038 | 0.052 | 0.1 | 1 | 2 |
| eif2s1b | 0.48126 | 0.209616279 | 0.235 | 0.15 | 1 | 2 |
| atf4a | 0.48135 | 0.203864539 | 0.174 | 0.1 | 1 | 2 |
| alg5 | 0.48239 | -0.16887787 | 0.061 | 0.1 | 1 | 2 |
| slc43a2a | 0.48281 | -0.17418743 | 0.078 | 0.125 | 1 | 2 |
| tefa | 0.48307 | -0.15203897 | 0.07 | 0.1 | 1 | 2 |
| snx10b | 0.48329 | 0.144039842 | 0.139 | 0.075 | 1 | 2 |
| xiap | 0.48394 | -0.15658375 | 0.035 | 0.075 | 1 | 2 |
| hk2 | 0.48451 | 0.106950438 | 0.07 | 0.025 | 1 | 2 |
| ndufa1 | 0.48522 | -0.17641474 | 0.174 | 0.225 | 1 | 2 |
| CABZ01073795.1 | 0.48711 | -0.14547971 | 0.017 | 0.05 | 1 | 2 |
| serf2 | 0.48763 | 0.128702072 | 0.409 | 0.375 | 1 | 2 |
| eif3f | 0.48937 | 0.162837777 | 0.609 | 0.5 | 1 | 2 |
| sem1 | 0.48988 | 0.160876419 | 0.287 | 0.2 | 1 | 2 |
| snrpf | 0.49085 | -0.13842016 | 0.165 | 0.2 | 1 | 2 |
| psmc2 | 0.49286 | 0.163744916 | 0.104 | 0.05 | 1 | 2 |
| osgn1 | 0.49356 | -0.15495117 | 0.035 | 0.075 | 1 | 2 |
| npepl1 | 0.49384 | -0.20632203 | 0.148 | 0.225 | 1 | 2 |
| tmem199 | 0.49391 | -0.13514645 | 0.017 | 0.05 | 1 | 2 |
| si:ch211-147a11.3 | 0.49407 | -0.10577937 | 0.017 | 0.05 | 1 | 2 |
| CR855311.1 | 0.49522 | -0.19975899 | 0.209 | 0.275 | 1 | 2 |
| XLOC-028370 | 0.49547 | 0.156636062 | 0.165 | 0.1 | 1 | 2 |
| ptges3b | 0.49581 | -0.18077098 | 0.122 | 0.2 | 1 | 2 |
| atp5fa1 | 0.49679 | 0.162649778 | 0.452 | 0.35 | 1 | 2 |
| snx2 | 0.49734 | -0.11471866 | 0.017 | 0.05 | 1 | 2 |
| XLOC-005449 | 0.49756 | 0.195753819 | 0.165 | 0.1 | 1 | 2 |
| psma3 | 0.49805 | 0.111776716 | 0.191 | 0.125 | 1 | 2 |
| aup1 | 0.49871 | -0.11970996 | 0.026 | 0.05 | 1 | 2 |
| mapk11 | 0.49876 | 0.120246228 | 0.07 | 0.025 | 1 | 2 |
| ddx21 | 0.49904 | -0.15522243 | 0.113 | 0.15 | 1 | 2 |
| ywhaqa | 0.49915 | -0.15811094 | 0.104 | 0.175 | 1 | 2 |
| prdx4 | 0.49927 | -0.1680944 | 0.104 | 0.15 | 1 | 2 |
| ufl1 | 0.49996 | -0.12321586 | 0.026 | 0.05 | 1 | 2 |
| sec11a | 0.50065 | 0.130618855 | 0.139 | 0.075 | 1 | 2 |
| atp6ap2 | 0.50087 | -0.13853036 | 0.165 | 0.2 | 1 | 2 |
| ctsh | 0.50243 | 0.114244209 | 0.27 | 0.2 | 1 | 2 |
| b2m | 0.50282 | 0.179798187 | 0.261 | 0.175 | 1 | 2 |
| zgc:194312 | 0.50318 | -0.12637025 | 0.017 | 0.05 | 1 | 2 |
| mfng | 0.5036 | -0.18177519 | 0.096 | 0.15 | 1 | 2 |
| hnrnpl | 0.50394 | -0.11645507 | 0.017 | 0.05 | 1 | 2 |
| xrn2 | 0.50436 | -0.13372879 | 0.026 | 0.05 | 1 | 2 |

| | | | | | | |
|---|---|---|---|---|---|---|
| sars | 0.50539 | 0.119594922 | 0.104 | 0.05 | 1 | 2 |
| galt | 0.50701 | 0.111284407 | 0.07 | 0.025 | 1 | 2 |
| wdr45 | 0.51185 | -0.13643449 | 0.017 | 0.05 | 1 | 2 |
| psmc1a | 0.51312 | -0.13689901 | 0.07 | 0.1 | 1 | 2 |
| sap18 | 0.51344 | 0.139304499 | 0.139 | 0.075 | 1 | 2 |
| ube2g2 | 0.51345 | 0.129017394 | 0.061 | 0.025 | 1 | 2 |
| ywhabl | 0.51438 | -0.17559177 | 0.304 | 0.375 | 1 | 2 |
| calm1a | 0.516 | 0.164571169 | 0.148 | 0.1 | 1 | 2 |
| eif3m | 0.51629 | 0.186571273 | 0.478 | 0.375 | 1 | 2 |
| parp8 | 0.51678 | -0.13574785 | 0.035 | 0.075 | 1 | 2 |
| gsdmeb | 0.51775 | 0.134599601 | 0.061 | 0.025 | 1 | 2 |
| fkbp3 | 0.518 | -0.14918563 | 0.061 | 0.1 | 1 | 2 |
| rab6a | 0.51808 | 0.1293617 | 0.07 | 0.025 | 1 | 2 |
| h2afy2 | 0.51821 | 0.105462129 | 0.07 | 0.025 | 1 | 2 |
| zfand5b | 0.51851 | 0.113719934 | 0.07 | 0.025 | 1 | 2 |
| brms1la | 0.51924 | -0.11522591 | 0.017 | 0.05 | 1 | 2 |
| dhdds | 0.51954 | 0.116469032 | 0.07 | 0.025 | 1 | 2 |
| etfb | 0.51981 | -0.19731304 | 0.104 | 0.175 | 1 | 2 |
| cops2 | 0.52025 | 0.110139912 | 0.07 | 0.025 | 1 | 2 |
| pfdn2 | 0.5207 | 0.136890406 | 0.07 | 0.025 | 1 | 2 |
| s100a10b | 0.52101 | -0.19253998 | 0.348 | 0.45 | 1 | 2 |
| lcp2a | 0.52114 | 0.130587932 | 0.07 | 0.025 | 1 | 2 |
| si:ch211-107o10.3 | 0.52184 | 0.113905673 | 0.07 | 0.025 | 1 | 2 |
| med6 | 0.52195 | 0.136767118 | 0.07 | 0.025 | 1 | 2 |
| ddx61 | 0.52214 | 0.181602957 | 0.139 | 0.075 | 1 | 2 |
| nenf | 0.52283 | 0.119302606 | 0.07 | 0.025 | 1 | 2 |
| rab44 | 0.52301 | 0.139706566 | 0.104 | 0.05 | 1 | 2 |
| maff | 0.52302 | -0.13831332 | 0.035 | 0.075 | 1 | 2 |
| tmem242 | 0.52335 | -0.12276785 | 0.017 | 0.05 | 1 | 2 |
| hspa5 | 0.52355 | -0.17689825 | 0.087 | 0.15 | 1 | 2 |
| khdrbs1a | 0.524 | -0.10022994 | 0.322 | 0.4 | 1 | 2 |
| gabarapa | 0.52417 | 0.158678333 | 0.322 | 0.275 | 1 | 2 |
| LEPROTL1 | 0.52467 | 0.117768516 | 0.07 | 0.025 | 1 | 2 |
| gnpda1 | 0.52494 | -0.11664937 | 0.2 | 0.225 | 1 | 2 |
| washc2c | 0.52594 | -0.11149918 | 0.017 | 0.05 | 1 | 2 |
| si:dkeyp-55f12.3 | 0.52614 | -0.13023556 | 0.017 | 0.05 | 1 | 2 |
| chmp2a | 0.5269 | -0.15850029 | 0.13 | 0.175 | 1 | 2 |
| si:ch211-262h13.3 | 0.52696 | -0.12678027 | 0.017 | 0.05 | 1 | 2 |
| irf5 | 0.5274 | 0.123828571 | 0.07 | 0.025 | 1 | 2 |
| selenbp1 | 0.52748 | 0.117180243 | 0.07 | 0.025 | 1 | 2 |
| lin37 | 0.52841 | 0.129080549 | 0.061 | 0.025 | 1 | 2 |
| stx11a | 0.52847 | 0.13823128 | 0.07 | 0.025 | 1 | 2 |
| tram1 | 0.52857 | -0.18656611 | 0.13 | 0.2 | 1 | 2 |
| caspa | 0.52876 | 0.123397579 | 0.07 | 0.025 | 1 | 2 |
| mars | 0.5288 | -0.13134727 | 0.017 | 0.05 | 1 | 2 |
| stxbp3 | 0.52918 | 0.111979759 | 0.07 | 0.025 | 1 | 2 |
| pam16 | 0.52971 | 0.1271279 | 0.07 | 0.025 | 1 | 2 |
| timm8b | 0.52985 | -0.15115558 | 0.035 | 0.075 | 1 | 2 |

| | | | | | | |
|---|---|---|---|---|---|---|
| psma2 | 0.53002 | 0.193994001 | 0.191 | 0.125 | 1 | 2 |
| ndufv1 | 0.53017 | -0.17106567 | 0.07 | 0.125 | 1 | 2 |
| CABZ01076275.1 | 0.53159 | -0.15872328 | 0.035 | 0.075 | 1 | 2 |
| id2a | 0.53178 | 0.142057006 | 0.07 | 0.025 | 1 | 2 |
| ak1 | 0.53193 | -0.11089718 | 0.061 | 0.075 | 1 | 2 |
| mtx2 | 0.53248 | -0.1346696 | 0.017 | 0.05 | 1 | 2 |
| riok3 | 0.53265 | -0.13074907 | 0.017 | 0.05 | 1 | 2 |
| rbm14b | 0.53273 | -0.12601816 | 0.017 | 0.05 | 1 | 2 |
| si:ch73-376l24.2 | 0.5329 | -0.17979585 | 0.07 | 0.125 | 1 | 2 |
| eed | 0.53314 | -0.10883306 | 0.017 | 0.05 | 1 | 2 |
| srp19 | 0.5333 | 0.136477154 | 0.07 | 0.025 | 1 | 2 |
| lxn | 0.5335 | -0.17482933 | 0.148 | 0.2 | 1 | 2 |
| birc2 | 0.53351 | 0.164618174 | 0.104 | 0.05 | 1 | 2 |
| CABZ01068358.1 | 0.53361 | 0.121575851 | 0.07 | 0.025 | 1 | 2 |
| cyth3a | 0.53471 | -0.12705423 | 0.017 | 0.05 | 1 | 2 |
| ddt | 0.53471 | 0.113684116 | 0.07 | 0.025 | 1 | 2 |
| fam49bb | 0.53574 | -0.14768339 | 0.087 | 0.15 | 1 | 2 |
| arih2 | 0.53587 | 0.129025314 | 0.061 | 0.025 | 1 | 2 |
| ppp3r1b | 0.53595 | -0.1499384 | 0.07 | 0.125 | 1 | 2 |
| cul5a | 0.536 | 0.125898146 | 0.07 | 0.025 | 1 | 2 |
| arf6a | 0.53779 | 0.163321321 | 0.139 | 0.075 | 1 | 2 |
| nolc1 | 0.54017 | -0.10233329 | 0.026 | 0.05 | 1 | 2 |
| zgc:110239 | 0.54029 | -0.11814526 | 0.2 | 0.225 | 1 | 2 |
| pabpn1 | 0.5405 | -0.11868873 | 0.113 | 0.175 | 1 | 2 |
| tsr3 | 0.54306 | -0.12379529 | 0.017 | 0.05 | 1 | 2 |
| ralbp1 | 0.54514 | -0.10126721 | 0.061 | 0.075 | 1 | 2 |
| trim35-28 | 0.54524 | -0.11324467 | 0.017 | 0.05 | 1 | 2 |
| ppm1g | 0.54546 | -0.15593956 | 0.07 | 0.125 | 1 | 2 |
| hspe1 | 0.54556 | -0.14676383 | 0.148 | 0.225 | 1 | 2 |
| med10 | 0.5457 | 0.148768558 | 0.104 | 0.05 | 1 | 2 |
| tcirg1b | 0.54654 | -0.16157315 | 0.052 | 0.1 | 1 | 2 |
| nocta | 0.54724 | -0.13386501 | 0.017 | 0.05 | 1 | 2 |
| lsm3 | 0.54948 | -0.13942599 | 0.07 | 0.1 | 1 | 2 |
| tspan14 | 0.55039 | 0.14043829 | 0.104 | 0.05 | 1 | 2 |
| mrps35 | 0.55091 | -0.11787052 | 0.017 | 0.05 | 1 | 2 |
| mapk14a | 0.55169 | 0.141641689 | 0.104 | 0.05 | 1 | 2 |
| elovl5 | 0.55186 | -0.1096721 | 0.017 | 0.05 | 1 | 2 |
| rps27.2 | 0.55206 | 0.136918537 | 0.722 | 0.65 | 1 | 2 |
| tia1l | 0.55249 | -0.14142505 | 0.052 | 0.1 | 1 | 2 |
| eepd1 | 0.55266 | -0.13176214 | 0.017 | 0.05 | 1 | 2 |
| ap1s2 | 0.553 | -0.1481755 | 0.087 | 0.15 | 1 | 2 |
| tmem9b | 0.55373 | -0.11859279 | 0.017 | 0.05 | 1 | 2 |
| mrpl27 | 0.55453 | -0.14002879 | 0.043 | 0.075 | 1 | 2 |
| nkap | 0.55482 | -0.13557564 | 0.07 | 0.125 | 1 | 2 |
| igf2bp1 | 0.55565 | -0.14541378 | 0.035 | 0.075 | 1 | 2 |
| imp4 | 0.55573 | 0.12197596 | 0.052 | 0.025 | 1 | 2 |
| f11r.1 | 0.55581 | 0.127959667 | 0.061 | 0.025 | 1 | 2 |
| igf2bp2b | 0.55628 | -0.10821597 | 0.017 | 0.05 | 1 | 2 |

| | | | | | | |
|---|---|---|---|---|---|---|
| si:ch211-217k17.7 | 0.55672 | -0.18242084 | 0.122 | 0.175 | 1 | 2 |
| stra6l | 0.55704 | -0.10303619 | 0.017 | 0.05 | 1 | 2 |
| si:zfos-2330d3.3 | 0.55709 | 0.206053193 | 0.165 | 0.1 | 1 | 2 |
| si:ch211-262i1.6 | 0.55776 | -0.13750725 | 0.035 | 0.075 | 1 | 2 |
| gga1 | 0.55913 | -0.13022423 | 0.017 | 0.05 | 1 | 2 |
| prrc1 | 0.55955 | -0.11359365 | 0.017 | 0.05 | 1 | 2 |
| si:dkey-16l2.16 | 0.56001 | -0.11581732 | 0.017 | 0.05 | 1 | 2 |
| rdh12l | 0.56029 | -0.11073881 | 0.052 | 0.1 | 1 | 2 |
| ubl7b | 0.56085 | 0.106879996 | 0.13 | 0.075 | 1 | 2 |
| dusp7 | 0.56161 | -0.10854649 | 0.035 | 0.075 | 1 | 2 |
| si:ch73-264p11.1 | 0.56254 | -0.10727713 | 0.017 | 0.05 | 1 | 2 |
| lcp1 | 0.56261 | -0.15899233 | 0.078 | 0.125 | 1 | 2 |
| COX3 | 0.56497 | 0.220264271 | 0.47 | 0.375 | 1 | 2 |
| smfn | 0.56552 | -0.13861195 | 0.052 | 0.1 | 1 | 2 |
| skp1 | 0.56692 | 0.112536454 | 0.243 | 0.175 | 1 | 2 |
| atp6v0d1 | 0.56693 | -0.16412738 | 0.104 | 0.15 | 1 | 2 |
| gnai1 | 0.56782 | -0.12177623 | 0.139 | 0.2 | 1 | 2 |
| ap3s1.1 | 0.56808 | -0.10712282 | 0.017 | 0.05 | 1 | 2 |
| ptpn9a | 0.56836 | -0.12455172 | 0.017 | 0.05 | 1 | 2 |
| mir21-1 | 0.56919 | -0.1711614 | 0.07 | 0.125 | 1 | 2 |
| LOC103909972 | 0.56938 | -0.14586195 | 0.07 | 0.125 | 1 | 2 |
| syncrip | 0.56954 | 0.167780016 | 0.226 | 0.15 | 1 | 2 |
| zgc:162396 | 0.5696 | -0.10670289 | 0.017 | 0.05 | 1 | 2 |
| tecrb | 0.56967 | -0.1217895 | 0.017 | 0.05 | 1 | 2 |
| mrpl11 | 0.57115 | -0.13229992 | 0.043 | 0.075 | 1 | 2 |
| irf9 | 0.57168 | -0.10864782 | 0.017 | 0.05 | 1 | 2 |
| zgc:77486 | 0.57176 | -0.15060463 | 0.087 | 0.125 | 1 | 2 |
| sdhaf3 | 0.57178 | -0.11734226 | 0.017 | 0.05 | 1 | 2 |
| bri3 | 0.57198 | -0.12929379 | 0.096 | 0.125 | 1 | 2 |
| si:dkey-11f4.14 | 0.57325 | 0.126891494 | 0.148 | 0.125 | 1 | 2 |
| etf1b | 0.57374 | -0.17557837 | 0.113 | 0.175 | 1 | 2 |
| aoc2 | 0.57447 | -0.12217063 | 0.026 | 0.05 | 1 | 2 |
| atp5po | 0.57469 | -0.18583003 | 0.226 | 0.3 | 1 | 2 |
| cst3 | 0.57578 | -0.1417872 | 0.052 | 0.1 | 1 | 2 |
| sept9b | 0.57624 | -0.12346333 | 0.026 | 0.05 | 1 | 2 |
| magt1 | 0.57697 | -0.13123846 | 0.113 | 0.175 | 1 | 2 |
| vamp3 | 0.57719 | 0.155053666 | 0.13 | 0.075 | 1 | 2 |
| mapk12a | 0.57779 | -0.14062857 | 0.07 | 0.125 | 1 | 2 |
| cox6a1 | 0.57842 | 0.12218875 | 0.461 | 0.375 | 1 | 2 |
| cxcl8b.1 | 0.57867 | 0.217441837 | 0.165 | 0.1 | 1 | 2 |
| plpp5 | 0.57948 | -0.108273 | 0.017 | 0.05 | 1 | 2 |
| mcm7 | 0.57957 | -0.10336974 | 0.017 | 0.05 | 1 | 2 |
| sec14l1 | 0.57979 | -0.10043915 | 0.017 | 0.05 | 1 | 2 |
| ino80c | 0.5809 | 0.111896338 | 0.061 | 0.025 | 1 | 2 |
| c1qa | 0.58146 | 0.166309861 | 0.47 | 0.4 | 1 | 2 |
| scaf1 | 0.58164 | -0.10236506 | 0.017 | 0.05 | 1 | 2 |
| timm8a | 0.58174 | 0.130723149 | 0.061 | 0.025 | 1 | 2 |
| naa15a | 0.58198 | -0.10804015 | 0.017 | 0.05 | 1 | 2 |

| | | | | | | |
|---|---|---|---|---|---|---|
| chek2 | 0.58288 | -0.10815236 | 0.017 | 0.05 | 1 | 2 |
| si:dkey-19a16.7 | 0.58311 | -0.10818835 | 0.017 | 0.05 | 1 | 2 |
| si:dkey-79d12.5 | 0.5836 | -0.11391574 | 0.017 | 0.05 | 1 | 2 |
| xrcc4 | 0.58417 | -0.10326086 | 0.017 | 0.05 | 1 | 2 |
| dicp1.17 | 0.58523 | 0.121260047 | 0.061 | 0.025 | 1 | 2 |
| pdcd6ip | 0.58617 | -0.14666188 | 0.052 | 0.1 | 1 | 2 |
| pfdn4 | 0.58734 | -0.11829686 | 0.174 | 0.2 | 1 | 2 |
| rnf115 | 0.58743 | 0.121582662 | 0.061 | 0.025 | 1 | 2 |
| mrpl34 | 0.58762 | -0.12247162 | 0.096 | 0.125 | 1 | 2 |
| si:dkey-7j14.6 | 0.58831 | 0.151154537 | 0.096 | 0.05 | 1 | 2 |
| sucla2 | 0.58954 | 0.100103046 | 0.096 | 0.05 | 1 | 2 |
| cmpk | 0.59067 | 0.170467994 | 0.157 | 0.1 | 1 | 2 |
| tomm6 | 0.59068 | -0.16151834 | 0.104 | 0.15 | 1 | 2 |
| sat1a.2 | 0.59214 | -0.15653605 | 0.383 | 0.475 | 1 | 2 |
| cyldl | 0.59462 | -0.14753544 | 0.035 | 0.075 | 1 | 2 |
| npm1a | 0.59533 | 0.115048169 | 0.243 | 0.175 | 1 | 2 |
| ndfip2 | 0.59661 | -0.10375937 | 0.035 | 0.075 | 1 | 2 |
| wdr43 | 0.59662 | -0.14232058 | 0.052 | 0.1 | 1 | 2 |
| tmed9 | 0.59811 | -0.13490712 | 0.043 | 0.075 | 1 | 2 |
| atp6v1c1a | 0.59869 | -0.14454182 | 0.052 | 0.1 | 1 | 2 |
| slc30a1a | 0.59968 | -0.12350978 | 0.035 | 0.075 | 1 | 2 |
| smarcd2 | 0.60011 | -0.10513977 | 0.026 | 0.05 | 1 | 2 |
| lrrc59 | 0.60011 | -0.12670558 | 0.07 | 0.1 | 1 | 2 |
| mtap | 0.60118 | 0.113625812 | 0.061 | 0.025 | 1 | 2 |
| dnase1l4.1 | 0.6024 | -0.12540861 | 0.035 | 0.075 | 1 | 2 |
| fam49al | 0.60338 | 0.122513078 | 0.061 | 0.025 | 1 | 2 |
| si:dkey-78l4.14 | 0.60398 | -0.12330777 | 0.035 | 0.075 | 1 | 2 |
| gtf2b | 0.60437 | -0.11498 | 0.052 | 0.075 | 1 | 2 |
| sik1 | 0.60453 | 0.103871976 | 0.061 | 0.025 | 1 | 2 |
| cap1 | 0.60484 | 0.106165624 | 0.53 | 0.45 | 1 | 2 |
| selenos | 0.60495 | -0.1222462 | 0.035 | 0.075 | 1 | 2 |
| atp2b2 | 0.60579 | -0.14583411 | 0.061 | 0.1 | 1 | 2 |
| chmp4ba | 0.60625 | -0.11328431 | 0.035 | 0.075 | 1 | 2 |
| cct5 | 0.60652 | 0.15537151 | 0.278 | 0.2 | 1 | 2 |
| ftsj3 | 0.60704 | -0.11074694 | 0.035 | 0.075 | 1 | 2 |
| inip | 0.60737 | -0.12826715 | 0.035 | 0.075 | 1 | 2 |
| vta1 | 0.60758 | -0.11580024 | 0.035 | 0.075 | 1 | 2 |
| tmem42a | 0.60774 | -0.11318249 | 0.035 | 0.075 | 1 | 2 |
| si:ch1073-392o20.2 | 0.60847 | 0.121689035 | 0.096 | 0.05 | 1 | 2 |
| mafbb | 0.60851 | -0.17832463 | 0.243 | 0.325 | 1 | 2 |
| cbr1l | 0.60862 | 0.149218381 | 0.191 | 0.125 | 1 | 2 |
| cx32.2 | 0.60974 | 0.1129892 | 0.183 | 0.125 | 1 | 2 |
| adcy7 | 0.61013 | 0.111053119 | 0.052 | 0.025 | 1 | 2 |
| dnm1l | 0.61187 | -0.1114441 | 0.026 | 0.05 | 1 | 2 |
| abi2b | 0.61201 | -0.11395633 | 0.104 | 0.125 | 1 | 2 |
| fcer1gl | 0.61215 | -0.1010474 | 0.783 | 0.85 | 1 | 2 |
| golga7 | 0.61247 | 0.118724133 | 0.096 | 0.05 | 1 | 2 |
| denr | 0.61332 | 0.141296057 | 0.13 | 0.075 | 1 | 2 |

| | | | | | | |
|---|---|---|---|---|---|---|
| BX649485.1 | 0.614 | -0.14244048 | 0.087 | 0.125 | 1 | 2 |
| si:ch211-74m13.3 | 0.61686 | -0.15895135 | 0.113 | 0.175 | 1 | 2 |
| exosc1 | 0.61722 | -0.10524863 | 0.026 | 0.05 | 1 | 2 |
| hdac1 | 0.61846 | -0.15319175 | 0.113 | 0.175 | 1 | 2 |
| mid1ip1b | 0.61951 | -0.10281519 | 0.035 | 0.05 | 1 | 2 |
| aip | 0.62029 | -0.12002191 | 0.043 | 0.075 | 1 | 2 |
| pfdn5 | 0.62034 | -0.12749029 | 0.148 | 0.175 | 1 | 2 |
| mapk3 | 0.62278 | 0.103193122 | 0.061 | 0.025 | 1 | 2 |
| si:ch211-283g2.2 | 0.62338 | 0.120695032 | 0.096 | 0.05 | 1 | 2 |
| ago2 | 0.6238 | -0.11664609 | 0.026 | 0.05 | 1 | 2 |
| blmh | 0.62425 | -0.11505984 | 0.078 | 0.1 | 1 | 2 |
| lig1 | 0.6243 | -0.11362958 | 0.026 | 0.05 | 1 | 2 |
| eif2b4 | 0.62722 | -0.11618127 | 0.026 | 0.05 | 1 | 2 |
| csnk1da | 0.62784 | 0.107085231 | 0.052 | 0.025 | 1 | 2 |
| ptprc | 0.62802 | 0.123930898 | 0.157 | 0.1 | 1 | 2 |
| kif5ba | 0.62921 | 0.11374127 | 0.096 | 0.05 | 1 | 2 |
| polr2k | 0.63213 | -0.10373378 | 0.026 | 0.05 | 1 | 2 |
| nckap5l | 0.63223 | -0.1111372 | 0.026 | 0.05 | 1 | 2 |
| srgn | 0.6339 | 0.151535436 | 0.348 | 0.3 | 1 | 2 |
| cd81a | 0.63426 | 0.144523491 | 0.165 | 0.125 | 1 | 2 |
| baxb | 0.63486 | 0.125995607 | 0.061 | 0.025 | 1 | 2 |
| XLOC-042595 | 0.63541 | 0.144163014 | 0.096 | 0.05 | 1 | 2 |
| tnrc6c1 | 0.6355 | 0.115342375 | 0.096 | 0.05 | 1 | 2 |
| psmd3 | 0.63869 | 0.104723578 | 0.061 | 0.025 | 1 | 2 |
| mapkapk2a | 0.63894 | -0.13536429 | 0.087 | 0.125 | 1 | 2 |
| supt16h | 0.63916 | 0.105039794 | 0.061 | 0.025 | 1 | 2 |
| abca1a | 0.63947 | -0.1224496 | 0.026 | 0.05 | 1 | 2 |
| vmp1 | 0.64012 | -0.13322529 | 0.061 | 0.1 | 1 | 2 |
| acbd3 | 0.64049 | 0.104371064 | 0.061 | 0.025 | 1 | 2 |
| ier2a | 0.64053 | -0.16378249 | 0.104 | 0.15 | 1 | 2 |
| ndufs7 | 0.64053 | 0.106900905 | 0.096 | 0.05 | 1 | 2 |
| ssuh2.4 | 0.64075 | 0.133961743 | 0.087 | 0.05 | 1 | 2 |
| calhm6 | 0.6409 | 0.112397439 | 0.096 | 0.05 | 1 | 2 |
| jak1 | 0.64159 | 0.133649878 | 0.096 | 0.05 | 1 | 2 |
| nrp1a | 0.64168 | 0.110256098 | 0.061 | 0.025 | 1 | 2 |
| si:dkey-251i10.1 | 0.64233 | -0.10579425 | 0.026 | 0.05 | 1 | 2 |
| plek | 0.64265 | 0.111851103 | 0.096 | 0.05 | 1 | 2 |
| glipr1a | 0.64277 | 0.123048518 | 0.096 | 0.05 | 1 | 2 |
| si:ch211-212k18.7 | 0.64278 | -0.13103539 | 0.33 | 0.375 | 1 | 2 |
| got2b | 0.64293 | 0.131156542 | 0.096 | 0.05 | 1 | 2 |
| vcp | 0.64302 | 0.131061706 | 0.096 | 0.05 | 1 | 2 |
| nudt5 | 0.64394 | -0.12951633 | 0.07 | 0.1 | 1 | 2 |
| ccl34b.8 | 0.64411 | 0.119172493 | 0.478 | 0.45 | 1 | 2 |
| mrpl24 | 0.64427 | -0.12382305 | 0.043 | 0.075 | 1 | 2 |
| cnbpa | 0.64478 | -0.12675678 | 0.07 | 0.1 | 1 | 2 |
| histh1l | 0.64692 | 0.16001772 | 0.157 | 0.1 | 1 | 2 |
| me1 | 0.64693 | -0.15633551 | 0.122 | 0.175 | 1 | 2 |
| mrps36 | 0.64713 | 0.118257176 | 0.087 | 0.05 | 1 | 2 |

| | | | | | | |
|---|---|---|---|---|---|---|
| grinab | 0.64726 | 0.102988103 | 0.148 | 0.1 | 1 | 2 |
| rab11a | 0.64795 | -0.15660439 | 0.078 | 0.125 | 1 | 2 |
| h1fx | 0.64829 | 0.125636902 | 0.183 | 0.125 | 1 | 2 |
| anapc11 | 0.64993 | -0.11608081 | 0.026 | 0.05 | 1 | 2 |
| anxa4 | 0.65116 | 0.160758242 | 0.183 | 0.125 | 1 | 2 |
| ube2d2 | 0.65191 | -0.13433596 | 0.183 | 0.25 | 1 | 2 |
| ubtf | 0.65348 | -0.10221301 | 0.026 | 0.05 | 1 | 2 |
| spi1a | 0.65463 | 0.132515606 | 0.348 | 0.275 | 1 | 2 |
| cpvl | 0.65507 | -0.14104052 | 0.087 | 0.125 | 1 | 2 |
| zgc:100829 | 0.65555 | -0.13113517 | 0.096 | 0.15 | 1 | 2 |
| skap2 | 0.65601 | -0.13123239 | 0.139 | 0.175 | 1 | 2 |
| ube2e1 | 0.65854 | -0.12812779 | 0.139 | 0.175 | 1 | 2 |
| vat1 | 0.65915 | -0.1579969 | 0.2 | 0.25 | 1 | 2 |
| zeb2a | 0.66017 | 0.101933558 | 0.052 | 0.025 | 1 | 2 |
| selenop | 0.66042 | -0.11373639 | 0.122 | 0.15 | 1 | 2 |
| si:ch211-1o7.3 | 0.66049 | 0.144129516 | 0.157 | 0.1 | 1 | 2 |
| rab8b | 0.6635 | -0.15673783 | 0.183 | 0.25 | 1 | 2 |
| tmem50a | 0.66359 | -0.11029522 | 0.026 | 0.05 | 1 | 2 |
| capn2a | 0.66478 | -0.10343227 | 0.026 | 0.05 | 1 | 2 |
| alas1 | 0.66618 | -0.14270549 | 0.252 | 0.325 | 1 | 2 |
| tma16 | 0.66676 | -0.11007018 | 0.026 | 0.05 | 1 | 2 |
| tuba1b | 0.66967 | -0.1305827 | 0.061 | 0.1 | 1 | 2 |
| cxcr4b | 0.67167 | 0.158076894 | 0.27 | 0.2 | 1 | 2 |
| cox7b | 0.67506 | -0.10478572 | 0.487 | 0.525 | 1 | 2 |
| fbxw11b | 0.67529 | -0.10595672 | 0.026 | 0.05 | 1 | 2 |
| eci1 | 0.67537 | -0.10393752 | 0.026 | 0.05 | 1 | 2 |
| tfe3a | 0.67717 | -0.10934091 | 0.078 | 0.1 | 1 | 2 |
| ftr50 | 0.67752 | -0.10216372 | 0.026 | 0.05 | 1 | 2 |
| nup37 | 0.67949 | -0.10505116 | 0.026 | 0.05 | 1 | 2 |
| rac2 | 0.68043 | 0.144839188 | 0.504 | 0.425 | 1 | 2 |
| eif3g | 0.68258 | -0.10664786 | 0.296 | 0.325 | 1 | 2 |
| taf7 | 0.68267 | -0.13426283 | 0.061 | 0.1 | 1 | 2 |
| si:ch73-248e21.5 | 0.68279 | -0.14199249 | 0.078 | 0.125 | 1 | 2 |
| actr2a | 0.68417 | 0.132924688 | 0.183 | 0.125 | 1 | 2 |
| CR753876.1 | 0.68437 | 0.153714054 | 0.122 | 0.075 | 1 | 2 |
| fosb | 0.68453 | -0.1431908 | 0.043 | 0.075 | 1 | 2 |
| stat3 | 0.68519 | -0.14273008 | 0.078 | 0.125 | 1 | 2 |
| hnrnpl2 | 0.68573 | -0.12021824 | 0.043 | 0.075 | 1 | 2 |
| ndufb9 | 0.68743 | 0.128356929 | 0.183 | 0.125 | 1 | 2 |
| tob1a | 0.68822 | -0.12588269 | 0.061 | 0.1 | 1 | 2 |
| nrros | 0.6891 | -0.12705394 | 0.061 | 0.1 | 1 | 2 |
| atp5meb | 0.68942 | -0.1398844 | 0.322 | 0.375 | 1 | 2 |
| banf1 | 0.6904 | 0.106949732 | 0.122 | 0.075 | 1 | 2 |
| lrrc15 | 0.69113 | -0.12070715 | 0.043 | 0.075 | 1 | 2 |
| hdlbpa | 0.69268 | -0.11540192 | 0.165 | 0.225 | 1 | 2 |
| metap2b | 0.69318 | -0.14021236 | 0.122 | 0.175 | 1 | 2 |
| NDUFB1 | 0.69339 | 0.112353982 | 0.209 | 0.15 | 1 | 2 |
| uck2a | 0.69411 | 0.100292301 | 0.052 | 0.025 | 1 | 2 |

| | | | | | | |
|---|---|---|---|---|---|---|
| snx12 | 0.69414 | -0.11017011 | 0.061 | 0.1 | 1 | 2 |
| sh3bgrl | 0.69443 | 0.124578948 | 0.122 | 0.075 | 1 | 2 |
| antxr2a | 0.69733 | -0.15339129 | 0.174 | 0.225 | 1 | 2 |
| si:dkey-7i4.24 | 0.69736 | 0.171804677 | 0.426 | 0.35 | 1 | 2 |
| h3f3c | 0.6989 | 0.136631264 | 0.217 | 0.175 | 1 | 2 |
| fam76b | 0.69927 | 0.111134636 | 0.052 | 0.025 | 1 | 2 |
| ddx27 | 0.69979 | -0.12266776 | 0.043 | 0.075 | 1 | 2 |
| rab5aa | 0.70015 | -0.1107436 | 0.043 | 0.075 | 1 | 2 |
| hikeshi | 0.7023 | -0.11691787 | 0.07 | 0.1 | 1 | 2 |
| si:ch211-222l21.1 | 0.70254 | 0.114691534 | 0.261 | 0.2 | 1 | 2 |
| per1a | 0.70271 | -0.10112855 | 0.026 | 0.05 | 1 | 2 |
| cox14 | 0.70354 | 0.10998935 | 0.087 | 0.05 | 1 | 2 |
| actr1 | 0.70476 | 0.103282944 | 0.052 | 0.025 | 1 | 2 |
| mrps18c | 0.70512 | 0.101834565 | 0.078 | 0.05 | 1 | 2 |
| gna12a | 0.70625 | -0.10684796 | 0.026 | 0.05 | 1 | 2 |
| tpmt.1 | 0.71082 | -0.10299276 | 0.052 | 0.075 | 1 | 2 |
| chmp4bb | 0.71108 | 0.106933677 | 0.104 | 0.075 | 1 | 2 |
| rcc2 | 0.71117 | 0.107217821 | 0.087 | 0.05 | 1 | 2 |
| ppp1caa | 0.71243 | 0.107844695 | 0.148 | 0.1 | 1 | 2 |
| ctss2.2 | 0.71243 | 0.136802211 | 0.522 | 0.45 | 1 | 2 |
| zgc:55558 | 0.71326 | -0.10245481 | 0.043 | 0.075 | 1 | 2 |
| ccdc88b | 0.71381 | -0.1016897 | 0.026 | 0.05 | 1 | 2 |
| XLOC-010899 | 0.71418 | -0.13173824 | 0.07 | 0.1 | 1 | 2 |
| psenen | 0.71437 | -0.12521541 | 0.061 | 0.1 | 1 | 2 |
| srsf6b | 0.71676 | -0.12169117 | 0.061 | 0.1 | 1 | 2 |
| sec13 | 0.71789 | -0.11253687 | 0.061 | 0.1 | 1 | 2 |
| ptp4a2b | 0.71889 | -0.13837989 | 0.13 | 0.175 | 1 | 2 |
| triqk | 0.71891 | 0.101335518 | 0.087 | 0.05 | 1 | 2 |
| ltb4r | 0.72237 | -0.10675317 | 0.026 | 0.05 | 1 | 2 |
| cox7c | 0.72353 | -0.10330569 | 0.487 | 0.525 | 1 | 2 |
| tradd | 0.72355 | -0.10851664 | 0.061 | 0.1 | 1 | 2 |
| nop10 | 0.72368 | 0.11893503 | 0.113 | 0.075 | 1 | 2 |
| stat2 | 0.7242 | -0.11084848 | 0.043 | 0.075 | 1 | 2 |
| xbp1 | 0.72428 | 0.137122458 | 0.383 | 0.325 | 1 | 2 |
| ccni | 0.72469 | 0.147348779 | 0.313 | 0.25 | 1 | 2 |
| ccng2 | 0.72523 | -0.1067167 | 0.078 | 0.1 | 1 | 2 |
| csrp1a | 0.72619 | 0.114542412 | 0.087 | 0.05 | 1 | 2 |
| myg1 | 0.72841 | 0.110491125 | 0.078 | 0.05 | 1 | 2 |
| sept2 | 0.72923 | 0.116428169 | 0.148 | 0.1 | 1 | 2 |
| rpz5 | 0.73344 | 0.112786192 | 0.13 | 0.1 | 1 | 2 |
| tln1 | 0.73385 | 0.118484146 | 0.148 | 0.1 | 1 | 2 |
| mbpb | 0.73428 | 0.111228283 | 0.087 | 0.05 | 1 | 2 |
| slc37a2 | 0.73439 | -0.10376925 | 0.078 | 0.1 | 1 | 2 |
| arf6b | 0.73494 | -0.12762108 | 0.13 | 0.175 | 1 | 2 |
| tnfaip3 | 0.73769 | -0.10093701 | 0.026 | 0.05 | 1 | 2 |
| rab10 | 0.73856 | -0.13181181 | 0.191 | 0.25 | 1 | 2 |
| ndufb2 | 0.74471 | -0.11077489 | 0.122 | 0.15 | 1 | 2 |
| cenpx | 0.74564 | -0.10509872 | 0.043 | 0.075 | 1 | 2 |

| | | | | | | |
|---|---|---|---|---|---|---|
| ugp2b | 0.74876 | -0.10894087 | 0.096 | 0.125 | 1 | 2 |
| apoeb | 0.74887 | -0.14460741 | 0.148 | 0.175 | 1 | 2 |
| vdac3 | 0.74981 | -0.13534358 | 0.2 | 0.25 | 1 | 2 |
| dbi | 0.75066 | 0.120306484 | 0.443 | 0.375 | 1 | 2 |
| rsl1d1 | 0.75069 | -0.12878217 | 0.13 | 0.175 | 1 | 2 |
| pa2g4a | 0.75171 | 0.109547623 | 0.113 | 0.075 | 1 | 2 |
| mrps25 | 0.75582 | -0.10345276 | 0.07 | 0.1 | 1 | 2 |
| si:dkey-112e7.2 | 0.75612 | -0.12915205 | 0.296 | 0.35 | 1 | 2 |
| clta | 0.75665 | -0.12981114 | 0.2 | 0.25 | 1 | 2 |
| anxa2a | 0.75952 | -0.13793928 | 0.13 | 0.175 | 1 | 2 |
| lmcd1 | 0.76424 | -0.11232368 | 0.07 | 0.1 | 1 | 2 |
| c1qbp | 0.76838 | 0.107332471 | 0.113 | 0.075 | 1 | 2 |
| egr1 | 0.77509 | -0.11950344 | 0.087 | 0.125 | 1 | 2 |
| tspan37 | 0.77872 | 0.101012843 | 0.113 | 0.075 | 1 | 2 |
| spi1b | 0.78108 | 0.126256897 | 0.278 | 0.225 | 1 | 2 |
| grb2b | 0.78124 | -0.10307202 | 0.122 | 0.15 | 1 | 2 |
| snrpa | 0.78199 | -0.10096415 | 0.07 | 0.1 | 1 | 2 |
| rhoaa | 0.78373 | -0.10269369 | 0.217 | 0.25 | 1 | 2 |
| eif3ha | 0.78613 | 0.116292844 | 0.243 | 0.2 | 1 | 2 |
| capgb | 0.78617 | 0.101701427 | 0.278 | 0.225 | 1 | 2 |
| dad1 | 0.78677 | 0.125827088 | 0.33 | 0.275 | 1 | 2 |
| sdr16c5b | 0.78742 | -0.10551346 | 0.087 | 0.125 | 1 | 2 |
| zgc:162944 | 0.78764 | 0.12515209 | 0.33 | 0.275 | 1 | 2 |
| CABZ01001434.1 | 0.79196 | -0.12066448 | 0.087 | 0.125 | 1 | 2 |
| LOC564660 | 0.79359 | -0.14575679 | 0.322 | 0.375 | 1 | 2 |
| ppib | 0.79521 | 0.108010168 | 0.461 | 0.4 | 1 | 2 |
| npc2 | 0.79711 | 0.117878013 | 0.478 | 0.425 | 1 | 2 |
| COX7A2 (1 of many) | 0.80302 | 0.1159279 | 0.33 | 0.275 | 1 | 2 |
| stk17b | 0.80731 | -0.10848791 | 0.113 | 0.15 | 1 | 2 |
| ccl39.1 | 0.81085 | 0.140692511 | 0.243 | 0.2 | 1 | 2 |
| ptp4a1 | 0.81494 | -0.10277378 | 0.139 | 0.175 | 1 | 2 |
| pdia3 | 0.81637 | 0.101355128 | 0.504 | 0.45 | 1 | 2 |
| zgc:63831 | 0.85692 | -0.10029067 | 0.139 | 0.175 | 1 | 2 |
| CR383676.1 | 0.88717 | -0.12066297 | 0.191 | 0.225 | 1 | 2 |
| gamt | 2.522929 | 0.626155234 | 0.227 | 0 | 0.5095561 | 3 |
| si:dkey-14d8.6 | 5.880944 | 0.763816895 | 0.307 | 0.02 | 1 | 3 |
| bhmt | 5.940917 | 0.687571358 | 0.28 | 0.02 | 1 | 3 |
| cpb1 | 8.849635 | 0.527475266 | 0.2 | 0 | 1 | 3 |
| eif4ebp3 | 0.0003 | 0.53739115 | 0.173 | 0 | 1 | 3 |
| rpl27a | 0.00054 | -0.27882118 | 0.453 | 0.51 | 1 | 3 |
| si:dkey-14d8.7 | 0.00056 | 0.381555687 | 0.16 | 0 | 1 | 3 |
| c6ast3 | 0.00102 | 0.32025434 | 0.147 | 0 | 1 | 3 |
| pla2g1b | 0.00102 | 0.356472304 | 0.147 | 0 | 1 | 3 |
| bckdk | 0.00115 | -0.40355087 | 0 | 0.122 | 1 | 3 |
| apoc1 | 0.00185 | 0.374711923 | 0.133 | 0 | 1 | 3 |
| si:ch211-240l19.6 | 0.00185 | 0.472543882 | 0.133 | 0 | 1 | 3 |
| erp27 | 0.00185 | 0.282914454 | 0.133 | 0 | 1 | 3 |
| si:ch211-246m6.4 | 0.00207 | -0.46986063 | 0.013 | 0.143 | 1 | 3 |

| | | | | | | |
|---|---|---|---|---|---|---|
| tmem263 | 0.00332 | -0.37464365 | 0 | 0.102 | 1 | 3 |
| baz1b | 0.00332 | -0.37359639 | 0 | 0.102 | 1 | 3 |
| fkbp4 | 0.00332 | -0.3765544 | 0 | 0.102 | 1 | 3 |
| nup50 | 0.00332 | -0.37596614 | 0 | 0.102 | 1 | 3 |
| apoda.2 | 0.00335 | 0.439093012 | 0.12 | 0 | 1 | 3 |
| khsrp | 0.00335 | 0.408951547 | 0.12 | 0 | 1 | 3 |
| si:ch211-240l19.8 | 0.00335 | 0.448611886 | 0.12 | 0 | 1 | 3 |
| zmat2 | 0.00335 | 0.381470299 | 0.12 | 0 | 1 | 3 |
| cel.1 | 0.00335 | 0.320021121 | 0.12 | 0 | 1 | 3 |
| mdh1aa | 0.0035 | 0.66541085 | 0.293 | 0.061 | 1 | 3 |
| ppiaa | 0.00546 | 0.203201454 | 0.987 | 0.837 | 1 | 3 |
| bzw1a | 0.00575 | -0.47877901 | 0.013 | 0.163 | 1 | 3 |
| zgc:153713 | 0.00606 | 0.374079151 | 0.107 | 0 | 1 | 3 |
| mrps7 | 0.00606 | 0.351775736 | 0.107 | 0 | 1 | 3 |
| rtraf | 0.00606 | 0.319769548 | 0.107 | 0 | 1 | 3 |
| ctrl | 0.00606 | 0.379973385 | 0.107 | 0 | 1 | 3 |
| lsm8 | 0.00606 | 0.358532332 | 0.107 | 0 | 1 | 3 |
| rpa2 | 0.00606 | 0.38664378 | 0.107 | 0 | 1 | 3 |
| mt-nd4 | 0.00637 | 0.509771154 | 0.267 | 0.061 | 1 | 3 |
| icn | 0.00769 | 0.716603835 | 0.24 | 0.061 | 1 | 3 |
| ppp4cb | 0.00962 | -0.2846802 | 0 | 0.082 | 1 | 3 |
| trappc2l | 0.00962 | -0.28378191 | 0 | 0.082 | 1 | 3 |
| parp2 | 0.00962 | -0.30662163 | 0 | 0.082 | 1 | 3 |
| idh3a | 0.00962 | -0.2906255 | 0 | 0.082 | 1 | 3 |
| fip1l1b | 0.00962 | -0.32353436 | 0 | 0.082 | 1 | 3 |
| cpa5 | 0.01095 | 0.32624421 | 0.093 | 0 | 1 | 3 |
| ela2 | 0.01095 | 0.39243001 | 0.093 | 0 | 1 | 3 |
| denr | 0.01095 | 0.283886349 | 0.093 | 0 | 1 | 3 |
| endouc | 0.01095 | 0.367931408 | 0.093 | 0 | 1 | 3 |
| fabp10a | 0.01095 | 0.263002957 | 0.093 | 0 | 1 | 3 |
| ndufs8a | 0.01095 | 0.28829875 | 0.093 | 0 | 1 | 3 |
| sf3a2 | 0.01095 | 0.336222883 | 0.093 | 0 | 1 | 3 |
| si:ch211-195b11.3 | 0.01095 | 0.218073055 | 0.093 | 0 | 1 | 3 |
| hnrnpk | 0.01095 | 0.348633423 | 0.093 | 0 | 1 | 3 |
| snrpd3l | 0.01095 | 0.293634533 | 0.093 | 0 | 1 | 3 |
| tomm70a | 0.01095 | 0.361493323 | 0.093 | 0 | 1 | 3 |
| cpa4 | 0.01332 | 0.439746688 | 0.173 | 0.02 | 1 | 3 |
| pde4ba | 0.01433 | -0.45204165 | 0.013 | 0.143 | 1 | 3 |
| mt-nd1 | 0.016 | 0.192416164 | 0.28 | 0.143 | 1 | 3 |
| dbi | 0.01732 | 0.241973645 | 0.36 | 0.204 | 1 | 3 |
| rps25 | 0.01903 | 0.19584834 | 0.973 | 0.837 | 1 | 3 |
| CR318588.1 | 0.01916 | -0.65429347 | 0.12 | 0.327 | 1 | 3 |
| rps3a | 0.01926 | 0.172162496 | 0.973 | 0.878 | 1 | 3 |
| cgref1 | 0.01975 | 0.288531586 | 0.08 | 0 | 1 | 3 |
| snrnp27 | 0.01975 | 0.28103528 | 0.08 | 0 | 1 | 3 |
| wu:fj16a03 | 0.01975 | 0.306018214 | 0.08 | 0 | 1 | 3 |
| atf1 | 0.01975 | 0.292008267 | 0.08 | 0 | 1 | 3 |
| ctrb1 | 0.01975 | 0.300737237 | 0.08 | 0 | 1 | 3 |

| | | | | | | |
|---|---|---|---|---|---|---|
| mgst1.2 | 0.01975 | 0.257435711 | 0.08 | 0 | 1 | 3 |
| noc2l | 0.01975 | 0.291124748 | 0.08 | 0 | 1 | 3 |
| polr2k | 0.01975 | 0.283320843 | 0.08 | 0 | 1 | 3 |
| scp2b | 0.01975 | 0.26921029 | 0.08 | 0 | 1 | 3 |
| si:ch211-286o17.1 | 0.01975 | 0.283206043 | 0.08 | 0 | 1 | 3 |
| si:dkey-180p18.9 | 0.01975 | 0.26695798 | 0.08 | 0 | 1 | 3 |
| fkbp11 | 0.01975 | 0.269581344 | 0.08 | 0 | 1 | 3 |
| ip6k2b | 0.01975 | 0.276143174 | 0.08 | 0 | 1 | 3 |
| cpa1 | 0.01975 | 0.268291282 | 0.08 | 0 | 1 | 3 |
| emc6 | 0.01975 | 0.245250409 | 0.08 | 0 | 1 | 3 |
| histh1l | 0.01975 | 0.307679977 | 0.08 | 0 | 1 | 3 |
| lamtor4 | 0.01975 | 0.237884549 | 0.08 | 0 | 1 | 3 |
| zgc:77262 | 0.01975 | 0.272106409 | 0.08 | 0 | 1 | 3 |
| cel.2 | 0.02009 | 0.380436675 | 0.16 | 0.02 | 1 | 3 |
| pfdn5 | 0.02391 | 0.46267288 | 0.16 | 0.02 | 1 | 3 |
| ybx1 | 0.02415 | 0.176560548 | 0.92 | 0.755 | 1 | 3 |
| psma2 | 0.02421 | 0.370054623 | 0.2 | 0.061 | 1 | 3 |
| si:ch1073-429i10.3.1 | 0.02505 | 0.383066781 | 0.147 | 0.02 | 1 | 3 |
| krt91 | 0.02789 | -0.41742264 | 0.04 | 0.184 | 1 | 3 |
| gclm | 0.028 | -0.22432199 | 0 | 0.061 | 1 | 3 |
| golga7 | 0.028 | -0.2491455 | 0 | 0.061 | 1 | 3 |
| nup58 | 0.028 | -0.2575439 | 0 | 0.061 | 1 | 3 |
| cebpb | 0.028 | -0.24697119 | 0 | 0.061 | 1 | 3 |
| dtnbp1a | 0.028 | -0.24629466 | 0 | 0.061 | 1 | 3 |
| rab18a | 0.028 | -0.20964266 | 0 | 0.061 | 1 | 3 |
| gpalpp1 | 0.028 | -0.24192158 | 0 | 0.061 | 1 | 3 |
| si:ch211-162e15.3 | 0.028 | -0.26053965 | 0 | 0.061 | 1 | 3 |
| etv4 | 0.028 | -0.22321289 | 0 | 0.061 | 1 | 3 |
| foxp4 | 0.028 | -0.22536943 | 0 | 0.061 | 1 | 3 |
| midn | 0.028 | -0.22516796 | 0 | 0.061 | 1 | 3 |
| mrpl48 | 0.028 | -0.22744092 | 0 | 0.061 | 1 | 3 |
| pitpnaa | 0.028 | -0.24389897 | 0 | 0.061 | 1 | 3 |
| nfkbie | 0.028 | -0.25722833 | 0 | 0.061 | 1 | 3 |
| pdcd10a | 0.028 | -0.24470219 | 0 | 0.061 | 1 | 3 |
| emc8 | 0.028 | -0.2453382 | 0 | 0.061 | 1 | 3 |
| acin1a | 0.028 | -0.25404009 | 0 | 0.061 | 1 | 3 |
| atp1a3b | 0.028 | -0.24282611 | 0 | 0.061 | 1 | 3 |
| rbp7b | 0.028 | -0.27172497 | 0 | 0.061 | 1 | 3 |
| u2af2a | 0.028 | -0.19655256 | 0 | 0.061 | 1 | 3 |
| ubl3a | 0.028 | -0.25037036 | 0 | 0.061 | 1 | 3 |
| ddb2 | 0.028 | -0.22134679 | 0 | 0.061 | 1 | 3 |
| dnajc12 | 0.028 | -0.2633399 | 0 | 0.061 | 1 | 3 |
| fam49a | 0.028 | -0.22060019 | 0 | 0.061 | 1 | 3 |
| fbxo28 | 0.028 | -0.2412183 | 0 | 0.061 | 1 | 3 |
| jam2a | 0.028 | -0.23103674 | 0 | 0.061 | 1 | 3 |
| phldb2b | 0.028 | -0.26275784 | 0 | 0.061 | 1 | 3 |
| pop5 | 0.028 | -0.23936727 | 0 | 0.061 | 1 | 3 |
| LOC101885938 | 0.028 | -0.23049444 | 0 | 0.061 | 1 | 3 |

| | | | | | | |
|---|---|---|---|---|---|---|
| akt2l | 0.028 | -0.20153853 | 0 | 0.061 | 1 | 3 |
| grwd1 | 0.028 | -0.26689768 | 0 | 0.061 | 1 | 3 |
| hspa12a | 0.028 | -0.21035587 | 0 | 0.061 | 1 | 3 |
| idh1 | 0.028 | -0.24910914 | 0 | 0.061 | 1 | 3 |
| mettl17 | 0.028 | -0.21707263 | 0 | 0.061 | 1 | 3 |
| prpf40a | 0.028 | -0.26047293 | 0 | 0.061 | 1 | 3 |
| si:ch211-284b7.3 | 0.028 | -0.22732509 | 0 | 0.061 | 1 | 3 |
| zgc:110626 | 0.028 | -0.24494116 | 0 | 0.061 | 1 | 3 |
| nap1l1 | 0.03369 | 0.439600741 | 0.333 | 0.143 | 1 | 3 |
| lgals2a | 0.03468 | 0.464572687 | 0.147 | 0.02 | 1 | 3 |
| hypk | 0.03537 | 0.427516148 | 0.307 | 0.122 | 1 | 3 |
| zgc:103697 | 0.03561 | -0.33122729 | 0.013 | 0.122 | 1 | 3 |
| pabpn1 | 0.0357 | -0.44773086 | 0.053 | 0.204 | 1 | 3 |
| gapdh | 0.03571 | 0.226473914 | 0.067 | 0 | 1 | 3 |
| LOC103909284 | 0.03571 | 0.225021888 | 0.067 | 0 | 1 | 3 |
| aebp1 | 0.03571 | 0.232484761 | 0.067 | 0 | 1 | 3 |
| amy2a | 0.03571 | 0.281999178 | 0.067 | 0 | 1 | 3 |
| cdk2ap2 | 0.03571 | 0.203656491 | 0.067 | 0 | 1 | 3 |
| dap | 0.03571 | 0.258956708 | 0.067 | 0 | 1 | 3 |
| eif1axa | 0.03571 | 0.260239849 | 0.067 | 0 | 1 | 3 |
| ela3l | 0.03571 | 0.296732709 | 0.067 | 0 | 1 | 3 |
| fhl1a | 0.03571 | 0.281754632 | 0.067 | 0 | 1 | 3 |
| glud1a | 0.03571 | 0.224688105 | 0.067 | 0 | 1 | 3 |
| hacd2 | 0.03571 | 0.243714179 | 0.067 | 0 | 1 | 3 |
| mbd3a | 0.03571 | 0.251292154 | 0.067 | 0 | 1 | 3 |
| myh11a | 0.03571 | 0.258146087 | 0.067 | 0 | 1 | 3 |
| ralbb | 0.03571 | 0.235279278 | 0.067 | 0 | 1 | 3 |
| si:dkey-67c22.2 | 0.03571 | 0.221399989 | 0.067 | 0 | 1 | 3 |
| slc25a20 | 0.03571 | 0.233736845 | 0.067 | 0 | 1 | 3 |
| aqp12 | 0.03571 | 0.224029706 | 0.067 | 0 | 1 | 3 |
| rrm2 | 0.03571 | 0.266074428 | 0.067 | 0 | 1 | 3 |
| ube2l3a | 0.03571 | 0.210875952 | 0.067 | 0 | 1 | 3 |
| vps29 | 0.03571 | 0.233439973 | 0.067 | 0 | 1 | 3 |
| sub1a | 0.03754 | 0.144995323 | 0.213 | 0.122 | 1 | 3 |
| ndufb8 | 0.03957 | 0.345199057 | 0.173 | 0.041 | 1 | 3 |
| rhoaa | 0.04016 | 0.299888067 | 0.16 | 0.041 | 1 | 3 |
| hint1 | 0.04596 | 0.339836702 | 0.187 | 0.061 | 1 | 3 |
| qkia | 0.04629 | 0.360320098 | 0.227 | 0.082 | 1 | 3 |
| pdia2 | 0.04763 | 0.368469043 | 0.133 | 0.02 | 1 | 3 |
| sparc | 0.04849 | 0.472673361 | 0.387 | 0.184 | 1 | 3 |
| rps27a | 0.04857 | 0.161698526 | 0.973 | 0.878 | 1 | 3 |
| sfpq | 0.04965 | -0.38862535 | 0.027 | 0.143 | 1 | 3 |
| eif2s3 | 0.05116 | 0.46907982 | 0.36 | 0.163 | 1 | 3 |
| anxa11b | 0.05117 | 0.454749957 | 0.213 | 0.061 | 1 | 3 |
| cox7c | 0.05456 | -0.39007154 | 0.253 | 0.388 | 1 | 3 |
| hspe1 | 0.05466 | -0.36991075 | 0.173 | 0.286 | 1 | 3 |
| zgc:193541 | 0.05548 | 0.245643513 | 0.347 | 0.204 | 1 | 3 |
| atp5if1a | 0.05584 | -0.46667777 | 0.107 | 0.265 | 1 | 3 |

| | | | | | | |
|---|---|---|---|---|---|---|
| trappc1 | 0.05629 | 0.332765697 | 0.133 | 0.02 | 1 | 3 |
| fthl27 | 0.05867 | -0.10782972 | 0.587 | 0.592 | 1 | 3 |
| ifi45 | 0.06024 | -0.37284142 | 0.053 | 0.122 | 1 | 3 |
| ptmaa | 0.06054 | -0.34890634 | 0.44 | 0.592 | 1 | 3 |
| rbp5 | 0.06349 | -0.4137267 | 0.093 | 0.245 | 1 | 3 |
| ywhabl | 0.06443 | 0.297691024 | 0.307 | 0.163 | 1 | 3 |
| timm9 | 0.06488 | 0.193684657 | 0.053 | 0 | 1 | 3 |
| LOC103910438 | 0.06488 | 0.190758282 | 0.053 | 0 | 1 | 3 |
| MCMDC2 | 0.06488 | 0.175583332 | 0.053 | 0 | 1 | 3 |
| MTERF4 | 0.06488 | 0.180583412 | 0.053 | 0 | 1 | 3 |
| ap3s1.1 | 0.06488 | 0.190772536 | 0.053 | 0 | 1 | 3 |
| asph | 0.06488 | 0.175780223 | 0.053 | 0 | 1 | 3 |
| cenpx | 0.06488 | 0.213405143 | 0.053 | 0 | 1 | 3 |
| chaf1a | 0.06488 | 0.192015308 | 0.053 | 0 | 1 | 3 |
| chchd1 | 0.06488 | 0.181634531 | 0.053 | 0 | 1 | 3 |
| cox14 | 0.06488 | 0.195927147 | 0.053 | 0 | 1 | 3 |
| cs | 0.06488 | 0.207285673 | 0.053 | 0 | 1 | 3 |
| ctsd | 0.06488 | 0.207965651 | 0.053 | 0 | 1 | 3 |
| ddx54 | 0.06488 | 0.187806695 | 0.053 | 0 | 1 | 3 |
| dtwd1 | 0.06488 | 0.181911015 | 0.053 | 0 | 1 | 3 |
| eif4a2 | 0.06488 | 0.20647205 | 0.053 | 0 | 1 | 3 |
| farsa | 0.06488 | 0.18350246 | 0.053 | 0 | 1 | 3 |
| gch2 | 0.06488 | 0.21264647 | 0.053 | 0 | 1 | 3 |
| got2b | 0.06488 | 0.198022354 | 0.053 | 0 | 1 | 3 |
| grb2b | 0.06488 | 0.201965338 | 0.053 | 0 | 1 | 3 |
| hif1ab | 0.06488 | 0.180383961 | 0.053 | 0 | 1 | 3 |
| ik | 0.06488 | 0.164131361 | 0.053 | 0 | 1 | 3 |
| ino80c | 0.06488 | 0.195522636 | 0.053 | 0 | 1 | 3 |
| larp7 | 0.06488 | 0.189827745 | 0.053 | 0 | 1 | 3 |
| luc7l | 0.06488 | 0.175960131 | 0.053 | 0 | 1 | 3 |
| meox1 | 0.06488 | 0.217418823 | 0.053 | 0 | 1 | 3 |
| mmgt1 | 0.06488 | 0.205648702 | 0.053 | 0 | 1 | 3 |
| mmp16a | 0.06488 | 0.192888944 | 0.053 | 0 | 1 | 3 |
| mmp2 | 0.06488 | 0.212538581 | 0.053 | 0 | 1 | 3 |
| mrps18b | 0.06488 | 0.200331351 | 0.053 | 0 | 1 | 3 |
| paics | 0.06488 | 0.202744529 | 0.053 | 0 | 1 | 3 |
| pak2a | 0.06488 | 0.223470791 | 0.053 | 0 | 1 | 3 |
| parp1 | 0.06488 | 0.191937908 | 0.053 | 0 | 1 | 3 |
| psmc6 | 0.06488 | 0.188709307 | 0.053 | 0 | 1 | 3 |
| psmd6 | 0.06488 | 0.196309738 | 0.053 | 0 | 1 | 3 |
| rab7a | 0.06488 | 0.202829342 | 0.053 | 0 | 1 | 3 |
| si:ch211-140m22.7 | 0.06488 | 0.197176651 | 0.053 | 0 | 1 | 3 |
| si:ch73-37h15.2 | 0.06488 | 0.224622982 | 0.053 | 0 | 1 | 3 |
| si:ch73-389b16.1 | 0.06488 | 0.239738956 | 0.053 | 0 | 1 | 3 |
| slc38a5b | 0.06488 | 0.191880558 | 0.053 | 0 | 1 | 3 |
| smim8 | 0.06488 | 0.169225642 | 0.053 | 0 | 1 | 3 |
| sumo3b | 0.06488 | 0.198464681 | 0.053 | 0 | 1 | 3 |
| sycn.2 | 0.06488 | 0.22671068 | 0.053 | 0 | 1 | 3 |

| | | | | | | |
|---|---|---|---|---|---|---|
| tmem243a | 0.06488 | 0.182074012 | 0.053 | 0 | 1 | 3 |
| vim | 0.06488 | 0.205628953 | 0.053 | 0 | 1 | 3 |
| waca | 0.06488 | 0.188922831 | 0.053 | 0 | 1 | 3 |
| wnt11r | 0.06488 | 0.187468739 | 0.053 | 0 | 1 | 3 |
| zcrb1 | 0.06488 | 0.178305252 | 0.053 | 0 | 1 | 3 |
| zgc:136461 | 0.06488 | 0.235939723 | 0.053 | 0 | 1 | 3 |
| zgc:92242 | 0.06488 | 0.21644128 | 0.053 | 0 | 1 | 3 |
| afp4 | 0.06488 | 0.166026831 | 0.053 | 0 | 1 | 3 |
| cox7a1 | 0.06488 | 0.197202302 | 0.053 | 0 | 1 | 3 |
| dpm2 | 0.06488 | 0.173777564 | 0.053 | 0 | 1 | 3 |
| lamb1a | 0.06488 | 0.177940747 | 0.053 | 0 | 1 | 3 |
| lamtor5 | 0.06488 | 0.18230922 | 0.053 | 0 | 1 | 3 |
| med8 | 0.06488 | 0.183753659 | 0.053 | 0 | 1 | 3 |
| nupr1b | 0.06488 | 0.175565815 | 0.053 | 0 | 1 | 3 |
| park7 | 0.06488 | 0.174770829 | 0.053 | 0 | 1 | 3 |
| slc30a4 | 0.06488 | 0.210711193 | 0.053 | 0 | 1 | 3 |
| tjp2b | 0.06488 | 0.196981183 | 0.053 | 0 | 1 | 3 |
| zgc:92137 | 0.06488 | 0.134919325 | 0.053 | 0 | 1 | 3 |
| ash2l | 0.06488 | 0.19816837 | 0.053 | 0 | 1 | 3 |
| clpp | 0.06488 | 0.201498679 | 0.053 | 0 | 1 | 3 |
| hmg20a | 0.06488 | 0.212965092 | 0.053 | 0 | 1 | 3 |
| mrps23 | 0.06488 | 0.189391648 | 0.053 | 0 | 1 | 3 |
| ncbp2 | 0.06488 | 0.201451948 | 0.053 | 0 | 1 | 3 |
| prss1 | 0.06488 | 0.228584576 | 0.053 | 0 | 1 | 3 |
| rtn1a | 0.06488 | 0.162236117 | 0.053 | 0 | 1 | 3 |
| sept9a | 0.06488 | 0.191833751 | 0.053 | 0 | 1 | 3 |
| yipf3 | 0.06488 | 0.200815515 | 0.053 | 0 | 1 | 3 |
| socs3a | 0.06688 | -0.30788724 | 0.053 | 0.163 | 1 | 3 |
| wdr83os | 0.06957 | 0.300969833 | 0.12 | 0.02 | 1 | 3 |
| eif3m | 0.06976 | -0.21856888 | 0.307 | 0.367 | 1 | 3 |
| zgc:92744 | 0.07027 | 0.432497242 | 0.373 | 0.184 | 1 | 3 |
| gpx1a | 0.07049 | 0.268426906 | 0.12 | 0.02 | 1 | 3 |
| psmb6 | 0.07225 | 0.234393883 | 0.24 | 0.122 | 1 | 3 |
| jpt1b | 0.0729 | -0.23286442 | 0.213 | 0.265 | 1 | 3 |
| tfa | 0.0736 | 0.290471977 | 0.12 | 0.02 | 1 | 3 |
| fabp11a | 0.07478 | -0.4799481 | 0.107 | 0.265 | 1 | 3 |
| CR318588.3 | 0.075 | -0.33639176 | 0.013 | 0.102 | 1 | 3 |
| myl9b | 0.07565 | 0.40547628 | 0.2 | 0.061 | 1 | 3 |
| cirbpb | 0.07848 | 0.275428627 | 0.813 | 0.633 | 1 | 3 |
| tpm1 | 0.07896 | 0.460948845 | 0.2 | 0.061 | 1 | 3 |
| rcc2 | 0.0798 | -0.29302941 | 0.013 | 0.102 | 1 | 3 |
| ssr1 | 0.08034 | 0.198494611 | 0.067 | 0.041 | 1 | 3 |
| eif4ebp2 | 0.08071 | 0.306009641 | 0.12 | 0.02 | 1 | 3 |
| cd74a | 0.081 | -0.12875469 | 0.013 | 0.061 | 1 | 3 |
| atp5f1b | 0.08723 | 0.165841785 | 0.36 | 0.245 | 1 | 3 |
| atp5f1c | 0.089 | 0.409203988 | 0.227 | 0.082 | 1 | 3 |
| st13 | 0.09074 | 0.217120306 | 0.093 | 0.02 | 1 | 3 |
| rps17 | 0.09512 | 0.137125181 | 0.907 | 0.776 | 1 | 3 |

| | | | | | | |
|---|---|---|---|---|---|---|
| eif3ba | 0.09734 | 0.310010242 | 0.333 | 0.184 | 1 | 3 |
| hnrnpabb | 0.09754 | -0.36010597 | 0.187 | 0.306 | 1 | 3 |
| rpl5a | 0.09867 | -0.22009119 | 0.64 | 0.735 | 1 | 3 |
| ppp2cb | 0.10243 | -0.33679621 | 0.04 | 0.143 | 1 | 3 |
| rps8a | 0.10305 | 0.136888564 | 0.973 | 0.898 | 1 | 3 |
| stau2 | 0.10747 | -0.35345914 | 0.04 | 0.143 | 1 | 3 |
| srsf7a | 0.10835 | 0.185580972 | 0.093 | 0.02 | 1 | 3 |
| tpm3 | 0.10895 | 0.321432899 | 0.307 | 0.163 | 1 | 3 |
| abcf1 | 0.10908 | -0.32049671 | 0.053 | 0.122 | 1 | 3 |
| gapdhs | 0.11223 | -0.29197548 | 0.2 | 0.286 | 1 | 3 |
| rpl12 | 0.11257 | 0.145674542 | 0.96 | 0.857 | 1 | 3 |
| mapre1b | 0.11302 | 0.198517689 | 0.093 | 0.02 | 1 | 3 |
| lypla2 | 0.11404 | -0.27647885 | 0.013 | 0.082 | 1 | 3 |
| rpl32 | 0.11472 | 0.106713482 | 0.973 | 0.939 | 1 | 3 |
| setb | 0.11482 | -0.20949588 | 0.253 | 0.306 | 1 | 3 |
| lsm4 | 0.11505 | 0.373300143 | 0.187 | 0.061 | 1 | 3 |
| hmgb3b | 0.11506 | 0.257811922 | 0.107 | 0.02 | 1 | 3 |
| NC-002333.4 | 0.11513 | 0.459197121 | 0.4 | 0.224 | 1 | 3 |
| tra2a | 0.11874 | 0.322232413 | 0.147 | 0.041 | 1 | 3 |
| smim15 | 0.11922 | 0.306966153 | 0.147 | 0.041 | 1 | 3 |
| rad21a | 0.1193 | -0.26488533 | 0.027 | 0.082 | 1 | 3 |
| ppiab | 0.12 | 0.226986313 | 0.8 | 0.633 | 1 | 3 |
| hmgb1a | 0.12136 | 0.405639259 | 0.24 | 0.102 | 1 | 3 |
| romo1 | 0.12215 | 0.367084375 | 0.373 | 0.204 | 1 | 3 |
| max | 0.12358 | 0.202348895 | 0.12 | 0.041 | 1 | 3 |
| rac2 | 0.12364 | -0.20681696 | 0.027 | 0.061 | 1 | 3 |
| nccrp1 | 0.12376 | 0.250593239 | 0.107 | 0.02 | 1 | 3 |
| smarce1 | 0.12761 | -0.24888628 | 0.093 | 0.143 | 1 | 3 |
| dad1 | 0.12794 | -0.34340285 | 0.107 | 0.204 | 1 | 3 |
| banf1 | 0.12975 | -0.32634332 | 0.08 | 0.163 | 1 | 3 |
| cfl1 | 0.12983 | -0.25402754 | 0.493 | 0.592 | 1 | 3 |
| hmgn3 | 0.13097 | -0.30713305 | 0.067 | 0.143 | 1 | 3 |
| rpl4 | 0.13334 | 0.211371088 | 0.853 | 0.714 | 1 | 3 |
| anp32e | 0.13336 | 0.332171645 | 0.147 | 0.041 | 1 | 3 |
| pik3r3a | 0.13612 | 0.251380422 | 0.107 | 0.02 | 1 | 3 |
| zgc:171480 | 0.13618 | -0.28877426 | 0.027 | 0.102 | 1 | 3 |
| cotl1 | 0.13623 | 0.26134189 | 0.133 | 0.041 | 1 | 3 |
| serbp1a | 0.1365 | 0.137243383 | 0.453 | 0.347 | 1 | 3 |
| cox6b2 | 0.13751 | -0.34332116 | 0.373 | 0.551 | 1 | 3 |
| lsm5 | 0.13803 | 0.236498778 | 0.16 | 0.061 | 1 | 3 |
| psmb2 | 0.13972 | -0.11135629 | 0.16 | 0.163 | 1 | 3 |
| dut | 0.14192 | 0.168920612 | 0.08 | 0.02 | 1 | 3 |
| dnaja2a | 0.14285 | 0.271216958 | 0.107 | 0.02 | 1 | 3 |
| sinhcafl | 0.14311 | -0.25953512 | 0.013 | 0.082 | 1 | 3 |
| pnrc2 | 0.14363 | 0.120195047 | 0.4 | 0.306 | 1 | 3 |
| casp8ap2 | 0.14438 | -0.2988307 | 0.013 | 0.082 | 1 | 3 |
| polr2d | 0.14709 | 0.299723538 | 0.107 | 0.02 | 1 | 3 |
| nop10 | 0.14767 | 0.273635947 | 0.107 | 0.02 | 1 | 3 |

| | | | | | | |
|---|---|---|---|---|---|---|
| cebpa | 0.14793 | -0.22213724 | 0.013 | 0.082 | 1 | 3 |
| irf2bp2a | 0.14865 | -0.24978492 | 0.013 | 0.082 | 1 | 3 |
| chmp4bb | 0.14905 | 0.292215802 | 0.107 | 0.02 | 1 | 3 |
| si:dkey-261h17.1 | 0.14972 | 0.332377699 | 0.107 | 0.02 | 1 | 3 |
| ptp4a2a | 0.15143 | -0.23141481 | 0.013 | 0.082 | 1 | 3 |
| xbp1 | 0.15148 | 0.34320124 | 0.173 | 0.061 | 1 | 3 |
| srsf5a | 0.15182 | -0.30272172 | 0.12 | 0.204 | 1 | 3 |
| pinx1 | 0.15194 | 0.271184493 | 0.133 | 0.041 | 1 | 3 |
| hmgb2b | 0.15357 | 0.311947414 | 0.4 | 0.245 | 1 | 3 |
| eif4e1c | 0.15636 | -0.26493543 | 0.013 | 0.082 | 1 | 3 |
| nedd8 | 0.15719 | 0.293628045 | 0.093 | 0.02 | 1 | 3 |
| sh3bgrl3 | 0.158 | 0.203634473 | 0.093 | 0.02 | 1 | 3 |
| snw1 | 0.15828 | 0.321780582 | 0.133 | 0.041 | 1 | 3 |
| mtdha | 0.15973 | 0.232518392 | 0.093 | 0.02 | 1 | 3 |
| fam162a | 0.15991 | -0.2683195 | 0.013 | 0.082 | 1 | 3 |
| pcna | 0.16156 | -0.18290718 | 0.12 | 0.143 | 1 | 3 |
| uchl3 | 0.16212 | -0.28324255 | 0.027 | 0.102 | 1 | 3 |
| rps27l | 0.16317 | 0.164884259 | 0.187 | 0.102 | 1 | 3 |
| snrpd2 | 0.16336 | 0.333004775 | 0.173 | 0.061 | 1 | 3 |
| nfyc | 0.16364 | -0.25238581 | 0.013 | 0.082 | 1 | 3 |
| ssr4 | 0.16431 | -0.21531506 | 0.24 | 0.286 | 1 | 3 |
| ptp4a2b | 0.16773 | -0.26021256 | 0.013 | 0.082 | 1 | 3 |
| slirp | 0.16785 | 0.231253421 | 0.093 | 0.02 | 1 | 3 |
| chd3 | 0.16922 | -0.2616209 | 0.053 | 0.143 | 1 | 3 |
| ppdpfb | 0.16956 | -0.10780287 | 0.4 | 0.408 | 1 | 3 |
| wdr46 | 0.17014 | -0.23993033 | 0.013 | 0.082 | 1 | 3 |
| aldh1a2 | 0.17175 | -0.33803824 | 0.067 | 0.143 | 1 | 3 |
| gpx4b | 0.17193 | -0.25187938 | 0.067 | 0.122 | 1 | 3 |
| selenoj | 0.17278 | -0.26291048 | 0.013 | 0.082 | 1 | 3 |
| mrps33 | 0.17319 | -0.25158957 | 0.013 | 0.082 | 1 | 3 |
| mcrip1 | 0.17335 | -0.26720415 | 0.013 | 0.082 | 1 | 3 |
| rpl7l1 | 0.17423 | -0.24808649 | 0.013 | 0.082 | 1 | 3 |
| nop53 | 0.17676 | 0.161947257 | 0.227 | 0.143 | 1 | 3 |
| reep5 | 0.17893 | -0.21992103 | 0.013 | 0.061 | 1 | 3 |
| si:dkey-222f8.3 | 0.17898 | -0.17344797 | 0.04 | 0.061 | 1 | 3 |
| rpa3 | 0.18059 | 0.146220549 | 0.08 | 0.02 | 1 | 3 |
| rad23b | 0.18108 | -0.30149811 | 0.053 | 0.122 | 1 | 3 |
| ccdc187 | 0.18163 | -0.13038312 | 0.053 | 0.061 | 1 | 3 |
| psmb5 | 0.1831 | 0.154540336 | 0.133 | 0.061 | 1 | 3 |
| mrps36 | 0.18593 | 0.243964465 | 0.093 | 0.02 | 1 | 3 |
| ppm1g | 0.18799 | 0.218817163 | 0.093 | 0.02 | 1 | 3 |
| si:zfos-169g10.3 | 0.1902 | 0.114520092 | 0.12 | 0.061 | 1 | 3 |
| gmfb | 0.19067 | -0.28863376 | 0.053 | 0.122 | 1 | 3 |
| mgst3a | 0.19285 | 0.225133758 | 0.12 | 0.041 | 1 | 3 |
| rps27.1 | 0.19409 | 0.137930999 | 0.947 | 0.857 | 1 | 3 |
| prdx6 | 0.19464 | 0.218566756 | 0.173 | 0.082 | 1 | 3 |
| uqcrb | 0.19558 | -0.2227722 | 0.013 | 0.061 | 1 | 3 |
| c1qbp | 0.19579 | 0.272208159 | 0.133 | 0.041 | 1 | 3 |

| | | | | | | |
|---|---|---|---|---|---|---|
| ube2l3b | 0.19616 | -0.28786535 | 0.027 | 0.102 | 1 | 3 |
| srsf2a | 0.19753 | 0.17340541 | 0.16 | 0.082 | 1 | 3 |
| bcas2 | 0.19764 | -0.24786171 | 0.04 | 0.102 | 1 | 3 |
| ppib | 0.19803 | 0.325110426 | 0.507 | 0.347 | 1 | 3 |
| ccdc12 | 0.19994 | 0.241882597 | 0.08 | 0.02 | 1 | 3 |
| calm2b | 0.20026 | 0.180504254 | 0.307 | 0.204 | 1 | 3 |
| fkbp1aa | 0.20043 | 0.275555796 | 0.48 | 0.327 | 1 | 3 |
| pa2g4b | 0.20358 | 0.246378273 | 0.093 | 0.02 | 1 | 3 |
| gas1a | 0.20418 | -0.26058197 | 0.027 | 0.102 | 1 | 3 |
| tmed10 | 0.20447 | -0.20803018 | 0.013 | 0.061 | 1 | 3 |
| hmgb2a | 0.20731 | 0.323201461 | 0.52 | 0.367 | 1 | 3 |
| eif5b | 0.20896 | 0.228606603 | 0.093 | 0.02 | 1 | 3 |
| ube2v1 | 0.21008 | 0.280711061 | 0.213 | 0.102 | 1 | 3 |
| rbm4.1 | 0.21032 | -0.25995059 | 0.027 | 0.102 | 1 | 3 |
| psmd3 | 0.21078 | -0.1816562 | 0.053 | 0.082 | 1 | 3 |
| cebpd | 0.21107 | -0.28702579 | 0.067 | 0.143 | 1 | 3 |
| atp6v1e1a | 0.21135 | -0.24818433 | 0.027 | 0.102 | 1 | 3 |
| trmt112 | 0.21245 | 0.216221319 | 0.12 | 0.041 | 1 | 3 |
| eif3ja | 0.21249 | -0.2984466 | 0.067 | 0.143 | 1 | 3 |
| atp5pb | 0.21426 | 0.271947913 | 0.333 | 0.204 | 1 | 3 |
| snrpd1 | 0.21432 | -0.21935959 | 0.093 | 0.143 | 1 | 3 |
| nme3 | 0.21541 | -0.25610783 | 0.027 | 0.102 | 1 | 3 |
| mdh2 | 0.21601 | 0.278125317 | 0.213 | 0.102 | 1 | 3 |
| eif1axb | 0.21613 | 0.286563317 | 0.187 | 0.082 | 1 | 3 |
| gstp1 | 0.21806 | 0.33407494 | 0.227 | 0.122 | 1 | 3 |
| atox1 | 0.21855 | -0.25949235 | 0.067 | 0.122 | 1 | 3 |
| aldh6a1 | 0.22023 | 0.252844784 | 0.093 | 0.02 | 1 | 3 |
| ost4 | 0.22039 | -0.32770237 | 0.067 | 0.163 | 1 | 3 |
| pef1 | 0.22099 | 0.251370972 | 0.093 | 0.02 | 1 | 3 |
| ndufb6 | 0.22158 | -0.27907078 | 0.067 | 0.163 | 1 | 3 |
| capzb | 0.22187 | 0.222297918 | 0.147 | 0.061 | 1 | 3 |
| dap1b | 0.22304 | -0.33300994 | 0.067 | 0.163 | 1 | 3 |
| top1l | 0.22324 | 0.16683867 | 0.08 | 0.02 | 1 | 3 |
| krt18b | 0.22485 | 0.130220539 | 0.213 | 0.143 | 1 | 3 |
| rpl7 | 0.22624 | 0.101880556 | 0.96 | 0.918 | 1 | 3 |
| rps5 | 0.22711 | 0.127477727 | 0.947 | 0.878 | 1 | 3 |
| sumo1 | 0.2281 | 0.157199733 | 0.107 | 0.041 | 1 | 3 |
| hnrnpa1b | 0.22892 | 0.125908337 | 0.12 | 0.061 | 1 | 3 |
| rps11 | 0.22988 | 0.149308165 | 0.907 | 0.816 | 1 | 3 |
| snrpe | 0.23073 | 0.156670127 | 0.227 | 0.143 | 1 | 3 |
| fhit | 0.23182 | -0.18995025 | 0.027 | 0.061 | 1 | 3 |
| tp53 | 0.23224 | -0.24720018 | 0.04 | 0.102 | 1 | 3 |
| si:ch73-281n10.2 | 0.23239 | 0.341121157 | 0.413 | 0.265 | 1 | 3 |
| arf2b | 0.23275 | -0.27685282 | 0.107 | 0.184 | 1 | 3 |
| krtcap2 | 0.23332 | 0.21925803 | 0.12 | 0.041 | 1 | 3 |
| prdx2 | 0.23463 | 0.17953391 | 0.533 | 0.408 | 1 | 3 |
| sumo2b | 0.23491 | -0.11175107 | 0.213 | 0.224 | 1 | 3 |
| atp5mc3a | 0.23552 | 0.199163759 | 0.213 | 0.122 | 1 | 3 |

| | | | | | | |
|---|---|---|---|---|---|---|
| timm10 | 0.23752 | -0.30467482 | 0.053 | 0.143 | 1 | 3 |
| osr1 | 0.23863 | -0.13950954 | 0.013 | 0.061 | 1 | 3 |
| glo1 | 0.23897 | -0.28112872 | 0.067 | 0.163 | 1 | 3 |
| adh5 | 0.24078 | 0.314433899 | 0.293 | 0.163 | 1 | 3 |
| ldhba | 0.24234 | 0.332434668 | 0.213 | 0.102 | 1 | 3 |
| nhp2 | 0.24292 | -0.23452672 | 0.067 | 0.122 | 1 | 3 |
| snrpf | 0.24375 | -0.33443639 | 0.213 | 0.347 | 1 | 3 |
| fbl | 0.24649 | 0.113728258 | 0.093 | 0.041 | 1 | 3 |
| h3f3d | 0.24732 | 0.260156869 | 0.493 | 0.347 | 1 | 3 |
| junba | 0.24863 | 0.223045685 | 0.08 | 0.02 | 1 | 3 |
| rps27.2 | 0.25025 | 0.170892666 | 0.68 | 0.551 | 1 | 3 |
| ucp2 | 0.25073 | 0.321069817 | 0.213 | 0.102 | 1 | 3 |
| eif3g | 0.25215 | 0.22265768 | 0.267 | 0.163 | 1 | 3 |
| ponzr1 | 0.25314 | -0.32791718 | 0.213 | 0.347 | 1 | 3 |
| zgc:92590 | 0.25366 | 0.207713149 | 0.08 | 0.02 | 1 | 3 |
| cd82a | 0.25925 | -0.2098149 | 0.013 | 0.061 | 1 | 3 |
| mettl5 | 0.26002 | -0.21107378 | 0.013 | 0.061 | 1 | 3 |
| atp5if1b | 0.26144 | 0.255304147 | 0.173 | 0.082 | 1 | 3 |
| h1fx | 0.26182 | 0.232193472 | 0.147 | 0.061 | 1 | 3 |
| morf4l1 | 0.26282 | -0.29115301 | 0.067 | 0.143 | 1 | 3 |
| eif2s2 | 0.26584 | 0.322666277 | 0.333 | 0.204 | 1 | 3 |
| cd63 | 0.26794 | 0.264230301 | 0.133 | 0.061 | 1 | 3 |
| stip1 | 0.26958 | 0.116661066 | 0.093 | 0.041 | 1 | 3 |
| ndufb4 | 0.26985 | 0.198443529 | 0.213 | 0.122 | 1 | 3 |
| psmc1a | 0.27006 | -0.10615096 | 0.053 | 0.061 | 1 | 3 |
| hp1bp3 | 0.27176 | 0.230322673 | 0.147 | 0.061 | 1 | 3 |
| fkbp1ab | 0.27179 | -0.17719852 | 0.013 | 0.061 | 1 | 3 |
| prmt1 | 0.27282 | 0.232175637 | 0.147 | 0.061 | 1 | 3 |
| h3f3a | 0.27295 | -0.22586874 | 0.24 | 0.306 | 1 | 3 |
| ppa1b | 0.27332 | 0.162132765 | 0.107 | 0.041 | 1 | 3 |
| eif3i | 0.27383 | -0.20136909 | 0.173 | 0.224 | 1 | 3 |
| eif4h | 0.27462 | 0.198718863 | 0.08 | 0.02 | 1 | 3 |
| uba1 | 0.27474 | 0.162914241 | 0.08 | 0.02 | 1 | 3 |
| ngdn | 0.27518 | 0.164013119 | 0.08 | 0.02 | 1 | 3 |
| pdgfrl | 0.2752 | -0.13740965 | 0.013 | 0.061 | 1 | 3 |
| ndufa8 | 0.27541 | 0.242622257 | 0.12 | 0.041 | 1 | 3 |
| krt94 | 0.27597 | -0.22875989 | 0.253 | 0.367 | 1 | 3 |
| mgst1.1 | 0.27904 | 0.146289292 | 0.067 | 0.02 | 1 | 3 |
| sms | 0.27923 | -0.21039258 | 0.013 | 0.061 | 1 | 3 |
| nutf2l | 0.28052 | 0.257874559 | 0.12 | 0.041 | 1 | 3 |
| gskip | 0.28193 | 0.182242428 | 0.08 | 0.02 | 1 | 3 |
| atp5mf | 0.28239 | 0.122118431 | 0.187 | 0.122 | 1 | 3 |
| taf15 | 0.28315 | 0.167670116 | 0.107 | 0.041 | 1 | 3 |
| ran | 0.28508 | 0.241928816 | 0.507 | 0.367 | 1 | 3 |
| zgc:91910 | 0.28727 | 0.150739992 | 0.147 | 0.082 | 1 | 3 |
| mvb12a | 0.28897 | 0.231484224 | 0.08 | 0.02 | 1 | 3 |
| rpa1 | 0.28958 | -0.14574279 | 0.013 | 0.061 | 1 | 3 |
| adssl | 0.29138 | 0.127284282 | 0.12 | 0.061 | 1 | 3 |

| | | | | | | |
|---|---|---|---|---|---|---|
| eif4e2rs1 | 0.29187 | -0.21995544 | 0.027 | 0.082 | 1 | 3 |
| cbx1a | 0.29263 | 0.12394381 | 0.067 | 0.02 | 1 | 3 |
| cfd | 0.29354 | -0.21200682 | 0.067 | 0.102 | 1 | 3 |
| tomm5 | 0.29453 | 0.123652004 | 0.213 | 0.143 | 1 | 3 |
| rbbp4 | 0.29503 | -0.24089639 | 0.027 | 0.082 | 1 | 3 |
| rbm28 | 0.29571 | 0.118497025 | 0.067 | 0.02 | 1 | 3 |
| zgc:66427 | 0.29703 | -0.1946178 | 0.013 | 0.061 | 1 | 3 |
| marcksb | 0.29743 | -0.19572055 | 0.04 | 0.082 | 1 | 3 |
| pcbp2 | 0.29767 | -0.26466469 | 0.04 | 0.102 | 1 | 3 |
| arf6a | 0.30059 | -0.14552031 | 0.04 | 0.061 | 1 | 3 |
| tagln | 0.3009 | -0.32477725 | 0.107 | 0.184 | 1 | 3 |
| stau1 | 0.30158 | -0.23070236 | 0.027 | 0.082 | 1 | 3 |
| ube2a | 0.30245 | -0.21828933 | 0.027 | 0.082 | 1 | 3 |
| cst14a.2 | 0.30324 | 0.295136275 | 0.173 | 0.082 | 1 | 3 |
| dctn2 | 0.30387 | 0.200700739 | 0.08 | 0.02 | 1 | 3 |
| si:dkey-112e7.2 | 0.30425 | 0.247845663 | 0.08 | 0.02 | 1 | 3 |
| osr2 | 0.30625 | -0.26066319 | 0.053 | 0.122 | 1 | 3 |
| mrps18a | 0.30694 | -0.18245483 | 0.013 | 0.061 | 1 | 3 |
| zgc:113208 | 0.30726 | 0.128342415 | 0.067 | 0.02 | 1 | 3 |
| smim13 | 0.30796 | -0.20571617 | 0.013 | 0.061 | 1 | 3 |
| rp9 | 0.30819 | -0.20694386 | 0.013 | 0.061 | 1 | 3 |
| zgc:175088 | 0.30978 | 0.189634771 | 0.08 | 0.02 | 1 | 3 |
| psmc5 | 0.31254 | -0.21143988 | 0.053 | 0.102 | 1 | 3 |
| lsm6 | 0.31374 | 0.262312929 | 0.147 | 0.061 | 1 | 3 |
| CABZ01068251.1 | 0.31385 | -0.14186661 | 0.04 | 0.061 | 1 | 3 |
| eif3ha | 0.31491 | 0.277786434 | 0.253 | 0.143 | 1 | 3 |
| commd2 | 0.31523 | -0.19467137 | 0.013 | 0.061 | 1 | 3 |
| arpc3 | 0.31612 | -0.2553177 | 0.133 | 0.204 | 1 | 3 |
| ranbp1 | 0.31629 | 0.279983326 | 0.227 | 0.122 | 1 | 3 |
| gstt1a | 0.31776 | 0.12841105 | 0.053 | 0.041 | 1 | 3 |
| hars | 0.31956 | -0.19620982 | 0.013 | 0.061 | 1 | 3 |
| csde1 | 0.31984 | 0.190742196 | 0.107 | 0.041 | 1 | 3 |
| si:dkey-188i13.9 | 0.32103 | -0.18843428 | 0.013 | 0.061 | 1 | 3 |
| calm1b | 0.32133 | -0.19428713 | 0.013 | 0.061 | 1 | 3 |
| si:ch211-68a17.7 | 0.32182 | 0.19161287 | 0.08 | 0.02 | 1 | 3 |
| ubap2b | 0.32315 | -0.15365392 | 0.04 | 0.061 | 1 | 3 |
| dhfr | 0.32409 | -0.19273395 | 0.013 | 0.061 | 1 | 3 |
| chmp2a | 0.32506 | 0.20369323 | 0.08 | 0.02 | 1 | 3 |
| ilk | 0.3251 | -0.19297584 | 0.013 | 0.061 | 1 | 3 |
| ing4 | 0.32567 | -0.20542935 | 0.013 | 0.061 | 1 | 3 |
| atp1a1a.1 | 0.32609 | -0.16512958 | 0.013 | 0.061 | 1 | 3 |
| psmd7 | 0.32611 | 0.196925324 | 0.08 | 0.02 | 1 | 3 |
| cox6b1 | 0.32611 | 0.206032449 | 0.08 | 0.02 | 1 | 3 |
| copz1 | 0.32635 | -0.25884144 | 0.067 | 0.143 | 1 | 3 |
| rab5c | 0.32639 | 0.222594215 | 0.08 | 0.02 | 1 | 3 |
| s100v2 | 0.32758 | -0.23693231 | 0.12 | 0.184 | 1 | 3 |
| cdk11b | 0.32861 | -0.18979552 | 0.013 | 0.061 | 1 | 3 |
| ak2 | 0.32911 | -0.27519822 | 0.107 | 0.204 | 1 | 3 |

| | | | | | | |
|---|---|---|---|---|---|---|
| mrpl54 | 0.32943 | 0.204494612 | 0.08 | 0.02 | 1 | 3 |
| CABZ01115113.1 | 0.32949 | 0.202733867 | 0.08 | 0.02 | 1 | 3 |
| psmd10 | 0.32968 | -0.18994817 | 0.013 | 0.061 | 1 | 3 |
| ufm1 | 0.32979 | -0.1685213 | 0.013 | 0.061 | 1 | 3 |
| stmn1a | 0.33011 | 0.203081726 | 0.08 | 0.02 | 1 | 3 |
| mrpl33 | 0.33219 | -0.19947297 | 0.013 | 0.061 | 1 | 3 |
| gdi2 | 0.33366 | 0.218507237 | 0.093 | 0.041 | 1 | 3 |
| arpc5a | 0.33571 | 0.127737144 | 0.067 | 0.02 | 1 | 3 |
| ca5a | 0.33601 | -0.1603216 | 0.013 | 0.061 | 1 | 3 |
| calr | 0.33798 | -0.16632121 | 0.013 | 0.061 | 1 | 3 |
| skp1 | 0.33924 | 0.116654953 | 0.253 | 0.184 | 1 | 3 |
| cct8 | 0.33982 | 0.116286496 | 0.227 | 0.163 | 1 | 3 |
| hnrnpl | 0.33995 | -0.21640734 | 0.027 | 0.082 | 1 | 3 |
| ywhae1 | 0.34069 | -0.18342744 | 0.12 | 0.163 | 1 | 3 |
| luc7l3 | 0.3408 | -0.17976974 | 0.013 | 0.061 | 1 | 3 |
| mak16 | 0.34128 | -0.21560961 | 0.027 | 0.082 | 1 | 3 |
| lsm14ab | 0.34216 | -0.18504985 | 0.013 | 0.061 | 1 | 3 |
| zgc:77056 | 0.34295 | -0.1647045 | 0.013 | 0.061 | 1 | 3 |
| prmt5 | 0.34511 | -0.17788115 | 0.027 | 0.061 | 1 | 3 |
| tln1 | 0.34629 | -0.17619366 | 0.013 | 0.061 | 1 | 3 |
| CDK2AP1 | 0.34664 | 0.122284187 | 0.067 | 0.02 | 1 | 3 |
| gnl3 | 0.34772 | -0.22930893 | 0.04 | 0.102 | 1 | 3 |
| pim1 | 0.34777 | -0.20362488 | 0.013 | 0.061 | 1 | 3 |
| cope | 0.34795 | -0.21195558 | 0.027 | 0.082 | 1 | 3 |
| homer3b | 0.34837 | -0.16983242 | 0.027 | 0.082 | 1 | 3 |
| ccdc124 | 0.34882 | -0.18395557 | 0.013 | 0.061 | 1 | 3 |
| si:ch73-238c9.1 | 0.3495 | -0.18423835 | 0.013 | 0.061 | 1 | 3 |
| elac1 | 0.34987 | -0.19905467 | 0.013 | 0.061 | 1 | 3 |
| cplx2l | 0.35004 | -0.16736354 | 0.013 | 0.061 | 1 | 3 |
| commd9 | 0.35032 | -0.18860234 | 0.013 | 0.061 | 1 | 3 |
| cdc37 | 0.35073 | -0.2012864 | 0.013 | 0.061 | 1 | 3 |
| prkar2aa | 0.35105 | -0.19832125 | 0.013 | 0.061 | 1 | 3 |
| inka1a | 0.35106 | -0.16660533 | 0.013 | 0.061 | 1 | 3 |
| taf10 | 0.35155 | -0.17625504 | 0.013 | 0.061 | 1 | 3 |
| nr2f1a | 0.35185 | -0.20196701 | 0.053 | 0.102 | 1 | 3 |
| sptlc2a | 0.3519 | -0.17811671 | 0.013 | 0.061 | 1 | 3 |
| magt1 | 0.35477 | -0.19815537 | 0.053 | 0.122 | 1 | 3 |
| syf2 | 0.35483 | 0.230393551 | 0.107 | 0.041 | 1 | 3 |
| syncrip | 0.35765 | -0.13677335 | 0.16 | 0.184 | 1 | 3 |
| sdhb | 0.35865 | -0.13911249 | 0.04 | 0.061 | 1 | 3 |
| anxa11a | 0.35987 | -0.15817072 | 0.053 | 0.082 | 1 | 3 |
| sae1 | 0.36242 | 0.125834341 | 0.067 | 0.02 | 1 | 3 |
| pfn2 | 0.3626 | -0.16143544 | 0.187 | 0.224 | 1 | 3 |
| ier2b | 0.36261 | -0.20262499 | 0.027 | 0.082 | 1 | 3 |
| aldoaa | 0.36267 | 0.249821069 | 0.147 | 0.082 | 1 | 3 |
| sec61b | 0.36316 | 0.193637085 | 0.373 | 0.265 | 1 | 3 |
| cops9 | 0.36519 | -0.204888 | 0.053 | 0.122 | 1 | 3 |
| lysmd2 | 0.36582 | -0.21302393 | 0.027 | 0.082 | 1 | 3 |

| | | | | | | |
|---|---|---|---|---|---|---|
| fkbp3 | 0.36717 | 0.194833824 | 0.133 | 0.061 | 1 | 3 |
| tspan36 | 0.36749 | 0.117561535 | 0.067 | 0.02 | 1 | 3 |
| atp5mc3b | 0.36956 | -0.12016691 | 0.533 | 0.571 | 1 | 3 |
| rap2c | 0.37042 | -0.13692768 | 0.04 | 0.061 | 1 | 3 |
| eif4a1a | 0.37156 | -0.22844981 | 0.12 | 0.184 | 1 | 3 |
| mcts1 | 0.37643 | 0.208619626 | 0.093 | 0.041 | 1 | 3 |
| ube2na | 0.37904 | -0.18837788 | 0.08 | 0.122 | 1 | 3 |
| sh3bgrl | 0.37935 | 0.105290326 | 0.067 | 0.02 | 1 | 3 |
| tuba8l3 | 0.38005 | 0.16263877 | 0.24 | 0.163 | 1 | 3 |
| mrpl57 | 0.38138 | -0.20823017 | 0.027 | 0.082 | 1 | 3 |
| ndufa11 | 0.38141 | 0.133704222 | 0.093 | 0.041 | 1 | 3 |
| llph | 0.38199 | -0.2249167 | 0.04 | 0.102 | 1 | 3 |
| psma6a | 0.38312 | -0.20650746 | 0.027 | 0.082 | 1 | 3 |
| atp6v0ca | 0.38488 | -0.1936256 | 0.027 | 0.082 | 1 | 3 |
| lrrc17 | 0.38565 | 0.127728118 | 0.067 | 0.02 | 1 | 3 |
| napab | 0.38734 | -0.2015238 | 0.027 | 0.082 | 1 | 3 |
| map1lc3a | 0.3875 | -0.24407662 | 0.053 | 0.122 | 1 | 3 |
| snu13b | 0.3883 | 0.149312765 | 0.213 | 0.143 | 1 | 3 |
| tomm20b | 0.3884 | 0.112364853 | 0.2 | 0.143 | 1 | 3 |
| col1a1a | 0.39103 | 0.239218611 | 0.107 | 0.041 | 1 | 3 |
| irf2bp2b | 0.39135 | -0.20036943 | 0.027 | 0.082 | 1 | 3 |
| lgals3a | 0.392 | -0.20538607 | 0.027 | 0.082 | 1 | 3 |
| add3a | 0.39289 | -0.21675244 | 0.053 | 0.122 | 1 | 3 |
| usf1l | 0.39343 | -0.11476037 | 0.067 | 0.082 | 1 | 3 |
| prelid1a | 0.39397 | 0.15889256 | 0.093 | 0.041 | 1 | 3 |
| ndufs5 | 0.3942 | 0.23717366 | 0.133 | 0.061 | 1 | 3 |
| edf1 | 0.3952 | 0.214422179 | 0.187 | 0.102 | 1 | 3 |
| cct4 | 0.3972 | -0.15358691 | 0.187 | 0.224 | 1 | 3 |
| zgc:153867 | 0.39894 | 0.250129277 | 0.467 | 0.347 | 1 | 3 |
| ctnnb1 | 0.4011 | -0.20145506 | 0.04 | 0.102 | 1 | 3 |
| hsf2 | 0.4015 | 0.129367596 | 0.067 | 0.02 | 1 | 3 |
| puf60b | 0.40183 | 0.145669714 | 0.067 | 0.02 | 1 | 3 |
| myl6 | 0.40218 | 0.233715908 | 0.16 | 0.082 | 1 | 3 |
| tob1a | 0.4024 | -0.13024585 | 0.04 | 0.061 | 1 | 3 |
| ywhabb | 0.40274 | -0.23585995 | 0.133 | 0.204 | 1 | 3 |
| ywhaqa | 0.4028 | 0.200481415 | 0.133 | 0.082 | 1 | 3 |
| si:ch211-196l7.4 | 0.40371 | 0.172146052 | 0.147 | 0.082 | 1 | 3 |
| capza1b | 0.4038 | -0.20909753 | 0.093 | 0.143 | 1 | 3 |
| zgc:195173 | 0.4041 | -0.18589374 | 0.04 | 0.082 | 1 | 3 |
| si:dkey-23i12.5 | 0.40472 | 0.186457614 | 0.067 | 0.02 | 1 | 3 |
| eef1db | 0.40479 | -0.25438442 | 0.213 | 0.306 | 1 | 3 |
| cox4i1 | 0.40541 | 0.25700386 | 0.28 | 0.184 | 1 | 3 |
| prdx3 | 0.40649 | -0.22184148 | 0.067 | 0.122 | 1 | 3 |
| uqcrh | 0.40653 | -0.17906933 | 0.12 | 0.163 | 1 | 3 |
| sdcbp2 | 0.40658 | -0.25370027 | 0.12 | 0.204 | 1 | 3 |
| gnai1 | 0.40751 | -0.16780629 | 0.027 | 0.061 | 1 | 3 |
| bccip | 0.40794 | 0.153470562 | 0.147 | 0.082 | 1 | 3 |
| akt2 | 0.40922 | 0.17282673 | 0.067 | 0.02 | 1 | 3 |

| | | | | | | |
|---|---|---|---|---|---|---|
| snrpa | 0.40952 | 0.194107808 | 0.093 | 0.041 | 1 | 3 |
| hmgn7 | 0.4127 | 0.114999888 | 0.427 | 0.347 | 1 | 3 |
| slc25a3b | 0.41393 | 0.144196727 | 0.44 | 0.347 | 1 | 3 |
| brk1 | 0.41424 | 0.232413964 | 0.16 | 0.082 | 1 | 3 |
| nanos1 | 0.41465 | 0.142311697 | 0.093 | 0.041 | 1 | 3 |
| zgc:110339 | 0.41494 | 0.193521875 | 0.107 | 0.082 | 1 | 3 |
| myef2 | 0.41702 | -0.15135475 | 0.027 | 0.061 | 1 | 3 |
| cct3 | 0.41898 | 0.197238175 | 0.227 | 0.143 | 1 | 3 |
| ppdpfa | 0.41967 | -0.25388032 | 0.173 | 0.265 | 1 | 3 |
| aimp1 | 0.41967 | -0.11275295 | 0.107 | 0.122 | 1 | 3 |
| cnp | 0.42037 | 0.161762225 | 0.067 | 0.02 | 1 | 3 |
| ywhaba | 0.42127 | 0.238484396 | 0.187 | 0.102 | 1 | 3 |
| fam168b | 0.42223 | 0.156310963 | 0.053 | 0.02 | 1 | 3 |
| uqcrc2a | 0.4229 | 0.120846353 | 0.067 | 0.02 | 1 | 3 |
| tmsb1 | 0.42538 | 0.113808881 | 0.053 | 0.041 | 1 | 3 |
| lamtor3 | 0.42546 | 0.15638189 | 0.067 | 0.02 | 1 | 3 |
| cdc42l | 0.42564 | -0.19395581 | 0.08 | 0.122 | 1 | 3 |
| eif4ebp1 | 0.4264 | 0.234732235 | 0.16 | 0.082 | 1 | 3 |
| arpc4l | 0.42823 | 0.147765988 | 0.093 | 0.041 | 1 | 3 |
| tfam | 0.4291 | 0.154638615 | 0.067 | 0.02 | 1 | 3 |
| si:dkey-188i13.10 | 0.42953 | -0.18391143 | 0.04 | 0.082 | 1 | 3 |
| tcea1 | 0.43192 | 0.138372894 | 0.067 | 0.02 | 1 | 3 |
| zgc:56493 | 0.43207 | 0.200135424 | 0.6 | 0.49 | 1 | 3 |
| b2ml | 0.43479 | -0.23849554 | 0.093 | 0.163 | 1 | 3 |
| serf2 | 0.43938 | -0.23593126 | 0.253 | 0.347 | 1 | 3 |
| arpc2 | 0.44375 | -0.20766808 | 0.147 | 0.204 | 1 | 3 |
| tcp1 | 0.445 | 0.182573999 | 0.2 | 0.122 | 1 | 3 |
| si:ch211-214p16.1 | 0.44636 | -0.10152443 | 0.053 | 0.061 | 1 | 3 |
| atp1b1a | 0.44638 | 0.19057782 | 0.093 | 0.041 | 1 | 3 |
| gstm.1 | 0.44704 | 0.168721142 | 0.333 | 0.245 | 1 | 3 |
| hnrnpa0l | 0.44812 | -0.1826201 | 0.44 | 0.551 | 1 | 3 |
| vdac1 | 0.44832 | -0.15795121 | 0.107 | 0.143 | 1 | 3 |
| srprb | 0.44891 | -0.16992367 | 0.027 | 0.061 | 1 | 3 |
| fth1a | 0.45036 | 0.218255933 | 0.52 | 0.408 | 1 | 3 |
| wasf2 | 0.45127 | 0.176867913 | 0.067 | 0.02 | 1 | 3 |
| arhgdia | 0.45136 | 0.213744101 | 0.16 | 0.102 | 1 | 3 |
| cox5aa | 0.45212 | 0.208342709 | 0.253 | 0.163 | 1 | 3 |
| rpf2 | 0.45381 | -0.17539322 | 0.04 | 0.082 | 1 | 3 |
| cpne1 | 0.45536 | 0.140878714 | 0.067 | 0.02 | 1 | 3 |
| sdhc | 0.45567 | 0.184216867 | 0.067 | 0.02 | 1 | 3 |
| prdx5 | 0.45742 | -0.23644162 | 0.107 | 0.184 | 1 | 3 |
| pkma | 0.45753 | -0.17912746 | 0.04 | 0.082 | 1 | 3 |
| nmt1a | 0.45802 | -0.13149601 | 0.04 | 0.061 | 1 | 3 |
| gstm.3 | 0.45912 | -0.2586842 | 0.347 | 0.449 | 1 | 3 |
| pmp22a | 0.45916 | -0.23575714 | 0.08 | 0.143 | 1 | 3 |
| gtf2f2a | 0.46245 | 0.146120201 | 0.067 | 0.02 | 1 | 3 |
| si:dkey-238c7.16 | 0.46351 | 0.175858494 | 0.067 | 0.02 | 1 | 3 |
| si:ch211-282b22.1 | 0.4637 | -0.18062872 | 0.04 | 0.082 | 1 | 3 |

| | | | | | | |
|---|---|---|---|---|---|---|
| oaz1a | 0.46821 | -0.22394732 | 0.28 | 0.367 | 1 | 3 |
| phb2a | 0.46889 | 0.14616955 | 0.067 | 0.02 | 1 | 3 |
| rtca | 0.46923 | 0.164959507 | 0.067 | 0.02 | 1 | 3 |
| spcs2 | 0.46928 | 0.162694157 | 0.067 | 0.02 | 1 | 3 |
| gabarapl2 | 0.4694 | 0.138695148 | 0.093 | 0.041 | 1 | 3 |
| gstm.2 | 0.46984 | -0.1678435 | 0.027 | 0.061 | 1 | 3 |
| cirbpa | 0.47098 | -0.10868157 | 0.52 | 0.551 | 1 | 3 |
| rap1b | 0.4736 | -0.15118588 | 0.093 | 0.122 | 1 | 3 |
| tuba8l4 | 0.47649 | -0.23456497 | 0.24 | 0.327 | 1 | 3 |
| creg2 | 0.47923 | 0.150402135 | 0.053 | 0.02 | 1 | 3 |
| sf3b5 | 0.48054 | -0.1915927 | 0.053 | 0.102 | 1 | 3 |
| uqcrfs1 | 0.48055 | -0.17912163 | 0.08 | 0.122 | 1 | 3 |
| si:ch211-222l21.1 | 0.48091 | -0.20364138 | 0.44 | 0.551 | 1 | 3 |
| ubc | 0.48361 | 0.152588025 | 0.347 | 0.265 | 1 | 3 |
| ptgr1 | 0.48364 | 0.105260118 | 0.053 | 0.02 | 1 | 3 |
| anxa2a | 0.48399 | -0.24146884 | 0.107 | 0.184 | 1 | 3 |
| sap30l | 0.48587 | -0.10549471 | 0.067 | 0.082 | 1 | 3 |
| lsm12b | 0.48948 | 0.117516086 | 0.08 | 0.061 | 1 | 3 |
| cct6a | 0.48975 | -0.21563682 | 0.08 | 0.143 | 1 | 3 |
| sap18 | 0.49019 | -0.1788216 | 0.04 | 0.082 | 1 | 3 |
| gars | 0.49037 | 0.159979278 | 0.093 | 0.041 | 1 | 3 |
| anp32b | 0.49048 | -0.20337621 | 0.067 | 0.122 | 1 | 3 |
| eif5a | 0.49056 | 0.216160838 | 0.453 | 0.347 | 1 | 3 |
| calm1a | 0.49155 | -0.22395837 | 0.08 | 0.143 | 1 | 3 |
| gosr2 | 0.49178 | 0.182091357 | 0.093 | 0.041 | 1 | 3 |
| capns1a | 0.49233 | 0.197887114 | 0.173 | 0.102 | 1 | 3 |
| hmgn6 | 0.49273 | 0.175792524 | 0.533 | 0.429 | 1 | 3 |
| smarca4a | 0.49316 | -0.17100203 | 0.027 | 0.061 | 1 | 3 |
| cnbpa | 0.49425 | -0.20788268 | 0.16 | 0.224 | 1 | 3 |
| cdkn2aipnl | 0.49438 | -0.14955362 | 0.067 | 0.102 | 1 | 3 |
| actb1 | 0.50107 | -0.19990158 | 0.44 | 0.531 | 1 | 3 |
| map1lc3b | 0.50149 | 0.172903812 | 0.093 | 0.041 | 1 | 3 |
| wls | 0.50517 | -0.12764766 | 0.04 | 0.061 | 1 | 3 |
| rwdd1 | 0.50727 | 0.11148334 | 0.107 | 0.061 | 1 | 3 |
| si:ch73-46j18.5 | 0.5082 | 0.156416647 | 0.28 | 0.204 | 1 | 3 |
| grcc10 | 0.50915 | 0.159542688 | 0.093 | 0.041 | 1 | 3 |
| rpf1 | 0.50921 | -0.16050591 | 0.04 | 0.082 | 1 | 3 |
| sarnp | 0.50948 | 0.202961089 | 0.12 | 0.061 | 1 | 3 |
| aldob | 0.5111 | 0.188339373 | 0.147 | 0.082 | 1 | 3 |
| snx1a | 0.5119 | 0.185215384 | 0.093 | 0.041 | 1 | 3 |
| NDUFB1 | 0.51293 | 0.110292234 | 0.133 | 0.082 | 1 | 3 |
| gtf2h5 | 0.51665 | 0.187854013 | 0.093 | 0.041 | 1 | 3 |
| si:dkey-88p24.11 | 0.51722 | -0.10201545 | 0.067 | 0.082 | 1 | 3 |
| col1a1b | 0.51963 | 0.171587005 | 0.093 | 0.041 | 1 | 3 |
| marcksl1a | 0.51992 | -0.20252203 | 0.253 | 0.347 | 1 | 3 |
| fkbp2 | 0.52138 | 0.166018871 | 0.08 | 0.041 | 1 | 3 |
| uqcrc1 | 0.52207 | 0.164110435 | 0.093 | 0.041 | 1 | 3 |
| eif1ad | 0.52218 | 0.167386691 | 0.093 | 0.041 | 1 | 3 |

| | | | | | | |
|---|---|---|---|---|---|---|
| rbmx | 0.5223 | 0.141173312 | 0.053 | 0.02 | 1 | 3 |
| mycbp | 0.52726 | -0.15423607 | 0.067 | 0.122 | 1 | 3 |
| psmb7 | 0.52938 | -0.16750322 | 0.213 | 0.265 | 1 | 3 |
| ptp4a1 | 0.52991 | 0.176832766 | 0.12 | 0.061 | 1 | 3 |
| cdc42 | 0.53129 | 0.189883619 | 0.147 | 0.082 | 1 | 3 |
| ddx39ab | 0.53174 | 0.179131513 | 0.12 | 0.061 | 1 | 3 |
| tmed2 | 0.53273 | -0.17359304 | 0.04 | 0.082 | 1 | 3 |
| ndufb7 | 0.53461 | 0.178234364 | 0.12 | 0.061 | 1 | 3 |
| nucks1b | 0.53474 | -0.17270751 | 0.04 | 0.082 | 1 | 3 |
| mrpl30 | 0.53497 | -0.13357533 | 0.053 | 0.082 | 1 | 3 |
| slc25a11 | 0.53658 | -0.16450254 | 0.04 | 0.082 | 1 | 3 |
| snrpb | 0.53686 | 0.162978055 | 0.213 | 0.143 | 1 | 3 |
| cox8a | 0.53733 | 0.169444071 | 0.12 | 0.061 | 1 | 3 |
| mt-nd3 | 0.5392 | 0.127921966 | 0.16 | 0.102 | 1 | 3 |
| eif3s10 | 0.5399 | 0.195515155 | 0.12 | 0.061 | 1 | 3 |
| mtpn | 0.54008 | 0.202072541 | 0.12 | 0.061 | 1 | 3 |
| ier3ip1 | 0.54034 | -0.16878396 | 0.04 | 0.082 | 1 | 3 |
| cdv3 | 0.5407 | -0.11997963 | 0.08 | 0.102 | 1 | 3 |
| psmb1 | 0.54262 | 0.10937514 | 0.107 | 0.061 | 1 | 3 |
| hnrnpl2 | 0.54342 | -0.16886404 | 0.04 | 0.082 | 1 | 3 |
| pno1 | 0.54349 | 0.135962061 | 0.133 | 0.082 | 1 | 3 |
| eif5 | 0.54359 | 0.112743278 | 0.133 | 0.082 | 1 | 3 |
| tmem88b | 0.54389 | -0.1595867 | 0.027 | 0.061 | 1 | 3 |
| txn2 | 0.5447 | -0.14172147 | 0.053 | 0.082 | 1 | 3 |
| khdrbs1a | 0.54869 | 0.158998248 | 0.48 | 0.388 | 1 | 3 |
| chd4a | 0.55145 | -0.15782799 | 0.027 | 0.061 | 1 | 3 |
| hibadhb | 0.55169 | 0.104517087 | 0.053 | 0.02 | 1 | 3 |
| uchl5 | 0.55369 | 0.101373574 | 0.053 | 0.02 | 1 | 3 |
| litaf | 0.55399 | -0.15410615 | 0.04 | 0.082 | 1 | 3 |
| rbm25b | 0.55576 | -0.10966671 | 0.027 | 0.061 | 1 | 3 |
| ap1s2 | 0.55583 | 0.114522239 | 0.08 | 0.041 | 1 | 3 |
| mt-cyb | 0.55756 | 0.155811923 | 0.307 | 0.224 | 1 | 3 |
| sec61a1 | 0.55934 | -0.11725187 | 0.12 | 0.143 | 1 | 3 |
| sept2 | 0.56035 | 0.163491458 | 0.187 | 0.122 | 1 | 3 |
| cnn2 | 0.56173 | 0.136920727 | 0.133 | 0.082 | 1 | 3 |
| arf5 | 0.56287 | -0.18742801 | 0.093 | 0.143 | 1 | 3 |
| mrps34 | 0.56532 | 0.101651895 | 0.053 | 0.02 | 1 | 3 |
| dars | 0.56657 | 0.114353761 | 0.053 | 0.02 | 1 | 3 |
| pfdn6 | 0.56932 | 0.130548538 | 0.133 | 0.082 | 1 | 3 |
| ndufv1 | 0.56965 | -0.14316826 | 0.027 | 0.061 | 1 | 3 |
| ptges3b | 0.57082 | -0.2092589 | 0.12 | 0.184 | 1 | 3 |
| tia1l | 0.5717 | -0.14442008 | 0.027 | 0.061 | 1 | 3 |
| fundc1 | 0.57284 | -0.120499 | 0.04 | 0.061 | 1 | 3 |
| ivd | 0.57313 | 0.107642841 | 0.08 | 0.041 | 1 | 3 |
| eno1a | 0.57414 | 0.163821454 | 0.08 | 0.041 | 1 | 3 |
| XLOC-035413 | 0.57752 | 0.141398093 | 0.053 | 0.02 | 1 | 3 |
| sub1b | 0.57782 | -0.15640704 | 0.067 | 0.102 | 1 | 3 |
| rpz5 | 0.57803 | -0.18801054 | 0.053 | 0.102 | 1 | 3 |

| | | | | | | |
|---|---|---|---|---|---|---|
| arl6ip1 | 0.57832 | -0.12682729 | 0.04 | 0.082 | 1 | 3 |
| sox4a | 0.57844 | -0.17018176 | 0.053 | 0.102 | 1 | 3 |
| hnrnpa0a | 0.57861 | 0.134487583 | 0.053 | 0.02 | 1 | 3 |
| ddx21 | 0.57878 | 0.119041295 | 0.107 | 0.061 | 1 | 3 |
| arl2 | 0.579 | -0.14989755 | 0.027 | 0.061 | 1 | 3 |
| ifi30 | 0.58152 | -0.16968541 | 0.053 | 0.102 | 1 | 3 |
| aldh9a1a.1 | 0.58206 | -0.18505279 | 0.147 | 0.204 | 1 | 3 |
| mrpl13 | 0.58303 | -0.15571799 | 0.027 | 0.061 | 1 | 3 |
| wdr1 | 0.58549 | -0.15943144 | 0.04 | 0.082 | 1 | 3 |
| ppa2 | 0.58581 | 0.115835014 | 0.053 | 0.02 | 1 | 3 |
| cfl1l | 0.58625 | -0.10461928 | 0.027 | 0.061 | 1 | 3 |
| hsd11b2 | 0.58685 | -0.10506753 | 0.027 | 0.061 | 1 | 3 |
| bcl2l10 | 0.58732 | -0.13706245 | 0.053 | 0.082 | 1 | 3 |
| gbp | 0.5878 | -0.15610785 | 0.04 | 0.082 | 1 | 3 |
| arpp19b | 0.59079 | 0.111720703 | 0.053 | 0.02 | 1 | 3 |
| ppp1cab | 0.59101 | 0.109040865 | 0.053 | 0.02 | 1 | 3 |
| zyx | 0.59101 | -0.1480937 | 0.027 | 0.061 | 1 | 3 |
| zgc:153675 | 0.5918 | 0.113271738 | 0.053 | 0.02 | 1 | 3 |
| dido1 | 0.59281 | 0.110910029 | 0.053 | 0.02 | 1 | 3 |
| sept7b | 0.59488 | 0.119727838 | 0.053 | 0.02 | 1 | 3 |
| actr1 | 0.59507 | 0.109409067 | 0.053 | 0.02 | 1 | 3 |
| rab11ba | 0.59636 | 0.102148465 | 0.08 | 0.041 | 1 | 3 |
| lamtor2 | 0.59674 | 0.118046314 | 0.107 | 0.061 | 1 | 3 |
| mrpl38 | 0.59764 | -0.14084309 | 0.027 | 0.061 | 1 | 3 |
| idh2 | 0.5982 | 0.182390418 | 0.213 | 0.143 | 1 | 3 |
| gemin2 | 0.59903 | -0.12158046 | 0.04 | 0.061 | 1 | 3 |
| pigbos1 | 0.59922 | -0.13889316 | 0.027 | 0.061 | 1 | 3 |
| ctsla | 0.6005 | 0.150198165 | 0.053 | 0.02 | 1 | 3 |
| nmi | 0.60289 | 0.116120798 | 0.053 | 0.02 | 1 | 3 |
| rpl5b | 0.60497 | 0.103148226 | 0.733 | 0.653 | 1 | 3 |
| ebag9 | 0.60538 | 0.134497148 | 0.053 | 0.02 | 1 | 3 |
| si:ch211-288g17.3 | 0.60682 | 0.193409408 | 0.213 | 0.143 | 1 | 3 |
| dnaja2b | 0.60944 | -0.13729132 | 0.027 | 0.061 | 1 | 3 |
| capza1a | 0.60963 | 0.187220647 | 0.16 | 0.102 | 1 | 3 |
| timm23a | 0.61065 | 0.112565881 | 0.053 | 0.02 | 1 | 3 |
| pdcd10b | 0.61145 | -0.14394885 | 0.027 | 0.061 | 1 | 3 |
| nop56 | 0.61159 | -0.16427011 | 0.08 | 0.122 | 1 | 3 |
| rhoua | 0.61248 | 0.103087863 | 0.053 | 0.02 | 1 | 3 |
| anxa5b | 0.61279 | 0.155341084 | 0.08 | 0.041 | 1 | 3 |
| gyg1a | 0.6129 | -0.12666277 | 0.04 | 0.082 | 1 | 3 |
| emc2 | 0.61464 | -0.1307822 | 0.04 | 0.082 | 1 | 3 |
| ier2a | 0.61624 | 0.174441969 | 0.107 | 0.061 | 1 | 3 |
| eif6 | 0.61644 | 0.126483946 | 0.107 | 0.061 | 1 | 3 |
| cxxc5a | 0.61679 | -0.13536328 | 0.04 | 0.082 | 1 | 3 |
| gtpbp4 | 0.61788 | -0.16537851 | 0.173 | 0.224 | 1 | 3 |
| tpm4a | 0.62024 | -0.19361949 | 0.12 | 0.184 | 1 | 3 |
| fen1 | 0.62041 | 0.103911573 | 0.053 | 0.02 | 1 | 3 |
| hnrpkl | 0.62058 | 0.108051759 | 0.08 | 0.041 | 1 | 3 |

| | | | | | | |
|---|---|---|---|---|---|---|
| calr3a | 0.62078 | -0.13474277 | 0.027 | 0.061 | 1 | 3 |
| sypl2b | 0.62096 | 0.113599042 | 0.053 | 0.02 | 1 | 3 |
| ciao2a | 0.62263 | -0.13721965 | 0.027 | 0.061 | 1 | 3 |
| polr3k | 0.6233 | 0.105270819 | 0.053 | 0.02 | 1 | 3 |
| klf6a | 0.62342 | 0.117332649 | 0.053 | 0.02 | 1 | 3 |
| tomm22 | 0.62405 | -0.14554074 | 0.04 | 0.082 | 1 | 3 |
| psme3 | 0.62651 | -0.12817634 | 0.053 | 0.082 | 1 | 3 |
| hdlbpa | 0.62657 | 0.160104022 | 0.08 | 0.041 | 1 | 3 |
| tmem107 | 0.62745 | 0.103204745 | 0.053 | 0.02 | 1 | 3 |
| pim3 | 0.62762 | -0.13566716 | 0.027 | 0.061 | 1 | 3 |
| atf4a | 0.62804 | -0.14252244 | 0.04 | 0.082 | 1 | 3 |
| ptmab | 0.62937 | -0.12141001 | 0.653 | 0.714 | 1 | 3 |
| zgc:77849 | 0.63076 | 0.121223493 | 0.067 | 0.041 | 1 | 3 |
| wu:fi37e09 | 0.63191 | 0.101534774 | 0.08 | 0.041 | 1 | 3 |
| TCIM (1 of many) | 0.63223 | 0.130820125 | 0.053 | 0.02 | 1 | 3 |
| wdr43 | 0.63287 | 0.122139887 | 0.053 | 0.02 | 1 | 3 |
| dctn3 | 0.63391 | 0.119530931 | 0.08 | 0.041 | 1 | 3 |
| ube2g1a | 0.63445 | -0.14750013 | 0.027 | 0.061 | 1 | 3 |
| zc3h15 | 0.63533 | 0.126798438 | 0.053 | 0.02 | 1 | 3 |
| dynll1 | 0.63589 | -0.16984672 | 0.267 | 0.327 | 1 | 3 |
| prps1a | 0.63642 | 0.110733046 | 0.053 | 0.02 | 1 | 3 |
| fam204a | 0.63651 | 0.108680945 | 0.053 | 0.02 | 1 | 3 |
| wdr12 | 0.63728 | -0.13946692 | 0.027 | 0.061 | 1 | 3 |
| coa5 | 0.63739 | 0.113924019 | 0.053 | 0.02 | 1 | 3 |
| mknk2b | 0.63753 | 0.107997895 | 0.053 | 0.02 | 1 | 3 |
| uqcrc2b | 0.63808 | 0.117049736 | 0.053 | 0.02 | 1 | 3 |
| nr2f2 | 0.63928 | 0.131316178 | 0.053 | 0.02 | 1 | 3 |
| ndufc2 | 0.63942 | 0.115818302 | 0.08 | 0.041 | 1 | 3 |
| CDPF1 | 0.64006 | 0.114147216 | 0.053 | 0.02 | 1 | 3 |
| anxa13 | 0.64014 | -0.1160212 | 0.04 | 0.061 | 1 | 3 |
| mrpl12 | 0.64029 | 0.113851952 | 0.053 | 0.02 | 1 | 3 |
| ube2e3 | 0.64065 | 0.162027787 | 0.16 | 0.102 | 1 | 3 |
| ndufa10 | 0.6408 | 0.124659728 | 0.053 | 0.02 | 1 | 3 |
| tnrc6c1 | 0.6413 | 0.123111676 | 0.053 | 0.02 | 1 | 3 |
| pdap1a | 0.6414 | 0.121823347 | 0.053 | 0.02 | 1 | 3 |
| chmp2bb | 0.64151 | 0.112503053 | 0.053 | 0.02 | 1 | 3 |
| XLOC-016094 | 0.64155 | -0.13331177 | 0.027 | 0.061 | 1 | 3 |
| gatc | 0.64164 | 0.116601103 | 0.053 | 0.02 | 1 | 3 |
| coa6 | 0.64169 | 0.124853545 | 0.053 | 0.02 | 1 | 3 |
| gnl3l | 0.64194 | 0.118527803 | 0.053 | 0.02 | 1 | 3 |
| mrps11 | 0.64203 | 0.113293827 | 0.053 | 0.02 | 1 | 3 |
| plscr3b | 0.64283 | -0.11908101 | 0.027 | 0.061 | 1 | 3 |
| dpp3 | 0.64372 | -0.13054972 | 0.027 | 0.061 | 1 | 3 |
| cmpk | 0.6439 | -0.13448667 | 0.027 | 0.061 | 1 | 3 |
| ppp6c | 0.64455 | -0.12088353 | 0.027 | 0.061 | 1 | 3 |
| nfixb | 0.64491 | 0.14433157 | 0.08 | 0.041 | 1 | 3 |
| gtf2b | 0.64538 | -0.12680879 | 0.027 | 0.061 | 1 | 3 |
| h2afx1 | 0.64541 | -0.17936792 | 0.107 | 0.163 | 1 | 3 |

| | | | | | | |
|---|---|---|---|---|---|---|
| si:dkey-159f12.2 | 0.64558 | -0.12016257 | 0.027 | 0.061 | 1 | 3 |
| myg1 | 0.6458 | 0.109867756 | 0.08 | 0.041 | 1 | 3 |
| atp5mea | 0.64667 | -0.11470286 | 0.04 | 0.061 | 1 | 3 |
| hnrnpa0b | 0.64757 | 0.180904159 | 0.32 | 0.245 | 1 | 3 |
| eci1 | 0.64849 | -0.10691639 | 0.04 | 0.061 | 1 | 3 |
| hmgn2 | 0.6515 | 0.161905382 | 0.573 | 0.49 | 1 | 3 |
| kmt5aa | 0.65216 | 0.12561984 | 0.08 | 0.041 | 1 | 3 |
| minos1 | 0.65773 | 0.113543089 | 0.08 | 0.041 | 1 | 3 |
| abracl | 0.65783 | 0.124459341 | 0.08 | 0.041 | 1 | 3 |
| cct7 | 0.65784 | 0.176638328 | 0.253 | 0.184 | 1 | 3 |
| postnb | 0.65873 | 0.150756071 | 0.08 | 0.041 | 1 | 3 |
| tomm20a | 0.65989 | -0.13533009 | 0.067 | 0.102 | 1 | 3 |
| ahcy | 0.66134 | 0.150852258 | 0.12 | 0.082 | 1 | 3 |
| bsg | 0.66195 | 0.132117144 | 0.08 | 0.041 | 1 | 3 |
| celf2 | 0.66371 | 0.146620895 | 0.08 | 0.041 | 1 | 3 |
| pa2g4a | 0.66423 | 0.14577136 | 0.133 | 0.082 | 1 | 3 |
| anp32a | 0.66423 | 0.161432445 | 0.133 | 0.082 | 1 | 3 |
| txnipa | 0.66447 | -0.15711067 | 0.08 | 0.122 | 1 | 3 |
| alas1 | 0.66482 | -0.10334583 | 0.04 | 0.061 | 1 | 3 |
| hnrnpaba | 0.66731 | 0.131907429 | 0.333 | 0.265 | 1 | 3 |
| fxyd1 | 0.67238 | 0.151602288 | 0.107 | 0.061 | 1 | 3 |
| eif5a2 | 0.67369 | -0.1498698 | 0.293 | 0.347 | 1 | 3 |
| itm2bb | 0.6779 | -0.12602306 | 0.053 | 0.082 | 1 | 3 |
| tmem167a | 0.68014 | -0.10894281 | 0.04 | 0.061 | 1 | 3 |
| chac1 | 0.68243 | 0.126708454 | 0.133 | 0.102 | 1 | 3 |
| tomm7 | 0.6835 | -0.1327214 | 0.067 | 0.102 | 1 | 3 |
| smarcd1 | 0.6852 | -0.10100391 | 0.04 | 0.061 | 1 | 3 |
| atf4b | 0.69385 | -0.13775575 | 0.067 | 0.102 | 1 | 3 |
| myl12.2 | 0.69759 | -0.12287744 | 0.053 | 0.082 | 1 | 3 |
| acadm | 0.70241 | -0.10039116 | 0.04 | 0.061 | 1 | 3 |
| ciarta | 0.71643 | -0.13362406 | 0.067 | 0.102 | 1 | 3 |
| cald1a | 0.72354 | 0.135578094 | 0.093 | 0.061 | 1 | 3 |
| lsm7 | 0.7305 | -0.11047658 | 0.08 | 0.102 | 1 | 3 |
| spred2b | 0.73951 | 0.107524317 | 0.12 | 0.082 | 1 | 3 |
| ccni | 0.74046 | 0.107707249 | 0.187 | 0.143 | 1 | 3 |
| rtn4a | 0.7412 | -0.14019293 | 0.08 | 0.122 | 1 | 3 |
| sem1 | 0.74336 | -0.11524114 | 0.133 | 0.163 | 1 | 3 |
| akt1s1 | 0.74399 | -0.10000548 | 0.04 | 0.061 | 1 | 3 |
| ctbp1 | 0.74604 | -0.11286869 | 0.053 | 0.082 | 1 | 3 |
| anapc16 | 0.74808 | -0.12764691 | 0.067 | 0.102 | 1 | 3 |
| pdia3 | 0.75958 | 0.13767401 | 0.147 | 0.102 | 1 | 3 |
| adh8a | 0.75992 | 0.122915806 | 0.067 | 0.041 | 1 | 3 |
| gnb1b | 0.76331 | -0.13428174 | 0.067 | 0.102 | 1 | 3 |
| smdt1b | 0.7679 | -0.11346247 | 0.053 | 0.082 | 1 | 3 |
| tuba8l | 0.76798 | -0.10483773 | 0.12 | 0.143 | 1 | 3 |
| eif3ea | 0.77201 | 0.12322493 | 0.28 | 0.224 | 1 | 3 |
| ywhaqb | 0.77601 | -0.1238997 | 0.067 | 0.102 | 1 | 3 |
| rac1a | 0.77802 | -0.11524375 | 0.053 | 0.082 | 1 | 3 |

| | | | | | | |
|---|---|---|---|---|---|---|
| etfb | 0.78081 | -0.10703872 | 0.053 | 0.082 | 1 | 3 |
| zfand3 | 0.78089 | -0.10099098 | 0.04 | 0.061 | 1 | 3 |
| gabarapa | 0.7817 | -0.11586239 | 0.093 | 0.122 | 1 | 3 |
| srsf9 | 0.78347 | -0.11191944 | 0.053 | 0.082 | 1 | 3 |
| elocb | 0.78561 | 0.116397151 | 0.12 | 0.082 | 1 | 3 |
| si:dkey-7j14.6 | 0.78628 | -0.10934615 | 0.053 | 0.082 | 1 | 3 |
| STMP1 | 0.78644 | 0.120331638 | 0.12 | 0.082 | 1 | 3 |
| itm2ba | 0.78784 | 0.12314722 | 0.12 | 0.082 | 1 | 3 |
| ssr2 | 0.78852 | 0.106190234 | 0.253 | 0.204 | 1 | 3 |
| mtbl | 0.79069 | 0.115929621 | 0.107 | 0.082 | 1 | 3 |
| sumo3a | 0.79291 | 0.132991278 | 0.253 | 0.204 | 1 | 3 |
| med31 | 0.79773 | -0.10988347 | 0.053 | 0.082 | 1 | 3 |
| ccl25b | 0.79894 | -0.12699383 | 0.053 | 0.082 | 1 | 3 |
| cyc1 | 0.80136 | 0.103011562 | 0.093 | 0.061 | 1 | 3 |
| atp5f1e | 0.80165 | 0.108380253 | 0.347 | 0.306 | 1 | 3 |
| rsl1d1 | 0.80342 | 0.10011811 | 0.093 | 0.061 | 1 | 3 |
| si:dkey-102m7.3 | 0.80532 | 0.105959227 | 0.093 | 0.061 | 1 | 3 |
| myl9a | 0.8057 | 0.102462231 | 0.067 | 0.041 | 1 | 3 |
| ndufs7 | 0.80827 | -0.11152729 | 0.053 | 0.082 | 1 | 3 |
| kctd5a | 0.80873 | 0.10854684 | 0.093 | 0.061 | 1 | 3 |
| ndufa5 | 0.80881 | 0.108372541 | 0.093 | 0.061 | 1 | 3 |
| eif4bb | 0.80923 | 0.100916514 | 0.093 | 0.061 | 1 | 3 |
| eif3d | 0.81173 | -0.11758219 | 0.253 | 0.306 | 1 | 3 |
| nupr1a | 0.81349 | -0.1105362 | 0.053 | 0.082 | 1 | 3 |
| cnbpb | 0.81504 | 0.101468373 | 0.16 | 0.122 | 1 | 3 |
| sf3b6 | 0.81512 | -0.10760244 | 0.16 | 0.204 | 1 | 3 |
| eif3k | 0.81525 | -0.11491258 | 0.187 | 0.224 | 1 | 3 |
| sat1a.2 | 0.82723 | 0.106586804 | 0.107 | 0.082 | 1 | 3 |
| rbm8a | 0.83675 | 0.108729506 | 0.227 | 0.184 | 1 | 3 |
| hsbp1b | 0.84252 | 0.101513362 | 0.16 | 0.122 | 1 | 3 |
| hist2h2l | 0.85082 | -0.10468094 | 0.093 | 0.122 | 1 | 3 |
| vat1 | 0.85303 | -0.10795496 | 0.093 | 0.122 | 1 | 3 |
| gstt1b | 0.85312 | -0.10056855 | 0.093 | 0.122 | 1 | 3 |
| mcl1b | 0.86481 | 0.105623014 | 0.133 | 0.102 | 1 | 3 |
| zgc:92744 | 2.927687 | 0.998715593 | 0.435 | 0 | 5.91305024 | 4 |
| c6ast3 | 1.541016 | 0.568464459 | 0.258 | 0 | 0.311239 | 4 |
| elof1 | 0.00025 | 0.556286776 | 0.194 | 0 | 1 | 4 |
| actb1 | 0.00085 | -0.12394155 | 1 | 1 | 1 | 4 |
| tpt1 | 0.00087 | 0.171901895 | 1 | 0.826 | 1 | 4 |
| pla2g1b | 0.00097 | 0.405634374 | 0.161 | 0 | 1 | 4 |
| cpb1 | 0.00097 | 0.408282258 | 0.161 | 0 | 1 | 4 |
| ctrl | 0.00097 | 0.572778148 | 0.161 | 0 | 1 | 4 |
| rps16 | 0.00113 | 0.313316525 | 0.968 | 0.783 | 1 | 4 |
| socs3a | 0.0019 | -0.57988822 | 0.016 | 0.217 | 1 | 4 |
| ucp2 | 0.00336 | 0.440652035 | 0.516 | 0.239 | 1 | 4 |
| si:dkey-14d8.6 | 0.00367 | 0.353098267 | 0.129 | 0 | 1 | 4 |
| cpa4 | 0.00367 | 0.301089326 | 0.129 | 0 | 1 | 4 |
| nfybb | 0.00367 | 0.380598229 | 0.129 | 0 | 1 | 4 |

| | | | | | | |
|---|---|---|---|---|---|---|
| CELA1 (1 of many) | 0.00367 | 0.56059365 | 0.129 | 0 | 1 | 4 |
| sqstm1 | 0.00367 | 0.357137546 | 0.129 | 0 | 1 | 4 |
| rpl22 | 0.00396 | 0.100536874 | 0.855 | 0.913 | 1 | 4 |
| spi1b | 0.00444 | -0.21355705 | 0.387 | 0.413 | 1 | 4 |
| ppp4cb | 0.00491 | -0.29242794 | 0 | 0.109 | 1 | 4 |
| faub | 0.00491 | -0.34950915 | 0 | 0.109 | 1 | 4 |
| idh3g | 0.00491 | -0.29958774 | 0 | 0.109 | 1 | 4 |
| ak1 | 0.00606 | 0.54200368 | 0.258 | 0.043 | 1 | 4 |
| amy2a | 0.00708 | 0.374993163 | 0.113 | 0 | 1 | 4 |
| atf1 | 0.00708 | 0.292771369 | 0.113 | 0 | 1 | 4 |
| cel.2 | 0.00708 | 0.245493018 | 0.113 | 0 | 1 | 4 |
| gamt | 0.00708 | 0.239754175 | 0.113 | 0 | 1 | 4 |
| mycbp | 0.00708 | 0.39622851 | 0.113 | 0 | 1 | 4 |
| prss59.1 | 0.00708 | 0.48095031 | 0.113 | 0 | 1 | 4 |
| spink4 | 0.00708 | 0.279314807 | 0.113 | 0 | 1 | 4 |
| zgc:152791 | 0.00708 | 0.333298601 | 0.113 | 0 | 1 | 4 |
| mt-cyb | 0.00728 | -0.27647626 | 0.145 | 0.348 | 1 | 4 |
| rhoaa | 0.00821 | 0.479055671 | 0.274 | 0.065 | 1 | 4 |
| rpl23a | 0.00858 | 0.240230268 | 0.806 | 0.761 | 1 | 4 |
| cpne1 | 0.009 | 0.394225189 | 0.226 | 0.043 | 1 | 4 |
| hnrnpa0b | 0.00908 | 0.544350144 | 0.355 | 0.109 | 1 | 4 |
| npsn | 0.01028 | 0.508765906 | 0.194 | 0.022 | 1 | 4 |
| serpinb1l3 | 0.01072 | 0.497986794 | 0.194 | 0.022 | 1 | 4 |
| uqcrfs1 | 0.01122 | 0.43388502 | 0.194 | 0.022 | 1 | 4 |
| hnrnpa0l | 0.01145 | -0.21010916 | 0.565 | 0.609 | 1 | 4 |
| fam49ba | 0.0116 | 0.479980834 | 0.532 | 0.261 | 1 | 4 |
| rpl19 | 0.01231 | 0.197948803 | 0.984 | 0.913 | 1 | 4 |
| nsfl1c | 0.01321 | -0.28734351 | 0 | 0.087 | 1 | 4 |
| si:dkey-9k7.3 | 0.01321 | -0.24517312 | 0 | 0.087 | 1 | 4 |
| nt5dc2 | 0.01321 | -0.30298172 | 0 | 0.087 | 1 | 4 |
| gmfg | 0.01321 | -0.2664037 | 0 | 0.087 | 1 | 4 |
| pdcl | 0.01321 | -0.326562 | 0 | 0.087 | 1 | 4 |
| dnph1 | 0.01321 | -0.26461277 | 0 | 0.087 | 1 | 4 |
| cdk2ap2 | 0.01321 | -0.32402661 | 0 | 0.087 | 1 | 4 |
| cpa5 | 0.01362 | 0.274446798 | 0.097 | 0 | 1 | 4 |
| erp27 | 0.01362 | 0.166050223 | 0.097 | 0 | 1 | 4 |
| fhit | 0.01362 | 0.276665145 | 0.097 | 0 | 1 | 4 |
| nkl.4 | 0.01362 | 0.468786741 | 0.097 | 0 | 1 | 4 |
| tnrc6b | 0.01362 | 0.283723982 | 0.097 | 0 | 1 | 4 |
| hsdl2 | 0.01362 | 0.286547404 | 0.097 | 0 | 1 | 4 |
| si:ch211-191j22.3 | 0.01362 | 0.27844537 | 0.097 | 0 | 1 | 4 |
| zgc:112160 | 0.01362 | 0.40582514 | 0.097 | 0 | 1 | 4 |
| ywhae1 | 0.01394 | -0.55950529 | 0.21 | 0.435 | 1 | 4 |
| rpl30 | 0.01638 | 0.242080472 | 0.935 | 0.826 | 1 | 4 |
| ccdc124 | 0.01745 | 0.380432963 | 0.242 | 0.065 | 1 | 4 |
| rpl27a | 0.01887 | -0.41455882 | 0.403 | 0.587 | 1 | 4 |
| mt-nd4 | 0.02071 | -0.55240301 | 0.113 | 0.326 | 1 | 4 |
| zgc:162730 | 0.02275 | -0.32845095 | 0.5 | 0.609 | 1 | 4 |

| | | | | | | |
|---|---|---|---|---|---|---|
| si:rp71-45k5.4 | 0.02304 | -0.39539027 | 0.016 | 0.152 | 1 | 4 |
| gapdhs | 0.02492 | 0.482585648 | 0.565 | 0.304 | 1 | 4 |
| calr3a | 0.02623 | 0.235106274 | 0.081 | 0 | 1 | 4 |
| ctrb1 | 0.02623 | 0.391983327 | 0.081 | 0 | 1 | 4 |
| dnajc5gb | 0.02623 | 0.254980087 | 0.081 | 0 | 1 | 4 |
| etfb | 0.02623 | 0.227494452 | 0.081 | 0 | 1 | 4 |
| fkbp5 | 0.02623 | 0.214617002 | 0.081 | 0 | 1 | 4 |
| gstm.1 | 0.02623 | 0.198867824 | 0.081 | 0 | 1 | 4 |
| hsf2 | 0.02623 | 0.227720915 | 0.081 | 0 | 1 | 4 |
| npdc1a | 0.02623 | 0.231068598 | 0.081 | 0 | 1 | 4 |
| prss1 | 0.02623 | 0.394231736 | 0.081 | 0 | 1 | 4 |
| si:dkey-53k12.1 | 0.02623 | 0.311433322 | 0.081 | 0 | 1 | 4 |
| si:zfos-1192g2.3 | 0.02623 | 0.226993844 | 0.081 | 0 | 1 | 4 |
| sycn.2 | 0.02623 | 0.231039959 | 0.081 | 0 | 1 | 4 |
| taf12 | 0.02623 | 0.230200632 | 0.081 | 0 | 1 | 4 |
| dock8 | 0.02623 | 0.268057084 | 0.081 | 0 | 1 | 4 |
| st13 | 0.02623 | 0.193803961 | 0.081 | 0 | 1 | 4 |
| zgc:136461 | 0.02623 | 0.338343836 | 0.081 | 0 | 1 | 4 |
| cpa1 | 0.02623 | 0.191097075 | 0.081 | 0 | 1 | 4 |
| emg1 | 0.02623 | 0.255166027 | 0.081 | 0 | 1 | 4 |
| lsm12b | 0.02623 | 0.244401159 | 0.081 | 0 | 1 | 4 |
| ndufab1a | 0.02623 | 0.198615348 | 0.081 | 0 | 1 | 4 |
| sumo1 | 0.02623 | 0.256391686 | 0.081 | 0 | 1 | 4 |
| birc5a | 0.02623 | 0.22283458 | 0.081 | 0 | 1 | 4 |
| gnb1b | 0.02623 | 0.222411741 | 0.081 | 0 | 1 | 4 |
| nudcd3 | 0.02623 | 0.221871377 | 0.081 | 0 | 1 | 4 |
| ophn1 | 0.02623 | 0.221101679 | 0.081 | 0 | 1 | 4 |
| si:ch211-240l19.5 | 0.02623 | 0.283092303 | 0.081 | 0 | 1 | 4 |
| tcea1 | 0.02623 | 0.218203905 | 0.081 | 0 | 1 | 4 |
| selenow1 | 0.02663 | 0.385525186 | 0.468 | 0.239 | 1 | 4 |
| dynlrb1 | 0.02686 | 0.348008099 | 0.161 | 0.022 | 1 | 4 |
| ncl | 0.02794 | -0.46993181 | 0.097 | 0.239 | 1 | 4 |
| cct2 | 0.02855 | 0.362028992 | 0.323 | 0.13 | 1 | 4 |
| RPL37A | 0.02862 | -0.39764717 | 0.016 | 0.13 | 1 | 4 |
| rpl35 | 0.03126 | 0.207410034 | 0.952 | 0.891 | 1 | 4 |
| DDX17 | 0.03566 | -0.20889621 | 0 | 0.065 | 1 | 4 |
| LOC108191465 | 0.03566 | -0.21492196 | 0 | 0.065 | 1 | 4 |
| ascc1 | 0.03566 | -0.19885203 | 0 | 0.065 | 1 | 4 |
| cherp | 0.03566 | -0.23022793 | 0 | 0.065 | 1 | 4 |
| prpf39 | 0.03566 | -0.1962178 | 0 | 0.065 | 1 | 4 |
| rab20 | 0.03566 | -0.23063773 | 0 | 0.065 | 1 | 4 |
| rpz2 | 0.03566 | -0.24105545 | 0 | 0.065 | 1 | 4 |
| tceanc2 | 0.03566 | -0.21424799 | 0 | 0.065 | 1 | 4 |
| vps35 | 0.03566 | -0.23588857 | 0 | 0.065 | 1 | 4 |
| BX322618.1 | 0.03566 | -0.21952857 | 0 | 0.065 | 1 | 4 |
| XLOC-011450 | 0.03566 | -0.18313601 | 0 | 0.065 | 1 | 4 |
| phf19 | 0.03566 | -0.20089372 | 0 | 0.065 | 1 | 4 |
| CABZ01044053.1 | 0.03566 | -0.22794657 | 0 | 0.065 | 1 | 4 |

| | | | | | | |
|---|---|---|---|---|---|---|
| CABZ01046432.1 | 0.03566 | -0.21055916 | 0 | 0.065 | 1 | 4 |
| LOC108190717 | 0.03566 | -0.239809 | 0 | 0.065 | 1 | 4 |
| XLOC-043760 | 0.03566 | -0.27817889 | 0 | 0.065 | 1 | 4 |
| erp44 | 0.03566 | -0.23990482 | 0 | 0.065 | 1 | 4 |
| pfkpa | 0.03566 | -0.230907 | 0 | 0.065 | 1 | 4 |
| pigm | 0.03566 | -0.23062017 | 0 | 0.065 | 1 | 4 |
| prpf31 | 0.03566 | -0.21091926 | 0 | 0.065 | 1 | 4 |
| si:ch211-105j21.9 | 0.03566 | -0.23872668 | 0 | 0.065 | 1 | 4 |
| trnau1apb | 0.03566 | -0.26311717 | 0 | 0.065 | 1 | 4 |
| ubtf | 0.03566 | -0.18054889 | 0 | 0.065 | 1 | 4 |
| zgc:92664 | 0.03566 | -0.18687269 | 0 | 0.065 | 1 | 4 |
| acox1 | 0.03566 | -0.22652368 | 0 | 0.065 | 1 | 4 |
| sh3bgrl3 | 0.03818 | 0.448119051 | 0.274 | 0.087 | 1 | 4 |
| atp5pb | 0.03936 | 0.425631998 | 0.306 | 0.109 | 1 | 4 |
| mrpl12 | 0.03952 | -0.40375158 | 0.016 | 0.13 | 1 | 4 |
| ndufb6 | 0.04057 | -0.41582136 | 0.032 | 0.174 | 1 | 4 |
| vaspa | 0.04146 | -0.29748061 | 0.565 | 0.696 | 1 | 4 |
| CR855389.1 | 0.04548 | 0.30051408 | 0.145 | 0.022 | 1 | 4 |
| rpl26 | 0.04563 | 0.136516146 | 1 | 0.978 | 1 | 4 |
| sumo3a | 0.0461 | 0.375488245 | 0.194 | 0.043 | 1 | 4 |
| ncalda | 0.05067 | 0.216861056 | 0.065 | 0 | 1 | 4 |
| XLOC-013964 | 0.05067 | 0.205287766 | 0.065 | 0 | 1 | 4 |
| jun | 0.05067 | 0.133736101 | 0.065 | 0 | 1 | 4 |
| ccdc9 | 0.05067 | 0.19900162 | 0.065 | 0 | 1 | 4 |
| ptena | 0.05067 | 0.227243845 | 0.065 | 0 | 1 | 4 |
| caap1 | 0.05067 | 0.196137588 | 0.065 | 0 | 1 | 4 |
| cab39l1 | 0.05067 | 0.177578483 | 0.065 | 0 | 1 | 4 |
| ccnb3 | 0.05067 | 0.16514168 | 0.065 | 0 | 1 | 4 |
| cebpb | 0.05067 | 0.181826684 | 0.065 | 0 | 1 | 4 |
| cel.1 | 0.05067 | 0.1975259 | 0.065 | 0 | 1 | 4 |
| chico | 0.05067 | 0.232924946 | 0.065 | 0 | 1 | 4 |
| cwc25 | 0.05067 | 0.200436947 | 0.065 | 0 | 1 | 4 |
| def6c | 0.05067 | 0.206542426 | 0.065 | 0 | 1 | 4 |
| dph5 | 0.05067 | 0.202949044 | 0.065 | 0 | 1 | 4 |
| gpr132b | 0.05067 | 0.226169 | 0.065 | 0 | 1 | 4 |
| knstrn | 0.05067 | 0.161348074 | 0.065 | 0 | 1 | 4 |
| lxn | 0.05067 | 0.152488463 | 0.065 | 0 | 1 | 4 |
| ndfip2 | 0.05067 | 0.195306092 | 0.065 | 0 | 1 | 4 |
| nfkbiaa | 0.05067 | 0.200881054 | 0.065 | 0 | 1 | 4 |
| prelid1a | 0.05067 | 0.182123569 | 0.065 | 0 | 1 | 4 |
| psmc5 | 0.05067 | 0.222025045 | 0.065 | 0 | 1 | 4 |
| si:ch211-269k10.2 | 0.05067 | 0.203898738 | 0.065 | 0 | 1 | 4 |
| smim4 | 0.05067 | 0.214800914 | 0.065 | 0 | 1 | 4 |
| spink2.3 | 0.05067 | 0.169162955 | 0.065 | 0 | 1 | 4 |
| spsb4b | 0.05067 | 0.166067064 | 0.065 | 0 | 1 | 4 |
| srsf4 | 0.05067 | 0.240572387 | 0.065 | 0 | 1 | 4 |
| tcta | 0.05067 | 0.17323757 | 0.065 | 0 | 1 | 4 |
| vps36 | 0.05067 | 0.188569043 | 0.065 | 0 | 1 | 4 |

| | | | | | | |
|---|---|---|---|---|---|---|
| agpat5 | 0.05067 | 0.217560671 | 0.065 | 0 | 1 | 4 |
| bach1a | 0.05067 | 0.228927371 | 0.065 | 0 | 1 | 4 |
| acot13 | 0.05067 | 0.196933803 | 0.065 | 0 | 1 | 4 |
| ccdc43 | 0.05067 | 0.19933579 | 0.065 | 0 | 1 | 4 |
| cd44b | 0.05067 | 0.226467754 | 0.065 | 0 | 1 | 4 |
| degs1 | 0.05067 | 0.189194314 | 0.065 | 0 | 1 | 4 |
| dnajc8 | 0.05067 | 0.159438857 | 0.065 | 0 | 1 | 4 |
| eif4ebp3 | 0.05067 | 0.167164139 | 0.065 | 0 | 1 | 4 |
| fdx1b | 0.05067 | 0.18405856 | 0.065 | 0 | 1 | 4 |
| gatc | 0.05067 | 0.197219892 | 0.065 | 0 | 1 | 4 |
| gmppaa | 0.05067 | 0.198284494 | 0.065 | 0 | 1 | 4 |
| mrpl19 | 0.05067 | 0.211134184 | 0.065 | 0 | 1 | 4 |
| nkl.3 | 0.05067 | 0.331723747 | 0.065 | 0 | 1 | 4 |
| pop7 | 0.05067 | 0.177846922 | 0.065 | 0 | 1 | 4 |
| ptges3a | 0.05067 | 0.17190626 | 0.065 | 0 | 1 | 4 |
| rbb4l | 0.05067 | 0.173097377 | 0.065 | 0 | 1 | 4 |
| rin3 | 0.05067 | 0.228130324 | 0.065 | 0 | 1 | 4 |
| si:dkey-14d8.7 | 0.05067 | 0.150327306 | 0.065 | 0 | 1 | 4 |
| si:dkey-7c18.24 | 0.05067 | 0.184692537 | 0.065 | 0 | 1 | 4 |
| slc38a5b | 0.05067 | 0.15695052 | 0.065 | 0 | 1 | 4 |
| slirp | 0.05067 | 0.179971081 | 0.065 | 0 | 1 | 4 |
| spc24 | 0.05067 | 0.200549412 | 0.065 | 0 | 1 | 4 |
| srsf10b | 0.05067 | 0.185184534 | 0.065 | 0 | 1 | 4 |
| srsf6b | 0.05067 | 0.204787724 | 0.065 | 0 | 1 | 4 |
| tacc3 | 0.05067 | 0.196930599 | 0.065 | 0 | 1 | 4 |
| thoc1 | 0.05067 | 0.1707623 | 0.065 | 0 | 1 | 4 |
| zbtb2b | 0.05067 | 0.194347172 | 0.065 | 0 | 1 | 4 |
| sec61b | 0.05168 | 0.277146482 | 0.435 | 0.239 | 1 | 4 |
| ier3ip1 | 0.05237 | -0.34087633 | 0.016 | 0.13 | 1 | 4 |
| yrk | 0.05324 | 0.467832117 | 0.323 | 0.13 | 1 | 4 |
| prmt1 | 0.05399 | 0.361239313 | 0.145 | 0.022 | 1 | 4 |
| ahcy | 0.05425 | -0.3603503 | 0.048 | 0.196 | 1 | 4 |
| arhgdig | 0.05425 | -0.408521 | 0.435 | 0.652 | 1 | 4 |
| ikzf2 | 0.05769 | 0.273509127 | 0.226 | 0.087 | 1 | 4 |
| sub1a | 0.05883 | -0.39386017 | 0.145 | 0.261 | 1 | 4 |
| mt-nd1 | 0.05971 | -0.21056622 | 0.048 | 0.152 | 1 | 4 |
| pgam1a | 0.06058 | 0.349526136 | 0.145 | 0.022 | 1 | 4 |
| ndufb8 | 0.06325 | 0.427502464 | 0.258 | 0.087 | 1 | 4 |
| LOC100536213 | 0.06444 | 0.429485448 | 0.565 | 0.348 | 1 | 4 |
| cox6b1 | 0.06524 | 0.33792283 | 0.242 | 0.087 | 1 | 4 |
| jpt2 | 0.06583 | -0.45239865 | 0.097 | 0.239 | 1 | 4 |
| CR753876.1 | 0.06806 | -0.30567828 | 0.29 | 0.37 | 1 | 4 |
| mt-atp6 | 0.07052 | -0.16201435 | 0.145 | 0.261 | 1 | 4 |
| eif4a1b | 0.07085 | 0.258112707 | 0.306 | 0.152 | 1 | 4 |
| gnb1a | 0.07156 | -0.40584274 | 0.113 | 0.239 | 1 | 4 |
| atp5mc3b | 0.07203 | 0.355689712 | 0.613 | 0.391 | 1 | 4 |
| tubb2b | 0.07346 | -0.48764882 | 0.161 | 0.326 | 1 | 4 |
| dad1 | 0.07416 | 0.231190627 | 0.161 | 0.043 | 1 | 4 |

| | | | | | | |
|---|---|---|---|---|---|---|
| tpd52l2b | 0.07453 | -0.33574444 | 0.032 | 0.152 | 1 | 4 |
| tomm5 | 0.07633 | 0.315569716 | 0.177 | 0.043 | 1 | 4 |
| sesn1 | 0.07636 | 0.352211519 | 0.242 | 0.087 | 1 | 4 |
| CR381673.1 | 0.07638 | -0.45853611 | 0.129 | 0.283 | 1 | 4 |
| sri | 0.07651 | -0.38664782 | 0.145 | 0.261 | 1 | 4 |
| rhoab | 0.07864 | -0.29114322 | 0.21 | 0.283 | 1 | 4 |
| arpc5a | 0.07912 | 0.260457854 | 0.29 | 0.152 | 1 | 4 |
| COX3 | 0.07968 | -0.2134868 | 0.258 | 0.413 | 1 | 4 |
| gpx4b | 0.08099 | 0.375300339 | 0.274 | 0.109 | 1 | 4 |
| csnk2b | 0.08142 | -0.42120994 | 0.129 | 0.304 | 1 | 4 |
| gtpbp4 | 0.08409 | 0.42251091 | 0.226 | 0.087 | 1 | 4 |
| si:ch211-220f21.3 | 0.08829 | 0.32273919 | 0.339 | 0.174 | 1 | 4 |
| rpl10a | 0.08963 | 0.155513115 | 0.839 | 0.804 | 1 | 4 |
| rpl11 | 0.08987 | 0.16116696 | 0.984 | 0.913 | 1 | 4 |
| taf9 | 0.0903 | 0.295534404 | 0.129 | 0.022 | 1 | 4 |
| zgc:162944 | 0.09038 | 0.257553139 | 0.403 | 0.239 | 1 | 4 |
| erh | 0.09078 | 0.191170883 | 0.355 | 0.217 | 1 | 4 |
| cdkn1bb | 0.09223 | 0.317139747 | 0.194 | 0.065 | 1 | 4 |
| srgn | 0.0934 | 0.402843265 | 0.387 | 0.196 | 1 | 4 |
| hint1 | 0.09349 | 0.278345499 | 0.161 | 0.043 | 1 | 4 |
| ndufb9 | 0.09449 | 0.302944954 | 0.258 | 0.109 | 1 | 4 |
| sf3b6 | 0.09623 | 0.234836887 | 0.177 | 0.065 | 1 | 4 |
| ppm1g | 0.09726 | -0.27741909 | 0.016 | 0.109 | 1 | 4 |
| npc2 | 0.09753 | 0.320556548 | 0.129 | 0.022 | 1 | 4 |
| fcer1gl | 0.0999 | 0.191751107 | 0.935 | 0.804 | 1 | 4 |
| sec11a | 0.10038 | 0.319767542 | 0.129 | 0.022 | 1 | 4 |
| taf15 | 0.10056 | -0.3205992 | 0.065 | 0.196 | 1 | 4 |
| si:zfos-169g10.3 | 0.10087 | 0.324739642 | 0.129 | 0.022 | 1 | 4 |
| aimp2 | 0.10096 | 0.193621803 | 0.113 | 0.022 | 1 | 4 |
| coro1a | 0.10211 | -0.17974911 | 0.839 | 0.935 | 1 | 4 |
| wdr1 | 0.10551 | -0.25336094 | 0.306 | 0.37 | 1 | 4 |
| sec61g | 0.106 | 0.188736968 | 0.742 | 0.565 | 1 | 4 |
| zgc:114188 | 0.10639 | 0.203520641 | 0.871 | 0.783 | 1 | 4 |
| dut | 0.10665 | -0.32509414 | 0.032 | 0.109 | 1 | 4 |
| btg1 | 0.10671 | 0.132687952 | 0.323 | 0.217 | 1 | 4 |
| rpl31 | 0.10687 | 0.139303344 | 0.935 | 0.804 | 1 | 4 |
| hck | 0.10871 | -0.33480201 | 0.032 | 0.109 | 1 | 4 |
| pgk1 | 0.10933 | -0.30248688 | 0.016 | 0.109 | 1 | 4 |
| snx5 | 0.10959 | -0.3136782 | 0.016 | 0.109 | 1 | 4 |
| cops4 | 0.11012 | 0.205101037 | 0.113 | 0.022 | 1 | 4 |
| rps14 | 0.11078 | 0.114311718 | 0.984 | 0.891 | 1 | 4 |
| smfn | 0.11101 | -0.31428844 | 0.016 | 0.109 | 1 | 4 |
| srsf5a | 0.11253 | 0.318736143 | 0.194 | 0.065 | 1 | 4 |
| hcls1 | 0.11358 | -0.25073815 | 0.339 | 0.413 | 1 | 4 |
| rpl28 | 0.11549 | 0.150993631 | 0.952 | 0.891 | 1 | 4 |
| ybx1 | 0.11568 | 0.163156103 | 0.806 | 0.652 | 1 | 4 |
| rps9 | 0.11862 | 0.163875163 | 0.919 | 0.87 | 1 | 4 |
| vps26c | 0.11865 | -0.3017181 | 0.032 | 0.13 | 1 | 4 |

| | | | | | | |
|---|---|---|---|---|---|---|
| hnrnpub | 0.12149 | -0.28743358 | 0.016 | 0.087 | 1 | 4 |
| rpl15 | 0.12279 | 0.171675476 | 0.952 | 0.848 | 1 | 4 |
| rnaset2l | 0.12779 | 0.397833652 | 0.258 | 0.109 | 1 | 4 |
| eif4ebp2 | 0.12858 | 0.239259646 | 0.306 | 0.174 | 1 | 4 |
| cox7a2a | 0.13137 | 0.295611034 | 0.274 | 0.13 | 1 | 4 |
| chac1 | 0.13174 | -0.28033145 | 0.016 | 0.087 | 1 | 4 |
| cdc42 | 0.13488 | 0.385194657 | 0.371 | 0.196 | 1 | 4 |
| arpc2 | 0.13741 | 0.253899663 | 0.806 | 0.652 | 1 | 4 |
| pxna | 0.14052 | 0.377000463 | 0.194 | 0.065 | 1 | 4 |
| mhc2dab | 0.14679 | 0.266462159 | 0.145 | 0.043 | 1 | 4 |
| oaz1b | 0.14862 | -0.28293891 | 0.274 | 0.37 | 1 | 4 |
| mbnl1 | 0.14906 | 0.257799669 | 0.21 | 0.087 | 1 | 4 |
| aldoab | 0.1491 | 0.239823836 | 0.113 | 0.022 | 1 | 4 |
| dbi | 0.15098 | -0.11162962 | 0.161 | 0.152 | 1 | 4 |
| alox5a | 0.15114 | -0.2742306 | 0.016 | 0.087 | 1 | 4 |
| mbpb | 0.15122 | 0.262946278 | 0.113 | 0.022 | 1 | 4 |
| cyc1 | 0.15265 | 0.172444615 | 0.21 | 0.109 | 1 | 4 |
| rpl23 | 0.15555 | 0.113406148 | 0.968 | 0.87 | 1 | 4 |
| ak2 | 0.15737 | 0.315658885 | 0.274 | 0.13 | 1 | 4 |
| nedd8 | 0.15861 | 0.249252941 | 0.113 | 0.022 | 1 | 4 |
| capgb | 0.16067 | 0.297300128 | 0.113 | 0.022 | 1 | 4 |
| cxcr4b | 0.16082 | 0.260846095 | 0.145 | 0.043 | 1 | 4 |
| abcf1 | 0.16342 | 0.293438252 | 0.113 | 0.022 | 1 | 4 |
| zgc:112356 | 0.16353 | 0.242762821 | 0.145 | 0.043 | 1 | 4 |
| ndufb3 | 0.16614 | -0.37554565 | 0.129 | 0.261 | 1 | 4 |
| zgc:153867 | 0.16789 | 0.245392396 | 0.79 | 0.63 | 1 | 4 |
| tefa | 0.16862 | -0.26029884 | 0.016 | 0.087 | 1 | 4 |
| cdc37 | 0.16944 | -0.25732655 | 0.016 | 0.087 | 1 | 4 |
| lsm6 | 0.17245 | -0.24127929 | 0.081 | 0.13 | 1 | 4 |
| eef1da | 0.17642 | 0.291661942 | 0.5 | 0.326 | 1 | 4 |
| btf3 | 0.17827 | -0.30432696 | 0.468 | 0.63 | 1 | 4 |
| fkbp1aa | 0.17862 | -0.32423515 | 0.065 | 0.152 | 1 | 4 |
| cyth4a | 0.17928 | -0.25688738 | 0.016 | 0.087 | 1 | 4 |
| rbm4.3 | 0.17954 | 0.252773996 | 0.226 | 0.109 | 1 | 4 |
| atp5f1d | 0.18404 | 0.278452843 | 0.21 | 0.087 | 1 | 4 |
| wipf1a | 0.18563 | -0.13992578 | 0.371 | 0.391 | 1 | 4 |
| eif5 | 0.18655 | 0.187769418 | 0.097 | 0.022 | 1 | 4 |
| fahd2a | 0.18818 | -0.23915536 | 0.016 | 0.087 | 1 | 4 |
| imp3 | 0.18864 | -0.16926542 | 0.032 | 0.109 | 1 | 4 |
| mt-nd3 | 0.18937 | -0.2651224 | 0.177 | 0.217 | 1 | 4 |
| banf1 | 0.19173 | -0.22735567 | 0.016 | 0.087 | 1 | 4 |
| nptna | 0.19186 | 0.299587034 | 0.145 | 0.043 | 1 | 4 |
| XLOC-043857 | 0.19233 | -0.28283578 | 0.016 | 0.087 | 1 | 4 |
| nusap1 | 0.1931 | 0.200665841 | 0.097 | 0.022 | 1 | 4 |
| rfc5 | 0.1934 | -0.22936657 | 0.016 | 0.087 | 1 | 4 |
| zfyve19 | 0.19489 | -0.23528629 | 0.016 | 0.087 | 1 | 4 |
| ndufa2 | 0.19549 | 0.286479225 | 0.145 | 0.043 | 1 | 4 |
| naca | 0.19571 | -0.17023589 | 0.758 | 0.891 | 1 | 4 |

| | | | | | | |
|---|---|---|---|---|---|---|
| arf6b | 0.19792 | 0.285365633 | 0.177 | 0.065 | 1 | 4 |
| chd4a | 0.19832 | -0.19992098 | 0.016 | 0.087 | 1 | 4 |
| snrpd2 | 0.20005 | -0.32200662 | 0.097 | 0.196 | 1 | 4 |
| nipsnap2 | 0.20034 | 0.264225374 | 0.097 | 0.022 | 1 | 4 |
| rplp0 | 0.20049 | 0.147986673 | 0.903 | 0.87 | 1 | 4 |
| itga1 | 0.20235 | 0.207285205 | 0.21 | 0.109 | 1 | 4 |
| LOC110437917 | 0.20329 | -0.27077701 | 0.016 | 0.087 | 1 | 4 |
| sept6 | 0.20423 | 0.173221841 | 0.355 | 0.239 | 1 | 4 |
| ndufs5 | 0.20487 | 0.174006441 | 0.097 | 0.022 | 1 | 4 |
| oaz1a | 0.2054 | 0.263041061 | 0.435 | 0.283 | 1 | 4 |
| hnrnpk | 0.20556 | 0.157458271 | 0.177 | 0.087 | 1 | 4 |
| ube2d2 | 0.20604 | 0.300106289 | 0.177 | 0.065 | 1 | 4 |
| pfdn4 | 0.20611 | 0.197020714 | 0.161 | 0.065 | 1 | 4 |
| lgals17 | 0.20763 | -0.21515383 | 0.016 | 0.087 | 1 | 4 |
| si:ch211-214p16.1 | 0.20837 | -0.21394549 | 0.71 | 0.848 | 1 | 4 |
| mtmr12 | 0.21032 | 0.187425602 | 0.097 | 0.022 | 1 | 4 |
| lcp1 | 0.21116 | -0.35204108 | 0.113 | 0.217 | 1 | 4 |
| NC-002333.4 | 0.21202 | 0.159618556 | 0.194 | 0.217 | 1 | 4 |
| eif1axb | 0.21595 | 0.24700209 | 0.226 | 0.109 | 1 | 4 |
| gdi2 | 0.21642 | -0.28135634 | 0.161 | 0.239 | 1 | 4 |
| CR318588.1 | 0.21852 | -0.17362917 | 0.242 | 0.261 | 1 | 4 |
| CU856520.1 | 0.21897 | -0.27500607 | 0.016 | 0.087 | 1 | 4 |
| snrnp27 | 0.21939 | -0.21293543 | 0.016 | 0.087 | 1 | 4 |
| rps8a | 0.21999 | 0.118729314 | 0.952 | 0.913 | 1 | 4 |
| nop56 | 0.22063 | 0.258316251 | 0.194 | 0.087 | 1 | 4 |
| cfl1 | 0.22146 | -0.10043066 | 0.903 | 0.935 | 1 | 4 |
| nutf2l | 0.22248 | -0.25138354 | 0.016 | 0.087 | 1 | 4 |
| ppp6r3 | 0.22272 | -0.25368479 | 0.016 | 0.087 | 1 | 4 |
| ptmaa | 0.22385 | 0.164568368 | 0.887 | 0.761 | 1 | 4 |
| mphosph8 | 0.22549 | 0.22472414 | 0.097 | 0.022 | 1 | 4 |
| trappc2l | 0.22558 | -0.24213551 | 0.032 | 0.087 | 1 | 4 |
| ubap2b | 0.22651 | 0.178981913 | 0.097 | 0.022 | 1 | 4 |
| trappc1 | 0.22842 | -0.27566856 | 0.048 | 0.13 | 1 | 4 |
| cd81a | 0.22849 | 0.216736099 | 0.097 | 0.022 | 1 | 4 |
| hprt1 | 0.23026 | -0.32790663 | 0.081 | 0.174 | 1 | 4 |
| ndufa5 | 0.23256 | 0.198431719 | 0.129 | 0.043 | 1 | 4 |
| snrpa | 0.23669 | 0.191494392 | 0.097 | 0.022 | 1 | 4 |
| commd7 | 0.23753 | 0.215410334 | 0.097 | 0.022 | 1 | 4 |
| def6a | 0.23876 | 0.215092489 | 0.097 | 0.022 | 1 | 4 |
| cbx5 | 0.23905 | 0.222210488 | 0.097 | 0.022 | 1 | 4 |
| gstr | 0.24002 | -0.28644094 | 0.048 | 0.13 | 1 | 4 |
| ppp3r1b | 0.24077 | 0.213143231 | 0.097 | 0.022 | 1 | 4 |
| mrpl51 | 0.24129 | 0.210154426 | 0.097 | 0.022 | 1 | 4 |
| anxa4 | 0.2418 | 0.229692747 | 0.097 | 0.022 | 1 | 4 |
| mid1ip1l | 0.24325 | 0.228385136 | 0.097 | 0.022 | 1 | 4 |
| mt-co2 | 0.24394 | -0.24015197 | 0.242 | 0.37 | 1 | 4 |
| reep5 | 0.24571 | 0.220895059 | 0.097 | 0.022 | 1 | 4 |
| si:dkey-222f2.7 | 0.24646 | 0.305865136 | 0.258 | 0.13 | 1 | 4 |

| | | | | | | |
|---|---|---|---|---|---|---|
| glo1 | 0.24769 | 0.311052586 | 0.258 | 0.13 | 1 | 4 |
| si:ch73-380l3.4 | 0.24805 | -0.15778633 | 0.113 | 0.13 | 1 | 4 |
| tspan14 | 0.2482 | 0.127923095 | 0.081 | 0.022 | 1 | 4 |
| lrrfip1a | 0.24979 | 0.106649004 | 0.129 | 0.065 | 1 | 4 |
| arf6a | 0.25013 | -0.25184696 | 0.129 | 0.196 | 1 | 4 |
| spink2.1 | 0.25049 | -0.14208712 | 0.081 | 0.087 | 1 | 4 |
| ndufa11 | 0.25105 | 0.210074475 | 0.097 | 0.022 | 1 | 4 |
| csnk1a1 | 0.2511 | -0.25547464 | 0.048 | 0.109 | 1 | 4 |
| LOC101884267 | 0.25142 | 0.258451048 | 0.097 | 0.022 | 1 | 4 |
| pcbp2 | 0.25165 | 0.205713906 | 0.097 | 0.022 | 1 | 4 |
| pik3ap1 | 0.25295 | 0.204445062 | 0.097 | 0.022 | 1 | 4 |
| dctn3 | 0.25482 | 0.239603861 | 0.097 | 0.022 | 1 | 4 |
| si:ch211-215e19.4 | 0.25544 | 0.14928714 | 0.081 | 0.022 | 1 | 4 |
| cenpx | 0.25602 | 0.154178363 | 0.113 | 0.043 | 1 | 4 |
| lyrm4 | 0.25865 | -0.25958045 | 0.032 | 0.109 | 1 | 4 |
| zgc:193541 | 0.25941 | 0.243370528 | 0.403 | 0.261 | 1 | 4 |
| rab5c | 0.25993 | 0.109659086 | 0.129 | 0.065 | 1 | 4 |
| sftpba | 0.26237 | -0.27265042 | 0.403 | 0.522 | 1 | 4 |
| rps26 | 0.2627 | 0.202116493 | 0.774 | 0.674 | 1 | 4 |
| fam49bb | 0.26282 | 0.285273778 | 0.129 | 0.043 | 1 | 4 |
| ndufa9a | 0.2653 | -0.21105407 | 0.048 | 0.087 | 1 | 4 |
| hmgb2b | 0.26808 | 0.160851398 | 0.323 | 0.217 | 1 | 4 |
| skp1 | 0.27011 | 0.178325211 | 0.226 | 0.13 | 1 | 4 |
| si:ch211-262e15.1 | 0.2708 | -0.26913335 | 0.048 | 0.13 | 1 | 4 |
| AL954142.1 | 0.27295 | -0.24055331 | 0.097 | 0.152 | 1 | 4 |
| khsrp | 0.27384 | 0.12897327 | 0.081 | 0.022 | 1 | 4 |
| il7r | 0.27632 | 0.121622396 | 0.081 | 0.022 | 1 | 4 |
| LOC103909115 | 0.28116 | 0.173475589 | 0.323 | 0.217 | 1 | 4 |
| cops8 | 0.28477 | 0.230627325 | 0.129 | 0.043 | 1 | 4 |
| psmb5 | 0.28605 | -0.20997323 | 0.016 | 0.065 | 1 | 4 |
| si:ch211-119e14.1 | 0.28724 | 0.220547011 | 0.129 | 0.043 | 1 | 4 |
| kctd5a | 0.28759 | 0.216992382 | 0.129 | 0.043 | 1 | 4 |
| ndufb10 | 0.28897 | -0.22611179 | 0.065 | 0.109 | 1 | 4 |
| si:ch1073-429i10.3.1 | 0.28958 | 0.205696582 | 0.306 | 0.196 | 1 | 4 |
| ccdc28a | 0.29065 | -0.125896 | 0.145 | 0.152 | 1 | 4 |
| rps27a | 0.29402 | 0.127578939 | 0.903 | 0.848 | 1 | 4 |
| arpc4 | 0.29486 | 0.20528132 | 0.71 | 0.565 | 1 | 4 |
| rbx1 | 0.29522 | 0.286001115 | 0.274 | 0.152 | 1 | 4 |
| ccng1 | 0.2954 | 0.270215893 | 0.274 | 0.152 | 1 | 4 |
| csk | 0.29711 | -0.22861157 | 0.274 | 0.348 | 1 | 4 |
| si:dkey-172o19.2 | 0.29727 | -0.26627626 | 0.081 | 0.152 | 1 | 4 |
| rac2 | 0.29929 | -0.17822832 | 0.581 | 0.717 | 1 | 4 |
| COX5B | 0.2993 | 0.291932553 | 0.403 | 0.261 | 1 | 4 |
| ube2nb | 0.30209 | 0.156131904 | 0.113 | 0.043 | 1 | 4 |
| atp1b1a | 0.30279 | -0.30142268 | 0.129 | 0.239 | 1 | 4 |
| selenof | 0.30513 | 0.109391953 | 0.226 | 0.152 | 1 | 4 |
| ebna1bp2 | 0.30678 | -0.19589924 | 0.016 | 0.065 | 1 | 4 |
| emd | 0.30701 | 0.131015459 | 0.081 | 0.022 | 1 | 4 |

| | | | | | | |
|---|---|---|---|---|---|---|
| hadh | 0.30751 | -0.20570876 | 0.032 | 0.087 | 1 | 4 |
| atp1a1a.1 | 0.30856 | -0.2609743 | 0.048 | 0.13 | 1 | 4 |
| pinx1 | 0.30898 | 0.134192748 | 0.081 | 0.022 | 1 | 4 |
| stmn1a | 0.31071 | -0.19828803 | 0.032 | 0.087 | 1 | 4 |
| vdac2 | 0.31077 | -0.22763096 | 0.258 | 0.326 | 1 | 4 |
| fcer1g | 0.31278 | 0.23837442 | 0.29 | 0.174 | 1 | 4 |
| mcl1a | 0.31293 | 0.190915547 | 0.726 | 0.587 | 1 | 4 |
| lamtor3 | 0.3169 | -0.21049753 | 0.016 | 0.065 | 1 | 4 |
| sh3kbp1 | 0.31695 | -0.21274421 | 0.081 | 0.13 | 1 | 4 |
| mgst1.2 | 0.31758 | 0.221255318 | 0.081 | 0.022 | 1 | 4 |
| actr2a | 0.31881 | -0.25642106 | 0.145 | 0.217 | 1 | 4 |
| seta | 0.31887 | 0.198351874 | 0.177 | 0.087 | 1 | 4 |
| elf1 | 0.3194 | -0.23442379 | 0.081 | 0.13 | 1 | 4 |
| celf2 | 0.31972 | -0.23842255 | 0.048 | 0.109 | 1 | 4 |
| prdx3 | 0.31985 | 0.120446731 | 0.113 | 0.043 | 1 | 4 |
| lamtor4 | 0.32009 | -0.18875176 | 0.016 | 0.065 | 1 | 4 |
| snrpb | 0.32088 | -0.24557885 | 0.21 | 0.283 | 1 | 4 |
| pggt1b | 0.3225 | -0.20474353 | 0.016 | 0.065 | 1 | 4 |
| tab1 | 0.32592 | 0.214013487 | 0.081 | 0.022 | 1 | 4 |
| wdr19 | 0.3274 | 0.217820695 | 0.081 | 0.022 | 1 | 4 |
| rps18 | 0.32749 | 0.106759027 | 0.968 | 0.913 | 1 | 4 |
| rps3 | 0.32789 | 0.129684329 | 0.887 | 0.826 | 1 | 4 |
| NDUFB1 | 0.32906 | -0.2881354 | 0.097 | 0.196 | 1 | 4 |
| snrpe | 0.3303 | -0.1950593 | 0.081 | 0.109 | 1 | 4 |
| ndufa8 | 0.3307 | 0.156783977 | 0.113 | 0.043 | 1 | 4 |
| alas1 | 0.33168 | 0.158060716 | 0.113 | 0.043 | 1 | 4 |
| atox1 | 0.33174 | 0.157331019 | 0.113 | 0.043 | 1 | 4 |
| cirbpa | 0.333 | 0.228856859 | 0.532 | 0.391 | 1 | 4 |
| eno3 | 0.33329 | -0.1722169 | 0.306 | 0.348 | 1 | 4 |
| gabarapa | 0.33466 | -0.23571645 | 0.129 | 0.196 | 1 | 4 |
| cops3 | 0.33613 | 0.205566594 | 0.081 | 0.022 | 1 | 4 |
| setb | 0.33643 | 0.136322703 | 0.306 | 0.217 | 1 | 4 |
| histh1l | 0.33823 | -0.29158169 | 0.129 | 0.217 | 1 | 4 |
| fdx1 | 0.33826 | 0.162565806 | 0.081 | 0.022 | 1 | 4 |
| rps10 | 0.34085 | 0.137259289 | 0.887 | 0.783 | 1 | 4 |
| zgc:100918 | 0.3422 | -0.20286105 | 0.016 | 0.065 | 1 | 4 |
| XLOC-025423 | 0.34298 | 0.133763488 | 0.258 | 0.174 | 1 | 4 |
| cnn2 | 0.34318 | 0.113151391 | 0.323 | 0.239 | 1 | 4 |
| cyb5a | 0.34352 | -0.1842991 | 0.016 | 0.065 | 1 | 4 |
| bod1 | 0.3462 | 0.182842736 | 0.081 | 0.022 | 1 | 4 |
| psmb10 | 0.34642 | 0.194400746 | 0.081 | 0.022 | 1 | 4 |
| cyfip1 | 0.34751 | 0.177107719 | 0.081 | 0.022 | 1 | 4 |
| hells | 0.34763 | -0.1997672 | 0.016 | 0.065 | 1 | 4 |
| cyb5r3 | 0.34914 | -0.21561072 | 0.032 | 0.087 | 1 | 4 |
| BOLA2 | 0.35157 | -0.19290618 | 0.032 | 0.087 | 1 | 4 |
| nop53 | 0.35424 | 0.104361329 | 0.177 | 0.109 | 1 | 4 |
| abracl | 0.35454 | 0.102953156 | 0.548 | 0.457 | 1 | 4 |
| gosr2 | 0.35971 | 0.193804862 | 0.081 | 0.022 | 1 | 4 |

| | | | | | | |
|---|---|---|---|---|---|---|
| pagr1 | 0.3606 | 0.190379283 | 0.081 | 0.022 | 1 | 4 |
| clta | 0.36098 | 0.150974789 | 0.113 | 0.043 | 1 | 4 |
| ube2v1 | 0.36147 | 0.195346639 | 0.226 | 0.13 | 1 | 4 |
| pycard | 0.3617 | 0.22679408 | 0.258 | 0.152 | 1 | 4 |
| mrps21 | 0.36304 | -0.192006 | 0.016 | 0.065 | 1 | 4 |
| abhd17c | 0.3648 | -0.17138677 | 0.016 | 0.065 | 1 | 4 |
| psme2 | 0.36662 | 0.15856778 | 0.081 | 0.022 | 1 | 4 |
| cmc1 | 0.36739 | 0.181596251 | 0.081 | 0.022 | 1 | 4 |
| BX323557.1 | 0.36795 | -0.17561849 | 0.016 | 0.065 | 1 | 4 |
| naa10 | 0.36818 | -0.11569192 | 0.161 | 0.174 | 1 | 4 |
| sept9b | 0.36944 | -0.15342659 | 0.065 | 0.087 | 1 | 4 |
| CR391910.2 | 0.36979 | -0.1814337 | 0.016 | 0.065 | 1 | 4 |
| cox8a | 0.37057 | 0.187667313 | 0.468 | 0.348 | 1 | 4 |
| ppp1r18 | 0.3718 | 0.176422103 | 0.194 | 0.109 | 1 | 4 |
| aip | 0.37229 | -0.17561199 | 0.016 | 0.065 | 1 | 4 |
| vamp8 | 0.37298 | 0.216353193 | 0.177 | 0.087 | 1 | 4 |
| gemin8 | 0.37365 | -0.12667862 | 0.016 | 0.065 | 1 | 4 |
| snrpd1 | 0.37414 | 0.24759043 | 0.145 | 0.065 | 1 | 4 |
| plscr3b | 0.37446 | -0.19869501 | 0.016 | 0.065 | 1 | 4 |
| ezrb | 0.375 | 0.186521549 | 0.081 | 0.022 | 1 | 4 |
| eif1b | 0.37547 | 0.268436298 | 0.339 | 0.217 | 1 | 4 |
| pdcd10a | 0.37553 | 0.187016884 | 0.081 | 0.022 | 1 | 4 |
| incenp | 0.37609 | -0.1713411 | 0.016 | 0.065 | 1 | 4 |
| ndufb5 | 0.37722 | -0.15855309 | 0.016 | 0.065 | 1 | 4 |
| dicp1.1 | 0.37724 | 0.246727895 | 0.177 | 0.087 | 1 | 4 |
| suclg1 | 0.37859 | 0.163085427 | 0.081 | 0.022 | 1 | 4 |
| atp6v1e1a | 0.37875 | 0.195397592 | 0.081 | 0.022 | 1 | 4 |
| tagapa | 0.37936 | 0.175414692 | 0.081 | 0.022 | 1 | 4 |
| mgst3a | 0.37967 | -0.13172808 | 0.016 | 0.065 | 1 | 4 |
| serf2 | 0.37989 | -0.24754056 | 0.274 | 0.37 | 1 | 4 |
| mdh2 | 0.37997 | 0.146317012 | 0.306 | 0.217 | 1 | 4 |
| eif4ebp1 | 0.3805 | 0.246895708 | 0.177 | 0.087 | 1 | 4 |
| kdm1a | 0.38154 | 0.144375912 | 0.081 | 0.022 | 1 | 4 |
| CT027744.2 | 0.38172 | 0.188686995 | 0.081 | 0.022 | 1 | 4 |
| ndufa7 | 0.38198 | -0.19146436 | 0.016 | 0.065 | 1 | 4 |
| gbp1 | 0.38267 | 0.175475938 | 0.081 | 0.022 | 1 | 4 |
| mhc2b | 0.38321 | -0.2464624 | 0.048 | 0.109 | 1 | 4 |
| atp6v0ca | 0.38329 | 0.18460239 | 0.081 | 0.022 | 1 | 4 |
| ndufa3 | 0.38488 | 0.147628226 | 0.081 | 0.022 | 1 | 4 |
| zdhhc23a | 0.38531 | -0.21211727 | 0.032 | 0.087 | 1 | 4 |
| mibp2 | 0.38585 | -0.18592078 | 0.565 | 0.652 | 1 | 4 |
| si:zfos-741a10.3 | 0.38718 | 0.25188701 | 0.177 | 0.087 | 1 | 4 |
| ptp4a2b | 0.38885 | 0.200238654 | 0.145 | 0.065 | 1 | 4 |
| PDCL3 | 0.38898 | -0.148487 | 0.016 | 0.065 | 1 | 4 |
| mcm3 | 0.39066 | -0.13855842 | 0.016 | 0.065 | 1 | 4 |
| ndufv3 | 0.39262 | -0.15127302 | 0.016 | 0.065 | 1 | 4 |
| serbp1a | 0.39287 | -0.13516589 | 0.403 | 0.435 | 1 | 4 |
| tpi1b | 0.39546 | 0.205175583 | 0.274 | 0.174 | 1 | 4 |

| | | | | | | |
|---|---|---|---|---|---|---|
| gspt1l | 0.39573 | -0.14825793 | 0.016 | 0.065 | 1 | 4 |
| map2k2a | 0.39575 | -0.19549134 | 0.016 | 0.065 | 1 | 4 |
| snrpf | 0.39599 | -0.20479701 | 0.161 | 0.217 | 1 | 4 |
| bxdc2 | 0.39624 | -0.16334412 | 0.016 | 0.065 | 1 | 4 |
| pabpn1 | 0.39638 | -0.24062215 | 0.048 | 0.109 | 1 | 4 |
| cd63 | 0.39793 | 0.187507757 | 0.371 | 0.261 | 1 | 4 |
| trappc5 | 0.39803 | -0.14821889 | 0.016 | 0.065 | 1 | 4 |
| vipas39 | 0.39804 | -0.19063835 | 0.016 | 0.065 | 1 | 4 |
| irf2bp2a | 0.39929 | 0.223352141 | 0.113 | 0.043 | 1 | 4 |
| sarnp | 0.39938 | -0.16475089 | 0.097 | 0.13 | 1 | 4 |
| ifi45 | 0.40115 | 0.105005357 | 0.145 | 0.087 | 1 | 4 |
| ddx61 | 0.40127 | -0.16844265 | 0.081 | 0.109 | 1 | 4 |
| ppp5c | 0.40177 | -0.14388555 | 0.016 | 0.065 | 1 | 4 |
| cox14 | 0.40198 | 0.110359353 | 0.097 | 0.043 | 1 | 4 |
| CABZ01063556.1 | 0.40226 | -0.15515376 | 0.016 | 0.065 | 1 | 4 |
| chmp7 | 0.40347 | -0.16720527 | 0.016 | 0.065 | 1 | 4 |
| nudt21 | 0.40347 | -0.19418043 | 0.016 | 0.065 | 1 | 4 |
| rwdd1 | 0.40548 | 0.230124047 | 0.145 | 0.065 | 1 | 4 |
| adssl | 0.40639 | 0.116507318 | 0.097 | 0.043 | 1 | 4 |
| rrm2 | 0.40721 | -0.2039369 | 0.032 | 0.087 | 1 | 4 |
| dbpa | 0.408 | -0.14497703 | 0.016 | 0.065 | 1 | 4 |
| prdx2 | 0.40847 | 0.18885412 | 0.323 | 0.217 | 1 | 4 |
| b2m | 0.4085 | 0.216855277 | 0.113 | 0.043 | 1 | 4 |
| CT573231.1 | 0.40852 | 0.20775025 | 0.113 | 0.043 | 1 | 4 |
| sat1a.2 | 0.40876 | 0.126829284 | 0.323 | 0.239 | 1 | 4 |
| CR847984.1 | 0.40949 | 0.215845588 | 0.113 | 0.043 | 1 | 4 |
| nfkbiab | 0.4096 | 0.12057556 | 0.097 | 0.043 | 1 | 4 |
| caprin1a | 0.41154 | -0.17530688 | 0.016 | 0.065 | 1 | 4 |
| pik3r3a | 0.41157 | -0.16896974 | 0.016 | 0.065 | 1 | 4 |
| ptp4a3 | 0.41168 | -0.17210691 | 0.016 | 0.065 | 1 | 4 |
| tufm | 0.41177 | -0.15728713 | 0.016 | 0.065 | 1 | 4 |
| eif5b | 0.41181 | -0.17090989 | 0.016 | 0.065 | 1 | 4 |
| timm10 | 0.41268 | -0.16125671 | 0.016 | 0.065 | 1 | 4 |
| pik3cb | 0.41396 | -0.16893006 | 0.016 | 0.065 | 1 | 4 |
| nup50 | 0.41447 | -0.15702581 | 0.016 | 0.065 | 1 | 4 |
| zgc:174945 | 0.41458 | -0.20550852 | 0.065 | 0.109 | 1 | 4 |
| atp5if1a | 0.41486 | -0.18506689 | 0.194 | 0.239 | 1 | 4 |
| mrpl37 | 0.41502 | -0.15519754 | 0.016 | 0.065 | 1 | 4 |
| eif3m | 0.41535 | 0.182084626 | 0.29 | 0.196 | 1 | 4 |
| pak2b | 0.41537 | -0.16999601 | 0.016 | 0.065 | 1 | 4 |
| hspa8 | 0.41539 | -0.10633999 | 0.79 | 0.826 | 1 | 4 |
| tln1 | 0.41566 | -0.2648245 | 0.177 | 0.283 | 1 | 4 |
| pa2g4a | 0.41598 | -0.17109166 | 0.016 | 0.065 | 1 | 4 |
| copa | 0.41661 | -0.17711466 | 0.016 | 0.065 | 1 | 4 |
| mvb12a | 0.41678 | -0.21486129 | 0.032 | 0.087 | 1 | 4 |
| tmem258 | 0.41688 | 0.139645236 | 0.21 | 0.13 | 1 | 4 |
| psmd14 | 0.4169 | -0.1510699 | 0.016 | 0.065 | 1 | 4 |
| si:ch211-262i1.6 | 0.41713 | -0.13440745 | 0.016 | 0.065 | 1 | 4 |

| | | | | | | |
|---|---|---|---|---|---|---|
| cdkn2aipnl | 0.41717 | -0.17951653 | 0.016 | 0.065 | 1 | 4 |
| tra2a | 0.41722 | -0.15962806 | 0.016 | 0.065 | 1 | 4 |
| dnajc1 | 0.41728 | -0.16700276 | 0.016 | 0.065 | 1 | 4 |
| nfkbie | 0.41732 | 0.144481178 | 0.177 | 0.109 | 1 | 4 |
| eomesa | 0.41764 | 0.255891733 | 0.29 | 0.196 | 1 | 4 |
| atp5if1b | 0.41985 | -0.10758458 | 0.145 | 0.152 | 1 | 4 |
| atp5f1e | 0.42156 | 0.126983616 | 0.484 | 0.391 | 1 | 4 |
| eif3ha | 0.42195 | 0.195811944 | 0.194 | 0.109 | 1 | 4 |
| pik3r1 | 0.42221 | -0.18850283 | 0.113 | 0.196 | 1 | 4 |
| vsir | 0.42534 | 0.102743733 | 0.097 | 0.043 | 1 | 4 |
| atp5f1b | 0.42544 | -0.19398303 | 0.355 | 0.413 | 1 | 4 |
| ahcyl1 | 0.42639 | -0.21466305 | 0.032 | 0.087 | 1 | 4 |
| tomm40 | 0.42798 | 0.108589554 | 0.097 | 0.043 | 1 | 4 |
| mcm5 | 0.42843 | -0.1382684 | 0.032 | 0.087 | 1 | 4 |
| cox6c | 0.42938 | 0.242630051 | 0.355 | 0.239 | 1 | 4 |
| zgc:56493 | 0.43286 | -0.13838753 | 0.629 | 0.739 | 1 | 4 |
| FO393424.2 | 0.43497 | -0.19712649 | 0.032 | 0.087 | 1 | 4 |
| rps27.2 | 0.43549 | 0.162064574 | 0.661 | 0.543 | 1 | 4 |
| stxbp2 | 0.43713 | -0.18903594 | 0.032 | 0.087 | 1 | 4 |
| sap30bp | 0.43848 | -0.18607939 | 0.032 | 0.087 | 1 | 4 |
| eif3d | 0.43888 | 0.217548282 | 0.226 | 0.13 | 1 | 4 |
| eloca | 0.43998 | 0.119381548 | 0.065 | 0.022 | 1 | 4 |
| ssr1 | 0.44111 | 0.110570088 | 0.097 | 0.043 | 1 | 4 |
| si:ch211-102c2.4 | 0.44196 | 0.241022862 | 0.484 | 0.37 | 1 | 4 |
| ccr9a | 0.44315 | -0.19739378 | 0.032 | 0.087 | 1 | 4 |
| mt-nd2 | 0.44558 | -0.23528013 | 0.113 | 0.196 | 1 | 4 |
| arhgap15 | 0.44936 | -0.16805433 | 0.032 | 0.087 | 1 | 4 |
| capzb | 0.45061 | -0.1629989 | 0.403 | 0.457 | 1 | 4 |
| odc1 | 0.45179 | -0.18817085 | 0.032 | 0.087 | 1 | 4 |
| si:dkey-112e7.2 | 0.45241 | 0.134078444 | 0.226 | 0.152 | 1 | 4 |
| bloc1s2 | 0.45354 | -0.15452957 | 0.032 | 0.087 | 1 | 4 |
| wasa | 0.45515 | 0.177595671 | 0.435 | 0.326 | 1 | 4 |
| ndufs3 | 0.45652 | -0.12581271 | 0.113 | 0.13 | 1 | 4 |
| rap1gap2a | 0.45652 | -0.17579236 | 0.048 | 0.087 | 1 | 4 |
| hmgn6 | 0.45661 | 0.221261699 | 0.323 | 0.217 | 1 | 4 |
| CABZ01109481.1 | 0.45785 | -0.20941801 | 0.032 | 0.087 | 1 | 4 |
| pdk2b | 0.45834 | -0.1735967 | 0.032 | 0.065 | 1 | 4 |
| cmpk | 0.45888 | -0.20514233 | 0.048 | 0.109 | 1 | 4 |
| ufc1 | 0.46118 | -0.14706432 | 0.065 | 0.087 | 1 | 4 |
| rpn1 | 0.4622 | -0.17926969 | 0.048 | 0.087 | 1 | 4 |
| adh5 | 0.4635 | 0.162165725 | 0.161 | 0.087 | 1 | 4 |
| pfdn5 | 0.46437 | -0.15461335 | 0.129 | 0.152 | 1 | 4 |
| CU207241.1 | 0.46499 | -0.16399864 | 0.032 | 0.087 | 1 | 4 |
| eif3i | 0.46606 | -0.19052533 | 0.21 | 0.261 | 1 | 4 |
| zgc:152774 | 0.46697 | -0.18284514 | 0.032 | 0.087 | 1 | 4 |
| max | 0.4682 | -0.17161728 | 0.032 | 0.087 | 1 | 4 |
| anxa11b | 0.46872 | -0.15337906 | 0.081 | 0.109 | 1 | 4 |
| mcm7 | 0.47205 | -0.16341846 | 0.032 | 0.087 | 1 | 4 |

| | | | | | | |
|---|---|---|---|---|---|---|
| ampd3b | 0.47279 | -0.18449923 | 0.032 | 0.087 | 1 | 4 |
| mif | 0.47333 | 0.204845754 | 0.194 | 0.109 | 1 | 4 |
| actn4 | 0.47334 | -0.17235444 | 0.065 | 0.13 | 1 | 4 |
| pabpc1a | 0.47472 | 0.124251285 | 0.645 | 0.543 | 1 | 4 |
| laptm4a | 0.47579 | -0.17834063 | 0.032 | 0.087 | 1 | 4 |
| chmp1a | 0.47602 | -0.15430773 | 0.032 | 0.087 | 1 | 4 |
| cct4 | 0.47685 | 0.165037167 | 0.161 | 0.087 | 1 | 4 |
| cox7c | 0.47694 | 0.22687312 | 0.371 | 0.261 | 1 | 4 |
| mrpl34 | 0.4775 | -0.14892777 | 0.032 | 0.087 | 1 | 4 |
| zgc:56304 | 0.47762 | -0.17550539 | 0.032 | 0.087 | 1 | 4 |
| LOC100535743 | 0.47764 | -0.18698607 | 0.032 | 0.087 | 1 | 4 |
| casp8 | 0.47846 | 0.101334247 | 0.065 | 0.022 | 1 | 4 |
| rhof | 0.47997 | 0.105552168 | 0.065 | 0.022 | 1 | 4 |
| fkbp2 | 0.47999 | -0.21760301 | 0.065 | 0.13 | 1 | 4 |
| tmem59 | 0.48051 | 0.19038833 | 0.097 | 0.043 | 1 | 4 |
| psmc4 | 0.48082 | 0.115038333 | 0.097 | 0.043 | 1 | 4 |
| limd2 | 0.48153 | -0.15725909 | 0.032 | 0.065 | 1 | 4 |
| denr | 0.48558 | -0.22413889 | 0.048 | 0.109 | 1 | 4 |
| ifi30 | 0.48707 | 0.110290041 | 0.145 | 0.087 | 1 | 4 |
| churc1 | 0.48719 | -0.16639769 | 0.065 | 0.13 | 1 | 4 |
| ldha | 0.48785 | -0.14912094 | 0.065 | 0.087 | 1 | 4 |
| ccdc167 | 0.48938 | 0.115757477 | 0.065 | 0.022 | 1 | 4 |
| pou2f2a | 0.48958 | 0.15738137 | 0.065 | 0.022 | 1 | 4 |
| psmb3 | 0.49055 | -0.17768295 | 0.113 | 0.152 | 1 | 4 |
| XLOC-023492 | 0.49082 | -0.20223924 | 0.048 | 0.109 | 1 | 4 |
| mat2b | 0.49145 | -0.12758788 | 0.048 | 0.065 | 1 | 4 |
| stk17b | 0.49209 | -0.1909078 | 0.226 | 0.283 | 1 | 4 |
| rab7a | 0.49283 | 0.108198784 | 0.065 | 0.022 | 1 | 4 |
| fabp3 | 0.49302 | -0.19464086 | 0.048 | 0.109 | 1 | 4 |
| mtpn | 0.49387 | -0.1381574 | 0.21 | 0.239 | 1 | 4 |
| sec61a1 | 0.4945 | 0.112567785 | 0.065 | 0.022 | 1 | 4 |
| ss18 | 0.49484 | 0.109115888 | 0.065 | 0.022 | 1 | 4 |
| mrps6 | 0.4959 | 0.106356559 | 0.065 | 0.022 | 1 | 4 |
| cry-dash | 0.49597 | 0.109202217 | 0.065 | 0.022 | 1 | 4 |
| clic1 | 0.49683 | 0.146520551 | 0.226 | 0.152 | 1 | 4 |
| cdc14b | 0.49748 | 0.104360395 | 0.065 | 0.022 | 1 | 4 |
| si:ch211-69g19.2 | 0.49832 | 0.117750018 | 0.065 | 0.022 | 1 | 4 |
| tmed2 | 0.50134 | -0.18494045 | 0.048 | 0.109 | 1 | 4 |
| XLOC-032002 | 0.50271 | -0.1915863 | 0.048 | 0.109 | 1 | 4 |
| tyms | 0.50288 | -0.20183073 | 0.048 | 0.109 | 1 | 4 |
| dpy30 | 0.50319 | -0.16208444 | 0.065 | 0.109 | 1 | 4 |
| efhd2 | 0.50335 | 0.154125079 | 0.129 | 0.065 | 1 | 4 |
| mt-co1 | 0.50383 | -0.17803904 | 0.306 | 0.413 | 1 | 4 |
| desi2 | 0.50585 | -0.16808105 | 0.032 | 0.065 | 1 | 4 |
| ptbp1a | 0.50609 | 0.154624825 | 0.065 | 0.022 | 1 | 4 |
| rab2a | 0.50609 | -0.18811662 | 0.065 | 0.13 | 1 | 4 |
| rps6ka1 | 0.50617 | 0.121579781 | 0.097 | 0.043 | 1 | 4 |
| syk | 0.5062 | 0.220997492 | 0.177 | 0.109 | 1 | 4 |

| | | | | | | |
|---|---|---|---|---|---|---|
| rpl22l1 | 0.5103 | 0.209362173 | 0.242 | 0.152 | 1 | 4 |
| tuba8l4 | 0.51049 | -0.17252795 | 0.258 | 0.304 | 1 | 4 |
| b4galt3 | 0.51267 | -0.17673091 | 0.065 | 0.109 | 1 | 4 |
| cdc20 | 0.51502 | 0.179324771 | 0.097 | 0.043 | 1 | 4 |
| jpt1b | 0.51576 | 0.146092123 | 0.065 | 0.022 | 1 | 4 |
| si:dkey-33c12.4 | 0.51675 | -0.20762453 | 0.097 | 0.152 | 1 | 4 |
| tnfaip8l2a | 0.51811 | 0.115021043 | 0.065 | 0.022 | 1 | 4 |
| aurkb | 0.51915 | 0.118750802 | 0.065 | 0.022 | 1 | 4 |
| si:ch211-217k17.7 | 0.52133 | 0.116723495 | 0.065 | 0.022 | 1 | 4 |
| calm1a | 0.52147 | 0.115298169 | 0.065 | 0.022 | 1 | 4 |
| chchd2 | 0.52207 | 0.148415404 | 0.065 | 0.022 | 1 | 4 |
| mrpl33 | 0.52278 | -0.16436373 | 0.048 | 0.087 | 1 | 4 |
| ppp3ccb | 0.52398 | -0.14374234 | 0.081 | 0.109 | 1 | 4 |
| krt18b | 0.52424 | 0.18273571 | 0.129 | 0.065 | 1 | 4 |
| lypla2 | 0.52542 | 0.14617553 | 0.065 | 0.022 | 1 | 4 |
| cers1 | 0.52582 | 0.178568924 | 0.097 | 0.043 | 1 | 4 |
| selenot1a | 0.52644 | 0.127730726 | 0.065 | 0.022 | 1 | 4 |
| jdp2b | 0.52679 | 0.102749984 | 0.065 | 0.022 | 1 | 4 |
| dbnlb | 0.52757 | 0.151847043 | 0.065 | 0.022 | 1 | 4 |
| arhgef3l | 0.52995 | -0.14945959 | 0.032 | 0.065 | 1 | 4 |
| fam162a | 0.53019 | 0.129771352 | 0.065 | 0.022 | 1 | 4 |
| CDPF1 | 0.53173 | 0.155381963 | 0.065 | 0.022 | 1 | 4 |
| ndufb2 | 0.53303 | 0.151297595 | 0.097 | 0.043 | 1 | 4 |
| dynlt3 | 0.53336 | 0.140565777 | 0.097 | 0.043 | 1 | 4 |
| park7 | 0.53349 | 0.144891112 | 0.065 | 0.022 | 1 | 4 |
| faua | 0.53405 | 0.125951136 | 0.823 | 0.761 | 1 | 4 |
| ppp1cb | 0.53523 | 0.112844608 | 0.065 | 0.022 | 1 | 4 |
| srsf3b | 0.53543 | 0.165924598 | 0.097 | 0.043 | 1 | 4 |
| idh2 | 0.5387 | -0.18098247 | 0.387 | 0.457 | 1 | 4 |
| usf1 | 0.54019 | 0.125680754 | 0.065 | 0.022 | 1 | 4 |
| ndufa12 | 0.54079 | -0.16934326 | 0.065 | 0.109 | 1 | 4 |
| ndufb7 | 0.54105 | -0.21302345 | 0.129 | 0.196 | 1 | 4 |
| eif2s2 | 0.54276 | 0.145007534 | 0.177 | 0.109 | 1 | 4 |
| ndufa1 | 0.54463 | -0.18460065 | 0.145 | 0.196 | 1 | 4 |
| hnrpkl | 0.54753 | 0.128192294 | 0.065 | 0.022 | 1 | 4 |
| ssr2 | 0.54801 | -0.19800006 | 0.113 | 0.174 | 1 | 4 |
| calm2a | 0.5488 | 0.116335558 | 0.065 | 0.022 | 1 | 4 |
| csf2rb | 0.54901 | 0.120003661 | 0.065 | 0.022 | 1 | 4 |
| aldh6a1 | 0.54939 | -0.14486004 | 0.032 | 0.065 | 1 | 4 |
| kmt5ab | 0.54992 | 0.120448317 | 0.065 | 0.022 | 1 | 4 |
| mrps16 | 0.54992 | 0.113262063 | 0.065 | 0.022 | 1 | 4 |
| cct7 | 0.55023 | -0.16561465 | 0.113 | 0.152 | 1 | 4 |
| cacybp | 0.55042 | 0.158888482 | 0.065 | 0.022 | 1 | 4 |
| spink2.2 | 0.55117 | 0.103140098 | 0.065 | 0.022 | 1 | 4 |
| ap2m1a | 0.55326 | 0.175931768 | 0.097 | 0.043 | 1 | 4 |
| polr2f | 0.5533 | 0.115549384 | 0.065 | 0.022 | 1 | 4 |
| smc1al | 0.55525 | 0.140543277 | 0.097 | 0.043 | 1 | 4 |
| sh3glb1b | 0.55542 | 0.125366682 | 0.065 | 0.022 | 1 | 4 |

| | | | | | | |
|---|---|---|---|---|---|---|
| ube2e3 | 0.55586 | -0.16300878 | 0.032 | 0.065 | 1 | 4 |
| ptprea | 0.55643 | 0.15115731 | 0.065 | 0.022 | 1 | 4 |
| si:dkeyp-44a8.2 | 0.55692 | 0.125119538 | 0.065 | 0.022 | 1 | 4 |
| kpnb1 | 0.55698 | 0.128615691 | 0.065 | 0.022 | 1 | 4 |
| atp5meb | 0.55729 | 0.173055865 | 0.306 | 0.217 | 1 | 4 |
| CABZ01088229.1 | 0.5576 | 0.15411215 | 0.065 | 0.022 | 1 | 4 |
| kmt5aa | 0.55796 | 0.131323183 | 0.065 | 0.022 | 1 | 4 |
| eef1g | 0.55811 | 0.117786399 | 0.806 | 0.717 | 1 | 4 |
| CABZ01073834.1 | 0.55822 | 0.134872854 | 0.065 | 0.022 | 1 | 4 |
| zbtb8os | 0.55846 | 0.133009713 | 0.065 | 0.022 | 1 | 4 |
| rbm22 | 0.55847 | 0.131645251 | 0.065 | 0.022 | 1 | 4 |
| hnrnpa0a | 0.5585 | 0.109562784 | 0.065 | 0.022 | 1 | 4 |
| rab44 | 0.55851 | 0.140750308 | 0.065 | 0.022 | 1 | 4 |
| actr2b | 0.55855 | 0.143222113 | 0.065 | 0.022 | 1 | 4 |
| srsf9 | 0.55856 | 0.120748743 | 0.065 | 0.022 | 1 | 4 |
| cnbpa | 0.55943 | -0.17261595 | 0.065 | 0.109 | 1 | 4 |
| csnk2a1 | 0.55946 | 0.169094482 | 0.097 | 0.043 | 1 | 4 |
| hmha1b | 0.55947 | -0.16032885 | 0.032 | 0.065 | 1 | 4 |
| gabarapb | 0.55987 | 0.1698324 | 0.339 | 0.283 | 1 | 4 |
| scinlb | 0.56022 | -0.20843299 | 0.113 | 0.174 | 1 | 4 |
| lin52 | 0.56222 | 0.14830914 | 0.097 | 0.043 | 1 | 4 |
| psenen | 0.5628 | -0.15377264 | 0.032 | 0.065 | 1 | 4 |
| tpx2 | 0.56302 | -0.11481443 | 0.048 | 0.065 | 1 | 4 |
| cdc42l | 0.56304 | -0.21242396 | 0.274 | 0.37 | 1 | 4 |
| u2af2b | 0.56363 | 0.142075127 | 0.097 | 0.043 | 1 | 4 |
| si:ch1073-340j19.1 | 0.56382 | -0.14217685 | 0.032 | 0.065 | 1 | 4 |
| alyref | 0.56417 | -0.14193816 | 0.032 | 0.065 | 1 | 4 |
| eif4a1a | 0.56626 | 0.158836547 | 0.177 | 0.109 | 1 | 4 |
| map1lc3b | 0.57096 | -0.1263254 | 0.048 | 0.065 | 1 | 4 |
| rps27l | 0.57239 | 0.145268956 | 0.081 | 0.043 | 1 | 4 |
| eef1b2 | 0.57418 | -0.11713887 | 0.742 | 0.804 | 1 | 4 |
| rpl5a | 0.57704 | 0.136542383 | 0.758 | 0.674 | 1 | 4 |
| arl1 | 0.58046 | -0.15948945 | 0.048 | 0.087 | 1 | 4 |
| smdt1b | 0.58265 | -0.17165433 | 0.081 | 0.13 | 1 | 4 |
| zgc:103700 | 0.585 | -0.1796686 | 0.129 | 0.174 | 1 | 4 |
| eif4h | 0.58556 | -0.14152717 | 0.032 | 0.065 | 1 | 4 |
| carhsp1 | 0.5866 | -0.15232295 | 0.032 | 0.065 | 1 | 4 |
| calm3b | 0.58668 | 0.162302576 | 0.371 | 0.283 | 1 | 4 |
| rpa2 | 0.58861 | -0.19540004 | 0.113 | 0.174 | 1 | 4 |
| idh1 | 0.58891 | -0.14732379 | 0.032 | 0.065 | 1 | 4 |
| sf3b5 | 0.59053 | -0.13779155 | 0.032 | 0.065 | 1 | 4 |
| brk1 | 0.59096 | 0.18667026 | 0.177 | 0.109 | 1 | 4 |
| ola1 | 0.597 | -0.10006503 | 0.161 | 0.174 | 1 | 4 |
| bcas2 | 0.59772 | -0.19145772 | 0.113 | 0.174 | 1 | 4 |
| mrtfaa | 0.59825 | -0.11547763 | 0.048 | 0.065 | 1 | 4 |
| pcmtd1 | 0.60097 | -0.11975979 | 0.065 | 0.087 | 1 | 4 |
| lamtor5 | 0.60164 | -0.17698382 | 0.081 | 0.13 | 1 | 4 |
| ube2v2 | 0.603 | -0.11619456 | 0.194 | 0.217 | 1 | 4 |

| | | | | | | |
|---|---|---|---|---|---|---|
| si:ch211-222l21.1 | 0.60305 | 0.125736271 | 0.242 | 0.174 | 1 | 4 |
| smndc1 | 0.60829 | -0.15589292 | 0.032 | 0.065 | 1 | 4 |
| ptmab | 0.61079 | 0.104966578 | 0.855 | 0.783 | 1 | 4 |
| drap1 | 0.61103 | 0.172658725 | 0.113 | 0.065 | 1 | 4 |
| fgfr1op2 | 0.61287 | -0.1502935 | 0.048 | 0.087 | 1 | 4 |
| dldh | 0.6144 | -0.15312699 | 0.048 | 0.087 | 1 | 4 |
| eif3c | 0.61793 | -0.17116443 | 0.145 | 0.217 | 1 | 4 |
| ccl35.1 | 0.62131 | 0.212241031 | 0.323 | 0.239 | 1 | 4 |
| ywhaba | 0.62379 | -0.12304006 | 0.29 | 0.326 | 1 | 4 |
| rsl24d1 | 0.62627 | 0.105693711 | 0.355 | 0.326 | 1 | 4 |
| TSTA3 (1 of many) | 0.6265 | -0.15224661 | 0.032 | 0.065 | 1 | 4 |
| psmb1 | 0.62812 | -0.143745 | 0.097 | 0.13 | 1 | 4 |
| ndufs8a | 0.62838 | -0.14325023 | 0.097 | 0.13 | 1 | 4 |
| tuba8l | 0.63147 | -0.21935249 | 0.129 | 0.196 | 1 | 4 |
| coa6 | 0.63324 | -0.12118194 | 0.065 | 0.087 | 1 | 4 |
| atf4b | 0.63475 | 0.148021906 | 0.081 | 0.043 | 1 | 4 |
| psmd4a | 0.63675 | -0.1417997 | 0.032 | 0.065 | 1 | 4 |
| pcna | 0.63715 | -0.11905239 | 0.081 | 0.13 | 1 | 4 |
| psma4 | 0.63793 | -0.13709967 | 0.048 | 0.087 | 1 | 4 |
| zmat2 | 0.64163 | 0.159647763 | 0.113 | 0.065 | 1 | 4 |
| rsl1d1 | 0.64321 | -0.142873 | 0.032 | 0.065 | 1 | 4 |
| pik3cd | 0.64457 | -0.14153484 | 0.032 | 0.065 | 1 | 4 |
| glrx | 0.64516 | 0.179012229 | 0.435 | 0.348 | 1 | 4 |
| mrps24 | 0.64533 | -0.13308395 | 0.032 | 0.065 | 1 | 4 |
| nop10 | 0.64715 | 0.152952695 | 0.081 | 0.043 | 1 | 4 |
| tbcb | 0.64752 | -0.13993516 | 0.048 | 0.087 | 1 | 4 |
| myd88 | 0.64769 | 0.168521098 | 0.145 | 0.087 | 1 | 4 |
| ssr4 | 0.65042 | -0.14772571 | 0.177 | 0.217 | 1 | 4 |
| tmed7 | 0.65176 | -0.10178029 | 0.065 | 0.109 | 1 | 4 |
| mdh1aa | 0.65432 | 0.135557232 | 0.242 | 0.174 | 1 | 4 |
| pin1 | 0.65586 | -0.16515546 | 0.081 | 0.13 | 1 | 4 |
| prdx5 | 0.65978 | 0.16285842 | 0.113 | 0.065 | 1 | 4 |
| tsc22d3 | 0.66014 | -0.1406491 | 0.097 | 0.152 | 1 | 4 |
| lbr | 0.66023 | -0.14164714 | 0.065 | 0.109 | 1 | 4 |
| stip1 | 0.66262 | -0.14388371 | 0.048 | 0.087 | 1 | 4 |
| hp1bp3 | 0.66279 | 0.113592758 | 0.113 | 0.065 | 1 | 4 |
| bzw1a | 0.67089 | -0.1175243 | 0.032 | 0.065 | 1 | 4 |
| srsf11 | 0.67092 | -0.12873682 | 0.032 | 0.065 | 1 | 4 |
| tegt | 0.67106 | 0.147603491 | 0.113 | 0.065 | 1 | 4 |
| dusp22b | 0.67143 | -0.11719691 | 0.065 | 0.087 | 1 | 4 |
| XLOC-026836 | 0.67195 | -0.15255258 | 0.113 | 0.152 | 1 | 4 |
| dazap2 | 0.67272 | -0.1262664 | 0.081 | 0.109 | 1 | 4 |
| crlf3 | 0.67287 | -0.14669122 | 0.097 | 0.152 | 1 | 4 |
| arf2b | 0.67746 | -0.12771764 | 0.032 | 0.065 | 1 | 4 |
| sdcbp2 | 0.67986 | -0.15448459 | 0.048 | 0.087 | 1 | 4 |
| prelid3b | 0.6807 | -0.12053324 | 0.065 | 0.087 | 1 | 4 |
| LOC110438870 | 0.68257 | -0.11697967 | 0.081 | 0.13 | 1 | 4 |
| nccrp1 | 0.68291 | -0.11077884 | 0.048 | 0.065 | 1 | 4 |

| | | | | | | |
|---|---|---|---|---|---|---|
| fosab | 0.6832 | -0.15345279 | 0.065 | 0.109 | 1 | 4 |
| ISCU (1 of many) | 0.68556 | -0.1112828 | 0.129 | 0.152 | 1 | 4 |
| COA4 | 0.68642 | 0.140729554 | 0.081 | 0.043 | 1 | 4 |
| u2af1 | 0.68708 | -0.14144817 | 0.065 | 0.109 | 1 | 4 |
| psma2 | 0.68764 | 0.125814367 | 0.113 | 0.065 | 1 | 4 |
| spcs3 | 0.68825 | 0.134843607 | 0.113 | 0.065 | 1 | 4 |
| si:busm1-266f07.2 | 0.68886 | 0.171647255 | 0.242 | 0.174 | 1 | 4 |
| si:dkey-78l4.14 | 0.6907 | -0.11937797 | 0.032 | 0.065 | 1 | 4 |
| marcksl1a | 0.69071 | -0.13345886 | 0.081 | 0.109 | 1 | 4 |
| si:ch211-214k5.3 | 0.69133 | 0.149597992 | 0.113 | 0.065 | 1 | 4 |
| cox6a1 | 0.69274 | 0.108197678 | 0.419 | 0.348 | 1 | 4 |
| gadd45bb | 0.69391 | -0.13584209 | 0.032 | 0.065 | 1 | 4 |
| phf5a | 0.69528 | -0.10120724 | 0.048 | 0.087 | 1 | 4 |
| mbd2 | 0.69636 | 0.135337492 | 0.081 | 0.043 | 1 | 4 |
| itm2ba | 0.69667 | -0.1331495 | 0.097 | 0.13 | 1 | 4 |
| c1qa | 0.6975 | 0.103629707 | 0.081 | 0.043 | 1 | 4 |
| rhoac | 0.69775 | -0.10652431 | 0.113 | 0.13 | 1 | 4 |
| rap1b | 0.69917 | -0.11740369 | 0.29 | 0.326 | 1 | 4 |
| h1fx | 0.70079 | 0.158474572 | 0.5 | 0.435 | 1 | 4 |
| twf1a | 0.70158 | 0.118525842 | 0.258 | 0.196 | 1 | 4 |
| LOC101882805 | 0.70197 | -0.15181949 | 0.081 | 0.13 | 1 | 4 |
| hypk | 0.70235 | -0.13905371 | 0.081 | 0.13 | 1 | 4 |
| pfdn2 | 0.70327 | 0.106890034 | 0.065 | 0.043 | 1 | 4 |
| h3f3d | 0.70357 | -0.1066312 | 0.403 | 0.435 | 1 | 4 |
| si:ch211-235e9.6 | 0.7056 | 0.128337844 | 0.081 | 0.043 | 1 | 4 |
| cox7b | 0.7069 | -0.11597007 | 0.355 | 0.391 | 1 | 4 |
| psma6b | 0.70711 | -0.13768746 | 0.065 | 0.109 | 1 | 4 |
| ube2na | 0.70784 | -0.14729545 | 0.113 | 0.152 | 1 | 4 |
| mfng | 0.70856 | -0.12534833 | 0.081 | 0.109 | 1 | 4 |
| itgb2 | 0.70925 | -0.13550318 | 0.161 | 0.196 | 1 | 4 |
| ppp2r3c | 0.70967 | -0.121885 | 0.032 | 0.065 | 1 | 4 |
| CU694197.1 | 0.71048 | -0.12421183 | 0.081 | 0.109 | 1 | 4 |
| ddx5 | 0.71157 | -0.13499809 | 0.032 | 0.065 | 1 | 4 |
| lrrc59 | 0.71222 | -0.13660553 | 0.048 | 0.087 | 1 | 4 |
| mrps26 | 0.71249 | -0.12375791 | 0.032 | 0.065 | 1 | 4 |
| esd | 0.71343 | -0.12969286 | 0.097 | 0.13 | 1 | 4 |
| selenoh | 0.71363 | -0.10362723 | 0.065 | 0.087 | 1 | 4 |
| hnrnpabb | 0.71421 | -0.11478354 | 0.129 | 0.152 | 1 | 4 |
| imp4 | 0.71605 | -0.13532939 | 0.032 | 0.065 | 1 | 4 |
| FERMT3 (1 of many) | 0.71695 | -0.12662293 | 0.065 | 0.109 | 1 | 4 |
| cct5 | 0.71708 | 0.127529246 | 0.21 | 0.152 | 1 | 4 |
| nek7 | 0.71748 | 0.111349553 | 0.065 | 0.043 | 1 | 4 |
| tomm7 | 0.71805 | -0.13646443 | 0.065 | 0.109 | 1 | 4 |
| trmt112 | 0.71852 | -0.11115862 | 0.048 | 0.087 | 1 | 4 |
| XLOC-000431 | 0.71991 | 0.115261821 | 0.081 | 0.043 | 1 | 4 |
| cks1b | 0.72039 | -0.12392518 | 0.048 | 0.087 | 1 | 4 |
| rtn4a | 0.72097 | -0.10229687 | 0.032 | 0.065 | 1 | 4 |
| calr3b | 0.72128 | -0.10520904 | 0.048 | 0.065 | 1 | 4 |

| | | | | | | |
|---|---|---|---|---|---|---|
| ube2l3b | 0.72143 | 0.131444116 | 0.161 | 0.109 | 1 | 4 |
| nap1l4b | 0.72172 | -0.12486126 | 0.032 | 0.065 | 1 | 4 |
| litaf | 0.72359 | 0.154820712 | 0.161 | 0.109 | 1 | 4 |
| snap23.1 | 0.72402 | 0.126648971 | 0.081 | 0.043 | 1 | 4 |
| lsm3 | 0.72405 | -0.10442868 | 0.048 | 0.087 | 1 | 4 |
| capns1b | 0.72446 | -0.12684364 | 0.081 | 0.109 | 1 | 4 |
| ppp2r5eb | 0.7255 | -0.12041932 | 0.048 | 0.087 | 1 | 4 |
| grasp | 0.72574 | 0.132746135 | 0.097 | 0.065 | 1 | 4 |
| cish | 0.72656 | -0.11847385 | 0.032 | 0.065 | 1 | 4 |
| chmp2a | 0.7269 | -0.12254692 | 0.032 | 0.065 | 1 | 4 |
| si:ch73-42p12.2 | 0.7271 | -0.11279515 | 0.032 | 0.065 | 1 | 4 |
| capza1b | 0.72728 | 0.137822071 | 0.452 | 0.391 | 1 | 4 |
| ube2g1a | 0.72787 | -0.12888948 | 0.048 | 0.087 | 1 | 4 |
| prkacba | 0.72812 | -0.10911509 | 0.048 | 0.087 | 1 | 4 |
| eef1db | 0.72855 | 0.141503544 | 0.161 | 0.109 | 1 | 4 |
| psmd11b | 0.72923 | -0.10652345 | 0.032 | 0.065 | 1 | 4 |
| si:ch73-27e22.7 | 0.72937 | -0.10977712 | 0.032 | 0.065 | 1 | 4 |
| LOC110438754 | 0.72937 | -0.11147101 | 0.032 | 0.065 | 1 | 4 |
| znf593 | 0.72938 | -0.10047313 | 0.032 | 0.065 | 1 | 4 |
| usp14 | 0.72941 | -0.11934881 | 0.032 | 0.065 | 1 | 4 |
| mtch2 | 0.73055 | 0.107637032 | 0.081 | 0.043 | 1 | 4 |
| fgl2a | 0.73226 | 0.111590418 | 0.081 | 0.043 | 1 | 4 |
| cyba | 0.74956 | -0.13077761 | 0.097 | 0.13 | 1 | 4 |
| lsm7 | 0.75225 | -0.11182687 | 0.081 | 0.109 | 1 | 4 |
| sub1b | 0.75528 | 0.112408808 | 0.113 | 0.087 | 1 | 4 |
| kctd12.2 | 0.76107 | 0.107165045 | 0.548 | 0.478 | 1 | 4 |
| hars | 0.76127 | 0.100908869 | 0.065 | 0.043 | 1 | 4 |
| atp5mf | 0.76446 | -0.14577735 | 0.145 | 0.196 | 1 | 4 |
| dtnbp1a | 0.76903 | -0.14645594 | 0.145 | 0.196 | 1 | 4 |
| zgc:64051 | 0.76984 | 0.135355104 | 0.371 | 0.304 | 1 | 4 |
| cops9 | 0.77081 | -0.11215765 | 0.065 | 0.087 | 1 | 4 |
| syncrip | 0.78252 | 0.112349969 | 0.129 | 0.087 | 1 | 4 |
| ube2e1 | 0.78399 | 0.128185344 | 0.129 | 0.087 | 1 | 4 |
| paip2b | 0.78535 | 0.108654376 | 0.129 | 0.087 | 1 | 4 |
| parvg | 0.7864 | -0.12951582 | 0.113 | 0.152 | 1 | 4 |
| mych | 0.79555 | -0.12477728 | 0.097 | 0.13 | 1 | 4 |
| ndufa4l | 0.80622 | 0.11561022 | 0.387 | 0.326 | 1 | 4 |
| plekhj1 | 0.81322 | 0.111253414 | 0.097 | 0.065 | 1 | 4 |
| ube2ib | 0.81825 | 0.10054323 | 0.113 | 0.087 | 1 | 4 |
| myh11a | 0.82719 | -0.11336882 | 0.113 | 0.152 | 1 | 4 |
| ppp1cab | 0.82898 | -0.11268247 | 0.097 | 0.13 | 1 | 4 |
| calm2b | 0.83074 | -0.12118598 | 0.274 | 0.326 | 1 | 4 |
| rbm8a | 0.83092 | 0.118782607 | 0.242 | 0.196 | 1 | 4 |
| tomm20b | 0.83448 | 0.108722406 | 0.145 | 0.109 | 1 | 4 |
| mob1a | 0.83897 | 0.116268337 | 0.242 | 0.196 | 1 | 4 |
| atp5mc3a | 0.84183 | 0.111499592 | 0.21 | 0.174 | 1 | 4 |
| ost4 | 0.84965 | 0.104029036 | 0.145 | 0.109 | 1 | 4 |
| gm2a | 0.87089 | -0.10585558 | 0.177 | 0.217 | 1 | 4 |

| | | | | | | |
|---|---|---|---|---|---|---|
| mt-cyb | 1.216888 | -1.09929074 | 0.206 | 0.871 | 2.45774894 | 5 |
| mt-co2 | 2.008338 | -0.92092935 | 0.381 | 0.968 | 0.0004056 | 5 |
| mt-nd2 | 6.984900 | -0.74221906 | 0.175 | 0.71 | 0.0141074 | 5 |
| mycbp | 8.358822 | 0.506478374 | 0.413 | 0 | 0.0168823 | 5 |
| rpl27a | 1.520131 | -0.369399 | 0.921 | 0.968 | 0.0307021 | 5 |
| mt-nd1 | 3.235662 | -0.81818686 | 0.238 | 0.774 | 0.0653507 | 5 |
| bzw1a | 1.629309 | -0.64129565 | 0.063 | 0.484 | 0.3290717 | 5 |
| LOC100536946 | 5.184002 | -0.68950001 | 0.556 | 0.806 | 1 | 5 |
| ccl34a.4 | 6.361764 | -0.56099983 | 0.556 | 0.71 | 1 | 5 |
| mt-nd4 | 0.00012 | -0.69572087 | 0.317 | 0.774 | 1 | 5 |
| samsn1a | 0.00014 | -0.59182825 | 0.429 | 0.806 | 1 | 5 |
| mt-nd3 | 0.00017 | -0.70748241 | 0.333 | 0.742 | 1 | 5 |
| cd40 | 0.00026 | 0.405655144 | 0.254 | 0 | 1 | 5 |
| ms4a17a.8 | 0.00031 | -0.328198 | 0 | 0.194 | 1 | 5 |
| snx4 | 0.00031 | -0.22753585 | 0 | 0.194 | 1 | 5 |
| lysmd3 | 0.00031 | -0.15924392 | 0 | 0.194 | 1 | 5 |
| smg7 | 0.00031 | -0.21687322 | 0 | 0.194 | 1 | 5 |
| apoc1 | 0.00031 | -0.55808576 | 0 | 0.194 | 1 | 5 |
| lsm6 | 0.0004 | 0.304077149 | 0.317 | 0.032 | 1 | 5 |
| zgc:110843 | 0.00049 | -0.40036831 | 0.032 | 0.323 | 1 | 5 |
| dennd4a | 0.00051 | -0.50393903 | 0.159 | 0.548 | 1 | 5 |
| BX649485.2 | 0.00062 | -0.51978138 | 0.683 | 0.774 | 1 | 5 |
| mt-atp6 | 0.00067 | -0.64523242 | 0.444 | 0.839 | 1 | 5 |
| tp53inp1 | 0.00073 | 0.3190091 | 0.222 | 0 | 1 | 5 |
| rgcc | 0.00111 | -0.21164612 | 0.032 | 0.194 | 1 | 5 |
| timm10b | 0.00114 | -0.20638204 | 0 | 0.161 | 1 | 5 |
| tnfrsf9a | 0.00114 | -0.32980725 | 0 | 0.161 | 1 | 5 |
| acvr1ba | 0.00114 | -0.23728544 | 0 | 0.161 | 1 | 5 |
| surf4 | 0.00114 | -0.21898258 | 0 | 0.161 | 1 | 5 |
| LOC103910163 | 0.00114 | -0.23110641 | 0 | 0.161 | 1 | 5 |
| afg1la | 0.00114 | -0.25891126 | 0 | 0.161 | 1 | 5 |
| glud1b | 0.00114 | -0.19490729 | 0 | 0.161 | 1 | 5 |
| rock1 | 0.00114 | -0.22398931 | 0 | 0.161 | 1 | 5 |
| LOC564660 | 0.0013 | -0.21015617 | 0.397 | 0.677 | 1 | 5 |
| larp4ab | 0.00138 | -0.44395444 | 0.127 | 0.419 | 1 | 5 |
| ccl39.2 | 0.00158 | -0.60255396 | 0.032 | 0.29 | 1 | 5 |
| fth1a | 0.00159 | -0.66883486 | 0.397 | 0.71 | 1 | 5 |
| slc35c2 | 0.00165 | -0.40430969 | 0.048 | 0.258 | 1 | 5 |
| CR753876.1 | 0.00184 | -0.37137117 | 0.016 | 0.226 | 1 | 5 |
| mpeg1.1 | 0.00197 | -0.23475619 | 0.079 | 0.29 | 1 | 5 |
| tmem189 | 0.00198 | -0.50547764 | 0.206 | 0.548 | 1 | 5 |
| tuba1b | 0.00199 | -0.46623587 | 0.175 | 0.516 | 1 | 5 |
| ccl34b.8 | 0.00205 | -0.44824369 | 0.619 | 0.935 | 1 | 5 |
| lima1a | 0.00228 | -0.50979741 | 0.143 | 0.484 | 1 | 5 |
| btg2 | 0.00266 | 0.568179222 | 0.619 | 0.29 | 1 | 5 |
| ndufa1 | 0.00302 | -0.44449046 | 0.492 | 0.806 | 1 | 5 |
| si:dkey-56i24.1 | 0.00333 | 0.299336696 | 0.175 | 0 | 1 | 5 |
| vrk2 | 0.00333 | 0.278254127 | 0.175 | 0 | 1 | 5 |

| | | | | | | |
|---|---|---|---|---|---|---|
| psmd11b | 0.00333 | 0.233596299 | 0.175 | 0 | 1 | 5 |
| illr3 | 0.00333 | 0.39727166 | 0.175 | 0 | 1 | 5 |
| si:dkey-58f10.11 | 0.00333 | 0.271901632 | 0.175 | 0 | 1 | 5 |
| si:ch211-165i18.2 | 0.00362 | -0.32483176 | 0.825 | 1 | 1 | 5 |
| si:dkeyp-27c8.2 | 0.00371 | -0.45551125 | 0.111 | 0.419 | 1 | 5 |
| COX3 | 0.00372 | -0.38941029 | 0.73 | 0.968 | 1 | 5 |
| cct5 | 0.0039 | 0.1579139 | 0.492 | 0.613 | 1 | 5 |
| pdss1 | 0.00394 | -0.28226192 | 0.032 | 0.258 | 1 | 5 |
| si:dkey-169i5.4 | 0.00414 | -0.16598326 | 0 | 0.129 | 1 | 5 |
| vars | 0.00414 | -0.13111168 | 0 | 0.129 | 1 | 5 |
| CU469362.1 | 0.00414 | -0.20779217 | 0 | 0.129 | 1 | 5 |
| armc1 | 0.00414 | -0.17448909 | 0 | 0.129 | 1 | 5 |
| cacul1 | 0.00414 | -0.14733262 | 0 | 0.129 | 1 | 5 |
| chmp3 | 0.00414 | -0.18339549 | 0 | 0.129 | 1 | 5 |
| rab18b | 0.00414 | -0.15591198 | 0 | 0.129 | 1 | 5 |
| rn7sk | 0.00414 | -0.14303878 | 0 | 0.129 | 1 | 5 |
| rnf19b | 0.00414 | -0.21264563 | 0 | 0.129 | 1 | 5 |
| si:ch211-137j23.7 | 0.00414 | -0.23619674 | 0 | 0.129 | 1 | 5 |
| CU929070.1 | 0.00414 | -0.14182114 | 0 | 0.129 | 1 | 5 |
| cdkn1a | 0.00414 | -0.12819439 | 0 | 0.129 | 1 | 5 |
| cgas | 0.00414 | -0.22183507 | 0 | 0.129 | 1 | 5 |
| glrx5 | 0.00414 | -0.10892979 | 0 | 0.129 | 1 | 5 |
| csf1ra | 0.00414 | -0.21098583 | 0 | 0.129 | 1 | 5 |
| id4 | 0.00414 | -0.18951792 | 0 | 0.129 | 1 | 5 |
| acss1 | 0.00414 | -0.18322416 | 0 | 0.129 | 1 | 5 |
| wbp2nl | 0.00432 | -0.46211983 | 0.302 | 0.613 | 1 | 5 |
| prdx2 | 0.00446 | -0.39245012 | 0.46 | 0.806 | 1 | 5 |
| icn | 0.00496 | 0.811496097 | 0.54 | 0.258 | 1 | 5 |
| zgc:193541 | 0.00502 | -0.40318432 | 0.683 | 0.903 | 1 | 5 |
| cxcr4b | 0.00533 | 0.47363473 | 0.857 | 0.613 | 1 | 5 |
| si:dkey-188i13.11 | 0.00548 | 0.225583877 | 0.159 | 0 | 1 | 5 |
| c1d | 0.00548 | 0.243136693 | 0.159 | 0 | 1 | 5 |
| btr31 | 0.00548 | 0.253956393 | 0.159 | 0 | 1 | 5 |
| adrb2a | 0.00548 | 0.263270776 | 0.159 | 0 | 1 | 5 |
| fam168a | 0.00548 | 0.208023828 | 0.159 | 0 | 1 | 5 |
| fbxl6 | 0.00548 | 0.269041001 | 0.159 | 0 | 1 | 5 |
| anp32e | 0.00548 | 0.184221871 | 0.159 | 0 | 1 | 5 |
| hadhaa | 0.00548 | 0.243612511 | 0.159 | 0 | 1 | 5 |
| ddx19 | 0.00548 | 0.234168794 | 0.159 | 0 | 1 | 5 |
| mgat3b | 0.00548 | 0.303931642 | 0.159 | 0 | 1 | 5 |
| thoc2 | 0.00548 | 0.278594415 | 0.159 | 0 | 1 | 5 |
| ap1s1 | 0.00571 | 0.384077237 | 0.333 | 0.065 | 1 | 5 |
| tbxas1 | 0.00603 | -0.2926826 | 0.016 | 0.194 | 1 | 5 |
| ponzr1 | 0.00662 | 0.381673723 | 0.825 | 0.71 | 1 | 5 |
| atp5fa1 | 0.00669 | 0.433592134 | 0.746 | 0.548 | 1 | 5 |
| acsl4a | 0.00726 | -0.42061577 | 0.079 | 0.323 | 1 | 5 |
| si:dkey-69o16.5 | 0.00739 | -0.27236792 | 0.222 | 0.516 | 1 | 5 |
| LOC100148254 | 0.00832 | -0.43201546 | 0.175 | 0.484 | 1 | 5 |

| | | | | | | |
|---|---|---|---|---|---|---|
| si:ch211-246m6.4 | 0.0085 | -0.44747633 | 0.079 | 0.29 | 1 | 5 |
| ndnl2 | 0.00897 | 0.205106411 | 0.143 | 0 | 1 | 5 |
| slc30a4 | 0.00897 | 0.188885433 | 0.143 | 0 | 1 | 5 |
| bola3 | 0.00897 | 0.224325757 | 0.143 | 0 | 1 | 5 |
| id1 | 0.00897 | 0.300090807 | 0.143 | 0 | 1 | 5 |
| ivd | 0.00897 | 0.18709591 | 0.143 | 0 | 1 | 5 |
| becn1 | 0.00897 | 0.193933245 | 0.143 | 0 | 1 | 5 |
| si:cabz01036006.1 | 0.00897 | 0.277668402 | 0.143 | 0 | 1 | 5 |
| etfa | 0.00921 | 0.3123782 | 0.317 | 0.065 | 1 | 5 |
| CR936442.1 | 0.00972 | 0.473089331 | 0.492 | 0.258 | 1 | 5 |
| tmem214 | 0.00985 | 0.315945386 | 0.254 | 0.032 | 1 | 5 |
| abhd17c | 0.00993 | -0.25423974 | 0.016 | 0.194 | 1 | 5 |
| wdr5 | 0.01059 | -0.25056457 | 0.016 | 0.194 | 1 | 5 |
| fitm2 | 0.01071 | -0.2539955 | 0.016 | 0.194 | 1 | 5 |
| hnrnpk | 0.01108 | 0.334806228 | 0.286 | 0.129 | 1 | 5 |
| ptp4a3 | 0.01125 | -0.21657289 | 0.016 | 0.194 | 1 | 5 |
| ndufa10 | 0.01137 | 0.387379789 | 0.333 | 0.129 | 1 | 5 |
| yipf2 | 0.01137 | -0.20484465 | 0.016 | 0.194 | 1 | 5 |
| si:ch211-233a24.2 | 0.01138 | -0.25508897 | 0.016 | 0.194 | 1 | 5 |
| pabpn1 | 0.01144 | -0.40266674 | 0.302 | 0.613 | 1 | 5 |
| mfsd11 | 0.01154 | -0.28725674 | 0.063 | 0.29 | 1 | 5 |
| mt-co1 | 0.01165 | -0.28587718 | 0.873 | 0.968 | 1 | 5 |
| psmd3 | 0.0119 | -0.37698109 | 0.19 | 0.452 | 1 | 5 |
| adi1 | 0.01199 | 0.251339748 | 0.206 | 0.129 | 1 | 5 |
| h1fx | 0.01227 | -0.55311227 | 0.238 | 0.484 | 1 | 5 |
| plin2 | 0.01229 | -0.52544118 | 0.175 | 0.452 | 1 | 5 |
| tuba8l | 0.0123 | -0.381968 | 0.19 | 0.484 | 1 | 5 |
| cebpb | 0.01238 | 0.235217027 | 0.952 | 0.903 | 1 | 5 |
| sarnp | 0.01275 | -0.22113168 | 0.254 | 0.516 | 1 | 5 |
| abce1 | 0.01283 | -0.24820549 | 0.079 | 0.323 | 1 | 5 |
| rgs19 | 0.01298 | 0.378303288 | 0.317 | 0.065 | 1 | 5 |
| hexb | 0.01311 | 0.482554811 | 0.397 | 0.129 | 1 | 5 |
| mafba | 0.01346 | -0.40939689 | 0.286 | 0.516 | 1 | 5 |
| rhoab | 0.01375 | -0.31446995 | 0.587 | 0.71 | 1 | 5 |
| map1lc3b | 0.01412 | -0.41398584 | 0.254 | 0.548 | 1 | 5 |
| ier5 | 0.01429 | 0.20671913 | 0.508 | 0.258 | 1 | 5 |
| bloc1s5 | 0.01464 | 0.18817714 | 0.127 | 0 | 1 | 5 |
| ftr67 | 0.01464 | 0.168515621 | 0.127 | 0 | 1 | 5 |
| gkap1 | 0.01464 | 0.168991243 | 0.127 | 0 | 1 | 5 |
| zranb2 | 0.01464 | 0.169678599 | 0.127 | 0 | 1 | 5 |
| cmc1 | 0.01464 | 0.172189032 | 0.127 | 0 | 1 | 5 |
| ctsf | 0.01464 | 0.151897411 | 0.127 | 0 | 1 | 5 |
| eif4e3 | 0.01464 | 0.199432039 | 0.127 | 0 | 1 | 5 |
| mapk12b | 0.01464 | 0.179893023 | 0.127 | 0 | 1 | 5 |
| sco1 | 0.01464 | 0.19493871 | 0.127 | 0 | 1 | 5 |
| FQ976913.1 | 0.01464 | 0.154181024 | 0.127 | 0 | 1 | 5 |
| gcsha | 0.01464 | 0.142719127 | 0.127 | 0 | 1 | 5 |
| samm50 | 0.01464 | 0.178610179 | 0.127 | 0 | 1 | 5 |

| | | | | | | |
|---|---|---|---|---|---|---|
| sept9b | 0.01464 | 0.178274871 | 0.127 | 0 | 1 | 5 |
| cdca4 | 0.01464 | 0.187966675 | 0.127 | 0 | 1 | 5 |
| bzw1b | 0.01475 | -0.38901435 | 0.619 | 0.806 | 1 | 5 |
| fam204a | 0.01477 | -0.25183173 | 0.032 | 0.226 | 1 | 5 |
| LOC101886751 | 0.01501 | -0.12851795 | 0 | 0.097 | 1 | 5 |
| cryzl1 | 0.01501 | -0.15105457 | 0 | 0.097 | 1 | 5 |
| hs6st1b | 0.01501 | -0.12108894 | 0 | 0.097 | 1 | 5 |
| ndc1 | 0.01501 | -0.10489715 | 0 | 0.097 | 1 | 5 |
| nrd1a | 0.01501 | -0.14212186 | 0 | 0.097 | 1 | 5 |
| pnp5b | 0.01501 | -0.16223389 | 0 | 0.097 | 1 | 5 |
| slc25a17 | 0.01501 | -0.16987231 | 0 | 0.097 | 1 | 5 |
| cdca8 | 0.01501 | -0.20056142 | 0 | 0.097 | 1 | 5 |
| gpbp1l1 | 0.01501 | -0.15003906 | 0 | 0.097 | 1 | 5 |
| lzic | 0.01501 | -0.11313992 | 0 | 0.097 | 1 | 5 |
| mibp | 0.01501 | -0.10494037 | 0 | 0.097 | 1 | 5 |
| BX908782.3 | 0.01501 | -0.28147986 | 0 | 0.097 | 1 | 5 |
| atxn10 | 0.01501 | -0.18094921 | 0 | 0.097 | 1 | 5 |
| golga3 | 0.01501 | -0.12858001 | 0 | 0.097 | 1 | 5 |
| hnrnpdl | 0.01501 | -0.1231636 | 0 | 0.097 | 1 | 5 |
| orc4 | 0.01501 | -0.10320725 | 0 | 0.097 | 1 | 5 |
| ctso | 0.01501 | -0.11510838 | 0 | 0.097 | 1 | 5 |
| pde4cb | 0.01501 | -0.19429808 | 0 | 0.097 | 1 | 5 |
| XLOC-032002 | 0.01501 | -0.12935128 | 0 | 0.097 | 1 | 5 |
| ascl1b | 0.01501 | -0.11489277 | 0 | 0.097 | 1 | 5 |
| ggact.2 | 0.01501 | -0.12568073 | 0 | 0.097 | 1 | 5 |
| itprip | 0.01501 | -0.14017652 | 0 | 0.097 | 1 | 5 |
| p2ry11 | 0.01501 | -0.17418602 | 0 | 0.097 | 1 | 5 |
| spint2 | 0.01501 | -0.18056686 | 0 | 0.097 | 1 | 5 |
| usp38 | 0.01501 | -0.15573035 | 0 | 0.097 | 1 | 5 |
| LOC110438144 | 0.01501 | -0.140141 | 0 | 0.097 | 1 | 5 |
| fbrs | 0.01501 | -0.15819438 | 0 | 0.097 | 1 | 5 |
| glyr1 | 0.01501 | -0.15626911 | 0 | 0.097 | 1 | 5 |
| lrch1 | 0.01501 | -0.12750576 | 0 | 0.097 | 1 | 5 |
| lrch4 | 0.01501 | -0.14246877 | 0 | 0.097 | 1 | 5 |
| mfsd12a | 0.01501 | -0.11791112 | 0 | 0.097 | 1 | 5 |
| mllt1a | 0.01501 | -0.1343735 | 0 | 0.097 | 1 | 5 |
| s1pr2 | 0.01501 | -0.11642099 | 0 | 0.097 | 1 | 5 |
| zgc:55262 | 0.01501 | -0.12626524 | 0 | 0.097 | 1 | 5 |
| XLOC-030587 | 0.01501 | -0.14679321 | 0 | 0.097 | 1 | 5 |
| mre11a | 0.01501 | -0.11224292 | 0 | 0.097 | 1 | 5 |
| tyms | 0.01501 | -0.1421804 | 0 | 0.097 | 1 | 5 |
| wu:fc33b09 | 0.01501 | -0.21100396 | 0 | 0.097 | 1 | 5 |
| znf865 | 0.01501 | -0.14315769 | 0 | 0.097 | 1 | 5 |
| caspbl | 0.01501 | -0.19296607 | 0 | 0.097 | 1 | 5 |
| BX908782.2 | 0.01501 | -0.16554001 | 0 | 0.097 | 1 | 5 |
| LOC100538052 | 0.01501 | -0.13948693 | 0 | 0.097 | 1 | 5 |
| klhl20 | 0.01501 | -0.10814623 | 0 | 0.097 | 1 | 5 |
| si:dkey-188i13.9 | 0.01501 | -0.1241531 | 0 | 0.097 | 1 | 5 |

| | | | | | | |
|---|---|---|---|---|---|---|
| slc25a37 | 0.01501 | -0.11297381 | 0 | 0.097 | 1 | 5 |
| rgs2 | 0.01549 | -0.13708963 | 0.206 | 0.387 | 1 | 5 |
| wdfy4 | 0.01575 | -0.32730329 | 0.048 | 0.258 | 1 | 5 |
| CR387933.1 | 0.01583 | -0.29389876 | 0.048 | 0.258 | 1 | 5 |
| tacc1 | 0.01641 | 0.316222552 | 0.238 | 0.032 | 1 | 5 |
| FP101887.1 | 0.01699 | -0.25812705 | 0.032 | 0.194 | 1 | 5 |
| zgc:64065 | 0.01722 | -0.41109507 | 0.54 | 0.806 | 1 | 5 |
| cox17 | 0.01735 | -0.2820615 | 0.476 | 0.774 | 1 | 5 |
| cndp2 | 0.01801 | -0.38080685 | 0.571 | 0.806 | 1 | 5 |
| itgae.2 | 0.01814 | -0.36556453 | 0.254 | 0.548 | 1 | 5 |
| stip1 | 0.01818 | -0.35242832 | 0.127 | 0.387 | 1 | 5 |
| faub | 0.01835 | 0.128315868 | 0.206 | 0.258 | 1 | 5 |
| ip6k2b | 0.01886 | 0.343983566 | 0.429 | 0.29 | 1 | 5 |
| h2afx1 | 0.01886 | 0.236491865 | 0.333 | 0.097 | 1 | 5 |
| fosl1a | 0.01919 | -0.38856069 | 0.111 | 0.323 | 1 | 5 |
| tubb2b | 0.01942 | -0.38339728 | 0.476 | 0.645 | 1 | 5 |
| pfdn6 | 0.01995 | 0.324934082 | 0.27 | 0.097 | 1 | 5 |
| eif3ha | 0.02087 | -0.21129161 | 0.571 | 0.548 | 1 | 5 |
| rpn1 | 0.02146 | 0.38067612 | 0.651 | 0.419 | 1 | 5 |
| dpm2 | 0.02153 | -0.29764509 | 0.111 | 0.355 | 1 | 5 |
| smim8 | 0.02185 | -0.15218178 | 0.016 | 0.161 | 1 | 5 |
| cwc25 | 0.02189 | 0.257109783 | 0.317 | 0.097 | 1 | 5 |
| clcn5b | 0.02211 | -0.40626456 | 0.048 | 0.226 | 1 | 5 |
| fabp3 | 0.02279 | -0.30849282 | 0.111 | 0.355 | 1 | 5 |
| zgc:123010 | 0.02282 | -0.20331979 | 0.016 | 0.161 | 1 | 5 |
| ier2a | 0.02336 | 0.442298443 | 0.746 | 0.484 | 1 | 5 |
| nipsnap3a | 0.02368 | -0.15982484 | 0.032 | 0.194 | 1 | 5 |
| babam2 | 0.02385 | 0.142175903 | 0.111 | 0 | 1 | 5 |
| cul2 | 0.02385 | 0.160348972 | 0.111 | 0 | 1 | 5 |
| fam45a | 0.02385 | 0.159755729 | 0.111 | 0 | 1 | 5 |
| fundc1 | 0.02385 | 0.140463105 | 0.111 | 0 | 1 | 5 |
| maip1 | 0.02385 | 0.112108805 | 0.111 | 0 | 1 | 5 |
| mgmt | 0.02385 | 0.122537874 | 0.111 | 0 | 1 | 5 |
| tapbpl | 0.02385 | 0.193731339 | 0.111 | 0 | 1 | 5 |
| tnfsf10 | 0.02385 | 0.213294671 | 0.111 | 0 | 1 | 5 |
| tram2 | 0.02385 | 0.210165348 | 0.111 | 0 | 1 | 5 |
| zgc:172090 | 0.02385 | 0.190075714 | 0.111 | 0 | 1 | 5 |
| NINJ1 | 0.02385 | 0.161306347 | 0.111 | 0 | 1 | 5 |
| ciao1 | 0.02385 | 0.147865461 | 0.111 | 0 | 1 | 5 |
| cmss1 | 0.02385 | 0.184294443 | 0.111 | 0 | 1 | 5 |
| coq4 | 0.02385 | 0.154717751 | 0.111 | 0 | 1 | 5 |
| dph5 | 0.02385 | 0.129401817 | 0.111 | 0 | 1 | 5 |
| dph7 | 0.02385 | 0.161821585 | 0.111 | 0 | 1 | 5 |
| mob1bb | 0.02385 | 0.156744174 | 0.111 | 0 | 1 | 5 |
| mrpl10 | 0.02385 | 0.141547675 | 0.111 | 0 | 1 | 5 |
| p4ha1b | 0.02385 | 0.156648182 | 0.111 | 0 | 1 | 5 |
| si:dkey-187a12.4 | 0.02385 | 0.207895755 | 0.111 | 0 | 1 | 5 |
| smg6 | 0.02385 | 0.180307816 | 0.111 | 0 | 1 | 5 |

| | | | | | | |
|---|---|---|---|---|---|---|
| ston2 | 0.02385 | 0.186848655 | 0.111 | 0 | 1 | 5 |
| tcta | 0.02385 | 0.160831655 | 0.111 | 0 | 1 | 5 |
| timm29 | 0.02385 | 0.162600636 | 0.111 | 0 | 1 | 5 |
| zcrb1 | 0.02385 | 0.152640137 | 0.111 | 0 | 1 | 5 |
| zgc:194312 | 0.02385 | 0.185320896 | 0.111 | 0 | 1 | 5 |
| prkacaa | 0.02385 | 0.187421226 | 0.111 | 0 | 1 | 5 |
| commd1 | 0.02457 | -0.17526446 | 0.016 | 0.161 | 1 | 5 |
| dnttip2 | 0.02463 | -0.19148709 | 0.016 | 0.161 | 1 | 5 |
| gnai1 | 0.0251 | 0.215576413 | 0.381 | 0.323 | 1 | 5 |
| ppp4cb | 0.02642 | -0.31155827 | 0.063 | 0.226 | 1 | 5 |
| pdia6 | 0.02647 | -0.44913739 | 0.286 | 0.516 | 1 | 5 |
| tuba8l4 | 0.02832 | -0.37001296 | 0.587 | 0.774 | 1 | 5 |
| PDCL3 | 0.02848 | -0.21518424 | 0.016 | 0.161 | 1 | 5 |
| snrpd1 | 0.02858 | 0.358696951 | 0.524 | 0.387 | 1 | 5 |
| snrpd2 | 0.02896 | 0.314486127 | 0.444 | 0.355 | 1 | 5 |
| txnipa | 0.02917 | 0.355618667 | 0.841 | 0.645 | 1 | 5 |
| iqgap1 | 0.02931 | 0.311973212 | 0.286 | 0.065 | 1 | 5 |
| ccnd1 | 0.0295 | -0.26065799 | 0.032 | 0.161 | 1 | 5 |
| rhocb | 0.02967 | -0.36054926 | 0.27 | 0.484 | 1 | 5 |
| zgc:174863 | 0.03001 | -0.19270298 | 0.254 | 0.484 | 1 | 5 |
| fam3c | 0.03029 | -0.15847254 | 0.016 | 0.161 | 1 | 5 |
| arfrp1 | 0.0304 | -0.21675075 | 0.016 | 0.161 | 1 | 5 |
| adam9 | 0.03042 | -0.24814346 | 0.032 | 0.194 | 1 | 5 |
| cxcl20 | 0.03066 | -0.30130136 | 0.032 | 0.129 | 1 | 5 |
| LOC103909796 | 0.03075 | -0.11722636 | 0.016 | 0.129 | 1 | 5 |
| XLOC-032112 | 0.03088 | -0.18657208 | 0.016 | 0.161 | 1 | 5 |
| ppp2r5ea | 0.03093 | -0.16646643 | 0.016 | 0.161 | 1 | 5 |
| ivns1abpa | 0.03101 | -0.24720943 | 0.016 | 0.161 | 1 | 5 |
| hmgcrb | 0.03116 | -0.22183436 | 0.016 | 0.161 | 1 | 5 |
| si:dkey-1h24.6 | 0.03129 | -0.21120822 | 0.016 | 0.161 | 1 | 5 |
| wipf1a | 0.03131 | -0.1801467 | 0.016 | 0.161 | 1 | 5 |
| eif4bb | 0.03154 | 0.292210432 | 0.381 | 0.129 | 1 | 5 |
| arl8a | 0.03182 | -0.27261213 | 0.079 | 0.29 | 1 | 5 |
| wdr1 | 0.03186 | 0.279124735 | 0.698 | 0.581 | 1 | 5 |
| pi4kb | 0.03189 | -0.21301396 | 0.016 | 0.161 | 1 | 5 |
| ehd1a | 0.03206 | -0.23470725 | 0.016 | 0.161 | 1 | 5 |
| si:dkey-171c9.3 | 0.03228 | -0.32251238 | 0.048 | 0.194 | 1 | 5 |
| dnase1l4.2 | 0.0323 | -0.22291918 | 0.016 | 0.161 | 1 | 5 |
| si:ch1073-392o20.2 | 0.03232 | -0.35330659 | 0.429 | 0.71 | 1 | 5 |
| ssr4 | 0.03233 | -0.33154308 | 0.746 | 0.839 | 1 | 5 |
| marcksl1b | 0.03239 | -0.254007 | 0.032 | 0.194 | 1 | 5 |
| ssr1 | 0.03244 | -0.39823219 | 0.365 | 0.581 | 1 | 5 |
| socs7 | 0.03267 | -0.22144385 | 0.016 | 0.161 | 1 | 5 |
| smim7 | 0.03301 | -0.26486009 | 0.079 | 0.29 | 1 | 5 |
| kdm6ba | 0.03304 | -0.31268786 | 0.079 | 0.29 | 1 | 5 |
| arrdc2 | 0.03307 | 0.37314431 | 0.317 | 0.097 | 1 | 5 |
| rpa2 | 0.0332 | -0.1725634 | 0.032 | 0.194 | 1 | 5 |
| pcnp | 0.03334 | -0.17398302 | 0.032 | 0.194 | 1 | 5 |

| | | | | | | |
|---|---|---|---|---|---|---|
| sec61g | 0.03439 | -0.19055992 | 0.905 | 1 | 1 | 5 |
| rab11fip1b | 0.03464 | -0.34601553 | 0.159 | 0.387 | 1 | 5 |
| spcs3 | 0.03502 | 0.394575858 | 0.54 | 0.29 | 1 | 5 |
| zgc:162730 | 0.03512 | 0.142726623 | 0.937 | 0.935 | 1 | 5 |
| ddx56 | 0.03541 | -0.23022365 | 0.032 | 0.194 | 1 | 5 |
| timm17a | 0.03559 | -0.14005041 | 0.127 | 0.323 | 1 | 5 |
| gale | 0.03566 | -0.19686026 | 0.032 | 0.194 | 1 | 5 |
| srp72 | 0.03571 | -0.16270499 | 0.159 | 0.161 | 1 | 5 |
| zfand5a | 0.03613 | -0.20713317 | 0.095 | 0.29 | 1 | 5 |
| tmem14ca | 0.03632 | -0.29549469 | 0.111 | 0.323 | 1 | 5 |
| klf2b | 0.03649 | 0.337096252 | 0.27 | 0.065 | 1 | 5 |
| gpr183a | 0.03755 | 0.393339538 | 0.524 | 0.258 | 1 | 5 |
| lpxn | 0.03857 | -0.29353649 | 0.048 | 0.226 | 1 | 5 |
| fam91a1 | 0.0388 | 0.160982907 | 0.095 | 0 | 1 | 5 |
| klf9 | 0.0388 | 0.107312008 | 0.095 | 0 | 1 | 5 |
| luzp1 | 0.0388 | 0.190548005 | 0.095 | 0 | 1 | 5 |
| myorg | 0.0388 | 0.171932551 | 0.095 | 0 | 1 | 5 |
| rab5b | 0.0388 | 0.136164932 | 0.095 | 0 | 1 | 5 |
| rnf40 | 0.0388 | 0.138126504 | 0.095 | 0 | 1 | 5 |
| si:dkey-172m14.1 | 0.0388 | 0.13575033 | 0.095 | 0 | 1 | 5 |
| slc25a51b | 0.0388 | 0.12221001 | 0.095 | 0 | 1 | 5 |
| slc45a4 | 0.0388 | 0.179193691 | 0.095 | 0 | 1 | 5 |
| ythdc1 | 0.0388 | 0.143868199 | 0.095 | 0 | 1 | 5 |
| znhit1 | 0.0388 | 0.129505191 | 0.095 | 0 | 1 | 5 |
| LO018176.1 | 0.0388 | 0.148675337 | 0.095 | 0 | 1 | 5 |
| abhd13 | 0.0388 | 0.114053363 | 0.095 | 0 | 1 | 5 |
| acacb | 0.0388 | 0.131500218 | 0.095 | 0 | 1 | 5 |
| acp5a | 0.0388 | 0.201454956 | 0.095 | 0 | 1 | 5 |
| arhgef15 | 0.0388 | 0.14975184 | 0.095 | 0 | 1 | 5 |
| ash1l | 0.0388 | 0.151711879 | 0.095 | 0 | 1 | 5 |
| crtc1a | 0.0388 | 0.144303641 | 0.095 | 0 | 1 | 5 |
| dctn5 | 0.0388 | 0.128638506 | 0.095 | 0 | 1 | 5 |
| ddx46 | 0.0388 | 0.14476018 | 0.095 | 0 | 1 | 5 |
| dnajc3b | 0.0388 | 0.164842775 | 0.095 | 0 | 1 | 5 |
| eaf2 | 0.0388 | 0.140976254 | 0.095 | 0 | 1 | 5 |
| fap | 0.0388 | 0.172508439 | 0.095 | 0 | 1 | 5 |
| golga2 | 0.0388 | 0.177220317 | 0.095 | 0 | 1 | 5 |
| hnrnpul1 | 0.0388 | 0.138148173 | 0.095 | 0 | 1 | 5 |
| idh3b | 0.0388 | 0.107234934 | 0.095 | 0 | 1 | 5 |
| mbtps2 | 0.0388 | 0.139453629 | 0.095 | 0 | 1 | 5 |
| mrpl40 | 0.0388 | 0.110573077 | 0.095 | 0 | 1 | 5 |
| pip4k2ca | 0.0388 | 0.159263391 | 0.095 | 0 | 1 | 5 |
| ppp1r10 | 0.0388 | 0.110515523 | 0.095 | 0 | 1 | 5 |
| rcc1 | 0.0388 | 0.113128651 | 0.095 | 0 | 1 | 5 |
| s100w | 0.0388 | 0.124337186 | 0.095 | 0 | 1 | 5 |
| slc30a6 | 0.0388 | 0.130872452 | 0.095 | 0 | 1 | 5 |
| snx14 | 0.0388 | 0.163886983 | 0.095 | 0 | 1 | 5 |
| tprkb | 0.0388 | 0.13777068 | 0.095 | 0 | 1 | 5 |

| | | | | | | |
|---|---|---|---|---|---|---|
| zgc:153759 | 0.0388 | 0.117766383 | 0.095 | 0 | 1 | 5 |
| aebp2 | 0.0388 | 0.119790419 | 0.095 | 0 | 1 | 5 |
| map4k5 | 0.0388 | 0.142118149 | 0.095 | 0 | 1 | 5 |
| tlr22 | 0.0388 | 0.147359254 | 0.095 | 0 | 1 | 5 |
| vdac3 | 0.03917 | -0.36409989 | 0.381 | 0.613 | 1 | 5 |
| si:ch211-158d24.4 | 0.03948 | -0.28816984 | 0.032 | 0.194 | 1 | 5 |
| larp4aa | 0.0395 | -0.19550183 | 0.032 | 0.194 | 1 | 5 |
| bahcc1b | 0.03979 | 0.347788661 | 0.222 | 0.065 | 1 | 5 |
| fam107b | 0.04001 | -0.27936483 | 0.048 | 0.226 | 1 | 5 |
| ctns | 0.04003 | -0.23082645 | 0.048 | 0.226 | 1 | 5 |
| lgals2a | 0.04048 | 0.107085637 | 0.952 | 0.968 | 1 | 5 |
| vsir | 0.04065 | 0.236699304 | 0.762 | 0.677 | 1 | 5 |
| dnajc19 | 0.04088 | -0.29877469 | 0.19 | 0.323 | 1 | 5 |
| parvg | 0.041 | 0.293867966 | 0.762 | 0.645 | 1 | 5 |
| dedd1 | 0.04117 | 0.353270899 | 0.381 | 0.194 | 1 | 5 |
| si:zfos-2326c3.2 | 0.04182 | -0.33178484 | 0.127 | 0.355 | 1 | 5 |
| ptmaa | 0.0424 | 0.211864186 | 0.873 | 0.871 | 1 | 5 |
| lxn | 0.04333 | 0.323102548 | 0.619 | 0.516 | 1 | 5 |
| il10ra | 0.04339 | 0.363468891 | 0.429 | 0.29 | 1 | 5 |
| txndc17 | 0.0438 | 0.220160149 | 0.19 | 0.032 | 1 | 5 |
| XLOC-042595 | 0.04431 | 0.398119488 | 0.556 | 0.323 | 1 | 5 |
| si:rp71-68n21.9 | 0.04461 | 0.329136289 | 0.317 | 0.097 | 1 | 5 |
| tsc22d2 | 0.04487 | -0.32778618 | 0.111 | 0.323 | 1 | 5 |
| zfyve21 | 0.04501 | -0.28883234 | 0.095 | 0.29 | 1 | 5 |
| smarca4a | 0.04504 | 0.23897536 | 0.206 | 0.032 | 1 | 5 |
| osbp | 0.04517 | 0.325444994 | 0.317 | 0.097 | 1 | 5 |
| dbi | 0.04612 | -0.24355235 | 0.683 | 0.903 | 1 | 5 |
| tent5ab | 0.04631 | 0.329247106 | 0.238 | 0.065 | 1 | 5 |
| ap5m1 | 0.0464 | -0.1956642 | 0.016 | 0.129 | 1 | 5 |
| tgm1l1 | 0.04704 | -0.33023143 | 0.095 | 0.29 | 1 | 5 |
| fthl27 | 0.04717 | -0.28435016 | 0.825 | 0.968 | 1 | 5 |
| si:ch211-194m7.3 | 0.04762 | 0.111014263 | 0.46 | 0.29 | 1 | 5 |
| LOC103910015 | 0.04861 | -0.16366201 | 0.063 | 0.226 | 1 | 5 |
| ccdc47 | 0.04906 | -0.14791166 | 0.111 | 0.29 | 1 | 5 |
| sh3kbp1 | 0.04946 | 0.171113781 | 0.175 | 0.032 | 1 | 5 |
| jac3 | 0.0495 | -0.20883135 | 0.444 | 0.677 | 1 | 5 |
| esd | 0.0495 | 0.311840377 | 0.27 | 0.129 | 1 | 5 |
| cited4a | 0.04997 | 0.297786745 | 0.206 | 0.065 | 1 | 5 |
| rras | 0.05083 | -0.26824112 | 0.079 | 0.226 | 1 | 5 |
| cox7c | 0.05205 | -0.23400608 | 0.81 | 0.935 | 1 | 5 |
| sh2b3 | 0.05223 | 0.179267309 | 0.302 | 0.29 | 1 | 5 |
| odc1 | 0.05259 | -0.42037813 | 0.238 | 0.452 | 1 | 5 |
| tmem199 | 0.05287 | -0.20397337 | 0.079 | 0.258 | 1 | 5 |
| selenoh | 0.05298 | -0.11511829 | 0.032 | 0.161 | 1 | 5 |
| cib1 | 0.05303 | -0.32457684 | 0.19 | 0.355 | 1 | 5 |
| gstt1a | 0.05305 | -0.32683878 | 0.159 | 0.387 | 1 | 5 |
| mrps14 | 0.05364 | 0.186820216 | 0.238 | 0.226 | 1 | 5 |
| tmem176l.1 | 0.05375 | -0.29456603 | 0.19 | 0.323 | 1 | 5 |

| | | | | | | |
|---|---|---|---|---|---|---|
| pdhx | 0.05375 | -0.20248304 | 0.032 | 0.161 | 1 | 5 |
| apoeb | 0.05404 | -0.5887652 | 0.048 | 0.194 | 1 | 5 |
| LOC110438280 | 0.05487 | -0.11833696 | 0 | 0.065 | 1 | 5 |
| XLOC-033403 | 0.05487 | -0.10755064 | 0 | 0.065 | 1 | 5 |
| brcc3 | 0.05487 | -0.13432165 | 0 | 0.065 | 1 | 5 |
| chd1l | 0.05487 | -0.1068493 | 0 | 0.065 | 1 | 5 |
| kdm5a | 0.05487 | -0.11310614 | 0 | 0.065 | 1 | 5 |
| ppa2 | 0.05487 | -0.10295033 | 0 | 0.065 | 1 | 5 |
| serpine2 | 0.05487 | -0.10505668 | 0 | 0.065 | 1 | 5 |
| si:ch211-22k7.9 | 0.05487 | -0.13731503 | 0 | 0.065 | 1 | 5 |
| si:ch211-285c6.5 | 0.05487 | -0.10016317 | 0 | 0.065 | 1 | 5 |
| slc44a1b | 0.05487 | -0.10254301 | 0 | 0.065 | 1 | 5 |
| trim36 | 0.05487 | -0.12613479 | 0 | 0.065 | 1 | 5 |
| trim44 | 0.05487 | -0.11040188 | 0 | 0.065 | 1 | 5 |
| ubxn6 | 0.05487 | -0.12204103 | 0 | 0.065 | 1 | 5 |
| usp37 | 0.05487 | -0.10618375 | 0 | 0.065 | 1 | 5 |
| utrn | 0.05487 | -0.12621425 | 0 | 0.065 | 1 | 5 |
| fam57a | 0.05487 | -0.10029078 | 0 | 0.065 | 1 | 5 |
| fastk | 0.05487 | -0.10775655 | 0 | 0.065 | 1 | 5 |
| si:ch1073-67j19.2 | 0.05487 | -0.10070336 | 0 | 0.065 | 1 | 5 |
| tesca | 0.05487 | -0.10851594 | 0 | 0.065 | 1 | 5 |
| zbtb44 | 0.05487 | -0.10359274 | 0 | 0.065 | 1 | 5 |
| cables2a | 0.05487 | -0.13076629 | 0 | 0.065 | 1 | 5 |
| cradd | 0.05487 | -0.10768327 | 0 | 0.065 | 1 | 5 |
| vav2 | 0.05487 | -0.11185647 | 0 | 0.065 | 1 | 5 |
| LOC103909099 | 0.05487 | -0.14024809 | 0 | 0.065 | 1 | 5 |
| LOC103910448 | 0.05487 | -0.17042514 | 0 | 0.065 | 1 | 5 |
| gda | 0.05487 | -0.22909523 | 0 | 0.065 | 1 | 5 |
| gpr184 | 0.05487 | -0.11623525 | 0 | 0.065 | 1 | 5 |
| ppp1r13ba | 0.05487 | -0.11685301 | 0 | 0.065 | 1 | 5 |
| slc9a8 | 0.05487 | -0.11919584 | 0 | 0.065 | 1 | 5 |
| spartb | 0.05487 | -0.10561555 | 0 | 0.065 | 1 | 5 |
| zfhx4 | 0.05487 | -0.12262378 | 0 | 0.065 | 1 | 5 |
| LOC108180204 | 0.05487 | -0.10538404 | 0 | 0.065 | 1 | 5 |
| ankmy2a | 0.05487 | -0.11068584 | 0 | 0.065 | 1 | 5 |
| b4galnt1a | 0.05487 | -0.11783298 | 0 | 0.065 | 1 | 5 |
| btbd7 | 0.05487 | -0.12468902 | 0 | 0.065 | 1 | 5 |
| chst12a | 0.05487 | -0.11062631 | 0 | 0.065 | 1 | 5 |
| gphna | 0.05487 | -0.11985766 | 0 | 0.065 | 1 | 5 |
| lrwd1 | 0.05487 | -0.1430859 | 0 | 0.065 | 1 | 5 |
| orc6 | 0.05487 | -0.13357348 | 0 | 0.065 | 1 | 5 |
| phf2 | 0.05487 | -0.11528953 | 0 | 0.065 | 1 | 5 |
| pole2 | 0.05487 | -0.11196334 | 0 | 0.065 | 1 | 5 |
| rab11fip4a | 0.05487 | -0.13961141 | 0 | 0.065 | 1 | 5 |
| retreg2 | 0.05487 | -0.10873278 | 0 | 0.065 | 1 | 5 |
| rhot1a | 0.05487 | -0.11671121 | 0 | 0.065 | 1 | 5 |
| si:ch211-107m4.1 | 0.05487 | -0.10125078 | 0 | 0.065 | 1 | 5 |
| si:ch211-156b7.4 | 0.05487 | -0.1135595 | 0 | 0.065 | 1 | 5 |

| | | | | | | |
|---|---|---|---|---|---|---|
| sin3b | 0.05487 | -0.13508303 | 0 | 0.065 | 1 | 5 |
| wdr20b | 0.05487 | -0.11314771 | 0 | 0.065 | 1 | 5 |
| zbtb34 | 0.05487 | -0.10546225 | 0 | 0.065 | 1 | 5 |
| znf148 | 0.05487 | -0.10690151 | 0 | 0.065 | 1 | 5 |
| si:ch211-213a13.2 | 0.05487 | -0.10380854 | 0 | 0.065 | 1 | 5 |
| traf3 | 0.05487 | -0.11735656 | 0 | 0.065 | 1 | 5 |
| rnf217 | 0.05487 | -0.14218731 | 0 | 0.065 | 1 | 5 |
| tmem63ba | 0.05487 | -0.12069754 | 0 | 0.065 | 1 | 5 |
| si:ch211-189a15.5 | 0.05487 | -0.1080824 | 0 | 0.065 | 1 | 5 |
| si:dkey-56e3.3 | 0.05487 | -0.11178087 | 0 | 0.065 | 1 | 5 |
| thada | 0.05487 | -0.12226423 | 0 | 0.065 | 1 | 5 |
| LOC101886030 | 0.05487 | -0.11235654 | 0 | 0.065 | 1 | 5 |
| arpc5lb | 0.05487 | -0.10190857 | 0 | 0.065 | 1 | 5 |
| dyrk2 | 0.05487 | -0.11768314 | 0 | 0.065 | 1 | 5 |
| il11b | 0.05487 | -0.10812405 | 0 | 0.065 | 1 | 5 |
| kat8 | 0.05487 | -0.11076771 | 0 | 0.065 | 1 | 5 |
| phactr4a | 0.05487 | -0.10389386 | 0 | 0.065 | 1 | 5 |
| si:dkey-85k7.12 | 0.05487 | -0.12482218 | 0 | 0.065 | 1 | 5 |
| slc39a10 | 0.05487 | -0.11351958 | 0 | 0.065 | 1 | 5 |
| snx10b | 0.05562 | 0.17706635 | 0.254 | 0.097 | 1 | 5 |
| ndufs8a | 0.05572 | 0.208982988 | 0.333 | 0.258 | 1 | 5 |
| nucb1 | 0.05592 | -0.18095766 | 0.111 | 0.29 | 1 | 5 |
| pdpk1b | 0.05593 | -0.19552877 | 0.063 | 0.226 | 1 | 5 |
| rhoca | 0.05656 | 0.320896212 | 0.365 | 0.194 | 1 | 5 |
| dnajc11a | 0.05685 | -0.21671884 | 0.032 | 0.161 | 1 | 5 |
| sdf2l1 | 0.05695 | -0.22648611 | 0.095 | 0.29 | 1 | 5 |
| BX539325.1 | 0.0575 | -0.16879947 | 0.016 | 0.129 | 1 | 5 |
| naa15a | 0.05751 | -0.11662063 | 0.048 | 0.194 | 1 | 5 |
| dusp6 | 0.05753 | 0.337310552 | 0.349 | 0.129 | 1 | 5 |
| LOC101882843 | 0.05754 | 0.272069488 | 0.19 | 0.032 | 1 | 5 |
| u2af1 | 0.05876 | 0.287844834 | 0.429 | 0.29 | 1 | 5 |
| zgc:153118 | 0.05999 | -0.32131991 | 0.794 | 0.903 | 1 | 5 |
| polr1d | 0.06001 | -0.22024878 | 0.095 | 0.29 | 1 | 5 |
| hsbp1b | 0.06059 | -0.30992632 | 0.302 | 0.516 | 1 | 5 |
| CABZ01045617.1 | 0.06107 | 0.345469832 | 0.286 | 0.097 | 1 | 5 |
| sap30l | 0.06111 | -0.24852831 | 0.27 | 0.516 | 1 | 5 |
| ncalda | 0.06147 | -0.27480521 | 0.079 | 0.258 | 1 | 5 |
| gyg1a | 0.06162 | 0.206990465 | 0.19 | 0.032 | 1 | 5 |
| prdm1a | 0.06174 | -0.21867232 | 0.016 | 0.129 | 1 | 5 |
| capns1b | 0.06186 | 0.250742653 | 0.254 | 0.065 | 1 | 5 |
| me1 | 0.06216 | -0.25416587 | 0.127 | 0.226 | 1 | 5 |
| eif3ja | 0.06265 | -0.21585643 | 0.397 | 0.452 | 1 | 5 |
| ywhaba | 0.0627 | -0.31052567 | 0.54 | 0.742 | 1 | 5 |
| pigx | 0.06321 | 0.1409412 | 0.079 | 0 | 1 | 5 |
| zdhhc20b | 0.06321 | 0.158460006 | 0.079 | 0 | 1 | 5 |
| gcdhb | 0.06321 | 0.138844689 | 0.079 | 0 | 1 | 5 |
| gtpbp10 | 0.06321 | 0.104814079 | 0.079 | 0 | 1 | 5 |
| nufip1 | 0.06321 | 0.1413338 | 0.079 | 0 | 1 | 5 |

| | | | | | | |
|---|---|---|---|---|---|---|
| CU639397.1 | 0.06321 | 0.160754602 | 0.079 | 0 | 1 | 5 |
| LOC101884921 | 0.06321 | 0.134641538 | 0.079 | 0 | 1 | 5 |
| XLOC-043861 | 0.06321 | 0.101343825 | 0.079 | 0 | 1 | 5 |
| acot13 | 0.06321 | 0.107228654 | 0.079 | 0 | 1 | 5 |
| ap1m3 | 0.06321 | 0.114398221 | 0.079 | 0 | 1 | 5 |
| ascc1 | 0.06321 | 0.102817314 | 0.079 | 0 | 1 | 5 |
| cmc4 | 0.06321 | 0.123795009 | 0.079 | 0 | 1 | 5 |
| crebrf | 0.06321 | 0.14146395 | 0.079 | 0 | 1 | 5 |
| def6a | 0.06321 | 0.159961675 | 0.079 | 0 | 1 | 5 |
| dhps | 0.06321 | 0.113328592 | 0.079 | 0 | 1 | 5 |
| fuom | 0.06321 | 0.120656244 | 0.079 | 0 | 1 | 5 |
| gpr65 | 0.06321 | 0.118074625 | 0.079 | 0 | 1 | 5 |
| jdp2a | 0.06321 | 0.137313962 | 0.079 | 0 | 1 | 5 |
| kdm5bb | 0.06321 | 0.111413084 | 0.079 | 0 | 1 | 5 |
| kdm6bb | 0.06321 | 0.116379644 | 0.079 | 0 | 1 | 5 |
| klhl11 | 0.06321 | 0.138513133 | 0.079 | 0 | 1 | 5 |
| kmt2a | 0.06321 | 0.155766167 | 0.079 | 0 | 1 | 5 |
| kmt2d | 0.06321 | 0.119580759 | 0.079 | 0 | 1 | 5 |
| mettl2a | 0.06321 | 0.12456654 | 0.079 | 0 | 1 | 5 |
| opa1 | 0.06321 | 0.132449538 | 0.079 | 0 | 1 | 5 |
| osgn1 | 0.06321 | 0.131732058 | 0.079 | 0 | 1 | 5 |
| pcsk7 | 0.06321 | 0.109262659 | 0.079 | 0 | 1 | 5 |
| pdrg1 | 0.06321 | 0.107619085 | 0.079 | 0 | 1 | 5 |
| ralgps1 | 0.06321 | 0.139861651 | 0.079 | 0 | 1 | 5 |
| scpp8 | 0.06321 | 0.142511857 | 0.079 | 0 | 1 | 5 |
| si:ch211-140b10.6 | 0.06321 | 0.110732984 | 0.079 | 0 | 1 | 5 |
| si:ch211-79k12.1 | 0.06321 | 0.146338494 | 0.079 | 0 | 1 | 5 |
| slc6a15 | 0.06321 | 0.112127099 | 0.079 | 0 | 1 | 5 |
| srsf1b | 0.06321 | 0.11829565 | 0.079 | 0 | 1 | 5 |
| syne2b | 0.06321 | 0.121617458 | 0.079 | 0 | 1 | 5 |
| tlk2 | 0.06321 | 0.112514985 | 0.079 | 0 | 1 | 5 |
| vdrb | 0.06321 | 0.12723427 | 0.079 | 0 | 1 | 5 |
| zbtb7b | 0.06321 | 0.163533317 | 0.079 | 0 | 1 | 5 |
| zdhhc17 | 0.06321 | 0.119874717 | 0.079 | 0 | 1 | 5 |
| cep89 | 0.06321 | 0.111215637 | 0.079 | 0 | 1 | 5 |
| dtx4b | 0.06321 | 0.142096194 | 0.079 | 0 | 1 | 5 |
| elp4 | 0.06321 | 0.101146355 | 0.079 | 0 | 1 | 5 |
| fut9b | 0.06321 | 0.125718712 | 0.079 | 0 | 1 | 5 |
| hace1 | 0.06321 | 0.104826865 | 0.079 | 0 | 1 | 5 |
| ints10 | 0.06321 | 0.14816916 | 0.079 | 0 | 1 | 5 |
| msl2b | 0.06321 | 0.108318954 | 0.079 | 0 | 1 | 5 |
| parp2 | 0.06321 | 0.116146949 | 0.079 | 0 | 1 | 5 |
| rab4a | 0.06321 | 0.111274254 | 0.079 | 0 | 1 | 5 |
| rufy3 | 0.06321 | 0.141375313 | 0.079 | 0 | 1 | 5 |
| si:ch73-236c18.8 | 0.06321 | 0.132071715 | 0.079 | 0 | 1 | 5 |
| tbc1d25 | 0.06321 | 0.100086363 | 0.079 | 0 | 1 | 5 |
| trnt1 | 0.06321 | 0.10441147 | 0.079 | 0 | 1 | 5 |
| zswim5 | 0.06321 | 0.123147734 | 0.079 | 0 | 1 | 5 |

| | | | | | | |
|---|---|---|---|---|---|---|
| XLOC-028370 | 0.0633 | 0.457361398 | 0.429 | 0.194 | 1 | 5 |
| nrros | 0.06345 | 0.365350248 | 0.476 | 0.258 | 1 | 5 |
| tnip1 | 0.06433 | -0.14991836 | 0.048 | 0.194 | 1 | 5 |
| ier2b | 0.06452 | 0.334297925 | 0.778 | 0.613 | 1 | 5 |
| eif6 | 0.06453 | -0.20762188 | 0.333 | 0.581 | 1 | 5 |
| uchl3 | 0.06457 | -0.21784551 | 0.048 | 0.194 | 1 | 5 |
| sult2st1 | 0.06477 | 0.201193601 | 0.159 | 0.032 | 1 | 5 |
| rpl27 | 0.06499 | 0.107473531 | 0.984 | 0.968 | 1 | 5 |
| si:ch211-199g17.2 | 0.06506 | 0.260074852 | 0.19 | 0.032 | 1 | 5 |
| arrb2b | 0.06589 | 0.332983509 | 0.381 | 0.194 | 1 | 5 |
| susd6 | 0.0666 | -0.28045284 | 0.063 | 0.226 | 1 | 5 |
| nono | 0.06743 | -0.28978482 | 0.127 | 0.323 | 1 | 5 |
| eif2s1a | 0.06766 | 0.2345455 | 0.175 | 0.032 | 1 | 5 |
| prmt5 | 0.06873 | -0.22368068 | 0.032 | 0.097 | 1 | 5 |
| sh3bgrl | 0.06879 | -0.31286579 | 0.143 | 0.355 | 1 | 5 |
| eif4a1a | 0.06881 | -0.34356225 | 0.46 | 0.645 | 1 | 5 |
| edem1 | 0.06883 | 0.17821448 | 0.19 | 0.032 | 1 | 5 |
| sdcbp2 | 0.069 | -0.27298876 | 0.571 | 0.806 | 1 | 5 |
| snu13b | 0.069 | -0.1693255 | 0.381 | 0.613 | 1 | 5 |
| si:ch211-74m13.3 | 0.06938 | -0.38333273 | 0.444 | 0.677 | 1 | 5 |
| smfn | 0.06943 | -0.25158464 | 0.111 | 0.29 | 1 | 5 |
| cope | 0.06987 | -0.31155721 | 0.222 | 0.452 | 1 | 5 |
| snx1a | 0.06996 | 0.31687395 | 0.492 | 0.323 | 1 | 5 |
| mast2 | 0.07034 | -0.20442791 | 0.016 | 0.129 | 1 | 5 |
| nufip2 | 0.07038 | -0.13965985 | 0.079 | 0.226 | 1 | 5 |
| si:ch73-111e15.1 | 0.07064 | -0.16618078 | 0.016 | 0.129 | 1 | 5 |
| sgta | 0.07075 | 0.159544012 | 0.19 | 0.032 | 1 | 5 |
| brdt | 0.07098 | 0.207011847 | 0.111 | 0.065 | 1 | 5 |
| kdelr2a | 0.07109 | -0.26639011 | 0.079 | 0.258 | 1 | 5 |
| thoc7 | 0.07132 | -0.26293966 | 0.063 | 0.226 | 1 | 5 |
| wdr18 | 0.07132 | -0.24338378 | 0.032 | 0.161 | 1 | 5 |
| zfp36l1a | 0.07187 | 0.303683125 | 0.619 | 0.516 | 1 | 5 |
| rab8b | 0.07206 | -0.27436176 | 0.429 | 0.677 | 1 | 5 |
| ccng2 | 0.07221 | 0.216615741 | 0.238 | 0.194 | 1 | 5 |
| si:ch73-41e3.7 | 0.07247 | -0.28463385 | 0.063 | 0.226 | 1 | 5 |
| slc6a6b | 0.07283 | 0.325928839 | 0.556 | 0.387 | 1 | 5 |
| hdlbpa | 0.07296 | -0.32165159 | 0.508 | 0.677 | 1 | 5 |
| sumo3a | 0.07314 | 0.26448125 | 0.317 | 0.194 | 1 | 5 |
| exosc1 | 0.07333 | -0.14682137 | 0.111 | 0.129 | 1 | 5 |
| ddx3xb | 0.07349 | -0.20816406 | 0.127 | 0.323 | 1 | 5 |
| rnf38 | 0.07353 | -0.13761929 | 0.048 | 0.194 | 1 | 5 |
| irak4 | 0.07357 | -0.26987316 | 0.127 | 0.29 | 1 | 5 |
| ndufab1b | 0.0736 | 0.30602819 | 0.349 | 0.161 | 1 | 5 |
| tbca | 0.07362 | -0.24715898 | 0.397 | 0.645 | 1 | 5 |
| smox | 0.07388 | 0.119471567 | 0.857 | 0.677 | 1 | 5 |
| cd99 | 0.07389 | -0.2808882 | 0.222 | 0.452 | 1 | 5 |
| NC-002333.4 | 0.0751 | -0.21216891 | 0.667 | 0.871 | 1 | 5 |
| rrbp1a | 0.07513 | 0.179402985 | 0.175 | 0.097 | 1 | 5 |

| | | | | | | |
|---|---|---|---|---|---|---|
| prpf6 | 0.07534 | 0.224665601 | 0.143 | 0.032 | 1 | 5 |
| sap18 | 0.07538 | -0.25055629 | 0.365 | 0.613 | 1 | 5 |
| si:ch211-264f5.2 | 0.07568 | -0.31375928 | 0.206 | 0.355 | 1 | 5 |
| hnrnpub | 0.07594 | -0.12633899 | 0.27 | 0.258 | 1 | 5 |
| si:dkeyp-120h9.1 | 0.07609 | -0.13889539 | 0.032 | 0.161 | 1 | 5 |
| cct2 | 0.0762 | 0.295473113 | 0.619 | 0.387 | 1 | 5 |
| si:ch211-14a17.11 | 0.07621 | 0.289259768 | 0.286 | 0.097 | 1 | 5 |
| si:dkey-42l23.4 | 0.07692 | -0.25524064 | 0.016 | 0.129 | 1 | 5 |
| zbtb16a | 0.07728 | 0.322386275 | 0.27 | 0.129 | 1 | 5 |
| txnl1 | 0.0777 | -0.11646985 | 0.016 | 0.129 | 1 | 5 |
| tm2d1 | 0.07866 | -0.16483875 | 0.016 | 0.129 | 1 | 5 |
| smarca2 | 0.07895 | -0.14621059 | 0.032 | 0.161 | 1 | 5 |
| polr1e | 0.07954 | -0.12499145 | 0.016 | 0.129 | 1 | 5 |
| azin1b | 0.08003 | -0.21065677 | 0.016 | 0.129 | 1 | 5 |
| MAP3K11 | 0.08022 | -0.26974153 | 0.048 | 0.194 | 1 | 5 |
| rnf168 | 0.08024 | -0.11919978 | 0.016 | 0.129 | 1 | 5 |
| stat2 | 0.08074 | -0.26144494 | 0.063 | 0.226 | 1 | 5 |
| serpinb1l1 | 0.08123 | -0.25375297 | 0.127 | 0.323 | 1 | 5 |
| fosab | 0.08192 | 0.294220816 | 0.873 | 0.742 | 1 | 5 |
| slc16a10 | 0.082 | -0.17862404 | 0.063 | 0.226 | 1 | 5 |
| elp5 | 0.0825 | -0.12622894 | 0.032 | 0.161 | 1 | 5 |
| LOC110437969 | 0.08315 | -0.23839562 | 0.032 | 0.161 | 1 | 5 |
| swap70b | 0.08324 | 0.20084662 | 0.27 | 0.097 | 1 | 5 |
| polr2j | 0.08341 | -0.1782315 | 0.016 | 0.129 | 1 | 5 |
| tomm20b | 0.08388 | 0.302036168 | 0.524 | 0.355 | 1 | 5 |
| sgk1 | 0.0839 | 0.145614016 | 0.206 | 0.065 | 1 | 5 |
| pitpnbl | 0.08394 | -0.17702578 | 0.016 | 0.129 | 1 | 5 |
| cat | 0.08415 | -0.15247149 | 0.016 | 0.129 | 1 | 5 |
| b3gnt3.4 | 0.08439 | -0.32674403 | 0.206 | 0.387 | 1 | 5 |
| rbx1 | 0.0847 | 0.275477722 | 0.619 | 0.516 | 1 | 5 |
| wu:fb16f03 | 0.08504 | -0.24009957 | 0.127 | 0.323 | 1 | 5 |
| scube1 | 0.08506 | -0.26708654 | 0.063 | 0.226 | 1 | 5 |
| gatad2ab | 0.08522 | -0.23006706 | 0.063 | 0.226 | 1 | 5 |
| pafah1b1b | 0.08553 | -0.13757657 | 0.032 | 0.161 | 1 | 5 |
| stau1 | 0.08556 | -0.17280832 | 0.016 | 0.129 | 1 | 5 |
| eci1 | 0.08577 | -0.16204919 | 0.032 | 0.161 | 1 | 5 |
| cox7b | 0.08596 | -0.19106706 | 0.825 | 0.968 | 1 | 5 |
| ms4a17a.6 | 0.08665 | 0.280629006 | 0.778 | 0.645 | 1 | 5 |
| mtbl | 0.08681 | -0.21808162 | 0.016 | 0.129 | 1 | 5 |
| tia1 | 0.08718 | -0.13114688 | 0.111 | 0.258 | 1 | 5 |
| bloc1s3 | 0.0872 | -0.13091364 | 0.016 | 0.129 | 1 | 5 |
| akap17a | 0.0872 | -0.13320039 | 0.016 | 0.129 | 1 | 5 |
| rnf11b | 0.08728 | -0.13349098 | 0.016 | 0.129 | 1 | 5 |
| rbm7 | 0.08743 | -0.18253659 | 0.032 | 0.161 | 1 | 5 |
| sc:d0202 | 0.08755 | -0.20827543 | 0.016 | 0.129 | 1 | 5 |
| tsc22d3 | 0.08763 | 0.357921013 | 0.492 | 0.323 | 1 | 5 |
| zc3h14 | 0.088 | -0.12442558 | 0.016 | 0.129 | 1 | 5 |
| LOC100006926 | 0.08804 | -0.19546063 | 0.032 | 0.161 | 1 | 5 |

| | | | | | | |
|---|---|---|---|---|---|---|
| uqcrc2a | 0.08813 | -0.11189191 | 0.048 | 0.194 | 1 | 5 |
| rnf146 | 0.08819 | -0.18616046 | 0.016 | 0.129 | 1 | 5 |
| zgc:174268 | 0.08827 | -0.1681638 | 0.063 | 0.097 | 1 | 5 |
| fdx1 | 0.08834 | -0.27032567 | 0.175 | 0.355 | 1 | 5 |
| rcc2 | 0.08907 | -0.15342741 | 0.143 | 0.323 | 1 | 5 |
| polr3h | 0.08909 | -0.19991806 | 0.048 | 0.194 | 1 | 5 |
| si:ch73-335l21.4 | 0.08993 | -0.2493286 | 0.048 | 0.194 | 1 | 5 |
| cxcl11.6 | 0.08996 | -0.40843217 | 0.048 | 0.194 | 1 | 5 |
| pdap1b | 0.09003 | -0.20860611 | 0.206 | 0.419 | 1 | 5 |
| rnf130 | 0.09052 | 0.298468556 | 0.286 | 0.097 | 1 | 5 |
| rwdd1 | 0.09062 | 0.21769228 | 0.333 | 0.29 | 1 | 5 |
| CU467861.1 | 0.09121 | -0.14867618 | 0.032 | 0.161 | 1 | 5 |
| abcf2a | 0.09172 | -0.16326095 | 0.159 | 0.323 | 1 | 5 |
| rdh12l | 0.0918 | -0.2135448 | 0.048 | 0.194 | 1 | 5 |
| plek | 0.092 | -0.28820055 | 0.222 | 0.355 | 1 | 5 |
| slc25a11 | 0.09203 | 0.266056096 | 0.27 | 0.097 | 1 | 5 |
| cdk12 | 0.09206 | -0.16546013 | 0.032 | 0.161 | 1 | 5 |
| BX901883.3 | 0.09243 | -0.17190936 | 0.032 | 0.161 | 1 | 5 |
| alas1 | 0.09251 | 0.311220856 | 0.603 | 0.452 | 1 | 5 |
| sprydd7a | 0.09263 | -0.18654308 | 0.048 | 0.194 | 1 | 5 |
| cnpy1 | 0.09266 | -0.27387589 | 0.159 | 0.355 | 1 | 5 |
| utp15 | 0.09296 | -0.19019312 | 0.048 | 0.194 | 1 | 5 |
| gstr | 0.09374 | -0.13026983 | 0.032 | 0.161 | 1 | 5 |
| CABZ01038697.1 | 0.0939 | -0.18937294 | 0.032 | 0.161 | 1 | 5 |
| trim8a | 0.09398 | -0.23551333 | 0.048 | 0.194 | 1 | 5 |
| spty2d1 | 0.09415 | -0.17755028 | 0.048 | 0.194 | 1 | 5 |
| LOC100536983 | 0.09424 | -0.23264567 | 0.048 | 0.194 | 1 | 5 |
| tmem165 | 0.09438 | 0.228996644 | 0.238 | 0.161 | 1 | 5 |
| ppp2r1ba | 0.09451 | -0.25809508 | 0.19 | 0.323 | 1 | 5 |
| mrpl34 | 0.09454 | -0.1780853 | 0.333 | 0.387 | 1 | 5 |
| mfge8b | 0.09475 | -0.1411824 | 0.032 | 0.161 | 1 | 5 |
| robo4 | 0.09555 | -0.21095007 | 0.032 | 0.161 | 1 | 5 |
| itm2bb | 0.09558 | 0.176465235 | 0.889 | 0.839 | 1 | 5 |
| ap2b1 | 0.09583 | 0.19489688 | 0.175 | 0.032 | 1 | 5 |
| coa6 | 0.09592 | -0.26436017 | 0.254 | 0.452 | 1 | 5 |
| zgc:152791 | 0.09634 | -0.21265694 | 0.032 | 0.161 | 1 | 5 |
| mfsd1 | 0.09684 | -0.20811749 | 0.175 | 0.355 | 1 | 5 |
| rab20 | 0.09702 | 0.274035698 | 0.27 | 0.097 | 1 | 5 |
| tmcc3 | 0.09723 | -0.19084444 | 0.032 | 0.161 | 1 | 5 |
| gnl2 | 0.09728 | -0.14671959 | 0.032 | 0.161 | 1 | 5 |
| zgc:112052 | 0.09748 | -0.20236374 | 0.063 | 0.194 | 1 | 5 |
| rpl8 | 0.09752 | 0.101026537 | 0.968 | 0.968 | 1 | 5 |
| naa38 | 0.09767 | -0.14754052 | 0.032 | 0.161 | 1 | 5 |
| dock11 | 0.09769 | -0.18840935 | 0.032 | 0.161 | 1 | 5 |
| kpna3 | 0.09793 | -0.25890115 | 0.095 | 0.258 | 1 | 5 |
| qkia | 0.09841 | 0.306870486 | 0.444 | 0.226 | 1 | 5 |
| CR855337.1 | 0.09875 | 0.299823039 | 0.19 | 0.065 | 1 | 5 |
| zgc:66388 | 0.09916 | 0.14955117 | 0.175 | 0.161 | 1 | 5 |

| | | | | | | |
|---|---|---|---|---|---|---|
| arpc3 | 0.0994 | 0.17545783 | 0.841 | 0.806 | 1 | 5 |
| si:dkeyp-68b7.12 | 0.10012 | 0.192986442 | 0.254 | 0.097 | 1 | 5 |
| sars | 0.10027 | -0.28820689 | 0.19 | 0.355 | 1 | 5 |
| mdh2 | 0.10203 | 0.239164195 | 0.635 | 0.548 | 1 | 5 |
| naa15b | 0.10336 | 0.120204738 | 0.063 | 0 | 1 | 5 |
| LOC101883468 | 0.10336 | 0.116817534 | 0.063 | 0 | 1 | 5 |
| LOC101885083 | 0.10336 | 0.108633444 | 0.063 | 0 | 1 | 5 |
| acvr2ab | 0.10336 | 0.123633998 | 0.063 | 0 | 1 | 5 |
| cebpg | 0.10336 | 0.11595604 | 0.063 | 0 | 1 | 5 |
| chd8 | 0.10336 | 0.104465517 | 0.063 | 0 | 1 | 5 |
| cnpy2 | 0.10336 | 0.112621476 | 0.063 | 0 | 1 | 5 |
| ctbs | 0.10336 | 0.128526465 | 0.063 | 0 | 1 | 5 |
| cul4b | 0.10336 | 0.100451859 | 0.063 | 0 | 1 | 5 |
| dis3l2 | 0.10336 | 0.102461065 | 0.063 | 0 | 1 | 5 |
| hddc2 | 0.10336 | 0.134562402 | 0.063 | 0 | 1 | 5 |
| mogs | 0.10336 | 0.105036291 | 0.063 | 0 | 1 | 5 |
| rpp25l | 0.10336 | 0.105951766 | 0.063 | 0 | 1 | 5 |
| si:ch211-171h4.6 | 0.10336 | 0.119419583 | 0.063 | 0 | 1 | 5 |
| si:dkey-217d24.6 | 0.10336 | 0.123078173 | 0.063 | 0 | 1 | 5 |
| si:dkeyp-104h9.5 | 0.10336 | 0.103687963 | 0.063 | 0 | 1 | 5 |
| slc38a5b | 0.10336 | 0.100328502 | 0.063 | 0 | 1 | 5 |
| tk2 | 0.10336 | 0.114258636 | 0.063 | 0 | 1 | 5 |
| tmem170a | 0.10336 | 0.109039536 | 0.063 | 0 | 1 | 5 |
| zmp:0000001132 | 0.10336 | 0.126135656 | 0.063 | 0 | 1 | 5 |
| znf704 | 0.10336 | 0.106265927 | 0.063 | 0 | 1 | 5 |
| CABZ01066694.1 | 0.10336 | 0.113498093 | 0.063 | 0 | 1 | 5 |
| adtrp1 | 0.10336 | 0.102902868 | 0.063 | 0 | 1 | 5 |
| card19 | 0.10336 | 0.106898919 | 0.063 | 0 | 1 | 5 |
| ciapin1 | 0.10336 | 0.120890633 | 0.063 | 0 | 1 | 5 |
| col12a1a | 0.10336 | 0.106141223 | 0.063 | 0 | 1 | 5 |
| dapk1 | 0.10336 | 0.105373043 | 0.063 | 0 | 1 | 5 |
| fzr1a | 0.10336 | 0.103932408 | 0.063 | 0 | 1 | 5 |
| gpatch3 | 0.10336 | 0.112663795 | 0.063 | 0 | 1 | 5 |
| il15 | 0.10336 | 0.104312262 | 0.063 | 0 | 1 | 5 |
| klf12b | 0.10336 | 0.10364898 | 0.063 | 0 | 1 | 5 |
| larp7 | 0.10336 | 0.127204681 | 0.063 | 0 | 1 | 5 |
| mapk8b | 0.10336 | 0.118735503 | 0.063 | 0 | 1 | 5 |
| nelfa | 0.10336 | 0.100906356 | 0.063 | 0 | 1 | 5 |
| pcyox1l | 0.10336 | 0.107118452 | 0.063 | 0 | 1 | 5 |
| pex14 | 0.10336 | 0.121850372 | 0.063 | 0 | 1 | 5 |
| ppip5k1b | 0.10336 | 0.103220516 | 0.063 | 0 | 1 | 5 |
| pxnb | 0.10336 | 0.105431251 | 0.063 | 0 | 1 | 5 |
| rab9b | 0.10336 | 0.129064264 | 0.063 | 0 | 1 | 5 |
| si:ch211-1i11.3 | 0.10336 | 0.114268716 | 0.063 | 0 | 1 | 5 |
| si:dkey-57k2.7 | 0.10336 | 0.110191364 | 0.063 | 0 | 1 | 5 |
| si:dkey-7i4.8 | 0.10336 | 0.126766057 | 0.063 | 0 | 1 | 5 |
| smc5 | 0.10336 | 0.122018324 | 0.063 | 0 | 1 | 5 |
| tatdn3 | 0.10336 | 0.117685774 | 0.063 | 0 | 1 | 5 |

| | | | | | | |
|---|---|---|---|---|---|---|
| ube2w | 0.10336 | 0.128627977 | 0.063 | 0 | 1 | 5 |
| xkr8.2 | 0.10336 | 0.107269654 | 0.063 | 0 | 1 | 5 |
| zgc:77086 | 0.10336 | 0.146000426 | 0.063 | 0 | 1 | 5 |
| im:7138535 | 0.10336 | 0.136110633 | 0.063 | 0 | 1 | 5 |
| mocs3 | 0.10336 | 0.102991123 | 0.063 | 0 | 1 | 5 |
| mtss1lb | 0.10336 | 0.132273475 | 0.063 | 0 | 1 | 5 |
| zgc:114130 | 0.10336 | 0.125013442 | 0.063 | 0 | 1 | 5 |
| XLOC-043789 | 0.10336 | 0.109033681 | 0.063 | 0 | 1 | 5 |
| proser3 | 0.10336 | 0.114061536 | 0.063 | 0 | 1 | 5 |
| CABZ01059406.1 | 0.10336 | 0.130096464 | 0.063 | 0 | 1 | 5 |
| XLOC-026010 | 0.10336 | 0.123907296 | 0.063 | 0 | 1 | 5 |
| ccdc130 | 0.10336 | 0.123311732 | 0.063 | 0 | 1 | 5 |
| enoph1 | 0.10336 | 0.118765744 | 0.063 | 0 | 1 | 5 |
| mmp13a | 0.10336 | 0.132008125 | 0.063 | 0 | 1 | 5 |
| pou2f1b | 0.10336 | 0.10081948 | 0.063 | 0 | 1 | 5 |
| riox1 | 0.10336 | 0.107845913 | 0.063 | 0 | 1 | 5 |
| si:ch211-256m1.8 | 0.10336 | 0.129996454 | 0.063 | 0 | 1 | 5 |
| zgc:153044 | 0.10336 | 0.137153556 | 0.063 | 0 | 1 | 5 |
| zgc:66475 | 0.10336 | 0.107246657 | 0.063 | 0 | 1 | 5 |
| znf1015 | 0.10336 | 0.136799264 | 0.063 | 0 | 1 | 5 |
| erap1a | 0.10336 | 0.103535549 | 0.063 | 0 | 1 | 5 |
| sulf2b | 0.10365 | 0.279451611 | 0.238 | 0.097 | 1 | 5 |
| sec61a1 | 0.10415 | -0.30749464 | 0.46 | 0.645 | 1 | 5 |
| XLOC-013964 | 0.10486 | 0.335386244 | 0.444 | 0.226 | 1 | 5 |
| srsf9 | 0.10524 | 0.188376789 | 0.302 | 0.258 | 1 | 5 |
| sf3a2 | 0.1057 | 0.132883833 | 0.302 | 0.323 | 1 | 5 |
| ptpmt1 | 0.106 | -0.11164118 | 0.063 | 0.194 | 1 | 5 |
| grcc10 | 0.10656 | -0.10480113 | 0.063 | 0.194 | 1 | 5 |
| map3k4 | 0.10765 | 0.245552702 | 0.159 | 0.032 | 1 | 5 |
| ankrd11 | 0.10869 | 0.292243986 | 0.365 | 0.161 | 1 | 5 |
| si:dkey-51e6.1 | 0.11012 | -0.26637071 | 0.111 | 0.258 | 1 | 5 |
| junba | 0.11028 | 0.239721975 | 0.873 | 0.774 | 1 | 5 |
| CU499336.2 | 0.11149 | -0.16853473 | 0.127 | 0.29 | 1 | 5 |
| csnk1g2b | 0.11209 | 0.226331805 | 0.19 | 0.065 | 1 | 5 |
| si:dkey-15j16.3 | 0.11279 | -0.13817539 | 0.048 | 0.161 | 1 | 5 |
| ssh2a | 0.11331 | -0.21911009 | 0.143 | 0.323 | 1 | 5 |
| p4hb | 0.11346 | -0.28161274 | 0.476 | 0.548 | 1 | 5 |
| paip2b | 0.11429 | 0.289940351 | 0.492 | 0.29 | 1 | 5 |
| ISCU (1 of many) | 0.1148 | 0.219533423 | 0.317 | 0.129 | 1 | 5 |
| cisd2 | 0.11482 | -0.22284562 | 0.063 | 0.194 | 1 | 5 |
| cmklr1 | 0.11531 | 0.354675976 | 0.651 | 0.452 | 1 | 5 |
| map2k6 | 0.11554 | 0.262618809 | 0.222 | 0.065 | 1 | 5 |
| pfn2 | 0.11609 | 0.171754235 | 0.27 | 0.258 | 1 | 5 |
| entpd2a.1 | 0.11614 | -0.33245202 | 0.206 | 0.387 | 1 | 5 |
| rab32a | 0.11628 | -0.1277478 | 0.175 | 0.323 | 1 | 5 |
| mbd2 | 0.11663 | -0.27618249 | 0.27 | 0.484 | 1 | 5 |
| nol12 | 0.11717 | -0.17885074 | 0.079 | 0.226 | 1 | 5 |
| pmm2 | 0.11781 | -0.24691481 | 0.127 | 0.29 | 1 | 5 |

| | | | | | | |
|---|---|---|---|---|---|---|
| capn2a | 0.11784 | -0.10711195 | 0.127 | 0.129 | 1 | 5 |
| higd1a | 0.11801 | -0.28089328 | 0.492 | 0.71 | 1 | 5 |
| si:ch73-376l24.2 | 0.11814 | 0.271809236 | 0.333 | 0.226 | 1 | 5 |
| PLBD2 | 0.11846 | -0.11267357 | 0.19 | 0.194 | 1 | 5 |
| rnaset2l | 0.11878 | 0.207421656 | 0.619 | 0.581 | 1 | 5 |
| CABZ01078737.1 | 0.11901 | -0.29497651 | 0.095 | 0.258 | 1 | 5 |
| dnajb9b | 0.11903 | -0.24389464 | 0.063 | 0.194 | 1 | 5 |
| hcls1 | 0.11927 | -0.30384907 | 0.365 | 0.581 | 1 | 5 |
| slc9a3r1a | 0.11984 | -0.23667939 | 0.079 | 0.194 | 1 | 5 |
| mknk2b | 0.12073 | 0.318591821 | 0.397 | 0.194 | 1 | 5 |
| malt1 | 0.12078 | 0.2266932 | 0.19 | 0.065 | 1 | 5 |
| ubr7 | 0.12224 | -0.1807569 | 0.048 | 0.161 | 1 | 5 |
| si:ch211-15b10.6 | 0.12265 | 0.2982486 | 0.492 | 0.323 | 1 | 5 |
| LO018340.1 | 0.12293 | -0.24511055 | 0.19 | 0.387 | 1 | 5 |
| si:dkey-42i9.4 | 0.12344 | 0.23826903 | 0.27 | 0.097 | 1 | 5 |
| mbnl1 | 0.12354 | 0.288505132 | 0.571 | 0.355 | 1 | 5 |
| tardbp | 0.12384 | 0.226387611 | 0.206 | 0.097 | 1 | 5 |
| ccdc88b | 0.12491 | -0.23432245 | 0.19 | 0.387 | 1 | 5 |
| setb | 0.12492 | -0.12645997 | 0.381 | 0.581 | 1 | 5 |
| coq9 | 0.12508 | 0.194350949 | 0.143 | 0.032 | 1 | 5 |
| si:ch211-235e9.6 | 0.12539 | 0.17796277 | 0.524 | 0.323 | 1 | 5 |
| tinagl1 | 0.12548 | -0.30681853 | 0.079 | 0.161 | 1 | 5 |
| slc43a2b | 0.12563 | -0.26163795 | 0.063 | 0.161 | 1 | 5 |
| ehmt1b | 0.12573 | -0.21679888 | 0.063 | 0.194 | 1 | 5 |
| sypl2a | 0.12594 | 0.206378593 | 0.159 | 0.032 | 1 | 5 |
| gnl3 | 0.12635 | -0.14307196 | 0.095 | 0.258 | 1 | 5 |
| namptb | 0.1269 | -0.17386627 | 0.032 | 0.129 | 1 | 5 |
| snx13 | 0.12692 | -0.2041687 | 0.079 | 0.226 | 1 | 5 |
| hint1 | 0.12695 | 0.245501767 | 0.397 | 0.194 | 1 | 5 |
| naca | 0.12699 | 0.154570377 | 0.984 | 0.935 | 1 | 5 |
| idh1 | 0.12705 | -0.2678352 | 0.222 | 0.387 | 1 | 5 |
| si:dkey-188i13.10 | 0.12716 | -0.27182019 | 0.19 | 0.355 | 1 | 5 |
| utp4 | 0.12725 | -0.19805222 | 0.079 | 0.226 | 1 | 5 |
| CABZ01080702.1 | 0.12789 | -0.26947751 | 0.508 | 0.645 | 1 | 5 |
| grap2b | 0.12791 | 0.245392399 | 0.683 | 0.581 | 1 | 5 |
| h2afy2 | 0.12792 | -0.20826326 | 0.19 | 0.387 | 1 | 5 |
| rnf44 | 0.12806 | 0.267437416 | 0.27 | 0.097 | 1 | 5 |
| cxcl8b.1 | 0.12834 | -0.43083394 | 0.397 | 0.613 | 1 | 5 |
| arpc4 | 0.12864 | 0.114494577 | 0.651 | 0.677 | 1 | 5 |
| ppil2 | 0.12895 | 0.156250822 | 0.111 | 0.032 | 1 | 5 |
| dpf2 | 0.12898 | -0.21633258 | 0.095 | 0.258 | 1 | 5 |
| dusp5 | 0.12923 | -0.28254751 | 0.19 | 0.387 | 1 | 5 |
| kif5bb | 0.13057 | -0.21005342 | 0.048 | 0.161 | 1 | 5 |
| acsl4b | 0.13082 | -0.18074242 | 0.381 | 0.419 | 1 | 5 |
| apbb1ip | 0.1318 | -0.14974562 | 0.111 | 0.258 | 1 | 5 |
| supt6h | 0.13184 | 0.175773319 | 0.206 | 0.161 | 1 | 5 |
| ndufb5 | 0.13191 | 0.115536852 | 0.444 | 0.258 | 1 | 5 |
| coro1a | 0.13274 | -0.20021873 | 0.794 | 0.935 | 1 | 5 |

| | | | | | | |
|---|---|---|---|---|---|---|
| CABZ01034698.2 | 0.13322 | -0.21085403 | 0.063 | 0.194 | 1 | 5 |
| tuba8l2 | 0.1343 | 0.288099047 | 0.587 | 0.452 | 1 | 5 |
| wu:fi37e09 | 0.13516 | -0.29006083 | 0.286 | 0.484 | 1 | 5 |
| socs1a | 0.13533 | -0.20014967 | 0.111 | 0.258 | 1 | 5 |
| lsm12b | 0.13634 | -0.25139232 | 0.397 | 0.613 | 1 | 5 |
| manf | 0.13714 | -0.13202559 | 0.286 | 0.452 | 1 | 5 |
| pdhb | 0.13726 | 0.192393139 | 0.159 | 0.032 | 1 | 5 |
| eny2 | 0.13734 | 0.227943855 | 0.349 | 0.161 | 1 | 5 |
| uqcrfs1 | 0.13742 | -0.26393927 | 0.397 | 0.548 | 1 | 5 |
| ppib | 0.13825 | -0.15903982 | 0.841 | 0.968 | 1 | 5 |
| nop53 | 0.13895 | -0.24639603 | 0.429 | 0.645 | 1 | 5 |
| mdm4 | 0.13933 | -0.16201249 | 0.032 | 0.129 | 1 | 5 |
| cfdp1 | 0.14027 | 0.189585178 | 0.143 | 0.032 | 1 | 5 |
| elmsan1b | 0.14039 | 0.229658662 | 0.206 | 0.065 | 1 | 5 |
| LOC103909115 | 0.14042 | 0.221896418 | 0.603 | 0.516 | 1 | 5 |
| btg1 | 0.14057 | 0.105520831 | 0.762 | 0.806 | 1 | 5 |
| cpsf4 | 0.14089 | -0.17625047 | 0.016 | 0.097 | 1 | 5 |
| capns1a | 0.14094 | 0.200389378 | 0.302 | 0.226 | 1 | 5 |
| copb1 | 0.14356 | -0.20815606 | 0.079 | 0.226 | 1 | 5 |
| si:ch211-197h24.9 | 0.1446 | 0.149567105 | 0.159 | 0.032 | 1 | 5 |
| chd4a | 0.14488 | 0.243900803 | 0.302 | 0.129 | 1 | 5 |
| tmsb1 | 0.14494 | 0.260952565 | 0.825 | 0.645 | 1 | 5 |
| frrs1b | 0.14574 | -0.1540633 | 0.032 | 0.129 | 1 | 5 |
| ywhae1 | 0.14616 | 0.260081924 | 0.508 | 0.419 | 1 | 5 |
| ddt | 0.1463 | 0.232770081 | 0.317 | 0.161 | 1 | 5 |
| tnfrsf18 | 0.14699 | -0.19770976 | 0.032 | 0.129 | 1 | 5 |
| CU929150.1 | 0.14699 | 0.253898832 | 0.365 | 0.194 | 1 | 5 |
| slc25a36a | 0.14723 | 0.172221488 | 0.159 | 0.032 | 1 | 5 |
| smndc1 | 0.14729 | 0.131893071 | 0.222 | 0.226 | 1 | 5 |
| cep85l | 0.14736 | -0.23859915 | 0.032 | 0.129 | 1 | 5 |
| usf1l | 0.14781 | -0.20058565 | 0.079 | 0.226 | 1 | 5 |
| histh1l | 0.14839 | -0.20976254 | 0.175 | 0.355 | 1 | 5 |
| ugt5b2 | 0.14851 | 0.272215355 | 0.238 | 0.097 | 1 | 5 |
| CR383676.1 | 0.14853 | 0.186144482 | 0.952 | 0.839 | 1 | 5 |
| fubp1 | 0.14877 | 0.265652836 | 0.286 | 0.129 | 1 | 5 |
| nktr | 0.14915 | -0.11644622 | 0.048 | 0.161 | 1 | 5 |
| egln1b | 0.14968 | -0.23060937 | 0.095 | 0.226 | 1 | 5 |
| ube2na | 0.15002 | 0.126754127 | 0.54 | 0.516 | 1 | 5 |
| tlr5b | 0.15008 | 0.20194704 | 0.143 | 0.032 | 1 | 5 |
| brd3b | 0.15022 | -0.17249503 | 0.048 | 0.161 | 1 | 5 |
| mydgf | 0.15124 | 0.180079639 | 0.143 | 0.032 | 1 | 5 |
| si:ch73-40a2.1 | 0.15205 | -0.26160703 | 0.079 | 0.194 | 1 | 5 |
| nrbf2b | 0.1522 | -0.11204267 | 0.063 | 0.065 | 1 | 5 |
| CABZ01020840.1 | 0.15225 | -0.20211781 | 0.127 | 0.29 | 1 | 5 |
| hectd3 | 0.15271 | -0.19236645 | 0.079 | 0.226 | 1 | 5 |
| LOC101882921 | 0.15331 | -0.22592939 | 0.032 | 0.129 | 1 | 5 |
| zgc:162339 | 0.15379 | 0.190794788 | 0.143 | 0.065 | 1 | 5 |
| eif3s10 | 0.15397 | -0.21950948 | 0.476 | 0.548 | 1 | 5 |

| | | | | | | |
|---|---|---|---|---|---|---|
| pin1 | 0.1541 | -0.18176528 | 0.19 | 0.258 | 1 | 5 |
| fam208b | 0.15477 | 0.195412308 | 0.206 | 0.065 | 1 | 5 |
| cox5aa | 0.15556 | -0.20450094 | 0.698 | 0.871 | 1 | 5 |
| sem1 | 0.15563 | 0.167053454 | 0.603 | 0.581 | 1 | 5 |
| prkcsh | 0.15566 | 0.268270899 | 0.302 | 0.161 | 1 | 5 |
| eloca | 0.1566 | 0.114829748 | 0.095 | 0.097 | 1 | 5 |
| si:dkey-9i23.15 | 0.15662 | 0.21726135 | 0.159 | 0.065 | 1 | 5 |
| zgc:103692 | 0.15672 | 0.15178989 | 0.111 | 0.032 | 1 | 5 |
| psenen | 0.15693 | 0.288395165 | 0.333 | 0.161 | 1 | 5 |
| ncf4 | 0.15717 | -0.26518165 | 0.111 | 0.258 | 1 | 5 |
| xcr1b.3 | 0.1572 | -0.23257786 | 0.063 | 0.194 | 1 | 5 |
| hbp1 | 0.15733 | 0.108431692 | 0.127 | 0.032 | 1 | 5 |
| zgc:113208 | 0.15754 | -0.15598533 | 0.063 | 0.194 | 1 | 5 |
| entpd1 | 0.15835 | -0.21336358 | 0.508 | 0.71 | 1 | 5 |
| txnl4a | 0.15868 | -0.16176513 | 0.159 | 0.323 | 1 | 5 |
| nop56 | 0.15885 | 0.275568341 | 0.556 | 0.355 | 1 | 5 |
| wee2 | 0.15939 | 0.251668704 | 0.302 | 0.129 | 1 | 5 |
| aclya | 0.15942 | 0.160265821 | 0.111 | 0.032 | 1 | 5 |
| zgc:136439 | 0.15943 | 0.13608453 | 0.19 | 0.065 | 1 | 5 |
| FO704821.1 | 0.15986 | -0.17458387 | 0.048 | 0.161 | 1 | 5 |
| alyref | 0.15988 | -0.26466874 | 0.254 | 0.452 | 1 | 5 |
| her9 | 0.16038 | 0.226536477 | 0.127 | 0.032 | 1 | 5 |
| eif4e1c | 0.16062 | -0.15915316 | 0.127 | 0.29 | 1 | 5 |
| mrps24 | 0.16083 | 0.207197403 | 0.143 | 0.032 | 1 | 5 |
| pbdc1 | 0.16084 | -0.15262007 | 0.365 | 0.387 | 1 | 5 |
| polr2f | 0.16139 | 0.145305969 | 0.27 | 0.258 | 1 | 5 |
| timm10 | 0.16141 | 0.199320889 | 0.159 | 0.065 | 1 | 5 |
| eif5a2 | 0.16155 | 0.266830781 | 0.762 | 0.645 | 1 | 5 |
| slc16a9a | 0.16179 | -0.28540604 | 0.159 | 0.29 | 1 | 5 |
| tpm4a | 0.1621 | -0.18901682 | 0.032 | 0.129 | 1 | 5 |
| tnfrsf11b | 0.16216 | -0.17957912 | 0.048 | 0.065 | 1 | 5 |
| celf2 | 0.16235 | 0.278354727 | 0.73 | 0.548 | 1 | 5 |
| klf2a | 0.16242 | 0.272128605 | 0.206 | 0.065 | 1 | 5 |
| casc3 | 0.16257 | -0.21910807 | 0.111 | 0.258 | 1 | 5 |
| ubap1 | 0.16281 | 0.221094537 | 0.143 | 0.032 | 1 | 5 |
| cfh | 0.1634 | -0.21676864 | 0.063 | 0.194 | 1 | 5 |
| commd4 | 0.16355 | 0.155557093 | 0.111 | 0.032 | 1 | 5 |
| arf2a | 0.16388 | -0.28835434 | 0.349 | 0.548 | 1 | 5 |
| CABZ01030278.1 | 0.16422 | -0.30421239 | 0.175 | 0.355 | 1 | 5 |
| rbbp4 | 0.16439 | -0.11976582 | 0.016 | 0.097 | 1 | 5 |
| rbpjb | 0.16457 | -0.17244534 | 0.238 | 0.29 | 1 | 5 |
| snrnp35 | 0.16491 | -0.13793201 | 0.016 | 0.097 | 1 | 5 |
| sdc4 | 0.16501 | -0.25735327 | 0.683 | 0.839 | 1 | 5 |
| cnbpb | 0.16572 | -0.19574641 | 0.397 | 0.452 | 1 | 5 |
| eef1da | 0.16597 | -0.16205004 | 0.857 | 0.968 | 1 | 5 |
| qrsl1 | 0.16602 | -0.17348063 | 0.016 | 0.097 | 1 | 5 |
| bcl7bb | 0.16656 | 0.21072984 | 0.19 | 0.065 | 1 | 5 |
| nup62l | 0.16662 | -0.20506385 | 0.111 | 0.258 | 1 | 5 |

| | | | | | | |
|---|---|---|---|---|---|---|
| naa10 | 0.16685 | 0.143101248 | 0.46 | 0.452 | 1 | 5 |
| spry4 | 0.16701 | -0.16101466 | 0.016 | 0.097 | 1 | 5 |
| si:ch1073-443f11.2 | 0.16703 | 0.222776612 | 0.19 | 0.065 | 1 | 5 |
| txn2 | 0.16708 | -0.21539541 | 0.27 | 0.355 | 1 | 5 |
| lypla2 | 0.16804 | 0.175522987 | 0.206 | 0.065 | 1 | 5 |
| cst3 | 0.16808 | -0.21681416 | 0.127 | 0.29 | 1 | 5 |
| vps29 | 0.16854 | 0.216883707 | 0.54 | 0.419 | 1 | 5 |
| plekho2 | 0.16864 | 0.160258074 | 0.238 | 0.194 | 1 | 5 |
| cdv3 | 0.16921 | -0.24805273 | 0.317 | 0.419 | 1 | 5 |
| gng12a | 0.16959 | -0.16700067 | 0.048 | 0.161 | 1 | 5 |
| stk24b | 0.16965 | -0.18250804 | 0.063 | 0.194 | 1 | 5 |
| trappc2l | 0.16996 | 0.217428639 | 0.254 | 0.097 | 1 | 5 |
| il1fma | 0.17 | -0.18027483 | 0.063 | 0.194 | 1 | 5 |
| sept2 | 0.17083 | -0.13773778 | 0.254 | 0.258 | 1 | 5 |
| atp5pf | 0.17089 | -0.20401324 | 0.619 | 0.806 | 1 | 5 |
| cebpa | 0.17113 | 0.155008377 | 0.651 | 0.645 | 1 | 5 |
| fbl | 0.17139 | -0.27854357 | 0.286 | 0.452 | 1 | 5 |
| CU659670.1 | 0.17162 | -0.18251785 | 0.048 | 0.161 | 1 | 5 |
| si:dkeyp-69e1.8 | 0.17164 | -0.15739246 | 0.032 | 0.129 | 1 | 5 |
| kras | 0.17196 | -0.16724752 | 0.063 | 0.194 | 1 | 5 |
| pcf11 | 0.172 | -0.11297833 | 0.238 | 0.387 | 1 | 5 |
| mef2d | 0.17231 | -0.21045176 | 0.063 | 0.194 | 1 | 5 |
| dffa | 0.17305 | -0.17039235 | 0.048 | 0.161 | 1 | 5 |
| serpine1 | 0.17317 | -0.28514952 | 0.063 | 0.194 | 1 | 5 |
| si:dkey-7i4.24 | 0.17321 | 0.360647659 | 0.476 | 0.419 | 1 | 5 |
| bpgm | 0.17373 | -0.17249988 | 0.016 | 0.097 | 1 | 5 |
| prdx6 | 0.17408 | -0.24962264 | 0.46 | 0.548 | 1 | 5 |
| dpy30 | 0.17445 | 0.205548328 | 0.286 | 0.161 | 1 | 5 |
| bccip | 0.17461 | -0.24561241 | 0.349 | 0.548 | 1 | 5 |
| arhgef4 | 0.17472 | -0.21913687 | 0.032 | 0.129 | 1 | 5 |
| si:dkey-262k9.4 | 0.17478 | -0.20582418 | 0.571 | 0.677 | 1 | 5 |
| adssl | 0.17547 | 0.124248289 | 0.175 | 0.065 | 1 | 5 |
| aars | 0.1756 | 0.239237916 | 0.302 | 0.161 | 1 | 5 |
| si:dkey-112e7.2 | 0.17651 | 0.114129961 | 0.667 | 0.677 | 1 | 5 |
| kdm5c | 0.17652 | -0.1815981 | 0.048 | 0.161 | 1 | 5 |
| cks1b | 0.17653 | -0.12557454 | 0.016 | 0.097 | 1 | 5 |
| FP102809.1 | 0.17674 | -0.14386392 | 0.016 | 0.097 | 1 | 5 |
| ube2e1 | 0.17716 | -0.24737174 | 0.206 | 0.355 | 1 | 5 |
| ddx39aa | 0.17739 | -0.15981651 | 0.27 | 0.452 | 1 | 5 |
| BX323596.2 | 0.17792 | -0.36525703 | 0.238 | 0.355 | 1 | 5 |
| dpp7 | 0.17811 | 0.264934377 | 0.683 | 0.548 | 1 | 5 |
| nt5dc2 | 0.17834 | 0.196736149 | 0.333 | 0.161 | 1 | 5 |
| cds1 | 0.17913 | -0.26167303 | 0.032 | 0.129 | 1 | 5 |
| snip1 | 0.17932 | -0.15899231 | 0.063 | 0.194 | 1 | 5 |
| casp9 | 0.17938 | -0.19707415 | 0.063 | 0.194 | 1 | 5 |
| ripor1.1 | 0.17938 | -0.2060233 | 0.063 | 0.194 | 1 | 5 |
| sept8a | 0.17961 | -0.20386611 | 0.159 | 0.323 | 1 | 5 |
| jarid2b | 0.17998 | -0.11721637 | 0.063 | 0.065 | 1 | 5 |

| | | | | | | |
|---|---|---|---|---|---|---|
| tmx1 | 0.18067 | 0.208536662 | 0.222 | 0.161 | 1 | 5 |
| arf5 | 0.1822 | 0.249269501 | 0.619 | 0.419 | 1 | 5 |
| tbc1d14 | 0.18249 | 0.166332536 | 0.127 | 0.032 | 1 | 5 |
| arhgap31 | 0.18254 | -0.12373516 | 0.016 | 0.097 | 1 | 5 |
| h2afy | 0.18266 | -0.21872299 | 0.048 | 0.161 | 1 | 5 |
| ostm1 | 0.18273 | 0.175758702 | 0.143 | 0.032 | 1 | 5 |
| tlr1 | 0.1829 | -0.16632259 | 0.032 | 0.129 | 1 | 5 |
| rps6ka5 | 0.18298 | 0.190753971 | 0.143 | 0.032 | 1 | 5 |
| dspa | 0.18301 | -0.15989378 | 0.111 | 0.161 | 1 | 5 |
| irf2a | 0.18309 | -0.11406253 | 0.159 | 0.29 | 1 | 5 |
| zfpl1 | 0.18327 | -0.11371857 | 0.016 | 0.097 | 1 | 5 |
| atp8a1 | 0.18333 | 0.167603406 | 0.127 | 0.065 | 1 | 5 |
| gskip | 0.18336 | 0.210203656 | 0.286 | 0.129 | 1 | 5 |
| emc3 | 0.18352 | -0.23045725 | 0.079 | 0.194 | 1 | 5 |
| selenow2a | 0.1842 | 0.189642162 | 0.143 | 0.032 | 1 | 5 |
| eif3d | 0.18497 | -0.16828975 | 0.587 | 0.774 | 1 | 5 |
| marveld1 | 0.1855 | -0.14511213 | 0.048 | 0.161 | 1 | 5 |
| lman2 | 0.18554 | 0.107394147 | 0.286 | 0.29 | 1 | 5 |
| zgc:92606 | 0.1857 | 0.150042679 | 0.143 | 0.032 | 1 | 5 |
| txndc15 | 0.1857 | -0.15763499 | 0.159 | 0.194 | 1 | 5 |
| mtss1 | 0.18648 | 0.21264628 | 0.127 | 0.032 | 1 | 5 |
| brd1a | 0.18717 | -0.12912068 | 0.016 | 0.097 | 1 | 5 |
| ing3 | 0.18721 | -0.18354408 | 0.032 | 0.129 | 1 | 5 |
| magoh | 0.18758 | 0.128923514 | 0.286 | 0.258 | 1 | 5 |
| rrp36 | 0.18837 | -0.15897472 | 0.048 | 0.161 | 1 | 5 |
| nucb2b | 0.18856 | -0.24229633 | 0.111 | 0.226 | 1 | 5 |
| gnb1b | 0.18867 | -0.23557671 | 0.508 | 0.645 | 1 | 5 |
| brk1 | 0.18881 | 0.228496872 | 0.349 | 0.226 | 1 | 5 |
| laptm5 | 0.18888 | 0.148751581 | 0.952 | 0.839 | 1 | 5 |
| tmem176l.3a | 0.18898 | 0.230223533 | 0.333 | 0.226 | 1 | 5 |
| ly6m5 | 0.1892 | -0.27940418 | 0.222 | 0.387 | 1 | 5 |
| si:ch1073-358c10.1 | 0.18939 | -0.12230697 | 0.048 | 0.065 | 1 | 5 |
| taf10 | 0.19011 | 0.136758133 | 0.095 | 0.032 | 1 | 5 |
| elk1 | 0.19028 | -0.1413861 | 0.032 | 0.129 | 1 | 5 |
| baz1a | 0.1908 | -0.20697147 | 0.159 | 0.323 | 1 | 5 |
| nfil3 | 0.1909 | -0.10608384 | 0.063 | 0.161 | 1 | 5 |
| tbl1xr1b | 0.19139 | -0.14617154 | 0.048 | 0.161 | 1 | 5 |
| igf2r | 0.19141 | -0.16082872 | 0.032 | 0.129 | 1 | 5 |
| spg21 | 0.19141 | 0.231391787 | 0.238 | 0.097 | 1 | 5 |
| mvb12ba | 0.19185 | -0.16662349 | 0.016 | 0.097 | 1 | 5 |
| mrps25 | 0.19185 | 0.185597359 | 0.206 | 0.097 | 1 | 5 |
| ctsl.1 | 0.19232 | 0.228771331 | 0.175 | 0.097 | 1 | 5 |
| wt1b | 0.19241 | -0.23452748 | 0.095 | 0.194 | 1 | 5 |
| cmip | 0.19276 | 0.214849937 | 0.254 | 0.161 | 1 | 5 |
| atp5pb | 0.19313 | 0.207178496 | 0.651 | 0.613 | 1 | 5 |
| LOC110437731 | 0.19322 | -0.22006812 | 0.778 | 0.903 | 1 | 5 |
| srsf10b | 0.19404 | 0.129780082 | 0.127 | 0.032 | 1 | 5 |
| rerea | 0.19425 | 0.156512346 | 0.127 | 0.032 | 1 | 5 |

| | | | | | | |
|---|---|---|---|---|---|---|
| slc2a3b | 0.19505 | -0.11866837 | 0.127 | 0.258 | 1 | 5 |
| chd1 | 0.19517 | -0.13188734 | 0.032 | 0.129 | 1 | 5 |
| snx17 | 0.19527 | 0.199403671 | 0.159 | 0.065 | 1 | 5 |
| blnk | 0.1959 | 0.222892005 | 0.159 | 0.065 | 1 | 5 |
| tomm34 | 0.19601 | -0.13387273 | 0.048 | 0.161 | 1 | 5 |
| spink2.1 | 0.19621 | -0.13738055 | 0.048 | 0.161 | 1 | 5 |
| prkcdb | 0.19666 | 0.186669545 | 0.143 | 0.032 | 1 | 5 |
| hmgb1a | 0.19674 | 0.160637935 | 0.317 | 0.29 | 1 | 5 |
| si:dkey-184a18.5 | 0.19729 | -0.10158886 | 0.032 | 0.129 | 1 | 5 |
| adar | 0.19741 | -0.17952617 | 0.016 | 0.097 | 1 | 5 |
| pxna | 0.19819 | 0.146760546 | 0.143 | 0.032 | 1 | 5 |
| hspa5 | 0.19842 | -0.31388386 | 0.46 | 0.613 | 1 | 5 |
| arpc1a | 0.19862 | -0.20237066 | 0.159 | 0.323 | 1 | 5 |
| mrpl22 | 0.19866 | -0.17424103 | 0.032 | 0.129 | 1 | 5 |
| myd88 | 0.19888 | -0.27271725 | 0.143 | 0.29 | 1 | 5 |
| actn4 | 0.1989 | -0.24304441 | 0.159 | 0.323 | 1 | 5 |
| mafbb | 0.1997 | -0.26252363 | 0.524 | 0.71 | 1 | 5 |
| thoc1 | 0.19985 | -0.20032717 | 0.079 | 0.194 | 1 | 5 |
| tars | 0.19988 | -0.21514999 | 0.111 | 0.258 | 1 | 5 |
| zmp:0000000936 | 0.20008 | 0.21724342 | 0.206 | 0.097 | 1 | 5 |
| mrps33 | 0.20038 | -0.13314187 | 0.048 | 0.161 | 1 | 5 |
| lrrc8da | 0.20046 | -0.15552083 | 0.032 | 0.129 | 1 | 5 |
| mtx2 | 0.20061 | -0.15056486 | 0.016 | 0.097 | 1 | 5 |
| glipr2 | 0.20065 | -0.25161303 | 0.159 | 0.323 | 1 | 5 |
| fam117ab | 0.20086 | 0.202405142 | 0.19 | 0.097 | 1 | 5 |
| ubc | 0.20098 | 0.134506414 | 0.825 | 0.839 | 1 | 5 |
| exosc4 | 0.20127 | -0.15838087 | 0.048 | 0.161 | 1 | 5 |
| pola1 | 0.20155 | -0.10314127 | 0.016 | 0.097 | 1 | 5 |
| mogat3b | 0.20167 | -0.1582407 | 0.016 | 0.097 | 1 | 5 |
| tcnbb | 0.20174 | 0.190342548 | 0.492 | 0.323 | 1 | 5 |
| emc1 | 0.20176 | -0.16594121 | 0.048 | 0.161 | 1 | 5 |
| CABZ01084575.1 | 0.20197 | -0.1582103 | 0.048 | 0.161 | 1 | 5 |
| ss18 | 0.20264 | -0.1196633 | 0.079 | 0.194 | 1 | 5 |
| nfs1 | 0.20274 | 0.151474503 | 0.127 | 0.032 | 1 | 5 |
| PQLC1 | 0.20278 | -0.13095994 | 0.016 | 0.097 | 1 | 5 |
| rit1 | 0.20279 | 0.141496111 | 0.143 | 0.032 | 1 | 5 |
| rbms1b | 0.20326 | -0.2158987 | 0.111 | 0.258 | 1 | 5 |
| rps6kb1a | 0.2034 | -0.14060776 | 0.048 | 0.161 | 1 | 5 |
| ccnl1a | 0.20352 | 0.145239739 | 0.143 | 0.032 | 1 | 5 |
| ppp1r12c | 0.20359 | -0.14593257 | 0.048 | 0.161 | 1 | 5 |
| uba5 | 0.20361 | -0.13240366 | 0.048 | 0.161 | 1 | 5 |
| slc39a1 | 0.20375 | -0.13739173 | 0.19 | 0.323 | 1 | 5 |
| ccr9a | 0.20418 | 0.275710064 | 0.365 | 0.194 | 1 | 5 |
| rcl1 | 0.20435 | -0.19341016 | 0.079 | 0.161 | 1 | 5 |
| slc38a5a | 0.20446 | -0.22467942 | 0.095 | 0.226 | 1 | 5 |
| rnaseh1 | 0.2045 | 0.163763567 | 0.127 | 0.032 | 1 | 5 |
| g6pc3 | 0.20483 | -0.14052285 | 0.032 | 0.129 | 1 | 5 |
| si:ch73-281k2.5 | 0.20486 | -0.16981238 | 0.048 | 0.129 | 1 | 5 |

| | | | | | | |
|---|---|---|---|---|---|---|
| exoc4 | 0.20496 | -0.14769512 | 0.016 | 0.097 | 1 | 5 |
| mid1ip1l | 0.20501 | 0.273800914 | 0.429 | 0.323 | 1 | 5 |
| tmem154 | 0.20523 | -0.20804242 | 0.111 | 0.258 | 1 | 5 |
| ube2nb | 0.20531 | 0.210973984 | 0.238 | 0.097 | 1 | 5 |
| cep131 | 0.20545 | -0.18405927 | 0.032 | 0.097 | 1 | 5 |
| sugt1 | 0.20556 | 0.154602715 | 0.238 | 0.194 | 1 | 5 |
| tmc8 | 0.20575 | 0.21757314 | 0.175 | 0.065 | 1 | 5 |
| hmgb2a | 0.20576 | -0.24810938 | 0.571 | 0.645 | 1 | 5 |
| ndufaf3 | 0.20578 | -0.15223584 | 0.032 | 0.129 | 1 | 5 |
| LOC101886114 | 0.2058 | -0.21314776 | 0.111 | 0.258 | 1 | 5 |
| ip6k2a | 0.20592 | 0.183336112 | 0.27 | 0.194 | 1 | 5 |
| nek9 | 0.20604 | -0.13544858 | 0.032 | 0.129 | 1 | 5 |
| si:dkeyp-87d8.8 | 0.20612 | -0.13009921 | 0.016 | 0.097 | 1 | 5 |
| capza1b | 0.20629 | -0.18362981 | 0.683 | 0.774 | 1 | 5 |
| kat5b | 0.20649 | -0.11461028 | 0.032 | 0.129 | 1 | 5 |
| cnot1 | 0.20664 | -0.22392614 | 0.127 | 0.258 | 1 | 5 |
| mcm3 | 0.207 | -0.15432147 | 0.032 | 0.129 | 1 | 5 |
| plekhg5b | 0.20723 | 0.177303997 | 0.127 | 0.032 | 1 | 5 |
| stat1a | 0.20782 | -0.22791053 | 0.175 | 0.258 | 1 | 5 |
| ube2j2 | 0.20809 | -0.1454942 | 0.032 | 0.129 | 1 | 5 |
| mbd4 | 0.20819 | 0.19468114 | 0.127 | 0.032 | 1 | 5 |
| dhx57 | 0.20829 | -0.14453738 | 0.016 | 0.097 | 1 | 5 |
| prkag1 | 0.2083 | -0.12482976 | 0.032 | 0.129 | 1 | 5 |
| elovl1b | 0.20844 | 0.215218019 | 0.571 | 0.484 | 1 | 5 |
| mcm7 | 0.20852 | -0.10888737 | 0.016 | 0.097 | 1 | 5 |
| aco1 | 0.20871 | -0.10142491 | 0.032 | 0.129 | 1 | 5 |
| CU074419.2 | 0.20892 | -0.11409585 | 0.016 | 0.097 | 1 | 5 |
| spice1 | 0.20901 | 0.21389349 | 0.19 | 0.065 | 1 | 5 |
| ptmab | 0.20907 | -0.17446852 | 0.81 | 0.935 | 1 | 5 |
| rprd2a | 0.20925 | -0.10946173 | 0.016 | 0.097 | 1 | 5 |
| osgep | 0.20927 | -0.12975341 | 0.095 | 0.226 | 1 | 5 |
| trappc11 | 0.20984 | -0.14592066 | 0.016 | 0.097 | 1 | 5 |
| dnase1l1 | 0.20987 | 0.230329377 | 0.429 | 0.355 | 1 | 5 |
| slc25a35 | 0.20988 | -0.12407749 | 0.016 | 0.097 | 1 | 5 |
| si:ch73-12o23.1 | 0.21 | -0.16825373 | 0.032 | 0.129 | 1 | 5 |
| nup93 | 0.2101 | -0.16599106 | 0.048 | 0.129 | 1 | 5 |
| wdr83os | 0.2101 | 0.25882715 | 0.349 | 0.226 | 1 | 5 |
| tescb | 0.21082 | 0.132629582 | 0.127 | 0.032 | 1 | 5 |
| cyp4f3 | 0.21097 | 0.236732602 | 0.238 | 0.097 | 1 | 5 |
| ptp4a1 | 0.21125 | -0.24274516 | 0.206 | 0.355 | 1 | 5 |
| XLOC-018182 | 0.21152 | -0.1494556 | 0.016 | 0.097 | 1 | 5 |
| si:ch1073-325m22.2 | 0.21156 | -0.14010579 | 0.714 | 0.871 | 1 | 5 |
| XLOC-004016 | 0.212 | -0.13366361 | 0.016 | 0.097 | 1 | 5 |
| si:dkeyp-80c12.4 | 0.21206 | -0.16283619 | 0.032 | 0.129 | 1 | 5 |
| lsm1 | 0.21216 | 0.101136429 | 0.079 | 0.065 | 1 | 5 |
| pgs1 | 0.21237 | -0.13817775 | 0.032 | 0.129 | 1 | 5 |
| lnpa | 0.21244 | -0.10982458 | 0.016 | 0.097 | 1 | 5 |
| pcmtd1 | 0.21263 | 0.165210802 | 0.127 | 0.032 | 1 | 5 |

| | | | | | | |
|---|---|---|---|---|---|---|
| eif4g2a | 0.21319 | -0.18792519 | 0.19 | 0.355 | 1 | 5 |
| ifi46 | 0.2132 | -0.22917842 | 0.556 | 0.71 | 1 | 5 |
| slc25a28 | 0.21382 | 0.186521856 | 0.175 | 0.097 | 1 | 5 |
| mrps21 | 0.21403 | -0.11637568 | 0.206 | 0.355 | 1 | 5 |
| srek1ip1 | 0.21413 | -0.14570143 | 0.032 | 0.129 | 1 | 5 |
| lpin2 | 0.2147 | -0.15158345 | 0.032 | 0.129 | 1 | 5 |
| si:dkey-11f4.14 | 0.21482 | 0.245587576 | 0.619 | 0.548 | 1 | 5 |
| paxip1 | 0.21483 | -0.12326699 | 0.016 | 0.097 | 1 | 5 |
| casp2 | 0.21553 | -0.13583591 | 0.032 | 0.129 | 1 | 5 |
| slc33a1 | 0.21587 | -0.14484555 | 0.016 | 0.097 | 1 | 5 |
| usp10 | 0.2159 | -0.16925172 | 0.143 | 0.29 | 1 | 5 |
| XLOC-022740 | 0.21621 | -0.12756668 | 0.016 | 0.097 | 1 | 5 |
| ptena | 0.21661 | 0.221977236 | 0.175 | 0.065 | 1 | 5 |
| si:dkey-100n23.5 | 0.21672 | -0.15221923 | 0.032 | 0.129 | 1 | 5 |
| rimkla | 0.21672 | -0.10627532 | 0.032 | 0.129 | 1 | 5 |
| GK3P | 0.21683 | -0.12980061 | 0.016 | 0.097 | 1 | 5 |
| si:dkey-222f2.7 | 0.21687 | -0.11687116 | 0.222 | 0.355 | 1 | 5 |
| ppp1r18 | 0.21711 | -0.15747501 | 0.175 | 0.323 | 1 | 5 |
| qtrt1 | 0.21725 | -0.1100926 | 0.016 | 0.097 | 1 | 5 |
| csf2rb | 0.21743 | -0.19565042 | 0.397 | 0.581 | 1 | 5 |
| bida | 0.21748 | -0.14146226 | 0.032 | 0.129 | 1 | 5 |
| CABZ01075242.1 | 0.21756 | -0.15984886 | 0.016 | 0.097 | 1 | 5 |
| BX957297.2 | 0.2189 | -0.11154972 | 0.016 | 0.097 | 1 | 5 |
| rbm8a | 0.21895 | 0.203283464 | 0.524 | 0.452 | 1 | 5 |
| BX640512.3 | 0.21923 | -0.16496827 | 0.032 | 0.129 | 1 | 5 |
| mgat1b | 0.21925 | -0.13115039 | 0.032 | 0.129 | 1 | 5 |
| pim2 | 0.21954 | 0.260480109 | 0.238 | 0.097 | 1 | 5 |
| slc16a6b | 0.21995 | -0.11893122 | 0.016 | 0.097 | 1 | 5 |
| si:dkey-32n7.4 | 0.21998 | -0.19388351 | 0.048 | 0.129 | 1 | 5 |
| galnt12 | 0.22 | -0.11357195 | 0.016 | 0.097 | 1 | 5 |
| si:dkey-23i12.5 | 0.2201 | -0.1469157 | 0.19 | 0.355 | 1 | 5 |
| nup210 | 0.22018 | -0.12020563 | 0.016 | 0.097 | 1 | 5 |
| fdps | 0.2203 | -0.10560328 | 0.016 | 0.097 | 1 | 5 |
| uqcrb | 0.22031 | -0.15423249 | 0.032 | 0.129 | 1 | 5 |
| fuca1.1 | 0.22037 | 0.206186949 | 0.238 | 0.161 | 1 | 5 |
| traf2b | 0.22038 | -0.16744453 | 0.016 | 0.097 | 1 | 5 |
| fnbp4 | 0.22048 | 0.186511015 | 0.127 | 0.032 | 1 | 5 |
| coq8a | 0.2207 | 0.123143813 | 0.222 | 0.097 | 1 | 5 |
| si:ch211-13c6.2 | 0.22085 | -0.12312472 | 0.032 | 0.129 | 1 | 5 |
| ube3a | 0.22091 | -0.11446895 | 0.032 | 0.129 | 1 | 5 |
| aqp7 | 0.22091 | -0.12748783 | 0.016 | 0.097 | 1 | 5 |
| carm1 | 0.22114 | -0.12775216 | 0.032 | 0.129 | 1 | 5 |
| si:ch211-197k17.3 | 0.22127 | -0.15542144 | 0.032 | 0.129 | 1 | 5 |
| apool | 0.2214 | -0.10558022 | 0.016 | 0.097 | 1 | 5 |
| prkacbb | 0.22142 | -0.12879314 | 0.016 | 0.097 | 1 | 5 |
| pla2g7 | 0.22143 | -0.10245535 | 0.016 | 0.097 | 1 | 5 |
| foxp1b | 0.22154 | -0.13687591 | 0.032 | 0.129 | 1 | 5 |
| dr1 | 0.22158 | -0.14088709 | 0.032 | 0.129 | 1 | 5 |

| | | | | | | |
|---|---|---|---|---|---|---|
| trim8b | 0.22165 | -0.17539983 | 0.032 | 0.129 | 1 | 5 |
| ndufaf5 | 0.2217 | -0.12725304 | 0.032 | 0.129 | 1 | 5 |
| arhgap42a | 0.22175 | -0.11870671 | 0.032 | 0.129 | 1 | 5 |
| dock10 | 0.22177 | -0.19045778 | 0.032 | 0.129 | 1 | 5 |
| AL732488.1 | 0.22181 | -0.17871988 | 0.032 | 0.129 | 1 | 5 |
| snphb | 0.22184 | -0.13614613 | 0.032 | 0.129 | 1 | 5 |
| stoml3b | 0.22224 | -0.10613777 | 0.175 | 0.29 | 1 | 5 |
| fam162a | 0.22253 | -0.21347632 | 0.175 | 0.29 | 1 | 5 |
| hspe1 | 0.22379 | -0.21459515 | 0.238 | 0.29 | 1 | 5 |
| si:dkeyp-23e4.3 | 0.22518 | 0.176512969 | 0.19 | 0.065 | 1 | 5 |
| xbp1 | 0.22562 | 0.155190417 | 0.937 | 0.903 | 1 | 5 |
| ndufa2 | 0.22571 | 0.233789723 | 0.524 | 0.355 | 1 | 5 |
| cops5 | 0.22638 | 0.170885842 | 0.175 | 0.097 | 1 | 5 |
| slc39a7 | 0.22643 | -0.17088872 | 0.127 | 0.194 | 1 | 5 |
| cnpy3 | 0.22702 | 0.252935433 | 0.27 | 0.129 | 1 | 5 |
| itpkcb | 0.22767 | 0.224225287 | 0.19 | 0.065 | 1 | 5 |
| tnfrsf21 | 0.22786 | 0.154833925 | 0.19 | 0.129 | 1 | 5 |
| gb:bc139872 | 0.22796 | 0.105450404 | 0.143 | 0.129 | 1 | 5 |
| kpnb1 | 0.22837 | -0.1141922 | 0.286 | 0.29 | 1 | 5 |
| larp1 | 0.22871 | 0.150465788 | 0.111 | 0.065 | 1 | 5 |
| slc37a2 | 0.22925 | 0.177861505 | 0.19 | 0.065 | 1 | 5 |
| si:dkeyp-97b10.3 | 0.22927 | 0.132574649 | 0.175 | 0.065 | 1 | 5 |
| znf827 | 0.22947 | 0.168016801 | 0.127 | 0.032 | 1 | 5 |
| psme2 | 0.22969 | 0.241170551 | 0.349 | 0.226 | 1 | 5 |
| cd276 | 0.23013 | -0.22858947 | 0.175 | 0.29 | 1 | 5 |
| dhrs9 | 0.23028 | 0.252285895 | 0.508 | 0.323 | 1 | 5 |
| snrnp200 | 0.2303 | -0.18137006 | 0.095 | 0.226 | 1 | 5 |
| fcf1 | 0.23097 | 0.186988028 | 0.19 | 0.065 | 1 | 5 |
| FAM83G | 0.23179 | -0.15369008 | 0.048 | 0.129 | 1 | 5 |
| tmem11 | 0.23181 | 0.131348894 | 0.19 | 0.065 | 1 | 5 |
| ilf3b | 0.23201 | -0.10746851 | 0.159 | 0.29 | 1 | 5 |
| slc12a7b | 0.23207 | 0.188238625 | 0.19 | 0.129 | 1 | 5 |
| eif3ba | 0.23277 | -0.23959968 | 0.556 | 0.71 | 1 | 5 |
| nbr1a | 0.23334 | 0.159694086 | 0.111 | 0.032 | 1 | 5 |
| gadd45ba | 0.23335 | 0.255446798 | 0.381 | 0.258 | 1 | 5 |
| ndufs3 | 0.23356 | 0.183559531 | 0.222 | 0.129 | 1 | 5 |
| si:dkey-222f8.3 | 0.23401 | 0.141839494 | 0.175 | 0.129 | 1 | 5 |
| acad9 | 0.23449 | -0.12049769 | 0.016 | 0.065 | 1 | 5 |
| optn | 0.23449 | -0.10184658 | 0.016 | 0.065 | 1 | 5 |
| eif3hb | 0.23449 | -0.1111156 | 0.016 | 0.065 | 1 | 5 |
| kitlga | 0.23449 | -0.10624248 | 0.016 | 0.065 | 1 | 5 |
| mrpl16 | 0.23449 | -0.11652475 | 0.016 | 0.065 | 1 | 5 |
| dus1l | 0.23449 | -0.13268477 | 0.016 | 0.065 | 1 | 5 |
| enah | 0.23449 | -0.11841709 | 0.016 | 0.065 | 1 | 5 |
| maml3 | 0.23449 | -0.119514 | 0.016 | 0.065 | 1 | 5 |
| nedd9 | 0.23449 | -0.12247076 | 0.016 | 0.065 | 1 | 5 |
| fbxl14a | 0.23449 | -0.12570284 | 0.016 | 0.065 | 1 | 5 |
| cdk11b | 0.23449 | -0.10819721 | 0.016 | 0.065 | 1 | 5 |

| | | | | | | |
|---|---|---|---|---|---|---|
| elk4 | 0.23449 | -0.10517072 | 0.016 | 0.065 | 1 | 5 |
| rinl | 0.23449 | -0.12209502 | 0.016 | 0.065 | 1 | 5 |
| CABZ01073795.1 | 0.23449 | -0.1123514 | 0.016 | 0.065 | 1 | 5 |
| CU984600.2 | 0.23449 | -0.10019735 | 0.016 | 0.065 | 1 | 5 |
| LOC103909107 | 0.23449 | -0.22594006 | 0.016 | 0.065 | 1 | 5 |
| ajuba | 0.23449 | -0.12925281 | 0.016 | 0.065 | 1 | 5 |
| antxr1c | 0.23449 | -0.1021094 | 0.016 | 0.065 | 1 | 5 |
| dyrk1b | 0.23449 | -0.10186006 | 0.016 | 0.065 | 1 | 5 |
| eif4e2 | 0.23449 | -0.11603253 | 0.016 | 0.065 | 1 | 5 |
| nptna | 0.23449 | -0.10806032 | 0.016 | 0.065 | 1 | 5 |
| ptpn11b | 0.23449 | -0.13842955 | 0.016 | 0.065 | 1 | 5 |
| vcanb | 0.23449 | -0.10400458 | 0.016 | 0.065 | 1 | 5 |
| xaf1 | 0.23463 | -0.24786163 | 0.175 | 0.323 | 1 | 5 |
| eif2s2 | 0.23472 | -0.2174433 | 0.635 | 0.742 | 1 | 5 |
| f3b | 0.23474 | -0.30327332 | 0.175 | 0.323 | 1 | 5 |
| cox6c | 0.23504 | -0.17547485 | 0.73 | 0.774 | 1 | 5 |
| zgc:56304 | 0.23516 | -0.14346031 | 0.175 | 0.323 | 1 | 5 |
| rars | 0.23584 | -0.22333521 | 0.175 | 0.29 | 1 | 5 |
| fgl2a | 0.23635 | -0.22738459 | 0.73 | 0.839 | 1 | 5 |
| si:ch211-171h4.5 | 0.23647 | 0.116844812 | 0.111 | 0.032 | 1 | 5 |
| commd3 | 0.23696 | -0.22682347 | 0.222 | 0.387 | 1 | 5 |
| snrnp70 | 0.23715 | -0.20092497 | 0.143 | 0.29 | 1 | 5 |
| sftpbb | 0.23723 | 0.222264207 | 0.778 | 0.677 | 1 | 5 |
| pdia3 | 0.2376 | -0.19708398 | 0.73 | 0.871 | 1 | 5 |
| gmps | 0.23762 | 0.165785251 | 0.111 | 0.032 | 1 | 5 |
| si:dkey-172h23.2 | 0.23768 | 0.210474247 | 0.286 | 0.161 | 1 | 5 |
| ets2 | 0.23851 | 0.242158122 | 0.571 | 0.387 | 1 | 5 |
| tpp1 | 0.23888 | 0.233983696 | 0.746 | 0.581 | 1 | 5 |
| mat2aa | 0.23912 | -0.22044122 | 0.111 | 0.226 | 1 | 5 |
| uri1 | 0.2392 | 0.130442303 | 0.143 | 0.129 | 1 | 5 |
| MCUR1 | 0.24068 | 0.120427174 | 0.127 | 0.032 | 1 | 5 |
| meaf6 | 0.24107 | -0.14300872 | 0.143 | 0.29 | 1 | 5 |
| api5 | 0.2413 | -0.14391684 | 0.095 | 0.226 | 1 | 5 |
| stx5a | 0.2416 | -0.19125518 | 0.095 | 0.226 | 1 | 5 |
| FO704661.1 | 0.24177 | 0.182350459 | 0.222 | 0.161 | 1 | 5 |
| ptgr1 | 0.24184 | -0.13706946 | 0.095 | 0.226 | 1 | 5 |
| ddx3xa | 0.24206 | 0.121325493 | 0.175 | 0.161 | 1 | 5 |
| syk | 0.2421 | 0.207545678 | 0.317 | 0.161 | 1 | 5 |
| mrpl12 | 0.24212 | 0.165531023 | 0.317 | 0.258 | 1 | 5 |
| gemin7 | 0.24319 | -0.18734603 | 0.127 | 0.258 | 1 | 5 |
| lrrc15 | 0.24325 | -0.24872841 | 0.206 | 0.355 | 1 | 5 |
| clk4a | 0.24334 | 0.176377753 | 0.175 | 0.065 | 1 | 5 |
| ptenb | 0.2438 | -0.15192581 | 0.127 | 0.258 | 1 | 5 |
| herpud2 | 0.2444 | 0.1144599 | 0.127 | 0.032 | 1 | 5 |
| MRPS12 | 0.2445 | -0.13722097 | 0.254 | 0.29 | 1 | 5 |
| si:dkey-56f14.7 | 0.24478 | 0.233395154 | 0.286 | 0.161 | 1 | 5 |
| cenpx | 0.24579 | 0.104746197 | 0.111 | 0.097 | 1 | 5 |
| ormdl2 | 0.24592 | 0.14617906 | 0.111 | 0.032 | 1 | 5 |

| | | | | | | |
|---|---|---|---|---|---|---|
| selenoj | 0.24615 | -0.11229008 | 0.063 | 0.097 | 1 | 5 |
| decr1 | 0.24728 | -0.14504943 | 0.079 | 0.194 | 1 | 5 |
| CABZ01026681.1 | 0.24748 | -0.21040902 | 0.143 | 0.29 | 1 | 5 |
| tbl3 | 0.24758 | 0.150433611 | 0.127 | 0.032 | 1 | 5 |
| si:ch211-212g7.6 | 0.24788 | -0.18147941 | 0.079 | 0.129 | 1 | 5 |
| lrrfip1a | 0.24792 | -0.1300129 | 0.19 | 0.323 | 1 | 5 |
| tm9sf2 | 0.24897 | -0.10391031 | 0.222 | 0.355 | 1 | 5 |
| elof1 | 0.24948 | 0.109991778 | 0.222 | 0.226 | 1 | 5 |
| mcm6 | 0.25081 | -0.13849335 | 0.063 | 0.097 | 1 | 5 |
| pwp2h | 0.25108 | -0.13006293 | 0.063 | 0.161 | 1 | 5 |
| gsk3b | 0.25121 | -0.19980084 | 0.048 | 0.129 | 1 | 5 |
| gorasp2 | 0.25189 | -0.17621027 | 0.111 | 0.226 | 1 | 5 |
| ptdss1a | 0.25209 | -0.14650306 | 0.048 | 0.129 | 1 | 5 |
| uprt | 0.25222 | -0.15071567 | 0.063 | 0.161 | 1 | 5 |
| fbxo7 | 0.25226 | 0.141419394 | 0.079 | 0.032 | 1 | 5 |
| yeats4 | 0.25251 | -0.1022815 | 0.079 | 0.194 | 1 | 5 |
| crlf3 | 0.25262 | -0.13284643 | 0.079 | 0.194 | 1 | 5 |
| hibadhb | 0.25281 | 0.114653751 | 0.175 | 0.065 | 1 | 5 |
| si:rp71-36a1.5 | 0.25291 | -0.17036462 | 0.095 | 0.161 | 1 | 5 |
| b3gnt2b | 0.25355 | -0.12795633 | 0.048 | 0.129 | 1 | 5 |
| ankrd10b | 0.254 | 0.183914021 | 0.175 | 0.065 | 1 | 5 |
| f5 | 0.25406 | -0.25790703 | 0.063 | 0.161 | 1 | 5 |
| tmem39a | 0.25463 | -0.19453028 | 0.079 | 0.194 | 1 | 5 |
| slc25a33 | 0.25503 | -0.18099144 | 0.063 | 0.161 | 1 | 5 |
| ganab | 0.25599 | 0.172300001 | 0.143 | 0.065 | 1 | 5 |
| rasal2 | 0.25673 | 0.169544033 | 0.111 | 0.032 | 1 | 5 |
| plrg1 | 0.25709 | 0.146771071 | 0.095 | 0.065 | 1 | 5 |
| rnf10 | 0.25744 | 0.224534425 | 0.429 | 0.258 | 1 | 5 |
| ndufa4l | 0.25754 | -0.15183879 | 0.857 | 0.903 | 1 | 5 |
| ndufs5 | 0.25764 | -0.21544875 | 0.286 | 0.452 | 1 | 5 |
| grpel2 | 0.25775 | 0.144171135 | 0.159 | 0.065 | 1 | 5 |
| psma5 | 0.259 | 0.218863579 | 0.381 | 0.29 | 1 | 5 |
| ndufa3 | 0.25942 | 0.178479347 | 0.429 | 0.258 | 1 | 5 |
| chd9 | 0.25959 | 0.173171079 | 0.175 | 0.097 | 1 | 5 |
| pdcd10b | 0.26007 | 0.182861401 | 0.206 | 0.097 | 1 | 5 |
| psmb10 | 0.26101 | -0.12145284 | 0.079 | 0.194 | 1 | 5 |
| RNF122 | 0.26286 | -0.19202996 | 0.079 | 0.194 | 1 | 5 |
| pygb | 0.26291 | 0.131810589 | 0.175 | 0.129 | 1 | 5 |
| zgc:173915 | 0.26327 | -0.14137226 | 0.063 | 0.161 | 1 | 5 |
| XLOC-041870 | 0.26349 | 0.153690331 | 0.143 | 0.097 | 1 | 5 |
| syap1 | 0.26357 | -0.16080855 | 0.079 | 0.194 | 1 | 5 |
| lrrc8aa | 0.26384 | 0.197017933 | 0.206 | 0.097 | 1 | 5 |
| sfpq | 0.26481 | -0.18882785 | 0.206 | 0.29 | 1 | 5 |
| acadm | 0.26497 | -0.18990648 | 0.159 | 0.29 | 1 | 5 |
| ccdc25 | 0.26507 | -0.20808367 | 0.286 | 0.419 | 1 | 5 |
| xiap | 0.2654 | -0.20664291 | 0.206 | 0.355 | 1 | 5 |
| CU019662.1 | 0.26541 | -0.18084927 | 0.063 | 0.161 | 1 | 5 |
| atp6v1f | 0.26555 | -0.20123464 | 0.111 | 0.226 | 1 | 5 |

| | | | | | | |
|---|---|---|---|---|---|---|
| taf12 | 0.26577 | 0.107303043 | 0.222 | 0.226 | 1 | 5 |
| glo1 | 0.26603 | -0.1744862 | 0.492 | 0.548 | 1 | 5 |
| spop | 0.26617 | 0.143836304 | 0.079 | 0.032 | 1 | 5 |
| si:ch211-107o10.3 | 0.26634 | -0.12841753 | 0.079 | 0.129 | 1 | 5 |
| ndufs4 | 0.26685 | -0.14856798 | 0.286 | 0.323 | 1 | 5 |
| ppp1r12a | 0.26734 | 0.14015696 | 0.175 | 0.065 | 1 | 5 |
| sar1b | 0.26741 | -0.15511419 | 0.175 | 0.323 | 1 | 5 |
| calr | 0.26747 | -0.16719847 | 0.127 | 0.258 | 1 | 5 |
| immt | 0.26808 | 0.109735904 | 0.127 | 0.032 | 1 | 5 |
| abcf1 | 0.26847 | 0.184220324 | 0.365 | 0.323 | 1 | 5 |
| timm23a | 0.26852 | 0.200056631 | 0.286 | 0.161 | 1 | 5 |
| cgrrf1 | 0.26873 | 0.171509683 | 0.127 | 0.032 | 1 | 5 |
| march5l | 0.26876 | 0.177698134 | 0.127 | 0.065 | 1 | 5 |
| cox6b1 | 0.27016 | 0.223578651 | 0.365 | 0.226 | 1 | 5 |
| ppm1aa | 0.27079 | 0.215337588 | 0.238 | 0.129 | 1 | 5 |
| entpd4 | 0.27108 | -0.11240521 | 0.063 | 0.161 | 1 | 5 |
| atp6v1c1b | 0.27308 | -0.19377928 | 0.063 | 0.161 | 1 | 5 |
| tfe3a | 0.27312 | -0.21777282 | 0.222 | 0.323 | 1 | 5 |
| cry1aa | 0.27348 | -0.16628875 | 0.079 | 0.194 | 1 | 5 |
| slc31a1 | 0.27536 | -0.15825561 | 0.095 | 0.161 | 1 | 5 |
| ppm1bb | 0.27582 | -0.13660836 | 0.079 | 0.129 | 1 | 5 |
| zgc:101583 | 0.27601 | -0.18004354 | 0.079 | 0.194 | 1 | 5 |
| ptp4a2b | 0.2761 | -0.14592779 | 0.333 | 0.355 | 1 | 5 |
| rnf220a | 0.27654 | 0.159799004 | 0.095 | 0.032 | 1 | 5 |
| tnfsf12 | 0.27658 | 0.155413582 | 0.175 | 0.129 | 1 | 5 |
| usp8 | 0.27672 | 0.167362548 | 0.127 | 0.032 | 1 | 5 |
| zgc:165573 | 0.27685 | -0.10054594 | 0.143 | 0.258 | 1 | 5 |
| si:ch73-248e21.7 | 0.27718 | 0.22798357 | 0.46 | 0.29 | 1 | 5 |
| zgc:175264 | 0.27786 | 0.138566027 | 0.127 | 0.032 | 1 | 5 |
| XLOC-000431 | 0.27905 | 0.185667811 | 0.825 | 0.677 | 1 | 5 |
| anxa4 | 0.27914 | 0.124645357 | 0.46 | 0.452 | 1 | 5 |
| atp5if1a | 0.27942 | -0.12029783 | 0.587 | 0.742 | 1 | 5 |
| ggcta | 0.27948 | 0.148365108 | 0.127 | 0.032 | 1 | 5 |
| tlr5a | 0.27962 | 0.2012952 | 0.111 | 0.032 | 1 | 5 |
| pik3r2 | 0.27993 | 0.130963626 | 0.127 | 0.032 | 1 | 5 |
| erh | 0.28043 | 0.212703992 | 0.683 | 0.548 | 1 | 5 |
| csk | 0.28077 | -0.16673012 | 0.127 | 0.258 | 1 | 5 |
| dlst | 0.28083 | 0.246137662 | 0.365 | 0.226 | 1 | 5 |
| cnot6a | 0.28096 | 0.145700301 | 0.127 | 0.032 | 1 | 5 |
| huwe1 | 0.281 | -0.1663035 | 0.079 | 0.194 | 1 | 5 |
| si:dkey-25e12.3 | 0.28157 | -0.22448789 | 0.175 | 0.323 | 1 | 5 |
| slc43a2a | 0.28186 | 0.276616035 | 0.476 | 0.323 | 1 | 5 |
| mfap1 | 0.28212 | 0.129401553 | 0.095 | 0.032 | 1 | 5 |
| grasp | 0.28214 | -0.21215193 | 0.175 | 0.323 | 1 | 5 |
| si:ch73-234b20.5 | 0.28228 | 0.220281773 | 0.429 | 0.29 | 1 | 5 |
| gpr132b | 0.28239 | -0.20480284 | 0.127 | 0.258 | 1 | 5 |
| dcp1b | 0.28244 | -0.17636071 | 0.032 | 0.097 | 1 | 5 |
| CABZ01109481.1 | 0.2827 | 0.135327892 | 0.127 | 0.032 | 1 | 5 |

| | | | | | | |
|---|---|---|---|---|---|---|
| tpst1l | 0.28286 | 0.126771834 | 0.127 | 0.032 | 1 | 5 |
| tnpo3 | 0.28299 | 0.128132647 | 0.127 | 0.032 | 1 | 5 |
| tmem147 | 0.28314 | -0.17456261 | 0.175 | 0.323 | 1 | 5 |
| atf6 | 0.28381 | -0.1773366 | 0.206 | 0.355 | 1 | 5 |
| krt18b | 0.28397 | -0.11213183 | 0.873 | 0.968 | 1 | 5 |
| strn3 | 0.28402 | 0.145327646 | 0.127 | 0.032 | 1 | 5 |
| cab39l1 | 0.28414 | -0.17743877 | 0.079 | 0.194 | 1 | 5 |
| pkma | 0.28449 | 0.170299749 | 0.714 | 0.645 | 1 | 5 |
| ccl39.6 | 0.28478 | 0.210422293 | 0.111 | 0.032 | 1 | 5 |
| cs | 0.28489 | 0.19277181 | 0.27 | 0.161 | 1 | 5 |
| gpatch11 | 0.28514 | 0.15533601 | 0.111 | 0.032 | 1 | 5 |
| zgc:103697 | 0.28539 | 0.129232163 | 0.127 | 0.065 | 1 | 5 |
| wls | 0.28633 | -0.18731928 | 0.286 | 0.452 | 1 | 5 |
| ubtd2 | 0.28664 | 0.174117321 | 0.111 | 0.032 | 1 | 5 |
| fam49ba | 0.28668 | -0.20127149 | 0.286 | 0.452 | 1 | 5 |
| bxdc2 | 0.28687 | -0.16305599 | 0.111 | 0.194 | 1 | 5 |
| cct3 | 0.2871 | -0.16301842 | 0.508 | 0.677 | 1 | 5 |
| mrpl43 | 0.28713 | 0.104605733 | 0.111 | 0.032 | 1 | 5 |
| fam49bb | 0.28769 | 0.223615437 | 0.333 | 0.194 | 1 | 5 |
| ddx61 | 0.28773 | 0.195650343 | 0.587 | 0.516 | 1 | 5 |
| ube4b | 0.28805 | -0.17294143 | 0.079 | 0.194 | 1 | 5 |
| emc7 | 0.28833 | -0.11316117 | 0.159 | 0.29 | 1 | 5 |
| snw1 | 0.28853 | -0.11565833 | 0.159 | 0.29 | 1 | 5 |
| stt3b | 0.28905 | -0.15824489 | 0.127 | 0.258 | 1 | 5 |
| afmid | 0.28908 | 0.177998051 | 0.222 | 0.129 | 1 | 5 |
| ilf2 | 0.28962 | 0.12083595 | 0.333 | 0.323 | 1 | 5 |
| gstk1 | 0.29004 | -0.18103489 | 0.127 | 0.226 | 1 | 5 |
| si:dkey-42l23.2 | 0.29018 | -0.1569943 | 0.079 | 0.194 | 1 | 5 |
| nsmce1 | 0.29073 | -0.12351154 | 0.063 | 0.129 | 1 | 5 |
| tmpob | 0.29113 | 0.175360857 | 0.159 | 0.065 | 1 | 5 |
| slc30a7 | 0.29123 | 0.13771709 | 0.286 | 0.258 | 1 | 5 |
| arid4a | 0.29183 | 0.169284834 | 0.159 | 0.065 | 1 | 5 |
| ell | 0.29204 | -0.15069018 | 0.079 | 0.194 | 1 | 5 |
| dnajc8 | 0.29212 | -0.10193093 | 0.079 | 0.194 | 1 | 5 |
| acbd3 | 0.29213 | -0.16623539 | 0.079 | 0.194 | 1 | 5 |
| jak1 | 0.29239 | 0.229804709 | 0.571 | 0.419 | 1 | 5 |
| zgc:77486 | 0.29247 | -0.16726858 | 0.238 | 0.387 | 1 | 5 |
| itgb2 | 0.29251 | 0.188596067 | 0.762 | 0.677 | 1 | 5 |
| si:ch211-210c8.7 | 0.29329 | 0.115130647 | 0.111 | 0.065 | 1 | 5 |
| nme7 | 0.29367 | -0.14255262 | 0.063 | 0.129 | 1 | 5 |
| yaf2 | 0.29416 | -0.11205613 | 0.048 | 0.097 | 1 | 5 |
| acsl1b | 0.29426 | 0.154159685 | 0.127 | 0.065 | 1 | 5 |
| kpnb3 | 0.29506 | -0.19061792 | 0.206 | 0.323 | 1 | 5 |
| snrnp27 | 0.29514 | 0.145905509 | 0.238 | 0.194 | 1 | 5 |
| ophn1 | 0.29646 | -0.17573331 | 0.063 | 0.129 | 1 | 5 |
| fam177a1 | 0.29766 | -0.20250849 | 0.159 | 0.29 | 1 | 5 |
| hk2 | 0.29827 | -0.19372367 | 0.127 | 0.258 | 1 | 5 |
| tmem59 | 0.29832 | 0.246427768 | 0.444 | 0.29 | 1 | 5 |

| | | | | | | |
|---|---|---|---|---|---|---|
| ube2q2 | 0.29863 | 0.146363778 | 0.159 | 0.097 | 1 | 5 |
| mych | 0.29864 | -0.20302253 | 0.206 | 0.355 | 1 | 5 |
| tmem134 | 0.29908 | 0.114104271 | 0.111 | 0.032 | 1 | 5 |
| bloc1s2 | 0.29987 | -0.18426731 | 0.127 | 0.258 | 1 | 5 |
| cipcb | 0.3001 | 0.132852099 | 0.127 | 0.097 | 1 | 5 |
| adrm1 | 0.30064 | -0.12903199 | 0.127 | 0.258 | 1 | 5 |
| trpc4apa | 0.30107 | -0.17536812 | 0.032 | 0.097 | 1 | 5 |
| casp8ap2 | 0.30118 | 0.170870432 | 0.143 | 0.065 | 1 | 5 |
| si:ch211-284b7.3 | 0.30165 | 0.11261383 | 0.111 | 0.032 | 1 | 5 |
| rab5ab | 0.3025 | -0.11735736 | 0.286 | 0.419 | 1 | 5 |
| elmo3 | 0.30273 | -0.16941218 | 0.063 | 0.161 | 1 | 5 |
| rnf185 | 0.30275 | -0.1925972 | 0.27 | 0.355 | 1 | 5 |
| sptlc2b | 0.30373 | 0.17825616 | 0.175 | 0.065 | 1 | 5 |
| rhbdf1a | 0.30387 | -0.11632219 | 0.27 | 0.29 | 1 | 5 |
| BX005183.3 | 0.30393 | 0.187702382 | 0.238 | 0.161 | 1 | 5 |
| rasgef1ba | 0.30428 | 0.208969098 | 0.175 | 0.065 | 1 | 5 |
| arl2 | 0.3044 | -0.1511922 | 0.048 | 0.129 | 1 | 5 |
| zgc:153713 | 0.30491 | 0.10895471 | 0.587 | 0.581 | 1 | 5 |
| taf13 | 0.30517 | 0.217583328 | 0.444 | 0.29 | 1 | 5 |
| furina | 0.30555 | 0.146609436 | 0.222 | 0.161 | 1 | 5 |
| rab2a | 0.30663 | -0.17251839 | 0.254 | 0.355 | 1 | 5 |
| zgc:91910 | 0.30681 | -0.17992557 | 0.413 | 0.581 | 1 | 5 |
| acin1a | 0.30705 | 0.219478002 | 0.349 | 0.258 | 1 | 5 |
| gnal | 0.3072 | -0.17277675 | 0.063 | 0.161 | 1 | 5 |
| vapb | 0.30724 | -0.17386816 | 0.111 | 0.194 | 1 | 5 |
| hsp90b1 | 0.30759 | -0.21291257 | 0.413 | 0.581 | 1 | 5 |
| mtdha | 0.30769 | -0.19486407 | 0.413 | 0.581 | 1 | 5 |
| ugp2b | 0.30781 | -0.2148655 | 0.206 | 0.355 | 1 | 5 |
| rrs1 | 0.30793 | -0.1861426 | 0.159 | 0.29 | 1 | 5 |
| mrpl54 | 0.30814 | -0.15947091 | 0.206 | 0.355 | 1 | 5 |
| nek7 | 0.30817 | 0.148270394 | 0.111 | 0.032 | 1 | 5 |
| zgc:112982 | 0.30867 | -0.15385382 | 0.032 | 0.097 | 1 | 5 |
| utp6 | 0.30887 | 0.131554329 | 0.111 | 0.065 | 1 | 5 |
| pdcd6ip | 0.30919 | -0.14418288 | 0.19 | 0.323 | 1 | 5 |
| pop7 | 0.30956 | 0.142015855 | 0.111 | 0.032 | 1 | 5 |
| srp68 | 0.31003 | -0.20962998 | 0.095 | 0.194 | 1 | 5 |
| atp5mea | 0.31018 | -0.17865617 | 0.206 | 0.355 | 1 | 5 |
| igf2b | 0.31072 | 0.156608644 | 0.206 | 0.097 | 1 | 5 |
| fgd4b | 0.31087 | -0.16769646 | 0.048 | 0.129 | 1 | 5 |
| ccnt2b | 0.31114 | 0.145617913 | 0.159 | 0.065 | 1 | 5 |
| psmd11a | 0.31162 | -0.15923926 | 0.111 | 0.226 | 1 | 5 |
| irf9 | 0.31205 | -0.17098949 | 0.063 | 0.161 | 1 | 5 |
| mcl1a | 0.31317 | 0.129871184 | 0.968 | 0.903 | 1 | 5 |
| mta2 | 0.31318 | -0.16790163 | 0.063 | 0.161 | 1 | 5 |
| naa50 | 0.31357 | -0.1790993 | 0.143 | 0.258 | 1 | 5 |
| zgc:64051 | 0.31387 | -0.17304698 | 0.746 | 0.871 | 1 | 5 |
| rnf144b | 0.31452 | 0.15639326 | 0.238 | 0.194 | 1 | 5 |
| zgc:101858 | 0.31499 | 0.160850657 | 0.159 | 0.065 | 1 | 5 |

| | | | | | | |
|---|---|---|---|---|---|---|
| edem3 | 0.31536 | -0.10077389 | 0.048 | 0.129 | 1 | 5 |
| smim20 | 0.31548 | 0.111879928 | 0.111 | 0.065 | 1 | 5 |
| col4a3bpb | 0.31556 | -0.11211142 | 0.095 | 0.129 | 1 | 5 |
| safb | 0.31575 | -0.11068807 | 0.111 | 0.226 | 1 | 5 |
| piezo1 | 0.31621 | -0.13344354 | 0.063 | 0.161 | 1 | 5 |
| ano6 | 0.31622 | -0.13685412 | 0.048 | 0.129 | 1 | 5 |
| chmp2bb | 0.31632 | -0.21441765 | 0.238 | 0.355 | 1 | 5 |
| zgc:175088 | 0.31649 | 0.167018455 | 0.222 | 0.129 | 1 | 5 |
| ranbp3b | 0.31719 | -0.12556673 | 0.032 | 0.097 | 1 | 5 |
| golt1bb | 0.31774 | -0.21389482 | 0.143 | 0.258 | 1 | 5 |
| si:ch1073-59l16.1 | 0.31833 | -0.11018437 | 0.048 | 0.129 | 1 | 5 |
| igf2bp2a | 0.31855 | 0.152366949 | 0.111 | 0.032 | 1 | 5 |
| il6r | 0.31914 | 0.114009152 | 0.127 | 0.097 | 1 | 5 |
| mesd | 0.31977 | -0.13715719 | 0.159 | 0.29 | 1 | 5 |
| si:dkey-237i9.1 | 0.31993 | 0.129466601 | 0.111 | 0.032 | 1 | 5 |
| polr2d | 0.31996 | -0.20034115 | 0.286 | 0.419 | 1 | 5 |
| lgals3bpb | 0.32063 | -0.24465629 | 0.635 | 0.774 | 1 | 5 |
| dctn3 | 0.32107 | 0.175233843 | 0.302 | 0.161 | 1 | 5 |
| ms4a17a.10 | 0.32176 | 0.209040225 | 0.825 | 0.742 | 1 | 5 |
| diaph2 | 0.32186 | -0.14499083 | 0.063 | 0.129 | 1 | 5 |
| XLOC-023492 | 0.32235 | -0.23469525 | 0.365 | 0.484 | 1 | 5 |
| tmem120a | 0.32242 | -0.11998605 | 0.063 | 0.161 | 1 | 5 |
| si:ch1073-314i13.4 | 0.3225 | -0.15512359 | 0.127 | 0.194 | 1 | 5 |
| lcp2a | 0.32261 | 0.149952126 | 0.254 | 0.129 | 1 | 5 |
| CR753886.1 | 0.32302 | -0.11464768 | 0.032 | 0.097 | 1 | 5 |
| ctsla | 0.32318 | -0.30688285 | 0.111 | 0.226 | 1 | 5 |
| numb | 0.32319 | -0.14576328 | 0.095 | 0.129 | 1 | 5 |
| mrps23 | 0.32359 | 0.135686186 | 0.111 | 0.065 | 1 | 5 |
| arl8bb | 0.32406 | 0.169127522 | 0.238 | 0.161 | 1 | 5 |
| fosb | 0.32424 | -0.13798709 | 0.095 | 0.194 | 1 | 5 |
| acadvl | 0.32443 | 0.193489273 | 0.333 | 0.194 | 1 | 5 |
| rhoaa | 0.32557 | -0.14574222 | 0.429 | 0.581 | 1 | 5 |
| fabp11a | 0.32571 | -0.11321417 | 0.143 | 0.129 | 1 | 5 |
| got2a | 0.32615 | 0.113046365 | 0.111 | 0.032 | 1 | 5 |
| lamp1a | 0.32665 | -0.15027216 | 0.429 | 0.581 | 1 | 5 |
| ddx6 | 0.32716 | 0.122582629 | 0.19 | 0.161 | 1 | 5 |
| hccsb | 0.32746 | -0.17891635 | 0.111 | 0.226 | 1 | 5 |
| hic1 | 0.32808 | 0.151535424 | 0.159 | 0.129 | 1 | 5 |
| tmem9b | 0.3281 | -0.14353913 | 0.111 | 0.226 | 1 | 5 |
| serpinb1l3 | 0.32878 | 0.116281017 | 0.381 | 0.355 | 1 | 5 |
| tob1b | 0.32882 | -0.22806611 | 0.286 | 0.419 | 1 | 5 |
| pnp5a | 0.32931 | -0.23701504 | 0.317 | 0.452 | 1 | 5 |
| si:dkey-52l18.4 | 0.32981 | -0.16130625 | 0.111 | 0.226 | 1 | 5 |
| sat1a.2 | 0.33054 | -0.16536836 | 0.746 | 0.839 | 1 | 5 |
| man2b1 | 0.33064 | -0.14063887 | 0.111 | 0.226 | 1 | 5 |
| mrpl48 | 0.33121 | -0.10035105 | 0.032 | 0.097 | 1 | 5 |
| srsf6a | 0.33137 | -0.14147586 | 0.095 | 0.161 | 1 | 5 |
| tmem263 | 0.3314 | -0.13339764 | 0.111 | 0.226 | 1 | 5 |

| | | | | | | |
|---|---|---|---|---|---|---|
| ap3d1 | 0.3321 | -0.1248718 | 0.063 | 0.161 | 1 | 5 |
| psma1 | 0.33294 | 0.191760461 | 0.683 | 0.548 | 1 | 5 |
| pfkfb4b | 0.33296 | 0.133774861 | 0.524 | 0.516 | 1 | 5 |
| pstpip2 | 0.33301 | 0.141714597 | 0.111 | 0.032 | 1 | 5 |
| si:dkeyp-87e7.4 | 0.33519 | 0.111871894 | 0.079 | 0.032 | 1 | 5 |
| rgs13 | 0.33531 | 0.105615954 | 0.381 | 0.258 | 1 | 5 |
| polr3k | 0.33538 | 0.136866895 | 0.19 | 0.097 | 1 | 5 |
| pim1 | 0.33547 | 0.22248241 | 0.635 | 0.484 | 1 | 5 |
| ruvbl1 | 0.33551 | -0.10618386 | 0.063 | 0.161 | 1 | 5 |
| asap1b | 0.33631 | 0.151797242 | 0.159 | 0.097 | 1 | 5 |
| cers2b | 0.33636 | -0.19575883 | 0.111 | 0.226 | 1 | 5 |
| ccl39.3 | 0.33659 | -0.26524503 | 0.159 | 0.29 | 1 | 5 |
| bbx | 0.33686 | -0.14313236 | 0.063 | 0.161 | 1 | 5 |
| il1b | 0.33689 | -0.14674045 | 0.063 | 0.161 | 1 | 5 |
| aktip | 0.33785 | 0.113334807 | 0.079 | 0.032 | 1 | 5 |
| LOC100538255 | 0.33799 | 0.205725523 | 0.238 | 0.194 | 1 | 5 |
| pnn | 0.33809 | 0.178362008 | 0.27 | 0.161 | 1 | 5 |
| nolc1 | 0.33812 | -0.19547783 | 0.127 | 0.226 | 1 | 5 |
| fmr1 | 0.33821 | -0.14193378 | 0.079 | 0.129 | 1 | 5 |
| fmnl1a | 0.33844 | 0.10561769 | 0.19 | 0.097 | 1 | 5 |
| pin4 | 0.3391 | -0.17652362 | 0.159 | 0.258 | 1 | 5 |
| galnt7 | 0.33957 | -0.13477819 | 0.111 | 0.226 | 1 | 5 |
| dnajc3a | 0.33964 | -0.15168713 | 0.111 | 0.226 | 1 | 5 |
| nt5c3a | 0.34011 | -0.12642583 | 0.063 | 0.161 | 1 | 5 |
| si:dkey-239h2.3 | 0.34015 | 0.183060986 | 0.254 | 0.161 | 1 | 5 |
| mrps9 | 0.34036 | 0.156810342 | 0.143 | 0.065 | 1 | 5 |
| LOC101883052 | 0.34039 | -0.13572159 | 0.063 | 0.161 | 1 | 5 |
| arl1 | 0.34066 | -0.20142646 | 0.19 | 0.323 | 1 | 5 |
| rpa1 | 0.34207 | -0.1144152 | 0.063 | 0.161 | 1 | 5 |
| llph | 0.34214 | -0.14053639 | 0.19 | 0.323 | 1 | 5 |
| trappc2 | 0.34233 | -0.14122491 | 0.063 | 0.161 | 1 | 5 |
| si:rp71-79p20.2 | 0.34242 | -0.12899295 | 0.063 | 0.161 | 1 | 5 |
| hif1ab | 0.34262 | -0.1809659 | 0.27 | 0.419 | 1 | 5 |
| mgrn1b | 0.34265 | -0.21814358 | 0.19 | 0.323 | 1 | 5 |
| srgap3 | 0.34308 | 0.180278003 | 0.206 | 0.129 | 1 | 5 |
| rbmx2 | 0.34315 | -0.11208923 | 0.063 | 0.161 | 1 | 5 |
| sox4b | 0.34355 | -0.10790623 | 0.111 | 0.129 | 1 | 5 |
| prex1 | 0.34366 | 0.166593495 | 0.111 | 0.032 | 1 | 5 |
| ctsa | 0.34367 | 0.188187485 | 0.73 | 0.581 | 1 | 5 |
| fut8a | 0.34434 | 0.140276307 | 0.111 | 0.032 | 1 | 5 |
| CU681842.1 | 0.34463 | 0.137763115 | 0.111 | 0.032 | 1 | 5 |
| atp5meb | 0.34475 | -0.21137701 | 0.508 | 0.645 | 1 | 5 |
| il16 | 0.34517 | 0.147736311 | 0.127 | 0.065 | 1 | 5 |
| ubb | 0.34582 | -0.15234168 | 0.556 | 0.71 | 1 | 5 |
| lonrf1l | 0.34608 | -0.13435593 | 0.032 | 0.097 | 1 | 5 |
| mrpl33 | 0.3461 | -0.20861039 | 0.349 | 0.484 | 1 | 5 |
| prpf39 | 0.34659 | -0.17275017 | 0.111 | 0.226 | 1 | 5 |
| nipsnap2 | 0.34673 | 0.138286107 | 0.19 | 0.097 | 1 | 5 |

| | | | | | | |
|---|---|---|---|---|---|---|
| hmha1b | 0.34743 | 0.12805039 | 0.54 | 0.516 | 1 | 5 |
| vps25 | 0.34779 | -0.16486119 | 0.111 | 0.226 | 1 | 5 |
| cox14 | 0.3482 | 0.181253434 | 0.286 | 0.194 | 1 | 5 |
| cabz01093075.1 | 0.34877 | -0.18341389 | 0.048 | 0.129 | 1 | 5 |
| puf60b | 0.34878 | 0.137070093 | 0.175 | 0.097 | 1 | 5 |
| klf7b | 0.34879 | 0.132248308 | 0.111 | 0.032 | 1 | 5 |
| mrpl42 | 0.34882 | 0.138279704 | 0.286 | 0.226 | 1 | 5 |
| gpr155a | 0.34891 | -0.1438253 | 0.063 | 0.129 | 1 | 5 |
| atp5md | 0.34919 | 0.216416946 | 0.683 | 0.548 | 1 | 5 |
| ggps1 | 0.34935 | -0.15862353 | 0.048 | 0.129 | 1 | 5 |
| hnrnpaba | 0.34958 | -0.16544331 | 0.556 | 0.613 | 1 | 5 |
| jac9 | 0.3496 | -0.19806097 | 0.333 | 0.484 | 1 | 5 |
| trim105 | 0.34978 | -0.1132411 | 0.048 | 0.129 | 1 | 5 |
| vps37a | 0.34998 | 0.143620164 | 0.127 | 0.065 | 1 | 5 |
| syf2 | 0.35008 | -0.14734507 | 0.19 | 0.323 | 1 | 5 |
| rtn4a | 0.35011 | -0.19639706 | 0.397 | 0.548 | 1 | 5 |
| BX005340.1 | 0.35066 | 0.177580715 | 0.254 | 0.129 | 1 | 5 |
| lcp1 | 0.35149 | -0.19809301 | 0.19 | 0.323 | 1 | 5 |
| atp13a2 | 0.35167 | -0.11905771 | 0.048 | 0.129 | 1 | 5 |
| ddx24 | 0.35173 | -0.16902821 | 0.111 | 0.226 | 1 | 5 |
| hif1an | 0.35196 | -0.14103166 | 0.032 | 0.097 | 1 | 5 |
| tmub1 | 0.3523 | 0.168462028 | 0.254 | 0.129 | 1 | 5 |
| ddx49 | 0.35258 | 0.136641386 | 0.111 | 0.032 | 1 | 5 |
| pih1d1 | 0.353 | -0.14627149 | 0.032 | 0.097 | 1 | 5 |
| si:ch211-198a12.6 | 0.35384 | 0.116951644 | 0.111 | 0.032 | 1 | 5 |
| selplg | 0.35472 | 0.213765025 | 0.254 | 0.129 | 1 | 5 |
| gng10 | 0.35533 | -0.15329764 | 0.667 | 0.806 | 1 | 5 |
| degs1 | 0.35564 | -0.16175467 | 0.333 | 0.484 | 1 | 5 |
| slc35a4 | 0.35611 | 0.140084401 | 0.159 | 0.097 | 1 | 5 |
| ndufa12 | 0.35703 | 0.104087022 | 0.365 | 0.355 | 1 | 5 |
| acin1b | 0.35709 | -0.17348798 | 0.111 | 0.226 | 1 | 5 |
| n4bp1 | 0.35786 | -0.10408851 | 0.095 | 0.194 | 1 | 5 |
| gemin2 | 0.35828 | -0.10961251 | 0.048 | 0.129 | 1 | 5 |
| kmt2bb | 0.3586 | 0.148569337 | 0.206 | 0.161 | 1 | 5 |
| dus3l | 0.35897 | 0.148461095 | 0.143 | 0.065 | 1 | 5 |
| atp5mf | 0.35921 | -0.19612398 | 0.603 | 0.71 | 1 | 5 |
| rbm25b | 0.35964 | -0.15377084 | 0.111 | 0.226 | 1 | 5 |
| lamtor5 | 0.36001 | -0.1912858 | 0.492 | 0.645 | 1 | 5 |
| si:zfos-169g10.3 | 0.36005 | 0.182295448 | 0.333 | 0.226 | 1 | 5 |
| nabp1a | 0.36063 | 0.129777412 | 0.143 | 0.065 | 1 | 5 |
| ube2v2 | 0.361 | 0.136759307 | 0.714 | 0.677 | 1 | 5 |
| znhit3 | 0.36119 | 0.114765165 | 0.111 | 0.032 | 1 | 5 |
| zgc:158343 | 0.36156 | -0.17781701 | 0.492 | 0.645 | 1 | 5 |
| mir21-1 | 0.36164 | 0.165929853 | 0.46 | 0.419 | 1 | 5 |
| gnpda1 | 0.36236 | 0.222246136 | 0.492 | 0.355 | 1 | 5 |
| kctd13 | 0.36278 | -0.11144118 | 0.048 | 0.129 | 1 | 5 |
| dse | 0.36285 | -0.11776711 | 0.175 | 0.29 | 1 | 5 |
| kpna1 | 0.36434 | -0.13821996 | 0.063 | 0.129 | 1 | 5 |

| | | | | | | |
|---|---|---|---|---|---|---|
| rraga | 0.36544 | 0.15508462 | 0.238 | 0.161 | 1 | 5 |
| si:dkey-73n10.1 | 0.3656 | 0.10126342 | 0.063 | 0.032 | 1 | 5 |
| isg20 | 0.36613 | -0.14063129 | 0.19 | 0.323 | 1 | 5 |
| sdhaf2 | 0.36626 | -0.12633584 | 0.032 | 0.097 | 1 | 5 |
| lsm5 | 0.36684 | 0.117995313 | 0.46 | 0.452 | 1 | 5 |
| ndufa11 | 0.36719 | -0.19401688 | 0.429 | 0.548 | 1 | 5 |
| map4k4 | 0.36789 | -0.10417571 | 0.048 | 0.129 | 1 | 5 |
| nap1l4b | 0.36836 | 0.107190689 | 0.111 | 0.032 | 1 | 5 |
| pxk | 0.36866 | 0.132163686 | 0.095 | 0.032 | 1 | 5 |
| cltca | 0.36871 | 0.153939318 | 0.397 | 0.258 | 1 | 5 |
| atxn1a | 0.36942 | -0.12472448 | 0.048 | 0.129 | 1 | 5 |
| pgrmc1 | 0.36952 | -0.12730777 | 0.159 | 0.194 | 1 | 5 |
| dedd | 0.36982 | 0.104361586 | 0.111 | 0.032 | 1 | 5 |
| psme1 | 0.36993 | 0.197093182 | 0.365 | 0.226 | 1 | 5 |
| fam126a | 0.37065 | -0.11580418 | 0.095 | 0.194 | 1 | 5 |
| LOC100150882 | 0.37106 | 0.229240005 | 0.19 | 0.097 | 1 | 5 |
| AL935174.2 | 0.37109 | -0.15822721 | 0.095 | 0.161 | 1 | 5 |
| psma6a | 0.37135 | -0.18674044 | 0.19 | 0.323 | 1 | 5 |
| si:zfos-943e10.1 | 0.37194 | 0.159925334 | 0.19 | 0.097 | 1 | 5 |
| pdcd11 | 0.37243 | 0.119148746 | 0.111 | 0.032 | 1 | 5 |
| slc9a7 | 0.37286 | 0.13305266 | 0.095 | 0.032 | 1 | 5 |
| rab7a | 0.37307 | 0.181399238 | 0.238 | 0.129 | 1 | 5 |
| baz1b | 0.37328 | -0.1543803 | 0.143 | 0.258 | 1 | 5 |
| psmd6 | 0.3733 | -0.1964819 | 0.206 | 0.29 | 1 | 5 |
| cd44b | 0.37346 | -0.13398828 | 0.032 | 0.097 | 1 | 5 |
| rexo4 | 0.3737 | 0.114307512 | 0.079 | 0.032 | 1 | 5 |
| galk1 | 0.37411 | -0.19379785 | 0.19 | 0.323 | 1 | 5 |
| clptm1 | 0.3747 | -0.12321867 | 0.048 | 0.129 | 1 | 5 |
| pde7a | 0.37487 | 0.135556384 | 0.111 | 0.032 | 1 | 5 |
| frg1 | 0.37487 | 0.134084839 | 0.095 | 0.032 | 1 | 5 |
| sesn1 | 0.37504 | 0.208946161 | 0.27 | 0.194 | 1 | 5 |
| si:dkey-180p18.9 | 0.37508 | 0.17411672 | 0.444 | 0.355 | 1 | 5 |
| FBXO48 | 0.3751 | -0.17986523 | 0.19 | 0.323 | 1 | 5 |
| cox7a2a | 0.37524 | -0.19642592 | 0.556 | 0.677 | 1 | 5 |
| notch2 | 0.37528 | -0.14967763 | 0.222 | 0.355 | 1 | 5 |
| arpin | 0.37531 | -0.15570933 | 0.143 | 0.258 | 1 | 5 |
| chmp6b | 0.37566 | -0.12730331 | 0.159 | 0.194 | 1 | 5 |
| adcy7 | 0.37635 | -0.19273618 | 0.19 | 0.323 | 1 | 5 |
| mob3a | 0.37831 | 0.108885906 | 0.317 | 0.29 | 1 | 5 |
| zgc:92360 | 0.37841 | -0.11409677 | 0.032 | 0.097 | 1 | 5 |
| dbn1 | 0.37842 | -0.19861267 | 0.286 | 0.419 | 1 | 5 |
| fam234a | 0.37899 | -0.16028722 | 0.048 | 0.129 | 1 | 5 |
| glipr1a | 0.37936 | 0.220670792 | 0.492 | 0.355 | 1 | 5 |
| epn2 | 0.37936 | 0.149809444 | 0.19 | 0.129 | 1 | 5 |
| ndrg3b | 0.37937 | 0.108183417 | 0.111 | 0.032 | 1 | 5 |
| tpp2 | 0.38018 | -0.13621505 | 0.048 | 0.129 | 1 | 5 |
| nucks1a | 0.3802 | -0.17981663 | 0.206 | 0.29 | 1 | 5 |
| tegt | 0.38049 | -0.17872709 | 0.524 | 0.613 | 1 | 5 |

| | | | | | | |
|---|---|---|---|---|---|---|
| tm2d2 | 0.38139 | -0.1003678 | 0.032 | 0.097 | 1 | 5 |
| LOC100007492 | 0.38246 | 0.104724057 | 0.095 | 0.032 | 1 | 5 |
| si:dkey-78l4.14 | 0.3825 | 0.171855116 | 0.206 | 0.097 | 1 | 5 |
| zdhhc18a | 0.38295 | 0.162038248 | 0.206 | 0.097 | 1 | 5 |
| zmiz2 | 0.38311 | -0.13820285 | 0.032 | 0.097 | 1 | 5 |
| atox1 | 0.38382 | -0.20783339 | 0.317 | 0.452 | 1 | 5 |
| aldh7a1 | 0.384 | 0.103894476 | 0.111 | 0.032 | 1 | 5 |
| itgb1a | 0.38402 | 0.148958226 | 0.111 | 0.032 | 1 | 5 |
| pfdn1 | 0.38429 | -0.11409351 | 0.19 | 0.226 | 1 | 5 |
| glmp | 0.38439 | 0.193000388 | 0.397 | 0.29 | 1 | 5 |
| kif21b | 0.38462 | 0.126380401 | 0.111 | 0.065 | 1 | 5 |
| anapc11 | 0.38497 | 0.14411905 | 0.222 | 0.161 | 1 | 5 |
| cant1a | 0.38507 | 0.114145474 | 0.111 | 0.065 | 1 | 5 |
| si:dkey-126g1.7 | 0.38535 | 0.104600712 | 0.111 | 0.032 | 1 | 5 |
| si:ch73-380l3.4 | 0.38553 | 0.19541194 | 0.206 | 0.097 | 1 | 5 |
| eif2b2 | 0.38561 | -0.10932424 | 0.095 | 0.194 | 1 | 5 |
| kri1 | 0.38563 | 0.165289839 | 0.206 | 0.097 | 1 | 5 |
| CABZ01084963.1 | 0.38633 | -0.10904987 | 0.048 | 0.129 | 1 | 5 |
| fbp1a | 0.38635 | 0.130753519 | 0.143 | 0.065 | 1 | 5 |
| sh3gl3a | 0.38649 | -0.14575083 | 0.143 | 0.258 | 1 | 5 |
| pdgfab | 0.38651 | -0.18330854 | 0.111 | 0.194 | 1 | 5 |
| nup107 | 0.387 | -0.12091277 | 0.048 | 0.129 | 1 | 5 |
| abracl | 0.38712 | -0.15188421 | 0.667 | 0.742 | 1 | 5 |
| reep1 | 0.38721 | -0.17665144 | 0.048 | 0.129 | 1 | 5 |
| timm9 | 0.38727 | -0.14339243 | 0.143 | 0.226 | 1 | 5 |
| si:ch211-212o1.2 | 0.38754 | 0.122204016 | 0.111 | 0.032 | 1 | 5 |
| hnrnpa1a | 0.38809 | -0.10891679 | 0.444 | 0.452 | 1 | 5 |
| aip | 0.38858 | 0.130072317 | 0.175 | 0.129 | 1 | 5 |
| arhgap22 | 0.38932 | -0.12733674 | 0.048 | 0.129 | 1 | 5 |
| si:dkey-66i24.9 | 0.38975 | 0.124570854 | 0.079 | 0.032 | 1 | 5 |
| ap3m2 | 0.38983 | -0.1824226 | 0.095 | 0.194 | 1 | 5 |
| pkig | 0.39012 | 0.110301923 | 0.159 | 0.097 | 1 | 5 |
| ak2 | 0.39028 | 0.118307067 | 0.587 | 0.581 | 1 | 5 |
| fas | 0.39033 | -0.13960853 | 0.079 | 0.129 | 1 | 5 |
| si:dkey-88p24.11 | 0.39037 | 0.108346528 | 0.143 | 0.065 | 1 | 5 |
| pdcd6 | 0.39071 | 0.171387948 | 0.222 | 0.129 | 1 | 5 |
| inpp5d | 0.39078 | 0.157925533 | 0.206 | 0.129 | 1 | 5 |
| PRRC2B | 0.39108 | -0.13150166 | 0.048 | 0.129 | 1 | 5 |
| si:ch211-276a23.5 | 0.39244 | -0.16246104 | 0.048 | 0.129 | 1 | 5 |
| atp6v0a1a | 0.39283 | -0.15236764 | 0.048 | 0.129 | 1 | 5 |
| tm9sf1 | 0.39302 | -0.16077762 | 0.095 | 0.194 | 1 | 5 |
| tfr1b | 0.39318 | -0.1563798 | 0.048 | 0.129 | 1 | 5 |
| atp5if1b | 0.39352 | -0.11870664 | 0.603 | 0.742 | 1 | 5 |
| arl6ip1 | 0.39364 | -0.11411633 | 0.841 | 0.935 | 1 | 5 |
| lyar | 0.39376 | 0.125450712 | 0.159 | 0.065 | 1 | 5 |
| lrrc59 | 0.3939 | 0.194294251 | 0.46 | 0.355 | 1 | 5 |
| tmed10 | 0.394 | 0.19031856 | 0.54 | 0.452 | 1 | 5 |
| gatd1 | 0.39401 | -0.1035347 | 0.032 | 0.097 | 1 | 5 |

| | | | | | | |
|---|---|---|---|---|---|---|
| ctss2.1 | 0.39413 | -0.23306885 | 0.381 | 0.484 | 1 | 5 |
| fbrsl1 | 0.39439 | -0.11539827 | 0.032 | 0.065 | 1 | 5 |
| csnk2b | 0.39482 | 0.103241689 | 0.46 | 0.452 | 1 | 5 |
| ndufa4 | 0.39486 | 0.121566452 | 0.095 | 0.032 | 1 | 5 |
| sf3b1 | 0.39492 | 0.15011217 | 0.159 | 0.065 | 1 | 5 |
| cdk7 | 0.39507 | -0.12253306 | 0.048 | 0.129 | 1 | 5 |
| surf2 | 0.39545 | 0.115559588 | 0.159 | 0.065 | 1 | 5 |
| si:dkey-222h21.2 | 0.39575 | 0.115424606 | 0.127 | 0.097 | 1 | 5 |
| GNB4 | 0.39622 | -0.10933948 | 0.095 | 0.194 | 1 | 5 |
| nubp2 | 0.39631 | -0.13791595 | 0.048 | 0.129 | 1 | 5 |
| socs1b | 0.39639 | -0.12916052 | 0.048 | 0.129 | 1 | 5 |
| gadd45ga | 0.39676 | -0.17961695 | 0.048 | 0.129 | 1 | 5 |
| rfc5 | 0.39677 | -0.10858926 | 0.048 | 0.129 | 1 | 5 |
| flot2a | 0.39703 | 0.127694936 | 0.143 | 0.065 | 1 | 5 |
| ngfrb | 0.39707 | -0.14391365 | 0.095 | 0.194 | 1 | 5 |
| slc40a1 | 0.39741 | 0.156473304 | 0.444 | 0.419 | 1 | 5 |
| map2k2a | 0.39745 | -0.11386834 | 0.095 | 0.194 | 1 | 5 |
| pdap1a | 0.39749 | -0.13746154 | 0.254 | 0.387 | 1 | 5 |
| cdk2ap2 | 0.39767 | -0.2107446 | 0.286 | 0.387 | 1 | 5 |
| tpcn2 | 0.39805 | -0.12394396 | 0.032 | 0.097 | 1 | 5 |
| gtf2f1 | 0.39933 | 0.111692894 | 0.159 | 0.065 | 1 | 5 |
| kars | 0.39944 | 0.115829919 | 0.365 | 0.323 | 1 | 5 |
| kctd5a | 0.39958 | -0.13842636 | 0.349 | 0.387 | 1 | 5 |
| surf1 | 0.3997 | 0.127589019 | 0.095 | 0.032 | 1 | 5 |
| mknk2a | 0.39971 | 0.172339691 | 0.175 | 0.097 | 1 | 5 |
| hiat1a | 0.39976 | -0.12208434 | 0.032 | 0.097 | 1 | 5 |
| zgc:158564 | 0.40001 | 0.141307592 | 0.159 | 0.065 | 1 | 5 |
| kdm5ba | 0.40011 | -0.12458769 | 0.127 | 0.161 | 1 | 5 |
| zgc:174904 | 0.40035 | -0.21162756 | 0.635 | 0.742 | 1 | 5 |
| erlec1 | 0.40038 | 0.138880339 | 0.159 | 0.065 | 1 | 5 |
| creb3l3l | 0.40045 | 0.107068492 | 0.238 | 0.226 | 1 | 5 |
| tmed5 | 0.40058 | -0.21180074 | 0.254 | 0.387 | 1 | 5 |
| metrnla | 0.40127 | -0.21718721 | 0.254 | 0.387 | 1 | 5 |
| hint2 | 0.40143 | -0.15997701 | 0.095 | 0.194 | 1 | 5 |
| pak2b | 0.40198 | 0.14099129 | 0.159 | 0.065 | 1 | 5 |
| ncf2 | 0.40206 | -0.17592033 | 0.238 | 0.323 | 1 | 5 |
| cd9b | 0.40208 | -0.25381712 | 0.222 | 0.355 | 1 | 5 |
| eef1b2 | 0.40305 | 0.124620941 | 0.937 | 0.871 | 1 | 5 |
| arhgef5 | 0.40353 | 0.130656188 | 0.381 | 0.258 | 1 | 5 |
| si:ch211-204c21.1 | 0.40413 | -0.21678379 | 0.381 | 0.516 | 1 | 5 |
| rarga | 0.40443 | 0.113706254 | 0.286 | 0.258 | 1 | 5 |
| arhgef12b | 0.40471 | 0.121323874 | 0.143 | 0.065 | 1 | 5 |
| srprb | 0.40519 | 0.198040769 | 0.46 | 0.323 | 1 | 5 |
| cycsb | 0.40524 | -0.16276929 | 0.683 | 0.806 | 1 | 5 |
| palm1b | 0.40563 | 0.101934232 | 0.095 | 0.032 | 1 | 5 |
| stimate | 0.40591 | 0.121305474 | 0.111 | 0.065 | 1 | 5 |
| thy1 | 0.40594 | 0.223829536 | 0.397 | 0.258 | 1 | 5 |
| zgc:136870 | 0.40596 | 0.19193141 | 0.397 | 0.258 | 1 | 5 |

| | | | | | | |
|---|---|---|---|---|---|---|
| nudcd3 | 0.40654 | 0.11220388 | 0.175 | 0.097 | 1 | 5 |
| cetn4 | 0.40678 | 0.101442517 | 0.095 | 0.065 | 1 | 5 |
| si:dkey-238c7.16 | 0.40682 | 0.202641494 | 0.27 | 0.161 | 1 | 5 |
| prss16 | 0.40684 | 0.191249181 | 0.302 | 0.226 | 1 | 5 |
| ctss2.2 | 0.40716 | -0.11030732 | 0.921 | 0.968 | 1 | 5 |
| echs1 | 0.40745 | 0.137180405 | 0.222 | 0.161 | 1 | 5 |
| si:ch211-198c19.3 | 0.40824 | -0.14545143 | 0.476 | 0.613 | 1 | 5 |
| nocta | 0.40851 | -0.15859027 | 0.143 | 0.258 | 1 | 5 |
| ms4a17a.9 | 0.40861 | 0.129325845 | 0.746 | 0.71 | 1 | 5 |
| parp8 | 0.41005 | 0.163653537 | 0.143 | 0.065 | 1 | 5 |
| tra2a | 0.41034 | -0.12824734 | 0.286 | 0.419 | 1 | 5 |
| yif1a | 0.41086 | -0.13201221 | 0.063 | 0.129 | 1 | 5 |
| rhebl1 | 0.41093 | -0.13218469 | 0.095 | 0.194 | 1 | 5 |
| fam13b | 0.41112 | 0.159009043 | 0.159 | 0.097 | 1 | 5 |
| fosl2 | 0.41168 | -0.1934063 | 0.54 | 0.677 | 1 | 5 |
| itpr1b | 0.41202 | 0.118636582 | 0.095 | 0.032 | 1 | 5 |
| per1b | 0.41211 | 0.136850593 | 0.159 | 0.097 | 1 | 5 |
| epg5 | 0.41268 | -0.12231456 | 0.032 | 0.097 | 1 | 5 |
| itprid2 | 0.41392 | 0.111233473 | 0.079 | 0.065 | 1 | 5 |
| atp6v0d1 | 0.41458 | -0.16271032 | 0.27 | 0.323 | 1 | 5 |
| carhsp1 | 0.41464 | -0.10942616 | 0.095 | 0.129 | 1 | 5 |
| ptpn22 | 0.41468 | -0.12581754 | 0.048 | 0.097 | 1 | 5 |
| CR855311.1 | 0.41535 | -0.17808381 | 0.698 | 0.774 | 1 | 5 |
| creg1 | 0.41585 | -0.11498341 | 0.095 | 0.194 | 1 | 5 |
| si:ch211-134c9.2 | 0.41735 | 0.118966931 | 0.095 | 0.032 | 1 | 5 |
| ddx39ab | 0.41753 | -0.18089077 | 0.27 | 0.387 | 1 | 5 |
| actr2a | 0.41766 | 0.154900408 | 0.476 | 0.419 | 1 | 5 |
| ell2 | 0.41847 | -0.1076107 | 0.095 | 0.194 | 1 | 5 |
| mid1ip1b | 0.41876 | -0.15546139 | 0.095 | 0.194 | 1 | 5 |
| fip1l1b | 0.41896 | -0.12309082 | 0.095 | 0.194 | 1 | 5 |
| waplb | 0.41969 | -0.10067854 | 0.032 | 0.097 | 1 | 5 |
| lifrb | 0.41978 | 0.204759305 | 0.19 | 0.097 | 1 | 5 |
| cpox | 0.42047 | -0.13674133 | 0.286 | 0.419 | 1 | 5 |
| cptp | 0.42108 | -0.14100884 | 0.095 | 0.194 | 1 | 5 |
| suclg1 | 0.42124 | -0.14208321 | 0.571 | 0.71 | 1 | 5 |
| usp14 | 0.42136 | 0.150002253 | 0.143 | 0.065 | 1 | 5 |
| acsl3b | 0.42152 | -0.15413608 | 0.032 | 0.097 | 1 | 5 |
| NPC2 (1 of many) | 0.42155 | -0.1262991 | 0.81 | 0.806 | 1 | 5 |
| cd2bp2 | 0.42163 | 0.10184553 | 0.079 | 0.032 | 1 | 5 |
| tmed2 | 0.42263 | -0.10369296 | 0.683 | 0.806 | 1 | 5 |
| msra | 0.42265 | 0.13365338 | 0.302 | 0.258 | 1 | 5 |
| klf11b | 0.42334 | 0.187099241 | 0.19 | 0.097 | 1 | 5 |
| tm9sf4 | 0.42392 | 0.136410291 | 0.175 | 0.097 | 1 | 5 |
| nucb2a | 0.42448 | 0.128169397 | 0.19 | 0.161 | 1 | 5 |
| pinx1 | 0.4246 | 0.10216753 | 0.444 | 0.323 | 1 | 5 |
| pip5k1aa | 0.42461 | -0.12358196 | 0.063 | 0.097 | 1 | 5 |
| ccl39.1 | 0.42476 | -0.24972634 | 0.238 | 0.29 | 1 | 5 |
| cul5a | 0.42586 | -0.1479966 | 0.095 | 0.194 | 1 | 5 |

| | | | | | | |
|---|---|---|---|---|---|---|
| si:dkey-17m8.1 | 0.4259 | -0.15199034 | 0.095 | 0.194 | 1 | 5 |
| drg1 | 0.4261 | -0.11680937 | 0.095 | 0.194 | 1 | 5 |
| gsdmeb | 0.42618 | -0.13363992 | 0.095 | 0.194 | 1 | 5 |
| sel1l | 0.4264 | 0.141454145 | 0.127 | 0.065 | 1 | 5 |
| rbks | 0.42649 | -0.11048708 | 0.032 | 0.097 | 1 | 5 |
| hif1al | 0.42702 | 0.117675496 | 0.19 | 0.161 | 1 | 5 |
| tufm | 0.42725 | -0.16496168 | 0.19 | 0.29 | 1 | 5 |
| gnai2b | 0.42843 | -0.11799458 | 0.127 | 0.226 | 1 | 5 |
| pnisr | 0.42889 | 0.110340146 | 0.159 | 0.129 | 1 | 5 |
| gar1 | 0.42912 | 0.168860697 | 0.27 | 0.161 | 1 | 5 |
| rap1b | 0.42941 | 0.165550576 | 0.778 | 0.677 | 1 | 5 |
| ephx2 | 0.42941 | -0.15536138 | 0.175 | 0.29 | 1 | 5 |
| clasrp | 0.43002 | 0.101509035 | 0.079 | 0.032 | 1 | 5 |
| asna1 | 0.43037 | -0.17984394 | 0.159 | 0.258 | 1 | 5 |
| si:ch211-222l21.1 | 0.43085 | -0.18732034 | 0.286 | 0.419 | 1 | 5 |
| psmb4 | 0.43194 | -0.18492972 | 0.413 | 0.548 | 1 | 5 |
| eno3 | 0.43199 | 0.176427534 | 0.73 | 0.645 | 1 | 5 |
| rabl3 | 0.43256 | 0.103714412 | 0.079 | 0.032 | 1 | 5 |
| fam32a | 0.43292 | 0.18932874 | 0.444 | 0.323 | 1 | 5 |
| vrk1 | 0.433 | 0.141290754 | 0.143 | 0.065 | 1 | 5 |
| slc25a3b | 0.43449 | 0.112373706 | 0.778 | 0.774 | 1 | 5 |
| ccr12b.2 | 0.43701 | -0.16889275 | 0.492 | 0.548 | 1 | 5 |
| stxbp3 | 0.4375 | 0.151851754 | 0.19 | 0.129 | 1 | 5 |
| fam172a | 0.43763 | 0.102212697 | 0.095 | 0.065 | 1 | 5 |
| lpcat4 | 0.43789 | -0.12294042 | 0.032 | 0.097 | 1 | 5 |
| rpf1 | 0.43793 | 0.160981262 | 0.254 | 0.161 | 1 | 5 |
| foxn2b | 0.43822 | -0.12870374 | 0.032 | 0.097 | 1 | 5 |
| eml3 | 0.43857 | 0.157950683 | 0.143 | 0.065 | 1 | 5 |
| mgst1.2 | 0.43871 | -0.12954 | 0.286 | 0.29 | 1 | 5 |
| si:ch211-147m6.1 | 0.43909 | -0.16642948 | 0.444 | 0.581 | 1 | 5 |
| bmt2 | 0.43921 | 0.149656836 | 0.143 | 0.065 | 1 | 5 |
| pcnt | 0.43936 | -0.12867342 | 0.048 | 0.097 | 1 | 5 |
| si:dkey-33c12.4 | 0.44016 | -0.17729203 | 0.302 | 0.419 | 1 | 5 |
| atp6v1ba | 0.44074 | 0.191003148 | 0.524 | 0.387 | 1 | 5 |
| dctn1a | 0.442 | -0.12735108 | 0.032 | 0.097 | 1 | 5 |
| nomo | 0.44257 | -0.16099957 | 0.111 | 0.194 | 1 | 5 |
| fuca2 | 0.44273 | 0.196155517 | 0.587 | 0.452 | 1 | 5 |
| kidins220b | 0.44276 | -0.10297668 | 0.032 | 0.097 | 1 | 5 |
| gnb1a | 0.44287 | 0.193803222 | 0.508 | 0.387 | 1 | 5 |
| lsm14ab | 0.4429 | 0.114632248 | 0.127 | 0.065 | 1 | 5 |
| mycbp2 | 0.44319 | 0.152985935 | 0.238 | 0.129 | 1 | 5 |
| rgs5b | 0.44355 | -0.21689851 | 0.175 | 0.29 | 1 | 5 |
| prrc1 | 0.44451 | -0.12864341 | 0.048 | 0.097 | 1 | 5 |
| CU570889.1 | 0.44503 | 0.160941546 | 0.127 | 0.065 | 1 | 5 |
| csrnp1b | 0.44505 | -0.18643345 | 0.381 | 0.516 | 1 | 5 |
| pnpla6 | 0.44515 | -0.11292903 | 0.032 | 0.097 | 1 | 5 |
| LOC110439414 | 0.44577 | -0.11862228 | 0.032 | 0.097 | 1 | 5 |
| tm9sf5 | 0.44597 | 0.112940498 | 0.095 | 0.032 | 1 | 5 |

| | | | | | | |
|---|---|---|---|---|---|---|
| CR318588.3 | 0.44622 | 0.174885372 | 0.127 | 0.065 | 1 | 5 |
| LOC108179139 | 0.4464 | -0.11365888 | 0.032 | 0.097 | 1 | 5 |
| abhd16a | 0.4465 | 0.107157918 | 0.143 | 0.129 | 1 | 5 |
| LOC101885445 | 0.44658 | -0.10114813 | 0.032 | 0.097 | 1 | 5 |
| npm3 | 0.44665 | 0.103148854 | 0.365 | 0.355 | 1 | 5 |
| abcc3 | 0.4472 | -0.15599692 | 0.175 | 0.29 | 1 | 5 |
| zbed4 | 0.44727 | -0.13695253 | 0.048 | 0.097 | 1 | 5 |
| baxa | 0.44807 | -0.15057715 | 0.127 | 0.226 | 1 | 5 |
| adipor1b | 0.44824 | 0.145329221 | 0.127 | 0.065 | 1 | 5 |
| bop1 | 0.44843 | -0.10327107 | 0.032 | 0.097 | 1 | 5 |
| lect2l | 0.44862 | 0.197601688 | 0.683 | 0.548 | 1 | 5 |
| rnf213b | 0.44879 | -0.14023408 | 0.032 | 0.097 | 1 | 5 |
| myo1cb | 0.44905 | -0.10427498 | 0.032 | 0.097 | 1 | 5 |
| si:ch211-136m16.8 | 0.44907 | 0.166919447 | 0.19 | 0.097 | 1 | 5 |
| lamp2 | 0.44907 | -0.15632378 | 0.317 | 0.452 | 1 | 5 |
| eno1a | 0.44934 | 0.17119153 | 0.492 | 0.355 | 1 | 5 |
| slc25a44a | 0.44937 | -0.10442844 | 0.032 | 0.097 | 1 | 5 |
| rbm4.3 | 0.44945 | 0.145788057 | 0.524 | 0.387 | 1 | 5 |
| mef2aa | 0.44965 | -0.11256589 | 0.159 | 0.258 | 1 | 5 |
| praf2 | 0.44967 | -0.1048002 | 0.032 | 0.097 | 1 | 5 |
| tspan3a | 0.44982 | -0.11058199 | 0.032 | 0.097 | 1 | 5 |
| anapc15 | 0.44988 | 0.112223843 | 0.079 | 0.032 | 1 | 5 |
| slc7a3a | 0.44991 | -0.11497413 | 0.095 | 0.129 | 1 | 5 |
| cpda | 0.44992 | -0.10282735 | 0.032 | 0.097 | 1 | 5 |
| fbxo34 | 0.45017 | -0.10115318 | 0.032 | 0.097 | 1 | 5 |
| tmem167b | 0.45029 | -0.10679742 | 0.032 | 0.097 | 1 | 5 |
| pfdn5 | 0.45036 | 0.110665968 | 0.381 | 0.355 | 1 | 5 |
| ndufs2 | 0.45042 | 0.16315066 | 0.302 | 0.194 | 1 | 5 |
| ubfd1 | 0.45049 | 0.133584505 | 0.206 | 0.129 | 1 | 5 |
| pxmp2 | 0.45057 | -0.10726647 | 0.032 | 0.097 | 1 | 5 |
| eepd1 | 0.451 | 0.137994326 | 0.159 | 0.097 | 1 | 5 |
| tpd52l2a | 0.45118 | 0.111101936 | 0.127 | 0.065 | 1 | 5 |
| ncoa4 | 0.45201 | 0.10263149 | 0.095 | 0.032 | 1 | 5 |
| arih1 | 0.45214 | -0.10526572 | 0.079 | 0.161 | 1 | 5 |
| LOC101886073 | 0.45259 | 0.157382504 | 0.238 | 0.161 | 1 | 5 |
| si:ch73-1a9.3 | 0.45321 | 0.174935664 | 0.651 | 0.516 | 1 | 5 |
| gadd45ab | 0.45438 | 0.111462766 | 0.143 | 0.065 | 1 | 5 |
| polr2a | 0.45506 | -0.16306966 | 0.111 | 0.194 | 1 | 5 |
| asah1b | 0.45511 | 0.159688591 | 0.587 | 0.452 | 1 | 5 |
| snrpg | 0.45572 | -0.14463152 | 0.46 | 0.516 | 1 | 5 |
| mcoln1a | 0.45604 | 0.112606428 | 0.095 | 0.032 | 1 | 5 |
| fhit | 0.45605 | -0.14925468 | 0.159 | 0.226 | 1 | 5 |
| abhd12 | 0.45664 | -0.16981823 | 0.127 | 0.226 | 1 | 5 |
| hmga1a | 0.45709 | 0.136651434 | 0.651 | 0.516 | 1 | 5 |
| si:ch211-102c2.4 | 0.45772 | -0.13789254 | 0.81 | 0.871 | 1 | 5 |
| waca | 0.45798 | 0.147414093 | 0.175 | 0.097 | 1 | 5 |
| ctdsplb | 0.45821 | -0.13580007 | 0.063 | 0.129 | 1 | 5 |
| CU929230.1 | 0.45855 | 0.105531162 | 0.095 | 0.032 | 1 | 5 |

| | | | | | | |
|---|---|---|---|---|---|---|
| aamp | 0.45888 | -0.11741152 | 0.095 | 0.161 | 1 | 5 |
| larp1b | 0.45892 | -0.15630345 | 0.063 | 0.129 | 1 | 5 |
| aldh2.2 | 0.45901 | 0.136074033 | 0.206 | 0.161 | 1 | 5 |
| cops3 | 0.45949 | -0.13066372 | 0.079 | 0.161 | 1 | 5 |
| ist1 | 0.45992 | -0.18319573 | 0.238 | 0.355 | 1 | 5 |
| zgc:100918 | 0.46012 | -0.11611359 | 0.127 | 0.226 | 1 | 5 |
| git2b | 0.46104 | -0.10584232 | 0.19 | 0.29 | 1 | 5 |
| XLOC-025423 | 0.46156 | 0.187043951 | 0.587 | 0.452 | 1 | 5 |
| bola1 | 0.46174 | -0.10758749 | 0.079 | 0.097 | 1 | 5 |
| prdx3 | 0.46268 | 0.138911612 | 0.333 | 0.258 | 1 | 5 |
| zgc:162025 | 0.4635 | -0.10062561 | 0.079 | 0.161 | 1 | 5 |
| atp6v0b | 0.46353 | -0.17252924 | 0.413 | 0.484 | 1 | 5 |
| chid1 | 0.46405 | 0.114000174 | 0.079 | 0.032 | 1 | 5 |
| atf7ip | 0.46481 | 0.129202036 | 0.095 | 0.032 | 1 | 5 |
| retsat | 0.46496 | 0.103271668 | 0.095 | 0.032 | 1 | 5 |
| LOC108191614 | 0.46507 | -0.15711661 | 0.095 | 0.161 | 1 | 5 |
| banf1 | 0.46513 | -0.17014734 | 0.27 | 0.387 | 1 | 5 |
| tbc1d22a | 0.46549 | 0.105837623 | 0.095 | 0.032 | 1 | 5 |
| si:busm1-71b9.3 | 0.46574 | 0.144557789 | 0.19 | 0.097 | 1 | 5 |
| gspt1l | 0.46579 | -0.16908715 | 0.27 | 0.387 | 1 | 5 |
| hikeshi | 0.46582 | -0.1535889 | 0.206 | 0.29 | 1 | 5 |
| eif4a1b | 0.46589 | 0.157545082 | 0.635 | 0.581 | 1 | 5 |
| mrpl18 | 0.46732 | -0.13815926 | 0.143 | 0.194 | 1 | 5 |
| blcap | 0.46848 | 0.167215936 | 0.508 | 0.387 | 1 | 5 |
| snrpa | 0.4687 | -0.10934018 | 0.127 | 0.226 | 1 | 5 |
| phf8 | 0.46881 | 0.111367202 | 0.079 | 0.032 | 1 | 5 |
| klf13 | 0.4706 | -0.105762 | 0.365 | 0.387 | 1 | 5 |
| mtch2 | 0.47071 | 0.13358362 | 0.27 | 0.161 | 1 | 5 |
| smarcd1 | 0.47168 | 0.126887984 | 0.143 | 0.065 | 1 | 5 |
| zgc:77650 | 0.47172 | -0.15565229 | 0.238 | 0.323 | 1 | 5 |
| gab1 | 0.47231 | -0.13321672 | 0.079 | 0.129 | 1 | 5 |
| nfil3-6 | 0.47273 | 0.187706987 | 0.651 | 0.548 | 1 | 5 |
| atp5f1c | 0.47323 | -0.10081596 | 0.619 | 0.742 | 1 | 5 |
| rab11a | 0.47349 | 0.174949245 | 0.381 | 0.258 | 1 | 5 |
| glod5 | 0.47365 | 0.125853323 | 0.175 | 0.097 | 1 | 5 |
| hnrnpr | 0.4742 | -0.10815559 | 0.127 | 0.226 | 1 | 5 |
| cyth1a | 0.47451 | 0.154327116 | 0.27 | 0.161 | 1 | 5 |
| fbxw2 | 0.47478 | 0.110609891 | 0.095 | 0.032 | 1 | 5 |
| ywhabb | 0.47513 | 0.172006166 | 0.603 | 0.484 | 1 | 5 |
| ppp1caa | 0.47549 | -0.15948951 | 0.365 | 0.484 | 1 | 5 |
| si:busm1-266f07.2 | 0.47586 | -0.12972225 | 0.81 | 0.903 | 1 | 5 |
| taf3 | 0.47602 | 0.151301145 | 0.143 | 0.065 | 1 | 5 |
| chmp1a | 0.47646 | 0.102620722 | 0.127 | 0.097 | 1 | 5 |
| sod2 | 0.47669 | 0.129644438 | 0.222 | 0.161 | 1 | 5 |
| spns1 | 0.47677 | -0.13670886 | 0.111 | 0.161 | 1 | 5 |
| rabgap1l | 0.47725 | -0.12651114 | 0.143 | 0.194 | 1 | 5 |
| supt4h1 | 0.47736 | -0.1394281 | 0.206 | 0.323 | 1 | 5 |
| srsf6b | 0.47754 | -0.14731115 | 0.159 | 0.258 | 1 | 5 |

| | | | | | | |
|---|---|---|---|---|---|---|
| znf576.2 | 0.47818 | 0.102678915 | 0.079 | 0.032 | 1 | 5 |
| rab10 | 0.47878 | -0.15418934 | 0.444 | 0.516 | 1 | 5 |
| hspa4b | 0.47899 | -0.12946873 | 0.175 | 0.226 | 1 | 5 |
| psmb7 | 0.47931 | -0.14146873 | 0.381 | 0.419 | 1 | 5 |
| coa5 | 0.47961 | 0.139438081 | 0.222 | 0.129 | 1 | 5 |
| plscr3b | 0.48059 | 0.150389255 | 0.317 | 0.258 | 1 | 5 |
| ssr2 | 0.48078 | -0.13731012 | 0.825 | 0.903 | 1 | 5 |
| rnf111 | 0.4808 | 0.118066725 | 0.095 | 0.032 | 1 | 5 |
| dnajc7 | 0.48095 | -0.13844868 | 0.111 | 0.194 | 1 | 5 |
| si:ch211-87m7.2 | 0.48132 | -0.16047901 | 0.127 | 0.226 | 1 | 5 |
| ncln | 0.48133 | 0.118632966 | 0.19 | 0.097 | 1 | 5 |
| serinc5 | 0.48185 | 0.163524674 | 0.127 | 0.065 | 1 | 5 |
| rnf121 | 0.48244 | -0.12789176 | 0.079 | 0.161 | 1 | 5 |
| ctnna1 | 0.48268 | -0.11658422 | 0.079 | 0.161 | 1 | 5 |
| nxt2 | 0.48288 | -0.12120176 | 0.127 | 0.226 | 1 | 5 |
| ptgir | 0.48298 | 0.106508272 | 0.095 | 0.032 | 1 | 5 |
| ehbp1l1a | 0.48355 | 0.10664685 | 0.095 | 0.065 | 1 | 5 |
| mt-nd5 | 0.48375 | -0.15213629 | 0.27 | 0.323 | 1 | 5 |
| atpaf2 | 0.48413 | 0.102339273 | 0.095 | 0.065 | 1 | 5 |
| ube2ib | 0.48441 | -0.17045644 | 0.254 | 0.355 | 1 | 5 |
| txn | 0.48471 | 0.182742019 | 0.698 | 0.581 | 1 | 5 |
| thumpd3 | 0.4848 | -0.11834861 | 0.079 | 0.161 | 1 | 5 |
| cd81a | 0.48519 | -0.17472117 | 0.508 | 0.613 | 1 | 5 |
| smdt1b | 0.48682 | 0.147635729 | 0.524 | 0.452 | 1 | 5 |
| ppp4r1 | 0.48882 | -0.13961123 | 0.063 | 0.129 | 1 | 5 |
| yy1a | 0.48937 | 0.106765746 | 0.222 | 0.129 | 1 | 5 |
| si:dkey-56m19.5 | 0.4897 | 0.167465162 | 0.27 | 0.161 | 1 | 5 |
| zgc:163080 | 0.49087 | 0.117067228 | 0.143 | 0.065 | 1 | 5 |
| svbp | 0.49122 | -0.11680319 | 0.079 | 0.129 | 1 | 5 |
| abi3a | 0.4919 | 0.138741714 | 0.175 | 0.097 | 1 | 5 |
| tnfrsf1a | 0.49228 | -0.14573683 | 0.349 | 0.419 | 1 | 5 |
| ppp2r2d | 0.49233 | -0.11680195 | 0.079 | 0.161 | 1 | 5 |
| dusp1 | 0.49298 | 0.142984275 | 0.524 | 0.484 | 1 | 5 |
| ptbp3 | 0.49302 | 0.13446729 | 0.19 | 0.129 | 1 | 5 |
| eif3k | 0.49343 | 0.131242303 | 0.556 | 0.516 | 1 | 5 |
| atf4a | 0.49356 | -0.16459049 | 0.492 | 0.581 | 1 | 5 |
| psma2 | 0.49373 | 0.10796904 | 0.413 | 0.387 | 1 | 5 |
| hipk3b | 0.49373 | 0.148555945 | 0.143 | 0.065 | 1 | 5 |
| lmf2b | 0.49392 | -0.13425584 | 0.095 | 0.161 | 1 | 5 |
| si:dkey-148a17.6 | 0.49448 | 0.135614354 | 0.19 | 0.129 | 1 | 5 |
| aldh9a1a.1 | 0.49464 | 0.139644234 | 0.46 | 0.419 | 1 | 5 |
| ipmkb | 0.49484 | -0.1196684 | 0.079 | 0.161 | 1 | 5 |
| vaspb | 0.49485 | 0.114546617 | 0.143 | 0.065 | 1 | 5 |
| cish | 0.495 | -0.12918255 | 0.079 | 0.161 | 1 | 5 |
| adnpa | 0.49549 | 0.125597522 | 0.143 | 0.065 | 1 | 5 |
| si:ch211-160o17.4 | 0.49651 | 0.101074604 | 0.079 | 0.032 | 1 | 5 |
| ndfip1l | 0.49702 | -0.14587098 | 0.238 | 0.355 | 1 | 5 |
| chmp2a | 0.49735 | -0.12234496 | 0.333 | 0.387 | 1 | 5 |

| | | | | | | |
|---|---|---|---|---|---|---|
| mrpl19 | 0.49777 | -0.12769366 | 0.079 | 0.161 | 1 | 5 |
| zgc:85858 | 0.49812 | 0.120027413 | 0.095 | 0.032 | 1 | 5 |
| vps36 | 0.4983 | -0.10781179 | 0.063 | 0.129 | 1 | 5 |
| apbb1 | 0.49839 | -0.10528007 | 0.032 | 0.065 | 1 | 5 |
| AL928650.1 | 0.49872 | -0.1791658 | 0.238 | 0.355 | 1 | 5 |
| tefb | 0.49884 | -0.11885847 | 0.079 | 0.161 | 1 | 5 |
| unc93b1 | 0.49893 | 0.169134812 | 0.206 | 0.129 | 1 | 5 |
| gemin8 | 0.49895 | -0.10483366 | 0.079 | 0.161 | 1 | 5 |
| aldh1a2 | 0.49924 | -0.10384228 | 0.032 | 0.065 | 1 | 5 |
| seta | 0.49938 | -0.16948475 | 0.476 | 0.581 | 1 | 5 |
| apmap | 0.49961 | -0.12934536 | 0.143 | 0.194 | 1 | 5 |
| DDX17 | 0.49973 | -0.1266679 | 0.079 | 0.161 | 1 | 5 |
| flna | 0.50079 | 0.146861925 | 0.444 | 0.323 | 1 | 5 |
| slc16a1b | 0.50081 | -0.11389666 | 0.079 | 0.161 | 1 | 5 |
| si:ch211-39i2.2 | 0.50134 | 0.170558481 | 0.508 | 0.419 | 1 | 5 |
| calm3a | 0.50261 | 0.115076173 | 0.143 | 0.065 | 1 | 5 |
| fam160b1 | 0.50265 | 0.109281824 | 0.095 | 0.032 | 1 | 5 |
| glula | 0.50291 | 0.134311455 | 0.095 | 0.065 | 1 | 5 |
| dtnbp1a | 0.50309 | 0.111007845 | 0.143 | 0.097 | 1 | 5 |
| plekho1b | 0.50376 | 0.181774464 | 0.508 | 0.387 | 1 | 5 |
| rbm47 | 0.50433 | 0.140170394 | 0.175 | 0.097 | 1 | 5 |
| ywhabl | 0.50458 | -0.11390867 | 0.603 | 0.645 | 1 | 5 |
| tmem106a | 0.50535 | -0.10814053 | 0.159 | 0.194 | 1 | 5 |
| arhgef1b | 0.50593 | 0.128710387 | 0.222 | 0.161 | 1 | 5 |
| bckdk | 0.50597 | -0.13713552 | 0.111 | 0.161 | 1 | 5 |
| rrp12 | 0.50598 | -0.12406264 | 0.063 | 0.129 | 1 | 5 |
| wdfy1 | 0.50602 | 0.10752732 | 0.143 | 0.065 | 1 | 5 |
| CHST13 | 0.50737 | -0.10006552 | 0.111 | 0.194 | 1 | 5 |
| tpi1b | 0.50744 | -0.13199412 | 0.587 | 0.71 | 1 | 5 |
| yipf4 | 0.50744 | 0.151240817 | 0.286 | 0.226 | 1 | 5 |
| cxcr3.2 | 0.50745 | 0.170191156 | 0.571 | 0.452 | 1 | 5 |
| zgc:175214 | 0.50754 | 0.119364664 | 0.143 | 0.065 | 1 | 5 |
| cox5ab | 0.5077 | 0.182376061 | 0.556 | 0.452 | 1 | 5 |
| ncoa3 | 0.50842 | -0.12001874 | 0.032 | 0.065 | 1 | 5 |
| il6st | 0.50924 | 0.138942193 | 0.175 | 0.097 | 1 | 5 |
| rnf169 | 0.50927 | 0.112099585 | 0.111 | 0.065 | 1 | 5 |
| polr1b | 0.50991 | -0.12298893 | 0.063 | 0.129 | 1 | 5 |
| mcf2l2 | 0.51055 | 0.11332068 | 0.095 | 0.032 | 1 | 5 |
| si:zfos-2330d3.3 | 0.51098 | -0.24164619 | 0.302 | 0.419 | 1 | 5 |
| scfd1 | 0.51101 | -0.10343601 | 0.079 | 0.129 | 1 | 5 |
| tax1bp1b | 0.51101 | 0.103439438 | 0.143 | 0.065 | 1 | 5 |
| si:ch73-158p21.2 | 0.51117 | 0.119243355 | 0.127 | 0.065 | 1 | 5 |
| CABZ01087560.1 | 0.51122 | 0.10023567 | 0.143 | 0.065 | 1 | 5 |
| srp54 | 0.51125 | 0.101701071 | 0.143 | 0.065 | 1 | 5 |
| ralab | 0.51138 | 0.107887221 | 0.095 | 0.032 | 1 | 5 |
| mapk3 | 0.51195 | 0.106107192 | 0.127 | 0.097 | 1 | 5 |
| vps26c | 0.51234 | 0.106603163 | 0.111 | 0.065 | 1 | 5 |
| csnk1db | 0.5129 | -0.11519243 | 0.159 | 0.258 | 1 | 5 |

| | | | | | | |
|---|---|---|---|---|---|---|
| top1l | 0.51378 | -0.14020926 | 0.27 | 0.387 | 1 | 5 |
| leng8 | 0.51493 | 0.107376879 | 0.063 | 0.032 | 1 | 5 |
| eif4g1a | 0.51567 | 0.169826542 | 0.333 | 0.226 | 1 | 5 |
| ubr4 | 0.51588 | 0.123935658 | 0.175 | 0.097 | 1 | 5 |
| mbpb | 0.51645 | 0.117732079 | 0.222 | 0.129 | 1 | 5 |
| rarab | 0.51676 | 0.105429016 | 0.063 | 0.032 | 1 | 5 |
| hmgn6 | 0.51728 | 0.118922689 | 0.571 | 0.548 | 1 | 5 |
| rab5c | 0.51747 | 0.111489074 | 0.54 | 0.419 | 1 | 5 |
| gadd45gip1 | 0.51749 | -0.13320696 | 0.159 | 0.258 | 1 | 5 |
| slc7a6os | 0.51769 | -0.11197285 | 0.095 | 0.129 | 1 | 5 |
| hdac1 | 0.5192 | -0.13995749 | 0.254 | 0.323 | 1 | 5 |
| rin3 | 0.52105 | -0.16719306 | 0.222 | 0.323 | 1 | 5 |
| stx11a | 0.52135 | 0.103408848 | 0.286 | 0.194 | 1 | 5 |
| clic1 | 0.52183 | 0.136311325 | 0.508 | 0.452 | 1 | 5 |
| gabarapb | 0.52203 | -0.12417757 | 0.698 | 0.806 | 1 | 5 |
| slc35f6 | 0.52256 | 0.185271015 | 0.476 | 0.355 | 1 | 5 |
| ackr3b | 0.52307 | -0.14302635 | 0.063 | 0.129 | 1 | 5 |
| calm3b | 0.52321 | -0.14584034 | 0.46 | 0.516 | 1 | 5 |
| AK6 | 0.52397 | 0.100641177 | 0.079 | 0.065 | 1 | 5 |
| exoc3 | 0.52401 | -0.10020409 | 0.032 | 0.065 | 1 | 5 |
| f11r.1 | 0.52408 | 0.146146781 | 0.317 | 0.258 | 1 | 5 |
| zgc:162698 | 0.52532 | 0.163786531 | 0.333 | 0.258 | 1 | 5 |
| zmynd11 | 0.52595 | -0.1179527 | 0.048 | 0.097 | 1 | 5 |
| nr4a1 | 0.52652 | 0.135364932 | 0.556 | 0.516 | 1 | 5 |
| ap2m1b | 0.52681 | -0.11920053 | 0.159 | 0.258 | 1 | 5 |
| rpl5b | 0.5272 | -0.10545027 | 0.873 | 0.935 | 1 | 5 |
| chd7 | 0.528 | -0.13336674 | 0.111 | 0.194 | 1 | 5 |
| impdh1b | 0.52806 | 0.110650274 | 0.127 | 0.065 | 1 | 5 |
| elavl1a | 0.52864 | -0.15202609 | 0.222 | 0.323 | 1 | 5 |
| ypel5 | 0.53014 | -0.12166708 | 0.095 | 0.161 | 1 | 5 |
| psmg2 | 0.53043 | 0.125917555 | 0.222 | 0.129 | 1 | 5 |
| atp5mc1 | 0.53111 | 0.160715801 | 0.603 | 0.484 | 1 | 5 |
| tmem209 | 0.53166 | -0.100892 | 0.095 | 0.129 | 1 | 5 |
| calr3a | 0.53262 | -0.14187123 | 0.524 | 0.645 | 1 | 5 |
| lsm8 | 0.5331 | 0.139504768 | 0.333 | 0.226 | 1 | 5 |
| SCARF2 | 0.53442 | -0.13840859 | 0.079 | 0.129 | 1 | 5 |
| alg2 | 0.53564 | 0.101925137 | 0.079 | 0.032 | 1 | 5 |
| atp6ap2 | 0.53615 | -0.16825613 | 0.476 | 0.581 | 1 | 5 |
| sh3bgrl3 | 0.53647 | -0.16455227 | 0.365 | 0.452 | 1 | 5 |
| atf4b | 0.53664 | -0.12586931 | 0.619 | 0.677 | 1 | 5 |
| fam53b | 0.53719 | 0.150446705 | 0.254 | 0.161 | 1 | 5 |
| dennd4b | 0.53759 | 0.105426199 | 0.111 | 0.065 | 1 | 5 |
| chchd7 | 0.53767 | -0.15536235 | 0.222 | 0.323 | 1 | 5 |
| dusp2 | 0.53785 | -0.19822277 | 0.286 | 0.387 | 1 | 5 |
| pcmt | 0.53851 | 0.124054999 | 0.127 | 0.065 | 1 | 5 |
| dnajc1 | 0.53919 | -0.1371387 | 0.333 | 0.452 | 1 | 5 |
| me2 | 0.53956 | -0.11225968 | 0.254 | 0.355 | 1 | 5 |
| tcima | 0.53957 | 0.140877285 | 0.667 | 0.548 | 1 | 5 |

| | | | | | | |
|---|---|---|---|---|---|---|
| vps28 | 0.53994 | 0.126776659 | 0.222 | 0.129 | 1 | 5 |
| lgals9l3 | 0.54031 | -0.15319664 | 0.492 | 0.613 | 1 | 5 |
| atp6v1d | 0.54073 | -0.15645712 | 0.27 | 0.355 | 1 | 5 |
| slc3a2b | 0.5408 | -0.14750661 | 0.333 | 0.452 | 1 | 5 |
| eif4ebp2 | 0.54175 | 0.138425877 | 0.508 | 0.387 | 1 | 5 |
| gpia | 0.54191 | 0.169988859 | 0.397 | 0.29 | 1 | 5 |
| mcm2 | 0.54242 | -0.12434742 | 0.063 | 0.129 | 1 | 5 |
| tab1 | 0.54269 | 0.116310118 | 0.175 | 0.097 | 1 | 5 |
| vps4b | 0.5427 | -0.15548473 | 0.222 | 0.323 | 1 | 5 |
| ero1b | 0.5438 | -0.11442098 | 0.111 | 0.194 | 1 | 5 |
| rsl1d1 | 0.54436 | 0.148615426 | 0.54 | 0.419 | 1 | 5 |
| triqk | 0.54457 | 0.15723973 | 0.333 | 0.226 | 1 | 5 |
| smap1 | 0.54497 | 0.110339386 | 0.143 | 0.097 | 1 | 5 |
| CABZ01066926.1 | 0.54556 | -0.11884942 | 0.048 | 0.097 | 1 | 5 |
| psmd13 | 0.54622 | 0.125823249 | 0.238 | 0.161 | 1 | 5 |
| cyp2k8 | 0.54645 | -0.11113273 | 0.111 | 0.194 | 1 | 5 |
| ywhaz | 0.54713 | 0.146692808 | 0.667 | 0.581 | 1 | 5 |
| ndufb8 | 0.54804 | 0.134292728 | 0.603 | 0.484 | 1 | 5 |
| med19a | 0.54911 | 0.135032449 | 0.206 | 0.129 | 1 | 5 |
| alox5ap | 0.54921 | -0.15318025 | 0.19 | 0.29 | 1 | 5 |
| rbm5 | 0.54957 | -0.11393577 | 0.111 | 0.194 | 1 | 5 |
| nop58 | 0.54983 | -0.1073868 | 0.381 | 0.387 | 1 | 5 |
| ptpn6 | 0.55052 | -0.1502905 | 0.524 | 0.613 | 1 | 5 |
| dad1 | 0.55103 | -0.10778017 | 0.762 | 0.806 | 1 | 5 |
| znf593 | 0.55119 | -0.12703758 | 0.19 | 0.29 | 1 | 5 |
| pomp | 0.55139 | 0.147485286 | 0.603 | 0.548 | 1 | 5 |
| gbp1 | 0.55353 | -0.13567622 | 0.127 | 0.194 | 1 | 5 |
| crip1 | 0.55383 | -0.1038574 | 0.889 | 0.935 | 1 | 5 |
| chmp5b | 0.55394 | 0.151522608 | 0.397 | 0.29 | 1 | 5 |
| cybb | 0.55401 | 0.147178331 | 0.333 | 0.226 | 1 | 5 |
| BX649485.1 | 0.55525 | -0.17068248 | 0.413 | 0.516 | 1 | 5 |
| serpinb14 | 0.55541 | -0.10685943 | 0.063 | 0.129 | 1 | 5 |
| glrx | 0.5555 | -0.12145573 | 0.794 | 0.871 | 1 | 5 |
| srsf1a | 0.55637 | -0.10561219 | 0.111 | 0.194 | 1 | 5 |
| bnip3lb | 0.55667 | -0.14134821 | 0.127 | 0.194 | 1 | 5 |
| rap2b | 0.55808 | -0.13915954 | 0.19 | 0.258 | 1 | 5 |
| dohh | 0.55816 | 0.10608292 | 0.175 | 0.097 | 1 | 5 |
| enosf1 | 0.55845 | 0.106355338 | 0.175 | 0.097 | 1 | 5 |
| rab5if | 0.56014 | -0.11120918 | 0.238 | 0.29 | 1 | 5 |
| XLOC-018981 | 0.56266 | -0.11757463 | 0.063 | 0.129 | 1 | 5 |
| si:ch73-37h15.2 | 0.56314 | 0.121605349 | 0.127 | 0.065 | 1 | 5 |
| mapk11 | 0.56376 | -0.12008672 | 0.222 | 0.323 | 1 | 5 |
| kdelr2b | 0.56407 | 0.122650611 | 0.254 | 0.161 | 1 | 5 |
| dub | 0.56436 | 0.148982644 | 0.254 | 0.161 | 1 | 5 |
| msna | 0.56495 | 0.125098765 | 0.841 | 0.774 | 1 | 5 |
| ndufv1 | 0.56552 | 0.132730581 | 0.159 | 0.097 | 1 | 5 |
| zgc:153184 | 0.56585 | -0.1083595 | 0.063 | 0.129 | 1 | 5 |
| si:ch73-343l4.8 | 0.56621 | 0.115493527 | 0.159 | 0.097 | 1 | 5 |

| | | | | | | |
|---|---|---|---|---|---|---|
| plp2 | 0.56652 | 0.134207112 | 0.317 | 0.258 | 1 | 5 |
| asb8 | 0.56751 | 0.104082202 | 0.127 | 0.065 | 1 | 5 |
| tomm40l | 0.56764 | -0.10443433 | 0.079 | 0.129 | 1 | 5 |
| kdm2ba | 0.56823 | -0.10725469 | 0.095 | 0.129 | 1 | 5 |
| polr2i | 0.56871 | 0.102787654 | 0.175 | 0.097 | 1 | 5 |
| zdhhc20a | 0.56891 | 0.106251542 | 0.111 | 0.065 | 1 | 5 |
| si:dkey-7j14.6 | 0.56907 | 0.150762735 | 0.619 | 0.548 | 1 | 5 |
| fbxo28 | 0.5706 | 0.106640052 | 0.175 | 0.097 | 1 | 5 |
| flot1b | 0.57085 | 0.111052308 | 0.206 | 0.129 | 1 | 5 |
| tnfrsf1b | 0.57164 | 0.129020549 | 0.159 | 0.097 | 1 | 5 |
| pa2g4b | 0.57251 | -0.1239905 | 0.238 | 0.323 | 1 | 5 |
| serp1 | 0.57301 | -0.15281062 | 0.46 | 0.516 | 1 | 5 |
| tradd | 0.5731 | 0.144346882 | 0.286 | 0.194 | 1 | 5 |
| trib3 | 0.57369 | -0.16229803 | 0.413 | 0.516 | 1 | 5 |
| arfgef1 | 0.57585 | -0.10720327 | 0.048 | 0.097 | 1 | 5 |
| taok3a | 0.57601 | 0.103526224 | 0.127 | 0.065 | 1 | 5 |
| nxpe3 | 0.57688 | 0.111424292 | 0.095 | 0.065 | 1 | 5 |
| ddx21 | 0.57751 | -0.14869354 | 0.381 | 0.484 | 1 | 5 |
| stk35 | 0.57757 | 0.109452065 | 0.206 | 0.161 | 1 | 5 |
| LOC110438237 | 0.5782 | 0.120999803 | 0.238 | 0.194 | 1 | 5 |
| asns | 0.57867 | -0.10946392 | 0.048 | 0.097 | 1 | 5 |
| tmem30aa | 0.57882 | -0.1500257 | 0.222 | 0.323 | 1 | 5 |
| parp16 | 0.57948 | 0.103344095 | 0.175 | 0.097 | 1 | 5 |
| tpm3 | 0.58139 | -0.11024148 | 0.587 | 0.613 | 1 | 5 |
| XLOC-028371 | 0.58155 | -0.14226404 | 0.063 | 0.129 | 1 | 5 |
| si:ch211-271d10.2 | 0.58201 | -0.11045474 | 0.063 | 0.129 | 1 | 5 |
| si:dkey-42i9.6 | 0.58232 | -0.1264091 | 0.222 | 0.323 | 1 | 5 |
| plekhj1 | 0.58278 | 0.103485164 | 0.111 | 0.065 | 1 | 5 |
| ube2al | 0.58303 | -0.13393809 | 0.222 | 0.323 | 1 | 5 |
| jagn1b | 0.58309 | 0.116113834 | 0.254 | 0.161 | 1 | 5 |
| rtf2 | 0.58316 | -0.10265772 | 0.095 | 0.161 | 1 | 5 |
| birc2 | 0.58325 | -0.1383871 | 0.286 | 0.355 | 1 | 5 |
| nudc | 0.58355 | -0.11388972 | 0.222 | 0.323 | 1 | 5 |
| fkbp3 | 0.5843 | -0.1057117 | 0.222 | 0.323 | 1 | 5 |
| hck | 0.58493 | -0.13442343 | 0.159 | 0.226 | 1 | 5 |
| rnf13 | 0.58579 | 0.113055867 | 0.111 | 0.065 | 1 | 5 |
| pdk2b | 0.58595 | 0.17006143 | 0.587 | 0.484 | 1 | 5 |
| uqcrq | 0.5865 | -0.12333543 | 0.73 | 0.774 | 1 | 5 |
| tm2d3 | 0.58658 | 0.118339205 | 0.175 | 0.097 | 1 | 5 |
| tomm20a | 0.58782 | -0.11553555 | 0.222 | 0.29 | 1 | 5 |
| eef1db | 0.58811 | 0.125875238 | 0.683 | 0.645 | 1 | 5 |
| BX324132.2 | 0.58863 | 0.128243427 | 0.159 | 0.097 | 1 | 5 |
| zgc:195175 | 0.58864 | 0.102246875 | 0.111 | 0.065 | 1 | 5 |
| sri | 0.58962 | -0.14731938 | 0.349 | 0.452 | 1 | 5 |
| ddost | 0.59002 | 0.121291115 | 0.397 | 0.29 | 1 | 5 |
| stag2a | 0.59003 | 0.101653887 | 0.079 | 0.032 | 1 | 5 |
| grk3 | 0.5906 | 0.104789106 | 0.175 | 0.097 | 1 | 5 |
| mmp14b | 0.59073 | 0.128375695 | 0.175 | 0.097 | 1 | 5 |

| | | | | | | |
|---|---|---|---|---|---|---|
| jun | 0.59073 | -0.16393546 | 0.349 | 0.452 | 1 | 5 |
| sowahd | 0.59084 | 0.133801429 | 0.206 | 0.129 | 1 | 5 |
| kctd12.2 | 0.59142 | 0.113803653 | 0.413 | 0.387 | 1 | 5 |
| tln1 | 0.59169 | -0.14946752 | 0.524 | 0.613 | 1 | 5 |
| pphln1 | 0.59426 | -0.1148751 | 0.079 | 0.129 | 1 | 5 |
| shoc2 | 0.59442 | -0.1037028 | 0.143 | 0.226 | 1 | 5 |
| prkd2 | 0.59471 | 0.102448146 | 0.063 | 0.032 | 1 | 5 |
| aldoaa | 0.59499 | -0.11935915 | 0.603 | 0.71 | 1 | 5 |
| fzd2 | 0.59664 | 0.104328923 | 0.127 | 0.065 | 1 | 5 |
| eif3jb | 0.59709 | -0.14836261 | 0.317 | 0.419 | 1 | 5 |
| wbp1lb | 0.59727 | 0.120625328 | 0.127 | 0.065 | 1 | 5 |
| dkc1 | 0.59885 | -0.14366263 | 0.302 | 0.387 | 1 | 5 |
| nop2 | 0.59932 | -0.13697705 | 0.175 | 0.258 | 1 | 5 |
| hspa9 | 0.60025 | -0.11857555 | 0.143 | 0.194 | 1 | 5 |
| cdc37 | 0.60131 | 0.107473716 | 0.079 | 0.032 | 1 | 5 |
| rasgrp4 | 0.60152 | -0.10314262 | 0.143 | 0.226 | 1 | 5 |
| arcn1b | 0.60178 | -0.10433351 | 0.079 | 0.129 | 1 | 5 |
| asf1bb | 0.60192 | -0.10137421 | 0.143 | 0.226 | 1 | 5 |
| si:dkey-10h3.2 | 0.60272 | 0.106463609 | 0.127 | 0.065 | 1 | 5 |
| zgc:161969 | 0.60291 | 0.108350246 | 0.127 | 0.065 | 1 | 5 |
| pho | 0.6033 | 0.13449588 | 0.222 | 0.161 | 1 | 5 |
| dnase2 | 0.60442 | 0.131735455 | 0.429 | 0.323 | 1 | 5 |
| prpf4ba | 0.60472 | 0.104340503 | 0.079 | 0.032 | 1 | 5 |
| sept6 | 0.60721 | -0.10631949 | 0.349 | 0.452 | 1 | 5 |
| zfand6 | 0.60797 | -0.11520565 | 0.143 | 0.194 | 1 | 5 |
| igf2bp2b | 0.60864 | -0.10413677 | 0.095 | 0.161 | 1 | 5 |
| MCMDC2 | 0.60886 | 0.10974116 | 0.159 | 0.097 | 1 | 5 |
| psmd7 | 0.60932 | 0.121076307 | 0.429 | 0.323 | 1 | 5 |
| clip1a | 0.60967 | -0.11095509 | 0.095 | 0.161 | 1 | 5 |
| akr7a3 | 0.61021 | 0.101106372 | 0.111 | 0.065 | 1 | 5 |
| zmp:0000001082 | 0.61227 | -0.1264473 | 0.286 | 0.387 | 1 | 5 |
| ddx23 | 0.61334 | -0.11659357 | 0.127 | 0.194 | 1 | 5 |
| lyrm4 | 0.61436 | 0.116169104 | 0.27 | 0.226 | 1 | 5 |
| selenof | 0.61457 | 0.124502595 | 0.698 | 0.645 | 1 | 5 |
| adam10a | 0.61649 | -0.13109169 | 0.286 | 0.387 | 1 | 5 |
| si:ch211-217k17.10 | 0.617 | -0.18558845 | 0.032 | 0.065 | 1 | 5 |
| ube2d2 | 0.617 | 0.134796252 | 0.508 | 0.419 | 1 | 5 |
| srsf2a | 0.61897 | -0.12295476 | 0.476 | 0.581 | 1 | 5 |
| smarca5 | 0.61952 | 0.10193017 | 0.095 | 0.065 | 1 | 5 |
| commd7 | 0.62084 | -0.13205956 | 0.397 | 0.484 | 1 | 5 |
| ewsr1b | 0.62162 | -0.13256305 | 0.238 | 0.323 | 1 | 5 |
| psmc1a | 0.6217 | -0.130738 | 0.127 | 0.194 | 1 | 5 |
| ube2l3b | 0.62178 | -0.12217473 | 0.397 | 0.452 | 1 | 5 |
| FP236812.2 | 0.62461 | 0.110295163 | 0.111 | 0.065 | 1 | 5 |
| derl2 | 0.62523 | 0.134870491 | 0.524 | 0.452 | 1 | 5 |
| uap1l1 | 0.62542 | 0.105728815 | 0.127 | 0.065 | 1 | 5 |
| adipor2 | 0.62604 | -0.1357351 | 0.317 | 0.419 | 1 | 5 |
| srsf5a | 0.62645 | -0.10452011 | 0.413 | 0.516 | 1 | 5 |

| | | | | | | |
|---|---|---|---|---|---|---|
| g3bp1 | 0.62712 | -0.14936612 | 0.302 | 0.387 | 1 | 5 |
| morc2 | 0.62738 | 0.112236719 | 0.127 | 0.065 | 1 | 5 |
| loxl2b | 0.62803 | 0.118995434 | 0.127 | 0.065 | 1 | 5 |
| csde1 | 0.62891 | -0.13199824 | 0.46 | 0.548 | 1 | 5 |
| vmp1 | 0.63067 | -0.16133444 | 0.317 | 0.387 | 1 | 5 |
| sdr16c5b | 0.63071 | 0.147795838 | 0.381 | 0.29 | 1 | 5 |
| eif3ea | 0.6319 | 0.107242646 | 0.651 | 0.548 | 1 | 5 |
| brms1la | 0.63205 | 0.112988358 | 0.159 | 0.097 | 1 | 5 |
| tnfb | 0.6322 | -0.15024606 | 0.175 | 0.258 | 1 | 5 |
| ciarta | 0.63272 | 0.109462297 | 0.159 | 0.097 | 1 | 5 |
| FP085398.1 | 0.63285 | -0.11838066 | 0.095 | 0.161 | 1 | 5 |
| eif3m | 0.63355 | 0.109327408 | 0.794 | 0.742 | 1 | 5 |
| gnptab | 0.63423 | -0.10473986 | 0.048 | 0.097 | 1 | 5 |
| grapa | 0.63429 | 0.141799572 | 0.317 | 0.226 | 1 | 5 |
| uck2a | 0.63668 | -0.12603615 | 0.159 | 0.226 | 1 | 5 |
| med6 | 0.63726 | 0.112963368 | 0.206 | 0.129 | 1 | 5 |
| morf4l1 | 0.63764 | 0.131538853 | 0.317 | 0.226 | 1 | 5 |
| LOC103911430 | 0.638 | 0.107985796 | 0.206 | 0.129 | 1 | 5 |
| CABZ01041494.1 | 0.6392 | -0.10229599 | 0.079 | 0.129 | 1 | 5 |
| mrc1a | 0.63978 | -0.13630904 | 0.175 | 0.258 | 1 | 5 |
| atp2b2 | 0.64022 | -0.13639116 | 0.492 | 0.581 | 1 | 5 |
| got2b | 0.64179 | 0.120433647 | 0.317 | 0.226 | 1 | 5 |
| metap2b | 0.64249 | 0.141415651 | 0.54 | 0.452 | 1 | 5 |
| ssuh2rs1 | 0.64332 | 0.108759756 | 0.206 | 0.129 | 1 | 5 |
| oxsr1a | 0.64403 | -0.10840217 | 0.127 | 0.194 | 1 | 5 |
| rap2c | 0.64409 | 0.115804605 | 0.206 | 0.129 | 1 | 5 |
| abca1b | 0.64436 | 0.123598266 | 0.365 | 0.323 | 1 | 5 |
| sypl2b | 0.64449 | -0.11233133 | 0.175 | 0.258 | 1 | 5 |
| clptm1l | 0.64469 | 0.121226313 | 0.254 | 0.194 | 1 | 5 |
| scarb2a | 0.64492 | -0.12114183 | 0.175 | 0.258 | 1 | 5 |
| tcea3 | 0.64492 | 0.107494398 | 0.206 | 0.129 | 1 | 5 |
| LOC101884451 | 0.6454 | -0.13743483 | 0.19 | 0.258 | 1 | 5 |
| zgc:66427 | 0.64603 | -0.11619708 | 0.175 | 0.258 | 1 | 5 |
| LOC100332329 | 0.64642 | -0.10528662 | 0.095 | 0.161 | 1 | 5 |
| hnrnpa0b | 0.64648 | -0.12081833 | 0.619 | 0.677 | 1 | 5 |
| elf2b | 0.64799 | -0.11116144 | 0.095 | 0.161 | 1 | 5 |
| UTP14C | 0.65005 | 0.102760069 | 0.159 | 0.097 | 1 | 5 |
| sde2 | 0.65162 | 0.102053041 | 0.063 | 0.032 | 1 | 5 |
| crfb4 | 0.65198 | 0.125719938 | 0.19 | 0.129 | 1 | 5 |
| copg2 | 0.65211 | -0.10366697 | 0.095 | 0.161 | 1 | 5 |
| si:dkey-67c22.2 | 0.65279 | 0.112449341 | 0.159 | 0.097 | 1 | 5 |
| idh3a | 0.65834 | -0.10300011 | 0.127 | 0.194 | 1 | 5 |
| XLOC-011085 | 0.65869 | 0.114752497 | 0.143 | 0.097 | 1 | 5 |
| zgc:63587 | 0.65985 | -0.11333157 | 0.048 | 0.097 | 1 | 5 |
| cd83 | 0.66266 | 0.144876975 | 0.349 | 0.258 | 1 | 5 |
| jdp2b | 0.66311 | 0.173024992 | 0.365 | 0.29 | 1 | 5 |
| smc1al | 0.66335 | -0.12731572 | 0.206 | 0.29 | 1 | 5 |
| eif4g2b | 0.66388 | 0.113053214 | 0.206 | 0.161 | 1 | 5 |

| | | | | | | |
|---|---|---|---|---|---|---|
| selenos | 0.66556 | -0.1192174 | 0.111 | 0.161 | 1 | 5 |
| skp1 | 0.66732 | 0.120841875 | 0.603 | 0.516 | 1 | 5 |
| wu:fb55g09 | 0.66931 | 0.139646752 | 0.413 | 0.323 | 1 | 5 |
| grn1 | 0.66999 | -0.11717421 | 0.841 | 0.871 | 1 | 5 |
| atp6ap1b | 0.67008 | -0.11861121 | 0.206 | 0.29 | 1 | 5 |
| chaf1a | 0.6705 | -0.10063611 | 0.048 | 0.097 | 1 | 5 |
| bcam | 0.67065 | 0.12274177 | 0.238 | 0.161 | 1 | 5 |
| pef1 | 0.67113 | -0.10604593 | 0.206 | 0.29 | 1 | 5 |
| baz2a | 0.67121 | -0.10947549 | 0.127 | 0.194 | 1 | 5 |
| sf3b5 | 0.67128 | -0.10840093 | 0.206 | 0.29 | 1 | 5 |
| abcc5 | 0.67262 | -0.13504574 | 0.27 | 0.355 | 1 | 5 |
| mxi1 | 0.67361 | -0.10089204 | 0.079 | 0.129 | 1 | 5 |
| clic2 | 0.67411 | 0.117583492 | 0.698 | 0.645 | 1 | 5 |
| si:ch211-9d9.8 | 0.68015 | 0.112234891 | 0.238 | 0.161 | 1 | 5 |
| trappc5 | 0.68036 | 0.102175165 | 0.222 | 0.161 | 1 | 5 |
| eif4ebp1 | 0.68036 | 0.126123016 | 0.683 | 0.613 | 1 | 5 |
| stk10 | 0.68111 | 0.115989522 | 0.175 | 0.129 | 1 | 5 |
| mrpl15 | 0.68209 | -0.10115988 | 0.127 | 0.194 | 1 | 5 |
| ehd4 | 0.68391 | -0.12482546 | 0.492 | 0.581 | 1 | 5 |
| SFMBT1 | 0.68451 | 0.109682589 | 0.143 | 0.097 | 1 | 5 |
| arhgef2 | 0.68577 | -0.10976805 | 0.079 | 0.129 | 1 | 5 |
| si:dkey-187e18.1 | 0.68825 | -0.10817148 | 0.175 | 0.226 | 1 | 5 |
| taf15 | 0.68849 | -0.11541794 | 0.238 | 0.323 | 1 | 5 |
| XLOC-010899 | 0.68913 | 0.148778491 | 0.476 | 0.387 | 1 | 5 |
| nfe2l2a | 0.68982 | -0.12824782 | 0.302 | 0.387 | 1 | 5 |
| LO018493.1 | 0.69164 | 0.102247708 | 0.587 | 0.548 | 1 | 5 |
| acp1 | 0.69308 | 0.105961465 | 0.222 | 0.161 | 1 | 5 |
| eif2s1b | 0.69309 | -0.11159761 | 0.54 | 0.613 | 1 | 5 |
| vat1 | 0.6959 | -0.14429403 | 0.444 | 0.516 | 1 | 5 |
| tomm6 | 0.69932 | -0.11681384 | 0.365 | 0.419 | 1 | 5 |
| h3f3c | 0.69965 | 0.101879119 | 0.444 | 0.355 | 1 | 5 |
| arfip1 | 0.70085 | 0.103233312 | 0.159 | 0.097 | 1 | 5 |
| cdkn1bb | 0.70165 | 0.124564007 | 0.317 | 0.258 | 1 | 5 |
| lpar5a | 0.70178 | 0.127147246 | 0.444 | 0.355 | 1 | 5 |
| BX004816.2 | 0.70332 | -0.11093322 | 0.079 | 0.129 | 1 | 5 |
| zgc:100846 | 0.70333 | -0.10036955 | 0.127 | 0.194 | 1 | 5 |
| mrpl20 | 0.7042 | 0.103340154 | 0.27 | 0.194 | 1 | 5 |
| antxr2a | 0.70701 | -0.12864099 | 0.587 | 0.645 | 1 | 5 |
| ptges3b | 0.70844 | -0.11474247 | 0.397 | 0.452 | 1 | 5 |
| pak2a | 0.70955 | -0.1017545 | 0.302 | 0.387 | 1 | 5 |
| ndufs7 | 0.71064 | 0.103713291 | 0.333 | 0.29 | 1 | 5 |
| LOC103909972 | 0.71636 | 0.107142275 | 0.286 | 0.226 | 1 | 5 |
| twf1a | 0.71683 | 0.120299817 | 0.46 | 0.387 | 1 | 5 |
| ddit3 | 0.71826 | 0.107663579 | 0.54 | 0.452 | 1 | 5 |
| pgam1a | 0.71908 | -0.10138878 | 0.397 | 0.484 | 1 | 5 |
| si:dkey-27i16.2 | 0.72083 | -0.13864245 | 0.397 | 0.484 | 1 | 5 |
| psmc2 | 0.72084 | 0.108351494 | 0.302 | 0.226 | 1 | 5 |
| si:zfos-741a10.3 | 0.72151 | -0.1093979 | 0.365 | 0.452 | 1 | 5 |

| | | | | | | |
|---|---|---|---|---|---|---|
| havcr2 | 0.72352 | -0.11033708 | 0.429 | 0.516 | 1 | 5 |
| slc8a4b | 0.72406 | -0.10315251 | 0.079 | 0.129 | 1 | 5 |
| jac2 | 0.72895 | -0.1395832 | 0.222 | 0.29 | 1 | 5 |
| ifi45 | 0.73 | -0.10629444 | 0.079 | 0.129 | 1 | 5 |
| gnai2a | 0.73026 | -0.10326267 | 0.492 | 0.548 | 1 | 5 |
| c1galt1c1 | 0.73112 | 0.116046709 | 0.19 | 0.129 | 1 | 5 |
| ppargc1b | 0.73214 | 0.108617275 | 0.222 | 0.161 | 1 | 5 |
| gpd1c | 0.73317 | -0.10967345 | 0.143 | 0.194 | 1 | 5 |
| adam8a | 0.73318 | -0.11271396 | 0.159 | 0.226 | 1 | 5 |
| si:ch211-152p11.4 | 0.73536 | 0.10459417 | 0.175 | 0.129 | 1 | 5 |
| nck1a | 0.73713 | 0.108572929 | 0.254 | 0.194 | 1 | 5 |
| serbp1b | 0.74002 | -0.10732925 | 0.19 | 0.258 | 1 | 5 |
| zc3h12b | 0.74026 | 0.109382544 | 0.19 | 0.129 | 1 | 5 |
| CR318588.1 | 0.74483 | 0.178122411 | 0.397 | 0.355 | 1 | 5 |
| ndufa7 | 0.74613 | -0.10101942 | 0.349 | 0.387 | 1 | 5 |
| mboat2a | 0.74823 | 0.101647009 | 0.222 | 0.161 | 1 | 5 |
| laptm4a | 0.74837 | 0.109413567 | 0.333 | 0.258 | 1 | 5 |
| uqcrh | 0.75599 | 0.10495463 | 0.651 | 0.581 | 1 | 5 |
| cnn2 | 0.75649 | -0.10565292 | 0.46 | 0.516 | 1 | 5 |
| cyfip1 | 0.75933 | -0.11267797 | 0.381 | 0.452 | 1 | 5 |
| ncor2 | 0.76198 | -0.10377032 | 0.27 | 0.323 | 1 | 5 |
| cmpk | 0.7675 | 0.100454379 | 0.365 | 0.29 | 1 | 5 |
| mrps18c | 0.76777 | -0.10363966 | 0.222 | 0.29 | 1 | 5 |
| CABZ01001434.1 | 0.76963 | -0.14486269 | 0.317 | 0.387 | 1 | 5 |
| rnf213a | 0.77009 | 0.119196351 | 0.397 | 0.323 | 1 | 5 |
| arpc5a | 0.77376 | 0.102801064 | 0.429 | 0.355 | 1 | 5 |
| FO907110.1 | 0.78554 | 0.111075102 | 0.302 | 0.258 | 1 | 5 |
| BX322618.1 | 0.81216 | -0.11028972 | 0.19 | 0.226 | 1 | 5 |
| si:ch211-283g2.3 | 0.82147 | -0.11672224 | 0.397 | 0.452 | 1 | 5 |
| ccl35.1 | 0.89402 | -0.10592964 | 0.365 | 0.387 | 1 | 5 |
| fabp11a | 4.814980 | -1.47864614 | 0.039 | 0.588 | 0.0097248 | 6 |
| tfa | 3.950768 | -1.19245722 | 0.013 | 0.471 | 0.0797937 | 6 |
| BX901920.1 | 0.00012 | -1.02027583 | 0.117 | 0.471 | 1 | 6 |
| foxp4 | 0.00014 | 0.983265615 | 0.597 | 0.118 | 1 | 6 |
| CR855311.1 | 0.00017 | -0.78046331 | 0.013 | 0.353 | 1 | 6 |
| vasna | 0.00021 | 0.787521445 | 0.39 | 0 | 1 | 6 |
| zgc:153704 | 0.00021 | 1.153309857 | 0.39 | 0 | 1 | 6 |
| rbp7b | 0.00027 | -0.47460091 | 0 | 0.235 | 1 | 6 |
| ms4a17a.8 | 0.00027 | -0.51037699 | 0 | 0.235 | 1 | 6 |
| nol8 | 0.00027 | -0.44267381 | 0 | 0.235 | 1 | 6 |
| mt-cyb | 0.0003 | -0.75474745 | 0.312 | 0.824 | 1 | 6 |
| ptena | 0.00053 | 0.600204887 | 0.351 | 0 | 1 | 6 |
| tob1a | 0.00072 | 0.626535491 | 0.338 | 0 | 1 | 6 |
| col12a1b | 0.00097 | 0.80310152 | 0.325 | 0 | 1 | 6 |
| fosab | 0.00101 | 0.944941967 | 0.688 | 0.294 | 1 | 6 |
| si:ch211-198c19.3 | 0.00105 | -0.89430772 | 0.156 | 0.529 | 1 | 6 |
| gpx1a | 0.00115 | -0.30340411 | 0.078 | 0.412 | 1 | 6 |
| XLOC-025423 | 0.00115 | -0.96158226 | 0.312 | 0.647 | 1 | 6 |

| | | | | | | |
|---|---|---|---|---|---|---|
| ms4a17a.14 | 0.00122 | -0.63846716 | 0.013 | 0.294 | 1 | 6 |
| nanos1 | 0.00131 | 0.646210567 | 0.312 | 0 | 1 | 6 |
| net1 | 0.00131 | 0.591409034 | 0.312 | 0 | 1 | 6 |
| ckbb | 0.00136 | -0.73240032 | 0.013 | 0.294 | 1 | 6 |
| XLOC-041870 | 0.00152 | -0.9757317 | 0.078 | 0.412 | 1 | 6 |
| nup98 | 0.00203 | -0.3384532 | 0 | 0.176 | 1 | 6 |
| CR936442.1 | 0.00203 | -0.20317049 | 0 | 0.176 | 1 | 6 |
| rap2ab | 0.00203 | -0.28386918 | 0 | 0.176 | 1 | 6 |
| tsg101b | 0.00203 | -0.28810706 | 0 | 0.176 | 1 | 6 |
| ccdc171 | 0.00203 | -0.3701637 | 0 | 0.176 | 1 | 6 |
| il16 | 0.00203 | -0.2001622 | 0 | 0.176 | 1 | 6 |
| klhl12 | 0.00203 | -0.34523749 | 0 | 0.176 | 1 | 6 |
| nek9 | 0.00203 | -0.35385815 | 0 | 0.176 | 1 | 6 |
| ugt1a7 | 0.00203 | -0.33028425 | 0 | 0.176 | 1 | 6 |
| ppp1caa | 0.00233 | 0.524174717 | 0.286 | 0 | 1 | 6 |
| pon3.2 | 0.00246 | -0.63544837 | 0.078 | 0.412 | 1 | 6 |
| dbi | 0.0025 | -0.67249561 | 0.455 | 0.647 | 1 | 6 |
| sf3b3 | 0.00257 | -0.58147012 | 0.039 | 0.353 | 1 | 6 |
| csnk2a1 | 0.00261 | -0.16103302 | 0.312 | 0.235 | 1 | 6 |
| lgals9l3 | 0.00278 | -0.67599328 | 0.182 | 0.471 | 1 | 6 |
| mxra8b | 0.0031 | 0.542686614 | 0.273 | 0 | 1 | 6 |
| rspo1 | 0.0031 | 0.711642392 | 0.273 | 0 | 1 | 6 |
| prdx3 | 0.00318 | -0.65803418 | 0.143 | 0.471 | 1 | 6 |
| krt18b | 0.00408 | -0.68284641 | 0.351 | 0.765 | 1 | 6 |
| ssu72 | 0.0041 | 0.468145774 | 0.26 | 0 | 1 | 6 |
| psmb3 | 0.0041 | 0.462156491 | 0.26 | 0 | 1 | 6 |
| rgs5a | 0.00441 | -0.40328203 | 0.39 | 0.765 | 1 | 6 |
| cxcl18b | 0.00451 | -0.48439025 | 0.195 | 0.294 | 1 | 6 |
| gnai3 | 0.00461 | -0.58175736 | 0.026 | 0.294 | 1 | 6 |
| jun | 0.005 | 0.675570194 | 0.792 | 0.471 | 1 | 6 |
| adob | 0.00532 | -0.55913306 | 0.013 | 0.235 | 1 | 6 |
| mt-co2 | 0.00536 | -0.70115693 | 0.403 | 0.824 | 1 | 6 |
| si:ch73-376l24.2 | 0.00541 | 0.522932368 | 0.247 | 0 | 1 | 6 |
| BX322530.1 | 0.00541 | 0.501170617 | 0.247 | 0 | 1 | 6 |
| rbfox1 | 0.00541 | 0.504390315 | 0.247 | 0 | 1 | 6 |
| BX322618.1 | 0.00548 | -0.86625084 | 0.195 | 0.588 | 1 | 6 |
| sfrp1a | 0.00576 | 0.822251978 | 0.416 | 0.059 | 1 | 6 |
| ube2v2 | 0.0059 | 0.651185454 | 0.481 | 0.176 | 1 | 6 |
| wbp2nl | 0.00591 | -0.52782204 | 0.052 | 0.353 | 1 | 6 |
| atp5if1a | 0.00656 | -0.26283636 | 0.442 | 0.412 | 1 | 6 |
| znf330 | 0.00671 | -0.60092326 | 0.078 | 0.353 | 1 | 6 |
| TMEM208 | 0.00678 | -0.42085587 | 0.013 | 0.235 | 1 | 6 |
| khdrbs1a | 0.00679 | 0.513526135 | 0.805 | 0.412 | 1 | 6 |
| naa20 | 0.00711 | 0.383359126 | 0.234 | 0 | 1 | 6 |
| plpp1a | 0.00711 | 0.479419224 | 0.234 | 0 | 1 | 6 |
| lyrm4 | 0.00725 | -0.37254354 | 0.104 | 0.176 | 1 | 6 |
| si:ch211-222l21.1 | 0.00772 | 0.463351144 | 0.714 | 0.588 | 1 | 6 |
| mt-atp6 | 0.0079 | -0.58730919 | 0.312 | 0.706 | 1 | 6 |

| | | | | | | |
|---|---|---|---|---|---|---|
| brd3b | 0.00933 | 0.429955068 | 0.221 | 0 | 1 | 6 |
| kdelr2a | 0.00933 | 0.450533261 | 0.221 | 0 | 1 | 6 |
| rai14 | 0.00933 | 0.399632373 | 0.221 | 0 | 1 | 6 |
| si:dkey-51e6.1 | 0.00933 | 0.422046088 | 0.221 | 0 | 1 | 6 |
| CR383676.1 | 0.00937 | 0.503143096 | 0.909 | 0.588 | 1 | 6 |
| ucp2 | 0.00992 | 0.6033193 | 0.403 | 0.059 | 1 | 6 |
| XLOC-005449 | 0.01011 | -0.74737856 | 0.13 | 0.412 | 1 | 6 |
| brd7 | 0.01012 | -0.36330238 | 0.195 | 0.294 | 1 | 6 |
| chchd3a | 0.01029 | -0.11238912 | 0.156 | 0.118 | 1 | 6 |
| naa50 | 0.01034 | -0.54867656 | 0.052 | 0.294 | 1 | 6 |
| acvrl1 | 0.01055 | -0.53046204 | 0.052 | 0.235 | 1 | 6 |
| rpl22l1 | 0.01096 | 0.381899956 | 0.818 | 0.471 | 1 | 6 |
| ier2b | 0.01182 | 0.65384026 | 0.416 | 0.118 | 1 | 6 |
| atp5mc3a | 0.01189 | -0.5217099 | 0.377 | 0.588 | 1 | 6 |
| irf2a | 0.0122 | 0.373751945 | 0.208 | 0 | 1 | 6 |
| stk35 | 0.0122 | 0.405919595 | 0.208 | 0 | 1 | 6 |
| ahr2 | 0.0122 | 0.404464651 | 0.208 | 0 | 1 | 6 |
| ifi45 | 0.01317 | -0.73287761 | 0.039 | 0.235 | 1 | 6 |
| ppdpfb | 0.01366 | 0.535133915 | 0.792 | 0.529 | 1 | 6 |
| ncl | 0.01379 | -0.40697164 | 0.571 | 0.706 | 1 | 6 |
| krt15 | 0.014 | 0.825096758 | 0.429 | 0.118 | 1 | 6 |
| rps18 | 0.01409 | 0.111771688 | 0.961 | 1 | 1 | 6 |
| XLOC-023492 | 0.01448 | -0.69722215 | 0.117 | 0.412 | 1 | 6 |
| crip1 | 0.01456 | -0.67135281 | 0.338 | 0.647 | 1 | 6 |
| CU914776.1 | 0.0147 | -0.29387088 | 0 | 0.118 | 1 | 6 |
| LOC101886002 | 0.0147 | -0.20681565 | 0 | 0.118 | 1 | 6 |
| TMEM19 | 0.0147 | -0.22016324 | 0 | 0.118 | 1 | 6 |
| XLOC-032112 | 0.0147 | -0.2052186 | 0 | 0.118 | 1 | 6 |
| ano10b | 0.0147 | -0.20371122 | 0 | 0.118 | 1 | 6 |
| fam160a2 | 0.0147 | -0.19347976 | 0 | 0.118 | 1 | 6 |
| gab1 | 0.0147 | -0.20496817 | 0 | 0.118 | 1 | 6 |
| isg15 | 0.0147 | -0.25119206 | 0 | 0.118 | 1 | 6 |
| porcnl | 0.0147 | -0.25505807 | 0 | 0.118 | 1 | 6 |
| prkacbb | 0.0147 | -0.18648858 | 0 | 0.118 | 1 | 6 |
| rnf139 | 0.0147 | -0.29613883 | 0 | 0.118 | 1 | 6 |
| si:ch211-212k18.7 | 0.0147 | -0.30938305 | 0 | 0.118 | 1 | 6 |
| si:ch211-255f4.9 | 0.0147 | -0.20790838 | 0 | 0.118 | 1 | 6 |
| si:ch73-352p4.5 | 0.0147 | -0.20242848 | 0 | 0.118 | 1 | 6 |
| si:dkey-188i13.9 | 0.0147 | -0.3331832 | 0 | 0.118 | 1 | 6 |
| vcpkmt | 0.0147 | -0.15717343 | 0 | 0.118 | 1 | 6 |
| zgc:162472 | 0.0147 | -0.1849281 | 0 | 0.118 | 1 | 6 |
| CABZ01084131.1 | 0.0147 | -0.2188969 | 0 | 0.118 | 1 | 6 |
| arl4cb | 0.0147 | -0.24335626 | 0 | 0.118 | 1 | 6 |
| atp1b4 | 0.0147 | -0.28124632 | 0 | 0.118 | 1 | 6 |
| b9d1 | 0.0147 | -0.22625421 | 0 | 0.118 | 1 | 6 |
| cdkn1ba | 0.0147 | -0.22235984 | 0 | 0.118 | 1 | 6 |
| cetn4 | 0.0147 | -0.2837129 | 0 | 0.118 | 1 | 6 |
| ercc6 | 0.0147 | -0.21225021 | 0 | 0.118 | 1 | 6 |

| | | | | | | |
|---|---|---|---|---|---|---|
| mak | 0.0147 | -0.25687205 | 0 | 0.118 | 1 | 6 |
| masp1 | 0.0147 | -0.25655578 | 0 | 0.118 | 1 | 6 |
| ppm1ba | 0.0147 | -0.21740495 | 0 | 0.118 | 1 | 6 |
| psmd2 | 0.0147 | -0.27099184 | 0 | 0.118 | 1 | 6 |
| ptpreb | 0.0147 | -0.26395183 | 0 | 0.118 | 1 | 6 |
| rc3h1a | 0.0147 | -0.21756462 | 0 | 0.118 | 1 | 6 |
| rragca | 0.0147 | -0.3024234 | 0 | 0.118 | 1 | 6 |
| si:ch211-245h14.1 | 0.0147 | -0.2322879 | 0 | 0.118 | 1 | 6 |
| zgc:195175 | 0.0147 | -0.17026872 | 0 | 0.118 | 1 | 6 |
| znf280d | 0.0147 | -0.21881652 | 0 | 0.118 | 1 | 6 |
| CABZ01088428.1 | 0.0147 | -0.1875344 | 0 | 0.118 | 1 | 6 |
| LOC110439449 | 0.0147 | -0.20933156 | 0 | 0.118 | 1 | 6 |
| arhgap29a | 0.0147 | -0.27070306 | 0 | 0.118 | 1 | 6 |
| calcrla | 0.0147 | -0.24211715 | 0 | 0.118 | 1 | 6 |
| coq4 | 0.0147 | -0.1448944 | 0 | 0.118 | 1 | 6 |
| gtf2h2 | 0.0147 | -0.20392205 | 0 | 0.118 | 1 | 6 |
| hdac12 | 0.0147 | -0.26995948 | 0 | 0.118 | 1 | 6 |
| laynb | 0.0147 | -0.18416196 | 0 | 0.118 | 1 | 6 |
| map7d3 | 0.0147 | -0.17062928 | 0 | 0.118 | 1 | 6 |
| med21 | 0.0147 | -0.26636871 | 0 | 0.118 | 1 | 6 |
| mrm3b | 0.0147 | -0.28222757 | 0 | 0.118 | 1 | 6 |
| pak1ip1 | 0.0147 | -0.30298873 | 0 | 0.118 | 1 | 6 |
| pcdh2ab7 | 0.0147 | -0.17023362 | 0 | 0.118 | 1 | 6 |
| rhoj | 0.0147 | -0.17368847 | 0 | 0.118 | 1 | 6 |
| selenou1b | 0.0147 | -0.25808052 | 0 | 0.118 | 1 | 6 |
| si:ch73-86n18.1 | 0.0147 | -0.16342744 | 0 | 0.118 | 1 | 6 |
| slc25a48 | 0.0147 | -0.28775154 | 0 | 0.118 | 1 | 6 |
| tfip11 | 0.0147 | -0.29246447 | 0 | 0.118 | 1 | 6 |
| tm4sf18 | 0.0147 | -0.17947595 | 0 | 0.118 | 1 | 6 |
| zgc:114104 | 0.0147 | -0.22783527 | 0 | 0.118 | 1 | 6 |
| angptl2b | 0.0147 | -0.26509898 | 0 | 0.118 | 1 | 6 |
| si:ch211-243j20.2 | 0.0147 | -0.23362029 | 0 | 0.118 | 1 | 6 |
| arsa | 0.0147 | -0.22836659 | 0 | 0.118 | 1 | 6 |
| gldc | 0.0147 | -0.26773405 | 0 | 0.118 | 1 | 6 |
| ivns1abpa | 0.0147 | -0.33060542 | 0 | 0.118 | 1 | 6 |
| kank4 | 0.0147 | -0.28844816 | 0 | 0.118 | 1 | 6 |
| pkmyt1 | 0.0147 | -0.21365185 | 0 | 0.118 | 1 | 6 |
| si:dkey-71l4.4 | 0.0147 | -0.30927832 | 0 | 0.118 | 1 | 6 |
| uhmk1 | 0.0147 | -0.33035091 | 0 | 0.118 | 1 | 6 |
| CABZ01078733.1 | 0.0147 | -0.2140546 | 0 | 0.118 | 1 | 6 |
| CT030188.1 | 0.0147 | -0.20039239 | 0 | 0.118 | 1 | 6 |
| XLOC-016167 | 0.0147 | -0.23922467 | 0 | 0.118 | 1 | 6 |
| amn1 | 0.0147 | -0.18011672 | 0 | 0.118 | 1 | 6 |
| caska | 0.0147 | -0.18928849 | 0 | 0.118 | 1 | 6 |
| cwf19l1 | 0.0147 | -0.21234816 | 0 | 0.118 | 1 | 6 |
| gbp1 | 0.0147 | -0.22842699 | 0 | 0.118 | 1 | 6 |
| oaz2a | 0.0147 | -0.39102796 | 0 | 0.118 | 1 | 6 |
| scp2a | 0.0147 | -0.30910655 | 0 | 0.118 | 1 | 6 |

| | | | | | | |
|---|---|---|---|---|---|---|
| sh3glb2a | 0.0147 | -0.2995819 | 0 | 0.118 | 1 | 6 |
| stx6 | 0.0147 | -0.21675583 | 0 | 0.118 | 1 | 6 |
| tmem86a | 0.0147 | -0.21592384 | 0 | 0.118 | 1 | 6 |
| zgc:92481 | 0.0147 | -0.23232453 | 0 | 0.118 | 1 | 6 |
| znf395a | 0.0147 | -0.27989218 | 0 | 0.118 | 1 | 6 |
| anxa5b | 0.01476 | -0.54423438 | 0.26 | 0.471 | 1 | 6 |
| mycb | 0.01505 | 0.621598559 | 0.364 | 0.059 | 1 | 6 |
| hmgb2a | 0.01562 | -0.52481185 | 0.481 | 0.647 | 1 | 6 |
| si:ch211-1a19.3 | 0.01585 | -0.42219439 | 0.026 | 0.235 | 1 | 6 |
| nfixb | 0.01591 | 0.668970948 | 0.571 | 0.235 | 1 | 6 |
| UBL4A | 0.01592 | 0.307672191 | 0.195 | 0 | 1 | 6 |
| cadm2a | 0.01592 | 0.486221074 | 0.195 | 0 | 1 | 6 |
| hoxc3a | 0.01592 | 0.410501831 | 0.195 | 0 | 1 | 6 |
| jmjd1cb | 0.01592 | 0.435153332 | 0.195 | 0 | 1 | 6 |
| podxl | 0.01592 | 0.542037532 | 0.195 | 0 | 1 | 6 |
| znf395b | 0.01592 | 0.39619422 | 0.195 | 0 | 1 | 6 |
| filip1l | 0.01592 | 0.359448563 | 0.195 | 0 | 1 | 6 |
| tp53 | 0.01592 | 0.343547198 | 0.195 | 0 | 1 | 6 |
| sept2 | 0.01671 | -0.32970216 | 0.364 | 0.412 | 1 | 6 |
| id1 | 0.01694 | 0.824216818 | 0.506 | 0.176 | 1 | 6 |
| atp5mf | 0.01739 | -0.49179719 | 0.13 | 0.412 | 1 | 6 |
| igf2b | 0.0177 | -0.61031178 | 0.234 | 0.588 | 1 | 6 |
| slc38a5b | 0.01796 | -0.30274989 | 0.377 | 0.412 | 1 | 6 |
| hnrnpr | 0.01801 | 0.195054041 | 0.221 | 0.059 | 1 | 6 |
| postnb | 0.019 | 0.684228764 | 0.701 | 0.353 | 1 | 6 |
| si:dkey-7j14.6 | 0.01911 | -0.11019208 | 0.558 | 0.471 | 1 | 6 |
| appa | 0.01984 | 0.579591325 | 0.312 | 0.059 | 1 | 6 |
| lxn | 0.01997 | -0.45154331 | 0.338 | 0.471 | 1 | 6 |
| cfd | 0.02005 | -0.60059471 | 0.442 | 0.588 | 1 | 6 |
| ablim1b | 0.02073 | 0.395414838 | 0.182 | 0 | 1 | 6 |
| dhrs3b | 0.02073 | 0.445145704 | 0.182 | 0 | 1 | 6 |
| CABZ01058650.1 | 0.02073 | 0.326899769 | 0.182 | 0 | 1 | 6 |
| inip | 0.02073 | 0.325248209 | 0.182 | 0 | 1 | 6 |
| nell2a | 0.02073 | 0.490458548 | 0.182 | 0 | 1 | 6 |
| rap1gap | 0.02073 | 0.427143291 | 0.182 | 0 | 1 | 6 |
| si:ch211-260e23.9 | 0.02073 | 0.290593401 | 0.182 | 0 | 1 | 6 |
| ubl3a | 0.02073 | 0.33735506 | 0.182 | 0 | 1 | 6 |
| CABZ01061591.1 | 0.0208 | -0.41490966 | 0.026 | 0.235 | 1 | 6 |
| pet100 | 0.02109 | -0.35065162 | 0.052 | 0.294 | 1 | 6 |
| uqcrq | 0.02165 | -0.26233432 | 0.299 | 0.294 | 1 | 6 |
| anp32b | 0.02193 | 0.207269894 | 0.416 | 0.176 | 1 | 6 |
| mt-nd1 | 0.0222 | -0.43608676 | 0.351 | 0.706 | 1 | 6 |
| pbdc1 | 0.02255 | 0.169940039 | 0.312 | 0.118 | 1 | 6 |
| cfap45 | 0.02287 | -0.4709661 | 0.026 | 0.235 | 1 | 6 |
| cycsb | 0.02308 | -0.37946711 | 0.182 | 0.294 | 1 | 6 |
| tspan9a | 0.02353 | -0.26298961 | 0.039 | 0.235 | 1 | 6 |
| BX908750.1 | 0.02367 | -0.58547519 | 0.052 | 0.294 | 1 | 6 |
| rpa2 | 0.02395 | -0.414413 | 0.026 | 0.235 | 1 | 6 |

| | | | | | | |
|---|---|---|---|---|---|---|
| TCIM (1 of many) | 0.02435 | -0.46787567 | 0.026 | 0.235 | 1 | 6 |
| fam133b | 0.02441 | 0.597571706 | 0.403 | 0.118 | 1 | 6 |
| klf6a | 0.02459 | 0.578023832 | 0.377 | 0.118 | 1 | 6 |
| cyp1c2 | 0.02516 | -0.59181075 | 0.195 | 0.529 | 1 | 6 |
| c1qtnf6a | 0.02531 | -0.53619876 | 0.104 | 0.294 | 1 | 6 |
| nop10 | 0.02569 | -0.43362428 | 0.299 | 0.471 | 1 | 6 |
| pcna | 0.02617 | -0.6874217 | 0.104 | 0.353 | 1 | 6 |
| gstt1b | 0.02637 | -0.36414439 | 0.299 | 0.412 | 1 | 6 |
| c1qtnf5 | 0.02637 | 0.194568396 | 0.364 | 0.176 | 1 | 6 |
| ccdc124 | 0.02693 | 0.346879312 | 0.169 | 0 | 1 | 6 |
| dag1 | 0.02693 | 0.342558083 | 0.169 | 0 | 1 | 6 |
| fam162a | 0.02693 | 0.295192675 | 0.169 | 0 | 1 | 6 |
| hoxd9a | 0.02693 | 0.335780347 | 0.169 | 0 | 1 | 6 |
| jund | 0.02693 | 0.348068091 | 0.169 | 0 | 1 | 6 |
| letm1 | 0.02693 | 0.285192355 | 0.169 | 0 | 1 | 6 |
| mef2aa | 0.02693 | 0.336624373 | 0.169 | 0 | 1 | 6 |
| mfap1 | 0.02693 | 0.313084328 | 0.169 | 0 | 1 | 6 |
| palld | 0.02693 | 0.354080326 | 0.169 | 0 | 1 | 6 |
| angpt2b | 0.02693 | 0.401626732 | 0.169 | 0 | 1 | 6 |
| atp6v1e1a | 0.02693 | 0.29219817 | 0.169 | 0 | 1 | 6 |
| prkar2aa | 0.02693 | 0.330721589 | 0.169 | 0 | 1 | 6 |
| srsf10b | 0.02693 | 0.31264527 | 0.169 | 0 | 1 | 6 |
| id2b | 0.02693 | 0.388167344 | 0.169 | 0 | 1 | 6 |
| zmynd11 | 0.02693 | 0.347014299 | 0.169 | 0 | 1 | 6 |
| hmgb3a | 0.02704 | 0.575462121 | 0.584 | 0.235 | 1 | 6 |
| btg2 | 0.02785 | 0.587965684 | 0.403 | 0.118 | 1 | 6 |
| taf15 | 0.02826 | -0.49255289 | 0.208 | 0.471 | 1 | 6 |
| ssr1 | 0.02838 | -0.44924265 | 0.39 | 0.529 | 1 | 6 |
| bcas2 | 0.02864 | -0.25006694 | 0.26 | 0.294 | 1 | 6 |
| col4a1 | 0.0301 | -0.60784107 | 0.234 | 0.471 | 1 | 6 |
| zgc:172106 | 0.03019 | -0.50146103 | 0.026 | 0.176 | 1 | 6 |
| nfyc | 0.03069 | -0.4728073 | 0.065 | 0.294 | 1 | 6 |
| fkbp9 | 0.03129 | -0.46411029 | 0.208 | 0.529 | 1 | 6 |
| cdkn1a | 0.03159 | -0.22909881 | 0.221 | 0.235 | 1 | 6 |
| picalma | 0.03165 | -0.27982604 | 0.013 | 0.176 | 1 | 6 |
| mvp | 0.0317 | -0.3231326 | 0.039 | 0.235 | 1 | 6 |
| gatd1 | 0.03178 | -0.35269459 | 0.013 | 0.176 | 1 | 6 |
| prdx4 | 0.03196 | -0.21497603 | 0.325 | 0.294 | 1 | 6 |
| bub3 | 0.03236 | -0.25218376 | 0.013 | 0.176 | 1 | 6 |
| synm | 0.03445 | -0.40727582 | 0.013 | 0.176 | 1 | 6 |
| cyb5r1 | 0.03492 | 0.306540346 | 0.156 | 0 | 1 | 6 |
| efna2b | 0.03492 | 0.310431977 | 0.156 | 0 | 1 | 6 |
| emid1 | 0.03492 | 0.418173526 | 0.156 | 0 | 1 | 6 |
| eya1 | 0.03492 | 0.320054918 | 0.156 | 0 | 1 | 6 |
| f3b | 0.03492 | 0.328023143 | 0.156 | 0 | 1 | 6 |
| frzb | 0.03492 | 0.389917257 | 0.156 | 0 | 1 | 6 |
| hif1al | 0.03492 | 0.311090298 | 0.156 | 0 | 1 | 6 |
| hsdl2 | 0.03492 | 0.284140201 | 0.156 | 0 | 1 | 6 |

| | | | | | | |
|---|---|---|---|---|---|---|
| ifrd1 | 0.03492 | 0.225160697 | 0.156 | 0 | 1 | 6 |
| itga9 | 0.03492 | 0.309209795 | 0.156 | 0 | 1 | 6 |
| ndrg2 | 0.03492 | 0.316179326 | 0.156 | 0 | 1 | 6 |
| pgam5 | 0.03492 | 0.259883924 | 0.156 | 0 | 1 | 6 |
| plod3 | 0.03492 | 0.293538712 | 0.156 | 0 | 1 | 6 |
| proza | 0.03492 | 0.333401713 | 0.156 | 0 | 1 | 6 |
| pudp | 0.03492 | 0.305049098 | 0.156 | 0 | 1 | 6 |
| si:dkey-19a16.7 | 0.03492 | 0.322188322 | 0.156 | 0 | 1 | 6 |
| svep1 | 0.03492 | 0.422583515 | 0.156 | 0 | 1 | 6 |
| txndc12 | 0.03492 | 0.278449514 | 0.156 | 0 | 1 | 6 |
| arf1 | 0.03492 | 0.311287631 | 0.156 | 0 | 1 | 6 |
| creb3l2 | 0.03492 | 0.308060563 | 0.156 | 0 | 1 | 6 |
| fank1 | 0.03498 | -0.5010934 | 0.065 | 0.294 | 1 | 6 |
| fth1a | 0.03518 | -0.51681837 | 0.364 | 0.529 | 1 | 6 |
| zgc:162509 | 0.03564 | -0.19026843 | 0.052 | 0.235 | 1 | 6 |
| strn4 | 0.03586 | -0.26939757 | 0.039 | 0.235 | 1 | 6 |
| lsm12b | 0.03591 | 0.237198107 | 0.325 | 0.118 | 1 | 6 |
| myh9b | 0.03704 | -0.33691854 | 0.013 | 0.176 | 1 | 6 |
| orc1 | 0.03728 | -0.3143507 | 0.013 | 0.176 | 1 | 6 |
| cirbpb | 0.03739 | 0.201066186 | 0.935 | 0.941 | 1 | 6 |
| ndufa9a | 0.03754 | -0.38306919 | 0.117 | 0.235 | 1 | 6 |
| slc29a1a | 0.03774 | -0.30710588 | 0.013 | 0.176 | 1 | 6 |
| sh3gl1b | 0.03793 | -0.42513894 | 0.091 | 0.353 | 1 | 6 |
| ctnnb2 | 0.0389 | 0.165655669 | 0.195 | 0.059 | 1 | 6 |
| rnf113a | 0.03933 | -0.29763482 | 0.013 | 0.176 | 1 | 6 |
| col4a2 | 0.03946 | -0.46174352 | 0.104 | 0.235 | 1 | 6 |
| LOC100537342 | 0.03953 | -0.23446845 | 0.013 | 0.176 | 1 | 6 |
| extl2 | 0.0397 | -0.25822913 | 0.013 | 0.176 | 1 | 6 |
| nucks1b | 0.03983 | 0.454052249 | 0.338 | 0.118 | 1 | 6 |
| casp8 | 0.04041 | -0.2550012 | 0.013 | 0.176 | 1 | 6 |
| acsl3b | 0.04069 | -0.28744824 | 0.013 | 0.176 | 1 | 6 |
| tomm20a | 0.04074 | -0.25765781 | 0.195 | 0.235 | 1 | 6 |
| gins2 | 0.04098 | -0.26392085 | 0.013 | 0.176 | 1 | 6 |
| egln1b | 0.04106 | -0.23612296 | 0.013 | 0.176 | 1 | 6 |
| rab1aa | 0.04113 | -0.22206069 | 0.013 | 0.176 | 1 | 6 |
| btg1 | 0.04193 | 0.248357531 | 0.468 | 0.235 | 1 | 6 |
| rpl27a | 0.04231 | -0.17354461 | 0.805 | 0.765 | 1 | 6 |
| egr1 | 0.04271 | 0.307573882 | 0.26 | 0.059 | 1 | 6 |
| ccdc126 | 0.04346 | -0.43167527 | 0.052 | 0.176 | 1 | 6 |
| cbx3a | 0.0439 | -0.34882662 | 0.312 | 0.412 | 1 | 6 |
| mrpl33 | 0.04409 | -0.24746002 | 0.156 | 0.176 | 1 | 6 |
| fn1b | 0.04411 | -0.44596942 | 0.455 | 0.588 | 1 | 6 |
| apbb2b | 0.04523 | 0.279136331 | 0.143 | 0 | 1 | 6 |
| cdca4 | 0.04523 | 0.287273545 | 0.143 | 0 | 1 | 6 |
| esr2b | 0.04523 | 0.325732964 | 0.143 | 0 | 1 | 6 |
| fgfr2 | 0.04523 | 0.335969881 | 0.143 | 0 | 1 | 6 |
| lpp | 0.04523 | 0.29036043 | 0.143 | 0 | 1 | 6 |
| pim3 | 0.04523 | 0.276894556 | 0.143 | 0 | 1 | 6 |

| | | | | | | |
|---|---|---|---|---|---|---|
| rab40b | 0.04523 | 0.28842734 | 0.143 | 0 | 1 | 6 |
| six2a | 0.04523 | 0.304370513 | 0.143 | 0 | 1 | 6 |
| ctdspla | 0.04523 | 0.293528268 | 0.143 | 0 | 1 | 6 |
| cyp2p6 | 0.04523 | 0.276892144 | 0.143 | 0 | 1 | 6 |
| ncam1a | 0.04523 | 0.312682606 | 0.143 | 0 | 1 | 6 |
| phf20b | 0.04523 | 0.267347403 | 0.143 | 0 | 1 | 6 |
| pik3r3b | 0.04523 | 0.330900375 | 0.143 | 0 | 1 | 6 |
| taf3 | 0.04523 | 0.320884533 | 0.143 | 0 | 1 | 6 |
| ugt1b5 | 0.04523 | 0.279053181 | 0.143 | 0 | 1 | 6 |
| waca | 0.04523 | 0.285842956 | 0.143 | 0 | 1 | 6 |
| aktip | 0.04523 | 0.262548632 | 0.143 | 0 | 1 | 6 |
| lmo4b | 0.04523 | 0.243861525 | 0.143 | 0 | 1 | 6 |
| mrpl4 | 0.04523 | 0.290246324 | 0.143 | 0 | 1 | 6 |
| usf1l | 0.04523 | 0.269046636 | 0.143 | 0 | 1 | 6 |
| wt1a | 0.04523 | 0.314154982 | 0.143 | 0 | 1 | 6 |
| rsu1 | 0.04523 | 0.301729986 | 0.143 | 0 | 1 | 6 |
| ubxn1 | 0.04523 | 0.242623607 | 0.143 | 0 | 1 | 6 |
| fbxl14a | 0.04523 | 0.288239563 | 0.143 | 0 | 1 | 6 |
| mbd3b | 0.04523 | 0.253173007 | 0.143 | 0 | 1 | 6 |
| oaz2b | 0.04523 | 0.310604541 | 0.143 | 0 | 1 | 6 |
| six1b | 0.04523 | 0.349031207 | 0.143 | 0 | 1 | 6 |
| cyc1 | 0.04652 | -0.40549194 | 0.273 | 0.412 | 1 | 6 |
| gadd45ab | 0.04795 | -0.41959169 | 0.026 | 0.176 | 1 | 6 |
| nudt17 | 0.04854 | -0.27108363 | 0.039 | 0.235 | 1 | 6 |
| ndufa1 | 0.04927 | 0.165557253 | 0.416 | 0.176 | 1 | 6 |
| fam129bb | 0.05044 | -0.31850098 | 0.039 | 0.235 | 1 | 6 |
| uchl3 | 0.05073 | -0.1672631 | 0.065 | 0.059 | 1 | 6 |
| slc25a3b | 0.0509 | -0.23557737 | 0.714 | 0.706 | 1 | 6 |
| tmem238a | 0.05117 | -0.1202284 | 0.078 | 0.059 | 1 | 6 |
| si:ch211-251b21.1 | 0.05334 | -0.1392596 | 0.156 | 0.118 | 1 | 6 |
| capza1b | 0.05417 | 0.483027525 | 0.416 | 0.176 | 1 | 6 |
| atp5f1d | 0.05459 | -0.39997448 | 0.429 | 0.529 | 1 | 6 |
| taf12 | 0.0554 | 0.439529211 | 0.273 | 0.059 | 1 | 6 |
| kctd5a | 0.05631 | 0.450745106 | 0.299 | 0.059 | 1 | 6 |
| myo1b | 0.05694 | -0.41661852 | 0.026 | 0.176 | 1 | 6 |
| eif3g | 0.05722 | 0.508105592 | 0.519 | 0.235 | 1 | 6 |
| id2a | 0.0573 | 0.535194618 | 0.312 | 0.059 | 1 | 6 |
| ddx51 | 0.05746 | -0.35995816 | 0.104 | 0.353 | 1 | 6 |
| psmd8 | 0.05763 | 0.484560652 | 0.273 | 0.059 | 1 | 6 |
| col6a2 | 0.05833 | 0.49760065 | 0.416 | 0.235 | 1 | 6 |
| si:dkey-222h21.2 | 0.05842 | -0.42284871 | 0.052 | 0.235 | 1 | 6 |
| dvl2 | 0.05851 | 0.216751 | 0.13 | 0 | 1 | 6 |
| lum | 0.05851 | 0.346368303 | 0.13 | 0 | 1 | 6 |
| ppfibp1a | 0.05851 | 0.310479578 | 0.13 | 0 | 1 | 6 |
| sh3bp5la | 0.05851 | 0.249676748 | 0.13 | 0 | 1 | 6 |
| ssbp4 | 0.05851 | 0.232659308 | 0.13 | 0 | 1 | 6 |
| taf7 | 0.05851 | 0.222399234 | 0.13 | 0 | 1 | 6 |
| wbp1 | 0.05851 | 0.257681444 | 0.13 | 0 | 1 | 6 |

| | | | | | | |
|---|---|---|---|---|---|---|
| camk2g1 | 0.05851 | 0.292083749 | 0.13 | 0 | 1 | 6 |
| daam1b | 0.05851 | 0.253537128 | 0.13 | 0 | 1 | 6 |
| eed | 0.05851 | 0.23937548 | 0.13 | 0 | 1 | 6 |
| gtf2a1 | 0.05851 | 0.234320055 | 0.13 | 0 | 1 | 6 |
| helz | 0.05851 | 0.251452656 | 0.13 | 0 | 1 | 6 |
| hoxa3a | 0.05851 | 0.302608257 | 0.13 | 0 | 1 | 6 |
| kri1 | 0.05851 | 0.212157188 | 0.13 | 0 | 1 | 6 |
| lmnb2 | 0.05851 | 0.246955358 | 0.13 | 0 | 1 | 6 |
| msi2b | 0.05851 | 0.248638696 | 0.13 | 0 | 1 | 6 |
| rnf220a | 0.05851 | 0.260659699 | 0.13 | 0 | 1 | 6 |
| sav1 | 0.05851 | 0.263128082 | 0.13 | 0 | 1 | 6 |
| ss18l2 | 0.05851 | 0.232647427 | 0.13 | 0 | 1 | 6 |
| stx2b | 0.05851 | 0.27326553 | 0.13 | 0 | 1 | 6 |
| mrpl11 | 0.05851 | 0.251007242 | 0.13 | 0 | 1 | 6 |
| polr2i | 0.05851 | 0.263162988 | 0.13 | 0 | 1 | 6 |
| KIF2A | 0.05851 | 0.273788332 | 0.13 | 0 | 1 | 6 |
| LEPROTL1 | 0.05851 | 0.238548468 | 0.13 | 0 | 1 | 6 |
| bida | 0.05851 | 0.217836026 | 0.13 | 0 | 1 | 6 |
| elovl1b | 0.05851 | 0.263327622 | 0.13 | 0 | 1 | 6 |
| fhl1b | 0.05851 | 0.303959387 | 0.13 | 0 | 1 | 6 |
| foxf1 | 0.05851 | 0.297846165 | 0.13 | 0 | 1 | 6 |
| gcga | 0.05851 | 0.258019258 | 0.13 | 0 | 1 | 6 |
| snx5 | 0.05851 | 0.252278922 | 0.13 | 0 | 1 | 6 |
| zranb2 | 0.05851 | 0.233436513 | 0.13 | 0 | 1 | 6 |
| CABZ01112656.1 | 0.05851 | 0.209851619 | 0.13 | 0 | 1 | 6 |
| wnt11r | 0.05851 | 0.306278132 | 0.13 | 0 | 1 | 6 |
| decr1 | 0.05851 | 0.233456487 | 0.13 | 0 | 1 | 6 |
| grb2b | 0.05851 | 0.252112332 | 0.13 | 0 | 1 | 6 |
| gtf2f1 | 0.05851 | 0.24532565 | 0.13 | 0 | 1 | 6 |
| hnrnpl | 0.05851 | 0.215258979 | 0.13 | 0 | 1 | 6 |
| lims1 | 0.05851 | 0.254478308 | 0.13 | 0 | 1 | 6 |
| mbnl1 | 0.05851 | 0.232609543 | 0.13 | 0 | 1 | 6 |
| prkacaa | 0.05851 | 0.282120057 | 0.13 | 0 | 1 | 6 |
| si:ch211-160j14.3 | 0.05851 | 0.281023555 | 0.13 | 0 | 1 | 6 |
| si:ch211-213a13.1 | 0.05851 | 0.282239767 | 0.13 | 0 | 1 | 6 |
| slc25a43 | 0.05851 | 0.252742279 | 0.13 | 0 | 1 | 6 |
| smoc1 | 0.05851 | 0.329815183 | 0.13 | 0 | 1 | 6 |
| sox6 | 0.05851 | 0.302753611 | 0.13 | 0 | 1 | 6 |
| zc4h2 | 0.05851 | 0.235714868 | 0.13 | 0 | 1 | 6 |
| zgc:153372 | 0.05851 | 0.266063671 | 0.13 | 0 | 1 | 6 |
| cdr2l | 0.05851 | 0.255186993 | 0.13 | 0 | 1 | 6 |
| frem2b | 0.05851 | 0.255563192 | 0.13 | 0 | 1 | 6 |
| plxna1a | 0.05851 | 0.266159621 | 0.13 | 0 | 1 | 6 |
| si:dkeyp-44a8.2 | 0.05851 | 0.245094614 | 0.13 | 0 | 1 | 6 |
| ube2al | 0.05851 | 0.208686914 | 0.13 | 0 | 1 | 6 |
| birc6-as2 | 0.05851 | 0.3265089 | 0.13 | 0 | 1 | 6 |
| usp10 | 0.05851 | 0.246557588 | 0.13 | 0 | 1 | 6 |
| dedd1 | 0.05858 | 0.545235715 | 0.442 | 0.176 | 1 | 6 |

| | | | | | | |
|---|---|---|---|---|---|---|
| zgc:101858 | 0.05901 | -0.39934567 | 0.052 | 0.235 | 1 | 6 |
| STMP1 | 0.05934 | -0.40804977 | 0.208 | 0.353 | 1 | 6 |
| ctdsp2 | 0.0613 | 0.346616928 | 0.273 | 0.059 | 1 | 6 |
| angptl1a | 0.06197 | 0.463534687 | 0.455 | 0.176 | 1 | 6 |
| XLOC-026836 | 0.06281 | -0.37175589 | 0.026 | 0.118 | 1 | 6 |
| pgk1 | 0.06322 | -0.27965057 | 0.078 | 0.118 | 1 | 6 |
| emc8 | 0.06325 | -0.38110879 | 0.052 | 0.235 | 1 | 6 |
| lgals3b | 0.06349 | 0.271954857 | 0.338 | 0.118 | 1 | 6 |
| CU695232.1 | 0.06362 | -0.20015359 | 0.013 | 0.118 | 1 | 6 |
| XLOC-043918 | 0.06362 | -0.17354887 | 0.013 | 0.118 | 1 | 6 |
| adcy6a | 0.06362 | -0.28838815 | 0.013 | 0.118 | 1 | 6 |
| amdhd2 | 0.06362 | -0.12338458 | 0.013 | 0.118 | 1 | 6 |
| baiap2b | 0.06362 | -0.13963394 | 0.013 | 0.118 | 1 | 6 |
| ep300a | 0.06362 | -0.30657657 | 0.013 | 0.118 | 1 | 6 |
| gpn3 | 0.06362 | -0.11828412 | 0.013 | 0.118 | 1 | 6 |
| kin | 0.06362 | -0.19072207 | 0.013 | 0.118 | 1 | 6 |
| rab6ba | 0.06362 | -0.25852949 | 0.013 | 0.118 | 1 | 6 |
| rab8a | 0.06362 | -0.12445426 | 0.013 | 0.118 | 1 | 6 |
| shroom4 | 0.06362 | -0.18753233 | 0.013 | 0.118 | 1 | 6 |
| znfx1 | 0.06362 | -0.22928386 | 0.013 | 0.118 | 1 | 6 |
| pnp5a | 0.06362 | -0.16393384 | 0.013 | 0.118 | 1 | 6 |
| rspry1 | 0.06362 | -0.25449642 | 0.013 | 0.118 | 1 | 6 |
| CABZ01117348.1 | 0.06362 | -0.13912826 | 0.013 | 0.118 | 1 | 6 |
| CU856344.1 | 0.06362 | -0.11699936 | 0.013 | 0.118 | 1 | 6 |
| acad8 | 0.06362 | -0.16205389 | 0.013 | 0.118 | 1 | 6 |
| enpp2 | 0.06362 | -0.13448367 | 0.013 | 0.118 | 1 | 6 |
| got1 | 0.06362 | -0.33848168 | 0.013 | 0.118 | 1 | 6 |
| gsdf | 0.06362 | -0.3257293 | 0.013 | 0.118 | 1 | 6 |
| mpv17l2 | 0.06362 | -0.20992446 | 0.013 | 0.118 | 1 | 6 |
| ndufaf4 | 0.06362 | -0.16469712 | 0.013 | 0.118 | 1 | 6 |
| neu3.5 | 0.06362 | -0.29971478 | 0.013 | 0.118 | 1 | 6 |
| patz1 | 0.06362 | -0.28077517 | 0.013 | 0.118 | 1 | 6 |
| pcnt | 0.06362 | -0.23393234 | 0.013 | 0.118 | 1 | 6 |
| ppp1r14aa | 0.06362 | -0.31383144 | 0.013 | 0.118 | 1 | 6 |
| si:dkey-58f10.11 | 0.06362 | -0.18873971 | 0.013 | 0.118 | 1 | 6 |
| smarcd2 | 0.06362 | -0.14393947 | 0.013 | 0.118 | 1 | 6 |
| sp100.1 | 0.06362 | -0.26110314 | 0.013 | 0.118 | 1 | 6 |
| taf2 | 0.06362 | -0.22182222 | 0.013 | 0.118 | 1 | 6 |
| znf292b | 0.06362 | -0.17351485 | 0.013 | 0.118 | 1 | 6 |
| ngfb | 0.06362 | -0.22682337 | 0.013 | 0.118 | 1 | 6 |
| pfkpa | 0.06362 | -0.13907348 | 0.013 | 0.118 | 1 | 6 |
| naa15b | 0.06362 | -0.17472848 | 0.013 | 0.118 | 1 | 6 |
| si:ch211-199m9.3 | 0.06362 | -0.29128363 | 0.013 | 0.118 | 1 | 6 |
| aldh3a2b | 0.06362 | -0.2827565 | 0.013 | 0.118 | 1 | 6 |
| cers2a | 0.06362 | -0.17010352 | 0.013 | 0.118 | 1 | 6 |
| cers2b | 0.06362 | -0.15232968 | 0.013 | 0.118 | 1 | 6 |
| chchd10 | 0.06362 | -0.22573985 | 0.013 | 0.118 | 1 | 6 |
| exd2 | 0.06362 | -0.18686498 | 0.013 | 0.118 | 1 | 6 |

| | | | | | | |
|---|---|---|---|---|---|---|
| galnt12 | 0.06362 | -0.16141195 | 0.013 | 0.118 | 1 | 6 |
| gart | 0.06362 | -0.1554344 | 0.013 | 0.118 | 1 | 6 |
| gm2a | 0.06362 | -0.12514226 | 0.013 | 0.118 | 1 | 6 |
| grm4 | 0.06362 | -0.16755504 | 0.013 | 0.118 | 1 | 6 |
| nipbla | 0.06362 | -0.15960328 | 0.013 | 0.118 | 1 | 6 |
| nup133 | 0.06362 | -0.25417171 | 0.013 | 0.118 | 1 | 6 |
| pde3a | 0.06362 | -0.34201004 | 0.013 | 0.118 | 1 | 6 |
| pla2g6 | 0.06362 | -0.2263667 | 0.013 | 0.118 | 1 | 6 |
| ptgdsb.1 | 0.06362 | -0.11469391 | 0.013 | 0.118 | 1 | 6 |
| scml2 | 0.06362 | -0.25414009 | 0.013 | 0.118 | 1 | 6 |
| si:ch211-87m7.2 | 0.06362 | -0.17393629 | 0.013 | 0.118 | 1 | 6 |
| ube2z | 0.06362 | -0.3051398 | 0.013 | 0.118 | 1 | 6 |
| usp6nl | 0.06362 | -0.21854355 | 0.013 | 0.118 | 1 | 6 |
| nudt3b | 0.06362 | -0.30645304 | 0.013 | 0.118 | 1 | 6 |
| rnf121 | 0.06362 | -0.16455114 | 0.013 | 0.118 | 1 | 6 |
| umps | 0.06362 | -0.17735494 | 0.013 | 0.118 | 1 | 6 |
| CABZ01066926.1 | 0.06362 | -0.21217116 | 0.013 | 0.118 | 1 | 6 |
| dhfr | 0.06362 | -0.34163913 | 0.013 | 0.118 | 1 | 6 |
| galnt11 | 0.06362 | -0.28535885 | 0.013 | 0.118 | 1 | 6 |
| mogs | 0.06362 | -0.17569695 | 0.013 | 0.118 | 1 | 6 |
| opa1 | 0.06362 | -0.21441314 | 0.013 | 0.118 | 1 | 6 |
| si:ch211-13c6.2 | 0.06362 | -0.2431793 | 0.013 | 0.118 | 1 | 6 |
| si:ch73-299h12.6 | 0.06362 | -0.17537189 | 0.013 | 0.118 | 1 | 6 |
| wdr83 | 0.06362 | -0.17003397 | 0.013 | 0.118 | 1 | 6 |
| zgc:110540 | 0.06362 | -0.16041937 | 0.013 | 0.118 | 1 | 6 |
| draxin | 0.06362 | -0.14772387 | 0.013 | 0.118 | 1 | 6 |
| kcnip1b | 0.06362 | -0.15095523 | 0.013 | 0.118 | 1 | 6 |
| mars | 0.06362 | -0.24550177 | 0.013 | 0.118 | 1 | 6 |
| pank4 | 0.06362 | -0.15381207 | 0.013 | 0.118 | 1 | 6 |
| pcdh10a | 0.06362 | -0.24878304 | 0.013 | 0.118 | 1 | 6 |
| pdhx | 0.06362 | -0.16762486 | 0.013 | 0.118 | 1 | 6 |
| srpx | 0.06362 | -0.15956821 | 0.013 | 0.118 | 1 | 6 |
| vps39 | 0.06362 | -0.25888752 | 0.013 | 0.118 | 1 | 6 |
| si:rp71-36a1.2 | 0.06362 | -0.29260093 | 0.013 | 0.118 | 1 | 6 |
| apeh | 0.06362 | -0.12779359 | 0.013 | 0.118 | 1 | 6 |
| c19h1orf109 | 0.06362 | -0.1714015 | 0.013 | 0.118 | 1 | 6 |
| col5a3a | 0.06362 | -0.16611982 | 0.013 | 0.118 | 1 | 6 |
| cuedc1b | 0.06362 | -0.2059319 | 0.013 | 0.118 | 1 | 6 |
| glb1l | 0.06362 | -0.21762363 | 0.013 | 0.118 | 1 | 6 |
| gpbar1 | 0.06362 | -0.23519748 | 0.013 | 0.118 | 1 | 6 |
| krt4 | 0.06362 | -0.24844682 | 0.013 | 0.118 | 1 | 6 |
| mapk9 | 0.06362 | -0.15475988 | 0.013 | 0.118 | 1 | 6 |
| rnf150b | 0.06362 | -0.30700449 | 0.013 | 0.118 | 1 | 6 |
| FBXO48 | 0.06392 | -0.20055317 | 0.052 | 0.059 | 1 | 6 |
| epha4b | 0.06454 | 0.519793539 | 0.26 | 0.059 | 1 | 6 |
| tuba8l4 | 0.06498 | -0.45470389 | 0.39 | 0.588 | 1 | 6 |
| tbp | 0.06507 | -0.39897228 | 0.026 | 0.176 | 1 | 6 |
| ccl25b | 0.06521 | -0.42584547 | 0.662 | 0.765 | 1 | 6 |

| | | | | | | |
|---|---|---|---|---|---|---|
| ms4a17a.11 | 0.06557 | 0.424100461 | 0.299 | 0.059 | 1 | 6 |
| si:ch211-132p1.3 | 0.06565 | -0.48238668 | 0.078 | 0.294 | 1 | 6 |
| hnrnpa1a | 0.06567 | -0.30210333 | 0.221 | 0.294 | 1 | 6 |
| gadd45ba | 0.06582 | 0.511281208 | 0.351 | 0.118 | 1 | 6 |
| fabp3 | 0.06635 | -0.24267367 | 0.234 | 0.471 | 1 | 6 |
| col6a1 | 0.06673 | 0.516949914 | 0.597 | 0.294 | 1 | 6 |
| csk | 0.0669 | 0.318277758 | 0.156 | 0.059 | 1 | 6 |
| zgc:158343 | 0.0674 | 0.436861019 | 0.299 | 0.059 | 1 | 6 |
| BX005183.3 | 0.06766 | -0.45096593 | 0.065 | 0.176 | 1 | 6 |
| ip6k2a | 0.06776 | -0.27113601 | 0.247 | 0.294 | 1 | 6 |
| zgc:162944 | 0.06814 | -0.47738882 | 0.312 | 0.588 | 1 | 6 |
| brdt | 0.06815 | -0.21683876 | 0.104 | 0.294 | 1 | 6 |
| sik2b | 0.0682 | -0.4204945 | 0.104 | 0.235 | 1 | 6 |
| nfyba | 0.07092 | 0.478001449 | 0.364 | 0.118 | 1 | 6 |
| rhag | 0.07142 | -0.40667241 | 0.065 | 0.176 | 1 | 6 |
| dbnlb | 0.07184 | -0.26816231 | 0.026 | 0.176 | 1 | 6 |
| snx3 | 0.0724 | -0.4640398 | 0.195 | 0.412 | 1 | 6 |
| mdkb | 0.07294 | -0.59904139 | 0.169 | 0.412 | 1 | 6 |
| crispld2 | 0.073 | -0.30253207 | 0.052 | 0.235 | 1 | 6 |
| tmod2 | 0.07325 | -0.27142431 | 0.052 | 0.235 | 1 | 6 |
| pfkfb4b | 0.07333 | 0.427517606 | 0.299 | 0.059 | 1 | 6 |
| ndufs2 | 0.0735 | -0.44812709 | 0.13 | 0.353 | 1 | 6 |
| dhrs3a | 0.07367 | -0.30415367 | 0.091 | 0.294 | 1 | 6 |
| chaf1a | 0.07426 | -0.40985923 | 0.052 | 0.235 | 1 | 6 |
| tagln2 | 0.07459 | -0.2632838 | 0.416 | 0.471 | 1 | 6 |
| akap8l | 0.07565 | 0.226033924 | 0.117 | 0 | 1 | 6 |
| ap2b1 | 0.07565 | 0.240929821 | 0.117 | 0 | 1 | 6 |
| coq9 | 0.07565 | 0.218950576 | 0.117 | 0 | 1 | 6 |
| fbxo25 | 0.07565 | 0.235461304 | 0.117 | 0 | 1 | 6 |
| fem1c | 0.07565 | 0.179917484 | 0.117 | 0 | 1 | 6 |
| fstb | 0.07565 | 0.284129184 | 0.117 | 0 | 1 | 6 |
| hspa12b | 0.07565 | 0.26708328 | 0.117 | 0 | 1 | 6 |
| ino80e | 0.07565 | 0.231658871 | 0.117 | 0 | 1 | 6 |
| plekha1a | 0.07565 | 0.209092744 | 0.117 | 0 | 1 | 6 |
| sccpdhb | 0.07565 | 0.196351866 | 0.117 | 0 | 1 | 6 |
| sh3pxd2aa | 0.07565 | 0.258337148 | 0.117 | 0 | 1 | 6 |
| si:dkey-69o16.5 | 0.07565 | 0.186731091 | 0.117 | 0 | 1 | 6 |
| smarca2 | 0.07565 | 0.227602696 | 0.117 | 0 | 1 | 6 |
| tcp11l2 | 0.07565 | 0.261931516 | 0.117 | 0 | 1 | 6 |
| tia1 | 0.07565 | 0.251666357 | 0.117 | 0 | 1 | 6 |
| zdhhc20a | 0.07565 | 0.247816391 | 0.117 | 0 | 1 | 6 |
| agmo | 0.07565 | 0.193858057 | 0.117 | 0 | 1 | 6 |
| cyfip1 | 0.07565 | 0.182912122 | 0.117 | 0 | 1 | 6 |
| ergic2 | 0.07565 | 0.212859564 | 0.117 | 0 | 1 | 6 |
| igfbp2b | 0.07565 | 0.240385842 | 0.117 | 0 | 1 | 6 |
| rexo4 | 0.07565 | 0.244430154 | 0.117 | 0 | 1 | 6 |
| sorbs1 | 0.07565 | 0.211457353 | 0.117 | 0 | 1 | 6 |
| zgc:123010 | 0.07565 | 0.252315669 | 0.117 | 0 | 1 | 6 |

| | | | | | | |
|---|---|---|---|---|---|---|
| cav1 | 0.07565 | 0.284781824 | 0.117 | 0 | 1 | 6 |
| gzf1 | 0.07565 | 0.204601499 | 0.117 | 0 | 1 | 6 |
| pigf | 0.07565 | 0.229846969 | 0.117 | 0 | 1 | 6 |
| pop5 | 0.07565 | 0.191423101 | 0.117 | 0 | 1 | 6 |
| si:dkey-17m8.1 | 0.07565 | 0.204653106 | 0.117 | 0 | 1 | 6 |
| srp54 | 0.07565 | 0.249032865 | 0.117 | 0 | 1 | 6 |
| zgc:154058 | 0.07565 | 0.216673459 | 0.117 | 0 | 1 | 6 |
| MZT1 | 0.07565 | 0.210521611 | 0.117 | 0 | 1 | 6 |
| acin1b | 0.07565 | 0.269368113 | 0.117 | 0 | 1 | 6 |
| aebp2 | 0.07565 | 0.210479809 | 0.117 | 0 | 1 | 6 |
| bmp1a | 0.07565 | 0.228970653 | 0.117 | 0 | 1 | 6 |
| cdh13 | 0.07565 | 0.25492199 | 0.117 | 0 | 1 | 6 |
| cib1 | 0.07565 | 0.200323438 | 0.117 | 0 | 1 | 6 |
| coch | 0.07565 | 0.380269398 | 0.117 | 0 | 1 | 6 |
| coro1ca | 0.07565 | 0.212380249 | 0.117 | 0 | 1 | 6 |
| crkl | 0.07565 | 0.23917024 | 0.117 | 0 | 1 | 6 |
| ctdspl2b | 0.07565 | 0.237388962 | 0.117 | 0 | 1 | 6 |
| eif2a | 0.07565 | 0.187009775 | 0.117 | 0 | 1 | 6 |
| hoxa10b | 0.07565 | 0.239582724 | 0.117 | 0 | 1 | 6 |
| il6st | 0.07565 | 0.245159162 | 0.117 | 0 | 1 | 6 |
| pdlim5a | 0.07565 | 0.285207914 | 0.117 | 0 | 1 | 6 |
| phf2 | 0.07565 | 0.244728233 | 0.117 | 0 | 1 | 6 |
| roraa | 0.07565 | 0.230178484 | 0.117 | 0 | 1 | 6 |
| si:dkey-159f12.2 | 0.07565 | 0.226114301 | 0.117 | 0 | 1 | 6 |
| yif1a | 0.07565 | 0.208377962 | 0.117 | 0 | 1 | 6 |
| eya4 | 0.07565 | 0.204953366 | 0.117 | 0 | 1 | 6 |
| fam49a | 0.07565 | 0.197473901 | 0.117 | 0 | 1 | 6 |
| fgfrl1a | 0.07565 | 0.231266088 | 0.117 | 0 | 1 | 6 |
| foxf2a | 0.07565 | 0.2550046 | 0.117 | 0 | 1 | 6 |
| selenon | 0.07565 | 0.24481079 | 0.117 | 0 | 1 | 6 |
| tox2 | 0.07565 | 0.206061168 | 0.117 | 0 | 1 | 6 |
| txn2 | 0.07565 | 0.174382098 | 0.117 | 0 | 1 | 6 |
| cd82a | 0.07588 | 0.286375466 | 0.325 | 0.118 | 1 | 6 |
| psme3 | 0.07609 | -0.36131595 | 0.078 | 0.294 | 1 | 6 |
| hsp90b1 | 0.07644 | -0.42626603 | 0.156 | 0.294 | 1 | 6 |
| osr1 | 0.07651 | 0.46266555 | 0.377 | 0.176 | 1 | 6 |
| si:dkey-78l4.14 | 0.07874 | -0.42038715 | 0.13 | 0.294 | 1 | 6 |
| ywhabl | 0.07898 | -0.34618306 | 0.455 | 0.588 | 1 | 6 |
| slc22a18 | 0.07919 | -0.30025023 | 0.026 | 0.176 | 1 | 6 |
| socs3a | 0.07947 | 0.563872647 | 0.584 | 0.294 | 1 | 6 |
| fblim1 | 0.0808 | 0.199403325 | 0.286 | 0.118 | 1 | 6 |
| serp1 | 0.08207 | -0.21238118 | 0.195 | 0.235 | 1 | 6 |
| agtr2 | 0.08208 | -0.31158333 | 0.13 | 0.176 | 1 | 6 |
| tril | 0.08317 | -0.45517848 | 0.143 | 0.353 | 1 | 6 |
| ifi30 | 0.08377 | 0.35504741 | 0.247 | 0.059 | 1 | 6 |
| erlin1 | 0.0839 | -0.32188273 | 0.039 | 0.118 | 1 | 6 |
| mrpl36 | 0.08415 | -0.42623137 | 0.143 | 0.353 | 1 | 6 |
| ppm1bb | 0.08436 | -0.26466981 | 0.026 | 0.176 | 1 | 6 |

| | | | | | | |
|---|---|---|---|---|---|---|
| tmem176 | 0.08441 | -0.45169149 | 0.312 | 0.529 | 1 | 6 |
| tnrc6c1 | 0.08458 | 0.446982778 | 0.455 | 0.176 | 1 | 6 |
| dut | 0.08533 | -0.43810421 | 0.039 | 0.176 | 1 | 6 |
| zgc:162025 | 0.08658 | -0.27222391 | 0.091 | 0.294 | 1 | 6 |
| pcolcea | 0.08685 | -0.38491215 | 0.584 | 0.765 | 1 | 6 |
| LO016987.2 | 0.08695 | -0.40131604 | 0.065 | 0.235 | 1 | 6 |
| mt-nd2 | 0.08699 | -0.37613214 | 0.416 | 0.706 | 1 | 6 |
| wu:fb55g09 | 0.087 | 0.498233851 | 0.481 | 0.235 | 1 | 6 |
| ewsr1b | 0.08712 | -0.39307729 | 0.104 | 0.294 | 1 | 6 |
| lrch3 | 0.08718 | -0.22006507 | 0.026 | 0.176 | 1 | 6 |
| mphosph8 | 0.08747 | 0.441252435 | 0.403 | 0.176 | 1 | 6 |
| b2ml | 0.08762 | 0.359396349 | 0.636 | 0.353 | 1 | 6 |
| adh5 | 0.08796 | 0.432663697 | 0.649 | 0.412 | 1 | 6 |
| mcl1b | 0.088 | 0.347060413 | 0.571 | 0.294 | 1 | 6 |
| gpatch11 | 0.08824 | -0.24519 | 0.052 | 0.235 | 1 | 6 |
| npm1a | 0.089 | -0.32624234 | 0.286 | 0.353 | 1 | 6 |
| arpc4l | 0.08963 | 0.3865178 | 0.286 | 0.059 | 1 | 6 |
| gramd4b | 0.09038 | -0.24983231 | 0.026 | 0.176 | 1 | 6 |
| wdr1 | 0.09038 | -0.35043558 | 0.117 | 0.235 | 1 | 6 |
| efnb1 | 0.09046 | 0.484180147 | 0.286 | 0.059 | 1 | 6 |
| tmem242 | 0.09094 | -0.20292588 | 0.026 | 0.176 | 1 | 6 |
| junba | 0.09163 | 0.499357023 | 0.623 | 0.353 | 1 | 6 |
| rasa3 | 0.09241 | -0.27506798 | 0.026 | 0.176 | 1 | 6 |
| prss23 | 0.09262 | -0.22996717 | 0.065 | 0.235 | 1 | 6 |
| pleca | 0.09269 | 0.454593668 | 0.247 | 0.059 | 1 | 6 |
| tma16 | 0.09404 | -0.29226126 | 0.052 | 0.235 | 1 | 6 |
| tufm | 0.09457 | -0.1492002 | 0.117 | 0.118 | 1 | 6 |
| flot1b | 0.0946 | -0.19194813 | 0.286 | 0.294 | 1 | 6 |
| rac1b | 0.09486 | -0.32466304 | 0.026 | 0.176 | 1 | 6 |
| arhgef5 | 0.095 | -0.29554851 | 0.052 | 0.118 | 1 | 6 |
| cnn3b | 0.09537 | -0.3595839 | 0.026 | 0.176 | 1 | 6 |
| xirp2a | 0.09551 | -0.34296693 | 0.052 | 0.235 | 1 | 6 |
| CU459094.2 | 0.09577 | -0.25718901 | 0.065 | 0.235 | 1 | 6 |
| tspan3a | 0.09594 | -0.29778563 | 0.052 | 0.235 | 1 | 6 |
| ghrb | 0.09603 | -0.36992766 | 0.052 | 0.176 | 1 | 6 |
| mdh1aa | 0.09624 | -0.21658078 | 0.494 | 0.529 | 1 | 6 |
| dtx4b | 0.09657 | -0.33740806 | 0.026 | 0.176 | 1 | 6 |
| atoh8 | 0.09721 | 0.477227009 | 0.351 | 0.118 | 1 | 6 |
| dhrs4 | 0.09751 | -0.20939277 | 0.026 | 0.176 | 1 | 6 |
| bicc1b | 0.09782 | 0.231588922 | 0.104 | 0 | 1 | 6 |
| CABZ01068251.1 | 0.09782 | 0.197667236 | 0.104 | 0 | 1 | 6 |
| FQ323119.1 | 0.09782 | 0.215274459 | 0.104 | 0 | 1 | 6 |
| aup1 | 0.09782 | 0.169264809 | 0.104 | 0 | 1 | 6 |
| caprin1a | 0.09782 | 0.216355672 | 0.104 | 0 | 1 | 6 |
| creb1a | 0.09782 | 0.192183423 | 0.104 | 0 | 1 | 6 |
| isl2b | 0.09782 | 0.256899542 | 0.104 | 0 | 1 | 6 |
| pgpep1 | 0.09782 | 0.173688605 | 0.104 | 0 | 1 | 6 |
| phb2b | 0.09782 | 0.171412664 | 0.104 | 0 | 1 | 6 |

| | | | | | | |
|---|---|---|---|---|---|---|
| rab18b | 0.09782 | 0.213965492 | 0.104 | 0 | 1 | 6 |
| si:dkey-266m15.6 | 0.09782 | 0.199567498 | 0.104 | 0 | 1 | 6 |
| stom | 0.09782 | 0.190704361 | 0.104 | 0 | 1 | 6 |
| adamts15b | 0.09782 | 0.248817837 | 0.104 | 0 | 1 | 6 |
| arl13b | 0.09782 | 0.199186804 | 0.104 | 0 | 1 | 6 |
| cdc34b | 0.09782 | 0.166969426 | 0.104 | 0 | 1 | 6 |
| cldn12 | 0.09782 | 0.179919314 | 0.104 | 0 | 1 | 6 |
| ctdsp1 | 0.09782 | 0.209345221 | 0.104 | 0 | 1 | 6 |
| eaf1 | 0.09782 | 0.183775412 | 0.104 | 0 | 1 | 6 |
| foxk2 | 0.09782 | 0.212744139 | 0.104 | 0 | 1 | 6 |
| glrx5 | 0.09782 | 0.184245803 | 0.104 | 0 | 1 | 6 |
| golgb1 | 0.09782 | 0.218126342 | 0.104 | 0 | 1 | 6 |
| hikeshi | 0.09782 | 0.159161745 | 0.104 | 0 | 1 | 6 |
| hoxa13b | 0.09782 | 0.270433053 | 0.104 | 0 | 1 | 6 |
| kpna1 | 0.09782 | 0.213627499 | 0.104 | 0 | 1 | 6 |
| lpar2a | 0.09782 | 0.197172509 | 0.104 | 0 | 1 | 6 |
| ltbp1 | 0.09782 | 0.210616328 | 0.104 | 0 | 1 | 6 |
| maff | 0.09782 | 0.215618307 | 0.104 | 0 | 1 | 6 |
| map3k3 | 0.09782 | 0.220923098 | 0.104 | 0 | 1 | 6 |
| mef2d | 0.09782 | 0.188892061 | 0.104 | 0 | 1 | 6 |
| nenf | 0.09782 | 0.202717568 | 0.104 | 0 | 1 | 6 |
| nol10 | 0.09782 | 0.154930772 | 0.104 | 0 | 1 | 6 |
| pdgfaa | 0.09782 | 0.234332902 | 0.104 | 0 | 1 | 6 |
| psmd12 | 0.09782 | 0.208417907 | 0.104 | 0 | 1 | 6 |
| ptx3a | 0.09782 | 0.290297783 | 0.104 | 0 | 1 | 6 |
| pxnb | 0.09782 | 0.206828329 | 0.104 | 0 | 1 | 6 |
| rnf170 | 0.09782 | 0.22205249 | 0.104 | 0 | 1 | 6 |
| rpz2 | 0.09782 | 0.228256531 | 0.104 | 0 | 1 | 6 |
| slc16a1b | 0.09782 | 0.205783282 | 0.104 | 0 | 1 | 6 |
| smn1 | 0.09782 | 0.17897903 | 0.104 | 0 | 1 | 6 |
| tcf7l1a | 0.09782 | 0.209215596 | 0.104 | 0 | 1 | 6 |
| tjp1b | 0.09782 | 0.224791005 | 0.104 | 0 | 1 | 6 |
| tmem134 | 0.09782 | 0.170190426 | 0.104 | 0 | 1 | 6 |
| zgc:163030 | 0.09782 | 0.194353852 | 0.104 | 0 | 1 | 6 |
| znf292a | 0.09782 | 0.17538328 | 0.104 | 0 | 1 | 6 |
| znf576.2 | 0.09782 | 0.200329899 | 0.104 | 0 | 1 | 6 |
| LOC110439840 | 0.09782 | 0.184417316 | 0.104 | 0 | 1 | 6 |
| TPGS1 | 0.09782 | 0.192448163 | 0.104 | 0 | 1 | 6 |
| UTP14C | 0.09782 | 0.167544482 | 0.104 | 0 | 1 | 6 |
| acbd3 | 0.09782 | 0.205089183 | 0.104 | 0 | 1 | 6 |
| arhgef4 | 0.09782 | 0.266279387 | 0.104 | 0 | 1 | 6 |
| bcl7bb | 0.09782 | 0.222585423 | 0.104 | 0 | 1 | 6 |
| cdh11 | 0.09782 | 0.237068648 | 0.104 | 0 | 1 | 6 |
| csnk1g2b | 0.09782 | 0.173396282 | 0.104 | 0 | 1 | 6 |
| cspg5a | 0.09782 | 0.217368709 | 0.104 | 0 | 1 | 6 |
| ctnna1 | 0.09782 | 0.225337633 | 0.104 | 0 | 1 | 6 |
| ech1 | 0.09782 | 0.232320583 | 0.104 | 0 | 1 | 6 |
| fam171a2a | 0.09782 | 0.187229968 | 0.104 | 0 | 1 | 6 |

| | | | | | | |
|---|---|---|---|---|---|---|
| fam204a | 0.09782 | 0.172855832 | 0.104 | 0 | 1 | 6 |
| fgfr1a | 0.09782 | 0.182034974 | 0.104 | 0 | 1 | 6 |
| fhl1a | 0.09782 | 0.269034083 | 0.104 | 0 | 1 | 6 |
| hars | 0.09782 | 0.218077 | 0.104 | 0 | 1 | 6 |
| hoxa11b | 0.09782 | 0.213343148 | 0.104 | 0 | 1 | 6 |
| impa1 | 0.09782 | 0.202644018 | 0.104 | 0 | 1 | 6 |
| ing4 | 0.09782 | 0.19962841 | 0.104 | 0 | 1 | 6 |
| mrpl17 | 0.09782 | 0.219850112 | 0.104 | 0 | 1 | 6 |
| nfya | 0.09782 | 0.213034958 | 0.104 | 0 | 1 | 6 |
| ociad1 | 0.09782 | 0.192963131 | 0.104 | 0 | 1 | 6 |
| pttg1ipb | 0.09782 | 0.209858239 | 0.104 | 0 | 1 | 6 |
| rcor1 | 0.09782 | 0.188175347 | 0.104 | 0 | 1 | 6 |
| rhogc | 0.09782 | 0.178126099 | 0.104 | 0 | 1 | 6 |
| si:ch211-237c6.4 | 0.09782 | 0.217346437 | 0.104 | 0 | 1 | 6 |
| si:dkey-46a10.3 | 0.09782 | 0.182456457 | 0.104 | 0 | 1 | 6 |
| srp72 | 0.09782 | 0.221315088 | 0.104 | 0 | 1 | 6 |
| u2af2a | 0.09782 | 0.180545894 | 0.104 | 0 | 1 | 6 |
| ube3a | 0.09782 | 0.226492472 | 0.104 | 0 | 1 | 6 |
| vstm4a | 0.09782 | 0.207768983 | 0.104 | 0 | 1 | 6 |
| znf410 | 0.09782 | 0.23061027 | 0.104 | 0 | 1 | 6 |
| barx1 | 0.09782 | 0.238929085 | 0.104 | 0 | 1 | 6 |
| calub | 0.09782 | 0.218594371 | 0.104 | 0 | 1 | 6 |
| LOC101883815 | 0.09782 | 0.197154126 | 0.104 | 0 | 1 | 6 |
| grb2a | 0.09782 | 0.21511492 | 0.104 | 0 | 1 | 6 |
| ntn1a | 0.09782 | 0.262650802 | 0.104 | 0 | 1 | 6 |
| phldb2b | 0.09782 | 0.2089095 | 0.104 | 0 | 1 | 6 |
| rps19bp1 | 0.09782 | 0.16855769 | 0.104 | 0 | 1 | 6 |
| zgc:153115 | 0.09782 | 0.22845109 | 0.104 | 0 | 1 | 6 |
| tnrc18 | 0.09782 | 0.186106754 | 0.104 | 0 | 1 | 6 |
| mmrn2a | 0.09794 | -0.30380535 | 0.026 | 0.176 | 1 | 6 |
| polr1e | 0.09806 | -0.26827788 | 0.026 | 0.176 | 1 | 6 |
| txnrd3 | 0.09875 | -0.33201002 | 0.091 | 0.294 | 1 | 6 |
| srsf3b | 0.099 | 0.425198826 | 0.416 | 0.176 | 1 | 6 |
| smim4 | 0.09902 | -0.11081938 | 0.078 | 0.059 | 1 | 6 |
| furina | 0.09904 | 0.122941493 | 0.169 | 0.059 | 1 | 6 |
| mrpl2 | 0.09931 | -0.2409512 | 0.026 | 0.176 | 1 | 6 |
| si:ch211-156b7.4 | 0.09939 | -0.3608981 | 0.065 | 0.235 | 1 | 6 |
| icn | 0.09957 | 0.60866036 | 0.506 | 0.235 | 1 | 6 |
| selenof | 0.09968 | 0.205192531 | 0.506 | 0.294 | 1 | 6 |
| gigyf2 | 0.09968 | -0.27500897 | 0.026 | 0.176 | 1 | 6 |
| hmgcl | 0.09987 | -0.24907261 | 0.026 | 0.176 | 1 | 6 |
| ndufb8 | 0.09988 | -0.18328842 | 0.338 | 0.353 | 1 | 6 |
| si:ch1073-340j19.1 | 0.10009 | -0.30838356 | 0.026 | 0.176 | 1 | 6 |
| prkd2 | 0.10043 | -0.24918801 | 0.026 | 0.176 | 1 | 6 |
| lsm6 | 0.10059 | 0.411398862 | 0.312 | 0.118 | 1 | 6 |
| gig2o | 0.10062 | -0.24457891 | 0.026 | 0.176 | 1 | 6 |
| col5a1 | 0.10075 | 0.344942085 | 0.792 | 0.529 | 1 | 6 |
| si:ch1073-429i10.3.1 | 0.10115 | 0.427171559 | 0.364 | 0.118 | 1 | 6 |

| | | | | | | |
|---|---|---|---|---|---|---|
| calm2b | 0.10161 | 0.313863245 | 0.494 | 0.235 | 1 | 6 |
| bxdc2 | 0.10167 | -0.30854495 | 0.169 | 0.235 | 1 | 6 |
| arpc5a | 0.10263 | -0.41054462 | 0.065 | 0.235 | 1 | 6 |
| wls | 0.10296 | 0.437380169 | 0.377 | 0.176 | 1 | 6 |
| hp1bp3 | 0.10409 | 0.348357503 | 0.39 | 0.235 | 1 | 6 |
| loxa | 0.10434 | 0.465443996 | 0.234 | 0.059 | 1 | 6 |
| aoc2 | 0.10485 | -0.53474631 | 0.325 | 0.529 | 1 | 6 |
| eif4e2 | 0.10585 | -0.2333783 | 0.078 | 0.118 | 1 | 6 |
| mt-nd3 | 0.10629 | -0.20645147 | 0.26 | 0.471 | 1 | 6 |
| lrp1aa | 0.10644 | -0.41301216 | 0.156 | 0.353 | 1 | 6 |
| bsg | 0.10663 | 0.441630607 | 0.597 | 0.353 | 1 | 6 |
| fosb | 0.10673 | 0.537397103 | 0.273 | 0.059 | 1 | 6 |
| hsbp1b | 0.10772 | 0.200710565 | 0.286 | 0.118 | 1 | 6 |
| psmd6 | 0.10784 | 0.349146116 | 0.26 | 0.059 | 1 | 6 |
| bhlhe41 | 0.10852 | 0.295957944 | 0.247 | 0.059 | 1 | 6 |
| rsl24d1 | 0.10921 | 0.416300784 | 0.688 | 0.471 | 1 | 6 |
| AREL1 | 0.1093 | -0.12325716 | 0 | 0.059 | 1 | 6 |
| ARHGEF17 | 0.1093 | -0.16232248 | 0 | 0.059 | 1 | 6 |
| BX571880.1 | 0.1093 | -0.12550843 | 0 | 0.059 | 1 | 6 |
| CABZ01059119.1 | 0.1093 | -0.16136413 | 0 | 0.059 | 1 | 6 |
| CABZ01118004.1 | 0.1093 | -0.10574404 | 0 | 0.059 | 1 | 6 |
| CU468920.1 | 0.1093 | -0.16307095 | 0 | 0.059 | 1 | 6 |
| CU639397.1 | 0.1093 | -0.10295488 | 0 | 0.059 | 1 | 6 |
| FP245456.1 | 0.1093 | -0.14738064 | 0 | 0.059 | 1 | 6 |
| LO018029.1 | 0.1093 | -0.15247612 | 0 | 0.059 | 1 | 6 |
| LOC100534805 | 0.1093 | -0.10604988 | 0 | 0.059 | 1 | 6 |
| LOC101884549 | 0.1093 | -0.10522323 | 0 | 0.059 | 1 | 6 |
| XLOC-005605 | 0.1093 | -0.1026225 | 0 | 0.059 | 1 | 6 |
| XLOC-019096 | 0.1093 | -0.12550843 | 0 | 0.059 | 1 | 6 |
| XLOC-026848 | 0.1093 | -0.1553288 | 0 | 0.059 | 1 | 6 |
| XLOC-043834 | 0.1093 | -0.15035497 | 0 | 0.059 | 1 | 6 |
| XLOC-044297 | 0.1093 | -0.15619618 | 0 | 0.059 | 1 | 6 |
| acad11 | 0.1093 | -0.15400196 | 0 | 0.059 | 1 | 6 |
| adam8b | 0.1093 | -0.10675827 | 0 | 0.059 | 1 | 6 |
| admb | 0.1093 | -0.20132301 | 0 | 0.059 | 1 | 6 |
| agla | 0.1093 | -0.15184456 | 0 | 0.059 | 1 | 6 |
| ago1 | 0.1093 | -0.16150497 | 0 | 0.059 | 1 | 6 |
| ap3b1a | 0.1093 | -0.10781591 | 0 | 0.059 | 1 | 6 |
| arhgef37 | 0.1093 | -0.14859887 | 0 | 0.059 | 1 | 6 |
| armc7 | 0.1093 | -0.10306865 | 0 | 0.059 | 1 | 6 |
| atp6ap1la | 0.1093 | -0.1553288 | 0 | 0.059 | 1 | 6 |
| bglapl | 0.1093 | -0.12550843 | 0 | 0.059 | 1 | 6 |
| brwd3 | 0.1093 | -0.10090007 | 0 | 0.059 | 1 | 6 |
| btbd11a | 0.1093 | -0.12550843 | 0 | 0.059 | 1 | 6 |
| ccser2a | 0.1093 | -0.16232444 | 0 | 0.059 | 1 | 6 |
| cenpl | 0.1093 | -0.22448041 | 0 | 0.059 | 1 | 6 |
| chst2b | 0.1093 | -0.1042135 | 0 | 0.059 | 1 | 6 |
| crebl2 | 0.1093 | -0.10515924 | 0 | 0.059 | 1 | 6 |

| | | | | | | |
|---|---|---|---|---|---|---|
| crtac1a | 0.1093 | -0.14896254 | 0 | 0.059 | 1 | 6 |
| cse1l | 0.1093 | -0.10643112 | 0 | 0.059 | 1 | 6 |
| cul1b | 0.1093 | -0.2235197 | 0 | 0.059 | 1 | 6 |
| cxcr3.3 | 0.1093 | -0.1199638 | 0 | 0.059 | 1 | 6 |
| dct | 0.1093 | -0.15041418 | 0 | 0.059 | 1 | 6 |
| ddx31 | 0.1093 | -0.10664621 | 0 | 0.059 | 1 | 6 |
| dgkza | 0.1093 | -0.16230103 | 0 | 0.059 | 1 | 6 |
| dnajc14 | 0.1093 | -0.11915137 | 0 | 0.059 | 1 | 6 |
| dpy19l1l | 0.1093 | -0.1247241 | 0 | 0.059 | 1 | 6 |
| eml3 | 0.1093 | -0.1076711 | 0 | 0.059 | 1 | 6 |
| ephx5 | 0.1093 | -0.12392235 | 0 | 0.059 | 1 | 6 |
| ext1c | 0.1093 | -0.10375248 | 0 | 0.059 | 1 | 6 |
| fam69aa | 0.1093 | -0.1030676 | 0 | 0.059 | 1 | 6 |
| fas | 0.1093 | -0.105825 | 0 | 0.059 | 1 | 6 |
| fgfr4 | 0.1093 | -0.12148853 | 0 | 0.059 | 1 | 6 |
| focad | 0.1093 | -0.10613863 | 0 | 0.059 | 1 | 6 |
| fthl30 | 0.1093 | -0.1553288 | 0 | 0.059 | 1 | 6 |
| fubp3 | 0.1093 | -0.10414164 | 0 | 0.059 | 1 | 6 |
| gatd3a | 0.1093 | -0.14068714 | 0 | 0.059 | 1 | 6 |
| haus5 | 0.1093 | -0.13947805 | 0 | 0.059 | 1 | 6 |
| hgd | 0.1093 | -0.12238313 | 0 | 0.059 | 1 | 6 |
| hmcn1 | 0.1093 | -0.16259643 | 0 | 0.059 | 1 | 6 |
| homer2 | 0.1093 | -0.10278307 | 0 | 0.059 | 1 | 6 |
| htra3a | 0.1093 | -0.10675827 | 0 | 0.059 | 1 | 6 |
| ifi35 | 0.1093 | -0.1479797 | 0 | 0.059 | 1 | 6 |
| ikzf4 | 0.1093 | -0.10623611 | 0 | 0.059 | 1 | 6 |
| irs2a | 0.1093 | -0.15035123 | 0 | 0.059 | 1 | 6 |
| jade2 | 0.1093 | -0.1241448 | 0 | 0.059 | 1 | 6 |
| kcnk1b | 0.1093 | -0.16307095 | 0 | 0.059 | 1 | 6 |
| kcnk5b | 0.1093 | -0.15360996 | 0 | 0.059 | 1 | 6 |
| kdsr | 0.1093 | -0.15406157 | 0 | 0.059 | 1 | 6 |
| klc1a | 0.1093 | -0.12550843 | 0 | 0.059 | 1 | 6 |
| lgals17 | 0.1093 | -0.16120335 | 0 | 0.059 | 1 | 6 |
| lnpk | 0.1093 | -0.10125471 | 0 | 0.059 | 1 | 6 |
| lrrc15 | 0.1093 | -0.15414163 | 0 | 0.059 | 1 | 6 |
| maco1b | 0.1093 | -0.12351873 | 0 | 0.059 | 1 | 6 |
| mafk | 0.1093 | -0.16307095 | 0 | 0.059 | 1 | 6 |
| mblac1 | 0.1093 | -0.14775676 | 0 | 0.059 | 1 | 6 |
| mboat2b | 0.1093 | -0.15985958 | 0 | 0.059 | 1 | 6 |
| mdm1 | 0.1093 | -0.10516954 | 0 | 0.059 | 1 | 6 |
| med1 | 0.1093 | -0.2205617 | 0 | 0.059 | 1 | 6 |
| megf6a | 0.1093 | -0.1054721 | 0 | 0.059 | 1 | 6 |
| mia3 | 0.1093 | -0.10385034 | 0 | 0.059 | 1 | 6 |
| mir22a | 0.1093 | -0.12434098 | 0 | 0.059 | 1 | 6 |
| mkrn4 | 0.1093 | -0.10346864 | 0 | 0.059 | 1 | 6 |
| mpzl2b | 0.1093 | -0.1053968 | 0 | 0.059 | 1 | 6 |
| nfil3-2 | 0.1093 | -0.16100367 | 0 | 0.059 | 1 | 6 |
| nfil3-5 | 0.1093 | -0.10911133 | 0 | 0.059 | 1 | 6 |

| | | | | | | |
|---|---|---|---|---|---|---|
| nhlrc2 | 0.1093 | -0.1397082 | 0 | 0.059 | 1 | 6 |
| nkx2.3 | 0.1093 | -0.1553288 | 0 | 0.059 | 1 | 6 |
| nlrc3l | 0.1093 | -0.10829595 | 0 | 0.059 | 1 | 6 |
| nrcama | 0.1093 | -0.12550843 | 0 | 0.059 | 1 | 6 |
| ntn1b | 0.1093 | -0.1056225 | 0 | 0.059 | 1 | 6 |
| ogfod1 | 0.1093 | -0.11808042 | 0 | 0.059 | 1 | 6 |
| pard6a | 0.1093 | -0.15453479 | 0 | 0.059 | 1 | 6 |
| pcgf1 | 0.1093 | -0.15298385 | 0 | 0.059 | 1 | 6 |
| pcp4a | 0.1093 | -0.10263824 | 0 | 0.059 | 1 | 6 |
| phf19 | 0.1093 | -0.15837797 | 0 | 0.059 | 1 | 6 |
| polr3d | 0.1093 | -0.10562981 | 0 | 0.059 | 1 | 6 |
| pon1 | 0.1093 | -0.12257183 | 0 | 0.059 | 1 | 6 |
| pparg | 0.1093 | -0.10120064 | 0 | 0.059 | 1 | 6 |
| ppfibp1b | 0.1093 | -0.15443266 | 0 | 0.059 | 1 | 6 |
| ppwd1 | 0.1093 | -0.11698934 | 0 | 0.059 | 1 | 6 |
| prf1.5 | 0.1093 | -0.16658208 | 0 | 0.059 | 1 | 6 |
| pygo1 | 0.1093 | -0.11915137 | 0 | 0.059 | 1 | 6 |
| rab39bb | 0.1093 | -0.10366638 | 0 | 0.059 | 1 | 6 |
| ralgapa2 | 0.1093 | -0.11975328 | 0 | 0.059 | 1 | 6 |
| raver1 | 0.1093 | -0.15198318 | 0 | 0.059 | 1 | 6 |
| rbsn | 0.1093 | -0.15501644 | 0 | 0.059 | 1 | 6 |
| relt | 0.1093 | -0.10185815 | 0 | 0.059 | 1 | 6 |
| riok3 | 0.1093 | -0.14991873 | 0 | 0.059 | 1 | 6 |
| rnf167 | 0.1093 | -0.15711282 | 0 | 0.059 | 1 | 6 |
| ruvbl2 | 0.1093 | -0.10372873 | 0 | 0.059 | 1 | 6 |
| sap130a | 0.1093 | -0.16223262 | 0 | 0.059 | 1 | 6 |
| sft2d2a | 0.1093 | -0.10237228 | 0 | 0.059 | 1 | 6 |
| si:cabz01007802.1 | 0.1093 | -0.16307095 | 0 | 0.059 | 1 | 6 |
| si:ch211-125m10.6 | 0.1093 | -0.10675827 | 0 | 0.059 | 1 | 6 |
| si:ch211-12e13.12 | 0.1093 | -0.10455358 | 0 | 0.059 | 1 | 6 |
| si:ch211-194p6.10 | 0.1093 | -0.10562262 | 0 | 0.059 | 1 | 6 |
| si:ch211-196f5.9 | 0.1093 | -0.12550843 | 0 | 0.059 | 1 | 6 |
| si:ch211-209a2.1 | 0.1093 | -0.1194708 | 0 | 0.059 | 1 | 6 |
| si:ch211-209l18.4 | 0.1093 | -0.10675827 | 0 | 0.059 | 1 | 6 |
| si:ch211-214k5.3 | 0.1093 | -0.10826941 | 0 | 0.059 | 1 | 6 |
| si:ch211-221j21.3 | 0.1093 | -0.10068831 | 0 | 0.059 | 1 | 6 |
| si:ch211-251f6.6 | 0.1093 | -0.17121979 | 0 | 0.059 | 1 | 6 |
| si:ch211-51c14.1 | 0.1093 | -0.10625175 | 0 | 0.059 | 1 | 6 |
| si:ch211-62a1.3 | 0.1093 | -0.12550843 | 0 | 0.059 | 1 | 6 |
| si:ch73-14h10.2 | 0.1093 | -0.10599121 | 0 | 0.059 | 1 | 6 |
| si:dkey-156j15.1 | 0.1093 | -0.10488643 | 0 | 0.059 | 1 | 6 |
| si:dkey-225k4.1 | 0.1093 | -0.10800607 | 0 | 0.059 | 1 | 6 |
| si:dkey-23i12.7 | 0.1093 | -0.10699025 | 0 | 0.059 | 1 | 6 |
| si:dkey-261e22.4 | 0.1093 | -0.11915137 | 0 | 0.059 | 1 | 6 |
| si:dkey-264f17.4 | 0.1093 | -0.16307095 | 0 | 0.059 | 1 | 6 |
| si:dkey-27i16.2 | 0.1093 | -0.11162599 | 0 | 0.059 | 1 | 6 |
| si:dkey-42p14.3 | 0.1093 | -0.10957547 | 0 | 0.059 | 1 | 6 |
| si:dkey-46i9.1 | 0.1093 | -0.14861316 | 0 | 0.059 | 1 | 6 |

| | | | | | | |
|---|---|---|---|---|---|---|
| si:dkey-93n13.2 | 0.1093 | -0.10618071 | 0 | 0.059 | 1 | 6 |
| si:dkeyp-53d3.3 | 0.1093 | -0.1507958 | 0 | 0.059 | 1 | 6 |
| si:dkeyp-97a10.2 | 0.1093 | -0.10793895 | 0 | 0.059 | 1 | 6 |
| skp2 | 0.1093 | -0.10671774 | 0 | 0.059 | 1 | 6 |
| slc25a18 | 0.1093 | -0.10729185 | 0 | 0.059 | 1 | 6 |
| slc25a32b | 0.1093 | -0.16307095 | 0 | 0.059 | 1 | 6 |
| sos1 | 0.1093 | -0.15242192 | 0 | 0.059 | 1 | 6 |
| ssbp2 | 0.1093 | -0.20693498 | 0 | 0.059 | 1 | 6 |
| stat1b | 0.1093 | -0.13764919 | 0 | 0.059 | 1 | 6 |
| steap4 | 0.1093 | -0.10797028 | 0 | 0.059 | 1 | 6 |
| stoml2 | 0.1093 | -0.14321726 | 0 | 0.059 | 1 | 6 |
| sun1 | 0.1093 | -0.12369805 | 0 | 0.059 | 1 | 6 |
| synpo2lb | 0.1093 | -0.15505725 | 0 | 0.059 | 1 | 6 |
| tbccd1 | 0.1093 | -0.12325716 | 0 | 0.059 | 1 | 6 |
| tmem184c | 0.1093 | -0.10520098 | 0 | 0.059 | 1 | 6 |
| tmf1 | 0.1093 | -0.12034151 | 0 | 0.059 | 1 | 6 |
| tnfsf13b | 0.1093 | -0.15280359 | 0 | 0.059 | 1 | 6 |
| tns3.2 | 0.1093 | -0.12007429 | 0 | 0.059 | 1 | 6 |
| tpp2 | 0.1093 | -0.15412956 | 0 | 0.059 | 1 | 6 |
| trmt61b | 0.1093 | -0.16013368 | 0 | 0.059 | 1 | 6 |
| tsen2 | 0.1093 | -0.1484242 | 0 | 0.059 | 1 | 6 |
| tspan37 | 0.1093 | -0.10335152 | 0 | 0.059 | 1 | 6 |
| ubr2 | 0.1093 | -0.10557055 | 0 | 0.059 | 1 | 6 |
| uhrf1 | 0.1093 | -0.10629083 | 0 | 0.059 | 1 | 6 |
| vav2 | 0.1093 | -0.15471874 | 0 | 0.059 | 1 | 6 |
| wasb | 0.1093 | -0.22391636 | 0 | 0.059 | 1 | 6 |
| wnk4b | 0.1093 | -0.12329939 | 0 | 0.059 | 1 | 6 |
| wu:fj29h11 | 0.1093 | -0.10530009 | 0 | 0.059 | 1 | 6 |
| xkr8.3 | 0.1093 | -0.10328997 | 0 | 0.059 | 1 | 6 |
| zfp64 | 0.1093 | -0.16060603 | 0 | 0.059 | 1 | 6 |
| zgc:112163 | 0.1093 | -0.10452694 | 0 | 0.059 | 1 | 6 |
| zgc:113298 | 0.1093 | -0.10675827 | 0 | 0.059 | 1 | 6 |
| zgc:152791 | 0.1093 | -0.18238282 | 0 | 0.059 | 1 | 6 |
| zgc:154046 | 0.1093 | -0.10770045 | 0 | 0.059 | 1 | 6 |
| zgc:92107 | 0.1093 | -0.10059027 | 0 | 0.059 | 1 | 6 |
| znf1153 | 0.1093 | -0.16135726 | 0 | 0.059 | 1 | 6 |
| znf1164 | 0.1093 | -0.10553853 | 0 | 0.059 | 1 | 6 |
| znf592 | 0.1093 | -0.15309981 | 0 | 0.059 | 1 | 6 |
| znf646 | 0.1093 | -0.10455972 | 0 | 0.059 | 1 | 6 |
| znfl2a | 0.1093 | -0.22667033 | 0 | 0.059 | 1 | 6 |
| CABZ01084575.1 | 0.1093 | -0.18579964 | 0 | 0.059 | 1 | 6 |
| CU570889.1 | 0.1093 | -0.19270714 | 0 | 0.059 | 1 | 6 |
| CU984600.2 | 0.1093 | -0.1914304 | 0 | 0.059 | 1 | 6 |
| FQ378018.1 | 0.1093 | -0.19190352 | 0 | 0.059 | 1 | 6 |
| LOC108182857 | 0.1093 | -0.1754975 | 0 | 0.059 | 1 | 6 |
| abl2 | 0.1093 | -0.19203352 | 0 | 0.059 | 1 | 6 |
| b3galnt2 | 0.1093 | -0.19170923 | 0 | 0.059 | 1 | 6 |
| ccnc | 0.1093 | -0.18933386 | 0 | 0.059 | 1 | 6 |

| | | | | | | |
|---|---|---|---|---|---|---|
| chmp5a | 0.1093 | -0.19137375 | 0 | 0.059 | 1 | 6 |
| rnf213b | 0.1093 | -0.19039589 | 0 | 0.059 | 1 | 6 |
| si:ch211-214p16.2 | 0.1093 | -0.14735865 | 0 | 0.059 | 1 | 6 |
| si:dkey-75a21.2 | 0.1093 | -0.19171839 | 0 | 0.059 | 1 | 6 |
| tmem110l | 0.1093 | -0.18608138 | 0 | 0.059 | 1 | 6 |
| zswim7 | 0.1093 | -0.17175088 | 0 | 0.059 | 1 | 6 |
| BX908782.3 | 0.1093 | -0.20091659 | 0 | 0.059 | 1 | 6 |
| CABZ01033180.1 | 0.1093 | -0.1186126 | 0 | 0.059 | 1 | 6 |
| CABZ01072487.2 | 0.1093 | -0.11857924 | 0 | 0.059 | 1 | 6 |
| CABZ01080568.1 | 0.1093 | -0.13033376 | 0 | 0.059 | 1 | 6 |
| CU464124.1 | 0.1093 | -0.17465066 | 0 | 0.059 | 1 | 6 |
| FRMD4A | 0.1093 | -0.17612378 | 0 | 0.059 | 1 | 6 |
| LOC107195253 | 0.1093 | -0.11495632 | 0 | 0.059 | 1 | 6 |
| XLOC-003453 | 0.1093 | -0.11455628 | 0 | 0.059 | 1 | 6 |
| XLOC-043761 | 0.1093 | -0.18697763 | 0 | 0.059 | 1 | 6 |
| abch1 | 0.1093 | -0.1918621 | 0 | 0.059 | 1 | 6 |
| abhd3 | 0.1093 | -0.11952501 | 0 | 0.059 | 1 | 6 |
| acta1b | 0.1093 | -0.18047476 | 0 | 0.059 | 1 | 6 |
| ank3a | 0.1093 | -0.12046864 | 0 | 0.059 | 1 | 6 |
| antxr1d | 0.1093 | -0.11570644 | 0 | 0.059 | 1 | 6 |
| arhgap36 | 0.1093 | -0.16215056 | 0 | 0.059 | 1 | 6 |
| arhgef7a | 0.1093 | -0.12021679 | 0 | 0.059 | 1 | 6 |
| atad2 | 0.1093 | -0.11773014 | 0 | 0.059 | 1 | 6 |
| atp10b | 0.1093 | -0.12046864 | 0 | 0.059 | 1 | 6 |
| avpr1ab | 0.1093 | -0.11709435 | 0 | 0.059 | 1 | 6 |
| bcl6ab | 0.1093 | -0.11852531 | 0 | 0.059 | 1 | 6 |
| bin2b | 0.1093 | -0.11628171 | 0 | 0.059 | 1 | 6 |
| castor1 | 0.1093 | -0.19128899 | 0 | 0.059 | 1 | 6 |
| cep162 | 0.1093 | -0.12046864 | 0 | 0.059 | 1 | 6 |
| chrnb2b | 0.1093 | -0.12046864 | 0 | 0.059 | 1 | 6 |
| clgn | 0.1093 | -0.16217678 | 0 | 0.059 | 1 | 6 |
| col18a1b | 0.1093 | -0.18337765 | 0 | 0.059 | 1 | 6 |
| cpped1 | 0.1093 | -0.11743125 | 0 | 0.059 | 1 | 6 |
| crabp1a | 0.1093 | -0.17344296 | 0 | 0.059 | 1 | 6 |
| cx39.4 | 0.1093 | -0.11939759 | 0 | 0.059 | 1 | 6 |
| dachb | 0.1093 | -0.19222563 | 0 | 0.059 | 1 | 6 |
| dazap1 | 0.1093 | -0.17248097 | 0 | 0.059 | 1 | 6 |
| dnajc16l | 0.1093 | -0.11812339 | 0 | 0.059 | 1 | 6 |
| epas1a | 0.1093 | -0.11974795 | 0 | 0.059 | 1 | 6 |
| eva1bb | 0.1093 | -0.19125746 | 0 | 0.059 | 1 | 6 |
| fam43a | 0.1093 | -0.11748893 | 0 | 0.059 | 1 | 6 |
| fancf | 0.1093 | -0.11633796 | 0 | 0.059 | 1 | 6 |
| fbxl18 | 0.1093 | -0.1189588 | 0 | 0.059 | 1 | 6 |
| fgf1b | 0.1093 | -0.18461928 | 0 | 0.059 | 1 | 6 |
| fig4a | 0.1093 | -0.190176 | 0 | 0.059 | 1 | 6 |
| ftr93 | 0.1093 | -0.11875454 | 0 | 0.059 | 1 | 6 |
| gemin4 | 0.1093 | -0.11851098 | 0 | 0.059 | 1 | 6 |
| gins4 | 0.1093 | -0.11569964 | 0 | 0.059 | 1 | 6 |

| | | | | | | |
|---|---|---|---|---|---|---|
| glg1b | 0.1093 | -0.11935452 | 0 | 0.059 | 1 | 6 |
| hey1 | 0.1093 | -0.19264778 | 0 | 0.059 | 1 | 6 |
| hey2 | 0.1093 | -0.16356855 | 0 | 0.059 | 1 | 6 |
| iqcb1 | 0.1093 | -0.11662809 | 0 | 0.059 | 1 | 6 |
| kif3a | 0.1093 | -0.16351128 | 0 | 0.059 | 1 | 6 |
| lgals1l1 | 0.1093 | -0.13060017 | 0 | 0.059 | 1 | 6 |
| lpcat1 | 0.1093 | -0.1169082 | 0 | 0.059 | 1 | 6 |
| map3k20 | 0.1093 | -0.11752473 | 0 | 0.059 | 1 | 6 |
| mat2al | 0.1093 | -0.16283754 | 0 | 0.059 | 1 | 6 |
| mcm8 | 0.1093 | -0.19124773 | 0 | 0.059 | 1 | 6 |
| med13a | 0.1093 | -0.13240315 | 0 | 0.059 | 1 | 6 |
| mtss1 | 0.1093 | -0.11796768 | 0 | 0.059 | 1 | 6 |
| ncaldb | 0.1093 | -0.18918981 | 0 | 0.059 | 1 | 6 |
| ndufa4l2a | 0.1093 | -0.11822093 | 0 | 0.059 | 1 | 6 |
| nek8 | 0.1093 | -0.11802733 | 0 | 0.059 | 1 | 6 |
| obscna | 0.1093 | -0.17234489 | 0 | 0.059 | 1 | 6 |
| pacrg | 0.1093 | -0.18911609 | 0 | 0.059 | 1 | 6 |
| palb2 | 0.1093 | -0.11804162 | 0 | 0.059 | 1 | 6 |
| pane1 | 0.1093 | -0.10547622 | 0 | 0.059 | 1 | 6 |
| pcdh1b | 0.1093 | -0.11964825 | 0 | 0.059 | 1 | 6 |
| pemt | 0.1093 | -0.13286621 | 0 | 0.059 | 1 | 6 |
| phldb1a | 0.1093 | -0.11938322 | 0 | 0.059 | 1 | 6 |
| pigo | 0.1093 | -0.11979649 | 0 | 0.059 | 1 | 6 |
| pmm2 | 0.1093 | -0.11686566 | 0 | 0.059 | 1 | 6 |
| ppargc1a | 0.1093 | -0.12046864 | 0 | 0.059 | 1 | 6 |
| prr14 | 0.1093 | -0.11926889 | 0 | 0.059 | 1 | 6 |
| qrsl1 | 0.1093 | -0.16272325 | 0 | 0.059 | 1 | 6 |
| rab24 | 0.1093 | -0.11892182 | 0 | 0.059 | 1 | 6 |
| retsatl | 0.1093 | -0.11948628 | 0 | 0.059 | 1 | 6 |
| sema6d | 0.1093 | -0.16407671 | 0 | 0.059 | 1 | 6 |
| sh3gl3a | 0.1093 | -0.1881379 | 0 | 0.059 | 1 | 6 |
| si:ch1073-268j14.1 | 0.1093 | -0.12046864 | 0 | 0.059 | 1 | 6 |
| si:ch1073-303k11.2 | 0.1093 | -0.1196331 | 0 | 0.059 | 1 | 6 |
| si:ch211-117c9.5 | 0.1093 | -0.1632032 | 0 | 0.059 | 1 | 6 |
| si:ch211-161h7.4 | 0.1093 | -0.13169956 | 0 | 0.059 | 1 | 6 |
| si:ch211-227m13.1 | 0.1093 | -0.12085806 | 0 | 0.059 | 1 | 6 |
| si:ch211-253h3.1 | 0.1093 | -0.11902932 | 0 | 0.059 | 1 | 6 |
| si:ch211-71m22.1 | 0.1093 | -0.18545312 | 0 | 0.059 | 1 | 6 |
| si:dkey-182i3.9 | 0.1093 | -0.12046864 | 0 | 0.059 | 1 | 6 |
| si:dkey-190j3.4 | 0.1093 | -0.11909399 | 0 | 0.059 | 1 | 6 |
| si:dkey-20d21.12 | 0.1093 | -0.18331781 | 0 | 0.059 | 1 | 6 |
| si:dkey-228a15.1 | 0.1093 | -0.11051639 | 0 | 0.059 | 1 | 6 |
| si:dkey-30k6.5 | 0.1093 | -0.1178204 | 0 | 0.059 | 1 | 6 |
| si:dkey-79d12.4 | 0.1093 | -0.11944158 | 0 | 0.059 | 1 | 6 |
| slc20a1a | 0.1093 | -0.11789687 | 0 | 0.059 | 1 | 6 |
| slc8a1a | 0.1093 | -0.12046864 | 0 | 0.059 | 1 | 6 |
| slx4 | 0.1093 | -0.12046864 | 0 | 0.059 | 1 | 6 |
| sorbs2a | 0.1093 | -0.18881782 | 0 | 0.059 | 1 | 6 |

| | | | | | | |
|---|---|---|---|---|---|---|
| sorbs2b | 0.1093 | -0.18047476 | 0 | 0.059 | 1 | 6 |
| srpk2 | 0.1093 | -0.19264778 | 0 | 0.059 | 1 | 6 |
| stxbp5b | 0.1093 | -0.12046864 | 0 | 0.059 | 1 | 6 |
| tcf4 | 0.1093 | -0.12027428 | 0 | 0.059 | 1 | 6 |
| terb1 | 0.1093 | -0.16912242 | 0 | 0.059 | 1 | 6 |
| tesca | 0.1093 | -0.18929939 | 0 | 0.059 | 1 | 6 |
| tmem120b | 0.1093 | -0.11574335 | 0 | 0.059 | 1 | 6 |
| tmem160 | 0.1093 | -0.11704728 | 0 | 0.059 | 1 | 6 |
| tmtops2a | 0.1093 | -0.1179893 | 0 | 0.059 | 1 | 6 |
| twnk | 0.1093 | -0.11817174 | 0 | 0.059 | 1 | 6 |
| ube2s | 0.1093 | -0.11854343 | 0 | 0.059 | 1 | 6 |
| ugt5b1 | 0.1093 | -0.11385103 | 0 | 0.059 | 1 | 6 |
| urod | 0.1093 | -0.10809079 | 0 | 0.059 | 1 | 6 |
| xrcc1 | 0.1093 | -0.11776932 | 0 | 0.059 | 1 | 6 |
| zbtb49 | 0.1093 | -0.19226111 | 0 | 0.059 | 1 | 6 |
| zdhhc7 | 0.1093 | -0.18881782 | 0 | 0.059 | 1 | 6 |
| zgc:101130 | 0.1093 | -0.11958119 | 0 | 0.059 | 1 | 6 |
| zgc:110224 | 0.1093 | -0.11556816 | 0 | 0.059 | 1 | 6 |
| zgc:112023 | 0.1093 | -0.1601316 | 0 | 0.059 | 1 | 6 |
| zgc:153157 | 0.1093 | -0.11718956 | 0 | 0.059 | 1 | 6 |
| zgc:172121 | 0.1093 | -0.17687535 | 0 | 0.059 | 1 | 6 |
| zgc:194629 | 0.1093 | -0.11343944 | 0 | 0.059 | 1 | 6 |
| zgc:63470 | 0.1093 | -0.18914633 | 0 | 0.059 | 1 | 6 |
| zgc:86839 | 0.1093 | -0.19138812 | 0 | 0.059 | 1 | 6 |
| CABZ01072487.1 | 0.1093 | -0.22739016 | 0 | 0.059 | 1 | 6 |
| CABZ01087502.1 | 0.1093 | -0.22644918 | 0 | 0.059 | 1 | 6 |
| CABZ01114105.1 | 0.1093 | -0.22710781 | 0 | 0.059 | 1 | 6 |
| CT583652.1 | 0.1093 | -0.22760775 | 0 | 0.059 | 1 | 6 |
| GTPBP8 | 0.1093 | -0.22760775 | 0 | 0.059 | 1 | 6 |
| LOC103910007 | 0.1093 | -0.22721338 | 0 | 0.059 | 1 | 6 |
| LOC110439953 | 0.1093 | -0.22760775 | 0 | 0.059 | 1 | 6 |
| USP34 | 0.1093 | -0.22667522 | 0 | 0.059 | 1 | 6 |
| XLOC-032002 | 0.1093 | -0.22496613 | 0 | 0.059 | 1 | 6 |
| XLOC-033378 | 0.1093 | -0.22485472 | 0 | 0.059 | 1 | 6 |
| acsbg1 | 0.1093 | -0.2263903 | 0 | 0.059 | 1 | 6 |
| ank2b | 0.1093 | -0.22731788 | 0 | 0.059 | 1 | 6 |
| atp11c | 0.1093 | -0.21692299 | 0 | 0.059 | 1 | 6 |
| carnmt1 | 0.1093 | -0.2282445 | 0 | 0.059 | 1 | 6 |
| ccdc191 | 0.1093 | -0.22502471 | 0 | 0.059 | 1 | 6 |
| cep70 | 0.1093 | -0.14754141 | 0 | 0.059 | 1 | 6 |
| dbnla | 0.1093 | -0.20025448 | 0 | 0.059 | 1 | 6 |
| ebf1b | 0.1093 | -0.27475405 | 0 | 0.059 | 1 | 6 |
| elmod2 | 0.1093 | -0.22585357 | 0 | 0.059 | 1 | 6 |
| lamtor1 | 0.1093 | -0.22720277 | 0 | 0.059 | 1 | 6 |
| nr1h3 | 0.1093 | -0.22689948 | 0 | 0.059 | 1 | 6 |
| oclna | 0.1093 | -0.216398 | 0 | 0.059 | 1 | 6 |
| poli | 0.1093 | -0.22760775 | 0 | 0.059 | 1 | 6 |
| renbp | 0.1093 | -0.19297673 | 0 | 0.059 | 1 | 6 |

| | | | | | | |
|---|---|---|---|---|---|---|
| sfxn2 | 0.1093 | -0.20045665 | 0 | 0.059 | 1 | 6 |
| si:dkey-190g11.3 | 0.1093 | -0.22760775 | 0 | 0.059 | 1 | 6 |
| si:dkeyp-13a3.10 | 0.1093 | -0.20829015 | 0 | 0.059 | 1 | 6 |
| ttll4 | 0.1093 | -0.21767181 | 0 | 0.059 | 1 | 6 |
| wars | 0.1093 | -0.21548001 | 0 | 0.059 | 1 | 6 |
| znf644b | 0.1093 | -0.22741819 | 0 | 0.059 | 1 | 6 |
| mfhas1 | 0.1093 | -0.11096068 | 0 | 0.059 | 1 | 6 |
| prx | 0.1093 | -0.15420758 | 0 | 0.059 | 1 | 6 |
| rbfox1l | 0.1093 | -0.10644504 | 0 | 0.059 | 1 | 6 |
| slc31a1 | 0.1093 | -0.10970811 | 0 | 0.059 | 1 | 6 |
| ednraa | 0.10951 | -0.36128217 | 0.234 | 0.353 | 1 | 6 |
| mrpl42 | 0.11045 | -0.14622375 | 0.182 | 0.176 | 1 | 6 |
| gsta.1 | 0.11059 | -0.31940649 | 0.26 | 0.353 | 1 | 6 |
| CABZ01080702.1 | 0.11119 | 0.133437775 | 0.325 | 0.353 | 1 | 6 |
| si:ch211-191i18.2 | 0.11121 | -0.35499627 | 0.104 | 0.176 | 1 | 6 |
| adh8a | 0.1118 | 0.158391294 | 0.208 | 0.235 | 1 | 6 |
| nid2a | 0.11269 | -0.44907065 | 0.312 | 0.529 | 1 | 6 |
| zgc:114188 | 0.11321 | 0.229043809 | 0.935 | 0.824 | 1 | 6 |
| csnk1db | 0.11449 | 0.381246041 | 0.299 | 0.118 | 1 | 6 |
| mrps14 | 0.11491 | 0.295575909 | 0.182 | 0.059 | 1 | 6 |
| rps16 | 0.11516 | 0.100358573 | 0.935 | 0.941 | 1 | 6 |
| sost | 0.11551 | 0.288349797 | 0.208 | 0.059 | 1 | 6 |
| gtpbp4 | 0.11552 | 0.249979089 | 0.468 | 0.235 | 1 | 6 |
| rpl7a | 0.11692 | 0.20002559 | 0.974 | 0.882 | 1 | 6 |
| tcp1 | 0.11708 | 0.386300491 | 0.455 | 0.294 | 1 | 6 |
| nop16 | 0.11781 | -0.33244125 | 0.078 | 0.235 | 1 | 6 |
| rpa3 | 0.11792 | -0.35403793 | 0.065 | 0.235 | 1 | 6 |
| arpc4 | 0.11814 | -0.28567477 | 0.195 | 0.294 | 1 | 6 |
| gbp | 0.11839 | 0.421845313 | 0.403 | 0.176 | 1 | 6 |
| baz2ba | 0.11961 | -0.41143642 | 0.104 | 0.294 | 1 | 6 |
| rab1ab | 0.11965 | 0.243421445 | 0.234 | 0.059 | 1 | 6 |
| hnrpkl | 0.11966 | 0.167611494 | 0.195 | 0.059 | 1 | 6 |
| h3f3c | 0.11971 | 0.317190179 | 0.857 | 0.647 | 1 | 6 |
| bet1l | 0.12029 | -0.12419846 | 0.065 | 0.059 | 1 | 6 |
| slc10a3 | 0.12064 | -0.27999569 | 0.039 | 0.118 | 1 | 6 |
| atp5po | 0.12065 | -0.1511237 | 0.39 | 0.353 | 1 | 6 |
| tsr2 | 0.12152 | -0.24304832 | 0.117 | 0.176 | 1 | 6 |
| zfhx3 | 0.12162 | 0.350444304 | 0.195 | 0.059 | 1 | 6 |
| slc30a7 | 0.12173 | -0.21204282 | 0.065 | 0.235 | 1 | 6 |
| txndc9 | 0.12173 | 0.40303131 | 0.312 | 0.118 | 1 | 6 |
| ppiaa | 0.12246 | 0.180241153 | 0.961 | 0.882 | 1 | 6 |
| rhoca | 0.12293 | 0.125749234 | 0.169 | 0.059 | 1 | 6 |
| CABZ01068366.1 | 0.12333 | -0.35601425 | 0.065 | 0.235 | 1 | 6 |
| si:ch73-111k22.2 | 0.12401 | -0.35098909 | 0.039 | 0.176 | 1 | 6 |
| dynll2b | 0.12428 | 0.167705348 | 0.195 | 0.059 | 1 | 6 |
| gpatch4 | 0.12439 | -0.34057843 | 0.117 | 0.235 | 1 | 6 |
| rps13 | 0.12481 | 0.176046931 | 0.974 | 0.882 | 1 | 6 |
| gtpbp1 | 0.12513 | -0.25574308 | 0.026 | 0.118 | 1 | 6 |

| | | | | | | |
|---|---|---|---|---|---|---|
| tmeff1b | 0.12526 | 0.357923934 | 0.273 | 0.118 | 1 | 6 |
| skp1 | 0.12561 | -0.26271307 | 0.468 | 0.529 | 1 | 6 |
| vim | 0.12604 | 0.405821019 | 0.351 | 0.118 | 1 | 6 |
| LOC110437917 | 0.12656 | 0.178013243 | 0.091 | 0 | 1 | 6 |
| ggcta | 0.12656 | 0.150863154 | 0.091 | 0 | 1 | 6 |
| hoxa11a | 0.12656 | 0.194882084 | 0.091 | 0 | 1 | 6 |
| marcksa | 0.12656 | 0.179688569 | 0.091 | 0 | 1 | 6 |
| neu3.2 | 0.12656 | 0.222598556 | 0.091 | 0 | 1 | 6 |
| pcnp | 0.12656 | 0.161116707 | 0.091 | 0 | 1 | 6 |
| per1b | 0.12656 | 0.19173687 | 0.091 | 0 | 1 | 6 |
| tpte | 0.12656 | 0.163766417 | 0.091 | 0 | 1 | 6 |
| yy1b | 0.12656 | 0.185751609 | 0.091 | 0 | 1 | 6 |
| zgc:112356 | 0.12656 | 0.185970643 | 0.091 | 0 | 1 | 6 |
| agrn | 0.12656 | 0.217130839 | 0.091 | 0 | 1 | 6 |
| akt2l | 0.12656 | 0.170237304 | 0.091 | 0 | 1 | 6 |
| auts2a | 0.12656 | 0.206105109 | 0.091 | 0 | 1 | 6 |
| b4galt7 | 0.12656 | 0.198712726 | 0.091 | 0 | 1 | 6 |
| ciao2b | 0.12656 | 0.19053028 | 0.091 | 0 | 1 | 6 |
| cyth1a | 0.12656 | 0.194543628 | 0.091 | 0 | 1 | 6 |
| fam3c | 0.12656 | 0.216620898 | 0.091 | 0 | 1 | 6 |
| kctd6b | 0.12656 | 0.171271089 | 0.091 | 0 | 1 | 6 |
| lysmd4 | 0.12656 | 0.184554973 | 0.091 | 0 | 1 | 6 |
| mllt10 | 0.12656 | 0.230167317 | 0.091 | 0 | 1 | 6 |
| naa40 | 0.12656 | 0.156743162 | 0.091 | 0 | 1 | 6 |
| otub1b | 0.12656 | 0.208555941 | 0.091 | 0 | 1 | 6 |
| phactr4a | 0.12656 | 0.220991645 | 0.091 | 0 | 1 | 6 |
| pi15b | 0.12656 | 0.21609721 | 0.091 | 0 | 1 | 6 |
| pigp | 0.12656 | 0.17949699 | 0.091 | 0 | 1 | 6 |
| polr3e | 0.12656 | 0.145748116 | 0.091 | 0 | 1 | 6 |
| ppig | 0.12656 | 0.1490188 | 0.091 | 0 | 1 | 6 |
| ppil3 | 0.12656 | 0.153726094 | 0.091 | 0 | 1 | 6 |
| prpf8 | 0.12656 | 0.154377625 | 0.091 | 0 | 1 | 6 |
| ptprda | 0.12656 | 0.206502968 | 0.091 | 0 | 1 | 6 |
| rangap1a | 0.12656 | 0.198889874 | 0.091 | 0 | 1 | 6 |
| rev3l | 0.12656 | 0.195570094 | 0.091 | 0 | 1 | 6 |
| rraga | 0.12656 | 0.191630322 | 0.091 | 0 | 1 | 6 |
| si:ch211-232b12.5 | 0.12656 | 0.176859891 | 0.091 | 0 | 1 | 6 |
| si:ch211-241b2.1 | 0.12656 | 0.175196392 | 0.091 | 0 | 1 | 6 |
| si:ch73-22o12.1 | 0.12656 | 0.194738347 | 0.091 | 0 | 1 | 6 |
| si:rp71-79p20.2 | 0.12656 | 0.237756108 | 0.091 | 0 | 1 | 6 |
| slmapb | 0.12656 | 0.171001609 | 0.091 | 0 | 1 | 6 |
| smg7 | 0.12656 | 0.184292046 | 0.091 | 0 | 1 | 6 |
| srek1ip1 | 0.12656 | 0.147233832 | 0.091 | 0 | 1 | 6 |
| srgap3 | 0.12656 | 0.15693004 | 0.091 | 0 | 1 | 6 |
| sssca1 | 0.12656 | 0.209256916 | 0.091 | 0 | 1 | 6 |
| supt16h | 0.12656 | 0.208695407 | 0.091 | 0 | 1 | 6 |
| surf4l | 0.12656 | 0.207817414 | 0.091 | 0 | 1 | 6 |
| urm1 | 0.12656 | 0.159889498 | 0.091 | 0 | 1 | 6 |

| | | | | | | |
|---|---|---|---|---|---|---|
| wwtr1 | 0.12656 | 0.174416306 | 0.091 | 0 | 1 | 6 |
| abcf2a | 0.12656 | 0.211000739 | 0.091 | 0 | 1 | 6 |
| bcl7a | 0.12656 | 0.206713783 | 0.091 | 0 | 1 | 6 |
| ciapin1 | 0.12656 | 0.142049431 | 0.091 | 0 | 1 | 6 |
| col4a6 | 0.12656 | 0.228298106 | 0.091 | 0 | 1 | 6 |
| cul4b | 0.12656 | 0.15944506 | 0.091 | 0 | 1 | 6 |
| cyp2ad2 | 0.12656 | 0.196853868 | 0.091 | 0 | 1 | 6 |
| desi1a | 0.12656 | 0.181925951 | 0.091 | 0 | 1 | 6 |
| dnajb1b | 0.12656 | 0.216098594 | 0.091 | 0 | 1 | 6 |
| dusp8a | 0.12656 | 0.202782862 | 0.091 | 0 | 1 | 6 |
| eif4eb | 0.12656 | 0.189179598 | 0.091 | 0 | 1 | 6 |
| elk1 | 0.12656 | 0.170037411 | 0.091 | 0 | 1 | 6 |
| gdf10a | 0.12656 | 0.207581525 | 0.091 | 0 | 1 | 6 |
| ggctb | 0.12656 | 0.188036249 | 0.091 | 0 | 1 | 6 |
| gnaia | 0.12656 | 0.210066737 | 0.091 | 0 | 1 | 6 |
| hoxa9b | 0.12656 | 0.168311342 | 0.091 | 0 | 1 | 6 |
| hoxc10a | 0.12656 | 0.174614829 | 0.091 | 0 | 1 | 6 |
| hoxd11a | 0.12656 | 0.198995739 | 0.091 | 0 | 1 | 6 |
| igf2bp1 | 0.12656 | 0.150394835 | 0.091 | 0 | 1 | 6 |
| irs1 | 0.12656 | 0.187303158 | 0.091 | 0 | 1 | 6 |
| jag1b | 0.12656 | 0.198951542 | 0.091 | 0 | 1 | 6 |
| katnb1 | 0.12656 | 0.170260027 | 0.091 | 0 | 1 | 6 |
| lzic | 0.12656 | 0.181295954 | 0.091 | 0 | 1 | 6 |
| mapk14b | 0.12656 | 0.160809489 | 0.091 | 0 | 1 | 6 |
| mrpl37 | 0.12656 | 0.138653507 | 0.091 | 0 | 1 | 6 |
| mrpl41 | 0.12656 | 0.188583849 | 0.091 | 0 | 1 | 6 |
| msi2a | 0.12656 | 0.169716425 | 0.091 | 0 | 1 | 6 |
| oxsr1b | 0.12656 | 0.184834546 | 0.091 | 0 | 1 | 6 |
| prpf38b | 0.12656 | 0.147422603 | 0.091 | 0 | 1 | 6 |
| psen1 | 0.12656 | 0.170562717 | 0.091 | 0 | 1 | 6 |
| quo | 0.12656 | 0.206641363 | 0.091 | 0 | 1 | 6 |
| ralbb | 0.12656 | 0.190998627 | 0.091 | 0 | 1 | 6 |
| rbm14a | 0.12656 | 0.221186529 | 0.091 | 0 | 1 | 6 |
| rbm34 | 0.12656 | 0.153818023 | 0.091 | 0 | 1 | 6 |
| rcl1 | 0.12656 | 0.14518809 | 0.091 | 0 | 1 | 6 |
| sc:d217 | 0.12656 | 0.180255625 | 0.091 | 0 | 1 | 6 |
| smad3a | 0.12656 | 0.196389676 | 0.091 | 0 | 1 | 6 |
| smad9 | 0.12656 | 0.168946422 | 0.091 | 0 | 1 | 6 |
| ss18 | 0.12656 | 0.181244979 | 0.091 | 0 | 1 | 6 |
| tmem178b | 0.12656 | 0.207430638 | 0.091 | 0 | 1 | 6 |
| tmem41aa | 0.12656 | 0.149938643 | 0.091 | 0 | 1 | 6 |
| wdr45 | 0.12656 | 0.231695782 | 0.091 | 0 | 1 | 6 |
| yju2 | 0.12656 | 0.184360516 | 0.091 | 0 | 1 | 6 |
| zgc:55733 | 0.12656 | 0.166776298 | 0.091 | 0 | 1 | 6 |
| tradd | 0.12656 | 0.179174539 | 0.091 | 0 | 1 | 6 |
| wdr46 | 0.12656 | 0.150298599 | 0.091 | 0 | 1 | 6 |
| ddx19 | 0.12656 | 0.170635098 | 0.091 | 0 | 1 | 6 |
| cachd1 | 0.12656 | 0.227856234 | 0.091 | 0 | 1 | 6 |

| | | | | | | |
|---|---|---|---|---|---|---|
| hexim1 | 0.12656 | 0.187587777 | 0.091 | 0 | 1 | 6 |
| map7d1a | 0.12656 | 0.179731041 | 0.091 | 0 | 1 | 6 |
| tnfsf12 | 0.12656 | 0.204379385 | 0.091 | 0 | 1 | 6 |
| ttc27 | 0.12656 | 0.206186212 | 0.091 | 0 | 1 | 6 |
| zgc:158564 | 0.12656 | 0.210603842 | 0.091 | 0 | 1 | 6 |
| gorasp2 | 0.12656 | 0.197240661 | 0.091 | 0 | 1 | 6 |
| nkd1 | 0.12656 | 0.191707934 | 0.091 | 0 | 1 | 6 |
| pdpk1b | 0.12656 | 0.176031373 | 0.091 | 0 | 1 | 6 |
| si:ch211-155e24.3 | 0.12656 | 0.150304803 | 0.091 | 0 | 1 | 6 |
| asip2b | 0.1269 | -0.34434711 | 0.338 | 0.588 | 1 | 6 |
| pa2g4b | 0.12808 | -0.18808392 | 0.169 | 0.353 | 1 | 6 |
| ier2a | 0.12828 | 0.489543904 | 0.558 | 0.294 | 1 | 6 |
| nrp1a | 0.12867 | -0.21250631 | 0.13 | 0.294 | 1 | 6 |
| atp5mc1 | 0.12882 | -0.27457257 | 0.286 | 0.529 | 1 | 6 |
| mgst3b | 0.12966 | 0.422277412 | 0.545 | 0.294 | 1 | 6 |
| rpl30 | 0.13007 | 0.189658938 | 0.948 | 0.882 | 1 | 6 |
| EML4 | 0.13037 | -0.14422226 | 0.052 | 0.176 | 1 | 6 |
| atf7ip | 0.13093 | 0.312147314 | 0.247 | 0.059 | 1 | 6 |
| nedd8 | 0.13283 | -0.1203307 | 0.13 | 0.118 | 1 | 6 |
| nucks1a | 0.13355 | 0.393478281 | 0.299 | 0.118 | 1 | 6 |
| trir | 0.13452 | 0.231302895 | 0.143 | 0.059 | 1 | 6 |
| cox7c | 0.13453 | -0.22842397 | 0.584 | 0.588 | 1 | 6 |
| glrx3 | 0.13472 | -0.1578686 | 0.13 | 0.294 | 1 | 6 |
| ndufa4l | 0.1349 | -0.22753425 | 0.636 | 0.647 | 1 | 6 |
| cygb1 | 0.13507 | -0.47410493 | 0.221 | 0.412 | 1 | 6 |
| gna13b | 0.13529 | -0.33521498 | 0.052 | 0.176 | 1 | 6 |
| CABZ01077217.1 | 0.13622 | -0.34248721 | 0.065 | 0.235 | 1 | 6 |
| ubc | 0.13828 | 0.257968215 | 0.818 | 0.588 | 1 | 6 |
| syf2 | 0.13832 | 0.368468441 | 0.299 | 0.118 | 1 | 6 |
| si:dkey-239h2.3 | 0.13877 | -0.34195991 | 0.221 | 0.353 | 1 | 6 |
| adipor2 | 0.13951 | -0.3405885 | 0.065 | 0.235 | 1 | 6 |
| med19a | 0.13972 | -0.27905003 | 0.065 | 0.235 | 1 | 6 |
| rbp5 | 0.1398 | -0.30986724 | 0.584 | 0.647 | 1 | 6 |
| ptpn1 | 0.14055 | -0.11944413 | 0.065 | 0.059 | 1 | 6 |
| terfa | 0.14138 | -0.25007889 | 0.065 | 0.118 | 1 | 6 |
| gpm6bb | 0.14149 | -0.34283311 | 0.065 | 0.235 | 1 | 6 |
| timp2a | 0.14151 | 0.458442202 | 0.325 | 0.118 | 1 | 6 |
| tspan36 | 0.14166 | -0.29209478 | 0.052 | 0.176 | 1 | 6 |
| yipf2 | 0.14366 | -0.1156579 | 0.065 | 0.059 | 1 | 6 |
| dync1i2a | 0.14382 | -0.19078036 | 0.091 | 0.118 | 1 | 6 |
| nfkbiab | 0.14407 | 0.30995707 | 0.338 | 0.118 | 1 | 6 |
| tmem119b | 0.14452 | 0.298074343 | 0.234 | 0.059 | 1 | 6 |
| si:ch211-284b7.3 | 0.14515 | -0.17883286 | 0.091 | 0.118 | 1 | 6 |
| prss35 | 0.14515 | 0.363014503 | 0.325 | 0.235 | 1 | 6 |
| fmnl2a | 0.1454 | -0.12269536 | 0.026 | 0.118 | 1 | 6 |
| pkp3b | 0.14598 | -0.40194425 | 0.143 | 0.353 | 1 | 6 |
| copz1 | 0.14629 | -0.23367648 | 0.182 | 0.235 | 1 | 6 |
| zc3h15 | 0.1467 | -0.2619063 | 0.065 | 0.235 | 1 | 6 |

| | | | | | | |
|---|---|---|---|---|---|---|
| nutf2l | 0.14708 | 0.155630614 | 0.273 | 0.118 | 1 | 6 |
| phf11 | 0.1473 | -0.11230247 | 0.065 | 0.059 | 1 | 6 |
| tnpo2 | 0.14774 | -0.18869408 | 0.039 | 0.176 | 1 | 6 |
| CABZ01034698.2 | 0.1479 | -0.35531602 | 0.104 | 0.294 | 1 | 6 |
| BX248082.1 | 0.14797 | -0.39593013 | 0.052 | 0.176 | 1 | 6 |
| hmgn6 | 0.149 | 0.29302078 | 0.818 | 0.588 | 1 | 6 |
| ran | 0.14927 | 0.329882687 | 0.805 | 0.588 | 1 | 6 |
| sdhdb | 0.1495 | -0.24631418 | 0.065 | 0.235 | 1 | 6 |
| gng5 | 0.15066 | -0.11804057 | 0.377 | 0.353 | 1 | 6 |
| rbp4 | 0.15086 | -0.42998587 | 0.442 | 0.588 | 1 | 6 |
| sdc2 | 0.15145 | 0.366630629 | 0.416 | 0.176 | 1 | 6 |
| snrnp25 | 0.15149 | -0.15460543 | 0.052 | 0.059 | 1 | 6 |
| hmgb1a | 0.15203 | 0.36853857 | 0.61 | 0.353 | 1 | 6 |
| prdx2 | 0.15236 | -0.3853756 | 0.481 | 0.647 | 1 | 6 |
| ccdc12 | 0.15333 | -0.23889786 | 0.065 | 0.235 | 1 | 6 |
| h2afy2 | 0.15392 | 0.359892794 | 0.468 | 0.294 | 1 | 6 |
| suclg2 | 0.15417 | -0.26822892 | 0.039 | 0.176 | 1 | 6 |
| rnf10 | 0.15458 | 0.344171652 | 0.351 | 0.176 | 1 | 6 |
| mex3b | 0.15472 | 0.263537479 | 0.299 | 0.118 | 1 | 6 |
| pfdn5 | 0.15493 | -0.20236475 | 0.208 | 0.235 | 1 | 6 |
| sparc | 0.15529 | 0.258826727 | 0.896 | 0.765 | 1 | 6 |
| vps26a | 0.15552 | -0.29419268 | 0.039 | 0.176 | 1 | 6 |
| bscl2 | 0.15653 | -0.17407398 | 0.039 | 0.176 | 1 | 6 |
| rad51 | 0.15656 | -0.19698805 | 0.039 | 0.059 | 1 | 6 |
| cox4i1 | 0.15675 | -0.22653573 | 0.662 | 0.882 | 1 | 6 |
| med27 | 0.15693 | -0.30426405 | 0.039 | 0.176 | 1 | 6 |
| rwdd1 | 0.15737 | 0.38218627 | 0.338 | 0.118 | 1 | 6 |
| nr2f2 | 0.15781 | -0.21912748 | 0.169 | 0.353 | 1 | 6 |
| commd8 | 0.15878 | -0.29941249 | 0.052 | 0.176 | 1 | 6 |
| adrm1 | 0.1588 | -0.36054192 | 0.091 | 0.235 | 1 | 6 |
| chd6 | 0.15899 | -0.14178442 | 0.182 | 0.176 | 1 | 6 |
| tomm40 | 0.15939 | -0.13539387 | 0.091 | 0.235 | 1 | 6 |
| eps15 | 0.15965 | -0.36531474 | 0.039 | 0.176 | 1 | 6 |
| spg7 | 0.15992 | -0.27859001 | 0.052 | 0.118 | 1 | 6 |
| hsd11b2 | 0.15999 | 0.327058756 | 0.714 | 0.471 | 1 | 6 |
| waplb | 0.16066 | -0.1155587 | 0.065 | 0.059 | 1 | 6 |
| ndufa8 | 0.16134 | 0.29191523 | 0.221 | 0.059 | 1 | 6 |
| ap2m1b | 0.16152 | -0.31845146 | 0.117 | 0.235 | 1 | 6 |
| ddx6 | 0.16165 | -0.29687463 | 0.104 | 0.294 | 1 | 6 |
| ccdc3b | 0.16168 | -0.27646543 | 0.117 | 0.294 | 1 | 6 |
| rps26 | 0.16179 | 0.14239382 | 0.909 | 0.882 | 1 | 6 |
| wu:fu71h07 | 0.16212 | -0.26547407 | 0.039 | 0.176 | 1 | 6 |
| hspe1 | 0.16214 | -0.31658321 | 0.234 | 0.294 | 1 | 6 |
| col1a2 | 0.16236 | 0.211757396 | 0.909 | 0.882 | 1 | 6 |
| rpap3 | 0.1628 | -0.14866992 | 0.052 | 0.059 | 1 | 6 |
| pik3ca | 0.1629 | -0.26966406 | 0.039 | 0.176 | 1 | 6 |
| sostdc1a | 0.16345 | 0.478406589 | 0.325 | 0.118 | 1 | 6 |
| vdac3 | 0.16371 | 0.360586767 | 0.545 | 0.294 | 1 | 6 |

| | | | | | | |
|---|---|---|---|---|---|---|
| idh2 | 0.16393 | 0.349646326 | 0.494 | 0.353 | 1 | 6 |
| arcn1a | 0.16401 | 0.141756667 | 0.078 | 0 | 1 | 6 |
| hnrnpul1l | 0.16401 | 0.152004131 | 0.078 | 0 | 1 | 6 |
| pdlim3a | 0.16401 | 0.163624858 | 0.078 | 0 | 1 | 6 |
| rnf145b | 0.16401 | 0.180924825 | 0.078 | 0 | 1 | 6 |
| abhd18 | 0.16401 | 0.123159325 | 0.078 | 0 | 1 | 6 |
| adhfe1 | 0.16401 | 0.168746486 | 0.078 | 0 | 1 | 6 |
| bud23 | 0.16401 | 0.125009093 | 0.078 | 0 | 1 | 6 |
| cldn11b | 0.16401 | 0.133201849 | 0.078 | 0 | 1 | 6 |
| eva1a | 0.16401 | 0.135730954 | 0.078 | 0 | 1 | 6 |
| klf7a | 0.16401 | 0.179339146 | 0.078 | 0 | 1 | 6 |
| n6amt1 | 0.16401 | 0.145284899 | 0.078 | 0 | 1 | 6 |
| ten1 | 0.16401 | 0.146549635 | 0.078 | 0 | 1 | 6 |
| tox | 0.16401 | 0.156374463 | 0.078 | 0 | 1 | 6 |
| uxt | 0.16401 | 0.169031321 | 0.078 | 0 | 1 | 6 |
| vps25 | 0.16401 | 0.132291612 | 0.078 | 0 | 1 | 6 |
| ZNF512B | 0.16401 | 0.212881417 | 0.078 | 0 | 1 | 6 |
| asb12a | 0.16401 | 0.147086922 | 0.078 | 0 | 1 | 6 |
| crispld1a | 0.16401 | 0.15887658 | 0.078 | 0 | 1 | 6 |
| herc2 | 0.16401 | 0.171419835 | 0.078 | 0 | 1 | 6 |
| morc2 | 0.16401 | 0.180312665 | 0.078 | 0 | 1 | 6 |
| nuak1b | 0.16401 | 0.12268841 | 0.078 | 0 | 1 | 6 |
| ppan | 0.16401 | 0.141908407 | 0.078 | 0 | 1 | 6 |
| rbm4.2 | 0.16401 | 0.143187439 | 0.078 | 0 | 1 | 6 |
| rgrb | 0.16401 | 0.106702614 | 0.078 | 0 | 1 | 6 |
| slc27a1a | 0.16401 | 0.163176587 | 0.078 | 0 | 1 | 6 |
| smad7 | 0.16401 | 0.146204094 | 0.078 | 0 | 1 | 6 |
| tdp1 | 0.16401 | 0.128231063 | 0.078 | 0 | 1 | 6 |
| tent4b | 0.16401 | 0.125726866 | 0.078 | 0 | 1 | 6 |
| BOLA2 | 0.16401 | 0.174281842 | 0.078 | 0 | 1 | 6 |
| BX470182.1 | 0.16401 | 0.19320537 | 0.078 | 0 | 1 | 6 |
| CABZ01005379.1 | 0.16401 | 0.161476273 | 0.078 | 0 | 1 | 6 |
| FQ976913.1 | 0.16401 | 0.151954872 | 0.078 | 0 | 1 | 6 |
| HACD4 | 0.16401 | 0.144356965 | 0.078 | 0 | 1 | 6 |
| RAMP1 | 0.16401 | 0.217039462 | 0.078 | 0 | 1 | 6 |
| XLOC-002129 | 0.16401 | 0.134554442 | 0.078 | 0 | 1 | 6 |
| XLOC-015238 | 0.16401 | 0.154813247 | 0.078 | 0 | 1 | 6 |
| abracl | 0.16401 | 0.153287478 | 0.078 | 0 | 1 | 6 |
| adipor1b | 0.16401 | 0.146767594 | 0.078 | 0 | 1 | 6 |
| antxr1c | 0.16401 | 0.147397065 | 0.078 | 0 | 1 | 6 |
| arpc5lb | 0.16401 | 0.144221323 | 0.078 | 0 | 1 | 6 |
| as3mt | 0.16401 | 0.127378911 | 0.078 | 0 | 1 | 6 |
| ash1l | 0.16401 | 0.149115256 | 0.078 | 0 | 1 | 6 |
| atp2b1a | 0.16401 | 0.166426148 | 0.078 | 0 | 1 | 6 |
| atp6v0cb | 0.16401 | 0.139403619 | 0.078 | 0 | 1 | 6 |
| baxa | 0.16401 | 0.141604822 | 0.078 | 0 | 1 | 6 |
| bbx | 0.16401 | 0.17961672 | 0.078 | 0 | 1 | 6 |
| bcl9 | 0.16401 | 0.180967407 | 0.078 | 0 | 1 | 6 |

| | | | | | | |
|---|---|---|---|---|---|---|
| cant1a | 0.16401 | 0.162364107 | 0.078 | 0 | 1 | 6 |
| cebpz | 0.16401 | 0.129064771 | 0.078 | 0 | 1 | 6 |
| chd2 | 0.16401 | 0.201394421 | 0.078 | 0 | 1 | 6 |
| chp2 | 0.16401 | 0.136409485 | 0.078 | 0 | 1 | 6 |
| chst14 | 0.16401 | 0.140812171 | 0.078 | 0 | 1 | 6 |
| cops5 | 0.16401 | 0.168405804 | 0.078 | 0 | 1 | 6 |
| cplane2 | 0.16401 | 0.160895707 | 0.078 | 0 | 1 | 6 |
| cpne3 | 0.16401 | 0.140570578 | 0.078 | 0 | 1 | 6 |
| crebbpb | 0.16401 | 0.145033108 | 0.078 | 0 | 1 | 6 |
| dazap2 | 0.16401 | 0.140583355 | 0.078 | 0 | 1 | 6 |
| ddx54 | 0.16401 | 0.170572046 | 0.078 | 0 | 1 | 6 |
| dnajc4 | 0.16401 | 0.159313345 | 0.078 | 0 | 1 | 6 |
| dpm1 | 0.16401 | 0.123634133 | 0.078 | 0 | 1 | 6 |
| dtnbp1a | 0.16401 | 0.123747123 | 0.078 | 0 | 1 | 6 |
| eif2b3 | 0.16401 | 0.187772917 | 0.078 | 0 | 1 | 6 |
| emc3 | 0.16401 | 0.155807485 | 0.078 | 0 | 1 | 6 |
| enoph1 | 0.16401 | 0.145163876 | 0.078 | 0 | 1 | 6 |
| faf1 | 0.16401 | 0.152133509 | 0.078 | 0 | 1 | 6 |
| fam122b | 0.16401 | 0.166577545 | 0.078 | 0 | 1 | 6 |
| fam210ab | 0.16401 | 0.153145045 | 0.078 | 0 | 1 | 6 |
| fdx1 | 0.16401 | 0.17949631 | 0.078 | 0 | 1 | 6 |
| flncb | 0.16401 | 0.172029374 | 0.078 | 0 | 1 | 6 |
| fosl2 | 0.16401 | 0.169312448 | 0.078 | 0 | 1 | 6 |
| foxo4 | 0.16401 | 0.138786228 | 0.078 | 0 | 1 | 6 |
| fzd1 | 0.16401 | 0.168116257 | 0.078 | 0 | 1 | 6 |
| ggnbp2 | 0.16401 | 0.183992222 | 0.078 | 0 | 1 | 6 |
| gigyf1a | 0.16401 | 0.157138939 | 0.078 | 0 | 1 | 6 |
| gnav1 | 0.16401 | 0.150320423 | 0.078 | 0 | 1 | 6 |
| grk4 | 0.16401 | 0.19058191 | 0.078 | 0 | 1 | 6 |
| hoxc11a | 0.16401 | 0.202176348 | 0.078 | 0 | 1 | 6 |
| hsbp1a | 0.16401 | 0.116142459 | 0.078 | 0 | 1 | 6 |
| ift74 | 0.16401 | 0.154868074 | 0.078 | 0 | 1 | 6 |
| igbp1 | 0.16401 | 0.151680526 | 0.078 | 0 | 1 | 6 |
| immp2l | 0.16401 | 0.16895302 | 0.078 | 0 | 1 | 6 |
| ing2 | 0.16401 | 0.170933321 | 0.078 | 0 | 1 | 6 |
| isl2a | 0.16401 | 0.178657996 | 0.078 | 0 | 1 | 6 |
| isy1 | 0.16401 | 0.153063065 | 0.078 | 0 | 1 | 6 |
| jam2b | 0.16401 | 0.146123632 | 0.078 | 0 | 1 | 6 |
| kirrel3l | 0.16401 | 0.166188934 | 0.078 | 0 | 1 | 6 |
| klf15 | 0.16401 | 0.162368802 | 0.078 | 0 | 1 | 6 |
| krr1 | 0.16401 | 0.1801229 | 0.078 | 0 | 1 | 6 |
| lats2 | 0.16401 | 0.136592397 | 0.078 | 0 | 1 | 6 |
| loxl3b | 0.16401 | 0.151713211 | 0.078 | 0 | 1 | 6 |
| lysmd3 | 0.16401 | 0.163663257 | 0.078 | 0 | 1 | 6 |
| m6pr | 0.16401 | 0.156019408 | 0.078 | 0 | 1 | 6 |
| mab21l2 | 0.16401 | 0.226367719 | 0.078 | 0 | 1 | 6 |
| macf1a | 0.16401 | 0.208000222 | 0.078 | 0 | 1 | 6 |
| maip1 | 0.16401 | 0.12726793 | 0.078 | 0 | 1 | 6 |

| | | | | | | |
|---|---|---|---|---|---|---|
| map2 | 0.16401 | 0.164016325 | 0.078 | 0 | 1 | 6 |
| mark3a | 0.16401 | 0.122968141 | 0.078 | 0 | 1 | 6 |
| mbnl3 | 0.16401 | 0.220368704 | 0.078 | 0 | 1 | 6 |
| mbtps1 | 0.16401 | 0.151163846 | 0.078 | 0 | 1 | 6 |
| mcrs1 | 0.16401 | 0.136081964 | 0.078 | 0 | 1 | 6 |
| me3 | 0.16401 | 0.14466532 | 0.078 | 0 | 1 | 6 |
| meis3 | 0.16401 | 0.136292759 | 0.078 | 0 | 1 | 6 |
| mkrn1 | 0.16401 | 0.161940653 | 0.078 | 0 | 1 | 6 |
| mlec | 0.16401 | 0.136428333 | 0.078 | 0 | 1 | 6 |
| mmp11b | 0.16401 | 0.160732229 | 0.078 | 0 | 1 | 6 |
| mta3 | 0.16401 | 0.165941398 | 0.078 | 0 | 1 | 6 |
| mtus1b | 0.16401 | 0.155275847 | 0.078 | 0 | 1 | 6 |
| myo18ab | 0.16401 | 0.175975624 | 0.078 | 0 | 1 | 6 |
| myo1cb | 0.16401 | 0.160399729 | 0.078 | 0 | 1 | 6 |
| nabp1a | 0.16401 | 0.152479406 | 0.078 | 0 | 1 | 6 |
| nap1l4b | 0.16401 | 0.140740542 | 0.078 | 0 | 1 | 6 |
| nfe2l2a | 0.16401 | 0.189476184 | 0.078 | 0 | 1 | 6 |
| nfe2l3 | 0.16401 | 0.151385505 | 0.078 | 0 | 1 | 6 |
| nmt2 | 0.16401 | 0.144408922 | 0.078 | 0 | 1 | 6 |
| nova2 | 0.16401 | 0.162550201 | 0.078 | 0 | 1 | 6 |
| npc1 | 0.16401 | 0.145222448 | 0.078 | 0 | 1 | 6 |
| nsd3 | 0.16401 | 0.196181502 | 0.078 | 0 | 1 | 6 |
| nup37 | 0.16401 | 0.151980044 | 0.078 | 0 | 1 | 6 |
| osbpl2a | 0.16401 | 0.161783133 | 0.078 | 0 | 1 | 6 |
| park7 | 0.16401 | 0.114285215 | 0.078 | 0 | 1 | 6 |
| pcmt | 0.16401 | 0.136313635 | 0.078 | 0 | 1 | 6 |
| phc2a | 0.16401 | 0.136008301 | 0.078 | 0 | 1 | 6 |
| phlda3 | 0.16401 | 0.151133969 | 0.078 | 0 | 1 | 6 |
| plat | 0.16401 | 0.156831046 | 0.078 | 0 | 1 | 6 |
| plecb | 0.16401 | 0.166844743 | 0.078 | 0 | 1 | 6 |
| plxna3 | 0.16401 | 0.167696266 | 0.078 | 0 | 1 | 6 |
| polr3glb | 0.16401 | 0.180132122 | 0.078 | 0 | 1 | 6 |
| pop7 | 0.16401 | 0.138023808 | 0.078 | 0 | 1 | 6 |
| pou2f1b | 0.16401 | 0.14415688 | 0.078 | 0 | 1 | 6 |
| ppm1da | 0.16401 | 0.168871116 | 0.078 | 0 | 1 | 6 |
| ptges3a | 0.16401 | 0.138031194 | 0.078 | 0 | 1 | 6 |
| ptgesl | 0.16401 | 0.127278998 | 0.078 | 0 | 1 | 6 |
| pwp2h | 0.16401 | 0.144756819 | 0.078 | 0 | 1 | 6 |
| qsox1 | 0.16401 | 0.156940951 | 0.078 | 0 | 1 | 6 |
| rab23 | 0.16401 | 0.145505297 | 0.078 | 0 | 1 | 6 |
| rab27a | 0.16401 | 0.188464744 | 0.078 | 0 | 1 | 6 |
| rae1 | 0.16401 | 0.189710337 | 0.078 | 0 | 1 | 6 |
| ranbp9 | 0.16401 | 0.154670148 | 0.078 | 0 | 1 | 6 |
| rassf2a | 0.16401 | 0.146348227 | 0.078 | 0 | 1 | 6 |
| rassf8a | 0.16401 | 0.147971088 | 0.078 | 0 | 1 | 6 |
| rbm12b | 0.16401 | 0.163773804 | 0.078 | 0 | 1 | 6 |
| rimkla | 0.16401 | 0.152729709 | 0.078 | 0 | 1 | 6 |
| riok2 | 0.16401 | 0.133846542 | 0.078 | 0 | 1 | 6 |

| | | | | | | |
|---|---|---|---|---|---|---|
| rnf11b | 0.16401 | 0.150987921 | 0.078 | 0 | 1 | 6 |
| rnf216 | 0.16401 | 0.137351381 | 0.078 | 0 | 1 | 6 |
| rpa1 | 0.16401 | 0.144434345 | 0.078 | 0 | 1 | 6 |
| rpp25l | 0.16401 | 0.151618845 | 0.078 | 0 | 1 | 6 |
| rpp38 | 0.16401 | 0.134200791 | 0.078 | 0 | 1 | 6 |
| rsrc2 | 0.16401 | 0.128167445 | 0.078 | 0 | 1 | 6 |
| sat2b | 0.16401 | 0.147593395 | 0.078 | 0 | 1 | 6 |
| serpinb1 | 0.16401 | 0.138132547 | 0.078 | 0 | 1 | 6 |
| si:ch211-160o17.4 | 0.16401 | 0.15880631 | 0.078 | 0 | 1 | 6 |
| si:ch211-161c3.5 | 0.16401 | 0.169933091 | 0.078 | 0 | 1 | 6 |
| si:ch211-195b15.8 | 0.16401 | 0.190764764 | 0.078 | 0 | 1 | 6 |
| si:ch211-195h23.3 | 0.16401 | 0.154247761 | 0.078 | 0 | 1 | 6 |
| si:ch211-206k20.5 | 0.16401 | 0.136968922 | 0.078 | 0 | 1 | 6 |
| si:ch211-71n6.4 | 0.16401 | 0.170042704 | 0.078 | 0 | 1 | 6 |
| si:dkey-12j5.1 | 0.16401 | 0.111110786 | 0.078 | 0 | 1 | 6 |
| slc16a5a | 0.16401 | 0.208897389 | 0.078 | 0 | 1 | 6 |
| slc44a1b | 0.16401 | 0.161075285 | 0.078 | 0 | 1 | 6 |
| smad5 | 0.16401 | 0.131754852 | 0.078 | 0 | 1 | 6 |
| srpra | 0.16401 | 0.114726845 | 0.078 | 0 | 1 | 6 |
| st7 | 0.16401 | 0.129948458 | 0.078 | 0 | 1 | 6 |
| stk17al | 0.16401 | 0.15017884 | 0.078 | 0 | 1 | 6 |
| styx | 0.16401 | 0.15535477 | 0.078 | 0 | 1 | 6 |
| tbc1d4 | 0.16401 | 0.184699064 | 0.078 | 0 | 1 | 6 |
| tcf7l2 | 0.16401 | 0.175294324 | 0.078 | 0 | 1 | 6 |
| tfcp2 | 0.16401 | 0.148453219 | 0.078 | 0 | 1 | 6 |
| tmed5 | 0.16401 | 0.15847274 | 0.078 | 0 | 1 | 6 |
| tmem251 | 0.16401 | 0.142155909 | 0.078 | 0 | 1 | 6 |
| triap1 | 0.16401 | 0.178339538 | 0.078 | 0 | 1 | 6 |
| trib2 | 0.16401 | 0.147661355 | 0.078 | 0 | 1 | 6 |
| tsr3 | 0.16401 | 0.142652565 | 0.078 | 0 | 1 | 6 |
| ttc26 | 0.16401 | 0.15934399 | 0.078 | 0 | 1 | 6 |
| ubr5 | 0.16401 | 0.129934353 | 0.078 | 0 | 1 | 6 |
| ubtfl | 0.16401 | 0.121963351 | 0.078 | 0 | 1 | 6 |
| utp6 | 0.16401 | 0.143310092 | 0.078 | 0 | 1 | 6 |
| vapal | 0.16401 | 0.161201937 | 0.078 | 0 | 1 | 6 |
| vclb | 0.16401 | 0.15803852 | 0.078 | 0 | 1 | 6 |
| vps26bl | 0.16401 | 0.161374059 | 0.078 | 0 | 1 | 6 |
| vrk1 | 0.16401 | 0.142959584 | 0.078 | 0 | 1 | 6 |
| wdr3 | 0.16401 | 0.148088151 | 0.078 | 0 | 1 | 6 |
| wnt7bb | 0.16401 | 0.148452876 | 0.078 | 0 | 1 | 6 |
| wscd2 | 0.16401 | 0.159294851 | 0.078 | 0 | 1 | 6 |
| ypel1 | 0.16401 | 0.159638794 | 0.078 | 0 | 1 | 6 |
| ythdf1 | 0.16401 | 0.141404179 | 0.078 | 0 | 1 | 6 |
| zc3h14 | 0.16401 | 0.128346379 | 0.078 | 0 | 1 | 6 |
| zgc:100829 | 0.16401 | 0.125822599 | 0.078 | 0 | 1 | 6 |
| zgc:100832 | 0.16401 | 0.158198559 | 0.078 | 0 | 1 | 6 |
| zgc:158640 | 0.16401 | 0.149402265 | 0.078 | 0 | 1 | 6 |
| zgc:92140 | 0.16401 | 0.150335027 | 0.078 | 0 | 1 | 6 |

| | | | | | | |
|---|---|---|---|---|---|---|
| zgc:92242 | 0.16401 | 0.154371388 | 0.078 | 0 | 1 | 6 |
| p3h3 | 0.16434 | -0.38899987 | 0.078 | 0.235 | 1 | 6 |
| si:dkey-85k7.7 | 0.16447 | -0.1606348 | 0.026 | 0.118 | 1 | 6 |
| ftsj3 | 0.1648 | -0.22096509 | 0.078 | 0.235 | 1 | 6 |
| tsr1 | 0.16491 | -0.34428015 | 0.039 | 0.176 | 1 | 6 |
| krt18a.1 | 0.16494 | 0.363474004 | 0.558 | 0.471 | 1 | 6 |
| tomm7 | 0.16516 | -0.32099714 | 0.26 | 0.412 | 1 | 6 |
| xbp1 | 0.16526 | 0.330121399 | 0.545 | 0.294 | 1 | 6 |
| pwwp2a | 0.16699 | -0.3506057 | 0.026 | 0.118 | 1 | 6 |
| ube2d2l | 0.1672 | -0.2624737 | 0.039 | 0.176 | 1 | 6 |
| manf | 0.16774 | -0.27548597 | 0.104 | 0.294 | 1 | 6 |
| cyp2n13 | 0.16775 | -0.36784951 | 0.104 | 0.294 | 1 | 6 |
| llph | 0.16794 | -0.15285046 | 0.078 | 0.235 | 1 | 6 |
| lgmn | 0.16909 | 0.177794597 | 0.195 | 0.059 | 1 | 6 |
| supt6h | 0.1694 | 0.171298771 | 0.182 | 0.059 | 1 | 6 |
| calm3b | 0.17007 | -0.35106652 | 0.377 | 0.529 | 1 | 6 |
| ier5l | 0.17056 | 0.228598236 | 0.195 | 0.059 | 1 | 6 |
| ube2d2 | 0.17093 | -0.35945471 | 0.286 | 0.471 | 1 | 6 |
| mef2cb | 0.17099 | -0.21272724 | 0.078 | 0.118 | 1 | 6 |
| gabarapl2 | 0.17216 | -0.15707408 | 0.234 | 0.235 | 1 | 6 |
| eif4e1c | 0.17273 | -0.25679968 | 0.039 | 0.176 | 1 | 6 |
| ube2q2 | 0.17295 | 0.259461238 | 0.169 | 0.059 | 1 | 6 |
| im:7152348 | 0.17532 | -0.2884629 | 0.026 | 0.118 | 1 | 6 |
| psmb2 | 0.17635 | -0.15959087 | 0.351 | 0.353 | 1 | 6 |
| zmpste24 | 0.17638 | -0.24353886 | 0.039 | 0.176 | 1 | 6 |
| phf8 | 0.17705 | -0.13121099 | 0.052 | 0.176 | 1 | 6 |
| ppm1g | 0.17737 | -0.26975109 | 0.143 | 0.235 | 1 | 6 |
| tuba8l2 | 0.17748 | -0.29616686 | 0.026 | 0.118 | 1 | 6 |
| zeb1b | 0.1779 | -0.27216138 | 0.039 | 0.176 | 1 | 6 |
| yaf2 | 0.17798 | 0.316888482 | 0.234 | 0.059 | 1 | 6 |
| rad54l | 0.17853 | -0.26261246 | 0.026 | 0.118 | 1 | 6 |
| sap18 | 0.17857 | 0.366654839 | 0.325 | 0.118 | 1 | 6 |
| ndufb6 | 0.17896 | -0.33932558 | 0.182 | 0.353 | 1 | 6 |
| tnfrsf19 | 0.17952 | 0.368086856 | 0.221 | 0.059 | 1 | 6 |
| elavl1a | 0.17956 | -0.31034382 | 0.078 | 0.235 | 1 | 6 |
| hells | 0.17969 | -0.40549019 | 0.104 | 0.235 | 1 | 6 |
| cyp27c1 | 0.18009 | 0.297650915 | 0.234 | 0.059 | 1 | 6 |
| coa1 | 0.18078 | -0.23925786 | 0.039 | 0.176 | 1 | 6 |
| si:ch73-1a9.3 | 0.18117 | 0.355232546 | 0.74 | 0.588 | 1 | 6 |
| tenm3 | 0.18127 | 0.393131614 | 0.481 | 0.294 | 1 | 6 |
| tprkb | 0.18182 | -0.22832451 | 0.039 | 0.176 | 1 | 6 |
| si:ch211-203d1.3 | 0.18187 | -0.35088999 | 0.026 | 0.118 | 1 | 6 |
| caskin1 | 0.18232 | -0.31867078 | 0.039 | 0.176 | 1 | 6 |
| prmt1 | 0.18237 | -0.34549098 | 0.273 | 0.412 | 1 | 6 |
| bysl | 0.1824 | -0.34252025 | 0.169 | 0.294 | 1 | 6 |
| si:ch211-114c12.2 | 0.18291 | -0.14358419 | 0.052 | 0.059 | 1 | 6 |
| cdca7b | 0.18292 | -0.25797384 | 0.039 | 0.176 | 1 | 6 |
| plxnb2a | 0.18297 | -0.29606943 | 0.039 | 0.176 | 1 | 6 |

| | | | | | | |
|---|---|---|---|---|---|---|
| rps25 | 0.18308 | 0.114526549 | 1 | 0.941 | 1 | 6 |
| ide | 0.18321 | -0.20984931 | 0.039 | 0.176 | 1 | 6 |
| mapre1b | 0.18334 | -0.27927097 | 0.169 | 0.294 | 1 | 6 |
| prps1b | 0.18336 | -0.29636046 | 0.039 | 0.176 | 1 | 6 |
| ndufaf3 | 0.1835 | -0.27901623 | 0.052 | 0.176 | 1 | 6 |
| atp5if1b | 0.18356 | -0.28566188 | 0.195 | 0.294 | 1 | 6 |
| spag9a | 0.1837 | -0.29857402 | 0.039 | 0.176 | 1 | 6 |
| ewsr1a | 0.18372 | 0.227769819 | 0.208 | 0.059 | 1 | 6 |
| clk2a | 0.18395 | -0.25123257 | 0.039 | 0.176 | 1 | 6 |
| cct6a | 0.18464 | 0.368050548 | 0.325 | 0.118 | 1 | 6 |
| eif4a3 | 0.18494 | -0.21483148 | 0.247 | 0.294 | 1 | 6 |
| si:ch211-214p13.9 | 0.1851 | -0.28992668 | 0.026 | 0.118 | 1 | 6 |
| mcm6 | 0.1851 | -0.24486453 | 0.117 | 0.176 | 1 | 6 |
| fam114a2 | 0.18563 | -0.32223163 | 0.052 | 0.176 | 1 | 6 |
| bhlhe40 | 0.18575 | 0.294432823 | 0.312 | 0.118 | 1 | 6 |
| si:ch211-196l7.4 | 0.18608 | 0.34481634 | 0.597 | 0.353 | 1 | 6 |
| naa10 | 0.18666 | 0.368355311 | 0.325 | 0.118 | 1 | 6 |
| hdac1 | 0.18671 | 0.110750844 | 0.377 | 0.235 | 1 | 6 |
| zgc:77849 | 0.18684 | 0.301068471 | 0.208 | 0.118 | 1 | 6 |
| h2afva | 0.1875 | 0.265121481 | 0.299 | 0.118 | 1 | 6 |
| rab7a | 0.18767 | -0.33261489 | 0.117 | 0.235 | 1 | 6 |
| sec24a | 0.18834 | -0.25687013 | 0.052 | 0.118 | 1 | 6 |
| adma | 0.189 | -0.14311303 | 0.052 | 0.059 | 1 | 6 |
| serf2 | 0.18916 | -0.30086105 | 0.662 | 0.824 | 1 | 6 |
| gsk3b | 0.1893 | -0.30244956 | 0.026 | 0.118 | 1 | 6 |
| uqcc2 | 0.18941 | -0.22093135 | 0.117 | 0.294 | 1 | 6 |
| eif6 | 0.18961 | -0.2809711 | 0.247 | 0.353 | 1 | 6 |
| pkig | 0.1918 | 0.333192445 | 0.234 | 0.059 | 1 | 6 |
| mrtfba | 0.19367 | -0.29168465 | 0.026 | 0.118 | 1 | 6 |
| mast2 | 0.19395 | -0.13768225 | 0.117 | 0.118 | 1 | 6 |
| atp2b2 | 0.19416 | 0.312920168 | 0.299 | 0.176 | 1 | 6 |
| BX284696.1 | 0.19425 | 0.369488743 | 0.234 | 0.059 | 1 | 6 |
| top1l | 0.19456 | 0.341718841 | 0.234 | 0.059 | 1 | 6 |
| nudcd3 | 0.19484 | -0.14615389 | 0.091 | 0.235 | 1 | 6 |
| ankrd49 | 0.19598 | -0.2787819 | 0.026 | 0.118 | 1 | 6 |
| smap1 | 0.1963 | -0.10251546 | 0.065 | 0.059 | 1 | 6 |
| dcn | 0.19659 | 0.244801643 | 0.792 | 0.706 | 1 | 6 |
| si:ch73-237c6.1 | 0.19659 | -0.33075229 | 0.052 | 0.176 | 1 | 6 |
| krt91 | 0.19703 | 0.370138418 | 0.208 | 0.059 | 1 | 6 |
| slc43a2a | 0.19754 | -0.33663189 | 0.117 | 0.294 | 1 | 6 |
| stx12 | 0.19756 | -0.29127465 | 0.078 | 0.176 | 1 | 6 |
| tshz3b | 0.1979 | -0.27418755 | 0.156 | 0.235 | 1 | 6 |
| si:ch211-262e15.1 | 0.19858 | 0.257237743 | 0.13 | 0.059 | 1 | 6 |
| prdx6 | 0.20033 | -0.32603059 | 0.208 | 0.353 | 1 | 6 |
| cad | 0.20161 | -0.28928451 | 0.026 | 0.118 | 1 | 6 |
| mif | 0.20193 | 0.368821725 | 0.442 | 0.235 | 1 | 6 |
| tomm22 | 0.20233 | 0.101235889 | 0.156 | 0.059 | 1 | 6 |
| rsl1d1 | 0.20294 | 0.343399588 | 0.312 | 0.118 | 1 | 6 |

| | | | | | | |
|---|---|---|---|---|---|---|
| purab | 0.20304 | 0.338293793 | 0.208 | 0.059 | 1 | 6 |
| smarcb1b | 0.20335 | 0.291422784 | 0.208 | 0.059 | 1 | 6 |
| s100u | 0.20419 | -0.26575955 | 0.117 | 0.294 | 1 | 6 |
| atp5pb | 0.20469 | 0.360678136 | 0.636 | 0.471 | 1 | 6 |
| snrpb2 | 0.20702 | -0.10515965 | 0.065 | 0.059 | 1 | 6 |
| apex1 | 0.2075 | -0.30316946 | 0.117 | 0.235 | 1 | 6 |
| hmgn3 | 0.20788 | -0.3780427 | 0.221 | 0.412 | 1 | 6 |
| ndfip1l | 0.20802 | 0.293366916 | 0.195 | 0.059 | 1 | 6 |
| fhl2a | 0.20862 | 0.193025476 | 0.143 | 0.118 | 1 | 6 |
| clic4 | 0.20969 | 0.144523275 | 0.247 | 0.118 | 1 | 6 |
| denr | 0.21035 | -0.28723193 | 0.195 | 0.294 | 1 | 6 |
| papola | 0.21079 | -0.13568362 | 0.052 | 0.059 | 1 | 6 |
| eif3c | 0.21086 | 0.368691086 | 0.351 | 0.176 | 1 | 6 |
| bnc2 | 0.21089 | 0.202209503 | 0.195 | 0.059 | 1 | 6 |
| ndufa7 | 0.21296 | -0.28084172 | 0.065 | 0.176 | 1 | 6 |
| gsdmeb | 0.213 | -0.24154465 | 0.052 | 0.118 | 1 | 6 |
| LOC103911404 | 0.21317 | 0.151049255 | 0.065 | 0 | 1 | 6 |
| ahi1 | 0.21317 | 0.153478718 | 0.065 | 0 | 1 | 6 |
| axin2 | 0.21317 | 0.125712505 | 0.065 | 0 | 1 | 6 |
| dusp1 | 0.21317 | 0.131947453 | 0.065 | 0 | 1 | 6 |
| gnal | 0.21317 | 0.123320418 | 0.065 | 0 | 1 | 6 |
| gorab | 0.21317 | 0.10725717 | 0.065 | 0 | 1 | 6 |
| gpc4 | 0.21317 | 0.162707777 | 0.065 | 0 | 1 | 6 |
| gpkow | 0.21317 | 0.14362627 | 0.065 | 0 | 1 | 6 |
| inhbab | 0.21317 | 0.134311624 | 0.065 | 0 | 1 | 6 |
| kpna4 | 0.21317 | 0.120057424 | 0.065 | 0 | 1 | 6 |
| rbmx2 | 0.21317 | 0.149930389 | 0.065 | 0 | 1 | 6 |
| reep3a | 0.21317 | 0.144166327 | 0.065 | 0 | 1 | 6 |
| rnf169 | 0.21317 | 0.124915719 | 0.065 | 0 | 1 | 6 |
| si:ch73-71c20.5 | 0.21317 | 0.162545416 | 0.065 | 0 | 1 | 6 |
| slc44a2 | 0.21317 | 0.138003845 | 0.065 | 0 | 1 | 6 |
| spag7 | 0.21317 | 0.115719127 | 0.065 | 0 | 1 | 6 |
| susd6 | 0.21317 | 0.110059084 | 0.065 | 0 | 1 | 6 |
| tcta | 0.21317 | 0.147563594 | 0.065 | 0 | 1 | 6 |
| tnfaip8l1 | 0.21317 | 0.113601484 | 0.065 | 0 | 1 | 6 |
| xpr1b | 0.21317 | 0.129384742 | 0.065 | 0 | 1 | 6 |
| zgc:162698 | 0.21317 | 0.119242808 | 0.065 | 0 | 1 | 6 |
| osgep | 0.21317 | 0.126773692 | 0.065 | 0 | 1 | 6 |
| si:dkey-151g10.3 | 0.21317 | 0.17637924 | 0.065 | 0 | 1 | 6 |
| tm2d1 | 0.21317 | 0.127315897 | 0.065 | 0 | 1 | 6 |
| tmx4 | 0.21317 | 0.178323865 | 0.065 | 0 | 1 | 6 |
| aspn | 0.21317 | 0.202736838 | 0.065 | 0 | 1 | 6 |
| efnb2b | 0.21317 | 0.152432595 | 0.065 | 0 | 1 | 6 |
| homer3b | 0.21317 | 0.125148377 | 0.065 | 0 | 1 | 6 |
| vkorc1 | 0.21317 | 0.144274954 | 0.065 | 0 | 1 | 6 |
| BX649300.2 | 0.21317 | 0.151145414 | 0.065 | 0 | 1 | 6 |
| CR751602.2 | 0.21317 | 0.129180214 | 0.065 | 0 | 1 | 6 |
| CT956002.2 | 0.21317 | 0.171803704 | 0.065 | 0 | 1 | 6 |

| | | | | | | |
|---|---|---|---|---|---|---|
| XLOC-005794 | 0.21317 | 0.140181447 | 0.065 | 0 | 1 | 6 |
| abhd4 | 0.21317 | 0.124137485 | 0.065 | 0 | 1 | 6 |
| acvr1l | 0.21317 | 0.12785967 | 0.065 | 0 | 1 | 6 |
| ankrd13c | 0.21317 | 0.121708119 | 0.065 | 0 | 1 | 6 |
| ankrd54 | 0.21317 | 0.104185355 | 0.065 | 0 | 1 | 6 |
| arhgef12b | 0.21317 | 0.155293151 | 0.065 | 0 | 1 | 6 |
| arl8ba | 0.21317 | 0.105909741 | 0.065 | 0 | 1 | 6 |
| bcas3 | 0.21317 | 0.125103016 | 0.065 | 0 | 1 | 6 |
| bmpr1ab | 0.21317 | 0.106069063 | 0.065 | 0 | 1 | 6 |
| ca8 | 0.21317 | 0.14328217 | 0.065 | 0 | 1 | 6 |
| cacna1c | 0.21317 | 0.163769831 | 0.065 | 0 | 1 | 6 |
| cd248b | 0.21317 | 0.196553684 | 0.065 | 0 | 1 | 6 |
| chd9 | 0.21317 | 0.115038966 | 0.065 | 0 | 1 | 6 |
| ches1 | 0.21317 | 0.12719336 | 0.065 | 0 | 1 | 6 |
| coil | 0.21317 | 0.132029292 | 0.065 | 0 | 1 | 6 |
| ctnnd1 | 0.21317 | 0.13181652 | 0.065 | 0 | 1 | 6 |
| ctsf | 0.21317 | 0.121151951 | 0.065 | 0 | 1 | 6 |
| cyp26a1 | 0.21317 | 0.155744968 | 0.065 | 0 | 1 | 6 |
| dek | 0.21317 | 0.123083974 | 0.065 | 0 | 1 | 6 |
| dnm3a | 0.21317 | 0.128477177 | 0.065 | 0 | 1 | 6 |
| fam84a | 0.21317 | 0.122606065 | 0.065 | 0 | 1 | 6 |
| fntb | 0.21317 | 0.151481485 | 0.065 | 0 | 1 | 6 |
| foxo1a | 0.21317 | 0.125378191 | 0.065 | 0 | 1 | 6 |
| fsta | 0.21317 | 0.154674878 | 0.065 | 0 | 1 | 6 |
| glod5 | 0.21317 | 0.109568812 | 0.065 | 0 | 1 | 6 |
| glrx2 | 0.21317 | 0.119755032 | 0.065 | 0 | 1 | 6 |
| gpr85 | 0.21317 | 0.133075562 | 0.065 | 0 | 1 | 6 |
| gtf2h3 | 0.21317 | 0.128906063 | 0.065 | 0 | 1 | 6 |
| hiat1b | 0.21317 | 0.140320043 | 0.065 | 0 | 1 | 6 |
| inpp5l | 0.21317 | 0.134436181 | 0.065 | 0 | 1 | 6 |
| klhl24b | 0.21317 | 0.125221112 | 0.065 | 0 | 1 | 6 |
| kxd1 | 0.21317 | 0.128742291 | 0.065 | 0 | 1 | 6 |
| lamtor5 | 0.21317 | 0.111583885 | 0.065 | 0 | 1 | 6 |
| lsp1 | 0.21317 | 0.122573204 | 0.065 | 0 | 1 | 6 |
| mcmbp | 0.21317 | 0.127329003 | 0.065 | 0 | 1 | 6 |
| mrpl48 | 0.21317 | 0.10030231 | 0.065 | 0 | 1 | 6 |
| mrps18b | 0.21317 | 0.104774952 | 0.065 | 0 | 1 | 6 |
| myoc | 0.21317 | 0.218887367 | 0.065 | 0 | 1 | 6 |
| ncdn | 0.21317 | 0.131028466 | 0.065 | 0 | 1 | 6 |
| nfil3 | 0.21317 | 0.134310672 | 0.065 | 0 | 1 | 6 |
| nfs1 | 0.21317 | 0.116123457 | 0.065 | 0 | 1 | 6 |
| nnt | 0.21317 | 0.136115094 | 0.065 | 0 | 1 | 6 |
| noc4l | 0.21317 | 0.128976954 | 0.065 | 0 | 1 | 6 |
| nsun5 | 0.21317 | 0.115440561 | 0.065 | 0 | 1 | 6 |
| pdk1 | 0.21317 | 0.156677721 | 0.065 | 0 | 1 | 6 |
| per3 | 0.21317 | 0.151552278 | 0.065 | 0 | 1 | 6 |
| ppp4ca | 0.21317 | 0.129454278 | 0.065 | 0 | 1 | 6 |
| pqbp1 | 0.21317 | 0.122255314 | 0.065 | 0 | 1 | 6 |

| | | | | | | |
|---|---|---|---|---|---|---|
| pum1 | 0.21317 | 0.151787537 | 0.065 | 0 | 1 | 6 |
| rab9b | 0.21317 | 0.116391212 | 0.065 | 0 | 1 | 6 |
| rasgef1bb | 0.21317 | 0.160198295 | 0.065 | 0 | 1 | 6 |
| rbm14b | 0.21317 | 0.129832289 | 0.065 | 0 | 1 | 6 |
| rc3h1b | 0.21317 | 0.110009622 | 0.065 | 0 | 1 | 6 |
| rgmd | 0.21317 | 0.117141355 | 0.065 | 0 | 1 | 6 |
| rorab | 0.21317 | 0.174183406 | 0.065 | 0 | 1 | 6 |
| sash1a | 0.21317 | 0.118902172 | 0.065 | 0 | 1 | 6 |
| scinla | 0.21317 | 0.144234721 | 0.065 | 0 | 1 | 6 |
| sfrp2 | 0.21317 | 0.219887445 | 0.065 | 0 | 1 | 6 |
| si:ch73-311h14.2 | 0.21317 | 0.125198074 | 0.065 | 0 | 1 | 6 |
| si:ch73-340n8.4 | 0.21317 | 0.187296627 | 0.065 | 0 | 1 | 6 |
| six1a | 0.21317 | 0.172709075 | 0.065 | 0 | 1 | 6 |
| sntb1 | 0.21317 | 0.13147288 | 0.065 | 0 | 1 | 6 |
| stag1b | 0.21317 | 0.123836727 | 0.065 | 0 | 1 | 6 |
| sv2 | 0.21317 | 0.1108168 | 0.065 | 0 | 1 | 6 |
| tdrd3 | 0.21317 | 0.122329467 | 0.065 | 0 | 1 | 6 |
| tead3b | 0.21317 | 0.161318732 | 0.065 | 0 | 1 | 6 |
| tmem203 | 0.21317 | 0.114398908 | 0.065 | 0 | 1 | 6 |
| tns2b | 0.21317 | 0.130575655 | 0.065 | 0 | 1 | 6 |
| ube2d4 | 0.21317 | 0.12449731 | 0.065 | 0 | 1 | 6 |
| ubr4 | 0.21317 | 0.138124106 | 0.065 | 0 | 1 | 6 |
| zeb1a | 0.21317 | 0.158452945 | 0.065 | 0 | 1 | 6 |
| zgc:77056 | 0.21317 | 0.116768061 | 0.065 | 0 | 1 | 6 |
| zgc:77929 | 0.21317 | 0.142848872 | 0.065 | 0 | 1 | 6 |
| zgc:85858 | 0.21317 | 0.134201898 | 0.065 | 0 | 1 | 6 |
| AL935174.5 | 0.21317 | 0.133219034 | 0.065 | 0 | 1 | 6 |
| ANKRD50 | 0.21317 | 0.149518583 | 0.065 | 0 | 1 | 6 |
| CABZ01055347.1 | 0.21317 | 0.117093806 | 0.065 | 0 | 1 | 6 |
| FAM53C | 0.21317 | 0.144223499 | 0.065 | 0 | 1 | 6 |
| LO018340.1 | 0.21317 | 0.149603882 | 0.065 | 0 | 1 | 6 |
| LOC405768 | 0.21317 | 0.107983463 | 0.065 | 0 | 1 | 6 |
| NFX1 | 0.21317 | 0.10427374 | 0.065 | 0 | 1 | 6 |
| abi2a | 0.21317 | 0.13060382 | 0.065 | 0 | 1 | 6 |
| aco1 | 0.21317 | 0.138030492 | 0.065 | 0 | 1 | 6 |
| adgrb1a | 0.21317 | 0.16629387 | 0.065 | 0 | 1 | 6 |
| ak3 | 0.21317 | 0.102535265 | 0.065 | 0 | 1 | 6 |
| alcama | 0.21317 | 0.167179197 | 0.065 | 0 | 1 | 6 |
| ankhd1 | 0.21317 | 0.139849225 | 0.065 | 0 | 1 | 6 |
| aph1b | 0.21317 | 0.127840349 | 0.065 | 0 | 1 | 6 |
| arl2 | 0.21317 | 0.111538671 | 0.065 | 0 | 1 | 6 |
| arl3 | 0.21317 | 0.135313977 | 0.065 | 0 | 1 | 6 |
| arpp19a | 0.21317 | 0.148343125 | 0.065 | 0 | 1 | 6 |
| bag5 | 0.21317 | 0.148193656 | 0.065 | 0 | 1 | 6 |
| cactin | 0.21317 | 0.165626309 | 0.065 | 0 | 1 | 6 |
| calb2a | 0.21317 | 0.175635318 | 0.065 | 0 | 1 | 6 |
| ccdc102a | 0.21317 | 0.176712516 | 0.065 | 0 | 1 | 6 |
| cdk5rap3 | 0.21317 | 0.13129788 | 0.065 | 0 | 1 | 6 |

| | | | | | | |
|---|---|---|---|---|---|---|
| cep290 | 0.21317 | 0.153846037 | 0.065 | 0 | 1 | 6 |
| chmp4ba | 0.21317 | 0.115547281 | 0.065 | 0 | 1 | 6 |
| cited4b | 0.21317 | 0.115308547 | 0.065 | 0 | 1 | 6 |
| ckap5 | 0.21317 | 0.119771711 | 0.065 | 0 | 1 | 6 |
| coq3 | 0.21317 | 0.123563683 | 0.065 | 0 | 1 | 6 |
| coro1cb | 0.21317 | 0.114888267 | 0.065 | 0 | 1 | 6 |
| crabp2b | 0.21317 | 0.104652539 | 0.065 | 0 | 1 | 6 |
| cyp27a7 | 0.21317 | 0.1072934 | 0.065 | 0 | 1 | 6 |
| dnajc11a | 0.21317 | 0.106062541 | 0.065 | 0 | 1 | 6 |
| dnmt3ba | 0.21317 | 0.110555503 | 0.065 | 0 | 1 | 6 |
| dok4 | 0.21317 | 0.14357554 | 0.065 | 0 | 1 | 6 |
| dync1li2 | 0.21317 | 0.128177935 | 0.065 | 0 | 1 | 6 |
| edem1 | 0.21317 | 0.119113803 | 0.065 | 0 | 1 | 6 |
| eif2b1 | 0.21317 | 0.111488261 | 0.065 | 0 | 1 | 6 |
| exosc7 | 0.21317 | 0.120111999 | 0.065 | 0 | 1 | 6 |
| fam171a1 | 0.21317 | 0.151625673 | 0.065 | 0 | 1 | 6 |
| fam198b | 0.21317 | 0.133514289 | 0.065 | 0 | 1 | 6 |
| farsa | 0.21317 | 0.115036322 | 0.065 | 0 | 1 | 6 |
| fhl3a | 0.21317 | 0.177341001 | 0.065 | 0 | 1 | 6 |
| fscn1a | 0.21317 | 0.11949118 | 0.065 | 0 | 1 | 6 |
| fzr1a | 0.21317 | 0.144077813 | 0.065 | 0 | 1 | 6 |
| g3bp2 | 0.21317 | 0.138741754 | 0.065 | 0 | 1 | 6 |
| gas1b | 0.21317 | 0.152301679 | 0.065 | 0 | 1 | 6 |
| gdf7 | 0.21317 | 0.141088432 | 0.065 | 0 | 1 | 6 |
| glmp | 0.21317 | 0.109167187 | 0.065 | 0 | 1 | 6 |
| gpn1 | 0.21317 | 0.117855265 | 0.065 | 0 | 1 | 6 |
| gse1 | 0.21317 | 0.124837405 | 0.065 | 0 | 1 | 6 |
| gusb | 0.21317 | 0.125971364 | 0.065 | 0 | 1 | 6 |
| hbs1l | 0.21317 | 0.10875728 | 0.065 | 0 | 1 | 6 |
| hdac4 | 0.21317 | 0.129747106 | 0.065 | 0 | 1 | 6 |
| hiat1a | 0.21317 | 0.114153962 | 0.065 | 0 | 1 | 6 |
| hmg20a | 0.21317 | 0.128980079 | 0.065 | 0 | 1 | 6 |
| hoxa9a | 0.21317 | 0.127055923 | 0.065 | 0 | 1 | 6 |
| htatip2 | 0.21317 | 0.105399542 | 0.065 | 0 | 1 | 6 |
| htatsf1 | 0.21317 | 0.12843661 | 0.065 | 0 | 1 | 6 |
| jam3b | 0.21317 | 0.150392803 | 0.065 | 0 | 1 | 6 |
| kcnq5a | 0.21317 | 0.154087006 | 0.065 | 0 | 1 | 6 |
| kctd10 | 0.21317 | 0.117576391 | 0.065 | 0 | 1 | 6 |
| kdm3b | 0.21317 | 0.114639641 | 0.065 | 0 | 1 | 6 |
| kif1bp | 0.21317 | 0.129072052 | 0.065 | 0 | 1 | 6 |
| kmt2e | 0.21317 | 0.111392192 | 0.065 | 0 | 1 | 6 |
| kremen1 | 0.21317 | 0.140579427 | 0.065 | 0 | 1 | 6 |
| lamb1b | 0.21317 | 0.135972151 | 0.065 | 0 | 1 | 6 |
| lhfpl6 | 0.21317 | 0.114334218 | 0.065 | 0 | 1 | 6 |
| lin52 | 0.21317 | 0.130287736 | 0.065 | 0 | 1 | 6 |
| maf1 | 0.21317 | 0.119628363 | 0.065 | 0 | 1 | 6 |
| mapkapk5 | 0.21317 | 0.143198163 | 0.065 | 0 | 1 | 6 |
| march6 | 0.21317 | 0.114010272 | 0.065 | 0 | 1 | 6 |

| | | | | | | |
|---|---|---|---|---|---|---|
| mboat2a | 0.21317 | 0.135391281 | 0.065 | 0 | 1 | 6 |
| mcm4 | 0.21317 | 0.102933832 | 0.065 | 0 | 1 | 6 |
| med15 | 0.21317 | 0.133519611 | 0.065 | 0 | 1 | 6 |
| med4 | 0.21317 | 0.149655217 | 0.065 | 0 | 1 | 6 |
| med8 | 0.21317 | 0.129924696 | 0.065 | 0 | 1 | 6 |
| mettl9 | 0.21317 | 0.149238882 | 0.065 | 0 | 1 | 6 |
| mmp11a | 0.21317 | 0.188025775 | 0.065 | 0 | 1 | 6 |
| mrps28 | 0.21317 | 0.130358464 | 0.065 | 0 | 1 | 6 |
| mtss1la | 0.21317 | 0.137502385 | 0.065 | 0 | 1 | 6 |
| myca | 0.21317 | 0.120236643 | 0.065 | 0 | 1 | 6 |
| nadkb | 0.21317 | 0.132725999 | 0.065 | 0 | 1 | 6 |
| nat10 | 0.21317 | 0.115266213 | 0.065 | 0 | 1 | 6 |
| nbr1b | 0.21317 | 0.136324534 | 0.065 | 0 | 1 | 6 |
| ncbp3 | 0.21317 | 0.106745734 | 0.065 | 0 | 1 | 6 |
| ncoa4 | 0.21317 | 0.128183258 | 0.065 | 0 | 1 | 6 |
| npl | 0.21317 | 0.132393239 | 0.065 | 0 | 1 | 6 |
| nsd2 | 0.21317 | 0.113449835 | 0.065 | 0 | 1 | 6 |
| nudt12 | 0.21317 | 0.13671099 | 0.065 | 0 | 1 | 6 |
| nupr1b | 0.21317 | 0.194995805 | 0.065 | 0 | 1 | 6 |
| ormdl2 | 0.21317 | 0.142820038 | 0.065 | 0 | 1 | 6 |
| pcmtl | 0.21317 | 0.136328379 | 0.065 | 0 | 1 | 6 |
| pdcd6ip | 0.21317 | 0.127573952 | 0.065 | 0 | 1 | 6 |
| per1a | 0.21317 | 0.117368746 | 0.065 | 0 | 1 | 6 |
| pex3 | 0.21317 | 0.159663214 | 0.065 | 0 | 1 | 6 |
| pgap2 | 0.21317 | 0.12932799 | 0.065 | 0 | 1 | 6 |
| phf21aa | 0.21317 | 0.152416984 | 0.065 | 0 | 1 | 6 |
| picalmb | 0.21317 | 0.1247272 | 0.065 | 0 | 1 | 6 |
| ppie | 0.21317 | 0.105948865 | 0.065 | 0 | 1 | 6 |
| prr12a | 0.21317 | 0.148630312 | 0.065 | 0 | 1 | 6 |
| prtfdc1 | 0.21317 | 0.112755311 | 0.065 | 0 | 1 | 6 |
| ptpn11a | 0.21317 | 0.107365311 | 0.065 | 0 | 1 | 6 |
| rab30 | 0.21317 | 0.139046267 | 0.065 | 0 | 1 | 6 |
| rab3db | 0.21317 | 0.150942904 | 0.065 | 0 | 1 | 6 |
| rasa1a | 0.21317 | 0.153750115 | 0.065 | 0 | 1 | 6 |
| rgs19 | 0.21317 | 0.164608695 | 0.065 | 0 | 1 | 6 |
| ror2 | 0.21317 | 0.140476252 | 0.065 | 0 | 1 | 6 |
| rxfp2a | 0.21317 | 0.133541481 | 0.065 | 0 | 1 | 6 |
| sbf2 | 0.21317 | 0.141234886 | 0.065 | 0 | 1 | 6 |
| sc:d0202 | 0.21317 | 0.174500279 | 0.065 | 0 | 1 | 6 |
| scaf11 | 0.21317 | 0.135926584 | 0.065 | 0 | 1 | 6 |
| scara5 | 0.21317 | 0.149789714 | 0.065 | 0 | 1 | 6 |
| sept9b | 0.21317 | 0.108420243 | 0.065 | 0 | 1 | 6 |
| serpinb14 | 0.21317 | 0.130680829 | 0.065 | 0 | 1 | 6 |
| si:ch211-216l23.2 | 0.21317 | 0.128416457 | 0.065 | 0 | 1 | 6 |
| si:ch211-256e16.11 | 0.21317 | 0.154067344 | 0.065 | 0 | 1 | 6 |
| si:ch211-59d15.9 | 0.21317 | 0.122940497 | 0.065 | 0 | 1 | 6 |
| si:ch73-257c13.2 | 0.21317 | 0.114402637 | 0.065 | 0 | 1 | 6 |
| si:ch73-390p7.2 | 0.21317 | 0.149868004 | 0.065 | 0 | 1 | 6 |

| | | | | | | |
|---|---|---|---|---|---|---|
| si:dkey-57k2.7 | 0.21317 | 0.128479999 | 0.065 | 0 | 1 | 6 |
| si:dkeyp-66d1.7 | 0.21317 | 0.114204051 | 0.065 | 0 | 1 | 6 |
| sipa1l2 | 0.21317 | 0.155897684 | 0.065 | 0 | 1 | 6 |
| slc37a2 | 0.21317 | 0.12930074 | 0.065 | 0 | 1 | 6 |
| slc39a9 | 0.21317 | 0.109277766 | 0.065 | 0 | 1 | 6 |
| snap23.1 | 0.21317 | 0.131780933 | 0.065 | 0 | 1 | 6 |
| snx27b | 0.21317 | 0.105334464 | 0.065 | 0 | 1 | 6 |
| spoplb | 0.21317 | 0.133688672 | 0.065 | 0 | 1 | 6 |
| srsf2b | 0.21317 | 0.126204789 | 0.065 | 0 | 1 | 6 |
| strada | 0.21317 | 0.104591751 | 0.065 | 0 | 1 | 6 |
| tbrg1 | 0.21317 | 0.122370724 | 0.065 | 0 | 1 | 6 |
| tcf20 | 0.21317 | 0.126818901 | 0.065 | 0 | 1 | 6 |
| tm2d2 | 0.21317 | 0.151803352 | 0.065 | 0 | 1 | 6 |
| tmtc1 | 0.21317 | 0.152950474 | 0.065 | 0 | 1 | 6 |
| tmtc4 | 0.21317 | 0.164021277 | 0.065 | 0 | 1 | 6 |
| tnfaip8l3 | 0.21317 | 0.151957083 | 0.065 | 0 | 1 | 6 |
| tubg1 | 0.21317 | 0.106210039 | 0.065 | 0 | 1 | 6 |
| tusc3 | 0.21317 | 0.133602628 | 0.065 | 0 | 1 | 6 |
| vhll | 0.21317 | 0.164428156 | 0.065 | 0 | 1 | 6 |
| vps26c | 0.21317 | 0.158285583 | 0.065 | 0 | 1 | 6 |
| wdr5 | 0.21317 | 0.12560044 | 0.065 | 0 | 1 | 6 |
| wu:fa19b12 | 0.21317 | 0.10391647 | 0.065 | 0 | 1 | 6 |
| zbtb2b | 0.21317 | 0.110958929 | 0.065 | 0 | 1 | 6 |
| zbtb33 | 0.21317 | 0.134286782 | 0.065 | 0 | 1 | 6 |
| zgc:162339 | 0.21317 | 0.156165001 | 0.065 | 0 | 1 | 6 |
| zgc:174263 | 0.21317 | 0.122456991 | 0.065 | 0 | 1 | 6 |
| zgc:55558 | 0.21317 | 0.135674288 | 0.065 | 0 | 1 | 6 |
| zgc:66448 | 0.21317 | 0.130834982 | 0.065 | 0 | 1 | 6 |
| znf1124 | 0.21317 | 0.11229121 | 0.065 | 0 | 1 | 6 |
| znf423 | 0.21317 | 0.128845986 | 0.065 | 0 | 1 | 6 |
| znf451 | 0.21317 | 0.148379215 | 0.065 | 0 | 1 | 6 |
| BX323087.1 | 0.21317 | 0.160692438 | 0.065 | 0 | 1 | 6 |
| CU929230.1 | 0.21317 | 0.162787767 | 0.065 | 0 | 1 | 6 |
| LO018102.1 | 0.21317 | 0.154401602 | 0.065 | 0 | 1 | 6 |
| aftpha | 0.21317 | 0.115407198 | 0.065 | 0 | 1 | 6 |
| ago2 | 0.21317 | 0.138106343 | 0.065 | 0 | 1 | 6 |
| bmpr2a | 0.21317 | 0.159635362 | 0.065 | 0 | 1 | 6 |
| cav2 | 0.21317 | 0.155779015 | 0.065 | 0 | 1 | 6 |
| dcaf13 | 0.21317 | 0.122082437 | 0.065 | 0 | 1 | 6 |
| dph3 | 0.21317 | 0.111325646 | 0.065 | 0 | 1 | 6 |
| ehd1a | 0.21317 | 0.103148318 | 0.065 | 0 | 1 | 6 |
| foxk1 | 0.21317 | 0.134252704 | 0.065 | 0 | 1 | 6 |
| ftsj1 | 0.21317 | 0.107575924 | 0.065 | 0 | 1 | 6 |
| gpd1l | 0.21317 | 0.153301362 | 0.065 | 0 | 1 | 6 |
| ikbkg | 0.21317 | 0.114760377 | 0.065 | 0 | 1 | 6 |
| irf2bp1 | 0.21317 | 0.150848909 | 0.065 | 0 | 1 | 6 |
| jmjd7 | 0.21317 | 0.126994896 | 0.065 | 0 | 1 | 6 |
| kdelc2 | 0.21317 | 0.119622066 | 0.065 | 0 | 1 | 6 |

| | | | | | | |
|---|---|---|---|---|---|---|
| luzp1 | 0.21317 | 0.115654155 | 0.065 | 0 | 1 | 6 |
| mettl7a | 0.21317 | 0.107748102 | 0.065 | 0 | 1 | 6 |
| mink1 | 0.21317 | 0.14928768 | 0.065 | 0 | 1 | 6 |
| mmadhc | 0.21317 | 0.129426496 | 0.065 | 0 | 1 | 6 |
| mrps31 | 0.21317 | 0.104088643 | 0.065 | 0 | 1 | 6 |
| myd88 | 0.21317 | 0.12495018 | 0.065 | 0 | 1 | 6 |
| pbrm1l | 0.21317 | 0.107546097 | 0.065 | 0 | 1 | 6 |
| pdik1l | 0.21317 | 0.147107676 | 0.065 | 0 | 1 | 6 |
| phactr2 | 0.21317 | 0.131965565 | 0.065 | 0 | 1 | 6 |
| plrg1 | 0.21317 | 0.106419897 | 0.065 | 0 | 1 | 6 |
| ppt2 | 0.21317 | 0.119007024 | 0.065 | 0 | 1 | 6 |
| ptk2ab | 0.21317 | 0.144326644 | 0.065 | 0 | 1 | 6 |
| sall1a | 0.21317 | 0.135226386 | 0.065 | 0 | 1 | 6 |
| scfd1 | 0.21317 | 0.113715577 | 0.065 | 0 | 1 | 6 |
| serpinh2 | 0.21317 | 0.138187091 | 0.065 | 0 | 1 | 6 |
| setd7 | 0.21317 | 0.111674349 | 0.065 | 0 | 1 | 6 |
| sfswap | 0.21317 | 0.116209423 | 0.065 | 0 | 1 | 6 |
| si:ch1073-322p19.1 | 0.21317 | 0.137322378 | 0.065 | 0 | 1 | 6 |
| si:ch1073-357b18.4 | 0.21317 | 0.129277211 | 0.065 | 0 | 1 | 6 |
| si:ch211-212g7.6 | 0.21317 | 0.128044704 | 0.065 | 0 | 1 | 6 |
| sigmar1 | 0.21317 | 0.126020439 | 0.065 | 0 | 1 | 6 |
| sos2 | 0.21317 | 0.117281013 | 0.065 | 0 | 1 | 6 |
| stx3a | 0.21317 | 0.127107164 | 0.065 | 0 | 1 | 6 |
| tipin | 0.21317 | 0.13102269 | 0.065 | 0 | 1 | 6 |
| top1 | 0.21317 | 0.142321326 | 0.065 | 0 | 1 | 6 |
| trdn | 0.21317 | 0.162531751 | 0.065 | 0 | 1 | 6 |
| zgc:113149 | 0.21317 | 0.128106533 | 0.065 | 0 | 1 | 6 |
| ikzf5 | 0.21317 | 0.14273526 | 0.065 | 0 | 1 | 6 |
| si:dkey-48p11.3 | 0.21317 | 0.120161178 | 0.065 | 0 | 1 | 6 |
| stk25b | 0.21317 | 0.126149956 | 0.065 | 0 | 1 | 6 |
| metap1 | 0.21333 | -0.33209541 | 0.117 | 0.235 | 1 | 6 |
| slit3 | 0.21396 | 0.218447859 | 0.195 | 0.059 | 1 | 6 |
| nop53 | 0.21573 | 0.364548829 | 0.519 | 0.294 | 1 | 6 |
| ifi46 | 0.21577 | -0.13561908 | 0.052 | 0.059 | 1 | 6 |
| si:dkey-28b4.7 | 0.21623 | -0.21174085 | 0.364 | 0.412 | 1 | 6 |
| LOC103909972 | 0.21678 | -0.12170218 | 0.117 | 0.118 | 1 | 6 |
| vdac1 | 0.21684 | 0.338430498 | 0.351 | 0.176 | 1 | 6 |
| surf2 | 0.21697 | -0.11038069 | 0.104 | 0.235 | 1 | 6 |
| cox7a2a | 0.21718 | -0.18650628 | 0.208 | 0.235 | 1 | 6 |
| gstt1a | 0.21727 | -0.2749381 | 0.117 | 0.176 | 1 | 6 |
| cpox | 0.21742 | -0.18692348 | 0.091 | 0.118 | 1 | 6 |
| il10rb | 0.21753 | -0.27268198 | 0.052 | 0.176 | 1 | 6 |
| midn | 0.21762 | 0.128538591 | 0.221 | 0.235 | 1 | 6 |
| notch2 | 0.21789 | 0.309274368 | 0.156 | 0.059 | 1 | 6 |
| phax | 0.21822 | -0.28814655 | 0.078 | 0.235 | 1 | 6 |
| anp32e | 0.21844 | -0.10114079 | 0.13 | 0.118 | 1 | 6 |
| mtpn | 0.21861 | -0.29756843 | 0.247 | 0.353 | 1 | 6 |
| ccng2 | 0.2191 | 0.254330741 | 0.169 | 0.059 | 1 | 6 |

| | | | | | | |
|---|---|---|---|---|---|---|
| FP102783.1 | 0.21914 | -0.33766247 | 0.078 | 0.235 | 1 | 6 |
| cdca7a | 0.21994 | -0.26089839 | 0.026 | 0.118 | 1 | 6 |
| brox | 0.22014 | -0.28359078 | 0.026 | 0.118 | 1 | 6 |
| znf318 | 0.22051 | -0.30395027 | 0.052 | 0.176 | 1 | 6 |
| arhgef1b | 0.22061 | 0.385169189 | 0.286 | 0.118 | 1 | 6 |
| LOC100333474 | 0.22161 | -0.28546227 | 0.026 | 0.118 | 1 | 6 |
| pdia3 | 0.22201 | -0.24321772 | 0.779 | 0.882 | 1 | 6 |
| ms4a17a.9 | 0.22211 | 0.354717531 | 0.325 | 0.176 | 1 | 6 |
| mgst1.1 | 0.22221 | 0.160059223 | 0.247 | 0.118 | 1 | 6 |
| cpne1 | 0.22229 | -0.31230246 | 0.182 | 0.294 | 1 | 6 |
| gtf2a2 | 0.22326 | -0.2474252 | 0.078 | 0.235 | 1 | 6 |
| arglu1a | 0.22476 | 0.288366196 | 0.234 | 0.118 | 1 | 6 |
| selenou1a | 0.225 | -0.2716187 | 0.052 | 0.176 | 1 | 6 |
| lig1 | 0.22535 | -0.28103639 | 0.143 | 0.235 | 1 | 6 |
| scube1 | 0.22559 | -0.12269344 | 0.026 | 0.118 | 1 | 6 |
| adam10a | 0.22576 | -0.3387762 | 0.117 | 0.294 | 1 | 6 |
| lrp10 | 0.22618 | -0.30054967 | 0.052 | 0.176 | 1 | 6 |
| syncrip | 0.22669 | -0.11336954 | 0.545 | 0.529 | 1 | 6 |
| colec12 | 0.2268 | -0.30064998 | 0.234 | 0.353 | 1 | 6 |
| rn7sk | 0.22697 | 0.230694557 | 0.195 | 0.059 | 1 | 6 |
| exosc5 | 0.22749 | -0.23563315 | 0.052 | 0.176 | 1 | 6 |
| cap1 | 0.22842 | -0.2823157 | 0.247 | 0.353 | 1 | 6 |
| psap | 0.22854 | 0.271928977 | 0.273 | 0.176 | 1 | 6 |
| adamts3 | 0.22914 | -0.13862439 | 0.026 | 0.118 | 1 | 6 |
| hdac9b | 0.22945 | -0.13442927 | 0.026 | 0.118 | 1 | 6 |
| tagln | 0.22949 | 0.405472948 | 0.636 | 0.412 | 1 | 6 |
| si:ch211-288g17.3 | 0.22979 | 0.310734539 | 0.519 | 0.294 | 1 | 6 |
| cd99l2 | 0.22987 | 0.322084749 | 0.221 | 0.059 | 1 | 6 |
| myg1 | 0.23 | 0.274675969 | 0.208 | 0.059 | 1 | 6 |
| cdh2 | 0.23004 | -0.11846932 | 0.325 | 0.294 | 1 | 6 |
| gli3 | 0.23053 | -0.32987401 | 0.091 | 0.235 | 1 | 6 |
| tmem165 | 0.23203 | -0.15760889 | 0.052 | 0.176 | 1 | 6 |
| LO018188.1 | 0.23207 | -0.29505357 | 0.312 | 0.529 | 1 | 6 |
| hgs | 0.23225 | 0.311904917 | 0.195 | 0.059 | 1 | 6 |
| spred2b | 0.23231 | 0.217319453 | 0.338 | 0.176 | 1 | 6 |
| ogt.1 | 0.23243 | 0.160543828 | 0.169 | 0.059 | 1 | 6 |
| sart1 | 0.23377 | 0.302652588 | 0.208 | 0.059 | 1 | 6 |
| tubb5 | 0.23406 | 0.326053949 | 0.195 | 0.059 | 1 | 6 |
| akt1s1 | 0.23442 | 0.2080748 | 0.182 | 0.118 | 1 | 6 |
| CU104709.1 | 0.23443 | -0.28526093 | 0.091 | 0.176 | 1 | 6 |
| CDPF1 | 0.23573 | -0.16563845 | 0.104 | 0.118 | 1 | 6 |
| sept9a | 0.23598 | 0.25536282 | 0.182 | 0.059 | 1 | 6 |
| mylka | 0.23638 | -0.30787449 | 0.078 | 0.176 | 1 | 6 |
| tsen34 | 0.23771 | -0.2689633 | 0.026 | 0.118 | 1 | 6 |
| LOC103910438 | 0.23774 | 0.228698975 | 0.247 | 0.176 | 1 | 6 |
| khdrbs1b | 0.23816 | 0.275235531 | 0.221 | 0.059 | 1 | 6 |
| prkab1b | 0.2393 | 0.308398548 | 0.286 | 0.176 | 1 | 6 |
| fermt2 | 0.23968 | -0.28463989 | 0.104 | 0.176 | 1 | 6 |

| | | | | | | |
|---|---|---|---|---|---|---|
| hm13 | 0.23971 | -0.31566014 | 0.13 | 0.294 | 1 | 6 |
| rnf8 | 0.23974 | -0.25515651 | 0.026 | 0.118 | 1 | 6 |
| pdap1b | 0.24057 | -0.13308834 | 0.052 | 0.176 | 1 | 6 |
| pole4 | 0.24183 | -0.25946009 | 0.182 | 0.294 | 1 | 6 |
| nme2b.1 | 0.24201 | -0.1430925 | 0.896 | 0.941 | 1 | 6 |
| LOC108190726 | 0.24238 | -0.29999405 | 0.052 | 0.176 | 1 | 6 |
| dlst | 0.24269 | -0.34057603 | 0.208 | 0.353 | 1 | 6 |
| srp9 | 0.24306 | -0.11723922 | 0.182 | 0.176 | 1 | 6 |
| gtf2f2a | 0.2435 | 0.259274294 | 0.169 | 0.059 | 1 | 6 |
| faub | 0.24355 | 0.164588852 | 0.117 | 0.059 | 1 | 6 |
| aldh2.2 | 0.24358 | -0.28467595 | 0.104 | 0.235 | 1 | 6 |
| trnt1 | 0.24358 | -0.13190775 | 0.052 | 0.059 | 1 | 6 |
| tuba8l | 0.24363 | -0.34910536 | 0.364 | 0.529 | 1 | 6 |
| FAT4 | 0.24412 | 0.145139366 | 0.169 | 0.059 | 1 | 6 |
| etv4 | 0.24461 | -0.22216924 | 0.052 | 0.176 | 1 | 6 |
| skia | 0.24502 | -0.25637634 | 0.052 | 0.176 | 1 | 6 |
| rpz5 | 0.24516 | 0.198824799 | 0.286 | 0.235 | 1 | 6 |
| zgc:86598 | 0.24608 | -0.32058286 | 0.182 | 0.294 | 1 | 6 |
| ttc3 | 0.2464 | -0.26314997 | 0.026 | 0.118 | 1 | 6 |
| emilin1a | 0.24668 | -0.15894603 | 0.208 | 0.235 | 1 | 6 |
| gigyf1b | 0.24695 | -0.30327333 | 0.117 | 0.235 | 1 | 6 |
| nhp2 | 0.24763 | -0.32153793 | 0.169 | 0.294 | 1 | 6 |
| cat | 0.24808 | -0.28362231 | 0.052 | 0.176 | 1 | 6 |
| zgc:85777 | 0.24888 | -0.22587599 | 0.026 | 0.118 | 1 | 6 |
| cct3 | 0.24905 | 0.347961469 | 0.468 | 0.294 | 1 | 6 |
| cenps | 0.24994 | -0.24451743 | 0.026 | 0.118 | 1 | 6 |
| pdgfrb | 0.25101 | -0.33942683 | 0.234 | 0.353 | 1 | 6 |
| psma5 | 0.25141 | 0.168979666 | 0.247 | 0.235 | 1 | 6 |
| tars | 0.25174 | -0.17348262 | 0.052 | 0.176 | 1 | 6 |
| mcm7 | 0.25195 | -0.22556104 | 0.117 | 0.176 | 1 | 6 |
| zgc:56676 | 0.25209 | -0.34811965 | 0.26 | 0.412 | 1 | 6 |
| syap1 | 0.25215 | -0.27524399 | 0.065 | 0.176 | 1 | 6 |
| snrpd2 | 0.25219 | -0.33688333 | 0.286 | 0.471 | 1 | 6 |
| XLOC-013964 | 0.25227 | -0.35048231 | 0.208 | 0.353 | 1 | 6 |
| sirt7 | 0.25285 | -0.21464425 | 0.13 | 0.294 | 1 | 6 |
| sel1l | 0.25429 | -0.10182611 | 0.026 | 0.118 | 1 | 6 |
| ilk | 0.25575 | 0.311198684 | 0.208 | 0.059 | 1 | 6 |
| rpl14 | 0.25611 | 0.182486484 | 0.883 | 0.824 | 1 | 6 |
| tmem107 | 0.25628 | -0.2150796 | 0.104 | 0.176 | 1 | 6 |
| spry4 | 0.25647 | 0.18637612 | 0.182 | 0.059 | 1 | 6 |
| atrx | 0.25682 | 0.241187536 | 0.494 | 0.294 | 1 | 6 |
| si:ch73-335l21.4 | 0.25713 | 0.367807148 | 0.208 | 0.059 | 1 | 6 |
| h2afx1 | 0.25731 | 0.372819499 | 0.519 | 0.353 | 1 | 6 |
| si:ch1073-459j12.1 | 0.25773 | 0.366312202 | 0.623 | 0.412 | 1 | 6 |
| mycbp | 0.25782 | 0.32646556 | 0.364 | 0.176 | 1 | 6 |
| dkk3b | 0.2583 | 0.347042752 | 0.377 | 0.176 | 1 | 6 |
| parvaa | 0.2597 | -0.1452263 | 0.104 | 0.118 | 1 | 6 |
| fam172a | 0.25983 | -0.24373237 | 0.052 | 0.176 | 1 | 6 |

| | | | | | | |
|---|---|---|---|---|---|---|
| pprc1 | 0.26002 | -0.1008841 | 0.026 | 0.118 | 1 | 6 |
| BX284638.1 | 0.26004 | -0.2177082 | 0.052 | 0.176 | 1 | 6 |
| sptlc2a | 0.26013 | -0.17674392 | 0.052 | 0.176 | 1 | 6 |
| exosc6 | 0.26081 | -0.24039038 | 0.052 | 0.176 | 1 | 6 |
| sprydd7a | 0.26089 | -0.24031319 | 0.039 | 0.118 | 1 | 6 |
| atf4a | 0.26115 | 0.324176535 | 0.195 | 0.059 | 1 | 6 |
| skib | 0.26124 | 0.268327908 | 0.195 | 0.059 | 1 | 6 |
| tspan18b | 0.26176 | -0.30195252 | 0.078 | 0.176 | 1 | 6 |
| slc35e1 | 0.26221 | -0.18476469 | 0.052 | 0.176 | 1 | 6 |
| med28 | 0.26272 | 0.216762636 | 0.143 | 0.059 | 1 | 6 |
| eif3ha | 0.26319 | 0.285073663 | 0.506 | 0.294 | 1 | 6 |
| plcg1 | 0.26327 | -0.25762419 | 0.026 | 0.118 | 1 | 6 |
| mt-nd5 | 0.26345 | 0.220835252 | 0.286 | 0.118 | 1 | 6 |
| eif2b4 | 0.26367 | -0.2583747 | 0.026 | 0.118 | 1 | 6 |
| thoc2 | 0.26481 | 0.296163596 | 0.208 | 0.059 | 1 | 6 |
| si:ch211-74m13.3 | 0.26521 | -0.12696596 | 0.052 | 0.059 | 1 | 6 |
| eral1 | 0.26529 | -0.22180629 | 0.026 | 0.118 | 1 | 6 |
| rbpms2b | 0.26565 | -0.35616941 | 0.169 | 0.353 | 1 | 6 |
| ap2s1 | 0.26584 | 0.243824416 | 0.208 | 0.059 | 1 | 6 |
| ypel3 | 0.26592 | -0.29211808 | 0.026 | 0.118 | 1 | 6 |
| si:ch211-167b20.8 | 0.26644 | -0.10622791 | 0.026 | 0.118 | 1 | 6 |
| si:ch211-276c2.4 | 0.26736 | -0.23345431 | 0.026 | 0.118 | 1 | 6 |
| si:ch211-191j22.3 | 0.26739 | 0.114618841 | 0.234 | 0.118 | 1 | 6 |
| cox10 | 0.26811 | -0.25469105 | 0.052 | 0.176 | 1 | 6 |
| timm8a | 0.26815 | -0.17481781 | 0.052 | 0.176 | 1 | 6 |
| polb | 0.26858 | -0.10047744 | 0.026 | 0.118 | 1 | 6 |
| inhbaa | 0.26942 | 0.278465695 | 0.143 | 0.059 | 1 | 6 |
| jdp2b | 0.26997 | 0.306147423 | 0.195 | 0.059 | 1 | 6 |
| bag2 | 0.27017 | -0.21890268 | 0.091 | 0.235 | 1 | 6 |
| nasp | 0.27026 | -0.22552568 | 0.052 | 0.176 | 1 | 6 |
| nr2f1a | 0.27034 | 0.214814224 | 0.234 | 0.176 | 1 | 6 |
| pde7a | 0.27085 | -0.16071535 | 0.039 | 0.059 | 1 | 6 |
| abce1 | 0.27107 | -0.19575443 | 0.078 | 0.118 | 1 | 6 |
| rad23b | 0.27201 | 0.343732955 | 0.364 | 0.176 | 1 | 6 |
| itpkcb | 0.27218 | -0.20882589 | 0.052 | 0.176 | 1 | 6 |
| CABZ01074363.1 | 0.27226 | -0.14094241 | 0.026 | 0.118 | 1 | 6 |
| fbln2 | 0.27318 | -0.23735798 | 0.052 | 0.176 | 1 | 6 |
| rpp30 | 0.27338 | -0.25486946 | 0.026 | 0.118 | 1 | 6 |
| gtf2e2 | 0.2749 | 0.226969176 | 0.208 | 0.059 | 1 | 6 |
| col27a1b | 0.27495 | -0.26498504 | 0.143 | 0.235 | 1 | 6 |
| lsm4 | 0.27507 | -0.21619062 | 0.338 | 0.412 | 1 | 6 |
| si:dkeyp-55f12.3 | 0.27508 | -0.2250988 | 0.052 | 0.176 | 1 | 6 |
| gabarapa | 0.27534 | 0.286272482 | 0.221 | 0.118 | 1 | 6 |
| si:dkey-33c12.4 | 0.27549 | 0.291636488 | 0.286 | 0.118 | 1 | 6 |
| bmper | 0.27579 | 0.183376915 | 0.39 | 0.235 | 1 | 6 |
| si:zfos-169g10.3 | 0.27593 | 0.262075864 | 0.338 | 0.235 | 1 | 6 |
| col6a4a | 0.27594 | -0.11589268 | 0.377 | 0.353 | 1 | 6 |
| fam32a | 0.27596 | 0.316708665 | 0.429 | 0.235 | 1 | 6 |

| | | | | | | |
|---|---|---|---|---|---|---|
| psenen | 0.276 | 0.26855209 | 0.208 | 0.059 | 1 | 6 |
| srpk1b | 0.27603 | -0.26441416 | 0.039 | 0.118 | 1 | 6 |
| nid1a | 0.27612 | 0.283273674 | 0.143 | 0.059 | 1 | 6 |
| tshz2 | 0.27645 | -0.23404962 | 0.052 | 0.176 | 1 | 6 |
| hnrnph1l | 0.27652 | -0.2213778 | 0.117 | 0.176 | 1 | 6 |
| aggf1 | 0.27658 | -0.16423187 | 0.039 | 0.059 | 1 | 6 |
| tmed10 | 0.27662 | 0.103613247 | 0.234 | 0.118 | 1 | 6 |
| fndc3ba | 0.27696 | -0.16360896 | 0.039 | 0.059 | 1 | 6 |
| si:ch211-105c13.3 | 0.2771 | 0.309221349 | 0.208 | 0.059 | 1 | 6 |
| dnlz | 0.27737 | -0.2390275 | 0.026 | 0.118 | 1 | 6 |
| txndc17 | 0.2777 | -0.27563546 | 0.091 | 0.235 | 1 | 6 |
| ssr4 | 0.27803 | 0.118081216 | 0.649 | 0.471 | 1 | 6 |
| zfp36l2 | 0.27805 | 0.312223829 | 0.247 | 0.118 | 1 | 6 |
| gfra1a | 0.27844 | 0.15475156 | 0.052 | 0 | 1 | 6 |
| rnasekb | 0.27844 | 0.105355266 | 0.052 | 0 | 1 | 6 |
| bmp16 | 0.27844 | 0.137152259 | 0.052 | 0 | 1 | 6 |
| cyp2v1 | 0.27844 | 0.105289321 | 0.052 | 0 | 1 | 6 |
| lnpep | 0.27844 | 0.108028314 | 0.052 | 0 | 1 | 6 |
| zdhhc8b | 0.27844 | 0.100713608 | 0.052 | 0 | 1 | 6 |
| LOC101887070 | 0.27844 | 0.108558731 | 0.052 | 0 | 1 | 6 |
| MCUR1 | 0.27844 | 0.100325129 | 0.052 | 0 | 1 | 6 |
| XLOC-034394 | 0.27844 | 0.154871701 | 0.052 | 0 | 1 | 6 |
| cald1b | 0.27844 | 0.188247438 | 0.052 | 0 | 1 | 6 |
| ek1 | 0.27844 | 0.122441924 | 0.052 | 0 | 1 | 6 |
| hagh | 0.27844 | 0.10351956 | 0.052 | 0 | 1 | 6 |
| ift52 | 0.27844 | 0.116161372 | 0.052 | 0 | 1 | 6 |
| itgb4 | 0.27844 | 0.116853971 | 0.052 | 0 | 1 | 6 |
| mat2b | 0.27844 | 0.106147591 | 0.052 | 0 | 1 | 6 |
| myo9aa | 0.27844 | 0.101943008 | 0.052 | 0 | 1 | 6 |
| nfkbie | 0.27844 | 0.105014946 | 0.052 | 0 | 1 | 6 |
| nudt18 | 0.27844 | 0.121903684 | 0.052 | 0 | 1 | 6 |
| ptpra | 0.27844 | 0.125043334 | 0.052 | 0 | 1 | 6 |
| rdh10b | 0.27844 | 0.144759859 | 0.052 | 0 | 1 | 6 |
| slc35b1 | 0.27844 | 0.111609721 | 0.052 | 0 | 1 | 6 |
| zswim8 | 0.27844 | 0.120872945 | 0.052 | 0 | 1 | 6 |
| borcs8 | 0.27844 | 0.120582138 | 0.052 | 0 | 1 | 6 |
| btd | 0.27844 | 0.11844544 | 0.052 | 0 | 1 | 6 |
| c1qtnf12 | 0.27844 | 0.124204031 | 0.052 | 0 | 1 | 6 |
| rem1 | 0.27844 | 0.112436188 | 0.052 | 0 | 1 | 6 |
| rnf14 | 0.27844 | 0.119085904 | 0.052 | 0 | 1 | 6 |
| si:dkey-283j8.1 | 0.27844 | 0.129397188 | 0.052 | 0 | 1 | 6 |
| slc30a4 | 0.27844 | 0.116763442 | 0.052 | 0 | 1 | 6 |
| CABZ01087514.1 | 0.27844 | 0.118208442 | 0.052 | 0 | 1 | 6 |
| PDE3B | 0.27844 | 0.123178371 | 0.052 | 0 | 1 | 6 |
| XLOC-042565 | 0.27844 | 0.102921355 | 0.052 | 0 | 1 | 6 |
| arglu1b | 0.27844 | 0.100574927 | 0.052 | 0 | 1 | 6 |
| c1galt1b | 0.27844 | 0.107022636 | 0.052 | 0 | 1 | 6 |
| casp3a | 0.27844 | 0.126562095 | 0.052 | 0 | 1 | 6 |

| | | | | | | |
|---|---|---|---|---|---|---|
| ccdc137 | 0.27844 | 0.10884132 | 0.052 | 0 | 1 | 6 |
| dock1 | 0.27844 | 0.116311596 | 0.052 | 0 | 1 | 6 |
| iah1 | 0.27844 | 0.110969437 | 0.052 | 0 | 1 | 6 |
| nck2a | 0.27844 | 0.119242925 | 0.052 | 0 | 1 | 6 |
| pkdccb | 0.27844 | 0.117355033 | 0.052 | 0 | 1 | 6 |
| si:ch211-106k21.5 | 0.27844 | 0.104453873 | 0.052 | 0 | 1 | 6 |
| st3gal4 | 0.27844 | 0.102556955 | 0.052 | 0 | 1 | 6 |
| utp11l | 0.27844 | 0.102429067 | 0.052 | 0 | 1 | 6 |
| wdr36 | 0.27844 | 0.101952084 | 0.052 | 0 | 1 | 6 |
| zfpm2b | 0.27844 | 0.114077206 | 0.052 | 0 | 1 | 6 |
| zgc:91976 | 0.27844 | 0.102574771 | 0.052 | 0 | 1 | 6 |
| BX571942.1 | 0.27844 | 0.111962501 | 0.052 | 0 | 1 | 6 |
| GNAZ | 0.27844 | 0.124614041 | 0.052 | 0 | 1 | 6 |
| HOOK3 | 0.27844 | 0.108913936 | 0.052 | 0 | 1 | 6 |
| LOC101885018 | 0.27844 | 0.117341036 | 0.052 | 0 | 1 | 6 |
| LOC108190702 | 0.27844 | 0.120001579 | 0.052 | 0 | 1 | 6 |
| LOC110438169 | 0.27844 | 0.108924548 | 0.052 | 0 | 1 | 6 |
| PLEKHH3 | 0.27844 | 0.109647215 | 0.052 | 0 | 1 | 6 |
| XLOC-005350 | 0.27844 | 0.117687112 | 0.052 | 0 | 1 | 6 |
| XLOC-007665 | 0.27844 | 0.102121945 | 0.052 | 0 | 1 | 6 |
| XLOC-024248 | 0.27844 | 0.115319971 | 0.052 | 0 | 1 | 6 |
| XLOC-032256 | 0.27844 | 0.115379096 | 0.052 | 0 | 1 | 6 |
| abca1b | 0.27844 | 0.115984606 | 0.052 | 0 | 1 | 6 |
| add3b | 0.27844 | 0.119691214 | 0.052 | 0 | 1 | 6 |
| asap1b | 0.27844 | 0.107741013 | 0.052 | 0 | 1 | 6 |
| asb7 | 0.27844 | 0.106299465 | 0.052 | 0 | 1 | 6 |
| bco2l | 0.27844 | 0.103698622 | 0.052 | 0 | 1 | 6 |
| bgna | 0.27844 | 0.173252234 | 0.052 | 0 | 1 | 6 |
| bloc1s2 | 0.27844 | 0.100384865 | 0.052 | 0 | 1 | 6 |
| bmt2 | 0.27844 | 0.114850309 | 0.052 | 0 | 1 | 6 |
| bnip2 | 0.27844 | 0.100152094 | 0.052 | 0 | 1 | 6 |
| btbd6b | 0.27844 | 0.133964894 | 0.052 | 0 | 1 | 6 |
| cacul1 | 0.27844 | 0.104152375 | 0.052 | 0 | 1 | 6 |
| cdk13 | 0.27844 | 0.100521123 | 0.052 | 0 | 1 | 6 |
| cdr2a | 0.27844 | 0.11263253 | 0.052 | 0 | 1 | 6 |
| cep19 | 0.27844 | 0.105093374 | 0.052 | 0 | 1 | 6 |
| cept1a | 0.27844 | 0.112479196 | 0.052 | 0 | 1 | 6 |
| cers5 | 0.27844 | 0.12089863 | 0.052 | 0 | 1 | 6 |
| cntrl | 0.27844 | 0.114553584 | 0.052 | 0 | 1 | 6 |
| cplx2 | 0.27844 | 0.14427711 | 0.052 | 0 | 1 | 6 |
| cpxm1a | 0.27844 | 0.128080659 | 0.052 | 0 | 1 | 6 |
| crtc3 | 0.27844 | 0.111550606 | 0.052 | 0 | 1 | 6 |
| cry-dash | 0.27844 | 0.122376206 | 0.052 | 0 | 1 | 6 |
| cry1ba | 0.27844 | 0.127866733 | 0.052 | 0 | 1 | 6 |
| cstf3 | 0.27844 | 0.104583674 | 0.052 | 0 | 1 | 6 |
| cybc1 | 0.27844 | 0.125871746 | 0.052 | 0 | 1 | 6 |
| dbpa | 0.27844 | 0.110921323 | 0.052 | 0 | 1 | 6 |
| dcun1d4 | 0.27844 | 0.103629743 | 0.052 | 0 | 1 | 6 |

| | | | | | | |
|---|---|---|---|---|---|---|
| dele1 | 0.27844 | 0.115889628 | 0.052 | 0 | 1 | 6 |
| dnm1l | 0.27844 | 0.104241561 | 0.052 | 0 | 1 | 6 |
| dpf3 | 0.27844 | 0.127980505 | 0.052 | 0 | 1 | 6 |
| dph1 | 0.27844 | 0.109838323 | 0.052 | 0 | 1 | 6 |
| dtd1 | 0.27844 | 0.101381639 | 0.052 | 0 | 1 | 6 |
| dync1i2b | 0.27844 | 0.114746139 | 0.052 | 0 | 1 | 6 |
| efna2a | 0.27844 | 0.108446091 | 0.052 | 0 | 1 | 6 |
| efna3b | 0.27844 | 0.120460394 | 0.052 | 0 | 1 | 6 |
| egfra | 0.27844 | 0.129991921 | 0.052 | 0 | 1 | 6 |
| emg1 | 0.27844 | 0.113207916 | 0.052 | 0 | 1 | 6 |
| entr1 | 0.27844 | 0.100542331 | 0.052 | 0 | 1 | 6 |
| epc2 | 0.27844 | 0.12911494 | 0.052 | 0 | 1 | 6 |
| epha4l | 0.27844 | 0.100881982 | 0.052 | 0 | 1 | 6 |
| epha7 | 0.27844 | 0.101976267 | 0.052 | 0 | 1 | 6 |
| eps15l1a | 0.27844 | 0.11462424 | 0.052 | 0 | 1 | 6 |
| errfi1a | 0.27844 | 0.104261084 | 0.052 | 0 | 1 | 6 |
| etv5a | 0.27844 | 0.104013283 | 0.052 | 0 | 1 | 6 |
| f11r.1 | 0.27844 | 0.106782396 | 0.052 | 0 | 1 | 6 |
| fam131bb | 0.27844 | 0.10311401 | 0.052 | 0 | 1 | 6 |
| fam173a | 0.27844 | 0.104065952 | 0.052 | 0 | 1 | 6 |
| fam181b | 0.27844 | 0.117529611 | 0.052 | 0 | 1 | 6 |
| fcho2 | 0.27844 | 0.125042744 | 0.052 | 0 | 1 | 6 |
| flrt2 | 0.27844 | 0.117869749 | 0.052 | 0 | 1 | 6 |
| fus | 0.27844 | 0.115160411 | 0.052 | 0 | 1 | 6 |
| fut8a | 0.27844 | 0.112618013 | 0.052 | 0 | 1 | 6 |
| gale | 0.27844 | 0.146248337 | 0.052 | 0 | 1 | 6 |
| gamt | 0.27844 | 0.136257545 | 0.052 | 0 | 1 | 6 |
| gbp3 | 0.27844 | 0.120976084 | 0.052 | 0 | 1 | 6 |
| glod4 | 0.27844 | 0.103935823 | 0.052 | 0 | 1 | 6 |
| gmds | 0.27844 | 0.107260961 | 0.052 | 0 | 1 | 6 |
| gng13b | 0.27844 | 0.105050745 | 0.052 | 0 | 1 | 6 |
| gosr1 | 0.27844 | 0.101204775 | 0.052 | 0 | 1 | 6 |
| gpr108 | 0.27844 | 0.106908325 | 0.052 | 0 | 1 | 6 |
| grin3bb | 0.27844 | 0.120029416 | 0.052 | 0 | 1 | 6 |
| hapln3 | 0.27844 | 0.108052156 | 0.052 | 0 | 1 | 6 |
| hmgxb4a | 0.27844 | 0.100243101 | 0.052 | 0 | 1 | 6 |
| hspa4a | 0.27844 | 0.115709101 | 0.052 | 0 | 1 | 6 |
| huwe1 | 0.27844 | 0.120594077 | 0.052 | 0 | 1 | 6 |
| igfbp2a | 0.27844 | 0.100067729 | 0.052 | 0 | 1 | 6 |
| igsf8 | 0.27844 | 0.127094002 | 0.052 | 0 | 1 | 6 |
| ilkap | 0.27844 | 0.101703539 | 0.052 | 0 | 1 | 6 |
| irak1bp1 | 0.27844 | 0.110504645 | 0.052 | 0 | 1 | 6 |
| kbtbd11 | 0.27844 | 0.124165063 | 0.052 | 0 | 1 | 6 |
| kctd3 | 0.27844 | 0.104887805 | 0.052 | 0 | 1 | 6 |
| kif13ba | 0.27844 | 0.125026008 | 0.052 | 0 | 1 | 6 |
| kmt2ba | 0.27844 | 0.110059164 | 0.052 | 0 | 1 | 6 |
| lin7c | 0.27844 | 0.105314929 | 0.052 | 0 | 1 | 6 |
| lzts1 | 0.27844 | 0.103153726 | 0.052 | 0 | 1 | 6 |

| | | | | | | |
|---|---|---|---|---|---|---|
| map3k14a | 0.27844 | 0.107184506 | 0.052 | 0 | 1 | 6 |
| mdc1 | 0.27844 | 0.102565059 | 0.052 | 0 | 1 | 6 |
| mical2a | 0.27844 | 0.114021178 | 0.052 | 0 | 1 | 6 |
| mlx | 0.27844 | 0.105428512 | 0.052 | 0 | 1 | 6 |
| mon2 | 0.27844 | 0.11737354 | 0.052 | 0 | 1 | 6 |
| mpped2a | 0.27844 | 0.116606876 | 0.052 | 0 | 1 | 6 |
| myofl | 0.27844 | 0.119841766 | 0.052 | 0 | 1 | 6 |
| ndufaf1 | 0.27844 | 0.102248708 | 0.052 | 0 | 1 | 6 |
| nf2b | 0.27844 | 0.109750607 | 0.052 | 0 | 1 | 6 |
| nkain1 | 0.27844 | 0.119007857 | 0.052 | 0 | 1 | 6 |
| ormdl1 | 0.27844 | 0.102557764 | 0.052 | 0 | 1 | 6 |
| pacsin2 | 0.27844 | 0.105143252 | 0.052 | 0 | 1 | 6 |
| pacsin3 | 0.27844 | 0.1011334 | 0.052 | 0 | 1 | 6 |
| parn | 0.27844 | 0.107140387 | 0.052 | 0 | 1 | 6 |
| pdcd7 | 0.27844 | 0.10118619 | 0.052 | 0 | 1 | 6 |
| pdgfab | 0.27844 | 0.106948898 | 0.052 | 0 | 1 | 6 |
| pdlim3b | 0.27844 | 0.118630356 | 0.052 | 0 | 1 | 6 |
| pdlim4 | 0.27844 | 0.100954516 | 0.052 | 0 | 1 | 6 |
| phf21ab | 0.27844 | 0.109261344 | 0.052 | 0 | 1 | 6 |
| phf23a | 0.27844 | 0.153150764 | 0.052 | 0 | 1 | 6 |
| pld1a | 0.27844 | 0.127260136 | 0.052 | 0 | 1 | 6 |
| plekha3 | 0.27844 | 0.132637959 | 0.052 | 0 | 1 | 6 |
| plekhg4 | 0.27844 | 0.113446669 | 0.052 | 0 | 1 | 6 |
| ppp2r1bb | 0.27844 | 0.102082171 | 0.052 | 0 | 1 | 6 |
| ppp2r5d | 0.27844 | 0.106141231 | 0.052 | 0 | 1 | 6 |
| prpsap2 | 0.27844 | 0.125536156 | 0.052 | 0 | 1 | 6 |
| ptger2b | 0.27844 | 0.113276301 | 0.052 | 0 | 1 | 6 |
| ptgs2a | 0.27844 | 0.117862623 | 0.052 | 0 | 1 | 6 |
| rab20 | 0.27844 | 0.143247439 | 0.052 | 0 | 1 | 6 |
| rabl6a | 0.27844 | 0.109458623 | 0.052 | 0 | 1 | 6 |
| rarab | 0.27844 | 0.11192351 | 0.052 | 0 | 1 | 6 |
| rest | 0.27844 | 0.117992051 | 0.052 | 0 | 1 | 6 |
| rfc3 | 0.27844 | 0.102937271 | 0.052 | 0 | 1 | 6 |
| rgs4 | 0.27844 | 0.136107251 | 0.052 | 0 | 1 | 6 |
| rhogd | 0.27844 | 0.106900145 | 0.052 | 0 | 1 | 6 |
| rilpl2 | 0.27844 | 0.11097205 | 0.052 | 0 | 1 | 6 |
| rnf24 | 0.27844 | 0.130386374 | 0.052 | 0 | 1 | 6 |
| sec22bb | 0.27844 | 0.101035618 | 0.052 | 0 | 1 | 6 |
| sema3b | 0.27844 | 0.101494196 | 0.052 | 0 | 1 | 6 |
| sema6bb | 0.27844 | 0.168328815 | 0.052 | 0 | 1 | 6 |
| setdb1a | 0.27844 | 0.110442022 | 0.052 | 0 | 1 | 6 |
| sh2b1 | 0.27844 | 0.106615346 | 0.052 | 0 | 1 | 6 |
| shisal1b | 0.27844 | 0.112320478 | 0.052 | 0 | 1 | 6 |
| si:ch1073-314i13.4 | 0.27844 | 0.108145701 | 0.052 | 0 | 1 | 6 |
| si:ch211-39k3.2 | 0.27844 | 0.100287592 | 0.052 | 0 | 1 | 6 |
| si:ch211-56a11.2 | 0.27844 | 0.115969994 | 0.052 | 0 | 1 | 6 |
| si:ch211-86h15.1 | 0.27844 | 0.107478896 | 0.052 | 0 | 1 | 6 |
| si:ch73-109d9.2 | 0.27844 | 0.126259843 | 0.052 | 0 | 1 | 6 |

| | | | | | | |
|---|---|---|---|---|---|---|
| si:ch73-221f6.4 | 0.27844 | 0.143244915 | 0.052 | 0 | 1 | 6 |
| si:ch73-347e22.4 | 0.27844 | 0.100288444 | 0.052 | 0 | 1 | 6 |
| si:dkey-182g1.2 | 0.27844 | 0.112813363 | 0.052 | 0 | 1 | 6 |
| si:dkey-184a18.5 | 0.27844 | 0.129174865 | 0.052 | 0 | 1 | 6 |
| si:dkey-222b8.1 | 0.27844 | 0.110367911 | 0.052 | 0 | 1 | 6 |
| si:dkey-238i5.2 | 0.27844 | 0.111771116 | 0.052 | 0 | 1 | 6 |
| si:dkey-78a14.4 | 0.27844 | 0.11294986 | 0.052 | 0 | 1 | 6 |
| si:dkeyp-69e1.8 | 0.27844 | 0.138343826 | 0.052 | 0 | 1 | 6 |
| six4b | 0.27844 | 0.119616783 | 0.052 | 0 | 1 | 6 |
| slc30a5 | 0.27844 | 0.108786852 | 0.052 | 0 | 1 | 6 |
| slc35a4 | 0.27844 | 0.133953374 | 0.052 | 0 | 1 | 6 |
| slc35b3 | 0.27844 | 0.118249484 | 0.052 | 0 | 1 | 6 |
| smad6a | 0.27844 | 0.124746298 | 0.052 | 0 | 1 | 6 |
| smad6b | 0.27844 | 0.127969782 | 0.052 | 0 | 1 | 6 |
| smim20 | 0.27844 | 0.109395195 | 0.052 | 0 | 1 | 6 |
| snapc3 | 0.27844 | 0.119898112 | 0.052 | 0 | 1 | 6 |
| snapc4 | 0.27844 | 0.107471189 | 0.052 | 0 | 1 | 6 |
| snx16 | 0.27844 | 0.104007063 | 0.052 | 0 | 1 | 6 |
| sox11b | 0.27844 | 0.105852793 | 0.052 | 0 | 1 | 6 |
| sox9a | 0.27844 | 0.153728401 | 0.052 | 0 | 1 | 6 |
| spon1a | 0.27844 | 0.109653652 | 0.052 | 0 | 1 | 6 |
| spopla | 0.27844 | 0.10156596 | 0.052 | 0 | 1 | 6 |
| stx4 | 0.27844 | 0.111782775 | 0.052 | 0 | 1 | 6 |
| sulf1 | 0.27844 | 0.111138186 | 0.052 | 0 | 1 | 6 |
| tanc2a | 0.27844 | 0.111216748 | 0.052 | 0 | 1 | 6 |
| tapt1a | 0.27844 | 0.122782577 | 0.052 | 0 | 1 | 6 |
| tbx2a | 0.27844 | 0.147254994 | 0.052 | 0 | 1 | 6 |
| tchp | 0.27844 | 0.103232708 | 0.052 | 0 | 1 | 6 |
| tfpia | 0.27844 | 0.133484501 | 0.052 | 0 | 1 | 6 |
| thap7 | 0.27844 | 0.101919199 | 0.052 | 0 | 1 | 6 |
| tmem198b | 0.27844 | 0.128478683 | 0.052 | 0 | 1 | 6 |
| tmem59l | 0.27844 | 0.106513112 | 0.052 | 0 | 1 | 6 |
| tnrc6c2 | 0.27844 | 0.134976164 | 0.052 | 0 | 1 | 6 |
| tp63 | 0.27844 | 0.129863773 | 0.052 | 0 | 1 | 6 |
| tprg1l | 0.27844 | 0.104273735 | 0.052 | 0 | 1 | 6 |
| traf7 | 0.27844 | 0.105128518 | 0.052 | 0 | 1 | 6 |
| trappc6b | 0.27844 | 0.125550391 | 0.052 | 0 | 1 | 6 |
| trappc8 | 0.27844 | 0.127517575 | 0.052 | 0 | 1 | 6 |
| trim44 | 0.27844 | 0.107561577 | 0.052 | 0 | 1 | 6 |
| twist1a | 0.27844 | 0.111537596 | 0.052 | 0 | 1 | 6 |
| ugt2a5 | 0.27844 | 0.106269941 | 0.052 | 0 | 1 | 6 |
| utp4 | 0.27844 | 0.101896219 | 0.052 | 0 | 1 | 6 |
| waslb | 0.27844 | 0.112005649 | 0.052 | 0 | 1 | 6 |
| wfdc2 | 0.27844 | 0.169197861 | 0.052 | 0 | 1 | 6 |
| wif1 | 0.27844 | 0.108103645 | 0.052 | 0 | 1 | 6 |
| wnt16 | 0.27844 | 0.103400022 | 0.052 | 0 | 1 | 6 |
| yjefn3 | 0.27844 | 0.10829148 | 0.052 | 0 | 1 | 6 |
| zbtb17 | 0.27844 | 0.101555911 | 0.052 | 0 | 1 | 6 |

| | | | | | | |
|---|---|---|---|---|---|---|
| zbtb46 | 0.27844 | 0.107777592 | 0.052 | 0 | 1 | 6 |
| zgc:110366 | 0.27844 | 0.113370318 | 0.052 | 0 | 1 | 6 |
| zgc:136308 | 0.27844 | 0.114039808 | 0.052 | 0 | 1 | 6 |
| znf385d | 0.27844 | 0.114739052 | 0.052 | 0 | 1 | 6 |
| znf618 | 0.27844 | 0.104314222 | 0.052 | 0 | 1 | 6 |
| znf654 | 0.27844 | 0.119183321 | 0.052 | 0 | 1 | 6 |
| znf687b | 0.27844 | 0.109426276 | 0.052 | 0 | 1 | 6 |
| crcp | 0.27844 | 0.107712887 | 0.052 | 0 | 1 | 6 |
| hbegfa | 0.27844 | 0.106416876 | 0.052 | 0 | 1 | 6 |
| plekhj1 | 0.27844 | 0.101625678 | 0.052 | 0 | 1 | 6 |
| bmp6 | 0.27844 | 0.137170929 | 0.052 | 0 | 1 | 6 |
| cdk4 | 0.27844 | 0.1145758 | 0.052 | 0 | 1 | 6 |
| ganab | 0.27844 | 0.108454604 | 0.052 | 0 | 1 | 6 |
| mettl24 | 0.27844 | 0.105923682 | 0.052 | 0 | 1 | 6 |
| si:ch1073-440b2.1 | 0.27844 | 0.107122753 | 0.052 | 0 | 1 | 6 |
| ube4b | 0.27844 | 0.1133207 | 0.052 | 0 | 1 | 6 |
| vps37a | 0.27844 | 0.124590351 | 0.052 | 0 | 1 | 6 |
| lmod1b | 0.27844 | 0.141502223 | 0.052 | 0 | 1 | 6 |
| rasl11b | 0.27844 | 0.112127328 | 0.052 | 0 | 1 | 6 |
| dap3 | 0.27844 | 0.101791358 | 0.052 | 0 | 1 | 6 |
| pias2 | 0.27844 | 0.117015036 | 0.052 | 0 | 1 | 6 |
| cnbpb | 0.27984 | -0.14570501 | 0.455 | 0.471 | 1 | 6 |
| mfap5 | 0.27987 | -0.31270859 | 0.403 | 0.529 | 1 | 6 |
| tmem161a | 0.27997 | -0.20562671 | 0.026 | 0.118 | 1 | 6 |
| cbx1b | 0.28036 | 0.137063761 | 0.182 | 0.176 | 1 | 6 |
| mdn1 | 0.28054 | -0.26005243 | 0.052 | 0.176 | 1 | 6 |
| rab2a | 0.28059 | 0.142058662 | 0.312 | 0.176 | 1 | 6 |
| eif3m | 0.28071 | 0.292796912 | 0.623 | 0.412 | 1 | 6 |
| ywhag1 | 0.2808 | -0.25131394 | 0.052 | 0.176 | 1 | 6 |
| chd7 | 0.28093 | -0.19092761 | 0.026 | 0.118 | 1 | 6 |
| pla2g7 | 0.28136 | -0.13355529 | 0.026 | 0.118 | 1 | 6 |
| actl6a | 0.28185 | -0.21849088 | 0.052 | 0.176 | 1 | 6 |
| p3h4 | 0.28201 | -0.25465533 | 0.052 | 0.118 | 1 | 6 |
| zgc:110158 | 0.28217 | 0.275442793 | 0.195 | 0.059 | 1 | 6 |
| sft2d1 | 0.28239 | -0.19329025 | 0.052 | 0.176 | 1 | 6 |
| ppid | 0.28254 | 0.238825793 | 0.208 | 0.059 | 1 | 6 |
| hmbox1a | 0.28267 | 0.250241712 | 0.195 | 0.059 | 1 | 6 |
| ppp2r1ba | 0.28304 | 0.25246007 | 0.208 | 0.059 | 1 | 6 |
| dnajc17 | 0.28316 | -0.13911821 | 0.026 | 0.118 | 1 | 6 |
| bnip3lb | 0.28351 | 0.223169066 | 0.208 | 0.059 | 1 | 6 |
| trip12 | 0.28352 | -0.2230205 | 0.052 | 0.176 | 1 | 6 |
| ier5 | 0.28358 | 0.360542395 | 0.351 | 0.176 | 1 | 6 |
| selenoj | 0.28378 | 0.192894775 | 0.13 | 0.059 | 1 | 6 |
| col11a1b | 0.28443 | 0.250069337 | 0.13 | 0.059 | 1 | 6 |
| gng12a | 0.28491 | -0.29843505 | 0.104 | 0.235 | 1 | 6 |
| idh1 | 0.28527 | -0.1589671 | 0.052 | 0.176 | 1 | 6 |
| cd82b | 0.28577 | -0.33500131 | 0.143 | 0.294 | 1 | 6 |
| st13 | 0.28601 | 0.155192591 | 0.247 | 0.118 | 1 | 6 |

| | | | | | | |
|---|---|---|---|---|---|---|
| exosc2 | 0.28628 | -0.21304131 | 0.026 | 0.118 | 1 | 6 |
| zfand4 | 0.28636 | -0.1259435 | 0.026 | 0.118 | 1 | 6 |
| tgfb3 | 0.28654 | 0.147806717 | 0.208 | 0.176 | 1 | 6 |
| gata5 | 0.28725 | 0.318297258 | 0.182 | 0.059 | 1 | 6 |
| zgc:113531 | 0.28784 | 0.120151403 | 0.299 | 0.176 | 1 | 6 |
| uqcr10 | 0.28803 | -0.18836274 | 0.312 | 0.353 | 1 | 6 |
| igfbp3 | 0.2883 | -0.12585453 | 0.026 | 0.118 | 1 | 6 |
| jade1 | 0.28857 | -0.20721045 | 0.026 | 0.118 | 1 | 6 |
| galns | 0.28928 | -0.1192238 | 0.026 | 0.118 | 1 | 6 |
| slc25a1a | 0.28928 | -0.19268138 | 0.026 | 0.118 | 1 | 6 |
| gpx4a | 0.28994 | -0.12039737 | 0.026 | 0.118 | 1 | 6 |
| pdcd5 | 0.29002 | 0.258030714 | 0.247 | 0.118 | 1 | 6 |
| LOC108180286 | 0.29038 | -0.22627214 | 0.026 | 0.118 | 1 | 6 |
| lsm5 | 0.29112 | 0.186052963 | 0.26 | 0.118 | 1 | 6 |
| epha3 | 0.29118 | 0.31184386 | 0.286 | 0.118 | 1 | 6 |
| trub2 | 0.29132 | -0.1387016 | 0.026 | 0.118 | 1 | 6 |
| got2b | 0.29136 | -0.20463832 | 0.065 | 0.118 | 1 | 6 |
| vdrb | 0.29181 | 0.118101835 | 0.221 | 0.235 | 1 | 6 |
| zgc:163080 | 0.29189 | -0.13239193 | 0.234 | 0.235 | 1 | 6 |
| CABZ01068367.1 | 0.29194 | -0.14334108 | 0.026 | 0.118 | 1 | 6 |
| kif5bb | 0.29326 | 0.24218375 | 0.156 | 0.059 | 1 | 6 |
| sec61a1 | 0.29355 | -0.18157163 | 0.299 | 0.294 | 1 | 6 |
| myl12.1 | 0.29378 | -0.15280214 | 0.649 | 0.647 | 1 | 6 |
| odc1 | 0.29395 | -0.19654532 | 0.039 | 0.118 | 1 | 6 |
| dbn1 | 0.29547 | 0.292676081 | 0.286 | 0.176 | 1 | 6 |
| znf593 | 0.29576 | -0.27291297 | 0.143 | 0.235 | 1 | 6 |
| DDX17 | 0.29626 | -0.15637468 | 0.039 | 0.059 | 1 | 6 |
| bud31 | 0.2966 | 0.297902025 | 0.325 | 0.176 | 1 | 6 |
| tmem106a | 0.29748 | -0.14562817 | 0.026 | 0.118 | 1 | 6 |
| eif4a1b | 0.29757 | 0.317136363 | 0.636 | 0.471 | 1 | 6 |
| psmd4b | 0.29821 | 0.277223929 | 0.234 | 0.118 | 1 | 6 |
| ginm1 | 0.2984 | 0.187574733 | 0.169 | 0.118 | 1 | 6 |
| smc1al | 0.29864 | 0.306740837 | 0.26 | 0.118 | 1 | 6 |
| cox5aa | 0.29909 | -0.32560557 | 0.377 | 0.529 | 1 | 6 |
| cacybp | 0.2991 | 0.20238188 | 0.182 | 0.059 | 1 | 6 |
| notch1b | 0.2995 | -0.31032739 | 0.026 | 0.118 | 1 | 6 |
| atxn1a | 0.29954 | 0.252082944 | 0.13 | 0.059 | 1 | 6 |
| tarbp2 | 0.30024 | -0.10237801 | 0.026 | 0.118 | 1 | 6 |
| cst3 | 0.3005 | -0.30821343 | 0.104 | 0.235 | 1 | 6 |
| XLOC-028370 | 0.3015 | -0.22936495 | 0.091 | 0.235 | 1 | 6 |
| ino80c | 0.30205 | 0.260180596 | 0.182 | 0.059 | 1 | 6 |
| parp3 | 0.30226 | -0.12552356 | 0.026 | 0.118 | 1 | 6 |
| asph | 0.30271 | -0.25476772 | 0.091 | 0.235 | 1 | 6 |
| chmp2a | 0.30274 | 0.185789765 | 0.234 | 0.176 | 1 | 6 |
| capza1a | 0.30276 | 0.241640862 | 0.351 | 0.176 | 1 | 6 |
| gpc1b | 0.30366 | -0.1430185 | 0.104 | 0.118 | 1 | 6 |
| spcs1 | 0.30388 | -0.26139721 | 0.416 | 0.529 | 1 | 6 |
| snai2 | 0.30389 | -0.2708361 | 0.091 | 0.235 | 1 | 6 |

| | | | | | | |
|---|---|---|---|---|---|---|
| tax1bp1b | 0.30408 | -0.26613183 | 0.091 | 0.235 | 1 | 6 |
| ppp1r14bb | 0.30454 | 0.224879927 | 0.156 | 0.059 | 1 | 6 |
| MDFI | 0.30474 | 0.298201626 | 0.247 | 0.118 | 1 | 6 |
| ahcy | 0.3051 | 0.198977594 | 0.143 | 0.059 | 1 | 6 |
| pdia6 | 0.30554 | -0.18431755 | 0.195 | 0.235 | 1 | 6 |
| sh3bgrl3 | 0.30609 | -0.25236572 | 0.13 | 0.235 | 1 | 6 |
| eci1 | 0.30642 | 0.237821239 | 0.182 | 0.059 | 1 | 6 |
| dicer1 | 0.30666 | -0.29213425 | 0.026 | 0.118 | 1 | 6 |
| cdc73 | 0.30701 | -0.25244643 | 0.039 | 0.118 | 1 | 6 |
| tsc1a | 0.30706 | -0.26122008 | 0.026 | 0.118 | 1 | 6 |
| supt4h1 | 0.30731 | -0.10442984 | 0.117 | 0.118 | 1 | 6 |
| grb10b | 0.30794 | 0.202786626 | 0.117 | 0.059 | 1 | 6 |
| ap3m2 | 0.3081 | -0.11275285 | 0.026 | 0.118 | 1 | 6 |
| atg9b | 0.30865 | -0.11758279 | 0.026 | 0.118 | 1 | 6 |
| si:dkey-57k2.6 | 0.309 | -0.14364377 | 0.026 | 0.118 | 1 | 6 |
| golph3 | 0.30938 | -0.23524173 | 0.065 | 0.176 | 1 | 6 |
| casp8ap2 | 0.30963 | -0.23918177 | 0.026 | 0.118 | 1 | 6 |
| sec61b | 0.30981 | 0.264541545 | 0.766 | 0.588 | 1 | 6 |
| hectd1 | 0.31059 | -0.26395252 | 0.026 | 0.118 | 1 | 6 |
| mcm5 | 0.31124 | -0.28044187 | 0.078 | 0.176 | 1 | 6 |
| p4ha2 | 0.31127 | -0.21942426 | 0.026 | 0.118 | 1 | 6 |
| plpp3 | 0.31201 | 0.360222048 | 0.403 | 0.235 | 1 | 6 |
| glula | 0.31254 | -0.30947422 | 0.182 | 0.353 | 1 | 6 |
| rtn4b | 0.31279 | 0.131797579 | 0.156 | 0.059 | 1 | 6 |
| baz2a | 0.31406 | -0.24019769 | 0.026 | 0.118 | 1 | 6 |
| tpcn3 | 0.31418 | -0.21430053 | 0.026 | 0.118 | 1 | 6 |
| exosc9 | 0.31448 | -0.14631868 | 0.026 | 0.118 | 1 | 6 |
| mcrip1 | 0.31449 | -0.26686391 | 0.104 | 0.235 | 1 | 6 |
| cyp2ad3 | 0.31451 | -0.22236962 | 0.247 | 0.294 | 1 | 6 |
| cmtm7 | 0.31488 | -0.25384933 | 0.104 | 0.235 | 1 | 6 |
| ssbp3b | 0.31514 | -0.116592 | 0.117 | 0.118 | 1 | 6 |
| fbln1 | 0.31524 | -0.14983495 | 0.026 | 0.118 | 1 | 6 |
| kars | 0.31532 | -0.2389726 | 0.273 | 0.353 | 1 | 6 |
| smo | 0.31541 | 0.206750864 | 0.182 | 0.059 | 1 | 6 |
| flot2a | 0.31553 | 0.203067844 | 0.26 | 0.118 | 1 | 6 |
| ak2 | 0.31564 | 0.268012643 | 0.429 | 0.235 | 1 | 6 |
| smim13 | 0.3163 | 0.258357777 | 0.195 | 0.059 | 1 | 6 |
| ctsa | 0.31677 | -0.25482814 | 0.13 | 0.235 | 1 | 6 |
| capn2b | 0.31699 | 0.122097353 | 0.065 | 0.059 | 1 | 6 |
| nol11 | 0.31703 | -0.19521937 | 0.026 | 0.118 | 1 | 6 |
| map2k4b | 0.31771 | -0.23202625 | 0.026 | 0.118 | 1 | 6 |
| CABZ01080375.1 | 0.31778 | -0.1033833 | 0.013 | 0.059 | 1 | 6 |
| abhd15a | 0.31778 | -0.12174907 | 0.013 | 0.059 | 1 | 6 |
| ccar2 | 0.31778 | -0.12155834 | 0.013 | 0.059 | 1 | 6 |
| fuom | 0.31778 | -0.17952114 | 0.013 | 0.059 | 1 | 6 |
| pcdh19 | 0.31778 | -0.12424074 | 0.013 | 0.059 | 1 | 6 |
| ppp2r2aa | 0.31778 | -0.12730058 | 0.013 | 0.059 | 1 | 6 |
| ptprz1b | 0.31778 | -0.14640575 | 0.013 | 0.059 | 1 | 6 |

| | | | | | | |
|---|---|---|---|---|---|---|
| si:dkey-264d12.5 | 0.31778 | -0.15982646 | 0.013 | 0.059 | 1 | 6 |
| FP101861.1 | 0.31778 | -0.19006479 | 0.013 | 0.059 | 1 | 6 |
| bokb | 0.31778 | -0.21945927 | 0.013 | 0.059 | 1 | 6 |
| cap2 | 0.31778 | -0.1721403 | 0.013 | 0.059 | 1 | 6 |
| gins3 | 0.31778 | -0.17761698 | 0.013 | 0.059 | 1 | 6 |
| rmdn3 | 0.31778 | -0.14846216 | 0.013 | 0.059 | 1 | 6 |
| zmym4 | 0.31778 | -0.19690574 | 0.013 | 0.059 | 1 | 6 |
| BX088707.1 | 0.31778 | -0.18652568 | 0.013 | 0.059 | 1 | 6 |
| adam15 | 0.31778 | -0.11006231 | 0.013 | 0.059 | 1 | 6 |
| arpc1b | 0.31778 | -0.19976498 | 0.013 | 0.059 | 1 | 6 |
| asxl1 | 0.31778 | -0.19366625 | 0.013 | 0.059 | 1 | 6 |
| edrf1 | 0.31778 | -0.10495799 | 0.013 | 0.059 | 1 | 6 |
| hipk3a | 0.31778 | -0.15077744 | 0.013 | 0.059 | 1 | 6 |
| hunk | 0.31778 | -0.19883549 | 0.013 | 0.059 | 1 | 6 |
| il4r.1 | 0.31778 | -0.12707387 | 0.013 | 0.059 | 1 | 6 |
| lrp5.1 | 0.31778 | -0.13001531 | 0.013 | 0.059 | 1 | 6 |
| ndst3 | 0.31778 | -0.10192166 | 0.013 | 0.059 | 1 | 6 |
| polh | 0.31778 | -0.14987297 | 0.013 | 0.059 | 1 | 6 |
| rabep2 | 0.31778 | -0.20532925 | 0.013 | 0.059 | 1 | 6 |
| rassf8b | 0.31778 | -0.17033628 | 0.013 | 0.059 | 1 | 6 |
| rgcc | 0.31778 | -0.20866431 | 0.013 | 0.059 | 1 | 6 |
| saal1 | 0.31778 | -0.16568139 | 0.013 | 0.059 | 1 | 6 |
| si:ch211-137i24.10 | 0.31778 | -0.13656595 | 0.013 | 0.059 | 1 | 6 |
| si:ch211-51h9.6 | 0.31778 | -0.10643221 | 0.013 | 0.059 | 1 | 6 |
| si:dkey-66i24.9 | 0.31778 | -0.14327755 | 0.013 | 0.059 | 1 | 6 |
| snrnp200 | 0.31778 | -0.12680048 | 0.013 | 0.059 | 1 | 6 |
| tada3l | 0.31778 | -0.19533984 | 0.013 | 0.059 | 1 | 6 |
| tceanc2 | 0.31778 | -0.16418446 | 0.013 | 0.059 | 1 | 6 |
| xpo1a | 0.31778 | -0.17135972 | 0.013 | 0.059 | 1 | 6 |
| yrk | 0.31778 | -0.17520549 | 0.013 | 0.059 | 1 | 6 |
| CABZ01029822.1 | 0.31778 | -0.13107699 | 0.013 | 0.059 | 1 | 6 |
| abca2 | 0.31778 | -0.10367104 | 0.013 | 0.059 | 1 | 6 |
| arhgap23a | 0.31778 | -0.11147673 | 0.013 | 0.059 | 1 | 6 |
| cd40 | 0.31778 | -0.12630007 | 0.013 | 0.059 | 1 | 6 |
| chl1b | 0.31778 | -0.13448435 | 0.013 | 0.059 | 1 | 6 |
| clstn3 | 0.31778 | -0.15664586 | 0.013 | 0.059 | 1 | 6 |
| dio1 | 0.31778 | -0.10739463 | 0.013 | 0.059 | 1 | 6 |
| elmo1 | 0.31778 | -0.12452292 | 0.013 | 0.059 | 1 | 6 |
| ephb2a | 0.31778 | -0.10349968 | 0.013 | 0.059 | 1 | 6 |
| gdf6a | 0.31778 | -0.10564142 | 0.013 | 0.059 | 1 | 6 |
| gnl3l | 0.31778 | -0.13068297 | 0.013 | 0.059 | 1 | 6 |
| hpca | 0.31778 | -0.16973958 | 0.013 | 0.059 | 1 | 6 |
| hsd20b2 | 0.31778 | -0.12514903 | 0.013 | 0.059 | 1 | 6 |
| ice2 | 0.31778 | -0.12505074 | 0.013 | 0.059 | 1 | 6 |
| itgb2 | 0.31778 | -0.10818521 | 0.013 | 0.059 | 1 | 6 |
| lmbrd2a | 0.31778 | -0.12889587 | 0.013 | 0.059 | 1 | 6 |
| lrata | 0.31778 | -0.17151343 | 0.013 | 0.059 | 1 | 6 |
| mettl14 | 0.31778 | -0.11791467 | 0.013 | 0.059 | 1 | 6 |

| | | | | | | |
|---|---|---|---|---|---|---|
| mroh1 | 0.31778 | -0.10334352 | 0.013 | 0.059 | 1 | 6 |
| nme7 | 0.31778 | -0.16530041 | 0.013 | 0.059 | 1 | 6 |
| nrarpa | 0.31778 | -0.13326614 | 0.013 | 0.059 | 1 | 6 |
| pbx2 | 0.31778 | -0.12375619 | 0.013 | 0.059 | 1 | 6 |
| pigb | 0.31778 | -0.10290135 | 0.013 | 0.059 | 1 | 6 |
| ppp5c | 0.31778 | -0.12047838 | 0.013 | 0.059 | 1 | 6 |
| rad23aa | 0.31778 | -0.10023045 | 0.013 | 0.059 | 1 | 6 |
| si:cabz01069012.2 | 0.31778 | -0.16604575 | 0.013 | 0.059 | 1 | 6 |
| si:ch211-11n16.2 | 0.31778 | -0.1023222 | 0.013 | 0.059 | 1 | 6 |
| si:dkey-220k22.1 | 0.31778 | -0.12602463 | 0.013 | 0.059 | 1 | 6 |
| si:dkeyp-19e1.3 | 0.31778 | -0.1226811 | 0.013 | 0.059 | 1 | 6 |
| socs6a | 0.31778 | -0.18113779 | 0.013 | 0.059 | 1 | 6 |
| stx18 | 0.31778 | -0.1307986 | 0.013 | 0.059 | 1 | 6 |
| tmem241 | 0.31778 | -0.12254335 | 0.013 | 0.059 | 1 | 6 |
| triob | 0.31778 | -0.13447067 | 0.013 | 0.059 | 1 | 6 |
| tspan5a | 0.31778 | -0.13566272 | 0.013 | 0.059 | 1 | 6 |
| washc4 | 0.31778 | -0.11126094 | 0.013 | 0.059 | 1 | 6 |
| zgc:162972 | 0.31778 | -0.10347901 | 0.013 | 0.059 | 1 | 6 |
| zgc:85722 | 0.31778 | -0.10574395 | 0.013 | 0.059 | 1 | 6 |
| XLOC-024446 | 0.31778 | -0.12837457 | 0.013 | 0.059 | 1 | 6 |
| XLOC-033137 | 0.31778 | -0.11915099 | 0.013 | 0.059 | 1 | 6 |
| ZNF462 | 0.31778 | -0.11405197 | 0.013 | 0.059 | 1 | 6 |
| adam17a | 0.31778 | -0.10572177 | 0.013 | 0.059 | 1 | 6 |
| bag4 | 0.31778 | -0.10834422 | 0.013 | 0.059 | 1 | 6 |
| enkur | 0.31778 | -0.11772241 | 0.013 | 0.059 | 1 | 6 |
| ftr56 | 0.31778 | -0.12608659 | 0.013 | 0.059 | 1 | 6 |
| fuk | 0.31778 | -0.14284743 | 0.013 | 0.059 | 1 | 6 |
| gde1 | 0.31778 | -0.10441525 | 0.013 | 0.059 | 1 | 6 |
| gmeb1 | 0.31778 | -0.1002607 | 0.013 | 0.059 | 1 | 6 |
| iffo2b | 0.31778 | -0.12889235 | 0.013 | 0.059 | 1 | 6 |
| lypla2 | 0.31778 | -0.12260837 | 0.013 | 0.059 | 1 | 6 |
| nkx3.3 | 0.31778 | -0.11985119 | 0.013 | 0.059 | 1 | 6 |
| nrp2b | 0.31778 | -0.17354726 | 0.013 | 0.059 | 1 | 6 |
| rogdi | 0.31778 | -0.16863649 | 0.013 | 0.059 | 1 | 6 |
| rwdd3 | 0.31778 | -0.12144067 | 0.013 | 0.059 | 1 | 6 |
| si:ch73-233k15.2 | 0.31778 | -0.15624433 | 0.013 | 0.059 | 1 | 6 |
| slc6a16a | 0.31778 | -0.12844587 | 0.013 | 0.059 | 1 | 6 |
| slmapa | 0.31778 | -0.12596326 | 0.013 | 0.059 | 1 | 6 |
| tmsb | 0.31778 | -0.21075791 | 0.013 | 0.059 | 1 | 6 |
| zgc:77262 | 0.31778 | -0.12210547 | 0.013 | 0.059 | 1 | 6 |
| dia1b | 0.31778 | -0.13883602 | 0.013 | 0.059 | 1 | 6 |
| CABZ01092982.1 | 0.31778 | -0.13611672 | 0.013 | 0.059 | 1 | 6 |
| atp1a1b | 0.31778 | -0.17197826 | 0.013 | 0.059 | 1 | 6 |
| gpr107 | 0.31778 | -0.10359403 | 0.013 | 0.059 | 1 | 6 |
| il13ra1 | 0.31778 | -0.1402174 | 0.013 | 0.059 | 1 | 6 |
| slit2 | 0.31778 | -0.10528654 | 0.013 | 0.059 | 1 | 6 |
| ssh2a | 0.31778 | -0.20454007 | 0.013 | 0.059 | 1 | 6 |
| CR751227.2 | 0.31778 | -0.20152116 | 0.013 | 0.059 | 1 | 6 |

| | | | | | | |
|---|---|---|---|---|---|---|
| LOC100330864 | 0.31778 | -0.10027602 | 0.013 | 0.059 | 1 | 6 |
| LOC110440142 | 0.31778 | -0.21027256 | 0.013 | 0.059 | 1 | 6 |
| LOC110440195 | 0.31778 | -0.19869806 | 0.013 | 0.059 | 1 | 6 |
| XLOC-005448 | 0.31778 | -0.15291879 | 0.013 | 0.059 | 1 | 6 |
| anapc1 | 0.31778 | -0.1237515 | 0.013 | 0.059 | 1 | 6 |
| cog6 | 0.31778 | -0.1940998 | 0.013 | 0.059 | 1 | 6 |
| col15a1b | 0.31778 | -0.12056183 | 0.013 | 0.059 | 1 | 6 |
| dact2 | 0.31778 | -0.12046819 | 0.013 | 0.059 | 1 | 6 |
| gtf3c5 | 0.31778 | -0.14046835 | 0.013 | 0.059 | 1 | 6 |
| lims2 | 0.31778 | -0.14235209 | 0.013 | 0.059 | 1 | 6 |
| mcf2l2 | 0.31778 | -0.19857325 | 0.013 | 0.059 | 1 | 6 |
| mrtfbb | 0.31778 | -0.19125806 | 0.013 | 0.059 | 1 | 6 |
| msx3 | 0.31778 | -0.16459428 | 0.013 | 0.059 | 1 | 6 |
| nup153 | 0.31778 | -0.11154067 | 0.013 | 0.059 | 1 | 6 |
| ostm1 | 0.31778 | -0.10184386 | 0.013 | 0.059 | 1 | 6 |
| plbd1 | 0.31778 | -0.17136109 | 0.013 | 0.059 | 1 | 6 |
| rfc1 | 0.31778 | -0.10409477 | 0.013 | 0.059 | 1 | 6 |
| sema5a | 0.31778 | -0.1250911 | 0.013 | 0.059 | 1 | 6 |
| si:dkey-163m14.2 | 0.31778 | -0.12981205 | 0.013 | 0.059 | 1 | 6 |
| slc30a9 | 0.31778 | -0.10228118 | 0.013 | 0.059 | 1 | 6 |
| tgfbr1a | 0.31778 | -0.13220666 | 0.013 | 0.059 | 1 | 6 |
| trpc6a | 0.31778 | -0.10435077 | 0.013 | 0.059 | 1 | 6 |
| ttll1 | 0.31778 | -0.2048607 | 0.013 | 0.059 | 1 | 6 |
| zfand1 | 0.31778 | -0.13749772 | 0.013 | 0.059 | 1 | 6 |
| zgc:101840 | 0.31778 | -0.18666094 | 0.013 | 0.059 | 1 | 6 |
| LOC101885536 | 0.31778 | -0.10044552 | 0.013 | 0.059 | 1 | 6 |
| RASA2 | 0.31778 | -0.20538972 | 0.013 | 0.059 | 1 | 6 |
| usp48 | 0.31778 | -0.12851666 | 0.013 | 0.059 | 1 | 6 |
| zgc:171967 | 0.31781 | -0.19690896 | 0.026 | 0.118 | 1 | 6 |
| spina | 0.31791 | 0.242454911 | 0.143 | 0.059 | 1 | 6 |
| ppp2r3c | 0.31797 | -0.17460312 | 0.026 | 0.118 | 1 | 6 |
| nedd8l | 0.31847 | -0.13139056 | 0.065 | 0.176 | 1 | 6 |
| MDFIC | 0.3191 | -0.13332244 | 0.052 | 0.059 | 1 | 6 |
| chtf8 | 0.31917 | -0.22266412 | 0.052 | 0.118 | 1 | 6 |
| sesn1 | 0.31919 | 0.240432835 | 0.273 | 0.118 | 1 | 6 |
| nudt8 | 0.32041 | -0.10627516 | 0.026 | 0.118 | 1 | 6 |
| zmiz1a | 0.32064 | 0.21475796 | 0.182 | 0.059 | 1 | 6 |
| zdhhc6 | 0.32075 | -0.14751311 | 0.039 | 0.059 | 1 | 6 |
| eif4a1a | 0.32107 | 0.133194973 | 0.442 | 0.294 | 1 | 6 |
| cpsf2 | 0.32111 | -0.13966554 | 0.026 | 0.118 | 1 | 6 |
| rnf175 | 0.32113 | -0.11469751 | 0.026 | 0.118 | 1 | 6 |
| lrp1ab | 0.3212 | 0.123557843 | 0.234 | 0.235 | 1 | 6 |
| tln1 | 0.32134 | 0.206311202 | 0.195 | 0.059 | 1 | 6 |
| asip1 | 0.32197 | -0.33070901 | 0.065 | 0.176 | 1 | 6 |
| hsp90ab1 | 0.32198 | 0.116776034 | 0.948 | 0.941 | 1 | 6 |
| march2 | 0.32203 | -0.26670155 | 0.078 | 0.176 | 1 | 6 |
| cldn5a | 0.32243 | -0.10357283 | 0.078 | 0.176 | 1 | 6 |
| ctsc | 0.32244 | -0.22650793 | 0.039 | 0.118 | 1 | 6 |

| | | | | | | |
|---|---|---|---|---|---|---|
| taok1a | 0.3226 | -0.11790087 | 0.052 | 0.059 | 1 | 6 |
| gabarapb | 0.32279 | 0.109622623 | 0.623 | 0.471 | 1 | 6 |
| pcdh7b | 0.32281 | -0.21758988 | 0.026 | 0.118 | 1 | 6 |
| tmem263 | 0.32323 | 0.272127009 | 0.169 | 0.059 | 1 | 6 |
| pcyt2 | 0.32327 | -0.18789836 | 0.026 | 0.118 | 1 | 6 |
| PLBD2 | 0.3238 | -0.16074858 | 0.026 | 0.118 | 1 | 6 |
| atxn2l | 0.32387 | -0.20225731 | 0.026 | 0.118 | 1 | 6 |
| psmb5 | 0.32405 | 0.266763745 | 0.195 | 0.059 | 1 | 6 |
| rpl13 | 0.32415 | 0.126047416 | 0.974 | 0.882 | 1 | 6 |
| mfap3l | 0.3242 | -0.11264446 | 0.026 | 0.118 | 1 | 6 |
| XLOC-037780 | 0.32589 | 0.258619013 | 0.156 | 0.059 | 1 | 6 |
| b3gnt2b | 0.32595 | -0.11237386 | 0.026 | 0.118 | 1 | 6 |
| zc3h6 | 0.32613 | -0.13493647 | 0.026 | 0.118 | 1 | 6 |
| lamc1 | 0.32626 | -0.17394277 | 0.026 | 0.118 | 1 | 6 |
| cep131 | 0.32632 | -0.2178737 | 0.026 | 0.118 | 1 | 6 |
| rca2.1 | 0.32642 | -0.13337832 | 0.065 | 0.176 | 1 | 6 |
| acp6 | 0.32647 | -0.20344832 | 0.026 | 0.118 | 1 | 6 |
| si:dkey-93m18.4 | 0.32687 | -0.18849675 | 0.026 | 0.118 | 1 | 6 |
| tob1b | 0.32708 | -0.29790124 | 0.182 | 0.353 | 1 | 6 |
| zgc:153993 | 0.32722 | -0.18257564 | 0.026 | 0.118 | 1 | 6 |
| pi15a | 0.3274 | 0.326133654 | 0.156 | 0.059 | 1 | 6 |
| ndnl2 | 0.32755 | -0.27404605 | 0.065 | 0.176 | 1 | 6 |
| tox4b | 0.32791 | -0.18176258 | 0.026 | 0.118 | 1 | 6 |
| atf5b | 0.32886 | -0.14551445 | 0.208 | 0.353 | 1 | 6 |
| gmfb | 0.32888 | -0.14039884 | 0.286 | 0.294 | 1 | 6 |
| MMP23B | 0.32948 | -0.29686809 | 0.169 | 0.294 | 1 | 6 |
| si:dkey-111k8.5 | 0.32954 | -0.17372876 | 0.026 | 0.118 | 1 | 6 |
| LOC101886030 | 0.32974 | -0.24784446 | 0.026 | 0.118 | 1 | 6 |
| jak1 | 0.32993 | -0.13939537 | 0.117 | 0.235 | 1 | 6 |
| ak4 | 0.32997 | -0.26070213 | 0.065 | 0.176 | 1 | 6 |
| serpinf1 | 0.33028 | 0.216690309 | 0.169 | 0.118 | 1 | 6 |
| CABZ01084963.1 | 0.33049 | -0.17502843 | 0.065 | 0.118 | 1 | 6 |
| cables2b | 0.33066 | -0.2479026 | 0.026 | 0.118 | 1 | 6 |
| ppic | 0.33093 | -0.19931096 | 0.026 | 0.118 | 1 | 6 |
| dph6 | 0.33127 | -0.13630165 | 0.026 | 0.118 | 1 | 6 |
| man2b1 | 0.33185 | -0.1236768 | 0.026 | 0.118 | 1 | 6 |
| si:ch211-152c8.4 | 0.332 | -0.19347197 | 0.026 | 0.118 | 1 | 6 |
| idh3g | 0.3327 | -0.15383775 | 0.026 | 0.118 | 1 | 6 |
| irf2bpl | 0.33288 | 0.239282433 | 0.143 | 0.059 | 1 | 6 |
| MTERF4 | 0.33303 | -0.14839098 | 0.026 | 0.118 | 1 | 6 |
| ptrh2 | 0.33322 | -0.22232408 | 0.052 | 0.118 | 1 | 6 |
| commd1 | 0.3338 | -0.18637243 | 0.026 | 0.118 | 1 | 6 |
| orc4 | 0.33402 | -0.13340666 | 0.026 | 0.118 | 1 | 6 |
| hsf2 | 0.33404 | 0.252124077 | 0.195 | 0.059 | 1 | 6 |
| XLOC-020079 | 0.33404 | -0.19757001 | 0.026 | 0.118 | 1 | 6 |
| pigc | 0.33418 | -0.16927156 | 0.026 | 0.118 | 1 | 6 |
| cux1b | 0.33428 | -0.15922505 | 0.026 | 0.118 | 1 | 6 |
| cdh6 | 0.33437 | -0.25567812 | 0.104 | 0.235 | 1 | 6 |

| | | | | | | |
|---|---|---|---|---|---|---|
| timm10b | 0.33438 | -0.17232297 | 0.026 | 0.118 | 1 | 6 |
| chpf2 | 0.33441 | -0.20474428 | 0.026 | 0.118 | 1 | 6 |
| LOC110438079 | 0.33481 | -0.13876962 | 0.026 | 0.118 | 1 | 6 |
| sptbn2 | 0.33511 | -0.15051992 | 0.026 | 0.118 | 1 | 6 |
| ugt5b4 | 0.33537 | -0.15790276 | 0.026 | 0.118 | 1 | 6 |
| LOC110439915 | 0.33538 | -0.17333802 | 0.026 | 0.118 | 1 | 6 |
| tubb2b | 0.3355 | -0.30363836 | 0.26 | 0.353 | 1 | 6 |
| mrpl3 | 0.33554 | -0.14290159 | 0.026 | 0.118 | 1 | 6 |
| babam2 | 0.33563 | -0.18754263 | 0.026 | 0.118 | 1 | 6 |
| trim24 | 0.33563 | -0.15299981 | 0.026 | 0.118 | 1 | 6 |
| CABZ01088274.1 | 0.33566 | -0.19824145 | 0.026 | 0.118 | 1 | 6 |
| vcam1b | 0.33578 | -0.20151219 | 0.026 | 0.118 | 1 | 6 |
| sema3fa | 0.33635 | 0.288830515 | 0.182 | 0.059 | 1 | 6 |
| arrdc3a | 0.33637 | -0.13165885 | 0.065 | 0.176 | 1 | 6 |
| nup58 | 0.33709 | -0.11829099 | 0.052 | 0.059 | 1 | 6 |
| mapre3b | 0.33737 | -0.19123948 | 0.026 | 0.118 | 1 | 6 |
| eif1axa | 0.33752 | -0.27752645 | 0.104 | 0.235 | 1 | 6 |
| twist3 | 0.33762 | -0.16567157 | 0.026 | 0.118 | 1 | 6 |
| rad18 | 0.3381 | -0.18640278 | 0.026 | 0.118 | 1 | 6 |
| mat2aa | 0.33812 | 0.209968286 | 0.13 | 0.059 | 1 | 6 |
| tomm40l | 0.3387 | -0.14665262 | 0.026 | 0.118 | 1 | 6 |
| mrpl22 | 0.33871 | -0.14337252 | 0.026 | 0.118 | 1 | 6 |
| invs | 0.33884 | -0.16205854 | 0.026 | 0.118 | 1 | 6 |
| ksr1b | 0.33898 | -0.20085459 | 0.026 | 0.118 | 1 | 6 |
| itsn1 | 0.33901 | -0.16593887 | 0.026 | 0.118 | 1 | 6 |
| si:dkeyp-121d4.3 | 0.33901 | -0.19907703 | 0.026 | 0.118 | 1 | 6 |
| igf2bp3 | 0.33901 | 0.272901953 | 0.182 | 0.059 | 1 | 6 |
| narfl | 0.33907 | -0.15529971 | 0.026 | 0.118 | 1 | 6 |
| si:ch73-375g18.1 | 0.33911 | -0.16752201 | 0.026 | 0.118 | 1 | 6 |
| pex19 | 0.33914 | -0.15377298 | 0.026 | 0.118 | 1 | 6 |
| otub1a | 0.33917 | -0.16234043 | 0.026 | 0.118 | 1 | 6 |
| ggt5a | 0.33919 | -0.17111016 | 0.026 | 0.118 | 1 | 6 |
| scp2b | 0.3397 | -0.25700928 | 0.078 | 0.176 | 1 | 6 |
| dab2ipb | 0.34005 | -0.12234389 | 0.052 | 0.059 | 1 | 6 |
| ywhah | 0.34019 | 0.272814759 | 0.26 | 0.118 | 1 | 6 |
| kdm4b | 0.34114 | 0.204852404 | 0.143 | 0.059 | 1 | 6 |
| arpc3 | 0.34115 | -0.1743662 | 0.351 | 0.353 | 1 | 6 |
| cxcl12a | 0.34132 | 0.168146749 | 0.519 | 0.353 | 1 | 6 |
| pepd | 0.34166 | -0.1826787 | 0.104 | 0.176 | 1 | 6 |
| rps10 | 0.34168 | -0.10666467 | 0.948 | 1 | 1 | 6 |
| inka1b | 0.34319 | 0.175803406 | 0.091 | 0.059 | 1 | 6 |
| tmem256 | 0.34334 | -0.28549428 | 0.104 | 0.235 | 1 | 6 |
| col1a1b | 0.34361 | 0.249052866 | 0.844 | 0.706 | 1 | 6 |
| mtrr | 0.34394 | 0.120233551 | 0.078 | 0.059 | 1 | 6 |
| inka1a | 0.34448 | -0.23251329 | 0.143 | 0.294 | 1 | 6 |
| wu:fb16f03 | 0.34455 | 0.207654333 | 0.156 | 0.118 | 1 | 6 |
| ppp1cb | 0.34471 | 0.211419151 | 0.156 | 0.059 | 1 | 6 |
| zbtb37 | 0.34504 | -0.17252338 | 0.104 | 0.235 | 1 | 6 |

| | | | | | | |
|---|---|---|---|---|---|---|
| eif3k | 0.34556 | -0.25075707 | 0.377 | 0.471 | 1 | 6 |
| tent5c | 0.3457 | -0.21909008 | 0.052 | 0.118 | 1 | 6 |
| sept15 | 0.34578 | -0.24040877 | 0.091 | 0.176 | 1 | 6 |
| rbms3 | 0.34582 | -0.31210033 | 0.143 | 0.294 | 1 | 6 |
| rps27.1 | 0.34604 | 0.167962882 | 0.935 | 0.824 | 1 | 6 |
| gstk1 | 0.34628 | -0.16916791 | 0.156 | 0.294 | 1 | 6 |
| psma3 | 0.3476 | 0.158254853 | 0.377 | 0.235 | 1 | 6 |
| nid1b | 0.34766 | -0.31156856 | 0.143 | 0.294 | 1 | 6 |
| rpn1 | 0.34791 | -0.21393033 | 0.143 | 0.294 | 1 | 6 |
| calr | 0.3481 | -0.28006601 | 0.143 | 0.294 | 1 | 6 |
| gdi2 | 0.34814 | 0.267385194 | 0.416 | 0.294 | 1 | 6 |
| LOC103910871 | 0.34834 | -0.24627543 | 0.091 | 0.176 | 1 | 6 |
| prdx5 | 0.34939 | 0.291343247 | 0.273 | 0.118 | 1 | 6 |
| alg5 | 0.34961 | -0.11023548 | 0.117 | 0.118 | 1 | 6 |
| rps26l | 0.34967 | 0.135912964 | 0.948 | 0.882 | 1 | 6 |
| bzw1a | 0.34981 | 0.217634113 | 0.273 | 0.118 | 1 | 6 |
| loxl1 | 0.35024 | 0.273912778 | 0.143 | 0.059 | 1 | 6 |
| pik3r3a | 0.35044 | 0.301087162 | 0.247 | 0.118 | 1 | 6 |
| BX005309.1 | 0.35047 | -0.27637368 | 0.065 | 0.176 | 1 | 6 |
| sh3pxd2b | 0.35073 | 0.252556404 | 0.182 | 0.059 | 1 | 6 |
| patl1 | 0.35127 | -0.11204762 | 0.078 | 0.176 | 1 | 6 |
| twistnb | 0.35179 | -0.11096687 | 0.117 | 0.118 | 1 | 6 |
| dnajc24 | 0.35232 | -0.21175869 | 0.039 | 0.118 | 1 | 6 |
| pxna | 0.35373 | 0.11155818 | 0.143 | 0.059 | 1 | 6 |
| sema3bl | 0.35457 | 0.17258733 | 0.13 | 0.118 | 1 | 6 |
| degs1 | 0.35553 | -0.14842555 | 0.208 | 0.235 | 1 | 6 |
| kmt2bb | 0.35556 | 0.256735392 | 0.182 | 0.059 | 1 | 6 |
| gnl3 | 0.35567 | -0.21145516 | 0.143 | 0.294 | 1 | 6 |
| mrps24 | 0.35591 | 0.197892686 | 0.117 | 0.059 | 1 | 6 |
| eno3 | 0.35605 | 0.153121194 | 0.26 | 0.118 | 1 | 6 |
| smim15 | 0.35612 | 0.273677742 | 0.312 | 0.176 | 1 | 6 |
| LOC797168 | 0.35639 | 0.169895374 | 0.091 | 0.059 | 1 | 6 |
| ddx18 | 0.35651 | -0.1545314 | 0.156 | 0.294 | 1 | 6 |
| rhoaa | 0.35712 | -0.1334256 | 0.338 | 0.353 | 1 | 6 |
| gpt2l | 0.35714 | -0.22125295 | 0.039 | 0.118 | 1 | 6 |
| si:dkey-226m8.10 | 0.35762 | -0.15372509 | 0.039 | 0.059 | 1 | 6 |
| fxyd1 | 0.35781 | 0.23729812 | 0.182 | 0.059 | 1 | 6 |
| sf3a2 | 0.35782 | 0.194136171 | 0.182 | 0.059 | 1 | 6 |
| cnot3a | 0.35803 | -0.23505282 | 0.039 | 0.118 | 1 | 6 |
| selenok | 0.35808 | -0.10745276 | 0.169 | 0.294 | 1 | 6 |
| bcar1 | 0.35857 | -0.1471125 | 0.039 | 0.059 | 1 | 6 |
| socs3b | 0.35869 | 0.204211818 | 0.195 | 0.118 | 1 | 6 |
| si:dkeyp-69b9.6 | 0.35907 | 0.221501664 | 0.13 | 0.059 | 1 | 6 |
| slco5a1b | 0.35921 | -0.12291505 | 0.039 | 0.118 | 1 | 6 |
| tpm4b | 0.35934 | 0.152960284 | 0.234 | 0.118 | 1 | 6 |
| si:dkey-11f4.14 | 0.35959 | 0.27793179 | 0.273 | 0.118 | 1 | 6 |
| ndufa10 | 0.35971 | 0.25293055 | 0.26 | 0.118 | 1 | 6 |
| atrxl | 0.35991 | -0.21525033 | 0.052 | 0.118 | 1 | 6 |

| | | | | | | |
|---|---|---|---|---|---|---|
| ctsba | 0.35992 | -0.12417711 | 0.169 | 0.294 | 1 | 6 |
| si:ch73-46j18.5 | 0.35998 | 0.200278328 | 0.857 | 0.765 | 1 | 6 |
| calm2a | 0.36003 | -0.21593458 | 0.247 | 0.412 | 1 | 6 |
| rnf7 | 0.36012 | 0.230250363 | 0.273 | 0.118 | 1 | 6 |
| gstr | 0.36032 | 0.217105935 | 0.273 | 0.118 | 1 | 6 |
| eif2s1b | 0.36066 | 0.277625435 | 0.416 | 0.235 | 1 | 6 |
| COX5B | 0.36072 | -0.14475795 | 0.545 | 0.529 | 1 | 6 |
| wnt5b | 0.36076 | 0.270719942 | 0.182 | 0.059 | 1 | 6 |
| tpst1 | 0.36222 | -0.15170089 | 0.065 | 0.176 | 1 | 6 |
| naxe | 0.36301 | -0.19448035 | 0.039 | 0.118 | 1 | 6 |
| id3 | 0.36366 | 0.33548089 | 0.39 | 0.235 | 1 | 6 |
| anp32a | 0.36392 | -0.17633617 | 0.377 | 0.412 | 1 | 6 |
| ssrp1a | 0.36433 | -0.11292629 | 0.052 | 0.059 | 1 | 6 |
| cebpd | 0.3644 | 0.302733365 | 0.481 | 0.294 | 1 | 6 |
| appbp2 | 0.36478 | -0.14543921 | 0.039 | 0.059 | 1 | 6 |
| ccnl1b | 0.36482 | 0.154987519 | 0.104 | 0.059 | 1 | 6 |
| adamts15a | 0.36514 | -0.24671205 | 0.273 | 0.353 | 1 | 6 |
| ipo7 | 0.36541 | 0.136632799 | 0.143 | 0.118 | 1 | 6 |
| AL732488.1 | 0.36544 | -0.2074941 | 0.065 | 0.118 | 1 | 6 |
| si:dkey-177p2.6 | 0.36645 | 0.195310401 | 0.247 | 0.176 | 1 | 6 |
| rtraf | 0.36672 | 0.224139784 | 0.195 | 0.118 | 1 | 6 |
| s100a10b | 0.36788 | -0.19937423 | 0.831 | 0.941 | 1 | 6 |
| klf11b | 0.37017 | 0.224170952 | 0.169 | 0.059 | 1 | 6 |
| gadd45gip1 | 0.37094 | -0.12875309 | 0.104 | 0.118 | 1 | 6 |
| pomp | 0.37139 | 0.195794506 | 0.468 | 0.412 | 1 | 6 |
| uba1 | 0.37164 | 0.220449615 | 0.182 | 0.059 | 1 | 6 |
| ctps1b | 0.3718 | -0.13587282 | 0.039 | 0.059 | 1 | 6 |
| dpp7 | 0.37189 | -0.23846494 | 0.156 | 0.235 | 1 | 6 |
| snx6 | 0.37191 | 0.164277044 | 0.104 | 0.059 | 1 | 6 |
| far1 | 0.37204 | -0.12111468 | 0.052 | 0.059 | 1 | 6 |
| psma2 | 0.37227 | 0.235627186 | 0.338 | 0.235 | 1 | 6 |
| rbfox2 | 0.37251 | -0.20648513 | 0.104 | 0.235 | 1 | 6 |
| tm9sf5 | 0.37274 | -0.19874758 | 0.039 | 0.118 | 1 | 6 |
| imp4 | 0.37296 | -0.20000617 | 0.065 | 0.118 | 1 | 6 |
| cebpa | 0.3733 | -0.35109364 | 0.221 | 0.353 | 1 | 6 |
| clns1a | 0.37365 | -0.21552379 | 0.104 | 0.176 | 1 | 6 |
| dnpep | 0.37392 | 0.247760817 | 0.182 | 0.059 | 1 | 6 |
| klf13 | 0.37447 | -0.20341737 | 0.156 | 0.294 | 1 | 6 |
| si:dkey-67c22.2 | 0.37508 | 0.211149328 | 0.26 | 0.176 | 1 | 6 |
| znf385a | 0.37579 | 0.141024285 | 0.221 | 0.118 | 1 | 6 |
| rps20 | 0.37619 | -0.10088348 | 0.922 | 1 | 1 | 6 |
| ssr3 | 0.37635 | -0.30775291 | 0.377 | 0.529 | 1 | 6 |
| s100v2 | 0.37653 | 0.284892255 | 0.571 | 0.412 | 1 | 6 |
| chd3 | 0.37694 | 0.25176183 | 0.506 | 0.412 | 1 | 6 |
| psmb4 | 0.37714 | -0.15411615 | 0.325 | 0.353 | 1 | 6 |
| wt1b | 0.37788 | 0.249422702 | 0.299 | 0.235 | 1 | 6 |
| zgc:111986 | 0.37817 | 0.279430202 | 0.364 | 0.235 | 1 | 6 |
| mrps21 | 0.37849 | -0.20892999 | 0.078 | 0.176 | 1 | 6 |

| | | | | | | |
|---|---|---|---|---|---|---|
| si:dkey-6n21.12 | 0.37854 | 0.1783973 | 0.117 | 0.059 | 1 | 6 |
| ccdc58 | 0.37879 | -0.11176888 | 0.078 | 0.176 | 1 | 6 |
| zfp36l1b | 0.37896 | 0.102586244 | 0.416 | 0.294 | 1 | 6 |
| aip | 0.38052 | 0.262136715 | 0.247 | 0.118 | 1 | 6 |
| CR749775.2 | 0.38111 | -0.20075405 | 0.039 | 0.118 | 1 | 6 |
| yipf4 | 0.38123 | -0.15182185 | 0.117 | 0.235 | 1 | 6 |
| ankrd11 | 0.3817 | 0.249868502 | 0.182 | 0.059 | 1 | 6 |
| eif3s6ip | 0.38233 | 0.225024889 | 0.312 | 0.235 | 1 | 6 |
| cdc42bpb | 0.38249 | -0.1998008 | 0.065 | 0.176 | 1 | 6 |
| fam168a | 0.38274 | 0.277475598 | 0.286 | 0.176 | 1 | 6 |
| sec62 | 0.38301 | 0.234391495 | 0.182 | 0.059 | 1 | 6 |
| srrm2 | 0.3841 | -0.14105294 | 0.117 | 0.235 | 1 | 6 |
| pim1 | 0.3843 | 0.251441132 | 0.169 | 0.059 | 1 | 6 |
| meaf6 | 0.3845 | 0.218781359 | 0.182 | 0.059 | 1 | 6 |
| ubtf | 0.38469 | 0.158051354 | 0.156 | 0.118 | 1 | 6 |
| chmp6b | 0.38528 | -0.17702102 | 0.065 | 0.176 | 1 | 6 |
| mt-co1 | 0.38536 | -0.30167992 | 0.571 | 0.706 | 1 | 6 |
| ube2nb | 0.38547 | -0.23729294 | 0.104 | 0.235 | 1 | 6 |
| chchd6b | 0.38562 | -0.1581659 | 0.065 | 0.176 | 1 | 6 |
| pdia4 | 0.38624 | -0.20586232 | 0.065 | 0.118 | 1 | 6 |
| sumo3a | 0.38634 | -0.17199076 | 0.532 | 0.588 | 1 | 6 |
| nkain4 | 0.38638 | -0.15511898 | 0.065 | 0.176 | 1 | 6 |
| smim29 | 0.38639 | -0.18135048 | 0.065 | 0.176 | 1 | 6 |
| pamr1 | 0.38672 | -0.21360296 | 0.039 | 0.118 | 1 | 6 |
| tmbim4 | 0.38684 | -0.12319196 | 0.078 | 0.176 | 1 | 6 |
| si:ch73-281n10.2 | 0.38697 | 0.237873349 | 0.753 | 0.588 | 1 | 6 |
| nck1b | 0.38729 | 0.258458108 | 0.169 | 0.059 | 1 | 6 |
| c1qbp | 0.38747 | -0.14249095 | 0.169 | 0.176 | 1 | 6 |
| mmgt1 | 0.38787 | -0.21631749 | 0.104 | 0.235 | 1 | 6 |
| smarcc2 | 0.38801 | 0.216438549 | 0.169 | 0.059 | 1 | 6 |
| trim105 | 0.38847 | -0.12698075 | 0.039 | 0.118 | 1 | 6 |
| mvb12a | 0.38849 | 0.204427586 | 0.13 | 0.059 | 1 | 6 |
| hirip3 | 0.38869 | 0.230934992 | 0.182 | 0.059 | 1 | 6 |
| actb1 | 0.3887 | -0.14364485 | 0.584 | 0.588 | 1 | 6 |
| gstp1 | 0.38984 | -0.29225548 | 0.299 | 0.471 | 1 | 6 |
| tspan18a | 0.39013 | 0.192019518 | 0.234 | 0.176 | 1 | 6 |
| anapc10 | 0.39068 | -0.16232641 | 0.065 | 0.176 | 1 | 6 |
| sirt1 | 0.39135 | -0.21911569 | 0.104 | 0.235 | 1 | 6 |
| atp5f1b | 0.39217 | -0.22805205 | 0.571 | 0.647 | 1 | 6 |
| ube2d1a | 0.3931 | -0.11817549 | 0.078 | 0.176 | 1 | 6 |
| ckap4 | 0.39334 | -0.23524702 | 0.26 | 0.353 | 1 | 6 |
| gstm.3 | 0.39343 | -0.32882173 | 0.351 | 0.529 | 1 | 6 |
| crfb4 | 0.39376 | -0.20318888 | 0.065 | 0.176 | 1 | 6 |
| s1pr5a | 0.39386 | -0.28077798 | 0.156 | 0.294 | 1 | 6 |
| si:ch211-212k18.8 | 0.39392 | -0.27955718 | 0.208 | 0.353 | 1 | 6 |
| gosr2 | 0.39464 | -0.22426103 | 0.065 | 0.176 | 1 | 6 |
| imp3 | 0.39503 | -0.2098566 | 0.065 | 0.176 | 1 | 6 |
| emilin1b | 0.3952 | -0.22956446 | 0.065 | 0.176 | 1 | 6 |

| | | | | | | |
|---|---|---|---|---|---|---|
| paxip1 | 0.39534 | -0.22080922 | 0.065 | 0.176 | 1 | 6 |
| hspa9 | 0.39552 | 0.157589637 | 0.104 | 0.059 | 1 | 6 |
| ap3s1.1 | 0.39616 | -0.17004476 | 0.117 | 0.176 | 1 | 6 |
| rsf1b.1 | 0.39638 | 0.240115525 | 0.221 | 0.118 | 1 | 6 |
| srsf6a | 0.39701 | -0.20990358 | 0.065 | 0.176 | 1 | 6 |
| h3f3d | 0.39723 | 0.201482767 | 0.805 | 0.647 | 1 | 6 |
| fbp1a | 0.39758 | -0.23125142 | 0.091 | 0.176 | 1 | 6 |
| thoc5 | 0.39891 | 0.121316403 | 0.091 | 0.059 | 1 | 6 |
| ccbe1 | 0.39915 | -0.17142406 | 0.091 | 0.118 | 1 | 6 |
| rps21 | 0.40005 | -0.10096566 | 0.935 | 1 | 1 | 6 |
| zgc:113208 | 0.40071 | -0.2148735 | 0.065 | 0.118 | 1 | 6 |
| zfand2a | 0.40132 | -0.10351708 | 0.052 | 0.059 | 1 | 6 |
| slc25a11 | 0.40189 | -0.23069318 | 0.091 | 0.176 | 1 | 6 |
| ccnl1a | 0.40311 | -0.14601635 | 0.039 | 0.059 | 1 | 6 |
| dctn2 | 0.40343 | -0.2389488 | 0.156 | 0.294 | 1 | 6 |
| xpo1b | 0.4044 | -0.1342701 | 0.039 | 0.059 | 1 | 6 |
| slc25a34 | 0.40495 | -0.2105079 | 0.052 | 0.118 | 1 | 6 |
| coa3a | 0.40534 | -0.22708712 | 0.117 | 0.176 | 1 | 6 |
| psmb7 | 0.4055 | -0.18530212 | 0.312 | 0.471 | 1 | 6 |
| rbm8a | 0.40563 | -0.14617744 | 0.455 | 0.471 | 1 | 6 |
| si:ch211-262i1.6 | 0.40591 | 0.105474823 | 0.143 | 0.059 | 1 | 6 |
| lysmd2 | 0.40609 | -0.10611001 | 0.078 | 0.176 | 1 | 6 |
| tmem231 | 0.40633 | 0.175012102 | 0.13 | 0.059 | 1 | 6 |
| lhfpl2a | 0.40685 | 0.223005909 | 0.143 | 0.059 | 1 | 6 |
| zgc:56493 | 0.40753 | -0.15835787 | 0.779 | 0.824 | 1 | 6 |
| entpd1 | 0.40809 | -0.24001448 | 0.247 | 0.412 | 1 | 6 |
| si:dkey-180p18.9 | 0.40926 | -0.25042876 | 0.208 | 0.294 | 1 | 6 |
| clic2 | 0.4097 | -0.16195869 | 0.039 | 0.118 | 1 | 6 |
| ccnd1 | 0.40975 | 0.216838228 | 0.195 | 0.118 | 1 | 6 |
| phb2a | 0.41018 | 0.125916603 | 0.078 | 0.059 | 1 | 6 |
| bckdk | 0.41073 | 0.227614663 | 0.156 | 0.059 | 1 | 6 |
| pomk | 0.41091 | -0.13612641 | 0.039 | 0.059 | 1 | 6 |
| si:dkey-276j7.1 | 0.41121 | -0.1523308 | 0.156 | 0.176 | 1 | 6 |
| fam208b | 0.41128 | 0.160408537 | 0.104 | 0.059 | 1 | 6 |
| mrps6 | 0.41142 | -0.14007143 | 0.143 | 0.176 | 1 | 6 |
| selenot1b | 0.41166 | 0.191252506 | 0.156 | 0.059 | 1 | 6 |
| psma1 | 0.41203 | 0.266676007 | 0.468 | 0.294 | 1 | 6 |
| histh1l | 0.41256 | 0.259230124 | 0.156 | 0.059 | 1 | 6 |
| dpt | 0.41261 | 0.425450325 | 0.234 | 0.118 | 1 | 6 |
| COX3 | 0.41274 | -0.28229991 | 0.597 | 0.706 | 1 | 6 |
| ube2w | 0.41287 | -0.11099777 | 0.039 | 0.118 | 1 | 6 |
| zgc:114200 | 0.4138 | -0.19898049 | 0.078 | 0.118 | 1 | 6 |
| ehd2b | 0.41412 | 0.148945778 | 0.143 | 0.118 | 1 | 6 |
| kirrel1a | 0.41421 | 0.262465013 | 0.169 | 0.059 | 1 | 6 |
| palm1b | 0.41422 | 0.248488207 | 0.156 | 0.059 | 1 | 6 |
| cox17 | 0.41475 | -0.27316452 | 0.078 | 0.176 | 1 | 6 |
| ppp2r5cb | 0.41503 | 0.142035217 | 0.091 | 0.059 | 1 | 6 |
| ctps1a | 0.41567 | -0.19821829 | 0.039 | 0.118 | 1 | 6 |

| | | | | | | |
|---|---|---|---|---|---|---|
| CABZ01049847.1 | 0.41703 | -0.12715253 | 0.039 | 0.118 | 1 | 6 |
| ptmab | 0.41707 | 0.149989876 | 0.857 | 0.824 | 1 | 6 |
| rdh10a | 0.41726 | 0.272605131 | 0.221 | 0.118 | 1 | 6 |
| ufm1 | 0.41808 | -0.11609818 | 0.078 | 0.176 | 1 | 6 |
| jagn1b | 0.41838 | 0.13942451 | 0.273 | 0.235 | 1 | 6 |
| ccdc32 | 0.41842 | -0.13276023 | 0.117 | 0.235 | 1 | 6 |
| tsku | 0.41847 | 0.108005983 | 0.182 | 0.176 | 1 | 6 |
| zgc:56106 | 0.41863 | -0.13059232 | 0.039 | 0.059 | 1 | 6 |
| klf11a | 0.4194 | 0.218323603 | 0.169 | 0.059 | 1 | 6 |
| tjp1a | 0.41942 | 0.175633429 | 0.182 | 0.118 | 1 | 6 |
| ephb3 | 0.41947 | 0.152688115 | 0.429 | 0.294 | 1 | 6 |
| rps11 | 0.41996 | 0.104597697 | 0.974 | 0.941 | 1 | 6 |
| usp14 | 0.42022 | -0.20996763 | 0.052 | 0.118 | 1 | 6 |
| ppt1 | 0.42121 | 0.142296619 | 0.364 | 0.235 | 1 | 6 |
| eif2s2 | 0.42127 | 0.139001485 | 0.558 | 0.412 | 1 | 6 |
| prkar1ab | 0.42138 | 0.199228485 | 0.13 | 0.059 | 1 | 6 |
| pafah1b3 | 0.42152 | -0.1306156 | 0.039 | 0.059 | 1 | 6 |
| rpe | 0.4217 | 0.220821742 | 0.169 | 0.059 | 1 | 6 |
| tefa | 0.42191 | 0.131332554 | 0.195 | 0.176 | 1 | 6 |
| CZQB01141835.1 | 0.42209 | 0.178882137 | 0.143 | 0.059 | 1 | 6 |
| kmt2cb | 0.42257 | 0.176998168 | 0.117 | 0.059 | 1 | 6 |
| metrnla | 0.42339 | 0.2280285 | 0.169 | 0.059 | 1 | 6 |
| ubb | 0.42427 | 0.162603481 | 0.506 | 0.353 | 1 | 6 |
| arl6ip4 | 0.42484 | 0.112746498 | 0.065 | 0.059 | 1 | 6 |
| exosc1 | 0.42538 | -0.18554973 | 0.039 | 0.118 | 1 | 6 |
| pam16 | 0.42592 | -0.21051313 | 0.156 | 0.294 | 1 | 6 |
| psmd10 | 0.42717 | -0.19062056 | 0.065 | 0.118 | 1 | 6 |
| selenot2 | 0.42737 | -0.18662012 | 0.104 | 0.176 | 1 | 6 |
| ptp4a1 | 0.42741 | 0.125840665 | 0.286 | 0.176 | 1 | 6 |
| psmc5 | 0.42742 | 0.133651678 | 0.299 | 0.176 | 1 | 6 |
| scrib | 0.4278 | -0.12202553 | 0.052 | 0.059 | 1 | 6 |
| carhsp1 | 0.4284 | -0.17946154 | 0.078 | 0.118 | 1 | 6 |
| cltb | 0.42849 | -0.14150256 | 0.039 | 0.059 | 1 | 6 |
| si:dkey-42i9.6 | 0.42983 | -0.1548939 | 0.117 | 0.235 | 1 | 6 |
| igf2a | 0.43053 | -0.16097398 | 0.052 | 0.118 | 1 | 6 |
| pkn2 | 0.43147 | -0.19281772 | 0.052 | 0.118 | 1 | 6 |
| hdlbpa | 0.43211 | 0.157234389 | 0.506 | 0.353 | 1 | 6 |
| ldlrad4a | 0.43256 | 0.157356094 | 0.091 | 0.059 | 1 | 6 |
| rasl12 | 0.43323 | -0.18219778 | 0.078 | 0.118 | 1 | 6 |
| MRPS12 | 0.43362 | 0.16423721 | 0.13 | 0.059 | 1 | 6 |
| SMIM10 | 0.43395 | -0.24142132 | 0.039 | 0.118 | 1 | 6 |
| adi1 | 0.43431 | 0.165600156 | 0.156 | 0.118 | 1 | 6 |
| fam102ab | 0.43504 | 0.229126381 | 0.143 | 0.059 | 1 | 6 |
| ube2na | 0.43527 | 0.120674725 | 0.221 | 0.118 | 1 | 6 |
| sept7b | 0.43544 | 0.177857295 | 0.117 | 0.059 | 1 | 6 |
| pcdh10b | 0.43586 | 0.132655525 | 0.078 | 0.059 | 1 | 6 |
| grpel2 | 0.43654 | 0.202572297 | 0.143 | 0.059 | 1 | 6 |
| cnn3a | 0.43655 | 0.248055195 | 0.416 | 0.294 | 1 | 6 |

| | | | | | | |
|---|---|---|---|---|---|---|
| cdc26 | 0.4382 | 0.237357425 | 0.286 | 0.176 | 1 | 6 |
| krt94 | 0.4391 | 0.288896098 | 0.662 | 0.529 | 1 | 6 |
| kmt5aa | 0.43952 | 0.139658468 | 0.234 | 0.118 | 1 | 6 |
| ppih | 0.43981 | -0.139043 | 0.091 | 0.118 | 1 | 6 |
| mmp16a | 0.44014 | -0.19115764 | 0.377 | 0.412 | 1 | 6 |
| hnrnpm | 0.4407 | 0.22897341 | 0.169 | 0.059 | 1 | 6 |
| dido1 | 0.44191 | 0.125704441 | 0.195 | 0.176 | 1 | 6 |
| si:rp71-36a1.3 | 0.44237 | -0.19387791 | 0.039 | 0.118 | 1 | 6 |
| aldh1a2 | 0.44262 | 0.308502401 | 0.325 | 0.176 | 1 | 6 |
| brk1 | 0.44315 | 0.124253899 | 0.312 | 0.294 | 1 | 6 |
| rbbp6 | 0.4432 | -0.26606572 | 0.156 | 0.294 | 1 | 6 |
| uba5 | 0.44402 | -0.21438243 | 0.078 | 0.176 | 1 | 6 |
| CR847566.1 | 0.44491 | 0.239733651 | 0.325 | 0.235 | 1 | 6 |
| ece2b | 0.44498 | -0.18302607 | 0.039 | 0.118 | 1 | 6 |
| LOC103909205 | 0.44546 | -0.1365888 | 0.039 | 0.059 | 1 | 6 |
| si:ch73-42p12.2 | 0.44552 | -0.10141138 | 0.234 | 0.235 | 1 | 6 |
| tbcb | 0.4468 | 0.101466196 | 0.13 | 0.118 | 1 | 6 |
| spon1b | 0.44703 | -0.12726073 | 0.182 | 0.294 | 1 | 6 |
| hmgn2 | 0.44716 | 0.113877765 | 0.584 | 0.647 | 1 | 6 |
| cnpy3 | 0.44753 | -0.17845579 | 0.052 | 0.118 | 1 | 6 |
| rcn1 | 0.44798 | -0.11432371 | 0.039 | 0.118 | 1 | 6 |
| tmem98 | 0.44801 | 0.157498399 | 0.104 | 0.059 | 1 | 6 |
| ube2e3 | 0.44887 | 0.232074788 | 0.403 | 0.294 | 1 | 6 |
| LOC101884451 | 0.44945 | -0.11875097 | 0.182 | 0.294 | 1 | 6 |
| nutf2 | 0.44986 | 0.214542418 | 0.169 | 0.059 | 1 | 6 |
| ddx23 | 0.45057 | -0.21959445 | 0.117 | 0.235 | 1 | 6 |
| prkcsh | 0.45104 | -0.2299901 | 0.117 | 0.235 | 1 | 6 |
| klf3 | 0.4512 | -0.21130514 | 0.052 | 0.118 | 1 | 6 |
| gadd45bb | 0.45141 | -0.12602493 | 0.039 | 0.059 | 1 | 6 |
| add3a | 0.45288 | 0.221416193 | 0.442 | 0.353 | 1 | 6 |
| pgrmc2 | 0.45317 | 0.126068182 | 0.091 | 0.059 | 1 | 6 |
| wdr82 | 0.45317 | -0.16336584 | 0.117 | 0.235 | 1 | 6 |
| oip5-as1 | 0.45394 | 0.244654511 | 0.221 | 0.118 | 1 | 6 |
| rnf2 | 0.45496 | 0.202244574 | 0.143 | 0.059 | 1 | 6 |
| psmb6 | 0.45532 | 0.197290013 | 0.351 | 0.294 | 1 | 6 |
| atp5f1e | 0.45595 | 0.203046188 | 0.636 | 0.471 | 1 | 6 |
| psmc6 | 0.45669 | 0.159956415 | 0.182 | 0.118 | 1 | 6 |
| nufip1 | 0.45706 | -0.141601 | 0.039 | 0.118 | 1 | 6 |
| nktr | 0.45727 | -0.12570759 | 0.039 | 0.118 | 1 | 6 |
| chmp1a | 0.45743 | -0.15943217 | 0.039 | 0.118 | 1 | 6 |
| caprin1b | 0.45824 | -0.25126607 | 0.325 | 0.471 | 1 | 6 |
| si:ch211-147a11.3 | 0.45911 | -0.18896437 | 0.065 | 0.118 | 1 | 6 |
| thg1l | 0.46071 | -0.16695775 | 0.052 | 0.118 | 1 | 6 |
| chka | 0.46165 | -0.20497735 | 0.039 | 0.118 | 1 | 6 |
| drap1 | 0.46214 | -0.16365412 | 0.169 | 0.294 | 1 | 6 |
| cyb5a | 0.4631 | -0.15578885 | 0.078 | 0.118 | 1 | 6 |
| csnk2b | 0.46333 | -0.17417505 | 0.312 | 0.353 | 1 | 6 |
| col12a1a | 0.46385 | 0.280986295 | 0.247 | 0.118 | 1 | 6 |

| | | | | | | |
|---|---|---|---|---|---|---|
| trim23 | 0.4646 | -0.12312176 | 0.039 | 0.059 | 1 | 6 |
| wnt4a | 0.46488 | 0.105548158 | 0.078 | 0.059 | 1 | 6 |
| timm9 | 0.46521 | 0.167897423 | 0.13 | 0.059 | 1 | 6 |
| pbx3b | 0.46565 | 0.225745933 | 0.208 | 0.118 | 1 | 6 |
| prickle2b | 0.46581 | -0.19648784 | 0.039 | 0.118 | 1 | 6 |
| hvcn1 | 0.46604 | 0.198308378 | 0.156 | 0.059 | 1 | 6 |
| luc7l | 0.46607 | -0.1541543 | 0.117 | 0.235 | 1 | 6 |
| si:dkey-9i23.16 | 0.46614 | 0.201186982 | 0.377 | 0.294 | 1 | 6 |
| psen2 | 0.46615 | -0.17467202 | 0.039 | 0.118 | 1 | 6 |
| eif4g2b | 0.46623 | 0.190439255 | 0.182 | 0.118 | 1 | 6 |
| aldh7a1 | 0.46624 | -0.2185062 | 0.117 | 0.235 | 1 | 6 |
| btf3 | 0.46646 | 0.190900171 | 0.792 | 0.647 | 1 | 6 |
| cavin2b | 0.46667 | 0.216384324 | 0.182 | 0.118 | 1 | 6 |
| abcf1 | 0.46777 | -0.16624209 | 0.169 | 0.294 | 1 | 6 |
| ap1s2 | 0.46783 | 0.204725695 | 0.247 | 0.118 | 1 | 6 |
| polr2k | 0.46784 | 0.249457778 | 0.221 | 0.118 | 1 | 6 |
| man1a2 | 0.46848 | -0.1281473 | 0.039 | 0.059 | 1 | 6 |
| atp6ap2 | 0.46992 | -0.13200656 | 0.078 | 0.176 | 1 | 6 |
| nipa2 | 0.47037 | -0.15399847 | 0.039 | 0.118 | 1 | 6 |
| mfap2 | 0.47046 | 0.226374068 | 0.688 | 0.529 | 1 | 6 |
| wbp11 | 0.47113 | -0.1821007 | 0.039 | 0.118 | 1 | 6 |
| flnb | 0.47197 | 0.201101347 | 0.117 | 0.059 | 1 | 6 |
| RPL37A | 0.47221 | -0.10295108 | 0.104 | 0.118 | 1 | 6 |
| uchl5 | 0.47286 | -0.22527979 | 0.078 | 0.176 | 1 | 6 |
| eef1akmt1 | 0.47361 | -0.1354229 | 0.039 | 0.118 | 1 | 6 |
| yipf1 | 0.47383 | -0.11776929 | 0.039 | 0.118 | 1 | 6 |
| si:ch73-37h15.2 | 0.47476 | 0.192275536 | 0.13 | 0.059 | 1 | 6 |
| zgc:195173 | 0.47515 | 0.134815596 | 0.377 | 0.353 | 1 | 6 |
| hcfc1b | 0.47538 | -0.15902335 | 0.039 | 0.118 | 1 | 6 |
| ugdh | 0.47593 | -0.17311934 | 0.039 | 0.118 | 1 | 6 |
| pbx1b | 0.476 | 0.190312806 | 0.13 | 0.059 | 1 | 6 |
| pak2a | 0.47604 | -0.1694566 | 0.039 | 0.118 | 1 | 6 |
| irf2bp2a | 0.47631 | 0.21759794 | 0.429 | 0.353 | 1 | 6 |
| clptm1 | 0.47654 | -0.19922687 | 0.039 | 0.118 | 1 | 6 |
| ndufa6 | 0.47833 | 0.210269834 | 0.338 | 0.294 | 1 | 6 |
| stip1 | 0.47853 | 0.210470124 | 0.156 | 0.059 | 1 | 6 |
| srsf9 | 0.4788 | 0.190623342 | 0.273 | 0.176 | 1 | 6 |
| cwc25 | 0.47918 | 0.217226245 | 0.208 | 0.118 | 1 | 6 |
| ube2l3b | 0.47967 | 0.179143275 | 0.312 | 0.176 | 1 | 6 |
| cnep1r1 | 0.47983 | 0.137919395 | 0.117 | 0.059 | 1 | 6 |
| pmaip1 | 0.48031 | -0.10110278 | 0.039 | 0.118 | 1 | 6 |
| si:ch73-103b11.2 | 0.48039 | -0.14279632 | 0.039 | 0.118 | 1 | 6 |
| ddt | 0.48129 | 0.182236081 | 0.195 | 0.118 | 1 | 6 |
| fzd2 | 0.48154 | 0.246165695 | 0.286 | 0.176 | 1 | 6 |
| zgc:91944 | 0.48186 | -0.13572112 | 0.078 | 0.176 | 1 | 6 |
| adcy9 | 0.48188 | -0.18087866 | 0.039 | 0.118 | 1 | 6 |
| lamtor4 | 0.4822 | 0.178387128 | 0.156 | 0.059 | 1 | 6 |
| hnrnpa1b | 0.48245 | -0.16878117 | 0.247 | 0.294 | 1 | 6 |

| | | | | | | |
|---|---|---|---|---|---|---|
| MED9 | 0.48286 | -0.21600026 | 0.052 | 0.118 | 1 | 6 |
| frg1 | 0.48323 | -0.17030999 | 0.078 | 0.176 | 1 | 6 |
| slc6a4a | 0.48408 | -0.14931822 | 0.039 | 0.118 | 1 | 6 |
| mrps23 | 0.48443 | -0.1482261 | 0.039 | 0.118 | 1 | 6 |
| ufc1 | 0.4845 | -0.13390457 | 0.039 | 0.118 | 1 | 6 |
| selenot1a | 0.48579 | -0.11317018 | 0.091 | 0.118 | 1 | 6 |
| LOC553316 | 0.48586 | -0.18971563 | 0.039 | 0.118 | 1 | 6 |
| gnb1b | 0.48586 | -0.23414147 | 0.377 | 0.529 | 1 | 6 |
| zc3h11a | 0.48674 | 0.143849689 | 0.091 | 0.059 | 1 | 6 |
| tns2a | 0.48701 | -0.14113493 | 0.13 | 0.235 | 1 | 6 |
| naxd | 0.48802 | -0.15811796 | 0.039 | 0.118 | 1 | 6 |
| cdc40 | 0.48866 | -0.22481365 | 0.078 | 0.176 | 1 | 6 |
| rex1bd | 0.48897 | 0.152531705 | 0.104 | 0.059 | 1 | 6 |
| rhoub | 0.48928 | -0.13385316 | 0.039 | 0.059 | 1 | 6 |
| ptp4a2b | 0.48937 | 0.194636727 | 0.156 | 0.059 | 1 | 6 |
| dpm3 | 0.48951 | -0.12682447 | 0.039 | 0.118 | 1 | 6 |
| khsrp | 0.49045 | -0.19732865 | 0.234 | 0.294 | 1 | 6 |
| acadm | 0.49058 | -0.2315763 | 0.182 | 0.294 | 1 | 6 |
| rhobtb4 | 0.49068 | -0.17087253 | 0.039 | 0.118 | 1 | 6 |
| rangrf | 0.49083 | -0.12492684 | 0.039 | 0.118 | 1 | 6 |
| dpf2 | 0.49169 | 0.103481873 | 0.052 | 0.059 | 1 | 6 |
| si:zfos-1192g2.3 | 0.49283 | 0.19194756 | 0.156 | 0.059 | 1 | 6 |
| adam17b | 0.49287 | -0.1120984 | 0.039 | 0.118 | 1 | 6 |
| rprd2b | 0.49306 | -0.13700348 | 0.078 | 0.176 | 1 | 6 |
| amotl2a | 0.49312 | -0.11259464 | 0.052 | 0.059 | 1 | 6 |
| BX649250.1 | 0.49386 | -0.12179595 | 0.039 | 0.118 | 1 | 6 |
| crim1 | 0.49399 | 0.136947642 | 0.143 | 0.059 | 1 | 6 |
| nfia | 0.49438 | -0.18902284 | 0.325 | 0.471 | 1 | 6 |
| peli1b | 0.49443 | -0.16504829 | 0.039 | 0.118 | 1 | 6 |
| actr10 | 0.49461 | -0.1264491 | 0.039 | 0.118 | 1 | 6 |
| mmp14a | 0.49473 | -0.20084664 | 0.156 | 0.235 | 1 | 6 |
| si:ch211-39i22.1 | 0.49492 | 0.272130303 | 0.442 | 0.294 | 1 | 6 |
| pbx1a | 0.4951 | 0.266471677 | 0.351 | 0.235 | 1 | 6 |
| tmem33 | 0.49526 | -0.14568696 | 0.039 | 0.118 | 1 | 6 |
| ccng1 | 0.49537 | -0.18108703 | 0.532 | 0.588 | 1 | 6 |
| top2b | 0.49629 | -0.13945184 | 0.13 | 0.235 | 1 | 6 |
| timm13 | 0.49664 | -0.20404048 | 0.052 | 0.118 | 1 | 6 |
| pomt1 | 0.49668 | -0.11991988 | 0.039 | 0.118 | 1 | 6 |
| alkal1 | 0.49678 | -0.12590391 | 0.039 | 0.118 | 1 | 6 |
| sumo3b | 0.49692 | -0.13136932 | 0.065 | 0.118 | 1 | 6 |
| NDUFAF8 | 0.49695 | -0.13883319 | 0.039 | 0.118 | 1 | 6 |
| wdr89 | 0.49699 | -0.14532364 | 0.039 | 0.118 | 1 | 6 |
| ddx61 | 0.49704 | 0.189104411 | 0.442 | 0.294 | 1 | 6 |
| srpk1a | 0.49718 | -0.18518405 | 0.052 | 0.118 | 1 | 6 |
| btf3l4 | 0.49734 | 0.17888886 | 0.234 | 0.118 | 1 | 6 |
| lrrfip1a | 0.49754 | -0.13398598 | 0.039 | 0.118 | 1 | 6 |
| twsg1a | 0.49774 | 0.140565376 | 0.104 | 0.059 | 1 | 6 |
| tuba1a | 0.49775 | -0.19160803 | 0.065 | 0.118 | 1 | 6 |

| | | | | | | |
|---|---|---|---|---|---|---|
| zgc:112982 | 0.49777 | -0.135508 | 0.039 | 0.118 | 1 | 6 |
| ruvbl1 | 0.49777 | -0.15362242 | 0.039 | 0.118 | 1 | 6 |
| si:ch211-197h24.6 | 0.49869 | -0.20155252 | 0.078 | 0.176 | 1 | 6 |
| cyp46a1.2 | 0.4987 | -0.13642908 | 0.039 | 0.118 | 1 | 6 |
| lgals2a | 0.49929 | -0.1626017 | 0.234 | 0.353 | 1 | 6 |
| hlx1 | 0.49942 | -0.20147319 | 0.052 | 0.118 | 1 | 6 |
| dap1b | 0.4997 | 0.157353859 | 0.364 | 0.235 | 1 | 6 |
| cct8 | 0.49981 | 0.172320495 | 0.519 | 0.471 | 1 | 6 |
| baz1a | 0.49998 | -0.13290275 | 0.039 | 0.118 | 1 | 6 |
| camk2n1a | 0.49999 | -0.1205322 | 0.039 | 0.118 | 1 | 6 |
| cdc14b | 0.50028 | 0.108865289 | 0.208 | 0.118 | 1 | 6 |
| psmd4a | 0.5005 | -0.139526 | 0.091 | 0.118 | 1 | 6 |
| clcn5a | 0.50096 | -0.14726974 | 0.039 | 0.118 | 1 | 6 |
| wasf3b | 0.50109 | -0.1130571 | 0.039 | 0.118 | 1 | 6 |
| selenop | 0.50154 | -0.19577584 | 0.078 | 0.176 | 1 | 6 |
| tmem9b | 0.50158 | 0.119919318 | 0.091 | 0.059 | 1 | 6 |
| dph2 | 0.50171 | -0.15826261 | 0.039 | 0.118 | 1 | 6 |
| hspa12a | 0.50176 | -0.13509666 | 0.039 | 0.118 | 1 | 6 |
| col1a1a | 0.50188 | 0.180533939 | 0.844 | 0.765 | 1 | 6 |
| hira | 0.50208 | -0.16405711 | 0.039 | 0.118 | 1 | 6 |
| ptprsa | 0.5021 | -0.19549886 | 0.078 | 0.176 | 1 | 6 |
| PYURF | 0.50223 | -0.18247243 | 0.078 | 0.176 | 1 | 6 |
| ddx27 | 0.50225 | 0.22635898 | 0.221 | 0.118 | 1 | 6 |
| cdc42se1 | 0.50233 | -0.15195586 | 0.039 | 0.118 | 1 | 6 |
| bach2b | 0.50239 | -0.17836256 | 0.039 | 0.118 | 1 | 6 |
| nkrf | 0.50253 | 0.167929052 | 0.104 | 0.059 | 1 | 6 |
| ddi2 | 0.50258 | -0.11671883 | 0.039 | 0.118 | 1 | 6 |
| larp1 | 0.5032 | -0.18904936 | 0.052 | 0.118 | 1 | 6 |
| dnph1 | 0.50321 | -0.12004287 | 0.039 | 0.118 | 1 | 6 |
| sh2b3 | 0.50326 | -0.15622734 | 0.039 | 0.118 | 1 | 6 |
| paplna | 0.50331 | -0.21652608 | 0.039 | 0.118 | 1 | 6 |
| trabd | 0.50333 | -0.1170128 | 0.039 | 0.118 | 1 | 6 |
| ywhae2 | 0.50336 | -0.17398807 | 0.039 | 0.118 | 1 | 6 |
| zgc:193742 | 0.50336 | -0.12521221 | 0.039 | 0.118 | 1 | 6 |
| h2afvb | 0.50351 | 0.173099041 | 0.623 | 0.471 | 1 | 6 |
| LOC101882592 | 0.50408 | -0.13549016 | 0.039 | 0.059 | 1 | 6 |
| mob4 | 0.50492 | 0.197183483 | 0.221 | 0.118 | 1 | 6 |
| sesn3 | 0.50551 | 0.16901865 | 0.117 | 0.059 | 1 | 6 |
| zgc:112437 | 0.50653 | 0.1876685 | 0.156 | 0.118 | 1 | 6 |
| strap | 0.50671 | -0.19531992 | 0.091 | 0.176 | 1 | 6 |
| tomm20b | 0.50712 | 0.160474699 | 0.351 | 0.294 | 1 | 6 |
| grwd1 | 0.50726 | -0.15946599 | 0.078 | 0.176 | 1 | 6 |
| tomm5 | 0.50753 | 0.186997925 | 0.442 | 0.294 | 1 | 6 |
| usp7 | 0.50763 | -0.17295591 | 0.052 | 0.118 | 1 | 6 |
| lmcd1 | 0.50774 | 0.175367646 | 0.143 | 0.059 | 1 | 6 |
| col2a1b | 0.50808 | 0.201061437 | 0.143 | 0.059 | 1 | 6 |
| rerea | 0.50866 | -0.15039116 | 0.091 | 0.118 | 1 | 6 |
| strn3 | 0.50946 | 0.167642565 | 0.104 | 0.059 | 1 | 6 |

| | | | | | | |
|---|---|---|---|---|---|---|
| cnot9 | 0.51004 | -0.17918678 | 0.078 | 0.176 | 1 | 6 |
| ndufab1b | 0.51096 | -0.2050746 | 0.208 | 0.294 | 1 | 6 |
| stag2b | 0.51107 | -0.17756522 | 0.078 | 0.176 | 1 | 6 |
| pole3 | 0.51127 | -0.19962836 | 0.091 | 0.176 | 1 | 6 |
| znf503 | 0.51132 | -0.16620142 | 0.078 | 0.176 | 1 | 6 |
| sntb2 | 0.51163 | 0.219034043 | 0.221 | 0.118 | 1 | 6 |
| tead1b | 0.5117 | 0.181692991 | 0.13 | 0.059 | 1 | 6 |
| pinx1 | 0.51248 | -0.24884609 | 0.247 | 0.353 | 1 | 6 |
| wu:fi37e09 | 0.5135 | 0.119747304 | 0.208 | 0.118 | 1 | 6 |
| gna11b | 0.5137 | -0.17012526 | 0.195 | 0.235 | 1 | 6 |
| slc25a51b | 0.514 | 0.125985645 | 0.091 | 0.059 | 1 | 6 |
| arpin | 0.51576 | -0.21705455 | 0.091 | 0.176 | 1 | 6 |
| zgc:153713 | 0.51621 | 0.233656877 | 0.338 | 0.235 | 1 | 6 |
| erbb2 | 0.51621 | 0.171265585 | 0.13 | 0.059 | 1 | 6 |
| sdhc | 0.51747 | -0.16675683 | 0.182 | 0.235 | 1 | 6 |
| si:ch211-194k22.8 | 0.51751 | 0.1925967 | 0.143 | 0.059 | 1 | 6 |
| rab14 | 0.5177 | 0.115262129 | 0.078 | 0.059 | 1 | 6 |
| stau2 | 0.51777 | 0.1127793 | 0.403 | 0.294 | 1 | 6 |
| ctgfa | 0.51778 | 0.240413059 | 0.156 | 0.059 | 1 | 6 |
| CABZ01078055.1 | 0.51781 | -0.20666655 | 0.052 | 0.118 | 1 | 6 |
| PCMTD2 (1 of many) | 0.51783 | -0.11766188 | 0.039 | 0.059 | 1 | 6 |
| gnb1a | 0.51873 | 0.23165355 | 0.273 | 0.176 | 1 | 6 |
| sdhb | 0.51892 | 0.165523418 | 0.156 | 0.059 | 1 | 6 |
| zgc:91910 | 0.51918 | -0.16845247 | 0.325 | 0.412 | 1 | 6 |
| mob1a | 0.51985 | 0.14034305 | 0.104 | 0.059 | 1 | 6 |
| LOC108179023 | 0.52022 | -0.11868304 | 0.039 | 0.059 | 1 | 6 |
| hnrnpk | 0.52027 | 0.208446439 | 0.234 | 0.118 | 1 | 6 |
| NC-002333.17 | 0.52087 | 0.111167746 | 0.143 | 0.059 | 1 | 6 |
| pfdn2 | 0.52172 | 0.164384165 | 0.182 | 0.118 | 1 | 6 |
| CABZ01045617.1 | 0.52179 | 0.261407797 | 0.299 | 0.176 | 1 | 6 |
| tmem47 | 0.52199 | 0.235132786 | 0.234 | 0.118 | 1 | 6 |
| srsf1b | 0.52207 | 0.160426532 | 0.156 | 0.059 | 1 | 6 |
| hpf1 | 0.52232 | -0.11929796 | 0.039 | 0.059 | 1 | 6 |
| zgc:158463 | 0.52233 | -0.19468469 | 0.221 | 0.353 | 1 | 6 |
| naca | 0.5225 | 0.115902905 | 0.909 | 0.882 | 1 | 6 |
| stmn1a | 0.52268 | -0.10329563 | 0.065 | 0.059 | 1 | 6 |
| si:ch211-204a13.2 | 0.523 | 0.155273233 | 0.156 | 0.118 | 1 | 6 |
| ddx3xa | 0.52317 | 0.191771466 | 0.143 | 0.059 | 1 | 6 |
| gapdhs | 0.52322 | -0.19271535 | 0.312 | 0.353 | 1 | 6 |
| sec11a | 0.5245 | 0.165171721 | 0.299 | 0.176 | 1 | 6 |
| fhl3b | 0.52474 | -0.14659278 | 0.143 | 0.235 | 1 | 6 |
| rab5c | 0.5251 | 0.174546934 | 0.195 | 0.118 | 1 | 6 |
| tbc1d10aa | 0.52523 | 0.186584707 | 0.156 | 0.059 | 1 | 6 |
| smndc1 | 0.52534 | -0.2259721 | 0.156 | 0.235 | 1 | 6 |
| gas1a | 0.52588 | 0.179180979 | 0.364 | 0.235 | 1 | 6 |
| sox4a | 0.52609 | 0.230743057 | 0.442 | 0.294 | 1 | 6 |
| sod1 | 0.5261 | 0.207745828 | 0.442 | 0.294 | 1 | 6 |
| arcn1b | 0.52632 | 0.168002812 | 0.156 | 0.059 | 1 | 6 |

| | | | | | | |
|---|---|---|---|---|---|---|
| psmd11a | 0.52659 | -0.12726308 | 0.078 | 0.118 | 1 | 6 |
| zfhx4 | 0.52703 | 0.180600482 | 0.143 | 0.059 | 1 | 6 |
| ndufa12 | 0.52717 | -0.19158154 | 0.221 | 0.294 | 1 | 6 |
| XLOC-011450 | 0.52734 | -0.14242348 | 0.078 | 0.118 | 1 | 6 |
| igflr1 | 0.52777 | -0.16806892 | 0.143 | 0.176 | 1 | 6 |
| klhl11 | 0.52858 | -0.12076056 | 0.039 | 0.059 | 1 | 6 |
| tcf3b | 0.52878 | 0.175912224 | 0.117 | 0.059 | 1 | 6 |
| bcl2a | 0.52935 | 0.122158981 | 0.078 | 0.059 | 1 | 6 |
| angpt1 | 0.52965 | -0.18487185 | 0.052 | 0.118 | 1 | 6 |
| XLOC-034538 | 0.53 | -0.18780591 | 0.026 | 0.059 | 1 | 6 |
| XLOC-042919 | 0.53 | -0.15261395 | 0.026 | 0.059 | 1 | 6 |
| adamts9 | 0.53 | -0.16306426 | 0.026 | 0.059 | 1 | 6 |
| cwf19l2 | 0.53 | -0.12360086 | 0.026 | 0.059 | 1 | 6 |
| pigm | 0.53 | -0.10196188 | 0.026 | 0.059 | 1 | 6 |
| rhpn2 | 0.53 | -0.11568643 | 0.026 | 0.059 | 1 | 6 |
| rspo2 | 0.53 | -0.14278167 | 0.026 | 0.059 | 1 | 6 |
| sh3bp5a | 0.53 | -0.1427066 | 0.026 | 0.059 | 1 | 6 |
| supt20 | 0.53 | -0.19807866 | 0.026 | 0.059 | 1 | 6 |
| tcerg1a | 0.53 | -0.16646829 | 0.026 | 0.059 | 1 | 6 |
| ube2r2 | 0.53 | -0.10548318 | 0.026 | 0.059 | 1 | 6 |
| AL954322.2 | 0.53 | -0.14433758 | 0.026 | 0.059 | 1 | 6 |
| GUCY1A2 | 0.53 | -0.14175251 | 0.026 | 0.059 | 1 | 6 |
| lyrm5a | 0.53 | -0.13113738 | 0.026 | 0.059 | 1 | 6 |
| mapk8b | 0.53 | -0.16777116 | 0.026 | 0.059 | 1 | 6 |
| ntpcr | 0.53 | -0.14464472 | 0.026 | 0.059 | 1 | 6 |
| palm2 | 0.53 | -0.14063797 | 0.026 | 0.059 | 1 | 6 |
| rab43 | 0.53 | -0.11607111 | 0.026 | 0.059 | 1 | 6 |
| rims2a | 0.53 | -0.11921371 | 0.026 | 0.059 | 1 | 6 |
| socs5b | 0.53 | -0.17652445 | 0.026 | 0.059 | 1 | 6 |
| syne1b | 0.53 | -0.15871689 | 0.026 | 0.059 | 1 | 6 |
| tbc1d12b | 0.53 | -0.12899155 | 0.026 | 0.059 | 1 | 6 |
| zgc:112496 | 0.53 | -0.14839677 | 0.026 | 0.059 | 1 | 6 |
| zgc:153293 | 0.53 | -0.15992428 | 0.026 | 0.059 | 1 | 6 |
| CABZ01074397.1 | 0.53 | -0.23188004 | 0.026 | 0.059 | 1 | 6 |
| dynll2a | 0.53 | -0.14363853 | 0.026 | 0.059 | 1 | 6 |
| eif4e2rs1 | 0.53 | -0.11081563 | 0.026 | 0.059 | 1 | 6 |
| etfdh | 0.53 | -0.13379833 | 0.026 | 0.059 | 1 | 6 |
| mhc1laa | 0.53 | -0.10729919 | 0.026 | 0.059 | 1 | 6 |
| naa35 | 0.53 | -0.10210896 | 0.026 | 0.059 | 1 | 6 |
| otud7b | 0.53 | -0.1547012 | 0.026 | 0.059 | 1 | 6 |
| rnf4 | 0.53 | -0.14624113 | 0.026 | 0.059 | 1 | 6 |
| upf1 | 0.53 | -0.10932515 | 0.026 | 0.059 | 1 | 6 |
| clmn | 0.53 | -0.11939721 | 0.026 | 0.059 | 1 | 6 |
| hapln1b | 0.53 | -0.10728152 | 0.026 | 0.059 | 1 | 6 |
| mynn | 0.53 | -0.17496549 | 0.026 | 0.059 | 1 | 6 |
| zbtb21 | 0.53 | -0.17511295 | 0.026 | 0.059 | 1 | 6 |
| CDK2AP1 | 0.53 | -0.14132341 | 0.026 | 0.059 | 1 | 6 |
| inafm2 | 0.53 | -0.12102496 | 0.026 | 0.059 | 1 | 6 |

| | | | | | | |
|---|---|---|---|---|---|---|
| abca5 | 0.53 | -0.17884881 | 0.026 | 0.059 | 1 | 6 |
| acsm3 | 0.53 | -0.13894251 | 0.026 | 0.059 | 1 | 6 |
| cdc42ep1b | 0.53 | -0.10692758 | 0.026 | 0.059 | 1 | 6 |
| fhit | 0.53 | -0.11228908 | 0.026 | 0.059 | 1 | 6 |
| herpud1 | 0.53 | -0.12678013 | 0.026 | 0.059 | 1 | 6 |
| mier1a | 0.53 | -0.13323097 | 0.026 | 0.059 | 1 | 6 |
| pelo | 0.53 | -0.11357084 | 0.026 | 0.059 | 1 | 6 |
| pola2 | 0.53 | -0.14646978 | 0.026 | 0.059 | 1 | 6 |
| sc:d189 | 0.53 | -0.10369605 | 0.026 | 0.059 | 1 | 6 |
| si:ch73-92e7.4 | 0.53 | -0.11111626 | 0.026 | 0.059 | 1 | 6 |
| CR753886.1 | 0.53 | -0.13420892 | 0.026 | 0.059 | 1 | 6 |
| XLOC-028371 | 0.53 | -0.13095782 | 0.026 | 0.059 | 1 | 6 |
| arhgef39 | 0.53 | -0.148796 | 0.026 | 0.059 | 1 | 6 |
| atad1b | 0.53 | -0.10183057 | 0.026 | 0.059 | 1 | 6 |
| ccnt2a | 0.53 | -0.17834437 | 0.026 | 0.059 | 1 | 6 |
| fopnl | 0.53 | -0.12531678 | 0.026 | 0.059 | 1 | 6 |
| hmox1a | 0.53 | -0.12810829 | 0.026 | 0.059 | 1 | 6 |
| kcnj8 | 0.53 | -0.10236659 | 0.026 | 0.059 | 1 | 6 |
| mosmob | 0.53 | -0.15775144 | 0.026 | 0.059 | 1 | 6 |
| mtor | 0.53 | -0.17400448 | 0.026 | 0.059 | 1 | 6 |
| si:dkey-6e2.2 | 0.53 | -0.17529762 | 0.026 | 0.059 | 1 | 6 |
| spty2d1 | 0.53 | -0.10867522 | 0.026 | 0.059 | 1 | 6 |
| swt1 | 0.53 | -0.14433934 | 0.026 | 0.059 | 1 | 6 |
| tsen54 | 0.53 | -0.10295378 | 0.026 | 0.059 | 1 | 6 |
| tyms | 0.53 | -0.14049355 | 0.026 | 0.059 | 1 | 6 |
| ube2f | 0.53 | -0.10028192 | 0.026 | 0.059 | 1 | 6 |
| zc3h7a | 0.53 | -0.1399319 | 0.026 | 0.059 | 1 | 6 |
| COLEC10 | 0.53 | -0.10467409 | 0.026 | 0.059 | 1 | 6 |
| LOC101883052 | 0.53 | -0.14067633 | 0.026 | 0.059 | 1 | 6 |
| cpsf3 | 0.53 | -0.11263501 | 0.026 | 0.059 | 1 | 6 |
| fkbp1b | 0.53 | -0.10889053 | 0.026 | 0.059 | 1 | 6 |
| hmga2 | 0.53 | -0.11516007 | 0.026 | 0.059 | 1 | 6 |
| mtif3 | 0.53 | -0.14404344 | 0.026 | 0.059 | 1 | 6 |
| nelfb | 0.53 | -0.11543935 | 0.026 | 0.059 | 1 | 6 |
| ptbp2a | 0.53 | -0.11479842 | 0.026 | 0.059 | 1 | 6 |
| rpz4 | 0.53 | -0.17240599 | 0.026 | 0.059 | 1 | 6 |
| setd3 | 0.53 | -0.1032184 | 0.026 | 0.059 | 1 | 6 |
| si:rp71-36a1.1 | 0.53 | -0.10442515 | 0.026 | 0.059 | 1 | 6 |
| slc25a25a | 0.53 | -0.13498317 | 0.026 | 0.059 | 1 | 6 |
| tnk2b | 0.53 | -0.14359544 | 0.026 | 0.059 | 1 | 6 |
| trappc10 | 0.53 | -0.13014823 | 0.026 | 0.059 | 1 | 6 |
| trappc2 | 0.53 | -0.12948631 | 0.026 | 0.059 | 1 | 6 |
| CR293507.1 | 0.53 | -0.1090776 | 0.026 | 0.059 | 1 | 6 |
| aqr | 0.53 | -0.11374458 | 0.026 | 0.059 | 1 | 6 |
| bmi1a | 0.53 | -0.11092534 | 0.026 | 0.059 | 1 | 6 |
| cpda | 0.53 | -0.10021634 | 0.026 | 0.059 | 1 | 6 |
| erf | 0.53 | -0.1112058 | 0.026 | 0.059 | 1 | 6 |
| nsun2 | 0.53 | -0.12184826 | 0.026 | 0.059 | 1 | 6 |

| | | | | | | |
|---|---|---|---|---|---|---|
| pold1 | 0.53 | -0.10026453 | 0.026 | 0.059 | 1 | 6 |
| rhoua | 0.53001 | -0.15246224 | 0.13 | 0.235 | 1 | 6 |
| map3k7 | 0.5306 | -0.16630807 | 0.052 | 0.118 | 1 | 6 |
| u2af1 | 0.53084 | 0.151475292 | 0.299 | 0.176 | 1 | 6 |
| mmp15b | 0.53103 | -0.11564999 | 0.091 | 0.118 | 1 | 6 |
| slc25a55a | 0.53187 | -0.15909455 | 0.078 | 0.118 | 1 | 6 |
| fam53b | 0.53188 | -0.13965343 | 0.091 | 0.118 | 1 | 6 |
| creb3l3l | 0.53189 | 0.193860147 | 0.208 | 0.118 | 1 | 6 |
| ctsz | 0.53199 | -0.2075696 | 0.104 | 0.176 | 1 | 6 |
| snx1a | 0.5323 | 0.166324303 | 0.299 | 0.176 | 1 | 6 |
| hist2h2l | 0.53235 | 0.194916931 | 0.351 | 0.294 | 1 | 6 |
| laptm4a | 0.53371 | 0.203871017 | 0.234 | 0.118 | 1 | 6 |
| atp10a | 0.53385 | 0.173524942 | 0.143 | 0.059 | 1 | 6 |
| ifitm1 | 0.53498 | -0.27917749 | 0.091 | 0.176 | 1 | 6 |
| eif1ad | 0.5351 | 0.224038469 | 0.234 | 0.118 | 1 | 6 |
| CU638870.1 | 0.53543 | 0.169583865 | 0.117 | 0.059 | 1 | 6 |
| slc3a2b | 0.53586 | -0.11539685 | 0.299 | 0.412 | 1 | 6 |
| inppl1a | 0.53608 | -0.18759644 | 0.052 | 0.118 | 1 | 6 |
| pnisr | 0.53623 | 0.11502516 | 0.13 | 0.118 | 1 | 6 |
| smad1 | 0.53704 | 0.124612717 | 0.104 | 0.059 | 1 | 6 |
| tmem198a | 0.53895 | -0.11743818 | 0.039 | 0.059 | 1 | 6 |
| ubr7 | 0.53945 | -0.15212541 | 0.117 | 0.176 | 1 | 6 |
| dkc1 | 0.53963 | -0.21035814 | 0.221 | 0.353 | 1 | 6 |
| tspan9b | 0.54045 | 0.119774349 | 0.143 | 0.118 | 1 | 6 |
| clcn3 | 0.54048 | -0.17465761 | 0.052 | 0.118 | 1 | 6 |
| erp44 | 0.54063 | -0.16603163 | 0.182 | 0.235 | 1 | 6 |
| tiprl | 0.54066 | -0.15046409 | 0.052 | 0.118 | 1 | 6 |
| trib3 | 0.54162 | 0.186467687 | 0.13 | 0.059 | 1 | 6 |
| nfkbib | 0.54186 | -0.21532 | 0.13 | 0.235 | 1 | 6 |
| si:dkey-112e7.2 | 0.54223 | 0.139181055 | 0.351 | 0.235 | 1 | 6 |
| kdm6bb | 0.54347 | 0.184018528 | 0.299 | 0.176 | 1 | 6 |
| eef1b2 | 0.54348 | -0.10103144 | 0.87 | 0.882 | 1 | 6 |
| rpl8 | 0.54352 | 0.10943971 | 0.948 | 0.882 | 1 | 6 |
| ccdc43 | 0.54413 | -0.11968254 | 0.039 | 0.059 | 1 | 6 |
| wdr26b | 0.54498 | -0.12703714 | 0.039 | 0.059 | 1 | 6 |
| sgce | 0.54505 | 0.208321977 | 0.221 | 0.118 | 1 | 6 |
| foxp1b | 0.54509 | 0.132772604 | 0.091 | 0.059 | 1 | 6 |
| ndufs7 | 0.54554 | -0.17331821 | 0.169 | 0.235 | 1 | 6 |
| emd | 0.54681 | -0.2180819 | 0.182 | 0.294 | 1 | 6 |
| rhbdf1a | 0.54824 | -0.1456435 | 0.091 | 0.118 | 1 | 6 |
| arih2 | 0.54911 | -0.16780917 | 0.065 | 0.118 | 1 | 6 |
| enpep | 0.54983 | 0.139423251 | 0.156 | 0.118 | 1 | 6 |
| cald1a | 0.55111 | 0.225384584 | 0.61 | 0.471 | 1 | 6 |
| chmp4bb | 0.55113 | -0.11229348 | 0.091 | 0.176 | 1 | 6 |
| eif5a2 | 0.55136 | 0.138509802 | 0.545 | 0.412 | 1 | 6 |
| zfp36l1a | 0.5521 | 0.193877448 | 0.351 | 0.294 | 1 | 6 |
| serinc1 | 0.55233 | 0.150699142 | 0.169 | 0.118 | 1 | 6 |
| adnpa | 0.55261 | 0.117729765 | 0.078 | 0.059 | 1 | 6 |

| | | | | | | |
|---|---|---|---|---|---|---|
| asap1a | 0.55294 | -0.16656241 | 0.052 | 0.118 | 1 | 6 |
| rbpjb | 0.55374 | -0.20396068 | 0.13 | 0.235 | 1 | 6 |
| lbh | 0.55433 | -0.18440735 | 0.13 | 0.176 | 1 | 6 |
| twist1b | 0.55468 | -0.1161125 | 0.039 | 0.059 | 1 | 6 |
| thbs2a | 0.55546 | 0.129532283 | 0.078 | 0.059 | 1 | 6 |
| pgam1a | 0.55616 | -0.17566325 | 0.13 | 0.235 | 1 | 6 |
| si:ch1073-325m22.2 | 0.55637 | -0.14204147 | 0.39 | 0.412 | 1 | 6 |
| ptmaa | 0.55708 | -0.19201622 | 0.649 | 0.706 | 1 | 6 |
| pdk2b | 0.55751 | -0.17012437 | 0.169 | 0.235 | 1 | 6 |
| edil3a | 0.55769 | -0.18536312 | 0.13 | 0.235 | 1 | 6 |
| cfl1 | 0.55849 | -0.12697614 | 0.688 | 0.706 | 1 | 6 |
| wdr45b | 0.55888 | -0.15337138 | 0.052 | 0.118 | 1 | 6 |
| cnbpa | 0.55895 | -0.21778303 | 0.364 | 0.471 | 1 | 6 |
| hspb1 | 0.5593 | 0.239370204 | 0.299 | 0.176 | 1 | 6 |
| zgc:77650 | 0.55945 | 0.167646157 | 0.117 | 0.059 | 1 | 6 |
| kat5b | 0.55962 | 0.103313627 | 0.13 | 0.059 | 1 | 6 |
| wbp1lb | 0.55964 | -0.1694586 | 0.065 | 0.118 | 1 | 6 |
| slu7 | 0.56125 | 0.121989103 | 0.091 | 0.059 | 1 | 6 |
| tpm4a | 0.56172 | 0.206505456 | 0.429 | 0.353 | 1 | 6 |
| creg2 | 0.56238 | -0.11207732 | 0.169 | 0.176 | 1 | 6 |
| wdr33 | 0.56273 | -0.16232836 | 0.052 | 0.118 | 1 | 6 |
| rbmx | 0.56342 | -0.18355958 | 0.117 | 0.176 | 1 | 6 |
| vipr2 | 0.56403 | -0.12619092 | 0.039 | 0.059 | 1 | 6 |
| triqk | 0.56428 | 0.137224157 | 0.078 | 0.059 | 1 | 6 |
| ankrd10b | 0.56509 | 0.122395104 | 0.078 | 0.059 | 1 | 6 |
| si:ch211-66k16.27 | 0.56617 | -0.19705612 | 0.273 | 0.353 | 1 | 6 |
| cox7b | 0.56677 | -0.17379725 | 0.442 | 0.471 | 1 | 6 |
| ik | 0.56696 | -0.10906869 | 0.091 | 0.176 | 1 | 6 |
| ppifb | 0.56707 | 0.146888727 | 0.143 | 0.059 | 1 | 6 |
| slc15a2 | 0.5674 | 0.109090574 | 0.078 | 0.059 | 1 | 6 |
| nars | 0.56768 | 0.128344982 | 0.143 | 0.059 | 1 | 6 |
| ube2e1 | 0.56779 | -0.16504372 | 0.234 | 0.294 | 1 | 6 |
| mrpl28 | 0.5679 | -0.14485732 | 0.13 | 0.235 | 1 | 6 |
| gar1 | 0.56864 | -0.18540034 | 0.104 | 0.176 | 1 | 6 |
| hint2 | 0.56866 | -0.10827582 | 0.039 | 0.059 | 1 | 6 |
| rab5ab | 0.56883 | 0.133052585 | 0.143 | 0.059 | 1 | 6 |
| si:ch211-286o17.1 | 0.56892 | -0.13041121 | 0.247 | 0.353 | 1 | 6 |
| aimp1 | 0.5693 | 0.152626483 | 0.247 | 0.176 | 1 | 6 |
| dock7 | 0.56969 | 0.184439519 | 0.143 | 0.059 | 1 | 6 |
| ppp2r2d | 0.57036 | 0.165669411 | 0.143 | 0.059 | 1 | 6 |
| acadvl | 0.57081 | -0.13527596 | 0.078 | 0.118 | 1 | 6 |
| magixa | 0.57091 | -0.10766881 | 0.052 | 0.118 | 1 | 6 |
| nme3 | 0.57096 | -0.13602255 | 0.325 | 0.353 | 1 | 6 |
| abcc5 | 0.57108 | 0.208376613 | 0.221 | 0.118 | 1 | 6 |
| rpl4 | 0.57127 | 0.131609459 | 0.883 | 0.824 | 1 | 6 |
| zgc:112255 | 0.57178 | -0.15529886 | 0.078 | 0.118 | 1 | 6 |
| krtcap2 | 0.57192 | -0.20673355 | 0.364 | 0.471 | 1 | 6 |
| med10 | 0.57227 | 0.12518617 | 0.208 | 0.118 | 1 | 6 |

| | | | | | | |
|---|---|---|---|---|---|---|
| pum2 | 0.57298 | 0.165849896 | 0.13 | 0.059 | 1 | 6 |
| si:ch211-22i13.2 | 0.57306 | -0.17498838 | 0.091 | 0.176 | 1 | 6 |
| psmd11b | 0.57355 | 0.148160348 | 0.13 | 0.059 | 1 | 6 |
| eif4bb | 0.57369 | -0.14381456 | 0.26 | 0.294 | 1 | 6 |
| atp5md | 0.57371 | -0.12508897 | 0.286 | 0.412 | 1 | 6 |
| lpar1 | 0.57421 | 0.141081606 | 0.208 | 0.118 | 1 | 6 |
| psmg3 | 0.57423 | 0.103831321 | 0.13 | 0.059 | 1 | 6 |
| puf60a | 0.57448 | 0.188514711 | 0.208 | 0.118 | 1 | 6 |
| samm50 | 0.57508 | -0.13316771 | 0.052 | 0.118 | 1 | 6 |
| map1lc3b | 0.57514 | 0.137502089 | 0.325 | 0.294 | 1 | 6 |
| ptp4a2a | 0.57524 | -0.11280384 | 0.247 | 0.353 | 1 | 6 |
| ppp4cb | 0.57679 | -0.15496291 | 0.13 | 0.235 | 1 | 6 |
| LOC108180269 | 0.57682 | -0.10147928 | 0.156 | 0.176 | 1 | 6 |
| sdc4 | 0.57705 | -0.14728022 | 0.234 | 0.353 | 1 | 6 |
| LOC101883507 | 0.57755 | -0.1684179 | 0.065 | 0.118 | 1 | 6 |
| si:dkey-56m19.5 | 0.578 | -0.14679812 | 0.247 | 0.353 | 1 | 6 |
| pcdh18a | 0.57814 | -0.12008466 | 0.039 | 0.059 | 1 | 6 |
| snrnp27 | 0.57985 | 0.153729983 | 0.143 | 0.059 | 1 | 6 |
| g3bp1 | 0.58188 | 0.128583983 | 0.143 | 0.059 | 1 | 6 |
| coa6 | 0.58226 | 0.116741952 | 0.143 | 0.059 | 1 | 6 |
| ecm2 | 0.58347 | -0.19419154 | 0.091 | 0.176 | 1 | 6 |
| rbx1 | 0.5836 | 0.199363936 | 0.429 | 0.294 | 1 | 6 |
| XLOC-043857 | 0.58361 | -0.1015237 | 0.091 | 0.176 | 1 | 6 |
| luzp2 | 0.58452 | 0.186763757 | 0.143 | 0.059 | 1 | 6 |
| raraa | 0.58484 | 0.185294585 | 0.13 | 0.059 | 1 | 6 |
| ndufs3 | 0.58534 | 0.133552017 | 0.143 | 0.059 | 1 | 6 |
| gskip | 0.58762 | 0.157521545 | 0.143 | 0.059 | 1 | 6 |
| ndufb5 | 0.58766 | -0.15739516 | 0.221 | 0.294 | 1 | 6 |
| hypk | 0.58855 | -0.16561529 | 0.338 | 0.471 | 1 | 6 |
| spcs3 | 0.58857 | -0.13707124 | 0.26 | 0.294 | 1 | 6 |
| thtpa | 0.58877 | -0.13626718 | 0.078 | 0.118 | 1 | 6 |
| immt | 0.58947 | 0.131541274 | 0.143 | 0.059 | 1 | 6 |
| sgta | 0.59045 | 0.162840527 | 0.13 | 0.059 | 1 | 6 |
| tmem88b | 0.59053 | 0.15881528 | 0.221 | 0.176 | 1 | 6 |
| pnn | 0.5906 | -0.10094589 | 0.221 | 0.235 | 1 | 6 |
| atp5meb | 0.59096 | 0.105580716 | 0.532 | 0.412 | 1 | 6 |
| selenbp1 | 0.59164 | 0.118248572 | 0.338 | 0.235 | 1 | 6 |
| adrb2a | 0.59224 | 0.105082408 | 0.065 | 0.059 | 1 | 6 |
| tmem88a | 0.59238 | 0.116180835 | 0.091 | 0.059 | 1 | 6 |
| arih1 | 0.59268 | -0.1433116 | 0.065 | 0.118 | 1 | 6 |
| smyd2a | 0.59306 | -0.14539584 | 0.052 | 0.118 | 1 | 6 |
| il11ra | 0.59331 | 0.130648795 | 0.13 | 0.059 | 1 | 6 |
| igfbp7 | 0.59334 | -0.24259489 | 0.143 | 0.235 | 1 | 6 |
| acta2 | 0.59348 | 0.115747417 | 0.195 | 0.118 | 1 | 6 |
| prickle1b | 0.59414 | 0.17872586 | 0.13 | 0.059 | 1 | 6 |
| polr3h | 0.59444 | -0.10215834 | 0.143 | 0.235 | 1 | 6 |
| aars | 0.59486 | 0.154191619 | 0.143 | 0.059 | 1 | 6 |
| sars | 0.59502 | -0.14258853 | 0.065 | 0.118 | 1 | 6 |

| | | | | | | |
|---|---|---|---|---|---|---|
| c1d | 0.59524 | 0.13347106 | 0.143 | 0.059 | 1 | 6 |
| szrd1 | 0.59599 | 0.138711874 | 0.143 | 0.059 | 1 | 6 |
| gemin8 | 0.59615 | 0.136498053 | 0.143 | 0.059 | 1 | 6 |
| rarga | 0.59622 | 0.177668943 | 0.221 | 0.118 | 1 | 6 |
| csad | 0.5964 | 0.11393456 | 0.104 | 0.059 | 1 | 6 |
| smu1a | 0.59748 | 0.147257273 | 0.13 | 0.059 | 1 | 6 |
| bcap31 | 0.59756 | 0.164733925 | 0.143 | 0.059 | 1 | 6 |
| ponzr1 | 0.59759 | -0.22249328 | 0.182 | 0.294 | 1 | 6 |
| kpnb1 | 0.59791 | -0.16806681 | 0.091 | 0.176 | 1 | 6 |
| blcap | 0.59839 | -0.11466454 | 0.195 | 0.235 | 1 | 6 |
| tpm1 | 0.59858 | -0.21483397 | 0.519 | 0.647 | 1 | 6 |
| atp6v0ca | 0.59883 | -0.12194818 | 0.208 | 0.235 | 1 | 6 |
| cstf2 | 0.59993 | 0.146145083 | 0.143 | 0.059 | 1 | 6 |
| bcl2l1 | 0.59993 | 0.161310443 | 0.143 | 0.059 | 1 | 6 |
| fbl | 0.60019 | -0.20416888 | 0.273 | 0.353 | 1 | 6 |
| si:ch73-138n13.1 | 0.60047 | 0.18420784 | 0.195 | 0.118 | 1 | 6 |
| mrpl30 | 0.60104 | 0.137007809 | 0.143 | 0.059 | 1 | 6 |
| aldh6a1 | 0.60109 | 0.134832311 | 0.221 | 0.118 | 1 | 6 |
| tmem167a | 0.6012 | -0.18551182 | 0.182 | 0.294 | 1 | 6 |
| rcc2 | 0.6013 | 0.186712077 | 0.208 | 0.118 | 1 | 6 |
| tmem131l | 0.60132 | -0.1934804 | 0.091 | 0.176 | 1 | 6 |
| actr1 | 0.60136 | 0.136548121 | 0.143 | 0.059 | 1 | 6 |
| ppp6c | 0.60144 | 0.152759806 | 0.143 | 0.059 | 1 | 6 |
| ndufv2 | 0.60155 | -0.1404375 | 0.091 | 0.176 | 1 | 6 |
| ezrb | 0.60159 | -0.14335649 | 0.065 | 0.118 | 1 | 6 |
| CU539058.1 | 0.60161 | 0.13869324 | 0.104 | 0.059 | 1 | 6 |
| npepl1 | 0.6018 | 0.153846891 | 0.143 | 0.059 | 1 | 6 |
| si:dkey-27b3.2 | 0.6018 | 0.138504249 | 0.143 | 0.059 | 1 | 6 |
| palm1a | 0.60215 | 0.100744795 | 0.078 | 0.059 | 1 | 6 |
| tmem59 | 0.6026 | 0.144939354 | 0.234 | 0.176 | 1 | 6 |
| tspan14 | 0.60284 | 0.181133615 | 0.221 | 0.118 | 1 | 6 |
| sult1st6 | 0.60337 | 0.165094039 | 0.13 | 0.059 | 1 | 6 |
| tmem183a | 0.60504 | -0.105872 | 0.039 | 0.059 | 1 | 6 |
| XLOC-035413 | 0.60698 | -0.11842673 | 0.091 | 0.176 | 1 | 6 |
| si:ch211-68a17.7 | 0.6074 | -0.1286042 | 0.052 | 0.118 | 1 | 6 |
| eif5 | 0.60892 | -0.19930874 | 0.273 | 0.353 | 1 | 6 |
| zgc:110239 | 0.60896 | 0.154998723 | 0.234 | 0.176 | 1 | 6 |
| znf703 | 0.61024 | -0.10920852 | 0.039 | 0.059 | 1 | 6 |
| wfdc1 | 0.61026 | 0.146941736 | 0.117 | 0.059 | 1 | 6 |
| brd3a | 0.61069 | 0.125706664 | 0.104 | 0.059 | 1 | 6 |
| sppl3 | 0.61226 | 0.116847827 | 0.104 | 0.059 | 1 | 6 |
| rfxap | 0.6129 | -0.12121946 | 0.143 | 0.176 | 1 | 6 |
| zfand3 | 0.61304 | 0.169649583 | 0.195 | 0.118 | 1 | 6 |
| si:ch211-217k17.7 | 0.61341 | -0.15119424 | 0.234 | 0.294 | 1 | 6 |
| hadhb | 0.61347 | -0.16479631 | 0.091 | 0.176 | 1 | 6 |
| cst14b.1 | 0.61495 | 0.155227406 | 0.169 | 0.118 | 1 | 6 |
| ube2l3a | 0.61499 | -0.13623761 | 0.091 | 0.176 | 1 | 6 |
| kank2 | 0.61521 | 0.164660093 | 0.117 | 0.059 | 1 | 6 |

| | | | | | | |
|---|---|---|---|---|---|---|
| cldn11a | 0.6161 | 0.135789354 | 0.208 | 0.118 | 1 | 6 |
| si:ch211-166a6.5 | 0.61706 | 0.139639156 | 0.104 | 0.059 | 1 | 6 |
| lrba | 0.61744 | -0.16757895 | 0.091 | 0.176 | 1 | 6 |
| cox7a2l | 0.61752 | -0.18109049 | 0.156 | 0.235 | 1 | 6 |
| myl9a | 0.61772 | 0.179530754 | 0.351 | 0.235 | 1 | 6 |
| ajuba | 0.61828 | -0.12336318 | 0.039 | 0.059 | 1 | 6 |
| loxl2b | 0.61841 | 0.217867206 | 0.208 | 0.118 | 1 | 6 |
| nmi | 0.6196 | 0.150760822 | 0.13 | 0.059 | 1 | 6 |
| LOC103909284 | 0.61986 | 0.180971375 | 0.351 | 0.235 | 1 | 6 |
| farsb | 0.62051 | -0.13280922 | 0.091 | 0.176 | 1 | 6 |
| ndufa3 | 0.62064 | 0.181579458 | 0.338 | 0.235 | 1 | 6 |
| dnajc5ga | 0.62131 | -0.13790267 | 0.065 | 0.118 | 1 | 6 |
| crebzf | 0.62181 | 0.149539938 | 0.273 | 0.235 | 1 | 6 |
| anxa13 | 0.62188 | -0.13318499 | 0.091 | 0.176 | 1 | 6 |
| tram1 | 0.62274 | -0.12731992 | 0.091 | 0.176 | 1 | 6 |
| col5a2a | 0.62286 | 0.208960654 | 0.481 | 0.353 | 1 | 6 |
| arhgap5 | 0.62419 | 0.16912561 | 0.117 | 0.059 | 1 | 6 |
| rbm5 | 0.62431 | 0.130791568 | 0.156 | 0.118 | 1 | 6 |
| romo1 | 0.62536 | -0.13218591 | 0.662 | 0.706 | 1 | 6 |
| tsen15 | 0.62574 | -0.14162104 | 0.052 | 0.118 | 1 | 6 |
| pabpc1a | 0.62673 | -0.10691034 | 0.844 | 0.882 | 1 | 6 |
| foxc1b | 0.62736 | -0.14666042 | 0.052 | 0.118 | 1 | 6 |
| hoxc6b | 0.62799 | 0.131068216 | 0.091 | 0.059 | 1 | 6 |
| sh3kbp1 | 0.62814 | -0.1249801 | 0.052 | 0.118 | 1 | 6 |
| lsm7 | 0.62837 | 0.135833642 | 0.221 | 0.176 | 1 | 6 |
| tbc1d15 | 0.62858 | -0.12861117 | 0.039 | 0.059 | 1 | 6 |
| ctbp2a | 0.6293 | 0.162678161 | 0.13 | 0.059 | 1 | 6 |
| ywhaqb | 0.63 | -0.20000919 | 0.247 | 0.353 | 1 | 6 |
| phip | 0.6303 | -0.13013987 | 0.052 | 0.118 | 1 | 6 |
| pdcd6 | 0.63062 | -0.16025473 | 0.143 | 0.235 | 1 | 6 |
| ptbp3 | 0.63066 | -0.1260972 | 0.078 | 0.118 | 1 | 6 |
| bms1 | 0.63096 | -0.13775116 | 0.052 | 0.118 | 1 | 6 |
| dachc | 0.63118 | -0.15788168 | 0.052 | 0.118 | 1 | 6 |
| tnrc6b | 0.63177 | -0.13246804 | 0.143 | 0.235 | 1 | 6 |
| cavin1b | 0.63235 | 0.103274213 | 0.117 | 0.059 | 1 | 6 |
| atp5l | 0.63305 | -0.20412271 | 0.416 | 0.529 | 1 | 6 |
| wwc1 | 0.63328 | 0.104260148 | 0.078 | 0.059 | 1 | 6 |
| pabpc1b | 0.63344 | -0.2061954 | 0.377 | 0.471 | 1 | 6 |
| ldlrad3 | 0.63371 | -0.16024456 | 0.052 | 0.118 | 1 | 6 |
| si:busm1-57f23.1 | 0.6342 | -0.18397641 | 0.052 | 0.118 | 1 | 6 |
| mcl1a | 0.63474 | 0.154959338 | 0.416 | 0.294 | 1 | 6 |
| efna5b | 0.63476 | 0.153420663 | 0.13 | 0.059 | 1 | 6 |
| eif4h | 0.63482 | 0.10159265 | 0.104 | 0.059 | 1 | 6 |
| ccni | 0.63496 | 0.201969618 | 0.519 | 0.412 | 1 | 6 |
| tspan4a | 0.63506 | 0.104527498 | 0.143 | 0.118 | 1 | 6 |
| rrp36 | 0.63515 | 0.105552908 | 0.13 | 0.059 | 1 | 6 |
| senp2 | 0.63579 | -0.14185646 | 0.052 | 0.118 | 1 | 6 |
| zgc:171480 | 0.63622 | 0.132482588 | 0.117 | 0.059 | 1 | 6 |

| | | | | | | |
|---|---|---|---|---|---|---|
| utrn | 0.63657 | -0.16884648 | 0.052 | 0.118 | 1 | 6 |
| mustn1a | 0.63822 | -0.10834705 | 0.052 | 0.118 | 1 | 6 |
| sox5 | 0.63879 | -0.11173383 | 0.052 | 0.118 | 1 | 6 |
| itga5 | 0.63883 | -0.10466035 | 0.052 | 0.118 | 1 | 6 |
| mrpl18 | 0.63897 | 0.128320266 | 0.13 | 0.059 | 1 | 6 |
| alcamb | 0.63899 | -0.17137192 | 0.052 | 0.118 | 1 | 6 |
| glrx | 0.63913 | -0.19197841 | 0.377 | 0.471 | 1 | 6 |
| arpp19b | 0.63914 | 0.111561921 | 0.208 | 0.176 | 1 | 6 |
| si:dkey-240h12.3 | 0.63942 | -0.10007663 | 0.052 | 0.118 | 1 | 6 |
| tia1l | 0.63977 | 0.151094395 | 0.117 | 0.059 | 1 | 6 |
| atp5pf | 0.63995 | -0.13490286 | 0.377 | 0.412 | 1 | 6 |
| atp1a1a.1 | 0.63996 | -0.17530127 | 0.169 | 0.235 | 1 | 6 |
| fkbp5 | 0.64016 | -0.12108765 | 0.208 | 0.235 | 1 | 6 |
| h2afx | 0.64025 | -0.11836517 | 0.052 | 0.118 | 1 | 6 |
| rsrp1 | 0.64025 | 0.159220909 | 0.169 | 0.118 | 1 | 6 |
| cope | 0.64055 | -0.13527889 | 0.247 | 0.294 | 1 | 6 |
| arrb2b | 0.64088 | -0.10729041 | 0.052 | 0.118 | 1 | 6 |
| si:ch211-261n11.5 | 0.64116 | -0.12929553 | 0.052 | 0.118 | 1 | 6 |
| dnaja2a | 0.64122 | 0.176778966 | 0.299 | 0.235 | 1 | 6 |
| dph5 | 0.64134 | -0.10835171 | 0.052 | 0.118 | 1 | 6 |
| atp1b1a | 0.64138 | 0.159279998 | 0.234 | 0.176 | 1 | 6 |
| fbxo11a | 0.64141 | 0.136938521 | 0.13 | 0.059 | 1 | 6 |
| ndufb4 | 0.64231 | 0.171608856 | 0.416 | 0.294 | 1 | 6 |
| fam50a | 0.64322 | 0.130175041 | 0.117 | 0.059 | 1 | 6 |
| birc6 | 0.64336 | -0.12929316 | 0.052 | 0.118 | 1 | 6 |
| tbl3 | 0.64411 | -0.10203478 | 0.052 | 0.118 | 1 | 6 |
| ptpn21 | 0.64429 | 0.14347683 | 0.117 | 0.059 | 1 | 6 |
| ak1 | 0.64509 | -0.10961722 | 0.247 | 0.353 | 1 | 6 |
| cry2 | 0.64544 | 0.139245367 | 0.13 | 0.059 | 1 | 6 |
| psd3l | 0.64557 | -0.1264689 | 0.052 | 0.118 | 1 | 6 |
| abi2b | 0.64561 | 0.115722506 | 0.104 | 0.059 | 1 | 6 |
| socs2 | 0.64577 | -0.10544033 | 0.039 | 0.059 | 1 | 6 |
| mgst3a | 0.64584 | 0.101055875 | 0.26 | 0.176 | 1 | 6 |
| her9 | 0.64599 | 0.128849403 | 0.104 | 0.059 | 1 | 6 |
| sec61a1l | 0.64606 | -0.11216908 | 0.052 | 0.118 | 1 | 6 |
| ylpm1 | 0.64616 | -0.13375019 | 0.052 | 0.118 | 1 | 6 |
| pdcd10a | 0.647 | -0.1146829 | 0.052 | 0.118 | 1 | 6 |
| fam192a | 0.64701 | 0.130665346 | 0.091 | 0.059 | 1 | 6 |
| rnf213a | 0.64761 | -0.14173952 | 0.065 | 0.118 | 1 | 6 |
| cd151l | 0.64774 | 0.163157404 | 0.195 | 0.118 | 1 | 6 |
| lama5 | 0.64797 | -0.12446461 | 0.052 | 0.118 | 1 | 6 |
| rpl5b | 0.64851 | -0.1000242 | 0.87 | 0.941 | 1 | 6 |
| ciao1 | 0.64881 | -0.11959179 | 0.052 | 0.118 | 1 | 6 |
| zgc:66433 | 0.64893 | -0.14029826 | 0.143 | 0.235 | 1 | 6 |
| sort1a | 0.64894 | 0.122241444 | 0.091 | 0.059 | 1 | 6 |
| hic1 | 0.64926 | 0.199363372 | 0.273 | 0.176 | 1 | 6 |
| mrpl38 | 0.64941 | -0.10217599 | 0.052 | 0.118 | 1 | 6 |
| qser1 | 0.64954 | -0.14765638 | 0.052 | 0.118 | 1 | 6 |

| | | | | | | |
|---|---|---|---|---|---|---|
| ist1 | 0.64972 | -0.10467093 | 0.052 | 0.118 | 1 | 6 |
| serpinh1a | 0.64972 | -0.12350488 | 0.052 | 0.118 | 1 | 6 |
| ift27 | 0.65 | 0.113715555 | 0.091 | 0.059 | 1 | 6 |
| stim1b | 0.65042 | 0.118833599 | 0.078 | 0.059 | 1 | 6 |
| BX323797.4 | 0.65081 | -0.12622036 | 0.052 | 0.118 | 1 | 6 |
| puf60b | 0.65089 | -0.13164888 | 0.052 | 0.118 | 1 | 6 |
| csnk1da | 0.65114 | -0.15075334 | 0.117 | 0.176 | 1 | 6 |
| anapc15 | 0.6512 | -0.16194394 | 0.052 | 0.118 | 1 | 6 |
| ctnnbl1 | 0.65132 | -0.10051366 | 0.052 | 0.118 | 1 | 6 |
| wdr44 | 0.65168 | -0.11286983 | 0.052 | 0.118 | 1 | 6 |
| mlf2 | 0.65185 | -0.13668477 | 0.13 | 0.176 | 1 | 6 |
| smek1 | 0.6524 | -0.13397689 | 0.052 | 0.118 | 1 | 6 |
| si:ch211-210c8.7 | 0.65244 | -0.13728193 | 0.052 | 0.118 | 1 | 6 |
| brd4 | 0.65308 | -0.12707935 | 0.052 | 0.118 | 1 | 6 |
| polr1d | 0.65316 | -0.12930796 | 0.065 | 0.118 | 1 | 6 |
| gstm.2 | 0.65322 | -0.10768109 | 0.052 | 0.118 | 1 | 6 |
| cnot2 | 0.65327 | -0.12293977 | 0.052 | 0.118 | 1 | 6 |
| timmdc1 | 0.65361 | -0.11272492 | 0.052 | 0.118 | 1 | 6 |
| ets1 | 0.65363 | -0.12559878 | 0.052 | 0.118 | 1 | 6 |
| usp44 | 0.65372 | -0.11496631 | 0.052 | 0.118 | 1 | 6 |
| dnajc15 | 0.65375 | -0.12323433 | 0.052 | 0.118 | 1 | 6 |
| zdhhc3a | 0.65388 | -0.11347423 | 0.052 | 0.118 | 1 | 6 |
| rrm2 | 0.65394 | -0.13021192 | 0.052 | 0.118 | 1 | 6 |
| tgfbr3 | 0.65395 | 0.166402078 | 0.117 | 0.059 | 1 | 6 |
| atp2a2b | 0.65395 | -0.14585608 | 0.052 | 0.118 | 1 | 6 |
| akirin1 | 0.65434 | 0.137572786 | 0.104 | 0.059 | 1 | 6 |
| csde1 | 0.65536 | 0.18473422 | 0.338 | 0.235 | 1 | 6 |
| ube2ib | 0.65567 | 0.152596525 | 0.351 | 0.294 | 1 | 6 |
| FQ323156.1 | 0.65591 | -0.13496655 | 0.078 | 0.118 | 1 | 6 |
| mre11a | 0.65591 | -0.12778174 | 0.065 | 0.118 | 1 | 6 |
| mylipb | 0.65679 | 0.15213642 | 0.195 | 0.118 | 1 | 6 |
| CABZ01076667.1 | 0.65708 | 0.13151708 | 0.208 | 0.118 | 1 | 6 |
| si:ch211-212k18.9 | 0.65766 | -0.16624425 | 0.351 | 0.471 | 1 | 6 |
| taf13 | 0.65851 | 0.139767058 | 0.338 | 0.294 | 1 | 6 |
| etfa | 0.65865 | 0.15912822 | 0.195 | 0.118 | 1 | 6 |
| vgll4b | 0.66094 | -0.1009376 | 0.091 | 0.118 | 1 | 6 |
| lman2 | 0.66098 | -0.17846563 | 0.299 | 0.412 | 1 | 6 |
| cpn1 | 0.66152 | 0.127412435 | 0.338 | 0.235 | 1 | 6 |
| fam69c | 0.66168 | 0.149672689 | 0.117 | 0.059 | 1 | 6 |
| rrp1 | 0.66203 | -0.15696561 | 0.143 | 0.235 | 1 | 6 |
| foxo3b | 0.66222 | -0.13297849 | 0.091 | 0.118 | 1 | 6 |
| reck | 0.66228 | 0.117341614 | 0.156 | 0.118 | 1 | 6 |
| dnaja2b | 0.66239 | -0.14160112 | 0.182 | 0.235 | 1 | 6 |
| commd3 | 0.66271 | 0.134839234 | 0.13 | 0.059 | 1 | 6 |
| dynll1 | 0.66318 | -0.19023214 | 0.481 | 0.588 | 1 | 6 |
| hspa5 | 0.66419 | 0.111725186 | 0.195 | 0.176 | 1 | 6 |
| smdt1b | 0.66469 | -0.16843423 | 0.325 | 0.412 | 1 | 6 |
| mt-nd4 | 0.66479 | -0.19755062 | 0.519 | 0.588 | 1 | 6 |

| | | | | | | |
|---|---|---|---|---|---|---|
| znf609a | 0.66487 | 0.120672306 | 0.091 | 0.059 | 1 | 6 |
| slc43a3b | 0.66532 | 0.135868327 | 0.117 | 0.059 | 1 | 6 |
| crk | 0.66562 | -0.15557746 | 0.143 | 0.235 | 1 | 6 |
| RSBN1 | 0.66576 | -0.12645376 | 0.039 | 0.059 | 1 | 6 |
| uqcrc2b | 0.66595 | 0.128577255 | 0.169 | 0.118 | 1 | 6 |
| atp5mea | 0.66635 | 0.100305556 | 0.143 | 0.118 | 1 | 6 |
| acp1 | 0.66672 | 0.115000422 | 0.13 | 0.059 | 1 | 6 |
| eef1da | 0.66689 | 0.137209051 | 0.805 | 0.706 | 1 | 6 |
| cmtm3 | 0.66747 | -0.12578291 | 0.065 | 0.118 | 1 | 6 |
| zgc:101569 | 0.66754 | -0.12165661 | 0.039 | 0.059 | 1 | 6 |
| cyb5r3 | 0.66781 | 0.125463704 | 0.104 | 0.059 | 1 | 6 |
| nucb2a | 0.668 | 0.182877787 | 0.26 | 0.176 | 1 | 6 |
| bsdc1 | 0.66828 | 0.142568081 | 0.208 | 0.118 | 1 | 6 |
| mcm3 | 0.669 | -0.15088947 | 0.078 | 0.118 | 1 | 6 |
| cercam | 0.66904 | 0.112541921 | 0.104 | 0.059 | 1 | 6 |
| ddx39aa | 0.6695 | -0.169654 | 0.234 | 0.294 | 1 | 6 |
| sypl2a | 0.66963 | 0.137749029 | 0.13 | 0.059 | 1 | 6 |
| sfxn4 | 0.66975 | 0.113926116 | 0.104 | 0.059 | 1 | 6 |
| crtap | 0.67024 | 0.189360755 | 0.273 | 0.176 | 1 | 6 |
| med11 | 0.67042 | -0.10912984 | 0.156 | 0.176 | 1 | 6 |
| pdlim1 | 0.67053 | 0.142687807 | 0.117 | 0.059 | 1 | 6 |
| fzd7b | 0.67089 | 0.157295154 | 0.208 | 0.118 | 1 | 6 |
| zgc:92744 | 0.67107 | 0.126292832 | 0.13 | 0.059 | 1 | 6 |
| fam89b | 0.6714 | 0.141534206 | 0.104 | 0.059 | 1 | 6 |
| ssr2 | 0.6719 | -0.14003073 | 0.558 | 0.588 | 1 | 6 |
| ndufab1a | 0.67194 | -0.10710578 | 0.208 | 0.235 | 1 | 6 |
| slc43a1a | 0.67227 | 0.137067494 | 0.13 | 0.059 | 1 | 6 |
| c1qtnf2 | 0.67382 | 0.130994575 | 0.13 | 0.059 | 1 | 6 |
| nop56 | 0.67406 | -0.15056078 | 0.247 | 0.294 | 1 | 6 |
| ebna1bp2 | 0.67434 | -0.14972999 | 0.13 | 0.176 | 1 | 6 |
| fam168b | 0.67466 | 0.130171228 | 0.13 | 0.059 | 1 | 6 |
| mrps16 | 0.67477 | -0.13365997 | 0.065 | 0.118 | 1 | 6 |
| isg20 | 0.67534 | 0.124947373 | 0.13 | 0.059 | 1 | 6 |
| rap1aa | 0.67607 | -0.16309054 | 0.169 | 0.235 | 1 | 6 |
| ddit3 | 0.67657 | 0.105672171 | 0.26 | 0.176 | 1 | 6 |
| cmasa | 0.67694 | 0.142416863 | 0.13 | 0.059 | 1 | 6 |
| arl5c | 0.67724 | 0.105390554 | 0.13 | 0.059 | 1 | 6 |
| actn4 | 0.67879 | 0.12169805 | 0.13 | 0.059 | 1 | 6 |
| zgc:77086 | 0.67893 | 0.163340678 | 0.195 | 0.118 | 1 | 6 |
| DST | 0.6791 | 0.132501528 | 0.117 | 0.059 | 1 | 6 |
| dohh | 0.67915 | 0.134276473 | 0.13 | 0.059 | 1 | 6 |
| eif4ea | 0.67965 | 0.13199514 | 0.104 | 0.059 | 1 | 6 |
| sod2 | 0.67981 | -0.15865434 | 0.104 | 0.176 | 1 | 6 |
| si:ch211-276i12.11 | 0.67983 | 0.133782852 | 0.13 | 0.059 | 1 | 6 |
| spcs2 | 0.68015 | -0.15820489 | 0.234 | 0.294 | 1 | 6 |
| pdap1a | 0.68022 | 0.114600594 | 0.13 | 0.059 | 1 | 6 |
| b2m | 0.68035 | -0.11030144 | 0.325 | 0.353 | 1 | 6 |
| abca1a | 0.68066 | 0.165195299 | 0.117 | 0.059 | 1 | 6 |

| | | | | | | |
|---|---|---|---|---|---|---|
| tspan7 | 0.68195 | 0.146900074 | 0.273 | 0.176 | 1 | 6 |
| cotl1 | 0.68241 | 0.122125549 | 0.104 | 0.059 | 1 | 6 |
| pfdn6 | 0.68283 | 0.140271446 | 0.221 | 0.176 | 1 | 6 |
| cpsf6 | 0.68352 | 0.172940633 | 0.195 | 0.118 | 1 | 6 |
| aldh1l1 | 0.68372 | -0.1098148 | 0.065 | 0.118 | 1 | 6 |
| fam120c | 0.68428 | 0.134906001 | 0.169 | 0.118 | 1 | 6 |
| cadm3 | 0.68597 | 0.102492675 | 0.078 | 0.059 | 1 | 6 |
| desi2 | 0.6861 | 0.125895319 | 0.104 | 0.059 | 1 | 6 |
| eif2s3 | 0.68737 | 0.163572019 | 0.558 | 0.471 | 1 | 6 |
| zgc:66388 | 0.68764 | -0.11853751 | 0.078 | 0.118 | 1 | 6 |
| tmem11 | 0.68843 | 0.122758599 | 0.117 | 0.059 | 1 | 6 |
| ctsk | 0.68956 | -0.14544528 | 0.195 | 0.235 | 1 | 6 |
| tmem106ba | 0.69002 | 0.139505436 | 0.104 | 0.059 | 1 | 6 |
| polr2c | 0.69017 | -0.12888442 | 0.065 | 0.118 | 1 | 6 |
| pdcd4a | 0.69087 | 0.12489668 | 0.117 | 0.059 | 1 | 6 |
| si:ch1073-291c23.2 | 0.69139 | -0.17057248 | 0.338 | 0.412 | 1 | 6 |
| znhit1 | 0.69145 | -0.1241925 | 0.065 | 0.118 | 1 | 6 |
| fam13b | 0.69167 | -0.14250165 | 0.078 | 0.118 | 1 | 6 |
| uba2 | 0.69196 | 0.102451479 | 0.143 | 0.118 | 1 | 6 |
| ppp3ccb | 0.69198 | -0.14949945 | 0.104 | 0.176 | 1 | 6 |
| zcchc24 | 0.69279 | -0.18042802 | 0.156 | 0.235 | 1 | 6 |
| casc4 | 0.69303 | 0.107557233 | 0.091 | 0.059 | 1 | 6 |
| icmt | 0.69373 | 0.113543907 | 0.091 | 0.059 | 1 | 6 |
| nat9 | 0.69378 | -0.13813522 | 0.117 | 0.176 | 1 | 6 |
| emc7 | 0.69485 | -0.12714401 | 0.078 | 0.118 | 1 | 6 |
| ssuh2rs1 | 0.69498 | -0.10280756 | 0.156 | 0.176 | 1 | 6 |
| trmt112 | 0.69554 | 0.134330732 | 0.286 | 0.235 | 1 | 6 |
| sat1a.2 | 0.69613 | 0.165864627 | 0.273 | 0.176 | 1 | 6 |
| vcp | 0.69736 | 0.154049779 | 0.273 | 0.176 | 1 | 6 |
| rabif | 0.69758 | 0.10751344 | 0.091 | 0.059 | 1 | 6 |
| CR382300.1 | 0.69892 | 0.135976877 | 0.117 | 0.059 | 1 | 6 |
| mibp2 | 0.69977 | 0.104690371 | 0.688 | 0.588 | 1 | 6 |
| ndufv1 | 0.70093 | -0.13037221 | 0.078 | 0.118 | 1 | 6 |
| slc20a2 | 0.70126 | 0.107147208 | 0.078 | 0.059 | 1 | 6 |
| slc25a24 | 0.70139 | 0.119238554 | 0.117 | 0.059 | 1 | 6 |
| sp4 | 0.70205 | -0.10218776 | 0.104 | 0.176 | 1 | 6 |
| yeats4 | 0.70248 | -0.12603649 | 0.065 | 0.118 | 1 | 6 |
| cux1a | 0.70306 | 0.170099928 | 0.195 | 0.118 | 1 | 6 |
| col6a3 | 0.70323 | 0.142477744 | 0.519 | 0.412 | 1 | 6 |
| zgc:153018 | 0.70374 | -0.11530417 | 0.078 | 0.118 | 1 | 6 |
| pitpnb | 0.70414 | 0.157583791 | 0.117 | 0.059 | 1 | 6 |
| h1f0 | 0.70577 | 0.151021657 | 0.169 | 0.118 | 1 | 6 |
| sulf2b | 0.70583 | 0.125296187 | 0.104 | 0.059 | 1 | 6 |
| rbms2a | 0.70642 | -0.10097177 | 0.104 | 0.118 | 1 | 6 |
| znf609b | 0.70664 | 0.125803835 | 0.104 | 0.059 | 1 | 6 |
| hacd2 | 0.70702 | -0.12460956 | 0.104 | 0.176 | 1 | 6 |
| cdkn1bb | 0.70771 | 0.13660501 | 0.221 | 0.176 | 1 | 6 |
| fundc2 | 0.70784 | -0.10425898 | 0.104 | 0.176 | 1 | 6 |

| | | | | | | |
|---|---|---|---|---|---|---|
| ehd1b | 0.70835 | 0.112961798 | 0.117 | 0.059 | 1 | 6 |
| tpm2 | 0.70859 | 0.175887536 | 0.117 | 0.059 | 1 | 6 |
| fat1a | 0.70972 | 0.116971921 | 0.117 | 0.059 | 1 | 6 |
| arhgdig | 0.70998 | 0.151418322 | 0.195 | 0.118 | 1 | 6 |
| zeb2a | 0.71035 | -0.10595267 | 0.156 | 0.235 | 1 | 6 |
| safb | 0.71135 | 0.125410092 | 0.104 | 0.059 | 1 | 6 |
| srsf5a | 0.71159 | 0.154827218 | 0.519 | 0.412 | 1 | 6 |
| nifk | 0.71193 | -0.16050602 | 0.169 | 0.235 | 1 | 6 |
| tmem126a | 0.71335 | -0.13663135 | 0.104 | 0.176 | 1 | 6 |
| ar | 0.7154 | -0.14423359 | 0.117 | 0.176 | 1 | 6 |
| edf1 | 0.71624 | 0.118282356 | 0.455 | 0.353 | 1 | 6 |
| itm2ba | 0.71626 | -0.11837212 | 0.545 | 0.588 | 1 | 6 |
| ciarta | 0.71632 | 0.120834465 | 0.39 | 0.294 | 1 | 6 |
| psmc3 | 0.71693 | 0.144218074 | 0.195 | 0.118 | 1 | 6 |
| zgc:100918 | 0.7183 | -0.13693544 | 0.104 | 0.176 | 1 | 6 |
| mxra8a | 0.71893 | -0.1555653 | 0.169 | 0.235 | 1 | 6 |
| ube2g2 | 0.71958 | -0.11749651 | 0.091 | 0.118 | 1 | 6 |
| polr2a | 0.71961 | 0.156555635 | 0.182 | 0.118 | 1 | 6 |
| si:dkey-261h17.1 | 0.71995 | 0.198902447 | 0.39 | 0.294 | 1 | 6 |
| wdfy1 | 0.72048 | 0.11866163 | 0.104 | 0.059 | 1 | 6 |
| lrrc8aa | 0.72142 | 0.143111332 | 0.104 | 0.059 | 1 | 6 |
| ntd5 | 0.7219 | -0.14964591 | 0.104 | 0.176 | 1 | 6 |
| cpvl | 0.72235 | 0.13558552 | 0.195 | 0.118 | 1 | 6 |
| timm29 | 0.72341 | 0.124399872 | 0.104 | 0.059 | 1 | 6 |
| itgb3b | 0.7235 | 0.122239713 | 0.091 | 0.059 | 1 | 6 |
| her6 | 0.7239 | -0.12910485 | 0.104 | 0.176 | 1 | 6 |
| cxcl8b.3 | 0.7243 | -0.1652059 | 0.117 | 0.176 | 1 | 6 |
| tspan31 | 0.72444 | -0.1381556 | 0.104 | 0.176 | 1 | 6 |
| pitpnbl | 0.72464 | -0.10653574 | 0.091 | 0.118 | 1 | 6 |
| gnai2a | 0.7249 | -0.14717189 | 0.156 | 0.235 | 1 | 6 |
| gkap1 | 0.72504 | 0.114022823 | 0.195 | 0.118 | 1 | 6 |
| hnrnpa0b | 0.7251 | -0.12068172 | 0.545 | 0.588 | 1 | 6 |
| hnrnpub | 0.72529 | -0.12212273 | 0.104 | 0.176 | 1 | 6 |
| dus3l | 0.72539 | -0.11250234 | 0.104 | 0.176 | 1 | 6 |
| gnai1 | 0.72548 | 0.133652198 | 0.117 | 0.059 | 1 | 6 |
| knop1 | 0.72567 | -0.12404547 | 0.104 | 0.176 | 1 | 6 |
| trnau1apb | 0.72625 | -0.10948749 | 0.143 | 0.176 | 1 | 6 |
| pard3bb | 0.72698 | 0.161568382 | 0.195 | 0.118 | 1 | 6 |
| cdc42l | 0.72833 | -0.11671872 | 0.247 | 0.294 | 1 | 6 |
| srrm1 | 0.72962 | 0.157839311 | 0.182 | 0.118 | 1 | 6 |
| mrps7 | 0.72965 | -0.13048584 | 0.078 | 0.118 | 1 | 6 |
| ugt2a7 | 0.72989 | -0.14435433 | 0.065 | 0.118 | 1 | 6 |
| serinc2 | 0.73073 | -0.13753088 | 0.312 | 0.412 | 1 | 6 |
| nap1l4a | 0.73178 | 0.105380335 | 0.104 | 0.059 | 1 | 6 |
| upf3b | 0.73301 | 0.117155433 | 0.117 | 0.059 | 1 | 6 |
| nip7 | 0.73327 | 0.111784644 | 0.104 | 0.059 | 1 | 6 |
| sfpq | 0.73416 | 0.168428991 | 0.26 | 0.176 | 1 | 6 |
| tnfsf10 | 0.73753 | 0.161762833 | 0.195 | 0.118 | 1 | 6 |

| | | | | | | |
|---|---|---|---|---|---|---|
| echs1 | 0.73853 | -0.15032556 | 0.156 | 0.235 | 1 | 6 |
| si:rp71-45k5.4 | 0.73883 | -0.16478626 | 0.325 | 0.412 | 1 | 6 |
| MYADM | 0.73892 | -0.12757745 | 0.065 | 0.118 | 1 | 6 |
| zgc:101016 | 0.73984 | -0.11170015 | 0.065 | 0.118 | 1 | 6 |
| cnot8 | 0.74015 | 0.113436856 | 0.117 | 0.059 | 1 | 6 |
| myl9b | 0.74099 | -0.10099738 | 0.558 | 0.588 | 1 | 6 |
| pfn2 | 0.74219 | 0.155449379 | 0.455 | 0.353 | 1 | 6 |
| st5 | 0.74239 | 0.129733023 | 0.117 | 0.059 | 1 | 6 |
| zc3h7bb | 0.743 | 0.12024011 | 0.117 | 0.059 | 1 | 6 |
| anapc16 | 0.7432 | 0.129772808 | 0.182 | 0.118 | 1 | 6 |
| loxl2a | 0.74368 | -0.1703845 | 0.156 | 0.235 | 1 | 6 |
| vmp1 | 0.7437 | -0.10031196 | 0.065 | 0.118 | 1 | 6 |
| api5 | 0.74398 | -0.10597555 | 0.143 | 0.176 | 1 | 6 |
| pdrg1 | 0.74427 | 0.121507227 | 0.117 | 0.059 | 1 | 6 |
| sdf2 | 0.74481 | 0.104546485 | 0.104 | 0.059 | 1 | 6 |
| marcksb | 0.74521 | -0.15174025 | 0.221 | 0.294 | 1 | 6 |
| ndufc2 | 0.74545 | 0.140170331 | 0.286 | 0.235 | 1 | 6 |
| anxa11b | 0.74608 | 0.126797636 | 0.403 | 0.353 | 1 | 6 |
| serbp1b | 0.74634 | -0.13647882 | 0.117 | 0.176 | 1 | 6 |
| ap2m1a | 0.74664 | 0.151430144 | 0.377 | 0.294 | 1 | 6 |
| atp6v0b | 0.74815 | -0.14567749 | 0.117 | 0.176 | 1 | 6 |
| ypel5 | 0.74933 | 0.112219624 | 0.104 | 0.059 | 1 | 6 |
| snrnp48 | 0.74974 | -0.1272232 | 0.065 | 0.118 | 1 | 6 |
| seh1l | 0.75052 | -0.10921591 | 0.065 | 0.118 | 1 | 6 |
| zmat5 | 0.75101 | -0.10796909 | 0.078 | 0.118 | 1 | 6 |
| lclat1 | 0.75275 | -0.10384234 | 0.065 | 0.118 | 1 | 6 |
| mrpl24 | 0.75306 | -0.10864105 | 0.078 | 0.118 | 1 | 6 |
| fuca1.1 | 0.75326 | -0.11801203 | 0.078 | 0.118 | 1 | 6 |
| zgc:162730 | 0.75338 | 0.176490759 | 0.416 | 0.353 | 1 | 6 |
| npc2 | 0.75372 | 0.113886797 | 0.221 | 0.176 | 1 | 6 |
| hbp1 | 0.75399 | 0.109266593 | 0.117 | 0.059 | 1 | 6 |
| uqcrc1 | 0.75401 | 0.11666151 | 0.26 | 0.176 | 1 | 6 |
| col16a1 | 0.75466 | 0.16565148 | 0.26 | 0.176 | 1 | 6 |
| COA4 | 0.75473 | 0.106118892 | 0.117 | 0.059 | 1 | 6 |
| XLOC-044204 | 0.75486 | 0.101268082 | 0.091 | 0.059 | 1 | 6 |
| ctnnbip1 | 0.75487 | 0.10784106 | 0.117 | 0.059 | 1 | 6 |
| zfand5a | 0.75493 | 0.102108202 | 0.117 | 0.059 | 1 | 6 |
| mapkapk2a | 0.75502 | 0.153987477 | 0.182 | 0.118 | 1 | 6 |
| irf2bp2b | 0.75529 | 0.134951729 | 0.325 | 0.235 | 1 | 6 |
| znf362a | 0.75621 | 0.107232729 | 0.117 | 0.059 | 1 | 6 |
| psip1b | 0.7568 | 0.153607273 | 0.26 | 0.176 | 1 | 6 |
| serpinh1b | 0.75796 | 0.148127925 | 0.26 | 0.176 | 1 | 6 |
| arl4d | 0.75824 | 0.142166041 | 0.117 | 0.059 | 1 | 6 |
| cd28 | 0.75827 | -0.11832851 | 0.065 | 0.118 | 1 | 6 |
| abi3bpb | 0.75881 | 0.136743006 | 0.117 | 0.059 | 1 | 6 |
| rbb4l | 0.75974 | 0.103033418 | 0.117 | 0.059 | 1 | 6 |
| prr12b | 0.75977 | 0.11109622 | 0.117 | 0.059 | 1 | 6 |
| copg2 | 0.7598 | 0.112014795 | 0.117 | 0.059 | 1 | 6 |

| | | | | | | |
|---|---|---|---|---|---|---|
| tmed9 | 0.76087 | -0.13099494 | 0.117 | 0.176 | 1 | 6 |
| lpin1 | 0.76292 | -0.10559473 | 0.065 | 0.118 | 1 | 6 |
| rrs1 | 0.76357 | -0.13884296 | 0.117 | 0.176 | 1 | 6 |
| psmc2 | 0.76552 | 0.145277215 | 0.234 | 0.176 | 1 | 6 |
| plxdc2 | 0.76573 | -0.11219316 | 0.091 | 0.118 | 1 | 6 |
| sf3b1 | 0.76576 | -0.11305323 | 0.065 | 0.118 | 1 | 6 |
| csnk1g2a | 0.76597 | -0.11156069 | 0.065 | 0.118 | 1 | 6 |
| zgc:56235 | 0.76633 | -0.10171723 | 0.065 | 0.118 | 1 | 6 |
| psme1 | 0.76663 | 0.119786224 | 0.156 | 0.118 | 1 | 6 |
| lasp1 | 0.76672 | 0.143109371 | 0.364 | 0.294 | 1 | 6 |
| pcbd1 | 0.76779 | -0.11617888 | 0.117 | 0.176 | 1 | 6 |
| BX005340.1 | 0.76854 | 0.100985431 | 0.091 | 0.059 | 1 | 6 |
| sdcbp2 | 0.76863 | -0.13947837 | 0.299 | 0.353 | 1 | 6 |
| vps29 | 0.7691 | -0.12346166 | 0.247 | 0.294 | 1 | 6 |
| mrpl44 | 0.76956 | 0.105697455 | 0.091 | 0.059 | 1 | 6 |
| marcksl1b | 0.77028 | -0.10268692 | 0.39 | 0.412 | 1 | 6 |
| usp36 | 0.77072 | 0.123699831 | 0.169 | 0.118 | 1 | 6 |
| ilf2 | 0.77258 | -0.15000355 | 0.221 | 0.294 | 1 | 6 |
| igfbp5b | 0.77375 | 0.111034697 | 0.429 | 0.412 | 1 | 6 |
| os9 | 0.77413 | -0.11656488 | 0.078 | 0.118 | 1 | 6 |
| cbr1l | 0.77553 | -0.11611307 | 0.195 | 0.235 | 1 | 6 |
| ythdc1 | 0.77561 | 0.100078114 | 0.104 | 0.059 | 1 | 6 |
| znf706 | 0.77774 | 0.112031928 | 0.182 | 0.118 | 1 | 6 |
| fubp1 | 0.77783 | 0.103702361 | 0.208 | 0.176 | 1 | 6 |
| commd7 | 0.77854 | 0.112678608 | 0.156 | 0.118 | 1 | 6 |
| si:ch211-195o20.7 | 0.7791 | -0.11654482 | 0.065 | 0.118 | 1 | 6 |
| tle2a | 0.78069 | -0.10710378 | 0.065 | 0.118 | 1 | 6 |
| cdc5l | 0.78069 | -0.10875616 | 0.065 | 0.118 | 1 | 6 |
| svila | 0.78087 | -0.10041281 | 0.065 | 0.118 | 1 | 6 |
| adamts2 | 0.78087 | -0.12231852 | 0.065 | 0.118 | 1 | 6 |
| cct2 | 0.78141 | -0.13037065 | 0.468 | 0.529 | 1 | 6 |
| wtap | 0.78313 | 0.107811548 | 0.104 | 0.059 | 1 | 6 |
| fbxl6 | 0.78469 | 0.126398564 | 0.182 | 0.118 | 1 | 6 |
| hipk3b | 0.78493 | 0.108354852 | 0.104 | 0.059 | 1 | 6 |
| ost4 | 0.78556 | -0.14033826 | 0.299 | 0.353 | 1 | 6 |
| anxa2a | 0.78561 | -0.15165274 | 0.13 | 0.176 | 1 | 6 |
| sdhaf4 | 0.78688 | -0.10254998 | 0.078 | 0.118 | 1 | 6 |
| vbp1 | 0.78948 | -0.12530571 | 0.117 | 0.176 | 1 | 6 |
| tbca | 0.79094 | 0.126481314 | 0.442 | 0.353 | 1 | 6 |
| glo1 | 0.79197 | 0.140487565 | 0.299 | 0.235 | 1 | 6 |
| ndrg3b | 0.79223 | -0.10744714 | 0.078 | 0.118 | 1 | 6 |
| sap30l | 0.79474 | -0.12120162 | 0.325 | 0.412 | 1 | 6 |
| rhoac | 0.79684 | 0.100792782 | 0.312 | 0.235 | 1 | 6 |
| tab1 | 0.79778 | 0.115228693 | 0.169 | 0.118 | 1 | 6 |
| nup62l | 0.79807 | 0.102719828 | 0.104 | 0.059 | 1 | 6 |
| kras | 0.79844 | 0.108095886 | 0.104 | 0.059 | 1 | 6 |
| hnrnpaba | 0.79943 | 0.111511833 | 0.727 | 0.647 | 1 | 6 |
| ube2d1b | 0.80065 | 0.110549482 | 0.182 | 0.118 | 1 | 6 |

| | | | | | | |
|---|---|---|---|---|---|---|
| hnrnpc | 0.80157 | 0.102957931 | 0.182 | 0.118 | 1 | 6 |
| cd276 | 0.80314 | 0.10411931 | 0.156 | 0.118 | 1 | 6 |
| rap1b | 0.80389 | 0.112478734 | 0.312 | 0.235 | 1 | 6 |
| sec13 | 0.80499 | 0.119417661 | 0.182 | 0.118 | 1 | 6 |
| pdgfra | 0.80512 | -0.11712991 | 0.221 | 0.294 | 1 | 6 |
| twf1a | 0.80605 | -0.10336656 | 0.117 | 0.176 | 1 | 6 |
| psmc4 | 0.80664 | -0.12267625 | 0.273 | 0.353 | 1 | 6 |
| fkbp10b | 0.80778 | -0.10030594 | 0.117 | 0.176 | 1 | 6 |
| tpbgb | 0.80787 | -0.11587443 | 0.078 | 0.118 | 1 | 6 |
| rnaseka | 0.80964 | -0.10709251 | 0.078 | 0.118 | 1 | 6 |
| mcm2 | 0.81045 | -0.10081455 | 0.117 | 0.176 | 1 | 6 |
| GCA | 0.81112 | -0.11362122 | 0.221 | 0.294 | 1 | 6 |
| add1 | 0.81236 | -0.11339767 | 0.169 | 0.235 | 1 | 6 |
| sept6 | 0.8133 | 0.111064072 | 0.169 | 0.118 | 1 | 6 |
| nop2 | 0.81383 | -0.11585065 | 0.117 | 0.176 | 1 | 6 |
| pak2b | 0.81459 | -0.116455 | 0.13 | 0.176 | 1 | 6 |
| ep300b | 0.8151 | 0.106933371 | 0.104 | 0.059 | 1 | 6 |
| gars | 0.81537 | -0.1004643 | 0.078 | 0.118 | 1 | 6 |
| hnrnpa0a | 0.81704 | 0.129604271 | 0.247 | 0.176 | 1 | 6 |
| ptgr1 | 0.81832 | -0.10662903 | 0.169 | 0.235 | 1 | 6 |
| bckdhbl | 0.81867 | -0.11022167 | 0.169 | 0.235 | 1 | 6 |
| esd | 0.81871 | -0.11406268 | 0.221 | 0.294 | 1 | 6 |
| rpia | 0.81901 | -0.10566004 | 0.078 | 0.118 | 1 | 6 |
| gng2 | 0.81911 | 0.10891928 | 0.104 | 0.059 | 1 | 6 |
| brd2a | 0.82004 | 0.111699193 | 0.156 | 0.118 | 1 | 6 |
| rbm39b | 0.82083 | 0.10309675 | 0.104 | 0.059 | 1 | 6 |
| msna | 0.8213 | 0.147284981 | 0.429 | 0.353 | 1 | 6 |
| tgfb1a | 0.82158 | 0.106459163 | 0.104 | 0.059 | 1 | 6 |
| ola1 | 0.8219 | 0.131546385 | 0.364 | 0.294 | 1 | 6 |
| rbbp4 | 0.82196 | -0.1155287 | 0.169 | 0.235 | 1 | 6 |
| phf14 | 0.822 | 0.106854397 | 0.104 | 0.059 | 1 | 6 |
| tbx2b | 0.82285 | -0.12912462 | 0.13 | 0.176 | 1 | 6 |
| hsd17b12b | 0.82384 | 0.101548991 | 0.104 | 0.059 | 1 | 6 |
| ptges | 0.82676 | 0.10306758 | 0.104 | 0.059 | 1 | 6 |
| mgst1.2 | 0.82755 | 0.107409209 | 0.208 | 0.176 | 1 | 6 |
| XLOC-042917 | 0.82842 | -0.1216591 | 0.078 | 0.118 | 1 | 6 |
| eif3ba | 0.82924 | 0.112061292 | 0.494 | 0.412 | 1 | 6 |
| pom121 | 0.83005 | -0.1059089 | 0.078 | 0.118 | 1 | 6 |
| acin1a | 0.83397 | 0.112784852 | 0.455 | 0.412 | 1 | 6 |
| acvr2ab | 0.8342 | -0.10419904 | 0.13 | 0.176 | 1 | 6 |
| lmnb1 | 0.83629 | -0.10538359 | 0.078 | 0.118 | 1 | 6 |
| tardbpl | 0.83792 | -0.12834173 | 0.234 | 0.294 | 1 | 6 |
| ctbp1 | 0.84082 | 0.110850615 | 0.234 | 0.176 | 1 | 6 |
| ccdc80 | 0.84583 | -0.10736818 | 0.078 | 0.118 | 1 | 6 |
| ece1 | 0.84635 | -0.12213146 | 0.13 | 0.176 | 1 | 6 |
| si:dkey-238c7.16 | 0.8485 | -0.12868805 | 0.286 | 0.353 | 1 | 6 |
| eva1ba | 0.84979 | -0.12737856 | 0.078 | 0.118 | 1 | 6 |
| map1lc3a | 0.84996 | 0.115208962 | 0.234 | 0.176 | 1 | 6 |

| | | | | | | |
|---|---|---|---|---|---|---|
| anxa1a | 0.8521 | -0.12380777 | 0.455 | 0.529 | 1 | 6 |
| lama4 | 0.85299 | -0.10838629 | 0.182 | 0.235 | 1 | 6 |
| myo1ca | 0.85353 | 0.108845163 | 0.169 | 0.118 | 1 | 6 |
| sarnp | 0.85687 | 0.109338822 | 0.299 | 0.235 | 1 | 6 |
| si:dkey-188i13.11 | 0.8601 | -0.1006005 | 0.26 | 0.294 | 1 | 6 |
| uqcrh | 0.86241 | -0.11149525 | 0.351 | 0.412 | 1 | 6 |
| mmp14b | 0.86251 | -0.11652292 | 0.234 | 0.294 | 1 | 6 |
| cyp2p7 | 0.86324 | -0.12279759 | 0.182 | 0.235 | 1 | 6 |
| sult6b1 | 0.86415 | 0.101257513 | 0.169 | 0.118 | 1 | 6 |
| meis2b | 0.86494 | 0.121668662 | 0.299 | 0.235 | 1 | 6 |
| mrpl54 | 0.86768 | -0.10052396 | 0.13 | 0.176 | 1 | 6 |
| yap1 | 0.87184 | 0.100231187 | 0.221 | 0.176 | 1 | 6 |
| svild | 0.87536 | -0.11522375 | 0.234 | 0.294 | 1 | 6 |
| dnmt3ab | 0.88691 | 0.104519254 | 0.416 | 0.353 | 1 | 6 |
| BX640522.2 | 0.91311 | 0.107865575 | 0.286 | 0.235 | 1 | 6 |
| mt-co2 | 4.982066 | -1.00618934 | 0.175 | 0.769 | 0.0100623 | 7 |
| mt-nd4 | 3.488486 | -0.46387961 | 0.111 | 0.5 | 0.7045697 | 7 |
| mt-cyb | 0.00011 | -0.85316675 | 0.063 | 0.462 | 1 | 7 |
| COX3 | 0.00012 | -0.82418675 | 0.286 | 0.769 | 1 | 7 |
| pnp5a | 0.00013 | -0.33370557 | 0 | 0.231 | 1 | 7 |
| mt-atp6 | 0.00021 | -0.6244179 | 0.175 | 0.615 | 1 | 7 |
| pnisr | 0.00027 | 0.512767316 | 0.286 | 0 | 1 | 7 |
| fbxo28 | 0.00036 | -0.49146002 | 0.032 | 0.346 | 1 | 7 |
| XLOC-043760 | 0.00038 | -0.6733403 | 0.063 | 0.423 | 1 | 7 |
| ddx49 | 0.00053 | -0.69876319 | 0.079 | 0.385 | 1 | 7 |
| GPR68 | 0.00056 | -0.2799045 | 0 | 0.192 | 1 | 7 |
| pak2b | 0.00056 | -0.29703738 | 0 | 0.192 | 1 | 7 |
| si:ch1073-394i4.1 | 0.00056 | -0.32639855 | 0 | 0.192 | 1 | 7 |
| bud31 | 0.00058 | 0.53799 | 0.333 | 0.077 | 1 | 7 |
| rpl27a | 0.00089 | -0.37486406 | 0.81 | 0.923 | 1 | 7 |
| mycbp | 0.00175 | 0.368450485 | 0.222 | 0 | 1 | 7 |
| skia | 0.0022 | -0.36250809 | 0.016 | 0.231 | 1 | 7 |
| seta | 0.00235 | 0.386616593 | 0.524 | 0.462 | 1 | 7 |
| gzm3 | 0.00238 | -0.23803855 | 0 | 0.154 | 1 | 7 |
| myh9b | 0.00238 | -0.30031994 | 0 | 0.154 | 1 | 7 |
| pcnt | 0.00238 | -0.24242231 | 0 | 0.154 | 1 | 7 |
| CU929418.1 | 0.00238 | -0.26896917 | 0 | 0.154 | 1 | 7 |
| tollip | 0.00238 | -0.21753241 | 0 | 0.154 | 1 | 7 |
| ube2j2 | 0.00238 | -0.24191047 | 0 | 0.154 | 1 | 7 |
| CABZ01029938.1 | 0.00238 | -0.29260011 | 0 | 0.154 | 1 | 7 |
| chst12a | 0.00238 | -0.30750999 | 0 | 0.154 | 1 | 7 |
| pnp5b | 0.00238 | -0.28311792 | 0 | 0.154 | 1 | 7 |
| tceb2 | 0.00238 | -0.23310861 | 0 | 0.154 | 1 | 7 |
| CABZ01045617.1 | 0.00273 | 0.365768674 | 0.206 | 0 | 1 | 7 |
| tfap4 | 0.00273 | 0.306424917 | 0.206 | 0 | 1 | 7 |
| mt-co1 | 0.00305 | -0.4992861 | 0.524 | 0.885 | 1 | 7 |
| LOC101882375 | 0.00426 | 0.429062051 | 0.19 | 0 | 1 | 7 |
| si:dkey-211g8.4 | 0.0043 | -0.40359172 | 0.063 | 0.346 | 1 | 7 |

| | | | | | | |
|---|---|---|---|---|---|---|
| med10 | 0.00457 | 0.223462637 | 0.302 | 0.077 | 1 | 7 |
| clic1 | 0.00479 | -0.44212265 | 0.492 | 0.846 | 1 | 7 |
| LOC566587 | 0.00554 | -0.24378668 | 0.032 | 0.231 | 1 | 7 |
| zgc:113411 | 0.00579 | -0.40029741 | 0.048 | 0.308 | 1 | 7 |
| coa5 | 0.0066 | 0.248547607 | 0.175 | 0 | 1 | 7 |
| ogdha | 0.0066 | 0.345613694 | 0.175 | 0 | 1 | 7 |
| si:ch211-162e15.3 | 0.0066 | 0.236056305 | 0.175 | 0 | 1 | 7 |
| mt-nd2 | 0.00672 | -0.19131001 | 0.079 | 0.269 | 1 | 7 |
| prkar1aa | 0.00722 | -0.2333166 | 0.048 | 0.231 | 1 | 7 |
| ddit3 | 0.0076 | -0.53590207 | 0.286 | 0.615 | 1 | 7 |
| sarnp | 0.00848 | -0.35267994 | 0.19 | 0.5 | 1 | 7 |
| LOC100330744 | 0.00996 | -0.21009047 | 0 | 0.115 | 1 | 7 |
| LOC101884253 | 0.00996 | -0.1986261 | 0 | 0.115 | 1 | 7 |
| LOC101885027 | 0.00996 | -0.26681441 | 0 | 0.115 | 1 | 7 |
| LOC103908877 | 0.00996 | -0.15166993 | 0 | 0.115 | 1 | 7 |
| XLOC-034235 | 0.00996 | -0.177818 | 0 | 0.115 | 1 | 7 |
| azin1b | 0.00996 | -0.29590326 | 0 | 0.115 | 1 | 7 |
| fam210b | 0.00996 | -0.16526016 | 0 | 0.115 | 1 | 7 |
| lpcat3 | 0.00996 | -0.23522103 | 0 | 0.115 | 1 | 7 |
| mat2b | 0.00996 | -0.14961191 | 0 | 0.115 | 1 | 7 |
| phf6 | 0.00996 | -0.15736976 | 0 | 0.115 | 1 | 7 |
| pir | 0.00996 | -0.19406723 | 0 | 0.115 | 1 | 7 |
| klf8 | 0.00996 | -0.20487655 | 0 | 0.115 | 1 | 7 |
| prox1b | 0.00996 | -0.25100829 | 0 | 0.115 | 1 | 7 |
| FP102158.1 | 0.00996 | -0.17403305 | 0 | 0.115 | 1 | 7 |
| LOC108190082 | 0.00996 | -0.20341187 | 0 | 0.115 | 1 | 7 |
| XLOC-032110 | 0.00996 | -0.24682198 | 0 | 0.115 | 1 | 7 |
| XLOC-032112 | 0.00996 | -0.1790029 | 0 | 0.115 | 1 | 7 |
| abl1 | 0.00996 | -0.15900918 | 0 | 0.115 | 1 | 7 |
| cyp3c1 | 0.00996 | -0.23834296 | 0 | 0.115 | 1 | 7 |
| evi5b | 0.00996 | -0.19799917 | 0 | 0.115 | 1 | 7 |
| fermt3b | 0.00996 | -0.22195972 | 0 | 0.115 | 1 | 7 |
| hnrnph3 | 0.00996 | -0.18665872 | 0 | 0.115 | 1 | 7 |
| iqgap2 | 0.00996 | -0.16476775 | 0 | 0.115 | 1 | 7 |
| per2 | 0.00996 | -0.19053654 | 0 | 0.115 | 1 | 7 |
| si:ch1073-228b5.2 | 0.00996 | -0.19181693 | 0 | 0.115 | 1 | 7 |
| si:dkey-42i9.6 | 0.00996 | -0.21925778 | 0 | 0.115 | 1 | 7 |
| ticam1 | 0.00996 | -0.20371548 | 0 | 0.115 | 1 | 7 |
| tmem101 | 0.00996 | -0.20392381 | 0 | 0.115 | 1 | 7 |
| usp48 | 0.00996 | -0.14201669 | 0 | 0.115 | 1 | 7 |
| vps26a | 0.00996 | -0.16440101 | 0 | 0.115 | 1 | 7 |
| zgc:153631 | 0.00996 | -0.18343332 | 0 | 0.115 | 1 | 7 |
| osbpl9 | 0.00996 | -0.16695769 | 0 | 0.115 | 1 | 7 |
| actl6a | 0.0102 | 0.269263725 | 0.159 | 0 | 1 | 7 |
| smarca4a | 0.0102 | 0.280016574 | 0.159 | 0 | 1 | 7 |
| dlgap5 | 0.0102 | 0.332500751 | 0.159 | 0 | 1 | 7 |
| hmgb1b | 0.0102 | 0.332127612 | 0.159 | 0 | 1 | 7 |
| kdm4b | 0.0102 | 0.260485842 | 0.159 | 0 | 1 | 7 |

| | | | | | | |
|---|---|---|---|---|---|---|
| sugt1 | 0.0102 | 0.262677795 | 0.159 | 0 | 1 | 7 |
| tnfb | 0.01021 | -0.64409638 | 0.238 | 0.462 | 1 | 7 |
| spcs3 | 0.01022 | 0.53687089 | 0.524 | 0.192 | 1 | 7 |
| cct3 | 0.01066 | 0.134754601 | 0.444 | 0.538 | 1 | 7 |
| dap1b | 0.01175 | -0.45598529 | 0.254 | 0.577 | 1 | 7 |
| XLOC-043761 | 0.01351 | -0.36346694 | 0.032 | 0.231 | 1 | 7 |
| CU972454.4 | 0.01366 | -0.33349384 | 0.016 | 0.192 | 1 | 7 |
| mob4 | 0.01372 | -0.22324952 | 0.079 | 0.308 | 1 | 7 |
| LOC103909972 | 0.01378 | -0.26680699 | 0.127 | 0.385 | 1 | 7 |
| trmt112 | 0.01381 | 0.108791156 | 0.19 | 0.269 | 1 | 7 |
| ruvbl1 | 0.01418 | 0.31045642 | 0.27 | 0.038 | 1 | 7 |
| sec61a1 | 0.0151 | -0.25771045 | 0.127 | 0.385 | 1 | 7 |
| lonrf1l | 0.01511 | -0.24139731 | 0.016 | 0.192 | 1 | 7 |
| mt-nd3 | 0.01533 | -0.16692066 | 0.143 | 0.385 | 1 | 7 |
| mrps35 | 0.0157 | 0.206254604 | 0.143 | 0 | 1 | 7 |
| cdkn3 | 0.0157 | 0.237442416 | 0.143 | 0 | 1 | 7 |
| cpsf3 | 0.0157 | 0.241494912 | 0.143 | 0 | 1 | 7 |
| itpka | 0.0157 | 0.268922609 | 0.143 | 0 | 1 | 7 |
| mrgbp | 0.0157 | 0.171509449 | 0.143 | 0 | 1 | 7 |
| phrf1 | 0.0157 | 0.251270358 | 0.143 | 0 | 1 | 7 |
| usp5 | 0.0157 | 0.22335555 | 0.143 | 0 | 1 | 7 |
| acadl | 0.0157 | 0.232880476 | 0.143 | 0 | 1 | 7 |
| birc5a | 0.0157 | 0.258218163 | 0.143 | 0 | 1 | 7 |
| cks1b | 0.0157 | 0.287735967 | 0.143 | 0 | 1 | 7 |
| slc25a43 | 0.01626 | -0.25177613 | 0.016 | 0.192 | 1 | 7 |
| si:dkey-260g12.1 | 0.01636 | -0.45429724 | 0.254 | 0.577 | 1 | 7 |
| tnfaip8l2a | 0.01748 | -0.35038627 | 0.048 | 0.269 | 1 | 7 |
| cox6a1 | 0.018 | 0.174261901 | 0.619 | 0.654 | 1 | 7 |
| irf4b | 0.0189 | -0.53176667 | 0.159 | 0.423 | 1 | 7 |
| acadvl | 0.0198 | 0.367916125 | 0.333 | 0.077 | 1 | 7 |
| btg2 | 0.02006 | -0.44716204 | 0.333 | 0.615 | 1 | 7 |
| ppt1 | 0.0204 | -0.31984109 | 0.349 | 0.654 | 1 | 7 |
| socs1a | 0.02058 | -0.49978859 | 0.159 | 0.385 | 1 | 7 |
| hcls1 | 0.02074 | -0.28567036 | 0.27 | 0.538 | 1 | 7 |
| fam208b | 0.02078 | -0.34905139 | 0.079 | 0.308 | 1 | 7 |
| bzw1a | 0.02097 | -0.45917061 | 0.127 | 0.385 | 1 | 7 |
| nutf2l | 0.02119 | 0.42788795 | 0.444 | 0.269 | 1 | 7 |
| irf2bp2a | 0.02173 | -0.33869476 | 0.349 | 0.654 | 1 | 7 |
| arpc1b | 0.02204 | -0.23180948 | 0.841 | 0.885 | 1 | 7 |
| pa2g4a | 0.02296 | 0.401563474 | 0.302 | 0.077 | 1 | 7 |
| gkap1 | 0.02411 | 0.179511246 | 0.127 | 0 | 1 | 7 |
| arih1 | 0.02411 | 0.258653293 | 0.127 | 0 | 1 | 7 |
| ash2l | 0.02411 | 0.255633403 | 0.127 | 0 | 1 | 7 |
| dynll2b | 0.02411 | 0.156636417 | 0.127 | 0 | 1 | 7 |
| hdgfl2 | 0.02411 | 0.249789523 | 0.127 | 0 | 1 | 7 |
| kif22 | 0.02411 | 0.240278624 | 0.127 | 0 | 1 | 7 |
| mphosph6 | 0.02411 | 0.162999329 | 0.127 | 0 | 1 | 7 |
| mrps34 | 0.02411 | 0.203557353 | 0.127 | 0 | 1 | 7 |

| | | | | | | |
|---|---|---|---|---|---|---|
| ncam1a | 0.02411 | 0.305342176 | 0.127 | 0 | 1 | 7 |
| polr1d | 0.02411 | 0.235103516 | 0.127 | 0 | 1 | 7 |
| rrm2 | 0.02411 | 0.430247495 | 0.127 | 0 | 1 | 7 |
| snx29 | 0.02411 | 0.291985005 | 0.127 | 0 | 1 | 7 |
| swi5 | 0.02411 | 0.252246569 | 0.127 | 0 | 1 | 7 |
| fus | 0.02411 | 0.236653815 | 0.127 | 0 | 1 | 7 |
| hdlbpa | 0.02411 | 0.221641958 | 0.127 | 0 | 1 | 7 |
| idh3b | 0.02411 | 0.210205963 | 0.127 | 0 | 1 | 7 |
| mgmt | 0.02411 | 0.193189446 | 0.127 | 0 | 1 | 7 |
| mrpl40 | 0.02411 | 0.225758617 | 0.127 | 0 | 1 | 7 |
| si:dkey-53k12.1 | 0.02411 | 0.220192575 | 0.127 | 0 | 1 | 7 |
| tcta | 0.02411 | 0.250166447 | 0.127 | 0 | 1 | 7 |
| tk2 | 0.02411 | 0.228403737 | 0.127 | 0 | 1 | 7 |
| ccdc167 | 0.02411 | 0.192682209 | 0.127 | 0 | 1 | 7 |
| cks2 | 0.02411 | 0.291069927 | 0.127 | 0 | 1 | 7 |
| paf1 | 0.02411 | 0.20308009 | 0.127 | 0 | 1 | 7 |
| zgc:112496 | 0.02411 | 0.187780093 | 0.127 | 0 | 1 | 7 |
| zgc:194981 | 0.02411 | 0.196548913 | 0.127 | 0 | 1 | 7 |
| zgc:66427 | 0.0245 | -0.378747 | 0.079 | 0.308 | 1 | 7 |
| dnaja2b | 0.02458 | 0.254701274 | 0.175 | 0.038 | 1 | 7 |
| socs3a | 0.02478 | -0.46590757 | 0.19 | 0.346 | 1 | 7 |
| si:ch211-226m16.2 | 0.02581 | -0.16621624 | 0.222 | 0.423 | 1 | 7 |
| naa15a | 0.02596 | 0.257719355 | 0.222 | 0.038 | 1 | 7 |
| ndufa8 | 0.02664 | 0.397421896 | 0.397 | 0.154 | 1 | 7 |
| snrpd1 | 0.02664 | 0.427472172 | 0.381 | 0.192 | 1 | 7 |
| pnrc2 | 0.02675 | -0.22074372 | 0.921 | 1 | 1 | 7 |
| BX120005.1 | 0.02678 | -0.28742758 | 0.206 | 0.462 | 1 | 7 |
| ndfip2 | 0.0288 | -0.14322397 | 0.111 | 0.269 | 1 | 7 |
| nr4a1 | 0.02919 | -0.44890275 | 0.524 | 0.769 | 1 | 7 |
| sik1 | 0.0299 | -0.26342827 | 0.222 | 0.462 | 1 | 7 |
| prkcbb | 0.03009 | -0.20833815 | 0.079 | 0.269 | 1 | 7 |
| zgc:162730 | 0.03109 | -0.36349213 | 0.619 | 0.885 | 1 | 7 |
| rps27l | 0.03153 | 0.404103536 | 0.381 | 0.115 | 1 | 7 |
| selenoh | 0.03215 | 0.216435724 | 0.175 | 0.154 | 1 | 7 |
| denr | 0.03465 | 0.33650193 | 0.206 | 0.038 | 1 | 7 |
| wasb | 0.03518 | -0.14934659 | 0.603 | 0.808 | 1 | 7 |
| adam19a | 0.03622 | -0.19888541 | 0.016 | 0.154 | 1 | 7 |
| XLOC-032002 | 0.03689 | -0.43901441 | 0.111 | 0.346 | 1 | 7 |
| mef2d | 0.03693 | -0.24700048 | 0.111 | 0.308 | 1 | 7 |
| inka1b | 0.03696 | 0.181554303 | 0.111 | 0 | 1 | 7 |
| rfc2 | 0.03696 | 0.183425144 | 0.111 | 0 | 1 | 7 |
| rcc1 | 0.03696 | 0.209091915 | 0.111 | 0 | 1 | 7 |
| aar2 | 0.03696 | 0.182961369 | 0.111 | 0 | 1 | 7 |
| mir181b-3 | 0.03696 | 0.24555016 | 0.111 | 0 | 1 | 7 |
| npm1b | 0.03696 | 0.173209111 | 0.111 | 0 | 1 | 7 |
| actr6 | 0.03696 | 0.164633692 | 0.111 | 0 | 1 | 7 |
| edaradd | 0.03696 | 0.223566665 | 0.111 | 0 | 1 | 7 |
| mrpl43 | 0.03696 | 0.186566518 | 0.111 | 0 | 1 | 7 |

| | | | | | | |
|---|---|---|---|---|---|---|
| rpia | 0.03696 | 0.129476636 | 0.111 | 0 | 1 | 7 |
| ubxn1 | 0.03696 | 0.191273469 | 0.111 | 0 | 1 | 7 |
| cars | 0.03696 | 0.203438084 | 0.111 | 0 | 1 | 7 |
| faf2 | 0.03696 | 0.184690736 | 0.111 | 0 | 1 | 7 |
| trub2 | 0.03696 | 0.208504956 | 0.111 | 0 | 1 | 7 |
| cbfa2t3 | 0.03696 | 0.304387185 | 0.111 | 0 | 1 | 7 |
| dnph1 | 0.03696 | 0.150891721 | 0.111 | 0 | 1 | 7 |
| med21 | 0.03696 | 0.237594522 | 0.111 | 0 | 1 | 7 |
| si:dkey-42i9.7 | 0.03696 | 0.204614426 | 0.111 | 0 | 1 | 7 |
| srp9 | 0.03696 | 0.17400155 | 0.111 | 0 | 1 | 7 |
| zranb2 | 0.03696 | 0.208663209 | 0.111 | 0 | 1 | 7 |
| ccl38a.4 | 0.03841 | -0.29697454 | 0.111 | 0.308 | 1 | 7 |
| ctsba | 0.03859 | -0.13039507 | 0.127 | 0.115 | 1 | 7 |
| pgrmc1 | 0.03887 | -0.26800199 | 0.016 | 0.154 | 1 | 7 |
| zfand5b | 0.03895 | -0.12512855 | 0.111 | 0.269 | 1 | 7 |
| CR381673.1 | 0.039 | -0.23360845 | 0.206 | 0.423 | 1 | 7 |
| rab8b | 0.04046 | -0.3074995 | 0.143 | 0.385 | 1 | 7 |
| rabgap1l | 0.04125 | -0.28963107 | 0.048 | 0.231 | 1 | 7 |
| si:dkeyp-113d7.10 | 0.0415 | -0.23514099 | 0.048 | 0.231 | 1 | 7 |
| rtf1 | 0.04153 | 0.24177017 | 0.206 | 0.038 | 1 | 7 |
| CT956002.2 | 0.04182 | -0.21238702 | 0 | 0.077 | 1 | 7 |
| aph1b | 0.04182 | -0.12720221 | 0 | 0.077 | 1 | 7 |
| bco2a | 0.04182 | -0.12833913 | 0 | 0.077 | 1 | 7 |
| lancl1 | 0.04182 | -0.10357251 | 0 | 0.077 | 1 | 7 |
| lztr1 | 0.04182 | -0.13948231 | 0 | 0.077 | 1 | 7 |
| ranbp10 | 0.04182 | -0.1269465 | 0 | 0.077 | 1 | 7 |
| sgk1 | 0.04182 | -0.21326714 | 0 | 0.077 | 1 | 7 |
| tmed8 | 0.04182 | -0.1290418 | 0 | 0.077 | 1 | 7 |
| wdfy3 | 0.04182 | -0.14368322 | 0 | 0.077 | 1 | 7 |
| LOC103909899 | 0.04182 | -0.14064524 | 0 | 0.077 | 1 | 7 |
| si:ch211-59h6.1 | 0.04182 | -0.14228128 | 0 | 0.077 | 1 | 7 |
| CABZ01074453.1 | 0.04182 | -0.14556482 | 0 | 0.077 | 1 | 7 |
| LOC101883507 | 0.04182 | -0.16209706 | 0 | 0.077 | 1 | 7 |
| XLOC-005726 | 0.04182 | -0.13335826 | 0 | 0.077 | 1 | 7 |
| drp2 | 0.04182 | -0.1056462 | 0 | 0.077 | 1 | 7 |
| erf | 0.04182 | -0.1407436 | 0 | 0.077 | 1 | 7 |
| fuca1.2 | 0.04182 | -0.16059794 | 0 | 0.077 | 1 | 7 |
| galnt12 | 0.04182 | -0.1240363 | 0 | 0.077 | 1 | 7 |
| gpbp1l1 | 0.04182 | -0.10564724 | 0 | 0.077 | 1 | 7 |
| hdac7b | 0.04182 | -0.17692939 | 0 | 0.077 | 1 | 7 |
| itsn1 | 0.04182 | -0.11614787 | 0 | 0.077 | 1 | 7 |
| lonp2 | 0.04182 | -0.17848864 | 0 | 0.077 | 1 | 7 |
| pnpla7a | 0.04182 | -0.20602405 | 0 | 0.077 | 1 | 7 |
| si:ch211-156p11.1 | 0.04182 | -0.20181978 | 0 | 0.077 | 1 | 7 |
| si:dkey-48p11.3 | 0.04182 | -0.15777574 | 0 | 0.077 | 1 | 7 |
| tbc1d17 | 0.04182 | -0.10621672 | 0 | 0.077 | 1 | 7 |
| zdhhc24 | 0.04182 | -0.11093623 | 0 | 0.077 | 1 | 7 |
| asah1b | 0.04182 | -0.16956525 | 0 | 0.077 | 1 | 7 |

| | | | | | | |
|---|---|---|---|---|---|---|
| bmpr2a | 0.04182 | -0.16405647 | 0 | 0.077 | 1 | 7 |
| cxcl32b.1 | 0.04182 | -0.14784937 | 0 | 0.077 | 1 | 7 |
| fbxo34 | 0.04182 | -0.1560572 | 0 | 0.077 | 1 | 7 |
| hopx | 0.04182 | -0.13343646 | 0 | 0.077 | 1 | 7 |
| il17rd | 0.04182 | -0.15280571 | 0 | 0.077 | 1 | 7 |
| mxd3 | 0.04182 | -0.16129435 | 0 | 0.077 | 1 | 7 |
| qrich1 | 0.04182 | -0.10404718 | 0 | 0.077 | 1 | 7 |
| sft2d2b | 0.04182 | -0.17412075 | 0 | 0.077 | 1 | 7 |
| si:ch211-244b2.3 | 0.04182 | -0.1262715 | 0 | 0.077 | 1 | 7 |
| si:ch73-141c7.1 | 0.04182 | -0.10338825 | 0 | 0.077 | 1 | 7 |
| si:dkey-79d12.4 | 0.04182 | -0.13890165 | 0 | 0.077 | 1 | 7 |
| tpst2 | 0.04182 | -0.1544733 | 0 | 0.077 | 1 | 7 |
| CR388029.1 | 0.04182 | -0.11344155 | 0 | 0.077 | 1 | 7 |
| LOC103908858 | 0.04182 | -0.11136136 | 0 | 0.077 | 1 | 7 |
| LOC103911652 | 0.04182 | -0.18205864 | 0 | 0.077 | 1 | 7 |
| LOC110439377 | 0.04182 | -0.13270873 | 0 | 0.077 | 1 | 7 |
| OSBPL8 | 0.04182 | -0.11484875 | 0 | 0.077 | 1 | 7 |
| XLOC-020099 | 0.04182 | -0.15980672 | 0 | 0.077 | 1 | 7 |
| acsf2 | 0.04182 | -0.14515192 | 0 | 0.077 | 1 | 7 |
| atp6v1d | 0.04182 | -0.11579493 | 0 | 0.077 | 1 | 7 |
| bnip3la | 0.04182 | -0.1039179 | 0 | 0.077 | 1 | 7 |
| c2cd5 | 0.04182 | -0.14824573 | 0 | 0.077 | 1 | 7 |
| colgalt1 | 0.04182 | -0.1485799 | 0 | 0.077 | 1 | 7 |
| dis3 | 0.04182 | -0.12653606 | 0 | 0.077 | 1 | 7 |
| dnase1l1l | 0.04182 | -0.13703991 | 0 | 0.077 | 1 | 7 |
| hdlbpb | 0.04182 | -0.11756489 | 0 | 0.077 | 1 | 7 |
| il2rga | 0.04182 | -0.12175502 | 0 | 0.077 | 1 | 7 |
| isoc1 | 0.04182 | -0.11327514 | 0 | 0.077 | 1 | 7 |
| kpna7 | 0.04182 | -0.17264233 | 0 | 0.077 | 1 | 7 |
| mtmr6 | 0.04182 | -0.11769042 | 0 | 0.077 | 1 | 7 |
| nod2 | 0.04182 | -0.15997487 | 0 | 0.077 | 1 | 7 |
| nucb2b | 0.04182 | -0.15769344 | 0 | 0.077 | 1 | 7 |
| ppp1r7 | 0.04182 | -0.10561032 | 0 | 0.077 | 1 | 7 |
| shc2 | 0.04182 | -0.20284304 | 0 | 0.077 | 1 | 7 |
| si:ch1073-385f13.3 | 0.04182 | -0.1619537 | 0 | 0.077 | 1 | 7 |
| si:ch73-174h16.4 | 0.04182 | -0.12826668 | 0 | 0.077 | 1 | 7 |
| si:dkey-207l24.2 | 0.04182 | -0.10086636 | 0 | 0.077 | 1 | 7 |
| slc12a9 | 0.04182 | -0.1363209 | 0 | 0.077 | 1 | 7 |
| slc35b4 | 0.04182 | -0.11967781 | 0 | 0.077 | 1 | 7 |
| slc3a2a | 0.04182 | -0.12015345 | 0 | 0.077 | 1 | 7 |
| soul3 | 0.04182 | -0.10866602 | 0 | 0.077 | 1 | 7 |
| tmem170b | 0.04182 | -0.14126757 | 0 | 0.077 | 1 | 7 |
| tti2 | 0.04182 | -0.12667212 | 0 | 0.077 | 1 | 7 |
| ublcp1 | 0.04182 | -0.10650089 | 0 | 0.077 | 1 | 7 |
| vipas39 | 0.04182 | -0.13485447 | 0 | 0.077 | 1 | 7 |
| yif1b | 0.04182 | -0.10622865 | 0 | 0.077 | 1 | 7 |
| zgc:101566 | 0.04182 | -0.1087951 | 0 | 0.077 | 1 | 7 |
| LO017877.2 | 0.04182 | -0.13621315 | 0 | 0.077 | 1 | 7 |

| | | | | | | |
|---|---|---|---|---|---|---|
| cd40 | 0.04182 | -0.13074839 | 0 | 0.077 | 1 | 7 |
| kdm3b | 0.04182 | -0.20855387 | 0 | 0.077 | 1 | 7 |
| lmbrd1 | 0.04182 | -0.14833285 | 0 | 0.077 | 1 | 7 |
| LOC100333482 | 0.04182 | -0.12917649 | 0 | 0.077 | 1 | 7 |
| LOC101882640 | 0.04182 | -0.13003683 | 0 | 0.077 | 1 | 7 |
| XLOC-043834 | 0.04182 | -0.1563657 | 0 | 0.077 | 1 | 7 |
| alcamb | 0.04182 | -0.11989633 | 0 | 0.077 | 1 | 7 |
| dhrs7 | 0.04182 | -0.17565272 | 0 | 0.077 | 1 | 7 |
| fam3a | 0.04182 | -0.12132316 | 0 | 0.077 | 1 | 7 |
| gdi1 | 0.04182 | -0.15967037 | 0 | 0.077 | 1 | 7 |
| ifng1 | 0.04182 | -0.13003683 | 0 | 0.077 | 1 | 7 |
| map7d3 | 0.04182 | -0.13914669 | 0 | 0.077 | 1 | 7 |
| mtbl | 0.04182 | -0.10518537 | 0 | 0.077 | 1 | 7 |
| nme3 | 0.04182 | -0.17740672 | 0 | 0.077 | 1 | 7 |
| optn | 0.04182 | -0.12451924 | 0 | 0.077 | 1 | 7 |
| rsad2 | 0.04182 | -0.15590721 | 0 | 0.077 | 1 | 7 |
| si:ch73-111k22.2 | 0.04182 | -0.17619312 | 0 | 0.077 | 1 | 7 |
| si:dkey-217d24.6 | 0.04182 | -0.14374254 | 0 | 0.077 | 1 | 7 |
| si:dkey-33c14.6 | 0.04182 | -0.13821751 | 0 | 0.077 | 1 | 7 |
| si:dkey-56e3.3 | 0.04182 | -0.19245088 | 0 | 0.077 | 1 | 7 |
| si:dkey-6e2.2 | 0.04182 | -0.22556751 | 0 | 0.077 | 1 | 7 |
| snx27a | 0.04182 | -0.12943985 | 0 | 0.077 | 1 | 7 |
| tfe3b | 0.04182 | -0.1260314 | 0 | 0.077 | 1 | 7 |
| tle3a | 0.04182 | -0.17978028 | 0 | 0.077 | 1 | 7 |
| wdr37 | 0.04182 | -0.12339587 | 0 | 0.077 | 1 | 7 |
| zgc:162898 | 0.04182 | -0.10153151 | 0 | 0.077 | 1 | 7 |
| zgc:174972 | 0.04182 | -0.11715805 | 0 | 0.077 | 1 | 7 |
| CABZ01066926.1 | 0.04182 | -0.10339105 | 0 | 0.077 | 1 | 7 |
| CU137717.1 | 0.04182 | -0.1262284 | 0 | 0.077 | 1 | 7 |
| CU467861.1 | 0.04182 | -0.14395868 | 0 | 0.077 | 1 | 7 |
| LOC101883097 | 0.04182 | -0.13185117 | 0 | 0.077 | 1 | 7 |
| XLOC-032256 | 0.04182 | -0.11930453 | 0 | 0.077 | 1 | 7 |
| arid3b | 0.04182 | -0.10623807 | 0 | 0.077 | 1 | 7 |
| armh3 | 0.04182 | -0.11673859 | 0 | 0.077 | 1 | 7 |
| brap | 0.04182 | -0.11912407 | 0 | 0.077 | 1 | 7 |
| chic1 | 0.04182 | -0.10065471 | 0 | 0.077 | 1 | 7 |
| fthl30 | 0.04182 | -0.12595348 | 0 | 0.077 | 1 | 7 |
| gpcpd1 | 0.04182 | -0.1342804 | 0 | 0.077 | 1 | 7 |
| hdac4 | 0.04182 | -0.12099899 | 0 | 0.077 | 1 | 7 |
| hexa | 0.04182 | -0.12681506 | 0 | 0.077 | 1 | 7 |
| igbp1 | 0.04182 | -0.12348089 | 0 | 0.077 | 1 | 7 |
| lactb | 0.04182 | -0.10701309 | 0 | 0.077 | 1 | 7 |
| lrrc3ca | 0.04182 | -0.11031929 | 0 | 0.077 | 1 | 7 |
| mettl13 | 0.04182 | -0.13713668 | 0 | 0.077 | 1 | 7 |
| nme7 | 0.04182 | -0.14335375 | 0 | 0.077 | 1 | 7 |
| otud4 | 0.04182 | -0.10746792 | 0 | 0.077 | 1 | 7 |
| ptdss1a | 0.04182 | -0.11613151 | 0 | 0.077 | 1 | 7 |
| raraa | 0.04182 | -0.13657267 | 0 | 0.077 | 1 | 7 |

| | | | | | | |
|---|---|---|---|---|---|---|
| rbm25a | 0.04182 | -0.11889219 | 0 | 0.077 | 1 | 7 |
| rgs16 | 0.04182 | -0.1374926 | 0 | 0.077 | 1 | 7 |
| rnf180 | 0.04182 | -0.13128882 | 0 | 0.077 | 1 | 7 |
| si:ch211-106f21.1 | 0.04182 | -0.12751923 | 0 | 0.077 | 1 | 7 |
| si:dkey-33c14.7 | 0.04182 | -0.13853091 | 0 | 0.077 | 1 | 7 |
| si:dkey-3h3.3 | 0.04182 | -0.13218023 | 0 | 0.077 | 1 | 7 |
| si:dkey-78l4.1 | 0.04182 | -0.20467347 | 0 | 0.077 | 1 | 7 |
| spi1b | 0.04182 | -0.1692244 | 0 | 0.077 | 1 | 7 |
| zgc:86609 | 0.04182 | -0.13645884 | 0 | 0.077 | 1 | 7 |
| ier2b | 0.0428 | -0.31640362 | 0.635 | 0.769 | 1 | 7 |
| h3f3d | 0.04327 | -0.30090518 | 0.714 | 0.923 | 1 | 7 |
| lyrm2 | 0.04345 | -0.29965147 | 0.048 | 0.192 | 1 | 7 |
| tgfbr2a | 0.04373 | -0.24178779 | 0.016 | 0.154 | 1 | 7 |
| prelid1b | 0.04417 | 0.223668737 | 0.19 | 0.038 | 1 | 7 |
| XLOC-013964 | 0.04435 | -0.22851109 | 0.857 | 1 | 1 | 7 |
| mpp1 | 0.04496 | -0.24020166 | 0.032 | 0.192 | 1 | 7 |
| abce1 | 0.04511 | 0.282411108 | 0.238 | 0.038 | 1 | 7 |
| fth1a | 0.04516 | -0.31187182 | 0.238 | 0.5 | 1 | 7 |
| uchl5 | 0.04551 | 0.295332666 | 0.19 | 0.038 | 1 | 7 |
| fam49bb | 0.0458 | -0.37734427 | 0.159 | 0.346 | 1 | 7 |
| nfu1 | 0.0459 | -0.27189722 | 0.048 | 0.231 | 1 | 7 |
| si:dkeyp-84f3.5 | 0.04659 | -0.12831089 | 0.016 | 0.115 | 1 | 7 |
| polr2l | 0.04663 | 0.170920201 | 0.27 | 0.269 | 1 | 7 |
| uqcrc1 | 0.0467 | 0.151317893 | 0.381 | 0.192 | 1 | 7 |
| zgc:110540 | 0.04682 | 0.161129765 | 0.079 | 0.077 | 1 | 7 |
| copa | 0.04699 | -0.24636834 | 0.048 | 0.231 | 1 | 7 |
| dnase1l4.2 | 0.04703 | -0.28130639 | 0.048 | 0.231 | 1 | 7 |
| mphosph8 | 0.04712 | 0.16614628 | 0.333 | 0.346 | 1 | 7 |
| ccl33.3 | 0.04784 | -0.33163515 | 0.048 | 0.192 | 1 | 7 |
| tmem176l.3a | 0.04797 | -0.32042465 | 0.048 | 0.231 | 1 | 7 |
| egr1 | 0.04804 | -0.36704334 | 0.238 | 0.5 | 1 | 7 |
| fam49ba | 0.04811 | -0.34336757 | 0.333 | 0.615 | 1 | 7 |
| tmem176l.1 | 0.04825 | -0.3325034 | 0.095 | 0.308 | 1 | 7 |
| fas | 0.04866 | -0.20960572 | 0.016 | 0.154 | 1 | 7 |
| aldh2.1 | 0.04871 | -0.12802888 | 0.016 | 0.154 | 1 | 7 |
| rpl5b | 0.04951 | -0.18315021 | 0.857 | 1 | 1 | 7 |
| sod2 | 0.05055 | 0.24641241 | 0.27 | 0.154 | 1 | 7 |
| ola1 | 0.05084 | 0.349247526 | 0.524 | 0.385 | 1 | 7 |
| pgam1a | 0.0514 | -0.13527716 | 0.492 | 0.692 | 1 | 7 |
| alad | 0.05172 | -0.20866392 | 0.016 | 0.154 | 1 | 7 |
| snx17 | 0.05177 | -0.21998956 | 0.032 | 0.192 | 1 | 7 |
| ostc | 0.05203 | -0.16768058 | 0.222 | 0.231 | 1 | 7 |
| il12rb2l | 0.05233 | -0.34284373 | 0.095 | 0.308 | 1 | 7 |
| arl5c | 0.05235 | -0.42010435 | 0.476 | 0.731 | 1 | 7 |
| ube2d4 | 0.05251 | -0.24255054 | 0.032 | 0.192 | 1 | 7 |
| dnttip2 | 0.05256 | -0.21273818 | 0.016 | 0.154 | 1 | 7 |
| cwf19l2 | 0.05275 | -0.23684254 | 0.016 | 0.154 | 1 | 7 |
| rsrp1 | 0.05282 | -0.30618801 | 0.19 | 0.308 | 1 | 7 |

| | | | | | | |
|---|---|---|---|---|---|---|
| cbx1b | 0.053 | -0.22175059 | 0.079 | 0.269 | 1 | 7 |
| cdk2ap2 | 0.05312 | 0.336453707 | 0.222 | 0.077 | 1 | 7 |
| golga7 | 0.0533 | 0.292512175 | 0.222 | 0.038 | 1 | 7 |
| zgc:165573 | 0.05352 | -0.35066872 | 0.095 | 0.308 | 1 | 7 |
| nfkbiaa | 0.05365 | -0.47244911 | 0.397 | 0.654 | 1 | 7 |
| rnaset2 | 0.05382 | -0.22259365 | 0.032 | 0.192 | 1 | 7 |
| tagapa | 0.05392 | -0.42022978 | 0.508 | 0.731 | 1 | 7 |
| mgat1a | 0.05396 | -0.23086969 | 0.032 | 0.192 | 1 | 7 |
| pdik1l | 0.05408 | -0.26986011 | 0.016 | 0.154 | 1 | 7 |
| szrd1 | 0.05431 | -0.23438823 | 0.032 | 0.192 | 1 | 7 |
| rfc4 | 0.05432 | 0.33082689 | 0.175 | 0.038 | 1 | 7 |
| nccrp1 | 0.05436 | -0.29111749 | 0.032 | 0.192 | 1 | 7 |
| adgre5b.3 | 0.0544 | -0.31909245 | 0.032 | 0.192 | 1 | 7 |
| tnrc6c1 | 0.05446 | -0.37890982 | 0.302 | 0.577 | 1 | 7 |
| urb1 | 0.05458 | -0.2399598 | 0.016 | 0.154 | 1 | 7 |
| csnk2a2a | 0.05474 | -0.26364038 | 0.016 | 0.154 | 1 | 7 |
| kdm6bb | 0.05494 | 0.420108996 | 0.302 | 0.115 | 1 | 7 |
| stat1a | 0.05498 | -0.28737342 | 0.127 | 0.346 | 1 | 7 |
| XLOC-000431 | 0.05535 | -0.21247011 | 0.952 | 1 | 1 | 7 |
| dusp22b | 0.05548 | -0.2334854 | 0.032 | 0.192 | 1 | 7 |
| timm23a | 0.05551 | 0.195653637 | 0.206 | 0.192 | 1 | 7 |
| slc35a4 | 0.05551 | -0.2614493 | 0.016 | 0.154 | 1 | 7 |
| LOC101886687 | 0.05552 | -0.25566912 | 0.016 | 0.154 | 1 | 7 |
| si:ch211-196c10.15 | 0.05554 | -0.26400567 | 0.032 | 0.192 | 1 | 7 |
| ptprc | 0.05592 | -0.20566988 | 0.508 | 0.538 | 1 | 7 |
| txnipa | 0.05601 | 0.379959708 | 0.524 | 0.385 | 1 | 7 |
| hnrnpaba | 0.05659 | 0.276469431 | 0.746 | 0.654 | 1 | 7 |
| capgb | 0.05659 | -0.43490663 | 0.429 | 0.692 | 1 | 7 |
| tsr3 | 0.05664 | 0.126764081 | 0.095 | 0 | 1 | 7 |
| aif1l | 0.05664 | 0.232776466 | 0.095 | 0 | 1 | 7 |
| bahcc1b | 0.05664 | 0.199256564 | 0.095 | 0 | 1 | 7 |
| bcl7a | 0.05664 | 0.181904989 | 0.095 | 0 | 1 | 7 |
| cetn3 | 0.05664 | 0.129176292 | 0.095 | 0 | 1 | 7 |
| ddx24 | 0.05664 | 0.158426846 | 0.095 | 0 | 1 | 7 |
| entpd4 | 0.05664 | 0.143348096 | 0.095 | 0 | 1 | 7 |
| exoc6 | 0.05664 | 0.162705871 | 0.095 | 0 | 1 | 7 |
| hacd2 | 0.05664 | 0.193806814 | 0.095 | 0 | 1 | 7 |
| myo1cb | 0.05664 | 0.204462916 | 0.095 | 0 | 1 | 7 |
| sel1l | 0.05664 | 0.202166878 | 0.095 | 0 | 1 | 7 |
| smg5 | 0.05664 | 0.208520149 | 0.095 | 0 | 1 | 7 |
| sulf2b | 0.05664 | 0.190741599 | 0.095 | 0 | 1 | 7 |
| tsr1 | 0.05664 | 0.141616093 | 0.095 | 0 | 1 | 7 |
| cd8b | 0.05664 | 0.295429212 | 0.095 | 0 | 1 | 7 |
| chd6 | 0.05664 | 0.172779063 | 0.095 | 0 | 1 | 7 |
| mrps2 | 0.05664 | 0.190431365 | 0.095 | 0 | 1 | 7 |
| amfra | 0.05664 | 0.193480704 | 0.095 | 0 | 1 | 7 |
| apeh | 0.05664 | 0.160701884 | 0.095 | 0 | 1 | 7 |
| apex1 | 0.05664 | 0.157398007 | 0.095 | 0 | 1 | 7 |

| | | | | | | |
|---|---|---|---|---|---|---|
| araf | 0.05664 | 0.193467823 | 0.095 | 0 | 1 | 7 |
| arid1aa | 0.05664 | 0.177151791 | 0.095 | 0 | 1 | 7 |
| cant1a | 0.05664 | 0.172135886 | 0.095 | 0 | 1 | 7 |
| cfdp1 | 0.05664 | 0.189493507 | 0.095 | 0 | 1 | 7 |
| dph5 | 0.05664 | 0.195998074 | 0.095 | 0 | 1 | 7 |
| gtf3aa | 0.05664 | 0.141651697 | 0.095 | 0 | 1 | 7 |
| hax1 | 0.05664 | 0.1499747 | 0.095 | 0 | 1 | 7 |
| kmt2ba | 0.05664 | 0.186877782 | 0.095 | 0 | 1 | 7 |
| mrpl51 | 0.05664 | 0.162771001 | 0.095 | 0 | 1 | 7 |
| nol11 | 0.05664 | 0.166995056 | 0.095 | 0 | 1 | 7 |
| pdia4 | 0.05664 | 0.154476647 | 0.095 | 0 | 1 | 7 |
| ppp1r2 | 0.05664 | 0.159390122 | 0.095 | 0 | 1 | 7 |
| psmc1a | 0.05664 | 0.147866479 | 0.095 | 0 | 1 | 7 |
| rnmt | 0.05664 | 0.153435985 | 0.095 | 0 | 1 | 7 |
| shmt1 | 0.05664 | 0.199638719 | 0.095 | 0 | 1 | 7 |
| si:ch211-286b5.5 | 0.05664 | 0.111098741 | 0.095 | 0 | 1 | 7 |
| slc43a2a | 0.05664 | 0.163486996 | 0.095 | 0 | 1 | 7 |
| slu7 | 0.05664 | 0.15480702 | 0.095 | 0 | 1 | 7 |
| spc25 | 0.05664 | 0.187538369 | 0.095 | 0 | 1 | 7 |
| srsf5b | 0.05664 | 0.15572498 | 0.095 | 0 | 1 | 7 |
| ssna1 | 0.05664 | 0.153844079 | 0.095 | 0 | 1 | 7 |
| supt5h | 0.05664 | 0.134201272 | 0.095 | 0 | 1 | 7 |
| tespa1 | 0.05664 | 0.224509816 | 0.095 | 0 | 1 | 7 |
| znf207b | 0.05664 | 0.169312668 | 0.095 | 0 | 1 | 7 |
| tfdp1b | 0.05664 | 0.200436563 | 0.095 | 0 | 1 | 7 |
| CABZ01113192.1 | 0.05664 | 0.135374691 | 0.095 | 0 | 1 | 7 |
| LOC108179023 | 0.05664 | 0.16111318 | 0.095 | 0 | 1 | 7 |
| cdc37 | 0.05664 | 0.154845111 | 0.095 | 0 | 1 | 7 |
| drg1 | 0.05664 | 0.125120627 | 0.095 | 0 | 1 | 7 |
| mrpl23 | 0.05664 | 0.169359875 | 0.095 | 0 | 1 | 7 |
| opa1 | 0.05664 | 0.16405092 | 0.095 | 0 | 1 | 7 |
| os9 | 0.05664 | 0.15844738 | 0.095 | 0 | 1 | 7 |
| parp3 | 0.05664 | 0.164694725 | 0.095 | 0 | 1 | 7 |
| pelp1 | 0.05664 | 0.155974877 | 0.095 | 0 | 1 | 7 |
| pip4k2aa | 0.05664 | 0.155476899 | 0.095 | 0 | 1 | 7 |
| ppp2r1bb | 0.05664 | 0.139742422 | 0.095 | 0 | 1 | 7 |
| seh1l | 0.05664 | 0.139630906 | 0.095 | 0 | 1 | 7 |
| sema4ab | 0.05664 | 0.202505491 | 0.095 | 0 | 1 | 7 |
| smarcd2 | 0.05664 | 0.191386648 | 0.095 | 0 | 1 | 7 |
| tbc1d10aa | 0.05664 | 0.171896816 | 0.095 | 0 | 1 | 7 |
| tox4a | 0.05664 | 0.211314066 | 0.095 | 0 | 1 | 7 |
| urod | 0.05664 | 0.135193366 | 0.095 | 0 | 1 | 7 |
| zgc:153115 | 0.05664 | 0.166810849 | 0.095 | 0 | 1 | 7 |
| lsr | 0.05664 | 0.303110049 | 0.095 | 0 | 1 | 7 |
| xpo6 | 0.0573 | -0.24728187 | 0.079 | 0.269 | 1 | 7 |
| lsm7 | 0.05857 | 0.282359405 | 0.413 | 0.346 | 1 | 7 |
| plk3 | 0.05858 | -0.40442162 | 0.206 | 0.462 | 1 | 7 |
| rbx1 | 0.05997 | 0.332850247 | 0.54 | 0.385 | 1 | 7 |

| | | | | | | |
|---|---|---|---|---|---|---|
| si:dkey-27i16.2 | 0.06026 | -0.29240594 | 0.429 | 0.692 | 1 | 7 |
| si:ch73-264p11.1 | 0.06145 | -0.42242212 | 0.111 | 0.308 | 1 | 7 |
| calm3b | 0.06487 | 0.129435311 | 0.413 | 0.462 | 1 | 7 |
| h3f3a | 0.06503 | 0.315406685 | 0.508 | 0.5 | 1 | 7 |
| sub1b | 0.06547 | 0.257902665 | 0.206 | 0.038 | 1 | 7 |
| si:dkeyp-44a8.2 | 0.06647 | -0.17910116 | 0.048 | 0.192 | 1 | 7 |
| zgc:56676 | 0.06668 | -0.20568163 | 0.27 | 0.5 | 1 | 7 |
| si:dkey-262k9.2 | 0.06805 | -0.27673243 | 0.063 | 0.231 | 1 | 7 |
| ebna1bp2 | 0.06834 | 0.241358332 | 0.19 | 0.115 | 1 | 7 |
| crebbpa | 0.0686 | -0.24179733 | 0.032 | 0.154 | 1 | 7 |
| fam107b | 0.06873 | -0.37425288 | 0.111 | 0.308 | 1 | 7 |
| psma5 | 0.06912 | 0.189391762 | 0.397 | 0.346 | 1 | 7 |
| pik3r3a | 0.06945 | 0.389949607 | 0.286 | 0.077 | 1 | 7 |
| atl3 | 0.0696 | -0.31730345 | 0.079 | 0.269 | 1 | 7 |
| lsm6 | 0.07016 | 0.239724158 | 0.286 | 0.231 | 1 | 7 |
| srgn | 0.07019 | -0.33046742 | 0.667 | 0.846 | 1 | 7 |
| TADA1 | 0.07041 | -0.17135098 | 0.016 | 0.115 | 1 | 7 |
| srrm2 | 0.07055 | 0.217473359 | 0.159 | 0.038 | 1 | 7 |
| lgals3a | 0.07169 | -0.17999282 | 0.222 | 0.423 | 1 | 7 |
| nipsnap2 | 0.07234 | 0.300863715 | 0.254 | 0.077 | 1 | 7 |
| ifrd1 | 0.07248 | -0.20236372 | 0.159 | 0.154 | 1 | 7 |
| srsf10b | 0.07252 | 0.275249104 | 0.206 | 0.115 | 1 | 7 |
| capns1a | 0.0726 | 0.32204493 | 0.254 | 0.077 | 1 | 7 |
| sdhc | 0.07292 | 0.224317805 | 0.302 | 0.231 | 1 | 7 |
| BX322618.1 | 0.07317 | 0.35838023 | 0.238 | 0.154 | 1 | 7 |
| atf4b | 0.07467 | -0.19221081 | 0.492 | 0.731 | 1 | 7 |
| chrac1 | 0.07577 | 0.247042918 | 0.159 | 0.077 | 1 | 7 |
| vamp3 | 0.07587 | -0.28940568 | 0.254 | 0.5 | 1 | 7 |
| aars | 0.07612 | 0.281745156 | 0.206 | 0.115 | 1 | 7 |
| rsf1b.1 | 0.07618 | 0.348360688 | 0.317 | 0.115 | 1 | 7 |
| trnau1apa | 0.07699 | -0.1574811 | 0.032 | 0.154 | 1 | 7 |
| cebpb | 0.07873 | -0.37390349 | 0.397 | 0.654 | 1 | 7 |
| atp5f1d | 0.07874 | -0.30013412 | 0.476 | 0.731 | 1 | 7 |
| fen1 | 0.07876 | 0.222626612 | 0.175 | 0.115 | 1 | 7 |
| nip7 | 0.07886 | 0.263990157 | 0.19 | 0.077 | 1 | 7 |
| myd88 | 0.07942 | -0.32147562 | 0.159 | 0.385 | 1 | 7 |
| abcf1 | 0.07947 | 0.334253535 | 0.286 | 0.154 | 1 | 7 |
| ssr3 | 0.07981 | -0.28539491 | 0.159 | 0.385 | 1 | 7 |
| lcorl | 0.08079 | -0.14032377 | 0.016 | 0.115 | 1 | 7 |
| ndufa6 | 0.08089 | 0.336985471 | 0.302 | 0.154 | 1 | 7 |
| golgb1 | 0.08107 | -0.21535296 | 0.095 | 0.269 | 1 | 7 |
| ube2v2 | 0.08142 | 0.264907582 | 0.54 | 0.462 | 1 | 7 |
| si:dkey-171c9.3 | 0.08225 | 0.288910056 | 0.286 | 0.192 | 1 | 7 |
| XLOC-014079 | 0.08248 | 0.460319326 | 0.333 | 0.192 | 1 | 7 |
| ywhaqa | 0.0829 | 0.135972076 | 0.476 | 0.5 | 1 | 7 |
| atp11c | 0.08376 | -0.15132921 | 0.016 | 0.115 | 1 | 7 |
| dusp2 | 0.08514 | -0.26669197 | 0.683 | 0.769 | 1 | 7 |
| kansl1b | 0.08543 | -0.212279 | 0.032 | 0.154 | 1 | 7 |

| | | | | | | |
|---|---|---|---|---|---|---|
| cpne1 | 0.08572 | -0.31936106 | 0.254 | 0.5 | 1 | 7 |
| timm17a | 0.08641 | 0.254000475 | 0.175 | 0.038 | 1 | 7 |
| slc6a6b | 0.08666 | -0.1431065 | 0.079 | 0.192 | 1 | 7 |
| LOC100332847 | 0.08693 | 0.159383255 | 0.079 | 0 | 1 | 7 |
| josd2 | 0.08693 | 0.166417973 | 0.079 | 0 | 1 | 7 |
| myo9bl | 0.08693 | 0.171707065 | 0.079 | 0 | 1 | 7 |
| nusap1 | 0.08693 | 0.193384773 | 0.079 | 0 | 1 | 7 |
| ptpn9a | 0.08693 | 0.157180739 | 0.079 | 0 | 1 | 7 |
| umps | 0.08693 | 0.141530852 | 0.079 | 0 | 1 | 7 |
| CABZ01088149.1 | 0.08693 | 0.107526477 | 0.079 | 0 | 1 | 7 |
| ankrd54 | 0.08693 | 0.110983 | 0.079 | 0 | 1 | 7 |
| ap3m2 | 0.08693 | 0.177402378 | 0.079 | 0 | 1 | 7 |
| cd164 | 0.08693 | 0.126777062 | 0.079 | 0 | 1 | 7 |
| cd28l | 0.08693 | 0.184661672 | 0.079 | 0 | 1 | 7 |
| cul3b | 0.08693 | 0.136579924 | 0.079 | 0 | 1 | 7 |
| cyba | 0.08693 | 0.208611196 | 0.079 | 0 | 1 | 7 |
| dot1l | 0.08693 | 0.17833757 | 0.079 | 0 | 1 | 7 |
| dr1 | 0.08693 | 0.12638358 | 0.079 | 0 | 1 | 7 |
| fam173a | 0.08693 | 0.151878586 | 0.079 | 0 | 1 | 7 |
| gfer | 0.08693 | 0.141595499 | 0.079 | 0 | 1 | 7 |
| hexim1 | 0.08693 | 0.127612719 | 0.079 | 0 | 1 | 7 |
| ipo4 | 0.08693 | 0.203162381 | 0.079 | 0 | 1 | 7 |
| irf2a | 0.08693 | 0.13863561 | 0.079 | 0 | 1 | 7 |
| klf12b | 0.08693 | 0.137932818 | 0.079 | 0 | 1 | 7 |
| lyrm9 | 0.08693 | 0.118170652 | 0.079 | 0 | 1 | 7 |
| mrpl52 | 0.08693 | 0.121704277 | 0.079 | 0 | 1 | 7 |
| mrps24 | 0.08693 | 0.137331834 | 0.079 | 0 | 1 | 7 |
| mydgf | 0.08693 | 0.128768601 | 0.079 | 0 | 1 | 7 |
| paqr3a | 0.08693 | 0.143518922 | 0.079 | 0 | 1 | 7 |
| pecr | 0.08693 | 0.14721616 | 0.079 | 0 | 1 | 7 |
| pmvk | 0.08693 | 0.156093045 | 0.079 | 0 | 1 | 7 |
| pptc7a | 0.08693 | 0.150823465 | 0.079 | 0 | 1 | 7 |
| prkag1 | 0.08693 | 0.122853347 | 0.079 | 0 | 1 | 7 |
| rab14l | 0.08693 | 0.141481175 | 0.079 | 0 | 1 | 7 |
| ranbp9 | 0.08693 | 0.166216781 | 0.079 | 0 | 1 | 7 |
| rcl1 | 0.08693 | 0.104754625 | 0.079 | 0 | 1 | 7 |
| sec22a | 0.08693 | 0.144670904 | 0.079 | 0 | 1 | 7 |
| si:ch1073-157b13.1 | 0.08693 | 0.129519352 | 0.079 | 0 | 1 | 7 |
| si:ch211-199g17.2 | 0.08693 | 0.205055859 | 0.079 | 0 | 1 | 7 |
| si:ch211-243a20.4 | 0.08693 | 0.177896582 | 0.079 | 0 | 1 | 7 |
| si:dkey-6i22.5 | 0.08693 | 0.151218013 | 0.079 | 0 | 1 | 7 |
| si:dkey-81j8.6 | 0.08693 | 0.140617141 | 0.079 | 0 | 1 | 7 |
| si:dkeyp-2e4.3 | 0.08693 | 0.147425121 | 0.079 | 0 | 1 | 7 |
| slx1b | 0.08693 | 0.156571208 | 0.079 | 0 | 1 | 7 |
| smc6 | 0.08693 | 0.112946886 | 0.079 | 0 | 1 | 7 |
| sorl1 | 0.08693 | 0.202050536 | 0.079 | 0 | 1 | 7 |
| srsf2b | 0.08693 | 0.117465475 | 0.079 | 0 | 1 | 7 |
| tfe3a | 0.08693 | 0.165243161 | 0.079 | 0 | 1 | 7 |

| | | | | | | |
|---|---|---|---|---|---|---|
| tmem131 | 0.08693 | 0.132855803 | 0.079 | 0 | 1 | 7 |
| trpc4apa | 0.08693 | 0.164425621 | 0.079 | 0 | 1 | 7 |
| uhrf1 | 0.08693 | 0.143915863 | 0.079 | 0 | 1 | 7 |
| usp4 | 0.08693 | 0.184963495 | 0.079 | 0 | 1 | 7 |
| usp8 | 0.08693 | 0.123931107 | 0.079 | 0 | 1 | 7 |
| vgll4b | 0.08693 | 0.150773866 | 0.079 | 0 | 1 | 7 |
| xrcc6 | 0.08693 | 0.114837103 | 0.079 | 0 | 1 | 7 |
| zbtb46 | 0.08693 | 0.145903993 | 0.079 | 0 | 1 | 7 |
| CDK2AP1 | 0.08693 | 0.137425616 | 0.079 | 0 | 1 | 7 |
| FAM53C | 0.08693 | 0.101552705 | 0.079 | 0 | 1 | 7 |
| LOC108185170 | 0.08693 | 0.183070422 | 0.079 | 0 | 1 | 7 |
| XLOC-043013 | 0.08693 | 0.207233118 | 0.079 | 0 | 1 | 7 |
| aaas | 0.08693 | 0.165319617 | 0.079 | 0 | 1 | 7 |
| ccdc90b | 0.08693 | 0.128853731 | 0.079 | 0 | 1 | 7 |
| coa1 | 0.08693 | 0.119288296 | 0.079 | 0 | 1 | 7 |
| cpsf1 | 0.08693 | 0.119675548 | 0.079 | 0 | 1 | 7 |
| gclm | 0.08693 | 0.10926627 | 0.079 | 0 | 1 | 7 |
| grinab | 0.08693 | 0.140079571 | 0.079 | 0 | 1 | 7 |
| lman1 | 0.08693 | 0.12270878 | 0.079 | 0 | 1 | 7 |
| med27 | 0.08693 | 0.140704197 | 0.079 | 0 | 1 | 7 |
| mettl1 | 0.08693 | 0.113879681 | 0.079 | 0 | 1 | 7 |
| mybbp1a | 0.08693 | 0.123251848 | 0.079 | 0 | 1 | 7 |
| nol9 | 0.08693 | 0.148330964 | 0.079 | 0 | 1 | 7 |
| pou2f1b | 0.08693 | 0.210762825 | 0.079 | 0 | 1 | 7 |
| prf1.9 | 0.08693 | 0.313873529 | 0.079 | 0 | 1 | 7 |
| si:ch211-281l24.3 | 0.08693 | 0.153646594 | 0.079 | 0 | 1 | 7 |
| slc25a26 | 0.08693 | 0.107153866 | 0.079 | 0 | 1 | 7 |
| slc29a1a | 0.08693 | 0.180084163 | 0.079 | 0 | 1 | 7 |
| tk1 | 0.08693 | 0.136461365 | 0.079 | 0 | 1 | 7 |
| trmt10a | 0.08693 | 0.149363272 | 0.079 | 0 | 1 | 7 |
| zdhhc4 | 0.08693 | 0.14830568 | 0.079 | 0 | 1 | 7 |
| zgc:153733 | 0.08693 | 0.161251196 | 0.079 | 0 | 1 | 7 |
| zgc:171967 | 0.08693 | 0.187427567 | 0.079 | 0 | 1 | 7 |
| ccdc93 | 0.08693 | 0.133203631 | 0.079 | 0 | 1 | 7 |
| itgb1bp1 | 0.08693 | 0.154218696 | 0.079 | 0 | 1 | 7 |
| lrpprc | 0.08693 | 0.130410423 | 0.079 | 0 | 1 | 7 |
| nefla | 0.08693 | 0.165136242 | 0.079 | 0 | 1 | 7 |
| arl9 | 0.08693 | 0.166996461 | 0.079 | 0 | 1 | 7 |
| klhdc10 | 0.08693 | 0.204284574 | 0.079 | 0 | 1 | 7 |
| larp4aa | 0.08693 | 0.125618368 | 0.079 | 0 | 1 | 7 |
| march5l | 0.08693 | 0.134301101 | 0.079 | 0 | 1 | 7 |
| raf1b | 0.08693 | 0.138078647 | 0.079 | 0 | 1 | 7 |
| rnf40 | 0.08693 | 0.15017017 | 0.079 | 0 | 1 | 7 |
| dop1b | 0.08693 | 0.151122647 | 0.079 | 0 | 1 | 7 |
| lto1 | 0.08693 | 0.164365428 | 0.079 | 0 | 1 | 7 |
| zgc:92907 | 0.08693 | 0.153162604 | 0.079 | 0 | 1 | 7 |
| XLOC-028370 | 0.08851 | 0.417623434 | 0.27 | 0.077 | 1 | 7 |
| zgc:66160 | 0.08894 | -0.14328745 | 0.048 | 0.192 | 1 | 7 |

| | | | | | | |
|---|---|---|---|---|---|---|
| nop56 | 0.08915 | -0.24438995 | 0.381 | 0.615 | 1 | 7 |
| qkia | 0.08957 | -0.35236514 | 0.143 | 0.346 | 1 | 7 |
| cabz01093075.1 | 0.08972 | -0.12290334 | 0.111 | 0.231 | 1 | 7 |
| si:ch73-308l14.2 | 0.09117 | 0.277501869 | 0.206 | 0.077 | 1 | 7 |
| march5 | 0.09164 | -0.19757684 | 0.095 | 0.269 | 1 | 7 |
| si:zfos-464b6.2 | 0.09174 | 0.246828348 | 0.206 | 0.038 | 1 | 7 |
| acp1 | 0.0933 | 0.248882466 | 0.19 | 0.038 | 1 | 7 |
| fosab | 0.09336 | -0.35798521 | 0.444 | 0.692 | 1 | 7 |
| ddx23 | 0.09343 | -0.19201496 | 0.063 | 0.231 | 1 | 7 |
| pkma | 0.09388 | -0.28145952 | 0.302 | 0.538 | 1 | 7 |
| dkc1 | 0.09496 | 0.256115482 | 0.302 | 0.231 | 1 | 7 |
| XLOC-043043 | 0.09503 | 0.178249928 | 0.492 | 0.538 | 1 | 7 |
| jpt1b | 0.09514 | -0.28788151 | 0.111 | 0.308 | 1 | 7 |
| mt-nd1 | 0.0955 | -0.28671754 | 0.079 | 0.231 | 1 | 7 |
| pho | 0.09599 | 0.230261817 | 0.206 | 0.154 | 1 | 7 |
| atp1a1a.1 | 0.09644 | -0.27295262 | 0.111 | 0.308 | 1 | 7 |
| cmtm6 | 0.09656 | -0.23547703 | 0.063 | 0.231 | 1 | 7 |
| ddx5 | 0.0976 | -0.30061915 | 0.524 | 0.654 | 1 | 7 |
| rae1 | 0.09802 | -0.23231963 | 0.063 | 0.231 | 1 | 7 |
| ip6k2a | 0.09828 | -0.20422743 | 0.095 | 0.269 | 1 | 7 |
| cpeb4b | 0.09909 | -0.17232549 | 0.016 | 0.115 | 1 | 7 |
| cox7a2a | 0.0991 | 0.336761817 | 0.556 | 0.308 | 1 | 7 |
| ythdf2 | 0.09939 | -0.28107442 | 0.032 | 0.154 | 1 | 7 |
| si:rp71-81e14.2 | 0.09956 | -0.40250747 | 0.222 | 0.423 | 1 | 7 |
| nudt9 | 0.10023 | -0.20041575 | 0.048 | 0.192 | 1 | 7 |
| wnk1a | 0.10042 | -0.28660283 | 0.048 | 0.154 | 1 | 7 |
| si:dkey-217l24.1 | 0.10072 | 0.305014147 | 0.175 | 0.038 | 1 | 7 |
| kdm2ba | 0.10212 | -0.19498693 | 0.079 | 0.231 | 1 | 7 |
| mhc2dab | 0.10313 | -0.15648915 | 0.063 | 0.192 | 1 | 7 |
| mycb | 0.10482 | -0.34039663 | 0.175 | 0.385 | 1 | 7 |
| FP101882.1 | 0.10496 | 0.303867347 | 0.222 | 0.077 | 1 | 7 |
| wdr82 | 0.10586 | 0.256097445 | 0.206 | 0.077 | 1 | 7 |
| rpf2 | 0.107 | -0.20194003 | 0.095 | 0.269 | 1 | 7 |
| u2af1 | 0.10727 | 0.211075867 | 0.397 | 0.346 | 1 | 7 |
| mavs | 0.10731 | -0.13761671 | 0.016 | 0.115 | 1 | 7 |
| pgpep1 | 0.10759 | -0.24007856 | 0.032 | 0.154 | 1 | 7 |
| cmah | 0.10772 | -0.30105801 | 0.063 | 0.192 | 1 | 7 |
| si:ch211-105j21.9 | 0.10886 | -0.37433901 | 0.206 | 0.423 | 1 | 7 |
| gng10 | 0.10935 | -0.25223393 | 0.032 | 0.154 | 1 | 7 |
| id2a | 0.10939 | 0.156888812 | 0.206 | 0.077 | 1 | 7 |
| lsm3 | 0.1095 | 0.258605919 | 0.27 | 0.192 | 1 | 7 |
| vapal | 0.10954 | -0.15499814 | 0.048 | 0.192 | 1 | 7 |
| lsm12a | 0.1097 | -0.22130426 | 0.048 | 0.192 | 1 | 7 |
| glipr1a | 0.10987 | 0.338064075 | 0.365 | 0.154 | 1 | 7 |
| lasp1 | 0.10992 | -0.28055291 | 0.492 | 0.731 | 1 | 7 |
| GCA | 0.11072 | -0.33011318 | 0.46 | 0.692 | 1 | 7 |
| cytip | 0.11157 | -0.34603524 | 0.175 | 0.346 | 1 | 7 |
| slc25a51b | 0.1116 | 0.185629898 | 0.127 | 0.038 | 1 | 7 |

| | | | | | | |
|---|---|---|---|---|---|---|
| usp47 | 0.11175 | -0.25728611 | 0.016 | 0.115 | 1 | 7 |
| mdh1aa | 0.11187 | 0.352676449 | 0.429 | 0.269 | 1 | 7 |
| adam10a | 0.11304 | -0.19170443 | 0.079 | 0.231 | 1 | 7 |
| gsto2 | 0.11352 | -0.2308657 | 0.048 | 0.192 | 1 | 7 |
| ccl34b.4 | 0.1144 | -0.30664884 | 0.159 | 0.192 | 1 | 7 |
| slc25a5 | 0.11451 | -0.11268805 | 0.952 | 1 | 1 | 7 |
| cct5 | 0.11452 | 0.149464422 | 0.556 | 0.577 | 1 | 7 |
| XLOC-023492 | 0.1146 | -0.39616544 | 0.254 | 0.462 | 1 | 7 |
| phb2a | 0.11464 | -0.11999689 | 0.016 | 0.115 | 1 | 7 |
| srp68 | 0.11475 | -0.24755641 | 0.032 | 0.154 | 1 | 7 |
| acp5b | 0.11479 | -0.15523445 | 0.032 | 0.154 | 1 | 7 |
| nansa | 0.11519 | -0.13988835 | 0.032 | 0.154 | 1 | 7 |
| cflara | 0.11551 | -0.24895391 | 0.016 | 0.115 | 1 | 7 |
| mef2b | 0.11594 | 0.397770361 | 0.19 | 0.038 | 1 | 7 |
| sf3b2 | 0.11609 | 0.279221916 | 0.254 | 0.077 | 1 | 7 |
| proser1 | 0.11713 | -0.22951132 | 0.032 | 0.154 | 1 | 7 |
| tax1bp1b | 0.11809 | -0.23582491 | 0.095 | 0.269 | 1 | 7 |
| si:ch211-260e23.9 | 0.11831 | 0.323852188 | 0.175 | 0.077 | 1 | 7 |
| nek6 | 0.11877 | -0.21131907 | 0.048 | 0.192 | 1 | 7 |
| tmem50a | 0.12008 | 0.306096458 | 0.238 | 0.077 | 1 | 7 |
| psmd6 | 0.12014 | 0.20752091 | 0.19 | 0.038 | 1 | 7 |
| mych | 0.12026 | -0.32851574 | 0.27 | 0.5 | 1 | 7 |
| edf1 | 0.12053 | 0.361298272 | 0.603 | 0.385 | 1 | 7 |
| cd4-1 | 0.12101 | 0.256479737 | 0.159 | 0.038 | 1 | 7 |
| COX5B | 0.12112 | 0.300443537 | 0.714 | 0.615 | 1 | 7 |
| adh5 | 0.12114 | 0.260467729 | 0.317 | 0.269 | 1 | 7 |
| kctd5a | 0.12122 | 0.313251981 | 0.524 | 0.385 | 1 | 7 |
| tnfrsf9a | 0.12147 | 0.300869611 | 0.19 | 0.038 | 1 | 7 |
| utp18 | 0.12165 | -0.2026208 | 0.048 | 0.192 | 1 | 7 |
| nkl.2 | 0.12184 | 0.384584488 | 0.238 | 0.192 | 1 | 7 |
| mlf2 | 0.12225 | -0.1993344 | 0.048 | 0.192 | 1 | 7 |
| inpp4ab | 0.12257 | -0.23736084 | 0.048 | 0.192 | 1 | 7 |
| dnajb1b | 0.12268 | -0.30122035 | 0.063 | 0.192 | 1 | 7 |
| ier2a | 0.12283 | -0.22238692 | 0.54 | 0.577 | 1 | 7 |
| hivep2a | 0.12316 | -0.2606834 | 0.032 | 0.115 | 1 | 7 |
| gstr | 0.1244 | -0.21302516 | 0.095 | 0.269 | 1 | 7 |
| ubap2b | 0.12456 | 0.235399786 | 0.19 | 0.038 | 1 | 7 |
| elmsan1b | 0.12491 | -0.20278611 | 0.032 | 0.154 | 1 | 7 |
| sdc4 | 0.12505 | -0.31917065 | 0.079 | 0.192 | 1 | 7 |
| si:ch73-234b20.5 | 0.12552 | -0.20942152 | 0.254 | 0.462 | 1 | 7 |
| LEPROTL1 | 0.12554 | -0.21367408 | 0.079 | 0.231 | 1 | 7 |
| commd1 | 0.12558 | -0.21387838 | 0.032 | 0.154 | 1 | 7 |
| wdr3 | 0.12624 | -0.31250495 | 0.159 | 0.346 | 1 | 7 |
| usp9 | 0.12648 | -0.24318656 | 0.095 | 0.269 | 1 | 7 |
| mlx | 0.12663 | -0.11178679 | 0.079 | 0.192 | 1 | 7 |
| srfa | 0.12676 | -0.1367374 | 0.048 | 0.154 | 1 | 7 |
| mrps18a | 0.12685 | 0.23984463 | 0.19 | 0.038 | 1 | 7 |
| taf11 | 0.12694 | 0.243437741 | 0.19 | 0.038 | 1 | 7 |

| | | | | | | |
|---|---|---|---|---|---|---|
| pde7a | 0.12708 | -0.10851092 | 0.095 | 0.231 | 1 | 7 |
| si:ch211-262e15.1 | 0.12739 | -0.26109516 | 0.111 | 0.269 | 1 | 7 |
| si:ch211-212k18.7 | 0.12757 | -0.25059577 | 0.095 | 0.269 | 1 | 7 |
| arl6ip1 | 0.12769 | 0.385337339 | 0.46 | 0.269 | 1 | 7 |
| csf2rb | 0.12778 | 0.331634305 | 0.175 | 0.038 | 1 | 7 |
| dock4b | 0.12805 | -0.14816525 | 0.016 | 0.115 | 1 | 7 |
| samsn1b | 0.12828 | -0.21451037 | 0.143 | 0.308 | 1 | 7 |
| psmc2 | 0.12861 | 0.259796015 | 0.254 | 0.077 | 1 | 7 |
| ttf1 | 0.12883 | -0.20821847 | 0.032 | 0.154 | 1 | 7 |
| tmem131l | 0.12883 | -0.2809045 | 0.095 | 0.269 | 1 | 7 |
| gzm3.3 | 0.12943 | -0.18767384 | 0.063 | 0.192 | 1 | 7 |
| snx12 | 0.12945 | -0.29448733 | 0.095 | 0.269 | 1 | 7 |
| ubtf | 0.12946 | -0.30044867 | 0.095 | 0.269 | 1 | 7 |
| elp2 | 0.13067 | -0.14565994 | 0.016 | 0.115 | 1 | 7 |
| map3k7 | 0.13106 | -0.16905845 | 0.016 | 0.115 | 1 | 7 |
| bnip2 | 0.13133 | -0.206907 | 0.032 | 0.154 | 1 | 7 |
| cnih1 | 0.13148 | -0.2122671 | 0.032 | 0.154 | 1 | 7 |
| gnb1a | 0.13188 | -0.26924179 | 0.349 | 0.577 | 1 | 7 |
| XLOC-038098 | 0.13191 | -0.13162184 | 0.016 | 0.115 | 1 | 7 |
| XLOC-039757 | 0.13234 | 0.283577503 | 0.365 | 0.308 | 1 | 7 |
| ppm1aa | 0.13285 | -0.29981343 | 0.095 | 0.231 | 1 | 7 |
| surf4l | 0.13295 | -0.22570316 | 0.016 | 0.115 | 1 | 7 |
| dhx38 | 0.13381 | -0.19515837 | 0.032 | 0.154 | 1 | 7 |
| BX001014.2 | 0.13395 | 0.15492135 | 0.063 | 0 | 1 | 7 |
| CR812481.1 | 0.13395 | 0.122578418 | 0.063 | 0 | 1 | 7 |
| CU914617.1 | 0.13395 | 0.16653515 | 0.063 | 0 | 1 | 7 |
| FAM72B | 0.13395 | 0.12697245 | 0.063 | 0 | 1 | 7 |
| LOC100004978 | 0.13395 | 0.135000273 | 0.063 | 0 | 1 | 7 |
| LOC570835 | 0.13395 | 0.121359982 | 0.063 | 0 | 1 | 7 |
| acad9 | 0.13395 | 0.122354364 | 0.063 | 0 | 1 | 7 |
| alox5a | 0.13395 | 0.163058459 | 0.063 | 0 | 1 | 7 |
| bscl2 | 0.13395 | 0.108672276 | 0.063 | 0 | 1 | 7 |
| col14a1b | 0.13395 | 0.124404451 | 0.063 | 0 | 1 | 7 |
| commd8 | 0.13395 | 0.108487375 | 0.063 | 0 | 1 | 7 |
| galnt6 | 0.13395 | 0.139236635 | 0.063 | 0 | 1 | 7 |
| gigyf1a | 0.13395 | 0.172146101 | 0.063 | 0 | 1 | 7 |
| haus4 | 0.13395 | 0.139234721 | 0.063 | 0 | 1 | 7 |
| ing3 | 0.13395 | 0.115918892 | 0.063 | 0 | 1 | 7 |
| klc1b | 0.13395 | 0.119636656 | 0.063 | 0 | 1 | 7 |
| lipea | 0.13395 | 0.104455428 | 0.063 | 0 | 1 | 7 |
| mcph1 | 0.13395 | 0.164115585 | 0.063 | 0 | 1 | 7 |
| mtmr8 | 0.13395 | 0.130516861 | 0.063 | 0 | 1 | 7 |
| mus81 | 0.13395 | 0.105495656 | 0.063 | 0 | 1 | 7 |
| mynn | 0.13395 | 0.156564284 | 0.063 | 0 | 1 | 7 |
| nfe2l2a | 0.13395 | 0.120962448 | 0.063 | 0 | 1 | 7 |
| nfil3-4 | 0.13395 | 0.118549206 | 0.063 | 0 | 1 | 7 |
| nudt1 | 0.13395 | 0.105607007 | 0.063 | 0 | 1 | 7 |
| racgap1 | 0.13395 | 0.119528104 | 0.063 | 0 | 1 | 7 |

| | | | | | | |
|---|---|---|---|---|---|---|
| rad54l | 0.13395 | 0.102536008 | 0.063 | 0 | 1 | 7 |
| si:ch1073-443f11.2 | 0.13395 | 0.121152327 | 0.063 | 0 | 1 | 7 |
| si:ch211-266i6.3 | 0.13395 | 0.101011129 | 0.063 | 0 | 1 | 7 |
| si:dkey-237i9.1 | 0.13395 | 0.100413663 | 0.063 | 0 | 1 | 7 |
| si:dkeyp-75h12.2 | 0.13395 | 0.119806947 | 0.063 | 0 | 1 | 7 |
| spc24 | 0.13395 | 0.125378962 | 0.063 | 0 | 1 | 7 |
| srek1ip1 | 0.13395 | 0.110205717 | 0.063 | 0 | 1 | 7 |
| tpx2 | 0.13395 | 0.114990582 | 0.063 | 0 | 1 | 7 |
| zfpl1 | 0.13395 | 0.120636194 | 0.063 | 0 | 1 | 7 |
| zgc:101783 | 0.13395 | 0.104094364 | 0.063 | 0 | 1 | 7 |
| zgc:110425 | 0.13395 | 0.121763757 | 0.063 | 0 | 1 | 7 |
| zgc:123105 | 0.13395 | 0.121236198 | 0.063 | 0 | 1 | 7 |
| znf576.2 | 0.13395 | 0.11650632 | 0.063 | 0 | 1 | 7 |
| hk1 | 0.13395 | 0.126864948 | 0.063 | 0 | 1 | 7 |
| nt5dc2 | 0.13395 | 0.138882783 | 0.063 | 0 | 1 | 7 |
| peak3 | 0.13395 | 0.146248284 | 0.063 | 0 | 1 | 7 |
| pigrl2.3 | 0.13395 | 0.127328149 | 0.063 | 0 | 1 | 7 |
| si:ch211-86h15.1 | 0.13395 | 0.126027895 | 0.063 | 0 | 1 | 7 |
| BX470189.1 | 0.13395 | 0.142465467 | 0.063 | 0 | 1 | 7 |
| FP236812.2 | 0.13395 | 0.131775583 | 0.063 | 0 | 1 | 7 |
| dffb | 0.13395 | 0.14465532 | 0.063 | 0 | 1 | 7 |
| zgc:158445 | 0.13395 | 0.189430381 | 0.063 | 0 | 1 | 7 |
| AL954696.1 | 0.13395 | 0.120418703 | 0.063 | 0 | 1 | 7 |
| CR318588.3 | 0.13395 | 0.159989856 | 0.063 | 0 | 1 | 7 |
| CU499330.1 | 0.13395 | 0.145517898 | 0.063 | 0 | 1 | 7 |
| LO017938.1 | 0.13395 | 0.101565113 | 0.063 | 0 | 1 | 7 |
| XLOC-041980 | 0.13395 | 0.152622433 | 0.063 | 0 | 1 | 7 |
| akip1 | 0.13395 | 0.116495424 | 0.063 | 0 | 1 | 7 |
| atad2 | 0.13395 | 0.112901063 | 0.063 | 0 | 1 | 7 |
| b3glcta | 0.13395 | 0.11403982 | 0.063 | 0 | 1 | 7 |
| bcar1 | 0.13395 | 0.138949898 | 0.063 | 0 | 1 | 7 |
| ccndbp1 | 0.13395 | 0.102101743 | 0.063 | 0 | 1 | 7 |
| ch25hl3 | 0.13395 | 0.110814574 | 0.063 | 0 | 1 | 7 |
| clec16a | 0.13395 | 0.137025162 | 0.063 | 0 | 1 | 7 |
| cul4b | 0.13395 | 0.121736099 | 0.063 | 0 | 1 | 7 |
| dapp1 | 0.13395 | 0.132573106 | 0.063 | 0 | 1 | 7 |
| dbt | 0.13395 | 0.108560106 | 0.063 | 0 | 1 | 7 |
| dcun1d5 | 0.13395 | 0.103692257 | 0.063 | 0 | 1 | 7 |
| desi1b | 0.13395 | 0.111054077 | 0.063 | 0 | 1 | 7 |
| dstyk | 0.13395 | 0.101015925 | 0.063 | 0 | 1 | 7 |
| dusp12 | 0.13395 | 0.107442231 | 0.063 | 0 | 1 | 7 |
| elp3 | 0.13395 | 0.11821342 | 0.063 | 0 | 1 | 7 |
| eml3 | 0.13395 | 0.125470503 | 0.063 | 0 | 1 | 7 |
| exosc7 | 0.13395 | 0.104165892 | 0.063 | 0 | 1 | 7 |
| fgfr1op | 0.13395 | 0.136579301 | 0.063 | 0 | 1 | 7 |
| ftsj1 | 0.13395 | 0.102942308 | 0.063 | 0 | 1 | 7 |
| gsk3ab | 0.13395 | 0.114588063 | 0.063 | 0 | 1 | 7 |
| hddc3 | 0.13395 | 0.100069611 | 0.063 | 0 | 1 | 7 |

| | | | | | | |
|---|---|---|---|---|---|---|
| hes6 | 0.13395 | 0.112397546 | 0.063 | 0 | 1 | 7 |
| im:7147486 | 0.13395 | 0.108548032 | 0.063 | 0 | 1 | 7 |
| kpna4 | 0.13395 | 0.114671283 | 0.063 | 0 | 1 | 7 |
| lsm14aa | 0.13395 | 0.156411462 | 0.063 | 0 | 1 | 7 |
| luc7l | 0.13395 | 0.110763309 | 0.063 | 0 | 1 | 7 |
| lxn | 0.13395 | 0.11246957 | 0.063 | 0 | 1 | 7 |
| marcksb | 0.13395 | 0.107480009 | 0.063 | 0 | 1 | 7 |
| mfn2 | 0.13395 | 0.119378074 | 0.063 | 0 | 1 | 7 |
| mfsd5 | 0.13395 | 0.106777786 | 0.063 | 0 | 1 | 7 |
| mob1ba | 0.13395 | 0.108630631 | 0.063 | 0 | 1 | 7 |
| necap2 | 0.13395 | 0.109345859 | 0.063 | 0 | 1 | 7 |
| nlk2 | 0.13395 | 0.127634103 | 0.063 | 0 | 1 | 7 |
| nsmaf | 0.13395 | 0.130569658 | 0.063 | 0 | 1 | 7 |
| parp12a | 0.13395 | 0.122473508 | 0.063 | 0 | 1 | 7 |
| pdpk1a | 0.13395 | 0.137198727 | 0.063 | 0 | 1 | 7 |
| pgrmc2 | 0.13395 | 0.135176112 | 0.063 | 0 | 1 | 7 |
| pou6f1 | 0.13395 | 0.14980671 | 0.063 | 0 | 1 | 7 |
| ppcdc | 0.13395 | 0.145663243 | 0.063 | 0 | 1 | 7 |
| prim1 | 0.13395 | 0.100351989 | 0.063 | 0 | 1 | 7 |
| psph | 0.13395 | 0.102991922 | 0.063 | 0 | 1 | 7 |
| pygb | 0.13395 | 0.12163681 | 0.063 | 0 | 1 | 7 |
| rbm45 | 0.13395 | 0.208240883 | 0.063 | 0 | 1 | 7 |
| rbms1b | 0.13395 | 0.190001293 | 0.063 | 0 | 1 | 7 |
| reps1 | 0.13395 | 0.105814282 | 0.063 | 0 | 1 | 7 |
| retsat | 0.13395 | 0.103655307 | 0.063 | 0 | 1 | 7 |
| serpinb14 | 0.13395 | 0.152533472 | 0.063 | 0 | 1 | 7 |
| sesn2 | 0.13395 | 0.107794519 | 0.063 | 0 | 1 | 7 |
| si:ch211-108d22.2 | 0.13395 | 0.122717331 | 0.063 | 0 | 1 | 7 |
| si:ch211-160o17.4 | 0.13395 | 0.141081642 | 0.063 | 0 | 1 | 7 |
| si:ch211-166g5.4 | 0.13395 | 0.159665199 | 0.063 | 0 | 1 | 7 |
| si:ch211-215e19.3 | 0.13395 | 0.124031229 | 0.063 | 0 | 1 | 7 |
| si:ch211-251p5.5 | 0.13395 | 0.100663573 | 0.063 | 0 | 1 | 7 |
| si:dkey-21c1.1 | 0.13395 | 0.107067255 | 0.063 | 0 | 1 | 7 |
| si:dkey-21e2.3 | 0.13395 | 0.140434069 | 0.063 | 0 | 1 | 7 |
| si:dkey-26c10.5 | 0.13395 | 0.100141542 | 0.063 | 0 | 1 | 7 |
| si:dkey-66i24.9 | 0.13395 | 0.101230565 | 0.063 | 0 | 1 | 7 |
| slbp | 0.13395 | 0.149462181 | 0.063 | 0 | 1 | 7 |
| slc6a9 | 0.13395 | 0.144233955 | 0.063 | 0 | 1 | 7 |
| tet1 | 0.13395 | 0.115676083 | 0.063 | 0 | 1 | 7 |
| tmem165 | 0.13395 | 0.10121248 | 0.063 | 0 | 1 | 7 |
| tmem184a | 0.13395 | 0.145495027 | 0.063 | 0 | 1 | 7 |
| tra2b | 0.13395 | 0.102914453 | 0.063 | 0 | 1 | 7 |
| trappc10 | 0.13395 | 0.157815096 | 0.063 | 0 | 1 | 7 |
| trim33l | 0.13395 | 0.151892004 | 0.063 | 0 | 1 | 7 |
| tubgcp5 | 0.13395 | 0.111863341 | 0.063 | 0 | 1 | 7 |
| unm-hu7910 | 0.13395 | 0.161710692 | 0.063 | 0 | 1 | 7 |
| vps11 | 0.13395 | 0.128542935 | 0.063 | 0 | 1 | 7 |
| wdr41 | 0.13395 | 0.130878423 | 0.063 | 0 | 1 | 7 |

| | | | | | | |
|---|---|---|---|---|---|---|
| zgc:152891 | 0.13395 | 0.121283234 | 0.063 | 0 | 1 | 7 |
| zgc:153901 | 0.13395 | 0.105534382 | 0.063 | 0 | 1 | 7 |
| zgc:77262 | 0.13395 | 0.11800333 | 0.063 | 0 | 1 | 7 |
| nkiras2 | 0.13395 | 0.128531295 | 0.063 | 0 | 1 | 7 |
| larp1b | 0.13434 | -0.20438374 | 0.063 | 0.154 | 1 | 7 |
| chchd2 | 0.13468 | 0.278508063 | 0.238 | 0.115 | 1 | 7 |
| mpnd | 0.13495 | -0.16625403 | 0.016 | 0.115 | 1 | 7 |
| cdca7b | 0.13533 | -0.18291099 | 0.032 | 0.154 | 1 | 7 |
| osgn1 | 0.13536 | -0.2690641 | 0.032 | 0.154 | 1 | 7 |
| pbx4 | 0.13588 | -0.22882878 | 0.032 | 0.154 | 1 | 7 |
| khsrp | 0.13589 | 0.281528837 | 0.365 | 0.269 | 1 | 7 |
| glmp | 0.13704 | -0.18107448 | 0.032 | 0.154 | 1 | 7 |
| psmb3 | 0.13799 | 0.186909946 | 0.397 | 0.346 | 1 | 7 |
| arpc5a | 0.13829 | 0.124416185 | 0.476 | 0.5 | 1 | 7 |
| calhm6 | 0.13863 | -0.13765954 | 0.016 | 0.115 | 1 | 7 |
| LOC101884451 | 0.139 | -0.33368542 | 0.127 | 0.308 | 1 | 7 |
| zgc:158803 | 0.13917 | 0.165946543 | 0.159 | 0.115 | 1 | 7 |
| epdl1 | 0.14082 | -0.15995934 | 0.016 | 0.115 | 1 | 7 |
| zgc:100918 | 0.14097 | -0.25370719 | 0.032 | 0.154 | 1 | 7 |
| uqcrh | 0.14108 | 0.138206532 | 0.46 | 0.5 | 1 | 7 |
| zgc:174863 | 0.14116 | -0.31421192 | 0.127 | 0.308 | 1 | 7 |
| clta | 0.14132 | -0.31491444 | 0.175 | 0.346 | 1 | 7 |
| znf217 | 0.14151 | -0.17367879 | 0.032 | 0.154 | 1 | 7 |
| eif3ba | 0.14177 | -0.17295099 | 0.651 | 0.846 | 1 | 7 |
| chtopa | 0.14184 | 0.23085691 | 0.19 | 0.115 | 1 | 7 |
| pdlim1 | 0.14229 | -0.20653567 | 0.032 | 0.154 | 1 | 7 |
| CU570894.1 | 0.14235 | -0.20429642 | 0.016 | 0.115 | 1 | 7 |
| si:dkey-262k9.4 | 0.14261 | -0.33272208 | 0.286 | 0.5 | 1 | 7 |
| wipf1a | 0.14302 | -0.28275045 | 0.254 | 0.462 | 1 | 7 |
| sptan1 | 0.14373 | -0.1716995 | 0.016 | 0.115 | 1 | 7 |
| hmgb3a | 0.14373 | -0.15169446 | 0.143 | 0.308 | 1 | 7 |
| gabarapb | 0.14496 | -0.34289897 | 0.397 | 0.615 | 1 | 7 |
| ciao2a | 0.14497 | -0.23039778 | 0.032 | 0.154 | 1 | 7 |
| csrnp1a | 0.1453 | -0.28327148 | 0.079 | 0.231 | 1 | 7 |
| elof1 | 0.14563 | -0.30566207 | 0.143 | 0.308 | 1 | 7 |
| rbm17 | 0.14582 | -0.18250349 | 0.032 | 0.154 | 1 | 7 |
| hnrnpa0l | 0.14595 | 0.103849709 | 0.857 | 0.885 | 1 | 7 |
| BX950187.1 | 0.14602 | -0.20574681 | 0.032 | 0.154 | 1 | 7 |
| si:ch211-165d12.4 | 0.14614 | -0.17370995 | 0.032 | 0.154 | 1 | 7 |
| r3hdm1 | 0.14615 | -0.158227 | 0.032 | 0.154 | 1 | 7 |
| ppp4ca | 0.14616 | -0.13074482 | 0.016 | 0.115 | 1 | 7 |
| st6galnac5b | 0.14649 | -0.19313315 | 0.032 | 0.154 | 1 | 7 |
| foxn2b | 0.14654 | -0.23615249 | 0.016 | 0.115 | 1 | 7 |
| tmem33 | 0.14664 | -0.18765822 | 0.032 | 0.154 | 1 | 7 |
| rap2b | 0.14666 | -0.2201925 | 0.032 | 0.154 | 1 | 7 |
| pigc | 0.14705 | -0.13117309 | 0.016 | 0.115 | 1 | 7 |
| cox6b1 | 0.14719 | 0.30582333 | 0.413 | 0.308 | 1 | 7 |
| marcksl1b | 0.14732 | -0.20950997 | 0.016 | 0.115 | 1 | 7 |

| | | | | | | |
|---|---|---|---|---|---|---|
| gtf2f2a | 0.14734 | 0.140524747 | 0.19 | 0.192 | 1 | 7 |
| eif4e1c | 0.14756 | 0.180024149 | 0.159 | 0.115 | 1 | 7 |
| SIPA1 | 0.1476 | -0.16292481 | 0.063 | 0.192 | 1 | 7 |
| upf3b | 0.14834 | -0.12268535 | 0.063 | 0.192 | 1 | 7 |
| erp44 | 0.1488 | 0.262603996 | 0.159 | 0.038 | 1 | 7 |
| mapkapk2a | 0.14886 | -0.26681361 | 0.127 | 0.308 | 1 | 7 |
| BX465227.1 | 0.14888 | -0.15165053 | 0.016 | 0.115 | 1 | 7 |
| mccc2 | 0.1493 | -0.14420973 | 0.016 | 0.115 | 1 | 7 |
| fbl | 0.14935 | 0.174195389 | 0.365 | 0.346 | 1 | 7 |
| klf11a | 0.14973 | -0.26799567 | 0.095 | 0.231 | 1 | 7 |
| gle1 | 0.15039 | -0.14387698 | 0.016 | 0.115 | 1 | 7 |
| uba2 | 0.15043 | 0.187618834 | 0.127 | 0.077 | 1 | 7 |
| chmp3 | 0.15045 | -0.16608954 | 0.016 | 0.115 | 1 | 7 |
| rb1 | 0.15073 | -0.14864384 | 0.016 | 0.115 | 1 | 7 |
| zgc:136864 | 0.15078 | -0.11382416 | 0.016 | 0.115 | 1 | 7 |
| ndufa11 | 0.15176 | 0.274068296 | 0.27 | 0.115 | 1 | 7 |
| cyb561a3a | 0.15232 | -0.18625487 | 0.016 | 0.115 | 1 | 7 |
| mrpl47 | 0.153 | 0.198285155 | 0.175 | 0.038 | 1 | 7 |
| nol7 | 0.15406 | -0.20041442 | 0.079 | 0.231 | 1 | 7 |
| si:ch211-167j9.5 | 0.15425 | 0.300952876 | 0.254 | 0.115 | 1 | 7 |
| samm50 | 0.15425 | -0.15237568 | 0.016 | 0.115 | 1 | 7 |
| tmem135 | 0.15442 | -0.136063 | 0.016 | 0.115 | 1 | 7 |
| si:dkey-56i24.1 | 0.15451 | 0.264165665 | 0.175 | 0.038 | 1 | 7 |
| hirip3 | 0.15472 | 0.181735011 | 0.127 | 0.038 | 1 | 7 |
| dbi | 0.15486 | -0.20251358 | 0.222 | 0.423 | 1 | 7 |
| FQ323156.1 | 0.15512 | -0.23047106 | 0.333 | 0.538 | 1 | 7 |
| pcyt1ab | 0.15524 | -0.16006442 | 0.016 | 0.115 | 1 | 7 |
| itm2bb | 0.15534 | -0.13400625 | 0.476 | 0.654 | 1 | 7 |
| fbxl14b | 0.15542 | -0.14428275 | 0.016 | 0.115 | 1 | 7 |
| dbnlb | 0.15576 | -0.22603816 | 0.048 | 0.154 | 1 | 7 |
| MRPS12 | 0.15604 | -0.23478414 | 0.111 | 0.231 | 1 | 7 |
| ube2q1 | 0.15609 | -0.13633944 | 0.016 | 0.115 | 1 | 7 |
| il10ra | 0.15621 | 0.218970511 | 0.317 | 0.154 | 1 | 7 |
| foxp1b | 0.15628 | -0.20380975 | 0.016 | 0.115 | 1 | 7 |
| cnp | 0.15651 | 0.173025074 | 0.317 | 0.154 | 1 | 7 |
| rpl39 | 0.15681 | -0.10366407 | 0.984 | 1 | 1 | 7 |
| ube2d3 | 0.15684 | -0.12353668 | 0.016 | 0.115 | 1 | 7 |
| efr3a | 0.15696 | -0.15942367 | 0.016 | 0.115 | 1 | 7 |
| trim2a | 0.15703 | -0.1610922 | 0.016 | 0.115 | 1 | 7 |
| ist1 | 0.15761 | -0.24978749 | 0.079 | 0.231 | 1 | 7 |
| si:ch211-127m7.2 | 0.15769 | -0.11538947 | 0.016 | 0.115 | 1 | 7 |
| ttc3 | 0.15777 | -0.17663065 | 0.016 | 0.115 | 1 | 7 |
| rps8a | 0.15798 | -0.12380578 | 0.905 | 1 | 1 | 7 |
| cog4 | 0.15811 | -0.16467384 | 0.016 | 0.115 | 1 | 7 |
| med13b | 0.15844 | -0.15945471 | 0.016 | 0.115 | 1 | 7 |
| rad21a | 0.15847 | 0.121311195 | 0.46 | 0.308 | 1 | 7 |
| msi2b | 0.15859 | -0.14222484 | 0.016 | 0.115 | 1 | 7 |
| glrx5 | 0.15863 | -0.24004253 | 0.032 | 0.115 | 1 | 7 |

| | | | | | | |
|---|---|---|---|---|---|---|
| wwp1 | 0.15881 | -0.16416301 | 0.016 | 0.115 | 1 | 7 |
| xpr1b | 0.15896 | -0.20107176 | 0.016 | 0.115 | 1 | 7 |
| spina | 0.15897 | -0.13165247 | 0.016 | 0.115 | 1 | 7 |
| cryz | 0.15956 | -0.13322251 | 0.016 | 0.115 | 1 | 7 |
| mdm1 | 0.15959 | -0.16564903 | 0.016 | 0.115 | 1 | 7 |
| srm | 0.15959 | -0.16834622 | 0.016 | 0.115 | 1 | 7 |
| rnf126 | 0.15971 | -0.13266638 | 0.016 | 0.115 | 1 | 7 |
| zc3h11a | 0.15973 | 0.254276266 | 0.159 | 0.038 | 1 | 7 |
| hmox2a | 0.15976 | -0.19008907 | 0.016 | 0.115 | 1 | 7 |
| ncbp3 | 0.15981 | 0.238733334 | 0.127 | 0.038 | 1 | 7 |
| casp2 | 0.15986 | -0.15535908 | 0.016 | 0.115 | 1 | 7 |
| maco1a | 0.15987 | -0.11712374 | 0.016 | 0.115 | 1 | 7 |
| echdc2 | 0.15988 | -0.17176357 | 0.016 | 0.115 | 1 | 7 |
| gpr65 | 0.15994 | -0.14155499 | 0.016 | 0.115 | 1 | 7 |
| sdc2 | 0.15995 | -0.16959095 | 0.016 | 0.115 | 1 | 7 |
| nxpe3 | 0.15996 | -0.19862407 | 0.016 | 0.115 | 1 | 7 |
| tbl3 | 0.15997 | -0.16089428 | 0.016 | 0.115 | 1 | 7 |
| FO834831.1 | 0.15998 | -0.15053129 | 0.016 | 0.115 | 1 | 7 |
| erbin | 0.16026 | 0.289294726 | 0.175 | 0.077 | 1 | 7 |
| psmd8 | 0.16026 | 0.206511162 | 0.159 | 0.038 | 1 | 7 |
| sh3bgrl | 0.16044 | 0.302892771 | 0.397 | 0.192 | 1 | 7 |
| selenot1a | 0.1612 | -0.24851525 | 0.159 | 0.346 | 1 | 7 |
| skap1 | 0.1615 | -0.29192975 | 0.159 | 0.346 | 1 | 7 |
| gpx4b | 0.16193 | -0.25332283 | 0.286 | 0.5 | 1 | 7 |
| cand1 | 0.16348 | -0.21846213 | 0.079 | 0.231 | 1 | 7 |
| atp5mc1 | 0.16411 | 0.334488115 | 0.413 | 0.231 | 1 | 7 |
| gtf2a2 | 0.16461 | -0.25974085 | 0.079 | 0.231 | 1 | 7 |
| calm1a | 0.16511 | -0.28395026 | 0.222 | 0.423 | 1 | 7 |
| brd2a | 0.1654 | -0.15281921 | 0.302 | 0.308 | 1 | 7 |
| ndufs2 | 0.16547 | -0.23835952 | 0.254 | 0.462 | 1 | 7 |
| gatd3a | 0.16619 | -0.22272941 | 0.079 | 0.231 | 1 | 7 |
| larp7 | 0.1662 | 0.158725957 | 0.127 | 0.115 | 1 | 7 |
| ube2l3a | 0.1671 | -0.25757957 | 0.111 | 0.269 | 1 | 7 |
| rgs3a | 0.16746 | -0.27771419 | 0.095 | 0.231 | 1 | 7 |
| ppp3r1b | 0.1677 | -0.28671414 | 0.143 | 0.308 | 1 | 7 |
| prelid1a | 0.16969 | -0.18803433 | 0.175 | 0.346 | 1 | 7 |
| ndufb5 | 0.16974 | 0.223075798 | 0.333 | 0.269 | 1 | 7 |
| unc50 | 0.1702 | -0.22717194 | 0.079 | 0.231 | 1 | 7 |
| ube2g1b | 0.17023 | 0.233932555 | 0.238 | 0.077 | 1 | 7 |
| smim15 | 0.17106 | 0.261193483 | 0.238 | 0.077 | 1 | 7 |
| dusp1 | 0.17115 | -0.14172483 | 0.27 | 0.423 | 1 | 7 |
| id3 | 0.17127 | -0.17284767 | 0.587 | 0.769 | 1 | 7 |
| si:ch211-93g21.1 | 0.17156 | -0.30789932 | 0.222 | 0.423 | 1 | 7 |
| bida | 0.17207 | -0.18014382 | 0.111 | 0.269 | 1 | 7 |
| eif2s1a | 0.17229 | 0.176956316 | 0.175 | 0.038 | 1 | 7 |
| tpm3 | 0.17307 | 0.15360135 | 0.492 | 0.5 | 1 | 7 |
| psmb2 | 0.17311 | 0.325463952 | 0.444 | 0.269 | 1 | 7 |
| ppp2r2d | 0.1732 | -0.15723263 | 0.095 | 0.231 | 1 | 7 |

| | | | | | | |
|---|---|---|---|---|---|---|
| sept12 | 0.17326 | -0.15879638 | 0.302 | 0.462 | 1 | 7 |
| capn2b | 0.17327 | -0.12332314 | 0.048 | 0.154 | 1 | 7 |
| eloca | 0.17392 | 0.149853239 | 0.111 | 0.077 | 1 | 7 |
| zgc:63882 | 0.17394 | 0.262118908 | 0.159 | 0.038 | 1 | 7 |
| si:ch73-248e21.7 | 0.17407 | -0.29681789 | 0.143 | 0.308 | 1 | 7 |
| rhoh | 0.17762 | -0.22379397 | 0.063 | 0.154 | 1 | 7 |
| gapdhs | 0.17843 | -0.28501421 | 0.587 | 0.731 | 1 | 7 |
| tdrd3 | 0.17857 | -0.22045847 | 0.032 | 0.115 | 1 | 7 |
| pet100 | 0.18023 | 0.284261807 | 0.222 | 0.077 | 1 | 7 |
| kdm2bb | 0.18041 | -0.12466822 | 0.016 | 0.077 | 1 | 7 |
| CU855821.1 | 0.18041 | -0.11559558 | 0.016 | 0.077 | 1 | 7 |
| adrb2a | 0.18041 | -0.11226606 | 0.016 | 0.077 | 1 | 7 |
| crk | 0.18041 | -0.11239836 | 0.016 | 0.077 | 1 | 7 |
| zzef1 | 0.18041 | -0.14259352 | 0.016 | 0.077 | 1 | 7 |
| cbx8a | 0.18041 | -0.11154836 | 0.016 | 0.077 | 1 | 7 |
| hgsnat | 0.18041 | -0.10023422 | 0.016 | 0.077 | 1 | 7 |
| mtr | 0.18041 | -0.12972405 | 0.016 | 0.077 | 1 | 7 |
| stk16 | 0.18041 | -0.16308855 | 0.016 | 0.077 | 1 | 7 |
| BX511129.1 | 0.18041 | -0.14916874 | 0.016 | 0.077 | 1 | 7 |
| ptprr | 0.18041 | -0.11822373 | 0.016 | 0.077 | 1 | 7 |
| si:ch211-210g13.5 | 0.18041 | -0.13253711 | 0.016 | 0.077 | 1 | 7 |
| kat6a | 0.18041 | -0.1272542 | 0.016 | 0.077 | 1 | 7 |
| dyrk3 | 0.18041 | -0.12098109 | 0.016 | 0.077 | 1 | 7 |
| elac2 | 0.18041 | -0.10930143 | 0.016 | 0.077 | 1 | 7 |
| insig1 | 0.18041 | -0.11089896 | 0.016 | 0.077 | 1 | 7 |
| ptpn11a | 0.18041 | -0.15097253 | 0.016 | 0.077 | 1 | 7 |
| si:dkey-93n13.2 | 0.18041 | -0.14905739 | 0.016 | 0.077 | 1 | 7 |
| zgc:103678 | 0.18041 | -0.12108315 | 0.016 | 0.077 | 1 | 7 |
| XLOC-002533 | 0.18041 | -0.11205552 | 0.016 | 0.077 | 1 | 7 |
| abhd10b | 0.18041 | -0.12172971 | 0.016 | 0.077 | 1 | 7 |
| ccl39.6 | 0.18041 | -0.14522587 | 0.016 | 0.077 | 1 | 7 |
| BX511021.2 | 0.18041 | -0.13344747 | 0.016 | 0.077 | 1 | 7 |
| CABZ01046432.1 | 0.18041 | -0.1003405 | 0.016 | 0.077 | 1 | 7 |
| CABZ01090373.1 | 0.18041 | -0.11524741 | 0.016 | 0.077 | 1 | 7 |
| CR388166.2 | 0.18041 | -0.11642517 | 0.016 | 0.077 | 1 | 7 |
| CU570889.1 | 0.18041 | -0.18349069 | 0.016 | 0.077 | 1 | 7 |
| LOC101883052 | 0.18041 | -0.13386993 | 0.016 | 0.077 | 1 | 7 |
| LOC101884753 | 0.18041 | -0.10466585 | 0.016 | 0.077 | 1 | 7 |
| LOC101887156 | 0.18041 | -0.1333016 | 0.016 | 0.077 | 1 | 7 |
| LOC110437825 | 0.18041 | -0.11722435 | 0.016 | 0.077 | 1 | 7 |
| ackr3b | 0.18041 | -0.10939759 | 0.016 | 0.077 | 1 | 7 |
| ada2a | 0.18041 | -0.14658927 | 0.016 | 0.077 | 1 | 7 |
| ankhd1 | 0.18041 | -0.15774489 | 0.016 | 0.077 | 1 | 7 |
| apoob | 0.18041 | -0.1167818 | 0.016 | 0.077 | 1 | 7 |
| arhgap42a | 0.18041 | -0.11809422 | 0.016 | 0.077 | 1 | 7 |
| atxn7l2b | 0.18041 | -0.11517622 | 0.016 | 0.077 | 1 | 7 |
| btr31 | 0.18041 | -0.11886505 | 0.016 | 0.077 | 1 | 7 |
| ccdc115 | 0.18041 | -0.15384313 | 0.016 | 0.077 | 1 | 7 |

| | | | | | | |
|---|---|---|---|---|---|---|
| ccdc174 | 0.18041 | -0.12360123 | 0.016 | 0.077 | 1 | 7 |
| cd226 | 0.18041 | -0.15728442 | 0.016 | 0.077 | 1 | 7 |
| cers5 | 0.18041 | -0.11056925 | 0.016 | 0.077 | 1 | 7 |
| cflarb | 0.18041 | -0.16058666 | 0.016 | 0.077 | 1 | 7 |
| clpp | 0.18041 | -0.11145955 | 0.016 | 0.077 | 1 | 7 |
| ctsk | 0.18041 | -0.19346609 | 0.016 | 0.077 | 1 | 7 |
| eloa | 0.18041 | -0.13616306 | 0.016 | 0.077 | 1 | 7 |
| foxp1a | 0.18041 | -0.11442137 | 0.016 | 0.077 | 1 | 7 |
| gpr174 | 0.18041 | -0.10764984 | 0.016 | 0.077 | 1 | 7 |
| gramd1a | 0.18041 | -0.18776659 | 0.016 | 0.077 | 1 | 7 |
| jak2b | 0.18041 | -0.1551095 | 0.016 | 0.077 | 1 | 7 |
| kdm4ab | 0.18041 | -0.1433078 | 0.016 | 0.077 | 1 | 7 |
| lgals9l3 | 0.18041 | -0.10809835 | 0.016 | 0.077 | 1 | 7 |
| me3 | 0.18041 | -0.14704479 | 0.016 | 0.077 | 1 | 7 |
| mllt1a | 0.18041 | -0.20807147 | 0.016 | 0.077 | 1 | 7 |
| nde1 | 0.18041 | -0.10909736 | 0.016 | 0.077 | 1 | 7 |
| nlrc3l1 | 0.18041 | -0.14998083 | 0.016 | 0.077 | 1 | 7 |
| pgbd4 | 0.18041 | -0.18838388 | 0.016 | 0.077 | 1 | 7 |
| polr2j | 0.18041 | -0.11372174 | 0.016 | 0.077 | 1 | 7 |
| ppp4r1l | 0.18041 | -0.1615451 | 0.016 | 0.077 | 1 | 7 |
| rab28 | 0.18041 | -0.10416419 | 0.016 | 0.077 | 1 | 7 |
| rangrf | 0.18041 | -0.18040578 | 0.016 | 0.077 | 1 | 7 |
| rras2 | 0.18041 | -0.15098821 | 0.016 | 0.077 | 1 | 7 |
| si:ch211-13c6.2 | 0.18041 | -0.12089575 | 0.016 | 0.077 | 1 | 7 |
| si:ch211-245h14.1 | 0.18041 | -0.1060926 | 0.016 | 0.077 | 1 | 7 |
| si:dkey-260j18.2 | 0.18041 | -0.1062003 | 0.016 | 0.077 | 1 | 7 |
| si:dkey-93h22.8 | 0.18041 | -0.11725746 | 0.016 | 0.077 | 1 | 7 |
| slc9a7 | 0.18041 | -0.10425441 | 0.016 | 0.077 | 1 | 7 |
| snx2 | 0.18041 | -0.12985917 | 0.016 | 0.077 | 1 | 7 |
| stat4 | 0.18041 | -0.10077376 | 0.016 | 0.077 | 1 | 7 |
| strada | 0.18041 | -0.10039157 | 0.016 | 0.077 | 1 | 7 |
| tnrc18 | 0.18041 | -0.1527447 | 0.016 | 0.077 | 1 | 7 |
| trabd | 0.18041 | -0.10029155 | 0.016 | 0.077 | 1 | 7 |
| ubac1 | 0.18041 | -0.10569796 | 0.016 | 0.077 | 1 | 7 |
| ugcg | 0.18041 | -0.12879978 | 0.016 | 0.077 | 1 | 7 |
| uggt1 | 0.18041 | -0.10136707 | 0.016 | 0.077 | 1 | 7 |
| vti1a | 0.18041 | -0.16840489 | 0.016 | 0.077 | 1 | 7 |
| zgc:110063 | 0.18041 | -0.13438758 | 0.016 | 0.077 | 1 | 7 |
| zgc:112023 | 0.18041 | -0.12849156 | 0.016 | 0.077 | 1 | 7 |
| zgc:152791 | 0.18041 | -0.15998773 | 0.016 | 0.077 | 1 | 7 |
| zgc:174680 | 0.18041 | -0.10625505 | 0.016 | 0.077 | 1 | 7 |
| zhx3 | 0.18041 | -0.1036092 | 0.016 | 0.077 | 1 | 7 |
| ccl38a.5 | 0.18052 | -0.51988128 | 0.206 | 0.385 | 1 | 7 |
| bag1 | 0.18139 | -0.17330071 | 0.063 | 0.192 | 1 | 7 |
| gmppb | 0.18354 | -0.2186722 | 0.063 | 0.192 | 1 | 7 |
| prmt1 | 0.1837 | 0.296019772 | 0.333 | 0.154 | 1 | 7 |
| scaf4b | 0.18432 | -0.12743147 | 0.048 | 0.154 | 1 | 7 |
| hsdl2 | 0.18471 | 0.132583443 | 0.143 | 0.038 | 1 | 7 |

| | | | | | | |
|---|---|---|---|---|---|---|
| usf1l | 0.18475 | -0.19166377 | 0.111 | 0.269 | 1 | 7 |
| ndufa4l | 0.185 | 0.221769371 | 0.698 | 0.654 | 1 | 7 |
| mrpl42 | 0.18511 | 0.216196151 | 0.19 | 0.077 | 1 | 7 |
| slc43a1a | 0.18575 | 0.205737142 | 0.143 | 0.038 | 1 | 7 |
| tap1 | 0.18602 | -0.18509868 | 0.063 | 0.192 | 1 | 7 |
| commd7 | 0.18645 | -0.23317244 | 0.079 | 0.192 | 1 | 7 |
| sap18 | 0.18852 | 0.172522564 | 0.302 | 0.269 | 1 | 7 |
| litaf | 0.18876 | -0.15286288 | 0.524 | 0.538 | 1 | 7 |
| nfyba | 0.18929 | 0.271542245 | 0.397 | 0.269 | 1 | 7 |
| rexo4 | 0.18937 | 0.169582597 | 0.127 | 0.038 | 1 | 7 |
| rnf10 | 0.1899 | -0.27866521 | 0.143 | 0.308 | 1 | 7 |
| jdp2b | 0.18997 | -0.12535963 | 0.254 | 0.231 | 1 | 7 |
| hmgb2b | 0.19008 | 0.41506449 | 0.429 | 0.231 | 1 | 7 |
| smdt1b | 0.19063 | 0.188476675 | 0.159 | 0.038 | 1 | 7 |
| dennd2da | 0.19072 | -0.19897033 | 0.048 | 0.154 | 1 | 7 |
| LOC103909115 | 0.19125 | -0.20305317 | 0.571 | 0.654 | 1 | 7 |
| capza1b | 0.1929 | -0.21137096 | 0.619 | 0.692 | 1 | 7 |
| mfap1 | 0.19314 | 0.220822972 | 0.127 | 0.038 | 1 | 7 |
| ptp4a1 | 0.19408 | 0.183731961 | 0.302 | 0.269 | 1 | 7 |
| trappc3 | 0.19418 | 0.211900728 | 0.143 | 0.077 | 1 | 7 |
| egr2a | 0.19424 | -0.23065337 | 0.063 | 0.192 | 1 | 7 |
| spcs1 | 0.19486 | 0.166759125 | 0.429 | 0.423 | 1 | 7 |
| COX7A2 (1 of many) | 0.19571 | 0.184462345 | 0.333 | 0.308 | 1 | 7 |
| pcnp | 0.19638 | -0.21689535 | 0.048 | 0.154 | 1 | 7 |
| myca | 0.19639 | -0.26368173 | 0.143 | 0.308 | 1 | 7 |
| csrp1a | 0.19655 | 0.315145107 | 0.222 | 0.077 | 1 | 7 |
| slc25a28 | 0.19705 | 0.111190438 | 0.159 | 0.154 | 1 | 7 |
| tradd | 0.19743 | -0.1527432 | 0.127 | 0.269 | 1 | 7 |
| dnajc8 | 0.19789 | 0.121938482 | 0.159 | 0.038 | 1 | 7 |
| zgc:63470 | 0.19804 | 0.163315105 | 0.111 | 0.077 | 1 | 7 |
| cct4 | 0.19842 | 0.293400772 | 0.571 | 0.423 | 1 | 7 |
| bola3 | 0.19856 | 0.15064912 | 0.095 | 0.038 | 1 | 7 |
| psmb5 | 0.19863 | 0.232562001 | 0.159 | 0.038 | 1 | 7 |
| anxa4 | 0.1988 | -0.29644186 | 0.095 | 0.231 | 1 | 7 |
| hypk | 0.1989 | 0.157769011 | 0.333 | 0.308 | 1 | 7 |
| itm2ba | 0.19913 | -0.25729795 | 0.333 | 0.538 | 1 | 7 |
| dazap2 | 0.19936 | -0.22288921 | 0.175 | 0.346 | 1 | 7 |
| si:ch211-51e12.7 | 0.20068 | 0.232949535 | 0.19 | 0.115 | 1 | 7 |
| kctd12.2 | 0.20107 | -0.26153315 | 0.111 | 0.269 | 1 | 7 |
| nop10 | 0.20305 | 0.270395332 | 0.349 | 0.231 | 1 | 7 |
| far1 | 0.20359 | -0.20173934 | 0.063 | 0.154 | 1 | 7 |
| sept7a | 0.20406 | 0.237170367 | 0.254 | 0.192 | 1 | 7 |
| tyms | 0.2042 | 0.237097283 | 0.127 | 0.038 | 1 | 7 |
| cmtm7 | 0.20424 | -0.26972853 | 0.111 | 0.231 | 1 | 7 |
| alkbh5 | 0.20432 | -0.20978127 | 0.032 | 0.115 | 1 | 7 |
| snrpa1 | 0.20438 | 0.200377185 | 0.238 | 0.154 | 1 | 7 |
| chchd3a | 0.20454 | -0.17588246 | 0.095 | 0.231 | 1 | 7 |
| higd1a | 0.20492 | -0.29735174 | 0.381 | 0.577 | 1 | 7 |

| | | | | | | |
|---|---|---|---|---|---|---|
| rangap1a | 0.20566 | -0.21132395 | 0.048 | 0.154 | 1 | 7 |
| grb2b | 0.20678 | -0.2108829 | 0.063 | 0.192 | 1 | 7 |
| ndufa12 | 0.20715 | 0.178783045 | 0.317 | 0.269 | 1 | 7 |
| tm9sf4 | 0.20852 | -0.20591849 | 0.048 | 0.154 | 1 | 7 |
| msna | 0.20944 | 0.115178277 | 0.73 | 0.731 | 1 | 7 |
| gtf2h5 | 0.21011 | -0.23369587 | 0.143 | 0.308 | 1 | 7 |
| srrt | 0.21083 | 0.192074414 | 0.143 | 0.038 | 1 | 7 |
| bin2b | 0.211 | 0.26216343 | 0.222 | 0.077 | 1 | 7 |
| lsm4 | 0.21166 | 0.300243333 | 0.286 | 0.154 | 1 | 7 |
| gsdmeb | 0.21174 | -0.2699311 | 0.127 | 0.231 | 1 | 7 |
| ankrd11 | 0.21176 | 0.259343878 | 0.286 | 0.192 | 1 | 7 |
| vta1 | 0.21249 | 0.221908235 | 0.159 | 0.038 | 1 | 7 |
| tmem14ca | 0.21269 | -0.11932617 | 0.159 | 0.154 | 1 | 7 |
| clint1a | 0.21284 | -0.23979569 | 0.048 | 0.154 | 1 | 7 |
| twist3 | 0.21286 | 0.256290212 | 0.111 | 0.038 | 1 | 7 |
| fam49a | 0.21292 | 0.291820391 | 0.302 | 0.154 | 1 | 7 |
| nkap | 0.21353 | 0.172529535 | 0.254 | 0.192 | 1 | 7 |
| larp6a | 0.21513 | -0.18083033 | 0.063 | 0.192 | 1 | 7 |
| zbtb20 | 0.2152 | 0.253702925 | 0.159 | 0.077 | 1 | 7 |
| prdx2 | 0.21526 | -0.2454656 | 0.413 | 0.615 | 1 | 7 |
| si:ch73-238c9.1 | 0.21635 | -0.17625682 | 0.063 | 0.192 | 1 | 7 |
| rps6ka1 | 0.21707 | 0.219298576 | 0.286 | 0.231 | 1 | 7 |
| rab5c | 0.21716 | -0.22448539 | 0.19 | 0.269 | 1 | 7 |
| rab11ba | 0.21753 | -0.26243621 | 0.19 | 0.346 | 1 | 7 |
| zgc:64051 | 0.2177 | -0.26114305 | 0.492 | 0.692 | 1 | 7 |
| eif4a1b | 0.2182 | -0.24087132 | 0.619 | 0.769 | 1 | 7 |
| nupr1a | 0.21866 | 0.249825893 | 0.175 | 0.077 | 1 | 7 |
| eif3ea | 0.2189 | -0.14641025 | 0.635 | 0.808 | 1 | 7 |
| p2ry10 | 0.21919 | -0.23229266 | 0.175 | 0.346 | 1 | 7 |
| march7 | 0.21939 | 0.19218784 | 0.127 | 0.038 | 1 | 7 |
| il16 | 0.21959 | -0.24818317 | 0.143 | 0.308 | 1 | 7 |
| gpx1a | 0.22007 | -0.15906478 | 0.222 | 0.385 | 1 | 7 |
| kif5bb | 0.22087 | -0.10866478 | 0.032 | 0.115 | 1 | 7 |
| nsd3 | 0.22158 | -0.11379804 | 0.032 | 0.115 | 1 | 7 |
| si:ch211-284f22.3 | 0.22173 | -0.11686507 | 0.048 | 0.154 | 1 | 7 |
| si:dkey-78l4.14 | 0.22201 | 0.205269182 | 0.159 | 0.038 | 1 | 7 |
| arhgap4b | 0.22412 | -0.16303272 | 0.127 | 0.269 | 1 | 7 |
| si:ch73-1a9.3 | 0.22426 | 0.25733381 | 0.651 | 0.615 | 1 | 7 |
| fnbp4 | 0.22469 | 0.266961932 | 0.143 | 0.038 | 1 | 7 |
| zgc:103700 | 0.22508 | -0.18620028 | 0.143 | 0.192 | 1 | 7 |
| snupn | 0.22516 | -0.21378021 | 0.048 | 0.154 | 1 | 7 |
| cry-dash | 0.22596 | 0.183500678 | 0.143 | 0.038 | 1 | 7 |
| si:dkey-21e2.12 | 0.22596 | 0.167271727 | 0.159 | 0.077 | 1 | 7 |
| smarcc1a | 0.226 | -0.18139614 | 0.032 | 0.115 | 1 | 7 |
| ormdl1 | 0.22631 | -0.14903609 | 0.048 | 0.154 | 1 | 7 |
| zgc:56095 | 0.22675 | -0.1502419 | 0.048 | 0.154 | 1 | 7 |
| sart1 | 0.22749 | -0.1758118 | 0.048 | 0.154 | 1 | 7 |
| srsf6b | 0.22769 | 0.272683415 | 0.349 | 0.192 | 1 | 7 |

| | | | | | | |
|---|---|---|---|---|---|---|
| txnrd3 | 0.22784 | -0.14671627 | 0.095 | 0.231 | 1 | 7 |
| sec11a | 0.22913 | 0.160088284 | 0.413 | 0.385 | 1 | 7 |
| mtap | 0.22935 | 0.248834192 | 0.206 | 0.077 | 1 | 7 |
| hnrnpr | 0.22944 | -0.22670585 | 0.175 | 0.346 | 1 | 7 |
| rgcc | 0.22994 | -0.25589716 | 0.048 | 0.115 | 1 | 7 |
| ywhaqb | 0.23049 | -0.25553589 | 0.238 | 0.423 | 1 | 7 |
| taf3 | 0.23068 | -0.1342494 | 0.048 | 0.154 | 1 | 7 |
| jun | 0.2307 | -0.33478629 | 0.095 | 0.192 | 1 | 7 |
| ctss2.1 | 0.23154 | -0.265539 | 0.206 | 0.385 | 1 | 7 |
| si:dkeyp-113d7.1 | 0.23177 | 0.192100855 | 0.159 | 0.038 | 1 | 7 |
| pole4 | 0.23203 | 0.175293786 | 0.159 | 0.077 | 1 | 7 |
| ccl32b.3 | 0.23211 | -0.1437232 | 0.175 | 0.308 | 1 | 7 |
| sh3bp5la | 0.23236 | -0.17139059 | 0.048 | 0.154 | 1 | 7 |
| arhgef3l | 0.23278 | -0.26529241 | 0.206 | 0.385 | 1 | 7 |
| actr2a | 0.23297 | 0.313386988 | 0.413 | 0.231 | 1 | 7 |
| eif4ea | 0.23357 | 0.15635116 | 0.079 | 0.038 | 1 | 7 |
| plcg2 | 0.23367 | 0.294070665 | 0.222 | 0.115 | 1 | 7 |
| rbm28 | 0.23385 | 0.231675409 | 0.222 | 0.115 | 1 | 7 |
| arfip1 | 0.23399 | -0.10965029 | 0.032 | 0.115 | 1 | 7 |
| midn | 0.23426 | -0.31502234 | 0.365 | 0.538 | 1 | 7 |
| uck2a | 0.23657 | 0.244535223 | 0.365 | 0.192 | 1 | 7 |
| rnaseh2b | 0.23712 | 0.118421625 | 0.143 | 0.038 | 1 | 7 |
| LOC103911430 | 0.23828 | 0.223045381 | 0.175 | 0.115 | 1 | 7 |
| klhdc4 | 0.23875 | -0.15834421 | 0.032 | 0.115 | 1 | 7 |
| nfkbiab | 0.23944 | -0.2819925 | 0.492 | 0.654 | 1 | 7 |
| mid1ip1l | 0.23946 | -0.24563867 | 0.222 | 0.346 | 1 | 7 |
| arf5 | 0.23949 | -0.29045004 | 0.254 | 0.423 | 1 | 7 |
| cwf19l1 | 0.23976 | -0.20239012 | 0.032 | 0.115 | 1 | 7 |
| hsd17b10 | 0.23978 | 0.152207293 | 0.143 | 0.077 | 1 | 7 |
| rhebl1 | 0.24028 | -0.20793959 | 0.079 | 0.192 | 1 | 7 |
| rsu1 | 0.24052 | -0.2392956 | 0.063 | 0.154 | 1 | 7 |
| pum2 | 0.24068 | -0.17651213 | 0.048 | 0.154 | 1 | 7 |
| pgam1b | 0.24117 | 0.216142269 | 0.127 | 0.038 | 1 | 7 |
| letm1 | 0.24281 | -0.15719062 | 0.048 | 0.154 | 1 | 7 |
| larp4ab | 0.24371 | -0.17874829 | 0.095 | 0.231 | 1 | 7 |
| ccng1 | 0.24371 | 0.306917871 | 0.365 | 0.231 | 1 | 7 |
| sec61g | 0.2445 | -0.2592767 | 0.635 | 0.769 | 1 | 7 |
| btg3 | 0.24495 | -0.18086757 | 0.159 | 0.231 | 1 | 7 |
| hnrnpc | 0.24507 | 0.179040238 | 0.143 | 0.038 | 1 | 7 |
| letm2 | 0.2451 | -0.22923769 | 0.079 | 0.192 | 1 | 7 |
| atrxl | 0.24527 | -0.18584185 | 0.048 | 0.154 | 1 | 7 |
| fabp3 | 0.24539 | -0.20042717 | 0.254 | 0.423 | 1 | 7 |
| golt1bb | 0.2459 | -0.16007855 | 0.048 | 0.154 | 1 | 7 |
| uri1 | 0.24625 | -0.11934146 | 0.048 | 0.154 | 1 | 7 |
| ddx3xa | 0.2469 | -0.2449534 | 0.143 | 0.269 | 1 | 7 |
| ncoa2 | 0.24752 | -0.11677567 | 0.032 | 0.115 | 1 | 7 |
| csnk2a1 | 0.24771 | -0.26753186 | 0.127 | 0.269 | 1 | 7 |
| ddx56 | 0.24827 | -0.14682175 | 0.079 | 0.115 | 1 | 7 |

| | | | | | | |
|---|---|---|---|---|---|---|
| fam193a | 0.24869 | -0.22917396 | 0.063 | 0.154 | 1 | 7 |
| rps6kb1b | 0.24987 | -0.13765656 | 0.095 | 0.115 | 1 | 7 |
| serpinb1l1 | 0.25012 | -0.18135721 | 0.127 | 0.269 | 1 | 7 |
| atp2a3 | 0.25031 | -0.17717143 | 0.127 | 0.269 | 1 | 7 |
| rhoab | 0.25036 | -0.24310091 | 0.508 | 0.692 | 1 | 7 |
| h2afy2 | 0.25088 | 0.26072665 | 0.238 | 0.115 | 1 | 7 |
| si:ch211-197h24.9 | 0.25166 | -0.17354559 | 0.095 | 0.231 | 1 | 7 |
| nedd9 | 0.25209 | -0.22750582 | 0.048 | 0.154 | 1 | 7 |
| pik3ap1 | 0.25237 | -0.14718943 | 0.079 | 0.192 | 1 | 7 |
| lsm12b | 0.25238 | -0.21363121 | 0.349 | 0.538 | 1 | 7 |
| ccdc106a | 0.25255 | -0.13327222 | 0.048 | 0.154 | 1 | 7 |
| nono | 0.25303 | -0.23657965 | 0.032 | 0.115 | 1 | 7 |
| ubap2l | 0.25359 | 0.189982514 | 0.19 | 0.115 | 1 | 7 |
| dpy30 | 0.25405 | 0.262800369 | 0.349 | 0.269 | 1 | 7 |
| plp2 | 0.25466 | -0.26990068 | 0.095 | 0.231 | 1 | 7 |
| acin1b | 0.255 | 0.22101686 | 0.143 | 0.038 | 1 | 7 |
| atf4a | 0.25613 | 0.164760463 | 0.286 | 0.269 | 1 | 7 |
| cct6a | 0.25702 | 0.122225651 | 0.429 | 0.423 | 1 | 7 |
| pdap1a | 0.25728 | 0.199651289 | 0.27 | 0.192 | 1 | 7 |
| det1 | 0.25733 | -0.15195446 | 0.032 | 0.115 | 1 | 7 |
| il2rgb | 0.25748 | -0.20006197 | 0.048 | 0.154 | 1 | 7 |
| twf1a | 0.25786 | -0.1902419 | 0.397 | 0.577 | 1 | 7 |
| rbb4l | 0.25806 | 0.144964907 | 0.206 | 0.077 | 1 | 7 |
| arrdc1b | 0.25819 | -0.1141912 | 0.032 | 0.115 | 1 | 7 |
| zgc:194224 | 0.25841 | 0.134917587 | 0.079 | 0.038 | 1 | 7 |
| zgc:113232 | 0.25897 | -0.22331739 | 0.048 | 0.154 | 1 | 7 |
| psmc5 | 0.25958 | 0.257899529 | 0.317 | 0.154 | 1 | 7 |
| spen | 0.25992 | -0.20256302 | 0.048 | 0.115 | 1 | 7 |
| mrpl3 | 0.26039 | 0.134364325 | 0.143 | 0.038 | 1 | 7 |
| rpz2 | 0.26072 | 0.160307346 | 0.19 | 0.154 | 1 | 7 |
| arglu1b | 0.26089 | -0.17803245 | 0.127 | 0.269 | 1 | 7 |
| aldoaa | 0.26128 | -0.26494043 | 0.429 | 0.615 | 1 | 7 |
| fmnl3 | 0.26217 | -0.18946595 | 0.032 | 0.115 | 1 | 7 |
| hyi | 0.26294 | -0.12230206 | 0.079 | 0.192 | 1 | 7 |
| si:ch73-281n10.2 | 0.26295 | 0.334796014 | 0.349 | 0.231 | 1 | 7 |
| CABZ01076758.1 | 0.2635 | -0.18060222 | 0.095 | 0.231 | 1 | 7 |
| gar1 | 0.26392 | 0.219600067 | 0.238 | 0.115 | 1 | 7 |
| mrps26 | 0.26404 | -0.12751818 | 0.032 | 0.115 | 1 | 7 |
| hgs | 0.26682 | -0.22765578 | 0.048 | 0.115 | 1 | 7 |
| snd1 | 0.26715 | -0.2260791 | 0.222 | 0.385 | 1 | 7 |
| ubfd1 | 0.26721 | 0.124738476 | 0.079 | 0.077 | 1 | 7 |
| itgav | 0.26727 | -0.1500131 | 0.048 | 0.154 | 1 | 7 |
| ets1 | 0.26766 | 0.159332803 | 0.175 | 0.154 | 1 | 7 |
| si:ch1073-322p19.1 | 0.26833 | -0.14737365 | 0.048 | 0.154 | 1 | 7 |
| sf3a2 | 0.2685 | 0.103028241 | 0.19 | 0.192 | 1 | 7 |
| degs1 | 0.26893 | -0.26761609 | 0.127 | 0.269 | 1 | 7 |
| LOC101885053 | 0.26955 | -0.18198168 | 0.048 | 0.154 | 1 | 7 |
| si:dkey-192g7.3 | 0.26958 | -0.13429163 | 0.032 | 0.115 | 1 | 7 |

| | | | | | | |
|---|---|---|---|---|---|---|
| pycard | 0.26958 | -0.18166199 | 0.27 | 0.423 | 1 | 7 |
| prkar1ab | 0.26973 | 0.196027199 | 0.159 | 0.115 | 1 | 7 |
| smg7 | 0.2699 | 0.20369919 | 0.127 | 0.038 | 1 | 7 |
| naxd | 0.27052 | 0.1852973 | 0.143 | 0.038 | 1 | 7 |
| eif3g | 0.27052 | 0.110580475 | 0.603 | 0.615 | 1 | 7 |
| junbb | 0.27068 | -0.16807319 | 0.032 | 0.115 | 1 | 7 |
| cbr1l | 0.27121 | -0.16926086 | 0.032 | 0.115 | 1 | 7 |
| cd74b | 0.27131 | -0.27160335 | 0.365 | 0.423 | 1 | 7 |
| rbm4.3 | 0.2715 | 0.279064654 | 0.54 | 0.385 | 1 | 7 |
| bpgm | 0.27193 | -0.10515128 | 0.063 | 0.154 | 1 | 7 |
| sf3b6 | 0.27197 | -0.14465526 | 0.302 | 0.462 | 1 | 7 |
| mfng | 0.27205 | 0.225261272 | 0.286 | 0.192 | 1 | 7 |
| map4k6 | 0.27221 | -0.10596547 | 0.127 | 0.231 | 1 | 7 |
| cd63 | 0.27295 | -0.14645385 | 0.651 | 0.654 | 1 | 7 |
| mob2b | 0.2734 | -0.17768805 | 0.048 | 0.154 | 1 | 7 |
| supt4h1 | 0.27353 | -0.1860368 | 0.127 | 0.269 | 1 | 7 |
| flot1b | 0.27431 | 0.11505532 | 0.159 | 0.154 | 1 | 7 |
| si:dkey-117m1.4 | 0.27461 | -0.19383909 | 0.032 | 0.115 | 1 | 7 |
| spcs2 | 0.2748 | -0.2363925 | 0.317 | 0.5 | 1 | 7 |
| srsf1a | 0.27486 | 0.216073034 | 0.143 | 0.038 | 1 | 7 |
| plekhj1 | 0.27494 | -0.17878174 | 0.048 | 0.154 | 1 | 7 |
| si:ch211-22i13.2 | 0.27508 | 0.224442609 | 0.206 | 0.077 | 1 | 7 |
| tmem42b | 0.27547 | -0.20300643 | 0.048 | 0.154 | 1 | 7 |
| LOC100007492 | 0.2761 | -0.20447953 | 0.048 | 0.154 | 1 | 7 |
| si:busm1-194e12.11 | 0.27611 | -0.19701191 | 0.048 | 0.154 | 1 | 7 |
| mrps21 | 0.27654 | 0.200410984 | 0.19 | 0.077 | 1 | 7 |
| plek | 0.2767 | -0.16975377 | 0.048 | 0.154 | 1 | 7 |
| smu1a | 0.27709 | -0.11903048 | 0.048 | 0.154 | 1 | 7 |
| dhx8 | 0.2774 | -0.17388509 | 0.048 | 0.154 | 1 | 7 |
| bhlhe41 | 0.27742 | -0.15094635 | 0.048 | 0.154 | 1 | 7 |
| gpn2 | 0.27748 | 0.107007376 | 0.111 | 0.115 | 1 | 7 |
| zgc:153993 | 0.27757 | -0.17516486 | 0.048 | 0.154 | 1 | 7 |
| DHDH | 0.27763 | -0.15887204 | 0.048 | 0.154 | 1 | 7 |
| pggt1b | 0.27768 | 0.174488023 | 0.127 | 0.038 | 1 | 7 |
| tcp11l2 | 0.27775 | -0.17220892 | 0.048 | 0.154 | 1 | 7 |
| lgals3b | 0.27791 | 0.147732615 | 0.238 | 0.115 | 1 | 7 |
| tmem59 | 0.2786 | 0.121812505 | 0.159 | 0.154 | 1 | 7 |
| snw1 | 0.27865 | 0.173589202 | 0.222 | 0.154 | 1 | 7 |
| gnb1b | 0.27912 | -0.1764819 | 0.492 | 0.538 | 1 | 7 |
| si:dkeyp-27c8.2 | 0.27981 | -0.17884062 | 0.048 | 0.115 | 1 | 7 |
| SMIM10 | 0.28071 | -0.16078817 | 0.032 | 0.077 | 1 | 7 |
| mrpl57 | 0.28094 | -0.21863153 | 0.127 | 0.231 | 1 | 7 |
| si:dkey-92i15.4 | 0.28171 | -0.11990375 | 0.032 | 0.115 | 1 | 7 |
| sco1 | 0.28277 | 0.187602445 | 0.143 | 0.038 | 1 | 7 |
| riok2 | 0.28324 | 0.165342339 | 0.127 | 0.038 | 1 | 7 |
| arhgef12a | 0.28435 | -0.11888189 | 0.032 | 0.115 | 1 | 7 |
| hsf2 | 0.28458 | -0.2184508 | 0.127 | 0.269 | 1 | 7 |
| ube2h | 0.28476 | -0.20620797 | 0.127 | 0.269 | 1 | 7 |

| | | | | | | |
|---|---|---|---|---|---|---|
| si:dkey-88p24.11 | 0.28495 | 0.157059507 | 0.143 | 0.077 | 1 | 7 |
| syap1 | 0.28507 | -0.12831734 | 0.079 | 0.192 | 1 | 7 |
| ponzr6 | 0.28546 | 0.169828558 | 0.079 | 0.038 | 1 | 7 |
| adssl | 0.28685 | 0.214827772 | 0.317 | 0.231 | 1 | 7 |
| mtmr10 | 0.28693 | -0.11477801 | 0.032 | 0.115 | 1 | 7 |
| prpf6 | 0.28794 | 0.212425327 | 0.206 | 0.077 | 1 | 7 |
| rab7a | 0.28826 | 0.149002681 | 0.143 | 0.038 | 1 | 7 |
| ankrd12 | 0.28865 | -0.18333335 | 0.127 | 0.269 | 1 | 7 |
| spg21 | 0.28893 | -0.22685411 | 0.127 | 0.269 | 1 | 7 |
| ccnt2b | 0.28945 | 0.165413341 | 0.127 | 0.038 | 1 | 7 |
| ccser2a | 0.28991 | -0.15819951 | 0.032 | 0.115 | 1 | 7 |
| arid4a | 0.29059 | 0.218469907 | 0.206 | 0.077 | 1 | 7 |
| dhps | 0.29062 | 0.110713514 | 0.079 | 0.038 | 1 | 7 |
| znf511 | 0.2909 | 0.112772235 | 0.063 | 0.038 | 1 | 7 |
| klhdc3 | 0.29098 | 0.105389465 | 0.111 | 0.115 | 1 | 7 |
| rag1 | 0.29104 | 0.222822661 | 0.175 | 0.077 | 1 | 7 |
| dut | 0.29213 | 0.194637203 | 0.175 | 0.154 | 1 | 7 |
| si:dkey-19e4.5 | 0.29219 | 0.186038583 | 0.143 | 0.038 | 1 | 7 |
| ywhabb | 0.29224 | 0.171781346 | 0.54 | 0.5 | 1 | 7 |
| ino80e | 0.29265 | 0.154978406 | 0.143 | 0.038 | 1 | 7 |
| grcc10 | 0.29302 | 0.232004156 | 0.222 | 0.115 | 1 | 7 |
| si:ch211-150o23.2 | 0.29385 | -0.15664126 | 0.079 | 0.192 | 1 | 7 |
| pef1 | 0.29405 | -0.19578192 | 0.27 | 0.346 | 1 | 7 |
| eef1db | 0.2942 | 0.179197592 | 0.19 | 0.115 | 1 | 7 |
| inpp5d | 0.29435 | -0.20831546 | 0.143 | 0.192 | 1 | 7 |
| psme1 | 0.29453 | 0.244665704 | 0.444 | 0.269 | 1 | 7 |
| lsm5 | 0.29585 | 0.239159826 | 0.413 | 0.269 | 1 | 7 |
| blcap | 0.29613 | -0.23335061 | 0.159 | 0.308 | 1 | 7 |
| ndufa2 | 0.2966 | 0.248101029 | 0.349 | 0.192 | 1 | 7 |
| scin | 0.29715 | -0.17390588 | 0.032 | 0.115 | 1 | 7 |
| apooa | 0.2972 | -0.16632187 | 0.063 | 0.154 | 1 | 7 |
| lcp2a | 0.29764 | -0.11420051 | 0.254 | 0.385 | 1 | 7 |
| eif2s2 | 0.29768 | 0.23153011 | 0.54 | 0.423 | 1 | 7 |
| BX248082.1 | 0.29774 | 0.168315367 | 0.127 | 0.038 | 1 | 7 |
| CABZ01111959.1 | 0.29856 | -0.10104684 | 0.095 | 0.192 | 1 | 7 |
| abi3a | 0.29932 | -0.22541603 | 0.19 | 0.346 | 1 | 7 |
| tes | 0.29988 | -0.23302854 | 0.159 | 0.308 | 1 | 7 |
| fbxl3a | 0.30045 | -0.14068697 | 0.079 | 0.192 | 1 | 7 |
| ube2e3 | 0.30056 | -0.10311573 | 0.254 | 0.385 | 1 | 7 |
| tusc2b | 0.30101 | -0.15072438 | 0.032 | 0.115 | 1 | 7 |
| sypl2b | 0.3011 | -0.24686169 | 0.159 | 0.308 | 1 | 7 |
| mir21-1 | 0.30128 | -0.2221448 | 0.079 | 0.154 | 1 | 7 |
| scgn | 0.30182 | -0.23520384 | 0.159 | 0.308 | 1 | 7 |
| kpnb3 | 0.30229 | 0.225250986 | 0.19 | 0.077 | 1 | 7 |
| nup50 | 0.3025 | 0.191815461 | 0.159 | 0.077 | 1 | 7 |
| snx1a | 0.30278 | -0.21434724 | 0.159 | 0.308 | 1 | 7 |
| nmi | 0.30282 | -0.22755117 | 0.159 | 0.308 | 1 | 7 |
| zgc:136605 | 0.30282 | -0.27724524 | 0.159 | 0.308 | 1 | 7 |

| | | | | | | |
|---|---|---|---|---|---|---|
| zmynd11 | 0.30282 | 0.161820256 | 0.127 | 0.038 | 1 | 7 |
| pigp | 0.30357 | -0.1386547 | 0.079 | 0.192 | 1 | 7 |
| agpat5 | 0.30403 | 0.208748138 | 0.143 | 0.038 | 1 | 7 |
| zfp36l1a | 0.30427 | -0.27181365 | 0.492 | 0.654 | 1 | 7 |
| fthl27 | 0.30435 | -0.30983519 | 0.492 | 0.615 | 1 | 7 |
| atp5meb | 0.3044 | 0.242929923 | 0.571 | 0.423 | 1 | 7 |
| mrpl28 | 0.30443 | 0.136674997 | 0.143 | 0.038 | 1 | 7 |
| ccar1 | 0.30458 | -0.13221873 | 0.032 | 0.077 | 1 | 7 |
| arf6a | 0.3051 | 0.268962351 | 0.397 | 0.231 | 1 | 7 |
| herpud1 | 0.30525 | 0.143146536 | 0.19 | 0.077 | 1 | 7 |
| si:dkey-286j15.3 | 0.30528 | -0.10006719 | 0.063 | 0.154 | 1 | 7 |
| naa10 | 0.30561 | 0.225852024 | 0.508 | 0.385 | 1 | 7 |
| acox1 | 0.3057 | 0.177848142 | 0.143 | 0.038 | 1 | 7 |
| tmed9 | 0.30587 | 0.230266393 | 0.27 | 0.154 | 1 | 7 |
| zswim5 | 0.30609 | 0.198156998 | 0.127 | 0.038 | 1 | 7 |
| rab5if | 0.30633 | 0.166448398 | 0.143 | 0.038 | 1 | 7 |
| med28 | 0.30663 | 0.161078205 | 0.143 | 0.038 | 1 | 7 |
| crfb4 | 0.30685 | 0.182311171 | 0.143 | 0.038 | 1 | 7 |
| pex26 | 0.30876 | -0.16024713 | 0.063 | 0.115 | 1 | 7 |
| ccdc6a | 0.30886 | -0.10936804 | 0.032 | 0.115 | 1 | 7 |
| slc7a3b | 0.30916 | -0.18410779 | 0.079 | 0.192 | 1 | 7 |
| aprt | 0.30953 | -0.13890222 | 0.079 | 0.192 | 1 | 7 |
| sdcbp2 | 0.30975 | -0.26160557 | 0.27 | 0.423 | 1 | 7 |
| psmd4a | 0.30979 | 0.166352691 | 0.127 | 0.038 | 1 | 7 |
| ost4 | 0.31037 | 0.209281447 | 0.27 | 0.154 | 1 | 7 |
| mau2 | 0.31078 | -0.16088052 | 0.111 | 0.231 | 1 | 7 |
| tspan33a | 0.31088 | -0.20193773 | 0.079 | 0.192 | 1 | 7 |
| pitpnc1b | 0.31114 | 0.183853172 | 0.127 | 0.038 | 1 | 7 |
| pes | 0.31122 | -0.10420545 | 0.079 | 0.192 | 1 | 7 |
| ncor2 | 0.31143 | -0.13765728 | 0.063 | 0.154 | 1 | 7 |
| srsf7a | 0.31164 | -0.15036917 | 0.381 | 0.538 | 1 | 7 |
| LOC100536213 | 0.31176 | -0.20749824 | 0.603 | 0.769 | 1 | 7 |
| snapc4 | 0.31179 | -0.14114321 | 0.032 | 0.115 | 1 | 7 |
| zbed4 | 0.31259 | -0.10838167 | 0.032 | 0.115 | 1 | 7 |
| guk1a | 0.31277 | 0.179525602 | 0.127 | 0.038 | 1 | 7 |
| si:dkey-102m7.3 | 0.31308 | 0.141764774 | 0.175 | 0.154 | 1 | 7 |
| oaz1a | 0.31338 | -0.1308343 | 0.746 | 0.885 | 1 | 7 |
| selenol | 0.3135 | -0.15559066 | 0.032 | 0.115 | 1 | 7 |
| rgs5b | 0.31375 | -0.10850096 | 0.127 | 0.231 | 1 | 7 |
| ctsd | 0.31422 | -0.1206147 | 0.032 | 0.115 | 1 | 7 |
| smg6 | 0.31518 | 0.221188934 | 0.19 | 0.077 | 1 | 7 |
| epc2 | 0.31556 | -0.1546923 | 0.032 | 0.115 | 1 | 7 |
| usp42 | 0.31565 | -0.14768772 | 0.032 | 0.115 | 1 | 7 |
| LOC100332221 | 0.31569 | -0.14569789 | 0.032 | 0.115 | 1 | 7 |
| psma8 | 0.31573 | 0.251576682 | 0.54 | 0.385 | 1 | 7 |
| yars | 0.31584 | 0.148758371 | 0.127 | 0.038 | 1 | 7 |
| ddi2 | 0.31625 | -0.20487051 | 0.063 | 0.154 | 1 | 7 |
| npepl1 | 0.31691 | -0.18496395 | 0.079 | 0.192 | 1 | 7 |

| | | | | | | |
|---|---|---|---|---|---|---|
| si:dkey-156n14.3 | 0.31854 | -0.12991701 | 0.032 | 0.115 | 1 | 7 |
| nkrf | 0.3191 | -0.22687456 | 0.079 | 0.192 | 1 | 7 |
| ppa1b | 0.32002 | 0.16873726 | 0.19 | 0.115 | 1 | 7 |
| chac1 | 0.32008 | -0.21539244 | 0.365 | 0.538 | 1 | 7 |
| ppid | 0.32199 | -0.1167764 | 0.143 | 0.269 | 1 | 7 |
| osbpl3b | 0.32248 | -0.13799254 | 0.032 | 0.115 | 1 | 7 |
| CU929070.1 | 0.3226 | -0.16203547 | 0.079 | 0.192 | 1 | 7 |
| ptmab | 0.32362 | -0.15325792 | 0.873 | 0.962 | 1 | 7 |
| stag2b | 0.3239 | -0.1660116 | 0.079 | 0.192 | 1 | 7 |
| nfatc2ip | 0.3246 | -0.16893152 | 0.032 | 0.115 | 1 | 7 |
| st3gal3a | 0.32474 | 0.153372464 | 0.127 | 0.038 | 1 | 7 |
| sptssa | 0.32533 | -0.16462172 | 0.079 | 0.192 | 1 | 7 |
| baz2a | 0.32542 | -0.13748623 | 0.095 | 0.115 | 1 | 7 |
| bcl2a | 0.32574 | -0.18493508 | 0.063 | 0.154 | 1 | 7 |
| odc1 | 0.32577 | -0.15040778 | 0.175 | 0.308 | 1 | 7 |
| rbbp4 | 0.32591 | 0.207631513 | 0.206 | 0.154 | 1 | 7 |
| cul1b | 0.32636 | -0.14135385 | 0.079 | 0.115 | 1 | 7 |
| psmd13 | 0.32642 | 0.204076284 | 0.19 | 0.077 | 1 | 7 |
| rnaseh2a | 0.32697 | 0.211324697 | 0.222 | 0.115 | 1 | 7 |
| junba | 0.32758 | -0.25819109 | 0.444 | 0.538 | 1 | 7 |
| fam3c | 0.32797 | -0.12280879 | 0.032 | 0.115 | 1 | 7 |
| ppp3r1a | 0.32804 | 0.133759526 | 0.159 | 0.154 | 1 | 7 |
| grk6 | 0.32893 | -0.18052249 | 0.032 | 0.115 | 1 | 7 |
| shrprbck1r | 0.32898 | -0.13390989 | 0.032 | 0.115 | 1 | 7 |
| cxcr4b | 0.32901 | -0.15226192 | 0.429 | 0.577 | 1 | 7 |
| bptf | 0.33001 | 0.249751917 | 0.238 | 0.115 | 1 | 7 |
| tmem167a | 0.33172 | 0.214936492 | 0.302 | 0.231 | 1 | 7 |
| ubqln4 | 0.33175 | -0.17237246 | 0.032 | 0.115 | 1 | 7 |
| sidt2 | 0.33186 | -0.17346918 | 0.032 | 0.115 | 1 | 7 |
| itsn2b | 0.33187 | -0.2175162 | 0.063 | 0.154 | 1 | 7 |
| gemin7 | 0.33215 | -0.12910058 | 0.111 | 0.231 | 1 | 7 |
| arhgap27l | 0.33241 | -0.16181172 | 0.032 | 0.115 | 1 | 7 |
| tbca | 0.33279 | 0.156383165 | 0.302 | 0.269 | 1 | 7 |
| tsen15 | 0.33309 | -0.10076807 | 0.032 | 0.115 | 1 | 7 |
| mark3b | 0.33322 | -0.18555434 | 0.032 | 0.115 | 1 | 7 |
| gpatch4 | 0.33326 | 0.132269994 | 0.19 | 0.077 | 1 | 7 |
| zfand2a | 0.33384 | -0.12468784 | 0.032 | 0.115 | 1 | 7 |
| si:dkey-199f5.8 | 0.334 | -0.14641019 | 0.032 | 0.115 | 1 | 7 |
| si:dkey-11f4.14 | 0.33443 | 0.281879659 | 0.333 | 0.192 | 1 | 7 |
| cited4b | 0.33479 | 0.191877357 | 0.19 | 0.115 | 1 | 7 |
| ddost | 0.33502 | 0.121367906 | 0.175 | 0.077 | 1 | 7 |
| chchd10 | 0.33513 | -0.14878524 | 0.079 | 0.192 | 1 | 7 |
| NDUFB1 | 0.33539 | -0.22476381 | 0.333 | 0.5 | 1 | 7 |
| XLOC-043861 | 0.33552 | 0.158351186 | 0.127 | 0.038 | 1 | 7 |
| mcm7 | 0.33567 | 0.165675554 | 0.143 | 0.077 | 1 | 7 |
| abca2 | 0.33582 | -0.10858558 | 0.063 | 0.154 | 1 | 7 |
| utp3 | 0.33585 | 0.108573899 | 0.127 | 0.115 | 1 | 7 |
| prkcha | 0.33635 | 0.147340788 | 0.079 | 0.038 | 1 | 7 |

| | | | | | | |
|---|---|---|---|---|---|---|
| ptprea | 0.33646 | -0.19107034 | 0.079 | 0.192 | 1 | 7 |
| ctdspl2b | 0.3365 | -0.12417238 | 0.032 | 0.115 | 1 | 7 |
| mid1ip1b | 0.33657 | -0.18411634 | 0.079 | 0.192 | 1 | 7 |
| sirt2 | 0.33683 | 0.203149818 | 0.127 | 0.038 | 1 | 7 |
| pinx1 | 0.33709 | 0.252150878 | 0.333 | 0.192 | 1 | 7 |
| lrba | 0.33744 | 0.168023989 | 0.111 | 0.038 | 1 | 7 |
| ift20 | 0.33769 | -0.17790576 | 0.111 | 0.192 | 1 | 7 |
| znf395b | 0.3379 | -0.21182299 | 0.079 | 0.154 | 1 | 7 |
| pgap2 | 0.33815 | -0.18269883 | 0.048 | 0.115 | 1 | 7 |
| traf3 | 0.3388 | -0.12194768 | 0.032 | 0.115 | 1 | 7 |
| ptp4a3 | 0.33885 | -0.13649494 | 0.032 | 0.115 | 1 | 7 |
| si:dkey-78l4.2 | 0.33939 | -0.37808321 | 0.032 | 0.115 | 1 | 7 |
| ubl3a | 0.3394 | -0.12982574 | 0.032 | 0.115 | 1 | 7 |
| cers1 | 0.33956 | -0.10102637 | 0.063 | 0.154 | 1 | 7 |
| znf609a | 0.33957 | -0.1831863 | 0.079 | 0.192 | 1 | 7 |
| med31 | 0.33958 | 0.19747496 | 0.175 | 0.077 | 1 | 7 |
| kat7b | 0.33976 | -0.15941365 | 0.048 | 0.115 | 1 | 7 |
| nucks1b | 0.3401 | 0.147150739 | 0.175 | 0.154 | 1 | 7 |
| cnbpa | 0.34032 | 0.195371374 | 0.381 | 0.308 | 1 | 7 |
| psmb10 | 0.34036 | 0.165209787 | 0.159 | 0.115 | 1 | 7 |
| faslg | 0.34074 | -0.16880804 | 0.175 | 0.308 | 1 | 7 |
| tma16 | 0.34094 | -0.1178444 | 0.032 | 0.115 | 1 | 7 |
| yipf6 | 0.34116 | -0.13555603 | 0.032 | 0.115 | 1 | 7 |
| cs | 0.34117 | 0.198224676 | 0.19 | 0.077 | 1 | 7 |
| thap4 | 0.34123 | -0.13707718 | 0.111 | 0.231 | 1 | 7 |
| stau1 | 0.3418 | -0.10586273 | 0.032 | 0.115 | 1 | 7 |
| mier2 | 0.34235 | -0.10244977 | 0.032 | 0.115 | 1 | 7 |
| zgc:153654 | 0.34274 | -0.14971006 | 0.032 | 0.115 | 1 | 7 |
| tbl2 | 0.34286 | -0.15874087 | 0.032 | 0.115 | 1 | 7 |
| adnpa | 0.34292 | -0.15017451 | 0.032 | 0.115 | 1 | 7 |
| lgals2a | 0.34298 | -0.28078298 | 0.032 | 0.115 | 1 | 7 |
| irf2bp2b | 0.34307 | -0.11358629 | 0.032 | 0.115 | 1 | 7 |
| tprn | 0.34307 | -0.14297321 | 0.032 | 0.115 | 1 | 7 |
| cdc27 | 0.34308 | -0.10481797 | 0.032 | 0.115 | 1 | 7 |
| st3gal4 | 0.34312 | -0.19442871 | 0.032 | 0.115 | 1 | 7 |
| si:ch211-214p16.2 | 0.34392 | 0.211821978 | 0.238 | 0.115 | 1 | 7 |
| sla2 | 0.34438 | -0.25900589 | 0.333 | 0.423 | 1 | 7 |
| c1d | 0.34551 | 0.100549258 | 0.095 | 0.077 | 1 | 7 |
| ifi45 | 0.34565 | -0.14519441 | 0.111 | 0.231 | 1 | 7 |
| tmem256 | 0.3467 | -0.17507855 | 0.111 | 0.192 | 1 | 7 |
| rgs18 | 0.34732 | -0.23703475 | 0.111 | 0.231 | 1 | 7 |
| CABZ01005379.1 | 0.34737 | 0.172525811 | 0.127 | 0.038 | 1 | 7 |
| creb1a | 0.34813 | 0.111778432 | 0.159 | 0.154 | 1 | 7 |
| rgs13 | 0.34921 | -0.24463606 | 0.492 | 0.654 | 1 | 7 |
| cdk7 | 0.34921 | 0.158530667 | 0.159 | 0.077 | 1 | 7 |
| si:ch211-230g15.5 | 0.34961 | -0.11380544 | 0.063 | 0.154 | 1 | 7 |
| ATP11B | 0.34967 | -0.18217967 | 0.095 | 0.192 | 1 | 7 |
| eno1a | 0.35088 | -0.12659646 | 0.365 | 0.5 | 1 | 7 |

| | | | | | | |
|---|---|---|---|---|---|---|
| rhoaa | 0.35103 | -0.21686116 | 0.302 | 0.385 | 1 | 7 |
| rasal2 | 0.35334 | 0.126192824 | 0.095 | 0.077 | 1 | 7 |
| yy1a | 0.3538 | 0.141138692 | 0.127 | 0.038 | 1 | 7 |
| psma4 | 0.35426 | -0.1279667 | 0.317 | 0.462 | 1 | 7 |
| pus7 | 0.35459 | 0.15379756 | 0.111 | 0.038 | 1 | 7 |
| si:dkey-53k12.2 | 0.35595 | -0.18984048 | 0.254 | 0.308 | 1 | 7 |
| naa25 | 0.35666 | -0.17557033 | 0.048 | 0.115 | 1 | 7 |
| ube4b | 0.35679 | -0.18826625 | 0.063 | 0.154 | 1 | 7 |
| chtf8 | 0.35682 | 0.173489238 | 0.127 | 0.038 | 1 | 7 |
| rps27.2 | 0.35686 | 0.129471689 | 0.841 | 0.808 | 1 | 7 |
| ppp6r2a | 0.35855 | -0.12350333 | 0.063 | 0.154 | 1 | 7 |
| BOLA2 | 0.35888 | 0.156217663 | 0.111 | 0.038 | 1 | 7 |
| fam162a | 0.35926 | 0.184956386 | 0.206 | 0.115 | 1 | 7 |
| zgc:195175 | 0.35936 | -0.21942425 | 0.095 | 0.192 | 1 | 7 |
| hectd1 | 0.35959 | 0.169989296 | 0.111 | 0.038 | 1 | 7 |
| wipi2 | 0.35994 | 0.156248037 | 0.127 | 0.038 | 1 | 7 |
| si:ch211-207n23.2 | 0.36004 | -0.32149879 | 0.175 | 0.308 | 1 | 7 |
| ppm1la | 0.36012 | 0.125008005 | 0.095 | 0.038 | 1 | 7 |
| ca2 | 0.36023 | 0.224578449 | 0.238 | 0.192 | 1 | 7 |
| tob1a | 0.36092 | -0.23748112 | 0.254 | 0.385 | 1 | 7 |
| casp3a | 0.36125 | 0.177778444 | 0.127 | 0.038 | 1 | 7 |
| zmp:0000001082 | 0.3616 | -0.23205879 | 0.127 | 0.231 | 1 | 7 |
| tmem18 | 0.36208 | 0.165037523 | 0.127 | 0.038 | 1 | 7 |
| ap4s1 | 0.36346 | 0.136995444 | 0.127 | 0.038 | 1 | 7 |
| btg1 | 0.36358 | 0.118888484 | 0.841 | 0.846 | 1 | 7 |
| hspd1 | 0.36366 | 0.191682913 | 0.19 | 0.077 | 1 | 7 |
| tfam | 0.36425 | -0.13143697 | 0.079 | 0.115 | 1 | 7 |
| nup98 | 0.36524 | -0.19890931 | 0.063 | 0.154 | 1 | 7 |
| ndfip1l | 0.36585 | 0.155860634 | 0.111 | 0.038 | 1 | 7 |
| mrpl30 | 0.36754 | -0.14506826 | 0.143 | 0.269 | 1 | 7 |
| vdac2 | 0.36853 | 0.142628852 | 0.651 | 0.654 | 1 | 7 |
| ptp4a2b | 0.36861 | -0.11186357 | 0.19 | 0.308 | 1 | 7 |
| rab4b | 0.3687 | 0.219213865 | 0.19 | 0.077 | 1 | 7 |
| tnfsf14 | 0.36876 | 0.192814271 | 0.159 | 0.077 | 1 | 7 |
| ssr1 | 0.37016 | -0.12141777 | 0.111 | 0.231 | 1 | 7 |
| htatip2 | 0.37021 | -0.10007569 | 0.063 | 0.154 | 1 | 7 |
| atp5f1b | 0.37026 | 0.232097069 | 0.603 | 0.538 | 1 | 7 |
| rrp15 | 0.3705 | -0.17101378 | 0.063 | 0.115 | 1 | 7 |
| txndc9 | 0.3706 | 0.193285409 | 0.317 | 0.192 | 1 | 7 |
| chmp5b | 0.37071 | -0.15206532 | 0.111 | 0.231 | 1 | 7 |
| si:dkey-78l4.4 | 0.37075 | 0.303613631 | 0.175 | 0.077 | 1 | 7 |
| mbpb | 0.37089 | -0.1314864 | 0.254 | 0.385 | 1 | 7 |
| phf3 | 0.3716 | -0.13070033 | 0.063 | 0.154 | 1 | 7 |
| ywhaba | 0.37251 | -0.23504908 | 0.508 | 0.654 | 1 | 7 |
| rnf2 | 0.37364 | 0.158044854 | 0.19 | 0.115 | 1 | 7 |
| scinlb | 0.37397 | -0.21494314 | 0.492 | 0.654 | 1 | 7 |
| NC-002333.4 | 0.37399 | 0.206004097 | 0.46 | 0.462 | 1 | 7 |
| thrap3b | 0.37407 | -0.21028105 | 0.27 | 0.423 | 1 | 7 |

| | | | | | | |
|---|---|---|---|---|---|---|
| mrps11 | 0.37415 | 0.125424965 | 0.143 | 0.115 | 1 | 7 |
| ccdc88b | 0.37535 | 0.114161661 | 0.095 | 0.077 | 1 | 7 |
| cndp2 | 0.3756 | 0.166704403 | 0.159 | 0.077 | 1 | 7 |
| TST | 0.37576 | 0.115453131 | 0.127 | 0.038 | 1 | 7 |
| XLOC-018981 | 0.37584 | 0.153503479 | 0.127 | 0.077 | 1 | 7 |
| tia1l | 0.37586 | -0.16175481 | 0.175 | 0.308 | 1 | 7 |
| pmm2 | 0.37613 | -0.15550692 | 0.127 | 0.192 | 1 | 7 |
| BX901920.1 | 0.37775 | -0.27391334 | 0.206 | 0.346 | 1 | 7 |
| smchd1 | 0.37832 | 0.141240765 | 0.095 | 0.038 | 1 | 7 |
| snrpg | 0.37838 | 0.236539705 | 0.381 | 0.269 | 1 | 7 |
| si:ch73-226l13.2 | 0.3786 | 0.203448725 | 0.143 | 0.115 | 1 | 7 |
| ccdc25 | 0.37899 | 0.133842587 | 0.238 | 0.192 | 1 | 7 |
| fmnl1a | 0.37915 | -0.24250116 | 0.143 | 0.269 | 1 | 7 |
| anp32e | 0.38133 | 0.148644029 | 0.27 | 0.231 | 1 | 7 |
| sesn1 | 0.38153 | 0.251408868 | 0.54 | 0.423 | 1 | 7 |
| akirin2 | 0.38166 | -0.15982783 | 0.143 | 0.269 | 1 | 7 |
| dnajb6a | 0.38206 | 0.152223153 | 0.175 | 0.077 | 1 | 7 |
| baz1b | 0.38221 | 0.228413243 | 0.238 | 0.115 | 1 | 7 |
| dnmt3ab | 0.38367 | -0.20917718 | 0.175 | 0.308 | 1 | 7 |
| mcm3 | 0.38373 | 0.167875081 | 0.159 | 0.077 | 1 | 7 |
| ndufb2 | 0.38397 | -0.13046313 | 0.143 | 0.269 | 1 | 7 |
| ccl36.1 | 0.38436 | 0.394646016 | 0.333 | 0.192 | 1 | 7 |
| si:ch211-218c6.8 | 0.38448 | 0.173794519 | 0.159 | 0.077 | 1 | 7 |
| orc6 | 0.3847 | 0.12093792 | 0.127 | 0.038 | 1 | 7 |
| cstf1 | 0.38488 | 0.141488425 | 0.127 | 0.038 | 1 | 7 |
| sltm | 0.38492 | 0.170528324 | 0.143 | 0.077 | 1 | 7 |
| fpr1 | 0.38556 | 0.137572466 | 0.111 | 0.077 | 1 | 7 |
| ndnl2 | 0.38559 | 0.15869761 | 0.127 | 0.038 | 1 | 7 |
| polr3k | 0.38653 | 0.114296832 | 0.111 | 0.077 | 1 | 7 |
| eif2s1b | 0.38665 | 0.230147527 | 0.571 | 0.423 | 1 | 7 |
| pak2a | 0.3867 | -0.22768671 | 0.206 | 0.346 | 1 | 7 |
| runx3 | 0.38703 | 0.236897889 | 0.286 | 0.154 | 1 | 7 |
| ccdc50 | 0.38802 | 0.186756057 | 0.159 | 0.115 | 1 | 7 |
| srpk1a | 0.38815 | 0.148191547 | 0.127 | 0.038 | 1 | 7 |
| polr1a | 0.38819 | 0.132732271 | 0.111 | 0.038 | 1 | 7 |
| eif3s6ip | 0.38837 | -0.17951245 | 0.381 | 0.538 | 1 | 7 |
| ddx6 | 0.38897 | -0.18707794 | 0.095 | 0.192 | 1 | 7 |
| phlda2 | 0.38917 | 0.190705536 | 0.127 | 0.038 | 1 | 7 |
| mrps18c | 0.38966 | 0.139709902 | 0.143 | 0.077 | 1 | 7 |
| adgrg1 | 0.38986 | 0.180955173 | 0.127 | 0.038 | 1 | 7 |
| pafah1b1a | 0.38997 | -0.18695892 | 0.143 | 0.269 | 1 | 7 |
| phf23a | 0.39108 | 0.138826959 | 0.111 | 0.038 | 1 | 7 |
| rbm19 | 0.39136 | 0.144284589 | 0.127 | 0.038 | 1 | 7 |
| caap1 | 0.39142 | 0.164029182 | 0.127 | 0.038 | 1 | 7 |
| XLOC-005448 | 0.39178 | 0.208317332 | 0.159 | 0.077 | 1 | 7 |
| nup62l | 0.39375 | -0.12126278 | 0.063 | 0.154 | 1 | 7 |
| prkrip1 | 0.39414 | 0.129215118 | 0.095 | 0.038 | 1 | 7 |
| hint1 | 0.39419 | 0.2218891 | 0.397 | 0.308 | 1 | 7 |

| | | | | | | |
|---|---|---|---|---|---|---|
| si:ch211-76m11.3 | 0.39428 | 0.169508874 | 0.111 | 0.038 | 1 | 7 |
| tktb | 0.3943 | 0.183224721 | 0.238 | 0.115 | 1 | 7 |
| rel | 0.39438 | -0.22001635 | 0.175 | 0.308 | 1 | 7 |
| desi2 | 0.39461 | 0.20849692 | 0.127 | 0.038 | 1 | 7 |
| rpl13a | 0.39463 | 0.111471498 | 0.952 | 0.885 | 1 | 7 |
| si:ch73-208g10.1 | 0.39502 | -0.23931869 | 0.159 | 0.269 | 1 | 7 |
| psmg1 | 0.39587 | 0.122299541 | 0.127 | 0.038 | 1 | 7 |
| gabarapa | 0.3963 | 0.132840854 | 0.206 | 0.192 | 1 | 7 |
| lrrfip1a | 0.39638 | -0.20233461 | 0.175 | 0.308 | 1 | 7 |
| rars | 0.39663 | 0.118115929 | 0.127 | 0.038 | 1 | 7 |
| gpalpp1 | 0.39675 | 0.173620158 | 0.111 | 0.038 | 1 | 7 |
| si:busm1-266f07.2 | 0.39816 | -0.29691182 | 0.175 | 0.269 | 1 | 7 |
| cd40lg | 0.39829 | -0.13623049 | 0.19 | 0.231 | 1 | 7 |
| eaf1 | 0.39857 | 0.130396915 | 0.127 | 0.038 | 1 | 7 |
| mrpl41 | 0.39875 | 0.112592918 | 0.127 | 0.038 | 1 | 7 |
| si:dkey-185m8.2 | 0.39943 | 0.160887938 | 0.317 | 0.192 | 1 | 7 |
| blk | 0.39957 | 0.166244921 | 0.127 | 0.038 | 1 | 7 |
| eif1ad | 0.39962 | 0.130610965 | 0.222 | 0.192 | 1 | 7 |
| lrrc47 | 0.39965 | 0.153264932 | 0.127 | 0.038 | 1 | 7 |
| dctn5 | 0.39968 | 0.147838636 | 0.127 | 0.038 | 1 | 7 |
| znf1035 | 0.3997 | 0.127825111 | 0.127 | 0.038 | 1 | 7 |
| nck1b | 0.3997 | 0.15176405 | 0.127 | 0.038 | 1 | 7 |
| dido1 | 0.40005 | 0.205785168 | 0.286 | 0.154 | 1 | 7 |
| znf865 | 0.40064 | 0.175354554 | 0.095 | 0.038 | 1 | 7 |
| dusp5 | 0.4011 | -0.21895972 | 0.095 | 0.192 | 1 | 7 |
| tmx1 | 0.40182 | 0.184642465 | 0.238 | 0.154 | 1 | 7 |
| rpa2 | 0.40289 | 0.185935871 | 0.143 | 0.077 | 1 | 7 |
| pbxip1b | 0.40353 | -0.15008623 | 0.063 | 0.154 | 1 | 7 |
| coil | 0.40371 | 0.121195803 | 0.079 | 0.038 | 1 | 7 |
| si:dkeyp-68b7.12 | 0.40407 | -0.15025894 | 0.222 | 0.346 | 1 | 7 |
| thoc1 | 0.40431 | -0.1102182 | 0.063 | 0.154 | 1 | 7 |
| krt8 | 0.40669 | 0.174442322 | 0.175 | 0.077 | 1 | 7 |
| ormdl2 | 0.40674 | 0.151354509 | 0.095 | 0.038 | 1 | 7 |
| supt3h | 0.40674 | 0.112925571 | 0.111 | 0.038 | 1 | 7 |
| calr | 0.40731 | 0.148226955 | 0.095 | 0.038 | 1 | 7 |
| ddx46 | 0.40952 | 0.145125901 | 0.127 | 0.077 | 1 | 7 |
| ywhae1 | 0.41098 | -0.16108584 | 0.429 | 0.577 | 1 | 7 |
| grapa | 0.41103 | -0.18059949 | 0.048 | 0.115 | 1 | 7 |
| dbpb | 0.41113 | 0.100986253 | 0.111 | 0.115 | 1 | 7 |
| sephs1 | 0.41157 | 0.116870636 | 0.095 | 0.038 | 1 | 7 |
| XLOC-026836 | 0.41173 | -0.2644168 | 0.397 | 0.538 | 1 | 7 |
| CABZ01102170.1 | 0.41189 | -0.18475015 | 0.048 | 0.115 | 1 | 7 |
| mak16 | 0.41259 | 0.131681593 | 0.111 | 0.038 | 1 | 7 |
| lck | 0.4132 | 0.122880526 | 0.206 | 0.115 | 1 | 7 |
| ik | 0.41326 | 0.120551361 | 0.095 | 0.038 | 1 | 7 |
| manf | 0.4134 | 0.135488435 | 0.127 | 0.077 | 1 | 7 |
| tmem209 | 0.41383 | -0.14070242 | 0.063 | 0.154 | 1 | 7 |
| si:dkey-11f4.20 | 0.41434 | -0.17033646 | 0.063 | 0.154 | 1 | 7 |

| | | | | | | |
|---|---|---|---|---|---|---|
| med17 | 0.41457 | -0.1277197 | 0.063 | 0.154 | 1 | 7 |
| ndufa1 | 0.41481 | -0.10131892 | 0.365 | 0.5 | 1 | 7 |
| sde2 | 0.41545 | 0.18307738 | 0.111 | 0.038 | 1 | 7 |
| syne1b | 0.41583 | 0.143738606 | 0.159 | 0.115 | 1 | 7 |
| si:ch211-209l18.2 | 0.41624 | -0.23700895 | 0.095 | 0.192 | 1 | 7 |
| ap2m1b | 0.41633 | -0.13945652 | 0.048 | 0.115 | 1 | 7 |
| sbf1 | 0.41653 | 0.162178299 | 0.111 | 0.038 | 1 | 7 |
| emc7 | 0.41702 | -0.22142582 | 0.127 | 0.231 | 1 | 7 |
| rchy1 | 0.41723 | -0.1407313 | 0.063 | 0.154 | 1 | 7 |
| zgc:152863 | 0.41756 | -0.13754552 | 0.159 | 0.269 | 1 | 7 |
| tomm5 | 0.41775 | 0.210156456 | 0.365 | 0.231 | 1 | 7 |
| arf1 | 0.41799 | 0.168519727 | 0.143 | 0.077 | 1 | 7 |
| faub | 0.41813 | -0.11182776 | 0.095 | 0.192 | 1 | 7 |
| znf704 | 0.41833 | -0.16506924 | 0.063 | 0.154 | 1 | 7 |
| b3gnt5a | 0.41871 | -0.22107643 | 0.222 | 0.346 | 1 | 7 |
| imp3 | 0.41996 | 0.118692158 | 0.095 | 0.038 | 1 | 7 |
| nlk1 | 0.42001 | -0.13267104 | 0.048 | 0.077 | 1 | 7 |
| sptlc2b | 0.42027 | -0.18307556 | 0.048 | 0.115 | 1 | 7 |
| atp5mc3a | 0.4205 | 0.201910175 | 0.429 | 0.346 | 1 | 7 |
| tsc22d2 | 0.42081 | -0.14343341 | 0.063 | 0.154 | 1 | 7 |
| ube2q2 | 0.4211 | -0.13642883 | 0.063 | 0.154 | 1 | 7 |
| ipcef1 | 0.42117 | -0.16550648 | 0.063 | 0.154 | 1 | 7 |
| zfr2 | 0.42173 | -0.15354338 | 0.063 | 0.154 | 1 | 7 |
| si:ch73-376l24.6 | 0.42248 | 0.145767151 | 0.175 | 0.077 | 1 | 7 |
| foxj3 | 0.42294 | 0.126962263 | 0.111 | 0.077 | 1 | 7 |
| si:ch211-68a17.7 | 0.42304 | 0.136338343 | 0.111 | 0.038 | 1 | 7 |
| myb | 0.42422 | 0.130740977 | 0.111 | 0.038 | 1 | 7 |
| zgc:55262 | 0.42429 | -0.16509196 | 0.063 | 0.154 | 1 | 7 |
| btbd10b | 0.42447 | 0.186632542 | 0.159 | 0.077 | 1 | 7 |
| pfdn4 | 0.42637 | 0.122790042 | 0.444 | 0.423 | 1 | 7 |
| psma6a | 0.42693 | 0.126471135 | 0.222 | 0.115 | 1 | 7 |
| med29 | 0.42697 | -0.13565019 | 0.111 | 0.154 | 1 | 7 |
| rasal3 | 0.42763 | -0.19656522 | 0.127 | 0.231 | 1 | 7 |
| tcf3b | 0.42797 | -0.14065292 | 0.063 | 0.154 | 1 | 7 |
| slka | 0.42846 | 0.142667383 | 0.095 | 0.038 | 1 | 7 |
| dctn3 | 0.42857 | -0.12730243 | 0.27 | 0.308 | 1 | 7 |
| ndufa3 | 0.42858 | -0.19052662 | 0.19 | 0.308 | 1 | 7 |
| CABZ01084963.1 | 0.42862 | -0.10683561 | 0.079 | 0.115 | 1 | 7 |
| knop1 | 0.42869 | 0.156987241 | 0.111 | 0.038 | 1 | 7 |
| ppifb | 0.42891 | -0.20676917 | 0.429 | 0.577 | 1 | 7 |
| gig2i | 0.42891 | -0.12914164 | 0.048 | 0.077 | 1 | 7 |
| blzf1 | 0.42925 | -0.1110964 | 0.032 | 0.077 | 1 | 7 |
| gnl3 | 0.42981 | 0.105876753 | 0.206 | 0.192 | 1 | 7 |
| si:ch211-66e2.3 | 0.43063 | -0.17867262 | 0.063 | 0.154 | 1 | 7 |
| eif1b | 0.43099 | -0.16357778 | 0.587 | 0.731 | 1 | 7 |
| jagn1b | 0.43125 | 0.157474269 | 0.143 | 0.077 | 1 | 7 |
| tcf7 | 0.43145 | -0.18195592 | 0.254 | 0.385 | 1 | 7 |
| ccl39.7 | 0.43216 | -0.27118275 | 0.063 | 0.154 | 1 | 7 |

| | | | | | | |
|---|---|---|---|---|---|---|
| tfr1b | 0.4323 | -0.12492895 | 0.063 | 0.154 | 1 | 7 |
| rin3 | 0.43274 | -0.16386874 | 0.063 | 0.154 | 1 | 7 |
| fntb | 0.43277 | -0.12701286 | 0.063 | 0.154 | 1 | 7 |
| LOC103909463 | 0.43279 | -0.13424702 | 0.095 | 0.192 | 1 | 7 |
| LO018102.1 | 0.43287 | -0.12627698 | 0.063 | 0.154 | 1 | 7 |
| spopla | 0.4332 | -0.12798976 | 0.063 | 0.154 | 1 | 7 |
| mrpl36 | 0.43337 | -0.11040693 | 0.159 | 0.269 | 1 | 7 |
| rbmx | 0.43499 | 0.175640578 | 0.175 | 0.077 | 1 | 7 |
| aga | 0.43512 | 0.131683522 | 0.095 | 0.038 | 1 | 7 |
| zgc:153409 | 0.43528 | 0.191399258 | 0.159 | 0.077 | 1 | 7 |
| atp5pd | 0.43604 | -0.15278312 | 0.429 | 0.577 | 1 | 7 |
| ap1s2 | 0.43652 | 0.134782475 | 0.175 | 0.077 | 1 | 7 |
| loxl2b | 0.43712 | -0.21653249 | 0.127 | 0.231 | 1 | 7 |
| pde12 | 0.43766 | 0.121411501 | 0.079 | 0.038 | 1 | 7 |
| CU694481.1 | 0.43781 | -0.16733549 | 0.048 | 0.115 | 1 | 7 |
| si:rp71-68n21.9 | 0.43817 | 0.178520082 | 0.302 | 0.231 | 1 | 7 |
| mrpl37 | 0.43891 | 0.136669687 | 0.095 | 0.038 | 1 | 7 |
| atg12 | 0.43916 | -0.12602528 | 0.032 | 0.077 | 1 | 7 |
| rbm39a | 0.43964 | 0.154938839 | 0.302 | 0.269 | 1 | 7 |
| gm2a | 0.43973 | -0.14192776 | 0.079 | 0.115 | 1 | 7 |
| mir223 | 0.4403 | 0.14732279 | 0.063 | 0.038 | 1 | 7 |
| si:ch211-246m6.4 | 0.44066 | -0.21905849 | 0.19 | 0.308 | 1 | 7 |
| tmpob | 0.44081 | 0.202402977 | 0.254 | 0.154 | 1 | 7 |
| cib1 | 0.44096 | -0.1719863 | 0.254 | 0.346 | 1 | 7 |
| psma6l | 0.44144 | -0.22344205 | 0.127 | 0.231 | 1 | 7 |
| vaspa | 0.44276 | -0.20855854 | 0.317 | 0.462 | 1 | 7 |
| npm1a | 0.44297 | 0.18245215 | 0.492 | 0.346 | 1 | 7 |
| actr1 | 0.443 | 0.126470951 | 0.127 | 0.077 | 1 | 7 |
| prkcsh | 0.44317 | 0.161410262 | 0.175 | 0.115 | 1 | 7 |
| dnaja2a | 0.44342 | 0.121276312 | 0.444 | 0.423 | 1 | 7 |
| zgc:162396 | 0.44347 | 0.105907034 | 0.111 | 0.077 | 1 | 7 |
| u2surp | 0.44364 | 0.193243303 | 0.206 | 0.115 | 1 | 7 |
| alyref | 0.44413 | 0.203375586 | 0.19 | 0.115 | 1 | 7 |
| ppdpfb | 0.44426 | -0.19784468 | 0.508 | 0.654 | 1 | 7 |
| mpc1 | 0.44499 | -0.11142262 | 0.048 | 0.115 | 1 | 7 |
| prpf39 | 0.4452 | -0.18502705 | 0.254 | 0.346 | 1 | 7 |
| kmt5aa | 0.44537 | -0.20040761 | 0.333 | 0.462 | 1 | 7 |
| smc1al | 0.4461 | -0.21218688 | 0.286 | 0.423 | 1 | 7 |
| lcp1 | 0.4464 | -0.20966363 | 0.206 | 0.308 | 1 | 7 |
| phf20l1 | 0.44656 | 0.128294795 | 0.111 | 0.038 | 1 | 7 |
| zdhhc5a | 0.4466 | -0.13538195 | 0.095 | 0.192 | 1 | 7 |
| prim2 | 0.44691 | 0.113373764 | 0.111 | 0.038 | 1 | 7 |
| rbpjb | 0.44734 | 0.10837428 | 0.159 | 0.077 | 1 | 7 |
| zyg11 | 0.44737 | 0.180783006 | 0.111 | 0.038 | 1 | 7 |
| csnk1g2b | 0.44751 | 0.103834615 | 0.127 | 0.115 | 1 | 7 |
| klf7b | 0.44821 | 0.171073539 | 0.127 | 0.077 | 1 | 7 |
| tufm | 0.44852 | 0.116714289 | 0.19 | 0.154 | 1 | 7 |
| pfdn6 | 0.44884 | 0.173544517 | 0.175 | 0.077 | 1 | 7 |

| | | | | | | |
|---|---|---|---|---|---|---|
| themis | 0.44944 | 0.185337918 | 0.159 | 0.077 | 1 | 7 |
| birc6 | 0.45014 | 0.116088307 | 0.095 | 0.077 | 1 | 7 |
| ncbp2 | 0.45036 | 0.11488088 | 0.206 | 0.192 | 1 | 7 |
| RPL37A | 0.451 | -0.12257748 | 0.095 | 0.192 | 1 | 7 |
| uqcrq | 0.45117 | 0.161257859 | 0.571 | 0.538 | 1 | 7 |
| lrpap1 | 0.45126 | -0.11899558 | 0.048 | 0.077 | 1 | 7 |
| smap1 | 0.45178 | -0.14596605 | 0.048 | 0.115 | 1 | 7 |
| CR293507.1 | 0.45179 | -0.19213179 | 0.079 | 0.154 | 1 | 7 |
| ndufs4 | 0.45235 | 0.10126584 | 0.175 | 0.154 | 1 | 7 |
| mgst3b | 0.45304 | -0.19131148 | 0.159 | 0.269 | 1 | 7 |
| b3gnt3.4 | 0.45305 | -0.14342732 | 0.048 | 0.115 | 1 | 7 |
| gtf2a1 | 0.45324 | 0.157415599 | 0.111 | 0.038 | 1 | 7 |
| nat10 | 0.45328 | 0.145694788 | 0.095 | 0.038 | 1 | 7 |
| creb5b | 0.45379 | 0.219654177 | 0.111 | 0.038 | 1 | 7 |
| wdr43 | 0.454 | -0.12754936 | 0.095 | 0.192 | 1 | 7 |
| syncripl | 0.45491 | 0.143231614 | 0.222 | 0.115 | 1 | 7 |
| stxbp2 | 0.45499 | 0.198086978 | 0.238 | 0.154 | 1 | 7 |
| gmip | 0.45504 | 0.148134007 | 0.095 | 0.038 | 1 | 7 |
| men1 | 0.45509 | 0.132588867 | 0.127 | 0.077 | 1 | 7 |
| zgc:113184 | 0.45572 | 0.128816454 | 0.111 | 0.038 | 1 | 7 |
| psmb4 | 0.45575 | 0.210093264 | 0.413 | 0.308 | 1 | 7 |
| zgc:122979 | 0.45615 | -0.1087081 | 0.048 | 0.115 | 1 | 7 |
| yeats4 | 0.45636 | 0.13701322 | 0.175 | 0.077 | 1 | 7 |
| si:ch73-42p12.2 | 0.45671 | 0.187550669 | 0.19 | 0.115 | 1 | 7 |
| rraga | 0.45689 | 0.152764523 | 0.111 | 0.038 | 1 | 7 |
| CR753876.1 | 0.45747 | -0.25821635 | 0.238 | 0.346 | 1 | 7 |
| vamp8 | 0.4575 | -0.14093431 | 0.159 | 0.269 | 1 | 7 |
| relb | 0.45751 | -0.13694835 | 0.095 | 0.192 | 1 | 7 |
| hmg20b | 0.45854 | 0.152210254 | 0.111 | 0.038 | 1 | 7 |
| gtf2e2 | 0.45867 | -0.12835598 | 0.095 | 0.192 | 1 | 7 |
| syncrip | 0.45939 | 0.143565547 | 0.571 | 0.538 | 1 | 7 |
| csnk1db | 0.45956 | -0.17455979 | 0.159 | 0.269 | 1 | 7 |
| crema | 0.45991 | 0.102029603 | 0.27 | 0.269 | 1 | 7 |
| rbm39b | 0.46095 | 0.128632915 | 0.143 | 0.115 | 1 | 7 |
| lamtor4 | 0.46111 | 0.135422136 | 0.111 | 0.038 | 1 | 7 |
| arid3a | 0.46181 | 0.147202688 | 0.111 | 0.038 | 1 | 7 |
| purba | 0.46185 | -0.11286826 | 0.095 | 0.192 | 1 | 7 |
| mrpl18 | 0.46189 | 0.160881548 | 0.19 | 0.115 | 1 | 7 |
| spink2.1 | 0.46426 | -0.1284695 | 0.079 | 0.154 | 1 | 7 |
| morc2 | 0.46486 | 0.112504099 | 0.095 | 0.077 | 1 | 7 |
| uprt | 0.4649 | -0.1526137 | 0.095 | 0.192 | 1 | 7 |
| si:ch73-248e21.5 | 0.46496 | -0.13649504 | 0.111 | 0.154 | 1 | 7 |
| tmc6a | 0.46543 | 0.152464848 | 0.079 | 0.038 | 1 | 7 |
| rassf2b | 0.46562 | -0.14093925 | 0.095 | 0.192 | 1 | 7 |
| mpp6a | 0.46575 | -0.11596501 | 0.032 | 0.077 | 1 | 7 |
| hdac1 | 0.46695 | -0.21069276 | 0.381 | 0.5 | 1 | 7 |
| fibpa | 0.467 | 0.127043744 | 0.095 | 0.038 | 1 | 7 |
| klf11b | 0.46706 | 0.121524443 | 0.143 | 0.115 | 1 | 7 |

| | | | | | | |
|---|---|---|---|---|---|---|
| arhgef1b | 0.46741 | 0.128435841 | 0.127 | 0.077 | 1 | 7 |
| ngly1 | 0.46742 | -0.13070648 | 0.048 | 0.115 | 1 | 7 |
| hnrnpd | 0.46774 | -0.13025662 | 0.111 | 0.154 | 1 | 7 |
| cd74a | 0.46897 | -0.29154734 | 0.27 | 0.385 | 1 | 7 |
| traf1 | 0.46924 | -0.14116857 | 0.127 | 0.231 | 1 | 7 |
| ciarta | 0.46964 | -0.20892395 | 0.222 | 0.346 | 1 | 7 |
| rab44 | 0.47054 | 0.211555526 | 0.222 | 0.115 | 1 | 7 |
| atp5f1e | 0.4706 | 0.170206585 | 0.778 | 0.654 | 1 | 7 |
| ginm1 | 0.47064 | 0.133796704 | 0.111 | 0.077 | 1 | 7 |
| nfkb2 | 0.471 | 0.159560245 | 0.143 | 0.077 | 1 | 7 |
| sept9b | 0.47185 | -0.20479324 | 0.254 | 0.385 | 1 | 7 |
| nr4a3 | 0.47227 | -0.18810933 | 0.079 | 0.154 | 1 | 7 |
| atp5if1b | 0.47233 | 0.196828475 | 0.524 | 0.462 | 1 | 7 |
| coro1a | 0.47298 | -0.10810915 | 0.921 | 0.962 | 1 | 7 |
| snrpc | 0.47321 | 0.175359848 | 0.317 | 0.192 | 1 | 7 |
| cenpx | 0.47375 | 0.158741112 | 0.159 | 0.115 | 1 | 7 |
| snrpb2 | 0.47378 | 0.162912818 | 0.175 | 0.115 | 1 | 7 |
| sucla2 | 0.47406 | -0.1154067 | 0.127 | 0.231 | 1 | 7 |
| nrbp1 | 0.47408 | 0.133634321 | 0.095 | 0.038 | 1 | 7 |
| ndufv2 | 0.47409 | 0.169579346 | 0.206 | 0.115 | 1 | 7 |
| cds2 | 0.47428 | -0.19292252 | 0.095 | 0.154 | 1 | 7 |
| ppp2r1ba | 0.47455 | -0.14520509 | 0.19 | 0.308 | 1 | 7 |
| snrpd3l | 0.47459 | 0.105567754 | 0.254 | 0.231 | 1 | 7 |
| mbnl1 | 0.47475 | 0.160136611 | 0.667 | 0.615 | 1 | 7 |
| gid8b | 0.47517 | 0.103894454 | 0.111 | 0.038 | 1 | 7 |
| rxrba | 0.47557 | -0.1245003 | 0.048 | 0.115 | 1 | 7 |
| ccl38.6 | 0.4756 | -0.23777187 | 0.365 | 0.5 | 1 | 7 |
| gigyf1b | 0.4764 | -0.15215219 | 0.048 | 0.115 | 1 | 7 |
| tspan37 | 0.47656 | 0.144248354 | 0.111 | 0.038 | 1 | 7 |
| g3bp2 | 0.47688 | 0.113753968 | 0.063 | 0.038 | 1 | 7 |
| uqcr10 | 0.47788 | 0.194539175 | 0.46 | 0.346 | 1 | 7 |
| vps26c | 0.4779 | 0.117915308 | 0.127 | 0.077 | 1 | 7 |
| tor1 | 0.4783 | -0.12740015 | 0.048 | 0.115 | 1 | 7 |
| cops5 | 0.47839 | 0.113254528 | 0.159 | 0.115 | 1 | 7 |
| lgals17 | 0.4784 | -0.12559875 | 0.032 | 0.077 | 1 | 7 |
| si:ch211-222l21.1 | 0.47846 | 0.115644814 | 0.794 | 0.808 | 1 | 7 |
| cdk2 | 0.47878 | 0.133578169 | 0.095 | 0.038 | 1 | 7 |
| nfya | 0.47909 | -0.15827017 | 0.063 | 0.115 | 1 | 7 |
| si:ch211-191j22.3 | 0.47934 | -0.19955611 | 0.238 | 0.346 | 1 | 7 |
| h2afvb | 0.4816 | 0.232388624 | 0.683 | 0.615 | 1 | 7 |
| fundc2 | 0.48289 | -0.10308249 | 0.127 | 0.154 | 1 | 7 |
| rpn1 | 0.48289 | 0.170622506 | 0.27 | 0.154 | 1 | 7 |
| si:zfos-169g10.3 | 0.48316 | -0.15329404 | 0.127 | 0.231 | 1 | 7 |
| rest | 0.48334 | -0.15549677 | 0.032 | 0.077 | 1 | 7 |
| sdhaf4 | 0.48381 | 0.147386966 | 0.111 | 0.038 | 1 | 7 |
| slirp | 0.48386 | 0.190802472 | 0.302 | 0.192 | 1 | 7 |
| tcf25 | 0.48434 | 0.103101288 | 0.111 | 0.038 | 1 | 7 |
| grasp | 0.4849 | 0.110793699 | 0.143 | 0.115 | 1 | 7 |

| | | | | | | |
|---|---|---|---|---|---|---|
| si:dkey-100n23.3 | 0.48526 | -0.10716898 | 0.032 | 0.077 | 1 | 7 |
| maff | 0.48576 | 0.119149978 | 0.111 | 0.038 | 1 | 7 |
| cnbpb | 0.48619 | 0.175791929 | 0.524 | 0.385 | 1 | 7 |
| CR786582.2 | 0.48673 | -0.15209356 | 0.206 | 0.269 | 1 | 7 |
| eif4h | 0.48679 | 0.174379892 | 0.19 | 0.115 | 1 | 7 |
| psmc3 | 0.48719 | 0.150906855 | 0.19 | 0.115 | 1 | 7 |
| il34 | 0.48719 | -0.18943239 | 0.19 | 0.308 | 1 | 7 |
| il10rb | 0.48732 | 0.148338613 | 0.095 | 0.038 | 1 | 7 |
| ssr4 | 0.48762 | -0.19542643 | 0.413 | 0.5 | 1 | 7 |
| hsd17b4 | 0.48791 | -0.13317713 | 0.032 | 0.077 | 1 | 7 |
| eif4ebp1 | 0.48823 | -0.16497719 | 0.222 | 0.346 | 1 | 7 |
| dynlt3 | 0.48826 | 0.121862108 | 0.254 | 0.154 | 1 | 7 |
| cd82b | 0.48849 | -0.17433656 | 0.079 | 0.154 | 1 | 7 |
| urm1 | 0.48851 | 0.140303334 | 0.111 | 0.038 | 1 | 7 |
| zgc:153293 | 0.48909 | -0.14014096 | 0.032 | 0.077 | 1 | 7 |
| vars | 0.48949 | -0.11746938 | 0.048 | 0.115 | 1 | 7 |
| anapc16 | 0.48969 | -0.14643746 | 0.19 | 0.231 | 1 | 7 |
| ISCU (1 of many) | 0.48974 | -0.17303509 | 0.222 | 0.346 | 1 | 7 |
| sh3d21 | 0.48984 | 0.152127178 | 0.127 | 0.077 | 1 | 7 |
| rassf1 | 0.49059 | 0.104880618 | 0.111 | 0.038 | 1 | 7 |
| lyn | 0.49087 | 0.156197498 | 0.111 | 0.038 | 1 | 7 |
| si:ch211-194k22.8 | 0.4912 | 0.151522116 | 0.095 | 0.038 | 1 | 7 |
| foxp3a | 0.49128 | -0.14346017 | 0.048 | 0.115 | 1 | 7 |
| si:ch211-197k17.3 | 0.49163 | 0.139900208 | 0.095 | 0.038 | 1 | 7 |
| rbm14a | 0.49181 | -0.17295344 | 0.127 | 0.231 | 1 | 7 |
| psmd7 | 0.49213 | -0.19087657 | 0.159 | 0.269 | 1 | 7 |
| bbx | 0.4922 | 0.147659363 | 0.127 | 0.077 | 1 | 7 |
| ctsa | 0.49296 | 0.120353531 | 0.111 | 0.038 | 1 | 7 |
| cldni | 0.49339 | -0.10585933 | 0.048 | 0.115 | 1 | 7 |
| itga4 | 0.49383 | -0.19466546 | 0.19 | 0.269 | 1 | 7 |
| mrps18b | 0.49503 | 0.170421296 | 0.143 | 0.077 | 1 | 7 |
| pola2 | 0.49523 | 0.120472219 | 0.095 | 0.038 | 1 | 7 |
| CU694197.1 | 0.49554 | -0.1904057 | 0.206 | 0.308 | 1 | 7 |
| smc5 | 0.49581 | 0.11309583 | 0.063 | 0.038 | 1 | 7 |
| mrps7 | 0.496 | 0.141301165 | 0.111 | 0.038 | 1 | 7 |
| prps1a | 0.4961 | -0.15132853 | 0.19 | 0.308 | 1 | 7 |
| gskip | 0.49641 | 0.163192175 | 0.19 | 0.115 | 1 | 7 |
| hnrnpl2 | 0.49701 | 0.172054793 | 0.238 | 0.154 | 1 | 7 |
| dusp6 | 0.49712 | -0.18016321 | 0.159 | 0.269 | 1 | 7 |
| CABZ01044053.1 | 0.49712 | 0.103797137 | 0.111 | 0.038 | 1 | 7 |
| dok2 | 0.49773 | -0.12447035 | 0.27 | 0.385 | 1 | 7 |
| RASSF5 | 0.49809 | 0.228393721 | 0.349 | 0.231 | 1 | 7 |
| fam133b | 0.49863 | 0.184496792 | 0.27 | 0.192 | 1 | 7 |
| rttn | 0.49883 | -0.17934328 | 0.032 | 0.077 | 1 | 7 |
| nmt1a | 0.49913 | -0.16390831 | 0.159 | 0.269 | 1 | 7 |
| zdhhc18b | 0.49921 | 0.125278397 | 0.095 | 0.038 | 1 | 7 |
| tmco1 | 0.49936 | -0.14293218 | 0.032 | 0.077 | 1 | 7 |
| tox | 0.49953 | 0.113244358 | 0.111 | 0.077 | 1 | 7 |

| | | | | | | |
|---|---|---|---|---|---|---|
| ndufb9 | 0.49953 | 0.16053893 | 0.397 | 0.269 | 1 | 7 |
| wdr46 | 0.49965 | 0.113034633 | 0.111 | 0.038 | 1 | 7 |
| smarca5 | 0.4997 | 0.113825618 | 0.111 | 0.038 | 1 | 7 |
| si:dkey-172h23.2 | 0.49999 | 0.13623552 | 0.111 | 0.038 | 1 | 7 |
| rnf185 | 0.50025 | -0.16092116 | 0.095 | 0.154 | 1 | 7 |
| myo1f | 0.50026 | -0.17084115 | 0.159 | 0.269 | 1 | 7 |
| usp36 | 0.50063 | 0.180237036 | 0.286 | 0.192 | 1 | 7 |
| MARK4 | 0.50134 | -0.14468434 | 0.048 | 0.077 | 1 | 7 |
| usp7 | 0.50149 | -0.17811075 | 0.111 | 0.192 | 1 | 7 |
| tuba1b | 0.50202 | 0.132941277 | 0.111 | 0.038 | 1 | 7 |
| si:ch211-212k18.4 | 0.50248 | 0.107989492 | 0.111 | 0.038 | 1 | 7 |
| diablob | 0.50323 | 0.136179218 | 0.095 | 0.038 | 1 | 7 |
| gmds | 0.50347 | 0.127905813 | 0.159 | 0.077 | 1 | 7 |
| trip12 | 0.50389 | 0.1348004 | 0.111 | 0.038 | 1 | 7 |
| pim1 | 0.50475 | -0.10038634 | 0.143 | 0.231 | 1 | 7 |
| zgc:153713 | 0.50514 | 0.128791522 | 0.46 | 0.423 | 1 | 7 |
| acaa2 | 0.50543 | 0.113132507 | 0.111 | 0.038 | 1 | 7 |
| cdc42l | 0.50558 | -0.17251545 | 0.619 | 0.731 | 1 | 7 |
| decr1 | 0.5061 | 0.102676194 | 0.111 | 0.038 | 1 | 7 |
| znf410 | 0.50693 | 0.107850813 | 0.111 | 0.038 | 1 | 7 |
| zgc:92818 | 0.50705 | -0.18018491 | 0.079 | 0.154 | 1 | 7 |
| mrps9 | 0.50758 | 0.114488755 | 0.095 | 0.038 | 1 | 7 |
| spty2d1 | 0.50764 | 0.139493531 | 0.111 | 0.038 | 1 | 7 |
| ifrd2 | 0.5082 | 0.112445167 | 0.111 | 0.038 | 1 | 7 |
| hnrnpabb | 0.50888 | 0.207372777 | 0.492 | 0.385 | 1 | 7 |
| mgaa | 0.50888 | 0.148085577 | 0.111 | 0.038 | 1 | 7 |
| mibp | 0.50895 | 0.153552116 | 0.095 | 0.038 | 1 | 7 |
| zgc:113208 | 0.50899 | -0.15295926 | 0.095 | 0.154 | 1 | 7 |
| mrpl53 | 0.5091 | 0.116719877 | 0.111 | 0.038 | 1 | 7 |
| scpep1 | 0.50927 | 0.114802769 | 0.111 | 0.038 | 1 | 7 |
| dnajb9b | 0.50938 | 0.123049352 | 0.079 | 0.038 | 1 | 7 |
| sft2d1 | 0.50946 | 0.131826242 | 0.111 | 0.038 | 1 | 7 |
| ezh2 | 0.5098 | 0.103355158 | 0.095 | 0.077 | 1 | 7 |
| CR392341.1 | 0.51002 | -0.12079238 | 0.349 | 0.462 | 1 | 7 |
| cox15 | 0.51035 | -0.11704385 | 0.048 | 0.115 | 1 | 7 |
| cmpk | 0.51113 | 0.179025467 | 0.206 | 0.115 | 1 | 7 |
| larp1 | 0.51155 | -0.1164133 | 0.048 | 0.115 | 1 | 7 |
| yipf3 | 0.51286 | 0.121613141 | 0.095 | 0.038 | 1 | 7 |
| capns1b | 0.51357 | 0.106930587 | 0.254 | 0.231 | 1 | 7 |
| BX005105.1 | 0.51389 | -0.20040726 | 0.206 | 0.308 | 1 | 7 |
| mpc2 | 0.51395 | 0.135757126 | 0.222 | 0.154 | 1 | 7 |
| susd6 | 0.51432 | -0.15322786 | 0.063 | 0.115 | 1 | 7 |
| tspan36 | 0.51478 | 0.102800399 | 0.095 | 0.038 | 1 | 7 |
| plekho2 | 0.51631 | -0.18517039 | 0.159 | 0.231 | 1 | 7 |
| ube2nb | 0.51638 | -0.17624025 | 0.222 | 0.308 | 1 | 7 |
| gtpbp4 | 0.51647 | -0.18634193 | 0.46 | 0.538 | 1 | 7 |
| prpf8 | 0.51655 | 0.109639698 | 0.19 | 0.154 | 1 | 7 |
| gnai2b | 0.51671 | 0.185526952 | 0.27 | 0.192 | 1 | 7 |

| | | | | | | |
|---|---|---|---|---|---|---|
| sdhb | 0.51686 | 0.179245976 | 0.238 | 0.154 | 1 | 7 |
| rnf220a | 0.51713 | 0.126442906 | 0.095 | 0.038 | 1 | 7 |
| stat2 | 0.51818 | 0.137539536 | 0.079 | 0.038 | 1 | 7 |
| cd28 | 0.5182 | 0.157845476 | 0.095 | 0.038 | 1 | 7 |
| LOC101886015 | 0.51878 | 0.144794665 | 0.095 | 0.038 | 1 | 7 |
| pdcl | 0.51888 | 0.172338175 | 0.159 | 0.077 | 1 | 7 |
| fkbp4 | 0.51997 | 0.115092428 | 0.095 | 0.077 | 1 | 7 |
| nbeal2 | 0.52021 | -0.12912721 | 0.048 | 0.115 | 1 | 7 |
| tp53 | 0.52024 | -0.14189336 | 0.27 | 0.385 | 1 | 7 |
| chmp1a | 0.52135 | 0.141148654 | 0.159 | 0.077 | 1 | 7 |
| FERMT3 (1 of many) | 0.52161 | 0.115869097 | 0.349 | 0.308 | 1 | 7 |
| stk17b | 0.52226 | 0.191323549 | 0.302 | 0.192 | 1 | 7 |
| ubtfl | 0.52237 | -0.13173446 | 0.063 | 0.115 | 1 | 7 |
| gse1 | 0.52256 | -0.16872265 | 0.127 | 0.192 | 1 | 7 |
| nr1d2b | 0.52353 | 0.129218902 | 0.111 | 0.077 | 1 | 7 |
| hspa4b | 0.52384 | 0.144589387 | 0.159 | 0.077 | 1 | 7 |
| srsf4 | 0.52458 | 0.122059402 | 0.238 | 0.192 | 1 | 7 |
| XLOC-011085 | 0.52515 | 0.194802563 | 0.222 | 0.154 | 1 | 7 |
| hnrnpa0a | 0.52539 | 0.147288712 | 0.159 | 0.077 | 1 | 7 |
| hspa9 | 0.52544 | 0.111558194 | 0.159 | 0.077 | 1 | 7 |
| irak3 | 0.52621 | -0.13223316 | 0.048 | 0.115 | 1 | 7 |
| tnfa | 0.52651 | -0.18478425 | 0.048 | 0.115 | 1 | 7 |
| cotl1 | 0.52738 | -0.15381355 | 0.778 | 0.846 | 1 | 7 |
| arhgef1a | 0.52779 | -0.10541559 | 0.048 | 0.115 | 1 | 7 |
| metap2b | 0.52831 | 0.183801354 | 0.381 | 0.308 | 1 | 7 |
| tpm1 | 0.52898 | -0.1919716 | 0.317 | 0.423 | 1 | 7 |
| selenof | 0.52903 | 0.204746561 | 0.46 | 0.346 | 1 | 7 |
| lamtor5 | 0.52952 | -0.1641147 | 0.111 | 0.192 | 1 | 7 |
| maip1 | 0.52999 | 0.112880744 | 0.095 | 0.038 | 1 | 7 |
| ncalda | 0.53015 | -0.12296537 | 0.079 | 0.154 | 1 | 7 |
| slc25a44b | 0.53041 | 0.112015223 | 0.079 | 0.038 | 1 | 7 |
| exosc1 | 0.53064 | -0.11717914 | 0.048 | 0.115 | 1 | 7 |
| kmt2cb | 0.53094 | -0.15760521 | 0.048 | 0.115 | 1 | 7 |
| otub1a | 0.53282 | 0.173823224 | 0.159 | 0.077 | 1 | 7 |
| gga1 | 0.53372 | -0.11885392 | 0.048 | 0.115 | 1 | 7 |
| gstm.2 | 0.53383 | -0.14060228 | 0.095 | 0.154 | 1 | 7 |
| zgc:171686 | 0.53614 | -0.18966571 | 0.063 | 0.115 | 1 | 7 |
| kat5b | 0.5381 | -0.11662249 | 0.048 | 0.115 | 1 | 7 |
| tomm70a | 0.53851 | 0.137305201 | 0.159 | 0.077 | 1 | 7 |
| pds5b | 0.54001 | 0.135893028 | 0.127 | 0.077 | 1 | 7 |
| mrpl22 | 0.54005 | -0.1084334 | 0.048 | 0.115 | 1 | 7 |
| BX649485.1 | 0.54023 | -0.16293647 | 0.048 | 0.115 | 1 | 7 |
| si:ch211-66k16.27 | 0.54037 | 0.155690491 | 0.254 | 0.154 | 1 | 7 |
| ptbp3 | 0.54062 | -0.19421525 | 0.143 | 0.231 | 1 | 7 |
| hmgn3 | 0.54101 | 0.207510468 | 0.333 | 0.231 | 1 | 7 |
| itm2ca | 0.54172 | -0.11299172 | 0.048 | 0.115 | 1 | 7 |
| thoc7 | 0.54199 | 0.125942869 | 0.095 | 0.038 | 1 | 7 |
| inip | 0.54248 | -0.14000913 | 0.143 | 0.231 | 1 | 7 |

| | | | | | | |
|---|---|---|---|---|---|---|
| edc3 | 0.54283 | -0.13502456 | 0.048 | 0.115 | 1 | 7 |
| vps36 | 0.54287 | 0.107331922 | 0.063 | 0.038 | 1 | 7 |
| tprg1 | 0.54288 | 0.125511224 | 0.111 | 0.077 | 1 | 7 |
| zgc:110269 | 0.54321 | 0.138414188 | 0.095 | 0.038 | 1 | 7 |
| ddx21 | 0.54373 | -0.18364407 | 0.429 | 0.538 | 1 | 7 |
| rabac1 | 0.54406 | -0.12418226 | 0.048 | 0.115 | 1 | 7 |
| dock8 | 0.54412 | 0.173481765 | 0.175 | 0.115 | 1 | 7 |
| ppp5c | 0.54427 | -0.14349117 | 0.048 | 0.115 | 1 | 7 |
| wdr26b | 0.54443 | -0.14914648 | 0.048 | 0.115 | 1 | 7 |
| map3k14a | 0.54446 | -0.11987441 | 0.048 | 0.115 | 1 | 7 |
| selenbp1 | 0.54448 | -0.11920701 | 0.048 | 0.115 | 1 | 7 |
| mcm6 | 0.54501 | 0.139462098 | 0.206 | 0.154 | 1 | 7 |
| akap9 | 0.54553 | -0.10776091 | 0.032 | 0.077 | 1 | 7 |
| nat9 | 0.54616 | -0.1384784 | 0.079 | 0.154 | 1 | 7 |
| dck | 0.5464 | 0.116305248 | 0.143 | 0.077 | 1 | 7 |
| tial1 | 0.5467 | 0.155761863 | 0.095 | 0.038 | 1 | 7 |
| gspt1l | 0.5477 | 0.134157703 | 0.159 | 0.077 | 1 | 7 |
| ptena | 0.54947 | 0.151875762 | 0.175 | 0.115 | 1 | 7 |
| btf3l4 | 0.55024 | 0.149979172 | 0.254 | 0.154 | 1 | 7 |
| hnrnpa0b | 0.55038 | 0.113626041 | 0.635 | 0.615 | 1 | 7 |
| msl2b | 0.55064 | 0.114118816 | 0.079 | 0.038 | 1 | 7 |
| crebzf | 0.55212 | 0.124655134 | 0.222 | 0.192 | 1 | 7 |
| CU639397.1 | 0.55253 | 0.129176113 | 0.095 | 0.038 | 1 | 7 |
| msl1b | 0.55262 | -0.13718216 | 0.032 | 0.077 | 1 | 7 |
| nuggc.1 | 0.55356 | -0.11028523 | 0.079 | 0.154 | 1 | 7 |
| arpp21 | 0.55415 | 0.23030184 | 0.159 | 0.077 | 1 | 7 |
| xpot | 0.55455 | -0.12297686 | 0.079 | 0.154 | 1 | 7 |
| fcer1g | 0.55472 | -0.18071753 | 0.111 | 0.192 | 1 | 7 |
| im:7142702 | 0.55529 | 0.177852792 | 0.159 | 0.115 | 1 | 7 |
| ncoa5 | 0.55549 | -0.12764119 | 0.079 | 0.154 | 1 | 7 |
| sec61b | 0.55631 | -0.16697873 | 0.413 | 0.538 | 1 | 7 |
| timm9 | 0.55682 | 0.103114827 | 0.159 | 0.077 | 1 | 7 |
| kpnb1 | 0.55693 | 0.16195163 | 0.254 | 0.154 | 1 | 7 |
| mafa | 0.55762 | 0.103730789 | 0.079 | 0.038 | 1 | 7 |
| si:dkey-183p4.10 | 0.5577 | 0.198660072 | 0.413 | 0.308 | 1 | 7 |
| myl12.1 | 0.55778 | -0.12511126 | 0.794 | 0.885 | 1 | 7 |
| mvp | 0.55796 | -0.12076173 | 0.048 | 0.077 | 1 | 7 |
| plekha1a | 0.55944 | -0.10536907 | 0.079 | 0.154 | 1 | 7 |
| si:ch211-196f2.6 | 0.56234 | 0.197936307 | 0.254 | 0.154 | 1 | 7 |
| stk35 | 0.56246 | 0.155325604 | 0.159 | 0.077 | 1 | 7 |
| si:ch211-102c2.4 | 0.5625 | -0.15711754 | 0.317 | 0.423 | 1 | 7 |
| nap1l4b | 0.56251 | 0.138353492 | 0.159 | 0.077 | 1 | 7 |
| antxr2a | 0.56279 | -0.11605518 | 0.206 | 0.308 | 1 | 7 |
| npc2 | 0.56285 | -0.15906127 | 0.27 | 0.385 | 1 | 7 |
| pa2g4b | 0.56359 | 0.168338437 | 0.317 | 0.231 | 1 | 7 |
| map3k3 | 0.56432 | 0.130813958 | 0.079 | 0.038 | 1 | 7 |
| snapc3 | 0.56508 | -0.1187137 | 0.048 | 0.077 | 1 | 7 |
| taf15 | 0.56552 | 0.162238723 | 0.254 | 0.154 | 1 | 7 |

| | | | | | | |
|---|---|---|---|---|---|---|
| si:dkey-159f12.2 | 0.56587 | 0.129145643 | 0.143 | 0.077 | 1 | 7 |
| CABZ01055347.1 | 0.56621 | -0.14595301 | 0.063 | 0.115 | 1 | 7 |
| ccdc82 | 0.56787 | -0.13463576 | 0.063 | 0.115 | 1 | 7 |
| ranbp1 | 0.56817 | -0.14483631 | 0.492 | 0.615 | 1 | 7 |
| wasa | 0.56846 | 0.104199513 | 0.476 | 0.462 | 1 | 7 |
| supt16h | 0.56895 | 0.126595433 | 0.143 | 0.077 | 1 | 7 |
| ggh | 0.56919 | 0.102074198 | 0.079 | 0.038 | 1 | 7 |
| nsfl1c | 0.56935 | 0.125782686 | 0.206 | 0.115 | 1 | 7 |
| xpo1a | 0.56959 | 0.132676615 | 0.095 | 0.038 | 1 | 7 |
| mplkip | 0.56968 | 0.102656523 | 0.095 | 0.038 | 1 | 7 |
| il2rb | 0.56975 | -0.1027772 | 0.111 | 0.192 | 1 | 7 |
| rcor1 | 0.57019 | 0.164261823 | 0.206 | 0.115 | 1 | 7 |
| ufc1 | 0.57093 | -0.15854333 | 0.143 | 0.231 | 1 | 7 |
| hcst | 0.57096 | 0.149894325 | 0.175 | 0.115 | 1 | 7 |
| ino80c | 0.57114 | -0.1098455 | 0.111 | 0.154 | 1 | 7 |
| si:ch211-153b23.7 | 0.57197 | 0.176512241 | 0.143 | 0.077 | 1 | 7 |
| chd2 | 0.5723 | 0.136801744 | 0.143 | 0.077 | 1 | 7 |
| sept7b | 0.57246 | 0.122143428 | 0.381 | 0.346 | 1 | 7 |
| cox7b | 0.5727 | 0.145395448 | 0.778 | 0.731 | 1 | 7 |
| rcc2 | 0.57275 | 0.155348444 | 0.333 | 0.269 | 1 | 7 |
| rhof | 0.57278 | -0.1252918 | 0.175 | 0.269 | 1 | 7 |
| slc38a5a | 0.57284 | -0.13953831 | 0.032 | 0.077 | 1 | 7 |
| rap1aa | 0.57297 | -0.12792219 | 0.238 | 0.346 | 1 | 7 |
| actb1 | 0.57343 | -0.1083825 | 0.889 | 0.923 | 1 | 7 |
| pik3cg | 0.57374 | -0.12210273 | 0.079 | 0.154 | 1 | 7 |
| adgrg3 | 0.57384 | 0.115407865 | 0.063 | 0.038 | 1 | 7 |
| gramd2aa | 0.57443 | -0.14585424 | 0.143 | 0.231 | 1 | 7 |
| mtfp1 | 0.57491 | -0.1313949 | 0.032 | 0.077 | 1 | 7 |
| rer1 | 0.57587 | 0.101920744 | 0.238 | 0.154 | 1 | 7 |
| atp5mea | 0.57646 | -0.16050088 | 0.238 | 0.346 | 1 | 7 |
| pitpnb | 0.57663 | -0.14711885 | 0.111 | 0.192 | 1 | 7 |
| bach1a | 0.57688 | -0.11122881 | 0.032 | 0.077 | 1 | 7 |
| bcl7bb | 0.57835 | -0.10711657 | 0.143 | 0.231 | 1 | 7 |
| elf1 | 0.57986 | 0.206165913 | 0.413 | 0.308 | 1 | 7 |
| nus1 | 0.58028 | -0.10850394 | 0.032 | 0.077 | 1 | 7 |
| ppp1r10 | 0.58113 | -0.11221199 | 0.111 | 0.192 | 1 | 7 |
| calm2a | 0.58115 | 0.160504313 | 0.206 | 0.154 | 1 | 7 |
| ciao1 | 0.58188 | 0.114645491 | 0.127 | 0.077 | 1 | 7 |
| LOC103909107 | 0.58229 | -0.16706599 | 0.079 | 0.154 | 1 | 7 |
| hps5 | 0.5825 | -0.12473003 | 0.032 | 0.077 | 1 | 7 |
| slc38a2 | 0.58274 | -0.1411533 | 0.079 | 0.115 | 1 | 7 |
| il7r | 0.58384 | 0.100430611 | 0.238 | 0.231 | 1 | 7 |
| GNB4 | 0.58593 | -0.10709674 | 0.032 | 0.077 | 1 | 7 |
| BX908750.1 | 0.58604 | -0.17175564 | 0.111 | 0.192 | 1 | 7 |
| meaf6 | 0.58643 | -0.13757949 | 0.143 | 0.231 | 1 | 7 |
| mrpl35 | 0.58689 | -0.12393301 | 0.079 | 0.154 | 1 | 7 |
| gnai2a | 0.58702 | -0.15400877 | 0.381 | 0.5 | 1 | 7 |
| cab39l1 | 0.58774 | -0.13310599 | 0.079 | 0.154 | 1 | 7 |

| | | | | | | |
|---|---|---|---|---|---|---|
| spint1a | 0.58841 | 0.195044344 | 0.222 | 0.154 | 1 | 7 |
| ide | 0.58871 | -0.11046779 | 0.032 | 0.077 | 1 | 7 |
| si:ch211-114c12.2 | 0.58886 | -0.14173751 | 0.063 | 0.115 | 1 | 7 |
| LOC101885675 | 0.58894 | -0.17216776 | 0.238 | 0.346 | 1 | 7 |
| fam32a | 0.58915 | 0.140694655 | 0.365 | 0.308 | 1 | 7 |
| ppp2cb | 0.58974 | -0.12788391 | 0.206 | 0.308 | 1 | 7 |
| zgc:77929 | 0.59071 | 0.115091621 | 0.127 | 0.077 | 1 | 7 |
| ccr2 | 0.591 | -0.15970002 | 0.397 | 0.462 | 1 | 7 |
| mef2aa | 0.59104 | -0.14426634 | 0.079 | 0.154 | 1 | 7 |
| vps26b | 0.59161 | -0.11741608 | 0.079 | 0.154 | 1 | 7 |
| lsm14ab | 0.59188 | -0.10889749 | 0.143 | 0.231 | 1 | 7 |
| mff | 0.59214 | 0.136903482 | 0.143 | 0.077 | 1 | 7 |
| ndufb7 | 0.59234 | 0.103246218 | 0.333 | 0.308 | 1 | 7 |
| ppp1cab | 0.59236 | -0.1523314 | 0.175 | 0.269 | 1 | 7 |
| nsd1a | 0.59245 | 0.147549701 | 0.143 | 0.077 | 1 | 7 |
| zfyve19 | 0.59249 | -0.11835556 | 0.032 | 0.077 | 1 | 7 |
| eif2s3 | 0.59333 | -0.1344915 | 0.619 | 0.731 | 1 | 7 |
| xrn2 | 0.59335 | -0.11133697 | 0.063 | 0.115 | 1 | 7 |
| ttyh2l | 0.5938 | -0.12861275 | 0.079 | 0.154 | 1 | 7 |
| ube2w | 0.59491 | 0.128921679 | 0.127 | 0.077 | 1 | 7 |
| cdv3 | 0.59498 | -0.1505145 | 0.27 | 0.346 | 1 | 7 |
| cdkn1bb | 0.59555 | 0.114057159 | 0.333 | 0.308 | 1 | 7 |
| slc5a3b | 0.59596 | -0.11678079 | 0.032 | 0.077 | 1 | 7 |
| farsa | 0.59673 | 0.112834128 | 0.095 | 0.038 | 1 | 7 |
| lrrc8aa | 0.59721 | -0.11990464 | 0.032 | 0.077 | 1 | 7 |
| hmgb3b | 0.59726 | 0.11368491 | 0.143 | 0.077 | 1 | 7 |
| nfkbie | 0.59732 | -0.16255113 | 0.317 | 0.385 | 1 | 7 |
| tpi1b | 0.5982 | -0.12897043 | 0.46 | 0.577 | 1 | 7 |
| dram2b | 0.59912 | -0.11063405 | 0.063 | 0.115 | 1 | 7 |
| btbd2b | 0.59923 | -0.11909028 | 0.032 | 0.077 | 1 | 7 |
| bcl2l10 | 0.60013 | -0.1438918 | 0.683 | 0.731 | 1 | 7 |
| fam49al | 0.60029 | 0.106307015 | 0.095 | 0.077 | 1 | 7 |
| man1b1b | 0.60071 | 0.137614977 | 0.079 | 0.038 | 1 | 7 |
| mrpl32 | 0.6008 | 0.150284718 | 0.238 | 0.154 | 1 | 7 |
| ncaldb | 0.60263 | -0.16850988 | 0.46 | 0.577 | 1 | 7 |
| dub | 0.6034 | 0.169816132 | 0.46 | 0.346 | 1 | 7 |
| fbxo9 | 0.60373 | 0.116874961 | 0.079 | 0.038 | 1 | 7 |
| dpf2 | 0.60398 | 0.124068494 | 0.095 | 0.038 | 1 | 7 |
| zgc:152774 | 0.60433 | 0.159549942 | 0.238 | 0.154 | 1 | 7 |
| atp6v1c1a | 0.60442 | -0.10726501 | 0.032 | 0.077 | 1 | 7 |
| wu:fi37e09 | 0.6045 | -0.1457158 | 0.143 | 0.231 | 1 | 7 |
| fkbp5 | 0.60487 | -0.13655474 | 0.111 | 0.192 | 1 | 7 |
| clk4b | 0.60525 | 0.106454616 | 0.095 | 0.038 | 1 | 7 |
| LOC103910438 | 0.60561 | -0.16288652 | 0.349 | 0.462 | 1 | 7 |
| si:ch1073-325m22.2 | 0.60566 | -0.12762558 | 0.46 | 0.577 | 1 | 7 |
| ets2 | 0.60597 | 0.111671121 | 0.111 | 0.077 | 1 | 7 |
| ctdsp1 | 0.6065 | -0.12668935 | 0.111 | 0.192 | 1 | 7 |
| mbd1b | 0.60677 | -0.11136125 | 0.032 | 0.077 | 1 | 7 |

| | | | | | | |
|---|---|---|---|---|---|---|
| tfg | 0.60735 | 0.16119973 | 0.175 | 0.115 | 1 | 7 |
| zgc:101016 | 0.6081 | 0.10411323 | 0.095 | 0.038 | 1 | 7 |
| ctnnb1 | 0.60831 | -0.126809 | 0.063 | 0.115 | 1 | 7 |
| parp8 | 0.60836 | -0.13120948 | 0.111 | 0.192 | 1 | 7 |
| spsb1 | 0.60854 | -0.10798055 | 0.032 | 0.077 | 1 | 7 |
| lpar5a | 0.60982 | -0.13722987 | 0.254 | 0.308 | 1 | 7 |
| flot2a | 0.61016 | 0.110401942 | 0.111 | 0.077 | 1 | 7 |
| egr2b | 0.61026 | -0.11836674 | 0.143 | 0.154 | 1 | 7 |
| lpar5b | 0.61055 | -0.13132404 | 0.143 | 0.231 | 1 | 7 |
| lbr | 0.6109 | -0.17463073 | 0.111 | 0.192 | 1 | 7 |
| spry2 | 0.61131 | -0.1202962 | 0.143 | 0.231 | 1 | 7 |
| malt1 | 0.61137 | 0.166915635 | 0.143 | 0.077 | 1 | 7 |
| phf8 | 0.61167 | 0.10612123 | 0.079 | 0.038 | 1 | 7 |
| anxa11a | 0.61226 | -0.111582 | 0.286 | 0.385 | 1 | 7 |
| mhc2b | 0.61275 | -0.17338017 | 0.175 | 0.269 | 1 | 7 |
| ndufc2 | 0.61278 | -0.14332116 | 0.175 | 0.231 | 1 | 7 |
| fignl1 | 0.61313 | 0.11115453 | 0.095 | 0.038 | 1 | 7 |
| CU207241.1 | 0.61393 | -0.15989201 | 0.143 | 0.231 | 1 | 7 |
| fut9b | 0.61579 | 0.102413978 | 0.095 | 0.038 | 1 | 7 |
| echs1 | 0.61643 | 0.125548175 | 0.19 | 0.115 | 1 | 7 |
| slc6a7 | 0.61737 | 0.124443162 | 0.095 | 0.038 | 1 | 7 |
| stx12 | 0.61739 | 0.105419607 | 0.095 | 0.038 | 1 | 7 |
| sumo1 | 0.61804 | 0.166119318 | 0.286 | 0.192 | 1 | 7 |
| tmem106a | 0.61873 | 0.161491249 | 0.143 | 0.077 | 1 | 7 |
| hcfc1b | 0.61895 | 0.129428785 | 0.095 | 0.038 | 1 | 7 |
| ucp2 | 0.61924 | -0.14453561 | 0.73 | 0.808 | 1 | 7 |
| nr4a2b | 0.61964 | -0.1466507 | 0.032 | 0.077 | 1 | 7 |
| nucks1a | 0.62073 | 0.158014323 | 0.333 | 0.231 | 1 | 7 |
| heatr5b | 0.62109 | -0.1091938 | 0.032 | 0.077 | 1 | 7 |
| tagapb | 0.62121 | -0.12643806 | 0.429 | 0.538 | 1 | 7 |
| sept8a | 0.62135 | 0.10443531 | 0.095 | 0.038 | 1 | 7 |
| eci2 | 0.62172 | -0.10007966 | 0.063 | 0.115 | 1 | 7 |
| roraa | 0.62198 | 0.101064041 | 0.095 | 0.038 | 1 | 7 |
| sox12 | 0.62281 | -0.14626463 | 0.063 | 0.115 | 1 | 7 |
| pik3c2b | 0.62344 | 0.108458475 | 0.063 | 0.038 | 1 | 7 |
| eno3 | 0.62396 | 0.178759029 | 0.524 | 0.423 | 1 | 7 |
| aldocb | 0.62416 | -0.11098947 | 0.254 | 0.346 | 1 | 7 |
| cyth4b | 0.62507 | -0.1001101 | 0.048 | 0.077 | 1 | 7 |
| mcl1b | 0.6258 | 0.117229289 | 0.095 | 0.077 | 1 | 7 |
| si:rp71-36a1.5 | 0.62671 | -0.10489752 | 0.032 | 0.077 | 1 | 7 |
| si:ch1073-220m6.1 | 0.62768 | -0.1042323 | 0.222 | 0.308 | 1 | 7 |
| pdlim5b | 0.62815 | -0.12401533 | 0.032 | 0.077 | 1 | 7 |
| cybc1 | 0.62843 | 0.106247229 | 0.095 | 0.038 | 1 | 7 |
| mtor | 0.62852 | 0.101646968 | 0.095 | 0.038 | 1 | 7 |
| chmp1b | 0.6287 | 0.145640831 | 0.333 | 0.269 | 1 | 7 |
| FP085398.1 | 0.62888 | -0.12568609 | 0.032 | 0.077 | 1 | 7 |
| irf7 | 0.62895 | -0.13670326 | 0.111 | 0.154 | 1 | 7 |
| nutf2 | 0.62951 | 0.106409545 | 0.143 | 0.077 | 1 | 7 |

| | | | | | | |
|---|---|---|---|---|---|---|
| ilf3b | 0.62956 | 0.112205877 | 0.222 | 0.192 | 1 | 7 |
| rhogb | 0.6297 | 0.172884525 | 0.365 | 0.269 | 1 | 7 |
| zgc:195282 | 0.6299 | 0.132260032 | 0.127 | 0.077 | 1 | 7 |
| kcmf1 | 0.63005 | -0.10053901 | 0.032 | 0.077 | 1 | 7 |
| tdh | 0.63025 | 0.130367187 | 0.159 | 0.115 | 1 | 7 |
| ptmaa | 0.63052 | -0.1230605 | 0.825 | 0.885 | 1 | 7 |
| sgta | 0.63136 | -0.15164959 | 0.143 | 0.192 | 1 | 7 |
| pfdn2 | 0.63232 | 0.118348254 | 0.238 | 0.192 | 1 | 7 |
| satb1b | 0.63259 | 0.163955781 | 0.143 | 0.077 | 1 | 7 |
| tmem39b | 0.63284 | 0.103079484 | 0.095 | 0.038 | 1 | 7 |
| rtn3 | 0.63378 | 0.108089279 | 0.079 | 0.038 | 1 | 7 |
| mrpl15 | 0.63394 | 0.109346453 | 0.095 | 0.038 | 1 | 7 |
| dpm2 | 0.63446 | 0.110892123 | 0.143 | 0.077 | 1 | 7 |
| fbxl6 | 0.63447 | -0.14823179 | 0.159 | 0.231 | 1 | 7 |
| brms1la | 0.63472 | 0.107949199 | 0.095 | 0.038 | 1 | 7 |
| slc20a1b | 0.63473 | 0.106659864 | 0.095 | 0.038 | 1 | 7 |
| ptpn4b | 0.63474 | 0.111048826 | 0.095 | 0.038 | 1 | 7 |
| timm10 | 0.63562 | 0.104966673 | 0.238 | 0.154 | 1 | 7 |
| dnmt1 | 0.6368 | 0.105920781 | 0.079 | 0.038 | 1 | 7 |
| npm3 | 0.63688 | 0.146274231 | 0.19 | 0.115 | 1 | 7 |
| ciapin1 | 0.6374 | 0.11184866 | 0.127 | 0.077 | 1 | 7 |
| arfgap2 | 0.63772 | 0.103200989 | 0.079 | 0.038 | 1 | 7 |
| chchd6b | 0.63889 | 0.110647771 | 0.127 | 0.077 | 1 | 7 |
| dtnbp1a | 0.63904 | -0.15369065 | 0.317 | 0.423 | 1 | 7 |
| gpr184 | 0.63922 | -0.12589919 | 0.095 | 0.154 | 1 | 7 |
| ppp1cb | 0.64264 | -0.13000076 | 0.159 | 0.231 | 1 | 7 |
| si:cabz01036006.1 | 0.64301 | -0.11004451 | 0.032 | 0.077 | 1 | 7 |
| ppp1caa | 0.64639 | -0.11275742 | 0.397 | 0.5 | 1 | 7 |
| tsr2 | 0.64658 | -0.12202354 | 0.127 | 0.192 | 1 | 7 |
| ilf2 | 0.64671 | 0.134996541 | 0.492 | 0.385 | 1 | 7 |
| vhl | 0.64779 | -0.10996384 | 0.032 | 0.077 | 1 | 7 |
| map2k2a | 0.64826 | 0.120494153 | 0.143 | 0.077 | 1 | 7 |
| BX649250.1 | 0.64849 | 0.103599239 | 0.111 | 0.077 | 1 | 7 |
| phf14 | 0.6516 | -0.10071473 | 0.032 | 0.077 | 1 | 7 |
| rbmx2 | 0.65178 | 0.115162699 | 0.079 | 0.038 | 1 | 7 |
| tmub2 | 0.65295 | -0.12596722 | 0.063 | 0.115 | 1 | 7 |
| c1qbp | 0.65405 | 0.102400022 | 0.222 | 0.192 | 1 | 7 |
| cdipt | 0.65493 | 0.133536922 | 0.238 | 0.154 | 1 | 7 |
| txndc12 | 0.65681 | -0.12297136 | 0.063 | 0.115 | 1 | 7 |
| ikzf1 | 0.66281 | 0.15723942 | 0.381 | 0.308 | 1 | 7 |
| cops2 | 0.66303 | 0.124060046 | 0.143 | 0.077 | 1 | 7 |
| atrx | 0.66345 | 0.137006828 | 0.413 | 0.346 | 1 | 7 |
| LOC100535806 | 0.66372 | 0.125746515 | 0.143 | 0.077 | 1 | 7 |
| trappc1 | 0.66391 | 0.104770024 | 0.222 | 0.192 | 1 | 7 |
| mcts1 | 0.66486 | 0.114675947 | 0.238 | 0.154 | 1 | 7 |
| pik3r1 | 0.66525 | -0.16118566 | 0.349 | 0.423 | 1 | 7 |
| zgc:111986 | 0.66566 | 0.136382287 | 0.238 | 0.154 | 1 | 7 |
| psmb7 | 0.66574 | 0.110331071 | 0.238 | 0.154 | 1 | 7 |

| | | | | | | |
|---|---|---|---|---|---|---|
| ssu72 | 0.66739 | -0.12961144 | 0.143 | 0.192 | 1 | 7 |
| pcna | 0.66767 | 0.170902039 | 0.317 | 0.269 | 1 | 7 |
| elf2b | 0.66769 | 0.144490765 | 0.19 | 0.115 | 1 | 7 |
| mrpl13 | 0.66786 | 0.109266389 | 0.127 | 0.077 | 1 | 7 |
| ifi46 | 0.6692 | -0.1115044 | 0.032 | 0.077 | 1 | 7 |
| pnn | 0.66929 | 0.112568563 | 0.19 | 0.115 | 1 | 7 |
| sdha | 0.66935 | 0.107318538 | 0.19 | 0.115 | 1 | 7 |
| si:ch73-223p23.2 | 0.66989 | -0.15199995 | 0.222 | 0.308 | 1 | 7 |
| hnrnpub | 0.67026 | 0.131562026 | 0.238 | 0.192 | 1 | 7 |
| mcm2 | 0.67129 | 0.128276561 | 0.19 | 0.115 | 1 | 7 |
| tmem30aa | 0.67191 | 0.155907889 | 0.19 | 0.115 | 1 | 7 |
| rad23b | 0.67341 | -0.14118434 | 0.254 | 0.346 | 1 | 7 |
| mrpl34 | 0.67462 | 0.135306528 | 0.222 | 0.154 | 1 | 7 |
| bcas2 | 0.67505 | 0.117787003 | 0.286 | 0.231 | 1 | 7 |
| srebf2 | 0.67544 | -0.10953186 | 0.032 | 0.077 | 1 | 7 |
| hspa14 | 0.67594 | 0.10331467 | 0.19 | 0.115 | 1 | 7 |
| gcnt7 | 0.67674 | 0.116598321 | 0.079 | 0.038 | 1 | 7 |
| mxd1 | 0.67682 | -0.10340571 | 0.032 | 0.077 | 1 | 7 |
| CR847906.1 | 0.67686 | -0.12104956 | 0.032 | 0.077 | 1 | 7 |
| ip6k2b | 0.67692 | -0.10159128 | 0.19 | 0.269 | 1 | 7 |
| armt1 | 0.67696 | -0.10552057 | 0.032 | 0.077 | 1 | 7 |
| stmn1a | 0.67713 | 0.110600589 | 0.127 | 0.115 | 1 | 7 |
| msrb1b | 0.67716 | -0.11588553 | 0.175 | 0.231 | 1 | 7 |
| snip1 | 0.67723 | 0.111193456 | 0.127 | 0.077 | 1 | 7 |
| ythdf1 | 0.67798 | 0.102922359 | 0.079 | 0.038 | 1 | 7 |
| prpf38b | 0.67918 | 0.103161042 | 0.159 | 0.115 | 1 | 7 |
| afap1l2 | 0.68073 | -0.10757681 | 0.048 | 0.077 | 1 | 7 |
| PDPR | 0.68156 | -0.10612051 | 0.063 | 0.115 | 1 | 7 |
| mcrip1 | 0.68288 | -0.12901096 | 0.143 | 0.192 | 1 | 7 |
| chd9 | 0.68378 | 0.115674626 | 0.19 | 0.154 | 1 | 7 |
| XLOC-033137 | 0.68732 | -0.15313016 | 0.159 | 0.231 | 1 | 7 |
| LOC110439931 | 0.68753 | 0.158439593 | 0.317 | 0.231 | 1 | 7 |
| idh2 | 0.68775 | 0.119818005 | 0.524 | 0.423 | 1 | 7 |
| rab5ab | 0.6886 | -0.14268673 | 0.127 | 0.192 | 1 | 7 |
| cox10 | 0.69097 | -0.10254108 | 0.063 | 0.115 | 1 | 7 |
| psma6b | 0.69123 | 0.125230233 | 0.27 | 0.192 | 1 | 7 |
| sept2 | 0.69157 | 0.143834615 | 0.587 | 0.5 | 1 | 7 |
| atp6ap2 | 0.69276 | -0.12008355 | 0.143 | 0.192 | 1 | 7 |
| mrpl12 | 0.69679 | 0.133818222 | 0.222 | 0.154 | 1 | 7 |
| stx11a | 0.6984 | 0.10118632 | 0.063 | 0.038 | 1 | 7 |
| ppp1r12c | 0.69869 | 0.101953398 | 0.143 | 0.115 | 1 | 7 |
| rfc3 | 0.69889 | 0.102021771 | 0.111 | 0.077 | 1 | 7 |
| afmid | 0.69977 | 0.102737547 | 0.127 | 0.077 | 1 | 7 |
| lyrm4 | 0.69977 | 0.109712938 | 0.206 | 0.154 | 1 | 7 |
| si:ch211-132p1.3 | 0.6998 | -0.14843755 | 0.333 | 0.423 | 1 | 7 |
| aip | 0.70277 | 0.114343957 | 0.111 | 0.077 | 1 | 7 |
| mcm5 | 0.70304 | 0.100762439 | 0.175 | 0.115 | 1 | 7 |
| pgk1 | 0.70512 | -0.1010377 | 0.19 | 0.269 | 1 | 7 |

| | | | | | | |
|---|---|---|---|---|---|---|
| cd83 | 0.7056 | -0.12080862 | 0.159 | 0.231 | 1 | 7 |
| cenpw | 0.70584 | 0.108203843 | 0.127 | 0.077 | 1 | 7 |
| lrp10 | 0.70678 | -0.12279737 | 0.095 | 0.154 | 1 | 7 |
| prdx3 | 0.70934 | -0.1136245 | 0.127 | 0.192 | 1 | 7 |
| dock11 | 0.71189 | 0.104296254 | 0.079 | 0.038 | 1 | 7 |
| tmem185 | 0.71303 | -0.12607507 | 0.063 | 0.115 | 1 | 7 |
| akap8l | 0.71327 | -0.11662681 | 0.159 | 0.231 | 1 | 7 |
| si:dkey-206d17.12 | 0.71413 | -0.10021724 | 0.127 | 0.192 | 1 | 7 |
| mapk3 | 0.71483 | -0.12321744 | 0.19 | 0.269 | 1 | 7 |
| CU499336.2 | 0.71777 | -0.11006775 | 0.095 | 0.154 | 1 | 7 |
| axin1 | 0.71851 | -0.11079937 | 0.063 | 0.115 | 1 | 7 |
| polr2f | 0.72017 | 0.123403839 | 0.238 | 0.192 | 1 | 7 |
| ccr9a | 0.7222 | -0.15020095 | 0.397 | 0.462 | 1 | 7 |
| si:ch211-184m13.4 | 0.72264 | -0.12021668 | 0.063 | 0.115 | 1 | 7 |
| map1lc3b | 0.72266 | -0.10836604 | 0.159 | 0.231 | 1 | 7 |
| arf6b | 0.72292 | 0.114550006 | 0.397 | 0.308 | 1 | 7 |
| lsm8 | 0.72298 | 0.108489004 | 0.27 | 0.192 | 1 | 7 |
| si:ch211-288g17.3 | 0.72506 | 0.140970289 | 0.349 | 0.308 | 1 | 7 |
| ptk2ba | 0.72615 | -0.14544388 | 0.175 | 0.231 | 1 | 7 |
| cpsf6 | 0.72635 | 0.127158406 | 0.27 | 0.192 | 1 | 7 |
| itpkb | 0.72671 | -0.10695657 | 0.063 | 0.115 | 1 | 7 |
| ppp1r18 | 0.72703 | -0.12355199 | 0.254 | 0.308 | 1 | 7 |
| anp32b | 0.72734 | 0.123484329 | 0.397 | 0.308 | 1 | 7 |
| XLOC-041870 | 0.72864 | -0.13073021 | 0.159 | 0.231 | 1 | 7 |
| rpl7l1 | 0.72894 | 0.107905653 | 0.222 | 0.154 | 1 | 7 |
| sh2d3ca | 0.72931 | 0.104586671 | 0.111 | 0.077 | 1 | 7 |
| mars | 0.73027 | -0.10389152 | 0.095 | 0.154 | 1 | 7 |
| ldha | 0.73214 | -0.11109417 | 0.159 | 0.231 | 1 | 7 |
| hist2h2l | 0.7324 | -0.10264901 | 0.159 | 0.231 | 1 | 7 |
| cltca | 0.73272 | -0.10186098 | 0.127 | 0.192 | 1 | 7 |
| cnpy3 | 0.73278 | 0.108722979 | 0.143 | 0.115 | 1 | 7 |
| hsp90b1 | 0.73312 | 0.118695757 | 0.127 | 0.077 | 1 | 7 |
| inpp5ka | 0.73442 | -0.11089807 | 0.079 | 0.115 | 1 | 7 |
| ehbp1l1b | 0.73461 | 0.105941991 | 0.063 | 0.038 | 1 | 7 |
| tubb2b | 0.73473 | 0.108172623 | 0.429 | 0.423 | 1 | 7 |
| pdcd10b | 0.73506 | 0.10279529 | 0.19 | 0.154 | 1 | 7 |
| phb | 0.73559 | -0.11070112 | 0.127 | 0.192 | 1 | 7 |
| bzw1b | 0.73607 | -0.11348543 | 0.302 | 0.346 | 1 | 7 |
| ago2 | 0.73725 | -0.10506737 | 0.048 | 0.077 | 1 | 7 |
| bcl11ba | 0.73788 | 0.132270028 | 0.302 | 0.269 | 1 | 7 |
| waca | 0.73949 | -0.10895359 | 0.127 | 0.192 | 1 | 7 |
| snrnp70 | 0.7399 | -0.10755746 | 0.127 | 0.192 | 1 | 7 |
| slc16a8 | 0.73999 | -0.1014085 | 0.127 | 0.192 | 1 | 7 |
| tox2 | 0.74036 | 0.124309614 | 0.175 | 0.115 | 1 | 7 |
| mrps15 | 0.74095 | 0.114506704 | 0.206 | 0.154 | 1 | 7 |
| pagr1 | 0.74223 | 0.111587403 | 0.127 | 0.077 | 1 | 7 |
| taf13 | 0.74258 | -0.12469182 | 0.302 | 0.385 | 1 | 7 |
| atp1b1a | 0.74716 | -0.11771609 | 0.349 | 0.423 | 1 | 7 |

| | | | | | | |
|---|---|---|---|---|---|---|
| naa40 | 0.74799 | 0.10531306 | 0.175 | 0.115 | 1 | 7 |
| serinc1 | 0.74928 | 0.106863758 | 0.079 | 0.038 | 1 | 7 |
| pde4ba | 0.74941 | -0.10814635 | 0.238 | 0.269 | 1 | 7 |
| rab10 | 0.75005 | 0.13364863 | 0.381 | 0.308 | 1 | 7 |
| LOC101884267 | 0.75033 | 0.118701369 | 0.079 | 0.038 | 1 | 7 |
| jak1 | 0.75072 | 0.118735112 | 0.222 | 0.154 | 1 | 7 |
| snrpa | 0.75101 | 0.101377152 | 0.222 | 0.154 | 1 | 7 |
| si:dkey-250d21.1 | 0.75112 | 0.106456074 | 0.159 | 0.115 | 1 | 7 |
| nrros | 0.75181 | 0.115645231 | 0.206 | 0.154 | 1 | 7 |
| tars | 0.75197 | 0.102163792 | 0.127 | 0.077 | 1 | 7 |
| eif4bb | 0.752 | 0.111553913 | 0.333 | 0.269 | 1 | 7 |
| foxo3b | 0.75512 | 0.102790609 | 0.127 | 0.077 | 1 | 7 |
| wdr83os | 0.7559 | 0.103067989 | 0.222 | 0.154 | 1 | 7 |
| nptna | 0.75662 | -0.12039964 | 0.079 | 0.115 | 1 | 7 |
| prkar2aa | 0.7581 | 0.126005604 | 0.254 | 0.192 | 1 | 7 |
| sri | 0.75915 | 0.115437175 | 0.349 | 0.269 | 1 | 7 |
| tomm20a | 0.76281 | -0.11908402 | 0.238 | 0.308 | 1 | 7 |
| khdrbs1b | 0.76311 | -0.10198636 | 0.143 | 0.192 | 1 | 7 |
| tenm3 | 0.76656 | 0.105872066 | 0.079 | 0.038 | 1 | 7 |
| grap2b | 0.76802 | -0.10777815 | 0.571 | 0.654 | 1 | 7 |
| plcg1 | 0.77034 | -0.12131083 | 0.143 | 0.192 | 1 | 7 |
| znrf1 | 0.77282 | 0.123820712 | 0.127 | 0.077 | 1 | 7 |
| yrk | 0.77359 | -0.12565451 | 0.175 | 0.231 | 1 | 7 |
| XLOC-005449 | 0.77602 | 0.140279043 | 0.302 | 0.231 | 1 | 7 |
| ralbp1 | 0.77859 | 0.101964014 | 0.159 | 0.115 | 1 | 7 |
| tefa | 0.77909 | -0.11665721 | 0.317 | 0.385 | 1 | 7 |
| tpst1l | 0.7796 | 0.106891366 | 0.127 | 0.077 | 1 | 7 |
| atp6v1e1a | 0.78528 | 0.114300683 | 0.159 | 0.115 | 1 | 7 |
| calr3b | 0.79122 | -0.13034835 | 0.238 | 0.308 | 1 | 7 |
| csnk1a1 | 0.79501 | -0.11737892 | 0.317 | 0.385 | 1 | 7 |
| ccnl1a | 0.79592 | 0.116468174 | 0.159 | 0.115 | 1 | 7 |
| capza1a | 0.80227 | 0.104255557 | 0.603 | 0.538 | 1 | 7 |
| sox13 | 0.80227 | -0.11442411 | 0.079 | 0.115 | 1 | 7 |
| anp32a | 0.80349 | 0.107748771 | 0.587 | 0.538 | 1 | 7 |
| hspe1 | 0.80429 | 0.100628993 | 0.206 | 0.154 | 1 | 7 |
| srsf3b | 0.80794 | -0.10786616 | 0.429 | 0.5 | 1 | 7 |
| dnm2a | 0.81307 | -0.10037116 | 0.111 | 0.154 | 1 | 7 |
| slc3a2b | 0.81452 | -0.11346432 | 0.254 | 0.308 | 1 | 7 |
| hnrnpa1a | 0.83276 | 0.10732234 | 0.333 | 0.269 | 1 | 7 |
| mknk2b | 0.83597 | -0.10623683 | 0.365 | 0.423 | 1 | 7 |
| zap70 | 0.83897 | 0.116922412 | 0.286 | 0.231 | 1 | 7 |
| tuba8l | 0.84831 | 0.113634799 | 0.397 | 0.346 | 1 | 7 |
| ccl35.1 | 0.85079 | -0.11009802 | 0.095 | 0.115 | 1 | 7 |
| sftpba | 0.87526 | -0.10277342 | 0.127 | 0.154 | 1 | 7 |
| CR318588.1 | 0.91456 | 0.118953408 | 0.19 | 0.154 | 1 | 7 |
| nupr1b | 5.895900 | 0.664285799 | 0.263 | 0 | 1 | 8 |
| tmem176 | 0.00016 | 0.617234248 | 0.237 | 0 | 1 | 8 |
| gamt | 0.00042 | 0.362472449 | 0.211 | 0 | 1 | 8 |

| | | | | | | |
|---|---|---|---|---|---|---|
| mmp2 | 0.00094 | -0.60580249 | 0 | 0.2 | 1 | 8 |
| zgc:92744 | 0.00104 | 0.650890257 | 0.395 | 0.067 | 1 | 8 |
| rpl7l1 | 0.00108 | 0.484886325 | 0.184 | 0 | 1 | 8 |
| ube2l3a | 0.00108 | 0.492274671 | 0.184 | 0 | 1 | 8 |
| si:dkey-14d8.6 | 0.00108 | 0.468818871 | 0.184 | 0 | 1 | 8 |
| rpl27a | 0.00139 | -0.4132217 | 0.632 | 0.778 | 1 | 8 |
| h2afva | 0.00199 | -0.50523263 | 0 | 0.178 | 1 | 8 |
| lmcd1 | 0.00276 | 0.439915581 | 0.158 | 0 | 1 | 8 |
| nedd8 | 0.00276 | 0.42794208 | 0.158 | 0 | 1 | 8 |
| pkig | 0.00276 | 0.43827944 | 0.158 | 0 | 1 | 8 |
| rpe | 0.00276 | 0.418958326 | 0.158 | 0 | 1 | 8 |
| rpl38 | 0.00589 | -0.20845211 | 0.921 | 1 | 1 | 8 |
| abracl | 0.00678 | 0.506026063 | 0.237 | 0.022 | 1 | 8 |
| ptp4a2b | 0.00701 | 0.384296566 | 0.132 | 0 | 1 | 8 |
| meis3 | 0.00701 | 0.37512764 | 0.132 | 0 | 1 | 8 |
| chd3 | 0.00701 | 0.488858483 | 0.132 | 0 | 1 | 8 |
| CABZ01072614.1 | 0.00701 | 0.402657574 | 0.132 | 0 | 1 | 8 |
| bhmt | 0.00701 | 0.332078683 | 0.132 | 0 | 1 | 8 |
| ctdspla | 0.00701 | 0.450192346 | 0.132 | 0 | 1 | 8 |
| noc4l | 0.00701 | 0.302927754 | 0.132 | 0 | 1 | 8 |
| sh3pxd2b | 0.00701 | 0.433767543 | 0.132 | 0 | 1 | 8 |
| med29 | 0.00701 | 0.433688349 | 0.132 | 0 | 1 | 8 |
| phc2a | 0.00701 | 0.400031744 | 0.132 | 0 | 1 | 8 |
| reep3a | 0.00701 | 0.387561298 | 0.132 | 0 | 1 | 8 |
| ugt1b5 | 0.00801 | 0.558063683 | 0.211 | 0.022 | 1 | 8 |
| rab10 | 0.00819 | 0.672093172 | 0.289 | 0.067 | 1 | 8 |
| col4a1 | 0.0088 | -0.42690394 | 0 | 0.133 | 1 | 8 |
| rpa3 | 0.0088 | -0.37208935 | 0 | 0.133 | 1 | 8 |
| dpp3 | 0.0088 | -0.40789627 | 0 | 0.133 | 1 | 8 |
| h3f3c | 0.00929 | 0.529200839 | 0.711 | 0.378 | 1 | 8 |
| ppdpfb | 0.01037 | 0.434054411 | 0.711 | 0.4 | 1 | 8 |
| chmp5b | 0.01405 | 0.503909524 | 0.263 | 0.044 | 1 | 8 |
| psmc4 | 0.01502 | 0.536681596 | 0.211 | 0.022 | 1 | 8 |
| dars | 0.01648 | 0.526094428 | 0.211 | 0.022 | 1 | 8 |
| psmc1a | 0.01686 | 0.4812972 | 0.211 | 0.022 | 1 | 8 |
| ap3m1 | 0.01769 | 0.369396696 | 0.105 | 0 | 1 | 8 |
| asip2b | 0.01769 | 0.365409092 | 0.105 | 0 | 1 | 8 |
| cebpg | 0.01769 | 0.32101634 | 0.105 | 0 | 1 | 8 |
| npdc1a | 0.01769 | 0.211252369 | 0.105 | 0 | 1 | 8 |
| polr2b | 0.01769 | 0.374924584 | 0.105 | 0 | 1 | 8 |
| ppm1aa | 0.01769 | 0.354947383 | 0.105 | 0 | 1 | 8 |
| rab3db | 0.01769 | 0.319939392 | 0.105 | 0 | 1 | 8 |
| rbp4 | 0.01769 | 0.360185619 | 0.105 | 0 | 1 | 8 |
| rnf44 | 0.01769 | 0.357230207 | 0.105 | 0 | 1 | 8 |
| snx17 | 0.01769 | 0.33488873 | 0.105 | 0 | 1 | 8 |
| spg21 | 0.01769 | 0.31930649 | 0.105 | 0 | 1 | 8 |
| zgc:114045 | 0.01769 | 0.340263083 | 0.105 | 0 | 1 | 8 |
| BX322530.1 | 0.01769 | 0.371526369 | 0.105 | 0 | 1 | 8 |

| | | | | | | |
|---|---|---|---|---|---|---|
| cyb5a | 0.01769 | 0.275932483 | 0.105 | 0 | 1 | 8 |
| ephb3 | 0.01769 | 0.420178908 | 0.105 | 0 | 1 | 8 |
| impa1 | 0.01769 | 0.266782686 | 0.105 | 0 | 1 | 8 |
| mrps36 | 0.01769 | 0.339702136 | 0.105 | 0 | 1 | 8 |
| myof | 0.01769 | 0.34605153 | 0.105 | 0 | 1 | 8 |
| si:dkey-67c22.2 | 0.01769 | 0.33541843 | 0.105 | 0 | 1 | 8 |
| srp9 | 0.01769 | 0.285909322 | 0.105 | 0 | 1 | 8 |
| mbd3b | 0.01769 | 0.287767453 | 0.105 | 0 | 1 | 8 |
| si:ch211-218c6.8 | 0.01769 | 0.306296625 | 0.105 | 0 | 1 | 8 |
| cd151l | 0.01769 | 0.323702765 | 0.105 | 0 | 1 | 8 |
| coro1ca | 0.01769 | 0.359131642 | 0.105 | 0 | 1 | 8 |
| eif4ebp3 | 0.01769 | 0.304601798 | 0.105 | 0 | 1 | 8 |
| gatd1 | 0.01769 | 0.292095173 | 0.105 | 0 | 1 | 8 |
| gtf2a1 | 0.01769 | 0.322092664 | 0.105 | 0 | 1 | 8 |
| hbp1 | 0.01769 | 0.305683426 | 0.105 | 0 | 1 | 8 |
| hsp90b1 | 0.01769 | 0.379937821 | 0.105 | 0 | 1 | 8 |
| mrpl20 | 0.01769 | 0.296499662 | 0.105 | 0 | 1 | 8 |
| rtn1a | 0.01769 | 0.342606795 | 0.105 | 0 | 1 | 8 |
| ap2b1 | 0.01835 | -0.32829562 | 0 | 0.111 | 1 | 8 |
| mcm2 | 0.01835 | -0.34573196 | 0 | 0.111 | 1 | 8 |
| sirt7 | 0.01835 | -0.34908763 | 0 | 0.111 | 1 | 8 |
| slc29a1b | 0.01835 | -0.36837433 | 0 | 0.111 | 1 | 8 |
| cenpx | 0.01835 | -0.38996577 | 0 | 0.111 | 1 | 8 |
| taf7 | 0.01835 | -0.32114609 | 0 | 0.111 | 1 | 8 |
| oaz1a | 0.02053 | 0.462508116 | 0.579 | 0.289 | 1 | 8 |
| mcl1b | 0.02255 | 0.559838934 | 0.395 | 0.133 | 1 | 8 |
| rpl10 | 0.02445 | -0.24247622 | 0.816 | 0.978 | 1 | 8 |
| rpl34 | 0.02459 | -0.17581067 | 0.868 | 1 | 1 | 8 |
| ccl25b | 0.02522 | 0.695557832 | 0.289 | 0.089 | 1 | 8 |
| rpsa | 0.02597 | -0.1871262 | 0.895 | 0.978 | 1 | 8 |
| rps29 | 0.02622 | -0.17654401 | 0.921 | 1 | 1 | 8 |
| add3a | 0.0266 | 0.527719476 | 0.237 | 0.044 | 1 | 8 |
| elof1 | 0.02765 | 0.383842309 | 0.184 | 0.022 | 1 | 8 |
| gabarapa | 0.02782 | 0.464866898 | 0.237 | 0.044 | 1 | 8 |
| itgav | 0.03146 | 0.490306778 | 0.237 | 0.044 | 1 | 8 |
| tagln | 0.03241 | 0.296084703 | 0.289 | 0.244 | 1 | 8 |
| gstm.3 | 0.03272 | -0.11665513 | 0.974 | 0.956 | 1 | 8 |
| adh8b | 0.03496 | 0.456813351 | 0.263 | 0.067 | 1 | 8 |
| arpc5a | 0.03539 | 0.41495883 | 0.184 | 0.022 | 1 | 8 |
| mknk2b | 0.03626 | 0.4350499 | 0.184 | 0.022 | 1 | 8 |
| sdcbp2 | 0.0363 | 0.431174908 | 0.184 | 0.022 | 1 | 8 |
| fam133b | 0.03669 | 0.566963893 | 0.342 | 0.111 | 1 | 8 |
| tnip1 | 0.03821 | -0.2729228 | 0 | 0.089 | 1 | 8 |
| CU459094.2 | 0.03821 | -0.24769447 | 0 | 0.089 | 1 | 8 |
| akt1s1 | 0.03821 | -0.26749988 | 0 | 0.089 | 1 | 8 |
| dhps | 0.03821 | -0.22791347 | 0 | 0.089 | 1 | 8 |
| eif4e2rs1 | 0.03821 | -0.29586315 | 0 | 0.089 | 1 | 8 |
| gtf2b | 0.03821 | -0.23981796 | 0 | 0.089 | 1 | 8 |

| | | | | | | |
|---|---|---|---|---|---|---|
| isg15 | 0.03821 | -0.32695511 | 0 | 0.089 | 1 | 8 |
| itgb1bp1 | 0.03821 | -0.31404147 | 0 | 0.089 | 1 | 8 |
| lmnb2 | 0.03821 | -0.28684672 | 0 | 0.089 | 1 | 8 |
| mtap | 0.03821 | -0.25949667 | 0 | 0.089 | 1 | 8 |
| mych | 0.03821 | -0.32249751 | 0 | 0.089 | 1 | 8 |
| ppig | 0.03821 | -0.29526159 | 0 | 0.089 | 1 | 8 |
| ppp1cb | 0.03821 | -0.2837156 | 0 | 0.089 | 1 | 8 |
| rbfox1 | 0.03821 | -0.3027924 | 0 | 0.089 | 1 | 8 |
| zgc:55943 | 0.03821 | -0.26867404 | 0 | 0.089 | 1 | 8 |
| hexim1 | 0.03821 | -0.28955169 | 0 | 0.089 | 1 | 8 |
| ifi46 | 0.03821 | -0.26197098 | 0 | 0.089 | 1 | 8 |
| CDPF1 | 0.03821 | -0.26267474 | 0 | 0.089 | 1 | 8 |
| cetn4 | 0.03821 | -0.31121327 | 0 | 0.089 | 1 | 8 |
| ckbb | 0.03821 | -0.32317659 | 0 | 0.089 | 1 | 8 |
| csrnp1b | 0.03821 | -0.26971442 | 0 | 0.089 | 1 | 8 |
| dkc1 | 0.03821 | -0.27963489 | 0 | 0.089 | 1 | 8 |
| fundc1 | 0.03821 | -0.29427046 | 0 | 0.089 | 1 | 8 |
| mob3a | 0.03821 | -0.23547308 | 0 | 0.089 | 1 | 8 |
| siah1 | 0.03821 | -0.27372489 | 0 | 0.089 | 1 | 8 |
| sult6b1 | 0.03821 | -0.23440379 | 0 | 0.089 | 1 | 8 |
| taf13 | 0.03981 | 0.474490737 | 0.263 | 0.067 | 1 | 8 |
| ndufv1 | 0.04008 | 0.294871837 | 0.158 | 0.022 | 1 | 8 |
| arhgdig | 0.0409 | 0.492240699 | 0.368 | 0.133 | 1 | 8 |
| bccip | 0.04109 | 0.424086244 | 0.263 | 0.067 | 1 | 8 |
| cth | 0.04471 | 0.209514743 | 0.079 | 0 | 1 | 8 |
| CR925817.1 | 0.04471 | 0.20558704 | 0.079 | 0 | 1 | 8 |
| acy1 | 0.04471 | 0.225084576 | 0.079 | 0 | 1 | 8 |
| arhgef4 | 0.04471 | 0.255246769 | 0.079 | 0 | 1 | 8 |
| ccdc32 | 0.04471 | 0.234446136 | 0.079 | 0 | 1 | 8 |
| cherp | 0.04471 | 0.261944092 | 0.079 | 0 | 1 | 8 |
| clic1 | 0.04471 | 0.267523278 | 0.079 | 0 | 1 | 8 |
| crlf3 | 0.04471 | 0.234419201 | 0.079 | 0 | 1 | 8 |
| emid1 | 0.04471 | 0.309237027 | 0.079 | 0 | 1 | 8 |
| exosc4 | 0.04471 | 0.22092485 | 0.079 | 0 | 1 | 8 |
| fam3a | 0.04471 | 0.29221911 | 0.079 | 0 | 1 | 8 |
| fam50a | 0.04471 | 0.198696987 | 0.079 | 0 | 1 | 8 |
| gtf2e2 | 0.04471 | 0.198106182 | 0.079 | 0 | 1 | 8 |
| inip | 0.04471 | 0.22854279 | 0.079 | 0 | 1 | 8 |
| klhdc3 | 0.04471 | 0.237627145 | 0.079 | 0 | 1 | 8 |
| lrrc17 | 0.04471 | 0.298279382 | 0.079 | 0 | 1 | 8 |
| mapkap1 | 0.04471 | 0.217442839 | 0.079 | 0 | 1 | 8 |
| mat1a | 0.04471 | 0.227400665 | 0.079 | 0 | 1 | 8 |
| mmp16a | 0.04471 | 0.30583377 | 0.079 | 0 | 1 | 8 |
| nagk | 0.04471 | 0.228508487 | 0.079 | 0 | 1 | 8 |
| ndfip1 | 0.04471 | 0.242980545 | 0.079 | 0 | 1 | 8 |
| nfkbiaa | 0.04471 | 0.249030571 | 0.079 | 0 | 1 | 8 |
| nmt2 | 0.04471 | 0.240785312 | 0.079 | 0 | 1 | 8 |
| nr2f6b | 0.04471 | 0.231128644 | 0.079 | 0 | 1 | 8 |

| | | | | | | |
|---|---|---|---|---|---|---|
| nudt9 | 0.04471 | 0.228910414 | 0.079 | 0 | 1 | 8 |
| nup188 | 0.04471 | 0.199439191 | 0.079 | 0 | 1 | 8 |
| paox1 | 0.04471 | 0.257804516 | 0.079 | 0 | 1 | 8 |
| pla2g4aa | 0.04471 | 0.267656498 | 0.079 | 0 | 1 | 8 |
| ppm1g | 0.04471 | 0.218350574 | 0.079 | 0 | 1 | 8 |
| ppm1la | 0.04471 | 0.206067668 | 0.079 | 0 | 1 | 8 |
| prpsap2 | 0.04471 | 0.209821254 | 0.079 | 0 | 1 | 8 |
| psat1 | 0.04471 | 0.218041416 | 0.079 | 0 | 1 | 8 |
| psmc3 | 0.04471 | 0.206773608 | 0.079 | 0 | 1 | 8 |
| ptbp1b | 0.04471 | 0.306225944 | 0.079 | 0 | 1 | 8 |
| scaf11 | 0.04471 | 0.240795508 | 0.079 | 0 | 1 | 8 |
| si:dkey-14d8.7 | 0.04471 | 0.177338107 | 0.079 | 0 | 1 | 8 |
| si:dkey-250d21.1 | 0.04471 | 0.283182244 | 0.079 | 0 | 1 | 8 |
| si:dkeyp-55f12.3 | 0.04471 | 0.22805869 | 0.079 | 0 | 1 | 8 |
| sirt2 | 0.04471 | 0.216320907 | 0.079 | 0 | 1 | 8 |
| srrm1 | 0.04471 | 0.279419696 | 0.079 | 0 | 1 | 8 |
| srsf3a | 0.04471 | 0.249669499 | 0.079 | 0 | 1 | 8 |
| stat6 | 0.04471 | 0.212361291 | 0.079 | 0 | 1 | 8 |
| taf10 | 0.04471 | 0.245291314 | 0.079 | 0 | 1 | 8 |
| ubxn1 | 0.04471 | 0.259166777 | 0.079 | 0 | 1 | 8 |
| usp8 | 0.04471 | 0.213573759 | 0.079 | 0 | 1 | 8 |
| wnt5b | 0.04471 | 0.259591478 | 0.079 | 0 | 1 | 8 |
| zgc:153284 | 0.04471 | 0.274722569 | 0.079 | 0 | 1 | 8 |
| zgc:154075 | 0.04471 | 0.210917841 | 0.079 | 0 | 1 | 8 |
| zgc:158343 | 0.04471 | 0.336947979 | 0.079 | 0 | 1 | 8 |
| cwc25 | 0.04471 | 0.244237557 | 0.079 | 0 | 1 | 8 |
| plekha1a | 0.04471 | 0.2502044 | 0.079 | 0 | 1 | 8 |
| zgc:92590 | 0.04471 | 0.204613426 | 0.079 | 0 | 1 | 8 |
| BX284638.1 | 0.04471 | 0.247846803 | 0.079 | 0 | 1 | 8 |
| RPL37A | 0.04471 | 0.202644986 | 0.079 | 0 | 1 | 8 |
| afp4 | 0.04471 | 0.17520301 | 0.079 | 0 | 1 | 8 |
| cdc73 | 0.04471 | 0.212081869 | 0.079 | 0 | 1 | 8 |
| cops4 | 0.04471 | 0.224822869 | 0.079 | 0 | 1 | 8 |
| ehd3 | 0.04471 | 0.293732247 | 0.079 | 0 | 1 | 8 |
| gnl3l | 0.04471 | 0.206289914 | 0.079 | 0 | 1 | 8 |
| gpc1b | 0.04471 | 0.242771264 | 0.079 | 0 | 1 | 8 |
| hmga2 | 0.04471 | 0.263513564 | 0.079 | 0 | 1 | 8 |
| igfbp7 | 0.04471 | 0.237961183 | 0.079 | 0 | 1 | 8 |
| mrps28 | 0.04471 | 0.248313466 | 0.079 | 0 | 1 | 8 |
| nacc2 | 0.04471 | 0.27578348 | 0.079 | 0 | 1 | 8 |
| phf5a | 0.04471 | 0.221734906 | 0.079 | 0 | 1 | 8 |
| puraa | 0.04471 | 0.258765833 | 0.079 | 0 | 1 | 8 |
| si:dkeyp-44a8.2 | 0.04471 | 0.223899686 | 0.079 | 0 | 1 | 8 |
| sorbs1 | 0.04471 | 0.247215399 | 0.079 | 0 | 1 | 8 |
| tmub1 | 0.04471 | 0.245397568 | 0.079 | 0 | 1 | 8 |
| usp10 | 0.04471 | 0.240783906 | 0.079 | 0 | 1 | 8 |
| aamdc | 0.04471 | 0.2377095 | 0.079 | 0 | 1 | 8 |
| adka | 0.04471 | 0.21369259 | 0.079 | 0 | 1 | 8 |

| | | | | | | |
|---|---|---|---|---|---|---|
| arpc5b | 0.04471 | 0.246429143 | 0.079 | 0 | 1 | 8 |
| capn7 | 0.04471 | 0.287472649 | 0.079 | 0 | 1 | 8 |
| commd4 | 0.04471 | 0.208287616 | 0.079 | 0 | 1 | 8 |
| commd5 | 0.04471 | 0.195450219 | 0.079 | 0 | 1 | 8 |
| ddb2 | 0.04471 | 0.224718201 | 0.079 | 0 | 1 | 8 |
| lrrc15 | 0.04471 | 0.239066689 | 0.079 | 0 | 1 | 8 |
| morc2 | 0.04471 | 0.220012499 | 0.079 | 0 | 1 | 8 |
| ndfip1l | 0.04471 | 0.288786149 | 0.079 | 0 | 1 | 8 |
| ndnf | 0.04471 | 0.239895447 | 0.079 | 0 | 1 | 8 |
| nthl1 | 0.04471 | 0.258873549 | 0.079 | 0 | 1 | 8 |
| si:dkey-72l14.4 | 0.04471 | 0.224016286 | 0.079 | 0 | 1 | 8 |
| vasna | 0.04471 | 0.29830104 | 0.079 | 0 | 1 | 8 |
| zgc:110626 | 0.04471 | 0.257068189 | 0.079 | 0 | 1 | 8 |
| znrf3 | 0.04471 | 0.283226768 | 0.079 | 0 | 1 | 8 |
| mylipb | 0.04471 | 0.267775704 | 0.079 | 0 | 1 | 8 |
| wt1b | 0.04525 | 0.516782958 | 0.263 | 0.067 | 1 | 8 |
| ndufb4 | 0.04558 | 0.462271241 | 0.263 | 0.067 | 1 | 8 |
| gstt1b | 0.04755 | -0.52683428 | 0.079 | 0.267 | 1 | 8 |
| tuba8l4 | 0.04759 | -0.20669147 | 0.263 | 0.267 | 1 | 8 |
| sumo3a | 0.04969 | 0.519031939 | 0.421 | 0.178 | 1 | 8 |
| id1 | 0.05246 | 0.510154983 | 0.579 | 0.333 | 1 | 8 |
| bmper | 0.05307 | 0.434275339 | 0.211 | 0.044 | 1 | 8 |
| ubc | 0.05411 | 0.39353551 | 0.684 | 0.422 | 1 | 8 |
| anxa5b | 0.05475 | 0.538482827 | 0.289 | 0.089 | 1 | 8 |
| gng5 | 0.05479 | 0.449675001 | 0.316 | 0.111 | 1 | 8 |
| yaf2 | 0.05709 | 0.386252976 | 0.132 | 0.022 | 1 | 8 |
| ppt1 | 0.05812 | -0.40481714 | 0.026 | 0.178 | 1 | 8 |
| rps24 | 0.05941 | -0.13250959 | 0.895 | 1 | 1 | 8 |
| cycsb | 0.0595 | 0.39405686 | 0.211 | 0.044 | 1 | 8 |
| si:zfos-1192g2.3 | 0.0599 | 0.402796698 | 0.158 | 0.022 | 1 | 8 |
| minos1 | 0.06049 | 0.358384516 | 0.158 | 0.022 | 1 | 8 |
| prelid3b | 0.06117 | -0.44317026 | 0.026 | 0.178 | 1 | 8 |
| prdx2 | 0.0613 | -0.27267965 | 0.395 | 0.467 | 1 | 8 |
| ctsla | 0.06346 | 0.390418007 | 0.158 | 0.022 | 1 | 8 |
| mapkapk2a | 0.06417 | 0.338644283 | 0.158 | 0.022 | 1 | 8 |
| marcksl1a | 0.06553 | 0.508865464 | 0.474 | 0.244 | 1 | 8 |
| snu13b | 0.06582 | -0.48589395 | 0.105 | 0.311 | 1 | 8 |
| scarb2a | 0.06809 | -0.41538862 | 0.026 | 0.133 | 1 | 8 |
| ranbp1 | 0.06878 | -0.34602166 | 0.184 | 0.267 | 1 | 8 |
| ptges3a | 0.07007 | 0.293087178 | 0.158 | 0.022 | 1 | 8 |
| wdr1 | 0.07255 | 0.403745228 | 0.158 | 0.022 | 1 | 8 |
| acin1a | 0.07299 | 0.439119753 | 0.158 | 0.022 | 1 | 8 |
| ifi45 | 0.07299 | -0.48642452 | 0.079 | 0.267 | 1 | 8 |
| rpl37 | 0.07317 | -0.13778849 | 0.921 | 0.978 | 1 | 8 |
| mtdha | 0.0737 | 0.348463265 | 0.158 | 0.022 | 1 | 8 |
| atp1b1a | 0.07452 | 0.427603974 | 0.158 | 0.022 | 1 | 8 |
| hoxc3a | 0.07545 | 0.377646728 | 0.132 | 0.022 | 1 | 8 |
| zfp36l1a | 0.07635 | -0.50474185 | 0.158 | 0.356 | 1 | 8 |

| | | | | | | |
|---|---|---|---|---|---|---|
| edf1 | 0.07951 | 0.403526022 | 0.421 | 0.2 | 1 | 8 |
| LOC101885938 | 0.07997 | -0.20842343 | 0 | 0.067 | 1 | 8 |
| XLOC-035413 | 0.07997 | -0.19784561 | 0 | 0.067 | 1 | 8 |
| anapc11 | 0.07997 | -0.14891543 | 0 | 0.067 | 1 | 8 |
| atoh8 | 0.07997 | -0.23084698 | 0 | 0.067 | 1 | 8 |
| carhsp1 | 0.07997 | -0.2107906 | 0 | 0.067 | 1 | 8 |
| ccdc137 | 0.07997 | -0.22080969 | 0 | 0.067 | 1 | 8 |
| chaf1a | 0.07997 | -0.1967189 | 0 | 0.067 | 1 | 8 |
| cyfip1 | 0.07997 | -0.20395461 | 0 | 0.067 | 1 | 8 |
| dhfr | 0.07997 | -0.23212504 | 0 | 0.067 | 1 | 8 |
| dhrs3a | 0.07997 | -0.1890613 | 0 | 0.067 | 1 | 8 |
| dhrs7 | 0.07997 | -0.19847822 | 0 | 0.067 | 1 | 8 |
| dnpep | 0.07997 | -0.2355422 | 0 | 0.067 | 1 | 8 |
| dock1 | 0.07997 | -0.25149169 | 0 | 0.067 | 1 | 8 |
| dr1 | 0.07997 | -0.27425386 | 0 | 0.067 | 1 | 8 |
| ebag9 | 0.07997 | -0.20078376 | 0 | 0.067 | 1 | 8 |
| elmo3 | 0.07997 | -0.20731585 | 0 | 0.067 | 1 | 8 |
| exosc6 | 0.07997 | -0.23791199 | 0 | 0.067 | 1 | 8 |
| fntb | 0.07997 | -0.24760433 | 0 | 0.067 | 1 | 8 |
| gnai1 | 0.07997 | -0.23228947 | 0 | 0.067 | 1 | 8 |
| hax1 | 0.07997 | -0.1785934 | 0 | 0.067 | 1 | 8 |
| junbb | 0.07997 | -0.22608825 | 0 | 0.067 | 1 | 8 |
| lgmn | 0.07997 | -0.1845872 | 0 | 0.067 | 1 | 8 |
| mid1ip1l | 0.07997 | -0.15939065 | 0 | 0.067 | 1 | 8 |
| mkrn1 | 0.07997 | -0.23771405 | 0 | 0.067 | 1 | 8 |
| ndufs8a | 0.07997 | -0.20998237 | 0 | 0.067 | 1 | 8 |
| nek7 | 0.07997 | -0.22530311 | 0 | 0.067 | 1 | 8 |
| patl1 | 0.07997 | -0.21927323 | 0 | 0.067 | 1 | 8 |
| pex5la | 0.07997 | -0.27464252 | 0 | 0.067 | 1 | 8 |
| phyhd1 | 0.07997 | -0.21001981 | 0 | 0.067 | 1 | 8 |
| polr2i | 0.07997 | -0.22838011 | 0 | 0.067 | 1 | 8 |
| ppp2r5d | 0.07997 | -0.17860736 | 0 | 0.067 | 1 | 8 |
| psme3 | 0.07997 | -0.19069131 | 0 | 0.067 | 1 | 8 |
| ptena | 0.07997 | -0.20882663 | 0 | 0.067 | 1 | 8 |
| purba | 0.07997 | -0.20418521 | 0 | 0.067 | 1 | 8 |
| ralbb | 0.07997 | -0.23972583 | 0 | 0.067 | 1 | 8 |
| sec13 | 0.07997 | -0.18263495 | 0 | 0.067 | 1 | 8 |
| si:ch211-191j22.3 | 0.07997 | -0.27474285 | 0 | 0.067 | 1 | 8 |
| si:ch211-206k20.5 | 0.07997 | -0.18000621 | 0 | 0.067 | 1 | 8 |
| si:dkey-78l4.14 | 0.07997 | -0.18587593 | 0 | 0.067 | 1 | 8 |
| snx30 | 0.07997 | -0.23037246 | 0 | 0.067 | 1 | 8 |
| spint1a | 0.07997 | -0.19985588 | 0 | 0.067 | 1 | 8 |
| spsb4a | 0.07997 | -0.23366456 | 0 | 0.067 | 1 | 8 |
| ss18 | 0.07997 | -0.24488798 | 0 | 0.067 | 1 | 8 |
| stxbp3 | 0.07997 | -0.21380207 | 0 | 0.067 | 1 | 8 |
| supt6h | 0.07997 | -0.23609912 | 0 | 0.067 | 1 | 8 |
| tbcb | 0.07997 | -0.21827225 | 0 | 0.067 | 1 | 8 |
| tmem59 | 0.07997 | -0.2011163 | 0 | 0.067 | 1 | 8 |

| | | | | | | |
|---|---|---|---|---|---|---|
| twist1a | 0.07997 | -0.16796504 | 0 | 0.067 | 1 | 8 |
| usp5 | 0.07997 | -0.23957578 | 0 | 0.067 | 1 | 8 |
| wdr45b | 0.07997 | -0.17443999 | 0 | 0.067 | 1 | 8 |
| zgc:194469 | 0.07997 | -0.22724589 | 0 | 0.067 | 1 | 8 |
| zgc:194981 | 0.07997 | -0.20685821 | 0 | 0.067 | 1 | 8 |
| zgc:55558 | 0.07997 | -0.21446766 | 0 | 0.067 | 1 | 8 |
| znf622 | 0.07997 | -0.24314953 | 0 | 0.067 | 1 | 8 |
| antxr2a | 0.07997 | -0.20935696 | 0 | 0.067 | 1 | 8 |
| ciapin1 | 0.07997 | -0.20792696 | 0 | 0.067 | 1 | 8 |
| lrrc1 | 0.07997 | -0.21160256 | 0 | 0.067 | 1 | 8 |
| mrpl28 | 0.07997 | -0.18095236 | 0 | 0.067 | 1 | 8 |
| prr13 | 0.07997 | -0.21155721 | 0 | 0.067 | 1 | 8 |
| rcc1 | 0.07997 | -0.21532198 | 0 | 0.067 | 1 | 8 |
| rfc2 | 0.07997 | -0.22410198 | 0 | 0.067 | 1 | 8 |
| rpa2 | 0.07997 | -0.25196324 | 0 | 0.067 | 1 | 8 |
| sf3a3 | 0.07997 | -0.18188774 | 0 | 0.067 | 1 | 8 |
| tgm1l1 | 0.07997 | -0.23348661 | 0 | 0.067 | 1 | 8 |
| ypel5 | 0.07997 | -0.20871374 | 0 | 0.067 | 1 | 8 |
| KIF2A | 0.07997 | -0.18810586 | 0 | 0.067 | 1 | 8 |
| abhd10a | 0.07997 | -0.2244357 | 0 | 0.067 | 1 | 8 |
| cdc40 | 0.07997 | -0.20097783 | 0 | 0.067 | 1 | 8 |
| ctsf | 0.07997 | -0.25488808 | 0 | 0.067 | 1 | 8 |
| dnajc8 | 0.07997 | -0.19776854 | 0 | 0.067 | 1 | 8 |
| emc6 | 0.07997 | -0.16781204 | 0 | 0.067 | 1 | 8 |
| faf2 | 0.07997 | -0.21086418 | 0 | 0.067 | 1 | 8 |
| irf2bpl | 0.07997 | -0.16183641 | 0 | 0.067 | 1 | 8 |
| isy1 | 0.07997 | -0.18133149 | 0 | 0.067 | 1 | 8 |
| lzic | 0.07997 | -0.13843562 | 0 | 0.067 | 1 | 8 |
| osbp | 0.07997 | -0.22489208 | 0 | 0.067 | 1 | 8 |
| pcid2 | 0.07997 | -0.20807038 | 0 | 0.067 | 1 | 8 |
| pdgfaa | 0.07997 | -0.18890723 | 0 | 0.067 | 1 | 8 |
| pik3ip1 | 0.07997 | -0.18912513 | 0 | 0.067 | 1 | 8 |
| polr2k | 0.07997 | -0.16629802 | 0 | 0.067 | 1 | 8 |
| ppan | 0.07997 | -0.22456193 | 0 | 0.067 | 1 | 8 |
| ppp2r3c | 0.07997 | -0.18239239 | 0 | 0.067 | 1 | 8 |
| rab7a | 0.07997 | -0.18890738 | 0 | 0.067 | 1 | 8 |
| rtf1 | 0.07997 | -0.21046817 | 0 | 0.067 | 1 | 8 |
| sephs1 | 0.07997 | -0.24850555 | 0 | 0.067 | 1 | 8 |
| smn1 | 0.07997 | -0.22559486 | 0 | 0.067 | 1 | 8 |
| snap23.1 | 0.07997 | -0.18380611 | 0 | 0.067 | 1 | 8 |
| snrpc | 0.07997 | -0.19481408 | 0 | 0.067 | 1 | 8 |
| socs2 | 0.07997 | -0.21174933 | 0 | 0.067 | 1 | 8 |
| tnrc6c2 | 0.07997 | -0.24855433 | 0 | 0.067 | 1 | 8 |
| trappc2l | 0.07997 | -0.24192855 | 0 | 0.067 | 1 | 8 |
| utrn | 0.07997 | -0.18451782 | 0 | 0.067 | 1 | 8 |
| zgc:193812 | 0.07997 | -0.22360582 | 0 | 0.067 | 1 | 8 |
| znf395a | 0.07997 | -0.16666387 | 0 | 0.067 | 1 | 8 |
| mphosph8 | 0.08045 | 0.451033332 | 0.237 | 0.089 | 1 | 8 |

| | | | | | | |
|---|---|---|---|---|---|---|
| c6 | 0.08106 | 0.515595897 | 0.211 | 0.067 | 1 | 8 |
| pcna | 0.08191 | -0.46905747 | 0.053 | 0.178 | 1 | 8 |
| NC-002333.4 | 0.08621 | 0.623685544 | 0.237 | 0.089 | 1 | 8 |
| rps20 | 0.08679 | -0.13706489 | 0.921 | 1 | 1 | 8 |
| cox7b | 0.08852 | 0.460513759 | 0.395 | 0.2 | 1 | 8 |
| arpc4 | 0.0891 | 0.304709898 | 0.184 | 0.044 | 1 | 8 |
| atp6v0ca | 0.08938 | 0.224061536 | 0.132 | 0.022 | 1 | 8 |
| ndufa4l | 0.08966 | 0.430666657 | 0.474 | 0.244 | 1 | 8 |
| xbp1 | 0.09177 | 0.394339956 | 0.184 | 0.044 | 1 | 8 |
| tax1bp3 | 0.09246 | 0.357051689 | 0.316 | 0.133 | 1 | 8 |
| calm3a | 0.09356 | 0.246583436 | 0.132 | 0.022 | 1 | 8 |
| snrpf | 0.09578 | -0.45654223 | 0.184 | 0.4 | 1 | 8 |
| rps27.1 | 0.09735 | -0.17658013 | 0.868 | 0.978 | 1 | 8 |
| cox5ab | 0.09844 | 0.354948233 | 0.289 | 0.111 | 1 | 8 |
| banf1 | 0.09911 | -0.43131379 | 0.026 | 0.156 | 1 | 8 |
| rbms2b | 0.10133 | 0.39644942 | 0.184 | 0.044 | 1 | 8 |
| aldh2.2 | 0.10164 | 0.417108346 | 0.184 | 0.044 | 1 | 8 |
| ppdpfa | 0.10366 | 0.132118101 | 0.211 | 0.244 | 1 | 8 |
| pfdn1 | 0.10372 | -0.4488329 | 0.105 | 0.289 | 1 | 8 |
| si:ch211-286b5.5 | 0.10551 | 0.35354275 | 0.289 | 0.111 | 1 | 8 |
| rac1a | 0.10602 | 0.368112827 | 0.132 | 0.022 | 1 | 8 |
| hmgb3a | 0.10856 | -0.47452839 | 0.132 | 0.311 | 1 | 8 |
| mibp2 | 0.10929 | 0.118160525 | 0.605 | 0.467 | 1 | 8 |
| eif5b | 0.1094 | 0.30234436 | 0.132 | 0.022 | 1 | 8 |
| atrx | 0.10977 | 0.340432177 | 0.184 | 0.044 | 1 | 8 |
| eif3s6ip | 0.11246 | -0.4276142 | 0.053 | 0.2 | 1 | 8 |
| lasp1 | 0.11293 | 0.423193695 | 0.368 | 0.178 | 1 | 8 |
| serbp1a | 0.11307 | 0.11511555 | 0.632 | 0.489 | 1 | 8 |
| meis1b | 0.11434 | 0.302900882 | 0.211 | 0.067 | 1 | 8 |
| zgc:101851 | 0.11457 | 0.12099471 | 0.053 | 0 | 1 | 8 |
| DDX17 | 0.11457 | 0.152293839 | 0.053 | 0 | 1 | 8 |
| abi1a | 0.11457 | 0.158774363 | 0.053 | 0 | 1 | 8 |
| apol1 | 0.11457 | 0.170462167 | 0.053 | 0 | 1 | 8 |
| arrb2b | 0.11457 | 0.180467994 | 0.053 | 0 | 1 | 8 |
| bet1l | 0.11457 | 0.191795352 | 0.053 | 0 | 1 | 8 |
| ctdsplb | 0.11457 | 0.180335872 | 0.053 | 0 | 1 | 8 |
| dnmbp | 0.11457 | 0.176167718 | 0.053 | 0 | 1 | 8 |
| dync1i2a | 0.11457 | 0.168988835 | 0.053 | 0 | 1 | 8 |
| gemin8 | 0.11457 | 0.158700583 | 0.053 | 0 | 1 | 8 |
| ghrb | 0.11457 | 0.163212415 | 0.053 | 0 | 1 | 8 |
| gpt2l | 0.11457 | 0.15442391 | 0.053 | 0 | 1 | 8 |
| gsk3b | 0.11457 | 0.17140971 | 0.053 | 0 | 1 | 8 |
| gtpbp1l | 0.11457 | 0.152560298 | 0.053 | 0 | 1 | 8 |
| hectd1 | 0.11457 | 0.226525053 | 0.053 | 0 | 1 | 8 |
| id2b | 0.11457 | 0.150220938 | 0.053 | 0 | 1 | 8 |
| kansl1b | 0.11457 | 0.169317241 | 0.053 | 0 | 1 | 8 |
| leo1 | 0.11457 | 0.169008956 | 0.053 | 0 | 1 | 8 |
| matr3l1.1 | 0.11457 | 0.135277425 | 0.053 | 0 | 1 | 8 |

| | | | | | | |
|---|---|---|---|---|---|---|
| mgrn1a | 0.11457 | 0.175677847 | 0.053 | 0 | 1 | 8 |
| pdcd7 | 0.11457 | 0.168504974 | 0.053 | 0 | 1 | 8 |
| plod1a | 0.11457 | 0.155273935 | 0.053 | 0 | 1 | 8 |
| psmd11b | 0.11457 | 0.159722962 | 0.053 | 0 | 1 | 8 |
| ptprda | 0.11457 | 0.173873162 | 0.053 | 0 | 1 | 8 |
| rbms1b | 0.11457 | 0.226916572 | 0.053 | 0 | 1 | 8 |
| rca2.1 | 0.11457 | 0.136582577 | 0.053 | 0 | 1 | 8 |
| sestd1 | 0.11457 | 0.184629514 | 0.053 | 0 | 1 | 8 |
| setd2 | 0.11457 | 0.173930225 | 0.053 | 0 | 1 | 8 |
| sf3b1 | 0.11457 | 0.179444616 | 0.053 | 0 | 1 | 8 |
| taf2 | 0.11457 | 0.173797661 | 0.053 | 0 | 1 | 8 |
| taf3 | 0.11457 | 0.229097593 | 0.053 | 0 | 1 | 8 |
| tgm2b | 0.11457 | 0.161763636 | 0.053 | 0 | 1 | 8 |
| tmx2b | 0.11457 | 0.12123717 | 0.053 | 0 | 1 | 8 |
| yeats4 | 0.11457 | 0.159134853 | 0.053 | 0 | 1 | 8 |
| zgc:110843 | 0.11457 | 0.176787925 | 0.053 | 0 | 1 | 8 |
| zgc:153631 | 0.11457 | 0.169962036 | 0.053 | 0 | 1 | 8 |
| BX005183.3 | 0.11457 | 0.139559641 | 0.053 | 0 | 1 | 8 |
| CABZ01068251.1 | 0.11457 | 0.130157907 | 0.053 | 0 | 1 | 8 |
| CABZ01084963.1 | 0.11457 | 0.155360469 | 0.053 | 0 | 1 | 8 |
| CABZ01110881.1 | 0.11457 | 0.191561798 | 0.053 | 0 | 1 | 8 |
| CR376737.1 | 0.11457 | 0.172627044 | 0.053 | 0 | 1 | 8 |
| CR847566.1 | 0.11457 | 0.211110631 | 0.053 | 0 | 1 | 8 |
| CU929259.1 | 0.11457 | 0.162518406 | 0.053 | 0 | 1 | 8 |
| EARS2 | 0.11457 | 0.18571432 | 0.053 | 0 | 1 | 8 |
| LOC796453 | 0.11457 | 0.156723627 | 0.053 | 0 | 1 | 8 |
| XLOC-005199 | 0.11457 | 0.19388145 | 0.053 | 0 | 1 | 8 |
| XLOC-016094 | 0.11457 | 0.231744828 | 0.053 | 0 | 1 | 8 |
| XLOC-034437 | 0.11457 | 0.188273861 | 0.053 | 0 | 1 | 8 |
| XLOC-037780 | 0.11457 | 0.214143731 | 0.053 | 0 | 1 | 8 |
| acsf2 | 0.11457 | 0.163989157 | 0.053 | 0 | 1 | 8 |
| adamts10 | 0.11457 | 0.209921164 | 0.053 | 0 | 1 | 8 |
| aebp1 | 0.11457 | 0.18417531 | 0.053 | 0 | 1 | 8 |
| aif1l | 0.11457 | 0.147181351 | 0.053 | 0 | 1 | 8 |
| aldocb | 0.11457 | 0.149262159 | 0.053 | 0 | 1 | 8 |
| alyref | 0.11457 | 0.155758775 | 0.053 | 0 | 1 | 8 |
| anapc15 | 0.11457 | 0.172133925 | 0.053 | 0 | 1 | 8 |
| ap1s3b | 0.11457 | 0.157886651 | 0.053 | 0 | 1 | 8 |
| appb | 0.11457 | 0.185326836 | 0.053 | 0 | 1 | 8 |
| asb12a | 0.11457 | 0.141193139 | 0.053 | 0 | 1 | 8 |
| atg3 | 0.11457 | 0.15376085 | 0.053 | 0 | 1 | 8 |
| bambib | 0.11457 | 0.169115822 | 0.053 | 0 | 1 | 8 |
| baz1b | 0.11457 | 0.228862998 | 0.053 | 0 | 1 | 8 |
| bckdhb | 0.11457 | 0.147675202 | 0.053 | 0 | 1 | 8 |
| bin1a | 0.11457 | 0.146694631 | 0.053 | 0 | 1 | 8 |
| bms1 | 0.11457 | 0.188009494 | 0.053 | 0 | 1 | 8 |
| brd3b | 0.11457 | 0.197157959 | 0.053 | 0 | 1 | 8 |
| btr04 | 0.11457 | 0.152496066 | 0.053 | 0 | 1 | 8 |

| | | | | | | |
|---|---|---|---|---|---|---|
| c1s | 0.11457 | 0.168377944 | 0.053 | 0 | 1 | 8 |
| caap1 | 0.11457 | 0.158317067 | 0.053 | 0 | 1 | 8 |
| cald1b | 0.11457 | 0.186696196 | 0.053 | 0 | 1 | 8 |
| cd9b | 0.11457 | 0.199032534 | 0.053 | 0 | 1 | 8 |
| cdk5rap3 | 0.11457 | 0.157539815 | 0.053 | 0 | 1 | 8 |
| cdkn1d | 0.11457 | 0.173001872 | 0.053 | 0 | 1 | 8 |
| clns1a | 0.11457 | 0.147257466 | 0.053 | 0 | 1 | 8 |
| cndp2 | 0.11457 | 0.169519147 | 0.053 | 0 | 1 | 8 |
| col12a1a | 0.11457 | 0.183406736 | 0.053 | 0 | 1 | 8 |
| col18a1a | 0.11457 | 0.1489923 | 0.053 | 0 | 1 | 8 |
| cpb1 | 0.11457 | 0.184441542 | 0.053 | 0 | 1 | 8 |
| crfb4 | 0.11457 | 0.184780664 | 0.053 | 0 | 1 | 8 |
| ctsz | 0.11457 | 0.17599853 | 0.053 | 0 | 1 | 8 |
| cygb2 | 0.11457 | 0.135790086 | 0.053 | 0 | 1 | 8 |
| cyp2p6 | 0.11457 | 0.155611077 | 0.053 | 0 | 1 | 8 |
| dap | 0.11457 | 0.121515813 | 0.053 | 0 | 1 | 8 |
| dcun1d5 | 0.11457 | 0.174135258 | 0.053 | 0 | 1 | 8 |
| dda1 | 0.11457 | 0.138508418 | 0.053 | 0 | 1 | 8 |
| dixdc1a | 0.11457 | 0.136852734 | 0.053 | 0 | 1 | 8 |
| dlat | 0.11457 | 0.155299762 | 0.053 | 0 | 1 | 8 |
| edil3a | 0.11457 | 0.182277573 | 0.053 | 0 | 1 | 8 |
| efnb2a | 0.11457 | 0.187917012 | 0.053 | 0 | 1 | 8 |
| ehbp1l1a | 0.11457 | 0.221537333 | 0.053 | 0 | 1 | 8 |
| emc7 | 0.11457 | 0.174593722 | 0.053 | 0 | 1 | 8 |
| enoph1 | 0.11457 | 0.146886457 | 0.053 | 0 | 1 | 8 |
| enpp5 | 0.11457 | 0.19176296 | 0.053 | 0 | 1 | 8 |
| entr1 | 0.11457 | 0.158430673 | 0.053 | 0 | 1 | 8 |
| eny2 | 0.11457 | 0.181258964 | 0.053 | 0 | 1 | 8 |
| erbin | 0.11457 | 0.184371739 | 0.053 | 0 | 1 | 8 |
| errfi1a | 0.11457 | 0.146939008 | 0.053 | 0 | 1 | 8 |
| fbrs | 0.11457 | 0.141353217 | 0.053 | 0 | 1 | 8 |
| fhl1a | 0.11457 | 0.137935789 | 0.053 | 0 | 1 | 8 |
| fkbp2 | 0.11457 | 0.144663988 | 0.053 | 0 | 1 | 8 |
| fnta | 0.11457 | 0.136879469 | 0.053 | 0 | 1 | 8 |
| fosb | 0.11457 | 0.214097988 | 0.053 | 0 | 1 | 8 |
| gadd45aa | 0.11457 | 0.125936753 | 0.053 | 0 | 1 | 8 |
| galnt1 | 0.11457 | 0.1943771 | 0.053 | 0 | 1 | 8 |
| gclm | 0.11457 | 0.175371287 | 0.053 | 0 | 1 | 8 |
| gemin2 | 0.11457 | 0.20126878 | 0.053 | 0 | 1 | 8 |
| glis2a | 0.11457 | 0.195440941 | 0.053 | 0 | 1 | 8 |
| grpel2 | 0.11457 | 0.158174855 | 0.053 | 0 | 1 | 8 |
| gsr | 0.11457 | 0.13464824 | 0.053 | 0 | 1 | 8 |
| herpud2 | 0.11457 | 0.148288454 | 0.053 | 0 | 1 | 8 |
| hsd11b1la | 0.11457 | 0.188033719 | 0.053 | 0 | 1 | 8 |
| hspa4b | 0.11457 | 0.151484554 | 0.053 | 0 | 1 | 8 |
| ifngr2 | 0.11457 | 0.164588742 | 0.053 | 0 | 1 | 8 |
| jpt2 | 0.11457 | 0.14413357 | 0.053 | 0 | 1 | 8 |
| kat7b | 0.11457 | 0.198472783 | 0.053 | 0 | 1 | 8 |

| | | | | | | |
|---|---|---|---|---|---|---|
| kdm3b | 0.11457 | 0.129114681 | 0.053 | 0 | 1 | 8 |
| kdm4b | 0.11457 | 0.153922858 | 0.053 | 0 | 1 | 8 |
| kmt2ba | 0.11457 | 0.215455935 | 0.053 | 0 | 1 | 8 |
| lama4 | 0.11457 | 0.14754074 | 0.053 | 0 | 1 | 8 |
| lamtor2 | 0.11457 | 0.182478066 | 0.053 | 0 | 1 | 8 |
| larp1 | 0.11457 | 0.160663022 | 0.053 | 0 | 1 | 8 |
| limch1b | 0.11457 | 0.142175797 | 0.053 | 0 | 1 | 8 |
| llgl1 | 0.11457 | 0.157710773 | 0.053 | 0 | 1 | 8 |
| lonrf1l | 0.11457 | 0.188326219 | 0.053 | 0 | 1 | 8 |
| lpgat1 | 0.11457 | 0.209724056 | 0.053 | 0 | 1 | 8 |
| lto1 | 0.11457 | 0.141464532 | 0.053 | 0 | 1 | 8 |
| mast2 | 0.11457 | 0.138396531 | 0.053 | 0 | 1 | 8 |
| mbd6 | 0.11457 | 0.180022125 | 0.053 | 0 | 1 | 8 |
| mbip | 0.11457 | 0.163468626 | 0.053 | 0 | 1 | 8 |
| mical2a | 0.11457 | 0.214775555 | 0.053 | 0 | 1 | 8 |
| mob4 | 0.11457 | 0.17179072 | 0.053 | 0 | 1 | 8 |
| mrpl15 | 0.11457 | 0.143245234 | 0.053 | 0 | 1 | 8 |
| mrpl9 | 0.11457 | 0.130615291 | 0.053 | 0 | 1 | 8 |
| mrps30 | 0.11457 | 0.202488812 | 0.053 | 0 | 1 | 8 |
| mtx2 | 0.11457 | 0.128002026 | 0.053 | 0 | 1 | 8 |
| mycn | 0.11457 | 0.170838968 | 0.053 | 0 | 1 | 8 |
| myd88 | 0.11457 | 0.207204509 | 0.053 | 0 | 1 | 8 |
| myh11a | 0.11457 | 0.188368727 | 0.053 | 0 | 1 | 8 |
| naprt | 0.11457 | 0.165461074 | 0.053 | 0 | 1 | 8 |
| nenf | 0.11457 | 0.154694653 | 0.053 | 0 | 1 | 8 |
| nfyal | 0.11457 | 0.143948854 | 0.053 | 0 | 1 | 8 |
| nkain1 | 0.11457 | 0.173248481 | 0.053 | 0 | 1 | 8 |
| nme4 | 0.11457 | 0.183009656 | 0.053 | 0 | 1 | 8 |
| nr2f6a | 0.11457 | 0.173727196 | 0.053 | 0 | 1 | 8 |
| nt5dc2 | 0.11457 | 0.18609107 | 0.053 | 0 | 1 | 8 |
| ogfod2 | 0.11457 | 0.177562048 | 0.053 | 0 | 1 | 8 |
| paip1 | 0.11457 | 0.164484948 | 0.053 | 0 | 1 | 8 |
| paqr6 | 0.11457 | 0.223668919 | 0.053 | 0 | 1 | 8 |
| pdap1a | 0.11457 | 0.118409132 | 0.053 | 0 | 1 | 8 |
| per1a | 0.11457 | 0.225812531 | 0.053 | 0 | 1 | 8 |
| pet100 | 0.11457 | 0.158152052 | 0.053 | 0 | 1 | 8 |
| pex26 | 0.11457 | 0.177179508 | 0.053 | 0 | 1 | 8 |
| pgd | 0.11457 | 0.165584997 | 0.053 | 0 | 1 | 8 |
| phf3 | 0.11457 | 0.197587689 | 0.053 | 0 | 1 | 8 |
| phlda3 | 0.11457 | 0.167817661 | 0.053 | 0 | 1 | 8 |
| pithd1 | 0.11457 | 0.159857504 | 0.053 | 0 | 1 | 8 |
| plcd3a | 0.11457 | 0.145992632 | 0.053 | 0 | 1 | 8 |
| ppa2 | 0.11457 | 0.140410348 | 0.053 | 0 | 1 | 8 |
| ppp1r13l | 0.11457 | 0.197575362 | 0.053 | 0 | 1 | 8 |
| ppp3r1b | 0.11457 | 0.168760596 | 0.053 | 0 | 1 | 8 |
| ppp4ca | 0.11457 | 0.144927148 | 0.053 | 0 | 1 | 8 |
| prickle1b | 0.11457 | 0.202961827 | 0.053 | 0 | 1 | 8 |
| prpf31 | 0.11457 | 0.115686242 | 0.053 | 0 | 1 | 8 |

| | | | | | | |
|---|---|---|---|---|---|---|
| psenen | 0.11457 | 0.163085344 | 0.053 | 0 | 1 | 8 |
| rab32a | 0.11457 | 0.147655139 | 0.053 | 0 | 1 | 8 |
| rab3il1 | 0.11457 | 0.174393655 | 0.053 | 0 | 1 | 8 |
| rab8b | 0.11457 | 0.224013931 | 0.053 | 0 | 1 | 8 |
| rae1 | 0.11457 | 0.178372066 | 0.053 | 0 | 1 | 8 |
| rap1gap | 0.11457 | 0.15197895 | 0.053 | 0 | 1 | 8 |
| raver2 | 0.11457 | 0.222190204 | 0.053 | 0 | 1 | 8 |
| rereb | 0.11457 | 0.160102247 | 0.053 | 0 | 1 | 8 |
| rhbdf1a | 0.11457 | 0.192088451 | 0.053 | 0 | 1 | 8 |
| rhoua | 0.11457 | 0.166257219 | 0.053 | 0 | 1 | 8 |
| rnf25 | 0.11457 | 0.126198706 | 0.053 | 0 | 1 | 8 |
| rprd2b | 0.11457 | 0.153645968 | 0.053 | 0 | 1 | 8 |
| rraga | 0.11457 | 0.202158205 | 0.053 | 0 | 1 | 8 |
| s100v1 | 0.11457 | 0.168768739 | 0.053 | 0 | 1 | 8 |
| sars | 0.11457 | 0.12938577 | 0.053 | 0 | 1 | 8 |
| sbf1 | 0.11457 | 0.17843876 | 0.053 | 0 | 1 | 8 |
| sec22ba | 0.11457 | 0.130298924 | 0.053 | 0 | 1 | 8 |
| sec24b | 0.11457 | 0.177740327 | 0.053 | 0 | 1 | 8 |
| selenot2 | 0.11457 | 0.188086004 | 0.053 | 0 | 1 | 8 |
| serpina1l | 0.11457 | 0.152971205 | 0.053 | 0 | 1 | 8 |
| sh2d5 | 0.11457 | 0.168075052 | 0.053 | 0 | 1 | 8 |
| sh3gl3a | 0.11457 | 0.166134424 | 0.053 | 0 | 1 | 8 |
| si:ch211-195h23.3 | 0.11457 | 0.134764113 | 0.053 | 0 | 1 | 8 |
| si:ch211-198a12.6 | 0.11457 | 0.182032701 | 0.053 | 0 | 1 | 8 |
| si:ch211-262e15.1 | 0.11457 | 0.148739238 | 0.053 | 0 | 1 | 8 |
| si:ch211-59p23.1 | 0.11457 | 0.146393871 | 0.053 | 0 | 1 | 8 |
| si:ch211-87m7.2 | 0.11457 | 0.202543323 | 0.053 | 0 | 1 | 8 |
| si:dkey-192l18.9 | 0.11457 | 0.18925383 | 0.053 | 0 | 1 | 8 |
| si:dkey-24l11.2 | 0.11457 | 0.119079814 | 0.053 | 0 | 1 | 8 |
| si:dkeyp-27c8.2 | 0.11457 | 0.123237322 | 0.053 | 0 | 1 | 8 |
| sinhcafl | 0.11457 | 0.134287608 | 0.053 | 0 | 1 | 8 |
| slc13a2 | 0.11457 | 0.147869034 | 0.053 | 0 | 1 | 8 |
| slc25a36b | 0.11457 | 0.202844977 | 0.053 | 0 | 1 | 8 |
| slit3 | 0.11457 | 0.189072162 | 0.053 | 0 | 1 | 8 |
| smim4 | 0.11457 | 0.196728827 | 0.053 | 0 | 1 | 8 |
| snx11 | 0.11457 | 0.123603038 | 0.053 | 0 | 1 | 8 |
| sox6 | 0.11457 | 0.179452544 | 0.053 | 0 | 1 | 8 |
| srsf5b | 0.11457 | 0.153421912 | 0.053 | 0 | 1 | 8 |
| stat5a | 0.11457 | 0.18792301 | 0.053 | 0 | 1 | 8 |
| suclg2 | 0.11457 | 0.127710024 | 0.053 | 0 | 1 | 8 |
| svild | 0.11457 | 0.157194459 | 0.053 | 0 | 1 | 8 |
| tax1bp1b | 0.11457 | 0.156028084 | 0.053 | 0 | 1 | 8 |
| thap7 | 0.11457 | 0.173992566 | 0.053 | 0 | 1 | 8 |
| tmem231 | 0.11457 | 0.197098102 | 0.053 | 0 | 1 | 8 |
| tp63 | 0.11457 | 0.166165371 | 0.053 | 0 | 1 | 8 |
| tspan5a | 0.11457 | 0.185069361 | 0.053 | 0 | 1 | 8 |
| tspan7 | 0.11457 | 0.170455287 | 0.053 | 0 | 1 | 8 |
| ubald1a | 0.11457 | 0.167920655 | 0.053 | 0 | 1 | 8 |

| | | | | | | |
|---|---|---|---|---|---|---|
| ube2r2 | 0.11457 | 0.187066344 | 0.053 | 0 | 1 | 8 |
| ubl7b | 0.11457 | 0.171988966 | 0.053 | 0 | 1 | 8 |
| uqcrc2a | 0.11457 | 0.140808294 | 0.053 | 0 | 1 | 8 |
| usf1l | 0.11457 | 0.137769271 | 0.053 | 0 | 1 | 8 |
| washc2c | 0.11457 | 0.170070879 | 0.053 | 0 | 1 | 8 |
| wdr26b | 0.11457 | 0.176083739 | 0.053 | 0 | 1 | 8 |
| wdr36 | 0.11457 | 0.198482048 | 0.053 | 0 | 1 | 8 |
| wdr45 | 0.11457 | 0.167410431 | 0.053 | 0 | 1 | 8 |
| wdr73 | 0.11457 | 0.170939155 | 0.053 | 0 | 1 | 8 |
| wdr83 | 0.11457 | 0.153051024 | 0.053 | 0 | 1 | 8 |
| wtap | 0.11457 | 0.167468228 | 0.053 | 0 | 1 | 8 |
| wu:fj16a03 | 0.11457 | 0.10042527 | 0.053 | 0 | 1 | 8 |
| zcrb1 | 0.11457 | 0.127452197 | 0.053 | 0 | 1 | 8 |
| zdhhc18b | 0.11457 | 0.173769519 | 0.053 | 0 | 1 | 8 |
| zfand2a | 0.11457 | 0.154432752 | 0.053 | 0 | 1 | 8 |
| zgc:136439 | 0.11457 | 0.165570185 | 0.053 | 0 | 1 | 8 |
| zgc:136493 | 0.11457 | 0.148720268 | 0.053 | 0 | 1 | 8 |
| zgc:158846 | 0.11457 | 0.142136516 | 0.053 | 0 | 1 | 8 |
| zgc:165573 | 0.11457 | 0.137692608 | 0.053 | 0 | 1 | 8 |
| zgc:56095 | 0.11457 | 0.17752071 | 0.053 | 0 | 1 | 8 |
| zgc:63831 | 0.11457 | 0.16764457 | 0.053 | 0 | 1 | 8 |
| zmynd19 | 0.11457 | 0.150173493 | 0.053 | 0 | 1 | 8 |
| znf576.2 | 0.11457 | 0.152444106 | 0.053 | 0 | 1 | 8 |
| zswim5 | 0.11457 | 0.202528323 | 0.053 | 0 | 1 | 8 |
| commd1 | 0.11457 | 0.193577358 | 0.053 | 0 | 1 | 8 |
| mak16 | 0.11457 | 0.153907544 | 0.053 | 0 | 1 | 8 |
| rras2 | 0.11457 | 0.194225065 | 0.053 | 0 | 1 | 8 |
| si:ch1073-143l10.2 | 0.11457 | 0.135546021 | 0.053 | 0 | 1 | 8 |
| si:ch211-51h9.7 | 0.11457 | 0.192587268 | 0.053 | 0 | 1 | 8 |
| si:dkey-69o16.5 | 0.11457 | 0.137918452 | 0.053 | 0 | 1 | 8 |
| sulf1 | 0.11457 | 0.195718994 | 0.053 | 0 | 1 | 8 |
| BX005254.1 | 0.11457 | 0.148470183 | 0.053 | 0 | 1 | 8 |
| BX901920.1 | 0.11457 | 0.23539437 | 0.053 | 0 | 1 | 8 |
| MMP23B | 0.11457 | 0.167212802 | 0.053 | 0 | 1 | 8 |
| PLEKHH3 | 0.11457 | 0.170955059 | 0.053 | 0 | 1 | 8 |
| actr6 | 0.11457 | 0.177822845 | 0.053 | 0 | 1 | 8 |
| akirin1 | 0.11457 | 0.179342029 | 0.053 | 0 | 1 | 8 |
| ankrd54 | 0.11457 | 0.220171521 | 0.053 | 0 | 1 | 8 |
| ap3s2 | 0.11457 | 0.158857739 | 0.053 | 0 | 1 | 8 |
| arfgap2 | 0.11457 | 0.157467778 | 0.053 | 0 | 1 | 8 |
| arglu1b | 0.11457 | 0.159031573 | 0.053 | 0 | 1 | 8 |
| casc3 | 0.11457 | 0.15055467 | 0.053 | 0 | 1 | 8 |
| ccdc102a | 0.11457 | 0.190365841 | 0.053 | 0 | 1 | 8 |
| ccdc43 | 0.11457 | 0.143002266 | 0.053 | 0 | 1 | 8 |
| cdk12 | 0.11457 | 0.179574358 | 0.053 | 0 | 1 | 8 |
| cdk2ap2 | 0.11457 | 0.118315002 | 0.053 | 0 | 1 | 8 |
| cdk9 | 0.11457 | 0.129631981 | 0.053 | 0 | 1 | 8 |
| chic2 | 0.11457 | 0.156667816 | 0.053 | 0 | 1 | 8 |

| | | | | | | |
|---|---|---|---|---|---|---|
| chmp3 | 0.11457 | 0.183440373 | 0.053 | 0 | 1 | 8 |
| cnot4a | 0.11457 | 0.149820565 | 0.053 | 0 | 1 | 8 |
| cox4i2 | 0.11457 | 0.177414529 | 0.053 | 0 | 1 | 8 |
| cpa4 | 0.11457 | 0.14530226 | 0.053 | 0 | 1 | 8 |
| cpne3 | 0.11457 | 0.162041071 | 0.053 | 0 | 1 | 8 |
| dapk1 | 0.11457 | 0.16705924 | 0.053 | 0 | 1 | 8 |
| dctd | 0.11457 | 0.20019711 | 0.053 | 0 | 1 | 8 |
| dhrs1 | 0.11457 | 0.183551714 | 0.053 | 0 | 1 | 8 |
| dok4 | 0.11457 | 0.156781055 | 0.053 | 0 | 1 | 8 |
| dph5 | 0.11457 | 0.158599965 | 0.053 | 0 | 1 | 8 |
| dsc2l | 0.11457 | 0.170764176 | 0.053 | 0 | 1 | 8 |
| eif4ba | 0.11457 | 0.148834601 | 0.053 | 0 | 1 | 8 |
| eri1 | 0.11457 | 0.157242016 | 0.053 | 0 | 1 | 8 |
| faf1 | 0.11457 | 0.157143601 | 0.053 | 0 | 1 | 8 |
| fdx1 | 0.11457 | 0.200594053 | 0.053 | 0 | 1 | 8 |
| fkbp10b | 0.11457 | 0.151392265 | 0.053 | 0 | 1 | 8 |
| foxp1b | 0.11457 | 0.20018736 | 0.053 | 0 | 1 | 8 |
| gale | 0.11457 | 0.178553386 | 0.053 | 0 | 1 | 8 |
| gart | 0.11457 | 0.158787852 | 0.053 | 0 | 1 | 8 |
| gmps | 0.11457 | 0.170995091 | 0.053 | 0 | 1 | 8 |
| gnai2a | 0.11457 | 0.17291886 | 0.053 | 0 | 1 | 8 |
| gorasp2 | 0.11457 | 0.169228772 | 0.053 | 0 | 1 | 8 |
| gpatch1 | 0.11457 | 0.149675204 | 0.053 | 0 | 1 | 8 |
| her6 | 0.11457 | 0.177483664 | 0.053 | 0 | 1 | 8 |
| hif1al | 0.11457 | 0.135890792 | 0.053 | 0 | 1 | 8 |
| hmg20b | 0.11457 | 0.130029552 | 0.053 | 0 | 1 | 8 |
| hoxb5b | 0.11457 | 0.166655097 | 0.053 | 0 | 1 | 8 |
| hoxb8a | 0.11457 | 0.178691862 | 0.053 | 0 | 1 | 8 |
| hprt1 | 0.11457 | 0.157251945 | 0.053 | 0 | 1 | 8 |
| ift52 | 0.11457 | 0.14785064 | 0.053 | 0 | 1 | 8 |
| igfbp1a | 0.11457 | 0.178714238 | 0.053 | 0 | 1 | 8 |
| itga6b | 0.11457 | 0.15520759 | 0.053 | 0 | 1 | 8 |
| kdm7aa | 0.11457 | 0.189146215 | 0.053 | 0 | 1 | 8 |
| laptm4a | 0.11457 | 0.142457584 | 0.053 | 0 | 1 | 8 |
| maip1 | 0.11457 | 0.133947871 | 0.053 | 0 | 1 | 8 |
| mks1 | 0.11457 | 0.135072194 | 0.053 | 0 | 1 | 8 |
| mmab | 0.11457 | 0.198464163 | 0.053 | 0 | 1 | 8 |
| mov10a | 0.11457 | 0.179315704 | 0.053 | 0 | 1 | 8 |
| mrpl4 | 0.11457 | 0.15739761 | 0.053 | 0 | 1 | 8 |
| mthfd1b | 0.11457 | 0.168046123 | 0.053 | 0 | 1 | 8 |
| mthfd2 | 0.11457 | 0.153355451 | 0.053 | 0 | 1 | 8 |
| mycbp2 | 0.11457 | 0.186519732 | 0.053 | 0 | 1 | 8 |
| nol6 | 0.11457 | 0.170350523 | 0.053 | 0 | 1 | 8 |
| nosip | 0.11457 | 0.170134685 | 0.053 | 0 | 1 | 8 |
| nr0b2a | 0.11457 | 0.185416964 | 0.053 | 0 | 1 | 8 |
| nudt17 | 0.11457 | 0.158578174 | 0.053 | 0 | 1 | 8 |
| pdcl | 0.11457 | 0.139068321 | 0.053 | 0 | 1 | 8 |
| pdzd11 | 0.11457 | 0.146840472 | 0.053 | 0 | 1 | 8 |

| | | | | | | |
|---|---|---|---|---|---|---|
| pi4k2a | 0.11457 | 0.189772724 | 0.053 | 0 | 1 | 8 |
| plp2 | 0.11457 | 0.210084702 | 0.053 | 0 | 1 | 8 |
| polr3k | 0.11457 | 0.134805518 | 0.053 | 0 | 1 | 8 |
| qser1 | 0.11457 | 0.139437734 | 0.053 | 0 | 1 | 8 |
| ralba | 0.11457 | 0.183720429 | 0.053 | 0 | 1 | 8 |
| rcan2 | 0.11457 | 0.169258093 | 0.053 | 0 | 1 | 8 |
| rergla | 0.11457 | 0.173934921 | 0.053 | 0 | 1 | 8 |
| rnf115 | 0.11457 | 0.15093767 | 0.053 | 0 | 1 | 8 |
| rpz | 0.11457 | 0.184648622 | 0.053 | 0 | 1 | 8 |
| rragcb | 0.11457 | 0.154762523 | 0.053 | 0 | 1 | 8 |
| scp2b | 0.11457 | 0.160954331 | 0.053 | 0 | 1 | 8 |
| si:ch211-110e21.3 | 0.11457 | 0.137037071 | 0.053 | 0 | 1 | 8 |
| si:ch211-89o9.6 | 0.11457 | 0.170760211 | 0.053 | 0 | 1 | 8 |
| si:dkey-17m8.1 | 0.11457 | 0.165856565 | 0.053 | 0 | 1 | 8 |
| slc11a2 | 0.11457 | 0.125760266 | 0.053 | 0 | 1 | 8 |
| slc25a1a | 0.11457 | 0.129824473 | 0.053 | 0 | 1 | 8 |
| snx7 | 0.11457 | 0.187684826 | 0.053 | 0 | 1 | 8 |
| ssbp3b | 0.11457 | 0.165123168 | 0.053 | 0 | 1 | 8 |
| sucla2 | 0.11457 | 0.149214323 | 0.053 | 0 | 1 | 8 |
| suds3 | 0.11457 | 0.164500466 | 0.053 | 0 | 1 | 8 |
| tmed9 | 0.11457 | 0.192490594 | 0.053 | 0 | 1 | 8 |
| traf6 | 0.11457 | 0.201830783 | 0.053 | 0 | 1 | 8 |
| ttpal | 0.11457 | 0.147695769 | 0.053 | 0 | 1 | 8 |
| txndc15 | 0.11457 | 0.166032219 | 0.053 | 0 | 1 | 8 |
| ubl3a | 0.11457 | 0.140661175 | 0.053 | 0 | 1 | 8 |
| ulk2 | 0.11457 | 0.149504884 | 0.053 | 0 | 1 | 8 |
| vps28 | 0.11457 | 0.182346693 | 0.053 | 0 | 1 | 8 |
| wu:fi36c02 | 0.11457 | 0.217768986 | 0.053 | 0 | 1 | 8 |
| zeb2a | 0.11457 | 0.170672123 | 0.053 | 0 | 1 | 8 |
| zgc:112970 | 0.11457 | 0.163890325 | 0.053 | 0 | 1 | 8 |
| zgc:114119 | 0.11457 | 0.150639734 | 0.053 | 0 | 1 | 8 |
| zgc:77739 | 0.11457 | 0.174498114 | 0.053 | 0 | 1 | 8 |
| zmp:0000001316 | 0.11457 | 0.190327973 | 0.053 | 0 | 1 | 8 |
| znf292b | 0.11457 | 0.213358051 | 0.053 | 0 | 1 | 8 |
| PDPR | 0.11457 | 0.182981831 | 0.053 | 0 | 1 | 8 |
| agpat5 | 0.11457 | 0.150122141 | 0.053 | 0 | 1 | 8 |
| ctrl | 0.11457 | 0.161794982 | 0.053 | 0 | 1 | 8 |
| mars | 0.11457 | 0.200607373 | 0.053 | 0 | 1 | 8 |
| ncln | 0.11457 | 0.143612177 | 0.053 | 0 | 1 | 8 |
| ube4b | 0.11457 | 0.148386938 | 0.053 | 0 | 1 | 8 |
| zgc:92606 | 0.11457 | 0.141552325 | 0.053 | 0 | 1 | 8 |
| lyrm4 | 0.11478 | 0.366493987 | 0.184 | 0.044 | 1 | 8 |
| ldhba | 0.11645 | -0.39916608 | 0.079 | 0.244 | 1 | 8 |
| fam168a | 0.11657 | 0.308566183 | 0.132 | 0.022 | 1 | 8 |
| si:ch211-213d14.2 | 0.11695 | -0.44110731 | 0.158 | 0.356 | 1 | 8 |
| timm9 | 0.11734 | 0.345609441 | 0.184 | 0.044 | 1 | 8 |
| pabpc1b | 0.11825 | 0.377274871 | 0.184 | 0.044 | 1 | 8 |
| rpl11 | 0.11842 | -0.1177063 | 0.921 | 1 | 1 | 8 |

| | | | | | | |
|---|---|---|---|---|---|---|
| rpl7a | 0.11893 | -0.16592721 | 0.868 | 0.978 | 1 | 8 |
| zgc:113531 | 0.11928 | 0.371499531 | 0.132 | 0.022 | 1 | 8 |
| zc3h7bb | 0.12062 | 0.310998319 | 0.132 | 0.022 | 1 | 8 |
| med28 | 0.12146 | 0.343495886 | 0.132 | 0.022 | 1 | 8 |
| tdh | 0.12272 | 0.264007642 | 0.132 | 0.022 | 1 | 8 |
| ctbp2a | 0.1229 | 0.348281558 | 0.132 | 0.022 | 1 | 8 |
| eif3k | 0.12292 | 0.434941113 | 0.342 | 0.156 | 1 | 8 |
| atp6v1e1a | 0.12552 | 0.333228324 | 0.132 | 0.022 | 1 | 8 |
| crtap | 0.12923 | -0.28094482 | 0.026 | 0.133 | 1 | 8 |
| sem1 | 0.12971 | 0.424270075 | 0.395 | 0.2 | 1 | 8 |
| paplna | 0.12979 | 0.381841893 | 0.132 | 0.022 | 1 | 8 |
| aoc2 | 0.13088 | 0.317692449 | 0.105 | 0.022 | 1 | 8 |
| cirbpa | 0.1316 | -0.20502594 | 0.632 | 0.689 | 1 | 8 |
| rpl15 | 0.13362 | -0.13097041 | 0.947 | 0.956 | 1 | 8 |
| arpc1b | 0.13383 | -0.41962211 | 0.079 | 0.222 | 1 | 8 |
| rpl36a | 0.13524 | -0.11034573 | 0.974 | 1 | 1 | 8 |
| CABZ01112656.1 | 0.13568 | 0.284957703 | 0.132 | 0.022 | 1 | 8 |
| npm1a | 0.13691 | 0.396983122 | 0.211 | 0.089 | 1 | 8 |
| spred2b | 0.13713 | 0.365702455 | 0.132 | 0.022 | 1 | 8 |
| rhoab | 0.13934 | -0.17884362 | 0.053 | 0.156 | 1 | 8 |
| ube2e3 | 0.13959 | 0.418841389 | 0.342 | 0.156 | 1 | 8 |
| hm13 | 0.14126 | 0.310398925 | 0.105 | 0.022 | 1 | 8 |
| sdha | 0.14182 | 0.19145214 | 0.105 | 0.022 | 1 | 8 |
| rpl10a | 0.14208 | -0.14561608 | 0.868 | 0.978 | 1 | 8 |
| strap | 0.1423 | 0.291787707 | 0.132 | 0.022 | 1 | 8 |
| si:ch211-105c13.3 | 0.14506 | 0.229605807 | 0.368 | 0.222 | 1 | 8 |
| akap8l | 0.14543 | 0.327620858 | 0.132 | 0.022 | 1 | 8 |
| tefa | 0.14619 | 0.325188185 | 0.132 | 0.022 | 1 | 8 |
| fn1b | 0.14766 | 0.435868449 | 0.395 | 0.2 | 1 | 8 |
| uba1 | 0.14812 | 0.33189764 | 0.132 | 0.022 | 1 | 8 |
| nifk | 0.14869 | 0.345348784 | 0.158 | 0.044 | 1 | 8 |
| anp32a | 0.15124 | 0.335204033 | 0.211 | 0.067 | 1 | 8 |
| lsm6 | 0.15272 | 0.351111492 | 0.211 | 0.067 | 1 | 8 |
| rhoac | 0.15449 | 0.242071474 | 0.237 | 0.111 | 1 | 8 |
| gata6 | 0.15654 | 0.340135939 | 0.105 | 0.022 | 1 | 8 |
| bag3 | 0.16367 | 0.325099458 | 0.105 | 0.022 | 1 | 8 |
| zfp36l1b | 0.16501 | -0.40803964 | 0.184 | 0.333 | 1 | 8 |
| wt1a | 0.16597 | -0.36725189 | 0.079 | 0.178 | 1 | 8 |
| prkab1b | 0.16741 | -0.35089123 | 0.026 | 0.133 | 1 | 8 |
| ywhae1 | 0.16796 | 0.296007559 | 0.368 | 0.2 | 1 | 8 |
| mt-atp6 | 0.16833 | 0.323582198 | 0.158 | 0.111 | 1 | 8 |
| CR318588.1 | 0.16841 | -0.48839863 | 0.132 | 0.244 | 1 | 8 |
| fam32a | 0.17086 | 0.284389698 | 0.316 | 0.156 | 1 | 8 |
| rpl29 | 0.17148 | -0.11238478 | 0.947 | 1 | 1 | 8 |
| rab11a | 0.17214 | 0.241175592 | 0.184 | 0.067 | 1 | 8 |
| med19a | 0.17221 | 0.255706819 | 0.158 | 0.044 | 1 | 8 |
| fblim1 | 0.17476 | 0.308415081 | 0.105 | 0.022 | 1 | 8 |
| tpm1 | 0.17634 | 0.386447106 | 0.237 | 0.089 | 1 | 8 |

| | | | | | | |
|---|---|---|---|---|---|---|
| trmt112 | 0.17705 | -0.29802677 | 0.026 | 0.133 | 1 | 8 |
| cst14a.2 | 0.17771 | 0.31121417 | 0.342 | 0.178 | 1 | 8 |
| ndufb3 | 0.17973 | 0.397610519 | 0.263 | 0.111 | 1 | 8 |
| cav2 | 0.18148 | -0.33914228 | 0.026 | 0.133 | 1 | 8 |
| hmgn6 | 0.18445 | 0.207340708 | 0.658 | 0.489 | 1 | 8 |
| rps27l | 0.1851 | 0.342418855 | 0.263 | 0.178 | 1 | 8 |
| igfbp5b | 0.18768 | 0.261820098 | 0.079 | 0.022 | 1 | 8 |
| rbpms2a | 0.18832 | 0.264533499 | 0.105 | 0.022 | 1 | 8 |
| atp5f1b | 0.18839 | 0.208233273 | 0.395 | 0.244 | 1 | 8 |
| wu:fb55g09 | 0.18884 | 0.374325737 | 0.263 | 0.111 | 1 | 8 |
| cxcl8b.1 | 0.189 | -0.3569234 | 0.053 | 0.178 | 1 | 8 |
| pfdn5 | 0.19061 | 0.303500464 | 0.158 | 0.044 | 1 | 8 |
| btg2 | 0.19419 | 0.330185899 | 0.132 | 0.044 | 1 | 8 |
| col1a1b | 0.19482 | -0.20096831 | 0.237 | 0.378 | 1 | 8 |
| cldn5a | 0.19496 | -0.40544936 | 0.132 | 0.289 | 1 | 8 |
| ehd2b | 0.19548 | 0.325371402 | 0.158 | 0.067 | 1 | 8 |
| sh3bgrl3 | 0.19549 | -0.34387975 | 0.053 | 0.178 | 1 | 8 |
| dnaja2b | 0.19711 | 0.261802009 | 0.158 | 0.044 | 1 | 8 |
| snrpd2 | 0.19866 | -0.35314965 | 0.158 | 0.267 | 1 | 8 |
| ms4a17a.11 | 0.19897 | 0.264383798 | 0.105 | 0.022 | 1 | 8 |
| nme2b.1 | 0.19942 | -0.13343689 | 0.921 | 0.978 | 1 | 8 |
| flna | 0.20035 | 0.353099808 | 0.158 | 0.044 | 1 | 8 |
| calr3a | 0.20235 | 0.312628405 | 0.158 | 0.044 | 1 | 8 |
| txnl4a | 0.20295 | 0.163718065 | 0.105 | 0.022 | 1 | 8 |
| krt18a.1 | 0.20314 | -0.30534598 | 0.579 | 0.733 | 1 | 8 |
| krt94 | 0.20329 | 0.27815876 | 0.789 | 0.644 | 1 | 8 |
| mtch2 | 0.20515 | 0.275576017 | 0.105 | 0.022 | 1 | 8 |
| glo1 | 0.20853 | 0.370318702 | 0.263 | 0.133 | 1 | 8 |
| dohh | 0.20885 | 0.257536345 | 0.105 | 0.022 | 1 | 8 |
| sult1st6 | 0.2097 | 0.258875708 | 0.105 | 0.022 | 1 | 8 |
| rbm8a | 0.21272 | 0.100123539 | 0.263 | 0.178 | 1 | 8 |
| nop53 | 0.21276 | 0.358626816 | 0.289 | 0.133 | 1 | 8 |
| rnf7 | 0.21282 | 0.302272399 | 0.158 | 0.044 | 1 | 8 |
| COX3 | 0.21413 | 0.355455148 | 0.184 | 0.133 | 1 | 8 |
| nip7 | 0.21854 | 0.200740242 | 0.105 | 0.022 | 1 | 8 |
| gsta.1 | 0.22045 | -0.34545702 | 0.053 | 0.156 | 1 | 8 |
| ybx1 | 0.22117 | -0.13742982 | 0.895 | 0.956 | 1 | 8 |
| vamp3 | 0.22178 | 0.242222488 | 0.289 | 0.156 | 1 | 8 |
| eif4a1a | 0.22542 | 0.242931835 | 0.289 | 0.156 | 1 | 8 |
| si:dkey-40c11.2 | 0.22588 | 0.20323473 | 0.105 | 0.022 | 1 | 8 |
| bysl | 0.22608 | 0.212893892 | 0.105 | 0.022 | 1 | 8 |
| ucp2 | 0.2276 | -0.34014661 | 0.158 | 0.267 | 1 | 8 |
| tmem263 | 0.22861 | 0.168547041 | 0.132 | 0.044 | 1 | 8 |
| atp5meb | 0.2296 | 0.38814207 | 0.342 | 0.2 | 1 | 8 |
| tpm3 | 0.23 | 0.347887251 | 0.421 | 0.244 | 1 | 8 |
| psma8 | 0.23096 | 0.348460595 | 0.368 | 0.2 | 1 | 8 |
| ubb | 0.23103 | 0.333234416 | 0.368 | 0.2 | 1 | 8 |
| cox14 | 0.23122 | 0.323061903 | 0.184 | 0.067 | 1 | 8 |

| | | | | | | |
|---|---|---|---|---|---|---|
| rpa1 | 0.23173 | -0.30163107 | 0.026 | 0.111 | 1 | 8 |
| rac3a | 0.23184 | -0.20437225 | 0.026 | 0.111 | 1 | 8 |
| pbx4 | 0.23228 | 0.185277413 | 0.132 | 0.044 | 1 | 8 |
| ucmaa | 0.2357 | -0.33494909 | 0.158 | 0.311 | 1 | 8 |
| nucks1a | 0.23614 | -0.17150104 | 0.158 | 0.267 | 1 | 8 |
| sept7a | 0.23707 | 0.232497863 | 0.105 | 0.022 | 1 | 8 |
| rwdd1 | 0.23739 | 0.233099958 | 0.263 | 0.133 | 1 | 8 |
| nhp2 | 0.23923 | 0.270811525 | 0.184 | 0.067 | 1 | 8 |
| LOC103909115 | 0.24141 | 0.208657366 | 0.105 | 0.022 | 1 | 8 |
| gadd45gip1 | 0.24436 | 0.189020989 | 0.105 | 0.022 | 1 | 8 |
| ciartb | 0.24475 | 0.248176669 | 0.105 | 0.022 | 1 | 8 |
| si:ch211-156j16.1 | 0.24482 | 0.316707764 | 0.184 | 0.111 | 1 | 8 |
| dbi | 0.24495 | 0.260345433 | 0.579 | 0.4 | 1 | 8 |
| rnf2 | 0.2457 | 0.190261796 | 0.105 | 0.022 | 1 | 8 |
| dynlrb1 | 0.24586 | 0.31687338 | 0.184 | 0.067 | 1 | 8 |
| prdx1 | 0.2473 | 0.286702398 | 0.105 | 0.022 | 1 | 8 |
| chmp4ba | 0.24938 | 0.202292266 | 0.105 | 0.022 | 1 | 8 |
| stxbp6 | 0.25417 | -0.27124584 | 0.026 | 0.111 | 1 | 8 |
| tsc22d3 | 0.25641 | 0.208479591 | 0.105 | 0.022 | 1 | 8 |
| cbx1b | 0.25739 | 0.255142164 | 0.105 | 0.022 | 1 | 8 |
| icn | 0.25758 | 0.212595238 | 0.842 | 0.689 | 1 | 8 |
| zgc:77849 | 0.25838 | 0.316043952 | 0.184 | 0.067 | 1 | 8 |
| si:ch211-235e9.6 | 0.25863 | 0.333418869 | 0.184 | 0.067 | 1 | 8 |
| ccnl1b | 0.25947 | 0.304606969 | 0.105 | 0.022 | 1 | 8 |
| srrm2 | 0.25982 | 0.265634368 | 0.105 | 0.022 | 1 | 8 |
| bxdc2 | 0.26005 | 0.19777824 | 0.105 | 0.022 | 1 | 8 |
| fthl27 | 0.26115 | 0.148401943 | 0.579 | 0.444 | 1 | 8 |
| eif3d | 0.26136 | -0.33115312 | 0.237 | 0.356 | 1 | 8 |
| rpl39 | 0.26208 | -0.11604995 | 0.895 | 0.978 | 1 | 8 |
| pabpc1a | 0.26232 | -0.17369625 | 0.605 | 0.667 | 1 | 8 |
| txn | 0.264 | 0.297405137 | 0.105 | 0.022 | 1 | 8 |
| skp1 | 0.26449 | -0.35931901 | 0.184 | 0.333 | 1 | 8 |
| gtf2h5 | 0.26477 | -0.28770936 | 0.026 | 0.111 | 1 | 8 |
| fxyd6l | 0.26514 | 0.337229513 | 0.132 | 0.044 | 1 | 8 |
| hmg20a | 0.26524 | 0.208039203 | 0.105 | 0.022 | 1 | 8 |
| lsm7 | 0.26667 | -0.27716833 | 0.105 | 0.178 | 1 | 8 |
| rtraf | 0.26769 | 0.246334622 | 0.105 | 0.022 | 1 | 8 |
| rps12 | 0.26849 | -0.13066372 | 0.895 | 0.933 | 1 | 8 |
| psmd4a | 0.2685 | 0.228433414 | 0.105 | 0.022 | 1 | 8 |
| rpl8 | 0.26855 | -0.11699742 | 0.921 | 0.978 | 1 | 8 |
| sdf2l1 | 0.2694 | -0.25173224 | 0.053 | 0.111 | 1 | 8 |
| ndufb7 | 0.26967 | 0.17009325 | 0.184 | 0.089 | 1 | 8 |
| fam3c | 0.27009 | 0.236933174 | 0.105 | 0.022 | 1 | 8 |
| chtopa | 0.27062 | 0.242668955 | 0.105 | 0.022 | 1 | 8 |
| psmb7 | 0.271 | 0.262919354 | 0.342 | 0.2 | 1 | 8 |
| loxl2a | 0.271 | 0.285645539 | 0.105 | 0.022 | 1 | 8 |
| rab11ba | 0.27136 | -0.32673711 | 0.053 | 0.156 | 1 | 8 |
| aldh7a1 | 0.27168 | 0.257911677 | 0.105 | 0.022 | 1 | 8 |

| | | | | | | |
|---|---|---|---|---|---|---|
| hsdl2 | 0.2718 | 0.209621525 | 0.105 | 0.022 | 1 | 8 |
| drg2 | 0.27194 | 0.220890573 | 0.105 | 0.022 | 1 | 8 |
| ssu72 | 0.2724 | 0.192764459 | 0.105 | 0.022 | 1 | 8 |
| glipr2l | 0.2725 | 0.22069043 | 0.105 | 0.022 | 1 | 8 |
| sec61g | 0.27309 | 0.265592858 | 0.684 | 0.511 | 1 | 8 |
| grinab | 0.27322 | 0.260294122 | 0.105 | 0.022 | 1 | 8 |
| sostdc1a | 0.27389 | -0.33476729 | 0.079 | 0.2 | 1 | 8 |
| tfa | 0.27453 | 0.314912975 | 0.105 | 0.022 | 1 | 8 |
| XLOC-025423 | 0.27475 | 0.246845126 | 0.079 | 0.022 | 1 | 8 |
| ccdc59 | 0.27482 | 0.248447094 | 0.105 | 0.022 | 1 | 8 |
| mex3b | 0.27509 | 0.234680972 | 0.105 | 0.022 | 1 | 8 |
| celf2 | 0.2766 | 0.209927474 | 0.132 | 0.044 | 1 | 8 |
| ddx24 | 0.27669 | 0.240003419 | 0.105 | 0.022 | 1 | 8 |
| cbx5 | 0.27691 | -0.33316455 | 0.079 | 0.2 | 1 | 8 |
| selenok | 0.27721 | 0.233584245 | 0.105 | 0.022 | 1 | 8 |
| eif2s1a | 0.27728 | 0.237965032 | 0.105 | 0.022 | 1 | 8 |
| ubap2b | 0.27733 | 0.23737143 | 0.105 | 0.022 | 1 | 8 |
| adam10a | 0.27762 | 0.252443828 | 0.105 | 0.022 | 1 | 8 |
| twistnb | 0.27773 | 0.22882282 | 0.079 | 0.022 | 1 | 8 |
| tubb2b | 0.27854 | -0.2919569 | 0.132 | 0.2 | 1 | 8 |
| polr2f | 0.27897 | -0.33103963 | 0.079 | 0.2 | 1 | 8 |
| erp44 | 0.27945 | -0.28348427 | 0.026 | 0.111 | 1 | 8 |
| paip2b | 0.27947 | -0.33388912 | 0.079 | 0.2 | 1 | 8 |
| coa6 | 0.27969 | 0.301194614 | 0.132 | 0.044 | 1 | 8 |
| igf2b | 0.28037 | -0.1830897 | 0.053 | 0.133 | 1 | 8 |
| pam16 | 0.28348 | 0.307672561 | 0.211 | 0.089 | 1 | 8 |
| zgc:162730 | 0.28502 | 0.101391851 | 0.316 | 0.222 | 1 | 8 |
| zgc:86598 | 0.28637 | 0.157755138 | 0.184 | 0.089 | 1 | 8 |
| wdr12 | 0.28724 | -0.27653837 | 0.026 | 0.111 | 1 | 8 |
| psmb4 | 0.28753 | 0.175451494 | 0.184 | 0.089 | 1 | 8 |
| ywhabb | 0.28771 | 0.32277091 | 0.421 | 0.267 | 1 | 8 |
| ssr1 | 0.29284 | 0.277906564 | 0.132 | 0.067 | 1 | 8 |
| LO018188.1 | 0.29349 | -0.28339524 | 0.026 | 0.111 | 1 | 8 |
| atp5mf | 0.29507 | -0.2503695 | 0.237 | 0.311 | 1 | 8 |
| mt-co2 | 0.29712 | 0.300765815 | 0.158 | 0.111 | 1 | 8 |
| znf385a | 0.29746 | -0.15804377 | 0.053 | 0.133 | 1 | 8 |
| rpl3 | 0.29896 | -0.11568271 | 0.921 | 0.978 | 1 | 8 |
| nutf2l | 0.30088 | -0.24942354 | 0.026 | 0.111 | 1 | 8 |
| igf2bp2a | 0.30244 | -0.27432495 | 0.026 | 0.111 | 1 | 8 |
| tomm22 | 0.30284 | -0.21701938 | 0.026 | 0.111 | 1 | 8 |
| ptbp1a | 0.30326 | -0.25613852 | 0.026 | 0.111 | 1 | 8 |
| si:dkey-112e7.2 | 0.30359 | -0.33014226 | 0.105 | 0.222 | 1 | 8 |
| rpl28 | 0.3046 | -0.10647354 | 0.921 | 0.978 | 1 | 8 |
| ap2m1b | 0.30524 | 0.197137123 | 0.132 | 0.044 | 1 | 8 |
| myl9a | 0.30629 | 0.281850785 | 0.289 | 0.156 | 1 | 8 |
| nfia | 0.30672 | 0.346924843 | 0.237 | 0.111 | 1 | 8 |
| nat9 | 0.30713 | 0.221331345 | 0.079 | 0.022 | 1 | 8 |
| psma3 | 0.30796 | 0.176322481 | 0.184 | 0.089 | 1 | 8 |

| | | | | | | |
|---|---|---|---|---|---|---|
| pkma | 0.30887 | 0.202062404 | 0.132 | 0.044 | 1 | 8 |
| zgc:175088 | 0.31006 | 0.234465437 | 0.132 | 0.044 | 1 | 8 |
| nop58 | 0.31161 | 0.106347317 | 0.184 | 0.111 | 1 | 8 |
| cct3 | 0.31176 | -0.31680395 | 0.105 | 0.222 | 1 | 8 |
| prmt1 | 0.31186 | 0.333341039 | 0.237 | 0.111 | 1 | 8 |
| cox6c | 0.3122 | 0.319636841 | 0.368 | 0.222 | 1 | 8 |
| rps28 | 0.3128 | -0.10415239 | 0.947 | 0.978 | 1 | 8 |
| cct2 | 0.31395 | 0.280499369 | 0.289 | 0.156 | 1 | 8 |
| ctsk | 0.31648 | 0.300546316 | 0.132 | 0.044 | 1 | 8 |
| si:dkey-28b4.7 | 0.31791 | 0.333622928 | 0.263 | 0.133 | 1 | 8 |
| CABZ01080702.1 | 0.31928 | 0.114628757 | 0.105 | 0.044 | 1 | 8 |
| rps15a | 0.31936 | -0.11719303 | 0.921 | 0.956 | 1 | 8 |
| jund | 0.32028 | 0.243668092 | 0.079 | 0.022 | 1 | 8 |
| cct6a | 0.32043 | -0.31552241 | 0.105 | 0.2 | 1 | 8 |
| khdrbs1a | 0.32117 | 0.147033758 | 0.395 | 0.378 | 1 | 8 |
| meaf6 | 0.32136 | -0.24043613 | 0.026 | 0.089 | 1 | 8 |
| gtf2f2a | 0.32253 | 0.191034737 | 0.132 | 0.044 | 1 | 8 |
| zgc:56493 | 0.32316 | 0.118022354 | 0.658 | 0.533 | 1 | 8 |
| hdlbpa | 0.32372 | -0.14937021 | 0.368 | 0.4 | 1 | 8 |
| klf2a | 0.32555 | 0.28132282 | 0.342 | 0.2 | 1 | 8 |
| ndufa1 | 0.32566 | -0.2936587 | 0.079 | 0.178 | 1 | 8 |
| sap30l | 0.32648 | 0.195172965 | 0.132 | 0.044 | 1 | 8 |
| psma4 | 0.32772 | 0.322242673 | 0.316 | 0.178 | 1 | 8 |
| sec61a1 | 0.32774 | 0.304653825 | 0.289 | 0.156 | 1 | 8 |
| rad23ab | 0.32819 | 0.245162504 | 0.079 | 0.022 | 1 | 8 |
| glrx | 0.32885 | 0.308110779 | 0.421 | 0.267 | 1 | 8 |
| mcl1a | 0.33118 | 0.291518595 | 0.289 | 0.156 | 1 | 8 |
| smim13 | 0.33354 | 0.229472483 | 0.132 | 0.044 | 1 | 8 |
| tuba8l2 | 0.33384 | 0.246040094 | 0.105 | 0.044 | 1 | 8 |
| gtpbp4 | 0.33887 | -0.31869856 | 0.132 | 0.244 | 1 | 8 |
| rack1 | 0.33917 | 0.152105283 | 0.921 | 0.822 | 1 | 8 |
| aldob | 0.34014 | 0.181926925 | 0.132 | 0.044 | 1 | 8 |
| mrpl36 | 0.34071 | 0.202000344 | 0.158 | 0.067 | 1 | 8 |
| rab1ab | 0.34099 | 0.216761332 | 0.132 | 0.044 | 1 | 8 |
| rpl19 | 0.34521 | -0.11166295 | 0.921 | 0.978 | 1 | 8 |
| gadd45ba | 0.34586 | 0.300931037 | 0.158 | 0.067 | 1 | 8 |
| nfkbie | 0.34631 | 0.202401064 | 0.132 | 0.044 | 1 | 8 |
| nfic | 0.34657 | 0.24601896 | 0.079 | 0.022 | 1 | 8 |
| si:ch73-42p12.2 | 0.34746 | 0.199319754 | 0.158 | 0.067 | 1 | 8 |
| cfd | 0.34764 | 0.238362278 | 0.184 | 0.156 | 1 | 8 |
| gkap1 | 0.34847 | 0.227259434 | 0.105 | 0.044 | 1 | 8 |
| elovl1b | 0.35125 | -0.21376755 | 0.132 | 0.244 | 1 | 8 |
| mt-nd1 | 0.35191 | 0.308479316 | 0.158 | 0.089 | 1 | 8 |
| psmd12 | 0.35194 | 0.208469049 | 0.132 | 0.044 | 1 | 8 |
| sod1 | 0.3542 | 0.17426905 | 0.342 | 0.222 | 1 | 8 |
| taf12 | 0.3547 | 0.210350079 | 0.132 | 0.044 | 1 | 8 |
| psmb6 | 0.35518 | 0.206860021 | 0.237 | 0.133 | 1 | 8 |
| pleca | 0.35608 | 0.239305384 | 0.079 | 0.022 | 1 | 8 |

| | | | | | | |
|---|---|---|---|---|---|---|
| kmt5aa | 0.35668 | 0.266075849 | 0.132 | 0.044 | 1 | 8 |
| srsf6b | 0.3567 | -0.13849044 | 0.026 | 0.089 | 1 | 8 |
| eif4ebp1 | 0.35718 | 0.312346581 | 0.211 | 0.111 | 1 | 8 |
| zgc:195173 | 0.35858 | 0.248983058 | 0.132 | 0.044 | 1 | 8 |
| rps14 | 0.35858 | -0.1135148 | 0.868 | 0.956 | 1 | 8 |
| bhlhe40 | 0.35959 | 0.255058398 | 0.132 | 0.044 | 1 | 8 |
| snrpa1 | 0.35995 | 0.225520613 | 0.132 | 0.044 | 1 | 8 |
| arpc1a | 0.36081 | -0.17947663 | 0.184 | 0.222 | 1 | 8 |
| im:7151449 | 0.36119 | 0.26380498 | 0.132 | 0.044 | 1 | 8 |
| fundc2 | 0.36169 | -0.12912321 | 0.026 | 0.089 | 1 | 8 |
| slc35c2 | 0.36186 | -0.23692732 | 0.026 | 0.089 | 1 | 8 |
| seta | 0.36294 | -0.30788594 | 0.105 | 0.2 | 1 | 8 |
| pdlim1 | 0.36608 | 0.115733674 | 0.079 | 0.022 | 1 | 8 |
| qkia | 0.367 | 0.312964198 | 0.263 | 0.156 | 1 | 8 |
| atf4a | 0.3685 | 0.236510832 | 0.158 | 0.067 | 1 | 8 |
| zgc:153867 | 0.36871 | 0.220151085 | 0.684 | 0.533 | 1 | 8 |
| rbm34 | 0.36916 | 0.214329152 | 0.079 | 0.022 | 1 | 8 |
| hnrnpa0a | 0.36929 | -0.26937025 | 0.053 | 0.133 | 1 | 8 |
| h1fx | 0.36939 | -0.28225319 | 0.079 | 0.156 | 1 | 8 |
| mfap5 | 0.3695 | -0.30437486 | 0.079 | 0.178 | 1 | 8 |
| s100v2 | 0.37049 | 0.300029984 | 0.474 | 0.333 | 1 | 8 |
| aurkaip1 | 0.37275 | 0.11640965 | 0.079 | 0.022 | 1 | 8 |
| pa2g4b | 0.37338 | 0.268708728 | 0.158 | 0.067 | 1 | 8 |
| rgs5a | 0.37344 | 0.139899547 | 0.079 | 0.022 | 1 | 8 |
| tob1a | 0.37803 | 0.282763029 | 0.158 | 0.067 | 1 | 8 |
| si:ch211-250c4.4 | 0.37892 | 0.233397716 | 0.079 | 0.022 | 1 | 8 |
| flot1b | 0.37956 | 0.281009173 | 0.158 | 0.067 | 1 | 8 |
| si:dkey-239h2.3 | 0.37983 | -0.25557896 | 0.105 | 0.178 | 1 | 8 |
| marcksl1b | 0.38108 | 0.28054506 | 0.158 | 0.067 | 1 | 8 |
| sox4a | 0.38303 | -0.27842253 | 0.053 | 0.133 | 1 | 8 |
| tp53 | 0.38436 | -0.22607048 | 0.026 | 0.089 | 1 | 8 |
| hist2h2l | 0.38501 | 0.107639789 | 0.132 | 0.133 | 1 | 8 |
| ptmaa | 0.3853 | -0.14431052 | 0.605 | 0.644 | 1 | 8 |
| si:dkey-188i13.11 | 0.38762 | 0.220134928 | 0.184 | 0.089 | 1 | 8 |
| mrps21 | 0.38826 | 0.256200958 | 0.132 | 0.067 | 1 | 8 |
| mt-co1 | 0.38946 | 0.35010016 | 0.184 | 0.089 | 1 | 8 |
| si:ch211-222l21.1 | 0.39019 | 0.196505852 | 0.632 | 0.489 | 1 | 8 |
| si:ch211-133n4.4 | 0.39037 | -0.2171596 | 0.026 | 0.089 | 1 | 8 |
| map1lc3a | 0.39042 | 0.107717334 | 0.158 | 0.089 | 1 | 8 |
| selenoh | 0.39106 | -0.30047917 | 0.079 | 0.178 | 1 | 8 |
| csnk2a1 | 0.39133 | 0.128706872 | 0.079 | 0.022 | 1 | 8 |
| ddx5 | 0.39165 | -0.28233642 | 0.079 | 0.156 | 1 | 8 |
| ndufa12 | 0.39235 | 0.232487938 | 0.158 | 0.067 | 1 | 8 |
| ndufb10 | 0.39272 | 0.293582886 | 0.211 | 0.111 | 1 | 8 |
| b2m | 0.39439 | -0.27575896 | 0.079 | 0.178 | 1 | 8 |
| echs1 | 0.39492 | -0.18265373 | 0.053 | 0.133 | 1 | 8 |
| nop10 | 0.3977 | -0.27387915 | 0.079 | 0.178 | 1 | 8 |
| eif2s2 | 0.39782 | -0.15665493 | 0.342 | 0.378 | 1 | 8 |

| | | | | | | |
|---|---|---|---|---|---|---|
| uqcr10 | 0.39793 | 0.200274995 | 0.368 | 0.244 | 1 | 8 |
| clta | 0.39905 | 0.28876646 | 0.211 | 0.111 | 1 | 8 |
| mrps2 | 0.3992 | 0.112901971 | 0.079 | 0.022 | 1 | 8 |
| sccpdha | 0.40154 | 0.198628632 | 0.079 | 0.022 | 1 | 8 |
| hsbp1a | 0.40178 | -0.29413949 | 0.105 | 0.2 | 1 | 8 |
| zgc:111986 | 0.40276 | -0.26925941 | 0.079 | 0.178 | 1 | 8 |
| zgc:163080 | 0.40318 | -0.15565407 | 0.026 | 0.089 | 1 | 8 |
| rnpc3 | 0.40391 | 0.223617208 | 0.079 | 0.022 | 1 | 8 |
| taf15 | 0.40418 | -0.2770995 | 0.053 | 0.133 | 1 | 8 |
| serbp1b | 0.40447 | 0.227298666 | 0.105 | 0.044 | 1 | 8 |
| zgc:153713 | 0.40474 | 0.250618624 | 0.158 | 0.067 | 1 | 8 |
| eif3g | 0.40585 | -0.28553554 | 0.184 | 0.289 | 1 | 8 |
| eif4a3 | 0.40694 | -0.25498521 | 0.053 | 0.133 | 1 | 8 |
| gbp1 | 0.40736 | 0.125367141 | 0.079 | 0.022 | 1 | 8 |
| ccdc124 | 0.4093 | 0.267736432 | 0.158 | 0.067 | 1 | 8 |
| tyms | 0.40945 | -0.15767409 | 0.026 | 0.089 | 1 | 8 |
| s100a10a | 0.40962 | -0.29181964 | 0.184 | 0.311 | 1 | 8 |
| sh3bgrl | 0.41034 | 0.105857155 | 0.211 | 0.133 | 1 | 8 |
| selenow1 | 0.41063 | 0.207876749 | 0.737 | 0.6 | 1 | 8 |
| gnb1b | 0.41171 | 0.237982527 | 0.158 | 0.067 | 1 | 8 |
| fbln5 | 0.41182 | 0.249115879 | 0.158 | 0.067 | 1 | 8 |
| hdac1 | 0.4121 | 0.259043011 | 0.158 | 0.067 | 1 | 8 |
| ube2na | 0.4121 | 0.244749135 | 0.158 | 0.067 | 1 | 8 |
| smarcd1 | 0.41362 | 0.187787885 | 0.079 | 0.022 | 1 | 8 |
| ptges3b | 0.4144 | 0.103956098 | 0.132 | 0.067 | 1 | 8 |
| si:ch211-241b2.1 | 0.4156 | 0.128559409 | 0.079 | 0.022 | 1 | 8 |
| eva1ba | 0.41693 | 0.139749803 | 0.079 | 0.022 | 1 | 8 |
| bnip3lb | 0.41714 | -0.21853072 | 0.026 | 0.089 | 1 | 8 |
| kctd5a | 0.41729 | -0.20443964 | 0.026 | 0.089 | 1 | 8 |
| fosab | 0.41775 | 0.280997019 | 0.211 | 0.156 | 1 | 8 |
| khsrp | 0.41782 | -0.26688553 | 0.053 | 0.133 | 1 | 8 |
| ap2m1a | 0.41965 | 0.118981497 | 0.184 | 0.111 | 1 | 8 |
| ilf3b | 0.42016 | -0.19069694 | 0.026 | 0.089 | 1 | 8 |
| tpd52l2a | 0.42033 | -0.15427825 | 0.026 | 0.089 | 1 | 8 |
| eif3ea | 0.4205 | -0.27726755 | 0.289 | 0.422 | 1 | 8 |
| epb41l5 | 0.42241 | -0.1114 | 0.053 | 0.111 | 1 | 8 |
| mycb | 0.4234 | 0.194664825 | 0.079 | 0.022 | 1 | 8 |
| litaf | 0.42501 | 0.249107868 | 0.184 | 0.089 | 1 | 8 |
| ciarta | 0.4251 | 0.209829651 | 0.211 | 0.111 | 1 | 8 |
| cdc26 | 0.42511 | 0.226076629 | 0.184 | 0.089 | 1 | 8 |
| ier2a | 0.42711 | -0.16451327 | 0.132 | 0.222 | 1 | 8 |
| prdx3 | 0.42757 | -0.25170555 | 0.079 | 0.156 | 1 | 8 |
| ddt | 0.42793 | 0.145421704 | 0.079 | 0.022 | 1 | 8 |
| bod1 | 0.42811 | 0.117978151 | 0.079 | 0.022 | 1 | 8 |
| si:ch211-196l7.4 | 0.42871 | 0.21093496 | 0.237 | 0.133 | 1 | 8 |
| lxn | 0.42953 | 0.125728738 | 0.316 | 0.222 | 1 | 8 |
| snrpd1 | 0.4296 | -0.27401487 | 0.132 | 0.222 | 1 | 8 |
| krt15 | 0.42998 | -0.13767431 | 0.816 | 0.911 | 1 | 8 |

| | | | | | | |
|---|---|---|---|---|---|---|
| gnl3 | 0.43009 | 0.102762991 | 0.132 | 0.067 | 1 | 8 |
| pdcd6ip | 0.43011 | 0.152026938 | 0.079 | 0.022 | 1 | 8 |
| hnrnpabb | 0.43071 | -0.21926902 | 0.184 | 0.244 | 1 | 8 |
| HACD4 | 0.43182 | 0.190236925 | 0.079 | 0.022 | 1 | 8 |
| ufc1 | 0.43255 | -0.20659826 | 0.026 | 0.089 | 1 | 8 |
| txndc17 | 0.43302 | 0.11944108 | 0.079 | 0.022 | 1 | 8 |
| ablim1b | 0.43368 | 0.162225012 | 0.079 | 0.022 | 1 | 8 |
| mrps34 | 0.43386 | 0.157645353 | 0.079 | 0.022 | 1 | 8 |
| zgc:77056 | 0.4342 | -0.23888291 | 0.053 | 0.133 | 1 | 8 |
| hmga1a | 0.43438 | 0.250286843 | 0.447 | 0.311 | 1 | 8 |
| serpine3 | 0.43494 | -0.21594755 | 0.026 | 0.089 | 1 | 8 |
| cs | 0.43573 | -0.1406164 | 0.026 | 0.089 | 1 | 8 |
| rp9 | 0.4358 | 0.14925811 | 0.079 | 0.022 | 1 | 8 |
| zgc:91910 | 0.43663 | -0.10406438 | 0.158 | 0.156 | 1 | 8 |
| tmem176l.1 | 0.4379 | 0.260266671 | 0.184 | 0.111 | 1 | 8 |
| pmp22a | 0.43799 | 0.291835096 | 0.211 | 0.111 | 1 | 8 |
| mrpl34 | 0.43811 | -0.2508594 | 0.053 | 0.133 | 1 | 8 |
| ciao2b | 0.43832 | -0.19419503 | 0.026 | 0.089 | 1 | 8 |
| brk1 | 0.43853 | 0.208145506 | 0.211 | 0.111 | 1 | 8 |
| col16a1 | 0.43928 | 0.154986688 | 0.079 | 0.022 | 1 | 8 |
| taf9 | 0.4414 | -0.16129887 | 0.026 | 0.089 | 1 | 8 |
| cstf1 | 0.44182 | 0.131118239 | 0.079 | 0.022 | 1 | 8 |
| psmd13 | 0.44183 | 0.150099608 | 0.132 | 0.111 | 1 | 8 |
| rbx1 | 0.44195 | 0.205822532 | 0.289 | 0.178 | 1 | 8 |
| vdac2 | 0.44212 | 0.274841925 | 0.368 | 0.244 | 1 | 8 |
| mgst3b | 0.44231 | -0.23121689 | 0.053 | 0.133 | 1 | 8 |
| caprin1b | 0.44304 | 0.238141063 | 0.184 | 0.089 | 1 | 8 |
| rmdn1 | 0.44308 | -0.20830599 | 0.053 | 0.133 | 1 | 8 |
| napab | 0.44332 | 0.186709783 | 0.079 | 0.022 | 1 | 8 |
| si:ch73-1a9.3 | 0.44343 | 0.207852861 | 0.605 | 0.511 | 1 | 8 |
| atp5pd | 0.44599 | 0.218669555 | 0.368 | 0.244 | 1 | 8 |
| leng1 | 0.44624 | 0.144107099 | 0.079 | 0.022 | 1 | 8 |
| ubfd1 | 0.44624 | 0.156793718 | 0.079 | 0.022 | 1 | 8 |
| hsd11b2 | 0.44649 | 0.231727882 | 0.395 | 0.267 | 1 | 8 |
| gabarapl2 | 0.44708 | 0.119499317 | 0.079 | 0.022 | 1 | 8 |
| spcs1 | 0.4475 | -0.24014113 | 0.053 | 0.133 | 1 | 8 |
| h1f0 | 0.4479 | -0.23091064 | 0.053 | 0.133 | 1 | 8 |
| med31 | 0.44837 | -0.21358204 | 0.026 | 0.089 | 1 | 8 |
| si:ch73-71c20.5 | 0.44917 | -0.19910562 | 0.026 | 0.089 | 1 | 8 |
| st13 | 0.4493 | -0.19934858 | 0.026 | 0.089 | 1 | 8 |
| fabp3 | 0.44938 | 0.215114055 | 0.105 | 0.067 | 1 | 8 |
| eif1b | 0.44948 | -0.24770667 | 0.289 | 0.422 | 1 | 8 |
| hibadhb | 0.45095 | 0.15444416 | 0.079 | 0.022 | 1 | 8 |
| btg1 | 0.45188 | -0.1168262 | 0.184 | 0.267 | 1 | 8 |
| zgc:162025 | 0.45196 | 0.167862584 | 0.079 | 0.022 | 1 | 8 |
| gnb1a | 0.45364 | -0.16244532 | 0.026 | 0.089 | 1 | 8 |
| cpn1 | 0.45416 | -0.14230513 | 0.026 | 0.089 | 1 | 8 |
| pigbos1 | 0.45584 | 0.200486991 | 0.079 | 0.022 | 1 | 8 |

| | | | | | | |
|---|---|---|---|---|---|---|
| creb3l3l | 0.45599 | 0.15912246 | 0.079 | 0.022 | 1 | 8 |
| nrp1a | 0.4564 | 0.206992798 | 0.079 | 0.022 | 1 | 8 |
| si:ch211-151g22.1 | 0.45658 | 0.232062158 | 0.079 | 0.022 | 1 | 8 |
| si:dkey-23i12.5 | 0.4577 | -0.19661854 | 0.026 | 0.089 | 1 | 8 |
| cebpd | 0.45779 | 0.171799822 | 0.474 | 0.356 | 1 | 8 |
| sec61b | 0.45836 | 0.234266085 | 0.579 | 0.444 | 1 | 8 |
| prss23 | 0.45844 | -0.19166329 | 0.026 | 0.089 | 1 | 8 |
| cbx1a | 0.45845 | 0.129587345 | 0.132 | 0.067 | 1 | 8 |
| gng7 | 0.45897 | -0.20149059 | 0.026 | 0.089 | 1 | 8 |
| rfc5 | 0.45912 | -0.20673134 | 0.026 | 0.089 | 1 | 8 |
| ebna1bp2 | 0.45932 | -0.19882525 | 0.026 | 0.089 | 1 | 8 |
| chka | 0.45944 | 0.195275103 | 0.079 | 0.044 | 1 | 8 |
| pinx1 | 0.45945 | 0.122818071 | 0.132 | 0.067 | 1 | 8 |
| fnbp1l | 0.45963 | -0.22364022 | 0.026 | 0.089 | 1 | 8 |
| si:ch211-282b22.1 | 0.46014 | 0.182986576 | 0.079 | 0.022 | 1 | 8 |
| mdka | 0.46151 | 0.237810113 | 0.105 | 0.044 | 1 | 8 |
| psmd7 | 0.46163 | -0.16520002 | 0.026 | 0.089 | 1 | 8 |
| wdr61 | 0.46266 | -0.16292716 | 0.026 | 0.089 | 1 | 8 |
| rbm4.2 | 0.46302 | -0.20346193 | 0.026 | 0.089 | 1 | 8 |
| cald1a | 0.46367 | 0.234721158 | 0.211 | 0.111 | 1 | 8 |
| psmb3 | 0.46384 | 0.248232853 | 0.211 | 0.111 | 1 | 8 |
| luc7l3 | 0.4644 | 0.102718524 | 0.105 | 0.044 | 1 | 8 |
| gapdhs | 0.46454 | 0.256872854 | 0.368 | 0.244 | 1 | 8 |
| si:ch73-376l24.2 | 0.46474 | 0.154006762 | 0.079 | 0.022 | 1 | 8 |
| rangap1a | 0.46487 | 0.139100123 | 0.079 | 0.022 | 1 | 8 |
| stx12 | 0.4653 | 0.16326702 | 0.079 | 0.022 | 1 | 8 |
| ash1l | 0.46572 | -0.17638045 | 0.026 | 0.089 | 1 | 8 |
| smim15 | 0.46573 | -0.17331696 | 0.079 | 0.111 | 1 | 8 |
| rps15 | 0.46577 | -0.10453723 | 0.895 | 0.956 | 1 | 8 |
| hspb1 | 0.46587 | 0.183023465 | 0.526 | 0.4 | 1 | 8 |
| mvb12a | 0.46718 | 0.100423191 | 0.053 | 0.022 | 1 | 8 |
| FP074902.2 | 0.46718 | 0.14015613 | 0.053 | 0.022 | 1 | 8 |
| map2k1 | 0.46718 | 0.109432353 | 0.053 | 0.022 | 1 | 8 |
| psma6l | 0.46718 | 0.134794954 | 0.053 | 0.022 | 1 | 8 |
| zgc:162509 | 0.46718 | 0.10352754 | 0.053 | 0.022 | 1 | 8 |
| LOC101882642 | 0.46718 | 0.119448896 | 0.053 | 0.022 | 1 | 8 |
| LOC103910438 | 0.46718 | 0.145243645 | 0.053 | 0.022 | 1 | 8 |
| arl6ip4 | 0.46718 | 0.101101965 | 0.053 | 0.022 | 1 | 8 |
| cavin2a | 0.46718 | 0.129627748 | 0.053 | 0.022 | 1 | 8 |
| cbsa | 0.46718 | 0.12449118 | 0.053 | 0.022 | 1 | 8 |
| cgnl1 | 0.46718 | 0.116146419 | 0.053 | 0.022 | 1 | 8 |
| cnot6a | 0.46718 | 0.157109458 | 0.053 | 0.022 | 1 | 8 |
| crnkl1 | 0.46718 | 0.133531567 | 0.053 | 0.022 | 1 | 8 |
| cry-dash | 0.46718 | 0.164201715 | 0.053 | 0.022 | 1 | 8 |
| cyr61 | 0.46718 | 0.192017979 | 0.053 | 0.022 | 1 | 8 |
| dbn1 | 0.46718 | 0.161180013 | 0.053 | 0.022 | 1 | 8 |
| evi5b | 0.46718 | 0.117245857 | 0.053 | 0.022 | 1 | 8 |
| fahd2a | 0.46718 | 0.133839807 | 0.053 | 0.022 | 1 | 8 |

| | | | | | | |
|---|---|---|---|---|---|---|
| fbxl5 | 0.46718 | 0.108733306 | 0.053 | 0.022 | 1 | 8 |
| fxr2 | 0.46718 | 0.136454889 | 0.053 | 0.022 | 1 | 8 |
| gusb | 0.46718 | 0.134758185 | 0.053 | 0.022 | 1 | 8 |
| hbegfa | 0.46718 | 0.133664571 | 0.053 | 0.022 | 1 | 8 |
| hpf1 | 0.46718 | 0.102208262 | 0.053 | 0.022 | 1 | 8 |
| iqgap1 | 0.46718 | 0.108274247 | 0.053 | 0.022 | 1 | 8 |
| kcnq5a | 0.46718 | 0.154707967 | 0.053 | 0.022 | 1 | 8 |
| kif5ba | 0.46718 | 0.101301514 | 0.053 | 0.022 | 1 | 8 |
| mao | 0.46718 | 0.158517516 | 0.053 | 0.022 | 1 | 8 |
| mmp14b | 0.46718 | 0.117013831 | 0.053 | 0.022 | 1 | 8 |
| mrpl2 | 0.46718 | 0.119640304 | 0.053 | 0.022 | 1 | 8 |
| net1 | 0.46718 | 0.148800116 | 0.053 | 0.022 | 1 | 8 |
| nfkbib | 0.46718 | 0.128356192 | 0.053 | 0.022 | 1 | 8 |
| nt5c3a | 0.46718 | 0.100149463 | 0.053 | 0.022 | 1 | 8 |
| pgrmc2 | 0.46718 | 0.124843403 | 0.053 | 0.022 | 1 | 8 |
| plpp3 | 0.46718 | 0.10712901 | 0.053 | 0.022 | 1 | 8 |
| psmc5 | 0.46718 | 0.117961063 | 0.053 | 0.022 | 1 | 8 |
| rbm22 | 0.46718 | 0.112071757 | 0.053 | 0.022 | 1 | 8 |
| rhogb | 0.46718 | 0.105418283 | 0.053 | 0.022 | 1 | 8 |
| rpp30 | 0.46718 | 0.111495911 | 0.053 | 0.022 | 1 | 8 |
| s1pr5a | 0.46718 | 0.10883471 | 0.053 | 0.022 | 1 | 8 |
| serpinb14 | 0.46718 | 0.110354966 | 0.053 | 0.022 | 1 | 8 |
| sh3pxd2aa | 0.46718 | 0.151096071 | 0.053 | 0.022 | 1 | 8 |
| si:ch211-137i24.10 | 0.46718 | 0.143234773 | 0.053 | 0.022 | 1 | 8 |
| si:ch211-219a4.3 | 0.46718 | 0.114616684 | 0.053 | 0.022 | 1 | 8 |
| slc37a2 | 0.46718 | 0.105472553 | 0.053 | 0.022 | 1 | 8 |
| spink2.1 | 0.46718 | 0.106655443 | 0.053 | 0.022 | 1 | 8 |
| timm8a | 0.46718 | 0.131754567 | 0.053 | 0.022 | 1 | 8 |
| tmem203 | 0.46718 | 0.104363904 | 0.053 | 0.022 | 1 | 8 |
| traf7 | 0.46718 | 0.122513808 | 0.053 | 0.022 | 1 | 8 |
| trim35-28 | 0.46718 | 0.104857559 | 0.053 | 0.022 | 1 | 8 |
| trip12 | 0.46718 | 0.118615272 | 0.053 | 0.022 | 1 | 8 |
| tubb6 | 0.46718 | 0.121619597 | 0.053 | 0.022 | 1 | 8 |
| ubap2l | 0.46718 | 0.12072363 | 0.053 | 0.022 | 1 | 8 |
| wu:fb18f06 | 0.46718 | 0.111703748 | 0.053 | 0.022 | 1 | 8 |
| CABZ01090890.1 | 0.46718 | 0.140522455 | 0.053 | 0.022 | 1 | 8 |
| CR753876.1 | 0.46718 | 0.1429176 | 0.053 | 0.022 | 1 | 8 |
| ahnak | 0.46718 | 0.16213873 | 0.053 | 0.022 | 1 | 8 |
| alkal2b | 0.46718 | 0.11172812 | 0.053 | 0.022 | 1 | 8 |
| ar | 0.46718 | 0.11565287 | 0.053 | 0.022 | 1 | 8 |
| arglu1a | 0.46718 | 0.1101822 | 0.053 | 0.022 | 1 | 8 |
| asph | 0.46718 | 0.13283025 | 0.053 | 0.022 | 1 | 8 |
| bgna | 0.46718 | 0.108422815 | 0.053 | 0.022 | 1 | 8 |
| ccbe1 | 0.46718 | 0.118157767 | 0.053 | 0.022 | 1 | 8 |
| cox17 | 0.46718 | 0.120955895 | 0.053 | 0.022 | 1 | 8 |
| csrnp1a | 0.46718 | 0.13246239 | 0.053 | 0.022 | 1 | 8 |
| efhd2 | 0.46718 | 0.134348937 | 0.053 | 0.022 | 1 | 8 |
| exoc6 | 0.46718 | 0.129628014 | 0.053 | 0.022 | 1 | 8 |

| | | | | | | |
|---|---|---|---|---|---|---|
| fer | 0.46718 | 0.154643854 | 0.053 | 0.022 | 1 | 8 |
| gstk1 | 0.46718 | 0.106584697 | 0.053 | 0.022 | 1 | 8 |
| march5l | 0.46718 | 0.109316207 | 0.053 | 0.022 | 1 | 8 |
| mark3a | 0.46718 | 0.114758554 | 0.053 | 0.022 | 1 | 8 |
| mpp6b | 0.46718 | 0.135373568 | 0.053 | 0.022 | 1 | 8 |
| palm1a | 0.46718 | 0.102493511 | 0.053 | 0.022 | 1 | 8 |
| pdk4 | 0.46718 | 0.115094631 | 0.053 | 0.022 | 1 | 8 |
| pdxka | 0.46718 | 0.159282844 | 0.053 | 0.022 | 1 | 8 |
| rbm39a | 0.46718 | 0.127747895 | 0.053 | 0.022 | 1 | 8 |
| si:dkey-247i3.6 | 0.46718 | 0.107643223 | 0.053 | 0.022 | 1 | 8 |
| slc25a32a | 0.46718 | 0.123550228 | 0.053 | 0.022 | 1 | 8 |
| surf6 | 0.46718 | 0.122305219 | 0.053 | 0.022 | 1 | 8 |
| tmem256 | 0.46718 | 0.116557204 | 0.053 | 0.022 | 1 | 8 |
| traf4a | 0.46718 | 0.115414261 | 0.053 | 0.022 | 1 | 8 |
| vapal | 0.46718 | 0.129796149 | 0.053 | 0.022 | 1 | 8 |
| zgc:175214 | 0.46718 | 0.117899427 | 0.053 | 0.022 | 1 | 8 |
| BX539336.1 | 0.46718 | 0.107346025 | 0.053 | 0.022 | 1 | 8 |
| dido1 | 0.46718 | 0.138783838 | 0.053 | 0.022 | 1 | 8 |
| fam207a | 0.46718 | 0.113127626 | 0.053 | 0.022 | 1 | 8 |
| ndufb5 | 0.46718 | 0.103050914 | 0.053 | 0.022 | 1 | 8 |
| rchy1 | 0.46718 | 0.123608437 | 0.053 | 0.022 | 1 | 8 |
| rtcb | 0.46718 | 0.131430122 | 0.053 | 0.022 | 1 | 8 |
| sf3b2 | 0.46718 | 0.100890426 | 0.053 | 0.022 | 1 | 8 |
| tmem107 | 0.46718 | 0.113652631 | 0.053 | 0.022 | 1 | 8 |
| dusp6 | 0.46718 | 0.110411371 | 0.053 | 0.022 | 1 | 8 |
| ttc26 | 0.46718 | 0.125523408 | 0.053 | 0.022 | 1 | 8 |
| upf3b | 0.46718 | 0.103394245 | 0.053 | 0.022 | 1 | 8 |
| tbk1 | 0.46718 | 0.13265813 | 0.053 | 0.022 | 1 | 8 |
| rex1bd | 0.46731 | 0.202407018 | 0.079 | 0.022 | 1 | 8 |
| katnb1 | 0.46753 | -0.19534843 | 0.026 | 0.089 | 1 | 8 |
| pcbd1 | 0.46762 | 0.142747836 | 0.079 | 0.022 | 1 | 8 |
| cxcl8b.3 | 0.46767 | -0.18643053 | 0.026 | 0.089 | 1 | 8 |
| atp5f1d | 0.46788 | 0.238688742 | 0.237 | 0.133 | 1 | 8 |
| cry2 | 0.46795 | 0.198919414 | 0.079 | 0.022 | 1 | 8 |
| ctdspl2a | 0.46809 | -0.21140448 | 0.026 | 0.089 | 1 | 8 |
| prdx4 | 0.46902 | -0.12941914 | 0.026 | 0.089 | 1 | 8 |
| ak2 | 0.46903 | 0.238194001 | 0.368 | 0.244 | 1 | 8 |
| srp72 | 0.46961 | -0.1832139 | 0.026 | 0.089 | 1 | 8 |
| rrs1 | 0.46963 | -0.1855843 | 0.026 | 0.089 | 1 | 8 |
| si:ch211-204c21.1 | 0.46987 | -0.21161921 | 0.026 | 0.089 | 1 | 8 |
| fbn2b | 0.47 | 0.18554696 | 0.079 | 0.022 | 1 | 8 |
| gsna | 0.47002 | 0.200271385 | 0.105 | 0.044 | 1 | 8 |
| max | 0.47011 | -0.17278132 | 0.026 | 0.089 | 1 | 8 |
| rab28 | 0.47029 | -0.18408661 | 0.026 | 0.089 | 1 | 8 |
| rcn2 | 0.47032 | 0.191378265 | 0.079 | 0.022 | 1 | 8 |
| bbip1 | 0.47032 | -0.17668399 | 0.026 | 0.089 | 1 | 8 |
| sdhb | 0.47044 | -0.20377811 | 0.026 | 0.089 | 1 | 8 |
| clic4 | 0.47049 | -0.13973672 | 0.158 | 0.178 | 1 | 8 |

| | | | | | | |
|---|---|---|---|---|---|---|
| rps25 | 0.47103 | -0.10233157 | 0.895 | 0.956 | 1 | 8 |
| adi1 | 0.47139 | -0.17408583 | 0.026 | 0.089 | 1 | 8 |
| cldn11a | 0.47154 | 0.231059145 | 0.105 | 0.044 | 1 | 8 |
| nucks1b | 0.47216 | 0.17240101 | 0.079 | 0.022 | 1 | 8 |
| arpp19a | 0.47246 | 0.171655882 | 0.079 | 0.022 | 1 | 8 |
| uqcc3 | 0.47275 | 0.143317196 | 0.079 | 0.022 | 1 | 8 |
| nars | 0.47278 | 0.196705586 | 0.079 | 0.022 | 1 | 8 |
| atp6v0d1 | 0.47293 | 0.173432044 | 0.079 | 0.022 | 1 | 8 |
| BNC1 | 0.47424 | 0.146624254 | 0.079 | 0.022 | 1 | 8 |
| gmppb | 0.47435 | 0.16250449 | 0.079 | 0.022 | 1 | 8 |
| dldh | 0.4748 | 0.148874623 | 0.079 | 0.022 | 1 | 8 |
| rhebl1 | 0.47488 | 0.21732839 | 0.079 | 0.022 | 1 | 8 |
| si:dkey-188i13.10 | 0.47529 | 0.236186379 | 0.237 | 0.133 | 1 | 8 |
| mapre1b | 0.47625 | 0.125255077 | 0.158 | 0.089 | 1 | 8 |
| nxn | 0.47639 | 0.133485875 | 0.079 | 0.022 | 1 | 8 |
| sumo2b | 0.47646 | 0.23401953 | 0.316 | 0.2 | 1 | 8 |
| dusp1 | 0.47669 | 0.166214437 | 0.079 | 0.022 | 1 | 8 |
| ndufb9 | 0.47722 | 0.254350782 | 0.237 | 0.156 | 1 | 8 |
| cdh2 | 0.47767 | 0.194360052 | 0.079 | 0.022 | 1 | 8 |
| nfil3-6 | 0.47799 | 0.214284613 | 0.105 | 0.044 | 1 | 8 |
| dcaf7 | 0.47811 | 0.203898245 | 0.105 | 0.044 | 1 | 8 |
| atp5fa1 | 0.47832 | 0.248451743 | 0.342 | 0.222 | 1 | 8 |
| prpsap1 | 0.47838 | 0.181868579 | 0.079 | 0.022 | 1 | 8 |
| yap1 | 0.47911 | 0.164611538 | 0.079 | 0.022 | 1 | 8 |
| ccdc126 | 0.47926 | 0.148657294 | 0.079 | 0.022 | 1 | 8 |
| zbtb4 | 0.47963 | 0.205415087 | 0.079 | 0.022 | 1 | 8 |
| cog4 | 0.48059 | 0.148239822 | 0.079 | 0.022 | 1 | 8 |
| itpkcb | 0.48065 | 0.166294597 | 0.079 | 0.022 | 1 | 8 |
| sec24a | 0.48086 | 0.176557939 | 0.079 | 0.022 | 1 | 8 |
| EML4 | 0.48111 | 0.1602073 | 0.079 | 0.022 | 1 | 8 |
| dkk3b | 0.48162 | 0.267264941 | 0.263 | 0.156 | 1 | 8 |
| midn | 0.48208 | 0.204987011 | 0.079 | 0.022 | 1 | 8 |
| arhgdia | 0.48247 | 0.256366051 | 0.289 | 0.178 | 1 | 8 |
| capza1b | 0.48281 | 0.126041065 | 0.237 | 0.156 | 1 | 8 |
| tmed7 | 0.48305 | 0.128275243 | 0.132 | 0.067 | 1 | 8 |
| zbtb10 | 0.48334 | 0.17651349 | 0.079 | 0.022 | 1 | 8 |
| tpd52l2b | 0.48347 | 0.161866545 | 0.079 | 0.022 | 1 | 8 |
| eloca | 0.48391 | 0.155454925 | 0.079 | 0.022 | 1 | 8 |
| fbxo9 | 0.48465 | 0.152188392 | 0.079 | 0.022 | 1 | 8 |
| capn1a | 0.48477 | 0.16238567 | 0.079 | 0.022 | 1 | 8 |
| acp1 | 0.48495 | 0.160328648 | 0.079 | 0.022 | 1 | 8 |
| ier5l | 0.48496 | 0.150654331 | 0.079 | 0.022 | 1 | 8 |
| fkbp3 | 0.48501 | 0.143125449 | 0.263 | 0.178 | 1 | 8 |
| suclg1 | 0.48512 | 0.141815814 | 0.079 | 0.022 | 1 | 8 |
| si:ch211-68a17.7 | 0.48552 | 0.17970952 | 0.079 | 0.022 | 1 | 8 |
| LOC101884451 | 0.48558 | 0.21026138 | 0.079 | 0.022 | 1 | 8 |
| mrps18a | 0.4857 | 0.208537615 | 0.105 | 0.044 | 1 | 8 |
| spry2 | 0.48573 | 0.187497643 | 0.079 | 0.022 | 1 | 8 |

| | | | | | | |
|---|---|---|---|---|---|---|
| zfand5b | 0.48595 | 0.179339994 | 0.079 | 0.022 | 1 | 8 |
| pum1 | 0.48597 | 0.148732328 | 0.079 | 0.022 | 1 | 8 |
| slc43a2a | 0.48598 | 0.196492444 | 0.079 | 0.022 | 1 | 8 |
| metap1 | 0.48619 | 0.155765229 | 0.079 | 0.022 | 1 | 8 |
| baz2a | 0.48644 | 0.166849999 | 0.079 | 0.022 | 1 | 8 |
| cfap45 | 0.48659 | 0.176284517 | 0.079 | 0.022 | 1 | 8 |
| rdh10a | 0.48666 | 0.207353499 | 0.079 | 0.022 | 1 | 8 |
| sub1a | 0.48723 | 0.133242221 | 0.237 | 0.156 | 1 | 8 |
| timm23a | 0.48793 | 0.124823911 | 0.105 | 0.044 | 1 | 8 |
| rhpn2 | 0.48874 | -0.19408705 | 0.053 | 0.089 | 1 | 8 |
| khdrbs1b | 0.48899 | 0.217921699 | 0.105 | 0.044 | 1 | 8 |
| col1a2 | 0.48902 | -0.20412856 | 0.447 | 0.511 | 1 | 8 |
| naa50 | 0.49006 | 0.130419959 | 0.105 | 0.044 | 1 | 8 |
| ddx39ab | 0.49022 | 0.112159848 | 0.105 | 0.044 | 1 | 8 |
| txn2 | 0.49094 | 0.184224811 | 0.079 | 0.044 | 1 | 8 |
| zgc:158463 | 0.49151 | 0.166404004 | 0.079 | 0.044 | 1 | 8 |
| klf6a | 0.49632 | -0.21289031 | 0.079 | 0.133 | 1 | 8 |
| b2ml | 0.50154 | -0.13765931 | 0.158 | 0.178 | 1 | 8 |
| gstm.1 | 0.50487 | -0.14376702 | 0.553 | 0.667 | 1 | 8 |
| timm10 | 0.50637 | -0.18898489 | 0.026 | 0.067 | 1 | 8 |
| si:rp71-45k5.4 | 0.50841 | 0.239260102 | 0.132 | 0.067 | 1 | 8 |
| hmgb1a | 0.5102 | 0.243528644 | 0.395 | 0.289 | 1 | 8 |
| tomm5 | 0.51049 | -0.26331268 | 0.211 | 0.311 | 1 | 8 |
| fam49a | 0.51153 | 0.125520739 | 0.079 | 0.067 | 1 | 8 |
| pfn2 | 0.51171 | 0.16473038 | 0.395 | 0.289 | 1 | 8 |
| podxl | 0.51207 | 0.162939547 | 0.395 | 0.289 | 1 | 8 |
| serpinb1l3 | 0.51297 | -0.19542018 | 0.053 | 0.111 | 1 | 8 |
| triqk | 0.51387 | -0.20188521 | 0.026 | 0.067 | 1 | 8 |
| ndufc2 | 0.514 | 0.191113565 | 0.105 | 0.044 | 1 | 8 |
| romo1 | 0.5146 | 0.177139131 | 0.421 | 0.311 | 1 | 8 |
| usp36 | 0.51508 | 0.136638009 | 0.105 | 0.044 | 1 | 8 |
| mif | 0.51803 | -0.20308924 | 0.184 | 0.289 | 1 | 8 |
| gstm.2 | 0.52004 | -0.23555342 | 0.079 | 0.156 | 1 | 8 |
| cirbpb | 0.52205 | 0.139104534 | 0.868 | 0.778 | 1 | 8 |
| slc25a5 | 0.52329 | 0.135052777 | 0.868 | 0.8 | 1 | 8 |
| stmn1a | 0.52367 | -0.23362841 | 0.053 | 0.111 | 1 | 8 |
| emc8 | 0.52382 | 0.136491105 | 0.105 | 0.044 | 1 | 8 |
| snrpg | 0.52702 | -0.25246638 | 0.105 | 0.178 | 1 | 8 |
| gstr | 0.52724 | -0.10678387 | 0.079 | 0.133 | 1 | 8 |
| rap1aa | 0.52738 | 0.150157895 | 0.105 | 0.044 | 1 | 8 |
| morf4l1 | 0.52865 | -0.22526659 | 0.079 | 0.156 | 1 | 8 |
| gar1 | 0.5289 | -0.13745253 | 0.053 | 0.067 | 1 | 8 |
| hypk | 0.52907 | -0.24328215 | 0.079 | 0.156 | 1 | 8 |
| ncbp2 | 0.53215 | 0.202321776 | 0.132 | 0.067 | 1 | 8 |
| ifi30 | 0.53218 | 0.153006353 | 0.105 | 0.044 | 1 | 8 |
| smarcb1a | 0.53361 | 0.175182427 | 0.105 | 0.044 | 1 | 8 |
| si:ch211-197h24.9 | 0.53364 | -0.17591066 | 0.079 | 0.156 | 1 | 8 |
| atp5mc1 | 0.53409 | -0.20363208 | 0.184 | 0.244 | 1 | 8 |

| | | | | | | |
|---|---|---|---|---|---|---|
| nap1l1 | 0.5341 | -0.23978791 | 0.184 | 0.289 | 1 | 8 |
| cox6a1 | 0.53602 | -0.20599699 | 0.184 | 0.244 | 1 | 8 |
| pomp | 0.53705 | -0.232625 | 0.184 | 0.267 | 1 | 8 |
| nsfl1c | 0.53709 | 0.150736997 | 0.132 | 0.067 | 1 | 8 |
| uqcrc1 | 0.53717 | -0.127698 | 0.053 | 0.067 | 1 | 8 |
| rbm4.1 | 0.53769 | 0.206237711 | 0.132 | 0.067 | 1 | 8 |
| slu7 | 0.53868 | -0.18161105 | 0.026 | 0.067 | 1 | 8 |
| marcksb | 0.53969 | 0.132167012 | 0.105 | 0.044 | 1 | 8 |
| tmem167a | 0.53976 | 0.158782566 | 0.105 | 0.044 | 1 | 8 |
| CR383676.1 | 0.54009 | 0.299016654 | 0.342 | 0.289 | 1 | 8 |
| cox4i1 | 0.54032 | 0.213541291 | 0.289 | 0.222 | 1 | 8 |
| rab18b | 0.54059 | 0.150149705 | 0.105 | 0.044 | 1 | 8 |
| arl4d | 0.54063 | 0.181023078 | 0.079 | 0.044 | 1 | 8 |
| cxadr | 0.54116 | 0.163874829 | 0.105 | 0.044 | 1 | 8 |
| hsp90ab1 | 0.54175 | -0.10006173 | 0.868 | 0.911 | 1 | 8 |
| ddx21 | 0.54253 | 0.142938265 | 0.158 | 0.133 | 1 | 8 |
| pfkfb4b | 0.54517 | 0.235937311 | 0.184 | 0.111 | 1 | 8 |
| tnfsf10 | 0.5459 | 0.205246361 | 0.105 | 0.044 | 1 | 8 |
| ackr3b | 0.54791 | -0.14998577 | 0.026 | 0.067 | 1 | 8 |
| foxp4 | 0.54797 | 0.170392202 | 0.132 | 0.067 | 1 | 8 |
| atp5mc3a | 0.54937 | -0.24356136 | 0.158 | 0.244 | 1 | 8 |
| aip | 0.55007 | 0.216099467 | 0.105 | 0.044 | 1 | 8 |
| PDCL3 | 0.555 | 0.165819362 | 0.079 | 0.044 | 1 | 8 |
| psmg3 | 0.55529 | 0.143403684 | 0.105 | 0.044 | 1 | 8 |
| ppp2r1ba | 0.55623 | -0.20446785 | 0.079 | 0.156 | 1 | 8 |
| arpc4l | 0.55691 | 0.164359862 | 0.237 | 0.156 | 1 | 8 |
| dctn3 | 0.55736 | 0.125535349 | 0.158 | 0.089 | 1 | 8 |
| si:dkeyp-75h12.5 | 0.55836 | 0.189479813 | 0.105 | 0.044 | 1 | 8 |
| ctnnb2 | 0.55886 | 0.188849309 | 0.105 | 0.044 | 1 | 8 |
| commd7 | 0.5606 | -0.17541627 | 0.053 | 0.089 | 1 | 8 |
| COX5B | 0.56141 | 0.237172195 | 0.316 | 0.222 | 1 | 8 |
| emc2 | 0.5623 | 0.146336874 | 0.132 | 0.067 | 1 | 8 |
| ssr4 | 0.56312 | 0.106055514 | 0.421 | 0.333 | 1 | 8 |
| nfkbiab | 0.56433 | 0.230846223 | 0.184 | 0.111 | 1 | 8 |
| mob1a | 0.56439 | 0.157670296 | 0.105 | 0.044 | 1 | 8 |
| rbpms2b | 0.56507 | 0.169735523 | 0.105 | 0.044 | 1 | 8 |
| dynll2b | 0.5654 | 0.178928567 | 0.105 | 0.044 | 1 | 8 |
| ndufv3 | 0.5655 | 0.175523693 | 0.105 | 0.044 | 1 | 8 |
| pfdn6 | 0.56568 | 0.194986928 | 0.105 | 0.044 | 1 | 8 |
| adrm1 | 0.5662 | 0.174484616 | 0.105 | 0.044 | 1 | 8 |
| rras | 0.56681 | 0.172979134 | 0.105 | 0.044 | 1 | 8 |
| dhdhl | 0.56683 | 0.183804747 | 0.105 | 0.044 | 1 | 8 |
| nr2f2 | 0.5672 | 0.182146387 | 0.105 | 0.044 | 1 | 8 |
| snx5 | 0.56814 | 0.168775645 | 0.105 | 0.044 | 1 | 8 |
| aamp | 0.56836 | 0.187075401 | 0.105 | 0.044 | 1 | 8 |
| snx3 | 0.56882 | 0.186028797 | 0.105 | 0.044 | 1 | 8 |
| ppil3 | 0.56902 | -0.16418668 | 0.026 | 0.067 | 1 | 8 |
| msna | 0.56949 | -0.15908781 | 0.053 | 0.111 | 1 | 8 |

| | | | | | | |
|---|---|---|---|---|---|---|
| atp5pb | 0.56993 | 0.128344355 | 0.474 | 0.378 | 1 | 8 |
| si:dkey-16p21.8 | 0.57059 | -0.19367554 | 0.105 | 0.156 | 1 | 8 |
| tegt | 0.57126 | 0.171855351 | 0.132 | 0.067 | 1 | 8 |
| dctn2 | 0.57184 | -0.17577757 | 0.026 | 0.067 | 1 | 8 |
| anxa2a | 0.57308 | 0.114687733 | 0.632 | 0.6 | 1 | 8 |
| mt-nd3 | 0.57336 | 0.152541444 | 0.079 | 0.044 | 1 | 8 |
| dad1 | 0.57775 | -0.14689436 | 0.184 | 0.267 | 1 | 8 |
| si:ch73-308l14.2 | 0.5787 | -0.18278363 | 0.026 | 0.067 | 1 | 8 |
| si:ch211-166a6.5 | 0.58145 | 0.195523448 | 0.132 | 0.067 | 1 | 8 |
| c1qbp | 0.5827 | -0.20871724 | 0.053 | 0.111 | 1 | 8 |
| hmgb2b | 0.58372 | -0.21190001 | 0.289 | 0.356 | 1 | 8 |
| mfap2 | 0.5845 | 0.208489173 | 0.132 | 0.067 | 1 | 8 |
| ckba | 0.58451 | 0.177463109 | 0.132 | 0.067 | 1 | 8 |
| crip1 | 0.58466 | -0.23421542 | 0.342 | 0.444 | 1 | 8 |
| lsm4 | 0.5848 | 0.15763575 | 0.158 | 0.089 | 1 | 8 |
| arpc3 | 0.58608 | 0.151749092 | 0.263 | 0.178 | 1 | 8 |
| ola1 | 0.58645 | -0.1223316 | 0.158 | 0.178 | 1 | 8 |
| eif2s1b | 0.58667 | -0.18631268 | 0.211 | 0.267 | 1 | 8 |
| mt-nd4 | 0.58706 | 0.237081523 | 0.158 | 0.089 | 1 | 8 |
| sf1 | 0.58756 | -0.19424323 | 0.053 | 0.111 | 1 | 8 |
| pin4 | 0.5885 | 0.159894538 | 0.079 | 0.044 | 1 | 8 |
| h3f3d | 0.58987 | 0.172569338 | 0.579 | 0.467 | 1 | 8 |
| irf2bp2a | 0.59019 | 0.201234126 | 0.132 | 0.067 | 1 | 8 |
| psmd1 | 0.5902 | -0.15019819 | 0.105 | 0.133 | 1 | 8 |
| rap2c | 0.59065 | -0.16845181 | 0.053 | 0.089 | 1 | 8 |
| puf60b | 0.59153 | -0.14065519 | 0.053 | 0.111 | 1 | 8 |
| col5a2a | 0.59161 | 0.207178722 | 0.132 | 0.067 | 1 | 8 |
| sdhdb | 0.59194 | 0.145229127 | 0.132 | 0.067 | 1 | 8 |
| tjp2b | 0.59233 | 0.190652257 | 0.132 | 0.067 | 1 | 8 |
| anp32b | 0.59319 | -0.16480947 | 0.053 | 0.089 | 1 | 8 |
| nop56 | 0.59333 | 0.210351261 | 0.158 | 0.089 | 1 | 8 |
| fkbp1aa | 0.59398 | 0.103004685 | 0.579 | 0.489 | 1 | 8 |
| uqcrh | 0.59554 | -0.22823684 | 0.158 | 0.244 | 1 | 8 |
| si:ch73-234b20.5 | 0.59823 | 0.216481819 | 0.184 | 0.111 | 1 | 8 |
| arf6b | 0.59891 | 0.167976512 | 0.158 | 0.089 | 1 | 8 |
| tmsb1 | 0.60184 | 0.188396909 | 0.658 | 0.556 | 1 | 8 |
| ccni | 0.60276 | 0.220644907 | 0.237 | 0.156 | 1 | 8 |
| atp5md | 0.60288 | 0.191046432 | 0.158 | 0.089 | 1 | 8 |
| hells | 0.60344 | -0.17777763 | 0.026 | 0.067 | 1 | 8 |
| hspe1 | 0.60463 | -0.19278551 | 0.158 | 0.244 | 1 | 8 |
| nutf2 | 0.60573 | -0.16252678 | 0.026 | 0.067 | 1 | 8 |
| zgc:100918 | 0.60653 | 0.183634558 | 0.132 | 0.067 | 1 | 8 |
| mt-nd2 | 0.6067 | 0.204789271 | 0.132 | 0.067 | 1 | 8 |
| ccng1 | 0.60765 | 0.202601203 | 0.368 | 0.289 | 1 | 8 |
| ndufb2 | 0.60774 | 0.206397684 | 0.158 | 0.089 | 1 | 8 |
| cul1a | 0.60776 | 0.10773526 | 0.053 | 0.044 | 1 | 8 |
| ISCU (1 of many) | 0.60871 | 0.161973117 | 0.132 | 0.067 | 1 | 8 |
| bloc1s6 | 0.60968 | 0.176722858 | 0.132 | 0.067 | 1 | 8 |

| | | | | | | |
|---|---|---|---|---|---|---|
| ndufa5 | 0.61036 | 0.163660066 | 0.158 | 0.089 | 1 | 8 |
| thrap3b | 0.61039 | -0.18747555 | 0.053 | 0.111 | 1 | 8 |
| cacybp | 0.61164 | -0.17948325 | 0.053 | 0.111 | 1 | 8 |
| rhoaa | 0.6129 | -0.15616942 | 0.184 | 0.222 | 1 | 8 |
| ttc38 | 0.61319 | -0.16455247 | 0.026 | 0.067 | 1 | 8 |
| ftr82 | 0.61379 | 0.122464617 | 0.132 | 0.111 | 1 | 8 |
| yy1a | 0.61409 | -0.17480399 | 0.026 | 0.067 | 1 | 8 |
| aldoaa | 0.61488 | 0.155215212 | 0.184 | 0.111 | 1 | 8 |
| cbr1l | 0.61583 | 0.16203298 | 0.184 | 0.111 | 1 | 8 |
| irf2bp2b | 0.61595 | 0.19441795 | 0.158 | 0.089 | 1 | 8 |
| ube2v2 | 0.61676 | 0.157558753 | 0.158 | 0.089 | 1 | 8 |
| slc38a5b | 0.61693 | -0.15630289 | 0.105 | 0.133 | 1 | 8 |
| dnajc1 | 0.61717 | -0.16004511 | 0.026 | 0.067 | 1 | 8 |
| tuba8l | 0.61786 | -0.20048928 | 0.158 | 0.244 | 1 | 8 |
| psme1 | 0.61797 | 0.215619134 | 0.237 | 0.156 | 1 | 8 |
| cnn2 | 0.61837 | -0.20976964 | 0.158 | 0.244 | 1 | 8 |
| serinc2 | 0.61838 | -0.1466117 | 0.026 | 0.067 | 1 | 8 |
| kars | 0.61854 | -0.19859215 | 0.158 | 0.244 | 1 | 8 |
| aimp2 | 0.61874 | -0.18462465 | 0.105 | 0.156 | 1 | 8 |
| sf3b6 | 0.61882 | -0.20103732 | 0.158 | 0.244 | 1 | 8 |
| eif4a1b | 0.62009 | 0.112961607 | 0.368 | 0.289 | 1 | 8 |
| zc3h15 | 0.6204 | -0.17125799 | 0.026 | 0.067 | 1 | 8 |
| sf3b5 | 0.62059 | -0.17741442 | 0.053 | 0.111 | 1 | 8 |
| cebpa | 0.62177 | 0.220022335 | 0.237 | 0.156 | 1 | 8 |
| aldh9a1a.1 | 0.62196 | -0.11553883 | 0.395 | 0.422 | 1 | 8 |
| creb1a | 0.62199 | -0.16744213 | 0.053 | 0.111 | 1 | 8 |
| ppp6c | 0.6226 | -0.1678883 | 0.053 | 0.111 | 1 | 8 |
| cox7a2l | 0.62273 | -0.16085842 | 0.026 | 0.067 | 1 | 8 |
| eef2b | 0.62317 | 0.126960182 | 0.816 | 0.733 | 1 | 8 |
| sept7b | 0.62498 | -0.16977639 | 0.053 | 0.111 | 1 | 8 |
| cpsf6 | 0.62509 | -0.16032524 | 0.053 | 0.111 | 1 | 8 |
| vcp | 0.62551 | -0.1670554 | 0.026 | 0.067 | 1 | 8 |
| tmed10 | 0.62569 | -0.17319068 | 0.053 | 0.111 | 1 | 8 |
| stau1 | 0.62582 | -0.14756023 | 0.053 | 0.111 | 1 | 8 |
| mrpl57 | 0.62582 | -0.16828695 | 0.053 | 0.111 | 1 | 8 |
| mrps14 | 0.62602 | -0.11744701 | 0.026 | 0.067 | 1 | 8 |
| ufm1 | 0.62683 | -0.11651771 | 0.053 | 0.067 | 1 | 8 |
| ube2l3b | 0.62748 | 0.182009568 | 0.158 | 0.089 | 1 | 8 |
| myl9b | 0.62834 | 0.182514893 | 0.158 | 0.089 | 1 | 8 |
| fam162a | 0.62902 | -0.1414254 | 0.026 | 0.067 | 1 | 8 |
| h2afx1 | 0.63027 | -0.15134618 | 0.184 | 0.222 | 1 | 8 |
| atp5po | 0.63052 | 0.188596865 | 0.316 | 0.222 | 1 | 8 |
| si:ch211-66k16.27 | 0.63083 | 0.183473257 | 0.158 | 0.089 | 1 | 8 |
| magoh | 0.6311 | -0.1515097 | 0.026 | 0.067 | 1 | 8 |
| golga7 | 0.63123 | -0.14047416 | 0.105 | 0.133 | 1 | 8 |
| ndufab1a | 0.63124 | 0.176460005 | 0.158 | 0.089 | 1 | 8 |
| gchfr | 0.63144 | 0.193351892 | 0.158 | 0.089 | 1 | 8 |
| si:ch73-46j18.5 | 0.63177 | 0.11132349 | 0.737 | 0.689 | 1 | 8 |

| | | | | | | |
|---|---|---|---|---|---|---|
| rsl1d1 | 0.63248 | 0.178496442 | 0.263 | 0.178 | 1 | 8 |
| psmc6 | 0.63402 | -0.1865267 | 0.079 | 0.133 | 1 | 8 |
| calm1a | 0.63409 | 0.180182862 | 0.263 | 0.178 | 1 | 8 |
| si:dkey-180p18.9 | 0.63714 | -0.14194453 | 0.079 | 0.111 | 1 | 8 |
| dnajb6a | 0.63866 | 0.101846919 | 0.053 | 0.044 | 1 | 8 |
| pim1 | 0.6394 | -0.11428735 | 0.053 | 0.067 | 1 | 8 |
| ptp4a1 | 0.63955 | -0.17496898 | 0.105 | 0.178 | 1 | 8 |
| capzb | 0.6397 | -0.15823107 | 0.184 | 0.222 | 1 | 8 |
| nfixb | 0.64011 | 0.179519445 | 0.132 | 0.089 | 1 | 8 |
| sept2 | 0.64024 | 0.184813708 | 0.263 | 0.178 | 1 | 8 |
| ilk | 0.6405 | 0.106044742 | 0.079 | 0.067 | 1 | 8 |
| rtn4a | 0.64199 | 0.17350045 | 0.211 | 0.133 | 1 | 8 |
| rfc4 | 0.64345 | -0.16103286 | 0.026 | 0.067 | 1 | 8 |
| vim | 0.64497 | 0.183101246 | 0.158 | 0.111 | 1 | 8 |
| chac1 | 0.64559 | 0.184733538 | 0.211 | 0.156 | 1 | 8 |
| rbm4.3 | 0.64614 | 0.1896529 | 0.211 | 0.133 | 1 | 8 |
| adh8a | 0.64686 | -0.17433275 | 0.184 | 0.222 | 1 | 8 |
| rps6kb1b | 0.64725 | -0.18922767 | 0.026 | 0.067 | 1 | 8 |
| mcts1 | 0.64895 | -0.10290536 | 0.026 | 0.067 | 1 | 8 |
| cops9 | 0.64926 | -0.16602161 | 0.053 | 0.089 | 1 | 8 |
| scn4ba | 0.64933 | 0.101843863 | 0.053 | 0.044 | 1 | 8 |
| ssr3 | 0.65096 | -0.19025526 | 0.263 | 0.356 | 1 | 8 |
| cdc37 | 0.65302 | -0.1343913 | 0.026 | 0.067 | 1 | 8 |
| u2af2b | 0.65305 | -0.12818533 | 0.026 | 0.067 | 1 | 8 |
| dut | 0.65309 | -0.16470661 | 0.026 | 0.067 | 1 | 8 |
| si:ch211-214j24.10 | 0.65838 | -0.17754942 | 0.105 | 0.156 | 1 | 8 |
| atp1a1a.1 | 0.65859 | -0.15935495 | 0.053 | 0.089 | 1 | 8 |
| seh1l | 0.65932 | -0.15187021 | 0.026 | 0.067 | 1 | 8 |
| u2af2a | 0.66022 | -0.15676767 | 0.053 | 0.089 | 1 | 8 |
| tktb | 0.66085 | -0.16021252 | 0.053 | 0.089 | 1 | 8 |
| afdna | 0.66108 | -0.13019475 | 0.026 | 0.067 | 1 | 8 |
| chmp1a | 0.66429 | -0.11984876 | 0.026 | 0.067 | 1 | 8 |
| sufu | 0.66464 | 0.144676249 | 0.079 | 0.044 | 1 | 8 |
| ppa1b | 0.6663 | -0.13141976 | 0.026 | 0.067 | 1 | 8 |
| mrpl3 | 0.66639 | -0.15057946 | 0.026 | 0.067 | 1 | 8 |
| calm3b | 0.66655 | -0.13821813 | 0.211 | 0.244 | 1 | 8 |
| zgc:110182 | 0.66683 | -0.19906182 | 0.184 | 0.267 | 1 | 8 |
| ube2nb | 0.66691 | -0.1108686 | 0.053 | 0.067 | 1 | 8 |
| pdcd5 | 0.66743 | -0.12330971 | 0.026 | 0.067 | 1 | 8 |
| psmg1 | 0.66769 | -0.11855317 | 0.026 | 0.067 | 1 | 8 |
| sec62 | 0.66779 | -0.12190901 | 0.026 | 0.067 | 1 | 8 |
| cct7 | 0.66878 | 0.165264632 | 0.158 | 0.111 | 1 | 8 |
| nr2f1a | 0.66959 | -0.15751698 | 0.053 | 0.089 | 1 | 8 |
| pole4 | 0.66982 | -0.12990464 | 0.026 | 0.067 | 1 | 8 |
| si:ch211-284b7.3 | 0.67 | -0.13109761 | 0.026 | 0.067 | 1 | 8 |
| cluap1 | 0.67014 | -0.10594543 | 0.026 | 0.067 | 1 | 8 |
| srsf5a | 0.67126 | 0.190954434 | 0.289 | 0.222 | 1 | 8 |
| rad23b | 0.6713 | -0.13737591 | 0.105 | 0.133 | 1 | 8 |

| | | | | | | |
|---|---|---|---|---|---|---|
| ran | 0.6728 | -0.14661921 | 0.579 | 0.667 | 1 | 8 |
| ddx18 | 0.67388 | -0.13427207 | 0.105 | 0.133 | 1 | 8 |
| mtpn | 0.67495 | -0.15403748 | 0.132 | 0.2 | 1 | 8 |
| polr2h | 0.67569 | -0.15236815 | 0.026 | 0.067 | 1 | 8 |
| phactr4a | 0.67575 | -0.12606633 | 0.026 | 0.067 | 1 | 8 |
| sec11a | 0.67593 | -0.10974082 | 0.053 | 0.067 | 1 | 8 |
| ubr7 | 0.6766 | -0.13770602 | 0.026 | 0.067 | 1 | 8 |
| calm2a | 0.67668 | -0.17739317 | 0.132 | 0.178 | 1 | 8 |
| ost4 | 0.67846 | -0.19670906 | 0.132 | 0.2 | 1 | 8 |
| slc43a3b | 0.67885 | -0.13286319 | 0.079 | 0.133 | 1 | 8 |
| ddx39b | 0.67976 | -0.13167253 | 0.026 | 0.067 | 1 | 8 |
| srsf1b | 0.67989 | -0.13429822 | 0.026 | 0.067 | 1 | 8 |
| lamb1b | 0.68009 | -0.13422165 | 0.026 | 0.067 | 1 | 8 |
| tcea2 | 0.68034 | -0.10173408 | 0.026 | 0.067 | 1 | 8 |
| nnt | 0.68188 | -0.11404552 | 0.026 | 0.067 | 1 | 8 |
| pbx1a | 0.68203 | -0.1327643 | 0.026 | 0.067 | 1 | 8 |
| mt-cyb | 0.68206 | 0.178626064 | 0.184 | 0.156 | 1 | 8 |
| il15 | 0.68212 | -0.10175602 | 0.026 | 0.067 | 1 | 8 |
| sh3gl1b | 0.68221 | -0.12662633 | 0.026 | 0.067 | 1 | 8 |
| uchl1 | 0.68277 | -0.11069182 | 0.026 | 0.067 | 1 | 8 |
| atg101 | 0.6832 | -0.12727706 | 0.026 | 0.067 | 1 | 8 |
| nol12 | 0.68344 | -0.14191234 | 0.026 | 0.067 | 1 | 8 |
| gstp1 | 0.68359 | 0.165445721 | 0.211 | 0.178 | 1 | 8 |
| cdc42se1 | 0.68373 | -0.10266479 | 0.026 | 0.067 | 1 | 8 |
| sart1 | 0.68442 | -0.1282848 | 0.026 | 0.067 | 1 | 8 |
| smim29 | 0.68447 | -0.11393317 | 0.026 | 0.067 | 1 | 8 |
| higd2a | 0.68458 | -0.10924279 | 0.026 | 0.067 | 1 | 8 |
| pcolcea | 0.68459 | -0.13635055 | 0.026 | 0.067 | 1 | 8 |
| pop5 | 0.68505 | -0.11264523 | 0.026 | 0.067 | 1 | 8 |
| itgb1a | 0.68563 | -0.15982902 | 0.026 | 0.067 | 1 | 8 |
| cdc42 | 0.68584 | -0.18547338 | 0.237 | 0.311 | 1 | 8 |
| exoc2 | 0.68585 | -0.12443844 | 0.026 | 0.067 | 1 | 8 |
| si:dkey-6n21.12 | 0.68593 | -0.13047441 | 0.026 | 0.067 | 1 | 8 |
| ube2g1b | 0.68608 | -0.12472546 | 0.026 | 0.067 | 1 | 8 |
| bud23 | 0.68622 | 0.149492515 | 0.079 | 0.044 | 1 | 8 |
| farsa | 0.68644 | -0.13765613 | 0.026 | 0.067 | 1 | 8 |
| hnrnpc | 0.68649 | -0.11434347 | 0.026 | 0.067 | 1 | 8 |
| drap1 | 0.68658 | -0.17078545 | 0.079 | 0.133 | 1 | 8 |
| uqcrfs1 | 0.68698 | -0.11412115 | 0.026 | 0.067 | 1 | 8 |
| nup50 | 0.68707 | -0.14854256 | 0.026 | 0.067 | 1 | 8 |
| CU459056.1 | 0.68724 | -0.11877477 | 0.026 | 0.067 | 1 | 8 |
| per1b | 0.68743 | -0.13530936 | 0.026 | 0.067 | 1 | 8 |
| irf9 | 0.68744 | -0.13449477 | 0.026 | 0.067 | 1 | 8 |
| mrpl17 | 0.68748 | -0.12656009 | 0.026 | 0.067 | 1 | 8 |
| si:dkey-9i23.16 | 0.68748 | -0.12033458 | 0.026 | 0.067 | 1 | 8 |
| syf2 | 0.68936 | -0.14591602 | 0.053 | 0.089 | 1 | 8 |
| hp1bp3 | 0.68997 | -0.16947741 | 0.079 | 0.133 | 1 | 8 |
| fxyd1 | 0.69054 | -0.19047726 | 0.132 | 0.2 | 1 | 8 |

| | | | | | | |
|---|---|---|---|---|---|---|
| abcf1 | 0.69331 | 0.144689584 | 0.105 | 0.067 | 1 | 8 |
| csnk1a1 | 0.69378 | 0.127993894 | 0.158 | 0.133 | 1 | 8 |
| ipo7 | 0.69415 | 0.139877473 | 0.079 | 0.044 | 1 | 8 |
| tspan4a | 0.69896 | -0.16731282 | 0.079 | 0.133 | 1 | 8 |
| snx6 | 0.70078 | 0.155223443 | 0.079 | 0.044 | 1 | 8 |
| mrpl12 | 0.70141 | -0.10431075 | 0.053 | 0.067 | 1 | 8 |
| cavin2b | 0.70178 | -0.10701538 | 0.289 | 0.311 | 1 | 8 |
| tnrc6c1 | 0.70363 | -0.13723323 | 0.079 | 0.133 | 1 | 8 |
| mpc1 | 0.70364 | -0.10594663 | 0.053 | 0.067 | 1 | 8 |
| rbbp4 | 0.70566 | -0.10460159 | 0.053 | 0.067 | 1 | 8 |
| cyp2ad3 | 0.70617 | 0.152558472 | 0.105 | 0.067 | 1 | 8 |
| si:ch211-132b12.7 | 0.70621 | 0.156027491 | 0.079 | 0.044 | 1 | 8 |
| knop1 | 0.70875 | 0.142948897 | 0.079 | 0.044 | 1 | 8 |
| si:dkey-7j14.6 | 0.71009 | -0.17408904 | 0.079 | 0.133 | 1 | 8 |
| ncl | 0.71164 | 0.176792453 | 0.395 | 0.311 | 1 | 8 |
| cebpb | 0.71189 | 0.1570745 | 0.158 | 0.133 | 1 | 8 |
| eif3ba | 0.71306 | -0.12814698 | 0.211 | 0.244 | 1 | 8 |
| capns1b | 0.71451 | -0.16916498 | 0.289 | 0.356 | 1 | 8 |
| pnrc2 | 0.71478 | 0.139032868 | 0.579 | 0.489 | 1 | 8 |
| tmem88b | 0.71955 | -0.14073537 | 0.079 | 0.133 | 1 | 8 |
| chmp2a | 0.72012 | -0.15302025 | 0.053 | 0.089 | 1 | 8 |
| cct5 | 0.72101 | 0.113807772 | 0.316 | 0.244 | 1 | 8 |
| acadm | 0.72103 | -0.13717555 | 0.079 | 0.111 | 1 | 8 |
| ywhaz | 0.72184 | 0.135343636 | 0.368 | 0.289 | 1 | 8 |
| rab5ab | 0.72234 | -0.11049714 | 0.053 | 0.067 | 1 | 8 |
| sf3a2 | 0.72548 | -0.14734489 | 0.079 | 0.133 | 1 | 8 |
| cox7a1 | 0.72577 | -0.13563 | 0.079 | 0.133 | 1 | 8 |
| rbp5 | 0.72705 | -0.15194226 | 0.079 | 0.111 | 1 | 8 |
| socs3a | 0.72786 | 0.137046252 | 0.395 | 0.356 | 1 | 8 |
| srsf3b | 0.72852 | 0.166653615 | 0.184 | 0.133 | 1 | 8 |
| tomm20a | 0.72872 | 0.156372686 | 0.105 | 0.067 | 1 | 8 |
| ube2d1b | 0.72947 | 0.100977557 | 0.211 | 0.156 | 1 | 8 |
| selenbp1 | 0.73112 | 0.166126094 | 0.211 | 0.156 | 1 | 8 |
| eif3ja | 0.73148 | -0.16787809 | 0.132 | 0.178 | 1 | 8 |
| pik3r3a | 0.7329 | 0.141327253 | 0.105 | 0.067 | 1 | 8 |
| psma1 | 0.73674 | -0.17162539 | 0.184 | 0.244 | 1 | 8 |
| pno1 | 0.7382 | -0.15040061 | 0.105 | 0.156 | 1 | 8 |
| hnrnpa0b | 0.74472 | 0.125758185 | 0.316 | 0.244 | 1 | 8 |
| npc2 | 0.74489 | 0.137493511 | 0.079 | 0.044 | 1 | 8 |
| anapc13 | 0.74645 | -0.13533299 | 0.053 | 0.089 | 1 | 8 |
| sri | 0.74788 | -0.17480291 | 0.158 | 0.222 | 1 | 8 |
| mdh2 | 0.74798 | -0.16282572 | 0.158 | 0.222 | 1 | 8 |
| si:ch211-288g17.3 | 0.74851 | 0.149024775 | 0.342 | 0.267 | 1 | 8 |
| vat1 | 0.75193 | -0.16917127 | 0.105 | 0.156 | 1 | 8 |
| txndc9 | 0.75686 | -0.13240212 | 0.079 | 0.111 | 1 | 8 |
| tpm4a | 0.75752 | 0.149681164 | 0.342 | 0.267 | 1 | 8 |
| cyp2n13 | 0.75771 | 0.124137986 | 0.079 | 0.044 | 1 | 8 |
| fgf24 | 0.75859 | -0.12069899 | 0.105 | 0.156 | 1 | 8 |

| | | | | | | |
|---|---|---|---|---|---|---|
| mrps10 | 0.76086 | 0.128007782 | 0.079 | 0.044 | 1 | 8 |
| col6a3 | 0.76313 | 0.103012657 | 0.105 | 0.067 | 1 | 8 |
| ppp1caa | 0.76366 | -0.1170897 | 0.105 | 0.156 | 1 | 8 |
| snrnp70 | 0.7641 | -0.13006981 | 0.053 | 0.089 | 1 | 8 |
| rap1b | 0.76491 | -0.111715 | 0.184 | 0.244 | 1 | 8 |
| palm1b | 0.76622 | -0.13974237 | 0.053 | 0.089 | 1 | 8 |
| psmd8 | 0.76656 | 0.132683915 | 0.105 | 0.067 | 1 | 8 |
| itm2ba | 0.7673 | 0.111858699 | 0.211 | 0.156 | 1 | 8 |
| tjp1a | 0.76783 | -0.14036712 | 0.053 | 0.089 | 1 | 8 |
| prdx5 | 0.768 | 0.108256544 | 0.211 | 0.156 | 1 | 8 |
| prdx6 | 0.7681 | 0.110188084 | 0.158 | 0.133 | 1 | 8 |
| cbx3a | 0.77 | 0.146797153 | 0.132 | 0.089 | 1 | 8 |
| zgc:162944 | 0.77025 | 0.124599587 | 0.289 | 0.222 | 1 | 8 |
| zgc:66388 | 0.77128 | 0.143233076 | 0.132 | 0.089 | 1 | 8 |
| atp5mea | 0.77156 | 0.117664955 | 0.184 | 0.156 | 1 | 8 |
| tgif1 | 0.77321 | -0.1424497 | 0.053 | 0.089 | 1 | 8 |
| rassf8b | 0.77372 | -0.12059029 | 0.053 | 0.089 | 1 | 8 |
| atp5f1c | 0.77496 | 0.153725447 | 0.237 | 0.178 | 1 | 8 |
| timm13 | 0.77506 | -0.12413657 | 0.053 | 0.089 | 1 | 8 |
| si:dkey-51e6.1 | 0.77733 | -0.13591199 | 0.132 | 0.178 | 1 | 8 |
| pitpnbl | 0.78341 | -0.11957711 | 0.053 | 0.089 | 1 | 8 |
| plscr3b | 0.78385 | 0.143131009 | 0.105 | 0.067 | 1 | 8 |
| pdgfrl | 0.78609 | 0.121216934 | 0.079 | 0.044 | 1 | 8 |
| sdhc | 0.78776 | 0.110297373 | 0.132 | 0.089 | 1 | 8 |
| churc1 | 0.78795 | -0.12574687 | 0.053 | 0.089 | 1 | 8 |
| lin7c | 0.78799 | 0.12516302 | 0.079 | 0.044 | 1 | 8 |
| srsf9 | 0.78847 | 0.126410908 | 0.079 | 0.044 | 1 | 8 |
| capza1a | 0.78885 | 0.123872138 | 0.237 | 0.178 | 1 | 8 |
| atp6v1e1b | 0.7912 | 0.122974723 | 0.079 | 0.044 | 1 | 8 |
| csnk1da | 0.79222 | 0.119469128 | 0.079 | 0.044 | 1 | 8 |
| si:ch73-376l24.3 | 0.79434 | -0.12461558 | 0.079 | 0.111 | 1 | 8 |
| ctdsp2 | 0.79505 | -0.10864472 | 0.132 | 0.156 | 1 | 8 |
| eif4a2 | 0.79506 | -0.1364021 | 0.053 | 0.089 | 1 | 8 |
| fkbp7 | 0.79592 | 0.14292986 | 0.184 | 0.133 | 1 | 8 |
| elf1 | 0.79612 | 0.12272066 | 0.079 | 0.044 | 1 | 8 |
| csnk2b | 0.79744 | -0.12264462 | 0.184 | 0.222 | 1 | 8 |
| gmfb | 0.79804 | 0.146979509 | 0.237 | 0.178 | 1 | 8 |
| si:ch1073-459j12.1 | 0.79908 | 0.149455855 | 0.237 | 0.178 | 1 | 8 |
| psmb1 | 0.79948 | 0.127745461 | 0.237 | 0.178 | 1 | 8 |
| anxa11b | 0.79953 | -0.1424533 | 0.132 | 0.178 | 1 | 8 |
| cnp | 0.80044 | 0.121651595 | 0.105 | 0.067 | 1 | 8 |
| cox7c | 0.80212 | -0.11841281 | 0.395 | 0.467 | 1 | 8 |
| lgals9l3 | 0.80226 | 0.107314874 | 0.158 | 0.111 | 1 | 8 |
| coa5 | 0.80333 | 0.130055404 | 0.105 | 0.067 | 1 | 8 |
| hsf2 | 0.80459 | -0.11801092 | 0.053 | 0.089 | 1 | 8 |
| vdac1 | 0.80539 | 0.13374404 | 0.184 | 0.133 | 1 | 8 |
| cyp27c1 | 0.80559 | 0.104684305 | 0.079 | 0.044 | 1 | 8 |
| arf2b | 0.80645 | 0.11837434 | 0.079 | 0.044 | 1 | 8 |

| | | | | | | |
|---|---|---|---|---|---|---|
| cyb5r1 | 0.80692 | 0.120597373 | 0.079 | 0.044 | 1 | 8 |
| si:ch1073-291c23.2 | 0.80729 | 0.100249237 | 0.079 | 0.044 | 1 | 8 |
| tia1l | 0.80792 | 0.106357753 | 0.079 | 0.044 | 1 | 8 |
| dnaja2a | 0.80966 | 0.136260086 | 0.184 | 0.133 | 1 | 8 |
| map2k6 | 0.81006 | -0.12469057 | 0.053 | 0.089 | 1 | 8 |
| mcm7 | 0.81161 | -0.10933772 | 0.053 | 0.089 | 1 | 8 |
| sntb2 | 0.81205 | -0.10945343 | 0.053 | 0.089 | 1 | 8 |
| entpd1 | 0.81244 | 0.122307205 | 0.158 | 0.111 | 1 | 8 |
| hnrnpa1a | 0.81327 | 0.118143818 | 0.158 | 0.111 | 1 | 8 |
| llph | 0.81405 | -0.11763442 | 0.079 | 0.111 | 1 | 8 |
| id2a | 0.81438 | 0.115047793 | 0.184 | 0.133 | 1 | 8 |
| cfl1l | 0.81516 | 0.115338522 | 0.105 | 0.067 | 1 | 8 |
| CU694197.1 | 0.81759 | 0.118098162 | 0.184 | 0.133 | 1 | 8 |
| ak1 | 0.81797 | -0.11278561 | 0.105 | 0.133 | 1 | 8 |
| arl6ip5b | 0.81811 | 0.120879508 | 0.184 | 0.133 | 1 | 8 |
| rab2a | 0.81958 | 0.107328026 | 0.105 | 0.067 | 1 | 8 |
| eif6 | 0.82017 | 0.100102804 | 0.132 | 0.089 | 1 | 8 |
| anapc16 | 0.82102 | 0.107433546 | 0.158 | 0.111 | 1 | 8 |
| tob1b | 0.82159 | 0.123739747 | 0.105 | 0.067 | 1 | 8 |
| gdi2 | 0.82193 | 0.10743678 | 0.158 | 0.111 | 1 | 8 |
| lamp2 | 0.8239 | 0.105658142 | 0.132 | 0.089 | 1 | 8 |
| cdc42l | 0.82741 | -0.12045607 | 0.079 | 0.111 | 1 | 8 |
| myl12.2 | 0.83161 | -0.11044746 | 0.079 | 0.111 | 1 | 8 |
| jun | 0.83969 | 0.107242204 | 0.368 | 0.333 | 1 | 8 |
| smarce1 | 0.84019 | -0.12512111 | 0.079 | 0.111 | 1 | 8 |
| sub1b | 0.84532 | -0.11133419 | 0.105 | 0.133 | 1 | 8 |
| zgc:193541 | 0.85952 | -0.11188787 | 0.342 | 0.4 | 1 | 8 |
| cct4 | 0.86403 | 0.112908803 | 0.316 | 0.267 | 1 | 8 |
| eif2s3 | 0.86836 | 0.114162627 | 0.289 | 0.244 | 1 | 8 |
| si:ch1073-429i10.3.1 | 0.86899 | -0.10925665 | 0.079 | 0.111 | 1 | 8 |
| si:ch211-246m6.4 | 0.87032 | -0.11452796 | 0.079 | 0.111 | 1 | 8 |
| zgc:65873 | 0.87683 | -0.10154268 | 0.079 | 0.111 | 1 | 8 |
| hmgb1b | 0.88155 | 0.113899496 | 0.263 | 0.222 | 1 | 8 |
| txnipa | 0.88251 | 0.103604042 | 0.211 | 0.178 | 1 | 8 |
| mt-atp6 | 5.912181 | -0.95967352 | 0.293 | 0.9 | 0.1194083 | 9 |
| hmgb2b | 1.039122 | 1.024724788 | 0.69 | 0.1 | 0.2098716 | 9 |
| sumo3a | 3.534224 | 0.890276245 | 0.621 | 0.1 | 0.7138073 | 9 |
| acsbg2 | 4.645169 | -0.75587116 | 0 | 0.3 | 0.9381849 | 9 |
| rpl8 | 0.00016 | 0.347760476 | 0.897 | 0.85 | 1 | 9 |
| rpl22l1 | 0.00024 | 0.820359331 | 0.828 | 0.35 | 1 | 9 |
| zgc:172339 | 0.00025 | -0.84786844 | 0 | 0.25 | 1 | 9 |
| cct7 | 0.00028 | 0.682576219 | 0.586 | 0.1 | 1 | 9 |
| rpl11 | 0.00028 | 0.5093185 | 0.966 | 0.8 | 1 | 9 |
| anapc16 | 0.00029 | 0.376312432 | 0.362 | 0.05 | 1 | 9 |
| txndc9 | 0.0003 | 0.496161165 | 0.345 | 0 | 1 | 9 |
| tex2 | 0.00037 | -0.64180071 | 0.034 | 0.4 | 1 | 9 |
| serbp1a | 0.00038 | 0.686228254 | 0.862 | 0.4 | 1 | 9 |
| dad1 | 0.00038 | -0.55357894 | 0.621 | 0.75 | 1 | 9 |

| | | | | | | |
|---|---|---|---|---|---|---|
| ost4 | 0.0004 | -0.7156116 | 0.362 | 0.6 | 1 | 9 |
| mt-co2 | 0.00046 | -0.87868234 | 0.362 | 0.85 | 1 | 9 |
| si:ch211-212k18.7 | 0.00047 | -0.55886594 | 0.293 | 0.45 | 1 | 9 |
| slirp | 0.00048 | 0.620226121 | 0.483 | 0.05 | 1 | 9 |
| sec61g | 0.00055 | -0.53903031 | 0.776 | 0.9 | 1 | 9 |
| ckbb | 0.00061 | -1.01344243 | 0.086 | 0.4 | 1 | 9 |
| id2b | 0.00063 | -0.72764171 | 0.017 | 0.3 | 1 | 9 |
| rpl22 | 0.00073 | 0.498959989 | 0.966 | 0.65 | 1 | 9 |
| atp2a3 | 0.00085 | -0.82248808 | 0.034 | 0.35 | 1 | 9 |
| tegt | 0.00086 | -0.41264054 | 0.621 | 0.65 | 1 | 9 |
| rps27.1 | 0.00096 | 0.390802593 | 0.914 | 0.8 | 1 | 9 |
| mt-nd2 | 0.00108 | -0.49706823 | 0.397 | 0.85 | 1 | 9 |
| rpf2 | 0.00108 | 0.494131981 | 0.293 | 0 | 1 | 9 |
| sf3b4 | 0.00108 | 0.441935811 | 0.293 | 0 | 1 | 9 |
| jagn1b | 0.00108 | 0.544411103 | 0.293 | 0 | 1 | 9 |
| LO017835.1 | 0.00127 | -0.37442149 | 0 | 0.2 | 1 | 9 |
| btk | 0.00127 | -0.58238559 | 0 | 0.2 | 1 | 9 |
| CR762483.1 | 0.00127 | -0.51454242 | 0 | 0.2 | 1 | 9 |
| XLOC-006268 | 0.00127 | -0.42029281 | 0 | 0.2 | 1 | 9 |
| cyp11c1 | 0.00127 | -0.45687921 | 0 | 0.2 | 1 | 9 |
| si:ch73-359m17.7 | 0.00127 | -0.67712666 | 0 | 0.2 | 1 | 9 |
| LOC100538255 | 0.00127 | -1.12522866 | 0 | 0.2 | 1 | 9 |
| grna | 0.00127 | -0.92435849 | 0 | 0.2 | 1 | 9 |
| myh11b | 0.00127 | -0.75042196 | 0 | 0.2 | 1 | 9 |
| si:dkeyp-93a5.3 | 0.00127 | -0.9013726 | 0 | 0.2 | 1 | 9 |
| ak2 | 0.00138 | 0.536798259 | 0.655 | 0.2 | 1 | 9 |
| rpl17 | 0.00142 | 0.444250466 | 0.983 | 0.75 | 1 | 9 |
| eef2b | 0.00162 | 0.424366767 | 0.948 | 0.6 | 1 | 9 |
| rpl7 | 0.00162 | 0.599892092 | 0.862 | 0.6 | 1 | 9 |
| fbl | 0.00164 | 0.512467123 | 0.276 | 0 | 1 | 9 |
| ubr7 | 0.00164 | 0.468873403 | 0.276 | 0 | 1 | 9 |
| eif5a | 0.00187 | 0.539948413 | 0.862 | 0.45 | 1 | 9 |
| eif3ha | 0.00197 | 0.55920406 | 0.569 | 0.15 | 1 | 9 |
| plcg2 | 0.00204 | -0.74685682 | 0.034 | 0.35 | 1 | 9 |
| pdia4 | 0.00213 | -0.54664495 | 0.155 | 0.3 | 1 | 9 |
| rgs5a | 0.00224 | -0.54238916 | 0.017 | 0.25 | 1 | 9 |
| magt1 | 0.00234 | 0.277950478 | 0.31 | 0.05 | 1 | 9 |
| h2afy2 | 0.00241 | 0.650237971 | 0.414 | 0.05 | 1 | 9 |
| rbm8a | 0.00245 | 0.456826761 | 0.655 | 0.25 | 1 | 9 |
| nme3 | 0.00247 | 0.435153388 | 0.259 | 0 | 1 | 9 |
| lmnb1 | 0.00247 | 0.532296546 | 0.259 | 0 | 1 | 9 |
| mrps11 | 0.00247 | 0.383090082 | 0.259 | 0 | 1 | 9 |
| smox | 0.00247 | 0.407487855 | 0.259 | 0 | 1 | 9 |
| rps3a | 0.00248 | 0.449114379 | 0.931 | 0.75 | 1 | 9 |
| si:ch211-288g17.3 | 0.00253 | 0.641827771 | 0.414 | 0.05 | 1 | 9 |
| COX5B | 0.0026 | 0.517216744 | 0.776 | 0.35 | 1 | 9 |
| rps16 | 0.00262 | 0.454477545 | 0.966 | 0.7 | 1 | 9 |
| rpl10 | 0.00266 | 0.496855718 | 0.948 | 0.65 | 1 | 9 |

| | | | | | | |
|---|---|---|---|---|---|---|
| calr3b | 0.00267 | -0.51103062 | 0.224 | 0.3 | 1 | 9 |
| alox5a | 0.00291 | -0.71279165 | 0.017 | 0.25 | 1 | 9 |
| rsl24d1 | 0.00316 | 0.592177624 | 0.828 | 0.45 | 1 | 9 |
| sdf2l1 | 0.00336 | -0.66303778 | 0.138 | 0.35 | 1 | 9 |
| mt-cyb | 0.00353 | -0.74345753 | 0.276 | 0.7 | 1 | 9 |
| rpl27 | 0.00357 | 0.43404806 | 0.948 | 0.75 | 1 | 9 |
| tuba8l | 0.00359 | -0.13486975 | 0.483 | 0.35 | 1 | 9 |
| myl9a | 0.00369 | -0.71151873 | 0.034 | 0.3 | 1 | 9 |
| mrpl12 | 0.0037 | 0.428374363 | 0.241 | 0 | 1 | 9 |
| ndrg2 | 0.0037 | 0.399491638 | 0.241 | 0 | 1 | 9 |
| rcl1 | 0.0037 | 0.353314074 | 0.241 | 0 | 1 | 9 |
| kdm6bb | 0.0037 | 0.418863934 | 0.241 | 0 | 1 | 9 |
| ftsj3 | 0.0037 | 0.382269217 | 0.241 | 0 | 1 | 9 |
| ilf2 | 0.00374 | 0.608882882 | 0.483 | 0.15 | 1 | 9 |
| rpl39 | 0.00391 | 0.359827484 | 0.948 | 0.85 | 1 | 9 |
| pabpn1 | 0.00399 | 0.607330394 | 0.379 | 0.05 | 1 | 9 |
| phb | 0.0041 | 0.16716738 | 0.276 | 0.05 | 1 | 9 |
| hspa5 | 0.00459 | -0.62204104 | 0.328 | 0.5 | 1 | 9 |
| seta | 0.00481 | 0.575096153 | 0.483 | 0.1 | 1 | 9 |
| srsf9 | 0.00483 | 0.115702045 | 0.345 | 0.15 | 1 | 9 |
| fabp3 | 0.00485 | 0.737315297 | 0.603 | 0.2 | 1 | 9 |
| eef1g | 0.00486 | 0.512061698 | 0.914 | 0.6 | 1 | 9 |
| tubb2b | 0.00489 | 0.219146765 | 0.655 | 0.35 | 1 | 9 |
| idh2 | 0.00504 | 0.56534065 | 0.552 | 0.15 | 1 | 9 |
| hsp90b1 | 0.00525 | -0.69186553 | 0.241 | 0.35 | 1 | 9 |
| cct6a | 0.00526 | 0.630482184 | 0.552 | 0.2 | 1 | 9 |
| vbp1 | 0.0055 | 0.379569867 | 0.224 | 0 | 1 | 9 |
| rab11ba | 0.0055 | 0.337195501 | 0.224 | 0 | 1 | 9 |
| zc4h2 | 0.0055 | 0.42017139 | 0.224 | 0 | 1 | 9 |
| tmpob | 0.0055 | 0.39084557 | 0.224 | 0 | 1 | 9 |
| si:dkey-206d17.12 | 0.00554 | -0.62432597 | 0.069 | 0.35 | 1 | 9 |
| ube2l3a | 0.00565 | -0.60320393 | 0.138 | 0.5 | 1 | 9 |
| jpt1b | 0.00565 | 0.626241072 | 0.466 | 0.1 | 1 | 9 |
| pbxip1b | 0.00565 | -0.65786704 | 0.069 | 0.3 | 1 | 9 |
| CABZ01056052.1 | 0.00574 | -0.67460506 | 0.017 | 0.2 | 1 | 9 |
| bik | 0.00596 | -0.53535521 | 0.103 | 0.45 | 1 | 9 |
| ppa1b | 0.00606 | 0.165941061 | 0.293 | 0.1 | 1 | 9 |
| ppp3ccb | 0.00615 | -0.4945331 | 0.259 | 0.4 | 1 | 9 |
| pabpc1a | 0.00625 | 0.30701585 | 0.914 | 0.6 | 1 | 9 |
| CU929070.1 | 0.00628 | 0.361776224 | 0.293 | 0.05 | 1 | 9 |
| ppp1r26 | 0.0063 | -0.24944169 | 0 | 0.15 | 1 | 9 |
| sfxn5b | 0.0063 | -0.39782271 | 0 | 0.15 | 1 | 9 |
| si:dkeyp-93a5.2 | 0.0063 | -0.53253296 | 0 | 0.15 | 1 | 9 |
| LOC100331140 | 0.0063 | -0.44036078 | 0 | 0.15 | 1 | 9 |
| LOC110438226 | 0.0063 | -0.49261673 | 0 | 0.15 | 1 | 9 |
| adgrf3a | 0.0063 | -0.4425953 | 0 | 0.15 | 1 | 9 |
| arfip2a | 0.0063 | -0.23682996 | 0 | 0.15 | 1 | 9 |
| ergic1 | 0.0063 | -0.25960772 | 0 | 0.15 | 1 | 9 |

| | | | | | | |
|---|---|---|---|---|---|---|
| grtp1a | 0.0063 | -0.40650451 | 0 | 0.15 | 1 | 9 |
| mon1bb | 0.0063 | -0.37578679 | 0 | 0.15 | 1 | 9 |
| prr15la | 0.0063 | -0.47321071 | 0 | 0.15 | 1 | 9 |
| si:ch1073-214b20.2 | 0.0063 | -0.34579517 | 0 | 0.15 | 1 | 9 |
| si:ch211-130h14.6 | 0.0063 | -0.28426583 | 0 | 0.15 | 1 | 9 |
| si:dkeyp-117b11.1 | 0.0063 | -0.4374906 | 0 | 0.15 | 1 | 9 |
| sorbs2a | 0.0063 | -0.24823864 | 0 | 0.15 | 1 | 9 |
| zgc:158412 | 0.0063 | -0.36840969 | 0 | 0.15 | 1 | 9 |
| zgc:162171 | 0.0063 | -0.24233956 | 0 | 0.15 | 1 | 9 |
| pik3cg | 0.0063 | -0.43483809 | 0 | 0.15 | 1 | 9 |
| rac2 | 0.0063 | -0.2854359 | 0 | 0.15 | 1 | 9 |
| LOC108190702 | 0.0063 | -0.39487918 | 0 | 0.15 | 1 | 9 |
| ccdc186 | 0.0063 | -0.28711534 | 0 | 0.15 | 1 | 9 |
| draxin | 0.0063 | -0.37194073 | 0 | 0.15 | 1 | 9 |
| fam57ba | 0.0063 | -0.33782785 | 0 | 0.15 | 1 | 9 |
| hipk2 | 0.0063 | -0.30768436 | 0 | 0.15 | 1 | 9 |
| znf1083 | 0.0063 | -0.25036302 | 0 | 0.15 | 1 | 9 |
| adamts10 | 0.0063 | -0.36971143 | 0 | 0.15 | 1 | 9 |
| arrdc2 | 0.00651 | -0.6675993 | 0.086 | 0.3 | 1 | 9 |
| h3f3d | 0.00678 | 0.459086603 | 0.862 | 0.5 | 1 | 9 |
| hmgb3b | 0.00678 | 0.509538943 | 0.379 | 0.05 | 1 | 9 |
| rpl4 | 0.00681 | 0.380025899 | 0.983 | 0.8 | 1 | 9 |
| hepacam2 | 0.00692 | -0.66473543 | 0.052 | 0.3 | 1 | 9 |
| traf4b | 0.00713 | -0.48888491 | 0.017 | 0.25 | 1 | 9 |
| hnrnpaba | 0.00714 | 0.611414377 | 0.759 | 0.4 | 1 | 9 |
| hmgn7 | 0.00715 | 0.565975051 | 0.741 | 0.35 | 1 | 9 |
| rpl28 | 0.00722 | 0.337763605 | 0.983 | 0.8 | 1 | 9 |
| hdlbpa | 0.00727 | -0.60258963 | 0.293 | 0.6 | 1 | 9 |
| hmga1a | 0.00737 | 0.652574875 | 0.724 | 0.4 | 1 | 9 |
| snrpd1 | 0.00759 | 0.618540862 | 0.466 | 0.1 | 1 | 9 |
| nutf2l | 0.00765 | 0.433954533 | 0.362 | 0.05 | 1 | 9 |
| rps9 | 0.00785 | 0.419242124 | 0.948 | 0.75 | 1 | 9 |
| zgc:77849 | 0.00797 | 0.419508782 | 0.431 | 0.1 | 1 | 9 |
| ccl19a.1 | 0.00803 | -0.71857454 | 0.069 | 0.25 | 1 | 9 |
| ifrd2 | 0.00814 | 0.32377118 | 0.207 | 0 | 1 | 9 |
| psenen | 0.00814 | 0.35063532 | 0.207 | 0 | 1 | 9 |
| chchd3a | 0.00814 | 0.317243669 | 0.207 | 0 | 1 | 9 |
| ctnna1 | 0.00814 | 0.303135611 | 0.207 | 0 | 1 | 9 |
| drg2 | 0.00814 | 0.319030525 | 0.207 | 0 | 1 | 9 |
| khdrbs1b | 0.00814 | 0.354791493 | 0.207 | 0 | 1 | 9 |
| marcksb | 0.00814 | 0.434778275 | 0.207 | 0 | 1 | 9 |
| mrpl2 | 0.00814 | 0.336851831 | 0.207 | 0 | 1 | 9 |
| ssb | 0.00814 | 0.352780307 | 0.207 | 0 | 1 | 9 |
| tufm | 0.00814 | 0.331414537 | 0.207 | 0 | 1 | 9 |
| ssr4 | 0.00824 | -0.24234799 | 0.621 | 0.55 | 1 | 9 |
| ndufab1b | 0.00844 | 0.454914411 | 0.362 | 0.05 | 1 | 9 |
| lyn | 0.00849 | -0.36988472 | 0.017 | 0.25 | 1 | 9 |
| hspe1 | 0.00855 | 0.379619741 | 0.414 | 0.1 | 1 | 9 |

| | | | | | | |
|---|---|---|---|---|---|---|
| rpl13a | 0.0086 | 0.36890152 | 0.966 | 0.8 | 1 | 9 |
| pdia6 | 0.00862 | -0.46434023 | 0.241 | 0.3 | 1 | 9 |
| dkc1 | 0.00863 | 0.401495677 | 0.328 | 0.05 | 1 | 9 |
| dag1 | 0.00917 | 0.258701562 | 0.259 | 0.05 | 1 | 9 |
| paip2b | 0.00922 | 0.515299093 | 0.586 | 0.2 | 1 | 9 |
| rab1ab | 0.00971 | -0.38900442 | 0.293 | 0.35 | 1 | 9 |
| ostf1 | 0.01009 | -0.52805894 | 0.034 | 0.25 | 1 | 9 |
| p2rx4b | 0.01033 | -0.5970444 | 0.034 | 0.25 | 1 | 9 |
| eif3c | 0.01036 | 0.20260371 | 0.466 | 0.2 | 1 | 9 |
| pfdn6 | 0.01051 | 0.150389746 | 0.293 | 0.1 | 1 | 9 |
| tuba1b | 0.01053 | -0.58318515 | 0.086 | 0.25 | 1 | 9 |
| gpx1b | 0.01084 | -0.69875165 | 0.052 | 0.3 | 1 | 9 |
| rps23 | 0.01109 | 0.381736906 | 0.966 | 0.75 | 1 | 9 |
| capza1a | 0.01122 | 0.336567224 | 0.466 | 0.15 | 1 | 9 |
| wbp2nl | 0.01166 | -0.52639833 | 0.207 | 0.4 | 1 | 9 |
| mbnl1 | 0.01175 | -0.56849627 | 0.121 | 0.35 | 1 | 9 |
| icn | 0.01193 | -0.72322002 | 0.431 | 0.8 | 1 | 9 |
| rn7sk | 0.01199 | 0.401355037 | 0.19 | 0 | 1 | 9 |
| cbx1a | 0.01199 | 0.394844323 | 0.19 | 0 | 1 | 9 |
| ccnl1b | 0.01199 | 0.295686094 | 0.19 | 0 | 1 | 9 |
| cd99l2 | 0.01199 | 0.314858052 | 0.19 | 0 | 1 | 9 |
| cnot9 | 0.01199 | 0.368491666 | 0.19 | 0 | 1 | 9 |
| gigyf1b | 0.01199 | 0.378558605 | 0.19 | 0 | 1 | 9 |
| lmnb2 | 0.01199 | 0.308425045 | 0.19 | 0 | 1 | 9 |
| rras | 0.01199 | 0.34833047 | 0.19 | 0 | 1 | 9 |
| rrp7a | 0.01199 | 0.289092374 | 0.19 | 0 | 1 | 9 |
| si:ch73-308l14.2 | 0.01199 | 0.334659493 | 0.19 | 0 | 1 | 9 |
| agrn | 0.01199 | 0.355307293 | 0.19 | 0 | 1 | 9 |
| ash2l | 0.01199 | 0.331183405 | 0.19 | 0 | 1 | 9 |
| bckdhb | 0.01199 | 0.336334813 | 0.19 | 0 | 1 | 9 |
| esd | 0.01199 | 0.288021939 | 0.19 | 0 | 1 | 9 |
| nfia | 0.01199 | 0.369411893 | 0.19 | 0 | 1 | 9 |
| nr1d2a | 0.01199 | 0.339660051 | 0.19 | 0 | 1 | 9 |
| phf10 | 0.01199 | 0.336056923 | 0.19 | 0 | 1 | 9 |
| rpa2 | 0.01199 | 0.257514309 | 0.19 | 0 | 1 | 9 |
| rtraf | 0.01199 | 0.354556978 | 0.19 | 0 | 1 | 9 |
| tmem33 | 0.01199 | 0.35515551 | 0.19 | 0 | 1 | 9 |
| uri1 | 0.01199 | 0.353519877 | 0.19 | 0 | 1 | 9 |
| dhx15 | 0.01199 | 0.303481408 | 0.19 | 0 | 1 | 9 |
| dph6 | 0.01199 | 0.320460209 | 0.19 | 0 | 1 | 9 |
| mcm5 | 0.01199 | 0.354241093 | 0.19 | 0 | 1 | 9 |
| ptgr1 | 0.01199 | 0.300362643 | 0.19 | 0 | 1 | 9 |
| rrp15 | 0.01199 | 0.329245074 | 0.19 | 0 | 1 | 9 |
| si:ch211-282b22.1 | 0.01199 | 0.293661695 | 0.19 | 0 | 1 | 9 |
| timm10 | 0.01199 | 0.281706071 | 0.19 | 0 | 1 | 9 |
| vwde | 0.01199 | 0.467973827 | 0.19 | 0 | 1 | 9 |
| mt-nd1 | 0.01206 | -0.68317281 | 0.293 | 0.65 | 1 | 9 |
| bysl | 0.01211 | 0.39602985 | 0.345 | 0.05 | 1 | 9 |

| | | | | | | |
|---|---|---|---|---|---|---|
| ube2g1a | 0.01212 | 0.107638789 | 0.19 | 0.05 | 1 | 9 |
| CABZ01072614.1 | 0.01233 | 0.636834708 | 0.448 | 0.2 | 1 | 9 |
| kctd5a | 0.01258 | -0.25361206 | 0.362 | 0.35 | 1 | 9 |
| lsm4 | 0.01271 | 0.548657082 | 0.431 | 0.15 | 1 | 9 |
| nop56 | 0.01275 | 0.57304451 | 0.466 | 0.15 | 1 | 9 |
| fybb | 0.01303 | -0.50494583 | 0.017 | 0.2 | 1 | 9 |
| atp5f1d | 0.01319 | 0.404055507 | 0.621 | 0.25 | 1 | 9 |
| rps15a | 0.01354 | 0.294457773 | 0.897 | 0.85 | 1 | 9 |
| serf2 | 0.01419 | 0.144433372 | 0.724 | 0.45 | 1 | 9 |
| ddx61 | 0.01426 | 0.421021734 | 0.552 | 0.2 | 1 | 9 |
| prdx2 | 0.01426 | 0.367306538 | 0.655 | 0.3 | 1 | 9 |
| si:busm1-57f23.1 | 0.01453 | 0.556729984 | 0.276 | 0.05 | 1 | 9 |
| rpl35 | 0.01458 | 0.403628847 | 0.897 | 0.75 | 1 | 9 |
| rpl3 | 0.01462 | 0.376919153 | 0.966 | 0.75 | 1 | 9 |
| abcc3 | 0.01513 | -0.59090534 | 0.034 | 0.25 | 1 | 9 |
| tnfrsf11b | 0.01514 | -0.5733179 | 0.017 | 0.2 | 1 | 9 |
| si:ch211-51e12.7 | 0.01545 | 0.561397747 | 0.397 | 0.1 | 1 | 9 |
| ostc | 0.01556 | -0.44749198 | 0.362 | 0.5 | 1 | 9 |
| cnpy1 | 0.0157 | -0.67324858 | 0.224 | 0.45 | 1 | 9 |
| selenoh | 0.01601 | 0.521058367 | 0.345 | 0.05 | 1 | 9 |
| ero1b | 0.01608 | -0.48810667 | 0.052 | 0.2 | 1 | 9 |
| ybx1 | 0.0161 | 0.240077907 | 0.983 | 0.9 | 1 | 9 |
| rtn4a | 0.01625 | 0.184309798 | 0.448 | 0.2 | 1 | 9 |
| rps7 | 0.01666 | 0.371515207 | 0.948 | 0.8 | 1 | 9 |
| ndfip1 | 0.01728 | -0.16281628 | 0.397 | 0.35 | 1 | 9 |
| ubc | 0.01737 | -0.18056891 | 0.862 | 0.8 | 1 | 9 |
| ccni | 0.01753 | -0.36585565 | 0.534 | 0.6 | 1 | 9 |
| gtf2f1 | 0.01759 | 0.256758399 | 0.172 | 0 | 1 | 9 |
| hsd17b10 | 0.01759 | 0.323635243 | 0.172 | 0 | 1 | 9 |
| bet1l | 0.01759 | 0.286329062 | 0.172 | 0 | 1 | 9 |
| bloc1s1 | 0.01759 | 0.273866736 | 0.172 | 0 | 1 | 9 |
| cbx1b | 0.01759 | 0.325574981 | 0.172 | 0 | 1 | 9 |
| ergic2 | 0.01759 | 0.277274918 | 0.172 | 0 | 1 | 9 |
| mrps17 | 0.01759 | 0.259604581 | 0.172 | 0 | 1 | 9 |
| ormdl1 | 0.01759 | 0.278694693 | 0.172 | 0 | 1 | 9 |
| pak2b | 0.01759 | 0.288282346 | 0.172 | 0 | 1 | 9 |
| rbb4l | 0.01759 | 0.262161415 | 0.172 | 0 | 1 | 9 |
| si:dkey-276j7.1 | 0.01759 | 0.343817684 | 0.172 | 0 | 1 | 9 |
| six1b | 0.01759 | 0.429333783 | 0.172 | 0 | 1 | 9 |
| tsen34 | 0.01759 | 0.26385921 | 0.172 | 0 | 1 | 9 |
| uqcrc2a | 0.01759 | 0.221242127 | 0.172 | 0 | 1 | 9 |
| ap3m1 | 0.01759 | 0.247325321 | 0.172 | 0 | 1 | 9 |
| arpc5b | 0.01759 | 0.309277035 | 0.172 | 0 | 1 | 9 |
| foxn3 | 0.01759 | 0.306306624 | 0.172 | 0 | 1 | 9 |
| gars | 0.01759 | 0.320594836 | 0.172 | 0 | 1 | 9 |
| gsta.1 | 0.01759 | 0.313678772 | 0.172 | 0 | 1 | 9 |
| gtf2e2 | 0.01759 | 0.254038744 | 0.172 | 0 | 1 | 9 |
| hnrnpub | 0.01759 | 0.312339074 | 0.172 | 0 | 1 | 9 |

| | | | | | | |
|---|---|---|---|---|---|---|
| inip | 0.01759 | 0.303644259 | 0.172 | 0 | 1 | 9 |
| npm1b | 0.01759 | 0.274685272 | 0.172 | 0 | 1 | 9 |
| si:dkey-88p24.11 | 0.01759 | 0.277004833 | 0.172 | 0 | 1 | 9 |
| slc22a18 | 0.01759 | 0.258007495 | 0.172 | 0 | 1 | 9 |
| u2af2a | 0.01759 | 0.325120862 | 0.172 | 0 | 1 | 9 |
| plecb | 0.01818 | -0.56884676 | 0.19 | 0.4 | 1 | 9 |
| rps21 | 0.01826 | 0.295885455 | 0.983 | 0.8 | 1 | 9 |
| rps4x | 0.01888 | 0.346244524 | 0.966 | 0.8 | 1 | 9 |
| si:zfos-169g10.3 | 0.01892 | 0.481157773 | 0.414 | 0.1 | 1 | 9 |
| mdkb | 0.01911 | -0.28859728 | 0.052 | 0.05 | 1 | 9 |
| pinx1 | 0.01944 | -0.13419091 | 0.207 | 0.15 | 1 | 9 |
| b2ml | 0.01944 | -0.34364796 | 0.621 | 0.7 | 1 | 9 |
| ywhabb | 0.01953 | -0.10942347 | 0.552 | 0.45 | 1 | 9 |
| hnrnph1l | 0.01968 | 0.393038669 | 0.328 | 0.05 | 1 | 9 |
| mt-co1 | 0.02006 | -0.51277312 | 0.672 | 0.9 | 1 | 9 |
| si:dkey-28d5.11 | 0.0204 | -0.48241494 | 0.034 | 0.15 | 1 | 9 |
| hmgb1b | 0.02059 | 0.619668124 | 0.414 | 0.15 | 1 | 9 |
| esrp2 | 0.02073 | -0.49031378 | 0.034 | 0.25 | 1 | 9 |
| paqr6 | 0.02094 | -0.45447153 | 0.052 | 0.3 | 1 | 9 |
| pou2f3 | 0.02133 | -0.51214742 | 0.052 | 0.3 | 1 | 9 |
| trmt112 | 0.02139 | 0.471389058 | 0.293 | 0.05 | 1 | 9 |
| LOC103910192 | 0.02143 | -0.86337343 | 0.017 | 0.2 | 1 | 9 |
| si:ch211-66k16.27 | 0.02166 | 0.40743622 | 0.31 | 0.05 | 1 | 9 |
| cd63 | 0.02192 | -0.13334164 | 0.569 | 0.45 | 1 | 9 |
| tomm5 | 0.02216 | 0.537760619 | 0.534 | 0.2 | 1 | 9 |
| acin1a | 0.0223 | 0.544865701 | 0.483 | 0.15 | 1 | 9 |
| snrpe | 0.02233 | 0.534494124 | 0.483 | 0.15 | 1 | 9 |
| tgif1 | 0.02236 | 0.291352986 | 0.276 | 0.05 | 1 | 9 |
| rpl36a | 0.02251 | 0.296116294 | 0.983 | 0.8 | 1 | 9 |
| sap18 | 0.02254 | 0.461379076 | 0.414 | 0.1 | 1 | 9 |
| cirbpb | 0.02267 | 0.404177562 | 0.914 | 0.65 | 1 | 9 |
| mrpl57 | 0.02268 | 0.237312004 | 0.276 | 0.05 | 1 | 9 |
| si:dkey-61f9.2 | 0.02279 | -0.51660769 | 0.034 | 0.15 | 1 | 9 |
| rps10 | 0.02281 | 0.29177957 | 0.983 | 0.8 | 1 | 9 |
| hypk | 0.0229 | 0.479808136 | 0.483 | 0.15 | 1 | 9 |
| rpl12 | 0.02295 | 0.337455329 | 0.948 | 0.8 | 1 | 9 |
| tpt1 | 0.02299 | 0.381962847 | 0.862 | 0.7 | 1 | 9 |
| vdac2 | 0.02318 | 0.192682464 | 0.776 | 0.5 | 1 | 9 |
| zgc:85858 | 0.02332 | -0.43757765 | 0.017 | 0.2 | 1 | 9 |
| tmed10 | 0.02349 | 0.132761032 | 0.431 | 0.2 | 1 | 9 |
| eif3d | 0.02362 | 0.280472049 | 0.603 | 0.3 | 1 | 9 |
| psmc1a | 0.02365 | 0.481874968 | 0.397 | 0.1 | 1 | 9 |
| slc35b4 | 0.0237 | -0.45922653 | 0.017 | 0.2 | 1 | 9 |
| BX855618.1 | 0.02452 | -0.80027049 | 0.017 | 0.2 | 1 | 9 |
| cox4i1 | 0.02453 | 0.458404644 | 0.741 | 0.4 | 1 | 9 |
| sox11a | 0.02496 | 0.616684323 | 0.31 | 0.05 | 1 | 9 |
| eml2 | 0.02546 | -0.45446407 | 0.103 | 0.2 | 1 | 9 |
| lims1 | 0.02557 | -0.4365486 | 0.017 | 0.2 | 1 | 9 |

| | | | | | | |
|---|---|---|---|---|---|---|
| XLOC-037780 | 0.02573 | 0.261334193 | 0.155 | 0 | 1 | 9 |
| gmps | 0.02573 | 0.247611611 | 0.155 | 0 | 1 | 9 |
| klhdc3 | 0.02573 | 0.256866168 | 0.155 | 0 | 1 | 9 |
| mat2b | 0.02573 | 0.265833108 | 0.155 | 0 | 1 | 9 |
| si:ch211-206k20.5 | 0.02573 | 0.282515548 | 0.155 | 0 | 1 | 9 |
| atoh8 | 0.02573 | 0.342564239 | 0.155 | 0 | 1 | 9 |
| brd7 | 0.02573 | 0.2793368 | 0.155 | 0 | 1 | 9 |
| chchd1 | 0.02573 | 0.315502212 | 0.155 | 0 | 1 | 9 |
| gatc | 0.02573 | 0.253050361 | 0.155 | 0 | 1 | 9 |
| hirip3 | 0.02573 | 0.28332413 | 0.155 | 0 | 1 | 9 |
| krtt1c19e | 0.02573 | 0.637975845 | 0.155 | 0 | 1 | 9 |
| msi2b | 0.02573 | 0.298063709 | 0.155 | 0 | 1 | 9 |
| nedd8l | 0.02573 | 0.26123926 | 0.155 | 0 | 1 | 9 |
| nrg1 | 0.02573 | 0.268158552 | 0.155 | 0 | 1 | 9 |
| oxa1l | 0.02573 | 0.235381991 | 0.155 | 0 | 1 | 9 |
| plekhf2 | 0.02573 | 0.299296058 | 0.155 | 0 | 1 | 9 |
| prmt5 | 0.02573 | 0.266464979 | 0.155 | 0 | 1 | 9 |
| rcor1 | 0.02573 | 0.257256359 | 0.155 | 0 | 1 | 9 |
| si:dkey-57k2.7 | 0.02573 | 0.300558774 | 0.155 | 0 | 1 | 9 |
| atg101 | 0.02573 | 0.305206521 | 0.155 | 0 | 1 | 9 |
| ccar1 | 0.02573 | 0.235309003 | 0.155 | 0 | 1 | 9 |
| cxxc5a | 0.02573 | 0.244428841 | 0.155 | 0 | 1 | 9 |
| idh3b | 0.02573 | 0.261008934 | 0.155 | 0 | 1 | 9 |
| lsm1 | 0.02573 | 0.216040373 | 0.155 | 0 | 1 | 9 |
| mat2ab | 0.02573 | 0.238839393 | 0.155 | 0 | 1 | 9 |
| mrps28 | 0.02573 | 0.255126744 | 0.155 | 0 | 1 | 9 |
| myod1 | 0.02573 | 0.374161311 | 0.155 | 0 | 1 | 9 |
| otub1a | 0.02573 | 0.260625492 | 0.155 | 0 | 1 | 9 |
| parp1 | 0.02573 | 0.273668702 | 0.155 | 0 | 1 | 9 |
| pdcd4a | 0.02573 | 0.251083538 | 0.155 | 0 | 1 | 9 |
| pes | 0.02573 | 0.247301952 | 0.155 | 0 | 1 | 9 |
| ppp2r1bb | 0.02573 | 0.266914662 | 0.155 | 0 | 1 | 9 |
| ptp4a2a | 0.02573 | 0.309976263 | 0.155 | 0 | 1 | 9 |
| rac1a | 0.02573 | 0.310437271 | 0.155 | 0 | 1 | 9 |
| rbfox2 | 0.02573 | 0.281999255 | 0.155 | 0 | 1 | 9 |
| rbm4.1 | 0.02573 | 0.281353159 | 0.155 | 0 | 1 | 9 |
| rnaseh2a | 0.02573 | 0.304440803 | 0.155 | 0 | 1 | 9 |
| scfd1 | 0.02573 | 0.282913862 | 0.155 | 0 | 1 | 9 |
| uck2a | 0.02573 | 0.251197835 | 0.155 | 0 | 1 | 9 |
| antxr2a | 0.02573 | 0.314975363 | 0.155 | 0 | 1 | 9 |
| ddb2 | 0.02573 | 0.297740444 | 0.155 | 0 | 1 | 9 |
| ddx47 | 0.02573 | 0.262916461 | 0.155 | 0 | 1 | 9 |
| gadd45ga | 0.02573 | 0.373007185 | 0.155 | 0 | 1 | 9 |
| megf10 | 0.02573 | 0.273396233 | 0.155 | 0 | 1 | 9 |
| si:rp71-1h20.9 | 0.02573 | 0.290226436 | 0.155 | 0 | 1 | 9 |
| gpatch4 | 0.02573 | 0.225808384 | 0.155 | 0 | 1 | 9 |
| polr2i | 0.02573 | 0.221544817 | 0.155 | 0 | 1 | 9 |
| ginm1 | 0.02573 | 0.281580291 | 0.155 | 0 | 1 | 9 |

| | | | | | | |
|---|---|---|---|---|---|---|
| cnot6a | 0.02595 | -0.55289526 | 0.069 | 0.3 | 1 | 9 |
| stmn1a | 0.02643 | 0.48831158 | 0.31 | 0.05 | 1 | 9 |
| srsf7a | 0.02643 | 0.491605202 | 0.397 | 0.1 | 1 | 9 |
| cldnh | 0.02643 | -0.35038188 | 0.172 | 0.2 | 1 | 9 |
| selenop | 0.02649 | -0.35127087 | 0.017 | 0.2 | 1 | 9 |
| rsrp1 | 0.02652 | -0.37383928 | 0.155 | 0.25 | 1 | 9 |
| furina | 0.02655 | -0.4637844 | 0.121 | 0.4 | 1 | 9 |
| cope | 0.02669 | -0.3590779 | 0.362 | 0.45 | 1 | 9 |
| zgc:55262 | 0.0268 | -0.36347494 | 0.034 | 0.25 | 1 | 9 |
| irf2bp2a | 0.0268 | 0.411771786 | 0.431 | 0.15 | 1 | 9 |
| tmed7 | 0.02688 | -0.10555565 | 0.586 | 0.5 | 1 | 9 |
| ccdc171 | 0.02704 | -0.44173865 | 0.034 | 0.25 | 1 | 9 |
| mrpl32 | 0.02726 | 0.382322731 | 0.31 | 0.05 | 1 | 9 |
| marcksl1b | 0.02732 | 0.617921194 | 0.534 | 0.2 | 1 | 9 |
| snrpd3l | 0.02767 | 0.476599984 | 0.397 | 0.1 | 1 | 9 |
| mdh2 | 0.02784 | 0.449255972 | 0.69 | 0.35 | 1 | 9 |
| si:ch211-217k17.7 | 0.02786 | 0.457819024 | 0.31 | 0.05 | 1 | 9 |
| CT573383.1 | 0.02794 | -0.40200991 | 0.069 | 0.25 | 1 | 9 |
| zgc:193726 | 0.028 | -1.04160132 | 0.017 | 0.2 | 1 | 9 |
| eif5a2 | 0.02801 | 0.465759806 | 0.69 | 0.35 | 1 | 9 |
| cs | 0.02812 | 0.466064982 | 0.31 | 0.05 | 1 | 9 |
| ppiaa | 0.02837 | 0.315930055 | 0.948 | 0.8 | 1 | 9 |
| erh | 0.02844 | 0.469576357 | 0.638 | 0.3 | 1 | 9 |
| lsm6 | 0.02848 | -0.12579486 | 0.362 | 0.3 | 1 | 9 |
| hnrnpa0b | 0.02943 | 0.455451175 | 0.638 | 0.3 | 1 | 9 |
| usp4 | 0.03015 | -0.46547676 | 0.034 | 0.2 | 1 | 9 |
| zgc:152774 | 0.03021 | -0.45560972 | 0.017 | 0.2 | 1 | 9 |
| cldn15a | 0.03026 | -0.55725265 | 0.017 | 0.2 | 1 | 9 |
| rbm4.3 | 0.03038 | 0.526396742 | 0.517 | 0.2 | 1 | 9 |
| si:ch73-106l15.4 | 0.03047 | -0.93104231 | 0.017 | 0.2 | 1 | 9 |
| brd2a | 0.03054 | 0.391330953 | 0.31 | 0.05 | 1 | 9 |
| ugdh | 0.03074 | -0.38076406 | 0.017 | 0.2 | 1 | 9 |
| rpl13 | 0.03075 | 0.288115228 | 0.948 | 0.9 | 1 | 9 |
| sem1 | 0.03086 | 0.493483134 | 0.621 | 0.3 | 1 | 9 |
| rsu1 | 0.03107 | -0.35941179 | 0.017 | 0.2 | 1 | 9 |
| CU137717.1 | 0.03107 | -0.15005256 | 0 | 0.1 | 1 | 9 |
| apcdd1l | 0.03107 | -0.20797835 | 0 | 0.1 | 1 | 9 |
| arsh | 0.03107 | -0.1449443 | 0 | 0.1 | 1 | 9 |
| bpifcl | 0.03107 | -0.11175555 | 0 | 0.1 | 1 | 9 |
| fzd10 | 0.03107 | -0.2954324 | 0 | 0.1 | 1 | 9 |
| hcfc1b | 0.03107 | -0.22493872 | 0 | 0.1 | 1 | 9 |
| myo5b | 0.03107 | -0.15176839 | 0 | 0.1 | 1 | 9 |
| pbxip1a | 0.03107 | -0.14377382 | 0 | 0.1 | 1 | 9 |
| pcdh7b | 0.03107 | -0.14050426 | 0 | 0.1 | 1 | 9 |
| s1pr1 | 0.03107 | -0.186771 | 0 | 0.1 | 1 | 9 |
| sh3bp4a | 0.03107 | -0.15313823 | 0 | 0.1 | 1 | 9 |
| slc35e3 | 0.03107 | -0.14968335 | 0 | 0.1 | 1 | 9 |
| tbce | 0.03107 | -0.19248772 | 0 | 0.1 | 1 | 9 |

| | | | | | | |
|---|---|---|---|---|---|---|
| ubash3ba | 0.03107 | -0.1787894 | 0 | 0.1 | 1 | 9 |
| il4r.2 | 0.03107 | -0.32497841 | 0 | 0.1 | 1 | 9 |
| mospd2 | 0.03107 | -0.19756461 | 0 | 0.1 | 1 | 9 |
| naprt | 0.03107 | -0.24348101 | 0 | 0.1 | 1 | 9 |
| ptp4a3 | 0.03107 | -0.23579707 | 0 | 0.1 | 1 | 9 |
| trim59 | 0.03107 | -0.32393611 | 0 | 0.1 | 1 | 9 |
| zgc:113090 | 0.03107 | -0.33935604 | 0 | 0.1 | 1 | 9 |
| CABZ01008730.1 | 0.03107 | -0.15622241 | 0 | 0.1 | 1 | 9 |
| CR847984.1 | 0.03107 | -0.17154804 | 0 | 0.1 | 1 | 9 |
| CU682640.1 | 0.03107 | -0.21246929 | 0 | 0.1 | 1 | 9 |
| CU984600.2 | 0.03107 | -0.15195999 | 0 | 0.1 | 1 | 9 |
| LOC100330744 | 0.03107 | -0.14281276 | 0 | 0.1 | 1 | 9 |
| XLOC-039166 | 0.03107 | -0.13704912 | 0 | 0.1 | 1 | 9 |
| ankle2 | 0.03107 | -0.15521473 | 0 | 0.1 | 1 | 9 |
| ano5b | 0.03107 | -0.17351936 | 0 | 0.1 | 1 | 9 |
| cacng6a | 0.03107 | -0.13704912 | 0 | 0.1 | 1 | 9 |
| gbp3 | 0.03107 | -0.16543382 | 0 | 0.1 | 1 | 9 |
| hps1 | 0.03107 | -0.16857725 | 0 | 0.1 | 1 | 9 |
| kpna5 | 0.03107 | -0.1576355 | 0 | 0.1 | 1 | 9 |
| lrrc28 | 0.03107 | -0.16045866 | 0 | 0.1 | 1 | 9 |
| mob2b | 0.03107 | -0.12728807 | 0 | 0.1 | 1 | 9 |
| nlrx1 | 0.03107 | -0.13985851 | 0 | 0.1 | 1 | 9 |
| ponzr10 | 0.03107 | -0.26665694 | 0 | 0.1 | 1 | 9 |
| si:ch211-225b10.3 | 0.03107 | -0.16786196 | 0 | 0.1 | 1 | 9 |
| si:ch73-196i15.3 | 0.03107 | -0.16786196 | 0 | 0.1 | 1 | 9 |
| si:ch73-340m8.2 | 0.03107 | -0.13039571 | 0 | 0.1 | 1 | 9 |
| si:ch73-97h19.2 | 0.03107 | -0.18686068 | 0 | 0.1 | 1 | 9 |
| si:dkeyp-115e12.7 | 0.03107 | -0.1422381 | 0 | 0.1 | 1 | 9 |
| si:dkeyp-71f10.5 | 0.03107 | -0.21494691 | 0 | 0.1 | 1 | 9 |
| smc6 | 0.03107 | -0.20898063 | 0 | 0.1 | 1 | 9 |
| smg1 | 0.03107 | -0.19271153 | 0 | 0.1 | 1 | 9 |
| ssh1a | 0.03107 | -0.15895534 | 0 | 0.1 | 1 | 9 |
| stk3 | 0.03107 | -0.15900665 | 0 | 0.1 | 1 | 9 |
| stx4 | 0.03107 | -0.15478719 | 0 | 0.1 | 1 | 9 |
| tmem56b | 0.03107 | -0.13111467 | 0 | 0.1 | 1 | 9 |
| zgc:172139 | 0.03107 | -0.15971526 | 0 | 0.1 | 1 | 9 |
| znfx1 | 0.03107 | -0.21847098 | 0 | 0.1 | 1 | 9 |
| LOC103911313 | 0.03107 | -0.2136415 | 0 | 0.1 | 1 | 9 |
| avil | 0.03107 | -0.32330592 | 0 | 0.1 | 1 | 9 |
| brpf3b | 0.03107 | -0.27673632 | 0 | 0.1 | 1 | 9 |
| coro2a | 0.03107 | -0.27721041 | 0 | 0.1 | 1 | 9 |
| crebl2 | 0.03107 | -0.17743696 | 0 | 0.1 | 1 | 9 |
| crp7 | 0.03107 | -0.48604732 | 0 | 0.1 | 1 | 9 |
| ftr66 | 0.03107 | -0.27801259 | 0 | 0.1 | 1 | 9 |
| gpatch3 | 0.03107 | -0.2119285 | 0 | 0.1 | 1 | 9 |
| gucd1 | 0.03107 | -0.2073254 | 0 | 0.1 | 1 | 9 |
| irak3 | 0.03107 | -0.21085013 | 0 | 0.1 | 1 | 9 |
| mios | 0.03107 | -0.26981927 | 0 | 0.1 | 1 | 9 |

| | | | | | | |
|---|---|---|---|---|---|---|
| neu1 | 0.03107 | -0.20920708 | 0 | 0.1 | 1 | 9 |
| rapgef5a | 0.03107 | -0.31479694 | 0 | 0.1 | 1 | 9 |
| serpinf2b | 0.03107 | -0.3203535 | 0 | 0.1 | 1 | 9 |
| si:ch211-282j17.8 | 0.03107 | -0.40511246 | 0 | 0.1 | 1 | 9 |
| si:dkey-88e18.2 | 0.03107 | -0.26691932 | 0 | 0.1 | 1 | 9 |
| spsb1 | 0.03107 | -0.27701012 | 0 | 0.1 | 1 | 9 |
| syvn1 | 0.03107 | -0.26600463 | 0 | 0.1 | 1 | 9 |
| zgc:110783 | 0.03107 | -0.16265088 | 0 | 0.1 | 1 | 9 |
| ano5a | 0.03107 | -0.2077613 | 0 | 0.1 | 1 | 9 |
| ano9a | 0.03107 | -0.20428512 | 0 | 0.1 | 1 | 9 |
| arid5b | 0.03107 | -0.20847835 | 0 | 0.1 | 1 | 9 |
| bmp16 | 0.03107 | -0.26494482 | 0 | 0.1 | 1 | 9 |
| chia.6 | 0.03107 | -0.34095768 | 0 | 0.1 | 1 | 9 |
| ints11 | 0.03107 | -0.19416208 | 0 | 0.1 | 1 | 9 |
| kcnh6a | 0.03107 | -0.241039 | 0 | 0.1 | 1 | 9 |
| nfkb2 | 0.03107 | -0.21174352 | 0 | 0.1 | 1 | 9 |
| rxrbb | 0.03107 | -0.24173278 | 0 | 0.1 | 1 | 9 |
| si:ch211-106j24.1 | 0.03107 | -0.24419185 | 0 | 0.1 | 1 | 9 |
| si:ch211-15j1.4 | 0.03107 | -0.20872059 | 0 | 0.1 | 1 | 9 |
| si:ch211-163l21.8 | 0.03107 | -0.23352233 | 0 | 0.1 | 1 | 9 |
| si:dkey-256h2.1 | 0.03107 | -0.17621002 | 0 | 0.1 | 1 | 9 |
| smpx | 0.03107 | -0.31221565 | 0 | 0.1 | 1 | 9 |
| tmem267 | 0.03107 | -0.20464102 | 0 | 0.1 | 1 | 9 |
| pfkla | 0.03107 | -0.14652832 | 0 | 0.1 | 1 | 9 |
| BX682550.1 | 0.03107 | -0.28806138 | 0 | 0.1 | 1 | 9 |
| FP102158.1 | 0.03107 | -0.15554807 | 0 | 0.1 | 1 | 9 |
| UNC13B | 0.03107 | -0.20095179 | 0 | 0.1 | 1 | 9 |
| fbxo21 | 0.03107 | -0.19578774 | 0 | 0.1 | 1 | 9 |
| fgf1b | 0.03107 | -0.25901348 | 0 | 0.1 | 1 | 9 |
| malt2 | 0.03107 | -0.21335404 | 0 | 0.1 | 1 | 9 |
| micu1 | 0.03107 | -0.19759236 | 0 | 0.1 | 1 | 9 |
| mnx2a | 0.03107 | -0.15691852 | 0 | 0.1 | 1 | 9 |
| ndnf | 0.03107 | -0.19427549 | 0 | 0.1 | 1 | 9 |
| osr1 | 0.03107 | -0.17131632 | 0 | 0.1 | 1 | 9 |
| pik3r5 | 0.03107 | -0.26988196 | 0 | 0.1 | 1 | 9 |
| prkn | 0.03107 | -0.19512903 | 0 | 0.1 | 1 | 9 |
| rbfa | 0.03107 | -0.15002582 | 0 | 0.1 | 1 | 9 |
| rwdd2b | 0.03107 | -0.20418427 | 0 | 0.1 | 1 | 9 |
| si:dkey-202e17.1 | 0.03107 | -0.15453402 | 0 | 0.1 | 1 | 9 |
| spire1a | 0.03107 | -0.2128327 | 0 | 0.1 | 1 | 9 |
| taf2 | 0.03107 | -0.24263674 | 0 | 0.1 | 1 | 9 |
| tor4ab | 0.03107 | -0.22363047 | 0 | 0.1 | 1 | 9 |
| tpbgb | 0.03107 | -0.20583897 | 0 | 0.1 | 1 | 9 |
| wdr19 | 0.03107 | -0.18610761 | 0 | 0.1 | 1 | 9 |
| zgc:158258 | 0.03107 | -0.29949376 | 0 | 0.1 | 1 | 9 |
| si:dkey-172k15.6 | 0.03107 | -0.16954393 | 0 | 0.1 | 1 | 9 |
| LOC100536643 | 0.03107 | -0.20109504 | 0 | 0.1 | 1 | 9 |
| LOC103909050 | 0.03107 | -0.32105888 | 0 | 0.1 | 1 | 9 |

| | | | | | | |
|---|---|---|---|---|---|---|
| acp2 | 0.03107 | -0.19850454 | 0 | 0.1 | 1 | 9 |
| arid6 | 0.03107 | -0.20109504 | 0 | 0.1 | 1 | 9 |
| cryba2b | 0.03107 | -0.19837805 | 0 | 0.1 | 1 | 9 |
| kif21b | 0.03107 | -0.24976035 | 0 | 0.1 | 1 | 9 |
| mcm3ap | 0.03107 | -0.1937756 | 0 | 0.1 | 1 | 9 |
| mindy2 | 0.03107 | -0.29967887 | 0 | 0.1 | 1 | 9 |
| CABZ01102240.1 | 0.03107 | -0.19429174 | 0 | 0.1 | 1 | 9 |
| CABZ01111959.1 | 0.03107 | -0.17032571 | 0 | 0.1 | 1 | 9 |
| LOC101882596 | 0.03107 | -0.17583377 | 0 | 0.1 | 1 | 9 |
| LOC110440019 | 0.03107 | -0.26447726 | 0 | 0.1 | 1 | 9 |
| dhrs9 | 0.03107 | -0.25008472 | 0 | 0.1 | 1 | 9 |
| kif13bb | 0.03107 | -0.30331829 | 0 | 0.1 | 1 | 9 |
| mboat2b | 0.03107 | -0.24126287 | 0 | 0.1 | 1 | 9 |
| stx7l | 0.03107 | -0.18973533 | 0 | 0.1 | 1 | 9 |
| traf1 | 0.03107 | -0.16188079 | 0 | 0.1 | 1 | 9 |
| wdr6 | 0.03107 | -0.18195507 | 0 | 0.1 | 1 | 9 |
| pip4k2ca | 0.03125 | -0.36497784 | 0.017 | 0.2 | 1 | 9 |
| nras | 0.03134 | -0.28192555 | 0.017 | 0.2 | 1 | 9 |
| rhobtb4 | 0.03151 | -0.315435 | 0.017 | 0.2 | 1 | 9 |
| ndufab1a | 0.03155 | 0.440413734 | 0.31 | 0.05 | 1 | 9 |
| tm9sf1 | 0.032 | -0.35090043 | 0.017 | 0.2 | 1 | 9 |
| ltc4s | 0.0325 | -0.5666293 | 0.034 | 0.2 | 1 | 9 |
| rpl19 | 0.03265 | 0.342098325 | 0.914 | 0.75 | 1 | 9 |
| rpl34 | 0.03301 | 0.354740838 | 0.931 | 0.75 | 1 | 9 |
| rpl18a | 0.03306 | 0.288289494 | 0.966 | 0.85 | 1 | 9 |
| pdcl | 0.03366 | -0.4165521 | 0.138 | 0.3 | 1 | 9 |
| ano9b | 0.03371 | -0.46357338 | 0.052 | 0.2 | 1 | 9 |
| chmp4bb | 0.0343 | -0.51097198 | 0.207 | 0.5 | 1 | 9 |
| tmsb | 0.03449 | 0.647655119 | 0.31 | 0.05 | 1 | 9 |
| eef1b2 | 0.03532 | 0.385287918 | 0.828 | 0.7 | 1 | 9 |
| zmat2 | 0.03541 | 0.357489305 | 0.379 | 0.1 | 1 | 9 |
| gata1a | 0.03558 | -0.43522782 | 0.017 | 0.15 | 1 | 9 |
| naca | 0.03566 | 0.390245427 | 0.897 | 0.7 | 1 | 9 |
| rack1 | 0.03585 | 0.29194351 | 0.948 | 0.85 | 1 | 9 |
| rps26l | 0.03587 | 0.301324476 | 0.966 | 0.8 | 1 | 9 |
| cct3 | 0.03612 | 0.317972096 | 0.603 | 0.3 | 1 | 9 |
| ssr1 | 0.03677 | -0.16999243 | 0.345 | 0.3 | 1 | 9 |
| eya1 | 0.03752 | 0.310387145 | 0.138 | 0 | 1 | 9 |
| supt3h | 0.03752 | 0.191057256 | 0.138 | 0 | 1 | 9 |
| asb8 | 0.03752 | 0.220420187 | 0.138 | 0 | 1 | 9 |
| casp3a | 0.03752 | 0.272026093 | 0.138 | 0 | 1 | 9 |
| mrps7 | 0.03752 | 0.2248534 | 0.138 | 0 | 1 | 9 |
| myf5 | 0.03752 | 0.372525808 | 0.138 | 0 | 1 | 9 |
| pik3ca | 0.03752 | 0.195696308 | 0.138 | 0 | 1 | 9 |
| pknox1.1 | 0.03752 | 0.215985302 | 0.138 | 0 | 1 | 9 |
| sh3glb1b | 0.03752 | 0.209921261 | 0.138 | 0 | 1 | 9 |
| si:ch73-238c9.1 | 0.03752 | 0.216840814 | 0.138 | 0 | 1 | 9 |
| thop1 | 0.03752 | 0.243520339 | 0.138 | 0 | 1 | 9 |

| | | | | | | |
|---|---|---|---|---|---|---|
| ubxn1 | 0.03752 | 0.218301607 | 0.138 | 0 | 1 | 9 |
| znhit3 | 0.03752 | 0.231949897 | 0.138 | 0 | 1 | 9 |
| chd6 | 0.03752 | 0.256302465 | 0.138 | 0 | 1 | 9 |
| hey1 | 0.03752 | 0.303017515 | 0.138 | 0 | 1 | 9 |
| pabpc4 | 0.03752 | 0.243026364 | 0.138 | 0 | 1 | 9 |
| pmpcb | 0.03752 | 0.240422234 | 0.138 | 0 | 1 | 9 |
| BX284638.1 | 0.03752 | 0.298096306 | 0.138 | 0 | 1 | 9 |
| cnot1 | 0.03752 | 0.220306838 | 0.138 | 0 | 1 | 9 |
| cux1a | 0.03752 | 0.21850726 | 0.138 | 0 | 1 | 9 |
| dda1 | 0.03752 | 0.191224317 | 0.138 | 0 | 1 | 9 |
| decr1 | 0.03752 | 0.242183576 | 0.138 | 0 | 1 | 9 |
| dhrs4 | 0.03752 | 0.201069735 | 0.138 | 0 | 1 | 9 |
| dnm1l | 0.03752 | 0.165397881 | 0.138 | 0 | 1 | 9 |
| drg1 | 0.03752 | 0.233533269 | 0.138 | 0 | 1 | 9 |
| dvl3a | 0.03752 | 0.265826082 | 0.138 | 0 | 1 | 9 |
| eif2a | 0.03752 | 0.243434698 | 0.138 | 0 | 1 | 9 |
| fosb | 0.03752 | 0.340414439 | 0.138 | 0 | 1 | 9 |
| hmg20b | 0.03752 | 0.267862139 | 0.138 | 0 | 1 | 9 |
| hspa14 | 0.03752 | 0.247936339 | 0.138 | 0 | 1 | 9 |
| kif5bb | 0.03752 | 0.264562279 | 0.138 | 0 | 1 | 9 |
| lysmd2 | 0.03752 | 0.232524609 | 0.138 | 0 | 1 | 9 |
| map7d1b | 0.03752 | 0.170920979 | 0.138 | 0 | 1 | 9 |
| mbd3b | 0.03752 | 0.229902885 | 0.138 | 0 | 1 | 9 |
| mcmbp | 0.03752 | 0.211220084 | 0.138 | 0 | 1 | 9 |
| myca | 0.03752 | 0.306326604 | 0.138 | 0 | 1 | 9 |
| napga | 0.03752 | 0.21104734 | 0.138 | 0 | 1 | 9 |
| ndfip2 | 0.03752 | 0.209810676 | 0.138 | 0 | 1 | 9 |
| nolc1 | 0.03752 | 0.237177805 | 0.138 | 0 | 1 | 9 |
| nutf2 | 0.03752 | 0.237242761 | 0.138 | 0 | 1 | 9 |
| polr3glb | 0.03752 | 0.286233413 | 0.138 | 0 | 1 | 9 |
| prkrip1 | 0.03752 | 0.273716238 | 0.138 | 0 | 1 | 9 |
| prr12b | 0.03752 | 0.270271801 | 0.138 | 0 | 1 | 9 |
| ptgesl | 0.03752 | 0.254280961 | 0.138 | 0 | 1 | 9 |
| pwp2h | 0.03752 | 0.214595921 | 0.138 | 0 | 1 | 9 |
| rap1ab | 0.03752 | 0.218014281 | 0.138 | 0 | 1 | 9 |
| rpa1 | 0.03752 | 0.253940816 | 0.138 | 0 | 1 | 9 |
| rpia | 0.03752 | 0.198754841 | 0.138 | 0 | 1 | 9 |
| rpp25l | 0.03752 | 0.247258567 | 0.138 | 0 | 1 | 9 |
| sec24a | 0.03752 | 0.237004343 | 0.138 | 0 | 1 | 9 |
| sept9b | 0.03752 | 0.210947712 | 0.138 | 0 | 1 | 9 |
| si:ch73-37h15.2 | 0.03752 | 0.247144718 | 0.138 | 0 | 1 | 9 |
| si:dkey-19e4.5 | 0.03752 | 0.209079931 | 0.138 | 0 | 1 | 9 |
| smad5 | 0.03752 | 0.206026285 | 0.138 | 0 | 1 | 9 |
| srp68 | 0.03752 | 0.230716926 | 0.138 | 0 | 1 | 9 |
| tbc1d23 | 0.03752 | 0.249783983 | 0.138 | 0 | 1 | 9 |
| tubb5 | 0.03752 | 0.346328954 | 0.138 | 0 | 1 | 9 |
| ube2h | 0.03752 | 0.246460394 | 0.138 | 0 | 1 | 9 |
| waslb | 0.03752 | 0.262339297 | 0.138 | 0 | 1 | 9 |

| | | | | | | |
|---|---|---|---|---|---|---|
| wrb | 0.03752 | 0.268900962 | 0.138 | 0 | 1 | 9 |
| zgc:101858 | 0.03752 | 0.195408079 | 0.138 | 0 | 1 | 9 |
| zgc:153665 | 0.03752 | 0.349521956 | 0.138 | 0 | 1 | 9 |
| zgc:165573 | 0.03752 | 0.252084546 | 0.138 | 0 | 1 | 9 |
| zgc:66160 | 0.03752 | 0.232619029 | 0.138 | 0 | 1 | 9 |
| znf207b | 0.03752 | 0.251153736 | 0.138 | 0 | 1 | 9 |
| zswim5 | 0.03752 | 0.266052939 | 0.138 | 0 | 1 | 9 |
| arl6ip1 | 0.03766 | -0.20269954 | 0.397 | 0.35 | 1 | 9 |
| tm9sf2 | 0.0378 | -0.5273608 | 0.121 | 0.35 | 1 | 9 |
| atp5f1c | 0.03794 | 0.527639532 | 0.586 | 0.3 | 1 | 9 |
| ccdc124 | 0.03816 | 0.480880781 | 0.379 | 0.1 | 1 | 9 |
| p4hb | 0.03849 | -0.49989474 | 0.276 | 0.45 | 1 | 9 |
| sdcbp2 | 0.03898 | -0.45171159 | 0.362 | 0.55 | 1 | 9 |
| ran | 0.03905 | 0.45886359 | 0.793 | 0.5 | 1 | 9 |
| gosr1 | 0.03922 | -0.11544866 | 0.138 | 0.1 | 1 | 9 |
| eif3m | 0.0393 | 0.459990279 | 0.672 | 0.35 | 1 | 9 |
| npc2 | 0.03945 | -0.41192484 | 0.362 | 0.45 | 1 | 9 |
| pcna | 0.04055 | 0.54627608 | 0.293 | 0.05 | 1 | 9 |
| ywhaba | 0.04057 | -0.17795797 | 0.552 | 0.5 | 1 | 9 |
| CABZ01079427.1 | 0.04101 | -0.45263268 | 0.172 | 0.4 | 1 | 9 |
| h2afx1 | 0.04138 | 0.292036521 | 0.638 | 0.35 | 1 | 9 |
| rps26 | 0.0416 | 0.199602581 | 0.897 | 0.9 | 1 | 9 |
| si:dkey-250d21.1 | 0.04161 | 0.402582881 | 0.293 | 0.05 | 1 | 9 |
| canx | 0.04182 | -0.43244684 | 0.276 | 0.4 | 1 | 9 |
| rpl15 | 0.04254 | 0.326455594 | 0.914 | 0.8 | 1 | 9 |
| zgc:92606 | 0.04283 | 0.178967383 | 0.241 | 0.05 | 1 | 9 |
| atp5fa1 | 0.04289 | 0.467550981 | 0.707 | 0.4 | 1 | 9 |
| faua | 0.0429 | 0.267090712 | 0.931 | 0.7 | 1 | 9 |
| selenot1b | 0.04301 | -0.17292428 | 0.172 | 0.15 | 1 | 9 |
| npm1a | 0.04306 | 0.479884973 | 0.448 | 0.15 | 1 | 9 |
| anp32b | 0.04342 | 0.382146824 | 0.379 | 0.1 | 1 | 9 |
| id1 | 0.04358 | 0.648682328 | 0.379 | 0.1 | 1 | 9 |
| ube2d2 | 0.04381 | 0.445453794 | 0.448 | 0.15 | 1 | 9 |
| hspa8 | 0.04398 | 0.276631924 | 0.966 | 0.85 | 1 | 9 |
| ndufa8 | 0.0443 | 0.416577228 | 0.448 | 0.15 | 1 | 9 |
| adi1 | 0.04557 | 0.256582349 | 0.259 | 0.05 | 1 | 9 |
| ube2v1 | 0.04557 | 0.157101232 | 0.414 | 0.2 | 1 | 9 |
| s100v1 | 0.04568 | -0.41930512 | 0.069 | 0.3 | 1 | 9 |
| myl9b | 0.04571 | -0.59520536 | 0.138 | 0.35 | 1 | 9 |
| hibadhb | 0.04647 | 0.204804864 | 0.224 | 0.05 | 1 | 9 |
| ralbp1 | 0.04659 | -0.49211889 | 0.19 | 0.4 | 1 | 9 |
| degs1 | 0.04686 | 0.393804898 | 0.293 | 0.05 | 1 | 9 |
| suco | 0.04691 | -0.4753924 | 0.052 | 0.2 | 1 | 9 |
| rpl7a | 0.04716 | 0.241927259 | 0.966 | 0.9 | 1 | 9 |
| atp2b2 | 0.04726 | 0.432270767 | 0.293 | 0.05 | 1 | 9 |
| COX3 | 0.04728 | -0.50368704 | 0.586 | 0.85 | 1 | 9 |
| prps1a | 0.04767 | 0.2290079 | 0.241 | 0.05 | 1 | 9 |
| tcp1 | 0.04778 | 0.487634314 | 0.517 | 0.25 | 1 | 9 |

| | | | | | | |
|---|---|---|---|---|---|---|
| si:dkey-28d5.12 | 0.04851 | -0.45008412 | 0.017 | 0.15 | 1 | 9 |
| tagln2 | 0.04892 | -0.53929426 | 0.276 | 0.45 | 1 | 9 |
| pfn2 | 0.04915 | 0.245113584 | 0.672 | 0.4 | 1 | 9 |
| selenof | 0.04918 | 0.119413719 | 0.621 | 0.4 | 1 | 9 |
| CABZ01090890.1 | 0.04937 | -0.41767775 | 0.155 | 0.35 | 1 | 9 |
| rpn1 | 0.04937 | -0.1956026 | 0.379 | 0.35 | 1 | 9 |
| actb2 | 0.04951 | -0.38366886 | 0.724 | 0.85 | 1 | 9 |
| psma4 | 0.04982 | -0.22797679 | 0.448 | 0.45 | 1 | 9 |
| ube2na | 0.05009 | -0.2555823 | 0.31 | 0.35 | 1 | 9 |
| si:dkey-77f5.10 | 0.05031 | -0.52603843 | 0.034 | 0.15 | 1 | 9 |
| egr1 | 0.0507 | -0.34036096 | 0.241 | 0.3 | 1 | 9 |
| atg13 | 0.05092 | -0.36535273 | 0.103 | 0.25 | 1 | 9 |
| metap2b | 0.05113 | 0.390019136 | 0.5 | 0.2 | 1 | 9 |
| rplp0 | 0.05183 | 0.296017156 | 0.914 | 0.85 | 1 | 9 |
| adgrg11 | 0.05197 | -0.47933013 | 0.052 | 0.15 | 1 | 9 |
| rab10 | 0.05203 | -0.12365209 | 0.466 | 0.4 | 1 | 9 |
| fam107b | 0.05286 | 0.114805716 | 0.19 | 0.05 | 1 | 9 |
| gnal | 0.05289 | -0.31160902 | 0.138 | 0.2 | 1 | 9 |
| LOC101886933 | 0.05301 | -0.45511941 | 0.034 | 0.2 | 1 | 9 |
| kansl1a | 0.05302 | -0.41692578 | 0.017 | 0.15 | 1 | 9 |
| calm1b | 0.05303 | -0.42838047 | 0.259 | 0.35 | 1 | 9 |
| psme3 | 0.05315 | 0.226155373 | 0.224 | 0.05 | 1 | 9 |
| wu:fi37e09 | 0.05342 | -0.43028938 | 0.138 | 0.3 | 1 | 9 |
| zgc:56525 | 0.05377 | -0.27607234 | 0.103 | 0.15 | 1 | 9 |
| cavin2a | 0.05449 | -0.45936507 | 0.121 | 0.3 | 1 | 9 |
| pfdn5 | 0.0546 | 0.42348063 | 0.431 | 0.15 | 1 | 9 |
| CABZ01063556.1 | 0.05463 | 0.193220181 | 0.121 | 0 | 1 | 9 |
| acbd3 | 0.05463 | 0.192664729 | 0.121 | 0 | 1 | 9 |
| ankrd52a | 0.05463 | 0.21893397 | 0.121 | 0 | 1 | 9 |
| atp1b2a | 0.05463 | 0.227571374 | 0.121 | 0 | 1 | 9 |
| atpaf1 | 0.05463 | 0.197566431 | 0.121 | 0 | 1 | 9 |
| baz1a | 0.05463 | 0.201580306 | 0.121 | 0 | 1 | 9 |
| ccdc32 | 0.05463 | 0.173540798 | 0.121 | 0 | 1 | 9 |
| cdk7 | 0.05463 | 0.225289615 | 0.121 | 0 | 1 | 9 |
| cdon | 0.05463 | 0.257076894 | 0.121 | 0 | 1 | 9 |
| celsr1a | 0.05463 | 0.227075504 | 0.121 | 0 | 1 | 9 |
| cfap36 | 0.05463 | 0.193609971 | 0.121 | 0 | 1 | 9 |
| chmp7 | 0.05463 | 0.203727525 | 0.121 | 0 | 1 | 9 |
| ciapin1 | 0.05463 | 0.182436608 | 0.121 | 0 | 1 | 9 |
| ddx19 | 0.05463 | 0.197417137 | 0.121 | 0 | 1 | 9 |
| ddx42 | 0.05463 | 0.173312532 | 0.121 | 0 | 1 | 9 |
| dnaja3a | 0.05463 | 0.183338587 | 0.121 | 0 | 1 | 9 |
| eed | 0.05463 | 0.225338863 | 0.121 | 0 | 1 | 9 |
| fkbp1b | 0.05463 | 0.202565977 | 0.121 | 0 | 1 | 9 |
| gnaia | 0.05463 | 0.190733751 | 0.121 | 0 | 1 | 9 |
| gnl3l | 0.05463 | 0.199733486 | 0.121 | 0 | 1 | 9 |
| grwd1 | 0.05463 | 0.227547798 | 0.121 | 0 | 1 | 9 |
| idh3g | 0.05463 | 0.197015171 | 0.121 | 0 | 1 | 9 |

| | | | | | | |
|---|---|---|---|---|---|---|
| mrpl10 | 0.05463 | 0.175084612 | 0.121 | 0 | 1 | 9 |
| mrpl17 | 0.05463 | 0.203799937 | 0.121 | 0 | 1 | 9 |
| mrto4 | 0.05463 | 0.187264347 | 0.121 | 0 | 1 | 9 |
| mylipa | 0.05463 | 0.174969845 | 0.121 | 0 | 1 | 9 |
| ndst2a | 0.05463 | 0.162512692 | 0.121 | 0 | 1 | 9 |
| pcmtd1 | 0.05463 | 0.242829452 | 0.121 | 0 | 1 | 9 |
| pdia5 | 0.05463 | 0.213823497 | 0.121 | 0 | 1 | 9 |
| phf20b | 0.05463 | 0.208535454 | 0.121 | 0 | 1 | 9 |
| pom121 | 0.05463 | 0.205471391 | 0.121 | 0 | 1 | 9 |
| ptpn11a | 0.05463 | 0.172836984 | 0.121 | 0 | 1 | 9 |
| sept9a | 0.05463 | 0.212243611 | 0.121 | 0 | 1 | 9 |
| si:ch211-137a8.4 | 0.05463 | 0.28314221 | 0.121 | 0 | 1 | 9 |
| si:ch211-197g15.6 | 0.05463 | 0.24431468 | 0.121 | 0 | 1 | 9 |
| si:ch211-271b14.1 | 0.05463 | 0.207818038 | 0.121 | 0 | 1 | 9 |
| smyd2a | 0.05463 | 0.193389097 | 0.121 | 0 | 1 | 9 |
| sox11b | 0.05463 | 0.335899433 | 0.121 | 0 | 1 | 9 |
| ssbp1 | 0.05463 | 0.196531668 | 0.121 | 0 | 1 | 9 |
| ssuh2rs1 | 0.05463 | 0.171490279 | 0.121 | 0 | 1 | 9 |
| tbx2b | 0.05463 | 0.257732665 | 0.121 | 0 | 1 | 9 |
| tfcp2 | 0.05463 | 0.237770263 | 0.121 | 0 | 1 | 9 |
| tmc6b | 0.05463 | 0.204904767 | 0.121 | 0 | 1 | 9 |
| tmem209 | 0.05463 | 0.193816712 | 0.121 | 0 | 1 | 9 |
| tspan12 | 0.05463 | 0.222962638 | 0.121 | 0 | 1 | 9 |
| ube2d1a | 0.05463 | 0.201214145 | 0.121 | 0 | 1 | 9 |
| zfr2 | 0.05463 | 0.233351383 | 0.121 | 0 | 1 | 9 |
| zgc:113426 | 0.05463 | 0.170962636 | 0.121 | 0 | 1 | 9 |
| zgc:162025 | 0.05463 | 0.231519512 | 0.121 | 0 | 1 | 9 |
| zgc:162193 | 0.05463 | 0.257593848 | 0.121 | 0 | 1 | 9 |
| znf395a | 0.05463 | 0.190894236 | 0.121 | 0 | 1 | 9 |
| CABZ01058650.1 | 0.05463 | 0.226698021 | 0.121 | 0 | 1 | 9 |
| ap1s2 | 0.05463 | 0.202921114 | 0.121 | 0 | 1 | 9 |
| b9d1 | 0.05463 | 0.183404588 | 0.121 | 0 | 1 | 9 |
| casp6l1 | 0.05463 | 0.243321719 | 0.121 | 0 | 1 | 9 |
| cbfa2t3 | 0.05463 | 0.1695722 | 0.121 | 0 | 1 | 9 |
| cd2ap | 0.05463 | 0.231894688 | 0.121 | 0 | 1 | 9 |
| cited4b | 0.05463 | 0.20285384 | 0.121 | 0 | 1 | 9 |
| dnmt3bb.1 | 0.05463 | 0.211575349 | 0.121 | 0 | 1 | 9 |
| eaf1 | 0.05463 | 0.212051344 | 0.121 | 0 | 1 | 9 |
| golga5 | 0.05463 | 0.185828999 | 0.121 | 0 | 1 | 9 |
| gstt1a | 0.05463 | 0.207155516 | 0.121 | 0 | 1 | 9 |
| her6 | 0.05463 | 0.325304103 | 0.121 | 0 | 1 | 9 |
| ift20 | 0.05463 | 0.219886243 | 0.121 | 0 | 1 | 9 |
| ift46 | 0.05463 | 0.210159266 | 0.121 | 0 | 1 | 9 |
| mbd1b | 0.05463 | 0.215746145 | 0.121 | 0 | 1 | 9 |
| mesd | 0.05463 | 0.231051045 | 0.121 | 0 | 1 | 9 |
| naa15b | 0.05463 | 0.210729185 | 0.121 | 0 | 1 | 9 |
| nanos1 | 0.05463 | 0.275688825 | 0.121 | 0 | 1 | 9 |
| nrarpa | 0.05463 | 0.253772226 | 0.121 | 0 | 1 | 9 |

| | | | | | | |
|---|---|---|---|---|---|---|
| parvaa | 0.05463 | 0.236372653 | 0.121 | 0 | 1 | 9 |
| pcca | 0.05463 | 0.204473649 | 0.121 | 0 | 1 | 9 |
| ppm1bb | 0.05463 | 0.191800245 | 0.121 | 0 | 1 | 9 |
| ppp1r7 | 0.05463 | 0.169905243 | 0.121 | 0 | 1 | 9 |
| ptpmt1 | 0.05463 | 0.200990171 | 0.121 | 0 | 1 | 9 |
| ptpn9a | 0.05463 | 0.227895018 | 0.121 | 0 | 1 | 9 |
| rnf4 | 0.05463 | 0.237646709 | 0.121 | 0 | 1 | 9 |
| sae1 | 0.05463 | 0.201152263 | 0.121 | 0 | 1 | 9 |
| sart3 | 0.05463 | 0.231309842 | 0.121 | 0 | 1 | 9 |
| si:ch211-13k12.2 | 0.05463 | 0.217562856 | 0.121 | 0 | 1 | 9 |
| si:ch211-195b11.3 | 0.05463 | 0.470982112 | 0.121 | 0 | 1 | 9 |
| si:dkey-286j15.3 | 0.05463 | 0.204990045 | 0.121 | 0 | 1 | 9 |
| si:dkey-61p9.7 | 0.05463 | 0.250003651 | 0.121 | 0 | 1 | 9 |
| stk10 | 0.05463 | 0.224450973 | 0.121 | 0 | 1 | 9 |
| tfr1b | 0.05463 | 0.209188124 | 0.121 | 0 | 1 | 9 |
| ubxn6 | 0.05463 | 0.204025697 | 0.121 | 0 | 1 | 9 |
| ugt1b5 | 0.05463 | 0.240672873 | 0.121 | 0 | 1 | 9 |
| zgc:77262 | 0.05463 | 0.190781556 | 0.121 | 0 | 1 | 9 |
| znf292a | 0.05463 | 0.252958771 | 0.121 | 0 | 1 | 9 |
| cdc34b | 0.05463 | 0.278478136 | 0.121 | 0 | 1 | 9 |
| dpm3 | 0.05463 | 0.181514156 | 0.121 | 0 | 1 | 9 |
| nek7 | 0.05463 | 0.234140894 | 0.121 | 0 | 1 | 9 |
| oip5-as1 | 0.05463 | 0.221472945 | 0.121 | 0 | 1 | 9 |
| csnk1g2a | 0.05463 | 0.204664575 | 0.121 | 0 | 1 | 9 |
| prpf4bb | 0.05463 | 0.285621026 | 0.121 | 0 | 1 | 9 |
| raraa | 0.05463 | 0.221106506 | 0.121 | 0 | 1 | 9 |
| stim1a | 0.0548 | -0.42946856 | 0.034 | 0.2 | 1 | 9 |
| pin4 | 0.05485 | 0.27210791 | 0.241 | 0.05 | 1 | 9 |
| rps27.2 | 0.05489 | 0.314808141 | 0.897 | 0.65 | 1 | 9 |
| rbbp4 | 0.05505 | 0.411422235 | 0.276 | 0.05 | 1 | 9 |
| mmgt1 | 0.05525 | 0.321569023 | 0.276 | 0.05 | 1 | 9 |
| anxa11a | 0.05591 | -0.43482949 | 0.293 | 0.45 | 1 | 9 |
| rap1aa | 0.0562 | 0.383317814 | 0.259 | 0.05 | 1 | 9 |
| gnpat | 0.05653 | -0.38419431 | 0.017 | 0.15 | 1 | 9 |
| calr3a | 0.05672 | -0.47056372 | 0.379 | 0.55 | 1 | 9 |
| glrx | 0.05677 | -0.40239477 | 0.517 | 0.7 | 1 | 9 |
| brk1 | 0.057 | 0.416554608 | 0.552 | 0.25 | 1 | 9 |
| sinhcafl | 0.05709 | 0.398970297 | 0.276 | 0.05 | 1 | 9 |
| gmppb | 0.05755 | -0.15507598 | 0.224 | 0.2 | 1 | 9 |
| zgc:112356 | 0.05769 | -0.10457655 | 0.086 | 0.05 | 1 | 9 |
| agr2 | 0.05817 | -0.65659172 | 0.052 | 0.25 | 1 | 9 |
| mapre1b | 0.05857 | 0.346198496 | 0.276 | 0.05 | 1 | 9 |
| selenok | 0.05872 | -0.44378945 | 0.241 | 0.45 | 1 | 9 |
| hsp90ab1 | 0.05894 | 0.342139584 | 0.914 | 0.75 | 1 | 9 |
| cbx3a | 0.05943 | 0.481570772 | 0.483 | 0.2 | 1 | 9 |
| uqcrc1 | 0.05946 | 0.404353505 | 0.362 | 0.1 | 1 | 9 |
| eif4a1b | 0.05963 | 0.395136916 | 0.741 | 0.45 | 1 | 9 |
| denr | 0.05968 | 0.390330975 | 0.362 | 0.1 | 1 | 9 |

| | | | | | | |
|---|---|---|---|---|---|---|
| cct5 | 0.0602 | 0.431012783 | 0.603 | 0.3 | 1 | 9 |
| rpl9 | 0.06033 | 0.346734668 | 0.914 | 0.7 | 1 | 9 |
| rpl30 | 0.06085 | 0.152237674 | 0.845 | 0.85 | 1 | 9 |
| eif4ebp3l | 0.06089 | 0.287861583 | 0.414 | 0.15 | 1 | 9 |
| si:ch211-222l21.1 | 0.06098 | 0.617692469 | 0.517 | 0.25 | 1 | 9 |
| sec61b | 0.06107 | -0.24705703 | 0.586 | 0.55 | 1 | 9 |
| ube2e3 | 0.06118 | 0.452861639 | 0.414 | 0.15 | 1 | 9 |
| eif3ba | 0.06165 | 0.476091901 | 0.534 | 0.3 | 1 | 9 |
| mtmr11 | 0.06172 | -0.34358067 | 0.017 | 0.15 | 1 | 9 |
| rfxap | 0.06212 | 0.368767419 | 0.276 | 0.05 | 1 | 9 |
| b3gnt2b | 0.0622 | -0.40062941 | 0.034 | 0.15 | 1 | 9 |
| pdlim1 | 0.06247 | -0.45556174 | 0.086 | 0.3 | 1 | 9 |
| chd3 | 0.0627 | 0.436876548 | 0.259 | 0.05 | 1 | 9 |
| aimp2 | 0.06286 | 0.271645074 | 0.328 | 0.1 | 1 | 9 |
| psmb7 | 0.06305 | 0.332185433 | 0.534 | 0.25 | 1 | 9 |
| lmo7b | 0.06325 | -0.42367599 | 0.017 | 0.15 | 1 | 9 |
| si:dkey-112e7.2 | 0.06336 | 0.206399614 | 0.431 | 0.2 | 1 | 9 |
| atp5mc3a | 0.06403 | 0.44044472 | 0.517 | 0.25 | 1 | 9 |
| dnajc1 | 0.06424 | -0.30183217 | 0.19 | 0.25 | 1 | 9 |
| asb12a | 0.06432 | -0.39494763 | 0.052 | 0.2 | 1 | 9 |
| atp5mea | 0.06455 | 0.329129077 | 0.276 | 0.05 | 1 | 9 |
| slc7a6os | 0.06476 | 0.178717258 | 0.207 | 0.05 | 1 | 9 |
| lig1 | 0.06483 | 0.419267637 | 0.276 | 0.05 | 1 | 9 |
| xkr8.2 | 0.06516 | -0.46848689 | 0.052 | 0.25 | 1 | 9 |
| tmem176l.3b | 0.06524 | -0.37554913 | 0.034 | 0.2 | 1 | 9 |
| mcl1a | 0.06526 | -0.21450722 | 0.552 | 0.5 | 1 | 9 |
| ggh | 0.06543 | -0.42209311 | 0.052 | 0.25 | 1 | 9 |
| smim19 | 0.06548 | -0.31835754 | 0.052 | 0.25 | 1 | 9 |
| eif1b | 0.06591 | 0.288379571 | 0.776 | 0.5 | 1 | 9 |
| top1l | 0.06594 | 0.236512867 | 0.31 | 0.1 | 1 | 9 |
| luc7l3 | 0.06632 | -0.35793333 | 0.034 | 0.2 | 1 | 9 |
| dlg3 | 0.06655 | -0.38649754 | 0.034 | 0.2 | 1 | 9 |
| epn1 | 0.06667 | -0.44990225 | 0.017 | 0.15 | 1 | 9 |
| gbp | 0.06675 | 0.307142419 | 0.397 | 0.15 | 1 | 9 |
| nap1l1 | 0.06686 | 0.398896027 | 0.483 | 0.2 | 1 | 9 |
| rbm39a | 0.06711 | 0.452061863 | 0.414 | 0.15 | 1 | 9 |
| arpc1a | 0.06713 | -0.44508468 | 0.293 | 0.55 | 1 | 9 |
| abhd16a | 0.06726 | -0.27254224 | 0.069 | 0.1 | 1 | 9 |
| lsm3 | 0.06768 | 0.213550026 | 0.224 | 0.05 | 1 | 9 |
| spry2 | 0.06774 | -0.50120748 | 0.224 | 0.35 | 1 | 9 |
| aph1b | 0.06782 | -0.4141115 | 0.086 | 0.3 | 1 | 9 |
| prcp | 0.06784 | -0.3724201 | 0.017 | 0.15 | 1 | 9 |
| csnk1g1 | 0.06786 | -0.30948082 | 0.017 | 0.15 | 1 | 9 |
| tmed1a | 0.06875 | -0.39581618 | 0.052 | 0.15 | 1 | 9 |
| actn1 | 0.06876 | -0.37751281 | 0.017 | 0.15 | 1 | 9 |
| si:dkeyp-110e4.6 | 0.06925 | -0.3721042 | 0.052 | 0.2 | 1 | 9 |
| junba | 0.0694 | -0.46145707 | 0.328 | 0.45 | 1 | 9 |
| copa | 0.06942 | -0.40265757 | 0.086 | 0.25 | 1 | 9 |

| | | | | | | |
|---|---|---|---|---|---|---|
| CR396595.1 | 0.0697 | -0.329111 | 0.017 | 0.15 | 1 | 9 |
| rnf141 | 0.06972 | -0.30025237 | 0.017 | 0.15 | 1 | 9 |
| ccl20a.3 | 0.06975 | -0.39844888 | 0.086 | 0.15 | 1 | 9 |
| cox8a | 0.06995 | 0.353715751 | 0.69 | 0.4 | 1 | 9 |
| pdia3 | 0.07029 | -0.28416697 | 0.638 | 0.65 | 1 | 9 |
| zgc:110158 | 0.07076 | -0.38063849 | 0.052 | 0.2 | 1 | 9 |
| CDPF1 | 0.07106 | 0.123584521 | 0.19 | 0.05 | 1 | 9 |
| pkp2 | 0.07126 | -0.16026348 | 0.017 | 0.15 | 1 | 9 |
| rpl18 | 0.07147 | 0.320822387 | 0.879 | 0.75 | 1 | 9 |
| timm13 | 0.07157 | 0.285158653 | 0.224 | 0.05 | 1 | 9 |
| tmbim4 | 0.07174 | -0.35849382 | 0.224 | 0.3 | 1 | 9 |
| si:ch73-22a13.3 | 0.07207 | -0.36337374 | 0.017 | 0.15 | 1 | 9 |
| dhrs13l1 | 0.07209 | -0.35341154 | 0.086 | 0.25 | 1 | 9 |
| rhogb | 0.07233 | -0.20590201 | 0.103 | 0.1 | 1 | 9 |
| hspd1 | 0.07327 | 0.365494828 | 0.259 | 0.05 | 1 | 9 |
| rnf7 | 0.07352 | 0.146547857 | 0.328 | 0.15 | 1 | 9 |
| ssh2a | 0.07362 | -0.45454372 | 0.086 | 0.25 | 1 | 9 |
| sap30l | 0.07384 | 0.357929046 | 0.379 | 0.25 | 1 | 9 |
| calm2b | 0.07427 | 0.363857442 | 0.776 | 0.5 | 1 | 9 |
| si:dkey-33c12.4 | 0.07443 | 0.268255278 | 0.379 | 0.15 | 1 | 9 |
| prdx6 | 0.07466 | -0.21068981 | 0.259 | 0.25 | 1 | 9 |
| eif4a1a | 0.07479 | 0.318204872 | 0.466 | 0.2 | 1 | 9 |
| cd82a | 0.07549 | -0.33113148 | 0.138 | 0.2 | 1 | 9 |
| eef1da | 0.07606 | 0.223667699 | 0.793 | 0.55 | 1 | 9 |
| slc25a55a | 0.07654 | -0.44223958 | 0.138 | 0.3 | 1 | 9 |
| tomm6 | 0.07655 | -0.10453832 | 0.293 | 0.25 | 1 | 9 |
| h2afva | 0.07683 | 0.286459346 | 0.328 | 0.1 | 1 | 9 |
| mocs3 | 0.07754 | -0.14282599 | 0.017 | 0.15 | 1 | 9 |
| psma8 | 0.07817 | 0.211485095 | 0.466 | 0.25 | 1 | 9 |
| crema | 0.07828 | -0.5106452 | 0.276 | 0.5 | 1 | 9 |
| fkbp1aa | 0.07878 | 0.216723325 | 0.793 | 0.55 | 1 | 9 |
| xbp1 | 0.0792 | -0.42366269 | 0.483 | 0.65 | 1 | 9 |
| BX322530.1 | 0.07951 | 0.195297426 | 0.103 | 0 | 1 | 9 |
| XLOC-018981 | 0.07951 | 0.19617695 | 0.103 | 0 | 1 | 9 |
| XLOC-040516 | 0.07951 | 0.284029981 | 0.103 | 0 | 1 | 9 |
| acads | 0.07951 | 0.169296327 | 0.103 | 0 | 1 | 9 |
| adnpa | 0.07951 | 0.24797671 | 0.103 | 0 | 1 | 9 |
| ap1m3 | 0.07951 | 0.197912051 | 0.103 | 0 | 1 | 9 |
| calua | 0.07951 | 0.179195717 | 0.103 | 0 | 1 | 9 |
| cdk12 | 0.07951 | 0.207434614 | 0.103 | 0 | 1 | 9 |
| dnajc4 | 0.07951 | 0.162591523 | 0.103 | 0 | 1 | 9 |
| dnph1 | 0.07951 | 0.115631335 | 0.103 | 0 | 1 | 9 |
| dph2 | 0.07951 | 0.167500416 | 0.103 | 0 | 1 | 9 |
| emc3 | 0.07951 | 0.185485589 | 0.103 | 0 | 1 | 9 |
| exosc5 | 0.07951 | 0.15995646 | 0.103 | 0 | 1 | 9 |
| fdps | 0.07951 | 0.181826577 | 0.103 | 0 | 1 | 9 |
| gabpb2a | 0.07951 | 0.181777308 | 0.103 | 0 | 1 | 9 |
| gtf2h4 | 0.07951 | 0.172939602 | 0.103 | 0 | 1 | 9 |

| | | | | | | |
|---|---|---|---|---|---|---|
| hes6 | 0.07951 | 0.203481348 | 0.103 | 0 | 1 | 9 |
| hmga2 | 0.07951 | 0.237214336 | 0.103 | 0 | 1 | 9 |
| hpcal1 | 0.07951 | 0.204241092 | 0.103 | 0 | 1 | 9 |
| inppl1a | 0.07951 | 0.14572164 | 0.103 | 0 | 1 | 9 |
| kdm6al | 0.07951 | 0.240645784 | 0.103 | 0 | 1 | 9 |
| lcmt1 | 0.07951 | 0.137985829 | 0.103 | 0 | 1 | 9 |
| mettl17 | 0.07951 | 0.171497809 | 0.103 | 0 | 1 | 9 |
| mib1 | 0.07951 | 0.203705407 | 0.103 | 0 | 1 | 9 |
| mrps18b | 0.07951 | 0.202775367 | 0.103 | 0 | 1 | 9 |
| nudt17 | 0.07951 | 0.212818352 | 0.103 | 0 | 1 | 9 |
| nup107 | 0.07951 | 0.200572022 | 0.103 | 0 | 1 | 9 |
| polb | 0.07951 | 0.223320385 | 0.103 | 0 | 1 | 9 |
| ppan | 0.07951 | 0.220160603 | 0.103 | 0 | 1 | 9 |
| prpsap2 | 0.07951 | 0.179849012 | 0.103 | 0 | 1 | 9 |
| rab4b | 0.07951 | 0.189624385 | 0.103 | 0 | 1 | 9 |
| rbl1 | 0.07951 | 0.203372244 | 0.103 | 0 | 1 | 9 |
| rhbdl2 | 0.07951 | 0.16127364 | 0.103 | 0 | 1 | 9 |
| scaf11 | 0.07951 | 0.130748197 | 0.103 | 0 | 1 | 9 |
| senp2 | 0.07951 | 0.169523344 | 0.103 | 0 | 1 | 9 |
| si:ch73-95l15.5 | 0.07951 | 0.225202353 | 0.103 | 0 | 1 | 9 |
| si:dkey-223p19.1 | 0.07951 | 0.254177359 | 0.103 | 0 | 1 | 9 |
| si:dkey-79d12.5 | 0.07951 | 0.212153369 | 0.103 | 0 | 1 | 9 |
| tecrb | 0.07951 | 0.189883348 | 0.103 | 0 | 1 | 9 |
| tex264a | 0.07951 | 0.194317334 | 0.103 | 0 | 1 | 9 |
| tgfbi | 0.07951 | 0.250202023 | 0.103 | 0 | 1 | 9 |
| thbs4b | 0.07951 | 0.290507584 | 0.103 | 0 | 1 | 9 |
| tmem106bb | 0.07951 | 0.137228531 | 0.103 | 0 | 1 | 9 |
| tmem243b | 0.07951 | 0.164147719 | 0.103 | 0 | 1 | 9 |
| zfr | 0.07951 | 0.168292927 | 0.103 | 0 | 1 | 9 |
| zgc:112023 | 0.07951 | 0.184788807 | 0.103 | 0 | 1 | 9 |
| LOC101885018 | 0.07951 | 0.221744124 | 0.103 | 0 | 1 | 9 |
| ctdsplb | 0.07951 | 0.186444266 | 0.103 | 0 | 1 | 9 |
| fam114a2 | 0.07951 | 0.179912653 | 0.103 | 0 | 1 | 9 |
| rpap1 | 0.07951 | 0.193476491 | 0.103 | 0 | 1 | 9 |
| zgc:162339 | 0.07951 | 0.196458697 | 0.103 | 0 | 1 | 9 |
| KIF2A | 0.07951 | 0.144484308 | 0.103 | 0 | 1 | 9 |
| LOC100535095 | 0.07951 | 0.226809721 | 0.103 | 0 | 1 | 9 |
| abcb7 | 0.07951 | 0.190198083 | 0.103 | 0 | 1 | 9 |
| abl1 | 0.07951 | 0.181295061 | 0.103 | 0 | 1 | 9 |
| acot13 | 0.07951 | 0.186831009 | 0.103 | 0 | 1 | 9 |
| adgrl2a | 0.07951 | 0.271887614 | 0.103 | 0 | 1 | 9 |
| afmid | 0.07951 | 0.119569313 | 0.103 | 0 | 1 | 9 |
| agap1 | 0.07951 | 0.224179078 | 0.103 | 0 | 1 | 9 |
| amfra | 0.07951 | 0.158039063 | 0.103 | 0 | 1 | 9 |
| angel2 | 0.07951 | 0.204800813 | 0.103 | 0 | 1 | 9 |
| ap4s1 | 0.07951 | 0.186625464 | 0.103 | 0 | 1 | 9 |
| apbb1 | 0.07951 | 0.170526261 | 0.103 | 0 | 1 | 9 |
| cadm3 | 0.07951 | 0.157049496 | 0.103 | 0 | 1 | 9 |

| | | | | | | |
|---|---|---|---|---|---|---|
| casc3 | 0.07951 | 0.153882551 | 0.103 | 0 | 1 | 9 |
| ccdc167 | 0.07951 | 0.18002315 | 0.103 | 0 | 1 | 9 |
| ccdc59 | 0.07951 | 0.184018349 | 0.103 | 0 | 1 | 9 |
| cdk11b | 0.07951 | 0.190929336 | 0.103 | 0 | 1 | 9 |
| cdk8 | 0.07951 | 0.190706613 | 0.103 | 0 | 1 | 9 |
| celf3a | 0.07951 | 0.230714544 | 0.103 | 0 | 1 | 9 |
| cetn3 | 0.07951 | 0.182182744 | 0.103 | 0 | 1 | 9 |
| cica | 0.07951 | 0.183447662 | 0.103 | 0 | 1 | 9 |
| clic4 | 0.07951 | 0.198391403 | 0.103 | 0 | 1 | 9 |
| commd2 | 0.07951 | 0.184048499 | 0.103 | 0 | 1 | 9 |
| crip2 | 0.07951 | 0.147376044 | 0.103 | 0 | 1 | 9 |
| crk | 0.07951 | 0.168747512 | 0.103 | 0 | 1 | 9 |
| crnkl1 | 0.07951 | 0.184344627 | 0.103 | 0 | 1 | 9 |
| cry-dash | 0.07951 | 0.18992668 | 0.103 | 0 | 1 | 9 |
| dap3 | 0.07951 | 0.188764266 | 0.103 | 0 | 1 | 9 |
| dbt | 0.07951 | 0.151745988 | 0.103 | 0 | 1 | 9 |
| dck | 0.07951 | 0.137608809 | 0.103 | 0 | 1 | 9 |
| ddx18 | 0.07951 | 0.163530918 | 0.103 | 0 | 1 | 9 |
| desma | 0.07951 | 0.22248698 | 0.103 | 0 | 1 | 9 |
| dnajb1b | 0.07951 | 0.158251329 | 0.103 | 0 | 1 | 9 |
| dpp7 | 0.07951 | 0.178067136 | 0.103 | 0 | 1 | 9 |
| exosc4 | 0.07951 | 0.193028526 | 0.103 | 0 | 1 | 9 |
| fdx1 | 0.07951 | 0.136089214 | 0.103 | 0 | 1 | 9 |
| fzd7a | 0.07951 | 0.192992034 | 0.103 | 0 | 1 | 9 |
| gli3 | 0.07951 | 0.226561872 | 0.103 | 0 | 1 | 9 |
| gng7 | 0.07951 | 0.179565549 | 0.103 | 0 | 1 | 9 |
| hsbp1a | 0.07951 | 0.23900995 | 0.103 | 0 | 1 | 9 |
| hsp90aa1.1 | 0.07951 | 0.238702885 | 0.103 | 0 | 1 | 9 |
| hspa9 | 0.07951 | 0.186278277 | 0.103 | 0 | 1 | 9 |
| irs1 | 0.07951 | 0.19293696 | 0.103 | 0 | 1 | 9 |
| lin52 | 0.07951 | 0.176566237 | 0.103 | 0 | 1 | 9 |
| lin54 | 0.07951 | 0.158820456 | 0.103 | 0 | 1 | 9 |
| lsm12a | 0.07951 | 0.125945127 | 0.103 | 0 | 1 | 9 |
| lypla2 | 0.07951 | 0.159958952 | 0.103 | 0 | 1 | 9 |
| lztfl1 | 0.07951 | 0.160460947 | 0.103 | 0 | 1 | 9 |
| mmaa | 0.07951 | 0.178360747 | 0.103 | 0 | 1 | 9 |
| mrpl45 | 0.07951 | 0.172958079 | 0.103 | 0 | 1 | 9 |
| mrps15 | 0.07951 | 0.205654764 | 0.103 | 0 | 1 | 9 |
| mrps23 | 0.07951 | 0.175478223 | 0.103 | 0 | 1 | 9 |
| mycn | 0.07951 | 0.200795018 | 0.103 | 0 | 1 | 9 |
| myog | 0.07951 | 0.364499724 | 0.103 | 0 | 1 | 9 |
| noc4l | 0.07951 | 0.136574039 | 0.103 | 0 | 1 | 9 |
| nrd1a | 0.07951 | 0.142968056 | 0.103 | 0 | 1 | 9 |
| nup43 | 0.07951 | 0.204247611 | 0.103 | 0 | 1 | 9 |
| parp3 | 0.07951 | 0.168018672 | 0.103 | 0 | 1 | 9 |
| phf20l1 | 0.07951 | 0.223912708 | 0.103 | 0 | 1 | 9 |
| pitpnbl | 0.07951 | 0.137932975 | 0.103 | 0 | 1 | 9 |
| poldip2 | 0.07951 | 0.136660842 | 0.103 | 0 | 1 | 9 |

| | | | | | | |
|---|---|---|---|---|---|---|
| polr1e | 0.07951 | 0.184971943 | 0.103 | 0 | 1 | 9 |
| polr2j | 0.07951 | 0.228357965 | 0.103 | 0 | 1 | 9 |
| ppp3cca | 0.07951 | 0.119700033 | 0.103 | 0 | 1 | 9 |
| prpf4 | 0.07951 | 0.139560965 | 0.103 | 0 | 1 | 9 |
| prss16 | 0.07951 | 0.195350987 | 0.103 | 0 | 1 | 9 |
| pus1 | 0.07951 | 0.18473474 | 0.103 | 0 | 1 | 9 |
| rab28 | 0.07951 | 0.171459783 | 0.103 | 0 | 1 | 9 |
| raf1b | 0.07951 | 0.126126476 | 0.103 | 0 | 1 | 9 |
| ralab | 0.07951 | 0.187592584 | 0.103 | 0 | 1 | 9 |
| rbm42 | 0.07951 | 0.180679123 | 0.103 | 0 | 1 | 9 |
| rfc2 | 0.07951 | 0.182585499 | 0.103 | 0 | 1 | 9 |
| rnd3a | 0.07951 | 0.175408464 | 0.103 | 0 | 1 | 9 |
| scamp2l | 0.07951 | 0.189554028 | 0.103 | 0 | 1 | 9 |
| serinc2l | 0.07951 | 0.15504993 | 0.103 | 0 | 1 | 9 |
| serpine2 | 0.07951 | 0.238344407 | 0.103 | 0 | 1 | 9 |
| sh3gl1a | 0.07951 | 0.192127033 | 0.103 | 0 | 1 | 9 |
| shmt1 | 0.07951 | 0.183024222 | 0.103 | 0 | 1 | 9 |
| si:ch211-173p18.3 | 0.07951 | 0.199245915 | 0.103 | 0 | 1 | 9 |
| si:ch211-197h24.9 | 0.07951 | 0.142289671 | 0.103 | 0 | 1 | 9 |
| si:ch211-286o17.1 | 0.07951 | 0.240835507 | 0.103 | 0 | 1 | 9 |
| si:ch73-147o17.1 | 0.07951 | 0.137482771 | 0.103 | 0 | 1 | 9 |
| six4b | 0.07951 | 0.182560575 | 0.103 | 0 | 1 | 9 |
| slc27a1a | 0.07951 | 0.151764103 | 0.103 | 0 | 1 | 9 |
| slc9a3r1a | 0.07951 | 0.196926603 | 0.103 | 0 | 1 | 9 |
| sox6 | 0.07951 | 0.179490988 | 0.103 | 0 | 1 | 9 |
| sppl2 | 0.07951 | 0.150911931 | 0.103 | 0 | 1 | 9 |
| stau1 | 0.07951 | 0.160943158 | 0.103 | 0 | 1 | 9 |
| stmn1b | 0.07951 | 0.239065474 | 0.103 | 0 | 1 | 9 |
| tada3l | 0.07951 | 0.166137517 | 0.103 | 0 | 1 | 9 |
| tdg.1 | 0.07951 | 0.169115581 | 0.103 | 0 | 1 | 9 |
| thumpd1 | 0.07951 | 0.160946946 | 0.103 | 0 | 1 | 9 |
| tmem106a | 0.07951 | 0.198983465 | 0.103 | 0 | 1 | 9 |
| tmx4 | 0.07951 | 0.183825943 | 0.103 | 0 | 1 | 9 |
| ube2d4 | 0.07951 | 0.164367251 | 0.103 | 0 | 1 | 9 |
| usp1 | 0.07951 | 0.165482828 | 0.103 | 0 | 1 | 9 |
| vamp5 | 0.07951 | 0.170185449 | 0.103 | 0 | 1 | 9 |
| wdr3 | 0.07951 | 0.179934942 | 0.103 | 0 | 1 | 9 |
| wu:fe05a04 | 0.07951 | 0.169814267 | 0.103 | 0 | 1 | 9 |
| zcrb1 | 0.07951 | 0.180839574 | 0.103 | 0 | 1 | 9 |
| zgc:92335 | 0.07951 | 0.153350284 | 0.103 | 0 | 1 | 9 |
| gal3st1a | 0.07951 | 0.217689862 | 0.103 | 0 | 1 | 9 |
| dus2 | 0.07951 | 0.172925377 | 0.103 | 0 | 1 | 9 |
| ipo9 | 0.07951 | 0.206429812 | 0.103 | 0 | 1 | 9 |
| mpped2a | 0.07951 | 0.167513074 | 0.103 | 0 | 1 | 9 |
| si:ch1073-406l10.2 | 0.07951 | 0.252711255 | 0.103 | 0 | 1 | 9 |
| timm44 | 0.07951 | 0.20705636 | 0.103 | 0 | 1 | 9 |
| si:dkey-16l2.16 | 0.07951 | 0.224118555 | 0.103 | 0 | 1 | 9 |
| drap1 | 0.07956 | 0.366752886 | 0.414 | 0.15 | 1 | 9 |

| | | | | | | |
|---|---|---|---|---|---|---|
| rps3 | 0.07975 | 0.306203326 | 0.897 | 0.8 | 1 | 9 |
| si:dkey-67c22.2 | 0.07985 | 0.400888406 | 0.345 | 0.1 | 1 | 9 |
| fam129ba | 0.07986 | -0.44662836 | 0.052 | 0.2 | 1 | 9 |
| mrpl30 | 0.07992 | 0.314890009 | 0.345 | 0.1 | 1 | 9 |
| slc30a7 | 0.08024 | 0.20211707 | 0.207 | 0.05 | 1 | 9 |
| got1 | 0.08052 | -0.10232362 | 0.086 | 0.05 | 1 | 9 |
| NC-002333.4 | 0.08087 | -0.22905558 | 0.845 | 0.85 | 1 | 9 |
| ndufa3 | 0.08101 | 0.442405133 | 0.517 | 0.25 | 1 | 9 |
| hars | 0.08126 | -0.1786568 | 0.172 | 0.15 | 1 | 9 |
| cldnc | 0.08233 | -0.53887432 | 0.086 | 0.25 | 1 | 9 |
| glud1b | 0.08233 | 0.299128029 | 0.259 | 0.05 | 1 | 9 |
| zgc:110339 | 0.08255 | 0.10798616 | 0.31 | 0.15 | 1 | 9 |
| atp1a1a.1 | 0.08281 | 0.395856405 | 0.259 | 0.05 | 1 | 9 |
| rps12 | 0.08299 | 0.274100165 | 0.914 | 0.85 | 1 | 9 |
| si:ch211-270g19.5 | 0.08311 | -0.56849968 | 0.034 | 0.2 | 1 | 9 |
| gbgt1l4 | 0.08318 | -0.43367393 | 0.034 | 0.2 | 1 | 9 |
| hyou1 | 0.08338 | -0.40016099 | 0.19 | 0.3 | 1 | 9 |
| flna | 0.08356 | -0.38678929 | 0.086 | 0.2 | 1 | 9 |
| krt18b | 0.08377 | 0.494849637 | 0.241 | 0.05 | 1 | 9 |
| si:ch73-281n10.2 | 0.08397 | 0.42264036 | 0.776 | 0.55 | 1 | 9 |
| cdh1 | 0.08414 | -0.48539997 | 0.293 | 0.5 | 1 | 9 |
| sf3b1 | 0.08443 | 0.343496058 | 0.259 | 0.05 | 1 | 9 |
| hsbp1b | 0.08456 | 0.41014157 | 0.448 | 0.2 | 1 | 9 |
| rps17 | 0.08459 | 0.288158072 | 0.862 | 0.75 | 1 | 9 |
| sdf4 | 0.08464 | -0.45532225 | 0.138 | 0.3 | 1 | 9 |
| wtap | 0.08494 | 0.244392153 | 0.241 | 0.05 | 1 | 9 |
| lrrc59 | 0.08502 | -0.29292194 | 0.155 | 0.25 | 1 | 9 |
| nop2 | 0.08568 | 0.39184266 | 0.241 | 0.05 | 1 | 9 |
| rca2.1 | 0.08572 | -0.38496055 | 0.103 | 0.25 | 1 | 9 |
| cox17 | 0.08574 | -0.1867834 | 0.293 | 0.3 | 1 | 9 |
| hnrnpd | 0.08579 | 0.349441956 | 0.224 | 0.05 | 1 | 9 |
| zgc:171497 | 0.08605 | -0.35072201 | 0.034 | 0.2 | 1 | 9 |
| lig3 | 0.08622 | -0.31541158 | 0.017 | 0.15 | 1 | 9 |
| paics | 0.08641 | 0.317810236 | 0.241 | 0.05 | 1 | 9 |
| h3f3c | 0.08662 | 0.339780941 | 0.724 | 0.45 | 1 | 9 |
| smc1al | 0.08681 | 0.327748372 | 0.397 | 0.15 | 1 | 9 |
| wdr43 | 0.08731 | 0.299758371 | 0.259 | 0.05 | 1 | 9 |
| cd3eap | 0.08758 | 0.323772042 | 0.259 | 0.05 | 1 | 9 |
| pxylp1 | 0.08777 | -0.33943484 | 0.017 | 0.15 | 1 | 9 |
| clcn3 | 0.08785 | -0.37774277 | 0.069 | 0.25 | 1 | 9 |
| eef1db | 0.08803 | 0.445464767 | 0.569 | 0.3 | 1 | 9 |
| xrn2 | 0.08806 | 0.321031473 | 0.259 | 0.05 | 1 | 9 |
| sptbn2 | 0.08829 | -0.46836174 | 0.086 | 0.25 | 1 | 9 |
| ctbp1 | 0.08876 | 0.371005105 | 0.259 | 0.05 | 1 | 9 |
| npm3 | 0.08894 | 0.362185203 | 0.241 | 0.05 | 1 | 9 |
| pbx3b | 0.089 | -0.30592605 | 0.034 | 0.2 | 1 | 9 |
| zgc:85789 | 0.08919 | -0.18870383 | 0.034 | 0.2 | 1 | 9 |
| alox12 | 0.08929 | -0.45610853 | 0.069 | 0.25 | 1 | 9 |

| | | | | | | |
|---|---|---|---|---|---|---|
| calcoco1a | 0.08931 | -0.29723648 | 0.052 | 0.2 | 1 | 9 |
| pstpip2 | 0.08953 | -0.22710407 | 0.034 | 0.2 | 1 | 9 |
| si:rp71-36a1.1 | 0.08954 | -0.32249757 | 0.034 | 0.2 | 1 | 9 |
| ddost | 0.08974 | -0.39519587 | 0.241 | 0.4 | 1 | 9 |
| rpl5b | 0.08978 | 0.295847355 | 0.879 | 0.65 | 1 | 9 |
| lsr | 0.09002 | -0.38085148 | 0.121 | 0.35 | 1 | 9 |
| CR318588.3 | 0.0904 | -0.61453501 | 0.052 | 0.2 | 1 | 9 |
| col4a1 | 0.09041 | 0.307956031 | 0.207 | 0.05 | 1 | 9 |
| si:ch211-71m22.1 | 0.09052 | -0.37033868 | 0.017 | 0.15 | 1 | 9 |
| ptbp3 | 0.09062 | -0.12795818 | 0.172 | 0.15 | 1 | 9 |
| entpd2a.1 | 0.09143 | -0.36471391 | 0.017 | 0.15 | 1 | 9 |
| pam | 0.09146 | -0.46447288 | 0.138 | 0.35 | 1 | 9 |
| rps6 | 0.09154 | 0.261769281 | 0.931 | 0.85 | 1 | 9 |
| ndufb10 | 0.09204 | 0.416797087 | 0.483 | 0.25 | 1 | 9 |
| ndufb7 | 0.09221 | 0.231230038 | 0.552 | 0.3 | 1 | 9 |
| eif4e2 | 0.09234 | -0.34325968 | 0.034 | 0.1 | 1 | 9 |
| pgrmc1 | 0.09259 | -0.22348874 | 0.328 | 0.35 | 1 | 9 |
| si:ch73-208h1.1 | 0.09266 | -0.47228825 | 0.017 | 0.15 | 1 | 9 |
| XLOC-015291 | 0.09271 | -0.32772629 | 0.017 | 0.15 | 1 | 9 |
| fam53b | 0.09343 | -0.16309328 | 0.224 | 0.2 | 1 | 9 |
| cdv3 | 0.0936 | 0.254163927 | 0.328 | 0.1 | 1 | 9 |
| gmfb | 0.09362 | 0.374000802 | 0.241 | 0.05 | 1 | 9 |
| atp5l | 0.09425 | 0.112325894 | 0.759 | 0.55 | 1 | 9 |
| srebf2 | 0.09436 | -0.33526005 | 0.052 | 0.15 | 1 | 9 |
| ier2b | 0.09547 | -0.33091857 | 0.466 | 0.55 | 1 | 9 |
| gosr2 | 0.09551 | -0.33579375 | 0.069 | 0.25 | 1 | 9 |
| ca2 | 0.0956 | 0.361931796 | 0.19 | 0.1 | 1 | 9 |
| ube2a | 0.09567 | 0.323684831 | 0.241 | 0.05 | 1 | 9 |
| rassf7b | 0.09571 | -0.45678537 | 0.103 | 0.3 | 1 | 9 |
| zgc:114188 | 0.09594 | 0.235271686 | 0.914 | 0.85 | 1 | 9 |
| rnasel4 | 0.09665 | -0.39568689 | 0.017 | 0.15 | 1 | 9 |
| s100w | 0.09692 | -0.34594318 | 0.224 | 0.3 | 1 | 9 |
| llph | 0.09708 | 0.197588382 | 0.259 | 0.1 | 1 | 9 |
| ube2nb | 0.09722 | 0.331359221 | 0.224 | 0.05 | 1 | 9 |
| mif | 0.09724 | 0.2987101 | 0.448 | 0.2 | 1 | 9 |
| gipc2 | 0.09734 | -0.40229793 | 0.155 | 0.3 | 1 | 9 |
| vdac3 | 0.09752 | 0.202071258 | 0.276 | 0.1 | 1 | 9 |
| cmtr2 | 0.09781 | -0.24266025 | 0.017 | 0.15 | 1 | 9 |
| cftr | 0.09781 | -0.50152477 | 0.017 | 0.15 | 1 | 9 |
| rhoca | 0.09855 | -0.39240614 | 0.172 | 0.3 | 1 | 9 |
| lrmp | 0.09895 | -0.33641235 | 0.017 | 0.15 | 1 | 9 |
| dap1b | 0.09927 | -0.16894746 | 0.379 | 0.35 | 1 | 9 |
| mir21-1 | 0.09931 | -0.20891716 | 0.017 | 0.15 | 1 | 9 |
| si:dkey-163m14.2 | 0.09966 | -0.39939077 | 0.034 | 0.15 | 1 | 9 |
| btf3l4 | 0.09989 | -0.2238047 | 0.276 | 0.3 | 1 | 9 |
| sf3b2 | 0.09992 | 0.345934456 | 0.328 | 0.1 | 1 | 9 |
| eif3s6ip | 0.09996 | 0.342838545 | 0.397 | 0.15 | 1 | 9 |
| romo1 | 0.10019 | 0.305854932 | 0.672 | 0.4 | 1 | 9 |

| | | | | | | |
|---|---|---|---|---|---|---|
| sema6bb | 0.1003 | -0.24750325 | 0.017 | 0.15 | 1 | 9 |
| gdi2 | 0.10043 | 0.144137412 | 0.448 | 0.25 | 1 | 9 |
| pcdh1b | 0.10123 | -0.34835606 | 0.086 | 0.25 | 1 | 9 |
| calcoco2 | 0.10132 | -0.35554828 | 0.069 | 0.25 | 1 | 9 |
| kmt2cb | 0.10148 | 0.274475869 | 0.224 | 0.05 | 1 | 9 |
| ccdc9 | 0.10164 | -0.35782443 | 0.069 | 0.25 | 1 | 9 |
| gnb1a | 0.10177 | 0.113717484 | 0.466 | 0.3 | 1 | 9 |
| anxa11b | 0.10183 | 0.390876003 | 0.31 | 0.1 | 1 | 9 |
| rps24 | 0.10187 | 0.259322281 | 0.948 | 0.8 | 1 | 9 |
| csrp2 | 0.10221 | -0.21451306 | 0.017 | 0.15 | 1 | 9 |
| mrpl15 | 0.10226 | -0.24918743 | 0.017 | 0.15 | 1 | 9 |
| hint1 | 0.10228 | 0.284680132 | 0.448 | 0.2 | 1 | 9 |
| relch | 0.10259 | -0.29160339 | 0.017 | 0.15 | 1 | 9 |
| prkacaa | 0.10285 | -0.24313155 | 0.017 | 0.15 | 1 | 9 |
| krt5 | 0.10286 | 0.428354513 | 0.31 | 0.25 | 1 | 9 |
| plxnb3 | 0.10289 | -0.24650697 | 0.017 | 0.15 | 1 | 9 |
| rtf2 | 0.10337 | -0.18572388 | 0.207 | 0.2 | 1 | 9 |
| jpt2 | 0.10364 | -0.39747109 | 0.276 | 0.45 | 1 | 9 |
| ptenb | 0.10365 | 0.329305801 | 0.207 | 0.05 | 1 | 9 |
| mhc1laa | 0.10395 | -0.24621901 | 0.017 | 0.15 | 1 | 9 |
| il4r.1 | 0.10423 | -0.32935028 | 0.034 | 0.15 | 1 | 9 |
| casp7 | 0.10464 | -0.18270784 | 0.017 | 0.15 | 1 | 9 |
| sept7b | 0.10476 | 0.215853186 | 0.224 | 0.05 | 1 | 9 |
| ywhae1 | 0.10484 | 0.368965795 | 0.517 | 0.25 | 1 | 9 |
| CU929145.1 | 0.10502 | -0.45111896 | 0.017 | 0.15 | 1 | 9 |
| tcea1 | 0.10508 | 0.324826825 | 0.241 | 0.05 | 1 | 9 |
| rpl21 | 0.10509 | 0.294010944 | 0.879 | 0.75 | 1 | 9 |
| krt94 | 0.10531 | -0.63270868 | 0.086 | 0.25 | 1 | 9 |
| naa10 | 0.10567 | 0.365569321 | 0.397 | 0.15 | 1 | 9 |
| bcl2l13 | 0.10582 | -0.42168144 | 0.138 | 0.25 | 1 | 9 |
| ube4b | 0.10589 | -0.13714686 | 0.121 | 0.1 | 1 | 9 |
| c9 | 0.10637 | -0.23219344 | 0.017 | 0.15 | 1 | 9 |
| gabarapa | 0.10649 | 0.285631742 | 0.362 | 0.15 | 1 | 9 |
| jtb | 0.10662 | -0.10758993 | 0.086 | 0.05 | 1 | 9 |
| noxo1a | 0.10676 | -0.26325479 | 0.017 | 0.15 | 1 | 9 |
| cobll1a | 0.10688 | -0.25841107 | 0.017 | 0.15 | 1 | 9 |
| enc1 | 0.1072 | -0.32637326 | 0.052 | 0.15 | 1 | 9 |
| il17ra1a | 0.10737 | -0.22098489 | 0.017 | 0.15 | 1 | 9 |
| CABZ01070852.1 | 0.10742 | -0.25946981 | 0.017 | 0.15 | 1 | 9 |
| ngdn | 0.10747 | 0.215477756 | 0.207 | 0.05 | 1 | 9 |
| syngr2a | 0.10781 | -0.28069499 | 0.103 | 0.15 | 1 | 9 |
| rps25 | 0.10791 | 0.258274357 | 0.931 | 0.8 | 1 | 9 |
| sft2d2b | 0.10794 | -0.22694235 | 0.017 | 0.15 | 1 | 9 |
| zgc:66484 | 0.108 | -0.19212 | 0.017 | 0.15 | 1 | 9 |
| larp4ab | 0.10833 | 0.300445981 | 0.241 | 0.05 | 1 | 9 |
| tyw1 | 0.10909 | -0.37302046 | 0.017 | 0.15 | 1 | 9 |
| sik3 | 0.10915 | -0.1853028 | 0.017 | 0.15 | 1 | 9 |
| si:ch211-195e19.1 | 0.10915 | -0.24864664 | 0.017 | 0.15 | 1 | 9 |

| | | | | | | |
|---|---|---|---|---|---|---|
| aco1 | 0.10933 | -0.22823262 | 0.017 | 0.15 | 1 | 9 |
| ptprjb.1 | 0.10948 | -0.40036244 | 0.052 | 0.2 | 1 | 9 |
| si:ch211-213a13.2 | 0.10956 | -0.24552976 | 0.017 | 0.15 | 1 | 9 |
| rplp1 | 0.10961 | 0.180000953 | 0.948 | 0.9 | 1 | 9 |
| calr | 0.10963 | -0.22133498 | 0.328 | 0.35 | 1 | 9 |
| si:dkey-58f10.11 | 0.10964 | -0.24535531 | 0.017 | 0.15 | 1 | 9 |
| nup214 | 0.10977 | -0.19418857 | 0.017 | 0.15 | 1 | 9 |
| AL590151.1 | 0.10987 | -0.23416758 | 0.017 | 0.15 | 1 | 9 |
| rdh12l | 0.10991 | -0.21947547 | 0.017 | 0.15 | 1 | 9 |
| LOC100537265 | 0.10991 | -0.2180766 | 0.017 | 0.15 | 1 | 9 |
| dtwd1 | 0.11046 | 0.303000424 | 0.224 | 0.05 | 1 | 9 |
| snu13b | 0.11056 | 0.380846178 | 0.379 | 0.15 | 1 | 9 |
| cnbpa | 0.1107 | 0.351419884 | 0.569 | 0.3 | 1 | 9 |
| calm2a | 0.11179 | -0.33563062 | 0.345 | 0.45 | 1 | 9 |
| ttc33 | 0.11235 | -0.29246044 | 0.052 | 0.15 | 1 | 9 |
| vamp8 | 0.113 | 0.36496885 | 0.241 | 0.1 | 1 | 9 |
| hnrnpabb | 0.11316 | 0.491286748 | 0.5 | 0.25 | 1 | 9 |
| cldn7b | 0.1141 | -0.38195235 | 0.172 | 0.4 | 1 | 9 |
| ranbp1 | 0.11413 | 0.335279063 | 0.431 | 0.3 | 1 | 9 |
| unc50 | 0.1143 | -0.30441633 | 0.069 | 0.25 | 1 | 9 |
| arl8ba | 0.11441 | -0.33733268 | 0.069 | 0.25 | 1 | 9 |
| chaf1a | 0.11466 | 0.376841081 | 0.241 | 0.05 | 1 | 9 |
| si:ch211-79k12.1 | 0.11514 | -0.44565576 | 0.19 | 0.3 | 1 | 9 |
| nupr1a | 0.11561 | -0.10975335 | 0.379 | 0.3 | 1 | 9 |
| epha3 | 0.11586 | 0.167943469 | 0.086 | 0 | 1 | 9 |
| gpr85 | 0.11586 | 0.190375966 | 0.086 | 0 | 1 | 9 |
| paf1 | 0.11586 | 0.161196335 | 0.086 | 0 | 1 | 9 |
| CU074419.2 | 0.11586 | 0.190576417 | 0.086 | 0 | 1 | 9 |
| skia | 0.11586 | 0.167531004 | 0.086 | 0 | 1 | 9 |
| strn4 | 0.11586 | 0.185137387 | 0.086 | 0 | 1 | 9 |
| zmp:0000000846 | 0.11586 | 0.229330162 | 0.086 | 0 | 1 | 9 |
| znf704 | 0.11586 | 0.181345583 | 0.086 | 0 | 1 | 9 |
| zgc:100920 | 0.11586 | 0.217552215 | 0.086 | 0 | 1 | 9 |
| ANKRD50 | 0.11586 | 0.184707321 | 0.086 | 0 | 1 | 9 |
| CABZ01113192.1 | 0.11586 | 0.16603378 | 0.086 | 0 | 1 | 9 |
| cdkn1ca | 0.11586 | 0.164558733 | 0.086 | 0 | 1 | 9 |
| cox20 | 0.11586 | 0.139120137 | 0.086 | 0 | 1 | 9 |
| fbxo33 | 0.11586 | 0.133463317 | 0.086 | 0 | 1 | 9 |
| fkbp7 | 0.11586 | 0.134564955 | 0.086 | 0 | 1 | 9 |
| fzd7b | 0.11586 | 0.128208354 | 0.086 | 0 | 1 | 9 |
| glg1a | 0.11586 | 0.128769791 | 0.086 | 0 | 1 | 9 |
| kpna6 | 0.11586 | 0.1348538 | 0.086 | 0 | 1 | 9 |
| ldb1b | 0.11586 | 0.139264164 | 0.086 | 0 | 1 | 9 |
| mcama | 0.11586 | 0.147799186 | 0.086 | 0 | 1 | 9 |
| mcamb | 0.11586 | 0.187589799 | 0.086 | 0 | 1 | 9 |
| mrpl38 | 0.11586 | 0.139767824 | 0.086 | 0 | 1 | 9 |
| pex5 | 0.11586 | 0.135870817 | 0.086 | 0 | 1 | 9 |
| phf23a | 0.11586 | 0.144802526 | 0.086 | 0 | 1 | 9 |

| | | | | | | |
|---|---|---|---|---|---|---|
| pigt | 0.11586 | 0.131056283 | 0.086 | 0 | 1 | 9 |
| si:ch211-276i12.11 | 0.11586 | 0.172423687 | 0.086 | 0 | 1 | 9 |
| smo | 0.11586 | 0.151010423 | 0.086 | 0 | 1 | 9 |
| tfb2m | 0.11586 | 0.133129736 | 0.086 | 0 | 1 | 9 |
| ttc1 | 0.11586 | 0.160459328 | 0.086 | 0 | 1 | 9 |
| ttll12 | 0.11586 | 0.138249876 | 0.086 | 0 | 1 | 9 |
| tvp23b | 0.11586 | 0.161594851 | 0.086 | 0 | 1 | 9 |
| CR387933.1 | 0.11586 | 0.125024094 | 0.086 | 0 | 1 | 9 |
| casp8ap2 | 0.11586 | 0.167973499 | 0.086 | 0 | 1 | 9 |
| cept1a | 0.11586 | 0.14417991 | 0.086 | 0 | 1 | 9 |
| gnas | 0.11586 | 0.153943492 | 0.086 | 0 | 1 | 9 |
| man2b2 | 0.11586 | 0.154353988 | 0.086 | 0 | 1 | 9 |
| miip | 0.11586 | 0.173551197 | 0.086 | 0 | 1 | 9 |
| nab1b | 0.11586 | 0.180707888 | 0.086 | 0 | 1 | 9 |
| nat8l | 0.11586 | 0.191469351 | 0.086 | 0 | 1 | 9 |
| ncbp3 | 0.11586 | 0.12221622 | 0.086 | 0 | 1 | 9 |
| nfatc2ip | 0.11586 | 0.161650167 | 0.086 | 0 | 1 | 9 |
| oxsm | 0.11586 | 0.1412612 | 0.086 | 0 | 1 | 9 |
| plxnb1a | 0.11586 | 0.168335895 | 0.086 | 0 | 1 | 9 |
| ptdss2 | 0.11586 | 0.169448658 | 0.086 | 0 | 1 | 9 |
| ramp2 | 0.11586 | 0.161757319 | 0.086 | 0 | 1 | 9 |
| rnf40 | 0.11586 | 0.139266808 | 0.086 | 0 | 1 | 9 |
| si:ch1073-291c23.2 | 0.11586 | 0.221397674 | 0.086 | 0 | 1 | 9 |
| sirt5 | 0.11586 | 0.144335617 | 0.086 | 0 | 1 | 9 |
| soga1 | 0.11586 | 0.193199312 | 0.086 | 0 | 1 | 9 |
| svbp | 0.11586 | 0.122440382 | 0.086 | 0 | 1 | 9 |
| tmco3 | 0.11586 | 0.172249033 | 0.086 | 0 | 1 | 9 |
| znf1124 | 0.11586 | 0.137976589 | 0.086 | 0 | 1 | 9 |
| CABZ01044053.1 | 0.11586 | 0.153471457 | 0.086 | 0 | 1 | 9 |
| CABZ01078600.1 | 0.11586 | 0.131389068 | 0.086 | 0 | 1 | 9 |
| CR318588.1 | 0.11586 | 0.383945548 | 0.086 | 0 | 1 | 9 |
| CT573344.1 | 0.11586 | 0.139402115 | 0.086 | 0 | 1 | 9 |
| HACD4 | 0.11586 | 0.165732241 | 0.086 | 0 | 1 | 9 |
| HOOK3 | 0.11586 | 0.184517334 | 0.086 | 0 | 1 | 9 |
| LO018148.1 | 0.11586 | 0.1695108 | 0.086 | 0 | 1 | 9 |
| NC-002333.17 | 0.11586 | 0.146250221 | 0.086 | 0 | 1 | 9 |
| SMIM10 | 0.11586 | 0.133670276 | 0.086 | 0 | 1 | 9 |
| aaas | 0.11586 | 0.131002882 | 0.086 | 0 | 1 | 9 |
| abcf2a | 0.11586 | 0.17574266 | 0.086 | 0 | 1 | 9 |
| adnpb | 0.11586 | 0.17293098 | 0.086 | 0 | 1 | 9 |
| agpat5 | 0.11586 | 0.126673402 | 0.086 | 0 | 1 | 9 |
| apool | 0.11586 | 0.119990597 | 0.086 | 0 | 1 | 9 |
| arl4ab | 0.11586 | 0.194271315 | 0.086 | 0 | 1 | 9 |
| arl4cb | 0.11586 | 0.179312498 | 0.086 | 0 | 1 | 9 |
| ascl1b | 0.11586 | 0.198698908 | 0.086 | 0 | 1 | 9 |
| asf1ba | 0.11586 | 0.148199335 | 0.086 | 0 | 1 | 9 |
| bahcc1b | 0.11586 | 0.157812775 | 0.086 | 0 | 1 | 9 |
| bmpr1ba | 0.11586 | 0.159673795 | 0.086 | 0 | 1 | 9 |

| | | | | | | |
|---|---|---|---|---|---|---|
| bod1 | 0.11586 | 0.18167383 | 0.086 | 0 | 1 | 9 |
| caspb | 0.11586 | 0.266823687 | 0.086 | 0 | 1 | 9 |
| cenpw | 0.11586 | 0.119392795 | 0.086 | 0 | 1 | 9 |
| chp1 | 0.11586 | 0.182727063 | 0.086 | 0 | 1 | 9 |
| cmc1 | 0.11586 | 0.127331307 | 0.086 | 0 | 1 | 9 |
| col4a2 | 0.11586 | 0.193009911 | 0.086 | 0 | 1 | 9 |
| coq10b | 0.11586 | 0.16557584 | 0.086 | 0 | 1 | 9 |
| cox15 | 0.11586 | 0.143918395 | 0.086 | 0 | 1 | 9 |
| crcp | 0.11586 | 0.138039777 | 0.086 | 0 | 1 | 9 |
| creb1b | 0.11586 | 0.181461849 | 0.086 | 0 | 1 | 9 |
| crybg1a | 0.11586 | 0.144989634 | 0.086 | 0 | 1 | 9 |
| ctps1b | 0.11586 | 0.136917787 | 0.086 | 0 | 1 | 9 |
| cul3a | 0.11586 | 0.137365829 | 0.086 | 0 | 1 | 9 |
| cxcl14 | 0.11586 | 0.193413341 | 0.086 | 0 | 1 | 9 |
| dgkaa | 0.11586 | 0.175523153 | 0.086 | 0 | 1 | 9 |
| dkk3b | 0.11586 | 0.25507258 | 0.086 | 0 | 1 | 9 |
| dnajc11a | 0.11586 | 0.134881548 | 0.086 | 0 | 1 | 9 |
| dnajc5ab | 0.11586 | 0.239602147 | 0.086 | 0 | 1 | 9 |
| dnajc5ga | 0.11586 | 0.11966065 | 0.086 | 0 | 1 | 9 |
| dvl2 | 0.11586 | 0.134903307 | 0.086 | 0 | 1 | 9 |
| ech1 | 0.11586 | 0.164287536 | 0.086 | 0 | 1 | 9 |
| eci2 | 0.11586 | 0.142111439 | 0.086 | 0 | 1 | 9 |
| eif4e3 | 0.11586 | 0.167466927 | 0.086 | 0 | 1 | 9 |
| fbxl14a | 0.11586 | 0.155257495 | 0.086 | 0 | 1 | 9 |
| fbxl14b | 0.11586 | 0.170812561 | 0.086 | 0 | 1 | 9 |
| gas6 | 0.11586 | 0.186028794 | 0.086 | 0 | 1 | 9 |
| gatd3a | 0.11586 | 0.185303657 | 0.086 | 0 | 1 | 9 |
| gde1 | 0.11586 | 0.110944265 | 0.086 | 0 | 1 | 9 |
| gfpt2 | 0.11586 | 0.214092633 | 0.086 | 0 | 1 | 9 |
| gnl1 | 0.11586 | 0.166193288 | 0.086 | 0 | 1 | 9 |
| golph3l | 0.11586 | 0.180644572 | 0.086 | 0 | 1 | 9 |
| gpc1b | 0.11586 | 0.161187536 | 0.086 | 0 | 1 | 9 |
| gusb | 0.11586 | 0.152263003 | 0.086 | 0 | 1 | 9 |
| hacd1 | 0.11586 | 0.136762863 | 0.086 | 0 | 1 | 9 |
| hgh1 | 0.11586 | 0.132821387 | 0.086 | 0 | 1 | 9 |
| hoxb2a | 0.11586 | 0.131110266 | 0.086 | 0 | 1 | 9 |
| ift27 | 0.11586 | 0.1308725 | 0.086 | 0 | 1 | 9 |
| inka1b | 0.11586 | 0.203158127 | 0.086 | 0 | 1 | 9 |
| ints12 | 0.11586 | 0.201340109 | 0.086 | 0 | 1 | 9 |
| larsb | 0.11586 | 0.132849905 | 0.086 | 0 | 1 | 9 |
| lmbr1 | 0.11586 | 0.119057141 | 0.086 | 0 | 1 | 9 |
| lrpprc | 0.11586 | 0.171501429 | 0.086 | 0 | 1 | 9 |
| mex3b | 0.11586 | 0.185980866 | 0.086 | 0 | 1 | 9 |
| mrgbp | 0.11586 | 0.142110262 | 0.086 | 0 | 1 | 9 |
| ms4a17a.10 | 0.11586 | 0.156911302 | 0.086 | 0 | 1 | 9 |
| msrb1a | 0.11586 | 0.154361 | 0.086 | 0 | 1 | 9 |
| mut | 0.11586 | 0.127837419 | 0.086 | 0 | 1 | 9 |
| myt1b | 0.11586 | 0.200264996 | 0.086 | 0 | 1 | 9 |

| | | | | | | |
|---|---|---|---|---|---|---|
| nfil3-5 | 0.11586 | 0.166483527 | 0.086 | 0 | 1 | 9 |
| nfkbib | 0.11586 | 0.139657464 | 0.086 | 0 | 1 | 9 |
| ngrn | 0.11586 | 0.120419714 | 0.086 | 0 | 1 | 9 |
| nup160 | 0.11586 | 0.109499052 | 0.086 | 0 | 1 | 9 |
| oser1 | 0.11586 | 0.106732291 | 0.086 | 0 | 1 | 9 |
| paip1 | 0.11586 | 0.133679337 | 0.086 | 0 | 1 | 9 |
| pax7a | 0.11586 | 0.258158639 | 0.086 | 0 | 1 | 9 |
| pcnp | 0.11586 | 0.172100367 | 0.086 | 0 | 1 | 9 |
| pitx3 | 0.11586 | 0.190411946 | 0.086 | 0 | 1 | 9 |
| plod2 | 0.11586 | 0.167592821 | 0.086 | 0 | 1 | 9 |
| plxna2 | 0.11586 | 0.119124905 | 0.086 | 0 | 1 | 9 |
| pola1 | 0.11586 | 0.196455635 | 0.086 | 0 | 1 | 9 |
| polr1c | 0.11586 | 0.124958054 | 0.086 | 0 | 1 | 9 |
| ponzr1 | 0.11586 | 0.184544378 | 0.086 | 0 | 1 | 9 |
| proza | 0.11586 | 0.132132047 | 0.086 | 0 | 1 | 9 |
| psmd9 | 0.11586 | 0.139792507 | 0.086 | 0 | 1 | 9 |
| rab1bb | 0.11586 | 0.161617836 | 0.086 | 0 | 1 | 9 |
| rab3da | 0.11586 | 0.198590318 | 0.086 | 0 | 1 | 9 |
| rbm6 | 0.11586 | 0.143160317 | 0.086 | 0 | 1 | 9 |
| rcc1 | 0.11586 | 0.155166294 | 0.086 | 0 | 1 | 9 |
| rit1 | 0.11586 | 0.167295719 | 0.086 | 0 | 1 | 9 |
| rnf146 | 0.11586 | 0.131365816 | 0.086 | 0 | 1 | 9 |
| rrm2 | 0.11586 | 0.234140817 | 0.086 | 0 | 1 | 9 |
| ruvbl2 | 0.11586 | 0.137062823 | 0.086 | 0 | 1 | 9 |
| scg3 | 0.11586 | 0.28501211 | 0.086 | 0 | 1 | 9 |
| si:ch211-212k18.4 | 0.11586 | 0.183080807 | 0.086 | 0 | 1 | 9 |
| si:ch211-86h15.1 | 0.11586 | 0.152431671 | 0.086 | 0 | 1 | 9 |
| si:dkey-19a16.2 | 0.11586 | 0.14841881 | 0.086 | 0 | 1 | 9 |
| si:dkey-30c15.2 | 0.11586 | 0.149723382 | 0.086 | 0 | 1 | 9 |
| siva1 | 0.11586 | 0.183325735 | 0.086 | 0 | 1 | 9 |
| slc25a51b | 0.11586 | 0.113071622 | 0.086 | 0 | 1 | 9 |
| slc30a5 | 0.11586 | 0.135057641 | 0.086 | 0 | 1 | 9 |
| smg6 | 0.11586 | 0.150262591 | 0.086 | 0 | 1 | 9 |
| snu13a | 0.11586 | 0.145539391 | 0.086 | 0 | 1 | 9 |
| spry1 | 0.11586 | 0.154301289 | 0.086 | 0 | 1 | 9 |
| srm | 0.11586 | 0.137493101 | 0.086 | 0 | 1 | 9 |
| stat2 | 0.11586 | 0.124699448 | 0.086 | 0 | 1 | 9 |
| stx5a | 0.11586 | 0.15281279 | 0.086 | 0 | 1 | 9 |
| sucla2 | 0.11586 | 0.157677908 | 0.086 | 0 | 1 | 9 |
| sumf2 | 0.11586 | 0.158480737 | 0.086 | 0 | 1 | 9 |
| ten1 | 0.11586 | 0.128416086 | 0.086 | 0 | 1 | 9 |
| tfpia | 0.11586 | 0.211356889 | 0.086 | 0 | 1 | 9 |
| tgm5l | 0.11586 | 0.297937072 | 0.086 | 0 | 1 | 9 |
| thumpd3 | 0.11586 | 0.123963011 | 0.086 | 0 | 1 | 9 |
| top2b | 0.11586 | 0.161815896 | 0.086 | 0 | 1 | 9 |
| tor1l2 | 0.11586 | 0.145571195 | 0.086 | 0 | 1 | 9 |
| tsr3 | 0.11586 | 0.153005837 | 0.086 | 0 | 1 | 9 |
| ube2j2 | 0.11586 | 0.142933211 | 0.086 | 0 | 1 | 9 |

| | | | | | | |
|---|---|---|---|---|---|---|
| ugt5b2 | 0.11586 | 0.155935787 | 0.086 | 0 | 1 | 9 |
| unga | 0.11586 | 0.169596737 | 0.086 | 0 | 1 | 9 |
| utp23 | 0.11586 | 0.12166437 | 0.086 | 0 | 1 | 9 |
| vps37a | 0.11586 | 0.136617073 | 0.086 | 0 | 1 | 9 |
| wdr12 | 0.11586 | 0.145055046 | 0.086 | 0 | 1 | 9 |
| wdr89 | 0.11586 | 0.16614978 | 0.086 | 0 | 1 | 9 |
| wu:fc21g02 | 0.11586 | 0.144407855 | 0.086 | 0 | 1 | 9 |
| zgc:110540 | 0.11586 | 0.17615114 | 0.086 | 0 | 1 | 9 |
| zgc:153901 | 0.11586 | 0.108253906 | 0.086 | 0 | 1 | 9 |
| zgc:91944 | 0.11586 | 0.183824269 | 0.086 | 0 | 1 | 9 |
| znf687b | 0.11586 | 0.139545757 | 0.086 | 0 | 1 | 9 |
| CR788248.1 | 0.11586 | 0.171290752 | 0.086 | 0 | 1 | 9 |
| FP101887.1 | 0.11586 | 0.108217725 | 0.086 | 0 | 1 | 9 |
| XLOC-034235 | 0.11586 | 0.134642209 | 0.086 | 0 | 1 | 9 |
| acvr1ba | 0.11586 | 0.154181995 | 0.086 | 0 | 1 | 9 |
| aftpha | 0.11586 | 0.131232141 | 0.086 | 0 | 1 | 9 |
| ahcyl1 | 0.11586 | 0.129182689 | 0.086 | 0 | 1 | 9 |
| ankrd54 | 0.11586 | 0.115143653 | 0.086 | 0 | 1 | 9 |
| arl6ip6 | 0.11586 | 0.150634829 | 0.086 | 0 | 1 | 9 |
| arvcfb | 0.11586 | 0.138663095 | 0.086 | 0 | 1 | 9 |
| atg16l1 | 0.11586 | 0.132419818 | 0.086 | 0 | 1 | 9 |
| atic | 0.11586 | 0.148158724 | 0.086 | 0 | 1 | 9 |
| atrip | 0.11586 | 0.163590055 | 0.086 | 0 | 1 | 9 |
| bckdk | 0.11586 | 0.15431777 | 0.086 | 0 | 1 | 9 |
| cand1 | 0.11586 | 0.123068131 | 0.086 | 0 | 1 | 9 |
| cluha | 0.11586 | 0.152263003 | 0.086 | 0 | 1 | 9 |
| cul1a | 0.11586 | 0.116942238 | 0.086 | 0 | 1 | 9 |
| dcbld2 | 0.11586 | 0.143070945 | 0.086 | 0 | 1 | 9 |
| decr2 | 0.11586 | 0.159415423 | 0.086 | 0 | 1 | 9 |
| eef2k | 0.11586 | 0.147623497 | 0.086 | 0 | 1 | 9 |
| eps8l1b | 0.11586 | 0.160292766 | 0.086 | 0 | 1 | 9 |
| erbb3b | 0.11586 | 0.180666164 | 0.086 | 0 | 1 | 9 |
| fam222bb | 0.11586 | 0.147622137 | 0.086 | 0 | 1 | 9 |
| fem1c | 0.11586 | 0.15679331 | 0.086 | 0 | 1 | 9 |
| gcga | 0.11586 | 0.327344522 | 0.086 | 0 | 1 | 9 |
| gna12a | 0.11586 | 0.158283191 | 0.086 | 0 | 1 | 9 |
| gpn2 | 0.11586 | 0.11582849 | 0.086 | 0 | 1 | 9 |
| herc2 | 0.11586 | 0.169202605 | 0.086 | 0 | 1 | 9 |
| homer3b | 0.11586 | 0.157259781 | 0.086 | 0 | 1 | 9 |
| hoxb3a | 0.11586 | 0.196249828 | 0.086 | 0 | 1 | 9 |
| ipcef1 | 0.11586 | 0.130297784 | 0.086 | 0 | 1 | 9 |
| lyrm7 | 0.11586 | 0.117096425 | 0.086 | 0 | 1 | 9 |
| mafba | 0.11586 | 0.223263125 | 0.086 | 0 | 1 | 9 |
| march5 | 0.11586 | 0.154247571 | 0.086 | 0 | 1 | 9 |
| mcee | 0.11586 | 0.128591889 | 0.086 | 0 | 1 | 9 |
| mical3a | 0.11586 | 0.138607271 | 0.086 | 0 | 1 | 9 |
| mta3 | 0.11586 | 0.157562038 | 0.086 | 0 | 1 | 9 |
| mtss1 | 0.11586 | 0.147144878 | 0.086 | 0 | 1 | 9 |

| | | | | | | |
|---|---|---|---|---|---|---|
| ncoa4 | 0.11586 | 0.123331787 | 0.086 | 0 | 1 | 9 |
| nkx3-1 | 0.11586 | 0.206674571 | 0.086 | 0 | 1 | 9 |
| nomo | 0.11586 | 0.16876562 | 0.086 | 0 | 1 | 9 |
| nsg2 | 0.11586 | 0.294136472 | 0.086 | 0 | 1 | 9 |
| ntmt1 | 0.11586 | 0.150278866 | 0.086 | 0 | 1 | 9 |
| nup58 | 0.11586 | 0.128614168 | 0.086 | 0 | 1 | 9 |
| palm1b | 0.11586 | 0.130824435 | 0.086 | 0 | 1 | 9 |
| pik3r2 | 0.11586 | 0.146613795 | 0.086 | 0 | 1 | 9 |
| plrg1 | 0.11586 | 0.152810314 | 0.086 | 0 | 1 | 9 |
| pls3 | 0.11586 | 0.147960682 | 0.086 | 0 | 1 | 9 |
| plxna3 | 0.11586 | 0.225491459 | 0.086 | 0 | 1 | 9 |
| pola2 | 0.11586 | 0.143893579 | 0.086 | 0 | 1 | 9 |
| pomk | 0.11586 | 0.113791511 | 0.086 | 0 | 1 | 9 |
| ppardb | 0.11586 | 0.171359627 | 0.086 | 0 | 1 | 9 |
| purbb | 0.11586 | 0.15612022 | 0.086 | 0 | 1 | 9 |
| rabl3 | 0.11586 | 0.152878674 | 0.086 | 0 | 1 | 9 |
| ralgps2 | 0.11586 | 0.137542098 | 0.086 | 0 | 1 | 9 |
| rassf1 | 0.11586 | 0.148851297 | 0.086 | 0 | 1 | 9 |
| rbbp5 | 0.11586 | 0.175916646 | 0.086 | 0 | 1 | 9 |
| relt | 0.11586 | 0.185537359 | 0.086 | 0 | 1 | 9 |
| rgmd | 0.11586 | 0.156534129 | 0.086 | 0 | 1 | 9 |
| rnf130 | 0.11586 | 0.169402133 | 0.086 | 0 | 1 | 9 |
| rnf216 | 0.11586 | 0.16130834 | 0.086 | 0 | 1 | 9 |
| rnf220a | 0.11586 | 0.11938481 | 0.086 | 0 | 1 | 9 |
| rrp36 | 0.11586 | 0.189796884 | 0.086 | 0 | 1 | 9 |
| samd13 | 0.11586 | 0.115478945 | 0.086 | 0 | 1 | 9 |
| si:dkey-175d9.2 | 0.11586 | 0.151917235 | 0.086 | 0 | 1 | 9 |
| si:rp71-17i16.6 | 0.11586 | 0.209833494 | 0.086 | 0 | 1 | 9 |
| slc30a1a | 0.11586 | 0.155739008 | 0.086 | 0 | 1 | 9 |
| slc48a1b | 0.11586 | 0.185409239 | 0.086 | 0 | 1 | 9 |
| suclg2 | 0.11586 | 0.143316506 | 0.086 | 0 | 1 | 9 |
| suv39h1b | 0.11586 | 0.141128721 | 0.086 | 0 | 1 | 9 |
| syngr2b | 0.11586 | 0.129891378 | 0.086 | 0 | 1 | 9 |
| tab2 | 0.11586 | 0.152846661 | 0.086 | 0 | 1 | 9 |
| taf5 | 0.11586 | 0.168892862 | 0.086 | 0 | 1 | 9 |
| tfb1m | 0.11586 | 0.129668181 | 0.086 | 0 | 1 | 9 |
| tjap1 | 0.11586 | 0.141006093 | 0.086 | 0 | 1 | 9 |
| tle2a | 0.11586 | 0.161961385 | 0.086 | 0 | 1 | 9 |
| ttn.2 | 0.11586 | 0.139738085 | 0.086 | 0 | 1 | 9 |
| uck2b | 0.11586 | 0.13575471 | 0.086 | 0 | 1 | 9 |
| vim | 0.11586 | 0.238128257 | 0.086 | 0 | 1 | 9 |
| vps4a | 0.11586 | 0.157650848 | 0.086 | 0 | 1 | 9 |
| wdfy2 | 0.11586 | 0.127045103 | 0.086 | 0 | 1 | 9 |
| zc3h14 | 0.11586 | 0.126191227 | 0.086 | 0 | 1 | 9 |
| zeb2a | 0.11586 | 0.176856999 | 0.086 | 0 | 1 | 9 |
| zgc:66427 | 0.11586 | 0.124341161 | 0.086 | 0 | 1 | 9 |
| zgc:77739 | 0.11586 | 0.132546105 | 0.086 | 0 | 1 | 9 |
| zgc:92818 | 0.11586 | 0.19372186 | 0.086 | 0 | 1 | 9 |

| | | | | | | |
|---|---|---|---|---|---|---|
| zgc:92907 | 0.11586 | 0.132466968 | 0.086 | 0 | 1 | 9 |
| zranb1b | 0.11586 | 0.121488806 | 0.086 | 0 | 1 | 9 |
| zrsr2 | 0.11586 | 0.133355249 | 0.086 | 0 | 1 | 9 |
| rnf2 | 0.11612 | 0.299770669 | 0.224 | 0.05 | 1 | 9 |
| actr10 | 0.11638 | 0.119224436 | 0.172 | 0.05 | 1 | 9 |
| nr1d2b | 0.11678 | -0.32635167 | 0.069 | 0.15 | 1 | 9 |
| BX664625.1 | 0.11685 | -0.33484014 | 0.034 | 0.1 | 1 | 9 |
| eif3ea | 0.11824 | 0.126902643 | 0.655 | 0.45 | 1 | 9 |
| tax1bp1b | 0.11836 | -0.40403768 | 0.259 | 0.5 | 1 | 9 |
| kras | 0.11872 | 0.359435387 | 0.241 | 0.05 | 1 | 9 |
| emc1 | 0.1189 | -0.35184413 | 0.069 | 0.2 | 1 | 9 |
| map1lc3b | 0.11903 | -0.24054633 | 0.224 | 0.25 | 1 | 9 |
| aktip | 0.11949 | 0.144512897 | 0.259 | 0.1 | 1 | 9 |
| eppk1 | 0.11951 | -0.31162004 | 0.052 | 0.15 | 1 | 9 |
| wu:fb16f03 | 0.11962 | -0.27793955 | 0.155 | 0.2 | 1 | 9 |
| eif2s3 | 0.1207 | 0.369117967 | 0.69 | 0.45 | 1 | 9 |
| gadd45gip1 | 0.12133 | 0.291223429 | 0.224 | 0.05 | 1 | 9 |
| mrpl34 | 0.12212 | 0.325622205 | 0.276 | 0.1 | 1 | 9 |
| celf2 | 0.12222 | 0.467412119 | 0.379 | 0.15 | 1 | 9 |
| naa20 | 0.12236 | 0.354501176 | 0.31 | 0.1 | 1 | 9 |
| add3a | 0.12275 | -0.26583841 | 0.138 | 0.2 | 1 | 9 |
| tm9sf3 | 0.12279 | 0.269225757 | 0.224 | 0.05 | 1 | 9 |
| CABZ01110881.1 | 0.12347 | 0.190721672 | 0.19 | 0.05 | 1 | 9 |
| srsf11 | 0.1235 | 0.225175497 | 0.293 | 0.1 | 1 | 9 |
| si:ch211-63p21.4 | 0.12387 | -0.43869093 | 0.034 | 0.15 | 1 | 9 |
| tjp1a | 0.12388 | -0.31019826 | 0.069 | 0.25 | 1 | 9 |
| c1d | 0.12427 | 0.301273027 | 0.224 | 0.05 | 1 | 9 |
| eef1a1l1 | 0.12451 | 0.154046913 | 1 | 0.95 | 1 | 9 |
| cdca4 | 0.12499 | -0.28957625 | 0.069 | 0.25 | 1 | 9 |
| becn1 | 0.12499 | -0.32914768 | 0.069 | 0.2 | 1 | 9 |
| ano10b | 0.1251 | -0.30537442 | 0.069 | 0.2 | 1 | 9 |
| pa2g4a | 0.12545 | 0.297276053 | 0.224 | 0.05 | 1 | 9 |
| ctdspla | 0.12559 | 0.230729547 | 0.207 | 0.05 | 1 | 9 |
| gtpbp4 | 0.12566 | 0.388993665 | 0.5 | 0.25 | 1 | 9 |
| psmd8 | 0.12567 | 0.14302845 | 0.241 | 0.1 | 1 | 9 |
| lgals2b | 0.12614 | -0.61808599 | 0.086 | 0.2 | 1 | 9 |
| sar1b | 0.12619 | -0.33061259 | 0.241 | 0.35 | 1 | 9 |
| mgst2 | 0.12651 | 0.304221994 | 0.19 | 0.05 | 1 | 9 |
| elovl1a | 0.12671 | 0.290168641 | 0.19 | 0.15 | 1 | 9 |
| cyb5a | 0.12692 | -0.33011873 | 0.103 | 0.25 | 1 | 9 |
| slc25a5 | 0.12711 | 0.255582774 | 0.948 | 0.85 | 1 | 9 |
| gstk1 | 0.12728 | -0.37088231 | 0.19 | 0.35 | 1 | 9 |
| psma5 | 0.12748 | 0.380979247 | 0.431 | 0.2 | 1 | 9 |
| ndufa9a | 0.12748 | 0.334743313 | 0.31 | 0.1 | 1 | 9 |
| thrb | 0.12779 | -0.2677525 | 0.034 | 0.15 | 1 | 9 |
| hrc | 0.12827 | 0.354662125 | 0.293 | 0.1 | 1 | 9 |
| CABZ01055365.1 | 0.12861 | -0.401994 | 0.034 | 0.15 | 1 | 9 |
| psmg3 | 0.12914 | 0.300536152 | 0.31 | 0.1 | 1 | 9 |

| | | | | | | |
|---|---|---|---|---|---|---|
| zfand5a | 0.12994 | 0.354049109 | 0.224 | 0.05 | 1 | 9 |
| rgs4 | 0.12996 | -0.11992894 | 0.069 | 0.05 | 1 | 9 |
| ctsf | 0.13028 | -0.28956115 | 0.207 | 0.25 | 1 | 9 |
| nhp2 | 0.13033 | 0.384812328 | 0.379 | 0.15 | 1 | 9 |
| ank3b | 0.13053 | -0.15522041 | 0.207 | 0.2 | 1 | 9 |
| h3f3a | 0.13088 | 0.422034123 | 0.603 | 0.35 | 1 | 9 |
| phlda2 | 0.13195 | 0.487470732 | 0.379 | 0.15 | 1 | 9 |
| psma6l | 0.1326 | -0.32868955 | 0.052 | 0.2 | 1 | 9 |
| si:ch73-42p12.2 | 0.13264 | -0.11426007 | 0.276 | 0.25 | 1 | 9 |
| cnp | 0.13296 | 0.260285979 | 0.293 | 0.1 | 1 | 9 |
| mt-nd3 | 0.13348 | -0.37447631 | 0.172 | 0.4 | 1 | 9 |
| m6pr | 0.13456 | -0.14240053 | 0.155 | 0.15 | 1 | 9 |
| mef2d | 0.13469 | -0.12355786 | 0.207 | 0.2 | 1 | 9 |
| CU655842.1 | 0.13494 | -0.16533327 | 0.017 | 0.1 | 1 | 9 |
| abhd12 | 0.13494 | -0.12505101 | 0.017 | 0.1 | 1 | 9 |
| arhgap24 | 0.13494 | -0.19773625 | 0.017 | 0.1 | 1 | 9 |
| asns | 0.13494 | -0.19604802 | 0.017 | 0.1 | 1 | 9 |
| atp11c | 0.13494 | -0.24238948 | 0.017 | 0.1 | 1 | 9 |
| camk2d1 | 0.13494 | -0.19988972 | 0.017 | 0.1 | 1 | 9 |
| ch25hl1.1 | 0.13494 | -0.18938888 | 0.017 | 0.1 | 1 | 9 |
| ches1 | 0.13494 | -0.18364923 | 0.017 | 0.1 | 1 | 9 |
| cog2 | 0.13494 | -0.17521935 | 0.017 | 0.1 | 1 | 9 |
| crfb16 | 0.13494 | -0.15750171 | 0.017 | 0.1 | 1 | 9 |
| dhx9 | 0.13494 | -0.11690742 | 0.017 | 0.1 | 1 | 9 |
| dnajc7 | 0.13494 | -0.14139359 | 0.017 | 0.1 | 1 | 9 |
| fdxr | 0.13494 | -0.25844136 | 0.017 | 0.1 | 1 | 9 |
| hes2.2 | 0.13494 | -0.27189321 | 0.017 | 0.1 | 1 | 9 |
| kbtbd2 | 0.13494 | -0.11463651 | 0.017 | 0.1 | 1 | 9 |
| mctp2b | 0.13494 | -0.22874837 | 0.017 | 0.1 | 1 | 9 |
| nr4a2b | 0.13494 | -0.22545665 | 0.017 | 0.1 | 1 | 9 |
| plekhb2 | 0.13494 | -0.21474726 | 0.017 | 0.1 | 1 | 9 |
| ppt2 | 0.13494 | -0.19802421 | 0.017 | 0.1 | 1 | 9 |
| rapgef2 | 0.13494 | -0.12888911 | 0.017 | 0.1 | 1 | 9 |
| si:ch211-285f17.1 | 0.13494 | -0.11744958 | 0.017 | 0.1 | 1 | 9 |
| sp1 | 0.13494 | -0.12365787 | 0.017 | 0.1 | 1 | 9 |
| spg21 | 0.13494 | -0.16805803 | 0.017 | 0.1 | 1 | 9 |
| tcerg1a | 0.13494 | -0.32167526 | 0.017 | 0.1 | 1 | 9 |
| tnfrsf11a | 0.13494 | -0.30434329 | 0.017 | 0.1 | 1 | 9 |
| tnfrsf1a | 0.13494 | -0.11476364 | 0.017 | 0.1 | 1 | 9 |
| txlng | 0.13494 | -0.1391944 | 0.017 | 0.1 | 1 | 9 |
| uba7 | 0.13494 | -0.21154843 | 0.017 | 0.1 | 1 | 9 |
| zgc:110182 | 0.13494 | -0.27146023 | 0.017 | 0.1 | 1 | 9 |
| zgc:162184 | 0.13494 | -0.16440851 | 0.017 | 0.1 | 1 | 9 |
| WDR7 | 0.13494 | -0.14746143 | 0.017 | 0.1 | 1 | 9 |
| XLOC-025423 | 0.13494 | -0.22420129 | 0.017 | 0.1 | 1 | 9 |
| abhd4 | 0.13494 | -0.12058889 | 0.017 | 0.1 | 1 | 9 |
| acoxl | 0.13494 | -0.21103805 | 0.017 | 0.1 | 1 | 9 |
| calml4a | 0.13494 | -0.10732488 | 0.017 | 0.1 | 1 | 9 |

| | | | | | | |
|---|---|---|---|---|---|---|
| gkup | 0.13494 | -0.16031945 | 0.017 | 0.1 | 1 | 9 |
| gng10 | 0.13494 | -0.2655029 | 0.017 | 0.1 | 1 | 9 |
| hsd20b2 | 0.13494 | -0.16653619 | 0.017 | 0.1 | 1 | 9 |
| igsf9a | 0.13494 | -0.13305596 | 0.017 | 0.1 | 1 | 9 |
| inavab | 0.13494 | -0.1286356 | 0.017 | 0.1 | 1 | 9 |
| ralgapa2 | 0.13494 | -0.1565132 | 0.017 | 0.1 | 1 | 9 |
| ranbp10 | 0.13494 | -0.12495564 | 0.017 | 0.1 | 1 | 9 |
| snx19b | 0.13494 | -0.18874255 | 0.017 | 0.1 | 1 | 9 |
| sp3a | 0.13494 | -0.21975973 | 0.017 | 0.1 | 1 | 9 |
| terf2ip | 0.13494 | -0.13421607 | 0.017 | 0.1 | 1 | 9 |
| tmbim1a | 0.13494 | -0.1564105 | 0.017 | 0.1 | 1 | 9 |
| bida | 0.13494 | -0.12195729 | 0.017 | 0.1 | 1 | 9 |
| cdx1a | 0.13494 | -0.18104457 | 0.017 | 0.1 | 1 | 9 |
| dr1 | 0.13494 | -0.13123485 | 0.017 | 0.1 | 1 | 9 |
| foxk1 | 0.13494 | -0.26201813 | 0.017 | 0.1 | 1 | 9 |
| lipea | 0.13494 | -0.11538483 | 0.017 | 0.1 | 1 | 9 |
| llgl2 | 0.13494 | -0.17094139 | 0.017 | 0.1 | 1 | 9 |
| pdx1 | 0.13494 | -0.11250157 | 0.017 | 0.1 | 1 | 9 |
| snap23.2 | 0.13494 | -0.31504688 | 0.017 | 0.1 | 1 | 9 |
| tfpt | 0.13494 | -0.13350548 | 0.017 | 0.1 | 1 | 9 |
| tle2c | 0.13494 | -0.16340386 | 0.017 | 0.1 | 1 | 9 |
| wdr44 | 0.13494 | -0.23834369 | 0.017 | 0.1 | 1 | 9 |
| CU639397.1 | 0.13494 | -0.17544483 | 0.017 | 0.1 | 1 | 9 |
| cdh17 | 0.13494 | -0.25005511 | 0.017 | 0.1 | 1 | 9 |
| clmn | 0.13494 | -0.18881661 | 0.017 | 0.1 | 1 | 9 |
| jak2a | 0.13494 | -0.17503841 | 0.017 | 0.1 | 1 | 9 |
| kif3a | 0.13494 | -0.2294343 | 0.017 | 0.1 | 1 | 9 |
| map3k12 | 0.13494 | -0.20483613 | 0.017 | 0.1 | 1 | 9 |
| march7 | 0.13494 | -0.11489816 | 0.017 | 0.1 | 1 | 9 |
| mink1 | 0.13494 | -0.12732736 | 0.017 | 0.1 | 1 | 9 |
| rdh10a | 0.13494 | -0.28877951 | 0.017 | 0.1 | 1 | 9 |
| sh3rf1 | 0.13494 | -0.16704045 | 0.017 | 0.1 | 1 | 9 |
| slc5a9 | 0.13494 | -0.20343478 | 0.017 | 0.1 | 1 | 9 |
| slc6a9 | 0.13494 | -0.25318171 | 0.017 | 0.1 | 1 | 9 |
| slc7a3b | 0.13494 | -0.18076267 | 0.017 | 0.1 | 1 | 9 |
| tbc1d4 | 0.13494 | -0.10866487 | 0.017 | 0.1 | 1 | 9 |
| unm-hu7910 | 0.13494 | -0.22762159 | 0.017 | 0.1 | 1 | 9 |
| vangl1 | 0.13494 | -0.16612033 | 0.017 | 0.1 | 1 | 9 |
| zgc:112982 | 0.13494 | -0.19030686 | 0.017 | 0.1 | 1 | 9 |
| zhx3 | 0.13494 | -0.13470151 | 0.017 | 0.1 | 1 | 9 |
| BX571884.1 | 0.13494 | -0.22549569 | 0.017 | 0.1 | 1 | 9 |
| LOC103910363 | 0.13494 | -0.11222178 | 0.017 | 0.1 | 1 | 9 |
| otud4 | 0.13494 | -0.11315948 | 0.017 | 0.1 | 1 | 9 |
| p3h2 | 0.13494 | -0.17905274 | 0.017 | 0.1 | 1 | 9 |
| scyl1 | 0.13494 | -0.18288537 | 0.017 | 0.1 | 1 | 9 |
| zdhhc17 | 0.13494 | -0.26052032 | 0.017 | 0.1 | 1 | 9 |
| zmynd8 | 0.13494 | -0.13853367 | 0.017 | 0.1 | 1 | 9 |
| CABZ01088428.1 | 0.13494 | -0.1418368 | 0.017 | 0.1 | 1 | 9 |

| | | | | | | |
|---|---|---|---|---|---|---|
| CU914776.2 | 0.13494 | -0.40806926 | 0.017 | 0.1 | 1 | 9 |
| SLC7A1 (1 of many) | 0.13494 | -0.26450667 | 0.017 | 0.1 | 1 | 9 |
| XLOC-040069 | 0.13494 | -0.11550788 | 0.017 | 0.1 | 1 | 9 |
| XLOC-042917 | 0.13494 | -0.24435755 | 0.017 | 0.1 | 1 | 9 |
| anpepb | 0.13494 | -0.17190079 | 0.017 | 0.1 | 1 | 9 |
| arhgap23a | 0.13494 | -0.16347718 | 0.017 | 0.1 | 1 | 9 |
| barx2 | 0.13494 | -0.13470028 | 0.017 | 0.1 | 1 | 9 |
| blnk | 0.13494 | -0.13310006 | 0.017 | 0.1 | 1 | 9 |
| bscl2 | 0.13494 | -0.18738102 | 0.017 | 0.1 | 1 | 9 |
| camsap3 | 0.13494 | -0.20229947 | 0.017 | 0.1 | 1 | 9 |
| clcn7 | 0.13494 | -0.20595717 | 0.017 | 0.1 | 1 | 9 |
| cobl | 0.13494 | -0.21064488 | 0.017 | 0.1 | 1 | 9 |
| crp6 | 0.13494 | -0.53910271 | 0.017 | 0.1 | 1 | 9 |
| crtap | 0.13494 | -0.22715178 | 0.017 | 0.1 | 1 | 9 |
| dapp1 | 0.13494 | -0.12907304 | 0.017 | 0.1 | 1 | 9 |
| eapp | 0.13494 | -0.17907993 | 0.017 | 0.1 | 1 | 9 |
| elf2a | 0.13494 | -0.16685954 | 0.017 | 0.1 | 1 | 9 |
| fbln5 | 0.13494 | -0.14980152 | 0.017 | 0.1 | 1 | 9 |
| frzb | 0.13494 | -0.16561559 | 0.017 | 0.1 | 1 | 9 |
| fto | 0.13494 | -0.13249793 | 0.017 | 0.1 | 1 | 9 |
| galnt11 | 0.13494 | -0.26036774 | 0.017 | 0.1 | 1 | 9 |
| gramd1a | 0.13494 | -0.25309298 | 0.017 | 0.1 | 1 | 9 |
| grm4 | 0.13494 | -0.15047575 | 0.017 | 0.1 | 1 | 9 |
| grsf1 | 0.13494 | -0.210559 | 0.017 | 0.1 | 1 | 9 |
| guk1a | 0.13494 | -0.16496186 | 0.017 | 0.1 | 1 | 9 |
| hdac6 | 0.13494 | -0.17797752 | 0.017 | 0.1 | 1 | 9 |
| herpud2 | 0.13494 | -0.1904286 | 0.017 | 0.1 | 1 | 9 |
| hmgcs1 | 0.13494 | -0.17746024 | 0.017 | 0.1 | 1 | 9 |
| il13ra2 | 0.13494 | -0.18038709 | 0.017 | 0.1 | 1 | 9 |
| intu | 0.13494 | -0.16309791 | 0.017 | 0.1 | 1 | 9 |
| iqsec2b | 0.13494 | -0.2395842 | 0.017 | 0.1 | 1 | 9 |
| itpr1a | 0.13494 | -0.26371161 | 0.017 | 0.1 | 1 | 9 |
| jade1 | 0.13494 | -0.11954965 | 0.017 | 0.1 | 1 | 9 |
| kat14 | 0.13494 | -0.17462963 | 0.017 | 0.1 | 1 | 9 |
| maco1b | 0.13494 | -0.22815305 | 0.017 | 0.1 | 1 | 9 |
| mgat2 | 0.13494 | -0.224624 | 0.017 | 0.1 | 1 | 9 |
| mul1a | 0.13494 | -0.15687183 | 0.017 | 0.1 | 1 | 9 |
| nrip1b | 0.13494 | -0.18440556 | 0.017 | 0.1 | 1 | 9 |
| pacsin2 | 0.13494 | -0.21778037 | 0.017 | 0.1 | 1 | 9 |
| pde12 | 0.13494 | -0.20246672 | 0.017 | 0.1 | 1 | 9 |
| pex26 | 0.13494 | -0.21057073 | 0.017 | 0.1 | 1 | 9 |
| piezo1 | 0.13494 | -0.13774089 | 0.017 | 0.1 | 1 | 9 |
| pip5k1aa | 0.13494 | -0.12404392 | 0.017 | 0.1 | 1 | 9 |
| pip5k1ab | 0.13494 | -0.13446309 | 0.017 | 0.1 | 1 | 9 |
| si:ch211-256e16.3 | 0.13494 | -0.26896307 | 0.017 | 0.1 | 1 | 9 |
| si:ch73-359m17.5 | 0.13494 | -0.60017404 | 0.017 | 0.1 | 1 | 9 |
| si:dkey-10c21.1 | 0.13494 | -0.19343332 | 0.017 | 0.1 | 1 | 9 |
| si:dkey-125i10.3 | 0.13494 | -0.2645538 | 0.017 | 0.1 | 1 | 9 |

| | | | | | | |
|---|---|---|---|---|---|---|
| si:dkey-222n6.2 | 0.13494 | -0.30401022 | 0.017 | 0.1 | 1 | 9 |
| si:dkey-6n21.12 | 0.13494 | -0.21033194 | 0.017 | 0.1 | 1 | 9 |
| slc12a9 | 0.13494 | -0.12563496 | 0.017 | 0.1 | 1 | 9 |
| slc39a11 | 0.13494 | -0.16463686 | 0.017 | 0.1 | 1 | 9 |
| sptlc3 | 0.13494 | -0.16021692 | 0.017 | 0.1 | 1 | 9 |
| stap2b | 0.13494 | -0.17879816 | 0.017 | 0.1 | 1 | 9 |
| stard3nl | 0.13494 | -0.19256788 | 0.017 | 0.1 | 1 | 9 |
| stk16 | 0.13494 | -0.11135835 | 0.017 | 0.1 | 1 | 9 |
| sult1st5 | 0.13494 | -0.11020012 | 0.017 | 0.1 | 1 | 9 |
| tacc1 | 0.13494 | -0.29713921 | 0.017 | 0.1 | 1 | 9 |
| tarbp2 | 0.13494 | -0.24026661 | 0.017 | 0.1 | 1 | 9 |
| ticam1 | 0.13494 | -0.25386937 | 0.017 | 0.1 | 1 | 9 |
| tmem183a | 0.13494 | -0.1122491 | 0.017 | 0.1 | 1 | 9 |
| tmem184a | 0.13494 | -0.1436033 | 0.017 | 0.1 | 1 | 9 |
| traf7 | 0.13494 | -0.16840519 | 0.017 | 0.1 | 1 | 9 |
| trioa | 0.13494 | -0.13819169 | 0.017 | 0.1 | 1 | 9 |
| ubr3 | 0.13494 | -0.16658085 | 0.017 | 0.1 | 1 | 9 |
| vil1 | 0.13494 | -0.22776917 | 0.017 | 0.1 | 1 | 9 |
| wdtc1 | 0.13494 | -0.10570962 | 0.017 | 0.1 | 1 | 9 |
| wrap73 | 0.13494 | -0.12514593 | 0.017 | 0.1 | 1 | 9 |
| mef2aa | 0.13498 | -0.14547444 | 0.172 | 0.15 | 1 | 9 |
| cdk5rap3 | 0.13507 | -0.31979614 | 0.052 | 0.15 | 1 | 9 |
| pde4ba | 0.13585 | -0.33036896 | 0.103 | 0.3 | 1 | 9 |
| smarca4a | 0.13638 | 0.302677187 | 0.31 | 0.1 | 1 | 9 |
| bzw1b | 0.1365 | 0.203912858 | 0.397 | 0.2 | 1 | 9 |
| hm13 | 0.13657 | 0.206276187 | 0.276 | 0.1 | 1 | 9 |
| spcs2 | 0.13662 | 0.225544492 | 0.466 | 0.25 | 1 | 9 |
| mrpl14 | 0.13663 | 0.267558411 | 0.207 | 0.05 | 1 | 9 |
| echs1 | 0.13673 | 0.314245894 | 0.31 | 0.1 | 1 | 9 |
| si:ch211-117c9.5 | 0.13682 | -0.34042812 | 0.034 | 0.15 | 1 | 9 |
| chd4b | 0.13729 | 0.210752822 | 0.397 | 0.2 | 1 | 9 |
| copb2 | 0.13748 | -0.37583987 | 0.103 | 0.25 | 1 | 9 |
| si:ch211-39i22.1 | 0.13816 | -0.2749834 | 0.345 | 0.4 | 1 | 9 |
| zgc:56676 | 0.13839 | 0.319630029 | 0.31 | 0.1 | 1 | 9 |
| sh3yl1 | 0.13847 | -0.25368484 | 0.086 | 0.25 | 1 | 9 |
| sparc | 0.13862 | 0.384077838 | 0.362 | 0.15 | 1 | 9 |
| dctn3 | 0.13894 | 0.308216504 | 0.31 | 0.1 | 1 | 9 |
| rtn3 | 0.13899 | -0.30263287 | 0.103 | 0.3 | 1 | 9 |
| pmp22a | 0.13907 | 0.475299749 | 0.31 | 0.1 | 1 | 9 |
| atp5pf | 0.13907 | 0.196106207 | 0.724 | 0.5 | 1 | 9 |
| ctnnb1 | 0.13932 | 0.37779093 | 0.379 | 0.15 | 1 | 9 |
| sec23b | 0.13945 | 0.233453898 | 0.207 | 0.2 | 1 | 9 |
| znf706 | 0.1397 | 0.350758121 | 0.379 | 0.15 | 1 | 9 |
| tbca | 0.13972 | 0.326490173 | 0.431 | 0.2 | 1 | 9 |
| ube2l3b | 0.13975 | 0.343983255 | 0.379 | 0.15 | 1 | 9 |
| LOC110439840 | 0.13989 | -0.30230972 | 0.069 | 0.2 | 1 | 9 |
| uba2 | 0.14033 | 0.229032158 | 0.207 | 0.05 | 1 | 9 |
| crip1 | 0.14152 | -0.47931807 | 0.207 | 0.4 | 1 | 9 |

| | | | | | | |
|---|---|---|---|---|---|---|
| cdc42bpb | 0.1419 | -0.24842278 | 0.086 | 0.15 | 1 | 9 |
| aqp8a.1 | 0.14311 | -0.43714791 | 0.052 | 0.15 | 1 | 9 |
| LEPROTL1 | 0.14348 | 0.13113573 | 0.293 | 0.15 | 1 | 9 |
| rps18 | 0.14369 | 0.222717626 | 0.914 | 0.85 | 1 | 9 |
| eif2s1a | 0.14411 | 0.292007436 | 0.207 | 0.05 | 1 | 9 |
| selenom | 0.14413 | 0.326569636 | 0.19 | 0.05 | 1 | 9 |
| anxa4 | 0.1442 | -0.34532821 | 0.379 | 0.4 | 1 | 9 |
| grb10a | 0.14628 | -0.31134399 | 0.052 | 0.2 | 1 | 9 |
| tp53rk | 0.14657 | -0.10964354 | 0.069 | 0.05 | 1 | 9 |
| metap1 | 0.14671 | 0.302430366 | 0.224 | 0.05 | 1 | 9 |
| sf1 | 0.14701 | -0.1693343 | 0.155 | 0.15 | 1 | 9 |
| syf2 | 0.14724 | 0.100164581 | 0.345 | 0.2 | 1 | 9 |
| usp24 | 0.14777 | -0.33030393 | 0.034 | 0.15 | 1 | 9 |
| eif2s2 | 0.14786 | 0.203888919 | 0.569 | 0.35 | 1 | 9 |
| cisd2 | 0.14838 | -0.31282736 | 0.103 | 0.2 | 1 | 9 |
| rgcc | 0.14839 | -0.30856798 | 0.052 | 0.2 | 1 | 9 |
| snx9b | 0.14841 | -0.30041362 | 0.052 | 0.2 | 1 | 9 |
| ube2ib | 0.14841 | 0.277099646 | 0.362 | 0.15 | 1 | 9 |
| wu:fb55g09 | 0.14864 | 0.359241134 | 0.362 | 0.15 | 1 | 9 |
| psph | 0.14865 | 0.102271708 | 0.172 | 0.05 | 1 | 9 |
| eif4a3 | 0.14883 | 0.283541882 | 0.345 | 0.25 | 1 | 9 |
| SBNO2 | 0.14897 | -0.16281304 | 0.052 | 0.05 | 1 | 9 |
| sub1b | 0.14942 | 0.135551386 | 0.431 | 0.25 | 1 | 9 |
| arhgef1b | 0.14965 | 0.272902347 | 0.345 | 0.15 | 1 | 9 |
| arpc1b | 0.15007 | -0.2463101 | 0.052 | 0.1 | 1 | 9 |
| tmem30b | 0.15025 | -0.27138884 | 0.121 | 0.15 | 1 | 9 |
| h1f0 | 0.15057 | 0.132804573 | 0.224 | 0.1 | 1 | 9 |
| snrpd2 | 0.15092 | 0.393617181 | 0.466 | 0.25 | 1 | 9 |
| iqgap1 | 0.15112 | -0.36829487 | 0.052 | 0.15 | 1 | 9 |
| mrpl36 | 0.15154 | 0.36208631 | 0.293 | 0.1 | 1 | 9 |
| rffl | 0.15281 | -0.37019385 | 0.052 | 0.2 | 1 | 9 |
| crfb4 | 0.15351 | -0.25381805 | 0.052 | 0.2 | 1 | 9 |
| clpp | 0.15357 | 0.204505331 | 0.207 | 0.05 | 1 | 9 |
| ctsla | 0.15377 | -0.14740668 | 0.466 | 0.4 | 1 | 9 |
| tbcb | 0.1539 | 0.134806742 | 0.224 | 0.1 | 1 | 9 |
| cotl1 | 0.15485 | -0.33402839 | 0.224 | 0.25 | 1 | 9 |
| bmpr1bb | 0.1549 | -0.26967319 | 0.052 | 0.1 | 1 | 9 |
| ptp4a1 | 0.15511 | 0.293429713 | 0.362 | 0.15 | 1 | 9 |
| pawr | 0.15551 | -0.30648892 | 0.052 | 0.2 | 1 | 9 |
| eif4a2 | 0.15573 | 0.247230107 | 0.224 | 0.05 | 1 | 9 |
| ucp2 | 0.15578 | -0.33411054 | 0.655 | 0.8 | 1 | 9 |
| DHDH | 0.15579 | -0.31036768 | 0.121 | 0.2 | 1 | 9 |
| btr25 | 0.156 | -0.21834753 | 0.052 | 0.2 | 1 | 9 |
| slc39a7 | 0.15627 | -0.36149968 | 0.172 | 0.3 | 1 | 9 |
| hnrnpc | 0.15657 | 0.2646335 | 0.224 | 0.05 | 1 | 9 |
| nasp | 0.15712 | 0.312079661 | 0.224 | 0.05 | 1 | 9 |
| tuba8l3 | 0.15752 | 0.406446962 | 0.224 | 0.05 | 1 | 9 |
| mcm7 | 0.15753 | 0.294019944 | 0.224 | 0.05 | 1 | 9 |

| | | | | | | |
|---|---|---|---|---|---|---|
| cbx5 | 0.15756 | 0.25445206 | 0.328 | 0.15 | 1 | 9 |
| hdac1 | 0.15794 | 0.318581561 | 0.431 | 0.2 | 1 | 9 |
| arf2a | 0.15802 | -0.26386904 | 0.293 | 0.35 | 1 | 9 |
| ndufa10 | 0.15813 | 0.322826274 | 0.293 | 0.1 | 1 | 9 |
| lpcat3 | 0.15826 | -0.22882331 | 0.052 | 0.2 | 1 | 9 |
| BX530721.1 | 0.15842 | -0.10146581 | 0 | 0.05 | 1 | 9 |
| CABZ01067348.1 | 0.15842 | -0.1285504 | 0 | 0.05 | 1 | 9 |
| CT025690.1 | 0.15842 | -0.1468488 | 0 | 0.05 | 1 | 9 |
| CU861477.1 | 0.15842 | -0.14750811 | 0 | 0.05 | 1 | 9 |
| LO018154.1 | 0.15842 | -0.10271442 | 0 | 0.05 | 1 | 9 |
| LO018197.1 | 0.15842 | -0.14704965 | 0 | 0.05 | 1 | 9 |
| LOC103909205 | 0.15842 | -0.10442989 | 0 | 0.05 | 1 | 9 |
| LOC108183588 | 0.15842 | -0.1255272 | 0 | 0.05 | 1 | 9 |
| LOC108190134 | 0.15842 | -0.1285504 | 0 | 0.05 | 1 | 9 |
| LOC110438633 | 0.15842 | -0.10112792 | 0 | 0.05 | 1 | 9 |
| XLOC-023329 | 0.15842 | -0.1285504 | 0 | 0.05 | 1 | 9 |
| XLOC-024455 | 0.15842 | -0.12590082 | 0 | 0.05 | 1 | 9 |
| XLOC-031367 | 0.15842 | -0.1479657 | 0 | 0.05 | 1 | 9 |
| aanat2 | 0.15842 | -0.14530171 | 0 | 0.05 | 1 | 9 |
| adamts1 | 0.15842 | -0.14679315 | 0 | 0.05 | 1 | 9 |
| adora2ab | 0.15842 | -0.12798402 | 0 | 0.05 | 1 | 9 |
| aldh4a1 | 0.15842 | -0.1279945 | 0 | 0.05 | 1 | 9 |
| arhgef10lb | 0.15842 | -0.18815964 | 0 | 0.05 | 1 | 9 |
| atg4c | 0.15842 | -0.14735171 | 0 | 0.05 | 1 | 9 |
| atp1b4 | 0.15842 | -0.28960652 | 0 | 0.05 | 1 | 9 |
| bbs7 | 0.15842 | -0.10362765 | 0 | 0.05 | 1 | 9 |
| bcan | 0.15842 | -0.1479657 | 0 | 0.05 | 1 | 9 |
| bgnb | 0.15842 | -0.18807269 | 0 | 0.05 | 1 | 9 |
| bmp1b | 0.15842 | -0.14725915 | 0 | 0.05 | 1 | 9 |
| ca4a | 0.15842 | -0.26315221 | 0 | 0.05 | 1 | 9 |
| caskin1 | 0.15842 | -0.18813363 | 0 | 0.05 | 1 | 9 |
| ccdc14 | 0.15842 | -0.14676407 | 0 | 0.05 | 1 | 9 |
| cdh13 | 0.15842 | -0.14651358 | 0 | 0.05 | 1 | 9 |
| cercam | 0.15842 | -0.10328063 | 0 | 0.05 | 1 | 9 |
| cers4a | 0.15842 | -0.14369408 | 0 | 0.05 | 1 | 9 |
| cldn5a | 0.15842 | -0.14690922 | 0 | 0.05 | 1 | 9 |
| cntn5 | 0.15842 | -0.14769092 | 0 | 0.05 | 1 | 9 |
| cpamd8 | 0.15842 | -0.10305203 | 0 | 0.05 | 1 | 9 |
| crip3 | 0.15842 | -0.1467833 | 0 | 0.05 | 1 | 9 |
| crot | 0.15842 | -0.14732709 | 0 | 0.05 | 1 | 9 |
| crtac1a | 0.15842 | -0.25331713 | 0 | 0.05 | 1 | 9 |
| cspg5a | 0.15842 | -0.1881195 | 0 | 0.05 | 1 | 9 |
| cspg5b | 0.15842 | -0.24316065 | 0 | 0.05 | 1 | 9 |
| cyp2x7 | 0.15842 | -0.14777714 | 0 | 0.05 | 1 | 9 |
| dennd1a | 0.15842 | -0.10398543 | 0 | 0.05 | 1 | 9 |
| ebp | 0.15842 | -0.14739697 | 0 | 0.05 | 1 | 9 |
| fads2 | 0.15842 | -0.18583066 | 0 | 0.05 | 1 | 9 |
| fbxl7 | 0.15842 | -0.14712289 | 0 | 0.05 | 1 | 9 |

| | | | | | | |
|---|---|---|---|---|---|---|
| fgf24 | 0.15842 | -0.10316634 | 0 | 0.05 | 1 | 9 |
| fkrp | 0.15842 | -0.1039712 | 0 | 0.05 | 1 | 9 |
| frmd4bb | 0.15842 | -0.12720764 | 0 | 0.05 | 1 | 9 |
| gnai2b | 0.15842 | -0.1461971 | 0 | 0.05 | 1 | 9 |
| gpam | 0.15842 | -0.14711748 | 0 | 0.05 | 1 | 9 |
| gpr39 | 0.15842 | -0.1285504 | 0 | 0.05 | 1 | 9 |
| gtf2a1l | 0.15842 | -0.126583 | 0 | 0.05 | 1 | 9 |
| hrh3 | 0.15842 | -0.1479657 | 0 | 0.05 | 1 | 9 |
| ilvbl | 0.15842 | -0.12792909 | 0 | 0.05 | 1 | 9 |
| jam3a | 0.15842 | -0.14622681 | 0 | 0.05 | 1 | 9 |
| ldb2b | 0.15842 | -0.14676345 | 0 | 0.05 | 1 | 9 |
| ldlrad2 | 0.15842 | -0.1479657 | 0 | 0.05 | 1 | 9 |
| lrrn2 | 0.15842 | -0.1479657 | 0 | 0.05 | 1 | 9 |
| map7d1a | 0.15842 | -0.10331765 | 0 | 0.05 | 1 | 9 |
| mettl16 | 0.15842 | -0.12783752 | 0 | 0.05 | 1 | 9 |
| mlc1 | 0.15842 | -0.18784892 | 0 | 0.05 | 1 | 9 |
| mon1a | 0.15842 | -0.14685851 | 0 | 0.05 | 1 | 9 |
| mpdu1a | 0.15842 | -0.1479657 | 0 | 0.05 | 1 | 9 |
| ms4a17a.7 | 0.15842 | -0.10442989 | 0 | 0.05 | 1 | 9 |
| ms4a17a.8 | 0.15842 | -0.10354259 | 0 | 0.05 | 1 | 9 |
| msc | 0.15842 | -0.10442989 | 0 | 0.05 | 1 | 9 |
| ndst1a | 0.15842 | -0.1886467 | 0 | 0.05 | 1 | 9 |
| ndufaf7 | 0.15842 | -0.10189317 | 0 | 0.05 | 1 | 9 |
| nid1b | 0.15842 | -0.14715967 | 0 | 0.05 | 1 | 9 |
| nog3 | 0.15842 | -0.14695041 | 0 | 0.05 | 1 | 9 |
| pak7 | 0.15842 | -0.10442989 | 0 | 0.05 | 1 | 9 |
| phospho2 | 0.15842 | -0.10324924 | 0 | 0.05 | 1 | 9 |
| pla2g12a | 0.15842 | -0.14741322 | 0 | 0.05 | 1 | 9 |
| plk2b | 0.15842 | -0.12604951 | 0 | 0.05 | 1 | 9 |
| polrmt | 0.15842 | -0.12766069 | 0 | 0.05 | 1 | 9 |
| primpol | 0.15842 | -0.10305566 | 0 | 0.05 | 1 | 9 |
| prmt9 | 0.15842 | -0.12710629 | 0 | 0.05 | 1 | 9 |
| ptgdsb.1 | 0.15842 | -0.29419745 | 0 | 0.05 | 1 | 9 |
| ptgdsb.2 | 0.15842 | -0.354267 | 0 | 0.05 | 1 | 9 |
| pxmp2 | 0.15842 | -0.14694257 | 0 | 0.05 | 1 | 9 |
| pyroxd1 | 0.15842 | -0.14650234 | 0 | 0.05 | 1 | 9 |
| rac3a | 0.15842 | -0.10239535 | 0 | 0.05 | 1 | 9 |
| rasgrp4 | 0.15842 | -0.16428212 | 0 | 0.05 | 1 | 9 |
| rh50 | 0.15842 | -0.1285504 | 0 | 0.05 | 1 | 9 |
| rhot1b | 0.15842 | -0.10442989 | 0 | 0.05 | 1 | 9 |
| ryr3 | 0.15842 | -0.12683688 | 0 | 0.05 | 1 | 9 |
| s100b | 0.15842 | -0.20724491 | 0 | 0.05 | 1 | 9 |
| sc:d189 | 0.15842 | -0.21280231 | 0 | 0.05 | 1 | 9 |
| senp3a | 0.15842 | -0.10273861 | 0 | 0.05 | 1 | 9 |
| sft2d3 | 0.15842 | -0.10265536 | 0 | 0.05 | 1 | 9 |
| si:ch211-106f21.1 | 0.15842 | -0.1479657 | 0 | 0.05 | 1 | 9 |
| si:ch211-149k12.3 | 0.15842 | -0.1285504 | 0 | 0.05 | 1 | 9 |
| si:ch211-1a19.3 | 0.15842 | -0.14729603 | 0 | 0.05 | 1 | 9 |

| | | | | | | |
|---|---|---|---|---|---|---|
| si:ch211-212k18.5 | 0.15842 | -0.14734684 | 0 | 0.05 | 1 | 9 |
| si:ch211-59d17.3 | 0.15842 | -0.18856327 | 0 | 0.05 | 1 | 9 |
| si:ch73-211e3.1 | 0.15842 | -0.12817944 | 0 | 0.05 | 1 | 9 |
| si:dkey-111k8.3 | 0.15842 | -0.10312679 | 0 | 0.05 | 1 | 9 |
| si:dkey-256i11.2 | 0.15842 | -0.10367524 | 0 | 0.05 | 1 | 9 |
| si:dkey-48p11.3 | 0.15842 | -0.10060674 | 0 | 0.05 | 1 | 9 |
| slc1a3b | 0.15842 | -0.24289525 | 0 | 0.05 | 1 | 9 |
| slc7a10b | 0.15842 | -0.1886467 | 0 | 0.05 | 1 | 9 |
| slc9a3r1b | 0.15842 | -0.12665615 | 0 | 0.05 | 1 | 9 |
| slco1d1 | 0.15842 | -0.1479657 | 0 | 0.05 | 1 | 9 |
| smug1 | 0.15842 | -0.14649864 | 0 | 0.05 | 1 | 9 |
| sqlea | 0.15842 | -0.1473012 | 0 | 0.05 | 1 | 9 |
| steap2 | 0.15842 | -0.1479657 | 0 | 0.05 | 1 | 9 |
| tbc1d16 | 0.15842 | -0.10283795 | 0 | 0.05 | 1 | 9 |
| tex261 | 0.15842 | -0.10030004 | 0 | 0.05 | 1 | 9 |
| tmem229b | 0.15842 | -0.10294606 | 0 | 0.05 | 1 | 9 |
| tor2a | 0.15842 | -0.14668157 | 0 | 0.05 | 1 | 9 |
| ttyh1 | 0.15842 | -0.1886467 | 0 | 0.05 | 1 | 9 |
| urb2 | 0.15842 | -0.10374658 | 0 | 0.05 | 1 | 9 |
| viml | 0.15842 | -0.13766502 | 0 | 0.05 | 1 | 9 |
| vwa2 | 0.15842 | -0.10442989 | 0 | 0.05 | 1 | 9 |
| wnt7aa | 0.15842 | -0.14761905 | 0 | 0.05 | 1 | 9 |
| zbtb12.2 | 0.15842 | -0.10306946 | 0 | 0.05 | 1 | 9 |
| zgc:112332 | 0.15842 | -0.18758036 | 0 | 0.05 | 1 | 9 |
| zgc:162962 | 0.15842 | -0.10285821 | 0 | 0.05 | 1 | 9 |
| znf1079 | 0.15842 | -0.10442989 | 0 | 0.05 | 1 | 9 |
| znf385d | 0.15842 | -0.10442989 | 0 | 0.05 | 1 | 9 |
| znf970 | 0.15842 | -0.14766848 | 0 | 0.05 | 1 | 9 |
| BX649398.2 | 0.15842 | -0.15219312 | 0 | 0.05 | 1 | 9 |
| CR352226.5 | 0.15842 | -0.1537679 | 0 | 0.05 | 1 | 9 |
| CR749168.2 | 0.15842 | -0.15318335 | 0 | 0.05 | 1 | 9 |
| CT573342.1 | 0.15842 | -0.14892062 | 0 | 0.05 | 1 | 9 |
| LO017771.1 | 0.15842 | -0.1547748 | 0 | 0.05 | 1 | 9 |
| LO017954.1 | 0.15842 | -0.1529804 | 0 | 0.05 | 1 | 9 |
| LO018608.1 | 0.15842 | -0.1547748 | 0 | 0.05 | 1 | 9 |
| LOC100537613 | 0.15842 | -0.15354273 | 0 | 0.05 | 1 | 9 |
| LOC103911430 | 0.15842 | -0.15022508 | 0 | 0.05 | 1 | 9 |
| LOC110438394 | 0.15842 | -0.15219312 | 0 | 0.05 | 1 | 9 |
| XLOC-011160 | 0.15842 | -0.10834691 | 0 | 0.05 | 1 | 9 |
| XLOC-039931 | 0.15842 | -0.15447934 | 0 | 0.05 | 1 | 9 |
| arih1l | 0.15842 | -0.15266256 | 0 | 0.05 | 1 | 9 |
| cbln11 | 0.15842 | -0.25207429 | 0 | 0.05 | 1 | 9 |
| chn2 | 0.15842 | -0.15219312 | 0 | 0.05 | 1 | 9 |
| cldn20 | 0.15842 | -0.1547748 | 0 | 0.05 | 1 | 9 |
| cmtm8a | 0.15842 | -0.1547748 | 0 | 0.05 | 1 | 9 |
| dph7 | 0.15842 | -0.15216886 | 0 | 0.05 | 1 | 9 |
| dtnbb | 0.15842 | -0.14800258 | 0 | 0.05 | 1 | 9 |
| efcab7 | 0.15842 | -0.15409011 | 0 | 0.05 | 1 | 9 |

| | | | | | | |
|---|---|---|---|---|---|---|
| fhl1a | 0.15842 | -0.10714104 | 0 | 0.05 | 1 | 9 |
| gata6 | 0.15842 | -0.15366256 | 0 | 0.05 | 1 | 9 |
| gramd4a | 0.15842 | -0.1547748 | 0 | 0.05 | 1 | 9 |
| hgfa | 0.15842 | -0.10765744 | 0 | 0.05 | 1 | 9 |
| hsd17b4 | 0.15842 | -0.15296566 | 0 | 0.05 | 1 | 9 |
| inpp5f | 0.15842 | -0.15419771 | 0 | 0.05 | 1 | 9 |
| irf1b | 0.15842 | -0.15416838 | 0 | 0.05 | 1 | 9 |
| lifrb | 0.15842 | -0.15261989 | 0 | 0.05 | 1 | 9 |
| lrp1aa | 0.15842 | -0.10776816 | 0 | 0.05 | 1 | 9 |
| map4k6 | 0.15842 | -0.15360896 | 0 | 0.05 | 1 | 9 |
| mgp | 0.15842 | -0.10699089 | 0 | 0.05 | 1 | 9 |
| mras | 0.15842 | -0.1546199 | 0 | 0.05 | 1 | 9 |
| mvda | 0.15842 | -0.15368428 | 0 | 0.05 | 1 | 9 |
| narf | 0.15842 | -0.1536124 | 0 | 0.05 | 1 | 9 |
| nxnl2 | 0.15842 | -0.15422868 | 0 | 0.05 | 1 | 9 |
| rbpms2b | 0.15842 | -0.15193337 | 0 | 0.05 | 1 | 9 |
| rgl3a | 0.15842 | -0.10566116 | 0 | 0.05 | 1 | 9 |
| rspo1 | 0.15842 | -0.15936465 | 0 | 0.05 | 1 | 9 |
| si:ch211-15d5.11 | 0.15842 | -0.1547748 | 0 | 0.05 | 1 | 9 |
| si:ch211-215p11.1 | 0.15842 | -0.1547748 | 0 | 0.05 | 1 | 9 |
| si:ch211-266o15.1 | 0.15842 | -0.14180992 | 0 | 0.05 | 1 | 9 |
| si:ch211-284e13.9 | 0.15842 | -0.15159205 | 0 | 0.05 | 1 | 9 |
| si:ch211-40k21.9 | 0.15842 | -0.14240395 | 0 | 0.05 | 1 | 9 |
| si:dkey-204l11.1 | 0.15842 | -0.1543553 | 0 | 0.05 | 1 | 9 |
| slc8a2a | 0.15842 | -0.14892062 | 0 | 0.05 | 1 | 9 |
| smtnl | 0.15842 | -0.15464032 | 0 | 0.05 | 1 | 9 |
| snx15 | 0.15842 | -0.14499081 | 0 | 0.05 | 1 | 9 |
| socs6a | 0.15842 | -0.15422147 | 0 | 0.05 | 1 | 9 |
| spns3 | 0.15842 | -0.15319548 | 0 | 0.05 | 1 | 9 |
| thbs3a | 0.15842 | -0.13378604 | 0 | 0.05 | 1 | 9 |
| tln2a | 0.15842 | -0.15404895 | 0 | 0.05 | 1 | 9 |
| zgc:163077 | 0.15842 | -0.15341072 | 0 | 0.05 | 1 | 9 |
| zmiz1a | 0.15842 | -0.14972723 | 0 | 0.05 | 1 | 9 |
| AL772362.1 | 0.15842 | -0.13889899 | 0 | 0.05 | 1 | 9 |
| BX908782.2 | 0.15842 | -0.18608097 | 0 | 0.05 | 1 | 9 |
| CR762407.1 | 0.15842 | -0.21242037 | 0 | 0.05 | 1 | 9 |
| FO834857.1 | 0.15842 | -0.14684653 | 0 | 0.05 | 1 | 9 |
| LOC100333048 | 0.15842 | -0.26284466 | 0 | 0.05 | 1 | 9 |
| LOC101883798 | 0.15842 | -0.22582696 | 0 | 0.05 | 1 | 9 |
| LOC101886880 | 0.15842 | -0.10825309 | 0 | 0.05 | 1 | 9 |
| MDFI | 0.15842 | -0.11363832 | 0 | 0.05 | 1 | 9 |
| adgrf3b | 0.15842 | -0.17466963 | 0 | 0.05 | 1 | 9 |
| agla | 0.15842 | -0.14458023 | 0 | 0.05 | 1 | 9 |
| calhm6 | 0.15842 | -0.13520709 | 0 | 0.05 | 1 | 9 |
| cdab | 0.15842 | -0.11225815 | 0 | 0.05 | 1 | 9 |
| cdh30 | 0.15842 | -0.13520709 | 0 | 0.05 | 1 | 9 |
| cep120 | 0.15842 | -0.13752773 | 0 | 0.05 | 1 | 9 |
| chn1 | 0.15842 | -0.13672141 | 0 | 0.05 | 1 | 9 |

| | | | | | | |
|---|---|---|---|---|---|---|
| clcn5a | 0.15842 | -0.14055662 | 0 | 0.05 | 1 | 9 |
| cldn34a | 0.15842 | -0.11630815 | 0 | 0.05 | 1 | 9 |
| cpa4 | 0.15842 | -0.20888617 | 0 | 0.05 | 1 | 9 |
| cxcr4a | 0.15842 | -0.114058 | 0 | 0.05 | 1 | 9 |
| cyb561d1 | 0.15842 | -0.13978655 | 0 | 0.05 | 1 | 9 |
| dip2cb | 0.15842 | -0.12863299 | 0 | 0.05 | 1 | 9 |
| dusp4 | 0.15842 | -0.18405922 | 0 | 0.05 | 1 | 9 |
| fam214b | 0.15842 | -0.14160031 | 0 | 0.05 | 1 | 9 |
| fat1b | 0.15842 | -0.13889899 | 0 | 0.05 | 1 | 9 |
| fep15 | 0.15842 | -0.17927766 | 0 | 0.05 | 1 | 9 |
| ikzf4 | 0.15842 | -0.22168553 | 0 | 0.05 | 1 | 9 |
| kazald2 | 0.15842 | -0.11662324 | 0 | 0.05 | 1 | 9 |
| lcn15 | 0.15842 | -0.11401461 | 0 | 0.05 | 1 | 9 |
| micall2b | 0.15842 | -0.14515246 | 0 | 0.05 | 1 | 9 |
| mpnd | 0.15842 | -0.10605623 | 0 | 0.05 | 1 | 9 |
| muc5d | 0.15842 | -0.11880625 | 0 | 0.05 | 1 | 9 |
| ovol1b | 0.15842 | -0.20718524 | 0 | 0.05 | 1 | 9 |
| pnpla7b | 0.15842 | -0.14229315 | 0 | 0.05 | 1 | 9 |
| scn4bb | 0.15842 | -0.11063978 | 0 | 0.05 | 1 | 9 |
| si:ch1073-385f13.3 | 0.15842 | -0.14115799 | 0 | 0.05 | 1 | 9 |
| si:ch211-193k19.1 | 0.15842 | -0.14242956 | 0 | 0.05 | 1 | 9 |
| si:ch211-39i2.2 | 0.15842 | -0.13683504 | 0 | 0.05 | 1 | 9 |
| si:ch73-158p21.2 | 0.15842 | -0.14542869 | 0 | 0.05 | 1 | 9 |
| si:ch73-252i11.1 | 0.15842 | -0.1376774 | 0 | 0.05 | 1 | 9 |
| si:dkey-21a6.5 | 0.15842 | -0.10688389 | 0 | 0.05 | 1 | 9 |
| si:dkey-28d5.13 | 0.15842 | -0.11214432 | 0 | 0.05 | 1 | 9 |
| si:dkey-56i24.1 | 0.15842 | -0.21242037 | 0 | 0.05 | 1 | 9 |
| si:dkeyp-82a1.4 | 0.15842 | -0.10825309 | 0 | 0.05 | 1 | 9 |
| slc14a2 | 0.15842 | -0.14008061 | 0 | 0.05 | 1 | 9 |
| snx20 | 0.15842 | -0.10344558 | 0 | 0.05 | 1 | 9 |
| supt7l | 0.15842 | -0.13443152 | 0 | 0.05 | 1 | 9 |
| sycn.1 | 0.15842 | -0.19379125 | 0 | 0.05 | 1 | 9 |
| tmem176l.2 | 0.15842 | -0.14268811 | 0 | 0.05 | 1 | 9 |
| tnfb | 0.15842 | -0.15866124 | 0 | 0.05 | 1 | 9 |
| zgc:173575 | 0.15842 | -0.11570074 | 0 | 0.05 | 1 | 9 |
| XLOC-034863 | 0.15842 | -0.11874426 | 0 | 0.05 | 1 | 9 |
| XLOC-043834 | 0.15842 | -0.16546806 | 0 | 0.05 | 1 | 9 |
| abhd3 | 0.15842 | -0.1963374 | 0 | 0.05 | 1 | 9 |
| homezb | 0.15842 | -0.13575593 | 0 | 0.05 | 1 | 9 |
| ndor1 | 0.15842 | -0.13507507 | 0 | 0.05 | 1 | 9 |
| pnpla3 | 0.15842 | -0.1882499 | 0 | 0.05 | 1 | 9 |
| prr14 | 0.15842 | -0.1954825 | 0 | 0.05 | 1 | 9 |
| ptpn6 | 0.15842 | -0.23756093 | 0 | 0.05 | 1 | 9 |
| si:ch211-169p10.1 | 0.15842 | -0.19651765 | 0 | 0.05 | 1 | 9 |
| si:ch211-91p5.3 | 0.15842 | -0.13575497 | 0 | 0.05 | 1 | 9 |
| si:dkey-58f10.12 | 0.15842 | -0.13626973 | 0 | 0.05 | 1 | 9 |
| wnt9b | 0.15842 | -0.1882499 | 0 | 0.05 | 1 | 9 |
| CABZ01056054.1 | 0.15842 | -0.13124228 | 0 | 0.05 | 1 | 9 |

| | | | | | | |
|---|---|---|---|---|---|---|
| CCDC69 | 0.15842 | -0.13615089 | 0 | 0.05 | 1 | 9 |
| GTPBP8 | 0.15842 | -0.1365531 | 0 | 0.05 | 1 | 9 |
| XLOC-009557 | 0.15842 | -0.13618656 | 0 | 0.05 | 1 | 9 |
| XLOC-040350 | 0.15842 | -0.1365531 | 0 | 0.05 | 1 | 9 |
| adgre10 | 0.15842 | -0.19984574 | 0 | 0.05 | 1 | 9 |
| arhgef11 | 0.15842 | -0.13600587 | 0 | 0.05 | 1 | 9 |
| bnip2 | 0.15842 | -0.13585157 | 0 | 0.05 | 1 | 9 |
| camk1db | 0.15842 | -0.13468646 | 0 | 0.05 | 1 | 9 |
| cbx8b | 0.15842 | -0.13492578 | 0 | 0.05 | 1 | 9 |
| dkk2 | 0.15842 | -0.1365531 | 0 | 0.05 | 1 | 9 |
| foxc1b | 0.15842 | -0.17592321 | 0 | 0.05 | 1 | 9 |
| gfod2 | 0.15842 | -0.13370669 | 0 | 0.05 | 1 | 9 |
| khk | 0.15842 | -0.13531157 | 0 | 0.05 | 1 | 9 |
| miga2 | 0.15842 | -0.1365531 | 0 | 0.05 | 1 | 9 |
| nbeal2 | 0.15842 | -0.13516349 | 0 | 0.05 | 1 | 9 |
| nemp2 | 0.15842 | -0.13201862 | 0 | 0.05 | 1 | 9 |
| pfkpb | 0.15842 | -0.13504904 | 0 | 0.05 | 1 | 9 |
| ptpn23a | 0.15842 | -0.13576697 | 0 | 0.05 | 1 | 9 |
| pts | 0.15842 | -0.1355324 | 0 | 0.05 | 1 | 9 |
| reps1 | 0.15842 | -0.13580404 | 0 | 0.05 | 1 | 9 |
| ryr1a | 0.15842 | -0.1365531 | 0 | 0.05 | 1 | 9 |
| serpinb1l1 | 0.15842 | -0.13459038 | 0 | 0.05 | 1 | 9 |
| si:dkey-86e18.1 | 0.15842 | -0.13282835 | 0 | 0.05 | 1 | 9 |
| si:dkeyp-13a3.10 | 0.15842 | -0.13330968 | 0 | 0.05 | 1 | 9 |
| si:rp71-36a1.3 | 0.15842 | -0.13595603 | 0 | 0.05 | 1 | 9 |
| vps33b | 0.15842 | -0.13529245 | 0 | 0.05 | 1 | 9 |
| zgc:174906 | 0.15842 | -0.13543624 | 0 | 0.05 | 1 | 9 |
| zgc:193807 | 0.15842 | -0.13518618 | 0 | 0.05 | 1 | 9 |
| zgc:85932 | 0.15842 | -0.1294133 | 0 | 0.05 | 1 | 9 |
| BX324142.1 | 0.15842 | -0.11761713 | 0 | 0.05 | 1 | 9 |
| CR354588.1 | 0.15842 | -0.11615851 | 0 | 0.05 | 1 | 9 |
| CR759845.1 | 0.15842 | -0.11130879 | 0 | 0.05 | 1 | 9 |
| LOC101884267 | 0.15842 | -0.17102447 | 0 | 0.05 | 1 | 9 |
| LOC110437760 | 0.15842 | -0.11574553 | 0 | 0.05 | 1 | 9 |
| XLOC-015268 | 0.15842 | -0.11761713 | 0 | 0.05 | 1 | 9 |
| XLOC-022621 | 0.15842 | -0.11684736 | 0 | 0.05 | 1 | 9 |
| agps | 0.15842 | -0.11593108 | 0 | 0.05 | 1 | 9 |
| bhlha15 | 0.15842 | -0.1528133 | 0 | 0.05 | 1 | 9 |
| cbln12 | 0.15842 | -0.11595299 | 0 | 0.05 | 1 | 9 |
| cxxc1b | 0.15842 | -0.13771151 | 0 | 0.05 | 1 | 9 |
| cyp2ae1 | 0.15842 | -0.11477899 | 0 | 0.05 | 1 | 9 |
| dnah9l | 0.15842 | -0.11761713 | 0 | 0.05 | 1 | 9 |
| dpf1 | 0.15842 | -0.11681106 | 0 | 0.05 | 1 | 9 |
| egln2 | 0.15842 | -0.11647482 | 0 | 0.05 | 1 | 9 |
| fam98b | 0.15842 | -0.11619287 | 0 | 0.05 | 1 | 9 |
| glb1 | 0.15842 | -0.11556757 | 0 | 0.05 | 1 | 9 |
| gpat4 | 0.15842 | -0.11502197 | 0 | 0.05 | 1 | 9 |
| hdr | 0.15842 | -0.15569002 | 0 | 0.05 | 1 | 9 |

| | | | | | | |
|---|---|---|---|---|---|---|
| jakmip1 | 0.15842 | -0.15415173 | 0 | 0.05 | 1 | 9 |
| kiss2 | 0.15842 | -0.11316229 | 0 | 0.05 | 1 | 9 |
| klhl12 | 0.15842 | -0.12182152 | 0 | 0.05 | 1 | 9 |
| kyat1 | 0.15842 | -0.13702518 | 0 | 0.05 | 1 | 9 |
| lipcb | 0.15842 | -0.11706846 | 0 | 0.05 | 1 | 9 |
| lpar3 | 0.15842 | -0.14072341 | 0 | 0.05 | 1 | 9 |
| magi3b | 0.15842 | -0.11761713 | 0 | 0.05 | 1 | 9 |
| mkxb | 0.15842 | -0.1150834 | 0 | 0.05 | 1 | 9 |
| nfrkb | 0.15842 | -0.11468596 | 0 | 0.05 | 1 | 9 |
| nit2 | 0.15842 | -0.11398631 | 0 | 0.05 | 1 | 9 |
| npr1a | 0.15842 | -0.20092582 | 0 | 0.05 | 1 | 9 |
| nuak1b | 0.15842 | -0.15625153 | 0 | 0.05 | 1 | 9 |
| nupl2 | 0.15842 | -0.15522744 | 0 | 0.05 | 1 | 9 |
| pacs1a | 0.15842 | -0.11638931 | 0 | 0.05 | 1 | 9 |
| pls1 | 0.15842 | -0.10769504 | 0 | 0.05 | 1 | 9 |
| rab21 | 0.15842 | -0.12840667 | 0 | 0.05 | 1 | 9 |
| rab34a | 0.15842 | -0.11492864 | 0 | 0.05 | 1 | 9 |
| rorcb | 0.15842 | -0.11402321 | 0 | 0.05 | 1 | 9 |
| si:ch211-119d14.3 | 0.15842 | -0.11642385 | 0 | 0.05 | 1 | 9 |
| si:ch211-171h4.6 | 0.15842 | -0.13750366 | 0 | 0.05 | 1 | 9 |
| si:dkey-23a13.5 | 0.15842 | -0.11684853 | 0 | 0.05 | 1 | 9 |
| si:dkey-65b12.6 | 0.15842 | -0.2052647 | 0 | 0.05 | 1 | 9 |
| srebf1 | 0.15842 | -0.1167152 | 0 | 0.05 | 1 | 9 |
| traf3ip2l | 0.15842 | -0.13052193 | 0 | 0.05 | 1 | 9 |
| ttc7a | 0.15842 | -0.11761713 | 0 | 0.05 | 1 | 9 |
| zgc:153654 | 0.15842 | -0.20831328 | 0 | 0.05 | 1 | 9 |
| zgc:154006 | 0.15842 | -0.11064381 | 0 | 0.05 | 1 | 9 |
| zgc:171686 | 0.15842 | -0.11593449 | 0 | 0.05 | 1 | 9 |
| znf1155 | 0.15842 | -0.11689902 | 0 | 0.05 | 1 | 9 |
| zc3h13 | 0.15897 | 0.169961867 | 0.19 | 0.05 | 1 | 9 |
| ndufs7 | 0.15952 | 0.210337231 | 0.345 | 0.15 | 1 | 9 |
| smg5 | 0.15986 | -0.10494899 | 0.069 | 0.05 | 1 | 9 |
| slc25a3b | 0.16021 | 0.376649115 | 0.707 | 0.5 | 1 | 9 |
| slc3a2b | 0.16043 | 0.336936618 | 0.431 | 0.2 | 1 | 9 |
| eif2s1b | 0.16062 | 0.291198842 | 0.414 | 0.2 | 1 | 9 |
| cct8 | 0.16118 | 0.32377721 | 0.431 | 0.2 | 1 | 9 |
| CABZ01092156.1 | 0.16136 | 0.147097187 | 0.241 | 0.1 | 1 | 9 |
| atp5mc3b | 0.16157 | 0.327143235 | 0.81 | 0.6 | 1 | 9 |
| si:ch211-140m22.7 | 0.16205 | -0.27039244 | 0.052 | 0.2 | 1 | 9 |
| tmem97 | 0.16263 | -0.30978355 | 0.052 | 0.15 | 1 | 9 |
| ccl20b | 0.16319 | -0.50211396 | 0.19 | 0.35 | 1 | 9 |
| eif5 | 0.16323 | 0.335719031 | 0.431 | 0.2 | 1 | 9 |
| baz2ba | 0.16333 | 0.296942271 | 0.224 | 0.1 | 1 | 9 |
| eif3i | 0.16343 | 0.343996542 | 0.586 | 0.4 | 1 | 9 |
| ip6k2b | 0.16391 | 0.380718378 | 0.362 | 0.15 | 1 | 9 |
| zgc:56304 | 0.16401 | 0.372490338 | 0.293 | 0.1 | 1 | 9 |
| si:dkey-42i9.4 | 0.16452 | 0.159200175 | 0.431 | 0.25 | 1 | 9 |
| nectin1b | 0.16456 | -0.31205167 | 0.19 | 0.3 | 1 | 9 |

| | | | | | | |
|---|---|---|---|---|---|---|
| pin1 | 0.16457 | 0.258152999 | 0.362 | 0.15 | 1 | 9 |
| rpl24 | 0.1646 | 0.212351181 | 0.897 | 0.85 | 1 | 9 |
| hsf2 | 0.16566 | 0.250966004 | 0.293 | 0.1 | 1 | 9 |
| otud7b | 0.16569 | -0.27044989 | 0.034 | 0.15 | 1 | 9 |
| rps27a | 0.16649 | 0.151034318 | 1 | 0.95 | 1 | 9 |
| rpsa | 0.16771 | 0.208164486 | 0.966 | 0.9 | 1 | 9 |
| pdcd10a | 0.16869 | -0.16082331 | 0.121 | 0.1 | 1 | 9 |
| cyp20a1 | 0.16932 | -0.12942689 | 0.121 | 0.1 | 1 | 9 |
| XLOC-030786 | 0.16947 | 0.146337136 | 0.069 | 0 | 1 | 9 |
| gpr107 | 0.16947 | 0.133647272 | 0.069 | 0 | 1 | 9 |
| itga6a | 0.16947 | 0.157843583 | 0.069 | 0 | 1 | 9 |
| lgals3bpa | 0.16947 | 0.183787613 | 0.069 | 0 | 1 | 9 |
| mapk12a | 0.16947 | 0.134745296 | 0.069 | 0 | 1 | 9 |
| nelfa | 0.16947 | 0.132120107 | 0.069 | 0 | 1 | 9 |
| ppfia3 | 0.16947 | 0.156167938 | 0.069 | 0 | 1 | 9 |
| pvalb8 | 0.16947 | 0.246106375 | 0.069 | 0 | 1 | 9 |
| rad1 | 0.16947 | 0.122387986 | 0.069 | 0 | 1 | 9 |
| ubiad1 | 0.16947 | 0.144107557 | 0.069 | 0 | 1 | 9 |
| usp2a | 0.16947 | 0.159038685 | 0.069 | 0 | 1 | 9 |
| AL935174.2 | 0.16947 | 0.135075729 | 0.069 | 0 | 1 | 9 |
| BX901957.1 | 0.16947 | 0.13056133 | 0.069 | 0 | 1 | 9 |
| CABZ01005379.1 | 0.16947 | 0.172251234 | 0.069 | 0 | 1 | 9 |
| CABZ01068366.1 | 0.16947 | 0.140616707 | 0.069 | 0 | 1 | 9 |
| CABZ01077217.1 | 0.16947 | 0.12146541 | 0.069 | 0 | 1 | 9 |
| CABZ01078261.1 | 0.16947 | 0.111388449 | 0.069 | 0 | 1 | 9 |
| CABZ01113954.1 | 0.16947 | 0.11538875 | 0.069 | 0 | 1 | 9 |
| CR356236.1 | 0.16947 | 0.158244757 | 0.069 | 0 | 1 | 9 |
| CR388166.2 | 0.16947 | 0.10111113 | 0.069 | 0 | 1 | 9 |
| CU467856.3 | 0.16947 | 0.112385314 | 0.069 | 0 | 1 | 9 |
| CU539058.1 | 0.16947 | 0.101276334 | 0.069 | 0 | 1 | 9 |
| EML4 | 0.16947 | 0.152909251 | 0.069 | 0 | 1 | 9 |
| FQ323119.1 | 0.16947 | 0.140322512 | 0.069 | 0 | 1 | 9 |
| LOC101883507 | 0.16947 | 0.102821355 | 0.069 | 0 | 1 | 9 |
| LOC103909439 | 0.16947 | 0.136093962 | 0.069 | 0 | 1 | 9 |
| LOC108182329 | 0.16947 | 0.119657411 | 0.069 | 0 | 1 | 9 |
| LOC110438518 | 0.16947 | 0.147892474 | 0.069 | 0 | 1 | 9 |
| LOC799574 | 0.16947 | 0.208894824 | 0.069 | 0 | 1 | 9 |
| XLOC-010899 | 0.16947 | 0.103369042 | 0.069 | 0 | 1 | 9 |
| XLOC-032256 | 0.16947 | 0.154596064 | 0.069 | 0 | 1 | 9 |
| acot8 | 0.16947 | 0.111130725 | 0.069 | 0 | 1 | 9 |
| acsl1b | 0.16947 | 0.119539967 | 0.069 | 0 | 1 | 9 |
| ankrd40 | 0.16947 | 0.137399096 | 0.069 | 0 | 1 | 9 |
| apbb2b | 0.16947 | 0.116794545 | 0.069 | 0 | 1 | 9 |
| apip | 0.16947 | 0.147248349 | 0.069 | 0 | 1 | 9 |
| apopt1 | 0.16947 | 0.111512848 | 0.069 | 0 | 1 | 9 |
| arhgap35b | 0.16947 | 0.127297149 | 0.069 | 0 | 1 | 9 |
| arhgef16 | 0.16947 | 0.10968586 | 0.069 | 0 | 1 | 9 |
| arl6ip5a | 0.16947 | 0.111177709 | 0.069 | 0 | 1 | 9 |

| | | | | | | |
|---|---|---|---|---|---|---|
| arnt2 | 0.16947 | 0.154858711 | 0.069 | 0 | 1 | 9 |
| asap1b | 0.16947 | 0.151544514 | 0.069 | 0 | 1 | 9 |
| atg3 | 0.16947 | 0.114253133 | 0.069 | 0 | 1 | 9 |
| atl2 | 0.16947 | 0.159121106 | 0.069 | 0 | 1 | 9 |
| atp6ap1a | 0.16947 | 0.112461745 | 0.069 | 0 | 1 | 9 |
| atp6v1c1b | 0.16947 | 0.128852427 | 0.069 | 0 | 1 | 9 |
| atp6v1f | 0.16947 | 0.105792719 | 0.069 | 0 | 1 | 9 |
| atrxl | 0.16947 | 0.106160388 | 0.069 | 0 | 1 | 9 |
| b4galt7 | 0.16947 | 0.123163643 | 0.069 | 0 | 1 | 9 |
| babam2 | 0.16947 | 0.129148608 | 0.069 | 0 | 1 | 9 |
| bap1 | 0.16947 | 0.123070415 | 0.069 | 0 | 1 | 9 |
| bckdhbl | 0.16947 | 0.155042054 | 0.069 | 0 | 1 | 9 |
| bcl2a | 0.16947 | 0.142191281 | 0.069 | 0 | 1 | 9 |
| bcs1l | 0.16947 | 0.10520898 | 0.069 | 0 | 1 | 9 |
| bin1a | 0.16947 | 0.122331475 | 0.069 | 0 | 1 | 9 |
| bnipl | 0.16947 | 0.128511203 | 0.069 | 0 | 1 | 9 |
| btbd7 | 0.16947 | 0.106419878 | 0.069 | 0 | 1 | 9 |
| calhm2 | 0.16947 | 0.133052226 | 0.069 | 0 | 1 | 9 |
| ccne1 | 0.16947 | 0.139496903 | 0.069 | 0 | 1 | 9 |
| cdh15 | 0.16947 | 0.117381361 | 0.069 | 0 | 1 | 9 |
| cds2 | 0.16947 | 0.106646373 | 0.069 | 0 | 1 | 9 |
| cemip2 | 0.16947 | 0.128054262 | 0.069 | 0 | 1 | 9 |
| chac2 | 0.16947 | 0.125794565 | 0.069 | 0 | 1 | 9 |
| chtf8 | 0.16947 | 0.159108161 | 0.069 | 0 | 1 | 9 |
| cisd1 | 0.16947 | 0.105544367 | 0.069 | 0 | 1 | 9 |
| clcn4 | 0.16947 | 0.10958102 | 0.069 | 0 | 1 | 9 |
| clstn1 | 0.16947 | 0.104017307 | 0.069 | 0 | 1 | 9 |
| commd8 | 0.16947 | 0.113759982 | 0.069 | 0 | 1 | 9 |
| coro1cb | 0.16947 | 0.163446404 | 0.069 | 0 | 1 | 9 |
| cpt1ab | 0.16947 | 0.100447557 | 0.069 | 0 | 1 | 9 |
| crata | 0.16947 | 0.150193555 | 0.069 | 0 | 1 | 9 |
| crls1 | 0.16947 | 0.125875818 | 0.069 | 0 | 1 | 9 |
| cspg4 | 0.16947 | 0.121844849 | 0.069 | 0 | 1 | 9 |
| ctdnep1a | 0.16947 | 0.105742897 | 0.069 | 0 | 1 | 9 |
| cth | 0.16947 | 0.126436916 | 0.069 | 0 | 1 | 9 |
| ctnna2 | 0.16947 | 0.149225807 | 0.069 | 0 | 1 | 9 |
| ctps1a | 0.16947 | 0.111897012 | 0.069 | 0 | 1 | 9 |
| cyhr1 | 0.16947 | 0.10542986 | 0.069 | 0 | 1 | 9 |
| cyp2n13 | 0.16947 | 0.114606842 | 0.069 | 0 | 1 | 9 |
| cyth1b | 0.16947 | 0.148666294 | 0.069 | 0 | 1 | 9 |
| cyyr1 | 0.16947 | 0.134184211 | 0.069 | 0 | 1 | 9 |
| daxx | 0.16947 | 0.141928602 | 0.069 | 0 | 1 | 9 |
| dclre1b | 0.16947 | 0.107051285 | 0.069 | 0 | 1 | 9 |
| dcp2 | 0.16947 | 0.138797721 | 0.069 | 0 | 1 | 9 |
| dcun1d2a | 0.16947 | 0.121415933 | 0.069 | 0 | 1 | 9 |
| ddx49 | 0.16947 | 0.107253541 | 0.069 | 0 | 1 | 9 |
| depdc7a | 0.16947 | 0.126913783 | 0.069 | 0 | 1 | 9 |
| dhfr | 0.16947 | 0.132815106 | 0.069 | 0 | 1 | 9 |

| | | | | | | |
|---|---|---|---|---|---|---|
| diaph2 | 0.16947 | 0.117484606 | 0.069 | 0 | 1 | 9 |
| dla | 0.16947 | 0.113276343 | 0.069 | 0 | 1 | 9 |
| dll4 | 0.16947 | 0.214229842 | 0.069 | 0 | 1 | 9 |
| dnajb12b | 0.16947 | 0.128631952 | 0.069 | 0 | 1 | 9 |
| dnajc2 | 0.16947 | 0.102105178 | 0.069 | 0 | 1 | 9 |
| dpcd | 0.16947 | 0.113958634 | 0.069 | 0 | 1 | 9 |
| dpf2 | 0.16947 | 0.110656402 | 0.069 | 0 | 1 | 9 |
| dus3l | 0.16947 | 0.109988563 | 0.069 | 0 | 1 | 9 |
| e2f3 | 0.16947 | 0.119253111 | 0.069 | 0 | 1 | 9 |
| eef1akmt2 | 0.16947 | 0.148062902 | 0.069 | 0 | 1 | 9 |
| eif4ebp3 | 0.16947 | 0.14208266 | 0.069 | 0 | 1 | 9 |
| elp6 | 0.16947 | 0.107360478 | 0.069 | 0 | 1 | 9 |
| emc9 | 0.16947 | 0.120519281 | 0.069 | 0 | 1 | 9 |
| eme1 | 0.16947 | 0.127811853 | 0.069 | 0 | 1 | 9 |
| epb41a | 0.16947 | 0.138790567 | 0.069 | 0 | 1 | 9 |
| etv5b | 0.16947 | 0.129613333 | 0.069 | 0 | 1 | 9 |
| ext1a | 0.16947 | 0.108198741 | 0.069 | 0 | 1 | 9 |
| fam89b | 0.16947 | 0.152924744 | 0.069 | 0 | 1 | 9 |
| fastkd1 | 0.16947 | 0.117765916 | 0.069 | 0 | 1 | 9 |
| fbxw11b | 0.16947 | 0.135403931 | 0.069 | 0 | 1 | 9 |
| fbxw4 | 0.16947 | 0.141063991 | 0.069 | 0 | 1 | 9 |
| fer | 0.16947 | 0.12621826 | 0.069 | 0 | 1 | 9 |
| flj11011l | 0.16947 | 0.110103187 | 0.069 | 0 | 1 | 9 |
| fn1b | 0.16947 | 0.159760653 | 0.069 | 0 | 1 | 9 |
| fopnl | 0.16947 | 0.121533766 | 0.069 | 0 | 1 | 9 |
| fryl | 0.16947 | 0.118810376 | 0.069 | 0 | 1 | 9 |
| gas2l1 | 0.16947 | 0.162670388 | 0.069 | 0 | 1 | 9 |
| gata3 | 0.16947 | 0.144133878 | 0.069 | 0 | 1 | 9 |
| gb:ai877918 | 0.16947 | 0.122392386 | 0.069 | 0 | 1 | 9 |
| gdpd5b | 0.16947 | 0.10483226 | 0.069 | 0 | 1 | 9 |
| gins4 | 0.16947 | 0.117337453 | 0.069 | 0 | 1 | 9 |
| gnsa | 0.16947 | 0.118255353 | 0.069 | 0 | 1 | 9 |
| gpkow | 0.16947 | 0.10153446 | 0.069 | 0 | 1 | 9 |
| grpel1 | 0.16947 | 0.111829211 | 0.069 | 0 | 1 | 9 |
| gtf2h3 | 0.16947 | 0.107579799 | 0.069 | 0 | 1 | 9 |
| hdx | 0.16947 | 0.135942423 | 0.069 | 0 | 1 | 9 |
| hoxb7a | 0.16947 | 0.11058594 | 0.069 | 0 | 1 | 9 |
| hs6st1b | 0.16947 | 0.134443045 | 0.069 | 0 | 1 | 9 |
| hsf1 | 0.16947 | 0.123669645 | 0.069 | 0 | 1 | 9 |
| hvcn1 | 0.16947 | 0.14368456 | 0.069 | 0 | 1 | 9 |
| ift52 | 0.16947 | 0.118550901 | 0.069 | 0 | 1 | 9 |
| ikzf5 | 0.16947 | 0.136401221 | 0.069 | 0 | 1 | 9 |
| ilf3a | 0.16947 | 0.123011951 | 0.069 | 0 | 1 | 9 |
| ilkap | 0.16947 | 0.13051281 | 0.069 | 0 | 1 | 9 |
| ino80 | 0.16947 | 0.118050954 | 0.069 | 0 | 1 | 9 |
| inpp5ka | 0.16947 | 0.112928185 | 0.069 | 0 | 1 | 9 |
| insm1a | 0.16947 | 0.243658018 | 0.069 | 0 | 1 | 9 |
| insm1b | 0.16947 | 0.143671843 | 0.069 | 0 | 1 | 9 |

| | | | | | | |
|---|---|---|---|---|---|---|
| isl1 | 0.16947 | 0.183018892 | 0.069 | 0 | 1 | 9 |
| ispd | 0.16947 | 0.134931941 | 0.069 | 0 | 1 | 9 |
| jam3b | 0.16947 | 0.12442704 | 0.069 | 0 | 1 | 9 |
| kank1a | 0.16947 | 0.137443488 | 0.069 | 0 | 1 | 9 |
| kdm2ab | 0.16947 | 0.109031401 | 0.069 | 0 | 1 | 9 |
| kif22 | 0.16947 | 0.154650111 | 0.069 | 0 | 1 | 9 |
| klf5l | 0.16947 | 0.12844889 | 0.069 | 0 | 1 | 9 |
| lactb | 0.16947 | 0.103472574 | 0.069 | 0 | 1 | 9 |
| lap3 | 0.16947 | 0.145886656 | 0.069 | 0 | 1 | 9 |
| lbh | 0.16947 | 0.101219571 | 0.069 | 0 | 1 | 9 |
| ldb1a | 0.16947 | 0.101162235 | 0.069 | 0 | 1 | 9 |
| lhfpl2a | 0.16947 | 0.116304818 | 0.069 | 0 | 1 | 9 |
| lmtk2 | 0.16947 | 0.153434299 | 0.069 | 0 | 1 | 9 |
| lrig1 | 0.16947 | 0.104120385 | 0.069 | 0 | 1 | 9 |
| ltv1 | 0.16947 | 0.116272065 | 0.069 | 0 | 1 | 9 |
| lzts2a | 0.16947 | 0.136629906 | 0.069 | 0 | 1 | 9 |
| mad1l1 | 0.16947 | 0.101817866 | 0.069 | 0 | 1 | 9 |
| mbtd1 | 0.16947 | 0.157814919 | 0.069 | 0 | 1 | 9 |
| med14 | 0.16947 | 0.105388948 | 0.069 | 0 | 1 | 9 |
| med27 | 0.16947 | 0.133843281 | 0.069 | 0 | 1 | 9 |
| mef2cb | 0.16947 | 0.105835641 | 0.069 | 0 | 1 | 9 |
| mgmt | 0.16947 | 0.147369544 | 0.069 | 0 | 1 | 9 |
| mid1ip1l | 0.16947 | 0.103903899 | 0.069 | 0 | 1 | 9 |
| mpp6a | 0.16947 | 0.10827553 | 0.069 | 0 | 1 | 9 |
| mpped2 | 0.16947 | 0.186352684 | 0.069 | 0 | 1 | 9 |
| mrpl21 | 0.16947 | 0.118751949 | 0.069 | 0 | 1 | 9 |
| mrps9 | 0.16947 | 0.10031562 | 0.069 | 0 | 1 | 9 |
| ms4a17a.14 | 0.16947 | 0.101425002 | 0.069 | 0 | 1 | 9 |
| ms4a17a.2 | 0.16947 | 0.107565164 | 0.069 | 0 | 1 | 9 |
| mxi1 | 0.16947 | 0.139571724 | 0.069 | 0 | 1 | 9 |
| myo10 | 0.16947 | 0.117483906 | 0.069 | 0 | 1 | 9 |
| naa30 | 0.16947 | 0.126071031 | 0.069 | 0 | 1 | 9 |
| nat16 | 0.16947 | 0.138485824 | 0.069 | 0 | 1 | 9 |
| ndufaf4 | 0.16947 | 0.11089407 | 0.069 | 0 | 1 | 9 |
| nf1b | 0.16947 | 0.105666286 | 0.069 | 0 | 1 | 9 |
| nfatc3a | 0.16947 | 0.137449923 | 0.069 | 0 | 1 | 9 |
| nle1 | 0.16947 | 0.106379265 | 0.069 | 0 | 1 | 9 |
| nlrc3l1 | 0.16947 | 0.11499424 | 0.069 | 0 | 1 | 9 |
| noc2l | 0.16947 | 0.124750582 | 0.069 | 0 | 1 | 9 |
| nol4lb | 0.16947 | 0.125350714 | 0.069 | 0 | 1 | 9 |
| nop14 | 0.16947 | 0.127179705 | 0.069 | 0 | 1 | 9 |
| nr1h3 | 0.16947 | 0.167896564 | 0.069 | 0 | 1 | 9 |
| nr2c2ap | 0.16947 | 0.103189932 | 0.069 | 0 | 1 | 9 |
| nr2f1a | 0.16947 | 0.165891938 | 0.069 | 0 | 1 | 9 |
| nsrp1 | 0.16947 | 0.114060756 | 0.069 | 0 | 1 | 9 |
| nt5c2a | 0.16947 | 0.111736534 | 0.069 | 0 | 1 | 9 |
| nvl | 0.16947 | 0.125402782 | 0.069 | 0 | 1 | 9 |
| osbp | 0.16947 | 0.133145066 | 0.069 | 0 | 1 | 9 |

| | | | | | | |
|---|---|---|---|---|---|---|
| otud6b | 0.16947 | 0.106228666 | 0.069 | 0 | 1 | 9 |
| pbx1a | 0.16947 | 0.122112199 | 0.069 | 0 | 1 | 9 |
| pdgfd | 0.16947 | 0.198823787 | 0.069 | 0 | 1 | 9 |
| pdgfrl | 0.16947 | 0.127589818 | 0.069 | 0 | 1 | 9 |
| pds5b | 0.16947 | 0.129247553 | 0.069 | 0 | 1 | 9 |
| pet100 | 0.16947 | 0.144748157 | 0.069 | 0 | 1 | 9 |
| pex19 | 0.16947 | 0.135118374 | 0.069 | 0 | 1 | 9 |
| phb2a | 0.16947 | 0.131156839 | 0.069 | 0 | 1 | 9 |
| phf21ab | 0.16947 | 0.14686113 | 0.069 | 0 | 1 | 9 |
| pigh | 0.16947 | 0.175311655 | 0.069 | 0 | 1 | 9 |
| pik3c2a | 0.16947 | 0.10634338 | 0.069 | 0 | 1 | 9 |
| pkp3b | 0.16947 | 0.122213265 | 0.069 | 0 | 1 | 9 |
| plxnb2a | 0.16947 | 0.151875474 | 0.069 | 0 | 1 | 9 |
| poldip3 | 0.16947 | 0.132575608 | 0.069 | 0 | 1 | 9 |
| pop4 | 0.16947 | 0.113100856 | 0.069 | 0 | 1 | 9 |
| ppat | 0.16947 | 0.126853064 | 0.069 | 0 | 1 | 9 |
| ppp2r3a | 0.16947 | 0.129759523 | 0.069 | 0 | 1 | 9 |
| ppp2r5cb | 0.16947 | 0.115701933 | 0.069 | 0 | 1 | 9 |
| psme4b | 0.16947 | 0.110811906 | 0.069 | 0 | 1 | 9 |
| ptar1 | 0.16947 | 0.128056993 | 0.069 | 0 | 1 | 9 |
| ptpa | 0.16947 | 0.151893657 | 0.069 | 0 | 1 | 9 |
| ptrh2 | 0.16947 | 0.117035555 | 0.069 | 0 | 1 | 9 |
| pus7 | 0.16947 | 0.115893785 | 0.069 | 0 | 1 | 9 |
| pxk | 0.16947 | 0.127436244 | 0.069 | 0 | 1 | 9 |
| qki2 | 0.16947 | 0.173298849 | 0.069 | 0 | 1 | 9 |
| qser1 | 0.16947 | 0.130477802 | 0.069 | 0 | 1 | 9 |
| rab24 | 0.16947 | 0.10706984 | 0.069 | 0 | 1 | 9 |
| ralba | 0.16947 | 0.132301246 | 0.069 | 0 | 1 | 9 |
| rbck1 | 0.16947 | 0.119677073 | 0.069 | 0 | 1 | 9 |
| rdh10b | 0.16947 | 0.122099068 | 0.069 | 0 | 1 | 9 |
| recql | 0.16947 | 0.133815903 | 0.069 | 0 | 1 | 9 |
| rhogc | 0.16947 | 0.112466684 | 0.069 | 0 | 1 | 9 |
| riox1 | 0.16947 | 0.162975733 | 0.069 | 0 | 1 | 9 |
| rnaseh2b | 0.16947 | 0.110307868 | 0.069 | 0 | 1 | 9 |
| rps6kal | 0.16947 | 0.100484235 | 0.069 | 0 | 1 | 9 |
| rrp12 | 0.16947 | 0.124465906 | 0.069 | 0 | 1 | 9 |
| samm50 | 0.16947 | 0.104587042 | 0.069 | 0 | 1 | 9 |
| sat1b | 0.16947 | 0.128605923 | 0.069 | 0 | 1 | 9 |
| scap | 0.16947 | 0.12365454 | 0.069 | 0 | 1 | 9 |
| scn4ab | 0.16947 | 0.281036559 | 0.069 | 0 | 1 | 9 |
| sdad1 | 0.16947 | 0.11356796 | 0.069 | 0 | 1 | 9 |
| sertad2a | 0.16947 | 0.14415544 | 0.069 | 0 | 1 | 9 |
| si:ch1073-322p19.1 | 0.16947 | 0.11438603 | 0.069 | 0 | 1 | 9 |
| si:ch211-161h7.4 | 0.16947 | 0.138005397 | 0.069 | 0 | 1 | 9 |
| si:ch211-212c13.10 | 0.16947 | 0.134547435 | 0.069 | 0 | 1 | 9 |
| si:ch211-229n2.6 | 0.16947 | 0.131541304 | 0.069 | 0 | 1 | 9 |
| si:ch211-239j9.1 | 0.16947 | 0.118866836 | 0.069 | 0 | 1 | 9 |
| si:ch211-275j6.5 | 0.16947 | 0.117233551 | 0.069 | 0 | 1 | 9 |

| | | | | | | |
|---|---|---|---|---|---|---|
| si:ch211-282j17.13 | 0.16947 | 0.112148226 | 0.069 | 0 | 1 | 9 |
| si:ch211-282j22.3 | 0.16947 | 0.116837469 | 0.069 | 0 | 1 | 9 |
| si:ch211-51c14.1 | 0.16947 | 0.115308716 | 0.069 | 0 | 1 | 9 |
| si:ch73-338o16.4 | 0.16947 | 0.13053391 | 0.069 | 0 | 1 | 9 |
| si:ch73-367f21.4 | 0.16947 | 0.139800832 | 0.069 | 0 | 1 | 9 |
| si:dkey-122a22.2 | 0.16947 | 0.129140381 | 0.069 | 0 | 1 | 9 |
| si:dkey-151g10.3 | 0.16947 | 0.101766597 | 0.069 | 0 | 1 | 9 |
| si:dkey-159f12.2 | 0.16947 | 0.135160621 | 0.069 | 0 | 1 | 9 |
| si:dkey-17o15.2 | 0.16947 | 0.100398234 | 0.069 | 0 | 1 | 9 |
| si:dkey-229e3.2 | 0.16947 | 0.137687054 | 0.069 | 0 | 1 | 9 |
| si:dkey-31f5.8 | 0.16947 | 0.110922408 | 0.069 | 0 | 1 | 9 |
| si:dkey-75a21.2 | 0.16947 | 0.131641253 | 0.069 | 0 | 1 | 9 |
| si:dkey-77f5.3 | 0.16947 | 0.122766727 | 0.069 | 0 | 1 | 9 |
| si:dkey-89b17.4 | 0.16947 | 0.146287061 | 0.069 | 0 | 1 | 9 |
| si:dkeyp-69e1.8 | 0.16947 | 0.139083615 | 0.069 | 0 | 1 | 9 |
| si:dkeyp-75h12.5 | 0.16947 | 0.11566295 | 0.069 | 0 | 1 | 9 |
| si:rp71-19m20.1 | 0.16947 | 0.125129631 | 0.069 | 0 | 1 | 9 |
| sil1 | 0.16947 | 0.104761382 | 0.069 | 0 | 1 | 9 |
| slc25a16 | 0.16947 | 0.138365475 | 0.069 | 0 | 1 | 9 |
| slc35b3 | 0.16947 | 0.131555655 | 0.069 | 0 | 1 | 9 |
| slc6a7 | 0.16947 | 0.127832094 | 0.069 | 0 | 1 | 9 |
| smad3a | 0.16947 | 0.134066625 | 0.069 | 0 | 1 | 9 |
| smarca5 | 0.16947 | 0.159670264 | 0.069 | 0 | 1 | 9 |
| smarcd2 | 0.16947 | 0.121326662 | 0.069 | 0 | 1 | 9 |
| snapin | 0.16947 | 0.14170206 | 0.069 | 0 | 1 | 9 |
| snf8 | 0.16947 | 0.102823791 | 0.069 | 0 | 1 | 9 |
| socs6b | 0.16947 | 0.12692806 | 0.069 | 0 | 1 | 9 |
| sparta | 0.16947 | 0.126658928 | 0.069 | 0 | 1 | 9 |
| sphk2 | 0.16947 | 0.11688381 | 0.069 | 0 | 1 | 9 |
| spty2d1 | 0.16947 | 0.137505954 | 0.069 | 0 | 1 | 9 |
| srbd1 | 0.16947 | 0.113498933 | 0.069 | 0 | 1 | 9 |
| srfbp1 | 0.16947 | 0.115179184 | 0.069 | 0 | 1 | 9 |
| srpk1b | 0.16947 | 0.111811129 | 0.069 | 0 | 1 | 9 |
| srsf2b | 0.16947 | 0.152816102 | 0.069 | 0 | 1 | 9 |
| ssna1 | 0.16947 | 0.119033344 | 0.069 | 0 | 1 | 9 |
| ssuh2.4 | 0.16947 | 0.143970568 | 0.069 | 0 | 1 | 9 |
| st3gal3a | 0.16947 | 0.136835806 | 0.069 | 0 | 1 | 9 |
| stam | 0.16947 | 0.133796868 | 0.069 | 0 | 1 | 9 |
| stard3 | 0.16947 | 0.121324585 | 0.069 | 0 | 1 | 9 |
| stat5a | 0.16947 | 0.140132926 | 0.069 | 0 | 1 | 9 |
| stk38l | 0.16947 | 0.146307172 | 0.069 | 0 | 1 | 9 |
| svild | 0.16947 | 0.158082771 | 0.069 | 0 | 1 | 9 |
| sypl2b | 0.16947 | 0.10049222 | 0.069 | 0 | 1 | 9 |
| tab3 | 0.16947 | 0.116368833 | 0.069 | 0 | 1 | 9 |
| taf10 | 0.16947 | 0.156052687 | 0.069 | 0 | 1 | 9 |
| taf6 | 0.16947 | 0.138023049 | 0.069 | 0 | 1 | 9 |
| tbc1d14 | 0.16947 | 0.107497077 | 0.069 | 0 | 1 | 9 |
| tcf4 | 0.16947 | 0.140046845 | 0.069 | 0 | 1 | 9 |

| | | | | | | |
|---|---|---|---|---|---|---|
| tgfbr3 | 0.16947 | 0.137772883 | 0.069 | 0 | 1 | 9 |
| thoc3 | 0.16947 | 0.136628913 | 0.069 | 0 | 1 | 9 |
| thraa | 0.16947 | 0.119132727 | 0.069 | 0 | 1 | 9 |
| tmem223 | 0.16947 | 0.125863848 | 0.069 | 0 | 1 | 9 |
| tmsb2 | 0.16947 | 0.188276482 | 0.069 | 0 | 1 | 9 |
| tns3.2 | 0.16947 | 0.159097691 | 0.069 | 0 | 1 | 9 |
| tp53inp2 | 0.16947 | 0.118518893 | 0.069 | 0 | 1 | 9 |
| tra2b | 0.16947 | 0.121246192 | 0.069 | 0 | 1 | 9 |
| trak1a | 0.16947 | 0.150289183 | 0.069 | 0 | 1 | 9 |
| trim8b | 0.16947 | 0.108557526 | 0.069 | 0 | 1 | 9 |
| tspan18b | 0.16947 | 0.111572305 | 0.069 | 0 | 1 | 9 |
| ttc9c | 0.16947 | 0.125249022 | 0.069 | 0 | 1 | 9 |
| ttll4 | 0.16947 | 0.122965541 | 0.069 | 0 | 1 | 9 |
| ube2c | 0.16947 | 0.140001121 | 0.069 | 0 | 1 | 9 |
| ube3c | 0.16947 | 0.138872349 | 0.069 | 0 | 1 | 9 |
| uchl1 | 0.16947 | 0.135722595 | 0.069 | 0 | 1 | 9 |
| uchl5 | 0.16947 | 0.106134887 | 0.069 | 0 | 1 | 9 |
| usf1 | 0.16947 | 0.115543667 | 0.069 | 0 | 1 | 9 |
| usp39 | 0.16947 | 0.102838645 | 0.069 | 0 | 1 | 9 |
| usp44 | 0.16947 | 0.102683073 | 0.069 | 0 | 1 | 9 |
| vrk1 | 0.16947 | 0.107439347 | 0.069 | 0 | 1 | 9 |
| wdr13 | 0.16947 | 0.132506566 | 0.069 | 0 | 1 | 9 |
| wdr18 | 0.16947 | 0.135276033 | 0.069 | 0 | 1 | 9 |
| wdr26b | 0.16947 | 0.110114817 | 0.069 | 0 | 1 | 9 |
| wdr45 | 0.16947 | 0.130715522 | 0.069 | 0 | 1 | 9 |
| wnk1a | 0.16947 | 0.147911305 | 0.069 | 0 | 1 | 9 |
| wu:fa19b12 | 0.16947 | 0.105472504 | 0.069 | 0 | 1 | 9 |
| wu:fb74b10 | 0.16947 | 0.103756519 | 0.069 | 0 | 1 | 9 |
| xpnpep1 | 0.16947 | 0.113358809 | 0.069 | 0 | 1 | 9 |
| xpr1b | 0.16947 | 0.121144654 | 0.069 | 0 | 1 | 9 |
| yif1b | 0.16947 | 0.144520461 | 0.069 | 0 | 1 | 9 |
| zdhhc24 | 0.16947 | 0.113667286 | 0.069 | 0 | 1 | 9 |
| zdhhc9 | 0.16947 | 0.137157919 | 0.069 | 0 | 1 | 9 |
| zeb2b | 0.16947 | 0.138908969 | 0.069 | 0 | 1 | 9 |
| zgc:153215 | 0.16947 | 0.122377483 | 0.069 | 0 | 1 | 9 |
| zgc:153409 | 0.16947 | 0.117994475 | 0.069 | 0 | 1 | 9 |
| zgc:162396 | 0.16947 | 0.129358221 | 0.069 | 0 | 1 | 9 |
| zgc:55733 | 0.16947 | 0.137684011 | 0.069 | 0 | 1 | 9 |
| zgc:63882 | 0.16947 | 0.11331496 | 0.069 | 0 | 1 | 9 |
| zgc:65894 | 0.16947 | 0.139852757 | 0.069 | 0 | 1 | 9 |
| zgc:92161 | 0.16947 | 0.126812315 | 0.069 | 0 | 1 | 9 |
| zmp:0000000881 | 0.16947 | 0.111614028 | 0.069 | 0 | 1 | 9 |
| znf1001 | 0.16947 | 0.113187763 | 0.069 | 0 | 1 | 9 |
| znf1024 | 0.16947 | 0.169625028 | 0.069 | 0 | 1 | 9 |
| znf280d | 0.16947 | 0.149211707 | 0.069 | 0 | 1 | 9 |
| znf367 | 0.16947 | 0.111218276 | 0.069 | 0 | 1 | 9 |
| znf598 | 0.16947 | 0.137974488 | 0.069 | 0 | 1 | 9 |
| cldn11b | 0.16947 | 0.123619466 | 0.069 | 0 | 1 | 9 |

| | | | | | | |
|---|---|---|---|---|---|---|
| cnot3b | 0.16947 | 0.141353178 | 0.069 | 0 | 1 | 9 |
| fzd8a | 0.16947 | 0.129608026 | 0.069 | 0 | 1 | 9 |
| msrb3 | 0.16947 | 0.113474027 | 0.069 | 0 | 1 | 9 |
| myd88 | 0.16947 | 0.125232182 | 0.069 | 0 | 1 | 9 |
| bcl9 | 0.16947 | 0.1319468 | 0.069 | 0 | 1 | 9 |
| ezh2 | 0.16947 | 0.130509352 | 0.069 | 0 | 1 | 9 |
| fam122b | 0.16947 | 0.14575482 | 0.069 | 0 | 1 | 9 |
| si:ch211-231f6.6 | 0.16947 | 0.153084389 | 0.069 | 0 | 1 | 9 |
| si:dkey-234h16.7 | 0.16947 | 0.138586962 | 0.069 | 0 | 1 | 9 |
| tent5bb | 0.16947 | 0.152204193 | 0.069 | 0 | 1 | 9 |
| zgc:64106 | 0.16947 | 0.104846153 | 0.069 | 0 | 1 | 9 |
| as3mt | 0.16947 | 0.121310551 | 0.069 | 0 | 1 | 9 |
| BX957362.1 | 0.16947 | 0.130153351 | 0.069 | 0 | 1 | 9 |
| ankrd13a | 0.16947 | 0.133229771 | 0.069 | 0 | 1 | 9 |
| asb7 | 0.16947 | 0.150836444 | 0.069 | 0 | 1 | 9 |
| ift57 | 0.16947 | 0.159212715 | 0.069 | 0 | 1 | 9 |
| CU856344.1 | 0.16947 | 0.130797923 | 0.069 | 0 | 1 | 9 |
| c1galt1c1 | 0.16947 | 0.129484747 | 0.069 | 0 | 1 | 9 |
| ccdc82 | 0.16947 | 0.152027003 | 0.069 | 0 | 1 | 9 |
| cilp | 0.16947 | 0.135794891 | 0.069 | 0 | 1 | 9 |
| cyb5b | 0.16947 | 0.142145958 | 0.069 | 0 | 1 | 9 |
| fam198b | 0.16947 | 0.183349266 | 0.069 | 0 | 1 | 9 |
| rpp40 | 0.16947 | 0.156209046 | 0.069 | 0 | 1 | 9 |
| sorbs1 | 0.16947 | 0.181590048 | 0.069 | 0 | 1 | 9 |
| tmem176 | 0.16947 | 0.206625509 | 0.069 | 0 | 1 | 9 |
| tril | 0.16947 | 0.171970689 | 0.069 | 0 | 1 | 9 |
| CU467961.1 | 0.16947 | 0.1243709 | 0.069 | 0 | 1 | 9 |
| LO018188.1 | 0.16947 | 0.186786105 | 0.069 | 0 | 1 | 9 |
| arl3 | 0.16947 | 0.123903017 | 0.069 | 0 | 1 | 9 |
| cdr2l | 0.16947 | 0.144541154 | 0.069 | 0 | 1 | 9 |
| ctbp2a | 0.16947 | 0.14381036 | 0.069 | 0 | 1 | 9 |
| fam208b | 0.16947 | 0.12681176 | 0.069 | 0 | 1 | 9 |
| itm2ca | 0.16947 | 0.121263533 | 0.069 | 0 | 1 | 9 |
| morc2 | 0.16947 | 0.116200726 | 0.069 | 0 | 1 | 9 |
| pcbp4 | 0.16947 | 0.137518405 | 0.069 | 0 | 1 | 9 |
| phkg2 | 0.16947 | 0.133992258 | 0.069 | 0 | 1 | 9 |
| slc25a10 | 0.16947 | 0.118495176 | 0.069 | 0 | 1 | 9 |
| srgap2 | 0.16947 | 0.168429369 | 0.069 | 0 | 1 | 9 |
| tspan7b | 0.16947 | 0.124768649 | 0.069 | 0 | 1 | 9 |
| ucp3 | 0.16947 | 0.191397843 | 0.069 | 0 | 1 | 9 |
| rbm25a | 0.16947 | 0.154775745 | 0.069 | 0 | 1 | 9 |
| zdhhc20a | 0.16947 | 0.176115835 | 0.069 | 0 | 1 | 9 |
| timm10b | 0.16947 | 0.138250488 | 0.069 | 0 | 1 | 9 |
| rps11 | 0.16959 | 0.210358716 | 0.931 | 0.85 | 1 | 9 |
| si:ch1073-429i10.3.1 | 0.16969 | 0.285463802 | 0.638 | 0.4 | 1 | 9 |
| cldnf | 0.16985 | -0.22790663 | 0.155 | 0.2 | 1 | 9 |
| sqstm1 | 0.1704 | -0.39095881 | 0.207 | 0.35 | 1 | 9 |
| rbm28 | 0.1706 | 0.357118519 | 0.293 | 0.1 | 1 | 9 |

| | | | | | | |
|---|---|---|---|---|---|---|
| copz1 | 0.17071 | 0.318157943 | 0.362 | 0.15 | 1 | 9 |
| ppt1 | 0.17079 | -0.28746308 | 0.362 | 0.45 | 1 | 9 |
| ogdha | 0.17088 | -0.18133624 | 0.172 | 0.2 | 1 | 9 |
| qkia | 0.17095 | 0.434932161 | 0.552 | 0.35 | 1 | 9 |
| wdr1 | 0.17214 | -0.35860707 | 0.207 | 0.4 | 1 | 9 |
| bola1 | 0.17232 | -0.19706126 | 0.121 | 0.15 | 1 | 9 |
| sqor | 0.17254 | -0.25229884 | 0.034 | 0.15 | 1 | 9 |
| mprip | 0.17273 | -0.2274061 | 0.052 | 0.2 | 1 | 9 |
| wbp1la | 0.17274 | -0.25026928 | 0.034 | 0.1 | 1 | 9 |
| gcsha | 0.17316 | 0.217462527 | 0.207 | 0.05 | 1 | 9 |
| LOC103909616 | 0.17419 | -0.35362879 | 0.034 | 0.15 | 1 | 9 |
| nop58 | 0.17442 | 0.28673807 | 0.345 | 0.15 | 1 | 9 |
| shq1 | 0.17557 | -0.29061987 | 0.034 | 0.15 | 1 | 9 |
| LOC103910438 | 0.17562 | 0.33802042 | 0.276 | 0.1 | 1 | 9 |
| lyrm4 | 0.17575 | -0.22656155 | 0.293 | 0.3 | 1 | 9 |
| slx1b | 0.17608 | 0.24992622 | 0.207 | 0.05 | 1 | 9 |
| si:ch211-213d14.2 | 0.17631 | -0.31822083 | 0.052 | 0.2 | 1 | 9 |
| fam76b | 0.17635 | -0.1339115 | 0.103 | 0.1 | 1 | 9 |
| zgc:113142 | 0.17642 | -0.44794897 | 0.052 | 0.2 | 1 | 9 |
| rrp8 | 0.17648 | 0.192054457 | 0.19 | 0.05 | 1 | 9 |
| dnajb11 | 0.177 | -0.21548238 | 0.155 | 0.2 | 1 | 9 |
| chmp1a | 0.1773 | 0.146979192 | 0.172 | 0.05 | 1 | 9 |
| rpl36 | 0.17736 | 0.178000654 | 0.948 | 0.8 | 1 | 9 |
| ecm1b | 0.17748 | 0.157403306 | 0.155 | 0.05 | 1 | 9 |
| sgk1 | 0.17767 | -0.36224574 | 0.155 | 0.3 | 1 | 9 |
| marcksl1a | 0.17774 | 0.445748214 | 0.414 | 0.25 | 1 | 9 |
| zgc:136864 | 0.17811 | -0.26005832 | 0.034 | 0.15 | 1 | 9 |
| ifi46 | 0.17837 | -0.30162355 | 0.069 | 0.2 | 1 | 9 |
| srsf3b | 0.17854 | 0.329026825 | 0.483 | 0.25 | 1 | 9 |
| si:ch211-15b10.6 | 0.17892 | -0.29376117 | 0.052 | 0.2 | 1 | 9 |
| psmd1 | 0.17897 | 0.167583881 | 0.379 | 0.2 | 1 | 9 |
| mt-nd5 | 0.17943 | 0.286144693 | 0.362 | 0.15 | 1 | 9 |
| rpl10a | 0.17947 | 0.245218755 | 0.931 | 0.8 | 1 | 9 |
| gdpd3a | 0.17962 | -0.38863707 | 0.103 | 0.25 | 1 | 9 |
| calm3b | 0.17983 | 0.137697665 | 0.638 | 0.45 | 1 | 9 |
| mpv17l2 | 0.17994 | -0.25039024 | 0.034 | 0.15 | 1 | 9 |
| sf3b5 | 0.18043 | 0.364978642 | 0.345 | 0.15 | 1 | 9 |
| rps5 | 0.18049 | 0.24873776 | 0.914 | 0.75 | 1 | 9 |
| lsm12b | 0.18059 | 0.217597353 | 0.569 | 0.35 | 1 | 9 |
| sec62 | 0.18061 | 0.137868976 | 0.172 | 0.05 | 1 | 9 |
| foxa2 | 0.18114 | -0.28085423 | 0.034 | 0.1 | 1 | 9 |
| hbs1l | 0.18143 | -0.28606335 | 0.103 | 0.2 | 1 | 9 |
| pfn2l | 0.18154 | 0.289607833 | 0.414 | 0.2 | 1 | 9 |
| LOC100332329 | 0.1824 | -0.28585394 | 0.069 | 0.2 | 1 | 9 |
| capza1b | 0.18272 | -0.2258655 | 0.31 | 0.35 | 1 | 9 |
| btg2 | 0.18283 | 0.259214598 | 0.517 | 0.3 | 1 | 9 |
| ppdpfb | 0.18331 | 0.35293959 | 0.759 | 0.6 | 1 | 9 |
| sat1a.2 | 0.18396 | -0.18016693 | 0.276 | 0.3 | 1 | 9 |

| | | | | | | |
|---|---|---|---|---|---|---|
| psma6a | 0.18463 | 0.134692794 | 0.293 | 0.15 | 1 | 9 |
| rpl6 | 0.18476 | 0.234166488 | 0.931 | 0.8 | 1 | 9 |
| rps2 | 0.18497 | 0.216833236 | 0.948 | 0.85 | 1 | 9 |
| BX284696.1 | 0.18539 | 0.190704869 | 0.31 | 0.15 | 1 | 9 |
| cald1a | 0.18581 | -0.38286924 | 0.121 | 0.3 | 1 | 9 |
| si:ch211-286b5.5 | 0.18633 | 0.167614134 | 0.483 | 0.3 | 1 | 9 |
| ap2s1 | 0.18653 | -0.24167404 | 0.207 | 0.25 | 1 | 9 |
| actb1 | 0.18669 | -0.36069475 | 0.621 | 0.8 | 1 | 9 |
| laptm4b | 0.18674 | -0.30758731 | 0.069 | 0.15 | 1 | 9 |
| FAR2 | 0.18725 | -0.25325096 | 0.069 | 0.1 | 1 | 9 |
| puf60a | 0.18792 | -0.27217993 | 0.172 | 0.25 | 1 | 9 |
| srsf1a | 0.18831 | 0.282292285 | 0.19 | 0.05 | 1 | 9 |
| sgta | 0.18864 | 0.255702372 | 0.207 | 0.05 | 1 | 9 |
| papss2b | 0.18868 | 0.265804081 | 0.172 | 0.05 | 1 | 9 |
| magoh | 0.18904 | 0.116901992 | 0.276 | 0.15 | 1 | 9 |
| fam49a | 0.18977 | -0.31239249 | 0.155 | 0.25 | 1 | 9 |
| ak1 | 0.19029 | -0.36297708 | 0.397 | 0.5 | 1 | 9 |
| eif4h | 0.19034 | -0.25376871 | 0.121 | 0.3 | 1 | 9 |
| cd81a | 0.19035 | 0.254928961 | 0.397 | 0.2 | 1 | 9 |
| hp1bp3 | 0.19038 | 0.363902726 | 0.414 | 0.2 | 1 | 9 |
| ms4a17a.5 | 0.19064 | 0.276140391 | 0.155 | 0.05 | 1 | 9 |
| gstm.3 | 0.19103 | 0.397047188 | 0.155 | 0.05 | 1 | 9 |
| mdh1aa | 0.19117 | 0.393019359 | 0.466 | 0.25 | 1 | 9 |
| rps20 | 0.19134 | 0.187210325 | 0.897 | 0.85 | 1 | 9 |
| rps8a | 0.19164 | 0.17200085 | 0.879 | 0.85 | 1 | 9 |
| hmgb3a | 0.19165 | 0.124579943 | 0.466 | 0.3 | 1 | 9 |
| sub1a | 0.19208 | 0.122594136 | 0.586 | 0.4 | 1 | 9 |
| col18a1b | 0.19253 | 0.143729413 | 0.155 | 0.05 | 1 | 9 |
| clk2a | 0.19253 | -0.24872047 | 0.034 | 0.15 | 1 | 9 |
| pdcd5 | 0.19259 | 0.325988849 | 0.345 | 0.15 | 1 | 9 |
| plekhf1 | 0.19272 | -0.28756624 | 0.034 | 0.15 | 1 | 9 |
| RASSF5 | 0.19283 | -0.31743347 | 0.052 | 0.15 | 1 | 9 |
| nme2b.1 | 0.19315 | 0.257518428 | 0.897 | 0.8 | 1 | 9 |
| taf9 | 0.1936 | 0.101415829 | 0.224 | 0.1 | 1 | 9 |
| zgc:158343 | 0.19363 | 0.178087808 | 0.241 | 0.1 | 1 | 9 |
| ufd1l | 0.19399 | 0.216344714 | 0.207 | 0.05 | 1 | 9 |
| mrpl19 | 0.19428 | 0.218607514 | 0.207 | 0.05 | 1 | 9 |
| tiparp | 0.19453 | 0.179736047 | 0.172 | 0.15 | 1 | 9 |
| mlec | 0.19483 | -0.20179978 | 0.121 | 0.15 | 1 | 9 |
| pttg1ipb | 0.19611 | -0.17613476 | 0.086 | 0.1 | 1 | 9 |
| eif3s10 | 0.19622 | 0.107864411 | 0.466 | 0.3 | 1 | 9 |
| mrps18c | 0.19625 | 0.13853385 | 0.172 | 0.05 | 1 | 9 |
| snrnp27 | 0.19644 | 0.258397169 | 0.207 | 0.05 | 1 | 9 |
| shmt2 | 0.19686 | -0.20803284 | 0.052 | 0.1 | 1 | 9 |
| pigf | 0.19696 | 0.193438642 | 0.19 | 0.05 | 1 | 9 |
| prss35 | 0.1977 | -0.13599934 | 0.052 | 0.05 | 1 | 9 |
| ptges3b | 0.19834 | 0.340450001 | 0.276 | 0.1 | 1 | 9 |
| il13ra1 | 0.19875 | -0.3037256 | 0.103 | 0.2 | 1 | 9 |

| | | | | | | |
|---|---|---|---|---|---|---|
| TMEM19 | 0.19884 | 0.125307013 | 0.172 | 0.05 | 1 | 9 |
| gcshb | 0.19894 | -0.34515843 | 0.103 | 0.2 | 1 | 9 |
| mbtps2 | 0.19933 | -0.27614567 | 0.052 | 0.15 | 1 | 9 |
| uqcrfs1 | 0.20036 | 0.346604174 | 0.397 | 0.2 | 1 | 9 |
| tnip1 | 0.20065 | -0.26417538 | 0.155 | 0.2 | 1 | 9 |
| nup62l | 0.20077 | 0.26474686 | 0.207 | 0.05 | 1 | 9 |
| si:ch73-335l21.4 | 0.20143 | 0.201264505 | 0.172 | 0.05 | 1 | 9 |
| cox8b | 0.20204 | -0.23573186 | 0.052 | 0.15 | 1 | 9 |
| atox1 | 0.20217 | -0.28669566 | 0.207 | 0.3 | 1 | 9 |
| adrb2a | 0.20218 | -0.29116367 | 0.034 | 0.1 | 1 | 9 |
| emc8 | 0.20253 | -0.21330076 | 0.103 | 0.15 | 1 | 9 |
| aars | 0.20254 | 0.234760323 | 0.19 | 0.05 | 1 | 9 |
| LOC103909284 | 0.20323 | 0.252273761 | 0.207 | 0.05 | 1 | 9 |
| zc3h7bb | 0.20341 | -0.26072779 | 0.086 | 0.25 | 1 | 9 |
| kdelr2a | 0.20345 | -0.36378687 | 0.103 | 0.25 | 1 | 9 |
| cep350 | 0.2036 | -0.2644332 | 0.052 | 0.1 | 1 | 9 |
| srp54 | 0.20385 | -0.29341948 | 0.121 | 0.2 | 1 | 9 |
| tob1b | 0.20385 | 0.375799419 | 0.345 | 0.2 | 1 | 9 |
| MED9 | 0.20417 | -0.311549 | 0.103 | 0.25 | 1 | 9 |
| brd1b | 0.20468 | -0.24721812 | 0.086 | 0.25 | 1 | 9 |
| fscn1a | 0.20471 | -0.20262757 | 0.121 | 0.15 | 1 | 9 |
| lrch3 | 0.20501 | -0.25338314 | 0.052 | 0.15 | 1 | 9 |
| ncl | 0.20518 | 0.37668365 | 0.534 | 0.35 | 1 | 9 |
| zgc:153867 | 0.2056 | -0.1682131 | 0.707 | 0.7 | 1 | 9 |
| knop1 | 0.20574 | 0.284220898 | 0.224 | 0.1 | 1 | 9 |
| tpm1 | 0.20593 | -0.34022893 | 0.103 | 0.2 | 1 | 9 |
| vapb | 0.20614 | 0.284793739 | 0.241 | 0.1 | 1 | 9 |
| mrpl27 | 0.20616 | 0.219467861 | 0.207 | 0.05 | 1 | 9 |
| cdca7a | 0.2062 | 0.276085054 | 0.207 | 0.05 | 1 | 9 |
| mdm2 | 0.20638 | -0.18303994 | 0.069 | 0.1 | 1 | 9 |
| si:ch211-68a17.7 | 0.20656 | 0.113996313 | 0.155 | 0.05 | 1 | 9 |
| zgc:92313 | 0.20661 | -0.17731196 | 0.276 | 0.3 | 1 | 9 |
| wsb1 | 0.20686 | -0.32272092 | 0.155 | 0.25 | 1 | 9 |
| rhbg | 0.20693 | 0.365054853 | 0.138 | 0.1 | 1 | 9 |
| tomm22 | 0.20739 | 0.274497555 | 0.241 | 0.1 | 1 | 9 |
| pbdc1 | 0.20821 | 0.325450984 | 0.345 | 0.15 | 1 | 9 |
| pias4a | 0.2083 | -0.18715232 | 0.121 | 0.15 | 1 | 9 |
| ctsd | 0.2083 | 0.162463063 | 0.31 | 0.15 | 1 | 9 |
| zbtb20 | 0.20835 | -0.32056806 | 0.121 | 0.3 | 1 | 9 |
| tusc3 | 0.20851 | -0.23990251 | 0.086 | 0.15 | 1 | 9 |
| cbr1l | 0.20856 | -0.31237949 | 0.103 | 0.25 | 1 | 9 |
| ctnnd1 | 0.20922 | -0.25354052 | 0.121 | 0.3 | 1 | 9 |
| tuft1a | 0.20954 | -0.32651846 | 0.069 | 0.2 | 1 | 9 |
| schip1 | 0.20994 | -0.18070683 | 0.034 | 0.15 | 1 | 9 |
| ube2v2 | 0.21061 | 0.326259549 | 0.552 | 0.35 | 1 | 9 |
| gsk3b | 0.21105 | -0.20077226 | 0.069 | 0.1 | 1 | 9 |
| vdac1 | 0.21123 | 0.288818819 | 0.241 | 0.1 | 1 | 9 |
| csk | 0.2115 | -0.33454123 | 0.103 | 0.25 | 1 | 9 |

| | | | | | | |
|---|---|---|---|---|---|---|
| atp5f1e | 0.21212 | 0.262037438 | 0.672 | 0.45 | 1 | 9 |
| pcbd1 | 0.21213 | 0.267762089 | 0.172 | 0.05 | 1 | 9 |
| pisd | 0.21229 | -0.29263728 | 0.069 | 0.2 | 1 | 9 |
| dctn5 | 0.2131 | -0.29114689 | 0.069 | 0.2 | 1 | 9 |
| ulk2 | 0.21322 | -0.28239446 | 0.121 | 0.2 | 1 | 9 |
| kdm1a | 0.21333 | 0.284743243 | 0.19 | 0.05 | 1 | 9 |
| CABZ01045617.1 | 0.21356 | 0.242649264 | 0.276 | 0.1 | 1 | 9 |
| camk2g2 | 0.21437 | -0.21479105 | 0.103 | 0.15 | 1 | 9 |
| zc3h15 | 0.21472 | 0.238343645 | 0.155 | 0.05 | 1 | 9 |
| ndufb9 | 0.21479 | 0.318709827 | 0.621 | 0.4 | 1 | 9 |
| si:dkey-102m7.3 | 0.21482 | 0.319896413 | 0.276 | 0.1 | 1 | 9 |
| fkbp5 | 0.21488 | -0.119729 | 0.328 | 0.3 | 1 | 9 |
| ndufb4 | 0.21671 | 0.323694195 | 0.466 | 0.25 | 1 | 9 |
| herpud1 | 0.21679 | -0.27284614 | 0.034 | 0.15 | 1 | 9 |
| zgc:153713 | 0.21705 | 0.296665431 | 0.379 | 0.2 | 1 | 9 |
| sdc4 | 0.21718 | -0.38496141 | 0.362 | 0.5 | 1 | 9 |
| tmem54b | 0.21747 | -0.25109343 | 0.034 | 0.15 | 1 | 9 |
| mta2 | 0.21806 | -0.2683928 | 0.103 | 0.2 | 1 | 9 |
| grpel2 | 0.21807 | 0.264183731 | 0.19 | 0.05 | 1 | 9 |
| c5 | 0.21831 | -0.26575936 | 0.034 | 0.15 | 1 | 9 |
| derl1 | 0.21842 | -0.31295448 | 0.052 | 0.15 | 1 | 9 |
| ppp2cb | 0.21886 | 0.121667689 | 0.517 | 0.35 | 1 | 9 |
| il17rc | 0.21891 | -0.28626508 | 0.034 | 0.15 | 1 | 9 |
| taf7 | 0.21931 | -0.32012757 | 0.155 | 0.3 | 1 | 9 |
| gna14 | 0.21944 | -0.28888198 | 0.052 | 0.15 | 1 | 9 |
| srrm2 | 0.2195 | 0.129381021 | 0.155 | 0.05 | 1 | 9 |
| nifk | 0.21962 | 0.285621157 | 0.19 | 0.05 | 1 | 9 |
| sec61a1 | 0.21997 | -0.33313645 | 0.241 | 0.4 | 1 | 9 |
| rbmx | 0.22035 | 0.271852886 | 0.345 | 0.15 | 1 | 9 |
| zmp:0000000906 | 0.22092 | 0.234460324 | 0.121 | 0.05 | 1 | 9 |
| si:dkey-208c12.2 | 0.22106 | -0.20014881 | 0.034 | 0.15 | 1 | 9 |
| ppp1cb | 0.22136 | 0.111062365 | 0.172 | 0.05 | 1 | 9 |
| fzd4 | 0.22163 | 0.221588595 | 0.19 | 0.05 | 1 | 9 |
| fxyd1 | 0.22217 | 0.132437473 | 0.397 | 0.25 | 1 | 9 |
| spcs3 | 0.2222 | 0.104624756 | 0.397 | 0.25 | 1 | 9 |
| socs3a | 0.22253 | -0.35224661 | 0.362 | 0.45 | 1 | 9 |
| yipf6 | 0.22307 | -0.35073177 | 0.121 | 0.25 | 1 | 9 |
| rab1ba | 0.22327 | -0.26265544 | 0.103 | 0.2 | 1 | 9 |
| atp1a1b | 0.22372 | -0.38521869 | 0.052 | 0.1 | 1 | 9 |
| fam3c | 0.22374 | 0.296640487 | 0.19 | 0.05 | 1 | 9 |
| soul2 | 0.22386 | 0.437571343 | 0.276 | 0.1 | 1 | 9 |
| cox6c | 0.224 | 0.355708415 | 0.603 | 0.4 | 1 | 9 |
| si:ch211-22i13.2 | 0.22412 | 0.209798915 | 0.276 | 0.1 | 1 | 9 |
| bcam | 0.22426 | 0.360382479 | 0.259 | 0.1 | 1 | 9 |
| ptena | 0.22439 | -0.17396767 | 0.241 | 0.25 | 1 | 9 |
| FO082781.1 | 0.22448 | 0.327567884 | 0.224 | 0.1 | 1 | 9 |
| esrp1 | 0.22471 | -0.21695202 | 0.034 | 0.15 | 1 | 9 |
| mcm6 | 0.22491 | 0.258461385 | 0.19 | 0.05 | 1 | 9 |

| | | | | | | |
|---|---|---|---|---|---|---|
| cltcb | 0.22507 | -0.29904872 | 0.086 | 0.2 | 1 | 9 |
| ddx27 | 0.22531 | 0.287233316 | 0.241 | 0.1 | 1 | 9 |
| sh3pxd2b | 0.22616 | -0.14822424 | 0.034 | 0.15 | 1 | 9 |
| wdfy1 | 0.22666 | 0.269307185 | 0.19 | 0.05 | 1 | 9 |
| oat | 0.22681 | -0.25974989 | 0.034 | 0.1 | 1 | 9 |
| cops9 | 0.22721 | 0.143900256 | 0.362 | 0.2 | 1 | 9 |
| eif3k | 0.22744 | 0.313324214 | 0.552 | 0.4 | 1 | 9 |
| napab | 0.2276 | 0.236223146 | 0.276 | 0.1 | 1 | 9 |
| sdhaf4 | 0.22785 | 0.199832868 | 0.19 | 0.05 | 1 | 9 |
| hsd17b14 | 0.22797 | -0.21123941 | 0.034 | 0.15 | 1 | 9 |
| ube2ia | 0.22807 | 0.268618742 | 0.276 | 0.1 | 1 | 9 |
| si:ch211-217k17.9 | 0.2282 | 0.392137902 | 0.172 | 0.1 | 1 | 9 |
| snx3 | 0.22837 | 0.272284162 | 0.276 | 0.1 | 1 | 9 |
| fam169aa | 0.22875 | -0.24193327 | 0.069 | 0.2 | 1 | 9 |
| snrpf | 0.2289 | 0.341694566 | 0.517 | 0.3 | 1 | 9 |
| ywhaz | 0.22891 | 0.345267708 | 0.603 | 0.4 | 1 | 9 |
| pitpnaa | 0.22923 | -0.2537001 | 0.155 | 0.25 | 1 | 9 |
| sox4a | 0.23048 | 0.375518852 | 0.276 | 0.1 | 1 | 9 |
| uqcrq | 0.2305 | 0.106655098 | 0.517 | 0.35 | 1 | 9 |
| tnfaip3 | 0.23055 | -0.26658776 | 0.034 | 0.15 | 1 | 9 |
| ogt.1 | 0.23081 | -0.26976798 | 0.241 | 0.35 | 1 | 9 |
| huwe1 | 0.23091 | 0.117986138 | 0.155 | 0.05 | 1 | 9 |
| map3k14a | 0.23104 | -0.23050238 | 0.034 | 0.15 | 1 | 9 |
| ptges3a | 0.23105 | 0.282057216 | 0.19 | 0.05 | 1 | 9 |
| eif2b2 | 0.23106 | -0.23006266 | 0.103 | 0.15 | 1 | 9 |
| aldh6a1 | 0.23107 | -0.1261747 | 0.224 | 0.2 | 1 | 9 |
| ddx3xa | 0.23144 | -0.28167724 | 0.086 | 0.2 | 1 | 9 |
| psmd3 | 0.23148 | -0.17028301 | 0.293 | 0.3 | 1 | 9 |
| eif3f | 0.2319 | 0.289118688 | 0.776 | 0.6 | 1 | 9 |
| fbrsl1 | 0.23226 | -0.26781804 | 0.034 | 0.15 | 1 | 9 |
| spopla | 0.23345 | -0.27105119 | 0.069 | 0.15 | 1 | 9 |
| ahcy | 0.2336 | 0.17693832 | 0.259 | 0.1 | 1 | 9 |
| tmod2 | 0.23378 | -0.33282034 | 0.155 | 0.3 | 1 | 9 |
| yif1a | 0.23399 | -0.27275018 | 0.086 | 0.15 | 1 | 9 |
| rcan1a | 0.23561 | -0.14695728 | 0.069 | 0.2 | 1 | 9 |
| patj | 0.23568 | -0.25909323 | 0.103 | 0.2 | 1 | 9 |
| sec13 | 0.23611 | -0.19947614 | 0.172 | 0.2 | 1 | 9 |
| trnau1apb | 0.2365 | 0.277963654 | 0.241 | 0.15 | 1 | 9 |
| efna5a | 0.23654 | -0.23639536 | 0.034 | 0.15 | 1 | 9 |
| rnf185 | 0.23672 | 0.194587079 | 0.259 | 0.1 | 1 | 9 |
| dcn | 0.23678 | -0.15763576 | 0.034 | 0.15 | 1 | 9 |
| tmem63ba | 0.23755 | -0.27200192 | 0.034 | 0.15 | 1 | 9 |
| sorl1 | 0.23787 | -0.30150048 | 0.069 | 0.2 | 1 | 9 |
| CZQB01141835.1 | 0.23798 | 0.284784254 | 0.19 | 0.05 | 1 | 9 |
| prrc2a | 0.23833 | -0.19405254 | 0.034 | 0.15 | 1 | 9 |
| tmem231 | 0.23834 | 0.112866469 | 0.155 | 0.05 | 1 | 9 |
| si:ch1073-325m22.2 | 0.23862 | 0.359944891 | 0.534 | 0.35 | 1 | 9 |
| smim4 | 0.23909 | 0.272757298 | 0.19 | 0.05 | 1 | 9 |

| | | | | | | |
|---|---|---|---|---|---|---|
| ndufb3 | 0.23921 | 0.112378804 | 0.569 | 0.4 | 1 | 9 |
| hells | 0.23999 | 0.303029201 | 0.19 | 0.05 | 1 | 9 |
| rps15 | 0.24019 | 0.175538549 | 0.897 | 0.85 | 1 | 9 |
| si:ch211-87m7.2 | 0.2404 | -0.28030385 | 0.086 | 0.2 | 1 | 9 |
| zgc:153317 | 0.24061 | -0.23428558 | 0.138 | 0.2 | 1 | 9 |
| ptpra | 0.24092 | -0.1538361 | 0.034 | 0.15 | 1 | 9 |
| skap2 | 0.24146 | -0.23667725 | 0.034 | 0.1 | 1 | 9 |
| naa15a | 0.24177 | 0.21658413 | 0.19 | 0.05 | 1 | 9 |
| zgc:110591 | 0.24184 | -0.22390471 | 0.034 | 0.15 | 1 | 9 |
| mbd6 | 0.24189 | -0.22693764 | 0.034 | 0.15 | 1 | 9 |
| fam210b | 0.2421 | -0.26521493 | 0.086 | 0.15 | 1 | 9 |
| rplp2l | 0.24241 | 0.20347314 | 0.931 | 0.85 | 1 | 9 |
| dyrk1ab | 0.24268 | -0.22631748 | 0.034 | 0.15 | 1 | 9 |
| ubap1 | 0.24293 | -0.18359184 | 0.034 | 0.15 | 1 | 9 |
| gabarapl2 | 0.24305 | -0.33739393 | 0.259 | 0.45 | 1 | 9 |
| cox6b2 | 0.24313 | 0.119170232 | 0.862 | 0.7 | 1 | 9 |
| parp12b | 0.24314 | -0.30231205 | 0.034 | 0.15 | 1 | 9 |
| arhgef18a | 0.24327 | 0.194451587 | 0.155 | 0.05 | 1 | 9 |
| spint2 | 0.24333 | -0.31997839 | 0.397 | 0.5 | 1 | 9 |
| gig2o | 0.24367 | -0.19316993 | 0.034 | 0.15 | 1 | 9 |
| lrp10 | 0.24385 | -0.20130541 | 0.034 | 0.15 | 1 | 9 |
| phf2 | 0.24396 | -0.17422443 | 0.034 | 0.15 | 1 | 9 |
| caprin1b | 0.24434 | 0.167522486 | 0.31 | 0.15 | 1 | 9 |
| si:dkey-112a7.4 | 0.24474 | 0.11861265 | 0.155 | 0.05 | 1 | 9 |
| nedd8 | 0.24488 | 0.245949455 | 0.241 | 0.1 | 1 | 9 |
| fthl28 | 0.24517 | -0.21312289 | 0.069 | 0.2 | 1 | 9 |
| slc7a10a | 0.24519 | -0.20037516 | 0.034 | 0.15 | 1 | 9 |
| dpm1 | 0.24521 | -0.32995073 | 0.138 | 0.25 | 1 | 9 |
| nbr1b | 0.24533 | 0.151021687 | 0.172 | 0.05 | 1 | 9 |
| vps13c | 0.24546 | -0.21982034 | 0.052 | 0.15 | 1 | 9 |
| noxa1 | 0.24555 | -0.21930234 | 0.034 | 0.15 | 1 | 9 |
| myl12.2 | 0.24556 | 0.19471292 | 0.259 | 0.1 | 1 | 9 |
| frg1 | 0.2456 | -0.20939672 | 0.034 | 0.15 | 1 | 9 |
| psap | 0.24588 | -0.27684868 | 0.345 | 0.4 | 1 | 9 |
| atp5meb | 0.24604 | 0.153499194 | 0.638 | 0.45 | 1 | 9 |
| pias2 | 0.24607 | -0.29046654 | 0.086 | 0.2 | 1 | 9 |
| ss18 | 0.24625 | 0.206211426 | 0.259 | 0.1 | 1 | 9 |
| LOC103910015 | 0.24661 | -0.17597873 | 0.034 | 0.15 | 1 | 9 |
| palm3 | 0.24675 | -0.19755663 | 0.034 | 0.15 | 1 | 9 |
| nsdhl | 0.24749 | -0.21920653 | 0.034 | 0.15 | 1 | 9 |
| si:dkey-56m19.5 | 0.2477 | 0.252748311 | 0.241 | 0.1 | 1 | 9 |
| gpa33b | 0.24773 | -0.25903085 | 0.034 | 0.15 | 1 | 9 |
| mccc1 | 0.24782 | -0.13484778 | 0.034 | 0.15 | 1 | 9 |
| rab18a | 0.24871 | 0.165952712 | 0.172 | 0.05 | 1 | 9 |
| ddx5 | 0.24886 | 0.350416573 | 0.5 | 0.3 | 1 | 9 |
| asf1bb | 0.24914 | 0.191421103 | 0.19 | 0.05 | 1 | 9 |
| stimate | 0.24978 | -0.23959532 | 0.069 | 0.2 | 1 | 9 |
| col18a1a | 0.24983 | 0.143718533 | 0.052 | 0 | 1 | 9 |

| | | | | | | |
|---|---|---|---|---|---|---|
| kdm2ba | 0.24983 | 0.129590588 | 0.052 | 0 | 1 | 9 |
| CABZ01074453.1 | 0.24983 | 0.111357696 | 0.052 | 0 | 1 | 9 |
| smim14 | 0.24983 | 0.102669602 | 0.052 | 0 | 1 | 9 |
| urm1 | 0.24983 | 0.100404497 | 0.052 | 0 | 1 | 9 |
| arsj | 0.24983 | 0.142502201 | 0.052 | 0 | 1 | 9 |
| thap7 | 0.24983 | 0.132955987 | 0.052 | 0 | 1 | 9 |
| FP102170.1 | 0.24983 | 0.107483166 | 0.052 | 0 | 1 | 9 |
| abraxas2 | 0.24983 | 0.117868043 | 0.052 | 0 | 1 | 9 |
| ccdc85al | 0.24983 | 0.125880467 | 0.052 | 0 | 1 | 9 |
| slc25a42 | 0.24983 | 0.141195426 | 0.052 | 0 | 1 | 9 |
| tcf7l1a | 0.24983 | 0.122618524 | 0.052 | 0 | 1 | 9 |
| zcchc24 | 0.24983 | 0.130047307 | 0.052 | 0 | 1 | 9 |
| zgc:66472 | 0.24983 | 0.115973153 | 0.052 | 0 | 1 | 9 |
| BX323557.1 | 0.24983 | 0.116585663 | 0.052 | 0 | 1 | 9 |
| CR848749.2 | 0.24983 | 0.132156271 | 0.052 | 0 | 1 | 9 |
| LOC100149931 | 0.24983 | 0.117075859 | 0.052 | 0 | 1 | 9 |
| LOC110438211 | 0.24983 | 0.190938189 | 0.052 | 0 | 1 | 9 |
| afap1l1a | 0.24983 | 0.130104826 | 0.052 | 0 | 1 | 9 |
| apol1 | 0.24983 | 0.101698044 | 0.052 | 0 | 1 | 9 |
| btbd6a | 0.24983 | 0.11223733 | 0.052 | 0 | 1 | 9 |
| celf4 | 0.24983 | 0.142025602 | 0.052 | 0 | 1 | 9 |
| chaf1b | 0.24983 | 0.107757945 | 0.052 | 0 | 1 | 9 |
| dachb | 0.24983 | 0.118567794 | 0.052 | 0 | 1 | 9 |
| dlx4b | 0.24983 | 0.121696841 | 0.052 | 0 | 1 | 9 |
| dpysl2b | 0.24983 | 0.125463282 | 0.052 | 0 | 1 | 9 |
| epdr1 | 0.24983 | 0.120776256 | 0.052 | 0 | 1 | 9 |
| flad1 | 0.24983 | 0.10488509 | 0.052 | 0 | 1 | 9 |
| fzd2 | 0.24983 | 0.11236963 | 0.052 | 0 | 1 | 9 |
| kctd10 | 0.24983 | 0.100360755 | 0.052 | 0 | 1 | 9 |
| map4l | 0.24983 | 0.122192063 | 0.052 | 0 | 1 | 9 |
| notch1b | 0.24983 | 0.101741765 | 0.052 | 0 | 1 | 9 |
| nrp2b | 0.24983 | 0.133894199 | 0.052 | 0 | 1 | 9 |
| nsun4 | 0.24983 | 0.133613692 | 0.052 | 0 | 1 | 9 |
| nudt1 | 0.24983 | 0.117459383 | 0.052 | 0 | 1 | 9 |
| pck2 | 0.24983 | 0.10059653 | 0.052 | 0 | 1 | 9 |
| rbpja | 0.24983 | 0.102780855 | 0.052 | 0 | 1 | 9 |
| rxraa | 0.24983 | 0.10156879 | 0.052 | 0 | 1 | 9 |
| senp7b | 0.24983 | 0.107526956 | 0.052 | 0 | 1 | 9 |
| serpinh1b | 0.24983 | 0.137228184 | 0.052 | 0 | 1 | 9 |
| si:ch211-276c2.4 | 0.24983 | 0.104978648 | 0.052 | 0 | 1 | 9 |
| si:ch211-67e16.4 | 0.24983 | 0.100716459 | 0.052 | 0 | 1 | 9 |
| si:ch73-314g15.3 | 0.24983 | 0.110308815 | 0.052 | 0 | 1 | 9 |
| si:dkey-73n10.1 | 0.24983 | 0.101630014 | 0.052 | 0 | 1 | 9 |
| si:dkey-82i20.2 | 0.24983 | 0.104687489 | 0.052 | 0 | 1 | 9 |
| slc15a2 | 0.24983 | 0.133047976 | 0.052 | 0 | 1 | 9 |
| smg9 | 0.24983 | 0.118590668 | 0.052 | 0 | 1 | 9 |
| spns2 | 0.24983 | 0.121160623 | 0.052 | 0 | 1 | 9 |
| tox4a | 0.24983 | 0.104646715 | 0.052 | 0 | 1 | 9 |

| | | | | | | |
|---|---|---|---|---|---|---|
| tp53i11b | 0.24983 | 0.134066963 | 0.052 | 0 | 1 | 9 |
| wrnip1 | 0.24983 | 0.106012419 | 0.052 | 0 | 1 | 9 |
| AL732567.1 | 0.24983 | 0.128936014 | 0.052 | 0 | 1 | 9 |
| BX322661.2 | 0.24983 | 0.128815794 | 0.052 | 0 | 1 | 9 |
| CABZ01029938.1 | 0.24983 | 0.109966523 | 0.052 | 0 | 1 | 9 |
| CABZ01068600.1 | 0.24983 | 0.106974591 | 0.052 | 0 | 1 | 9 |
| CT027702.1 | 0.24983 | 0.101310704 | 0.052 | 0 | 1 | 9 |
| CT027980.1 | 0.24983 | 0.142663083 | 0.052 | 0 | 1 | 9 |
| LOC100333241 | 0.24983 | 0.101537108 | 0.052 | 0 | 1 | 9 |
| LOC103911115 | 0.24983 | 0.131163153 | 0.052 | 0 | 1 | 9 |
| LOC108181504 | 0.24983 | 0.127972604 | 0.052 | 0 | 1 | 9 |
| LOC108191465 | 0.24983 | 0.11082745 | 0.052 | 0 | 1 | 9 |
| TPGS1 | 0.24983 | 0.117657775 | 0.052 | 0 | 1 | 9 |
| XLOC-018182 | 0.24983 | 0.123169646 | 0.052 | 0 | 1 | 9 |
| aatkb | 0.24983 | 0.152557756 | 0.052 | 0 | 1 | 9 |
| abca1b | 0.24983 | 0.114249008 | 0.052 | 0 | 1 | 9 |
| ackr4b | 0.24983 | 0.166210988 | 0.052 | 0 | 1 | 9 |
| ankrd50l | 0.24983 | 0.105727636 | 0.052 | 0 | 1 | 9 |
| atp13a1 | 0.24983 | 0.103203254 | 0.052 | 0 | 1 | 9 |
| atp1b1b | 0.24983 | 0.214219963 | 0.052 | 0 | 1 | 9 |
| cadm4 | 0.24983 | 0.149111305 | 0.052 | 0 | 1 | 9 |
| capsla | 0.24983 | 0.193121228 | 0.052 | 0 | 1 | 9 |
| ccne2 | 0.24983 | 0.116912797 | 0.052 | 0 | 1 | 9 |
| cdkn2aip | 0.24983 | 0.106229639 | 0.052 | 0 | 1 | 9 |
| chga | 0.24983 | 0.168439335 | 0.052 | 0 | 1 | 9 |
| chrna5 | 0.24983 | 0.141912771 | 0.052 | 0 | 1 | 9 |
| chuk | 0.24983 | 0.115070456 | 0.052 | 0 | 1 | 9 |
| cks2 | 0.24983 | 0.119504403 | 0.052 | 0 | 1 | 9 |
| cpeb2 | 0.24983 | 0.152378656 | 0.052 | 0 | 1 | 9 |
| dbr1 | 0.24983 | 0.102275696 | 0.052 | 0 | 1 | 9 |
| dctn1a | 0.24983 | 0.110850949 | 0.052 | 0 | 1 | 9 |
| diablob | 0.24983 | 0.103775855 | 0.052 | 0 | 1 | 9 |
| dusp14 | 0.24983 | 0.107152398 | 0.052 | 0 | 1 | 9 |
| ebf3a | 0.24983 | 0.144724538 | 0.052 | 0 | 1 | 9 |
| efemp2a | 0.24983 | 0.137112245 | 0.052 | 0 | 1 | 9 |
| elp2 | 0.24983 | 0.125898429 | 0.052 | 0 | 1 | 9 |
| ercc1 | 0.24983 | 0.118214344 | 0.052 | 0 | 1 | 9 |
| errfi1a | 0.24983 | 0.106878907 | 0.052 | 0 | 1 | 9 |
| esyt2a | 0.24983 | 0.101057725 | 0.052 | 0 | 1 | 9 |
| fev | 0.24983 | 0.170046473 | 0.052 | 0 | 1 | 9 |
| ftr98 | 0.24983 | 0.120009753 | 0.052 | 0 | 1 | 9 |
| gbgt1l3 | 0.24983 | 0.138064316 | 0.052 | 0 | 1 | 9 |
| gemin6 | 0.24983 | 0.112627778 | 0.052 | 0 | 1 | 9 |
| gng2 | 0.24983 | 0.120845175 | 0.052 | 0 | 1 | 9 |
| gpr137 | 0.24983 | 0.107270746 | 0.052 | 0 | 1 | 9 |
| gspt1 | 0.24983 | 0.103600988 | 0.052 | 0 | 1 | 9 |
| gxylt2 | 0.24983 | 0.102108293 | 0.052 | 0 | 1 | 9 |
| h6pd | 0.24983 | 0.1078533 | 0.052 | 0 | 1 | 9 |

| | | | | | | |
|---|---|---|---|---|---|---|
| hand2 | 0.24983 | 0.250548776 | 0.052 | 0 | 1 | 9 |
| herc3 | 0.24983 | 0.134404786 | 0.052 | 0 | 1 | 9 |
| hoxb5b | 0.24983 | 0.196791713 | 0.052 | 0 | 1 | 9 |
| idi1 | 0.24983 | 0.126828949 | 0.052 | 0 | 1 | 9 |
| im:7147486 | 0.24983 | 0.127162755 | 0.052 | 0 | 1 | 9 |
| ints2 | 0.24983 | 0.133728269 | 0.052 | 0 | 1 | 9 |
| ints8 | 0.24983 | 0.10711588 | 0.052 | 0 | 1 | 9 |
| iqsec1b | 0.24983 | 0.101604556 | 0.052 | 0 | 1 | 9 |
| itgb5 | 0.24983 | 0.106309346 | 0.052 | 0 | 1 | 9 |
| itsn2a | 0.24983 | 0.112969665 | 0.052 | 0 | 1 | 9 |
| jade3 | 0.24983 | 0.108854689 | 0.052 | 0 | 1 | 9 |
| lamb2l | 0.24983 | 0.103212085 | 0.052 | 0 | 1 | 9 |
| ldlrad4a | 0.24983 | 0.117333914 | 0.052 | 0 | 1 | 9 |
| mad2l2 | 0.24983 | 0.110744215 | 0.052 | 0 | 1 | 9 |
| marveld1 | 0.24983 | 0.12755748 | 0.052 | 0 | 1 | 9 |
| men1 | 0.24983 | 0.115088925 | 0.052 | 0 | 1 | 9 |
| nemp1 | 0.24983 | 0.106437823 | 0.052 | 0 | 1 | 9 |
| neurod1 | 0.24983 | 0.187344712 | 0.052 | 0 | 1 | 9 |
| nol9 | 0.24983 | 0.113273792 | 0.052 | 0 | 1 | 9 |
| pard6b | 0.24983 | 0.10739022 | 0.052 | 0 | 1 | 9 |
| pcdh20.1 | 0.24983 | 0.100343681 | 0.052 | 0 | 1 | 9 |
| phf3 | 0.24983 | 0.110727896 | 0.052 | 0 | 1 | 9 |
| phox2bb | 0.24983 | 0.214151375 | 0.052 | 0 | 1 | 9 |
| pitrm1 | 0.24983 | 0.116317862 | 0.052 | 0 | 1 | 9 |
| praf2 | 0.24983 | 0.116515617 | 0.052 | 0 | 1 | 9 |
| psat1 | 0.24983 | 0.113645323 | 0.052 | 0 | 1 | 9 |
| rabep1 | 0.24983 | 0.111219385 | 0.052 | 0 | 1 | 9 |
| rbfox1 | 0.24983 | 0.110942686 | 0.052 | 0 | 1 | 9 |
| rcan3 | 0.24983 | 0.122037897 | 0.052 | 0 | 1 | 9 |
| rexo1 | 0.24983 | 0.105785384 | 0.052 | 0 | 1 | 9 |
| rgmb | 0.24983 | 0.104773219 | 0.052 | 0 | 1 | 9 |
| rnaset2 | 0.24983 | 0.104073919 | 0.052 | 0 | 1 | 9 |
| rps6kb1a | 0.24983 | 0.102965564 | 0.052 | 0 | 1 | 9 |
| ryk | 0.24983 | 0.107848684 | 0.052 | 0 | 1 | 9 |
| sap130a | 0.24983 | 0.108033004 | 0.052 | 0 | 1 | 9 |
| sema3ab | 0.24983 | 0.122351121 | 0.052 | 0 | 1 | 9 |
| serpina10a | 0.24983 | 0.100177965 | 0.052 | 0 | 1 | 9 |
| sgcg | 0.24983 | 0.116408425 | 0.052 | 0 | 1 | 9 |
| si:ch211-194b1.1 | 0.24983 | 0.112959224 | 0.052 | 0 | 1 | 9 |
| si:ch211-200p22.4 | 0.24983 | 0.108154314 | 0.052 | 0 | 1 | 9 |
| si:ch211-250e5.2 | 0.24983 | 0.106722012 | 0.052 | 0 | 1 | 9 |
| si:ch211-71n6.4 | 0.24983 | 0.10818743 | 0.052 | 0 | 1 | 9 |
| si:ch73-109d9.3 | 0.24983 | 0.124223831 | 0.052 | 0 | 1 | 9 |
| si:dkey-102c8.3 | 0.24983 | 0.100488666 | 0.052 | 0 | 1 | 9 |
| si:dkey-262g12.10 | 0.24983 | 0.102227432 | 0.052 | 0 | 1 | 9 |
| si:dkey-26m3.3 | 0.24983 | 0.107208962 | 0.052 | 0 | 1 | 9 |
| si:rp71-77l1.1 | 0.24983 | 0.156947447 | 0.052 | 0 | 1 | 9 |
| skib | 0.24983 | 0.110073423 | 0.052 | 0 | 1 | 9 |

| | | | | | | |
|---|---|---|---|---|---|---|
| slc16a1b | 0.24983 | 0.109738153 | 0.052 | 0 | 1 | 9 |
| slc6a16b | 0.24983 | 0.12086429 | 0.052 | 0 | 1 | 9 |
| snip1 | 0.24983 | 0.103123618 | 0.052 | 0 | 1 | 9 |
| socs3b | 0.24983 | 0.112569191 | 0.052 | 0 | 1 | 9 |
| sp3b | 0.24983 | 0.107545619 | 0.052 | 0 | 1 | 9 |
| stard8 | 0.24983 | 0.127257812 | 0.052 | 0 | 1 | 9 |
| sulf1 | 0.24983 | 0.12118401 | 0.052 | 0 | 1 | 9 |
| suz12b | 0.24983 | 0.100223277 | 0.052 | 0 | 1 | 9 |
| tapbp.2 | 0.24983 | 0.119428067 | 0.052 | 0 | 1 | 9 |
| tbx15 | 0.24983 | 0.101349838 | 0.052 | 0 | 1 | 9 |
| tgm1 | 0.24983 | 0.122157573 | 0.052 | 0 | 1 | 9 |
| tmem154 | 0.24983 | 0.1197896 | 0.052 | 0 | 1 | 9 |
| tom1l2a | 0.24983 | 0.104392442 | 0.052 | 0 | 1 | 9 |
| tp53i11a | 0.24983 | 0.114265664 | 0.052 | 0 | 1 | 9 |
| trappc2 | 0.24983 | 0.100841293 | 0.052 | 0 | 1 | 9 |
| trim2a | 0.24983 | 0.100410158 | 0.052 | 0 | 1 | 9 |
| trub2 | 0.24983 | 0.113680939 | 0.052 | 0 | 1 | 9 |
| ttbk2a | 0.24983 | 0.125022662 | 0.052 | 0 | 1 | 9 |
| ttpal | 0.24983 | 0.114326965 | 0.052 | 0 | 1 | 9 |
| vgll2a | 0.24983 | 0.113533351 | 0.052 | 0 | 1 | 9 |
| zbtb16b | 0.24983 | 0.104222073 | 0.052 | 0 | 1 | 9 |
| zeb1a | 0.24983 | 0.121476409 | 0.052 | 0 | 1 | 9 |
| zgc:113336 | 0.24983 | 0.12489679 | 0.052 | 0 | 1 | 9 |
| zgc:175214 | 0.24983 | 0.147230583 | 0.052 | 0 | 1 | 9 |
| zgc:55558 | 0.24983 | 0.119055922 | 0.052 | 0 | 1 | 9 |
| zgc:56556 | 0.24983 | 0.108539229 | 0.052 | 0 | 1 | 9 |
| zgc:65895 | 0.24983 | 0.125826858 | 0.052 | 0 | 1 | 9 |
| zmp:0000000624 | 0.24983 | 0.101537787 | 0.052 | 0 | 1 | 9 |
| znf385a | 0.24983 | 0.102142341 | 0.052 | 0 | 1 | 9 |
| LOC110438361 | 0.24983 | 0.11862105 | 0.052 | 0 | 1 | 9 |
| adgra3 | 0.24983 | 0.10878123 | 0.052 | 0 | 1 | 9 |
| adgrb1a | 0.24983 | 0.105979864 | 0.052 | 0 | 1 | 9 |
| akap17a | 0.24983 | 0.104990081 | 0.052 | 0 | 1 | 9 |
| akt3b | 0.24983 | 0.109273577 | 0.052 | 0 | 1 | 9 |
| aldh1a2 | 0.24983 | 0.171039773 | 0.052 | 0 | 1 | 9 |
| b9d2 | 0.24983 | 0.107354572 | 0.052 | 0 | 1 | 9 |
| ccdc80 | 0.24983 | 0.116898378 | 0.052 | 0 | 1 | 9 |
| cks1b | 0.24983 | 0.119286314 | 0.052 | 0 | 1 | 9 |
| clu | 0.24983 | 0.13349835 | 0.052 | 0 | 1 | 9 |
| col4a5 | 0.24983 | 0.102683424 | 0.052 | 0 | 1 | 9 |
| crim1 | 0.24983 | 0.118417209 | 0.052 | 0 | 1 | 9 |
| cxl34b.11 | 0.24983 | 0.122287863 | 0.052 | 0 | 1 | 9 |
| dhrs12 | 0.24983 | 0.102206764 | 0.052 | 0 | 1 | 9 |
| dscaml1 | 0.24983 | 0.11851782 | 0.052 | 0 | 1 | 9 |
| enox2 | 0.24983 | 0.127411306 | 0.052 | 0 | 1 | 9 |
| ext2 | 0.24983 | 0.109017505 | 0.052 | 0 | 1 | 9 |
| fbrs | 0.24983 | 0.108247325 | 0.052 | 0 | 1 | 9 |
| fgfrl1a | 0.24983 | 0.104334086 | 0.052 | 0 | 1 | 9 |

| | | | | | | |
|---|---|---|---|---|---|---|
| fnbp1l | 0.24983 | 0.112293653 | 0.052 | 0 | 1 | 9 |
| foxd3 | 0.24983 | 0.221873797 | 0.052 | 0 | 1 | 9 |
| frmd6 | 0.24983 | 0.102031299 | 0.052 | 0 | 1 | 9 |
| galnt2 | 0.24983 | 0.116111142 | 0.052 | 0 | 1 | 9 |
| gamt | 0.24983 | 0.100339668 | 0.052 | 0 | 1 | 9 |
| gfra3 | 0.24983 | 0.125824211 | 0.052 | 0 | 1 | 9 |
| hdac8 | 0.24983 | 0.103257164 | 0.052 | 0 | 1 | 9 |
| im:7157749 | 0.24983 | 0.111257634 | 0.052 | 0 | 1 | 9 |
| itgb4 | 0.24983 | 0.138352307 | 0.052 | 0 | 1 | 9 |
| june | 0.24983 | 0.122801038 | 0.052 | 0 | 1 | 9 |
| kat6a | 0.24983 | 0.103529807 | 0.052 | 0 | 1 | 9 |
| laynb | 0.24983 | 0.10921403 | 0.052 | 0 | 1 | 9 |
| napbb | 0.24983 | 0.114116135 | 0.052 | 0 | 1 | 9 |
| nck2a | 0.24983 | 0.114146422 | 0.052 | 0 | 1 | 9 |
| negr1 | 0.24983 | 0.118676742 | 0.052 | 0 | 1 | 9 |
| nek1 | 0.24983 | 0.102797378 | 0.052 | 0 | 1 | 9 |
| nsmce1 | 0.24983 | 0.112449963 | 0.052 | 0 | 1 | 9 |
| orc5 | 0.24983 | 0.102530932 | 0.052 | 0 | 1 | 9 |
| pax2a | 0.24983 | 0.121164437 | 0.052 | 0 | 1 | 9 |
| pigr | 0.24983 | 0.105503289 | 0.052 | 0 | 1 | 9 |
| ptn | 0.24983 | 0.130712706 | 0.052 | 0 | 1 | 9 |
| rab3il1 | 0.24983 | 0.117945613 | 0.052 | 0 | 1 | 9 |
| ribc1 | 0.24983 | 0.129097381 | 0.052 | 0 | 1 | 9 |
| scml2 | 0.24983 | 0.109889556 | 0.052 | 0 | 1 | 9 |
| si:ch1073-416d2.4 | 0.24983 | 0.134674464 | 0.052 | 0 | 1 | 9 |
| si:ch211-212k18.8 | 0.24983 | 0.12328904 | 0.052 | 0 | 1 | 9 |
| si:dkey-14d8.6 | 0.24983 | 0.133844756 | 0.052 | 0 | 1 | 9 |
| si:dkey-222f2.1 | 0.24983 | 0.139103508 | 0.052 | 0 | 1 | 9 |
| snap25a | 0.24983 | 0.10577127 | 0.052 | 0 | 1 | 9 |
| stx2a | 0.24983 | 0.11618404 | 0.052 | 0 | 1 | 9 |
| tmem119b | 0.24983 | 0.119686621 | 0.052 | 0 | 1 | 9 |
| trap1 | 0.24983 | 0.105527833 | 0.052 | 0 | 1 | 9 |
| ube2ql1 | 0.24983 | 0.119013147 | 0.052 | 0 | 1 | 9 |
| zeb1b | 0.24983 | 0.108822975 | 0.052 | 0 | 1 | 9 |
| zgc:123010 | 0.24983 | 0.104227312 | 0.052 | 0 | 1 | 9 |
| zgc:55461 | 0.24983 | 0.17717236 | 0.052 | 0 | 1 | 9 |
| zgc:64022 | 0.24983 | 0.100421191 | 0.052 | 0 | 1 | 9 |
| CR376783.1 | 0.24983 | 0.113879484 | 0.052 | 0 | 1 | 9 |
| FBXO46 | 0.24983 | 0.108735381 | 0.052 | 0 | 1 | 9 |
| adm2a | 0.24983 | 0.107720466 | 0.052 | 0 | 1 | 9 |
| arxa | 0.24983 | 0.103747744 | 0.052 | 0 | 1 | 9 |
| cdh11 | 0.24983 | 0.142556439 | 0.052 | 0 | 1 | 9 |
| cntnap2a | 0.24983 | 0.112340628 | 0.052 | 0 | 1 | 9 |
| dnal4b | 0.24983 | 0.119318637 | 0.052 | 0 | 1 | 9 |
| dpysl3 | 0.24983 | 0.157887036 | 0.052 | 0 | 1 | 9 |
| ece1 | 0.24983 | 0.114450703 | 0.052 | 0 | 1 | 9 |
| gng3 | 0.24983 | 0.135487696 | 0.052 | 0 | 1 | 9 |
| hk1 | 0.24983 | 0.103780937 | 0.052 | 0 | 1 | 9 |

| | | | | | | |
|---|---|---|---|---|---|---|
| igf2b | 0.24983 | 0.105421774 | 0.052 | 0 | 1 | 9 |
| itgb1b.1 | 0.24983 | 0.121706844 | 0.052 | 0 | 1 | 9 |
| kcnk5b | 0.24983 | 0.114266523 | 0.052 | 0 | 1 | 9 |
| kdelr3 | 0.24983 | 0.105350602 | 0.052 | 0 | 1 | 9 |
| l3mbtl2 | 0.24983 | 0.111505684 | 0.052 | 0 | 1 | 9 |
| nid2a | 0.24983 | 0.117581661 | 0.052 | 0 | 1 | 9 |
| otulina | 0.24983 | 0.100566319 | 0.052 | 0 | 1 | 9 |
| pdk2a | 0.24983 | 0.105975768 | 0.052 | 0 | 1 | 9 |
| si:ch211-197k17.3 | 0.24983 | 0.106682965 | 0.052 | 0 | 1 | 9 |
| si:dkey-153k10.9 | 0.24983 | 0.106808336 | 0.052 | 0 | 1 | 9 |
| six4a | 0.24983 | 0.111397669 | 0.052 | 0 | 1 | 9 |
| syt1a | 0.24983 | 0.129410142 | 0.052 | 0 | 1 | 9 |
| tada2b | 0.24983 | 0.104864772 | 0.052 | 0 | 1 | 9 |
| tango2 | 0.24983 | 0.118949674 | 0.052 | 0 | 1 | 9 |
| tmem59l | 0.24983 | 0.134294359 | 0.052 | 0 | 1 | 9 |
| zgc:171435 | 0.24983 | 0.109404148 | 0.052 | 0 | 1 | 9 |
| zgc:173548 | 0.24983 | 0.122162438 | 0.052 | 0 | 1 | 9 |
| znf1104 | 0.24983 | 0.128622964 | 0.052 | 0 | 1 | 9 |
| zswim8 | 0.24983 | 0.117545654 | 0.052 | 0 | 1 | 9 |
| frmd8 | 0.25019 | -0.22319954 | 0.034 | 0.1 | 1 | 9 |
| bloc1s6 | 0.25095 | -0.26989831 | 0.224 | 0.35 | 1 | 9 |
| si:ch211-105c13.3 | 0.25141 | -0.30178772 | 0.069 | 0.2 | 1 | 9 |
| myl6 | 0.25285 | 0.245179087 | 0.19 | 0.05 | 1 | 9 |
| si:ch73-52f15.5 | 0.25412 | -0.2076509 | 0.172 | 0.2 | 1 | 9 |
| h2afvb | 0.25533 | 0.246803335 | 0.707 | 0.5 | 1 | 9 |
| ndufa11 | 0.25538 | -0.21929841 | 0.362 | 0.4 | 1 | 9 |
| hspa4b | 0.25543 | 0.214475752 | 0.19 | 0.05 | 1 | 9 |
| si:ch211-260e23.9 | 0.25688 | 0.287012925 | 0.293 | 0.15 | 1 | 9 |
| krt96 | 0.25692 | 0.27145307 | 0.138 | 0.05 | 1 | 9 |
| gemin2 | 0.25826 | 0.125281045 | 0.172 | 0.05 | 1 | 9 |
| tspan31 | 0.25832 | 0.126570944 | 0.155 | 0.05 | 1 | 9 |
| angptl4 | 0.25844 | 0.29460119 | 0.19 | 0.05 | 1 | 9 |
| tmed2 | 0.25848 | 0.294658298 | 0.483 | 0.35 | 1 | 9 |
| hnrnpa1b | 0.25901 | 0.247833473 | 0.448 | 0.25 | 1 | 9 |
| btf3 | 0.25906 | 0.238981688 | 0.862 | 0.75 | 1 | 9 |
| anxa3a | 0.25927 | 0.278510713 | 0.207 | 0.1 | 1 | 9 |
| serinc2 | 0.25943 | 0.248143007 | 0.328 | 0.15 | 1 | 9 |
| tmem45b | 0.25954 | -0.28954297 | 0.155 | 0.3 | 1 | 9 |
| golga7 | 0.25977 | 0.249630434 | 0.328 | 0.15 | 1 | 9 |
| si:ch1073-345a8.1 | 0.26012 | -0.23648451 | 0.138 | 0.2 | 1 | 9 |
| trappc1 | 0.26015 | -0.15700591 | 0.345 | 0.35 | 1 | 9 |
| cx30.3 | 0.26032 | -0.36872027 | 0.069 | 0.2 | 1 | 9 |
| pole3 | 0.26054 | -0.17443819 | 0.086 | 0.1 | 1 | 9 |
| taf15 | 0.26121 | 0.294911621 | 0.31 | 0.15 | 1 | 9 |
| mrpl28 | 0.26148 | 0.177373896 | 0.19 | 0.05 | 1 | 9 |
| UBL4A | 0.26153 | 0.225076282 | 0.19 | 0.05 | 1 | 9 |
| rgs2 | 0.26175 | 0.339508658 | 0.224 | 0.15 | 1 | 9 |
| pcdh11 | 0.26183 | -0.1212764 | 0.052 | 0.05 | 1 | 9 |

| | | | | | | |
|---|---|---|---|---|---|---|
| dusp8a | 0.26207 | -0.22753628 | 0.052 | 0.1 | 1 | 9 |
| hopx | 0.26215 | -0.33690652 | 0.172 | 0.35 | 1 | 9 |
| wwp1 | 0.26219 | -0.31346842 | 0.052 | 0.15 | 1 | 9 |
| hunk | 0.2623 | 0.264536132 | 0.19 | 0.05 | 1 | 9 |
| ptprfb | 0.26246 | 0.275662808 | 0.259 | 0.1 | 1 | 9 |
| dtx4b | 0.26282 | -0.19214341 | 0.069 | 0.1 | 1 | 9 |
| wipi2 | 0.26364 | -0.24717726 | 0.103 | 0.2 | 1 | 9 |
| banf1 | 0.26448 | 0.320727462 | 0.414 | 0.25 | 1 | 9 |
| fen1 | 0.26458 | 0.261306754 | 0.19 | 0.05 | 1 | 9 |
| rbx1 | 0.26463 | 0.206879517 | 0.603 | 0.4 | 1 | 9 |
| mcm4 | 0.26499 | 0.233826013 | 0.19 | 0.05 | 1 | 9 |
| rps13 | 0.26517 | 0.183900173 | 0.914 | 0.85 | 1 | 9 |
| cav2 | 0.26609 | -0.35663196 | 0.121 | 0.25 | 1 | 9 |
| aco2 | 0.26611 | 0.211356113 | 0.19 | 0.05 | 1 | 9 |
| si:dkey-96g2.1 | 0.26622 | -0.57425601 | 0.069 | 0.2 | 1 | 9 |
| elf1 | 0.26661 | -0.27853507 | 0.138 | 0.3 | 1 | 9 |
| eif4bb | 0.26663 | 0.291526133 | 0.31 | 0.15 | 1 | 9 |
| atp5pd | 0.26679 | 0.208507789 | 0.655 | 0.45 | 1 | 9 |
| eif1ad | 0.2671 | 0.19424577 | 0.31 | 0.15 | 1 | 9 |
| gemin8 | 0.26802 | 0.236791138 | 0.155 | 0.05 | 1 | 9 |
| trim25 | 0.26909 | -0.15093621 | 0.086 | 0.1 | 1 | 9 |
| rps14 | 0.26915 | 0.180582697 | 0.948 | 0.85 | 1 | 9 |
| tmem167a | 0.27032 | 0.25078791 | 0.241 | 0.1 | 1 | 9 |
| stat1a | 0.2706 | -0.34393017 | 0.19 | 0.35 | 1 | 9 |
| tial1 | 0.27086 | -0.16165726 | 0.086 | 0.1 | 1 | 9 |
| sart1 | 0.27173 | 0.128237419 | 0.241 | 0.1 | 1 | 9 |
| stk26 | 0.27213 | -0.22743003 | 0.069 | 0.15 | 1 | 9 |
| cyc1 | 0.27247 | 0.144156801 | 0.414 | 0.25 | 1 | 9 |
| map1lc3a | 0.27327 | 0.201357238 | 0.259 | 0.1 | 1 | 9 |
| rpp14 | 0.27441 | 0.245004362 | 0.155 | 0.05 | 1 | 9 |
| gcnt4a | 0.27451 | 0.280569549 | 0.138 | 0.05 | 1 | 9 |
| myg1 | 0.2746 | 0.23384284 | 0.207 | 0.1 | 1 | 9 |
| ankrd11 | 0.27493 | 0.243857787 | 0.328 | 0.15 | 1 | 9 |
| aldh7a1 | 0.27507 | -0.23238719 | 0.172 | 0.25 | 1 | 9 |
| wdr83os | 0.27522 | 0.209600952 | 0.259 | 0.1 | 1 | 9 |
| sf3b6 | 0.27546 | 0.302045927 | 0.5 | 0.3 | 1 | 9 |
| abhd2a | 0.27552 | -0.20687421 | 0.069 | 0.2 | 1 | 9 |
| glula | 0.27563 | 0.161754992 | 0.155 | 0.05 | 1 | 9 |
| si:zfos-1192g2.3 | 0.27587 | 0.17138835 | 0.241 | 0.1 | 1 | 9 |
| hmgcrb | 0.27595 | 0.165107537 | 0.086 | 0.05 | 1 | 9 |
| tmem131 | 0.2764 | -0.27922567 | 0.069 | 0.15 | 1 | 9 |
| si:dkey-11f4.14 | 0.27691 | -0.28199299 | 0.172 | 0.35 | 1 | 9 |
| abce1 | 0.27749 | 0.104902443 | 0.224 | 0.1 | 1 | 9 |
| dusp5 | 0.27756 | -0.1774874 | 0.069 | 0.2 | 1 | 9 |
| lgals3a | 0.27765 | 0.260136518 | 0.328 | 0.15 | 1 | 9 |
| notch2 | 0.27801 | -0.18507932 | 0.069 | 0.1 | 1 | 9 |
| ehf | 0.27809 | -0.30881731 | 0.069 | 0.2 | 1 | 9 |
| mcm3 | 0.27819 | 0.15144112 | 0.155 | 0.05 | 1 | 9 |

| | | | | | | |
|---|---|---|---|---|---|---|
| kpnb1 | 0.27861 | 0.159999775 | 0.241 | 0.1 | 1 | 9 |
| smarce1 | 0.27947 | 0.286748569 | 0.448 | 0.25 | 1 | 9 |
| COX7A2 (1 of many) | 0.27974 | 0.159284624 | 0.362 | 0.2 | 1 | 9 |
| tspan37 | 0.28008 | -0.23184055 | 0.069 | 0.15 | 1 | 9 |
| metrnla | 0.28052 | -0.23003364 | 0.155 | 0.2 | 1 | 9 |
| tspan35 | 0.28064 | -0.31741987 | 0.052 | 0.15 | 1 | 9 |
| ggnbp2 | 0.28122 | -0.24728702 | 0.069 | 0.2 | 1 | 9 |
| mvp | 0.282 | -0.32380817 | 0.155 | 0.3 | 1 | 9 |
| vamp3 | 0.28206 | 0.175404119 | 0.362 | 0.2 | 1 | 9 |
| ier5 | 0.28288 | -0.25994621 | 0.328 | 0.4 | 1 | 9 |
| pfkfb4b | 0.28342 | 0.277043877 | 0.328 | 0.15 | 1 | 9 |
| CABZ01088109.1 | 0.28381 | -0.20172523 | 0.052 | 0.1 | 1 | 9 |
| CU694197.1 | 0.28387 | 0.265337806 | 0.328 | 0.15 | 1 | 9 |
| tox | 0.28414 | -0.26580561 | 0.086 | 0.15 | 1 | 9 |
| selenow1 | 0.28423 | 0.101331184 | 0.81 | 0.65 | 1 | 9 |
| XLOC-004556 | 0.28429 | -0.28656636 | 0.069 | 0.2 | 1 | 9 |
| slc25a24 | 0.28434 | 0.216695713 | 0.172 | 0.05 | 1 | 9 |
| rpl23 | 0.28462 | 0.243631722 | 0.879 | 0.75 | 1 | 9 |
| prox1a | 0.28522 | -0.26040616 | 0.069 | 0.2 | 1 | 9 |
| fam57a | 0.28529 | -0.21665481 | 0.034 | 0.1 | 1 | 9 |
| cyfip1 | 0.28541 | -0.19282384 | 0.103 | 0.15 | 1 | 9 |
| myl12.1 | 0.28553 | -0.14702876 | 0.655 | 0.65 | 1 | 9 |
| MARK4 | 0.28557 | -0.24868609 | 0.052 | 0.15 | 1 | 9 |
| vangl2 | 0.28645 | -0.21996598 | 0.103 | 0.15 | 1 | 9 |
| lgals3b | 0.28835 | 0.260783538 | 0.31 | 0.15 | 1 | 9 |
| atp5po | 0.28843 | 0.283494591 | 0.5 | 0.3 | 1 | 9 |
| LOC100330250 | 0.28915 | -0.23461729 | 0.052 | 0.1 | 1 | 9 |
| patl1 | 0.28948 | -0.10096555 | 0.052 | 0.15 | 1 | 9 |
| bbox1 | 0.28958 | -0.25556929 | 0.103 | 0.25 | 1 | 9 |
| fstl1b | 0.29009 | 0.351275981 | 0.259 | 0.1 | 1 | 9 |
| cyp27a7 | 0.29013 | -0.2188967 | 0.052 | 0.15 | 1 | 9 |
| rcc2 | 0.29015 | 0.30274261 | 0.259 | 0.1 | 1 | 9 |
| lpar2b | 0.2906 | -0.19175474 | 0.069 | 0.2 | 1 | 9 |
| zgc:100868 | 0.29096 | -0.2181984 | 0.103 | 0.15 | 1 | 9 |
| odc1 | 0.29152 | -0.2552365 | 0.172 | 0.25 | 1 | 9 |
| si:dkeyp-72e1.6 | 0.29161 | -0.23936738 | 0.069 | 0.2 | 1 | 9 |
| yipf3 | 0.29181 | -0.21937645 | 0.069 | 0.2 | 1 | 9 |
| hebp2 | 0.29181 | -0.179254 | 0.069 | 0.2 | 1 | 9 |
| hnrnpl2 | 0.29181 | 0.256964453 | 0.259 | 0.1 | 1 | 9 |
| f2rl1.2 | 0.29191 | -0.25463869 | 0.086 | 0.15 | 1 | 9 |
| foxp1a | 0.29201 | -0.26949534 | 0.069 | 0.2 | 1 | 9 |
| cttn | 0.29234 | -0.1840119 | 0.069 | 0.2 | 1 | 9 |
| mrpl37 | 0.29266 | 0.132968125 | 0.155 | 0.05 | 1 | 9 |
| cnot2 | 0.29267 | -0.20622486 | 0.069 | 0.2 | 1 | 9 |
| atp5mf | 0.29333 | -0.2769241 | 0.483 | 0.6 | 1 | 9 |
| vps4b | 0.29339 | -0.26115145 | 0.276 | 0.35 | 1 | 9 |
| smndc1 | 0.29368 | 0.208785849 | 0.379 | 0.2 | 1 | 9 |
| anp32e | 0.2939 | 0.154021508 | 0.293 | 0.15 | 1 | 9 |

| | | | | | | |
|---|---|---|---|---|---|---|
| krtcap2 | 0.29391 | -0.22311815 | 0.448 | 0.5 | 1 | 9 |
| ciarta | 0.29391 | 0.21130534 | 0.31 | 0.15 | 1 | 9 |
| pdlim2 | 0.29407 | -0.21912804 | 0.069 | 0.2 | 1 | 9 |
| prss23 | 0.29412 | -0.12045924 | 0.052 | 0.05 | 1 | 9 |
| dynll2a | 0.29479 | -0.23853875 | 0.052 | 0.15 | 1 | 9 |
| fntb | 0.29497 | 0.142696768 | 0.155 | 0.05 | 1 | 9 |
| sypl2a | 0.29498 | -0.27923873 | 0.138 | 0.3 | 1 | 9 |
| ssrp1a | 0.29546 | 0.143514434 | 0.172 | 0.05 | 1 | 9 |
| ppp1r10 | 0.29558 | -0.27370416 | 0.103 | 0.25 | 1 | 9 |
| slc43a1a | 0.29569 | 0.248534215 | 0.172 | 0.05 | 1 | 9 |
| eno1a | 0.29569 | -0.11227414 | 0.121 | 0.1 | 1 | 9 |
| plpp1a | 0.29833 | -0.27902293 | 0.069 | 0.15 | 1 | 9 |
| si:dkeyp-113d7.1 | 0.29879 | 0.23975455 | 0.155 | 0.05 | 1 | 9 |
| cflara | 0.29881 | -0.24529497 | 0.103 | 0.25 | 1 | 9 |
| mfap1 | 0.29908 | 0.153177583 | 0.172 | 0.05 | 1 | 9 |
| copz2 | 0.29913 | -0.25712693 | 0.052 | 0.15 | 1 | 9 |
| dlx3b | 0.2998 | 0.106778485 | 0.138 | 0.15 | 1 | 9 |
| pitpnb | 0.29983 | 0.144909389 | 0.155 | 0.05 | 1 | 9 |
| ppp1r14bb | 0.30031 | 0.185793206 | 0.241 | 0.1 | 1 | 9 |
| ube2g1b | 0.30157 | 0.222394111 | 0.155 | 0.05 | 1 | 9 |
| palm2 | 0.30162 | -0.2557048 | 0.052 | 0.15 | 1 | 9 |
| usp9 | 0.30232 | -0.10136868 | 0.086 | 0.2 | 1 | 9 |
| rpl32 | 0.30268 | 0.128430835 | 0.931 | 0.9 | 1 | 9 |
| dhx38 | 0.30311 | -0.25409265 | 0.069 | 0.15 | 1 | 9 |
| rsf1b.1 | 0.3032 | 0.213160993 | 0.172 | 0.05 | 1 | 9 |
| cst14a.2 | 0.30397 | 0.127689198 | 0.448 | 0.3 | 1 | 9 |
| rpa3 | 0.30402 | 0.283743227 | 0.172 | 0.05 | 1 | 9 |
| utp3 | 0.30443 | 0.228817381 | 0.172 | 0.05 | 1 | 9 |
| zfyve27 | 0.30479 | -0.27453881 | 0.069 | 0.15 | 1 | 9 |
| pagr1 | 0.3051 | 0.245825157 | 0.172 | 0.05 | 1 | 9 |
| ildr1a | 0.30511 | -0.23351959 | 0.069 | 0.15 | 1 | 9 |
| cnn2 | 0.30537 | -0.30618783 | 0.207 | 0.35 | 1 | 9 |
| arhgdig | 0.30544 | -0.28229673 | 0.172 | 0.25 | 1 | 9 |
| apex1 | 0.3056 | 0.193775841 | 0.241 | 0.1 | 1 | 9 |
| qars | 0.30611 | 0.205577764 | 0.121 | 0.05 | 1 | 9 |
| cox7a2a | 0.30668 | 0.314821353 | 0.431 | 0.25 | 1 | 9 |
| tmed5 | 0.30705 | -0.22867713 | 0.121 | 0.2 | 1 | 9 |
| surf2 | 0.30727 | 0.122237193 | 0.155 | 0.05 | 1 | 9 |
| rpl23a | 0.30728 | 0.159546772 | 0.948 | 0.9 | 1 | 9 |
| pycard | 0.30747 | -0.21265212 | 0.259 | 0.3 | 1 | 9 |
| rpl26 | 0.30831 | 0.13397363 | 0.983 | 0.95 | 1 | 9 |
| plpp5 | 0.30845 | -0.29351392 | 0.103 | 0.2 | 1 | 9 |
| prkcba | 0.30888 | -0.23358954 | 0.052 | 0.1 | 1 | 9 |
| selenot1a | 0.3095 | -0.197307 | 0.224 | 0.25 | 1 | 9 |
| CABZ01076667.1 | 0.30957 | 0.213571112 | 0.172 | 0.05 | 1 | 9 |
| fam50a | 0.30978 | 0.251053243 | 0.224 | 0.1 | 1 | 9 |
| tmsb4x | 0.31008 | -0.20949884 | 0.586 | 0.6 | 1 | 9 |
| macc1 | 0.31046 | -0.1121461 | 0.052 | 0.15 | 1 | 9 |

| | | | | | | |
|---|---|---|---|---|---|---|
| ISCU (1 of many) | 0.31049 | 0.29468925 | 0.31 | 0.15 | 1 | 9 |
| uqcrc2b | 0.31051 | -0.27189432 | 0.155 | 0.25 | 1 | 9 |
| lamp1a | 0.31077 | -0.28918138 | 0.138 | 0.25 | 1 | 9 |
| tpm3 | 0.31118 | -0.16056478 | 0.483 | 0.5 | 1 | 9 |
| ppp1caa | 0.3113 | -0.2240356 | 0.259 | 0.35 | 1 | 9 |
| st13 | 0.31179 | 0.270968864 | 0.293 | 0.15 | 1 | 9 |
| khdrbs1a | 0.31296 | 0.245983194 | 0.776 | 0.6 | 1 | 9 |
| tefb | 0.31336 | -0.21854534 | 0.052 | 0.15 | 1 | 9 |
| nkap | 0.31367 | 0.238035188 | 0.259 | 0.15 | 1 | 9 |
| akirin2 | 0.31369 | 0.121017251 | 0.138 | 0.05 | 1 | 9 |
| jam2a | 0.31478 | 0.294928852 | 0.172 | 0.05 | 1 | 9 |
| midn | 0.31537 | -0.3320183 | 0.31 | 0.45 | 1 | 9 |
| wdr33 | 0.31568 | 0.243279055 | 0.241 | 0.1 | 1 | 9 |
| fam129aa | 0.3159 | -0.2609915 | 0.052 | 0.15 | 1 | 9 |
| ptbp1a | 0.316 | -0.26280887 | 0.155 | 0.3 | 1 | 9 |
| histh1l | 0.31626 | -0.30206974 | 0.155 | 0.3 | 1 | 9 |
| tdh | 0.3164 | -0.33130457 | 0.293 | 0.45 | 1 | 9 |
| ada | 0.31642 | -0.18393505 | 0.034 | 0.1 | 1 | 9 |
| slc35f6 | 0.31646 | 0.145202618 | 0.155 | 0.05 | 1 | 9 |
| dusp1 | 0.31746 | -0.12430762 | 0.121 | 0.25 | 1 | 9 |
| zgc:175264 | 0.3175 | 0.144883404 | 0.155 | 0.05 | 1 | 9 |
| pdcd4b | 0.31769 | -0.11092825 | 0.052 | 0.05 | 1 | 9 |
| anxa13 | 0.31775 | 0.109229449 | 0.138 | 0.05 | 1 | 9 |
| atrx | 0.31784 | 0.21599427 | 0.362 | 0.2 | 1 | 9 |
| zgc:113425 | 0.31824 | -0.24802726 | 0.052 | 0.15 | 1 | 9 |
| rhoaa | 0.31924 | 0.196163452 | 0.483 | 0.3 | 1 | 9 |
| psma2 | 0.31954 | 0.254303389 | 0.379 | 0.2 | 1 | 9 |
| ndufc2 | 0.31967 | 0.278935985 | 0.466 | 0.3 | 1 | 9 |
| rpz2 | 0.32047 | -0.155855 | 0.086 | 0.1 | 1 | 9 |
| dlst | 0.32067 | 0.108672008 | 0.328 | 0.2 | 1 | 9 |
| akap12b | 0.32315 | 0.278051727 | 0.138 | 0.05 | 1 | 9 |
| nucks1b | 0.32316 | 0.128096926 | 0.207 | 0.2 | 1 | 9 |
| zgc:195173 | 0.32444 | 0.201635103 | 0.172 | 0.05 | 1 | 9 |
| mapkapk2a | 0.32507 | 0.149914537 | 0.293 | 0.15 | 1 | 9 |
| atp5if1b | 0.32541 | -0.29296036 | 0.448 | 0.6 | 1 | 9 |
| prkci | 0.32555 | -0.15444951 | 0.121 | 0.15 | 1 | 9 |
| prrg2 | 0.32564 | -0.2855808 | 0.241 | 0.4 | 1 | 9 |
| gnai2a | 0.32617 | 0.112976351 | 0.259 | 0.15 | 1 | 9 |
| rangap1a | 0.3263 | 0.220695981 | 0.155 | 0.05 | 1 | 9 |
| csnk1g2b | 0.32659 | 0.175510439 | 0.172 | 0.05 | 1 | 9 |
| adam17a | 0.32663 | -0.12964344 | 0.086 | 0.1 | 1 | 9 |
| eif4ebp2 | 0.32962 | 0.274693619 | 0.345 | 0.2 | 1 | 9 |
| rbm14b | 0.3298 | 0.206077916 | 0.121 | 0.05 | 1 | 9 |
| stat3 | 0.33123 | -0.25812605 | 0.052 | 0.15 | 1 | 9 |
| LOC100331080 | 0.33164 | 0.290412966 | 0.138 | 0.05 | 1 | 9 |
| relb | 0.33204 | -0.22191709 | 0.052 | 0.15 | 1 | 9 |
| dync1i2a | 0.33215 | -0.20367725 | 0.034 | 0.1 | 1 | 9 |
| arl4d | 0.33226 | 0.237461176 | 0.155 | 0.05 | 1 | 9 |

| | | | | | | |
|---|---|---|---|---|---|---|
| adh5 | 0.33243 | 0.20282794 | 0.534 | 0.35 | 1 | 9 |
| gid8a | 0.33252 | -0.26703088 | 0.052 | 0.15 | 1 | 9 |
| rab3ip | 0.33288 | -0.22074454 | 0.069 | 0.15 | 1 | 9 |
| tsc22d2 | 0.3337 | -0.24552934 | 0.103 | 0.2 | 1 | 9 |
| aqp3a | 0.3344 | -0.30406058 | 0.103 | 0.2 | 1 | 9 |
| tmem115 | 0.33462 | -0.22925681 | 0.052 | 0.15 | 1 | 9 |
| si:ch73-22o12.1 | 0.33465 | -0.27838738 | 0.103 | 0.2 | 1 | 9 |
| ap1b1 | 0.33554 | -0.20371045 | 0.103 | 0.15 | 1 | 9 |
| ncor1 | 0.33648 | 0.150472364 | 0.172 | 0.05 | 1 | 9 |
| cdc40 | 0.33683 | 0.198403036 | 0.172 | 0.05 | 1 | 9 |
| exosc6 | 0.33728 | 0.141976574 | 0.155 | 0.05 | 1 | 9 |
| sumo3b | 0.33775 | 0.233092523 | 0.172 | 0.05 | 1 | 9 |
| krt97 | 0.3378 | 0.34034681 | 0.172 | 0.05 | 1 | 9 |
| hnrnpa1a | 0.33784 | 0.14637954 | 0.345 | 0.2 | 1 | 9 |
| coa5 | 0.33796 | 0.180975076 | 0.172 | 0.05 | 1 | 9 |
| idh1 | 0.33848 | -0.10578992 | 0.155 | 0.15 | 1 | 9 |
| dhx16 | 0.33867 | 0.142427507 | 0.103 | 0.1 | 1 | 9 |
| arpp19b | 0.3393 | 0.223702065 | 0.172 | 0.05 | 1 | 9 |
| rpl7l1 | 0.33941 | 0.207191403 | 0.172 | 0.05 | 1 | 9 |
| ppm1g | 0.33944 | 0.222217002 | 0.172 | 0.05 | 1 | 9 |
| rspry1 | 0.33947 | -0.17845907 | 0.103 | 0.15 | 1 | 9 |
| nav2b | 0.34018 | -0.21276201 | 0.052 | 0.15 | 1 | 9 |
| glrx3 | 0.34088 | 0.196230158 | 0.31 | 0.15 | 1 | 9 |
| cdkn2aipnl | 0.3409 | 0.200449462 | 0.241 | 0.1 | 1 | 9 |
| serbp1b | 0.34138 | 0.225871068 | 0.207 | 0.1 | 1 | 9 |
| nop53 | 0.34147 | 0.221338578 | 0.638 | 0.45 | 1 | 9 |
| acbd7 | 0.34182 | -0.24855913 | 0.052 | 0.1 | 1 | 9 |
| surf6 | 0.3419 | 0.230588586 | 0.138 | 0.05 | 1 | 9 |
| fthl27 | 0.34245 | 0.192454557 | 0.724 | 0.55 | 1 | 9 |
| lto1 | 0.34268 | -0.22044982 | 0.034 | 0.1 | 1 | 9 |
| si:rp71-45k5.4 | 0.34395 | 0.180799797 | 0.414 | 0.25 | 1 | 9 |
| sirt7 | 0.34397 | 0.19556037 | 0.138 | 0.05 | 1 | 9 |
| mrpl24 | 0.34439 | 0.188839293 | 0.155 | 0.05 | 1 | 9 |
| si:ch73-91k6.2 | 0.34506 | -0.27036137 | 0.155 | 0.25 | 1 | 9 |
| eps8l1a | 0.34583 | -0.23394162 | 0.155 | 0.25 | 1 | 9 |
| chmp2bb | 0.34624 | -0.15235416 | 0.241 | 0.25 | 1 | 9 |
| bcap31 | 0.34624 | -0.10479913 | 0.155 | 0.15 | 1 | 9 |
| ddx39b | 0.34684 | -0.15909603 | 0.069 | 0.1 | 1 | 9 |
| tmem263 | 0.34767 | -0.18746212 | 0.155 | 0.2 | 1 | 9 |
| fkbp9 | 0.34768 | 0.166921049 | 0.155 | 0.05 | 1 | 9 |
| gpm6ab | 0.34849 | 0.124594388 | 0.121 | 0.05 | 1 | 9 |
| bcl7bb | 0.34871 | -0.11298102 | 0.052 | 0.05 | 1 | 9 |
| psmd7 | 0.34902 | 0.206681618 | 0.31 | 0.15 | 1 | 9 |
| arl8bb | 0.34932 | -0.21952021 | 0.052 | 0.1 | 1 | 9 |
| tmem50a | 0.34998 | -0.10321443 | 0.362 | 0.35 | 1 | 9 |
| march6 | 0.35082 | -0.18361886 | 0.052 | 0.1 | 1 | 9 |
| pam16 | 0.35136 | 0.115381637 | 0.155 | 0.05 | 1 | 9 |
| mrpl11 | 0.35147 | 0.198455076 | 0.241 | 0.1 | 1 | 9 |

| | | | | | | |
|---|---|---|---|---|---|---|
| cops8 | 0.35246 | 0.225307735 | 0.155 | 0.05 | 1 | 9 |
| bloc1s5 | 0.35292 | -0.21357869 | 0.052 | 0.1 | 1 | 9 |
| si:dkey-46a10.3 | 0.35304 | -0.12913448 | 0.103 | 0.1 | 1 | 9 |
| nol10 | 0.35324 | 0.236347823 | 0.138 | 0.05 | 1 | 9 |
| lsm14ab | 0.35368 | -0.17205691 | 0.155 | 0.3 | 1 | 9 |
| znf207a | 0.3537 | 0.221158515 | 0.155 | 0.05 | 1 | 9 |
| pgpep1 | 0.35379 | -0.18914868 | 0.172 | 0.2 | 1 | 9 |
| ing4 | 0.35424 | 0.245639081 | 0.207 | 0.1 | 1 | 9 |
| bloc1s2 | 0.3547 | -0.21535585 | 0.086 | 0.15 | 1 | 9 |
| psmb1 | 0.35495 | 0.241340695 | 0.534 | 0.35 | 1 | 9 |
| gabarapb | 0.35506 | 0.124127217 | 0.552 | 0.4 | 1 | 9 |
| abcb5 | 0.35772 | 0.244549467 | 0.207 | 0.15 | 1 | 9 |
| dnase1l4.1 | 0.35798 | -0.17656374 | 0.052 | 0.15 | 1 | 9 |
| ddx39ab | 0.35821 | 0.182735291 | 0.362 | 0.2 | 1 | 9 |
| setb | 0.35828 | 0.253251361 | 0.534 | 0.35 | 1 | 9 |
| pef1 | 0.35832 | -0.11285231 | 0.155 | 0.15 | 1 | 9 |
| si:ch73-376l24.2 | 0.35837 | 0.175059476 | 0.155 | 0.05 | 1 | 9 |
| CYTH2 | 0.3585 | -0.27654755 | 0.086 | 0.15 | 1 | 9 |
| gng12a | 0.35863 | -0.22611966 | 0.052 | 0.15 | 1 | 9 |
| lamtor4 | 0.35865 | 0.202490846 | 0.155 | 0.05 | 1 | 9 |
| blcap | 0.35884 | -0.10568852 | 0.259 | 0.25 | 1 | 9 |
| lpcat2 | 0.35889 | -0.18644424 | 0.034 | 0.1 | 1 | 9 |
| wbp1lb | 0.35947 | -0.23446871 | 0.086 | 0.2 | 1 | 9 |
| zgc:92744 | 0.35973 | -0.29539858 | 0.19 | 0.25 | 1 | 9 |
| hnrnpk | 0.36058 | 0.199962013 | 0.362 | 0.2 | 1 | 9 |
| atxn2l | 0.3606 | -0.20866876 | 0.069 | 0.15 | 1 | 9 |
| kmt5aa | 0.3613 | 0.228401483 | 0.241 | 0.1 | 1 | 9 |
| tpd52l2a | 0.36214 | -0.12058912 | 0.19 | 0.2 | 1 | 9 |
| cebpb | 0.36339 | -0.10800775 | 0.103 | 0.1 | 1 | 9 |
| znf622 | 0.36531 | -0.19651299 | 0.052 | 0.15 | 1 | 9 |
| si:ch211-162e15.3 | 0.36568 | 0.156343476 | 0.155 | 0.05 | 1 | 9 |
| b3gnt2l | 0.36685 | -0.17704073 | 0.121 | 0.15 | 1 | 9 |
| selenou1a | 0.3673 | 0.207865425 | 0.155 | 0.05 | 1 | 9 |
| sox8b | 0.36971 | -0.16339542 | 0.069 | 0.1 | 1 | 9 |
| gtf2a2 | 0.37032 | -0.13235823 | 0.155 | 0.15 | 1 | 9 |
| si:ch211-284b7.3 | 0.37092 | 0.169253366 | 0.138 | 0.05 | 1 | 9 |
| zgc:92481 | 0.37093 | -0.22759745 | 0.155 | 0.3 | 1 | 9 |
| sipa1l2 | 0.37113 | -0.13715915 | 0.086 | 0.1 | 1 | 9 |
| taf3 | 0.37165 | -0.194734 | 0.052 | 0.15 | 1 | 9 |
| rps19 | 0.37202 | 0.158959894 | 0.948 | 0.85 | 1 | 9 |
| ncoa5 | 0.37266 | 0.23226391 | 0.155 | 0.05 | 1 | 9 |
| rbbp8l | 0.37287 | -0.20348671 | 0.052 | 0.15 | 1 | 9 |
| si:ch211-243g18.2 | 0.3745 | -0.27432804 | 0.034 | 0.1 | 1 | 9 |
| atp6v1g1 | 0.37456 | 0.289793504 | 0.293 | 0.15 | 1 | 9 |
| arrb2b | 0.37519 | -0.1554871 | 0.172 | 0.2 | 1 | 9 |
| mrps35 | 0.37597 | 0.18913255 | 0.155 | 0.05 | 1 | 9 |
| cox5ab | 0.3767 | 0.284917156 | 0.466 | 0.3 | 1 | 9 |
| polr3h | 0.37765 | 0.135286391 | 0.155 | 0.05 | 1 | 9 |

| | | | | | | |
|---|---|---|---|---|---|---|
| slc25a33 | 0.37848 | 0.224955292 | 0.138 | 0.05 | 1 | 9 |
| zmp:0000001114 | 0.37871 | 0.185981567 | 0.155 | 0.05 | 1 | 9 |
| cdkn1bb | 0.37875 | 0.284738409 | 0.293 | 0.15 | 1 | 9 |
| XLOC-040108 | 0.37932 | -0.22138577 | 0.034 | 0.1 | 1 | 9 |
| nap1l4b | 0.37944 | -0.18686744 | 0.034 | 0.1 | 1 | 9 |
| eif4e1c | 0.37961 | 0.151540511 | 0.155 | 0.05 | 1 | 9 |
| snx27b | 0.37973 | -0.20995736 | 0.052 | 0.15 | 1 | 9 |
| si:dkey-172m14.1 | 0.38027 | -0.16327218 | 0.052 | 0.1 | 1 | 9 |
| stx12 | 0.38096 | -0.19527896 | 0.086 | 0.15 | 1 | 9 |
| timm8b | 0.38131 | 0.206451451 | 0.138 | 0.05 | 1 | 9 |
| yipf4 | 0.38137 | 0.246689905 | 0.362 | 0.2 | 1 | 9 |
| snx12 | 0.38149 | -0.2316582 | 0.069 | 0.15 | 1 | 9 |
| foxp1b | 0.38209 | 0.157556517 | 0.224 | 0.1 | 1 | 9 |
| itsn2b | 0.38215 | -0.22440346 | 0.086 | 0.15 | 1 | 9 |
| nop10 | 0.38282 | 0.284212797 | 0.345 | 0.2 | 1 | 9 |
| zdhhc20b | 0.38314 | -0.12979933 | 0.052 | 0.15 | 1 | 9 |
| si:dkey-188i13.10 | 0.38349 | -0.25690948 | 0.086 | 0.2 | 1 | 9 |
| BX548028.1 | 0.38393 | -0.16077318 | 0.069 | 0.1 | 1 | 9 |
| UTP14C | 0.38395 | 0.134010362 | 0.155 | 0.05 | 1 | 9 |
| efnb1 | 0.38465 | 0.217144196 | 0.138 | 0.05 | 1 | 9 |
| syt11a | 0.3847 | -0.29744349 | 0.069 | 0.15 | 1 | 9 |
| sfxn4 | 0.38479 | 0.212614533 | 0.155 | 0.05 | 1 | 9 |
| nop16 | 0.38499 | 0.203088677 | 0.155 | 0.05 | 1 | 9 |
| pnp5a | 0.38518 | 0.270465989 | 0.19 | 0.1 | 1 | 9 |
| cox10 | 0.38528 | 0.180310237 | 0.121 | 0.05 | 1 | 9 |
| btbd10b | 0.38548 | -0.28917792 | 0.276 | 0.4 | 1 | 9 |
| LOC796453 | 0.38586 | -0.17301405 | 0.034 | 0.1 | 1 | 9 |
| prmt1 | 0.38652 | 0.282316934 | 0.362 | 0.25 | 1 | 9 |
| rgma | 0.38694 | -0.13804214 | 0.138 | 0.15 | 1 | 9 |
| bnip3lb | 0.38716 | -0.11122732 | 0.155 | 0.15 | 1 | 9 |
| tmeff1b | 0.38742 | 0.19274891 | 0.155 | 0.05 | 1 | 9 |
| syncripl | 0.38745 | 0.18671146 | 0.224 | 0.1 | 1 | 9 |
| rnaseka | 0.38753 | -0.27022759 | 0.259 | 0.4 | 1 | 9 |
| peli1b | 0.38761 | -0.15793307 | 0.052 | 0.15 | 1 | 9 |
| tmem198b | 0.38767 | -0.18714145 | 0.052 | 0.1 | 1 | 9 |
| ep300b | 0.38769 | 0.228360505 | 0.121 | 0.05 | 1 | 9 |
| rbm5 | 0.38801 | -0.29062572 | 0.207 | 0.35 | 1 | 9 |
| cdc42ep5 | 0.38809 | 0.190804168 | 0.155 | 0.05 | 1 | 9 |
| polr2h | 0.38826 | 0.143172035 | 0.155 | 0.05 | 1 | 9 |
| si:dkey-247k7.2 | 0.38868 | -0.41594197 | 0.224 | 0.35 | 1 | 9 |
| phf8 | 0.38887 | 0.159465718 | 0.155 | 0.05 | 1 | 9 |
| lrrc17 | 0.38887 | 0.195812534 | 0.241 | 0.2 | 1 | 9 |
| gnpda1 | 0.38926 | -0.18913264 | 0.034 | 0.1 | 1 | 9 |
| grinab | 0.39002 | -0.22376403 | 0.207 | 0.3 | 1 | 9 |
| DDX17 | 0.39002 | -0.22098999 | 0.086 | 0.2 | 1 | 9 |
| pycr1b | 0.39026 | -0.15959915 | 0.052 | 0.15 | 1 | 9 |
| triqk | 0.39075 | -0.18246866 | 0.052 | 0.1 | 1 | 9 |
| lgals9l3 | 0.3908 | -0.20092033 | 0.034 | 0.1 | 1 | 9 |

| | | | | | | |
|---|---|---|---|---|---|---|
| zgc:101744 | 0.3908 | -0.1841816 | 0.052 | 0.15 | 1 | 9 |
| lamp2 | 0.39097 | 0.145763746 | 0.155 | 0.05 | 1 | 9 |
| tom1 | 0.39142 | -0.20091365 | 0.034 | 0.1 | 1 | 9 |
| cxcl20 | 0.39159 | -0.18388469 | 0.052 | 0.15 | 1 | 9 |
| perp | 0.39172 | -0.23299016 | 0.414 | 0.5 | 1 | 9 |
| si:ch1073-314i13.4 | 0.39207 | 0.206031378 | 0.155 | 0.05 | 1 | 9 |
| smim7 | 0.39235 | -0.2187664 | 0.19 | 0.25 | 1 | 9 |
| psmb4 | 0.39256 | -0.25990954 | 0.431 | 0.55 | 1 | 9 |
| zgc:92380 | 0.3926 | -0.36597533 | 0.121 | 0.25 | 1 | 9 |
| AL929237.1 | 0.39271 | -0.20900485 | 0.121 | 0.15 | 1 | 9 |
| tnks1bp1 | 0.39348 | -0.32305958 | 0.328 | 0.45 | 1 | 9 |
| ntan1 | 0.39354 | 0.184108468 | 0.155 | 0.05 | 1 | 9 |
| rab6a | 0.39362 | -0.15815532 | 0.069 | 0.1 | 1 | 9 |
| sirt6 | 0.39406 | -0.18709843 | 0.034 | 0.1 | 1 | 9 |
| slc35b2 | 0.3947 | -0.17270691 | 0.052 | 0.15 | 1 | 9 |
| rab42b | 0.39494 | -0.19313558 | 0.034 | 0.1 | 1 | 9 |
| adgrf8 | 0.39517 | -0.17547903 | 0.052 | 0.15 | 1 | 9 |
| si:ch73-61d6.3 | 0.39559 | -0.16807994 | 0.052 | 0.15 | 1 | 9 |
| hspa4a | 0.39582 | -0.16730624 | 0.103 | 0.15 | 1 | 9 |
| arl4aa | 0.39625 | -0.21812518 | 0.034 | 0.1 | 1 | 9 |
| cct4 | 0.39626 | 0.259764181 | 0.5 | 0.35 | 1 | 9 |
| nbeal1 | 0.39744 | -0.21296899 | 0.052 | 0.15 | 1 | 9 |
| pgam1a | 0.39774 | 0.120661669 | 0.155 | 0.05 | 1 | 9 |
| si:ch211-137i24.10 | 0.39785 | -0.35985263 | 0.086 | 0.2 | 1 | 9 |
| atf1 | 0.39882 | 0.230131426 | 0.259 | 0.15 | 1 | 9 |
| spint1a | 0.39946 | -0.28393322 | 0.259 | 0.4 | 1 | 9 |
| dffa | 0.39986 | -0.19015092 | 0.052 | 0.15 | 1 | 9 |
| gtf2f2a | 0.40099 | 0.175887564 | 0.293 | 0.15 | 1 | 9 |
| si:ch211-262i1.6 | 0.40117 | -0.21126992 | 0.103 | 0.15 | 1 | 9 |
| lipia | 0.40219 | -0.16095463 | 0.086 | 0.2 | 1 | 9 |
| sp4 | 0.40264 | -0.19001663 | 0.052 | 0.15 | 1 | 9 |
| fbxo11b | 0.40275 | -0.17445745 | 0.052 | 0.15 | 1 | 9 |
| tspan7 | 0.40278 | 0.135795223 | 0.207 | 0.1 | 1 | 9 |
| si:ch211-262h13.3 | 0.40283 | 0.17165884 | 0.155 | 0.05 | 1 | 9 |
| nosip | 0.40324 | 0.102528341 | 0.138 | 0.05 | 1 | 9 |
| zgc:92242 | 0.40366 | 0.120297887 | 0.31 | 0.2 | 1 | 9 |
| max | 0.40375 | -0.16165919 | 0.259 | 0.3 | 1 | 9 |
| hnrnpl | 0.40429 | 0.200514821 | 0.155 | 0.05 | 1 | 9 |
| rab32a | 0.40432 | -0.15138712 | 0.052 | 0.15 | 1 | 9 |
| pdzk1ip1 | 0.4047 | -0.25054046 | 0.052 | 0.15 | 1 | 9 |
| cdkn1a | 0.40514 | 0.127345243 | 0.207 | 0.1 | 1 | 9 |
| tomm70a | 0.40539 | 0.197649513 | 0.224 | 0.1 | 1 | 9 |
| arpc4 | 0.40556 | 0.206199894 | 0.207 | 0.1 | 1 | 9 |
| trmt61a | 0.40564 | 0.201385144 | 0.138 | 0.05 | 1 | 9 |
| ifngr1 | 0.40608 | -0.13382565 | 0.052 | 0.15 | 1 | 9 |
| cmc2 | 0.40647 | 0.187614053 | 0.121 | 0.05 | 1 | 9 |
| cish | 0.40686 | -0.12004885 | 0.052 | 0.15 | 1 | 9 |
| ing3 | 0.40725 | -0.11329905 | 0.052 | 0.15 | 1 | 9 |

| | | | | | | |
|---|---|---|---|---|---|---|
| acaa2 | 0.40738 | 0.232269533 | 0.224 | 0.1 | 1 | 9 |
| pho | 0.40822 | 0.120794099 | 0.259 | 0.15 | 1 | 9 |
| s1pr2 | 0.40823 | -0.17239746 | 0.052 | 0.15 | 1 | 9 |
| tbl3 | 0.40849 | -0.18130951 | 0.069 | 0.1 | 1 | 9 |
| wnt11r | 0.40858 | -0.21158006 | 0.052 | 0.15 | 1 | 9 |
| rbms1a | 0.40887 | -0.18320082 | 0.052 | 0.15 | 1 | 9 |
| clk4a | 0.40893 | -0.22886205 | 0.121 | 0.25 | 1 | 9 |
| copg2 | 0.40904 | -0.22322847 | 0.155 | 0.25 | 1 | 9 |
| hsd17b12a | 0.40907 | 0.114831275 | 0.103 | 0.1 | 1 | 9 |
| etfb | 0.40909 | 0.22984789 | 0.293 | 0.15 | 1 | 9 |
| ddah2 | 0.40914 | -0.18376448 | 0.052 | 0.15 | 1 | 9 |
| sptan1 | 0.4093 | -0.15183235 | 0.121 | 0.15 | 1 | 9 |
| cdc42 | 0.40964 | -0.22004818 | 0.241 | 0.4 | 1 | 9 |
| clic2 | 0.41011 | -0.2361996 | 0.052 | 0.15 | 1 | 9 |
| hnrnpul1 | 0.41013 | 0.194171626 | 0.155 | 0.05 | 1 | 9 |
| ctnnbip1 | 0.41042 | -0.16409774 | 0.086 | 0.2 | 1 | 9 |
| riok1 | 0.41042 | 0.207295195 | 0.138 | 0.05 | 1 | 9 |
| nfil3-6 | 0.41044 | -0.2042513 | 0.103 | 0.15 | 1 | 9 |
| eif4g2b | 0.41064 | -0.12939925 | 0.052 | 0.15 | 1 | 9 |
| MZT1 | 0.41114 | 0.147473373 | 0.155 | 0.05 | 1 | 9 |
| fbxo28 | 0.41129 | 0.189679302 | 0.155 | 0.05 | 1 | 9 |
| asah1b | 0.41181 | 0.114474103 | 0.138 | 0.05 | 1 | 9 |
| b3gnt3.4 | 0.41203 | -0.20835109 | 0.103 | 0.15 | 1 | 9 |
| polr2f | 0.41244 | 0.164113766 | 0.397 | 0.25 | 1 | 9 |
| echdc2 | 0.41252 | -0.1362281 | 0.052 | 0.15 | 1 | 9 |
| arl6ip5b | 0.41268 | 0.234961229 | 0.293 | 0.15 | 1 | 9 |
| polr3k | 0.41281 | -0.14698567 | 0.121 | 0.25 | 1 | 9 |
| ctso | 0.41293 | -0.17923657 | 0.052 | 0.15 | 1 | 9 |
| emg1 | 0.41298 | -0.15249962 | 0.052 | 0.15 | 1 | 9 |
| gpm6bb | 0.413 | -0.21901877 | 0.052 | 0.15 | 1 | 9 |
| gspt1l | 0.41303 | 0.177772115 | 0.155 | 0.05 | 1 | 9 |
| plp2 | 0.41307 | -0.22540778 | 0.121 | 0.25 | 1 | 9 |
| hmgb2a | 0.41308 | 0.206765849 | 0.759 | 0.6 | 1 | 9 |
| msra | 0.41332 | 0.187636865 | 0.138 | 0.05 | 1 | 9 |
| gbp1 | 0.41351 | -0.10901757 | 0.052 | 0.05 | 1 | 9 |
| egfra | 0.41372 | -0.20056569 | 0.052 | 0.15 | 1 | 9 |
| lxn | 0.41373 | -0.256038 | 0.224 | 0.35 | 1 | 9 |
| leng1 | 0.41375 | -0.17348217 | 0.052 | 0.15 | 1 | 9 |
| ppfibp1b | 0.41385 | -0.19205553 | 0.052 | 0.15 | 1 | 9 |
| fuom | 0.41397 | -0.16719209 | 0.052 | 0.15 | 1 | 9 |
| dip2ca | 0.41402 | -0.16697847 | 0.052 | 0.15 | 1 | 9 |
| hsh2d | 0.41419 | -0.17936598 | 0.052 | 0.15 | 1 | 9 |
| elovl6l | 0.41432 | -0.22230099 | 0.086 | 0.2 | 1 | 9 |
| phc2a | 0.41438 | 0.161274192 | 0.155 | 0.05 | 1 | 9 |
| gprc5c | 0.41455 | -0.18251707 | 0.086 | 0.2 | 1 | 9 |
| elof1 | 0.41475 | 0.166100381 | 0.276 | 0.15 | 1 | 9 |
| wdr61 | 0.41484 | 0.130171381 | 0.155 | 0.05 | 1 | 9 |
| hspg2 | 0.41515 | 0.224212915 | 0.155 | 0.05 | 1 | 9 |

| | | | | | | |
|---|---|---|---|---|---|---|
| aes | 0.41536 | 0.216510238 | 0.155 | 0.05 | 1 | 9 |
| sf3a3 | 0.41544 | 0.159165483 | 0.155 | 0.05 | 1 | 9 |
| atp2b1a | 0.41545 | -0.20458815 | 0.086 | 0.15 | 1 | 9 |
| mrps16 | 0.41566 | 0.178923236 | 0.155 | 0.05 | 1 | 9 |
| ppargc1b | 0.41594 | -0.22704493 | 0.086 | 0.2 | 1 | 9 |
| btg3 | 0.41604 | 0.167074573 | 0.155 | 0.05 | 1 | 9 |
| mccc2 | 0.41612 | 0.12647235 | 0.155 | 0.05 | 1 | 9 |
| zgc:194981 | 0.41619 | 0.16360984 | 0.155 | 0.05 | 1 | 9 |
| tmem47 | 0.41632 | -0.1006463 | 0.052 | 0.05 | 1 | 9 |
| zgc:193505 | 0.4165 | 0.233965509 | 0.276 | 0.25 | 1 | 9 |
| epdl1 | 0.41681 | -0.23845442 | 0.086 | 0.2 | 1 | 9 |
| si:dkey-16p21.8 | 0.41716 | 0.218216815 | 0.207 | 0.15 | 1 | 9 |
| nck1b | 0.41723 | 0.170337071 | 0.155 | 0.05 | 1 | 9 |
| sult6b1 | 0.41735 | 0.169658917 | 0.224 | 0.1 | 1 | 9 |
| cast | 0.41741 | -0.17632962 | 0.172 | 0.3 | 1 | 9 |
| xirp2a | 0.41802 | -0.19733088 | 0.086 | 0.2 | 1 | 9 |
| flvcr1 | 0.41829 | -0.19049807 | 0.034 | 0.1 | 1 | 9 |
| PDE3B | 0.41839 | -0.19737987 | 0.086 | 0.15 | 1 | 9 |
| glrx5 | 0.41905 | 0.174726079 | 0.155 | 0.05 | 1 | 9 |
| cyba | 0.41958 | -0.16769791 | 0.034 | 0.1 | 1 | 9 |
| dcun1d5 | 0.41976 | 0.12947294 | 0.086 | 0.05 | 1 | 9 |
| ap3d1 | 0.41977 | -0.16888126 | 0.034 | 0.1 | 1 | 9 |
| lta4h | 0.41979 | -0.18861788 | 0.034 | 0.1 | 1 | 9 |
| got2b | 0.42035 | 0.16658908 | 0.155 | 0.05 | 1 | 9 |
| nxt2 | 0.42036 | 0.163644899 | 0.155 | 0.05 | 1 | 9 |
| pprc1 | 0.42048 | 0.217054642 | 0.155 | 0.05 | 1 | 9 |
| akirin1 | 0.42118 | 0.19541862 | 0.293 | 0.15 | 1 | 9 |
| homer2 | 0.42211 | -0.23521285 | 0.103 | 0.2 | 1 | 9 |
| creld2 | 0.42244 | -0.22208152 | 0.052 | 0.1 | 1 | 9 |
| mknk2b | 0.42267 | 0.232236081 | 0.293 | 0.15 | 1 | 9 |
| pgam5 | 0.42274 | 0.174567231 | 0.121 | 0.05 | 1 | 9 |
| tmco1 | 0.42294 | -0.21702312 | 0.086 | 0.2 | 1 | 9 |
| tcf7l2 | 0.4235 | 0.182637125 | 0.103 | 0.05 | 1 | 9 |
| ebna1bp2 | 0.42369 | 0.144216816 | 0.207 | 0.1 | 1 | 9 |
| os9 | 0.42372 | 0.150079628 | 0.155 | 0.1 | 1 | 9 |
| ppp4r3b | 0.42407 | 0.172057402 | 0.155 | 0.05 | 1 | 9 |
| prkd2 | 0.42425 | -0.2139109 | 0.052 | 0.1 | 1 | 9 |
| gtf2h5 | 0.42443 | 0.154691814 | 0.155 | 0.05 | 1 | 9 |
| srprb | 0.42451 | -0.15369387 | 0.276 | 0.3 | 1 | 9 |
| farsb | 0.42475 | 0.167915423 | 0.155 | 0.05 | 1 | 9 |
| syncrip | 0.42501 | 0.22713558 | 0.569 | 0.4 | 1 | 9 |
| si:ch211-156b7.4 | 0.42565 | 0.185988705 | 0.155 | 0.05 | 1 | 9 |
| eif2b4 | 0.42568 | 0.148564908 | 0.155 | 0.05 | 1 | 9 |
| mrps10 | 0.42574 | 0.145501829 | 0.155 | 0.05 | 1 | 9 |
| bxdc2 | 0.42579 | 0.195860532 | 0.155 | 0.05 | 1 | 9 |
| gsr | 0.42596 | -0.18328037 | 0.086 | 0.2 | 1 | 9 |
| ptk7a | 0.42599 | 0.19301689 | 0.155 | 0.05 | 1 | 9 |
| ak3 | 0.42603 | -0.17346759 | 0.052 | 0.1 | 1 | 9 |

| | | | | | | |
|---|---|---|---|---|---|---|
| cox6b1 | 0.42608 | -0.24644135 | 0.086 | 0.2 | 1 | 9 |
| CDK2AP1 | 0.42609 | 0.185213027 | 0.155 | 0.05 | 1 | 9 |
| chchd2 | 0.42615 | 0.217563717 | 0.31 | 0.2 | 1 | 9 |
| vps28 | 0.42663 | 0.158258005 | 0.397 | 0.25 | 1 | 9 |
| psmc1b | 0.42677 | 0.105415939 | 0.138 | 0.05 | 1 | 9 |
| nudt21 | 0.42698 | 0.224500113 | 0.224 | 0.1 | 1 | 9 |
| gpatch8 | 0.42727 | -0.27268106 | 0.207 | 0.35 | 1 | 9 |
| dnajc8 | 0.42821 | 0.125716803 | 0.207 | 0.1 | 1 | 9 |
| mt-nd4 | 0.42843 | -0.27549501 | 0.552 | 0.7 | 1 | 9 |
| zbtb37 | 0.42855 | -0.19965319 | 0.086 | 0.15 | 1 | 9 |
| gsto2 | 0.4293 | 0.242300597 | 0.276 | 0.15 | 1 | 9 |
| arhgap29a | 0.4293 | -0.16034162 | 0.034 | 0.1 | 1 | 9 |
| xrcc6 | 0.42945 | -0.19385019 | 0.086 | 0.15 | 1 | 9 |
| lpar2a | 0.4299 | -0.18358436 | 0.034 | 0.1 | 1 | 9 |
| gadd45ba | 0.42999 | -0.19979198 | 0.397 | 0.45 | 1 | 9 |
| trappc3 | 0.43052 | -0.1271253 | 0.086 | 0.1 | 1 | 9 |
| qsox1 | 0.43101 | -0.17029189 | 0.052 | 0.1 | 1 | 9 |
| litaf | 0.43122 | -0.26965389 | 0.224 | 0.35 | 1 | 9 |
| zgc:100829 | 0.43144 | 0.197206344 | 0.138 | 0.05 | 1 | 9 |
| si:ch73-1a9.3 | 0.43149 | 0.249091949 | 0.707 | 0.6 | 1 | 9 |
| vkorc1l1 | 0.43165 | -0.13501092 | 0.19 | 0.2 | 1 | 9 |
| mob2a | 0.43165 | 0.183136918 | 0.103 | 0.05 | 1 | 9 |
| rnf38 | 0.43168 | -0.14594141 | 0.069 | 0.1 | 1 | 9 |
| hcfc1a | 0.43191 | 0.183760593 | 0.224 | 0.1 | 1 | 9 |
| vmp1 | 0.43194 | -0.19385529 | 0.052 | 0.1 | 1 | 9 |
| cyldb | 0.4323 | 0.212637923 | 0.172 | 0.1 | 1 | 9 |
| fam168b | 0.43231 | 0.193070325 | 0.224 | 0.1 | 1 | 9 |
| mrpl52 | 0.43269 | 0.186455032 | 0.224 | 0.1 | 1 | 9 |
| arl14ep | 0.43275 | 0.193500883 | 0.138 | 0.05 | 1 | 9 |
| capns1a | 0.43338 | -0.23333702 | 0.379 | 0.45 | 1 | 9 |
| ebag9 | 0.43411 | -0.18074828 | 0.207 | 0.25 | 1 | 9 |
| map2k2a | 0.43446 | 0.11464461 | 0.138 | 0.05 | 1 | 9 |
| nqo1 | 0.43451 | 0.273779487 | 0.207 | 0.1 | 1 | 9 |
| dnajc19 | 0.43502 | 0.2261369 | 0.276 | 0.15 | 1 | 9 |
| isca2 | 0.43507 | -0.1481279 | 0.069 | 0.1 | 1 | 9 |
| ponzr6 | 0.43511 | -0.21168088 | 0.034 | 0.1 | 1 | 9 |
| nkain1 | 0.43541 | 0.186081299 | 0.138 | 0.05 | 1 | 9 |
| ap2a1 | 0.43544 | -0.13758848 | 0.121 | 0.15 | 1 | 9 |
| tcta | 0.43582 | 0.210175567 | 0.224 | 0.1 | 1 | 9 |
| pof1b | 0.43607 | -0.22580549 | 0.207 | 0.35 | 1 | 9 |
| igf2bp2b | 0.43639 | 0.202980484 | 0.207 | 0.1 | 1 | 9 |
| papola | 0.43687 | -0.19892179 | 0.172 | 0.25 | 1 | 9 |
| dbnla | 0.43693 | 0.150210254 | 0.138 | 0.1 | 1 | 9 |
| rpe | 0.43695 | 0.183467011 | 0.19 | 0.1 | 1 | 9 |
| ssu72 | 0.437 | -0.21386864 | 0.121 | 0.2 | 1 | 9 |
| dnajb6a | 0.43716 | 0.215899157 | 0.207 | 0.1 | 1 | 9 |
| ccnd1 | 0.43718 | -0.14163463 | 0.276 | 0.3 | 1 | 9 |
| s100u | 0.43779 | 0.226783101 | 0.224 | 0.1 | 1 | 9 |

| | | | | | | |
|---|---|---|---|---|---|---|
| si:ch211-198a12.6 | 0.4379 | 0.18965768 | 0.224 | 0.1 | 1 | 9 |
| arfgap3 | 0.43833 | 0.13796276 | 0.207 | 0.1 | 1 | 9 |
| surf4l | 0.43868 | -0.20744009 | 0.19 | 0.25 | 1 | 9 |
| mtpn | 0.43887 | 0.120651544 | 0.328 | 0.2 | 1 | 9 |
| zyg11 | 0.43924 | 0.207248522 | 0.138 | 0.05 | 1 | 9 |
| cpvl | 0.43952 | -0.17262209 | 0.034 | 0.1 | 1 | 9 |
| nup50 | 0.43982 | 0.201021732 | 0.224 | 0.1 | 1 | 9 |
| reep3b | 0.44051 | -0.17567431 | 0.259 | 0.3 | 1 | 9 |
| tia1l | 0.44069 | 0.163673104 | 0.224 | 0.1 | 1 | 9 |
| lasp1 | 0.44109 | -0.13419247 | 0.328 | 0.35 | 1 | 9 |
| aldh9a1a.1 | 0.44197 | 0.205766378 | 0.345 | 0.2 | 1 | 9 |
| atp6v1h | 0.44245 | 0.227506666 | 0.19 | 0.1 | 1 | 9 |
| fat1a | 0.44282 | -0.11276979 | 0.086 | 0.1 | 1 | 9 |
| zgc:163080 | 0.4431 | -0.25165549 | 0.224 | 0.35 | 1 | 9 |
| cbx8a | 0.44352 | -0.17367211 | 0.034 | 0.1 | 1 | 9 |
| gnptg | 0.44378 | -0.13993877 | 0.138 | 0.15 | 1 | 9 |
| tmem39a | 0.44412 | -0.17159632 | 0.069 | 0.1 | 1 | 9 |
| tspan3a | 0.44523 | -0.24177007 | 0.19 | 0.3 | 1 | 9 |
| snx27a | 0.44547 | -0.18980279 | 0.034 | 0.1 | 1 | 9 |
| nfkbiaa | 0.44591 | -0.20614759 | 0.207 | 0.25 | 1 | 9 |
| si:ch73-78o10.1 | 0.44612 | -0.1581125 | 0.069 | 0.1 | 1 | 9 |
| arf6a | 0.4467 | -0.19657438 | 0.103 | 0.15 | 1 | 9 |
| atp5mc1 | 0.44685 | 0.133394657 | 0.552 | 0.4 | 1 | 9 |
| rps29 | 0.44738 | 0.12198079 | 0.966 | 0.9 | 1 | 9 |
| atg12 | 0.44785 | 0.173641331 | 0.121 | 0.05 | 1 | 9 |
| csnk2b | 0.4479 | 0.235192485 | 0.431 | 0.3 | 1 | 9 |
| hiat1a | 0.44832 | -0.18260517 | 0.069 | 0.15 | 1 | 9 |
| pdgfaa | 0.44902 | 0.15628597 | 0.138 | 0.05 | 1 | 9 |
| eloca | 0.44922 | -0.19853756 | 0.069 | 0.15 | 1 | 9 |
| utp25 | 0.44927 | -0.11236187 | 0.086 | 0.1 | 1 | 9 |
| pgk1 | 0.44951 | -0.24017986 | 0.103 | 0.2 | 1 | 9 |
| ubl7b | 0.44971 | -0.15118981 | 0.155 | 0.2 | 1 | 9 |
| vta1 | 0.44995 | -0.11605914 | 0.138 | 0.15 | 1 | 9 |
| tp53bp2b | 0.45013 | -0.16157675 | 0.069 | 0.1 | 1 | 9 |
| nip7 | 0.45031 | 0.238414198 | 0.259 | 0.15 | 1 | 9 |
| map2k6 | 0.45122 | -0.18530695 | 0.155 | 0.2 | 1 | 9 |
| mibp2 | 0.45133 | 0.193865429 | 0.828 | 0.7 | 1 | 9 |
| chkb | 0.45174 | -0.14182718 | 0.069 | 0.15 | 1 | 9 |
| ccnt2b | 0.45242 | 0.204560201 | 0.207 | 0.1 | 1 | 9 |
| entpd1 | 0.45256 | -0.18058409 | 0.052 | 0.1 | 1 | 9 |
| padi2 | 0.45308 | -0.16547293 | 0.103 | 0.15 | 1 | 9 |
| mrps36 | 0.45324 | 0.21911989 | 0.276 | 0.15 | 1 | 9 |
| snx5 | 0.45354 | -0.13638459 | 0.069 | 0.1 | 1 | 9 |
| zgc:113279 | 0.45356 | 0.2334804 | 0.121 | 0.05 | 1 | 9 |
| prdx1 | 0.45386 | 0.151931589 | 0.138 | 0.05 | 1 | 9 |
| exoc7 | 0.45391 | -0.11305299 | 0.086 | 0.1 | 1 | 9 |
| psmd11b | 0.45403 | -0.14483784 | 0.172 | 0.2 | 1 | 9 |
| tmem120a | 0.45476 | -0.12981089 | 0.069 | 0.1 | 1 | 9 |

| | | | | | | |
|---|---|---|---|---|---|---|
| u2af2b | 0.45612 | 0.230577164 | 0.345 | 0.2 | 1 | 9 |
| cldnb | 0.45726 | -0.31628131 | 0.379 | 0.5 | 1 | 9 |
| fkbp14 | 0.45787 | -0.15559663 | 0.034 | 0.1 | 1 | 9 |
| deptor | 0.45888 | -0.18528649 | 0.069 | 0.15 | 1 | 9 |
| atp8b1 | 0.46015 | -0.17553437 | 0.069 | 0.1 | 1 | 9 |
| sox13 | 0.46027 | -0.20970834 | 0.052 | 0.1 | 1 | 9 |
| FP017215.1 | 0.46065 | 0.146556468 | 0.19 | 0.15 | 1 | 9 |
| commd3 | 0.46085 | -0.1660319 | 0.207 | 0.25 | 1 | 9 |
| fosab | 0.46136 | -0.2229279 | 0.638 | 0.7 | 1 | 9 |
| wdr82 | 0.46148 | 0.11479009 | 0.069 | 0.05 | 1 | 9 |
| znf654 | 0.46183 | -0.16002465 | 0.052 | 0.1 | 1 | 9 |
| net1 | 0.46184 | 0.228034782 | 0.345 | 0.2 | 1 | 9 |
| LOC110438237 | 0.46188 | -0.16472898 | 0.034 | 0.1 | 1 | 9 |
| snrpb | 0.46281 | 0.247628685 | 0.552 | 0.4 | 1 | 9 |
| nmt1a | 0.46438 | -0.19632625 | 0.224 | 0.3 | 1 | 9 |
| mrpl54 | 0.46448 | 0.18320642 | 0.276 | 0.2 | 1 | 9 |
| sf3b3 | 0.46499 | -0.14856279 | 0.069 | 0.15 | 1 | 9 |
| ccdc22 | 0.4652 | -0.1658078 | 0.052 | 0.1 | 1 | 9 |
| si:dkey-28b4.7 | 0.46526 | 0.243617651 | 0.448 | 0.3 | 1 | 9 |
| zgc:66475 | 0.46614 | -0.20325542 | 0.034 | 0.1 | 1 | 9 |
| timp2a | 0.46622 | 0.15433013 | 0.207 | 0.1 | 1 | 9 |
| si:ch211-253h3.1 | 0.46754 | -0.20251789 | 0.034 | 0.1 | 1 | 9 |
| smap1 | 0.46786 | -0.13189364 | 0.086 | 0.1 | 1 | 9 |
| gas1a | 0.46793 | 0.225386802 | 0.121 | 0.05 | 1 | 9 |
| gpatch11 | 0.46884 | -0.18070839 | 0.052 | 0.1 | 1 | 9 |
| actn4 | 0.46899 | 0.227202461 | 0.328 | 0.2 | 1 | 9 |
| sort1b | 0.46905 | -0.21002566 | 0.069 | 0.15 | 1 | 9 |
| kat7b | 0.47011 | 0.161265465 | 0.121 | 0.05 | 1 | 9 |
| lnx2b | 0.47033 | -0.21109019 | 0.103 | 0.2 | 1 | 9 |
| rnf10 | 0.4704 | -0.15000603 | 0.31 | 0.35 | 1 | 9 |
| stk35l | 0.47055 | -0.11182964 | 0.138 | 0.15 | 1 | 9 |
| rnf25 | 0.47058 | -0.18475766 | 0.052 | 0.1 | 1 | 9 |
| sts | 0.47072 | 0.211750027 | 0.138 | 0.05 | 1 | 9 |
| si:ch211-251b21.1 | 0.47077 | -0.26049494 | 0.052 | 0.1 | 1 | 9 |
| yaf2 | 0.47098 | 0.177655693 | 0.138 | 0.05 | 1 | 9 |
| BX957297.2 | 0.4711 | -0.11632515 | 0.017 | 0.05 | 1 | 9 |
| EARS2 | 0.4711 | -0.10875362 | 0.017 | 0.05 | 1 | 9 |
| XLOC-002129 | 0.4711 | -0.16062528 | 0.017 | 0.05 | 1 | 9 |
| XLOC-009550 | 0.4711 | -0.12019313 | 0.017 | 0.05 | 1 | 9 |
| XLOC-021534 | 0.4711 | -0.12293217 | 0.017 | 0.05 | 1 | 9 |
| XLOC-027844 | 0.4711 | -0.10813236 | 0.017 | 0.05 | 1 | 9 |
| abi1b | 0.4711 | -0.10243212 | 0.017 | 0.05 | 1 | 9 |
| agk | 0.4711 | -0.11834306 | 0.017 | 0.05 | 1 | 9 |
| alg6 | 0.4711 | -0.12721511 | 0.017 | 0.05 | 1 | 9 |
| ankib1b | 0.4711 | -0.10883917 | 0.017 | 0.05 | 1 | 9 |
| apc | 0.4711 | -0.10930847 | 0.017 | 0.05 | 1 | 9 |
| cachd1 | 0.4711 | -0.12114105 | 0.017 | 0.05 | 1 | 9 |
| cdip1 | 0.4711 | -0.1301803 | 0.017 | 0.05 | 1 | 9 |

| | | | | | | |
|---|---|---|---|---|---|---|
| cdk4 | 0.4711 | -0.18197352 | 0.017 | 0.05 | 1 | 9 |
| cdyl | 0.4711 | -0.17512649 | 0.017 | 0.05 | 1 | 9 |
| cox5b2 | 0.4711 | -0.19176337 | 0.017 | 0.05 | 1 | 9 |
| cox6a2 | 0.4711 | -0.15986872 | 0.017 | 0.05 | 1 | 9 |
| dbpa | 0.4711 | -0.1055375 | 0.017 | 0.05 | 1 | 9 |
| ddx20 | 0.4711 | -0.13697082 | 0.017 | 0.05 | 1 | 9 |
| dtd2 | 0.4711 | -0.13449441 | 0.017 | 0.05 | 1 | 9 |
| ftr50 | 0.4711 | -0.14277174 | 0.017 | 0.05 | 1 | 9 |
| galk1 | 0.4711 | -0.10807769 | 0.017 | 0.05 | 1 | 9 |
| galnt6 | 0.4711 | -0.11041167 | 0.017 | 0.05 | 1 | 9 |
| gna15.1 | 0.4711 | -0.10541412 | 0.017 | 0.05 | 1 | 9 |
| hmbsb | 0.4711 | -0.12418144 | 0.017 | 0.05 | 1 | 9 |
| hs3st1l2 | 0.4711 | -0.12194343 | 0.017 | 0.05 | 1 | 9 |
| itgb1bp1 | 0.4711 | -0.13779358 | 0.017 | 0.05 | 1 | 9 |
| kirrel3l | 0.4711 | -0.16629815 | 0.017 | 0.05 | 1 | 9 |
| klf12b | 0.4711 | -0.16097043 | 0.017 | 0.05 | 1 | 9 |
| l3hypdh | 0.4711 | -0.1734095 | 0.017 | 0.05 | 1 | 9 |
| lamb1b | 0.4711 | -0.1693706 | 0.017 | 0.05 | 1 | 9 |
| mhc1lba | 0.4711 | -0.14783293 | 0.017 | 0.05 | 1 | 9 |
| nkd1 | 0.4711 | -0.14393362 | 0.017 | 0.05 | 1 | 9 |
| nudt18 | 0.4711 | -0.11014926 | 0.017 | 0.05 | 1 | 9 |
| olfml2ba | 0.4711 | -0.21328316 | 0.017 | 0.05 | 1 | 9 |
| pde6d | 0.4711 | -0.11900121 | 0.017 | 0.05 | 1 | 9 |
| pim2 | 0.4711 | -0.18562352 | 0.017 | 0.05 | 1 | 9 |
| plcd3a | 0.4711 | -0.10978902 | 0.017 | 0.05 | 1 | 9 |
| rasgef1bb | 0.4711 | -0.14592275 | 0.017 | 0.05 | 1 | 9 |
| rbl2 | 0.4711 | -0.12639476 | 0.017 | 0.05 | 1 | 9 |
| rgl3b | 0.4711 | -0.13698016 | 0.017 | 0.05 | 1 | 9 |
| scaper | 0.4711 | -0.13682009 | 0.017 | 0.05 | 1 | 9 |
| scp2a | 0.4711 | -0.11631749 | 0.017 | 0.05 | 1 | 9 |
| scube3 | 0.4711 | -0.11380676 | 0.017 | 0.05 | 1 | 9 |
| sec63 | 0.4711 | -0.13670581 | 0.017 | 0.05 | 1 | 9 |
| setd1ba | 0.4711 | -0.10285247 | 0.017 | 0.05 | 1 | 9 |
| si:ch211-79k12.2 | 0.4711 | -0.10281106 | 0.017 | 0.05 | 1 | 9 |
| si:ch211-79l17.1 | 0.4711 | -0.10739696 | 0.017 | 0.05 | 1 | 9 |
| si:dkey-197j19.6 | 0.4711 | -0.20803232 | 0.017 | 0.05 | 1 | 9 |
| si:zfos-80g12.1 | 0.4711 | -0.10249388 | 0.017 | 0.05 | 1 | 9 |
| slbp | 0.4711 | -0.11550265 | 0.017 | 0.05 | 1 | 9 |
| slc4a4a | 0.4711 | -0.21585343 | 0.017 | 0.05 | 1 | 9 |
| smpd3 | 0.4711 | -0.15757164 | 0.017 | 0.05 | 1 | 9 |
| tap1 | 0.4711 | -0.12359995 | 0.017 | 0.05 | 1 | 9 |
| tapt1a | 0.4711 | -0.10629408 | 0.017 | 0.05 | 1 | 9 |
| thnsl2 | 0.4711 | -0.12431671 | 0.017 | 0.05 | 1 | 9 |
| tmem145 | 0.4711 | -0.12990414 | 0.017 | 0.05 | 1 | 9 |
| tyk2 | 0.4711 | -0.1523662 | 0.017 | 0.05 | 1 | 9 |
| ube2r2 | 0.4711 | -0.108658 | 0.017 | 0.05 | 1 | 9 |
| vps35l | 0.4711 | -0.13046345 | 0.017 | 0.05 | 1 | 9 |
| vsir | 0.4711 | -0.12719986 | 0.017 | 0.05 | 1 | 9 |

| | | | | | | |
|---|---|---|---|---|---|---|
| vwa8 | 0.4711 | -0.12620777 | 0.017 | 0.05 | 1 | 9 |
| wbp1 | 0.4711 | -0.10757253 | 0.017 | 0.05 | 1 | 9 |
| wnk1b | 0.4711 | -0.12662864 | 0.017 | 0.05 | 1 | 9 |
| zbtb46 | 0.4711 | -0.13209103 | 0.017 | 0.05 | 1 | 9 |
| zgc:113423 | 0.4711 | -0.10475123 | 0.017 | 0.05 | 1 | 9 |
| zgc:136493 | 0.4711 | -0.20260792 | 0.017 | 0.05 | 1 | 9 |
| zgc:174890 | 0.4711 | -0.13182672 | 0.017 | 0.05 | 1 | 9 |
| zgc:77880 | 0.4711 | -0.10483523 | 0.017 | 0.05 | 1 | 9 |
| BX548015.2 | 0.4711 | -0.1256465 | 0.017 | 0.05 | 1 | 9 |
| CABZ01073795.1 | 0.4711 | -0.10798912 | 0.017 | 0.05 | 1 | 9 |
| CU469577.1 | 0.4711 | -0.10152296 | 0.017 | 0.05 | 1 | 9 |
| LOC100334439 | 0.4711 | -0.11770723 | 0.017 | 0.05 | 1 | 9 |
| LOC556776 | 0.4711 | -0.11404128 | 0.017 | 0.05 | 1 | 9 |
| XLOC-020187 | 0.4711 | -0.13000878 | 0.017 | 0.05 | 1 | 9 |
| XLOC-042200 | 0.4711 | -0.10524065 | 0.017 | 0.05 | 1 | 9 |
| ablim1a | 0.4711 | -0.12263856 | 0.017 | 0.05 | 1 | 9 |
| acot7 | 0.4711 | -0.12286442 | 0.017 | 0.05 | 1 | 9 |
| atmin | 0.4711 | -0.12450883 | 0.017 | 0.05 | 1 | 9 |
| casp2 | 0.4711 | -0.11051429 | 0.017 | 0.05 | 1 | 9 |
| ccr6a | 0.4711 | -0.10204613 | 0.017 | 0.05 | 1 | 9 |
| cdc42ep1b | 0.4711 | -0.10001443 | 0.017 | 0.05 | 1 | 9 |
| cep57l1 | 0.4711 | -0.11211197 | 0.017 | 0.05 | 1 | 9 |
| chd1l | 0.4711 | -0.105531 | 0.017 | 0.05 | 1 | 9 |
| cipca | 0.4711 | -0.11805585 | 0.017 | 0.05 | 1 | 9 |
| coq8b | 0.4711 | -0.1049665 | 0.017 | 0.05 | 1 | 9 |
| cx43 | 0.4711 | -0.29688517 | 0.017 | 0.05 | 1 | 9 |
| dennd5a | 0.4711 | -0.12522123 | 0.017 | 0.05 | 1 | 9 |
| dmxl2 | 0.4711 | -0.1032088 | 0.017 | 0.05 | 1 | 9 |
| efna3a | 0.4711 | -0.13268556 | 0.017 | 0.05 | 1 | 9 |
| egr2a | 0.4711 | -0.10297763 | 0.017 | 0.05 | 1 | 9 |
| fyco1a | 0.4711 | -0.10234917 | 0.017 | 0.05 | 1 | 9 |
| ggctb | 0.4711 | -0.12421981 | 0.017 | 0.05 | 1 | 9 |
| lrrc4ba | 0.4711 | -0.10878939 | 0.017 | 0.05 | 1 | 9 |
| mbd5 | 0.4711 | -0.12318082 | 0.017 | 0.05 | 1 | 9 |
| mettl27 | 0.4711 | -0.10930569 | 0.017 | 0.05 | 1 | 9 |
| pax7b | 0.4711 | -0.10717062 | 0.017 | 0.05 | 1 | 9 |
| pax9 | 0.4711 | -0.11450522 | 0.017 | 0.05 | 1 | 9 |
| pcnt | 0.4711 | -0.11099642 | 0.017 | 0.05 | 1 | 9 |
| pign | 0.4711 | -0.15194251 | 0.017 | 0.05 | 1 | 9 |
| plaa | 0.4711 | -0.1039957 | 0.017 | 0.05 | 1 | 9 |
| plppr1 | 0.4711 | -0.11141934 | 0.017 | 0.05 | 1 | 9 |
| ptdss1a | 0.4711 | -0.11258005 | 0.017 | 0.05 | 1 | 9 |
| rab39bb | 0.4711 | -0.10679735 | 0.017 | 0.05 | 1 | 9 |
| ranbp3a | 0.4711 | -0.11095837 | 0.017 | 0.05 | 1 | 9 |
| rnf166 | 0.4711 | -0.10836468 | 0.017 | 0.05 | 1 | 9 |
| sepsecs | 0.4711 | -0.11018022 | 0.017 | 0.05 | 1 | 9 |
| sgsh | 0.4711 | -0.12838437 | 0.017 | 0.05 | 1 | 9 |
| si:ch1073-303k11.2 | 0.4711 | -0.18291146 | 0.017 | 0.05 | 1 | 9 |

| | | | | | | |
|---|---|---|---|---|---|---|
| si:ch211-113e8.11 | 0.4711 | -0.12284758 | 0.017 | 0.05 | 1 | 9 |
| si:ch211-261n11.7 | 0.4711 | -0.10374395 | 0.017 | 0.05 | 1 | 9 |
| si:ch73-144d13.7 | 0.4711 | -0.12780044 | 0.017 | 0.05 | 1 | 9 |
| si:ch73-281f12.4 | 0.4711 | -0.10021242 | 0.017 | 0.05 | 1 | 9 |
| snx17 | 0.4711 | -0.11765131 | 0.017 | 0.05 | 1 | 9 |
| thrap3a | 0.4711 | -0.10882277 | 0.017 | 0.05 | 1 | 9 |
| tmem234 | 0.4711 | -0.1217945 | 0.017 | 0.05 | 1 | 9 |
| unc45a | 0.4711 | -0.10913725 | 0.017 | 0.05 | 1 | 9 |
| wu:fc34e06 | 0.4711 | -0.12336335 | 0.017 | 0.05 | 1 | 9 |
| zfand1 | 0.4711 | -0.10258327 | 0.017 | 0.05 | 1 | 9 |
| zgc:101016 | 0.4711 | -0.11942581 | 0.017 | 0.05 | 1 | 9 |
| zgc:112998 | 0.4711 | -0.10016225 | 0.017 | 0.05 | 1 | 9 |
| zgc:174972 | 0.4711 | -0.13539542 | 0.017 | 0.05 | 1 | 9 |
| znf1007 | 0.4711 | -0.11357033 | 0.017 | 0.05 | 1 | 9 |
| CABZ01028771.1 | 0.4711 | -0.1178951 | 0.017 | 0.05 | 1 | 9 |
| bhmt | 0.4711 | -0.10765826 | 0.017 | 0.05 | 1 | 9 |
| blzf1 | 0.4711 | -0.11748418 | 0.017 | 0.05 | 1 | 9 |
| si:ch211-122f10.4 | 0.4711 | -0.20478993 | 0.017 | 0.05 | 1 | 9 |
| si:ch211-240l19.5 | 0.4711 | -0.16472993 | 0.017 | 0.05 | 1 | 9 |
| sorbs3 | 0.4711 | -0.10328801 | 0.017 | 0.05 | 1 | 9 |
| spryd7b | 0.4711 | -0.11251671 | 0.017 | 0.05 | 1 | 9 |
| trim32 | 0.4711 | -0.11272164 | 0.017 | 0.05 | 1 | 9 |
| CABZ01050249.1 | 0.4711 | -0.11864971 | 0.017 | 0.05 | 1 | 9 |
| RNF14 (1 of many) | 0.4711 | -0.10856346 | 0.017 | 0.05 | 1 | 9 |
| cxcl18a.1 | 0.4711 | -0.16166084 | 0.017 | 0.05 | 1 | 9 |
| fig4a | 0.4711 | -0.14195274 | 0.017 | 0.05 | 1 | 9 |
| fnbp1a | 0.4711 | -0.15644145 | 0.017 | 0.05 | 1 | 9 |
| glceb | 0.4711 | -0.10784017 | 0.017 | 0.05 | 1 | 9 |
| mfsd6a | 0.4711 | -0.13001152 | 0.017 | 0.05 | 1 | 9 |
| pip5k1ba | 0.4711 | -0.11683975 | 0.017 | 0.05 | 1 | 9 |
| rap1gap | 0.4711 | -0.11544569 | 0.017 | 0.05 | 1 | 9 |
| riok3 | 0.4711 | -0.12334911 | 0.017 | 0.05 | 1 | 9 |
| si:ch211-106k21.5 | 0.4711 | -0.10266086 | 0.017 | 0.05 | 1 | 9 |
| slc25a19 | 0.4711 | -0.19238514 | 0.017 | 0.05 | 1 | 9 |
| slc38a6 | 0.4711 | -0.10826061 | 0.017 | 0.05 | 1 | 9 |
| tnfsf13b | 0.4711 | -0.16444115 | 0.017 | 0.05 | 1 | 9 |
| trappc9 | 0.4711 | -0.15830289 | 0.017 | 0.05 | 1 | 9 |
| xrcc4 | 0.4711 | -0.19169616 | 0.017 | 0.05 | 1 | 9 |
| tsen54 | 0.4711 | -0.12548592 | 0.017 | 0.05 | 1 | 9 |
| vasna | 0.4711 | -0.11244501 | 0.017 | 0.05 | 1 | 9 |
| LO018176.1 | 0.47115 | -0.15774972 | 0.034 | 0.1 | 1 | 9 |
| eif6 | 0.47148 | 0.145759265 | 0.328 | 0.2 | 1 | 9 |
| zfpm2b | 0.4717 | -0.10192161 | 0.034 | 0.1 | 1 | 9 |
| hmgn6 | 0.47229 | 0.235306284 | 0.672 | 0.55 | 1 | 9 |
| cfl1 | 0.47231 | 0.149980751 | 0.69 | 0.55 | 1 | 9 |
| amacr | 0.47243 | -0.13380058 | 0.034 | 0.1 | 1 | 9 |
| si:ch211-191j22.3 | 0.47418 | -0.18155409 | 0.276 | 0.3 | 1 | 9 |
| kpnb3 | 0.47448 | -0.17667592 | 0.103 | 0.15 | 1 | 9 |

| | | | | | | |
|---|---|---|---|---|---|---|
| capn2l | 0.47576 | -0.22046195 | 0.069 | 0.15 | 1 | 9 |
| faub | 0.47697 | -0.1298616 | 0.086 | 0.1 | 1 | 9 |
| fam120c | 0.47705 | 0.195507159 | 0.259 | 0.15 | 1 | 9 |
| skp1 | 0.47759 | 0.161314291 | 0.552 | 0.4 | 1 | 9 |
| FO904970.1 | 0.47795 | -0.10111008 | 0.052 | 0.05 | 1 | 9 |
| ube2q2 | 0.47839 | 0.166535442 | 0.121 | 0.05 | 1 | 9 |
| si:dkey-27n6.4 | 0.47867 | -0.17326363 | 0.034 | 0.1 | 1 | 9 |
| tex10 | 0.47993 | -0.16161339 | 0.052 | 0.1 | 1 | 9 |
| gfm1 | 0.48003 | -0.17747929 | 0.034 | 0.1 | 1 | 9 |
| chchd10 | 0.4807 | 0.126692619 | 0.207 | 0.1 | 1 | 9 |
| slc9a8 | 0.48089 | 0.154936702 | 0.138 | 0.05 | 1 | 9 |
| anxa1b | 0.48098 | 0.133342251 | 0.224 | 0.15 | 1 | 9 |
| mrpl48 | 0.48134 | -0.21127472 | 0.138 | 0.25 | 1 | 9 |
| nr3c1 | 0.48136 | -0.14467638 | 0.172 | 0.2 | 1 | 9 |
| nansb | 0.48158 | -0.26367321 | 0.069 | 0.15 | 1 | 9 |
| selenol | 0.48174 | -0.1688211 | 0.086 | 0.15 | 1 | 9 |
| hmgn2 | 0.48182 | 0.233077018 | 0.69 | 0.65 | 1 | 9 |
| wu:fu71h07 | 0.48271 | -0.15676323 | 0.052 | 0.1 | 1 | 9 |
| si:dkey-222b8.1 | 0.48274 | -0.17838746 | 0.034 | 0.1 | 1 | 9 |
| arl8 | 0.48312 | 0.195862822 | 0.138 | 0.05 | 1 | 9 |
| hexa | 0.48343 | -0.15917281 | 0.034 | 0.1 | 1 | 9 |
| tmem79b | 0.48464 | 0.147785792 | 0.103 | 0.05 | 1 | 9 |
| ndufs2 | 0.48571 | 0.15469073 | 0.328 | 0.2 | 1 | 9 |
| ctsk | 0.486 | -0.11580004 | 0.103 | 0.2 | 1 | 9 |
| tax1bp3 | 0.48714 | -0.24020131 | 0.224 | 0.35 | 1 | 9 |
| cmpk | 0.48871 | 0.197836548 | 0.276 | 0.15 | 1 | 9 |
| supt16h | 0.48893 | -0.22498016 | 0.103 | 0.2 | 1 | 9 |
| arfgef1 | 0.48901 | -0.16741405 | 0.034 | 0.1 | 1 | 9 |
| cbln17 | 0.48924 | -0.15228357 | 0.034 | 0.1 | 1 | 9 |
| dido1 | 0.48944 | 0.214374862 | 0.31 | 0.2 | 1 | 9 |
| csrnp1b | 0.48945 | -0.18450573 | 0.069 | 0.15 | 1 | 9 |
| hnrpkl | 0.49046 | 0.134528396 | 0.259 | 0.15 | 1 | 9 |
| tssc4 | 0.49078 | 0.111076116 | 0.138 | 0.05 | 1 | 9 |
| f3b | 0.49081 | -0.20651898 | 0.138 | 0.2 | 1 | 9 |
| pdxdc1 | 0.49087 | -0.18183014 | 0.086 | 0.15 | 1 | 9 |
| sec11a | 0.49103 | 0.217589091 | 0.328 | 0.25 | 1 | 9 |
| sdha | 0.49142 | 0.155456462 | 0.138 | 0.05 | 1 | 9 |
| dnajc15 | 0.49151 | 0.119064855 | 0.138 | 0.05 | 1 | 9 |
| zgc:112496 | 0.49158 | 0.122779734 | 0.138 | 0.05 | 1 | 9 |
| arhgap1 | 0.49201 | -0.16344109 | 0.034 | 0.1 | 1 | 9 |
| XLOC-005982 | 0.49238 | -0.19805956 | 0.052 | 0.1 | 1 | 9 |
| tpp1 | 0.49274 | 0.144595459 | 0.207 | 0.1 | 1 | 9 |
| hadhb | 0.49279 | 0.147502959 | 0.138 | 0.05 | 1 | 9 |
| rnf175 | 0.4929 | 0.15516452 | 0.121 | 0.05 | 1 | 9 |
| si:ch211-22k7.9 | 0.49345 | -0.16038132 | 0.034 | 0.1 | 1 | 9 |
| cyb5r3 | 0.49383 | 0.156495734 | 0.207 | 0.1 | 1 | 9 |
| rasa3 | 0.49418 | -0.19571027 | 0.034 | 0.1 | 1 | 9 |
| CABZ01085658.1 | 0.49421 | -0.17110335 | 0.034 | 0.1 | 1 | 9 |

| | | | | | | |
|---|---|---|---|---|---|---|
| ttc5 | 0.49449 | -0.12768669 | 0.034 | 0.1 | 1 | 9 |
| FQ323156.1 | 0.49491 | 0.181015019 | 0.172 | 0.1 | 1 | 9 |
| ppp3r1b | 0.49606 | -0.20037302 | 0.172 | 0.3 | 1 | 9 |
| emc6 | 0.49623 | 0.170887893 | 0.276 | 0.15 | 1 | 9 |
| zgc:153932 | 0.49687 | 0.172001987 | 0.103 | 0.05 | 1 | 9 |
| bpnt1 | 0.49713 | -0.14322088 | 0.069 | 0.1 | 1 | 9 |
| sdr16c5b | 0.49733 | -0.19505944 | 0.052 | 0.1 | 1 | 9 |
| irf6 | 0.4975 | -0.17576313 | 0.034 | 0.1 | 1 | 9 |
| agr1 | 0.49821 | 0.239204971 | 0.207 | 0.1 | 1 | 9 |
| cbfb | 0.4984 | -0.15377413 | 0.034 | 0.1 | 1 | 9 |
| pfdn4 | 0.49847 | 0.198390924 | 0.448 | 0.3 | 1 | 9 |
| emc7 | 0.49847 | -0.19418709 | 0.19 | 0.25 | 1 | 9 |
| rab25a | 0.49878 | 0.154554588 | 0.121 | 0.05 | 1 | 9 |
| tram1 | 0.49887 | -0.19887657 | 0.121 | 0.2 | 1 | 9 |
| anxa1c | 0.4991 | -0.16752741 | 0.241 | 0.25 | 1 | 9 |
| rsl1d1 | 0.49921 | 0.231029498 | 0.362 | 0.25 | 1 | 9 |
| pbx2 | 0.49934 | 0.150762627 | 0.138 | 0.05 | 1 | 9 |
| dnaja2b | 0.49939 | 0.162378863 | 0.155 | 0.1 | 1 | 9 |
| thoc2 | 0.4998 | 0.179862377 | 0.207 | 0.1 | 1 | 9 |
| ubr5 | 0.50003 | 0.115876998 | 0.138 | 0.05 | 1 | 9 |
| ythdf2 | 0.50017 | 0.114341619 | 0.138 | 0.05 | 1 | 9 |
| hgs | 0.50088 | -0.1431984 | 0.121 | 0.15 | 1 | 9 |
| nudt5 | 0.50115 | 0.129437109 | 0.138 | 0.05 | 1 | 9 |
| golt1ba | 0.50155 | 0.123180077 | 0.138 | 0.05 | 1 | 9 |
| BX511129.1 | 0.50194 | 0.151843766 | 0.138 | 0.05 | 1 | 9 |
| clptm1l | 0.50225 | -0.18596149 | 0.069 | 0.15 | 1 | 9 |
| hif1ab | 0.50288 | -0.12047431 | 0.19 | 0.3 | 1 | 9 |
| cox5aa | 0.5032 | -0.12497442 | 0.569 | 0.55 | 1 | 9 |
| psmd13 | 0.5034 | -0.10058302 | 0.362 | 0.35 | 1 | 9 |
| fam168a | 0.50355 | 0.199104546 | 0.276 | 0.15 | 1 | 9 |
| prdx4 | 0.50387 | -0.12813068 | 0.414 | 0.4 | 1 | 9 |
| ufc1 | 0.50397 | 0.152684302 | 0.138 | 0.05 | 1 | 9 |
| XLOC-041945 | 0.50437 | 0.19268078 | 0.138 | 0.05 | 1 | 9 |
| cd151l | 0.50448 | 0.150721859 | 0.207 | 0.1 | 1 | 9 |
| pop5 | 0.50472 | 0.120479605 | 0.138 | 0.05 | 1 | 9 |
| higd1a | 0.50535 | -0.23887801 | 0.414 | 0.55 | 1 | 9 |
| tomm34 | 0.50541 | -0.15487494 | 0.034 | 0.1 | 1 | 9 |
| ccnt1 | 0.50569 | -0.14350651 | 0.034 | 0.1 | 1 | 9 |
| ndufa7 | 0.50584 | 0.169388324 | 0.207 | 0.1 | 1 | 9 |
| mych | 0.50604 | -0.13413699 | 0.138 | 0.15 | 1 | 9 |
| mpc1 | 0.50643 | 0.108965578 | 0.31 | 0.2 | 1 | 9 |
| cmah | 0.50646 | -0.19893069 | 0.121 | 0.2 | 1 | 9 |
| med15 | 0.50667 | -0.11389872 | 0.034 | 0.1 | 1 | 9 |
| picalma | 0.50712 | -0.13162517 | 0.069 | 0.1 | 1 | 9 |
| farp2 | 0.50736 | -0.15342915 | 0.052 | 0.1 | 1 | 9 |
| grcc10 | 0.50757 | 0.159267327 | 0.138 | 0.05 | 1 | 9 |
| tmem39b | 0.50796 | 0.150825111 | 0.138 | 0.05 | 1 | 9 |
| ppp3r1a | 0.50887 | -0.12612047 | 0.172 | 0.2 | 1 | 9 |

| | | | | | | |
|---|---|---|---|---|---|---|
| snx7 | 0.50896 | 0.146907631 | 0.103 | 0.05 | 1 | 9 |
| znf576.2 | 0.50927 | 0.158898787 | 0.121 | 0.05 | 1 | 9 |
| stk17al | 0.50984 | -0.16239593 | 0.138 | 0.25 | 1 | 9 |
| mid1ip1a | 0.51074 | -0.11003328 | 0.259 | 0.35 | 1 | 9 |
| gpia | 0.5111 | 0.185886855 | 0.207 | 0.1 | 1 | 9 |
| ptbp1b | 0.51151 | -0.15789757 | 0.034 | 0.1 | 1 | 9 |
| CU855930.1 | 0.51187 | -0.1422005 | 0.034 | 0.1 | 1 | 9 |
| ppp1r9bb | 0.51214 | -0.16124984 | 0.034 | 0.1 | 1 | 9 |
| mgat5 | 0.51217 | -0.18648141 | 0.052 | 0.1 | 1 | 9 |
| mbd4 | 0.51234 | -0.1555829 | 0.052 | 0.1 | 1 | 9 |
| tenm3 | 0.5124 | 0.167432306 | 0.138 | 0.05 | 1 | 9 |
| zgc:162698 | 0.51317 | 0.174823415 | 0.172 | 0.1 | 1 | 9 |
| zgc:113921 | 0.51321 | -0.13485425 | 0.034 | 0.1 | 1 | 9 |
| gtf2a1 | 0.51337 | 0.143930406 | 0.138 | 0.05 | 1 | 9 |
| ccl25b | 0.51362 | -0.21165845 | 0.034 | 0.1 | 1 | 9 |
| abcc5 | 0.51471 | 0.191198971 | 0.138 | 0.05 | 1 | 9 |
| itm2bb | 0.51473 | -0.16890719 | 0.414 | 0.45 | 1 | 9 |
| dicer1 | 0.51532 | 0.132262194 | 0.138 | 0.05 | 1 | 9 |
| clint1a | 0.51558 | 0.153416982 | 0.207 | 0.1 | 1 | 9 |
| sel1l | 0.51593 | -0.19459467 | 0.052 | 0.1 | 1 | 9 |
| mydgf | 0.51598 | -0.11660648 | 0.103 | 0.2 | 1 | 9 |
| txndc17 | 0.51632 | 0.154389994 | 0.138 | 0.05 | 1 | 9 |
| ppid | 0.51671 | 0.157666105 | 0.172 | 0.1 | 1 | 9 |
| prpf38b | 0.51687 | 0.178272925 | 0.207 | 0.15 | 1 | 9 |
| cbwd | 0.51687 | -0.16018102 | 0.086 | 0.15 | 1 | 9 |
| coil | 0.51739 | 0.135874593 | 0.138 | 0.05 | 1 | 9 |
| hddc3 | 0.51782 | 0.11997114 | 0.103 | 0.05 | 1 | 9 |
| baz1b | 0.51813 | 0.109173202 | 0.19 | 0.1 | 1 | 9 |
| cx43.4 | 0.51847 | 0.193991965 | 0.121 | 0.05 | 1 | 9 |
| prxl2b | 0.51873 | -0.13084996 | 0.069 | 0.15 | 1 | 9 |
| fahd2a | 0.51889 | -0.16097211 | 0.121 | 0.15 | 1 | 9 |
| rnf170 | 0.51914 | -0.15841501 | 0.069 | 0.15 | 1 | 9 |
| CABZ01068251.1 | 0.51968 | -0.10417845 | 0.086 | 0.1 | 1 | 9 |
| mlx | 0.51982 | 0.132100086 | 0.138 | 0.05 | 1 | 9 |
| slc2a1b | 0.52005 | 0.140710657 | 0.103 | 0.05 | 1 | 9 |
| psmd4b | 0.5201 | 0.101290751 | 0.19 | 0.1 | 1 | 9 |
| ino80e | 0.52014 | 0.140876331 | 0.138 | 0.05 | 1 | 9 |
| ide | 0.5204 | 0.179776504 | 0.138 | 0.05 | 1 | 9 |
| si:ch211-25g7.5 | 0.52078 | -0.19117167 | 0.138 | 0.25 | 1 | 9 |
| si:ch211-133n4.4 | 0.52083 | 0.216824298 | 0.207 | 0.1 | 1 | 9 |
| anxa5b | 0.52083 | -0.23551242 | 0.138 | 0.25 | 1 | 9 |
| CABZ01112656.1 | 0.521 | -0.17166887 | 0.086 | 0.15 | 1 | 9 |
| ndufv3 | 0.521 | 0.184704009 | 0.19 | 0.1 | 1 | 9 |
| ddx51 | 0.52172 | 0.178972428 | 0.138 | 0.05 | 1 | 9 |
| zgc:162939 | 0.52183 | 0.150518339 | 0.138 | 0.05 | 1 | 9 |
| csrnp1a | 0.52228 | -0.19221279 | 0.103 | 0.2 | 1 | 9 |
| rps27l | 0.52235 | 0.117823229 | 0.379 | 0.25 | 1 | 9 |
| tead3b | 0.52298 | 0.166389953 | 0.207 | 0.1 | 1 | 9 |

| | | | | | | |
|---|---|---|---|---|---|---|
| mrtfba | 0.52309 | -0.13968 | 0.034 | 0.1 | 1 | 9 |
| ptpn2b | 0.52333 | -0.15264527 | 0.034 | 0.1 | 1 | 9 |
| coq9 | 0.52338 | 0.101557873 | 0.121 | 0.05 | 1 | 9 |
| CABZ01071848.1 | 0.52339 | -0.1381572 | 0.069 | 0.1 | 1 | 9 |
| LOC100333218 | 0.5234 | -0.17881454 | 0.052 | 0.1 | 1 | 9 |
| si:dkey-187e18.1 | 0.52381 | 0.172607267 | 0.138 | 0.05 | 1 | 9 |
| supt4h1 | 0.52455 | 0.166333955 | 0.207 | 0.1 | 1 | 9 |
| sod1 | 0.52491 | -0.20277542 | 0.534 | 0.6 | 1 | 9 |
| mapk3 | 0.52552 | 0.215208771 | 0.207 | 0.1 | 1 | 9 |
| nfe2l3 | 0.52561 | 0.15253408 | 0.138 | 0.05 | 1 | 9 |
| nsa2 | 0.52569 | 0.162236553 | 0.69 | 0.55 | 1 | 9 |
| cops5 | 0.52576 | 0.159076124 | 0.138 | 0.05 | 1 | 9 |
| rbm22 | 0.52582 | 0.128140197 | 0.138 | 0.05 | 1 | 9 |
| pdap1a | 0.52587 | 0.158906826 | 0.138 | 0.05 | 1 | 9 |
| npepl1 | 0.52588 | 0.154698427 | 0.138 | 0.05 | 1 | 9 |
| si:ch211-157c3.4 | 0.52638 | 0.128627836 | 0.172 | 0.1 | 1 | 9 |
| robo1 | 0.52638 | 0.162529944 | 0.121 | 0.05 | 1 | 9 |
| lyrm2 | 0.52641 | -0.18102476 | 0.138 | 0.25 | 1 | 9 |
| creb3l3l | 0.5272 | 0.163639401 | 0.155 | 0.1 | 1 | 9 |
| tmem214 | 0.52755 | -0.11842052 | 0.138 | 0.15 | 1 | 9 |
| tars | 0.52785 | 0.174602579 | 0.207 | 0.1 | 1 | 9 |
| nubp2 | 0.52787 | -0.10475738 | 0.086 | 0.1 | 1 | 9 |
| fuca2 | 0.52827 | -0.2150151 | 0.19 | 0.25 | 1 | 9 |
| snrnp70 | 0.52832 | 0.206095734 | 0.19 | 0.1 | 1 | 9 |
| tnksa | 0.52834 | -0.11470948 | 0.034 | 0.1 | 1 | 9 |
| macf1a | 0.52863 | 0.191182923 | 0.207 | 0.1 | 1 | 9 |
| hadh | 0.52952 | -0.18277351 | 0.19 | 0.25 | 1 | 9 |
| ndufs5 | 0.52961 | 0.135410436 | 0.483 | 0.35 | 1 | 9 |
| pole4 | 0.52961 | 0.17358834 | 0.207 | 0.1 | 1 | 9 |
| ska2 | 0.53012 | -0.13477936 | 0.121 | 0.15 | 1 | 9 |
| rbm39b | 0.53018 | -0.16466376 | 0.103 | 0.2 | 1 | 9 |
| susd6 | 0.53022 | -0.1786444 | 0.138 | 0.25 | 1 | 9 |
| tmem11 | 0.53077 | 0.160166857 | 0.207 | 0.1 | 1 | 9 |
| ypel5 | 0.53167 | -0.19875209 | 0.155 | 0.25 | 1 | 9 |
| dctn2 | 0.53234 | -0.12392564 | 0.224 | 0.25 | 1 | 9 |
| rhebl1 | 0.53235 | -0.19489546 | 0.103 | 0.2 | 1 | 9 |
| srsf10b | 0.53243 | 0.187488335 | 0.19 | 0.1 | 1 | 9 |
| tollip | 0.53248 | -0.19323284 | 0.069 | 0.15 | 1 | 9 |
| pafah1b3 | 0.53297 | -0.12427953 | 0.103 | 0.2 | 1 | 9 |
| nol12 | 0.53335 | -0.175642 | 0.069 | 0.15 | 1 | 9 |
| ddx10 | 0.53355 | -0.14587421 | 0.034 | 0.1 | 1 | 9 |
| ddx23 | 0.53363 | -0.17077464 | 0.086 | 0.15 | 1 | 9 |
| epas1a | 0.53408 | -0.19190985 | 0.052 | 0.1 | 1 | 9 |
| arfip1 | 0.53495 | -0.21371118 | 0.155 | 0.25 | 1 | 9 |
| zgc:198329 | 0.535 | -0.15307666 | 0.034 | 0.1 | 1 | 9 |
| taf1a | 0.53506 | -0.15153122 | 0.052 | 0.1 | 1 | 9 |
| vegfab | 0.53524 | -0.18011591 | 0.069 | 0.15 | 1 | 9 |
| acbd4 | 0.5354 | -0.16221012 | 0.034 | 0.1 | 1 | 9 |

| | | | | | | |
|---|---|---|---|---|---|---|
| snap23.1 | 0.53557 | -0.1493402 | 0.034 | 0.1 | 1 | 9 |
| slc35b1 | 0.53563 | -0.17562361 | 0.052 | 0.1 | 1 | 9 |
| fkbp3 | 0.53569 | 0.168865644 | 0.328 | 0.2 | 1 | 9 |
| sos1 | 0.536 | -0.13616719 | 0.034 | 0.1 | 1 | 9 |
| pde4d | 0.53641 | -0.15128621 | 0.052 | 0.1 | 1 | 9 |
| b2m | 0.53649 | -0.13997225 | 0.483 | 0.5 | 1 | 9 |
| kat2b | 0.53653 | -0.15933568 | 0.034 | 0.1 | 1 | 9 |
| pigb | 0.53664 | -0.10250163 | 0.086 | 0.1 | 1 | 9 |
| optn | 0.53667 | -0.16799732 | 0.034 | 0.1 | 1 | 9 |
| ifrd1 | 0.53726 | 0.190817586 | 0.19 | 0.1 | 1 | 9 |
| krt1-19d | 0.5373 | 0.132119899 | 0.19 | 0.2 | 1 | 9 |
| msmo1 | 0.53763 | -0.14657546 | 0.052 | 0.1 | 1 | 9 |
| cry1aa | 0.53778 | -0.16016571 | 0.103 | 0.2 | 1 | 9 |
| wdr21 | 0.53824 | 0.123707133 | 0.069 | 0.05 | 1 | 9 |
| n4bp1 | 0.53897 | -0.14942327 | 0.034 | 0.1 | 1 | 9 |
| kalrna | 0.53899 | -0.14601258 | 0.034 | 0.1 | 1 | 9 |
| ankrd46a | 0.53916 | -0.13931733 | 0.069 | 0.1 | 1 | 9 |
| fip1l1b | 0.53922 | 0.119682489 | 0.121 | 0.1 | 1 | 9 |
| pcmt | 0.53926 | 0.140278444 | 0.121 | 0.05 | 1 | 9 |
| nufip2 | 0.5393 | -0.15542342 | 0.034 | 0.1 | 1 | 9 |
| gramd4b | 0.53961 | -0.13920564 | 0.034 | 0.1 | 1 | 9 |
| dynll2b | 0.54006 | -0.17335482 | 0.19 | 0.25 | 1 | 9 |
| rhoab | 0.54057 | 0.125548971 | 0.31 | 0.2 | 1 | 9 |
| tmppe | 0.54079 | -0.14640401 | 0.034 | 0.1 | 1 | 9 |
| tp53bp2a | 0.54114 | -0.12781188 | 0.034 | 0.1 | 1 | 9 |
| XLOC-043857 | 0.54115 | -0.18715899 | 0.155 | 0.25 | 1 | 9 |
| chd1 | 0.54116 | 0.124314625 | 0.069 | 0.05 | 1 | 9 |
| r3hdm2 | 0.54141 | -0.1513879 | 0.052 | 0.1 | 1 | 9 |
| XLOC-000946 | 0.54209 | -0.1782564 | 0.069 | 0.15 | 1 | 9 |
| tmprss4a | 0.54218 | -0.19464724 | 0.052 | 0.1 | 1 | 9 |
| tanc1b | 0.54234 | -0.13912604 | 0.034 | 0.1 | 1 | 9 |
| cttnbp2 | 0.54254 | -0.14850489 | 0.034 | 0.1 | 1 | 9 |
| ccdc47 | 0.54351 | -0.18020651 | 0.19 | 0.25 | 1 | 9 |
| CU693379.1 | 0.54378 | -0.20969127 | 0.069 | 0.15 | 1 | 9 |
| brms1la | 0.54465 | 0.109488111 | 0.121 | 0.05 | 1 | 9 |
| gng5 | 0.54469 | 0.148045161 | 0.483 | 0.35 | 1 | 9 |
| zgc:91890 | 0.54487 | -0.13869758 | 0.052 | 0.1 | 1 | 9 |
| cgnb | 0.5449 | -0.16888328 | 0.086 | 0.15 | 1 | 9 |
| grk4 | 0.54517 | -0.16998814 | 0.052 | 0.1 | 1 | 9 |
| s100v2 | 0.54528 | -0.24305814 | 0.466 | 0.6 | 1 | 9 |
| irf2bp2b | 0.54561 | 0.148709448 | 0.259 | 0.15 | 1 | 9 |
| ivns1abpb | 0.54666 | 0.147654861 | 0.121 | 0.05 | 1 | 9 |
| calub | 0.54674 | -0.10408647 | 0.034 | 0.1 | 1 | 9 |
| ppiab | 0.54732 | -0.11526695 | 0.897 | 0.95 | 1 | 9 |
| cbx6b | 0.54734 | -0.14723628 | 0.034 | 0.1 | 1 | 9 |
| mpc2 | 0.5475 | -0.20324548 | 0.276 | 0.4 | 1 | 9 |
| fbxl6 | 0.54797 | 0.122550594 | 0.19 | 0.1 | 1 | 9 |
| srd5a2a | 0.54836 | -0.18516941 | 0.138 | 0.2 | 1 | 9 |

| | | | | | | |
|---|---|---|---|---|---|---|
| foxo3b | 0.54893 | 0.150517277 | 0.121 | 0.05 | 1 | 9 |
| sumo1 | 0.54906 | -0.20350743 | 0.224 | 0.35 | 1 | 9 |
| maff | 0.54917 | -0.10596383 | 0.069 | 0.15 | 1 | 9 |
| ripor1.1 | 0.54922 | -0.10892063 | 0.034 | 0.1 | 1 | 9 |
| tardbpl | 0.54928 | 0.11839167 | 0.31 | 0.2 | 1 | 9 |
| hoxc3a | 0.55019 | -0.1695616 | 0.069 | 0.15 | 1 | 9 |
| si:dkey-202l22.6 | 0.55033 | -0.11888814 | 0.069 | 0.15 | 1 | 9 |
| cstf1 | 0.55044 | -0.11110035 | 0.034 | 0.1 | 1 | 9 |
| itgb1a | 0.55049 | -0.11472528 | 0.034 | 0.1 | 1 | 9 |
| psip1b | 0.55069 | 0.163514953 | 0.259 | 0.15 | 1 | 9 |
| fis1 | 0.55079 | 0.13621735 | 0.121 | 0.05 | 1 | 9 |
| nkd2a | 0.55095 | -0.1436769 | 0.034 | 0.1 | 1 | 9 |
| vcp | 0.55118 | -0.11223956 | 0.293 | 0.3 | 1 | 9 |
| sptlc1 | 0.55174 | -0.13480281 | 0.034 | 0.1 | 1 | 9 |
| si:ch73-236c18.3 | 0.5518 | -0.16904338 | 0.034 | 0.1 | 1 | 9 |
| zgc:193541 | 0.55182 | 0.148908318 | 0.638 | 0.5 | 1 | 9 |
| hivep2a | 0.55267 | -0.17018115 | 0.034 | 0.1 | 1 | 9 |
| si:ch211-153b23.5 | 0.55278 | -0.22615603 | 0.034 | 0.1 | 1 | 9 |
| tead1b | 0.55278 | 0.156806165 | 0.121 | 0.05 | 1 | 9 |
| morf4l1 | 0.55348 | 0.192031285 | 0.345 | 0.25 | 1 | 9 |
| zdhhc18a | 0.55373 | -0.1000364 | 0.034 | 0.1 | 1 | 9 |
| si:ch73-237c6.1 | 0.55385 | -0.15427645 | 0.069 | 0.1 | 1 | 9 |
| yju2 | 0.55396 | 0.136386729 | 0.121 | 0.05 | 1 | 9 |
| rars | 0.5546 | 0.129972261 | 0.086 | 0.05 | 1 | 9 |
| ergic3 | 0.55548 | -0.13286271 | 0.034 | 0.1 | 1 | 9 |
| mif4gda | 0.55573 | -0.11790595 | 0.034 | 0.1 | 1 | 9 |
| arhgap35a | 0.55621 | -0.18844766 | 0.086 | 0.15 | 1 | 9 |
| tnfsf10 | 0.55678 | -0.18866193 | 0.034 | 0.1 | 1 | 9 |
| dusp12 | 0.55753 | -0.11862651 | 0.034 | 0.1 | 1 | 9 |
| serinc1 | 0.5576 | -0.10268681 | 0.086 | 0.1 | 1 | 9 |
| mcts1 | 0.55769 | 0.191384994 | 0.259 | 0.15 | 1 | 9 |
| zgc:174931 | 0.55778 | -0.13469831 | 0.034 | 0.1 | 1 | 9 |
| crebbpa | 0.55807 | 0.17126657 | 0.19 | 0.1 | 1 | 9 |
| trim16 | 0.55814 | -0.13907179 | 0.034 | 0.1 | 1 | 9 |
| tmem8a | 0.55846 | -0.15796163 | 0.034 | 0.1 | 1 | 9 |
| sytl4 | 0.55871 | -0.14134906 | 0.034 | 0.1 | 1 | 9 |
| CABZ01078055.1 | 0.55881 | -0.10046572 | 0.086 | 0.1 | 1 | 9 |
| kmt5b | 0.55904 | -0.12008815 | 0.034 | 0.1 | 1 | 9 |
| rab8b | 0.55918 | -0.153738 | 0.034 | 0.1 | 1 | 9 |
| acat1 | 0.55927 | 0.120901756 | 0.121 | 0.05 | 1 | 9 |
| tmem242 | 0.55931 | -0.12321767 | 0.034 | 0.1 | 1 | 9 |
| gsk3ab | 0.55932 | -0.12799006 | 0.069 | 0.1 | 1 | 9 |
| LOC100333482 | 0.55974 | -0.1346199 | 0.034 | 0.1 | 1 | 9 |
| mylka | 0.55977 | -0.12474833 | 0.034 | 0.1 | 1 | 9 |
| snrpa1 | 0.5598 | -0.12354529 | 0.121 | 0.15 | 1 | 9 |
| mbnl3 | 0.56007 | -0.19720016 | 0.155 | 0.25 | 1 | 9 |
| prrc2c | 0.56016 | -0.12275533 | 0.069 | 0.15 | 1 | 9 |
| uba5 | 0.56056 | -0.15364586 | 0.034 | 0.1 | 1 | 9 |

| | | | | | | |
|---|---|---|---|---|---|---|
| ldhba | 0.56097 | 0.155799328 | 0.586 | 0.45 | 1 | 9 |
| capgb | 0.56111 | -0.16413782 | 0.086 | 0.1 | 1 | 9 |
| cinp | 0.56111 | 0.127377204 | 0.121 | 0.05 | 1 | 9 |
| ufsp2 | 0.56155 | 0.131761909 | 0.121 | 0.05 | 1 | 9 |
| zfp36l2 | 0.56166 | -0.16105123 | 0.155 | 0.2 | 1 | 9 |
| sars | 0.56171 | 0.181131841 | 0.259 | 0.15 | 1 | 9 |
| vps16 | 0.5618 | -0.13643539 | 0.034 | 0.1 | 1 | 9 |
| yap1 | 0.56185 | 0.145448441 | 0.103 | 0.05 | 1 | 9 |
| si:ch211-226m16.2 | 0.56228 | -0.14911681 | 0.086 | 0.15 | 1 | 9 |
| si:dkey-111e8.5 | 0.56249 | -0.13700444 | 0.034 | 0.1 | 1 | 9 |
| zdhhc5a | 0.56267 | -0.1630359 | 0.052 | 0.1 | 1 | 9 |
| pou2f1b | 0.56289 | -0.14451405 | 0.103 | 0.15 | 1 | 9 |
| zgc:153759 | 0.56323 | -0.12662314 | 0.034 | 0.1 | 1 | 9 |
| dazap1 | 0.56335 | -0.14496656 | 0.034 | 0.1 | 1 | 9 |
| rpf1 | 0.5634 | 0.149718921 | 0.121 | 0.05 | 1 | 9 |
| plekha1a | 0.56406 | -0.11460294 | 0.034 | 0.1 | 1 | 9 |
| epcam | 0.56419 | -0.21969452 | 0.5 | 0.6 | 1 | 9 |
| fmnl2a | 0.56462 | -0.1124909 | 0.034 | 0.1 | 1 | 9 |
| LOC101886114 | 0.56489 | -0.19244806 | 0.034 | 0.1 | 1 | 9 |
| hadhaa | 0.56513 | -0.1394933 | 0.052 | 0.1 | 1 | 9 |
| cdkn1ba | 0.56522 | -0.11997105 | 0.034 | 0.1 | 1 | 9 |
| wsb2 | 0.56537 | -0.16684184 | 0.103 | 0.2 | 1 | 9 |
| CABZ01086877.1 | 0.56538 | -0.15659796 | 0.052 | 0.1 | 1 | 9 |
| gmnn | 0.56565 | -0.13413833 | 0.121 | 0.15 | 1 | 9 |
| ptgs2a | 0.566 | -0.14334223 | 0.034 | 0.1 | 1 | 9 |
| uba52 | 0.56604 | 0.113135723 | 0.897 | 0.8 | 1 | 9 |
| lpar1 | 0.56636 | 0.212799337 | 0.19 | 0.1 | 1 | 9 |
| dynlrb1 | 0.56641 | 0.190793185 | 0.379 | 0.25 | 1 | 9 |
| si:dkey-11o15.7 | 0.56665 | -0.11331772 | 0.034 | 0.1 | 1 | 9 |
| retreg2 | 0.56688 | -0.11554079 | 0.034 | 0.1 | 1 | 9 |
| tbl1xr1a | 0.56723 | 0.159957345 | 0.241 | 0.15 | 1 | 9 |
| flcn | 0.56747 | -0.12753526 | 0.069 | 0.1 | 1 | 9 |
| wdr37 | 0.56759 | -0.12114745 | 0.034 | 0.1 | 1 | 9 |
| ldha | 0.56766 | -0.12134342 | 0.034 | 0.1 | 1 | 9 |
| BX640522.2 | 0.56784 | -0.13379432 | 0.034 | 0.1 | 1 | 9 |
| rfc3 | 0.56815 | -0.11604815 | 0.034 | 0.1 | 1 | 9 |
| jun | 0.56819 | -0.20175487 | 0.569 | 0.7 | 1 | 9 |
| tfap2a | 0.56829 | -0.19157699 | 0.069 | 0.15 | 1 | 9 |
| kifc3 | 0.56878 | -0.11413825 | 0.034 | 0.1 | 1 | 9 |
| dhdds | 0.56886 | -0.12000914 | 0.034 | 0.1 | 1 | 9 |
| ikzf2 | 0.56928 | -0.11376431 | 0.034 | 0.1 | 1 | 9 |
| zgc:153018 | 0.56939 | -0.1230499 | 0.052 | 0.1 | 1 | 9 |
| myo1ea | 0.5694 | -0.13008507 | 0.034 | 0.1 | 1 | 9 |
| rassf3 | 0.56956 | -0.10430186 | 0.034 | 0.1 | 1 | 9 |
| hmgcra | 0.56959 | -0.20850664 | 0.138 | 0.2 | 1 | 9 |
| eif2ak3 | 0.56965 | -0.12415484 | 0.034 | 0.1 | 1 | 9 |
| igf2r | 0.5698 | -0.12474005 | 0.034 | 0.1 | 1 | 9 |
| thrab | 0.56984 | -0.12862113 | 0.034 | 0.1 | 1 | 9 |

| | | | | | | |
|---|---|---|---|---|---|---|
| podxl | 0.56995 | -0.1625489 | 0.034 | 0.1 | 1 | 9 |
| atp5md | 0.57007 | 0.194706419 | 0.534 | 0.4 | 1 | 9 |
| fxyd6l | 0.57021 | 0.201060303 | 0.241 | 0.15 | 1 | 9 |
| stox2a | 0.5715 | -0.12721092 | 0.069 | 0.15 | 1 | 9 |
| acsl1a | 0.57166 | 0.101518264 | 0.121 | 0.05 | 1 | 9 |
| lsm14aa | 0.57232 | -0.12548408 | 0.069 | 0.1 | 1 | 9 |
| emd | 0.57239 | -0.18723395 | 0.224 | 0.3 | 1 | 9 |
| col5a1 | 0.57274 | 0.154418069 | 0.138 | 0.1 | 1 | 9 |
| nedd4l | 0.57309 | -0.11858158 | 0.069 | 0.15 | 1 | 9 |
| wfdc2 | 0.57327 | 0.199155376 | 0.103 | 0.05 | 1 | 9 |
| srsf6b | 0.57401 | -0.14193636 | 0.207 | 0.25 | 1 | 9 |
| tmem134 | 0.57401 | 0.159565629 | 0.121 | 0.05 | 1 | 9 |
| capn2b | 0.57458 | 0.115455672 | 0.121 | 0.05 | 1 | 9 |
| sgce | 0.57481 | 0.143703351 | 0.103 | 0.05 | 1 | 9 |
| rnf183 | 0.57499 | -0.18968763 | 0.086 | 0.15 | 1 | 9 |
| rgs12b | 0.57501 | -0.13008474 | 0.069 | 0.15 | 1 | 9 |
| chd4a | 0.57514 | 0.213270806 | 0.31 | 0.2 | 1 | 9 |
| alg5 | 0.57599 | -0.17454537 | 0.138 | 0.2 | 1 | 9 |
| itprid2 | 0.57603 | -0.10181618 | 0.069 | 0.15 | 1 | 9 |
| eny2 | 0.57621 | 0.196247831 | 0.241 | 0.15 | 1 | 9 |
| ezrb | 0.5765 | -0.17166335 | 0.276 | 0.35 | 1 | 9 |
| rfc4 | 0.57701 | 0.15386007 | 0.121 | 0.05 | 1 | 9 |
| ppig | 0.57722 | 0.120798159 | 0.19 | 0.1 | 1 | 9 |
| mvb12ba | 0.57774 | -0.16492649 | 0.069 | 0.15 | 1 | 9 |
| plcb3 | 0.57775 | -0.12691788 | 0.069 | 0.15 | 1 | 9 |
| casz1 | 0.57781 | -0.22107503 | 0.121 | 0.2 | 1 | 9 |
| desi2 | 0.57787 | -0.15778598 | 0.069 | 0.15 | 1 | 9 |
| dyrk3 | 0.57893 | -0.11389026 | 0.069 | 0.15 | 1 | 9 |
| coq3 | 0.57949 | -0.12208084 | 0.069 | 0.1 | 1 | 9 |
| dsc2l | 0.57983 | -0.1969116 | 0.155 | 0.25 | 1 | 9 |
| CABZ01027551.1 | 0.57992 | -0.17948308 | 0.155 | 0.2 | 1 | 9 |
| taok1b | 0.58099 | 0.168648054 | 0.155 | 0.1 | 1 | 9 |
| ctsz | 0.58214 | 0.105251647 | 0.241 | 0.15 | 1 | 9 |
| spink2.1 | 0.58226 | -0.14622101 | 0.069 | 0.15 | 1 | 9 |
| col6a1 | 0.58296 | 0.187308806 | 0.121 | 0.05 | 1 | 9 |
| arl1 | 0.58313 | -0.19271217 | 0.19 | 0.3 | 1 | 9 |
| zbtb22a | 0.58348 | -0.1569935 | 0.052 | 0.1 | 1 | 9 |
| pqlc3 | 0.58362 | -0.18827631 | 0.069 | 0.15 | 1 | 9 |
| CU914622.2 | 0.58379 | -0.22526688 | 0.069 | 0.15 | 1 | 9 |
| esf1 | 0.58379 | -0.11905498 | 0.069 | 0.15 | 1 | 9 |
| zgc:123295 | 0.58427 | -0.1537246 | 0.069 | 0.15 | 1 | 9 |
| lgals2a | 0.58457 | 0.193819541 | 0.103 | 0.05 | 1 | 9 |
| crtc3 | 0.58472 | 0.154331483 | 0.121 | 0.05 | 1 | 9 |
| tasp1 | 0.58594 | -0.15380176 | 0.103 | 0.15 | 1 | 9 |
| taf11 | 0.58647 | -0.14671702 | 0.103 | 0.15 | 1 | 9 |
| bptf | 0.58706 | 0.108125518 | 0.121 | 0.05 | 1 | 9 |
| rad23b | 0.58725 | 0.114585922 | 0.466 | 0.35 | 1 | 9 |
| cltca | 0.58826 | -0.20217311 | 0.155 | 0.25 | 1 | 9 |

| | | | | | | |
|---|---|---|---|---|---|---|
| brd3b | 0.58842 | 0.181111486 | 0.19 | 0.1 | 1 | 9 |
| bag4 | 0.58965 | -0.11602139 | 0.069 | 0.1 | 1 | 9 |
| zgc:153916 | 0.58984 | -0.14464246 | 0.052 | 0.1 | 1 | 9 |
| ap2b1 | 0.58998 | 0.103055287 | 0.121 | 0.1 | 1 | 9 |
| TSTA3 (1 of many) | 0.59058 | -0.14077723 | 0.086 | 0.15 | 1 | 9 |
| dctn4 | 0.59082 | 0.132516594 | 0.121 | 0.05 | 1 | 9 |
| gnpnat1 | 0.59094 | 0.135655708 | 0.103 | 0.05 | 1 | 9 |
| LOC110439500 | 0.59111 | 0.11436474 | 0.069 | 0.05 | 1 | 9 |
| sox4b | 0.59196 | 0.163368347 | 0.103 | 0.05 | 1 | 9 |
| tent5ba | 0.59249 | 0.131095231 | 0.121 | 0.05 | 1 | 9 |
| CABZ01055347.1 | 0.59284 | -0.15077543 | 0.155 | 0.2 | 1 | 9 |
| sssca1 | 0.59319 | 0.119716123 | 0.103 | 0.05 | 1 | 9 |
| tmem126a | 0.59343 | -0.100302 | 0.086 | 0.1 | 1 | 9 |
| csnk1db | 0.59345 | 0.112097037 | 0.241 | 0.15 | 1 | 9 |
| tspan9a | 0.59372 | 0.203891319 | 0.155 | 0.1 | 1 | 9 |
| aldob | 0.59391 | -0.24566888 | 0.19 | 0.25 | 1 | 9 |
| rtf1 | 0.5964 | -0.17112023 | 0.224 | 0.3 | 1 | 9 |
| zgc:77929 | 0.59668 | 0.117675496 | 0.121 | 0.05 | 1 | 9 |
| sarnp | 0.59723 | -0.17293669 | 0.293 | 0.35 | 1 | 9 |
| CABZ01039859.1 | 0.59736 | 0.135167889 | 0.103 | 0.05 | 1 | 9 |
| zgc:194392 | 0.59805 | -0.12993024 | 0.052 | 0.1 | 1 | 9 |
| tfg | 0.59868 | -0.15253162 | 0.19 | 0.3 | 1 | 9 |
| si:ch211-222k6.3 | 0.59942 | 0.170676258 | 0.19 | 0.1 | 1 | 9 |
| nckap1 | 0.6006 | -0.13817819 | 0.121 | 0.15 | 1 | 9 |
| srpk1a | 0.6007 | -0.10697004 | 0.19 | 0.2 | 1 | 9 |
| pgap2 | 0.60204 | 0.100722702 | 0.069 | 0.05 | 1 | 9 |
| psmd4a | 0.60254 | 0.129412324 | 0.19 | 0.1 | 1 | 9 |
| zfp36l1a | 0.60254 | 0.14375079 | 0.259 | 0.2 | 1 | 9 |
| cd9b | 0.60313 | -0.20619168 | 0.379 | 0.45 | 1 | 9 |
| sc5d | 0.60373 | -0.11973134 | 0.069 | 0.1 | 1 | 9 |
| si:dkey-71l4.4 | 0.60449 | 0.121329183 | 0.086 | 0.05 | 1 | 9 |
| serpinb1 | 0.60477 | -0.14711423 | 0.086 | 0.15 | 1 | 9 |
| srsf5b | 0.60493 | -0.15483284 | 0.103 | 0.15 | 1 | 9 |
| rmdn1 | 0.60495 | 0.167730426 | 0.241 | 0.15 | 1 | 9 |
| etfa | 0.60509 | 0.142007796 | 0.31 | 0.2 | 1 | 9 |
| cct2 | 0.60514 | 0.124596273 | 0.569 | 0.45 | 1 | 9 |
| phactr4a | 0.60516 | -0.19429844 | 0.121 | 0.2 | 1 | 9 |
| vma21 | 0.60518 | 0.10273883 | 0.241 | 0.15 | 1 | 9 |
| zgc:154054 | 0.60531 | -0.17683279 | 0.138 | 0.2 | 1 | 9 |
| tmed9 | 0.60547 | -0.10333519 | 0.19 | 0.2 | 1 | 9 |
| nfic | 0.60555 | 0.130411592 | 0.121 | 0.05 | 1 | 9 |
| plekhj1 | 0.60612 | -0.13964913 | 0.103 | 0.15 | 1 | 9 |
| gkap1 | 0.60619 | 0.134476265 | 0.19 | 0.1 | 1 | 9 |
| anapc13 | 0.60742 | -0.16174434 | 0.052 | 0.1 | 1 | 9 |
| zgc:162730 | 0.60743 | -0.13149778 | 0.5 | 0.5 | 1 | 9 |
| pbx4 | 0.60817 | 0.138072993 | 0.103 | 0.05 | 1 | 9 |
| ftr82 | 0.60879 | -0.1997945 | 0.121 | 0.2 | 1 | 9 |
| mphosph6 | 0.60909 | 0.121421857 | 0.121 | 0.05 | 1 | 9 |

| | | | | | | |
|---|---|---|---|---|---|---|
| march2 | 0.60918 | -0.13842339 | 0.052 | 0.1 | 1 | 9 |
| tomm7 | 0.60935 | 0.126738492 | 0.414 | 0.3 | 1 | 9 |
| tlcd1 | 0.60988 | 0.145614172 | 0.155 | 0.1 | 1 | 9 |
| ccdc90b | 0.61131 | 0.125017246 | 0.121 | 0.05 | 1 | 9 |
| zfhx3 | 0.61158 | 0.105228704 | 0.121 | 0.1 | 1 | 9 |
| mrpl43 | 0.61292 | 0.102626054 | 0.121 | 0.05 | 1 | 9 |
| nmi | 0.61302 | -0.14363554 | 0.207 | 0.25 | 1 | 9 |
| usp10 | 0.61319 | 0.12644923 | 0.19 | 0.1 | 1 | 9 |
| arl5a | 0.61389 | 0.123306162 | 0.086 | 0.05 | 1 | 9 |
| ube2ka | 0.61411 | -0.13008019 | 0.052 | 0.1 | 1 | 9 |
| ovol1a | 0.61497 | 0.141600907 | 0.103 | 0.05 | 1 | 9 |
| acin1b | 0.61538 | 0.155358439 | 0.19 | 0.1 | 1 | 9 |
| kdm4b | 0.61564 | 0.134403211 | 0.121 | 0.05 | 1 | 9 |
| taok2a | 0.61626 | -0.14137476 | 0.052 | 0.1 | 1 | 9 |
| blmh | 0.61651 | 0.151114758 | 0.19 | 0.1 | 1 | 9 |
| gstr | 0.61734 | -0.11544983 | 0.19 | 0.2 | 1 | 9 |
| etf1b | 0.61734 | 0.165260102 | 0.31 | 0.2 | 1 | 9 |
| srrm1 | 0.6176 | 0.189050588 | 0.241 | 0.15 | 1 | 9 |
| dtd1 | 0.61779 | 0.133279426 | 0.121 | 0.05 | 1 | 9 |
| scpep1 | 0.61847 | 0.119194107 | 0.121 | 0.05 | 1 | 9 |
| rhpn2 | 0.61978 | 0.13758383 | 0.138 | 0.1 | 1 | 9 |
| rrp1 | 0.61979 | 0.10870528 | 0.241 | 0.15 | 1 | 9 |
| bhlhe41 | 0.62038 | -0.14937531 | 0.121 | 0.15 | 1 | 9 |
| mrps31 | 0.62059 | 0.117393673 | 0.121 | 0.05 | 1 | 9 |
| six2a | 0.62065 | 0.137989735 | 0.121 | 0.05 | 1 | 9 |
| cobll1b | 0.62071 | -0.10077542 | 0.086 | 0.1 | 1 | 9 |
| rabac1 | 0.62085 | -0.16734392 | 0.138 | 0.2 | 1 | 9 |
| ano6 | 0.6216 | 0.112418034 | 0.069 | 0.05 | 1 | 9 |
| ktn1 | 0.62175 | 0.102338261 | 0.121 | 0.05 | 1 | 9 |
| ranbp3b | 0.62215 | 0.101417393 | 0.121 | 0.05 | 1 | 9 |
| alcama | 0.62277 | 0.158410596 | 0.121 | 0.05 | 1 | 9 |
| zgc:111983 | 0.62318 | 0.1291344 | 0.207 | 0.15 | 1 | 9 |
| myclb | 0.62386 | 0.115549606 | 0.172 | 0.15 | 1 | 9 |
| eaf2 | 0.62419 | 0.122511008 | 0.121 | 0.05 | 1 | 9 |
| mid1ip1b | 0.6242 | 0.200887453 | 0.276 | 0.2 | 1 | 9 |
| LOC101882729 | 0.62441 | 0.204982341 | 0.19 | 0.1 | 1 | 9 |
| prpf8 | 0.62489 | 0.131609857 | 0.121 | 0.05 | 1 | 9 |
| stxbp2 | 0.62507 | 0.137173801 | 0.19 | 0.1 | 1 | 9 |
| myzap | 0.62515 | 0.135437871 | 0.103 | 0.05 | 1 | 9 |
| ufm1 | 0.62534 | 0.113914916 | 0.19 | 0.1 | 1 | 9 |
| ip6k2a | 0.62582 | 0.160152018 | 0.19 | 0.1 | 1 | 9 |
| rimkla | 0.62587 | 0.116064443 | 0.121 | 0.05 | 1 | 9 |
| sbf1 | 0.62653 | 0.111707637 | 0.069 | 0.05 | 1 | 9 |
| pno1 | 0.62673 | 0.159485374 | 0.31 | 0.2 | 1 | 9 |
| flot1b | 0.62693 | 0.120574426 | 0.155 | 0.1 | 1 | 9 |
| mmp16a | 0.62771 | 0.162806182 | 0.103 | 0.05 | 1 | 9 |
| sulf2b | 0.62789 | 0.161899208 | 0.121 | 0.05 | 1 | 9 |
| hoxb6a | 0.62793 | 0.10841895 | 0.121 | 0.05 | 1 | 9 |

| | | | | | | |
|---|---|---|---|---|---|---|
| map2k4a | 0.62842 | -0.12714104 | 0.121 | 0.15 | 1 | 9 |
| vsig8b | 0.62854 | 0.148480697 | 0.121 | 0.05 | 1 | 9 |
| exosc2 | 0.62857 | 0.112517147 | 0.121 | 0.05 | 1 | 9 |
| sdhaf2 | 0.62929 | -0.15563213 | 0.052 | 0.1 | 1 | 9 |
| letm1 | 0.62931 | 0.117628773 | 0.121 | 0.05 | 1 | 9 |
| ndufaf2 | 0.62986 | 0.137785621 | 0.121 | 0.05 | 1 | 9 |
| hmgb1a | 0.63014 | 0.193840166 | 0.534 | 0.45 | 1 | 9 |
| ola1 | 0.63034 | 0.164900529 | 0.466 | 0.4 | 1 | 9 |
| uevld | 0.63054 | -0.14024369 | 0.052 | 0.1 | 1 | 9 |
| tln1 | 0.63151 | 0.151767773 | 0.121 | 0.05 | 1 | 9 |
| crebzf | 0.63195 | -0.11386629 | 0.207 | 0.3 | 1 | 9 |
| cox7b | 0.63207 | 0.152127191 | 0.759 | 0.65 | 1 | 9 |
| ubap2a | 0.63284 | -0.17661432 | 0.086 | 0.15 | 1 | 9 |
| cldna | 0.63301 | -0.14386046 | 0.069 | 0.1 | 1 | 9 |
| osgep | 0.63307 | 0.111854085 | 0.121 | 0.05 | 1 | 9 |
| cdc14b | 0.63311 | 0.120231127 | 0.121 | 0.05 | 1 | 9 |
| med19b | 0.63319 | 0.12456749 | 0.103 | 0.05 | 1 | 9 |
| akr1a1b | 0.63337 | 0.112116643 | 0.121 | 0.05 | 1 | 9 |
| faf2 | 0.6337 | 0.116357519 | 0.121 | 0.05 | 1 | 9 |
| plpp3 | 0.63391 | -0.16402001 | 0.052 | 0.1 | 1 | 9 |
| postnb | 0.63503 | 0.225843777 | 0.121 | 0.05 | 1 | 9 |
| rbms2a | 0.63515 | 0.101155478 | 0.121 | 0.05 | 1 | 9 |
| ube2d2l | 0.63532 | 0.111438531 | 0.086 | 0.05 | 1 | 9 |
| lrrfip1a | 0.63539 | 0.123890627 | 0.121 | 0.05 | 1 | 9 |
| timm17a | 0.63546 | -0.15703306 | 0.155 | 0.25 | 1 | 9 |
| btr12 | 0.63547 | 0.127395757 | 0.121 | 0.05 | 1 | 9 |
| mmp2 | 0.63554 | 0.164242526 | 0.103 | 0.05 | 1 | 9 |
| ctgfa | 0.63571 | 0.144586593 | 0.121 | 0.05 | 1 | 9 |
| kdf1b | 0.63586 | -0.1566684 | 0.052 | 0.1 | 1 | 9 |
| si:ch211-13c6.2 | 0.6359 | 0.131221482 | 0.121 | 0.05 | 1 | 9 |
| CU459056.1 | 0.63594 | 0.131275909 | 0.121 | 0.05 | 1 | 9 |
| aifm1 | 0.63618 | 0.101561445 | 0.121 | 0.05 | 1 | 9 |
| cfdp1 | 0.63624 | -0.17348791 | 0.086 | 0.15 | 1 | 9 |
| atp6v0cb | 0.63643 | 0.144086972 | 0.121 | 0.05 | 1 | 9 |
| tmed1b | 0.63763 | 0.107996932 | 0.069 | 0.05 | 1 | 9 |
| chmp5b | 0.63786 | 0.151383564 | 0.362 | 0.25 | 1 | 9 |
| fermt2 | 0.63893 | 0.132624415 | 0.086 | 0.05 | 1 | 9 |
| cox7a1 | 0.63959 | 0.111078684 | 0.103 | 0.05 | 1 | 9 |
| atp1b3a | 0.64066 | 0.131594389 | 0.086 | 0.05 | 1 | 9 |
| ndufaf3 | 0.64066 | -0.11462586 | 0.069 | 0.1 | 1 | 9 |
| zgc:111986 | 0.64245 | -0.16581074 | 0.414 | 0.5 | 1 | 9 |
| ifi30 | 0.64295 | -0.2048755 | 0.31 | 0.4 | 1 | 9 |
| ndufv2 | 0.64386 | 0.170730452 | 0.241 | 0.15 | 1 | 9 |
| cap1 | 0.64458 | -0.15304449 | 0.31 | 0.35 | 1 | 9 |
| lrba | 0.64466 | -0.14155228 | 0.103 | 0.15 | 1 | 9 |
| ndufs6 | 0.64609 | 0.174203481 | 0.414 | 0.3 | 1 | 9 |
| hexb | 0.6464 | 0.140435639 | 0.155 | 0.1 | 1 | 9 |
| trim35-28 | 0.6465 | -0.12694778 | 0.086 | 0.15 | 1 | 9 |

| | | | | | | |
|---|---|---|---|---|---|---|
| LOC407643 | 0.6469 | 0.121955162 | 0.103 | 0.05 | 1 | 9 |
| prpf3 | 0.64734 | -0.13700542 | 0.052 | 0.1 | 1 | 9 |
| rassf10a | 0.64748 | -0.10220352 | 0.138 | 0.15 | 1 | 9 |
| mgrn1a | 0.64797 | 0.131729931 | 0.103 | 0.05 | 1 | 9 |
| mapk14a | 0.64798 | 0.100288695 | 0.172 | 0.1 | 1 | 9 |
| rpl35a | 0.64905 | 0.130294102 | 0.879 | 0.8 | 1 | 9 |
| si:dkey-29d8.3 | 0.64952 | -0.14063463 | 0.052 | 0.1 | 1 | 9 |
| phf5a | 0.64971 | 0.109893912 | 0.276 | 0.25 | 1 | 9 |
| mapk14b | 0.65159 | -0.12741354 | 0.052 | 0.1 | 1 | 9 |
| MCMDC2 | 0.65166 | 0.129114801 | 0.103 | 0.05 | 1 | 9 |
| nfybb | 0.65198 | 0.116588279 | 0.103 | 0.05 | 1 | 9 |
| selenow2b | 0.65204 | 0.128315717 | 0.241 | 0.15 | 1 | 9 |
| cul4b | 0.65223 | -0.14708418 | 0.086 | 0.15 | 1 | 9 |
| spns1 | 0.65317 | -0.15782278 | 0.086 | 0.15 | 1 | 9 |
| purab | 0.65317 | -0.11838327 | 0.172 | 0.2 | 1 | 9 |
| mcrs1 | 0.65321 | -0.11839503 | 0.103 | 0.15 | 1 | 9 |
| mrpl42 | 0.65322 | 0.140163852 | 0.207 | 0.15 | 1 | 9 |
| elf2b | 0.65453 | -0.11992231 | 0.069 | 0.1 | 1 | 9 |
| dnttip2 | 0.65457 | -0.13400171 | 0.052 | 0.1 | 1 | 9 |
| tm4sf4 | 0.65523 | -0.27551443 | 0.086 | 0.15 | 1 | 9 |
| ftr83 | 0.6556 | -0.12138126 | 0.052 | 0.1 | 1 | 9 |
| gpa33a | 0.65576 | -0.16092097 | 0.121 | 0.2 | 1 | 9 |
| dynll1 | 0.65581 | 0.183255995 | 0.5 | 0.45 | 1 | 9 |
| xiap | 0.65619 | -0.14921628 | 0.259 | 0.3 | 1 | 9 |
| sik1 | 0.65668 | -0.1397172 | 0.121 | 0.15 | 1 | 9 |
| zbtb10 | 0.65725 | -0.14024019 | 0.155 | 0.2 | 1 | 9 |
| dhrs13a.3 | 0.6579 | -0.11889588 | 0.069 | 0.1 | 1 | 9 |
| pi15a | 0.65841 | 0.127526802 | 0.069 | 0.05 | 1 | 9 |
| si:ch211-59d15.9 | 0.65867 | -0.13282535 | 0.052 | 0.1 | 1 | 9 |
| mir203a | 0.65909 | -0.12442563 | 0.052 | 0.1 | 1 | 9 |
| eif4ebp1 | 0.66005 | 0.151136968 | 0.414 | 0.3 | 1 | 9 |
| slc25a43 | 0.66017 | -0.16408989 | 0.052 | 0.1 | 1 | 9 |
| atf7ip | 0.66023 | 0.156366573 | 0.224 | 0.15 | 1 | 9 |
| bcl6aa | 0.66121 | 0.108621564 | 0.086 | 0.05 | 1 | 9 |
| nfu1 | 0.66127 | 0.121026942 | 0.103 | 0.05 | 1 | 9 |
| LOC110438610 | 0.66203 | 0.119349015 | 0.103 | 0.05 | 1 | 9 |
| emc2 | 0.66252 | 0.159928762 | 0.241 | 0.15 | 1 | 9 |
| vaspb | 0.66297 | -0.13975111 | 0.207 | 0.25 | 1 | 9 |
| scrn3 | 0.66338 | 0.109261196 | 0.086 | 0.05 | 1 | 9 |
| prkab1b | 0.6638 | -0.16021315 | 0.224 | 0.3 | 1 | 9 |
| marcksa | 0.66384 | 0.124988269 | 0.103 | 0.05 | 1 | 9 |
| cpne1 | 0.66416 | -0.16594777 | 0.224 | 0.3 | 1 | 9 |
| pelp1 | 0.66426 | -0.15830586 | 0.121 | 0.2 | 1 | 9 |
| elmsan1b | 0.66431 | -0.11535487 | 0.052 | 0.1 | 1 | 9 |
| hoxc8a | 0.66441 | -0.13773895 | 0.103 | 0.15 | 1 | 9 |
| tmem176l.3a | 0.66468 | -0.12305673 | 0.224 | 0.25 | 1 | 9 |
| si:ch211-98n17.5 | 0.66493 | 0.157240262 | 0.103 | 0.05 | 1 | 9 |
| csnk1da | 0.66618 | -0.1020819 | 0.241 | 0.25 | 1 | 9 |

| | | | | | | |
|---|---|---|---|---|---|---|
| acvr2ab | 0.66648 | 0.138005028 | 0.103 | 0.05 | 1 | 9 |
| cav1 | 0.66762 | -0.14906627 | 0.138 | 0.2 | 1 | 9 |
| setd1a | 0.66793 | -0.1150063 | 0.069 | 0.1 | 1 | 9 |
| ngfrb | 0.66881 | 0.106514108 | 0.069 | 0.05 | 1 | 9 |
| zgc:65873 | 0.66886 | -0.13600137 | 0.121 | 0.2 | 1 | 9 |
| nucks1a | 0.66915 | 0.118929776 | 0.293 | 0.2 | 1 | 9 |
| zyx | 0.66999 | -0.13790044 | 0.086 | 0.15 | 1 | 9 |
| traf4a | 0.67007 | -0.11552768 | 0.052 | 0.1 | 1 | 9 |
| adrm1 | 0.67014 | 0.148069782 | 0.172 | 0.1 | 1 | 9 |
| slc38a5a | 0.67015 | -0.11303174 | 0.069 | 0.1 | 1 | 9 |
| tfam | 0.6704 | 0.156044566 | 0.172 | 0.1 | 1 | 9 |
| pa2g4b | 0.67042 | 0.130764675 | 0.241 | 0.2 | 1 | 9 |
| ehmt1a | 0.67104 | 0.101986094 | 0.086 | 0.05 | 1 | 9 |
| brd8 | 0.67174 | 0.116424531 | 0.103 | 0.05 | 1 | 9 |
| med6 | 0.67193 | 0.102600192 | 0.103 | 0.05 | 1 | 9 |
| si:ch73-347e22.8 | 0.67322 | -0.11723358 | 0.121 | 0.2 | 1 | 9 |
| arid3a | 0.67404 | 0.10913812 | 0.103 | 0.05 | 1 | 9 |
| tp53inp1 | 0.67444 | 0.181100003 | 0.276 | 0.2 | 1 | 9 |
| ppifb | 0.67508 | -0.17633461 | 0.328 | 0.4 | 1 | 9 |
| zfand5b | 0.67533 | 0.153946869 | 0.293 | 0.2 | 1 | 9 |
| me3 | 0.67585 | 0.126864176 | 0.086 | 0.05 | 1 | 9 |
| slc37a2 | 0.67591 | -0.13736983 | 0.103 | 0.15 | 1 | 9 |
| SEMA4F | 0.67644 | 0.1048647 | 0.086 | 0.05 | 1 | 9 |
| zdhhc6 | 0.67663 | 0.112119254 | 0.086 | 0.05 | 1 | 9 |
| aamp | 0.67699 | 0.12570066 | 0.241 | 0.15 | 1 | 9 |
| vps35 | 0.67752 | 0.108967074 | 0.103 | 0.05 | 1 | 9 |
| tcnbb | 0.67755 | 0.126897236 | 0.155 | 0.15 | 1 | 9 |
| ubxn4 | 0.67764 | 0.125495167 | 0.103 | 0.05 | 1 | 9 |
| mrps24 | 0.67781 | -0.11164746 | 0.121 | 0.15 | 1 | 9 |
| coa3a | 0.67791 | 0.129347548 | 0.241 | 0.15 | 1 | 9 |
| tmem147 | 0.67809 | 0.136503789 | 0.172 | 0.1 | 1 | 9 |
| zgc:153293 | 0.67824 | 0.100124103 | 0.086 | 0.05 | 1 | 9 |
| sh3bp5la | 0.67904 | -0.13226084 | 0.052 | 0.1 | 1 | 9 |
| wapla | 0.68027 | -0.10788108 | 0.069 | 0.1 | 1 | 9 |
| ppp2r1ba | 0.68033 | 0.134688628 | 0.241 | 0.15 | 1 | 9 |
| cx34.4 | 0.68044 | 0.11640991 | 0.086 | 0.05 | 1 | 9 |
| twistnb | 0.68048 | 0.146251681 | 0.241 | 0.15 | 1 | 9 |
| si:dkey-16l2.20 | 0.68051 | 0.104709464 | 0.121 | 0.1 | 1 | 9 |
| tomm20a | 0.68081 | 0.160511952 | 0.345 | 0.25 | 1 | 9 |
| ndufa2 | 0.6809 | 0.166152656 | 0.328 | 0.25 | 1 | 9 |
| ndufv1 | 0.68138 | 0.148637409 | 0.224 | 0.15 | 1 | 9 |
| erlec1 | 0.68143 | 0.102001379 | 0.172 | 0.1 | 1 | 9 |
| srsf6a | 0.68223 | 0.153870623 | 0.172 | 0.1 | 1 | 9 |
| tpm4a | 0.68244 | -0.15996322 | 0.155 | 0.2 | 1 | 9 |
| ptpn1 | 0.68311 | 0.119805738 | 0.103 | 0.05 | 1 | 9 |
| si:dkey-238k10.2 | 0.68321 | 0.112020363 | 0.103 | 0.05 | 1 | 9 |
| ddx21 | 0.68322 | 0.104987233 | 0.397 | 0.3 | 1 | 9 |
| ndufa4l | 0.68358 | 0.126985402 | 0.707 | 0.6 | 1 | 9 |

| | | | | | | |
|---|---|---|---|---|---|---|
| tcp11l2 | 0.68373 | 0.10583579 | 0.138 | 0.1 | 1 | 9 |
| tcf12 | 0.68385 | 0.153936235 | 0.172 | 0.1 | 1 | 9 |
| onecut1 | 0.68476 | -0.14263556 | 0.052 | 0.1 | 1 | 9 |
| stk35 | 0.68484 | 0.154294144 | 0.259 | 0.2 | 1 | 9 |
| tns1b | 0.68504 | 0.103218214 | 0.069 | 0.05 | 1 | 9 |
| kctd12.2 | 0.68521 | -0.12300988 | 0.052 | 0.1 | 1 | 9 |
| col1a1b | 0.68555 | 0.18278598 | 0.155 | 0.1 | 1 | 9 |
| ell | 0.68563 | 0.12643775 | 0.155 | 0.1 | 1 | 9 |
| st3gal7 | 0.68601 | -0.11280303 | 0.069 | 0.1 | 1 | 9 |
| fam174b | 0.68782 | 0.14140063 | 0.103 | 0.05 | 1 | 9 |
| hlfa | 0.68782 | -0.1317 | 0.086 | 0.15 | 1 | 9 |
| lygl1 | 0.68792 | -0.11562862 | 0.19 | 0.2 | 1 | 9 |
| mgll | 0.68939 | -0.12681054 | 0.121 | 0.2 | 1 | 9 |
| adipor2 | 0.6894 | -0.16078759 | 0.172 | 0.25 | 1 | 9 |
| dbpb | 0.69049 | -0.12680882 | 0.155 | 0.2 | 1 | 9 |
| s100a10b | 0.69059 | -0.21898148 | 0.414 | 0.5 | 1 | 9 |
| thoc7 | 0.6922 | 0.11504508 | 0.103 | 0.05 | 1 | 9 |
| vdrb | 0.69234 | 0.14059285 | 0.207 | 0.15 | 1 | 9 |
| aldh2.2 | 0.69254 | -0.14452746 | 0.086 | 0.15 | 1 | 9 |
| si:ch211-197h24.6 | 0.69265 | -0.11284581 | 0.121 | 0.15 | 1 | 9 |
| nr2f2 | 0.69323 | 0.117018793 | 0.103 | 0.05 | 1 | 9 |
| ptmab | 0.69329 | 0.139827856 | 0.793 | 0.7 | 1 | 9 |
| pptc7b | 0.69333 | 0.111655999 | 0.103 | 0.05 | 1 | 9 |
| appbp2 | 0.69355 | -0.1382147 | 0.086 | 0.15 | 1 | 9 |
| krt8 | 0.6941 | -0.17680511 | 0.569 | 0.65 | 1 | 9 |
| si:ch211-248a14.8 | 0.6942 | -0.12160045 | 0.052 | 0.1 | 1 | 9 |
| dnlz | 0.69444 | -0.11698478 | 0.052 | 0.1 | 1 | 9 |
| tmem238a | 0.69477 | -0.15646057 | 0.138 | 0.2 | 1 | 9 |
| yars | 0.69513 | -0.11878298 | 0.052 | 0.1 | 1 | 9 |
| pwwp2b | 0.69536 | -0.12491397 | 0.052 | 0.1 | 1 | 9 |
| nucb2b | 0.69593 | -0.17939871 | 0.121 | 0.2 | 1 | 9 |
| uba1 | 0.69605 | 0.155773105 | 0.276 | 0.2 | 1 | 9 |
| trip12 | 0.69673 | -0.12012735 | 0.052 | 0.1 | 1 | 9 |
| slka | 0.69678 | 0.112733379 | 0.103 | 0.05 | 1 | 9 |
| hsdl2 | 0.69708 | -0.11479661 | 0.121 | 0.2 | 1 | 9 |
| klf13 | 0.69811 | -0.11914961 | 0.052 | 0.1 | 1 | 9 |
| krt15 | 0.69814 | -0.14450274 | 0.31 | 0.3 | 1 | 9 |
| znf609b | 0.6983 | 0.127328197 | 0.138 | 0.1 | 1 | 9 |
| supt6h | 0.69935 | 0.109958991 | 0.103 | 0.05 | 1 | 9 |
| smc3 | 0.6994 | 0.112966845 | 0.103 | 0.05 | 1 | 9 |
| aga | 0.69946 | 0.12215435 | 0.138 | 0.1 | 1 | 9 |
| ube2al | 0.69948 | -0.134707 | 0.086 | 0.15 | 1 | 9 |
| tnksb | 0.69979 | 0.100423904 | 0.103 | 0.05 | 1 | 9 |
| nt5c2b | 0.69997 | -0.13832214 | 0.138 | 0.2 | 1 | 9 |
| zgc:114119 | 0.70016 | -0.12964992 | 0.052 | 0.1 | 1 | 9 |
| BX004785.3 | 0.70028 | -0.12295763 | 0.069 | 0.1 | 1 | 9 |
| tmem18 | 0.7005 | -0.15659014 | 0.172 | 0.25 | 1 | 9 |
| prpf19 | 0.70087 | 0.12221957 | 0.103 | 0.05 | 1 | 9 |

| | | | | | | |
|---|---|---|---|---|---|---|
| mrpl20 | 0.70104 | 0.125962145 | 0.293 | 0.2 | 1 | 9 |
| cryz | 0.70151 | 0.129218344 | 0.086 | 0.05 | 1 | 9 |
| stxbp6 | 0.703 | -0.12788129 | 0.103 | 0.15 | 1 | 9 |
| imp4 | 0.70307 | -0.10453665 | 0.052 | 0.1 | 1 | 9 |
| LO016987.2 | 0.70309 | 0.112558788 | 0.103 | 0.05 | 1 | 9 |
| zgc:174935 | 0.70416 | -0.15048541 | 0.155 | 0.2 | 1 | 9 |
| tmem131l | 0.70478 | -0.13639397 | 0.086 | 0.15 | 1 | 9 |
| tardbp | 0.70485 | 0.133584534 | 0.293 | 0.2 | 1 | 9 |
| mtmr8 | 0.70507 | 0.108213622 | 0.103 | 0.05 | 1 | 9 |
| lin37 | 0.70584 | -0.11849733 | 0.052 | 0.1 | 1 | 9 |
| grk5 | 0.70657 | -0.11506627 | 0.052 | 0.1 | 1 | 9 |
| cwc25 | 0.70658 | 0.110644272 | 0.172 | 0.1 | 1 | 9 |
| tmem192 | 0.7073 | -0.12046424 | 0.052 | 0.1 | 1 | 9 |
| nfe2l1b | 0.70771 | -0.11628431 | 0.069 | 0.1 | 1 | 9 |
| dhx8 | 0.70803 | 0.101997544 | 0.103 | 0.05 | 1 | 9 |
| ypel1 | 0.70862 | 0.12660069 | 0.086 | 0.05 | 1 | 9 |
| tmub2 | 0.70942 | 0.11071468 | 0.103 | 0.05 | 1 | 9 |
| rab5ab | 0.70961 | 0.151670292 | 0.224 | 0.15 | 1 | 9 |
| khsrp | 0.71013 | 0.135818178 | 0.293 | 0.2 | 1 | 9 |
| arrdc1a | 0.71193 | -0.14159275 | 0.155 | 0.2 | 1 | 9 |
| rtca | 0.71258 | 0.111499711 | 0.138 | 0.1 | 1 | 9 |
| psmb2 | 0.71302 | 0.118163484 | 0.397 | 0.3 | 1 | 9 |
| PYURF | 0.71395 | 0.110089681 | 0.103 | 0.05 | 1 | 9 |
| LOC100001846 | 0.71418 | -0.12864868 | 0.052 | 0.1 | 1 | 9 |
| mpzl1l | 0.71418 | -0.1104865 | 0.086 | 0.15 | 1 | 9 |
| cldn23a | 0.71431 | -0.11723274 | 0.121 | 0.15 | 1 | 9 |
| taf1 | 0.71446 | 0.105789636 | 0.086 | 0.05 | 1 | 9 |
| rerea | 0.71484 | 0.118995732 | 0.155 | 0.1 | 1 | 9 |
| pbrm1l | 0.71495 | -0.12061477 | 0.052 | 0.1 | 1 | 9 |
| exosc8 | 0.71553 | -0.11085827 | 0.069 | 0.1 | 1 | 9 |
| cmss1 | 0.71797 | 0.100857632 | 0.086 | 0.05 | 1 | 9 |
| LOC100333235 | 0.71801 | 0.126102931 | 0.172 | 0.1 | 1 | 9 |
| impdh1b | 0.71807 | 0.110749327 | 0.086 | 0.05 | 1 | 9 |
| ddx3xb | 0.7182 | 0.101397761 | 0.172 | 0.1 | 1 | 9 |
| ociad1 | 0.71826 | 0.100170567 | 0.172 | 0.1 | 1 | 9 |
| slc20a2 | 0.71921 | 0.113340947 | 0.086 | 0.05 | 1 | 9 |
| sephs2 | 0.71931 | 0.137274188 | 0.224 | 0.15 | 1 | 9 |
| tsr2 | 0.71934 | 0.10275382 | 0.172 | 0.1 | 1 | 9 |
| coa7 | 0.72036 | 0.120397366 | 0.103 | 0.05 | 1 | 9 |
| styk1b | 0.72129 | 0.132412117 | 0.103 | 0.05 | 1 | 9 |
| plpp2b | 0.72174 | -0.11597267 | 0.086 | 0.15 | 1 | 9 |
| hmbox1a | 0.72197 | 0.118587367 | 0.172 | 0.15 | 1 | 9 |
| GGT7 | 0.72272 | 0.113933463 | 0.103 | 0.05 | 1 | 9 |
| alcamb | 0.72413 | 0.160317832 | 0.172 | 0.1 | 1 | 9 |
| rexo4 | 0.72443 | 0.102416536 | 0.172 | 0.1 | 1 | 9 |
| dlat | 0.72453 | -0.11584534 | 0.121 | 0.15 | 1 | 9 |
| lcorl | 0.72477 | 0.100898301 | 0.103 | 0.05 | 1 | 9 |
| cipcb | 0.72493 | 0.103688878 | 0.103 | 0.05 | 1 | 9 |

| | | | | | | |
|---|---|---|---|---|---|---|
| pdhb | 0.72504 | -0.12310528 | 0.103 | 0.15 | 1 | 9 |
| pdcd10b | 0.72514 | 0.125440649 | 0.172 | 0.1 | 1 | 9 |
| zranb2 | 0.7256 | 0.135800046 | 0.172 | 0.1 | 1 | 9 |
| mrps33 | 0.72566 | 0.106859211 | 0.172 | 0.1 | 1 | 9 |
| LOC108190726 | 0.72569 | 0.110038871 | 0.103 | 0.05 | 1 | 9 |
| ppp2r5d | 0.72641 | 0.100430313 | 0.103 | 0.05 | 1 | 9 |
| fgl2a | 0.72646 | -0.10984682 | 0.052 | 0.1 | 1 | 9 |
| si:ch211-221f10.2 | 0.7269 | -0.12101637 | 0.086 | 0.15 | 1 | 9 |
| ssbp3b | 0.7272 | -0.10407826 | 0.086 | 0.15 | 1 | 9 |
| lamtor5 | 0.72727 | 0.142193032 | 0.155 | 0.1 | 1 | 9 |
| ms4a17a.11 | 0.72734 | 0.127797076 | 0.103 | 0.05 | 1 | 9 |
| ap3s2 | 0.72766 | -0.10658506 | 0.086 | 0.15 | 1 | 9 |
| pak2a | 0.728 | -0.11911842 | 0.086 | 0.15 | 1 | 9 |
| flii | 0.72802 | 0.101592867 | 0.086 | 0.05 | 1 | 9 |
| g3bp1 | 0.72835 | -0.15465391 | 0.138 | 0.2 | 1 | 9 |
| babam1 | 0.72858 | -0.10450782 | 0.069 | 0.1 | 1 | 9 |
| cnbpb | 0.72872 | 0.116689109 | 0.552 | 0.45 | 1 | 9 |
| pleca | 0.72917 | -0.12539323 | 0.259 | 0.3 | 1 | 9 |
| mrpl47 | 0.72991 | -0.1411484 | 0.103 | 0.15 | 1 | 9 |
| selenow2a | 0.7303 | -0.11396597 | 0.086 | 0.15 | 1 | 9 |
| ugp2b | 0.73064 | 0.112106186 | 0.086 | 0.05 | 1 | 9 |
| celf1 | 0.73163 | 0.11146552 | 0.103 | 0.05 | 1 | 9 |
| rnf144b | 0.73177 | -0.11150484 | 0.086 | 0.15 | 1 | 9 |
| eif1axa | 0.73189 | -0.10486347 | 0.086 | 0.15 | 1 | 9 |
| atl3 | 0.73226 | 0.126006975 | 0.103 | 0.05 | 1 | 9 |
| spsb4b | 0.73255 | 0.118434022 | 0.103 | 0.05 | 1 | 9 |
| metrn | 0.73353 | 0.112041454 | 0.086 | 0.05 | 1 | 9 |
| rap1b | 0.73359 | -0.13858565 | 0.241 | 0.3 | 1 | 9 |
| rbm38 | 0.73401 | 0.104897326 | 0.103 | 0.05 | 1 | 9 |
| swt1 | 0.73468 | -0.1167406 | 0.086 | 0.15 | 1 | 9 |
| cdc37 | 0.73539 | 0.10484463 | 0.103 | 0.05 | 1 | 9 |
| rabgap1l | 0.73709 | -0.10807643 | 0.052 | 0.1 | 1 | 9 |
| ap1g1 | 0.73768 | -0.12173618 | 0.086 | 0.15 | 1 | 9 |
| ankrd12 | 0.73771 | -0.11369881 | 0.086 | 0.15 | 1 | 9 |
| CU660013.4 | 0.73815 | -0.1192594 | 0.086 | 0.15 | 1 | 9 |
| helz | 0.73885 | -0.14096035 | 0.138 | 0.2 | 1 | 9 |
| bccip | 0.73919 | 0.114183358 | 0.448 | 0.35 | 1 | 9 |
| her15.1 | 0.73969 | -0.12807584 | 0.069 | 0.1 | 1 | 9 |
| ppp2r5eb | 0.73986 | 0.115151155 | 0.224 | 0.15 | 1 | 9 |
| tspan11 | 0.74139 | -0.11307114 | 0.069 | 0.1 | 1 | 9 |
| col5a2a | 0.74165 | 0.127879023 | 0.086 | 0.05 | 1 | 9 |
| arf5 | 0.7417 | 0.125358916 | 0.448 | 0.35 | 1 | 9 |
| calm3a | 0.74206 | -0.12496836 | 0.103 | 0.15 | 1 | 9 |
| il34 | 0.74272 | -0.11045066 | 0.052 | 0.1 | 1 | 9 |
| pmp22b | 0.74341 | 0.117042735 | 0.086 | 0.05 | 1 | 9 |
| zgc:194627 | 0.74354 | -0.13122059 | 0.052 | 0.1 | 1 | 9 |
| si:dkey-7c18.24 | 0.7444 | 0.161929849 | 0.207 | 0.15 | 1 | 9 |
| tor1 | 0.74446 | 0.119742752 | 0.103 | 0.05 | 1 | 9 |

| | | | | | | |
|---|---|---|---|---|---|---|
| CABZ01088149.1 | 0.74584 | -0.1014172 | 0.052 | 0.1 | 1 | 9 |
| rnf14 | 0.7471 | -0.10245309 | 0.069 | 0.1 | 1 | 9 |
| psmc2 | 0.74736 | 0.122056328 | 0.224 | 0.15 | 1 | 9 |
| notch3 | 0.74829 | 0.15536687 | 0.259 | 0.2 | 1 | 9 |
| ier2a | 0.74832 | -0.13760987 | 0.466 | 0.5 | 1 | 9 |
| per1a | 0.74841 | 0.114837857 | 0.103 | 0.05 | 1 | 9 |
| NDUFB1 | 0.74912 | -0.10691988 | 0.466 | 0.5 | 1 | 9 |
| elf3 | 0.74959 | -0.11937503 | 0.172 | 0.2 | 1 | 9 |
| si:dkey-221j11.3 | 0.74999 | 0.13369728 | 0.138 | 0.1 | 1 | 9 |
| sap30bp | 0.75194 | -0.13206242 | 0.138 | 0.2 | 1 | 9 |
| CABZ01034698.2 | 0.75196 | 0.107152717 | 0.103 | 0.05 | 1 | 9 |
| col6a3 | 0.75232 | 0.161432046 | 0.103 | 0.05 | 1 | 9 |
| osbpl7 | 0.75232 | 0.117949217 | 0.103 | 0.05 | 1 | 9 |
| hic2 | 0.75248 | -0.10442506 | 0.052 | 0.1 | 1 | 9 |
| fbxl3a | 0.75288 | 0.105687213 | 0.086 | 0.05 | 1 | 9 |
| gatad2b | 0.75445 | -0.13935467 | 0.172 | 0.25 | 1 | 9 |
| kars | 0.75447 | 0.114971477 | 0.241 | 0.2 | 1 | 9 |
| fkbp4 | 0.756 | 0.140463145 | 0.224 | 0.15 | 1 | 9 |
| lmo7a | 0.75807 | -0.13763166 | 0.172 | 0.25 | 1 | 9 |
| lamtor2 | 0.75919 | -0.13421597 | 0.155 | 0.2 | 1 | 9 |
| yy1a | 0.75955 | 0.110901499 | 0.207 | 0.15 | 1 | 9 |
| entpd8 | 0.75971 | -0.11288028 | 0.052 | 0.1 | 1 | 9 |
| tmem176l.4 | 0.75979 | 0.113659142 | 0.328 | 0.25 | 1 | 9 |
| znf770 | 0.7601 | -0.11336948 | 0.052 | 0.1 | 1 | 9 |
| bcl2l1 | 0.76032 | -0.14306274 | 0.103 | 0.15 | 1 | 9 |
| cep131 | 0.7622 | -0.10533911 | 0.052 | 0.1 | 1 | 9 |
| igf2a | 0.76232 | -0.11305295 | 0.052 | 0.1 | 1 | 9 |
| elovl1b | 0.76257 | 0.109472022 | 0.172 | 0.15 | 1 | 9 |
| sash1a | 0.76315 | -0.11490195 | 0.052 | 0.1 | 1 | 9 |
| nfixb | 0.76334 | 0.10658706 | 0.172 | 0.15 | 1 | 9 |
| tjp2b | 0.76381 | -0.11831883 | 0.224 | 0.25 | 1 | 9 |
| h1fx | 0.76408 | -0.11646235 | 0.293 | 0.3 | 1 | 9 |
| igfbp7 | 0.76476 | 0.147845644 | 0.069 | 0.05 | 1 | 9 |
| etnk1 | 0.7652 | 0.130174743 | 0.138 | 0.1 | 1 | 9 |
| dedd1 | 0.76608 | 0.149730852 | 0.431 | 0.35 | 1 | 9 |
| srp72 | 0.76712 | 0.114802133 | 0.155 | 0.1 | 1 | 9 |
| wipf2a | 0.7673 | -0.1220515 | 0.138 | 0.2 | 1 | 9 |
| armc9 | 0.76735 | -0.15624146 | 0.034 | 0.05 | 1 | 9 |
| NMNAT3 | 0.76735 | -0.10230223 | 0.034 | 0.05 | 1 | 9 |
| crabp1a | 0.76735 | -0.17716275 | 0.034 | 0.05 | 1 | 9 |
| LO017965.1 | 0.76735 | -0.12979129 | 0.034 | 0.05 | 1 | 9 |
| cxcl8b.1 | 0.76735 | -0.1371422 | 0.034 | 0.05 | 1 | 9 |
| znf653 | 0.76735 | -0.12692419 | 0.034 | 0.05 | 1 | 9 |
| e2f4 | 0.76735 | -0.14670971 | 0.034 | 0.05 | 1 | 9 |
| nr5a2 | 0.76735 | -0.11646988 | 0.034 | 0.05 | 1 | 9 |
| si:ch211-195d17.2 | 0.76735 | -0.15075601 | 0.034 | 0.05 | 1 | 9 |
| b3gnt5a | 0.76735 | -0.12180544 | 0.034 | 0.05 | 1 | 9 |
| si:ch211-230g15.5 | 0.76735 | -0.1002895 | 0.034 | 0.05 | 1 | 9 |

| | | | | | | |
|---|---|---|---|---|---|---|
| zgc:172053 | 0.76735 | 0.133890694 | 0.034 | 0.05 | 1 | 9 |
| GNAZ | 0.76735 | -0.11058032 | 0.034 | 0.05 | 1 | 9 |
| LOC799656 | 0.76735 | -0.11521961 | 0.034 | 0.05 | 1 | 9 |
| RALGDS | 0.76735 | -0.13357858 | 0.034 | 0.05 | 1 | 9 |
| dusp2 | 0.76735 | -0.10100299 | 0.034 | 0.05 | 1 | 9 |
| elovl2 | 0.76735 | -0.11480991 | 0.034 | 0.05 | 1 | 9 |
| faxdc2 | 0.76735 | -0.19380368 | 0.034 | 0.05 | 1 | 9 |
| fxn | 0.76735 | -0.16271837 | 0.034 | 0.05 | 1 | 9 |
| grhl2b | 0.76735 | -0.12600442 | 0.034 | 0.05 | 1 | 9 |
| hacd3 | 0.76735 | -0.17395446 | 0.034 | 0.05 | 1 | 9 |
| heatr5b | 0.76735 | -0.13868067 | 0.034 | 0.05 | 1 | 9 |
| igf1rb | 0.76735 | -0.10046126 | 0.034 | 0.05 | 1 | 9 |
| insig1 | 0.76735 | -0.13738357 | 0.034 | 0.05 | 1 | 9 |
| ptpn4a | 0.76735 | -0.14349756 | 0.034 | 0.05 | 1 | 9 |
| rybpa | 0.76735 | -0.13499458 | 0.034 | 0.05 | 1 | 9 |
| si:ch211-153b23.7 | 0.76735 | -0.10767793 | 0.034 | 0.05 | 1 | 9 |
| slc35e4 | 0.76735 | -0.12318375 | 0.034 | 0.05 | 1 | 9 |
| slc44a2 | 0.76735 | -0.10823896 | 0.034 | 0.05 | 1 | 9 |
| uap1l1 | 0.76735 | -0.10486177 | 0.034 | 0.05 | 1 | 9 |
| utp6 | 0.76735 | -0.13472246 | 0.034 | 0.05 | 1 | 9 |
| brf1b | 0.76735 | -0.16038616 | 0.034 | 0.05 | 1 | 9 |
| esrra | 0.76735 | -0.10414424 | 0.034 | 0.05 | 1 | 9 |
| smc4 | 0.76735 | -0.1287198 | 0.034 | 0.05 | 1 | 9 |
| wdhd1 | 0.76735 | -0.10125045 | 0.034 | 0.05 | 1 | 9 |
| zgc:100846 | 0.76735 | -0.1428851 | 0.034 | 0.05 | 1 | 9 |
| ubr4 | 0.7677 | -0.10934624 | 0.069 | 0.1 | 1 | 9 |
| sox3 | 0.76797 | -0.11682342 | 0.052 | 0.1 | 1 | 9 |
| lmf2b | 0.76819 | 0.105329666 | 0.172 | 0.15 | 1 | 9 |
| btg1 | 0.76834 | 0.113019785 | 0.69 | 0.6 | 1 | 9 |
| bbx | 0.76842 | -0.11576224 | 0.138 | 0.2 | 1 | 9 |
| hikeshi | 0.76913 | 0.119231942 | 0.224 | 0.15 | 1 | 9 |
| timmdc1 | 0.76914 | 0.100171839 | 0.086 | 0.05 | 1 | 9 |
| kpna4 | 0.77064 | 0.112899322 | 0.155 | 0.1 | 1 | 9 |
| dstyk | 0.77178 | 0.108043907 | 0.155 | 0.1 | 1 | 9 |
| ccdc28a | 0.77205 | -0.11269849 | 0.224 | 0.25 | 1 | 9 |
| glod4 | 0.77342 | -0.10353938 | 0.069 | 0.1 | 1 | 9 |
| cntfr | 0.77535 | 0.160986264 | 0.155 | 0.1 | 1 | 9 |
| gmds | 0.77537 | -0.13027409 | 0.103 | 0.15 | 1 | 9 |
| ndufb5 | 0.77538 | -0.13954211 | 0.362 | 0.45 | 1 | 9 |
| tmem176l.1 | 0.77666 | -0.11153111 | 0.172 | 0.2 | 1 | 9 |
| naa40 | 0.78284 | -0.11842311 | 0.103 | 0.15 | 1 | 9 |
| smarcd1 | 0.78333 | 0.122314931 | 0.276 | 0.2 | 1 | 9 |
| abracl | 0.78437 | -0.10700274 | 0.276 | 0.3 | 1 | 9 |
| igf2bp1 | 0.78678 | -0.11341298 | 0.069 | 0.1 | 1 | 9 |
| dspa | 0.78789 | 0.121890604 | 0.241 | 0.2 | 1 | 9 |
| rad21a | 0.78813 | 0.104770706 | 0.328 | 0.25 | 1 | 9 |
| adssl | 0.78854 | 0.103779512 | 0.276 | 0.2 | 1 | 9 |
| si:dkey-177p2.6 | 0.78857 | 0.126489693 | 0.276 | 0.2 | 1 | 9 |

| | | | | | | |
|---|---|---|---|---|---|---|
| eif4g2a | 0.78905 | 0.103059128 | 0.138 | 0.1 | 1 | 9 |
| fundc1 | 0.78958 | -0.1056667 | 0.138 | 0.2 | 1 | 9 |
| stx11b.1 | 0.79138 | 0.104504324 | 0.155 | 0.1 | 1 | 9 |
| prkar2aa | 0.79221 | 0.114347984 | 0.276 | 0.2 | 1 | 9 |
| tuba8l2 | 0.79225 | 0.100377198 | 0.328 | 0.25 | 1 | 9 |
| htatip2 | 0.79248 | 0.11134273 | 0.19 | 0.15 | 1 | 9 |
| ube2e2 | 0.79469 | 0.122884993 | 0.155 | 0.1 | 1 | 9 |
| rnf128a | 0.79558 | -0.10917085 | 0.224 | 0.3 | 1 | 9 |
| col6a2 | 0.79615 | 0.111788441 | 0.086 | 0.05 | 1 | 9 |
| tmem59 | 0.79764 | -0.11300294 | 0.414 | 0.45 | 1 | 9 |
| rbms1b | 0.80076 | 0.107391877 | 0.086 | 0.05 | 1 | 9 |
| mgst3a | 0.80122 | 0.124067197 | 0.31 | 0.25 | 1 | 9 |
| prr13 | 0.80212 | -0.12513572 | 0.155 | 0.2 | 1 | 9 |
| prelid3b | 0.8027 | 0.10095125 | 0.379 | 0.3 | 1 | 9 |
| lsm7 | 0.80321 | 0.106058089 | 0.328 | 0.25 | 1 | 9 |
| sccpdhb | 0.8037 | -0.10028367 | 0.155 | 0.2 | 1 | 9 |
| wu:fi04e12 | 0.80388 | -0.11933399 | 0.138 | 0.2 | 1 | 9 |
| bcas2 | 0.8044 | -0.12143018 | 0.19 | 0.25 | 1 | 9 |
| mat2aa | 0.80603 | 0.114263252 | 0.328 | 0.25 | 1 | 9 |
| rab2a | 0.8101 | 0.108271959 | 0.379 | 0.3 | 1 | 9 |
| tmx1 | 0.81027 | -0.10077281 | 0.121 | 0.15 | 1 | 9 |
| mpzl2b | 0.81035 | -0.11789676 | 0.172 | 0.2 | 1 | 9 |
| pkig | 0.8105 | -0.1087079 | 0.103 | 0.15 | 1 | 9 |
| asph | 0.81068 | -0.10964713 | 0.138 | 0.2 | 1 | 9 |
| brd3a | 0.81092 | -0.10611022 | 0.138 | 0.2 | 1 | 9 |
| afdna | 0.81135 | 0.118320323 | 0.086 | 0.05 | 1 | 9 |
| med29 | 0.81263 | 0.10898577 | 0.138 | 0.1 | 1 | 9 |
| icn2 | 0.81491 | -0.11372442 | 0.276 | 0.3 | 1 | 9 |
| cdc42l | 0.81541 | 0.115905732 | 0.431 | 0.35 | 1 | 9 |
| zgc:113625 | 0.8155 | -0.1106116 | 0.069 | 0.1 | 1 | 9 |
| uqcc2 | 0.81577 | 0.104604965 | 0.155 | 0.1 | 1 | 9 |
| tmsb1 | 0.81792 | -0.18581954 | 0.293 | 0.35 | 1 | 9 |
| rab18b | 0.81928 | -0.11537051 | 0.103 | 0.15 | 1 | 9 |
| klf3 | 0.81974 | 0.113267548 | 0.155 | 0.1 | 1 | 9 |
| si:ch73-204p21.2 | 0.82054 | 0.117992601 | 0.155 | 0.1 | 1 | 9 |
| ythdc1 | 0.82092 | 0.109119103 | 0.155 | 0.1 | 1 | 9 |
| polr2a | 0.82269 | -0.12023778 | 0.103 | 0.15 | 1 | 9 |
| stard14 | 0.82334 | -0.14041216 | 0.207 | 0.25 | 1 | 9 |
| bzw1a | 0.82471 | -0.11314061 | 0.276 | 0.35 | 1 | 9 |
| sdhc | 0.82864 | -0.12493329 | 0.293 | 0.35 | 1 | 9 |
| chtopa | 0.83268 | 0.110962579 | 0.207 | 0.15 | 1 | 9 |
| dhrs13a.2 | 0.8389 | 0.145033324 | 0.207 | 0.15 | 1 | 9 |
| CR854839.1 | 0.84029 | 0.112141693 | 0.138 | 0.1 | 1 | 9 |
| si:ch211-207n23.2 | 0.84958 | 0.156476915 | 0.19 | 0.15 | 1 | 9 |
| sigmar1 | 0.85122 | -0.10037187 | 0.103 | 0.15 | 1 | 9 |
| rhcgl1 | 0.85154 | 0.156961422 | 0.207 | 0.15 | 1 | 9 |
| znf185 | 0.8537 | 0.135920938 | 0.259 | 0.2 | 1 | 9 |
| fbp1a | 0.85443 | -0.10503693 | 0.172 | 0.2 | 1 | 9 |

| | | | | | | |
|---|---|---|---|---|---|---|
| f11r.1 | 0.85485 | -0.11306982 | 0.379 | 0.45 | 1 | 9 |
| dynlt3 | 0.8558 | 0.104158259 | 0.31 | 0.25 | 1 | 9 |
| adipor1b | 0.85591 | -0.10073293 | 0.103 | 0.15 | 1 | 9 |
| klf7b | 0.85824 | 0.103598324 | 0.086 | 0.05 | 1 | 9 |
| ponzr2 | 0.85975 | -0.10617626 | 0.103 | 0.15 | 1 | 9 |
| ctsba | 0.86208 | 0.102094069 | 0.293 | 0.25 | 1 | 9 |
| rab11al | 0.87412 | -0.10032813 | 0.069 | 0.1 | 1 | 9 |
| si:dkeyp-121d4.3 | 0.87617 | -0.10102836 | 0.069 | 0.1 | 1 | 9 |
| cxadr | 0.87962 | -0.10546567 | 0.155 | 0.2 | 1 | 9 |
| si:dkey-248g15.3 | 0.91104 | 0.110364656 | 0.172 | 0.15 | 1 | 9 |
| gamt | 0.00017 | 0.742522199 | 0.275 | 0 | 1 | 10 |
| bhmt | 0.00039 | 0.655383036 | 0.25 | 0 | 1 | 10 |
| cnp | 0.0004 | -0.62941123 | 0 | 0.235 | 1 | 10 |
| si:dkey-14d8.6 | 0.00085 | 0.576657575 | 0.225 | 0 | 1 | 10 |
| nupr1b | 0.00085 | 0.645302868 | 0.225 | 0 | 1 | 10 |
| trmt112 | 0.00125 | -0.71282353 | 0.025 | 0.324 | 1 | 10 |
| cpb1 | 0.00184 | 0.612895242 | 0.2 | 0 | 1 | 10 |
| chchd6b | 0.00273 | -0.62119635 | 0 | 0.176 | 1 | 10 |
| cdh1 | 0.00273 | -0.54185871 | 0 | 0.176 | 1 | 10 |
| c6ast4 | 0.00395 | 0.51184685 | 0.175 | 0 | 1 | 10 |
| ecrg4a | 0.00666 | -0.47735164 | 0.175 | 0.471 | 1 | 10 |
| smad1 | 0.00698 | -0.43854916 | 0 | 0.147 | 1 | 10 |
| sox9a | 0.00698 | -0.45820439 | 0 | 0.147 | 1 | 10 |
| ccdc51 | 0.00698 | -0.40597849 | 0 | 0.147 | 1 | 10 |
| timm23a | 0.00698 | -0.50442665 | 0 | 0.147 | 1 | 10 |
| wdr82 | 0.00698 | -0.430198 | 0 | 0.147 | 1 | 10 |
| snrpb2 | 0.00698 | -0.38184302 | 0 | 0.147 | 1 | 10 |
| vps35 | 0.00698 | -0.4568007 | 0 | 0.147 | 1 | 10 |
| agrn | 0.00698 | -0.48684755 | 0 | 0.147 | 1 | 10 |
| aldh1a2 | 0.00718 | -0.4720317 | 0.625 | 0.882 | 1 | 10 |
| cpa4 | 0.00839 | 0.436797578 | 0.15 | 0 | 1 | 10 |
| eci1 | 0.00839 | 0.425298166 | 0.15 | 0 | 1 | 10 |
| si:dkey-14d8.7 | 0.00839 | 0.430825185 | 0.15 | 0 | 1 | 10 |
| psmb3 | 0.0085 | -0.69299065 | 0.125 | 0.441 | 1 | 10 |
| sec61b | 0.00861 | 0.61184674 | 0.7 | 0.353 | 1 | 10 |
| psmd14 | 0.01334 | -0.57938178 | 0.05 | 0.294 | 1 | 10 |
| lsm6 | 0.01354 | -0.65990675 | 0.05 | 0.294 | 1 | 10 |
| rbm4.3 | 0.01451 | -0.60411642 | 0.05 | 0.294 | 1 | 10 |
| zgc:92744 | 0.01726 | 0.351160374 | 0.8 | 0.706 | 1 | 10 |
| jcada | 0.0177 | -0.39443909 | 0 | 0.118 | 1 | 10 |
| marcksb | 0.0177 | -0.39048749 | 0 | 0.118 | 1 | 10 |
| ap1s1 | 0.0177 | -0.3446361 | 0 | 0.118 | 1 | 10 |
| cpsf6 | 0.0177 | -0.36406079 | 0 | 0.118 | 1 | 10 |
| mcm7 | 0.0177 | -0.39044587 | 0 | 0.118 | 1 | 10 |
| arpin | 0.0177 | -0.31016503 | 0 | 0.118 | 1 | 10 |
| cetn3 | 0.0177 | -0.33879239 | 0 | 0.118 | 1 | 10 |
| lin52 | 0.0177 | -0.38921717 | 0 | 0.118 | 1 | 10 |
| ndufs2 | 0.0177 | -0.38058485 | 0 | 0.118 | 1 | 10 |

| | | | | | | |
|---|---|---|---|---|---|---|
| pdlim1 | 0.0177 | -0.37572343 | 0 | 0.118 | 1 | 10 |
| si:dkey-239h2.3 | 0.0177 | -0.32225747 | 0 | 0.118 | 1 | 10 |
| vaspb | 0.0177 | -0.31076008 | 0 | 0.118 | 1 | 10 |
| zgc:136864 | 0.0177 | -0.39954663 | 0 | 0.118 | 1 | 10 |
| arrb2b | 0.0177 | -0.37355651 | 0 | 0.118 | 1 | 10 |
| ccl19a.1 | 0.0177 | -0.3897976 | 0 | 0.118 | 1 | 10 |
| chrac1 | 0.0177 | -0.31551106 | 0 | 0.118 | 1 | 10 |
| kif22 | 0.0177 | -0.33987546 | 0 | 0.118 | 1 | 10 |
| pik3r3a | 0.0177 | -0.4033989 | 0 | 0.118 | 1 | 10 |
| sdha | 0.0177 | -0.33178159 | 0 | 0.118 | 1 | 10 |
| smad2 | 0.0177 | -0.36839675 | 0 | 0.118 | 1 | 10 |
| tia1 | 0.0177 | -0.34859704 | 0 | 0.118 | 1 | 10 |
| dr1 | 0.0177 | -0.31042545 | 0 | 0.118 | 1 | 10 |
| idh3b | 0.0177 | -0.36404125 | 0 | 0.118 | 1 | 10 |
| namptb | 0.0177 | -0.37208814 | 0 | 0.118 | 1 | 10 |
| ncoa5 | 0.0177 | -0.41772458 | 0 | 0.118 | 1 | 10 |
| psip1b | 0.0177 | -0.36115229 | 0 | 0.118 | 1 | 10 |
| rcor1 | 0.0177 | -0.38909556 | 0 | 0.118 | 1 | 10 |
| sec13 | 0.0177 | -0.3642446 | 0 | 0.118 | 1 | 10 |
| timm29 | 0.0177 | -0.37230441 | 0 | 0.118 | 1 | 10 |
| top2a | 0.0177 | -0.38879428 | 0 | 0.118 | 1 | 10 |
| ubqln4 | 0.0177 | -0.36146649 | 0 | 0.118 | 1 | 10 |
| tnrc6b | 0.0177 | -0.3992945 | 0 | 0.118 | 1 | 10 |
| zgc:114181 | 0.0177 | -0.37834341 | 0 | 0.118 | 1 | 10 |
| apoc1 | 0.01771 | 0.310569014 | 0.125 | 0 | 1 | 10 |
| pitpnbl | 0.01771 | 0.368205185 | 0.125 | 0 | 1 | 10 |
| si:ch211-195b11.3 | 0.01771 | 0.276433629 | 0.125 | 0 | 1 | 10 |
| c6ast3 | 0.01771 | 0.417324338 | 0.125 | 0 | 1 | 10 |
| rps6kb1b | 0.01771 | 0.380673611 | 0.125 | 0 | 1 | 10 |
| arl6ip4 | 0.01771 | 0.397670106 | 0.125 | 0 | 1 | 10 |
| mob4 | 0.01771 | 0.363178372 | 0.125 | 0 | 1 | 10 |
| prkab1a | 0.01771 | 0.39568091 | 0.125 | 0 | 1 | 10 |
| tpi1b | 0.01771 | 0.389438122 | 0.125 | 0 | 1 | 10 |
| trappc4 | 0.01771 | 0.41340124 | 0.125 | 0 | 1 | 10 |
| h3f3d | 0.02863 | -0.38914724 | 0.575 | 0.853 | 1 | 10 |
| prdx5 | 0.03017 | -0.58378887 | 0.05 | 0.265 | 1 | 10 |
| rps17 | 0.03117 | 0.306754377 | 0.925 | 0.765 | 1 | 10 |
| etfa | 0.03119 | -0.45171626 | 0.025 | 0.206 | 1 | 10 |
| sap30bp | 0.0314 | -0.46309631 | 0.025 | 0.206 | 1 | 10 |
| mgst3b | 0.03274 | 0.535381401 | 0.225 | 0.029 | 1 | 10 |
| rpl7 | 0.03337 | 0.23781614 | 0.925 | 0.853 | 1 | 10 |
| psmc4 | 0.03467 | -0.50030989 | 0.025 | 0.206 | 1 | 10 |
| abce1 | 0.03532 | -0.49207538 | 0.025 | 0.206 | 1 | 10 |
| cct6a | 0.03693 | -0.5324382 | 0.075 | 0.294 | 1 | 10 |
| rpl18a | 0.0372 | 0.160199318 | 0.925 | 0.912 | 1 | 10 |
| apoda.2 | 0.03725 | 0.275550453 | 0.1 | 0 | 1 | 10 |
| cpa5 | 0.03725 | 0.410733212 | 0.1 | 0 | 1 | 10 |
| ddx51 | 0.03725 | 0.309259168 | 0.1 | 0 | 1 | 10 |

| | | | | | | |
|---|---|---|---|---|---|---|
| eif4ebp3 | 0.03725 | 0.345341969 | 0.1 | 0 | 1 | 10 |
| fdps | 0.03725 | 0.326216763 | 0.1 | 0 | 1 | 10 |
| gatm | 0.03725 | 0.354363566 | 0.1 | 0 | 1 | 10 |
| lsm12a | 0.03725 | 0.340372511 | 0.1 | 0 | 1 | 10 |
| rrp9 | 0.03725 | 0.362266939 | 0.1 | 0 | 1 | 10 |
| smad5 | 0.03725 | 0.34161483 | 0.1 | 0 | 1 | 10 |
| tmem243b | 0.03725 | 0.307228818 | 0.1 | 0 | 1 | 10 |
| tradd | 0.03725 | 0.34859208 | 0.1 | 0 | 1 | 10 |
| zyx | 0.03725 | 0.367670078 | 0.1 | 0 | 1 | 10 |
| camlg | 0.03725 | 0.288687474 | 0.1 | 0 | 1 | 10 |
| frrs1b | 0.03725 | 0.32398689 | 0.1 | 0 | 1 | 10 |
| gpx4a | 0.03725 | 0.33201739 | 0.1 | 0 | 1 | 10 |
| kdelr2a | 0.03725 | 0.35409408 | 0.1 | 0 | 1 | 10 |
| lamp2 | 0.03725 | 0.345145293 | 0.1 | 0 | 1 | 10 |
| larp7 | 0.03725 | 0.323530881 | 0.1 | 0 | 1 | 10 |
| leg1.1 | 0.03725 | 0.325368766 | 0.1 | 0 | 1 | 10 |
| scp2b | 0.03725 | 0.319842996 | 0.1 | 0 | 1 | 10 |
| smim29 | 0.03725 | 0.330239619 | 0.1 | 0 | 1 | 10 |
| tmsb1 | 0.03725 | 0.36587537 | 0.1 | 0 | 1 | 10 |
| tomm70a | 0.03725 | 0.298592593 | 0.1 | 0 | 1 | 10 |
| actr2a | 0.03725 | 0.37309275 | 0.1 | 0 | 1 | 10 |
| eif4a1a | 0.03949 | -0.51956163 | 0.175 | 0.441 | 1 | 10 |
| cap1 | 0.04217 | -0.28330797 | 0.075 | 0.235 | 1 | 10 |
| cox20 | 0.04486 | -0.25127988 | 0 | 0.088 | 1 | 10 |
| eef1akmt1 | 0.04486 | -0.2868907 | 0 | 0.088 | 1 | 10 |
| AL935174.5 | 0.04486 | -0.264132 | 0 | 0.088 | 1 | 10 |
| ace | 0.04486 | -0.29769483 | 0 | 0.088 | 1 | 10 |
| creb3l3l | 0.04486 | -0.22993922 | 0 | 0.088 | 1 | 10 |
| ddx23 | 0.04486 | -0.31544891 | 0 | 0.088 | 1 | 10 |
| dync1li2 | 0.04486 | -0.25297299 | 0 | 0.088 | 1 | 10 |
| ece2b | 0.04486 | -0.28410372 | 0 | 0.088 | 1 | 10 |
| eif2b1 | 0.04486 | -0.2863338 | 0 | 0.088 | 1 | 10 |
| f11r.1 | 0.04486 | -0.33178979 | 0 | 0.088 | 1 | 10 |
| fn1b | 0.04486 | -0.27781394 | 0 | 0.088 | 1 | 10 |
| golt1bb | 0.04486 | -0.24944045 | 0 | 0.088 | 1 | 10 |
| hdac4 | 0.04486 | -0.31211242 | 0 | 0.088 | 1 | 10 |
| igf2bp2a | 0.04486 | -0.25769633 | 0 | 0.088 | 1 | 10 |
| ip6k2b | 0.04486 | -0.266981 | 0 | 0.088 | 1 | 10 |
| mdka | 0.04486 | -0.24994651 | 0 | 0.088 | 1 | 10 |
| nudt1 | 0.04486 | -0.32658253 | 0 | 0.088 | 1 | 10 |
| paics | 0.04486 | -0.32422571 | 0 | 0.088 | 1 | 10 |
| pole4 | 0.04486 | -0.28621133 | 0 | 0.088 | 1 | 10 |
| psat1 | 0.04486 | -0.25120639 | 0 | 0.088 | 1 | 10 |
| rbm22 | 0.04486 | -0.27482491 | 0 | 0.088 | 1 | 10 |
| selenot1a | 0.04486 | -0.25411466 | 0 | 0.088 | 1 | 10 |
| sema3ab | 0.04486 | -0.33571277 | 0 | 0.088 | 1 | 10 |
| si:ch211-162i8.5 | 0.04486 | -0.27882679 | 0 | 0.088 | 1 | 10 |
| si:dkey-27b3.2 | 0.04486 | -0.26409345 | 0 | 0.088 | 1 | 10 |

| | | | | | | |
|---|---|---|---|---|---|---|
| smoc2 | 0.04486 | -0.31388148 | 0 | 0.088 | 1 | 10 |
| stat3 | 0.04486 | -0.26893511 | 0 | 0.088 | 1 | 10 |
| tdp1 | 0.04486 | -0.32997557 | 0 | 0.088 | 1 | 10 |
| tfr1a | 0.04486 | -0.30205867 | 0 | 0.088 | 1 | 10 |
| tra2b | 0.04486 | -0.26345886 | 0 | 0.088 | 1 | 10 |
| trappc5 | 0.04486 | -0.288373 | 0 | 0.088 | 1 | 10 |
| ugp2b | 0.04486 | -0.25494499 | 0 | 0.088 | 1 | 10 |
| zgc:114200 | 0.04486 | -0.31032389 | 0 | 0.088 | 1 | 10 |
| CABZ01068251.1 | 0.04486 | -0.25616624 | 0 | 0.088 | 1 | 10 |
| CDPF1 | 0.04486 | -0.24743693 | 0 | 0.088 | 1 | 10 |
| CU104709.1 | 0.04486 | -0.30237219 | 0 | 0.088 | 1 | 10 |
| CZQB01141835.1 | 0.04486 | -0.23053635 | 0 | 0.088 | 1 | 10 |
| aldh3a2b | 0.04486 | -0.24904296 | 0 | 0.088 | 1 | 10 |
| anapc11 | 0.04486 | -0.32006557 | 0 | 0.088 | 1 | 10 |
| atp1a3b | 0.04486 | -0.27544741 | 0 | 0.088 | 1 | 10 |
| atpaf1 | 0.04486 | -0.22762216 | 0 | 0.088 | 1 | 10 |
| bcat2 | 0.04486 | -0.23818961 | 0 | 0.088 | 1 | 10 |
| camk2g1 | 0.04486 | -0.28615924 | 0 | 0.088 | 1 | 10 |
| cavin1b | 0.04486 | -0.27559434 | 0 | 0.088 | 1 | 10 |
| cdk1 | 0.04486 | -0.32218723 | 0 | 0.088 | 1 | 10 |
| cox17 | 0.04486 | -0.24979094 | 0 | 0.088 | 1 | 10 |
| cox6a2 | 0.04486 | -0.33391003 | 0 | 0.088 | 1 | 10 |
| ddx19 | 0.04486 | -0.25433354 | 0 | 0.088 | 1 | 10 |
| dusp7 | 0.04486 | -0.29961901 | 0 | 0.088 | 1 | 10 |
| erlin1 | 0.04486 | -0.23093068 | 0 | 0.088 | 1 | 10 |
| gmppb | 0.04486 | -0.26617391 | 0 | 0.088 | 1 | 10 |
| gpr108 | 0.04486 | -0.25559174 | 0 | 0.088 | 1 | 10 |
| hbegfb | 0.04486 | -0.21513996 | 0 | 0.088 | 1 | 10 |
| hikeshi | 0.04486 | -0.30436879 | 0 | 0.088 | 1 | 10 |
| hirip3 | 0.04486 | -0.34108474 | 0 | 0.088 | 1 | 10 |
| hmgcl | 0.04486 | -0.26556704 | 0 | 0.088 | 1 | 10 |
| hnrnpdl | 0.04486 | -0.20073761 | 0 | 0.088 | 1 | 10 |
| kif13ba | 0.04486 | -0.29577404 | 0 | 0.088 | 1 | 10 |
| lrpap1 | 0.04486 | -0.24642445 | 0 | 0.088 | 1 | 10 |
| mark2a | 0.04486 | -0.23926896 | 0 | 0.088 | 1 | 10 |
| mbnl1 | 0.04486 | -0.2760442 | 0 | 0.088 | 1 | 10 |
| mef2cb | 0.04486 | -0.26534938 | 0 | 0.088 | 1 | 10 |
| mibp | 0.04486 | -0.29636857 | 0 | 0.088 | 1 | 10 |
| mrps33 | 0.04486 | -0.21132509 | 0 | 0.088 | 1 | 10 |
| n4bp2l2 | 0.04486 | -0.27128736 | 0 | 0.088 | 1 | 10 |
| nasp | 0.04486 | -0.25847802 | 0 | 0.088 | 1 | 10 |
| nfil3 | 0.04486 | -0.28451538 | 0 | 0.088 | 1 | 10 |
| nop58 | 0.04486 | -0.26757086 | 0 | 0.088 | 1 | 10 |
| nudt8 | 0.04486 | -0.24415617 | 0 | 0.088 | 1 | 10 |
| nxt2 | 0.04486 | -0.23632613 | 0 | 0.088 | 1 | 10 |
| pbxip1b | 0.04486 | -0.34975489 | 0 | 0.088 | 1 | 10 |
| pdcd6ip | 0.04486 | -0.26549514 | 0 | 0.088 | 1 | 10 |
| pfdn2 | 0.04486 | -0.2678441 | 0 | 0.088 | 1 | 10 |

| | | | | | | |
|---|---|---|---|---|---|---|
| ppp1r1b | 0.04486 | -0.28031619 | 0 | 0.088 | 1 | 10 |
| ppp1r7 | 0.04486 | -0.23442228 | 0 | 0.088 | 1 | 10 |
| psmd10 | 0.04486 | -0.23946426 | 0 | 0.088 | 1 | 10 |
| pycr1b | 0.04486 | -0.23918236 | 0 | 0.088 | 1 | 10 |
| rbm28 | 0.04486 | -0.34180063 | 0 | 0.088 | 1 | 10 |
| rnls | 0.04486 | -0.21018492 | 0 | 0.088 | 1 | 10 |
| sdf2l1 | 0.04486 | -0.28079905 | 0 | 0.088 | 1 | 10 |
| serpinb1l3 | 0.04486 | -0.27301603 | 0 | 0.088 | 1 | 10 |
| si:ch211-266i6.3 | 0.04486 | -0.22368334 | 0 | 0.088 | 1 | 10 |
| si:dkey-6n21.12 | 0.04486 | -0.26756122 | 0 | 0.088 | 1 | 10 |
| si:dkeyp-113d7.1 | 0.04486 | -0.27365748 | 0 | 0.088 | 1 | 10 |
| snx7 | 0.04486 | -0.32884199 | 0 | 0.088 | 1 | 10 |
| tars | 0.04486 | -0.28269911 | 0 | 0.088 | 1 | 10 |
| ttc5 | 0.04486 | -0.23872896 | 0 | 0.088 | 1 | 10 |
| ube2c | 0.04486 | -0.31585008 | 0 | 0.088 | 1 | 10 |
| usp8 | 0.04486 | -0.25546036 | 0 | 0.088 | 1 | 10 |
| wdr43 | 0.04486 | -0.28828647 | 0 | 0.088 | 1 | 10 |
| wnk1b | 0.04486 | -0.27272017 | 0 | 0.088 | 1 | 10 |
| surf4l | 0.04486 | -0.31129309 | 0 | 0.088 | 1 | 10 |
| cops2 | 0.04486 | -0.2870299 | 0 | 0.088 | 1 | 10 |
| gfer | 0.04486 | -0.22009371 | 0 | 0.088 | 1 | 10 |
| mcm4 | 0.04486 | -0.31775548 | 0 | 0.088 | 1 | 10 |
| nosip | 0.04486 | -0.25714142 | 0 | 0.088 | 1 | 10 |
| skib | 0.04486 | -0.26779143 | 0 | 0.088 | 1 | 10 |
| srsf1a | 0.04486 | -0.23848055 | 0 | 0.088 | 1 | 10 |
| syt11a | 0.04486 | -0.31517703 | 0 | 0.088 | 1 | 10 |
| snrpb | 0.04495 | -0.56400696 | 0.2 | 0.471 | 1 | 10 |
| mrpl11 | 0.04852 | -0.48374244 | 0.025 | 0.176 | 1 | 10 |
| psmb5 | 0.0491 | -0.48629932 | 0.1 | 0.324 | 1 | 10 |
| arpc1a | 0.05476 | -0.53102009 | 0.1 | 0.324 | 1 | 10 |
| NDUFB1 | 0.05704 | -0.48778338 | 0.05 | 0.235 | 1 | 10 |
| jam2a | 0.05806 | 0.541286724 | 0.575 | 0.353 | 1 | 10 |
| magoh | 0.05897 | -0.43110318 | 0.025 | 0.176 | 1 | 10 |
| nedd8 | 0.05944 | -0.5003235 | 0.05 | 0.235 | 1 | 10 |
| gdi2 | 0.05988 | -0.4835034 | 0.05 | 0.235 | 1 | 10 |
| rplp0 | 0.06086 | 0.244145891 | 0.95 | 0.765 | 1 | 10 |
| caprin1b | 0.06132 | -0.41099409 | 0.075 | 0.265 | 1 | 10 |
| mt-nd4 | 0.06185 | -0.44661 | 0.2 | 0.441 | 1 | 10 |
| rps25 | 0.06217 | -0.26963719 | 0.75 | 0.941 | 1 | 10 |
| ldhba | 0.06459 | -0.53746631 | 0.15 | 0.382 | 1 | 10 |
| hnrnpk | 0.06476 | -0.49427728 | 0.125 | 0.353 | 1 | 10 |
| ilf3b | 0.06504 | 0.536103267 | 0.25 | 0.059 | 1 | 10 |
| rpl34 | 0.06538 | 0.216628487 | 0.975 | 0.824 | 1 | 10 |
| psmd11a | 0.06734 | -0.3656401 | 0.025 | 0.176 | 1 | 10 |
| chac1 | 0.06948 | -0.52918577 | 0.15 | 0.382 | 1 | 10 |
| canx | 0.07154 | -0.43271174 | 0.025 | 0.176 | 1 | 10 |
| cx43 | 0.07213 | 0.581023225 | 0.35 | 0.147 | 1 | 10 |
| csde1 | 0.0738 | -0.44024287 | 0.025 | 0.176 | 1 | 10 |

| | | | | | | |
|---|---|---|---|---|---|---|
| glulb | 0.07383 | -0.53406727 | 0.175 | 0.412 | 1 | 10 |
| ywhabb | 0.07457 | 0.547150416 | 0.275 | 0.088 | 1 | 10 |
| hmgb1b | 0.07711 | -0.52520795 | 0.25 | 0.471 | 1 | 10 |
| CABZ01058650.1 | 0.07863 | 0.206901493 | 0.075 | 0 | 1 | 10 |
| COLEC10 | 0.07863 | 0.292682825 | 0.075 | 0 | 1 | 10 |
| CR388143.1 | 0.07863 | 0.194217705 | 0.075 | 0 | 1 | 10 |
| LO018102.1 | 0.07863 | 0.219366633 | 0.075 | 0 | 1 | 10 |
| LOC553316 | 0.07863 | 0.260146469 | 0.075 | 0 | 1 | 10 |
| XLOC-014336 | 0.07863 | 0.216321186 | 0.075 | 0 | 1 | 10 |
| arl8ba | 0.07863 | 0.237086792 | 0.075 | 0 | 1 | 10 |
| atp6ap2 | 0.07863 | 0.223969991 | 0.075 | 0 | 1 | 10 |
| bckdha | 0.07863 | 0.23576274 | 0.075 | 0 | 1 | 10 |
| ccdc47 | 0.07863 | 0.287988053 | 0.075 | 0 | 1 | 10 |
| cdca4 | 0.07863 | 0.220424197 | 0.075 | 0 | 1 | 10 |
| cdh2 | 0.07863 | 0.235322909 | 0.075 | 0 | 1 | 10 |
| commd3 | 0.07863 | 0.192347094 | 0.075 | 0 | 1 | 10 |
| cops3 | 0.07863 | 0.249734966 | 0.075 | 0 | 1 | 10 |
| cse1l | 0.07863 | 0.244609344 | 0.075 | 0 | 1 | 10 |
| ctrl | 0.07863 | 0.319455398 | 0.075 | 0 | 1 | 10 |
| ctsd | 0.07863 | 0.275944948 | 0.075 | 0 | 1 | 10 |
| cyb5r3 | 0.07863 | 0.289007419 | 0.075 | 0 | 1 | 10 |
| dap | 0.07863 | 0.264485876 | 0.075 | 0 | 1 | 10 |
| dctn4 | 0.07863 | 0.233381527 | 0.075 | 0 | 1 | 10 |
| dnaja2b | 0.07863 | 0.238729225 | 0.075 | 0 | 1 | 10 |
| elavl4 | 0.07863 | 0.274241732 | 0.075 | 0 | 1 | 10 |
| fabp10a | 0.07863 | 0.128486801 | 0.075 | 0 | 1 | 10 |
| glod5 | 0.07863 | 0.196468105 | 0.075 | 0 | 1 | 10 |
| hddc3 | 0.07863 | 0.219352271 | 0.075 | 0 | 1 | 10 |
| hsd17b1 | 0.07863 | 0.28084636 | 0.075 | 0 | 1 | 10 |
| igf2bp2b | 0.07863 | 0.198069735 | 0.075 | 0 | 1 | 10 |
| krt18b | 0.07863 | 0.200367128 | 0.075 | 0 | 1 | 10 |
| lxn | 0.07863 | 0.20831162 | 0.075 | 0 | 1 | 10 |
| map3k1 | 0.07863 | 0.266156499 | 0.075 | 0 | 1 | 10 |
| marcksa | 0.07863 | 0.271300689 | 0.075 | 0 | 1 | 10 |
| mat2b | 0.07863 | 0.208796074 | 0.075 | 0 | 1 | 10 |
| mov10b.1 | 0.07863 | 0.225238746 | 0.075 | 0 | 1 | 10 |
| mrpl22 | 0.07863 | 0.248958296 | 0.075 | 0 | 1 | 10 |
| mrpl43 | 0.07863 | 0.223951976 | 0.075 | 0 | 1 | 10 |
| nelfe | 0.07863 | 0.262192563 | 0.075 | 0 | 1 | 10 |
| nras | 0.07863 | 0.200636751 | 0.075 | 0 | 1 | 10 |
| nsd1a | 0.07863 | 0.282331584 | 0.075 | 0 | 1 | 10 |
| ociad2 | 0.07863 | 0.243076248 | 0.075 | 0 | 1 | 10 |
| pdcd4a | 0.07863 | 0.208458067 | 0.075 | 0 | 1 | 10 |
| pgm1 | 0.07863 | 0.206311512 | 0.075 | 0 | 1 | 10 |
| ppa2 | 0.07863 | 0.238720313 | 0.075 | 0 | 1 | 10 |
| ppil3 | 0.07863 | 0.278860193 | 0.075 | 0 | 1 | 10 |
| ppm1da | 0.07863 | 0.270204676 | 0.075 | 0 | 1 | 10 |
| ppp2r1ba | 0.07863 | 0.284034594 | 0.075 | 0 | 1 | 10 |

| | | | | | | |
|---|---|---|---|---|---|---|
| prss59.1 | 0.07863 | 0.34076175 | 0.075 | 0 | 1 | 10 |
| ptrhd1 | 0.07863 | 0.261863756 | 0.075 | 0 | 1 | 10 |
| pwwp2b | 0.07863 | 0.210804857 | 0.075 | 0 | 1 | 10 |
| rab1aa | 0.07863 | 0.204677181 | 0.075 | 0 | 1 | 10 |
| rdh8a | 0.07863 | 0.225952398 | 0.075 | 0 | 1 | 10 |
| rnf2 | 0.07863 | 0.277728019 | 0.075 | 0 | 1 | 10 |
| rsrp1 | 0.07863 | 0.236953246 | 0.075 | 0 | 1 | 10 |
| serpina1 | 0.07863 | 0.273642585 | 0.075 | 0 | 1 | 10 |
| serpinf1 | 0.07863 | 0.227906467 | 0.075 | 0 | 1 | 10 |
| si:ch211-214j24.10 | 0.07863 | 0.198699904 | 0.075 | 0 | 1 | 10 |
| si:ch211-222k6.3 | 0.07863 | 0.190945957 | 0.075 | 0 | 1 | 10 |
| si:ch211-67e16.4 | 0.07863 | 0.25899196 | 0.075 | 0 | 1 | 10 |
| si:ch73-308l14.2 | 0.07863 | 0.278004952 | 0.075 | 0 | 1 | 10 |
| si:dkey-204f11.64 | 0.07863 | 0.222111497 | 0.075 | 0 | 1 | 10 |
| slc20a2 | 0.07863 | 0.28917772 | 0.075 | 0 | 1 | 10 |
| sox11b | 0.07863 | 0.216165271 | 0.075 | 0 | 1 | 10 |
| stk16 | 0.07863 | 0.247602083 | 0.075 | 0 | 1 | 10 |
| tatdn1 | 0.07863 | 0.220590593 | 0.075 | 0 | 1 | 10 |
| ubl7b | 0.07863 | 0.211656741 | 0.075 | 0 | 1 | 10 |
| utp3 | 0.07863 | 0.241131363 | 0.075 | 0 | 1 | 10 |
| xrcc5 | 0.07863 | 0.212796569 | 0.075 | 0 | 1 | 10 |
| zdhhc3a | 0.07863 | 0.253702228 | 0.075 | 0 | 1 | 10 |
| zgc:123258 | 0.07863 | 0.263661403 | 0.075 | 0 | 1 | 10 |
| zgc:162939 | 0.07863 | 0.265647368 | 0.075 | 0 | 1 | 10 |
| zgc:55262 | 0.07863 | 0.22157021 | 0.075 | 0 | 1 | 10 |
| CABZ01085658.1 | 0.07863 | 0.295871507 | 0.075 | 0 | 1 | 10 |
| cops8 | 0.07863 | 0.235055613 | 0.075 | 0 | 1 | 10 |
| cryzl1 | 0.07863 | 0.224277668 | 0.075 | 0 | 1 | 10 |
| dtd1 | 0.07863 | 0.281764311 | 0.075 | 0 | 1 | 10 |
| eif4g1a | 0.07863 | 0.299461069 | 0.075 | 0 | 1 | 10 |
| erp27 | 0.07863 | 0.194636518 | 0.075 | 0 | 1 | 10 |
| lgmn | 0.07863 | 0.269619131 | 0.075 | 0 | 1 | 10 |
| lmnb2 | 0.07863 | 0.243813634 | 0.075 | 0 | 1 | 10 |
| lrrc20 | 0.07863 | 0.27082507 | 0.075 | 0 | 1 | 10 |
| nolc1 | 0.07863 | 0.242149182 | 0.075 | 0 | 1 | 10 |
| pcdh1b | 0.07863 | 0.266596703 | 0.075 | 0 | 1 | 10 |
| picalmb | 0.07863 | 0.281955148 | 0.075 | 0 | 1 | 10 |
| rtraf | 0.07863 | 0.271379967 | 0.075 | 0 | 1 | 10 |
| sdf2 | 0.07863 | 0.255136677 | 0.075 | 0 | 1 | 10 |
| si:ch211-244b2.4 | 0.07863 | 0.234408849 | 0.075 | 0 | 1 | 10 |
| si:dkey-159f12.2 | 0.07863 | 0.236634481 | 0.075 | 0 | 1 | 10 |
| smg7 | 0.07863 | 0.293585044 | 0.075 | 0 | 1 | 10 |
| stx3a | 0.07863 | 0.258854191 | 0.075 | 0 | 1 | 10 |
| yipf4 | 0.07863 | 0.240834984 | 0.075 | 0 | 1 | 10 |
| zgc:103678 | 0.07863 | 0.247048998 | 0.075 | 0 | 1 | 10 |
| zgc:92590 | 0.07863 | 0.288693726 | 0.075 | 0 | 1 | 10 |
| odc1 | 0.07915 | -0.18820778 | 0.225 | 0.382 | 1 | 10 |
| ppdpfa | 0.08014 | 0.540666467 | 0.35 | 0.176 | 1 | 10 |

| | | | | | | |
|---|---|---|---|---|---|---|
| rpl26 | 0.08364 | 0.143118807 | 0.975 | 0.941 | 1 | 10 |
| il15 | 0.0876 | -0.36135022 | 0.05 | 0.206 | 1 | 10 |
| calm2b | 0.08772 | -0.36636004 | 0.35 | 0.588 | 1 | 10 |
| cdh4 | 0.09003 | -0.49548748 | 0.05 | 0.206 | 1 | 10 |
| rsl24d1 | 0.09452 | 0.506265855 | 0.5 | 0.265 | 1 | 10 |
| hsbp1a | 0.09743 | -0.42072643 | 0.025 | 0.147 | 1 | 10 |
| tomm6 | 0.10003 | -0.28003865 | 0.175 | 0.353 | 1 | 10 |
| tmem134 | 0.10088 | 0.428900594 | 0.175 | 0.029 | 1 | 10 |
| top1l | 0.10106 | -0.46427447 | 0.1 | 0.294 | 1 | 10 |
| ifi30 | 0.10144 | -0.43986744 | 0.15 | 0.353 | 1 | 10 |
| sat1a.2 | 0.10159 | -0.48096576 | 0.1 | 0.294 | 1 | 10 |
| ptmab | 0.10159 | 0.340347568 | 0.8 | 0.618 | 1 | 10 |
| ran | 0.10181 | 0.202688585 | 0.75 | 0.559 | 1 | 10 |
| mycbp | 0.10183 | -0.20020249 | 0.05 | 0.176 | 1 | 10 |
| mafba | 0.10214 | 0.411006582 | 0.175 | 0.029 | 1 | 10 |
| si:dkey-78l4.14 | 0.10253 | -0.26906041 | 0.025 | 0.147 | 1 | 10 |
| pinx1 | 0.10262 | -0.35229916 | 0.05 | 0.206 | 1 | 10 |
| hsd11b2 | 0.10359 | 0.405615591 | 0.175 | 0.029 | 1 | 10 |
| si:ch211-197h24.9 | 0.1047 | 0.398450385 | 0.175 | 0.029 | 1 | 10 |
| s100v2 | 0.1097 | -0.47826068 | 0.125 | 0.324 | 1 | 10 |
| aldh9a1a.1 | 0.11104 | 0.540059797 | 0.3 | 0.118 | 1 | 10 |
| churc1 | 0.11416 | -0.39615579 | 0.05 | 0.206 | 1 | 10 |
| rps5 | 0.11435 | 0.164002505 | 0.975 | 0.912 | 1 | 10 |
| supt4h1 | 0.11448 | -0.38433336 | 0.025 | 0.147 | 1 | 10 |
| LOC100536647 | 0.11505 | -0.18770109 | 0 | 0.059 | 1 | 10 |
| atp11a | 0.11505 | -0.19355899 | 0 | 0.059 | 1 | 10 |
| b4galt6 | 0.11505 | -0.20585888 | 0 | 0.059 | 1 | 10 |
| bmp1a | 0.11505 | -0.19177166 | 0 | 0.059 | 1 | 10 |
| brpf3b | 0.11505 | -0.1802364 | 0 | 0.059 | 1 | 10 |
| cep131 | 0.11505 | -0.23998092 | 0 | 0.059 | 1 | 10 |
| col14a1a | 0.11505 | -0.17276026 | 0 | 0.059 | 1 | 10 |
| col5a2a | 0.11505 | -0.22724845 | 0 | 0.059 | 1 | 10 |
| crfb2 | 0.11505 | -0.22730302 | 0 | 0.059 | 1 | 10 |
| dnajc12 | 0.11505 | -0.21374276 | 0 | 0.059 | 1 | 10 |
| fam129bb | 0.11505 | -0.20239563 | 0 | 0.059 | 1 | 10 |
| klhl24b | 0.11505 | -0.16546028 | 0 | 0.059 | 1 | 10 |
| ncapd2 | 0.11505 | -0.17322062 | 0 | 0.059 | 1 | 10 |
| nlrc3 | 0.11505 | -0.19291697 | 0 | 0.059 | 1 | 10 |
| nts | 0.11505 | -0.22346129 | 0 | 0.059 | 1 | 10 |
| nubp2 | 0.11505 | -0.21344409 | 0 | 0.059 | 1 | 10 |
| nup43 | 0.11505 | -0.16589624 | 0 | 0.059 | 1 | 10 |
| pecr | 0.11505 | -0.20990541 | 0 | 0.059 | 1 | 10 |
| pik3r3b | 0.11505 | -0.22373192 | 0 | 0.059 | 1 | 10 |
| ppp1r2 | 0.11505 | -0.22188766 | 0 | 0.059 | 1 | 10 |
| psmd9 | 0.11505 | -0.17105412 | 0 | 0.059 | 1 | 10 |
| rab3il1 | 0.11505 | -0.22604061 | 0 | 0.059 | 1 | 10 |
| ralgps2 | 0.11505 | -0.20126323 | 0 | 0.059 | 1 | 10 |
| rangap1a | 0.11505 | -0.17211984 | 0 | 0.059 | 1 | 10 |

| | | | | | | |
|---|---|---|---|---|---|---|
| rangap1b | 0.11505 | -0.20084066 | 0 | 0.059 | 1 | 10 |
| rbm45 | 0.11505 | -0.18210916 | 0 | 0.059 | 1 | 10 |
| rnf213b | 0.11505 | -0.16783541 | 0 | 0.059 | 1 | 10 |
| rnf41 | 0.11505 | -0.19341388 | 0 | 0.059 | 1 | 10 |
| rps6ka4 | 0.11505 | -0.20511821 | 0 | 0.059 | 1 | 10 |
| selenoj | 0.11505 | -0.17135426 | 0 | 0.059 | 1 | 10 |
| si:ch211-225b11.1 | 0.11505 | -0.17215323 | 0 | 0.059 | 1 | 10 |
| si:dkey-22o12.2 | 0.11505 | -0.21030092 | 0 | 0.059 | 1 | 10 |
| sik1 | 0.11505 | -0.14661491 | 0 | 0.059 | 1 | 10 |
| snx25 | 0.11505 | -0.18760027 | 0 | 0.059 | 1 | 10 |
| taok1a | 0.11505 | -0.14874926 | 0 | 0.059 | 1 | 10 |
| tbc1d16 | 0.11505 | -0.21741195 | 0 | 0.059 | 1 | 10 |
| tbk1 | 0.11505 | -0.17185109 | 0 | 0.059 | 1 | 10 |
| tex10 | 0.11505 | -0.15799262 | 0 | 0.059 | 1 | 10 |
| tmem254 | 0.11505 | -0.20709911 | 0 | 0.059 | 1 | 10 |
| tprkb | 0.11505 | -0.20075176 | 0 | 0.059 | 1 | 10 |
| tsc22d1 | 0.11505 | -0.2194642 | 0 | 0.059 | 1 | 10 |
| ube2h | 0.11505 | -0.18665427 | 0 | 0.059 | 1 | 10 |
| wipi1 | 0.11505 | -0.15210107 | 0 | 0.059 | 1 | 10 |
| wwp1 | 0.11505 | -0.2334627 | 0 | 0.059 | 1 | 10 |
| xrn2 | 0.11505 | -0.20063025 | 0 | 0.059 | 1 | 10 |
| BX088707.1 | 0.11505 | -0.21264844 | 0 | 0.059 | 1 | 10 |
| CR293507.1 | 0.11505 | -0.19485517 | 0 | 0.059 | 1 | 10 |
| acad9 | 0.11505 | -0.19367414 | 0 | 0.059 | 1 | 10 |
| aifm2 | 0.11505 | -0.19639226 | 0 | 0.059 | 1 | 10 |
| arhgef2 | 0.11505 | -0.20103351 | 0 | 0.059 | 1 | 10 |
| atp6v1d | 0.11505 | -0.20161467 | 0 | 0.059 | 1 | 10 |
| bbs7 | 0.11505 | -0.16632062 | 0 | 0.059 | 1 | 10 |
| btbd10a | 0.11505 | -0.19696939 | 0 | 0.059 | 1 | 10 |
| ccna2 | 0.11505 | -0.16238206 | 0 | 0.059 | 1 | 10 |
| clic1 | 0.11505 | -0.19791202 | 0 | 0.059 | 1 | 10 |
| coil | 0.11505 | -0.1332031 | 0 | 0.059 | 1 | 10 |
| cst14b.1 | 0.11505 | -0.13332197 | 0 | 0.059 | 1 | 10 |
| ctdnep1a | 0.11505 | -0.18370211 | 0 | 0.059 | 1 | 10 |
| ctr9 | 0.11505 | -0.13276242 | 0 | 0.059 | 1 | 10 |
| degs1 | 0.11505 | -0.20859137 | 0 | 0.059 | 1 | 10 |
| dnaja1 | 0.11505 | -0.15351835 | 0 | 0.059 | 1 | 10 |
| dtymk | 0.11505 | -0.18834247 | 0 | 0.059 | 1 | 10 |
| eif4eb | 0.11505 | -0.13426377 | 0 | 0.059 | 1 | 10 |
| enoph1 | 0.11505 | -0.13257454 | 0 | 0.059 | 1 | 10 |
| ephb3 | 0.11505 | -0.22850796 | 0 | 0.059 | 1 | 10 |
| erf | 0.11505 | -0.1734984 | 0 | 0.059 | 1 | 10 |
| fthl31 | 0.11505 | -0.13135219 | 0 | 0.059 | 1 | 10 |
| gabpa | 0.11505 | -0.15356252 | 0 | 0.059 | 1 | 10 |
| gpn2 | 0.11505 | -0.12960619 | 0 | 0.059 | 1 | 10 |
| gsna | 0.11505 | -0.15789975 | 0 | 0.059 | 1 | 10 |
| hcfc2 | 0.11505 | -0.15409734 | 0 | 0.059 | 1 | 10 |
| hdhd2 | 0.11505 | -0.13573703 | 0 | 0.059 | 1 | 10 |

| | | | | | | |
|---|---|---|---|---|---|---|
| idh3a | 0.11505 | -0.19248836 | 0 | 0.059 | 1 | 10 |
| ireb2 | 0.11505 | -0.19398642 | 0 | 0.059 | 1 | 10 |
| kdelr2b | 0.11505 | -0.2088861 | 0 | 0.059 | 1 | 10 |
| lama5 | 0.11505 | -0.21065385 | 0 | 0.059 | 1 | 10 |
| lims1 | 0.11505 | -0.23084498 | 0 | 0.059 | 1 | 10 |
| lmf2b | 0.11505 | -0.21018761 | 0 | 0.059 | 1 | 10 |
| mapk12a | 0.11505 | -0.18365276 | 0 | 0.059 | 1 | 10 |
| mcmbp | 0.11505 | -0.23576353 | 0 | 0.059 | 1 | 10 |
| med27 | 0.11505 | -0.13780423 | 0 | 0.059 | 1 | 10 |
| mef2d | 0.11505 | -0.1911107 | 0 | 0.059 | 1 | 10 |
| mesd | 0.11505 | -0.22740132 | 0 | 0.059 | 1 | 10 |
| mettl5 | 0.11505 | -0.2150795 | 0 | 0.059 | 1 | 10 |
| mpzl1l | 0.11505 | -0.15506109 | 0 | 0.059 | 1 | 10 |
| myclb | 0.11505 | -0.15917885 | 0 | 0.059 | 1 | 10 |
| napga | 0.11505 | -0.15759833 | 0 | 0.059 | 1 | 10 |
| nipblb | 0.11505 | -0.22044563 | 0 | 0.059 | 1 | 10 |
| nol4lb | 0.11505 | -0.13774192 | 0 | 0.059 | 1 | 10 |
| parp2 | 0.11505 | -0.24686098 | 0 | 0.059 | 1 | 10 |
| pelo | 0.11505 | -0.14077814 | 0 | 0.059 | 1 | 10 |
| ppfibp1a | 0.11505 | -0.16912387 | 0 | 0.059 | 1 | 10 |
| ppp6r2a | 0.11505 | -0.17330444 | 0 | 0.059 | 1 | 10 |
| pum1 | 0.11505 | -0.1361213 | 0 | 0.059 | 1 | 10 |
| pxnb | 0.11505 | -0.2369163 | 0 | 0.059 | 1 | 10 |
| rab11fip1b | 0.11505 | -0.20766855 | 0 | 0.059 | 1 | 10 |
| racgap1 | 0.11505 | -0.16987735 | 0 | 0.059 | 1 | 10 |
| ranbp10 | 0.11505 | -0.17712378 | 0 | 0.059 | 1 | 10 |
| rtcb | 0.11505 | -0.16792761 | 0 | 0.059 | 1 | 10 |
| sec63 | 0.11505 | -0.18228646 | 0 | 0.059 | 1 | 10 |
| selenot2 | 0.11505 | -0.19607074 | 0 | 0.059 | 1 | 10 |
| serpinh1a | 0.11505 | -0.18904199 | 0 | 0.059 | 1 | 10 |
| sesn2 | 0.11505 | -0.20865645 | 0 | 0.059 | 1 | 10 |
| sf3b1 | 0.11505 | -0.19941727 | 0 | 0.059 | 1 | 10 |
| si:ch211-150o23.2 | 0.11505 | -0.19895602 | 0 | 0.059 | 1 | 10 |
| si:ch211-194b1.1 | 0.11505 | -0.16732015 | 0 | 0.059 | 1 | 10 |
| si:dkeyp-121d4.3 | 0.11505 | -0.18794147 | 0 | 0.059 | 1 | 10 |
| sipa1l2 | 0.11505 | -0.18190582 | 0 | 0.059 | 1 | 10 |
| slc27a2a | 0.11505 | -0.21744393 | 0 | 0.059 | 1 | 10 |
| smc4 | 0.11505 | -0.16476391 | 0 | 0.059 | 1 | 10 |
| socs1a | 0.11505 | -0.13262454 | 0 | 0.059 | 1 | 10 |
| stip1 | 0.11505 | -0.24159233 | 0 | 0.059 | 1 | 10 |
| tent4b | 0.11505 | -0.19055467 | 0 | 0.059 | 1 | 10 |
| tiprl | 0.11505 | -0.13051253 | 0 | 0.059 | 1 | 10 |
| tnip1 | 0.11505 | -0.19008158 | 0 | 0.059 | 1 | 10 |
| tor4aa | 0.11505 | -0.19830405 | 0 | 0.059 | 1 | 10 |
| traf7 | 0.11505 | -0.18884194 | 0 | 0.059 | 1 | 10 |
| tubb5 | 0.11505 | -0.17687277 | 0 | 0.059 | 1 | 10 |
| ufsp2 | 0.11505 | -0.13752315 | 0 | 0.059 | 1 | 10 |
| vrk1 | 0.11505 | -0.20158795 | 0 | 0.059 | 1 | 10 |

| | | | | | | |
|---|---|---|---|---|---|---|
| xpot | 0.11505 | -0.17380955 | 0 | 0.059 | 1 | 10 |
| zfr | 0.11505 | -0.15459929 | 0 | 0.059 | 1 | 10 |
| zfyve9a | 0.11505 | -0.22443082 | 0 | 0.059 | 1 | 10 |
| zgc:100918 | 0.11505 | -0.19950603 | 0 | 0.059 | 1 | 10 |
| zgc:110783 | 0.11505 | -0.21417414 | 0 | 0.059 | 1 | 10 |
| zgc:153115 | 0.11505 | -0.12804668 | 0 | 0.059 | 1 | 10 |
| zgc:56235 | 0.11505 | -0.1956151 | 0 | 0.059 | 1 | 10 |
| zmp:0000000912 | 0.11505 | -0.14968346 | 0 | 0.059 | 1 | 10 |
| adamts9 | 0.11505 | -0.18652118 | 0 | 0.059 | 1 | 10 |
| ap2a1 | 0.11505 | -0.19139821 | 0 | 0.059 | 1 | 10 |
| arid1b | 0.11505 | -0.15294135 | 0 | 0.059 | 1 | 10 |
| ccnd1 | 0.11505 | -0.17398115 | 0 | 0.059 | 1 | 10 |
| chfr | 0.11505 | -0.20409637 | 0 | 0.059 | 1 | 10 |
| cops5 | 0.11505 | -0.16465474 | 0 | 0.059 | 1 | 10 |
| cptp | 0.11505 | -0.1804518 | 0 | 0.059 | 1 | 10 |
| gsk3b | 0.11505 | -0.18684366 | 0 | 0.059 | 1 | 10 |
| hnrnpul1l | 0.11505 | -0.18648295 | 0 | 0.059 | 1 | 10 |
| htatip2 | 0.11505 | -0.18717377 | 0 | 0.059 | 1 | 10 |
| kif2c | 0.11505 | -0.16607698 | 0 | 0.059 | 1 | 10 |
| klhl20 | 0.11505 | -0.18562628 | 0 | 0.059 | 1 | 10 |
| med15 | 0.11505 | -0.17784213 | 0 | 0.059 | 1 | 10 |
| miga1 | 0.11505 | -0.12616876 | 0 | 0.059 | 1 | 10 |
| nckap5l | 0.11505 | -0.23133877 | 0 | 0.059 | 1 | 10 |
| ndc80 | 0.11505 | -0.14913322 | 0 | 0.059 | 1 | 10 |
| nr1d2b | 0.11505 | -0.14676304 | 0 | 0.059 | 1 | 10 |
| nusap1 | 0.11505 | -0.17165357 | 0 | 0.059 | 1 | 10 |
| phf19 | 0.11505 | -0.17268341 | 0 | 0.059 | 1 | 10 |
| sh2b3 | 0.11505 | -0.21346416 | 0 | 0.059 | 1 | 10 |
| si:dkey-112a7.4 | 0.11505 | -0.22247631 | 0 | 0.059 | 1 | 10 |
| stub1 | 0.11505 | -0.21338335 | 0 | 0.059 | 1 | 10 |
| thap12a | 0.11505 | -0.12645014 | 0 | 0.059 | 1 | 10 |
| tmed1b | 0.11505 | -0.19582514 | 0 | 0.059 | 1 | 10 |
| trnau1apb | 0.11505 | -0.16041251 | 0 | 0.059 | 1 | 10 |
| ube2w | 0.11505 | -0.15888615 | 0 | 0.059 | 1 | 10 |
| urod | 0.11505 | -0.18409908 | 0 | 0.059 | 1 | 10 |
| zc3h14 | 0.11505 | -0.19697271 | 0 | 0.059 | 1 | 10 |
| arih1 | 0.11505 | -0.22022612 | 0 | 0.059 | 1 | 10 |
| arrb2a | 0.11505 | -0.17468575 | 0 | 0.059 | 1 | 10 |
| commd8 | 0.11505 | -0.18227861 | 0 | 0.059 | 1 | 10 |
| cttn | 0.11505 | -0.20108841 | 0 | 0.059 | 1 | 10 |
| cyba | 0.11505 | -0.1771446 | 0 | 0.059 | 1 | 10 |
| ddx20 | 0.11505 | -0.16223693 | 0 | 0.059 | 1 | 10 |
| dnajc2 | 0.11505 | -0.16131381 | 0 | 0.059 | 1 | 10 |
| fkbp1ab | 0.11505 | -0.19867934 | 0 | 0.059 | 1 | 10 |
| fkbp5 | 0.11505 | -0.17371661 | 0 | 0.059 | 1 | 10 |
| g3bp2 | 0.11505 | -0.23128444 | 0 | 0.059 | 1 | 10 |
| get4 | 0.11505 | -0.18537082 | 0 | 0.059 | 1 | 10 |
| ggcta | 0.11505 | -0.20884606 | 0 | 0.059 | 1 | 10 |

| | | | | | | |
|---|---|---|---|---|---|---|
| hadhab | 0.11505 | -0.20129252 | 0 | 0.059 | 1 | 10 |
| jkamp | 0.11505 | -0.23210593 | 0 | 0.059 | 1 | 10 |
| mplkip | 0.11505 | -0.16238062 | 0 | 0.059 | 1 | 10 |
| msh4 | 0.11505 | -0.16446362 | 0 | 0.059 | 1 | 10 |
| ncs1a | 0.11505 | -0.17458062 | 0 | 0.059 | 1 | 10 |
| pdgfc | 0.11505 | -0.21335278 | 0 | 0.059 | 1 | 10 |
| ppp1r10 | 0.11505 | -0.21144767 | 0 | 0.059 | 1 | 10 |
| rpusd2 | 0.11505 | -0.19414619 | 0 | 0.059 | 1 | 10 |
| si:ch211-271b14.1 | 0.11505 | -0.16161397 | 0 | 0.059 | 1 | 10 |
| si:dkey-165n16.1 | 0.11505 | -0.18520488 | 0 | 0.059 | 1 | 10 |
| stag1b | 0.11505 | -0.20039017 | 0 | 0.059 | 1 | 10 |
| trappc11 | 0.11505 | -0.23355254 | 0 | 0.059 | 1 | 10 |
| ube2d3 | 0.11505 | -0.20927566 | 0 | 0.059 | 1 | 10 |
| zcrb1 | 0.11505 | -0.19460872 | 0 | 0.059 | 1 | 10 |
| zgc:55461 | 0.11505 | -0.16589372 | 0 | 0.059 | 1 | 10 |
| znf839 | 0.11505 | -0.17523864 | 0 | 0.059 | 1 | 10 |
| CR855311.1 | 0.11505 | -0.17276738 | 0 | 0.059 | 1 | 10 |
| abcd3a | 0.11505 | -0.20601601 | 0 | 0.059 | 1 | 10 |
| afg3l2 | 0.11505 | -0.19371963 | 0 | 0.059 | 1 | 10 |
| ankrd40 | 0.11505 | -0.19134929 | 0 | 0.059 | 1 | 10 |
| arhgap1 | 0.11505 | -0.1929777 | 0 | 0.059 | 1 | 10 |
| aurka | 0.11505 | -0.17850152 | 0 | 0.059 | 1 | 10 |
| chad | 0.11505 | -0.21442209 | 0 | 0.059 | 1 | 10 |
| chordc1b | 0.11505 | -0.17390459 | 0 | 0.059 | 1 | 10 |
| col4a5 | 0.11505 | -0.21817212 | 0 | 0.059 | 1 | 10 |
| hectd1 | 0.11505 | -0.21376369 | 0 | 0.059 | 1 | 10 |
| irs1 | 0.11505 | -0.19229836 | 0 | 0.059 | 1 | 10 |
| limk2 | 0.11505 | -0.20853833 | 0 | 0.059 | 1 | 10 |
| mrpl23 | 0.11505 | -0.22708091 | 0 | 0.059 | 1 | 10 |
| mthfr | 0.11505 | -0.18368701 | 0 | 0.059 | 1 | 10 |
| myof | 0.11505 | -0.1934042 | 0 | 0.059 | 1 | 10 |
| sat1a.1 | 0.11505 | -0.22739251 | 0 | 0.059 | 1 | 10 |
| slc7a5 | 0.11505 | -0.24219131 | 0 | 0.059 | 1 | 10 |
| smtnl1 | 0.11505 | -0.21840861 | 0 | 0.059 | 1 | 10 |
| sostdc1a | 0.11505 | -0.22068928 | 0 | 0.059 | 1 | 10 |
| ssbp3b | 0.11505 | -0.22334815 | 0 | 0.059 | 1 | 10 |
| tfap4 | 0.11505 | -0.21424336 | 0 | 0.059 | 1 | 10 |
| usp3 | 0.11505 | -0.20577678 | 0 | 0.059 | 1 | 10 |
| vamp4 | 0.11505 | -0.19090309 | 0 | 0.059 | 1 | 10 |
| xpnpep3 | 0.11505 | -0.18072918 | 0 | 0.059 | 1 | 10 |
| zgc:194224 | 0.11505 | -0.20128108 | 0 | 0.059 | 1 | 10 |
| znf865 | 0.11505 | -0.19861795 | 0 | 0.059 | 1 | 10 |
| FQ323156.1 | 0.11505 | -0.17459469 | 0 | 0.059 | 1 | 10 |
| NFX1 | 0.11505 | -0.19610602 | 0 | 0.059 | 1 | 10 |
| aifm1 | 0.11505 | -0.17893705 | 0 | 0.059 | 1 | 10 |
| atp6v0d1 | 0.11505 | -0.19967045 | 0 | 0.059 | 1 | 10 |
| bend3 | 0.11505 | -0.23508236 | 0 | 0.059 | 1 | 10 |
| cbx4 | 0.11505 | -0.18443153 | 0 | 0.059 | 1 | 10 |

| | | | | | | |
|---|---|---|---|---|---|---|
| celf1 | 0.11505 | -0.18323315 | 0 | 0.059 | 1 | 10 |
| cuedc1b | 0.11505 | -0.16326951 | 0 | 0.059 | 1 | 10 |
| ddx47 | 0.11505 | -0.19610091 | 0 | 0.059 | 1 | 10 |
| ddx56 | 0.11505 | -0.17110097 | 0 | 0.059 | 1 | 10 |
| dnajc5aa | 0.11505 | -0.22971545 | 0 | 0.059 | 1 | 10 |
| elp5 | 0.11505 | -0.17645795 | 0 | 0.059 | 1 | 10 |
| fam13b | 0.11505 | -0.22217402 | 0 | 0.059 | 1 | 10 |
| fam210ab | 0.11505 | -0.19023258 | 0 | 0.059 | 1 | 10 |
| fbxl7 | 0.11505 | -0.22097749 | 0 | 0.059 | 1 | 10 |
| fuom | 0.11505 | -0.17079009 | 0 | 0.059 | 1 | 10 |
| gpatch4 | 0.11505 | -0.14976355 | 0 | 0.059 | 1 | 10 |
| hiat1b | 0.11505 | -0.22012574 | 0 | 0.059 | 1 | 10 |
| hmbsb | 0.11505 | -0.16186944 | 0 | 0.059 | 1 | 10 |
| hspa14 | 0.11505 | -0.17153081 | 0 | 0.059 | 1 | 10 |
| ift22 | 0.11505 | -0.15068403 | 0 | 0.059 | 1 | 10 |
| junbb | 0.11505 | -0.146952 | 0 | 0.059 | 1 | 10 |
| kdm5c | 0.11505 | -0.14654387 | 0 | 0.059 | 1 | 10 |
| mak16 | 0.11505 | -0.17314276 | 0 | 0.059 | 1 | 10 |
| mospd1 | 0.11505 | -0.14940646 | 0 | 0.059 | 1 | 10 |
| mus81 | 0.11505 | -0.19490911 | 0 | 0.059 | 1 | 10 |
| nck1a | 0.11505 | -0.18831408 | 0 | 0.059 | 1 | 10 |
| neu3.2 | 0.11505 | -0.21743612 | 0 | 0.059 | 1 | 10 |
| nipsnap3a | 0.11505 | -0.18184069 | 0 | 0.059 | 1 | 10 |
| polb | 0.11505 | -0.19740187 | 0 | 0.059 | 1 | 10 |
| ptbp2b | 0.11505 | -0.1831951 | 0 | 0.059 | 1 | 10 |
| rab18b | 0.11505 | -0.20211818 | 0 | 0.059 | 1 | 10 |
| rmnd5b | 0.11505 | -0.15361499 | 0 | 0.059 | 1 | 10 |
| rnaseh2c | 0.11505 | -0.18748822 | 0 | 0.059 | 1 | 10 |
| si:ch211-107m4.1 | 0.11505 | -0.20035842 | 0 | 0.059 | 1 | 10 |
| si:ch211-147a11.3 | 0.11505 | -0.219745 | 0 | 0.059 | 1 | 10 |
| si:dkey-100n23.5 | 0.11505 | -0.19138117 | 0 | 0.059 | 1 | 10 |
| si:dkeyp-55f12.3 | 0.11505 | -0.14962617 | 0 | 0.059 | 1 | 10 |
| smarcc1a | 0.11505 | -0.18854652 | 0 | 0.059 | 1 | 10 |
| stt3b | 0.11505 | -0.19966812 | 0 | 0.059 | 1 | 10 |
| tbl1xr1b | 0.11505 | -0.18085077 | 0 | 0.059 | 1 | 10 |
| tnpo2 | 0.11505 | -0.18358736 | 0 | 0.059 | 1 | 10 |
| tor1 | 0.11505 | -0.19338337 | 0 | 0.059 | 1 | 10 |
| uhrf1 | 0.11505 | -0.17993146 | 0 | 0.059 | 1 | 10 |
| wdr26b | 0.11505 | -0.16191678 | 0 | 0.059 | 1 | 10 |
| zc3h12b | 0.11505 | -0.18678928 | 0 | 0.059 | 1 | 10 |
| znf1124 | 0.11505 | -0.18593387 | 0 | 0.059 | 1 | 10 |
| ddx39b | 0.11505 | -0.16000701 | 0 | 0.059 | 1 | 10 |
| CABZ01113192.1 | 0.11505 | -0.21066838 | 0 | 0.059 | 1 | 10 |
| CR450785.1 | 0.11505 | -0.23047772 | 0 | 0.059 | 1 | 10 |
| CU137717.1 | 0.11505 | -0.18329375 | 0 | 0.059 | 1 | 10 |
| FP074902.2 | 0.11505 | -0.15702857 | 0 | 0.059 | 1 | 10 |
| XLOC-005199 | 0.11505 | -0.20979949 | 0 | 0.059 | 1 | 10 |
| XLOC-025423 | 0.11505 | -0.23612026 | 0 | 0.059 | 1 | 10 |

| | | | | | | |
|---|---|---|---|---|---|---|
| XLOC-030237 | 0.11505 | -0.16339465 | 0 | 0.059 | 1 | 10 |
| abracl | 0.11505 | -0.18160142 | 0 | 0.059 | 1 | 10 |
| arid2 | 0.11505 | -0.17790782 | 0 | 0.059 | 1 | 10 |
| arid4a | 0.11505 | -0.20750239 | 0 | 0.059 | 1 | 10 |
| atxn3 | 0.11505 | -0.2001941 | 0 | 0.059 | 1 | 10 |
| cat | 0.11505 | -0.23865112 | 0 | 0.059 | 1 | 10 |
| cks2 | 0.11505 | -0.16478602 | 0 | 0.059 | 1 | 10 |
| coq5 | 0.11505 | -0.19709571 | 0 | 0.059 | 1 | 10 |
| crip1 | 0.11505 | -0.19670977 | 0 | 0.059 | 1 | 10 |
| cst3 | 0.11505 | -0.18249721 | 0 | 0.059 | 1 | 10 |
| dicer1 | 0.11505 | -0.24940066 | 0 | 0.059 | 1 | 10 |
| dimt1l | 0.11505 | -0.22496693 | 0 | 0.059 | 1 | 10 |
| dnmt3bb.3 | 0.11505 | -0.17507748 | 0 | 0.059 | 1 | 10 |
| elf2b | 0.11505 | -0.17795725 | 0 | 0.059 | 1 | 10 |
| eloa | 0.11505 | -0.14008326 | 0 | 0.059 | 1 | 10 |
| eps15l1a | 0.11505 | -0.16572144 | 0 | 0.059 | 1 | 10 |
| etnk1 | 0.11505 | -0.20231757 | 0 | 0.059 | 1 | 10 |
| evi5b | 0.11505 | -0.20154601 | 0 | 0.059 | 1 | 10 |
| hddc2 | 0.11505 | -0.19608638 | 0 | 0.059 | 1 | 10 |
| hmbsa | 0.11505 | -0.17203978 | 0 | 0.059 | 1 | 10 |
| id2a | 0.11505 | -0.22407125 | 0 | 0.059 | 1 | 10 |
| jade3 | 0.11505 | -0.19653512 | 0 | 0.059 | 1 | 10 |
| jak1 | 0.11505 | -0.18375094 | 0 | 0.059 | 1 | 10 |
| kmt5ab | 0.11505 | -0.1660749 | 0 | 0.059 | 1 | 10 |
| lamb2l | 0.11505 | -0.20657526 | 0 | 0.059 | 1 | 10 |
| lpcat3 | 0.11505 | -0.18512864 | 0 | 0.059 | 1 | 10 |
| lrrfip2 | 0.11505 | -0.21043114 | 0 | 0.059 | 1 | 10 |
| luc7l3 | 0.11505 | -0.21504761 | 0 | 0.059 | 1 | 10 |
| mcm2 | 0.11505 | -0.1767269 | 0 | 0.059 | 1 | 10 |
| morc2 | 0.11505 | -0.13879153 | 0 | 0.059 | 1 | 10 |
| mtrf1 | 0.11505 | -0.14599107 | 0 | 0.059 | 1 | 10 |
| mul1a | 0.11505 | -0.21300486 | 0 | 0.059 | 1 | 10 |
| naa15b | 0.11505 | -0.14889589 | 0 | 0.059 | 1 | 10 |
| nek7 | 0.11505 | -0.19694505 | 0 | 0.059 | 1 | 10 |
| npepps | 0.11505 | -0.17205822 | 0 | 0.059 | 1 | 10 |
| nrf1 | 0.11505 | -0.17318463 | 0 | 0.059 | 1 | 10 |
| nsun2 | 0.11505 | -0.2109726 | 0 | 0.059 | 1 | 10 |
| pggt1b | 0.11505 | -0.17072048 | 0 | 0.059 | 1 | 10 |
| piezo2a.2 | 0.11505 | -0.17049794 | 0 | 0.059 | 1 | 10 |
| plk1 | 0.11505 | -0.18162644 | 0 | 0.059 | 1 | 10 |
| plpp5 | 0.11505 | -0.16657595 | 0 | 0.059 | 1 | 10 |
| ppp2r5ea | 0.11505 | -0.18542407 | 0 | 0.059 | 1 | 10 |
| prkaa1 | 0.11505 | -0.14304247 | 0 | 0.059 | 1 | 10 |
| prpf31 | 0.11505 | -0.16979431 | 0 | 0.059 | 1 | 10 |
| psph | 0.11505 | -0.19318774 | 0 | 0.059 | 1 | 10 |
| rab8a | 0.11505 | -0.20582074 | 0 | 0.059 | 1 | 10 |
| rabggtb | 0.11505 | -0.143053 | 0 | 0.059 | 1 | 10 |
| rfxap | 0.11505 | -0.22413984 | 0 | 0.059 | 1 | 10 |

| | | | | | | |
|---|---|---|---|---|---|---|
| rn7sk | 0.11505 | -0.27610267 | 0 | 0.059 | 1 | 10 |
| rnaseh2b | 0.11505 | -0.17618905 | 0 | 0.059 | 1 | 10 |
| rnf167 | 0.11505 | -0.23263252 | 0 | 0.059 | 1 | 10 |
| scarb2a | 0.11505 | -0.24431717 | 0 | 0.059 | 1 | 10 |
| sdhaf4 | 0.11505 | -0.16561901 | 0 | 0.059 | 1 | 10 |
| si:ch211-149b19.4 | 0.11505 | -0.16982834 | 0 | 0.059 | 1 | 10 |
| si:ch211-255f4.5 | 0.11505 | -0.2193749 | 0 | 0.059 | 1 | 10 |
| si:ch211-69g19.2 | 0.11505 | -0.18206128 | 0 | 0.059 | 1 | 10 |
| si:dkey-260j18.2 | 0.11505 | -0.17567001 | 0 | 0.059 | 1 | 10 |
| si:dkeyp-38g8.5 | 0.11505 | -0.14384303 | 0 | 0.059 | 1 | 10 |
| slc7a3a | 0.11505 | -0.26934466 | 0 | 0.059 | 1 | 10 |
| stk17b | 0.11505 | -0.2003004 | 0 | 0.059 | 1 | 10 |
| stk35l | 0.11505 | -0.18282118 | 0 | 0.059 | 1 | 10 |
| tbl1x | 0.11505 | -0.1951626 | 0 | 0.059 | 1 | 10 |
| tnfrsf19 | 0.11505 | -0.15677778 | 0 | 0.059 | 1 | 10 |
| uap1l1 | 0.11505 | -0.14338088 | 0 | 0.059 | 1 | 10 |
| vps25 | 0.11505 | -0.17448715 | 0 | 0.059 | 1 | 10 |
| wu:fc33b09 | 0.11505 | -0.19185276 | 0 | 0.059 | 1 | 10 |
| xpo7 | 0.11505 | -0.23228012 | 0 | 0.059 | 1 | 10 |
| xrcc4 | 0.11505 | -0.177987 | 0 | 0.059 | 1 | 10 |
| zc3h4 | 0.11505 | -0.19976861 | 0 | 0.059 | 1 | 10 |
| zc3h6 | 0.11505 | -0.18378517 | 0 | 0.059 | 1 | 10 |
| zgc:162472 | 0.11505 | -0.18575961 | 0 | 0.059 | 1 | 10 |
| alcamb | 0.11522 | 0.46481325 | 0.225 | 0.059 | 1 | 10 |
| clpp | 0.11692 | 0.424261141 | 0.15 | 0.029 | 1 | 10 |
| thoc5 | 0.11838 | -0.27374005 | 0.025 | 0.147 | 1 | 10 |
| rps29 | 0.11971 | -0.14296451 | 0.925 | 0.971 | 1 | 10 |
| mtap | 0.12101 | -0.32296025 | 0.025 | 0.147 | 1 | 10 |
| snrpe | 0.12142 | 0.350210814 | 0.25 | 0.088 | 1 | 10 |
| psma5 | 0.12159 | -0.45960527 | 0.125 | 0.294 | 1 | 10 |
| hypk | 0.12386 | -0.40806804 | 0.175 | 0.382 | 1 | 10 |
| ywhabl | 0.12684 | -0.38811562 | 0.275 | 0.5 | 1 | 10 |
| cnih1 | 0.13009 | 0.401264212 | 0.15 | 0.029 | 1 | 10 |
| rpl11 | 0.1302 | 0.109984484 | 1 | 1 | 1 | 10 |
| vdac2 | 0.13325 | -0.21301984 | 0.25 | 0.412 | 1 | 10 |
| eif4e1c | 0.13346 | -0.29335651 | 0.025 | 0.147 | 1 | 10 |
| baz2ba | 0.13677 | -0.30126795 | 0.025 | 0.147 | 1 | 10 |
| rpl18 | 0.14068 | 0.114600833 | 0.95 | 0.941 | 1 | 10 |
| zfp36l1a | 0.14092 | -0.38115887 | 0.025 | 0.147 | 1 | 10 |
| ndufb10 | 0.14185 | 0.356780285 | 0.15 | 0.029 | 1 | 10 |
| ndnl2 | 0.14236 | -0.32707403 | 0.025 | 0.147 | 1 | 10 |
| capza1b | 0.14269 | 0.415989445 | 0.15 | 0.029 | 1 | 10 |
| pa2g4b | 0.14284 | -0.32779205 | 0.025 | 0.147 | 1 | 10 |
| h2afva | 0.14291 | -0.46017392 | 0.075 | 0.235 | 1 | 10 |
| ufm1 | 0.14327 | -0.34063744 | 0.025 | 0.147 | 1 | 10 |
| fam204a | 0.14329 | -0.34788964 | 0.025 | 0.147 | 1 | 10 |
| ing4 | 0.14438 | -0.3232215 | 0.025 | 0.147 | 1 | 10 |
| emx2 | 0.1455 | 0.417248828 | 0.25 | 0.088 | 1 | 10 |

| | | | | | | |
|---|---|---|---|---|---|---|
| eif2s2 | 0.14594 | -0.22607057 | 0.325 | 0.5 | 1 | 10 |
| mrpl54 | 0.14624 | -0.30210777 | 0.025 | 0.147 | 1 | 10 |
| acp1 | 0.14699 | -0.31041043 | 0.025 | 0.147 | 1 | 10 |
| arrdc3a | 0.14711 | 0.299629178 | 0.15 | 0.029 | 1 | 10 |
| arpp19a | 0.14774 | -0.30539197 | 0.025 | 0.147 | 1 | 10 |
| smndc1 | 0.1484 | -0.38593203 | 0.075 | 0.235 | 1 | 10 |
| pi15a | 0.14843 | 0.222664412 | 0.225 | 0.206 | 1 | 10 |
| CR318588.3 | 0.1489 | -0.30532012 | 0.325 | 0.529 | 1 | 10 |
| sec61g | 0.14935 | -0.26231429 | 0.525 | 0.735 | 1 | 10 |
| dynll2b | 0.15006 | -0.45144413 | 0.075 | 0.235 | 1 | 10 |
| eif1axb | 0.15204 | -0.29626726 | 0.225 | 0.412 | 1 | 10 |
| myl12.2 | 0.15239 | 0.42483149 | 0.15 | 0.029 | 1 | 10 |
| rpl32 | 0.15281 | 0.110929475 | 0.975 | 0.971 | 1 | 10 |
| rps27.2 | 0.1529 | -0.23885096 | 0.7 | 0.882 | 1 | 10 |
| snrpd1 | 0.1536 | -0.32285387 | 0.125 | 0.294 | 1 | 10 |
| tax1bp1b | 0.15553 | -0.29590962 | 0.05 | 0.176 | 1 | 10 |
| chchd10 | 0.1565 | 0.360112691 | 0.15 | 0.029 | 1 | 10 |
| pde7a | 0.15872 | 0.367853512 | 0.15 | 0.029 | 1 | 10 |
| copz1 | 0.1588 | -0.3006815 | 0.05 | 0.176 | 1 | 10 |
| anp32a | 0.15927 | -0.28511992 | 0.075 | 0.206 | 1 | 10 |
| rps21 | 0.16382 | -0.12578736 | 0.925 | 1 | 1 | 10 |
| atp5f1e | 0.16403 | -0.32108921 | 0.375 | 0.588 | 1 | 10 |
| glrx | 0.16524 | -0.36312676 | 0.125 | 0.294 | 1 | 10 |
| cyc1 | 0.16682 | -0.28612538 | 0.075 | 0.206 | 1 | 10 |
| actb2 | 0.16696 | 0.246191684 | 0.9 | 0.794 | 1 | 10 |
| krt8 | 0.16721 | 0.342632475 | 0.575 | 0.441 | 1 | 10 |
| amy2a | 0.16842 | 0.151525312 | 0.05 | 0 | 1 | 10 |
| atf7b | 0.16842 | 0.181728541 | 0.05 | 0 | 1 | 10 |
| bicra | 0.16842 | 0.186867798 | 0.05 | 0 | 1 | 10 |
| leng8 | 0.16842 | 0.18338759 | 0.05 | 0 | 1 | 10 |
| si:ch211-236l14.4 | 0.16842 | 0.189062039 | 0.05 | 0 | 1 | 10 |
| stk24b | 0.16842 | 0.194528852 | 0.05 | 0 | 1 | 10 |
| CABZ01102240.1 | 0.16842 | 0.145538262 | 0.05 | 0 | 1 | 10 |
| CABZ01110881.1 | 0.16842 | 0.200046036 | 0.05 | 0 | 1 | 10 |
| CU459094.2 | 0.16842 | 0.157756718 | 0.05 | 0 | 1 | 10 |
| CU639397.1 | 0.16842 | 0.169710579 | 0.05 | 0 | 1 | 10 |
| LOC108179036 | 0.16842 | 0.170141053 | 0.05 | 0 | 1 | 10 |
| TMEM19 | 0.16842 | 0.144781425 | 0.05 | 0 | 1 | 10 |
| XLOC-040582 | 0.16842 | 0.174726 | 0.05 | 0 | 1 | 10 |
| XLOC-042292 | 0.16842 | 0.190089578 | 0.05 | 0 | 1 | 10 |
| ahcyl2 | 0.16842 | 0.14060831 | 0.05 | 0 | 1 | 10 |
| ahsg2 | 0.16842 | 0.191075011 | 0.05 | 0 | 1 | 10 |
| aldh16a1 | 0.16842 | 0.169214656 | 0.05 | 0 | 1 | 10 |
| amh | 0.16842 | 0.182881672 | 0.05 | 0 | 1 | 10 |
| bag4 | 0.16842 | 0.179329062 | 0.05 | 0 | 1 | 10 |
| bcl6aa | 0.16842 | 0.171905952 | 0.05 | 0 | 1 | 10 |
| btr25 | 0.16842 | 0.160643459 | 0.05 | 0 | 1 | 10 |
| cd2ap | 0.16842 | 0.182086736 | 0.05 | 0 | 1 | 10 |

| | | | | | | |
|---|---|---|---|---|---|---|
| cgref1 | 0.16842 | 0.189620736 | 0.05 | 0 | 1 | 10 |
| chchd7 | 0.16842 | 0.146422792 | 0.05 | 0 | 1 | 10 |
| clstn1 | 0.16842 | 0.191070559 | 0.05 | 0 | 1 | 10 |
| crebzf | 0.16842 | 0.163241697 | 0.05 | 0 | 1 | 10 |
| csrp2 | 0.16842 | 0.192631811 | 0.05 | 0 | 1 | 10 |
| dock9b | 0.16842 | 0.172569471 | 0.05 | 0 | 1 | 10 |
| fastkd2 | 0.16842 | 0.195701286 | 0.05 | 0 | 1 | 10 |
| ftr36 | 0.16842 | 0.196981225 | 0.05 | 0 | 1 | 10 |
| fxyd1 | 0.16842 | 0.128974271 | 0.05 | 0 | 1 | 10 |
| hivep1 | 0.16842 | 0.165188292 | 0.05 | 0 | 1 | 10 |
| hmg20b | 0.16842 | 0.172929949 | 0.05 | 0 | 1 | 10 |
| ikzf4 | 0.16842 | 0.19127744 | 0.05 | 0 | 1 | 10 |
| larp1 | 0.16842 | 0.141238074 | 0.05 | 0 | 1 | 10 |
| lcorl | 0.16842 | 0.148820519 | 0.05 | 0 | 1 | 10 |
| manbal | 0.16842 | 0.146237787 | 0.05 | 0 | 1 | 10 |
| map2k4b | 0.16842 | 0.152248165 | 0.05 | 0 | 1 | 10 |
| mgrn1a | 0.16842 | 0.172009085 | 0.05 | 0 | 1 | 10 |
| msantd1 | 0.16842 | 0.182007703 | 0.05 | 0 | 1 | 10 |
| nr2f6b | 0.16842 | 0.138486233 | 0.05 | 0 | 1 | 10 |
| pdhx | 0.16842 | 0.189177645 | 0.05 | 0 | 1 | 10 |
| pmpcb | 0.16842 | 0.139567033 | 0.05 | 0 | 1 | 10 |
| prkcaa | 0.16842 | 0.144423069 | 0.05 | 0 | 1 | 10 |
| prss1 | 0.16842 | 0.251522372 | 0.05 | 0 | 1 | 10 |
| pxna | 0.16842 | 0.185340592 | 0.05 | 0 | 1 | 10 |
| r3hdm1 | 0.16842 | 0.203789983 | 0.05 | 0 | 1 | 10 |
| rbpja | 0.16842 | 0.177671685 | 0.05 | 0 | 1 | 10 |
| rraga | 0.16842 | 0.178881372 | 0.05 | 0 | 1 | 10 |
| sesn3 | 0.16842 | 0.134049506 | 0.05 | 0 | 1 | 10 |
| si:ch211-166g5.4 | 0.16842 | 0.169624102 | 0.05 | 0 | 1 | 10 |
| si:ch211-244b2.3 | 0.16842 | 0.156452222 | 0.05 | 0 | 1 | 10 |
| si:ch73-223f5.1 | 0.16842 | 0.181562573 | 0.05 | 0 | 1 | 10 |
| si:ch73-238c9.1 | 0.16842 | 0.1965816 | 0.05 | 0 | 1 | 10 |
| si:dkey-19a16.2 | 0.16842 | 0.146556035 | 0.05 | 0 | 1 | 10 |
| si:dkey-71l4.4 | 0.16842 | 0.203060686 | 0.05 | 0 | 1 | 10 |
| stau1 | 0.16842 | 0.124822651 | 0.05 | 0 | 1 | 10 |
| tcf12 | 0.16842 | 0.153340276 | 0.05 | 0 | 1 | 10 |
| tgfbrap1 | 0.16842 | 0.172131064 | 0.05 | 0 | 1 | 10 |
| tmem177 | 0.16842 | 0.176458753 | 0.05 | 0 | 1 | 10 |
| tnksa | 0.16842 | 0.148890866 | 0.05 | 0 | 1 | 10 |
| tp53bp2a | 0.16842 | 0.187550982 | 0.05 | 0 | 1 | 10 |
| tram2 | 0.16842 | 0.154564712 | 0.05 | 0 | 1 | 10 |
| utp4 | 0.16842 | 0.160029774 | 0.05 | 0 | 1 | 10 |
| vps26c | 0.16842 | 0.160335096 | 0.05 | 0 | 1 | 10 |
| wdr55 | 0.16842 | 0.153014121 | 0.05 | 0 | 1 | 10 |
| zgc:112982 | 0.16842 | 0.139599748 | 0.05 | 0 | 1 | 10 |
| zgc:153215 | 0.16842 | 0.146026843 | 0.05 | 0 | 1 | 10 |
| zgc:172128 | 0.16842 | 0.148482066 | 0.05 | 0 | 1 | 10 |
| FQ377903.3 | 0.16842 | 0.129270873 | 0.05 | 0 | 1 | 10 |

| | | | | | | |
|---|---|---|---|---|---|---|
| XLOC-043857 | 0.16842 | 0.126845263 | 0.05 | 0 | 1 | 10 |
| acbd3 | 0.16842 | 0.170536577 | 0.05 | 0 | 1 | 10 |
| add1 | 0.16842 | 0.154845106 | 0.05 | 0 | 1 | 10 |
| alas1 | 0.16842 | 0.191311726 | 0.05 | 0 | 1 | 10 |
| apoa1b | 0.16842 | 0.127529855 | 0.05 | 0 | 1 | 10 |
| arcn1b | 0.16842 | 0.112447175 | 0.05 | 0 | 1 | 10 |
| arhgap42a | 0.16842 | 0.146090885 | 0.05 | 0 | 1 | 10 |
| atf6 | 0.16842 | 0.1859504 | 0.05 | 0 | 1 | 10 |
| atp23 | 0.16842 | 0.18389787 | 0.05 | 0 | 1 | 10 |
| aven | 0.16842 | 0.179406935 | 0.05 | 0 | 1 | 10 |
| cdc26 | 0.16842 | 0.179144555 | 0.05 | 0 | 1 | 10 |
| cdk8 | 0.16842 | 0.195519477 | 0.05 | 0 | 1 | 10 |
| cmtm6 | 0.16842 | 0.191248216 | 0.05 | 0 | 1 | 10 |
| cnga1a | 0.16842 | 0.172010469 | 0.05 | 0 | 1 | 10 |
| col1a1a | 0.16842 | 0.185346822 | 0.05 | 0 | 1 | 10 |
| commd1 | 0.16842 | 0.184390935 | 0.05 | 0 | 1 | 10 |
| crim1 | 0.16842 | 0.141355706 | 0.05 | 0 | 1 | 10 |
| csnk1g2a | 0.16842 | 0.195801849 | 0.05 | 0 | 1 | 10 |
| cux1a | 0.16842 | 0.136823511 | 0.05 | 0 | 1 | 10 |
| dph3 | 0.16842 | 0.153593918 | 0.05 | 0 | 1 | 10 |
| fis1 | 0.16842 | 0.175484948 | 0.05 | 0 | 1 | 10 |
| hadhaa | 0.16842 | 0.198679542 | 0.05 | 0 | 1 | 10 |
| hnrnpd | 0.16842 | 0.193451848 | 0.05 | 0 | 1 | 10 |
| igbp1 | 0.16842 | 0.183491797 | 0.05 | 0 | 1 | 10 |
| ilkap | 0.16842 | 0.176778613 | 0.05 | 0 | 1 | 10 |
| ints8 | 0.16842 | 0.142980361 | 0.05 | 0 | 1 | 10 |
| maip1 | 0.16842 | 0.133777921 | 0.05 | 0 | 1 | 10 |
| mbpb | 0.16842 | 0.162902213 | 0.05 | 0 | 1 | 10 |
| mgst1.1 | 0.16842 | 0.187243645 | 0.05 | 0 | 1 | 10 |
| micu2 | 0.16842 | 0.179477316 | 0.05 | 0 | 1 | 10 |
| mmadhc | 0.16842 | 0.160502615 | 0.05 | 0 | 1 | 10 |
| mob2a | 0.16842 | 0.199331494 | 0.05 | 0 | 1 | 10 |
| mrpl51 | 0.16842 | 0.17992498 | 0.05 | 0 | 1 | 10 |
| nck1b | 0.16842 | 0.178068545 | 0.05 | 0 | 1 | 10 |
| ntmt1 | 0.16842 | 0.154986482 | 0.05 | 0 | 1 | 10 |
| patz1 | 0.16842 | 0.160927414 | 0.05 | 0 | 1 | 10 |
| plpp2b | 0.16842 | 0.191832407 | 0.05 | 0 | 1 | 10 |
| plpp3 | 0.16842 | 0.185938377 | 0.05 | 0 | 1 | 10 |
| pold3 | 0.16842 | 0.143837437 | 0.05 | 0 | 1 | 10 |
| prkag2a | 0.16842 | 0.151325947 | 0.05 | 0 | 1 | 10 |
| pwp2h | 0.16842 | 0.164306935 | 0.05 | 0 | 1 | 10 |
| rarres3l | 0.16842 | 0.113949026 | 0.05 | 0 | 1 | 10 |
| rbfox1 | 0.16842 | 0.190506968 | 0.05 | 0 | 1 | 10 |
| sec23b | 0.16842 | 0.112060666 | 0.05 | 0 | 1 | 10 |
| si:ch211-51h9.7 | 0.16842 | 0.162495828 | 0.05 | 0 | 1 | 10 |
| si:dkey-102m7.3 | 0.16842 | 0.169728693 | 0.05 | 0 | 1 | 10 |
| si:dkey-56m19.5 | 0.16842 | 0.189265086 | 0.05 | 0 | 1 | 10 |
| slc44a2 | 0.16842 | 0.206244137 | 0.05 | 0 | 1 | 10 |

| | | | | | | |
|---|---|---|---|---|---|---|
| smurf2 | 0.16842 | 0.182194995 | 0.05 | 0 | 1 | 10 |
| tbcelb | 0.16842 | 0.192166473 | 0.05 | 0 | 1 | 10 |
| tcf3a | 0.16842 | 0.184498123 | 0.05 | 0 | 1 | 10 |
| trabd | 0.16842 | 0.114105509 | 0.05 | 0 | 1 | 10 |
| ubr3 | 0.16842 | 0.130418759 | 0.05 | 0 | 1 | 10 |
| wipi2 | 0.16842 | 0.161949814 | 0.05 | 0 | 1 | 10 |
| zcchc8 | 0.16842 | 0.18927973 | 0.05 | 0 | 1 | 10 |
| zgc:103499 | 0.16842 | 0.19564304 | 0.05 | 0 | 1 | 10 |
| zgc:110239 | 0.16842 | 0.190986649 | 0.05 | 0 | 1 | 10 |
| zgc:112160 | 0.16842 | 0.176472999 | 0.05 | 0 | 1 | 10 |
| zgc:162816 | 0.16842 | 0.176544845 | 0.05 | 0 | 1 | 10 |
| arl9 | 0.16842 | 0.183242978 | 0.05 | 0 | 1 | 10 |
| rnf169 | 0.16842 | 0.193087934 | 0.05 | 0 | 1 | 10 |
| rnf216 | 0.16842 | 0.176738539 | 0.05 | 0 | 1 | 10 |
| rprd2b | 0.16842 | 0.196580421 | 0.05 | 0 | 1 | 10 |
| BX649300.2 | 0.16842 | 0.171995696 | 0.05 | 0 | 1 | 10 |
| BX908782.1 | 0.16842 | 0.161301057 | 0.05 | 0 | 1 | 10 |
| CELA1 (1 of many) | 0.16842 | 0.119518172 | 0.05 | 0 | 1 | 10 |
| TPGS1 | 0.16842 | 0.169025574 | 0.05 | 0 | 1 | 10 |
| ajuba | 0.16842 | 0.189706893 | 0.05 | 0 | 1 | 10 |
| apoa2 | 0.16842 | 0.172935799 | 0.05 | 0 | 1 | 10 |
| brf1b | 0.16842 | 0.16710765 | 0.05 | 0 | 1 | 10 |
| chchd4a | 0.16842 | 0.160061016 | 0.05 | 0 | 1 | 10 |
| chd7 | 0.16842 | 0.163022326 | 0.05 | 0 | 1 | 10 |
| cipcb | 0.16842 | 0.191243819 | 0.05 | 0 | 1 | 10 |
| cplx2l | 0.16842 | 0.186784591 | 0.05 | 0 | 1 | 10 |
| creb1a | 0.16842 | 0.154117223 | 0.05 | 0 | 1 | 10 |
| creb3l2 | 0.16842 | 0.16334919 | 0.05 | 0 | 1 | 10 |
| creg1 | 0.16842 | 0.182294577 | 0.05 | 0 | 1 | 10 |
| cyb5r1 | 0.16842 | 0.140057176 | 0.05 | 0 | 1 | 10 |
| dab2ipa | 0.16842 | 0.19326322 | 0.05 | 0 | 1 | 10 |
| dnmt3bb.1 | 0.16842 | 0.171230847 | 0.05 | 0 | 1 | 10 |
| dyrk1ab | 0.16842 | 0.162456081 | 0.05 | 0 | 1 | 10 |
| ech1 | 0.16842 | 0.150147355 | 0.05 | 0 | 1 | 10 |
| fam102ab | 0.16842 | 0.178254309 | 0.05 | 0 | 1 | 10 |
| fer | 0.16842 | 0.152844421 | 0.05 | 0 | 1 | 10 |
| fip1l1b | 0.16842 | 0.193307142 | 0.05 | 0 | 1 | 10 |
| fshr | 0.16842 | 0.168226466 | 0.05 | 0 | 1 | 10 |
| gbe1b | 0.16842 | 0.17171149 | 0.05 | 0 | 1 | 10 |
| gcc1 | 0.16842 | 0.189803812 | 0.05 | 0 | 1 | 10 |
| gid8b | 0.16842 | 0.158401009 | 0.05 | 0 | 1 | 10 |
| gsk3ab | 0.16842 | 0.168208722 | 0.05 | 0 | 1 | 10 |
| gtf2a2 | 0.16842 | 0.20850944 | 0.05 | 0 | 1 | 10 |
| hmcn2 | 0.16842 | 0.188219071 | 0.05 | 0 | 1 | 10 |
| htt | 0.16842 | 0.166876033 | 0.05 | 0 | 1 | 10 |
| hyal2b | 0.16842 | 0.182296703 | 0.05 | 0 | 1 | 10 |
| kctd12.1 | 0.16842 | 0.195385979 | 0.05 | 0 | 1 | 10 |
| lcmt1 | 0.16842 | 0.161301935 | 0.05 | 0 | 1 | 10 |

| | | | | | | |
|---|---|---|---|---|---|---|
| limch1b | 0.16842 | 0.160849873 | 0.05 | 0 | 1 | 10 |
| lman2 | 0.16842 | 0.180970108 | 0.05 | 0 | 1 | 10 |
| lsm14aa | 0.16842 | 0.200798967 | 0.05 | 0 | 1 | 10 |
| lysmd3 | 0.16842 | 0.126129392 | 0.05 | 0 | 1 | 10 |
| morc3a | 0.16842 | 0.160534602 | 0.05 | 0 | 1 | 10 |
| mtor | 0.16842 | 0.163647533 | 0.05 | 0 | 1 | 10 |
| ndufaf5 | 0.16842 | 0.177283582 | 0.05 | 0 | 1 | 10 |
| ndufv2 | 0.16842 | 0.169975588 | 0.05 | 0 | 1 | 10 |
| ngly1 | 0.16842 | 0.193936634 | 0.05 | 0 | 1 | 10 |
| nipal3 | 0.16842 | 0.187774277 | 0.05 | 0 | 1 | 10 |
| ociad1 | 0.16842 | 0.166121904 | 0.05 | 0 | 1 | 10 |
| otud3 | 0.16842 | 0.157059517 | 0.05 | 0 | 1 | 10 |
| parp1 | 0.16842 | 0.154588588 | 0.05 | 0 | 1 | 10 |
| pdhb | 0.16842 | 0.150480988 | 0.05 | 0 | 1 | 10 |
| phf3 | 0.16842 | 0.153146182 | 0.05 | 0 | 1 | 10 |
| pi4kb | 0.16842 | 0.191023674 | 0.05 | 0 | 1 | 10 |
| ppp4r2a | 0.16842 | 0.14167683 | 0.05 | 0 | 1 | 10 |
| prkcda | 0.16842 | 0.161823188 | 0.05 | 0 | 1 | 10 |
| prpf38a | 0.16842 | 0.172208011 | 0.05 | 0 | 1 | 10 |
| prxl2b | 0.16842 | 0.19670795 | 0.05 | 0 | 1 | 10 |
| ptpn2a | 0.16842 | 0.195581486 | 0.05 | 0 | 1 | 10 |
| pus1 | 0.16842 | 0.165054401 | 0.05 | 0 | 1 | 10 |
| relb | 0.16842 | 0.189611388 | 0.05 | 0 | 1 | 10 |
| rheb | 0.16842 | 0.136675476 | 0.05 | 0 | 1 | 10 |
| rock2a | 0.16842 | 0.163386998 | 0.05 | 0 | 1 | 10 |
| sdf4 | 0.16842 | 0.151418636 | 0.05 | 0 | 1 | 10 |
| setd3 | 0.16842 | 0.18141908 | 0.05 | 0 | 1 | 10 |
| si:ch211-113p18.3 | 0.16842 | 0.176212567 | 0.05 | 0 | 1 | 10 |
| si:ch211-214p16.1 | 0.16842 | 0.150260474 | 0.05 | 0 | 1 | 10 |
| si:ch211-76m11.3 | 0.16842 | 0.177632389 | 0.05 | 0 | 1 | 10 |
| si:ch73-44m9.5 | 0.16842 | 0.153304031 | 0.05 | 0 | 1 | 10 |
| slc25a16 | 0.16842 | 0.162935081 | 0.05 | 0 | 1 | 10 |
| slc27a1a | 0.16842 | 0.182960531 | 0.05 | 0 | 1 | 10 |
| slc40a1 | 0.16842 | 0.187736248 | 0.05 | 0 | 1 | 10 |
| sqor | 0.16842 | 0.169560806 | 0.05 | 0 | 1 | 10 |
| sra1 | 0.16842 | 0.177087083 | 0.05 | 0 | 1 | 10 |
| srp9 | 0.16842 | 0.143736424 | 0.05 | 0 | 1 | 10 |
| srsf10b | 0.16842 | 0.187464138 | 0.05 | 0 | 1 | 10 |
| strip1 | 0.16842 | 0.14165016 | 0.05 | 0 | 1 | 10 |
| suz12b | 0.16842 | 0.208633522 | 0.05 | 0 | 1 | 10 |
| taf11 | 0.16842 | 0.170773185 | 0.05 | 0 | 1 | 10 |
| tarbp2 | 0.16842 | 0.19132064 | 0.05 | 0 | 1 | 10 |
| tbx18 | 0.16842 | 0.187774277 | 0.05 | 0 | 1 | 10 |
| tcf7l1a | 0.16842 | 0.176368787 | 0.05 | 0 | 1 | 10 |
| tes | 0.16842 | 0.192813123 | 0.05 | 0 | 1 | 10 |
| tm9sf1 | 0.16842 | 0.142452386 | 0.05 | 0 | 1 | 10 |
| tmeff1b | 0.16842 | 0.196426915 | 0.05 | 0 | 1 | 10 |
| trim35-28 | 0.16842 | 0.211667744 | 0.05 | 0 | 1 | 10 |

| | | | | | | |
|---|---|---|---|---|---|---|
| trioa | 0.16842 | 0.165225993 | 0.05 | 0 | 1 | 10 |
| tsc1b | 0.16842 | 0.183261181 | 0.05 | 0 | 1 | 10 |
| uchl3 | 0.16842 | 0.194264931 | 0.05 | 0 | 1 | 10 |
| vps26b | 0.16842 | 0.141519339 | 0.05 | 0 | 1 | 10 |
| wu:fb74b10 | 0.16842 | 0.173732894 | 0.05 | 0 | 1 | 10 |
| zfr2 | 0.16842 | 0.178995472 | 0.05 | 0 | 1 | 10 |
| zgc:110269 | 0.16842 | 0.210061029 | 0.05 | 0 | 1 | 10 |
| zgc:66433 | 0.16842 | 0.187614925 | 0.05 | 0 | 1 | 10 |
| BX569787.1 | 0.16842 | 0.182099169 | 0.05 | 0 | 1 | 10 |
| CR751602.2 | 0.16842 | 0.187367719 | 0.05 | 0 | 1 | 10 |
| CR759918.1 | 0.16842 | 0.156918704 | 0.05 | 0 | 1 | 10 |
| FO082877.1 | 0.16842 | 0.182531618 | 0.05 | 0 | 1 | 10 |
| FP101887.1 | 0.16842 | 0.11874069 | 0.05 | 0 | 1 | 10 |
| LOC100534805 | 0.16842 | 0.183227024 | 0.05 | 0 | 1 | 10 |
| LOC101885027 | 0.16842 | 0.186890629 | 0.05 | 0 | 1 | 10 |
| RSPH1 | 0.16842 | 0.155613988 | 0.05 | 0 | 1 | 10 |
| acox1 | 0.16842 | 0.138854483 | 0.05 | 0 | 1 | 10 |
| adka | 0.16842 | 0.130099102 | 0.05 | 0 | 1 | 10 |
| aes | 0.16842 | 0.136688203 | 0.05 | 0 | 1 | 10 |
| agrp | 0.16842 | 0.150746557 | 0.05 | 0 | 1 | 10 |
| akirin1 | 0.16842 | 0.182362428 | 0.05 | 0 | 1 | 10 |
| aktip | 0.16842 | 0.10793777 | 0.05 | 0 | 1 | 10 |
| ankrd9 | 0.16842 | 0.200012695 | 0.05 | 0 | 1 | 10 |
| arrdc2 | 0.16842 | 0.150154507 | 0.05 | 0 | 1 | 10 |
| atpv0e2 | 0.16842 | 0.207154078 | 0.05 | 0 | 1 | 10 |
| bmpr2a | 0.16842 | 0.18205336 | 0.05 | 0 | 1 | 10 |
| bub3 | 0.16842 | 0.138428478 | 0.05 | 0 | 1 | 10 |
| ccdc32 | 0.16842 | 0.16341602 | 0.05 | 0 | 1 | 10 |
| ccdc3b | 0.16842 | 0.121321787 | 0.05 | 0 | 1 | 10 |
| cdk5rap3 | 0.16842 | 0.148729794 | 0.05 | 0 | 1 | 10 |
| cetn4 | 0.16842 | 0.184961243 | 0.05 | 0 | 1 | 10 |
| chchd6a | 0.16842 | 0.12979041 | 0.05 | 0 | 1 | 10 |
| creg2 | 0.16842 | 0.139362115 | 0.05 | 0 | 1 | 10 |
| ctsf | 0.16842 | 0.180854 | 0.05 | 0 | 1 | 10 |
| ddx6 | 0.16842 | 0.163617937 | 0.05 | 0 | 1 | 10 |
| dhrs13b | 0.16842 | 0.199792374 | 0.05 | 0 | 1 | 10 |
| dph6 | 0.16842 | 0.149632826 | 0.05 | 0 | 1 | 10 |
| dvl2 | 0.16842 | 0.157147847 | 0.05 | 0 | 1 | 10 |
| elovl8b | 0.16842 | 0.154593097 | 0.05 | 0 | 1 | 10 |
| ext1c | 0.16842 | 0.136607313 | 0.05 | 0 | 1 | 10 |
| fbxl18 | 0.16842 | 0.178123719 | 0.05 | 0 | 1 | 10 |
| fbxw11a | 0.16842 | 0.164613712 | 0.05 | 0 | 1 | 10 |
| ftr63 | 0.16842 | 0.144534603 | 0.05 | 0 | 1 | 10 |
| gng2 | 0.16842 | 0.183937741 | 0.05 | 0 | 1 | 10 |
| hacd2 | 0.16842 | 0.188137538 | 0.05 | 0 | 1 | 10 |
| isoc1 | 0.16842 | 0.150401796 | 0.05 | 0 | 1 | 10 |
| kank4 | 0.16842 | 0.161884786 | 0.05 | 0 | 1 | 10 |
| kif3b | 0.16842 | 0.167017966 | 0.05 | 0 | 1 | 10 |

| | | | | | | |
|---|---|---|---|---|---|---|
| kita | 0.16842 | 0.140809096 | 0.05 | 0 | 1 | 10 |
| klhl8 | 0.16842 | 0.125250719 | 0.05 | 0 | 1 | 10 |
| kpna4 | 0.16842 | 0.136118687 | 0.05 | 0 | 1 | 10 |
| lhfpl2b | 0.16842 | 0.107937963 | 0.05 | 0 | 1 | 10 |
| lman1 | 0.16842 | 0.14253603 | 0.05 | 0 | 1 | 10 |
| mapk3 | 0.16842 | 0.153031726 | 0.05 | 0 | 1 | 10 |
| midn | 0.16842 | 0.166486182 | 0.05 | 0 | 1 | 10 |
| mvb12a | 0.16842 | 0.164299179 | 0.05 | 0 | 1 | 10 |
| ncalda | 0.16842 | 0.161028323 | 0.05 | 0 | 1 | 10 |
| nit2 | 0.16842 | 0.113651738 | 0.05 | 0 | 1 | 10 |
| nme6 | 0.16842 | 0.184635534 | 0.05 | 0 | 1 | 10 |
| nt5c2b | 0.16842 | 0.183985884 | 0.05 | 0 | 1 | 10 |
| pam | 0.16842 | 0.181930456 | 0.05 | 0 | 1 | 10 |
| pcsk5b | 0.16842 | 0.166602967 | 0.05 | 0 | 1 | 10 |
| pde6d | 0.16842 | 0.185604563 | 0.05 | 0 | 1 | 10 |
| pdgfab | 0.16842 | 0.184615666 | 0.05 | 0 | 1 | 10 |
| phb2a | 0.16842 | 0.148442997 | 0.05 | 0 | 1 | 10 |
| pip4p1a | 0.16842 | 0.118948303 | 0.05 | 0 | 1 | 10 |
| pla2g4aa | 0.16842 | 0.171705111 | 0.05 | 0 | 1 | 10 |
| ptk7a | 0.16842 | 0.176493239 | 0.05 | 0 | 1 | 10 |
| ptprc | 0.16842 | 0.168353506 | 0.05 | 0 | 1 | 10 |
| pttg1ipb | 0.16842 | 0.156052412 | 0.05 | 0 | 1 | 10 |
| puraa | 0.16842 | 0.195390668 | 0.05 | 0 | 1 | 10 |
| rab40b | 0.16842 | 0.187105959 | 0.05 | 0 | 1 | 10 |
| rabl2 | 0.16842 | 0.122352408 | 0.05 | 0 | 1 | 10 |
| rabl6b | 0.16842 | 0.197284758 | 0.05 | 0 | 1 | 10 |
| rasa1b | 0.16842 | 0.152105983 | 0.05 | 0 | 1 | 10 |
| sbds | 0.16842 | 0.164106317 | 0.05 | 0 | 1 | 10 |
| sdad1 | 0.16842 | 0.177565644 | 0.05 | 0 | 1 | 10 |
| si:ch211-193c2.2 | 0.16842 | 0.189944579 | 0.05 | 0 | 1 | 10 |
| si:ch211-225h24.2 | 0.16842 | 0.121321787 | 0.05 | 0 | 1 | 10 |
| si:ch211-240l19.8 | 0.16842 | 0.222169398 | 0.05 | 0 | 1 | 10 |
| si:ch73-141c7.1 | 0.16842 | 0.201044076 | 0.05 | 0 | 1 | 10 |
| si:dkey-12e7.1 | 0.16842 | 0.139276517 | 0.05 | 0 | 1 | 10 |
| slc25a28 | 0.16842 | 0.165811218 | 0.05 | 0 | 1 | 10 |
| slc4a11 | 0.16842 | 0.207495328 | 0.05 | 0 | 1 | 10 |
| spata5l1 | 0.16842 | 0.11576623 | 0.05 | 0 | 1 | 10 |
| supt3h | 0.16842 | 0.105409186 | 0.05 | 0 | 1 | 10 |
| taf6l | 0.16842 | 0.110896342 | 0.05 | 0 | 1 | 10 |
| tfdp1b | 0.16842 | 0.132302446 | 0.05 | 0 | 1 | 10 |
| tm2d2 | 0.16842 | 0.180861414 | 0.05 | 0 | 1 | 10 |
| tmem203 | 0.16842 | 0.224096074 | 0.05 | 0 | 1 | 10 |
| vps18 | 0.16842 | 0.179767626 | 0.05 | 0 | 1 | 10 |
| wapla | 0.16842 | 0.157029208 | 0.05 | 0 | 1 | 10 |
| wdfy1 | 0.16842 | 0.157827931 | 0.05 | 0 | 1 | 10 |
| wdr32 | 0.16842 | 0.137102044 | 0.05 | 0 | 1 | 10 |
| zfpm2b | 0.16842 | 0.154044262 | 0.05 | 0 | 1 | 10 |
| zgc:101577 | 0.16842 | 0.153470054 | 0.05 | 0 | 1 | 10 |

| | | | | | | |
|---|---|---|---|---|---|---|
| zgc:153521 | 0.16842 | 0.115564799 | 0.05 | 0 | 1 | 10 |
| znf451 | 0.16842 | 0.116097659 | 0.05 | 0 | 1 | 10 |
| flj11011l | 0.16842 | 0.166688287 | 0.05 | 0 | 1 | 10 |
| hira | 0.16842 | 0.201433753 | 0.05 | 0 | 1 | 10 |
| rab11bb | 0.16842 | 0.159700492 | 0.05 | 0 | 1 | 10 |
| hspa4b | 0.17114 | 0.289498948 | 0.15 | 0.029 | 1 | 10 |
| rps16 | 0.172 | 0.144060943 | 0.95 | 0.912 | 1 | 10 |
| hnrnpa1b | 0.17355 | -0.27411545 | 0.1 | 0.235 | 1 | 10 |
| cyp17a1 | 0.17415 | 0.385671341 | 0.325 | 0.147 | 1 | 10 |
| ddx5 | 0.17489 | -0.4123846 | 0.125 | 0.265 | 1 | 10 |
| uqcc2 | 0.17548 | -0.27747365 | 0.05 | 0.176 | 1 | 10 |
| phc2a | 0.17561 | -0.40108291 | 0.1 | 0.265 | 1 | 10 |
| ndufa1 | 0.17568 | -0.4437801 | 0.125 | 0.294 | 1 | 10 |
| mrpl36 | 0.17644 | 0.365981072 | 0.2 | 0.059 | 1 | 10 |
| CABZ01115113.1 | 0.17711 | -0.43577879 | 0.05 | 0.176 | 1 | 10 |
| slc38a5b | 0.17739 | 0.381475074 | 0.2 | 0.059 | 1 | 10 |
| ssu72 | 0.17796 | 0.345222112 | 0.15 | 0.029 | 1 | 10 |
| myh10 | 0.17856 | 0.366902587 | 0.15 | 0.029 | 1 | 10 |
| gmfb | 0.17956 | 0.421489632 | 0.2 | 0.059 | 1 | 10 |
| zgc:55733 | 0.18109 | -0.35668213 | 0.025 | 0.118 | 1 | 10 |
| ctnnbl1 | 0.18451 | -0.35190672 | 0.025 | 0.118 | 1 | 10 |
| hells | 0.18835 | -0.36606771 | 0.025 | 0.118 | 1 | 10 |
| rps27a | 0.19101 | -0.11628515 | 0.925 | 1 | 1 | 10 |
| rpn1 | 0.19179 | 0.424490575 | 0.25 | 0.118 | 1 | 10 |
| gstm.3 | 0.19198 | 0.215286516 | 0.775 | 0.706 | 1 | 10 |
| rpl29 | 0.19394 | -0.11425411 | 0.925 | 1 | 1 | 10 |
| tmed9 | 0.19429 | -0.38833583 | 0.05 | 0.176 | 1 | 10 |
| pomp | 0.19567 | -0.18245227 | 0.35 | 0.5 | 1 | 10 |
| dynlrb1 | 0.20082 | -0.39044301 | 0.15 | 0.324 | 1 | 10 |
| marcksl1b | 0.20241 | 0.260322673 | 0.4 | 0.353 | 1 | 10 |
| fam107b | 0.20324 | 0.4413421 | 0.3 | 0.147 | 1 | 10 |
| si:ch211-156b7.4 | 0.2048 | -0.33710064 | 0.025 | 0.118 | 1 | 10 |
| hmga1a | 0.20489 | -0.30336267 | 0.275 | 0.353 | 1 | 10 |
| nop10 | 0.20602 | -0.36511853 | 0.05 | 0.176 | 1 | 10 |
| mtpn | 0.20603 | -0.37440479 | 0.175 | 0.353 | 1 | 10 |
| tomm20a | 0.20728 | -0.37349518 | 0.15 | 0.324 | 1 | 10 |
| zgc:66388 | 0.20826 | 0.365955581 | 0.125 | 0.029 | 1 | 10 |
| adi1 | 0.20965 | -0.32368223 | 0.025 | 0.118 | 1 | 10 |
| jagn1b | 0.21018 | -0.24421572 | 0.025 | 0.118 | 1 | 10 |
| pspc1 | 0.21102 | -0.32640549 | 0.025 | 0.118 | 1 | 10 |
| tuba8l4 | 0.21156 | -0.39322608 | 0.25 | 0.382 | 1 | 10 |
| sucla2 | 0.21234 | -0.33688138 | 0.05 | 0.176 | 1 | 10 |
| hnrnpl2 | 0.21558 | -0.27636896 | 0.075 | 0.206 | 1 | 10 |
| hspa8 | 0.21728 | -0.15009889 | 0.725 | 0.765 | 1 | 10 |
| hnrnpa0a | 0.21793 | -0.31649312 | 0.025 | 0.118 | 1 | 10 |
| zgc:193541 | 0.2197 | -0.35422522 | 0.375 | 0.529 | 1 | 10 |
| rap1b | 0.22304 | 0.385453055 | 0.175 | 0.059 | 1 | 10 |
| selenom | 0.22436 | -0.22641195 | 0.025 | 0.118 | 1 | 10 |

| | | | | | | |
|---|---|---|---|---|---|---|
| dldh | 0.22511 | -0.24831818 | 0.025 | 0.118 | 1 | 10 |
| pfdn6 | 0.22644 | 0.381555007 | 0.225 | 0.088 | 1 | 10 |
| uba52 | 0.22807 | 0.160803647 | 0.85 | 0.794 | 1 | 10 |
| ddx42 | 0.22887 | -0.3379897 | 0.025 | 0.118 | 1 | 10 |
| atf4b | 0.229 | 0.428093293 | 0.375 | 0.206 | 1 | 10 |
| rpl10a | 0.22918 | 0.107855864 | 1 | 0.971 | 1 | 10 |
| cdc14b | 0.22976 | -0.2063327 | 0.025 | 0.118 | 1 | 10 |
| tpd52l2a | 0.22979 | -0.2053453 | 0.025 | 0.118 | 1 | 10 |
| vat1 | 0.23319 | -0.34820008 | 0.025 | 0.118 | 1 | 10 |
| hnrnph1l | 0.23326 | -0.29656186 | 0.025 | 0.118 | 1 | 10 |
| ppp1cbl | 0.23872 | -0.28484608 | 0.025 | 0.118 | 1 | 10 |
| rps2 | 0.23899 | 0.10098587 | 1 | 0.941 | 1 | 10 |
| cox7a2l | 0.2393 | 0.274312311 | 0.175 | 0.059 | 1 | 10 |
| hmgn6 | 0.24004 | 0.328968568 | 0.625 | 0.441 | 1 | 10 |
| h2afx1 | 0.24067 | -0.26621559 | 0.25 | 0.412 | 1 | 10 |
| slirp | 0.2413 | -0.36712328 | 0.075 | 0.206 | 1 | 10 |
| si:ch211-222l21.1 | 0.24321 | -0.21730084 | 0.575 | 0.647 | 1 | 10 |
| si:busm1-266f07.2 | 0.24369 | -0.30822606 | 0.025 | 0.088 | 1 | 10 |
| med11 | 0.24653 | -0.32618528 | 0.025 | 0.118 | 1 | 10 |
| rad23b | 0.24662 | -0.41959053 | 0.15 | 0.294 | 1 | 10 |
| ppp2r5eb | 0.24977 | 0.308417711 | 0.125 | 0.029 | 1 | 10 |
| mob3a | 0.25067 | -0.3487491 | 0.075 | 0.206 | 1 | 10 |
| pcna | 0.25084 | -0.40725678 | 0.075 | 0.206 | 1 | 10 |
| ldlrap1a | 0.25328 | 0.335723673 | 0.125 | 0.029 | 1 | 10 |
| gtf2h4 | 0.2533 | -0.29925109 | 0.025 | 0.118 | 1 | 10 |
| ppid | 0.25333 | -0.20355119 | 0.025 | 0.118 | 1 | 10 |
| si:ch211-235e9.6 | 0.25366 | -0.35601665 | 0.075 | 0.206 | 1 | 10 |
| zgc:152791 | 0.25378 | -0.32747385 | 0.075 | 0.206 | 1 | 10 |
| ccdc90b | 0.25395 | -0.22020793 | 0.025 | 0.118 | 1 | 10 |
| sox4a | 0.25398 | 0.417867859 | 0.25 | 0.118 | 1 | 10 |
| parp3 | 0.25469 | -0.23789654 | 0.025 | 0.118 | 1 | 10 |
| sri | 0.25505 | -0.35301779 | 0.075 | 0.206 | 1 | 10 |
| CR383676.1 | 0.25584 | 0.645625142 | 0.225 | 0.088 | 1 | 10 |
| si:ch211-66k16.27 | 0.25662 | -0.32865026 | 0.075 | 0.206 | 1 | 10 |
| srsf7a | 0.25776 | -0.32610174 | 0.075 | 0.206 | 1 | 10 |
| clic4 | 0.25875 | -0.33139275 | 0.025 | 0.118 | 1 | 10 |
| tp53inp1 | 0.25878 | -0.24479449 | 0.05 | 0.147 | 1 | 10 |
| srsf2a | 0.2605 | -0.2802808 | 0.025 | 0.118 | 1 | 10 |
| znf207b | 0.26154 | -0.23922498 | 0.025 | 0.118 | 1 | 10 |
| gstm.1 | 0.26231 | 0.253723303 | 0.525 | 0.441 | 1 | 10 |
| zgc:56493 | 0.2631 | 0.337192631 | 0.575 | 0.412 | 1 | 10 |
| rpl27a | 0.26331 | -0.31388922 | 0.5 | 0.647 | 1 | 10 |
| ndufb3 | 0.26416 | 0.291465381 | 0.275 | 0.206 | 1 | 10 |
| anp32b | 0.26568 | -0.38076157 | 0.1 | 0.235 | 1 | 10 |
| u2af2b | 0.26682 | -0.26948302 | 0.025 | 0.118 | 1 | 10 |
| rab1ab | 0.26688 | -0.2617993 | 0.025 | 0.118 | 1 | 10 |
| sdhc | 0.26774 | 0.236417899 | 0.125 | 0.029 | 1 | 10 |
| prdx4 | 0.26794 | 0.380091779 | 0.225 | 0.118 | 1 | 10 |

| | | | | | | |
|---|---|---|---|---|---|---|
| isg15 | 0.26803 | 0.333592459 | 0.175 | 0.088 | 1 | 10 |
| psma2 | 0.27032 | -0.27559955 | 0.125 | 0.265 | 1 | 10 |
| cox14 | 0.27061 | -0.33428185 | 0.05 | 0.147 | 1 | 10 |
| nudt9 | 0.27153 | -0.28726171 | 0.025 | 0.118 | 1 | 10 |
| rhoab | 0.27176 | -0.2411975 | 0.025 | 0.118 | 1 | 10 |
| psmd1 | 0.2722 | 0.311924252 | 0.125 | 0.029 | 1 | 10 |
| mt-nd5 | 0.27232 | -0.27425844 | 0.025 | 0.118 | 1 | 10 |
| rsu1 | 0.27253 | 0.241226221 | 0.125 | 0.029 | 1 | 10 |
| cnpy1 | 0.27321 | -0.21953783 | 0.025 | 0.118 | 1 | 10 |
| carm1 | 0.27356 | -0.28341569 | 0.025 | 0.118 | 1 | 10 |
| prdx1 | 0.27408 | -0.32461033 | 0.025 | 0.118 | 1 | 10 |
| zgc:194246 | 0.27451 | -0.26165067 | 0.025 | 0.118 | 1 | 10 |
| ipo7 | 0.27471 | -0.28724421 | 0.025 | 0.118 | 1 | 10 |
| igf3 | 0.27531 | 0.375216948 | 0.375 | 0.206 | 1 | 10 |
| ddx3xb | 0.27616 | 0.309398993 | 0.125 | 0.029 | 1 | 10 |
| smfn | 0.27671 | -0.254426 | 0.025 | 0.118 | 1 | 10 |
| zgc:123010 | 0.27684 | -0.29050884 | 0.025 | 0.118 | 1 | 10 |
| fgd5a | 0.27715 | -0.29417645 | 0.025 | 0.118 | 1 | 10 |
| sfpq | 0.27783 | -0.28941625 | 0.025 | 0.118 | 1 | 10 |
| si:ch211-207d6.2 | 0.27795 | 0.39526816 | 0.175 | 0.059 | 1 | 10 |
| arf1 | 0.27829 | -0.26355976 | 0.025 | 0.118 | 1 | 10 |
| acads | 0.27839 | -0.29821675 | 0.025 | 0.118 | 1 | 10 |
| nup62l | 0.27845 | -0.2710632 | 0.025 | 0.118 | 1 | 10 |
| ctdsp2 | 0.27847 | -0.29921482 | 0.025 | 0.118 | 1 | 10 |
| zgc:174863 | 0.2785 | -0.32929887 | 0.025 | 0.118 | 1 | 10 |
| CABZ01080702.1 | 0.27856 | -0.29790716 | 0.025 | 0.118 | 1 | 10 |
| ubald2 | 0.27856 | -0.30177026 | 0.025 | 0.118 | 1 | 10 |
| calm3b | 0.27863 | -0.14318587 | 0.425 | 0.559 | 1 | 10 |
| mrpl52 | 0.27936 | 0.251137018 | 0.125 | 0.029 | 1 | 10 |
| serpinb1 | 0.27973 | 0.254707273 | 0.125 | 0.088 | 1 | 10 |
| smarce1 | 0.28127 | -0.33256723 | 0.1 | 0.235 | 1 | 10 |
| ndufs6 | 0.28313 | 0.336841965 | 0.125 | 0.029 | 1 | 10 |
| snrnp27 | 0.28411 | -0.3587614 | 0.1 | 0.235 | 1 | 10 |
| mgmt | 0.28686 | 0.272689572 | 0.125 | 0.029 | 1 | 10 |
| tmed10 | 0.2886 | -0.3101958 | 0.1 | 0.235 | 1 | 10 |
| ctsla | 0.29114 | -0.19518874 | 0.1 | 0.206 | 1 | 10 |
| tpd52l2b | 0.29223 | 0.288434045 | 0.125 | 0.029 | 1 | 10 |
| arf6b | 0.2928 | 0.347496161 | 0.175 | 0.059 | 1 | 10 |
| chmp1a | 0.29283 | 0.326610836 | 0.125 | 0.029 | 1 | 10 |
| tmed7 | 0.29299 | -0.21959911 | 0.05 | 0.147 | 1 | 10 |
| eif4ebp1 | 0.29365 | 0.33476374 | 0.5 | 0.353 | 1 | 10 |
| tktb | 0.29368 | 0.320712547 | 0.175 | 0.059 | 1 | 10 |
| eny2 | 0.29383 | 0.348895014 | 0.175 | 0.059 | 1 | 10 |
| slc16a10 | 0.29401 | 0.327693213 | 0.125 | 0.029 | 1 | 10 |
| vps4b | 0.29457 | 0.265068991 | 0.125 | 0.029 | 1 | 10 |
| mmp2 | 0.29507 | -0.36055266 | 0.25 | 0.382 | 1 | 10 |
| efhd1 | 0.29563 | 0.296624648 | 0.125 | 0.029 | 1 | 10 |
| twf1a | 0.29608 | -0.20527238 | 0.05 | 0.147 | 1 | 10 |

| | | | | | | |
|---|---|---|---|---|---|---|
| atp6v1e1a | 0.29621 | 0.280172624 | 0.125 | 0.029 | 1 | 10 |
| bcap31 | 0.29661 | 0.318525887 | 0.125 | 0.029 | 1 | 10 |
| taf9 | 0.29664 | -0.30339832 | 0.05 | 0.147 | 1 | 10 |
| mt-co1 | 0.29868 | -0.36707212 | 0.425 | 0.588 | 1 | 10 |
| rbm8a | 0.29915 | 0.371104784 | 0.4 | 0.235 | 1 | 10 |
| pabpc1a | 0.30179 | -0.26975884 | 0.5 | 0.676 | 1 | 10 |
| cox4i1 | 0.30191 | -0.28983992 | 0.3 | 0.471 | 1 | 10 |
| elof1 | 0.30405 | -0.28947156 | 0.15 | 0.294 | 1 | 10 |
| bzw1b | 0.30565 | -0.29819737 | 0.175 | 0.324 | 1 | 10 |
| crip2 | 0.30846 | -0.28716743 | 0.025 | 0.088 | 1 | 10 |
| sf3b6 | 0.3117 | -0.30025361 | 0.15 | 0.294 | 1 | 10 |
| atp5mc3b | 0.31317 | -0.28797045 | 0.475 | 0.647 | 1 | 10 |
| rap1aa | 0.3139 | -0.20118182 | 0.1 | 0.206 | 1 | 10 |
| pfdn1 | 0.31397 | -0.32391633 | 0.05 | 0.147 | 1 | 10 |
| ciarta | 0.3167 | -0.31182815 | 0.05 | 0.147 | 1 | 10 |
| rpa2 | 0.31708 | -0.24175093 | 0.05 | 0.147 | 1 | 10 |
| tegt | 0.31719 | 0.375865848 | 0.275 | 0.147 | 1 | 10 |
| ecm1b | 0.31757 | 0.358960752 | 0.2 | 0.088 | 1 | 10 |
| CABZ01087560.1 | 0.31809 | -0.10392412 | 0.025 | 0.088 | 1 | 10 |
| socs3a | 0.31809 | -0.35798127 | 0.175 | 0.324 | 1 | 10 |
| eif4g2a | 0.31833 | -0.23535969 | 0.05 | 0.147 | 1 | 10 |
| eif3ha | 0.32125 | 0.343674235 | 0.375 | 0.235 | 1 | 10 |
| gstt1b | 0.32232 | -0.12058854 | 0.25 | 0.353 | 1 | 10 |
| edem1 | 0.32425 | -0.22694107 | 0.075 | 0.176 | 1 | 10 |
| mt-atp6 | 0.32449 | -0.22242515 | 0.375 | 0.529 | 1 | 10 |
| gapdhs | 0.32454 | -0.33530419 | 0.15 | 0.294 | 1 | 10 |
| tomm7 | 0.32893 | 0.30991627 | 0.175 | 0.088 | 1 | 10 |
| syf2 | 0.33203 | -0.22188653 | 0.05 | 0.147 | 1 | 10 |
| ifi45 | 0.33233 | -0.28757313 | 0.475 | 0.588 | 1 | 10 |
| eif4a3 | 0.33249 | -0.32048824 | 0.175 | 0.324 | 1 | 10 |
| uqcrq | 0.33276 | 0.150744759 | 0.175 | 0.176 | 1 | 10 |
| erh | 0.33289 | -0.33637445 | 0.175 | 0.324 | 1 | 10 |
| rpl6 | 0.33453 | 0.135619718 | 0.875 | 0.824 | 1 | 10 |
| tomm40 | 0.3347 | -0.27639865 | 0.025 | 0.088 | 1 | 10 |
| snrpg | 0.33642 | -0.32526109 | 0.2 | 0.353 | 1 | 10 |
| atp5fa1 | 0.33684 | -0.32882676 | 0.225 | 0.382 | 1 | 10 |
| max | 0.33692 | -0.24143875 | 0.05 | 0.147 | 1 | 10 |
| STMP1 | 0.33754 | 0.359122416 | 0.3 | 0.176 | 1 | 10 |
| ppdpfb | 0.33877 | 0.314692404 | 0.575 | 0.412 | 1 | 10 |
| mgst3a | 0.34065 | -0.29808797 | 0.05 | 0.118 | 1 | 10 |
| hnrnpaba | 0.34134 | -0.24423313 | 0.425 | 0.588 | 1 | 10 |
| ptges3a | 0.34187 | -0.32002636 | 0.05 | 0.147 | 1 | 10 |
| ppifb | 0.34221 | -0.30092562 | 0.05 | 0.147 | 1 | 10 |
| ndufb9 | 0.34382 | 0.326429945 | 0.275 | 0.176 | 1 | 10 |
| mt-co2 | 0.34394 | -0.38154246 | 0.375 | 0.529 | 1 | 10 |
| sub1b | 0.34733 | -0.17841755 | 0.2 | 0.324 | 1 | 10 |
| timm13 | 0.34796 | -0.25113981 | 0.05 | 0.147 | 1 | 10 |
| lyrm9 | 0.34839 | -0.24707274 | 0.05 | 0.147 | 1 | 10 |

| | | | | | | |
|---|---|---|---|---|---|---|
| capzb | 0.35038 | 0.296497026 | 0.225 | 0.147 | 1 | 10 |
| si:ch73-281n10.2 | 0.35154 | 0.265871562 | 0.575 | 0.412 | 1 | 10 |
| txn | 0.35188 | 0.284845786 | 0.1 | 0.029 | 1 | 10 |
| pa2g4a | 0.35294 | -0.28006174 | 0.05 | 0.147 | 1 | 10 |
| rab6a | 0.35454 | 0.272002761 | 0.1 | 0.029 | 1 | 10 |
| zc3h15 | 0.35519 | -0.14850079 | 0.025 | 0.088 | 1 | 10 |
| ndufb5 | 0.35558 | 0.327136108 | 0.2 | 0.088 | 1 | 10 |
| chmp6b | 0.35612 | -0.27099728 | 0.05 | 0.147 | 1 | 10 |
| ola1 | 0.35619 | 0.147798406 | 0.275 | 0.176 | 1 | 10 |
| pdia5 | 0.35691 | -0.26677177 | 0.05 | 0.147 | 1 | 10 |
| naa10 | 0.35706 | -0.2577823 | 0.05 | 0.147 | 1 | 10 |
| mrps6 | 0.35763 | -0.29541326 | 0.05 | 0.147 | 1 | 10 |
| polr3k | 0.35993 | -0.2769479 | 0.05 | 0.147 | 1 | 10 |
| si:ch211-284b7.3 | 0.35999 | -0.25123221 | 0.05 | 0.147 | 1 | 10 |
| atrx | 0.36 | -0.26293155 | 0.05 | 0.147 | 1 | 10 |
| fkbp7 | 0.36007 | -0.28850944 | 0.05 | 0.147 | 1 | 10 |
| tuba8l3 | 0.3605 | -0.23390661 | 0.35 | 0.5 | 1 | 10 |
| idh1 | 0.36173 | 0.266787526 | 0.1 | 0.029 | 1 | 10 |
| atp1a1a.1 | 0.36278 | 0.279213259 | 0.1 | 0.029 | 1 | 10 |
| phf5a | 0.36335 | 0.303617994 | 0.15 | 0.059 | 1 | 10 |
| ube2e3 | 0.36371 | 0.254037658 | 0.2 | 0.088 | 1 | 10 |
| calr3a | 0.36608 | -0.33022947 | 0.125 | 0.235 | 1 | 10 |
| prdx6 | 0.36649 | 0.282223594 | 0.2 | 0.088 | 1 | 10 |
| serf2 | 0.36675 | 0.328843111 | 0.325 | 0.206 | 1 | 10 |
| rhoaa | 0.37334 | 0.321255812 | 0.225 | 0.118 | 1 | 10 |
| lsm12b | 0.37386 | 0.169000859 | 0.225 | 0.206 | 1 | 10 |
| magt1 | 0.37426 | 0.276842891 | 0.1 | 0.029 | 1 | 10 |
| txnl4a | 0.37624 | 0.295292168 | 0.15 | 0.059 | 1 | 10 |
| xbp1 | 0.37897 | 0.141538974 | 0.225 | 0.235 | 1 | 10 |
| sub1a | 0.37919 | 0.117021699 | 0.325 | 0.324 | 1 | 10 |
| paip2b | 0.38149 | 0.321836052 | 0.35 | 0.206 | 1 | 10 |
| si:dkey-28b4.7 | 0.38185 | 0.266955297 | 0.275 | 0.147 | 1 | 10 |
| sdhdb | 0.38353 | 0.264565784 | 0.1 | 0.029 | 1 | 10 |
| glo1 | 0.38524 | 0.253194468 | 0.1 | 0.029 | 1 | 10 |
| ap1b1 | 0.38546 | -0.11684405 | 0.05 | 0.118 | 1 | 10 |
| man2c1 | 0.38665 | 0.300797912 | 0.2 | 0.088 | 1 | 10 |
| tmsb | 0.38677 | 0.202242752 | 0.175 | 0.147 | 1 | 10 |
| exosc2 | 0.38889 | -0.23774178 | 0.025 | 0.088 | 1 | 10 |
| serbp1b | 0.38974 | 0.280214354 | 0.15 | 0.059 | 1 | 10 |
| irf2bp2b | 0.39065 | 0.304441832 | 0.2 | 0.088 | 1 | 10 |
| atf1 | 0.39086 | 0.246324734 | 0.1 | 0.029 | 1 | 10 |
| anxa13l | 0.39128 | 0.322213201 | 0.2 | 0.088 | 1 | 10 |
| NC-002333.4 | 0.39137 | -0.31150934 | 0.4 | 0.559 | 1 | 10 |
| COX3 | 0.39225 | -0.3201055 | 0.475 | 0.618 | 1 | 10 |
| si:ch211-117c9.5 | 0.39325 | -0.26210388 | 0.075 | 0.176 | 1 | 10 |
| sptssa | 0.39394 | 0.274001131 | 0.125 | 0.059 | 1 | 10 |
| CABZ01044746.1 | 0.39501 | 0.291807072 | 0.15 | 0.059 | 1 | 10 |
| eif5a2 | 0.39645 | -0.23357115 | 0.325 | 0.471 | 1 | 10 |

| | | | | | | |
|---|---|---|---|---|---|---|
| ost4 | 0.39713 | -0.32868865 | 0.15 | 0.265 | 1 | 10 |
| trir | 0.39786 | 0.178055349 | 0.1 | 0.029 | 1 | 10 |
| sumo3b | 0.40084 | -0.11991652 | 0.025 | 0.088 | 1 | 10 |
| eif3s10 | 0.40117 | -0.26984853 | 0.075 | 0.176 | 1 | 10 |
| si:ch73-376l24.3 | 0.40187 | -0.27200971 | 0.05 | 0.118 | 1 | 10 |
| cebpg | 0.40202 | -0.110699 | 0.025 | 0.088 | 1 | 10 |
| rbfox2 | 0.40234 | -0.231846 | 0.025 | 0.088 | 1 | 10 |
| sf3b5 | 0.40277 | -0.21165029 | 0.175 | 0.294 | 1 | 10 |
| polr2b | 0.40288 | -0.13231035 | 0.025 | 0.088 | 1 | 10 |
| pop5 | 0.40369 | 0.177923596 | 0.1 | 0.029 | 1 | 10 |
| mrpl33 | 0.40375 | -0.10656354 | 0.05 | 0.118 | 1 | 10 |
| nudc | 0.40609 | -0.28573255 | 0.1 | 0.176 | 1 | 10 |
| ndufb2 | 0.40611 | 0.227201767 | 0.15 | 0.059 | 1 | 10 |
| ablim1b | 0.40639 | -0.25682794 | 0.025 | 0.088 | 1 | 10 |
| emc7 | 0.40743 | -0.11760076 | 0.025 | 0.088 | 1 | 10 |
| COX5B | 0.40796 | 0.223881202 | 0.45 | 0.382 | 1 | 10 |
| tmem59 | 0.40834 | 0.291404038 | 0.15 | 0.059 | 1 | 10 |
| mrpl28 | 0.4084 | -0.25512583 | 0.025 | 0.088 | 1 | 10 |
| mrps11 | 0.40928 | -0.2839796 | 0.075 | 0.176 | 1 | 10 |
| npc2 | 0.40967 | -0.28068024 | 0.075 | 0.176 | 1 | 10 |
| tmem50a | 0.41014 | 0.233722106 | 0.1 | 0.029 | 1 | 10 |
| ndufs7 | 0.41087 | -0.24382065 | 0.075 | 0.176 | 1 | 10 |
| seta | 0.41113 | -0.33254482 | 0.175 | 0.294 | 1 | 10 |
| atp6v1ba | 0.41132 | -0.22465243 | 0.025 | 0.088 | 1 | 10 |
| spred2b | 0.41175 | -0.29804345 | 0.075 | 0.176 | 1 | 10 |
| acadm | 0.41194 | -0.27282986 | 0.075 | 0.176 | 1 | 10 |
| chmp4bb | 0.41213 | -0.23956186 | 0.075 | 0.176 | 1 | 10 |
| atp6v0ca | 0.41255 | -0.29665746 | 0.075 | 0.176 | 1 | 10 |
| psmb2 | 0.41268 | -0.32913006 | 0.125 | 0.235 | 1 | 10 |
| rpl9 | 0.41311 | 0.130298434 | 0.95 | 0.882 | 1 | 10 |
| pgd | 0.4132 | 0.294964002 | 0.15 | 0.059 | 1 | 10 |
| timm8a | 0.4133 | 0.254957017 | 0.1 | 0.029 | 1 | 10 |
| prkar2aa | 0.41373 | -0.24398476 | 0.125 | 0.235 | 1 | 10 |
| dhps | 0.41419 | -0.22777092 | 0.025 | 0.088 | 1 | 10 |
| ranbp3b | 0.41448 | -0.22545184 | 0.025 | 0.088 | 1 | 10 |
| cnep1r1 | 0.41554 | 0.256295525 | 0.1 | 0.029 | 1 | 10 |
| magi1b | 0.41603 | 0.182088388 | 0.1 | 0.029 | 1 | 10 |
| fn1a | 0.41695 | 0.233036901 | 0.125 | 0.088 | 1 | 10 |
| snx5 | 0.41721 | -0.24977076 | 0.025 | 0.088 | 1 | 10 |
| smim7 | 0.41944 | -0.13137843 | 0.025 | 0.088 | 1 | 10 |
| ndufa8 | 0.41987 | -0.31358148 | 0.1 | 0.206 | 1 | 10 |
| aip | 0.42006 | -0.26769062 | 0.025 | 0.088 | 1 | 10 |
| ptenb | 0.42293 | -0.24336871 | 0.025 | 0.088 | 1 | 10 |
| rpl22l1 | 0.42328 | -0.13763956 | 0.575 | 0.706 | 1 | 10 |
| ndp | 0.42598 | -0.21729723 | 0.025 | 0.088 | 1 | 10 |
| aldh6a1 | 0.42608 | -0.25633077 | 0.025 | 0.088 | 1 | 10 |
| rab1bb | 0.42736 | -0.10874669 | 0.025 | 0.088 | 1 | 10 |
| pfn2 | 0.42847 | 0.286363852 | 0.525 | 0.382 | 1 | 10 |

| | | | | | | |
|---|---|---|---|---|---|---|
| bckdhb | 0.42891 | -0.16359347 | 0.025 | 0.088 | 1 | 10 |
| rab11ba | 0.42899 | 0.233157942 | 0.225 | 0.118 | 1 | 10 |
| ppp3r1b | 0.43004 | 0.277702309 | 0.15 | 0.059 | 1 | 10 |
| bms1 | 0.43021 | -0.12691286 | 0.025 | 0.088 | 1 | 10 |
| mt-nd3 | 0.43109 | -0.23905937 | 0.2 | 0.324 | 1 | 10 |
| zgc:92140 | 0.43145 | 0.230255033 | 0.15 | 0.059 | 1 | 10 |
| metap1 | 0.43156 | -0.29042212 | 0.075 | 0.147 | 1 | 10 |
| ppiab | 0.43212 | -0.11298228 | 0.75 | 0.765 | 1 | 10 |
| rdh10a | 0.4327 | 0.339848294 | 0.225 | 0.118 | 1 | 10 |
| si:ch211-213a13.1 | 0.43401 | -0.15751386 | 0.025 | 0.088 | 1 | 10 |
| cmpk | 0.4341 | 0.22705134 | 0.15 | 0.059 | 1 | 10 |
| zgc:65873 | 0.43415 | -0.24826876 | 0.025 | 0.088 | 1 | 10 |
| myg1 | 0.43433 | 0.263352425 | 0.15 | 0.059 | 1 | 10 |
| LOC103909972 | 0.43524 | 0.227520549 | 0.1 | 0.029 | 1 | 10 |
| lsr | 0.43539 | -0.13164321 | 0.05 | 0.118 | 1 | 10 |
| sod1 | 0.43593 | -0.26364396 | 0.325 | 0.471 | 1 | 10 |
| ube2d2 | 0.43595 | -0.22938685 | 0.175 | 0.294 | 1 | 10 |
| cnbpa | 0.43599 | -0.24366636 | 0.125 | 0.235 | 1 | 10 |
| mtx2 | 0.43605 | 0.261693248 | 0.15 | 0.059 | 1 | 10 |
| fbxw2 | 0.43625 | -0.14755497 | 0.025 | 0.088 | 1 | 10 |
| cicb | 0.43746 | -0.24382227 | 0.025 | 0.088 | 1 | 10 |
| wu:fb55g09 | 0.43756 | -0.29686256 | 0.1 | 0.206 | 1 | 10 |
| ppm1g | 0.43768 | 0.240475001 | 0.15 | 0.059 | 1 | 10 |
| ndufs4 | 0.43774 | 0.280379241 | 0.15 | 0.059 | 1 | 10 |
| si:ch211-15d5.12 | 0.43845 | -0.13329374 | 0.025 | 0.088 | 1 | 10 |
| fundc1 | 0.44031 | -0.26995523 | 0.1 | 0.206 | 1 | 10 |
| ppp1r14bb | 0.44041 | 0.193019202 | 0.1 | 0.029 | 1 | 10 |
| ube2d1b | 0.44082 | 0.286544017 | 0.2 | 0.118 | 1 | 10 |
| psmg2 | 0.44153 | -0.14986001 | 0.025 | 0.088 | 1 | 10 |
| actb1 | 0.44198 | -0.23973342 | 0.5 | 0.647 | 1 | 10 |
| uri1 | 0.442 | -0.2051206 | 0.025 | 0.088 | 1 | 10 |
| lyrm7 | 0.44214 | 0.236958413 | 0.075 | 0.029 | 1 | 10 |
| mapre1a | 0.4424 | -0.21319258 | 0.025 | 0.088 | 1 | 10 |
| cbx3a | 0.4425 | -0.23774102 | 0.175 | 0.294 | 1 | 10 |
| pdcd6 | 0.44262 | 0.262661906 | 0.1 | 0.029 | 1 | 10 |
| mut | 0.44268 | -0.15311892 | 0.025 | 0.088 | 1 | 10 |
| yy1a | 0.44334 | -0.27883426 | 0.1 | 0.206 | 1 | 10 |
| psmc6 | 0.44369 | -0.26715589 | 0.1 | 0.206 | 1 | 10 |
| rps26l | 0.44373 | 0.150120238 | 0.9 | 0.824 | 1 | 10 |
| ppp2r1bb | 0.44416 | 0.207571015 | 0.1 | 0.029 | 1 | 10 |
| rpl5a | 0.44431 | 0.206551582 | 0.75 | 0.618 | 1 | 10 |
| edf1 | 0.44456 | 0.253773895 | 0.35 | 0.265 | 1 | 10 |
| uprt | 0.44469 | 0.189573171 | 0.1 | 0.029 | 1 | 10 |
| mrpl34 | 0.44495 | -0.12343961 | 0.075 | 0.147 | 1 | 10 |
| nkap | 0.44496 | 0.227493484 | 0.1 | 0.029 | 1 | 10 |
| ilk | 0.44559 | -0.15491269 | 0.025 | 0.088 | 1 | 10 |
| arhgef4 | 0.44604 | -0.2476532 | 0.025 | 0.088 | 1 | 10 |
| srprb | 0.44644 | 0.222479124 | 0.1 | 0.029 | 1 | 10 |

| | | | | | | |
|---|---|---|---|---|---|---|
| eno1a | 0.44671 | 0.198491925 | 0.1 | 0.029 | 1 | 10 |
| fez1 | 0.44778 | -0.26124571 | 0.125 | 0.235 | 1 | 10 |
| rbbp4 | 0.44781 | -0.12902255 | 0.025 | 0.088 | 1 | 10 |
| ube2g2 | 0.44809 | 0.189209095 | 0.1 | 0.029 | 1 | 10 |
| cbr4 | 0.4481 | 0.221473885 | 0.1 | 0.029 | 1 | 10 |
| lsm4 | 0.44816 | -0.25417567 | 0.125 | 0.235 | 1 | 10 |
| lsm8 | 0.4486 | -0.1303412 | 0.025 | 0.088 | 1 | 10 |
| psma4 | 0.44862 | -0.13253691 | 0.225 | 0.324 | 1 | 10 |
| ccdc25 | 0.44933 | 0.236946875 | 0.1 | 0.029 | 1 | 10 |
| prpf19 | 0.44956 | -0.19674451 | 0.025 | 0.088 | 1 | 10 |
| akt1s1 | 0.45007 | 0.206609841 | 0.1 | 0.029 | 1 | 10 |
| si:dkey-188i13.11 | 0.4504 | -0.27253232 | 0.3 | 0.441 | 1 | 10 |
| actl6a | 0.45115 | 0.210513318 | 0.1 | 0.029 | 1 | 10 |
| btr04 | 0.4518 | -0.15760493 | 0.025 | 0.088 | 1 | 10 |
| hadh | 0.45296 | 0.204710878 | 0.1 | 0.029 | 1 | 10 |
| serinc1 | 0.45324 | 0.248564269 | 0.1 | 0.029 | 1 | 10 |
| kras | 0.45407 | -0.15917363 | 0.025 | 0.088 | 1 | 10 |
| wdr1 | 0.45567 | 0.223820916 | 0.1 | 0.029 | 1 | 10 |
| XLOC-035413 | 0.45623 | 0.238143493 | 0.1 | 0.029 | 1 | 10 |
| b2ml | 0.45691 | -0.27179908 | 0.125 | 0.235 | 1 | 10 |
| ralbp1 | 0.45714 | -0.17221946 | 0.025 | 0.088 | 1 | 10 |
| vkorc1l1 | 0.45762 | -0.14348819 | 0.025 | 0.088 | 1 | 10 |
| dpp3 | 0.45794 | 0.313600203 | 0.25 | 0.147 | 1 | 10 |
| uqcrc2b | 0.45903 | -0.16871392 | 0.025 | 0.088 | 1 | 10 |
| rnf185 | 0.45908 | 0.226163863 | 0.1 | 0.029 | 1 | 10 |
| acaa2 | 0.45921 | -0.15697952 | 0.025 | 0.088 | 1 | 10 |
| mrpl37 | 0.45929 | -0.14791292 | 0.025 | 0.088 | 1 | 10 |
| ccdc97 | 0.45939 | -0.14709058 | 0.025 | 0.088 | 1 | 10 |
| si:ch211-68a17.7 | 0.45964 | -0.2431115 | 0.025 | 0.088 | 1 | 10 |
| si:ch73-109d9.2 | 0.45972 | 0.223070626 | 0.1 | 0.029 | 1 | 10 |
| srsf6b | 0.4612 | -0.18675496 | 0.025 | 0.088 | 1 | 10 |
| fkbp3 | 0.46159 | -0.27442085 | 0.125 | 0.235 | 1 | 10 |
| eif4ebp2 | 0.46188 | 0.243726788 | 0.275 | 0.206 | 1 | 10 |
| rnf7 | 0.46219 | 0.198988527 | 0.175 | 0.088 | 1 | 10 |
| pgpep1 | 0.46225 | -0.21872434 | 0.025 | 0.088 | 1 | 10 |
| krtcap2 | 0.46239 | -0.26028807 | 0.125 | 0.235 | 1 | 10 |
| snx9b | 0.4624 | 0.237677799 | 0.1 | 0.029 | 1 | 10 |
| zgc:171480 | 0.46241 | -0.16479053 | 0.025 | 0.088 | 1 | 10 |
| suclg1 | 0.46245 | -0.1072844 | 0.075 | 0.147 | 1 | 10 |
| arfgap2 | 0.46245 | 0.203821172 | 0.1 | 0.029 | 1 | 10 |
| mcts1 | 0.46255 | -0.26117273 | 0.125 | 0.235 | 1 | 10 |
| si:ch211-133n4.4 | 0.46258 | -0.2017691 | 0.025 | 0.088 | 1 | 10 |
| fam168a | 0.46263 | 0.250782829 | 0.15 | 0.088 | 1 | 10 |
| si:ch211-39i22.1 | 0.46277 | 0.234561809 | 0.1 | 0.029 | 1 | 10 |
| cd74b | 0.46288 | -0.2867015 | 0.125 | 0.235 | 1 | 10 |
| srsf9 | 0.46313 | -0.20693937 | 0.025 | 0.088 | 1 | 10 |
| lasp1 | 0.46319 | -0.25827517 | 0.125 | 0.235 | 1 | 10 |
| syncrip | 0.46321 | -0.28922902 | 0.125 | 0.235 | 1 | 10 |

| | | | | | | |
|---|---|---|---|---|---|---|
| bxdc2 | 0.46357 | -0.1728134 | 0.025 | 0.088 | 1 | 10 |
| zgc:77849 | 0.46373 | 0.237984102 | 0.1 | 0.029 | 1 | 10 |
| atp5if1b | 0.46374 | -0.23875518 | 0.15 | 0.265 | 1 | 10 |
| ube2d1a | 0.46439 | 0.220364983 | 0.1 | 0.029 | 1 | 10 |
| rer1 | 0.46467 | 0.217367196 | 0.1 | 0.029 | 1 | 10 |
| si:dkey-46a10.3 | 0.46472 | 0.191000165 | 0.1 | 0.029 | 1 | 10 |
| sh3pxd2b | 0.46476 | -0.16884406 | 0.025 | 0.088 | 1 | 10 |
| hif1ab | 0.46478 | 0.237610348 | 0.1 | 0.029 | 1 | 10 |
| gspt1l | 0.46484 | -0.17145417 | 0.025 | 0.088 | 1 | 10 |
| cwc27 | 0.46485 | -0.22662018 | 0.025 | 0.088 | 1 | 10 |
| sinhcafl | 0.46547 | -0.18196859 | 0.025 | 0.088 | 1 | 10 |
| nol10 | 0.46562 | -0.20341899 | 0.025 | 0.088 | 1 | 10 |
| lamc1 | 0.46611 | -0.16929032 | 0.025 | 0.088 | 1 | 10 |
| UBL4A | 0.4664 | -0.17595082 | 0.025 | 0.088 | 1 | 10 |
| swap70b | 0.46689 | -0.12610207 | 0.025 | 0.088 | 1 | 10 |
| caprin1a | 0.46698 | -0.16300864 | 0.025 | 0.088 | 1 | 10 |
| snrnp40 | 0.46788 | -0.1538769 | 0.025 | 0.088 | 1 | 10 |
| psma8 | 0.46857 | -0.25349298 | 0.2 | 0.324 | 1 | 10 |
| tango2 | 0.46905 | -0.15576166 | 0.025 | 0.088 | 1 | 10 |
| SMIM10 | 0.46936 | -0.1827237 | 0.025 | 0.088 | 1 | 10 |
| bola1 | 0.46966 | -0.13016252 | 0.025 | 0.059 | 1 | 10 |
| btc | 0.46966 | -0.12036801 | 0.025 | 0.059 | 1 | 10 |
| kansl1b | 0.46966 | -0.10072661 | 0.025 | 0.059 | 1 | 10 |
| svild | 0.46966 | -0.14680566 | 0.025 | 0.059 | 1 | 10 |
| orc4 | 0.46966 | -0.1353581 | 0.025 | 0.059 | 1 | 10 |
| srek1ip1 | 0.46966 | -0.14496762 | 0.025 | 0.059 | 1 | 10 |
| ches1 | 0.46966 | -0.14320627 | 0.025 | 0.059 | 1 | 10 |
| hnrnpc | 0.46966 | -0.10132418 | 0.025 | 0.059 | 1 | 10 |
| med29 | 0.46966 | -0.10061355 | 0.025 | 0.059 | 1 | 10 |
| ndrg1a | 0.46966 | -0.13493572 | 0.025 | 0.059 | 1 | 10 |
| phf6 | 0.46966 | -0.11100966 | 0.025 | 0.059 | 1 | 10 |
| rbm12b | 0.46966 | -0.10349511 | 0.025 | 0.059 | 1 | 10 |
| scaf11 | 0.46966 | -0.11138422 | 0.025 | 0.059 | 1 | 10 |
| snw1 | 0.46966 | -0.10893553 | 0.025 | 0.059 | 1 | 10 |
| CU929259.1 | 0.46966 | -0.15282408 | 0.025 | 0.059 | 1 | 10 |
| adgrl2a | 0.46966 | -0.13233206 | 0.025 | 0.059 | 1 | 10 |
| aff2 | 0.46966 | -0.12935499 | 0.025 | 0.059 | 1 | 10 |
| aldh7a1 | 0.46966 | -0.12267734 | 0.025 | 0.059 | 1 | 10 |
| ap3s2 | 0.46966 | -0.13447742 | 0.025 | 0.059 | 1 | 10 |
| bud23 | 0.46966 | -0.17164288 | 0.025 | 0.059 | 1 | 10 |
| cbr1 | 0.46966 | -0.13490978 | 0.025 | 0.059 | 1 | 10 |
| cdca7a | 0.46966 | -0.16358785 | 0.025 | 0.059 | 1 | 10 |
| chaf1a | 0.46966 | -0.13725595 | 0.025 | 0.059 | 1 | 10 |
| col18a1a | 0.46966 | -0.10613482 | 0.025 | 0.059 | 1 | 10 |
| copb2 | 0.46966 | -0.12744276 | 0.025 | 0.059 | 1 | 10 |
| crybg1a | 0.46966 | -0.15857384 | 0.025 | 0.059 | 1 | 10 |
| ets2 | 0.46966 | -0.11959262 | 0.025 | 0.059 | 1 | 10 |
| fam207a | 0.46966 | -0.10460162 | 0.025 | 0.059 | 1 | 10 |

| | | | | | | |
|---|---|---|---|---|---|---|
| foxp1a | 0.46966 | -0.12878928 | 0.025 | 0.059 | 1 | 10 |
| ftr80 | 0.46966 | -0.12617565 | 0.025 | 0.059 | 1 | 10 |
| her9 | 0.46966 | -0.10236267 | 0.025 | 0.059 | 1 | 10 |
| homer3b | 0.46966 | -0.12826551 | 0.025 | 0.059 | 1 | 10 |
| leng1 | 0.46966 | -0.10335947 | 0.025 | 0.059 | 1 | 10 |
| lysmd2 | 0.46966 | -0.10574039 | 0.025 | 0.059 | 1 | 10 |
| mccc2 | 0.46966 | -0.13871359 | 0.025 | 0.059 | 1 | 10 |
| ncoa3 | 0.46966 | -0.12815411 | 0.025 | 0.059 | 1 | 10 |
| net1 | 0.46966 | -0.14136037 | 0.025 | 0.059 | 1 | 10 |
| rabgap1l | 0.46966 | -0.10793647 | 0.025 | 0.059 | 1 | 10 |
| rpp25l | 0.46966 | -0.12961911 | 0.025 | 0.059 | 1 | 10 |
| selenot1b | 0.46966 | -0.1791206 | 0.025 | 0.059 | 1 | 10 |
| sf3b2 | 0.46966 | -0.10163034 | 0.025 | 0.059 | 1 | 10 |
| si:ch211-264f5.6 | 0.46966 | -0.12358171 | 0.025 | 0.059 | 1 | 10 |
| si:ch73-375g18.1 | 0.46966 | -0.13870665 | 0.025 | 0.059 | 1 | 10 |
| si:ch73-71c20.5 | 0.46966 | -0.14967178 | 0.025 | 0.059 | 1 | 10 |
| smek1 | 0.46966 | -0.11346234 | 0.025 | 0.059 | 1 | 10 |
| snd1 | 0.46966 | -0.12057371 | 0.025 | 0.059 | 1 | 10 |
| srrt | 0.46966 | -0.11917965 | 0.025 | 0.059 | 1 | 10 |
| syvn1 | 0.46966 | -0.10675381 | 0.025 | 0.059 | 1 | 10 |
| tbpl1 | 0.46966 | -0.14804945 | 0.025 | 0.059 | 1 | 10 |
| tdh | 0.46966 | -0.12605003 | 0.025 | 0.059 | 1 | 10 |
| usp5 | 0.46966 | -0.12756702 | 0.025 | 0.059 | 1 | 10 |
| waplb | 0.46966 | -0.1325413 | 0.025 | 0.059 | 1 | 10 |
| zgc:103697 | 0.46966 | -0.12835579 | 0.025 | 0.059 | 1 | 10 |
| aclya | 0.46966 | -0.11842829 | 0.025 | 0.059 | 1 | 10 |
| chmp3 | 0.46966 | -0.14945995 | 0.025 | 0.059 | 1 | 10 |
| fnbp4 | 0.46966 | -0.11927741 | 0.025 | 0.059 | 1 | 10 |
| gclc | 0.46966 | -0.15255994 | 0.025 | 0.059 | 1 | 10 |
| gtf2h3 | 0.46966 | -0.10690822 | 0.025 | 0.059 | 1 | 10 |
| hs2st1b | 0.46966 | -0.14215094 | 0.025 | 0.059 | 1 | 10 |
| mbd2 | 0.46966 | -0.12938676 | 0.025 | 0.059 | 1 | 10 |
| naa15a | 0.46966 | -0.15980408 | 0.025 | 0.059 | 1 | 10 |
| nup93 | 0.46966 | -0.14372301 | 0.025 | 0.059 | 1 | 10 |
| rab1ba | 0.46966 | -0.16933859 | 0.025 | 0.059 | 1 | 10 |
| rfc4 | 0.46966 | -0.13766754 | 0.025 | 0.059 | 1 | 10 |
| rtn4b | 0.46966 | -0.10113946 | 0.025 | 0.059 | 1 | 10 |
| seh1l | 0.46966 | -0.11846957 | 0.025 | 0.059 | 1 | 10 |
| si:dkey-177p2.6 | 0.46966 | -0.13308681 | 0.025 | 0.059 | 1 | 10 |
| tbcb | 0.46966 | -0.12731287 | 0.025 | 0.059 | 1 | 10 |
| tfam | 0.46966 | -0.13780189 | 0.025 | 0.059 | 1 | 10 |
| uba2 | 0.46966 | -0.13085218 | 0.025 | 0.059 | 1 | 10 |
| zbtb10 | 0.46966 | -0.12339118 | 0.025 | 0.059 | 1 | 10 |
| acbd6 | 0.46966 | -0.11505571 | 0.025 | 0.059 | 1 | 10 |
| birc5a | 0.46966 | -0.11773027 | 0.025 | 0.059 | 1 | 10 |
| ccnd2b | 0.46966 | -0.14450153 | 0.025 | 0.059 | 1 | 10 |
| celf2 | 0.46966 | -0.10794835 | 0.025 | 0.059 | 1 | 10 |
| crk | 0.46966 | -0.10516365 | 0.025 | 0.059 | 1 | 10 |

| | | | | | | |
|---|---|---|---|---|---|---|
| cstf2 | 0.46966 | -0.12528486 | 0.025 | 0.059 | 1 | 10 |
| dnpep | 0.46966 | -0.10395502 | 0.025 | 0.059 | 1 | 10 |
| gid8a | 0.46966 | -0.10014228 | 0.025 | 0.059 | 1 | 10 |
| gtf2f1 | 0.46966 | -0.14348886 | 0.025 | 0.059 | 1 | 10 |
| lamb1a | 0.46966 | -0.13863906 | 0.025 | 0.059 | 1 | 10 |
| mettl1 | 0.46966 | -0.1020959 | 0.025 | 0.059 | 1 | 10 |
| mrto4 | 0.46966 | -0.12122897 | 0.025 | 0.059 | 1 | 10 |
| pbk | 0.46966 | -0.12318918 | 0.025 | 0.059 | 1 | 10 |
| rbpms2b | 0.46966 | -0.10526935 | 0.025 | 0.059 | 1 | 10 |
| rpe | 0.46966 | -0.11353486 | 0.025 | 0.059 | 1 | 10 |
| rtf2 | 0.46966 | -0.13389652 | 0.025 | 0.059 | 1 | 10 |
| sec24d | 0.46966 | -0.10541731 | 0.025 | 0.059 | 1 | 10 |
| si:ch211-241b2.1 | 0.46966 | -0.1232222 | 0.025 | 0.059 | 1 | 10 |
| si:ch73-111k22.2 | 0.46966 | -0.13253334 | 0.025 | 0.059 | 1 | 10 |
| slc2a1b | 0.46966 | -0.11807826 | 0.025 | 0.059 | 1 | 10 |
| slc30a1a | 0.46966 | -0.11636712 | 0.025 | 0.059 | 1 | 10 |
| smim14 | 0.46966 | -0.11845499 | 0.025 | 0.059 | 1 | 10 |
| snrnp25 | 0.46966 | -0.12420503 | 0.025 | 0.059 | 1 | 10 |
| tfg | 0.46966 | -0.10899322 | 0.025 | 0.059 | 1 | 10 |
| tnfaip6 | 0.46966 | -0.10488135 | 0.025 | 0.059 | 1 | 10 |
| trip12 | 0.46966 | -0.1049683 | 0.025 | 0.059 | 1 | 10 |
| CABZ01092156.1 | 0.46966 | -0.10345609 | 0.025 | 0.059 | 1 | 10 |
| XLOC-018328 | 0.46966 | -0.10004966 | 0.025 | 0.059 | 1 | 10 |
| adsl | 0.46966 | -0.14037962 | 0.025 | 0.059 | 1 | 10 |
| ap2b1 | 0.46966 | -0.10747801 | 0.025 | 0.059 | 1 | 10 |
| cd2bp2 | 0.46966 | -0.16143427 | 0.025 | 0.059 | 1 | 10 |
| cd82b | 0.46966 | -0.11227731 | 0.025 | 0.059 | 1 | 10 |
| chchd1 | 0.46966 | -0.14234035 | 0.025 | 0.059 | 1 | 10 |
| chd2 | 0.46966 | -0.12842519 | 0.025 | 0.059 | 1 | 10 |
| cib1 | 0.46966 | -0.15351077 | 0.025 | 0.059 | 1 | 10 |
| copg2 | 0.46966 | -0.11391502 | 0.025 | 0.059 | 1 | 10 |
| dmac2l | 0.46966 | -0.12518071 | 0.025 | 0.059 | 1 | 10 |
| dnajc1 | 0.46966 | -0.10370559 | 0.025 | 0.059 | 1 | 10 |
| dvl3a | 0.46966 | -0.11518098 | 0.025 | 0.059 | 1 | 10 |
| eri1 | 0.46966 | -0.11136948 | 0.025 | 0.059 | 1 | 10 |
| errfi1a | 0.46966 | -0.12304351 | 0.025 | 0.059 | 1 | 10 |
| fdx2 | 0.46966 | -0.14146151 | 0.025 | 0.059 | 1 | 10 |
| foxo1a | 0.46966 | -0.14840563 | 0.025 | 0.059 | 1 | 10 |
| gtf2b | 0.46966 | -0.10633122 | 0.025 | 0.059 | 1 | 10 |
| ik | 0.46966 | -0.11577244 | 0.025 | 0.059 | 1 | 10 |
| mfap2 | 0.46966 | -0.14554875 | 0.025 | 0.059 | 1 | 10 |
| mrps35 | 0.46966 | -0.10019286 | 0.025 | 0.059 | 1 | 10 |
| nat15 | 0.46966 | -0.12225383 | 0.025 | 0.059 | 1 | 10 |
| patj | 0.46966 | -0.15647638 | 0.025 | 0.059 | 1 | 10 |
| pdcl | 0.46966 | -0.12643199 | 0.025 | 0.059 | 1 | 10 |
| plgrkt | 0.46966 | -0.11830172 | 0.025 | 0.059 | 1 | 10 |
| ppp1r37 | 0.46966 | -0.14671401 | 0.025 | 0.059 | 1 | 10 |
| pudp | 0.46966 | -0.1244696 | 0.025 | 0.059 | 1 | 10 |

| | | | | | | |
|---|---|---|---|---|---|---|
| rbm4.2 | 0.46966 | -0.11679745 | 0.025 | 0.059 | 1 | 10 |
| rpf2 | 0.46966 | -0.16067823 | 0.025 | 0.059 | 1 | 10 |
| si:ch211-156j16.1 | 0.46966 | -0.12371657 | 0.025 | 0.059 | 1 | 10 |
| si:ch211-22i13.2 | 0.46966 | -0.12539259 | 0.025 | 0.059 | 1 | 10 |
| si:ch73-61d6.3 | 0.46966 | -0.12478591 | 0.025 | 0.059 | 1 | 10 |
| srsf1b | 0.46966 | -0.14733359 | 0.025 | 0.059 | 1 | 10 |
| stk11 | 0.46966 | -0.11238668 | 0.025 | 0.059 | 1 | 10 |
| thtpa | 0.46966 | -0.12353156 | 0.025 | 0.059 | 1 | 10 |
| trappc3 | 0.46966 | -0.11708885 | 0.025 | 0.059 | 1 | 10 |
| ubr7 | 0.46966 | -0.14026195 | 0.025 | 0.059 | 1 | 10 |
| zgc:162200 | 0.46966 | -0.13081608 | 0.025 | 0.059 | 1 | 10 |
| znf536 | 0.46966 | -0.16918835 | 0.025 | 0.059 | 1 | 10 |
| znf740b | 0.46966 | -0.11355583 | 0.025 | 0.059 | 1 | 10 |
| snrnp70 | 0.46966 | -0.12383205 | 0.025 | 0.059 | 1 | 10 |
| trappc1 | 0.46972 | 0.297814409 | 0.275 | 0.176 | 1 | 10 |
| nudcd3 | 0.47038 | -0.22670974 | 0.025 | 0.088 | 1 | 10 |
| mov10a | 0.47135 | -0.22144683 | 0.025 | 0.088 | 1 | 10 |
| sh3bgrl | 0.47205 | -0.18472418 | 0.025 | 0.088 | 1 | 10 |
| inha | 0.47251 | -0.18111397 | 0.025 | 0.088 | 1 | 10 |
| col1a1b | 0.47286 | 0.133679032 | 0.125 | 0.059 | 1 | 10 |
| eif4bb | 0.47326 | -0.22794295 | 0.025 | 0.088 | 1 | 10 |
| sfxn4 | 0.47326 | -0.19480264 | 0.025 | 0.088 | 1 | 10 |
| wdr3 | 0.47347 | -0.18676063 | 0.025 | 0.088 | 1 | 10 |
| ssr1 | 0.47422 | -0.11567474 | 0.1 | 0.176 | 1 | 10 |
| rwdd1 | 0.4744 | -0.28946652 | 0.15 | 0.265 | 1 | 10 |
| arl8a | 0.47459 | -0.22236829 | 0.025 | 0.088 | 1 | 10 |
| dnajb1b | 0.47495 | -0.12636683 | 0.025 | 0.088 | 1 | 10 |
| glula | 0.47505 | 0.212636645 | 0.125 | 0.088 | 1 | 10 |
| LOC103909284 | 0.47567 | -0.17262721 | 0.025 | 0.088 | 1 | 10 |
| s100u | 0.47616 | -0.22465775 | 0.025 | 0.088 | 1 | 10 |
| dynll1 | 0.47665 | -0.27863222 | 0.225 | 0.353 | 1 | 10 |
| puf60a | 0.47674 | -0.22405524 | 0.025 | 0.088 | 1 | 10 |
| sap18 | 0.47689 | 0.279663935 | 0.2 | 0.118 | 1 | 10 |
| mphosph6 | 0.47703 | -0.21536718 | 0.025 | 0.088 | 1 | 10 |
| csnk1a1 | 0.4771 | 0.286279297 | 0.275 | 0.176 | 1 | 10 |
| sdc4 | 0.47726 | 0.138685274 | 0.125 | 0.118 | 1 | 10 |
| gps2 | 0.47727 | -0.16826775 | 0.025 | 0.088 | 1 | 10 |
| crema | 0.47801 | -0.13221242 | 0.125 | 0.206 | 1 | 10 |
| eif3g | 0.47846 | -0.28343378 | 0.175 | 0.294 | 1 | 10 |
| manf | 0.47847 | -0.19616109 | 0.025 | 0.088 | 1 | 10 |
| htra1b | 0.47902 | -0.15504999 | 0.025 | 0.088 | 1 | 10 |
| zgc:101851 | 0.47926 | -0.23170667 | 0.025 | 0.088 | 1 | 10 |
| h3f3a | 0.47927 | -0.2924132 | 0.2 | 0.324 | 1 | 10 |
| cope | 0.47929 | -0.27256636 | 0.175 | 0.294 | 1 | 10 |
| krt15 | 0.48029 | 0.20418474 | 0.1 | 0.088 | 1 | 10 |
| cltca | 0.48043 | -0.20513111 | 0.025 | 0.088 | 1 | 10 |
| plxna4 | 0.48051 | -0.17119148 | 0.025 | 0.088 | 1 | 10 |
| alad | 0.48095 | -0.17449215 | 0.025 | 0.088 | 1 | 10 |

| | | | | | | |
|---|---|---|---|---|---|---|
| prpf40a | 0.48133 | -0.18413043 | 0.025 | 0.088 | 1 | 10 |
| hgh1 | 0.48155 | -0.19252226 | 0.025 | 0.088 | 1 | 10 |
| smc3 | 0.48185 | -0.19421743 | 0.025 | 0.088 | 1 | 10 |
| dhx15 | 0.48191 | -0.20134869 | 0.025 | 0.088 | 1 | 10 |
| cdkn1a | 0.48194 | -0.23763979 | 0.025 | 0.088 | 1 | 10 |
| pafah1b1b | 0.48208 | -0.14115861 | 0.025 | 0.088 | 1 | 10 |
| reep5 | 0.48221 | 0.291982476 | 0.175 | 0.088 | 1 | 10 |
| atf7ip | 0.48242 | -0.2343549 | 0.025 | 0.088 | 1 | 10 |
| asph | 0.48358 | -0.22612922 | 0.025 | 0.088 | 1 | 10 |
| fblim1 | 0.48363 | -0.19122469 | 0.025 | 0.088 | 1 | 10 |
| tubb2b | 0.48371 | -0.19156125 | 0.2 | 0.235 | 1 | 10 |
| si:dkey-250d21.1 | 0.48383 | -0.18772432 | 0.025 | 0.088 | 1 | 10 |
| pim1 | 0.48421 | -0.16133063 | 0.05 | 0.118 | 1 | 10 |
| LOC108190056 | 0.48446 | -0.20555564 | 0.025 | 0.088 | 1 | 10 |
| suclg2 | 0.48491 | -0.18104044 | 0.025 | 0.088 | 1 | 10 |
| pwp1 | 0.48491 | -0.23338335 | 0.025 | 0.088 | 1 | 10 |
| eif1b | 0.48516 | 0.197401578 | 0.45 | 0.382 | 1 | 10 |
| tprb | 0.48529 | -0.18489046 | 0.025 | 0.088 | 1 | 10 |
| prrc2a | 0.4855 | -0.1898875 | 0.025 | 0.088 | 1 | 10 |
| med22 | 0.48606 | -0.18655994 | 0.025 | 0.088 | 1 | 10 |
| corin | 0.48626 | -0.23489466 | 0.025 | 0.088 | 1 | 10 |
| polr1e | 0.4863 | -0.17677387 | 0.025 | 0.088 | 1 | 10 |
| rabl3 | 0.48638 | -0.20206641 | 0.025 | 0.088 | 1 | 10 |
| nupr1a | 0.48641 | -0.15826123 | 0.05 | 0.118 | 1 | 10 |
| got2b | 0.48658 | -0.19822813 | 0.025 | 0.088 | 1 | 10 |
| cfap36 | 0.48699 | -0.19253054 | 0.025 | 0.088 | 1 | 10 |
| si:ch211-85n16.3 | 0.48704 | -0.19127237 | 0.025 | 0.088 | 1 | 10 |
| msi2a | 0.48705 | -0.21228454 | 0.025 | 0.088 | 1 | 10 |
| pnn | 0.48732 | -0.21674507 | 0.025 | 0.088 | 1 | 10 |
| atic | 0.48736 | -0.16596751 | 0.025 | 0.088 | 1 | 10 |
| tab1 | 0.48749 | -0.19079194 | 0.025 | 0.088 | 1 | 10 |
| pgp | 0.48754 | -0.2099837 | 0.025 | 0.088 | 1 | 10 |
| birc2 | 0.48757 | -0.17231063 | 0.025 | 0.088 | 1 | 10 |
| cyp19a1a | 0.4876 | 0.216082615 | 0.075 | 0.029 | 1 | 10 |
| wls | 0.48786 | -0.23648729 | 0.025 | 0.088 | 1 | 10 |
| sars | 0.4882 | -0.19894797 | 0.025 | 0.088 | 1 | 10 |
| metrn | 0.48823 | -0.22125564 | 0.025 | 0.088 | 1 | 10 |
| parvaa | 0.48831 | -0.16674512 | 0.025 | 0.088 | 1 | 10 |
| pdap1a | 0.48832 | -0.16322821 | 0.025 | 0.088 | 1 | 10 |
| col4a1 | 0.48835 | -0.2516881 | 0.025 | 0.088 | 1 | 10 |
| tspo | 0.4884 | -0.21300942 | 0.025 | 0.088 | 1 | 10 |
| ap3d1 | 0.48843 | -0.23064398 | 0.025 | 0.088 | 1 | 10 |
| hm13 | 0.48843 | -0.21801067 | 0.025 | 0.088 | 1 | 10 |
| sirt7 | 0.48843 | -0.21371459 | 0.025 | 0.088 | 1 | 10 |
| zfyve21 | 0.48843 | -0.17785329 | 0.025 | 0.088 | 1 | 10 |
| pgam1a | 0.48843 | -0.17035219 | 0.025 | 0.088 | 1 | 10 |
| map1lc3b | 0.49033 | 0.134488074 | 0.125 | 0.118 | 1 | 10 |
| si:ch73-42p12.2 | 0.49178 | -0.12976231 | 0.05 | 0.118 | 1 | 10 |

| | | | | | | |
|---|---|---|---|---|---|---|
| bysl | 0.49358 | -0.23898353 | 0.075 | 0.147 | 1 | 10 |
| p4hb | 0.49365 | -0.16210879 | 0.075 | 0.147 | 1 | 10 |
| baz1a | 0.4941 | -0.26016203 | 0.05 | 0.118 | 1 | 10 |
| srsf5a | 0.49696 | 0.159301213 | 0.3 | 0.206 | 1 | 10 |
| si:ch211-212k18.7 | 0.49858 | 0.255843864 | 0.125 | 0.059 | 1 | 10 |
| tcea1 | 0.50187 | -0.26670464 | 0.1 | 0.176 | 1 | 10 |
| ap2m1a | 0.50497 | 0.102092939 | 0.45 | 0.441 | 1 | 10 |
| nup50 | 0.50581 | -0.14474811 | 0.05 | 0.118 | 1 | 10 |
| prkab1b | 0.50699 | 0.203052503 | 0.225 | 0.176 | 1 | 10 |
| bccip | 0.50784 | -0.26248369 | 0.075 | 0.147 | 1 | 10 |
| cct2 | 0.50877 | -0.27925446 | 0.225 | 0.324 | 1 | 10 |
| ppp2cb | 0.5096 | 0.241620221 | 0.25 | 0.176 | 1 | 10 |
| psme1 | 0.51213 | 0.262994182 | 0.325 | 0.206 | 1 | 10 |
| vdac3 | 0.51221 | 0.277305602 | 0.25 | 0.147 | 1 | 10 |
| cd81a | 0.51221 | -0.28490953 | 0.1 | 0.176 | 1 | 10 |
| snrpf | 0.51349 | -0.10026647 | 0.3 | 0.382 | 1 | 10 |
| zgc:100829 | 0.51429 | 0.27605896 | 0.175 | 0.088 | 1 | 10 |
| knop1 | 0.51579 | -0.19332329 | 0.05 | 0.118 | 1 | 10 |
| wwtr1 | 0.52027 | -0.17310515 | 0.05 | 0.118 | 1 | 10 |
| msna | 0.5206 | 0.26817547 | 0.175 | 0.088 | 1 | 10 |
| pfkfb4b | 0.5219 | -0.22979358 | 0.175 | 0.235 | 1 | 10 |
| mrps21 | 0.52283 | -0.17597758 | 0.05 | 0.118 | 1 | 10 |
| mrpl14 | 0.5233 | -0.15066483 | 0.075 | 0.147 | 1 | 10 |
| tomm20b | 0.52341 | -0.2603133 | 0.125 | 0.206 | 1 | 10 |
| lsm7 | 0.52359 | 0.26431278 | 0.225 | 0.147 | 1 | 10 |
| nid1b | 0.52655 | -0.25146194 | 0.05 | 0.118 | 1 | 10 |
| gstm.2 | 0.52715 | 0.162673543 | 0.075 | 0.059 | 1 | 10 |
| si:ch211-196l7.4 | 0.52728 | 0.139497933 | 0.125 | 0.059 | 1 | 10 |
| romo1 | 0.52735 | 0.240887207 | 0.425 | 0.324 | 1 | 10 |
| mapkapk2a | 0.52781 | 0.25710666 | 0.125 | 0.059 | 1 | 10 |
| fgfr3 | 0.52864 | 0.151647435 | 0.125 | 0.059 | 1 | 10 |
| si:dkey-26c10.5 | 0.52902 | -0.27330779 | 0.075 | 0.147 | 1 | 10 |
| aldh2.2 | 0.52988 | -0.15172144 | 0.05 | 0.118 | 1 | 10 |
| rhoca | 0.52995 | 0.241127964 | 0.125 | 0.059 | 1 | 10 |
| bsg | 0.53032 | 0.275273172 | 0.175 | 0.088 | 1 | 10 |
| sumo2b | 0.53079 | 0.194428102 | 0.35 | 0.294 | 1 | 10 |
| csnk1g2b | 0.53141 | -0.19971114 | 0.05 | 0.088 | 1 | 10 |
| adh8b | 0.53215 | 0.206285936 | 0.075 | 0.029 | 1 | 10 |
| ttll12 | 0.53716 | 0.273042961 | 0.15 | 0.088 | 1 | 10 |
| g3bp1 | 0.53719 | -0.12840171 | 0.05 | 0.118 | 1 | 10 |
| cox7c | 0.53792 | -0.226575 | 0.4 | 0.529 | 1 | 10 |
| gar1 | 0.53813 | -0.1578484 | 0.05 | 0.118 | 1 | 10 |
| jpt2 | 0.53813 | 0.23107163 | 0.175 | 0.088 | 1 | 10 |
| rpl38 | 0.53856 | -0.10032326 | 0.875 | 0.912 | 1 | 10 |
| apex1 | 0.53881 | -0.23593798 | 0.05 | 0.118 | 1 | 10 |
| dnajc8 | 0.53939 | -0.22653758 | 0.05 | 0.118 | 1 | 10 |
| si:ch211-1a19.3 | 0.53939 | -0.17276422 | 0.05 | 0.118 | 1 | 10 |
| arrdc3b | 0.5412 | -0.14530707 | 0.05 | 0.118 | 1 | 10 |

| | | | | | | |
|---|---|---|---|---|---|---|
| tubb4b | 0.54154 | -0.24936514 | 0.375 | 0.5 | 1 | 10 |
| anxa2a | 0.54199 | 0.295241459 | 0.15 | 0.088 | 1 | 10 |
| pkma | 0.54211 | -0.22410898 | 0.05 | 0.118 | 1 | 10 |
| gstr | 0.54307 | 0.228817599 | 0.175 | 0.088 | 1 | 10 |
| ptgr1 | 0.54502 | 0.231755387 | 0.175 | 0.088 | 1 | 10 |
| pagr1 | 0.54784 | 0.142057119 | 0.125 | 0.059 | 1 | 10 |
| cd3eap | 0.54893 | -0.21913123 | 0.05 | 0.118 | 1 | 10 |
| ppt1 | 0.54912 | 0.255535255 | 0.125 | 0.059 | 1 | 10 |
| cox5aa | 0.55079 | -0.23111135 | 0.375 | 0.5 | 1 | 10 |
| bambia | 0.55485 | 0.267264102 | 0.2 | 0.118 | 1 | 10 |
| banf1 | 0.55541 | -0.25406506 | 0.1 | 0.176 | 1 | 10 |
| jpt1b | 0.55655 | 0.257415948 | 0.35 | 0.235 | 1 | 10 |
| arpp19b | 0.55674 | -0.20430586 | 0.05 | 0.118 | 1 | 10 |
| si:ch73-46j18.5 | 0.55706 | 0.199911635 | 0.325 | 0.265 | 1 | 10 |
| si:dkey-19e4.5 | 0.55791 | -0.15782711 | 0.05 | 0.118 | 1 | 10 |
| hsp90b1 | 0.55834 | 0.201159986 | 0.1 | 0.059 | 1 | 10 |
| dcaf7 | 0.55865 | -0.15415216 | 0.05 | 0.118 | 1 | 10 |
| srsf3b | 0.55938 | 0.200289079 | 0.225 | 0.176 | 1 | 10 |
| psap | 0.55989 | 0.22539021 | 0.15 | 0.088 | 1 | 10 |
| angptl4 | 0.56001 | -0.20860663 | 0.05 | 0.118 | 1 | 10 |
| gtf2h5 | 0.56012 | -0.22806577 | 0.05 | 0.118 | 1 | 10 |
| fabp3 | 0.56012 | -0.14652498 | 0.125 | 0.206 | 1 | 10 |
| blmh | 0.56044 | -0.2220948 | 0.05 | 0.118 | 1 | 10 |
| cops7a | 0.56112 | -0.23507758 | 0.05 | 0.118 | 1 | 10 |
| cox5ab | 0.56171 | 0.105197951 | 0.3 | 0.294 | 1 | 10 |
| chchd2 | 0.56218 | -0.22728421 | 0.075 | 0.147 | 1 | 10 |
| si:dkey-44g23.5 | 0.56268 | -0.17891133 | 0.05 | 0.118 | 1 | 10 |
| ndufs3 | 0.56335 | -0.20034334 | 0.05 | 0.118 | 1 | 10 |
| tomm5 | 0.5636 | 0.14273153 | 0.225 | 0.206 | 1 | 10 |
| mrps16 | 0.56406 | -0.19542743 | 0.05 | 0.118 | 1 | 10 |
| uqcrc1 | 0.56494 | -0.19874568 | 0.05 | 0.118 | 1 | 10 |
| psmg1 | 0.56564 | -0.19199306 | 0.05 | 0.118 | 1 | 10 |
| snx12 | 0.5669 | -0.21349818 | 0.05 | 0.118 | 1 | 10 |
| znf330 | 0.56694 | 0.136039292 | 0.125 | 0.059 | 1 | 10 |
| ndufa7 | 0.5672 | -0.20115005 | 0.05 | 0.118 | 1 | 10 |
| si:ch211-195o20.7 | 0.56807 | -0.17537131 | 0.05 | 0.118 | 1 | 10 |
| aplp2 | 0.56808 | -0.20980249 | 0.05 | 0.118 | 1 | 10 |
| ISCU (1 of many) | 0.56834 | 0.234749594 | 0.125 | 0.059 | 1 | 10 |
| hexb | 0.5686 | -0.2217367 | 0.05 | 0.118 | 1 | 10 |
| gstp1 | 0.56865 | 0.121639344 | 0.25 | 0.235 | 1 | 10 |
| rbp5 | 0.56869 | -0.20501862 | 0.05 | 0.118 | 1 | 10 |
| CABZ01088149.1 | 0.56933 | -0.19627342 | 0.05 | 0.118 | 1 | 10 |
| rpl28 | 0.56956 | -0.10279843 | 0.875 | 0.941 | 1 | 10 |
| gskip | 0.56957 | -0.17431579 | 0.05 | 0.118 | 1 | 10 |
| MZT1 | 0.57068 | -0.19372724 | 0.05 | 0.118 | 1 | 10 |
| cnn3a | 0.57078 | -0.1800213 | 0.05 | 0.118 | 1 | 10 |
| chd4a | 0.57091 | -0.14666157 | 0.075 | 0.147 | 1 | 10 |
| pde4ba | 0.57154 | -0.25217168 | 0.175 | 0.265 | 1 | 10 |

| | | | | | | |
|---|---|---|---|---|---|---|
| atp5pd | 0.57168 | 0.251483601 | 0.375 | 0.265 | 1 | 10 |
| adam10a | 0.57405 | 0.249410973 | 0.125 | 0.059 | 1 | 10 |
| mta3 | 0.57595 | 0.189562422 | 0.075 | 0.029 | 1 | 10 |
| akap8l | 0.57602 | 0.152075915 | 0.075 | 0.059 | 1 | 10 |
| selenoh | 0.57632 | -0.17339516 | 0.1 | 0.176 | 1 | 10 |
| med30 | 0.57755 | 0.202382055 | 0.075 | 0.029 | 1 | 10 |
| ndufv3 | 0.57848 | -0.15667245 | 0.1 | 0.176 | 1 | 10 |
| fosab | 0.5801 | -0.17514458 | 0.075 | 0.147 | 1 | 10 |
| rps27l | 0.5806 | -0.16471302 | 0.175 | 0.265 | 1 | 10 |
| aldoaa | 0.58123 | 0.25717211 | 0.2 | 0.118 | 1 | 10 |
| brd7 | 0.58155 | -0.21554215 | 0.075 | 0.147 | 1 | 10 |
| mrpl27 | 0.5817 | 0.19121912 | 0.075 | 0.029 | 1 | 10 |
| wsb1 | 0.58357 | 0.240544569 | 0.125 | 0.059 | 1 | 10 |
| gabarapb | 0.58386 | 0.179616111 | 0.375 | 0.324 | 1 | 10 |
| tia1l | 0.585 | 0.181076829 | 0.075 | 0.029 | 1 | 10 |
| fkbp2 | 0.58553 | -0.18838431 | 0.1 | 0.176 | 1 | 10 |
| grcc10 | 0.58584 | -0.15612382 | 0.1 | 0.176 | 1 | 10 |
| ddt | 0.58596 | 0.195875476 | 0.125 | 0.059 | 1 | 10 |
| ctnna1 | 0.58713 | 0.200936862 | 0.075 | 0.029 | 1 | 10 |
| phlda1 | 0.59022 | 0.182300508 | 0.125 | 0.088 | 1 | 10 |
| calm1b | 0.59034 | 0.222606903 | 0.125 | 0.059 | 1 | 10 |
| timm8b | 0.59215 | -0.17382608 | 0.075 | 0.147 | 1 | 10 |
| wbp2nl | 0.59234 | -0.18438037 | 0.075 | 0.147 | 1 | 10 |
| eif2s1a | 0.5925 | 0.212944544 | 0.125 | 0.059 | 1 | 10 |
| lamtor5 | 0.59293 | -0.20655584 | 0.1 | 0.147 | 1 | 10 |
| hmgn7 | 0.59298 | 0.225352543 | 0.375 | 0.294 | 1 | 10 |
| znf1104 | 0.59332 | 0.1980951 | 0.075 | 0.029 | 1 | 10 |
| hnrnpa0l | 0.59367 | -0.18089343 | 0.6 | 0.706 | 1 | 10 |
| tmub1 | 0.59404 | 0.183170161 | 0.075 | 0.029 | 1 | 10 |
| scaf1 | 0.5941 | 0.190290804 | 0.075 | 0.029 | 1 | 10 |
| fam162a | 0.59416 | 0.185286761 | 0.075 | 0.029 | 1 | 10 |
| tpt1 | 0.59508 | -0.13869803 | 0.725 | 0.824 | 1 | 10 |
| snx6 | 0.5952 | -0.16458446 | 0.075 | 0.147 | 1 | 10 |
| taf12 | 0.59602 | -0.16386333 | 0.125 | 0.206 | 1 | 10 |
| farsa | 0.59639 | 0.190709687 | 0.125 | 0.059 | 1 | 10 |
| flot1b | 0.59703 | -0.22065504 | 0.2 | 0.265 | 1 | 10 |
| rpl5b | 0.59821 | -0.1567864 | 0.625 | 0.735 | 1 | 10 |
| tsc22d3 | 0.6002 | -0.18707335 | 0.075 | 0.147 | 1 | 10 |
| h1f0 | 0.60036 | -0.24789254 | 0.1 | 0.176 | 1 | 10 |
| mt-nd2 | 0.60063 | -0.20435416 | 0.25 | 0.353 | 1 | 10 |
| cirbpb | 0.60067 | -0.1598264 | 0.65 | 0.735 | 1 | 10 |
| zmat2 | 0.60128 | 0.14332295 | 0.125 | 0.059 | 1 | 10 |
| lsm5 | 0.60151 | -0.17923223 | 0.075 | 0.147 | 1 | 10 |
| ilf2 | 0.60164 | 0.242016383 | 0.225 | 0.147 | 1 | 10 |
| psme2 | 0.60172 | 0.136632259 | 0.225 | 0.206 | 1 | 10 |
| si:rp71-45k5.4 | 0.60181 | 0.194774382 | 0.2 | 0.118 | 1 | 10 |
| atp5pf | 0.60218 | -0.22575373 | 0.3 | 0.412 | 1 | 10 |
| bcas2 | 0.60319 | 0.153326946 | 0.125 | 0.059 | 1 | 10 |

| | | | | | | |
|---|---|---|---|---|---|---|
| eif4h | 0.60354 | -0.23087847 | 0.1 | 0.176 | 1 | 10 |
| etv4 | 0.60444 | -0.19655628 | 0.075 | 0.147 | 1 | 10 |
| dut | 0.60459 | -0.19382813 | 0.075 | 0.147 | 1 | 10 |
| zgc:110339 | 0.60494 | 0.212416762 | 0.125 | 0.059 | 1 | 10 |
| ankrd11 | 0.60497 | -0.19555469 | 0.05 | 0.088 | 1 | 10 |
| pam16 | 0.60576 | -0.18188071 | 0.05 | 0.088 | 1 | 10 |
| emd | 0.60667 | 0.181687114 | 0.125 | 0.088 | 1 | 10 |
| eef1g | 0.60673 | 0.134836059 | 0.8 | 0.735 | 1 | 10 |
| maff | 0.60741 | 0.177337786 | 0.075 | 0.029 | 1 | 10 |
| mbnl3 | 0.6078 | -0.21918203 | 0.075 | 0.147 | 1 | 10 |
| anxa5b | 0.60786 | 0.16682607 | 0.125 | 0.059 | 1 | 10 |
| dnajb6a | 0.60863 | 0.189593265 | 0.075 | 0.029 | 1 | 10 |
| ap3s1.1 | 0.60871 | 0.174436447 | 0.075 | 0.029 | 1 | 10 |
| mcl1b | 0.60962 | -0.1886973 | 0.075 | 0.147 | 1 | 10 |
| bzw1a | 0.61021 | 0.223996287 | 0.125 | 0.059 | 1 | 10 |
| cdv3 | 0.61026 | -0.19402227 | 0.075 | 0.147 | 1 | 10 |
| uchl1 | 0.61112 | -0.18680464 | 0.075 | 0.147 | 1 | 10 |
| rasl11a | 0.61245 | 0.189859478 | 0.125 | 0.059 | 1 | 10 |
| gng5 | 0.61272 | 0.229783506 | 0.325 | 0.235 | 1 | 10 |
| csnk1da | 0.61322 | 0.149403375 | 0.075 | 0.059 | 1 | 10 |
| ftr82 | 0.61405 | 0.182303575 | 0.125 | 0.059 | 1 | 10 |
| perp | 0.61463 | 0.198061323 | 0.125 | 0.059 | 1 | 10 |
| fam133b | 0.61467 | 0.184763103 | 0.125 | 0.059 | 1 | 10 |
| cnbpb | 0.61467 | 0.213972513 | 0.2 | 0.118 | 1 | 10 |
| hars | 0.61534 | -0.1886315 | 0.1 | 0.176 | 1 | 10 |
| rsf1b.1 | 0.61625 | 0.179566466 | 0.075 | 0.029 | 1 | 10 |
| zgc:92818 | 0.61648 | 0.198998856 | 0.125 | 0.059 | 1 | 10 |
| uqcrh | 0.61726 | 0.196614629 | 0.125 | 0.059 | 1 | 10 |
| snu13b | 0.61748 | -0.16813588 | 0.125 | 0.206 | 1 | 10 |
| elocb | 0.61805 | 0.230786612 | 0.2 | 0.118 | 1 | 10 |
| nfybb | 0.61841 | 0.184399909 | 0.125 | 0.088 | 1 | 10 |
| LOC103910903 | 0.61845 | 0.227282807 | 0.2 | 0.118 | 1 | 10 |
| rab5c | 0.61851 | -0.14584529 | 0.125 | 0.147 | 1 | 10 |
| gpx1a | 0.62123 | -0.2213365 | 0.1 | 0.176 | 1 | 10 |
| si:ch211-197h24.6 | 0.62155 | 0.174512817 | 0.075 | 0.029 | 1 | 10 |
| cct4 | 0.62168 | -0.21186634 | 0.25 | 0.324 | 1 | 10 |
| stx12 | 0.62265 | 0.183915081 | 0.075 | 0.029 | 1 | 10 |
| npm1a | 0.6228 | -0.24599682 | 0.2 | 0.294 | 1 | 10 |
| cct5 | 0.6231 | -0.22744079 | 0.25 | 0.353 | 1 | 10 |
| snrpd2 | 0.62462 | -0.20062261 | 0.225 | 0.324 | 1 | 10 |
| btf3 | 0.62521 | 0.170495675 | 0.575 | 0.5 | 1 | 10 |
| LEPROTL1 | 0.62528 | -0.17935238 | 0.125 | 0.206 | 1 | 10 |
| srp19 | 0.62636 | 0.187078788 | 0.125 | 0.088 | 1 | 10 |
| rpz5 | 0.62665 | -0.12167203 | 0.55 | 0.647 | 1 | 10 |
| metap2b | 0.62678 | 0.210242184 | 0.2 | 0.118 | 1 | 10 |
| cisd2 | 0.62923 | 0.184220018 | 0.1 | 0.059 | 1 | 10 |
| rpl14 | 0.62968 | 0.147556026 | 0.775 | 0.676 | 1 | 10 |
| fkbp1b | 0.63011 | -0.21107391 | 0.1 | 0.176 | 1 | 10 |

| | | | | | | |
|---|---|---|---|---|---|---|
| psma6a | 0.63022 | -0.17895437 | 0.1 | 0.176 | 1 | 10 |
| eif5 | 0.63174 | -0.18386931 | 0.1 | 0.176 | 1 | 10 |
| pdha1a | 0.63221 | -0.21430358 | 0.1 | 0.176 | 1 | 10 |
| ndufb6 | 0.63286 | 0.209090804 | 0.175 | 0.118 | 1 | 10 |
| cops4 | 0.63331 | -0.19748871 | 0.1 | 0.176 | 1 | 10 |
| mknk2b | 0.63376 | 0.141876114 | 0.15 | 0.088 | 1 | 10 |
| gtf2f2a | 0.63411 | 0.170831864 | 0.075 | 0.029 | 1 | 10 |
| fkbp4 | 0.63498 | 0.10424526 | 0.075 | 0.029 | 1 | 10 |
| sec11a | 0.63648 | 0.111802206 | 0.075 | 0.029 | 1 | 10 |
| itm2ba | 0.63768 | -0.21988794 | 0.25 | 0.324 | 1 | 10 |
| ndufb8 | 0.63826 | -0.22071485 | 0.2 | 0.294 | 1 | 10 |
| cygb1 | 0.63842 | 0.182709916 | 0.075 | 0.029 | 1 | 10 |
| strap | 0.63849 | 0.1715496 | 0.1 | 0.059 | 1 | 10 |
| ufc1 | 0.6415 | -0.17750198 | 0.05 | 0.088 | 1 | 10 |
| nme3 | 0.64184 | -0.22040463 | 0.15 | 0.235 | 1 | 10 |
| ccdc187 | 0.64218 | 0.222377113 | 0.35 | 0.265 | 1 | 10 |
| pnp5a | 0.64218 | -0.20202319 | 0.125 | 0.176 | 1 | 10 |
| polr2d | 0.64513 | -0.19185152 | 0.15 | 0.235 | 1 | 10 |
| emc8 | 0.64544 | 0.16552448 | 0.075 | 0.029 | 1 | 10 |
| ssr2 | 0.64732 | -0.20360861 | 0.15 | 0.235 | 1 | 10 |
| cct3 | 0.64759 | -0.20143941 | 0.125 | 0.176 | 1 | 10 |
| si:ch211-288g17.3 | 0.64769 | -0.23271059 | 0.175 | 0.265 | 1 | 10 |
| pbdc1 | 0.64941 | -0.19015339 | 0.1 | 0.147 | 1 | 10 |
| atg4c | 0.64949 | 0.112786172 | 0.075 | 0.029 | 1 | 10 |
| smu1a | 0.65074 | 0.132743136 | 0.075 | 0.029 | 1 | 10 |
| surf2 | 0.65184 | 0.113601881 | 0.075 | 0.029 | 1 | 10 |
| apol1 | 0.65203 | 0.162236552 | 0.075 | 0.029 | 1 | 10 |
| mxa | 0.65223 | 0.144411209 | 0.075 | 0.029 | 1 | 10 |
| tp53 | 0.65282 | 0.178568432 | 0.25 | 0.206 | 1 | 10 |
| mrpl39 | 0.65285 | 0.105757408 | 0.075 | 0.029 | 1 | 10 |
| naa38 | 0.65297 | 0.173903246 | 0.075 | 0.029 | 1 | 10 |
| dctn5 | 0.65393 | 0.155634522 | 0.075 | 0.029 | 1 | 10 |
| gpx1b | 0.65405 | 0.250564869 | 0.15 | 0.088 | 1 | 10 |
| ppp1cab | 0.65577 | 0.170108116 | 0.125 | 0.088 | 1 | 10 |
| golga7 | 0.65608 | 0.122161729 | 0.225 | 0.206 | 1 | 10 |
| ccdc85b | 0.65608 | 0.159723688 | 0.075 | 0.029 | 1 | 10 |
| grpel2 | 0.65694 | 0.173704449 | 0.075 | 0.029 | 1 | 10 |
| tmem88b | 0.65707 | 0.136975725 | 0.2 | 0.176 | 1 | 10 |
| dcn | 0.65718 | 0.10803943 | 0.05 | 0.029 | 1 | 10 |
| dspa | 0.65718 | 0.123508833 | 0.05 | 0.029 | 1 | 10 |
| slc39a7 | 0.65718 | 0.108478639 | 0.05 | 0.029 | 1 | 10 |
| slka | 0.65718 | 0.114539018 | 0.05 | 0.029 | 1 | 10 |
| zbtb20 | 0.65718 | 0.118843055 | 0.05 | 0.029 | 1 | 10 |
| nfia | 0.65718 | 0.107403873 | 0.05 | 0.029 | 1 | 10 |
| serinc5 | 0.65718 | 0.123488322 | 0.05 | 0.029 | 1 | 10 |
| sh3glb1b | 0.65718 | 0.100353898 | 0.05 | 0.029 | 1 | 10 |
| sprydb7b | 0.65718 | 0.128831082 | 0.05 | 0.029 | 1 | 10 |
| txnl1 | 0.65718 | 0.117709528 | 0.05 | 0.029 | 1 | 10 |

| | | | | | | |
|---|---|---|---|---|---|---|
| wtap | 0.65718 | 0.103882547 | 0.05 | 0.029 | 1 | 10 |
| zfpm1 | 0.65718 | 0.102431503 | 0.05 | 0.029 | 1 | 10 |
| asna1 | 0.65718 | 0.114121664 | 0.05 | 0.029 | 1 | 10 |
| calm3a | 0.65718 | 0.111325859 | 0.05 | 0.029 | 1 | 10 |
| cdc73 | 0.65718 | 0.13359328 | 0.05 | 0.029 | 1 | 10 |
| gsta.1 | 0.65718 | 0.120680828 | 0.05 | 0.029 | 1 | 10 |
| man1a2 | 0.65718 | 0.101382704 | 0.05 | 0.029 | 1 | 10 |
| mrpl42 | 0.65718 | 0.10046561 | 0.05 | 0.029 | 1 | 10 |
| nub1 | 0.65718 | 0.142048739 | 0.05 | 0.029 | 1 | 10 |
| papolg | 0.65718 | 0.121675395 | 0.05 | 0.029 | 1 | 10 |
| pgls | 0.65718 | 0.112599131 | 0.05 | 0.029 | 1 | 10 |
| pmp22a | 0.65718 | 0.137837797 | 0.05 | 0.029 | 1 | 10 |
| rnf8 | 0.65718 | 0.127372125 | 0.05 | 0.029 | 1 | 10 |
| slc9a6a | 0.65718 | 0.112353248 | 0.05 | 0.029 | 1 | 10 |
| sod2 | 0.65718 | 0.127527249 | 0.05 | 0.029 | 1 | 10 |
| xrcc6 | 0.65718 | 0.122676435 | 0.05 | 0.029 | 1 | 10 |
| CABZ01112656.1 | 0.65718 | 0.126391821 | 0.05 | 0.029 | 1 | 10 |
| sf1 | 0.65718 | 0.10631584 | 0.05 | 0.029 | 1 | 10 |
| sf3a3 | 0.65718 | 0.111322195 | 0.05 | 0.029 | 1 | 10 |
| si:ch73-335l21.4 | 0.65718 | 0.105413974 | 0.05 | 0.029 | 1 | 10 |
| si:dkey-51e6.1 | 0.65718 | 0.111272755 | 0.05 | 0.029 | 1 | 10 |
| tufm | 0.65718 | 0.1380139 | 0.05 | 0.029 | 1 | 10 |
| cnot8 | 0.65718 | 0.12026808 | 0.05 | 0.029 | 1 | 10 |
| gsto2 | 0.65718 | 0.12855242 | 0.05 | 0.029 | 1 | 10 |
| her6 | 0.65718 | 0.134684659 | 0.05 | 0.029 | 1 | 10 |
| lzic | 0.65718 | 0.137195837 | 0.05 | 0.029 | 1 | 10 |
| osr2 | 0.65718 | 0.103290139 | 0.05 | 0.029 | 1 | 10 |
| rbm14a | 0.65718 | 0.131270188 | 0.05 | 0.029 | 1 | 10 |
| si:ch1073-440b2.1 | 0.65718 | 0.103454614 | 0.05 | 0.029 | 1 | 10 |
| smox | 0.65718 | 0.100107693 | 0.05 | 0.029 | 1 | 10 |
| ubap2a | 0.65718 | 0.10946208 | 0.05 | 0.029 | 1 | 10 |
| wdr75 | 0.65718 | 0.124811807 | 0.05 | 0.029 | 1 | 10 |
| ccdc167 | 0.65718 | 0.109243837 | 0.05 | 0.029 | 1 | 10 |
| cstf1 | 0.65718 | 0.144388451 | 0.05 | 0.029 | 1 | 10 |
| gdi1 | 0.65718 | 0.132440957 | 0.05 | 0.029 | 1 | 10 |
| gpia | 0.65718 | 0.104472235 | 0.05 | 0.029 | 1 | 10 |
| immp2l | 0.65718 | 0.120773428 | 0.05 | 0.029 | 1 | 10 |
| si:ch73-234b20.5 | 0.65718 | 0.130098614 | 0.05 | 0.029 | 1 | 10 |
| si:dkey-185e18.7 | 0.65718 | 0.145744356 | 0.05 | 0.029 | 1 | 10 |
| u2af2a | 0.65718 | 0.116757763 | 0.05 | 0.029 | 1 | 10 |
| zgc:85789 | 0.65718 | 0.107946604 | 0.05 | 0.029 | 1 | 10 |
| prpf6 | 0.65746 | 0.167041742 | 0.075 | 0.029 | 1 | 10 |
| rbm42 | 0.65756 | 0.158692669 | 0.075 | 0.029 | 1 | 10 |
| nudcd2 | 0.65812 | 0.122856762 | 0.1 | 0.088 | 1 | 10 |
| ubxn7 | 0.65859 | 0.115364022 | 0.075 | 0.029 | 1 | 10 |
| DHDH | 0.65919 | 0.16772542 | 0.075 | 0.029 | 1 | 10 |
| sept2 | 0.65963 | -0.19356153 | 0.075 | 0.118 | 1 | 10 |
| zgc:162698 | 0.66005 | 0.175169625 | 0.075 | 0.029 | 1 | 10 |

| | | | | | | |
|---|---|---|---|---|---|---|
| zgc:77086 | 0.66197 | 0.132661318 | 0.075 | 0.029 | 1 | 10 |
| cndp2 | 0.66359 | 0.151307823 | 0.075 | 0.029 | 1 | 10 |
| chac2 | 0.66474 | 0.170809339 | 0.075 | 0.029 | 1 | 10 |
| id2b | 0.66551 | 0.12084015 | 0.1 | 0.088 | 1 | 10 |
| zgc:91910 | 0.66563 | 0.124937817 | 0.225 | 0.206 | 1 | 10 |
| zgc:92242 | 0.66629 | 0.123936207 | 0.075 | 0.029 | 1 | 10 |
| adrm1 | 0.6665 | 0.119174797 | 0.075 | 0.029 | 1 | 10 |
| lgals2a | 0.66665 | 0.16299085 | 0.075 | 0.029 | 1 | 10 |
| nectin1b | 0.667 | 0.204483933 | 0.1 | 0.059 | 1 | 10 |
| dgcr8 | 0.66719 | 0.123481562 | 0.075 | 0.029 | 1 | 10 |
| ormdl1 | 0.66725 | 0.134519826 | 0.075 | 0.029 | 1 | 10 |
| slc25a20 | 0.66761 | 0.127510389 | 0.075 | 0.029 | 1 | 10 |
| psma1 | 0.66779 | -0.21198378 | 0.3 | 0.382 | 1 | 10 |
| foxl2a | 0.66837 | 0.177780031 | 0.1 | 0.059 | 1 | 10 |
| polr2c | 0.66856 | 0.123167302 | 0.075 | 0.029 | 1 | 10 |
| smarcd1 | 0.66898 | 0.133812252 | 0.075 | 0.029 | 1 | 10 |
| hpf1 | 0.669 | 0.157825586 | 0.075 | 0.029 | 1 | 10 |
| drap1 | 0.66913 | 0.175530317 | 0.1 | 0.059 | 1 | 10 |
| hmgn2 | 0.67056 | -0.18359252 | 0.55 | 0.647 | 1 | 10 |
| nars | 0.67096 | 0.137734181 | 0.075 | 0.029 | 1 | 10 |
| itga3b | 0.67163 | 0.148408099 | 0.075 | 0.029 | 1 | 10 |
| si:dkey-42i9.4 | 0.67267 | 0.148805885 | 0.175 | 0.147 | 1 | 10 |
| si:ch1073-325m22.2 | 0.67304 | 0.215239205 | 0.225 | 0.147 | 1 | 10 |
| aimp1 | 0.67326 | 0.13036407 | 0.075 | 0.029 | 1 | 10 |
| timm9 | 0.67336 | -0.18518931 | 0.075 | 0.118 | 1 | 10 |
| spsb4a | 0.67376 | 0.150501428 | 0.075 | 0.029 | 1 | 10 |
| igf2bp3 | 0.67452 | 0.145206181 | 0.075 | 0.029 | 1 | 10 |
| pgk1 | 0.6759 | 0.137288635 | 0.075 | 0.029 | 1 | 10 |
| ppih | 0.67615 | 0.146394877 | 0.075 | 0.029 | 1 | 10 |
| pak2a | 0.6764 | 0.161745924 | 0.075 | 0.029 | 1 | 10 |
| sh3gl3b | 0.67679 | 0.146733953 | 0.075 | 0.029 | 1 | 10 |
| brd3a | 0.67683 | -0.1847432 | 0.05 | 0.088 | 1 | 10 |
| anxa4 | 0.67686 | 0.111887535 | 0.125 | 0.118 | 1 | 10 |
| si:ch211-191j22.3 | 0.67724 | 0.195071452 | 0.175 | 0.118 | 1 | 10 |
| ifi46 | 0.67754 | 0.204341389 | 0.45 | 0.353 | 1 | 10 |
| cox5b2 | 0.67775 | 0.16076801 | 0.075 | 0.029 | 1 | 10 |
| psmd12 | 0.67781 | 0.172586673 | 0.075 | 0.029 | 1 | 10 |
| ypel5 | 0.67782 | 0.159187064 | 0.075 | 0.029 | 1 | 10 |
| gig2e | 0.67809 | 0.172667975 | 0.075 | 0.029 | 1 | 10 |
| bcl2l10 | 0.67831 | 0.157948658 | 0.075 | 0.029 | 1 | 10 |
| kif5bb | 0.67837 | 0.144147195 | 0.075 | 0.029 | 1 | 10 |
| srp72 | 0.67871 | 0.176656511 | 0.075 | 0.029 | 1 | 10 |
| igflr1 | 0.67969 | 0.137641509 | 0.075 | 0.029 | 1 | 10 |
| flot2a | 0.67997 | 0.150830166 | 0.075 | 0.029 | 1 | 10 |
| inip | 0.68006 | 0.128081578 | 0.075 | 0.029 | 1 | 10 |
| derl2 | 0.68015 | 0.125702408 | 0.075 | 0.029 | 1 | 10 |
| gng10 | 0.68016 | 0.162356751 | 0.075 | 0.029 | 1 | 10 |
| bod1 | 0.68045 | 0.134252863 | 0.075 | 0.029 | 1 | 10 |

| | | | | | | |
|---|---|---|---|---|---|---|
| cald1a | 0.68085 | 0.138803649 | 0.075 | 0.029 | 1 | 10 |
| cwc25 | 0.68126 | 0.163216065 | 0.075 | 0.029 | 1 | 10 |
| anxa11a | 0.68128 | 0.114318441 | 0.075 | 0.029 | 1 | 10 |
| urm1 | 0.6813 | 0.144216002 | 0.075 | 0.029 | 1 | 10 |
| gcsha | 0.68139 | 0.13262838 | 0.075 | 0.029 | 1 | 10 |
| cplx2 | 0.6815 | 0.145958734 | 0.075 | 0.029 | 1 | 10 |
| suds3 | 0.68165 | 0.106448818 | 0.075 | 0.029 | 1 | 10 |
| pip4k2ca | 0.68175 | 0.130033407 | 0.075 | 0.029 | 1 | 10 |
| flna | 0.68197 | 0.166856239 | 0.075 | 0.029 | 1 | 10 |
| atg13 | 0.68198 | 0.145611787 | 0.075 | 0.029 | 1 | 10 |
| mff | 0.68206 | 0.164459024 | 0.075 | 0.029 | 1 | 10 |
| eif1axa | 0.68216 | 0.12709738 | 0.075 | 0.029 | 1 | 10 |
| srp54 | 0.68216 | 0.124277746 | 0.075 | 0.029 | 1 | 10 |
| dhdds | 0.68223 | 0.13078943 | 0.075 | 0.029 | 1 | 10 |
| imp3 | 0.68223 | 0.166354431 | 0.075 | 0.029 | 1 | 10 |
| dpm1 | 0.68228 | 0.171349068 | 0.075 | 0.029 | 1 | 10 |
| pdcd10b | 0.68235 | 0.145356397 | 0.15 | 0.088 | 1 | 10 |
| hnrnpa0b | 0.68253 | -0.18608962 | 0.4 | 0.5 | 1 | 10 |
| txnrd3 | 0.68314 | -0.1655551 | 0.05 | 0.088 | 1 | 10 |
| lsm1 | 0.68722 | 0.176883016 | 0.1 | 0.059 | 1 | 10 |
| mrps14 | 0.69027 | -0.17822908 | 0.075 | 0.118 | 1 | 10 |
| psma6b | 0.69575 | -0.18258795 | 0.125 | 0.176 | 1 | 10 |
| sugt1 | 0.69586 | -0.15546696 | 0.05 | 0.088 | 1 | 10 |
| wdr83os | 0.69634 | 0.122631721 | 0.1 | 0.088 | 1 | 10 |
| rrm1 | 0.69893 | -0.15647317 | 0.05 | 0.088 | 1 | 10 |
| eif3jb | 0.69983 | 0.207934519 | 0.15 | 0.088 | 1 | 10 |
| ube2v1 | 0.70515 | -0.10102626 | 0.275 | 0.294 | 1 | 10 |
| skp1 | 0.70663 | -0.19812019 | 0.175 | 0.235 | 1 | 10 |
| npm3 | 0.70698 | -0.16821646 | 0.075 | 0.118 | 1 | 10 |
| ubxn1 | 0.70713 | -0.13656275 | 0.125 | 0.147 | 1 | 10 |
| cebpd | 0.70779 | 0.187429631 | 0.225 | 0.176 | 1 | 10 |
| esf1 | 0.70817 | -0.16533231 | 0.05 | 0.088 | 1 | 10 |
| atp5mea | 0.70832 | 0.130126203 | 0.2 | 0.176 | 1 | 10 |
| rsl1d1 | 0.70884 | 0.140045776 | 0.15 | 0.088 | 1 | 10 |
| mif | 0.7091 | -0.19066898 | 0.175 | 0.235 | 1 | 10 |
| podxl | 0.71155 | 0.208342491 | 0.15 | 0.088 | 1 | 10 |
| rpz | 0.71237 | -0.16668626 | 0.05 | 0.088 | 1 | 10 |
| nap1l1 | 0.71247 | -0.1904896 | 0.2 | 0.265 | 1 | 10 |
| si:ch211-246m6.4 | 0.71303 | -0.13018471 | 0.1 | 0.118 | 1 | 10 |
| dnajc11a | 0.71398 | 0.167357728 | 0.1 | 0.059 | 1 | 10 |
| khdrbs1a | 0.7148 | -0.13059206 | 0.525 | 0.618 | 1 | 10 |
| atp5if1a | 0.71541 | -0.1296872 | 0.275 | 0.353 | 1 | 10 |
| tpm1 | 0.7156 | 0.152154497 | 0.15 | 0.088 | 1 | 10 |
| ar | 0.71561 | 0.180146428 | 0.15 | 0.088 | 1 | 10 |
| psmc1a | 0.71577 | 0.114452853 | 0.1 | 0.088 | 1 | 10 |
| ndufs8a | 0.7159 | -0.17897394 | 0.1 | 0.147 | 1 | 10 |
| mbd3a | 0.71847 | 0.163356765 | 0.1 | 0.059 | 1 | 10 |
| abcf1 | 0.71912 | -0.15431903 | 0.25 | 0.294 | 1 | 10 |

| | | | | | | |
|---|---|---|---|---|---|---|
| ier3ip1 | 0.71959 | 0.124841606 | 0.2 | 0.176 | 1 | 10 |
| cd63 | 0.71965 | 0.163153377 | 0.25 | 0.176 | 1 | 10 |
| pgam5 | 0.72076 | 0.111365194 | 0.1 | 0.088 | 1 | 10 |
| thoc3 | 0.72126 | 0.123816657 | 0.075 | 0.059 | 1 | 10 |
| shc1 | 0.72339 | -0.1606085 | 0.05 | 0.088 | 1 | 10 |
| fam177a1 | 0.72497 | 0.166692711 | 0.1 | 0.059 | 1 | 10 |
| XLOC-005605 | 0.72711 | 0.174893088 | 0.325 | 0.265 | 1 | 10 |
| psmb1 | 0.72915 | 0.179113459 | 0.35 | 0.265 | 1 | 10 |
| commd7 | 0.73072 | -0.15192068 | 0.05 | 0.088 | 1 | 10 |
| ywhah | 0.73307 | -0.16381664 | 0.075 | 0.118 | 1 | 10 |
| nanos1 | 0.73358 | 0.172158409 | 0.1 | 0.059 | 1 | 10 |
| hdgfl2 | 0.73469 | -0.11967658 | 0.075 | 0.088 | 1 | 10 |
| agpat3 | 0.73536 | -0.11450912 | 0.075 | 0.088 | 1 | 10 |
| eif6 | 0.73681 | -0.17121457 | 0.075 | 0.118 | 1 | 10 |
| trappc2l | 0.73726 | 0.16590998 | 0.1 | 0.059 | 1 | 10 |
| arl6ip1 | 0.73761 | 0.154009171 | 0.15 | 0.118 | 1 | 10 |
| vma21 | 0.73823 | 0.156273893 | 0.1 | 0.059 | 1 | 10 |
| zc3h7bb | 0.73911 | -0.1539377 | 0.05 | 0.088 | 1 | 10 |
| arpc5a | 0.74026 | 0.160333232 | 0.1 | 0.059 | 1 | 10 |
| vapal | 0.74078 | -0.15517663 | 0.05 | 0.088 | 1 | 10 |
| ucp2 | 0.74166 | 0.17057655 | 0.425 | 0.353 | 1 | 10 |
| rhocb | 0.74183 | 0.151186467 | 0.15 | 0.118 | 1 | 10 |
| krt94 | 0.74209 | 0.154657608 | 0.475 | 0.412 | 1 | 10 |
| mvp | 0.74355 | 0.123287446 | 0.075 | 0.059 | 1 | 10 |
| adh5 | 0.74394 | 0.182092693 | 0.25 | 0.176 | 1 | 10 |
| capns1a | 0.74487 | 0.124856823 | 0.175 | 0.118 | 1 | 10 |
| nr2f1a | 0.74555 | 0.193093743 | 0.275 | 0.206 | 1 | 10 |
| atp5mc3a | 0.74744 | -0.14744744 | 0.3 | 0.382 | 1 | 10 |
| eif2s3 | 0.74982 | 0.172297495 | 0.325 | 0.265 | 1 | 10 |
| aars | 0.75055 | -0.13566923 | 0.05 | 0.088 | 1 | 10 |
| zgc:193742 | 0.75153 | 0.11927822 | 0.075 | 0.059 | 1 | 10 |
| anapc16 | 0.75309 | 0.161293294 | 0.1 | 0.059 | 1 | 10 |
| si:ch1073-429i10.3.1 | 0.75366 | 0.175330462 | 0.3 | 0.235 | 1 | 10 |
| eif3ba | 0.75647 | -0.1840799 | 0.225 | 0.294 | 1 | 10 |
| ssr4 | 0.75712 | -0.16574269 | 0.3 | 0.382 | 1 | 10 |
| id1 | 0.75799 | 0.199200392 | 0.275 | 0.206 | 1 | 10 |
| gabarapl2 | 0.75805 | 0.156477938 | 0.125 | 0.088 | 1 | 10 |
| pnrc2 | 0.75882 | -0.10068256 | 0.475 | 0.5 | 1 | 10 |
| gbp | 0.75898 | 0.143638644 | 0.1 | 0.059 | 1 | 10 |
| ube2v2 | 0.76356 | 0.161668969 | 0.375 | 0.294 | 1 | 10 |
| txn2 | 0.76618 | 0.120071215 | 0.075 | 0.059 | 1 | 10 |
| znf706 | 0.76622 | -0.17206261 | 0.175 | 0.235 | 1 | 10 |
| atp1b3b | 0.76653 | -0.18302079 | 0.15 | 0.206 | 1 | 10 |
| psmd8 | 0.7668 | -0.15373248 | 0.075 | 0.118 | 1 | 10 |
| snrpc | 0.76718 | 0.15468984 | 0.125 | 0.088 | 1 | 10 |
| psmb4 | 0.76748 | -0.16535582 | 0.275 | 0.353 | 1 | 10 |
| eif1ad | 0.76764 | 0.136364021 | 0.1 | 0.059 | 1 | 10 |
| sar1b | 0.76802 | -0.15114066 | 0.1 | 0.147 | 1 | 10 |

| | | | | | | |
|---|---|---|---|---|---|---|
| spcs1 | 0.7722 | 0.177058809 | 0.175 | 0.118 | 1 | 10 |
| crls1 | 0.77241 | -0.12307532 | 0.05 | 0.088 | 1 | 10 |
| lypla2 | 0.7734 | 0.155587845 | 0.1 | 0.059 | 1 | 10 |
| plscr3b | 0.77413 | 0.1111539 | 0.1 | 0.059 | 1 | 10 |
| nucks1b | 0.77449 | 0.169450774 | 0.175 | 0.118 | 1 | 10 |
| lamtor4 | 0.77518 | 0.133629147 | 0.175 | 0.118 | 1 | 10 |
| coa3a | 0.77529 | -0.10352286 | 0.075 | 0.118 | 1 | 10 |
| glrx3 | 0.77588 | -0.16081893 | 0.1 | 0.147 | 1 | 10 |
| dusp14 | 0.7763 | 0.161312335 | 0.1 | 0.059 | 1 | 10 |
| appb | 0.77823 | -0.16118101 | 0.05 | 0.088 | 1 | 10 |
| cox7a1 | 0.77843 | 0.147289462 | 0.1 | 0.059 | 1 | 10 |
| esd | 0.77861 | 0.145223967 | 0.175 | 0.118 | 1 | 10 |
| timm10 | 0.779 | -0.12421372 | 0.05 | 0.088 | 1 | 10 |
| acaa1 | 0.77914 | 0.158579139 | 0.1 | 0.059 | 1 | 10 |
| eif4ebp3l | 0.78014 | 0.139698039 | 0.175 | 0.118 | 1 | 10 |
| psmc2 | 0.78017 | -0.12133537 | 0.05 | 0.088 | 1 | 10 |
| zgc:56095 | 0.78024 | -0.15849714 | 0.075 | 0.118 | 1 | 10 |
| myl6 | 0.78027 | -0.15696911 | 0.1 | 0.147 | 1 | 10 |
| park7 | 0.78056 | -0.15484007 | 0.125 | 0.176 | 1 | 10 |
| lgals3b | 0.78179 | 0.171419134 | 0.1 | 0.059 | 1 | 10 |
| tpm4a | 0.78211 | -0.16323057 | 0.125 | 0.176 | 1 | 10 |
| atp6v1e1b | 0.78241 | -0.14935207 | 0.05 | 0.088 | 1 | 10 |
| rab11a | 0.78288 | 0.147131781 | 0.175 | 0.118 | 1 | 10 |
| h3f3c | 0.78329 | -0.15577515 | 0.325 | 0.382 | 1 | 10 |
| selenbp1 | 0.78329 | -0.15426188 | 0.05 | 0.088 | 1 | 10 |
| eif5a | 0.78397 | 0.138039608 | 0.575 | 0.5 | 1 | 10 |
| sparc | 0.78469 | 0.136537766 | 0.525 | 0.471 | 1 | 10 |
| arpc3 | 0.78503 | 0.165503371 | 0.275 | 0.206 | 1 | 10 |
| rexo4 | 0.78522 | 0.146534945 | 0.1 | 0.059 | 1 | 10 |
| arpc1b | 0.78525 | 0.142832961 | 0.15 | 0.118 | 1 | 10 |
| impdh1b | 0.78587 | 0.112257536 | 0.075 | 0.059 | 1 | 10 |
| zgc:111986 | 0.78705 | 0.102863699 | 0.1 | 0.088 | 1 | 10 |
| ash2l | 0.78787 | -0.1444835 | 0.05 | 0.088 | 1 | 10 |
| tcp1 | 0.78875 | -0.12901896 | 0.2 | 0.265 | 1 | 10 |
| etfb | 0.78895 | -0.11162065 | 0.05 | 0.088 | 1 | 10 |
| tmem176l.1 | 0.78897 | 0.160282678 | 0.225 | 0.176 | 1 | 10 |
| rbm19 | 0.78969 | 0.146523705 | 0.1 | 0.059 | 1 | 10 |
| ccnd2a | 0.79075 | -0.14880806 | 0.05 | 0.088 | 1 | 10 |
| cbx8a | 0.79087 | 0.138502445 | 0.1 | 0.059 | 1 | 10 |
| glrx5 | 0.79097 | 0.170647495 | 0.2 | 0.147 | 1 | 10 |
| tjp2b | 0.79163 | 0.101557651 | 0.1 | 0.059 | 1 | 10 |
| zgc:77056 | 0.7925 | -0.13865951 | 0.05 | 0.088 | 1 | 10 |
| hmgb3a | 0.79254 | 0.144231747 | 0.4 | 0.324 | 1 | 10 |
| smdt1b | 0.79268 | -0.16706701 | 0.15 | 0.206 | 1 | 10 |
| btbd10b | 0.79269 | 0.102176892 | 0.1 | 0.059 | 1 | 10 |
| lrp1aa | 0.79379 | 0.155861304 | 0.1 | 0.059 | 1 | 10 |
| cotl1 | 0.79436 | 0.128651023 | 0.1 | 0.059 | 1 | 10 |
| rbb4l | 0.7947 | -0.12660196 | 0.05 | 0.088 | 1 | 10 |

| | | | | | | |
|---|---|---|---|---|---|---|
| fuca1.1 | 0.79575 | 0.126998126 | 0.1 | 0.059 | 1 | 10 |
| COA4 | 0.7959 | -0.13628035 | 0.05 | 0.088 | 1 | 10 |
| ak1 | 0.79627 | -0.12147954 | 0.05 | 0.088 | 1 | 10 |
| nsa2 | 0.7974 | 0.114798571 | 0.3 | 0.235 | 1 | 10 |
| hnrnpul1 | 0.79927 | 0.102491823 | 0.1 | 0.059 | 1 | 10 |
| dnmt3ab | 0.79968 | 0.110755474 | 0.075 | 0.059 | 1 | 10 |
| mrpl13 | 0.79988 | -0.11047238 | 0.05 | 0.088 | 1 | 10 |
| hspd1 | 0.79989 | 0.100717811 | 0.2 | 0.147 | 1 | 10 |
| klf6a | 0.79992 | 0.159469426 | 0.1 | 0.059 | 1 | 10 |
| pip5k1ab | 0.79992 | -0.14155263 | 0.05 | 0.088 | 1 | 10 |
| crtap | 0.80016 | -0.10316293 | 0.05 | 0.088 | 1 | 10 |
| tspan9b | 0.8005 | 0.151060911 | 0.1 | 0.059 | 1 | 10 |
| ncl | 0.80144 | 0.120131177 | 0.3 | 0.265 | 1 | 10 |
| otud5a | 0.80153 | -0.12390158 | 0.05 | 0.088 | 1 | 10 |
| siva1 | 0.80209 | -0.10132905 | 0.05 | 0.088 | 1 | 10 |
| pgam1b | 0.80221 | 0.145363381 | 0.1 | 0.059 | 1 | 10 |
| rbm4.1 | 0.8025 | 0.117063962 | 0.1 | 0.059 | 1 | 10 |
| uba1 | 0.80256 | -0.10097174 | 0.1 | 0.147 | 1 | 10 |
| rrm2 | 0.80265 | -0.15221665 | 0.05 | 0.088 | 1 | 10 |
| cd82a | 0.80273 | -0.13429365 | 0.05 | 0.088 | 1 | 10 |
| trappc6bl | 0.80337 | 0.118424991 | 0.1 | 0.059 | 1 | 10 |
| pcmt | 0.80369 | -0.11994265 | 0.05 | 0.088 | 1 | 10 |
| usf1l | 0.80383 | -0.15400084 | 0.075 | 0.118 | 1 | 10 |
| txndc17 | 0.80431 | -0.11581659 | 0.05 | 0.088 | 1 | 10 |
| notch2 | 0.8044 | -0.11535031 | 0.05 | 0.088 | 1 | 10 |
| si:dkey-286j15.3 | 0.80457 | 0.128336959 | 0.1 | 0.059 | 1 | 10 |
| pole3 | 0.80472 | -0.12151133 | 0.05 | 0.088 | 1 | 10 |
| polr2eb | 0.80489 | -0.11331685 | 0.125 | 0.176 | 1 | 10 |
| emc10 | 0.80491 | 0.120979752 | 0.1 | 0.059 | 1 | 10 |
| pin1 | 0.80506 | -0.16108742 | 0.175 | 0.235 | 1 | 10 |
| rrp7a | 0.80523 | -0.10093935 | 0.05 | 0.088 | 1 | 10 |
| ndufa11 | 0.80529 | 0.124432381 | 0.1 | 0.059 | 1 | 10 |
| CABZ01068366.1 | 0.80571 | 0.116867981 | 0.1 | 0.059 | 1 | 10 |
| b2m | 0.80572 | 0.126492258 | 0.425 | 0.353 | 1 | 10 |
| ncbp2 | 0.80614 | -0.15666933 | 0.075 | 0.118 | 1 | 10 |
| oaz1b | 0.80654 | 0.1363646 | 0.5 | 0.441 | 1 | 10 |
| enc1 | 0.80674 | -0.13866568 | 0.05 | 0.088 | 1 | 10 |
| commd4 | 0.80684 | -0.12615879 | 0.05 | 0.088 | 1 | 10 |
| ddx24 | 0.80696 | 0.113000597 | 0.1 | 0.059 | 1 | 10 |
| mrpl12 | 0.80705 | 0.113541975 | 0.1 | 0.059 | 1 | 10 |
| arl1 | 0.80705 | -0.12359339 | 0.05 | 0.088 | 1 | 10 |
| sept6 | 0.80717 | 0.105882149 | 0.075 | 0.059 | 1 | 10 |
| ccdc12 | 0.80744 | 0.104623895 | 0.1 | 0.059 | 1 | 10 |
| si:dkey-42i9.7 | 0.80768 | -0.13468783 | 0.05 | 0.088 | 1 | 10 |
| si:dkeyp-27c8.2 | 0.80785 | -0.11590397 | 0.05 | 0.088 | 1 | 10 |
| tnrc6c1 | 0.80799 | -0.1234793 | 0.05 | 0.088 | 1 | 10 |
| ewsr1a | 0.80804 | -0.14067132 | 0.1 | 0.147 | 1 | 10 |
| stat2 | 0.80815 | 0.123562544 | 0.1 | 0.059 | 1 | 10 |

| | | | | | | |
|---|---|---|---|---|---|---|
| pfn1 | 0.80836 | 0.12783975 | 0.1 | 0.059 | 1 | 10 |
| si:dkey-67c22.2 | 0.8084 | -0.11517889 | 0.05 | 0.088 | 1 | 10 |
| irf2bp2a | 0.80876 | 0.132101169 | 0.1 | 0.059 | 1 | 10 |
| ywhaqb | 0.80886 | -0.14982229 | 0.175 | 0.235 | 1 | 10 |
| tpp1 | 0.80919 | -0.13406288 | 0.05 | 0.088 | 1 | 10 |
| zgc:77650 | 0.80955 | -0.13801601 | 0.075 | 0.118 | 1 | 10 |
| nfs1 | 0.80956 | 0.119304674 | 0.1 | 0.059 | 1 | 10 |
| si:ch211-218c6.8 | 0.8096 | 0.127767499 | 0.1 | 0.059 | 1 | 10 |
| CABZ01068367.1 | 0.8097 | -0.13099925 | 0.05 | 0.088 | 1 | 10 |
| tgif1 | 0.81007 | 0.156545431 | 0.2 | 0.147 | 1 | 10 |
| ube2l3a | 0.81081 | -0.10922706 | 0.05 | 0.088 | 1 | 10 |
| si:ch211-140m22.7 | 0.81095 | -0.11033526 | 0.05 | 0.088 | 1 | 10 |
| si:dkey-187e18.1 | 0.81096 | -0.13195982 | 0.05 | 0.088 | 1 | 10 |
| smarcb1a | 0.81097 | -0.10264967 | 0.05 | 0.088 | 1 | 10 |
| st5 | 0.81104 | -0.11509784 | 0.05 | 0.088 | 1 | 10 |
| nudt5 | 0.81104 | -0.11779324 | 0.05 | 0.088 | 1 | 10 |
| prkci | 0.81104 | 0.106482014 | 0.075 | 0.059 | 1 | 10 |
| zranb2 | 0.81104 | -0.10261866 | 0.05 | 0.088 | 1 | 10 |
| rgrb | 0.81105 | -0.12345918 | 0.05 | 0.088 | 1 | 10 |
| uck2a | 0.81117 | -0.13484458 | 0.075 | 0.118 | 1 | 10 |
| btf3l4 | 0.81127 | -0.14591484 | 0.175 | 0.235 | 1 | 10 |
| hadhb | 0.81205 | -0.10344182 | 0.075 | 0.118 | 1 | 10 |
| zgc:162944 | 0.81274 | 0.131049361 | 0.4 | 0.353 | 1 | 10 |
| arpc4 | 0.81294 | -0.14204604 | 0.1 | 0.147 | 1 | 10 |
| nsfl1c | 0.81338 | -0.10521898 | 0.075 | 0.118 | 1 | 10 |
| ddx39ab | 0.81539 | -0.10761947 | 0.1 | 0.147 | 1 | 10 |
| smc1al | 0.81625 | 0.119799187 | 0.2 | 0.147 | 1 | 10 |
| denr | 0.81681 | -0.14540581 | 0.1 | 0.147 | 1 | 10 |
| srsf11 | 0.81681 | -0.11200424 | 0.1 | 0.147 | 1 | 10 |
| idh2 | 0.817 | -0.14426986 | 0.15 | 0.206 | 1 | 10 |
| mcrip1 | 0.81756 | -0.12462724 | 0.125 | 0.176 | 1 | 10 |
| fth1a | 0.81853 | 0.13635508 | 0.4 | 0.353 | 1 | 10 |
| mtch2 | 0.81904 | -0.13695332 | 0.075 | 0.118 | 1 | 10 |
| mlf2 | 0.82029 | -0.10516599 | 0.075 | 0.118 | 1 | 10 |
| ptges3b | 0.8204 | 0.140712893 | 0.3 | 0.235 | 1 | 10 |
| si:dkey-33c12.4 | 0.8209 | -0.13462178 | 0.15 | 0.206 | 1 | 10 |
| si:ch211-132p1.3 | 0.82123 | -0.13488225 | 0.075 | 0.118 | 1 | 10 |
| aimp2 | 0.82212 | -0.13464029 | 0.1 | 0.147 | 1 | 10 |
| zgc:162509 | 0.82227 | -0.10778921 | 0.075 | 0.118 | 1 | 10 |
| polr2f | 0.82302 | -0.12780567 | 0.075 | 0.118 | 1 | 10 |
| psmc3 | 0.82312 | -0.10396404 | 0.075 | 0.118 | 1 | 10 |
| stau2 | 0.82361 | 0.131133798 | 0.125 | 0.088 | 1 | 10 |
| ndufab1a | 0.82383 | -0.11743221 | 0.125 | 0.176 | 1 | 10 |
| rap2c | 0.82396 | -0.12073976 | 0.1 | 0.147 | 1 | 10 |
| cd74a | 0.82433 | -0.11278214 | 0.075 | 0.118 | 1 | 10 |
| syap1 | 0.82456 | -0.1227004 | 0.075 | 0.118 | 1 | 10 |
| selenok | 0.82459 | -0.12071955 | 0.075 | 0.118 | 1 | 10 |
| polr2h | 0.82461 | -0.11604537 | 0.075 | 0.118 | 1 | 10 |

| | | | | | | |
|---|---|---|---|---|---|---|
| gnai1 | 0.82461 | -0.12467107 | 0.075 | 0.118 | 1 | 10 |
| ctnnbip1 | 0.82469 | -0.12070252 | 0.075 | 0.118 | 1 | 10 |
| mpzl2b | 0.82485 | -0.13624864 | 0.2 | 0.235 | 1 | 10 |
| zgc:175088 | 0.82612 | -0.13181042 | 0.125 | 0.176 | 1 | 10 |
| pno1 | 0.82623 | -0.12818677 | 0.125 | 0.176 | 1 | 10 |
| nmt1a | 0.82693 | -0.1443025 | 0.1 | 0.147 | 1 | 10 |
| cox6b1 | 0.82812 | 0.140225462 | 0.125 | 0.088 | 1 | 10 |
| selenof | 0.82864 | -0.12005851 | 0.4 | 0.471 | 1 | 10 |
| gkap1 | 0.83013 | -0.10730357 | 0.1 | 0.118 | 1 | 10 |
| lyrm4 | 0.83183 | 0.11034015 | 0.075 | 0.059 | 1 | 10 |
| si:dkey-188i13.9 | 0.8353 | -0.10683237 | 0.1 | 0.118 | 1 | 10 |
| ap1s2 | 0.84725 | 0.137367423 | 0.225 | 0.176 | 1 | 10 |
| naxe | 0.84927 | 0.131743958 | 0.125 | 0.088 | 1 | 10 |
| dpy30 | 0.85181 | -0.1219568 | 0.225 | 0.265 | 1 | 10 |
| eif3d | 0.8534 | 0.102247369 | 0.225 | 0.176 | 1 | 10 |
| cycsb | 0.85348 | 0.134648445 | 0.225 | 0.176 | 1 | 10 |
| cox6c | 0.86302 | 0.130041016 | 0.35 | 0.294 | 1 | 10 |
| gnb1a | 0.86899 | 0.120272479 | 0.125 | 0.088 | 1 | 10 |
| tmem258 | 0.87338 | 0.120516104 | 0.275 | 0.235 | 1 | 10 |
| hsd3b7 | 0.87389 | 0.116732396 | 0.225 | 0.176 | 1 | 10 |
| prdx3 | 0.87629 | -0.11256158 | 0.175 | 0.206 | 1 | 10 |
| si:dkey-6n6.1 | 0.87648 | 0.1228327 | 0.125 | 0.088 | 1 | 10 |
| sdcbp2 | 0.87652 | 0.11526714 | 0.35 | 0.294 | 1 | 10 |
| pgrmc1 | 0.87737 | 0.12631388 | 0.25 | 0.206 | 1 | 10 |
| cops9 | 0.87851 | 0.119937663 | 0.125 | 0.088 | 1 | 10 |
| atp5pb | 0.88574 | 0.103377416 | 0.425 | 0.382 | 1 | 10 |
| vamp3 | 0.88604 | -0.10025752 | 0.125 | 0.147 | 1 | 10 |
| sumo3a | 0.88896 | -0.10205008 | 0.425 | 0.471 | 1 | 10 |
| eif3m | 0.89667 | -0.11127382 | 0.275 | 0.324 | 1 | 10 |
| mdh1aa | 0.89984 | -0.1082219 | 0.2 | 0.235 | 1 | 10 |
| tuba8l | 0.90326 | -0.11751292 | 0.225 | 0.265 | 1 | 10 |
| gnb1b | 0.91377 | -0.10119418 | 0.225 | 0.265 | 1 | 10 |
| fth1a | 7.642068 | -0.92978848 | 0.278 | 0.857 | 0.0154347 | 11 |
| krt91 | 0.00049 | -0.30143095 | 0.667 | 0.607 | 1 | 11 |
| exosc6 | 0.00065 | -0.37117003 | 0 | 0.25 | 1 | 11 |
| hmgb1b | 0.00092 | 0.698342132 | 0.556 | 0.25 | 1 | 11 |
| cebpa | 0.00103 | -0.65303477 | 0.333 | 0.536 | 1 | 11 |
| CU104709.1 | 0.00183 | -0.33737848 | 0 | 0.214 | 1 | 11 |
| XLOC-016167 | 0.00183 | -0.47296876 | 0 | 0.214 | 1 | 11 |
| eif4ea | 0.00183 | -0.26134138 | 0 | 0.214 | 1 | 11 |
| fabp3 | 0.00183 | -0.35647254 | 0 | 0.214 | 1 | 11 |
| ms4a17a.8 | 0.00183 | -0.38002353 | 0 | 0.214 | 1 | 11 |
| setd5 | 0.00183 | -0.34908533 | 0 | 0.214 | 1 | 11 |
| dpy30 | 0.00185 | 0.488915077 | 0.5 | 0.107 | 1 | 11 |
| abcf1 | 0.00195 | 0.456664924 | 0.361 | 0.25 | 1 | 11 |
| tmtc1 | 0.00213 | 0.317506346 | 0.278 | 0.036 | 1 | 11 |
| nfyba | 0.00234 | 0.562771192 | 0.444 | 0.071 | 1 | 11 |
| bhlhe41 | 0.0025 | 0.348307972 | 0.222 | 0 | 1 | 11 |

| | | | | | | |
|---|---|---|---|---|---|---|
| ppt2 | 0.0025 | 0.393547502 | 0.222 | 0 | 1 | 11 |
| sfpq | 0.00266 | 0.585738471 | 0.361 | 0.036 | 1 | 11 |
| pgrmc2 | 0.0028 | 0.399146423 | 0.333 | 0.036 | 1 | 11 |
| gpx1a | 0.00282 | -0.57590031 | 0.194 | 0.607 | 1 | 11 |
| CR847566.1 | 0.00296 | 0.710360649 | 0.639 | 0.286 | 1 | 11 |
| cbr1l | 0.00337 | 0.453024958 | 0.361 | 0.036 | 1 | 11 |
| itga6b | 0.0043 | -0.47558942 | 0.056 | 0.357 | 1 | 11 |
| atp6v0d1 | 0.00506 | -0.21401382 | 0 | 0.179 | 1 | 11 |
| bcl7a | 0.00506 | -0.28426975 | 0 | 0.179 | 1 | 11 |
| btbd2a | 0.00506 | -0.21120796 | 0 | 0.179 | 1 | 11 |
| lrrc15 | 0.00506 | -0.27083469 | 0 | 0.179 | 1 | 11 |
| rnf38 | 0.00506 | -0.28051815 | 0 | 0.179 | 1 | 11 |
| scamp2 | 0.00506 | -0.22997971 | 0 | 0.179 | 1 | 11 |
| tipin | 0.00506 | -0.19222431 | 0 | 0.179 | 1 | 11 |
| zgc:112185 | 0.00506 | -0.26172227 | 0 | 0.179 | 1 | 11 |
| zgc:162509 | 0.00506 | -0.22145202 | 0 | 0.179 | 1 | 11 |
| acvr2ab | 0.00506 | -0.27113348 | 0 | 0.179 | 1 | 11 |
| atxn3 | 0.00506 | -0.25894664 | 0 | 0.179 | 1 | 11 |
| ca5a | 0.00506 | -0.27939192 | 0 | 0.179 | 1 | 11 |
| glg1a | 0.00506 | -0.29805414 | 0 | 0.179 | 1 | 11 |
| smim4 | 0.00506 | -0.23045666 | 0 | 0.179 | 1 | 11 |
| tfa | 0.00506 | -0.51513765 | 0 | 0.179 | 1 | 11 |
| tspan37 | 0.00506 | -0.26750565 | 0 | 0.179 | 1 | 11 |
| tspo | 0.00506 | -0.22225129 | 0 | 0.179 | 1 | 11 |
| utp18 | 0.00506 | -0.22949971 | 0 | 0.179 | 1 | 11 |
| LOC553316 | 0.0052 | 0.370968871 | 0.194 | 0 | 1 | 11 |
| smc5 | 0.0052 | 0.359936843 | 0.194 | 0 | 1 | 11 |
| kdm3b | 0.0052 | 0.284857269 | 0.194 | 0 | 1 | 11 |
| tfdp1a | 0.0052 | 0.31858111 | 0.194 | 0 | 1 | 11 |
| uxt | 0.0052 | 0.309819039 | 0.194 | 0 | 1 | 11 |
| epn2 | 0.0052 | 0.365189708 | 0.194 | 0 | 1 | 11 |
| mt-nd3 | 0.0054 | -0.58206369 | 0.278 | 0.679 | 1 | 11 |
| atp10a | 0.00612 | -0.48943044 | 0.222 | 0.607 | 1 | 11 |
| wbp2nl | 0.00626 | -0.52509152 | 0.139 | 0.5 | 1 | 11 |
| histh1l | 0.00648 | -0.53494851 | 0.083 | 0.321 | 1 | 11 |
| epha3 | 0.00749 | 0.434554443 | 0.389 | 0.107 | 1 | 11 |
| ythdf1 | 0.00753 | -0.44385517 | 0.028 | 0.286 | 1 | 11 |
| rpl27a | 0.00769 | -0.35024046 | 0.778 | 0.893 | 1 | 11 |
| foxo3b | 0.00823 | 0.549693652 | 0.389 | 0.071 | 1 | 11 |
| krt4 | 0.00895 | -0.38496456 | 0.806 | 0.893 | 1 | 11 |
| uchl1 | 0.00958 | -0.16055944 | 0.028 | 0.25 | 1 | 11 |
| lsm6 | 0.00982 | 0.598485852 | 0.472 | 0.143 | 1 | 11 |
| thoc2 | 0.01034 | -0.43309247 | 0.083 | 0.393 | 1 | 11 |
| dhrs13a.3 | 0.01067 | 0.245696658 | 0.167 | 0 | 1 | 11 |
| ing2 | 0.01067 | 0.299141921 | 0.167 | 0 | 1 | 11 |
| pafah1b3 | 0.01067 | 0.258773359 | 0.167 | 0 | 1 | 11 |
| polr3glb | 0.01067 | 0.257138076 | 0.167 | 0 | 1 | 11 |
| vps26a | 0.01067 | 0.301914603 | 0.167 | 0 | 1 | 11 |

| | | | | | | |
|---|---|---|---|---|---|---|
| plrg1 | 0.01067 | 0.266553204 | 0.167 | 0 | 1 | 11 |
| plxna3 | 0.01067 | 0.288837502 | 0.167 | 0 | 1 | 11 |
| si:ch211-271b14.1 | 0.01067 | 0.240438331 | 0.167 | 0 | 1 | 11 |
| si:dkey-101k6.5 | 0.01067 | 0.26456065 | 0.167 | 0 | 1 | 11 |
| zeb1b | 0.01067 | 0.33900425 | 0.167 | 0 | 1 | 11 |
| zgc:153018 | 0.01067 | 0.310120983 | 0.167 | 0 | 1 | 11 |
| spon1b | 0.01087 | 0.335800246 | 0.25 | 0.036 | 1 | 11 |
| slit3 | 0.01151 | -0.16457348 | 0.111 | 0.321 | 1 | 11 |
| tma7 | 0.01215 | 0.525638311 | 0.778 | 0.5 | 1 | 11 |
| mycbp | 0.01283 | 0.536666622 | 0.639 | 0.286 | 1 | 11 |
| XLOC-043834 | 0.01381 | -0.15426574 | 0 | 0.143 | 1 | 11 |
| nipa2 | 0.01381 | -0.20155657 | 0 | 0.143 | 1 | 11 |
| sbno1 | 0.01381 | -0.27147334 | 0 | 0.143 | 1 | 11 |
| si:ch1073-392o20.2 | 0.01381 | -0.26242916 | 0 | 0.143 | 1 | 11 |
| atg3 | 0.01381 | -0.22265513 | 0 | 0.143 | 1 | 11 |
| cacna1c | 0.01381 | -0.32882989 | 0 | 0.143 | 1 | 11 |
| cmc2 | 0.01381 | -0.24844186 | 0 | 0.143 | 1 | 11 |
| edem3 | 0.01381 | -0.23722273 | 0 | 0.143 | 1 | 11 |
| nono | 0.01381 | -0.17983455 | 0 | 0.143 | 1 | 11 |
| pag1 | 0.01381 | -0.29828679 | 0 | 0.143 | 1 | 11 |
| ppp4r3b | 0.01381 | -0.21639986 | 0 | 0.143 | 1 | 11 |
| sec24a | 0.01381 | -0.23580997 | 0 | 0.143 | 1 | 11 |
| sh2b1 | 0.01381 | -0.2312306 | 0 | 0.143 | 1 | 11 |
| skap2 | 0.01381 | -0.14252278 | 0 | 0.143 | 1 | 11 |
| slc35b1 | 0.01381 | -0.16643548 | 0 | 0.143 | 1 | 11 |
| chrdl2 | 0.01381 | -0.2179147 | 0 | 0.143 | 1 | 11 |
| MCMDC2 | 0.01381 | -0.21090486 | 0 | 0.143 | 1 | 11 |
| SMIM10 | 0.01381 | -0.17401178 | 0 | 0.143 | 1 | 11 |
| arih1l | 0.01381 | -0.21687789 | 0 | 0.143 | 1 | 11 |
| atl2 | 0.01381 | -0.21702636 | 0 | 0.143 | 1 | 11 |
| farsa | 0.01381 | -0.20110399 | 0 | 0.143 | 1 | 11 |
| syngr2a | 0.01381 | -0.15690334 | 0 | 0.143 | 1 | 11 |
| alas1 | 0.01381 | -0.17419459 | 0 | 0.143 | 1 | 11 |
| ccnt1 | 0.01381 | -0.22730466 | 0 | 0.143 | 1 | 11 |
| xkrx | 0.01381 | -0.16760801 | 0 | 0.143 | 1 | 11 |
| zc3h7a | 0.01381 | -0.19403297 | 0 | 0.143 | 1 | 11 |
| pcsk5b | 0.01381 | -0.22506792 | 0 | 0.143 | 1 | 11 |
| pygo2 | 0.01381 | -0.2538062 | 0 | 0.143 | 1 | 11 |
| trim8b | 0.01381 | -0.25018389 | 0 | 0.143 | 1 | 11 |
| blzf1 | 0.01381 | -0.23082554 | 0 | 0.143 | 1 | 11 |
| kpnb1 | 0.01399 | -0.37418751 | 0.028 | 0.25 | 1 | 11 |
| nt5e | 0.01594 | -0.44654383 | 0.056 | 0.321 | 1 | 11 |
| CABZ01045617.1 | 0.01677 | 0.529227593 | 0.472 | 0.179 | 1 | 11 |
| maco1a | 0.01729 | -0.34331241 | 0.056 | 0.321 | 1 | 11 |
| CABZ01090890.1 | 0.0175 | -0.47110097 | 0.056 | 0.286 | 1 | 11 |
| prdx2 | 0.01754 | -0.50216176 | 0.5 | 0.786 | 1 | 11 |
| elovl1b | 0.01872 | -0.51054338 | 0.333 | 0.679 | 1 | 11 |
| nat16 | 0.0192 | -0.47423508 | 0.028 | 0.25 | 1 | 11 |

| | | | | | | |
|---|---|---|---|---|---|---|
| zfand3 | 0.01972 | -0.30459032 | 0.167 | 0.464 | 1 | 11 |
| srsf11 | 0.02097 | 0.437679582 | 0.333 | 0.107 | 1 | 11 |
| tmem231 | 0.02104 | 0.407088785 | 0.278 | 0.036 | 1 | 11 |
| wnt4a | 0.02145 | 0.409343583 | 0.278 | 0.036 | 1 | 11 |
| bptf | 0.02174 | 0.205524652 | 0.139 | 0 | 1 | 11 |
| crlf3 | 0.02174 | 0.211311383 | 0.139 | 0 | 1 | 11 |
| ddx54 | 0.02174 | 0.250851959 | 0.139 | 0 | 1 | 11 |
| dnajc12 | 0.02174 | 0.204785896 | 0.139 | 0 | 1 | 11 |
| fuca1.1 | 0.02174 | 0.213050834 | 0.139 | 0 | 1 | 11 |
| lbr | 0.02174 | 0.212602498 | 0.139 | 0 | 1 | 11 |
| lsp1 | 0.02174 | 0.226220065 | 0.139 | 0 | 1 | 11 |
| myo10 | 0.02174 | 0.223725319 | 0.139 | 0 | 1 | 11 |
| otud7b | 0.02174 | 0.237478945 | 0.139 | 0 | 1 | 11 |
| rab34a | 0.02174 | 0.251864377 | 0.139 | 0 | 1 | 11 |
| si:ch73-71c20.5 | 0.02174 | 0.234706353 | 0.139 | 0 | 1 | 11 |
| tmem251 | 0.02174 | 0.280930282 | 0.139 | 0 | 1 | 11 |
| zbtb44 | 0.02174 | 0.242200135 | 0.139 | 0 | 1 | 11 |
| rrp7a | 0.02174 | 0.223467625 | 0.139 | 0 | 1 | 11 |
| syt5b | 0.02174 | 0.268020889 | 0.139 | 0 | 1 | 11 |
| tmem107 | 0.02174 | 0.175510308 | 0.139 | 0 | 1 | 11 |
| ptpn23a | 0.02174 | 0.249891735 | 0.139 | 0 | 1 | 11 |
| ttl | 0.02174 | 0.22326857 | 0.139 | 0 | 1 | 11 |
| med28 | 0.0218 | -0.3037447 | 0.028 | 0.25 | 1 | 11 |
| creg1 | 0.02233 | -0.14129824 | 0.028 | 0.214 | 1 | 11 |
| slc25a26 | 0.02346 | -0.3745215 | 0.028 | 0.25 | 1 | 11 |
| scarb2a | 0.02404 | -0.43900083 | 0.306 | 0.429 | 1 | 11 |
| eif5a | 0.0244 | 0.301948288 | 0.917 | 0.643 | 1 | 11 |
| epb41l4a | 0.02451 | -0.44983992 | 0.083 | 0.357 | 1 | 11 |
| psma5 | 0.02526 | 0.417013606 | 0.5 | 0.321 | 1 | 11 |
| dad1 | 0.02558 | -0.4866506 | 0.444 | 0.75 | 1 | 11 |
| eif4ebp3l | 0.02726 | -0.4816218 | 0.5 | 0.679 | 1 | 11 |
| ccdc80 | 0.02787 | -0.49245603 | 0.194 | 0.5 | 1 | 11 |
| pwp1 | 0.02819 | -0.27251513 | 0.056 | 0.286 | 1 | 11 |
| atp5md | 0.02889 | -0.46687826 | 0.222 | 0.536 | 1 | 11 |
| dbi | 0.02925 | -0.50209175 | 0.389 | 0.679 | 1 | 11 |
| mmp11b | 0.02962 | -0.34418541 | 0.056 | 0.286 | 1 | 11 |
| babam1 | 0.0302 | -0.19398376 | 0.028 | 0.214 | 1 | 11 |
| uqcrc2b | 0.03078 | -0.17249785 | 0.056 | 0.25 | 1 | 11 |
| ppiab | 0.03111 | 0.110709916 | 1 | 0.857 | 1 | 11 |
| h2afx1 | 0.03163 | 0.522021221 | 0.611 | 0.357 | 1 | 11 |
| lsm7 | 0.0322 | 0.513338199 | 0.444 | 0.179 | 1 | 11 |
| lsm4 | 0.03231 | 0.17934581 | 0.389 | 0.429 | 1 | 11 |
| ppiaa | 0.0327 | 0.109652899 | 0.944 | 0.964 | 1 | 11 |
| mycb | 0.03336 | -0.46121028 | 0.222 | 0.536 | 1 | 11 |
| ar | 0.03387 | -0.41156452 | 0.056 | 0.286 | 1 | 11 |
| pdgfab | 0.03395 | 0.38272852 | 0.25 | 0.036 | 1 | 11 |
| p4hb | 0.03413 | -0.4304382 | 0.333 | 0.429 | 1 | 11 |
| ntan1 | 0.03415 | -0.44790092 | 0.056 | 0.286 | 1 | 11 |

| | | | | | | |
|---|---|---|---|---|---|---|
| s100a10b | 0.03442 | -0.15700116 | 0.972 | 1 | 1 | 11 |
| naa50 | 0.03468 | -0.43926062 | 0.083 | 0.321 | 1 | 11 |
| arrdc2 | 0.03535 | 0.319264077 | 0.25 | 0.036 | 1 | 11 |
| twf1a | 0.0359 | -0.27561909 | 0.056 | 0.286 | 1 | 11 |
| itm2ba | 0.03685 | -0.35705248 | 0.611 | 0.714 | 1 | 11 |
| BX936439.1 | 0.03742 | -0.1836747 | 0 | 0.107 | 1 | 11 |
| LOC108179036 | 0.03742 | -0.16097561 | 0 | 0.107 | 1 | 11 |
| atf7b | 0.03742 | -0.14538682 | 0 | 0.107 | 1 | 11 |
| chchd6a | 0.03742 | -0.10405264 | 0 | 0.107 | 1 | 11 |
| enosf1 | 0.03742 | -0.13699499 | 0 | 0.107 | 1 | 11 |
| eps8l3b | 0.03742 | -0.19450855 | 0 | 0.107 | 1 | 11 |
| gcsha | 0.03742 | -0.18779739 | 0 | 0.107 | 1 | 11 |
| gss | 0.03742 | -0.13789514 | 0 | 0.107 | 1 | 11 |
| large1 | 0.03742 | -0.18065708 | 0 | 0.107 | 1 | 11 |
| med29 | 0.03742 | -0.17164177 | 0 | 0.107 | 1 | 11 |
| mlx | 0.03742 | -0.13352422 | 0 | 0.107 | 1 | 11 |
| mthfd1b | 0.03742 | -0.21639775 | 0 | 0.107 | 1 | 11 |
| neu3.5 | 0.03742 | -0.1695246 | 0 | 0.107 | 1 | 11 |
| pdik1l | 0.03742 | -0.2147151 | 0 | 0.107 | 1 | 11 |
| pold1 | 0.03742 | -0.21022841 | 0 | 0.107 | 1 | 11 |
| ppp1r16a | 0.03742 | -0.21942343 | 0 | 0.107 | 1 | 11 |
| prrx1a | 0.03742 | -0.2528933 | 0 | 0.107 | 1 | 11 |
| pter | 0.03742 | -0.20277118 | 0 | 0.107 | 1 | 11 |
| purg | 0.03742 | -0.17546862 | 0 | 0.107 | 1 | 11 |
| rock1 | 0.03742 | -0.19795898 | 0 | 0.107 | 1 | 11 |
| si:ch1073-184j22.1 | 0.03742 | -0.16509832 | 0 | 0.107 | 1 | 11 |
| si:ch211-107n13.1 | 0.03742 | -0.15826847 | 0 | 0.107 | 1 | 11 |
| si:ch211-85n16.3 | 0.03742 | -0.11234793 | 0 | 0.107 | 1 | 11 |
| si:ch211-85n16.4 | 0.03742 | -0.18253432 | 0 | 0.107 | 1 | 11 |
| si:ch73-221f6.4 | 0.03742 | -0.14565911 | 0 | 0.107 | 1 | 11 |
| si:dkey-23k10.5 | 0.03742 | -0.20430638 | 0 | 0.107 | 1 | 11 |
| slc25a36a | 0.03742 | -0.24181667 | 0 | 0.107 | 1 | 11 |
| slc43a1a | 0.03742 | -0.16537932 | 0 | 0.107 | 1 | 11 |
| slco5a1b | 0.03742 | -0.2250912 | 0 | 0.107 | 1 | 11 |
| smurf2 | 0.03742 | -0.15357859 | 0 | 0.107 | 1 | 11 |
| tm9sf5 | 0.03742 | -0.13003615 | 0 | 0.107 | 1 | 11 |
| tmem259 | 0.03742 | -0.14087255 | 0 | 0.107 | 1 | 11 |
| ube2d4 | 0.03742 | -0.18318448 | 0 | 0.107 | 1 | 11 |
| zgc:113411 | 0.03742 | -0.18665376 | 0 | 0.107 | 1 | 11 |
| zmynd19 | 0.03742 | -0.22606417 | 0 | 0.107 | 1 | 11 |
| CU137717.1 | 0.03742 | -0.16121408 | 0 | 0.107 | 1 | 11 |
| arhgap32a | 0.03742 | -0.1855075 | 0 | 0.107 | 1 | 11 |
| dnajc5ga | 0.03742 | -0.1089434 | 0 | 0.107 | 1 | 11 |
| gstt1a | 0.03742 | -0.24772542 | 0 | 0.107 | 1 | 11 |
| hps4 | 0.03742 | -0.15900902 | 0 | 0.107 | 1 | 11 |
| ifih1 | 0.03742 | -0.15547032 | 0 | 0.107 | 1 | 11 |
| lsm14aa | 0.03742 | -0.22224025 | 0 | 0.107 | 1 | 11 |
| lyrm2 | 0.03742 | -0.15956932 | 0 | 0.107 | 1 | 11 |

| | | | | | | |
|---|---|---|---|---|---|---|
| neil1 | 0.03742 | -0.16615694 | 0 | 0.107 | 1 | 11 |
| nf1b | 0.03742 | -0.19538873 | 0 | 0.107 | 1 | 11 |
| nhsl1a | 0.03742 | -0.20134798 | 0 | 0.107 | 1 | 11 |
| prr5a | 0.03742 | -0.19333168 | 0 | 0.107 | 1 | 11 |
| rarab | 0.03742 | -0.15213842 | 0 | 0.107 | 1 | 11 |
| si:ch211-256e16.11 | 0.03742 | -0.20574979 | 0 | 0.107 | 1 | 11 |
| si:dkey-72l14.4 | 0.03742 | -0.12454701 | 0 | 0.107 | 1 | 11 |
| si:zfos-1714f5.3 | 0.03742 | -0.22240448 | 0 | 0.107 | 1 | 11 |
| synm | 0.03742 | -0.1536909 | 0 | 0.107 | 1 | 11 |
| tmem39b | 0.03742 | -0.12992711 | 0 | 0.107 | 1 | 11 |
| ugt5c2 | 0.03742 | -0.13846832 | 0 | 0.107 | 1 | 11 |
| zcchc7 | 0.03742 | -0.12777595 | 0 | 0.107 | 1 | 11 |
| znf740b | 0.03742 | -0.19006417 | 0 | 0.107 | 1 | 11 |
| SCCPDH (1 of many) | 0.03742 | -0.18948089 | 0 | 0.107 | 1 | 11 |
| ago4 | 0.03742 | -0.19130263 | 0 | 0.107 | 1 | 11 |
| ank1a | 0.03742 | -0.25758513 | 0 | 0.107 | 1 | 11 |
| aph1b | 0.03742 | -0.27876336 | 0 | 0.107 | 1 | 11 |
| atg10 | 0.03742 | -0.18971971 | 0 | 0.107 | 1 | 11 |
| auts2b | 0.03742 | -0.17543367 | 0 | 0.107 | 1 | 11 |
| ccnh | 0.03742 | -0.10405141 | 0 | 0.107 | 1 | 11 |
| cdip1 | 0.03742 | -0.20883669 | 0 | 0.107 | 1 | 11 |
| cnppd1 | 0.03742 | -0.11879751 | 0 | 0.107 | 1 | 11 |
| cplx2l | 0.03742 | -0.26902505 | 0 | 0.107 | 1 | 11 |
| def8 | 0.03742 | -0.19145666 | 0 | 0.107 | 1 | 11 |
| irak1 | 0.03742 | -0.15689558 | 0 | 0.107 | 1 | 11 |
| kdm6ba | 0.03742 | -0.26735969 | 0 | 0.107 | 1 | 11 |
| kif1b | 0.03742 | -0.19045024 | 0 | 0.107 | 1 | 11 |
| klf12a | 0.03742 | -0.13855827 | 0 | 0.107 | 1 | 11 |
| malt3 | 0.03742 | -0.20083953 | 0 | 0.107 | 1 | 11 |
| mboat7 | 0.03742 | -0.16293401 | 0 | 0.107 | 1 | 11 |
| mib1 | 0.03742 | -0.15362316 | 0 | 0.107 | 1 | 11 |
| msl2b | 0.03742 | -0.29160559 | 0 | 0.107 | 1 | 11 |
| nol11 | 0.03742 | -0.11085499 | 0 | 0.107 | 1 | 11 |
| nudt17 | 0.03742 | -0.15183032 | 0 | 0.107 | 1 | 11 |
| nus1 | 0.03742 | -0.1890091 | 0 | 0.107 | 1 | 11 |
| ormdl3 | 0.03742 | -0.16661442 | 0 | 0.107 | 1 | 11 |
| pdxp | 0.03742 | -0.22291269 | 0 | 0.107 | 1 | 11 |
| pex5la | 0.03742 | -0.20984036 | 0 | 0.107 | 1 | 11 |
| rabl3 | 0.03742 | -0.11454538 | 0 | 0.107 | 1 | 11 |
| rbm42 | 0.03742 | -0.17920689 | 0 | 0.107 | 1 | 11 |
| rnf25 | 0.03742 | -0.18319072 | 0 | 0.107 | 1 | 11 |
| rrbp1b | 0.03742 | -0.1418885 | 0 | 0.107 | 1 | 11 |
| sema6e | 0.03742 | -0.22135457 | 0 | 0.107 | 1 | 11 |
| sgcb | 0.03742 | -0.1792174 | 0 | 0.107 | 1 | 11 |
| si:ch1073-314i13.4 | 0.03742 | -0.22393795 | 0 | 0.107 | 1 | 11 |
| sik3 | 0.03742 | -0.25103665 | 0 | 0.107 | 1 | 11 |
| slc9a3r1a | 0.03742 | -0.1848402 | 0 | 0.107 | 1 | 11 |
| smchd1 | 0.03742 | -0.14907382 | 0 | 0.107 | 1 | 11 |

| | | | | | | |
|---|---|---|---|---|---|---|
| smyd2a | 0.03742 | -0.10595164 | 0 | 0.107 | 1 | 11 |
| snapc1b | 0.03742 | -0.12721487 | 0 | 0.107 | 1 | 11 |
| tanc1b | 0.03742 | -0.18349232 | 0 | 0.107 | 1 | 11 |
| tnpo1 | 0.03742 | -0.20001806 | 0 | 0.107 | 1 | 11 |
| tspan5a | 0.03742 | -0.15534161 | 0 | 0.107 | 1 | 11 |
| twsg1a | 0.03742 | -0.21571823 | 0 | 0.107 | 1 | 11 |
| vps39 | 0.03742 | -0.22761109 | 0 | 0.107 | 1 | 11 |
| zc3h10 | 0.03742 | -0.16050403 | 0 | 0.107 | 1 | 11 |
| zgc:154055 | 0.03742 | -0.1538769 | 0 | 0.107 | 1 | 11 |
| dsc2l | 0.03759 | -0.14708471 | 0.111 | 0.286 | 1 | 11 |
| cygb1 | 0.0379 | -0.45991274 | 0.528 | 0.714 | 1 | 11 |
| hmgn6 | 0.03824 | 0.248506887 | 0.833 | 0.786 | 1 | 11 |
| phc2a | 0.03977 | 0.352671141 | 0.25 | 0.036 | 1 | 11 |
| csnk1db | 0.03997 | -0.29476971 | 0.194 | 0.464 | 1 | 11 |
| tob1a | 0.04018 | 0.449142495 | 0.528 | 0.321 | 1 | 11 |
| psmd7 | 0.04189 | 0.188964246 | 0.222 | 0.25 | 1 | 11 |
| nova2 | 0.04346 | 0.478932225 | 0.25 | 0.071 | 1 | 11 |
| XLOC-011107 | 0.0441 | 0.180436599 | 0.111 | 0 | 1 | 11 |
| XLOC-016546 | 0.0441 | 0.215659165 | 0.111 | 0 | 1 | 11 |
| adsl | 0.0441 | 0.138344772 | 0.111 | 0 | 1 | 11 |
| ash1l | 0.0441 | 0.224639863 | 0.111 | 0 | 1 | 11 |
| bach2b | 0.0441 | 0.17870863 | 0.111 | 0 | 1 | 11 |
| ccdc82 | 0.0441 | 0.226396136 | 0.111 | 0 | 1 | 11 |
| cttnbp2nlb | 0.0441 | 0.243408954 | 0.111 | 0 | 1 | 11 |
| ddx42 | 0.0441 | 0.181385893 | 0.111 | 0 | 1 | 11 |
| dynll2a | 0.0441 | 0.173961275 | 0.111 | 0 | 1 | 11 |
| keap1b | 0.0441 | 0.246296622 | 0.111 | 0 | 1 | 11 |
| nkain1 | 0.0441 | 0.197976687 | 0.111 | 0 | 1 | 11 |
| oga | 0.0441 | 0.149511157 | 0.111 | 0 | 1 | 11 |
| prdm2b | 0.0441 | 0.209681071 | 0.111 | 0 | 1 | 11 |
| ralba | 0.0441 | 0.252942007 | 0.111 | 0 | 1 | 11 |
| rbbp5 | 0.0441 | 0.199897592 | 0.111 | 0 | 1 | 11 |
| rnf4 | 0.0441 | 0.160421847 | 0.111 | 0 | 1 | 11 |
| si:ch211-156b7.4 | 0.0441 | 0.208984583 | 0.111 | 0 | 1 | 11 |
| siva1 | 0.0441 | 0.186763343 | 0.111 | 0 | 1 | 11 |
| smap1 | 0.0441 | 0.248276896 | 0.111 | 0 | 1 | 11 |
| snrnp200 | 0.0441 | 0.23378001 | 0.111 | 0 | 1 | 11 |
| snrpb2 | 0.0441 | 0.156138201 | 0.111 | 0 | 1 | 11 |
| stag1b | 0.0441 | 0.180564328 | 0.111 | 0 | 1 | 11 |
| tkfc | 0.0441 | 0.152554711 | 0.111 | 0 | 1 | 11 |
| trim3b | 0.0441 | 0.194976761 | 0.111 | 0 | 1 | 11 |
| ubald1b | 0.0441 | 0.160233372 | 0.111 | 0 | 1 | 11 |
| ugt5b2 | 0.0441 | 0.182502163 | 0.111 | 0 | 1 | 11 |
| zhx2a | 0.0441 | 0.232017625 | 0.111 | 0 | 1 | 11 |
| arid1b | 0.0441 | 0.18516388 | 0.111 | 0 | 1 | 11 |
| cep19 | 0.0441 | 0.166373587 | 0.111 | 0 | 1 | 11 |
| cfl1l | 0.0441 | 0.119863326 | 0.111 | 0 | 1 | 11 |
| cipca | 0.0441 | 0.251774553 | 0.111 | 0 | 1 | 11 |

| | | | | | | |
|---|---|---|---|---|---|---|
| creb3l2 | 0.0441 | 0.212318387 | 0.111 | 0 | 1 | 11 |
| dnajc3a | 0.0441 | 0.212708529 | 0.111 | 0 | 1 | 11 |
| elp3 | 0.0441 | 0.235340992 | 0.111 | 0 | 1 | 11 |
| grk4 | 0.0441 | 0.172898139 | 0.111 | 0 | 1 | 11 |
| gtf2a2 | 0.0441 | 0.182223235 | 0.111 | 0 | 1 | 11 |
| nudt22 | 0.0441 | 0.200779405 | 0.111 | 0 | 1 | 11 |
| parp8 | 0.0441 | 0.225806321 | 0.111 | 0 | 1 | 11 |
| pcsk7 | 0.0441 | 0.255041069 | 0.111 | 0 | 1 | 11 |
| si:ch211-191a24.4 | 0.0441 | 0.204517964 | 0.111 | 0 | 1 | 11 |
| tcf20 | 0.0441 | 0.242449465 | 0.111 | 0 | 1 | 11 |
| AK6 | 0.0441 | 0.164221968 | 0.111 | 0 | 1 | 11 |
| TIMM21 | 0.0441 | 0.177268152 | 0.111 | 0 | 1 | 11 |
| bin2b | 0.0441 | 0.143523651 | 0.111 | 0 | 1 | 11 |
| gpatch1 | 0.0441 | 0.236432052 | 0.111 | 0 | 1 | 11 |
| luc7l | 0.0441 | 0.161340947 | 0.111 | 0 | 1 | 11 |
| mosmob | 0.0441 | 0.210411743 | 0.111 | 0 | 1 | 11 |
| mrps2 | 0.0441 | 0.183568739 | 0.111 | 0 | 1 | 11 |
| slc38a10 | 0.0441 | 0.166650513 | 0.111 | 0 | 1 | 11 |
| wnt7aa | 0.0441 | 0.198146957 | 0.111 | 0 | 1 | 11 |
| aftpha | 0.0441 | 0.205177699 | 0.111 | 0 | 1 | 11 |
| arpc5lb | 0.0441 | 0.166273085 | 0.111 | 0 | 1 | 11 |
| calm1b | 0.0441 | 0.164749874 | 0.111 | 0 | 1 | 11 |
| col4a5 | 0.0441 | 0.186082585 | 0.111 | 0 | 1 | 11 |
| cutc | 0.0441 | 0.205485304 | 0.111 | 0 | 1 | 11 |
| dcp1b | 0.0441 | 0.214214825 | 0.111 | 0 | 1 | 11 |
| fam162a | 0.0441 | 0.159858662 | 0.111 | 0 | 1 | 11 |
| gch1 | 0.0441 | 0.228102054 | 0.111 | 0 | 1 | 11 |
| gnaia | 0.0441 | 0.205792554 | 0.111 | 0 | 1 | 11 |
| idh3b | 0.0441 | 0.143225114 | 0.111 | 0 | 1 | 11 |
| lsm1 | 0.0441 | 0.188857742 | 0.111 | 0 | 1 | 11 |
| mrpl44 | 0.0441 | 0.131149767 | 0.111 | 0 | 1 | 11 |
| npm1b | 0.0441 | 0.162671789 | 0.111 | 0 | 1 | 11 |
| osbpl2a | 0.0441 | 0.215036431 | 0.111 | 0 | 1 | 11 |
| pcid2 | 0.0441 | 0.17636076 | 0.111 | 0 | 1 | 11 |
| si:ch73-386h18.1 | 0.0441 | 0.23615953 | 0.111 | 0 | 1 | 11 |
| si:dkey-27b3.2 | 0.0441 | 0.209132516 | 0.111 | 0 | 1 | 11 |
| slco5a1a | 0.0441 | 0.175955262 | 0.111 | 0 | 1 | 11 |
| spc25 | 0.0441 | 0.192824849 | 0.111 | 0 | 1 | 11 |
| ythdf3 | 0.0441 | 0.232908451 | 0.111 | 0 | 1 | 11 |
| zbtb41 | 0.0441 | 0.218674601 | 0.111 | 0 | 1 | 11 |
| zgc:66472 | 0.0441 | 0.16915715 | 0.111 | 0 | 1 | 11 |
| znf507 | 0.0441 | 0.237219362 | 0.111 | 0 | 1 | 11 |
| axin1 | 0.0441 | 0.183236329 | 0.111 | 0 | 1 | 11 |
| nup160 | 0.0441 | 0.194124317 | 0.111 | 0 | 1 | 11 |
| palld | 0.0441 | 0.207000746 | 0.111 | 0 | 1 | 11 |
| aar2 | 0.0441 | 0.19791032 | 0.111 | 0 | 1 | 11 |
| asmtl | 0.0441 | 0.157807228 | 0.111 | 0 | 1 | 11 |
| cwf19l2 | 0.0441 | 0.177525649 | 0.111 | 0 | 1 | 11 |

| | | | | | | |
|---|---|---|---|---|---|---|
| fam214a | 0.0441 | 0.173312915 | 0.111 | 0 | 1 | 11 |
| pgm1 | 0.0441 | 0.182860952 | 0.111 | 0 | 1 | 11 |
| si:ch1073-157b13.1 | 0.0441 | 0.161962642 | 0.111 | 0 | 1 | 11 |
| tmem42a | 0.0441 | 0.218789134 | 0.111 | 0 | 1 | 11 |
| eml3 | 0.0441 | 0.202336053 | 0.111 | 0 | 1 | 11 |
| as3mt | 0.0441 | 0.206972228 | 0.111 | 0 | 1 | 11 |
| thy1 | 0.04464 | -0.50920473 | 0.083 | 0.286 | 1 | 11 |
| frzb | 0.04488 | 0.576235858 | 0.556 | 0.25 | 1 | 11 |
| ssr1 | 0.04541 | -0.25505447 | 0.278 | 0.571 | 1 | 11 |
| ndufa9a | 0.04681 | -0.14082728 | 0.083 | 0.286 | 1 | 11 |
| klf11b | 0.04727 | 0.335211895 | 0.306 | 0.071 | 1 | 11 |
| cox5ab | 0.04799 | -0.43447245 | 0.25 | 0.464 | 1 | 11 |
| hoxc10a | 0.0481 | 0.294543612 | 0.222 | 0.036 | 1 | 11 |
| si:ch211-217k17.7 | 0.04856 | -0.4104434 | 0.25 | 0.536 | 1 | 11 |
| frs2b | 0.04874 | -0.36079809 | 0.028 | 0.214 | 1 | 11 |
| eif1axb | 0.04895 | -0.42955247 | 0.361 | 0.607 | 1 | 11 |
| gsna | 0.04968 | -0.15290353 | 0.028 | 0.179 | 1 | 11 |
| slx1b | 0.04969 | 0.187573677 | 0.194 | 0.036 | 1 | 11 |
| plekha1a | 0.04974 | -0.33742364 | 0.028 | 0.214 | 1 | 11 |
| tmem47 | 0.04989 | -0.29623808 | 0.028 | 0.214 | 1 | 11 |
| sumo1 | 0.05019 | 0.381591618 | 0.444 | 0.286 | 1 | 11 |
| cyb5r3 | 0.05041 | -0.30797645 | 0.028 | 0.214 | 1 | 11 |
| waca | 0.05047 | 0.38897489 | 0.222 | 0.036 | 1 | 11 |
| tmed9 | 0.05072 | -0.34362969 | 0.083 | 0.321 | 1 | 11 |
| b3glcta | 0.05083 | -0.27887447 | 0.028 | 0.214 | 1 | 11 |
| taf12 | 0.05201 | -0.30117655 | 0.25 | 0.536 | 1 | 11 |
| jdp2b | 0.05222 | 0.501972636 | 0.361 | 0.107 | 1 | 11 |
| emd | 0.05268 | 0.432957807 | 0.333 | 0.143 | 1 | 11 |
| parp3 | 0.05315 | -0.22314929 | 0.028 | 0.214 | 1 | 11 |
| igf2bp3 | 0.05325 | -0.25247623 | 0.028 | 0.214 | 1 | 11 |
| CR383676.1 | 0.05326 | 0.268790086 | 0.972 | 0.786 | 1 | 11 |
| slc25a11 | 0.05328 | 0.361923331 | 0.361 | 0.107 | 1 | 11 |
| sypl1 | 0.0533 | -0.29401476 | 0.028 | 0.214 | 1 | 11 |
| sav1 | 0.05345 | -0.27442235 | 0.028 | 0.214 | 1 | 11 |
| arl2 | 0.05366 | -0.25310458 | 0.028 | 0.214 | 1 | 11 |
| si:ch211-262e15.1 | 0.05381 | 0.426090412 | 0.333 | 0.107 | 1 | 11 |
| si:ch211-276i12.11 | 0.05392 | -0.19747084 | 0.056 | 0.25 | 1 | 11 |
| si:ch211-213a13.1 | 0.05404 | -0.31559996 | 0.222 | 0.5 | 1 | 11 |
| sirt7 | 0.05685 | 0.318216987 | 0.222 | 0.036 | 1 | 11 |
| itgb4 | 0.05864 | -0.42909381 | 0.111 | 0.357 | 1 | 11 |
| sub1b | 0.05877 | 0.389808891 | 0.5 | 0.214 | 1 | 11 |
| fam168b | 0.05877 | 0.355802973 | 0.222 | 0.036 | 1 | 11 |
| FP016056.1 | 0.05983 | -0.42032631 | 0.111 | 0.357 | 1 | 11 |
| churc1 | 0.05984 | 0.351535699 | 0.278 | 0.071 | 1 | 11 |
| dag1 | 0.05996 | -0.2596262 | 0.222 | 0.464 | 1 | 11 |
| si:dkey-266m15.6 | 0.05996 | 0.234319632 | 0.222 | 0.036 | 1 | 11 |
| si:ch211-222l21.1 | 0.06151 | 0.303102891 | 0.944 | 0.75 | 1 | 11 |
| si:ch73-22o12.1 | 0.06256 | -0.41147464 | 0.139 | 0.393 | 1 | 11 |

| | | | | | | |
|---|---|---|---|---|---|---|
| cd82a | 0.06458 | 0.362421686 | 0.222 | 0.036 | 1 | 11 |
| selenoh | 0.06498 | 0.408216803 | 0.389 | 0.143 | 1 | 11 |
| lpl | 0.06521 | -0.40361613 | 0.056 | 0.25 | 1 | 11 |
| hoxb3a | 0.06612 | 0.401240763 | 0.278 | 0.071 | 1 | 11 |
| xrn2 | 0.06691 | -0.17307227 | 0.028 | 0.179 | 1 | 11 |
| creb1a | 0.06696 | 0.365041979 | 0.306 | 0.107 | 1 | 11 |
| khsrp | 0.06793 | -0.38397356 | 0.194 | 0.464 | 1 | 11 |
| akap12b | 0.06806 | 0.432781858 | 0.472 | 0.286 | 1 | 11 |
| rras | 0.06821 | -0.35376353 | 0.167 | 0.429 | 1 | 11 |
| mafba | 0.06836 | 0.430732647 | 0.417 | 0.179 | 1 | 11 |
| smndc1 | 0.0693 | 0.40370247 | 0.361 | 0.143 | 1 | 11 |
| aoc2 | 0.06965 | -0.56619437 | 0.361 | 0.607 | 1 | 11 |
| ube2al | 0.07077 | 0.301457522 | 0.278 | 0.071 | 1 | 11 |
| sltm | 0.07136 | 0.364343761 | 0.278 | 0.071 | 1 | 11 |
| oaz2b | 0.07258 | 0.332600132 | 0.389 | 0.143 | 1 | 11 |
| si:dkey-204f11.64 | 0.07276 | -0.19482096 | 0.028 | 0.179 | 1 | 11 |
| hmgb1a | 0.07306 | 0.294664162 | 0.75 | 0.714 | 1 | 11 |
| matr3l1.1 | 0.07312 | 0.207106353 | 0.25 | 0.071 | 1 | 11 |
| kif13ba | 0.07328 | 0.29712476 | 0.222 | 0.036 | 1 | 11 |
| prpf40a | 0.07362 | -0.24333765 | 0.056 | 0.25 | 1 | 11 |
| mmp16a | 0.07379 | 0.429585698 | 0.389 | 0.143 | 1 | 11 |
| cebpg | 0.07387 | 0.370512952 | 0.306 | 0.107 | 1 | 11 |
| ube2g1a | 0.0739 | 0.255670462 | 0.222 | 0.036 | 1 | 11 |
| rbpms2b | 0.07444 | 0.330568076 | 0.361 | 0.25 | 1 | 11 |
| lclat1 | 0.0754 | -0.33971012 | 0.056 | 0.25 | 1 | 11 |
| hmgn2 | 0.07591 | 0.39057393 | 0.528 | 0.536 | 1 | 11 |
| gadd45ab | 0.07635 | -0.42888021 | 0.083 | 0.286 | 1 | 11 |
| arf2b | 0.07669 | -0.12773891 | 0.194 | 0.393 | 1 | 11 |
| si:dkey-16p21.8 | 0.07708 | -0.27487965 | 0.056 | 0.25 | 1 | 11 |
| mgst1.1 | 0.07723 | 0.355150249 | 0.444 | 0.321 | 1 | 11 |
| atp5pf | 0.07748 | -0.46238252 | 0.278 | 0.464 | 1 | 11 |
| calm3b | 0.07751 | 0.368196648 | 0.444 | 0.321 | 1 | 11 |
| cops4 | 0.07793 | -0.10255048 | 0.083 | 0.25 | 1 | 11 |
| ciartb | 0.07971 | -0.34910755 | 0.056 | 0.25 | 1 | 11 |
| tnfsf10l | 0.08011 | 0.178450869 | 0.5 | 0.5 | 1 | 11 |
| hmgb2b | 0.08036 | 0.371799456 | 0.583 | 0.536 | 1 | 11 |
| tjp2b | 0.08063 | -0.20945669 | 0.306 | 0.536 | 1 | 11 |
| tnrc6c1 | 0.0814 | 0.421358471 | 0.444 | 0.214 | 1 | 11 |
| rnd3a | 0.08148 | 0.274078038 | 0.278 | 0.071 | 1 | 11 |
| smo | 0.08154 | -0.33466095 | 0.056 | 0.25 | 1 | 11 |
| si:dkey-42i9.4 | 0.08191 | 0.366940619 | 0.389 | 0.143 | 1 | 11 |
| slc25a28 | 0.08214 | -0.32964601 | 0.056 | 0.25 | 1 | 11 |
| snrpd3l | 0.08216 | 0.187005574 | 0.194 | 0.036 | 1 | 11 |
| kirrel1b | 0.08236 | -0.21263081 | 0.028 | 0.179 | 1 | 11 |
| ubald1a | 0.08452 | -0.20898545 | 0.056 | 0.214 | 1 | 11 |
| neu3.2 | 0.08531 | -0.16872923 | 0.056 | 0.214 | 1 | 11 |
| h2afvb | 0.08545 | 0.4010364 | 0.833 | 0.643 | 1 | 11 |
| itga5 | 0.08561 | -0.41113018 | 0.139 | 0.357 | 1 | 11 |

| | | | | | | |
|---|---|---|---|---|---|---|
| cab39l | 0.08597 | -0.27136709 | 0.083 | 0.286 | 1 | 11 |
| twistnb | 0.08651 | -0.21882365 | 0.028 | 0.179 | 1 | 11 |
| plpp3 | 0.08665 | -0.41692803 | 0.111 | 0.321 | 1 | 11 |
| tp53 | 0.08756 | 0.331864043 | 0.333 | 0.107 | 1 | 11 |
| dchs1b | 0.08809 | 0.381176472 | 0.333 | 0.107 | 1 | 11 |
| rrbp1a | 0.0888 | 0.349217781 | 0.278 | 0.071 | 1 | 11 |
| apbb1 | 0.08962 | 0.116388182 | 0.083 | 0 | 1 | 11 |
| hoxd10a | 0.08962 | 0.158243053 | 0.083 | 0 | 1 | 11 |
| slc37a4b | 0.08962 | 0.143192983 | 0.083 | 0 | 1 | 11 |
| CABZ01033394.1 | 0.08962 | 0.173200796 | 0.083 | 0 | 1 | 11 |
| CABZ01090373.1 | 0.08962 | 0.132255023 | 0.083 | 0 | 1 | 11 |
| CABZ01113599.1 | 0.08962 | 0.153569526 | 0.083 | 0 | 1 | 11 |
| CT025690.1 | 0.08962 | 0.129731521 | 0.083 | 0 | 1 | 11 |
| FQ323156.1 | 0.08962 | 0.146883764 | 0.083 | 0 | 1 | 11 |
| LOC101885376 | 0.08962 | 0.143865764 | 0.083 | 0 | 1 | 11 |
| MAP3K11 | 0.08962 | 0.181192376 | 0.083 | 0 | 1 | 11 |
| MTERF4 | 0.08962 | 0.146185686 | 0.083 | 0 | 1 | 11 |
| UTP14C | 0.08962 | 0.108740945 | 0.083 | 0 | 1 | 11 |
| XLOC-004016 | 0.08962 | 0.166638483 | 0.083 | 0 | 1 | 11 |
| XLOC-026010 | 0.08962 | 0.132311074 | 0.083 | 0 | 1 | 11 |
| XLOC-035413 | 0.08962 | 0.147456305 | 0.083 | 0 | 1 | 11 |
| abcb7 | 0.08962 | 0.153695763 | 0.083 | 0 | 1 | 11 |
| adrb2a | 0.08962 | 0.157902751 | 0.083 | 0 | 1 | 11 |
| agbl5 | 0.08962 | 0.118761762 | 0.083 | 0 | 1 | 11 |
| amd1 | 0.08962 | 0.165084903 | 0.083 | 0 | 1 | 11 |
| anxa3b | 0.08962 | 0.112308752 | 0.083 | 0 | 1 | 11 |
| ap3s2 | 0.08962 | 0.155836096 | 0.083 | 0 | 1 | 11 |
| arl6ip6 | 0.08962 | 0.143990604 | 0.083 | 0 | 1 | 11 |
| arntl1b | 0.08962 | 0.195433364 | 0.083 | 0 | 1 | 11 |
| atad5a | 0.08962 | 0.154446339 | 0.083 | 0 | 1 | 11 |
| bahd1 | 0.08962 | 0.177787827 | 0.083 | 0 | 1 | 11 |
| bbox1 | 0.08962 | 0.194730944 | 0.083 | 0 | 1 | 11 |
| c1galt1c1 | 0.08962 | 0.149409734 | 0.083 | 0 | 1 | 11 |
| cactin | 0.08962 | 0.151280708 | 0.083 | 0 | 1 | 11 |
| ccdc61 | 0.08962 | 0.191657061 | 0.083 | 0 | 1 | 11 |
| chac2 | 0.08962 | 0.166398545 | 0.083 | 0 | 1 | 11 |
| clec16a | 0.08962 | 0.134309936 | 0.083 | 0 | 1 | 11 |
| cmss1 | 0.08962 | 0.184533989 | 0.083 | 0 | 1 | 11 |
| cyb561a3a | 0.08962 | 0.155294548 | 0.083 | 0 | 1 | 11 |
| dffa | 0.08962 | 0.142025929 | 0.083 | 0 | 1 | 11 |
| dguok | 0.08962 | 0.127307832 | 0.083 | 0 | 1 | 11 |
| diaph2 | 0.08962 | 0.144637021 | 0.083 | 0 | 1 | 11 |
| dip2ca | 0.08962 | 0.201620278 | 0.083 | 0 | 1 | 11 |
| dlg3 | 0.08962 | 0.165455594 | 0.083 | 0 | 1 | 11 |
| efs | 0.08962 | 0.147730646 | 0.083 | 0 | 1 | 11 |
| eral1 | 0.08962 | 0.138610107 | 0.083 | 0 | 1 | 11 |
| eya3 | 0.08962 | 0.176694518 | 0.083 | 0 | 1 | 11 |
| fkrp | 0.08962 | 0.183526411 | 0.083 | 0 | 1 | 11 |

| | | | | | | |
|---|---|---|---|---|---|---|
| foxp1a | 0.08962 | 0.1544012 | 0.083 | 0 | 1 | 11 |
| gab2 | 0.08962 | 0.113440373 | 0.083 | 0 | 1 | 11 |
| gmppaa | 0.08962 | 0.152358272 | 0.083 | 0 | 1 | 11 |
| hsd17b12b | 0.08962 | 0.139566737 | 0.083 | 0 | 1 | 11 |
| hsf1 | 0.08962 | 0.156612648 | 0.083 | 0 | 1 | 11 |
| igf1ra | 0.08962 | 0.148355097 | 0.083 | 0 | 1 | 11 |
| impa2 | 0.08962 | 0.13160923 | 0.083 | 0 | 1 | 11 |
| impdh1b | 0.08962 | 0.147904667 | 0.083 | 0 | 1 | 11 |
| lrig1 | 0.08962 | 0.196654584 | 0.083 | 0 | 1 | 11 |
| lyar | 0.08962 | 0.13324549 | 0.083 | 0 | 1 | 11 |
| map3k14a | 0.08962 | 0.117174009 | 0.083 | 0 | 1 | 11 |
| mapk4 | 0.08962 | 0.151655695 | 0.083 | 0 | 1 | 11 |
| mdc1 | 0.08962 | 0.139345769 | 0.083 | 0 | 1 | 11 |
| mmrn2a | 0.08962 | 0.220177653 | 0.083 | 0 | 1 | 11 |
| mocs3 | 0.08962 | 0.169995165 | 0.083 | 0 | 1 | 11 |
| mterf2 | 0.08962 | 0.170531125 | 0.083 | 0 | 1 | 11 |
| ndc1 | 0.08962 | 0.122895496 | 0.083 | 0 | 1 | 11 |
| ndufaf6 | 0.08962 | 0.160418047 | 0.083 | 0 | 1 | 11 |
| nek7 | 0.08962 | 0.163446753 | 0.083 | 0 | 1 | 11 |
| nop14 | 0.08962 | 0.119239648 | 0.083 | 0 | 1 | 11 |
| nup50 | 0.08962 | 0.136350193 | 0.083 | 0 | 1 | 11 |
| paqr3a | 0.08962 | 0.167211837 | 0.083 | 0 | 1 | 11 |
| pde4ca | 0.08962 | 0.148526416 | 0.083 | 0 | 1 | 11 |
| pex14 | 0.08962 | 0.146559538 | 0.083 | 0 | 1 | 11 |
| phrf1 | 0.08962 | 0.155289809 | 0.083 | 0 | 1 | 11 |
| pip4k2aa | 0.08962 | 0.128956404 | 0.083 | 0 | 1 | 11 |
| pitpnaa | 0.08962 | 0.137199491 | 0.083 | 0 | 1 | 11 |
| pitx1 | 0.08962 | 0.146479166 | 0.083 | 0 | 1 | 11 |
| plekho1b | 0.08962 | 0.208376111 | 0.083 | 0 | 1 | 11 |
| pold2 | 0.08962 | 0.204202317 | 0.083 | 0 | 1 | 11 |
| pomt1 | 0.08962 | 0.159089195 | 0.083 | 0 | 1 | 11 |
| ppa1b | 0.08962 | 0.160418732 | 0.083 | 0 | 1 | 11 |
| pros1 | 0.08962 | 0.149187968 | 0.083 | 0 | 1 | 11 |
| prr12a | 0.08962 | 0.131696664 | 0.083 | 0 | 1 | 11 |
| rab28 | 0.08962 | 0.120088967 | 0.083 | 0 | 1 | 11 |
| rbm12b | 0.08962 | 0.12109311 | 0.083 | 0 | 1 | 11 |
| rell1 | 0.08962 | 0.180981952 | 0.083 | 0 | 1 | 11 |
| rereb | 0.08962 | 0.155353007 | 0.083 | 0 | 1 | 11 |
| rpz | 0.08962 | 0.169045207 | 0.083 | 0 | 1 | 11 |
| scrib | 0.08962 | 0.17064097 | 0.083 | 0 | 1 | 11 |
| selenon | 0.08962 | 0.175744237 | 0.083 | 0 | 1 | 11 |
| selenow2a | 0.08962 | 0.102081005 | 0.083 | 0 | 1 | 11 |
| senp7b | 0.08962 | 0.180145652 | 0.083 | 0 | 1 | 11 |
| sh3bp4a | 0.08962 | 0.128605509 | 0.083 | 0 | 1 | 11 |
| si:ch211-106j24.1 | 0.08962 | 0.143196938 | 0.083 | 0 | 1 | 11 |
| si:ch211-11n16.2 | 0.08962 | 0.211640227 | 0.083 | 0 | 1 | 11 |
| si:ch211-233a24.2 | 0.08962 | 0.159788016 | 0.083 | 0 | 1 | 11 |
| si:dkey-6i22.5 | 0.08962 | 0.125770553 | 0.083 | 0 | 1 | 11 |

| | | | | | | |
|---|---|---|---|---|---|---|
| slc12a2 | 0.08962 | 0.205151929 | 0.083 | 0 | 1 | 11 |
| spag7 | 0.08962 | 0.160836797 | 0.083 | 0 | 1 | 11 |
| spg7 | 0.08962 | 0.120085072 | 0.083 | 0 | 1 | 11 |
| stam | 0.08962 | 0.136246502 | 0.083 | 0 | 1 | 11 |
| stambpa | 0.08962 | 0.104478483 | 0.083 | 0 | 1 | 11 |
| stx2b | 0.08962 | 0.15332549 | 0.083 | 0 | 1 | 11 |
| swt1 | 0.08962 | 0.115660044 | 0.083 | 0 | 1 | 11 |
| thrb | 0.08962 | 0.136636298 | 0.083 | 0 | 1 | 11 |
| timm50 | 0.08962 | 0.120147906 | 0.083 | 0 | 1 | 11 |
| tmx2b | 0.08962 | 0.128719048 | 0.083 | 0 | 1 | 11 |
| tom1 | 0.08962 | 0.149191008 | 0.083 | 0 | 1 | 11 |
| trap1 | 0.08962 | 0.160762585 | 0.083 | 0 | 1 | 11 |
| txnl4b | 0.08962 | 0.140730669 | 0.083 | 0 | 1 | 11 |
| usp47 | 0.08962 | 0.203874118 | 0.083 | 0 | 1 | 11 |
| utp6 | 0.08962 | 0.120485007 | 0.083 | 0 | 1 | 11 |
| wdr74 | 0.08962 | 0.105902605 | 0.083 | 0 | 1 | 11 |
| zer1 | 0.08962 | 0.117473927 | 0.083 | 0 | 1 | 11 |
| zgc:101744 | 0.08962 | 0.138771079 | 0.083 | 0 | 1 | 11 |
| zgc:101851 | 0.08962 | 0.15186809 | 0.083 | 0 | 1 | 11 |
| zgc:153990 | 0.08962 | 0.110624889 | 0.083 | 0 | 1 | 11 |
| zgc:171971 | 0.08962 | 0.171653514 | 0.083 | 0 | 1 | 11 |
| zgc:63587 | 0.08962 | 0.163416972 | 0.083 | 0 | 1 | 11 |
| zgc:77058 | 0.08962 | 0.133441793 | 0.083 | 0 | 1 | 11 |
| zgc:91860 | 0.08962 | 0.12456923 | 0.083 | 0 | 1 | 11 |
| cog2 | 0.08962 | 0.11798723 | 0.083 | 0 | 1 | 11 |
| cxxc1a | 0.08962 | 0.132543199 | 0.083 | 0 | 1 | 11 |
| dctpp1 | 0.08962 | 0.116758875 | 0.083 | 0 | 1 | 11 |
| dis3l2 | 0.08962 | 0.129241454 | 0.083 | 0 | 1 | 11 |
| dnajc24 | 0.08962 | 0.129357242 | 0.083 | 0 | 1 | 11 |
| efr3bb | 0.08962 | 0.179217607 | 0.083 | 0 | 1 | 11 |
| eif2b1 | 0.08962 | 0.124632008 | 0.083 | 0 | 1 | 11 |
| elovl6 | 0.08962 | 0.152994576 | 0.083 | 0 | 1 | 11 |
| fam91a1 | 0.08962 | 0.12364946 | 0.083 | 0 | 1 | 11 |
| kin | 0.08962 | 0.162770599 | 0.083 | 0 | 1 | 11 |
| klhl18 | 0.08962 | 0.128408206 | 0.083 | 0 | 1 | 11 |
| kmt2d | 0.08962 | 0.150843172 | 0.083 | 0 | 1 | 11 |
| mmadhc | 0.08962 | 0.138443092 | 0.083 | 0 | 1 | 11 |
| mthfd2 | 0.08962 | 0.10469316 | 0.083 | 0 | 1 | 11 |
| ncaph2 | 0.08962 | 0.167589023 | 0.083 | 0 | 1 | 11 |
| nfyal | 0.08962 | 0.155059576 | 0.083 | 0 | 1 | 11 |
| nr1h3 | 0.08962 | 0.113600627 | 0.083 | 0 | 1 | 11 |
| nrde2 | 0.08962 | 0.163719953 | 0.083 | 0 | 1 | 11 |
| nup210 | 0.08962 | 0.137826645 | 0.083 | 0 | 1 | 11 |
| parvab | 0.08962 | 0.155880904 | 0.083 | 0 | 1 | 11 |
| pole2 | 0.08962 | 0.133552884 | 0.083 | 0 | 1 | 11 |
| rabgef1 | 0.08962 | 0.13145587 | 0.083 | 0 | 1 | 11 |
| serpinh1a | 0.08962 | 0.191190935 | 0.083 | 0 | 1 | 11 |
| setd3 | 0.08962 | 0.115709689 | 0.083 | 0 | 1 | 11 |

| | | | | | | |
|---|---|---|---|---|---|---|
| si:ch211-221f10.2 | 0.08962 | 0.134013312 | 0.083 | 0 | 1 | 11 |
| si:dkey-22o12.2 | 0.08962 | 0.159579937 | 0.083 | 0 | 1 | 11 |
| tanc2a | 0.08962 | 0.157408997 | 0.083 | 0 | 1 | 11 |
| tcf7l2 | 0.08962 | 0.12249571 | 0.083 | 0 | 1 | 11 |
| tnfsf10l4 | 0.08962 | 0.15804298 | 0.083 | 0 | 1 | 11 |
| xpc | 0.08962 | 0.171044851 | 0.083 | 0 | 1 | 11 |
| zcchc24 | 0.08962 | 0.131636923 | 0.083 | 0 | 1 | 11 |
| zgc:158846 | 0.08962 | 0.160306808 | 0.083 | 0 | 1 | 11 |
| zpr1 | 0.08962 | 0.121026038 | 0.083 | 0 | 1 | 11 |
| DDX17 | 0.08962 | 0.135797447 | 0.083 | 0 | 1 | 11 |
| LOC110437709 | 0.08962 | 0.200616043 | 0.083 | 0 | 1 | 11 |
| akr1a1b | 0.08962 | 0.114768162 | 0.083 | 0 | 1 | 11 |
| arl2bp | 0.08962 | 0.131090531 | 0.083 | 0 | 1 | 11 |
| bmpr1aa | 0.08962 | 0.171722961 | 0.083 | 0 | 1 | 11 |
| cenps | 0.08962 | 0.152156105 | 0.083 | 0 | 1 | 11 |
| ckap5 | 0.08962 | 0.135226129 | 0.083 | 0 | 1 | 11 |
| clcn7 | 0.08962 | 0.136266778 | 0.083 | 0 | 1 | 11 |
| dtd1 | 0.08962 | 0.138414389 | 0.083 | 0 | 1 | 11 |
| epas1b | 0.08962 | 0.174622769 | 0.083 | 0 | 1 | 11 |
| hira | 0.08962 | 0.136525681 | 0.083 | 0 | 1 | 11 |
| irs1 | 0.08962 | 0.137790356 | 0.083 | 0 | 1 | 11 |
| itm2cb | 0.08962 | 0.111464962 | 0.083 | 0 | 1 | 11 |
| kctd13 | 0.08962 | 0.118916623 | 0.083 | 0 | 1 | 11 |
| lhfpl6 | 0.08962 | 0.127456145 | 0.083 | 0 | 1 | 11 |
| maff | 0.08962 | 0.208191079 | 0.083 | 0 | 1 | 11 |
| med25 | 0.08962 | 0.141628604 | 0.083 | 0 | 1 | 11 |
| metrnlb | 0.08962 | 0.155864219 | 0.083 | 0 | 1 | 11 |
| mon1bb | 0.08962 | 0.151371445 | 0.083 | 0 | 1 | 11 |
| mrpl19 | 0.08962 | 0.162044636 | 0.083 | 0 | 1 | 11 |
| mrps23 | 0.08962 | 0.136394993 | 0.083 | 0 | 1 | 11 |
| notch1b | 0.08962 | 0.117583473 | 0.083 | 0 | 1 | 11 |
| ppp2r5cb | 0.08962 | 0.150372712 | 0.083 | 0 | 1 | 11 |
| raf1a | 0.08962 | 0.126221032 | 0.083 | 0 | 1 | 11 |
| retreg3 | 0.08962 | 0.167519493 | 0.083 | 0 | 1 | 11 |
| rfc3 | 0.08962 | 0.166813051 | 0.083 | 0 | 1 | 11 |
| sat1b | 0.08962 | 0.151429607 | 0.083 | 0 | 1 | 11 |
| si:ch211-157b11.12 | 0.08962 | 0.15424208 | 0.083 | 0 | 1 | 11 |
| si:ch73-238c9.1 | 0.08962 | 0.148384616 | 0.083 | 0 | 1 | 11 |
| snu13a | 0.08962 | 0.116768528 | 0.083 | 0 | 1 | 11 |
| stard8 | 0.08962 | 0.145629546 | 0.083 | 0 | 1 | 11 |
| tatdn1 | 0.08962 | 0.106540844 | 0.083 | 0 | 1 | 11 |
| tefm | 0.08962 | 0.106965658 | 0.083 | 0 | 1 | 11 |
| timm10b | 0.08962 | 0.140854475 | 0.083 | 0 | 1 | 11 |
| tmem230a | 0.08962 | 0.195307098 | 0.083 | 0 | 1 | 11 |
| tns3.2 | 0.08962 | 0.208428879 | 0.083 | 0 | 1 | 11 |
| trim47 | 0.08962 | 0.169654853 | 0.083 | 0 | 1 | 11 |
| zdhhc18b | 0.08962 | 0.139339304 | 0.083 | 0 | 1 | 11 |
| zgc:100846 | 0.08962 | 0.117366531 | 0.083 | 0 | 1 | 11 |

| | | | | | | |
|---|---|---|---|---|---|---|
| zgc:152830 | 0.08962 | 0.102787442 | 0.083 | 0 | 1 | 11 |
| zgc:153631 | 0.08962 | 0.160128379 | 0.083 | 0 | 1 | 11 |
| zgc:174288 | 0.08962 | 0.129387128 | 0.083 | 0 | 1 | 11 |
| BX005254.1 | 0.08962 | 0.116381848 | 0.083 | 0 | 1 | 11 |
| CR382370.2 | 0.08962 | 0.188432871 | 0.083 | 0 | 1 | 11 |
| arid4a | 0.08962 | 0.155066684 | 0.083 | 0 | 1 | 11 |
| flad1 | 0.08962 | 0.126432164 | 0.083 | 0 | 1 | 11 |
| ngrn | 0.08962 | 0.135155296 | 0.083 | 0 | 1 | 11 |
| relch | 0.08962 | 0.13058869 | 0.083 | 0 | 1 | 11 |
| rgs3a | 0.08962 | 0.169027012 | 0.083 | 0 | 1 | 11 |
| si:ch211-259g3.4 | 0.08962 | 0.1584793 | 0.083 | 0 | 1 | 11 |
| smarcc1a | 0.08962 | 0.122369363 | 0.083 | 0 | 1 | 11 |
| tada2b | 0.08962 | 0.109695808 | 0.083 | 0 | 1 | 11 |
| tmem161b | 0.08962 | 0.13454378 | 0.083 | 0 | 1 | 11 |
| nasp | 0.08965 | 0.115107184 | 0.167 | 0.036 | 1 | 11 |
| vdac3 | 0.0901 | -0.41179634 | 0.333 | 0.571 | 1 | 11 |
| asf1bb | 0.09013 | -0.26938479 | 0.028 | 0.179 | 1 | 11 |
| map2k1 | 0.0914 | -0.14948877 | 0.167 | 0.357 | 1 | 11 |
| nolc1 | 0.09159 | 0.285996235 | 0.333 | 0.107 | 1 | 11 |
| si:ch211-286b5.5 | 0.09165 | -0.21228472 | 0.333 | 0.571 | 1 | 11 |
| atp6v1aa | 0.0927 | -0.12649547 | 0.028 | 0.179 | 1 | 11 |
| fam32a | 0.09336 | 0.272103438 | 0.5 | 0.429 | 1 | 11 |
| stimate | 0.09338 | -0.17709469 | 0.028 | 0.179 | 1 | 11 |
| cav1 | 0.09413 | 0.178210366 | 0.889 | 0.679 | 1 | 11 |
| sf1 | 0.09421 | -0.31570921 | 0.083 | 0.286 | 1 | 11 |
| prkab1b | 0.09589 | 0.348275488 | 0.472 | 0.214 | 1 | 11 |
| adrm1 | 0.0966 | -0.24249733 | 0.083 | 0.286 | 1 | 11 |
| ube2d2l | 0.09678 | -0.19664898 | 0.083 | 0.286 | 1 | 11 |
| znf1035 | 0.09835 | -0.23376641 | 0.028 | 0.179 | 1 | 11 |
| fibinb | 0.09836 | 0.434655398 | 0.417 | 0.214 | 1 | 11 |
| ednraa | 0.09874 | -0.30909637 | 0.083 | 0.286 | 1 | 11 |
| cd81a | 0.099 | -0.20953815 | 0.861 | 0.964 | 1 | 11 |
| XLOC-010899 | 0.10021 | -0.25926 | 0.111 | 0.321 | 1 | 11 |
| cdca4 | 0.10029 | -0.28112896 | 0.083 | 0.286 | 1 | 11 |
| vps4b | 0.10047 | -0.15330094 | 0.056 | 0.214 | 1 | 11 |
| suclg1 | 0.10059 | -0.31411887 | 0.167 | 0.357 | 1 | 11 |
| tm9sf2 | 0.10076 | -0.26825212 | 0.083 | 0.286 | 1 | 11 |
| nuak1b | 0.101 | -0.30342217 | 0.028 | 0.179 | 1 | 11 |
| tagln | 0.10142 | 0.323386498 | 0.722 | 0.464 | 1 | 11 |
| dhdhl | 0.10177 | 0.251625203 | 0.194 | 0.036 | 1 | 11 |
| CU457819.1 | 0.10213 | -0.11869952 | 0 | 0.071 | 1 | 11 |
| CU929391.2 | 0.10213 | -0.17105809 | 0 | 0.071 | 1 | 11 |
| XLOC-018082 | 0.10213 | -0.17207516 | 0 | 0.071 | 1 | 11 |
| ankrd10b | 0.10213 | -0.15369117 | 0 | 0.071 | 1 | 11 |
| atraid | 0.10213 | -0.13765307 | 0 | 0.071 | 1 | 11 |
| casp2 | 0.10213 | -0.11407036 | 0 | 0.071 | 1 | 11 |
| dcun1d4 | 0.10213 | -0.11022356 | 0 | 0.071 | 1 | 11 |
| dlg1l | 0.10213 | -0.12764073 | 0 | 0.071 | 1 | 11 |

| | | | | | | |
|---|---|---|---|---|---|---|
| dnajb2 | 0.10213 | -0.11281575 | 0 | 0.071 | 1 | 11 |
| ephb4a | 0.10213 | -0.14023922 | 0 | 0.071 | 1 | 11 |
| gphnb | 0.10213 | -0.10693451 | 0 | 0.071 | 1 | 11 |
| maco1b | 0.10213 | -0.16068744 | 0 | 0.071 | 1 | 11 |
| mfsd5 | 0.10213 | -0.17577641 | 0 | 0.071 | 1 | 11 |
| mindy3 | 0.10213 | -0.10274235 | 0 | 0.071 | 1 | 11 |
| mn1b | 0.10213 | -0.15240013 | 0 | 0.071 | 1 | 11 |
| nipbla | 0.10213 | -0.11318938 | 0 | 0.071 | 1 | 11 |
| nqo1 | 0.10213 | -0.15776326 | 0 | 0.071 | 1 | 11 |
| pde4ba | 0.10213 | -0.15444245 | 0 | 0.071 | 1 | 11 |
| ppp1r3cb | 0.10213 | -0.10022314 | 0 | 0.071 | 1 | 11 |
| si:ch211-266k8.6 | 0.10213 | -0.10908338 | 0 | 0.071 | 1 | 11 |
| si:ch73-193c12.2 | 0.10213 | -0.11491392 | 0 | 0.071 | 1 | 11 |
| si:dkey-193c22.1 | 0.10213 | -0.11072082 | 0 | 0.071 | 1 | 11 |
| si:dkey-235d18.5 | 0.10213 | -0.15612976 | 0 | 0.071 | 1 | 11 |
| si:dkey-246i14.3 | 0.10213 | -0.10597902 | 0 | 0.071 | 1 | 11 |
| si:dkeyp-2e4.3 | 0.10213 | -0.14330314 | 0 | 0.071 | 1 | 11 |
| si:dkeyp-53e4.4 | 0.10213 | -0.11636144 | 0 | 0.071 | 1 | 11 |
| sidt2 | 0.10213 | -0.10084388 | 0 | 0.071 | 1 | 11 |
| slc1a5 | 0.10213 | -0.14751651 | 0 | 0.071 | 1 | 11 |
| syt11a | 0.10213 | -0.15994101 | 0 | 0.071 | 1 | 11 |
| tgfbrap1 | 0.10213 | -0.16495012 | 0 | 0.071 | 1 | 11 |
| tshz1 | 0.10213 | -0.11875774 | 0 | 0.071 | 1 | 11 |
| usp31 | 0.10213 | -0.11988022 | 0 | 0.071 | 1 | 11 |
| zgc:153039 | 0.10213 | -0.15730987 | 0 | 0.071 | 1 | 11 |
| BX957317.2 | 0.10213 | -0.10989158 | 0 | 0.071 | 1 | 11 |
| CABZ01088274.1 | 0.10213 | -0.10159215 | 0 | 0.071 | 1 | 11 |
| CHST13 | 0.10213 | -0.15085847 | 0 | 0.071 | 1 | 11 |
| CT027638.1 | 0.10213 | -0.12626608 | 0 | 0.071 | 1 | 11 |
| CU633762.1 | 0.10213 | -0.145939 | 0 | 0.071 | 1 | 11 |
| CU695223.1 | 0.10213 | -0.19421261 | 0 | 0.071 | 1 | 11 |
| GAN | 0.10213 | -0.11044107 | 0 | 0.071 | 1 | 11 |
| LOC100000748 | 0.10213 | -0.11790158 | 0 | 0.071 | 1 | 11 |
| LOC101884680 | 0.10213 | -0.16653425 | 0 | 0.071 | 1 | 11 |
| LOC101884921 | 0.10213 | -0.13264011 | 0 | 0.071 | 1 | 11 |
| LOC103908766 | 0.10213 | -0.12854546 | 0 | 0.071 | 1 | 11 |
| LOC108190785 | 0.10213 | -0.15334955 | 0 | 0.071 | 1 | 11 |
| LOC110437758 | 0.10213 | -0.11510163 | 0 | 0.071 | 1 | 11 |
| LOC110438055 | 0.10213 | -0.13911312 | 0 | 0.071 | 1 | 11 |
| LOC110439998 | 0.10213 | -0.17723744 | 0 | 0.071 | 1 | 11 |
| LOC110440195 | 0.10213 | -0.11678495 | 0 | 0.071 | 1 | 11 |
| XLOC-006330 | 0.10213 | -0.1269594 | 0 | 0.071 | 1 | 11 |
| XLOC-008119 | 0.10213 | -0.14123496 | 0 | 0.071 | 1 | 11 |
| XLOC-044048 | 0.10213 | -0.21119993 | 0 | 0.071 | 1 | 11 |
| alcamb | 0.10213 | -0.12177694 | 0 | 0.071 | 1 | 11 |
| alkal2a | 0.10213 | -0.14378814 | 0 | 0.071 | 1 | 11 |
| ankfy1 | 0.10213 | -0.10658467 | 0 | 0.071 | 1 | 11 |
| apoob | 0.10213 | -0.2013076 | 0 | 0.071 | 1 | 11 |

| | | | | | | |
|---|---|---|---|---|---|---|
| asap2a | 0.10213 | -0.13219498 | 0 | 0.071 | 1 | 11 |
| bicral | 0.10213 | -0.23194251 | 0 | 0.071 | 1 | 11 |
| casp8 | 0.10213 | -0.1187114 | 0 | 0.071 | 1 | 11 |
| casp8ap2 | 0.10213 | -0.11014423 | 0 | 0.071 | 1 | 11 |
| ccr8.1 | 0.10213 | -0.11774881 | 0 | 0.071 | 1 | 11 |
| cdh18a | 0.10213 | -0.14020432 | 0 | 0.071 | 1 | 11 |
| cdkn1d | 0.10213 | -0.10281201 | 0 | 0.071 | 1 | 11 |
| cenpo | 0.10213 | -0.12876476 | 0 | 0.071 | 1 | 11 |
| cerk | 0.10213 | -0.10706634 | 0 | 0.071 | 1 | 11 |
| chpf2 | 0.10213 | -0.10655643 | 0 | 0.071 | 1 | 11 |
| clint1b | 0.10213 | -0.12186456 | 0 | 0.071 | 1 | 11 |
| cluha | 0.10213 | -0.10427811 | 0 | 0.071 | 1 | 11 |
| cnot10 | 0.10213 | -0.12743396 | 0 | 0.071 | 1 | 11 |
| creld1b | 0.10213 | -0.15406742 | 0 | 0.071 | 1 | 11 |
| cst14b.1 | 0.10213 | -0.11162549 | 0 | 0.071 | 1 | 11 |
| cul5a | 0.10213 | -0.15697133 | 0 | 0.071 | 1 | 11 |
| ddx10 | 0.10213 | -0.13800695 | 0 | 0.071 | 1 | 11 |
| dhrs11b | 0.10213 | -0.14677205 | 0 | 0.071 | 1 | 11 |
| dia1a | 0.10213 | -0.13091501 | 0 | 0.071 | 1 | 11 |
| disp1 | 0.10213 | -0.10998041 | 0 | 0.071 | 1 | 11 |
| exoc3l1 | 0.10213 | -0.16550287 | 0 | 0.071 | 1 | 11 |
| fbxl17 | 0.10213 | -0.17001147 | 0 | 0.071 | 1 | 11 |
| fen1 | 0.10213 | -0.15763726 | 0 | 0.071 | 1 | 11 |
| galt | 0.10213 | -0.13666025 | 0 | 0.071 | 1 | 11 |
| glsa | 0.10213 | -0.13957628 | 0 | 0.071 | 1 | 11 |
| gpat3 | 0.10213 | -0.14944893 | 0 | 0.071 | 1 | 11 |
| gtf3aa | 0.10213 | -0.12829631 | 0 | 0.071 | 1 | 11 |
| gtpbp10 | 0.10213 | -0.13947366 | 0 | 0.071 | 1 | 11 |
| helb | 0.10213 | -0.15469752 | 0 | 0.071 | 1 | 11 |
| hibch | 0.10213 | -0.10734512 | 0 | 0.071 | 1 | 11 |
| hivep2a | 0.10213 | -0.10822738 | 0 | 0.071 | 1 | 11 |
| hsbp1l1 | 0.10213 | -0.12981512 | 0 | 0.071 | 1 | 11 |
| ifngr1l | 0.10213 | -0.12354689 | 0 | 0.071 | 1 | 11 |
| intu | 0.10213 | -0.11814025 | 0 | 0.071 | 1 | 11 |
| itsn2b | 0.10213 | -0.12121048 | 0 | 0.071 | 1 | 11 |
| katna1 | 0.10213 | -0.11444152 | 0 | 0.071 | 1 | 11 |
| map2k7 | 0.10213 | -0.15521371 | 0 | 0.071 | 1 | 11 |
| mapk8a | 0.10213 | -0.14351319 | 0 | 0.071 | 1 | 11 |
| mccc1 | 0.10213 | -0.10622805 | 0 | 0.071 | 1 | 11 |
| mettl3 | 0.10213 | -0.14019609 | 0 | 0.071 | 1 | 11 |
| mgarp | 0.10213 | -0.12550893 | 0 | 0.071 | 1 | 11 |
| mif4gda | 0.10213 | -0.13985873 | 0 | 0.071 | 1 | 11 |
| mindy2 | 0.10213 | -0.1097651 | 0 | 0.071 | 1 | 11 |
| ms4a17a.14 | 0.10213 | -0.14301341 | 0 | 0.071 | 1 | 11 |
| mta1 | 0.10213 | -0.11208857 | 0 | 0.071 | 1 | 11 |
| mtx1a | 0.10213 | -0.11321427 | 0 | 0.071 | 1 | 11 |
| myo10l3 | 0.10213 | -0.19616601 | 0 | 0.071 | 1 | 11 |
| nck2b | 0.10213 | -0.12565992 | 0 | 0.071 | 1 | 11 |

| | | | | | | |
|---|---|---|---|---|---|---|
| nfxl1 | 0.10213 | -0.15193737 | 0 | 0.071 | 1 | 11 |
| nlrc3 | 0.10213 | -0.10648702 | 0 | 0.071 | 1 | 11 |
| nt5c2a | 0.10213 | -0.13766301 | 0 | 0.071 | 1 | 11 |
| numb | 0.10213 | -0.1161791 | 0 | 0.071 | 1 | 11 |
| pgghg | 0.10213 | -0.10525143 | 0 | 0.071 | 1 | 11 |
| pi15b | 0.10213 | -0.15647489 | 0 | 0.071 | 1 | 11 |
| pi4kaa | 0.10213 | -0.15446403 | 0 | 0.071 | 1 | 11 |
| pitpnc1b | 0.10213 | -0.13385316 | 0 | 0.071 | 1 | 11 |
| plekha1b | 0.10213 | -0.15550085 | 0 | 0.071 | 1 | 11 |
| pltp | 0.10213 | -0.12526983 | 0 | 0.071 | 1 | 11 |
| prpf4bb | 0.10213 | -0.10319395 | 0 | 0.071 | 1 | 11 |
| rac1l | 0.10213 | -0.10143733 | 0 | 0.071 | 1 | 11 |
| rassf8a | 0.10213 | -0.11102164 | 0 | 0.071 | 1 | 11 |
| rbm48 | 0.10213 | -0.13228346 | 0 | 0.071 | 1 | 11 |
| rcbtb2 | 0.10213 | -0.18715806 | 0 | 0.071 | 1 | 11 |
| rem1 | 0.10213 | -0.11113111 | 0 | 0.071 | 1 | 11 |
| rmnd1 | 0.10213 | -0.13310566 | 0 | 0.071 | 1 | 11 |
| rnf141 | 0.10213 | -0.13416853 | 0 | 0.071 | 1 | 11 |
| sart3 | 0.10213 | -0.11315868 | 0 | 0.071 | 1 | 11 |
| si:ch211-128m15.3 | 0.10213 | -0.19166812 | 0 | 0.071 | 1 | 11 |
| si:ch211-194p6.10 | 0.10213 | -0.11078135 | 0 | 0.071 | 1 | 11 |
| si:ch211-195b13.1 | 0.10213 | -0.17741454 | 0 | 0.071 | 1 | 11 |
| si:ch211-212o1.2 | 0.10213 | -0.11127255 | 0 | 0.071 | 1 | 11 |
| si:ch211-245h14.1 | 0.10213 | -0.10658294 | 0 | 0.071 | 1 | 11 |
| si:ch211-59h6.1 | 0.10213 | -0.12851341 | 0 | 0.071 | 1 | 11 |
| si:ch73-141c7.1 | 0.10213 | -0.1130119 | 0 | 0.071 | 1 | 11 |
| si:ch73-266f23.1 | 0.10213 | -0.1266796 | 0 | 0.071 | 1 | 11 |
| si:ch73-92e7.4 | 0.10213 | -0.11566689 | 0 | 0.071 | 1 | 11 |
| si:dkey-27b3.4 | 0.10213 | -0.12583451 | 0 | 0.071 | 1 | 11 |
| si:dkey-91m11.5 | 0.10213 | -0.11602098 | 0 | 0.071 | 1 | 11 |
| si:dkeyp-38g8.5 | 0.10213 | -0.13530238 | 0 | 0.071 | 1 | 11 |
| si:dkeyp-66d1.7 | 0.10213 | -0.11760208 | 0 | 0.071 | 1 | 11 |
| si:zfos-464b6.2 | 0.10213 | -0.11899362 | 0 | 0.071 | 1 | 11 |
| sirt4 | 0.10213 | -0.11000212 | 0 | 0.071 | 1 | 11 |
| slc7a8a | 0.10213 | -0.11022171 | 0 | 0.071 | 1 | 11 |
| smad7 | 0.10213 | -0.13772205 | 0 | 0.071 | 1 | 11 |
| snupn | 0.10213 | -0.12325797 | 0 | 0.071 | 1 | 11 |
| sowahcb | 0.10213 | -0.13117738 | 0 | 0.071 | 1 | 11 |
| stam2 | 0.10213 | -0.1143903 | 0 | 0.071 | 1 | 11 |
| stambpb | 0.10213 | -0.14158594 | 0 | 0.071 | 1 | 11 |
| stk10 | 0.10213 | -0.10512239 | 0 | 0.071 | 1 | 11 |
| sult2st1 | 0.10213 | -0.11486436 | 0 | 0.071 | 1 | 11 |
| sympk | 0.10213 | -0.10071894 | 0 | 0.071 | 1 | 11 |
| tbc1d10b | 0.10213 | -0.14369577 | 0 | 0.071 | 1 | 11 |
| tbxa2r | 0.10213 | -0.11500179 | 0 | 0.071 | 1 | 11 |
| tcima | 0.10213 | -0.13626668 | 0 | 0.071 | 1 | 11 |
| thbs1b | 0.10213 | -0.11066191 | 0 | 0.071 | 1 | 11 |
| timeless | 0.10213 | -0.15049634 | 0 | 0.071 | 1 | 11 |

| | | | | | | |
|---|---|---|---|---|---|---|
| timm17b | 0.10213 | -0.10343795 | 0 | 0.071 | 1 | 11 |
| tmed4 | 0.10213 | -0.10320625 | 0 | 0.071 | 1 | 11 |
| tmod4 | 0.10213 | -0.12384826 | 0 | 0.071 | 1 | 11 |
| tor2a | 0.10213 | -0.14044784 | 0 | 0.071 | 1 | 11 |
| tox4b | 0.10213 | -0.12825643 | 0 | 0.071 | 1 | 11 |
| tsen15 | 0.10213 | -0.10823493 | 0 | 0.071 | 1 | 11 |
| ube2t | 0.10213 | -0.14335693 | 0 | 0.071 | 1 | 11 |
| vav2 | 0.10213 | -0.11056876 | 0 | 0.071 | 1 | 11 |
| wdr20a | 0.10213 | -0.15003813 | 0 | 0.071 | 1 | 11 |
| zcchc2 | 0.10213 | -0.16072907 | 0 | 0.071 | 1 | 11 |
| zgc:110626 | 0.10213 | -0.18258681 | 0 | 0.071 | 1 | 11 |
| zgc:110843 | 0.10213 | -0.14605932 | 0 | 0.071 | 1 | 11 |
| zgc:154093 | 0.10213 | -0.1409941 | 0 | 0.071 | 1 | 11 |
| zgc:171857 | 0.10213 | -0.10139566 | 0 | 0.071 | 1 | 11 |
| zgc:173575 | 0.10213 | -0.11145644 | 0 | 0.071 | 1 | 11 |
| zgc:92818 | 0.10213 | -0.12021635 | 0 | 0.071 | 1 | 11 |
| zmp:0000000711 | 0.10213 | -0.12092817 | 0 | 0.071 | 1 | 11 |
| zmp:0000001316 | 0.10213 | -0.11683075 | 0 | 0.071 | 1 | 11 |
| CABZ01059408.1 | 0.10213 | -0.10760025 | 0 | 0.071 | 1 | 11 |
| GNB4 | 0.10213 | -0.10243665 | 0 | 0.071 | 1 | 11 |
| LOC110439489 | 0.10213 | -0.24829476 | 0 | 0.071 | 1 | 11 |
| badb | 0.10213 | -0.10176426 | 0 | 0.071 | 1 | 11 |
| bbs1 | 0.10213 | -0.15480711 | 0 | 0.071 | 1 | 11 |
| frem2b | 0.10213 | -0.10701271 | 0 | 0.071 | 1 | 11 |
| fsd1l | 0.10213 | -0.10949694 | 0 | 0.071 | 1 | 11 |
| gnas | 0.10213 | -0.10479613 | 0 | 0.071 | 1 | 11 |
| mios | 0.10213 | -0.13093178 | 0 | 0.071 | 1 | 11 |
| mpdu1b | 0.10213 | -0.12109778 | 0 | 0.071 | 1 | 11 |
| sh3bp5a | 0.10213 | -0.21027212 | 0 | 0.071 | 1 | 11 |
| zgc:153012 | 0.10213 | -0.10920013 | 0 | 0.071 | 1 | 11 |
| epha2a | 0.10213 | -0.14877228 | 0 | 0.071 | 1 | 11 |
| fpgt | 0.10213 | -0.11804816 | 0 | 0.071 | 1 | 11 |
| irak4 | 0.10213 | -0.10873248 | 0 | 0.071 | 1 | 11 |
| lonrf1 | 0.10213 | -0.12310342 | 0 | 0.071 | 1 | 11 |
| mxra7 | 0.10213 | -0.14885412 | 0 | 0.071 | 1 | 11 |
| odr4 | 0.10213 | -0.11227752 | 0 | 0.071 | 1 | 11 |
| tspan2a | 0.10213 | -0.11804001 | 0 | 0.071 | 1 | 11 |
| CU024870.1 | 0.10213 | -0.15831409 | 0 | 0.071 | 1 | 11 |
| XLOC-013497 | 0.10213 | -0.13610907 | 0 | 0.071 | 1 | 11 |
| acad8 | 0.10213 | -0.12171033 | 0 | 0.071 | 1 | 11 |
| asphd2 | 0.10213 | -0.15237116 | 0 | 0.071 | 1 | 11 |
| b4galt1l | 0.10213 | -0.20680688 | 0 | 0.071 | 1 | 11 |
| ccz1 | 0.10213 | -0.16475014 | 0 | 0.071 | 1 | 11 |
| cpa6 | 0.10213 | -0.15661718 | 0 | 0.071 | 1 | 11 |
| elmo1 | 0.10213 | -0.11199853 | 0 | 0.071 | 1 | 11 |
| frem2a | 0.10213 | -0.11133861 | 0 | 0.071 | 1 | 11 |
| gcat | 0.10213 | -0.11725621 | 0 | 0.071 | 1 | 11 |
| itln1l | 0.10213 | -0.14333469 | 0 | 0.071 | 1 | 11 |

| | | | | | | |
|---|---|---|---|---|---|---|
| mul1a | 0.10213 | -0.20831403 | 0 | 0.071 | 1 | 11 |
| naa25 | 0.10213 | -0.16586055 | 0 | 0.071 | 1 | 11 |
| nfatc1 | 0.10213 | -0.11554891 | 0 | 0.071 | 1 | 11 |
| olfml2ba | 0.10213 | -0.1018374 | 0 | 0.071 | 1 | 11 |
| preb | 0.10213 | -0.13143666 | 0 | 0.071 | 1 | 11 |
| rptor | 0.10213 | -0.16879767 | 0 | 0.071 | 1 | 11 |
| si:ch73-182a11.2 | 0.10213 | -0.11772722 | 0 | 0.071 | 1 | 11 |
| uros | 0.10213 | -0.14858592 | 0 | 0.071 | 1 | 11 |
| wu:fc27b11 | 0.10213 | -0.15105847 | 0 | 0.071 | 1 | 11 |
| zgc:162939 | 0.10213 | -0.1112816 | 0 | 0.071 | 1 | 11 |
| zgc:92360 | 0.10213 | -0.14924129 | 0 | 0.071 | 1 | 11 |
| zmp:0000000912 | 0.10213 | -0.14069043 | 0 | 0.071 | 1 | 11 |
| c1galt1a | 0.10213 | -0.11102448 | 0 | 0.071 | 1 | 11 |
| katnb1 | 0.10213 | -0.11796146 | 0 | 0.071 | 1 | 11 |
| mpv17l2 | 0.10213 | -0.11288998 | 0 | 0.071 | 1 | 11 |
| si:dkey-10o6.2 | 0.10213 | -0.14174888 | 0 | 0.071 | 1 | 11 |
| slc35b3 | 0.10213 | -0.12577926 | 0 | 0.071 | 1 | 11 |
| tomm34 | 0.10213 | -0.10371917 | 0 | 0.071 | 1 | 11 |
| ucmaa | 0.10219 | -0.50606979 | 0.25 | 0.464 | 1 | 11 |
| CABZ01072614.1 | 0.10253 | 0.411753836 | 0.639 | 0.429 | 1 | 11 |
| cox7a2l | 0.10268 | -0.34669754 | 0.139 | 0.357 | 1 | 11 |
| gsta.1 | 0.10427 | 0.376994544 | 0.444 | 0.25 | 1 | 11 |
| tgm5l | 0.10439 | -0.37496208 | 0.056 | 0.143 | 1 | 11 |
| aebp1 | 0.10444 | 0.392579822 | 0.417 | 0.179 | 1 | 11 |
| magoh | 0.1045 | 0.333275926 | 0.278 | 0.107 | 1 | 11 |
| ccdc32 | 0.10549 | -0.23667345 | 0.028 | 0.179 | 1 | 11 |
| dhrs12 | 0.10584 | 0.154671579 | 0.167 | 0.036 | 1 | 11 |
| mt-co2 | 0.10584 | -0.3655948 | 0.5 | 0.75 | 1 | 11 |
| pfn2 | 0.10586 | 0.268105307 | 0.694 | 0.643 | 1 | 11 |
| ints1 | 0.10588 | -0.33357875 | 0.028 | 0.179 | 1 | 11 |
| si:ch211-276i12.9 | 0.10591 | 0.341767444 | 0.194 | 0.036 | 1 | 11 |
| fap | 0.10632 | 0.281694725 | 0.25 | 0.071 | 1 | 11 |
| psmb3 | 0.10637 | -0.36845407 | 0.25 | 0.464 | 1 | 11 |
| acot15 | 0.10663 | -0.25813145 | 0.028 | 0.179 | 1 | 11 |
| cercam | 0.10673 | -0.23674351 | 0.028 | 0.179 | 1 | 11 |
| rnf44 | 0.10673 | 0.36577418 | 0.278 | 0.107 | 1 | 11 |
| ppp1r3ca | 0.10747 | -0.19369944 | 0.139 | 0.143 | 1 | 11 |
| si:dkey-69o16.5 | 0.10772 | -0.31905071 | 0.083 | 0.25 | 1 | 11 |
| cav2 | 0.10781 | -0.41975534 | 0.194 | 0.393 | 1 | 11 |
| bicc1b | 0.10827 | -0.20209657 | 0.111 | 0.286 | 1 | 11 |
| polr2d | 0.10868 | -0.25147513 | 0.139 | 0.357 | 1 | 11 |
| ndufa6 | 0.10881 | -0.2806302 | 0.278 | 0.536 | 1 | 11 |
| angptl1a | 0.10919 | -0.40445521 | 0.278 | 0.536 | 1 | 11 |
| ruvbl1 | 0.10955 | -0.24106248 | 0.028 | 0.179 | 1 | 11 |
| LO018340.1 | 0.10963 | -0.3494591 | 0.028 | 0.179 | 1 | 11 |
| ptges3b | 0.10974 | 0.256833483 | 0.222 | 0.214 | 1 | 11 |
| mfap1 | 0.10984 | 0.312598006 | 0.222 | 0.071 | 1 | 11 |
| stxbp2 | 0.10994 | -0.24321905 | 0.028 | 0.179 | 1 | 11 |

| | | | | | | |
|---|---|---|---|---|---|---|
| stx12 | 0.11009 | -0.23920064 | 0.028 | 0.179 | 1 | 11 |
| srpra | 0.11029 | 0.282477304 | 0.194 | 0.036 | 1 | 11 |
| mat2aa | 0.11106 | 0.269207044 | 0.194 | 0.036 | 1 | 11 |
| kank4 | 0.11111 | -0.25488183 | 0.028 | 0.179 | 1 | 11 |
| c7a | 0.11172 | 0.408173919 | 0.5 | 0.25 | 1 | 11 |
| prickle2b | 0.11177 | -0.21152099 | 0.028 | 0.179 | 1 | 11 |
| csnk1g2b | 0.11193 | 0.278513366 | 0.306 | 0.107 | 1 | 11 |
| junbb | 0.11226 | -0.33807268 | 0.111 | 0.321 | 1 | 11 |
| laptm4a | 0.11233 | -0.2933098 | 0.25 | 0.5 | 1 | 11 |
| meis2b | 0.11259 | 0.216041576 | 0.306 | 0.286 | 1 | 11 |
| notch2 | 0.11301 | 0.385084962 | 0.472 | 0.25 | 1 | 11 |
| psmg2 | 0.11323 | -0.23235523 | 0.028 | 0.179 | 1 | 11 |
| mtpn | 0.11418 | -0.35289634 | 0.306 | 0.536 | 1 | 11 |
| RASSF5 | 0.11451 | -0.28288594 | 0.028 | 0.179 | 1 | 11 |
| mpp1 | 0.11461 | -0.21378248 | 0.028 | 0.179 | 1 | 11 |
| si:dkeyp-120h9.1 | 0.11472 | -0.20784736 | 0.028 | 0.179 | 1 | 11 |
| glsb | 0.11478 | -0.24111714 | 0.028 | 0.179 | 1 | 11 |
| pigbos1 | 0.11479 | -0.21204327 | 0.028 | 0.179 | 1 | 11 |
| mrps7 | 0.11482 | -0.20010433 | 0.028 | 0.179 | 1 | 11 |
| vapal | 0.11488 | -0.25149903 | 0.028 | 0.179 | 1 | 11 |
| fbxw4 | 0.11494 | -0.2628539 | 0.028 | 0.179 | 1 | 11 |
| blmh | 0.11511 | -0.20719691 | 0.028 | 0.179 | 1 | 11 |
| si:ch211-151g22.1 | 0.1152 | -0.21136723 | 0.028 | 0.179 | 1 | 11 |
| si:dkey-74k8.3 | 0.1152 | -0.20221655 | 0.028 | 0.179 | 1 | 11 |
| smarcd1 | 0.11526 | -0.2104751 | 0.028 | 0.179 | 1 | 11 |
| pdlim5a | 0.11529 | -0.28494681 | 0.028 | 0.179 | 1 | 11 |
| btg3 | 0.11529 | -0.17793093 | 0.028 | 0.179 | 1 | 11 |
| CR855311.1 | 0.11533 | -0.41053006 | 0.028 | 0.143 | 1 | 11 |
| wasf2 | 0.11585 | -0.27588913 | 0.111 | 0.321 | 1 | 11 |
| coa1 | 0.11619 | -0.2642213 | 0.028 | 0.143 | 1 | 11 |
| BX569787.1 | 0.11653 | 0.121357874 | 0.417 | 0.25 | 1 | 11 |
| rprd1b | 0.11676 | -0.2237449 | 0.056 | 0.214 | 1 | 11 |
| tefb | 0.1181 | 0.378916336 | 0.25 | 0.071 | 1 | 11 |
| slc22a31 | 0.11822 | -0.18218688 | 0.056 | 0.214 | 1 | 11 |
| bin3 | 0.11879 | -0.14529354 | 0.028 | 0.143 | 1 | 11 |
| chd9 | 0.11906 | 0.324903409 | 0.194 | 0.036 | 1 | 11 |
| arf2a | 0.11926 | -0.35739328 | 0.167 | 0.393 | 1 | 11 |
| tmpob | 0.11927 | -0.34625766 | 0.056 | 0.214 | 1 | 11 |
| serpinb14 | 0.12133 | 0.273535193 | 0.194 | 0.036 | 1 | 11 |
| sinhcafl | 0.12198 | -0.23957717 | 0.056 | 0.214 | 1 | 11 |
| slc6a4a | 0.122 | -0.33754183 | 0.056 | 0.214 | 1 | 11 |
| edil3a | 0.12226 | -0.3118415 | 0.139 | 0.357 | 1 | 11 |
| rab14 | 0.1223 | -0.29008323 | 0.139 | 0.357 | 1 | 11 |
| arvcfb | 0.12307 | -0.39300397 | 0.111 | 0.286 | 1 | 11 |
| vdac1 | 0.12319 | 0.189060569 | 0.444 | 0.429 | 1 | 11 |
| si:ch211-204a13.2 | 0.12428 | 0.331484823 | 0.278 | 0.107 | 1 | 11 |
| si:dkey-52l18.4 | 0.12485 | 0.243676808 | 0.194 | 0.036 | 1 | 11 |
| ckba | 0.12519 | -0.24747272 | 0.167 | 0.393 | 1 | 11 |

| | | | | | | |
|---|---|---|---|---|---|---|
| hectd3 | 0.12689 | 0.248893958 | 0.194 | 0.036 | 1 | 11 |
| pmp22b | 0.12709 | -0.31397039 | 0.167 | 0.393 | 1 | 11 |
| mesd | 0.12769 | 0.299215686 | 0.306 | 0.107 | 1 | 11 |
| ginm1 | 0.1277 | 0.36474997 | 0.389 | 0.179 | 1 | 11 |
| si:dkey-188i13.11 | 0.12831 | 0.364683819 | 0.444 | 0.25 | 1 | 11 |
| si:ch211-66k16.27 | 0.1284 | 0.29543947 | 0.306 | 0.107 | 1 | 11 |
| aldh1a2 | 0.12904 | -0.1951244 | 0.806 | 0.964 | 1 | 11 |
| ywhae1 | 0.12915 | -0.21047049 | 0.444 | 0.679 | 1 | 11 |
| tgfbr2b | 0.12926 | -0.35005652 | 0.056 | 0.214 | 1 | 11 |
| ier5l | 0.13131 | 0.267272644 | 0.194 | 0.036 | 1 | 11 |
| fbxl6 | 0.1319 | 0.335369611 | 0.361 | 0.143 | 1 | 11 |
| ppp6c | 0.132 | 0.27082983 | 0.361 | 0.143 | 1 | 11 |
| asip2b | 0.13208 | 0.477135537 | 0.361 | 0.143 | 1 | 11 |
| mt-cyb | 0.13317 | -0.32924721 | 0.444 | 0.679 | 1 | 11 |
| atrxl | 0.1339 | -0.13641083 | 0.028 | 0.143 | 1 | 11 |
| sdc2 | 0.13457 | 0.118199103 | 0.25 | 0.286 | 1 | 11 |
| si:ch211-66e2.3 | 0.13483 | 0.125093784 | 0.583 | 0.607 | 1 | 11 |
| llph | 0.13537 | -0.10893928 | 0.167 | 0.321 | 1 | 11 |
| kmt5aa | 0.13555 | 0.301656453 | 0.25 | 0.071 | 1 | 11 |
| ivns1abpa | 0.13607 | -0.18573461 | 0.083 | 0.25 | 1 | 11 |
| gadd45ba | 0.1367 | 0.277451477 | 0.611 | 0.536 | 1 | 11 |
| ube2e1 | 0.13785 | 0.176814361 | 0.333 | 0.321 | 1 | 11 |
| ncor1 | 0.13793 | -0.30559973 | 0.083 | 0.25 | 1 | 11 |
| negr1 | 0.13839 | -0.36213757 | 0.028 | 0.143 | 1 | 11 |
| itm2bb | 0.13874 | 0.25047318 | 0.25 | 0.071 | 1 | 11 |
| mgst1.2 | 0.13879 | -0.11641941 | 0.444 | 0.357 | 1 | 11 |
| cyp27a1.4 | 0.13909 | -0.12181428 | 0.028 | 0.143 | 1 | 11 |
| rac3b | 0.14019 | 0.320160703 | 0.167 | 0.036 | 1 | 11 |
| si:busm1-57f23.1 | 0.14036 | -0.27742671 | 0.056 | 0.214 | 1 | 11 |
| zgc:163080 | 0.1404 | 0.251909062 | 0.25 | 0.071 | 1 | 11 |
| capza1a | 0.14163 | -0.3482266 | 0.389 | 0.571 | 1 | 11 |
| suds3 | 0.14252 | 0.314635534 | 0.194 | 0.071 | 1 | 11 |
| cbx1b | 0.14298 | -0.18030517 | 0.083 | 0.25 | 1 | 11 |
| flnb | 0.14316 | 0.220631242 | 0.472 | 0.429 | 1 | 11 |
| psmb10 | 0.14334 | 0.290940242 | 0.25 | 0.071 | 1 | 11 |
| tnrc6c2 | 0.14465 | -0.25237756 | 0.056 | 0.214 | 1 | 11 |
| zeb2a | 0.14506 | 0.112385617 | 0.194 | 0.071 | 1 | 11 |
| abca1a | 0.14568 | -0.18001185 | 0.056 | 0.214 | 1 | 11 |
| tsr1 | 0.14628 | -0.22511858 | 0.111 | 0.286 | 1 | 11 |
| nufip2 | 0.14649 | -0.26215119 | 0.056 | 0.214 | 1 | 11 |
| cbx7a | 0.14687 | -0.24326973 | 0.083 | 0.25 | 1 | 11 |
| pus7 | 0.14718 | -0.28361038 | 0.028 | 0.143 | 1 | 11 |
| bhlhe40 | 0.14788 | 0.384301902 | 0.556 | 0.321 | 1 | 11 |
| tmem88b | 0.14817 | 0.349441072 | 0.722 | 0.536 | 1 | 11 |
| pkig | 0.14825 | 0.279527222 | 0.25 | 0.071 | 1 | 11 |
| sem1 | 0.14837 | 0.291221025 | 0.667 | 0.571 | 1 | 11 |
| rpz2 | 0.1489 | -0.35221785 | 0.083 | 0.214 | 1 | 11 |
| ftr82 | 0.14925 | 0.364803313 | 0.583 | 0.393 | 1 | 11 |

| | | | | | | |
|---|---|---|---|---|---|---|
| endouc | 0.14948 | -0.16165701 | 0.694 | 0.679 | 1 | 11 |
| glis2a | 0.14993 | -0.12456049 | 0.111 | 0.25 | 1 | 11 |
| abl1 | 0.15069 | -0.10152756 | 0.083 | 0.214 | 1 | 11 |
| pno1 | 0.1509 | -0.15324173 | 0.194 | 0.393 | 1 | 11 |
| ckap4 | 0.15094 | -0.3420555 | 0.306 | 0.536 | 1 | 11 |
| slc43a3b | 0.15288 | -0.33784243 | 0.444 | 0.679 | 1 | 11 |
| myl6 | 0.15291 | 0.130717559 | 0.639 | 0.679 | 1 | 11 |
| psmb1 | 0.15303 | 0.366092524 | 0.556 | 0.357 | 1 | 11 |
| jak1 | 0.15345 | -0.10098522 | 0.083 | 0.214 | 1 | 11 |
| hmgn7 | 0.15364 | 0.286286027 | 0.833 | 0.679 | 1 | 11 |
| hbp1 | 0.15393 | -0.31471862 | 0.083 | 0.25 | 1 | 11 |
| cebpb | 0.154 | 0.406174598 | 0.611 | 0.429 | 1 | 11 |
| irf2 | 0.15403 | -0.30187487 | 0.083 | 0.25 | 1 | 11 |
| cxcl8b.1 | 0.15405 | -0.43306284 | 0.278 | 0.429 | 1 | 11 |
| eef1b2 | 0.15508 | -0.13993051 | 0.806 | 0.786 | 1 | 11 |
| olfml3b | 0.15537 | 0.223115469 | 0.528 | 0.464 | 1 | 11 |
| ythdc1 | 0.15596 | 0.304257877 | 0.167 | 0.036 | 1 | 11 |
| aurkaip1 | 0.1576 | -0.15087702 | 0.056 | 0.214 | 1 | 11 |
| ndufs6 | 0.15853 | 0.237960481 | 0.361 | 0.286 | 1 | 11 |
| psmc5 | 0.15855 | 0.287022865 | 0.306 | 0.214 | 1 | 11 |
| rab8b | 0.16003 | -0.22899568 | 0.056 | 0.214 | 1 | 11 |
| DHDH | 0.16084 | 0.127619504 | 0.222 | 0.25 | 1 | 11 |
| crk | 0.16095 | -0.23682273 | 0.056 | 0.214 | 1 | 11 |
| gatc | 0.1613 | -0.23007135 | 0.056 | 0.214 | 1 | 11 |
| XLOC-043857 | 0.16135 | -0.18477124 | 0.056 | 0.214 | 1 | 11 |
| chic2 | 0.16148 | -0.22958551 | 0.056 | 0.214 | 1 | 11 |
| ppp3r1b | 0.16154 | -0.28192742 | 0.056 | 0.214 | 1 | 11 |
| plxnb2a | 0.16156 | -0.2220289 | 0.111 | 0.286 | 1 | 11 |
| snrpf | 0.1618 | 0.288506814 | 0.667 | 0.429 | 1 | 11 |
| slc7a1 | 0.16208 | -0.33188999 | 0.111 | 0.286 | 1 | 11 |
| lsm5 | 0.16255 | 0.318988651 | 0.389 | 0.179 | 1 | 11 |
| ube2l3a | 0.16262 | -0.31134915 | 0.167 | 0.357 | 1 | 11 |
| eif4a1a | 0.16297 | -0.26795515 | 0.306 | 0.536 | 1 | 11 |
| si:ch211-288g17.3 | 0.16342 | 0.296391959 | 0.611 | 0.536 | 1 | 11 |
| rhoca | 0.16459 | -0.17860559 | 0.139 | 0.321 | 1 | 11 |
| zcchc17 | 0.166 | -0.3162214 | 0.056 | 0.179 | 1 | 11 |
| eny2 | 0.16608 | -0.17586604 | 0.139 | 0.321 | 1 | 11 |
| NC-002333.4 | 0.16614 | 0.298395512 | 0.889 | 0.714 | 1 | 11 |
| nr3c1 | 0.16665 | -0.30681287 | 0.111 | 0.286 | 1 | 11 |
| arl14ep | 0.16694 | -0.11901392 | 0.056 | 0.179 | 1 | 11 |
| krt8 | 0.16709 | -0.1227615 | 0.944 | 1 | 1 | 11 |
| pabpc1a | 0.16781 | -0.2574303 | 0.722 | 0.893 | 1 | 11 |
| glo1 | 0.16787 | 0.289706271 | 0.583 | 0.429 | 1 | 11 |
| pitpnbl | 0.16801 | -0.1767196 | 0.111 | 0.286 | 1 | 11 |
| tfg | 0.16959 | -0.11510157 | 0.139 | 0.286 | 1 | 11 |
| cmpk | 0.17107 | 0.279591773 | 0.222 | 0.071 | 1 | 11 |
| mcts1 | 0.17132 | 0.284987969 | 0.306 | 0.214 | 1 | 11 |
| cnot9 | 0.17137 | -0.24447842 | 0.083 | 0.25 | 1 | 11 |

| | | | | | | |
|---|---|---|---|---|---|---|
| LO018148.1 | 0.1717 | 0.358713048 | 0.222 | 0.071 | 1 | 11 |
| anxa2a | 0.17182 | -0.12354711 | 0.778 | 0.75 | 1 | 11 |
| mdh2 | 0.17265 | -0.12293842 | 0.472 | 0.429 | 1 | 11 |
| asip1 | 0.173 | -0.32901608 | 0.028 | 0.107 | 1 | 11 |
| pfdn6 | 0.17347 | 0.273182424 | 0.306 | 0.179 | 1 | 11 |
| reep3a | 0.17386 | 0.324301913 | 0.389 | 0.179 | 1 | 11 |
| efnb1 | 0.17433 | 0.338647111 | 0.278 | 0.107 | 1 | 11 |
| gstt1b | 0.17433 | -0.31475193 | 0.306 | 0.536 | 1 | 11 |
| mbnl2 | 0.17441 | -0.12763048 | 0.056 | 0.179 | 1 | 11 |
| ndufs4 | 0.17445 | -0.24395259 | 0.167 | 0.214 | 1 | 11 |
| si:ch211-226m16.2 | 0.17509 | 0.135216225 | 0.083 | 0.107 | 1 | 11 |
| gnb1a | 0.17603 | 0.179942851 | 0.306 | 0.286 | 1 | 11 |
| psma2 | 0.17638 | 0.275821158 | 0.389 | 0.179 | 1 | 11 |
| bgnb | 0.17744 | -0.10526417 | 0.056 | 0.179 | 1 | 11 |
| gsdf | 0.17832 | -0.29881767 | 0.083 | 0.179 | 1 | 11 |
| dlst | 0.17847 | -0.35680245 | 0.194 | 0.393 | 1 | 11 |
| mat2ab | 0.17876 | 0.246149461 | 0.25 | 0.143 | 1 | 11 |
| sesn1 | 0.17899 | 0.222107482 | 0.556 | 0.536 | 1 | 11 |
| sumo3a | 0.17963 | 0.175353648 | 0.556 | 0.571 | 1 | 11 |
| ptmab | 0.17984 | 0.222620957 | 0.917 | 0.821 | 1 | 11 |
| ostc | 0.18176 | 0.239583781 | 0.389 | 0.321 | 1 | 11 |
| capns1b | 0.18195 | -0.32287782 | 0.472 | 0.643 | 1 | 11 |
| sec61g | 0.18198 | -0.29535425 | 0.639 | 0.75 | 1 | 11 |
| gpr155a | 0.18206 | -0.35316254 | 0.111 | 0.286 | 1 | 11 |
| cul3a | 0.18257 | 0.24893893 | 0.167 | 0.036 | 1 | 11 |
| ppp2r2d | 0.18309 | 0.138122649 | 0.194 | 0.071 | 1 | 11 |
| paplna | 0.1839 | -0.26211805 | 0.417 | 0.643 | 1 | 11 |
| s100u | 0.18429 | 0.33432545 | 0.306 | 0.143 | 1 | 11 |
| CABZ01078607.1 | 0.18454 | 0.12246301 | 0.056 | 0 | 1 | 11 |
| CU138547.1 | 0.18454 | 0.108593055 | 0.056 | 0 | 1 | 11 |
| CU929418.1 | 0.18454 | 0.135907294 | 0.056 | 0 | 1 | 11 |
| LOC100535543 | 0.18454 | 0.114250768 | 0.056 | 0 | 1 | 11 |
| LOC101885646 | 0.18454 | 0.12637658 | 0.056 | 0 | 1 | 11 |
| LOC110438754 | 0.18454 | 0.118325961 | 0.056 | 0 | 1 | 11 |
| NDUFAF8 | 0.18454 | 0.114142972 | 0.056 | 0 | 1 | 11 |
| WDR70 | 0.18454 | 0.112713996 | 0.056 | 0 | 1 | 11 |
| XLOC-006824 | 0.18454 | 0.105137674 | 0.056 | 0 | 1 | 11 |
| XLOC-024750 | 0.18454 | 0.129627521 | 0.056 | 0 | 1 | 11 |
| XLOC-029591 | 0.18454 | 0.103364801 | 0.056 | 0 | 1 | 11 |
| abhd15a | 0.18454 | 0.140801312 | 0.056 | 0 | 1 | 11 |
| adgrd1 | 0.18454 | 0.11406604 | 0.056 | 0 | 1 | 11 |
| adgrl3.1 | 0.18454 | 0.122371652 | 0.056 | 0 | 1 | 11 |
| aldh9a1b | 0.18454 | 0.104318838 | 0.056 | 0 | 1 | 11 |
| ankmy2a | 0.18454 | 0.100999711 | 0.056 | 0 | 1 | 11 |
| ankrd10a | 0.18454 | 0.101326816 | 0.056 | 0 | 1 | 11 |
| arhgap12b | 0.18454 | 0.123451012 | 0.056 | 0 | 1 | 11 |
| asb12a | 0.18454 | 0.109000358 | 0.056 | 0 | 1 | 11 |
| ascl1b | 0.18454 | 0.115848824 | 0.056 | 0 | 1 | 11 |

| | | | | | | |
|---|---|---|---|---|---|---|
| atm | 0.18454 | 0.141828104 | 0.056 | 0 | 1 | 11 |
| atp1b2a | 0.18454 | 0.101919345 | 0.056 | 0 | 1 | 11 |
| bambib | 0.18454 | 0.132518169 | 0.056 | 0 | 1 | 11 |
| bmp4 | 0.18454 | 0.11349924 | 0.056 | 0 | 1 | 11 |
| ccdc57 | 0.18454 | 0.110380879 | 0.056 | 0 | 1 | 11 |
| ccl20b | 0.18454 | 0.118817929 | 0.056 | 0 | 1 | 11 |
| cdca8 | 0.18454 | 0.12060441 | 0.056 | 0 | 1 | 11 |
| clec11a | 0.18454 | 0.140839365 | 0.056 | 0 | 1 | 11 |
| cnot4b | 0.18454 | 0.135636554 | 0.056 | 0 | 1 | 11 |
| cog7 | 0.18454 | 0.100579792 | 0.056 | 0 | 1 | 11 |
| ctps1b | 0.18454 | 0.10891763 | 0.056 | 0 | 1 | 11 |
| dbt | 0.18454 | 0.104927313 | 0.056 | 0 | 1 | 11 |
| ddx56 | 0.18454 | 0.128593093 | 0.056 | 0 | 1 | 11 |
| dixdc1b | 0.18454 | 0.103001961 | 0.056 | 0 | 1 | 11 |
| dlc | 0.18454 | 0.12555893 | 0.056 | 0 | 1 | 11 |
| dnajc18 | 0.18454 | 0.119370895 | 0.056 | 0 | 1 | 11 |
| dpf2 | 0.18454 | 0.126812253 | 0.056 | 0 | 1 | 11 |
| dpf3 | 0.18454 | 0.100100642 | 0.056 | 0 | 1 | 11 |
| drosha | 0.18454 | 0.101919345 | 0.056 | 0 | 1 | 11 |
| eefsec | 0.18454 | 0.128096773 | 0.056 | 0 | 1 | 11 |
| exoc7 | 0.18454 | 0.163391474 | 0.056 | 0 | 1 | 11 |
| fbxo22 | 0.18454 | 0.1271426 | 0.056 | 0 | 1 | 11 |
| fgf10a | 0.18454 | 0.114272905 | 0.056 | 0 | 1 | 11 |
| fryb | 0.18454 | 0.160220704 | 0.056 | 0 | 1 | 11 |
| gcdhb | 0.18454 | 0.112980434 | 0.056 | 0 | 1 | 11 |
| gmds | 0.18454 | 0.103991588 | 0.056 | 0 | 1 | 11 |
| golim4a.1 | 0.18454 | 0.101080696 | 0.056 | 0 | 1 | 11 |
| gorab | 0.18454 | 0.135080286 | 0.056 | 0 | 1 | 11 |
| grk1b | 0.18454 | 0.105005186 | 0.056 | 0 | 1 | 11 |
| gstt2 | 0.18454 | 0.129411428 | 0.056 | 0 | 1 | 11 |
| gtdc1 | 0.18454 | 0.127441277 | 0.056 | 0 | 1 | 11 |
| hic1 | 0.18454 | 0.105869783 | 0.056 | 0 | 1 | 11 |
| hoxa11a | 0.18454 | 0.106442886 | 0.056 | 0 | 1 | 11 |
| iah1 | 0.18454 | 0.108813744 | 0.056 | 0 | 1 | 11 |
| igdcc3 | 0.18454 | 0.136719127 | 0.056 | 0 | 1 | 11 |
| im:7147486 | 0.18454 | 0.121464071 | 0.056 | 0 | 1 | 11 |
| iqca1 | 0.18454 | 0.10909846 | 0.056 | 0 | 1 | 11 |
| irf2a | 0.18454 | 0.1132691 | 0.056 | 0 | 1 | 11 |
| kiaa1958 | 0.18454 | 0.100535907 | 0.056 | 0 | 1 | 11 |
| kif20a | 0.18454 | 0.132446785 | 0.056 | 0 | 1 | 11 |
| kmt5b | 0.18454 | 0.144280368 | 0.056 | 0 | 1 | 11 |
| lmbr1 | 0.18454 | 0.117669771 | 0.056 | 0 | 1 | 11 |
| lonp2 | 0.18454 | 0.103129496 | 0.056 | 0 | 1 | 11 |
| lrrc38a | 0.18454 | 0.10011356 | 0.056 | 0 | 1 | 11 |
| lyst | 0.18454 | 0.124187873 | 0.056 | 0 | 1 | 11 |
| madd | 0.18454 | 0.106154346 | 0.056 | 0 | 1 | 11 |
| magi2a | 0.18454 | 0.180103792 | 0.056 | 0 | 1 | 11 |
| mepce | 0.18454 | 0.116022059 | 0.056 | 0 | 1 | 11 |

| | | | | | | |
|---|---|---|---|---|---|---|
| mief1 | 0.18454 | 0.133675621 | 0.056 | 0 | 1 | 11 |
| mrpl40 | 0.18454 | 0.117276419 | 0.056 | 0 | 1 | 11 |
| mtfr2 | 0.18454 | 0.126944199 | 0.056 | 0 | 1 | 11 |
| myca | 0.18454 | 0.105593038 | 0.056 | 0 | 1 | 11 |
| nckap5l | 0.18454 | 0.122632333 | 0.056 | 0 | 1 | 11 |
| nfybb | 0.18454 | 0.124945248 | 0.056 | 0 | 1 | 11 |
| ngfrb | 0.18454 | 0.106400322 | 0.056 | 0 | 1 | 11 |
| nipal3 | 0.18454 | 0.100100642 | 0.056 | 0 | 1 | 11 |
| nthl1 | 0.18454 | 0.120166773 | 0.056 | 0 | 1 | 11 |
| pacsin3 | 0.18454 | 0.108706684 | 0.056 | 0 | 1 | 11 |
| pan3 | 0.18454 | 0.165139461 | 0.056 | 0 | 1 | 11 |
| pde3a | 0.18454 | 0.123798218 | 0.056 | 0 | 1 | 11 |
| pdss1 | 0.18454 | 0.103761508 | 0.056 | 0 | 1 | 11 |
| pex12 | 0.18454 | 0.107454364 | 0.056 | 0 | 1 | 11 |
| pigb | 0.18454 | 0.133990685 | 0.056 | 0 | 1 | 11 |
| ppp1r3da | 0.18454 | 0.14076124 | 0.056 | 0 | 1 | 11 |
| ppp2cab | 0.18454 | 0.124550073 | 0.056 | 0 | 1 | 11 |
| ppp2r2aa | 0.18454 | 0.109013138 | 0.056 | 0 | 1 | 11 |
| ptgs2a | 0.18454 | 0.146362929 | 0.056 | 0 | 1 | 11 |
| ptprq | 0.18454 | 0.174859471 | 0.056 | 0 | 1 | 11 |
| pycard | 0.18454 | 0.126353891 | 0.056 | 0 | 1 | 11 |
| r3hdm2 | 0.18454 | 0.120732292 | 0.056 | 0 | 1 | 11 |
| rab11fip1a | 0.18454 | 0.115634181 | 0.056 | 0 | 1 | 11 |
| rapgef1a | 0.18454 | 0.117771458 | 0.056 | 0 | 1 | 11 |
| rb1cc1 | 0.18454 | 0.108210472 | 0.056 | 0 | 1 | 11 |
| rgs12b | 0.18454 | 0.117613024 | 0.056 | 0 | 1 | 11 |
| riox2 | 0.18454 | 0.134971065 | 0.056 | 0 | 1 | 11 |
| rrm2b | 0.18454 | 0.125736863 | 0.056 | 0 | 1 | 11 |
| samd13 | 0.18454 | 0.105264588 | 0.056 | 0 | 1 | 11 |
| scly | 0.18454 | 0.103622907 | 0.056 | 0 | 1 | 11 |
| sema7a | 0.18454 | 0.118635272 | 0.056 | 0 | 1 | 11 |
| setdb1b | 0.18454 | 0.130110585 | 0.056 | 0 | 1 | 11 |
| si:ch211-214b16.4 | 0.18454 | 0.117247277 | 0.056 | 0 | 1 | 11 |
| si:ch211-214c7.4 | 0.18454 | 0.114142429 | 0.056 | 0 | 1 | 11 |
| si:ch211-225h24.2 | 0.18454 | 0.10295113 | 0.056 | 0 | 1 | 11 |
| si:ch211-261d7.6 | 0.18454 | 0.111718706 | 0.056 | 0 | 1 | 11 |
| si:ch211-266i6.3 | 0.18454 | 0.137627161 | 0.056 | 0 | 1 | 11 |
| si:ch211-42i9.8 | 0.18454 | 0.104398668 | 0.056 | 0 | 1 | 11 |
| si:ch73-358j7.2 | 0.18454 | 0.107946875 | 0.056 | 0 | 1 | 11 |
| si:dkey-163m14.2 | 0.18454 | 0.129306239 | 0.056 | 0 | 1 | 11 |
| si:dkey-170l10.1 | 0.18454 | 0.106505759 | 0.056 | 0 | 1 | 11 |
| si:dkey-30k6.5 | 0.18454 | 0.105137674 | 0.056 | 0 | 1 | 11 |
| si:dkey-56e3.3 | 0.18454 | 0.100269286 | 0.056 | 0 | 1 | 11 |
| si:dkey-66i24.7 | 0.18454 | 0.102550344 | 0.056 | 0 | 1 | 11 |
| si:dkey-9i23.15 | 0.18454 | 0.105925678 | 0.056 | 0 | 1 | 11 |
| skiv2l | 0.18454 | 0.104957906 | 0.056 | 0 | 1 | 11 |
| slc10a7 | 0.18454 | 0.130475431 | 0.056 | 0 | 1 | 11 |
| slc11a2 | 0.18454 | 0.116766993 | 0.056 | 0 | 1 | 11 |

| | | | | | | |
|---|---|---|---|---|---|---|
| slc9a6a | 0.18454 | 0.104957906 | 0.056 | 0 | 1 | 11 |
| slka | 0.18454 | 0.100535907 | 0.056 | 0 | 1 | 11 |
| smim12 | 0.18454 | 0.134609713 | 0.056 | 0 | 1 | 11 |
| sntb1 | 0.18454 | 0.122295039 | 0.056 | 0 | 1 | 11 |
| sp3a | 0.18454 | 0.110443165 | 0.056 | 0 | 1 | 11 |
| spc24 | 0.18454 | 0.101270746 | 0.056 | 0 | 1 | 11 |
| stk38b | 0.18454 | 0.100506757 | 0.056 | 0 | 1 | 11 |
| taf2 | 0.18454 | 0.110198376 | 0.056 | 0 | 1 | 11 |
| tbl1xr1b | 0.18454 | 0.108875579 | 0.056 | 0 | 1 | 11 |
| tex2 | 0.18454 | 0.127629684 | 0.056 | 0 | 1 | 11 |
| tjp1b | 0.18454 | 0.117554724 | 0.056 | 0 | 1 | 11 |
| tmem119b | 0.18454 | 0.103622907 | 0.056 | 0 | 1 | 11 |
| tmem132e | 0.18454 | 0.166458853 | 0.056 | 0 | 1 | 11 |
| tmem234 | 0.18454 | 0.101324891 | 0.056 | 0 | 1 | 11 |
| tmem26a | 0.18454 | 0.145501037 | 0.056 | 0 | 1 | 11 |
| tp53inp2 | 0.18454 | 0.136300179 | 0.056 | 0 | 1 | 11 |
| tppp2 | 0.18454 | 0.137325432 | 0.056 | 0 | 1 | 11 |
| tshz3a | 0.18454 | 0.100735751 | 0.056 | 0 | 1 | 11 |
| ttyh3b | 0.18454 | 0.127245689 | 0.056 | 0 | 1 | 11 |
| vps18 | 0.18454 | 0.10738323 | 0.056 | 0 | 1 | 11 |
| xpo1b | 0.18454 | 0.116602962 | 0.056 | 0 | 1 | 11 |
| xrcc3 | 0.18454 | 0.116740939 | 0.056 | 0 | 1 | 11 |
| zdhhc14 | 0.18454 | 0.13501457 | 0.056 | 0 | 1 | 11 |
| zgc:113452 | 0.18454 | 0.103561374 | 0.056 | 0 | 1 | 11 |
| zgc:153704 | 0.18454 | 0.110281497 | 0.056 | 0 | 1 | 11 |
| zgc:153901 | 0.18454 | 0.109181773 | 0.056 | 0 | 1 | 11 |
| zgc:162971 | 0.18454 | 0.161693384 | 0.056 | 0 | 1 | 11 |
| zgc:172139 | 0.18454 | 0.157731832 | 0.056 | 0 | 1 | 11 |
| zgc:194281 | 0.18454 | 0.109060138 | 0.056 | 0 | 1 | 11 |
| zgc:55781 | 0.18454 | 0.101727261 | 0.056 | 0 | 1 | 11 |
| znf1094 | 0.18454 | 0.12706414 | 0.056 | 0 | 1 | 11 |
| znf1154 | 0.18454 | 0.121987146 | 0.056 | 0 | 1 | 11 |
| znf1157 | 0.18454 | 0.12538314 | 0.056 | 0 | 1 | 11 |
| znf654 | 0.18454 | 0.10208194 | 0.056 | 0 | 1 | 11 |
| znf704 | 0.18454 | 0.133153352 | 0.056 | 0 | 1 | 11 |
| AL954696.1 | 0.18454 | 0.131254014 | 0.056 | 0 | 1 | 11 |
| CABZ01064859.1 | 0.18454 | 0.154368418 | 0.056 | 0 | 1 | 11 |
| CR450780.2 | 0.18454 | 0.120876477 | 0.056 | 0 | 1 | 11 |
| CU655961.1 | 0.18454 | 0.111734231 | 0.056 | 0 | 1 | 11 |
| LO018029.1 | 0.18454 | 0.129534145 | 0.056 | 0 | 1 | 11 |
| RAMP1 | 0.18454 | 0.104152139 | 0.056 | 0 | 1 | 11 |
| XLOC-001102 | 0.18454 | 0.100873898 | 0.056 | 0 | 1 | 11 |
| ache | 0.18454 | 0.158703681 | 0.056 | 0 | 1 | 11 |
| adgre5a | 0.18454 | 0.104738654 | 0.056 | 0 | 1 | 11 |
| adgre5b.3 | 0.18454 | 0.130892782 | 0.056 | 0 | 1 | 11 |
| arhgef1a | 0.18454 | 0.123066799 | 0.056 | 0 | 1 | 11 |
| bcl3 | 0.18454 | 0.117534291 | 0.056 | 0 | 1 | 11 |
| cbr1 | 0.18454 | 0.109174017 | 0.056 | 0 | 1 | 11 |

| | | | | | | |
|---|---|---|---|---|---|---|
| ccne1 | 0.18454 | 0.104804136 | 0.056 | 0 | 1 | 11 |
| cdk5rap2 | 0.18454 | 0.120199778 | 0.056 | 0 | 1 | 11 |
| cmtr1 | 0.18454 | 0.140133017 | 0.056 | 0 | 1 | 11 |
| ctso | 0.18454 | 0.107978237 | 0.056 | 0 | 1 | 11 |
| dhps | 0.18454 | 0.12474218 | 0.056 | 0 | 1 | 11 |
| dyrk1b | 0.18454 | 0.14568306 | 0.056 | 0 | 1 | 11 |
| elavl1b | 0.18454 | 0.104698037 | 0.056 | 0 | 1 | 11 |
| epg5 | 0.18454 | 0.118343565 | 0.056 | 0 | 1 | 11 |
| ercc5 | 0.18454 | 0.12247573 | 0.056 | 0 | 1 | 11 |
| errfi1a | 0.18454 | 0.113544928 | 0.056 | 0 | 1 | 11 |
| fndc3a | 0.18454 | 0.126183636 | 0.056 | 0 | 1 | 11 |
| fnip1 | 0.18454 | 0.149144478 | 0.056 | 0 | 1 | 11 |
| fopnl | 0.18454 | 0.114820615 | 0.056 | 0 | 1 | 11 |
| gb:ai877918 | 0.18454 | 0.116386498 | 0.056 | 0 | 1 | 11 |
| igsf8 | 0.18454 | 0.128964847 | 0.056 | 0 | 1 | 11 |
| kcnd3 | 0.18454 | 0.100800556 | 0.056 | 0 | 1 | 11 |
| kdm4ab | 0.18454 | 0.113092167 | 0.056 | 0 | 1 | 11 |
| kdm7aa | 0.18454 | 0.110519794 | 0.056 | 0 | 1 | 11 |
| klhl2 | 0.18454 | 0.124782059 | 0.056 | 0 | 1 | 11 |
| larsb | 0.18454 | 0.137818506 | 0.056 | 0 | 1 | 11 |
| lmtk2 | 0.18454 | 0.128964847 | 0.056 | 0 | 1 | 11 |
| lpgat1 | 0.18454 | 0.11615165 | 0.056 | 0 | 1 | 11 |
| map3k20 | 0.18454 | 0.127376384 | 0.056 | 0 | 1 | 11 |
| mapk8b | 0.18454 | 0.103728529 | 0.056 | 0 | 1 | 11 |
| mki67 | 0.18454 | 0.120090088 | 0.056 | 0 | 1 | 11 |
| morc3a | 0.18454 | 0.140715583 | 0.056 | 0 | 1 | 11 |
| mpp6b | 0.18454 | 0.165491603 | 0.056 | 0 | 1 | 11 |
| naa35 | 0.18454 | 0.109016042 | 0.056 | 0 | 1 | 11 |
| nisch | 0.18454 | 0.104556149 | 0.056 | 0 | 1 | 11 |
| nrarpa | 0.18454 | 0.115213426 | 0.056 | 0 | 1 | 11 |
| patj | 0.18454 | 0.119175042 | 0.056 | 0 | 1 | 11 |
| pde8a | 0.18454 | 0.104804136 | 0.056 | 0 | 1 | 11 |
| ppfia1 | 0.18454 | 0.167843054 | 0.056 | 0 | 1 | 11 |
| rbpjb | 0.18454 | 0.139782871 | 0.056 | 0 | 1 | 11 |
| rnf20 | 0.18454 | 0.118271435 | 0.056 | 0 | 1 | 11 |
| rpusd4 | 0.18454 | 0.127376384 | 0.056 | 0 | 1 | 11 |
| scocb | 0.18454 | 0.10034968 | 0.056 | 0 | 1 | 11 |
| sema5bb | 0.18454 | 0.11975456 | 0.056 | 0 | 1 | 11 |
| setd1ba | 0.18454 | 0.108817777 | 0.056 | 0 | 1 | 11 |
| si:ch211-262i1.5 | 0.18454 | 0.116640132 | 0.056 | 0 | 1 | 11 |
| si:dkey-20i10.7 | 0.18454 | 0.125577123 | 0.056 | 0 | 1 | 11 |
| si:dkey-253d23.3 | 0.18454 | 0.101646637 | 0.056 | 0 | 1 | 11 |
| si:dkey-3d4.3 | 0.18454 | 0.14055049 | 0.056 | 0 | 1 | 11 |
| si:dkey-82o10.4 | 0.18454 | 0.122962758 | 0.056 | 0 | 1 | 11 |
| sort1b | 0.18454 | 0.101307341 | 0.056 | 0 | 1 | 11 |
| sox4b | 0.18454 | 0.121772448 | 0.056 | 0 | 1 | 11 |
| srebf2 | 0.18454 | 0.132731472 | 0.056 | 0 | 1 | 11 |
| stk17a | 0.18454 | 0.110063575 | 0.056 | 0 | 1 | 11 |

| | | | | | | |
|---|---|---|---|---|---|---|
| tbx2a | 0.18454 | 0.100614175 | 0.056 | 0 | 1 | 11 |
| tigarb | 0.18454 | 0.121854776 | 0.056 | 0 | 1 | 11 |
| ube3c | 0.18454 | 0.167822669 | 0.056 | 0 | 1 | 11 |
| ubiad1 | 0.18454 | 0.101038036 | 0.056 | 0 | 1 | 11 |
| ubr4 | 0.18454 | 0.118023083 | 0.056 | 0 | 1 | 11 |
| usp33 | 0.18454 | 0.100333179 | 0.056 | 0 | 1 | 11 |
| vps8 | 0.18454 | 0.107656324 | 0.056 | 0 | 1 | 11 |
| zdhhc9 | 0.18454 | 0.121106405 | 0.056 | 0 | 1 | 11 |
| zgc:63831 | 0.18454 | 0.119446754 | 0.056 | 0 | 1 | 11 |
| zgc:76872 | 0.18454 | 0.120012607 | 0.056 | 0 | 1 | 11 |
| zmp:0000000652 | 0.18454 | 0.127364271 | 0.056 | 0 | 1 | 11 |
| znf385d | 0.18454 | 0.100800556 | 0.056 | 0 | 1 | 11 |
| BX323458.1 | 0.18454 | 0.123448913 | 0.056 | 0 | 1 | 11 |
| si:dkeyp-110c7.4 | 0.18454 | 0.132353029 | 0.056 | 0 | 1 | 11 |
| AL732567.1 | 0.18454 | 0.109931888 | 0.056 | 0 | 1 | 11 |
| gpr157 | 0.18454 | 0.132051204 | 0.056 | 0 | 1 | 11 |
| mrpl38 | 0.18454 | 0.118022315 | 0.056 | 0 | 1 | 11 |
| msto1 | 0.18454 | 0.114384378 | 0.056 | 0 | 1 | 11 |
| mycn | 0.18454 | 0.112535583 | 0.056 | 0 | 1 | 11 |
| ppie | 0.18454 | 0.100303728 | 0.056 | 0 | 1 | 11 |
| prkrip1 | 0.18454 | 0.103422314 | 0.056 | 0 | 1 | 11 |
| si:ch211-281l24.3 | 0.18454 | 0.107448468 | 0.056 | 0 | 1 | 11 |
| si:dkey-256h2.1 | 0.18454 | 0.101574975 | 0.056 | 0 | 1 | 11 |
| sox18 | 0.18454 | 0.117909308 | 0.056 | 0 | 1 | 11 |
| ttc5 | 0.18454 | 0.107698356 | 0.056 | 0 | 1 | 11 |
| c1qtnf1 | 0.18454 | 0.104087821 | 0.056 | 0 | 1 | 11 |
| dock11 | 0.18454 | 0.103007716 | 0.056 | 0 | 1 | 11 |
| jag2b | 0.18454 | 0.139702441 | 0.056 | 0 | 1 | 11 |
| pacrg | 0.18454 | 0.102643089 | 0.056 | 0 | 1 | 11 |
| pik3r4 | 0.18454 | 0.102680552 | 0.056 | 0 | 1 | 11 |
| sema3e | 0.18454 | 0.102643089 | 0.056 | 0 | 1 | 11 |
| si:dkey-247i3.6 | 0.18454 | 0.121513458 | 0.056 | 0 | 1 | 11 |
| tmem192 | 0.18454 | 0.108507156 | 0.056 | 0 | 1 | 11 |
| wu:fj29h11 | 0.18454 | 0.102643089 | 0.056 | 0 | 1 | 11 |
| mast2 | 0.18496 | -0.3142115 | 0.083 | 0.214 | 1 | 11 |
| foxk1 | 0.18509 | 0.179717397 | 0.167 | 0.036 | 1 | 11 |
| bysl | 0.18526 | -0.21268584 | 0.25 | 0.464 | 1 | 11 |
| ranbp9 | 0.18531 | 0.280243476 | 0.278 | 0.107 | 1 | 11 |
| col1a2 | 0.18567 | -0.12171087 | 0.972 | 1 | 1 | 11 |
| cdc42se1 | 0.18603 | -0.10184865 | 0.139 | 0.286 | 1 | 11 |
| psen1 | 0.18616 | 0.261698722 | 0.167 | 0.036 | 1 | 11 |
| gorasp2 | 0.18647 | -0.24248424 | 0.083 | 0.25 | 1 | 11 |
| psmb6 | 0.1869 | -0.17368135 | 0.306 | 0.5 | 1 | 11 |
| si:rp71-45k5.4 | 0.18703 | 0.141210682 | 0.306 | 0.321 | 1 | 11 |
| rbms2a | 0.1872 | -0.22366191 | 0.139 | 0.321 | 1 | 11 |
| sephs2 | 0.18726 | -0.14309671 | 0.028 | 0.143 | 1 | 11 |
| rwdd1 | 0.18774 | -0.20567193 | 0.194 | 0.393 | 1 | 11 |
| rhocb | 0.18776 | -0.24747444 | 0.25 | 0.464 | 1 | 11 |

| | | | | | | |
|---|---|---|---|---|---|---|
| zgc:163030 | 0.18834 | 0.239993458 | 0.167 | 0.036 | 1 | 11 |
| jpt1b | 0.18851 | 0.228023824 | 0.75 | 0.536 | 1 | 11 |
| podxl | 0.18862 | -0.13728054 | 0.917 | 1 | 1 | 11 |
| ttf1 | 0.18895 | -0.2405388 | 0.028 | 0.143 | 1 | 11 |
| gpc4 | 0.18907 | -0.31257542 | 0.139 | 0.321 | 1 | 11 |
| hnrnpl2 | 0.19004 | -0.26192178 | 0.083 | 0.25 | 1 | 11 |
| si:ch73-91k6.2 | 0.19022 | 0.307124664 | 0.222 | 0.071 | 1 | 11 |
| mrpl34 | 0.19032 | -0.27050448 | 0.111 | 0.286 | 1 | 11 |
| add1 | 0.19037 | 0.301220908 | 0.306 | 0.214 | 1 | 11 |
| zgc:77486 | 0.19089 | -0.23165292 | 0.056 | 0.179 | 1 | 11 |
| cdv3 | 0.19138 | -0.21925606 | 0.222 | 0.429 | 1 | 11 |
| nfil3-6 | 0.19238 | 0.150310067 | 0.333 | 0.357 | 1 | 11 |
| sephs1 | 0.19291 | -0.22897922 | 0.028 | 0.143 | 1 | 11 |
| anxa1a | 0.19331 | -0.17423844 | 0.917 | 0.964 | 1 | 11 |
| snx17 | 0.19349 | 0.356767981 | 0.417 | 0.214 | 1 | 11 |
| fkbp5 | 0.19425 | -0.30035759 | 0.306 | 0.429 | 1 | 11 |
| hmga1a | 0.19462 | 0.343962172 | 0.611 | 0.393 | 1 | 11 |
| ugt1b5 | 0.19466 | 0.31940818 | 0.611 | 0.429 | 1 | 11 |
| mtss1 | 0.19554 | -0.13150752 | 0.028 | 0.143 | 1 | 11 |
| znf609a | 0.19618 | 0.326684546 | 0.222 | 0.107 | 1 | 11 |
| setdb1a | 0.1965 | 0.115699312 | 0.139 | 0.036 | 1 | 11 |
| vclb | 0.19657 | 0.240314747 | 0.167 | 0.036 | 1 | 11 |
| cd40 | 0.19701 | -0.25918427 | 0.028 | 0.143 | 1 | 11 |
| gtf2b | 0.19728 | 0.264926349 | 0.139 | 0.036 | 1 | 11 |
| tktb | 0.1977 | 0.180044348 | 0.222 | 0.071 | 1 | 11 |
| slc43a2b | 0.19792 | -0.26349332 | 0.028 | 0.143 | 1 | 11 |
| si:dkey-23i12.5 | 0.19829 | 0.273781535 | 0.333 | 0.143 | 1 | 11 |
| srgap3 | 0.19854 | -0.14972099 | 0.028 | 0.143 | 1 | 11 |
| akt2 | 0.1995 | -0.25840509 | 0.028 | 0.143 | 1 | 11 |
| mrpl45 | 0.19952 | -0.13164069 | 0.028 | 0.143 | 1 | 11 |
| tdrd3 | 0.20049 | -0.13147814 | 0.028 | 0.143 | 1 | 11 |
| ext1a | 0.20104 | -0.2725469 | 0.028 | 0.143 | 1 | 11 |
| si:dkey-122a22.2 | 0.20124 | -0.14351129 | 0.028 | 0.143 | 1 | 11 |
| epha4l | 0.20263 | 0.345447518 | 0.417 | 0.214 | 1 | 11 |
| si:ch73-269m14.2 | 0.20312 | 0.225174348 | 0.167 | 0.036 | 1 | 11 |
| adamts8a | 0.20384 | -0.28745982 | 0.028 | 0.143 | 1 | 11 |
| bcl9 | 0.20408 | -0.19585044 | 0.028 | 0.143 | 1 | 11 |
| itgb1b.1 | 0.20451 | -0.14651565 | 0.028 | 0.143 | 1 | 11 |
| calm1a | 0.20474 | -0.18304082 | 0.333 | 0.536 | 1 | 11 |
| rpl4 | 0.2052 | -0.12355342 | 0.917 | 1 | 1 | 11 |
| palm1b | 0.20524 | -0.2729668 | 0.222 | 0.429 | 1 | 11 |
| xaf1 | 0.20535 | -0.13829598 | 0.056 | 0.179 | 1 | 11 |
| plekhf2 | 0.20536 | -0.11259082 | 0.056 | 0.179 | 1 | 11 |
| cpsf3 | 0.2057 | -0.17806617 | 0.028 | 0.143 | 1 | 11 |
| zgc:92140 | 0.2061 | 0.273357963 | 0.222 | 0.071 | 1 | 11 |
| tufm | 0.20652 | -0.23563507 | 0.111 | 0.286 | 1 | 11 |
| hadhb | 0.20664 | 0.231274433 | 0.139 | 0.036 | 1 | 11 |
| tpst1l | 0.20734 | 0.255822285 | 0.167 | 0.036 | 1 | 11 |

| | | | | | | |
|---|---|---|---|---|---|---|
| pik3r3a | 0.20735 | -0.31178805 | 0.167 | 0.321 | 1 | 11 |
| ppme1 | 0.20799 | -0.28531131 | 0.111 | 0.286 | 1 | 11 |
| bmper | 0.20813 | 0.208043718 | 0.667 | 0.464 | 1 | 11 |
| copz1 | 0.20935 | -0.23742382 | 0.222 | 0.429 | 1 | 11 |
| mff | 0.2101 | 0.191692393 | 0.167 | 0.036 | 1 | 11 |
| parvaa | 0.21062 | 0.223108334 | 0.222 | 0.071 | 1 | 11 |
| cd151 | 0.21074 | -0.28299409 | 0.222 | 0.429 | 1 | 11 |
| idh2 | 0.21097 | -0.32966353 | 0.417 | 0.571 | 1 | 11 |
| cyb5r1 | 0.21102 | -0.21289572 | 0.139 | 0.321 | 1 | 11 |
| exosc4 | 0.21118 | 0.254923633 | 0.167 | 0.036 | 1 | 11 |
| smim20 | 0.21125 | 0.112460293 | 0.139 | 0.036 | 1 | 11 |
| arl6ip1 | 0.21199 | -0.24308798 | 0.194 | 0.393 | 1 | 11 |
| sept10 | 0.21242 | -0.27851556 | 0.167 | 0.357 | 1 | 11 |
| ccdc187 | 0.21255 | -0.14555236 | 0.028 | 0.143 | 1 | 11 |
| hagh | 0.21259 | -0.16603169 | 0.028 | 0.143 | 1 | 11 |
| sh3gl3a | 0.21266 | -0.18294866 | 0.028 | 0.143 | 1 | 11 |
| galnt7 | 0.21348 | 0.253595217 | 0.167 | 0.036 | 1 | 11 |
| atp5mc1 | 0.2138 | 0.113164709 | 0.278 | 0.143 | 1 | 11 |
| frs2a | 0.21437 | -0.12606575 | 0.028 | 0.143 | 1 | 11 |
| tnpo2 | 0.21458 | -0.16757015 | 0.028 | 0.143 | 1 | 11 |
| surf4 | 0.21499 | -0.21727919 | 0.028 | 0.143 | 1 | 11 |
| ankhd1 | 0.2152 | 0.214884197 | 0.167 | 0.036 | 1 | 11 |
| cand1 | 0.21521 | 0.229420195 | 0.167 | 0.036 | 1 | 11 |
| rab5c | 0.21531 | -0.24407587 | 0.167 | 0.357 | 1 | 11 |
| si:ch211-236l14.4 | 0.21543 | 0.226938905 | 0.167 | 0.036 | 1 | 11 |
| atp6v1e1b | 0.2156 | 0.171429651 | 0.167 | 0.036 | 1 | 11 |
| si:ch211-107o10.3 | 0.21561 | -0.19351457 | 0.028 | 0.143 | 1 | 11 |
| ier3ip1 | 0.21593 | -0.23459622 | 0.167 | 0.357 | 1 | 11 |
| pak2b | 0.21605 | 0.273572639 | 0.167 | 0.071 | 1 | 11 |
| rnaseh1 | 0.21616 | 0.212305915 | 0.167 | 0.036 | 1 | 11 |
| yes1 | 0.21679 | 0.22155275 | 0.167 | 0.036 | 1 | 11 |
| cavin2b | 0.21721 | -0.24428564 | 0.778 | 0.893 | 1 | 11 |
| dhrs13l1 | 0.2175 | 0.194192926 | 0.306 | 0.286 | 1 | 11 |
| pik3r1 | 0.21795 | 0.271641921 | 0.278 | 0.143 | 1 | 11 |
| gng5 | 0.21819 | -0.23969025 | 0.389 | 0.607 | 1 | 11 |
| pdcd4a | 0.21844 | -0.10731984 | 0.056 | 0.179 | 1 | 11 |
| supt6h | 0.21852 | 0.259097917 | 0.222 | 0.071 | 1 | 11 |
| tmed7 | 0.2187 | -0.23257687 | 0.389 | 0.607 | 1 | 11 |
| rpl14 | 0.21877 | -0.12079287 | 0.917 | 1 | 1 | 11 |
| gstm.2 | 0.21951 | 0.179266078 | 0.167 | 0.143 | 1 | 11 |
| rps27l | 0.22013 | 0.256157166 | 0.611 | 0.393 | 1 | 11 |
| si:dkey-199f5.8 | 0.22019 | 0.292055005 | 0.194 | 0.071 | 1 | 11 |
| zgc:162952 | 0.22057 | -0.18652977 | 0.028 | 0.143 | 1 | 11 |
| lamtor3 | 0.22066 | -0.17428257 | 0.028 | 0.143 | 1 | 11 |
| myo1ca | 0.22069 | -0.13096786 | 0.111 | 0.25 | 1 | 11 |
| mrps28 | 0.22076 | -0.24833618 | 0.028 | 0.143 | 1 | 11 |
| zgc:163098 | 0.221 | -0.18529271 | 0.028 | 0.143 | 1 | 11 |
| BX548028.1 | 0.22123 | -0.2891271 | 0.056 | 0.179 | 1 | 11 |

| | | | | | | |
|---|---|---|---|---|---|---|
| CABZ01068251.1 | 0.22186 | 0.214514524 | 0.222 | 0.071 | 1 | 11 |
| BX957297.2 | 0.22225 | -0.28571707 | 0.056 | 0.179 | 1 | 11 |
| styx | 0.22274 | -0.15788336 | 0.028 | 0.143 | 1 | 11 |
| noc3l | 0.2228 | -0.12055055 | 0.028 | 0.143 | 1 | 11 |
| MDFI | 0.22286 | -0.18168947 | 0.028 | 0.143 | 1 | 11 |
| emg1 | 0.22356 | -0.11616206 | 0.028 | 0.143 | 1 | 11 |
| tmem170a | 0.2239 | -0.14931433 | 0.028 | 0.143 | 1 | 11 |
| si:dkey-253d23.4 | 0.22394 | -0.13329589 | 0.056 | 0.179 | 1 | 11 |
| cnn3a | 0.22398 | 0.311608715 | 0.333 | 0.179 | 1 | 11 |
| hdgfl2 | 0.22436 | -0.20235492 | 0.028 | 0.143 | 1 | 11 |
| zgc:162698 | 0.22467 | -0.21275857 | 0.028 | 0.143 | 1 | 11 |
| CABZ01110881.1 | 0.22511 | -0.15112227 | 0.028 | 0.143 | 1 | 11 |
| usp43b | 0.2252 | 0.296231837 | 0.278 | 0.107 | 1 | 11 |
| c1qtnf12 | 0.22608 | -0.11744431 | 0.056 | 0.179 | 1 | 11 |
| clptm1l | 0.2263 | -0.20263047 | 0.028 | 0.143 | 1 | 11 |
| rpl7l1 | 0.22667 | -0.3262875 | 0.167 | 0.321 | 1 | 11 |
| taf15 | 0.22727 | 0.269580336 | 0.333 | 0.214 | 1 | 11 |
| atg101 | 0.22736 | -0.22741269 | 0.028 | 0.143 | 1 | 11 |
| itpa | 0.22753 | -0.15714683 | 0.028 | 0.143 | 1 | 11 |
| pola1 | 0.22762 | -0.21568028 | 0.028 | 0.143 | 1 | 11 |
| ptpn2b | 0.22788 | -0.20069217 | 0.028 | 0.143 | 1 | 11 |
| arfrp1 | 0.2284 | -0.18364605 | 0.028 | 0.143 | 1 | 11 |
| cgna | 0.22846 | 0.292416463 | 0.167 | 0.071 | 1 | 11 |
| ube4b | 0.22847 | -0.12978568 | 0.028 | 0.143 | 1 | 11 |
| zmat2 | 0.22855 | 0.131623382 | 0.306 | 0.321 | 1 | 11 |
| psma8 | 0.2286 | 0.213475766 | 0.611 | 0.571 | 1 | 11 |
| ehd1a | 0.22865 | -0.17826122 | 0.028 | 0.143 | 1 | 11 |
| pmaip1 | 0.22866 | -0.14692307 | 0.028 | 0.143 | 1 | 11 |
| pigp | 0.22891 | -0.15559693 | 0.028 | 0.143 | 1 | 11 |
| map3k4 | 0.22929 | -0.18945426 | 0.028 | 0.143 | 1 | 11 |
| fbp1a | 0.22976 | -0.18588707 | 0.028 | 0.143 | 1 | 11 |
| mak16 | 0.2301 | -0.19108474 | 0.028 | 0.143 | 1 | 11 |
| prpf38a | 0.23016 | -0.1767559 | 0.028 | 0.143 | 1 | 11 |
| tomm40l | 0.23017 | -0.16282814 | 0.028 | 0.143 | 1 | 11 |
| ndufb2 | 0.23052 | 0.292361912 | 0.528 | 0.357 | 1 | 11 |
| nudt8 | 0.23059 | -0.21034983 | 0.028 | 0.143 | 1 | 11 |
| prpsap1 | 0.23062 | -0.17857274 | 0.028 | 0.143 | 1 | 11 |
| hsd11b1la | 0.23066 | -0.18582298 | 0.028 | 0.143 | 1 | 11 |
| prps1a | 0.23067 | 0.156471297 | 0.194 | 0.071 | 1 | 11 |
| si:dkey-42i9.6 | 0.23097 | -0.14763734 | 0.028 | 0.143 | 1 | 11 |
| UBE2O | 0.23106 | -0.19071696 | 0.028 | 0.143 | 1 | 11 |
| ptgesl | 0.23126 | -0.13274327 | 0.028 | 0.143 | 1 | 11 |
| susd6 | 0.2313 | -0.15264487 | 0.028 | 0.143 | 1 | 11 |
| crkl | 0.2315 | -0.21210643 | 0.028 | 0.143 | 1 | 11 |
| pcnp | 0.2316 | -0.16976842 | 0.028 | 0.143 | 1 | 11 |
| zgc:91944 | 0.23182 | -0.18792539 | 0.028 | 0.143 | 1 | 11 |
| eed | 0.23188 | -0.15124243 | 0.028 | 0.143 | 1 | 11 |
| raraa | 0.23193 | -0.16542009 | 0.028 | 0.143 | 1 | 11 |

| | | | | | | |
|---|---|---|---|---|---|---|
| spast | 0.23198 | -0.15508143 | 0.028 | 0.143 | 1 | 11 |
| klf12b | 0.23212 | -0.16748091 | 0.028 | 0.143 | 1 | 11 |
| plekha6 | 0.2322 | 0.273273878 | 0.139 | 0.036 | 1 | 11 |
| MCUR1 | 0.23229 | -0.16007594 | 0.028 | 0.143 | 1 | 11 |
| il13ra1 | 0.23231 | -0.1363895 | 0.028 | 0.143 | 1 | 11 |
| kmt2a | 0.23231 | -0.1880078 | 0.028 | 0.143 | 1 | 11 |
| gadd45bb | 0.23233 | -0.23753446 | 0.083 | 0.214 | 1 | 11 |
| nup93 | 0.23236 | -0.16554151 | 0.028 | 0.143 | 1 | 11 |
| HOOK3 | 0.23238 | -0.16572307 | 0.028 | 0.143 | 1 | 11 |
| nt5dc2 | 0.23247 | -0.21281703 | 0.028 | 0.143 | 1 | 11 |
| bag1 | 0.23251 | -0.15763846 | 0.028 | 0.143 | 1 | 11 |
| tpp1 | 0.23252 | -0.18080652 | 0.028 | 0.143 | 1 | 11 |
| dsg2l | 0.23253 | -0.16154995 | 0.028 | 0.143 | 1 | 11 |
| rcn3 | 0.23253 | -0.16129783 | 0.028 | 0.143 | 1 | 11 |
| ubtf | 0.23263 | 0.287626673 | 0.361 | 0.179 | 1 | 11 |
| metap1 | 0.23331 | -0.11655998 | 0.083 | 0.214 | 1 | 11 |
| snrpd1 | 0.23411 | 0.247535269 | 0.361 | 0.286 | 1 | 11 |
| rnf13 | 0.2343 | 0.210203023 | 0.25 | 0.214 | 1 | 11 |
| lnx2b | 0.23521 | -0.25429069 | 0.056 | 0.179 | 1 | 11 |
| tpm1 | 0.23578 | -0.26808114 | 0.361 | 0.571 | 1 | 11 |
| hoxa9b | 0.23613 | 0.17744712 | 0.194 | 0.071 | 1 | 11 |
| ctbp1 | 0.23632 | 0.193207104 | 0.333 | 0.321 | 1 | 11 |
| ubb | 0.23674 | 0.209688192 | 0.667 | 0.464 | 1 | 11 |
| qki2 | 0.23811 | -0.14668152 | 0.056 | 0.179 | 1 | 11 |
| pdap1a | 0.23833 | 0.158576273 | 0.139 | 0.143 | 1 | 11 |
| rhoaa | 0.24034 | 0.175187134 | 0.528 | 0.5 | 1 | 11 |
| s100a10a | 0.24151 | -0.34097199 | 0.444 | 0.643 | 1 | 11 |
| pnn | 0.24156 | 0.287904196 | 0.278 | 0.179 | 1 | 11 |
| si:zfos-169g10.3 | 0.24192 | -0.32537475 | 0.25 | 0.393 | 1 | 11 |
| phactr4a | 0.24272 | 0.27744035 | 0.361 | 0.25 | 1 | 11 |
| crim1 | 0.24278 | 0.320854063 | 0.306 | 0.143 | 1 | 11 |
| lpar1 | 0.24305 | -0.31233247 | 0.139 | 0.286 | 1 | 11 |
| tead3b | 0.24309 | -0.25316017 | 0.056 | 0.179 | 1 | 11 |
| riox1 | 0.24384 | 0.193800905 | 0.139 | 0.036 | 1 | 11 |
| ucp2 | 0.24412 | -0.24638321 | 0.361 | 0.571 | 1 | 11 |
| chchd3a | 0.24485 | -0.12812709 | 0.083 | 0.214 | 1 | 11 |
| prkar2aa | 0.24565 | -0.11483736 | 0.111 | 0.25 | 1 | 11 |
| marcksl1a | 0.2465 | 0.213027901 | 0.806 | 0.75 | 1 | 11 |
| chd3 | 0.24655 | 0.301931143 | 0.639 | 0.464 | 1 | 11 |
| tmed2 | 0.24738 | 0.120932894 | 0.389 | 0.429 | 1 | 11 |
| uri1 | 0.24947 | 0.27690305 | 0.306 | 0.143 | 1 | 11 |
| flot2a | 0.2495 | 0.287550669 | 0.417 | 0.25 | 1 | 11 |
| slc35f6 | 0.25024 | -0.12657206 | 0.056 | 0.179 | 1 | 11 |
| otub1a | 0.25106 | -0.21944572 | 0.056 | 0.179 | 1 | 11 |
| znhit1 | 0.25125 | 0.117032804 | 0.139 | 0.036 | 1 | 11 |
| sap18 | 0.25126 | 0.286298655 | 0.444 | 0.25 | 1 | 11 |
| rcc2 | 0.25133 | 0.248560487 | 0.194 | 0.071 | 1 | 11 |
| cpne1 | 0.25187 | -0.10260596 | 0.25 | 0.393 | 1 | 11 |

| | | | | | | |
|---|---|---|---|---|---|---|
| tomm20a | 0.25246 | 0.31133676 | 0.472 | 0.286 | 1 | 11 |
| si:ch211-214j24.10 | 0.25339 | 0.259435999 | 0.472 | 0.357 | 1 | 11 |
| si:ch211-14a17.11 | 0.25424 | 0.225193865 | 0.194 | 0.071 | 1 | 11 |
| si:ch211-218c6.8 | 0.25495 | 0.215108719 | 0.194 | 0.143 | 1 | 11 |
| ciao2a | 0.25512 | 0.258418561 | 0.222 | 0.107 | 1 | 11 |
| nccrp1 | 0.25531 | -0.25551122 | 0.278 | 0.357 | 1 | 11 |
| fam204a | 0.25538 | 0.120232592 | 0.167 | 0.179 | 1 | 11 |
| ran | 0.25541 | 0.277575172 | 0.75 | 0.571 | 1 | 11 |
| dnase1l1 | 0.25581 | -0.23830282 | 0.056 | 0.179 | 1 | 11 |
| kri1 | 0.25591 | 0.215524219 | 0.139 | 0.036 | 1 | 11 |
| smg6 | 0.25609 | 0.26937639 | 0.139 | 0.036 | 1 | 11 |
| etfa | 0.25628 | 0.158119939 | 0.333 | 0.321 | 1 | 11 |
| emilin1a | 0.25649 | -0.30955264 | 0.333 | 0.536 | 1 | 11 |
| pcf11 | 0.25693 | -0.14003447 | 0.111 | 0.25 | 1 | 11 |
| arf1 | 0.25701 | 0.27594253 | 0.306 | 0.143 | 1 | 11 |
| srpx | 0.25774 | -0.32188024 | 0.083 | 0.214 | 1 | 11 |
| loxl2a | 0.2581 | 0.239881866 | 0.444 | 0.357 | 1 | 11 |
| si:dkey-17m8.1 | 0.25856 | 0.245041231 | 0.194 | 0.071 | 1 | 11 |
| cldnc | 0.25959 | 0.166728318 | 0.806 | 0.643 | 1 | 11 |
| si:dkey-9i23.16 | 0.2601 | 0.166214676 | 0.417 | 0.393 | 1 | 11 |
| vangl2 | 0.26028 | -0.15626688 | 0.083 | 0.214 | 1 | 11 |
| ilf3b | 0.26032 | 0.194726382 | 0.222 | 0.179 | 1 | 11 |
| si:ch73-281n10.2 | 0.26046 | 0.179561741 | 0.611 | 0.643 | 1 | 11 |
| ddx27 | 0.26057 | 0.262890607 | 0.25 | 0.143 | 1 | 11 |
| myl12.2 | 0.26103 | 0.151121771 | 0.25 | 0.25 | 1 | 11 |
| cux1a | 0.26123 | 0.301412173 | 0.333 | 0.214 | 1 | 11 |
| hdac4 | 0.26143 | -0.15936359 | 0.083 | 0.214 | 1 | 11 |
| gcga | 0.26145 | 0.32333573 | 0.528 | 0.357 | 1 | 11 |
| lamp2 | 0.26182 | -0.29370477 | 0.306 | 0.5 | 1 | 11 |
| piezo2a.2 | 0.26261 | 0.286228905 | 0.25 | 0.107 | 1 | 11 |
| plscr3b | 0.26298 | 0.261859352 | 0.389 | 0.286 | 1 | 11 |
| CU459094.2 | 0.26301 | -0.20011667 | 0.167 | 0.321 | 1 | 11 |
| hlx1 | 0.26359 | -0.26677958 | 0.111 | 0.179 | 1 | 11 |
| vim | 0.26448 | -0.29940974 | 0.333 | 0.536 | 1 | 11 |
| arl5a | 0.26495 | 0.217883789 | 0.139 | 0.036 | 1 | 11 |
| cnp | 0.26534 | 0.118285253 | 0.361 | 0.214 | 1 | 11 |
| slco2b1 | 0.26548 | 0.14263293 | 0.472 | 0.321 | 1 | 11 |
| ube2d1a | 0.26588 | 0.210791999 | 0.139 | 0.036 | 1 | 11 |
| hoxc9a | 0.26589 | 0.300502315 | 0.25 | 0.107 | 1 | 11 |
| cttn | 0.2677 | -0.13835145 | 0.083 | 0.214 | 1 | 11 |
| eaf1 | 0.26777 | -0.20739696 | 0.056 | 0.179 | 1 | 11 |
| sgce | 0.26786 | -0.16643675 | 0.111 | 0.25 | 1 | 11 |
| cyr61 | 0.26806 | 0.262162597 | 0.25 | 0.107 | 1 | 11 |
| dnmbp | 0.26844 | -0.17197804 | 0.083 | 0.214 | 1 | 11 |
| chac1 | 0.26892 | -0.27207355 | 0.306 | 0.5 | 1 | 11 |
| tomm6 | 0.26906 | 0.157028754 | 0.333 | 0.321 | 1 | 11 |
| wdr75 | 0.26918 | -0.22705294 | 0.056 | 0.179 | 1 | 11 |
| hoxa3a | 0.26919 | 0.225434609 | 0.139 | 0.036 | 1 | 11 |

| | | | | | | |
|---|---|---|---|---|---|---|
| tmem11 | 0.26938 | 0.25883104 | 0.194 | 0.107 | 1 | 11 |
| FP102018.1 | 0.27037 | 0.221637411 | 0.167 | 0.107 | 1 | 11 |
| znf330 | 0.27061 | -0.26714656 | 0.083 | 0.214 | 1 | 11 |
| baz2ba | 0.27063 | 0.2343597 | 0.222 | 0.107 | 1 | 11 |
| hdac1 | 0.27196 | 0.285473349 | 0.472 | 0.321 | 1 | 11 |
| rpz5 | 0.27199 | -0.28256084 | 0.444 | 0.607 | 1 | 11 |
| tspan4a | 0.2729 | -0.25501412 | 0.306 | 0.393 | 1 | 11 |
| atp1a1a.1 | 0.27327 | 0.211734327 | 0.139 | 0.036 | 1 | 11 |
| cyp2p10 | 0.27339 | 0.210180166 | 0.139 | 0.036 | 1 | 11 |
| zgc:113208 | 0.2735 | 0.227148374 | 0.139 | 0.036 | 1 | 11 |
| prdx4 | 0.27417 | 0.220646898 | 0.306 | 0.214 | 1 | 11 |
| mid1ip1a | 0.27534 | -0.27713533 | 0.056 | 0.179 | 1 | 11 |
| hsd11b2 | 0.27589 | 0.229878058 | 0.889 | 0.75 | 1 | 11 |
| srsf5a | 0.27595 | 0.152196615 | 0.583 | 0.571 | 1 | 11 |
| col2a1b | 0.27602 | -0.16607098 | 0.056 | 0.179 | 1 | 11 |
| per3 | 0.27655 | 0.210263811 | 0.306 | 0.143 | 1 | 11 |
| usp10 | 0.27659 | 0.286242184 | 0.25 | 0.107 | 1 | 11 |
| prpf6 | 0.2772 | 0.227547322 | 0.194 | 0.107 | 1 | 11 |
| eif4g2a | 0.27781 | -0.16959774 | 0.056 | 0.179 | 1 | 11 |
| mob3a | 0.27827 | 0.271843981 | 0.25 | 0.107 | 1 | 11 |
| snrpd2 | 0.27904 | 0.232496846 | 0.306 | 0.143 | 1 | 11 |
| zmynd8 | 0.27946 | 0.240337307 | 0.139 | 0.036 | 1 | 11 |
| arf6a | 0.28024 | 0.20908124 | 0.139 | 0.036 | 1 | 11 |
| psmd11a | 0.28037 | 0.224113086 | 0.194 | 0.071 | 1 | 11 |
| aldh2.2 | 0.28049 | -0.13327883 | 0.472 | 0.464 | 1 | 11 |
| man1b1b | 0.28053 | 0.243376765 | 0.167 | 0.107 | 1 | 11 |
| fkbp4 | 0.28073 | 0.234814639 | 0.25 | 0.107 | 1 | 11 |
| si:dkey-222f8.3 | 0.28079 | 0.201923741 | 0.139 | 0.036 | 1 | 11 |
| cdc73 | 0.28109 | -0.22066764 | 0.056 | 0.179 | 1 | 11 |
| sfrp1a | 0.28132 | 0.253517949 | 0.778 | 0.607 | 1 | 11 |
| slc29a1b | 0.28167 | -0.21213471 | 0.583 | 0.607 | 1 | 11 |
| dnajc5aa | 0.28204 | 0.216930276 | 0.139 | 0.036 | 1 | 11 |
| zgc:77929 | 0.28252 | -0.16932208 | 0.056 | 0.179 | 1 | 11 |
| nudt21 | 0.28281 | 0.123921743 | 0.194 | 0.214 | 1 | 11 |
| aqp8a.1 | 0.28285 | -0.37450505 | 0.056 | 0.179 | 1 | 11 |
| spaca4l | 0.28295 | -0.25019693 | 0.056 | 0.179 | 1 | 11 |
| zhx3 | 0.28345 | -0.16065433 | 0.056 | 0.179 | 1 | 11 |
| ei24 | 0.28379 | -0.19975631 | 0.056 | 0.179 | 1 | 11 |
| zgc:91910 | 0.28385 | -0.1181028 | 0.306 | 0.464 | 1 | 11 |
| sf3b3 | 0.28411 | -0.16010339 | 0.056 | 0.179 | 1 | 11 |
| mcm4 | 0.28425 | 0.205993967 | 0.111 | 0.036 | 1 | 11 |
| adam19a | 0.28436 | -0.19509339 | 0.056 | 0.179 | 1 | 11 |
| tardbpl | 0.28442 | 0.20889943 | 0.278 | 0.214 | 1 | 11 |
| cops8 | 0.28462 | -0.21844913 | 0.056 | 0.179 | 1 | 11 |
| ms4a17a.11 | 0.2852 | 0.249588693 | 0.25 | 0.107 | 1 | 11 |
| mrpl11 | 0.2853 | 0.215346621 | 0.167 | 0.071 | 1 | 11 |
| pcolcea | 0.28538 | -0.33882507 | 0.056 | 0.179 | 1 | 11 |
| mych | 0.28554 | -0.26155024 | 0.333 | 0.464 | 1 | 11 |

| | | | | | | |
|---|---|---|---|---|---|---|
| bccip | 0.28556 | -0.25935427 | 0.278 | 0.464 | 1 | 11 |
| spns2 | 0.2862 | -0.2044361 | 0.056 | 0.179 | 1 | 11 |
| mfap3l | 0.28691 | 0.213475527 | 0.139 | 0.036 | 1 | 11 |
| pxnb | 0.28705 | -0.20528407 | 0.056 | 0.179 | 1 | 11 |
| derl2 | 0.28782 | -0.23213287 | 0.083 | 0.107 | 1 | 11 |
| cpsf6 | 0.28793 | -0.18486859 | 0.139 | 0.286 | 1 | 11 |
| gabarapa | 0.28813 | -0.30395619 | 0.278 | 0.429 | 1 | 11 |
| LOC101886030 | 0.28861 | 0.276370103 | 0.25 | 0.107 | 1 | 11 |
| ppm1la | 0.2901 | -0.18071321 | 0.056 | 0.179 | 1 | 11 |
| smco4 | 0.29124 | -0.15237328 | 0.056 | 0.179 | 1 | 11 |
| cops2 | 0.29252 | -0.16911262 | 0.111 | 0.25 | 1 | 11 |
| znf131 | 0.29252 | 0.216370562 | 0.139 | 0.036 | 1 | 11 |
| snx27a | 0.29287 | 0.22047507 | 0.194 | 0.071 | 1 | 11 |
| BX511129.1 | 0.29295 | -0.20177392 | 0.056 | 0.179 | 1 | 11 |
| hspa12a | 0.29303 | -0.19548864 | 0.056 | 0.179 | 1 | 11 |
| socs3a | 0.29317 | -0.17382756 | 0.694 | 0.857 | 1 | 11 |
| fabp11a | 0.29477 | -0.30591563 | 0.083 | 0.214 | 1 | 11 |
| col6a1 | 0.29538 | -0.13661477 | 0.528 | 0.536 | 1 | 11 |
| heatr3 | 0.29573 | -0.2077017 | 0.056 | 0.179 | 1 | 11 |
| fam129bb | 0.2959 | -0.20295021 | 0.056 | 0.179 | 1 | 11 |
| tcf4 | 0.29594 | -0.18486854 | 0.056 | 0.179 | 1 | 11 |
| ccdc28a | 0.29596 | -0.14331852 | 0.056 | 0.179 | 1 | 11 |
| rnf220a | 0.29599 | -0.27408512 | 0.056 | 0.179 | 1 | 11 |
| hnrnpa1b | 0.29606 | -0.26093504 | 0.278 | 0.464 | 1 | 11 |
| sec11a | 0.29618 | 0.279067802 | 0.389 | 0.214 | 1 | 11 |
| zswim5 | 0.2962 | 0.219710531 | 0.111 | 0.036 | 1 | 11 |
| si:ch73-376l24.6 | 0.29663 | -0.14157533 | 0.111 | 0.25 | 1 | 11 |
| cavin2a | 0.29703 | 0.254033097 | 0.167 | 0.071 | 1 | 11 |
| itga10 | 0.29756 | -0.24912798 | 0.056 | 0.143 | 1 | 11 |
| si:ch211-140m22.7 | 0.29767 | 0.236221324 | 0.306 | 0.143 | 1 | 11 |
| s100v2 | 0.29768 | -0.17165296 | 0.611 | 0.786 | 1 | 11 |
| pleca | 0.29774 | 0.279992474 | 0.639 | 0.536 | 1 | 11 |
| ciarta | 0.29781 | -0.28014811 | 0.278 | 0.464 | 1 | 11 |
| surf2 | 0.29832 | 0.177439781 | 0.139 | 0.036 | 1 | 11 |
| rnf213a | 0.29859 | -0.18086082 | 0.139 | 0.286 | 1 | 11 |
| smarcc2 | 0.29867 | 0.155170897 | 0.139 | 0.036 | 1 | 11 |
| gnb1b | 0.29974 | 0.282589524 | 0.444 | 0.321 | 1 | 11 |
| macf1a | 0.29974 | 0.264478345 | 0.306 | 0.143 | 1 | 11 |
| osgep | 0.30019 | 0.180086598 | 0.139 | 0.036 | 1 | 11 |
| fkbp7 | 0.30029 | -0.23676485 | 0.278 | 0.464 | 1 | 11 |
| nedd8l | 0.30029 | 0.202942013 | 0.167 | 0.071 | 1 | 11 |
| proza | 0.30044 | -0.19901205 | 0.083 | 0.214 | 1 | 11 |
| eif3m | 0.30079 | -0.22422108 | 0.611 | 0.714 | 1 | 11 |
| tmem185 | 0.30088 | 0.206337914 | 0.139 | 0.036 | 1 | 11 |
| btf3l4 | 0.30104 | -0.24848976 | 0.194 | 0.357 | 1 | 11 |
| gbp1 | 0.30116 | -0.17881574 | 0.083 | 0.214 | 1 | 11 |
| rbm19 | 0.30125 | 0.251484914 | 0.222 | 0.107 | 1 | 11 |
| tab1 | 0.30171 | 0.218339735 | 0.194 | 0.071 | 1 | 11 |

| | | | | | | |
|---|---|---|---|---|---|---|
| sod2 | 0.30181 | 0.104356072 | 0.139 | 0.036 | 1 | 11 |
| kras | 0.30274 | 0.19669153 | 0.139 | 0.036 | 1 | 11 |
| ampd2b | 0.30282 | 0.220166249 | 0.139 | 0.036 | 1 | 11 |
| hnrpkl | 0.30311 | -0.16328167 | 0.167 | 0.321 | 1 | 11 |
| rpsa | 0.30319 | -0.11331198 | 1 | 1 | 1 | 11 |
| gars | 0.30347 | -0.23947196 | 0.139 | 0.286 | 1 | 11 |
| sept6 | 0.30348 | -0.27619737 | 0.139 | 0.286 | 1 | 11 |
| psmc1a | 0.30354 | -0.10085925 | 0.222 | 0.357 | 1 | 11 |
| clk2a | 0.30406 | 0.211204696 | 0.194 | 0.071 | 1 | 11 |
| gng2 | 0.3045 | 0.194609757 | 0.139 | 0.036 | 1 | 11 |
| jam2b | 0.30567 | -0.11899249 | 0.556 | 0.714 | 1 | 11 |
| ppfibp1a | 0.30585 | 0.273304466 | 0.389 | 0.214 | 1 | 11 |
| pcsk9 | 0.30611 | -0.29900909 | 0.111 | 0.25 | 1 | 11 |
| nars | 0.30726 | 0.274431814 | 0.222 | 0.107 | 1 | 11 |
| map3k1 | 0.30731 | -0.22448078 | 0.083 | 0.143 | 1 | 11 |
| fgf24 | 0.30747 | -0.23826755 | 0.222 | 0.393 | 1 | 11 |
| sirt5 | 0.30793 | 0.224929034 | 0.194 | 0.071 | 1 | 11 |
| nfkbiab | 0.308 | 0.249769885 | 0.472 | 0.286 | 1 | 11 |
| gtf2h4 | 0.30822 | 0.212819586 | 0.111 | 0.036 | 1 | 11 |
| nanos1 | 0.3085 | 0.312019766 | 0.472 | 0.286 | 1 | 11 |
| vps28 | 0.30995 | -0.16833987 | 0.139 | 0.286 | 1 | 11 |
| cald1a | 0.31016 | 0.241370021 | 0.75 | 0.607 | 1 | 11 |
| serbp1a | 0.31042 | 0.139760849 | 0.861 | 0.714 | 1 | 11 |
| smad3a | 0.31045 | 0.259681053 | 0.194 | 0.071 | 1 | 11 |
| lancl1 | 0.31072 | 0.198860336 | 0.139 | 0.036 | 1 | 11 |
| cnn3b | 0.31098 | 0.251166813 | 0.167 | 0.071 | 1 | 11 |
| tnfrsf1a | 0.31113 | -0.19531341 | 0.167 | 0.321 | 1 | 11 |
| id3 | 0.31209 | 0.261651025 | 0.194 | 0.107 | 1 | 11 |
| si:ch211-106k21.5 | 0.31266 | 0.127908421 | 0.111 | 0.107 | 1 | 11 |
| sra1 | 0.31281 | -0.25988054 | 0.028 | 0.107 | 1 | 11 |
| cyp1c1 | 0.31321 | -0.19872645 | 0.111 | 0.25 | 1 | 11 |
| strn | 0.31321 | -0.19954112 | 0.056 | 0.107 | 1 | 11 |
| hmbox1a | 0.31366 | -0.27230849 | 0.111 | 0.25 | 1 | 11 |
| dbn1 | 0.3138 | -0.25975327 | 0.361 | 0.464 | 1 | 11 |
| nupr1a | 0.31418 | -0.17890435 | 0.278 | 0.286 | 1 | 11 |
| ifi45 | 0.31449 | 0.361390672 | 0.306 | 0.179 | 1 | 11 |
| srp68 | 0.31466 | 0.218680105 | 0.194 | 0.071 | 1 | 11 |
| top1l | 0.31481 | 0.261817592 | 0.278 | 0.143 | 1 | 11 |
| dram2b | 0.31535 | 0.207762751 | 0.139 | 0.036 | 1 | 11 |
| celf2 | 0.31601 | 0.22774465 | 0.694 | 0.607 | 1 | 11 |
| ewsr1a | 0.31611 | 0.141432951 | 0.139 | 0.036 | 1 | 11 |
| rexo4 | 0.31666 | 0.127162102 | 0.139 | 0.036 | 1 | 11 |
| pfdn4 | 0.31708 | 0.245164369 | 0.444 | 0.321 | 1 | 11 |
| wipf2a | 0.31709 | -0.21272096 | 0.083 | 0.214 | 1 | 11 |
| LOC100536647 | 0.31768 | 0.110156671 | 0.167 | 0.179 | 1 | 11 |
| col16a1 | 0.31793 | -0.21882784 | 0.222 | 0.286 | 1 | 11 |
| rrp8 | 0.3181 | -0.19869206 | 0.083 | 0.214 | 1 | 11 |
| yipf5 | 0.31813 | -0.20452328 | 0.083 | 0.214 | 1 | 11 |

| | | | | | | |
|---|---|---|---|---|---|---|
| ivns1abpb | 0.31846 | 0.164417426 | 0.111 | 0.036 | 1 | 11 |
| pcbd1 | 0.31869 | -0.11508242 | 0.111 | 0.25 | 1 | 11 |
| caprin1a | 0.31871 | 0.253478063 | 0.194 | 0.071 | 1 | 11 |
| si:ch211-63p21.1 | 0.31914 | 0.283840274 | 0.389 | 0.214 | 1 | 11 |
| krtcap2 | 0.3195 | -0.10420818 | 0.417 | 0.393 | 1 | 11 |
| ythdf2 | 0.31973 | 0.211934325 | 0.139 | 0.071 | 1 | 11 |
| bsdc1 | 0.31983 | -0.15559048 | 0.111 | 0.25 | 1 | 11 |
| smek1 | 0.32017 | 0.224557482 | 0.194 | 0.071 | 1 | 11 |
| psme2 | 0.32038 | -0.21378581 | 0.25 | 0.429 | 1 | 11 |
| mir21-1 | 0.32043 | 0.254451936 | 0.222 | 0.107 | 1 | 11 |
| ccdc6a | 0.32048 | -0.20853354 | 0.111 | 0.25 | 1 | 11 |
| ylpm1 | 0.32084 | 0.176060899 | 0.139 | 0.036 | 1 | 11 |
| ufd1l | 0.32086 | -0.21502437 | 0.111 | 0.25 | 1 | 11 |
| fam50a | 0.32108 | -0.11650681 | 0.083 | 0.214 | 1 | 11 |
| gpr78a | 0.32185 | -0.24398597 | 0.083 | 0.214 | 1 | 11 |
| gcdha | 0.32205 | -0.15301695 | 0.083 | 0.214 | 1 | 11 |
| prps1b | 0.32236 | 0.232336026 | 0.167 | 0.071 | 1 | 11 |
| tsn | 0.32272 | 0.202344238 | 0.111 | 0.036 | 1 | 11 |
| mkrn1 | 0.32329 | -0.24650871 | 0.167 | 0.286 | 1 | 11 |
| slc38a5b | 0.32374 | -0.24781389 | 0.5 | 0.679 | 1 | 11 |
| cnbpb | 0.32375 | -0.25890291 | 0.528 | 0.643 | 1 | 11 |
| cs | 0.32389 | -0.24256206 | 0.111 | 0.25 | 1 | 11 |
| appb | 0.3242 | 0.267037301 | 0.278 | 0.143 | 1 | 11 |
| mrps33 | 0.32421 | 0.135467206 | 0.139 | 0.036 | 1 | 11 |
| kdm4b | 0.32425 | 0.22774525 | 0.25 | 0.107 | 1 | 11 |
| ppid | 0.32428 | 0.160084496 | 0.139 | 0.036 | 1 | 11 |
| kpnb3 | 0.32515 | -0.22309895 | 0.083 | 0.214 | 1 | 11 |
| zgc:101858 | 0.32534 | -0.16621399 | 0.083 | 0.214 | 1 | 11 |
| znf503 | 0.32548 | 0.130153638 | 0.139 | 0.143 | 1 | 11 |
| btf3 | 0.32563 | 0.167906356 | 0.833 | 0.786 | 1 | 11 |
| taf10 | 0.32625 | -0.15935955 | 0.083 | 0.214 | 1 | 11 |
| gulp1a | 0.32633 | -0.12054359 | 0.194 | 0.321 | 1 | 11 |
| itgb1a | 0.32706 | -0.2750965 | 0.111 | 0.25 | 1 | 11 |
| nutf2 | 0.32725 | -0.16146721 | 0.139 | 0.286 | 1 | 11 |
| fubp1 | 0.32736 | 0.166025994 | 0.194 | 0.179 | 1 | 11 |
| bcap31 | 0.32777 | 0.226568455 | 0.25 | 0.143 | 1 | 11 |
| cops9 | 0.32789 | 0.100764031 | 0.361 | 0.393 | 1 | 11 |
| zgc:66388 | 0.32806 | -0.22312663 | 0.083 | 0.214 | 1 | 11 |
| s1pr5a | 0.3284 | -0.20050029 | 0.111 | 0.25 | 1 | 11 |
| neo1a | 0.32868 | -0.18794008 | 0.083 | 0.214 | 1 | 11 |
| znf706 | 0.32948 | -0.28033655 | 0.278 | 0.429 | 1 | 11 |
| zgc:56095 | 0.32958 | 0.154510633 | 0.25 | 0.107 | 1 | 11 |
| slc25a51b | 0.33014 | -0.19123815 | 0.083 | 0.214 | 1 | 11 |
| atf5b | 0.33015 | -0.18957751 | 0.083 | 0.214 | 1 | 11 |
| cdk2ap2 | 0.33015 | -0.20810488 | 0.083 | 0.214 | 1 | 11 |
| CABZ01080702.1 | 0.3303 | -0.27394584 | 0.472 | 0.643 | 1 | 11 |
| ncam1a | 0.33038 | -0.10899349 | 0.111 | 0.214 | 1 | 11 |
| si:dkey-177p2.6 | 0.33057 | -0.29745771 | 0.111 | 0.214 | 1 | 11 |

| | | | | | | |
|---|---|---|---|---|---|---|
| oip5-as1 | 0.33064 | 0.235053902 | 0.194 | 0.071 | 1 | 11 |
| ubr7 | 0.3308 | -0.20198519 | 0.111 | 0.25 | 1 | 11 |
| cox7c | 0.33108 | 0.155766879 | 0.778 | 0.75 | 1 | 11 |
| tm9sf4 | 0.33138 | -0.20551295 | 0.111 | 0.25 | 1 | 11 |
| zgc:56676 | 0.33149 | -0.2602588 | 0.139 | 0.286 | 1 | 11 |
| si:ch211-191j22.3 | 0.33176 | -0.21087172 | 0.194 | 0.357 | 1 | 11 |
| ric8b | 0.33183 | -0.10497395 | 0.083 | 0.071 | 1 | 11 |
| arcn1a | 0.33195 | 0.161119825 | 0.139 | 0.036 | 1 | 11 |
| sv2 | 0.33211 | 0.155568069 | 0.139 | 0.036 | 1 | 11 |
| zc4h2 | 0.33218 | 0.182145888 | 0.139 | 0.036 | 1 | 11 |
| sugt1 | 0.33239 | 0.165032522 | 0.139 | 0.036 | 1 | 11 |
| hmgb3a | 0.33269 | 0.308183913 | 0.444 | 0.286 | 1 | 11 |
| eif4g1a | 0.33381 | -0.22252089 | 0.222 | 0.393 | 1 | 11 |
| gnl3 | 0.33409 | 0.189702894 | 0.194 | 0.143 | 1 | 11 |
| ints11 | 0.33424 | 0.167953498 | 0.139 | 0.036 | 1 | 11 |
| kdelr2a | 0.33434 | 0.134594831 | 0.194 | 0.179 | 1 | 11 |
| rasgef1bb | 0.33444 | 0.172147439 | 0.139 | 0.036 | 1 | 11 |
| cyfip1 | 0.33457 | 0.20952574 | 0.111 | 0.036 | 1 | 11 |
| rps6ka5 | 0.33464 | -0.27693994 | 0.056 | 0.143 | 1 | 11 |
| trim35-28 | 0.33494 | 0.193700086 | 0.139 | 0.036 | 1 | 11 |
| kitlga | 0.33545 | -0.26847221 | 0.083 | 0.179 | 1 | 11 |
| kank1a | 0.33554 | -0.10340748 | 0.139 | 0.25 | 1 | 11 |
| rap1b | 0.33608 | -0.27534501 | 0.417 | 0.571 | 1 | 11 |
| rab5aa | 0.33619 | 0.258419419 | 0.333 | 0.214 | 1 | 11 |
| aplp2 | 0.33692 | -0.17966482 | 0.111 | 0.25 | 1 | 11 |
| cmip | 0.33808 | -0.10364193 | 0.083 | 0.071 | 1 | 11 |
| ssbp4 | 0.33814 | 0.226597542 | 0.194 | 0.071 | 1 | 11 |
| phldb2a | 0.33867 | 0.199587146 | 0.139 | 0.036 | 1 | 11 |
| pgam5 | 0.33883 | 0.19673695 | 0.194 | 0.107 | 1 | 11 |
| txnl4a | 0.33933 | 0.146523785 | 0.111 | 0.107 | 1 | 11 |
| bet1l | 0.3396 | 0.189006727 | 0.139 | 0.036 | 1 | 11 |
| brd7 | 0.3396 | -0.24558239 | 0.167 | 0.321 | 1 | 11 |
| raver2 | 0.33983 | 0.184739084 | 0.139 | 0.036 | 1 | 11 |
| drg1 | 0.33995 | 0.141347511 | 0.139 | 0.036 | 1 | 11 |
| selenol | 0.3405 | 0.135844089 | 0.139 | 0.036 | 1 | 11 |
| fbxo34 | 0.34061 | 0.176684772 | 0.139 | 0.036 | 1 | 11 |
| myo5ab | 0.34076 | 0.146173942 | 0.139 | 0.036 | 1 | 11 |
| cdk4 | 0.34093 | -0.19409426 | 0.111 | 0.25 | 1 | 11 |
| ror2 | 0.3413 | 0.179604512 | 0.139 | 0.036 | 1 | 11 |
| kmt2cb | 0.3416 | 0.244272253 | 0.222 | 0.107 | 1 | 11 |
| sik1 | 0.34169 | 0.210719543 | 0.194 | 0.071 | 1 | 11 |
| pigk | 0.3417 | -0.1001092 | 0.028 | 0.107 | 1 | 11 |
| syncripl | 0.34181 | 0.183201716 | 0.194 | 0.071 | 1 | 11 |
| nutf2l | 0.3424 | 0.149166167 | 0.139 | 0.036 | 1 | 11 |
| uqcc3 | 0.34274 | 0.205445201 | 0.194 | 0.071 | 1 | 11 |
| hbs1l | 0.34297 | 0.15721194 | 0.139 | 0.036 | 1 | 11 |
| sufu | 0.343 | 0.166568359 | 0.139 | 0.036 | 1 | 11 |
| ak2 | 0.34355 | -0.21713791 | 0.444 | 0.536 | 1 | 11 |

| | | | | | | |
|---|---|---|---|---|---|---|
| hopx | 0.34371 | 0.224607284 | 0.194 | 0.071 | 1 | 11 |
| si:dkey-118j18.2 | 0.34372 | -0.22390284 | 0.083 | 0.179 | 1 | 11 |
| sh3pxd2aa | 0.34416 | 0.16041998 | 0.167 | 0.143 | 1 | 11 |
| chmp2a | 0.34432 | -0.18526976 | 0.111 | 0.25 | 1 | 11 |
| CABZ01084963.1 | 0.34444 | 0.216649581 | 0.167 | 0.071 | 1 | 11 |
| degs1 | 0.34475 | -0.21444918 | 0.167 | 0.321 | 1 | 11 |
| zfand5a | 0.34485 | -0.18239174 | 0.139 | 0.286 | 1 | 11 |
| ivd | 0.34494 | -0.18507705 | 0.167 | 0.321 | 1 | 11 |
| fundc2 | 0.34547 | -0.1988393 | 0.111 | 0.25 | 1 | 11 |
| fbl | 0.34551 | -0.23032792 | 0.139 | 0.286 | 1 | 11 |
| gpc1b | 0.34596 | -0.24624541 | 0.139 | 0.25 | 1 | 11 |
| srrm1 | 0.34616 | -0.20410605 | 0.139 | 0.286 | 1 | 11 |
| cst3 | 0.3464 | 0.195781709 | 0.222 | 0.179 | 1 | 11 |
| pfkfb4b | 0.34689 | 0.192430728 | 0.528 | 0.357 | 1 | 11 |
| jun | 0.34721 | 0.163508744 | 0.889 | 0.75 | 1 | 11 |
| ankrd11 | 0.3473 | 0.197740851 | 0.333 | 0.179 | 1 | 11 |
| lurap1 | 0.34733 | -0.10580575 | 0.333 | 0.464 | 1 | 11 |
| chrac1 | 0.34781 | 0.221148928 | 0.194 | 0.071 | 1 | 11 |
| id2a | 0.35005 | -0.30205111 | 0.278 | 0.429 | 1 | 11 |
| zfand5b | 0.35064 | 0.180269399 | 0.194 | 0.071 | 1 | 11 |
| bsg | 0.3508 | 0.202245539 | 0.694 | 0.607 | 1 | 11 |
| nup37 | 0.35096 | 0.163637801 | 0.194 | 0.071 | 1 | 11 |
| gnal | 0.35129 | 0.195036181 | 0.194 | 0.071 | 1 | 11 |
| adipor1b | 0.35154 | 0.188138266 | 0.111 | 0.036 | 1 | 11 |
| flii | 0.35157 | -0.20822477 | 0.139 | 0.286 | 1 | 11 |
| gchfr | 0.35182 | -0.24470622 | 0.139 | 0.286 | 1 | 11 |
| itgb2 | 0.35208 | -0.16890524 | 0.028 | 0.107 | 1 | 11 |
| hmcn1 | 0.3521 | -0.1122987 | 0.028 | 0.107 | 1 | 11 |
| mrpl12 | 0.35218 | -0.18182405 | 0.139 | 0.286 | 1 | 11 |
| dtwd1 | 0.35246 | 0.20457549 | 0.111 | 0.036 | 1 | 11 |
| msh6 | 0.35246 | -0.24062742 | 0.028 | 0.107 | 1 | 11 |
| rpa2 | 0.35268 | 0.190152973 | 0.194 | 0.071 | 1 | 11 |
| fstl3 | 0.35281 | -0.22885968 | 0.056 | 0.143 | 1 | 11 |
| mgst3a | 0.35354 | -0.21231046 | 0.139 | 0.286 | 1 | 11 |
| clpp | 0.35367 | -0.2443247 | 0.111 | 0.214 | 1 | 11 |
| atf7ip | 0.35418 | 0.244694739 | 0.222 | 0.107 | 1 | 11 |
| rps26 | 0.35564 | 0.131462191 | 0.944 | 0.893 | 1 | 11 |
| usp39 | 0.3573 | -0.18733762 | 0.028 | 0.107 | 1 | 11 |
| nrg2a | 0.35735 | -0.10392125 | 0.028 | 0.107 | 1 | 11 |
| si:ch1073-170o4.1 | 0.35804 | 0.214160219 | 0.111 | 0.036 | 1 | 11 |
| BX322530.1 | 0.35932 | 0.15073162 | 0.306 | 0.286 | 1 | 11 |
| mpzl1l | 0.35969 | -0.10356625 | 0.056 | 0.143 | 1 | 11 |
| yaf2 | 0.35999 | -0.20928093 | 0.111 | 0.179 | 1 | 11 |
| cct3 | 0.3602 | 0.226631454 | 0.528 | 0.429 | 1 | 11 |
| zgc:174895 | 0.3604 | -0.21207288 | 0.028 | 0.107 | 1 | 11 |
| BX465862.1 | 0.36062 | 0.208975016 | 0.111 | 0.036 | 1 | 11 |
| trit1 | 0.36065 | 0.200442781 | 0.111 | 0.036 | 1 | 11 |
| seta | 0.36069 | 0.24387792 | 0.556 | 0.429 | 1 | 11 |

| | | | | | | |
|---|---|---|---|---|---|---|
| cnpy1 | 0.36109 | 0.17976235 | 0.222 | 0.143 | 1 | 11 |
| mrpl20 | 0.36131 | 0.230841986 | 0.194 | 0.107 | 1 | 11 |
| net1 | 0.36223 | 0.272577082 | 0.278 | 0.143 | 1 | 11 |
| nme3 | 0.3623 | 0.175645178 | 0.306 | 0.25 | 1 | 11 |
| adamts10 | 0.36342 | -0.15259288 | 0.028 | 0.107 | 1 | 11 |
| klf2a | 0.36347 | -0.23137367 | 0.583 | 0.75 | 1 | 11 |
| ctgfa | 0.36355 | 0.124172915 | 0.25 | 0.143 | 1 | 11 |
| si:dkey-33c12.4 | 0.36368 | 0.16541669 | 0.306 | 0.286 | 1 | 11 |
| lcorl | 0.36586 | -0.13980618 | 0.139 | 0.143 | 1 | 11 |
| irf2bp2a | 0.36634 | 0.210570163 | 0.444 | 0.357 | 1 | 11 |
| zc3h18 | 0.36723 | -0.10289763 | 0.028 | 0.107 | 1 | 11 |
| sms | 0.36724 | -0.16909159 | 0.028 | 0.107 | 1 | 11 |
| psmc4 | 0.36781 | 0.245432766 | 0.306 | 0.179 | 1 | 11 |
| ece1 | 0.36815 | 0.242987343 | 0.167 | 0.071 | 1 | 11 |
| epb41l5 | 0.36847 | 0.208724671 | 0.417 | 0.357 | 1 | 11 |
| RTKN | 0.36854 | -0.11270916 | 0.028 | 0.107 | 1 | 11 |
| selenof | 0.36876 | 0.138448769 | 0.694 | 0.679 | 1 | 11 |
| nucks1a | 0.36999 | 0.22983079 | 0.333 | 0.179 | 1 | 11 |
| psmd1 | 0.37023 | -0.12095871 | 0.222 | 0.357 | 1 | 11 |
| pkma | 0.37078 | 0.201094496 | 0.278 | 0.179 | 1 | 11 |
| prim2 | 0.37157 | -0.21008346 | 0.028 | 0.107 | 1 | 11 |
| foxp4 | 0.37186 | 0.282328162 | 0.5 | 0.357 | 1 | 11 |
| b9d1 | 0.37244 | -0.15146882 | 0.028 | 0.107 | 1 | 11 |
| adh8a | 0.37253 | 0.111544863 | 0.5 | 0.357 | 1 | 11 |
| dhrs4 | 0.37282 | -0.21723929 | 0.111 | 0.214 | 1 | 11 |
| elavl4 | 0.37336 | -0.22189702 | 0.028 | 0.107 | 1 | 11 |
| si:ch211-282b22.1 | 0.37351 | 0.204372473 | 0.222 | 0.107 | 1 | 11 |
| wdr93 | 0.37359 | -0.19083662 | 0.028 | 0.107 | 1 | 11 |
| si:ch211-1a19.3 | 0.37392 | -0.28965688 | 0.306 | 0.429 | 1 | 11 |
| zgc:153293 | 0.37402 | 0.177404667 | 0.111 | 0.071 | 1 | 11 |
| cd3eap | 0.37477 | 0.228598084 | 0.278 | 0.143 | 1 | 11 |
| vkorc1 | 0.37487 | 0.124826249 | 0.111 | 0.107 | 1 | 11 |
| si:ch1073-406l10.2 | 0.37535 | 0.191477596 | 0.111 | 0.036 | 1 | 11 |
| pcdh10b | 0.37578 | 0.277221466 | 0.389 | 0.25 | 1 | 11 |
| tpm4a | 0.37614 | 0.194185076 | 0.667 | 0.607 | 1 | 11 |
| lgals3b | 0.37671 | -0.24470568 | 0.472 | 0.643 | 1 | 11 |
| hspa5 | 0.37676 | 0.28508256 | 0.278 | 0.143 | 1 | 11 |
| utp25 | 0.37692 | 0.201715849 | 0.111 | 0.036 | 1 | 11 |
| crip3 | 0.37702 | -0.1382422 | 0.028 | 0.107 | 1 | 11 |
| pacsin2 | 0.37769 | -0.1505727 | 0.028 | 0.107 | 1 | 11 |
| CU539058.1 | 0.37839 | 0.241416577 | 0.194 | 0.107 | 1 | 11 |
| sept15 | 0.37952 | 0.220532682 | 0.111 | 0.036 | 1 | 11 |
| fam107b | 0.37957 | -0.2079765 | 0.278 | 0.357 | 1 | 11 |
| dlg5a | 0.38007 | -0.20285658 | 0.028 | 0.107 | 1 | 11 |
| txndc15 | 0.38013 | 0.114482455 | 0.222 | 0.107 | 1 | 11 |
| elmsan1b | 0.38033 | -0.17287221 | 0.028 | 0.107 | 1 | 11 |
| smim15 | 0.38049 | -0.15684733 | 0.306 | 0.464 | 1 | 11 |
| dstyk | 0.38058 | 0.121621139 | 0.167 | 0.071 | 1 | 11 |

| | | | | | | |
|---|---|---|---|---|---|---|
| mmp15a | 0.38065 | 0.214938302 | 0.139 | 0.071 | 1 | 11 |
| myh9b | 0.38075 | -0.18147737 | 0.028 | 0.107 | 1 | 11 |
| arl9 | 0.38201 | -0.20527216 | 0.028 | 0.107 | 1 | 11 |
| gm2a | 0.38206 | -0.18109603 | 0.056 | 0.143 | 1 | 11 |
| lims1 | 0.38276 | 0.180821491 | 0.25 | 0.179 | 1 | 11 |
| clcn5a | 0.38289 | -0.19827729 | 0.028 | 0.107 | 1 | 11 |
| vdac2 | 0.38309 | -0.14833265 | 0.472 | 0.643 | 1 | 11 |
| ptpn21 | 0.38311 | -0.13148587 | 0.222 | 0.357 | 1 | 11 |
| col12a1a | 0.3832 | 0.23211133 | 0.5 | 0.429 | 1 | 11 |
| nipblb | 0.38327 | 0.186715986 | 0.111 | 0.036 | 1 | 11 |
| hsdl2 | 0.38335 | -0.25142787 | 0.139 | 0.214 | 1 | 11 |
| atg5 | 0.38476 | -0.21230552 | 0.028 | 0.107 | 1 | 11 |
| bpnt1 | 0.38577 | -0.11334074 | 0.028 | 0.107 | 1 | 11 |
| snw1 | 0.38654 | 0.236884039 | 0.278 | 0.179 | 1 | 11 |
| selenoj | 0.38673 | 0.199344848 | 0.194 | 0.107 | 1 | 11 |
| ubap2b | 0.38686 | 0.179043719 | 0.194 | 0.107 | 1 | 11 |
| actn4 | 0.38802 | 0.201409948 | 0.25 | 0.179 | 1 | 11 |
| LOC103911473 | 0.3894 | -0.19711197 | 0.028 | 0.107 | 1 | 11 |
| myo1cb | 0.3897 | 0.193973929 | 0.417 | 0.357 | 1 | 11 |
| ube2a | 0.38984 | 0.198586164 | 0.194 | 0.107 | 1 | 11 |
| eif6 | 0.39166 | 0.161926354 | 0.222 | 0.214 | 1 | 11 |
| XLOC-023492 | 0.39296 | -0.17282452 | 0.056 | 0.107 | 1 | 11 |
| gng7 | 0.39326 | 0.138613505 | 0.306 | 0.179 | 1 | 11 |
| gabarapb | 0.39373 | -0.20367967 | 0.472 | 0.643 | 1 | 11 |
| cldn5a | 0.39413 | 0.203031105 | 0.806 | 0.714 | 1 | 11 |
| oaz1b | 0.39426 | -0.20188693 | 0.639 | 0.714 | 1 | 11 |
| rdh10a | 0.39459 | -0.1934202 | 0.472 | 0.536 | 1 | 11 |
| nr2f1a | 0.39574 | 0.198073648 | 0.278 | 0.143 | 1 | 11 |
| hibadhb | 0.39579 | 0.197320403 | 0.167 | 0.071 | 1 | 11 |
| ptprk | 0.39603 | -0.25769999 | 0.306 | 0.429 | 1 | 11 |
| igf2bp2b | 0.39668 | 0.210619574 | 0.139 | 0.071 | 1 | 11 |
| eif2s2 | 0.39671 | -0.2063078 | 0.611 | 0.679 | 1 | 11 |
| cpox | 0.39676 | -0.14390073 | 0.056 | 0.071 | 1 | 11 |
| txnrd3 | 0.3969 | -0.18800201 | 0.056 | 0.143 | 1 | 11 |
| tvp23b | 0.39727 | -0.10082581 | 0.028 | 0.107 | 1 | 11 |
| rab14l | 0.39777 | -0.15107193 | 0.028 | 0.107 | 1 | 11 |
| march5l | 0.3996 | 0.170541926 | 0.139 | 0.107 | 1 | 11 |
| fbxo11b | 0.40011 | -0.17338317 | 0.028 | 0.107 | 1 | 11 |
| rab2a | 0.4005 | -0.25275306 | 0.306 | 0.464 | 1 | 11 |
| adhfe1 | 0.4006 | -0.10097524 | 0.028 | 0.107 | 1 | 11 |
| si:dkeyp-82a1.6 | 0.40068 | -0.17920116 | 0.028 | 0.107 | 1 | 11 |
| ccdc137 | 0.40121 | 0.18151802 | 0.083 | 0.036 | 1 | 11 |
| tns1b | 0.40124 | -0.13935497 | 0.167 | 0.286 | 1 | 11 |
| si:rp71-36a1.2 | 0.40215 | -0.10734254 | 0.083 | 0.179 | 1 | 11 |
| sh2b3 | 0.40228 | -0.14770028 | 0.028 | 0.107 | 1 | 11 |
| hsbp1a | 0.40305 | -0.19327933 | 0.306 | 0.464 | 1 | 11 |
| tgfbr3 | 0.40336 | 0.124616642 | 0.167 | 0.071 | 1 | 11 |
| ube2ia | 0.40347 | 0.109258936 | 0.278 | 0.286 | 1 | 11 |

| | | | | | | |
|---|---|---|---|---|---|---|
| cdk12 | 0.40373 | -0.17426147 | 0.028 | 0.107 | 1 | 11 |
| commd1 | 0.40374 | -0.11559756 | 0.028 | 0.107 | 1 | 11 |
| cfap45 | 0.40378 | -0.23706086 | 0.111 | 0.214 | 1 | 11 |
| esd | 0.40384 | 0.195846093 | 0.306 | 0.214 | 1 | 11 |
| si:ch211-39a7.1 | 0.40395 | -0.17116253 | 0.028 | 0.107 | 1 | 11 |
| pmp22a | 0.40424 | 0.138778826 | 0.861 | 0.821 | 1 | 11 |
| dnaja2a | 0.40427 | -0.2094677 | 0.333 | 0.5 | 1 | 11 |
| aldoaa | 0.40454 | -0.25119651 | 0.333 | 0.464 | 1 | 11 |
| si:ch211-253b8.5 | 0.40462 | 0.198839428 | 0.194 | 0.107 | 1 | 11 |
| nfkbiaa | 0.40582 | 0.248070403 | 0.222 | 0.107 | 1 | 11 |
| cdc40 | 0.40623 | -0.18503727 | 0.056 | 0.107 | 1 | 11 |
| hnrnpl | 0.40726 | 0.115097739 | 0.167 | 0.071 | 1 | 11 |
| si:ch211-196c10.15 | 0.4074 | -0.16999712 | 0.028 | 0.107 | 1 | 11 |
| XLOC-000946 | 0.40763 | 0.115300626 | 0.222 | 0.214 | 1 | 11 |
| pofut2 | 0.40765 | -0.10906438 | 0.028 | 0.107 | 1 | 11 |
| si:ch211-194c3.5 | 0.40795 | -0.11085878 | 0.028 | 0.107 | 1 | 11 |
| mrps11 | 0.4083 | 0.169797278 | 0.167 | 0.107 | 1 | 11 |
| igf2bp2a | 0.40844 | -0.10227297 | 0.083 | 0.179 | 1 | 11 |
| spsb3a | 0.40864 | -0.1704934 | 0.028 | 0.107 | 1 | 11 |
| smu1a | 0.40882 | -0.19387801 | 0.056 | 0.143 | 1 | 11 |
| cbln2b | 0.4091 | -0.11574805 | 0.028 | 0.107 | 1 | 11 |
| men1 | 0.4092 | -0.17921702 | 0.028 | 0.107 | 1 | 11 |
| tnikb | 0.40925 | 0.21700569 | 0.194 | 0.107 | 1 | 11 |
| shroom2a | 0.41032 | -0.15179139 | 0.028 | 0.107 | 1 | 11 |
| galnt1 | 0.41034 | -0.15236366 | 0.028 | 0.107 | 1 | 11 |
| entpd1 | 0.41044 | -0.15685424 | 0.611 | 0.643 | 1 | 11 |
| calcrla | 0.41094 | -0.10550531 | 0.028 | 0.107 | 1 | 11 |
| wu:fb55g09 | 0.41134 | 0.22459047 | 0.694 | 0.536 | 1 | 11 |
| rbbp4 | 0.41138 | 0.107061225 | 0.222 | 0.107 | 1 | 11 |
| cxadr | 0.41169 | 0.245879596 | 0.444 | 0.286 | 1 | 11 |
| zgc:56304 | 0.413 | 0.187658918 | 0.167 | 0.107 | 1 | 11 |
| arl6ip5b | 0.41315 | -0.18642846 | 0.306 | 0.464 | 1 | 11 |
| commd5 | 0.41333 | -0.15586427 | 0.028 | 0.107 | 1 | 11 |
| cetn4 | 0.41399 | -0.16821097 | 0.028 | 0.107 | 1 | 11 |
| zbtb46 | 0.41423 | 0.202930396 | 0.167 | 0.071 | 1 | 11 |
| mrpl46 | 0.41436 | 0.137938728 | 0.111 | 0.036 | 1 | 11 |
| got2a | 0.41437 | -0.15394328 | 0.028 | 0.107 | 1 | 11 |
| lims2 | 0.41456 | 0.151915811 | 0.111 | 0.036 | 1 | 11 |
| aldh3a2a | 0.41459 | 0.177877019 | 0.111 | 0.036 | 1 | 11 |
| msna | 0.4152 | 0.222091115 | 0.722 | 0.571 | 1 | 11 |
| znf451 | 0.41529 | 0.197195771 | 0.111 | 0.036 | 1 | 11 |
| adma | 0.41534 | -0.1082322 | 0.028 | 0.107 | 1 | 11 |
| cntf | 0.4155 | -0.14523362 | 0.028 | 0.107 | 1 | 11 |
| fzd1 | 0.4163 | -0.12318451 | 0.028 | 0.107 | 1 | 11 |
| MDFIC | 0.41645 | -0.17521787 | 0.028 | 0.107 | 1 | 11 |
| gtpbp1 | 0.41672 | -0.16192768 | 0.028 | 0.107 | 1 | 11 |
| wdr26b | 0.41681 | 0.111088633 | 0.111 | 0.036 | 1 | 11 |
| pwwp2b | 0.41694 | 0.181786781 | 0.111 | 0.036 | 1 | 11 |

| | | | | | | |
|---|---|---|---|---|---|---|
| eif4e3 | 0.41711 | -0.15758936 | 0.028 | 0.107 | 1 | 11 |
| ndufs2 | 0.41756 | -0.23000177 | 0.167 | 0.286 | 1 | 11 |
| cratb | 0.41785 | 0.142016898 | 0.083 | 0.071 | 1 | 11 |
| stat2 | 0.418 | 0.115981973 | 0.111 | 0.036 | 1 | 11 |
| arf6b | 0.41812 | 0.167547046 | 0.167 | 0.071 | 1 | 11 |
| spata13 | 0.41813 | -0.15528 | 0.028 | 0.107 | 1 | 11 |
| cyp2p6 | 0.41828 | -0.27439996 | 0.139 | 0.25 | 1 | 11 |
| fzd2 | 0.41842 | -0.15948514 | 0.278 | 0.429 | 1 | 11 |
| LOC100334706 | 0.41842 | 0.182673222 | 0.278 | 0.143 | 1 | 11 |
| sulf1 | 0.41848 | -0.24920336 | 0.083 | 0.179 | 1 | 11 |
| cd2bp2 | 0.41872 | 0.178254468 | 0.167 | 0.071 | 1 | 11 |
| egln1b | 0.41874 | -0.15268504 | 0.028 | 0.107 | 1 | 11 |
| kcnq5a | 0.41887 | 0.247451364 | 0.278 | 0.143 | 1 | 11 |
| u2af1 | 0.41905 | 0.178437411 | 0.333 | 0.286 | 1 | 11 |
| fam83hb | 0.4191 | -0.16849112 | 0.028 | 0.107 | 1 | 11 |
| pbx4 | 0.41924 | 0.237675611 | 0.389 | 0.25 | 1 | 11 |
| erp44 | 0.41926 | -0.18332849 | 0.25 | 0.321 | 1 | 11 |
| crtap | 0.41931 | 0.206542119 | 0.25 | 0.179 | 1 | 11 |
| chsy1 | 0.41937 | -0.16304152 | 0.028 | 0.107 | 1 | 11 |
| abcf2a | 0.41955 | -0.148689 | 0.028 | 0.107 | 1 | 11 |
| snx14 | 0.41963 | 0.21396159 | 0.139 | 0.071 | 1 | 11 |
| mical2a | 0.42002 | -0.12732791 | 0.111 | 0.214 | 1 | 11 |
| LOC101885758 | 0.42013 | -0.13627694 | 0.028 | 0.107 | 1 | 11 |
| seh1l | 0.42069 | 0.181160725 | 0.111 | 0.036 | 1 | 11 |
| echs1 | 0.42079 | 0.102751127 | 0.167 | 0.071 | 1 | 11 |
| nol7 | 0.42079 | 0.194332656 | 0.167 | 0.071 | 1 | 11 |
| tmem35 | 0.42082 | -0.20708941 | 0.028 | 0.107 | 1 | 11 |
| nedd8 | 0.42103 | 0.184216603 | 0.194 | 0.143 | 1 | 11 |
| polr2h | 0.42106 | 0.131815322 | 0.083 | 0.071 | 1 | 11 |
| cdc14ab | 0.42108 | -0.15126331 | 0.028 | 0.107 | 1 | 11 |
| adssl | 0.42114 | 0.19629211 | 0.278 | 0.143 | 1 | 11 |
| rbm47 | 0.42115 | 0.180067543 | 0.111 | 0.036 | 1 | 11 |
| dnmt3ab | 0.42124 | -0.15526825 | 0.25 | 0.393 | 1 | 11 |
| sesn3 | 0.42129 | -0.13338573 | 0.028 | 0.107 | 1 | 11 |
| ncor2 | 0.42137 | 0.231672465 | 0.389 | 0.25 | 1 | 11 |
| kbtbd2 | 0.42165 | -0.16449364 | 0.028 | 0.107 | 1 | 11 |
| nop53 | 0.42188 | 0.233896862 | 0.583 | 0.464 | 1 | 11 |
| ubl3b | 0.42224 | -0.13269978 | 0.028 | 0.107 | 1 | 11 |
| zgc:110783 | 0.42233 | -0.14003669 | 0.028 | 0.107 | 1 | 11 |
| kdm6a | 0.42238 | -0.11697953 | 0.028 | 0.071 | 1 | 11 |
| LOC101886337 | 0.42238 | -0.10200068 | 0.028 | 0.071 | 1 | 11 |
| LRRC75A | 0.42238 | -0.13138547 | 0.028 | 0.071 | 1 | 11 |
| chd7 | 0.42238 | -0.10423378 | 0.028 | 0.071 | 1 | 11 |
| cyp27a7 | 0.42238 | -0.1331658 | 0.028 | 0.071 | 1 | 11 |
| hmga1b | 0.42238 | -0.17291679 | 0.028 | 0.071 | 1 | 11 |
| june | 0.42238 | -0.1255009 | 0.028 | 0.071 | 1 | 11 |
| kif1bp | 0.42238 | -0.11078016 | 0.028 | 0.071 | 1 | 11 |
| maf1 | 0.42238 | -0.11453405 | 0.028 | 0.071 | 1 | 11 |

| | | | | | | |
|---|---|---|---|---|---|---|
| mertka | 0.42238 | -0.15449414 | 0.028 | 0.071 | 1 | 11 |
| mrm2 | 0.42238 | -0.1075454 | 0.028 | 0.071 | 1 | 11 |
| prrg4 | 0.42238 | -0.11819527 | 0.028 | 0.071 | 1 | 11 |
| ptbp2b | 0.42238 | -0.10958006 | 0.028 | 0.071 | 1 | 11 |
| scaf11 | 0.42238 | -0.14191978 | 0.028 | 0.071 | 1 | 11 |
| si:rp71-1g18.1 | 0.42238 | -0.1133473 | 0.028 | 0.071 | 1 | 11 |
| tchp | 0.42238 | -0.11230707 | 0.028 | 0.071 | 1 | 11 |
| tmem51a | 0.42238 | -0.10675363 | 0.028 | 0.071 | 1 | 11 |
| znf367 | 0.42238 | -0.16141742 | 0.028 | 0.071 | 1 | 11 |
| CABZ01029938.1 | 0.42238 | -0.12524878 | 0.028 | 0.071 | 1 | 11 |
| acox3 | 0.42238 | -0.11213756 | 0.028 | 0.071 | 1 | 11 |
| adgrl2a | 0.42238 | -0.11333457 | 0.028 | 0.071 | 1 | 11 |
| amn1 | 0.42238 | -0.13292435 | 0.028 | 0.071 | 1 | 11 |
| anxa6 | 0.42238 | -0.11810388 | 0.028 | 0.071 | 1 | 11 |
| b3galt1a | 0.42238 | -0.11146273 | 0.028 | 0.071 | 1 | 11 |
| bag2 | 0.42238 | -0.11689905 | 0.028 | 0.071 | 1 | 11 |
| capn12 | 0.42238 | -0.16261934 | 0.028 | 0.071 | 1 | 11 |
| clns1a | 0.42238 | -0.1107473 | 0.028 | 0.071 | 1 | 11 |
| dctd | 0.42238 | -0.15061376 | 0.028 | 0.071 | 1 | 11 |
| heatr5b | 0.42238 | -0.10019821 | 0.028 | 0.071 | 1 | 11 |
| il11ra | 0.42238 | -0.10214598 | 0.028 | 0.071 | 1 | 11 |
| ncoa4 | 0.42238 | -0.18997322 | 0.028 | 0.071 | 1 | 11 |
| pkp3a | 0.42238 | -0.12153511 | 0.028 | 0.071 | 1 | 11 |
| si:ch73-211e3.1 | 0.42238 | -0.12609816 | 0.028 | 0.071 | 1 | 11 |
| smarca1 | 0.42238 | -0.12942635 | 0.028 | 0.071 | 1 | 11 |
| tmem107l | 0.42238 | -0.10394391 | 0.028 | 0.071 | 1 | 11 |
| tmem56a | 0.42238 | -0.13041743 | 0.028 | 0.071 | 1 | 11 |
| zgc:162255 | 0.42238 | -0.10037516 | 0.028 | 0.071 | 1 | 11 |
| F7 (1 of many) | 0.42238 | -0.12393793 | 0.028 | 0.071 | 1 | 11 |
| FP101914.1 | 0.42238 | -0.12994926 | 0.028 | 0.071 | 1 | 11 |
| LOC103911404 | 0.42238 | -0.11018144 | 0.028 | 0.071 | 1 | 11 |
| bop1 | 0.42238 | -0.10469394 | 0.028 | 0.071 | 1 | 11 |
| ccdc3b | 0.42238 | -0.10130306 | 0.028 | 0.071 | 1 | 11 |
| dpysl3 | 0.42238 | -0.1078969 | 0.028 | 0.071 | 1 | 11 |
| exoc5 | 0.42238 | -0.13558787 | 0.028 | 0.071 | 1 | 11 |
| hapln3 | 0.42238 | -0.10821922 | 0.028 | 0.071 | 1 | 11 |
| hcst | 0.42238 | -0.12932179 | 0.028 | 0.071 | 1 | 11 |
| lmo1 | 0.42238 | -0.11562648 | 0.028 | 0.071 | 1 | 11 |
| phf10 | 0.42238 | -0.10727223 | 0.028 | 0.071 | 1 | 11 |
| sash1a | 0.42238 | -0.10650625 | 0.028 | 0.071 | 1 | 11 |
| syvn1 | 0.42238 | -0.14893698 | 0.028 | 0.071 | 1 | 11 |
| tet3 | 0.42238 | -0.10247066 | 0.028 | 0.071 | 1 | 11 |
| diablob | 0.42238 | -0.10646098 | 0.028 | 0.071 | 1 | 11 |
| mcu | 0.42238 | -0.11108577 | 0.028 | 0.071 | 1 | 11 |
| nr1d1 | 0.42238 | -0.12424384 | 0.028 | 0.071 | 1 | 11 |
| entpd6 | 0.42239 | -0.12016543 | 0.028 | 0.107 | 1 | 11 |
| harbi1 | 0.42241 | -0.1013035 | 0.028 | 0.107 | 1 | 11 |
| ppap2d | 0.42249 | 0.179597881 | 0.111 | 0.036 | 1 | 11 |

| | | | | | | |
|---|---|---|---|---|---|---|
| CABZ01058046.1 | 0.42319 | -0.12045374 | 0.028 | 0.107 | 1 | 11 |
| nudt12 | 0.42348 | -0.14580576 | 0.028 | 0.107 | 1 | 11 |
| zgc:66475 | 0.42362 | -0.15530364 | 0.028 | 0.107 | 1 | 11 |
| sh3bgrl | 0.4237 | -0.23682079 | 0.278 | 0.429 | 1 | 11 |
| surf4l | 0.42375 | -0.11349076 | 0.028 | 0.107 | 1 | 11 |
| arrb2b | 0.42407 | -0.17778679 | 0.083 | 0.179 | 1 | 11 |
| nrp2a | 0.42424 | -0.16346639 | 0.028 | 0.107 | 1 | 11 |
| rpl23 | 0.42436 | 0.111486318 | 0.972 | 0.929 | 1 | 11 |
| trnt1 | 0.4244 | -0.13123064 | 0.028 | 0.107 | 1 | 11 |
| cnbpa | 0.4244 | 0.199369739 | 0.444 | 0.286 | 1 | 11 |
| cdh2 | 0.42475 | -0.19701193 | 0.556 | 0.714 | 1 | 11 |
| ttpa | 0.42503 | -0.14225613 | 0.028 | 0.107 | 1 | 11 |
| glrx3 | 0.42511 | -0.19313356 | 0.056 | 0.143 | 1 | 11 |
| si:ch73-109i22.2 | 0.42511 | -0.14469697 | 0.028 | 0.107 | 1 | 11 |
| mrpl51 | 0.4253 | 0.154425586 | 0.139 | 0.071 | 1 | 11 |
| si:dkey-125i10.3 | 0.42539 | 0.17672199 | 0.111 | 0.036 | 1 | 11 |
| cul4a | 0.42551 | -0.12974574 | 0.028 | 0.107 | 1 | 11 |
| h2afva | 0.42554 | -0.11561172 | 0.222 | 0.357 | 1 | 11 |
| pmpcb | 0.42555 | -0.12957819 | 0.028 | 0.107 | 1 | 11 |
| trim24 | 0.42573 | -0.14222902 | 0.028 | 0.107 | 1 | 11 |
| tmem106ba | 0.42574 | 0.232832562 | 0.222 | 0.107 | 1 | 11 |
| pigx | 0.42587 | 0.171802353 | 0.111 | 0.036 | 1 | 11 |
| aldh6a1 | 0.4259 | 0.203694304 | 0.167 | 0.071 | 1 | 11 |
| tiparp | 0.4261 | -0.10584353 | 0.028 | 0.107 | 1 | 11 |
| apbb2b | 0.42626 | -0.15324927 | 0.028 | 0.107 | 1 | 11 |
| zgc:195173 | 0.42642 | 0.232078905 | 0.639 | 0.5 | 1 | 11 |
| si:ch73-100l22.3 | 0.42642 | -0.10028187 | 0.028 | 0.107 | 1 | 11 |
| gnl2 | 0.4268 | -0.11369727 | 0.028 | 0.107 | 1 | 11 |
| timp2a | 0.42687 | -0.24801908 | 0.25 | 0.393 | 1 | 11 |
| fam219b | 0.42718 | -0.12148185 | 0.028 | 0.107 | 1 | 11 |
| lgals9l3 | 0.42723 | 0.155317198 | 0.333 | 0.286 | 1 | 11 |
| zgc:66160 | 0.42728 | -0.15468815 | 0.028 | 0.107 | 1 | 11 |
| tent4b | 0.42742 | 0.152517795 | 0.139 | 0.107 | 1 | 11 |
| rab1ab | 0.42779 | -0.23426329 | 0.278 | 0.429 | 1 | 11 |
| zgc:174888 | 0.4279 | -0.14484397 | 0.028 | 0.107 | 1 | 11 |
| impa1 | 0.42791 | -0.1331314 | 0.028 | 0.107 | 1 | 11 |
| g3bp1 | 0.42817 | -0.2150495 | 0.056 | 0.143 | 1 | 11 |
| sec24b | 0.42821 | -0.10036727 | 0.056 | 0.143 | 1 | 11 |
| dbpa | 0.42839 | -0.13506998 | 0.028 | 0.107 | 1 | 11 |
| tmem176l.1 | 0.42892 | -0.20732761 | 0.667 | 0.786 | 1 | 11 |
| washc2c | 0.42935 | 0.182034901 | 0.139 | 0.071 | 1 | 11 |
| dpcd | 0.42937 | 0.131966846 | 0.111 | 0.107 | 1 | 11 |
| si:zfos-1011f11.2 | 0.42949 | -0.10101196 | 0.028 | 0.107 | 1 | 11 |
| col5a2a | 0.42951 | -0.15943332 | 0.639 | 0.786 | 1 | 11 |
| si:dkey-7j14.6 | 0.42954 | 0.115316915 | 0.667 | 0.536 | 1 | 11 |
| CABZ01085139.1 | 0.4298 | -0.13955442 | 0.028 | 0.107 | 1 | 11 |
| fbxo9 | 0.42982 | 0.125991317 | 0.222 | 0.107 | 1 | 11 |
| rc3h1b | 0.42989 | -0.12020323 | 0.028 | 0.107 | 1 | 11 |

| | | | | | | |
|---|---|---|---|---|---|---|
| casc4 | 0.43013 | 0.100884342 | 0.111 | 0.036 | 1 | 11 |
| hprt1 | 0.4302 | -0.11693957 | 0.028 | 0.107 | 1 | 11 |
| gsto2 | 0.43026 | -0.24713487 | 0.111 | 0.214 | 1 | 11 |
| dctn4 | 0.43045 | -0.11008936 | 0.028 | 0.107 | 1 | 11 |
| gna12a | 0.43051 | -0.13349709 | 0.194 | 0.321 | 1 | 11 |
| si:ch73-217n20.1 | 0.43067 | -0.18612681 | 0.028 | 0.107 | 1 | 11 |
| vgll4a | 0.43071 | -0.13596549 | 0.028 | 0.107 | 1 | 11 |
| colgalt1 | 0.43099 | -0.12924557 | 0.028 | 0.107 | 1 | 11 |
| sorl1 | 0.43101 | 0.173342502 | 0.111 | 0.036 | 1 | 11 |
| mxd1 | 0.43104 | -0.13045908 | 0.028 | 0.107 | 1 | 11 |
| ggctb | 0.43115 | -0.10538651 | 0.028 | 0.107 | 1 | 11 |
| znf710b | 0.43138 | -0.1261032 | 0.056 | 0.143 | 1 | 11 |
| prr13 | 0.43146 | -0.10000121 | 0.028 | 0.107 | 1 | 11 |
| mettl6 | 0.43165 | -0.10003159 | 0.028 | 0.107 | 1 | 11 |
| ubald2 | 0.43185 | -0.10902385 | 0.028 | 0.107 | 1 | 11 |
| rad21a | 0.43198 | 0.100218447 | 0.278 | 0.286 | 1 | 11 |
| cyp20a1 | 0.43202 | -0.19137177 | 0.083 | 0.179 | 1 | 11 |
| pum3 | 0.43219 | -0.20676997 | 0.139 | 0.214 | 1 | 11 |
| sar1aa | 0.43226 | -0.10950007 | 0.028 | 0.107 | 1 | 11 |
| sec22a | 0.43258 | -0.12406767 | 0.028 | 0.107 | 1 | 11 |
| eif3s10 | 0.43259 | 0.195112069 | 0.556 | 0.393 | 1 | 11 |
| hif1an | 0.43261 | -0.12150983 | 0.028 | 0.107 | 1 | 11 |
| hpf1 | 0.43265 | -0.12224281 | 0.028 | 0.107 | 1 | 11 |
| nlrc3l | 0.43281 | -0.11102649 | 0.028 | 0.107 | 1 | 11 |
| hes6 | 0.43297 | -0.11242317 | 0.028 | 0.107 | 1 | 11 |
| pkp4 | 0.43299 | -0.13652551 | 0.028 | 0.107 | 1 | 11 |
| mphosph10 | 0.43311 | -0.17344261 | 0.028 | 0.107 | 1 | 11 |
| sec31a | 0.43312 | -0.11168541 | 0.028 | 0.107 | 1 | 11 |
| mrpl35 | 0.43316 | -0.1125596 | 0.028 | 0.107 | 1 | 11 |
| btbd10a | 0.43327 | -0.11707304 | 0.028 | 0.107 | 1 | 11 |
| palm1a | 0.43329 | -0.1174802 | 0.028 | 0.107 | 1 | 11 |
| ier5 | 0.43332 | 0.221763628 | 0.333 | 0.214 | 1 | 11 |
| cad | 0.43333 | -0.12983988 | 0.028 | 0.107 | 1 | 11 |
| lman1 | 0.43336 | -0.10005544 | 0.028 | 0.107 | 1 | 11 |
| rnf168 | 0.43336 | -0.10789041 | 0.028 | 0.107 | 1 | 11 |
| eftud2 | 0.43336 | -0.10030228 | 0.028 | 0.107 | 1 | 11 |
| marveld2a | 0.43341 | -0.10472703 | 0.028 | 0.107 | 1 | 11 |
| mettl9 | 0.43344 | -0.11565243 | 0.028 | 0.107 | 1 | 11 |
| si:cabz01071911.3 | 0.43348 | -0.10743175 | 0.028 | 0.107 | 1 | 11 |
| bbip1 | 0.4335 | -0.15232987 | 0.028 | 0.107 | 1 | 11 |
| klf3 | 0.43352 | -0.19307909 | 0.056 | 0.143 | 1 | 11 |
| zgc:113184 | 0.43357 | -0.12441694 | 0.028 | 0.107 | 1 | 11 |
| lysmd4 | 0.43359 | -0.14629009 | 0.028 | 0.107 | 1 | 11 |
| epc2 | 0.43365 | -0.12951626 | 0.028 | 0.107 | 1 | 11 |
| gnsb | 0.43367 | -0.11855444 | 0.028 | 0.107 | 1 | 11 |
| pdpk1b | 0.43376 | -0.11474451 | 0.028 | 0.107 | 1 | 11 |
| rfx7b | 0.43377 | -0.10792074 | 0.028 | 0.107 | 1 | 11 |
| sat2b | 0.43383 | -0.11022401 | 0.028 | 0.107 | 1 | 11 |

| | | | | | | |
|---|---|---|---|---|---|---|
| vegfd | 0.43387 | -0.11257502 | 0.056 | 0.143 | 1 | 11 |
| cyp46a1.2 | 0.43391 | -0.10978464 | 0.028 | 0.107 | 1 | 11 |
| dis3 | 0.43395 | -0.13379804 | 0.028 | 0.107 | 1 | 11 |
| arl3 | 0.43404 | -0.11257644 | 0.028 | 0.107 | 1 | 11 |
| pcnx4 | 0.43404 | -0.11988899 | 0.028 | 0.107 | 1 | 11 |
| cwf19l1 | 0.43405 | -0.10927623 | 0.028 | 0.107 | 1 | 11 |
| alg2 | 0.43407 | -0.12434144 | 0.028 | 0.107 | 1 | 11 |
| si:dkey-172m14.1 | 0.43408 | -0.10494876 | 0.028 | 0.107 | 1 | 11 |
| chchd1 | 0.4341 | -0.10097844 | 0.028 | 0.107 | 1 | 11 |
| pnpla6 | 0.43412 | -0.11380708 | 0.028 | 0.107 | 1 | 11 |
| rcan2 | 0.43416 | -0.10603136 | 0.028 | 0.107 | 1 | 11 |
| LOC405768 | 0.43417 | -0.11296503 | 0.028 | 0.107 | 1 | 11 |
| meaf6 | 0.43524 | -0.15870402 | 0.278 | 0.321 | 1 | 11 |
| zgc:136439 | 0.43559 | 0.174555523 | 0.111 | 0.036 | 1 | 11 |
| slc39a7 | 0.43573 | 0.165366125 | 0.111 | 0.071 | 1 | 11 |
| lhfpl2a | 0.43601 | -0.10216237 | 0.083 | 0.179 | 1 | 11 |
| si:ch211-136m16.8 | 0.43625 | 0.172197007 | 0.111 | 0.036 | 1 | 11 |
| ndufb5 | 0.43675 | 0.178846222 | 0.361 | 0.214 | 1 | 11 |
| TTC28 | 0.43681 | 0.203343181 | 0.167 | 0.071 | 1 | 11 |
| fat1a | 0.43681 | 0.174301632 | 0.222 | 0.179 | 1 | 11 |
| nansa | 0.43695 | 0.13762072 | 0.111 | 0.036 | 1 | 11 |
| med1 | 0.43701 | 0.186821016 | 0.111 | 0.036 | 1 | 11 |
| oxr1a | 0.43743 | 0.179011386 | 0.083 | 0.036 | 1 | 11 |
| kpna3 | 0.43768 | -0.10912517 | 0.056 | 0.143 | 1 | 11 |
| sgta | 0.43836 | -0.17312717 | 0.278 | 0.321 | 1 | 11 |
| gata5 | 0.43839 | 0.225944947 | 0.222 | 0.107 | 1 | 11 |
| spcs1 | 0.43859 | 0.204532987 | 0.528 | 0.464 | 1 | 11 |
| ehd1b | 0.43874 | -0.21118005 | 0.389 | 0.464 | 1 | 11 |
| dedd1 | 0.43933 | 0.189800131 | 0.528 | 0.464 | 1 | 11 |
| dr1 | 0.43976 | 0.16666935 | 0.111 | 0.036 | 1 | 11 |
| rsl24d1 | 0.43976 | 0.159106876 | 0.639 | 0.607 | 1 | 11 |
| rbm8a | 0.43997 | 0.130337533 | 0.389 | 0.393 | 1 | 11 |
| rac1a | 0.44008 | 0.141861568 | 0.306 | 0.179 | 1 | 11 |
| calr3a | 0.44023 | 0.131678711 | 0.25 | 0.25 | 1 | 11 |
| lrig2 | 0.44026 | 0.184536486 | 0.111 | 0.036 | 1 | 11 |
| map1lc3b | 0.44217 | -0.22447197 | 0.361 | 0.464 | 1 | 11 |
| ago2 | 0.44235 | 0.180312431 | 0.111 | 0.036 | 1 | 11 |
| golga7 | 0.44271 | -0.18411788 | 0.278 | 0.429 | 1 | 11 |
| tnfsf10 | 0.44317 | 0.238332438 | 0.583 | 0.429 | 1 | 11 |
| hells | 0.4433 | 0.10389458 | 0.167 | 0.071 | 1 | 11 |
| pdhx | 0.44331 | 0.140174097 | 0.111 | 0.036 | 1 | 11 |
| uba2 | 0.44362 | 0.160400575 | 0.222 | 0.107 | 1 | 11 |
| osr1 | 0.44378 | 0.17390674 | 0.167 | 0.107 | 1 | 11 |
| lama4 | 0.44389 | -0.13031713 | 0.361 | 0.5 | 1 | 11 |
| hoxc6b | 0.44393 | 0.21723026 | 0.167 | 0.071 | 1 | 11 |
| txndc12 | 0.44488 | -0.12517888 | 0.167 | 0.286 | 1 | 11 |
| eif4a2 | 0.44529 | -0.15256913 | 0.222 | 0.357 | 1 | 11 |
| baiap2a | 0.44564 | 0.164601766 | 0.111 | 0.036 | 1 | 11 |

| | | | | | | |
|---|---|---|---|---|---|---|
| cul1b | 0.44569 | -0.20374946 | 0.083 | 0.179 | 1 | 11 |
| gtf2f1 | 0.44624 | 0.219532874 | 0.25 | 0.143 | 1 | 11 |
| smdt1b | 0.44651 | 0.227684307 | 0.556 | 0.429 | 1 | 11 |
| herc2 | 0.44732 | -0.10077951 | 0.083 | 0.179 | 1 | 11 |
| btbd10b | 0.44741 | 0.183590164 | 0.139 | 0.071 | 1 | 11 |
| zgc:162944 | 0.4475 | -0.11175884 | 0.361 | 0.5 | 1 | 11 |
| coa3a | 0.44781 | -0.19786057 | 0.111 | 0.214 | 1 | 11 |
| mbnl1 | 0.44783 | 0.180463934 | 0.361 | 0.286 | 1 | 11 |
| dusp6 | 0.44825 | -0.12364801 | 0.056 | 0.143 | 1 | 11 |
| msrb1b | 0.44843 | 0.121698594 | 0.111 | 0.036 | 1 | 11 |
| mmp14b | 0.44932 | 0.195779079 | 0.333 | 0.25 | 1 | 11 |
| smarce1 | 0.44964 | -0.23025295 | 0.222 | 0.357 | 1 | 11 |
| efna1b | 0.44967 | -0.23795396 | 0.278 | 0.429 | 1 | 11 |
| mrpl36 | 0.44985 | -0.18094864 | 0.278 | 0.429 | 1 | 11 |
| csnk1a1 | 0.44999 | 0.21079976 | 0.389 | 0.25 | 1 | 11 |
| brd9 | 0.4507 | -0.17611519 | 0.056 | 0.143 | 1 | 11 |
| glrx5 | 0.45088 | 0.116679694 | 0.222 | 0.107 | 1 | 11 |
| ywhaz | 0.45091 | -0.19381725 | 0.611 | 0.75 | 1 | 11 |
| jmjd1cb | 0.45182 | 0.202108672 | 0.25 | 0.143 | 1 | 11 |
| pdlim3b | 0.45213 | 0.129387141 | 0.25 | 0.143 | 1 | 11 |
| ephb3 | 0.45228 | 0.131029408 | 0.306 | 0.286 | 1 | 11 |
| pef1 | 0.45296 | 0.108865337 | 0.111 | 0.107 | 1 | 11 |
| stau1 | 0.45314 | 0.213992894 | 0.306 | 0.214 | 1 | 11 |
| clic4 | 0.45347 | 0.206076172 | 0.611 | 0.5 | 1 | 11 |
| gosr2 | 0.45351 | 0.149143919 | 0.167 | 0.071 | 1 | 11 |
| ext2 | 0.45355 | -0.11419769 | 0.056 | 0.143 | 1 | 11 |
| kif5ba | 0.45378 | -0.1108175 | 0.111 | 0.214 | 1 | 11 |
| klf8 | 0.45388 | -0.19202963 | 0.056 | 0.143 | 1 | 11 |
| zfyve21 | 0.4539 | -0.12578232 | 0.167 | 0.286 | 1 | 11 |
| grb2b | 0.45427 | 0.170483972 | 0.111 | 0.036 | 1 | 11 |
| tomm40 | 0.45438 | 0.136699282 | 0.111 | 0.036 | 1 | 11 |
| ndufb3 | 0.45488 | 0.213660969 | 0.556 | 0.464 | 1 | 11 |
| tra2a | 0.45494 | -0.23219465 | 0.333 | 0.429 | 1 | 11 |
| tcf25 | 0.45527 | 0.238158169 | 0.194 | 0.107 | 1 | 11 |
| b3gnt5a | 0.45545 | 0.206895302 | 0.222 | 0.107 | 1 | 11 |
| uqcc2 | 0.45563 | -0.1077337 | 0.139 | 0.25 | 1 | 11 |
| pawr | 0.45708 | -0.21629126 | 0.111 | 0.214 | 1 | 11 |
| sostdc1a | 0.45743 | 0.25228289 | 0.583 | 0.429 | 1 | 11 |
| zgc:103697 | 0.4576 | 0.122275109 | 0.111 | 0.036 | 1 | 11 |
| actr1 | 0.45775 | -0.13481087 | 0.056 | 0.143 | 1 | 11 |
| bag3 | 0.458 | -0.20860026 | 0.111 | 0.214 | 1 | 11 |
| surf6 | 0.45813 | 0.142246227 | 0.111 | 0.036 | 1 | 11 |
| atp2b2 | 0.45817 | 0.142493002 | 0.361 | 0.321 | 1 | 11 |
| si:ch211-225b7.5 | 0.45856 | 0.181192019 | 0.139 | 0.071 | 1 | 11 |
| wasla | 0.45866 | 0.133198321 | 0.167 | 0.071 | 1 | 11 |
| zgc:193812 | 0.4587 | 0.175118707 | 0.167 | 0.071 | 1 | 11 |
| lmnb2 | 0.45882 | 0.146279669 | 0.222 | 0.107 | 1 | 11 |
| snx9b | 0.45901 | 0.196718206 | 0.167 | 0.071 | 1 | 11 |

| | | | | | | |
|---|---|---|---|---|---|---|
| scn2b | 0.46036 | 0.170848969 | 0.111 | 0.036 | 1 | 11 |
| magixa | 0.46047 | 0.13506165 | 0.111 | 0.107 | 1 | 11 |
| hnrnpc | 0.46115 | -0.22855589 | 0.111 | 0.214 | 1 | 11 |
| tspan9b | 0.46125 | -0.14826743 | 0.194 | 0.321 | 1 | 11 |
| rpl22l1 | 0.46155 | -0.18902712 | 0.694 | 0.821 | 1 | 11 |
| mier1a | 0.46174 | 0.146096772 | 0.083 | 0.071 | 1 | 11 |
| bloc1s6 | 0.46183 | -0.13758372 | 0.139 | 0.179 | 1 | 11 |
| tyw1 | 0.46195 | -0.17944508 | 0.056 | 0.107 | 1 | 11 |
| ep300b | 0.46197 | 0.155158665 | 0.111 | 0.036 | 1 | 11 |
| sri | 0.46214 | 0.192221522 | 0.278 | 0.214 | 1 | 11 |
| pex26 | 0.46227 | -0.15863659 | 0.056 | 0.107 | 1 | 11 |
| rrp12 | 0.46271 | -0.17585595 | 0.056 | 0.143 | 1 | 11 |
| trappc2l | 0.46298 | 0.168373999 | 0.222 | 0.107 | 1 | 11 |
| rps6kb1b | 0.4631 | -0.13098763 | 0.167 | 0.286 | 1 | 11 |
| tspan7 | 0.46329 | 0.190993171 | 0.389 | 0.321 | 1 | 11 |
| ube2ka | 0.46376 | -0.1159178 | 0.083 | 0.179 | 1 | 11 |
| si:ch211-232b12.5 | 0.46418 | -0.17692909 | 0.056 | 0.143 | 1 | 11 |
| carhsp1 | 0.4646 | 0.159447771 | 0.167 | 0.071 | 1 | 11 |
| ralgapa1 | 0.46516 | 0.151488854 | 0.111 | 0.036 | 1 | 11 |
| agrn | 0.46523 | -0.23897171 | 0.139 | 0.25 | 1 | 11 |
| rac1b | 0.46559 | 0.159360735 | 0.111 | 0.036 | 1 | 11 |
| zgc:112356 | 0.46583 | -0.11206113 | 0.139 | 0.143 | 1 | 11 |
| si:ch211-166g5.4 | 0.4661 | -0.16856383 | 0.056 | 0.143 | 1 | 11 |
| ap3b1a | 0.46615 | 0.157106375 | 0.111 | 0.036 | 1 | 11 |
| eif2b2 | 0.46627 | 0.177358251 | 0.194 | 0.107 | 1 | 11 |
| st5 | 0.46638 | 0.150699266 | 0.111 | 0.036 | 1 | 11 |
| copb2 | 0.46645 | -0.20966669 | 0.056 | 0.143 | 1 | 11 |
| dnajb1b | 0.46698 | 0.135127439 | 0.167 | 0.071 | 1 | 11 |
| rhobtb4 | 0.46711 | -0.18284977 | 0.083 | 0.179 | 1 | 11 |
| pdk2a | 0.46715 | 0.179788942 | 0.111 | 0.036 | 1 | 11 |
| lamp1a | 0.46721 | 0.162956373 | 0.222 | 0.107 | 1 | 11 |
| lrp1aa | 0.46722 | -0.17102322 | 0.222 | 0.357 | 1 | 11 |
| ska2 | 0.46741 | 0.153852443 | 0.167 | 0.071 | 1 | 11 |
| eef2b | 0.46767 | -0.16207169 | 0.833 | 0.893 | 1 | 11 |
| aco2 | 0.4682 | 0.125302065 | 0.083 | 0.071 | 1 | 11 |
| nfkbib | 0.46834 | 0.167361005 | 0.222 | 0.107 | 1 | 11 |
| CABZ01086029.1 | 0.46841 | 0.190922723 | 0.222 | 0.107 | 1 | 11 |
| CABZ01064670.1 | 0.46869 | 0.156132115 | 0.111 | 0.036 | 1 | 11 |
| cwc27 | 0.46878 | -0.15684052 | 0.056 | 0.143 | 1 | 11 |
| yipf6 | 0.46918 | -0.19256885 | 0.111 | 0.214 | 1 | 11 |
| impact | 0.46935 | -0.1855034 | 0.056 | 0.143 | 1 | 11 |
| marcksl1b | 0.4695 | 0.26280543 | 0.444 | 0.321 | 1 | 11 |
| tln1 | 0.47067 | 0.17222013 | 0.333 | 0.286 | 1 | 11 |
| vamp3 | 0.47074 | 0.209790892 | 0.472 | 0.393 | 1 | 11 |
| sdcbp2 | 0.47078 | 0.177354146 | 0.444 | 0.393 | 1 | 11 |
| gpr146 | 0.47096 | 0.185307837 | 0.167 | 0.071 | 1 | 11 |
| rab18a | 0.47099 | -0.10067044 | 0.083 | 0.179 | 1 | 11 |
| snx6 | 0.4717 | 0.126909222 | 0.167 | 0.071 | 1 | 11 |

| | | | | | | |
|---|---|---|---|---|---|---|
| hspb8 | 0.47212 | -0.23023921 | 0.111 | 0.214 | 1 | 11 |
| npm1a | 0.47221 | 0.159849739 | 0.389 | 0.25 | 1 | 11 |
| sf3b6 | 0.47239 | 0.198689521 | 0.556 | 0.429 | 1 | 11 |
| timp2b | 0.47249 | 0.14199457 | 0.111 | 0.036 | 1 | 11 |
| dapk3 | 0.47278 | 0.188967779 | 0.194 | 0.107 | 1 | 11 |
| eif3ha | 0.47314 | 0.158438697 | 0.472 | 0.321 | 1 | 11 |
| ccng1 | 0.47333 | 0.221906369 | 0.528 | 0.393 | 1 | 11 |
| copb1 | 0.4735 | 0.181198462 | 0.167 | 0.071 | 1 | 11 |
| pvrl2l | 0.47379 | 0.182024487 | 0.139 | 0.071 | 1 | 11 |
| ak3 | 0.4739 | -0.11447063 | 0.056 | 0.143 | 1 | 11 |
| vps26c | 0.47394 | 0.199729297 | 0.167 | 0.071 | 1 | 11 |
| sf3b2 | 0.47424 | 0.168648539 | 0.25 | 0.179 | 1 | 11 |
| abl2 | 0.47433 | 0.151507679 | 0.111 | 0.036 | 1 | 11 |
| ube2l3b | 0.47487 | -0.19670664 | 0.25 | 0.393 | 1 | 11 |
| tmem205 | 0.47513 | 0.150711332 | 0.111 | 0.036 | 1 | 11 |
| schip1 | 0.47565 | 0.174720568 | 0.167 | 0.071 | 1 | 11 |
| lrrc58b | 0.47583 | -0.2340418 | 0.111 | 0.214 | 1 | 11 |
| LOC103909284 | 0.47595 | -0.17262102 | 0.389 | 0.536 | 1 | 11 |
| mibp2 | 0.47614 | -0.12469079 | 0.611 | 0.75 | 1 | 11 |
| mvp | 0.47677 | -0.14113629 | 0.194 | 0.321 | 1 | 11 |
| rbb4l | 0.47693 | 0.133996645 | 0.139 | 0.107 | 1 | 11 |
| vipr2 | 0.47698 | 0.155222916 | 0.167 | 0.071 | 1 | 11 |
| fthl27 | 0.47704 | -0.18619093 | 0.694 | 0.786 | 1 | 11 |
| banf1 | 0.47716 | 0.149975427 | 0.222 | 0.179 | 1 | 11 |
| psmd11b | 0.47717 | -0.10759444 | 0.083 | 0.179 | 1 | 11 |
| FP101887.1 | 0.47759 | -0.13739997 | 0.083 | 0.107 | 1 | 11 |
| tmod2 | 0.47763 | -0.22845095 | 0.139 | 0.25 | 1 | 11 |
| rpl30 | 0.47814 | -0.11167911 | 0.889 | 0.964 | 1 | 11 |
| nmt1a | 0.47837 | 0.197406548 | 0.167 | 0.071 | 1 | 11 |
| kdelr2b | 0.47861 | -0.21170127 | 0.306 | 0.429 | 1 | 11 |
| zgc:114045 | 0.4789 | -0.12084413 | 0.056 | 0.143 | 1 | 11 |
| hnrnpd | 0.47945 | 0.178788143 | 0.167 | 0.071 | 1 | 11 |
| MZT1 | 0.47968 | 0.130461648 | 0.111 | 0.036 | 1 | 11 |
| fis1 | 0.47973 | -0.19589264 | 0.083 | 0.179 | 1 | 11 |
| aifm2 | 0.47974 | -0.10115909 | 0.056 | 0.143 | 1 | 11 |
| mllt10 | 0.47988 | 0.134808494 | 0.167 | 0.071 | 1 | 11 |
| BX005309.1 | 0.47999 | 0.18511324 | 0.167 | 0.071 | 1 | 11 |
| arhgap5 | 0.48014 | 0.130251855 | 0.167 | 0.071 | 1 | 11 |
| COA4 | 0.48029 | -0.18612144 | 0.139 | 0.214 | 1 | 11 |
| pbx1a | 0.48062 | 0.244026336 | 0.333 | 0.214 | 1 | 11 |
| zgc:56525 | 0.4808 | -0.15486022 | 0.056 | 0.143 | 1 | 11 |
| ppic | 0.48089 | 0.149239059 | 0.111 | 0.036 | 1 | 11 |
| slc2a1b | 0.48116 | 0.15248802 | 0.111 | 0.036 | 1 | 11 |
| lin52 | 0.48135 | 0.191363156 | 0.25 | 0.143 | 1 | 11 |
| akap1b | 0.48142 | 0.153223216 | 0.111 | 0.036 | 1 | 11 |
| pagr1 | 0.4815 | 0.144496808 | 0.111 | 0.036 | 1 | 11 |
| cox6c | 0.48209 | -0.21628708 | 0.444 | 0.571 | 1 | 11 |
| cyba | 0.4821 | -0.12659356 | 0.056 | 0.143 | 1 | 11 |

| | | | | | | |
|---|---|---|---|---|---|---|
| khdrbs1b | 0.48211 | -0.13373083 | 0.167 | 0.286 | 1 | 11 |
| ywhah | 0.48211 | -0.14629066 | 0.194 | 0.321 | 1 | 11 |
| scyl1 | 0.48242 | 0.137656097 | 0.111 | 0.036 | 1 | 11 |
| thtpa | 0.48297 | -0.16313081 | 0.056 | 0.143 | 1 | 11 |
| rbm26 | 0.48318 | 0.140148147 | 0.111 | 0.036 | 1 | 11 |
| tmem258 | 0.48366 | -0.14779511 | 0.611 | 0.643 | 1 | 11 |
| ndufb6 | 0.48374 | -0.13917528 | 0.25 | 0.286 | 1 | 11 |
| acsf2 | 0.48399 | -0.19299141 | 0.056 | 0.107 | 1 | 11 |
| stmn1a | 0.48416 | 0.234048131 | 0.139 | 0.071 | 1 | 11 |
| vgll4b | 0.48454 | 0.157875121 | 0.111 | 0.036 | 1 | 11 |
| b2m | 0.48487 | 0.161087852 | 0.611 | 0.464 | 1 | 11 |
| tcf3b | 0.48511 | 0.159889796 | 0.111 | 0.036 | 1 | 11 |
| zfhx3 | 0.48513 | 0.142218081 | 0.111 | 0.036 | 1 | 11 |
| bcl2l1 | 0.48523 | -0.21428563 | 0.056 | 0.143 | 1 | 11 |
| selenos | 0.48535 | 0.173784128 | 0.111 | 0.036 | 1 | 11 |
| fibpa | 0.48546 | 0.142324877 | 0.111 | 0.036 | 1 | 11 |
| p3h2 | 0.48558 | -0.13051825 | 0.083 | 0.179 | 1 | 11 |
| aldh9a1a.1 | 0.48578 | -0.17790634 | 0.694 | 0.786 | 1 | 11 |
| nfe2l2a | 0.48594 | 0.151576631 | 0.111 | 0.036 | 1 | 11 |
| ppp4r2a | 0.48647 | 0.147562442 | 0.111 | 0.036 | 1 | 11 |
| pof1b | 0.48687 | -0.10092135 | 0.056 | 0.143 | 1 | 11 |
| fam172a | 0.4871 | -0.13813419 | 0.056 | 0.143 | 1 | 11 |
| ndufab1b | 0.48717 | 0.19055563 | 0.361 | 0.286 | 1 | 11 |
| r3hdm1 | 0.48722 | 0.142487964 | 0.111 | 0.036 | 1 | 11 |
| si:ch211-76m11.3 | 0.48728 | 0.103669494 | 0.111 | 0.036 | 1 | 11 |
| nosip | 0.48779 | 0.143386786 | 0.167 | 0.071 | 1 | 11 |
| lmbr1l | 0.48782 | 0.149915065 | 0.167 | 0.071 | 1 | 11 |
| tmem59 | 0.48792 | -0.19024096 | 0.222 | 0.357 | 1 | 11 |
| rbm4.3 | 0.48834 | 0.217968282 | 0.556 | 0.429 | 1 | 11 |
| htra3b | 0.48857 | -0.15254715 | 0.056 | 0.143 | 1 | 11 |
| ccdc167 | 0.48873 | 0.143350423 | 0.083 | 0.036 | 1 | 11 |
| rgs19 | 0.48895 | -0.12753916 | 0.056 | 0.143 | 1 | 11 |
| si:ch211-108c6.2 | 0.48903 | 0.154907217 | 0.111 | 0.036 | 1 | 11 |
| mbpb | 0.48926 | -0.11465978 | 0.083 | 0.179 | 1 | 11 |
| mrpl39 | 0.48952 | 0.128859984 | 0.111 | 0.036 | 1 | 11 |
| znf395a | 0.48959 | -0.12129145 | 0.056 | 0.143 | 1 | 11 |
| glrx | 0.48996 | 0.190741927 | 0.5 | 0.429 | 1 | 11 |
| ciao1 | 0.48998 | -0.13153785 | 0.056 | 0.143 | 1 | 11 |
| npat | 0.49003 | -0.13255478 | 0.056 | 0.143 | 1 | 11 |
| CU929070.1 | 0.49016 | 0.180513723 | 0.194 | 0.143 | 1 | 11 |
| rpe | 0.49019 | 0.137426954 | 0.222 | 0.179 | 1 | 11 |
| cnot1 | 0.49028 | 0.164250748 | 0.167 | 0.071 | 1 | 11 |
| vcp | 0.49075 | -0.16778319 | 0.222 | 0.357 | 1 | 11 |
| cul4b | 0.49092 | -0.16560121 | 0.056 | 0.143 | 1 | 11 |
| foxn3 | 0.49118 | -0.17295289 | 0.056 | 0.143 | 1 | 11 |
| si:rp71-36a1.1 | 0.49154 | -0.14672345 | 0.056 | 0.143 | 1 | 11 |
| znf384l | 0.49156 | -0.14898576 | 0.056 | 0.143 | 1 | 11 |
| prpf19 | 0.4916 | -0.10138789 | 0.083 | 0.179 | 1 | 11 |

| | | | | | | |
|---|---|---|---|---|---|---|
| rbm14b | 0.49181 | -0.10219902 | 0.056 | 0.143 | 1 | 11 |
| slc43a2a | 0.49192 | 0.211281382 | 0.167 | 0.071 | 1 | 11 |
| ctdspla | 0.49212 | -0.22067098 | 0.139 | 0.25 | 1 | 11 |
| mao | 0.49221 | -0.14169914 | 0.056 | 0.143 | 1 | 11 |
| ormdl1 | 0.49228 | -0.11031489 | 0.083 | 0.179 | 1 | 11 |
| zgc:55262 | 0.49257 | -0.11196501 | 0.083 | 0.179 | 1 | 11 |
| pwp2h | 0.4929 | 0.117544769 | 0.111 | 0.036 | 1 | 11 |
| anxa4 | 0.493 | -0.18924517 | 0.389 | 0.536 | 1 | 11 |
| strn3 | 0.49406 | 0.170914981 | 0.111 | 0.036 | 1 | 11 |
| utp23 | 0.49412 | -0.15073656 | 0.056 | 0.143 | 1 | 11 |
| psmd3 | 0.49436 | 0.151138668 | 0.222 | 0.179 | 1 | 11 |
| usp7 | 0.49439 | -0.18584817 | 0.083 | 0.179 | 1 | 11 |
| cnot2 | 0.49449 | -0.18534577 | 0.167 | 0.214 | 1 | 11 |
| tns2a | 0.49486 | 0.13428211 | 0.167 | 0.143 | 1 | 11 |
| atoh8 | 0.49513 | -0.13049549 | 0.056 | 0.143 | 1 | 11 |
| lrp10 | 0.49517 | -0.14155547 | 0.056 | 0.143 | 1 | 11 |
| cmtm7 | 0.49537 | 0.108582056 | 0.111 | 0.036 | 1 | 11 |
| snrpe | 0.49538 | 0.171927996 | 0.25 | 0.143 | 1 | 11 |
| rab1aa | 0.49553 | 0.163146958 | 0.111 | 0.036 | 1 | 11 |
| frg1 | 0.49562 | -0.10657852 | 0.056 | 0.143 | 1 | 11 |
| BX284696.1 | 0.49567 | 0.184107972 | 0.333 | 0.25 | 1 | 11 |
| rfxap | 0.49568 | -0.16507817 | 0.056 | 0.143 | 1 | 11 |
| lin7c | 0.49581 | 0.105761618 | 0.111 | 0.036 | 1 | 11 |
| atad1a | 0.49604 | -0.10745651 | 0.056 | 0.143 | 1 | 11 |
| cdca7a | 0.4961 | 0.13049536 | 0.111 | 0.036 | 1 | 11 |
| rab18b | 0.49612 | -0.13810758 | 0.056 | 0.143 | 1 | 11 |
| pcbp2 | 0.49617 | 0.134840276 | 0.25 | 0.25 | 1 | 11 |
| zgc:158343 | 0.49636 | 0.203367963 | 0.389 | 0.25 | 1 | 11 |
| ppp3ccb | 0.49648 | 0.163506851 | 0.167 | 0.071 | 1 | 11 |
| parp12b | 0.49671 | -0.13911185 | 0.056 | 0.143 | 1 | 11 |
| prpsap2 | 0.49683 | -0.10998549 | 0.056 | 0.143 | 1 | 11 |
| ehbp1l1a | 0.49694 | -0.12780196 | 0.056 | 0.143 | 1 | 11 |
| prelid3b | 0.49707 | 0.162761589 | 0.194 | 0.107 | 1 | 11 |
| virma | 0.49712 | 0.135980003 | 0.111 | 0.036 | 1 | 11 |
| ncoa5 | 0.49724 | -0.131387 | 0.056 | 0.143 | 1 | 11 |
| zgc:153993 | 0.49735 | -0.13510538 | 0.056 | 0.143 | 1 | 11 |
| aes | 0.49775 | 0.123998912 | 0.111 | 0.036 | 1 | 11 |
| aida | 0.49776 | -0.18907937 | 0.194 | 0.321 | 1 | 11 |
| gnl1 | 0.49789 | 0.110887622 | 0.111 | 0.107 | 1 | 11 |
| get4 | 0.49792 | -0.12913588 | 0.083 | 0.179 | 1 | 11 |
| PDPR | 0.49831 | -0.13964717 | 0.056 | 0.143 | 1 | 11 |
| TMEM19 | 0.49843 | 0.172618306 | 0.139 | 0.071 | 1 | 11 |
| pcolce2b | 0.49856 | -0.12931973 | 0.056 | 0.143 | 1 | 11 |
| dclk1a | 0.49859 | -0.13618778 | 0.056 | 0.143 | 1 | 11 |
| psmd10 | 0.49878 | 0.139557557 | 0.111 | 0.036 | 1 | 11 |
| zranb2 | 0.49881 | 0.182845464 | 0.306 | 0.179 | 1 | 11 |
| perp | 0.49885 | 0.160485358 | 0.194 | 0.143 | 1 | 11 |
| epn1 | 0.49896 | 0.10251608 | 0.111 | 0.036 | 1 | 11 |

| | | | | | | |
|---|---|---|---|---|---|---|
| enpep | 0.49943 | 0.174226031 | 0.167 | 0.107 | 1 | 11 |
| lrtm2a | 0.4996 | -0.13472243 | 0.139 | 0.25 | 1 | 11 |
| ndufa5 | 0.49991 | -0.18041418 | 0.389 | 0.536 | 1 | 11 |
| elocb | 0.50049 | 0.203611708 | 0.389 | 0.286 | 1 | 11 |
| tcf12 | 0.50072 | 0.175998032 | 0.306 | 0.179 | 1 | 11 |
| ntd5 | 0.50089 | -0.14582821 | 0.139 | 0.25 | 1 | 11 |
| hspe1 | 0.5009 | 0.151413415 | 0.306 | 0.179 | 1 | 11 |
| tril | 0.5009 | 0.217248525 | 0.306 | 0.179 | 1 | 11 |
| slc37a2 | 0.50102 | -0.1565258 | 0.056 | 0.107 | 1 | 11 |
| mtx2 | 0.50133 | 0.113432535 | 0.111 | 0.036 | 1 | 11 |
| march7 | 0.50153 | 0.107027011 | 0.111 | 0.036 | 1 | 11 |
| praf2 | 0.50164 | 0.133176167 | 0.111 | 0.036 | 1 | 11 |
| thumpd1 | 0.50181 | 0.128664351 | 0.111 | 0.036 | 1 | 11 |
| sp3b | 0.50189 | 0.167212951 | 0.111 | 0.036 | 1 | 11 |
| zgc:100918 | 0.50189 | 0.143917066 | 0.167 | 0.071 | 1 | 11 |
| marcksb | 0.50202 | -0.13045618 | 0.111 | 0.214 | 1 | 11 |
| hexim1 | 0.50204 | -0.12030422 | 0.111 | 0.214 | 1 | 11 |
| gata2a | 0.50219 | 0.154730738 | 0.167 | 0.071 | 1 | 11 |
| nfya | 0.50231 | -0.19344813 | 0.083 | 0.143 | 1 | 11 |
| vps26b | 0.50236 | 0.122839878 | 0.111 | 0.036 | 1 | 11 |
| ywhae2 | 0.50242 | -0.11807897 | 0.083 | 0.179 | 1 | 11 |
| nt5c3a | 0.50256 | 0.146698783 | 0.167 | 0.071 | 1 | 11 |
| ssb | 0.50256 | 0.146323321 | 0.167 | 0.071 | 1 | 11 |
| idh1 | 0.50267 | -0.18682462 | 0.111 | 0.214 | 1 | 11 |
| ywhabl | 0.50269 | 0.181139006 | 0.667 | 0.571 | 1 | 11 |
| CABZ01088109.1 | 0.5035 | 0.14270738 | 0.167 | 0.071 | 1 | 11 |
| si:dkey-28b4.7 | 0.5035 | -0.15258755 | 0.5 | 0.643 | 1 | 11 |
| rnf167 | 0.50358 | 0.176745769 | 0.111 | 0.036 | 1 | 11 |
| phldb2b | 0.50367 | 0.131263984 | 0.111 | 0.036 | 1 | 11 |
| mrps14 | 0.50396 | -0.11186806 | 0.083 | 0.179 | 1 | 11 |
| arcn1b | 0.50421 | 0.178036348 | 0.167 | 0.071 | 1 | 11 |
| zgc:112982 | 0.50429 | 0.176116431 | 0.167 | 0.107 | 1 | 11 |
| pkmyt1 | 0.50481 | 0.15932827 | 0.083 | 0.036 | 1 | 11 |
| mrto4 | 0.50485 | 0.158854597 | 0.167 | 0.071 | 1 | 11 |
| cox7b | 0.50493 | -0.22067312 | 0.472 | 0.607 | 1 | 11 |
| mylipb | 0.5053 | 0.17179418 | 0.167 | 0.071 | 1 | 11 |
| atp6v1g1 | 0.50548 | -0.16466325 | 0.083 | 0.179 | 1 | 11 |
| eef1db | 0.50548 | 0.180901165 | 0.556 | 0.5 | 1 | 11 |
| dnaja2b | 0.50558 | 0.160375809 | 0.167 | 0.071 | 1 | 11 |
| maip1 | 0.50573 | 0.119269148 | 0.167 | 0.071 | 1 | 11 |
| dcaf7 | 0.50581 | 0.158394362 | 0.167 | 0.071 | 1 | 11 |
| cry1aa | 0.50603 | 0.149228907 | 0.167 | 0.071 | 1 | 11 |
| pdgfrb | 0.50604 | 0.171697367 | 0.167 | 0.071 | 1 | 11 |
| cdc42l | 0.50631 | 0.127474074 | 0.389 | 0.357 | 1 | 11 |
| enc1 | 0.50632 | 0.1310898 | 0.25 | 0.143 | 1 | 11 |
| porb | 0.5066 | 0.117980952 | 0.111 | 0.036 | 1 | 11 |
| mnta | 0.50671 | 0.157138171 | 0.111 | 0.036 | 1 | 11 |
| mapkapk2a | 0.50756 | 0.168575854 | 0.306 | 0.25 | 1 | 11 |

| | | | | | | |
|---|---|---|---|---|---|---|
| ccdc59 | 0.50779 | 0.166438677 | 0.111 | 0.071 | 1 | 11 |
| idh3a | 0.50825 | -0.12086323 | 0.111 | 0.214 | 1 | 11 |
| pum1 | 0.50844 | 0.157427867 | 0.139 | 0.071 | 1 | 11 |
| rnaseka | 0.5085 | 0.180945843 | 0.194 | 0.107 | 1 | 11 |
| dhcr7 | 0.50857 | 0.1066196 | 0.111 | 0.036 | 1 | 11 |
| ptgs1 | 0.50858 | -0.18910198 | 0.194 | 0.321 | 1 | 11 |
| ano10b | 0.5087 | 0.126057835 | 0.111 | 0.036 | 1 | 11 |
| tmem165 | 0.50893 | -0.11699988 | 0.083 | 0.179 | 1 | 11 |
| LOC103909840 | 0.50938 | 0.109439874 | 0.111 | 0.036 | 1 | 11 |
| ppp1caa | 0.50967 | -0.20262035 | 0.25 | 0.357 | 1 | 11 |
| spg21 | 0.50983 | 0.109983511 | 0.111 | 0.036 | 1 | 11 |
| zfpm2b | 0.50994 | 0.10285834 | 0.111 | 0.036 | 1 | 11 |
| usp9 | 0.51031 | -0.20953965 | 0.222 | 0.286 | 1 | 11 |
| slc3a2b | 0.51033 | -0.19332335 | 0.194 | 0.321 | 1 | 11 |
| cox17 | 0.51036 | 0.176239858 | 0.194 | 0.107 | 1 | 11 |
| snrnp70 | 0.51048 | -0.18831697 | 0.139 | 0.25 | 1 | 11 |
| rab11ba | 0.51094 | -0.15896299 | 0.167 | 0.286 | 1 | 11 |
| pgam1a | 0.51109 | 0.153442876 | 0.222 | 0.143 | 1 | 11 |
| ercc3 | 0.51114 | 0.111285257 | 0.111 | 0.036 | 1 | 11 |
| MRPS12 | 0.51118 | -0.18347192 | 0.139 | 0.25 | 1 | 11 |
| si:ch211-170d8.8 | 0.51124 | -0.20711065 | 0.111 | 0.214 | 1 | 11 |
| si:dkey-276j7.1 | 0.51137 | 0.195937325 | 0.25 | 0.143 | 1 | 11 |
| rab3gap1 | 0.51155 | -0.12218731 | 0.083 | 0.179 | 1 | 11 |
| selenou1a | 0.51163 | -0.11623896 | 0.111 | 0.214 | 1 | 11 |
| cyyr1 | 0.51173 | -0.19607951 | 0.083 | 0.179 | 1 | 11 |
| usf1l | 0.51205 | -0.14262608 | 0.056 | 0.107 | 1 | 11 |
| rbm39b | 0.51238 | -0.20296968 | 0.139 | 0.25 | 1 | 11 |
| pard6b | 0.51243 | 0.155849801 | 0.111 | 0.036 | 1 | 11 |
| lrrfip2 | 0.5127 | 0.112132135 | 0.111 | 0.036 | 1 | 11 |
| myd88 | 0.51283 | 0.158574676 | 0.111 | 0.036 | 1 | 11 |
| slc20a2 | 0.51299 | 0.118617674 | 0.111 | 0.036 | 1 | 11 |
| rtca | 0.513 | 0.14334521 | 0.111 | 0.036 | 1 | 11 |
| frya | 0.51305 | -0.14355646 | 0.139 | 0.25 | 1 | 11 |
| anos1b | 0.51317 | 0.125261096 | 0.111 | 0.036 | 1 | 11 |
| il6st | 0.51326 | 0.211253446 | 0.278 | 0.179 | 1 | 11 |
| glod4 | 0.51343 | 0.113289101 | 0.083 | 0.071 | 1 | 11 |
| si:dkey-283j8.1 | 0.51362 | -0.19052551 | 0.083 | 0.179 | 1 | 11 |
| zfr | 0.51383 | 0.10984129 | 0.111 | 0.036 | 1 | 11 |
| eif1ad | 0.51413 | 0.207689315 | 0.278 | 0.179 | 1 | 11 |
| amfra | 0.51424 | -0.13633057 | 0.083 | 0.179 | 1 | 11 |
| uqcr10 | 0.51427 | 0.193730424 | 0.667 | 0.536 | 1 | 11 |
| slc36a4 | 0.51429 | -0.12673178 | 0.056 | 0.071 | 1 | 11 |
| bloc1s4 | 0.51435 | 0.118435562 | 0.111 | 0.036 | 1 | 11 |
| si:dkey-78l4.14 | 0.51459 | -0.20231527 | 0.111 | 0.214 | 1 | 11 |
| immp2l | 0.51461 | 0.155161215 | 0.083 | 0.036 | 1 | 11 |
| cgnl1 | 0.51505 | -0.16920848 | 0.417 | 0.464 | 1 | 11 |
| arhgap35a | 0.51514 | 0.119967896 | 0.111 | 0.036 | 1 | 11 |
| si:ch211-241b2.1 | 0.51523 | 0.104857399 | 0.111 | 0.107 | 1 | 11 |

| | | | | | | |
|---|---|---|---|---|---|---|
| mrpl16 | 0.51556 | 0.105218337 | 0.111 | 0.036 | 1 | 11 |
| si:dkey-266m15.5 | 0.51566 | 0.108179929 | 0.111 | 0.036 | 1 | 11 |
| kif7 | 0.51571 | 0.14990482 | 0.111 | 0.036 | 1 | 11 |
| ufm1 | 0.51576 | -0.12603431 | 0.083 | 0.179 | 1 | 11 |
| chd4b | 0.51599 | 0.154153409 | 0.222 | 0.179 | 1 | 11 |
| dlat | 0.51623 | 0.110422355 | 0.111 | 0.036 | 1 | 11 |
| FQ311908.1 | 0.5163 | 0.117395308 | 0.111 | 0.036 | 1 | 11 |
| safb | 0.51715 | 0.141283845 | 0.111 | 0.036 | 1 | 11 |
| fam151b | 0.51717 | 0.137216215 | 0.111 | 0.036 | 1 | 11 |
| lgals2a | 0.51736 | 0.190391361 | 0.111 | 0.036 | 1 | 11 |
| egfra | 0.51765 | 0.139518002 | 0.111 | 0.036 | 1 | 11 |
| far1 | 0.51766 | 0.164510364 | 0.139 | 0.071 | 1 | 11 |
| vps41 | 0.51793 | 0.15102857 | 0.111 | 0.036 | 1 | 11 |
| abhd17c | 0.51795 | 0.13480767 | 0.111 | 0.036 | 1 | 11 |
| kmt2e | 0.51795 | 0.1228483 | 0.111 | 0.036 | 1 | 11 |
| col18a1a | 0.51811 | 0.148924727 | 0.111 | 0.036 | 1 | 11 |
| plcd3a | 0.51826 | 0.175624228 | 0.167 | 0.107 | 1 | 11 |
| alox12 | 0.51861 | -0.16316497 | 0.083 | 0.179 | 1 | 11 |
| adnpa | 0.5187 | -0.14818502 | 0.083 | 0.179 | 1 | 11 |
| hmgb3b | 0.51873 | -0.20002304 | 0.139 | 0.214 | 1 | 11 |
| pdia5 | 0.51898 | -0.1275213 | 0.111 | 0.214 | 1 | 11 |
| arhgap31 | 0.51912 | -0.14748395 | 0.083 | 0.179 | 1 | 11 |
| tmem108 | 0.51923 | -0.19161865 | 0.083 | 0.179 | 1 | 11 |
| cd99 | 0.51947 | -0.15463307 | 0.167 | 0.286 | 1 | 11 |
| dohh | 0.51954 | -0.12228246 | 0.111 | 0.214 | 1 | 11 |
| mtap | 0.51964 | 0.157504238 | 0.194 | 0.107 | 1 | 11 |
| ap2s1 | 0.51995 | -0.11572637 | 0.083 | 0.179 | 1 | 11 |
| park7 | 0.52066 | 0.158204865 | 0.194 | 0.107 | 1 | 11 |
| utp3 | 0.52097 | -0.14158142 | 0.083 | 0.179 | 1 | 11 |
| lrrc17 | 0.52102 | -0.11587688 | 0.278 | 0.393 | 1 | 11 |
| micall1a | 0.52132 | -0.17953391 | 0.111 | 0.214 | 1 | 11 |
| foxp1b | 0.52148 | 0.22413706 | 0.25 | 0.143 | 1 | 11 |
| eif2s1a | 0.52155 | -0.17410456 | 0.111 | 0.214 | 1 | 11 |
| ralbp1 | 0.52181 | -0.15076869 | 0.139 | 0.25 | 1 | 11 |
| si:ch211-212k18.4 | 0.52214 | -0.1346225 | 0.083 | 0.179 | 1 | 11 |
| camk2g1 | 0.5222 | -0.12411049 | 0.083 | 0.179 | 1 | 11 |
| srsf6b | 0.52318 | -0.16160302 | 0.167 | 0.286 | 1 | 11 |
| selenot2 | 0.52322 | -0.1528175 | 0.167 | 0.286 | 1 | 11 |
| ccnt2b | 0.52383 | -0.12815061 | 0.083 | 0.179 | 1 | 11 |
| tmem30ab | 0.52391 | -0.13298476 | 0.083 | 0.179 | 1 | 11 |
| zgc:171679 | 0.52392 | -0.14561563 | 0.083 | 0.179 | 1 | 11 |
| pcyox1 | 0.5241 | -0.16355418 | 0.083 | 0.179 | 1 | 11 |
| selenot1a | 0.52466 | -0.11217945 | 0.083 | 0.179 | 1 | 11 |
| ankrd12 | 0.52467 | -0.12530885 | 0.083 | 0.179 | 1 | 11 |
| stom | 0.52487 | -0.16954395 | 0.083 | 0.143 | 1 | 11 |
| rap2b | 0.52497 | -0.15746196 | 0.083 | 0.179 | 1 | 11 |
| anxa11b | 0.52502 | -0.19365385 | 0.333 | 0.429 | 1 | 11 |
| cadm4 | 0.52516 | -0.16490672 | 0.083 | 0.179 | 1 | 11 |

| | | | | | | |
|---|---|---|---|---|---|---|
| marcksa | 0.52519 | -0.1430119 | 0.083 | 0.179 | 1 | 11 |
| col4a3bpb | 0.52521 | -0.13451027 | 0.083 | 0.179 | 1 | 11 |
| bckdhb | 0.52521 | -0.12765977 | 0.083 | 0.179 | 1 | 11 |
| polr2i | 0.52531 | 0.141797722 | 0.222 | 0.143 | 1 | 11 |
| znf385a | 0.52552 | -0.20166956 | 0.361 | 0.5 | 1 | 11 |
| cox14 | 0.52593 | 0.166415982 | 0.278 | 0.179 | 1 | 11 |
| cpxm1a | 0.52602 | 0.1279088 | 0.194 | 0.179 | 1 | 11 |
| polr1d | 0.52635 | 0.142261909 | 0.083 | 0.036 | 1 | 11 |
| lgmn | 0.5264 | -0.13371437 | 0.139 | 0.25 | 1 | 11 |
| si:ch211-74m13.3 | 0.52701 | -0.14683052 | 0.139 | 0.25 | 1 | 11 |
| si:dkey-88p24.11 | 0.52765 | -0.13345406 | 0.111 | 0.214 | 1 | 11 |
| col6a4a | 0.52809 | -0.12917512 | 0.722 | 0.75 | 1 | 11 |
| cenpx | 0.52819 | -0.10306189 | 0.111 | 0.107 | 1 | 11 |
| cnn2 | 0.52884 | 0.212928517 | 0.556 | 0.429 | 1 | 11 |
| rrp1 | 0.52953 | 0.199010377 | 0.333 | 0.214 | 1 | 11 |
| pam | 0.52966 | 0.158638947 | 0.139 | 0.071 | 1 | 11 |
| acbd3 | 0.53 | 0.114679683 | 0.111 | 0.107 | 1 | 11 |
| nr2f2 | 0.53073 | -0.17211086 | 0.139 | 0.25 | 1 | 11 |
| tcea1 | 0.53104 | 0.168569961 | 0.167 | 0.107 | 1 | 11 |
| srrm2 | 0.53187 | -0.15649139 | 0.139 | 0.25 | 1 | 11 |
| ten1 | 0.5323 | -0.15341471 | 0.111 | 0.214 | 1 | 11 |
| tmem9b | 0.5329 | 0.15019093 | 0.167 | 0.107 | 1 | 11 |
| sf3a3 | 0.53292 | -0.14655918 | 0.111 | 0.214 | 1 | 11 |
| chd4a | 0.53306 | -0.10003681 | 0.25 | 0.357 | 1 | 11 |
| b2ml | 0.53368 | 0.111786262 | 0.889 | 0.786 | 1 | 11 |
| ak4 | 0.53369 | -0.18093121 | 0.111 | 0.214 | 1 | 11 |
| si:ch211-204c21.1 | 0.53589 | 0.173197065 | 0.222 | 0.179 | 1 | 11 |
| stmn4 | 0.53614 | 0.109880505 | 0.083 | 0.071 | 1 | 11 |
| ap1m1 | 0.53626 | -0.19731747 | 0.167 | 0.25 | 1 | 11 |
| crip1 | 0.53638 | -0.22145062 | 0.556 | 0.643 | 1 | 11 |
| XLOC-016094 | 0.53641 | 0.161381483 | 0.25 | 0.214 | 1 | 11 |
| zbtb4 | 0.53696 | 0.147394788 | 0.111 | 0.071 | 1 | 11 |
| cdk11b | 0.53698 | 0.120326678 | 0.083 | 0.071 | 1 | 11 |
| trappc5 | 0.53734 | 0.159618951 | 0.139 | 0.071 | 1 | 11 |
| ypel5 | 0.53759 | -0.16821373 | 0.139 | 0.214 | 1 | 11 |
| erbin | 0.53781 | 0.162886124 | 0.167 | 0.107 | 1 | 11 |
| ddx21 | 0.53785 | -0.18694819 | 0.278 | 0.357 | 1 | 11 |
| rpl5a | 0.53816 | -0.12941266 | 0.833 | 0.893 | 1 | 11 |
| si:ch211-198c19.3 | 0.5383 | -0.17898829 | 0.667 | 0.75 | 1 | 11 |
| pbx2 | 0.53899 | 0.153501059 | 0.083 | 0.036 | 1 | 11 |
| rnf7 | 0.53946 | -0.13485183 | 0.333 | 0.357 | 1 | 11 |
| ncoa6 | 0.54053 | 0.160316739 | 0.083 | 0.036 | 1 | 11 |
| ctdsp2 | 0.54072 | 0.176402873 | 0.417 | 0.286 | 1 | 11 |
| chmp6b | 0.54093 | 0.15305966 | 0.25 | 0.179 | 1 | 11 |
| brms1la | 0.54174 | -0.13546556 | 0.056 | 0.071 | 1 | 11 |
| c1galt1b | 0.54183 | 0.106999844 | 0.111 | 0.107 | 1 | 11 |
| igf2b | 0.54194 | -0.16476397 | 0.333 | 0.464 | 1 | 11 |
| meis1b | 0.54267 | -0.17372713 | 0.639 | 0.75 | 1 | 11 |

| | | | | | | |
|---|---|---|---|---|---|---|
| ndufaf1 | 0.54315 | -0.15079387 | 0.083 | 0.107 | 1 | 11 |
| taf9 | 0.54391 | 0.140218187 | 0.139 | 0.107 | 1 | 11 |
| eno1a | 0.54391 | -0.1790887 | 0.333 | 0.464 | 1 | 11 |
| f3b | 0.54427 | -0.21447848 | 0.306 | 0.393 | 1 | 11 |
| sf3a1 | 0.54559 | 0.148643379 | 0.139 | 0.071 | 1 | 11 |
| antxr2a | 0.54591 | 0.129048202 | 0.361 | 0.25 | 1 | 11 |
| wu:fb16f03 | 0.54607 | 0.131812187 | 0.361 | 0.321 | 1 | 11 |
| cab39l1 | 0.54615 | -0.13190137 | 0.056 | 0.071 | 1 | 11 |
| si:ch73-37h15.2 | 0.54743 | 0.166678779 | 0.139 | 0.071 | 1 | 11 |
| actn1 | 0.54746 | 0.115586114 | 0.194 | 0.179 | 1 | 11 |
| znf703 | 0.54799 | 0.12899081 | 0.083 | 0.071 | 1 | 11 |
| col27a1b | 0.54817 | 0.114503558 | 0.333 | 0.321 | 1 | 11 |
| gstr | 0.54909 | -0.17123561 | 0.306 | 0.429 | 1 | 11 |
| mrpl28 | 0.54928 | -0.13799475 | 0.056 | 0.107 | 1 | 11 |
| pa2g4b | 0.54959 | -0.11001211 | 0.278 | 0.393 | 1 | 11 |
| nkap | 0.54985 | 0.215657283 | 0.306 | 0.214 | 1 | 11 |
| ewsr1b | 0.55044 | 0.107028819 | 0.278 | 0.179 | 1 | 11 |
| fsta | 0.5507 | 0.168866404 | 0.333 | 0.214 | 1 | 11 |
| vat1 | 0.55104 | -0.16648687 | 0.472 | 0.607 | 1 | 11 |
| setb | 0.55194 | 0.171598893 | 0.667 | 0.536 | 1 | 11 |
| mt-atp6 | 0.55229 | -0.20331661 | 0.583 | 0.714 | 1 | 11 |
| eif4eb | 0.55233 | 0.148387019 | 0.083 | 0.036 | 1 | 11 |
| CU633999.2 | 0.55264 | 0.150557452 | 0.083 | 0.036 | 1 | 11 |
| atp5fa1 | 0.55286 | -0.14922707 | 0.583 | 0.714 | 1 | 11 |
| creb3l3l | 0.55356 | 0.11196655 | 0.278 | 0.179 | 1 | 11 |
| ddx39ab | 0.55429 | 0.162768716 | 0.139 | 0.071 | 1 | 11 |
| rsu1 | 0.55499 | -0.1735621 | 0.361 | 0.429 | 1 | 11 |
| sco1 | 0.55586 | 0.121562166 | 0.083 | 0.036 | 1 | 11 |
| siah2l | 0.55593 | 0.175867887 | 0.139 | 0.071 | 1 | 11 |
| mrpl53 | 0.55661 | -0.15287662 | 0.111 | 0.179 | 1 | 11 |
| ackr3b | 0.55707 | -0.2062259 | 0.333 | 0.429 | 1 | 11 |
| psmb4 | 0.55708 | -0.10838682 | 0.444 | 0.571 | 1 | 11 |
| zgc:175088 | 0.5572 | 0.194697265 | 0.278 | 0.179 | 1 | 11 |
| os9 | 0.55756 | -0.20013545 | 0.194 | 0.286 | 1 | 11 |
| hm13 | 0.55827 | -0.21319877 | 0.25 | 0.321 | 1 | 11 |
| eif3d | 0.55952 | -0.19230061 | 0.417 | 0.5 | 1 | 11 |
| pnisr | 0.5597 | 0.159711382 | 0.222 | 0.179 | 1 | 11 |
| rbp5 | 0.56037 | 0.201481038 | 0.444 | 0.321 | 1 | 11 |
| XLOC-011085 | 0.56096 | 0.112169621 | 0.083 | 0.071 | 1 | 11 |
| purab | 0.56127 | 0.191013222 | 0.278 | 0.179 | 1 | 11 |
| rp9 | 0.56151 | -0.107998 | 0.139 | 0.143 | 1 | 11 |
| wdr1 | 0.5618 | 0.162373424 | 0.25 | 0.143 | 1 | 11 |
| trib2 | 0.56184 | -0.11924069 | 0.056 | 0.071 | 1 | 11 |
| hint2 | 0.56261 | 0.157785345 | 0.25 | 0.143 | 1 | 11 |
| brd2a | 0.56297 | 0.194838146 | 0.333 | 0.25 | 1 | 11 |
| arpin | 0.56304 | -0.18318037 | 0.111 | 0.179 | 1 | 11 |
| brd3b | 0.56322 | 0.156184275 | 0.306 | 0.25 | 1 | 11 |
| phf5a | 0.5633 | 0.167614935 | 0.25 | 0.143 | 1 | 11 |

| | | | | | | |
|---|---|---|---|---|---|---|
| rnf185 | 0.56378 | 0.158428176 | 0.194 | 0.107 | 1 | 11 |
| mrpl30 | 0.56402 | 0.117687625 | 0.25 | 0.143 | 1 | 11 |
| atf1 | 0.56434 | 0.144682532 | 0.194 | 0.107 | 1 | 11 |
| si:rp71-1k13.6 | 0.56469 | -0.14743442 | 0.056 | 0.107 | 1 | 11 |
| ola1 | 0.56511 | 0.195655017 | 0.472 | 0.357 | 1 | 11 |
| emc7 | 0.56558 | -0.1031441 | 0.167 | 0.179 | 1 | 11 |
| rab8a | 0.56578 | 0.138907893 | 0.083 | 0.036 | 1 | 11 |
| si:ch211-213d14.2 | 0.56579 | -0.15312838 | 0.25 | 0.286 | 1 | 11 |
| tefa | 0.56623 | 0.191727993 | 0.333 | 0.25 | 1 | 11 |
| fbxo7 | 0.56677 | 0.140166975 | 0.083 | 0.036 | 1 | 11 |
| cipcb | 0.5669 | -0.14811093 | 0.083 | 0.107 | 1 | 11 |
| CABZ01087560.1 | 0.56695 | 0.152284812 | 0.194 | 0.143 | 1 | 11 |
| tpst1 | 0.56813 | 0.131739869 | 0.25 | 0.214 | 1 | 11 |
| klhdc3 | 0.56829 | 0.162425144 | 0.139 | 0.071 | 1 | 11 |
| api5 | 0.56836 | 0.158118693 | 0.167 | 0.107 | 1 | 11 |
| atp5f1b | 0.5688 | 0.114759929 | 0.583 | 0.571 | 1 | 11 |
| zfp36l1b | 0.56897 | -0.1830297 | 0.556 | 0.679 | 1 | 11 |
| aip | 0.56906 | -0.18549243 | 0.222 | 0.286 | 1 | 11 |
| XLOC-041269 | 0.57085 | 0.134724947 | 0.278 | 0.179 | 1 | 11 |
| tbca | 0.57198 | 0.108707831 | 0.556 | 0.429 | 1 | 11 |
| sntb2 | 0.57206 | 0.175478473 | 0.222 | 0.143 | 1 | 11 |
| snd1 | 0.57285 | -0.14423761 | 0.167 | 0.214 | 1 | 11 |
| mid1ip1b | 0.57371 | -0.15051175 | 0.056 | 0.107 | 1 | 11 |
| znf609b | 0.57397 | 0.112508567 | 0.167 | 0.143 | 1 | 11 |
| dhx15 | 0.57431 | 0.163328405 | 0.194 | 0.107 | 1 | 11 |
| slc25a44b | 0.57475 | 0.125814948 | 0.139 | 0.107 | 1 | 11 |
| ctsf | 0.57496 | 0.156239577 | 0.389 | 0.321 | 1 | 11 |
| nid1a | 0.5752 | 0.135491677 | 0.139 | 0.107 | 1 | 11 |
| irf2bpl | 0.57531 | 0.190336578 | 0.306 | 0.214 | 1 | 11 |
| fxr2 | 0.57598 | 0.136336325 | 0.083 | 0.036 | 1 | 11 |
| fgd1 | 0.57644 | 0.160379338 | 0.139 | 0.071 | 1 | 11 |
| cbx5 | 0.57662 | -0.19328025 | 0.389 | 0.5 | 1 | 11 |
| lasp1 | 0.57683 | -0.20148386 | 0.306 | 0.429 | 1 | 11 |
| btr09 | 0.57707 | 0.134112945 | 0.083 | 0.036 | 1 | 11 |
| ndufc2 | 0.57946 | 0.19487173 | 0.444 | 0.321 | 1 | 11 |
| tjp1a | 0.5797 | 0.104666724 | 0.444 | 0.429 | 1 | 11 |
| rab3db | 0.58118 | 0.194257204 | 0.278 | 0.179 | 1 | 11 |
| naa20 | 0.58155 | 0.11443669 | 0.194 | 0.179 | 1 | 11 |
| ddx24 | 0.5818 | 0.149351561 | 0.278 | 0.214 | 1 | 11 |
| si:ch211-156j16.1 | 0.58214 | 0.103090536 | 0.75 | 0.75 | 1 | 11 |
| rbm4.2 | 0.58227 | 0.142182139 | 0.167 | 0.107 | 1 | 11 |
| atp5mea | 0.58265 | 0.111228778 | 0.389 | 0.393 | 1 | 11 |
| pdia3 | 0.58273 | -0.16804337 | 0.639 | 0.75 | 1 | 11 |
| leng8 | 0.58309 | 0.157935044 | 0.167 | 0.107 | 1 | 11 |
| chmp2bb | 0.58314 | 0.132080967 | 0.139 | 0.107 | 1 | 11 |
| junba | 0.58358 | 0.170157552 | 0.694 | 0.607 | 1 | 11 |
| ccdc25 | 0.58389 | 0.17193636 | 0.278 | 0.179 | 1 | 11 |
| piezo1 | 0.58407 | 0.100601741 | 0.111 | 0.107 | 1 | 11 |

| | | | | | | |
|---|---|---|---|---|---|---|
| rbpms2a | 0.5844 | 0.129899044 | 0.139 | 0.107 | 1 | 11 |
| eif3ja | 0.5845 | -0.16861275 | 0.306 | 0.429 | 1 | 11 |
| mgst3b | 0.58596 | -0.15699045 | 0.556 | 0.679 | 1 | 11 |
| selenok | 0.58611 | -0.1273649 | 0.222 | 0.25 | 1 | 11 |
| prrc2a | 0.58713 | 0.136358589 | 0.083 | 0.036 | 1 | 11 |
| spcs3 | 0.58753 | 0.153333357 | 0.306 | 0.25 | 1 | 11 |
| hsp90b1 | 0.58803 | -0.12225243 | 0.167 | 0.179 | 1 | 11 |
| tmem256 | 0.5884 | 0.140523319 | 0.194 | 0.107 | 1 | 11 |
| gabpa | 0.58912 | 0.129621033 | 0.111 | 0.071 | 1 | 11 |
| thbs1a | 0.58959 | 0.140654234 | 0.083 | 0.036 | 1 | 11 |
| ctdsplb | 0.59051 | 0.164893073 | 0.139 | 0.071 | 1 | 11 |
| cybrd1 | 0.59104 | 0.11722595 | 0.167 | 0.143 | 1 | 11 |
| zgc:86598 | 0.59123 | 0.170112359 | 0.333 | 0.25 | 1 | 11 |
| lrp1ab | 0.59193 | -0.19371243 | 0.278 | 0.393 | 1 | 11 |
| si:ch73-61d6.3 | 0.59221 | -0.10597667 | 0.139 | 0.143 | 1 | 11 |
| pfdn5 | 0.59249 | 0.167417966 | 0.278 | 0.179 | 1 | 11 |
| snrpa1 | 0.5929 | 0.138876816 | 0.139 | 0.071 | 1 | 11 |
| chd2 | 0.59313 | 0.135327598 | 0.083 | 0.036 | 1 | 11 |
| acox1 | 0.59341 | -0.12985985 | 0.056 | 0.071 | 1 | 11 |
| slc27a1a | 0.59438 | -0.15033129 | 0.083 | 0.143 | 1 | 11 |
| cdc123 | 0.59463 | 0.12972139 | 0.111 | 0.071 | 1 | 11 |
| sars | 0.59506 | 0.105965676 | 0.083 | 0.071 | 1 | 11 |
| trub1 | 0.5959 | 0.117970372 | 0.083 | 0.036 | 1 | 11 |
| cyp2ad3 | 0.59637 | 0.170244528 | 0.583 | 0.5 | 1 | 11 |
| nelfe | 0.59639 | 0.135521491 | 0.083 | 0.036 | 1 | 11 |
| zgc:161969 | 0.59662 | 0.159307361 | 0.139 | 0.071 | 1 | 11 |
| tuba8l | 0.5971 | -0.16600519 | 0.444 | 0.571 | 1 | 11 |
| tmed10 | 0.5974 | -0.18143478 | 0.25 | 0.357 | 1 | 11 |
| col6a2 | 0.59741 | 0.159074056 | 0.556 | 0.429 | 1 | 11 |
| ptges | 0.59753 | -0.20871774 | 0.139 | 0.214 | 1 | 11 |
| qkia | 0.59781 | 0.140629472 | 0.75 | 0.679 | 1 | 11 |
| slc35e1 | 0.5987 | -0.12877354 | 0.056 | 0.071 | 1 | 11 |
| ppp2r1bb | 0.6 | 0.131486424 | 0.139 | 0.107 | 1 | 11 |
| atp5pb | 0.6002 | -0.15055294 | 0.556 | 0.679 | 1 | 11 |
| FP074902.2 | 0.6004 | 0.125633708 | 0.139 | 0.107 | 1 | 11 |
| aadac | 0.6005 | -0.19144157 | 0.056 | 0.107 | 1 | 11 |
| pinx1 | 0.60053 | -0.19894251 | 0.194 | 0.286 | 1 | 11 |
| ndfip1l | 0.60063 | -0.1546204 | 0.278 | 0.393 | 1 | 11 |
| rpa1 | 0.60116 | -0.14565059 | 0.083 | 0.143 | 1 | 11 |
| magt1 | 0.60161 | 0.122923143 | 0.417 | 0.393 | 1 | 11 |
| fbln5 | 0.60173 | -0.12401924 | 0.389 | 0.393 | 1 | 11 |
| spty2d1 | 0.60258 | 0.131437581 | 0.083 | 0.036 | 1 | 11 |
| gatad2ab | 0.60271 | 0.121881299 | 0.083 | 0.036 | 1 | 11 |
| dync1li2 | 0.60281 | -0.11142571 | 0.056 | 0.071 | 1 | 11 |
| si:ch211-195h23.3 | 0.60362 | 0.146155683 | 0.194 | 0.107 | 1 | 11 |
| ndufb8 | 0.60362 | 0.11166078 | 0.472 | 0.464 | 1 | 11 |
| BX284638.1 | 0.60372 | 0.104598538 | 0.083 | 0.071 | 1 | 11 |
| ociad1 | 0.60403 | 0.136495144 | 0.139 | 0.071 | 1 | 11 |

| | | | | | | |
|---|---|---|---|---|---|---|
| mrps18a | 0.60513 | 0.14019372 | 0.139 | 0.071 | 1 | 11 |
| si:dkey-239h2.3 | 0.60546 | -0.15908071 | 0.278 | 0.321 | 1 | 11 |
| tsc22d3 | 0.60555 | -0.14620852 | 0.333 | 0.393 | 1 | 11 |
| hnrnpa0l | 0.60556 | 0.122022579 | 0.833 | 0.786 | 1 | 11 |
| gps2 | 0.60561 | -0.14450364 | 0.056 | 0.107 | 1 | 11 |
| si:ch73-86n18.1 | 0.60568 | -0.1009336 | 0.056 | 0.071 | 1 | 11 |
| wipi1 | 0.60619 | -0.16695989 | 0.056 | 0.107 | 1 | 11 |
| atp6v0ca | 0.60718 | 0.111698513 | 0.278 | 0.179 | 1 | 11 |
| vta1 | 0.60744 | 0.123700649 | 0.278 | 0.179 | 1 | 11 |
| si:ch73-340n8.4 | 0.6076 | 0.126584882 | 0.139 | 0.107 | 1 | 11 |
| col4a2 | 0.60794 | 0.151776009 | 0.194 | 0.107 | 1 | 11 |
| fam168a | 0.60838 | 0.179702007 | 0.417 | 0.321 | 1 | 11 |
| tm2d3 | 0.60857 | 0.14103538 | 0.167 | 0.107 | 1 | 11 |
| litaf | 0.60872 | 0.104353177 | 0.306 | 0.286 | 1 | 11 |
| llgl1 | 0.60994 | 0.181164699 | 0.194 | 0.107 | 1 | 11 |
| fzd4 | 0.61012 | 0.210601051 | 0.278 | 0.179 | 1 | 11 |
| ap2a1 | 0.6104 | 0.159048538 | 0.139 | 0.071 | 1 | 11 |
| ube2ib | 0.61066 | -0.1414081 | 0.278 | 0.393 | 1 | 11 |
| brd4 | 0.61166 | 0.119378023 | 0.194 | 0.107 | 1 | 11 |
| serinc2l | 0.61182 | -0.181398 | 0.083 | 0.143 | 1 | 11 |
| spns1 | 0.61184 | -0.12159573 | 0.083 | 0.107 | 1 | 11 |
| aktip | 0.61261 | 0.114438019 | 0.194 | 0.107 | 1 | 11 |
| polr2f | 0.61388 | 0.129217512 | 0.25 | 0.214 | 1 | 11 |
| tmem98 | 0.61419 | 0.139275175 | 0.5 | 0.464 | 1 | 11 |
| cxcl8b.3 | 0.61432 | -0.14997551 | 0.222 | 0.25 | 1 | 11 |
| tlk1b | 0.61454 | 0.128580647 | 0.083 | 0.036 | 1 | 11 |
| huwe1 | 0.61463 | 0.141295195 | 0.222 | 0.179 | 1 | 11 |
| wu:fi37e09 | 0.61589 | -0.12049302 | 0.25 | 0.286 | 1 | 11 |
| glulb | 0.61606 | -0.13337269 | 0.111 | 0.143 | 1 | 11 |
| si:dkey-46a10.3 | 0.61621 | 0.124099374 | 0.111 | 0.071 | 1 | 11 |
| serpina10a | 0.61699 | 0.170324219 | 0.194 | 0.107 | 1 | 11 |
| lamb1a | 0.6176 | 0.125893651 | 0.389 | 0.357 | 1 | 11 |
| hoxc3a | 0.61782 | 0.18588084 | 0.333 | 0.25 | 1 | 11 |
| si:ch211-108d22.2 | 0.61795 | 0.133943799 | 0.083 | 0.036 | 1 | 11 |
| npc1 | 0.61824 | 0.106445006 | 0.083 | 0.071 | 1 | 11 |
| march6 | 0.61841 | 0.133469636 | 0.083 | 0.036 | 1 | 11 |
| aimp1 | 0.61873 | -0.15738964 | 0.167 | 0.214 | 1 | 11 |
| tomm70a | 0.61891 | 0.117323678 | 0.278 | 0.179 | 1 | 11 |
| ppp2r5d | 0.61907 | 0.181543232 | 0.194 | 0.107 | 1 | 11 |
| yeats4 | 0.6192 | 0.107033218 | 0.194 | 0.107 | 1 | 11 |
| bckdk | 0.61941 | -0.13919026 | 0.111 | 0.179 | 1 | 11 |
| efna1a | 0.61963 | 0.151800374 | 0.194 | 0.107 | 1 | 11 |
| si:ch211-260e23.9 | 0.61971 | 0.129165648 | 0.139 | 0.071 | 1 | 11 |
| si:ch211-126j24.1 | 0.62069 | 0.134741442 | 0.083 | 0.036 | 1 | 11 |
| psmg3 | 0.62161 | -0.1388799 | 0.278 | 0.393 | 1 | 11 |
| rrp15 | 0.6217 | 0.160668447 | 0.167 | 0.107 | 1 | 11 |
| ipmkb | 0.62173 | 0.143180453 | 0.111 | 0.071 | 1 | 11 |
| hipk3a | 0.62203 | 0.135513762 | 0.083 | 0.036 | 1 | 11 |

| | | | | | | |
|---|---|---|---|---|---|---|
| ATP11B | 0.62288 | 0.117208367 | 0.167 | 0.143 | 1 | 11 |
| cd63 | 0.62302 | 0.141076893 | 0.722 | 0.607 | 1 | 11 |
| pdcd10b | 0.62302 | 0.138575375 | 0.139 | 0.071 | 1 | 11 |
| asb8 | 0.62305 | -0.11377136 | 0.056 | 0.107 | 1 | 11 |
| ptprsa | 0.6232 | 0.1152 | 0.333 | 0.321 | 1 | 11 |
| XLOC-000431 | 0.62329 | -0.19311547 | 0.25 | 0.357 | 1 | 11 |
| npr2 | 0.62352 | 0.163468806 | 0.222 | 0.143 | 1 | 11 |
| sult1st1 | 0.62416 | -0.14343669 | 0.083 | 0.143 | 1 | 11 |
| wdr45b | 0.6242 | -0.1218827 | 0.111 | 0.143 | 1 | 11 |
| cyp2n13 | 0.62443 | -0.16253776 | 0.278 | 0.393 | 1 | 11 |
| ndufs5 | 0.62471 | -0.11389939 | 0.278 | 0.393 | 1 | 11 |
| eno3 | 0.62472 | -0.1423014 | 0.278 | 0.393 | 1 | 11 |
| sdha | 0.62506 | 0.120165759 | 0.194 | 0.107 | 1 | 11 |
| arhgef1b | 0.62552 | -0.17098194 | 0.194 | 0.286 | 1 | 11 |
| smim7 | 0.6256 | 0.129909409 | 0.194 | 0.107 | 1 | 11 |
| hnrnph1l | 0.62562 | 0.107996715 | 0.278 | 0.25 | 1 | 11 |
| sppl3 | 0.62565 | 0.15457709 | 0.194 | 0.107 | 1 | 11 |
| kdm6bb | 0.62571 | 0.101220995 | 0.389 | 0.286 | 1 | 11 |
| krt18a.1 | 0.62582 | -0.11218608 | 0.806 | 0.893 | 1 | 11 |
| ywhag1 | 0.62622 | 0.123013282 | 0.194 | 0.107 | 1 | 11 |
| ddit3 | 0.6265 | 0.134769375 | 0.194 | 0.107 | 1 | 11 |
| dpm3 | 0.62664 | 0.130523163 | 0.083 | 0.036 | 1 | 11 |
| ebag9 | 0.62667 | 0.104554948 | 0.194 | 0.107 | 1 | 11 |
| copa | 0.6269 | 0.128711512 | 0.139 | 0.107 | 1 | 11 |
| glud1a | 0.62721 | -0.14104563 | 0.389 | 0.429 | 1 | 11 |
| cul3b | 0.6276 | 0.116692477 | 0.083 | 0.036 | 1 | 11 |
| afap1 | 0.62801 | 0.126476374 | 0.111 | 0.071 | 1 | 11 |
| arpc4l | 0.6285 | 0.162827362 | 0.472 | 0.393 | 1 | 11 |
| mrps18c | 0.62885 | 0.125950397 | 0.167 | 0.143 | 1 | 11 |
| arpp19b | 0.62902 | 0.113580395 | 0.111 | 0.071 | 1 | 11 |
| cnot3a | 0.6293 | 0.126101934 | 0.083 | 0.036 | 1 | 11 |
| akt2l | 0.62969 | -0.12824929 | 0.111 | 0.143 | 1 | 11 |
| ntn1a | 0.62981 | 0.157418347 | 0.139 | 0.071 | 1 | 11 |
| smim29 | 0.62991 | 0.107502007 | 0.083 | 0.036 | 1 | 11 |
| pudp | 0.62999 | 0.131112783 | 0.139 | 0.107 | 1 | 11 |
| rprd1a | 0.63043 | 0.127669848 | 0.083 | 0.036 | 1 | 11 |
| atp6v1ba | 0.6308 | 0.107772092 | 0.083 | 0.036 | 1 | 11 |
| dda1 | 0.63082 | 0.149890608 | 0.167 | 0.107 | 1 | 11 |
| aldh18a1 | 0.63109 | 0.132242882 | 0.083 | 0.036 | 1 | 11 |
| ppm1g | 0.63131 | 0.110691173 | 0.139 | 0.107 | 1 | 11 |
| anp32a | 0.63212 | 0.17605162 | 0.472 | 0.357 | 1 | 11 |
| zbtb10 | 0.6324 | -0.14877083 | 0.083 | 0.143 | 1 | 11 |
| zgc:162344 | 0.63245 | 0.111036549 | 0.083 | 0.071 | 1 | 11 |
| rpl7 | 0.63251 | 0.113393992 | 0.917 | 0.857 | 1 | 11 |
| mcmbp | 0.6327 | 0.121882172 | 0.083 | 0.036 | 1 | 11 |
| CU694197.1 | 0.63295 | 0.141747304 | 0.361 | 0.25 | 1 | 11 |
| MED9 | 0.63376 | 0.119555908 | 0.111 | 0.071 | 1 | 11 |
| gps1 | 0.63382 | -0.1372369 | 0.056 | 0.107 | 1 | 11 |

| | | | | | | |
|---|---|---|---|---|---|---|
| fam171a2a | 0.63435 | -0.13218374 | 0.25 | 0.357 | 1 | 11 |
| cct7 | 0.63456 | -0.1020425 | 0.389 | 0.5 | 1 | 11 |
| sept12 | 0.63465 | -0.14025234 | 0.083 | 0.107 | 1 | 11 |
| trioa | 0.63509 | 0.129922064 | 0.083 | 0.036 | 1 | 11 |
| ddr2l | 0.63515 | 0.127280075 | 0.083 | 0.036 | 1 | 11 |
| sestd1 | 0.63525 | 0.129349759 | 0.083 | 0.036 | 1 | 11 |
| si:ch211-255f4.5 | 0.63552 | 0.108309056 | 0.083 | 0.036 | 1 | 11 |
| ralbb | 0.63625 | 0.123541504 | 0.111 | 0.071 | 1 | 11 |
| edf1 | 0.63636 | -0.11233633 | 0.528 | 0.643 | 1 | 11 |
| mxra8a | 0.63656 | -0.1152224 | 0.222 | 0.321 | 1 | 11 |
| gpatch8 | 0.63699 | 0.169105656 | 0.222 | 0.143 | 1 | 11 |
| yipf4 | 0.63709 | 0.150285547 | 0.278 | 0.214 | 1 | 11 |
| idh3g | 0.63731 | 0.124333475 | 0.083 | 0.036 | 1 | 11 |
| si:ch211-15b10.6 | 0.63752 | 0.135512868 | 0.111 | 0.071 | 1 | 11 |
| chka | 0.63755 | -0.14533192 | 0.139 | 0.214 | 1 | 11 |
| ranbp3b | 0.6382 | 0.133268751 | 0.111 | 0.071 | 1 | 11 |
| auts2a | 0.63844 | 0.159027216 | 0.139 | 0.071 | 1 | 11 |
| nfia | 0.63882 | 0.152080959 | 0.583 | 0.464 | 1 | 11 |
| her9 | 0.63926 | 0.162874844 | 0.278 | 0.179 | 1 | 11 |
| cul1a | 0.63985 | 0.139209472 | 0.139 | 0.071 | 1 | 11 |
| appa | 0.63992 | 0.175016243 | 0.278 | 0.179 | 1 | 11 |
| serinc5 | 0.64052 | 0.137271486 | 0.194 | 0.143 | 1 | 11 |
| sv2bb | 0.64096 | 0.12523716 | 0.111 | 0.071 | 1 | 11 |
| ndufa7 | 0.64098 | 0.145710997 | 0.194 | 0.143 | 1 | 11 |
| ms4a17a.9 | 0.64187 | 0.126251744 | 0.278 | 0.179 | 1 | 11 |
| gsk3aa | 0.64205 | 0.103173224 | 0.083 | 0.071 | 1 | 11 |
| si:ch211-166a6.5 | 0.64219 | -0.14482609 | 0.25 | 0.357 | 1 | 11 |
| pfn2l | 0.6424 | -0.12192966 | 0.528 | 0.643 | 1 | 11 |
| sec62 | 0.64261 | 0.157637036 | 0.222 | 0.179 | 1 | 11 |
| sdhc | 0.64276 | 0.121013881 | 0.194 | 0.143 | 1 | 11 |
| fosab | 0.64307 | 0.121506198 | 0.75 | 0.714 | 1 | 11 |
| slc25a43 | 0.64314 | 0.12430565 | 0.139 | 0.071 | 1 | 11 |
| pou2f1b | 0.64322 | 0.122380762 | 0.083 | 0.036 | 1 | 11 |
| plekhg4 | 0.64364 | 0.128725559 | 0.083 | 0.036 | 1 | 11 |
| med10 | 0.64375 | 0.101522085 | 0.389 | 0.286 | 1 | 11 |
| slmapb | 0.64381 | 0.160892042 | 0.278 | 0.179 | 1 | 11 |
| hipk3b | 0.64386 | 0.123473123 | 0.139 | 0.107 | 1 | 11 |
| XLOC-032256 | 0.64419 | 0.124618624 | 0.083 | 0.036 | 1 | 11 |
| map2k4a | 0.64431 | 0.140350553 | 0.139 | 0.071 | 1 | 11 |
| pkn1a | 0.6452 | 0.132533436 | 0.083 | 0.036 | 1 | 11 |
| LOC108180269 | 0.6452 | 0.148662958 | 0.139 | 0.071 | 1 | 11 |
| tmem263 | 0.64551 | 0.115580121 | 0.083 | 0.036 | 1 | 11 |
| sdc4 | 0.64595 | -0.16135529 | 0.389 | 0.464 | 1 | 11 |
| hs2st1b | 0.64614 | 0.118851656 | 0.083 | 0.036 | 1 | 11 |
| fbxl7 | 0.64672 | 0.13022456 | 0.139 | 0.071 | 1 | 11 |
| nr1d2b | 0.64685 | -0.16029246 | 0.111 | 0.179 | 1 | 11 |
| mettl23 | 0.64688 | 0.116728834 | 0.083 | 0.036 | 1 | 11 |
| fndc3ba | 0.64744 | -0.15393123 | 0.139 | 0.214 | 1 | 11 |

| | | | | | | |
|---|---|---|---|---|---|---|
| jupa | 0.64821 | -0.15625316 | 0.333 | 0.393 | 1 | 11 |
| mrps24 | 0.64844 | -0.12878567 | 0.083 | 0.143 | 1 | 11 |
| zgc:110591 | 0.64844 | 0.114570873 | 0.083 | 0.036 | 1 | 11 |
| anxa11a | 0.64878 | -0.14935844 | 0.25 | 0.357 | 1 | 11 |
| ctcf | 0.65038 | 0.127052837 | 0.139 | 0.071 | 1 | 11 |
| si:dkey-112a7.4 | 0.65071 | -0.12471128 | 0.056 | 0.107 | 1 | 11 |
| rchy1 | 0.65074 | 0.1227835 | 0.139 | 0.071 | 1 | 11 |
| apopt1 | 0.65078 | 0.106728323 | 0.083 | 0.036 | 1 | 11 |
| arglu1a | 0.65115 | -0.15196555 | 0.194 | 0.25 | 1 | 11 |
| lgals17 | 0.65138 | 0.117358657 | 0.083 | 0.036 | 1 | 11 |
| ap3s1.1 | 0.6519 | -0.11413834 | 0.167 | 0.179 | 1 | 11 |
| coq6 | 0.65199 | 0.119031684 | 0.083 | 0.036 | 1 | 11 |
| CR388029.1 | 0.65226 | 0.121309311 | 0.083 | 0.036 | 1 | 11 |
| cyp1c2 | 0.65248 | -0.1518637 | 0.139 | 0.179 | 1 | 11 |
| stag2b | 0.65274 | 0.130273047 | 0.139 | 0.071 | 1 | 11 |
| reck | 0.65377 | -0.11756304 | 0.111 | 0.143 | 1 | 11 |
| mrps15 | 0.65465 | 0.109608186 | 0.083 | 0.036 | 1 | 11 |
| acvr1ba | 0.65487 | 0.131400965 | 0.167 | 0.107 | 1 | 11 |
| spop | 0.65496 | 0.110259546 | 0.083 | 0.036 | 1 | 11 |
| XLOC-037780 | 0.65504 | 0.1241916 | 0.139 | 0.107 | 1 | 11 |
| cdkal1 | 0.65512 | 0.116293887 | 0.083 | 0.036 | 1 | 11 |
| hand2 | 0.65524 | -0.16413769 | 0.083 | 0.143 | 1 | 11 |
| lamc1 | 0.65531 | 0.137223585 | 0.222 | 0.179 | 1 | 11 |
| mknk2b | 0.65536 | 0.150483636 | 0.611 | 0.5 | 1 | 11 |
| mmp14a | 0.65612 | -0.14500379 | 0.167 | 0.25 | 1 | 11 |
| ppig | 0.65643 | 0.108733264 | 0.139 | 0.071 | 1 | 11 |
| eef1da | 0.65648 | 0.116765568 | 0.75 | 0.714 | 1 | 11 |
| rprd2b | 0.65694 | 0.12911656 | 0.139 | 0.107 | 1 | 11 |
| bms1 | 0.65747 | 0.105862942 | 0.083 | 0.036 | 1 | 11 |
| LOC101885018 | 0.65834 | 0.120945907 | 0.083 | 0.036 | 1 | 11 |
| prdm11 | 0.65851 | 0.114638198 | 0.083 | 0.036 | 1 | 11 |
| jam3b | 0.65874 | -0.12920335 | 0.083 | 0.107 | 1 | 11 |
| si:dkey-40c11.2 | 0.65882 | 0.146638165 | 0.222 | 0.143 | 1 | 11 |
| sfswap | 0.65885 | -0.13347025 | 0.056 | 0.107 | 1 | 11 |
| rbfox1 | 0.65894 | -0.16191749 | 0.056 | 0.107 | 1 | 11 |
| nf2b | 0.66001 | 0.12904001 | 0.194 | 0.143 | 1 | 11 |
| parp1 | 0.66243 | 0.136468394 | 0.139 | 0.071 | 1 | 11 |
| rabac1 | 0.66245 | 0.127174414 | 0.139 | 0.071 | 1 | 11 |
| rtf2 | 0.66267 | -0.11778332 | 0.139 | 0.179 | 1 | 11 |
| mmp2 | 0.66342 | 0.125024706 | 0.833 | 0.75 | 1 | 11 |
| klhl13 | 0.66351 | 0.101586318 | 0.083 | 0.036 | 1 | 11 |
| zgc:114181 | 0.66354 | 0.134764535 | 0.194 | 0.143 | 1 | 11 |
| srpk1a | 0.66383 | 0.102767812 | 0.083 | 0.036 | 1 | 11 |
| ubc | 0.66395 | 0.117841688 | 0.861 | 0.821 | 1 | 11 |
| pkp3b | 0.66408 | -0.10451973 | 0.5 | 0.607 | 1 | 11 |
| ncalda | 0.66465 | 0.107076748 | 0.083 | 0.036 | 1 | 11 |
| sec24c | 0.66492 | 0.117205926 | 0.083 | 0.036 | 1 | 11 |
| sarnp | 0.66537 | -0.15333078 | 0.194 | 0.286 | 1 | 11 |

| | | | | | | |
|---|---|---|---|---|---|---|
| rbm7 | 0.6666 | 0.115427698 | 0.083 | 0.036 | 1 | 11 |
| hist2h2l | 0.66662 | -0.14345668 | 0.389 | 0.429 | 1 | 11 |
| ywhabb | 0.66696 | -0.15364619 | 0.5 | 0.607 | 1 | 11 |
| per2 | 0.66726 | 0.110630212 | 0.083 | 0.036 | 1 | 11 |
| ubtd2 | 0.66742 | 0.106674166 | 0.139 | 0.071 | 1 | 11 |
| erlec1 | 0.66754 | 0.139845268 | 0.167 | 0.107 | 1 | 11 |
| grnb | 0.66781 | 0.105335068 | 0.139 | 0.071 | 1 | 11 |
| sh3pxd2b | 0.66842 | -0.11368582 | 0.222 | 0.321 | 1 | 11 |
| gsk3b | 0.66849 | -0.15043682 | 0.056 | 0.107 | 1 | 11 |
| zyx | 0.66879 | 0.11077532 | 0.361 | 0.321 | 1 | 11 |
| stard7 | 0.66906 | 0.103423109 | 0.083 | 0.036 | 1 | 11 |
| abhd14b | 0.66938 | 0.108101202 | 0.083 | 0.036 | 1 | 11 |
| ndufa11 | 0.66999 | -0.11967194 | 0.222 | 0.321 | 1 | 11 |
| arid2 | 0.67004 | -0.13260233 | 0.083 | 0.143 | 1 | 11 |
| tax1bp3 | 0.67009 | 0.164817624 | 0.389 | 0.286 | 1 | 11 |
| taf13 | 0.67068 | 0.106028182 | 0.417 | 0.393 | 1 | 11 |
| lamtor4 | 0.67093 | 0.138478575 | 0.167 | 0.107 | 1 | 11 |
| mrpl23 | 0.671 | 0.116270897 | 0.083 | 0.036 | 1 | 11 |
| mt-atp8 | 0.67118 | 0.112701278 | 0.083 | 0.036 | 1 | 11 |
| sgms1 | 0.67187 | -0.11315095 | 0.056 | 0.071 | 1 | 11 |
| caprin1b | 0.67218 | -0.15958006 | 0.333 | 0.429 | 1 | 11 |
| mapkap1 | 0.67247 | 0.129106224 | 0.083 | 0.036 | 1 | 11 |
| si:dkey-19e4.5 | 0.67256 | 0.110942574 | 0.083 | 0.036 | 1 | 11 |
| cnot3b | 0.67284 | -0.13159218 | 0.083 | 0.143 | 1 | 11 |
| tomm20b | 0.67319 | 0.146039588 | 0.389 | 0.321 | 1 | 11 |
| ost4 | 0.67399 | -0.13370054 | 0.389 | 0.5 | 1 | 11 |
| LO018188.1 | 0.67402 | 0.152328933 | 0.389 | 0.286 | 1 | 11 |
| zfyve27 | 0.67464 | 0.118006635 | 0.139 | 0.071 | 1 | 11 |
| nfyc | 0.67505 | 0.11661945 | 0.083 | 0.036 | 1 | 11 |
| lyrm9 | 0.67507 | -0.13958607 | 0.056 | 0.107 | 1 | 11 |
| jak2a | 0.67523 | 0.102455345 | 0.083 | 0.036 | 1 | 11 |
| nckap1 | 0.6762 | -0.13915131 | 0.139 | 0.214 | 1 | 11 |
| adamts2 | 0.67668 | 0.149937572 | 0.194 | 0.143 | 1 | 11 |
| wdr12 | 0.6768 | 0.108428385 | 0.139 | 0.107 | 1 | 11 |
| tsku | 0.67703 | 0.129971323 | 0.139 | 0.071 | 1 | 11 |
| atf6 | 0.6776 | 0.128266484 | 0.083 | 0.036 | 1 | 11 |
| coro1ca | 0.67821 | 0.164989078 | 0.333 | 0.25 | 1 | 11 |
| naa10 | 0.67833 | 0.114236362 | 0.361 | 0.321 | 1 | 11 |
| rab5ab | 0.67854 | 0.104572941 | 0.306 | 0.214 | 1 | 11 |
| mapk3 | 0.67855 | 0.135590845 | 0.139 | 0.071 | 1 | 11 |
| sh3gl1b | 0.67866 | 0.128585401 | 0.194 | 0.143 | 1 | 11 |
| sbf1 | 0.67918 | 0.132358031 | 0.111 | 0.071 | 1 | 11 |
| akirin2 | 0.67939 | 0.112842129 | 0.139 | 0.071 | 1 | 11 |
| sdf2 | 0.6795 | 0.11256574 | 0.083 | 0.036 | 1 | 11 |
| ndrg3a | 0.67951 | 0.115020997 | 0.083 | 0.036 | 1 | 11 |
| ninl | 0.68054 | 0.125540604 | 0.111 | 0.071 | 1 | 11 |
| si:dkeyp-19e1.3 | 0.6809 | 0.102062857 | 0.083 | 0.036 | 1 | 11 |
| kat6a | 0.68107 | 0.114461573 | 0.139 | 0.071 | 1 | 11 |

| | | | | | | |
|---|---|---|---|---|---|---|
| cndp2 | 0.68108 | -0.12207729 | 0.111 | 0.143 | 1 | 11 |
| mycbp2 | 0.68113 | 0.14470816 | 0.139 | 0.071 | 1 | 11 |
| cemip | 0.68117 | -0.16237883 | 0.056 | 0.107 | 1 | 11 |
| cfdp1 | 0.68173 | 0.134109247 | 0.167 | 0.107 | 1 | 11 |
| hcfc1a | 0.68174 | 0.103113202 | 0.083 | 0.036 | 1 | 11 |
| polr2c | 0.68175 | 0.103648274 | 0.139 | 0.071 | 1 | 11 |
| trir | 0.68237 | 0.107686263 | 0.139 | 0.071 | 1 | 11 |
| mrgbp | 0.68242 | -0.15522513 | 0.111 | 0.179 | 1 | 11 |
| LOC110439416 | 0.68251 | 0.133939886 | 0.139 | 0.071 | 1 | 11 |
| zgc:100832 | 0.68266 | 0.114003951 | 0.083 | 0.036 | 1 | 11 |
| si:ch211-218g4.2 | 0.68275 | 0.113879234 | 0.139 | 0.071 | 1 | 11 |
| sod1 | 0.6835 | -0.10598015 | 0.5 | 0.607 | 1 | 11 |
| p3h3 | 0.68381 | 0.130947615 | 0.111 | 0.071 | 1 | 11 |
| pcmtd1 | 0.68393 | -0.12228136 | 0.056 | 0.107 | 1 | 11 |
| pcca | 0.68403 | 0.115500291 | 0.139 | 0.071 | 1 | 11 |
| mrps6 | 0.68412 | 0.10395517 | 0.111 | 0.071 | 1 | 11 |
| nfic | 0.68474 | -0.1676088 | 0.194 | 0.286 | 1 | 11 |
| pdgfaa | 0.68484 | 0.113605577 | 0.139 | 0.071 | 1 | 11 |
| atp2a2b | 0.68491 | 0.105887029 | 0.139 | 0.071 | 1 | 11 |
| nr1d2a | 0.68503 | -0.13555683 | 0.278 | 0.321 | 1 | 11 |
| aga | 0.68512 | 0.102760093 | 0.139 | 0.071 | 1 | 11 |
| rgl1 | 0.68522 | 0.110312873 | 0.139 | 0.071 | 1 | 11 |
| nap1l1 | 0.68529 | -0.14139661 | 0.5 | 0.607 | 1 | 11 |
| nfkbie | 0.68532 | -0.11786519 | 0.083 | 0.143 | 1 | 11 |
| bnip3lb | 0.68538 | -0.13810483 | 0.167 | 0.214 | 1 | 11 |
| fzr1a | 0.68584 | 0.122865749 | 0.083 | 0.036 | 1 | 11 |
| ppp2cb | 0.68778 | 0.146827386 | 0.389 | 0.286 | 1 | 11 |
| eif4ebp1 | 0.68785 | 0.130787587 | 0.417 | 0.357 | 1 | 11 |
| nck1a | 0.688 | -0.10912924 | 0.056 | 0.107 | 1 | 11 |
| noc2l | 0.68865 | -0.13627309 | 0.083 | 0.143 | 1 | 11 |
| drap1 | 0.68867 | -0.12001705 | 0.306 | 0.321 | 1 | 11 |
| smad3b | 0.68868 | 0.103621543 | 0.083 | 0.036 | 1 | 11 |
| CABZ01083943.1 | 0.68893 | -0.13823314 | 0.056 | 0.107 | 1 | 11 |
| naa30 | 0.68927 | -0.11653636 | 0.083 | 0.107 | 1 | 11 |
| med13a | 0.69012 | 0.101733292 | 0.083 | 0.036 | 1 | 11 |
| urb1 | 0.69023 | 0.103058812 | 0.083 | 0.036 | 1 | 11 |
| rnd3b | 0.69077 | -0.1527848 | 0.361 | 0.464 | 1 | 11 |
| dpm2 | 0.69099 | -0.12138697 | 0.083 | 0.143 | 1 | 11 |
| itga1 | 0.69125 | 0.156790413 | 0.222 | 0.143 | 1 | 11 |
| tia1l | 0.69193 | -0.14385279 | 0.167 | 0.25 | 1 | 11 |
| thoc1 | 0.69194 | -0.11261812 | 0.056 | 0.107 | 1 | 11 |
| iqgap1 | 0.69206 | -0.11439527 | 0.194 | 0.286 | 1 | 11 |
| ing4 | 0.69225 | 0.10239499 | 0.167 | 0.143 | 1 | 11 |
| eif2a | 0.69244 | 0.104810568 | 0.083 | 0.036 | 1 | 11 |
| hnrnpa0b | 0.69264 | -0.13384014 | 0.5 | 0.607 | 1 | 11 |
| adam17a | 0.69276 | 0.100499305 | 0.083 | 0.036 | 1 | 11 |
| cyp2ad6 | 0.69292 | -0.13525638 | 0.056 | 0.107 | 1 | 11 |
| pcm1 | 0.69364 | 0.102726317 | 0.111 | 0.071 | 1 | 11 |

| | | | | | | |
|---|---|---|---|---|---|---|
| nid1b | 0.69369 | -0.12480558 | 0.194 | 0.286 | 1 | 11 |
| selenot1b | 0.69372 | -0.10034608 | 0.083 | 0.107 | 1 | 11 |
| rbmx2 | 0.69417 | 0.109429374 | 0.083 | 0.036 | 1 | 11 |
| fam3c | 0.69466 | -0.13731547 | 0.194 | 0.286 | 1 | 11 |
| caska | 0.69471 | 0.122007617 | 0.111 | 0.071 | 1 | 11 |
| ddx51 | 0.69485 | -0.12515369 | 0.056 | 0.107 | 1 | 11 |
| fam234a | 0.69632 | 0.106454165 | 0.083 | 0.036 | 1 | 11 |
| rbx1 | 0.6973 | 0.160192737 | 0.472 | 0.393 | 1 | 11 |
| srsf6a | 0.69731 | -0.13175984 | 0.111 | 0.179 | 1 | 11 |
| rsf1b.1 | 0.69743 | 0.130517169 | 0.222 | 0.179 | 1 | 11 |
| PCP4 | 0.69763 | 0.10182487 | 0.083 | 0.036 | 1 | 11 |
| blcap | 0.69817 | -0.13952212 | 0.278 | 0.321 | 1 | 11 |
| ube2e2 | 0.69879 | 0.132588654 | 0.278 | 0.214 | 1 | 11 |
| zgc:136864 | 0.69947 | -0.10452108 | 0.056 | 0.107 | 1 | 11 |
| snx7 | 0.69962 | 0.154531367 | 0.167 | 0.107 | 1 | 11 |
| si:dkey-240h12.3 | 0.6998 | -0.13068785 | 0.083 | 0.143 | 1 | 11 |
| aamp | 0.70011 | 0.144587174 | 0.194 | 0.143 | 1 | 11 |
| LOC101886015 | 0.70065 | 0.107078834 | 0.083 | 0.036 | 1 | 11 |
| sema3d | 0.70086 | 0.132789506 | 0.306 | 0.214 | 1 | 11 |
| cdr2l | 0.70088 | -0.15207776 | 0.139 | 0.214 | 1 | 11 |
| sucla2 | 0.7012 | -0.10516355 | 0.083 | 0.143 | 1 | 11 |
| foxj2 | 0.70175 | -0.12995978 | 0.083 | 0.143 | 1 | 11 |
| map2k6 | 0.70205 | 0.149930795 | 0.417 | 0.321 | 1 | 11 |
| usp1 | 0.70205 | 0.117956722 | 0.167 | 0.107 | 1 | 11 |
| vasna | 0.70211 | 0.156677896 | 0.306 | 0.214 | 1 | 11 |
| nid2a | 0.70228 | 0.121217827 | 0.306 | 0.214 | 1 | 11 |
| ehd2b | 0.70254 | -0.15806496 | 0.472 | 0.571 | 1 | 11 |
| srfbp1 | 0.70288 | 0.119373143 | 0.111 | 0.071 | 1 | 11 |
| LO016987.2 | 0.70308 | -0.12563017 | 0.056 | 0.107 | 1 | 11 |
| FO744854.1 | 0.70376 | -0.12421843 | 0.056 | 0.107 | 1 | 11 |
| dspa | 0.70401 | 0.163028449 | 0.25 | 0.179 | 1 | 11 |
| ubap2l | 0.70431 | 0.123475509 | 0.222 | 0.143 | 1 | 11 |
| rasgrp3 | 0.70599 | 0.118062796 | 0.083 | 0.036 | 1 | 11 |
| sept7a | 0.70601 | 0.123586422 | 0.194 | 0.143 | 1 | 11 |
| pabpn1 | 0.70648 | 0.118736304 | 0.639 | 0.536 | 1 | 11 |
| hacd2 | 0.70651 | 0.119494329 | 0.083 | 0.036 | 1 | 11 |
| pbx1b | 0.70733 | 0.106397918 | 0.111 | 0.071 | 1 | 11 |
| plxna1a | 0.70736 | 0.135589201 | 0.194 | 0.143 | 1 | 11 |
| atp5f1e | 0.70791 | 0.13931567 | 0.694 | 0.607 | 1 | 11 |
| ppp6r3 | 0.70895 | -0.12202364 | 0.167 | 0.25 | 1 | 11 |
| anxa13 | 0.70907 | -0.14077437 | 0.139 | 0.214 | 1 | 11 |
| h2afy2 | 0.70954 | 0.117772436 | 0.5 | 0.464 | 1 | 11 |
| mrps36 | 0.70956 | 0.103894255 | 0.306 | 0.214 | 1 | 11 |
| hnrnpaba | 0.70974 | -0.11014505 | 0.694 | 0.786 | 1 | 11 |
| tspan14 | 0.71015 | -0.14688747 | 0.278 | 0.357 | 1 | 11 |
| dnpep | 0.71064 | 0.129272213 | 0.167 | 0.107 | 1 | 11 |
| dhrs9 | 0.7108 | -0.11075868 | 0.139 | 0.179 | 1 | 11 |
| c1d | 0.7109 | -0.10366494 | 0.167 | 0.25 | 1 | 11 |

| | | | | | | |
|---|---|---|---|---|---|---|
| ehd3 | 0.71098 | -0.13270163 | 0.111 | 0.179 | 1 | 11 |
| si:ch211-51e12.7 | 0.71106 | 0.144640382 | 0.306 | 0.214 | 1 | 11 |
| daam1b | 0.71117 | 0.14593933 | 0.222 | 0.143 | 1 | 11 |
| znf395b | 0.71123 | -0.12506691 | 0.139 | 0.214 | 1 | 11 |
| ube2c | 0.71131 | 0.111785207 | 0.083 | 0.036 | 1 | 11 |
| hmgcrb | 0.71147 | -0.10652423 | 0.056 | 0.107 | 1 | 11 |
| XLOC-014336 | 0.71172 | 0.104356119 | 0.222 | 0.143 | 1 | 11 |
| bckdha | 0.71184 | 0.120937359 | 0.056 | 0.036 | 1 | 11 |
| fnta | 0.71184 | 0.108640209 | 0.056 | 0.036 | 1 | 11 |
| ogfrl2 | 0.71184 | -0.11060027 | 0.056 | 0.036 | 1 | 11 |
| usp14 | 0.71184 | 0.101570074 | 0.056 | 0.036 | 1 | 11 |
| rnf111 | 0.71184 | 0.100716724 | 0.056 | 0.036 | 1 | 11 |
| rwdd3 | 0.71184 | 0.104745888 | 0.056 | 0.036 | 1 | 11 |
| shroom4 | 0.71184 | 0.102693095 | 0.056 | 0.036 | 1 | 11 |
| zgc:194627 | 0.71184 | 0.15963167 | 0.056 | 0.036 | 1 | 11 |
| hnrnpul1l | 0.71204 | 0.113179139 | 0.222 | 0.143 | 1 | 11 |
| si:ch211-150o23.3 | 0.71251 | -0.10713557 | 0.056 | 0.107 | 1 | 11 |
| adam17b | 0.71272 | -0.11220232 | 0.083 | 0.143 | 1 | 11 |
| skilb | 0.71272 | 0.110413406 | 0.083 | 0.036 | 1 | 11 |
| actr2a | 0.71307 | 0.110940812 | 0.222 | 0.143 | 1 | 11 |
| malt1 | 0.71308 | 0.109008668 | 0.111 | 0.071 | 1 | 11 |
| fstl1b | 0.7139 | -0.10577004 | 0.806 | 0.857 | 1 | 11 |
| mtr | 0.71421 | 0.102320364 | 0.083 | 0.036 | 1 | 11 |
| ap1s2 | 0.71531 | -0.11900649 | 0.167 | 0.25 | 1 | 11 |
| mrps18b | 0.71534 | 0.11109041 | 0.167 | 0.143 | 1 | 11 |
| bahcc1b | 0.71581 | 0.115612252 | 0.111 | 0.071 | 1 | 11 |
| copg2 | 0.71617 | 0.135195276 | 0.167 | 0.107 | 1 | 11 |
| pcna | 0.71637 | 0.147255054 | 0.222 | 0.143 | 1 | 11 |
| si:dkeyp-44a8.2 | 0.7165 | 0.148866788 | 0.167 | 0.107 | 1 | 11 |
| ddx55 | 0.7167 | 0.107382775 | 0.111 | 0.071 | 1 | 11 |
| ghrb | 0.71683 | 0.124555246 | 0.222 | 0.143 | 1 | 11 |
| ssu72 | 0.71739 | 0.126118214 | 0.222 | 0.143 | 1 | 11 |
| nub1 | 0.71765 | 0.101391529 | 0.083 | 0.036 | 1 | 11 |
| ptenb | 0.71786 | -0.11395206 | 0.083 | 0.143 | 1 | 11 |
| rbms2b | 0.71828 | 0.121977876 | 0.222 | 0.143 | 1 | 11 |
| dldh | 0.71835 | 0.110552624 | 0.222 | 0.143 | 1 | 11 |
| ctsa | 0.71881 | -0.12143324 | 0.139 | 0.214 | 1 | 11 |
| vaspb | 0.71899 | -0.1253103 | 0.111 | 0.179 | 1 | 11 |
| cflara | 0.7191 | -0.13841945 | 0.111 | 0.179 | 1 | 11 |
| arhgdia | 0.71928 | 0.116255885 | 0.5 | 0.464 | 1 | 11 |
| zgc:171497 | 0.71969 | -0.11690458 | 0.056 | 0.107 | 1 | 11 |
| exosc1 | 0.71972 | -0.10880018 | 0.111 | 0.179 | 1 | 11 |
| supt5h | 0.71991 | 0.13121174 | 0.167 | 0.107 | 1 | 11 |
| GNAZ | 0.72016 | 0.12326785 | 0.167 | 0.107 | 1 | 11 |
| odc1 | 0.72053 | 0.104871674 | 0.083 | 0.036 | 1 | 11 |
| ppp2r5ea | 0.72062 | -0.16249393 | 0.083 | 0.143 | 1 | 11 |
| pdcd6ip | 0.7208 | -0.13273423 | 0.139 | 0.214 | 1 | 11 |
| ube2g2 | 0.7211 | -0.10913369 | 0.056 | 0.107 | 1 | 11 |

| | | | | | | |
|---|---|---|---|---|---|---|
| sema4bb | 0.72111 | -0.1191972 | 0.056 | 0.107 | 1 | 11 |
| sdhb | 0.72129 | -0.11078413 | 0.139 | 0.214 | 1 | 11 |
| grb10b | 0.7218 | -0.12937518 | 0.167 | 0.214 | 1 | 11 |
| mtor | 0.72198 | -0.1076821 | 0.083 | 0.143 | 1 | 11 |
| pik3r2 | 0.72215 | 0.113374643 | 0.389 | 0.357 | 1 | 11 |
| h2afx | 0.72218 | 0.113587799 | 0.111 | 0.071 | 1 | 11 |
| il15l | 0.72265 | 0.131232981 | 0.083 | 0.036 | 1 | 11 |
| c4b | 0.72282 | -0.14146546 | 0.083 | 0.143 | 1 | 11 |
| fam20b | 0.72285 | -0.10674948 | 0.056 | 0.107 | 1 | 11 |
| alg5 | 0.72292 | -0.11056507 | 0.111 | 0.179 | 1 | 11 |
| trmt10a | 0.72305 | 0.101868097 | 0.083 | 0.036 | 1 | 11 |
| wfdc1 | 0.72315 | 0.10267696 | 0.083 | 0.036 | 1 | 11 |
| si:ch211-133n4.4 | 0.72322 | -0.11377197 | 0.333 | 0.429 | 1 | 11 |
| rnf121 | 0.72427 | -0.10188874 | 0.083 | 0.143 | 1 | 11 |
| dock7 | 0.72546 | 0.114547483 | 0.083 | 0.036 | 1 | 11 |
| stox1 | 0.72606 | -0.11936709 | 0.056 | 0.107 | 1 | 11 |
| luzp1 | 0.72608 | -0.11716325 | 0.056 | 0.107 | 1 | 11 |
| si:ch1073-325m22.2 | 0.72636 | -0.15079111 | 0.444 | 0.5 | 1 | 11 |
| ube2h | 0.72687 | -0.11787604 | 0.139 | 0.214 | 1 | 11 |
| si:dkey-7j14.5 | 0.72728 | -0.14218322 | 0.111 | 0.179 | 1 | 11 |
| FP101882.1 | 0.72843 | 0.130102457 | 0.111 | 0.071 | 1 | 11 |
| waplb | 0.72856 | 0.125168042 | 0.083 | 0.036 | 1 | 11 |
| aars | 0.72872 | 0.117489996 | 0.167 | 0.107 | 1 | 11 |
| ccnd2b | 0.72872 | -0.1471548 | 0.111 | 0.179 | 1 | 11 |
| tpd52l2b | 0.72906 | 0.118476697 | 0.278 | 0.214 | 1 | 11 |
| txndc5 | 0.72981 | -0.1249704 | 0.139 | 0.214 | 1 | 11 |
| ube2r2 | 0.73004 | -0.10292887 | 0.056 | 0.107 | 1 | 11 |
| agtr2 | 0.73045 | -0.11572934 | 0.139 | 0.214 | 1 | 11 |
| prrc2c | 0.73047 | -0.10354016 | 0.083 | 0.143 | 1 | 11 |
| stk17al | 0.731 | 0.115304062 | 0.083 | 0.036 | 1 | 11 |
| foxc1b | 0.73133 | 0.102668978 | 0.083 | 0.036 | 1 | 11 |
| ywhaqa | 0.73218 | -0.11859163 | 0.472 | 0.571 | 1 | 11 |
| ptbp1a | 0.73232 | 0.110813488 | 0.333 | 0.286 | 1 | 11 |
| si:dkeyp-75h12.5 | 0.73236 | 0.132895753 | 0.278 | 0.214 | 1 | 11 |
| eif3k | 0.73277 | -0.11088407 | 0.472 | 0.571 | 1 | 11 |
| slmapa | 0.73334 | -0.10010105 | 0.056 | 0.107 | 1 | 11 |
| cbx3a | 0.73366 | 0.125276562 | 0.361 | 0.321 | 1 | 11 |
| cx43.4 | 0.73408 | 0.118441828 | 0.194 | 0.143 | 1 | 11 |
| gtf2a1 | 0.73425 | -0.12918567 | 0.111 | 0.179 | 1 | 11 |
| trip11 | 0.73447 | -0.10327598 | 0.056 | 0.107 | 1 | 11 |
| pgd | 0.73547 | -0.11050027 | 0.111 | 0.179 | 1 | 11 |
| pdhb | 0.73551 | -0.11360817 | 0.111 | 0.179 | 1 | 11 |
| timm17a | 0.73554 | 0.113352183 | 0.167 | 0.107 | 1 | 11 |
| mettl5 | 0.73583 | -0.12456671 | 0.056 | 0.107 | 1 | 11 |
| ghra | 0.73623 | 0.130871959 | 0.417 | 0.321 | 1 | 11 |
| bin1a | 0.73833 | -0.10846424 | 0.056 | 0.107 | 1 | 11 |
| si:ch211-194k22.8 | 0.73863 | 0.10827713 | 0.139 | 0.107 | 1 | 11 |
| arhgef4 | 0.73962 | 0.105468928 | 0.111 | 0.071 | 1 | 11 |

| | | | | | | |
|---|---|---|---|---|---|---|
| si:dkey-56m19.5 | 0.74018 | 0.131387397 | 0.444 | 0.393 | 1 | 11 |
| si:ch1073-459j12.1 | 0.74033 | 0.11612889 | 0.778 | 0.714 | 1 | 11 |
| tm9sf1 | 0.74102 | 0.123324805 | 0.111 | 0.071 | 1 | 11 |
| tpbgb | 0.74192 | -0.11584164 | 0.083 | 0.143 | 1 | 11 |
| vipr1a | 0.74237 | -0.10276768 | 0.111 | 0.179 | 1 | 11 |
| si:dkey-51e6.1 | 0.74244 | 0.138110411 | 0.389 | 0.321 | 1 | 11 |
| ptmaa | 0.74267 | 0.103163136 | 0.861 | 0.821 | 1 | 11 |
| slc25a55a | 0.7432 | 0.110404153 | 0.25 | 0.214 | 1 | 11 |
| march5 | 0.74351 | 0.116693357 | 0.167 | 0.107 | 1 | 11 |
| grinab | 0.74395 | -0.14810578 | 0.278 | 0.357 | 1 | 11 |
| ttc4 | 0.74423 | 0.1118957 | 0.167 | 0.107 | 1 | 11 |
| dnajb6a | 0.74581 | -0.10163613 | 0.111 | 0.179 | 1 | 11 |
| meis1a | 0.7459 | -0.1045303 | 0.111 | 0.143 | 1 | 11 |
| EML4 | 0.74652 | -0.10881044 | 0.111 | 0.179 | 1 | 11 |
| fbxw11b | 0.74658 | -0.12328291 | 0.111 | 0.179 | 1 | 11 |
| mboat2a | 0.74832 | 0.134971195 | 0.167 | 0.107 | 1 | 11 |
| CABZ01078261.1 | 0.74856 | -0.10437517 | 0.056 | 0.107 | 1 | 11 |
| foxo4 | 0.74922 | -0.10167886 | 0.056 | 0.107 | 1 | 11 |
| tars | 0.74931 | -0.10521585 | 0.083 | 0.143 | 1 | 11 |
| pank2 | 0.74953 | -0.11463495 | 0.083 | 0.143 | 1 | 11 |
| COX5B | 0.74958 | -0.1269459 | 0.444 | 0.5 | 1 | 11 |
| hmgb2a | 0.74989 | 0.102266431 | 0.556 | 0.571 | 1 | 11 |
| tgm2b | 0.75007 | 0.131826502 | 0.25 | 0.179 | 1 | 11 |
| zgc:85777 | 0.75007 | -0.11043178 | 0.083 | 0.143 | 1 | 11 |
| MYADM | 0.75132 | 0.100881292 | 0.111 | 0.071 | 1 | 11 |
| smoc1 | 0.75137 | -0.11033371 | 0.056 | 0.107 | 1 | 11 |
| sema3bl | 0.75177 | -0.13299161 | 0.083 | 0.143 | 1 | 11 |
| timm8b | 0.75279 | -0.12190738 | 0.139 | 0.179 | 1 | 11 |
| lin7a | 0.75282 | -0.10996073 | 0.056 | 0.107 | 1 | 11 |
| akirin1 | 0.75355 | 0.112909479 | 0.167 | 0.107 | 1 | 11 |
| eci1 | 0.75356 | 0.102358407 | 0.139 | 0.107 | 1 | 11 |
| flrt3 | 0.75369 | -0.10148999 | 0.083 | 0.143 | 1 | 11 |
| serpinb1l3 | 0.75375 | -0.12148786 | 0.194 | 0.25 | 1 | 11 |
| BX640522.2 | 0.75413 | -0.1660807 | 0.472 | 0.536 | 1 | 11 |
| ndrg1a | 0.75444 | -0.10891475 | 0.083 | 0.143 | 1 | 11 |
| arglu1b | 0.75484 | -0.11828931 | 0.083 | 0.143 | 1 | 11 |
| emc1 | 0.75485 | -0.1176361 | 0.083 | 0.143 | 1 | 11 |
| fosb | 0.7549 | 0.155496443 | 0.333 | 0.25 | 1 | 11 |
| cd151l | 0.75582 | -0.11346528 | 0.306 | 0.357 | 1 | 11 |
| ctsk | 0.75587 | -0.10485544 | 0.361 | 0.393 | 1 | 11 |
| twist1a | 0.75591 | -0.10796365 | 0.056 | 0.071 | 1 | 11 |
| ispd | 0.75593 | 0.129437904 | 0.111 | 0.071 | 1 | 11 |
| ap1s3b | 0.75682 | -0.10552374 | 0.139 | 0.179 | 1 | 11 |
| CABZ01068358.1 | 0.7574 | 0.10213802 | 0.167 | 0.143 | 1 | 11 |
| selenop | 0.75797 | 0.102376203 | 0.167 | 0.107 | 1 | 11 |
| dap1b | 0.75847 | -0.13772596 | 0.444 | 0.536 | 1 | 11 |
| mrpl33 | 0.75873 | -0.13257741 | 0.222 | 0.286 | 1 | 11 |
| dhrs3a | 0.75881 | 0.130536809 | 0.25 | 0.179 | 1 | 11 |

| | | | | | | |
|---|---|---|---|---|---|---|
| eif1b | 0.76028 | 0.11786215 | 0.694 | 0.607 | 1 | 11 |
| psmc6 | 0.76262 | -0.11972969 | 0.25 | 0.321 | 1 | 11 |
| ssr4 | 0.76359 | -0.11466717 | 0.667 | 0.714 | 1 | 11 |
| atp6v1e1a | 0.76359 | -0.11852328 | 0.111 | 0.143 | 1 | 11 |
| tenm3 | 0.76384 | 0.135058011 | 0.556 | 0.464 | 1 | 11 |
| ctnnb1 | 0.76498 | 0.129594926 | 0.528 | 0.464 | 1 | 11 |
| eif5b | 0.76508 | -0.12997607 | 0.306 | 0.393 | 1 | 11 |
| BX548015.2 | 0.76618 | 0.105991155 | 0.111 | 0.071 | 1 | 11 |
| tomm5 | 0.76635 | 0.130629421 | 0.583 | 0.5 | 1 | 11 |
| si:dkey-16l2.20 | 0.76808 | 0.125914071 | 0.333 | 0.25 | 1 | 11 |
| naa15a | 0.76865 | 0.100822196 | 0.111 | 0.071 | 1 | 11 |
| lsm12b | 0.76881 | -0.10678669 | 0.444 | 0.536 | 1 | 11 |
| nmt2 | 0.76958 | 0.101938009 | 0.111 | 0.071 | 1 | 11 |
| gpm6ab | 0.76999 | 0.11250661 | 0.167 | 0.143 | 1 | 11 |
| mrtfab | 0.77035 | -0.13798261 | 0.194 | 0.25 | 1 | 11 |
| cish | 0.77126 | -0.10735962 | 0.111 | 0.143 | 1 | 11 |
| thrab | 0.77269 | 0.134032773 | 0.111 | 0.071 | 1 | 11 |
| ldlrad3 | 0.77313 | 0.116780354 | 0.111 | 0.071 | 1 | 11 |
| wls | 0.77354 | -0.13230797 | 0.389 | 0.464 | 1 | 11 |
| si:ch211-39i22.1 | 0.77495 | 0.116245816 | 0.5 | 0.464 | 1 | 11 |
| rab13 | 0.77501 | 0.121646054 | 0.167 | 0.107 | 1 | 11 |
| hnrnpr | 0.7756 | 0.100570713 | 0.111 | 0.071 | 1 | 11 |
| klf6a | 0.77636 | -0.12251964 | 0.528 | 0.571 | 1 | 11 |
| ddt | 0.77805 | 0.104785632 | 0.278 | 0.214 | 1 | 11 |
| zc3h6 | 0.77903 | 0.104676164 | 0.111 | 0.071 | 1 | 11 |
| lrrc1 | 0.7794 | 0.107279978 | 0.111 | 0.071 | 1 | 11 |
| sccpdha | 0.77946 | 0.138953987 | 0.167 | 0.107 | 1 | 11 |
| taf7 | 0.78066 | 0.110552772 | 0.25 | 0.179 | 1 | 11 |
| add3a | 0.78073 | 0.134185203 | 0.472 | 0.393 | 1 | 11 |
| si:ch1073-340j19.1 | 0.7837 | 0.102383352 | 0.111 | 0.071 | 1 | 11 |
| FO082781.1 | 0.78406 | 0.107806488 | 0.167 | 0.107 | 1 | 11 |
| ywhaqb | 0.78634 | -0.12917978 | 0.444 | 0.5 | 1 | 11 |
| tgm1l1 | 0.78649 | 0.118223617 | 0.194 | 0.143 | 1 | 11 |
| nup62l | 0.78683 | 0.126610575 | 0.194 | 0.143 | 1 | 11 |
| gkap1 | 0.78724 | 0.103224134 | 0.25 | 0.179 | 1 | 11 |
| ldlrad4a | 0.78737 | 0.102096154 | 0.167 | 0.107 | 1 | 11 |
| phyh | 0.78762 | 0.105777659 | 0.25 | 0.179 | 1 | 11 |
| sptan1 | 0.78786 | -0.1179751 | 0.194 | 0.25 | 1 | 11 |
| hecw2b | 0.79363 | -0.11631513 | 0.139 | 0.179 | 1 | 11 |
| erap1a | 0.79383 | 0.104358066 | 0.111 | 0.071 | 1 | 11 |
| fstb | 0.79412 | -0.11764887 | 0.278 | 0.321 | 1 | 11 |
| eva1ba | 0.79491 | -0.10446863 | 0.222 | 0.25 | 1 | 11 |
| anapc16 | 0.79711 | -0.12271763 | 0.25 | 0.321 | 1 | 11 |
| cpn1 | 0.79813 | -0.12030043 | 0.472 | 0.536 | 1 | 11 |
| cox7a2a | 0.79841 | -0.11809862 | 0.222 | 0.286 | 1 | 11 |
| metap2b | 0.79853 | -0.11173078 | 0.389 | 0.429 | 1 | 11 |
| fam69c | 0.80095 | -0.11041993 | 0.167 | 0.214 | 1 | 11 |
| sept7b | 0.80548 | -0.1136131 | 0.222 | 0.286 | 1 | 11 |

| | | | | | | |
|---|---|---|---|---|---|---|
| atp5po | 0.80693 | -0.11479389 | 0.389 | 0.429 | 1 | 11 |
| eif3f | 0.80892 | 0.109242451 | 0.722 | 0.679 | 1 | 11 |
| si:dkey-19a16.7 | 0.80895 | -0.10830996 | 0.167 | 0.214 | 1 | 11 |
| rbp4 | 0.81031 | 0.101400941 | 0.444 | 0.429 | 1 | 11 |
| golgb1 | 0.81203 | -0.1284029 | 0.194 | 0.25 | 1 | 11 |
| mt-nd2 | 0.81219 | -0.12267044 | 0.639 | 0.679 | 1 | 11 |
| mphosph6 | 0.81291 | -0.11161345 | 0.111 | 0.143 | 1 | 11 |
| usp25 | 0.81416 | -0.10118802 | 0.083 | 0.107 | 1 | 11 |
| psma6b | 0.81486 | 0.102203323 | 0.361 | 0.286 | 1 | 11 |
| zgc:113531 | 0.81496 | 0.113752015 | 0.361 | 0.286 | 1 | 11 |
| kars | 0.81505 | 0.106886597 | 0.306 | 0.25 | 1 | 11 |
| socs2 | 0.81737 | -0.10020178 | 0.083 | 0.107 | 1 | 11 |
| kdm1a | 0.81752 | 0.115966758 | 0.306 | 0.25 | 1 | 11 |
| arl4d | 0.81934 | -0.11215273 | 0.25 | 0.321 | 1 | 11 |
| ranbp1 | 0.81946 | 0.111262928 | 0.5 | 0.429 | 1 | 11 |
| utrn | 0.81963 | -0.12829808 | 0.194 | 0.25 | 1 | 11 |
| igfbp5b | 0.82017 | -0.12261258 | 0.25 | 0.321 | 1 | 11 |
| CABZ01092746.1 | 0.82118 | 0.117962882 | 0.444 | 0.393 | 1 | 11 |
| csnk2a1 | 0.82138 | -0.11405051 | 0.306 | 0.357 | 1 | 11 |
| kmt2bb | 0.82337 | 0.122455522 | 0.417 | 0.357 | 1 | 11 |
| CU914487.1 | 0.82391 | -0.10046251 | 0.083 | 0.107 | 1 | 11 |
| fnbp1l | 0.82604 | 0.105137035 | 0.167 | 0.143 | 1 | 11 |
| pttg1ipb | 0.82755 | 0.103012011 | 0.111 | 0.071 | 1 | 11 |
| pho | 0.83628 | -0.10867668 | 0.167 | 0.214 | 1 | 11 |
| tgfb3 | 0.83689 | 0.111527683 | 0.278 | 0.214 | 1 | 11 |
| hsbp1b | 0.83696 | 0.101489906 | 0.444 | 0.393 | 1 | 11 |
| fgl2a | 0.83946 | 0.104595103 | 0.472 | 0.429 | 1 | 11 |
| ube2e3 | 0.83963 | 0.103083703 | 0.5 | 0.429 | 1 | 11 |
| bzw1a | 0.84048 | -0.10939072 | 0.194 | 0.25 | 1 | 11 |
| clk4a | 0.8449 | 0.109932652 | 0.194 | 0.143 | 1 | 11 |
| ifi30 | 0.84914 | -0.10776522 | 0.278 | 0.321 | 1 | 11 |
| zmiz1a | 0.85022 | 0.105434083 | 0.222 | 0.179 | 1 | 11 |
| ppp1cb | 0.85317 | 0.108628713 | 0.306 | 0.25 | 1 | 11 |
| skib | 0.85594 | -0.10447551 | 0.167 | 0.214 | 1 | 11 |
| esr2a | 0.85649 | 0.100770817 | 0.222 | 0.179 | 1 | 11 |
| fam133b | 0.86144 | -0.10694659 | 0.306 | 0.357 | 1 | 11 |
| gas1a | 0.86318 | 0.112903715 | 0.417 | 0.357 | 1 | 11 |
| zgc:154058 | 0.865 | 0.104383534 | 0.222 | 0.179 | 1 | 11 |
| rhoua | 0.8664 | -0.10779835 | 0.194 | 0.25 | 1 | 11 |
| rspo3 | 0.86888 | 0.105679353 | 0.222 | 0.179 | 1 | 11 |
| h3f3a | 0.87016 | 0.110974578 | 0.444 | 0.393 | 1 | 11 |
| ms4a17a.8 | 0.00021 | -0.80296572 | 0 | 0.296 | 1 | 12 |
| tuba8l | 0.00028 | -0.97305742 | 0.143 | 0.63 | 1 | 12 |
| fabp11a | 0.00187 | -0.78158404 | 0.429 | 0.852 | 1 | 12 |
| calm3b | 0.002 | -0.83056774 | 0.229 | 0.667 | 1 | 12 |
| emc6 | 0.00255 | 0.608424733 | 0.229 | 0 | 1 | 12 |
| emc8 | 0.00488 | -0.5739064 | 0 | 0.185 | 1 | 12 |
| calr | 0.00488 | -0.47520948 | 0 | 0.185 | 1 | 12 |

| | | | | | | |
|---|---|---|---|---|---|---|
| si:dkey-262k9.2 | 0.00488 | -0.61474522 | 0 | 0.185 | 1 | 12 |
| copb1 | 0.00488 | -0.49773571 | 0 | 0.185 | 1 | 12 |
| hdac1 | 0.00488 | -0.504523 | 0 | 0.185 | 1 | 12 |
| ivns1abpa | 0.00529 | 0.59530408 | 0.2 | 0 | 1 | 12 |
| pfkfb4b | 0.00529 | 0.562217587 | 0.2 | 0 | 1 | 12 |
| si:dkeyp-44a8.2 | 0.00529 | 0.485743233 | 0.2 | 0 | 1 | 12 |
| srgn | 0.00529 | 0.619421868 | 0.2 | 0 | 1 | 12 |
| npdc1a | 0.00529 | 0.563795548 | 0.2 | 0 | 1 | 12 |
| eef1b2 | 0.00642 | 0.25483353 | 0.857 | 0.556 | 1 | 12 |
| prdx2 | 0.00957 | 0.730116688 | 0.457 | 0.111 | 1 | 12 |
| rpl3 | 0.00971 | 0.338066514 | 0.971 | 0.852 | 1 | 12 |
| rwdd1 | 0.01085 | 0.397214801 | 0.171 | 0 | 1 | 12 |
| timp2a | 0.01085 | 0.45526026 | 0.171 | 0 | 1 | 12 |
| cwc25 | 0.01085 | 0.542072889 | 0.171 | 0 | 1 | 12 |
| mlf2 | 0.01085 | 0.435262329 | 0.171 | 0 | 1 | 12 |
| tnrc6c1 | 0.01085 | 0.403920621 | 0.171 | 0 | 1 | 12 |
| eif4ebp3 | 0.01085 | 0.455840855 | 0.171 | 0 | 1 | 12 |
| si:dkey-14d8.6 | 0.01085 | 0.411189924 | 0.171 | 0 | 1 | 12 |
| zranb2 | 0.01085 | 0.475426625 | 0.171 | 0 | 1 | 12 |
| rpl28 | 0.01295 | 0.298255444 | 1 | 0.815 | 1 | 12 |
| slc25a55a | 0.01344 | -0.35544124 | 0 | 0.148 | 1 | 12 |
| rbbp4 | 0.01344 | -0.47254952 | 0 | 0.148 | 1 | 12 |
| acvrl1 | 0.01344 | -0.4063827 | 0 | 0.148 | 1 | 12 |
| cyr61l2 | 0.01344 | -0.47169352 | 0 | 0.148 | 1 | 12 |
| plgrkt | 0.01344 | -0.44904719 | 0 | 0.148 | 1 | 12 |
| sdr39u1 | 0.01344 | -0.39623543 | 0 | 0.148 | 1 | 12 |
| zgc:56304 | 0.01344 | -0.52656014 | 0 | 0.148 | 1 | 12 |
| XLOC-000946 | 0.01344 | -0.39868718 | 0 | 0.148 | 1 | 12 |
| arpp19b | 0.01344 | -0.44774321 | 0 | 0.148 | 1 | 12 |
| asf1ba | 0.01344 | -0.47781152 | 0 | 0.148 | 1 | 12 |
| cflara | 0.01344 | -0.39143148 | 0 | 0.148 | 1 | 12 |
| dnajc9 | 0.01344 | -0.48388055 | 0 | 0.148 | 1 | 12 |
| nasp | 0.01344 | -0.43704962 | 0 | 0.148 | 1 | 12 |
| si:dkey-276j7.1 | 0.01344 | -0.37673903 | 0 | 0.148 | 1 | 12 |
| smfn | 0.01344 | -0.4365059 | 0 | 0.148 | 1 | 12 |
| smo | 0.01344 | -0.3874905 | 0 | 0.148 | 1 | 12 |
| zgc:113531 | 0.01344 | -0.32378985 | 0 | 0.148 | 1 | 12 |
| zgc:56525 | 0.01344 | -0.37009245 | 0 | 0.148 | 1 | 12 |
| si:ch211-169p10.1 | 0.01344 | -0.3711137 | 0 | 0.148 | 1 | 12 |
| dhx15 | 0.01344 | -0.43770678 | 0 | 0.148 | 1 | 12 |
| nono | 0.01344 | -0.37965963 | 0 | 0.148 | 1 | 12 |
| tubb2b | 0.01626 | -0.78817895 | 0.229 | 0.481 | 1 | 12 |
| gamt | 0.02207 | 0.425037849 | 0.143 | 0 | 1 | 12 |
| LOC100535806 | 0.02207 | 0.387352835 | 0.143 | 0 | 1 | 12 |
| atoh8 | 0.02207 | 0.380808414 | 0.143 | 0 | 1 | 12 |
| cyp2ad3 | 0.02207 | 0.481824486 | 0.143 | 0 | 1 | 12 |
| arpc5b | 0.02207 | 0.382607104 | 0.143 | 0 | 1 | 12 |
| bhmt | 0.02207 | 0.31590224 | 0.143 | 0 | 1 | 12 |

| | | | | | | |
|---|---|---|---|---|---|---|
| dll4 | 0.02207 | 0.345688662 | 0.143 | 0 | 1 | 12 |
| idh1 | 0.02207 | 0.363534673 | 0.143 | 0 | 1 | 12 |
| ik | 0.02207 | 0.394197565 | 0.143 | 0 | 1 | 12 |
| midn | 0.02207 | 0.350154181 | 0.143 | 0 | 1 | 12 |
| nfkbie | 0.02207 | 0.350514664 | 0.143 | 0 | 1 | 12 |
| nupr1b | 0.02207 | 0.476160196 | 0.143 | 0 | 1 | 12 |
| otub1b | 0.02207 | 0.386450053 | 0.143 | 0 | 1 | 12 |
| pdk2b | 0.02207 | 0.327054848 | 0.143 | 0 | 1 | 12 |
| ptprb | 0.02207 | 0.394564169 | 0.143 | 0 | 1 | 12 |
| ralab | 0.02207 | 0.365429573 | 0.143 | 0 | 1 | 12 |
| sept7a | 0.02207 | 0.322748991 | 0.143 | 0 | 1 | 12 |
| si:dkey-112a7.4 | 0.02207 | 0.34814294 | 0.143 | 0 | 1 | 12 |
| u2surp | 0.02207 | 0.476977667 | 0.143 | 0 | 1 | 12 |
| ash1l | 0.02207 | 0.411613599 | 0.143 | 0 | 1 | 12 |
| exosc6 | 0.02207 | 0.414640703 | 0.143 | 0 | 1 | 12 |
| cdkn1bb | 0.02207 | 0.407121228 | 0.143 | 0 | 1 | 12 |
| tia1 | 0.02207 | 0.339932583 | 0.143 | 0 | 1 | 12 |
| cox14 | 0.02207 | 0.373012026 | 0.143 | 0 | 1 | 12 |
| dbi | 0.02423 | -0.70814182 | 0.2 | 0.519 | 1 | 12 |
| rpl14 | 0.02568 | 0.217220726 | 0.943 | 0.704 | 1 | 12 |
| si:dkey-126g1.9 | 0.02688 | -0.57321989 | 0.143 | 0.444 | 1 | 12 |
| b2ml | 0.02992 | 0.690610906 | 0.4 | 0.111 | 1 | 12 |
| taf15 | 0.03139 | -0.63008668 | 0.029 | 0.222 | 1 | 12 |
| plk2b | 0.03169 | 0.655968851 | 0.257 | 0.037 | 1 | 12 |
| ppm1g | 0.03375 | -0.60532461 | 0.029 | 0.222 | 1 | 12 |
| zgc:92744 | 0.03659 | 0.626363498 | 0.314 | 0.074 | 1 | 12 |
| ccdc9 | 0.03671 | -0.28828864 | 0 | 0.111 | 1 | 12 |
| cetn3 | 0.03671 | -0.34241908 | 0 | 0.111 | 1 | 12 |
| cyldl | 0.03671 | -0.33555169 | 0 | 0.111 | 1 | 12 |
| dck | 0.03671 | -0.35198674 | 0 | 0.111 | 1 | 12 |
| letm1 | 0.03671 | -0.27757255 | 0 | 0.111 | 1 | 12 |
| rab6a | 0.03671 | -0.29827196 | 0 | 0.111 | 1 | 12 |
| rif1 | 0.03671 | -0.2689151 | 0 | 0.111 | 1 | 12 |
| tnrc6b | 0.03671 | -0.3205838 | 0 | 0.111 | 1 | 12 |
| CR762483.1 | 0.03671 | -0.31224496 | 0 | 0.111 | 1 | 12 |
| brd9 | 0.03671 | -0.2965273 | 0 | 0.111 | 1 | 12 |
| commd3 | 0.03671 | -0.40161229 | 0 | 0.111 | 1 | 12 |
| dctd | 0.03671 | -0.36919177 | 0 | 0.111 | 1 | 12 |
| dnph1 | 0.03671 | -0.35462203 | 0 | 0.111 | 1 | 12 |
| foxp1a | 0.03671 | -0.19791205 | 0 | 0.111 | 1 | 12 |
| hadh | 0.03671 | -0.32125535 | 0 | 0.111 | 1 | 12 |
| igfbp1a | 0.03671 | -0.39869111 | 0 | 0.111 | 1 | 12 |
| kdm6bb | 0.03671 | -0.24145153 | 0 | 0.111 | 1 | 12 |
| ltv1 | 0.03671 | -0.40435918 | 0 | 0.111 | 1 | 12 |
| mrpl51 | 0.03671 | -0.25569551 | 0 | 0.111 | 1 | 12 |
| mrpl52 | 0.03671 | -0.29143438 | 0 | 0.111 | 1 | 12 |
| ppifb | 0.03671 | -0.40124644 | 0 | 0.111 | 1 | 12 |
| ppip5k2 | 0.03671 | -0.27571615 | 0 | 0.111 | 1 | 12 |

| | | | | | | |
|---|---|---|---|---|---|---|
| ppp1r9ba | 0.03671 | -0.31945783 | 0 | 0.111 | 1 | 12 |
| ppp2r1ba | 0.03671 | -0.19684936 | 0 | 0.111 | 1 | 12 |
| prdx6 | 0.03671 | -0.31566612 | 0 | 0.111 | 1 | 12 |
| rnf213a | 0.03671 | -0.32712226 | 0 | 0.111 | 1 | 12 |
| taok2a | 0.03671 | -0.31609207 | 0 | 0.111 | 1 | 12 |
| alad | 0.03671 | -0.31665694 | 0 | 0.111 | 1 | 12 |
| c2cd2 | 0.03671 | -0.29213456 | 0 | 0.111 | 1 | 12 |
| c2cd5 | 0.03671 | -0.33596162 | 0 | 0.111 | 1 | 12 |
| cdc25b | 0.03671 | -0.26895388 | 0 | 0.111 | 1 | 12 |
| chchd10 | 0.03671 | -0.35137281 | 0 | 0.111 | 1 | 12 |
| chkb | 0.03671 | -0.37898826 | 0 | 0.111 | 1 | 12 |
| chmp4bb | 0.03671 | -0.27064956 | 0 | 0.111 | 1 | 12 |
| cnpy1 | 0.03671 | -0.27288293 | 0 | 0.111 | 1 | 12 |
| eif4g3b | 0.03671 | -0.38746425 | 0 | 0.111 | 1 | 12 |
| hspa9 | 0.03671 | -0.24157798 | 0 | 0.111 | 1 | 12 |
| mrps18c | 0.03671 | -0.29733756 | 0 | 0.111 | 1 | 12 |
| noc4l | 0.03671 | -0.29597716 | 0 | 0.111 | 1 | 12 |
| nup93 | 0.03671 | -0.32210382 | 0 | 0.111 | 1 | 12 |
| pgfb | 0.03671 | -0.3203891 | 0 | 0.111 | 1 | 12 |
| prkrip1 | 0.03671 | -0.25623538 | 0 | 0.111 | 1 | 12 |
| rnaseh2c | 0.03671 | -0.38304704 | 0 | 0.111 | 1 | 12 |
| rnf185 | 0.03671 | -0.281775 | 0 | 0.111 | 1 | 12 |
| rrm1 | 0.03671 | -0.3336505 | 0 | 0.111 | 1 | 12 |
| sh3glb2b | 0.03671 | -0.34245052 | 0 | 0.111 | 1 | 12 |
| si:ch73-61d6.3 | 0.03671 | -0.2974283 | 0 | 0.111 | 1 | 12 |
| slc25a46 | 0.03671 | -0.38955306 | 0 | 0.111 | 1 | 12 |
| slc31a1 | 0.03671 | -0.25114727 | 0 | 0.111 | 1 | 12 |
| smu1a | 0.03671 | -0.29944722 | 0 | 0.111 | 1 | 12 |
| tox2 | 0.03671 | -0.35509694 | 0 | 0.111 | 1 | 12 |
| ufm1 | 0.03671 | -0.28076769 | 0 | 0.111 | 1 | 12 |
| zgc:174906 | 0.03671 | -0.24018667 | 0 | 0.111 | 1 | 12 |
| epas1a | 0.03671 | -0.29921713 | 0 | 0.111 | 1 | 12 |
| kcnc4 | 0.03671 | -0.37400524 | 0 | 0.111 | 1 | 12 |
| lrch3 | 0.03671 | -0.27337332 | 0 | 0.111 | 1 | 12 |
| smim12 | 0.03671 | -0.36613882 | 0 | 0.111 | 1 | 12 |
| tcf3b | 0.03671 | -0.24681699 | 0 | 0.111 | 1 | 12 |
| tgfbi | 0.03671 | -0.30589865 | 0 | 0.111 | 1 | 12 |
| kat7b | 0.03671 | -0.36720267 | 0 | 0.111 | 1 | 12 |
| ppp2r2d | 0.03671 | -0.40231686 | 0 | 0.111 | 1 | 12 |
| utp4 | 0.03671 | -0.30409243 | 0 | 0.111 | 1 | 12 |
| fcf1 | 0.03671 | -0.38948145 | 0 | 0.111 | 1 | 12 |
| rbp2a | 0.03678 | -0.72443523 | 0.171 | 0.296 | 1 | 12 |
| cox6b2 | 0.03705 | 0.568565115 | 0.657 | 0.333 | 1 | 12 |
| rpl27 | 0.03888 | 0.414105083 | 0.886 | 0.667 | 1 | 12 |
| si:dkey-112e7.2 | 0.03919 | -0.63181002 | 0.086 | 0.333 | 1 | 12 |
| vwa1 | 0.04007 | 0.685270213 | 0.257 | 0.037 | 1 | 12 |
| nop56 | 0.04196 | 0.529187828 | 0.257 | 0.037 | 1 | 12 |
| serf2 | 0.04263 | 0.514408068 | 0.571 | 0.259 | 1 | 12 |

| | | | | | | |
|---|---|---|---|---|---|---|
| chka | 0.04468 | 0.324890217 | 0.114 | 0 | 1 | 12 |
| dlc | 0.04468 | 0.311028946 | 0.114 | 0 | 1 | 12 |
| emc2 | 0.04468 | 0.332375493 | 0.114 | 0 | 1 | 12 |
| gpc4 | 0.04468 | 0.351773317 | 0.114 | 0 | 1 | 12 |
| mak16 | 0.04468 | 0.381611377 | 0.114 | 0 | 1 | 12 |
| mfap1 | 0.04468 | 0.297684363 | 0.114 | 0 | 1 | 12 |
| mtch2 | 0.04468 | 0.361460769 | 0.114 | 0 | 1 | 12 |
| phf11 | 0.04468 | 0.33138235 | 0.114 | 0 | 1 | 12 |
| rab11fip1b | 0.04468 | 0.330532768 | 0.114 | 0 | 1 | 12 |
| sdad1 | 0.04468 | 0.357084932 | 0.114 | 0 | 1 | 12 |
| smg7 | 0.04468 | 0.315635235 | 0.114 | 0 | 1 | 12 |
| smim13 | 0.04468 | 0.283315035 | 0.114 | 0 | 1 | 12 |
| smim29 | 0.04468 | 0.31733896 | 0.114 | 0 | 1 | 12 |
| stk24b | 0.04468 | 0.329248206 | 0.114 | 0 | 1 | 12 |
| stom | 0.04468 | 0.2761301 | 0.114 | 0 | 1 | 12 |
| taz | 0.04468 | 0.240693418 | 0.114 | 0 | 1 | 12 |
| tia1l | 0.04468 | 0.24941899 | 0.114 | 0 | 1 | 12 |
| ube2h | 0.04468 | 0.236980033 | 0.114 | 0 | 1 | 12 |
| zcchc24 | 0.04468 | 0.285991513 | 0.114 | 0 | 1 | 12 |
| zgc:110626 | 0.04468 | 0.285156277 | 0.114 | 0 | 1 | 12 |
| aldob | 0.04468 | 0.322256357 | 0.114 | 0 | 1 | 12 |
| arrdc2 | 0.04468 | 0.283369389 | 0.114 | 0 | 1 | 12 |
| dcaf7 | 0.04468 | 0.352308323 | 0.114 | 0 | 1 | 12 |
| dctn1a | 0.04468 | 0.328491477 | 0.114 | 0 | 1 | 12 |
| kras | 0.04468 | 0.278631902 | 0.114 | 0 | 1 | 12 |
| polr2k | 0.04468 | 0.304652625 | 0.114 | 0 | 1 | 12 |
| prkar2aa | 0.04468 | 0.255735864 | 0.114 | 0 | 1 | 12 |
| proza | 0.04468 | 0.285245652 | 0.114 | 0 | 1 | 12 |
| rca2.2 | 0.04468 | 0.340097601 | 0.114 | 0 | 1 | 12 |
| rhag | 0.04468 | 0.286087629 | 0.114 | 0 | 1 | 12 |
| si:ch211-213a13.1 | 0.04468 | 0.243899125 | 0.114 | 0 | 1 | 12 |
| zfand6 | 0.04468 | 0.348512463 | 0.114 | 0 | 1 | 12 |
| zgc:92107 | 0.04468 | 0.291346007 | 0.114 | 0 | 1 | 12 |
| BX569787.1 | 0.04468 | 0.282911114 | 0.114 | 0 | 1 | 12 |
| fxyd6l | 0.04468 | 0.373401201 | 0.114 | 0 | 1 | 12 |
| stau2 | 0.04468 | 0.276263667 | 0.114 | 0 | 1 | 12 |
| suds3 | 0.04468 | 0.31785454 | 0.114 | 0 | 1 | 12 |
| CT033796.1 | 0.04468 | 0.291902683 | 0.114 | 0 | 1 | 12 |
| cpa4 | 0.04468 | 0.318158931 | 0.114 | 0 | 1 | 12 |
| dnajc19 | 0.04468 | 0.293891529 | 0.114 | 0 | 1 | 12 |
| fbxl6 | 0.04468 | 0.303112375 | 0.114 | 0 | 1 | 12 |
| mrpl37 | 0.04468 | 0.267806434 | 0.114 | 0 | 1 | 12 |
| rrs1 | 0.04468 | 0.342655417 | 0.114 | 0 | 1 | 12 |
| si:ch211-212g7.6 | 0.04468 | 0.336627253 | 0.114 | 0 | 1 | 12 |
| zgc:92818 | 0.04468 | 0.301988019 | 0.114 | 0 | 1 | 12 |
| sc:d189 | 0.04468 | 0.393197277 | 0.114 | 0 | 1 | 12 |
| arih2 | 0.04468 | 0.292042727 | 0.114 | 0 | 1 | 12 |
| ccz1 | 0.04468 | 0.338015752 | 0.114 | 0 | 1 | 12 |

| | | | | | | |
|---|---|---|---|---|---|---|
| isoc2 | 0.04468 | 0.320237304 | 0.114 | 0 | 1 | 12 |
| st13 | 0.04468 | 0.31670183 | 0.114 | 0 | 1 | 12 |
| uri1 | 0.04468 | 0.273655665 | 0.114 | 0 | 1 | 12 |
| wu:fj16a03 | 0.04468 | 0.371908298 | 0.114 | 0 | 1 | 12 |
| rps27a | 0.04576 | 0.150863228 | 1 | 0.852 | 1 | 12 |
| rps14 | 0.04842 | -0.10136069 | 0.857 | 1 | 1 | 12 |
| si:dkey-78l4.14 | 0.04889 | -0.54206212 | 0.029 | 0.222 | 1 | 12 |
| nsfl1c | 0.04948 | -0.54761499 | 0.029 | 0.222 | 1 | 12 |
| ms4a17a.9 | 0.05048 | 0.615951119 | 0.314 | 0.074 | 1 | 12 |
| atp5f1e | 0.05233 | -0.61927904 | 0.286 | 0.556 | 1 | 12 |
| hmgn2 | 0.05314 | -0.40835267 | 0.629 | 0.889 | 1 | 12 |
| uqcrfs1 | 0.05568 | 0.405351549 | 0.229 | 0.037 | 1 | 12 |
| hsdl2 | 0.06478 | -0.54097745 | 0.057 | 0.259 | 1 | 12 |
| fscn1a | 0.07115 | -0.58251052 | 0.057 | 0.259 | 1 | 12 |
| CR383676.1 | 0.07126 | 0.784881286 | 0.543 | 0.259 | 1 | 12 |
| psmd8 | 0.07243 | -0.53433706 | 0.029 | 0.185 | 1 | 12 |
| pora | 0.07473 | 0.438890611 | 0.229 | 0.037 | 1 | 12 |
| lmo4a | 0.07602 | 0.557872274 | 0.286 | 0.074 | 1 | 12 |
| tmem198b | 0.07644 | -0.52753072 | 0.029 | 0.185 | 1 | 12 |
| rbp7b | 0.07713 | -0.68551099 | 0.057 | 0.259 | 1 | 12 |
| tagln2 | 0.07714 | 0.401427211 | 0.686 | 0.407 | 1 | 12 |
| ctnnb1 | 0.08155 | 0.452613237 | 0.343 | 0.111 | 1 | 12 |
| grb10a | 0.08215 | 0.581877059 | 0.286 | 0.074 | 1 | 12 |
| rab10 | 0.08528 | 0.510303358 | 0.4 | 0.148 | 1 | 12 |
| atp5md | 0.08776 | -0.58247382 | 0.2 | 0.444 | 1 | 12 |
| irf2bp2a | 0.08819 | 0.553709764 | 0.343 | 0.111 | 1 | 12 |
| cox7c | 0.08911 | -0.32923486 | 0.543 | 0.63 | 1 | 12 |
| eif3ba | 0.09056 | 0.519047085 | 0.343 | 0.111 | 1 | 12 |
| BX284638.1 | 0.09058 | 0.25809807 | 0.086 | 0 | 1 | 12 |
| CR735102.1 | 0.09058 | 0.231181186 | 0.086 | 0 | 1 | 12 |
| CR855337.1 | 0.09058 | 0.223041201 | 0.086 | 0 | 1 | 12 |
| DHDH | 0.09058 | 0.22061852 | 0.086 | 0 | 1 | 12 |
| LOC101882770 | 0.09058 | 0.259231735 | 0.086 | 0 | 1 | 12 |
| SEMA4F | 0.09058 | 0.309611195 | 0.086 | 0 | 1 | 12 |
| XLOC-035413 | 0.09058 | 0.247741108 | 0.086 | 0 | 1 | 12 |
| adora2b | 0.09058 | 0.273663783 | 0.086 | 0 | 1 | 12 |
| akt2 | 0.09058 | 0.253313951 | 0.086 | 0 | 1 | 12 |
| arhgap5 | 0.09058 | 0.21474373 | 0.086 | 0 | 1 | 12 |
| atg3 | 0.09058 | 0.265091657 | 0.086 | 0 | 1 | 12 |
| bnip4 | 0.09058 | 0.252905727 | 0.086 | 0 | 1 | 12 |
| borcs8 | 0.09058 | 0.226379171 | 0.086 | 0 | 1 | 12 |
| ccnq | 0.09058 | 0.228213036 | 0.086 | 0 | 1 | 12 |
| cetn4 | 0.09058 | 0.217253103 | 0.086 | 0 | 1 | 12 |
| chrac1 | 0.09058 | 0.238925158 | 0.086 | 0 | 1 | 12 |
| cog4 | 0.09058 | 0.189333918 | 0.086 | 0 | 1 | 12 |
| crema | 0.09058 | 0.249050042 | 0.086 | 0 | 1 | 12 |
| crybg1a | 0.09058 | 0.19026845 | 0.086 | 0 | 1 | 12 |
| dennd2db | 0.09058 | 0.272194014 | 0.086 | 0 | 1 | 12 |

| | | | | | | |
|---|---|---|---|---|---|---|
| dnmbp | 0.09058 | 0.230248649 | 0.086 | 0 | 1 | 12 |
| draxin | 0.09058 | 0.248098332 | 0.086 | 0 | 1 | 12 |
| dtnbp1a | 0.09058 | 0.22821996 | 0.086 | 0 | 1 | 12 |
| etnk1 | 0.09058 | 0.24766262 | 0.086 | 0 | 1 | 12 |
| fam208b | 0.09058 | 0.220205039 | 0.086 | 0 | 1 | 12 |
| gatad2ab | 0.09058 | 0.214646444 | 0.086 | 0 | 1 | 12 |
| got1 | 0.09058 | 0.199547949 | 0.086 | 0 | 1 | 12 |
| gps2 | 0.09058 | 0.226442817 | 0.086 | 0 | 1 | 12 |
| gsto2 | 0.09058 | 0.263731761 | 0.086 | 0 | 1 | 12 |
| gtpbp10 | 0.09058 | 0.247795239 | 0.086 | 0 | 1 | 12 |
| hadhaa | 0.09058 | 0.229084508 | 0.086 | 0 | 1 | 12 |
| insra | 0.09058 | 0.264075007 | 0.086 | 0 | 1 | 12 |
| lamtor2 | 0.09058 | 0.237423995 | 0.086 | 0 | 1 | 12 |
| lamtor4 | 0.09058 | 0.176191669 | 0.086 | 0 | 1 | 12 |
| lmcd1 | 0.09058 | 0.188695451 | 0.086 | 0 | 1 | 12 |
| lrch1 | 0.09058 | 0.23210087 | 0.086 | 0 | 1 | 12 |
| mgat3b | 0.09058 | 0.2834793 | 0.086 | 0 | 1 | 12 |
| mrpl23 | 0.09058 | 0.270755882 | 0.086 | 0 | 1 | 12 |
| ncbp2 | 0.09058 | 0.238188296 | 0.086 | 0 | 1 | 12 |
| nfe2l3 | 0.09058 | 0.223806293 | 0.086 | 0 | 1 | 12 |
| nol12 | 0.09058 | 0.241709343 | 0.086 | 0 | 1 | 12 |
| os9 | 0.09058 | 0.263737433 | 0.086 | 0 | 1 | 12 |
| pmaip1 | 0.09058 | 0.170420997 | 0.086 | 0 | 1 | 12 |
| polr2i | 0.09058 | 0.252991432 | 0.086 | 0 | 1 | 12 |
| ppp5c | 0.09058 | 0.267597522 | 0.086 | 0 | 1 | 12 |
| ptpn23a | 0.09058 | 0.257672797 | 0.086 | 0 | 1 | 12 |
| rab1ba | 0.09058 | 0.258589571 | 0.086 | 0 | 1 | 12 |
| ralgapa2 | 0.09058 | 0.239794664 | 0.086 | 0 | 1 | 12 |
| rassf2a | 0.09058 | 0.237437717 | 0.086 | 0 | 1 | 12 |
| rbm28 | 0.09058 | 0.234990082 | 0.086 | 0 | 1 | 12 |
| rftn2 | 0.09058 | 0.24392183 | 0.086 | 0 | 1 | 12 |
| sephs2 | 0.09058 | 0.26271081 | 0.086 | 0 | 1 | 12 |
| si:ch211-140m22.7 | 0.09058 | 0.160926482 | 0.086 | 0 | 1 | 12 |
| si:ch211-240l19.6 | 0.09058 | 0.386243177 | 0.086 | 0 | 1 | 12 |
| si:ch73-37h15.2 | 0.09058 | 0.255393147 | 0.086 | 0 | 1 | 12 |
| si:dkey-14d8.7 | 0.09058 | 0.135583594 | 0.086 | 0 | 1 | 12 |
| si:dkey-6i22.5 | 0.09058 | 0.238420026 | 0.086 | 0 | 1 | 12 |
| slc16a9a | 0.09058 | 0.223709005 | 0.086 | 0 | 1 | 12 |
| slc25a24 | 0.09058 | 0.211665017 | 0.086 | 0 | 1 | 12 |
| slc25a28 | 0.09058 | 0.231207739 | 0.086 | 0 | 1 | 12 |
| slc30a4 | 0.09058 | 0.265974454 | 0.086 | 0 | 1 | 12 |
| spty2d1 | 0.09058 | 0.241269078 | 0.086 | 0 | 1 | 12 |
| srgap1b | 0.09058 | 0.245199798 | 0.086 | 0 | 1 | 12 |
| sult6b1 | 0.09058 | 0.254824054 | 0.086 | 0 | 1 | 12 |
| surf6 | 0.09058 | 0.277897537 | 0.086 | 0 | 1 | 12 |
| tarbp2 | 0.09058 | 0.23847986 | 0.086 | 0 | 1 | 12 |
| tssc4 | 0.09058 | 0.239929444 | 0.086 | 0 | 1 | 12 |
| twistnb | 0.09058 | 0.270153443 | 0.086 | 0 | 1 | 12 |

| | | | | | | |
|---|---|---|---|---|---|---|
| waca | 0.09058 | 0.221608407 | 0.086 | 0 | 1 | 12 |
| waslb | 0.09058 | 0.250931058 | 0.086 | 0 | 1 | 12 |
| zgc:101783 | 0.09058 | 0.292386094 | 0.086 | 0 | 1 | 12 |
| zgc:162339 | 0.09058 | 0.280512036 | 0.086 | 0 | 1 | 12 |
| zyg11 | 0.09058 | 0.176710646 | 0.086 | 0 | 1 | 12 |
| PAQR9 | 0.09058 | 0.219538293 | 0.086 | 0 | 1 | 12 |
| caskin1 | 0.09058 | 0.238441569 | 0.086 | 0 | 1 | 12 |
| dap | 0.09058 | 0.235583722 | 0.086 | 0 | 1 | 12 |
| dnajc2 | 0.09058 | 0.211664919 | 0.086 | 0 | 1 | 12 |
| erp44 | 0.09058 | 0.280505749 | 0.086 | 0 | 1 | 12 |
| grb2b | 0.09058 | 0.245360472 | 0.086 | 0 | 1 | 12 |
| il6st | 0.09058 | 0.23930956 | 0.086 | 0 | 1 | 12 |
| kdm1a | 0.09058 | 0.288114502 | 0.086 | 0 | 1 | 12 |
| lrrfip2 | 0.09058 | 0.237761241 | 0.086 | 0 | 1 | 12 |
| lxn | 0.09058 | 0.235788237 | 0.086 | 0 | 1 | 12 |
| map3k20 | 0.09058 | 0.294954943 | 0.086 | 0 | 1 | 12 |
| ncoa4 | 0.09058 | 0.234655793 | 0.086 | 0 | 1 | 12 |
| nmi | 0.09058 | 0.219567821 | 0.086 | 0 | 1 | 12 |
| nucb2b | 0.09058 | 0.272653963 | 0.086 | 0 | 1 | 12 |
| rnf126 | 0.09058 | 0.292479414 | 0.086 | 0 | 1 | 12 |
| si:ch211-132g1.3 | 0.09058 | 0.280002466 | 0.086 | 0 | 1 | 12 |
| tll1 | 0.09058 | 0.246405858 | 0.086 | 0 | 1 | 12 |
| zgc:153018 | 0.09058 | 0.292545859 | 0.086 | 0 | 1 | 12 |
| amfra | 0.09058 | 0.176928713 | 0.086 | 0 | 1 | 12 |
| bin2b | 0.09058 | 0.171779791 | 0.086 | 0 | 1 | 12 |
| ctdsp1 | 0.09058 | 0.20143327 | 0.086 | 0 | 1 | 12 |
| edem1 | 0.09058 | 0.214056194 | 0.086 | 0 | 1 | 12 |
| naxe | 0.09058 | 0.260005277 | 0.086 | 0 | 1 | 12 |
| nkd1 | 0.09058 | 0.276905557 | 0.086 | 0 | 1 | 12 |
| nrp2b | 0.09058 | 0.213965082 | 0.086 | 0 | 1 | 12 |
| si:ch211-107n13.1 | 0.09058 | 0.189141301 | 0.086 | 0 | 1 | 12 |
| si:ch211-225h24.2 | 0.09058 | 0.278719182 | 0.086 | 0 | 1 | 12 |
| timeless | 0.09058 | 0.165699522 | 0.086 | 0 | 1 | 12 |
| zfr | 0.09058 | 0.195354425 | 0.086 | 0 | 1 | 12 |
| ap1s2 | 0.09058 | 0.222777686 | 0.086 | 0 | 1 | 12 |
| arrdc3b | 0.09058 | 0.241679536 | 0.086 | 0 | 1 | 12 |
| bahcc1a | 0.09058 | 0.254305703 | 0.086 | 0 | 1 | 12 |
| birc2 | 0.09058 | 0.202177808 | 0.086 | 0 | 1 | 12 |
| castor2 | 0.09058 | 0.202852929 | 0.086 | 0 | 1 | 12 |
| cry-dash | 0.09058 | 0.250463842 | 0.086 | 0 | 1 | 12 |
| dcun1d5 | 0.09058 | 0.191108192 | 0.086 | 0 | 1 | 12 |
| eva1ba | 0.09058 | 0.254531815 | 0.086 | 0 | 1 | 12 |
| fbxo28 | 0.09058 | 0.166067951 | 0.086 | 0 | 1 | 12 |
| fthl31 | 0.09058 | 0.267132876 | 0.086 | 0 | 1 | 12 |
| glulb | 0.09058 | 0.301838753 | 0.086 | 0 | 1 | 12 |
| luc7l | 0.09058 | 0.183233781 | 0.086 | 0 | 1 | 12 |
| march6 | 0.09058 | 0.2055363 | 0.086 | 0 | 1 | 12 |
| nlrc3l1 | 0.09058 | 0.264310493 | 0.086 | 0 | 1 | 12 |

| | | | | | | |
|---|---|---|---|---|---|---|
| pik3ip1 | 0.09058 | 0.22634024 | 0.086 | 0 | 1 | 12 |
| rbms2a | 0.09058 | 0.215760752 | 0.086 | 0 | 1 | 12 |
| rnf169 | 0.09058 | 0.216329395 | 0.086 | 0 | 1 | 12 |
| scpep1 | 0.09058 | 0.176551819 | 0.086 | 0 | 1 | 12 |
| selenow2a | 0.09058 | 0.252239858 | 0.086 | 0 | 1 | 12 |
| si:ch211-51h9.7 | 0.09058 | 0.197061708 | 0.086 | 0 | 1 | 12 |
| tbk1 | 0.09058 | 0.18235021 | 0.086 | 0 | 1 | 12 |
| thtpa | 0.09058 | 0.289633735 | 0.086 | 0 | 1 | 12 |
| tsc22d1 | 0.09058 | 0.175824101 | 0.086 | 0 | 1 | 12 |
| zgc:92140 | 0.09058 | 0.205722839 | 0.086 | 0 | 1 | 12 |
| CABZ01044053.1 | 0.09058 | 0.257077922 | 0.086 | 0 | 1 | 12 |
| ctdspla | 0.09058 | 0.247534203 | 0.086 | 0 | 1 | 12 |
| f11r.1 | 0.09058 | 0.194930837 | 0.086 | 0 | 1 | 12 |
| heg1 | 0.09058 | 0.264045394 | 0.086 | 0 | 1 | 12 |
| hhex | 0.09058 | 0.199957365 | 0.086 | 0 | 1 | 12 |
| map3k3 | 0.09058 | 0.2499363 | 0.086 | 0 | 1 | 12 |
| meox2b | 0.09058 | 0.288712878 | 0.086 | 0 | 1 | 12 |
| pgpep1 | 0.09058 | 0.265821533 | 0.086 | 0 | 1 | 12 |
| pkn1a | 0.09058 | 0.267481139 | 0.086 | 0 | 1 | 12 |
| rsrp1 | 0.09058 | 0.212727698 | 0.086 | 0 | 1 | 12 |
| si:ch211-225b11.1 | 0.09058 | 0.209732458 | 0.086 | 0 | 1 | 12 |
| sorbs1 | 0.09058 | 0.263343318 | 0.086 | 0 | 1 | 12 |
| atl3 | 0.09058 | 0.26099967 | 0.086 | 0 | 1 | 12 |
| clk2b | 0.09058 | 0.233925879 | 0.086 | 0 | 1 | 12 |
| dbpb | 0.09058 | 0.203955634 | 0.086 | 0 | 1 | 12 |
| ddx51 | 0.09058 | 0.221918475 | 0.086 | 0 | 1 | 12 |
| ide | 0.09058 | 0.213096169 | 0.086 | 0 | 1 | 12 |
| mir21-1 | 0.09058 | 0.276276387 | 0.086 | 0 | 1 | 12 |
| mrpl45 | 0.09058 | 0.299367011 | 0.086 | 0 | 1 | 12 |
| puraa | 0.09058 | 0.20346214 | 0.086 | 0 | 1 | 12 |
| rras | 0.09058 | 0.286312901 | 0.086 | 0 | 1 | 12 |
| zgc:162939 | 0.09058 | 0.258201561 | 0.086 | 0 | 1 | 12 |
| kbtbd8 | 0.09058 | 0.284408828 | 0.086 | 0 | 1 | 12 |
| ubr2 | 0.09058 | 0.266368834 | 0.086 | 0 | 1 | 12 |
| COX7A2 (1 of many) | 0.09145 | 0.535507911 | 0.257 | 0.074 | 1 | 12 |
| elovl1b | 0.09173 | 0.476719379 | 0.286 | 0.074 | 1 | 12 |
| lsm12b | 0.09283 | 0.496084499 | 0.286 | 0.074 | 1 | 12 |
| cox4i1 | 0.09783 | 0.387449125 | 0.6 | 0.333 | 1 | 12 |
| rad21a | 0.09786 | -0.4933119 | 0.086 | 0.296 | 1 | 12 |
| stat1a | 0.09852 | -0.3821248 | 0.029 | 0.185 | 1 | 12 |
| rpl35a | 0.10014 | 0.158983575 | 0.971 | 0.815 | 1 | 12 |
| FQ976913.1 | 0.1009 | -0.20437405 | 0 | 0.074 | 1 | 12 |
| atad2 | 0.1009 | -0.23088891 | 0 | 0.074 | 1 | 12 |
| atp6v1ba | 0.1009 | -0.17066925 | 0 | 0.074 | 1 | 12 |
| bap1 | 0.1009 | -0.25238449 | 0 | 0.074 | 1 | 12 |
| bmp2k | 0.1009 | -0.2178408 | 0 | 0.074 | 1 | 12 |
| cand1 | 0.1009 | -0.20370666 | 0 | 0.074 | 1 | 12 |
| celsr1b | 0.1009 | -0.20453178 | 0 | 0.074 | 1 | 12 |

| | | | | | | |
|---|---|---|---|---|---|---|
| clgn | 0.1009 | -0.26082678 | 0 | 0.074 | 1 | 12 |
| crnkl1 | 0.1009 | -0.25793667 | 0 | 0.074 | 1 | 12 |
| dpcd | 0.1009 | -0.24074653 | 0 | 0.074 | 1 | 12 |
| ei24 | 0.1009 | -0.21069396 | 0 | 0.074 | 1 | 12 |
| esco2 | 0.1009 | -0.22875395 | 0 | 0.074 | 1 | 12 |
| furina | 0.1009 | -0.1872342 | 0 | 0.074 | 1 | 12 |
| gdf7 | 0.1009 | -0.20453178 | 0 | 0.074 | 1 | 12 |
| guk1a | 0.1009 | -0.21625956 | 0 | 0.074 | 1 | 12 |
| hoxb7a | 0.1009 | -0.23766643 | 0 | 0.074 | 1 | 12 |
| marcksa | 0.1009 | -0.20408856 | 0 | 0.074 | 1 | 12 |
| mfap3l | 0.1009 | -0.2339701 | 0 | 0.074 | 1 | 12 |
| mgat4b | 0.1009 | -0.22071443 | 0 | 0.074 | 1 | 12 |
| mmp15a | 0.1009 | -0.21613203 | 0 | 0.074 | 1 | 12 |
| oafb | 0.1009 | -0.17853737 | 0 | 0.074 | 1 | 12 |
| pigt | 0.1009 | -0.21082638 | 0 | 0.074 | 1 | 12 |
| plcd1a | 0.1009 | -0.23448738 | 0 | 0.074 | 1 | 12 |
| ppp3ccb | 0.1009 | -0.18654161 | 0 | 0.074 | 1 | 12 |
| rnf11a | 0.1009 | -0.2185019 | 0 | 0.074 | 1 | 12 |
| samm50l | 0.1009 | -0.22381516 | 0 | 0.074 | 1 | 12 |
| snrnp200 | 0.1009 | -0.18787509 | 0 | 0.074 | 1 | 12 |
| srsf1a | 0.1009 | -0.2337391 | 0 | 0.074 | 1 | 12 |
| tmem150ab | 0.1009 | -0.21019033 | 0 | 0.074 | 1 | 12 |
| tmem254 | 0.1009 | -0.17290844 | 0 | 0.074 | 1 | 12 |
| tpst1 | 0.1009 | -0.18652575 | 0 | 0.074 | 1 | 12 |
| usp9 | 0.1009 | -0.20459544 | 0 | 0.074 | 1 | 12 |
| zfr2 | 0.1009 | -0.2053746 | 0 | 0.074 | 1 | 12 |
| znf207b | 0.1009 | -0.202547 | 0 | 0.074 | 1 | 12 |
| abhd17ab | 0.1009 | -0.22450971 | 0 | 0.074 | 1 | 12 |
| acvr1ba | 0.1009 | -0.19177808 | 0 | 0.074 | 1 | 12 |
| ambra1a | 0.1009 | -0.25037765 | 0 | 0.074 | 1 | 12 |
| amot | 0.1009 | -0.22729249 | 0 | 0.074 | 1 | 12 |
| asna1 | 0.1009 | -0.19013088 | 0 | 0.074 | 1 | 12 |
| birc5a | 0.1009 | -0.21504052 | 0 | 0.074 | 1 | 12 |
| clpp | 0.1009 | -0.1763925 | 0 | 0.074 | 1 | 12 |
| dedd | 0.1009 | -0.23945263 | 0 | 0.074 | 1 | 12 |
| dhrs11a | 0.1009 | -0.2458986 | 0 | 0.074 | 1 | 12 |
| dkk3b | 0.1009 | -0.29024016 | 0 | 0.074 | 1 | 12 |
| emc3 | 0.1009 | -0.2311821 | 0 | 0.074 | 1 | 12 |
| hdac4 | 0.1009 | -0.24859577 | 0 | 0.074 | 1 | 12 |
| idh3b | 0.1009 | -0.22610501 | 0 | 0.074 | 1 | 12 |
| il4r.1 | 0.1009 | -0.24753899 | 0 | 0.074 | 1 | 12 |
| ing4 | 0.1009 | -0.30784574 | 0 | 0.074 | 1 | 12 |
| marveld2a | 0.1009 | -0.18611476 | 0 | 0.074 | 1 | 12 |
| mxd4 | 0.1009 | -0.18542835 | 0 | 0.074 | 1 | 12 |
| mxi1 | 0.1009 | -0.23945441 | 0 | 0.074 | 1 | 12 |
| nr1d2b | 0.1009 | -0.18545886 | 0 | 0.074 | 1 | 12 |
| pbx1b | 0.1009 | -0.26292642 | 0 | 0.074 | 1 | 12 |
| polr1d | 0.1009 | -0.24074352 | 0 | 0.074 | 1 | 12 |

| | | | | | | |
|---|---|---|---|---|---|---|
| rasal2 | 0.1009 | -0.23790401 | 0 | 0.074 | 1 | 12 |
| rbm17 | 0.1009 | -0.27079564 | 0 | 0.074 | 1 | 12 |
| rmnd5b | 0.1009 | -0.22937335 | 0 | 0.074 | 1 | 12 |
| schip1 | 0.1009 | -0.18684562 | 0 | 0.074 | 1 | 12 |
| sgcd | 0.1009 | -0.21613884 | 0 | 0.074 | 1 | 12 |
| si:ch73-238c9.1 | 0.1009 | -0.18146858 | 0 | 0.074 | 1 | 12 |
| tbl1xr1a | 0.1009 | -0.30254264 | 0 | 0.074 | 1 | 12 |
| tead3b | 0.1009 | -0.26405672 | 0 | 0.074 | 1 | 12 |
| uck2b | 0.1009 | -0.1816106 | 0 | 0.074 | 1 | 12 |
| uxt | 0.1009 | -0.27181541 | 0 | 0.074 | 1 | 12 |
| vkorc1l1 | 0.1009 | -0.20650538 | 0 | 0.074 | 1 | 12 |
| wu:fu71h07 | 0.1009 | -0.23451661 | 0 | 0.074 | 1 | 12 |
| BX908782.2 | 0.1009 | -0.25601797 | 0 | 0.074 | 1 | 12 |
| CABZ01076667.1 | 0.1009 | -0.2758893 | 0 | 0.074 | 1 | 12 |
| DOCK9 | 0.1009 | -0.19171674 | 0 | 0.074 | 1 | 12 |
| LOC101882826 | 0.1009 | -0.13922516 | 0 | 0.074 | 1 | 12 |
| LOC103910363 | 0.1009 | -0.18144759 | 0 | 0.074 | 1 | 12 |
| acvr2ab | 0.1009 | -0.18594455 | 0 | 0.074 | 1 | 12 |
| adam17b | 0.1009 | -0.1895949 | 0 | 0.074 | 1 | 12 |
| arnt2 | 0.1009 | -0.18947116 | 0 | 0.074 | 1 | 12 |
| ca4a | 0.1009 | -0.22953415 | 0 | 0.074 | 1 | 12 |
| calcrla | 0.1009 | -0.20233554 | 0 | 0.074 | 1 | 12 |
| calm1b | 0.1009 | -0.1661347 | 0 | 0.074 | 1 | 12 |
| cers2a | 0.1009 | -0.13737449 | 0 | 0.074 | 1 | 12 |
| cfap20 | 0.1009 | -0.13481742 | 0 | 0.074 | 1 | 12 |
| ddx11 | 0.1009 | -0.1636508 | 0 | 0.074 | 1 | 12 |
| ddx54 | 0.1009 | -0.13532491 | 0 | 0.074 | 1 | 12 |
| fam193a | 0.1009 | -0.13611865 | 0 | 0.074 | 1 | 12 |
| fen1 | 0.1009 | -0.20712629 | 0 | 0.074 | 1 | 12 |
| gmps | 0.1009 | -0.21972074 | 0 | 0.074 | 1 | 12 |
| gtf3c3 | 0.1009 | -0.21901445 | 0 | 0.074 | 1 | 12 |
| gxylt2 | 0.1009 | -0.19773038 | 0 | 0.074 | 1 | 12 |
| hspa14 | 0.1009 | -0.22216618 | 0 | 0.074 | 1 | 12 |
| ilf3a | 0.1009 | -0.20917156 | 0 | 0.074 | 1 | 12 |
| kbtbd11 | 0.1009 | -0.19422341 | 0 | 0.074 | 1 | 12 |
| kif11 | 0.1009 | -0.22418713 | 0 | 0.074 | 1 | 12 |
| kif22 | 0.1009 | -0.26392089 | 0 | 0.074 | 1 | 12 |
| klf12b | 0.1009 | -0.13709406 | 0 | 0.074 | 1 | 12 |
| larp6a | 0.1009 | -0.23066509 | 0 | 0.074 | 1 | 12 |
| mad2l1 | 0.1009 | -0.12263855 | 0 | 0.074 | 1 | 12 |
| mcm4 | 0.1009 | -0.21854266 | 0 | 0.074 | 1 | 12 |
| mrpl32 | 0.1009 | -0.27744583 | 0 | 0.074 | 1 | 12 |
| msh6 | 0.1009 | -0.20292563 | 0 | 0.074 | 1 | 12 |
| mxra8a | 0.1009 | -0.19425786 | 0 | 0.074 | 1 | 12 |
| nhsl1b | 0.1009 | -0.1658444 | 0 | 0.074 | 1 | 12 |
| nup210 | 0.1009 | -0.16557197 | 0 | 0.074 | 1 | 12 |
| otud5b | 0.1009 | -0.22284138 | 0 | 0.074 | 1 | 12 |
| pcdh1b | 0.1009 | -0.13802897 | 0 | 0.074 | 1 | 12 |

| | | | | | | |
|---|---|---|---|---|---|---|
| ppm1aa | 0.1009 | -0.28050103 | 0 | 0.074 | 1 | 12 |
| prc1a | 0.1009 | -0.21015851 | 0 | 0.074 | 1 | 12 |
| ptpn4a | 0.1009 | -0.2379809 | 0 | 0.074 | 1 | 12 |
| rb1 | 0.1009 | -0.16442671 | 0 | 0.074 | 1 | 12 |
| rfc4 | 0.1009 | -0.25681398 | 0 | 0.074 | 1 | 12 |
| rnd1a | 0.1009 | -0.19494556 | 0 | 0.074 | 1 | 12 |
| sbf1 | 0.1009 | -0.19434905 | 0 | 0.074 | 1 | 12 |
| senp7b | 0.1009 | -0.1825723 | 0 | 0.074 | 1 | 12 |
| si:ch211-156b7.4 | 0.1009 | -0.20460328 | 0 | 0.074 | 1 | 12 |
| si:ch211-156j22.4 | 0.1009 | -0.26611672 | 0 | 0.074 | 1 | 12 |
| si:dkey-102f14.5 | 0.1009 | -0.21291564 | 0 | 0.074 | 1 | 12 |
| sptlc1 | 0.1009 | -0.22601323 | 0 | 0.074 | 1 | 12 |
| ssbp3a | 0.1009 | -0.13473575 | 0 | 0.074 | 1 | 12 |
| st3gal8 | 0.1009 | -0.21131702 | 0 | 0.074 | 1 | 12 |
| stk17al | 0.1009 | -0.25105842 | 0 | 0.074 | 1 | 12 |
| svbp | 0.1009 | -0.22311439 | 0 | 0.074 | 1 | 12 |
| tgm1l1 | 0.1009 | -0.27594554 | 0 | 0.074 | 1 | 12 |
| tmem63ba | 0.1009 | -0.18620011 | 0 | 0.074 | 1 | 12 |
| tprb | 0.1009 | -0.26507623 | 0 | 0.074 | 1 | 12 |
| traf2b | 0.1009 | -0.16583825 | 0 | 0.074 | 1 | 12 |
| ttc9c | 0.1009 | -0.21407483 | 0 | 0.074 | 1 | 12 |
| tusc2b | 0.1009 | -0.13523632 | 0 | 0.074 | 1 | 12 |
| vrk1 | 0.1009 | -0.16118442 | 0 | 0.074 | 1 | 12 |
| vrk3 | 0.1009 | -0.26281801 | 0 | 0.074 | 1 | 12 |
| wu:fc17b08 | 0.1009 | -0.19082727 | 0 | 0.074 | 1 | 12 |
| znf704 | 0.1009 | -0.13468522 | 0 | 0.074 | 1 | 12 |
| amdhd2 | 0.1009 | -0.2868969 | 0 | 0.074 | 1 | 12 |
| carm1 | 0.1009 | -0.28482915 | 0 | 0.074 | 1 | 12 |
| ing5b | 0.1009 | -0.26747823 | 0 | 0.074 | 1 | 12 |
| marveld1 | 0.1009 | -0.27578039 | 0 | 0.074 | 1 | 12 |
| rflnb | 0.1009 | -0.27556837 | 0 | 0.074 | 1 | 12 |
| amotl2a | 0.1009 | -0.19997856 | 0 | 0.074 | 1 | 12 |
| cse1l | 0.1009 | -0.19705477 | 0 | 0.074 | 1 | 12 |
| dcbld1 | 0.1009 | -0.20087577 | 0 | 0.074 | 1 | 12 |
| dhx37 | 0.1009 | -0.19642578 | 0 | 0.074 | 1 | 12 |
| mcf2l2 | 0.1009 | -0.19605335 | 0 | 0.074 | 1 | 12 |
| pole3 | 0.1009 | -0.19352479 | 0 | 0.074 | 1 | 12 |
| rnaseka | 0.1009 | -0.26380143 | 0 | 0.074 | 1 | 12 |
| tspan4a | 0.1009 | -0.21359205 | 0 | 0.074 | 1 | 12 |
| ube2r2 | 0.1009 | -0.19668265 | 0 | 0.074 | 1 | 12 |
| ATP11B | 0.1009 | -0.22985356 | 0 | 0.074 | 1 | 12 |
| CR847936.1 | 0.1009 | -0.19481447 | 0 | 0.074 | 1 | 12 |
| CU929070.1 | 0.1009 | -0.20044767 | 0 | 0.074 | 1 | 12 |
| LOC100537613 | 0.1009 | -0.1992926 | 0 | 0.074 | 1 | 12 |
| aclya | 0.1009 | -0.2427777 | 0 | 0.074 | 1 | 12 |
| akt1s1 | 0.1009 | -0.28601171 | 0 | 0.074 | 1 | 12 |
| ankrd54 | 0.1009 | -0.20125252 | 0 | 0.074 | 1 | 12 |
| ascc1 | 0.1009 | -0.20725876 | 0 | 0.074 | 1 | 12 |

| | | | | | | |
|---|---|---|---|---|---|---|
| b9d2 | 0.1009 | -0.21049168 | 0 | 0.074 | 1 | 12 |
| bahd1 | 0.1009 | -0.25315234 | 0 | 0.074 | 1 | 12 |
| ccnb1 | 0.1009 | -0.25871709 | 0 | 0.074 | 1 | 12 |
| cpne2 | 0.1009 | -0.25982933 | 0 | 0.074 | 1 | 12 |
| ctdspl2b | 0.1009 | -0.2175175 | 0 | 0.074 | 1 | 12 |
| cyyr1 | 0.1009 | -0.25648138 | 0 | 0.074 | 1 | 12 |
| gga3a | 0.1009 | -0.2441236 | 0 | 0.074 | 1 | 12 |
| grap2a | 0.1009 | -0.24413535 | 0 | 0.074 | 1 | 12 |
| gsk3b | 0.1009 | -0.21227476 | 0 | 0.074 | 1 | 12 |
| gtf2h4 | 0.1009 | -0.19075402 | 0 | 0.074 | 1 | 12 |
| hnrnpc | 0.1009 | -0.31497393 | 0 | 0.074 | 1 | 12 |
| kdm6al | 0.1009 | -0.21414364 | 0 | 0.074 | 1 | 12 |
| llph | 0.1009 | -0.1941487 | 0 | 0.074 | 1 | 12 |
| mrps9 | 0.1009 | -0.21046075 | 0 | 0.074 | 1 | 12 |
| mtfp1 | 0.1009 | -0.22779584 | 0 | 0.074 | 1 | 12 |
| pold1 | 0.1009 | -0.22363258 | 0 | 0.074 | 1 | 12 |
| prpf19 | 0.1009 | -0.2497489 | 0 | 0.074 | 1 | 12 |
| prrg4 | 0.1009 | -0.26728824 | 0 | 0.074 | 1 | 12 |
| rab3db | 0.1009 | -0.191346 | 0 | 0.074 | 1 | 12 |
| rnd1b | 0.1009 | -0.23145538 | 0 | 0.074 | 1 | 12 |
| sco2 | 0.1009 | -0.24990476 | 0 | 0.074 | 1 | 12 |
| serpina10a | 0.1009 | -0.28366314 | 0 | 0.074 | 1 | 12 |
| si:ch211-243j20.2 | 0.1009 | -0.26541825 | 0 | 0.074 | 1 | 12 |
| si:zfos-1192g2.3 | 0.1009 | -0.17207884 | 0 | 0.074 | 1 | 12 |
| slc6a9 | 0.1009 | -0.25068179 | 0 | 0.074 | 1 | 12 |
| srd5a2a | 0.1009 | -0.24903205 | 0 | 0.074 | 1 | 12 |
| tecrb | 0.1009 | -0.25898528 | 0 | 0.074 | 1 | 12 |
| tmem11 | 0.1009 | -0.28333953 | 0 | 0.074 | 1 | 12 |
| tnfsf10l4 | 0.1009 | -0.29654852 | 0 | 0.074 | 1 | 12 |
| zgc:171967 | 0.1009 | -0.178836 | 0 | 0.074 | 1 | 12 |
| zgc:77739 | 0.1009 | -0.1904366 | 0 | 0.074 | 1 | 12 |
| znf865 | 0.1009 | -0.17700086 | 0 | 0.074 | 1 | 12 |
| GNB4 | 0.1009 | -0.27635256 | 0 | 0.074 | 1 | 12 |
| agpat4 | 0.1009 | -0.24870966 | 0 | 0.074 | 1 | 12 |
| apmap | 0.1009 | -0.23864439 | 0 | 0.074 | 1 | 12 |
| edc4 | 0.1009 | -0.233236 | 0 | 0.074 | 1 | 12 |
| gatad1 | 0.1009 | -0.29664611 | 0 | 0.074 | 1 | 12 |
| gins2 | 0.1009 | -0.23850975 | 0 | 0.074 | 1 | 12 |
| jag2b | 0.1009 | -0.22529753 | 0 | 0.074 | 1 | 12 |
| kaznb | 0.1009 | -0.23438557 | 0 | 0.074 | 1 | 12 |
| mia3 | 0.1009 | -0.24247264 | 0 | 0.074 | 1 | 12 |
| mpped2a | 0.1009 | -0.25962963 | 0 | 0.074 | 1 | 12 |
| mprip | 0.1009 | -0.22529803 | 0 | 0.074 | 1 | 12 |
| ndfip2 | 0.1009 | -0.22426726 | 0 | 0.074 | 1 | 12 |
| nup98 | 0.1009 | -0.24567032 | 0 | 0.074 | 1 | 12 |
| orc2 | 0.1009 | -0.27480809 | 0 | 0.074 | 1 | 12 |
| si:ch211-198a12.6 | 0.1009 | -0.25723522 | 0 | 0.074 | 1 | 12 |
| si:ch211-261d7.3 | 0.1009 | -0.24003935 | 0 | 0.074 | 1 | 12 |

| | | | | | | |
|---|---|---|---|---|---|---|
| slc12a2 | 0.1009 | -0.22583436 | 0 | 0.074 | 1 | 12 |
| tmem26a | 0.1009 | -0.23154016 | 0 | 0.074 | 1 | 12 |
| zeb1b | 0.1009 | -0.24120734 | 0 | 0.074 | 1 | 12 |
| zgc:152791 | 0.1009 | -0.27659897 | 0 | 0.074 | 1 | 12 |
| KIF2A | 0.1009 | -0.22434058 | 0 | 0.074 | 1 | 12 |
| adipor1a | 0.1009 | -0.18825428 | 0 | 0.074 | 1 | 12 |
| apooa | 0.1009 | -0.18767718 | 0 | 0.074 | 1 | 12 |
| cenpw | 0.1009 | -0.16932486 | 0 | 0.074 | 1 | 12 |
| clk2a | 0.1009 | -0.17466366 | 0 | 0.074 | 1 | 12 |
| dnase1l4.2 | 0.1009 | -0.29792241 | 0 | 0.074 | 1 | 12 |
| gmeb1 | 0.1009 | -0.20102242 | 0 | 0.074 | 1 | 12 |
| mettl9 | 0.1009 | -0.17300725 | 0 | 0.074 | 1 | 12 |
| nle1 | 0.1009 | -0.28222622 | 0 | 0.074 | 1 | 12 |
| osgep | 0.1009 | -0.17047028 | 0 | 0.074 | 1 | 12 |
| polr3k | 0.1009 | -0.19124255 | 0 | 0.074 | 1 | 12 |
| si:ch211-221j21.3 | 0.1009 | -0.30749289 | 0 | 0.074 | 1 | 12 |
| slc39a7 | 0.1009 | -0.17395548 | 0 | 0.074 | 1 | 12 |
| sufu | 0.1009 | -0.2693676 | 0 | 0.074 | 1 | 12 |
| trappc2l | 0.1009 | -0.17700166 | 0 | 0.074 | 1 | 12 |
| tubg1 | 0.1009 | -0.18966413 | 0 | 0.074 | 1 | 12 |
| vegfab | 0.1009 | -0.26986969 | 0 | 0.074 | 1 | 12 |
| LOC103911473 | 0.1009 | -0.20161732 | 0 | 0.074 | 1 | 12 |
| XLOC-018182 | 0.1009 | -0.14033422 | 0 | 0.074 | 1 | 12 |
| arfgap2 | 0.1009 | -0.10008846 | 0 | 0.074 | 1 | 12 |
| cactin | 0.1009 | -0.24509142 | 0 | 0.074 | 1 | 12 |
| chmp1a | 0.1009 | -0.2643355 | 0 | 0.074 | 1 | 12 |
| cmip | 0.1009 | -0.20692822 | 0 | 0.074 | 1 | 12 |
| cux1a | 0.1009 | -0.23061153 | 0 | 0.074 | 1 | 12 |
| ebf1b | 0.1009 | -0.19818981 | 0 | 0.074 | 1 | 12 |
| esyt2a | 0.1009 | -0.14898996 | 0 | 0.074 | 1 | 12 |
| ftr50 | 0.1009 | -0.14755 | 0 | 0.074 | 1 | 12 |
| hax1 | 0.1009 | -0.26071405 | 0 | 0.074 | 1 | 12 |
| hcfc1a | 0.1009 | -0.23469138 | 0 | 0.074 | 1 | 12 |
| lap3 | 0.1009 | -0.20518361 | 0 | 0.074 | 1 | 12 |
| leo1 | 0.1009 | -0.20233751 | 0 | 0.074 | 1 | 12 |
| luzp1 | 0.1009 | -0.20196677 | 0 | 0.074 | 1 | 12 |
| map4k5 | 0.1009 | -0.1871327 | 0 | 0.074 | 1 | 12 |
| mapk1 | 0.1009 | -0.18784106 | 0 | 0.074 | 1 | 12 |
| ndufaf2 | 0.1009 | -0.14596384 | 0 | 0.074 | 1 | 12 |
| ppp4r3b | 0.1009 | -0.20586767 | 0 | 0.074 | 1 | 12 |
| rybpa | 0.1009 | -0.17423757 | 0 | 0.074 | 1 | 12 |
| samd4b | 0.1009 | -0.18984881 | 0 | 0.074 | 1 | 12 |
| si:dkey-17o15.2 | 0.1009 | -0.14618038 | 0 | 0.074 | 1 | 12 |
| slc35a4 | 0.1009 | -0.2027808 | 0 | 0.074 | 1 | 12 |
| st3gal3a | 0.1009 | -0.14834008 | 0 | 0.074 | 1 | 12 |
| tpmt.1 | 0.1009 | -0.26585877 | 0 | 0.074 | 1 | 12 |
| utrn | 0.1009 | -0.14885697 | 0 | 0.074 | 1 | 12 |
| vwa8 | 0.1009 | -0.14918013 | 0 | 0.074 | 1 | 12 |

| | | | | | | |
|---|---|---|---|---|---|---|
| cdca4 | 0.1009 | -0.12518308 | 0 | 0.074 | 1 | 12 |
| cttnbp2nlb | 0.1009 | -0.18870684 | 0 | 0.074 | 1 | 12 |
| dcun1d1 | 0.1009 | -0.15956447 | 0 | 0.074 | 1 | 12 |
| fam76b | 0.1009 | -0.30454941 | 0 | 0.074 | 1 | 12 |
| hm13 | 0.1009 | -0.11993653 | 0 | 0.074 | 1 | 12 |
| ncapd2 | 0.1009 | -0.20184079 | 0 | 0.074 | 1 | 12 |
| ncaph | 0.1009 | -0.20346519 | 0 | 0.074 | 1 | 12 |
| nsmce2 | 0.1009 | -0.19363737 | 0 | 0.074 | 1 | 12 |
| p3h1 | 0.1009 | -0.12289144 | 0 | 0.074 | 1 | 12 |
| pdia4 | 0.1009 | -0.12410045 | 0 | 0.074 | 1 | 12 |
| polb | 0.1009 | -0.12259124 | 0 | 0.074 | 1 | 12 |
| rca2.1 | 0.1009 | -0.20061717 | 0 | 0.074 | 1 | 12 |
| slc16a9b | 0.1009 | -0.12296116 | 0 | 0.074 | 1 | 12 |
| slkb | 0.1009 | -0.12462278 | 0 | 0.074 | 1 | 12 |
| tom1l2a | 0.1009 | -0.12362722 | 0 | 0.074 | 1 | 12 |
| ube2g1b | 0.1009 | -0.28300435 | 0 | 0.074 | 1 | 12 |
| ube4b | 0.1009 | -0.15074244 | 0 | 0.074 | 1 | 12 |
| ubl7b | 0.1009 | -0.14668318 | 0 | 0.074 | 1 | 12 |
| usp12a | 0.1009 | -0.12378728 | 0 | 0.074 | 1 | 12 |
| wdr44 | 0.1009 | -0.12199492 | 0 | 0.074 | 1 | 12 |
| wdr61 | 0.1009 | -0.19142627 | 0 | 0.074 | 1 | 12 |
| zbtb10 | 0.1009 | -0.2020061 | 0 | 0.074 | 1 | 12 |
| zdhhc6 | 0.1009 | -0.12293798 | 0 | 0.074 | 1 | 12 |
| zgc:66443 | 0.1009 | -0.23266826 | 0 | 0.074 | 1 | 12 |
| zgpat | 0.1009 | -0.20081887 | 0 | 0.074 | 1 | 12 |
| znf1035 | 0.1009 | -0.12520589 | 0 | 0.074 | 1 | 12 |
| znf609b | 0.1009 | -0.16748361 | 0 | 0.074 | 1 | 12 |
| CABZ01074307.1 | 0.1009 | -0.17585329 | 0 | 0.074 | 1 | 12 |
| cdc14b | 0.1009 | -0.16249673 | 0 | 0.074 | 1 | 12 |
| exoc4 | 0.1009 | -0.1656924 | 0 | 0.074 | 1 | 12 |
| fam135a | 0.1009 | -0.16390488 | 0 | 0.074 | 1 | 12 |
| notch1a | 0.1009 | -0.17716056 | 0 | 0.074 | 1 | 12 |
| ntrk2a | 0.1009 | -0.16437682 | 0 | 0.074 | 1 | 12 |
| pafah1b1b | 0.1009 | -0.20548384 | 0 | 0.074 | 1 | 12 |
| srp9 | 0.1009 | -0.16240824 | 0 | 0.074 | 1 | 12 |
| tmed1b | 0.1009 | -0.16518137 | 0 | 0.074 | 1 | 12 |
| tmtc2a | 0.1009 | -0.14497089 | 0 | 0.074 | 1 | 12 |
| wu:fi46h04 | 0.1009 | -0.16515477 | 0 | 0.074 | 1 | 12 |
| ciao2b | 0.10219 | -0.4770368 | 0.029 | 0.185 | 1 | 12 |
| tcima | 0.10264 | 0.308373576 | 0.343 | 0.148 | 1 | 12 |
| npm3 | 0.10306 | -0.35580258 | 0.029 | 0.185 | 1 | 12 |
| snrpd1 | 0.10408 | -0.44073367 | 0.143 | 0.37 | 1 | 12 |
| hnrnpl | 0.10501 | -0.36710358 | 0.029 | 0.185 | 1 | 12 |
| sinhcafl | 0.10601 | -0.41383279 | 0.029 | 0.185 | 1 | 12 |
| ifi45 | 0.10619 | -0.59849312 | 0.057 | 0.222 | 1 | 12 |
| dut | 0.10846 | -0.53293824 | 0.114 | 0.333 | 1 | 12 |
| selenou1a | 0.11022 | -0.43714471 | 0.029 | 0.185 | 1 | 12 |
| taf12 | 0.11097 | -0.39660497 | 0.029 | 0.185 | 1 | 12 |

| | | | | | | |
|---|---|---|---|---|---|---|
| ilk | 0.11107 | -0.45118367 | 0.029 | 0.185 | 1 | 12 |
| mt-nd3 | 0.1116 | -0.52470399 | 0.171 | 0.407 | 1 | 12 |
| etf1b | 0.11162 | -0.45049275 | 0.029 | 0.185 | 1 | 12 |
| stmn1a | 0.11173 | -0.39672297 | 0.029 | 0.185 | 1 | 12 |
| grapa | 0.11188 | -0.45812961 | 0.029 | 0.185 | 1 | 12 |
| arf1 | 0.11203 | -0.34090397 | 0.029 | 0.185 | 1 | 12 |
| zgc:158343 | 0.1123 | 0.481523301 | 0.2 | 0.037 | 1 | 12 |
| krt18a.1 | 0.11247 | 0.394495477 | 0.771 | 0.519 | 1 | 12 |
| tuba8l4 | 0.11556 | -0.3396306 | 0.543 | 0.63 | 1 | 12 |
| lman2 | 0.116 | 0.375438418 | 0.257 | 0.074 | 1 | 12 |
| qkia | 0.11657 | 0.506690264 | 0.429 | 0.185 | 1 | 12 |
| cfd | 0.11827 | 0.638358093 | 0.229 | 0.111 | 1 | 12 |
| snx1a | 0.1191 | -0.53071127 | 0.057 | 0.222 | 1 | 12 |
| she | 0.12156 | 0.475845403 | 0.514 | 0.259 | 1 | 12 |
| zmat2 | 0.12302 | 0.449536761 | 0.2 | 0.037 | 1 | 12 |
| ISCU (1 of many) | 0.1241 | 0.307569265 | 0.2 | 0.037 | 1 | 12 |
| hnrnpa0b | 0.12646 | -0.1393324 | 0.6 | 0.556 | 1 | 12 |
| uba52 | 0.12679 | 0.154489368 | 0.886 | 0.704 | 1 | 12 |
| lgals3b | 0.12804 | 0.384012125 | 0.257 | 0.074 | 1 | 12 |
| cavin2b | 0.12919 | 0.450883143 | 0.371 | 0.148 | 1 | 12 |
| rpl22l1 | 0.1319 | -0.17202316 | 0.686 | 0.667 | 1 | 12 |
| ndufb9 | 0.13249 | 0.405395661 | 0.2 | 0.037 | 1 | 12 |
| fblim1 | 0.13283 | 0.395983492 | 0.2 | 0.037 | 1 | 12 |
| snrpc | 0.13318 | -0.52326625 | 0.086 | 0.259 | 1 | 12 |
| tmsb1 | 0.13343 | 0.600285569 | 0.371 | 0.148 | 1 | 12 |
| btg2 | 0.13533 | 0.526706766 | 0.229 | 0.074 | 1 | 12 |
| edf1 | 0.13829 | 0.532593509 | 0.314 | 0.148 | 1 | 12 |
| rhocb | 0.14061 | -0.45394166 | 0.4 | 0.556 | 1 | 12 |
| max | 0.14073 | 0.389887426 | 0.257 | 0.074 | 1 | 12 |
| si:ch73-42p12.2 | 0.14097 | 0.463695421 | 0.257 | 0.074 | 1 | 12 |
| eif3m | 0.1444 | 0.215450005 | 0.486 | 0.296 | 1 | 12 |
| anxa1a | 0.14523 | 0.506895446 | 0.257 | 0.074 | 1 | 12 |
| zfp36l2 | 0.14596 | 0.486785807 | 0.314 | 0.111 | 1 | 12 |
| cpne1 | 0.14609 | -0.36783253 | 0.057 | 0.222 | 1 | 12 |
| ndufs6 | 0.1463 | 0.466152274 | 0.314 | 0.111 | 1 | 12 |
| anxa13 | 0.14703 | 0.242781952 | 0.171 | 0.037 | 1 | 12 |
| ddx21 | 0.1474 | 0.420461081 | 0.257 | 0.074 | 1 | 12 |
| spcs1 | 0.14894 | 0.471543901 | 0.314 | 0.111 | 1 | 12 |
| nsa2 | 0.15071 | 0.454308394 | 0.457 | 0.222 | 1 | 12 |
| fkbp1aa | 0.15509 | 0.177704187 | 0.886 | 0.704 | 1 | 12 |
| actb1 | 0.1555 | -0.30782862 | 0.714 | 0.889 | 1 | 12 |
| selenow1 | 0.15683 | 0.491731422 | 0.486 | 0.259 | 1 | 12 |
| s100a10b | 0.16209 | 0.279895964 | 0.457 | 0.259 | 1 | 12 |
| ndufb5 | 0.16844 | 0.483680947 | 0.229 | 0.074 | 1 | 12 |
| sub1a | 0.16971 | -0.52288471 | 0.143 | 0.296 | 1 | 12 |
| rpl27a | 0.16981 | -0.40207871 | 0.486 | 0.704 | 1 | 12 |
| arl6ip1 | 0.17003 | 0.462120498 | 0.286 | 0.111 | 1 | 12 |
| hnrnpaba | 0.17118 | -0.48443406 | 0.286 | 0.444 | 1 | 12 |

| | | | | | | |
|---|---|---|---|---|---|---|
| h3f3a | 0.17124 | -0.29485002 | 0.343 | 0.37 | 1 | 12 |
| rps27l | 0.17464 | 0.471253607 | 0.4 | 0.185 | 1 | 12 |
| ap2s1 | 0.17593 | -0.42936485 | 0.086 | 0.259 | 1 | 12 |
| b2m | 0.17729 | 0.452672517 | 0.4 | 0.185 | 1 | 12 |
| timm9 | 0.17736 | 0.405238965 | 0.171 | 0.037 | 1 | 12 |
| kdr | 0.17737 | -0.3994389 | 0.086 | 0.259 | 1 | 12 |
| hmga1a | 0.17951 | -0.48511894 | 0.314 | 0.519 | 1 | 12 |
| ckba | 0.18051 | 0.458580455 | 0.4 | 0.185 | 1 | 12 |
| rpa3 | 0.18088 | -0.50606992 | 0.057 | 0.185 | 1 | 12 |
| rps21 | 0.18149 | 0.19425107 | 0.971 | 0.926 | 1 | 12 |
| psme3 | 0.18268 | -0.40588085 | 0.086 | 0.259 | 1 | 12 |
| sap30l | 0.18417 | -0.41706991 | 0.086 | 0.259 | 1 | 12 |
| pcna | 0.18587 | -0.52212918 | 0.143 | 0.333 | 1 | 12 |
| agmo | 0.18597 | 0.177813116 | 0.057 | 0 | 1 | 12 |
| celf1 | 0.18597 | 0.178785719 | 0.057 | 0 | 1 | 12 |
| ppa1b | 0.18597 | 0.176273258 | 0.057 | 0 | 1 | 12 |
| si:dkey-12j5.1 | 0.18597 | 0.17811926 | 0.057 | 0 | 1 | 12 |
| usp5 | 0.18597 | 0.188264161 | 0.057 | 0 | 1 | 12 |
| LOC110438055 | 0.18597 | 0.164856141 | 0.057 | 0 | 1 | 12 |
| PLEKHH3 | 0.18597 | 0.171683733 | 0.057 | 0 | 1 | 12 |
| aco2 | 0.18597 | 0.178224792 | 0.057 | 0 | 1 | 12 |
| ell | 0.18597 | 0.165623176 | 0.057 | 0 | 1 | 12 |
| fbxo33 | 0.18597 | 0.209032858 | 0.057 | 0 | 1 | 12 |
| si:ch73-390p7.2 | 0.18597 | 0.200097126 | 0.057 | 0 | 1 | 12 |
| si:dkey-48p11.3 | 0.18597 | 0.156212537 | 0.057 | 0 | 1 | 12 |
| smg6 | 0.18597 | 0.163434111 | 0.057 | 0 | 1 | 12 |
| tardbpl | 0.18597 | 0.176579226 | 0.057 | 0 | 1 | 12 |
| JMY | 0.18597 | 0.154364243 | 0.057 | 0 | 1 | 12 |
| ankrd40 | 0.18597 | 0.173327957 | 0.057 | 0 | 1 | 12 |
| crtap | 0.18597 | 0.19521778 | 0.057 | 0 | 1 | 12 |
| ddx27 | 0.18597 | 0.206233136 | 0.057 | 0 | 1 | 12 |
| e2f4 | 0.18597 | 0.18959992 | 0.057 | 0 | 1 | 12 |
| ero1b | 0.18597 | 0.156877355 | 0.057 | 0 | 1 | 12 |
| hmg20b | 0.18597 | 0.180670157 | 0.057 | 0 | 1 | 12 |
| hyal1 | 0.18597 | 0.185261602 | 0.057 | 0 | 1 | 12 |
| josd2 | 0.18597 | 0.122595649 | 0.057 | 0 | 1 | 12 |
| map3k4 | 0.18597 | 0.15477299 | 0.057 | 0 | 1 | 12 |
| nadkb | 0.18597 | 0.112327997 | 0.057 | 0 | 1 | 12 |
| pxnb | 0.18597 | 0.189730309 | 0.057 | 0 | 1 | 12 |
| si:ch211-196c10.15 | 0.18597 | 0.189730309 | 0.057 | 0 | 1 | 12 |
| smg1 | 0.18597 | 0.143228895 | 0.057 | 0 | 1 | 12 |
| srgap3 | 0.18597 | 0.193784088 | 0.057 | 0 | 1 | 12 |
| tbl3 | 0.18597 | 0.195129765 | 0.057 | 0 | 1 | 12 |
| tmem169a | 0.18597 | 0.182918277 | 0.057 | 0 | 1 | 12 |
| tpm2 | 0.18597 | 0.191382808 | 0.057 | 0 | 1 | 12 |
| zbtb16a | 0.18597 | 0.121136504 | 0.057 | 0 | 1 | 12 |
| zgc:153115 | 0.18597 | 0.177481451 | 0.057 | 0 | 1 | 12 |
| BX470189.1 | 0.18597 | 0.166694368 | 0.057 | 0 | 1 | 12 |

| | | | | | | |
|---|---|---|---|---|---|---|
| FP074902.2 | 0.18597 | 0.154876378 | 0.057 | 0 | 1 | 12 |
| LOC101886679 | 0.18597 | 0.151488103 | 0.057 | 0 | 1 | 12 |
| aamp | 0.18597 | 0.175573898 | 0.057 | 0 | 1 | 12 |
| akirin1 | 0.18597 | 0.193579707 | 0.057 | 0 | 1 | 12 |
| aldh2.1 | 0.18597 | 0.210368984 | 0.057 | 0 | 1 | 12 |
| anapc13 | 0.18597 | 0.193666446 | 0.057 | 0 | 1 | 12 |
| atg12 | 0.18597 | 0.152523701 | 0.057 | 0 | 1 | 12 |
| cavin2a | 0.18597 | 0.184011839 | 0.057 | 0 | 1 | 12 |
| cep170aa | 0.18597 | 0.203483446 | 0.057 | 0 | 1 | 12 |
| chd4b | 0.18597 | 0.147677217 | 0.057 | 0 | 1 | 12 |
| col1a1b | 0.18597 | 0.117784404 | 0.057 | 0 | 1 | 12 |
| cxcl18a.1 | 0.18597 | 0.177092007 | 0.057 | 0 | 1 | 12 |
| entr1 | 0.18597 | 0.130311938 | 0.057 | 0 | 1 | 12 |
| fam167b | 0.18597 | 0.166116913 | 0.057 | 0 | 1 | 12 |
| fam207a | 0.18597 | 0.167135563 | 0.057 | 0 | 1 | 12 |
| fnbp4 | 0.18597 | 0.176122565 | 0.057 | 0 | 1 | 12 |
| foxk1 | 0.18597 | 0.158797988 | 0.057 | 0 | 1 | 12 |
| gnb2 | 0.18597 | 0.137367543 | 0.057 | 0 | 1 | 12 |
| grb2a | 0.18597 | 0.155015361 | 0.057 | 0 | 1 | 12 |
| kctd12.2 | 0.18597 | 0.130891713 | 0.057 | 0 | 1 | 12 |
| klf9 | 0.18597 | 0.16974622 | 0.057 | 0 | 1 | 12 |
| maff | 0.18597 | 0.19456412 | 0.057 | 0 | 1 | 12 |
| med12 | 0.18597 | 0.185620068 | 0.057 | 0 | 1 | 12 |
| mgll | 0.18597 | 0.195623109 | 0.057 | 0 | 1 | 12 |
| mphosph6 | 0.18597 | 0.154924781 | 0.057 | 0 | 1 | 12 |
| mta2 | 0.18597 | 0.210615275 | 0.057 | 0 | 1 | 12 |
| nansa | 0.18597 | 0.186820657 | 0.057 | 0 | 1 | 12 |
| ndrg1a | 0.18597 | 0.2064825 | 0.057 | 0 | 1 | 12 |
| ndufaf3 | 0.18597 | 0.137771775 | 0.057 | 0 | 1 | 12 |
| nek6 | 0.18597 | 0.147745332 | 0.057 | 0 | 1 | 12 |
| nudt3b | 0.18597 | 0.180604334 | 0.057 | 0 | 1 | 12 |
| osgin2 | 0.18597 | 0.171953266 | 0.057 | 0 | 1 | 12 |
| psip1a | 0.18597 | 0.199248708 | 0.057 | 0 | 1 | 12 |
| rbfox2 | 0.18597 | 0.162945789 | 0.057 | 0 | 1 | 12 |
| rxrga | 0.18597 | 0.195648858 | 0.057 | 0 | 1 | 12 |
| sap30bp | 0.18597 | 0.13993538 | 0.057 | 0 | 1 | 12 |
| si:ch211-130m23.2 | 0.18597 | 0.151488103 | 0.057 | 0 | 1 | 12 |
| si:dkey-165a24.9 | 0.18597 | 0.21984758 | 0.057 | 0 | 1 | 12 |
| si:dkey-229e3.2 | 0.18597 | 0.19384636 | 0.057 | 0 | 1 | 12 |
| si:zfos-80g12.1 | 0.18597 | 0.135708074 | 0.057 | 0 | 1 | 12 |
| slc12a4 | 0.18597 | 0.207793914 | 0.057 | 0 | 1 | 12 |
| slc7a8a | 0.18597 | 0.113310093 | 0.057 | 0 | 1 | 12 |
| smad9 | 0.18597 | 0.148954846 | 0.057 | 0 | 1 | 12 |
| taf11 | 0.18597 | 0.150077866 | 0.057 | 0 | 1 | 12 |
| tnfrsf19 | 0.18597 | 0.101051388 | 0.057 | 0 | 1 | 12 |
| ubap1 | 0.18597 | 0.138832709 | 0.057 | 0 | 1 | 12 |
| ubfd1 | 0.18597 | 0.12470666 | 0.057 | 0 | 1 | 12 |
| uqcc1 | 0.18597 | 0.198605418 | 0.057 | 0 | 1 | 12 |

| | | | | | | |
|---|---|---|---|---|---|---|
| vps26b | 0.18597 | 0.184769311 | 0.057 | 0 | 1 | 12 |
| wwc3 | 0.18597 | 0.166116913 | 0.057 | 0 | 1 | 12 |
| yaf2 | 0.18597 | 0.195232413 | 0.057 | 0 | 1 | 12 |
| zgc:113294 | 0.18597 | 0.155957151 | 0.057 | 0 | 1 | 12 |
| zgc:154093 | 0.18597 | 0.14300333 | 0.057 | 0 | 1 | 12 |
| zgc:66474 | 0.18597 | 0.134246323 | 0.057 | 0 | 1 | 12 |
| znrf1 | 0.18597 | 0.188244069 | 0.057 | 0 | 1 | 12 |
| BX936439.1 | 0.18597 | 0.190885983 | 0.057 | 0 | 1 | 12 |
| CABZ01021592.1 | 0.18597 | 0.246750878 | 0.057 | 0 | 1 | 12 |
| CR382370.2 | 0.18597 | 0.144379848 | 0.057 | 0 | 1 | 12 |
| CU570894.1 | 0.18597 | 0.121920536 | 0.057 | 0 | 1 | 12 |
| CU655842.1 | 0.18597 | 0.178368631 | 0.057 | 0 | 1 | 12 |
| CU929259.1 | 0.18597 | 0.166401735 | 0.057 | 0 | 1 | 12 |
| CZQB01141835.1 | 0.18597 | 0.155481353 | 0.057 | 0 | 1 | 12 |
| FO082877.1 | 0.18597 | 0.171959945 | 0.057 | 0 | 1 | 12 |
| FP101914.1 | 0.18597 | 0.146516711 | 0.057 | 0 | 1 | 12 |
| FP102809.1 | 0.18597 | 0.106041416 | 0.057 | 0 | 1 | 12 |
| LOC101883052 | 0.18597 | 0.179086094 | 0.057 | 0 | 1 | 12 |
| LOC101883815 | 0.18597 | 0.140100976 | 0.057 | 0 | 1 | 12 |
| LOC101885376 | 0.18597 | 0.188415576 | 0.057 | 0 | 1 | 12 |
| LOC101885907 | 0.18597 | 0.206985423 | 0.057 | 0 | 1 | 12 |
| LOC108191614 | 0.18597 | 0.139457182 | 0.057 | 0 | 1 | 12 |
| MCMDC2 | 0.18597 | 0.211791615 | 0.057 | 0 | 1 | 12 |
| RASA2 | 0.18597 | 0.177642312 | 0.057 | 0 | 1 | 12 |
| XLOC-037780 | 0.18597 | 0.165651918 | 0.057 | 0 | 1 | 12 |
| XLOC-043857 | 0.18597 | 0.167267744 | 0.057 | 0 | 1 | 12 |
| acads | 0.18597 | 0.138121668 | 0.057 | 0 | 1 | 12 |
| acadvl | 0.18597 | 0.181264314 | 0.057 | 0 | 1 | 12 |
| ackr3b | 0.18597 | 0.17251308 | 0.057 | 0 | 1 | 12 |
| ada2a | 0.18597 | 0.142180088 | 0.057 | 0 | 1 | 12 |
| adma | 0.18597 | 0.164334198 | 0.057 | 0 | 1 | 12 |
| adra2da | 0.18597 | 0.205135811 | 0.057 | 0 | 1 | 12 |
| ap3s2 | 0.18597 | 0.175957248 | 0.057 | 0 | 1 | 12 |
| apoc1 | 0.18597 | 0.133999399 | 0.057 | 0 | 1 | 12 |
| apoda.2 | 0.18597 | 0.111903783 | 0.057 | 0 | 1 | 12 |
| aqp12 | 0.18597 | 0.194855357 | 0.057 | 0 | 1 | 12 |
| arcn1b | 0.18597 | 0.180251108 | 0.057 | 0 | 1 | 12 |
| arl14ep | 0.18597 | 0.229324135 | 0.057 | 0 | 1 | 12 |
| asnsd1 | 0.18597 | 0.176352181 | 0.057 | 0 | 1 | 12 |
| atp2b1a | 0.18597 | 0.131582841 | 0.057 | 0 | 1 | 12 |
| atp6v1aa | 0.18597 | 0.178600103 | 0.057 | 0 | 1 | 12 |
| bace2 | 0.18597 | 0.15679581 | 0.057 | 0 | 1 | 12 |
| bambib | 0.18597 | 0.201314548 | 0.057 | 0 | 1 | 12 |
| bcl2l1 | 0.18597 | 0.165965063 | 0.057 | 0 | 1 | 12 |
| bloc1s3 | 0.18597 | 0.122867755 | 0.057 | 0 | 1 | 12 |
| c6ast3 | 0.18597 | 0.145964523 | 0.057 | 0 | 1 | 12 |
| cacul1 | 0.18597 | 0.125503368 | 0.057 | 0 | 1 | 12 |
| camk2g2 | 0.18597 | 0.144568847 | 0.057 | 0 | 1 | 12 |

| | | | | | | |
|---|---|---|---|---|---|---|
| cat | 0.18597 | 0.17105291 | 0.057 | 0 | 1 | 12 |
| ccdc28a | 0.18597 | 0.176950177 | 0.057 | 0 | 1 | 12 |
| ccnt2b | 0.18597 | 0.213800695 | 0.057 | 0 | 1 | 12 |
| cd2ap | 0.18597 | 0.106978941 | 0.057 | 0 | 1 | 12 |
| cd99 | 0.18597 | 0.131244134 | 0.057 | 0 | 1 | 12 |
| cdk6 | 0.18597 | 0.174977868 | 0.057 | 0 | 1 | 12 |
| cdr2l | 0.18597 | 0.140788681 | 0.057 | 0 | 1 | 12 |
| cel.2 | 0.18597 | 0.118906187 | 0.057 | 0 | 1 | 12 |
| cep131 | 0.18597 | 0.164657158 | 0.057 | 0 | 1 | 12 |
| chchd2 | 0.18597 | 0.183333622 | 0.057 | 0 | 1 | 12 |
| chchd4a | 0.18597 | 0.150372948 | 0.057 | 0 | 1 | 12 |
| chchd7 | 0.18597 | 0.166680778 | 0.057 | 0 | 1 | 12 |
| chic2 | 0.18597 | 0.167558471 | 0.057 | 0 | 1 | 12 |
| chmp4ba | 0.18597 | 0.166492427 | 0.057 | 0 | 1 | 12 |
| ciartb | 0.18597 | 0.134165051 | 0.057 | 0 | 1 | 12 |
| cmtm3 | 0.18597 | 0.13249566 | 0.057 | 0 | 1 | 12 |
| cmtr1 | 0.18597 | 0.186771117 | 0.057 | 0 | 1 | 12 |
| cnot2 | 0.18597 | 0.162041965 | 0.057 | 0 | 1 | 12 |
| cnppd1 | 0.18597 | 0.228676795 | 0.057 | 0 | 1 | 12 |
| commd5 | 0.18597 | 0.174207945 | 0.057 | 0 | 1 | 12 |
| commd6 | 0.18597 | 0.139074561 | 0.057 | 0 | 1 | 12 |
| cops7a | 0.18597 | 0.121668727 | 0.057 | 0 | 1 | 12 |
| coq10b | 0.18597 | 0.211691749 | 0.057 | 0 | 1 | 12 |
| coq7 | 0.18597 | 0.106526899 | 0.057 | 0 | 1 | 12 |
| cpa1 | 0.18597 | 0.195002879 | 0.057 | 0 | 1 | 12 |
| csrp2 | 0.18597 | 0.160064692 | 0.057 | 0 | 1 | 12 |
| cst3 | 0.18597 | 0.141397297 | 0.057 | 0 | 1 | 12 |
| ctdsplb | 0.18597 | 0.128168944 | 0.057 | 0 | 1 | 12 |
| ctr9 | 0.18597 | 0.178795337 | 0.057 | 0 | 1 | 12 |
| cul3b | 0.18597 | 0.15503426 | 0.057 | 0 | 1 | 12 |
| cxcl20 | 0.18597 | 0.125729038 | 0.057 | 0 | 1 | 12 |
| cyp2ad6 | 0.18597 | 0.199742352 | 0.057 | 0 | 1 | 12 |
| dact2 | 0.18597 | 0.110025542 | 0.057 | 0 | 1 | 12 |
| dgkza | 0.18597 | 0.151911037 | 0.057 | 0 | 1 | 12 |
| dhrs12 | 0.18597 | 0.131412542 | 0.057 | 0 | 1 | 12 |
| diablob | 0.18597 | 0.196411439 | 0.057 | 0 | 1 | 12 |
| dnajc12 | 0.18597 | 0.145923788 | 0.057 | 0 | 1 | 12 |
| dnajc4 | 0.18597 | 0.164386466 | 0.057 | 0 | 1 | 12 |
| dok1b | 0.18597 | 0.154102867 | 0.057 | 0 | 1 | 12 |
| dr1 | 0.18597 | 0.197382521 | 0.057 | 0 | 1 | 12 |
| eef2a.1 | 0.18597 | 0.206057936 | 0.057 | 0 | 1 | 12 |
| efnb2a | 0.18597 | 0.19070247 | 0.057 | 0 | 1 | 12 |
| elf2a | 0.18597 | 0.136488913 | 0.057 | 0 | 1 | 12 |
| eloa | 0.18597 | 0.110025542 | 0.057 | 0 | 1 | 12 |
| eprs | 0.18597 | 0.138788855 | 0.057 | 0 | 1 | 12 |
| esf1 | 0.18597 | 0.136136995 | 0.057 | 0 | 1 | 12 |
| etv5a | 0.18597 | 0.19582772 | 0.057 | 0 | 1 | 12 |
| ewsr1a | 0.18597 | 0.162021382 | 0.057 | 0 | 1 | 12 |

| | | | | | | |
|---|---|---|---|---|---|---|
| fabp10a | 0.18597 | 0.120029463 | 0.057 | 0 | 1 | 12 |
| fam110a | 0.18597 | 0.127699832 | 0.057 | 0 | 1 | 12 |
| fam114a1 | 0.18597 | 0.209549147 | 0.057 | 0 | 1 | 12 |
| fam114a2 | 0.18597 | 0.169229746 | 0.057 | 0 | 1 | 12 |
| fam204a | 0.18597 | 0.187377609 | 0.057 | 0 | 1 | 12 |
| fas | 0.18597 | 0.132515016 | 0.057 | 0 | 1 | 12 |
| faxdc2 | 0.18597 | 0.164278983 | 0.057 | 0 | 1 | 12 |
| fbxo34 | 0.18597 | 0.149105286 | 0.057 | 0 | 1 | 12 |
| fkbp11 | 0.18597 | 0.141569194 | 0.057 | 0 | 1 | 12 |
| fkbp4 | 0.18597 | 0.16622027 | 0.057 | 0 | 1 | 12 |
| fmnl3 | 0.18597 | 0.201293079 | 0.057 | 0 | 1 | 12 |
| fndc3bb | 0.18597 | 0.170961856 | 0.057 | 0 | 1 | 12 |
| frmd4ba | 0.18597 | 0.12668667 | 0.057 | 0 | 1 | 12 |
| fryl | 0.18597 | 0.15207885 | 0.057 | 0 | 1 | 12 |
| fut8a | 0.18597 | 0.104659891 | 0.057 | 0 | 1 | 12 |
| galm | 0.18597 | 0.1636544 | 0.057 | 0 | 1 | 12 |
| gnas | 0.18597 | 0.136424601 | 0.057 | 0 | 1 | 12 |
| gng2 | 0.18597 | 0.173654949 | 0.057 | 0 | 1 | 12 |
| gnl3l | 0.18597 | 0.155526963 | 0.057 | 0 | 1 | 12 |
| gnsb | 0.18597 | 0.122505747 | 0.057 | 0 | 1 | 12 |
| h1f0 | 0.18597 | 0.132759368 | 0.057 | 0 | 1 | 12 |
| hccsb | 0.18597 | 0.225302891 | 0.057 | 0 | 1 | 12 |
| hk2 | 0.18597 | 0.181862889 | 0.057 | 0 | 1 | 12 |
| hoxb6a | 0.18597 | 0.129805923 | 0.057 | 0 | 1 | 12 |
| hoxb8a | 0.18597 | 0.109846536 | 0.057 | 0 | 1 | 12 |
| hyi | 0.18597 | 0.204236101 | 0.057 | 0 | 1 | 12 |
| id2b | 0.18597 | 0.137319286 | 0.057 | 0 | 1 | 12 |
| igfbp3 | 0.18597 | 0.123440114 | 0.057 | 0 | 1 | 12 |
| igflr1 | 0.18597 | 0.127463478 | 0.057 | 0 | 1 | 12 |
| il10rb | 0.18597 | 0.189355131 | 0.057 | 0 | 1 | 12 |
| impdh1b | 0.18597 | 0.173053404 | 0.057 | 0 | 1 | 12 |
| intu | 0.18597 | 0.186719197 | 0.057 | 0 | 1 | 12 |
| jade1 | 0.18597 | 0.120007279 | 0.057 | 0 | 1 | 12 |
| kctd1 | 0.18597 | 0.220427882 | 0.057 | 0 | 1 | 12 |
| kdm7ab | 0.18597 | 0.143274812 | 0.057 | 0 | 1 | 12 |
| kif20a | 0.18597 | 0.182572814 | 0.057 | 0 | 1 | 12 |
| kirrel3l | 0.18597 | 0.168652254 | 0.057 | 0 | 1 | 12 |
| klf12a | 0.18597 | 0.177484723 | 0.057 | 0 | 1 | 12 |
| kmt2ba | 0.18597 | 0.153701592 | 0.057 | 0 | 1 | 12 |
| kmt2e | 0.18597 | 0.174078795 | 0.057 | 0 | 1 | 12 |
| ktn1 | 0.18597 | 0.131315275 | 0.057 | 0 | 1 | 12 |
| lama4 | 0.18597 | 0.159172389 | 0.057 | 0 | 1 | 12 |
| leng8 | 0.18597 | 0.192319522 | 0.057 | 0 | 1 | 12 |
| letmd1 | 0.18597 | 0.187273248 | 0.057 | 0 | 1 | 12 |
| lonp2 | 0.18597 | 0.183586665 | 0.057 | 0 | 1 | 12 |
| loxl3b | 0.18597 | 0.156573338 | 0.057 | 0 | 1 | 12 |
| lpar1 | 0.18597 | 0.15943392 | 0.057 | 0 | 1 | 12 |
| maip1 | 0.18597 | 0.215093167 | 0.057 | 0 | 1 | 12 |

| | | | | | | |
|---|---|---|---|---|---|---|
| manbal | 0.18597 | 0.161949905 | 0.057 | 0 | 1 | 12 |
| manf | 0.18597 | 0.16633882 | 0.057 | 0 | 1 | 12 |
| map1sa | 0.18597 | 0.219663034 | 0.057 | 0 | 1 | 12 |
| mapkapk2a | 0.18597 | 0.119525756 | 0.057 | 0 | 1 | 12 |
| march2 | 0.18597 | 0.123591133 | 0.057 | 0 | 1 | 12 |
| micall1a | 0.18597 | 0.141117759 | 0.057 | 0 | 1 | 12 |
| mnta | 0.18597 | 0.203394193 | 0.057 | 0 | 1 | 12 |
| mphosph10 | 0.18597 | 0.140682254 | 0.057 | 0 | 1 | 12 |
| mrpl44 | 0.18597 | 0.168044235 | 0.057 | 0 | 1 | 12 |
| mrps16 | 0.18597 | 0.180831839 | 0.057 | 0 | 1 | 12 |
| nacc1a | 0.18597 | 0.144599262 | 0.057 | 0 | 1 | 12 |
| ncor1 | 0.18597 | 0.199436342 | 0.057 | 0 | 1 | 12 |
| ncor2 | 0.18597 | 0.161173832 | 0.057 | 0 | 1 | 12 |
| ndufaf1 | 0.18597 | 0.123464659 | 0.057 | 0 | 1 | 12 |
| nedd9 | 0.18597 | 0.198669978 | 0.057 | 0 | 1 | 12 |
| neurl4 | 0.18597 | 0.110025542 | 0.057 | 0 | 1 | 12 |
| nipblb | 0.18597 | 0.1452448 | 0.057 | 0 | 1 | 12 |
| nkd3 | 0.18597 | 0.174041664 | 0.057 | 0 | 1 | 12 |
| nol7 | 0.18597 | 0.131751679 | 0.057 | 0 | 1 | 12 |
| nt5dc2 | 0.18597 | 0.162518759 | 0.057 | 0 | 1 | 12 |
| nup43 | 0.18597 | 0.143295936 | 0.057 | 0 | 1 | 12 |
| opa1 | 0.18597 | 0.189015139 | 0.057 | 0 | 1 | 12 |
| parga | 0.18597 | 0.125467367 | 0.057 | 0 | 1 | 12 |
| pbrm1l | 0.18597 | 0.17706452 | 0.057 | 0 | 1 | 12 |
| pbx2 | 0.18597 | 0.122163904 | 0.057 | 0 | 1 | 12 |
| pbx4 | 0.18597 | 0.143172009 | 0.057 | 0 | 1 | 12 |
| pcyox1 | 0.18597 | 0.134997848 | 0.057 | 0 | 1 | 12 |
| pdf | 0.18597 | 0.17070505 | 0.057 | 0 | 1 | 12 |
| pdgfaa | 0.18597 | 0.189546695 | 0.057 | 0 | 1 | 12 |
| pdia5 | 0.18597 | 0.120032501 | 0.057 | 0 | 1 | 12 |
| pepd | 0.18597 | 0.192435151 | 0.057 | 0 | 1 | 12 |
| per2 | 0.18597 | 0.161161778 | 0.057 | 0 | 1 | 12 |
| pigbos1 | 0.18597 | 0.180839201 | 0.057 | 0 | 1 | 12 |
| pip4k2aa | 0.18597 | 0.101447893 | 0.057 | 0 | 1 | 12 |
| plekhb2 | 0.18597 | 0.171482723 | 0.057 | 0 | 1 | 12 |
| plxna1a | 0.18597 | 0.170256194 | 0.057 | 0 | 1 | 12 |
| plxnb3 | 0.18597 | 0.153727709 | 0.057 | 0 | 1 | 12 |
| polr1e | 0.18597 | 0.158603363 | 0.057 | 0 | 1 | 12 |
| ppat | 0.18597 | 0.201063036 | 0.057 | 0 | 1 | 12 |
| pphln1 | 0.18597 | 0.148631556 | 0.057 | 0 | 1 | 12 |
| ppp2r2ab | 0.18597 | 0.152913913 | 0.057 | 0 | 1 | 12 |
| ppp3cca | 0.18597 | 0.180026118 | 0.057 | 0 | 1 | 12 |
| prpf38a | 0.18597 | 0.155378178 | 0.057 | 0 | 1 | 12 |
| prpsap1 | 0.18597 | 0.172619854 | 0.057 | 0 | 1 | 12 |
| psme4b | 0.18597 | 0.143685817 | 0.057 | 0 | 1 | 12 |
| psmg1 | 0.18597 | 0.164293834 | 0.057 | 0 | 1 | 12 |
| ptbp1b | 0.18597 | 0.145238536 | 0.057 | 0 | 1 | 12 |
| ptcd1 | 0.18597 | 0.143581001 | 0.057 | 0 | 1 | 12 |

| | | | | | | |
|---|---|---|---|---|---|---|
| ptgr1 | 0.18597 | 0.15666097 | 0.057 | 0 | 1 | 12 |
| ptrh2 | 0.18597 | 0.118266867 | 0.057 | 0 | 1 | 12 |
| pum1 | 0.18597 | 0.156173897 | 0.057 | 0 | 1 | 12 |
| purbb | 0.18597 | 0.135651164 | 0.057 | 0 | 1 | 12 |
| pxk | 0.18597 | 0.134995594 | 0.057 | 0 | 1 | 12 |
| rab4b | 0.18597 | 0.128707298 | 0.057 | 0 | 1 | 12 |
| rac2 | 0.18597 | 0.183201066 | 0.057 | 0 | 1 | 12 |
| rad23ab | 0.18597 | 0.160834959 | 0.057 | 0 | 1 | 12 |
| ranbp9 | 0.18597 | 0.155400732 | 0.057 | 0 | 1 | 12 |
| rapgef3 | 0.18597 | 0.131100808 | 0.057 | 0 | 1 | 12 |
| rapgef5a | 0.18597 | 0.192026717 | 0.057 | 0 | 1 | 12 |
| rassf8a | 0.18597 | 0.142844308 | 0.057 | 0 | 1 | 12 |
| rbfa | 0.18597 | 0.182702505 | 0.057 | 0 | 1 | 12 |
| rbm14a | 0.18597 | 0.143649176 | 0.057 | 0 | 1 | 12 |
| rbms2b | 0.18597 | 0.192764071 | 0.057 | 0 | 1 | 12 |
| rcbtb1 | 0.18597 | 0.153581703 | 0.057 | 0 | 1 | 12 |
| relb | 0.18597 | 0.157854354 | 0.057 | 0 | 1 | 12 |
| rex1bd | 0.18597 | 0.173290691 | 0.057 | 0 | 1 | 12 |
| rilpl2 | 0.18597 | 0.121988586 | 0.057 | 0 | 1 | 12 |
| rnf38 | 0.18597 | 0.174977338 | 0.057 | 0 | 1 | 12 |
| rps6ka3a | 0.18597 | 0.130682501 | 0.057 | 0 | 1 | 12 |
| rrp15 | 0.18597 | 0.163546896 | 0.057 | 0 | 1 | 12 |
| rrp36 | 0.18597 | 0.160809242 | 0.057 | 0 | 1 | 12 |
| sccpdhb | 0.18597 | 0.123477833 | 0.057 | 0 | 1 | 12 |
| sdc2 | 0.18597 | 0.148540168 | 0.057 | 0 | 1 | 12 |
| sdr16c5a | 0.18597 | 0.17238778 | 0.057 | 0 | 1 | 12 |
| sec14l1 | 0.18597 | 0.182376763 | 0.057 | 0 | 1 | 12 |
| selenol | 0.18597 | 0.141112875 | 0.057 | 0 | 1 | 12 |
| senp2 | 0.18597 | 0.19904613 | 0.057 | 0 | 1 | 12 |
| serpinb1l3 | 0.18597 | 0.154590325 | 0.057 | 0 | 1 | 12 |
| sesn3 | 0.18597 | 0.147404323 | 0.057 | 0 | 1 | 12 |
| sfxn3 | 0.18597 | 0.106900241 | 0.057 | 0 | 1 | 12 |
| sgta | 0.18597 | 0.137978387 | 0.057 | 0 | 1 | 12 |
| sh3glb1b | 0.18597 | 0.153073098 | 0.057 | 0 | 1 | 12 |
| sh3pxd2aa | 0.18597 | 0.168557412 | 0.057 | 0 | 1 | 12 |
| si:ch1073-268j14.1 | 0.18597 | 0.193356128 | 0.057 | 0 | 1 | 12 |
| si:ch211-108c17.2 | 0.18597 | 0.105062073 | 0.057 | 0 | 1 | 12 |
| si:ch211-117c9.5 | 0.18597 | 0.138558512 | 0.057 | 0 | 1 | 12 |
| si:ch211-132p1.3 | 0.18597 | 0.169782608 | 0.057 | 0 | 1 | 12 |
| si:ch211-160o17.2 | 0.18597 | 0.131268673 | 0.057 | 0 | 1 | 12 |
| si:ch211-160o17.4 | 0.18597 | 0.115122585 | 0.057 | 0 | 1 | 12 |
| si:ch211-210c8.7 | 0.18597 | 0.19034029 | 0.057 | 0 | 1 | 12 |
| si:ch211-282j22.3 | 0.18597 | 0.18905878 | 0.057 | 0 | 1 | 12 |
| si:ch73-248e21.1 | 0.18597 | 0.137053919 | 0.057 | 0 | 1 | 12 |
| si:ch73-380l3.2 | 0.18597 | 0.193630565 | 0.057 | 0 | 1 | 12 |
| si:dkey-16l2.16 | 0.18597 | 0.190615853 | 0.057 | 0 | 1 | 12 |
| si:dkey-17m8.1 | 0.18597 | 0.20001939 | 0.057 | 0 | 1 | 12 |
| si:dkey-56i24.1 | 0.18597 | 0.198563261 | 0.057 | 0 | 1 | 12 |

| | | | | | | |
|---|---|---|---|---|---|---|
| si:dkey-72l14.4 | 0.18597 | 0.133525588 | 0.057 | 0 | 1 | 12 |
| si:dkey-91m11.5 | 0.18597 | 0.115941512 | 0.057 | 0 | 1 | 12 |
| si:dkeyp-113d7.1 | 0.18597 | 0.165611819 | 0.057 | 0 | 1 | 12 |
| si:dkeyp-82a1.6 | 0.18597 | 0.158856536 | 0.057 | 0 | 1 | 12 |
| sin3ab | 0.18597 | 0.129971186 | 0.057 | 0 | 1 | 12 |
| sirt7 | 0.18597 | 0.166118585 | 0.057 | 0 | 1 | 12 |
| skia | 0.18597 | 0.14289514 | 0.057 | 0 | 1 | 12 |
| slc12a7b | 0.18597 | 0.182324707 | 0.057 | 0 | 1 | 12 |
| slc25a18 | 0.18597 | 0.210599422 | 0.057 | 0 | 1 | 12 |
| slc38a2 | 0.18597 | 0.144877992 | 0.057 | 0 | 1 | 12 |
| slc48a1a | 0.18597 | 0.119507366 | 0.057 | 0 | 1 | 12 |
| sltm | 0.18597 | 0.11648858 | 0.057 | 0 | 1 | 12 |
| smc5 | 0.18597 | 0.115439261 | 0.057 | 0 | 1 | 12 |
| snai1a | 0.18597 | 0.134975466 | 0.057 | 0 | 1 | 12 |
| snx7 | 0.18597 | 0.176755448 | 0.057 | 0 | 1 | 12 |
| sox13 | 0.18597 | 0.212720128 | 0.057 | 0 | 1 | 12 |
| sox4b | 0.18597 | 0.181744104 | 0.057 | 0 | 1 | 12 |
| spopla | 0.18597 | 0.178661328 | 0.057 | 0 | 1 | 12 |
| srebf2 | 0.18597 | 0.160095029 | 0.057 | 0 | 1 | 12 |
| srpra | 0.18597 | 0.148947262 | 0.057 | 0 | 1 | 12 |
| srsf3a | 0.18597 | 0.151964119 | 0.057 | 0 | 1 | 12 |
| stard3 | 0.18597 | 0.14557958 | 0.057 | 0 | 1 | 12 |
| strn3 | 0.18597 | 0.171960711 | 0.057 | 0 | 1 | 12 |
| strn4 | 0.18597 | 0.153325691 | 0.057 | 0 | 1 | 12 |
| surf2 | 0.18597 | 0.169360165 | 0.057 | 0 | 1 | 12 |
| tab2 | 0.18597 | 0.222741903 | 0.057 | 0 | 1 | 12 |
| taf1b | 0.18597 | 0.183011187 | 0.057 | 0 | 1 | 12 |
| taf6l | 0.18597 | 0.179883943 | 0.057 | 0 | 1 | 12 |
| tbc1d16 | 0.18597 | 0.167131115 | 0.057 | 0 | 1 | 12 |
| tcea2 | 0.18597 | 0.106872066 | 0.057 | 0 | 1 | 12 |
| tefa | 0.18597 | 0.155394943 | 0.057 | 0 | 1 | 12 |
| tfap4 | 0.18597 | 0.145063457 | 0.057 | 0 | 1 | 12 |
| tfpt | 0.18597 | 0.130921079 | 0.057 | 0 | 1 | 12 |
| thoc7 | 0.18597 | 0.152319305 | 0.057 | 0 | 1 | 12 |
| thrap3b | 0.18597 | 0.177070421 | 0.057 | 0 | 1 | 12 |
| tmem185 | 0.18597 | 0.135418627 | 0.057 | 0 | 1 | 12 |
| tmem51a | 0.18597 | 0.167027125 | 0.057 | 0 | 1 | 12 |
| tmub2 | 0.18597 | 0.149407311 | 0.057 | 0 | 1 | 12 |
| tmx4 | 0.18597 | 0.141412059 | 0.057 | 0 | 1 | 12 |
| tnfaip3 | 0.18597 | 0.18121497 | 0.057 | 0 | 1 | 12 |
| tnfsf10 | 0.18597 | 0.163182555 | 0.057 | 0 | 1 | 12 |
| tnksb | 0.18597 | 0.171015926 | 0.057 | 0 | 1 | 12 |
| trim105 | 0.18597 | 0.142214823 | 0.057 | 0 | 1 | 12 |
| trip10a | 0.18597 | 0.217235476 | 0.057 | 0 | 1 | 12 |
| tspan17 | 0.18597 | 0.167858092 | 0.057 | 0 | 1 | 12 |
| tspan31 | 0.18597 | 0.145717791 | 0.057 | 0 | 1 | 12 |
| tspan9a | 0.18597 | 0.148461364 | 0.057 | 0 | 1 | 12 |
| ttc13 | 0.18597 | 0.185390701 | 0.057 | 0 | 1 | 12 |

| | | | | | | |
|---|---|---|---|---|---|---|
| ttpa | 0.18597 | 0.120647305 | 0.057 | 0 | 1 | 12 |
| ttyh2 | 0.18597 | 0.164117675 | 0.057 | 0 | 1 | 12 |
| ube2d3 | 0.18597 | 0.13835313 | 0.057 | 0 | 1 | 12 |
| ube2d4 | 0.18597 | 0.116248494 | 0.057 | 0 | 1 | 12 |
| upf3b | 0.18597 | 0.176598263 | 0.057 | 0 | 1 | 12 |
| uqcrc2a | 0.18597 | 0.20305021 | 0.057 | 0 | 1 | 12 |
| vash2 | 0.18597 | 0.189030187 | 0.057 | 0 | 1 | 12 |
| vps16 | 0.18597 | 0.123942427 | 0.057 | 0 | 1 | 12 |
| vps33b | 0.18597 | 0.110092961 | 0.057 | 0 | 1 | 12 |
| vps8 | 0.18597 | 0.168821964 | 0.057 | 0 | 1 | 12 |
| wasla | 0.18597 | 0.189028398 | 0.057 | 0 | 1 | 12 |
| wu:fc21g02 | 0.18597 | 0.188415576 | 0.057 | 0 | 1 | 12 |
| wwp1 | 0.18597 | 0.213047517 | 0.057 | 0 | 1 | 12 |
| xkr8.3 | 0.18597 | 0.170599944 | 0.057 | 0 | 1 | 12 |
| xpot | 0.18597 | 0.151915386 | 0.057 | 0 | 1 | 12 |
| zc3h11a | 0.18597 | 0.168880165 | 0.057 | 0 | 1 | 12 |
| zdhhc8b | 0.18597 | 0.179502261 | 0.057 | 0 | 1 | 12 |
| zgc:101679 | 0.18597 | 0.186580051 | 0.057 | 0 | 1 | 12 |
| zgc:101858 | 0.18597 | 0.16485841 | 0.057 | 0 | 1 | 12 |
| zgc:136439 | 0.18597 | 0.178659744 | 0.057 | 0 | 1 | 12 |
| zgc:158412 | 0.18597 | 0.127583484 | 0.057 | 0 | 1 | 12 |
| zgc:162472 | 0.18597 | 0.166992242 | 0.057 | 0 | 1 | 12 |
| zgc:163077 | 0.18597 | 0.102190352 | 0.057 | 0 | 1 | 12 |
| zgc:171480 | 0.18597 | 0.17205263 | 0.057 | 0 | 1 | 12 |
| zgc:193812 | 0.18597 | 0.133459074 | 0.057 | 0 | 1 | 12 |
| zgc:63882 | 0.18597 | 0.167414642 | 0.057 | 0 | 1 | 12 |
| zmiz1a | 0.18597 | 0.147815882 | 0.057 | 0 | 1 | 12 |
| znf1015 | 0.18597 | 0.175085017 | 0.057 | 0 | 1 | 12 |
| znf367 | 0.18597 | 0.138354786 | 0.057 | 0 | 1 | 12 |
| znfx1 | 0.18597 | 0.111080284 | 0.057 | 0 | 1 | 12 |
| zyx | 0.18597 | 0.133723591 | 0.057 | 0 | 1 | 12 |
| CR848667.1 | 0.18597 | 0.171344264 | 0.057 | 0 | 1 | 12 |
| LO018102.1 | 0.18597 | 0.200677703 | 0.057 | 0 | 1 | 12 |
| adh8b | 0.18597 | 0.14992527 | 0.057 | 0 | 1 | 12 |
| aga | 0.18597 | 0.178693171 | 0.057 | 0 | 1 | 12 |
| bin3 | 0.18597 | 0.167643056 | 0.057 | 0 | 1 | 12 |
| bola1 | 0.18597 | 0.158013335 | 0.057 | 0 | 1 | 12 |
| chd2 | 0.18597 | 0.172636361 | 0.057 | 0 | 1 | 12 |
| ckap5 | 0.18597 | 0.172043175 | 0.057 | 0 | 1 | 12 |
| cttn | 0.18597 | 0.219660751 | 0.057 | 0 | 1 | 12 |
| fkbp14 | 0.18597 | 0.173668989 | 0.057 | 0 | 1 | 12 |
| gchfr | 0.18597 | 0.177239282 | 0.057 | 0 | 1 | 12 |
| gig2j | 0.18597 | 0.163069919 | 0.057 | 0 | 1 | 12 |
| itpkcb | 0.18597 | 0.198021226 | 0.057 | 0 | 1 | 12 |
| mmp2 | 0.18597 | 0.137381062 | 0.057 | 0 | 1 | 12 |
| mrpl22 | 0.18597 | 0.138830302 | 0.057 | 0 | 1 | 12 |
| nolc1 | 0.18597 | 0.195427578 | 0.057 | 0 | 1 | 12 |
| pdia2 | 0.18597 | 0.148367997 | 0.057 | 0 | 1 | 12 |

| | | | | | | |
|---|---|---|---|---|---|---|
| phf8 | 0.18597 | 0.150483061 | 0.057 | 0 | 1 | 12 |
| pir | 0.18597 | 0.135728267 | 0.057 | 0 | 1 | 12 |
| pla2g1b | 0.18597 | 0.107177214 | 0.057 | 0 | 1 | 12 |
| psmd10 | 0.18597 | 0.188915788 | 0.057 | 0 | 1 | 12 |
| raph1b | 0.18597 | 0.187770454 | 0.057 | 0 | 1 | 12 |
| sestd1 | 0.18597 | 0.174698954 | 0.057 | 0 | 1 | 12 |
| smurf2 | 0.18597 | 0.175427189 | 0.057 | 0 | 1 | 12 |
| trim13 | 0.18597 | 0.175565896 | 0.057 | 0 | 1 | 12 |
| uap1l1 | 0.18597 | 0.155863355 | 0.057 | 0 | 1 | 12 |
| yipf5 | 0.18597 | 0.197380663 | 0.057 | 0 | 1 | 12 |
| zc4h2 | 0.18597 | 0.176955012 | 0.057 | 0 | 1 | 12 |
| atg10 | 0.18597 | 0.1647385 | 0.057 | 0 | 1 | 12 |
| kctd6b | 0.18597 | 0.216353257 | 0.057 | 0 | 1 | 12 |
| si:ch73-44m9.5 | 0.18597 | 0.139252483 | 0.057 | 0 | 1 | 12 |
| myef2 | 0.18597 | 0.206434832 | 0.057 | 0 | 1 | 12 |
| dcun1d2b | 0.18597 | 0.151808905 | 0.057 | 0 | 1 | 12 |
| wdr3 | 0.18597 | 0.156024657 | 0.057 | 0 | 1 | 12 |
| myl12.1 | 0.18729 | -0.36560561 | 0.514 | 0.667 | 1 | 12 |
| sarnp | 0.18766 | 0.405386194 | 0.171 | 0.037 | 1 | 12 |
| sf3b1 | 0.18772 | 0.392560919 | 0.171 | 0.037 | 1 | 12 |
| ppdpfa | 0.18862 | 0.426490219 | 0.171 | 0.037 | 1 | 12 |
| sypl2a | 0.18872 | 0.300142997 | 0.229 | 0.074 | 1 | 12 |
| snrpb | 0.19092 | 0.231782583 | 0.4 | 0.222 | 1 | 12 |
| golt1bb | 0.19241 | -0.34071704 | 0.029 | 0.148 | 1 | 12 |
| ehd3 | 0.19434 | 0.473268928 | 0.171 | 0.037 | 1 | 12 |
| hars | 0.19482 | -0.43623354 | 0.029 | 0.148 | 1 | 12 |
| pfn2 | 0.19585 | -0.43233991 | 0.4 | 0.593 | 1 | 12 |
| sft2d1 | 0.19646 | 0.312494594 | 0.171 | 0.037 | 1 | 12 |
| krtcap2 | 0.19723 | -0.43024687 | 0.114 | 0.296 | 1 | 12 |
| ranbp1 | 0.19858 | -0.44318964 | 0.114 | 0.296 | 1 | 12 |
| rpl39 | 0.19952 | 0.130452122 | 0.943 | 0.963 | 1 | 12 |
| hmgb2a | 0.19973 | -0.36725074 | 0.6 | 0.741 | 1 | 12 |
| banf1 | 0.20009 | -0.44877947 | 0.114 | 0.296 | 1 | 12 |
| eif6 | 0.2009 | -0.4396439 | 0.029 | 0.148 | 1 | 12 |
| add1 | 0.20415 | 0.323701004 | 0.171 | 0.037 | 1 | 12 |
| hint1 | 0.20524 | -0.47402061 | 0.2 | 0.333 | 1 | 12 |
| arf2a | 0.20569 | -0.46041049 | 0.086 | 0.222 | 1 | 12 |
| rpl21 | 0.20636 | 0.228257508 | 0.943 | 0.815 | 1 | 12 |
| mcl1b | 0.20711 | 0.417823148 | 0.343 | 0.148 | 1 | 12 |
| fam174b | 0.20764 | 0.441983686 | 0.286 | 0.111 | 1 | 12 |
| rpl36 | 0.20772 | 0.144353329 | 0.971 | 0.963 | 1 | 12 |
| ndufa8 | 0.21059 | 0.24562674 | 0.171 | 0.037 | 1 | 12 |
| celf2 | 0.21184 | 0.353519365 | 0.171 | 0.037 | 1 | 12 |
| cx43 | 0.21272 | 0.412558284 | 0.171 | 0.037 | 1 | 12 |
| tmem50a | 0.21295 | 0.429470584 | 0.229 | 0.074 | 1 | 12 |
| cyb5a | 0.21301 | 0.303953394 | 0.171 | 0.037 | 1 | 12 |
| zgc:110591 | 0.21325 | 0.354838089 | 0.171 | 0.037 | 1 | 12 |
| zgc:86598 | 0.21387 | -0.37544283 | 0.029 | 0.148 | 1 | 12 |

| | | | | | | |
|---|---|---|---|---|---|---|
| wdr45 | 0.21408 | -0.39463132 | 0.029 | 0.148 | 1 | 12 |
| id2a | 0.21532 | 0.365042393 | 0.171 | 0.037 | 1 | 12 |
| naa20 | 0.21542 | -0.37031135 | 0.029 | 0.148 | 1 | 12 |
| esd | 0.21616 | 0.396847848 | 0.171 | 0.037 | 1 | 12 |
| si:zfos-464b6.2 | 0.21622 | 0.291376662 | 0.171 | 0.037 | 1 | 12 |
| loxl1 | 0.21773 | 0.408792275 | 0.286 | 0.111 | 1 | 12 |
| mesd | 0.21829 | -0.34613209 | 0.029 | 0.148 | 1 | 12 |
| drg2 | 0.21898 | -0.23381731 | 0.029 | 0.148 | 1 | 12 |
| spry4 | 0.21932 | 0.355579393 | 0.171 | 0.037 | 1 | 12 |
| gpr182 | 0.21948 | 0.361790983 | 0.171 | 0.037 | 1 | 12 |
| slc44a2 | 0.21967 | 0.313632504 | 0.171 | 0.037 | 1 | 12 |
| gabarapb | 0.21984 | 0.476043613 | 0.371 | 0.185 | 1 | 12 |
| rrm2 | 0.22041 | -0.3135337 | 0.029 | 0.148 | 1 | 12 |
| ywhabb | 0.2209 | 0.387533734 | 0.514 | 0.296 | 1 | 12 |
| ppib | 0.22097 | 0.375981613 | 0.629 | 0.407 | 1 | 12 |
| selenos | 0.22189 | -0.3257582 | 0.029 | 0.148 | 1 | 12 |
| si:ch211-162e15.3 | 0.22233 | -0.32803078 | 0.029 | 0.148 | 1 | 12 |
| si:dkey-250d21.1 | 0.22245 | -0.33572274 | 0.029 | 0.148 | 1 | 12 |
| gtf2a2 | 0.2234 | -0.39145841 | 0.029 | 0.148 | 1 | 12 |
| CABZ01084131.1 | 0.22381 | -0.27701613 | 0.029 | 0.148 | 1 | 12 |
| fth1a | 0.22397 | -0.40819837 | 0.371 | 0.519 | 1 | 12 |
| hnrnpub | 0.22473 | -0.33073472 | 0.029 | 0.148 | 1 | 12 |
| tbcb | 0.22496 | -0.3873263 | 0.029 | 0.148 | 1 | 12 |
| noc2l | 0.22552 | -0.28304722 | 0.029 | 0.148 | 1 | 12 |
| ccdc32 | 0.22564 | -0.39157252 | 0.029 | 0.148 | 1 | 12 |
| mrpl53 | 0.22569 | -0.33471056 | 0.029 | 0.148 | 1 | 12 |
| rbx1 | 0.22643 | 0.386103198 | 0.429 | 0.222 | 1 | 12 |
| snapin | 0.22662 | -0.31539908 | 0.029 | 0.148 | 1 | 12 |
| gmds | 0.22668 | -0.34477066 | 0.029 | 0.148 | 1 | 12 |
| si:ch73-46j18.5 | 0.22674 | 0.400376782 | 0.429 | 0.222 | 1 | 12 |
| si:dkey-102m7.3 | 0.22692 | -0.3128114 | 0.029 | 0.148 | 1 | 12 |
| gng12a | 0.22709 | -0.30644901 | 0.029 | 0.148 | 1 | 12 |
| usp7 | 0.22711 | -0.28847222 | 0.029 | 0.148 | 1 | 12 |
| srsf5b | 0.22739 | -0.25188557 | 0.029 | 0.148 | 1 | 12 |
| WDR7 | 0.22746 | -0.25653307 | 0.029 | 0.148 | 1 | 12 |
| tmco1 | 0.22756 | -0.26082367 | 0.029 | 0.148 | 1 | 12 |
| clec14a | 0.2278 | 0.411008994 | 0.286 | 0.111 | 1 | 12 |
| CABZ01080702.1 | 0.22784 | -0.32473694 | 0.029 | 0.148 | 1 | 12 |
| zgc:66433 | 0.22795 | -0.354704 | 0.029 | 0.148 | 1 | 12 |
| emp2 | 0.22796 | -0.34008272 | 0.029 | 0.148 | 1 | 12 |
| psma4 | 0.23024 | -0.43965033 | 0.257 | 0.407 | 1 | 12 |
| atp5pf | 0.23255 | -0.32177379 | 0.371 | 0.444 | 1 | 12 |
| lsm4 | 0.23347 | 0.356963627 | 0.229 | 0.074 | 1 | 12 |
| eif2s3 | 0.23444 | 0.25684359 | 0.4 | 0.222 | 1 | 12 |
| rasgef1bb | 0.23512 | 0.361234444 | 0.229 | 0.074 | 1 | 12 |
| lyrm4 | 0.23512 | -0.47618471 | 0.057 | 0.185 | 1 | 12 |
| si:rp71-45k5.4 | 0.2362 | 0.372270403 | 0.229 | 0.074 | 1 | 12 |
| emd | 0.23774 | 0.359842537 | 0.229 | 0.074 | 1 | 12 |

| | | | | | | |
|---|---|---|---|---|---|---|
| sox18 | 0.23778 | 0.381468782 | 0.229 | 0.074 | 1 | 12 |
| dctn3 | 0.23788 | 0.363881492 | 0.229 | 0.074 | 1 | 12 |
| c1qtnf6a | 0.24157 | -0.27784501 | 0.057 | 0.185 | 1 | 12 |
| actn4 | 0.24453 | -0.43952883 | 0.114 | 0.259 | 1 | 12 |
| rsl24d1 | 0.24546 | 0.3247792 | 0.457 | 0.259 | 1 | 12 |
| gnb1a | 0.24666 | 0.275979174 | 0.314 | 0.148 | 1 | 12 |
| ebna1bp2 | 0.24718 | 0.18155698 | 0.2 | 0.074 | 1 | 12 |
| cope | 0.25029 | -0.34543591 | 0.057 | 0.185 | 1 | 12 |
| psap | 0.2528 | 0.258032038 | 0.314 | 0.148 | 1 | 12 |
| rps10 | 0.2545 | 0.184355541 | 0.971 | 0.889 | 1 | 12 |
| ehd2b | 0.25527 | -0.40222583 | 0.057 | 0.185 | 1 | 12 |
| CR318588.1 | 0.25666 | -0.31520144 | 0.257 | 0.444 | 1 | 12 |
| eif1axa | 0.25899 | -0.24842215 | 0.086 | 0.222 | 1 | 12 |
| txn | 0.25961 | -0.26335923 | 0.171 | 0.333 | 1 | 12 |
| sar1b | 0.26424 | -0.36467273 | 0.057 | 0.185 | 1 | 12 |
| rpl12 | 0.26576 | 0.237937443 | 0.914 | 0.778 | 1 | 12 |
| anp32b | 0.27248 | -0.43656182 | 0.086 | 0.222 | 1 | 12 |
| hspb1 | 0.27292 | 0.318322783 | 0.314 | 0.148 | 1 | 12 |
| adam8a | 0.27465 | -0.39510713 | 0.057 | 0.185 | 1 | 12 |
| cox6a1 | 0.27648 | 0.279322637 | 0.4 | 0.222 | 1 | 12 |
| cldn5b | 0.27691 | 0.25360414 | 0.514 | 0.333 | 1 | 12 |
| si:ch73-234b20.5 | 0.27715 | 0.225195004 | 0.2 | 0.074 | 1 | 12 |
| glula | 0.27786 | 0.441526575 | 0.314 | 0.148 | 1 | 12 |
| zgc:114188 | 0.27909 | 0.181020684 | 0.943 | 0.815 | 1 | 12 |
| mob4 | 0.2796 | -0.32295677 | 0.057 | 0.185 | 1 | 12 |
| dynlt3 | 0.28175 | -0.43990432 | 0.143 | 0.259 | 1 | 12 |
| snu13b | 0.28204 | 0.175294545 | 0.286 | 0.148 | 1 | 12 |
| ssr1 | 0.28399 | 0.304798124 | 0.343 | 0.296 | 1 | 12 |
| eif3ha | 0.28428 | 0.284530075 | 0.314 | 0.148 | 1 | 12 |
| cox8b | 0.28576 | -0.41764969 | 0.171 | 0.259 | 1 | 12 |
| ifitm1 | 0.28578 | 0.504672184 | 0.143 | 0.037 | 1 | 12 |
| cnbpb | 0.28613 | 0.415159026 | 0.314 | 0.148 | 1 | 12 |
| syngr2a | 0.28731 | -0.30356304 | 0.057 | 0.185 | 1 | 12 |
| plscr3b | 0.28768 | -0.32603517 | 0.057 | 0.185 | 1 | 12 |
| MYADM | 0.28809 | -0.36536396 | 0.057 | 0.185 | 1 | 12 |
| u2af2b | 0.2889 | -0.32473425 | 0.057 | 0.185 | 1 | 12 |
| fli1b | 0.28896 | -0.32851682 | 0.057 | 0.185 | 1 | 12 |
| si:dkey-9i23.16 | 0.28927 | -0.32524839 | 0.057 | 0.185 | 1 | 12 |
| khsrp | 0.28935 | -0.32506342 | 0.057 | 0.185 | 1 | 12 |
| arpin | 0.29001 | 0.340639343 | 0.143 | 0.037 | 1 | 12 |
| snx12 | 0.29006 | -0.42344458 | 0.114 | 0.259 | 1 | 12 |
| naa50 | 0.29023 | 0.168409111 | 0.143 | 0.037 | 1 | 12 |
| reep3a | 0.29085 | 0.185809282 | 0.143 | 0.037 | 1 | 12 |
| ube2na | 0.29176 | 0.404512045 | 0.2 | 0.074 | 1 | 12 |
| ahcy | 0.29199 | 0.173458201 | 0.2 | 0.074 | 1 | 12 |
| nap1l1 | 0.29267 | -0.39438977 | 0.257 | 0.444 | 1 | 12 |
| cib1 | 0.29275 | 0.229654451 | 0.143 | 0.037 | 1 | 12 |
| hmgb3b | 0.29286 | 0.33289997 | 0.143 | 0.037 | 1 | 12 |

| | | | | | | |
|---|---|---|---|---|---|---|
| si:ch211-152c2.3 | 0.29334 | -0.3289552 | 0.171 | 0.333 | 1 | 12 |
| timm13 | 0.29468 | 0.329447681 | 0.143 | 0.037 | 1 | 12 |
| nhp2 | 0.29531 | 0.321063445 | 0.314 | 0.148 | 1 | 12 |
| zgc:77849 | 0.29873 | 0.327924779 | 0.143 | 0.037 | 1 | 12 |
| rpl24 | 0.30054 | 0.152042015 | 0.943 | 0.926 | 1 | 12 |
| si:dkeyp-97a10.2 | 0.30065 | -0.39942273 | 0.229 | 0.407 | 1 | 12 |
| nucks1b | 0.3013 | -0.36855006 | 0.029 | 0.111 | 1 | 12 |
| cd81a | 0.30345 | -0.39976917 | 0.343 | 0.519 | 1 | 12 |
| thoc2 | 0.30799 | 0.363628609 | 0.143 | 0.037 | 1 | 12 |
| myo1cb | 0.30979 | -0.38581908 | 0.086 | 0.222 | 1 | 12 |
| fstl1b | 0.31018 | -0.39947829 | 0.229 | 0.407 | 1 | 12 |
| si:ch73-376l24.2 | 0.31046 | 0.299814709 | 0.143 | 0.037 | 1 | 12 |
| si:ch211-191j22.3 | 0.31083 | 0.302554584 | 0.143 | 0.037 | 1 | 12 |
| hmgb2b | 0.31315 | -0.35514329 | 0.4 | 0.593 | 1 | 12 |
| clic1 | 0.31373 | -0.36278385 | 0.086 | 0.222 | 1 | 12 |
| cyth1b | 0.31435 | -0.34390204 | 0.029 | 0.111 | 1 | 12 |
| caprin1b | 0.31502 | 0.324069876 | 0.143 | 0.037 | 1 | 12 |
| gnb1b | 0.31531 | 0.283183977 | 0.257 | 0.111 | 1 | 12 |
| paip2b | 0.31777 | -0.33535025 | 0.257 | 0.333 | 1 | 12 |
| erh | 0.31883 | 0.410971373 | 0.343 | 0.185 | 1 | 12 |
| psmb1 | 0.31901 | -0.35567211 | 0.229 | 0.407 | 1 | 12 |
| atp5if1a | 0.32039 | 0.253614977 | 0.343 | 0.185 | 1 | 12 |
| ehd1b | 0.32142 | -0.34350815 | 0.086 | 0.222 | 1 | 12 |
| blcap | 0.32295 | 0.255202152 | 0.2 | 0.074 | 1 | 12 |
| pros1 | 0.32687 | 0.366825652 | 0.2 | 0.074 | 1 | 12 |
| calm2b | 0.32695 | -0.34714766 | 0.343 | 0.444 | 1 | 12 |
| hsbp1b | 0.32709 | -0.35665426 | 0.114 | 0.259 | 1 | 12 |
| CU694197.1 | 0.32714 | 0.377265085 | 0.257 | 0.111 | 1 | 12 |
| ralba | 0.32753 | 0.250202757 | 0.143 | 0.037 | 1 | 12 |
| romo1 | 0.32803 | 0.35434571 | 0.514 | 0.333 | 1 | 12 |
| zgc:162944 | 0.32809 | 0.379483724 | 0.257 | 0.111 | 1 | 12 |
| psma6l | 0.32845 | 0.257749492 | 0.143 | 0.037 | 1 | 12 |
| pdcd6 | 0.32874 | 0.230045662 | 0.2 | 0.074 | 1 | 12 |
| pmp22a | 0.33114 | -0.34213771 | 0.2 | 0.37 | 1 | 12 |
| pim3 | 0.33193 | 0.258951562 | 0.143 | 0.037 | 1 | 12 |
| sap18 | 0.33249 | 0.342233195 | 0.257 | 0.111 | 1 | 12 |
| krt8 | 0.33359 | 0.187339247 | 0.857 | 0.704 | 1 | 12 |
| ap1b1 | 0.33544 | 0.26955619 | 0.143 | 0.037 | 1 | 12 |
| afap1l1b | 0.33609 | -0.35906913 | 0.114 | 0.259 | 1 | 12 |
| tmed2 | 0.33612 | 0.333964903 | 0.257 | 0.111 | 1 | 12 |
| znf706 | 0.33647 | 0.21732292 | 0.143 | 0.037 | 1 | 12 |
| tnfrsfa | 0.33772 | 0.282212967 | 0.143 | 0.037 | 1 | 12 |
| ubc | 0.33774 | 0.322428219 | 0.657 | 0.481 | 1 | 12 |
| arhgdig | 0.33794 | 0.24563593 | 0.143 | 0.037 | 1 | 12 |
| svild | 0.33798 | -0.38879113 | 0.143 | 0.296 | 1 | 12 |
| glrx | 0.33799 | -0.34415657 | 0.171 | 0.333 | 1 | 12 |
| ptp4a1 | 0.3394 | 0.247022486 | 0.143 | 0.037 | 1 | 12 |
| psmd3 | 0.3397 | 0.240729289 | 0.2 | 0.074 | 1 | 12 |

| | | | | | | |
|---|---|---|---|---|---|---|
| srrm1 | 0.33981 | 0.248258868 | 0.143 | 0.037 | 1 | 12 |
| sh2b3 | 0.34037 | 0.33714584 | 0.2 | 0.074 | 1 | 12 |
| dusp6 | 0.34104 | 0.266123917 | 0.2 | 0.074 | 1 | 12 |
| prkab1b | 0.34143 | 0.265926496 | 0.143 | 0.037 | 1 | 12 |
| eif5a2 | 0.3416 | 0.223978488 | 0.543 | 0.37 | 1 | 12 |
| sh3bgrl3 | 0.34239 | 0.263558168 | 0.143 | 0.037 | 1 | 12 |
| oaz2b | 0.34381 | 0.346852829 | 0.2 | 0.074 | 1 | 12 |
| sept15 | 0.3444 | 0.237013544 | 0.143 | 0.037 | 1 | 12 |
| ndufab1a | 0.34522 | 0.277580477 | 0.143 | 0.037 | 1 | 12 |
| NDUFB1 | 0.34549 | 0.347756203 | 0.2 | 0.074 | 1 | 12 |
| si:ch73-91k6.2 | 0.34652 | 0.254836576 | 0.143 | 0.037 | 1 | 12 |
| tor1 | 0.34669 | 0.259700458 | 0.143 | 0.037 | 1 | 12 |
| ddah2 | 0.34736 | 0.251788745 | 0.143 | 0.037 | 1 | 12 |
| XLOC-016094 | 0.34737 | 0.277404593 | 0.143 | 0.037 | 1 | 12 |
| ccdc85b | 0.34773 | 0.274318519 | 0.2 | 0.074 | 1 | 12 |
| ube2e3 | 0.34778 | 0.263738177 | 0.2 | 0.074 | 1 | 12 |
| nck1b | 0.35067 | 0.338421547 | 0.2 | 0.074 | 1 | 12 |
| vcp | 0.35083 | 0.341571505 | 0.2 | 0.074 | 1 | 12 |
| cdc42 | 0.35132 | 0.342526286 | 0.514 | 0.333 | 1 | 12 |
| psmg2 | 0.35269 | -0.35968117 | 0.057 | 0.148 | 1 | 12 |
| cavin1b | 0.35276 | -0.42856421 | 0.114 | 0.222 | 1 | 12 |
| jupa | 0.35404 | 0.288294447 | 0.2 | 0.074 | 1 | 12 |
| ndufb7 | 0.35431 | -0.38351767 | 0.229 | 0.37 | 1 | 12 |
| kdelr2b | 0.3561 | 0.335807521 | 0.2 | 0.074 | 1 | 12 |
| zgc:110339 | 0.35704 | 0.287403498 | 0.2 | 0.074 | 1 | 12 |
| aldh9a1a.1 | 0.35732 | 0.289807132 | 0.2 | 0.074 | 1 | 12 |
| bri3 | 0.35908 | -0.27776595 | 0.029 | 0.111 | 1 | 12 |
| icn | 0.35971 | 0.202424559 | 0.114 | 0.037 | 1 | 12 |
| ostf1 | 0.36052 | -0.27294902 | 0.029 | 0.111 | 1 | 12 |
| ddx23 | 0.36249 | -0.38208824 | 0.086 | 0.185 | 1 | 12 |
| atp5meb | 0.36418 | 0.36793907 | 0.371 | 0.222 | 1 | 12 |
| nme3 | 0.36705 | -0.27957998 | 0.029 | 0.111 | 1 | 12 |
| phb2a | 0.36718 | -0.29338709 | 0.029 | 0.111 | 1 | 12 |
| kdm3b | 0.36982 | -0.27266359 | 0.029 | 0.111 | 1 | 12 |
| ormdl1 | 0.37164 | -0.2669728 | 0.029 | 0.111 | 1 | 12 |
| cox17 | 0.37193 | 0.342544856 | 0.171 | 0.074 | 1 | 12 |
| ybx1 | 0.37221 | 0.167669564 | 0.943 | 0.852 | 1 | 12 |
| plvapb | 0.37496 | 0.360845296 | 0.429 | 0.259 | 1 | 12 |
| atf4b | 0.37537 | 0.335204168 | 0.343 | 0.185 | 1 | 12 |
| plekhg5a | 0.37563 | 0.369936806 | 0.229 | 0.111 | 1 | 12 |
| cnot1 | 0.37598 | -0.26476468 | 0.029 | 0.111 | 1 | 12 |
| sart3 | 0.3771 | -0.27353006 | 0.029 | 0.111 | 1 | 12 |
| brd3b | 0.3796 | -0.30907989 | 0.029 | 0.111 | 1 | 12 |
| khdrbs1b | 0.38027 | -0.25588318 | 0.029 | 0.111 | 1 | 12 |
| eef1g | 0.38098 | 0.186571126 | 0.8 | 0.741 | 1 | 12 |
| arhgdia | 0.38369 | -0.33063976 | 0.314 | 0.407 | 1 | 12 |
| lyve1b | 0.38399 | -0.40130101 | 0.171 | 0.296 | 1 | 12 |
| hvcn1 | 0.38479 | 0.341657334 | 0.114 | 0.037 | 1 | 12 |

| | | | | | | |
|---|---|---|---|---|---|---|
| pprc1 | 0.38729 | -0.14213432 | 0.029 | 0.111 | 1 | 12 |
| magt1 | 0.38955 | -0.15809802 | 0.029 | 0.111 | 1 | 12 |
| rad23b | 0.3898 | -0.35053748 | 0.143 | 0.222 | 1 | 12 |
| naca | 0.38989 | -0.19249965 | 0.8 | 0.889 | 1 | 12 |
| psma2 | 0.3906 | 0.342053631 | 0.229 | 0.111 | 1 | 12 |
| COX3 | 0.39179 | -0.13640885 | 0.429 | 0.556 | 1 | 12 |
| pdcd6ip | 0.39273 | -0.24211646 | 0.029 | 0.111 | 1 | 12 |
| gcnt4a | 0.39281 | -0.32539285 | 0.029 | 0.111 | 1 | 12 |
| rbbp6 | 0.39353 | -0.2329357 | 0.029 | 0.111 | 1 | 12 |
| ahi1 | 0.39424 | -0.142918 | 0.029 | 0.111 | 1 | 12 |
| abca1a | 0.39489 | -0.21523541 | 0.029 | 0.111 | 1 | 12 |
| rpl4 | 0.39618 | 0.262476023 | 0.743 | 0.593 | 1 | 12 |
| rps8a | 0.39654 | 0.147880835 | 0.943 | 0.889 | 1 | 12 |
| meox2a | 0.39722 | 0.312462036 | 0.114 | 0.037 | 1 | 12 |
| ndufv2 | 0.39724 | -0.29775485 | 0.029 | 0.111 | 1 | 12 |
| psmd7 | 0.39728 | 0.138783136 | 0.171 | 0.074 | 1 | 12 |
| me3 | 0.39732 | -0.38349116 | 0.086 | 0.185 | 1 | 12 |
| si:ch211-282b22.1 | 0.39822 | -0.31323714 | 0.029 | 0.111 | 1 | 12 |
| arf5 | 0.39977 | -0.36382388 | 0.086 | 0.148 | 1 | 12 |
| parp12b | 0.40023 | -0.21780066 | 0.029 | 0.111 | 1 | 12 |
| atp5l | 0.40129 | 0.203404098 | 0.571 | 0.407 | 1 | 12 |
| mrpl33 | 0.40295 | -0.33865374 | 0.057 | 0.148 | 1 | 12 |
| cct6a | 0.40311 | 0.361021586 | 0.314 | 0.185 | 1 | 12 |
| selenoj | 0.40316 | -0.24460876 | 0.029 | 0.111 | 1 | 12 |
| fkbp3 | 0.40351 | 0.355868147 | 0.286 | 0.148 | 1 | 12 |
| si:ch211-288d18.1 | 0.40459 | -0.20692996 | 0.029 | 0.111 | 1 | 12 |
| si:ch211-212k18.4 | 0.40471 | -0.28148265 | 0.029 | 0.111 | 1 | 12 |
| psmd11a | 0.40539 | -0.28506323 | 0.029 | 0.111 | 1 | 12 |
| zgc:153713 | 0.40586 | 0.256422432 | 0.286 | 0.148 | 1 | 12 |
| cltcb | 0.40665 | -0.30289194 | 0.029 | 0.111 | 1 | 12 |
| sdc4 | 0.40676 | -0.28287284 | 0.029 | 0.111 | 1 | 12 |
| atp5mc3b | 0.40686 | 0.171170913 | 0.714 | 0.556 | 1 | 12 |
| agpat3 | 0.40773 | -0.1678145 | 0.029 | 0.111 | 1 | 12 |
| cirbpa | 0.40828 | -0.20599402 | 0.6 | 0.667 | 1 | 12 |
| spred2a | 0.4086 | -0.29401126 | 0.029 | 0.111 | 1 | 12 |
| aip | 0.4089 | 0.341257855 | 0.286 | 0.148 | 1 | 12 |
| slc37a2 | 0.40893 | -0.1397979 | 0.029 | 0.111 | 1 | 12 |
| mmadhc | 0.40897 | -0.18452248 | 0.029 | 0.111 | 1 | 12 |
| slc35f6 | 0.40906 | -0.2695068 | 0.029 | 0.111 | 1 | 12 |
| arpc1b | 0.40959 | 0.152548978 | 0.229 | 0.111 | 1 | 12 |
| fam89b | 0.41009 | -0.19166278 | 0.029 | 0.111 | 1 | 12 |
| aoc2 | 0.41088 | 0.420624946 | 0.257 | 0.148 | 1 | 12 |
| h2afy2 | 0.41219 | -0.34892771 | 0.086 | 0.185 | 1 | 12 |
| slc16a5a | 0.41256 | -0.26585569 | 0.029 | 0.111 | 1 | 12 |
| ccnd2a | 0.4132 | 0.323923412 | 0.171 | 0.074 | 1 | 12 |
| pitpnaa | 0.41373 | -0.28313931 | 0.029 | 0.111 | 1 | 12 |
| per1b | 0.41421 | -0.14729436 | 0.029 | 0.111 | 1 | 12 |
| rxrba | 0.41694 | -0.25420823 | 0.029 | 0.111 | 1 | 12 |

| | | | | | | |
|---|---|---|---|---|---|---|
| slc43a2a | 0.41716 | -0.18490659 | 0.029 | 0.111 | 1 | 12 |
| SCARF2 | 0.41793 | -0.16063045 | 0.029 | 0.074 | 1 | 12 |
| itgb1a | 0.41793 | -0.11823894 | 0.029 | 0.074 | 1 | 12 |
| ppp4r2b | 0.41793 | -0.13787003 | 0.029 | 0.074 | 1 | 12 |
| scaf4a | 0.41793 | -0.15626493 | 0.029 | 0.074 | 1 | 12 |
| slc22a18 | 0.41793 | -0.11770568 | 0.029 | 0.074 | 1 | 12 |
| taf3 | 0.41793 | -0.11798924 | 0.029 | 0.074 | 1 | 12 |
| CABZ01110881.1 | 0.41793 | -0.15280602 | 0.029 | 0.074 | 1 | 12 |
| bcl2a | 0.41793 | -0.18085832 | 0.029 | 0.074 | 1 | 12 |
| eif4g2b | 0.41793 | -0.19649806 | 0.029 | 0.074 | 1 | 12 |
| fdps | 0.41793 | -0.1806987 | 0.029 | 0.074 | 1 | 12 |
| fn1a | 0.41793 | -0.11767903 | 0.029 | 0.074 | 1 | 12 |
| hnrnpua | 0.41793 | -0.195925 | 0.029 | 0.074 | 1 | 12 |
| inppl1a | 0.41793 | -0.11925944 | 0.029 | 0.074 | 1 | 12 |
| mob3a | 0.41793 | -0.18122295 | 0.029 | 0.074 | 1 | 12 |
| ncln | 0.41793 | -0.20509847 | 0.029 | 0.074 | 1 | 12 |
| pole4 | 0.41793 | -0.12514331 | 0.029 | 0.074 | 1 | 12 |
| rgs5b | 0.41793 | -0.13393113 | 0.029 | 0.074 | 1 | 12 |
| si:ch211-125a15.1 | 0.41793 | -0.1899214 | 0.029 | 0.074 | 1 | 12 |
| tomm40 | 0.41793 | -0.11394437 | 0.029 | 0.074 | 1 | 12 |
| abhd17b | 0.41793 | -0.14533339 | 0.029 | 0.074 | 1 | 12 |
| anxa2a | 0.41793 | -0.21292112 | 0.029 | 0.074 | 1 | 12 |
| apln | 0.41793 | -0.28967665 | 0.029 | 0.074 | 1 | 12 |
| arhgap20b | 0.41793 | -0.12135371 | 0.029 | 0.074 | 1 | 12 |
| arpp19a | 0.41793 | -0.12496883 | 0.029 | 0.074 | 1 | 12 |
| asb8 | 0.41793 | -0.1764029 | 0.029 | 0.074 | 1 | 12 |
| cdkal1 | 0.41793 | -0.22656048 | 0.029 | 0.074 | 1 | 12 |
| col18a1a | 0.41793 | -0.20665467 | 0.029 | 0.074 | 1 | 12 |
| cops8 | 0.41793 | -0.12795317 | 0.029 | 0.074 | 1 | 12 |
| creg2 | 0.41793 | -0.16630183 | 0.029 | 0.074 | 1 | 12 |
| cyp2v1 | 0.41793 | -0.10111591 | 0.029 | 0.074 | 1 | 12 |
| dab2ipb | 0.41793 | -0.14045196 | 0.029 | 0.074 | 1 | 12 |
| dapk3 | 0.41793 | -0.18407718 | 0.029 | 0.074 | 1 | 12 |
| ddb1 | 0.41793 | -0.16339386 | 0.029 | 0.074 | 1 | 12 |
| dicer1 | 0.41793 | -0.13774287 | 0.029 | 0.074 | 1 | 12 |
| dram2b | 0.41793 | -0.14087089 | 0.029 | 0.074 | 1 | 12 |
| dync1li1 | 0.41793 | -0.14044242 | 0.029 | 0.074 | 1 | 12 |
| eci1 | 0.41793 | -0.1921468 | 0.029 | 0.074 | 1 | 12 |
| gatd3a | 0.41793 | -0.21124234 | 0.029 | 0.074 | 1 | 12 |
| gnpat | 0.41793 | -0.21479636 | 0.029 | 0.074 | 1 | 12 |
| gnptg | 0.41793 | -0.13739311 | 0.029 | 0.074 | 1 | 12 |
| golga5 | 0.41793 | -0.12570833 | 0.029 | 0.074 | 1 | 12 |
| gosr1 | 0.41793 | -0.14657957 | 0.029 | 0.074 | 1 | 12 |
| heatr3 | 0.41793 | -0.11015878 | 0.029 | 0.074 | 1 | 12 |
| kansl3 | 0.41793 | -0.2057459 | 0.029 | 0.074 | 1 | 12 |
| lyrm5a | 0.41793 | -0.1374687 | 0.029 | 0.074 | 1 | 12 |
| lysmd2 | 0.41793 | -0.17395737 | 0.029 | 0.074 | 1 | 12 |
| map7d3 | 0.41793 | -0.15155266 | 0.029 | 0.074 | 1 | 12 |

| | | | | | | |
|---|---|---|---|---|---|---|
| mapre2 | 0.41793 | -0.177121 | 0.029 | 0.074 | 1 | 12 |
| mbnl1 | 0.41793 | -0.11935756 | 0.029 | 0.074 | 1 | 12 |
| metrnlb | 0.41793 | -0.10848558 | 0.029 | 0.074 | 1 | 12 |
| mgat1b | 0.41793 | -0.11444868 | 0.029 | 0.074 | 1 | 12 |
| mpzl1l | 0.41793 | -0.17959887 | 0.029 | 0.074 | 1 | 12 |
| mrpl11 | 0.41793 | -0.18925513 | 0.029 | 0.074 | 1 | 12 |
| mrpl47 | 0.41793 | -0.14470139 | 0.029 | 0.074 | 1 | 12 |
| mvp | 0.41793 | -0.17842446 | 0.029 | 0.074 | 1 | 12 |
| mylipb | 0.41793 | -0.18191531 | 0.029 | 0.074 | 1 | 12 |
| ncalda | 0.41793 | -0.16102207 | 0.029 | 0.074 | 1 | 12 |
| nr3c1 | 0.41793 | -0.14875661 | 0.029 | 0.074 | 1 | 12 |
| pard6gb | 0.41793 | -0.14459605 | 0.029 | 0.074 | 1 | 12 |
| parp1 | 0.41793 | -0.20894336 | 0.029 | 0.074 | 1 | 12 |
| pcyt2 | 0.41793 | -0.1091322 | 0.029 | 0.074 | 1 | 12 |
| polr3h | 0.41793 | -0.1799226 | 0.029 | 0.074 | 1 | 12 |
| ppp6r2a | 0.41793 | -0.12377206 | 0.029 | 0.074 | 1 | 12 |
| prox1a | 0.41793 | -0.15178242 | 0.029 | 0.074 | 1 | 12 |
| prrc2a | 0.41793 | -0.2167771 | 0.029 | 0.074 | 1 | 12 |
| psat1 | 0.41793 | -0.10840129 | 0.029 | 0.074 | 1 | 12 |
| rock2a | 0.41793 | -0.1815977 | 0.029 | 0.074 | 1 | 12 |
| scinla | 0.41793 | -0.16915734 | 0.029 | 0.074 | 1 | 12 |
| sec13 | 0.41793 | -0.18168568 | 0.029 | 0.074 | 1 | 12 |
| si:dkey-100n23.5 | 0.41793 | -0.1600145 | 0.029 | 0.074 | 1 | 12 |
| si:dkey-163m14.2 | 0.41793 | -0.15074972 | 0.029 | 0.074 | 1 | 12 |
| si:dkey-177p2.6 | 0.41793 | -0.12891081 | 0.029 | 0.074 | 1 | 12 |
| si:dkey-67c22.2 | 0.41793 | -0.11430976 | 0.029 | 0.074 | 1 | 12 |
| smc2 | 0.41793 | -0.16249738 | 0.029 | 0.074 | 1 | 12 |
| snapc3 | 0.41793 | -0.11096917 | 0.029 | 0.074 | 1 | 12 |
| spns2 | 0.41793 | -0.12201164 | 0.029 | 0.074 | 1 | 12 |
| srsf6a | 0.41793 | -0.19036711 | 0.029 | 0.074 | 1 | 12 |
| tmem141 | 0.41793 | -0.13664428 | 0.029 | 0.074 | 1 | 12 |
| tmem150aa | 0.41793 | -0.14366566 | 0.029 | 0.074 | 1 | 12 |
| tmem165 | 0.41793 | -0.17327511 | 0.029 | 0.074 | 1 | 12 |
| tmod2 | 0.41793 | -0.16896684 | 0.029 | 0.074 | 1 | 12 |
| tpp1 | 0.41793 | -0.14036114 | 0.029 | 0.074 | 1 | 12 |
| tsen34 | 0.41793 | -0.24078702 | 0.029 | 0.074 | 1 | 12 |
| ubap2l | 0.41793 | -0.16425823 | 0.029 | 0.074 | 1 | 12 |
| ube2c | 0.41793 | -0.15715947 | 0.029 | 0.074 | 1 | 12 |
| ubxn1 | 0.41793 | -0.1409142 | 0.029 | 0.074 | 1 | 12 |
| uqcc3 | 0.41793 | -0.11274701 | 0.029 | 0.074 | 1 | 12 |
| usp4 | 0.41793 | -0.17658055 | 0.029 | 0.074 | 1 | 12 |
| vmp1 | 0.41793 | -0.15123059 | 0.029 | 0.074 | 1 | 12 |
| wdr46 | 0.41793 | -0.12250656 | 0.029 | 0.074 | 1 | 12 |
| wdr75 | 0.41793 | -0.17979669 | 0.029 | 0.074 | 1 | 12 |
| yy1a | 0.41793 | -0.13149095 | 0.029 | 0.074 | 1 | 12 |
| zgc:158803 | 0.41793 | -0.12634227 | 0.029 | 0.074 | 1 | 12 |
| zgc:162025 | 0.41793 | -0.11895715 | 0.029 | 0.074 | 1 | 12 |
| CABZ01074453.1 | 0.41793 | -0.12812252 | 0.029 | 0.074 | 1 | 12 |

| | | | | | | |
|---|---|---|---|---|---|---|
| MRPS12 | 0.41793 | -0.21532193 | 0.029 | 0.074 | 1 | 12 |
| PAXBP1 | 0.41793 | -0.13409025 | 0.029 | 0.074 | 1 | 12 |
| TPGS1 | 0.41793 | -0.11781431 | 0.029 | 0.074 | 1 | 12 |
| abcf2a | 0.41793 | -0.15709218 | 0.029 | 0.074 | 1 | 12 |
| abhd16a | 0.41793 | -0.14925501 | 0.029 | 0.074 | 1 | 12 |
| adipor2 | 0.41793 | -0.12112545 | 0.029 | 0.074 | 1 | 12 |
| aldoab | 0.41793 | -0.13927523 | 0.029 | 0.074 | 1 | 12 |
| anapc11 | 0.41793 | -0.14791335 | 0.029 | 0.074 | 1 | 12 |
| apc | 0.41793 | -0.13337461 | 0.029 | 0.074 | 1 | 12 |
| aph1b | 0.41793 | -0.18316441 | 0.029 | 0.074 | 1 | 12 |
| arhgap29a | 0.41793 | -0.11755175 | 0.029 | 0.074 | 1 | 12 |
| arhgap29b | 0.41793 | -0.12021883 | 0.029 | 0.074 | 1 | 12 |
| b3gnt3.4 | 0.41793 | -0.11840699 | 0.029 | 0.074 | 1 | 12 |
| bxdc2 | 0.41793 | -0.1454398 | 0.029 | 0.074 | 1 | 12 |
| cdk2 | 0.41793 | -0.12264559 | 0.029 | 0.074 | 1 | 12 |
| chaf1a | 0.41793 | -0.11314586 | 0.029 | 0.074 | 1 | 12 |
| cpsf6 | 0.41793 | -0.10153516 | 0.029 | 0.074 | 1 | 12 |
| cul3a | 0.41793 | -0.15917426 | 0.029 | 0.074 | 1 | 12 |
| daam1a | 0.41793 | -0.1694715 | 0.029 | 0.074 | 1 | 12 |
| dbn1 | 0.41793 | -0.13595785 | 0.029 | 0.074 | 1 | 12 |
| dbt | 0.41793 | -0.16642786 | 0.029 | 0.074 | 1 | 12 |
| ddt | 0.41793 | -0.1056335 | 0.029 | 0.074 | 1 | 12 |
| dnmt1 | 0.41793 | -0.10159688 | 0.029 | 0.074 | 1 | 12 |
| dusp8a | 0.41793 | -0.16421107 | 0.029 | 0.074 | 1 | 12 |
| elk4 | 0.41793 | -0.13115546 | 0.029 | 0.074 | 1 | 12 |
| erlec1 | 0.41793 | -0.16115167 | 0.029 | 0.074 | 1 | 12 |
| fam13b | 0.41793 | -0.16124826 | 0.029 | 0.074 | 1 | 12 |
| fbxw2 | 0.41793 | -0.18739848 | 0.029 | 0.074 | 1 | 12 |
| gabarapl2 | 0.41793 | -0.16269716 | 0.029 | 0.074 | 1 | 12 |
| gcn1 | 0.41793 | -0.14741091 | 0.029 | 0.074 | 1 | 12 |
| gnpda1 | 0.41793 | -0.19351348 | 0.029 | 0.074 | 1 | 12 |
| helz | 0.41793 | -0.13467295 | 0.029 | 0.074 | 1 | 12 |
| isg15 | 0.41793 | -0.20352799 | 0.029 | 0.074 | 1 | 12 |
| kpna1 | 0.41793 | -0.23640946 | 0.029 | 0.074 | 1 | 12 |
| mkrn4 | 0.41793 | -0.22437288 | 0.029 | 0.074 | 1 | 12 |
| mrpl16 | 0.41793 | -0.10091608 | 0.029 | 0.074 | 1 | 12 |
| nme2b.2 | 0.41793 | -0.17574538 | 0.029 | 0.074 | 1 | 12 |
| pgd | 0.41793 | -0.12381049 | 0.029 | 0.074 | 1 | 12 |
| phactr2 | 0.41793 | -0.17109727 | 0.029 | 0.074 | 1 | 12 |
| plp2 | 0.41793 | -0.12452343 | 0.029 | 0.074 | 1 | 12 |
| pls3 | 0.41793 | -0.10522965 | 0.029 | 0.074 | 1 | 12 |
| ppme1 | 0.41793 | -0.10788838 | 0.029 | 0.074 | 1 | 12 |
| ppp2r5ea | 0.41793 | -0.1393568 | 0.029 | 0.074 | 1 | 12 |
| psmd11b | 0.41793 | -0.20662027 | 0.029 | 0.074 | 1 | 12 |
| ptp4a2b | 0.41793 | -0.10341435 | 0.029 | 0.074 | 1 | 12 |
| pwp2h | 0.41793 | -0.16151308 | 0.029 | 0.074 | 1 | 12 |
| rab7a | 0.41793 | -0.11126745 | 0.029 | 0.074 | 1 | 12 |
| rbpms2b | 0.41793 | -0.12038165 | 0.029 | 0.074 | 1 | 12 |

| | | | | | | |
|---|---|---|---|---|---|---|
| rrp8 | 0.41793 | -0.18879285 | 0.029 | 0.074 | 1 | 12 |
| ruvbl1 | 0.41793 | -0.13579329 | 0.029 | 0.074 | 1 | 12 |
| sart1 | 0.41793 | -0.25363957 | 0.029 | 0.074 | 1 | 12 |
| selenot1a | 0.41793 | -0.10763476 | 0.029 | 0.074 | 1 | 12 |
| sema3ga | 0.41793 | -0.18423908 | 0.029 | 0.074 | 1 | 12 |
| serpinh1a | 0.41793 | -0.15931708 | 0.029 | 0.074 | 1 | 12 |
| si:ch211-286b5.5 | 0.41793 | -0.10531342 | 0.029 | 0.074 | 1 | 12 |
| si:ch73-130a3.4 | 0.41793 | -0.10892961 | 0.029 | 0.074 | 1 | 12 |
| smco4 | 0.41793 | -0.15735489 | 0.029 | 0.074 | 1 | 12 |
| sox4a | 0.41793 | -0.15101266 | 0.029 | 0.074 | 1 | 12 |
| spag7 | 0.41793 | -0.11731358 | 0.029 | 0.074 | 1 | 12 |
| stk25b | 0.41793 | -0.11706821 | 0.029 | 0.074 | 1 | 12 |
| taf7 | 0.41793 | -0.20395741 | 0.029 | 0.074 | 1 | 12 |
| uqcrb | 0.41793 | -0.21368478 | 0.029 | 0.074 | 1 | 12 |
| usp10 | 0.41793 | -0.11842085 | 0.029 | 0.074 | 1 | 12 |
| ythdf1 | 0.41793 | -0.10178474 | 0.029 | 0.074 | 1 | 12 |
| zgc:77650 | 0.41793 | -0.19549552 | 0.029 | 0.074 | 1 | 12 |
| CABZ01068366.1 | 0.41793 | -0.10606343 | 0.029 | 0.074 | 1 | 12 |
| abcc5 | 0.41793 | -0.17451172 | 0.029 | 0.074 | 1 | 12 |
| bms1 | 0.41793 | -0.13054346 | 0.029 | 0.074 | 1 | 12 |
| camk2g1 | 0.41793 | -0.20412364 | 0.029 | 0.074 | 1 | 12 |
| cbx1b | 0.41793 | -0.219093 | 0.029 | 0.074 | 1 | 12 |
| ccdc137 | 0.41793 | -0.11645203 | 0.029 | 0.074 | 1 | 12 |
| dachd | 0.41793 | -0.20382327 | 0.029 | 0.074 | 1 | 12 |
| dhrs13l1 | 0.41793 | -0.1781862 | 0.029 | 0.074 | 1 | 12 |
| dusp12 | 0.41793 | -0.17348513 | 0.029 | 0.074 | 1 | 12 |
| fam210aa | 0.41793 | -0.161364 | 0.029 | 0.074 | 1 | 12 |
| fhl1b | 0.41793 | -0.17486932 | 0.029 | 0.074 | 1 | 12 |
| lpl | 0.41793 | -0.21085902 | 0.029 | 0.074 | 1 | 12 |
| mrps10 | 0.41793 | -0.15196745 | 0.029 | 0.074 | 1 | 12 |
| mrps21 | 0.41793 | -0.15712879 | 0.029 | 0.074 | 1 | 12 |
| plekha3 | 0.41793 | -0.17564853 | 0.029 | 0.074 | 1 | 12 |
| rcc1 | 0.41793 | -0.14388175 | 0.029 | 0.074 | 1 | 12 |
| sh3gl3a | 0.41793 | -0.10844152 | 0.029 | 0.074 | 1 | 12 |
| si:ch211-133n4.4 | 0.41793 | -0.1128388 | 0.029 | 0.074 | 1 | 12 |
| si:ch211-225b7.5 | 0.41793 | -0.16952748 | 0.029 | 0.074 | 1 | 12 |
| si:ch211-85n16.3 | 0.41793 | -0.13698403 | 0.029 | 0.074 | 1 | 12 |
| si:dkey-230p4.1 | 0.41793 | -0.16417352 | 0.029 | 0.074 | 1 | 12 |
| smarcad1a | 0.41793 | -0.16517845 | 0.029 | 0.074 | 1 | 12 |
| srp72 | 0.41793 | -0.14179378 | 0.029 | 0.074 | 1 | 12 |
| ss18 | 0.41793 | -0.19167722 | 0.029 | 0.074 | 1 | 12 |
| stx5a | 0.41793 | -0.18191608 | 0.029 | 0.074 | 1 | 12 |
| tns3.2 | 0.41793 | -0.21952088 | 0.029 | 0.074 | 1 | 12 |
| zgc:77056 | 0.41793 | -0.13409526 | 0.029 | 0.074 | 1 | 12 |
| TST | 0.41793 | -0.15879292 | 0.029 | 0.074 | 1 | 12 |
| exosc8 | 0.41793 | -0.18695608 | 0.029 | 0.074 | 1 | 12 |
| igf2bp1 | 0.41793 | -0.12989487 | 0.029 | 0.074 | 1 | 12 |
| rbms1a | 0.41793 | -0.12246762 | 0.029 | 0.074 | 1 | 12 |

| | | | | | | |
|---|---|---|---|---|---|---|
| si:dkeyp-27c8.2 | 0.41793 | -0.12371582 | 0.029 | 0.074 | 1 | 12 |
| snrnp70 | 0.41793 | -0.14055958 | 0.029 | 0.074 | 1 | 12 |
| tmem234 | 0.41793 | -0.10186088 | 0.029 | 0.074 | 1 | 12 |
| abi1a | 0.41793 | -0.10465772 | 0.029 | 0.074 | 1 | 12 |
| faf1 | 0.41793 | -0.19271864 | 0.029 | 0.074 | 1 | 12 |
| gdi1 | 0.41793 | -0.16120097 | 0.029 | 0.074 | 1 | 12 |
| as3mt | 0.41793 | -0.18113836 | 0.029 | 0.074 | 1 | 12 |
| pcdh17 | 0.41793 | -0.11013281 | 0.029 | 0.074 | 1 | 12 |
| mmp9 | 0.41849 | -0.30816995 | 0.029 | 0.111 | 1 | 12 |
| cyth1a | 0.41879 | -0.16180273 | 0.029 | 0.111 | 1 | 12 |
| syncripl | 0.41902 | -0.20119369 | 0.029 | 0.111 | 1 | 12 |
| jund | 0.41934 | -0.3028214 | 0.029 | 0.111 | 1 | 12 |
| si:ch73-376l24.3 | 0.41987 | 0.320489931 | 0.171 | 0.074 | 1 | 12 |
| hspe1 | 0.41991 | 0.35081734 | 0.171 | 0.074 | 1 | 12 |
| cct5 | 0.42082 | -0.31847545 | 0.286 | 0.444 | 1 | 12 |
| rmdn1 | 0.42084 | -0.27124335 | 0.029 | 0.111 | 1 | 12 |
| ndufa1 | 0.4209 | -0.33432279 | 0.171 | 0.296 | 1 | 12 |
| kmt2cb | 0.42118 | -0.19739583 | 0.029 | 0.111 | 1 | 12 |
| LOC108180269 | 0.42127 | -0.14425618 | 0.029 | 0.111 | 1 | 12 |
| myl6 | 0.42136 | 0.109104128 | 0.371 | 0.259 | 1 | 12 |
| lhx6 | 0.42138 | -0.25736556 | 0.029 | 0.111 | 1 | 12 |
| myg1 | 0.42257 | -0.24402484 | 0.029 | 0.111 | 1 | 12 |
| caskb | 0.42277 | -0.20824171 | 0.029 | 0.111 | 1 | 12 |
| dab2 | 0.42322 | 0.185803694 | 0.229 | 0.111 | 1 | 12 |
| btbd10b | 0.42334 | -0.22532939 | 0.029 | 0.111 | 1 | 12 |
| h2afx1 | 0.42348 | 0.277179026 | 0.286 | 0.148 | 1 | 12 |
| zgc:163030 | 0.42351 | -0.25275238 | 0.029 | 0.111 | 1 | 12 |
| mydgf | 0.4245 | -0.2032491 | 0.029 | 0.111 | 1 | 12 |
| ctdsp2 | 0.42457 | 0.293846719 | 0.286 | 0.148 | 1 | 12 |
| aurkb | 0.42484 | -0.24255395 | 0.029 | 0.111 | 1 | 12 |
| tspan18a | 0.425 | -0.21938259 | 0.029 | 0.111 | 1 | 12 |
| tsr2 | 0.42502 | -0.23059313 | 0.029 | 0.111 | 1 | 12 |
| ube2l3a | 0.42561 | -0.19669245 | 0.029 | 0.111 | 1 | 12 |
| kars | 0.42576 | -0.2115864 | 0.029 | 0.111 | 1 | 12 |
| slc25a20 | 0.42622 | -0.22856722 | 0.029 | 0.111 | 1 | 12 |
| mpc2 | 0.42623 | -0.19582572 | 0.029 | 0.111 | 1 | 12 |
| klf3 | 0.42631 | 0.30792199 | 0.114 | 0.037 | 1 | 12 |
| entpd6 | 0.42633 | -0.23223772 | 0.029 | 0.111 | 1 | 12 |
| dldh | 0.42641 | -0.24167108 | 0.029 | 0.111 | 1 | 12 |
| ngdn | 0.42647 | -0.20553199 | 0.029 | 0.111 | 1 | 12 |
| ndnl2 | 0.42651 | -0.27548169 | 0.029 | 0.111 | 1 | 12 |
| sms | 0.42709 | -0.2308689 | 0.029 | 0.111 | 1 | 12 |
| mbnl2 | 0.42715 | -0.23042615 | 0.029 | 0.111 | 1 | 12 |
| supt16h | 0.42722 | -0.19184691 | 0.029 | 0.111 | 1 | 12 |
| dda1 | 0.42733 | -0.26052699 | 0.029 | 0.111 | 1 | 12 |
| wu:fc33b09 | 0.42734 | -0.2159057 | 0.029 | 0.111 | 1 | 12 |
| smc3 | 0.42752 | -0.25466319 | 0.029 | 0.111 | 1 | 12 |
| clic4 | 0.42764 | -0.25183089 | 0.029 | 0.111 | 1 | 12 |

| | | | | | | |
|---|---|---|---|---|---|---|
| fosb | 0.42783 | -0.24041018 | 0.029 | 0.111 | 1 | 12 |
| tnrc18 | 0.42787 | -0.23033424 | 0.029 | 0.111 | 1 | 12 |
| zgc:110182 | 0.42787 | -0.26236737 | 0.029 | 0.111 | 1 | 12 |
| snapc1b | 0.42797 | -0.18095743 | 0.029 | 0.111 | 1 | 12 |
| tie1 | 0.42823 | 0.347967952 | 0.229 | 0.111 | 1 | 12 |
| p4hb | 0.42841 | -0.219954 | 0.029 | 0.111 | 1 | 12 |
| rpl23 | 0.42842 | 0.103251038 | 0.971 | 0.889 | 1 | 12 |
| rassf9 | 0.4285 | -0.23504713 | 0.029 | 0.111 | 1 | 12 |
| hdgfl2 | 0.42853 | -0.23835323 | 0.029 | 0.111 | 1 | 12 |
| lbr | 0.42855 | -0.25063523 | 0.029 | 0.111 | 1 | 12 |
| ext2 | 0.42856 | -0.20824823 | 0.029 | 0.111 | 1 | 12 |
| c1d | 0.42857 | -0.20693406 | 0.029 | 0.111 | 1 | 12 |
| mzt2b | 0.42858 | -0.2369876 | 0.029 | 0.111 | 1 | 12 |
| cdc42bpb | 0.42862 | -0.21028248 | 0.029 | 0.111 | 1 | 12 |
| rbmx | 0.42862 | -0.18601741 | 0.029 | 0.111 | 1 | 12 |
| si:ch211-166a6.5 | 0.42868 | -0.23724349 | 0.029 | 0.111 | 1 | 12 |
| sema6a | 0.42874 | -0.21126806 | 0.029 | 0.111 | 1 | 12 |
| gnaia | 0.42876 | -0.2473239 | 0.029 | 0.111 | 1 | 12 |
| dnajb12a | 0.42877 | -0.23281319 | 0.029 | 0.111 | 1 | 12 |
| smim4 | 0.42878 | -0.2324149 | 0.029 | 0.111 | 1 | 12 |
| id1 | 0.43154 | 0.315143894 | 0.457 | 0.296 | 1 | 12 |
| kpna4 | 0.43159 | -0.28969377 | 0.057 | 0.148 | 1 | 12 |
| zgc:110239 | 0.43223 | -0.31152044 | 0.086 | 0.185 | 1 | 12 |
| rpa2 | 0.43292 | -0.33548481 | 0.057 | 0.148 | 1 | 12 |
| dars | 0.43406 | -0.29548055 | 0.057 | 0.148 | 1 | 12 |
| ube2ib | 0.43597 | 0.286306368 | 0.114 | 0.037 | 1 | 12 |
| srsf2a | 0.43625 | -0.34557637 | 0.2 | 0.333 | 1 | 12 |
| sept2 | 0.4365 | -0.24753206 | 0.286 | 0.333 | 1 | 12 |
| fam177a1 | 0.43678 | 0.131349452 | 0.114 | 0.037 | 1 | 12 |
| rgl2 | 0.4375 | -0.27726161 | 0.057 | 0.148 | 1 | 12 |
| sf3b6 | 0.43828 | 0.219149788 | 0.229 | 0.111 | 1 | 12 |
| creb3l3l | 0.43846 | 0.287317931 | 0.114 | 0.037 | 1 | 12 |
| sem1 | 0.44008 | -0.24737938 | 0.257 | 0.296 | 1 | 12 |
| wipi2 | 0.44125 | 0.101998656 | 0.114 | 0.037 | 1 | 12 |
| vclb | 0.44166 | 0.291130011 | 0.114 | 0.037 | 1 | 12 |
| slc30a7 | 0.44191 | 0.134324543 | 0.114 | 0.037 | 1 | 12 |
| fam133b | 0.44323 | -0.37235228 | 0.114 | 0.222 | 1 | 12 |
| tegt | 0.44381 | 0.321039756 | 0.371 | 0.222 | 1 | 12 |
| CU929150.1 | 0.44407 | -0.39536121 | 0.086 | 0.185 | 1 | 12 |
| slc9a3r1b | 0.44439 | -0.31285905 | 0.086 | 0.185 | 1 | 12 |
| CABZ01076275.1 | 0.44512 | -0.36353884 | 0.057 | 0.148 | 1 | 12 |
| ak2 | 0.44521 | 0.356062696 | 0.314 | 0.185 | 1 | 12 |
| srprb | 0.44569 | 0.300754102 | 0.171 | 0.074 | 1 | 12 |
| chac1 | 0.44662 | 0.323702184 | 0.171 | 0.074 | 1 | 12 |
| mrps33 | 0.44711 | 0.1753239 | 0.114 | 0.037 | 1 | 12 |
| fam120c | 0.4486 | -0.29697612 | 0.086 | 0.185 | 1 | 12 |
| snap23.1 | 0.45078 | -0.34138616 | 0.114 | 0.222 | 1 | 12 |
| srsf3b | 0.45094 | -0.23104423 | 0.2 | 0.333 | 1 | 12 |

| | | | | | | |
|---|---|---|---|---|---|---|
| hif1al | 0.45096 | 0.237353202 | 0.229 | 0.111 | 1 | 12 |
| sh3kbp1 | 0.45117 | 0.268852141 | 0.171 | 0.074 | 1 | 12 |
| pitpnb | 0.45155 | 0.12856457 | 0.114 | 0.037 | 1 | 12 |
| jpt1b | 0.45203 | 0.335340466 | 0.4 | 0.259 | 1 | 12 |
| nop16 | 0.45222 | 0.261823478 | 0.114 | 0.037 | 1 | 12 |
| LOC101882264 | 0.45259 | -0.31598641 | 0.057 | 0.148 | 1 | 12 |
| wdr1 | 0.45439 | -0.32273408 | 0.086 | 0.185 | 1 | 12 |
| fzd4 | 0.45537 | 0.324361196 | 0.229 | 0.111 | 1 | 12 |
| mef2aa | 0.45538 | -0.35154176 | 0.114 | 0.222 | 1 | 12 |
| ndufab1b | 0.45596 | 0.241336657 | 0.229 | 0.111 | 1 | 12 |
| gemin7 | 0.45718 | 0.263127094 | 0.114 | 0.037 | 1 | 12 |
| dpy30 | 0.45763 | -0.29263695 | 0.086 | 0.185 | 1 | 12 |
| vamp5 | 0.45928 | -0.3250005 | 0.114 | 0.222 | 1 | 12 |
| ptp4a2a | 0.45984 | 0.280332617 | 0.114 | 0.037 | 1 | 12 |
| uqcr10 | 0.46057 | -0.33462689 | 0.2 | 0.333 | 1 | 12 |
| plekhg5b | 0.46107 | 0.270539808 | 0.114 | 0.037 | 1 | 12 |
| aida | 0.46141 | 0.125651224 | 0.114 | 0.037 | 1 | 12 |
| mrto4 | 0.46259 | -0.15285202 | 0.057 | 0.148 | 1 | 12 |
| si:ch211-284f22.3 | 0.46281 | 0.310697356 | 0.229 | 0.111 | 1 | 12 |
| CABZ01072614.1 | 0.46301 | 0.306859877 | 0.171 | 0.074 | 1 | 12 |
| stx3a | 0.4632 | -0.1727122 | 0.057 | 0.148 | 1 | 12 |
| sele | 0.46441 | 0.314072028 | 0.229 | 0.111 | 1 | 12 |
| etv2 | 0.46475 | 0.318519682 | 0.229 | 0.111 | 1 | 12 |
| snrpg | 0.46489 | -0.2180454 | 0.171 | 0.296 | 1 | 12 |
| zgc:65873 | 0.46503 | -0.31132583 | 0.086 | 0.185 | 1 | 12 |
| mcm7 | 0.46579 | -0.3119115 | 0.086 | 0.185 | 1 | 12 |
| nop53 | 0.46611 | 0.322392859 | 0.314 | 0.185 | 1 | 12 |
| bcl6b | 0.46635 | 0.235646146 | 0.229 | 0.111 | 1 | 12 |
| rtca | 0.46672 | 0.115340355 | 0.114 | 0.037 | 1 | 12 |
| lrrfip1a | 0.46837 | 0.162414287 | 0.114 | 0.037 | 1 | 12 |
| ralgapa1 | 0.46905 | 0.256661186 | 0.114 | 0.037 | 1 | 12 |
| si:ch211-217k17.7 | 0.46925 | 0.183498173 | 0.171 | 0.074 | 1 | 12 |
| brd2a | 0.46943 | 0.250337385 | 0.114 | 0.037 | 1 | 12 |
| hells | 0.47003 | -0.26665433 | 0.057 | 0.148 | 1 | 12 |
| mrpl15 | 0.47242 | -0.28790122 | 0.057 | 0.148 | 1 | 12 |
| anxa11a | 0.47265 | -0.30445078 | 0.114 | 0.222 | 1 | 12 |
| jpt2 | 0.47266 | -0.32373032 | 0.114 | 0.222 | 1 | 12 |
| dnase1l1 | 0.47281 | -0.23987847 | 0.057 | 0.148 | 1 | 12 |
| parvab | 0.47312 | -0.33912906 | 0.171 | 0.259 | 1 | 12 |
| psmg3 | 0.47345 | 0.257555007 | 0.229 | 0.111 | 1 | 12 |
| ralbp1 | 0.47401 | 0.285754919 | 0.114 | 0.037 | 1 | 12 |
| ccdc187 | 0.47403 | 0.10133506 | 0.114 | 0.037 | 1 | 12 |
| commd1 | 0.47405 | 0.263345337 | 0.114 | 0.037 | 1 | 12 |
| aldoaa | 0.47416 | 0.308128165 | 0.229 | 0.111 | 1 | 12 |
| ccdc124 | 0.4748 | 0.203698238 | 0.171 | 0.074 | 1 | 12 |
| cycsb | 0.475 | -0.18590203 | 0.057 | 0.148 | 1 | 12 |
| mef2d | 0.47574 | 0.241445193 | 0.114 | 0.037 | 1 | 12 |
| rps6 | 0.4758 | 0.160744973 | 0.914 | 0.852 | 1 | 12 |

| | | | | | | |
|---|---|---|---|---|---|---|
| dysf | 0.47617 | 0.3052274 | 0.143 | 0.074 | 1 | 12 |
| sdf2l1 | 0.47656 | 0.279764568 | 0.114 | 0.037 | 1 | 12 |
| acadl | 0.47695 | -0.19808322 | 0.057 | 0.148 | 1 | 12 |
| rab2a | 0.47726 | -0.30732769 | 0.143 | 0.259 | 1 | 12 |
| rpl34 | 0.47765 | 0.153413985 | 0.914 | 0.852 | 1 | 12 |
| zgc:194627 | 0.47771 | -0.32312365 | 0.057 | 0.148 | 1 | 12 |
| h2afvb | 0.4779 | -0.31237633 | 0.229 | 0.37 | 1 | 12 |
| gtf2b | 0.478 | -0.21048075 | 0.057 | 0.148 | 1 | 12 |
| FO082781.1 | 0.47881 | -0.21853689 | 0.057 | 0.148 | 1 | 12 |
| supt6h | 0.47922 | 0.24214485 | 0.114 | 0.037 | 1 | 12 |
| wwtr1 | 0.47936 | -0.21275298 | 0.057 | 0.148 | 1 | 12 |
| grinab | 0.48004 | 0.205823003 | 0.171 | 0.074 | 1 | 12 |
| rpl18 | 0.48035 | 0.111089184 | 0.971 | 0.926 | 1 | 12 |
| med10 | 0.4804 | -0.19677844 | 0.114 | 0.222 | 1 | 12 |
| rhogc | 0.48044 | 0.151651171 | 0.114 | 0.037 | 1 | 12 |
| lgi2a | 0.48047 | -0.24140288 | 0.057 | 0.148 | 1 | 12 |
| ap1s3b | 0.48077 | 0.248704537 | 0.114 | 0.037 | 1 | 12 |
| tgfbr2a | 0.48092 | -0.19396177 | 0.057 | 0.148 | 1 | 12 |
| bzw1b | 0.48112 | 0.191540246 | 0.171 | 0.074 | 1 | 12 |
| cbx3a | 0.48138 | -0.31964487 | 0.171 | 0.296 | 1 | 12 |
| wu:fi37e09 | 0.48203 | 0.282673109 | 0.171 | 0.074 | 1 | 12 |
| adi1 | 0.48221 | -0.18397331 | 0.057 | 0.148 | 1 | 12 |
| cox7a2l | 0.48265 | -0.30115807 | 0.143 | 0.259 | 1 | 12 |
| ccng1 | 0.48278 | 0.32429001 | 0.314 | 0.185 | 1 | 12 |
| clta | 0.4828 | -0.24696747 | 0.171 | 0.296 | 1 | 12 |
| mgst1.2 | 0.48288 | 0.285823722 | 0.171 | 0.074 | 1 | 12 |
| rpl5a | 0.48331 | -0.11541437 | 0.771 | 0.778 | 1 | 12 |
| rpp14 | 0.48343 | 0.157959897 | 0.114 | 0.037 | 1 | 12 |
| ppp6c | 0.48371 | 0.173673457 | 0.171 | 0.074 | 1 | 12 |
| ccm2l | 0.48388 | -0.28274212 | 0.086 | 0.185 | 1 | 12 |
| rab11a | 0.48419 | -0.32026809 | 0.143 | 0.259 | 1 | 12 |
| cd81b | 0.48434 | 0.201157429 | 0.257 | 0.148 | 1 | 12 |
| ppp3r1b | 0.48449 | 0.226413777 | 0.114 | 0.037 | 1 | 12 |
| cdv3 | 0.48469 | -0.19642409 | 0.143 | 0.259 | 1 | 12 |
| reep1 | 0.48518 | 0.284377366 | 0.171 | 0.074 | 1 | 12 |
| tmem59l | 0.48544 | -0.25830207 | 0.057 | 0.148 | 1 | 12 |
| vma21 | 0.48578 | -0.30916566 | 0.086 | 0.148 | 1 | 12 |
| tmem18 | 0.48607 | -0.22145864 | 0.057 | 0.148 | 1 | 12 |
| MZT1 | 0.48659 | -0.21758887 | 0.057 | 0.148 | 1 | 12 |
| hist2h2l | 0.48674 | -0.21233177 | 0.057 | 0.148 | 1 | 12 |
| glo1 | 0.48707 | 0.296207158 | 0.171 | 0.074 | 1 | 12 |
| sema3ab | 0.48726 | -0.27057487 | 0.057 | 0.148 | 1 | 12 |
| clint1a | 0.48809 | -0.19322589 | 0.057 | 0.148 | 1 | 12 |
| hnrnpa1a | 0.48854 | -0.3086932 | 0.114 | 0.222 | 1 | 12 |
| adgrl4 | 0.48855 | -0.19715527 | 0.057 | 0.148 | 1 | 12 |
| map4k2 | 0.48904 | -0.19109973 | 0.057 | 0.148 | 1 | 12 |
| mcm5 | 0.48945 | -0.29471562 | 0.057 | 0.148 | 1 | 12 |
| wdr83os | 0.48981 | 0.315194527 | 0.257 | 0.148 | 1 | 12 |

| | | | | | | |
|---|---|---|---|---|---|---|
| ndufs2 | 0.48993 | -0.22368833 | 0.057 | 0.148 | 1 | 12 |
| emg1 | 0.4903 | -0.22528081 | 0.057 | 0.148 | 1 | 12 |
| sptlc2a | 0.49036 | 0.172368309 | 0.114 | 0.037 | 1 | 12 |
| rps4x | 0.49039 | 0.116045017 | 0.943 | 0.852 | 1 | 12 |
| psmc2 | 0.49047 | -0.2745248 | 0.057 | 0.148 | 1 | 12 |
| txn2 | 0.49063 | -0.29458977 | 0.086 | 0.185 | 1 | 12 |
| gata2a | 0.49091 | -0.19763311 | 0.057 | 0.148 | 1 | 12 |
| gkap1 | 0.49092 | 0.181561995 | 0.171 | 0.074 | 1 | 12 |
| rpl7a | 0.49098 | 0.114947645 | 0.943 | 0.852 | 1 | 12 |
| LEPROTL1 | 0.49111 | -0.2092 | 0.057 | 0.148 | 1 | 12 |
| tgif1 | 0.49121 | -0.19258688 | 0.057 | 0.148 | 1 | 12 |
| rbpjb | 0.49122 | 0.224130904 | 0.114 | 0.037 | 1 | 12 |
| gspt1l | 0.49122 | -0.20584808 | 0.057 | 0.148 | 1 | 12 |
| emc7 | 0.49122 | -0.29572417 | 0.114 | 0.222 | 1 | 12 |
| uchl3 | 0.49126 | -0.24181076 | 0.057 | 0.148 | 1 | 12 |
| ivd | 0.49128 | -0.25358084 | 0.057 | 0.148 | 1 | 12 |
| gnpda2 | 0.49134 | -0.24226643 | 0.057 | 0.148 | 1 | 12 |
| cldn5a | 0.49135 | -0.19406468 | 0.057 | 0.148 | 1 | 12 |
| baz2ba | 0.49146 | 0.261472172 | 0.114 | 0.037 | 1 | 12 |
| tjp2b | 0.49279 | 0.253656383 | 0.114 | 0.037 | 1 | 12 |
| brd7 | 0.4933 | 0.137149731 | 0.114 | 0.037 | 1 | 12 |
| si:ch211-212k18.7 | 0.4939 | 0.212791598 | 0.114 | 0.037 | 1 | 12 |
| ptbp1a | 0.49413 | 0.252711433 | 0.114 | 0.037 | 1 | 12 |
| vat1 | 0.49474 | 0.258326466 | 0.629 | 0.481 | 1 | 12 |
| prelid1a | 0.49531 | 0.227942818 | 0.114 | 0.037 | 1 | 12 |
| si:ch211-14a17.11 | 0.49573 | 0.142647356 | 0.114 | 0.037 | 1 | 12 |
| hsd3b7 | 0.49632 | 0.22880833 | 0.114 | 0.037 | 1 | 12 |
| oxsr1a | 0.49707 | 0.183371918 | 0.114 | 0.037 | 1 | 12 |
| pip4k2ca | 0.49785 | 0.145804829 | 0.114 | 0.037 | 1 | 12 |
| atp5mc3a | 0.49804 | -0.30079225 | 0.171 | 0.296 | 1 | 12 |
| suclg1 | 0.49825 | 0.262655801 | 0.171 | 0.074 | 1 | 12 |
| naa10 | 0.49962 | -0.20648375 | 0.143 | 0.148 | 1 | 12 |
| rnf10 | 0.50006 | 0.123964983 | 0.2 | 0.111 | 1 | 12 |
| sub1b | 0.5001 | -0.18487926 | 0.114 | 0.222 | 1 | 12 |
| twist3 | 0.50038 | 0.197776144 | 0.114 | 0.037 | 1 | 12 |
| txnl4a | 0.50046 | 0.135594464 | 0.114 | 0.037 | 1 | 12 |
| arhgap31 | 0.50058 | 0.230265228 | 0.171 | 0.074 | 1 | 12 |
| pgam5 | 0.50121 | 0.183115309 | 0.114 | 0.037 | 1 | 12 |
| vps36 | 0.50158 | 0.171908971 | 0.114 | 0.037 | 1 | 12 |
| sfxn4 | 0.50222 | 0.212056621 | 0.114 | 0.037 | 1 | 12 |
| klf7b | 0.50296 | 0.182662863 | 0.114 | 0.037 | 1 | 12 |
| gna11b | 0.50345 | 0.195743905 | 0.114 | 0.037 | 1 | 12 |
| ypel5 | 0.50349 | 0.215154567 | 0.114 | 0.037 | 1 | 12 |
| twf1a | 0.50426 | 0.225447407 | 0.171 | 0.074 | 1 | 12 |
| ube2d1b | 0.50428 | 0.228060353 | 0.114 | 0.037 | 1 | 12 |
| cops5 | 0.50461 | 0.235086554 | 0.114 | 0.037 | 1 | 12 |
| nenf | 0.50488 | 0.214251052 | 0.114 | 0.037 | 1 | 12 |
| prdx1 | 0.50508 | -0.22547892 | 0.086 | 0.185 | 1 | 12 |

| | | | | | | |
|---|---|---|---|---|---|---|
| nckap1 | 0.50528 | 0.178565966 | 0.171 | 0.074 | 1 | 12 |
| stard8 | 0.50536 | 0.230528304 | 0.114 | 0.037 | 1 | 12 |
| mycn | 0.50596 | -0.30558164 | 0.086 | 0.185 | 1 | 12 |
| rhoac | 0.50608 | 0.294672446 | 0.314 | 0.185 | 1 | 12 |
| prpf38b | 0.50633 | 0.208995306 | 0.114 | 0.037 | 1 | 12 |
| cotl1 | 0.50695 | 0.258323544 | 0.314 | 0.185 | 1 | 12 |
| rab5c | 0.50704 | 0.28496037 | 0.314 | 0.185 | 1 | 12 |
| cald1a | 0.50746 | 0.256685753 | 0.314 | 0.185 | 1 | 12 |
| fbxo9 | 0.50787 | 0.19105874 | 0.114 | 0.037 | 1 | 12 |
| tmem88a | 0.50792 | 0.264274892 | 0.314 | 0.185 | 1 | 12 |
| si:ch73-334d15.2 | 0.50795 | 0.259914929 | 0.314 | 0.185 | 1 | 12 |
| psma5 | 0.50823 | 0.262495704 | 0.314 | 0.185 | 1 | 12 |
| fam171a2a | 0.50867 | 0.234654347 | 0.171 | 0.074 | 1 | 12 |
| ywhaqa | 0.5089 | -0.2558611 | 0.171 | 0.296 | 1 | 12 |
| atp1a1a.1 | 0.50966 | 0.199483678 | 0.114 | 0.037 | 1 | 12 |
| uck2a | 0.50998 | 0.247195021 | 0.171 | 0.074 | 1 | 12 |
| etfb | 0.51 | 0.229651223 | 0.171 | 0.074 | 1 | 12 |
| degs1 | 0.51002 | 0.189129953 | 0.114 | 0.037 | 1 | 12 |
| tanc2a | 0.51028 | 0.20275122 | 0.114 | 0.037 | 1 | 12 |
| selenop | 0.51029 | -0.30288347 | 0.114 | 0.222 | 1 | 12 |
| ppp2r5eb | 0.5107 | 0.198974973 | 0.114 | 0.037 | 1 | 12 |
| tomm6 | 0.51084 | 0.260470024 | 0.171 | 0.074 | 1 | 12 |
| chd3 | 0.51089 | 0.229891242 | 0.171 | 0.074 | 1 | 12 |
| calcrlb | 0.51109 | 0.259323658 | 0.171 | 0.074 | 1 | 12 |
| ddx39ab | 0.51114 | -0.28548043 | 0.114 | 0.222 | 1 | 12 |
| atp5pb | 0.5113 | -0.21329966 | 0.4 | 0.444 | 1 | 12 |
| supt4h1 | 0.51157 | 0.226587796 | 0.171 | 0.074 | 1 | 12 |
| cox4i2 | 0.51236 | 0.255290842 | 0.114 | 0.037 | 1 | 12 |
| si:ch1073-325m22.2 | 0.51438 | -0.28464783 | 0.371 | 0.481 | 1 | 12 |
| btg1 | 0.51451 | 0.223176355 | 0.429 | 0.37 | 1 | 12 |
| canx | 0.51479 | -0.24416074 | 0.086 | 0.185 | 1 | 12 |
| mgst3a | 0.51485 | -0.26545309 | 0.086 | 0.185 | 1 | 12 |
| si:dkey-28n18.9 | 0.51513 | 0.18039288 | 0.114 | 0.037 | 1 | 12 |
| sult1st1 | 0.51559 | -0.25936528 | 0.086 | 0.185 | 1 | 12 |
| eif3ja | 0.51604 | -0.28657267 | 0.086 | 0.185 | 1 | 12 |
| cyth3a | 0.51606 | 0.187060242 | 0.114 | 0.037 | 1 | 12 |
| eif4h | 0.5162 | -0.23437271 | 0.086 | 0.185 | 1 | 12 |
| fam129ba | 0.51751 | 0.218577177 | 0.114 | 0.037 | 1 | 12 |
| cxcl12a | 0.51764 | -0.35379772 | 0.257 | 0.37 | 1 | 12 |
| pin1 | 0.51769 | -0.2961254 | 0.114 | 0.222 | 1 | 12 |
| hmga2 | 0.51798 | 0.168881248 | 0.114 | 0.037 | 1 | 12 |
| pdlim1 | 0.51889 | -0.25565696 | 0.143 | 0.259 | 1 | 12 |
| rasip1 | 0.51911 | -0.27514566 | 0.143 | 0.259 | 1 | 12 |
| dusp1 | 0.51946 | 0.194360648 | 0.114 | 0.037 | 1 | 12 |
| emid1 | 0.51969 | 0.190086651 | 0.114 | 0.037 | 1 | 12 |
| stab1 | 0.5198 | -0.26463712 | 0.143 | 0.259 | 1 | 12 |
| rtn3 | 0.51981 | 0.158191397 | 0.114 | 0.037 | 1 | 12 |
| psma6b | 0.52 | -0.28864295 | 0.114 | 0.222 | 1 | 12 |

| | | | | | | |
|---|---|---|---|---|---|---|
| ckap4 | 0.52009 | 0.197323294 | 0.114 | 0.037 | 1 | 12 |
| pafah1b3 | 0.52046 | 0.197157152 | 0.114 | 0.037 | 1 | 12 |
| rbms3 | 0.52061 | 0.217913947 | 0.114 | 0.037 | 1 | 12 |
| zgc:162730 | 0.52066 | 0.171298165 | 0.114 | 0.037 | 1 | 12 |
| cdc26 | 0.52071 | 0.156999277 | 0.114 | 0.037 | 1 | 12 |
| pno1 | 0.52078 | 0.255183854 | 0.114 | 0.037 | 1 | 12 |
| magoh | 0.52089 | 0.246408366 | 0.114 | 0.037 | 1 | 12 |
| si:ch211-250c4.4 | 0.52103 | 0.168193765 | 0.114 | 0.037 | 1 | 12 |
| igf2a | 0.52134 | 0.204444622 | 0.114 | 0.037 | 1 | 12 |
| fam3c | 0.5214 | 0.177534828 | 0.114 | 0.037 | 1 | 12 |
| zgc:77086 | 0.52149 | 0.191729893 | 0.114 | 0.037 | 1 | 12 |
| ponzr1 | 0.52161 | 0.192599105 | 0.657 | 0.593 | 1 | 12 |
| hspa8 | 0.52187 | 0.128334104 | 0.886 | 0.778 | 1 | 12 |
| prpf39 | 0.52191 | 0.216855595 | 0.114 | 0.037 | 1 | 12 |
| mbd3a | 0.52199 | 0.237518045 | 0.114 | 0.037 | 1 | 12 |
| sulf2b | 0.52227 | 0.220332893 | 0.114 | 0.037 | 1 | 12 |
| rps27.1 | 0.52229 | 0.12766905 | 0.886 | 0.778 | 1 | 12 |
| im:7151449 | 0.5223 | 0.200379651 | 0.114 | 0.037 | 1 | 12 |
| gpr183a | 0.52243 | 0.231319802 | 0.114 | 0.037 | 1 | 12 |
| zgc:113184 | 0.5225 | 0.228904168 | 0.114 | 0.037 | 1 | 12 |
| zgc:174863 | 0.5225 | 0.163927543 | 0.114 | 0.037 | 1 | 12 |
| pdap1a | 0.52258 | 0.221069352 | 0.114 | 0.037 | 1 | 12 |
| hp1bp3 | 0.5226 | 0.238480195 | 0.114 | 0.037 | 1 | 12 |
| lsm7 | 0.52309 | -0.27968112 | 0.114 | 0.222 | 1 | 12 |
| ube2l3b | 0.52721 | 0.293626581 | 0.143 | 0.074 | 1 | 12 |
| socs3a | 0.52791 | 0.142991436 | 0.286 | 0.185 | 1 | 12 |
| pfn2l | 0.53138 | -0.19397427 | 0.314 | 0.444 | 1 | 12 |
| timm23a | 0.5344 | -0.29550732 | 0.057 | 0.111 | 1 | 12 |
| CT030188.1 | 0.53534 | -0.31301517 | 0.143 | 0.222 | 1 | 12 |
| mt-nd4 | 0.536 | -0.11753666 | 0.343 | 0.333 | 1 | 12 |
| mrps24 | 0.53623 | -0.27911501 | 0.057 | 0.111 | 1 | 12 |
| morf4l1 | 0.53696 | 0.134080014 | 0.2 | 0.111 | 1 | 12 |
| laptm4a | 0.53796 | 0.120388329 | 0.2 | 0.111 | 1 | 12 |
| hnrpkl | 0.5438 | 0.275690158 | 0.143 | 0.074 | 1 | 12 |
| krt15 | 0.54461 | 0.377038735 | 0.286 | 0.185 | 1 | 12 |
| ndufa5 | 0.54546 | -0.28717627 | 0.171 | 0.259 | 1 | 12 |
| snrpf | 0.54562 | 0.21828374 | 0.429 | 0.296 | 1 | 12 |
| atp1b1a | 0.55086 | 0.291806001 | 0.429 | 0.296 | 1 | 12 |
| mycbp | 0.55091 | 0.276286671 | 0.257 | 0.185 | 1 | 12 |
| si:dkey-7j14.6 | 0.55161 | 0.211268941 | 0.343 | 0.222 | 1 | 12 |
| exoc6 | 0.5558 | 0.131840803 | 0.2 | 0.111 | 1 | 12 |
| crip1 | 0.55585 | -0.21027754 | 0.314 | 0.444 | 1 | 12 |
| ndufa4l | 0.5559 | 0.290507542 | 0.343 | 0.222 | 1 | 12 |
| hnrnpabb | 0.56075 | -0.26944639 | 0.343 | 0.444 | 1 | 12 |
| eif5a | 0.56088 | 0.247722133 | 0.571 | 0.444 | 1 | 12 |
| tpm3 | 0.56128 | 0.136797306 | 0.486 | 0.37 | 1 | 12 |
| capns1b | 0.56139 | 0.258482529 | 0.257 | 0.148 | 1 | 12 |
| cdh6 | 0.56388 | -0.21383431 | 0.257 | 0.296 | 1 | 12 |

| | | | | | | |
|---|---|---|---|---|---|---|
| ldb2a | 0.56482 | 0.238436428 | 0.257 | 0.148 | 1 | 12 |
| atp5mc1 | 0.56597 | -0.29943381 | 0.229 | 0.333 | 1 | 12 |
| atp5po | 0.56713 | 0.242868552 | 0.257 | 0.148 | 1 | 12 |
| eif3k | 0.56827 | 0.252130472 | 0.257 | 0.148 | 1 | 12 |
| aqp1a.1 | 0.56896 | -0.3105399 | 0.086 | 0.148 | 1 | 12 |
| dap1b | 0.5708 | 0.255867549 | 0.257 | 0.148 | 1 | 12 |
| egfl7 | 0.57106 | 0.260833626 | 0.257 | 0.148 | 1 | 12 |
| ndufa6 | 0.57485 | 0.224553998 | 0.343 | 0.222 | 1 | 12 |
| mdka | 0.57881 | -0.19394165 | 0.2 | 0.222 | 1 | 12 |
| fbl | 0.58202 | -0.2892512 | 0.143 | 0.222 | 1 | 12 |
| ran | 0.58219 | -0.22778278 | 0.514 | 0.63 | 1 | 12 |
| smim15 | 0.5824 | -0.27607462 | 0.114 | 0.185 | 1 | 12 |
| pabpc1a | 0.58287 | -0.11061557 | 0.657 | 0.667 | 1 | 12 |
| aimp1 | 0.58346 | 0.275863512 | 0.143 | 0.074 | 1 | 12 |
| mdh1aa | 0.58396 | 0.253447877 | 0.2 | 0.148 | 1 | 12 |
| rbm8a | 0.58529 | -0.28415627 | 0.143 | 0.222 | 1 | 12 |
| junba | 0.58666 | -0.20617411 | 0.229 | 0.259 | 1 | 12 |
| rps9 | 0.58998 | -0.11330917 | 0.886 | 0.926 | 1 | 12 |
| nova2 | 0.59287 | -0.119978 | 0.171 | 0.259 | 1 | 12 |
| hadhb | 0.59331 | 0.119021394 | 0.143 | 0.074 | 1 | 12 |
| bccip | 0.59398 | 0.260049038 | 0.143 | 0.074 | 1 | 12 |
| csnk2a1 | 0.59425 | 0.171799629 | 0.2 | 0.111 | 1 | 12 |
| kank3 | 0.59509 | 0.257197238 | 0.143 | 0.074 | 1 | 12 |
| taf9 | 0.59554 | 0.241491304 | 0.143 | 0.074 | 1 | 12 |
| snrnp27 | 0.59803 | 0.237500083 | 0.143 | 0.074 | 1 | 12 |
| nfkbiab | 0.59824 | 0.275491907 | 0.286 | 0.185 | 1 | 12 |
| mcl1a | 0.59843 | 0.108235481 | 0.314 | 0.222 | 1 | 12 |
| serbp1a | 0.59897 | 0.228059488 | 0.571 | 0.444 | 1 | 12 |
| dynlrb1 | 0.60182 | 0.242451117 | 0.286 | 0.222 | 1 | 12 |
| cox5aa | 0.60347 | -0.26126599 | 0.229 | 0.296 | 1 | 12 |
| mrps14 | 0.60441 | -0.25875973 | 0.086 | 0.148 | 1 | 12 |
| lhfpl2a | 0.60642 | 0.109361455 | 0.143 | 0.074 | 1 | 12 |
| iqsec1b | 0.60653 | -0.26823797 | 0.057 | 0.111 | 1 | 12 |
| si:ch211-33e4.2 | 0.60711 | 0.168021966 | 0.286 | 0.185 | 1 | 12 |
| golga7 | 0.609 | 0.261318824 | 0.2 | 0.111 | 1 | 12 |
| zfyve21 | 0.60947 | -0.26885407 | 0.057 | 0.111 | 1 | 12 |
| cap1 | 0.61047 | -0.25967913 | 0.143 | 0.222 | 1 | 12 |
| rab11bb | 0.61293 | 0.183783984 | 0.286 | 0.185 | 1 | 12 |
| fzd7b | 0.61325 | 0.229805146 | 0.086 | 0.037 | 1 | 12 |
| pfdn2 | 0.61372 | 0.110225769 | 0.143 | 0.074 | 1 | 12 |
| actr2a | 0.61524 | 0.117459588 | 0.143 | 0.074 | 1 | 12 |
| tm4sf18 | 0.61825 | 0.24891324 | 0.2 | 0.111 | 1 | 12 |
| clic2 | 0.61826 | -0.24684453 | 0.171 | 0.222 | 1 | 12 |
| pfdn1 | 0.61882 | -0.26637861 | 0.114 | 0.185 | 1 | 12 |
| syncrip | 0.61941 | -0.20342661 | 0.257 | 0.296 | 1 | 12 |
| cnn2 | 0.62019 | 0.187043897 | 0.371 | 0.259 | 1 | 12 |
| ndufb8 | 0.62104 | 0.259720922 | 0.286 | 0.185 | 1 | 12 |
| ndufb3 | 0.62438 | 0.11156537 | 0.314 | 0.222 | 1 | 12 |

| | | | | | | |
|---|---|---|---|---|---|---|
| ifi30 | 0.62444 | 0.196539685 | 0.2 | 0.111 | 1 | 12 |
| uqcrq | 0.6247 | -0.25911545 | 0.2 | 0.296 | 1 | 12 |
| arf2b | 0.62484 | 0.222816024 | 0.143 | 0.074 | 1 | 12 |
| pnn | 0.62506 | 0.23473117 | 0.143 | 0.074 | 1 | 12 |
| pomp | 0.62535 | -0.11857518 | 0.371 | 0.37 | 1 | 12 |
| reep5 | 0.62557 | -0.25465315 | 0.143 | 0.222 | 1 | 12 |
| crk | 0.62726 | -0.25955931 | 0.086 | 0.148 | 1 | 12 |
| jak1 | 0.62757 | 0.196052602 | 0.2 | 0.111 | 1 | 12 |
| atp5pd | 0.6278 | 0.104732882 | 0.314 | 0.222 | 1 | 12 |
| aplp2 | 0.6295 | 0.21669556 | 0.2 | 0.111 | 1 | 12 |
| ucp2 | 0.62995 | 0.235384643 | 0.371 | 0.296 | 1 | 12 |
| eif4a1a | 0.62997 | 0.206295819 | 0.2 | 0.111 | 1 | 12 |
| ppp1cab | 0.63149 | -0.25048014 | 0.114 | 0.185 | 1 | 12 |
| ldhba | 0.63156 | -0.18402006 | 0.171 | 0.185 | 1 | 12 |
| plpp1a | 0.63265 | 0.214819153 | 0.2 | 0.111 | 1 | 12 |
| STMP1 | 0.63288 | 0.20121769 | 0.2 | 0.111 | 1 | 12 |
| psmd14 | 0.63323 | -0.245393 | 0.057 | 0.111 | 1 | 12 |
| cyc1 | 0.63326 | 0.193683873 | 0.2 | 0.111 | 1 | 12 |
| GCA | 0.6333 | -0.23322898 | 0.257 | 0.37 | 1 | 12 |
| flot1b | 0.63361 | 0.210348487 | 0.2 | 0.111 | 1 | 12 |
| si:ch211-145b13.6 | 0.63399 | 0.190191845 | 0.371 | 0.333 | 1 | 12 |
| caprin1a | 0.63417 | 0.214536839 | 0.086 | 0.037 | 1 | 12 |
| tomm70a | 0.63422 | 0.228891983 | 0.143 | 0.074 | 1 | 12 |
| ccl38.1 | 0.63497 | -0.2954558 | 0.229 | 0.333 | 1 | 12 |
| si:ch211-51e12.7 | 0.63499 | 0.216282667 | 0.143 | 0.111 | 1 | 12 |
| arpc4l | 0.6355 | 0.254489014 | 0.314 | 0.222 | 1 | 12 |
| cmpk | 0.63631 | -0.2624321 | 0.086 | 0.148 | 1 | 12 |
| csrp1a | 0.63633 | 0.247157464 | 0.286 | 0.185 | 1 | 12 |
| si:dkey-118j18.2 | 0.63756 | 0.21145047 | 0.086 | 0.037 | 1 | 12 |
| zgc:162509 | 0.63801 | 0.128844447 | 0.143 | 0.074 | 1 | 12 |
| foxp1b | 0.63876 | 0.138370509 | 0.143 | 0.074 | 1 | 12 |
| LO018340.1 | 0.6393 | 0.216782805 | 0.086 | 0.037 | 1 | 12 |
| s1pr5a | 0.63933 | 0.210546229 | 0.086 | 0.037 | 1 | 12 |
| eif2s2 | 0.64009 | -0.17265648 | 0.314 | 0.333 | 1 | 12 |
| LOC101884451 | 0.64047 | -0.18906955 | 0.229 | 0.333 | 1 | 12 |
| crebzf | 0.64348 | 0.227312785 | 0.143 | 0.074 | 1 | 12 |
| ilf3b | 0.64409 | 0.224058874 | 0.114 | 0.074 | 1 | 12 |
| slc40a1 | 0.64495 | 0.242833615 | 0.143 | 0.074 | 1 | 12 |
| rab14 | 0.64557 | -0.23163261 | 0.143 | 0.185 | 1 | 12 |
| brd4 | 0.64611 | 0.222633574 | 0.086 | 0.037 | 1 | 12 |
| ier2a | 0.6465 | 0.211455773 | 0.286 | 0.185 | 1 | 12 |
| acsbg2 | 0.64706 | -0.25174334 | 0.086 | 0.148 | 1 | 12 |
| fam53b | 0.64743 | 0.20455217 | 0.086 | 0.037 | 1 | 12 |
| mt-co1 | 0.64798 | -0.14326308 | 0.457 | 0.481 | 1 | 12 |
| fis1 | 0.64821 | -0.20682603 | 0.057 | 0.074 | 1 | 12 |
| tm9sf2 | 0.64892 | -0.22908984 | 0.057 | 0.111 | 1 | 12 |
| rtn4a | 0.64946 | 0.200678815 | 0.086 | 0.037 | 1 | 12 |
| cyfip1 | 0.64978 | -0.2443476 | 0.171 | 0.259 | 1 | 12 |

| | | | | | | |
|---|---|---|---|---|---|---|
| LOC110438237 | 0.65334 | -0.22926 | 0.057 | 0.111 | 1 | 12 |
| ctsf | 0.65438 | 0.124677533 | 0.143 | 0.074 | 1 | 12 |
| oaz2a | 0.65612 | -0.27447266 | 0.086 | 0.148 | 1 | 12 |
| appb | 0.65622 | 0.19943918 | 0.086 | 0.037 | 1 | 12 |
| gadd45ba | 0.65649 | 0.241410176 | 0.257 | 0.185 | 1 | 12 |
| hnrnpa1b | 0.65664 | -0.24521942 | 0.143 | 0.222 | 1 | 12 |
| ctnnd1 | 0.65745 | -0.23319765 | 0.057 | 0.111 | 1 | 12 |
| cox8a | 0.65746 | -0.20358071 | 0.229 | 0.333 | 1 | 12 |
| zgc:153311 | 0.65998 | 0.196297955 | 0.086 | 0.037 | 1 | 12 |
| flt4 | 0.66181 | -0.10768887 | 0.229 | 0.222 | 1 | 12 |
| fam43a | 0.66223 | 0.205948775 | 0.086 | 0.037 | 1 | 12 |
| bnip3lb | 0.66274 | 0.200641473 | 0.086 | 0.037 | 1 | 12 |
| btf3 | 0.66305 | 0.113763873 | 0.657 | 0.556 | 1 | 12 |
| ctnnb2 | 0.6631 | -0.25312052 | 0.143 | 0.222 | 1 | 12 |
| spryd7a | 0.66327 | 0.191319848 | 0.086 | 0.037 | 1 | 12 |
| mrpl34 | 0.66367 | 0.188342304 | 0.086 | 0.037 | 1 | 12 |
| med21 | 0.66504 | 0.189986832 | 0.086 | 0.037 | 1 | 12 |
| ccni | 0.66595 | -0.18072713 | 0.2 | 0.296 | 1 | 12 |
| ugt1b5 | 0.667 | 0.140531115 | 0.143 | 0.074 | 1 | 12 |
| rarga | 0.66821 | 0.122622657 | 0.143 | 0.074 | 1 | 12 |
| alas1 | 0.6689 | 0.183427399 | 0.086 | 0.037 | 1 | 12 |
| chd6 | 0.669 | 0.187775365 | 0.086 | 0.037 | 1 | 12 |
| tmed9 | 0.66902 | -0.15623168 | 0.143 | 0.148 | 1 | 12 |
| rps6ka3b | 0.67027 | 0.212215405 | 0.143 | 0.074 | 1 | 12 |
| fam32a | 0.67078 | 0.130106826 | 0.229 | 0.148 | 1 | 12 |
| antxr2a | 0.67193 | -0.23148132 | 0.2 | 0.296 | 1 | 12 |
| metrnla | 0.67193 | 0.195684406 | 0.086 | 0.037 | 1 | 12 |
| med29 | 0.67237 | 0.183172531 | 0.086 | 0.037 | 1 | 12 |
| knop1 | 0.67396 | -0.23485728 | 0.086 | 0.148 | 1 | 12 |
| ppp1caa | 0.67482 | -0.25006806 | 0.086 | 0.148 | 1 | 12 |
| slc25a5 | 0.67516 | -0.1069654 | 0.886 | 0.926 | 1 | 12 |
| wu:fb16f03 | 0.67567 | -0.22264295 | 0.086 | 0.148 | 1 | 12 |
| tpt1 | 0.67638 | -0.10106027 | 0.857 | 0.926 | 1 | 12 |
| anp32a | 0.67642 | -0.22795486 | 0.2 | 0.296 | 1 | 12 |
| tmem47 | 0.67651 | 0.205556547 | 0.086 | 0.037 | 1 | 12 |
| mrps11 | 0.67695 | 0.189354538 | 0.143 | 0.074 | 1 | 12 |
| arl6ip5b | 0.67711 | 0.157835707 | 0.143 | 0.074 | 1 | 12 |
| znf516 | 0.6776 | 0.208860438 | 0.086 | 0.037 | 1 | 12 |
| atp2b2 | 0.67855 | 0.18153082 | 0.086 | 0.037 | 1 | 12 |
| prdx5 | 0.67872 | -0.15044987 | 0.143 | 0.148 | 1 | 12 |
| gstt1b | 0.67991 | -0.23170371 | 0.2 | 0.259 | 1 | 12 |
| pkig | 0.67995 | 0.220292131 | 0.143 | 0.074 | 1 | 12 |
| jun | 0.68041 | -0.24628694 | 0.171 | 0.259 | 1 | 12 |
| sox7 | 0.68093 | 0.22773684 | 0.229 | 0.148 | 1 | 12 |
| pacsin2 | 0.68154 | 0.199639988 | 0.086 | 0.037 | 1 | 12 |
| eno3 | 0.68163 | -0.11548244 | 0.114 | 0.185 | 1 | 12 |
| akap12b | 0.68259 | -0.2148976 | 0.2 | 0.296 | 1 | 12 |
| ndufa10 | 0.68281 | -0.20783536 | 0.057 | 0.111 | 1 | 12 |

| | | | | | | |
|---|---|---|---|---|---|---|
| pafah1b1a | 0.68293 | -0.21294689 | 0.057 | 0.111 | 1 | 12 |
| prps1a | 0.68315 | 0.152533763 | 0.143 | 0.074 | 1 | 12 |
| map1lc3b | 0.6833 | 0.145040686 | 0.143 | 0.074 | 1 | 12 |
| pfn1 | 0.68351 | 0.220216179 | 0.314 | 0.222 | 1 | 12 |
| pcf11 | 0.684 | 0.171775378 | 0.086 | 0.037 | 1 | 12 |
| scamp2 | 0.6849 | -0.20580845 | 0.057 | 0.111 | 1 | 12 |
| rsrc2 | 0.68554 | 0.183696479 | 0.086 | 0.037 | 1 | 12 |
| eif5b | 0.68554 | -0.22079957 | 0.143 | 0.185 | 1 | 12 |
| arf6b | 0.68587 | 0.190973607 | 0.143 | 0.074 | 1 | 12 |
| kdm4b | 0.68607 | 0.196637881 | 0.086 | 0.037 | 1 | 12 |
| ier5 | 0.68691 | 0.169446069 | 0.143 | 0.074 | 1 | 12 |
| arf6a | 0.68734 | 0.161865888 | 0.143 | 0.074 | 1 | 12 |
| copg2 | 0.68742 | -0.18238846 | 0.057 | 0.111 | 1 | 12 |
| pfdn6 | 0.68759 | 0.208114195 | 0.143 | 0.074 | 1 | 12 |
| tsc22d3 | 0.68776 | 0.139360736 | 0.143 | 0.074 | 1 | 12 |
| faub | 0.68781 | 0.10685623 | 0.086 | 0.037 | 1 | 12 |
| bambia | 0.68804 | 0.201595667 | 0.086 | 0.037 | 1 | 12 |
| smdt1b | 0.68843 | -0.14780156 | 0.171 | 0.259 | 1 | 12 |
| sfpq | 0.68856 | 0.166313954 | 0.086 | 0.037 | 1 | 12 |
| denr | 0.68929 | 0.168971935 | 0.143 | 0.074 | 1 | 12 |
| pitpnbl | 0.68961 | 0.18185872 | 0.143 | 0.074 | 1 | 12 |
| tns2a | 0.68986 | 0.165732675 | 0.143 | 0.074 | 1 | 12 |
| arglu1a | 0.6899 | -0.20290296 | 0.114 | 0.185 | 1 | 12 |
| anks1b | 0.68998 | 0.181688802 | 0.086 | 0.037 | 1 | 12 |
| eif1ad | 0.69007 | 0.182526356 | 0.143 | 0.074 | 1 | 12 |
| BX470259.2 | 0.69031 | 0.179054552 | 0.086 | 0.037 | 1 | 12 |
| mob1a | 0.69033 | 0.16302987 | 0.143 | 0.074 | 1 | 12 |
| tcp1 | 0.69063 | 0.172759666 | 0.143 | 0.074 | 1 | 12 |
| pkma | 0.69085 | 0.15830492 | 0.143 | 0.074 | 1 | 12 |
| sars | 0.69124 | -0.22280521 | 0.057 | 0.111 | 1 | 12 |
| CU104709.1 | 0.69248 | 0.194033822 | 0.086 | 0.037 | 1 | 12 |
| zgc:92606 | 0.6927 | 0.186029968 | 0.086 | 0.037 | 1 | 12 |
| LOC100535225 | 0.69338 | -0.15795816 | 0.143 | 0.222 | 1 | 12 |
| cops4 | 0.69392 | 0.163656945 | 0.086 | 0.037 | 1 | 12 |
| BX548028.1 | 0.69398 | 0.168233002 | 0.086 | 0.037 | 1 | 12 |
| XLOC-039817 | 0.69494 | 0.186944588 | 0.086 | 0.037 | 1 | 12 |
| pabpn1 | 0.6959 | -0.12109917 | 0.114 | 0.185 | 1 | 12 |
| krt94 | 0.69596 | 0.19041622 | 0.514 | 0.407 | 1 | 12 |
| myl12.2 | 0.69741 | -0.23250869 | 0.114 | 0.185 | 1 | 12 |
| swap70b | 0.69744 | 0.184993969 | 0.086 | 0.037 | 1 | 12 |
| gsta.1 | 0.69812 | 0.157912988 | 0.086 | 0.037 | 1 | 12 |
| sbds | 0.69861 | 0.153306166 | 0.086 | 0.037 | 1 | 12 |
| atp5mea | 0.69961 | 0.147000641 | 0.114 | 0.111 | 1 | 12 |
| rac1b | 0.69976 | -0.19359603 | 0.057 | 0.111 | 1 | 12 |
| ssrp1a | 0.70083 | -0.19956537 | 0.057 | 0.111 | 1 | 12 |
| cdc5l | 0.70148 | 0.167468772 | 0.086 | 0.037 | 1 | 12 |
| tfe3a | 0.7015 | 0.172581983 | 0.086 | 0.037 | 1 | 12 |
| mdh2 | 0.70306 | -0.19071813 | 0.171 | 0.259 | 1 | 12 |

| | | | | | | |
|---|---|---|---|---|---|---|
| sf3b4 | 0.70335 | 0.163448366 | 0.086 | 0.037 | 1 | 12 |
| si:ch211-288g17.3 | 0.70369 | -0.2087816 | 0.171 | 0.259 | 1 | 12 |
| mrpl24 | 0.704 | 0.162024273 | 0.086 | 0.037 | 1 | 12 |
| tmem30ab | 0.7044 | 0.176887496 | 0.086 | 0.037 | 1 | 12 |
| prelid3b | 0.70448 | -0.14320576 | 0.143 | 0.148 | 1 | 12 |
| mycb | 0.70596 | 0.147205628 | 0.086 | 0.037 | 1 | 12 |
| prcp | 0.70605 | 0.119084262 | 0.086 | 0.037 | 1 | 12 |
| appa | 0.70632 | -0.1467756 | 0.143 | 0.222 | 1 | 12 |
| mrpl54 | 0.70702 | 0.103295538 | 0.086 | 0.037 | 1 | 12 |
| nop10 | 0.70734 | 0.201318489 | 0.114 | 0.074 | 1 | 12 |
| rnd3a | 0.70739 | -0.2081335 | 0.057 | 0.111 | 1 | 12 |
| bzw1a | 0.70825 | -0.21672605 | 0.143 | 0.222 | 1 | 12 |
| hnrnpl2 | 0.70863 | 0.211846109 | 0.229 | 0.148 | 1 | 12 |
| itga5 | 0.7088 | 0.158685413 | 0.086 | 0.037 | 1 | 12 |
| taok1a | 0.709 | 0.100502083 | 0.086 | 0.037 | 1 | 12 |
| gng10 | 0.70906 | -0.23000173 | 0.114 | 0.185 | 1 | 12 |
| tp53 | 0.70963 | -0.14411873 | 0.143 | 0.222 | 1 | 12 |
| sdhb | 0.71037 | 0.103203206 | 0.086 | 0.037 | 1 | 12 |
| gnai2a | 0.71043 | 0.21093531 | 0.314 | 0.222 | 1 | 12 |
| rassf1 | 0.71069 | 0.139680122 | 0.086 | 0.037 | 1 | 12 |
| tktb | 0.7109 | -0.15943416 | 0.057 | 0.111 | 1 | 12 |
| gmppb | 0.71117 | -0.21364708 | 0.057 | 0.111 | 1 | 12 |
| vdrb | 0.71132 | 0.111703803 | 0.086 | 0.037 | 1 | 12 |
| si:dkey-23i12.5 | 0.71138 | 0.149083291 | 0.086 | 0.037 | 1 | 12 |
| acaa2 | 0.71151 | 0.222314726 | 0.171 | 0.111 | 1 | 12 |
| entpd1 | 0.7119 | 0.169218344 | 0.086 | 0.037 | 1 | 12 |
| minos1 | 0.71203 | -0.20699961 | 0.086 | 0.148 | 1 | 12 |
| ece1 | 0.71212 | 0.167981382 | 0.086 | 0.037 | 1 | 12 |
| sept10 | 0.71232 | -0.19242182 | 0.086 | 0.148 | 1 | 12 |
| il13ra1 | 0.7124 | 0.217717981 | 0.171 | 0.111 | 1 | 12 |
| tfa | 0.7125 | 0.144441143 | 0.086 | 0.037 | 1 | 12 |
| arhgef12b | 0.71314 | -0.18783353 | 0.057 | 0.111 | 1 | 12 |
| adam10a | 0.71332 | -0.19557183 | 0.114 | 0.185 | 1 | 12 |
| fdx2 | 0.71341 | 0.140589602 | 0.086 | 0.037 | 1 | 12 |
| drap1 | 0.71398 | -0.20408602 | 0.086 | 0.148 | 1 | 12 |
| hnrnph1l | 0.71418 | 0.110724102 | 0.086 | 0.037 | 1 | 12 |
| rps27.2 | 0.71484 | 0.161784149 | 0.657 | 0.556 | 1 | 12 |
| si:ch1073-291c23.2 | 0.71484 | 0.151614722 | 0.086 | 0.037 | 1 | 12 |
| spred2b | 0.71488 | -0.16859809 | 0.143 | 0.222 | 1 | 12 |
| mcu | 0.71636 | 0.159418281 | 0.086 | 0.037 | 1 | 12 |
| eif4a1b | 0.71649 | -0.20677724 | 0.314 | 0.407 | 1 | 12 |
| si:ch211-207d6.2 | 0.71679 | -0.20758976 | 0.057 | 0.111 | 1 | 12 |
| CABZ01058650.1 | 0.71684 | 0.120498824 | 0.057 | 0.037 | 1 | 12 |
| acp1 | 0.71684 | 0.103589296 | 0.057 | 0.037 | 1 | 12 |
| agtr2 | 0.71684 | 0.107245461 | 0.057 | 0.037 | 1 | 12 |
| dync1li2 | 0.71684 | 0.148201787 | 0.057 | 0.037 | 1 | 12 |
| loxl2a | 0.71684 | 0.110825868 | 0.057 | 0.037 | 1 | 12 |
| notch1b | 0.71684 | 0.149125044 | 0.057 | 0.037 | 1 | 12 |

| | | | | | | |
|---|---|---|---|---|---|---|
| pam16 | 0.71684 | 0.15658458 | 0.057 | 0.037 | 1 | 12 |
| pawr | 0.71684 | 0.137230989 | 0.057 | 0.037 | 1 | 12 |
| pdk2a | 0.71684 | 0.127191004 | 0.057 | 0.037 | 1 | 12 |
| sh3bp5a | 0.71684 | 0.136911718 | 0.057 | 0.037 | 1 | 12 |
| stim1a | 0.71684 | 0.126514136 | 0.057 | 0.037 | 1 | 12 |
| tent4b | 0.71684 | 0.109310023 | 0.057 | 0.037 | 1 | 12 |
| tmem263 | 0.71684 | 0.103625931 | 0.057 | 0.037 | 1 | 12 |
| tspan7 | 0.71684 | 0.154651141 | 0.057 | 0.037 | 1 | 12 |
| zc3h15 | 0.71684 | 0.101664802 | 0.057 | 0.037 | 1 | 12 |
| copb2 | 0.71684 | 0.1598018 | 0.057 | 0.037 | 1 | 12 |
| fam102ab | 0.71684 | 0.148978381 | 0.057 | 0.037 | 1 | 12 |
| gstk1 | 0.71684 | 0.105076434 | 0.057 | 0.037 | 1 | 12 |
| smarca4a | 0.71684 | 0.128377162 | 0.057 | 0.037 | 1 | 12 |
| vps26a | 0.71684 | 0.135176924 | 0.057 | 0.037 | 1 | 12 |
| laptm4b | 0.71684 | 0.117974549 | 0.057 | 0.037 | 1 | 12 |
| mettl26 | 0.71684 | 0.143542675 | 0.057 | 0.037 | 1 | 12 |
| nr2f1a | 0.71684 | 0.102877297 | 0.057 | 0.037 | 1 | 12 |
| rerea | 0.71684 | 0.12173399 | 0.057 | 0.037 | 1 | 12 |
| sel1l | 0.71684 | 0.150041296 | 0.057 | 0.037 | 1 | 12 |
| stx4 | 0.71684 | 0.125663253 | 0.057 | 0.037 | 1 | 12 |
| tp53i11b | 0.71684 | 0.113382947 | 0.057 | 0.037 | 1 | 12 |
| LOC101886114 | 0.71684 | 0.110444845 | 0.057 | 0.037 | 1 | 12 |
| arl8ba | 0.71684 | 0.114346703 | 0.057 | 0.037 | 1 | 12 |
| enpep | 0.71684 | 0.119761247 | 0.057 | 0.037 | 1 | 12 |
| hddc2 | 0.71684 | 0.107001531 | 0.057 | 0.037 | 1 | 12 |
| kif5ba | 0.71684 | 0.147189293 | 0.057 | 0.037 | 1 | 12 |
| med30 | 0.71684 | 0.106987571 | 0.057 | 0.037 | 1 | 12 |
| metap1 | 0.71684 | 0.118503827 | 0.057 | 0.037 | 1 | 12 |
| nr2f6a | 0.71684 | 0.101568086 | 0.057 | 0.037 | 1 | 12 |
| nrarpb | 0.71684 | 0.100843594 | 0.057 | 0.037 | 1 | 12 |
| plekhf2 | 0.71684 | 0.123886968 | 0.057 | 0.037 | 1 | 12 |
| ranbp3b | 0.71684 | 0.12521382 | 0.057 | 0.037 | 1 | 12 |
| surf4 | 0.71684 | 0.117561312 | 0.057 | 0.037 | 1 | 12 |
| trib3 | 0.71684 | 0.141156494 | 0.057 | 0.037 | 1 | 12 |
| tuba1b | 0.71684 | 0.1083367 | 0.057 | 0.037 | 1 | 12 |
| ubr4 | 0.71684 | 0.121166847 | 0.057 | 0.037 | 1 | 12 |
| zgc:55558 | 0.71684 | 0.115404358 | 0.057 | 0.037 | 1 | 12 |
| abhd17c | 0.71684 | 0.102961339 | 0.057 | 0.037 | 1 | 12 |
| ccdc58 | 0.71684 | 0.106204401 | 0.057 | 0.037 | 1 | 12 |
| mrpl13 | 0.71684 | 0.105124566 | 0.057 | 0.037 | 1 | 12 |
| pwwp2a | 0.71684 | 0.117328703 | 0.057 | 0.037 | 1 | 12 |
| trir | 0.71684 | 0.128970839 | 0.057 | 0.037 | 1 | 12 |
| atg9b | 0.71684 | 0.11181784 | 0.057 | 0.037 | 1 | 12 |
| mlx | 0.71684 | 0.111918319 | 0.057 | 0.037 | 1 | 12 |
| phf14 | 0.71684 | 0.121355483 | 0.057 | 0.037 | 1 | 12 |
| si:dkey-147f3.8 | 0.71684 | 0.114210804 | 0.057 | 0.037 | 1 | 12 |
| epb41l4a | 0.71684 | 0.138923544 | 0.057 | 0.037 | 1 | 12 |
| mcm2 | 0.71684 | 0.131033924 | 0.057 | 0.037 | 1 | 12 |

| | | | | | | |
|---|---|---|---|---|---|---|
| pdcd5 | 0.71684 | 0.101448628 | 0.057 | 0.037 | 1 | 12 |
| rprd2b | 0.71684 | 0.111711427 | 0.057 | 0.037 | 1 | 12 |
| bri3bp | 0.71684 | 0.127102258 | 0.057 | 0.037 | 1 | 12 |
| scaf1 | 0.71688 | 0.133668478 | 0.086 | 0.037 | 1 | 12 |
| smad1 | 0.71695 | -0.19820296 | 0.086 | 0.148 | 1 | 12 |
| ddx61 | 0.71755 | -0.19136988 | 0.086 | 0.111 | 1 | 12 |
| snrpe | 0.71762 | 0.196840216 | 0.229 | 0.148 | 1 | 12 |
| rhoaa | 0.71769 | 0.208694737 | 0.314 | 0.222 | 1 | 12 |
| nudcd3 | 0.7183 | 0.158013191 | 0.086 | 0.037 | 1 | 12 |
| tars | 0.71851 | -0.19569855 | 0.057 | 0.111 | 1 | 12 |
| selenof | 0.71914 | 0.137326992 | 0.429 | 0.333 | 1 | 12 |
| mgst3b | 0.71923 | -0.16549157 | 0.2 | 0.222 | 1 | 12 |
| foxo1a | 0.71934 | 0.123229227 | 0.086 | 0.037 | 1 | 12 |
| cnbpa | 0.71936 | 0.203129153 | 0.114 | 0.074 | 1 | 12 |
| ahnak | 0.71941 | 0.184993064 | 0.314 | 0.222 | 1 | 12 |
| ehd4 | 0.72016 | 0.186184611 | 0.229 | 0.148 | 1 | 12 |
| cxxc5a | 0.7208 | 0.139763937 | 0.086 | 0.037 | 1 | 12 |
| cnn3a | 0.72088 | 0.209661961 | 0.229 | 0.148 | 1 | 12 |
| safb | 0.72094 | 0.139972692 | 0.086 | 0.037 | 1 | 12 |
| epha2a | 0.72137 | 0.14095877 | 0.086 | 0.037 | 1 | 12 |
| bhlhe40 | 0.72138 | 0.149088999 | 0.086 | 0.037 | 1 | 12 |
| nid1b | 0.72187 | 0.184171663 | 0.086 | 0.037 | 1 | 12 |
| llgl2 | 0.72204 | -0.17848525 | 0.057 | 0.111 | 1 | 12 |
| eif4a2 | 0.72216 | 0.136983746 | 0.086 | 0.037 | 1 | 12 |
| ppp1cb | 0.72222 | -0.15652414 | 0.057 | 0.111 | 1 | 12 |
| CABZ01078055.1 | 0.72247 | 0.181048371 | 0.086 | 0.037 | 1 | 12 |
| ilf2 | 0.7227 | -0.1913809 | 0.143 | 0.222 | 1 | 12 |
| trmt112 | 0.72271 | -0.18876777 | 0.143 | 0.222 | 1 | 12 |
| rapgef2 | 0.72275 | 0.121956729 | 0.086 | 0.037 | 1 | 12 |
| si:ch73-1a9.3 | 0.72294 | -0.17561111 | 0.486 | 0.556 | 1 | 12 |
| psma8 | 0.72319 | -0.20764726 | 0.314 | 0.407 | 1 | 12 |
| snx3 | 0.72328 | -0.15240603 | 0.114 | 0.185 | 1 | 12 |
| spast | 0.72344 | 0.105310353 | 0.086 | 0.037 | 1 | 12 |
| tpm4a | 0.72431 | 0.201592702 | 0.457 | 0.37 | 1 | 12 |
| hmgb3a | 0.72455 | -0.19731041 | 0.086 | 0.111 | 1 | 12 |
| ube2e1 | 0.7246 | 0.174604289 | 0.229 | 0.148 | 1 | 12 |
| prickle1b | 0.72467 | 0.172919717 | 0.086 | 0.037 | 1 | 12 |
| aplnrb | 0.72484 | 0.149290158 | 0.086 | 0.037 | 1 | 12 |
| copz1 | 0.72495 | -0.14081864 | 0.086 | 0.148 | 1 | 12 |
| si:dkey-199f5.8 | 0.7251 | 0.119634626 | 0.086 | 0.037 | 1 | 12 |
| uqcrc1 | 0.72516 | 0.173600526 | 0.229 | 0.148 | 1 | 12 |
| metap2b | 0.72525 | 0.203495249 | 0.229 | 0.148 | 1 | 12 |
| zgc:56676 | 0.72527 | 0.184046424 | 0.229 | 0.148 | 1 | 12 |
| csde1 | 0.72549 | 0.18632246 | 0.229 | 0.148 | 1 | 12 |
| xpr1b | 0.72599 | 0.115731382 | 0.086 | 0.037 | 1 | 12 |
| srsf6b | 0.72606 | -0.17208692 | 0.057 | 0.111 | 1 | 12 |
| palm1a | 0.72617 | -0.11532824 | 0.057 | 0.111 | 1 | 12 |
| pdhb | 0.72631 | 0.105105849 | 0.086 | 0.037 | 1 | 12 |

| | | | | | | |
|---|---|---|---|---|---|---|
| pdcd10a | 0.72657 | -0.12674386 | 0.086 | 0.148 | 1 | 12 |
| phlda2 | 0.72674 | -0.21797441 | 0.114 | 0.185 | 1 | 12 |
| ifi46 | 0.72679 | 0.130223843 | 0.086 | 0.037 | 1 | 12 |
| jam3b | 0.72703 | -0.20795126 | 0.086 | 0.148 | 1 | 12 |
| ifrd1 | 0.72728 | 0.103419177 | 0.086 | 0.037 | 1 | 12 |
| zbtb20 | 0.72729 | 0.12865427 | 0.086 | 0.037 | 1 | 12 |
| rfxap | 0.72747 | -0.18639627 | 0.057 | 0.111 | 1 | 12 |
| snx9b | 0.72801 | -0.20425656 | 0.143 | 0.185 | 1 | 12 |
| snrpd2 | 0.72817 | -0.21282183 | 0.171 | 0.222 | 1 | 12 |
| rgl1 | 0.72825 | -0.20587107 | 0.114 | 0.185 | 1 | 12 |
| pdha1a | 0.72828 | -0.20888949 | 0.057 | 0.111 | 1 | 12 |
| LOC103909972 | 0.72832 | 0.101725876 | 0.086 | 0.037 | 1 | 12 |
| tmem176l.3a | 0.72834 | 0.140358949 | 0.086 | 0.037 | 1 | 12 |
| prpf8 | 0.72861 | 0.115315964 | 0.086 | 0.037 | 1 | 12 |
| slirp | 0.72866 | -0.15723188 | 0.114 | 0.185 | 1 | 12 |
| fosab | 0.72896 | -0.16332394 | 0.114 | 0.185 | 1 | 12 |
| tob1a | 0.72898 | 0.213138515 | 0.171 | 0.111 | 1 | 12 |
| arpc4 | 0.72916 | 0.133640819 | 0.086 | 0.037 | 1 | 12 |
| polr2gl | 0.72916 | 0.14408557 | 0.086 | 0.037 | 1 | 12 |
| mapre1b | 0.72939 | -0.20192842 | 0.114 | 0.148 | 1 | 12 |
| capn2a | 0.72987 | 0.115030075 | 0.086 | 0.037 | 1 | 12 |
| hspa5 | 0.72995 | 0.166068914 | 0.086 | 0.037 | 1 | 12 |
| tcf12 | 0.73006 | -0.19871196 | 0.114 | 0.185 | 1 | 12 |
| mrpl30 | 0.7302 | 0.149449204 | 0.086 | 0.037 | 1 | 12 |
| ufc1 | 0.73026 | 0.194266074 | 0.086 | 0.037 | 1 | 12 |
| tle2a | 0.73051 | 0.112860455 | 0.086 | 0.037 | 1 | 12 |
| pon3.2 | 0.73065 | 0.148175714 | 0.086 | 0.037 | 1 | 12 |
| zmynd11 | 0.73113 | 0.123438635 | 0.086 | 0.037 | 1 | 12 |
| yipf4 | 0.73148 | -0.18043317 | 0.114 | 0.185 | 1 | 12 |
| rbm19 | 0.73166 | 0.160652482 | 0.086 | 0.037 | 1 | 12 |
| polr2f | 0.73174 | -0.18370634 | 0.114 | 0.185 | 1 | 12 |
| mif | 0.73184 | -0.15235498 | 0.114 | 0.185 | 1 | 12 |
| LOC103910438 | 0.73206 | 0.120289131 | 0.086 | 0.037 | 1 | 12 |
| cd63 | 0.73243 | 0.19416444 | 0.571 | 0.481 | 1 | 12 |
| lzts2a | 0.73248 | 0.116390924 | 0.086 | 0.037 | 1 | 12 |
| tomm20a | 0.73258 | 0.216909063 | 0.171 | 0.111 | 1 | 12 |
| gnl2 | 0.73271 | 0.173365119 | 0.086 | 0.037 | 1 | 12 |
| gadd45gip1 | 0.73295 | -0.18799989 | 0.057 | 0.111 | 1 | 12 |
| parvaa | 0.733 | 0.124677778 | 0.086 | 0.037 | 1 | 12 |
| col15a1a | 0.73309 | 0.133867269 | 0.086 | 0.037 | 1 | 12 |
| eif4a3 | 0.73335 | -0.20438621 | 0.086 | 0.148 | 1 | 12 |
| yap1 | 0.7334 | 0.126339279 | 0.086 | 0.037 | 1 | 12 |
| ccdc82 | 0.73346 | 0.113483772 | 0.086 | 0.037 | 1 | 12 |
| nrarpa | 0.73361 | -0.18799005 | 0.057 | 0.111 | 1 | 12 |
| si:ch211-262i1.6 | 0.73361 | 0.163293805 | 0.086 | 0.037 | 1 | 12 |
| rab1ab | 0.73363 | -0.15878103 | 0.114 | 0.185 | 1 | 12 |
| nr2f2 | 0.73379 | 0.138182927 | 0.086 | 0.037 | 1 | 12 |
| prdx3 | 0.734 | 0.145400218 | 0.086 | 0.037 | 1 | 12 |

| | | | | | | |
|---|---|---|---|---|---|---|
| si:dkey-239h2.3 | 0.73401 | 0.15170122 | 0.086 | 0.037 | 1 | 12 |
| galns | 0.73412 | 0.135974134 | 0.086 | 0.037 | 1 | 12 |
| rpz5 | 0.73434 | 0.132898602 | 0.086 | 0.037 | 1 | 12 |
| btbd10a | 0.73437 | 0.148846039 | 0.086 | 0.037 | 1 | 12 |
| LOC100333111 | 0.73437 | 0.107485716 | 0.086 | 0.037 | 1 | 12 |
| dido1 | 0.73478 | 0.115387951 | 0.086 | 0.037 | 1 | 12 |
| osbpl10b | 0.73478 | 0.101346937 | 0.086 | 0.037 | 1 | 12 |
| pho | 0.73498 | 0.123180542 | 0.086 | 0.037 | 1 | 12 |
| znf410 | 0.73521 | 0.146801652 | 0.086 | 0.037 | 1 | 12 |
| si:ch211-227m13.1 | 0.73527 | 0.143750685 | 0.086 | 0.037 | 1 | 12 |
| si:ch211-262e15.1 | 0.73529 | 0.139566732 | 0.086 | 0.037 | 1 | 12 |
| rbms1b | 0.73538 | 0.110582947 | 0.086 | 0.037 | 1 | 12 |
| ogdha | 0.73541 | 0.139074848 | 0.086 | 0.037 | 1 | 12 |
| brd3a | 0.73547 | 0.160275299 | 0.086 | 0.037 | 1 | 12 |
| irf2bp2b | 0.73548 | 0.144108112 | 0.086 | 0.037 | 1 | 12 |
| ildr2 | 0.73553 | 0.109746379 | 0.086 | 0.037 | 1 | 12 |
| rnf7 | 0.7356 | 0.127559966 | 0.086 | 0.037 | 1 | 12 |
| nutf2l | 0.73561 | -0.15441347 | 0.057 | 0.111 | 1 | 12 |
| aimp2 | 0.73563 | 0.16785742 | 0.086 | 0.037 | 1 | 12 |
| rfc5 | 0.73566 | -0.12874549 | 0.057 | 0.111 | 1 | 12 |
| gnl3 | 0.7357 | 0.128883235 | 0.086 | 0.037 | 1 | 12 |
| pik3r3a | 0.73571 | 0.105267742 | 0.086 | 0.037 | 1 | 12 |
| plpp3 | 0.73624 | -0.17780016 | 0.057 | 0.111 | 1 | 12 |
| selenok | 0.73663 | -0.13674327 | 0.057 | 0.111 | 1 | 12 |
| trappc1 | 0.73698 | -0.14962737 | 0.114 | 0.185 | 1 | 12 |
| ier3ip1 | 0.73707 | -0.16832351 | 0.114 | 0.185 | 1 | 12 |
| ube2nb | 0.73708 | -0.1739985 | 0.114 | 0.185 | 1 | 12 |
| rgcc | 0.73709 | 0.219530948 | 0.371 | 0.296 | 1 | 12 |
| pde4ba | 0.73718 | -0.10423542 | 0.086 | 0.148 | 1 | 12 |
| faua | 0.73753 | 0.108905195 | 0.886 | 0.852 | 1 | 12 |
| ndufa7 | 0.73759 | -0.13905003 | 0.086 | 0.148 | 1 | 12 |
| lgals9l3 | 0.73919 | -0.20413765 | 0.171 | 0.222 | 1 | 12 |
| si:ch73-138n13.1 | 0.73939 | -0.17216638 | 0.086 | 0.148 | 1 | 12 |
| git1 | 0.7402 | -0.15561889 | 0.057 | 0.111 | 1 | 12 |
| CU459056.1 | 0.74024 | -0.14742646 | 0.057 | 0.111 | 1 | 12 |
| srp19 | 0.7404 | -0.13000741 | 0.057 | 0.111 | 1 | 12 |
| zc3h7bb | 0.74042 | -0.18613341 | 0.057 | 0.111 | 1 | 12 |
| pak2a | 0.74056 | -0.16315856 | 0.057 | 0.111 | 1 | 12 |
| xbp1 | 0.74067 | 0.158835379 | 0.2 | 0.185 | 1 | 12 |
| mphosph8 | 0.74083 | -0.12941756 | 0.086 | 0.148 | 1 | 12 |
| fam168a | 0.74117 | -0.164407 | 0.057 | 0.111 | 1 | 12 |
| shank3b | 0.74147 | -0.17260575 | 0.057 | 0.111 | 1 | 12 |
| palm2 | 0.74153 | 0.110298327 | 0.171 | 0.111 | 1 | 12 |
| slc25a3b | 0.74173 | 0.16773682 | 0.543 | 0.444 | 1 | 12 |
| si:dkey-19b23.8 | 0.74182 | -0.16368935 | 0.057 | 0.111 | 1 | 12 |
| ssr4 | 0.74187 | 0.155512464 | 0.429 | 0.333 | 1 | 12 |
| abce1 | 0.7427 | -0.19319216 | 0.057 | 0.111 | 1 | 12 |
| arpc2 | 0.74279 | 0.118213743 | 0.343 | 0.259 | 1 | 12 |

| | | | | | | |
|---|---|---|---|---|---|---|
| nfyba | 0.74331 | -0.19193901 | 0.086 | 0.148 | 1 | 12 |
| med11 | 0.74348 | -0.17642722 | 0.057 | 0.074 | 1 | 12 |
| mych | 0.74356 | -0.13271256 | 0.057 | 0.111 | 1 | 12 |
| ms4a17a.11 | 0.74448 | 0.211910681 | 0.257 | 0.185 | 1 | 12 |
| baz1b | 0.74462 | -0.12412113 | 0.057 | 0.111 | 1 | 12 |
| sptssa | 0.74486 | -0.15422843 | 0.057 | 0.111 | 1 | 12 |
| slc11a2 | 0.74487 | -0.16691764 | 0.057 | 0.111 | 1 | 12 |
| pgam1a | 0.74514 | -0.14986591 | 0.086 | 0.148 | 1 | 12 |
| psmc6 | 0.74522 | -0.13775247 | 0.057 | 0.111 | 1 | 12 |
| nup62l | 0.74537 | -0.15461169 | 0.057 | 0.111 | 1 | 12 |
| CABZ01115113.1 | 0.74537 | -0.14117436 | 0.057 | 0.111 | 1 | 12 |
| tango2 | 0.74567 | -0.10504831 | 0.057 | 0.111 | 1 | 12 |
| pgrmc2 | 0.74577 | -0.11972443 | 0.057 | 0.111 | 1 | 12 |
| szrd1 | 0.74608 | -0.14555958 | 0.057 | 0.111 | 1 | 12 |
| rab5aa | 0.74608 | -0.14144316 | 0.057 | 0.111 | 1 | 12 |
| g3bp1 | 0.74636 | 0.109842241 | 0.171 | 0.111 | 1 | 12 |
| fkbp5 | 0.74638 | -0.12893187 | 0.057 | 0.111 | 1 | 12 |
| rps19bp1 | 0.74639 | -0.11848705 | 0.057 | 0.111 | 1 | 12 |
| sf3b5 | 0.74659 | -0.13341401 | 0.057 | 0.111 | 1 | 12 |
| scarb2a | 0.74661 | -0.14680465 | 0.086 | 0.148 | 1 | 12 |
| egr1 | 0.74662 | -0.12823916 | 0.057 | 0.111 | 1 | 12 |
| trappc4 | 0.74682 | -0.13837096 | 0.057 | 0.111 | 1 | 12 |
| mrpl17 | 0.74685 | -0.15027207 | 0.057 | 0.111 | 1 | 12 |
| myh9a | 0.74686 | -0.1420464 | 0.057 | 0.111 | 1 | 12 |
| chd4a | 0.74689 | -0.14697188 | 0.057 | 0.111 | 1 | 12 |
| si:ch211-218c6.8 | 0.74689 | -0.17174743 | 0.057 | 0.111 | 1 | 12 |
| purba | 0.74692 | -0.15495846 | 0.057 | 0.111 | 1 | 12 |
| derl2 | 0.74695 | -0.13146537 | 0.057 | 0.111 | 1 | 12 |
| ddx3xb | 0.74702 | -0.16026884 | 0.086 | 0.148 | 1 | 12 |
| flna | 0.74704 | -0.15669793 | 0.086 | 0.148 | 1 | 12 |
| mapk3 | 0.74727 | -0.16051707 | 0.086 | 0.148 | 1 | 12 |
| prmt1 | 0.74736 | -0.13503094 | 0.086 | 0.148 | 1 | 12 |
| atf5b | 0.74826 | -0.11788877 | 0.114 | 0.111 | 1 | 12 |
| calm2a | 0.74901 | -0.19344453 | 0.4 | 0.481 | 1 | 12 |
| emc10 | 0.75054 | 0.155311298 | 0.086 | 0.074 | 1 | 12 |
| pin4 | 0.75094 | 0.211606833 | 0.171 | 0.111 | 1 | 12 |
| si:ch211-204c21.1 | 0.75154 | 0.111382744 | 0.257 | 0.185 | 1 | 12 |
| hmgb1a | 0.75432 | -0.19627247 | 0.257 | 0.333 | 1 | 12 |
| eif4ebp1 | 0.75607 | -0.15393962 | 0.171 | 0.185 | 1 | 12 |
| si:ch211-14k19.8 | 0.75614 | -0.11757766 | 0.114 | 0.111 | 1 | 12 |
| rbm4.2 | 0.75744 | 0.177281247 | 0.114 | 0.074 | 1 | 12 |
| nos1apa | 0.75938 | 0.107784717 | 0.171 | 0.111 | 1 | 12 |
| cdc42l | 0.76015 | 0.132260189 | 0.314 | 0.296 | 1 | 12 |
| snx5 | 0.76068 | 0.194543324 | 0.171 | 0.111 | 1 | 12 |
| ywhaba | 0.7615 | -0.11665431 | 0.286 | 0.296 | 1 | 12 |
| rbm39b | 0.76209 | 0.10519522 | 0.171 | 0.111 | 1 | 12 |
| mt-nd1 | 0.76315 | 0.121460223 | 0.457 | 0.37 | 1 | 12 |
| polr2l | 0.76713 | 0.11258113 | 0.257 | 0.185 | 1 | 12 |

| | | | | | | |
|---|---|---|---|---|---|---|
| tspan14 | 0.76781 | 0.108746399 | 0.171 | 0.111 | 1 | 12 |
| stip1 | 0.76916 | -0.11394171 | 0.114 | 0.111 | 1 | 12 |
| setb | 0.76947 | -0.1714907 | 0.286 | 0.333 | 1 | 12 |
| perp | 0.76948 | 0.19265435 | 0.2 | 0.148 | 1 | 12 |
| ushbp1 | 0.77217 | 0.180135307 | 0.171 | 0.111 | 1 | 12 |
| gmfb | 0.77272 | -0.19953529 | 0.171 | 0.222 | 1 | 12 |
| cct3 | 0.77414 | -0.13861782 | 0.343 | 0.37 | 1 | 12 |
| cct7 | 0.77419 | -0.18909911 | 0.143 | 0.185 | 1 | 12 |
| atp5fa1 | 0.77426 | 0.156165735 | 0.343 | 0.259 | 1 | 12 |
| trnau1apb | 0.77488 | 0.157318352 | 0.171 | 0.148 | 1 | 12 |
| kctd5a | 0.77629 | 0.171984718 | 0.343 | 0.259 | 1 | 12 |
| capza1a | 0.77687 | 0.163460674 | 0.343 | 0.259 | 1 | 12 |
| eef1db | 0.77864 | -0.17372876 | 0.314 | 0.37 | 1 | 12 |
| carhsp1 | 0.77962 | -0.16859703 | 0.286 | 0.37 | 1 | 12 |
| ephb4a | 0.7804 | 0.173895892 | 0.171 | 0.111 | 1 | 12 |
| idh2 | 0.78177 | -0.19079846 | 0.229 | 0.296 | 1 | 12 |
| hmgn6 | 0.78186 | 0.138367182 | 0.571 | 0.519 | 1 | 12 |
| rpl7l1 | 0.78272 | 0.124973512 | 0.114 | 0.111 | 1 | 12 |
| si:ch211-132b12.7 | 0.78342 | 0.1674422 | 0.114 | 0.074 | 1 | 12 |
| mmp14b | 0.78393 | -0.1870589 | 0.257 | 0.333 | 1 | 12 |
| arap3 | 0.78687 | -0.15681561 | 0.057 | 0.074 | 1 | 12 |
| mrpl36 | 0.78733 | -0.18316825 | 0.143 | 0.185 | 1 | 12 |
| nedd8 | 0.78752 | 0.128758206 | 0.171 | 0.111 | 1 | 12 |
| pecam1 | 0.78764 | 0.150288163 | 0.257 | 0.185 | 1 | 12 |
| si:dkey-33c12.4 | 0.78821 | 0.120509448 | 0.171 | 0.111 | 1 | 12 |
| ncl | 0.79034 | -0.13973312 | 0.343 | 0.37 | 1 | 12 |
| h3f3d | 0.79085 | 0.102590349 | 0.6 | 0.519 | 1 | 12 |
| psmb4 | 0.79098 | -0.17994798 | 0.229 | 0.296 | 1 | 12 |
| zfp36l1a | 0.79136 | 0.145951733 | 0.257 | 0.185 | 1 | 12 |
| bud31 | 0.79205 | 0.185749019 | 0.2 | 0.148 | 1 | 12 |
| klf6a | 0.79268 | 0.159729575 | 0.257 | 0.185 | 1 | 12 |
| dnpep | 0.79323 | -0.15499333 | 0.057 | 0.074 | 1 | 12 |
| zgc:193541 | 0.79354 | -0.16781799 | 0.314 | 0.37 | 1 | 12 |
| atp5f1c | 0.79463 | 0.173222787 | 0.257 | 0.185 | 1 | 12 |
| arrb2b | 0.79601 | 0.159613906 | 0.114 | 0.074 | 1 | 12 |
| sdhc | 0.79616 | 0.141946245 | 0.086 | 0.074 | 1 | 12 |
| tek | 0.79629 | 0.146157268 | 0.114 | 0.074 | 1 | 12 |
| tmed7 | 0.79632 | 0.125346979 | 0.171 | 0.111 | 1 | 12 |
| ndufs5 | 0.79665 | 0.143196264 | 0.171 | 0.111 | 1 | 12 |
| tpte | 0.79704 | 0.158122112 | 0.171 | 0.111 | 1 | 12 |
| capns1a | 0.79713 | 0.136544742 | 0.171 | 0.111 | 1 | 12 |
| dusp5 | 0.79773 | 0.164414513 | 0.114 | 0.074 | 1 | 12 |
| aldh2.2 | 0.79836 | 0.166598038 | 0.114 | 0.074 | 1 | 12 |
| sec61b | 0.80196 | 0.171011635 | 0.371 | 0.296 | 1 | 12 |
| rhoca | 0.80439 | -0.1643345 | 0.286 | 0.333 | 1 | 12 |
| ptbp3 | 0.80499 | 0.148154716 | 0.114 | 0.074 | 1 | 12 |
| vaspb | 0.80555 | -0.14836512 | 0.057 | 0.074 | 1 | 12 |
| gskip | 0.80656 | 0.170594119 | 0.143 | 0.111 | 1 | 12 |

| | | | | | | |
|---|---|---|---|---|---|---|
| ostc | 0.80994 | -0.16876412 | 0.229 | 0.296 | 1 | 12 |
| rcc2 | 0.8112 | -0.15617227 | 0.086 | 0.111 | 1 | 12 |
| ubr7 | 0.81138 | -0.17083196 | 0.114 | 0.148 | 1 | 12 |
| zgc:56095 | 0.81322 | -0.10594 | 0.114 | 0.111 | 1 | 12 |
| cs | 0.8162 | -0.16973214 | 0.143 | 0.185 | 1 | 12 |
| atp6v0ca | 0.8168 | -0.16828947 | 0.171 | 0.222 | 1 | 12 |
| gabarapa | 0.81773 | -0.17356443 | 0.171 | 0.222 | 1 | 12 |
| zgc:195173 | 0.81791 | 0.173888174 | 0.371 | 0.296 | 1 | 12 |
| mmgt1 | 0.81791 | -0.16726152 | 0.114 | 0.148 | 1 | 12 |
| gbp | 0.81807 | -0.15798737 | 0.086 | 0.111 | 1 | 12 |
| ywhaz | 0.81853 | 0.161172417 | 0.371 | 0.296 | 1 | 12 |
| sumo2b | 0.81925 | -0.16285024 | 0.314 | 0.37 | 1 | 12 |
| col4a1 | 0.82051 | -0.19492604 | 0.2 | 0.259 | 1 | 12 |
| ankrd11 | 0.82095 | 0.153091344 | 0.143 | 0.111 | 1 | 12 |
| ctsh | 0.82254 | -0.17733555 | 0.171 | 0.222 | 1 | 12 |
| si:ch211-66e2.3 | 0.82425 | 0.145256955 | 0.371 | 0.296 | 1 | 12 |
| krt18b | 0.82514 | 0.103497179 | 0.286 | 0.222 | 1 | 12 |
| sumo3a | 0.82649 | -0.12059855 | 0.229 | 0.296 | 1 | 12 |
| sept7b | 0.82699 | 0.102006002 | 0.286 | 0.222 | 1 | 12 |
| akr1a1b | 0.82723 | 0.110158245 | 0.114 | 0.111 | 1 | 12 |
| cav1 | 0.82844 | -0.18409556 | 0.171 | 0.222 | 1 | 12 |
| skp1 | 0.8287 | -0.12507784 | 0.229 | 0.296 | 1 | 12 |
| ccdc12 | 0.82895 | 0.129896433 | 0.086 | 0.074 | 1 | 12 |
| ppdpfb | 0.8293 | 0.145977522 | 0.543 | 0.481 | 1 | 12 |
| ftr82 | 0.82962 | -0.11296605 | 0.143 | 0.148 | 1 | 12 |
| zgc:55943 | 0.83356 | -0.15138024 | 0.086 | 0.111 | 1 | 12 |
| sesn1 | 0.83464 | -0.15702573 | 0.114 | 0.148 | 1 | 12 |
| fabp3 | 0.83585 | 0.135127916 | 0.114 | 0.074 | 1 | 12 |
| ap2m1a | 0.83645 | 0.157481324 | 0.314 | 0.259 | 1 | 12 |
| iscub | 0.83694 | 0.14279301 | 0.171 | 0.148 | 1 | 12 |
| ndfip1 | 0.83704 | 0.15280286 | 0.2 | 0.148 | 1 | 12 |
| ubqln4 | 0.83842 | 0.150796485 | 0.114 | 0.074 | 1 | 12 |
| cox6b1 | 0.83948 | -0.11273002 | 0.143 | 0.148 | 1 | 12 |
| smarcd1 | 0.83953 | -0.16452308 | 0.114 | 0.148 | 1 | 12 |
| zfand5b | 0.8397 | -0.13800086 | 0.057 | 0.074 | 1 | 12 |
| oaz1a | 0.83972 | -0.15019249 | 0.286 | 0.333 | 1 | 12 |
| zgc:158463 | 0.83987 | 0.148326809 | 0.114 | 0.074 | 1 | 12 |
| gas6 | 0.83996 | 0.15046106 | 0.2 | 0.148 | 1 | 12 |
| anapc16 | 0.84074 | 0.14324811 | 0.114 | 0.074 | 1 | 12 |
| wasf3b | 0.84111 | -0.13126504 | 0.2 | 0.222 | 1 | 12 |
| atp5f1d | 0.84113 | -0.15267674 | 0.2 | 0.259 | 1 | 12 |
| lmo2 | 0.84128 | -0.16367416 | 0.143 | 0.185 | 1 | 12 |
| sntb2 | 0.84141 | -0.1521651 | 0.114 | 0.148 | 1 | 12 |
| lsm3 | 0.8416 | 0.14008172 | 0.114 | 0.074 | 1 | 12 |
| gapdhs | 0.84173 | -0.14126448 | 0.371 | 0.444 | 1 | 12 |
| txnipa | 0.84187 | 0.109697254 | 0.314 | 0.296 | 1 | 12 |
| sos1 | 0.84323 | 0.145909542 | 0.114 | 0.074 | 1 | 12 |
| spcs3 | 0.84327 | -0.1531802 | 0.086 | 0.111 | 1 | 12 |

| | | | | | | |
|---|---|---|---|---|---|---|
| nfia | 0.847 | 0.142887933 | 0.114 | 0.074 | 1 | 12 |
| arhgef1a | 0.84966 | 0.140249587 | 0.114 | 0.074 | 1 | 12 |
| syap1 | 0.84994 | 0.137099679 | 0.114 | 0.074 | 1 | 12 |
| zgc:194246 | 0.85013 | 0.128539051 | 0.114 | 0.074 | 1 | 12 |
| echs1 | 0.85242 | -0.15066756 | 0.114 | 0.148 | 1 | 12 |
| mtss1la | 0.85338 | 0.126235448 | 0.114 | 0.074 | 1 | 12 |
| bace1 | 0.85358 | 0.139136082 | 0.114 | 0.074 | 1 | 12 |
| flt1 | 0.85554 | -0.1360444 | 0.057 | 0.074 | 1 | 12 |
| exoc3l2a | 0.85677 | 0.161864003 | 0.229 | 0.185 | 1 | 12 |
| psmd1 | 0.8568 | -0.15155863 | 0.171 | 0.222 | 1 | 12 |
| s100v2 | 0.85691 | 0.127013425 | 0.114 | 0.074 | 1 | 12 |
| eif3s6ip | 0.8579 | 0.142424581 | 0.2 | 0.148 | 1 | 12 |
| ndufb4 | 0.85892 | -0.1281827 | 0.2 | 0.259 | 1 | 12 |
| tomm5 | 0.8592 | -0.12706281 | 0.2 | 0.259 | 1 | 12 |
| akap8l | 0.85947 | 0.12622208 | 0.114 | 0.074 | 1 | 12 |
| atox1 | 0.86178 | 0.112639363 | 0.2 | 0.185 | 1 | 12 |
| ppp2cb | 0.86186 | 0.108191257 | 0.2 | 0.148 | 1 | 12 |
| si:dkey-188i13.10 | 0.86238 | 0.131582496 | 0.114 | 0.074 | 1 | 12 |
| rnf44 | 0.8626 | 0.114632246 | 0.114 | 0.074 | 1 | 12 |
| cxcl18b | 0.86263 | 0.1185462 | 0.086 | 0.074 | 1 | 12 |
| bsg | 0.86274 | -0.14116962 | 0.343 | 0.407 | 1 | 12 |
| nudc | 0.86278 | -0.15903499 | 0.143 | 0.185 | 1 | 12 |
| ndufc2 | 0.86347 | -0.12650711 | 0.2 | 0.222 | 1 | 12 |
| tm2d3 | 0.86493 | 0.115250031 | 0.086 | 0.074 | 1 | 12 |
| cryl1 | 0.86494 | 0.11517034 | 0.114 | 0.074 | 1 | 12 |
| cox5ab | 0.8651 | -0.14019486 | 0.171 | 0.222 | 1 | 12 |
| mtpn | 0.86516 | -0.14673087 | 0.143 | 0.185 | 1 | 12 |
| snx2 | 0.86539 | 0.117756223 | 0.114 | 0.074 | 1 | 12 |
| smarce1 | 0.86558 | 0.12497246 | 0.2 | 0.148 | 1 | 12 |
| mcfd2 | 0.86672 | 0.120226795 | 0.114 | 0.074 | 1 | 12 |
| gstp1 | 0.86706 | 0.109995203 | 0.114 | 0.074 | 1 | 12 |
| tmem147 | 0.86731 | 0.123522544 | 0.114 | 0.074 | 1 | 12 |
| selenbp1 | 0.86773 | 0.12572196 | 0.2 | 0.148 | 1 | 12 |
| eppk1 | 0.86775 | 0.11438518 | 0.086 | 0.074 | 1 | 12 |
| eif3c | 0.86777 | 0.132558063 | 0.2 | 0.148 | 1 | 12 |
| pak4 | 0.86861 | -0.12929564 | 0.057 | 0.074 | 1 | 12 |
| rac1a | 0.86889 | -0.14979873 | 0.143 | 0.185 | 1 | 12 |
| si:ch211-214j24.10 | 0.8701 | -0.12640298 | 0.057 | 0.074 | 1 | 12 |
| taf13 | 0.87046 | -0.13396928 | 0.229 | 0.259 | 1 | 12 |
| ptmab | 0.87053 | -0.10075791 | 0.657 | 0.704 | 1 | 12 |
| cd151l | 0.87068 | 0.140456718 | 0.143 | 0.111 | 1 | 12 |
| nck1a | 0.87081 | -0.1376885 | 0.086 | 0.111 | 1 | 12 |
| ndufa3 | 0.87141 | 0.145301444 | 0.229 | 0.185 | 1 | 12 |
| si:ch211-114c12.2 | 0.87144 | 0.115087034 | 0.086 | 0.074 | 1 | 12 |
| cbx1a | 0.87177 | 0.129181734 | 0.143 | 0.111 | 1 | 12 |
| atf6 | 0.87303 | -0.12589438 | 0.057 | 0.074 | 1 | 12 |
| tbc1d4 | 0.87304 | -0.13240466 | 0.086 | 0.111 | 1 | 12 |
| HACD4 | 0.87353 | -0.12713556 | 0.057 | 0.074 | 1 | 12 |

| | | | | | | |
|---|---|---|---|---|---|---|
| ost4 | 0.87363 | 0.134887381 | 0.257 | 0.222 | 1 | 12 |
| arfrp1 | 0.87481 | -0.12568366 | 0.057 | 0.074 | 1 | 12 |
| naga | 0.875 | -0.13804317 | 0.086 | 0.111 | 1 | 12 |
| dhdhl | 0.87569 | 0.116807157 | 0.086 | 0.074 | 1 | 12 |
| dlst | 0.87608 | -0.13268328 | 0.086 | 0.111 | 1 | 12 |
| pcbp2 | 0.87686 | 0.139538833 | 0.229 | 0.185 | 1 | 12 |
| lrrc20 | 0.87728 | -0.13162918 | 0.057 | 0.074 | 1 | 12 |
| anxa4 | 0.87839 | -0.10539008 | 0.171 | 0.185 | 1 | 12 |
| ywhah | 0.87866 | -0.11703442 | 0.057 | 0.074 | 1 | 12 |
| eif3g | 0.8797 | -0.13592901 | 0.143 | 0.185 | 1 | 12 |
| hmgb1b | 0.87978 | -0.12710907 | 0.171 | 0.222 | 1 | 12 |
| cd276 | 0.88037 | -0.12433877 | 0.171 | 0.222 | 1 | 12 |
| si:dkey-261h17.1 | 0.88057 | -0.14249398 | 0.171 | 0.222 | 1 | 12 |
| ppt1 | 0.88082 | -0.128217 | 0.114 | 0.148 | 1 | 12 |
| si:ch211-196l7.4 | 0.88086 | 0.137188623 | 0.229 | 0.185 | 1 | 12 |
| mrpl57 | 0.88088 | -0.11819227 | 0.057 | 0.074 | 1 | 12 |
| tra2a | 0.88131 | -0.12560742 | 0.086 | 0.111 | 1 | 12 |
| fkbp2 | 0.8822 | -0.10907342 | 0.171 | 0.222 | 1 | 12 |
| si:ch1073-429i10.3.1 | 0.8824 | -0.12111292 | 0.171 | 0.222 | 1 | 12 |
| crip2 | 0.88369 | -0.12796778 | 0.114 | 0.148 | 1 | 12 |
| vps25 | 0.886 | -0.11751534 | 0.057 | 0.074 | 1 | 12 |
| mgat4a | 0.88704 | -0.12404004 | 0.057 | 0.074 | 1 | 12 |
| ammecr1 | 0.88831 | -0.11348222 | 0.086 | 0.111 | 1 | 12 |
| mef2cb | 0.88937 | -0.12563233 | 0.114 | 0.148 | 1 | 12 |
| rpa1 | 0.89094 | -0.11958109 | 0.057 | 0.074 | 1 | 12 |
| h3f3c | 0.89288 | 0.106960454 | 0.571 | 0.519 | 1 | 12 |
| ednraa | 0.89315 | 0.123444162 | 0.314 | 0.259 | 1 | 12 |
| oip5-as1 | 0.89329 | 0.105760485 | 0.086 | 0.074 | 1 | 12 |
| tmem256 | 0.8934 | -0.11946761 | 0.057 | 0.074 | 1 | 12 |
| dazap2 | 0.89344 | -0.11990867 | 0.057 | 0.074 | 1 | 12 |
| chmp2a | 0.89385 | -0.12827964 | 0.143 | 0.185 | 1 | 12 |
| esama | 0.89394 | -0.12140683 | 0.086 | 0.111 | 1 | 12 |
| calm1a | 0.89471 | -0.13398977 | 0.143 | 0.185 | 1 | 12 |
| psmc1a | 0.89482 | -0.11535891 | 0.086 | 0.111 | 1 | 12 |
| cox7a2a | 0.89568 | -0.12444318 | 0.143 | 0.185 | 1 | 12 |
| cops3 | 0.89594 | 0.106917278 | 0.086 | 0.074 | 1 | 12 |
| copa | 0.8967 | -0.11732026 | 0.057 | 0.074 | 1 | 12 |
| hnrnpd | 0.89712 | -0.11783644 | 0.057 | 0.074 | 1 | 12 |
| smap1 | 0.89873 | -0.12069934 | 0.057 | 0.074 | 1 | 12 |
| tpd52l2b | 0.89941 | -0.11849539 | 0.086 | 0.111 | 1 | 12 |
| zgc:163080 | 0.90053 | -0.11385784 | 0.057 | 0.074 | 1 | 12 |
| rap2b | 0.90183 | -0.10443277 | 0.057 | 0.074 | 1 | 12 |
| gemin8 | 0.90314 | -0.11402486 | 0.057 | 0.074 | 1 | 12 |
| anxa5b | 0.90319 | -0.11820224 | 0.086 | 0.111 | 1 | 12 |
| ddx39aa | 0.90332 | -0.11559038 | 0.143 | 0.185 | 1 | 12 |
| csnk1db | 0.90438 | -0.1108077 | 0.114 | 0.148 | 1 | 12 |
| serinc2 | 0.90443 | -0.1039242 | 0.143 | 0.185 | 1 | 12 |
| podxl | 0.90505 | -0.12419385 | 0.086 | 0.111 | 1 | 12 |

| | | | | | | |
|---|---|---|---|---|---|---|
| h2afva | 0.90531 | 0.114722173 | 0.229 | 0.185 | 1 | 12 |
| si:dkey-28b4.7 | 0.90692 | 0.109406942 | 0.143 | 0.111 | 1 | 12 |
| etfa | 0.90784 | -0.12082582 | 0.086 | 0.111 | 1 | 12 |
| zc3h13 | 0.90942 | -0.1043557 | 0.057 | 0.074 | 1 | 12 |
| wasf2 | 0.90968 | -0.10558774 | 0.057 | 0.074 | 1 | 12 |
| smox | 0.9098 | -0.10314195 | 0.057 | 0.074 | 1 | 12 |
| eif2s1b | 0.90996 | -0.11484202 | 0.114 | 0.148 | 1 | 12 |
| rbm4.3 | 0.91107 | 0.118397797 | 0.229 | 0.185 | 1 | 12 |
| piezo2a.2 | 0.91126 | 0.116293442 | 0.143 | 0.111 | 1 | 12 |
| ptena | 0.91161 | -0.10656724 | 0.086 | 0.111 | 1 | 12 |
| psmc4 | 0.91168 | -0.11177012 | 0.086 | 0.111 | 1 | 12 |
| golph3 | 0.91221 | -0.1079307 | 0.057 | 0.074 | 1 | 12 |
| BX572619.1 | 0.91326 | -0.10611335 | 0.057 | 0.074 | 1 | 12 |
| si:ch211-248e11.2 | 0.91729 | -0.12085085 | 0.114 | 0.148 | 1 | 12 |
| abi2b | 0.91731 | -0.10275619 | 0.057 | 0.074 | 1 | 12 |
| glud1a | 0.9175 | -0.10672341 | 0.114 | 0.148 | 1 | 12 |
| gosr2 | 0.91757 | -0.10768148 | 0.086 | 0.111 | 1 | 12 |
| znf423 | 0.91812 | -0.10938716 | 0.114 | 0.148 | 1 | 12 |
| ube2g2 | 0.92029 | -0.10449847 | 0.086 | 0.111 | 1 | 12 |
| vps35 | 0.92063 | -0.10811281 | 0.086 | 0.111 | 1 | 12 |
| snd1 | 0.92073 | -0.10193814 | 0.086 | 0.111 | 1 | 12 |
| dnajc1 | 0.92244 | -0.10025762 | 0.114 | 0.148 | 1 | 12 |
| si:ch211-156j16.1 | 0.92281 | 0.109447428 | 0.343 | 0.296 | 1 | 12 |
| eef1da | 0.92427 | -0.10032067 | 0.371 | 0.407 | 1 | 12 |
| tcea1 | 0.92787 | 0.103964981 | 0.143 | 0.111 | 1 | 12 |
| ctnnbip1 | 0.92979 | -0.10183147 | 0.057 | 0.074 | 1 | 12 |
| zgc:175088 | 0.93031 | -0.10158698 | 0.057 | 0.074 | 1 | 12 |
| rbm4.1 | 0.93224 | -0.10177648 | 0.057 | 0.074 | 1 | 12 |
| CABZ01030107.1 | 0.93224 | -0.10246853 | 0.086 | 0.111 | 1 | 12 |
| fam32a | 9.151278 | -0.6869272 | 0.118 | 0.8 | 1 | 13 |
| fhl1b | 0.00013 | -0.41296815 | 0 | 0.4 | 1 | 13 |
| mbd3a | 0.00013 | -0.3006286 | 0 | 0.4 | 1 | 13 |
| serinc1 | 0.00013 | -0.43379482 | 0 | 0.4 | 1 | 13 |
| LOC110437920 | 0.00014 | -0.75010477 | 0.216 | 0.9 | 1 | 13 |
| cdipt | 0.00027 | -0.49458402 | 0.176 | 0.8 | 1 | 13 |
| scinlb | 0.00029 | -0.31370757 | 0.451 | 1 | 1 | 13 |
| itm2ba | 0.00041 | -0.41534437 | 0.275 | 0.9 | 1 | 13 |
| zgc:153867 | 0.00051 | -0.18500182 | 0.549 | 1 | 1 | 13 |
| mgst1.2 | 0.00065 | -0.53657744 | 0.412 | 1 | 1 | 13 |
| selenow1 | 0.00075 | -0.3760988 | 0.471 | 1 | 1 | 13 |
| ndufa1 | 0.00079 | -0.76483239 | 0.294 | 0.9 | 1 | 13 |
| si:rp71-68n21.9 | 0.00082 | -0.58380886 | 0.118 | 0.7 | 1 | 13 |
| cap1 | 0.00111 | -0.22863462 | 0.549 | 1 | 1 | 13 |
| spink2.2 | 0.00122 | -0.16280076 | 1 | 1 | 1 | 13 |
| ier2a | 0.00125 | -0.21035524 | 0.275 | 0.8 | 1 | 13 |
| eif4h | 0.00125 | -0.30474048 | 0 | 0.3 | 1 | 13 |
| mpc1 | 0.00125 | -0.24319796 | 0 | 0.3 | 1 | 13 |
| cat | 0.00125 | -0.2983524 | 0 | 0.3 | 1 | 13 |

| | | | | | | |
|---|---|---|---|---|---|---|
| hmox2a | 0.00125 | -0.35370912 | 0 | 0.3 | 1 | 13 |
| kansl1b | 0.00125 | -0.26100791 | 0 | 0.3 | 1 | 13 |
| kdm4b | 0.00125 | -0.25791359 | 0 | 0.3 | 1 | 13 |
| nfatc3b | 0.00125 | -0.27646393 | 0 | 0.3 | 1 | 13 |
| rsf1b.1 | 0.00125 | -0.29093452 | 0 | 0.3 | 1 | 13 |
| acadl | 0.00125 | -0.27899572 | 0 | 0.3 | 1 | 13 |
| borcs7 | 0.00125 | -0.36134452 | 0 | 0.3 | 1 | 13 |
| bpnt1 | 0.00125 | -0.32643501 | 0 | 0.3 | 1 | 13 |
| brms1la | 0.00125 | -0.20037279 | 0 | 0.3 | 1 | 13 |
| rbm33a | 0.00125 | -0.29317751 | 0 | 0.3 | 1 | 13 |
| sars | 0.00125 | -0.22890806 | 0 | 0.3 | 1 | 13 |
| si:dkey-58f10.11 | 0.00125 | -0.3253439 | 0 | 0.3 | 1 | 13 |
| timm44 | 0.00125 | -0.35127041 | 0 | 0.3 | 1 | 13 |
| si:ch211-102c2.4 | 0.00138 | -0.27521011 | 0.392 | 0.9 | 1 | 13 |
| eif4bb | 0.00152 | -0.26223758 | 0.098 | 0.6 | 1 | 13 |
| zgc:113208 | 0.00159 | -0.50032207 | 0.039 | 0.5 | 1 | 13 |
| dynlt3 | 0.00177 | -0.52525074 | 0.137 | 0.7 | 1 | 13 |
| si:dkey-203a12.5 | 0.00178 | 0.362238455 | 0.922 | 1 | 1 | 13 |
| timp4.2 | 0.00183 | -0.70991404 | 0.471 | 1 | 1 | 13 |
| eif5 | 0.00211 | -0.19929175 | 0.196 | 0.7 | 1 | 13 |
| cox5ab | 0.00222 | -0.32981872 | 0.373 | 0.9 | 1 | 13 |
| ndufb9 | 0.00242 | -0.39055751 | 0.157 | 0.7 | 1 | 13 |
| tegt | 0.00245 | -0.34277391 | 0.176 | 0.7 | 1 | 13 |
| zgc:162730 | 0.0025 | -0.17052097 | 0.451 | 0.9 | 1 | 13 |
| btf3l4 | 0.00251 | -0.27783757 | 0.098 | 0.6 | 1 | 13 |
| aldoaa | 0.00251 | -0.46456057 | 0.353 | 0.9 | 1 | 13 |
| glrx | 0.0026 | -0.38959751 | 0.255 | 0.8 | 1 | 13 |
| tomm6 | 0.00303 | -0.44256601 | 0.098 | 0.6 | 1 | 13 |
| ccni | 0.00312 | -0.43256475 | 0.353 | 0.9 | 1 | 13 |
| psmc5 | 0.00345 | -0.47116081 | 0.098 | 0.6 | 1 | 13 |
| psmb4 | 0.00353 | -0.29300372 | 0.294 | 0.8 | 1 | 13 |
| DIPK2B | 0.00364 | -0.44721865 | 0.02 | 0.4 | 1 | 13 |
| hprt1 | 0.00369 | -0.49272669 | 0.02 | 0.4 | 1 | 13 |
| eaf1 | 0.00375 | -0.40502136 | 0.02 | 0.4 | 1 | 13 |
| hmgn3 | 0.0038 | -0.47201146 | 0.353 | 0.9 | 1 | 13 |
| xylt1 | 0.004 | -0.44759522 | 0.02 | 0.4 | 1 | 13 |
| prdx2 | 0.00405 | -0.49432351 | 0.02 | 0.4 | 1 | 13 |
| os9 | 0.00406 | -0.42354315 | 0.176 | 0.7 | 1 | 13 |
| FO203432.1 | 0.00422 | -0.40998034 | 0.843 | 1 | 1 | 13 |
| si:dkey-203a12.4 | 0.00462 | 1.438124694 | 0.549 | 0.1 | 1 | 13 |
| ak2 | 0.00471 | -0.37648076 | 0.392 | 0.9 | 1 | 13 |
| psma2 | 0.00476 | -0.13264595 | 0.137 | 0.6 | 1 | 13 |
| si:ch1073-443f11.2 | 0.00539 | -0.32227282 | 0.412 | 0.9 | 1 | 13 |
| tm2d3 | 0.00577 | -0.3615413 | 0.078 | 0.5 | 1 | 13 |
| tnfaip8l2a | 0.00632 | -0.53842716 | 0.118 | 0.6 | 1 | 13 |
| gnai2a | 0.00683 | -0.43221545 | 0.196 | 0.7 | 1 | 13 |
| STMP1 | 0.00713 | -0.14638674 | 0.137 | 0.6 | 1 | 13 |
| cdaa | 0.0073 | -0.45266795 | 0.392 | 0.9 | 1 | 13 |

| | | | | | | |
|---|---|---|---|---|---|---|
| tmed2 | 0.00746 | -0.16843703 | 0.373 | 0.8 | 1 | 13 |
| tmem39a | 0.00796 | -0.28917226 | 0.078 | 0.5 | 1 | 13 |
| myo18ab | 0.00857 | -0.33836861 | 0.039 | 0.4 | 1 | 13 |
| plp2 | 0.00893 | -0.42329279 | 0.137 | 0.6 | 1 | 13 |
| tob1a | 0.00913 | -0.25762558 | 0.078 | 0.5 | 1 | 13 |
| hectd1 | 0.00919 | -0.29593193 | 0.078 | 0.5 | 1 | 13 |
| tspan14 | 0.00926 | -0.43724354 | 0.216 | 0.7 | 1 | 13 |
| tagln2 | 0.00934 | -0.3480319 | 0.451 | 0.9 | 1 | 13 |
| si:dkey-203a12.2 | 0.00945 | -0.1210046 | 1 | 1 | 1 | 13 |
| cnp | 0.00948 | -0.32301951 | 0.314 | 0.8 | 1 | 13 |
| rasal3 | 0.00967 | -0.42027713 | 0.157 | 0.6 | 1 | 13 |
| pdcd5 | 0.00975 | -0.32031442 | 0.078 | 0.5 | 1 | 13 |
| mbnl3 | 0.01008 | -0.43552013 | 0.078 | 0.5 | 1 | 13 |
| lcp1 | 0.01014 | -0.39982786 | 0.078 | 0.5 | 1 | 13 |
| stxbp2 | 0.01019 | -0.36062865 | 0.137 | 0.6 | 1 | 13 |
| gcnt3 | 0.01042 | -0.49222379 | 0.078 | 0.5 | 1 | 13 |
| rab35b | 0.01074 | -0.27081003 | 0.039 | 0.4 | 1 | 13 |
| srpra | 0.01079 | -0.11876137 | 0.098 | 0.5 | 1 | 13 |
| eno3 | 0.01096 | -0.39355022 | 0.333 | 0.8 | 1 | 13 |
| sri | 0.01114 | -0.45574142 | 0.137 | 0.6 | 1 | 13 |
| CABZ01085658.1 | 0.01115 | -0.15265806 | 0 | 0.2 | 1 | 13 |
| anapc10 | 0.01115 | -0.15019856 | 0 | 0.2 | 1 | 13 |
| ano5b | 0.01115 | -0.24641856 | 0 | 0.2 | 1 | 13 |
| ap3d1 | 0.01115 | -0.21773805 | 0 | 0.2 | 1 | 13 |
| becn1 | 0.01115 | -0.14124241 | 0 | 0.2 | 1 | 13 |
| dgcr2 | 0.01115 | -0.17262394 | 0 | 0.2 | 1 | 13 |
| glsa | 0.01115 | -0.24509201 | 0 | 0.2 | 1 | 13 |
| gpr174 | 0.01115 | -0.2291235 | 0 | 0.2 | 1 | 13 |
| hmg20b | 0.01115 | -0.11970054 | 0 | 0.2 | 1 | 13 |
| il10rb | 0.01115 | -0.22797006 | 0 | 0.2 | 1 | 13 |
| il12rb2l | 0.01115 | -0.26085583 | 0 | 0.2 | 1 | 13 |
| naa15a | 0.01115 | -0.24670182 | 0 | 0.2 | 1 | 13 |
| nudt19 | 0.01115 | -0.23265004 | 0 | 0.2 | 1 | 13 |
| osbpl1a | 0.01115 | -0.24667461 | 0 | 0.2 | 1 | 13 |
| puraa | 0.01115 | -0.26539755 | 0 | 0.2 | 1 | 13 |
| reep1 | 0.01115 | -0.29533469 | 0 | 0.2 | 1 | 13 |
| rfxap | 0.01115 | -0.15133215 | 0 | 0.2 | 1 | 13 |
| rprd1b | 0.01115 | -0.16641398 | 0 | 0.2 | 1 | 13 |
| sec31b | 0.01115 | -0.27066738 | 0 | 0.2 | 1 | 13 |
| si:ch211-171h4.6 | 0.01115 | -0.27194128 | 0 | 0.2 | 1 | 13 |
| si:ch211-194k22.8 | 0.01115 | -0.13481328 | 0 | 0.2 | 1 | 13 |
| si:dkey-7i4.1 | 0.01115 | -0.18718588 | 0 | 0.2 | 1 | 13 |
| si:dkeyp-113d7.1 | 0.01115 | -0.18383675 | 0 | 0.2 | 1 | 13 |
| slc6a7 | 0.01115 | -0.30150236 | 0 | 0.2 | 1 | 13 |
| strn3 | 0.01115 | -0.19606374 | 0 | 0.2 | 1 | 13 |
| tfdp2 | 0.01115 | -0.13437398 | 0 | 0.2 | 1 | 13 |
| tmem106c | 0.01115 | -0.17261303 | 0 | 0.2 | 1 | 13 |
| tpp1 | 0.01115 | -0.18364383 | 0 | 0.2 | 1 | 13 |

| | | | | | | |
|---|---|---|---|---|---|---|
| upf1 | 0.01115 | -0.29321767 | 0 | 0.2 | 1 | 13 |
| wipf2a | 0.01115 | -0.19459606 | 0 | 0.2 | 1 | 13 |
| zgc:112982 | 0.01115 | -0.15253162 | 0 | 0.2 | 1 | 13 |
| zgc:173575 | 0.01115 | -0.19857266 | 0 | 0.2 | 1 | 13 |
| AL772300.1 | 0.01115 | -0.21404767 | 0 | 0.2 | 1 | 13 |
| CABZ01064670.1 | 0.01115 | -0.2630775 | 0 | 0.2 | 1 | 13 |
| XLOC-005199 | 0.01115 | -0.19829309 | 0 | 0.2 | 1 | 13 |
| abhd14a | 0.01115 | -0.2599482 | 0 | 0.2 | 1 | 13 |
| arpc1a | 0.01115 | -0.14635273 | 0 | 0.2 | 1 | 13 |
| atp2b2 | 0.01115 | -0.26676315 | 0 | 0.2 | 1 | 13 |
| ccnd1 | 0.01115 | -0.11834662 | 0 | 0.2 | 1 | 13 |
| cdkn2aipnl | 0.01115 | -0.24114119 | 0 | 0.2 | 1 | 13 |
| csnk1g1 | 0.01115 | -0.23248977 | 0 | 0.2 | 1 | 13 |
| cst3 | 0.01115 | -0.27256945 | 0 | 0.2 | 1 | 13 |
| deptor | 0.01115 | -0.24100979 | 0 | 0.2 | 1 | 13 |
| dhdhl | 0.01115 | -0.11634634 | 0 | 0.2 | 1 | 13 |
| eed | 0.01115 | -0.15059927 | 0 | 0.2 | 1 | 13 |
| fbxw2 | 0.01115 | -0.26612968 | 0 | 0.2 | 1 | 13 |
| fnip1 | 0.01115 | -0.26003862 | 0 | 0.2 | 1 | 13 |
| gna11b | 0.01115 | -0.25957801 | 0 | 0.2 | 1 | 13 |
| hexim1 | 0.01115 | -0.13208346 | 0 | 0.2 | 1 | 13 |
| map7d3 | 0.01115 | -0.21520451 | 0 | 0.2 | 1 | 13 |
| p2ry13 | 0.01115 | -0.28154623 | 0 | 0.2 | 1 | 13 |
| pard6b | 0.01115 | -0.34294269 | 0 | 0.2 | 1 | 13 |
| pde8a | 0.01115 | -0.16013501 | 0 | 0.2 | 1 | 13 |
| prpf38b | 0.01115 | -0.25207636 | 0 | 0.2 | 1 | 13 |
| si:ch211-198a12.6 | 0.01115 | -0.25307178 | 0 | 0.2 | 1 | 13 |
| smfn | 0.01115 | -0.15689796 | 0 | 0.2 | 1 | 13 |
| sp2 | 0.01115 | -0.24939899 | 0 | 0.2 | 1 | 13 |
| sybl1 | 0.01115 | -0.23528182 | 0 | 0.2 | 1 | 13 |
| sypl2a | 0.01115 | -0.11237973 | 0 | 0.2 | 1 | 13 |
| tespa1 | 0.01115 | -0.26443499 | 0 | 0.2 | 1 | 13 |
| trit1 | 0.01115 | -0.24250747 | 0 | 0.2 | 1 | 13 |
| txnl4a | 0.01115 | -0.1405008 | 0 | 0.2 | 1 | 13 |
| usp4 | 0.01115 | -0.19335175 | 0 | 0.2 | 1 | 13 |
| utp11l | 0.01115 | -0.10390481 | 0 | 0.2 | 1 | 13 |
| zc3h11a | 0.01115 | -0.23101485 | 0 | 0.2 | 1 | 13 |
| LOC108191614 | 0.01115 | -0.12065488 | 0 | 0.2 | 1 | 13 |
| XLOC-005905 | 0.01115 | -0.15271487 | 0 | 0.2 | 1 | 13 |
| b3gnt3.4 | 0.01115 | -0.36348219 | 0 | 0.2 | 1 | 13 |
| cab39l | 0.01115 | -0.2765293 | 0 | 0.2 | 1 | 13 |
| chd7 | 0.01115 | -0.17064177 | 0 | 0.2 | 1 | 13 |
| cops2 | 0.01115 | -0.12969013 | 0 | 0.2 | 1 | 13 |
| dtd2 | 0.01115 | -0.20709301 | 0 | 0.2 | 1 | 13 |
| glud1a | 0.01115 | -0.13552359 | 0 | 0.2 | 1 | 13 |
| lclat1 | 0.01115 | -0.20570412 | 0 | 0.2 | 1 | 13 |
| myb | 0.01115 | -0.19523321 | 0 | 0.2 | 1 | 13 |
| nup93 | 0.01115 | -0.19312405 | 0 | 0.2 | 1 | 13 |

| | | | | | | |
|---|---|---|---|---|---|---|
| prpf4 | 0.01115 | -0.1152456 | 0 | 0.2 | 1 | 13 |
| rap2c | 0.01115 | -0.1111996 | 0 | 0.2 | 1 | 13 |
| rchy1 | 0.01115 | -0.10211264 | 0 | 0.2 | 1 | 13 |
| si:ch211-282b22.1 | 0.01115 | -0.17744594 | 0 | 0.2 | 1 | 13 |
| si:zfos-2326c3.2 | 0.01115 | -0.21087776 | 0 | 0.2 | 1 | 13 |
| slc25a25b | 0.01115 | -0.15636101 | 0 | 0.2 | 1 | 13 |
| smim29 | 0.01115 | -0.1770066 | 0 | 0.2 | 1 | 13 |
| snx4 | 0.01115 | -0.19098692 | 0 | 0.2 | 1 | 13 |
| suclg2 | 0.01115 | -0.10453948 | 0 | 0.2 | 1 | 13 |
| uhrf1bp1l | 0.01115 | -0.18792931 | 0 | 0.2 | 1 | 13 |
| unm-sa1261 | 0.01115 | -0.26923211 | 0 | 0.2 | 1 | 13 |
| urod | 0.01115 | -0.15379811 | 0 | 0.2 | 1 | 13 |
| lta4h | 0.01115 | -0.59573804 | 0.294 | 0.8 | 1 | 13 |
| ssr4 | 0.01126 | -0.14695545 | 0.686 | 1 | 1 | 13 |
| wdr1 | 0.01134 | 0.121269862 | 0.549 | 0.8 | 1 | 13 |
| si:ch1073-429i10.3.1 | 0.01146 | -0.31756118 | 0.235 | 0.7 | 1 | 13 |
| mgst3b | 0.01186 | -0.36896424 | 0.314 | 0.8 | 1 | 13 |
| prkchb | 0.01198 | -0.51666578 | 0.039 | 0.4 | 1 | 13 |
| mrps11 | 0.01209 | -0.31883985 | 0.039 | 0.4 | 1 | 13 |
| piezo1 | 0.01209 | -0.45467381 | 0.039 | 0.4 | 1 | 13 |
| rflnb | 0.01225 | -0.46433568 | 0.039 | 0.4 | 1 | 13 |
| ndufa11 | 0.01239 | -0.15897305 | 0.039 | 0.4 | 1 | 13 |
| tmem9b | 0.01252 | -0.50423158 | 0.039 | 0.4 | 1 | 13 |
| pqlc3 | 0.01255 | -0.32559255 | 0.039 | 0.4 | 1 | 13 |
| atp6ap1b | 0.01257 | -0.44391484 | 0.039 | 0.4 | 1 | 13 |
| nop56 | 0.0126 | -0.14298344 | 0.118 | 0.5 | 1 | 13 |
| mtdha | 0.01275 | -0.36496177 | 0.314 | 0.8 | 1 | 13 |
| sat1a.2 | 0.01292 | -0.10028326 | 0.412 | 0.8 | 1 | 13 |
| gng2 | 0.01342 | -0.48799482 | 0.216 | 0.7 | 1 | 13 |
| cst14b.2 | 0.01346 | -0.60089426 | 0.588 | 1 | 1 | 13 |
| si:dkey-118j18.2 | 0.01348 | -0.13681385 | 0.216 | 0.6 | 1 | 13 |
| pdia4 | 0.01354 | -0.33394152 | 0.373 | 0.8 | 1 | 13 |
| ptprr | 0.01358 | -0.34737459 | 0.176 | 0.6 | 1 | 13 |
| atp5pf | 0.01365 | -0.26735698 | 0.451 | 0.9 | 1 | 13 |
| b2m | 0.01393 | -0.11679707 | 0.176 | 0.6 | 1 | 13 |
| tram1 | 0.01443 | -0.28288648 | 0.373 | 0.8 | 1 | 13 |
| arpc2 | 0.01493 | -0.3386137 | 0.608 | 1 | 1 | 13 |
| pik3cd | 0.01494 | -0.31404231 | 0.098 | 0.5 | 1 | 13 |
| chmp2a | 0.01495 | -0.38199863 | 0.098 | 0.5 | 1 | 13 |
| hnrnph1l | 0.01501 | -0.134781 | 0.059 | 0.4 | 1 | 13 |
| bida | 0.01506 | -0.15039916 | 0.02 | 0.3 | 1 | 13 |
| abracl | 0.01592 | -0.42019016 | 0.333 | 0.8 | 1 | 13 |
| rab27a | 0.01601 | -0.38893246 | 0.098 | 0.5 | 1 | 13 |
| ldhba | 0.01616 | -0.50561127 | 0.314 | 0.8 | 1 | 13 |
| si:dkeyp-68b7.12 | 0.01625 | -0.21624927 | 0.196 | 0.6 | 1 | 13 |
| hcls1 | 0.01683 | -0.47651394 | 0.235 | 0.7 | 1 | 13 |
| trappc2 | 0.01688 | -0.24813284 | 0.02 | 0.3 | 1 | 13 |
| hipk3a | 0.01738 | -0.27555554 | 0.02 | 0.3 | 1 | 13 |

| | | | | | | |
|---|---|---|---|---|---|---|
| cox7c | 0.0177 | -0.37249449 | 0.608 | 1 | 1 | 13 |
| ncbp2 | 0.0181 | -0.38544483 | 0.157 | 0.6 | 1 | 13 |
| mt-cyb | 0.0183 | -0.30856218 | 0.353 | 0.7 | 1 | 13 |
| ube2l3a | 0.01895 | -0.46220747 | 0.098 | 0.5 | 1 | 13 |
| gars | 0.01918 | -0.17092686 | 0.02 | 0.3 | 1 | 13 |
| cxcl8b.1 | 0.01923 | 0.652391081 | 0.294 | 0 | 1 | 13 |
| viml | 0.01933 | -0.43884714 | 0.353 | 0.8 | 1 | 13 |
| ran | 0.01968 | 0.155276931 | 0.608 | 0.8 | 1 | 13 |
| ndufa7 | 0.0201 | -0.32156286 | 0.098 | 0.5 | 1 | 13 |
| tomm5 | 0.0201 | -0.63041405 | 0.118 | 0.5 | 1 | 13 |
| si:dkey-112a7.4 | 0.02015 | -0.10194333 | 0.137 | 0.5 | 1 | 13 |
| si:ch211-212k18.7 | 0.02018 | -0.37230553 | 0.098 | 0.5 | 1 | 13 |
| crip1 | 0.02042 | -0.29015665 | 0.647 | 1 | 1 | 13 |
| CT030188.1 | 0.02062 | -0.37925866 | 0.471 | 0.9 | 1 | 13 |
| serp1 | 0.02066 | -0.55007231 | 0.235 | 0.7 | 1 | 13 |
| tpt1 | 0.0207 | -0.35500097 | 0.627 | 1 | 1 | 13 |
| spcs3 | 0.0211 | -0.37144867 | 0.353 | 0.8 | 1 | 13 |
| fubp1 | 0.02214 | -0.23279376 | 0.078 | 0.4 | 1 | 13 |
| cox8a | 0.0223 | -0.31107328 | 0.647 | 1 | 1 | 13 |
| cox6c | 0.02253 | -0.38379257 | 0.353 | 0.8 | 1 | 13 |
| trir | 0.0229 | -0.24345536 | 0.059 | 0.4 | 1 | 13 |
| arpc4 | 0.02292 | -0.21013873 | 0.529 | 0.9 | 1 | 13 |
| hnrnpaba | 0.02348 | 0.118742552 | 0.392 | 0.7 | 1 | 13 |
| cirbpb | 0.02355 | 0.184410227 | 0.686 | 0.9 | 1 | 13 |
| nutf2 | 0.02367 | -0.12067117 | 0.02 | 0.3 | 1 | 13 |
| rhof | 0.02382 | -0.26657034 | 0.118 | 0.5 | 1 | 13 |
| sod1 | 0.02432 | -0.15491931 | 0.314 | 0.7 | 1 | 13 |
| st7 | 0.02451 | -0.23365895 | 0.02 | 0.3 | 1 | 13 |
| calm3b | 0.02462 | 0.485218 | 0.451 | 0.5 | 1 | 13 |
| naa38 | 0.0248 | -0.35418167 | 0.02 | 0.3 | 1 | 13 |
| cfl1l | 0.02508 | 0.478115076 | 0.275 | 0 | 1 | 13 |
| pla2g4aa | 0.02508 | 0.525862868 | 0.275 | 0 | 1 | 13 |
| elovl1b | 0.02508 | 0.490679671 | 0.275 | 0 | 1 | 13 |
| jagn1b | 0.02512 | -0.36514488 | 0.059 | 0.4 | 1 | 13 |
| agpat5 | 0.02512 | -0.35568682 | 0.02 | 0.3 | 1 | 13 |
| hadhb | 0.02543 | -0.28564763 | 0.059 | 0.4 | 1 | 13 |
| mrps16 | 0.02568 | -0.22059898 | 0.02 | 0.3 | 1 | 13 |
| keap1b | 0.02571 | -0.27369037 | 0.02 | 0.3 | 1 | 13 |
| LOC110438722 | 0.02579 | -0.3261268 | 0.49 | 0.9 | 1 | 13 |
| CABZ01026681.1 | 0.02581 | -0.28121171 | 0.02 | 0.3 | 1 | 13 |
| mrpl28 | 0.02589 | -0.28769767 | 0.02 | 0.3 | 1 | 13 |
| fip1l1b | 0.02598 | -0.35574202 | 0.02 | 0.3 | 1 | 13 |
| vaspb | 0.02609 | -0.28464679 | 0.059 | 0.4 | 1 | 13 |
| fermt3b | 0.02631 | -0.33931552 | 0.02 | 0.3 | 1 | 13 |
| si:dkey-10c21.1 | 0.02636 | -0.26895365 | 0.02 | 0.3 | 1 | 13 |
| gata2a | 0.02642 | -0.41422654 | 0.471 | 0.9 | 1 | 13 |
| calm1b | 0.0265 | 0.140963498 | 0.725 | 0.9 | 1 | 13 |
| CABZ01074307.1 | 0.0265 | -0.26932466 | 0.02 | 0.3 | 1 | 13 |

| | | | | | | |
|---|---|---|---|---|---|---|
| napab | 0.02652 | -0.28995903 | 0.059 | 0.4 | 1 | 13 |
| kdm5c | 0.02667 | -0.26909927 | 0.02 | 0.3 | 1 | 13 |
| slc2a3b | 0.02667 | -0.36564323 | 0.118 | 0.5 | 1 | 13 |
| txndc5 | 0.02671 | -0.21234932 | 0.02 | 0.3 | 1 | 13 |
| cpsf6 | 0.02673 | -0.18960876 | 0.02 | 0.3 | 1 | 13 |
| tagapb | 0.02673 | -0.44594791 | 0.176 | 0.6 | 1 | 13 |
| si:ch211-140m22.7 | 0.02673 | -0.14079279 | 0.02 | 0.3 | 1 | 13 |
| tsen34 | 0.02674 | -0.27801069 | 0.02 | 0.3 | 1 | 13 |
| xrcc6 | 0.02675 | -0.24548929 | 0.02 | 0.3 | 1 | 13 |
| hibadhb | 0.02675 | -0.28393119 | 0.02 | 0.3 | 1 | 13 |
| fam91a1 | 0.02675 | -0.2820854 | 0.02 | 0.3 | 1 | 13 |
| si:ch1073-67j19.1 | 0.02697 | -0.97691639 | 0.176 | 0.6 | 1 | 13 |
| cyba | 0.02712 | -0.45298822 | 0.471 | 0.9 | 1 | 13 |
| cox6b2 | 0.02738 | 0.162507799 | 0.471 | 0.7 | 1 | 13 |
| peak3 | 0.02743 | -0.2786598 | 0.078 | 0.4 | 1 | 13 |
| tmed7 | 0.02762 | -0.30933324 | 0.392 | 0.8 | 1 | 13 |
| nme2b.1 | 0.02797 | 0.183769123 | 0.784 | 0.9 | 1 | 13 |
| mt-nd1 | 0.02805 | -0.40830307 | 0.294 | 0.7 | 1 | 13 |
| calm3a | 0.02824 | -0.13416654 | 0.137 | 0.5 | 1 | 13 |
| kdelr3 | 0.02838 | -0.44337863 | 0.059 | 0.4 | 1 | 13 |
| cdv3 | 0.02844 | -0.32791201 | 0.059 | 0.4 | 1 | 13 |
| mpdu1b | 0.02847 | -0.38635978 | 0.059 | 0.4 | 1 | 13 |
| BOLA2 | 0.02859 | -0.28834719 | 0.059 | 0.4 | 1 | 13 |
| glo1 | 0.02901 | -0.11540084 | 0.333 | 0.7 | 1 | 13 |
| cfhl1 | 0.02909 | -0.24271003 | 0.137 | 0.5 | 1 | 13 |
| fybb | 0.02909 | -0.3248623 | 0.275 | 0.7 | 1 | 13 |
| CABZ01072950.1 | 0.02931 | -0.25073113 | 0.216 | 0.6 | 1 | 13 |
| hmgn7 | 0.02953 | -0.47667332 | 0.353 | 0.8 | 1 | 13 |
| spcs1 | 0.02957 | -0.36560446 | 0.49 | 0.9 | 1 | 13 |
| kctd5a | 0.02958 | -0.12046981 | 0.098 | 0.4 | 1 | 13 |
| lgals3b | 0.02976 | -0.10175269 | 0.137 | 0.5 | 1 | 13 |
| cmpk | 0.03024 | -0.29261022 | 0.196 | 0.6 | 1 | 13 |
| spcs2 | 0.03037 | -0.1902483 | 0.529 | 0.9 | 1 | 13 |
| adgre13 | 0.03053 | -0.37336639 | 0.275 | 0.7 | 1 | 13 |
| uqcrfs1 | 0.03114 | -0.25125338 | 0.118 | 0.5 | 1 | 13 |
| ewsr1b | 0.03146 | -0.25692507 | 0.137 | 0.5 | 1 | 13 |
| si:dkey-262k9.4 | 0.03226 | -0.27612152 | 0.216 | 0.6 | 1 | 13 |
| si:ch211-210b2.4 | 0.03256 | 0.560771035 | 0.255 | 0 | 1 | 13 |
| pycard | 0.03349 | -0.23674028 | 0.431 | 0.8 | 1 | 13 |
| mt-nd2 | 0.03493 | -0.21931938 | 0.275 | 0.6 | 1 | 13 |
| mid1ip1l | 0.03501 | -0.2788377 | 0.078 | 0.4 | 1 | 13 |
| rab1ab | 0.03515 | -0.29220411 | 0.294 | 0.7 | 1 | 13 |
| capzb | 0.03535 | -0.11268063 | 0.588 | 0.9 | 1 | 13 |
| rgs3a | 0.03553 | -0.22641801 | 0.078 | 0.4 | 1 | 13 |
| arpc5b | 0.03718 | -0.31588831 | 0.294 | 0.7 | 1 | 13 |
| clstn1 | 0.03743 | -0.33505258 | 0.216 | 0.6 | 1 | 13 |
| pnrc2 | 0.03858 | 0.243088199 | 0.686 | 0.8 | 1 | 13 |
| pdap1b | 0.03947 | -0.14924991 | 0.078 | 0.4 | 1 | 13 |

| | | | | | | |
|---|---|---|---|---|---|---|
| elmo1 | 0.04124 | -0.28254366 | 0.216 | 0.6 | 1 | 13 |
| wasb | 0.04197 | -0.39847611 | 0.392 | 0.8 | 1 | 13 |
| pfdn4 | 0.04207 | -0.18535688 | 0.137 | 0.5 | 1 | 13 |
| CT573383.1 | 0.0421 | 0.436998367 | 0.235 | 0 | 1 | 13 |
| dpy30 | 0.0421 | 0.386166527 | 0.235 | 0 | 1 | 13 |
| si:ch211-270n8.1 | 0.0421 | 0.459337868 | 0.235 | 0 | 1 | 13 |
| bud31 | 0.0421 | 0.421238869 | 0.235 | 0 | 1 | 13 |
| sdf2l1 | 0.04221 | -0.25339388 | 0.235 | 0.6 | 1 | 13 |
| sec11a | 0.0423 | -0.1748132 | 0.569 | 0.9 | 1 | 13 |
| aldh6a1 | 0.04243 | -0.15473915 | 0.137 | 0.5 | 1 | 13 |
| CABZ01058647.1 | 0.04251 | -0.68995083 | 0.216 | 0.6 | 1 | 13 |
| psmb6 | 0.04265 | -0.21281473 | 0.216 | 0.6 | 1 | 13 |
| ufc1 | 0.0443 | -0.15946879 | 0.098 | 0.4 | 1 | 13 |
| denr | 0.04431 | -0.2447705 | 0.137 | 0.5 | 1 | 13 |
| eif3ha | 0.04442 | -0.1246417 | 0.235 | 0.6 | 1 | 13 |
| cltca | 0.04451 | -0.29597602 | 0.216 | 0.6 | 1 | 13 |
| rhoac | 0.04469 | -0.29668846 | 0.216 | 0.6 | 1 | 13 |
| LOC101883382 | 0.0451 | -0.39914453 | 0.51 | 0.9 | 1 | 13 |
| triobpa | 0.04543 | -0.16883911 | 0.098 | 0.4 | 1 | 13 |
| mir223 | 0.04584 | -0.40935354 | 0.392 | 0.8 | 1 | 13 |
| arf2a | 0.04629 | -0.25969019 | 0.078 | 0.4 | 1 | 13 |
| srprb | 0.04649 | -0.10084461 | 0.255 | 0.6 | 1 | 13 |
| spink2.5 | 0.04662 | -0.39469361 | 0.765 | 0.8 | 1 | 13 |
| psma1 | 0.04712 | -0.17486223 | 0.333 | 0.7 | 1 | 13 |
| rpl6 | 0.04715 | -0.44406468 | 0.706 | 1 | 1 | 13 |
| abi3a | 0.04717 | -0.32655546 | 0.216 | 0.6 | 1 | 13 |
| arfgap2 | 0.04723 | -0.25958304 | 0.078 | 0.4 | 1 | 13 |
| tmem258 | 0.04798 | -0.19166721 | 0.569 | 0.9 | 1 | 13 |
| ca6 | 0.04807 | 0.533016005 | 0.451 | 0.1 | 1 | 13 |
| cited4a | 0.04822 | -0.23039625 | 0.02 | 0.2 | 1 | 13 |
| mir21-1 | 0.04822 | -0.27887122 | 0.02 | 0.2 | 1 | 13 |
| mntb | 0.04822 | -0.20928454 | 0.02 | 0.2 | 1 | 13 |
| pde7a | 0.04822 | -0.17959988 | 0.02 | 0.2 | 1 | 13 |
| si:ch73-71c20.5 | 0.04822 | -0.14240722 | 0.02 | 0.2 | 1 | 13 |
| si:dkey-253i9.4 | 0.04822 | -0.20388269 | 0.02 | 0.2 | 1 | 13 |
| rab33ba | 0.04822 | -0.17992859 | 0.02 | 0.2 | 1 | 13 |
| CMKLR1 (1 of many) | 0.04822 | -0.29214749 | 0.02 | 0.2 | 1 | 13 |
| XLOC-014336 | 0.04822 | -0.12529814 | 0.02 | 0.2 | 1 | 13 |
| abcf2a | 0.04822 | -0.20561899 | 0.02 | 0.2 | 1 | 13 |
| ap3m2 | 0.04822 | -0.14085224 | 0.02 | 0.2 | 1 | 13 |
| bag1 | 0.04822 | -0.14165039 | 0.02 | 0.2 | 1 | 13 |
| brwd1 | 0.04822 | -0.22530235 | 0.02 | 0.2 | 1 | 13 |
| bxdc2 | 0.04822 | -0.27759463 | 0.02 | 0.2 | 1 | 13 |
| cdc5l | 0.04822 | -0.21591781 | 0.02 | 0.2 | 1 | 13 |
| cep135 | 0.04822 | -0.22934303 | 0.02 | 0.2 | 1 | 13 |
| cept1b | 0.04822 | -0.31948442 | 0.02 | 0.2 | 1 | 13 |
| coa1 | 0.04822 | -0.14447446 | 0.02 | 0.2 | 1 | 13 |
| cog2 | 0.04822 | -0.10773625 | 0.02 | 0.2 | 1 | 13 |

| | | | | | | |
|---|---|---|---|---|---|---|
| cog4 | 0.04822 | -0.1622915 | 0.02 | 0.2 | 1 | 13 |
| ctcf | 0.04822 | -0.14863416 | 0.02 | 0.2 | 1 | 13 |
| ctnnb2 | 0.04822 | -0.12086705 | 0.02 | 0.2 | 1 | 13 |
| cwc25 | 0.04822 | -0.14429912 | 0.02 | 0.2 | 1 | 13 |
| dok1b | 0.04822 | -0.29216073 | 0.02 | 0.2 | 1 | 13 |
| fam84b | 0.04822 | -0.17337936 | 0.02 | 0.2 | 1 | 13 |
| glrx3 | 0.04822 | -0.27692127 | 0.02 | 0.2 | 1 | 13 |
| grk6 | 0.04822 | -0.17421702 | 0.02 | 0.2 | 1 | 13 |
| hsbp1a | 0.04822 | -0.10329618 | 0.02 | 0.2 | 1 | 13 |
| lrrc47 | 0.04822 | -0.22067935 | 0.02 | 0.2 | 1 | 13 |
| luc7l3 | 0.04822 | -0.28217806 | 0.02 | 0.2 | 1 | 13 |
| mrpl47 | 0.04822 | -0.16516431 | 0.02 | 0.2 | 1 | 13 |
| mrps31 | 0.04822 | -0.19641266 | 0.02 | 0.2 | 1 | 13 |
| mtch2 | 0.04822 | -0.18061816 | 0.02 | 0.2 | 1 | 13 |
| pdcd6 | 0.04822 | -0.28114214 | 0.02 | 0.2 | 1 | 13 |
| pgm2 | 0.04822 | -0.23251122 | 0.02 | 0.2 | 1 | 13 |
| phf20l1 | 0.04822 | -0.24133034 | 0.02 | 0.2 | 1 | 13 |
| pigp | 0.04822 | -0.16216658 | 0.02 | 0.2 | 1 | 13 |
| pmaip1 | 0.04822 | -0.24886639 | 0.02 | 0.2 | 1 | 13 |
| ppig | 0.04822 | -0.37038642 | 0.02 | 0.2 | 1 | 13 |
| ppip5k1b | 0.04822 | -0.13653388 | 0.02 | 0.2 | 1 | 13 |
| prkdc | 0.04822 | -0.15528829 | 0.02 | 0.2 | 1 | 13 |
| psmd11b | 0.04822 | -0.1477168 | 0.02 | 0.2 | 1 | 13 |
| rprd2b | 0.04822 | -0.16602901 | 0.02 | 0.2 | 1 | 13 |
| rps6ka3a | 0.04822 | -0.23544179 | 0.02 | 0.2 | 1 | 13 |
| si:ch211-15b10.6 | 0.04822 | -0.25863036 | 0.02 | 0.2 | 1 | 13 |
| si:ch73-170d6.3 | 0.04822 | -0.12394424 | 0.02 | 0.2 | 1 | 13 |
| si:ch73-368j24.10.1 | 0.04822 | -0.30344758 | 0.02 | 0.2 | 1 | 13 |
| si:dkey-261j15.2 | 0.04822 | -0.28268024 | 0.02 | 0.2 | 1 | 13 |
| si:dkey-5n18.1 | 0.04822 | -0.26130951 | 0.02 | 0.2 | 1 | 13 |
| slc9a8 | 0.04822 | -0.2645574 | 0.02 | 0.2 | 1 | 13 |
| sp4 | 0.04822 | -0.11657604 | 0.02 | 0.2 | 1 | 13 |
| srpk1a | 0.04822 | -0.26184441 | 0.02 | 0.2 | 1 | 13 |
| thtpa | 0.04822 | -0.20924261 | 0.02 | 0.2 | 1 | 13 |
| tmbim1a | 0.04822 | -0.23355729 | 0.02 | 0.2 | 1 | 13 |
| trappc11 | 0.04822 | -0.18226292 | 0.02 | 0.2 | 1 | 13 |
| vps13a | 0.04822 | -0.20753855 | 0.02 | 0.2 | 1 | 13 |
| yars | 0.04822 | -0.18397577 | 0.02 | 0.2 | 1 | 13 |
| zc3h12b | 0.04822 | -0.1462926 | 0.02 | 0.2 | 1 | 13 |
| zcchc17 | 0.04822 | -0.20992619 | 0.02 | 0.2 | 1 | 13 |
| BX649485.2 | 0.04822 | -0.10406281 | 0.02 | 0.2 | 1 | 13 |
| CU571315.1 | 0.04822 | -0.19305234 | 0.02 | 0.2 | 1 | 13 |
| CU681842.1 | 0.04822 | -0.17167086 | 0.02 | 0.2 | 1 | 13 |
| LOC100002469 | 0.04822 | -0.25994136 | 0.02 | 0.2 | 1 | 13 |
| MAP3K11 | 0.04822 | -0.2060582 | 0.02 | 0.2 | 1 | 13 |
| akap17a | 0.04822 | -0.11481979 | 0.02 | 0.2 | 1 | 13 |
| arhgef18b | 0.04822 | -0.15556722 | 0.02 | 0.2 | 1 | 13 |
| atp6v0d1 | 0.04822 | -0.27520854 | 0.02 | 0.2 | 1 | 13 |

| | | | | | | |
|---|---|---|---|---|---|---|
| bckdk | 0.04822 | -0.11787079 | 0.02 | 0.2 | 1 | 13 |
| bnip2 | 0.04822 | -0.23145732 | 0.02 | 0.2 | 1 | 13 |
| capn2b | 0.04822 | -0.13107979 | 0.02 | 0.2 | 1 | 13 |
| cds1 | 0.04822 | -0.16390105 | 0.02 | 0.2 | 1 | 13 |
| cndp2 | 0.04822 | -0.23130371 | 0.02 | 0.2 | 1 | 13 |
| crnkl1 | 0.04822 | -0.16206621 | 0.02 | 0.2 | 1 | 13 |
| cyb561a3a | 0.04822 | -0.21662968 | 0.02 | 0.2 | 1 | 13 |
| cyfip1 | 0.04822 | -0.21549966 | 0.02 | 0.2 | 1 | 13 |
| dcun1d5 | 0.04822 | -0.11855487 | 0.02 | 0.2 | 1 | 13 |
| dido1 | 0.04822 | -0.2384054 | 0.02 | 0.2 | 1 | 13 |
| dnase1l1l | 0.04822 | -0.18103865 | 0.02 | 0.2 | 1 | 13 |
| eif2b4 | 0.04822 | -0.10543457 | 0.02 | 0.2 | 1 | 13 |
| gpt2 | 0.04822 | -0.22831865 | 0.02 | 0.2 | 1 | 13 |
| grwd1 | 0.04822 | -0.19239391 | 0.02 | 0.2 | 1 | 13 |
| mroh1 | 0.04822 | -0.17809972 | 0.02 | 0.2 | 1 | 13 |
| nap1l4b | 0.04822 | -0.11517739 | 0.02 | 0.2 | 1 | 13 |
| ndufa4 | 0.04822 | -0.26892152 | 0.02 | 0.2 | 1 | 13 |
| nrd1b | 0.04822 | -0.15891418 | 0.02 | 0.2 | 1 | 13 |
| pes | 0.04822 | -0.10704885 | 0.02 | 0.2 | 1 | 13 |
| pet100 | 0.04822 | -0.12378176 | 0.02 | 0.2 | 1 | 13 |
| plaa | 0.04822 | -0.11144332 | 0.02 | 0.2 | 1 | 13 |
| ppp2r2d | 0.04822 | -0.12107064 | 0.02 | 0.2 | 1 | 13 |
| ppp2r3c | 0.04822 | -0.12297161 | 0.02 | 0.2 | 1 | 13 |
| rab39bb | 0.04822 | -0.15755372 | 0.02 | 0.2 | 1 | 13 |
| rhebl1 | 0.04822 | -0.27475313 | 0.02 | 0.2 | 1 | 13 |
| rtn4ip1 | 0.04822 | -0.15898321 | 0.02 | 0.2 | 1 | 13 |
| sh3bp2 | 0.04822 | -0.25414986 | 0.02 | 0.2 | 1 | 13 |
| si:ch211-132p1.3 | 0.04822 | -0.16097316 | 0.02 | 0.2 | 1 | 13 |
| si:dkey-165a24.10 | 0.04822 | -0.17682428 | 0.02 | 0.2 | 1 | 13 |
| si:dkey-42i9.4 | 0.04822 | -0.1526171 | 0.02 | 0.2 | 1 | 13 |
| sirt1 | 0.04822 | -0.18988595 | 0.02 | 0.2 | 1 | 13 |
| stimate | 0.04822 | -0.20596043 | 0.02 | 0.2 | 1 | 13 |
| sypl2b | 0.04822 | -0.21749453 | 0.02 | 0.2 | 1 | 13 |
| tradd | 0.04822 | -0.20983089 | 0.02 | 0.2 | 1 | 13 |
| ubr7 | 0.04822 | -0.21547348 | 0.02 | 0.2 | 1 | 13 |
| uqcc3 | 0.04822 | -0.1819259 | 0.02 | 0.2 | 1 | 13 |
| vps33b | 0.04822 | -0.16926893 | 0.02 | 0.2 | 1 | 13 |
| zc3h13 | 0.04822 | -0.12605874 | 0.02 | 0.2 | 1 | 13 |
| zgc:174268 | 0.04822 | -0.12653374 | 0.02 | 0.2 | 1 | 13 |
| zgc:55558 | 0.04822 | -0.12374203 | 0.02 | 0.2 | 1 | 13 |
| zgc:56525 | 0.04822 | -0.20438526 | 0.02 | 0.2 | 1 | 13 |
| atp5mf | 0.04853 | -0.36467294 | 0.294 | 0.7 | 1 | 13 |
| hnrpkl | 0.04864 | -0.22809517 | 0.078 | 0.4 | 1 | 13 |
| naa10 | 0.04875 | -0.21108793 | 0.137 | 0.5 | 1 | 13 |
| ppdpfb | 0.04951 | -0.56112281 | 0.569 | 0.9 | 1 | 13 |
| qsox1 | 0.05048 | -0.17826096 | 0.353 | 0.7 | 1 | 13 |
| sfxn3 | 0.05082 | -0.1676419 | 0.039 | 0.3 | 1 | 13 |
| rsl1d1 | 0.05082 | -0.21481042 | 0.078 | 0.4 | 1 | 13 |

| | | | | | | |
|---|---|---|---|---|---|---|
| alox5ap | 0.05084 | -0.37935346 | 0.216 | 0.6 | 1 | 13 |
| cnbpb | 0.05087 | -0.20833187 | 0.078 | 0.4 | 1 | 13 |
| arglu1a | 0.05161 | -0.17617115 | 0.098 | 0.4 | 1 | 13 |
| myca | 0.05167 | -0.13605069 | 0.059 | 0.3 | 1 | 13 |
| tmed10 | 0.05169 | 0.297555218 | 0.471 | 0.6 | 1 | 13 |
| zgc:193541 | 0.05169 | -0.32460402 | 0.314 | 0.7 | 1 | 13 |
| eef1db | 0.05185 | -0.21645864 | 0.216 | 0.6 | 1 | 13 |
| itm2cb | 0.05186 | -0.38016654 | 0.078 | 0.4 | 1 | 13 |
| tomm7 | 0.05202 | -0.29387207 | 0.137 | 0.5 | 1 | 13 |
| uba1 | 0.05203 | -0.15810387 | 0.176 | 0.5 | 1 | 13 |
| ube2v2 | 0.05214 | 0.112984558 | 0.333 | 0.6 | 1 | 13 |
| lsm4 | 0.05245 | -0.29405246 | 0.216 | 0.6 | 1 | 13 |
| etnk1 | 0.05249 | -0.37716608 | 0.078 | 0.4 | 1 | 13 |
| nfkbie | 0.05283 | -0.36908592 | 0.078 | 0.4 | 1 | 13 |
| otub1a | 0.05286 | -0.11673422 | 0.098 | 0.4 | 1 | 13 |
| id3 | 0.05305 | -0.28801458 | 0.078 | 0.4 | 1 | 13 |
| usp5 | 0.05315 | -0.31617518 | 0.078 | 0.4 | 1 | 13 |
| slc20a1b | 0.05329 | -0.24032597 | 0.039 | 0.3 | 1 | 13 |
| rgcc | 0.05335 | -0.17228144 | 0.451 | 0.8 | 1 | 13 |
| rpn1 | 0.05343 | -0.34044992 | 0.412 | 0.8 | 1 | 13 |
| echdc2 | 0.05344 | -0.24825023 | 0.078 | 0.4 | 1 | 13 |
| tuba8l4 | 0.05349 | -0.10829474 | 0.373 | 0.7 | 1 | 13 |
| rspry1 | 0.05426 | 0.282288516 | 0.216 | 0 | 1 | 13 |
| psma3 | 0.05511 | -0.26700606 | 0.235 | 0.6 | 1 | 13 |
| cox7b | 0.05514 | -0.44698143 | 0.392 | 0.8 | 1 | 13 |
| mgst3a | 0.05598 | -0.13890335 | 0.118 | 0.4 | 1 | 13 |
| khdrbs1a | 0.05712 | 0.194833663 | 0.51 | 0.7 | 1 | 13 |
| lgmn | 0.05732 | -0.23734748 | 0.039 | 0.3 | 1 | 13 |
| XLOC-008119 | 0.05743 | -0.30588117 | 0.039 | 0.3 | 1 | 13 |
| mrps7 | 0.05764 | -0.28469287 | 0.039 | 0.3 | 1 | 13 |
| yipf4 | 0.05783 | -0.13925559 | 0.098 | 0.4 | 1 | 13 |
| CABZ01111959.1 | 0.05828 | -0.12232977 | 0.059 | 0.3 | 1 | 13 |
| si:ch211-276a23.5 | 0.05829 | -0.27686362 | 0.333 | 0.7 | 1 | 13 |
| pigrl2.3 | 0.05854 | -0.296089 | 0.157 | 0.5 | 1 | 13 |
| mrpl11 | 0.05978 | -0.24196811 | 0.039 | 0.3 | 1 | 13 |
| rap1b | 0.06067 | -0.28131034 | 0.333 | 0.7 | 1 | 13 |
| scgn | 0.0609 | -0.28367349 | 0.039 | 0.3 | 1 | 13 |
| ost4 | 0.06094 | -0.28971534 | 0.549 | 0.9 | 1 | 13 |
| gadd45ga | 0.06188 | -0.24374826 | 0.039 | 0.3 | 1 | 13 |
| LOC103911430 | 0.06223 | -0.24055195 | 0.176 | 0.5 | 1 | 13 |
| LOC100536213 | 0.06229 | -0.33741017 | 0.314 | 0.7 | 1 | 13 |
| psmb10 | 0.06233 | -0.20361097 | 0.039 | 0.3 | 1 | 13 |
| bloc1s2 | 0.06283 | -0.40700114 | 0.039 | 0.3 | 1 | 13 |
| si:ch211-196l7.4 | 0.06291 | -0.10947054 | 0.275 | 0.6 | 1 | 13 |
| XLOC-035413 | 0.06291 | -0.27148533 | 0.039 | 0.3 | 1 | 13 |
| copb2 | 0.0631 | -0.14945478 | 0.098 | 0.4 | 1 | 13 |
| cdk5rap3 | 0.06317 | -0.21794832 | 0.039 | 0.3 | 1 | 13 |
| nasp | 0.06324 | -0.26454236 | 0.039 | 0.3 | 1 | 13 |

| | | | | | | |
|---|---|---|---|---|---|---|
| nfil3-6 | 0.06332 | -0.1932728 | 0.255 | 0.6 | 1 | 13 |
| grap2b | 0.06337 | -0.15151044 | 0.627 | 0.9 | 1 | 13 |
| vps28 | 0.06358 | -0.2627433 | 0.039 | 0.3 | 1 | 13 |
| cdkn1bb | 0.06365 | -0.23641821 | 0.157 | 0.5 | 1 | 13 |
| galk1 | 0.06372 | -0.21640018 | 0.039 | 0.3 | 1 | 13 |
| ostf1 | 0.06396 | -0.30671575 | 0.235 | 0.6 | 1 | 13 |
| npm1b | 0.06427 | -0.18641324 | 0.039 | 0.3 | 1 | 13 |
| zgc:174863 | 0.06427 | -0.23427569 | 0.039 | 0.3 | 1 | 13 |
| khsrp | 0.06476 | -0.26960087 | 0.039 | 0.3 | 1 | 13 |
| csnk1da | 0.06505 | -0.26856775 | 0.039 | 0.3 | 1 | 13 |
| pa2g4a | 0.06509 | -0.26961393 | 0.039 | 0.3 | 1 | 13 |
| pdlim5b | 0.06523 | -0.23184952 | 0.039 | 0.3 | 1 | 13 |
| ncln | 0.06523 | -0.16966986 | 0.039 | 0.3 | 1 | 13 |
| ywhaz | 0.06526 | 0.123315176 | 0.412 | 0.6 | 1 | 13 |
| atg12 | 0.06548 | -0.22509183 | 0.039 | 0.3 | 1 | 13 |
| cd81a | 0.06572 | -0.14617047 | 0.765 | 1 | 1 | 13 |
| rhbdf1a | 0.06599 | -0.29502179 | 0.039 | 0.3 | 1 | 13 |
| ino80c | 0.06616 | -0.2252821 | 0.039 | 0.3 | 1 | 13 |
| rps27.2 | 0.06625 | -0.19341642 | 0.569 | 0.9 | 1 | 13 |
| slc10a7 | 0.06627 | -0.29555818 | 0.039 | 0.3 | 1 | 13 |
| ociad1 | 0.06631 | -0.14830636 | 0.039 | 0.3 | 1 | 13 |
| snrpa | 0.06652 | -0.17068893 | 0.039 | 0.3 | 1 | 13 |
| naxd | 0.06661 | -0.21352653 | 0.039 | 0.3 | 1 | 13 |
| UBL4A | 0.06663 | -0.22024752 | 0.039 | 0.3 | 1 | 13 |
| nucks1b | 0.06684 | -0.17310601 | 0.039 | 0.3 | 1 | 13 |
| ada2a | 0.06699 | -0.26153894 | 0.039 | 0.3 | 1 | 13 |
| si:ch211-215e19.3 | 0.06713 | -0.39043885 | 0.039 | 0.3 | 1 | 13 |
| rpl7l1 | 0.0672 | -0.22675989 | 0.039 | 0.3 | 1 | 13 |
| sf3a1 | 0.06726 | -0.20079541 | 0.039 | 0.3 | 1 | 13 |
| oaz1b | 0.0673 | -0.21488076 | 0.569 | 0.9 | 1 | 13 |
| tmem39b | 0.06741 | -0.31578622 | 0.039 | 0.3 | 1 | 13 |
| tmem11 | 0.06742 | -0.18300915 | 0.039 | 0.3 | 1 | 13 |
| sptan1 | 0.06744 | -0.28301648 | 0.039 | 0.3 | 1 | 13 |
| si:dkey-180p18.9 | 0.06796 | -0.19274046 | 0.098 | 0.4 | 1 | 13 |
| vps26c | 0.068 | -0.15784019 | 0.118 | 0.4 | 1 | 13 |
| ier2b | 0.06875 | -0.16636607 | 0.471 | 0.8 | 1 | 13 |
| cops4 | 0.06971 | 0.330042963 | 0.196 | 0 | 1 | 13 |
| si:dkey-73n10.1 | 0.06971 | 0.40666445 | 0.196 | 0 | 1 | 13 |
| mycbp | 0.06971 | 0.371058513 | 0.196 | 0 | 1 | 13 |
| svila | 0.06971 | 0.253220368 | 0.196 | 0 | 1 | 13 |
| pho | 0.07042 | -0.27810112 | 0.098 | 0.4 | 1 | 13 |
| mibp2 | 0.07058 | -0.10020609 | 0.471 | 0.8 | 1 | 13 |
| tpst2 | 0.07097 | -0.27455145 | 0.098 | 0.4 | 1 | 13 |
| ywhaba | 0.07143 | -0.24597155 | 0.451 | 0.8 | 1 | 13 |
| CABZ01058650.1 | 0.07162 | -0.24812528 | 0.098 | 0.4 | 1 | 13 |
| erh | 0.0719 | -0.28006694 | 0.333 | 0.7 | 1 | 13 |
| spink2.3 | 0.07227 | -0.15921432 | 1 | 1 | 1 | 13 |
| si:dkey-102g19.3 | 0.07275 | 0.33442248 | 0.667 | 0.7 | 1 | 13 |

| | | | | | | |
|---|---|---|---|---|---|---|
| spink2.4 | 0.07276 | -0.10728152 | 0.882 | 0.8 | 1 | 13 |
| snrpd2 | 0.07393 | -0.25646814 | 0.157 | 0.5 | 1 | 13 |
| BX284696.1 | 0.07414 | -0.18053742 | 0.059 | 0.3 | 1 | 13 |
| ola1 | 0.07517 | 0.459582437 | 0.373 | 0.4 | 1 | 13 |
| psme2 | 0.07651 | -0.11962251 | 0.196 | 0.5 | 1 | 13 |
| btk | 0.07654 | -0.32223128 | 0.235 | 0.6 | 1 | 13 |
| syf2 | 0.07798 | -0.19201289 | 0.098 | 0.4 | 1 | 13 |
| sult5a1 | 0.07817 | 0.275542981 | 0.392 | 0.5 | 1 | 13 |
| skp1 | 0.07869 | -0.31933213 | 0.157 | 0.5 | 1 | 13 |
| atp5mc1 | 0.08007 | -0.19958219 | 0.196 | 0.5 | 1 | 13 |
| si:ch211-288g17.3 | 0.08033 | -0.10330052 | 0.216 | 0.5 | 1 | 13 |
| def6c | 0.08067 | -0.34256806 | 0.157 | 0.5 | 1 | 13 |
| si:dkey-203a12.7 | 0.08154 | -0.34341153 | 0.902 | 1 | 1 | 13 |
| arf1 | 0.08206 | 0.330733507 | 0.294 | 0.4 | 1 | 13 |
| si:ch211-218c6.8 | 0.08331 | -0.46500244 | 0.098 | 0.4 | 1 | 13 |
| creg1 | 0.0842 | -0.16427579 | 0.118 | 0.4 | 1 | 13 |
| eef1g | 0.08497 | -0.16283398 | 0.608 | 0.9 | 1 | 13 |
| fnbp4 | 0.08526 | -0.14434805 | 0.059 | 0.3 | 1 | 13 |
| krt8 | 0.08555 | 0.650747331 | 0.373 | 0.1 | 1 | 13 |
| ucp2 | 0.08574 | -0.19212315 | 0.471 | 0.8 | 1 | 13 |
| plpp1a | 0.08575 | -0.32027961 | 0.098 | 0.4 | 1 | 13 |
| psmc1a | 0.08578 | -0.33026004 | 0.098 | 0.4 | 1 | 13 |
| arpc3 | 0.08616 | -0.13505113 | 0.765 | 1 | 1 | 13 |
| magt1 | 0.0863 | -0.37060973 | 0.235 | 0.6 | 1 | 13 |
| psmd6 | 0.08675 | -0.21559621 | 0.098 | 0.4 | 1 | 13 |
| selenot2 | 0.08798 | -0.1883712 | 0.059 | 0.3 | 1 | 13 |
| supt4h1 | 0.08813 | -0.36337881 | 0.098 | 0.4 | 1 | 13 |
| si:ch211-226m16.2 | 0.08856 | -0.28659477 | 0.255 | 0.6 | 1 | 13 |
| si:ch1073-376c22.1 | 0.08886 | -0.34835743 | 0.765 | 1 | 1 | 13 |
| wu:fi37e09 | 0.08908 | -0.3668436 | 0.098 | 0.4 | 1 | 13 |
| sc:d0284 | 0.08917 | -0.37123376 | 0.098 | 0.4 | 1 | 13 |
| scfd1 | 0.08922 | -0.39867992 | 0.098 | 0.4 | 1 | 13 |
| dync1i2a | 0.08932 | -0.15441318 | 0.059 | 0.3 | 1 | 13 |
| ap1g1 | 0.08934 | 0.346582054 | 0.176 | 0 | 1 | 13 |
| mia3 | 0.08934 | 0.38529609 | 0.176 | 0 | 1 | 13 |
| ddx3xb | 0.08934 | 0.395942803 | 0.176 | 0 | 1 | 13 |
| hspa4b | 0.08934 | 0.319193258 | 0.176 | 0 | 1 | 13 |
| si:dkey-126g1.7 | 0.08934 | 0.359689549 | 0.176 | 0 | 1 | 13 |
| ccdc124 | 0.0894 | -0.20376592 | 0.176 | 0.5 | 1 | 13 |
| tmem106a | 0.08949 | -0.38946317 | 0.098 | 0.4 | 1 | 13 |
| trappc5 | 0.0902 | -0.15803319 | 0.059 | 0.3 | 1 | 13 |
| cebpb | 0.09021 | -0.21740218 | 0.745 | 1 | 1 | 13 |
| sumo2b | 0.09072 | -0.33012295 | 0.333 | 0.7 | 1 | 13 |
| ndufs7 | 0.09144 | -0.22961369 | 0.176 | 0.5 | 1 | 13 |
| dynlrb1 | 0.09237 | -0.1800552 | 0.255 | 0.6 | 1 | 13 |
| banf1 | 0.09248 | -0.1258288 | 0.196 | 0.5 | 1 | 13 |
| mt-atp6 | 0.09496 | -0.20410784 | 0.529 | 0.8 | 1 | 13 |
| lasp1 | 0.0951 | -0.20541689 | 0.49 | 0.8 | 1 | 13 |

| | | | | | | |
|---|---|---|---|---|---|---|
| ndufa4l | 0.0953 | 0.10963245 | 0.49 | 0.7 | 1 | 13 |
| carhsp1 | 0.09559 | -0.28160544 | 0.176 | 0.5 | 1 | 13 |
| ubald2 | 0.09579 | -0.24038176 | 0.059 | 0.3 | 1 | 13 |
| h3f3d | 0.09698 | -0.35118737 | 0.333 | 0.7 | 1 | 13 |
| BX284638.1 | 0.09717 | -0.11402652 | 0 | 0.1 | 1 | 13 |
| BX511061.1 | 0.09717 | -0.14662326 | 0 | 0.1 | 1 | 13 |
| CABZ01090361.1 | 0.09717 | -0.10957892 | 0 | 0.1 | 1 | 13 |
| CABZ01113830.1 | 0.09717 | -0.12173106 | 0 | 0.1 | 1 | 13 |
| CR318588.1 | 0.09717 | -0.29516496 | 0 | 0.1 | 1 | 13 |
| CU660013.4 | 0.09717 | -0.14109473 | 0 | 0.1 | 1 | 13 |
| LOC101885853 | 0.09717 | -0.12173106 | 0 | 0.1 | 1 | 13 |
| LOC110437818 | 0.09717 | -0.12173106 | 0 | 0.1 | 1 | 13 |
| NPAS3 | 0.09717 | -0.1112179 | 0 | 0.1 | 1 | 13 |
| NPC2 (1 of many) | 0.09717 | -0.1475301 | 0 | 0.1 | 1 | 13 |
| aak1b | 0.09717 | -0.12173106 | 0 | 0.1 | 1 | 13 |
| abca1a | 0.09717 | -0.14869581 | 0 | 0.1 | 1 | 13 |
| abi2a | 0.09717 | -0.1647753 | 0 | 0.1 | 1 | 13 |
| adam8b | 0.09717 | -0.15419284 | 0 | 0.1 | 1 | 13 |
| adarb1b | 0.09717 | -0.12173106 | 0 | 0.1 | 1 | 13 |
| alpk2 | 0.09717 | -0.1112179 | 0 | 0.1 | 1 | 13 |
| bco2a | 0.09717 | -0.11718459 | 0 | 0.1 | 1 | 13 |
| bloc1s3 | 0.09717 | -0.14135853 | 0 | 0.1 | 1 | 13 |
| borcs5 | 0.09717 | -0.11820653 | 0 | 0.1 | 1 | 13 |
| c2cd2 | 0.09717 | -0.14489229 | 0 | 0.1 | 1 | 13 |
| calcoco1a | 0.09717 | -0.10466199 | 0 | 0.1 | 1 | 13 |
| casc1 | 0.09717 | -0.15419284 | 0 | 0.1 | 1 | 13 |
| cgas | 0.09717 | -0.14820252 | 0 | 0.1 | 1 | 13 |
| clybl | 0.09717 | -0.12290539 | 0 | 0.1 | 1 | 13 |
| cmc4 | 0.09717 | -0.13207894 | 0 | 0.1 | 1 | 13 |
| cnot6l | 0.09717 | -0.11782458 | 0 | 0.1 | 1 | 13 |
| cox10 | 0.09717 | -0.14389312 | 0 | 0.1 | 1 | 13 |
| crebl2 | 0.09717 | -0.10112825 | 0 | 0.1 | 1 | 13 |
| cryl1 | 0.09717 | -0.13052351 | 0 | 0.1 | 1 | 13 |
| dbn1 | 0.09717 | -0.1405938 | 0 | 0.1 | 1 | 13 |
| dnajc15 | 0.09717 | -0.1026378 | 0 | 0.1 | 1 | 13 |
| dnmt3bb.1 | 0.09717 | -0.10856908 | 0 | 0.1 | 1 | 13 |
| eprs | 0.09717 | -0.1367833 | 0 | 0.1 | 1 | 13 |
| ergic1 | 0.09717 | -0.10890514 | 0 | 0.1 | 1 | 13 |
| etv4 | 0.09717 | -0.11515132 | 0 | 0.1 | 1 | 13 |
| fap | 0.09717 | -0.14892346 | 0 | 0.1 | 1 | 13 |
| fbxl5 | 0.09717 | -0.10829561 | 0 | 0.1 | 1 | 13 |
| fig4a | 0.09717 | -0.15154524 | 0 | 0.1 | 1 | 13 |
| foxo1a | 0.09717 | -0.15031265 | 0 | 0.1 | 1 | 13 |
| fthl28 | 0.09717 | -0.11669715 | 0 | 0.1 | 1 | 13 |
| fuom | 0.09717 | -0.10458545 | 0 | 0.1 | 1 | 13 |
| galk2 | 0.09717 | -0.11642524 | 0 | 0.1 | 1 | 13 |
| glb1 | 0.09717 | -0.15067911 | 0 | 0.1 | 1 | 13 |
| havcr2 | 0.09717 | -0.15188844 | 0 | 0.1 | 1 | 13 |

| | | | | | | |
|---|---|---|---|---|---|---|
| hccsb | 0.09717 | -0.14024611 | 0 | 0.1 | 1 | 13 |
| hmgb3a | 0.09717 | -0.13539314 | 0 | 0.1 | 1 | 13 |
| hnrnph1 | 0.09717 | -0.13763859 | 0 | 0.1 | 1 | 13 |
| ifi46 | 0.09717 | -0.10271027 | 0 | 0.1 | 1 | 13 |
| ift140 | 0.09717 | -0.1112179 | 0 | 0.1 | 1 | 13 |
| kynu | 0.09717 | -0.14677527 | 0 | 0.1 | 1 | 13 |
| lrch4 | 0.09717 | -0.11896461 | 0 | 0.1 | 1 | 13 |
| lrrtm1 | 0.09717 | -0.14223534 | 0 | 0.1 | 1 | 13 |
| ly6m5 | 0.09717 | -0.15377466 | 0 | 0.1 | 1 | 13 |
| lygl1 | 0.09717 | -0.21102591 | 0 | 0.1 | 1 | 13 |
| lypd6 | 0.09717 | -0.1112179 | 0 | 0.1 | 1 | 13 |
| lysmd2 | 0.09717 | -0.13919195 | 0 | 0.1 | 1 | 13 |
| matn3b | 0.09717 | -0.14223534 | 0 | 0.1 | 1 | 13 |
| mcm7 | 0.09717 | -0.18787615 | 0 | 0.1 | 1 | 13 |
| mfap4 | 0.09717 | -0.28863384 | 0 | 0.1 | 1 | 13 |
| mical1 | 0.09717 | -0.14202991 | 0 | 0.1 | 1 | 13 |
| mlh1 | 0.09717 | -0.13816967 | 0 | 0.1 | 1 | 13 |
| mllt1a | 0.09717 | -0.11917132 | 0 | 0.1 | 1 | 13 |
| mrpl16 | 0.09717 | -0.10190826 | 0 | 0.1 | 1 | 13 |
| mrpl58 | 0.09717 | -0.14065334 | 0 | 0.1 | 1 | 13 |
| mtfp1 | 0.09717 | -0.13862382 | 0 | 0.1 | 1 | 13 |
| necap2 | 0.09717 | -0.14468333 | 0 | 0.1 | 1 | 13 |
| nktr | 0.09717 | -0.14354792 | 0 | 0.1 | 1 | 13 |
| nsun4 | 0.09717 | -0.12506627 | 0 | 0.1 | 1 | 13 |
| nudcd1 | 0.09717 | -0.11138713 | 0 | 0.1 | 1 | 13 |
| nudt9 | 0.09717 | -0.1069343 | 0 | 0.1 | 1 | 13 |
| nxt2 | 0.09717 | -0.10087474 | 0 | 0.1 | 1 | 13 |
| pbx3b | 0.09717 | -0.19882109 | 0 | 0.1 | 1 | 13 |
| pcnp | 0.09717 | -0.13142047 | 0 | 0.1 | 1 | 13 |
| pcsk9 | 0.09717 | -0.11402652 | 0 | 0.1 | 1 | 13 |
| polr3gla | 0.09717 | -0.10994948 | 0 | 0.1 | 1 | 13 |
| praf2 | 0.09717 | -0.1111596 | 0 | 0.1 | 1 | 13 |
| prkx | 0.09717 | -0.14748732 | 0 | 0.1 | 1 | 13 |
| psip1b | 0.09717 | -0.10516736 | 0 | 0.1 | 1 | 13 |
| ranbp3b | 0.09717 | -0.13400249 | 0 | 0.1 | 1 | 13 |
| rfxank | 0.09717 | -0.15057485 | 0 | 0.1 | 1 | 13 |
| rgmd | 0.09717 | -0.11187651 | 0 | 0.1 | 1 | 13 |
| rgs5b | 0.09717 | -0.15068532 | 0 | 0.1 | 1 | 13 |
| rlf | 0.09717 | -0.14646082 | 0 | 0.1 | 1 | 13 |
| rpz5 | 0.09717 | -0.16768821 | 0 | 0.1 | 1 | 13 |
| runx1t1 | 0.09717 | -0.11457689 | 0 | 0.1 | 1 | 13 |
| scaf4b | 0.09717 | -0.11422824 | 0 | 0.1 | 1 | 13 |
| sec22c | 0.09717 | -0.14994578 | 0 | 0.1 | 1 | 13 |
| si:ch211-191i18.2 | 0.09717 | -0.14158101 | 0 | 0.1 | 1 | 13 |
| si:ch211-234h8.7 | 0.09717 | -0.1450647 | 0 | 0.1 | 1 | 13 |
| si:dkey-287g12.6 | 0.09717 | -0.10219127 | 0 | 0.1 | 1 | 13 |
| si:dkey-3k24.5 | 0.09717 | -0.14963869 | 0 | 0.1 | 1 | 13 |
| si:dkey-66a8.7 | 0.09717 | -0.11566023 | 0 | 0.1 | 1 | 13 |

| | | | | | | |
|---|---|---|---|---|---|---|
| si:rp71-1f1.4 | 0.09717 | -0.17531672 | 0 | 0.1 | 1 | 13 |
| slc16a13 | 0.09717 | -0.13932381 | 0 | 0.1 | 1 | 13 |
| slc2a12 | 0.09717 | -0.1113658 | 0 | 0.1 | 1 | 13 |
| slc48a1a | 0.09717 | -0.14223534 | 0 | 0.1 | 1 | 13 |
| smarcd1 | 0.09717 | -0.11804788 | 0 | 0.1 | 1 | 13 |
| snx27b | 0.09717 | -0.11946834 | 0 | 0.1 | 1 | 13 |
| socs6a | 0.09717 | -0.11237112 | 0 | 0.1 | 1 | 13 |
| taok2a | 0.09717 | -0.11972564 | 0 | 0.1 | 1 | 13 |
| thraa | 0.09717 | -0.10633581 | 0 | 0.1 | 1 | 13 |
| tmem241 | 0.09717 | -0.14097099 | 0 | 0.1 | 1 | 13 |
| trmu | 0.09717 | -0.15419284 | 0 | 0.1 | 1 | 13 |
| tsg101a | 0.09717 | -0.10364657 | 0 | 0.1 | 1 | 13 |
| ttyh3a | 0.09717 | -0.15289104 | 0 | 0.1 | 1 | 13 |
| tut4 | 0.09717 | -0.14223534 | 0 | 0.1 | 1 | 13 |
| vps16 | 0.09717 | -0.13919389 | 0 | 0.1 | 1 | 13 |
| vps26a | 0.09717 | -0.13260919 | 0 | 0.1 | 1 | 13 |
| wt1b | 0.09717 | -0.10248701 | 0 | 0.1 | 1 | 13 |
| wu:fu71h07 | 0.09717 | -0.14356124 | 0 | 0.1 | 1 | 13 |
| ylpm1 | 0.09717 | -0.10314926 | 0 | 0.1 | 1 | 13 |
| zdhhc3a | 0.09717 | -0.10552212 | 0 | 0.1 | 1 | 13 |
| zeb1a | 0.09717 | -0.11129843 | 0 | 0.1 | 1 | 13 |
| zgc:101559 | 0.09717 | -0.10243616 | 0 | 0.1 | 1 | 13 |
| zgc:162193 | 0.09717 | -0.15251219 | 0 | 0.1 | 1 | 13 |
| zgc:173915 | 0.09717 | -0.11816954 | 0 | 0.1 | 1 | 13 |
| zgc:65895 | 0.09717 | -0.11425777 | 0 | 0.1 | 1 | 13 |
| zmp:0000001228 | 0.09717 | -0.10250837 | 0 | 0.1 | 1 | 13 |
| znf1155 | 0.09717 | -0.1112179 | 0 | 0.1 | 1 | 13 |
| znf385c | 0.09717 | -0.12173106 | 0 | 0.1 | 1 | 13 |
| znf628 | 0.09717 | -0.15419284 | 0 | 0.1 | 1 | 13 |
| BX000363.1 | 0.09717 | -0.11732409 | 0 | 0.1 | 1 | 13 |
| CR450808.1 | 0.09717 | -0.12006938 | 0 | 0.1 | 1 | 13 |
| CU571081.1 | 0.09717 | -0.12355558 | 0 | 0.1 | 1 | 13 |
| XLOC-015481 | 0.09717 | -0.1258768 | 0 | 0.1 | 1 | 13 |
| chfr | 0.09717 | -0.10409413 | 0 | 0.1 | 1 | 13 |
| dclre1b | 0.09717 | -0.11409555 | 0 | 0.1 | 1 | 13 |
| e2f4 | 0.09717 | -0.10493893 | 0 | 0.1 | 1 | 13 |
| flot2b | 0.09717 | -0.11902022 | 0 | 0.1 | 1 | 13 |
| itcha | 0.09717 | -0.1112759 | 0 | 0.1 | 1 | 13 |
| jade2 | 0.09717 | -0.11515466 | 0 | 0.1 | 1 | 13 |
| klf2a | 0.09717 | -0.12000208 | 0 | 0.1 | 1 | 13 |
| map3k5 | 0.09717 | -0.1258768 | 0 | 0.1 | 1 | 13 |
| orai1b | 0.09717 | -0.11702793 | 0 | 0.1 | 1 | 13 |
| p2ry11 | 0.09717 | -0.1258768 | 0 | 0.1 | 1 | 13 |
| plekhm3 | 0.09717 | -0.1258768 | 0 | 0.1 | 1 | 13 |
| psip1a | 0.09717 | -0.1058563 | 0 | 0.1 | 1 | 13 |
| rad54l2 | 0.09717 | -0.11396702 | 0 | 0.1 | 1 | 13 |
| rfx2 | 0.09717 | -0.10316051 | 0 | 0.1 | 1 | 13 |
| sec24c | 0.09717 | -0.11480154 | 0 | 0.1 | 1 | 13 |

| | | | | | | |
|---|---|---|---|---|---|---|
| sfmbt2 | 0.09717 | -0.1258768 | 0 | 0.1 | 1 | 13 |
| siae | 0.09717 | -0.11654475 | 0 | 0.1 | 1 | 13 |
| slc15a4 | 0.09717 | -0.11840473 | 0 | 0.1 | 1 | 13 |
| vps25 | 0.09717 | -0.11260782 | 0 | 0.1 | 1 | 13 |
| zgc:172128 | 0.09717 | -0.10878395 | 0 | 0.1 | 1 | 13 |
| BX901883.3 | 0.09717 | -0.12162492 | 0 | 0.1 | 1 | 13 |
| BX957297.2 | 0.09717 | -0.11699463 | 0 | 0.1 | 1 | 13 |
| CU856539.1 | 0.09717 | -0.11149778 | 0 | 0.1 | 1 | 13 |
| CU856539.2 | 0.09717 | -0.17690661 | 0 | 0.1 | 1 | 13 |
| EIF2AK4 | 0.09717 | -0.10234987 | 0 | 0.1 | 1 | 13 |
| LOC100535806 | 0.09717 | -0.11315192 | 0 | 0.1 | 1 | 13 |
| LOC101882111 | 0.09717 | -0.12162492 | 0 | 0.1 | 1 | 13 |
| LOC108182673 | 0.09717 | -0.12162492 | 0 | 0.1 | 1 | 13 |
| XLOC-018082 | 0.09717 | -0.11456112 | 0 | 0.1 | 1 | 13 |
| atf7b | 0.09717 | -0.11544614 | 0 | 0.1 | 1 | 13 |
| atp2a2a | 0.09717 | -0.11446278 | 0 | 0.1 | 1 | 13 |
| cacnb4b | 0.09717 | -0.11111721 | 0 | 0.1 | 1 | 13 |
| chpf2 | 0.09717 | -0.10896256 | 0 | 0.1 | 1 | 13 |
| cpsf4 | 0.09717 | -0.10907843 | 0 | 0.1 | 1 | 13 |
| csnk2a2a | 0.09717 | -0.11673112 | 0 | 0.1 | 1 | 13 |
| dhx57 | 0.09717 | -0.10676422 | 0 | 0.1 | 1 | 13 |
| elmo3 | 0.09717 | -0.11947245 | 0 | 0.1 | 1 | 13 |
| fam208ab | 0.09717 | -0.10972703 | 0 | 0.1 | 1 | 13 |
| irf1b | 0.09717 | -0.10207996 | 0 | 0.1 | 1 | 13 |
| mybl2b | 0.09717 | -0.10869773 | 0 | 0.1 | 1 | 13 |
| ncapd3 | 0.09717 | -0.1040967 | 0 | 0.1 | 1 | 13 |
| ndc1 | 0.09717 | -0.15115983 | 0 | 0.1 | 1 | 13 |
| paxip1 | 0.09717 | -0.11922359 | 0 | 0.1 | 1 | 13 |
| pcif1 | 0.09717 | -0.11506486 | 0 | 0.1 | 1 | 13 |
| pik3r5 | 0.09717 | -0.11643474 | 0 | 0.1 | 1 | 13 |
| psmd2 | 0.09717 | -0.10452946 | 0 | 0.1 | 1 | 13 |
| rab8b | 0.09717 | -0.11640794 | 0 | 0.1 | 1 | 13 |
| recql4 | 0.09717 | -0.10783959 | 0 | 0.1 | 1 | 13 |
| samd1b | 0.09717 | -0.11190806 | 0 | 0.1 | 1 | 13 |
| scly | 0.09717 | -0.11042622 | 0 | 0.1 | 1 | 13 |
| shrprbck1r | 0.09717 | -0.10617882 | 0 | 0.1 | 1 | 13 |
| si:ch211-156b7.4 | 0.09717 | -0.20788 | 0 | 0.1 | 1 | 13 |
| si:dkey-9i23.15 | 0.09717 | -0.10152371 | 0 | 0.1 | 1 | 13 |
| si:rp71-36a1.3 | 0.09717 | -0.1142041 | 0 | 0.1 | 1 | 13 |
| smurf1 | 0.09717 | -0.11277556 | 0 | 0.1 | 1 | 13 |
| ssbp3a | 0.09717 | -0.10732704 | 0 | 0.1 | 1 | 13 |
| ttc17 | 0.09717 | -0.12162492 | 0 | 0.1 | 1 | 13 |
| utp25 | 0.09717 | -0.11035151 | 0 | 0.1 | 1 | 13 |
| wdr37 | 0.09717 | -0.10931825 | 0 | 0.1 | 1 | 13 |
| AL954142.1 | 0.09717 | -0.10452902 | 0 | 0.1 | 1 | 13 |
| CR383680.1 | 0.09717 | -0.11073201 | 0 | 0.1 | 1 | 13 |
| RF00560 | 0.09717 | -0.17677205 | 0 | 0.1 | 1 | 13 |
| XLOC-021534 | 0.09717 | -0.19773178 | 0 | 0.1 | 1 | 13 |

| | | | | | | |
|---|---|---|---|---|---|---|
| abhd17b | 0.09717 | -0.19022932 | 0 | 0.1 | 1 | 13 |
| alg8 | 0.09717 | -0.10466137 | 0 | 0.1 | 1 | 13 |
| arid1b | 0.09717 | -0.18904027 | 0 | 0.1 | 1 | 13 |
| arrdc3b | 0.09717 | -0.16026194 | 0 | 0.1 | 1 | 13 |
| arsa | 0.09717 | -0.18747294 | 0 | 0.1 | 1 | 13 |
| cabp1b | 0.09717 | -0.11073201 | 0 | 0.1 | 1 | 13 |
| camkmt | 0.09717 | -0.10599311 | 0 | 0.1 | 1 | 13 |
| ccdc171 | 0.09717 | -0.15533531 | 0 | 0.1 | 1 | 13 |
| cerk | 0.09717 | -0.18895743 | 0 | 0.1 | 1 | 13 |
| chchd10 | 0.09717 | -0.14060338 | 0 | 0.1 | 1 | 13 |
| clpxa | 0.09717 | -0.17454375 | 0 | 0.1 | 1 | 13 |
| coq8b | 0.09717 | -0.10095819 | 0 | 0.1 | 1 | 13 |
| cyth4a | 0.09717 | -0.10299546 | 0 | 0.1 | 1 | 13 |
| dkk3b | 0.09717 | -0.12800654 | 0 | 0.1 | 1 | 13 |
| dse | 0.09717 | -0.19530824 | 0 | 0.1 | 1 | 13 |
| ebna1bp2 | 0.09717 | -0.14979002 | 0 | 0.1 | 1 | 13 |
| flad1 | 0.09717 | -0.18342454 | 0 | 0.1 | 1 | 13 |
| fmnl2a | 0.09717 | -0.16941975 | 0 | 0.1 | 1 | 13 |
| galnt2 | 0.09717 | -0.1918216 | 0 | 0.1 | 1 | 13 |
| gpr155a | 0.09717 | -0.10906064 | 0 | 0.1 | 1 | 13 |
| homezb | 0.09717 | -0.17995417 | 0 | 0.1 | 1 | 13 |
| irak1 | 0.09717 | -0.10375901 | 0 | 0.1 | 1 | 13 |
| itpr2 | 0.09717 | -0.19773178 | 0 | 0.1 | 1 | 13 |
| lrmda | 0.09717 | -0.11073201 | 0 | 0.1 | 1 | 13 |
| mafbb | 0.09717 | -0.19519384 | 0 | 0.1 | 1 | 13 |
| mplkip | 0.09717 | -0.17432557 | 0 | 0.1 | 1 | 13 |
| mrm3a | 0.09717 | -0.18443211 | 0 | 0.1 | 1 | 13 |
| nr2f6a | 0.09717 | -0.19369526 | 0 | 0.1 | 1 | 13 |
| nr4a2b | 0.09717 | -0.15256988 | 0 | 0.1 | 1 | 13 |
| ormdl2 | 0.09717 | -0.17945895 | 0 | 0.1 | 1 | 13 |
| osbpl5 | 0.09717 | -0.10374567 | 0 | 0.1 | 1 | 13 |
| oxct1a | 0.09717 | -0.10189713 | 0 | 0.1 | 1 | 13 |
| pam | 0.09717 | -0.18211816 | 0 | 0.1 | 1 | 13 |
| paox1 | 0.09717 | -0.10493759 | 0 | 0.1 | 1 | 13 |
| pbrm1l | 0.09717 | -0.10487023 | 0 | 0.1 | 1 | 13 |
| ppp3cca | 0.09717 | -0.10560453 | 0 | 0.1 | 1 | 13 |
| pros1 | 0.09717 | -0.18397479 | 0 | 0.1 | 1 | 13 |
| pspc1 | 0.09717 | -0.1834577 | 0 | 0.1 | 1 | 13 |
| rbm25a | 0.09717 | -0.17530347 | 0 | 0.1 | 1 | 13 |
| rxraa | 0.09717 | -0.19046219 | 0 | 0.1 | 1 | 13 |
| si:ch1073-224n8.1 | 0.09717 | -0.10793642 | 0 | 0.1 | 1 | 13 |
| si:ch211-113e8.11 | 0.09717 | -0.15526352 | 0 | 0.1 | 1 | 13 |
| si:ch211-132g1.3 | 0.09717 | -0.19773178 | 0 | 0.1 | 1 | 13 |
| si:ch211-195b13.1 | 0.09717 | -0.10483254 | 0 | 0.1 | 1 | 13 |
| si:ch73-109i22.2 | 0.09717 | -0.11073201 | 0 | 0.1 | 1 | 13 |
| si:dkey-111e8.5 | 0.09717 | -0.16941975 | 0 | 0.1 | 1 | 13 |
| slc16a10 | 0.09717 | -0.16746161 | 0 | 0.1 | 1 | 13 |
| spata6 | 0.09717 | -0.10081135 | 0 | 0.1 | 1 | 13 |

| | | | | | | |
|---|---|---|---|---|---|---|
| sumo3b | 0.09717 | -0.13211041 | 0 | 0.1 | 1 | 13 |
| syt11a | 0.09717 | -0.10188622 | 0 | 0.1 | 1 | 13 |
| tm2d1 | 0.09717 | -0.10286482 | 0 | 0.1 | 1 | 13 |
| tnfrsf1b | 0.09717 | -0.19725154 | 0 | 0.1 | 1 | 13 |
| traf6 | 0.09717 | -0.10375012 | 0 | 0.1 | 1 | 13 |
| trpm5 | 0.09717 | -0.10081135 | 0 | 0.1 | 1 | 13 |
| ttf1 | 0.09717 | -0.17807536 | 0 | 0.1 | 1 | 13 |
| tyw3 | 0.09717 | -0.18659859 | 0 | 0.1 | 1 | 13 |
| uap1l1 | 0.09717 | -0.17586904 | 0 | 0.1 | 1 | 13 |
| ubl7a | 0.09717 | -0.1833125 | 0 | 0.1 | 1 | 13 |
| upp1 | 0.09717 | -0.10756255 | 0 | 0.1 | 1 | 13 |
| uxs1 | 0.09717 | -0.15464542 | 0 | 0.1 | 1 | 13 |
| wu:fi46h04 | 0.09717 | -0.18296201 | 0 | 0.1 | 1 | 13 |
| zak | 0.09717 | -0.19499049 | 0 | 0.1 | 1 | 13 |
| zgc:194281 | 0.09717 | -0.10863875 | 0 | 0.1 | 1 | 13 |
| zhx2a | 0.09717 | -0.10016598 | 0 | 0.1 | 1 | 13 |
| LOC100332847 | 0.09717 | -0.11117781 | 0 | 0.1 | 1 | 13 |
| ndufaf1 | 0.09717 | -0.10021277 | 0 | 0.1 | 1 | 13 |
| rnpc3 | 0.09717 | -0.10722442 | 0 | 0.1 | 1 | 13 |
| timp4.1 | 0.09717 | -0.13269997 | 0 | 0.1 | 1 | 13 |
| CABZ01077220.2 | 0.09717 | -0.18717692 | 0 | 0.1 | 1 | 13 |
| CABZ01078737.1 | 0.09717 | -0.20825545 | 0 | 0.1 | 1 | 13 |
| LOC101886078 | 0.09717 | -0.2085699 | 0 | 0.1 | 1 | 13 |
| LOC103911004 | 0.09717 | -0.20129902 | 0 | 0.1 | 1 | 13 |
| arhgap11a | 0.09717 | -0.20567051 | 0 | 0.1 | 1 | 13 |
| atg9b | 0.09717 | -0.14281954 | 0 | 0.1 | 1 | 13 |
| cbr4 | 0.09717 | -0.17476838 | 0 | 0.1 | 1 | 13 |
| cisd1 | 0.09717 | -0.19835314 | 0 | 0.1 | 1 | 13 |
| ctnnbl1 | 0.09717 | -0.12561877 | 0 | 0.1 | 1 | 13 |
| emx2 | 0.09717 | -0.1076602 | 0 | 0.1 | 1 | 13 |
| entpd6 | 0.09717 | -0.20286095 | 0 | 0.1 | 1 | 13 |
| erlin1 | 0.09717 | -0.14542837 | 0 | 0.1 | 1 | 13 |
| exoc7 | 0.09717 | -0.27495307 | 0 | 0.1 | 1 | 13 |
| fam193a | 0.09717 | -0.13370523 | 0 | 0.1 | 1 | 13 |
| gon4l | 0.09717 | -0.19995561 | 0 | 0.1 | 1 | 13 |
| kcnq5a | 0.09717 | -0.2085699 | 0 | 0.1 | 1 | 13 |
| kdm5a | 0.09717 | -0.20315838 | 0 | 0.1 | 1 | 13 |
| mcmbp | 0.09717 | -0.19609663 | 0 | 0.1 | 1 | 13 |
| mettl1 | 0.09717 | -0.18995832 | 0 | 0.1 | 1 | 13 |
| pank2 | 0.09717 | -0.12585414 | 0 | 0.1 | 1 | 13 |
| pde10a | 0.09717 | -0.2085699 | 0 | 0.1 | 1 | 13 |
| phf21ab | 0.09717 | -0.20631843 | 0 | 0.1 | 1 | 13 |
| pik3c2b | 0.09717 | -0.19868369 | 0 | 0.1 | 1 | 13 |
| pip4p1a | 0.09717 | -0.19946188 | 0 | 0.1 | 1 | 13 |
| pyroxd1 | 0.09717 | -0.1617431 | 0 | 0.1 | 1 | 13 |
| rabgap1 | 0.09717 | -0.20050933 | 0 | 0.1 | 1 | 13 |
| ranbp10 | 0.09717 | -0.20420288 | 0 | 0.1 | 1 | 13 |
| rpf2 | 0.09717 | -0.1954005 | 0 | 0.1 | 1 | 13 |

| | | | | | | |
|---|---|---|---|---|---|---|
| si:ch211-91p5.3 | 0.09717 | -0.19914056 | 0 | 0.1 | 1 | 13 |
| si:dkey-206f10.1 | 0.09717 | -0.19502333 | 0 | 0.1 | 1 | 13 |
| slc38a10 | 0.09717 | -0.20635518 | 0 | 0.1 | 1 | 13 |
| slc44a1b | 0.09717 | -0.20058059 | 0 | 0.1 | 1 | 13 |
| snx2 | 0.09717 | -0.20589426 | 0 | 0.1 | 1 | 13 |
| tdrd3 | 0.09717 | -0.19888264 | 0 | 0.1 | 1 | 13 |
| tmem94 | 0.09717 | -0.2067703 | 0 | 0.1 | 1 | 13 |
| tmx3a | 0.09717 | -0.17968186 | 0 | 0.1 | 1 | 13 |
| zfpm1 | 0.09717 | -0.1651167 | 0 | 0.1 | 1 | 13 |
| zgc:171679 | 0.09717 | -0.20570795 | 0 | 0.1 | 1 | 13 |
| znf1059 | 0.09717 | -0.19836726 | 0 | 0.1 | 1 | 13 |
| SHKBP1 | 0.09717 | -0.16705889 | 0 | 0.1 | 1 | 13 |
| SMIM10 | 0.09717 | -0.15459067 | 0 | 0.1 | 1 | 13 |
| acap2 | 0.09717 | -0.17631439 | 0 | 0.1 | 1 | 13 |
| alkbh4 | 0.09717 | -0.15193144 | 0 | 0.1 | 1 | 13 |
| arhgap25 | 0.09717 | -0.17765774 | 0 | 0.1 | 1 | 13 |
| igsf5b | 0.09717 | -0.15060145 | 0 | 0.1 | 1 | 13 |
| im:7147486 | 0.09717 | -0.15207603 | 0 | 0.1 | 1 | 13 |
| med20 | 0.09717 | -0.13394116 | 0 | 0.1 | 1 | 13 |
| nfil3-2 | 0.09717 | -0.16490222 | 0 | 0.1 | 1 | 13 |
| nthl1 | 0.09717 | -0.17663728 | 0 | 0.1 | 1 | 13 |
| ppic | 0.09717 | -0.17598625 | 0 | 0.1 | 1 | 13 |
| prpsap1 | 0.09717 | -0.13319028 | 0 | 0.1 | 1 | 13 |
| slc25a37 | 0.09717 | -0.17015244 | 0 | 0.1 | 1 | 13 |
| tcp11l1 | 0.09717 | -0.16360308 | 0 | 0.1 | 1 | 13 |
| tmem154 | 0.09717 | -0.17321488 | 0 | 0.1 | 1 | 13 |
| uvssa | 0.09717 | -0.16562739 | 0 | 0.1 | 1 | 13 |
| wdsub1 | 0.09717 | -0.14174172 | 0 | 0.1 | 1 | 13 |
| zbtb43 | 0.09717 | -0.17546667 | 0 | 0.1 | 1 | 13 |
| zgc:112023 | 0.09717 | -0.17015913 | 0 | 0.1 | 1 | 13 |
| sept7a | 0.09788 | -0.27196088 | 0.059 | 0.3 | 1 | 13 |
| mt-nd4 | 0.09843 | 0.234260312 | 0.529 | 0.7 | 1 | 13 |
| eif3d | 0.09945 | 0.329433002 | 0.294 | 0.4 | 1 | 13 |
| vta1 | 0.09965 | -0.13405782 | 0.059 | 0.3 | 1 | 13 |
| lxn | 0.0999 | -0.10439351 | 0.431 | 0.7 | 1 | 13 |
| grb2b | 0.10054 | -0.25745121 | 0.176 | 0.5 | 1 | 13 |
| rgs19 | 0.10133 | -0.30012626 | 0.176 | 0.5 | 1 | 13 |
| CABZ01112049.1 | 0.1022 | 0.141624787 | 0.961 | 1 | 1 | 13 |
| prdx4 | 0.10223 | -0.12644567 | 0.51 | 0.8 | 1 | 13 |
| cirbpa | 0.10312 | 0.258849747 | 0.706 | 0.8 | 1 | 13 |
| ppp1r10 | 0.1032 | -0.23615215 | 0.059 | 0.3 | 1 | 13 |
| fkbp1aa | 0.10323 | -0.19583645 | 0.373 | 0.7 | 1 | 13 |
| slc44a4 | 0.10554 | -0.22702851 | 0.118 | 0.4 | 1 | 13 |
| rsl24d1 | 0.10593 | -0.2355591 | 0.275 | 0.6 | 1 | 13 |
| pitpnb | 0.10623 | -0.1521754 | 0.059 | 0.3 | 1 | 13 |
| ptbp3 | 0.10639 | -0.20591904 | 0.059 | 0.3 | 1 | 13 |
| usp1 | 0.10804 | -0.15279541 | 0.059 | 0.3 | 1 | 13 |
| capgb | 0.10816 | -0.30379307 | 0.588 | 0.9 | 1 | 13 |

| | | | | | | |
|---|---|---|---|---|---|---|
| CABZ01079427.1 | 0.10826 | -0.13969575 | 0.059 | 0.3 | 1 | 13 |
| sec62 | 0.10917 | -0.23550918 | 0.118 | 0.4 | 1 | 13 |
| eif3f | 0.11077 | -0.11475676 | 0.412 | 0.7 | 1 | 13 |
| si:dkey-27h10.2 | 0.11083 | 0.317239126 | 0.471 | 0.5 | 1 | 13 |
| adam8a | 0.11112 | 0.350127772 | 0.588 | 0.6 | 1 | 13 |
| zgc:162698 | 0.1129 | -0.25569774 | 0.059 | 0.3 | 1 | 13 |
| gadd45ba | 0.11318 | -0.59116606 | 0.098 | 0.3 | 1 | 13 |
| mrpl52 | 0.11322 | -0.1540119 | 0.118 | 0.4 | 1 | 13 |
| si:ch211-26b3.4 | 0.11346 | -0.19105998 | 0.059 | 0.3 | 1 | 13 |
| snd1 | 0.11373 | 0.259993301 | 0.471 | 0.6 | 1 | 13 |
| rgs2 | 0.11396 | -0.27559287 | 0.118 | 0.4 | 1 | 13 |
| dnajc19 | 0.11408 | -0.19496339 | 0.059 | 0.3 | 1 | 13 |
| agap2 | 0.11429 | 0.367279733 | 0.157 | 0 | 1 | 13 |
| anp32b | 0.11429 | 0.235247867 | 0.157 | 0 | 1 | 13 |
| arhgef1b | 0.11429 | 0.322780303 | 0.157 | 0 | 1 | 13 |
| asb12a | 0.11429 | 0.259958831 | 0.157 | 0 | 1 | 13 |
| bsg | 0.11429 | 0.293115926 | 0.157 | 0 | 1 | 13 |
| camk2g1 | 0.11429 | 0.223032032 | 0.157 | 0 | 1 | 13 |
| itpk1b | 0.11429 | 0.304507699 | 0.157 | 0 | 1 | 13 |
| lmf2b | 0.11429 | 0.305510466 | 0.157 | 0 | 1 | 13 |
| lsm6 | 0.11429 | 0.234063184 | 0.157 | 0 | 1 | 13 |
| mfng | 0.11429 | 0.319598126 | 0.157 | 0 | 1 | 13 |
| mrpl33 | 0.11429 | 0.19426367 | 0.157 | 0 | 1 | 13 |
| si:dkey-261l7.2 | 0.11429 | 0.340504732 | 0.157 | 0 | 1 | 13 |
| arhgef12a | 0.11429 | 0.300147323 | 0.157 | 0 | 1 | 13 |
| eif5b | 0.11429 | 0.330410506 | 0.157 | 0 | 1 | 13 |
| gmip | 0.11429 | 0.35241744 | 0.157 | 0 | 1 | 13 |
| dnajc10 | 0.11429 | 0.324447722 | 0.157 | 0 | 1 | 13 |
| mapkapk2a | 0.11429 | 0.322626197 | 0.157 | 0 | 1 | 13 |
| ddx46 | 0.11434 | -0.16659281 | 0.059 | 0.3 | 1 | 13 |
| si:ch211-262i1.6 | 0.11585 | -0.14477303 | 0.118 | 0.4 | 1 | 13 |
| csrp1a | 0.116 | -0.36446209 | 0.255 | 0.6 | 1 | 13 |
| vps13c | 0.11647 | -0.21349092 | 0.059 | 0.3 | 1 | 13 |
| cxcr2 | 0.11661 | -0.4348123 | 0.176 | 0.5 | 1 | 13 |
| dgkzb | 0.11666 | -0.18575429 | 0.059 | 0.3 | 1 | 13 |
| tom1l2a | 0.11683 | -0.12591384 | 0.078 | 0.3 | 1 | 13 |
| gmfg | 0.11716 | -0.12457095 | 0.137 | 0.4 | 1 | 13 |
| cfbl | 0.1173 | 0.185554998 | 0.98 | 1 | 1 | 13 |
| mif | 0.11748 | 0.18908631 | 0.314 | 0.5 | 1 | 13 |
| sart1 | 0.11753 | 0.11146871 | 0.098 | 0.3 | 1 | 13 |
| atp1a3a | 0.11771 | -0.12526085 | 0.314 | 0.6 | 1 | 13 |
| ndufb8 | 0.11834 | 0.102898177 | 0.275 | 0.5 | 1 | 13 |
| mrpl18 | 0.11855 | -0.13599968 | 0.059 | 0.3 | 1 | 13 |
| abo | 0.11905 | -0.21539231 | 0.059 | 0.3 | 1 | 13 |
| snrpd3l | 0.1194 | -0.13794784 | 0.059 | 0.3 | 1 | 13 |
| cct8 | 0.12007 | -0.11168225 | 0.137 | 0.4 | 1 | 13 |
| elof1 | 0.12031 | -0.15296058 | 0.059 | 0.3 | 1 | 13 |
| hnrnpd | 0.12041 | -0.32812507 | 0.059 | 0.3 | 1 | 13 |

| | | | | | | |
|---|---|---|---|---|---|---|
| csnk1g2a | 0.12047 | -0.21582025 | 0.059 | 0.3 | 1 | 13 |
| chchd3a | 0.12077 | -0.18638827 | 0.059 | 0.3 | 1 | 13 |
| ddost | 0.12089 | -0.14953468 | 0.412 | 0.7 | 1 | 13 |
| polr2i | 0.12099 | -0.14313785 | 0.059 | 0.3 | 1 | 13 |
| pgp | 0.12117 | -0.14895704 | 0.059 | 0.3 | 1 | 13 |
| rpl27a | 0.12157 | -0.48889176 | 0.569 | 0.8 | 1 | 13 |
| slc39a7 | 0.12179 | -0.17834176 | 0.196 | 0.5 | 1 | 13 |
| arhgap10 | 0.1219 | -0.20591703 | 0.059 | 0.3 | 1 | 13 |
| si:zfos-169g10.3 | 0.1223 | -0.19913147 | 0.196 | 0.5 | 1 | 13 |
| si:dkey-112e7.2 | 0.12235 | -0.22203845 | 0.118 | 0.4 | 1 | 13 |
| slc25a10 | 0.12258 | -0.15444886 | 0.059 | 0.3 | 1 | 13 |
| nop53 | 0.12272 | -0.21855267 | 0.118 | 0.4 | 1 | 13 |
| ncalda | 0.12283 | -0.344809 | 0.059 | 0.3 | 1 | 13 |
| reep5 | 0.12351 | -0.29565776 | 0.275 | 0.6 | 1 | 13 |
| slc38a2 | 0.12386 | -0.26119703 | 0.059 | 0.3 | 1 | 13 |
| zfand6 | 0.12393 | -0.18034838 | 0.059 | 0.3 | 1 | 13 |
| eif3m | 0.12395 | -0.17982833 | 0.392 | 0.7 | 1 | 13 |
| sorl1 | 0.12422 | -0.28136422 | 0.059 | 0.3 | 1 | 13 |
| march5l | 0.1249 | -0.16630708 | 0.059 | 0.3 | 1 | 13 |
| dctd | 0.12519 | -0.21633878 | 0.059 | 0.3 | 1 | 13 |
| rgl2 | 0.12546 | -0.16711969 | 0.078 | 0.3 | 1 | 13 |
| kif21b | 0.12546 | -0.33387417 | 0.059 | 0.3 | 1 | 13 |
| ndufb5 | 0.12577 | -0.12653314 | 0.059 | 0.3 | 1 | 13 |
| sucla2 | 0.12579 | -0.28347597 | 0.059 | 0.3 | 1 | 13 |
| rac2 | 0.12602 | -0.15878261 | 0.784 | 1 | 1 | 13 |
| ca2 | 0.12614 | -0.25215599 | 0.059 | 0.3 | 1 | 13 |
| FP236734.1 | 0.12657 | -0.32406749 | 0.059 | 0.3 | 1 | 13 |
| mcm4 | 0.1268 | -0.22976853 | 0.059 | 0.3 | 1 | 13 |
| zbtb10 | 0.12742 | -0.36541588 | 0.059 | 0.3 | 1 | 13 |
| dctn3 | 0.12746 | -0.2493169 | 0.059 | 0.3 | 1 | 13 |
| pkn1b | 0.12747 | -0.33689862 | 0.059 | 0.3 | 1 | 13 |
| ypel5 | 0.1275 | -0.22172653 | 0.059 | 0.3 | 1 | 13 |
| laptm5 | 0.1275 | -0.1327942 | 0.549 | 0.8 | 1 | 13 |
| wdr83os | 0.12753 | -0.310873 | 0.059 | 0.3 | 1 | 13 |
| zbtb8os | 0.12753 | -0.22206308 | 0.059 | 0.3 | 1 | 13 |
| rps23 | 0.12856 | -0.40462071 | 0.588 | 0.9 | 1 | 13 |
| h2afvb | 0.12936 | 0.364976233 | 0.549 | 0.5 | 1 | 13 |
| prdx6 | 0.12965 | -0.24822169 | 0.118 | 0.4 | 1 | 13 |
| tbca | 0.12995 | -0.20589784 | 0.118 | 0.4 | 1 | 13 |
| bri3bp | 0.13083 | -0.21701633 | 0.137 | 0.4 | 1 | 13 |
| ralbp1 | 0.13167 | -0.17018127 | 0.137 | 0.4 | 1 | 13 |
| tma7 | 0.13319 | -0.23955354 | 0.373 | 0.7 | 1 | 13 |
| rps9 | 0.13385 | -0.28218433 | 0.608 | 0.9 | 1 | 13 |
| erbin | 0.13525 | -0.1337131 | 0.078 | 0.3 | 1 | 13 |
| tmed5 | 0.13525 | -0.30863744 | 0.118 | 0.4 | 1 | 13 |
| rasgrp4 | 0.13544 | -0.17445993 | 0.137 | 0.4 | 1 | 13 |
| anapc16 | 0.13655 | 0.100160994 | 0.137 | 0.3 | 1 | 13 |
| hmgn2 | 0.13686 | -0.23185229 | 0.627 | 0.9 | 1 | 13 |

| | | | | | | |
|---|---|---|---|---|---|---|
| camk2g2 | 0.13787 | -0.1962831 | 0.216 | 0.5 | 1 | 13 |
| csnk2b | 0.1381 | -0.16974273 | 0.196 | 0.5 | 1 | 13 |
| ndufb6 | 0.13853 | -0.14030631 | 0.137 | 0.4 | 1 | 13 |
| hnrnpa0a | 0.1386 | -0.13448322 | 0.137 | 0.4 | 1 | 13 |
| mmp25b | 0.13863 | -0.14913306 | 0.078 | 0.3 | 1 | 13 |
| hdlbpa | 0.14022 | 0.174951062 | 0.451 | 0.6 | 1 | 13 |
| mlx | 0.14066 | -0.14273378 | 0.137 | 0.4 | 1 | 13 |
| rps2 | 0.14072 | -0.1085176 | 0.569 | 0.8 | 1 | 13 |
| CR381673.1 | 0.14158 | -0.17799991 | 0.137 | 0.4 | 1 | 13 |
| samsn1b | 0.14175 | -0.10335387 | 0.176 | 0.4 | 1 | 13 |
| gabarapa | 0.14216 | -0.23305792 | 0.196 | 0.5 | 1 | 13 |
| alyref | 0.14238 | -0.16235919 | 0.137 | 0.4 | 1 | 13 |
| rpl37 | 0.14244 | -0.42461589 | 0.627 | 0.8 | 1 | 13 |
| selenof | 0.14363 | 0.29768359 | 0.686 | 0.7 | 1 | 13 |
| cd7al | 0.14567 | -0.12344263 | 0.333 | 0.6 | 1 | 13 |
| klf7b | 0.14607 | 0.22870149 | 0.137 | 0 | 1 | 13 |
| pik3r1 | 0.14607 | 0.186423284 | 0.137 | 0 | 1 | 13 |
| gm2a | 0.14607 | 0.19441115 | 0.137 | 0 | 1 | 13 |
| itga1 | 0.14607 | 0.220843087 | 0.137 | 0 | 1 | 13 |
| usp8 | 0.14607 | 0.213717747 | 0.137 | 0 | 1 | 13 |
| zc3h7bb | 0.14607 | 0.238733765 | 0.137 | 0 | 1 | 13 |
| adcy7 | 0.14607 | 0.21566062 | 0.137 | 0 | 1 | 13 |
| asph | 0.14607 | 0.207042375 | 0.137 | 0 | 1 | 13 |
| slmapa | 0.14607 | 0.179689601 | 0.137 | 0 | 1 | 13 |
| tnrc6a | 0.14607 | 0.212977286 | 0.137 | 0 | 1 | 13 |
| ubap2b | 0.14607 | 0.19306542 | 0.137 | 0 | 1 | 13 |
| atp2b1a | 0.14607 | 0.288111466 | 0.137 | 0 | 1 | 13 |
| camkk1a | 0.14607 | 0.325552554 | 0.137 | 0 | 1 | 13 |
| dock11 | 0.14607 | 0.21209885 | 0.137 | 0 | 1 | 13 |
| hk2 | 0.14607 | 0.304704425 | 0.137 | 0 | 1 | 13 |
| march7 | 0.14607 | 0.265235131 | 0.137 | 0 | 1 | 13 |
| rnaset2 | 0.14607 | 0.252009881 | 0.137 | 0 | 1 | 13 |
| rps27l | 0.14607 | 0.212663783 | 0.137 | 0 | 1 | 13 |
| sf3b1 | 0.14607 | 0.266818379 | 0.137 | 0 | 1 | 13 |
| trim25 | 0.14607 | 0.233016313 | 0.137 | 0 | 1 | 13 |
| ilk | 0.14607 | 0.26819286 | 0.137 | 0 | 1 | 13 |
| rffl | 0.14607 | 0.169303282 | 0.137 | 0 | 1 | 13 |
| sh3kbp1 | 0.1477 | -0.24051773 | 0.294 | 0.6 | 1 | 13 |
| btg1 | 0.14794 | -0.46667502 | 0.275 | 0.6 | 1 | 13 |
| mt-co2 | 0.14812 | -0.22388956 | 0.451 | 0.7 | 1 | 13 |
| tprg1 | 0.1489 | -0.19118167 | 0.137 | 0.4 | 1 | 13 |
| arpc1b | 0.14934 | -0.10073463 | 0.824 | 1 | 1 | 13 |
| rps13 | 0.14974 | -0.19969722 | 0.784 | 1 | 1 | 13 |
| tubb4b | 0.14993 | 0.626759776 | 0.49 | 0.3 | 1 | 13 |
| dbi | 0.15014 | -0.38280243 | 0.608 | 0.9 | 1 | 13 |
| lcp2a | 0.15014 | -0.21075435 | 0.137 | 0.4 | 1 | 13 |
| slc6a6b | 0.15015 | 0.145880842 | 0.373 | 0.5 | 1 | 13 |
| uqcr10 | 0.1507 | 0.386554596 | 0.255 | 0.3 | 1 | 13 |

| | | | | | | |
|---|---|---|---|---|---|---|
| lman1 | 0.1518 | -0.10678676 | 0.078 | 0.3 | 1 | 13 |
| rab11fip1b | 0.15358 | -0.19368368 | 0.216 | 0.5 | 1 | 13 |
| rpl8 | 0.1536 | -0.22561745 | 0.627 | 0.9 | 1 | 13 |
| txnipa | 0.15372 | -0.21257071 | 0.392 | 0.7 | 1 | 13 |
| ube2v1 | 0.15416 | -0.14829137 | 0.314 | 0.6 | 1 | 13 |
| sh3bgrl3 | 0.15541 | -0.128813 | 0.235 | 0.5 | 1 | 13 |
| atp5if1b | 0.15726 | -0.40783121 | 0.373 | 0.7 | 1 | 13 |
| CT027744.2 | 0.15729 | -0.1619518 | 0.078 | 0.3 | 1 | 13 |
| psma6b | 0.15732 | -0.19862048 | 0.137 | 0.4 | 1 | 13 |
| atp5f1e | 0.15771 | -0.2637894 | 0.49 | 0.8 | 1 | 13 |
| tpm1 | 0.15774 | -0.15528174 | 0.216 | 0.5 | 1 | 13 |
| si:dkey-276j7.1 | 0.1582 | -0.10120664 | 0.078 | 0.3 | 1 | 13 |
| fkbp3 | 0.16027 | 0.153930119 | 0.216 | 0.4 | 1 | 13 |
| irf2 | 0.1609 | -0.11024688 | 0.157 | 0.4 | 1 | 13 |
| ssr1 | 0.16169 | 0.288320497 | 0.529 | 0.6 | 1 | 13 |
| actn1 | 0.1617 | -0.14513334 | 0.235 | 0.5 | 1 | 13 |
| prdx5 | 0.16208 | 0.182823412 | 0.373 | 0.5 | 1 | 13 |
| atp5meb | 0.16358 | -0.16698539 | 0.235 | 0.5 | 1 | 13 |
| sh3glb1b | 0.1639 | -0.17903714 | 0.078 | 0.3 | 1 | 13 |
| arpc4l | 0.16433 | -0.1377699 | 0.333 | 0.6 | 1 | 13 |
| lpgat1 | 0.16503 | -0.13952287 | 0.078 | 0.3 | 1 | 13 |
| stim1a | 0.16627 | -0.24012434 | 0.137 | 0.4 | 1 | 13 |
| stk17b | 0.16663 | 0.134122735 | 0.353 | 0.5 | 1 | 13 |
| zbtb20 | 0.16712 | -0.24827672 | 0.314 | 0.6 | 1 | 13 |
| kpnb3 | 0.1674 | -0.22197668 | 0.078 | 0.3 | 1 | 13 |
| si:dkey-67c22.2 | 0.16812 | -0.15927564 | 0.078 | 0.3 | 1 | 13 |
| rrbp1a | 0.1688 | 0.525418871 | 0.294 | 0.1 | 1 | 13 |
| si:ch211-260e23.9 | 0.17001 | -0.10589336 | 0.039 | 0.2 | 1 | 13 |
| arhgap15 | 0.1712 | -0.15416394 | 0.078 | 0.3 | 1 | 13 |
| ier5 | 0.17326 | -0.10021716 | 0.176 | 0.4 | 1 | 13 |
| si:ch73-234b20.5 | 0.17503 | -0.31568923 | 0.294 | 0.6 | 1 | 13 |
| mlf2 | 0.17556 | -0.15093598 | 0.078 | 0.3 | 1 | 13 |
| cox6a2 | 0.17718 | -0.18433004 | 0.137 | 0.4 | 1 | 13 |
| LOC558761 | 0.17731 | -0.2337292 | 0.078 | 0.3 | 1 | 13 |
| mrpl20 | 0.17756 | -0.13728514 | 0.078 | 0.3 | 1 | 13 |
| tpd52l2b | 0.17852 | 0.161348256 | 0.157 | 0.3 | 1 | 13 |
| aldh18a1 | 0.17894 | -0.14403315 | 0.078 | 0.3 | 1 | 13 |
| si:dkeyp-66d1.7 | 0.17914 | 0.504198361 | 0.451 | 0.3 | 1 | 13 |
| vps4b | 0.17968 | -0.11588105 | 0.157 | 0.4 | 1 | 13 |
| si:dkey-78l4.14 | 0.18056 | -0.12253439 | 0.078 | 0.3 | 1 | 13 |
| snap23.1 | 0.18064 | -0.24026304 | 0.137 | 0.4 | 1 | 13 |
| nus1 | 0.18194 | -0.21976404 | 0.078 | 0.3 | 1 | 13 |
| higd2a | 0.18208 | -0.1991049 | 0.137 | 0.4 | 1 | 13 |
| si:ch211-214p16.2 | 0.18237 | -0.26712608 | 0.824 | 1 | 1 | 13 |
| vcp | 0.18315 | -0.22322615 | 0.216 | 0.5 | 1 | 13 |
| snrpf | 0.18454 | -0.29724498 | 0.294 | 0.6 | 1 | 13 |
| tmx1 | 0.1846 | -0.26933347 | 0.216 | 0.5 | 1 | 13 |
| plxnc1 | 0.18534 | 0.301909114 | 0.294 | 0.3 | 1 | 13 |

| | | | | | | |
|---|---|---|---|---|---|---|
| scn4bb | 0.18668 | 0.2167808 | 0.118 | 0 | 1 | 13 |
| si:ch73-359m17.5 | 0.18668 | 0.191804504 | 0.118 | 0 | 1 | 13 |
| ddx3xa | 0.18668 | 0.211167692 | 0.118 | 0 | 1 | 13 |
| zgc:194312 | 0.18668 | 0.161549506 | 0.118 | 0 | 1 | 13 |
| FQ976913.1 | 0.18668 | 0.152109397 | 0.118 | 0 | 1 | 13 |
| SLA (1 of many) | 0.18668 | 0.24006983 | 0.118 | 0 | 1 | 13 |
| arl2 | 0.18668 | 0.194211837 | 0.118 | 0 | 1 | 13 |
| cltb | 0.18668 | 0.23855581 | 0.118 | 0 | 1 | 13 |
| cmbl | 0.18668 | 0.130137316 | 0.118 | 0 | 1 | 13 |
| cnot1 | 0.18668 | 0.282750446 | 0.118 | 0 | 1 | 13 |
| dicer1 | 0.18668 | 0.190669772 | 0.118 | 0 | 1 | 13 |
| dnaja2b | 0.18668 | 0.268022678 | 0.118 | 0 | 1 | 13 |
| fzr1a | 0.18668 | 0.253628443 | 0.118 | 0 | 1 | 13 |
| gsk3b | 0.18668 | 0.303282803 | 0.118 | 0 | 1 | 13 |
| hspd1 | 0.18668 | 0.248122844 | 0.118 | 0 | 1 | 13 |
| itpkb | 0.18668 | 0.205284615 | 0.118 | 0 | 1 | 13 |
| kcnk5b | 0.18668 | 0.321044496 | 0.118 | 0 | 1 | 13 |
| kpna2 | 0.18668 | 0.22144652 | 0.118 | 0 | 1 | 13 |
| lpar5b | 0.18668 | 0.298558729 | 0.118 | 0 | 1 | 13 |
| mycbp2 | 0.18668 | 0.164944247 | 0.118 | 0 | 1 | 13 |
| nfil3 | 0.18668 | 0.183701452 | 0.118 | 0 | 1 | 13 |
| pcmt | 0.18668 | 0.195417103 | 0.118 | 0 | 1 | 13 |
| pdcl | 0.18668 | 0.247917124 | 0.118 | 0 | 1 | 13 |
| scarb2a | 0.18668 | 0.308403465 | 0.118 | 0 | 1 | 13 |
| srrm2 | 0.18668 | 0.152148332 | 0.118 | 0 | 1 | 13 |
| tpd52 | 0.18668 | 0.205570624 | 0.118 | 0 | 1 | 13 |
| ubtd2 | 0.18668 | 0.231553341 | 0.118 | 0 | 1 | 13 |
| ugcg | 0.18668 | 0.213201642 | 0.118 | 0 | 1 | 13 |
| unm-hu7910 | 0.18668 | 0.265866849 | 0.118 | 0 | 1 | 13 |
| zgc:92606 | 0.18668 | 0.216233704 | 0.118 | 0 | 1 | 13 |
| dpm1 | 0.18668 | 0.157542522 | 0.118 | 0 | 1 | 13 |
| kmt2bb | 0.18668 | 0.2222537 | 0.118 | 0 | 1 | 13 |
| luc7l | 0.18668 | 0.218901597 | 0.118 | 0 | 1 | 13 |
| rab1bb | 0.18668 | 0.241890668 | 0.118 | 0 | 1 | 13 |
| sf3a3 | 0.18668 | 0.222425396 | 0.118 | 0 | 1 | 13 |
| bysl | 0.18668 | 0.219290219 | 0.118 | 0 | 1 | 13 |
| cox7a2a | 0.18712 | -0.19930777 | 0.314 | 0.6 | 1 | 13 |
| u2surp | 0.18859 | -0.25662937 | 0.078 | 0.3 | 1 | 13 |
| p2ry10 | 0.18903 | -0.3637591 | 0.137 | 0.4 | 1 | 13 |
| hectd3 | 0.18912 | -0.17223943 | 0.078 | 0.3 | 1 | 13 |
| wipf1a | 0.18951 | -0.29512159 | 0.137 | 0.4 | 1 | 13 |
| ndufb7 | 0.19075 | -0.27960564 | 0.294 | 0.6 | 1 | 13 |
| CABZ01115113.1 | 0.19226 | -0.28030301 | 0.137 | 0.4 | 1 | 13 |
| psmb7 | 0.19234 | -0.28909342 | 0.137 | 0.4 | 1 | 13 |
| chac1 | 0.19244 | 0.227334136 | 0.392 | 0.5 | 1 | 13 |
| got2b | 0.19274 | -0.14882077 | 0.078 | 0.3 | 1 | 13 |
| nhp2 | 0.19291 | -0.30715683 | 0.137 | 0.4 | 1 | 13 |
| nmi | 0.19305 | -0.30595834 | 0.137 | 0.4 | 1 | 13 |

| | | | | | | |
|---|---|---|---|---|---|---|
| ches1 | 0.19351 | -0.15712337 | 0.039 | 0.2 | 1 | 13 |
| dub | 0.19393 | -0.30762013 | 0.216 | 0.5 | 1 | 13 |
| chchd7 | 0.19426 | -0.19106538 | 0.078 | 0.3 | 1 | 13 |
| mrps25 | 0.19476 | -0.19316472 | 0.078 | 0.3 | 1 | 13 |
| erp44 | 0.19496 | -0.28515206 | 0.216 | 0.5 | 1 | 13 |
| madd | 0.19519 | -0.10644652 | 0.039 | 0.2 | 1 | 13 |
| selp | 0.19528 | -0.26233933 | 0.078 | 0.3 | 1 | 13 |
| churc1 | 0.19615 | -0.19797383 | 0.078 | 0.3 | 1 | 13 |
| psmb3 | 0.19645 | -0.14942223 | 0.157 | 0.4 | 1 | 13 |
| mob4 | 0.19749 | -0.15703591 | 0.078 | 0.3 | 1 | 13 |
| tnpo3 | 0.1978 | -0.11482357 | 0.039 | 0.2 | 1 | 13 |
| vdac2 | 0.1983 | -0.11810599 | 0.235 | 0.5 | 1 | 13 |
| capza1a | 0.19833 | 0.347083119 | 0.392 | 0.4 | 1 | 13 |
| psma6a | 0.19919 | -0.14747055 | 0.078 | 0.3 | 1 | 13 |
| lig1 | 0.19942 | -0.29118686 | 0.078 | 0.3 | 1 | 13 |
| rpl15 | 0.2003 | -0.40332268 | 0.647 | 0.9 | 1 | 13 |
| si:dkeyp-87d8.8 | 0.20057 | -0.10315113 | 0.039 | 0.2 | 1 | 13 |
| dlst | 0.20117 | -0.25595054 | 0.078 | 0.3 | 1 | 13 |
| lrrfip1a | 0.20181 | -0.22549569 | 0.314 | 0.6 | 1 | 13 |
| ccdc167 | 0.20181 | -0.22929875 | 0.078 | 0.3 | 1 | 13 |
| nop10 | 0.20182 | -0.22575005 | 0.078 | 0.3 | 1 | 13 |
| nucks1a | 0.20239 | -0.28273953 | 0.078 | 0.3 | 1 | 13 |
| rab3ab | 0.20241 | -0.25932794 | 0.078 | 0.3 | 1 | 13 |
| clic1 | 0.20268 | -0.23696485 | 0.412 | 0.7 | 1 | 13 |
| sar1b | 0.20322 | -0.25427888 | 0.078 | 0.3 | 1 | 13 |
| rin3 | 0.20379 | -0.29000337 | 0.078 | 0.3 | 1 | 13 |
| arhgef3l | 0.20399 | -0.28562022 | 0.078 | 0.3 | 1 | 13 |
| mxi1 | 0.20408 | -0.29075347 | 0.078 | 0.3 | 1 | 13 |
| cbfa2t3 | 0.20448 | 0.447656048 | 0.275 | 0.2 | 1 | 13 |
| ogt.1 | 0.20452 | -0.2059466 | 0.078 | 0.3 | 1 | 13 |
| rpl10a | 0.20498 | -0.16129423 | 0.569 | 0.8 | 1 | 13 |
| tpm3 | 0.20817 | 0.113982492 | 0.588 | 0.7 | 1 | 13 |
| ctss1 | 0.20967 | -0.18732689 | 0.157 | 0.4 | 1 | 13 |
| zgc:63831 | 0.21105 | -0.13126274 | 0.353 | 0.6 | 1 | 13 |
| pgd | 0.21204 | 0.467609185 | 0.353 | 0.1 | 1 | 13 |
| camk1b | 0.21238 | -0.11315568 | 0.176 | 0.4 | 1 | 13 |
| actb2 | 0.2128 | 0.100105819 | 0.98 | 1 | 1 | 13 |
| git2b | 0.21436 | -0.10769903 | 0.039 | 0.2 | 1 | 13 |
| anks1b | 0.2144 | -0.13268424 | 0.157 | 0.4 | 1 | 13 |
| atp6v1g1 | 0.21701 | -0.14541028 | 0.235 | 0.5 | 1 | 13 |
| atp2a3 | 0.21777 | -0.13532758 | 0.353 | 0.6 | 1 | 13 |
| limd2 | 0.21851 | -0.16069797 | 0.451 | 0.7 | 1 | 13 |
| rps4x | 0.21919 | -0.1320659 | 0.686 | 0.9 | 1 | 13 |
| hnrnpa0b | 0.2224 | -0.11926559 | 0.353 | 0.6 | 1 | 13 |
| anxa11a | 0.22247 | -0.19524121 | 0.157 | 0.4 | 1 | 13 |
| skap2 | 0.2231 | -0.20152742 | 0.235 | 0.5 | 1 | 13 |
| hmga1a | 0.22314 | 0.324503083 | 0.451 | 0.5 | 1 | 13 |
| pacsin2 | 0.22355 | -0.13646987 | 0.098 | 0.3 | 1 | 13 |

| | | | | | | |
|---|---|---|---|---|---|---|
| ptges3b | 0.22561 | -0.14950925 | 0.157 | 0.4 | 1 | 13 |
| sept2 | 0.22574 | -0.1388884 | 0.098 | 0.3 | 1 | 13 |
| sf3b6 | 0.22638 | -0.1695124 | 0.157 | 0.4 | 1 | 13 |
| arf5 | 0.22806 | -0.27962068 | 0.314 | 0.6 | 1 | 13 |
| ccng2 | 0.22946 | -0.17410008 | 0.039 | 0.2 | 1 | 13 |
| fmr1 | 0.22981 | -0.10662646 | 0.039 | 0.2 | 1 | 13 |
| smox | 0.23212 | -0.22486038 | 0.157 | 0.4 | 1 | 13 |
| fth1a | 0.23255 | -0.110721 | 0.373 | 0.6 | 1 | 13 |
| atp5md | 0.23262 | 0.242028477 | 0.412 | 0.5 | 1 | 13 |
| romo1 | 0.23343 | 0.1983806 | 0.314 | 0.4 | 1 | 13 |
| rps6 | 0.23458 | -0.15422984 | 0.686 | 0.9 | 1 | 13 |
| si:dkey-79d12.5 | 0.23494 | -0.16419546 | 0.039 | 0.2 | 1 | 13 |
| ccdc32 | 0.23514 | -0.16464447 | 0.039 | 0.2 | 1 | 13 |
| ap1ar | 0.23527 | -0.1249239 | 0.039 | 0.2 | 1 | 13 |
| apbb1ip | 0.2354 | -0.14696732 | 0.039 | 0.2 | 1 | 13 |
| ostc | 0.23587 | -0.31145805 | 0.412 | 0.7 | 1 | 13 |
| timm23a | 0.23656 | 0.10185495 | 0.059 | 0.2 | 1 | 13 |
| foxj3 | 0.2372 | -0.10244549 | 0.039 | 0.2 | 1 | 13 |
| si:ch211-194m7.8 | 0.23762 | 0.193017127 | 1 | 1 | 1 | 13 |
| hnrnpa1b | 0.23792 | -0.24444925 | 0.157 | 0.4 | 1 | 13 |
| glyr1 | 0.23817 | -0.12035263 | 0.098 | 0.3 | 1 | 13 |
| LOC101884098 | 0.2389 | 0.167041475 | 0.098 | 0 | 1 | 13 |
| TMEM19 | 0.2389 | 0.1593252 | 0.098 | 0 | 1 | 13 |
| abca2 | 0.2389 | 0.248960304 | 0.098 | 0 | 1 | 13 |
| acsbg2 | 0.2389 | 0.18671576 | 0.098 | 0 | 1 | 13 |
| agla | 0.2389 | 0.16547911 | 0.098 | 0 | 1 | 13 |
| ap2b1 | 0.2389 | 0.175448775 | 0.098 | 0 | 1 | 13 |
| arih1 | 0.2389 | 0.156694562 | 0.098 | 0 | 1 | 13 |
| atg5 | 0.2389 | 0.236352258 | 0.098 | 0 | 1 | 13 |
| atp6v1h | 0.2389 | 0.214622226 | 0.098 | 0 | 1 | 13 |
| bahcc1b | 0.2389 | 0.227003254 | 0.098 | 0 | 1 | 13 |
| brd1b | 0.2389 | 0.126217427 | 0.098 | 0 | 1 | 13 |
| c1d | 0.2389 | 0.125030884 | 0.098 | 0 | 1 | 13 |
| cacybp | 0.2389 | 0.184321791 | 0.098 | 0 | 1 | 13 |
| caprin1a | 0.2389 | 0.205830614 | 0.098 | 0 | 1 | 13 |
| cept1a | 0.2389 | 0.169882493 | 0.098 | 0 | 1 | 13 |
| cherp | 0.2389 | 0.212632276 | 0.098 | 0 | 1 | 13 |
| crcp | 0.2389 | 0.190567841 | 0.098 | 0 | 1 | 13 |
| crebbpb | 0.2389 | 0.195564618 | 0.098 | 0 | 1 | 13 |
| eif4e1c | 0.2389 | 0.166553804 | 0.098 | 0 | 1 | 13 |
| fhit | 0.2389 | 0.148790959 | 0.098 | 0 | 1 | 13 |
| ggps1 | 0.2389 | 0.161387849 | 0.098 | 0 | 1 | 13 |
| ginm1 | 0.2389 | 0.20615712 | 0.098 | 0 | 1 | 13 |
| gsto2 | 0.2389 | 0.21732631 | 0.098 | 0 | 1 | 13 |
| gucd1 | 0.2389 | 0.175243164 | 0.098 | 0 | 1 | 13 |
| hcfc1b | 0.2389 | 0.158475038 | 0.098 | 0 | 1 | 13 |
| il4r.1 | 0.2389 | 0.215893518 | 0.098 | 0 | 1 | 13 |
| impa1 | 0.2389 | 0.184861329 | 0.098 | 0 | 1 | 13 |

| | | | | | | |
|---|---|---|---|---|---|---|
| lrba | 0.2389 | 0.149962306 | 0.098 | 0 | 1 | 13 |
| mast2 | 0.2389 | 0.213343 | 0.098 | 0 | 1 | 13 |
| mknk1 | 0.2389 | 0.144929652 | 0.098 | 0 | 1 | 13 |
| mmp28 | 0.2389 | 0.177402818 | 0.098 | 0 | 1 | 13 |
| npm3 | 0.2389 | 0.183837875 | 0.098 | 0 | 1 | 13 |
| nutf2l | 0.2389 | 0.135208081 | 0.098 | 0 | 1 | 13 |
| pcm1 | 0.2389 | 0.191680207 | 0.098 | 0 | 1 | 13 |
| pfdn5 | 0.2389 | 0.15045594 | 0.098 | 0 | 1 | 13 |
| plscr3b | 0.2389 | 0.265851819 | 0.098 | 0 | 1 | 13 |
| prkar1ab | 0.2389 | 0.134369271 | 0.098 | 0 | 1 | 13 |
| prmt1 | 0.2389 | 0.114870046 | 0.098 | 0 | 1 | 13 |
| rbm7 | 0.2389 | 0.14695817 | 0.098 | 0 | 1 | 13 |
| rnf121 | 0.2389 | 0.112578874 | 0.098 | 0 | 1 | 13 |
| sept7b | 0.2389 | 0.157693783 | 0.098 | 0 | 1 | 13 |
| shoc2 | 0.2389 | 0.189350723 | 0.098 | 0 | 1 | 13 |
| si:ch211-119e14.1 | 0.2389 | 0.189229675 | 0.098 | 0 | 1 | 13 |
| si:ch211-11k18.4 | 0.2389 | 0.185939972 | 0.098 | 0 | 1 | 13 |
| si:ch211-162e15.3 | 0.2389 | 0.136313402 | 0.098 | 0 | 1 | 13 |
| si:ch211-212d10.2 | 0.2389 | 0.160882393 | 0.098 | 0 | 1 | 13 |
| si:dkey-159f12.2 | 0.2389 | 0.191061572 | 0.098 | 0 | 1 | 13 |
| si:dkey-46a10.3 | 0.2389 | 0.168276687 | 0.098 | 0 | 1 | 13 |
| si:dkeyp-80c12.4 | 0.2389 | 0.194864599 | 0.098 | 0 | 1 | 13 |
| snrnp27 | 0.2389 | 0.14397272 | 0.098 | 0 | 1 | 13 |
| szrd1 | 0.2389 | 0.171873588 | 0.098 | 0 | 1 | 13 |
| tm9sf1 | 0.2389 | 0.227508381 | 0.098 | 0 | 1 | 13 |
| tmub1 | 0.2389 | 0.162682267 | 0.098 | 0 | 1 | 13 |
| tob1b | 0.2389 | 0.217743584 | 0.098 | 0 | 1 | 13 |
| tph1b | 0.2389 | 0.232938588 | 0.098 | 0 | 1 | 13 |
| tspan36 | 0.2389 | 0.115268869 | 0.098 | 0 | 1 | 13 |
| txndc12 | 0.2389 | 0.147276286 | 0.098 | 0 | 1 | 13 |
| ubxn4 | 0.2389 | 0.162056793 | 0.098 | 0 | 1 | 13 |
| uck1 | 0.2389 | 0.203991065 | 0.098 | 0 | 1 | 13 |
| usf1 | 0.2389 | 0.107255827 | 0.098 | 0 | 1 | 13 |
| vbp1 | 0.2389 | 0.125932016 | 0.098 | 0 | 1 | 13 |
| wnk1b | 0.2389 | 0.166628043 | 0.098 | 0 | 1 | 13 |
| zfand5a | 0.2389 | 0.178587007 | 0.098 | 0 | 1 | 13 |
| zgc:158564 | 0.2389 | 0.13182087 | 0.098 | 0 | 1 | 13 |
| LOC110438407 | 0.2389 | 0.225482592 | 0.098 | 0 | 1 | 13 |
| acot13 | 0.2389 | 0.16968099 | 0.098 | 0 | 1 | 13 |
| gigyf1b | 0.2389 | 0.170040539 | 0.098 | 0 | 1 | 13 |
| pno1 | 0.2389 | 0.164316288 | 0.098 | 0 | 1 | 13 |
| tmem106bb | 0.2389 | 0.177390824 | 0.098 | 0 | 1 | 13 |
| ttc7b | 0.2389 | 0.264212633 | 0.098 | 0 | 1 | 13 |
| vdac1 | 0.2389 | 0.15959766 | 0.098 | 0 | 1 | 13 |
| ctdsp1 | 0.2389 | 0.223844489 | 0.098 | 0 | 1 | 13 |
| sept6 | 0.23948 | 0.163955722 | 0.294 | 0.4 | 1 | 13 |
| cct4 | 0.23976 | -0.16237193 | 0.176 | 0.4 | 1 | 13 |
| usp36 | 0.24092 | -0.12447465 | 0.098 | 0.3 | 1 | 13 |

| | | | | | | |
|---|---|---|---|---|---|---|
| zgc:77486 | 0.24324 | -0.17561276 | 0.039 | 0.2 | 1 | 13 |
| irf2bp2a | 0.24327 | -0.15686424 | 0.255 | 0.5 | 1 | 13 |
| LOC103910871 | 0.24482 | -0.12935671 | 0.039 | 0.2 | 1 | 13 |
| proser1 | 0.24496 | -0.12878099 | 0.039 | 0.2 | 1 | 13 |
| wasa | 0.24503 | -0.10570691 | 0.373 | 0.6 | 1 | 13 |
| plcg2 | 0.24547 | -0.25237669 | 0.157 | 0.4 | 1 | 13 |
| shc1 | 0.24556 | -0.2296456 | 0.039 | 0.2 | 1 | 13 |
| tm9sf2 | 0.24579 | -0.25546081 | 0.235 | 0.5 | 1 | 13 |
| snrpe | 0.2458 | 0.158563453 | 0.176 | 0.3 | 1 | 13 |
| ddx39ab | 0.24618 | 0.140754504 | 0.176 | 0.3 | 1 | 13 |
| cox6a1 | 0.2463 | -0.19285026 | 0.431 | 0.7 | 1 | 13 |
| unc93b1 | 0.24724 | -0.16603346 | 0.039 | 0.2 | 1 | 13 |
| nedd9 | 0.25037 | -0.14023301 | 0.039 | 0.2 | 1 | 13 |
| myh11a | 0.25072 | 0.18972675 | 0.667 | 0.7 | 1 | 13 |
| bin2b | 0.25094 | -0.28823067 | 0.157 | 0.4 | 1 | 13 |
| yeats4 | 0.25185 | -0.25248534 | 0.039 | 0.2 | 1 | 13 |
| ranbp1 | 0.25186 | 0.334588153 | 0.275 | 0.3 | 1 | 13 |
| rpl18 | 0.25345 | -0.12295109 | 0.706 | 0.9 | 1 | 13 |
| etf1b | 0.25362 | -0.10599175 | 0.098 | 0.3 | 1 | 13 |
| pcca | 0.25387 | -0.12242165 | 0.098 | 0.3 | 1 | 13 |
| syt14b | 0.25388 | -0.11949878 | 0.039 | 0.2 | 1 | 13 |
| gfi1ab | 0.25394 | -0.27148276 | 0.235 | 0.5 | 1 | 13 |
| eif4ebp1 | 0.25399 | -0.16058969 | 0.451 | 0.7 | 1 | 13 |
| fthl27 | 0.25458 | -0.35294978 | 0.157 | 0.4 | 1 | 13 |
| wu:fc33b09 | 0.25518 | -0.26614018 | 0.039 | 0.2 | 1 | 13 |
| taf12 | 0.25567 | -0.10207647 | 0.039 | 0.2 | 1 | 13 |
| serpinb1l3 | 0.25587 | -0.14118284 | 0.176 | 0.4 | 1 | 13 |
| trappc9 | 0.25626 | -0.11851368 | 0.039 | 0.2 | 1 | 13 |
| CABZ01087514.1 | 0.25701 | -0.16980632 | 0.039 | 0.2 | 1 | 13 |
| ppp1r13bb | 0.25736 | -0.13098168 | 0.039 | 0.2 | 1 | 13 |
| rnf175 | 0.25738 | -0.12605566 | 0.039 | 0.2 | 1 | 13 |
| sik1 | 0.25805 | -0.10925889 | 0.039 | 0.2 | 1 | 13 |
| itm2bb | 0.25821 | -0.14525422 | 0.451 | 0.7 | 1 | 13 |
| birc6 | 0.25915 | -0.17384954 | 0.039 | 0.2 | 1 | 13 |
| letm1 | 0.25949 | -0.19823019 | 0.098 | 0.3 | 1 | 13 |
| zmynd11 | 0.25951 | -0.13312131 | 0.039 | 0.2 | 1 | 13 |
| vat1 | 0.26011 | 0.411369252 | 0.255 | 0.2 | 1 | 13 |
| smu1a | 0.26045 | -0.1220841 | 0.039 | 0.2 | 1 | 13 |
| traf7 | 0.261 | -0.17383956 | 0.098 | 0.3 | 1 | 13 |
| igf2bp2b | 0.26159 | -0.15791575 | 0.039 | 0.2 | 1 | 13 |
| add1 | 0.26176 | -0.12783896 | 0.039 | 0.2 | 1 | 13 |
| b2ml | 0.26185 | 0.114794291 | 0.373 | 0.5 | 1 | 13 |
| hm13 | 0.26283 | -0.12605383 | 0.275 | 0.5 | 1 | 13 |
| arhgap27 | 0.26365 | -0.12571615 | 0.039 | 0.2 | 1 | 13 |
| aph1b | 0.2647 | -0.13770489 | 0.039 | 0.2 | 1 | 13 |
| ptpn6 | 0.26477 | 0.138566873 | 0.49 | 0.6 | 1 | 13 |
| psmd12 | 0.26524 | -0.13116627 | 0.039 | 0.2 | 1 | 13 |
| arhgap36 | 0.26555 | -0.17050053 | 0.039 | 0.2 | 1 | 13 |

| | | | | | | |
|---|---|---|---|---|---|---|
| cep170aa | 0.26667 | -0.21383939 | 0.039 | 0.2 | 1 | 13 |
| nudt21 | 0.26699 | -0.13735916 | 0.039 | 0.2 | 1 | 13 |
| MZT1 | 0.26712 | -0.10694323 | 0.039 | 0.2 | 1 | 13 |
| tgfbr2a | 0.26731 | -0.12045825 | 0.039 | 0.2 | 1 | 13 |
| cops5 | 0.26755 | -0.13475697 | 0.039 | 0.2 | 1 | 13 |
| wipf1b | 0.26793 | -0.29678582 | 0.039 | 0.2 | 1 | 13 |
| heatr5a | 0.26848 | -0.30619213 | 0.039 | 0.2 | 1 | 13 |
| sec31a | 0.26861 | -0.10065256 | 0.039 | 0.2 | 1 | 13 |
| slc43a1a | 0.26882 | -0.19888743 | 0.039 | 0.2 | 1 | 13 |
| fdxr | 0.26901 | -0.12177083 | 0.039 | 0.2 | 1 | 13 |
| BX649411.1 | 0.26904 | -0.20562678 | 0.039 | 0.2 | 1 | 13 |
| zfr | 0.26941 | -0.16771111 | 0.039 | 0.2 | 1 | 13 |
| ptpn22 | 0.26943 | -0.17315305 | 0.098 | 0.3 | 1 | 13 |
| spice1 | 0.26964 | -0.16501992 | 0.039 | 0.2 | 1 | 13 |
| ulk1a | 0.27008 | -0.29767097 | 0.039 | 0.2 | 1 | 13 |
| pbxip1b | 0.27033 | -0.20959728 | 0.098 | 0.3 | 1 | 13 |
| zgc:85858 | 0.27057 | -0.14212281 | 0.039 | 0.2 | 1 | 13 |
| ppp1r14bb | 0.27074 | -0.10156717 | 0.039 | 0.2 | 1 | 13 |
| sec63 | 0.27078 | -0.17176137 | 0.039 | 0.2 | 1 | 13 |
| fosb | 0.27099 | -0.22014617 | 0.039 | 0.2 | 1 | 13 |
| abcc3 | 0.27147 | -0.19544287 | 0.039 | 0.2 | 1 | 13 |
| dctn5 | 0.27157 | -0.1501889 | 0.039 | 0.2 | 1 | 13 |
| eif2b1 | 0.27207 | -0.2071584 | 0.039 | 0.2 | 1 | 13 |
| mbd2 | 0.27211 | -0.22698444 | 0.039 | 0.2 | 1 | 13 |
| arf2b | 0.27264 | -0.18912821 | 0.176 | 0.4 | 1 | 13 |
| mgat4a | 0.27296 | -0.13047676 | 0.039 | 0.2 | 1 | 13 |
| asah1b | 0.27318 | -0.22828011 | 0.039 | 0.2 | 1 | 13 |
| ctsh | 0.27347 | -0.18241849 | 0.039 | 0.2 | 1 | 13 |
| dgkza | 0.27351 | -0.22031734 | 0.039 | 0.2 | 1 | 13 |
| mpzl2b | 0.27359 | 0.108978771 | 0.078 | 0.2 | 1 | 13 |
| tor1 | 0.27366 | -0.26093834 | 0.039 | 0.2 | 1 | 13 |
| nabp1a | 0.27371 | -0.20648919 | 0.039 | 0.2 | 1 | 13 |
| ikbkg | 0.27402 | -0.18276796 | 0.039 | 0.2 | 1 | 13 |
| tbk1 | 0.27406 | -0.24591656 | 0.039 | 0.2 | 1 | 13 |
| stk10 | 0.27418 | -0.16221401 | 0.039 | 0.2 | 1 | 13 |
| fam177a1 | 0.27419 | -0.13679234 | 0.039 | 0.2 | 1 | 13 |
| tmem56b | 0.27434 | -0.2058409 | 0.039 | 0.2 | 1 | 13 |
| nup153 | 0.27449 | -0.13152073 | 0.039 | 0.2 | 1 | 13 |
| glmp | 0.27453 | -0.2056628 | 0.039 | 0.2 | 1 | 13 |
| psmd3 | 0.27475 | -0.12837564 | 0.039 | 0.2 | 1 | 13 |
| srfa | 0.27483 | -0.16982479 | 0.039 | 0.2 | 1 | 13 |
| rbm14a | 0.27485 | -0.14548518 | 0.039 | 0.2 | 1 | 13 |
| pdzd8 | 0.27487 | -0.18729923 | 0.039 | 0.2 | 1 | 13 |
| bsdc1 | 0.27499 | -0.13549057 | 0.039 | 0.2 | 1 | 13 |
| uqcc1 | 0.2751 | -0.11925193 | 0.039 | 0.2 | 1 | 13 |
| tnfrsf9a | 0.27521 | -0.26770269 | 0.039 | 0.2 | 1 | 13 |
| hnrnpua | 0.27542 | -0.15726059 | 0.039 | 0.2 | 1 | 13 |
| cstf3 | 0.27551 | -0.19191097 | 0.039 | 0.2 | 1 | 13 |

| | | | | | | |
|---|---|---|---|---|---|---|
| mapk14a | 0.27558 | -0.23996761 | 0.039 | 0.2 | 1 | 13 |
| smim19 | 0.27568 | -0.15407889 | 0.039 | 0.2 | 1 | 13 |
| rell1 | 0.27569 | -0.13041174 | 0.039 | 0.2 | 1 | 13 |
| stub1 | 0.27579 | -0.20622552 | 0.039 | 0.2 | 1 | 13 |
| grna | 0.27585 | -0.18793278 | 0.039 | 0.2 | 1 | 13 |
| si:dkey-100n23.5 | 0.27587 | -0.20069763 | 0.039 | 0.2 | 1 | 13 |
| timmdc1 | 0.27587 | -0.19925245 | 0.039 | 0.2 | 1 | 13 |
| hadh | 0.27587 | -0.14174933 | 0.039 | 0.2 | 1 | 13 |
| LOC799656 | 0.27591 | -0.2506093 | 0.039 | 0.2 | 1 | 13 |
| emc3 | 0.27602 | -0.2036095 | 0.039 | 0.2 | 1 | 13 |
| phip | 0.27613 | -0.22662074 | 0.039 | 0.2 | 1 | 13 |
| pcgf5b | 0.27615 | -0.23603563 | 0.039 | 0.2 | 1 | 13 |
| ccdc47 | 0.27617 | -0.1639934 | 0.039 | 0.2 | 1 | 13 |
| MRPS12 | 0.27618 | -0.23780503 | 0.039 | 0.2 | 1 | 13 |
| wdr44 | 0.27619 | -0.12804938 | 0.039 | 0.2 | 1 | 13 |
| sgta | 0.2762 | -0.16108006 | 0.039 | 0.2 | 1 | 13 |
| decr1 | 0.27622 | -0.22759292 | 0.039 | 0.2 | 1 | 13 |
| zfr2 | 0.27622 | -0.14863237 | 0.039 | 0.2 | 1 | 13 |
| si:dkeyp-82a1.6 | 0.27624 | -0.24062722 | 0.039 | 0.2 | 1 | 13 |
| cfi | 0.27739 | 0.165287851 | 0.98 | 1 | 1 | 13 |
| sh2d3ca | 0.2777 | -0.11821646 | 0.118 | 0.3 | 1 | 13 |
| rbm4.2 | 0.27984 | -0.32696299 | 0.098 | 0.3 | 1 | 13 |
| gstp1 | 0.2803 | -0.16566061 | 0.098 | 0.3 | 1 | 13 |
| ssr3 | 0.28069 | -0.15092873 | 0.843 | 1 | 1 | 13 |
| BX936461.1 | 0.28074 | -0.10366224 | 0.02 | 0.1 | 1 | 13 |
| EML4 | 0.28074 | -0.1142002 | 0.02 | 0.1 | 1 | 13 |
| LOC100150882 | 0.28074 | -0.18090553 | 0.02 | 0.1 | 1 | 13 |
| ankrd10a | 0.28074 | -0.12877491 | 0.02 | 0.1 | 1 | 13 |
| atp8a1 | 0.28074 | -0.12685647 | 0.02 | 0.1 | 1 | 13 |
| baz2a | 0.28074 | -0.1387631 | 0.02 | 0.1 | 1 | 13 |
| cacul1 | 0.28074 | -0.11540329 | 0.02 | 0.1 | 1 | 13 |
| cmc2 | 0.28074 | -0.12512351 | 0.02 | 0.1 | 1 | 13 |
| commd7 | 0.28074 | -0.11035293 | 0.02 | 0.1 | 1 | 13 |
| dbr1 | 0.28074 | -0.16069396 | 0.02 | 0.1 | 1 | 13 |
| dcaf17 | 0.28074 | -0.12304757 | 0.02 | 0.1 | 1 | 13 |
| dffb | 0.28074 | -0.12128624 | 0.02 | 0.1 | 1 | 13 |
| exosc6 | 0.28074 | -0.15784299 | 0.02 | 0.1 | 1 | 13 |
| fcho2 | 0.28074 | -0.11406711 | 0.02 | 0.1 | 1 | 13 |
| g6pc3 | 0.28074 | -0.15783938 | 0.02 | 0.1 | 1 | 13 |
| gadd45aa | 0.28074 | -0.1784078 | 0.02 | 0.1 | 1 | 13 |
| glmna | 0.28074 | -0.12396857 | 0.02 | 0.1 | 1 | 13 |
| gtf2f2a | 0.28074 | -0.10789679 | 0.02 | 0.1 | 1 | 13 |
| gtf2h3 | 0.28074 | -0.10823951 | 0.02 | 0.1 | 1 | 13 |
| helz | 0.28074 | -0.10331384 | 0.02 | 0.1 | 1 | 13 |
| irf2bp1 | 0.28074 | -0.10632831 | 0.02 | 0.1 | 1 | 13 |
| jak2b | 0.28074 | -0.10587759 | 0.02 | 0.1 | 1 | 13 |
| kansl3 | 0.28074 | -0.10172043 | 0.02 | 0.1 | 1 | 13 |
| lims1 | 0.28074 | -0.14636895 | 0.02 | 0.1 | 1 | 13 |

| | | | | | | |
|---|---|---|---|---|---|---|
| mgrn1b | 0.28074 | -0.11997871 | 0.02 | 0.1 | 1 | 13 |
| mia2 | 0.28074 | -0.12671408 | 0.02 | 0.1 | 1 | 13 |
| mob2b | 0.28074 | -0.10272938 | 0.02 | 0.1 | 1 | 13 |
| mrps23 | 0.28074 | -0.16609015 | 0.02 | 0.1 | 1 | 13 |
| mvb12ba | 0.28074 | -0.13167608 | 0.02 | 0.1 | 1 | 13 |
| naga | 0.28074 | -0.13296533 | 0.02 | 0.1 | 1 | 13 |
| pfkpa | 0.28074 | -0.10251022 | 0.02 | 0.1 | 1 | 13 |
| pigb | 0.28074 | -0.14037128 | 0.02 | 0.1 | 1 | 13 |
| pik3cg | 0.28074 | -0.12119495 | 0.02 | 0.1 | 1 | 13 |
| plcg1 | 0.28074 | -0.13539062 | 0.02 | 0.1 | 1 | 13 |
| pmpca | 0.28074 | -0.1298666 | 0.02 | 0.1 | 1 | 13 |
| ppm1ba | 0.28074 | -0.13369928 | 0.02 | 0.1 | 1 | 13 |
| ptp4a2a | 0.28074 | -0.11340644 | 0.02 | 0.1 | 1 | 13 |
| rock2a | 0.28074 | -0.16028642 | 0.02 | 0.1 | 1 | 13 |
| si:ch211-107o10.3 | 0.28074 | -0.11292817 | 0.02 | 0.1 | 1 | 13 |
| si:ch211-196c10.15 | 0.28074 | -0.11732987 | 0.02 | 0.1 | 1 | 13 |
| si:ch211-221j21.3 | 0.28074 | -0.11375004 | 0.02 | 0.1 | 1 | 13 |
| si:ch211-223a10.1 | 0.28074 | -0.10740051 | 0.02 | 0.1 | 1 | 13 |
| sirt2 | 0.28074 | -0.11574904 | 0.02 | 0.1 | 1 | 13 |
| slain2 | 0.28074 | -0.12298082 | 0.02 | 0.1 | 1 | 13 |
| slc3a2a | 0.28074 | -0.11937341 | 0.02 | 0.1 | 1 | 13 |
| smpd2b | 0.28074 | -0.11527472 | 0.02 | 0.1 | 1 | 13 |
| stam2 | 0.28074 | -0.1010116 | 0.02 | 0.1 | 1 | 13 |
| strbp | 0.28074 | -0.13954708 | 0.02 | 0.1 | 1 | 13 |
| sugp1 | 0.28074 | -0.10674493 | 0.02 | 0.1 | 1 | 13 |
| tmed1b | 0.28074 | -0.16837151 | 0.02 | 0.1 | 1 | 13 |
| tmem129 | 0.28074 | -0.10096372 | 0.02 | 0.1 | 1 | 13 |
| trim8a | 0.28074 | -0.13359205 | 0.02 | 0.1 | 1 | 13 |
| ube2f | 0.28074 | -0.13501245 | 0.02 | 0.1 | 1 | 13 |
| unkl | 0.28074 | -0.11065727 | 0.02 | 0.1 | 1 | 13 |
| wdr41 | 0.28074 | -0.10065054 | 0.02 | 0.1 | 1 | 13 |
| zbtb49 | 0.28074 | -0.10379203 | 0.02 | 0.1 | 1 | 13 |
| zdhhc14 | 0.28074 | -0.17000855 | 0.02 | 0.1 | 1 | 13 |
| zgc:100918 | 0.28074 | -0.10493203 | 0.02 | 0.1 | 1 | 13 |
| zgc:193742 | 0.28074 | -0.12055165 | 0.02 | 0.1 | 1 | 13 |
| GFOD1 | 0.28074 | -0.18037995 | 0.02 | 0.1 | 1 | 13 |
| abl1 | 0.28074 | -0.17693899 | 0.02 | 0.1 | 1 | 13 |
| acaa2 | 0.28074 | -0.13067892 | 0.02 | 0.1 | 1 | 13 |
| acbd6 | 0.28074 | -0.100599 | 0.02 | 0.1 | 1 | 13 |
| adtrp1 | 0.28074 | -0.16889866 | 0.02 | 0.1 | 1 | 13 |
| alg3 | 0.28074 | -0.11421862 | 0.02 | 0.1 | 1 | 13 |
| asmtl | 0.28074 | -0.10421579 | 0.02 | 0.1 | 1 | 13 |
| atg4c | 0.28074 | -0.12692766 | 0.02 | 0.1 | 1 | 13 |
| brd1a | 0.28074 | -0.12323208 | 0.02 | 0.1 | 1 | 13 |
| cd74a | 0.28074 | -0.10968665 | 0.02 | 0.1 | 1 | 13 |
| cdca5 | 0.28074 | -0.14499468 | 0.02 | 0.1 | 1 | 13 |
| chmp4ba | 0.28074 | -0.16097891 | 0.02 | 0.1 | 1 | 13 |
| clasp1a | 0.28074 | -0.24931473 | 0.02 | 0.1 | 1 | 13 |

| | | | | | | |
|---|---|---|---|---|---|---|
| cog6 | 0.28074 | -0.18153604 | 0.02 | 0.1 | 1 | 13 |
| cox11 | 0.28074 | -0.14965303 | 0.02 | 0.1 | 1 | 13 |
| ctss2.1 | 0.28074 | -0.11915537 | 0.02 | 0.1 | 1 | 13 |
| dmap1 | 0.28074 | -0.16887375 | 0.02 | 0.1 | 1 | 13 |
| epn1 | 0.28074 | -0.16811439 | 0.02 | 0.1 | 1 | 13 |
| faah2a | 0.28074 | -0.12910633 | 0.02 | 0.1 | 1 | 13 |
| fxr1 | 0.28074 | -0.13667633 | 0.02 | 0.1 | 1 | 13 |
| galt | 0.28074 | -0.16604089 | 0.02 | 0.1 | 1 | 13 |
| hyal2b | 0.28074 | -0.18317649 | 0.02 | 0.1 | 1 | 13 |
| ifi45 | 0.28074 | -0.16023799 | 0.02 | 0.1 | 1 | 13 |
| ints4 | 0.28074 | -0.17749833 | 0.02 | 0.1 | 1 | 13 |
| klhdc8b | 0.28074 | -0.16943737 | 0.02 | 0.1 | 1 | 13 |
| kmt2ba | 0.28074 | -0.13991943 | 0.02 | 0.1 | 1 | 13 |
| lamtor3 | 0.28074 | -0.15661745 | 0.02 | 0.1 | 1 | 13 |
| map1lc3b | 0.28074 | -0.18511566 | 0.02 | 0.1 | 1 | 13 |
| mrpl19 | 0.28074 | -0.16315694 | 0.02 | 0.1 | 1 | 13 |
| mrpl22 | 0.28074 | -0.16149646 | 0.02 | 0.1 | 1 | 13 |
| npc1 | 0.28074 | -0.16940339 | 0.02 | 0.1 | 1 | 13 |
| nt5c2b | 0.28074 | -0.18078948 | 0.02 | 0.1 | 1 | 13 |
| olfml2ba | 0.28074 | -0.15083389 | 0.02 | 0.1 | 1 | 13 |
| phf12a | 0.28074 | -0.15426197 | 0.02 | 0.1 | 1 | 13 |
| ppp2r5d | 0.28074 | -0.13471706 | 0.02 | 0.1 | 1 | 13 |
| ralgapa2 | 0.28074 | -0.15362624 | 0.02 | 0.1 | 1 | 13 |
| rap2aa | 0.28074 | -0.17964452 | 0.02 | 0.1 | 1 | 13 |
| reck | 0.28074 | -0.1111173 | 0.02 | 0.1 | 1 | 13 |
| ripor2 | 0.28074 | -0.10826969 | 0.02 | 0.1 | 1 | 13 |
| rnf6 | 0.28074 | -0.13945686 | 0.02 | 0.1 | 1 | 13 |
| rorb | 0.28074 | -0.15836688 | 0.02 | 0.1 | 1 | 13 |
| rrp12 | 0.28074 | -0.1714796 | 0.02 | 0.1 | 1 | 13 |
| si:ch211-106e7.2 | 0.28074 | -0.13858229 | 0.02 | 0.1 | 1 | 13 |
| si:ch211-59h6.1 | 0.28074 | -0.18290448 | 0.02 | 0.1 | 1 | 13 |
| si:dkey-188i13.10 | 0.28074 | -0.14239585 | 0.02 | 0.1 | 1 | 13 |
| si:dkey-192k22.2 | 0.28074 | -0.18027548 | 0.02 | 0.1 | 1 | 13 |
| si:dkey-66i24.7 | 0.28074 | -0.17125013 | 0.02 | 0.1 | 1 | 13 |
| si:dkeyp-44a8.2 | 0.28074 | -0.19222089 | 0.02 | 0.1 | 1 | 13 |
| sirt6 | 0.28074 | -0.14428233 | 0.02 | 0.1 | 1 | 13 |
| slc4a4b | 0.28074 | -0.15554966 | 0.02 | 0.1 | 1 | 13 |
| snx30 | 0.28074 | -0.16012115 | 0.02 | 0.1 | 1 | 13 |
| srrt | 0.28074 | -0.14653184 | 0.02 | 0.1 | 1 | 13 |
| stk35 | 0.28074 | -0.16228744 | 0.02 | 0.1 | 1 | 13 |
| sts | 0.28074 | -0.1758275 | 0.02 | 0.1 | 1 | 13 |
| tm9sf5 | 0.28074 | -0.13059177 | 0.02 | 0.1 | 1 | 13 |
| tmem110l | 0.28074 | -0.14056048 | 0.02 | 0.1 | 1 | 13 |
| tmem222a | 0.28074 | -0.12832124 | 0.02 | 0.1 | 1 | 13 |
| trmt10b | 0.28074 | -0.1762825 | 0.02 | 0.1 | 1 | 13 |
| trpc4apb | 0.28074 | -0.12397892 | 0.02 | 0.1 | 1 | 13 |
| vrk3 | 0.28074 | -0.15194814 | 0.02 | 0.1 | 1 | 13 |
| wdr21 | 0.28074 | -0.11439453 | 0.02 | 0.1 | 1 | 13 |

| | | | | | | |
|---|---|---|---|---|---|---|
| zbtb1 | 0.28074 | -0.1795155 | 0.02 | 0.1 | 1 | 13 |
| zdhhc23b | 0.28074 | -0.17982564 | 0.02 | 0.1 | 1 | 13 |
| zgc:158398 | 0.28074 | -0.1037059 | 0.02 | 0.1 | 1 | 13 |
| zgc:66443 | 0.28074 | -0.15040798 | 0.02 | 0.1 | 1 | 13 |
| zmp:0000001132 | 0.28074 | -0.10557687 | 0.02 | 0.1 | 1 | 13 |
| kpnb1 | 0.28119 | -0.18155759 | 0.098 | 0.3 | 1 | 13 |
| srgn | 0.28124 | 0.125395018 | 0.843 | 0.9 | 1 | 13 |
| LOC100151196 | 0.28417 | -0.21730096 | 0.098 | 0.3 | 1 | 13 |
| ap3s1.1 | 0.28649 | -0.17900917 | 0.176 | 0.4 | 1 | 13 |
| stx12 | 0.28703 | 0.115725042 | 0.098 | 0.2 | 1 | 13 |
| pcna | 0.28874 | -0.28977028 | 0.098 | 0.3 | 1 | 13 |
| cfh | 0.28888 | -0.25182124 | 0.549 | 0.8 | 1 | 13 |
| rgs13 | 0.28904 | -0.18446651 | 0.176 | 0.4 | 1 | 13 |
| tefa | 0.28918 | -0.22578286 | 0.098 | 0.3 | 1 | 13 |
| phf23a | 0.28978 | -0.10757292 | 0.059 | 0.2 | 1 | 13 |
| adh5 | 0.29099 | -0.14432879 | 0.098 | 0.3 | 1 | 13 |
| tspan31 | 0.29228 | -0.23339078 | 0.098 | 0.3 | 1 | 13 |
| si:dkeyp-19e1.3 | 0.29272 | -0.28576927 | 0.098 | 0.3 | 1 | 13 |
| azin1b | 0.2937 | -0.19528238 | 0.098 | 0.3 | 1 | 13 |
| gpr108 | 0.29376 | -0.26756467 | 0.098 | 0.3 | 1 | 13 |
| rbm28 | 0.29464 | -0.18964327 | 0.098 | 0.3 | 1 | 13 |
| tmco1 | 0.29471 | -0.30187312 | 0.098 | 0.3 | 1 | 13 |
| klf6a | 0.29738 | -0.19062725 | 0.176 | 0.4 | 1 | 13 |
| copz1 | 0.29906 | -0.25519335 | 0.255 | 0.5 | 1 | 13 |
| prkcbb | 0.29991 | -0.14027787 | 0.275 | 0.5 | 1 | 13 |
| ndufc2 | 0.30283 | 0.399294144 | 0.255 | 0.2 | 1 | 13 |
| snx3 | 0.30494 | -0.17006394 | 0.176 | 0.4 | 1 | 13 |
| LOC110438970 | 0.30641 | 0.145562621 | 0.98 | 1 | 1 | 13 |
| agtrap | 0.30673 | 0.177562933 | 0.078 | 0 | 1 | 13 |
| ash2l | 0.30673 | 0.117537594 | 0.078 | 0 | 1 | 13 |
| dusp5 | 0.30673 | 0.212046025 | 0.078 | 0 | 1 | 13 |
| eif6 | 0.30673 | 0.160204286 | 0.078 | 0 | 1 | 13 |
| kidins220b | 0.30673 | 0.177064348 | 0.078 | 0 | 1 | 13 |
| mpp1 | 0.30673 | 0.206440876 | 0.078 | 0 | 1 | 13 |
| pik3ca | 0.30673 | 0.126183907 | 0.078 | 0 | 1 | 13 |
| prkacab | 0.30673 | 0.166394163 | 0.078 | 0 | 1 | 13 |
| slc7a11 | 0.30673 | 0.224963669 | 0.078 | 0 | 1 | 13 |
| socs3b | 0.30673 | 0.182488413 | 0.078 | 0 | 1 | 13 |
| sv2 | 0.30673 | 0.119597131 | 0.078 | 0 | 1 | 13 |
| vps9d1 | 0.30673 | 0.113528871 | 0.078 | 0 | 1 | 13 |
| FAR2 | 0.30673 | 0.131251449 | 0.078 | 0 | 1 | 13 |
| adam17a | 0.30673 | 0.172039511 | 0.078 | 0 | 1 | 13 |
| atf1 | 0.30673 | 0.103554579 | 0.078 | 0 | 1 | 13 |
| calhm6 | 0.30673 | 0.140591425 | 0.078 | 0 | 1 | 13 |
| ccnt2b | 0.30673 | 0.123514331 | 0.078 | 0 | 1 | 13 |
| cdk1 | 0.30673 | 0.180343499 | 0.078 | 0 | 1 | 13 |
| golga5 | 0.30673 | 0.112508872 | 0.078 | 0 | 1 | 13 |
| ing4 | 0.30673 | 0.110762626 | 0.078 | 0 | 1 | 13 |

| | | | | | | |
|---|---|---|---|---|---|---|
| kif5ba | 0.30673 | 0.129288039 | 0.078 | 0 | 1 | 13 |
| lyrm2 | 0.30673 | 0.139009518 | 0.078 | 0 | 1 | 13 |
| mtfr1l | 0.30673 | 0.15321133 | 0.078 | 0 | 1 | 13 |
| runx3 | 0.30673 | 0.14972003 | 0.078 | 0 | 1 | 13 |
| sbds | 0.30673 | 0.114585301 | 0.078 | 0 | 1 | 13 |
| si:dkey-21c1.1 | 0.30673 | 0.167089007 | 0.078 | 0 | 1 | 13 |
| si:dkey-33c9.6 | 0.30673 | 0.173427816 | 0.078 | 0 | 1 | 13 |
| si:dkeyp-69e1.8 | 0.30673 | 0.137402777 | 0.078 | 0 | 1 | 13 |
| ssuh2.3 | 0.30673 | 0.194138081 | 0.078 | 0 | 1 | 13 |
| tefb | 0.30673 | 0.132174785 | 0.078 | 0 | 1 | 13 |
| ubl3a | 0.30673 | 0.128929245 | 0.078 | 0 | 1 | 13 |
| uchl5 | 0.30673 | 0.120381562 | 0.078 | 0 | 1 | 13 |
| uckl1b | 0.30673 | 0.164137198 | 0.078 | 0 | 1 | 13 |
| utrn | 0.30673 | 0.116876172 | 0.078 | 0 | 1 | 13 |
| wwp2 | 0.30673 | 0.136666759 | 0.078 | 0 | 1 | 13 |
| zgc:158803 | 0.30673 | 0.154806583 | 0.078 | 0 | 1 | 13 |
| chd9 | 0.30673 | 0.176931154 | 0.078 | 0 | 1 | 13 |
| ranbp9 | 0.30673 | 0.155504572 | 0.078 | 0 | 1 | 13 |
| rasgef1ba | 0.30673 | 0.177093571 | 0.078 | 0 | 1 | 13 |
| rpa3 | 0.30673 | 0.212494468 | 0.078 | 0 | 1 | 13 |
| arih2 | 0.30673 | 0.148126222 | 0.078 | 0 | 1 | 13 |
| cdk5rap1 | 0.30673 | 0.180840879 | 0.078 | 0 | 1 | 13 |
| creld2 | 0.30673 | 0.181752709 | 0.078 | 0 | 1 | 13 |
| cul4b | 0.30673 | 0.120098517 | 0.078 | 0 | 1 | 13 |
| ddb1 | 0.30673 | 0.168842254 | 0.078 | 0 | 1 | 13 |
| exosc9 | 0.30673 | 0.142829633 | 0.078 | 0 | 1 | 13 |
| glsb | 0.30673 | 0.160696615 | 0.078 | 0 | 1 | 13 |
| leng8 | 0.30673 | 0.180444938 | 0.078 | 0 | 1 | 13 |
| lmod1a | 0.30673 | 0.124187066 | 0.078 | 0 | 1 | 13 |
| magoh | 0.30673 | 0.151631657 | 0.078 | 0 | 1 | 13 |
| mak16 | 0.30673 | 0.143667742 | 0.078 | 0 | 1 | 13 |
| ovca2 | 0.30673 | 0.145708542 | 0.078 | 0 | 1 | 13 |
| pkig | 0.30673 | 0.160927613 | 0.078 | 0 | 1 | 13 |
| plcb3 | 0.30673 | 0.216581247 | 0.078 | 0 | 1 | 13 |
| psmd4b | 0.30673 | 0.202299272 | 0.078 | 0 | 1 | 13 |
| rps6kb1a | 0.30673 | 0.166751669 | 0.078 | 0 | 1 | 13 |
| si:ch73-376l24.6 | 0.30673 | 0.154885214 | 0.078 | 0 | 1 | 13 |
| sqstm1 | 0.30673 | 0.155208432 | 0.078 | 0 | 1 | 13 |
| swi5 | 0.30673 | 0.122713987 | 0.078 | 0 | 1 | 13 |
| tmem30ab | 0.30673 | 0.136375431 | 0.078 | 0 | 1 | 13 |
| tmpob | 0.30673 | 0.1741528 | 0.078 | 0 | 1 | 13 |
| tomm22 | 0.30673 | 0.135484135 | 0.078 | 0 | 1 | 13 |
| trappc6bl | 0.30673 | 0.107972426 | 0.078 | 0 | 1 | 13 |
| AL831768.1 | 0.30673 | 0.17068958 | 0.078 | 0 | 1 | 13 |
| CABZ01029938.1 | 0.30673 | 0.198492866 | 0.078 | 0 | 1 | 13 |
| CABZ01088229.1 | 0.30673 | 0.190182792 | 0.078 | 0 | 1 | 13 |
| CYTH2 | 0.30673 | 0.105110036 | 0.078 | 0 | 1 | 13 |
| LOC100329806 | 0.30673 | 0.176173512 | 0.078 | 0 | 1 | 13 |

| | | | | | | |
|---|---|---|---|---|---|---|
| XLOC-042229 | 0.30673 | 0.12998316 | 0.078 | 0 | 1 | 13 |
| aldh2.1 | 0.30673 | 0.178406045 | 0.078 | 0 | 1 | 13 |
| atp6v1ba | 0.30673 | 0.12376698 | 0.078 | 0 | 1 | 13 |
| ccz1 | 0.30673 | 0.171768266 | 0.078 | 0 | 1 | 13 |
| cdc14b | 0.30673 | 0.170516216 | 0.078 | 0 | 1 | 13 |
| commd2 | 0.30673 | 0.151906417 | 0.078 | 0 | 1 | 13 |
| cyp3c1 | 0.30673 | 0.165145764 | 0.078 | 0 | 1 | 13 |
| edc4 | 0.30673 | 0.15649457 | 0.078 | 0 | 1 | 13 |
| ehbp1l1a | 0.30673 | 0.114299338 | 0.078 | 0 | 1 | 13 |
| fam45a | 0.30673 | 0.142115564 | 0.078 | 0 | 1 | 13 |
| fndc3ba | 0.30673 | 0.124776689 | 0.078 | 0 | 1 | 13 |
| get4 | 0.30673 | 0.135595728 | 0.078 | 0 | 1 | 13 |
| jak3 | 0.30673 | 0.142252089 | 0.078 | 0 | 1 | 13 |
| klf12a | 0.30673 | 0.183206838 | 0.078 | 0 | 1 | 13 |
| larp1 | 0.30673 | 0.183128648 | 0.078 | 0 | 1 | 13 |
| manea | 0.30673 | 0.152446869 | 0.078 | 0 | 1 | 13 |
| mat2aa | 0.30673 | 0.155092882 | 0.078 | 0 | 1 | 13 |
| per1b | 0.30673 | 0.105946844 | 0.078 | 0 | 1 | 13 |
| pik3r3a | 0.30673 | 0.195132585 | 0.078 | 0 | 1 | 13 |
| pnp5a | 0.30673 | 0.116927938 | 0.078 | 0 | 1 | 13 |
| polr3glb | 0.30673 | 0.128470791 | 0.078 | 0 | 1 | 13 |
| rrm2 | 0.30673 | 0.129736589 | 0.078 | 0 | 1 | 13 |
| sept9b | 0.30673 | 0.112571195 | 0.078 | 0 | 1 | 13 |
| tfe3b | 0.30673 | 0.153640848 | 0.078 | 0 | 1 | 13 |
| tiam2a | 0.30673 | 0.130690769 | 0.078 | 0 | 1 | 13 |
| wdr43 | 0.30673 | 0.19492271 | 0.078 | 0 | 1 | 13 |
| zbtb2b | 0.30673 | 0.105679067 | 0.078 | 0 | 1 | 13 |
| CR855389.1 | 0.30673 | 0.132292795 | 0.078 | 0 | 1 | 13 |
| LO017872.1 | 0.30673 | 0.175320307 | 0.078 | 0 | 1 | 13 |
| XLOC-010289 | 0.30673 | 0.130758675 | 0.078 | 0 | 1 | 13 |
| XLOC-018981 | 0.30673 | 0.20584289 | 0.078 | 0 | 1 | 13 |
| abcb7 | 0.30673 | 0.102412432 | 0.078 | 0 | 1 | 13 |
| actr1 | 0.30673 | 0.141553498 | 0.078 | 0 | 1 | 13 |
| agpat3 | 0.30673 | 0.138904448 | 0.078 | 0 | 1 | 13 |
| apeh | 0.30673 | 0.127645208 | 0.078 | 0 | 1 | 13 |
| bbs1 | 0.30673 | 0.16216849 | 0.078 | 0 | 1 | 13 |
| birc5a | 0.30673 | 0.151554661 | 0.078 | 0 | 1 | 13 |
| blmh | 0.30673 | 0.10517566 | 0.078 | 0 | 1 | 13 |
| cmtm6 | 0.30673 | 0.125641452 | 0.078 | 0 | 1 | 13 |
| coq9 | 0.30673 | 0.147508515 | 0.078 | 0 | 1 | 13 |
| csnk1g2b | 0.30673 | 0.126020093 | 0.078 | 0 | 1 | 13 |
| dhx36 | 0.30673 | 0.177146101 | 0.078 | 0 | 1 | 13 |
| diablob | 0.30673 | 0.104002411 | 0.078 | 0 | 1 | 13 |
| dynll2b | 0.30673 | 0.105913497 | 0.078 | 0 | 1 | 13 |
| fahd2a | 0.30673 | 0.100157888 | 0.078 | 0 | 1 | 13 |
| gb:bc139872 | 0.30673 | 0.122739105 | 0.078 | 0 | 1 | 13 |
| gra | 0.30673 | 0.129773132 | 0.078 | 0 | 1 | 13 |
| gtf2a2 | 0.30673 | 0.197496934 | 0.078 | 0 | 1 | 13 |

| | | | | | | |
|---|---|---|---|---|---|---|
| guk1a | 0.30673 | 0.165342183 | 0.078 | 0 | 1 | 13 |
| gys1 | 0.30673 | 0.199474391 | 0.078 | 0 | 1 | 13 |
| hnrnpc | 0.30673 | 0.107375027 | 0.078 | 0 | 1 | 13 |
| hnrnpul1 | 0.30673 | 0.138647775 | 0.078 | 0 | 1 | 13 |
| htatsf1 | 0.30673 | 0.163490905 | 0.078 | 0 | 1 | 13 |
| itsn1 | 0.30673 | 0.122030975 | 0.078 | 0 | 1 | 13 |
| kif22 | 0.30673 | 0.116147023 | 0.078 | 0 | 1 | 13 |
| kifc1 | 0.30673 | 0.209553742 | 0.078 | 0 | 1 | 13 |
| lbh | 0.30673 | 0.122207778 | 0.078 | 0 | 1 | 13 |
| lpar2b | 0.30673 | 0.131850141 | 0.078 | 0 | 1 | 13 |
| lsm1 | 0.30673 | 0.123774078 | 0.078 | 0 | 1 | 13 |
| map3k4 | 0.30673 | 0.135986604 | 0.078 | 0 | 1 | 13 |
| mzt2b | 0.30673 | 0.128150797 | 0.078 | 0 | 1 | 13 |
| nucb2a | 0.30673 | 0.177585388 | 0.078 | 0 | 1 | 13 |
| otud5a | 0.30673 | 0.179351445 | 0.078 | 0 | 1 | 13 |
| polr3k | 0.30673 | 0.122814359 | 0.078 | 0 | 1 | 13 |
| prkar2aa | 0.30673 | 0.126891558 | 0.078 | 0 | 1 | 13 |
| prkcba | 0.30673 | 0.193372237 | 0.078 | 0 | 1 | 13 |
| psmg2 | 0.30673 | 0.114256954 | 0.078 | 0 | 1 | 13 |
| sardh | 0.30673 | 0.133138769 | 0.078 | 0 | 1 | 13 |
| si:ch211-209f23.6 | 0.30673 | 0.203074456 | 0.078 | 0 | 1 | 13 |
| si:ch211-271d10.2 | 0.30673 | 0.137187535 | 0.078 | 0 | 1 | 13 |
| si:dkey-187e18.1 | 0.30673 | 0.135052223 | 0.078 | 0 | 1 | 13 |
| snx27a | 0.30673 | 0.187692209 | 0.078 | 0 | 1 | 13 |
| snx6 | 0.30673 | 0.104099924 | 0.078 | 0 | 1 | 13 |
| tagapa | 0.30673 | 0.127710804 | 0.078 | 0 | 1 | 13 |
| tank | 0.30673 | 0.144191718 | 0.078 | 0 | 1 | 13 |
| tmem131 | 0.30673 | 0.118909602 | 0.078 | 0 | 1 | 13 |
| tmem222b | 0.30673 | 0.147527022 | 0.078 | 0 | 1 | 13 |
| tspy | 0.30673 | 0.12154168 | 0.078 | 0 | 1 | 13 |
| tuba8l2 | 0.30673 | 0.142631851 | 0.078 | 0 | 1 | 13 |
| ugt5b1 | 0.30673 | 0.133622581 | 0.078 | 0 | 1 | 13 |
| vkorc1l1 | 0.30673 | 0.111310265 | 0.078 | 0 | 1 | 13 |
| vps41 | 0.30673 | 0.163025869 | 0.078 | 0 | 1 | 13 |
| wee2 | 0.30673 | 0.11181145 | 0.078 | 0 | 1 | 13 |
| zgc:77086 | 0.30673 | 0.140999914 | 0.078 | 0 | 1 | 13 |
| atp13a1 | 0.30673 | 0.113303513 | 0.078 | 0 | 1 | 13 |
| atp6v1aa | 0.30673 | 0.176253739 | 0.078 | 0 | 1 | 13 |
| dctn4 | 0.30673 | 0.139564729 | 0.078 | 0 | 1 | 13 |
| e2f8 | 0.30673 | 0.121948935 | 0.078 | 0 | 1 | 13 |
| tgm1l1 | 0.30673 | 0.233628522 | 0.078 | 0 | 1 | 13 |
| top2a | 0.30673 | 0.161563032 | 0.078 | 0 | 1 | 13 |
| uap1 | 0.30673 | 0.131907915 | 0.078 | 0 | 1 | 13 |
| limk1a | 0.30673 | 0.173322383 | 0.078 | 0 | 1 | 13 |
| eif5a2 | 0.30805 | 0.386285485 | 0.314 | 0.1 | 1 | 13 |
| spink2.1 | 0.31083 | 0.199091103 | 0.98 | 0.9 | 1 | 13 |
| gnao1a | 0.31177 | -0.11614823 | 0.118 | 0.3 | 1 | 13 |
| syk | 0.31242 | 0.275522556 | 0.392 | 0.4 | 1 | 13 |

| | | | | | | |
|---|---|---|---|---|---|---|
| mt-nd3 | 0.31309 | -0.30992373 | 0.176 | 0.4 | 1 | 13 |
| slc7a3a | 0.31316 | -0.27223627 | 0.176 | 0.4 | 1 | 13 |
| tpmt.1 | 0.31458 | -0.24449541 | 0.176 | 0.4 | 1 | 13 |
| tln1 | 0.31572 | -0.17834057 | 0.373 | 0.6 | 1 | 13 |
| rogdi | 0.31603 | -0.11642401 | 0.196 | 0.4 | 1 | 13 |
| rps7 | 0.3176 | -0.30936724 | 0.686 | 0.9 | 1 | 13 |
| rps26 | 0.3182 | -0.31024183 | 0.706 | 0.9 | 1 | 13 |
| mknk2b | 0.31917 | 0.422069109 | 0.235 | 0.1 | 1 | 13 |
| XLOC-000431 | 0.31943 | -0.25764782 | 0.176 | 0.4 | 1 | 13 |
| si:ch73-359m17.2 | 0.31998 | 0.165221736 | 1 | 1 | 1 | 13 |
| ywhae1 | 0.3206 | 0.245861421 | 0.216 | 0.3 | 1 | 13 |
| gapdhs | 0.3219 | -0.21733055 | 0.686 | 0.9 | 1 | 13 |
| midn | 0.32295 | 0.330003065 | 0.216 | 0.2 | 1 | 13 |
| cycsb | 0.32312 | -0.10148482 | 0.196 | 0.4 | 1 | 13 |
| csnk2a1 | 0.32333 | -0.12485682 | 0.118 | 0.3 | 1 | 13 |
| LOC101883999 | 0.32333 | -0.11792582 | 0.118 | 0.3 | 1 | 13 |
| atp5f1c | 0.32565 | 0.149513519 | 0.294 | 0.4 | 1 | 13 |
| rpl32 | 0.32715 | -0.25624417 | 0.686 | 0.9 | 1 | 13 |
| cox4i1 | 0.32893 | 0.166205571 | 0.51 | 0.6 | 1 | 13 |
| fosab | 0.33015 | -0.1678838 | 0.275 | 0.5 | 1 | 13 |
| myh9a | 0.33127 | 0.197754988 | 0.333 | 0.4 | 1 | 13 |
| ndufa6 | 0.3328 | 0.34513409 | 0.294 | 0.1 | 1 | 13 |
| plekhj1 | 0.33485 | -0.1357152 | 0.118 | 0.3 | 1 | 13 |
| tmem263 | 0.33744 | -0.109262 | 0.118 | 0.3 | 1 | 13 |
| rps14 | 0.33774 | -0.13417647 | 0.725 | 0.9 | 1 | 13 |
| itgae.2 | 0.34194 | -0.15426695 | 0.196 | 0.4 | 1 | 13 |
| slc5a3b | 0.34287 | -0.13957958 | 0.059 | 0.2 | 1 | 13 |
| ccr9a | 0.34453 | 0.329541135 | 0.529 | 0.5 | 1 | 13 |
| cdkn3 | 0.34509 | 0.14790616 | 0.196 | 0.3 | 1 | 13 |
| eef1da | 0.34514 | -0.25528024 | 0.451 | 0.7 | 1 | 13 |
| hsp90ab1 | 0.34649 | -0.11814021 | 0.706 | 0.9 | 1 | 13 |
| c3a.4 | 0.34944 | 0.202869496 | 0.784 | 0.8 | 1 | 13 |
| rpl26 | 0.35068 | -0.30035835 | 0.745 | 0.9 | 1 | 13 |
| aspm | 0.35095 | 0.190709423 | 0.137 | 0.2 | 1 | 13 |
| actn4 | 0.35115 | 0.338685013 | 0.235 | 0.2 | 1 | 13 |
| irf2bp2b | 0.35128 | 0.198303356 | 0.137 | 0.2 | 1 | 13 |
| twf1a | 0.35212 | 0.410507023 | 0.431 | 0.2 | 1 | 13 |
| zmp:0000001082 | 0.35317 | -0.21261493 | 0.275 | 0.5 | 1 | 13 |
| CR392341.1 | 0.35326 | -0.16892203 | 0.118 | 0.3 | 1 | 13 |
| hif1ab | 0.35382 | -0.11361962 | 0.216 | 0.4 | 1 | 13 |
| max | 0.35389 | 0.123904135 | 0.196 | 0.3 | 1 | 13 |
| bcap31 | 0.35511 | 0.124204127 | 0.098 | 0.2 | 1 | 13 |
| si:ch73-233k15.2 | 0.35523 | 0.132483596 | 0.098 | 0.2 | 1 | 13 |
| snx1a | 0.35712 | -0.1008647 | 0.118 | 0.3 | 1 | 13 |
| naca | 0.35714 | -0.28901971 | 0.627 | 0.7 | 1 | 13 |
| si:ch73-1a9.3 | 0.35987 | 0.130277563 | 0.196 | 0.3 | 1 | 13 |
| srsf7a | 0.36322 | -0.15420788 | 0.196 | 0.4 | 1 | 13 |
| pef1 | 0.3633 | -0.11322315 | 0.118 | 0.3 | 1 | 13 |

| | | | | | | |
|---|---|---|---|---|---|---|
| rps26l | 0.36518 | -0.13853271 | 0.725 | 0.9 | 1 | 13 |
| rps15 | 0.36551 | -0.20759546 | 0.706 | 0.9 | 1 | 13 |
| osmr | 0.36741 | 0.407479465 | 0.294 | 0.1 | 1 | 13 |
| mt-co1 | 0.36766 | 0.119250031 | 0.824 | 0.9 | 1 | 13 |
| slc25a3b | 0.36876 | -0.19112681 | 0.471 | 0.7 | 1 | 13 |
| gnb1b | 0.372 | 0.212551756 | 0.255 | 0.3 | 1 | 13 |
| hipk3b | 0.37207 | -0.13474631 | 0.059 | 0.2 | 1 | 13 |
| ptmab | 0.37292 | -0.17772131 | 0.706 | 0.9 | 1 | 13 |
| smim15 | 0.3769 | -0.11480844 | 0.118 | 0.3 | 1 | 13 |
| egr1 | 0.37718 | 0.224587487 | 0.275 | 0.3 | 1 | 13 |
| mef2aa | 0.37752 | -0.18298018 | 0.118 | 0.3 | 1 | 13 |
| eif3s6ip | 0.3791 | -0.13864641 | 0.118 | 0.3 | 1 | 13 |
| tuba8l | 0.38159 | 0.313054431 | 0.176 | 0.2 | 1 | 13 |
| gyg1a | 0.38273 | 0.405692925 | 0.412 | 0.2 | 1 | 13 |
| sult6b1 | 0.38359 | -0.14656955 | 0.118 | 0.3 | 1 | 13 |
| klf2b | 0.38382 | -0.22447738 | 0.118 | 0.3 | 1 | 13 |
| CABZ01076667.1 | 0.38461 | -0.13894496 | 0.118 | 0.3 | 1 | 13 |
| si:ch211-222l21.1 | 0.38586 | -0.1336549 | 0.235 | 0.4 | 1 | 13 |
| mdh1aa | 0.38722 | -0.18987325 | 0.118 | 0.3 | 1 | 13 |
| srsf4 | 0.38831 | -0.25416084 | 0.118 | 0.3 | 1 | 13 |
| zfpl1 | 0.38868 | -0.14381157 | 0.059 | 0.2 | 1 | 13 |
| zgc:174904 | 0.38912 | -0.34834853 | 0.137 | 0.3 | 1 | 13 |
| hmgb2b | 0.39034 | 0.32526293 | 0.294 | 0.1 | 1 | 13 |
| eif4ebp2 | 0.39111 | 0.329523159 | 0.49 | 0.4 | 1 | 13 |
| LOC101885769 | 0.39146 | -0.22694787 | 0.118 | 0.3 | 1 | 13 |
| rpsa | 0.39173 | 0.257786745 | 0.882 | 0.8 | 1 | 13 |
| furina | 0.39312 | 0.26483327 | 0.314 | 0.3 | 1 | 13 |
| ponzr1 | 0.39315 | -0.30959443 | 0.059 | 0.2 | 1 | 13 |
| ets2 | 0.39333 | -0.24675919 | 0.118 | 0.3 | 1 | 13 |
| kpna3 | 0.39341 | -0.108098 | 0.059 | 0.2 | 1 | 13 |
| COX7A2 (1 of many) | 0.39356 | -0.21408249 | 0.118 | 0.3 | 1 | 13 |
| rab44 | 0.39464 | -0.28481503 | 0.275 | 0.5 | 1 | 13 |
| mydgf | 0.39572 | 0.124401082 | 0.216 | 0.3 | 1 | 13 |
| XLOC-012680 | 0.39637 | 0.132293277 | 0.059 | 0 | 1 | 13 |
| arl9 | 0.39637 | 0.106756189 | 0.059 | 0 | 1 | 13 |
| arrb2a | 0.39637 | 0.132176424 | 0.059 | 0 | 1 | 13 |
| ccnl1b | 0.39637 | 0.159079268 | 0.059 | 0 | 1 | 13 |
| cd83 | 0.39637 | 0.159494048 | 0.059 | 0 | 1 | 13 |
| chp1 | 0.39637 | 0.142235252 | 0.059 | 0 | 1 | 13 |
| cluha | 0.39637 | 0.127176993 | 0.059 | 0 | 1 | 13 |
| dbt | 0.39637 | 0.10125799 | 0.059 | 0 | 1 | 13 |
| ddx27 | 0.39637 | 0.100682695 | 0.059 | 0 | 1 | 13 |
| dusp2 | 0.39637 | 0.127891545 | 0.059 | 0 | 1 | 13 |
| eri1 | 0.39637 | 0.122398775 | 0.059 | 0 | 1 | 13 |
| inpp5l | 0.39637 | 0.132487764 | 0.059 | 0 | 1 | 13 |
| kif11 | 0.39637 | 0.108686687 | 0.059 | 0 | 1 | 13 |
| lpxn | 0.39637 | 0.124499675 | 0.059 | 0 | 1 | 13 |
| mafgb | 0.39637 | 0.122258023 | 0.059 | 0 | 1 | 13 |

| | | | | | | |
|---|---|---|---|---|---|---|
| micall1a | 0.39637 | 0.104556664 | 0.059 | 0 | 1 | 13 |
| ndufaf5 | 0.39637 | 0.113294597 | 0.059 | 0 | 1 | 13 |
| pde6h | 0.39637 | 0.119332943 | 0.059 | 0 | 1 | 13 |
| prpf8 | 0.39637 | 0.145360951 | 0.059 | 0 | 1 | 13 |
| rab8a | 0.39637 | 0.16014666 | 0.059 | 0 | 1 | 13 |
| reps1 | 0.39637 | 0.165828555 | 0.059 | 0 | 1 | 13 |
| rnf19a | 0.39637 | 0.149183944 | 0.059 | 0 | 1 | 13 |
| si:ch211-68a17.7 | 0.39637 | 0.10561953 | 0.059 | 0 | 1 | 13 |
| si:ch73-37h15.2 | 0.39637 | 0.1462011 | 0.059 | 0 | 1 | 13 |
| si:dkeyp-104h9.5 | 0.39637 | 0.157615626 | 0.059 | 0 | 1 | 13 |
| slc11a2 | 0.39637 | 0.115237949 | 0.059 | 0 | 1 | 13 |
| spen | 0.39637 | 0.132444976 | 0.059 | 0 | 1 | 13 |
| stoml1 | 0.39637 | 0.161619624 | 0.059 | 0 | 1 | 13 |
| tnfsf11 | 0.39637 | 0.137433018 | 0.059 | 0 | 1 | 13 |
| tuba1b | 0.39637 | 0.126373298 | 0.059 | 0 | 1 | 13 |
| usp11 | 0.39637 | 0.129424195 | 0.059 | 0 | 1 | 13 |
| vma21 | 0.39637 | 0.104326255 | 0.059 | 0 | 1 | 13 |
| wdr5 | 0.39637 | 0.125143229 | 0.059 | 0 | 1 | 13 |
| zgc:113184 | 0.39637 | 0.136298728 | 0.059 | 0 | 1 | 13 |
| arf4a | 0.39637 | 0.121616269 | 0.059 | 0 | 1 | 13 |
| ccdc58 | 0.39637 | 0.138308553 | 0.059 | 0 | 1 | 13 |
| ccr6a | 0.39637 | 0.113621272 | 0.059 | 0 | 1 | 13 |
| hsd17b12b | 0.39637 | 0.128985576 | 0.059 | 0 | 1 | 13 |
| irak3 | 0.39637 | 0.131756124 | 0.059 | 0 | 1 | 13 |
| mcrip1 | 0.39637 | 0.110952339 | 0.059 | 0 | 1 | 13 |
| pcf11 | 0.39637 | 0.157196052 | 0.059 | 0 | 1 | 13 |
| soul4 | 0.39637 | 0.141609064 | 0.059 | 0 | 1 | 13 |
| ssrp1a | 0.39637 | 0.102060698 | 0.059 | 0 | 1 | 13 |
| tmtc3 | 0.39637 | 0.141492293 | 0.059 | 0 | 1 | 13 |
| tnfaip3 | 0.39637 | 0.126184611 | 0.059 | 0 | 1 | 13 |
| tra2b | 0.39637 | 0.104667041 | 0.059 | 0 | 1 | 13 |
| ubash3bb | 0.39637 | 0.113468048 | 0.059 | 0 | 1 | 13 |
| ube2c | 0.39637 | 0.136285234 | 0.059 | 0 | 1 | 13 |
| CU207221.1 | 0.39637 | 0.146639239 | 0.059 | 0 | 1 | 13 |
| LOC101883815 | 0.39637 | 0.144219458 | 0.059 | 0 | 1 | 13 |
| MCMDC2 | 0.39637 | 0.109700722 | 0.059 | 0 | 1 | 13 |
| SBNO2 | 0.39637 | 0.116944925 | 0.059 | 0 | 1 | 13 |
| aaas | 0.39637 | 0.101831719 | 0.059 | 0 | 1 | 13 |
| acads | 0.39637 | 0.130491658 | 0.059 | 0 | 1 | 13 |
| aktip | 0.39637 | 0.106562435 | 0.059 | 0 | 1 | 13 |
| ankrd13a | 0.39637 | 0.125462412 | 0.059 | 0 | 1 | 13 |
| ap1b1 | 0.39637 | 0.103416937 | 0.059 | 0 | 1 | 13 |
| arl8bb | 0.39637 | 0.10264159 | 0.059 | 0 | 1 | 13 |
| atp9b | 0.39637 | 0.155495587 | 0.059 | 0 | 1 | 13 |
| aurkb | 0.39637 | 0.10338426 | 0.059 | 0 | 1 | 13 |
| b4galt6 | 0.39637 | 0.151941711 | 0.059 | 0 | 1 | 13 |
| bcl2l1 | 0.39637 | 0.11599685 | 0.059 | 0 | 1 | 13 |
| bnip3lb | 0.39637 | 0.119310422 | 0.059 | 0 | 1 | 13 |

| | | | | | | |
|---|---|---|---|---|---|---|
| ccdc6a | 0.39637 | 0.109116848 | 0.059 | 0 | 1 | 13 |
| cdc27 | 0.39637 | 0.126286085 | 0.059 | 0 | 1 | 13 |
| cfap20 | 0.39637 | 0.113890759 | 0.059 | 0 | 1 | 13 |
| chuk | 0.39637 | 0.170024287 | 0.059 | 0 | 1 | 13 |
| csrnp1b | 0.39637 | 0.169745142 | 0.059 | 0 | 1 | 13 |
| ctdspl2a | 0.39637 | 0.112172716 | 0.059 | 0 | 1 | 13 |
| cxcr3.3 | 0.39637 | 0.11180763 | 0.059 | 0 | 1 | 13 |
| ddx18 | 0.39637 | 0.14631167 | 0.059 | 0 | 1 | 13 |
| ddx47 | 0.39637 | 0.129890139 | 0.059 | 0 | 1 | 13 |
| dis3l2 | 0.39637 | 0.129378002 | 0.059 | 0 | 1 | 13 |
| dlgap5 | 0.39637 | 0.117142688 | 0.059 | 0 | 1 | 13 |
| dnase2 | 0.39637 | 0.111455463 | 0.059 | 0 | 1 | 13 |
| dolpp1 | 0.39637 | 0.102597567 | 0.059 | 0 | 1 | 13 |
| edem3 | 0.39637 | 0.10601246 | 0.059 | 0 | 1 | 13 |
| eif1ad | 0.39637 | 0.124081941 | 0.059 | 0 | 1 | 13 |
| eif4e3 | 0.39637 | 0.143842037 | 0.059 | 0 | 1 | 13 |
| fam151b | 0.39637 | 0.122183009 | 0.059 | 0 | 1 | 13 |
| flot2a | 0.39637 | 0.125295494 | 0.059 | 0 | 1 | 13 |
| foxo4 | 0.39637 | 0.101531307 | 0.059 | 0 | 1 | 13 |
| gcc1 | 0.39637 | 0.126480375 | 0.059 | 0 | 1 | 13 |
| glod5 | 0.39637 | 0.119860351 | 0.059 | 0 | 1 | 13 |
| gmppab | 0.39637 | 0.116643581 | 0.059 | 0 | 1 | 13 |
| gpr146 | 0.39637 | 0.111962789 | 0.059 | 0 | 1 | 13 |
| grk3 | 0.39637 | 0.128081888 | 0.059 | 0 | 1 | 13 |
| inpp4aa | 0.39637 | 0.129983321 | 0.059 | 0 | 1 | 13 |
| itgb7 | 0.39637 | 0.10983999 | 0.059 | 0 | 1 | 13 |
| mad1l1 | 0.39637 | 0.146855625 | 0.059 | 0 | 1 | 13 |
| mad2l1 | 0.39637 | 0.133050769 | 0.059 | 0 | 1 | 13 |
| mapre2 | 0.39637 | 0.112707277 | 0.059 | 0 | 1 | 13 |
| med21 | 0.39637 | 0.107362637 | 0.059 | 0 | 1 | 13 |
| megf8l | 0.39637 | 0.121017313 | 0.059 | 0 | 1 | 13 |
| mest | 0.39637 | 0.116247738 | 0.059 | 0 | 1 | 13 |
| mff | 0.39637 | 0.100559998 | 0.059 | 0 | 1 | 13 |
| mrpl45 | 0.39637 | 0.140113908 | 0.059 | 0 | 1 | 13 |
| naa15b | 0.39637 | 0.124715588 | 0.059 | 0 | 1 | 13 |
| nip7 | 0.39637 | 0.102749154 | 0.059 | 0 | 1 | 13 |
| nrbp2a | 0.39637 | 0.160435539 | 0.059 | 0 | 1 | 13 |
| nrip1a | 0.39637 | 0.160213579 | 0.059 | 0 | 1 | 13 |
| nt5c2a | 0.39637 | 0.16236687 | 0.059 | 0 | 1 | 13 |
| nup98 | 0.39637 | 0.107702296 | 0.059 | 0 | 1 | 13 |
| ogdha | 0.39637 | 0.105191559 | 0.059 | 0 | 1 | 13 |
| osgn1 | 0.39637 | 0.127555574 | 0.059 | 0 | 1 | 13 |
| parpbp | 0.39637 | 0.110936054 | 0.059 | 0 | 1 | 13 |
| pdpk1b | 0.39637 | 0.11461261 | 0.059 | 0 | 1 | 13 |
| ppp2r5ea | 0.39637 | 0.11599116 | 0.059 | 0 | 1 | 13 |
| prdx1 | 0.39637 | 0.161434303 | 0.059 | 0 | 1 | 13 |
| ptgis | 0.39637 | 0.11369858 | 0.059 | 0 | 1 | 13 |
| rab11bb | 0.39637 | 0.134284656 | 0.059 | 0 | 1 | 13 |

| | | | | | | |
|---|---|---|---|---|---|---|
| rab3gap1 | 0.39637 | 0.105537509 | 0.059 | 0 | 1 | 13 |
| rasgrp3 | 0.39637 | 0.169267646 | 0.059 | 0 | 1 | 13 |
| rnf213a | 0.39637 | 0.136745923 | 0.059 | 0 | 1 | 13 |
| rnpepl1 | 0.39637 | 0.13541278 | 0.059 | 0 | 1 | 13 |
| sec22bb | 0.39637 | 0.106043495 | 0.059 | 0 | 1 | 13 |
| si:ch211-284b7.3 | 0.39637 | 0.102844057 | 0.059 | 0 | 1 | 13 |
| si:ch211-39i2.2 | 0.39637 | 0.153478548 | 0.059 | 0 | 1 | 13 |
| si:dkey-105i14.1 | 0.39637 | 0.117967891 | 0.059 | 0 | 1 | 13 |
| si:dkeyp-121d4.3 | 0.39637 | 0.132779984 | 0.059 | 0 | 1 | 13 |
| sipa1l3 | 0.39637 | 0.132233552 | 0.059 | 0 | 1 | 13 |
| slc7a3b | 0.39637 | 0.111023012 | 0.059 | 0 | 1 | 13 |
| sox5 | 0.39637 | 0.124204521 | 0.059 | 0 | 1 | 13 |
| srcap | 0.39637 | 0.110046728 | 0.059 | 0 | 1 | 13 |
| stambpb | 0.39637 | 0.125199556 | 0.059 | 0 | 1 | 13 |
| suds3 | 0.39637 | 0.105601469 | 0.059 | 0 | 1 | 13 |
| supt6h | 0.39637 | 0.102339727 | 0.059 | 0 | 1 | 13 |
| taz | 0.39637 | 0.1115508 | 0.059 | 0 | 1 | 13 |
| tbc1d10c | 0.39637 | 0.123658608 | 0.059 | 0 | 1 | 13 |
| timm10b | 0.39637 | 0.145756379 | 0.059 | 0 | 1 | 13 |
| tmem33 | 0.39637 | 0.106708201 | 0.059 | 0 | 1 | 13 |
| tmx4 | 0.39637 | 0.128188899 | 0.059 | 0 | 1 | 13 |
| ube2a | 0.39637 | 0.112811287 | 0.059 | 0 | 1 | 13 |
| upf3b | 0.39637 | 0.1214394 | 0.059 | 0 | 1 | 13 |
| usp44 | 0.39637 | 0.153783121 | 0.059 | 0 | 1 | 13 |
| zbtb7b | 0.39637 | 0.113808671 | 0.059 | 0 | 1 | 13 |
| zgc:163080 | 0.39637 | 0.106645242 | 0.059 | 0 | 1 | 13 |
| zmpste24 | 0.39637 | 0.146495477 | 0.059 | 0 | 1 | 13 |
| RANBP2 | 0.39637 | 0.159131055 | 0.059 | 0 | 1 | 13 |
| arfgap3 | 0.39637 | 0.151662623 | 0.059 | 0 | 1 | 13 |
| babam2 | 0.39637 | 0.135359216 | 0.059 | 0 | 1 | 13 |
| dennd4a | 0.39637 | 0.118787623 | 0.059 | 0 | 1 | 13 |
| ftsj1 | 0.39637 | 0.128400585 | 0.059 | 0 | 1 | 13 |
| gemin7 | 0.39637 | 0.139714731 | 0.059 | 0 | 1 | 13 |
| lpcat3 | 0.39637 | 0.101819594 | 0.059 | 0 | 1 | 13 |
| mea1 | 0.39637 | 0.100498607 | 0.059 | 0 | 1 | 13 |
| pcsk7 | 0.39637 | 0.100682574 | 0.059 | 0 | 1 | 13 |
| NC-002333.17 | 0.39637 | 0.138452151 | 0.059 | 0 | 1 | 13 |
| slc12a2 | 0.39637 | 0.150276953 | 0.059 | 0 | 1 | 13 |
| srek1ip1 | 0.39637 | 0.124415469 | 0.059 | 0 | 1 | 13 |
| XLOC-032002 | 0.39637 | 0.134022416 | 0.059 | 0 | 1 | 13 |
| b3gnt2l | 0.39637 | 0.130678789 | 0.059 | 0 | 1 | 13 |
| med30 | 0.39637 | 0.177230112 | 0.059 | 0 | 1 | 13 |
| rab1ba | 0.39637 | 0.158744143 | 0.059 | 0 | 1 | 13 |
| si:ch211-286b5.5 | 0.39637 | 0.116741946 | 0.059 | 0 | 1 | 13 |
| tmc8 | 0.39637 | 0.18835166 | 0.059 | 0 | 1 | 13 |
| wnk1a | 0.39637 | 0.124315216 | 0.059 | 0 | 1 | 13 |
| CABZ01033394.1 | 0.39637 | 0.160310753 | 0.059 | 0 | 1 | 13 |
| TADA1 | 0.39637 | 0.128045832 | 0.059 | 0 | 1 | 13 |

| | | | | | | |
|---|---|---|---|---|---|---|
| anxa1a | 0.39637 | 0.109406904 | 0.059 | 0 | 1 | 13 |
| fosl2 | 0.39637 | 0.186692362 | 0.059 | 0 | 1 | 13 |
| hspe1 | 0.39637 | 0.136080294 | 0.059 | 0 | 1 | 13 |
| irf5 | 0.39637 | 0.111201754 | 0.059 | 0 | 1 | 13 |
| mboat2a | 0.39637 | 0.131320502 | 0.059 | 0 | 1 | 13 |
| mrps26 | 0.39637 | 0.141929172 | 0.059 | 0 | 1 | 13 |
| pgap2 | 0.39637 | 0.117475443 | 0.059 | 0 | 1 | 13 |
| pon1 | 0.39637 | 0.113535221 | 0.059 | 0 | 1 | 13 |
| rab41 | 0.39637 | 0.122410879 | 0.059 | 0 | 1 | 13 |
| ralba | 0.39637 | 0.123109054 | 0.059 | 0 | 1 | 13 |
| rbl1 | 0.39637 | 0.153268204 | 0.059 | 0 | 1 | 13 |
| senp2 | 0.39637 | 0.112572399 | 0.059 | 0 | 1 | 13 |
| serpina10a | 0.39637 | 0.134358481 | 0.059 | 0 | 1 | 13 |
| si:dkey-19e4.5 | 0.39637 | 0.113248571 | 0.059 | 0 | 1 | 13 |
| si:dkeyp-61b2.1 | 0.39637 | 0.129347902 | 0.059 | 0 | 1 | 13 |
| spns1 | 0.39637 | 0.117063174 | 0.059 | 0 | 1 | 13 |
| tmem192 | 0.39637 | 0.10987962 | 0.059 | 0 | 1 | 13 |
| ttc39c | 0.39637 | 0.112470965 | 0.059 | 0 | 1 | 13 |
| zgc:110626 | 0.39637 | 0.116402964 | 0.059 | 0 | 1 | 13 |
| zgc:153409 | 0.39637 | 0.134587198 | 0.059 | 0 | 1 | 13 |
| zgc:153733 | 0.39637 | 0.171863005 | 0.059 | 0 | 1 | 13 |
| bloc1s6 | 0.39637 | 0.127983947 | 0.059 | 0 | 1 | 13 |
| kmt2d | 0.39637 | 0.106689917 | 0.059 | 0 | 1 | 13 |
| mnta | 0.39637 | 0.15694724 | 0.059 | 0 | 1 | 13 |
| pafah1b1b | 0.39637 | 0.179117767 | 0.059 | 0 | 1 | 13 |
| ppcs | 0.39637 | 0.121059769 | 0.059 | 0 | 1 | 13 |
| pphln1 | 0.39637 | 0.10975701 | 0.059 | 0 | 1 | 13 |
| psmd11a | 0.39637 | 0.148816622 | 0.059 | 0 | 1 | 13 |
| rnf2 | 0.39637 | 0.162367897 | 0.059 | 0 | 1 | 13 |
| spina | 0.39637 | 0.182347273 | 0.059 | 0 | 1 | 13 |
| vps37b | 0.39637 | 0.100974914 | 0.059 | 0 | 1 | 13 |
| FP085398.1 | 0.39637 | 0.110239196 | 0.059 | 0 | 1 | 13 |
| XLOC-000946 | 0.39637 | 0.117808334 | 0.059 | 0 | 1 | 13 |
| aldh3a2b | 0.39637 | 0.154294626 | 0.059 | 0 | 1 | 13 |
| rhogd | 0.39637 | 0.122528932 | 0.059 | 0 | 1 | 13 |
| smg7 | 0.39637 | 0.12741274 | 0.059 | 0 | 1 | 13 |
| tp53inp1 | 0.39637 | 0.157089176 | 0.059 | 0 | 1 | 13 |
| zgc:174906 | 0.39637 | 0.128535406 | 0.059 | 0 | 1 | 13 |
| CR848841.1 | 0.39706 | 0.103308287 | 0.196 | 0.3 | 1 | 13 |
| rp2 | 0.39727 | -0.12651078 | 0.059 | 0.2 | 1 | 13 |
| atp6v1e1a | 0.3973 | -0.26726532 | 0.059 | 0.2 | 1 | 13 |
| rpl13 | 0.39773 | 0.292200161 | 0.824 | 0.7 | 1 | 13 |
| flot1b | 0.39844 | -0.10671764 | 0.059 | 0.2 | 1 | 13 |
| unc50 | 0.39864 | -0.13508922 | 0.059 | 0.2 | 1 | 13 |
| rpl5b | 0.40403 | -0.32044689 | 0.588 | 0.7 | 1 | 13 |
| BX511132.1 | 0.40555 | -0.16255245 | 0.216 | 0.4 | 1 | 13 |
| lsm7 | 0.4059 | 0.332586026 | 0.196 | 0.1 | 1 | 13 |
| smarca2 | 0.40709 | -0.1202207 | 0.059 | 0.2 | 1 | 13 |

| | | | | | | |
|---|---|---|---|---|---|---|
| aclya | 0.40726 | 0.210687495 | 0.137 | 0.2 | 1 | 13 |
| spopla | 0.40744 | -0.17065946 | 0.059 | 0.2 | 1 | 13 |
| mrps9 | 0.40793 | -0.12076468 | 0.059 | 0.2 | 1 | 13 |
| celf2 | 0.40848 | 0.198498095 | 0.471 | 0.5 | 1 | 13 |
| gpr183b | 0.4099 | -0.14021898 | 0.137 | 0.3 | 1 | 13 |
| rer1 | 0.40992 | -0.22998986 | 0.196 | 0.4 | 1 | 13 |
| dock8 | 0.41019 | -0.25521852 | 0.196 | 0.4 | 1 | 13 |
| ndfip2 | 0.41035 | 0.359884589 | 0.196 | 0.1 | 1 | 13 |
| GCA | 0.4108 | -0.12655701 | 0.314 | 0.5 | 1 | 13 |
| rab5aa | 0.41151 | 0.122732367 | 0.314 | 0.4 | 1 | 13 |
| phf8 | 0.41189 | -0.2446682 | 0.059 | 0.2 | 1 | 13 |
| btg3 | 0.41212 | -0.19411668 | 0.059 | 0.2 | 1 | 13 |
| ncbp3 | 0.41349 | 0.103085322 | 0.098 | 0.2 | 1 | 13 |
| gpalpp1 | 0.41351 | -0.10523255 | 0.059 | 0.2 | 1 | 13 |
| spi1b | 0.41353 | 0.290691979 | 0.451 | 0.4 | 1 | 13 |
| gadd45bb | 0.41359 | -0.27530756 | 0.059 | 0.2 | 1 | 13 |
| atad1b | 0.41426 | -0.11483958 | 0.059 | 0.2 | 1 | 13 |
| eif3c | 0.41451 | 0.127451362 | 0.294 | 0.4 | 1 | 13 |
| usp33 | 0.41515 | -0.22332701 | 0.059 | 0.2 | 1 | 13 |
| uggt1 | 0.41649 | -0.11390877 | 0.059 | 0.2 | 1 | 13 |
| cdc42 | 0.41709 | 0.116584915 | 0.412 | 0.5 | 1 | 13 |
| knop1 | 0.41711 | -0.10135834 | 0.059 | 0.2 | 1 | 13 |
| rwdd1 | 0.41766 | -0.12350682 | 0.059 | 0.2 | 1 | 13 |
| dars | 0.41769 | -0.15553585 | 0.059 | 0.2 | 1 | 13 |
| plecb | 0.41789 | -0.29934659 | 0.059 | 0.2 | 1 | 13 |
| polr2d | 0.4191 | -0.1523437 | 0.059 | 0.2 | 1 | 13 |
| si:ch73-238c9.1 | 0.41994 | -0.12427534 | 0.059 | 0.2 | 1 | 13 |
| foxg1c | 0.42011 | -0.19544212 | 0.059 | 0.2 | 1 | 13 |
| cd63 | 0.4202 | 0.106189133 | 0.882 | 0.9 | 1 | 13 |
| nmt1a | 0.42036 | -0.15667199 | 0.059 | 0.2 | 1 | 13 |
| meaf6 | 0.42041 | -0.12351494 | 0.059 | 0.2 | 1 | 13 |
| fundc2 | 0.42084 | 0.132222536 | 0.118 | 0.2 | 1 | 13 |
| nop58 | 0.42085 | -0.16936145 | 0.059 | 0.2 | 1 | 13 |
| tufm | 0.4209 | -0.2077705 | 0.059 | 0.2 | 1 | 13 |
| rae1 | 0.42114 | -0.17102157 | 0.059 | 0.2 | 1 | 13 |
| mrpl12 | 0.42119 | -0.11525042 | 0.059 | 0.2 | 1 | 13 |
| mrps18b | 0.42129 | -0.2071737 | 0.059 | 0.2 | 1 | 13 |
| ssh2b | 0.4214 | -0.15422357 | 0.059 | 0.2 | 1 | 13 |
| ddt | 0.42155 | -0.10927147 | 0.059 | 0.2 | 1 | 13 |
| si:ch73-248e21.7 | 0.42456 | -0.11328827 | 0.314 | 0.5 | 1 | 13 |
| cnpy1 | 0.42601 | -0.19673 | 0.392 | 0.6 | 1 | 13 |
| rpl36 | 0.42641 | -0.1672385 | 0.784 | 0.8 | 1 | 13 |
| eif4a3 | 0.42687 | 0.170721523 | 0.137 | 0.2 | 1 | 13 |
| capza1b | 0.42719 | -0.17094119 | 0.392 | 0.6 | 1 | 13 |
| gnai1 | 0.42847 | -0.13345284 | 0.216 | 0.4 | 1 | 13 |
| mvp | 0.43246 | 0.134058745 | 0.118 | 0.2 | 1 | 13 |
| slc30a7 | 0.43265 | 0.223454097 | 0.157 | 0.2 | 1 | 13 |
| ube2ia | 0.43559 | 0.154030394 | 0.118 | 0.2 | 1 | 13 |

| | | | | | | |
|---|---|---|---|---|---|---|
| tcp1 | 0.43699 | 0.116141408 | 0.294 | 0.4 | 1 | 13 |
| gorasp2 | 0.43796 | 0.190743777 | 0.157 | 0.2 | 1 | 13 |
| rpl29 | 0.43802 | -0.25384452 | 0.784 | 0.9 | 1 | 13 |
| fkbp5 | 0.43807 | -0.11366551 | 0.137 | 0.3 | 1 | 13 |
| tmem214 | 0.443 | 0.138791205 | 0.118 | 0.2 | 1 | 13 |
| spns3 | 0.44326 | 0.384232075 | 0.353 | 0.2 | 1 | 13 |
| pim1 | 0.44368 | 0.381096825 | 0.216 | 0.1 | 1 | 13 |
| gabarapb | 0.44392 | -0.13355658 | 0.216 | 0.4 | 1 | 13 |
| tmed9 | 0.44413 | 0.335829319 | 0.373 | 0.3 | 1 | 13 |
| atp1b1a | 0.4461 | -0.1571644 | 0.216 | 0.4 | 1 | 13 |
| copb1 | 0.44629 | -0.11312401 | 0.137 | 0.3 | 1 | 13 |
| rbx1 | 0.44633 | 0.197955042 | 0.255 | 0.3 | 1 | 13 |
| moxd1 | 0.44787 | -0.17834246 | 0.137 | 0.3 | 1 | 13 |
| serf2 | 0.44976 | -0.1218926 | 0.627 | 0.8 | 1 | 13 |
| h6pd | 0.45033 | 0.172154042 | 0.235 | 0.3 | 1 | 13 |
| rpl11 | 0.45198 | -0.23726258 | 0.725 | 0.9 | 1 | 13 |
| cyth1a | 0.45333 | -0.17204722 | 0.216 | 0.4 | 1 | 13 |
| nme2b.2 | 0.45362 | -0.12007137 | 0.216 | 0.4 | 1 | 13 |
| yrk | 0.45523 | 0.329990142 | 0.392 | 0.3 | 1 | 13 |
| si:ch73-91k6.2 | 0.45539 | 0.242839393 | 0.196 | 0.2 | 1 | 13 |
| slc25a5 | 0.45613 | -0.10400806 | 0.627 | 0.8 | 1 | 13 |
| brk1 | 0.45795 | -0.12124257 | 0.137 | 0.3 | 1 | 13 |
| myo1f | 0.46284 | 0.124751178 | 0.333 | 0.4 | 1 | 13 |
| mlec | 0.46558 | -0.16300626 | 0.137 | 0.3 | 1 | 13 |
| rpl23a | 0.46584 | -0.28998036 | 0.667 | 0.8 | 1 | 13 |
| rpl19 | 0.46754 | -0.22948506 | 0.608 | 0.8 | 1 | 13 |
| krtcap2 | 0.46771 | -0.16045848 | 0.51 | 0.7 | 1 | 13 |
| rpl17 | 0.46797 | -0.22410854 | 0.608 | 0.8 | 1 | 13 |
| rpl28 | 0.46914 | -0.25705403 | 0.745 | 0.9 | 1 | 13 |
| srrm1 | 0.46938 | 0.316502134 | 0.196 | 0.1 | 1 | 13 |
| rpl27 | 0.4698 | -0.12852083 | 0.745 | 0.9 | 1 | 13 |
| metap2b | 0.47047 | 0.260293683 | 0.216 | 0.2 | 1 | 13 |
| sept12 | 0.47116 | -0.20304492 | 0.137 | 0.3 | 1 | 13 |
| abcd3a | 0.47157 | -0.12199768 | 0.039 | 0.1 | 1 | 13 |
| itprip | 0.47157 | -0.11953421 | 0.039 | 0.1 | 1 | 13 |
| atrxl | 0.47157 | -0.10676999 | 0.039 | 0.1 | 1 | 13 |
| hbxb | 0.47157 | -0.1143428 | 0.039 | 0.1 | 1 | 13 |
| cebpg | 0.47157 | -0.14929828 | 0.039 | 0.1 | 1 | 13 |
| ciartb | 0.47157 | -0.13016469 | 0.039 | 0.1 | 1 | 13 |
| fem1c | 0.47157 | -0.13017498 | 0.039 | 0.1 | 1 | 13 |
| sf3b3 | 0.47157 | -0.10464675 | 0.039 | 0.1 | 1 | 13 |
| si:dkeyp-113d7.10 | 0.47157 | -0.12164692 | 0.039 | 0.1 | 1 | 13 |
| tspan3a | 0.47157 | -0.1183751 | 0.039 | 0.1 | 1 | 13 |
| zc3h15 | 0.47157 | -0.1015324 | 0.039 | 0.1 | 1 | 13 |
| cant1a | 0.47157 | -0.1203859 | 0.039 | 0.1 | 1 | 13 |
| pggt1b | 0.47157 | -0.12023182 | 0.039 | 0.1 | 1 | 13 |
| sh2b3 | 0.47157 | -0.11486504 | 0.039 | 0.1 | 1 | 13 |
| sppl3 | 0.47157 | -0.10909626 | 0.039 | 0.1 | 1 | 13 |

| | | | | | | |
|---|---|---|---|---|---|---|
| abhd15a | 0.47157 | -0.12887235 | 0.039 | 0.1 | 1 | 13 |
| emc1 | 0.47157 | -0.15870876 | 0.039 | 0.1 | 1 | 13 |
| fam78ab | 0.47157 | -0.13161598 | 0.039 | 0.1 | 1 | 13 |
| gskip | 0.47157 | -0.12685318 | 0.039 | 0.1 | 1 | 13 |
| hgsnat | 0.47157 | -0.13075504 | 0.039 | 0.1 | 1 | 13 |
| pgam5 | 0.47157 | -0.10474901 | 0.039 | 0.1 | 1 | 13 |
| rhbdd1 | 0.47157 | -0.12347512 | 0.039 | 0.1 | 1 | 13 |
| tmed4 | 0.47157 | -0.11906605 | 0.039 | 0.1 | 1 | 13 |
| zfp36l1a | 0.47157 | -0.13927371 | 0.039 | 0.1 | 1 | 13 |
| CABZ01086293.1 | 0.47157 | -0.14221593 | 0.039 | 0.1 | 1 | 13 |
| ddx23 | 0.47157 | -0.10881106 | 0.039 | 0.1 | 1 | 13 |
| dut | 0.47157 | -0.18519675 | 0.039 | 0.1 | 1 | 13 |
| gpr107 | 0.47157 | -0.12765202 | 0.039 | 0.1 | 1 | 13 |
| gramd4b | 0.47157 | -0.1056581 | 0.039 | 0.1 | 1 | 13 |
| ip6k2a | 0.47157 | -0.14519499 | 0.039 | 0.1 | 1 | 13 |
| mark2b | 0.47157 | -0.10714215 | 0.039 | 0.1 | 1 | 13 |
| nat15 | 0.47157 | -0.13360566 | 0.039 | 0.1 | 1 | 13 |
| ndufs8a | 0.47157 | -0.13705797 | 0.039 | 0.1 | 1 | 13 |
| tbc1d15 | 0.47157 | -0.16974677 | 0.039 | 0.1 | 1 | 13 |
| tcnbb | 0.47157 | -0.11496946 | 0.039 | 0.1 | 1 | 13 |
| zgc:165656 | 0.47157 | -0.11294675 | 0.039 | 0.1 | 1 | 13 |
| myo9bl | 0.47157 | -0.13786832 | 0.039 | 0.1 | 1 | 13 |
| wdr45b | 0.47157 | -0.10722271 | 0.039 | 0.1 | 1 | 13 |
| arrb2b | 0.47401 | 0.139023973 | 0.333 | 0.4 | 1 | 13 |
| csde1 | 0.47436 | -0.12119352 | 0.137 | 0.3 | 1 | 13 |
| mapk11 | 0.47451 | -0.16241733 | 0.137 | 0.3 | 1 | 13 |
| NDUFB1 | 0.47755 | 0.22038914 | 0.373 | 0.4 | 1 | 13 |
| lrp10 | 0.47901 | -0.13295133 | 0.078 | 0.2 | 1 | 13 |
| btf3 | 0.48066 | 0.250630197 | 0.627 | 0.6 | 1 | 13 |
| hmgb3b | 0.4809 | 0.181279793 | 0.137 | 0.2 | 1 | 13 |
| ank3b | 0.4824 | 0.349612973 | 0.235 | 0.1 | 1 | 13 |
| ier3ip1 | 0.48334 | 0.223399358 | 0.275 | 0.3 | 1 | 13 |
| ybx1 | 0.4844 | -0.23283457 | 0.745 | 0.9 | 1 | 13 |
| uqcrc1 | 0.48492 | 0.124428779 | 0.118 | 0.2 | 1 | 13 |
| canx | 0.4853 | 0.350096313 | 0.314 | 0.2 | 1 | 13 |
| rpl38 | 0.48798 | -0.2362434 | 0.745 | 0.9 | 1 | 13 |
| hp1bp3 | 0.48922 | -0.22433543 | 0.137 | 0.3 | 1 | 13 |
| mtpn | 0.49122 | -0.1354883 | 0.137 | 0.3 | 1 | 13 |
| capns1a | 0.49142 | 0.129897279 | 0.118 | 0.2 | 1 | 13 |
| cyfip2 | 0.49477 | -0.25855704 | 0.137 | 0.3 | 1 | 13 |
| srsf2a | 0.49495 | -0.21079333 | 0.216 | 0.4 | 1 | 13 |
| seta | 0.4958 | -0.20875546 | 0.137 | 0.3 | 1 | 13 |
| raf1b | 0.49581 | -0.18497948 | 0.137 | 0.3 | 1 | 13 |
| elocb | 0.50111 | 0.30688331 | 0.275 | 0.2 | 1 | 13 |
| rps28 | 0.50121 | -0.10403077 | 0.765 | 0.9 | 1 | 13 |
| rps21 | 0.50345 | -0.23793042 | 0.784 | 0.9 | 1 | 13 |
| hnrnpa0l | 0.50591 | 0.208343919 | 0.49 | 0.5 | 1 | 13 |
| xiap | 0.50802 | -0.10043639 | 0.078 | 0.2 | 1 | 13 |

| | | | | | | |
|---|---|---|---|---|---|---|
| LOC108179162 | 0.51265 | -0.1524362 | 0.745 | 0.9 | 1 | 13 |
| fam49bb | 0.51378 | 0.258824459 | 0.235 | 0.2 | 1 | 13 |
| srp9 | 0.51481 | 0.149671785 | 0.118 | 0.2 | 1 | 13 |
| ckap5 | 0.51487 | 0.284662883 | 0.137 | 0.1 | 1 | 13 |
| pcbp2 | 0.5164 | -0.21006892 | 0.078 | 0.2 | 1 | 13 |
| herpud1 | 0.51867 | 0.125927405 | 0.235 | 0.3 | 1 | 13 |
| rpl36a | 0.51989 | -0.17989409 | 0.745 | 0.9 | 1 | 13 |
| hnrnpabb | 0.5218 | 0.319285808 | 0.235 | 0.1 | 1 | 13 |
| si:dkey-203a12.9 | 0.52244 | 0.356528629 | 0.373 | 0.2 | 1 | 13 |
| CR855337.1 | 0.52327 | -0.11606509 | 0.078 | 0.2 | 1 | 13 |
| eif3i | 0.52403 | 0.227570661 | 0.294 | 0.3 | 1 | 13 |
| dnajc1 | 0.52721 | 0.15893692 | 0.373 | 0.4 | 1 | 13 |
| cd74b | 0.52767 | 0.11265598 | 0.235 | 0.3 | 1 | 13 |
| tmem189 | 0.52947 | -0.10196202 | 0.078 | 0.2 | 1 | 13 |
| cep85l | 0.53322 | -0.10768215 | 0.078 | 0.2 | 1 | 13 |
| dnase1l1 | 0.53844 | -0.11774195 | 0.078 | 0.2 | 1 | 13 |
| mctp2b | 0.53844 | 0.322356229 | 0.235 | 0.1 | 1 | 13 |
| stat1a | 0.53965 | 0.265737832 | 0.235 | 0.2 | 1 | 13 |
| slc44a2 | 0.54004 | -0.16654927 | 0.078 | 0.2 | 1 | 13 |
| map4k6 | 0.54456 | -0.11715672 | 0.078 | 0.2 | 1 | 13 |
| ubr4 | 0.54473 | -0.11751675 | 0.078 | 0.2 | 1 | 13 |
| lypla2 | 0.54522 | 0.144654286 | 0.157 | 0.2 | 1 | 13 |
| si:dkey-151g10.6 | 0.54663 | -0.15238894 | 0.765 | 0.8 | 1 | 13 |
| ptgir | 0.54681 | -0.16343212 | 0.078 | 0.2 | 1 | 13 |
| LEPROTL1 | 0.54725 | -0.11978363 | 0.078 | 0.2 | 1 | 13 |
| zmp:0000000521 | 0.54738 | -0.15371117 | 0.078 | 0.2 | 1 | 13 |
| mrtfab | 0.54791 | 0.309051983 | 0.235 | 0.1 | 1 | 13 |
| tdh | 0.54803 | -0.12803245 | 0.078 | 0.2 | 1 | 13 |
| esyt2a | 0.54812 | -0.12081541 | 0.078 | 0.2 | 1 | 13 |
| zgc:77112 | 0.54882 | 0.280516798 | 0.196 | 0.1 | 1 | 13 |
| pdlim1 | 0.55163 | 0.106898035 | 0.235 | 0.3 | 1 | 13 |
| u2af1 | 0.55259 | 0.163352879 | 0.157 | 0.2 | 1 | 13 |
| rad23b | 0.55274 | 0.115583437 | 0.235 | 0.3 | 1 | 13 |
| sft2d1 | 0.55324 | -0.1168756 | 0.078 | 0.2 | 1 | 13 |
| rgs14a | 0.55439 | 0.255859135 | 0.235 | 0.1 | 1 | 13 |
| rhoaa | 0.55442 | 0.250256554 | 0.235 | 0.2 | 1 | 13 |
| LOC100148254 | 0.55491 | 0.155258039 | 0.275 | 0.3 | 1 | 13 |
| maf1 | 0.5556 | -0.12576785 | 0.078 | 0.2 | 1 | 13 |
| manf | 0.55571 | -0.16167624 | 0.333 | 0.5 | 1 | 13 |
| CU075735.1 | 0.55583 | -0.16685139 | 0.235 | 0.4 | 1 | 13 |
| rnaseka | 0.5564 | -0.25158579 | 0.157 | 0.3 | 1 | 13 |
| pbdc1 | 0.5576 | -0.15627074 | 0.078 | 0.2 | 1 | 13 |
| nr3c1 | 0.55784 | -0.14434834 | 0.078 | 0.2 | 1 | 13 |
| rab5if | 0.55945 | -0.12672238 | 0.078 | 0.2 | 1 | 13 |
| klf13 | 0.55974 | -0.15525888 | 0.078 | 0.2 | 1 | 13 |
| psmb5 | 0.55975 | -0.13106446 | 0.078 | 0.2 | 1 | 13 |
| lnpa | 0.56024 | -0.1847457 | 0.078 | 0.2 | 1 | 13 |
| bri3 | 0.56028 | -0.15854892 | 0.078 | 0.2 | 1 | 13 |

| | | | | | | |
|---|---|---|---|---|---|---|
| rab11fip4b | 0.56087 | -0.14947448 | 0.078 | 0.2 | 1 | 13 |
| si:ch73-269m14.2 | 0.56102 | -0.16479293 | 0.078 | 0.2 | 1 | 13 |
| mphosph8 | 0.56111 | 0.275715128 | 0.196 | 0.1 | 1 | 13 |
| pdha1a | 0.56121 | -0.12616829 | 0.078 | 0.2 | 1 | 13 |
| si:ch73-62l21.1 | 0.56123 | -0.1413641 | 0.078 | 0.2 | 1 | 13 |
| prkcdb | 0.56255 | -0.19653317 | 0.078 | 0.2 | 1 | 13 |
| pin1 | 0.56274 | -0.11628771 | 0.078 | 0.2 | 1 | 13 |
| ralbb | 0.56311 | -0.13244111 | 0.078 | 0.2 | 1 | 13 |
| fem1b | 0.5632 | -0.21417059 | 0.078 | 0.2 | 1 | 13 |
| commd9 | 0.5632 | -0.15607637 | 0.078 | 0.2 | 1 | 13 |
| mogs | 0.56324 | -0.16843693 | 0.078 | 0.2 | 1 | 13 |
| arl5c | 0.56326 | -0.15824603 | 0.157 | 0.3 | 1 | 13 |
| sort1b | 0.56331 | -0.14716613 | 0.078 | 0.2 | 1 | 13 |
| mcm5 | 0.56341 | -0.18128409 | 0.078 | 0.2 | 1 | 13 |
| ilf3a | 0.56343 | -0.17328847 | 0.078 | 0.2 | 1 | 13 |
| cox8b | 0.56352 | 0.139821017 | 0.059 | 0.1 | 1 | 13 |
| rhoab | 0.56591 | 0.107754197 | 0.431 | 0.5 | 1 | 13 |
| lsm3 | 0.57016 | 0.158310541 | 0.157 | 0.2 | 1 | 13 |
| myofl | 0.57579 | 0.137570995 | 0.353 | 0.4 | 1 | 13 |
| calm2b | 0.57773 | 0.305141629 | 0.275 | 0.2 | 1 | 13 |
| si:ch73-208g10.1 | 0.58053 | 0.161850426 | 0.275 | 0.3 | 1 | 13 |
| sh3bgrl | 0.58174 | 0.247721443 | 0.333 | 0.3 | 1 | 13 |
| syncrip | 0.5844 | 0.249590325 | 0.314 | 0.3 | 1 | 13 |
| si:ch211-213a13.1 | 0.58454 | 0.234717796 | 0.235 | 0.1 | 1 | 13 |
| snrpd1 | 0.58645 | 0.205231858 | 0.196 | 0.2 | 1 | 13 |
| ISCU (1 of many) | 0.58767 | 0.236215016 | 0.235 | 0.2 | 1 | 13 |
| fli1b | 0.58777 | 0.263738189 | 0.216 | 0.1 | 1 | 13 |
| psme1 | 0.58803 | 0.255135964 | 0.235 | 0.2 | 1 | 13 |
| zgc:77650 | 0.5895 | 0.240474841 | 0.235 | 0.2 | 1 | 13 |
| slc9a3r1a | 0.59076 | 0.291285863 | 0.294 | 0.2 | 1 | 13 |
| rps25 | 0.59103 | -0.20249978 | 0.784 | 0.9 | 1 | 13 |
| si:dkey-163m14.2 | 0.59125 | 0.277865554 | 0.176 | 0.1 | 1 | 13 |
| hsbp1b | 0.59154 | -0.1959486 | 0.157 | 0.3 | 1 | 13 |
| gpx1a | 0.59274 | 0.185124749 | 0.196 | 0.2 | 1 | 13 |
| smim7 | 0.59291 | -0.12294627 | 0.157 | 0.3 | 1 | 13 |
| parp8 | 0.593 | 0.291338391 | 0.176 | 0.1 | 1 | 13 |
| mcfd2 | 0.59474 | -0.19388513 | 0.157 | 0.3 | 1 | 13 |
| eif4g1a | 0.59498 | 0.231153848 | 0.216 | 0.2 | 1 | 13 |
| sec22ba | 0.59503 | 0.108928866 | 0.235 | 0.3 | 1 | 13 |
| dedd1 | 0.59616 | 0.255532528 | 0.118 | 0.1 | 1 | 13 |
| ppp3ccb | 0.59654 | 0.226075879 | 0.235 | 0.1 | 1 | 13 |
| jpt2 | 0.59685 | 0.246931796 | 0.235 | 0.1 | 1 | 13 |
| hikeshi | 0.59703 | 0.136600912 | 0.157 | 0.2 | 1 | 13 |
| ppiaa | 0.59784 | -0.16236622 | 0.765 | 0.9 | 1 | 13 |
| cst14b.1 | 0.59857 | 0.175771466 | 0.196 | 0.2 | 1 | 13 |
| pafah1b1a | 0.59962 | 0.173462606 | 0.157 | 0.2 | 1 | 13 |
| hmha1b | 0.60062 | -0.12539098 | 0.255 | 0.4 | 1 | 13 |
| agr2 | 0.60415 | 0.142081839 | 0.941 | 0.9 | 1 | 13 |

| | | | | | | |
|---|---|---|---|---|---|---|
| cd59 | 0.60428 | 0.124546577 | 0.255 | 0.3 | 1 | 13 |
| alox5a | 0.60478 | 0.173617072 | 0.294 | 0.3 | 1 | 13 |
| sf1 | 0.60726 | 0.241106489 | 0.137 | 0.1 | 1 | 13 |
| ppp1r12a | 0.60813 | 0.245723559 | 0.157 | 0.1 | 1 | 13 |
| ubl3b | 0.60821 | 0.264346095 | 0.176 | 0.1 | 1 | 13 |
| pfkfb4b | 0.60909 | 0.175947685 | 0.176 | 0.2 | 1 | 13 |
| cyb5r3 | 0.60927 | 0.139388693 | 0.157 | 0.2 | 1 | 13 |
| lpcat2 | 0.61087 | 0.241895785 | 0.255 | 0.2 | 1 | 13 |
| anxa4 | 0.61156 | 0.279180062 | 0.196 | 0.1 | 1 | 13 |
| LOC101886015 | 0.61251 | 0.150043192 | 0.078 | 0.1 | 1 | 13 |
| dnajb9b | 0.61282 | 0.273470913 | 0.157 | 0.1 | 1 | 13 |
| si:ch73-281n10.2 | 0.61321 | 0.31309587 | 0.314 | 0.2 | 1 | 13 |
| pdia6 | 0.61321 | -0.23212227 | 0.333 | 0.5 | 1 | 13 |
| ube2na | 0.61342 | -0.1074481 | 0.176 | 0.3 | 1 | 13 |
| calr3b | 0.61546 | -0.10634934 | 0.373 | 0.5 | 1 | 13 |
| stx5al | 0.61586 | 0.125379766 | 0.059 | 0.1 | 1 | 13 |
| hmgn6 | 0.61849 | -0.1144593 | 0.451 | 0.6 | 1 | 13 |
| unc13d | 0.61944 | 0.139974165 | 0.275 | 0.3 | 1 | 13 |
| rpl10 | 0.61953 | -0.23506395 | 0.686 | 0.8 | 1 | 13 |
| hyou1 | 0.6202 | 0.116382949 | 0.255 | 0.3 | 1 | 13 |
| tmub2 | 0.62021 | 0.161332789 | 0.078 | 0.1 | 1 | 13 |
| mt-nd5 | 0.62026 | 0.106092492 | 0.157 | 0.2 | 1 | 13 |
| pitpnaa | 0.62035 | 0.249057156 | 0.157 | 0.1 | 1 | 13 |
| tardbpl | 0.62064 | 0.222013329 | 0.176 | 0.1 | 1 | 13 |
| XLOC-016094 | 0.62305 | 0.247291654 | 0.137 | 0.1 | 1 | 13 |
| syvn1 | 0.62337 | 0.193199417 | 0.196 | 0.2 | 1 | 13 |
| eef1b2 | 0.62346 | -0.18504248 | 0.647 | 0.8 | 1 | 13 |
| samsn1a | 0.62356 | 0.282089463 | 0.216 | 0.1 | 1 | 13 |
| psma4 | 0.62378 | 0.237641574 | 0.216 | 0.2 | 1 | 13 |
| zgc:172053 | 0.62539 | 0.205966547 | 0.216 | 0.2 | 1 | 13 |
| btg2 | 0.62544 | -0.26781294 | 0.098 | 0.2 | 1 | 13 |
| sdc4 | 0.62633 | 0.250161109 | 0.137 | 0.1 | 1 | 13 |
| acat2 | 0.62657 | 0.155997202 | 0.078 | 0.1 | 1 | 13 |
| vwa11 | 0.62815 | 0.285140909 | 0.176 | 0.1 | 1 | 13 |
| tspan2b | 0.63012 | 0.222468822 | 0.216 | 0.2 | 1 | 13 |
| rps5 | 0.6317 | -0.12123257 | 0.667 | 0.8 | 1 | 13 |
| fmnl1a | 0.63403 | 0.142862274 | 0.275 | 0.3 | 1 | 13 |
| FERMT3 (1 of many) | 0.63437 | -0.11771907 | 0.176 | 0.3 | 1 | 13 |
| snx17 | 0.63879 | 0.24194099 | 0.176 | 0.1 | 1 | 13 |
| faub | 0.63963 | 0.110525859 | 0.137 | 0.2 | 1 | 13 |
| ero1b | 0.64413 | 0.22235717 | 0.216 | 0.1 | 1 | 13 |
| si:dkey-11f4.14 | 0.64539 | 0.242011542 | 0.118 | 0.1 | 1 | 13 |
| rps15a | 0.6466 | -0.22138814 | 0.667 | 0.8 | 1 | 13 |
| ptprea | 0.64882 | -0.21264475 | 0.098 | 0.2 | 1 | 13 |
| ppp3r1a | 0.64921 | 0.250168996 | 0.196 | 0.1 | 1 | 13 |
| eif3ea | 0.65026 | 0.264535328 | 0.314 | 0.2 | 1 | 13 |
| ppm1aa | 0.6517 | 0.231088298 | 0.196 | 0.1 | 1 | 13 |
| anp32e | 0.65209 | 0.163784989 | 0.176 | 0.2 | 1 | 13 |

| | | | | | | |
|---|---|---|---|---|---|---|
| glipr1a | 0.65384 | 0.200282395 | 0.549 | 0.5 | 1 | 13 |
| rps19 | 0.65482 | -0.15908327 | 0.667 | 0.8 | 1 | 13 |
| itpa | 0.65583 | 0.121542814 | 0.059 | 0.1 | 1 | 13 |
| hck | 0.65696 | -0.12666907 | 0.176 | 0.3 | 1 | 13 |
| tmed8 | 0.65825 | -0.10755 | 0.098 | 0.2 | 1 | 13 |
| zgc:175264 | 0.66082 | -0.12142346 | 0.098 | 0.2 | 1 | 13 |
| h2afx1 | 0.6612 | 0.24076841 | 0.157 | 0.1 | 1 | 13 |
| dnajc3b | 0.66158 | 0.253921455 | 0.157 | 0.1 | 1 | 13 |
| ip6k2b | 0.66193 | 0.116357044 | 0.157 | 0.2 | 1 | 13 |
| derl2 | 0.66297 | 0.239097575 | 0.176 | 0.1 | 1 | 13 |
| itpripl2 | 0.66319 | 0.159241841 | 0.078 | 0.1 | 1 | 13 |
| atp5po | 0.66383 | 0.178253082 | 0.294 | 0.3 | 1 | 13 |
| psmc3 | 0.66415 | -0.10727379 | 0.176 | 0.3 | 1 | 13 |
| ndufa3 | 0.66532 | 0.23448418 | 0.176 | 0.1 | 1 | 13 |
| ptmaa | 0.66743 | -0.16300721 | 0.804 | 0.9 | 1 | 13 |
| dhrs13a.1 | 0.66754 | 0.208912223 | 0.235 | 0.2 | 1 | 13 |
| ppifb | 0.66792 | 0.193681732 | 0.216 | 0.1 | 1 | 13 |
| si:ch211-66k16.27 | 0.6682 | 0.233799671 | 0.118 | 0.1 | 1 | 13 |
| ndufab1b | 0.66839 | 0.100565193 | 0.137 | 0.2 | 1 | 13 |
| cmtm7 | 0.66847 | 0.161018052 | 0.216 | 0.1 | 1 | 13 |
| mrps33 | 0.6715 | 0.190841753 | 0.098 | 0.1 | 1 | 13 |
| syngr2a | 0.67213 | -0.15946148 | 0.353 | 0.5 | 1 | 13 |
| smarce1 | 0.67219 | 0.128834982 | 0.157 | 0.2 | 1 | 13 |
| ptk2ba | 0.67243 | -0.10398472 | 0.098 | 0.2 | 1 | 13 |
| eif4g2a | 0.67261 | -0.20202993 | 0.098 | 0.2 | 1 | 13 |
| fam120c | 0.67287 | 0.250129627 | 0.157 | 0.1 | 1 | 13 |
| safb | 0.6737 | 0.113957677 | 0.059 | 0.1 | 1 | 13 |
| rps20 | 0.67376 | -0.15335285 | 0.784 | 0.9 | 1 | 13 |
| zgc:92360 | 0.67421 | -0.11447907 | 0.098 | 0.2 | 1 | 13 |
| rpl18a | 0.67431 | -0.10567819 | 0.588 | 0.7 | 1 | 13 |
| tes | 0.67449 | 0.144779561 | 0.275 | 0.3 | 1 | 13 |
| arhgap4b | 0.67555 | 0.186679414 | 0.196 | 0.2 | 1 | 13 |
| txndc15 | 0.67711 | 0.133037391 | 0.157 | 0.2 | 1 | 13 |
| si:ch211-199g17.2 | 0.67763 | 0.140800916 | 0.078 | 0.1 | 1 | 13 |
| akap8l | 0.678 | 0.211691591 | 0.157 | 0.1 | 1 | 13 |
| hnrnpul1l | 0.68029 | -0.10646904 | 0.098 | 0.2 | 1 | 13 |
| tacc1 | 0.68127 | 0.238306109 | 0.176 | 0.1 | 1 | 13 |
| vsir | 0.6823 | -0.1667599 | 0.176 | 0.3 | 1 | 13 |
| kmt5aa | 0.68394 | 0.207523481 | 0.235 | 0.2 | 1 | 13 |
| si:ch211-203d1.3 | 0.68444 | -0.15480201 | 0.098 | 0.2 | 1 | 13 |
| cse1l | 0.68445 | 0.164683974 | 0.078 | 0.1 | 1 | 13 |
| si:dkey-21o19.2 | 0.68484 | -0.1578849 | 0.176 | 0.3 | 1 | 13 |
| zgc:110591 | 0.68522 | -0.13846855 | 0.098 | 0.2 | 1 | 13 |
| brd2a | 0.6858 | 0.220912724 | 0.137 | 0.1 | 1 | 13 |
| atp2c1 | 0.68641 | -0.15076863 | 0.098 | 0.2 | 1 | 13 |
| m6pr | 0.68873 | -0.13209794 | 0.098 | 0.2 | 1 | 13 |
| stat5a | 0.68925 | -0.12108686 | 0.098 | 0.2 | 1 | 13 |
| ptp4a2b | 0.68965 | 0.194492757 | 0.196 | 0.2 | 1 | 13 |

| | | | | | | |
|---|---|---|---|---|---|---|
| rps6ka1 | 0.68991 | 0.112279481 | 0.157 | 0.2 | 1 | 13 |
| dhrs9 | 0.69012 | -0.14454081 | 0.098 | 0.2 | 1 | 13 |
| idh1 | 0.69088 | -0.12704657 | 0.098 | 0.2 | 1 | 13 |
| sel1l | 0.69115 | -0.16293362 | 0.098 | 0.2 | 1 | 13 |
| sdhb | 0.6912 | 0.167966595 | 0.196 | 0.2 | 1 | 13 |
| stt3b | 0.69145 | 0.23589809 | 0.137 | 0.1 | 1 | 13 |
| usp9 | 0.69152 | -0.15903405 | 0.098 | 0.2 | 1 | 13 |
| rpl30 | 0.69167 | -0.17286293 | 0.667 | 0.8 | 1 | 13 |
| tollip | 0.69167 | -0.13181796 | 0.098 | 0.2 | 1 | 13 |
| uso1 | 0.69188 | -0.16827053 | 0.098 | 0.2 | 1 | 13 |
| vamp8 | 0.69192 | -0.14686044 | 0.098 | 0.2 | 1 | 13 |
| rrbp1b | 0.69208 | -0.14388672 | 0.176 | 0.3 | 1 | 13 |
| rab2a | 0.69276 | 0.174714896 | 0.196 | 0.2 | 1 | 13 |
| sf3b2 | 0.69613 | 0.203500603 | 0.137 | 0.1 | 1 | 13 |
| rbm8a | 0.69773 | 0.251382244 | 0.294 | 0.2 | 1 | 13 |
| psmb2 | 0.69992 | 0.123597247 | 0.078 | 0.1 | 1 | 13 |
| gnaia | 0.70097 | 0.21694161 | 0.137 | 0.1 | 1 | 13 |
| hspa5 | 0.70274 | 0.276650101 | 0.608 | 0.5 | 1 | 13 |
| cyb5a | 0.70515 | 0.219178365 | 0.157 | 0.1 | 1 | 13 |
| npm1a | 0.7057 | 0.25973404 | 0.157 | 0.1 | 1 | 13 |
| sumo3a | 0.70592 | 0.246961283 | 0.275 | 0.2 | 1 | 13 |
| snrpg | 0.70818 | 0.196947788 | 0.176 | 0.1 | 1 | 13 |
| rpl24 | 0.71019 | -0.19138111 | 0.686 | 0.8 | 1 | 13 |
| sec61a1l | 0.71159 | 0.152249816 | 0.294 | 0.3 | 1 | 13 |
| gdi2 | 0.71258 | 0.194624884 | 0.235 | 0.2 | 1 | 13 |
| ptbp1a | 0.71443 | 0.203932218 | 0.137 | 0.1 | 1 | 13 |
| gpia | 0.7164 | 0.203036342 | 0.137 | 0.1 | 1 | 13 |
| hypk | 0.7172 | 0.157658809 | 0.196 | 0.2 | 1 | 13 |
| hk1 | 0.71749 | 0.149116889 | 0.196 | 0.2 | 1 | 13 |
| hnrnpk | 0.71877 | 0.199409291 | 0.235 | 0.2 | 1 | 13 |
| rab5c | 0.71915 | 0.239753745 | 0.275 | 0.2 | 1 | 13 |
| gcat | 0.71925 | 0.138278201 | 0.078 | 0.1 | 1 | 13 |
| hmgb1a | 0.71951 | 0.192322106 | 0.314 | 0.3 | 1 | 13 |
| chd4a | 0.72335 | 0.129925563 | 0.078 | 0.1 | 1 | 13 |
| seh1l | 0.72556 | 0.100551875 | 0.059 | 0.1 | 1 | 13 |
| ankrd52a | 0.72594 | 0.139384803 | 0.078 | 0.1 | 1 | 13 |
| illr4 | 0.72648 | 0.180627224 | 0.216 | 0.2 | 1 | 13 |
| crlf3 | 0.72776 | 0.179978505 | 0.216 | 0.2 | 1 | 13 |
| cct7 | 0.72831 | 0.227320998 | 0.176 | 0.1 | 1 | 13 |
| fkbp2 | 0.7285 | 0.209767164 | 0.255 | 0.2 | 1 | 13 |
| fam162a | 0.72951 | 0.152647571 | 0.196 | 0.1 | 1 | 13 |
| lrrc59 | 0.73162 | 0.169344514 | 0.314 | 0.3 | 1 | 13 |
| ube2h | 0.73368 | -0.17978556 | 0.118 | 0.2 | 1 | 13 |
| znf706 | 0.73515 | 0.144033664 | 0.196 | 0.1 | 1 | 13 |
| puf60b | 0.73641 | 0.167093624 | 0.098 | 0.1 | 1 | 13 |
| calr | 0.73828 | -0.2334785 | 0.294 | 0.4 | 1 | 13 |
| npc2 | 0.74176 | 0.149760897 | 0.196 | 0.2 | 1 | 13 |
| rbbp6 | 0.7418 | 0.184530616 | 0.137 | 0.1 | 1 | 13 |

| | | | | | | |
|---|---|---|---|---|---|---|
| sap30l | 0.74199 | 0.177775707 | 0.216 | 0.2 | 1 | 13 |
| minos1 | 0.7446 | 0.165936891 | 0.196 | 0.2 | 1 | 13 |
| syap1 | 0.74516 | 0.182217932 | 0.137 | 0.1 | 1 | 13 |
| itgav | 0.74573 | 0.228374751 | 0.255 | 0.2 | 1 | 13 |
| si:ch211-22i13.2 | 0.74664 | 0.152949558 | 0.098 | 0.1 | 1 | 13 |
| LOC110438237 | 0.74749 | 0.179805427 | 0.235 | 0.2 | 1 | 13 |
| rps17 | 0.74803 | -0.1680852 | 0.706 | 0.8 | 1 | 13 |
| fuca2 | 0.75045 | 0.159773248 | 0.196 | 0.1 | 1 | 13 |
| lgals3a | 0.75088 | 0.195025758 | 0.157 | 0.1 | 1 | 13 |
| rhocb | 0.75113 | 0.191113807 | 0.176 | 0.1 | 1 | 13 |
| kcnk1a | 0.75173 | 0.191339682 | 0.137 | 0.1 | 1 | 13 |
| nono | 0.75265 | 0.1564386 | 0.098 | 0.1 | 1 | 13 |
| BX323555.3 | 0.75373 | -0.19310376 | 0.118 | 0.2 | 1 | 13 |
| cited4b | 0.75384 | 0.183551982 | 0.157 | 0.1 | 1 | 13 |
| ankrd44 | 0.75462 | 0.187716277 | 0.137 | 0.1 | 1 | 13 |
| si:ch211-262e15.1 | 0.75494 | 0.127288846 | 0.078 | 0.1 | 1 | 13 |
| pygb | 0.75514 | -0.16663922 | 0.373 | 0.5 | 1 | 13 |
| zfand3 | 0.75708 | 0.101560118 | 0.059 | 0.1 | 1 | 13 |
| si:dkey-177p2.6 | 0.76004 | 0.177151567 | 0.118 | 0.1 | 1 | 13 |
| pigt | 0.76028 | 0.144389448 | 0.098 | 0.1 | 1 | 13 |
| ppp2cb | 0.76228 | 0.16027134 | 0.176 | 0.1 | 1 | 13 |
| mettl5 | 0.76304 | 0.180553741 | 0.176 | 0.1 | 1 | 13 |
| anapc15 | 0.76485 | 0.179838448 | 0.118 | 0.1 | 1 | 13 |
| mki67 | 0.76599 | 0.111265388 | 0.078 | 0.1 | 1 | 13 |
| polr2b | 0.76876 | 0.121659544 | 0.078 | 0.1 | 1 | 13 |
| si:zfos-464b6.2 | 0.76906 | -0.15986049 | 0.196 | 0.3 | 1 | 13 |
| dnajc8 | 0.7699 | 0.114665751 | 0.078 | 0.1 | 1 | 13 |
| ap1m1 | 0.77039 | 0.115263875 | 0.176 | 0.2 | 1 | 13 |
| tm9sf3 | 0.77089 | 0.149433791 | 0.196 | 0.2 | 1 | 13 |
| txndc9 | 0.77378 | 0.160970783 | 0.157 | 0.1 | 1 | 13 |
| pnisr | 0.77422 | 0.163023777 | 0.157 | 0.1 | 1 | 13 |
| fdx1 | 0.77565 | 0.106395048 | 0.078 | 0.1 | 1 | 13 |
| rassf2a | 0.77627 | 0.139806456 | 0.098 | 0.1 | 1 | 13 |
| fundc1 | 0.7776 | 0.187694445 | 0.157 | 0.1 | 1 | 13 |
| mapk3 | 0.77991 | -0.16932758 | 0.118 | 0.2 | 1 | 13 |
| rps3a | 0.78351 | -0.13405619 | 0.49 | 0.6 | 1 | 13 |
| elf2b | 0.78467 | -0.10770129 | 0.118 | 0.2 | 1 | 13 |
| sptbn2 | 0.78799 | 0.117668069 | 0.176 | 0.2 | 1 | 13 |
| mief1 | 0.78828 | 0.106911234 | 0.078 | 0.1 | 1 | 13 |
| asna1 | 0.7885 | 0.100072238 | 0.078 | 0.1 | 1 | 13 |
| hars | 0.789 | 0.162985732 | 0.098 | 0.1 | 1 | 13 |
| grpel2 | 0.78916 | 0.165732412 | 0.118 | 0.1 | 1 | 13 |
| rtn4a | 0.78931 | 0.103522874 | 0.176 | 0.2 | 1 | 13 |
| prpf19 | 0.79069 | 0.101054935 | 0.078 | 0.1 | 1 | 13 |
| tfg | 0.79219 | 0.176263157 | 0.137 | 0.1 | 1 | 13 |
| cox5aa | 0.79362 | 0.174114788 | 0.431 | 0.4 | 1 | 13 |
| CABZ01079663.1 | 0.79378 | 0.153490957 | 0.118 | 0.1 | 1 | 13 |
| mef2d | 0.79529 | 0.127896983 | 0.098 | 0.1 | 1 | 13 |

| | | | | | | |
|---|---|---|---|---|---|---|
| top1l | 0.79572 | 0.111898858 | 0.078 | 0.1 | 1 | 13 |
| thoc5 | 0.79882 | 0.155556294 | 0.118 | 0.1 | 1 | 13 |
| cds2 | 0.79887 | 0.117526835 | 0.196 | 0.2 | 1 | 13 |
| pdia5 | 0.79991 | -0.14230208 | 0.118 | 0.2 | 1 | 13 |
| mrpl57 | 0.80051 | -0.10824359 | 0.118 | 0.2 | 1 | 13 |
| bhlhe40 | 0.80154 | -0.18569877 | 0.137 | 0.2 | 1 | 13 |
| bccip | 0.80254 | 0.161044015 | 0.137 | 0.1 | 1 | 13 |
| ap2m1a | 0.80269 | 0.104420459 | 0.275 | 0.3 | 1 | 13 |
| tardbp | 0.80323 | 0.174947301 | 0.157 | 0.1 | 1 | 13 |
| jdp2b | 0.80517 | 0.159446513 | 0.118 | 0.1 | 1 | 13 |
| cipcb | 0.80593 | 0.144684327 | 0.098 | 0.1 | 1 | 13 |
| adssl | 0.80754 | 0.14361394 | 0.118 | 0.1 | 1 | 13 |
| eif3ba | 0.80758 | -0.14311813 | 0.294 | 0.4 | 1 | 13 |
| svild | 0.80911 | 0.158777274 | 0.137 | 0.1 | 1 | 13 |
| si:ch211-105j21.9 | 0.8112 | 0.171516882 | 0.157 | 0.1 | 1 | 13 |
| map2k6 | 0.81121 | 0.141791037 | 0.216 | 0.2 | 1 | 13 |
| rplp2l | 0.81455 | -0.11124328 | 0.765 | 0.8 | 1 | 13 |
| zgc:85932 | 0.81567 | 0.141118896 | 0.118 | 0.1 | 1 | 13 |
| dnaja2a | 0.81699 | 0.149636008 | 0.176 | 0.1 | 1 | 13 |
| pxdc1b | 0.81722 | 0.14249501 | 0.118 | 0.1 | 1 | 13 |
| si:ch211-241f5.3 | 0.81879 | 0.142508822 | 0.176 | 0.1 | 1 | 13 |
| snx5 | 0.81905 | 0.156584419 | 0.176 | 0.1 | 1 | 13 |
| LOC101884451 | 0.81945 | 0.149506897 | 0.118 | 0.1 | 1 | 13 |
| blcap | 0.81987 | 0.157410367 | 0.176 | 0.1 | 1 | 13 |
| txnrd3 | 0.81999 | 0.15148544 | 0.118 | 0.1 | 1 | 13 |
| si:dkey-206d17.12 | 0.82053 | 0.156082612 | 0.118 | 0.1 | 1 | 13 |
| gtf2b | 0.82056 | 0.151805685 | 0.137 | 0.1 | 1 | 13 |
| eif3ja | 0.82059 | 0.125242464 | 0.176 | 0.1 | 1 | 13 |
| mcts1 | 0.82106 | -0.14389365 | 0.059 | 0.1 | 1 | 13 |
| aqp3a | 0.8218 | 0.141008353 | 0.098 | 0.1 | 1 | 13 |
| ncaldb | 0.82198 | 0.11810806 | 0.176 | 0.1 | 1 | 13 |
| si:dkey-199f5.8 | 0.82254 | 0.164881569 | 0.137 | 0.1 | 1 | 13 |
| zgc:77849 | 0.82263 | 0.148609805 | 0.137 | 0.1 | 1 | 13 |
| NC-002333.4 | 0.82619 | -0.11423725 | 0.314 | 0.4 | 1 | 13 |
| pgam1b | 0.82832 | 0.149428408 | 0.118 | 0.1 | 1 | 13 |
| mrps21 | 0.82849 | 0.1380381 | 0.098 | 0.1 | 1 | 13 |
| acbd3 | 0.83009 | 0.119338339 | 0.098 | 0.1 | 1 | 13 |
| ctsd | 0.8309 | -0.14656859 | 0.275 | 0.3 | 1 | 13 |
| mrps14 | 0.83132 | -0.12973192 | 0.059 | 0.1 | 1 | 13 |
| si:ch73-42p12.2 | 0.83133 | 0.158661936 | 0.157 | 0.1 | 1 | 13 |
| inpp5d | 0.83306 | 0.119099707 | 0.196 | 0.2 | 1 | 13 |
| creb3l3l | 0.83376 | 0.140241726 | 0.118 | 0.1 | 1 | 13 |
| stk40 | 0.83421 | 0.150150196 | 0.118 | 0.1 | 1 | 13 |
| psenen | 0.83511 | 0.106298098 | 0.196 | 0.2 | 1 | 13 |
| rps12 | 0.83546 | -0.1313369 | 0.745 | 0.8 | 1 | 13 |
| rpl13a | 0.83912 | -0.13771898 | 0.725 | 0.8 | 1 | 13 |
| ndufv2 | 0.84113 | 0.124152225 | 0.098 | 0.1 | 1 | 13 |
| wu:fb55g09 | 0.84241 | 0.133993534 | 0.118 | 0.1 | 1 | 13 |

| | | | | | | |
|---|---|---|---|---|---|---|
| aars | 0.84478 | 0.159008112 | 0.137 | 0.1 | 1 | 13 |
| rusc1 | 0.84596 | -0.13607505 | 0.059 | 0.1 | 1 | 13 |
| zgc:63587 | 0.84682 | 0.119571106 | 0.098 | 0.1 | 1 | 13 |
| cltcb | 0.8476 | 0.143318228 | 0.157 | 0.1 | 1 | 13 |
| zmat2 | 0.84897 | 0.14155497 | 0.118 | 0.1 | 1 | 13 |
| polr2l | 0.8498 | 0.120119077 | 0.098 | 0.1 | 1 | 13 |
| ndufa8 | 0.85361 | 0.143484478 | 0.216 | 0.2 | 1 | 13 |
| cdc42l | 0.85414 | 0.118341462 | 0.412 | 0.4 | 1 | 13 |
| march8 | 0.85465 | 0.130556555 | 0.157 | 0.1 | 1 | 13 |
| zmp:0000000650 | 0.8562 | 0.127030243 | 0.118 | 0.1 | 1 | 13 |
| CABZ01117603.1 | 0.85635 | -0.16847785 | 0.059 | 0.1 | 1 | 13 |
| slc38a5a | 0.85913 | -0.12995472 | 0.059 | 0.1 | 1 | 13 |
| ube2e3 | 0.85952 | 0.119719357 | 0.098 | 0.1 | 1 | 13 |
| gmds | 0.86199 | -0.15487063 | 0.235 | 0.3 | 1 | 13 |
| lin37 | 0.8635 | -0.12041783 | 0.059 | 0.1 | 1 | 13 |
| tm9sf4 | 0.86415 | 0.103573544 | 0.098 | 0.1 | 1 | 13 |
| rps24 | 0.86558 | -0.11886026 | 0.725 | 0.8 | 1 | 13 |
| arhgef1a | 0.86609 | 0.104799065 | 0.098 | 0.1 | 1 | 13 |
| odc1 | 0.86622 | 0.154996331 | 0.137 | 0.1 | 1 | 13 |
| trappc10 | 0.86802 | -0.10203575 | 0.098 | 0.1 | 1 | 13 |
| pfdn2 | 0.86892 | 0.116683389 | 0.118 | 0.1 | 1 | 13 |
| gabarapl2 | 0.87149 | 0.131794228 | 0.118 | 0.1 | 1 | 13 |
| rps3 | 0.87223 | -0.12183363 | 0.745 | 0.8 | 1 | 13 |
| rpl35a | 0.87379 | -0.11532709 | 0.725 | 0.8 | 1 | 13 |
| smndc1 | 0.87461 | 0.130364901 | 0.157 | 0.1 | 1 | 13 |
| ece2b | 0.87468 | -0.12926236 | 0.078 | 0.1 | 1 | 13 |
| fam173a | 0.87599 | 0.132860595 | 0.118 | 0.1 | 1 | 13 |
| marcksl1a | 0.87618 | 0.12801311 | 0.275 | 0.2 | 1 | 13 |
| LO017700.1 | 0.87628 | 0.14448105 | 0.137 | 0.1 | 1 | 13 |
| eif3hb | 0.87673 | 0.112161285 | 0.098 | 0.1 | 1 | 13 |
| lamtor5 | 0.87705 | 0.124555156 | 0.137 | 0.1 | 1 | 13 |
| mboat7 | 0.87991 | 0.133833204 | 0.118 | 0.1 | 1 | 13 |
| emc6 | 0.87991 | 0.125699352 | 0.118 | 0.1 | 1 | 13 |
| zgc:136439 | 0.88039 | -0.11252102 | 0.059 | 0.1 | 1 | 13 |
| rps6ka5 | 0.88129 | -0.11100688 | 0.059 | 0.1 | 1 | 13 |
| copa | 0.88142 | -0.12358241 | 0.137 | 0.2 | 1 | 13 |
| hic1 | 0.88322 | 0.125748885 | 0.137 | 0.1 | 1 | 13 |
| hexb | 0.88335 | 0.114123315 | 0.137 | 0.1 | 1 | 13 |
| LOC101885065 | 0.88378 | -0.10812216 | 0.137 | 0.2 | 1 | 13 |
| cul3b | 0.88491 | -0.10699183 | 0.059 | 0.1 | 1 | 13 |
| amfra | 0.88769 | 0.11647641 | 0.118 | 0.1 | 1 | 13 |
| mphosph6 | 0.88845 | -0.10264982 | 0.059 | 0.1 | 1 | 13 |
| btbd10b | 0.88925 | 0.100165103 | 0.098 | 0.1 | 1 | 13 |
| XLOC-033403 | 0.89613 | -0.10135658 | 0.059 | 0.1 | 1 | 13 |
| cox17 | 0.8963 | 0.126648655 | 0.333 | 0.3 | 1 | 13 |
| pmm2 | 0.8977 | 0.100233336 | 0.137 | 0.1 | 1 | 13 |
| pum1 | 0.89867 | 0.11419679 | 0.118 | 0.1 | 1 | 13 |
| adam10a | 0.90092 | 0.13384327 | 0.255 | 0.2 | 1 | 13 |

| | | | | | | |
|---|---|---|---|---|---|---|
| eif2a | 0.90156 | 0.104126586 | 0.118 | 0.1 | 1 | 13 |
| ap1s2 | 0.90201 | 0.105060934 | 0.118 | 0.1 | 1 | 13 |
| hnrnpm | 0.90225 | 0.107406073 | 0.118 | 0.1 | 1 | 13 |
| lamtor2 | 0.90334 | 0.104366593 | 0.118 | 0.1 | 1 | 13 |
| nap1l1 | 0.90414 | 0.112281214 | 0.216 | 0.2 | 1 | 13 |
| CABZ01075242.1 | 0.90692 | 0.106943081 | 0.137 | 0.1 | 1 | 13 |
| zyg11 | 0.91236 | 0.122299795 | 0.137 | 0.1 | 1 | 13 |
| ktn1 | 0.91314 | 0.104235044 | 0.118 | 0.1 | 1 | 13 |
| ugt1a7 | 0.91348 | 0.116336376 | 0.118 | 0.1 | 1 | 13 |
| XLOC-011085 | 0.91621 | 0.106863047 | 0.118 | 0.1 | 1 | 13 |
| caspa | 0.91778 | -0.1072414 | 0.157 | 0.2 | 1 | 13 |
| si:dkey-262k9.2 | 0.91875 | 0.108458544 | 0.118 | 0.1 | 1 | 13 |
| cpne1 | 0.92519 | 0.102498893 | 0.137 | 0.1 | 1 | 13 |
| kdm1a | 0.92703 | 0.108021388 | 0.118 | 0.1 | 1 | 13 |
| ms4a17a.8 | 6.011683 | -0.71979296 | 0 | 0.35 | 1 | 14 |
| itm2ba | 0.00016 | -0.79331723 | 0.3 | 0.85 | 1 | 14 |
| gpx4a | 0.00025 | -0.4990056 | 0 | 0.3 | 1 | 14 |
| timp2a | 0.00036 | -0.64452372 | 0.475 | 0.95 | 1 | 14 |
| krt18a.1 | 0.00042 | 0.872066033 | 0.625 | 0.25 | 1 | 14 |
| rpl24 | 0.00047 | 0.417175863 | 1 | 0.75 | 1 | 14 |
| adh8a | 0.00066 | -0.87102255 | 0.35 | 0.85 | 1 | 14 |
| XLOC-032002 | 0.00079 | -0.80534841 | 0.025 | 0.4 | 1 | 14 |
| triqk | 0.00101 | -0.46820337 | 0 | 0.25 | 1 | 14 |
| sppl3 | 0.00101 | -0.57721343 | 0 | 0.25 | 1 | 14 |
| atp6ap1b | 0.00101 | -0.464135 | 0 | 0.25 | 1 | 14 |
| fbln1 | 0.00101 | -0.60844959 | 0 | 0.25 | 1 | 14 |
| thbs3a | 0.00101 | -0.60337151 | 0 | 0.25 | 1 | 14 |
| tmem63bb | 0.00101 | -0.50090698 | 0 | 0.25 | 1 | 14 |
| xbp1 | 0.00117 | -0.71132516 | 0.425 | 0.85 | 1 | 14 |
| rps7 | 0.00142 | 0.21429381 | 0.85 | 0.85 | 1 | 14 |
| rpl36a | 0.00152 | 0.23880854 | 1 | 0.95 | 1 | 14 |
| mt-co2 | 0.00156 | -0.53514792 | 0.65 | 1 | 1 | 14 |
| rps10 | 0.00205 | 0.335274029 | 1 | 0.9 | 1 | 14 |
| rps13 | 0.00207 | 0.309477447 | 1 | 0.9 | 1 | 14 |
| tmem176l.1 | 0.00227 | -0.59673487 | 0.475 | 0.75 | 1 | 14 |
| LOC108190717 | 0.00231 | -0.67062261 | 0.025 | 0.35 | 1 | 14 |
| vwde | 0.00244 | 0.903670585 | 0.275 | 0 | 1 | 14 |
| lsm6 | 0.00244 | 0.512733563 | 0.275 | 0 | 1 | 14 |
| cirbpb | 0.00263 | 0.464358042 | 0.975 | 0.75 | 1 | 14 |
| anxa1a | 0.0028 | -0.53104262 | 0.325 | 0.75 | 1 | 14 |
| col1a2 | 0.00298 | -0.21272004 | 1 | 1 | 1 | 14 |
| rpl17 | 0.00323 | 0.254962031 | 0.95 | 0.95 | 1 | 14 |
| rpl35a | 0.00367 | 0.350044383 | 1 | 0.75 | 1 | 14 |
| trabd2b | 0.00375 | -0.55880434 | 0.05 | 0.4 | 1 | 14 |
| galnt12 | 0.00388 | -0.39770852 | 0 | 0.2 | 1 | 14 |
| cers2a | 0.00388 | -0.37927391 | 0 | 0.2 | 1 | 14 |
| fam114a2 | 0.00388 | -0.34112298 | 0 | 0.2 | 1 | 14 |
| gadd45gip1 | 0.00388 | -0.32119607 | 0 | 0.2 | 1 | 14 |

| | | | | | | |
|---|---|---|---|---|---|---|
| kif1bp | 0.00388 | -0.38509636 | 0 | 0.2 | 1 | 14 |
| rcn3 | 0.00388 | -0.43609296 | 0 | 0.2 | 1 | 14 |
| sc:d217 | 0.00388 | -0.40287234 | 0 | 0.2 | 1 | 14 |
| tnfrsf1a | 0.00388 | -0.48452613 | 0 | 0.2 | 1 | 14 |
| XLOC-033137 | 0.00388 | -0.49837724 | 0 | 0.2 | 1 | 14 |
| cstf1 | 0.00388 | -0.33016827 | 0 | 0.2 | 1 | 14 |
| edem1 | 0.00388 | -0.40567249 | 0 | 0.2 | 1 | 14 |
| emilin2a | 0.00388 | -0.47990284 | 0 | 0.2 | 1 | 14 |
| gpx1a | 0.00388 | -0.42545814 | 0 | 0.2 | 1 | 14 |
| stk25b | 0.00388 | -0.36991786 | 0 | 0.2 | 1 | 14 |
| tent5c | 0.00388 | -0.37834849 | 0 | 0.2 | 1 | 14 |
| thrap3a | 0.00388 | -0.44059248 | 0 | 0.2 | 1 | 14 |
| vmp1 | 0.00388 | -0.38148197 | 0 | 0.2 | 1 | 14 |
| mgat1a | 0.00388 | -0.37071951 | 0 | 0.2 | 1 | 14 |
| castor2 | 0.00423 | 0.532422646 | 0.25 | 0 | 1 | 14 |
| sf1 | 0.00423 | 0.533978346 | 0.25 | 0 | 1 | 14 |
| sost | 0.00423 | 0.726137985 | 0.25 | 0 | 1 | 14 |
| sub1b | 0.00423 | 0.564644494 | 0.25 | 0 | 1 | 14 |
| ndufa12 | 0.00423 | 0.531460799 | 0.25 | 0 | 1 | 14 |
| epha3 | 0.00423 | 0.616599452 | 0.25 | 0 | 1 | 14 |
| ppp3ccb | 0.00447 | -0.47641658 | 0.05 | 0.35 | 1 | 14 |
| XLOC-026836 | 0.0045 | -0.95311461 | 0.1 | 0.45 | 1 | 14 |
| faua | 0.00462 | 0.360306434 | 0.95 | 0.9 | 1 | 14 |
| fkbp7 | 0.00479 | -0.67047107 | 0.25 | 0.55 | 1 | 14 |
| col2a1b | 0.0053 | -0.66969916 | 0.225 | 0.65 | 1 | 14 |
| lman2 | 0.00564 | -0.60049079 | 0.15 | 0.55 | 1 | 14 |
| mt-nd3 | 0.00565 | -0.49741064 | 0.5 | 0.9 | 1 | 14 |
| rpl36 | 0.00567 | 0.323846683 | 1 | 0.8 | 1 | 14 |
| si:ch73-1a9.3 | 0.006 | 0.356869429 | 0.7 | 0.65 | 1 | 14 |
| si:ch211-51e12.7 | 0.0063 | 0.140853483 | 0.275 | 0.35 | 1 | 14 |
| aoc2 | 0.0068 | -0.49862994 | 0.6 | 0.75 | 1 | 14 |
| CR753876.1 | 0.00692 | -0.734732 | 0.125 | 0.5 | 1 | 14 |
| hdac1 | 0.007 | 0.675146744 | 0.35 | 0.05 | 1 | 14 |
| snrpe | 0.00711 | 0.619058214 | 0.35 | 0.05 | 1 | 14 |
| ntn1b | 0.00726 | 0.564747973 | 0.225 | 0 | 1 | 14 |
| CR855375.3 | 0.00726 | 0.569007588 | 0.225 | 0 | 1 | 14 |
| hspe1 | 0.00726 | 0.358103763 | 0.225 | 0 | 1 | 14 |
| icn | 0.00726 | 0.508593711 | 0.225 | 0 | 1 | 14 |
| tmem26b | 0.00726 | 0.560447947 | 0.225 | 0 | 1 | 14 |
| arglu1a | 0.00726 | 0.523370146 | 0.225 | 0 | 1 | 14 |
| si:ch211-262e15.1 | 0.00726 | 0.498283539 | 0.225 | 0 | 1 | 14 |
| hoxa9b | 0.00726 | 0.518997501 | 0.225 | 0 | 1 | 14 |
| cldn11a | 0.00752 | -0.48776513 | 0.275 | 0.65 | 1 | 14 |
| tmem167a | 0.00784 | -0.62022648 | 0.1 | 0.45 | 1 | 14 |
| mt-atp6 | 0.00796 | -0.49629923 | 0.6 | 0.95 | 1 | 14 |
| htra1a | 0.00832 | -0.77158919 | 0.125 | 0.5 | 1 | 14 |
| mfap5 | 0.00836 | -0.60304132 | 0.5 | 0.85 | 1 | 14 |
| cav1 | 0.00854 | -0.75092198 | 0.175 | 0.55 | 1 | 14 |

| | | | | | | |
|---|---|---|---|---|---|---|
| rplp2l | 0.0086 | 0.395915812 | 0.975 | 0.8 | 1 | 14 |
| rps14 | 0.00871 | 0.220476334 | 0.95 | 0.95 | 1 | 14 |
| ccl25b | 0.00876 | -0.24123078 | 0.25 | 0.5 | 1 | 14 |
| rps27a | 0.00887 | 0.287631884 | 1 | 0.95 | 1 | 14 |
| zgc:195173 | 0.00901 | -0.6645328 | 0.05 | 0.35 | 1 | 14 |
| ptgdsb.2 | 0.00948 | -0.5420224 | 0.025 | 0.3 | 1 | 14 |
| hnrnpabb | 0.00983 | 0.280723895 | 0.525 | 0.55 | 1 | 14 |
| cxcl12a | 0.01034 | -0.72120241 | 0.1 | 0.45 | 1 | 14 |
| cnih1 | 0.01038 | -0.56839168 | 0.05 | 0.35 | 1 | 14 |
| hsp90ab1 | 0.01041 | 0.37393995 | 0.925 | 0.8 | 1 | 14 |
| ppt1 | 0.01053 | -0.55749681 | 0.175 | 0.55 | 1 | 14 |
| palld | 0.0107 | -0.45535368 | 0.125 | 0.45 | 1 | 14 |
| h2afvb | 0.01089 | 0.59642009 | 0.65 | 0.45 | 1 | 14 |
| ccdc80 | 0.01115 | -0.64606836 | 0.05 | 0.35 | 1 | 14 |
| arl6ip5b | 0.0112 | -0.59869909 | 0.25 | 0.65 | 1 | 14 |
| CABZ01085139.1 | 0.0116 | -0.62841363 | 0.05 | 0.35 | 1 | 14 |
| anxa2a | 0.01174 | -0.54914664 | 0.05 | 0.35 | 1 | 14 |
| tor1 | 0.01176 | -0.52189464 | 0.05 | 0.35 | 1 | 14 |
| RAMP1 | 0.01235 | 0.475858891 | 0.2 | 0 | 1 | 14 |
| abca1a | 0.01235 | 0.437993576 | 0.2 | 0 | 1 | 14 |
| fgfrl1a | 0.01235 | 0.465736561 | 0.2 | 0 | 1 | 14 |
| fkbp5 | 0.01235 | 0.393690066 | 0.2 | 0 | 1 | 14 |
| rbm7 | 0.01235 | 0.420754935 | 0.2 | 0 | 1 | 14 |
| lrp1aa | 0.01308 | -0.18163895 | 0.45 | 0.7 | 1 | 14 |
| lrrc17 | 0.01354 | 0.694844088 | 0.55 | 0.25 | 1 | 14 |
| h3f3d | 0.01417 | 0.398720406 | 0.875 | 0.75 | 1 | 14 |
| lgals2a | 0.01445 | -0.71272369 | 0.325 | 0.65 | 1 | 14 |
| XLOC-043761 | 0.01463 | -0.33685554 | 0 | 0.15 | 1 | 14 |
| atad1b | 0.01463 | -0.38082024 | 0 | 0.15 | 1 | 14 |
| gnl2 | 0.01463 | -0.2989541 | 0 | 0.15 | 1 | 14 |
| zgc:162939 | 0.01463 | -0.30870747 | 0 | 0.15 | 1 | 14 |
| zgc:171497 | 0.01463 | -0.33127968 | 0 | 0.15 | 1 | 14 |
| limk2 | 0.01463 | -0.32357421 | 0 | 0.15 | 1 | 14 |
| lmbr1l | 0.01463 | -0.30885863 | 0 | 0.15 | 1 | 14 |
| slc22a15 | 0.01463 | -0.27980012 | 0 | 0.15 | 1 | 14 |
| spryd7a | 0.01463 | -0.26543004 | 0 | 0.15 | 1 | 14 |
| xkr8.3 | 0.01463 | -0.366678 | 0 | 0.15 | 1 | 14 |
| ano10a | 0.01463 | -0.32038077 | 0 | 0.15 | 1 | 14 |
| cbx7a | 0.01463 | -0.44218386 | 0 | 0.15 | 1 | 14 |
| dcun1d4 | 0.01463 | -0.27401851 | 0 | 0.15 | 1 | 14 |
| galns | 0.01463 | -0.31138334 | 0 | 0.15 | 1 | 14 |
| hectd2 | 0.01463 | -0.3224344 | 0 | 0.15 | 1 | 14 |
| id2a | 0.01463 | -0.30931144 | 0 | 0.15 | 1 | 14 |
| kcnh7 | 0.01463 | -0.28384953 | 0 | 0.15 | 1 | 14 |
| mrpl37 | 0.01463 | -0.26749011 | 0 | 0.15 | 1 | 14 |
| mrpl40 | 0.01463 | -0.21846811 | 0 | 0.15 | 1 | 14 |
| nrp2a | 0.01463 | -0.31815547 | 0 | 0.15 | 1 | 14 |
| rras2 | 0.01463 | -0.2489494 | 0 | 0.15 | 1 | 14 |

| | | | | | | |
|---|---|---|---|---|---|---|
| snf8 | 0.01463 | -0.29347543 | 0 | 0.15 | 1 | 14 |
| srebf2 | 0.01463 | -0.31640488 | 0 | 0.15 | 1 | 14 |
| supt3h | 0.01463 | -0.3474633 | 0 | 0.15 | 1 | 14 |
| tmem241 | 0.01463 | -0.26104489 | 0 | 0.15 | 1 | 14 |
| trub1 | 0.01463 | -0.27366848 | 0 | 0.15 | 1 | 14 |
| zdhhc20a | 0.01463 | -0.29933896 | 0 | 0.15 | 1 | 14 |
| CU660013.4 | 0.01463 | -0.33371441 | 0 | 0.15 | 1 | 14 |
| ascc1 | 0.01463 | -0.21737261 | 0 | 0.15 | 1 | 14 |
| bco2l | 0.01463 | -0.27062378 | 0 | 0.15 | 1 | 14 |
| hnrnpdl | 0.01463 | -0.37014951 | 0 | 0.15 | 1 | 14 |
| sema3ab | 0.01463 | -0.32790022 | 0 | 0.15 | 1 | 14 |
| st3gal3a | 0.01463 | -0.2574753 | 0 | 0.15 | 1 | 14 |
| LOC103911361 | 0.01463 | -0.29668725 | 0 | 0.15 | 1 | 14 |
| arid2 | 0.01463 | -0.2853627 | 0 | 0.15 | 1 | 14 |
| sec13 | 0.01463 | -0.33100228 | 0 | 0.15 | 1 | 14 |
| dlg1l | 0.01463 | -0.27121531 | 0 | 0.15 | 1 | 14 |
| dyrk2 | 0.01463 | -0.31130383 | 0 | 0.15 | 1 | 14 |
| mier3a | 0.01463 | -0.32628944 | 0 | 0.15 | 1 | 14 |
| mprip | 0.01463 | -0.28187283 | 0 | 0.15 | 1 | 14 |
| si:ch211-204c21.1 | 0.01463 | -0.29828903 | 0 | 0.15 | 1 | 14 |
| tdg.1 | 0.01463 | -0.29465357 | 0 | 0.15 | 1 | 14 |
| asah1a | 0.01463 | -0.28000207 | 0 | 0.15 | 1 | 14 |
| dcbld2 | 0.01463 | -0.34109972 | 0 | 0.15 | 1 | 14 |
| fgf16 | 0.01463 | -0.38140962 | 0 | 0.15 | 1 | 14 |
| phlda3 | 0.01463 | -0.25679064 | 0 | 0.15 | 1 | 14 |
| rpusd2 | 0.01463 | -0.30616035 | 0 | 0.15 | 1 | 14 |
| si:dkey-199f5.8 | 0.01463 | -0.31435497 | 0 | 0.15 | 1 | 14 |
| tiam2a | 0.01463 | -0.26718239 | 0 | 0.15 | 1 | 14 |
| vps39 | 0.01463 | -0.38056662 | 0 | 0.15 | 1 | 14 |
| zgc:101016 | 0.01463 | -0.31465284 | 0 | 0.15 | 1 | 14 |
| eci2 | 0.01463 | -0.24828608 | 0 | 0.15 | 1 | 14 |
| furinb | 0.01463 | -0.33485784 | 0 | 0.15 | 1 | 14 |
| med7 | 0.01463 | -0.27337404 | 0 | 0.15 | 1 | 14 |
| plvapb | 0.01463 | -0.47877484 | 0 | 0.15 | 1 | 14 |
| tmem178b | 0.01463 | -0.35534796 | 0 | 0.15 | 1 | 14 |
| traf7 | 0.01463 | -0.30661481 | 0 | 0.15 | 1 | 14 |
| tuba8l2 | 0.01463 | -0.17336107 | 0 | 0.15 | 1 | 14 |
| znf1104 | 0.01463 | -0.29608168 | 0 | 0.15 | 1 | 14 |
| LOC110438375 | 0.01463 | -0.32362242 | 0 | 0.15 | 1 | 14 |
| psma8 | 0.01509 | 0.223716667 | 0.275 | 0.3 | 1 | 14 |
| adamts15a | 0.01516 | -0.50220197 | 0.275 | 0.65 | 1 | 14 |
| CR383676.1 | 0.01589 | -0.11727421 | 0.975 | 1 | 1 | 14 |
| tln1 | 0.0169 | -0.59176465 | 0.125 | 0.45 | 1 | 14 |
| net1 | 0.01696 | 0.546313812 | 0.35 | 0.05 | 1 | 14 |
| rpl26 | 0.01696 | 0.284125803 | 1 | 0.85 | 1 | 14 |
| glis2a | 0.01709 | -0.5512221 | 0.15 | 0.5 | 1 | 14 |
| serinc2 | 0.01726 | -0.59837948 | 0.15 | 0.5 | 1 | 14 |
| rpl11 | 0.01738 | 0.305105918 | 0.975 | 0.85 | 1 | 14 |

| | | | | | | |
|---|---|---|---|---|---|---|
| creb3l3l | 0.01785 | -0.48720726 | 0.125 | 0.45 | 1 | 14 |
| zgc:114188 | 0.01856 | 0.237387804 | 0.975 | 0.9 | 1 | 14 |
| ddx5 | 0.01914 | -0.5886315 | 0.325 | 0.7 | 1 | 14 |
| ccl39.7 | 0.01993 | -0.60131482 | 0.075 | 0.35 | 1 | 14 |
| pdlim1 | 0.02002 | -0.44918887 | 0.075 | 0.35 | 1 | 14 |
| lpl | 0.0203 | 0.58472754 | 0.35 | 0.05 | 1 | 14 |
| rpl37 | 0.02041 | 0.283686022 | 0.975 | 0.85 | 1 | 14 |
| si:ch211-15b10.6 | 0.02086 | 0.367971471 | 0.175 | 0 | 1 | 14 |
| ahcy | 0.02086 | 0.349980505 | 0.175 | 0 | 1 | 14 |
| ccdc12 | 0.02086 | 0.376302908 | 0.175 | 0 | 1 | 14 |
| ext2 | 0.02086 | 0.373515439 | 0.175 | 0 | 1 | 14 |
| loxl2b | 0.02086 | 0.478643094 | 0.175 | 0 | 1 | 14 |
| ptprsa | 0.02086 | 0.427281737 | 0.175 | 0 | 1 | 14 |
| atp5if1b | 0.02086 | 0.358347318 | 0.175 | 0 | 1 | 14 |
| sec63 | 0.02086 | 0.391445097 | 0.175 | 0 | 1 | 14 |
| usp42 | 0.02086 | 0.383844704 | 0.175 | 0 | 1 | 14 |
| znf706 | 0.02086 | 0.342050443 | 0.175 | 0 | 1 | 14 |
| mgst1.1 | 0.02124 | -0.44874958 | 0.25 | 0.6 | 1 | 14 |
| sema3d | 0.02218 | -0.23374629 | 0.2 | 0.45 | 1 | 14 |
| rps17 | 0.02255 | 0.314797064 | 0.775 | 0.7 | 1 | 14 |
| med15 | 0.02278 | -0.38304962 | 0.05 | 0.3 | 1 | 14 |
| fzd2 | 0.02284 | -0.5397014 | 0.125 | 0.45 | 1 | 14 |
| magt1 | 0.02302 | -0.50985871 | 0.125 | 0.45 | 1 | 14 |
| ctnnb1 | 0.02321 | 0.492322431 | 0.425 | 0.25 | 1 | 14 |
| hexb | 0.02333 | -0.34727897 | 0.025 | 0.25 | 1 | 14 |
| rpl9 | 0.02399 | 0.117440272 | 0.95 | 1 | 1 | 14 |
| dap1b | 0.02411 | -0.42013756 | 0.15 | 0.45 | 1 | 14 |
| dbn1 | 0.02427 | 0.691080043 | 0.475 | 0.15 | 1 | 14 |
| XLOC-000946 | 0.02428 | -0.38024666 | 0.05 | 0.3 | 1 | 14 |
| si:dkey-12l12.1 | 0.02437 | -0.62671521 | 0.3 | 0.65 | 1 | 14 |
| tnc | 0.02447 | 0.589842352 | 0.275 | 0.1 | 1 | 14 |
| mmp2 | 0.02497 | -0.37130284 | 0.725 | 0.9 | 1 | 14 |
| cfd | 0.02523 | -0.54854014 | 0.3 | 0.45 | 1 | 14 |
| abhd17c | 0.02535 | -0.43021107 | 0.025 | 0.25 | 1 | 14 |
| napab | 0.02565 | -0.27389334 | 0.1 | 0.35 | 1 | 14 |
| pcsk5b | 0.02573 | -0.49814823 | 0.025 | 0.25 | 1 | 14 |
| mt-nd2 | 0.02599 | -0.14880677 | 0.7 | 0.95 | 1 | 14 |
| cd81a | 0.02647 | -0.24709493 | 0.775 | 1 | 1 | 14 |
| golt1bb | 0.02718 | -0.56522555 | 0.125 | 0.4 | 1 | 14 |
| plpp5 | 0.0273 | -0.49253796 | 0.025 | 0.25 | 1 | 14 |
| cd28 | 0.0275 | -0.40548181 | 0.025 | 0.25 | 1 | 14 |
| oaz1b | 0.02768 | 0.242274352 | 0.5 | 0.5 | 1 | 14 |
| tgfb3 | 0.02784 | -0.58224796 | 0.15 | 0.4 | 1 | 14 |
| slit3 | 0.02787 | -0.4555128 | 0.225 | 0.55 | 1 | 14 |
| XLOC-000431 | 0.02827 | 0.801459301 | 0.45 | 0.15 | 1 | 14 |
| ppp1cb | 0.02849 | -0.47771505 | 0.025 | 0.25 | 1 | 14 |
| sept10 | 0.0285 | -0.50468021 | 0.1 | 0.4 | 1 | 14 |
| plekho2 | 0.02896 | -0.48081076 | 0.025 | 0.25 | 1 | 14 |

| | | | | | | |
|---|---|---|---|---|---|---|
| efna5a | 0.02927 | -0.51229124 | 0.025 | 0.25 | 1 | 14 |
| lasp1 | 0.0293 | -0.30623529 | 0.2 | 0.45 | 1 | 14 |
| sox18 | 0.02933 | -0.43965573 | 0.05 | 0.3 | 1 | 14 |
| prss23 | 0.02936 | -0.49087673 | 0.075 | 0.35 | 1 | 14 |
| CU929237.1 | 0.02976 | -0.54065516 | 0.4 | 0.75 | 1 | 14 |
| sptlc1 | 0.02979 | -0.45857342 | 0.025 | 0.25 | 1 | 14 |
| FP102052.2 | 0.0298 | -0.44118464 | 0.025 | 0.25 | 1 | 14 |
| sumo2b | 0.02986 | 0.444450074 | 0.55 | 0.4 | 1 | 14 |
| wbp2nl | 0.03016 | -0.48574672 | 0.05 | 0.3 | 1 | 14 |
| nucks1a | 0.03043 | 0.36966657 | 0.4 | 0.3 | 1 | 14 |
| vkorc1 | 0.03055 | -0.50767965 | 0.05 | 0.3 | 1 | 14 |
| rps23 | 0.03076 | 0.267257067 | 0.9 | 0.85 | 1 | 14 |
| sparc | 0.03114 | -0.25343431 | 0.9 | 1 | 1 | 14 |
| dad1 | 0.03125 | -0.45293463 | 0.35 | 0.7 | 1 | 14 |
| cdca4 | 0.03131 | -0.48905159 | 0.075 | 0.35 | 1 | 14 |
| loxl3b | 0.03134 | -0.56821102 | 0.05 | 0.3 | 1 | 14 |
| c1qtnf6a | 0.03247 | -0.53157694 | 0.05 | 0.3 | 1 | 14 |
| anxa13 | 0.03279 | -0.53534852 | 0.05 | 0.3 | 1 | 14 |
| pora | 0.03325 | -0.47137186 | 0.05 | 0.3 | 1 | 14 |
| zgc:162612 | 0.03374 | -0.48562405 | 0.05 | 0.3 | 1 | 14 |
| cct2 | 0.03378 | 0.523764815 | 0.4 | 0.15 | 1 | 14 |
| tuba8l | 0.03439 | 0.538355378 | 0.575 | 0.3 | 1 | 14 |
| COA4 | 0.03502 | 0.29686075 | 0.15 | 0 | 1 | 14 |
| aspn | 0.03502 | 0.410977547 | 0.15 | 0 | 1 | 14 |
| cpvl | 0.03502 | 0.362589911 | 0.15 | 0 | 1 | 14 |
| igsf10 | 0.03502 | 0.360423588 | 0.15 | 0 | 1 | 14 |
| kansl1a | 0.03502 | 0.346805177 | 0.15 | 0 | 1 | 14 |
| lsm8 | 0.03502 | 0.33010415 | 0.15 | 0 | 1 | 14 |
| mfap1 | 0.03502 | 0.330864383 | 0.15 | 0 | 1 | 14 |
| nccrp1 | 0.03502 | 0.284565201 | 0.15 | 0 | 1 | 14 |
| p4ha1b | 0.03502 | 0.361371137 | 0.15 | 0 | 1 | 14 |
| papola | 0.03502 | 0.358971138 | 0.15 | 0 | 1 | 14 |
| pttg1ipb | 0.03502 | 0.302076012 | 0.15 | 0 | 1 | 14 |
| rin3 | 0.03502 | 0.403801361 | 0.15 | 0 | 1 | 14 |
| scp2b | 0.03502 | 0.316289795 | 0.15 | 0 | 1 | 14 |
| stk35l | 0.03502 | 0.315938922 | 0.15 | 0 | 1 | 14 |
| stxbp6 | 0.03502 | 0.302155793 | 0.15 | 0 | 1 | 14 |
| tardbp | 0.03502 | 0.320634366 | 0.15 | 0 | 1 | 14 |
| tmem131l | 0.03502 | 0.353576638 | 0.15 | 0 | 1 | 14 |
| txndc12 | 0.03502 | 0.38576458 | 0.15 | 0 | 1 | 14 |
| znf292a | 0.03502 | 0.36705273 | 0.15 | 0 | 1 | 14 |
| cox17 | 0.03502 | 0.224947391 | 0.15 | 0 | 1 | 14 |
| cpxm1b | 0.03502 | 0.365483745 | 0.15 | 0 | 1 | 14 |
| fnbp1l | 0.03502 | 0.305137193 | 0.15 | 0 | 1 | 14 |
| gna13b | 0.03502 | 0.312200979 | 0.15 | 0 | 1 | 14 |
| lmnb1 | 0.03502 | 0.324588974 | 0.15 | 0 | 1 | 14 |
| med30 | 0.03502 | 0.287660119 | 0.15 | 0 | 1 | 14 |
| meox2b | 0.03502 | 0.354107517 | 0.15 | 0 | 1 | 14 |

| | | | | | | |
|---|---|---|---|---|---|---|
| mrps6 | 0.03502 | 0.306842665 | 0.15 | 0 | 1 | 14 |
| nuak1b | 0.03502 | 0.326525641 | 0.15 | 0 | 1 | 14 |
| rcc2 | 0.03502 | 0.393837986 | 0.15 | 0 | 1 | 14 |
| tshz2 | 0.03502 | 0.333090949 | 0.15 | 0 | 1 | 14 |
| vdrb | 0.03502 | 0.41527446 | 0.15 | 0 | 1 | 14 |
| si:dkey-102m7.3 | 0.03502 | 0.333070881 | 0.15 | 0 | 1 | 14 |
| snrpd3l | 0.03502 | 0.32966672 | 0.15 | 0 | 1 | 14 |
| col16a1 | 0.03523 | -0.48550019 | 0.35 | 0.7 | 1 | 14 |
| cd151 | 0.03524 | -0.49432581 | 0.15 | 0.45 | 1 | 14 |
| crip1 | 0.03592 | 0.53060262 | 0.5 | 0.4 | 1 | 14 |
| rps8a | 0.03665 | 0.280325686 | 0.925 | 0.85 | 1 | 14 |
| tpm4b | 0.037 | -0.6076354 | 0.25 | 0.55 | 1 | 14 |
| erh | 0.03718 | 0.256507797 | 0.35 | 0.35 | 1 | 14 |
| sumo3a | 0.03741 | 0.133724619 | 0.425 | 0.5 | 1 | 14 |
| hmgb2b | 0.03769 | 0.381108744 | 0.525 | 0.5 | 1 | 14 |
| cap1 | 0.0386 | -0.48344521 | 0.1 | 0.25 | 1 | 14 |
| tmem50a | 0.03861 | -0.43458977 | 0.15 | 0.45 | 1 | 14 |
| si:ch211-243a20.3 | 0.03883 | -0.4499454 | 0.625 | 0.9 | 1 | 14 |
| mdka | 0.03915 | 0.610288543 | 0.4 | 0.15 | 1 | 14 |
| fkbp3 | 0.04026 | 0.513543924 | 0.4 | 0.1 | 1 | 14 |
| si:dkey-151g10.6 | 0.04048 | 0.213273977 | 1 | 0.85 | 1 | 14 |
| galnt7 | 0.04119 | -0.35131211 | 0.075 | 0.3 | 1 | 14 |
| jund | 0.04346 | -0.3541414 | 0.1 | 0.35 | 1 | 14 |
| npm1a | 0.04368 | 0.439182937 | 0.3 | 0.05 | 1 | 14 |
| rab11a | 0.04377 | -0.51890518 | 0.15 | 0.45 | 1 | 14 |
| pomp | 0.04384 | 0.373950088 | 0.575 | 0.45 | 1 | 14 |
| hsd3b7 | 0.0439 | -0.47354809 | 0.15 | 0.45 | 1 | 14 |
| CABZ01074363.1 | 0.04431 | -0.49520287 | 0.125 | 0.4 | 1 | 14 |
| igfbp5b | 0.04433 | -0.5209442 | 0.275 | 0.6 | 1 | 14 |
| hnrnpa0l | 0.0445 | 0.378396352 | 0.825 | 0.65 | 1 | 14 |
| emx2 | 0.04523 | -0.51834383 | 0.225 | 0.55 | 1 | 14 |
| ppfibp1a | 0.04577 | -0.46907284 | 0.15 | 0.45 | 1 | 14 |
| rps9 | 0.04622 | 0.325304153 | 0.95 | 0.8 | 1 | 14 |
| sar1b | 0.04673 | -0.49616701 | 0.15 | 0.45 | 1 | 14 |
| slc43a3b | 0.04686 | -0.40196915 | 0.15 | 0.25 | 1 | 14 |
| hmgn7 | 0.04745 | 0.37628818 | 0.625 | 0.5 | 1 | 14 |
| rasgef1bb | 0.0478 | -0.34257856 | 0.075 | 0.3 | 1 | 14 |
| smu1a | 0.04845 | -0.22477866 | 0.05 | 0.25 | 1 | 14 |
| rps24 | 0.04923 | 0.27473154 | 0.95 | 0.85 | 1 | 14 |
| rbx1 | 0.04946 | 0.410426949 | 0.425 | 0.3 | 1 | 14 |
| cyp2n13 | 0.05055 | -0.52311827 | 0.025 | 0.2 | 1 | 14 |
| uqcrq | 0.05059 | 0.473907478 | 0.55 | 0.25 | 1 | 14 |
| si:dkey-177p2.6 | 0.05065 | 0.46993636 | 0.3 | 0.05 | 1 | 14 |
| angptl1a | 0.05101 | 0.723499016 | 0.3 | 0.05 | 1 | 14 |
| cntfr | 0.05158 | -0.12033895 | 0.4 | 0.6 | 1 | 14 |
| rpl27 | 0.05163 | 0.227018435 | 0.925 | 0.9 | 1 | 14 |
| ubc | 0.05173 | 0.439959152 | 0.75 | 0.55 | 1 | 14 |
| fkbp2 | 0.05174 | 0.497436086 | 0.3 | 0.1 | 1 | 14 |

| | | | | | | |
|---|---|---|---|---|---|---|
| gpatch8 | 0.05374 | -0.36006442 | 0.1 | 0.35 | 1 | 14 |
| chad | 0.05405 | -0.61251606 | 0.125 | 0.4 | 1 | 14 |
| cald1b | 0.05409 | -0.52302046 | 0.025 | 0.2 | 1 | 14 |
| gsna | 0.05427 | -0.17371223 | 0.05 | 0.2 | 1 | 14 |
| fabp3 | 0.05471 | -0.36056986 | 0.125 | 0.4 | 1 | 14 |
| CABZ01088271.1 | 0.05471 | -0.20280411 | 0 | 0.1 | 1 | 14 |
| CMKLR1 (1 of many) | 0.05471 | -0.26089785 | 0 | 0.1 | 1 | 14 |
| LOC103910007 | 0.05471 | -0.28580055 | 0 | 0.1 | 1 | 14 |
| XLOC-043788 | 0.05471 | -0.22197332 | 0 | 0.1 | 1 | 14 |
| abat | 0.05471 | -0.19544025 | 0 | 0.1 | 1 | 14 |
| acin1b | 0.05471 | -0.18962597 | 0 | 0.1 | 1 | 14 |
| acsl4a | 0.05471 | -0.22681849 | 0 | 0.1 | 1 | 14 |
| ak3 | 0.05471 | -0.23410774 | 0 | 0.1 | 1 | 14 |
| alkbh5 | 0.05471 | -0.17825804 | 0 | 0.1 | 1 | 14 |
| arhgap24 | 0.05471 | -0.20560706 | 0 | 0.1 | 1 | 14 |
| ccl38.1 | 0.05471 | -0.2405392 | 0 | 0.1 | 1 | 14 |
| chic2 | 0.05471 | -0.21660801 | 0 | 0.1 | 1 | 14 |
| coa5 | 0.05471 | -0.21269327 | 0 | 0.1 | 1 | 14 |
| col24a1 | 0.05471 | -0.24412516 | 0 | 0.1 | 1 | 14 |
| csnk1e | 0.05471 | -0.25349785 | 0 | 0.1 | 1 | 14 |
| dennd4b | 0.05471 | -0.13956076 | 0 | 0.1 | 1 | 14 |
| dlgap4b | 0.05471 | -0.15819189 | 0 | 0.1 | 1 | 14 |
| dynlt1 | 0.05471 | -0.24735706 | 0 | 0.1 | 1 | 14 |
| fads2 | 0.05471 | -0.15990684 | 0 | 0.1 | 1 | 14 |
| gemin4 | 0.05471 | -0.22859285 | 0 | 0.1 | 1 | 14 |
| glyr1 | 0.05471 | -0.23080045 | 0 | 0.1 | 1 | 14 |
| gspt1 | 0.05471 | -0.24800348 | 0 | 0.1 | 1 | 14 |
| gss | 0.05471 | -0.22633073 | 0 | 0.1 | 1 | 14 |
| her9 | 0.05471 | -0.26535802 | 0 | 0.1 | 1 | 14 |
| hpcal1 | 0.05471 | -0.20474118 | 0 | 0.1 | 1 | 14 |
| ildr2 | 0.05471 | -0.29031004 | 0 | 0.1 | 1 | 14 |
| kdm5ba | 0.05471 | -0.2744164 | 0 | 0.1 | 1 | 14 |
| lancl2 | 0.05471 | -0.15622902 | 0 | 0.1 | 1 | 14 |
| lias | 0.05471 | -0.20980535 | 0 | 0.1 | 1 | 14 |
| limd2 | 0.05471 | -0.2255889 | 0 | 0.1 | 1 | 14 |
| llgl1 | 0.05471 | -0.2434376 | 0 | 0.1 | 1 | 14 |
| lmod1b | 0.05471 | -0.2735354 | 0 | 0.1 | 1 | 14 |
| lrrc4.1 | 0.05471 | -0.17627276 | 0 | 0.1 | 1 | 14 |
| ltbp1 | 0.05471 | -0.27497669 | 0 | 0.1 | 1 | 14 |
| metrnlb | 0.05471 | -0.19038554 | 0 | 0.1 | 1 | 14 |
| mrps18b | 0.05471 | -0.17283624 | 0 | 0.1 | 1 | 14 |
| msl1a | 0.05471 | -0.1525625 | 0 | 0.1 | 1 | 14 |
| narfl | 0.05471 | -0.16241468 | 0 | 0.1 | 1 | 14 |
| olfm2a | 0.05471 | -0.20737706 | 0 | 0.1 | 1 | 14 |
| parga | 0.05471 | -0.18917019 | 0 | 0.1 | 1 | 14 |
| pi15a | 0.05471 | -0.21309896 | 0 | 0.1 | 1 | 14 |
| prkcba | 0.05471 | -0.29495154 | 0 | 0.1 | 1 | 14 |
| rab32a | 0.05471 | -0.22192477 | 0 | 0.1 | 1 | 14 |

| | | | | | | |
|---|---|---|---|---|---|---|
| rnf170 | 0.05471 | -0.18059156 | 0 | 0.1 | 1 | 14 |
| rnf25 | 0.05471 | -0.29057588 | 0 | 0.1 | 1 | 14 |
| rufy2 | 0.05471 | -0.15735521 | 0 | 0.1 | 1 | 14 |
| s100w | 0.05471 | -0.25376035 | 0 | 0.1 | 1 | 14 |
| sdk1a | 0.05471 | -0.25220512 | 0 | 0.1 | 1 | 14 |
| sec23a | 0.05471 | -0.19185883 | 0 | 0.1 | 1 | 14 |
| setd7 | 0.05471 | -0.15767378 | 0 | 0.1 | 1 | 14 |
| she | 0.05471 | -0.15780826 | 0 | 0.1 | 1 | 14 |
| si:ch211-132g1.3 | 0.05471 | -0.16667337 | 0 | 0.1 | 1 | 14 |
| si:dkeyp-38g8.5 | 0.05471 | -0.19415623 | 0 | 0.1 | 1 | 14 |
| siae | 0.05471 | -0.31019781 | 0 | 0.1 | 1 | 14 |
| sipa1l2 | 0.05471 | -0.31014226 | 0 | 0.1 | 1 | 14 |
| stim1a | 0.05471 | -0.15219616 | 0 | 0.1 | 1 | 14 |
| tapt1a | 0.05471 | -0.19015144 | 0 | 0.1 | 1 | 14 |
| tbc1d15 | 0.05471 | -0.15600789 | 0 | 0.1 | 1 | 14 |
| thap1 | 0.05471 | -0.16436578 | 0 | 0.1 | 1 | 14 |
| tmem184c | 0.05471 | -0.20688739 | 0 | 0.1 | 1 | 14 |
| xrcc3 | 0.05471 | -0.14360827 | 0 | 0.1 | 1 | 14 |
| yy1b | 0.05471 | -0.18613388 | 0 | 0.1 | 1 | 14 |
| zgc:56235 | 0.05471 | -0.25477401 | 0 | 0.1 | 1 | 14 |
| CR847953.1 | 0.05471 | -0.17514065 | 0 | 0.1 | 1 | 14 |
| LOC570526 | 0.05471 | -0.23512043 | 0 | 0.1 | 1 | 14 |
| OSBPL8 | 0.05471 | -0.20382879 | 0 | 0.1 | 1 | 14 |
| abhd10a | 0.05471 | -0.23793298 | 0 | 0.1 | 1 | 14 |
| adam15 | 0.05471 | -0.19957012 | 0 | 0.1 | 1 | 14 |
| aldh3a2b | 0.05471 | -0.23012637 | 0 | 0.1 | 1 | 14 |
| ankrd10b | 0.05471 | -0.16277384 | 0 | 0.1 | 1 | 14 |
| ankrd46a | 0.05471 | -0.26867199 | 0 | 0.1 | 1 | 14 |
| arpp19a | 0.05471 | -0.2453042 | 0 | 0.1 | 1 | 14 |
| atp6v1aa | 0.05471 | -0.19684207 | 0 | 0.1 | 1 | 14 |
| bco1 | 0.05471 | -0.21584519 | 0 | 0.1 | 1 | 14 |
| c1qtnf12 | 0.05471 | -0.30877667 | 0 | 0.1 | 1 | 14 |
| ccdc80l2 | 0.05471 | -0.23512043 | 0 | 0.1 | 1 | 14 |
| chka | 0.05471 | -0.18460585 | 0 | 0.1 | 1 | 14 |
| cnot6l | 0.05471 | -0.19917434 | 0 | 0.1 | 1 | 14 |
| coro2ba | 0.05471 | -0.22987797 | 0 | 0.1 | 1 | 14 |
| dact3a | 0.05471 | -0.24730501 | 0 | 0.1 | 1 | 14 |
| ehbp1 | 0.05471 | -0.22586394 | 0 | 0.1 | 1 | 14 |
| eif2a | 0.05471 | -0.20204965 | 0 | 0.1 | 1 | 14 |
| elmo3 | 0.05471 | -0.21193758 | 0 | 0.1 | 1 | 14 |
| ep300b | 0.05471 | -0.27450068 | 0 | 0.1 | 1 | 14 |
| fundc1 | 0.05471 | -0.10383831 | 0 | 0.1 | 1 | 14 |
| gde1 | 0.05471 | -0.15010618 | 0 | 0.1 | 1 | 14 |
| hdhd2 | 0.05471 | -0.1842585 | 0 | 0.1 | 1 | 14 |
| igfbp2a | 0.05471 | -0.23020667 | 0 | 0.1 | 1 | 14 |
| igfbp3 | 0.05471 | -0.21477233 | 0 | 0.1 | 1 | 14 |
| il12rb2l | 0.05471 | -0.21574722 | 0 | 0.1 | 1 | 14 |
| lmna | 0.05471 | -0.20846728 | 0 | 0.1 | 1 | 14 |

| | | | | | | |
|---|---|---|---|---|---|---|
| lonp1 | 0.05471 | -0.21728481 | 0 | 0.1 | 1 | 14 |
| m6pr | 0.05471 | -0.232419 | 0 | 0.1 | 1 | 14 |
| mbtps2 | 0.05471 | -0.23079074 | 0 | 0.1 | 1 | 14 |
| micu3a | 0.05471 | -0.16191319 | 0 | 0.1 | 1 | 14 |
| mrpl14 | 0.05471 | -0.16111939 | 0 | 0.1 | 1 | 14 |
| mthfs | 0.05471 | -0.18593216 | 0 | 0.1 | 1 | 14 |
| mtx2 | 0.05471 | -0.10849967 | 0 | 0.1 | 1 | 14 |
| myh9b | 0.05471 | -0.21788205 | 0 | 0.1 | 1 | 14 |
| natd1 | 0.05471 | -0.21202632 | 0 | 0.1 | 1 | 14 |
| nbr1b | 0.05471 | -0.20848872 | 0 | 0.1 | 1 | 14 |
| nfkb2 | 0.05471 | -0.21112284 | 0 | 0.1 | 1 | 14 |
| npl | 0.05471 | -0.21990282 | 0 | 0.1 | 1 | 14 |
| nudt7 | 0.05471 | -0.15026911 | 0 | 0.1 | 1 | 14 |
| pargl | 0.05471 | -0.1838766 | 0 | 0.1 | 1 | 14 |
| pigbos1 | 0.05471 | -0.18846597 | 0 | 0.1 | 1 | 14 |
| ppp1cbl | 0.05471 | -0.1931103 | 0 | 0.1 | 1 | 14 |
| prcc | 0.05471 | -0.22950255 | 0 | 0.1 | 1 | 14 |
| ptprea | 0.05471 | -0.13924859 | 0 | 0.1 | 1 | 14 |
| rab24 | 0.05471 | -0.23460296 | 0 | 0.1 | 1 | 14 |
| rasgrp3 | 0.05471 | -0.23830419 | 0 | 0.1 | 1 | 14 |
| rogdi | 0.05471 | -0.15447371 | 0 | 0.1 | 1 | 14 |
| rpp14 | 0.05471 | -0.21143386 | 0 | 0.1 | 1 | 14 |
| serpine2 | 0.05471 | -0.26916805 | 0 | 0.1 | 1 | 14 |
| shmt2 | 0.05471 | -0.15313477 | 0 | 0.1 | 1 | 14 |
| si:ch211-207e14.4 | 0.05471 | -0.25251951 | 0 | 0.1 | 1 | 14 |
| si:ch73-147o17.1 | 0.05471 | -0.19529154 | 0 | 0.1 | 1 | 14 |
| si:ch73-334d15.1 | 0.05471 | -0.16605813 | 0 | 0.1 | 1 | 14 |
| si:ch73-61d6.3 | 0.05471 | -0.24477319 | 0 | 0.1 | 1 | 14 |
| si:dkey-10o6.2 | 0.05471 | -0.19866011 | 0 | 0.1 | 1 | 14 |
| si:dkey-122a22.2 | 0.05471 | -0.20459326 | 0 | 0.1 | 1 | 14 |
| si:dkey-222b8.1 | 0.05471 | -0.19585725 | 0 | 0.1 | 1 | 14 |
| si:dkey-28e7.3 | 0.05471 | -0.24785996 | 0 | 0.1 | 1 | 14 |
| si:dkey-52l18.4 | 0.05471 | -0.24923154 | 0 | 0.1 | 1 | 14 |
| smarca4a | 0.05471 | -0.16095473 | 0 | 0.1 | 1 | 14 |
| smg8 | 0.05471 | -0.19827784 | 0 | 0.1 | 1 | 14 |
| tmcc1b | 0.05471 | -0.26421364 | 0 | 0.1 | 1 | 14 |
| tmem161a | 0.05471 | -0.20755898 | 0 | 0.1 | 1 | 14 |
| tmem43 | 0.05471 | -0.22983837 | 0 | 0.1 | 1 | 14 |
| tnfaip8l3 | 0.05471 | -0.19866524 | 0 | 0.1 | 1 | 14 |
| trabd | 0.05471 | -0.21724895 | 0 | 0.1 | 1 | 14 |
| twsg1b | 0.05471 | -0.15482642 | 0 | 0.1 | 1 | 14 |
| vps13c | 0.05471 | -0.28253135 | 0 | 0.1 | 1 | 14 |
| wipf1b | 0.05471 | -0.23006888 | 0 | 0.1 | 1 | 14 |
| yipf1 | 0.05471 | -0.16391259 | 0 | 0.1 | 1 | 14 |
| zbtb11 | 0.05471 | -0.17074263 | 0 | 0.1 | 1 | 14 |
| zcchc7 | 0.05471 | -0.1640308 | 0 | 0.1 | 1 | 14 |
| zgc:100832 | 0.05471 | -0.24651731 | 0 | 0.1 | 1 | 14 |
| AL929210.2 | 0.05471 | -0.17106818 | 0 | 0.1 | 1 | 14 |

| | | | | | | |
|---|---|---|---|---|---|---|
| CR848833.1 | 0.05471 | -0.21763232 | 0 | 0.1 | 1 | 14 |
| FP101861.1 | 0.05471 | -0.17969518 | 0 | 0.1 | 1 | 14 |
| LOC103911404 | 0.05471 | -0.1969722 | 0 | 0.1 | 1 | 14 |
| XLOC-014336 | 0.05471 | -0.22926536 | 0 | 0.1 | 1 | 14 |
| abca1b | 0.05471 | -0.18346867 | 0 | 0.1 | 1 | 14 |
| acvrl1 | 0.05471 | -0.21550846 | 0 | 0.1 | 1 | 14 |
| adam12 | 0.05471 | -0.24510746 | 0 | 0.1 | 1 | 14 |
| arl4aa | 0.05471 | -0.19495606 | 0 | 0.1 | 1 | 14 |
| atp7a | 0.05471 | -0.191105 | 0 | 0.1 | 1 | 14 |
| clcf1 | 0.05471 | -0.34626505 | 0 | 0.1 | 1 | 14 |
| cog6 | 0.05471 | -0.20328538 | 0 | 0.1 | 1 | 14 |
| cul4a | 0.05471 | -0.30838288 | 0 | 0.1 | 1 | 14 |
| ddx1 | 0.05471 | -0.20320256 | 0 | 0.1 | 1 | 14 |
| ehd1a | 0.05471 | -0.1764954 | 0 | 0.1 | 1 | 14 |
| evi5b | 0.05471 | -0.21812742 | 0 | 0.1 | 1 | 14 |
| fam193b | 0.05471 | -0.22042856 | 0 | 0.1 | 1 | 14 |
| golph3 | 0.05471 | -0.20815893 | 0 | 0.1 | 1 | 14 |
| hmgcrb | 0.05471 | -0.19468505 | 0 | 0.1 | 1 | 14 |
| leo1 | 0.05471 | -0.20786536 | 0 | 0.1 | 1 | 14 |
| lix1l | 0.05471 | -0.23934251 | 0 | 0.1 | 1 | 14 |
| lmtk2 | 0.05471 | -0.22033167 | 0 | 0.1 | 1 | 14 |
| lrrc59 | 0.05471 | -0.13530507 | 0 | 0.1 | 1 | 14 |
| man2b2 | 0.05471 | -0.25891462 | 0 | 0.1 | 1 | 14 |
| mrpl3 | 0.05471 | -0.23005736 | 0 | 0.1 | 1 | 14 |
| msh6 | 0.05471 | -0.16403791 | 0 | 0.1 | 1 | 14 |
| nfyal | 0.05471 | -0.22818699 | 0 | 0.1 | 1 | 14 |
| parp8 | 0.05471 | -0.28762966 | 0 | 0.1 | 1 | 14 |
| pdcd7 | 0.05471 | -0.1746672 | 0 | 0.1 | 1 | 14 |
| pom121 | 0.05471 | -0.25355701 | 0 | 0.1 | 1 | 14 |
| psmb10 | 0.05471 | -0.19026933 | 0 | 0.1 | 1 | 14 |
| qrich1 | 0.05471 | -0.25529729 | 0 | 0.1 | 1 | 14 |
| rangrf | 0.05471 | -0.22256878 | 0 | 0.1 | 1 | 14 |
| retreg2 | 0.05471 | -0.23093328 | 0 | 0.1 | 1 | 14 |
| rhbdd3 | 0.05471 | -0.18520145 | 0 | 0.1 | 1 | 14 |
| sf3a1 | 0.05471 | -0.18287126 | 0 | 0.1 | 1 | 14 |
| sfswap | 0.05471 | -0.27052623 | 0 | 0.1 | 1 | 14 |
| si:ch1073-228b5.2 | 0.05471 | -0.18807217 | 0 | 0.1 | 1 | 14 |
| si:ch211-114c12.2 | 0.05471 | -0.26722207 | 0 | 0.1 | 1 | 14 |
| si:ch211-81a5.1 | 0.05471 | -0.20087503 | 0 | 0.1 | 1 | 14 |
| si:ch73-340n8.4 | 0.05471 | -0.17765438 | 0 | 0.1 | 1 | 14 |
| si:dkey-286j15.1 | 0.05471 | -0.18074748 | 0 | 0.1 | 1 | 14 |
| si:dkeyp-69e1.8 | 0.05471 | -0.28065403 | 0 | 0.1 | 1 | 14 |
| slc6a4a | 0.05471 | -0.40399918 | 0 | 0.1 | 1 | 14 |
| sntb1 | 0.05471 | -0.19507707 | 0 | 0.1 | 1 | 14 |
| spina | 0.05471 | -0.21315478 | 0 | 0.1 | 1 | 14 |
| tctex1d2 | 0.05471 | -0.19246604 | 0 | 0.1 | 1 | 14 |
| tmem170a | 0.05471 | -0.21532242 | 0 | 0.1 | 1 | 14 |
| tyw3 | 0.05471 | -0.20867119 | 0 | 0.1 | 1 | 14 |

| | | | | | | |
|---|---|---|---|---|---|---|
| ythdf3 | 0.05471 | -0.18361642 | 0 | 0.1 | 1 | 14 |
| BX530097.1 | 0.05471 | -0.1729684 | 0 | 0.1 | 1 | 14 |
| CU539058.1 | 0.05471 | -0.2047003 | 0 | 0.1 | 1 | 14 |
| FNDC10 | 0.05471 | -0.20741215 | 0 | 0.1 | 1 | 14 |
| LO017938.1 | 0.05471 | -0.27963427 | 0 | 0.1 | 1 | 14 |
| XLOC-040020 | 0.05471 | -0.30825801 | 0 | 0.1 | 1 | 14 |
| aak1b | 0.05471 | -0.19944483 | 0 | 0.1 | 1 | 14 |
| abhd13 | 0.05471 | -0.1815668 | 0 | 0.1 | 1 | 14 |
| acad9 | 0.05471 | -0.1665383 | 0 | 0.1 | 1 | 14 |
| ap1b1 | 0.05471 | -0.17655187 | 0 | 0.1 | 1 | 14 |
| arrdc3a | 0.05471 | -0.2023946 | 0 | 0.1 | 1 | 14 |
| atp6v0a2b | 0.05471 | -0.20083906 | 0 | 0.1 | 1 | 14 |
| braf | 0.05471 | -0.18513368 | 0 | 0.1 | 1 | 14 |
| chp1 | 0.05471 | -0.20059535 | 0 | 0.1 | 1 | 14 |
| cpsf2 | 0.05471 | -0.19333281 | 0 | 0.1 | 1 | 14 |
| dbt | 0.05471 | -0.19454931 | 0 | 0.1 | 1 | 14 |
| dele1 | 0.05471 | -0.15025906 | 0 | 0.1 | 1 | 14 |
| dennd5b | 0.05471 | -0.23485632 | 0 | 0.1 | 1 | 14 |
| dip2ba | 0.05471 | -0.20248276 | 0 | 0.1 | 1 | 14 |
| fndc3a | 0.05471 | -0.21561156 | 0 | 0.1 | 1 | 14 |
| ifi45 | 0.05471 | -0.17694941 | 0 | 0.1 | 1 | 14 |
| il34 | 0.05471 | -0.24564366 | 0 | 0.1 | 1 | 14 |
| ism2a | 0.05471 | -0.30207308 | 0 | 0.1 | 1 | 14 |
| ivns1abpa | 0.05471 | -0.24814367 | 0 | 0.1 | 1 | 14 |
| kdelc1 | 0.05471 | -0.23264073 | 0 | 0.1 | 1 | 14 |
| lipg | 0.05471 | -0.19713193 | 0 | 0.1 | 1 | 14 |
| magi1b | 0.05471 | -0.21075014 | 0 | 0.1 | 1 | 14 |
| mul1a | 0.05471 | -0.16662507 | 0 | 0.1 | 1 | 14 |
| mvb12ba | 0.05471 | -0.1874974 | 0 | 0.1 | 1 | 14 |
| plcd1b | 0.05471 | -0.2674474 | 0 | 0.1 | 1 | 14 |
| ppm1k | 0.05471 | -0.24208122 | 0 | 0.1 | 1 | 14 |
| ptpn11b | 0.05471 | -0.25037667 | 0 | 0.1 | 1 | 14 |
| rassf2a | 0.05471 | -0.24432943 | 0 | 0.1 | 1 | 14 |
| rbpms2b | 0.05471 | -0.30602739 | 0 | 0.1 | 1 | 14 |
| sema4ab | 0.05471 | -0.26187133 | 0 | 0.1 | 1 | 14 |
| shdb | 0.05471 | -0.18140608 | 0 | 0.1 | 1 | 14 |
| si:dkeyp-110e4.6 | 0.05471 | -0.22965386 | 0 | 0.1 | 1 | 14 |
| sin3b | 0.05471 | -0.18813907 | 0 | 0.1 | 1 | 14 |
| slc20a1b | 0.05471 | -0.17046674 | 0 | 0.1 | 1 | 14 |
| slc6a22.1 | 0.05471 | -0.20093223 | 0 | 0.1 | 1 | 14 |
| spty2d1 | 0.05471 | -0.17256608 | 0 | 0.1 | 1 | 14 |
| stx18 | 0.05471 | -0.17528282 | 0 | 0.1 | 1 | 14 |
| tmem184ba | 0.05471 | -0.20383792 | 0 | 0.1 | 1 | 14 |
| tubgcp3 | 0.05471 | -0.19067559 | 0 | 0.1 | 1 | 14 |
| vhl | 0.05471 | -0.27091715 | 0 | 0.1 | 1 | 14 |
| zc3h11a | 0.05471 | -0.20548417 | 0 | 0.1 | 1 | 14 |
| zgc:92481 | 0.05471 | -0.19791343 | 0 | 0.1 | 1 | 14 |
| ARHGAP22 (1 of many) | 0.05471 | -0.22865177 | 0 | 0.1 | 1 | 14 |

| | | | | | | |
|---|---|---|---|---|---|---|
| CABZ01078055.1 | 0.05471 | -0.18270663 | 0 | 0.1 | 1 | 14 |
| LOC101886114 | 0.05471 | -0.18476552 | 0 | 0.1 | 1 | 14 |
| ccng2 | 0.05471 | -0.2744635 | 0 | 0.1 | 1 | 14 |
| chid1 | 0.05471 | -0.17903455 | 0 | 0.1 | 1 | 14 |
| col7a1 | 0.05471 | -0.19488475 | 0 | 0.1 | 1 | 14 |
| cratb | 0.05471 | -0.17531783 | 0 | 0.1 | 1 | 14 |
| ephx2 | 0.05471 | -0.13014667 | 0 | 0.1 | 1 | 14 |
| epn1 | 0.05471 | -0.16553406 | 0 | 0.1 | 1 | 14 |
| kank3 | 0.05471 | -0.21404226 | 0 | 0.1 | 1 | 14 |
| lrp3 | 0.05471 | -0.21259589 | 0 | 0.1 | 1 | 14 |
| neo1b | 0.05471 | -0.18747893 | 0 | 0.1 | 1 | 14 |
| nfybb | 0.05471 | -0.11945985 | 0 | 0.1 | 1 | 14 |
| rnpepl1 | 0.05471 | -0.20699735 | 0 | 0.1 | 1 | 14 |
| si:ch73-27e22.1 | 0.05471 | -0.18487921 | 0 | 0.1 | 1 | 14 |
| si:dkey-192l18.9 | 0.05471 | -0.19074351 | 0 | 0.1 | 1 | 14 |
| sin3ab | 0.05471 | -0.2439668 | 0 | 0.1 | 1 | 14 |
| tanc1b | 0.05471 | -0.21885708 | 0 | 0.1 | 1 | 14 |
| dennd4a | 0.05471 | -0.18239762 | 0 | 0.1 | 1 | 14 |
| fbxw4 | 0.05471 | -0.17981153 | 0 | 0.1 | 1 | 14 |
| gnal2 | 0.05471 | -0.2370164 | 0 | 0.1 | 1 | 14 |
| nfkbie | 0.05471 | -0.1728388 | 0 | 0.1 | 1 | 14 |
| riok1 | 0.05471 | -0.20935995 | 0 | 0.1 | 1 | 14 |
| LOC103909115 | 0.05471 | -0.27943765 | 0 | 0.1 | 1 | 14 |
| btbd2a | 0.05471 | -0.27329192 | 0 | 0.1 | 1 | 14 |
| cables1 | 0.05471 | -0.20222279 | 0 | 0.1 | 1 | 14 |
| cdc27 | 0.05471 | -0.18570899 | 0 | 0.1 | 1 | 14 |
| epn2 | 0.05471 | -0.23790427 | 0 | 0.1 | 1 | 14 |
| ggt5a | 0.05471 | -0.23090507 | 0 | 0.1 | 1 | 14 |
| heg1 | 0.05471 | -0.29237946 | 0 | 0.1 | 1 | 14 |
| hoxb9a | 0.05471 | -0.19707453 | 0 | 0.1 | 1 | 14 |
| lsg1 | 0.05471 | -0.15662061 | 0 | 0.1 | 1 | 14 |
| mid1ip1a | 0.05471 | -0.32594136 | 0 | 0.1 | 1 | 14 |
| mipep | 0.05471 | -0.23620092 | 0 | 0.1 | 1 | 14 |
| naa30 | 0.05471 | -0.18868888 | 0 | 0.1 | 1 | 14 |
| ppp3r1a | 0.05471 | -0.19175907 | 0 | 0.1 | 1 | 14 |
| prrg2 | 0.05471 | -0.20782944 | 0 | 0.1 | 1 | 14 |
| sap130a | 0.05471 | -0.2077619 | 0 | 0.1 | 1 | 14 |
| sgk1 | 0.05471 | -0.20675674 | 0 | 0.1 | 1 | 14 |
| shroom4 | 0.05471 | -0.25928293 | 0 | 0.1 | 1 | 14 |
| smg5 | 0.05471 | -0.19218291 | 0 | 0.1 | 1 | 14 |
| wnk1b | 0.05471 | -0.2721192 | 0 | 0.1 | 1 | 14 |
| zbtb10 | 0.05471 | -0.23692513 | 0 | 0.1 | 1 | 14 |
| zdhhc24 | 0.05471 | -0.1960556 | 0 | 0.1 | 1 | 14 |
| clstn1 | 0.05471 | -0.24471061 | 0 | 0.1 | 1 | 14 |
| igfbp5a | 0.05471 | -0.22589686 | 0 | 0.1 | 1 | 14 |
| rab10 | 0.05512 | -0.49081598 | 0.325 | 0.65 | 1 | 14 |
| phf6 | 0.05583 | -0.26813971 | 0.025 | 0.2 | 1 | 14 |
| scara3 | 0.05645 | -0.55351441 | 0.125 | 0.4 | 1 | 14 |

| | | | | | | |
|---|---|---|---|---|---|---|
| pleca | 0.05712 | -0.40142824 | 0.175 | 0.45 | 1 | 14 |
| mmp23bb | 0.05726 | -0.30514951 | 0.025 | 0.2 | 1 | 14 |
| cdh11 | 0.05766 | -0.32014256 | 0.375 | 0.45 | 1 | 14 |
| efna1b | 0.05822 | -0.42441639 | 0.1 | 0.35 | 1 | 14 |
| jcada | 0.05824 | -0.52927595 | 0.05 | 0.25 | 1 | 14 |
| cilp2 | 0.05859 | 0.359015159 | 0.125 | 0 | 1 | 14 |
| fam162a | 0.05859 | 0.281208974 | 0.125 | 0 | 1 | 14 |
| gpia | 0.05859 | 0.207619035 | 0.125 | 0 | 1 | 14 |
| nutf2 | 0.05859 | 0.283205425 | 0.125 | 0 | 1 | 14 |
| rnf40 | 0.05859 | 0.256492337 | 0.125 | 0 | 1 | 14 |
| si:ch73-211l13.2 | 0.05859 | 0.294008425 | 0.125 | 0 | 1 | 14 |
| ktn1 | 0.05859 | 0.259660765 | 0.125 | 0 | 1 | 14 |
| map3k7 | 0.05859 | 0.300934484 | 0.125 | 0 | 1 | 14 |
| mcm4 | 0.05859 | 0.266392902 | 0.125 | 0 | 1 | 14 |
| mylipb | 0.05859 | 0.315292277 | 0.125 | 0 | 1 | 14 |
| si:dkey-9i23.15 | 0.05859 | 0.285674431 | 0.125 | 0 | 1 | 14 |
| ube2d4 | 0.05859 | 0.260418356 | 0.125 | 0 | 1 | 14 |
| wt1a | 0.05859 | 0.333432566 | 0.125 | 0 | 1 | 14 |
| CABZ01088109.1 | 0.05859 | 0.33392574 | 0.125 | 0 | 1 | 14 |
| EIF2AK4 | 0.05859 | 0.278978693 | 0.125 | 0 | 1 | 14 |
| PYURF | 0.05859 | 0.253382714 | 0.125 | 0 | 1 | 14 |
| ccl44 | 0.05859 | 0.409082134 | 0.125 | 0 | 1 | 14 |
| clip1a | 0.05859 | 0.270231438 | 0.125 | 0 | 1 | 14 |
| eif1axa | 0.05859 | 0.252291871 | 0.125 | 0 | 1 | 14 |
| elavl1b | 0.05859 | 0.330725541 | 0.125 | 0 | 1 | 14 |
| fgd1 | 0.05859 | 0.331548866 | 0.125 | 0 | 1 | 14 |
| lrrfip2 | 0.05859 | 0.262442269 | 0.125 | 0 | 1 | 14 |
| per2 | 0.05859 | 0.227443145 | 0.125 | 0 | 1 | 14 |
| pigx | 0.05859 | 0.305728402 | 0.125 | 0 | 1 | 14 |
| rasl11b | 0.05859 | 0.268742028 | 0.125 | 0 | 1 | 14 |
| rnaseh2a | 0.05859 | 0.253383747 | 0.125 | 0 | 1 | 14 |
| rybpb | 0.05859 | 0.324748527 | 0.125 | 0 | 1 | 14 |
| setbp1 | 0.05859 | 0.307049383 | 0.125 | 0 | 1 | 14 |
| smad1 | 0.05859 | 0.331796962 | 0.125 | 0 | 1 | 14 |
| tmtc3 | 0.05859 | 0.30482789 | 0.125 | 0 | 1 | 14 |
| trappc6bl | 0.05859 | 0.25613369 | 0.125 | 0 | 1 | 14 |
| ufsp2 | 0.05859 | 0.268769967 | 0.125 | 0 | 1 | 14 |
| PAXBP1 | 0.05859 | 0.245638645 | 0.125 | 0 | 1 | 14 |
| clcn5a | 0.05859 | 0.302030935 | 0.125 | 0 | 1 | 14 |
| cnot1 | 0.05859 | 0.264670827 | 0.125 | 0 | 1 | 14 |
| dhrs7b | 0.05859 | 0.337912354 | 0.125 | 0 | 1 | 14 |
| fbxl14a | 0.05859 | 0.259899385 | 0.125 | 0 | 1 | 14 |
| gigyf1a | 0.05859 | 0.230707083 | 0.125 | 0 | 1 | 14 |
| herpud2 | 0.05859 | 0.238772552 | 0.125 | 0 | 1 | 14 |
| igf2bp1 | 0.05859 | 0.242754323 | 0.125 | 0 | 1 | 14 |
| mcm6 | 0.05859 | 0.305069103 | 0.125 | 0 | 1 | 14 |
| med13a | 0.05859 | 0.31356236 | 0.125 | 0 | 1 | 14 |
| med20 | 0.05859 | 0.294981627 | 0.125 | 0 | 1 | 14 |

| | | | | | | |
|---|---|---|---|---|---|---|
| mrpl18 | 0.05859 | 0.211718006 | 0.125 | 0 | 1 | 14 |
| nt5e | 0.05859 | 0.275869863 | 0.125 | 0 | 1 | 14 |
| pam | 0.05859 | 0.307793497 | 0.125 | 0 | 1 | 14 |
| rab34a | 0.05859 | 0.325268079 | 0.125 | 0 | 1 | 14 |
| rab7a | 0.05859 | 0.276408312 | 0.125 | 0 | 1 | 14 |
| schip1 | 0.05859 | 0.29451821 | 0.125 | 0 | 1 | 14 |
| tspan31 | 0.05859 | 0.259913206 | 0.125 | 0 | 1 | 14 |
| ube2w | 0.05859 | 0.320427332 | 0.125 | 0 | 1 | 14 |
| usf1 | 0.05859 | 0.263482652 | 0.125 | 0 | 1 | 14 |
| waca | 0.05859 | 0.317266574 | 0.125 | 0 | 1 | 14 |
| eci1 | 0.05859 | 0.302724771 | 0.125 | 0 | 1 | 14 |
| got2b | 0.05859 | 0.329571775 | 0.125 | 0 | 1 | 14 |
| hells | 0.05859 | 0.354191327 | 0.125 | 0 | 1 | 14 |
| naa38 | 0.05859 | 0.28077583 | 0.125 | 0 | 1 | 14 |
| srpra | 0.05859 | 0.25197765 | 0.125 | 0 | 1 | 14 |
| TIMM21 | 0.05859 | 0.325165475 | 0.125 | 0 | 1 | 14 |
| aars | 0.05859 | 0.214852462 | 0.125 | 0 | 1 | 14 |
| asip2b | 0.05859 | 0.303353246 | 0.125 | 0 | 1 | 14 |
| dapk3 | 0.05859 | 0.335799524 | 0.125 | 0 | 1 | 14 |
| entpd6 | 0.05859 | 0.273728766 | 0.125 | 0 | 1 | 14 |
| gxylt2 | 0.05859 | 0.328785831 | 0.125 | 0 | 1 | 14 |
| prrx1b | 0.05859 | 0.311689246 | 0.125 | 0 | 1 | 14 |
| ripor3 | 0.05859 | 0.311131971 | 0.125 | 0 | 1 | 14 |
| ssbp1 | 0.05859 | 0.273710901 | 0.125 | 0 | 1 | 14 |
| sumf2 | 0.05859 | 0.279962269 | 0.125 | 0 | 1 | 14 |
| vps26b | 0.05859 | 0.282997113 | 0.125 | 0 | 1 | 14 |
| col6a4a | 0.05936 | -0.40715718 | 0.3 | 0.4 | 1 | 14 |
| myh10 | 0.05948 | -0.47856287 | 0.125 | 0.4 | 1 | 14 |
| fam20a | 0.05964 | -0.37512508 | 0.025 | 0.2 | 1 | 14 |
| rhoca | 0.05986 | 0.46298526 | 0.25 | 0.05 | 1 | 14 |
| ist1 | 0.06011 | -0.4395763 | 0.025 | 0.2 | 1 | 14 |
| fosb | 0.06102 | 0.172444734 | 0.25 | 0.3 | 1 | 14 |
| chrna5 | 0.06174 | -0.61952736 | 0.05 | 0.25 | 1 | 14 |
| zgc:111986 | 0.06248 | 0.39109806 | 0.35 | 0.25 | 1 | 14 |
| ywhaqb | 0.06249 | -0.48856234 | 0.2 | 0.5 | 1 | 14 |
| si:ch1073-340j19.1 | 0.06253 | -0.27366162 | 0.025 | 0.2 | 1 | 14 |
| slc25a3b | 0.06269 | -0.30138565 | 0.35 | 0.65 | 1 | 14 |
| ebag9 | 0.06388 | -0.29116825 | 0.05 | 0.25 | 1 | 14 |
| rpl5b | 0.06413 | 0.346139302 | 0.9 | 0.7 | 1 | 14 |
| si:ch211-222l21.1 | 0.06432 | 0.36237503 | 0.675 | 0.6 | 1 | 14 |
| polr2k | 0.0644 | -0.27633253 | 0.05 | 0.25 | 1 | 14 |
| mmp14a | 0.0645 | -0.39177215 | 0.3 | 0.6 | 1 | 14 |
| cnp | 0.06526 | -0.30587696 | 0.075 | 0.3 | 1 | 14 |
| atp1a1a.1 | 0.06587 | -0.30773354 | 0.05 | 0.25 | 1 | 14 |
| si:ch211-251b21.1 | 0.06622 | 0.582248555 | 0.35 | 0.1 | 1 | 14 |
| brd3b | 0.06704 | 0.537111539 | 0.35 | 0.1 | 1 | 14 |
| hhipl2 | 0.06765 | -0.40652753 | 0.075 | 0.3 | 1 | 14 |
| rps16 | 0.06794 | 0.235150877 | 0.975 | 0.85 | 1 | 14 |

| | | | | | | |
|---|---|---|---|---|---|---|
| nme2b.1 | 0.06864 | 0.190961018 | 0.925 | 0.9 | 1 | 14 |
| mmp14b | 0.06877 | -0.51914695 | 0.3 | 0.6 | 1 | 14 |
| filip1l | 0.0688 | 0.427630291 | 0.275 | 0.05 | 1 | 14 |
| cyhr1 | 0.06899 | -0.30233626 | 0.025 | 0.2 | 1 | 14 |
| adam10a | 0.06918 | -0.39770803 | 0.075 | 0.3 | 1 | 14 |
| hspg2 | 0.0693 | -0.47388517 | 0.1 | 0.35 | 1 | 14 |
| pabpc1b | 0.07088 | -0.27267311 | 0.3 | 0.55 | 1 | 14 |
| ccni | 0.07109 | -0.29226198 | 0.2 | 0.45 | 1 | 14 |
| cpe | 0.07123 | -0.36444497 | 0.125 | 0.35 | 1 | 14 |
| rpl39 | 0.07202 | 0.211195981 | 0.95 | 0.9 | 1 | 14 |
| dnajc8 | 0.07226 | -0.41531538 | 0.1 | 0.35 | 1 | 14 |
| gadd45ba | 0.0723 | -0.51055804 | 0.325 | 0.6 | 1 | 14 |
| sod1 | 0.07241 | 0.506178744 | 0.55 | 0.25 | 1 | 14 |
| cst14a.2 | 0.07247 | -0.29718609 | 0.325 | 0.6 | 1 | 14 |
| atf4b | 0.07267 | 0.220051695 | 0.475 | 0.45 | 1 | 14 |
| rsu1 | 0.07282 | -0.45901895 | 0.1 | 0.35 | 1 | 14 |
| podxl | 0.0731 | -0.39910925 | 0.025 | 0.2 | 1 | 14 |
| pgrmc2 | 0.07337 | -0.22163143 | 0.1 | 0.3 | 1 | 14 |
| rpl12 | 0.07344 | 0.325908627 | 0.925 | 0.7 | 1 | 14 |
| gnas | 0.07392 | -0.3230205 | 0.025 | 0.2 | 1 | 14 |
| srsf1b | 0.07395 | -0.31028838 | 0.05 | 0.25 | 1 | 14 |
| smox | 0.07405 | -0.24684093 | 0.025 | 0.2 | 1 | 14 |
| plpp3 | 0.07427 | -0.21534085 | 0.55 | 0.8 | 1 | 14 |
| spns1 | 0.07449 | -0.29678297 | 0.025 | 0.2 | 1 | 14 |
| LOC108191614 | 0.07641 | -0.25340633 | 0.025 | 0.2 | 1 | 14 |
| col1a1a | 0.07689 | -0.17188127 | 1 | 1 | 1 | 14 |
| nipblb | 0.07719 | -0.33737269 | 0.05 | 0.25 | 1 | 14 |
| cd82a | 0.07745 | -0.52079945 | 0.075 | 0.3 | 1 | 14 |
| rplp1 | 0.07756 | 0.24615323 | 0.975 | 0.85 | 1 | 14 |
| mcl1b | 0.07762 | 0.511870908 | 0.575 | 0.35 | 1 | 14 |
| map2k6 | 0.07827 | -0.43282346 | 0.075 | 0.3 | 1 | 14 |
| cyr61 | 0.07844 | -0.43576051 | 0.525 | 0.8 | 1 | 14 |
| tpte | 0.07866 | -0.47742217 | 0.075 | 0.25 | 1 | 14 |
| CABZ01090890.1 | 0.07869 | -0.21382005 | 0.025 | 0.15 | 1 | 14 |
| snrpf | 0.07871 | 0.4733932 | 0.45 | 0.25 | 1 | 14 |
| sema6bb | 0.07943 | -0.44985329 | 0.025 | 0.2 | 1 | 14 |
| zgc:86598 | 0.0795 | -0.35739896 | 0.075 | 0.3 | 1 | 14 |
| si:ch211-160o17.4 | 0.08088 | -0.35687398 | 0.05 | 0.25 | 1 | 14 |
| fam198a | 0.08127 | -0.46570143 | 0.075 | 0.3 | 1 | 14 |
| npc2 | 0.08156 | -0.45905393 | 0.175 | 0.45 | 1 | 14 |
| rack1 | 0.08162 | 0.299508458 | 0.925 | 0.7 | 1 | 14 |
| CZQB01141835.1 | 0.08214 | -0.300879 | 0.025 | 0.2 | 1 | 14 |
| si:ch211-59h6.1 | 0.08228 | -0.3636185 | 0.025 | 0.2 | 1 | 14 |
| ppiaa | 0.08237 | 0.190294405 | 0.975 | 0.9 | 1 | 14 |
| bida | 0.08252 | -0.28939918 | 0.025 | 0.2 | 1 | 14 |
| dnttip2 | 0.08257 | -0.33168394 | 0.025 | 0.2 | 1 | 14 |
| ches1 | 0.08262 | -0.37571018 | 0.05 | 0.25 | 1 | 14 |
| ctsh | 0.08267 | -0.2921078 | 0.025 | 0.2 | 1 | 14 |

| | | | | | | |
|---|---|---|---|---|---|---|
| CU571074.1 | 0.08267 | -0.35517431 | 0.025 | 0.2 | 1 | 14 |
| prmt1 | 0.0828 | 0.405995272 | 0.275 | 0.05 | 1 | 14 |
| ube2g1b | 0.08281 | -0.3685299 | 0.025 | 0.2 | 1 | 14 |
| mgst3a | 0.08304 | 0.528126297 | 0.35 | 0.15 | 1 | 14 |
| atf6 | 0.08311 | -0.38386072 | 0.025 | 0.2 | 1 | 14 |
| jam3b | 0.08321 | -0.38237232 | 0.025 | 0.2 | 1 | 14 |
| hmg20a | 0.08337 | -0.29645323 | 0.05 | 0.25 | 1 | 14 |
| tbl2 | 0.08344 | -0.39206827 | 0.025 | 0.2 | 1 | 14 |
| dnajc1 | 0.08347 | -0.35317729 | 0.025 | 0.2 | 1 | 14 |
| upf1 | 0.08349 | -0.35195804 | 0.025 | 0.2 | 1 | 14 |
| psmg1 | 0.08368 | -0.26326805 | 0.025 | 0.2 | 1 | 14 |
| abhd15a | 0.08377 | -0.47726295 | 0.075 | 0.3 | 1 | 14 |
| fbxl5 | 0.08387 | -0.27635696 | 0.025 | 0.2 | 1 | 14 |
| tns3.2 | 0.08396 | -0.31127994 | 0.025 | 0.2 | 1 | 14 |
| spcs2 | 0.08405 | -0.46338057 | 0.175 | 0.45 | 1 | 14 |
| eaf1 | 0.08406 | -0.32160536 | 0.025 | 0.2 | 1 | 14 |
| lrrc58b | 0.08428 | -0.33120244 | 0.025 | 0.2 | 1 | 14 |
| mrps35 | 0.08428 | -0.31306277 | 0.025 | 0.2 | 1 | 14 |
| calub | 0.08442 | -0.36827684 | 0.075 | 0.3 | 1 | 14 |
| prnprs3 | 0.08459 | -0.34082049 | 0.025 | 0.2 | 1 | 14 |
| slc43a1a | 0.08461 | -0.31943458 | 0.025 | 0.2 | 1 | 14 |
| ppp6r2a | 0.08462 | -0.30621584 | 0.025 | 0.2 | 1 | 14 |
| taf3 | 0.08465 | -0.35674953 | 0.025 | 0.2 | 1 | 14 |
| march7 | 0.08467 | -0.32084083 | 0.025 | 0.2 | 1 | 14 |
| tmem243b | 0.08468 | -0.2814223 | 0.025 | 0.2 | 1 | 14 |
| mrps14 | 0.08469 | -0.30017339 | 0.025 | 0.2 | 1 | 14 |
| manbal | 0.08472 | -0.29952637 | 0.025 | 0.2 | 1 | 14 |
| tgm1l1 | 0.08473 | -0.41794679 | 0.025 | 0.2 | 1 | 14 |
| gclc | 0.08477 | -0.29460487 | 0.025 | 0.2 | 1 | 14 |
| spg7 | 0.08477 | -0.28739404 | 0.025 | 0.2 | 1 | 14 |
| calr3a | 0.08495 | -0.42831962 | 0.3 | 0.45 | 1 | 14 |
| psmb2 | 0.0854 | 0.442796668 | 0.35 | 0.1 | 1 | 14 |
| slc6a2 | 0.08544 | -0.3750854 | 0.05 | 0.25 | 1 | 14 |
| rps21 | 0.08583 | 0.250941715 | 0.95 | 0.85 | 1 | 14 |
| tmed7 | 0.08605 | -0.45829418 | 0.3 | 0.55 | 1 | 14 |
| zfhx3 | 0.08605 | 0.511583936 | 0.425 | 0.15 | 1 | 14 |
| rplp0 | 0.08635 | 0.303799626 | 0.95 | 0.85 | 1 | 14 |
| si:dkey-17m8.1 | 0.08691 | -0.38698134 | 0.05 | 0.25 | 1 | 14 |
| ctsz | 0.08709 | -0.38465406 | 0.05 | 0.25 | 1 | 14 |
| rnasekb | 0.08749 | -0.39627917 | 0.05 | 0.25 | 1 | 14 |
| llph | 0.08832 | -0.30701793 | 0.05 | 0.25 | 1 | 14 |
| abcc5 | 0.08852 | -0.22658827 | 0.05 | 0.2 | 1 | 14 |
| rpl34 | 0.08855 | 0.221868974 | 0.975 | 0.85 | 1 | 14 |
| cox8a | 0.08866 | 0.450897711 | 0.475 | 0.3 | 1 | 14 |
| rbbp4 | 0.08888 | 0.349529773 | 0.225 | 0.15 | 1 | 14 |
| ndufs6 | 0.08893 | 0.449396615 | 0.25 | 0.05 | 1 | 14 |
| puf60b | 0.08981 | -0.39530011 | 0.05 | 0.25 | 1 | 14 |
| pdk2b | 0.08988 | -0.40638458 | 0.05 | 0.25 | 1 | 14 |

| | | | | | | |
|---|---|---|---|---|---|---|
| esyt2a | 0.08996 | -0.40275148 | 0.05 | 0.25 | 1 | 14 |
| sptan1 | 0.09 | -0.38920593 | 0.05 | 0.25 | 1 | 14 |
| ptx3a | 0.09005 | -0.42462714 | 0.05 | 0.25 | 1 | 14 |
| tktb | 0.09037 | -0.39484634 | 0.05 | 0.25 | 1 | 14 |
| psmb3 | 0.09043 | 0.462939662 | 0.325 | 0.1 | 1 | 14 |
| ik | 0.09078 | -0.32911531 | 0.05 | 0.25 | 1 | 14 |
| si:zfos-169g10.3 | 0.09096 | -0.32692395 | 0.15 | 0.4 | 1 | 14 |
| snrpd2 | 0.0929 | 0.362300828 | 0.2 | 0.05 | 1 | 14 |
| prkag1 | 0.09336 | -0.1846246 | 0.025 | 0.15 | 1 | 14 |
| pin1 | 0.09401 | -0.3100873 | 0.125 | 0.35 | 1 | 14 |
| polr2l | 0.09493 | 0.350927543 | 0.25 | 0.15 | 1 | 14 |
| rsrp1 | 0.09667 | -0.2967885 | 0.175 | 0.4 | 1 | 14 |
| eif3m | 0.09687 | 0.162172177 | 0.525 | 0.55 | 1 | 14 |
| mt-nd1 | 0.09709 | -0.17761062 | 0.575 | 0.8 | 1 | 14 |
| h2afx1 | 0.09737 | 0.404733666 | 0.725 | 0.55 | 1 | 14 |
| cirbpa | 0.09739 | 0.348411284 | 0.8 | 0.65 | 1 | 14 |
| dstyk | 0.09746 | -0.17750333 | 0.1 | 0.25 | 1 | 14 |
| snrpb | 0.09772 | 0.240391909 | 0.425 | 0.4 | 1 | 14 |
| cd28l | 0.09783 | -0.47209278 | 0.15 | 0.4 | 1 | 14 |
| smn1 | 0.09798 | 0.160190691 | 0.1 | 0 | 1 | 14 |
| hipk3b | 0.09798 | 0.247345382 | 0.1 | 0 | 1 | 14 |
| kpna1 | 0.09798 | 0.227604695 | 0.1 | 0 | 1 | 14 |
| ncln | 0.09798 | 0.182028367 | 0.1 | 0 | 1 | 14 |
| pin4 | 0.09798 | 0.204365467 | 0.1 | 0 | 1 | 14 |
| stat6 | 0.09798 | 0.256908528 | 0.1 | 0 | 1 | 14 |
| supt5h | 0.09798 | 0.236795475 | 0.1 | 0 | 1 | 14 |
| DDX17 | 0.09798 | 0.287605916 | 0.1 | 0 | 1 | 14 |
| UTP14C | 0.09798 | 0.229567247 | 0.1 | 0 | 1 | 14 |
| XLOC-041870 | 0.09798 | 0.291369008 | 0.1 | 0 | 1 | 14 |
| acsf2 | 0.09798 | 0.238610217 | 0.1 | 0 | 1 | 14 |
| akt2l | 0.09798 | 0.216656933 | 0.1 | 0 | 1 | 14 |
| anks3 | 0.09798 | 0.217128756 | 0.1 | 0 | 1 | 14 |
| baz2ba | 0.09798 | 0.281948633 | 0.1 | 0 | 1 | 14 |
| bloc1s1 | 0.09798 | 0.199576345 | 0.1 | 0 | 1 | 14 |
| cdc14b | 0.09798 | 0.273824921 | 0.1 | 0 | 1 | 14 |
| cdc37 | 0.09798 | 0.190478958 | 0.1 | 0 | 1 | 14 |
| commd4 | 0.09798 | 0.261669093 | 0.1 | 0 | 1 | 14 |
| cxadr | 0.09798 | 0.25002612 | 0.1 | 0 | 1 | 14 |
| desi2 | 0.09798 | 0.215991933 | 0.1 | 0 | 1 | 14 |
| ebf1a | 0.09798 | 0.234741537 | 0.1 | 0 | 1 | 14 |
| fam13b | 0.09798 | 0.204691582 | 0.1 | 0 | 1 | 14 |
| gars | 0.09798 | 0.250439158 | 0.1 | 0 | 1 | 14 |
| gsc | 0.09798 | 0.195136783 | 0.1 | 0 | 1 | 14 |
| gse1 | 0.09798 | 0.210143461 | 0.1 | 0 | 1 | 14 |
| hoxa10b | 0.09798 | 0.269581673 | 0.1 | 0 | 1 | 14 |
| hsdl2 | 0.09798 | 0.226644175 | 0.1 | 0 | 1 | 14 |
| htra1b | 0.09798 | 0.226593147 | 0.1 | 0 | 1 | 14 |
| idh1 | 0.09798 | 0.197289164 | 0.1 | 0 | 1 | 14 |

| | | | | | | |
|---|---|---|---|---|---|---|
| irf9 | 0.09798 | 0.255889035 | 0.1 | 0 | 1 | 14 |
| klf11b | 0.09798 | 0.263290111 | 0.1 | 0 | 1 | 14 |
| lrrc57 | 0.09798 | 0.226269511 | 0.1 | 0 | 1 | 14 |
| lztfl1 | 0.09798 | 0.224661873 | 0.1 | 0 | 1 | 14 |
| mapk3 | 0.09798 | 0.197779867 | 0.1 | 0 | 1 | 14 |
| mlx | 0.09798 | 0.252807858 | 0.1 | 0 | 1 | 14 |
| mpped2 | 0.09798 | 0.241424182 | 0.1 | 0 | 1 | 14 |
| mrpl20 | 0.09798 | 0.194008591 | 0.1 | 0 | 1 | 14 |
| mrpl23 | 0.09798 | 0.248865789 | 0.1 | 0 | 1 | 14 |
| mycn | 0.09798 | 0.257828185 | 0.1 | 0 | 1 | 14 |
| nob1 | 0.09798 | 0.199066896 | 0.1 | 0 | 1 | 14 |
| nufip1 | 0.09798 | 0.216859087 | 0.1 | 0 | 1 | 14 |
| pdcd4a | 0.09798 | 0.205519786 | 0.1 | 0 | 1 | 14 |
| phf23a | 0.09798 | 0.266182459 | 0.1 | 0 | 1 | 14 |
| rab18b | 0.09798 | 0.222930982 | 0.1 | 0 | 1 | 14 |
| rnaseh2c | 0.09798 | 0.154783823 | 0.1 | 0 | 1 | 14 |
| shisa4 | 0.09798 | 0.216414545 | 0.1 | 0 | 1 | 14 |
| si:ch73-111k22.2 | 0.09798 | 0.234564302 | 0.1 | 0 | 1 | 14 |
| si:ch73-1a9.4 | 0.09798 | 0.254018482 | 0.1 | 0 | 1 | 14 |
| si:ch73-71c20.5 | 0.09798 | 0.200636658 | 0.1 | 0 | 1 | 14 |
| si:dkey-187e18.1 | 0.09798 | 0.205883098 | 0.1 | 0 | 1 | 14 |
| stip1 | 0.09798 | 0.220656743 | 0.1 | 0 | 1 | 14 |
| suds3 | 0.09798 | 0.239065544 | 0.1 | 0 | 1 | 14 |
| tbl3 | 0.09798 | 0.215070448 | 0.1 | 0 | 1 | 14 |
| trim25 | 0.09798 | 0.219493654 | 0.1 | 0 | 1 | 14 |
| tuba1a | 0.09798 | 0.227003766 | 0.1 | 0 | 1 | 14 |
| ube2ka | 0.09798 | 0.190342752 | 0.1 | 0 | 1 | 14 |
| uchl3 | 0.09798 | 0.191021338 | 0.1 | 0 | 1 | 14 |
| vgll4a | 0.09798 | 0.259571284 | 0.1 | 0 | 1 | 14 |
| wt1b | 0.09798 | 0.246455966 | 0.1 | 0 | 1 | 14 |
| adamts10 | 0.09798 | 0.224816498 | 0.1 | 0 | 1 | 14 |
| arid4a | 0.09798 | 0.235822003 | 0.1 | 0 | 1 | 14 |
| baxa | 0.09798 | 0.199692871 | 0.1 | 0 | 1 | 14 |
| bcl2l10 | 0.09798 | 0.206138644 | 0.1 | 0 | 1 | 14 |
| cdc123 | 0.09798 | 0.206360223 | 0.1 | 0 | 1 | 14 |
| creb1b | 0.09798 | 0.292868663 | 0.1 | 0 | 1 | 14 |
| cyth1a | 0.09798 | 0.216255356 | 0.1 | 0 | 1 | 14 |
| diaph2 | 0.09798 | 0.227513545 | 0.1 | 0 | 1 | 14 |
| exosc2 | 0.09798 | 0.167619342 | 0.1 | 0 | 1 | 14 |
| glceb | 0.09798 | 0.232728701 | 0.1 | 0 | 1 | 14 |
| hmgb3b | 0.09798 | 0.250788235 | 0.1 | 0 | 1 | 14 |
| hmox2b | 0.09798 | 0.229380471 | 0.1 | 0 | 1 | 14 |
| lsm14aa | 0.09798 | 0.167290623 | 0.1 | 0 | 1 | 14 |
| nhsl1a | 0.09798 | 0.260694538 | 0.1 | 0 | 1 | 14 |
| nol12 | 0.09798 | 0.209498435 | 0.1 | 0 | 1 | 14 |
| ophn1 | 0.09798 | 0.196944961 | 0.1 | 0 | 1 | 14 |
| pbxip1a | 0.09798 | 0.220418097 | 0.1 | 0 | 1 | 14 |
| sco1 | 0.09798 | 0.228716917 | 0.1 | 0 | 1 | 14 |

| | | | | | | |
|---|---|---|---|---|---|---|
| smad3a | 0.09798 | 0.257514374 | 0.1 | 0 | 1 | 14 |
| srp9 | 0.09798 | 0.186788819 | 0.1 | 0 | 1 | 14 |
| srpk1a | 0.09798 | 0.208544209 | 0.1 | 0 | 1 | 14 |
| stam2 | 0.09798 | 0.218757751 | 0.1 | 0 | 1 | 14 |
| stxbp2 | 0.09798 | 0.164946245 | 0.1 | 0 | 1 | 14 |
| tab2 | 0.09798 | 0.230011775 | 0.1 | 0 | 1 | 14 |
| taok1b | 0.09798 | 0.218057127 | 0.1 | 0 | 1 | 14 |
| tbx3a | 0.09798 | 0.236295648 | 0.1 | 0 | 1 | 14 |
| tom1l2b | 0.09798 | 0.215577646 | 0.1 | 0 | 1 | 14 |
| tusc2a | 0.09798 | 0.298656717 | 0.1 | 0 | 1 | 14 |
| wdr82 | 0.09798 | 0.201872298 | 0.1 | 0 | 1 | 14 |
| zgc:162025 | 0.09798 | 0.201806643 | 0.1 | 0 | 1 | 14 |
| znf410 | 0.09798 | 0.252826719 | 0.1 | 0 | 1 | 14 |
| eef1e1 | 0.09798 | 0.216799396 | 0.1 | 0 | 1 | 14 |
| rnd3a | 0.09798 | 0.252146998 | 0.1 | 0 | 1 | 14 |
| selenbp1 | 0.09798 | 0.244858869 | 0.1 | 0 | 1 | 14 |
| suclg2 | 0.09798 | 0.185545853 | 0.1 | 0 | 1 | 14 |
| ptrh2 | 0.09798 | 0.178557354 | 0.1 | 0 | 1 | 14 |
| BX663516.2 | 0.09798 | 0.284884734 | 0.1 | 0 | 1 | 14 |
| arl8ba | 0.09798 | 0.183039311 | 0.1 | 0 | 1 | 14 |
| atp6v1ba | 0.09798 | 0.284433391 | 0.1 | 0 | 1 | 14 |
| cant1a | 0.09798 | 0.206547917 | 0.1 | 0 | 1 | 14 |
| carm1 | 0.09798 | 0.253006727 | 0.1 | 0 | 1 | 14 |
| cd3eap | 0.09798 | 0.229609596 | 0.1 | 0 | 1 | 14 |
| cyp1d1 | 0.09798 | 0.202075284 | 0.1 | 0 | 1 | 14 |
| dnajb14 | 0.09798 | 0.231106242 | 0.1 | 0 | 1 | 14 |
| elp5 | 0.09798 | 0.232155319 | 0.1 | 0 | 1 | 14 |
| etnk1 | 0.09798 | 0.281687879 | 0.1 | 0 | 1 | 14 |
| fndc3bb | 0.09798 | 0.254984841 | 0.1 | 0 | 1 | 14 |
| frs2b | 0.09798 | 0.221282136 | 0.1 | 0 | 1 | 14 |
| harbi1 | 0.09798 | 0.25109518 | 0.1 | 0 | 1 | 14 |
| hspa4a | 0.09798 | 0.233230751 | 0.1 | 0 | 1 | 14 |
| ift20 | 0.09798 | 0.201227063 | 0.1 | 0 | 1 | 14 |
| itga10 | 0.09798 | 0.259417767 | 0.1 | 0 | 1 | 14 |
| megf6a | 0.09798 | 0.289948601 | 0.1 | 0 | 1 | 14 |
| mrps34 | 0.09798 | 0.254899375 | 0.1 | 0 | 1 | 14 |
| nedd4l | 0.09798 | 0.264515486 | 0.1 | 0 | 1 | 14 |
| noc3l | 0.09798 | 0.235235613 | 0.1 | 0 | 1 | 14 |
| nsun2 | 0.09798 | 0.208934265 | 0.1 | 0 | 1 | 14 |
| pigt | 0.09798 | 0.197448149 | 0.1 | 0 | 1 | 14 |
| pitx1 | 0.09798 | 0.234523375 | 0.1 | 0 | 1 | 14 |
| rnaseh1 | 0.09798 | 0.195650092 | 0.1 | 0 | 1 | 14 |
| scaf4a | 0.09798 | 0.24982759 | 0.1 | 0 | 1 | 14 |
| sema5a | 0.09798 | 0.250551983 | 0.1 | 0 | 1 | 14 |
| smpd5 | 0.09798 | 0.18617563 | 0.1 | 0 | 1 | 14 |
| tfdp1a | 0.09798 | 0.25857043 | 0.1 | 0 | 1 | 14 |
| trit1 | 0.09798 | 0.263724042 | 0.1 | 0 | 1 | 14 |
| ubxn1 | 0.09798 | 0.246294975 | 0.1 | 0 | 1 | 14 |

| | | | | | | |
|---|---|---|---|---|---|---|
| zgc:153115 | 0.09798 | 0.210687007 | 0.1 | 0 | 1 | 14 |
| fzd7a | 0.09802 | -0.23354203 | 0.2 | 0.4 | 1 | 14 |
| psmb5 | 0.09841 | -0.16847236 | 0.125 | 0.3 | 1 | 14 |
| rhoac | 0.09889 | 0.295118204 | 0.3 | 0.25 | 1 | 14 |
| rpl13 | 0.09962 | 0.214655168 | 0.975 | 0.9 | 1 | 14 |
| krtcap2 | 0.0999 | -0.46810144 | 0.225 | 0.5 | 1 | 14 |
| endouc | 0.10001 | -0.43378367 | 0.525 | 0.8 | 1 | 14 |
| meis1a | 0.10007 | 0.467156054 | 0.25 | 0.05 | 1 | 14 |
| zfp36l2 | 0.10025 | 0.417312409 | 0.325 | 0.1 | 1 | 14 |
| csrp2 | 0.10031 | -0.12947993 | 0.05 | 0.2 | 1 | 14 |
| si:dkey-33c12.4 | 0.10072 | 0.232466861 | 0.225 | 0.2 | 1 | 14 |
| ptbp1a | 0.10276 | -0.28275917 | 0.1 | 0.3 | 1 | 14 |
| tnfsf12 | 0.10333 | -0.40469544 | 0.15 | 0.4 | 1 | 14 |
| ebf3a | 0.10428 | -0.172549 | 0.175 | 0.35 | 1 | 14 |
| tmed9 | 0.10445 | -0.32752107 | 0.2 | 0.45 | 1 | 14 |
| rpl3 | 0.10486 | 0.169031667 | 0.9 | 0.9 | 1 | 14 |
| si:dkey-112e7.2 | 0.10643 | 0.370828765 | 0.25 | 0.05 | 1 | 14 |
| tnmd | 0.10645 | -0.49714885 | 0.3 | 0.5 | 1 | 14 |
| chd6 | 0.10819 | -0.43906639 | 0.15 | 0.4 | 1 | 14 |
| rps27l | 0.10846 | 0.395313969 | 0.3 | 0.1 | 1 | 14 |
| si:dkey-42i9.6 | 0.10928 | -0.43795703 | 0.1 | 0.3 | 1 | 14 |
| psmd12 | 0.10962 | 0.293118043 | 0.225 | 0.05 | 1 | 14 |
| rps12 | 0.10992 | 0.26557182 | 0.95 | 0.85 | 1 | 14 |
| hnrnpa1b | 0.11084 | 0.414393942 | 0.25 | 0.05 | 1 | 14 |
| eif4g1a | 0.11117 | 0.382103498 | 0.25 | 0.05 | 1 | 14 |
| marcksl1a | 0.11231 | -0.28426558 | 0.775 | 0.9 | 1 | 14 |
| btg3 | 0.11268 | 0.390190649 | 0.25 | 0.05 | 1 | 14 |
| calm2a | 0.11281 | -0.40476651 | 0.275 | 0.55 | 1 | 14 |
| zgc:56493 | 0.11312 | 0.267421631 | 0.625 | 0.6 | 1 | 14 |
| pfn2 | 0.1137 | -0.19881912 | 0.375 | 0.6 | 1 | 14 |
| tomm7 | 0.11419 | 0.177440876 | 0.25 | 0.25 | 1 | 14 |
| cast | 0.11487 | -0.21061369 | 0.1 | 0.25 | 1 | 14 |
| rpl23 | 0.11528 | 0.250729091 | 0.95 | 0.8 | 1 | 14 |
| tssc4 | 0.11563 | -0.17080563 | 0.025 | 0.15 | 1 | 14 |
| lsm5 | 0.11595 | 0.136958441 | 0.2 | 0.25 | 1 | 14 |
| hsd11b2 | 0.11633 | -0.40594521 | 0.6 | 0.75 | 1 | 14 |
| mcm2 | 0.11727 | 0.403907332 | 0.225 | 0.05 | 1 | 14 |
| csnk1a1 | 0.11758 | 0.438861824 | 0.325 | 0.1 | 1 | 14 |
| rps2 | 0.11773 | 0.210374401 | 0.975 | 0.9 | 1 | 14 |
| rpl10 | 0.11782 | 0.14644617 | 0.925 | 0.95 | 1 | 14 |
| XLOC-010899 | 0.11813 | 0.399678887 | 0.25 | 0.05 | 1 | 14 |
| fstl1b | 0.11859 | -0.23273988 | 0.875 | 0.95 | 1 | 14 |
| rps19 | 0.11911 | 0.16387732 | 1 | 0.9 | 1 | 14 |
| psma5 | 0.11951 | -0.20323561 | 0.15 | 0.35 | 1 | 14 |
| trmt112 | 0.11976 | 0.169640936 | 0.125 | 0.15 | 1 | 14 |
| yap1 | 0.11977 | -0.22075354 | 0.025 | 0.15 | 1 | 14 |
| atp5f1b | 0.11986 | -0.37926238 | 0.425 | 0.7 | 1 | 14 |
| tnfsf10l | 0.12018 | 0.478470987 | 0.25 | 0.05 | 1 | 14 |

| | | | | | | |
|---|---|---|---|---|---|---|
| srsf5b | 0.1207 | -0.19695808 | 0.025 | 0.15 | 1 | 14 |
| anp32b | 0.12104 | 0.353377188 | 0.2 | 0.05 | 1 | 14 |
| mt-nd4 | 0.12124 | -0.20178649 | 0.75 | 0.95 | 1 | 14 |
| fhl1b | 0.12156 | 0.401894531 | 0.225 | 0.05 | 1 | 14 |
| pik3r3a | 0.12242 | 0.428382138 | 0.275 | 0.1 | 1 | 14 |
| cox4i1 | 0.12256 | -0.31416778 | 0.425 | 0.7 | 1 | 14 |
| sec11a | 0.12361 | -0.36014556 | 0.325 | 0.6 | 1 | 14 |
| nfyba | 0.12397 | 0.410033488 | 0.25 | 0.05 | 1 | 14 |
| twist1a | 0.12474 | -0.42013847 | 0.475 | 0.7 | 1 | 14 |
| gmps | 0.12494 | -0.1953129 | 0.05 | 0.2 | 1 | 14 |
| si:dkey-188i13.11 | 0.12538 | 0.385672182 | 0.225 | 0.05 | 1 | 14 |
| XLOC-013964 | 0.12649 | 0.542676889 | 0.325 | 0.1 | 1 | 14 |
| eif5a | 0.12698 | -0.12695746 | 0.55 | 0.75 | 1 | 14 |
| vwa1 | 0.12877 | -0.4472369 | 0.2 | 0.45 | 1 | 14 |
| rbm4.3 | 0.12963 | 0.153429069 | 0.3 | 0.3 | 1 | 14 |
| rpl32 | 0.12992 | 0.154198605 | 0.925 | 0.9 | 1 | 14 |
| srp19 | 0.13001 | -0.20601708 | 0.05 | 0.2 | 1 | 14 |
| LOC100537342 | 0.13044 | 0.478131218 | 0.425 | 0.2 | 1 | 14 |
| atp2b2 | 0.13086 | -0.41378049 | 0.125 | 0.35 | 1 | 14 |
| ncbp2 | 0.13159 | 0.39165041 | 0.2 | 0.05 | 1 | 14 |
| cox6b2 | 0.13195 | 0.254845211 | 0.75 | 0.7 | 1 | 14 |
| lsp1 | 0.13195 | 0.46629461 | 0.275 | 0.1 | 1 | 14 |
| mfap2 | 0.1329 | -0.16750808 | 0.875 | 0.9 | 1 | 14 |
| mkxa | 0.13347 | -0.21156295 | 0.075 | 0.2 | 1 | 14 |
| ptmaa | 0.13401 | 0.406859856 | 0.575 | 0.45 | 1 | 14 |
| atp6ap2 | 0.13421 | -0.15672158 | 0.025 | 0.15 | 1 | 14 |
| pfkfb4b | 0.13531 | 0.470299599 | 0.2 | 0.05 | 1 | 14 |
| ginm1 | 0.13591 | -0.40767328 | 0.125 | 0.35 | 1 | 14 |
| pho | 0.13705 | -0.2464608 | 0.2 | 0.4 | 1 | 14 |
| ube2l3b | 0.13714 | -0.13261724 | 0.15 | 0.3 | 1 | 14 |
| tcp1 | 0.13727 | 0.420868475 | 0.375 | 0.2 | 1 | 14 |
| herpud1 | 0.13747 | -0.34408712 | 0.05 | 0.15 | 1 | 14 |
| SCARF2 | 0.13747 | -0.25472495 | 0.15 | 0.35 | 1 | 14 |
| socs3b | 0.13803 | -0.47456243 | 0.225 | 0.4 | 1 | 14 |
| pepd | 0.13815 | -0.23977522 | 0.05 | 0.2 | 1 | 14 |
| hmgb1a | 0.14 | -0.28682061 | 0.55 | 0.8 | 1 | 14 |
| si:ch211-133n4.4 | 0.14007 | -0.2753362 | 0.275 | 0.5 | 1 | 14 |
| si:ch1073-325m22.2 | 0.14019 | 0.3461045 | 0.275 | 0.1 | 1 | 14 |
| snrpg | 0.14023 | 0.343327111 | 0.275 | 0.15 | 1 | 14 |
| reck | 0.14069 | -0.33420291 | 0.1 | 0.3 | 1 | 14 |
| rps3 | 0.14105 | 0.2644364 | 0.925 | 0.75 | 1 | 14 |
| hsbp1b | 0.14247 | -0.41518171 | 0.2 | 0.4 | 1 | 14 |
| arpc1a | 0.14301 | -0.41654435 | 0.1 | 0.3 | 1 | 14 |
| flna | 0.14505 | -0.18159637 | 0.35 | 0.55 | 1 | 14 |
| rps26 | 0.14528 | 0.218854429 | 0.975 | 0.85 | 1 | 14 |
| rpl19 | 0.14554 | 0.233876736 | 0.9 | 0.8 | 1 | 14 |
| adamts15b | 0.14604 | -0.48667653 | 0.1 | 0.3 | 1 | 14 |
| rhoab | 0.14607 | -0.41875127 | 0.4 | 0.65 | 1 | 14 |

| | | | | | | |
|---|---|---|---|---|---|---|
| sri | 0.14623 | 0.354878137 | 0.275 | 0.15 | 1 | 14 |
| fam171a1 | 0.14714 | -0.22579797 | 0.025 | 0.15 | 1 | 14 |
| map4k5 | 0.14721 | -0.16749317 | 0.025 | 0.15 | 1 | 14 |
| tcf25 | 0.14735 | -0.24725326 | 0.075 | 0.25 | 1 | 14 |
| rpl13a | 0.14792 | 0.209957619 | 0.975 | 0.85 | 1 | 14 |
| cav2 | 0.14822 | -0.29924921 | 0.275 | 0.5 | 1 | 14 |
| calr | 0.14824 | -0.42171296 | 0.25 | 0.5 | 1 | 14 |
| pef1 | 0.14829 | -0.39000925 | 0.15 | 0.35 | 1 | 14 |
| tspan9b | 0.14933 | -0.3846761 | 0.1 | 0.3 | 1 | 14 |
| nup188 | 0.14959 | -0.18992387 | 0.025 | 0.15 | 1 | 14 |
| ap3s1.1 | 0.14972 | -0.22098103 | 0.05 | 0.2 | 1 | 14 |
| hif1ab | 0.1498 | -0.17624791 | 0.175 | 0.35 | 1 | 14 |
| cxcl18b | 0.15015 | -0.47163252 | 0.175 | 0.4 | 1 | 14 |
| hnrnpr | 0.15091 | -0.10341594 | 0.125 | 0.25 | 1 | 14 |
| aqp3a | 0.15133 | -0.60990192 | 0.25 | 0.45 | 1 | 14 |
| nop10 | 0.15244 | -0.40731037 | 0.15 | 0.35 | 1 | 14 |
| anxa11a | 0.15279 | -0.3125806 | 0.1 | 0.3 | 1 | 14 |
| rpl23a | 0.15367 | 0.246907339 | 0.875 | 0.8 | 1 | 14 |
| aktip | 0.1544 | -0.26896238 | 0.075 | 0.25 | 1 | 14 |
| dhrs3a | 0.15461 | -0.4814934 | 0.15 | 0.35 | 1 | 14 |
| psmg3 | 0.15584 | -0.16170607 | 0.1 | 0.25 | 1 | 14 |
| slain2 | 0.1559 | -0.19511693 | 0.025 | 0.15 | 1 | 14 |
| sh3bgrl3 | 0.15625 | -0.33449367 | 0.1 | 0.3 | 1 | 14 |
| sept7b | 0.1565 | -0.32242766 | 0.1 | 0.3 | 1 | 14 |
| rbp4 | 0.15675 | 0.498285059 | 0.5 | 0.25 | 1 | 14 |
| si:dkey-11f4.14 | 0.15693 | 0.345942979 | 0.3 | 0.1 | 1 | 14 |
| si:ch211-212k18.9 | 0.15746 | -0.33939961 | 0.175 | 0.4 | 1 | 14 |
| bccip | 0.15762 | -0.1804954 | 0.175 | 0.35 | 1 | 14 |
| si:ch73-376l24.3 | 0.15807 | -0.32299532 | 0.225 | 0.45 | 1 | 14 |
| taf15 | 0.15809 | 0.287660913 | 0.225 | 0.15 | 1 | 14 |
| NC-002333.4 | 0.15869 | 0.174558436 | 0.9 | 0.9 | 1 | 14 |
| coro1ca | 0.15886 | -0.38684731 | 0.025 | 0.15 | 1 | 14 |
| hspa8 | 0.15919 | 0.362865688 | 0.775 | 0.6 | 1 | 14 |
| rdh10a | 0.15952 | -0.46470477 | 0.05 | 0.2 | 1 | 14 |
| emc8 | 0.15972 | -0.15488837 | 0.025 | 0.15 | 1 | 14 |
| parvab | 0.15994 | -0.18941354 | 0.025 | 0.15 | 1 | 14 |
| rpl31 | 0.1606 | 0.197715237 | 0.975 | 0.85 | 1 | 14 |
| mrc2 | 0.16117 | -0.40341025 | 0.1 | 0.3 | 1 | 14 |
| ran | 0.1616 | 0.241968363 | 0.6 | 0.55 | 1 | 14 |
| chpfa | 0.16192 | -0.22789705 | 0.025 | 0.15 | 1 | 14 |
| fam168b | 0.16217 | -0.32775214 | 0.1 | 0.3 | 1 | 14 |
| ndufc2 | 0.16345 | -0.24438345 | 0.325 | 0.55 | 1 | 14 |
| rpl35 | 0.16346 | 0.188878881 | 0.925 | 0.85 | 1 | 14 |
| ucp2 | 0.16379 | -0.11579167 | 0.275 | 0.45 | 1 | 14 |
| ccdc3b | 0.16386 | -0.40645203 | 0.1 | 0.3 | 1 | 14 |
| man2b1 | 0.16391 | -0.42282501 | 0.075 | 0.25 | 1 | 14 |
| tmem88a | 0.16402 | -0.39937638 | 0.1 | 0.3 | 1 | 14 |
| hyou1 | 0.16459 | 0.159921631 | 0.075 | 0 | 1 | 14 |

| | | | | | | |
|---|---|---|---|---|---|---|
| nav2b | 0.16459 | 0.234176848 | 0.075 | 0 | 1 | 14 |
| si:ch211-241b2.1 | 0.16459 | 0.149278137 | 0.075 | 0 | 1 | 14 |
| BX537342.1 | 0.16459 | 0.16303244 | 0.075 | 0 | 1 | 14 |
| LOC100333572 | 0.16459 | 0.213367718 | 0.075 | 0 | 1 | 14 |
| atoh8 | 0.16459 | 0.172986606 | 0.075 | 0 | 1 | 14 |
| lrig2 | 0.16459 | 0.200209671 | 0.075 | 0 | 1 | 14 |
| wipi2 | 0.16459 | 0.173128483 | 0.075 | 0 | 1 | 14 |
| CABZ01110881.1 | 0.16459 | 0.199024544 | 0.075 | 0 | 1 | 14 |
| XLOC-042595 | 0.16459 | 0.261115911 | 0.075 | 0 | 1 | 14 |
| adam19a | 0.16459 | 0.165930031 | 0.075 | 0 | 1 | 14 |
| arl4d | 0.16459 | 0.183152494 | 0.075 | 0 | 1 | 14 |
| arvcfb | 0.16459 | 0.202041394 | 0.075 | 0 | 1 | 14 |
| bckdha | 0.16459 | 0.199709198 | 0.075 | 0 | 1 | 14 |
| bmpr1aa | 0.16459 | 0.240346145 | 0.075 | 0 | 1 | 14 |
| bmt2 | 0.16459 | 0.171905514 | 0.075 | 0 | 1 | 14 |
| cebpa | 0.16459 | 0.166081371 | 0.075 | 0 | 1 | 14 |
| cemip2 | 0.16459 | 0.220436583 | 0.075 | 0 | 1 | 14 |
| cilp | 0.16459 | 0.196003272 | 0.075 | 0 | 1 | 14 |
| coa6 | 0.16459 | 0.170824986 | 0.075 | 0 | 1 | 14 |
| cpsf1 | 0.16459 | 0.220561111 | 0.075 | 0 | 1 | 14 |
| cry1ba | 0.16459 | 0.245913197 | 0.075 | 0 | 1 | 14 |
| cul1b | 0.16459 | 0.140344304 | 0.075 | 0 | 1 | 14 |
| dhrs13a.3 | 0.16459 | 0.189891744 | 0.075 | 0 | 1 | 14 |
| eif2b1 | 0.16459 | 0.159193744 | 0.075 | 0 | 1 | 14 |
| eif4eb | 0.16459 | 0.225767727 | 0.075 | 0 | 1 | 14 |
| eif4g2b | 0.16459 | 0.155813447 | 0.075 | 0 | 1 | 14 |
| elovl6 | 0.16459 | 0.190045519 | 0.075 | 0 | 1 | 14 |
| emc9 | 0.16459 | 0.150452268 | 0.075 | 0 | 1 | 14 |
| ephb3a | 0.16459 | 0.21030332 | 0.075 | 0 | 1 | 14 |
| eya1 | 0.16459 | 0.229916001 | 0.075 | 0 | 1 | 14 |
| fgf12b | 0.16459 | 0.146797102 | 0.075 | 0 | 1 | 14 |
| fgfr1op2 | 0.16459 | 0.188047896 | 0.075 | 0 | 1 | 14 |
| fnbp4 | 0.16459 | 0.185423221 | 0.075 | 0 | 1 | 14 |
| gadd45bb | 0.16459 | 0.171523731 | 0.075 | 0 | 1 | 14 |
| gpatch3 | 0.16459 | 0.167480109 | 0.075 | 0 | 1 | 14 |
| gyg1a | 0.16459 | 0.207007134 | 0.075 | 0 | 1 | 14 |
| ing5a | 0.16459 | 0.159726318 | 0.075 | 0 | 1 | 14 |
| lsm1 | 0.16459 | 0.153520321 | 0.075 | 0 | 1 | 14 |
| map1sa | 0.16459 | 0.183917355 | 0.075 | 0 | 1 | 14 |
| mapkapk3 | 0.16459 | 0.219983694 | 0.075 | 0 | 1 | 14 |
| med12 | 0.16459 | 0.192289752 | 0.075 | 0 | 1 | 14 |
| mogs | 0.16459 | 0.166918558 | 0.075 | 0 | 1 | 14 |
| ms4a17a.7 | 0.16459 | 0.190076769 | 0.075 | 0 | 1 | 14 |
| ntan1 | 0.16459 | 0.181560105 | 0.075 | 0 | 1 | 14 |
| nudt3b | 0.16459 | 0.181026487 | 0.075 | 0 | 1 | 14 |
| pdik1l | 0.16459 | 0.181872354 | 0.075 | 0 | 1 | 14 |
| pkma | 0.16459 | 0.188723764 | 0.075 | 0 | 1 | 14 |
| rbm25a | 0.16459 | 0.213259813 | 0.075 | 0 | 1 | 14 |

| | | | | | | |
|---|---|---|---|---|---|---|
| si:ch211-151g22.1 | 0.16459 | 0.163252816 | 0.075 | 0 | 1 | 14 |
| si:ch211-226m16.2 | 0.16459 | 0.156669523 | 0.075 | 0 | 1 | 14 |
| si:ch211-276c2.4 | 0.16459 | 0.1367571 | 0.075 | 0 | 1 | 14 |
| si:ch211-284b7.3 | 0.16459 | 0.145776825 | 0.075 | 0 | 1 | 14 |
| srgap1b | 0.16459 | 0.246494566 | 0.075 | 0 | 1 | 14 |
| stmn1b | 0.16459 | 0.115333136 | 0.075 | 0 | 1 | 14 |
| timmdc1 | 0.16459 | 0.168602761 | 0.075 | 0 | 1 | 14 |
| tprb | 0.16459 | 0.197218052 | 0.075 | 0 | 1 | 14 |
| wdr75 | 0.16459 | 0.179729302 | 0.075 | 0 | 1 | 14 |
| zgc:100920 | 0.16459 | 0.17683401 | 0.075 | 0 | 1 | 14 |
| znhit3 | 0.16459 | 0.187711131 | 0.075 | 0 | 1 | 14 |
| ndnl2 | 0.16459 | 0.177095484 | 0.075 | 0 | 1 | 14 |
| rcn1 | 0.16459 | 0.177730871 | 0.075 | 0 | 1 | 14 |
| tmem206 | 0.16459 | 0.167031873 | 0.075 | 0 | 1 | 14 |
| BX957317.2 | 0.16459 | 0.165193548 | 0.075 | 0 | 1 | 14 |
| CABZ01038697.1 | 0.16459 | 0.144871232 | 0.075 | 0 | 1 | 14 |
| CU467639.1 | 0.16459 | 0.14002627 | 0.075 | 0 | 1 | 14 |
| CU638870.1 | 0.16459 | 0.159546947 | 0.075 | 0 | 1 | 14 |
| FO904870.1 | 0.16459 | 0.166245951 | 0.075 | 0 | 1 | 14 |
| LO018148.1 | 0.16459 | 0.173930223 | 0.075 | 0 | 1 | 14 |
| LOC100330186 | 0.16459 | 0.167308923 | 0.075 | 0 | 1 | 14 |
| LOC101883507 | 0.16459 | 0.167171616 | 0.075 | 0 | 1 | 14 |
| LOC110438169 | 0.16459 | 0.19440193 | 0.075 | 0 | 1 | 14 |
| adamts5 | 0.16459 | 0.189268632 | 0.075 | 0 | 1 | 14 |
| aida | 0.16459 | 0.154282239 | 0.075 | 0 | 1 | 14 |
| amt | 0.16459 | 0.116713617 | 0.075 | 0 | 1 | 14 |
| asna1 | 0.16459 | 0.15264292 | 0.075 | 0 | 1 | 14 |
| atl2 | 0.16459 | 0.140858112 | 0.075 | 0 | 1 | 14 |
| bmp5 | 0.16459 | 0.201439742 | 0.075 | 0 | 1 | 14 |
| brd7 | 0.16459 | 0.170384173 | 0.075 | 0 | 1 | 14 |
| bscl2 | 0.16459 | 0.156559266 | 0.075 | 0 | 1 | 14 |
| ca10a | 0.16459 | 0.185722486 | 0.075 | 0 | 1 | 14 |
| cdc6 | 0.16459 | 0.188870872 | 0.075 | 0 | 1 | 14 |
| cebpz | 0.16459 | 0.172181814 | 0.075 | 0 | 1 | 14 |
| chd9 | 0.16459 | 0.167214648 | 0.075 | 0 | 1 | 14 |
| cherp | 0.16459 | 0.22633293 | 0.075 | 0 | 1 | 14 |
| chst14 | 0.16459 | 0.212172357 | 0.075 | 0 | 1 | 14 |
| ciapin1 | 0.16459 | 0.146784742 | 0.075 | 0 | 1 | 14 |
| ckap5 | 0.16459 | 0.196922923 | 0.075 | 0 | 1 | 14 |
| clptm1l | 0.16459 | 0.176895181 | 0.075 | 0 | 1 | 14 |
| clpxa | 0.16459 | 0.203373949 | 0.075 | 0 | 1 | 14 |
| coa1 | 0.16459 | 0.150329989 | 0.075 | 0 | 1 | 14 |
| cpox | 0.16459 | 0.135280595 | 0.075 | 0 | 1 | 14 |
| cryl1 | 0.16459 | 0.141890543 | 0.075 | 0 | 1 | 14 |
| csrp1a | 0.16459 | 0.159441948 | 0.075 | 0 | 1 | 14 |
| ctnnbl1 | 0.16459 | 0.191077264 | 0.075 | 0 | 1 | 14 |
| ctps1b | 0.16459 | 0.144313914 | 0.075 | 0 | 1 | 14 |
| cul3a | 0.16459 | 0.162237 | 0.075 | 0 | 1 | 14 |

| | | | | | | |
|---|---|---|---|---|---|---|
| cyp26a1 | 0.16459 | 0.225319933 | 0.075 | 0 | 1 | 14 |
| dhx9 | 0.16459 | 0.169636914 | 0.075 | 0 | 1 | 14 |
| dimt1l | 0.16459 | 0.165330777 | 0.075 | 0 | 1 | 14 |
| dnajb4 | 0.16459 | 0.144181878 | 0.075 | 0 | 1 | 14 |
| dph2 | 0.16459 | 0.222514127 | 0.075 | 0 | 1 | 14 |
| eif2b3 | 0.16459 | 0.136848476 | 0.075 | 0 | 1 | 14 |
| enc1 | 0.16459 | 0.190872473 | 0.075 | 0 | 1 | 14 |
| fam102ab | 0.16459 | 0.145012446 | 0.075 | 0 | 1 | 14 |
| fank1 | 0.16459 | 0.204971884 | 0.075 | 0 | 1 | 14 |
| fen1 | 0.16459 | 0.18548582 | 0.075 | 0 | 1 | 14 |
| fgfr1op | 0.16459 | 0.153058432 | 0.075 | 0 | 1 | 14 |
| fnta | 0.16459 | 0.144627266 | 0.075 | 0 | 1 | 14 |
| gapvd1 | 0.16459 | 0.195600748 | 0.075 | 0 | 1 | 14 |
| gng3 | 0.16459 | 0.179802195 | 0.075 | 0 | 1 | 14 |
| grem1b | 0.16459 | 0.239022452 | 0.075 | 0 | 1 | 14 |
| gsk3b | 0.16459 | 0.195307187 | 0.075 | 0 | 1 | 14 |
| huwe1 | 0.16459 | 0.213189445 | 0.075 | 0 | 1 | 14 |
| ift27 | 0.16459 | 0.17089817 | 0.075 | 0 | 1 | 14 |
| ing2 | 0.16459 | 0.1852207 | 0.075 | 0 | 1 | 14 |
| itpka | 0.16459 | 0.170300327 | 0.075 | 0 | 1 | 14 |
| kazald3 | 0.16459 | 0.143997647 | 0.075 | 0 | 1 | 14 |
| kdm5c | 0.16459 | 0.161662602 | 0.075 | 0 | 1 | 14 |
| kmt2d | 0.16459 | 0.226040995 | 0.075 | 0 | 1 | 14 |
| kmt5b | 0.16459 | 0.199214506 | 0.075 | 0 | 1 | 14 |
| lamtor3 | 0.16459 | 0.166652517 | 0.075 | 0 | 1 | 14 |
| mdc1 | 0.16459 | 0.225444369 | 0.075 | 0 | 1 | 14 |
| mfap3l | 0.16459 | 0.136286191 | 0.075 | 0 | 1 | 14 |
| mier1a | 0.16459 | 0.178456162 | 0.075 | 0 | 1 | 14 |
| mrpl16 | 0.16459 | 0.105224104 | 0.075 | 0 | 1 | 14 |
| mrps28 | 0.16459 | 0.181224408 | 0.075 | 0 | 1 | 14 |
| myoc | 0.16459 | 0.241978337 | 0.075 | 0 | 1 | 14 |
| ndufaf5 | 0.16459 | 0.178591341 | 0.075 | 0 | 1 | 14 |
| nedd8l | 0.16459 | 0.149624133 | 0.075 | 0 | 1 | 14 |
| nfkbib | 0.16459 | 0.158049971 | 0.075 | 0 | 1 | 14 |
| nsd3 | 0.16459 | 0.208799104 | 0.075 | 0 | 1 | 14 |
| nufip2 | 0.16459 | 0.192777781 | 0.075 | 0 | 1 | 14 |
| nutf2l | 0.16459 | 0.124645909 | 0.075 | 0 | 1 | 14 |
| pamr1 | 0.16459 | 0.183427991 | 0.075 | 0 | 1 | 14 |
| pggt1b | 0.16459 | 0.134708423 | 0.075 | 0 | 1 | 14 |
| phc2b | 0.16459 | 0.130188426 | 0.075 | 0 | 1 | 14 |
| phgdh | 0.16459 | 0.15590609 | 0.075 | 0 | 1 | 14 |
| pop5 | 0.16459 | 0.150587549 | 0.075 | 0 | 1 | 14 |
| ppil2 | 0.16459 | 0.17321773 | 0.075 | 0 | 1 | 14 |
| prdm11 | 0.16459 | 0.197523753 | 0.075 | 0 | 1 | 14 |
| ptpn9a | 0.16459 | 0.140153092 | 0.075 | 0 | 1 | 14 |
| ptprfb | 0.16459 | 0.1718708 | 0.075 | 0 | 1 | 14 |
| rae1 | 0.16459 | 0.150415326 | 0.075 | 0 | 1 | 14 |
| rpa2 | 0.16459 | 0.157078588 | 0.075 | 0 | 1 | 14 |

| | | | | | | |
|---|---|---|---|---|---|---|
| si:ch211-210g13.5 | 0.16459 | 0.258899231 | 0.075 | 0 | 1 | 14 |
| si:ch211-276i12.11 | 0.16459 | 0.178469992 | 0.075 | 0 | 1 | 14 |
| si:dkeyp-66d1.7 | 0.16459 | 0.161023349 | 0.075 | 0 | 1 | 14 |
| slc9a6a | 0.16459 | 0.177947972 | 0.075 | 0 | 1 | 14 |
| slco5a1a | 0.16459 | 0.177847588 | 0.075 | 0 | 1 | 14 |
| smarcb1a | 0.16459 | 0.172528276 | 0.075 | 0 | 1 | 14 |
| smek1 | 0.16459 | 0.151294868 | 0.075 | 0 | 1 | 14 |
| srsf1a | 0.16459 | 0.180709839 | 0.075 | 0 | 1 | 14 |
| ssrp1a | 0.16459 | 0.146337332 | 0.075 | 0 | 1 | 14 |
| stx5a | 0.16459 | 0.197434905 | 0.075 | 0 | 1 | 14 |
| sugt1 | 0.16459 | 0.173769851 | 0.075 | 0 | 1 | 14 |
| tab3 | 0.16459 | 0.106775084 | 0.075 | 0 | 1 | 14 |
| tasp1 | 0.16459 | 0.203932321 | 0.075 | 0 | 1 | 14 |
| tent4b | 0.16459 | 0.168607656 | 0.075 | 0 | 1 | 14 |
| thoc3 | 0.16459 | 0.16401093 | 0.075 | 0 | 1 | 14 |
| tipin | 0.16459 | 0.175704201 | 0.075 | 0 | 1 | 14 |
| tle2a | 0.16459 | 0.171569315 | 0.075 | 0 | 1 | 14 |
| tmem131 | 0.16459 | 0.178853286 | 0.075 | 0 | 1 | 14 |
| tmem42a | 0.16459 | 0.159499129 | 0.075 | 0 | 1 | 14 |
| trappc6b | 0.16459 | 0.147526993 | 0.075 | 0 | 1 | 14 |
| tusc3 | 0.16459 | 0.134137243 | 0.075 | 0 | 1 | 14 |
| ubald1a | 0.16459 | 0.166786556 | 0.075 | 0 | 1 | 14 |
| uri1 | 0.16459 | 0.172569481 | 0.075 | 0 | 1 | 14 |
| usp8 | 0.16459 | 0.19962415 | 0.075 | 0 | 1 | 14 |
| uxt | 0.16459 | 0.165399192 | 0.075 | 0 | 1 | 14 |
| vps37b | 0.16459 | 0.157728158 | 0.075 | 0 | 1 | 14 |
| waplb | 0.16459 | 0.160934377 | 0.075 | 0 | 1 | 14 |
| wdr5 | 0.16459 | 0.145758335 | 0.075 | 0 | 1 | 14 |
| zbtb33 | 0.16459 | 0.145148719 | 0.075 | 0 | 1 | 14 |
| zgc:110796 | 0.16459 | 0.174900071 | 0.075 | 0 | 1 | 14 |
| zgc:171727 | 0.16459 | 0.173981729 | 0.075 | 0 | 1 | 14 |
| zgc:56556 | 0.16459 | 0.180820222 | 0.075 | 0 | 1 | 14 |
| znf281a | 0.16459 | 0.213183277 | 0.075 | 0 | 1 | 14 |
| bnip4 | 0.16459 | 0.156388687 | 0.075 | 0 | 1 | 14 |
| bola3 | 0.16459 | 0.193414354 | 0.075 | 0 | 1 | 14 |
| cldn12 | 0.16459 | 0.169405362 | 0.075 | 0 | 1 | 14 |
| clockb | 0.16459 | 0.212569751 | 0.075 | 0 | 1 | 14 |
| hint2 | 0.16459 | 0.170462755 | 0.075 | 0 | 1 | 14 |
| inavab | 0.16459 | 0.187219049 | 0.075 | 0 | 1 | 14 |
| jkamp | 0.16459 | 0.167002828 | 0.075 | 0 | 1 | 14 |
| lamp1b | 0.16459 | 0.151955052 | 0.075 | 0 | 1 | 14 |
| msl2b | 0.16459 | 0.192713423 | 0.075 | 0 | 1 | 14 |
| purbb | 0.16459 | 0.240924323 | 0.075 | 0 | 1 | 14 |
| rfc4 | 0.16459 | 0.212737092 | 0.075 | 0 | 1 | 14 |
| si:ch211-106k21.5 | 0.16459 | 0.181780463 | 0.075 | 0 | 1 | 14 |
| svbp | 0.16459 | 0.156237657 | 0.075 | 0 | 1 | 14 |
| tet3 | 0.16459 | 0.223734961 | 0.075 | 0 | 1 | 14 |
| timm44 | 0.16459 | 0.155388734 | 0.075 | 0 | 1 | 14 |

| | | | | | | |
|---|---|---|---|---|---|---|
| trappc3 | 0.16459 | 0.156666056 | 0.075 | 0 | 1 | 14 |
| utp25 | 0.16459 | 0.145020243 | 0.075 | 0 | 1 | 14 |
| vipr2 | 0.16459 | 0.239768519 | 0.075 | 0 | 1 | 14 |
| yme1l1a | 0.16459 | 0.203467791 | 0.075 | 0 | 1 | 14 |
| zgc:194981 | 0.16459 | 0.183107383 | 0.075 | 0 | 1 | 14 |
| znf576.1 | 0.16459 | 0.148224039 | 0.075 | 0 | 1 | 14 |
| LOC101884549 | 0.16459 | 0.136268602 | 0.075 | 0 | 1 | 14 |
| atxn7 | 0.16459 | 0.167319497 | 0.075 | 0 | 1 | 14 |
| bcl7bb | 0.16459 | 0.151382739 | 0.075 | 0 | 1 | 14 |
| cmbl | 0.16459 | 0.14338543 | 0.075 | 0 | 1 | 14 |
| desi1a | 0.16459 | 0.1869331 | 0.075 | 0 | 1 | 14 |
| fam208b | 0.16459 | 0.142973621 | 0.075 | 0 | 1 | 14 |
| im:7136021 | 0.16459 | 0.167864023 | 0.075 | 0 | 1 | 14 |
| lonrf1l | 0.16459 | 0.177473952 | 0.075 | 0 | 1 | 14 |
| myca | 0.16459 | 0.217502986 | 0.075 | 0 | 1 | 14 |
| pdpk1b | 0.16459 | 0.178246591 | 0.075 | 0 | 1 | 14 |
| polb | 0.16459 | 0.145936414 | 0.075 | 0 | 1 | 14 |
| ppm1bb | 0.16459 | 0.165072534 | 0.075 | 0 | 1 | 14 |
| rcc1 | 0.16459 | 0.137108687 | 0.075 | 0 | 1 | 14 |
| tmed3 | 0.16459 | 0.132515439 | 0.075 | 0 | 1 | 14 |
| tmem150ab | 0.16459 | 0.197349024 | 0.075 | 0 | 1 | 14 |
| tpbga | 0.16459 | 0.170038077 | 0.075 | 0 | 1 | 14 |
| trappc8 | 0.16459 | 0.198645545 | 0.075 | 0 | 1 | 14 |
| ube2d2l | 0.16459 | 0.144777762 | 0.075 | 0 | 1 | 14 |
| vezf1a | 0.16459 | 0.208327934 | 0.075 | 0 | 1 | 14 |
| wdr3 | 0.16459 | 0.161116599 | 0.075 | 0 | 1 | 14 |
| znf423 | 0.16459 | 0.158343272 | 0.075 | 0 | 1 | 14 |
| CABZ01049847.1 | 0.16459 | 0.180650923 | 0.075 | 0 | 1 | 14 |
| abce1 | 0.16459 | 0.122708753 | 0.075 | 0 | 1 | 14 |
| adma | 0.16459 | 0.151449867 | 0.075 | 0 | 1 | 14 |
| bap1 | 0.16459 | 0.19862592 | 0.075 | 0 | 1 | 14 |
| bckdk | 0.16459 | 0.196065494 | 0.075 | 0 | 1 | 14 |
| ccdc106a | 0.16459 | 0.161140278 | 0.075 | 0 | 1 | 14 |
| cfap298 | 0.16459 | 0.156491796 | 0.075 | 0 | 1 | 14 |
| chmp4ba | 0.16459 | 0.145938514 | 0.075 | 0 | 1 | 14 |
| ctr9 | 0.16459 | 0.174335732 | 0.075 | 0 | 1 | 14 |
| dhdds | 0.16459 | 0.159260245 | 0.075 | 0 | 1 | 14 |
| iqcc | 0.16459 | 0.168444558 | 0.075 | 0 | 1 | 14 |
| lrba | 0.16459 | 0.242774002 | 0.075 | 0 | 1 | 14 |
| lysmd3 | 0.16459 | 0.142219259 | 0.075 | 0 | 1 | 14 |
| mrps2 | 0.16459 | 0.135338645 | 0.075 | 0 | 1 | 14 |
| phactr4b | 0.16459 | 0.189934118 | 0.075 | 0 | 1 | 14 |
| ppa1b | 0.16459 | 0.128307778 | 0.075 | 0 | 1 | 14 |
| ppp1r7 | 0.16459 | 0.154932189 | 0.075 | 0 | 1 | 14 |
| rassf8a | 0.16459 | 0.212339529 | 0.075 | 0 | 1 | 14 |
| rex1bd | 0.16459 | 0.176815362 | 0.075 | 0 | 1 | 14 |
| selenos | 0.16459 | 0.138137545 | 0.075 | 0 | 1 | 14 |
| shmt1 | 0.16459 | 0.164095255 | 0.075 | 0 | 1 | 14 |

| | | | | | | |
|---|---|---|---|---|---|---|
| si:ch1073-322p19.1 | 0.16459 | 0.201640971 | 0.075 | 0 | 1 | 14 |
| si:ch211-156b7.4 | 0.16459 | 0.142452362 | 0.075 | 0 | 1 | 14 |
| si:ch73-380l3.2 | 0.16459 | 0.206161401 | 0.075 | 0 | 1 | 14 |
| sorbs1 | 0.16459 | 0.206356098 | 0.075 | 0 | 1 | 14 |
| tanc2a | 0.16459 | 0.213936518 | 0.075 | 0 | 1 | 14 |
| tpd52l2a | 0.16459 | 0.10404308 | 0.075 | 0 | 1 | 14 |
| tubb6 | 0.16459 | 0.161652994 | 0.075 | 0 | 1 | 14 |
| xrcc6 | 0.16459 | 0.147010267 | 0.075 | 0 | 1 | 14 |
| yars | 0.16459 | 0.187192197 | 0.075 | 0 | 1 | 14 |
| zgc:101569 | 0.16459 | 0.148434844 | 0.075 | 0 | 1 | 14 |
| zgc:171971 | 0.16459 | 0.167672682 | 0.075 | 0 | 1 | 14 |
| derl2 | 0.16459 | 0.169619728 | 0.075 | 0 | 1 | 14 |
| clic4 | 0.16471 | -0.38815487 | 0.1 | 0.3 | 1 | 14 |
| CR847994.1 | 0.16479 | -0.19471887 | 0.025 | 0.15 | 1 | 14 |
| taf13 | 0.16496 | 0.366120986 | 0.325 | 0.2 | 1 | 14 |
| XLOC-043760 | 0.16538 | -0.22987071 | 0.025 | 0.15 | 1 | 14 |
| ppp4cb | 0.16539 | -0.21229154 | 0.05 | 0.2 | 1 | 14 |
| khdrbs1a | 0.1655 | 0.190455868 | 0.775 | 0.75 | 1 | 14 |
| ugp2b | 0.16557 | -0.20252911 | 0.05 | 0.2 | 1 | 14 |
| chaf1a | 0.16578 | 0.165237051 | 0.125 | 0.15 | 1 | 14 |
| selenot2 | 0.16615 | -0.36949545 | 0.05 | 0.2 | 1 | 14 |
| tspan18b | 0.16634 | -0.30121277 | 0.075 | 0.25 | 1 | 14 |
| ggh | 0.16669 | -0.21431101 | 0.025 | 0.15 | 1 | 14 |
| rps29 | 0.16672 | 0.191383352 | 0.975 | 0.85 | 1 | 14 |
| uba52 | 0.16708 | 0.229645533 | 0.95 | 0.8 | 1 | 14 |
| fth1a | 0.16757 | -0.31100773 | 0.35 | 0.6 | 1 | 14 |
| trib3 | 0.16765 | -0.34710208 | 0.05 | 0.2 | 1 | 14 |
| rpl14 | 0.1682 | 0.197694812 | 0.975 | 0.85 | 1 | 14 |
| tpst1 | 0.16822 | 0.403147476 | 0.275 | 0.1 | 1 | 14 |
| hspa12a | 0.16866 | 0.393267886 | 0.2 | 0.05 | 1 | 14 |
| atp5pb | 0.1688 | 0.287326017 | 0.625 | 0.55 | 1 | 14 |
| coch | 0.16942 | -0.45743827 | 0.175 | 0.4 | 1 | 14 |
| hccsa.1 | 0.16989 | -0.10221312 | 0.025 | 0.15 | 1 | 14 |
| COX3 | 0.17023 | -0.19293029 | 0.9 | 1 | 1 | 14 |
| si:dkey-56m19.5 | 0.17099 | 0.401627545 | 0.175 | 0.05 | 1 | 14 |
| daam1a | 0.17133 | -0.33678545 | 0.025 | 0.15 | 1 | 14 |
| rps3a | 0.1714 | 0.158141382 | 0.9 | 0.9 | 1 | 14 |
| col6a3 | 0.17164 | -0.35351138 | 0.6 | 0.8 | 1 | 14 |
| agrn | 0.17259 | -0.40832722 | 0.175 | 0.4 | 1 | 14 |
| si:dkey-118j18.2 | 0.17297 | -0.35375212 | 0.075 | 0.25 | 1 | 14 |
| osr1 | 0.17339 | -0.4297452 | 0.075 | 0.25 | 1 | 14 |
| cotl1 | 0.17364 | -0.29946832 | 0.075 | 0.25 | 1 | 14 |
| trir | 0.17417 | -0.20878783 | 0.05 | 0.2 | 1 | 14 |
| cdkn1bb | 0.17452 | 0.129644926 | 0.275 | 0.3 | 1 | 14 |
| lrp5.1 | 0.17563 | -0.30645799 | 0.05 | 0.2 | 1 | 14 |
| rrbp1a | 0.17589 | -0.36301225 | 0.15 | 0.3 | 1 | 14 |
| csad | 0.17626 | -0.22290507 | 0.075 | 0.25 | 1 | 14 |
| pde4ba | 0.17671 | -0.44344008 | 0.075 | 0.25 | 1 | 14 |

| | | | | | | |
|---|---|---|---|---|---|---|
| eif5 | 0.17726 | -0.18782915 | 0.175 | 0.35 | 1 | 14 |
| nfkbiab | 0.17776 | -0.35663333 | 0.175 | 0.4 | 1 | 14 |
| mt-cyb | 0.17781 | -0.31261777 | 0.625 | 0.85 | 1 | 14 |
| slc22a18 | 0.17839 | -0.33438961 | 0.075 | 0.25 | 1 | 14 |
| selenot1b | 0.17844 | -0.36808053 | 0.075 | 0.25 | 1 | 14 |
| perp | 0.17985 | -0.12155217 | 0.075 | 0.2 | 1 | 14 |
| arl14ep | 0.18004 | -0.18860789 | 0.05 | 0.2 | 1 | 14 |
| rpl18a | 0.18005 | 0.185248003 | 0.975 | 0.9 | 1 | 14 |
| smc1al | 0.1802 | 0.350565603 | 0.225 | 0.05 | 1 | 14 |
| plxdc2 | 0.18031 | 0.236414234 | 0.2 | 0.15 | 1 | 14 |
| cald1a | 0.18064 | -0.18756768 | 0.45 | 0.65 | 1 | 14 |
| hnrnpa0a | 0.18115 | 0.223470986 | 0.225 | 0.2 | 1 | 14 |
| si:ch211-227m13.1 | 0.18128 | -0.37069535 | 0.05 | 0.2 | 1 | 14 |
| purba | 0.18169 | -0.32669927 | 0.025 | 0.15 | 1 | 14 |
| arf6b | 0.18219 | -0.30295935 | 0.075 | 0.25 | 1 | 14 |
| fam107b | 0.18223 | -0.42961697 | 0.225 | 0.4 | 1 | 14 |
| tmem115 | 0.18253 | -0.31867005 | 0.075 | 0.25 | 1 | 14 |
| CABZ01113812.1 | 0.18266 | -0.17841404 | 0.025 | 0.15 | 1 | 14 |
| psma6b | 0.18285 | 0.173339378 | 0.175 | 0.05 | 1 | 14 |
| meis1b | 0.18327 | -0.39103036 | 0.35 | 0.6 | 1 | 14 |
| fbln5 | 0.18352 | -0.39391516 | 0.35 | 0.6 | 1 | 14 |
| copz1 | 0.18354 | -0.23998485 | 0.075 | 0.25 | 1 | 14 |
| enpp1 | 0.1836 | -0.3787276 | 0.075 | 0.25 | 1 | 14 |
| metap2b | 0.18372 | 0.258032691 | 0.25 | 0.2 | 1 | 14 |
| si:dkey-7j14.6 | 0.1839 | -0.31709449 | 0.5 | 0.65 | 1 | 14 |
| ube2nb | 0.18396 | -0.22259347 | 0.05 | 0.2 | 1 | 14 |
| txnipa | 0.18403 | -0.32735305 | 0.4 | 0.65 | 1 | 14 |
| fbl | 0.1842 | 0.328735925 | 0.325 | 0.2 | 1 | 14 |
| pafah1b1b | 0.18465 | -0.20903107 | 0.025 | 0.15 | 1 | 14 |
| cox7b | 0.18507 | 0.370157906 | 0.475 | 0.25 | 1 | 14 |
| edil3a | 0.18531 | -0.40137822 | 0.075 | 0.25 | 1 | 14 |
| palm1a | 0.18541 | -0.15409666 | 0.325 | 0.5 | 1 | 14 |
| rps26l | 0.18546 | 0.220248473 | 0.95 | 0.8 | 1 | 14 |
| scfd1 | 0.1856 | -0.30465044 | 0.05 | 0.2 | 1 | 14 |
| mcrs1 | 0.18637 | -0.25939429 | 0.025 | 0.15 | 1 | 14 |
| zgc:77486 | 0.1865 | -0.29796708 | 0.075 | 0.25 | 1 | 14 |
| gkap1 | 0.18699 | 0.346997971 | 0.2 | 0.05 | 1 | 14 |
| si:ch211-178n15.1 | 0.18711 | -0.26701726 | 0.025 | 0.15 | 1 | 14 |
| cd63 | 0.18714 | -0.23905503 | 0.575 | 0.8 | 1 | 14 |
| eif4e2rs1 | 0.18729 | -0.18423183 | 0.025 | 0.15 | 1 | 14 |
| palm1b | 0.18745 | -0.33563734 | 0.075 | 0.25 | 1 | 14 |
| nop53 | 0.18786 | 0.249263282 | 0.475 | 0.4 | 1 | 14 |
| isg20 | 0.18868 | 0.212033527 | 0.125 | 0.1 | 1 | 14 |
| selenoo1 | 0.18871 | -0.21168468 | 0.025 | 0.15 | 1 | 14 |
| PRRC2B | 0.18872 | -0.31239578 | 0.025 | 0.15 | 1 | 14 |
| ugcg | 0.18911 | -0.20871857 | 0.025 | 0.15 | 1 | 14 |
| srsf9 | 0.18927 | -0.21913767 | 0.25 | 0.45 | 1 | 14 |
| sept6 | 0.18953 | -0.33923255 | 0.075 | 0.25 | 1 | 14 |

| | | | | | | |
|---|---|---|---|---|---|---|
| rab41 | 0.18994 | -0.19578362 | 0.025 | 0.15 | 1 | 14 |
| srrm2 | 0.19012 | -0.28510039 | 0.025 | 0.15 | 1 | 14 |
| rnf10 | 0.19099 | 0.306500388 | 0.3 | 0.2 | 1 | 14 |
| ndfip2 | 0.19114 | -0.30609211 | 0.075 | 0.25 | 1 | 14 |
| lbr | 0.19136 | -0.29077084 | 0.05 | 0.2 | 1 | 14 |
| cd276 | 0.19138 | -0.31040267 | 0.075 | 0.25 | 1 | 14 |
| pcsk6 | 0.19151 | 0.388339551 | 0.2 | 0.05 | 1 | 14 |
| psma4 | 0.19176 | 0.232268949 | 0.35 | 0.3 | 1 | 14 |
| map1lc3a | 0.19195 | -0.36001331 | 0.075 | 0.25 | 1 | 14 |
| rpl29 | 0.19196 | 0.202803387 | 0.9 | 0.8 | 1 | 14 |
| ccdc32 | 0.19246 | -0.24755385 | 0.05 | 0.2 | 1 | 14 |
| mvp | 0.19255 | -0.3687404 | 0.075 | 0.25 | 1 | 14 |
| nid1a | 0.19258 | -0.34394539 | 0.075 | 0.25 | 1 | 14 |
| tspan4a | 0.19315 | -0.33201788 | 0.05 | 0.2 | 1 | 14 |
| LO018102.1 | 0.1932 | -0.21158543 | 0.025 | 0.15 | 1 | 14 |
| psmb7 | 0.19381 | 0.186271766 | 0.375 | 0.35 | 1 | 14 |
| vamp3 | 0.1941 | -0.1842827 | 0.025 | 0.15 | 1 | 14 |
| LOC405768 | 0.19411 | -0.27227307 | 0.05 | 0.2 | 1 | 14 |
| pmp22a | 0.19418 | -0.11937869 | 0.8 | 0.95 | 1 | 14 |
| ndfip1l | 0.19432 | -0.33419184 | 0.15 | 0.35 | 1 | 14 |
| ackr3b | 0.19502 | -0.4398195 | 0.175 | 0.35 | 1 | 14 |
| rpl22l1 | 0.19524 | 0.371661624 | 0.7 | 0.5 | 1 | 14 |
| hmbox1a | 0.1954 | -0.18646835 | 0.275 | 0.45 | 1 | 14 |
| capns1b | 0.1962 | -0.37875384 | 0.225 | 0.45 | 1 | 14 |
| golt1ba | 0.19622 | -0.29296886 | 0.025 | 0.15 | 1 | 14 |
| kdelr3 | 0.19643 | -0.27270265 | 0.05 | 0.2 | 1 | 14 |
| paip2b | 0.19647 | 0.136092277 | 0.45 | 0.45 | 1 | 14 |
| XLOC-015238 | 0.19676 | -0.32018542 | 0.025 | 0.15 | 1 | 14 |
| eif4a1a | 0.19686 | 0.355458455 | 0.425 | 0.25 | 1 | 14 |
| serpinh2 | 0.19713 | -0.15837718 | 0.075 | 0.2 | 1 | 14 |
| ube2d1b | 0.19716 | 0.200013711 | 0.175 | 0.15 | 1 | 14 |
| zfhx4 | 0.19868 | 0.440013681 | 0.425 | 0.2 | 1 | 14 |
| krt4 | 0.20024 | -0.43712296 | 0.25 | 0.45 | 1 | 14 |
| efna5b | 0.20115 | -0.39099825 | 0.15 | 0.35 | 1 | 14 |
| zbtb47b | 0.20116 | -0.26603614 | 0.025 | 0.15 | 1 | 14 |
| tmem251 | 0.202 | -0.22906061 | 0.025 | 0.15 | 1 | 14 |
| dpm2 | 0.20203 | -0.21110568 | 0.025 | 0.15 | 1 | 14 |
| il4r.1 | 0.20205 | -0.32453722 | 0.05 | 0.2 | 1 | 14 |
| tmem165 | 0.20261 | -0.14993757 | 0.15 | 0.3 | 1 | 14 |
| rpl7a | 0.20295 | 0.208289814 | 0.95 | 0.8 | 1 | 14 |
| LOC110439449 | 0.20314 | -0.26411261 | 0.025 | 0.15 | 1 | 14 |
| rtca | 0.20335 | -0.16631936 | 0.025 | 0.15 | 1 | 14 |
| ap2m1b | 0.20343 | 0.329272901 | 0.175 | 0.05 | 1 | 14 |
| cavin2a | 0.20357 | 0.107525321 | 0.125 | 0.15 | 1 | 14 |
| lamp2 | 0.20415 | -0.38766373 | 0.225 | 0.45 | 1 | 14 |
| actr1 | 0.20446 | -0.34738536 | 0.05 | 0.2 | 1 | 14 |
| dnpep | 0.20447 | -0.27168236 | 0.025 | 0.15 | 1 | 14 |
| nfixb | 0.20448 | -0.15942577 | 0.425 | 0.6 | 1 | 14 |

| | | | | | | |
|---|---|---|---|---|---|---|
| reep3b | 0.2048 | -0.31214925 | 0.05 | 0.2 | 1 | 14 |
| ndufb6 | 0.20482 | -0.16302555 | 0.1 | 0.25 | 1 | 14 |
| slc39a6 | 0.20496 | -0.21257079 | 0.025 | 0.15 | 1 | 14 |
| rasa1b | 0.20508 | -0.3116274 | 0.025 | 0.15 | 1 | 14 |
| sufu | 0.20554 | -0.28273727 | 0.05 | 0.2 | 1 | 14 |
| itgb4 | 0.20562 | -0.3252795 | 0.05 | 0.2 | 1 | 14 |
| supt16h | 0.20572 | -0.26192309 | 0.05 | 0.2 | 1 | 14 |
| npepps | 0.20586 | -0.28887845 | 0.025 | 0.15 | 1 | 14 |
| rnaseka | 0.20615 | -0.23000787 | 0.025 | 0.15 | 1 | 14 |
| emilin1b | 0.20617 | -0.33577846 | 0.15 | 0.35 | 1 | 14 |
| kpnb1 | 0.20647 | -0.33384665 | 0.05 | 0.2 | 1 | 14 |
| ap2m1a | 0.20651 | 0.215337976 | 0.325 | 0.3 | 1 | 14 |
| il17rd | 0.20672 | -0.3037094 | 0.05 | 0.2 | 1 | 14 |
| ccdc25 | 0.20688 | -0.1709406 | 0.15 | 0.3 | 1 | 14 |
| ufd1l | 0.20736 | -0.28473556 | 0.05 | 0.2 | 1 | 14 |
| ptbp2b | 0.20748 | -0.20179486 | 0.025 | 0.15 | 1 | 14 |
| b4galt6 | 0.2075 | -0.3954767 | 0.1 | 0.25 | 1 | 14 |
| crebbpb | 0.20762 | -0.29943874 | 0.05 | 0.2 | 1 | 14 |
| casp8 | 0.20771 | -0.27410366 | 0.025 | 0.15 | 1 | 14 |
| hist2h2l | 0.20776 | 0.375780299 | 0.425 | 0.2 | 1 | 14 |
| ddr1 | 0.2079 | -0.31936379 | 0.05 | 0.2 | 1 | 14 |
| sdr16c5b | 0.20807 | -0.31222102 | 0.025 | 0.15 | 1 | 14 |
| nol11 | 0.20836 | -0.23518155 | 0.025 | 0.15 | 1 | 14 |
| CDPF1 | 0.20869 | -0.19874787 | 0.025 | 0.15 | 1 | 14 |
| pank2 | 0.20876 | -0.30492615 | 0.05 | 0.2 | 1 | 14 |
| dldh | 0.20886 | -0.15922587 | 0.1 | 0.25 | 1 | 14 |
| CABZ01068358.1 | 0.20943 | -0.26288622 | 0.025 | 0.15 | 1 | 14 |
| usp5 | 0.20956 | -0.27382496 | 0.025 | 0.15 | 1 | 14 |
| tm2d3 | 0.20989 | -0.29907547 | 0.05 | 0.2 | 1 | 14 |
| lactb | 0.2099 | -0.17888012 | 0.025 | 0.15 | 1 | 14 |
| ntd5 | 0.20992 | -0.36438977 | 0.025 | 0.15 | 1 | 14 |
| plpp1a | 0.21029 | -0.30052444 | 0.025 | 0.15 | 1 | 14 |
| rpap3 | 0.21044 | -0.23588738 | 0.025 | 0.15 | 1 | 14 |
| brd1b | 0.21058 | -0.26569224 | 0.025 | 0.15 | 1 | 14 |
| AL590149.1 | 0.21124 | -0.10554527 | 0 | 0.05 | 1 | 14 |
| AL954741.2 | 0.21124 | -0.1628688 | 0 | 0.05 | 1 | 14 |
| BX855614.1 | 0.21124 | -0.12616904 | 0 | 0.05 | 1 | 14 |
| CABZ01064670.1 | 0.21124 | -0.15772298 | 0 | 0.05 | 1 | 14 |
| CABZ01064859.2 | 0.21124 | -0.10554527 | 0 | 0.05 | 1 | 14 |
| CABZ01073097.1 | 0.21124 | -0.12703576 | 0 | 0.05 | 1 | 14 |
| CABZ01084963.1 | 0.21124 | -0.11494264 | 0 | 0.05 | 1 | 14 |
| CR354588.1 | 0.21124 | -0.12181401 | 0 | 0.05 | 1 | 14 |
| CT955967.1 | 0.21124 | -0.10078319 | 0 | 0.05 | 1 | 14 |
| CU062487.1 | 0.21124 | -0.12228033 | 0 | 0.05 | 1 | 14 |
| DST | 0.21124 | -0.16251215 | 0 | 0.05 | 1 | 14 |
| FO203432.1 | 0.21124 | -0.11779768 | 0 | 0.05 | 1 | 14 |
| FQ378018.1 | 0.21124 | -0.14397037 | 0 | 0.05 | 1 | 14 |
| LO018474.1 | 0.21124 | -0.16073801 | 0 | 0.05 | 1 | 14 |

| | | | | | | |
|---|---|---|---|---|---|---|
| LOC100535208 | 0.21124 | -0.12669088 | 0 | 0.05 | 1 | 14 |
| LOC101882492 | 0.21124 | -0.10210235 | 0 | 0.05 | 1 | 14 |
| LOC101882607 | 0.21124 | -0.10078319 | 0 | 0.05 | 1 | 14 |
| LOC101885776 | 0.21124 | -0.13228314 | 0 | 0.05 | 1 | 14 |
| LOC108179274 | 0.21124 | -0.12808228 | 0 | 0.05 | 1 | 14 |
| LOC108190763 | 0.21124 | -0.12228033 | 0 | 0.05 | 1 | 14 |
| LOC110437711 | 0.21124 | -0.10554527 | 0 | 0.05 | 1 | 14 |
| LOC110437926 | 0.21124 | -0.10011078 | 0 | 0.05 | 1 | 14 |
| LOC110439758 | 0.21124 | -0.10078319 | 0 | 0.05 | 1 | 14 |
| NFX1 | 0.21124 | -0.10066569 | 0 | 0.05 | 1 | 14 |
| NPAS3 | 0.21124 | -0.12710386 | 0 | 0.05 | 1 | 14 |
| SPAG9 | 0.21124 | -0.10053885 | 0 | 0.05 | 1 | 14 |
| XLOC-004016 | 0.21124 | -0.11589312 | 0 | 0.05 | 1 | 14 |
| XLOC-005350 | 0.21124 | -0.10078319 | 0 | 0.05 | 1 | 14 |
| XLOC-006268 | 0.21124 | -0.14447757 | 0 | 0.05 | 1 | 14 |
| XLOC-016167 | 0.21124 | -0.12477302 | 0 | 0.05 | 1 | 14 |
| XLOC-026835 | 0.21124 | -0.17377028 | 0 | 0.05 | 1 | 14 |
| XLOC-029041 | 0.21124 | -0.10078319 | 0 | 0.05 | 1 | 14 |
| XLOC-039167 | 0.21124 | -0.10554527 | 0 | 0.05 | 1 | 14 |
| ZBTB26 | 0.21124 | -0.16066776 | 0 | 0.05 | 1 | 14 |
| abi2a | 0.21124 | -0.14202918 | 0 | 0.05 | 1 | 14 |
| acot18 | 0.21124 | -0.10006315 | 0 | 0.05 | 1 | 14 |
| adcyap1b | 0.21124 | -0.18530559 | 0 | 0.05 | 1 | 14 |
| agtr2 | 0.21124 | -0.14455393 | 0 | 0.05 | 1 | 14 |
| ahi1 | 0.21124 | -0.12644627 | 0 | 0.05 | 1 | 14 |
| alg10 | 0.21124 | -0.11211761 | 0 | 0.05 | 1 | 14 |
| alg6 | 0.21124 | -0.12391482 | 0 | 0.05 | 1 | 14 |
| ap5b1 | 0.21124 | -0.10007535 | 0 | 0.05 | 1 | 14 |
| b4galt1l | 0.21124 | -0.15934531 | 0 | 0.05 | 1 | 14 |
| baiap2l2b | 0.21124 | -0.12808228 | 0 | 0.05 | 1 | 14 |
| bcl10 | 0.21124 | -0.12038638 | 0 | 0.05 | 1 | 14 |
| c6h1orf226 | 0.21124 | -0.14488838 | 0 | 0.05 | 1 | 14 |
| calb2b | 0.21124 | -0.13247574 | 0 | 0.05 | 1 | 14 |
| calhm5.2 | 0.21124 | -0.10266322 | 0 | 0.05 | 1 | 14 |
| capn15 | 0.21124 | -0.10261294 | 0 | 0.05 | 1 | 14 |
| ccdc93 | 0.21124 | -0.10171024 | 0 | 0.05 | 1 | 14 |
| cdkn1ba | 0.21124 | -0.12789955 | 0 | 0.05 | 1 | 14 |
| cep295 | 0.21124 | -0.12655806 | 0 | 0.05 | 1 | 14 |
| chp2 | 0.21124 | -0.14209975 | 0 | 0.05 | 1 | 14 |
| clcn3 | 0.21124 | -0.1442623 | 0 | 0.05 | 1 | 14 |
| clcn7 | 0.21124 | -0.10009467 | 0 | 0.05 | 1 | 14 |
| cldn15lb | 0.21124 | -0.14741336 | 0 | 0.05 | 1 | 14 |
| clec3bb | 0.21124 | -0.12574057 | 0 | 0.05 | 1 | 14 |
| crata | 0.21124 | -0.16230024 | 0 | 0.05 | 1 | 14 |
| crip3 | 0.21124 | -0.10390133 | 0 | 0.05 | 1 | 14 |
| csnk2a2b | 0.21124 | -0.10676704 | 0 | 0.05 | 1 | 14 |
| cyp2p10 | 0.21124 | -0.14224869 | 0 | 0.05 | 1 | 14 |
| dhrsx | 0.21124 | -0.12357616 | 0 | 0.05 | 1 | 14 |

| | | | | | | |
|---|---|---|---|---|---|---|
| dio2 | 0.21124 | -0.10059315 | 0 | 0.05 | 1 | 14 |
| dnajb1a | 0.21124 | -0.12325889 | 0 | 0.05 | 1 | 14 |
| dnajc28 | 0.21124 | -0.10407155 | 0 | 0.05 | 1 | 14 |
| dpt | 0.21124 | -0.14454689 | 0 | 0.05 | 1 | 14 |
| eif2ak3 | 0.21124 | -0.12350596 | 0 | 0.05 | 1 | 14 |
| epb41l4b | 0.21124 | -0.10187476 | 0 | 0.05 | 1 | 14 |
| fam214a | 0.21124 | -0.12484002 | 0 | 0.05 | 1 | 14 |
| faxcb | 0.21124 | -0.12614707 | 0 | 0.05 | 1 | 14 |
| fbxo32 | 0.21124 | -0.16199158 | 0 | 0.05 | 1 | 14 |
| fut10 | 0.21124 | -0.10078319 | 0 | 0.05 | 1 | 14 |
| gak | 0.21124 | -0.10303419 | 0 | 0.05 | 1 | 14 |
| gas2l3 | 0.21124 | -0.10078319 | 0 | 0.05 | 1 | 14 |
| gcc1 | 0.21124 | -0.16108678 | 0 | 0.05 | 1 | 14 |
| ggctb | 0.21124 | -0.15996362 | 0 | 0.05 | 1 | 14 |
| gigyf2 | 0.21124 | -0.10209491 | 0 | 0.05 | 1 | 14 |
| gli1 | 0.21124 | -0.10366042 | 0 | 0.05 | 1 | 14 |
| guk1a | 0.21124 | -0.16018507 | 0 | 0.05 | 1 | 14 |
| hlfb | 0.21124 | -0.12808228 | 0 | 0.05 | 1 | 14 |
| ildr1b | 0.21124 | -0.1183527 | 0 | 0.05 | 1 | 14 |
| im:7151449 | 0.21124 | -0.18525468 | 0 | 0.05 | 1 | 14 |
| irf7 | 0.21124 | -0.12636197 | 0 | 0.05 | 1 | 14 |
| jph1b | 0.21124 | -0.1567499 | 0 | 0.05 | 1 | 14 |
| kdm7ab | 0.21124 | -0.10198392 | 0 | 0.05 | 1 | 14 |
| larp6a | 0.21124 | -0.13078865 | 0 | 0.05 | 1 | 14 |
| lgals1l1 | 0.21124 | -0.12885389 | 0 | 0.05 | 1 | 14 |
| lrrc1 | 0.21124 | -0.12703373 | 0 | 0.05 | 1 | 14 |
| lrrtm2 | 0.21124 | -0.13231094 | 0 | 0.05 | 1 | 14 |
| lrtm2a | 0.21124 | -0.1439844 | 0 | 0.05 | 1 | 14 |
| lye | 0.21124 | -0.15495652 | 0 | 0.05 | 1 | 14 |
| map3k12 | 0.21124 | -0.10554527 | 0 | 0.05 | 1 | 14 |
| mast2 | 0.21124 | -0.12617463 | 0 | 0.05 | 1 | 14 |
| mdm2 | 0.21124 | -0.12004472 | 0 | 0.05 | 1 | 14 |
| mettl27 | 0.21124 | -0.10127348 | 0 | 0.05 | 1 | 14 |
| mfsd1 | 0.21124 | -0.10078319 | 0 | 0.05 | 1 | 14 |
| mid2 | 0.21124 | -0.12808228 | 0 | 0.05 | 1 | 14 |
| mmrn2a | 0.21124 | -0.14470597 | 0 | 0.05 | 1 | 14 |
| mpi | 0.21124 | -0.1199035 | 0 | 0.05 | 1 | 14 |
| mrps26 | 0.21124 | -0.12135054 | 0 | 0.05 | 1 | 14 |
| msantd4 | 0.21124 | -0.10034478 | 0 | 0.05 | 1 | 14 |
| nacc1b | 0.21124 | -0.16213575 | 0 | 0.05 | 1 | 14 |
| ndrg4 | 0.21124 | -0.14081077 | 0 | 0.05 | 1 | 14 |
| neu3.5 | 0.21124 | -0.13243978 | 0 | 0.05 | 1 | 14 |
| nle1 | 0.21124 | -0.11985832 | 0 | 0.05 | 1 | 14 |
| npm1b | 0.21124 | -0.14165073 | 0 | 0.05 | 1 | 14 |
| nsfb | 0.21124 | -0.12571889 | 0 | 0.05 | 1 | 14 |
| nup205 | 0.21124 | -0.12508713 | 0 | 0.05 | 1 | 14 |
| osbp | 0.21124 | -0.14379339 | 0 | 0.05 | 1 | 14 |
| osbpl3b | 0.21124 | -0.10879259 | 0 | 0.05 | 1 | 14 |

| | | | | | | |
|---|---|---|---|---|---|---|
| oxsm | 0.21124 | -0.15888053 | 0 | 0.05 | 1 | 14 |
| pak4 | 0.21124 | -0.10241683 | 0 | 0.05 | 1 | 14 |
| parla | 0.21124 | -0.12537852 | 0 | 0.05 | 1 | 14 |
| pbrm1 | 0.21124 | -0.11889586 | 0 | 0.05 | 1 | 14 |
| pcdh1g14 | 0.21124 | -0.10078319 | 0 | 0.05 | 1 | 14 |
| pcyt1aa | 0.21124 | -0.10258734 | 0 | 0.05 | 1 | 14 |
| phospho2 | 0.21124 | -0.12416037 | 0 | 0.05 | 1 | 14 |
| phpt1 | 0.21124 | -0.11574888 | 0 | 0.05 | 1 | 14 |
| pip5k1aa | 0.21124 | -0.14272854 | 0 | 0.05 | 1 | 14 |
| pon1 | 0.21124 | -0.14359796 | 0 | 0.05 | 1 | 14 |
| ppm1lb | 0.21124 | -0.14244197 | 0 | 0.05 | 1 | 14 |
| prss12 | 0.21124 | -0.14431519 | 0 | 0.05 | 1 | 14 |
| rapgef5b | 0.21124 | -0.10132837 | 0 | 0.05 | 1 | 14 |
| rbm33a | 0.21124 | -0.10336626 | 0 | 0.05 | 1 | 14 |
| rbm48 | 0.21124 | -0.16619004 | 0 | 0.05 | 1 | 14 |
| rfxank | 0.21124 | -0.12600397 | 0 | 0.05 | 1 | 14 |
| rgn | 0.21124 | -0.11760037 | 0 | 0.05 | 1 | 14 |
| rictora | 0.21124 | -0.12627005 | 0 | 0.05 | 1 | 14 |
| rnf11a | 0.21124 | -0.16115218 | 0 | 0.05 | 1 | 14 |
| rnf220a | 0.21124 | -0.10199682 | 0 | 0.05 | 1 | 14 |
| setmar | 0.21124 | -0.16074421 | 0 | 0.05 | 1 | 14 |
| si:ch211-127i16.2 | 0.21124 | -0.16161521 | 0 | 0.05 | 1 | 14 |
| si:ch211-181d7.3 | 0.21124 | -0.14171382 | 0 | 0.05 | 1 | 14 |
| si:ch211-203d1.3 | 0.21124 | -0.16108388 | 0 | 0.05 | 1 | 14 |
| si:ch211-207d6.2 | 0.21124 | -0.1610282 | 0 | 0.05 | 1 | 14 |
| si:ch211-254p10.2 | 0.21124 | -0.10554527 | 0 | 0.05 | 1 | 14 |
| si:ch73-109i22.2 | 0.21124 | -0.10078319 | 0 | 0.05 | 1 | 14 |
| si:ch73-390p7.2 | 0.21124 | -0.10001541 | 0 | 0.05 | 1 | 14 |
| si:dkey-110g7.8 | 0.21124 | -0.12004016 | 0 | 0.05 | 1 | 14 |
| si:dkey-112e17.1 | 0.21124 | -0.13365306 | 0 | 0.05 | 1 | 14 |
| si:dkey-184a18.5 | 0.21124 | -0.11909824 | 0 | 0.05 | 1 | 14 |
| si:dkey-206f10.1 | 0.21124 | -0.13890213 | 0 | 0.05 | 1 | 14 |
| si:dkey-23i12.7 | 0.21124 | -0.10472739 | 0 | 0.05 | 1 | 14 |
| si:dkey-264d12.5 | 0.21124 | -0.14446653 | 0 | 0.05 | 1 | 14 |
| si:dkey-60d5.3 | 0.21124 | -0.12750025 | 0 | 0.05 | 1 | 14 |
| si:dkeyp-80c12.7 | 0.21124 | -0.12474666 | 0 | 0.05 | 1 | 14 |
| sinhcaf | 0.21124 | -0.12427438 | 0 | 0.05 | 1 | 14 |
| slc25a16 | 0.21124 | -0.10879003 | 0 | 0.05 | 1 | 14 |
| slc29a3 | 0.21124 | -0.12488554 | 0 | 0.05 | 1 | 14 |
| slc2a12 | 0.21124 | -0.10287914 | 0 | 0.05 | 1 | 14 |
| slc38a3a | 0.21124 | -0.13228314 | 0 | 0.05 | 1 | 14 |
| snx24 | 0.21124 | -0.10407252 | 0 | 0.05 | 1 | 14 |
| sort1b | 0.21124 | -0.12394719 | 0 | 0.05 | 1 | 14 |
| soul5 | 0.21124 | -0.14476591 | 0 | 0.05 | 1 | 14 |
| spns2 | 0.21124 | -0.15021901 | 0 | 0.05 | 1 | 14 |
| srxn1 | 0.21124 | -0.12808228 | 0 | 0.05 | 1 | 14 |
| taf8 | 0.21124 | -0.13841935 | 0 | 0.05 | 1 | 14 |
| tbc1d16 | 0.21124 | -0.10234048 | 0 | 0.05 | 1 | 14 |

| | | | | | | |
|---|---|---|---|---|---|---|
| tbc1d17 | 0.21124 | -0.1440031 | 0 | 0.05 | 1 | 14 |
| thbs1b | 0.21124 | -0.12082896 | 0 | 0.05 | 1 | 14 |
| tmem141 | 0.21124 | -0.14190822 | 0 | 0.05 | 1 | 14 |
| tmem186 | 0.21124 | -0.12693327 | 0 | 0.05 | 1 | 14 |
| tmem230b | 0.21124 | -0.10554527 | 0 | 0.05 | 1 | 14 |
| tor2a | 0.21124 | -0.12633816 | 0 | 0.05 | 1 | 14 |
| trdn | 0.21124 | -0.10554527 | 0 | 0.05 | 1 | 14 |
| trim16 | 0.21124 | -0.20321922 | 0 | 0.05 | 1 | 14 |
| ugt2a7 | 0.21124 | -0.10142872 | 0 | 0.05 | 1 | 14 |
| ugt5c2 | 0.21124 | -0.12134346 | 0 | 0.05 | 1 | 14 |
| unk | 0.21124 | -0.14442004 | 0 | 0.05 | 1 | 14 |
| ywhag2 | 0.21124 | -0.11876471 | 0 | 0.05 | 1 | 14 |
| zdhhc1 | 0.21124 | -0.10345368 | 0 | 0.05 | 1 | 14 |
| zdhhc16b | 0.21124 | -0.10353106 | 0 | 0.05 | 1 | 14 |
| zgc:101765 | 0.21124 | -0.14409769 | 0 | 0.05 | 1 | 14 |
| zgc:110006 | 0.21124 | -0.1262481 | 0 | 0.05 | 1 | 14 |
| zgc:153681 | 0.21124 | -0.14542647 | 0 | 0.05 | 1 | 14 |
| zgc:162509 | 0.21124 | -0.14074602 | 0 | 0.05 | 1 | 14 |
| zgc:194398 | 0.21124 | -0.10078319 | 0 | 0.05 | 1 | 14 |
| zgc:86896 | 0.21124 | -0.10078319 | 0 | 0.05 | 1 | 14 |
| znf1001 | 0.21124 | -0.16188094 | 0 | 0.05 | 1 | 14 |
| znf1041 | 0.21124 | -0.10078319 | 0 | 0.05 | 1 | 14 |
| CABZ01034698.2 | 0.21124 | -0.12361865 | 0 | 0.05 | 1 | 14 |
| CU457778.3 | 0.21124 | -0.12467599 | 0 | 0.05 | 1 | 14 |
| ERC1 | 0.21124 | -0.11864629 | 0 | 0.05 | 1 | 14 |
| FP074902.2 | 0.21124 | -0.11417575 | 0 | 0.05 | 1 | 14 |
| GNAZ | 0.21124 | -0.15841258 | 0 | 0.05 | 1 | 14 |
| LOC101885444 | 0.21124 | -0.11834868 | 0 | 0.05 | 1 | 14 |
| LOC110438291 | 0.21124 | -0.12304167 | 0 | 0.05 | 1 | 14 |
| MYO1D | 0.21124 | -0.11751372 | 0 | 0.05 | 1 | 14 |
| XLOC-014378 | 0.21124 | -0.12004852 | 0 | 0.05 | 1 | 14 |
| XLOC-029130 | 0.21124 | -0.12004852 | 0 | 0.05 | 1 | 14 |
| acap3b | 0.21124 | -0.1182614 | 0 | 0.05 | 1 | 14 |
| ankrd49 | 0.21124 | -0.11346633 | 0 | 0.05 | 1 | 14 |
| apaf1 | 0.21124 | -0.10811296 | 0 | 0.05 | 1 | 14 |
| aqp11 | 0.21124 | -0.11755021 | 0 | 0.05 | 1 | 14 |
| atr | 0.21124 | -0.10811296 | 0 | 0.05 | 1 | 14 |
| azin1b | 0.21124 | -0.12144536 | 0 | 0.05 | 1 | 14 |
| baz1a | 0.21124 | -0.10536135 | 0 | 0.05 | 1 | 14 |
| btbd6a | 0.21124 | -0.11730274 | 0 | 0.05 | 1 | 14 |
| cby1 | 0.21124 | -0.12004852 | 0 | 0.05 | 1 | 14 |
| chuk | 0.21124 | -0.11738183 | 0 | 0.05 | 1 | 14 |
| crbn | 0.21124 | -0.11559547 | 0 | 0.05 | 1 | 14 |
| dbr1 | 0.21124 | -0.11632254 | 0 | 0.05 | 1 | 14 |
| dhodh | 0.21124 | -0.12175849 | 0 | 0.05 | 1 | 14 |
| dusp4 | 0.21124 | -0.10683578 | 0 | 0.05 | 1 | 14 |
| ecpas | 0.21124 | -0.11751522 | 0 | 0.05 | 1 | 14 |
| eya3 | 0.21124 | -0.12224272 | 0 | 0.05 | 1 | 14 |

| | | | | | | |
|---|---|---|---|---|---|---|
| fam198b | 0.21124 | -0.11738645 | 0 | 0.05 | 1 | 14 |
| fam57a | 0.21124 | -0.11824728 | 0 | 0.05 | 1 | 14 |
| gpr1 | 0.21124 | -0.12004852 | 0 | 0.05 | 1 | 14 |
| grin3ba | 0.21124 | -0.11891989 | 0 | 0.05 | 1 | 14 |
| grip1 | 0.21124 | -0.11822524 | 0 | 0.05 | 1 | 14 |
| haus6 | 0.21124 | -0.11329158 | 0 | 0.05 | 1 | 14 |
| hnrnph1 | 0.21124 | -0.10987348 | 0 | 0.05 | 1 | 14 |
| ints6l | 0.21124 | -0.11891989 | 0 | 0.05 | 1 | 14 |
| kpna6 | 0.21124 | -0.11947665 | 0 | 0.05 | 1 | 14 |
| lats1 | 0.21124 | -0.12228459 | 0 | 0.05 | 1 | 14 |
| lrp12 | 0.21124 | -0.11904526 | 0 | 0.05 | 1 | 14 |
| mbd5 | 0.21124 | -0.1179725 | 0 | 0.05 | 1 | 14 |
| mbd6 | 0.21124 | -0.11893161 | 0 | 0.05 | 1 | 14 |
| mllt1a | 0.21124 | -0.12422449 | 0 | 0.05 | 1 | 14 |
| nadsyn1.1 | 0.21124 | -0.11333233 | 0 | 0.05 | 1 | 14 |
| nisch | 0.21124 | -0.1183283 | 0 | 0.05 | 1 | 14 |
| nme6 | 0.21124 | -0.11177497 | 0 | 0.05 | 1 | 14 |
| nme7 | 0.21124 | -0.12089469 | 0 | 0.05 | 1 | 14 |
| pemt | 0.21124 | -0.10118929 | 0 | 0.05 | 1 | 14 |
| pgls | 0.21124 | -0.11045941 | 0 | 0.05 | 1 | 14 |
| phf12b | 0.21124 | -0.11823494 | 0 | 0.05 | 1 | 14 |
| plekhh2 | 0.21124 | -0.12004852 | 0 | 0.05 | 1 | 14 |
| pnpla7a | 0.21124 | -0.12255274 | 0 | 0.05 | 1 | 14 |
| ppp2caa | 0.21124 | -0.12004852 | 0 | 0.05 | 1 | 14 |
| ppp2r2aa | 0.21124 | -0.11757624 | 0 | 0.05 | 1 | 14 |
| pptc7a | 0.21124 | -0.11858323 | 0 | 0.05 | 1 | 14 |
| ptcd3 | 0.21124 | -0.11373473 | 0 | 0.05 | 1 | 14 |
| pygo2 | 0.21124 | -0.16329651 | 0 | 0.05 | 1 | 14 |
| rc3h1b | 0.21124 | -0.12136459 | 0 | 0.05 | 1 | 14 |
| recql | 0.21124 | -0.11756345 | 0 | 0.05 | 1 | 14 |
| rfx1a | 0.21124 | -0.12176571 | 0 | 0.05 | 1 | 14 |
| samm50l | 0.21124 | -0.11685963 | 0 | 0.05 | 1 | 14 |
| si:ch211-108d22.2 | 0.21124 | -0.11784721 | 0 | 0.05 | 1 | 14 |
| si:ch211-241f5.3 | 0.21124 | -0.1200268 | 0 | 0.05 | 1 | 14 |
| si:ch211-259g3.4 | 0.21124 | -0.12229615 | 0 | 0.05 | 1 | 14 |
| si:ch211-63p21.1 | 0.21124 | -0.11919743 | 0 | 0.05 | 1 | 14 |
| si:ch73-234b20.5 | 0.21124 | -0.11484746 | 0 | 0.05 | 1 | 14 |
| si:dkey-111k8.5 | 0.21124 | -0.11936841 | 0 | 0.05 | 1 | 14 |
| si:dkey-178e17.3 | 0.21124 | -0.10735952 | 0 | 0.05 | 1 | 14 |
| slc1a5 | 0.21124 | -0.12208814 | 0 | 0.05 | 1 | 14 |
| smad6b | 0.21124 | -0.12246217 | 0 | 0.05 | 1 | 14 |
| snx4 | 0.21124 | -0.12148391 | 0 | 0.05 | 1 | 14 |
| srpk3 | 0.21124 | -0.12467599 | 0 | 0.05 | 1 | 14 |
| stk39 | 0.21124 | -0.12004852 | 0 | 0.05 | 1 | 14 |
| tbc1d23 | 0.21124 | -0.11770673 | 0 | 0.05 | 1 | 14 |
| tbc1d7 | 0.21124 | -0.11965455 | 0 | 0.05 | 1 | 14 |
| tgm5l | 0.21124 | -0.11924615 | 0 | 0.05 | 1 | 14 |
| tmem132e | 0.21124 | -0.12467599 | 0 | 0.05 | 1 | 14 |

| | | | | | | |
|---|---|---|---|---|---|---|
| tmem160 | 0.21124 | -0.11984498 | 0 | 0.05 | 1 | 14 |
| tmem201 | 0.21124 | -0.11090925 | 0 | 0.05 | 1 | 14 |
| tmpoa | 0.21124 | -0.12077429 | 0 | 0.05 | 1 | 14 |
| tpgs2 | 0.21124 | -0.12138473 | 0 | 0.05 | 1 | 14 |
| trim9 | 0.21124 | -0.10548108 | 0 | 0.05 | 1 | 14 |
| washc2c | 0.21124 | -0.11639677 | 0 | 0.05 | 1 | 14 |
| wipf2b | 0.21124 | -0.11783327 | 0 | 0.05 | 1 | 14 |
| wu:fb74b10 | 0.21124 | -0.10138104 | 0 | 0.05 | 1 | 14 |
| zbtb18 | 0.21124 | -0.12210692 | 0 | 0.05 | 1 | 14 |
| zbtb2a | 0.21124 | -0.11435987 | 0 | 0.05 | 1 | 14 |
| zdhhc7 | 0.21124 | -0.11891989 | 0 | 0.05 | 1 | 14 |
| zgc:113625 | 0.21124 | -0.12467599 | 0 | 0.05 | 1 | 14 |
| zgc:92907 | 0.21124 | -0.11588007 | 0 | 0.05 | 1 | 14 |
| znf1152 | 0.21124 | -0.12277993 | 0 | 0.05 | 1 | 14 |
| BX248324.1 | 0.21124 | -0.10746141 | 0 | 0.05 | 1 | 14 |
| CABZ01057057.1 | 0.21124 | -0.16815969 | 0 | 0.05 | 1 | 14 |
| CABZ01080056.1 | 0.21124 | -0.1565098 | 0 | 0.05 | 1 | 14 |
| GID4 | 0.21124 | -0.1178218 | 0 | 0.05 | 1 | 14 |
| XLOC-017060 | 0.21124 | -0.16815969 | 0 | 0.05 | 1 | 14 |
| XLOC-027844 | 0.21124 | -0.1182416 | 0 | 0.05 | 1 | 14 |
| artnb | 0.21124 | -0.11846219 | 0 | 0.05 | 1 | 14 |
| asns | 0.21124 | -0.16622148 | 0 | 0.05 | 1 | 14 |
| btr31 | 0.21124 | -0.15564546 | 0 | 0.05 | 1 | 14 |
| cad | 0.21124 | -0.15380909 | 0 | 0.05 | 1 | 14 |
| cbfa2t2 | 0.21124 | -0.11770563 | 0 | 0.05 | 1 | 14 |
| chpf2 | 0.21124 | -0.11822998 | 0 | 0.05 | 1 | 14 |
| col4a2 | 0.21124 | -0.16779996 | 0 | 0.05 | 1 | 14 |
| cop1 | 0.21124 | -0.10801303 | 0 | 0.05 | 1 | 14 |
| csf1b | 0.21124 | -0.19667642 | 0 | 0.05 | 1 | 14 |
| cx40.8 | 0.21124 | -0.16815969 | 0 | 0.05 | 1 | 14 |
| cyp2aa7 | 0.21124 | -0.11856479 | 0 | 0.05 | 1 | 14 |
| dgkzb | 0.21124 | -0.16179014 | 0 | 0.05 | 1 | 14 |
| dlg5a | 0.21124 | -0.11807538 | 0 | 0.05 | 1 | 14 |
| fstl1a | 0.21124 | -0.15516582 | 0 | 0.05 | 1 | 14 |
| ftr67 | 0.21124 | -0.16658199 | 0 | 0.05 | 1 | 14 |
| habp2 | 0.21124 | -0.15611643 | 0 | 0.05 | 1 | 14 |
| hcar1-4 | 0.21124 | -0.11690351 | 0 | 0.05 | 1 | 14 |
| hmgcl | 0.21124 | -0.10457002 | 0 | 0.05 | 1 | 14 |
| hs2st1b | 0.21124 | -0.11601953 | 0 | 0.05 | 1 | 14 |
| ice1 | 0.21124 | -0.14834829 | 0 | 0.05 | 1 | 14 |
| ift74 | 0.21124 | -0.11268795 | 0 | 0.05 | 1 | 14 |
| itm2cb | 0.21124 | -0.10711449 | 0 | 0.05 | 1 | 14 |
| klhl24b | 0.21124 | -0.1152733 | 0 | 0.05 | 1 | 14 |
| lmo2 | 0.21124 | -0.11087386 | 0 | 0.05 | 1 | 14 |
| lpin1 | 0.21124 | -0.11646496 | 0 | 0.05 | 1 | 14 |
| lrp1bb | 0.21124 | -0.1565098 | 0 | 0.05 | 1 | 14 |
| lurap1 | 0.21124 | -0.11854526 | 0 | 0.05 | 1 | 14 |
| mettl23 | 0.21124 | -0.11291631 | 0 | 0.05 | 1 | 14 |

| | | | | | | |
|---|---|---|---|---|---|---|
| mogat3b | 0.21124 | -0.11369558 | 0 | 0.05 | 1 | 14 |
| ms4a17a.5 | 0.21124 | -0.10871428 | 0 | 0.05 | 1 | 14 |
| myo10l3 | 0.21124 | -0.1185524 | 0 | 0.05 | 1 | 14 |
| naga | 0.21124 | -0.11437437 | 0 | 0.05 | 1 | 14 |
| nbn | 0.21124 | -0.11300463 | 0 | 0.05 | 1 | 14 |
| pank4 | 0.21124 | -0.11607824 | 0 | 0.05 | 1 | 14 |
| pds5b | 0.21124 | -0.1181719 | 0 | 0.05 | 1 | 14 |
| pex2 | 0.21124 | -0.1560503 | 0 | 0.05 | 1 | 14 |
| pomt2 | 0.21124 | -0.15554431 | 0 | 0.05 | 1 | 14 |
| porcnl | 0.21124 | -0.15627786 | 0 | 0.05 | 1 | 14 |
| ppp1r12c | 0.21124 | -0.11239491 | 0 | 0.05 | 1 | 14 |
| qdprb1 | 0.21124 | -0.16339927 | 0 | 0.05 | 1 | 14 |
| saa | 0.21124 | -0.19742693 | 0 | 0.05 | 1 | 14 |
| sccpdha | 0.21124 | -0.15513225 | 0 | 0.05 | 1 | 14 |
| si:dkey-192g7.3 | 0.21124 | -0.19656488 | 0 | 0.05 | 1 | 14 |
| si:dkey-48p11.3 | 0.21124 | -0.10984513 | 0 | 0.05 | 1 | 14 |
| si:dkey-76k16.5 | 0.21124 | -0.11732003 | 0 | 0.05 | 1 | 14 |
| si:dkey-87k14.1 | 0.21124 | -0.11937387 | 0 | 0.05 | 1 | 14 |
| slc23a2 | 0.21124 | -0.11937387 | 0 | 0.05 | 1 | 14 |
| slc30a5 | 0.21124 | -0.16689601 | 0 | 0.05 | 1 | 14 |
| slc39a3 | 0.21124 | -0.11831281 | 0 | 0.05 | 1 | 14 |
| slc8a1a | 0.21124 | -0.1565098 | 0 | 0.05 | 1 | 14 |
| slit1b | 0.21124 | -0.1504312 | 0 | 0.05 | 1 | 14 |
| sox12 | 0.21124 | -0.11937387 | 0 | 0.05 | 1 | 14 |
| ston2 | 0.21124 | -0.16632372 | 0 | 0.05 | 1 | 14 |
| tap1 | 0.21124 | -0.16624253 | 0 | 0.05 | 1 | 14 |
| them4 | 0.21124 | -0.16267682 | 0 | 0.05 | 1 | 14 |
| tmem101 | 0.21124 | -0.15144639 | 0 | 0.05 | 1 | 14 |
| tmem69 | 0.21124 | -0.11386059 | 0 | 0.05 | 1 | 14 |
| trim13 | 0.21124 | -0.15823657 | 0 | 0.05 | 1 | 14 |
| tshz3b | 0.21124 | -0.16800185 | 0 | 0.05 | 1 | 14 |
| ttyh3a | 0.21124 | -0.16798294 | 0 | 0.05 | 1 | 14 |
| uggt1 | 0.21124 | -0.1119247 | 0 | 0.05 | 1 | 14 |
| wash1 | 0.21124 | -0.11582565 | 0 | 0.05 | 1 | 14 |
| wdr73 | 0.21124 | -0.11645381 | 0 | 0.05 | 1 | 14 |
| xpr1b | 0.21124 | -0.15379148 | 0 | 0.05 | 1 | 14 |
| ykt6 | 0.21124 | -0.11714621 | 0 | 0.05 | 1 | 14 |
| zgc:110063 | 0.21124 | -0.15596714 | 0 | 0.05 | 1 | 14 |
| zgc:123238 | 0.21124 | -0.11215041 | 0 | 0.05 | 1 | 14 |
| zgc:171500 | 0.21124 | -0.15622917 | 0 | 0.05 | 1 | 14 |
| zgc:91890 | 0.21124 | -0.11703301 | 0 | 0.05 | 1 | 14 |
| znf1174 | 0.21124 | -0.11313834 | 0 | 0.05 | 1 | 14 |
| ATE1 | 0.21124 | -0.16606865 | 0 | 0.05 | 1 | 14 |
| BX927219.1 | 0.21124 | -0.105466 | 0 | 0.05 | 1 | 14 |
| CABZ01072309.1 | 0.21124 | -0.15825992 | 0 | 0.05 | 1 | 14 |
| CABZ01084575.1 | 0.21124 | -0.16034497 | 0 | 0.05 | 1 | 14 |
| CR383676.2 | 0.21124 | -0.11668797 | 0 | 0.05 | 1 | 14 |
| CR855311.1 | 0.21124 | -0.16312921 | 0 | 0.05 | 1 | 14 |

| | | | | | | |
|---|---|---|---|---|---|---|
| CT027638.1 | 0.21124 | -0.1181992 | 0 | 0.05 | 1 | 14 |
| CU024870.1 | 0.21124 | -0.12628761 | 0 | 0.05 | 1 | 14 |
| CU467856.1 | 0.21124 | -0.16291189 | 0 | 0.05 | 1 | 14 |
| CU856516.1 | 0.21124 | -0.10160706 | 0 | 0.05 | 1 | 14 |
| LOC103910009 | 0.21124 | -0.12680787 | 0 | 0.05 | 1 | 14 |
| LOC796314 | 0.21124 | -0.16242401 | 0 | 0.05 | 1 | 14 |
| SNTA1 | 0.21124 | -0.15825992 | 0 | 0.05 | 1 | 14 |
| XLOC-001115 | 0.21124 | -0.16458193 | 0 | 0.05 | 1 | 14 |
| XLOC-005905 | 0.21124 | -0.10350789 | 0 | 0.05 | 1 | 14 |
| abcc1 | 0.21124 | -0.12445369 | 0 | 0.05 | 1 | 14 |
| adamts3 | 0.21124 | -0.10463246 | 0 | 0.05 | 1 | 14 |
| adnpa | 0.21124 | -0.12246287 | 0 | 0.05 | 1 | 14 |
| akap7 | 0.21124 | -0.15468077 | 0 | 0.05 | 1 | 14 |
| ankle2 | 0.21124 | -0.12447341 | 0 | 0.05 | 1 | 14 |
| antxr1a | 0.21124 | -0.13593105 | 0 | 0.05 | 1 | 14 |
| atp13a1 | 0.21124 | -0.11833871 | 0 | 0.05 | 1 | 14 |
| atp8a1 | 0.21124 | -0.10596596 | 0 | 0.05 | 1 | 14 |
| bco2a | 0.21124 | -0.15807346 | 0 | 0.05 | 1 | 14 |
| bmp7a | 0.21124 | -0.16606865 | 0 | 0.05 | 1 | 14 |
| brox | 0.21124 | -0.16458131 | 0 | 0.05 | 1 | 14 |
| cacna2d1a | 0.21124 | -0.12577759 | 0 | 0.05 | 1 | 14 |
| cbx6b | 0.21124 | -0.15761647 | 0 | 0.05 | 1 | 14 |
| ccdc125 | 0.21124 | -0.10417746 | 0 | 0.05 | 1 | 14 |
| cdhr5b | 0.21124 | -0.15825992 | 0 | 0.05 | 1 | 14 |
| cep57 | 0.21124 | -0.16507633 | 0 | 0.05 | 1 | 14 |
| ckmt2b | 0.21124 | -0.15216955 | 0 | 0.05 | 1 | 14 |
| cog2 | 0.21124 | -0.15818546 | 0 | 0.05 | 1 | 14 |
| cpne4b | 0.21124 | -0.15993194 | 0 | 0.05 | 1 | 14 |
| crebrf | 0.21124 | -0.15806102 | 0 | 0.05 | 1 | 14 |
| cyp2u1 | 0.21124 | -0.12680787 | 0 | 0.05 | 1 | 14 |
| dennd5a | 0.21124 | -0.16497083 | 0 | 0.05 | 1 | 14 |
| desi1b | 0.21124 | -0.13155649 | 0 | 0.05 | 1 | 14 |
| dhx57 | 0.21124 | -0.11913805 | 0 | 0.05 | 1 | 14 |
| dnaja1 | 0.21124 | -0.12423696 | 0 | 0.05 | 1 | 14 |
| dnajc16 | 0.21124 | -0.15216955 | 0 | 0.05 | 1 | 14 |
| dub | 0.21124 | -0.11861546 | 0 | 0.05 | 1 | 14 |
| dusp7 | 0.21124 | -0.1559127 | 0 | 0.05 | 1 | 14 |
| epb41l5 | 0.21124 | -0.1573552 | 0 | 0.05 | 1 | 14 |
| epg5 | 0.21124 | -0.14372104 | 0 | 0.05 | 1 | 14 |
| farp2 | 0.21124 | -0.10507026 | 0 | 0.05 | 1 | 14 |
| fbxo10 | 0.21124 | -0.1070716 | 0 | 0.05 | 1 | 14 |
| fbxo11b | 0.21124 | -0.1037715 | 0 | 0.05 | 1 | 14 |
| foxred2 | 0.21124 | -0.10411862 | 0 | 0.05 | 1 | 14 |
| fryl | 0.21124 | -0.10290678 | 0 | 0.05 | 1 | 14 |
| gcat | 0.21124 | -0.16091752 | 0 | 0.05 | 1 | 14 |
| gkup | 0.21124 | -0.10237922 | 0 | 0.05 | 1 | 14 |
| glmna | 0.21124 | -0.12107674 | 0 | 0.05 | 1 | 14 |
| gpr146 | 0.21124 | -0.10618664 | 0 | 0.05 | 1 | 14 |

| | | | | | | |
|---|---|---|---|---|---|---|
| gpr157 | 0.21124 | -0.12550358 | 0 | 0.05 | 1 | 14 |
| gramd1bb | 0.21124 | -0.10160706 | 0 | 0.05 | 1 | 14 |
| hectd3 | 0.21124 | -0.12427143 | 0 | 0.05 | 1 | 14 |
| hephl1a | 0.21124 | -0.12590172 | 0 | 0.05 | 1 | 14 |
| herc1 | 0.21124 | -0.11912523 | 0 | 0.05 | 1 | 14 |
| igfbp2b | 0.21124 | -0.1264204 | 0 | 0.05 | 1 | 14 |
| il16 | 0.21124 | -0.10295136 | 0 | 0.05 | 1 | 14 |
| insrb | 0.21124 | -0.16606865 | 0 | 0.05 | 1 | 14 |
| itsn2b | 0.21124 | -0.12405576 | 0 | 0.05 | 1 | 14 |
| ivns1abpb | 0.21124 | -0.15004137 | 0 | 0.05 | 1 | 14 |
| jade3 | 0.21124 | -0.10583777 | 0 | 0.05 | 1 | 14 |
| kansl3 | 0.21124 | -0.16562315 | 0 | 0.05 | 1 | 14 |
| klhl29 | 0.21124 | -0.1070716 | 0 | 0.05 | 1 | 14 |
| limd1a | 0.21124 | -0.13673625 | 0 | 0.05 | 1 | 14 |
| lipt2 | 0.21124 | -0.15556098 | 0 | 0.05 | 1 | 14 |
| mafa | 0.21124 | -0.16305188 | 0 | 0.05 | 1 | 14 |
| magi3a | 0.21124 | -0.10654637 | 0 | 0.05 | 1 | 14 |
| mettl24 | 0.21124 | -0.13706035 | 0 | 0.05 | 1 | 14 |
| mfsd4b | 0.21124 | -0.16606865 | 0 | 0.05 | 1 | 14 |
| mgll | 0.21124 | -0.13566113 | 0 | 0.05 | 1 | 14 |
| mib2 | 0.21124 | -0.15825992 | 0 | 0.05 | 1 | 14 |
| mindy3 | 0.21124 | -0.15390852 | 0 | 0.05 | 1 | 14 |
| mmrn2b | 0.21124 | -0.10160706 | 0 | 0.05 | 1 | 14 |
| mpp7a | 0.21124 | -0.12680787 | 0 | 0.05 | 1 | 14 |
| mxf | 0.21124 | -0.14541657 | 0 | 0.05 | 1 | 14 |
| mylkb | 0.21124 | -0.15825992 | 0 | 0.05 | 1 | 14 |
| n4bp2 | 0.21124 | -0.12217014 | 0 | 0.05 | 1 | 14 |
| nbas | 0.21124 | -0.12359536 | 0 | 0.05 | 1 | 14 |
| ncoa6 | 0.21124 | -0.15743423 | 0 | 0.05 | 1 | 14 |
| nlrc3l | 0.21124 | -0.10599461 | 0 | 0.05 | 1 | 14 |
| nmt1a | 0.21124 | -0.10250327 | 0 | 0.05 | 1 | 14 |
| nog2 | 0.21124 | -0.10169528 | 0 | 0.05 | 1 | 14 |
| pard3ab | 0.21124 | -0.1070716 | 0 | 0.05 | 1 | 14 |
| pard6b | 0.21124 | -0.16606865 | 0 | 0.05 | 1 | 14 |
| pde4d | 0.21124 | -0.15265421 | 0 | 0.05 | 1 | 14 |
| phactr3a | 0.21124 | -0.10160706 | 0 | 0.05 | 1 | 14 |
| phyhipla | 0.21124 | -0.12680787 | 0 | 0.05 | 1 | 14 |
| pid1 | 0.21124 | -0.15776903 | 0 | 0.05 | 1 | 14 |
| pip4k2ab | 0.21124 | -0.11875579 | 0 | 0.05 | 1 | 14 |
| pkdcca | 0.21124 | -0.1070716 | 0 | 0.05 | 1 | 14 |
| ppfibp2b | 0.21124 | -0.1070716 | 0 | 0.05 | 1 | 14 |
| psda | 0.21124 | -0.1070716 | 0 | 0.05 | 1 | 14 |
| ptger1b | 0.21124 | -0.13722795 | 0 | 0.05 | 1 | 14 |
| pxnb | 0.21124 | -0.15760569 | 0 | 0.05 | 1 | 14 |
| rab11fip4a | 0.21124 | -0.15825992 | 0 | 0.05 | 1 | 14 |
| rarres3 | 0.21124 | -0.1048832 | 0 | 0.05 | 1 | 14 |
| rbm15 | 0.21124 | -0.10547704 | 0 | 0.05 | 1 | 14 |
| rdh14a | 0.21124 | -0.10094391 | 0 | 0.05 | 1 | 14 |

| | | | | | | |
|---|---|---|---|---|---|---|
| rev3l | 0.21124 | -0.1566327 | 0 | 0.05 | 1 | 14 |
| rfk | 0.21124 | -0.10461687 | 0 | 0.05 | 1 | 14 |
| senp3b | 0.21124 | -0.10299272 | 0 | 0.05 | 1 | 14 |
| senp7b | 0.21124 | -0.15375379 | 0 | 0.05 | 1 | 14 |
| sfmbt2 | 0.21124 | -0.15825992 | 0 | 0.05 | 1 | 14 |
| si:cabz01071911.3 | 0.21124 | -0.11913561 | 0 | 0.05 | 1 | 14 |
| si:ch1073-157b13.1 | 0.21124 | -0.10077786 | 0 | 0.05 | 1 | 14 |
| si:ch1073-174d20.2 | 0.21124 | -0.16503467 | 0 | 0.05 | 1 | 14 |
| si:ch211-107m4.1 | 0.21124 | -0.10160706 | 0 | 0.05 | 1 | 14 |
| si:ch211-113p18.3 | 0.21124 | -0.12528817 | 0 | 0.05 | 1 | 14 |
| si:ch211-191c10.1 | 0.21124 | -0.12432361 | 0 | 0.05 | 1 | 14 |
| si:ch211-195e19.1 | 0.21124 | -0.15765679 | 0 | 0.05 | 1 | 14 |
| si:ch211-215a10.4 | 0.21124 | -0.11599269 | 0 | 0.05 | 1 | 14 |
| si:ch211-246b8.5 | 0.21124 | -0.1070716 | 0 | 0.05 | 1 | 14 |
| si:ch211-255f4.9 | 0.21124 | -0.16560261 | 0 | 0.05 | 1 | 14 |
| si:ch211-260e23.9 | 0.21124 | -0.10289754 | 0 | 0.05 | 1 | 14 |
| si:ch73-248e21.1 | 0.21124 | -0.15338647 | 0 | 0.05 | 1 | 14 |
| si:ch73-44m9.1 | 0.21124 | -0.12553192 | 0 | 0.05 | 1 | 14 |
| si:dkey-101k6.5 | 0.21124 | -0.10672353 | 0 | 0.05 | 1 | 14 |
| si:dkey-14k9.3 | 0.21124 | -0.16606865 | 0 | 0.05 | 1 | 14 |
| si:dkey-235d18.5 | 0.21124 | -0.15801276 | 0 | 0.05 | 1 | 14 |
| si:dkey-31c13.1 | 0.21124 | -0.16322398 | 0 | 0.05 | 1 | 14 |
| si:dkey-33c14.6 | 0.21124 | -0.15216955 | 0 | 0.05 | 1 | 14 |
| si:dkeyp-2e4.6 | 0.21124 | -0.10758229 | 0 | 0.05 | 1 | 14 |
| si:dkeyp-53d3.3 | 0.21124 | -0.10124139 | 0 | 0.05 | 1 | 14 |
| six4b | 0.21124 | -0.16606865 | 0 | 0.05 | 1 | 14 |
| slc27a6 | 0.21124 | -0.15758396 | 0 | 0.05 | 1 | 14 |
| slc4a10a | 0.21124 | -0.1653447 | 0 | 0.05 | 1 | 14 |
| slx4ip | 0.21124 | -0.1070716 | 0 | 0.05 | 1 | 14 |
| sobpa | 0.21124 | -0.15777072 | 0 | 0.05 | 1 | 14 |
| stk24b | 0.21124 | -0.1638088 | 0 | 0.05 | 1 | 14 |
| suz12a | 0.21124 | -0.15624448 | 0 | 0.05 | 1 | 14 |
| tanc1a | 0.21124 | -0.15734161 | 0 | 0.05 | 1 | 14 |
| tbc1d12b | 0.21124 | -0.15777921 | 0 | 0.05 | 1 | 14 |
| tent5ba | 0.21124 | -0.14315208 | 0 | 0.05 | 1 | 14 |
| tex2 | 0.21124 | -0.14446722 | 0 | 0.05 | 1 | 14 |
| timp2b | 0.21124 | -0.15496913 | 0 | 0.05 | 1 | 14 |
| tmem245 | 0.21124 | -0.10626182 | 0 | 0.05 | 1 | 14 |
| tmem97 | 0.21124 | -0.12006644 | 0 | 0.05 | 1 | 14 |
| tnksa | 0.21124 | -0.15603904 | 0 | 0.05 | 1 | 14 |
| traf2b | 0.21124 | -0.15648092 | 0 | 0.05 | 1 | 14 |
| traf3ip1 | 0.21124 | -0.12354271 | 0 | 0.05 | 1 | 14 |
| tshz3a | 0.21124 | -0.15783606 | 0 | 0.05 | 1 | 14 |
| ttc39c | 0.21124 | -0.12357319 | 0 | 0.05 | 1 | 14 |
| tyw1 | 0.21124 | -0.12338609 | 0 | 0.05 | 1 | 14 |
| usp24 | 0.21124 | -0.125626 | 0 | 0.05 | 1 | 14 |
| vstm4b | 0.21124 | -0.1070716 | 0 | 0.05 | 1 | 14 |
| wdhd1 | 0.21124 | -0.15522976 | 0 | 0.05 | 1 | 14 |

| | | | | | | |
|---|---|---|---|---|---|---|
| wtip | 0.21124 | -0.12429603 | 0 | 0.05 | 1 | 14 |
| zfand4 | 0.21124 | -0.12680787 | 0 | 0.05 | 1 | 14 |
| zgc:77439 | 0.21124 | -0.15588926 | 0 | 0.05 | 1 | 14 |
| znf219 | 0.21124 | -0.15825992 | 0 | 0.05 | 1 | 14 |
| znf292b | 0.21124 | -0.12486071 | 0 | 0.05 | 1 | 14 |
| zswim8 | 0.21124 | -0.12470843 | 0 | 0.05 | 1 | 14 |
| sptlc2b | 0.21136 | -0.26507475 | 0.05 | 0.2 | 1 | 14 |
| spag7 | 0.21137 | -0.29263258 | 0.05 | 0.2 | 1 | 14 |
| sephs2 | 0.21148 | -0.24801972 | 0.05 | 0.2 | 1 | 14 |
| ror2 | 0.2117 | -0.26871246 | 0.025 | 0.15 | 1 | 14 |
| ap1g1 | 0.21173 | -0.29272678 | 0.05 | 0.2 | 1 | 14 |
| zfp36l1a | 0.21173 | 0.431312254 | 0.35 | 0.15 | 1 | 14 |
| ccdc97 | 0.2124 | -0.33197632 | 0.05 | 0.2 | 1 | 14 |
| rpa3 | 0.21248 | 0.279692936 | 0.175 | 0.05 | 1 | 14 |
| rpl5a | 0.21262 | 0.151013619 | 0.875 | 0.7 | 1 | 14 |
| si:ch211-225b7.5 | 0.21271 | -0.28809189 | 0.025 | 0.15 | 1 | 14 |
| prelid1a | 0.21285 | -0.28221351 | 0.025 | 0.15 | 1 | 14 |
| si:ch211-212k18.7 | 0.21314 | -0.20311362 | 0.025 | 0.15 | 1 | 14 |
| usf2 | 0.21315 | -0.2893379 | 0.025 | 0.15 | 1 | 14 |
| tcima | 0.21341 | -0.2311859 | 0.025 | 0.15 | 1 | 14 |
| fuca2 | 0.21353 | -0.27722532 | 0.025 | 0.15 | 1 | 14 |
| ncoa3 | 0.21366 | -0.22893576 | 0.025 | 0.15 | 1 | 14 |
| c7a | 0.21372 | -0.36006268 | 0.05 | 0.2 | 1 | 14 |
| cflara | 0.21378 | -0.31203277 | 0.05 | 0.2 | 1 | 14 |
| CABZ01078261.1 | 0.2138 | -0.32001441 | 0.05 | 0.2 | 1 | 14 |
| notum1a | 0.21402 | -0.30021069 | 0.025 | 0.15 | 1 | 14 |
| tmub2 | 0.21413 | -0.33605476 | 0.05 | 0.2 | 1 | 14 |
| jdp2b | 0.21417 | 0.357876609 | 0.15 | 0.05 | 1 | 14 |
| plp2 | 0.21438 | -0.3032311 | 0.05 | 0.2 | 1 | 14 |
| kdm5a | 0.21438 | -0.2293277 | 0.025 | 0.15 | 1 | 14 |
| tpp1 | 0.21439 | -0.26779838 | 0.025 | 0.15 | 1 | 14 |
| TTC28 | 0.21492 | -0.30534168 | 0.05 | 0.2 | 1 | 14 |
| ndrg3a | 0.21497 | -0.28803063 | 0.025 | 0.15 | 1 | 14 |
| f11r.1 | 0.21501 | -0.29682801 | 0.025 | 0.15 | 1 | 14 |
| ifngr2 | 0.21502 | -0.29351042 | 0.05 | 0.2 | 1 | 14 |
| fbxl7 | 0.21508 | -0.28421382 | 0.025 | 0.15 | 1 | 14 |
| nifk | 0.21513 | 0.281988964 | 0.125 | 0.05 | 1 | 14 |
| dcp2 | 0.21539 | -0.22020412 | 0.025 | 0.15 | 1 | 14 |
| eppk1 | 0.21563 | -0.22154179 | 0.025 | 0.15 | 1 | 14 |
| osbpl9 | 0.21578 | -0.29581232 | 0.025 | 0.15 | 1 | 14 |
| trmt13 | 0.2158 | -0.18994442 | 0.025 | 0.15 | 1 | 14 |
| elk1 | 0.21582 | -0.24159092 | 0.025 | 0.15 | 1 | 14 |
| CU929418.1 | 0.21601 | -0.23748893 | 0.025 | 0.15 | 1 | 14 |
| txn2 | 0.21628 | -0.22700018 | 0.025 | 0.15 | 1 | 14 |
| kdm2ba | 0.21638 | -0.27934075 | 0.025 | 0.15 | 1 | 14 |
| trappc2l | 0.21673 | -0.19451169 | 0.025 | 0.15 | 1 | 14 |
| XLOC-035413 | 0.2171 | -0.21720324 | 0.125 | 0.3 | 1 | 14 |
| fgl2a | 0.21739 | -0.32462563 | 0.025 | 0.15 | 1 | 14 |

| | | | | | | |
|---|---|---|---|---|---|---|
| selenou1a | 0.21751 | -0.20466091 | 0.025 | 0.15 | 1 | 14 |
| mtdhb | 0.21754 | -0.26725346 | 0.025 | 0.15 | 1 | 14 |
| ADGRL3 | 0.21755 | -0.24510447 | 0.025 | 0.15 | 1 | 14 |
| myl10 | 0.21767 | -0.21812924 | 0.025 | 0.15 | 1 | 14 |
| rab4b | 0.21771 | -0.25298424 | 0.025 | 0.15 | 1 | 14 |
| zfand6 | 0.21797 | -0.23431026 | 0.025 | 0.15 | 1 | 14 |
| dvl2 | 0.21805 | -0.25455901 | 0.025 | 0.15 | 1 | 14 |
| grwd1 | 0.21807 | -0.2404382 | 0.025 | 0.15 | 1 | 14 |
| nop16 | 0.21809 | -0.24885512 | 0.025 | 0.15 | 1 | 14 |
| arcn1a | 0.21809 | -0.20811616 | 0.025 | 0.15 | 1 | 14 |
| tgfbr2a | 0.21812 | -0.35143974 | 0.025 | 0.15 | 1 | 14 |
| znf395b | 0.21813 | -0.20839844 | 0.025 | 0.15 | 1 | 14 |
| ssu72 | 0.21815 | -0.22001009 | 0.025 | 0.15 | 1 | 14 |
| becn1 | 0.21818 | -0.21344872 | 0.025 | 0.15 | 1 | 14 |
| zgc:66443 | 0.21822 | -0.20342118 | 0.025 | 0.15 | 1 | 14 |
| bcap31 | 0.21825 | -0.26904297 | 0.025 | 0.15 | 1 | 14 |
| thoc1 | 0.21828 | -0.20198621 | 0.025 | 0.15 | 1 | 14 |
| nudt5 | 0.21835 | -0.18544034 | 0.025 | 0.15 | 1 | 14 |
| ubald1b | 0.21837 | -0.21240898 | 0.025 | 0.15 | 1 | 14 |
| nrf1 | 0.2184 | -0.22821194 | 0.025 | 0.15 | 1 | 14 |
| mxra8b | 0.21848 | -0.22293218 | 0.025 | 0.15 | 1 | 14 |
| cox5ab | 0.21851 | -0.28370218 | 0.1 | 0.2 | 1 | 14 |
| bms1 | 0.21851 | -0.20345012 | 0.025 | 0.15 | 1 | 14 |
| gemin8 | 0.21852 | -0.20962501 | 0.025 | 0.15 | 1 | 14 |
| ikbkg | 0.21854 | -0.19839616 | 0.025 | 0.15 | 1 | 14 |
| arpc4 | 0.22102 | -0.28208932 | 0.15 | 0.35 | 1 | 14 |
| si:dkey-239h2.3 | 0.22224 | 0.326029644 | 0.35 | 0.15 | 1 | 14 |
| itga11a | 0.22325 | -0.39911753 | 0.125 | 0.3 | 1 | 14 |
| psap | 0.22362 | -0.34001843 | 0.15 | 0.35 | 1 | 14 |
| abcf1 | 0.22383 | 0.23449701 | 0.225 | 0.2 | 1 | 14 |
| vcp | 0.22471 | -0.24989602 | 0.2 | 0.4 | 1 | 14 |
| rsl1d1 | 0.2248 | 0.395441481 | 0.35 | 0.15 | 1 | 14 |
| psmd7 | 0.22685 | 0.333460223 | 0.2 | 0.05 | 1 | 14 |
| lsm4 | 0.22733 | 0.285268136 | 0.275 | 0.2 | 1 | 14 |
| sec23ip | 0.2283 | -0.12451185 | 0.05 | 0.15 | 1 | 14 |
| rpl15 | 0.22878 | 0.231597522 | 0.925 | 0.8 | 1 | 14 |
| slc25a5 | 0.22887 | 0.21733956 | 0.95 | 0.85 | 1 | 14 |
| cox6a1 | 0.22936 | -0.17211917 | 0.5 | 0.5 | 1 | 14 |
| tgif1 | 0.2321 | 0.331731734 | 0.175 | 0.05 | 1 | 14 |
| si:ch211-217k17.7 | 0.23229 | 0.293784487 | 0.2 | 0.05 | 1 | 14 |
| sept7a | 0.23249 | 0.37303111 | 0.325 | 0.15 | 1 | 14 |
| rab13 | 0.23525 | 0.347810603 | 0.2 | 0.05 | 1 | 14 |
| dnajc12 | 0.23525 | -0.1026336 | 0.025 | 0.1 | 1 | 14 |
| hiat1a | 0.23525 | -0.15925435 | 0.025 | 0.1 | 1 | 14 |
| nkain4 | 0.23525 | -0.20023306 | 0.025 | 0.1 | 1 | 14 |
| nup58 | 0.23525 | -0.1183574 | 0.025 | 0.1 | 1 | 14 |
| rab11fip1a | 0.23525 | -0.22840593 | 0.025 | 0.1 | 1 | 14 |
| tram1 | 0.23525 | -0.19624937 | 0.025 | 0.1 | 1 | 14 |

| | | | | | | |
|---|---|---|---|---|---|---|
| zgc:153215 | 0.23525 | -0.16453284 | 0.025 | 0.1 | 1 | 14 |
| CABZ01084131.1 | 0.23525 | -0.22260896 | 0.025 | 0.1 | 1 | 14 |
| cdh13 | 0.23525 | -0.10222638 | 0.025 | 0.1 | 1 | 14 |
| cnn3b | 0.23525 | -0.14809358 | 0.025 | 0.1 | 1 | 14 |
| ghrb | 0.23525 | -0.13163162 | 0.025 | 0.1 | 1 | 14 |
| inip | 0.23525 | -0.11359157 | 0.025 | 0.1 | 1 | 14 |
| si:dkey-19e4.5 | 0.23525 | -0.12188809 | 0.025 | 0.1 | 1 | 14 |
| sypl1 | 0.23525 | -0.23062274 | 0.025 | 0.1 | 1 | 14 |
| sypl2b | 0.23525 | -0.2206856 | 0.025 | 0.1 | 1 | 14 |
| tma16 | 0.23525 | -0.10942609 | 0.025 | 0.1 | 1 | 14 |
| tsc22d2 | 0.23525 | -0.20699694 | 0.025 | 0.1 | 1 | 14 |
| zfx | 0.23525 | -0.19410691 | 0.025 | 0.1 | 1 | 14 |
| b3gnt5a | 0.23525 | -0.14822779 | 0.025 | 0.1 | 1 | 14 |
| f3b | 0.23525 | -0.16612527 | 0.025 | 0.1 | 1 | 14 |
| leng1 | 0.23525 | -0.13707489 | 0.025 | 0.1 | 1 | 14 |
| lyrm5a | 0.23525 | -0.2135249 | 0.025 | 0.1 | 1 | 14 |
| ndufaf3 | 0.23525 | -0.10332777 | 0.025 | 0.1 | 1 | 14 |
| pdlim5a | 0.23525 | -0.16003127 | 0.025 | 0.1 | 1 | 14 |
| ptgs1 | 0.23525 | -0.14702793 | 0.025 | 0.1 | 1 | 14 |
| si:ch73-25f10.6 | 0.23525 | -0.11162386 | 0.025 | 0.1 | 1 | 14 |
| slc35a1 | 0.23525 | -0.12896404 | 0.025 | 0.1 | 1 | 14 |
| tspan35 | 0.23525 | -0.19180513 | 0.025 | 0.1 | 1 | 14 |
| CCDC69 | 0.23525 | -0.15503552 | 0.025 | 0.1 | 1 | 14 |
| ank3b | 0.23525 | -0.12681376 | 0.025 | 0.1 | 1 | 14 |
| calcr | 0.23525 | -0.18438553 | 0.025 | 0.1 | 1 | 14 |
| jdp2a | 0.23525 | -0.12797321 | 0.025 | 0.1 | 1 | 14 |
| junbb | 0.23525 | -0.15315047 | 0.025 | 0.1 | 1 | 14 |
| lzic | 0.23525 | -0.1633924 | 0.025 | 0.1 | 1 | 14 |
| n6amt1 | 0.23525 | -0.11417622 | 0.025 | 0.1 | 1 | 14 |
| nfatc3a | 0.23525 | -0.20174211 | 0.025 | 0.1 | 1 | 14 |
| phf14 | 0.23525 | -0.10568961 | 0.025 | 0.1 | 1 | 14 |
| prpf18 | 0.23525 | -0.11914796 | 0.025 | 0.1 | 1 | 14 |
| si:ch73-237c6.1 | 0.23525 | -0.1392062 | 0.025 | 0.1 | 1 | 14 |
| si:dkey-31f5.8 | 0.23525 | -0.18117656 | 0.025 | 0.1 | 1 | 14 |
| stk17b | 0.23525 | -0.19887335 | 0.025 | 0.1 | 1 | 14 |
| tcf7l1a | 0.23525 | -0.16339947 | 0.025 | 0.1 | 1 | 14 |
| uggt2 | 0.23525 | -0.21823244 | 0.025 | 0.1 | 1 | 14 |
| vldlr | 0.23525 | -0.15382118 | 0.025 | 0.1 | 1 | 14 |
| yipf3 | 0.23525 | -0.14147907 | 0.025 | 0.1 | 1 | 14 |
| zgc:162255 | 0.23525 | -0.19125609 | 0.025 | 0.1 | 1 | 14 |
| BX548028.1 | 0.23525 | -0.21278423 | 0.025 | 0.1 | 1 | 14 |
| CABZ01117603.1 | 0.23525 | -0.12541679 | 0.025 | 0.1 | 1 | 14 |
| FO704797.1 | 0.23525 | -0.2169268 | 0.025 | 0.1 | 1 | 14 |
| LOC108179036 | 0.23525 | -0.11307918 | 0.025 | 0.1 | 1 | 14 |
| XLOC-005605 | 0.23525 | -0.2472356 | 0.025 | 0.1 | 1 | 14 |
| arid5b | 0.23525 | -0.13254002 | 0.025 | 0.1 | 1 | 14 |
| bicd2 | 0.23525 | -0.10111472 | 0.025 | 0.1 | 1 | 14 |
| cstf3 | 0.23525 | -0.16507848 | 0.025 | 0.1 | 1 | 14 |

| | | | | | | |
|---|---|---|---|---|---|---|
| cttn | 0.23525 | -0.14160738 | 0.025 | 0.1 | 1 | 14 |
| glsb | 0.23525 | -0.14582039 | 0.025 | 0.1 | 1 | 14 |
| got2a | 0.23525 | -0.11380269 | 0.025 | 0.1 | 1 | 14 |
| hlfa | 0.23525 | -0.16397187 | 0.025 | 0.1 | 1 | 14 |
| hmcn2 | 0.23525 | -0.19184171 | 0.025 | 0.1 | 1 | 14 |
| lamb1b | 0.23525 | -0.22084725 | 0.025 | 0.1 | 1 | 14 |
| mapk14b | 0.23525 | -0.17840115 | 0.025 | 0.1 | 1 | 14 |
| mbd1a | 0.23525 | -0.1136202 | 0.025 | 0.1 | 1 | 14 |
| mrpl41 | 0.23525 | -0.1057874 | 0.025 | 0.1 | 1 | 14 |
| papss2b | 0.23525 | -0.10901743 | 0.025 | 0.1 | 1 | 14 |
| phactr4a | 0.23525 | -0.14023576 | 0.025 | 0.1 | 1 | 14 |
| raver2 | 0.23525 | -0.15395934 | 0.025 | 0.1 | 1 | 14 |
| rtcb | 0.23525 | -0.11739842 | 0.025 | 0.1 | 1 | 14 |
| sft2d1 | 0.23525 | -0.16147234 | 0.025 | 0.1 | 1 | 14 |
| si:dkeyp-27c8.2 | 0.23525 | -0.11534485 | 0.025 | 0.1 | 1 | 14 |
| si:rp71-36a1.2 | 0.23525 | -0.18968708 | 0.025 | 0.1 | 1 | 14 |
| slc30a6 | 0.23525 | -0.17353382 | 0.025 | 0.1 | 1 | 14 |
| slx1b | 0.23525 | -0.13686373 | 0.025 | 0.1 | 1 | 14 |
| spata2 | 0.23525 | -0.1067842 | 0.025 | 0.1 | 1 | 14 |
| zgc:101851 | 0.23525 | -0.11558778 | 0.025 | 0.1 | 1 | 14 |
| zgc:65895 | 0.23525 | -0.12179922 | 0.025 | 0.1 | 1 | 14 |
| CU467856.3 | 0.23525 | -0.1654369 | 0.025 | 0.1 | 1 | 14 |
| aagab | 0.23525 | -0.12562944 | 0.025 | 0.1 | 1 | 14 |
| aebp2 | 0.23525 | -0.20104392 | 0.025 | 0.1 | 1 | 14 |
| brinp3a.1 | 0.23525 | -0.23456611 | 0.025 | 0.1 | 1 | 14 |
| ccdc82 | 0.23525 | -0.14027742 | 0.025 | 0.1 | 1 | 14 |
| ech1 | 0.23525 | -0.21316245 | 0.025 | 0.1 | 1 | 14 |
| erap1a | 0.23525 | -0.2110853 | 0.025 | 0.1 | 1 | 14 |
| nfe2l3 | 0.23525 | -0.17512155 | 0.025 | 0.1 | 1 | 14 |
| nup107 | 0.23525 | -0.15909002 | 0.025 | 0.1 | 1 | 14 |
| rbl1 | 0.23525 | -0.18823248 | 0.025 | 0.1 | 1 | 14 |
| rgs5a | 0.23525 | -0.11205754 | 0.025 | 0.1 | 1 | 14 |
| rnf11b | 0.23525 | -0.14798985 | 0.025 | 0.1 | 1 | 14 |
| sirt2 | 0.23525 | -0.14457181 | 0.025 | 0.1 | 1 | 14 |
| ugt5e1 | 0.23525 | -0.16977984 | 0.025 | 0.1 | 1 | 14 |
| uso1 | 0.23525 | -0.22570203 | 0.025 | 0.1 | 1 | 14 |
| zgc:110591 | 0.23525 | -0.12522549 | 0.025 | 0.1 | 1 | 14 |
| znf703 | 0.23525 | -0.1500851 | 0.025 | 0.1 | 1 | 14 |
| AL935174.2 | 0.23525 | -0.15404926 | 0.025 | 0.1 | 1 | 14 |
| BX005183.3 | 0.23525 | -0.15380094 | 0.025 | 0.1 | 1 | 14 |
| CU138547.1 | 0.23525 | -0.15707629 | 0.025 | 0.1 | 1 | 14 |
| acsm3 | 0.23525 | -0.20191461 | 0.025 | 0.1 | 1 | 14 |
| aff2 | 0.23525 | -0.1797735 | 0.025 | 0.1 | 1 | 14 |
| atic | 0.23525 | -0.16433437 | 0.025 | 0.1 | 1 | 14 |
| c1d | 0.23525 | -0.18051436 | 0.025 | 0.1 | 1 | 14 |
| capn12 | 0.23525 | -0.16322881 | 0.025 | 0.1 | 1 | 14 |
| ccnd2b | 0.23525 | -0.10580074 | 0.025 | 0.1 | 1 | 14 |
| ccser2a | 0.23525 | -0.15308896 | 0.025 | 0.1 | 1 | 14 |

| | | | | | | |
|---|---|---|---|---|---|---|
| chchd6b | 0.23525 | -0.12992146 | 0.025 | 0.1 | 1 | 14 |
| cndp2 | 0.23525 | -0.12297354 | 0.025 | 0.1 | 1 | 14 |
| crema | 0.23525 | -0.12968556 | 0.025 | 0.1 | 1 | 14 |
| csnk1g2b | 0.23525 | -0.17441119 | 0.025 | 0.1 | 1 | 14 |
| ctns | 0.23525 | -0.13672457 | 0.025 | 0.1 | 1 | 14 |
| ddx51 | 0.23525 | -0.16905037 | 0.025 | 0.1 | 1 | 14 |
| dennd6aa | 0.23525 | -0.14301773 | 0.025 | 0.1 | 1 | 14 |
| dnm2a | 0.23525 | -0.1075978 | 0.025 | 0.1 | 1 | 14 |
| dync1i2b | 0.23525 | -0.16837577 | 0.025 | 0.1 | 1 | 14 |
| ezrb | 0.23525 | -0.12308725 | 0.025 | 0.1 | 1 | 14 |
| fam169aa | 0.23525 | -0.10843227 | 0.025 | 0.1 | 1 | 14 |
| fgd4a | 0.23525 | -0.21051443 | 0.025 | 0.1 | 1 | 14 |
| ggnbp2 | 0.23525 | -0.15686386 | 0.025 | 0.1 | 1 | 14 |
| gtpbp10 | 0.23525 | -0.16520307 | 0.025 | 0.1 | 1 | 14 |
| ipo9 | 0.23525 | -0.11480943 | 0.025 | 0.1 | 1 | 14 |
| irak1 | 0.23525 | -0.19953282 | 0.025 | 0.1 | 1 | 14 |
| map2k1 | 0.23525 | -0.17476487 | 0.025 | 0.1 | 1 | 14 |
| marveld2a | 0.23525 | -0.11099181 | 0.025 | 0.1 | 1 | 14 |
| mib1 | 0.23525 | -0.17680489 | 0.025 | 0.1 | 1 | 14 |
| micall1a | 0.23525 | -0.18551947 | 0.025 | 0.1 | 1 | 14 |
| mob2a | 0.23525 | -0.20647235 | 0.025 | 0.1 | 1 | 14 |
| mon2 | 0.23525 | -0.1799327 | 0.025 | 0.1 | 1 | 14 |
| mtch2 | 0.23525 | -0.21642127 | 0.025 | 0.1 | 1 | 14 |
| nubpl | 0.23525 | -0.14685541 | 0.025 | 0.1 | 1 | 14 |
| orai1a | 0.23525 | -0.1182496 | 0.025 | 0.1 | 1 | 14 |
| pacsin2 | 0.23525 | -0.20079323 | 0.025 | 0.1 | 1 | 14 |
| paqr6 | 0.23525 | -0.13981429 | 0.025 | 0.1 | 1 | 14 |
| pcdh1b | 0.23525 | -0.18773904 | 0.025 | 0.1 | 1 | 14 |
| pgm3 | 0.23525 | -0.11101879 | 0.025 | 0.1 | 1 | 14 |
| polr2eb | 0.23525 | -0.11022382 | 0.025 | 0.1 | 1 | 14 |
| pomt1 | 0.23525 | -0.11993059 | 0.025 | 0.1 | 1 | 14 |
| ppardb | 0.23525 | -0.1365417 | 0.025 | 0.1 | 1 | 14 |
| ppm1da | 0.23525 | -0.17085988 | 0.025 | 0.1 | 1 | 14 |
| pus1 | 0.23525 | -0.13539236 | 0.025 | 0.1 | 1 | 14 |
| rarab | 0.23525 | -0.1574057 | 0.025 | 0.1 | 1 | 14 |
| reep3a | 0.23525 | -0.11513229 | 0.025 | 0.1 | 1 | 14 |
| rgs19 | 0.23525 | -0.24518613 | 0.025 | 0.1 | 1 | 14 |
| rnf215 | 0.23525 | -0.13784059 | 0.025 | 0.1 | 1 | 14 |
| sel1l | 0.23525 | -0.10113058 | 0.025 | 0.1 | 1 | 14 |
| si:ch211-198a12.6 | 0.23525 | -0.17350793 | 0.025 | 0.1 | 1 | 14 |
| si:ch211-51h9.7 | 0.23525 | -0.15115136 | 0.025 | 0.1 | 1 | 14 |
| skilb | 0.23525 | -0.23607858 | 0.025 | 0.1 | 1 | 14 |
| spag9a | 0.23525 | -0.13528862 | 0.025 | 0.1 | 1 | 14 |
| sqstm1 | 0.23525 | -0.14712682 | 0.025 | 0.1 | 1 | 14 |
| srp54 | 0.23525 | -0.16535083 | 0.025 | 0.1 | 1 | 14 |
| sv2 | 0.23525 | -0.12323475 | 0.025 | 0.1 | 1 | 14 |
| tcf3b | 0.23525 | -0.13747957 | 0.025 | 0.1 | 1 | 14 |
| tjp1b | 0.23525 | -0.19114415 | 0.025 | 0.1 | 1 | 14 |

| | | | | | | |
|---|---|---|---|---|---|---|
| tpp2 | 0.23525 | -0.13618659 | 0.025 | 0.1 | 1 | 14 |
| trpv4 | 0.23525 | -0.12485428 | 0.025 | 0.1 | 1 | 14 |
| yif1b | 0.23525 | -0.13447048 | 0.025 | 0.1 | 1 | 14 |
| zeb1b | 0.23525 | -0.12814705 | 0.025 | 0.1 | 1 | 14 |
| CU929230.1 | 0.23525 | -0.15752284 | 0.025 | 0.1 | 1 | 14 |
| XLOC-009815 | 0.23525 | -0.23399468 | 0.025 | 0.1 | 1 | 14 |
| arhgap20b | 0.23525 | -0.14394047 | 0.025 | 0.1 | 1 | 14 |
| arl8 | 0.23525 | -0.16857752 | 0.025 | 0.1 | 1 | 14 |
| cd2ap | 0.23525 | -0.16350795 | 0.025 | 0.1 | 1 | 14 |
| chmp3 | 0.23525 | -0.10371109 | 0.025 | 0.1 | 1 | 14 |
| cnpy3 | 0.23525 | -0.15615639 | 0.025 | 0.1 | 1 | 14 |
| epha4a | 0.23525 | -0.13057819 | 0.025 | 0.1 | 1 | 14 |
| esr2b | 0.23525 | -0.10778122 | 0.025 | 0.1 | 1 | 14 |
| faf1 | 0.23525 | -0.14013377 | 0.025 | 0.1 | 1 | 14 |
| gpc1b | 0.23525 | -0.10177181 | 0.025 | 0.1 | 1 | 14 |
| hmgxb4a | 0.23525 | -0.18960538 | 0.025 | 0.1 | 1 | 14 |
| lnpa | 0.23525 | -0.10726525 | 0.025 | 0.1 | 1 | 14 |
| mapk9 | 0.23525 | -0.14420321 | 0.025 | 0.1 | 1 | 14 |
| mpzl1l | 0.23525 | -0.21638094 | 0.025 | 0.1 | 1 | 14 |
| nudt8 | 0.23525 | -0.1371706 | 0.025 | 0.1 | 1 | 14 |
| parp16 | 0.23525 | -0.10294836 | 0.025 | 0.1 | 1 | 14 |
| prkar1aa | 0.23525 | -0.16704525 | 0.025 | 0.1 | 1 | 14 |
| prrc2a | 0.23525 | -0.21609276 | 0.025 | 0.1 | 1 | 14 |
| psen2 | 0.23525 | -0.13420435 | 0.025 | 0.1 | 1 | 14 |
| saraf | 0.23525 | -0.13302877 | 0.025 | 0.1 | 1 | 14 |
| si:ch211-194c3.5 | 0.23525 | -0.20532408 | 0.025 | 0.1 | 1 | 14 |
| si:ch211-67e16.4 | 0.23525 | -0.15917174 | 0.025 | 0.1 | 1 | 14 |
| si:ch73-91k6.2 | 0.23525 | -0.15910052 | 0.025 | 0.1 | 1 | 14 |
| si:dkey-79d12.5 | 0.23525 | -0.12177677 | 0.025 | 0.1 | 1 | 14 |
| si:dkeyp-68b7.7 | 0.23525 | -0.15995176 | 0.025 | 0.1 | 1 | 14 |
| si:zfos-128g4.1 | 0.23525 | -0.11730682 | 0.025 | 0.1 | 1 | 14 |
| sigmar1 | 0.23525 | -0.10087017 | 0.025 | 0.1 | 1 | 14 |
| surf1 | 0.23525 | -0.17696334 | 0.025 | 0.1 | 1 | 14 |
| vrk2 | 0.23525 | -0.22190694 | 0.025 | 0.1 | 1 | 14 |
| wbp11 | 0.23525 | -0.12869446 | 0.025 | 0.1 | 1 | 14 |
| zgc:171480 | 0.23525 | -0.15373331 | 0.025 | 0.1 | 1 | 14 |
| XLOC-037780 | 0.23525 | -0.11646068 | 0.025 | 0.1 | 1 | 14 |
| XLOC-042919 | 0.23525 | -0.14872303 | 0.025 | 0.1 | 1 | 14 |
| arf4a | 0.23525 | -0.16464839 | 0.025 | 0.1 | 1 | 14 |
| dse | 0.23525 | -0.17937843 | 0.025 | 0.1 | 1 | 14 |
| fem1c | 0.23525 | -0.19969987 | 0.025 | 0.1 | 1 | 14 |
| hiat1b | 0.23525 | -0.17418709 | 0.025 | 0.1 | 1 | 14 |
| ispd | 0.23525 | -0.13394637 | 0.025 | 0.1 | 1 | 14 |
| mrps9 | 0.23525 | -0.12482072 | 0.025 | 0.1 | 1 | 14 |
| nf1a | 0.23525 | -0.14219651 | 0.025 | 0.1 | 1 | 14 |
| polr3glb | 0.23525 | -0.15657317 | 0.025 | 0.1 | 1 | 14 |
| prelp | 0.23525 | -0.14022555 | 0.025 | 0.1 | 1 | 14 |
| si:ch211-235i11.5 | 0.23525 | -0.12183514 | 0.025 | 0.1 | 1 | 14 |

| | | | | | | |
|---|---|---|---|---|---|---|
| tmem242 | 0.23525 | -0.19192316 | 0.025 | 0.1 | 1 | 14 |
| tmie | 0.23525 | -0.19063673 | 0.025 | 0.1 | 1 | 14 |
| PLEKHH3 | 0.23525 | -0.21597622 | 0.025 | 0.1 | 1 | 14 |
| acacb | 0.23525 | -0.15739587 | 0.025 | 0.1 | 1 | 14 |
| actr10 | 0.23525 | -0.18482742 | 0.025 | 0.1 | 1 | 14 |
| bmi1b | 0.23525 | -0.14337192 | 0.025 | 0.1 | 1 | 14 |
| cdr2a | 0.23525 | -0.10292246 | 0.025 | 0.1 | 1 | 14 |
| chd8 | 0.23525 | -0.15771333 | 0.025 | 0.1 | 1 | 14 |
| csnk1g1 | 0.23525 | -0.11415503 | 0.025 | 0.1 | 1 | 14 |
| dap3 | 0.23525 | -0.16499044 | 0.025 | 0.1 | 1 | 14 |
| ddb2 | 0.23525 | -0.1326707 | 0.025 | 0.1 | 1 | 14 |
| dnm3a | 0.23525 | -0.15540151 | 0.025 | 0.1 | 1 | 14 |
| lyar | 0.23525 | -0.1011078 | 0.025 | 0.1 | 1 | 14 |
| nnt | 0.23525 | -0.17242463 | 0.025 | 0.1 | 1 | 14 |
| nup43 | 0.23525 | -0.1835899 | 0.025 | 0.1 | 1 | 14 |
| ppat | 0.23525 | -0.14007863 | 0.025 | 0.1 | 1 | 14 |
| ppic | 0.23525 | -0.21420618 | 0.025 | 0.1 | 1 | 14 |
| prickle2b | 0.23525 | -0.15590845 | 0.025 | 0.1 | 1 | 14 |
| senp2 | 0.23525 | -0.27964635 | 0.025 | 0.1 | 1 | 14 |
| slc43a2b | 0.23525 | -0.17427345 | 0.025 | 0.1 | 1 | 14 |
| tm9sf1 | 0.23525 | -0.15559882 | 0.025 | 0.1 | 1 | 14 |
| smoc2 | 0.23525 | -0.10444488 | 0.025 | 0.1 | 1 | 14 |
| CABZ01075242.1 | 0.23525 | -0.16832266 | 0.025 | 0.1 | 1 | 14 |
| LOC101885758 | 0.23525 | -0.26363932 | 0.025 | 0.1 | 1 | 14 |
| LOC108191970 | 0.23525 | -0.16330869 | 0.025 | 0.1 | 1 | 14 |
| adssl | 0.23525 | -0.11007735 | 0.025 | 0.1 | 1 | 14 |
| agpat2 | 0.23525 | -0.26435664 | 0.025 | 0.1 | 1 | 14 |
| copz2 | 0.23525 | -0.16874812 | 0.025 | 0.1 | 1 | 14 |
| cyp3c1 | 0.23525 | -0.20522007 | 0.025 | 0.1 | 1 | 14 |
| emg1 | 0.23525 | -0.14053798 | 0.025 | 0.1 | 1 | 14 |
| faxdc2 | 0.23525 | -0.2121717 | 0.025 | 0.1 | 1 | 14 |
| fhl2a | 0.23525 | -0.13883084 | 0.025 | 0.1 | 1 | 14 |
| gsr | 0.23525 | -0.1128703 | 0.025 | 0.1 | 1 | 14 |
| hs6st2 | 0.23525 | -0.15426789 | 0.025 | 0.1 | 1 | 14 |
| klf7a | 0.23525 | -0.15973304 | 0.025 | 0.1 | 1 | 14 |
| mau2 | 0.23525 | -0.13070551 | 0.025 | 0.1 | 1 | 14 |
| mrps17 | 0.23525 | -0.18728645 | 0.025 | 0.1 | 1 | 14 |
| rbm12b | 0.23525 | -0.10344757 | 0.025 | 0.1 | 1 | 14 |
| samd10a | 0.23525 | -0.18367053 | 0.025 | 0.1 | 1 | 14 |
| sat1b | 0.23525 | -0.11732101 | 0.025 | 0.1 | 1 | 14 |
| scara5 | 0.23525 | -0.17579284 | 0.025 | 0.1 | 1 | 14 |
| setx | 0.23525 | -0.14183445 | 0.025 | 0.1 | 1 | 14 |
| sh3kbp1 | 0.23525 | -0.12860736 | 0.025 | 0.1 | 1 | 14 |
| si:ch211-173p18.3 | 0.23525 | -0.18422229 | 0.025 | 0.1 | 1 | 14 |
| stard7 | 0.23525 | -0.1630709 | 0.025 | 0.1 | 1 | 14 |
| tle3a | 0.23525 | -0.12973863 | 0.025 | 0.1 | 1 | 14 |
| trim23 | 0.23525 | -0.12266181 | 0.025 | 0.1 | 1 | 14 |
| xpot | 0.23525 | -0.21147122 | 0.025 | 0.1 | 1 | 14 |

| | | | | | | |
|---|---|---|---|---|---|---|
| zeb1a | 0.23525 | -0.13881313 | 0.025 | 0.1 | 1 | 14 |
| rbfox2 | 0.2369 | -0.28478164 | 0.125 | 0.3 | 1 | 14 |
| pfdn5 | 0.23763 | 0.232082683 | 0.2 | 0.15 | 1 | 14 |
| tubb2b | 0.23951 | 0.39372041 | 0.325 | 0.15 | 1 | 14 |
| caprin1b | 0.24083 | 0.346753386 | 0.25 | 0.1 | 1 | 14 |
| ptgr1 | 0.24188 | 0.114520787 | 0.225 | 0.25 | 1 | 14 |
| hmga2 | 0.24196 | 0.327050382 | 0.2 | 0.05 | 1 | 14 |
| eny2 | 0.24264 | 0.311278159 | 0.2 | 0.05 | 1 | 14 |
| ier3ip1 | 0.24285 | -0.20420484 | 0.175 | 0.35 | 1 | 14 |
| jmjd1cb | 0.24292 | 0.325619195 | 0.175 | 0.05 | 1 | 14 |
| pigr | 0.2431 | -0.30886411 | 0.2 | 0.4 | 1 | 14 |
| id1 | 0.24461 | 0.411889428 | 0.375 | 0.25 | 1 | 14 |
| eif3d | 0.24498 | -0.13552436 | 0.475 | 0.65 | 1 | 14 |
| seta | 0.24528 | 0.149400512 | 0.4 | 0.4 | 1 | 14 |
| dedd1 | 0.24561 | 0.245012971 | 0.475 | 0.4 | 1 | 14 |
| zgc:110339 | 0.24579 | 0.313005683 | 0.2 | 0.05 | 1 | 14 |
| prpf19 | 0.24582 | -0.15708008 | 0.075 | 0.2 | 1 | 14 |
| calm1a | 0.24597 | 0.37300094 | 0.35 | 0.2 | 1 | 14 |
| lbh | 0.2463 | 0.249093674 | 0.125 | 0.05 | 1 | 14 |
| inka1a | 0.24684 | 0.353903257 | 0.2 | 0.05 | 1 | 14 |
| tgfbi | 0.24698 | 0.277125902 | 0.175 | 0.1 | 1 | 14 |
| CABZ01005379.1 | 0.24786 | 0.232047495 | 0.125 | 0.1 | 1 | 14 |
| arhgdia | 0.25 | -0.24644592 | 0.25 | 0.45 | 1 | 14 |
| ndufa5 | 0.25074 | 0.366273659 | 0.3 | 0.15 | 1 | 14 |
| prkab1b | 0.25107 | 0.305183324 | 0.25 | 0.15 | 1 | 14 |
| gpatch4 | 0.25386 | -0.13758968 | 0.075 | 0.2 | 1 | 14 |
| csnk2b | 0.25518 | 0.250353264 | 0.325 | 0.25 | 1 | 14 |
| ndufs4 | 0.25582 | 0.218519636 | 0.175 | 0.15 | 1 | 14 |
| dctn3 | 0.25608 | 0.300156114 | 0.2 | 0.05 | 1 | 14 |
| ms4a17a.9 | 0.2562 | 0.260700914 | 0.15 | 0.1 | 1 | 14 |
| ctnna1 | 0.25621 | -0.14229608 | 0.075 | 0.2 | 1 | 14 |
| si:rp71-45k5.4 | 0.25666 | -0.14475785 | 0.2 | 0.35 | 1 | 14 |
| si:ch73-281n10.2 | 0.25688 | 0.3753046 | 0.6 | 0.4 | 1 | 14 |
| rps28 | 0.25692 | 0.103947148 | 0.975 | 0.95 | 1 | 14 |
| atp5mc1 | 0.25738 | 0.248339137 | 0.225 | 0.2 | 1 | 14 |
| ssr1 | 0.25846 | -0.34837041 | 0.425 | 0.6 | 1 | 14 |
| rnf7 | 0.25888 | -0.1056672 | 0.175 | 0.3 | 1 | 14 |
| jpt1b | 0.25904 | 0.313963125 | 0.65 | 0.5 | 1 | 14 |
| ddx61 | 0.25964 | 0.273157329 | 0.45 | 0.25 | 1 | 14 |
| prss35 | 0.25998 | -0.21301245 | 0.25 | 0.4 | 1 | 14 |
| fam69c | 0.26012 | 0.138050736 | 0.3 | 0.3 | 1 | 14 |
| fkbp10b | 0.26118 | -0.24569657 | 0.1 | 0.25 | 1 | 14 |
| ndufa10 | 0.26124 | -0.34510277 | 0.125 | 0.3 | 1 | 14 |
| yif1a | 0.26173 | -0.18835518 | 0.1 | 0.25 | 1 | 14 |
| hspa5 | 0.26183 | -0.37945408 | 0.2 | 0.4 | 1 | 14 |
| pdap1a | 0.26183 | -0.25994526 | 0.125 | 0.3 | 1 | 14 |
| zgc:113531 | 0.26405 | -0.38370254 | 0.125 | 0.3 | 1 | 14 |
| ube2d2 | 0.26436 | -0.3323842 | 0.2 | 0.4 | 1 | 14 |

| | | | | | | |
|---|---|---|---|---|---|---|
| krt91 | 0.26442 | 0.11706014 | 0.4 | 0.45 | 1 | 14 |
| si:dkeyp-113d7.1 | 0.26551 | -0.21744031 | 0.1 | 0.25 | 1 | 14 |
| ifngr1l | 0.26561 | -0.27617626 | 0.05 | 0.15 | 1 | 14 |
| rpl28 | 0.26571 | 0.10681402 | 0.95 | 0.95 | 1 | 14 |
| emd | 0.26626 | 0.281818842 | 0.2 | 0.1 | 1 | 14 |
| cyc1 | 0.26634 | 0.145972762 | 0.25 | 0.25 | 1 | 14 |
| btf3l4 | 0.26676 | -0.31449284 | 0.2 | 0.4 | 1 | 14 |
| rbp5 | 0.26682 | -0.32485037 | 0.475 | 0.65 | 1 | 14 |
| ramp2 | 0.26792 | -0.37260112 | 0.075 | 0.2 | 1 | 14 |
| tmem30aa | 0.26795 | -0.32223963 | 0.125 | 0.3 | 1 | 14 |
| atp5mc3b | 0.26799 | -0.11827565 | 0.7 | 0.65 | 1 | 14 |
| cnn3a | 0.26871 | 0.381612416 | 0.4 | 0.2 | 1 | 14 |
| ywhabb | 0.26931 | -0.16753813 | 0.475 | 0.65 | 1 | 14 |
| mcfd2 | 0.26988 | 0.107036394 | 0.175 | 0.2 | 1 | 14 |
| fmnl3 | 0.27048 | -0.15869221 | 0.05 | 0.15 | 1 | 14 |
| FQ323119.1 | 0.27082 | -0.34682366 | 0.075 | 0.2 | 1 | 14 |
| eif4e2 | 0.27143 | -0.22970387 | 0.1 | 0.25 | 1 | 14 |
| bcl2l1 | 0.27162 | -0.29338061 | 0.125 | 0.3 | 1 | 14 |
| rps11 | 0.27233 | 0.208330926 | 0.925 | 0.85 | 1 | 14 |
| ilf2 | 0.27287 | 0.251644436 | 0.275 | 0.2 | 1 | 14 |
| thbs2a | 0.27508 | -0.35745929 | 0.125 | 0.3 | 1 | 14 |
| fpr1 | 0.27521 | -0.34098592 | 0.125 | 0.3 | 1 | 14 |
| anos1b | 0.27522 | -0.32535895 | 0.125 | 0.3 | 1 | 14 |
| ahnak | 0.27607 | -0.21882225 | 0.5 | 0.7 | 1 | 14 |
| sgcb | 0.27615 | -0.3495173 | 0.1 | 0.25 | 1 | 14 |
| eif3k | 0.27652 | 0.319330545 | 0.375 | 0.25 | 1 | 14 |
| wu:fi37e09 | 0.27996 | -0.31830583 | 0.225 | 0.35 | 1 | 14 |
| cmtm6 | 0.28013 | -0.19737423 | 0.075 | 0.2 | 1 | 14 |
| ptgr2 | 0.28039 | 0.13540546 | 0.05 | 0 | 1 | 14 |
| pum2 | 0.28039 | 0.133142535 | 0.05 | 0 | 1 | 14 |
| tbl1xr1b | 0.28039 | 0.153817 | 0.05 | 0 | 1 | 14 |
| JMY | 0.28039 | 0.147581443 | 0.05 | 0 | 1 | 14 |
| LOC101886030 | 0.28039 | 0.124518069 | 0.05 | 0 | 1 | 14 |
| LOC571912 | 0.28039 | 0.121175627 | 0.05 | 0 | 1 | 14 |
| myo10 | 0.28039 | 0.116975176 | 0.05 | 0 | 1 | 14 |
| pex14 | 0.28039 | 0.11535086 | 0.05 | 0 | 1 | 14 |
| utp15 | 0.28039 | 0.114910014 | 0.05 | 0 | 1 | 14 |
| LOC103910921 | 0.28039 | 0.151759734 | 0.05 | 0 | 1 | 14 |
| LOC110439948 | 0.28039 | 0.16514428 | 0.05 | 0 | 1 | 14 |
| SFMBT1 | 0.28039 | 0.128600323 | 0.05 | 0 | 1 | 14 |
| atat1 | 0.28039 | 0.157078533 | 0.05 | 0 | 1 | 14 |
| cited4b | 0.28039 | 0.142150699 | 0.05 | 0 | 1 | 14 |
| dis3l2 | 0.28039 | 0.141255297 | 0.05 | 0 | 1 | 14 |
| ercc5 | 0.28039 | 0.175451068 | 0.05 | 0 | 1 | 14 |
| faub | 0.28039 | 0.126070161 | 0.05 | 0 | 1 | 14 |
| fmoda | 0.28039 | 0.183106794 | 0.05 | 0 | 1 | 14 |
| focad | 0.28039 | 0.104693565 | 0.05 | 0 | 1 | 14 |
| gcdha | 0.28039 | 0.132097551 | 0.05 | 0 | 1 | 14 |

| | | | | | | |
|---|---|---|---|---|---|---|
| gnsb | 0.28039 | 0.140416851 | 0.05 | 0 | 1 | 14 |
| gusb | 0.28039 | 0.12910752 | 0.05 | 0 | 1 | 14 |
| hsf1 | 0.28039 | 0.156612665 | 0.05 | 0 | 1 | 14 |
| kat8 | 0.28039 | 0.128273807 | 0.05 | 0 | 1 | 14 |
| lrrc39 | 0.28039 | 0.159876646 | 0.05 | 0 | 1 | 14 |
| nbr1a | 0.28039 | 0.136665941 | 0.05 | 0 | 1 | 14 |
| ndufv1 | 0.28039 | 0.150479384 | 0.05 | 0 | 1 | 14 |
| pcyt1ab | 0.28039 | 0.117249766 | 0.05 | 0 | 1 | 14 |
| pias4a | 0.28039 | 0.105760025 | 0.05 | 0 | 1 | 14 |
| pola1 | 0.28039 | 0.106766672 | 0.05 | 0 | 1 | 14 |
| ppcdc | 0.28039 | 0.102938005 | 0.05 | 0 | 1 | 14 |
| psme4b | 0.28039 | 0.121127155 | 0.05 | 0 | 1 | 14 |
| sdhaf3 | 0.28039 | 0.129465427 | 0.05 | 0 | 1 | 14 |
| sec23b | 0.28039 | 0.141463181 | 0.05 | 0 | 1 | 14 |
| si:ch211-261d7.3 | 0.28039 | 0.102281744 | 0.05 | 0 | 1 | 14 |
| si:dkey-185e18.7 | 0.28039 | 0.107567239 | 0.05 | 0 | 1 | 14 |
| stk38b | 0.28039 | 0.142579004 | 0.05 | 0 | 1 | 14 |
| tbc1d14 | 0.28039 | 0.122510281 | 0.05 | 0 | 1 | 14 |
| tmem63ba | 0.28039 | 0.116657813 | 0.05 | 0 | 1 | 14 |
| tspan5a | 0.28039 | 0.122178183 | 0.05 | 0 | 1 | 14 |
| ubxn6 | 0.28039 | 0.140460772 | 0.05 | 0 | 1 | 14 |
| usp45 | 0.28039 | 0.107567239 | 0.05 | 0 | 1 | 14 |
| vps13a | 0.28039 | 0.145059374 | 0.05 | 0 | 1 | 14 |
| BX001023.1 | 0.28039 | 0.112462032 | 0.05 | 0 | 1 | 14 |
| XLOC-032526 | 0.28039 | 0.151040174 | 0.05 | 0 | 1 | 14 |
| bag3 | 0.28039 | 0.11270434 | 0.05 | 0 | 1 | 14 |
| bahcc1b | 0.28039 | 0.140087446 | 0.05 | 0 | 1 | 14 |
| card19 | 0.28039 | 0.108551196 | 0.05 | 0 | 1 | 14 |
| clybl | 0.28039 | 0.106971566 | 0.05 | 0 | 1 | 14 |
| ift140 | 0.28039 | 0.112511151 | 0.05 | 0 | 1 | 14 |
| kmt5c | 0.28039 | 0.111793254 | 0.05 | 0 | 1 | 14 |
| pld1a | 0.28039 | 0.129373639 | 0.05 | 0 | 1 | 14 |
| rab23 | 0.28039 | 0.114078079 | 0.05 | 0 | 1 | 14 |
| rap2aa | 0.28039 | 0.114958179 | 0.05 | 0 | 1 | 14 |
| rit1 | 0.28039 | 0.129175987 | 0.05 | 0 | 1 | 14 |
| rngtt | 0.28039 | 0.113792974 | 0.05 | 0 | 1 | 14 |
| si:ch211-167b20.8 | 0.28039 | 0.115934261 | 0.05 | 0 | 1 | 14 |
| si:ch73-233k15.2 | 0.28039 | 0.126832613 | 0.05 | 0 | 1 | 14 |
| tep1 | 0.28039 | 0.10603798 | 0.05 | 0 | 1 | 14 |
| zgc:158640 | 0.28039 | 0.128220455 | 0.05 | 0 | 1 | 14 |
| znf280d | 0.28039 | 0.100814825 | 0.05 | 0 | 1 | 14 |
| BX322618.1 | 0.28039 | 0.145261441 | 0.05 | 0 | 1 | 14 |
| BX465862.1 | 0.28039 | 0.109636094 | 0.05 | 0 | 1 | 14 |
| BX544877.1 | 0.28039 | 0.159187179 | 0.05 | 0 | 1 | 14 |
| CABZ01117348.1 | 0.28039 | 0.123725032 | 0.05 | 0 | 1 | 14 |
| CR936442.1 | 0.28039 | 0.143611443 | 0.05 | 0 | 1 | 14 |
| DXO | 0.28039 | 0.138919279 | 0.05 | 0 | 1 | 14 |
| LOC101882111 | 0.28039 | 0.120232838 | 0.05 | 0 | 1 | 14 |

| | | | | | | |
|---|---|---|---|---|---|---|
| LOC101886390 | 0.28039 | 0.135520514 | 0.05 | 0 | 1 | 14 |
| LOC103910363 | 0.28039 | 0.101320556 | 0.05 | 0 | 1 | 14 |
| PCMTD2 (1 of many) | 0.28039 | 0.135119422 | 0.05 | 0 | 1 | 14 |
| XLOC-007541 | 0.28039 | 0.14491029 | 0.05 | 0 | 1 | 14 |
| XLOC-030786 | 0.28039 | 0.124032543 | 0.05 | 0 | 1 | 14 |
| abcc4 | 0.28039 | 0.116101979 | 0.05 | 0 | 1 | 14 |
| acox1 | 0.28039 | 0.114876524 | 0.05 | 0 | 1 | 14 |
| adgrd1 | 0.28039 | 0.112246571 | 0.05 | 0 | 1 | 14 |
| adipor2 | 0.28039 | 0.123830918 | 0.05 | 0 | 1 | 14 |
| adtrp1 | 0.28039 | 0.160392786 | 0.05 | 0 | 1 | 14 |
| agfg1b | 0.28039 | 0.121479128 | 0.05 | 0 | 1 | 14 |
| aggf1 | 0.28039 | 0.109707356 | 0.05 | 0 | 1 | 14 |
| angptl7 | 0.28039 | 0.185206694 | 0.05 | 0 | 1 | 14 |
| ap2a1 | 0.28039 | 0.130407965 | 0.05 | 0 | 1 | 14 |
| apoob | 0.28039 | 0.102373095 | 0.05 | 0 | 1 | 14 |
| araf | 0.28039 | 0.117004622 | 0.05 | 0 | 1 | 14 |
| arhgef40 | 0.28039 | 0.101914579 | 0.05 | 0 | 1 | 14 |
| asap1b | 0.28039 | 0.13644911 | 0.05 | 0 | 1 | 14 |
| atg9b | 0.28039 | 0.151121469 | 0.05 | 0 | 1 | 14 |
| atpv0e2 | 0.28039 | 0.109789651 | 0.05 | 0 | 1 | 14 |
| atxn1b | 0.28039 | 0.124179302 | 0.05 | 0 | 1 | 14 |
| b9d2 | 0.28039 | 0.117508375 | 0.05 | 0 | 1 | 14 |
| babam2 | 0.28039 | 0.100304573 | 0.05 | 0 | 1 | 14 |
| bach1a | 0.28039 | 0.107060858 | 0.05 | 0 | 1 | 14 |
| bdh1 | 0.28039 | 0.121417382 | 0.05 | 0 | 1 | 14 |
| bend5 | 0.28039 | 0.151260408 | 0.05 | 0 | 1 | 14 |
| bri3 | 0.28039 | 0.160075108 | 0.05 | 0 | 1 | 14 |
| c2cd2 | 0.28039 | 0.112406963 | 0.05 | 0 | 1 | 14 |
| casc3 | 0.28039 | 0.119480379 | 0.05 | 0 | 1 | 14 |
| cavin1b | 0.28039 | 0.150550478 | 0.05 | 0 | 1 | 14 |
| cbx4 | 0.28039 | 0.10916334 | 0.05 | 0 | 1 | 14 |
| cbx6a | 0.28039 | 0.125030666 | 0.05 | 0 | 1 | 14 |
| ccdc77 | 0.28039 | 0.112406963 | 0.05 | 0 | 1 | 14 |
| ccnl1b | 0.28039 | 0.118873121 | 0.05 | 0 | 1 | 14 |
| cd302 | 0.28039 | 0.140837417 | 0.05 | 0 | 1 | 14 |
| cd36 | 0.28039 | 0.130519599 | 0.05 | 0 | 1 | 14 |
| cdk8 | 0.28039 | 0.114575597 | 0.05 | 0 | 1 | 14 |
| cep120 | 0.28039 | 0.123571068 | 0.05 | 0 | 1 | 14 |
| chchd7 | 0.28039 | 0.123932831 | 0.05 | 0 | 1 | 14 |
| chek2 | 0.28039 | 0.165126961 | 0.05 | 0 | 1 | 14 |
| chl1b | 0.28039 | 0.153022899 | 0.05 | 0 | 1 | 14 |
| chst10 | 0.28039 | 0.107889379 | 0.05 | 0 | 1 | 14 |
| cisd2 | 0.28039 | 0.160884906 | 0.05 | 0 | 1 | 14 |
| clip2 | 0.28039 | 0.105861656 | 0.05 | 0 | 1 | 14 |
| crebbpa | 0.28039 | 0.135582023 | 0.05 | 0 | 1 | 14 |
| crnkl1 | 0.28039 | 0.142281558 | 0.05 | 0 | 1 | 14 |
| ctso | 0.28039 | 0.116362436 | 0.05 | 0 | 1 | 14 |
| cylda | 0.28039 | 0.119237627 | 0.05 | 0 | 1 | 14 |

| | | | | | | |
|---|---|---|---|---|---|---|
| dcaf13 | 0.28039 | 0.119498742 | 0.05 | 0 | 1 | 14 |
| dclre1b | 0.28039 | 0.157397086 | 0.05 | 0 | 1 | 14 |
| dhrs11a | 0.28039 | 0.1455615 | 0.05 | 0 | 1 | 14 |
| dhx16 | 0.28039 | 0.108660888 | 0.05 | 0 | 1 | 14 |
| dio3b | 0.28039 | 0.138989752 | 0.05 | 0 | 1 | 14 |
| dnajb6a | 0.28039 | 0.116560207 | 0.05 | 0 | 1 | 14 |
| dram2b | 0.28039 | 0.108659005 | 0.05 | 0 | 1 | 14 |
| dtymk | 0.28039 | 0.129909955 | 0.05 | 0 | 1 | 14 |
| dyrk1ab | 0.28039 | 0.163756216 | 0.05 | 0 | 1 | 14 |
| eef2a.1 | 0.28039 | 0.125330496 | 0.05 | 0 | 1 | 14 |
| ehmt1b | 0.28039 | 0.107129349 | 0.05 | 0 | 1 | 14 |
| elp6 | 0.28039 | 0.127784959 | 0.05 | 0 | 1 | 14 |
| ephb2b | 0.28039 | 0.124475682 | 0.05 | 0 | 1 | 14 |
| eps15l1a | 0.28039 | 0.155027158 | 0.05 | 0 | 1 | 14 |
| fam129bb | 0.28039 | 0.101014164 | 0.05 | 0 | 1 | 14 |
| fbxo44 | 0.28039 | 0.101739207 | 0.05 | 0 | 1 | 14 |
| ftr06 | 0.28039 | 0.150974101 | 0.05 | 0 | 1 | 14 |
| gab1 | 0.28039 | 0.139786899 | 0.05 | 0 | 1 | 14 |
| gas8 | 0.28039 | 0.107634096 | 0.05 | 0 | 1 | 14 |
| gmds | 0.28039 | 0.143065533 | 0.05 | 0 | 1 | 14 |
| gmeb1 | 0.28039 | 0.15969198 | 0.05 | 0 | 1 | 14 |
| gpkow | 0.28039 | 0.10065902 | 0.05 | 0 | 1 | 14 |
| gripap1 | 0.28039 | 0.106148615 | 0.05 | 0 | 1 | 14 |
| gsk3ab | 0.28039 | 0.123571068 | 0.05 | 0 | 1 | 14 |
| gtf2ird1 | 0.28039 | 0.111664655 | 0.05 | 0 | 1 | 14 |
| hip1 | 0.28039 | 0.158172901 | 0.05 | 0 | 1 | 14 |
| hlx1 | 0.28039 | 0.123442385 | 0.05 | 0 | 1 | 14 |
| hoxa11b | 0.28039 | 0.126920259 | 0.05 | 0 | 1 | 14 |
| hs3st3b1b | 0.28039 | 0.134552292 | 0.05 | 0 | 1 | 14 |
| hsbp1a | 0.28039 | 0.12274205 | 0.05 | 0 | 1 | 14 |
| htatsf1 | 0.28039 | 0.117982732 | 0.05 | 0 | 1 | 14 |
| igf2bp2a | 0.28039 | 0.108916153 | 0.05 | 0 | 1 | 14 |
| igf2bp2b | 0.28039 | 0.133075217 | 0.05 | 0 | 1 | 14 |
| imp4 | 0.28039 | 0.119875841 | 0.05 | 0 | 1 | 14 |
| ireb2 | 0.28039 | 0.129393897 | 0.05 | 0 | 1 | 14 |
| itpkb | 0.28039 | 0.109052306 | 0.05 | 0 | 1 | 14 |
| itprip | 0.28039 | 0.126405434 | 0.05 | 0 | 1 | 14 |
| kat7b | 0.28039 | 0.158390901 | 0.05 | 0 | 1 | 14 |
| kita | 0.28039 | 0.111032254 | 0.05 | 0 | 1 | 14 |
| klhl20 | 0.28039 | 0.118630991 | 0.05 | 0 | 1 | 14 |
| krr1 | 0.28039 | 0.113011283 | 0.05 | 0 | 1 | 14 |
| lig3 | 0.28039 | 0.125908897 | 0.05 | 0 | 1 | 14 |
| lipf | 0.28039 | 0.102784775 | 0.05 | 0 | 1 | 14 |
| lpgat1 | 0.28039 | 0.148417734 | 0.05 | 0 | 1 | 14 |
| lrrc40 | 0.28039 | 0.131750712 | 0.05 | 0 | 1 | 14 |
| lto1 | 0.28039 | 0.11532309 | 0.05 | 0 | 1 | 14 |
| lztr1 | 0.28039 | 0.115952936 | 0.05 | 0 | 1 | 14 |
| maml3 | 0.28039 | 0.104075877 | 0.05 | 0 | 1 | 14 |

| | | | | | | |
|---|---|---|---|---|---|---|
| mao | 0.28039 | 0.140089605 | 0.05 | 0 | 1 | 14 |
| mapkap1 | 0.28039 | 0.148729638 | 0.05 | 0 | 1 | 14 |
| mark3b | 0.28039 | 0.134449693 | 0.05 | 0 | 1 | 14 |
| med1 | 0.28039 | 0.112467047 | 0.05 | 0 | 1 | 14 |
| mgat4a | 0.28039 | 0.132195226 | 0.05 | 0 | 1 | 14 |
| mical3b | 0.28039 | 0.122357234 | 0.05 | 0 | 1 | 14 |
| mier1b | 0.28039 | 0.107889379 | 0.05 | 0 | 1 | 14 |
| mplkip | 0.28039 | 0.109562353 | 0.05 | 0 | 1 | 14 |
| mrpl22 | 0.28039 | 0.133162897 | 0.05 | 0 | 1 | 14 |
| mrpl51 | 0.28039 | 0.108597983 | 0.05 | 0 | 1 | 14 |
| mxd3 | 0.28039 | 0.116193008 | 0.05 | 0 | 1 | 14 |
| myef2 | 0.28039 | 0.152317464 | 0.05 | 0 | 1 | 14 |
| naglu | 0.28039 | 0.126359219 | 0.05 | 0 | 1 | 14 |
| ncalda | 0.28039 | 0.104111529 | 0.05 | 0 | 1 | 14 |
| ncam1a | 0.28039 | 0.129861543 | 0.05 | 0 | 1 | 14 |
| ndp | 0.28039 | 0.1120376 | 0.05 | 0 | 1 | 14 |
| neurl1b | 0.28039 | 0.126199835 | 0.05 | 0 | 1 | 14 |
| nfatc2ip | 0.28039 | 0.119138018 | 0.05 | 0 | 1 | 14 |
| nlrc3l1 | 0.28039 | 0.155558809 | 0.05 | 0 | 1 | 14 |
| nsmce1 | 0.28039 | 0.102036009 | 0.05 | 0 | 1 | 14 |
| nup153 | 0.28039 | 0.111045679 | 0.05 | 0 | 1 | 14 |
| nvl | 0.28039 | 0.117218089 | 0.05 | 0 | 1 | 14 |
| ogdha | 0.28039 | 0.119961542 | 0.05 | 0 | 1 | 14 |
| pbx1b | 0.28039 | 0.103594861 | 0.05 | 0 | 1 | 14 |
| pcsk7 | 0.28039 | 0.107363956 | 0.05 | 0 | 1 | 14 |
| pde6d | 0.28039 | 0.10251145 | 0.05 | 0 | 1 | 14 |
| pex5 | 0.28039 | 0.135091331 | 0.05 | 0 | 1 | 14 |
| phf21ab | 0.28039 | 0.115693853 | 0.05 | 0 | 1 | 14 |
| pip5k1ab | 0.28039 | 0.117313372 | 0.05 | 0 | 1 | 14 |
| pja2 | 0.28039 | 0.119597668 | 0.05 | 0 | 1 | 14 |
| plekha1b | 0.28039 | 0.157391992 | 0.05 | 0 | 1 | 14 |
| plekhh1 | 0.28039 | 0.126733957 | 0.05 | 0 | 1 | 14 |
| pm20d1.2 | 0.28039 | 0.126810711 | 0.05 | 0 | 1 | 14 |
| polg2 | 0.28039 | 0.127942537 | 0.05 | 0 | 1 | 14 |
| ppargc1b | 0.28039 | 0.150685461 | 0.05 | 0 | 1 | 14 |
| ppfia1 | 0.28039 | 0.113513242 | 0.05 | 0 | 1 | 14 |
| ppip5k1a | 0.28039 | 0.155369647 | 0.05 | 0 | 1 | 14 |
| prkrip1 | 0.28039 | 0.13892146 | 0.05 | 0 | 1 | 14 |
| prp | 0.28039 | 0.167206664 | 0.05 | 0 | 1 | 14 |
| prpf4 | 0.28039 | 0.110924662 | 0.05 | 0 | 1 | 14 |
| ptges | 0.28039 | 0.152938692 | 0.05 | 0 | 1 | 14 |
| ptpn23a | 0.28039 | 0.119254917 | 0.05 | 0 | 1 | 14 |
| pus7 | 0.28039 | 0.11567104 | 0.05 | 0 | 1 | 14 |
| pyroxd2 | 0.28039 | 0.103805861 | 0.05 | 0 | 1 | 14 |
| rab3gap1 | 0.28039 | 0.126673093 | 0.05 | 0 | 1 | 14 |
| rab40c | 0.28039 | 0.143648255 | 0.05 | 0 | 1 | 14 |
| rac3a | 0.28039 | 0.111984578 | 0.05 | 0 | 1 | 14 |
| rad54l2 | 0.28039 | 0.146590274 | 0.05 | 0 | 1 | 14 |

| | | | | | | |
|---|---|---|---|---|---|---|
| ralgapa1 | 0.28039 | 0.131486705 | 0.05 | 0 | 1 | 14 |
| raph1b | 0.28039 | 0.110636916 | 0.05 | 0 | 1 | 14 |
| rbpjb | 0.28039 | 0.105365407 | 0.05 | 0 | 1 | 14 |
| relch | 0.28039 | 0.121841382 | 0.05 | 0 | 1 | 14 |
| rem1 | 0.28039 | 0.106934517 | 0.05 | 0 | 1 | 14 |
| rnf168 | 0.28039 | 0.109466029 | 0.05 | 0 | 1 | 14 |
| rock1 | 0.28039 | 0.1506071 | 0.05 | 0 | 1 | 14 |
| rps6ka3a | 0.28039 | 0.103089807 | 0.05 | 0 | 1 | 14 |
| rpusd4 | 0.28039 | 0.111623344 | 0.05 | 0 | 1 | 14 |
| rusc2 | 0.28039 | 0.14103119 | 0.05 | 0 | 1 | 14 |
| rxraa | 0.28039 | 0.117306063 | 0.05 | 0 | 1 | 14 |
| sccpdhb | 0.28039 | 0.103660431 | 0.05 | 0 | 1 | 14 |
| scxa | 0.28039 | 0.188690555 | 0.05 | 0 | 1 | 14 |
| scyl3 | 0.28039 | 0.109483818 | 0.05 | 0 | 1 | 14 |
| sesn3 | 0.28039 | 0.113198067 | 0.05 | 0 | 1 | 14 |
| setd2 | 0.28039 | 0.111239057 | 0.05 | 0 | 1 | 14 |
| sh3rf1 | 0.28039 | 0.12931741 | 0.05 | 0 | 1 | 14 |
| si:ch211-140b10.6 | 0.28039 | 0.104802949 | 0.05 | 0 | 1 | 14 |
| si:ch211-154o6.2 | 0.28039 | 0.137741007 | 0.05 | 0 | 1 | 14 |
| si:ch211-43f4.1 | 0.28039 | 0.162136738 | 0.05 | 0 | 1 | 14 |
| si:ch211-57i17.1 | 0.28039 | 0.116776503 | 0.05 | 0 | 1 | 14 |
| si:ch73-299h12.6 | 0.28039 | 0.104428482 | 0.05 | 0 | 1 | 14 |
| si:ch73-367f21.4 | 0.28039 | 0.10916334 | 0.05 | 0 | 1 | 14 |
| si:ch73-376l24.6 | 0.28039 | 0.141124458 | 0.05 | 0 | 1 | 14 |
| si:dkey-154b15.1 | 0.28039 | 0.147227559 | 0.05 | 0 | 1 | 14 |
| si:dkey-156n14.3 | 0.28039 | 0.154381046 | 0.05 | 0 | 1 | 14 |
| si:dkey-172m14.1 | 0.28039 | 0.143438646 | 0.05 | 0 | 1 | 14 |
| si:dkey-253d23.3 | 0.28039 | 0.110731395 | 0.05 | 0 | 1 | 14 |
| si:dkey-25g12.4 | 0.28039 | 0.155732239 | 0.05 | 0 | 1 | 14 |
| si:dkey-286j15.3 | 0.28039 | 0.146819199 | 0.05 | 0 | 1 | 14 |
| sik2b | 0.28039 | 0.128052305 | 0.05 | 0 | 1 | 14 |
| six2a | 0.28039 | 0.143385786 | 0.05 | 0 | 1 | 14 |
| slc16a5a | 0.28039 | 0.103716913 | 0.05 | 0 | 1 | 14 |
| slc27a1a | 0.28039 | 0.130645559 | 0.05 | 0 | 1 | 14 |
| slc7a10a | 0.28039 | 0.133295759 | 0.05 | 0 | 1 | 14 |
| smarca5 | 0.28039 | 0.101914579 | 0.05 | 0 | 1 | 14 |
| smtnl1 | 0.28039 | 0.120251539 | 0.05 | 0 | 1 | 14 |
| snx27a | 0.28039 | 0.147616593 | 0.05 | 0 | 1 | 14 |
| snx27b | 0.28039 | 0.157811166 | 0.05 | 0 | 1 | 14 |
| snx5 | 0.28039 | 0.1013109 | 0.05 | 0 | 1 | 14 |
| srd5a2a | 0.28039 | 0.123069208 | 0.05 | 0 | 1 | 14 |
| srsf2b | 0.28039 | 0.150968538 | 0.05 | 0 | 1 | 14 |
| stard3nl | 0.28039 | 0.109228176 | 0.05 | 0 | 1 | 14 |
| stk11 | 0.28039 | 0.131416204 | 0.05 | 0 | 1 | 14 |
| stxbp3 | 0.28039 | 0.108204632 | 0.05 | 0 | 1 | 14 |
| suco | 0.28039 | 0.12344694 | 0.05 | 0 | 1 | 14 |
| susd6 | 0.28039 | 0.126467814 | 0.05 | 0 | 1 | 14 |
| tbc1d4 | 0.28039 | 0.129683015 | 0.05 | 0 | 1 | 14 |

| | | | | | | |
|---|---|---|---|---|---|---|
| tbrg1 | 0.28039 | 0.137119724 | 0.05 | 0 | 1 | 14 |
| tfdp2 | 0.28039 | 0.128705595 | 0.05 | 0 | 1 | 14 |
| tmem106a | 0.28039 | 0.106476847 | 0.05 | 0 | 1 | 14 |
| tmem185 | 0.28039 | 0.165181453 | 0.05 | 0 | 1 | 14 |
| tmem218 | 0.28039 | 0.156929348 | 0.05 | 0 | 1 | 14 |
| tmem35 | 0.28039 | 0.123486728 | 0.05 | 0 | 1 | 14 |
| tnfaip1 | 0.28039 | 0.130038213 | 0.05 | 0 | 1 | 14 |
| tnfrsf19 | 0.28039 | 0.10580005 | 0.05 | 0 | 1 | 14 |
| tox4a | 0.28039 | 0.105087554 | 0.05 | 0 | 1 | 14 |
| tpm2 | 0.28039 | 0.132272564 | 0.05 | 0 | 1 | 14 |
| tradd | 0.28039 | 0.107545598 | 0.05 | 0 | 1 | 14 |
| trappc10 | 0.28039 | 0.129430524 | 0.05 | 0 | 1 | 14 |
| tsnaxip1 | 0.28039 | 0.114696438 | 0.05 | 0 | 1 | 14 |
| ttc14 | 0.28039 | 0.110268272 | 0.05 | 0 | 1 | 14 |
| ttc37 | 0.28039 | 0.14103119 | 0.05 | 0 | 1 | 14 |
| unc50 | 0.28039 | 0.141759968 | 0.05 | 0 | 1 | 14 |
| uqcc1 | 0.28039 | 0.101582693 | 0.05 | 0 | 1 | 14 |
| usp19 | 0.28039 | 0.137570079 | 0.05 | 0 | 1 | 14 |
| usp49 | 0.28039 | 0.106711563 | 0.05 | 0 | 1 | 14 |
| vti1b | 0.28039 | 0.107787646 | 0.05 | 0 | 1 | 14 |
| wdr12 | 0.28039 | 0.109086956 | 0.05 | 0 | 1 | 14 |
| wee1 | 0.28039 | 0.139085162 | 0.05 | 0 | 1 | 14 |
| wu:fj29h11 | 0.28039 | 0.131319799 | 0.05 | 0 | 1 | 14 |
| zgc:101577 | 0.28039 | 0.123725032 | 0.05 | 0 | 1 | 14 |
| zgc:110366 | 0.28039 | 0.130154529 | 0.05 | 0 | 1 | 14 |
| zgc:110626 | 0.28039 | 0.161004565 | 0.05 | 0 | 1 | 14 |
| zgc:113149 | 0.28039 | 0.138233831 | 0.05 | 0 | 1 | 14 |
| zgc:113210 | 0.28039 | 0.114346596 | 0.05 | 0 | 1 | 14 |
| zgc:114200 | 0.28039 | 0.108554177 | 0.05 | 0 | 1 | 14 |
| znf1079 | 0.28039 | 0.110956647 | 0.05 | 0 | 1 | 14 |
| znf1155 | 0.28039 | 0.105766288 | 0.05 | 0 | 1 | 14 |
| ACKR2 | 0.28039 | 0.109741708 | 0.05 | 0 | 1 | 14 |
| BX323087.1 | 0.28039 | 0.128861954 | 0.05 | 0 | 1 | 14 |
| CABZ01068367.1 | 0.28039 | 0.109096041 | 0.05 | 0 | 1 | 14 |
| CR376737.1 | 0.28039 | 0.131352994 | 0.05 | 0 | 1 | 14 |
| LOC101882640 | 0.28039 | 0.165375801 | 0.05 | 0 | 1 | 14 |
| XLOC-038035 | 0.28039 | 0.155040264 | 0.05 | 0 | 1 | 14 |
| admb | 0.28039 | 0.208984007 | 0.05 | 0 | 1 | 14 |
| agpat4 | 0.28039 | 0.140306495 | 0.05 | 0 | 1 | 14 |
| alg2 | 0.28039 | 0.113357098 | 0.05 | 0 | 1 | 14 |
| caska | 0.28039 | 0.106998353 | 0.05 | 0 | 1 | 14 |
| cetn3 | 0.28039 | 0.133205085 | 0.05 | 0 | 1 | 14 |
| chst2b | 0.28039 | 0.129640795 | 0.05 | 0 | 1 | 14 |
| cldn17 | 0.28039 | 0.106998353 | 0.05 | 0 | 1 | 14 |
| cmc4 | 0.28039 | 0.112068332 | 0.05 | 0 | 1 | 14 |
| cnih4 | 0.28039 | 0.140624251 | 0.05 | 0 | 1 | 14 |
| dcun1d1 | 0.28039 | 0.13922368 | 0.05 | 0 | 1 | 14 |
| dlgap4a | 0.28039 | 0.124095636 | 0.05 | 0 | 1 | 14 |

| | | | | | | |
|---|---|---|---|---|---|---|
| gpr155a | 0.28039 | 0.136369771 | 0.05 | 0 | 1 | 14 |
| gramd1ba | 0.28039 | 0.100096623 | 0.05 | 0 | 1 | 14 |
| hapln1a | 0.28039 | 0.220618008 | 0.05 | 0 | 1 | 14 |
| hgfb | 0.28039 | 0.100329141 | 0.05 | 0 | 1 | 14 |
| hnrnpul1l | 0.28039 | 0.112908098 | 0.05 | 0 | 1 | 14 |
| ldhd | 0.28039 | 0.140483668 | 0.05 | 0 | 1 | 14 |
| lhfpl3 | 0.28039 | 0.103812417 | 0.05 | 0 | 1 | 14 |
| mcmbp | 0.28039 | 0.103329059 | 0.05 | 0 | 1 | 14 |
| mia3 | 0.28039 | 0.14274104 | 0.05 | 0 | 1 | 14 |
| morc2 | 0.28039 | 0.118505071 | 0.05 | 0 | 1 | 14 |
| mpp5a | 0.28039 | 0.13931194 | 0.05 | 0 | 1 | 14 |
| mrpl46 | 0.28039 | 0.113701546 | 0.05 | 0 | 1 | 14 |
| mtfp1 | 0.28039 | 0.132501249 | 0.05 | 0 | 1 | 14 |
| nova1 | 0.28039 | 0.134725628 | 0.05 | 0 | 1 | 14 |
| nsd1b | 0.28039 | 0.112647693 | 0.05 | 0 | 1 | 14 |
| oxsr1b | 0.28039 | 0.102917129 | 0.05 | 0 | 1 | 14 |
| psd2 | 0.28039 | 0.184767365 | 0.05 | 0 | 1 | 14 |
| rasa1a | 0.28039 | 0.150863507 | 0.05 | 0 | 1 | 14 |
| rexo1 | 0.28039 | 0.121205358 | 0.05 | 0 | 1 | 14 |
| si:ch73-347e22.4 | 0.28039 | 0.100329141 | 0.05 | 0 | 1 | 14 |
| si:dkey-253d23.4 | 0.28039 | 0.127566984 | 0.05 | 0 | 1 | 14 |
| stx10 | 0.28039 | 0.103812417 | 0.05 | 0 | 1 | 14 |
| tbx2a | 0.28039 | 0.117595177 | 0.05 | 0 | 1 | 14 |
| thoc5 | 0.28039 | 0.158762681 | 0.05 | 0 | 1 | 14 |
| trpm7 | 0.28039 | 0.122468847 | 0.05 | 0 | 1 | 14 |
| wrap53 | 0.28039 | 0.107024698 | 0.05 | 0 | 1 | 14 |
| xrcc4 | 0.28039 | 0.119150743 | 0.05 | 0 | 1 | 14 |
| znf1015 | 0.28039 | 0.169247361 | 0.05 | 0 | 1 | 14 |
| zpr1 | 0.28039 | 0.1161713 | 0.05 | 0 | 1 | 14 |
| CABZ01080375.1 | 0.28039 | 0.104521855 | 0.05 | 0 | 1 | 14 |
| CABZ01118270.1 | 0.28039 | 0.105443673 | 0.05 | 0 | 1 | 14 |
| adamts17 | 0.28039 | 0.10638916 | 0.05 | 0 | 1 | 14 |
| arhgap29b | 0.28039 | 0.116677879 | 0.05 | 0 | 1 | 14 |
| asb6 | 0.28039 | 0.12096319 | 0.05 | 0 | 1 | 14 |
| bud23 | 0.28039 | 0.101074501 | 0.05 | 0 | 1 | 14 |
| cdc37l1 | 0.28039 | 0.10638916 | 0.05 | 0 | 1 | 14 |
| clasrp | 0.28039 | 0.118477117 | 0.05 | 0 | 1 | 14 |
| ddx47 | 0.28039 | 0.107243426 | 0.05 | 0 | 1 | 14 |
| dgkza | 0.28039 | 0.116422472 | 0.05 | 0 | 1 | 14 |
| dusp3a | 0.28039 | 0.119253069 | 0.05 | 0 | 1 | 14 |
| efnb3a | 0.28039 | 0.143865075 | 0.05 | 0 | 1 | 14 |
| eprs | 0.28039 | 0.123170015 | 0.05 | 0 | 1 | 14 |
| fam98a | 0.28039 | 0.102261638 | 0.05 | 0 | 1 | 14 |
| fbxl20 | 0.28039 | 0.135508531 | 0.05 | 0 | 1 | 14 |
| gadd45ab | 0.28039 | 0.111729832 | 0.05 | 0 | 1 | 14 |
| hcfc2 | 0.28039 | 0.100217979 | 0.05 | 0 | 1 | 14 |
| ipo13 | 0.28039 | 0.123103755 | 0.05 | 0 | 1 | 14 |
| kmt2a | 0.28039 | 0.123666601 | 0.05 | 0 | 1 | 14 |

| | | | | | | |
|---|---|---|---|---|---|---|
| lcmt1 | 0.28039 | 0.10722797 | 0.05 | 0 | 1 | 14 |
| nat14 | 0.28039 | 0.114678058 | 0.05 | 0 | 1 | 14 |
| plekhg2 | 0.28039 | 0.132214594 | 0.05 | 0 | 1 | 14 |
| ptpn5 | 0.28039 | 0.109826808 | 0.05 | 0 | 1 | 14 |
| shc1 | 0.28039 | 0.107719164 | 0.05 | 0 | 1 | 14 |
| si:ch211-152c8.4 | 0.28039 | 0.104403153 | 0.05 | 0 | 1 | 14 |
| si:ch211-194b1.1 | 0.28039 | 0.105723647 | 0.05 | 0 | 1 | 14 |
| srebf1 | 0.28039 | 0.119150415 | 0.05 | 0 | 1 | 14 |
| syvn1 | 0.28039 | 0.122230842 | 0.05 | 0 | 1 | 14 |
| tet1 | 0.28039 | 0.118262885 | 0.05 | 0 | 1 | 14 |
| trip12 | 0.28039 | 0.119339126 | 0.05 | 0 | 1 | 14 |
| ube2z | 0.28039 | 0.119104089 | 0.05 | 0 | 1 | 14 |
| wdr37 | 0.28039 | 0.10638916 | 0.05 | 0 | 1 | 14 |
| CABZ01077218.1 | 0.28039 | 0.129166242 | 0.05 | 0 | 1 | 14 |
| CABZ01113599.1 | 0.28039 | 0.119846371 | 0.05 | 0 | 1 | 14 |
| CR388029.1 | 0.28039 | 0.163563887 | 0.05 | 0 | 1 | 14 |
| LO018197.1 | 0.28039 | 0.108339421 | 0.05 | 0 | 1 | 14 |
| LOC100535806 | 0.28039 | 0.129482619 | 0.05 | 0 | 1 | 14 |
| LOC110437782 | 0.28039 | 0.116514457 | 0.05 | 0 | 1 | 14 |
| LOC110440195 | 0.28039 | 0.179434639 | 0.05 | 0 | 1 | 14 |
| MTERF4 | 0.28039 | 0.121420974 | 0.05 | 0 | 1 | 14 |
| TMEM19 | 0.28039 | 0.14540944 | 0.05 | 0 | 1 | 14 |
| TSTD1 | 0.28039 | 0.106944828 | 0.05 | 0 | 1 | 14 |
| XLOC-001102 | 0.28039 | 0.140789678 | 0.05 | 0 | 1 | 14 |
| aasdhppt | 0.28039 | 0.1430086 | 0.05 | 0 | 1 | 14 |
| abhd16a | 0.28039 | 0.121375617 | 0.05 | 0 | 1 | 14 |
| actr2a | 0.28039 | 0.134993558 | 0.05 | 0 | 1 | 14 |
| ada | 0.28039 | 0.143625634 | 0.05 | 0 | 1 | 14 |
| adcy9 | 0.28039 | 0.145178854 | 0.05 | 0 | 1 | 14 |
| ahctf1 | 0.28039 | 0.116596773 | 0.05 | 0 | 1 | 14 |
| ahcyl2 | 0.28039 | 0.131026604 | 0.05 | 0 | 1 | 14 |
| als2a | 0.28039 | 0.14523504 | 0.05 | 0 | 1 | 14 |
| amd1 | 0.28039 | 0.123857224 | 0.05 | 0 | 1 | 14 |
| ankib1a | 0.28039 | 0.172274393 | 0.05 | 0 | 1 | 14 |
| apc | 0.28039 | 0.117616347 | 0.05 | 0 | 1 | 14 |
| arhgap35b | 0.28039 | 0.124548974 | 0.05 | 0 | 1 | 14 |
| arntl1b | 0.28039 | 0.104847143 | 0.05 | 0 | 1 | 14 |
| b3glcta | 0.28039 | 0.154482509 | 0.05 | 0 | 1 | 14 |
| bach1b | 0.28039 | 0.167162561 | 0.05 | 0 | 1 | 14 |
| bach2b | 0.28039 | 0.12560576 | 0.05 | 0 | 1 | 14 |
| bcat2 | 0.28039 | 0.113010467 | 0.05 | 0 | 1 | 14 |
| camk2g2 | 0.28039 | 0.108779434 | 0.05 | 0 | 1 | 14 |
| capn7 | 0.28039 | 0.146663101 | 0.05 | 0 | 1 | 14 |
| cars | 0.28039 | 0.105883657 | 0.05 | 0 | 1 | 14 |
| cbx2 | 0.28039 | 0.119389948 | 0.05 | 0 | 1 | 14 |
| cc2d1b | 0.28039 | 0.108692683 | 0.05 | 0 | 1 | 14 |
| ccdc85ca | 0.28039 | 0.170345439 | 0.05 | 0 | 1 | 14 |
| ccl20b | 0.28039 | 0.13333001 | 0.05 | 0 | 1 | 14 |

| | | | | | | |
|---|---|---|---|---|---|---|
| ccnf | 0.28039 | 0.124346254 | 0.05 | 0 | 1 | 14 |
| cdkn1d | 0.28039 | 0.109478916 | 0.05 | 0 | 1 | 14 |
| cep19 | 0.28039 | 0.123566924 | 0.05 | 0 | 1 | 14 |
| chaf1b | 0.28039 | 0.106134327 | 0.05 | 0 | 1 | 14 |
| cited2 | 0.28039 | 0.128104219 | 0.05 | 0 | 1 | 14 |
| clcn6 | 0.28039 | 0.123962298 | 0.05 | 0 | 1 | 14 |
| coq8b | 0.28039 | 0.133345959 | 0.05 | 0 | 1 | 14 |
| cpxm1a | 0.28039 | 0.135565543 | 0.05 | 0 | 1 | 14 |
| cuedc1b | 0.28039 | 0.179434639 | 0.05 | 0 | 1 | 14 |
| cul5a | 0.28039 | 0.127376604 | 0.05 | 0 | 1 | 14 |
| cwf19l1 | 0.28039 | 0.113992242 | 0.05 | 0 | 1 | 14 |
| depdc5 | 0.28039 | 0.133331639 | 0.05 | 0 | 1 | 14 |
| dlg3 | 0.28039 | 0.117938712 | 0.05 | 0 | 1 | 14 |
| dnajb12b | 0.28039 | 0.128001634 | 0.05 | 0 | 1 | 14 |
| dvl3a | 0.28039 | 0.120921547 | 0.05 | 0 | 1 | 14 |
| ebpl | 0.28039 | 0.12671504 | 0.05 | 0 | 1 | 14 |
| eda | 0.28039 | 0.135954142 | 0.05 | 0 | 1 | 14 |
| eif4ba | 0.28039 | 0.12979536 | 0.05 | 0 | 1 | 14 |
| elac1 | 0.28039 | 0.115538768 | 0.05 | 0 | 1 | 14 |
| enc2 | 0.28039 | 0.185116319 | 0.05 | 0 | 1 | 14 |
| entr1 | 0.28039 | 0.111362886 | 0.05 | 0 | 1 | 14 |
| exosc7 | 0.28039 | 0.122207898 | 0.05 | 0 | 1 | 14 |
| fbxo33 | 0.28039 | 0.129596419 | 0.05 | 0 | 1 | 14 |
| fdps | 0.28039 | 0.158700574 | 0.05 | 0 | 1 | 14 |
| fgf10a | 0.28039 | 0.110011305 | 0.05 | 0 | 1 | 14 |
| fitm2 | 0.28039 | 0.148191284 | 0.05 | 0 | 1 | 14 |
| gnpda2 | 0.28039 | 0.112963883 | 0.05 | 0 | 1 | 14 |
| golim4a.1 | 0.28039 | 0.134068441 | 0.05 | 0 | 1 | 14 |
| gpm6bb | 0.28039 | 0.161191221 | 0.05 | 0 | 1 | 14 |
| gpr63 | 0.28039 | 0.103929459 | 0.05 | 0 | 1 | 14 |
| gpt2l | 0.28039 | 0.174592554 | 0.05 | 0 | 1 | 14 |
| grtp1a | 0.28039 | 0.142506335 | 0.05 | 0 | 1 | 14 |
| gys1 | 0.28039 | 0.126296152 | 0.05 | 0 | 1 | 14 |
| hipk3a | 0.28039 | 0.121806391 | 0.05 | 0 | 1 | 14 |
| hoxd9a | 0.28039 | 0.128313431 | 0.05 | 0 | 1 | 14 |
| il13ra2 | 0.28039 | 0.124269692 | 0.05 | 0 | 1 | 14 |
| im:7140055 | 0.28039 | 0.107990649 | 0.05 | 0 | 1 | 14 |
| ing3 | 0.28039 | 0.129146869 | 0.05 | 0 | 1 | 14 |
| ints2 | 0.28039 | 0.104913028 | 0.05 | 0 | 1 | 14 |
| itga4 | 0.28039 | 0.105593711 | 0.05 | 0 | 1 | 14 |
| kirrel1b | 0.28039 | 0.152713031 | 0.05 | 0 | 1 | 14 |
| lpin2 | 0.28039 | 0.120644383 | 0.05 | 0 | 1 | 14 |
| lyrm2 | 0.28039 | 0.121369551 | 0.05 | 0 | 1 | 14 |
| maco1a | 0.28039 | 0.147541206 | 0.05 | 0 | 1 | 14 |
| map3k4 | 0.28039 | 0.117864472 | 0.05 | 0 | 1 | 14 |
| mark4a | 0.28039 | 0.121762997 | 0.05 | 0 | 1 | 14 |
| matn4 | 0.28039 | 0.16426462 | 0.05 | 0 | 1 | 14 |
| mfhas1 | 0.28039 | 0.104215099 | 0.05 | 0 | 1 | 14 |

| | | | | | | |
|---|---|---|---|---|---|---|
| mink1 | 0.28039 | 0.114845341 | 0.05 | 0 | 1 | 14 |
| mri1 | 0.28039 | 0.127678806 | 0.05 | 0 | 1 | 14 |
| mta2 | 0.28039 | 0.141187747 | 0.05 | 0 | 1 | 14 |
| narf | 0.28039 | 0.120281721 | 0.05 | 0 | 1 | 14 |
| nr2f6a | 0.28039 | 0.17514556 | 0.05 | 0 | 1 | 14 |
| nt5dc2 | 0.28039 | 0.101130692 | 0.05 | 0 | 1 | 14 |
| olfm1a | 0.28039 | 0.181200106 | 0.05 | 0 | 1 | 14 |
| osbpl2a | 0.28039 | 0.10170248 | 0.05 | 0 | 1 | 14 |
| palm3 | 0.28039 | 0.119856855 | 0.05 | 0 | 1 | 14 |
| paqr3a | 0.28039 | 0.129254147 | 0.05 | 0 | 1 | 14 |
| pcmtd1 | 0.28039 | 0.130864178 | 0.05 | 0 | 1 | 14 |
| pde11al | 0.28039 | 0.140664021 | 0.05 | 0 | 1 | 14 |
| pde12 | 0.28039 | 0.115538768 | 0.05 | 0 | 1 | 14 |
| pdgfc | 0.28039 | 0.165172267 | 0.05 | 0 | 1 | 14 |
| pgap3 | 0.28039 | 0.146494603 | 0.05 | 0 | 1 | 14 |
| pgm1 | 0.28039 | 0.143960441 | 0.05 | 0 | 1 | 14 |
| piezo2a.2 | 0.28039 | 0.101544973 | 0.05 | 0 | 1 | 14 |
| pkd2 | 0.28039 | 0.127255175 | 0.05 | 0 | 1 | 14 |
| pla2g6 | 0.28039 | 0.135466585 | 0.05 | 0 | 1 | 14 |
| pla2g7 | 0.28039 | 0.106419099 | 0.05 | 0 | 1 | 14 |
| plod2 | 0.28039 | 0.162117346 | 0.05 | 0 | 1 | 14 |
| plpp7 | 0.28039 | 0.149722725 | 0.05 | 0 | 1 | 14 |
| pole | 0.28039 | 0.118216986 | 0.05 | 0 | 1 | 14 |
| ppp2r5cb | 0.28039 | 0.130681593 | 0.05 | 0 | 1 | 14 |
| prdm9 | 0.28039 | 0.112649328 | 0.05 | 0 | 1 | 14 |
| ptgdsb.1 | 0.28039 | 0.140366849 | 0.05 | 0 | 1 | 14 |
| pycard | 0.28039 | 0.108202452 | 0.05 | 0 | 1 | 14 |
| qser1 | 0.28039 | 0.123993429 | 0.05 | 0 | 1 | 14 |
| r3hdm1 | 0.28039 | 0.153762638 | 0.05 | 0 | 1 | 14 |
| rab28 | 0.28039 | 0.11835202 | 0.05 | 0 | 1 | 14 |
| rbm45 | 0.28039 | 0.130842194 | 0.05 | 0 | 1 | 14 |
| rpp21 | 0.28039 | 0.113238865 | 0.05 | 0 | 1 | 14 |
| rprd2b | 0.28039 | 0.164420614 | 0.05 | 0 | 1 | 14 |
| rrn3 | 0.28039 | 0.158710804 | 0.05 | 0 | 1 | 14 |
| samd1a | 0.28039 | 0.135766785 | 0.05 | 0 | 1 | 14 |
| sardh | 0.28039 | 0.122810232 | 0.05 | 0 | 1 | 14 |
| sgk3 | 0.28039 | 0.114523437 | 0.05 | 0 | 1 | 14 |
| si:ch211-150o23.3 | 0.28039 | 0.122971739 | 0.05 | 0 | 1 | 14 |
| si:ch211-218g4.2 | 0.28039 | 0.153943215 | 0.05 | 0 | 1 | 14 |
| si:ch211-251p5.5 | 0.28039 | 0.131681004 | 0.05 | 0 | 1 | 14 |
| si:ch211-275j6.5 | 0.28039 | 0.175707921 | 0.05 | 0 | 1 | 14 |
| si:ch73-22o12.1 | 0.28039 | 0.132511305 | 0.05 | 0 | 1 | 14 |
| si:ch73-335l21.1 | 0.28039 | 0.112477984 | 0.05 | 0 | 1 | 14 |
| si:dkey-229b18.3 | 0.28039 | 0.168647854 | 0.05 | 0 | 1 | 14 |
| si:dkey-266m15.6 | 0.28039 | 0.144222327 | 0.05 | 0 | 1 | 14 |
| slc10a3 | 0.28039 | 0.13382258 | 0.05 | 0 | 1 | 14 |
| slc25a22a | 0.28039 | 0.117330695 | 0.05 | 0 | 1 | 14 |
| snapc1b | 0.28039 | 0.119803454 | 0.05 | 0 | 1 | 14 |

| | | | | | | |
|---|---|---|---|---|---|---|
| sncgb | 0.28039 | 0.10906819 | 0.05 | 0 | 1 | 14 |
| snrka | 0.28039 | 0.100060452 | 0.05 | 0 | 1 | 14 |
| snrnp35 | 0.28039 | 0.103805724 | 0.05 | 0 | 1 | 14 |
| snx19b | 0.28039 | 0.161374384 | 0.05 | 0 | 1 | 14 |
| sort1a | 0.28039 | 0.129991148 | 0.05 | 0 | 1 | 14 |
| sra1 | 0.28039 | 0.104052356 | 0.05 | 0 | 1 | 14 |
| stox2b | 0.28039 | 0.139935179 | 0.05 | 0 | 1 | 14 |
| terfa | 0.28039 | 0.152778591 | 0.05 | 0 | 1 | 14 |
| thrb | 0.28039 | 0.193403874 | 0.05 | 0 | 1 | 14 |
| thumpd1 | 0.28039 | 0.111572012 | 0.05 | 0 | 1 | 14 |
| tm2d2 | 0.28039 | 0.124083756 | 0.05 | 0 | 1 | 14 |
| tmem33 | 0.28039 | 0.127310792 | 0.05 | 0 | 1 | 14 |
| tmem39b | 0.28039 | 0.11546209 | 0.05 | 0 | 1 | 14 |
| tmem56b | 0.28039 | 0.113142616 | 0.05 | 0 | 1 | 14 |
| tnip1 | 0.28039 | 0.127265772 | 0.05 | 0 | 1 | 14 |
| tp53i11a | 0.28039 | 0.131893969 | 0.05 | 0 | 1 | 14 |
| ttc13 | 0.28039 | 0.134749142 | 0.05 | 0 | 1 | 14 |
| txlna | 0.28039 | 0.100107603 | 0.05 | 0 | 1 | 14 |
| ube2t | 0.28039 | 0.104182725 | 0.05 | 0 | 1 | 14 |
| ube3b | 0.28039 | 0.130810369 | 0.05 | 0 | 1 | 14 |
| ubxn8 | 0.28039 | 0.141625778 | 0.05 | 0 | 1 | 14 |
| usp33 | 0.28039 | 0.12799758 | 0.05 | 0 | 1 | 14 |
| uvssa | 0.28039 | 0.157560458 | 0.05 | 0 | 1 | 14 |
| vps13d | 0.28039 | 0.103924659 | 0.05 | 0 | 1 | 14 |
| vti1a | 0.28039 | 0.128944841 | 0.05 | 0 | 1 | 14 |
| wdr55 | 0.28039 | 0.107011373 | 0.05 | 0 | 1 | 14 |
| zbtb22a | 0.28039 | 0.117548258 | 0.05 | 0 | 1 | 14 |
| zc2hc1c | 0.28039 | 0.12706404 | 0.05 | 0 | 1 | 14 |
| zfyve9a | 0.28039 | 0.125716485 | 0.05 | 0 | 1 | 14 |
| zgc:174895 | 0.28039 | 0.148257397 | 0.05 | 0 | 1 | 14 |
| zgc:194281 | 0.28039 | 0.146663101 | 0.05 | 0 | 1 | 14 |
| zmynd11 | 0.28039 | 0.128268717 | 0.05 | 0 | 1 | 14 |
| tfpia | 0.28047 | -0.26838666 | 0.175 | 0.35 | 1 | 14 |
| pdia3 | 0.28136 | -0.22648377 | 0.6 | 0.8 | 1 | 14 |
| XLOC-025423 | 0.28148 | -0.38382958 | 0.2 | 0.3 | 1 | 14 |
| crfb4 | 0.28327 | -0.11597504 | 0.05 | 0.15 | 1 | 14 |
| sox11a | 0.28517 | 0.401200137 | 0.375 | 0.2 | 1 | 14 |
| gspt1l | 0.28547 | 0.291472393 | 0.175 | 0.05 | 1 | 14 |
| rbmx | 0.28677 | 0.267071419 | 0.15 | 0.05 | 1 | 14 |
| caskb | 0.28693 | 0.32827553 | 0.175 | 0.05 | 1 | 14 |
| CR847566.1 | 0.28766 | 0.151704205 | 0.4 | 0.4 | 1 | 14 |
| rab18a | 0.28768 | 0.281354135 | 0.15 | 0.05 | 1 | 14 |
| mcm3 | 0.28881 | 0.238227924 | 0.15 | 0.1 | 1 | 14 |
| ip6k2a | 0.28952 | 0.300105673 | 0.175 | 0.05 | 1 | 14 |
| mgmt | 0.29037 | 0.156672259 | 0.1 | 0.1 | 1 | 14 |
| fzd1 | 0.29085 | -0.22906787 | 0.1 | 0.15 | 1 | 14 |
| ponzr1 | 0.29122 | 0.246487171 | 0.175 | 0.05 | 1 | 14 |
| loxl1 | 0.29124 | -0.2878028 | 0.3 | 0.5 | 1 | 14 |

| | | | | | | |
|---|---|---|---|---|---|---|
| ube2v2 | 0.29245 | 0.159889524 | 0.375 | 0.35 | 1 | 14 |
| metrnla | 0.29281 | -0.33439908 | 0.475 | 0.65 | 1 | 14 |
| rpz5 | 0.2939 | -0.2297577 | 0.15 | 0.3 | 1 | 14 |
| si:ch211-51c14.1 | 0.29487 | -0.19381941 | 0.075 | 0.2 | 1 | 14 |
| crhbp | 0.29605 | -0.35308433 | 0.1 | 0.25 | 1 | 14 |
| tnrc6c1 | 0.29645 | -0.18035096 | 0.375 | 0.55 | 1 | 14 |
| eif4a1b | 0.29694 | 0.341905448 | 0.575 | 0.4 | 1 | 14 |
| arih1 | 0.29708 | -0.14742439 | 0.075 | 0.2 | 1 | 14 |
| si:ch73-42p12.2 | 0.29821 | 0.215207411 | 0.2 | 0.15 | 1 | 14 |
| glsl | 0.29867 | -0.1498721 | 0.05 | 0.15 | 1 | 14 |
| ciao2b | 0.29893 | -0.1398664 | 0.05 | 0.15 | 1 | 14 |
| larp4ab | 0.29899 | -0.23475361 | 0.1 | 0.25 | 1 | 14 |
| arl2 | 0.29911 | 0.258558045 | 0.175 | 0.05 | 1 | 14 |
| tmed5 | 0.29927 | -0.26905007 | 0.075 | 0.15 | 1 | 14 |
| myl9b | 0.29962 | 0.361858491 | 0.525 | 0.35 | 1 | 14 |
| rtn4a | 0.29986 | -0.323006 | 0.25 | 0.45 | 1 | 14 |
| med10 | 0.30008 | -0.31052125 | 0.1 | 0.25 | 1 | 14 |
| nfil3-6 | 0.30018 | 0.32910316 | 0.175 | 0.05 | 1 | 14 |
| stt3b | 0.3002 | -0.27688715 | 0.05 | 0.15 | 1 | 14 |
| cyp1c1 | 0.30029 | -0.16903858 | 0.05 | 0.15 | 1 | 14 |
| rer1 | 0.30108 | -0.32843738 | 0.225 | 0.4 | 1 | 14 |
| rab8b | 0.30226 | 0.312385966 | 0.15 | 0.05 | 1 | 14 |
| romo1 | 0.30273 | 0.199660852 | 0.525 | 0.5 | 1 | 14 |
| litaf | 0.30335 | -0.19018625 | 0.075 | 0.2 | 1 | 14 |
| tbl1xr1a | 0.30391 | -0.22686403 | 0.075 | 0.2 | 1 | 14 |
| aip | 0.30406 | 0.243628611 | 0.2 | 0.15 | 1 | 14 |
| FP101882.1 | 0.30497 | -0.37351749 | 0.1 | 0.25 | 1 | 14 |
| ss18l2 | 0.30591 | 0.282956803 | 0.15 | 0.05 | 1 | 14 |
| dcaf7 | 0.30633 | 0.246970001 | 0.175 | 0.1 | 1 | 14 |
| kdm5bb | 0.30689 | 0.323423588 | 0.175 | 0.05 | 1 | 14 |
| cyb5a | 0.30708 | -0.2725835 | 0.1 | 0.25 | 1 | 14 |
| tax1bp3 | 0.30783 | 0.266421353 | 0.25 | 0.15 | 1 | 14 |
| stat3 | 0.30789 | 0.119738666 | 0.125 | 0.05 | 1 | 14 |
| grinab | 0.30827 | -0.27917043 | 0.175 | 0.35 | 1 | 14 |
| mdh2 | 0.30936 | -0.10126015 | 0.3 | 0.45 | 1 | 14 |
| psmb1 | 0.30942 | 0.34655907 | 0.425 | 0.25 | 1 | 14 |
| ppie | 0.30992 | -0.13138431 | 0.05 | 0.15 | 1 | 14 |
| cux1a | 0.31172 | -0.12062482 | 0.175 | 0.3 | 1 | 14 |
| eif2s2 | 0.31175 | 0.208675094 | 0.6 | 0.55 | 1 | 14 |
| RPL37A | 0.31199 | -0.29123896 | 0.05 | 0.15 | 1 | 14 |
| lum | 0.31201 | -0.26799879 | 0.55 | 0.75 | 1 | 14 |
| dolpp1 | 0.31328 | -0.11200078 | 0.05 | 0.15 | 1 | 14 |
| ing4 | 0.31344 | 0.298087641 | 0.175 | 0.05 | 1 | 14 |
| plpp2b | 0.31347 | -0.29573416 | 0.1 | 0.25 | 1 | 14 |
| tmem258 | 0.31472 | -0.11279236 | 0.5 | 0.65 | 1 | 14 |
| tmeff2a | 0.3148 | 0.300442437 | 0.25 | 0.1 | 1 | 14 |
| hnrnpaba | 0.31567 | 0.132682019 | 0.6 | 0.6 | 1 | 14 |
| gpc4 | 0.31628 | -0.19550746 | 0.075 | 0.2 | 1 | 14 |

| | | | | | | |
|---|---|---|---|---|---|---|
| psmc5 | 0.31653 | -0.22643941 | 0.225 | 0.4 | 1 | 14 |
| arf5 | 0.317 | -0.32862361 | 0.325 | 0.5 | 1 | 14 |
| tm9sf2 | 0.31755 | -0.31479256 | 0.1 | 0.25 | 1 | 14 |
| calua | 0.31768 | -0.35220402 | 0.175 | 0.35 | 1 | 14 |
| CT030188.1 | 0.318 | -0.24377652 | 0.1 | 0.25 | 1 | 14 |
| gabarapa | 0.31898 | 0.277929958 | 0.275 | 0.2 | 1 | 14 |
| otud5b | 0.31927 | 0.289695495 | 0.175 | 0.05 | 1 | 14 |
| rps20 | 0.31941 | 0.181771535 | 0.95 | 0.85 | 1 | 14 |
| ecrg4a | 0.31979 | -0.35220427 | 0.375 | 0.45 | 1 | 14 |
| aebp1 | 0.3205 | 0.319849036 | 0.3 | 0.15 | 1 | 14 |
| ndufab1b | 0.3205 | 0.244196174 | 0.175 | 0.05 | 1 | 14 |
| tob1a | 0.32062 | -0.25434005 | 0.225 | 0.4 | 1 | 14 |
| dpy30 | 0.32138 | 0.118652623 | 0.225 | 0.25 | 1 | 14 |
| cpsf6 | 0.32195 | 0.278621818 | 0.175 | 0.05 | 1 | 14 |
| col6a2 | 0.3239 | 0.132906587 | 0.5 | 0.5 | 1 | 14 |
| sdcbp2 | 0.32428 | -0.28970274 | 0.175 | 0.35 | 1 | 14 |
| fam133b | 0.32461 | 0.209844372 | 0.25 | 0.2 | 1 | 14 |
| atp1b3a | 0.32545 | -0.13221206 | 0.05 | 0.15 | 1 | 14 |
| rps25 | 0.32547 | 0.157717042 | 0.95 | 0.9 | 1 | 14 |
| h2afva | 0.32579 | 0.339096043 | 0.325 | 0.2 | 1 | 14 |
| ugt1b5 | 0.32657 | -0.31833196 | 0.1 | 0.25 | 1 | 14 |
| eif3c | 0.3271 | -0.24447711 | 0.175 | 0.35 | 1 | 14 |
| safb | 0.3272 | 0.154415946 | 0.1 | 0.1 | 1 | 14 |
| pcna | 0.32727 | 0.353067798 | 0.325 | 0.2 | 1 | 14 |
| fkbp14 | 0.32778 | -0.27538609 | 0.1 | 0.25 | 1 | 14 |
| blcap | 0.3278 | 0.235755469 | 0.15 | 0.1 | 1 | 14 |
| gstp1 | 0.32865 | -0.22778247 | 0.225 | 0.4 | 1 | 14 |
| anxa4 | 0.32876 | 0.321704745 | 0.325 | 0.15 | 1 | 14 |
| pfdn4 | 0.32901 | -0.28774251 | 0.1 | 0.25 | 1 | 14 |
| ppp1r10 | 0.32902 | 0.151996553 | 0.15 | 0.05 | 1 | 14 |
| ppm1la | 0.32954 | -0.26645737 | 0.1 | 0.25 | 1 | 14 |
| vat1 | 0.32982 | -0.27698929 | 0.5 | 0.7 | 1 | 14 |
| rpz2 | 0.32994 | 0.272784645 | 0.175 | 0.05 | 1 | 14 |
| atp1b1a | 0.32999 | -0.23168578 | 0.175 | 0.35 | 1 | 14 |
| mcm7 | 0.33017 | 0.273867655 | 0.15 | 0.05 | 1 | 14 |
| rasl12 | 0.33079 | -0.31850253 | 0.1 | 0.25 | 1 | 14 |
| eef1g | 0.33086 | 0.151303275 | 0.925 | 0.9 | 1 | 14 |
| max | 0.33105 | 0.182799973 | 0.175 | 0.15 | 1 | 14 |
| dock1 | 0.3311 | -0.14832536 | 0.05 | 0.15 | 1 | 14 |
| ptena | 0.33133 | 0.281655229 | 0.2 | 0.1 | 1 | 14 |
| iqgap1 | 0.33157 | 0.348207237 | 0.175 | 0.05 | 1 | 14 |
| btg2 | 0.33163 | 0.389318004 | 0.375 | 0.2 | 1 | 14 |
| hmgn6 | 0.33211 | 0.135215265 | 0.675 | 0.65 | 1 | 14 |
| eif3s6ip | 0.33216 | 0.326035592 | 0.25 | 0.1 | 1 | 14 |
| cyp2ad3 | 0.33235 | -0.23576217 | 0.075 | 0.2 | 1 | 14 |
| eif1axb | 0.33257 | -0.19445716 | 0.325 | 0.5 | 1 | 14 |
| emc7 | 0.33278 | -0.15460792 | 0.125 | 0.25 | 1 | 14 |
| nap1l1 | 0.33345 | 0.107652613 | 0.475 | 0.5 | 1 | 14 |

| | | | | | | |
|---|---|---|---|---|---|---|
| si:busm1-57f23.1 | 0.33396 | -0.22562721 | 0.075 | 0.2 | 1 | 14 |
| emc2 | 0.33462 | -0.28947407 | 0.175 | 0.35 | 1 | 14 |
| atp5l | 0.33496 | 0.302168085 | 0.575 | 0.4 | 1 | 14 |
| rbm5 | 0.33556 | -0.14151651 | 0.05 | 0.15 | 1 | 14 |
| ttf1 | 0.33653 | 0.288575443 | 0.175 | 0.05 | 1 | 14 |
| lamp1a | 0.3379 | -0.15903321 | 0.125 | 0.25 | 1 | 14 |
| serbp1b | 0.33823 | -0.19157466 | 0.075 | 0.2 | 1 | 14 |
| ndufb4 | 0.33998 | 0.104166126 | 0.375 | 0.4 | 1 | 14 |
| sat1a.2 | 0.34011 | 0.272257549 | 0.225 | 0.1 | 1 | 14 |
| hmgn3 | 0.34015 | 0.239400365 | 0.2 | 0.15 | 1 | 14 |
| smarce1 | 0.34071 | 0.287536925 | 0.25 | 0.1 | 1 | 14 |
| angptl4 | 0.34099 | 0.294490669 | 0.25 | 0.15 | 1 | 14 |
| BX323797.4 | 0.3416 | -0.10086472 | 0.1 | 0.2 | 1 | 14 |
| serf2 | 0.34266 | -0.12942324 | 0.65 | 0.65 | 1 | 14 |
| tpm1 | 0.34491 | -0.36392307 | 0.25 | 0.4 | 1 | 14 |
| spry4 | 0.34493 | 0.243985945 | 0.175 | 0.05 | 1 | 14 |
| lbx2 | 0.34565 | -0.27569049 | 0.05 | 0.15 | 1 | 14 |
| CABZ01055347.1 | 0.34668 | -0.14227868 | 0.05 | 0.15 | 1 | 14 |
| c6 | 0.34683 | -0.39302766 | 0.2 | 0.35 | 1 | 14 |
| grm4 | 0.34701 | 0.331562537 | 0.25 | 0.1 | 1 | 14 |
| bnc2 | 0.34712 | -0.240458 | 0.15 | 0.3 | 1 | 14 |
| ldhba | 0.34792 | 0.323049244 | 0.35 | 0.2 | 1 | 14 |
| slc38a10 | 0.34804 | -0.16287468 | 0.05 | 0.15 | 1 | 14 |
| vdac3 | 0.34856 | 0.303676077 | 0.425 | 0.3 | 1 | 14 |
| srsf10b | 0.34932 | 0.211455721 | 0.125 | 0.05 | 1 | 14 |
| dars | 0.34992 | -0.14419485 | 0.05 | 0.15 | 1 | 14 |
| gng5 | 0.35028 | -0.20774603 | 0.325 | 0.5 | 1 | 14 |
| ppp6r3 | 0.35028 | -0.30902515 | 0.075 | 0.2 | 1 | 14 |
| pa2g4b | 0.35035 | 0.250692793 | 0.25 | 0.1 | 1 | 14 |
| ube2a | 0.35078 | 0.264819623 | 0.175 | 0.05 | 1 | 14 |
| ubl3a | 0.35104 | -0.1272507 | 0.05 | 0.15 | 1 | 14 |
| xrn2 | 0.35122 | -0.28914172 | 0.075 | 0.2 | 1 | 14 |
| bloc1s2 | 0.35182 | 0.202037853 | 0.1 | 0.05 | 1 | 14 |
| aifm2 | 0.35187 | 0.253085669 | 0.175 | 0.05 | 1 | 14 |
| CU694197.1 | 0.35205 | 0.287350257 | 0.325 | 0.2 | 1 | 14 |
| col21a1 | 0.35235 | 0.130254208 | 0.25 | 0.25 | 1 | 14 |
| si:ch211-162e15.3 | 0.3524 | 0.252533354 | 0.175 | 0.05 | 1 | 14 |
| dido1 | 0.35244 | 0.300017637 | 0.25 | 0.1 | 1 | 14 |
| actn1 | 0.35266 | -0.16715278 | 0.25 | 0.4 | 1 | 14 |
| tagln | 0.3537 | -0.43036514 | 0.2 | 0.35 | 1 | 14 |
| vbp1 | 0.35372 | 0.1697892 | 0.175 | 0.15 | 1 | 14 |
| tomm20a | 0.35393 | 0.192976321 | 0.25 | 0.2 | 1 | 14 |
| psma1 | 0.35424 | 0.1656112 | 0.425 | 0.4 | 1 | 14 |
| nip7 | 0.35428 | 0.258946485 | 0.2 | 0.1 | 1 | 14 |
| CABZ01045617.1 | 0.35492 | 0.208959249 | 0.15 | 0.1 | 1 | 14 |
| marcksb | 0.35542 | 0.230932812 | 0.175 | 0.05 | 1 | 14 |
| serpinf1 | 0.3557 | -0.2510347 | 0.35 | 0.45 | 1 | 14 |
| lhfpl6 | 0.35593 | -0.16499874 | 0.35 | 0.5 | 1 | 14 |

| | | | | | | |
|---|---|---|---|---|---|---|
| tmtc4 | 0.35715 | -0.13532683 | 0.05 | 0.15 | 1 | 14 |
| zgc:153704 | 0.3572 | 0.396107007 | 0.3 | 0.15 | 1 | 14 |
| nkrf | 0.35796 | -0.29216895 | 0.075 | 0.2 | 1 | 14 |
| psmc3 | 0.35847 | 0.268803585 | 0.25 | 0.1 | 1 | 14 |
| loxl2a | 0.35856 | 0.310310687 | 0.25 | 0.1 | 1 | 14 |
| capza1a | 0.35874 | -0.20764816 | 0.325 | 0.5 | 1 | 14 |
| mex3b | 0.35877 | 0.311960245 | 0.25 | 0.1 | 1 | 14 |
| csnk1da | 0.35879 | 0.27572271 | 0.15 | 0.05 | 1 | 14 |
| eif4a2 | 0.35912 | 0.268522916 | 0.15 | 0.05 | 1 | 14 |
| sf3b2 | 0.35981 | 0.263978102 | 0.175 | 0.05 | 1 | 14 |
| mgst1.2 | 0.35996 | -0.2974819 | 0.425 | 0.6 | 1 | 14 |
| styx | 0.36024 | 0.246668488 | 0.175 | 0.05 | 1 | 14 |
| smarcb1b | 0.36027 | 0.26526236 | 0.175 | 0.05 | 1 | 14 |
| usp14 | 0.36076 | -0.24548415 | 0.05 | 0.15 | 1 | 14 |
| pcyox1 | 0.36117 | 0.303789354 | 0.225 | 0.1 | 1 | 14 |
| tmem198b | 0.36157 | -0.19943822 | 0.075 | 0.2 | 1 | 14 |
| lhfpl2a | 0.36179 | 0.272582156 | 0.15 | 0.05 | 1 | 14 |
| scarb2a | 0.36188 | -0.27529534 | 0.05 | 0.15 | 1 | 14 |
| ncor1 | 0.36212 | 0.195729843 | 0.125 | 0.1 | 1 | 14 |
| dynlrb1 | 0.3634 | 0.269181433 | 0.375 | 0.2 | 1 | 14 |
| dkk3b | 0.36354 | 0.332542847 | 0.3 | 0.15 | 1 | 14 |
| adh5 | 0.36453 | 0.326151282 | 0.45 | 0.3 | 1 | 14 |
| brms1la | 0.36482 | 0.250700131 | 0.15 | 0.05 | 1 | 14 |
| cdk9 | 0.36517 | -0.28851053 | 0.075 | 0.2 | 1 | 14 |
| pitpnc1b | 0.36573 | -0.26461401 | 0.075 | 0.2 | 1 | 14 |
| col12a1a | 0.36647 | 0.335952821 | 0.35 | 0.25 | 1 | 14 |
| BX322530.1 | 0.3666 | -0.25242426 | 0.15 | 0.3 | 1 | 14 |
| scpep1 | 0.36725 | -0.18322204 | 0.075 | 0.2 | 1 | 14 |
| sf3b6 | 0.36801 | 0.314633187 | 0.3 | 0.15 | 1 | 14 |
| crim1 | 0.36826 | 0.285171126 | 0.15 | 0.05 | 1 | 14 |
| h3f3a | 0.36911 | 0.272601568 | 0.45 | 0.35 | 1 | 14 |
| babam1 | 0.36961 | -0.16330838 | 0.05 | 0.15 | 1 | 14 |
| si:ch211-250c4.4 | 0.36981 | -0.31924382 | 0.15 | 0.3 | 1 | 14 |
| thbs2b | 0.37003 | 0.283981116 | 0.35 | 0.25 | 1 | 14 |
| mphosph8 | 0.37024 | 0.183916462 | 0.175 | 0.15 | 1 | 14 |
| tax1bp1b | 0.37049 | -0.11975573 | 0.1 | 0.2 | 1 | 14 |
| edf1 | 0.37101 | 0.297012715 | 0.375 | 0.2 | 1 | 14 |
| zgc:171679 | 0.37107 | -0.22673593 | 0.075 | 0.2 | 1 | 14 |
| irf2bp1 | 0.37107 | -0.14385465 | 0.05 | 0.15 | 1 | 14 |
| mapkapk2a | 0.37169 | -0.27265408 | 0.075 | 0.2 | 1 | 14 |
| gli3 | 0.37334 | -0.23664863 | 0.15 | 0.3 | 1 | 14 |
| rbms2a | 0.37347 | 0.180023629 | 0.15 | 0.05 | 1 | 14 |
| susd5 | 0.37369 | -0.28146273 | 0.075 | 0.2 | 1 | 14 |
| rab6a | 0.37374 | -0.27110532 | 0.075 | 0.2 | 1 | 14 |
| uqcrc2a | 0.37449 | 0.188817438 | 0.1 | 0.05 | 1 | 14 |
| tma7 | 0.37515 | 0.299889132 | 0.575 | 0.4 | 1 | 14 |
| bzw1b | 0.3759 | -0.11861325 | 0.175 | 0.3 | 1 | 14 |
| cish | 0.376 | -0.11836686 | 0.1 | 0.2 | 1 | 14 |

| | | | | | | |
|---|---|---|---|---|---|---|
| znf704 | 0.37615 | 0.244541434 | 0.125 | 0.05 | 1 | 14 |
| myh9a | 0.377 | -0.27560843 | 0.05 | 0.15 | 1 | 14 |
| tm9sf3 | 0.37767 | -0.24889318 | 0.075 | 0.2 | 1 | 14 |
| si:ch211-198c19.3 | 0.37793 | 0.166104059 | 0.275 | 0.25 | 1 | 14 |
| cwf19l2 | 0.37927 | -0.13835949 | 0.05 | 0.15 | 1 | 14 |
| acat1 | 0.37961 | -0.22064576 | 0.075 | 0.2 | 1 | 14 |
| pdgfrl | 0.38016 | -0.2591556 | 0.575 | 0.75 | 1 | 14 |
| zgc:100918 | 0.38032 | -0.24093203 | 0.075 | 0.2 | 1 | 14 |
| abracl | 0.38105 | 0.11100849 | 0.15 | 0.05 | 1 | 14 |
| lgals9l3 | 0.38112 | 0.211835403 | 0.25 | 0.2 | 1 | 14 |
| sdf2 | 0.38119 | 0.217723737 | 0.15 | 0.05 | 1 | 14 |
| ap1m1 | 0.38153 | -0.14744681 | 0.05 | 0.15 | 1 | 14 |
| myl9a | 0.3821 | -0.23963996 | 0.325 | 0.5 | 1 | 14 |
| dbi | 0.38233 | -0.17413826 | 0.275 | 0.45 | 1 | 14 |
| tmem39a | 0.38282 | -0.15157538 | 0.05 | 0.15 | 1 | 14 |
| cd2bp2 | 0.38311 | -0.14031041 | 0.05 | 0.15 | 1 | 14 |
| ptpn13 | 0.3833 | -0.15958023 | 0.05 | 0.15 | 1 | 14 |
| alg11 | 0.3833 | -0.13050326 | 0.05 | 0.15 | 1 | 14 |
| calr3b | 0.38344 | 0.317717273 | 0.35 | 0.2 | 1 | 14 |
| ndufa1 | 0.38512 | -0.26932034 | 0.3 | 0.45 | 1 | 14 |
| map1lc3b | 0.38521 | 0.228416627 | 0.325 | 0.25 | 1 | 14 |
| ddx42 | 0.38523 | -0.26505226 | 0.075 | 0.2 | 1 | 14 |
| ccz1 | 0.38524 | -0.19926937 | 0.075 | 0.2 | 1 | 14 |
| dctn2 | 0.38535 | -0.14194215 | 0.05 | 0.15 | 1 | 14 |
| LOC100536647 | 0.38568 | -0.24507424 | 0.075 | 0.2 | 1 | 14 |
| tagln2 | 0.38595 | -0.24882526 | 0.275 | 0.45 | 1 | 14 |
| col5a2a | 0.386 | 0.182123659 | 0.825 | 0.75 | 1 | 14 |
| LOC103910871 | 0.38631 | 0.287525309 | 0.15 | 0.05 | 1 | 14 |
| uqcrc1 | 0.38667 | -0.27909207 | 0.075 | 0.2 | 1 | 14 |
| prdx6 | 0.3871 | -0.17907091 | 0.075 | 0.2 | 1 | 14 |
| sec61a1l | 0.38724 | -0.2715327 | 0.075 | 0.2 | 1 | 14 |
| tspan7 | 0.38731 | -0.303393 | 0.25 | 0.4 | 1 | 14 |
| BX284696.1 | 0.38734 | -0.18315759 | 0.125 | 0.25 | 1 | 14 |
| mknk2b | 0.38801 | 0.203100249 | 0.15 | 0.1 | 1 | 14 |
| vasna | 0.38852 | 0.132599097 | 0.25 | 0.25 | 1 | 14 |
| lsm3 | 0.38869 | -0.10427126 | 0.225 | 0.35 | 1 | 14 |
| pprc1 | 0.38873 | -0.14876941 | 0.05 | 0.15 | 1 | 14 |
| vps4b | 0.38876 | -0.25224929 | 0.075 | 0.2 | 1 | 14 |
| rps6 | 0.38898 | 0.146201275 | 0.925 | 0.8 | 1 | 14 |
| gpx8 | 0.38909 | -0.17070676 | 0.05 | 0.15 | 1 | 14 |
| pdia4 | 0.38922 | -0.24564592 | 0.075 | 0.2 | 1 | 14 |
| igf1 | 0.38938 | -0.28275796 | 0.075 | 0.2 | 1 | 14 |
| mid1ip1b | 0.3895 | -0.23392763 | 0.075 | 0.2 | 1 | 14 |
| vps28 | 0.38968 | -0.22368986 | 0.075 | 0.2 | 1 | 14 |
| itga9 | 0.3899 | -0.2958843 | 0.075 | 0.2 | 1 | 14 |
| calm3a | 0.38996 | -0.24148486 | 0.075 | 0.2 | 1 | 14 |
| klf11a | 0.39007 | -0.22159944 | 0.075 | 0.2 | 1 | 14 |
| plxna1a | 0.39012 | -0.25517054 | 0.075 | 0.2 | 1 | 14 |

| | | | | | | |
|---|---|---|---|---|---|---|
| twf1a | 0.39017 | -0.14265907 | 0.05 | 0.15 | 1 | 14 |
| eif3ja | 0.39074 | 0.168467227 | 0.275 | 0.25 | 1 | 14 |
| phlda2 | 0.39093 | -0.19347338 | 0.05 | 0.15 | 1 | 14 |
| psmd14 | 0.39105 | 0.127680861 | 0.15 | 0.15 | 1 | 14 |
| si:dkey-28b4.7 | 0.39196 | 0.292801644 | 0.25 | 0.15 | 1 | 14 |
| fndc3ba | 0.39216 | -0.16067895 | 0.05 | 0.15 | 1 | 14 |
| faf2 | 0.39219 | -0.25196798 | 0.05 | 0.15 | 1 | 14 |
| grcc10 | 0.39296 | -0.2726297 | 0.15 | 0.3 | 1 | 14 |
| stard13b | 0.39392 | -0.29072932 | 0.05 | 0.15 | 1 | 14 |
| adrm1 | 0.39444 | -0.15333459 | 0.05 | 0.15 | 1 | 14 |
| CABZ01072614.1 | 0.39636 | -0.10645405 | 0.15 | 0.25 | 1 | 14 |
| spcs3 | 0.39649 | -0.22535446 | 0.15 | 0.3 | 1 | 14 |
| cmtm4 | 0.39652 | -0.24097441 | 0.05 | 0.15 | 1 | 14 |
| BX539336.1 | 0.39705 | -0.27468073 | 0.05 | 0.15 | 1 | 14 |
| fibinb | 0.39793 | 0.19803305 | 0.4 | 0.35 | 1 | 14 |
| hbp1 | 0.39809 | 0.243299087 | 0.125 | 0.05 | 1 | 14 |
| ndufb10 | 0.39895 | 0.117358667 | 0.25 | 0.25 | 1 | 14 |
| arhgap31 | 0.39984 | -0.1373002 | 0.05 | 0.15 | 1 | 14 |
| rpl21 | 0.39995 | 0.188305917 | 0.9 | 0.85 | 1 | 14 |
| foxc1b | 0.40004 | 0.183626385 | 0.15 | 0.05 | 1 | 14 |
| dpm3 | 0.40004 | -0.25097451 | 0.05 | 0.15 | 1 | 14 |
| eif3i | 0.4001 | 0.19095064 | 0.35 | 0.3 | 1 | 14 |
| esf1 | 0.40067 | 0.22084662 | 0.15 | 0.1 | 1 | 14 |
| hoxc6b | 0.40161 | -0.17017918 | 0.05 | 0.15 | 1 | 14 |
| abi3bpb | 0.40206 | -0.31703074 | 0.275 | 0.45 | 1 | 14 |
| pbx3b | 0.40287 | -0.24356136 | 0.575 | 0.75 | 1 | 14 |
| slc37a2 | 0.40324 | -0.15689475 | 0.05 | 0.15 | 1 | 14 |
| taf7 | 0.40337 | -0.13017936 | 0.05 | 0.15 | 1 | 14 |
| ncl | 0.40365 | 0.261990528 | 0.45 | 0.35 | 1 | 14 |
| ldlrad4a | 0.40427 | -0.26336024 | 0.05 | 0.15 | 1 | 14 |
| pnisr | 0.40509 | 0.259871631 | 0.175 | 0.1 | 1 | 14 |
| canx | 0.40538 | -0.24077579 | 0.15 | 0.3 | 1 | 14 |
| si:dkey-16l2.16 | 0.40557 | -0.24860421 | 0.15 | 0.3 | 1 | 14 |
| crkl | 0.40562 | -0.24578095 | 0.05 | 0.15 | 1 | 14 |
| CT956002.2 | 0.40602 | -0.16853226 | 0.05 | 0.15 | 1 | 14 |
| hmgb3a | 0.40711 | 0.287098554 | 0.625 | 0.45 | 1 | 14 |
| slc15a2 | 0.40716 | -0.24889818 | 0.05 | 0.15 | 1 | 14 |
| cops2 | 0.40754 | 0.244595521 | 0.15 | 0.05 | 1 | 14 |
| srsf6a | 0.40758 | -0.24051215 | 0.05 | 0.15 | 1 | 14 |
| adgra2 | 0.40802 | -0.27907091 | 0.15 | 0.3 | 1 | 14 |
| dusp1 | 0.4084 | -0.28809188 | 0.05 | 0.15 | 1 | 14 |
| gna11b | 0.40843 | -0.28836221 | 0.15 | 0.3 | 1 | 14 |
| si:ch211-276i12.9 | 0.40918 | 0.229174593 | 0.15 | 0.05 | 1 | 14 |
| myo1cb | 0.40924 | -0.17418843 | 0.05 | 0.15 | 1 | 14 |
| zgc:91910 | 0.40949 | -0.21057139 | 0.425 | 0.6 | 1 | 14 |
| s100u | 0.40961 | -0.17900911 | 0.125 | 0.25 | 1 | 14 |
| tbca | 0.41013 | 0.189496864 | 0.35 | 0.3 | 1 | 14 |
| si:ch211-14a17.11 | 0.41096 | -0.12597716 | 0.05 | 0.15 | 1 | 14 |

| | | | | | | |
|---|---|---|---|---|---|---|
| sf3a3 | 0.41143 | -0.10238938 | 0.05 | 0.15 | 1 | 14 |
| polr1c | 0.41154 | 0.253630467 | 0.15 | 0.05 | 1 | 14 |
| pabpn1 | 0.41181 | 0.238495346 | 0.45 | 0.35 | 1 | 14 |
| rbfox1 | 0.41293 | 0.166226218 | 0.325 | 0.3 | 1 | 14 |
| col4a1 | 0.41367 | 0.346577592 | 0.3 | 0.15 | 1 | 14 |
| atp5po | 0.41441 | -0.22149468 | 0.325 | 0.5 | 1 | 14 |
| itgb3b | 0.41442 | -0.15868879 | 0.125 | 0.15 | 1 | 14 |
| capza1b | 0.41452 | 0.287684202 | 0.375 | 0.25 | 1 | 14 |
| ppp1r26 | 0.41463 | -0.23648076 | 0.05 | 0.15 | 1 | 14 |
| rnf213a | 0.41531 | 0.23038846 | 0.125 | 0.05 | 1 | 14 |
| akap12b | 0.41628 | 0.265956971 | 0.525 | 0.35 | 1 | 14 |
| ccng1 | 0.41638 | 0.195964412 | 0.4 | 0.35 | 1 | 14 |
| tubb5 | 0.41747 | -0.16979106 | 0.175 | 0.3 | 1 | 14 |
| ten1 | 0.41752 | 0.223276222 | 0.125 | 0.05 | 1 | 14 |
| glrx | 0.4178 | -0.145755 | 0.35 | 0.5 | 1 | 14 |
| dpm1 | 0.41816 | 0.168310127 | 0.075 | 0.05 | 1 | 14 |
| tomm5 | 0.41849 | 0.152771722 | 0.375 | 0.35 | 1 | 14 |
| mob4 | 0.41917 | 0.113529111 | 0.1 | 0.1 | 1 | 14 |
| atp5f1c | 0.41946 | 0.195138965 | 0.35 | 0.3 | 1 | 14 |
| cycsb | 0.41994 | 0.114027986 | 0.275 | 0.3 | 1 | 14 |
| kmt2bb | 0.42016 | 0.279927408 | 0.3 | 0.2 | 1 | 14 |
| mir21-1 | 0.42075 | -0.27176431 | 0.05 | 0.15 | 1 | 14 |
| aldh2.1 | 0.42195 | 0.217598812 | 0.125 | 0.05 | 1 | 14 |
| nipsnap2 | 0.42266 | 0.184369998 | 0.175 | 0.15 | 1 | 14 |
| ewsr1a | 0.42272 | -0.14755405 | 0.05 | 0.15 | 1 | 14 |
| efcc1 | 0.4232 | 0.239155525 | 0.225 | 0.15 | 1 | 14 |
| ogt.1 | 0.42336 | -0.12894317 | 0.1 | 0.2 | 1 | 14 |
| qsox1 | 0.42378 | -0.22002345 | 0.25 | 0.4 | 1 | 14 |
| si:ch211-286o17.1 | 0.42424 | 0.31051452 | 0.3 | 0.15 | 1 | 14 |
| ndufa6 | 0.42431 | 0.154060981 | 0.35 | 0.2 | 1 | 14 |
| lamc3 | 0.42433 | 0.24411049 | 0.15 | 0.05 | 1 | 14 |
| rras | 0.42446 | -0.165411 | 0.05 | 0.15 | 1 | 14 |
| ythdc1 | 0.42452 | -0.18663025 | 0.05 | 0.15 | 1 | 14 |
| higd1a | 0.42463 | -0.23556161 | 0.375 | 0.55 | 1 | 14 |
| otub1b | 0.42501 | 0.250530059 | 0.125 | 0.05 | 1 | 14 |
| gpr108 | 0.42503 | -0.17985382 | 0.075 | 0.1 | 1 | 14 |
| ndfip1 | 0.42599 | -0.2711712 | 0.325 | 0.5 | 1 | 14 |
| ism1 | 0.426 | -0.23321594 | 0.25 | 0.4 | 1 | 14 |
| cct5 | 0.42686 | 0.175579224 | 0.325 | 0.3 | 1 | 14 |
| apcdd1l | 0.42686 | 0.174971145 | 0.325 | 0.3 | 1 | 14 |
| FAT4 | 0.42703 | 0.258451388 | 0.175 | 0.1 | 1 | 14 |
| hbegfa | 0.42715 | 0.234498627 | 0.2 | 0.15 | 1 | 14 |
| pdcl | 0.42718 | 0.193959986 | 0.1 | 0.05 | 1 | 14 |
| anks1b | 0.42757 | -0.22192636 | 0.05 | 0.15 | 1 | 14 |
| mtr | 0.42799 | -0.2026592 | 0.05 | 0.15 | 1 | 14 |
| hsp70.2 | 0.42819 | -0.17596094 | 0.05 | 0.15 | 1 | 14 |
| ctsla | 0.4282 | -0.20805432 | 0.725 | 0.85 | 1 | 14 |
| crabp1a | 0.42851 | -0.18782548 | 0.3 | 0.45 | 1 | 14 |

| | | | | | | |
|---|---|---|---|---|---|---|
| naa10 | 0.42859 | 0.248967043 | 0.3 | 0.2 | 1 | 14 |
| sesn1 | 0.429 | -0.24091643 | 0.05 | 0.15 | 1 | 14 |
| ehd2b | 0.42918 | 0.279679445 | 0.3 | 0.2 | 1 | 14 |
| clta | 0.42995 | -0.2321572 | 0.05 | 0.15 | 1 | 14 |
| vdac1 | 0.43008 | 0.256981302 | 0.3 | 0.2 | 1 | 14 |
| dhrs3b | 0.43077 | -0.27807359 | 0.15 | 0.25 | 1 | 14 |
| dicer1 | 0.43089 | 0.250146685 | 0.15 | 0.05 | 1 | 14 |
| uqcrc2b | 0.4314 | 0.242861003 | 0.2 | 0.1 | 1 | 14 |
| wdr43 | 0.43159 | 0.206672586 | 0.15 | 0.05 | 1 | 14 |
| sdhb | 0.43174 | -0.13522522 | 0.05 | 0.15 | 1 | 14 |
| psmc1b | 0.43205 | -0.202957 | 0.05 | 0.15 | 1 | 14 |
| pcolce2b | 0.43218 | -0.2542805 | 0.55 | 0.7 | 1 | 14 |
| eef1a1l1 | 0.43249 | 0.11935439 | 0.975 | 0.9 | 1 | 14 |
| kpna4 | 0.43297 | -0.22954395 | 0.05 | 0.15 | 1 | 14 |
| asb12a | 0.43317 | -0.19831016 | 0.05 | 0.15 | 1 | 14 |
| sars | 0.43414 | -0.14918484 | 0.05 | 0.15 | 1 | 14 |
| kcmf1 | 0.43454 | -0.18581073 | 0.05 | 0.15 | 1 | 14 |
| znf385d | 0.43454 | 0.225368189 | 0.125 | 0.05 | 1 | 14 |
| zfyve21 | 0.4346 | -0.1629887 | 0.125 | 0.25 | 1 | 14 |
| nsfl1c | 0.43486 | 0.22564751 | 0.175 | 0.1 | 1 | 14 |
| bsg | 0.43513 | -0.26174756 | 0.55 | 0.7 | 1 | 14 |
| sox4a | 0.4354 | 0.210374173 | 0.3 | 0.25 | 1 | 14 |
| git2b | 0.43622 | 0.172340011 | 0.075 | 0.05 | 1 | 14 |
| actn4 | 0.43644 | 0.303345743 | 0.2 | 0.1 | 1 | 14 |
| igbp1 | 0.43674 | -0.1634456 | 0.05 | 0.15 | 1 | 14 |
| nt5c3a | 0.43706 | -0.2269021 | 0.05 | 0.15 | 1 | 14 |
| olfml3b | 0.43761 | -0.21085328 | 0.475 | 0.55 | 1 | 14 |
| rps18 | 0.43785 | 0.1341653 | 0.95 | 0.85 | 1 | 14 |
| si:ch73-138n13.1 | 0.43786 | -0.2181776 | 0.05 | 0.15 | 1 | 14 |
| lhfpl2b | 0.43827 | -0.22410545 | 0.05 | 0.15 | 1 | 14 |
| surf2 | 0.43844 | -0.16465307 | 0.05 | 0.15 | 1 | 14 |
| pitpnbl | 0.43877 | 0.227621282 | 0.15 | 0.05 | 1 | 14 |
| sema3aa | 0.43958 | -0.22727567 | 0.05 | 0.15 | 1 | 14 |
| bmpr2a | 0.43959 | -0.21180219 | 0.05 | 0.15 | 1 | 14 |
| nkain1 | 0.44018 | -0.14384625 | 0.05 | 0.15 | 1 | 14 |
| mafba | 0.44059 | -0.2813391 | 0.175 | 0.3 | 1 | 14 |
| ptpra | 0.4407 | -0.19744745 | 0.05 | 0.15 | 1 | 14 |
| mrps36 | 0.44117 | 0.11834529 | 0.15 | 0.15 | 1 | 14 |
| si:ch211-108c17.2 | 0.44153 | -0.2305883 | 0.05 | 0.15 | 1 | 14 |
| stag2b | 0.44198 | 0.233285779 | 0.15 | 0.05 | 1 | 14 |
| mrpl36 | 0.44286 | 0.253552247 | 0.2 | 0.1 | 1 | 14 |
| itga1 | 0.4429 | -0.2073362 | 0.05 | 0.15 | 1 | 14 |
| si:ch211-194k22.8 | 0.44302 | -0.23400266 | 0.05 | 0.15 | 1 | 14 |
| gchfr | 0.4432 | -0.22070089 | 0.05 | 0.15 | 1 | 14 |
| tnfrsfa | 0.44342 | -0.1636832 | 0.05 | 0.15 | 1 | 14 |
| vcam1b | 0.44373 | 0.254014858 | 0.125 | 0.05 | 1 | 14 |
| ep400 | 0.44399 | -0.19893667 | 0.05 | 0.15 | 1 | 14 |
| pgpep1 | 0.44422 | -0.1950022 | 0.05 | 0.15 | 1 | 14 |

| | | | | | | |
|---|---|---|---|---|---|---|
| zgc:158564 | 0.44424 | 0.121759515 | 0.15 | 0.15 | 1 | 14 |
| rbpms2a | 0.4448 | -0.21341782 | 0.05 | 0.15 | 1 | 14 |
| galnt11 | 0.44514 | -0.20808947 | 0.05 | 0.15 | 1 | 14 |
| bcor | 0.44515 | -0.2472764 | 0.05 | 0.15 | 1 | 14 |
| gnai1 | 0.44525 | -0.16770265 | 0.05 | 0.15 | 1 | 14 |
| mt-co1 | 0.44549 | -0.15622098 | 0.85 | 0.95 | 1 | 14 |
| rab11ba | 0.44568 | 0.227864158 | 0.15 | 0.05 | 1 | 14 |
| rbm39b | 0.44576 | -0.16471857 | 0.05 | 0.15 | 1 | 14 |
| XLOC-037556 | 0.44585 | -0.22216713 | 0.05 | 0.15 | 1 | 14 |
| eif4e1c | 0.44599 | -0.11831497 | 0.05 | 0.15 | 1 | 14 |
| chrdl2 | 0.44611 | -0.21112847 | 0.05 | 0.15 | 1 | 14 |
| sde2 | 0.44612 | -0.15774988 | 0.05 | 0.15 | 1 | 14 |
| srp72 | 0.44694 | -0.1778349 | 0.05 | 0.15 | 1 | 14 |
| decr1 | 0.44727 | 0.22767578 | 0.175 | 0.1 | 1 | 14 |
| rapgef2 | 0.44759 | -0.19318361 | 0.05 | 0.15 | 1 | 14 |
| ednraa | 0.44761 | -0.22570372 | 0.05 | 0.15 | 1 | 14 |
| tmem189 | 0.44773 | -0.17647072 | 0.05 | 0.15 | 1 | 14 |
| pcmt | 0.44794 | -0.16670542 | 0.05 | 0.15 | 1 | 14 |
| atrnl1b | 0.44804 | -0.19420242 | 0.05 | 0.15 | 1 | 14 |
| adka | 0.44811 | -0.16497267 | 0.05 | 0.15 | 1 | 14 |
| kif13ba | 0.44822 | -0.20855974 | 0.05 | 0.15 | 1 | 14 |
| nrp2b | 0.4483 | -0.21324441 | 0.05 | 0.15 | 1 | 14 |
| dspa | 0.4485 | -0.20380327 | 0.05 | 0.15 | 1 | 14 |
| rab1bb | 0.44851 | -0.16788021 | 0.05 | 0.15 | 1 | 14 |
| abcf2a | 0.44853 | -0.16916828 | 0.05 | 0.15 | 1 | 14 |
| fam20b | 0.44853 | -0.17506205 | 0.05 | 0.15 | 1 | 14 |
| larp1 | 0.44854 | -0.18910592 | 0.05 | 0.15 | 1 | 14 |
| sptlc2a | 0.44854 | -0.17193448 | 0.05 | 0.15 | 1 | 14 |
| fblim1 | 0.44945 | 0.214714291 | 0.15 | 0.05 | 1 | 14 |
| pak2b | 0.44975 | 0.15823554 | 0.175 | 0.15 | 1 | 14 |
| uqcrh | 0.44989 | 0.15786128 | 0.375 | 0.35 | 1 | 14 |
| sh3pxd2b | 0.45043 | 0.241065118 | 0.15 | 0.05 | 1 | 14 |
| pdia6 | 0.45059 | -0.27381049 | 0.2 | 0.35 | 1 | 14 |
| commd5 | 0.45203 | 0.227298414 | 0.125 | 0.05 | 1 | 14 |
| ddx3xb | 0.45235 | 0.238135942 | 0.15 | 0.05 | 1 | 14 |
| si:dkey-16l2.20 | 0.45336 | -0.17518101 | 0.175 | 0.3 | 1 | 14 |
| capns1a | 0.45444 | 0.18443906 | 0.35 | 0.3 | 1 | 14 |
| ube2ia | 0.45465 | 0.165701406 | 0.175 | 0.15 | 1 | 14 |
| flot2a | 0.45472 | -0.27450425 | 0.3 | 0.45 | 1 | 14 |
| si:dkey-250d21.1 | 0.45514 | 0.238182508 | 0.15 | 0.05 | 1 | 14 |
| tmem176 | 0.4562 | 0.219192066 | 0.375 | 0.3 | 1 | 14 |
| irf2bp2a | 0.45682 | -0.1323824 | 0.275 | 0.4 | 1 | 14 |
| gskip | 0.45703 | 0.263119916 | 0.225 | 0.1 | 1 | 14 |
| tmf1 | 0.45783 | 0.221459152 | 0.125 | 0.05 | 1 | 14 |
| laptm4a | 0.45785 | -0.21732447 | 0.25 | 0.4 | 1 | 14 |
| morf4l1 | 0.45845 | -0.22902886 | 0.2 | 0.35 | 1 | 14 |
| pabpc1a | 0.46004 | 0.18868539 | 0.825 | 0.75 | 1 | 14 |
| aldh9a1a.1 | 0.46029 | -0.11546462 | 0.275 | 0.4 | 1 | 14 |

| | | | | | | |
|---|---|---|---|---|---|---|
| smndc1 | 0.46096 | -0.24053135 | 0.2 | 0.35 | 1 | 14 |
| bbip1 | 0.4611 | -0.22645313 | 0.1 | 0.2 | 1 | 14 |
| commd7 | 0.4617 | 0.15985298 | 0.125 | 0.1 | 1 | 14 |
| kank4 | 0.46214 | -0.26639567 | 0.075 | 0.15 | 1 | 14 |
| dlst | 0.46342 | 0.183740494 | 0.25 | 0.2 | 1 | 14 |
| appb | 0.46519 | 0.301254223 | 0.225 | 0.1 | 1 | 14 |
| CABZ01058650.1 | 0.46526 | -0.20532774 | 0.125 | 0.25 | 1 | 14 |
| pbx1a | 0.46613 | 0.270533155 | 0.225 | 0.1 | 1 | 14 |
| tpd52l2b | 0.46643 | -0.16170452 | 0.175 | 0.3 | 1 | 14 |
| pafah1b1a | 0.46669 | 0.223251582 | 0.225 | 0.15 | 1 | 14 |
| hnrnpa1a | 0.46729 | 0.264881923 | 0.225 | 0.1 | 1 | 14 |
| puf60a | 0.46754 | -0.17312072 | 0.1 | 0.2 | 1 | 14 |
| eef1b2 | 0.46851 | 0.179606252 | 0.9 | 0.8 | 1 | 14 |
| mdh1aa | 0.46877 | 0.258053797 | 0.325 | 0.25 | 1 | 14 |
| lig1 | 0.46965 | 0.247033057 | 0.15 | 0.05 | 1 | 14 |
| tmeff1b | 0.46988 | 0.244892035 | 0.35 | 0.2 | 1 | 14 |
| impad1 | 0.47043 | 0.224966012 | 0.15 | 0.05 | 1 | 14 |
| snrpc | 0.47045 | 0.162554794 | 0.15 | 0.05 | 1 | 14 |
| selenoh | 0.47058 | 0.258514512 | 0.3 | 0.2 | 1 | 14 |
| rbm4.1 | 0.47059 | -0.1815523 | 0.175 | 0.3 | 1 | 14 |
| ylpm1 | 0.47108 | -0.19858537 | 0.125 | 0.25 | 1 | 14 |
| olfml3a | 0.47151 | 0.229308117 | 0.15 | 0.05 | 1 | 14 |
| si:ch211-210c8.7 | 0.47174 | 0.219326447 | 0.125 | 0.05 | 1 | 14 |
| rpp25l | 0.47197 | 0.207623603 | 0.1 | 0.05 | 1 | 14 |
| plod1a | 0.47234 | 0.260103051 | 0.225 | 0.1 | 1 | 14 |
| rbm28 | 0.47244 | 0.226491088 | 0.225 | 0.1 | 1 | 14 |
| ndufb5 | 0.47276 | 0.210593837 | 0.325 | 0.25 | 1 | 14 |
| jun | 0.4729 | -0.20995764 | 0.65 | 0.8 | 1 | 14 |
| si:ch211-288g17.3 | 0.47318 | 0.253151126 | 0.4 | 0.3 | 1 | 14 |
| hspa4b | 0.47422 | 0.171931118 | 0.15 | 0.05 | 1 | 14 |
| tmed2 | 0.47453 | 0.113931742 | 0.35 | 0.35 | 1 | 14 |
| sgta | 0.47461 | 0.244270358 | 0.225 | 0.15 | 1 | 14 |
| klf6a | 0.47541 | -0.14300109 | 0.525 | 0.55 | 1 | 14 |
| pphln1 | 0.47565 | 0.1664278 | 0.125 | 0.1 | 1 | 14 |
| selenoj | 0.4761 | 0.16919897 | 0.125 | 0.05 | 1 | 14 |
| cdh2 | 0.47639 | -0.23795956 | 0.125 | 0.25 | 1 | 14 |
| ndufa9a | 0.47648 | -0.19034698 | 0.125 | 0.25 | 1 | 14 |
| acadm | 0.47652 | 0.1825675 | 0.35 | 0.3 | 1 | 14 |
| ctsba | 0.47668 | -0.1287921 | 0.225 | 0.35 | 1 | 14 |
| mcrip1 | 0.47686 | 0.191230481 | 0.15 | 0.05 | 1 | 14 |
| cmpk | 0.47811 | 0.18851214 | 0.125 | 0.05 | 1 | 14 |
| srsf11 | 0.47821 | 0.188369406 | 0.25 | 0.2 | 1 | 14 |
| vps26a | 0.4783 | 0.206580144 | 0.125 | 0.05 | 1 | 14 |
| znf1035 | 0.47934 | 0.106560642 | 0.2 | 0.2 | 1 | 14 |
| ppfibp1b | 0.47956 | -0.20750874 | 0.125 | 0.25 | 1 | 14 |
| psmc4 | 0.48017 | -0.19283973 | 0.125 | 0.25 | 1 | 14 |
| pfdn6 | 0.48058 | 0.230545007 | 0.15 | 0.05 | 1 | 14 |
| wu:fu71h07 | 0.48068 | -0.23351327 | 0.075 | 0.15 | 1 | 14 |

| | | | | | | |
|---|---|---|---|---|---|---|
| eif4bb | 0.48104 | 0.213674351 | 0.15 | 0.1 | 1 | 14 |
| serinc1 | 0.48113 | 0.103360045 | 0.25 | 0.25 | 1 | 14 |
| siah1 | 0.48116 | 0.195756039 | 0.15 | 0.05 | 1 | 14 |
| cox4i2 | 0.48133 | 0.235480323 | 0.15 | 0.05 | 1 | 14 |
| rnf185 | 0.48151 | -0.23595192 | 0.125 | 0.25 | 1 | 14 |
| fn1a | 0.48189 | -0.24209058 | 0.3 | 0.45 | 1 | 14 |
| neo1a | 0.48292 | 0.195224355 | 0.15 | 0.05 | 1 | 14 |
| gmfb | 0.48381 | 0.179132116 | 0.15 | 0.1 | 1 | 14 |
| slc25a33 | 0.48398 | -0.22977891 | 0.1 | 0.2 | 1 | 14 |
| cdv3 | 0.48431 | -0.24441292 | 0.225 | 0.35 | 1 | 14 |
| arhgdig | 0.48446 | 0.175535505 | 0.15 | 0.05 | 1 | 14 |
| cct4 | 0.48452 | -0.12561322 | 0.275 | 0.4 | 1 | 14 |
| phax | 0.48471 | 0.150498785 | 0.075 | 0.05 | 1 | 14 |
| pitx2 | 0.48529 | 0.291469183 | 0.275 | 0.15 | 1 | 14 |
| btbd7 | 0.48565 | 0.20769033 | 0.15 | 0.05 | 1 | 14 |
| XLOC-042917 | 0.48569 | -0.17768926 | 0.225 | 0.35 | 1 | 14 |
| si:ch211-218c6.8 | 0.48571 | 0.233563017 | 0.125 | 0.05 | 1 | 14 |
| hmgn2 | 0.48579 | 0.206747801 | 0.775 | 0.75 | 1 | 14 |
| hnrnpub | 0.48588 | 0.216207465 | 0.15 | 0.05 | 1 | 14 |
| ptenb | 0.48588 | 0.22833185 | 0.15 | 0.05 | 1 | 14 |
| fcf1 | 0.48592 | 0.2072189 | 0.15 | 0.05 | 1 | 14 |
| actl6a | 0.48611 | 0.135230799 | 0.1 | 0.1 | 1 | 14 |
| mif | 0.48647 | -0.16121921 | 0.4 | 0.45 | 1 | 14 |
| foxo4 | 0.48685 | 0.217227764 | 0.1 | 0.05 | 1 | 14 |
| smim7 | 0.48691 | -0.13927441 | 0.1 | 0.2 | 1 | 14 |
| scamp2 | 0.48702 | 0.221625891 | 0.15 | 0.05 | 1 | 14 |
| zc3h7bb | 0.48703 | 0.212176646 | 0.15 | 0.05 | 1 | 14 |
| cdc42bpb | 0.48703 | 0.210534047 | 0.15 | 0.05 | 1 | 14 |
| si:dkey-180p18.9 | 0.48732 | -0.10213371 | 0.25 | 0.25 | 1 | 14 |
| XLOC-040108 | 0.48822 | -0.11440687 | 0.1 | 0.1 | 1 | 14 |
| atp5if1a | 0.48837 | 0.25862589 | 0.5 | 0.35 | 1 | 14 |
| znf362b | 0.4887 | 0.198244159 | 0.1 | 0.05 | 1 | 14 |
| hmga1a | 0.49006 | 0.226799638 | 0.45 | 0.35 | 1 | 14 |
| leng8 | 0.49055 | -0.23697365 | 0.125 | 0.25 | 1 | 14 |
| elovl1b | 0.49058 | -0.22102762 | 0.125 | 0.25 | 1 | 14 |
| tnfsf10 | 0.49059 | -0.25175899 | 0.125 | 0.25 | 1 | 14 |
| efemp2a | 0.49104 | -0.1377794 | 0.15 | 0.25 | 1 | 14 |
| si:ch73-376l24.2 | 0.49122 | -0.18643977 | 0.175 | 0.3 | 1 | 14 |
| ndufab1a | 0.49223 | 0.113922308 | 0.15 | 0.15 | 1 | 14 |
| calcoco1a | 0.49289 | 0.117497008 | 0.125 | 0.05 | 1 | 14 |
| tada3l | 0.49349 | 0.166641129 | 0.125 | 0.05 | 1 | 14 |
| si:ch211-71n6.4 | 0.49424 | -0.13075796 | 0.15 | 0.25 | 1 | 14 |
| pbx4 | 0.4943 | 0.104894484 | 0.2 | 0.2 | 1 | 14 |
| wdr83os | 0.49643 | 0.253862873 | 0.275 | 0.15 | 1 | 14 |
| zfand5a | 0.49724 | 0.208336651 | 0.15 | 0.1 | 1 | 14 |
| eif4g2a | 0.49762 | -0.13047288 | 0.1 | 0.2 | 1 | 14 |
| ghra | 0.49809 | -0.23660007 | 0.075 | 0.15 | 1 | 14 |
| ptk2aa | 0.49845 | -0.23762271 | 0.075 | 0.15 | 1 | 14 |

| | | | | | | |
|---|---|---|---|---|---|---|
| pam16 | 0.49946 | 0.245327813 | 0.2 | 0.1 | 1 | 14 |
| parvaa | 0.50062 | 0.245798907 | 0.25 | 0.15 | 1 | 14 |
| si:ch211-140m22.7 | 0.50069 | 0.232831821 | 0.175 | 0.1 | 1 | 14 |
| palm2 | 0.50135 | 0.160008248 | 0.125 | 0.1 | 1 | 14 |
| rtn3 | 0.50154 | 0.104203611 | 0.15 | 0.15 | 1 | 14 |
| wnt5b | 0.50163 | 0.252296758 | 0.225 | 0.15 | 1 | 14 |
| wls | 0.50192 | -0.15256381 | 0.275 | 0.4 | 1 | 14 |
| cnbpb | 0.50276 | -0.12385526 | 0.325 | 0.45 | 1 | 14 |
| magixa | 0.50302 | -0.23853486 | 0.075 | 0.15 | 1 | 14 |
| hectd1 | 0.50305 | 0.184024659 | 0.15 | 0.1 | 1 | 14 |
| amotl2b | 0.50315 | -0.24923889 | 0.1 | 0.2 | 1 | 14 |
| yipf6 | 0.50457 | -0.15308916 | 0.075 | 0.1 | 1 | 14 |
| si:dkey-282h22.5 | 0.50562 | 0.115605299 | 0.1 | 0.1 | 1 | 14 |
| mir214a | 0.50604 | 0.263295983 | 0.2 | 0.1 | 1 | 14 |
| dcun1d5 | 0.5065 | 0.152022737 | 0.075 | 0.05 | 1 | 14 |
| hexim1 | 0.50731 | -0.14864032 | 0.075 | 0.1 | 1 | 14 |
| puraa | 0.50744 | 0.13319302 | 0.25 | 0.15 | 1 | 14 |
| med19a | 0.50755 | 0.205489925 | 0.125 | 0.05 | 1 | 14 |
| plod3 | 0.50823 | -0.12546595 | 0.2 | 0.3 | 1 | 14 |
| mycb | 0.50838 | -0.10198683 | 0.25 | 0.35 | 1 | 14 |
| kctd5a | 0.50863 | 0.238186265 | 0.275 | 0.2 | 1 | 14 |
| mrps11 | 0.50863 | 0.165330597 | 0.15 | 0.1 | 1 | 14 |
| FP102018.1 | 0.50892 | -0.23799529 | 0.3 | 0.45 | 1 | 14 |
| cpda | 0.50968 | 0.194831268 | 0.1 | 0.05 | 1 | 14 |
| acin1a | 0.50994 | 0.182193371 | 0.45 | 0.4 | 1 | 14 |
| ddx39ab | 0.51069 | 0.185240673 | 0.1 | 0.05 | 1 | 14 |
| si:ch211-235e9.6 | 0.5108 | -0.11627301 | 0.15 | 0.25 | 1 | 14 |
| tra2a | 0.5114 | 0.145379494 | 0.225 | 0.2 | 1 | 14 |
| gas1a | 0.5119 | 0.2289305 | 0.45 | 0.35 | 1 | 14 |
| dnaja2a | 0.51248 | 0.110113154 | 0.2 | 0.2 | 1 | 14 |
| anp32a | 0.51281 | -0.13839237 | 0.275 | 0.4 | 1 | 14 |
| tram2 | 0.51296 | -0.14502454 | 0.1 | 0.2 | 1 | 14 |
| tsc22d3 | 0.51298 | 0.104594753 | 0.2 | 0.2 | 1 | 14 |
| pmp22b | 0.51392 | -0.24773007 | 0.1 | 0.2 | 1 | 14 |
| krt8 | 0.51428 | 0.135199157 | 0.775 | 0.75 | 1 | 14 |
| COX5B | 0.51491 | 0.229495053 | 0.375 | 0.3 | 1 | 14 |
| iqsec1b | 0.515 | 0.239071971 | 0.125 | 0.05 | 1 | 14 |
| rhocb | 0.5159 | 0.18849635 | 0.275 | 0.15 | 1 | 14 |
| crtap | 0.51626 | -0.19677445 | 0.2 | 0.25 | 1 | 14 |
| gorab | 0.51668 | 0.164894527 | 0.075 | 0.05 | 1 | 14 |
| ywhah | 0.51717 | -0.23415569 | 0.25 | 0.35 | 1 | 14 |
| bcas2 | 0.51733 | 0.242334074 | 0.275 | 0.15 | 1 | 14 |
| dnajc9 | 0.5183 | 0.193280037 | 0.1 | 0.05 | 1 | 14 |
| ythdf2 | 0.51831 | 0.178585472 | 0.1 | 0.05 | 1 | 14 |
| gtf2a2 | 0.51874 | -0.12728935 | 0.1 | 0.2 | 1 | 14 |
| efna2a | 0.52092 | 0.197985673 | 0.275 | 0.15 | 1 | 14 |
| zgc:193812 | 0.52151 | 0.203020462 | 0.125 | 0.05 | 1 | 14 |
| kdm6ba | 0.52175 | 0.133442978 | 0.125 | 0.05 | 1 | 14 |

| | | | | | | |
|---|---|---|---|---|---|---|
| mbnl3 | 0.52201 | 0.12565871 | 0.125 | 0.05 | 1 | 14 |
| b2ml | 0.52206 | -0.12717832 | 0.425 | 0.55 | 1 | 14 |
| tsku | 0.52331 | -0.27048935 | 0.2 | 0.3 | 1 | 14 |
| rab1ab | 0.52363 | 0.24454701 | 0.325 | 0.2 | 1 | 14 |
| ier2b | 0.52413 | -0.13213911 | 0.475 | 0.6 | 1 | 14 |
| efna1a | 0.5247 | -0.10215668 | 0.075 | 0.15 | 1 | 14 |
| gstt1b | 0.52524 | 0.247513877 | 0.425 | 0.3 | 1 | 14 |
| unm-hu7910 | 0.52549 | -0.10807624 | 0.075 | 0.15 | 1 | 14 |
| arhgef1b | 0.52601 | 0.141434596 | 0.125 | 0.05 | 1 | 14 |
| ehd1b | 0.52758 | 0.180181295 | 0.125 | 0.05 | 1 | 14 |
| yipf4 | 0.52889 | 0.168512205 | 0.2 | 0.1 | 1 | 14 |
| aamp | 0.52932 | 0.177686144 | 0.125 | 0.05 | 1 | 14 |
| smad2 | 0.52998 | -0.16665247 | 0.1 | 0.2 | 1 | 14 |
| serbp1a | 0.53159 | 0.197069003 | 0.775 | 0.65 | 1 | 14 |
| zgc:173548 | 0.53178 | 0.167793241 | 0.075 | 0.05 | 1 | 14 |
| aif1l | 0.53179 | 0.200010075 | 0.125 | 0.05 | 1 | 14 |
| tmsb | 0.53326 | -0.19897169 | 0.1 | 0.2 | 1 | 14 |
| fbxo9 | 0.5333 | -0.19922771 | 0.075 | 0.15 | 1 | 14 |
| tpt1 | 0.53382 | 0.115118268 | 0.925 | 0.9 | 1 | 14 |
| arpc5b | 0.53406 | 0.20980133 | 0.125 | 0.05 | 1 | 14 |
| gng7 | 0.53421 | -0.24557111 | 0.275 | 0.4 | 1 | 14 |
| drap1 | 0.53496 | -0.22701434 | 0.175 | 0.3 | 1 | 14 |
| tjp1a | 0.53531 | 0.105957893 | 0.175 | 0.1 | 1 | 14 |
| lypla2 | 0.53545 | -0.1196064 | 0.1 | 0.2 | 1 | 14 |
| si:ch211-214j24.10 | 0.53587 | -0.15119883 | 0.1 | 0.2 | 1 | 14 |
| sap18 | 0.53613 | 0.100444582 | 0.2 | 0.2 | 1 | 14 |
| cry1aa | 0.53752 | 0.204329058 | 0.125 | 0.05 | 1 | 14 |
| MCMDC2 | 0.53796 | 0.180246156 | 0.125 | 0.05 | 1 | 14 |
| znf609a | 0.53966 | 0.185769746 | 0.1 | 0.05 | 1 | 14 |
| ndufa4l | 0.53999 | -0.13037987 | 0.725 | 0.85 | 1 | 14 |
| gulp1a | 0.54092 | -0.13433131 | 0.425 | 0.55 | 1 | 14 |
| timm8b | 0.54182 | 0.199200464 | 0.125 | 0.05 | 1 | 14 |
| actb1 | 0.54184 | -0.19648818 | 0.7 | 0.8 | 1 | 14 |
| prkar2aa | 0.54358 | 0.198756817 | 0.175 | 0.1 | 1 | 14 |
| glrx3 | 0.54387 | 0.186052643 | 0.15 | 0.1 | 1 | 14 |
| meox1 | 0.54498 | 0.243235988 | 0.175 | 0.1 | 1 | 14 |
| syncrip | 0.54552 | 0.218391358 | 0.6 | 0.45 | 1 | 14 |
| ywhaba | 0.54573 | 0.231657289 | 0.3 | 0.2 | 1 | 14 |
| myo1ca | 0.54574 | -0.14265735 | 0.15 | 0.25 | 1 | 14 |
| selenot1a | 0.54637 | -0.18434825 | 0.1 | 0.15 | 1 | 14 |
| ccl19a.1 | 0.54665 | 0.306266172 | 0.225 | 0.15 | 1 | 14 |
| ppp2r1ba | 0.54689 | 0.180464346 | 0.2 | 0.1 | 1 | 14 |
| mych | 0.54752 | 0.154366017 | 0.175 | 0.15 | 1 | 14 |
| ctnnbip1 | 0.54816 | 0.189200947 | 0.125 | 0.05 | 1 | 14 |
| clns1a | 0.5482 | 0.15340208 | 0.1 | 0.05 | 1 | 14 |
| supt6h | 0.54856 | -0.25144792 | 0.1 | 0.2 | 1 | 14 |
| atp5mf | 0.54862 | -0.19256813 | 0.25 | 0.3 | 1 | 14 |
| snrpa | 0.54882 | 0.252738442 | 0.2 | 0.1 | 1 | 14 |

| | | | | | | |
|---|---|---|---|---|---|---|
| rnd3b | 0.54895 | 0.167990008 | 0.425 | 0.3 | 1 | 14 |
| igf2b | 0.5499 | 0.100594867 | 0.2 | 0.2 | 1 | 14 |
| mcm5 | 0.55006 | -0.16492849 | 0.1 | 0.2 | 1 | 14 |
| zmat2 | 0.55046 | -0.12505353 | 0.15 | 0.25 | 1 | 14 |
| cnpy4 | 0.55068 | 0.126734651 | 0.275 | 0.25 | 1 | 14 |
| scyl1 | 0.55198 | 0.200674666 | 0.1 | 0.05 | 1 | 14 |
| pon3.2 | 0.55208 | 0.186849254 | 0.1 | 0.05 | 1 | 14 |
| nckap1 | 0.55294 | -0.23844264 | 0.1 | 0.2 | 1 | 14 |
| CABZ01087560.1 | 0.55311 | -0.22706595 | 0.1 | 0.2 | 1 | 14 |
| LOC108190726 | 0.55379 | 0.18516189 | 0.1 | 0.05 | 1 | 14 |
| tmed10 | 0.5538 | -0.20303116 | 0.175 | 0.3 | 1 | 14 |
| mrpl12 | 0.55388 | -0.20025288 | 0.175 | 0.3 | 1 | 14 |
| cebpg | 0.55455 | 0.183235649 | 0.125 | 0.05 | 1 | 14 |
| brd3a | 0.55457 | 0.220450579 | 0.175 | 0.1 | 1 | 14 |
| nme3 | 0.55559 | 0.214043996 | 0.2 | 0.1 | 1 | 14 |
| skib | 0.55637 | 0.179217211 | 0.15 | 0.1 | 1 | 14 |
| sarnp | 0.55644 | 0.131864795 | 0.175 | 0.15 | 1 | 14 |
| zgc:85858 | 0.55695 | 0.207400703 | 0.125 | 0.05 | 1 | 14 |
| LOC108182329 | 0.55741 | 0.145575464 | 0.125 | 0.05 | 1 | 14 |
| utp23 | 0.55767 | 0.171324385 | 0.1 | 0.05 | 1 | 14 |
| tgfbr2b | 0.55771 | -0.18838062 | 0.1 | 0.2 | 1 | 14 |
| rrp15 | 0.55777 | 0.185315923 | 0.15 | 0.1 | 1 | 14 |
| sltm | 0.55787 | -0.14495813 | 0.15 | 0.25 | 1 | 14 |
| anapc11 | 0.55842 | 0.208108833 | 0.175 | 0.1 | 1 | 14 |
| ola1 | 0.55853 | 0.244880995 | 0.3 | 0.2 | 1 | 14 |
| skp1 | 0.55939 | 0.242517097 | 0.375 | 0.25 | 1 | 14 |
| zgc:158463 | 0.5606 | -0.15306938 | 0.225 | 0.35 | 1 | 14 |
| LOC108180269 | 0.56113 | 0.252073403 | 0.2 | 0.1 | 1 | 14 |
| rrp1 | 0.56139 | 0.189537065 | 0.2 | 0.1 | 1 | 14 |
| si:ch211-195b15.8 | 0.56186 | 0.207393227 | 0.125 | 0.05 | 1 | 14 |
| cnppd1 | 0.56188 | 0.142281159 | 0.075 | 0.05 | 1 | 14 |
| slc20a2 | 0.56225 | 0.21463576 | 0.125 | 0.05 | 1 | 14 |
| pik3r3b | 0.56258 | 0.102660964 | 0.1 | 0.1 | 1 | 14 |
| mrps25 | 0.56418 | 0.146189885 | 0.125 | 0.05 | 1 | 14 |
| hprt1 | 0.56454 | -0.1427877 | 0.1 | 0.2 | 1 | 14 |
| nt5c2b | 0.56468 | 0.144533872 | 0.075 | 0.05 | 1 | 14 |
| col18a1a | 0.5652 | -0.22656617 | 0.1 | 0.2 | 1 | 14 |
| alg5 | 0.56551 | -0.15842924 | 0.1 | 0.2 | 1 | 14 |
| taf9 | 0.56578 | -0.12676843 | 0.15 | 0.25 | 1 | 14 |
| s1pr2 | 0.56665 | -0.15533667 | 0.15 | 0.25 | 1 | 14 |
| mpc1 | 0.56696 | -0.14032719 | 0.1 | 0.2 | 1 | 14 |
| srfa | 0.56759 | 0.160588498 | 0.075 | 0.05 | 1 | 14 |
| CR293507.1 | 0.56769 | 0.14076502 | 0.125 | 0.1 | 1 | 14 |
| actr6 | 0.56857 | 0.178205106 | 0.1 | 0.05 | 1 | 14 |
| creb3l2 | 0.5689 | -0.21111165 | 0.1 | 0.2 | 1 | 14 |
| tia1l | 0.5696 | -0.16681606 | 0.1 | 0.2 | 1 | 14 |
| tufm | 0.57016 | 0.158574741 | 0.1 | 0.05 | 1 | 14 |
| kitlga | 0.57033 | -0.18180358 | 0.1 | 0.2 | 1 | 14 |

| | | | | | | |
|---|---|---|---|---|---|---|
| zgc:153293 | 0.57034 | 0.168803127 | 0.1 | 0.05 | 1 | 14 |
| laptm4b | 0.57063 | 0.141346184 | 0.125 | 0.05 | 1 | 14 |
| ddx23 | 0.57074 | 0.126181529 | 0.175 | 0.15 | 1 | 14 |
| trappc5 | 0.57145 | 0.181841949 | 0.125 | 0.05 | 1 | 14 |
| hadhaa | 0.57176 | 0.138729948 | 0.1 | 0.05 | 1 | 14 |
| praf2 | 0.57224 | -0.19716403 | 0.1 | 0.2 | 1 | 14 |
| hm13 | 0.57252 | -0.18912708 | 0.2 | 0.25 | 1 | 14 |
| si:dkeyp-120h9.1 | 0.57276 | 0.177487416 | 0.125 | 0.05 | 1 | 14 |
| proza | 0.57338 | 0.218474385 | 0.2 | 0.1 | 1 | 14 |
| immt | 0.57346 | 0.217435147 | 0.2 | 0.1 | 1 | 14 |
| brd2a | 0.57365 | 0.214278519 | 0.275 | 0.2 | 1 | 14 |
| cd99l2 | 0.5737 | 0.23992908 | 0.325 | 0.2 | 1 | 14 |
| zgc:77849 | 0.57397 | 0.183654041 | 0.125 | 0.05 | 1 | 14 |
| lrp5 | 0.57549 | -0.21401591 | 0.1 | 0.2 | 1 | 14 |
| nova2 | 0.57584 | 0.118237676 | 0.175 | 0.1 | 1 | 14 |
| chd4a | 0.57619 | -0.14694816 | 0.15 | 0.25 | 1 | 14 |
| rhogb | 0.57677 | -0.18831073 | 0.1 | 0.2 | 1 | 14 |
| emc6 | 0.57685 | -0.18247169 | 0.1 | 0.2 | 1 | 14 |
| rpia | 0.57751 | -0.17845272 | 0.1 | 0.2 | 1 | 14 |
| ddah2 | 0.57754 | -0.19987963 | 0.1 | 0.2 | 1 | 14 |
| ecm1b | 0.57799 | -0.22649864 | 0.475 | 0.6 | 1 | 14 |
| mlf2 | 0.57809 | -0.17841137 | 0.1 | 0.2 | 1 | 14 |
| myg1 | 0.5783 | 0.170631191 | 0.1 | 0.05 | 1 | 14 |
| s100a10b | 0.5784 | -0.18553944 | 0.675 | 0.8 | 1 | 14 |
| pex3 | 0.57843 | 0.13380957 | 0.075 | 0.05 | 1 | 14 |
| hopx | 0.57851 | -0.18053612 | 0.1 | 0.2 | 1 | 14 |
| pole4 | 0.57858 | -0.18042815 | 0.1 | 0.2 | 1 | 14 |
| ulk2 | 0.57899 | -0.20305188 | 0.1 | 0.2 | 1 | 14 |
| churc1 | 0.57903 | 0.191208419 | 0.125 | 0.05 | 1 | 14 |
| tomm70a | 0.57927 | -0.2134533 | 0.075 | 0.15 | 1 | 14 |
| cct8 | 0.57937 | 0.222363491 | 0.35 | 0.25 | 1 | 14 |
| slc39a7 | 0.57966 | -0.17354038 | 0.1 | 0.2 | 1 | 14 |
| btbd10b | 0.5797 | 0.188177248 | 0.125 | 0.05 | 1 | 14 |
| spg21 | 0.57995 | -0.15188322 | 0.1 | 0.2 | 1 | 14 |
| fah | 0.58021 | 0.130697456 | 0.125 | 0.05 | 1 | 14 |
| inhbaa | 0.58028 | -0.21288757 | 0.1 | 0.15 | 1 | 14 |
| nkd1 | 0.58041 | -0.18048185 | 0.05 | 0.1 | 1 | 14 |
| btf3 | 0.58059 | -0.17012427 | 0.7 | 0.8 | 1 | 14 |
| ap1ar | 0.58256 | 0.187467947 | 0.125 | 0.05 | 1 | 14 |
| pls3 | 0.58278 | 0.174357549 | 0.1 | 0.05 | 1 | 14 |
| yipf5 | 0.5831 | 0.226755372 | 0.2 | 0.1 | 1 | 14 |
| wtap | 0.58317 | -0.18331034 | 0.075 | 0.15 | 1 | 14 |
| zcchc24 | 0.58426 | 0.183762264 | 0.125 | 0.05 | 1 | 14 |
| eef1db | 0.58449 | 0.232752658 | 0.3 | 0.2 | 1 | 14 |
| npm3 | 0.58549 | 0.167692965 | 0.2 | 0.1 | 1 | 14 |
| timm9 | 0.58579 | 0.136820513 | 0.125 | 0.1 | 1 | 14 |
| crk | 0.586 | -0.17463232 | 0.1 | 0.15 | 1 | 14 |
| rbbp6 | 0.58601 | 0.209678287 | 0.225 | 0.15 | 1 | 14 |

| | | | | | | |
|---|---|---|---|---|---|---|
| hnrnph1l | 0.5867 | 0.157921642 | 0.15 | 0.1 | 1 | 14 |
| slc25a24 | 0.58685 | -0.22292098 | 0.075 | 0.15 | 1 | 14 |
| lxn | 0.58731 | 0.14649008 | 0.275 | 0.25 | 1 | 14 |
| sec61b | 0.58744 | 0.193978318 | 0.725 | 0.6 | 1 | 14 |
| rfxap | 0.58801 | -0.18067982 | 0.05 | 0.1 | 1 | 14 |
| egr1 | 0.58805 | 0.146445797 | 0.125 | 0.05 | 1 | 14 |
| bckdhbl | 0.58821 | -0.18932334 | 0.1 | 0.15 | 1 | 14 |
| tox | 0.58825 | 0.141434755 | 0.125 | 0.1 | 1 | 14 |
| stau2 | 0.58871 | -0.21416144 | 0.15 | 0.25 | 1 | 14 |
| alas1 | 0.58905 | -0.17465079 | 0.05 | 0.1 | 1 | 14 |
| eml2 | 0.59026 | -0.17840732 | 0.05 | 0.1 | 1 | 14 |
| kctd12.2 | 0.59047 | -0.12820272 | 0.2 | 0.3 | 1 | 14 |
| cdc26 | 0.59047 | 0.168872857 | 0.15 | 0.1 | 1 | 14 |
| birc2 | 0.59098 | 0.16997551 | 0.125 | 0.05 | 1 | 14 |
| serp1 | 0.59111 | -0.17214757 | 0.225 | 0.3 | 1 | 14 |
| svep1 | 0.59211 | 0.130647782 | 0.175 | 0.15 | 1 | 14 |
| kdm2ab | 0.59228 | 0.176776676 | 0.1 | 0.05 | 1 | 14 |
| ccnl1a | 0.59253 | -0.13111697 | 0.075 | 0.15 | 1 | 14 |
| glo1 | 0.59319 | -0.22151508 | 0.35 | 0.45 | 1 | 14 |
| rbms2b | 0.5932 | 0.19185911 | 0.175 | 0.1 | 1 | 14 |
| pdcd6ip | 0.59423 | -0.20733247 | 0.075 | 0.15 | 1 | 14 |
| march2 | 0.59443 | 0.177271052 | 0.125 | 0.05 | 1 | 14 |
| polr3f | 0.59482 | 0.161962168 | 0.125 | 0.05 | 1 | 14 |
| tpbgb | 0.5957 | -0.1100015 | 0.075 | 0.15 | 1 | 14 |
| fam177a1 | 0.59584 | 0.173626288 | 0.125 | 0.05 | 1 | 14 |
| sfrp1a | 0.59675 | -0.22148793 | 0.45 | 0.55 | 1 | 14 |
| cox7c | 0.59696 | -0.15606978 | 0.525 | 0.55 | 1 | 14 |
| ndufs2 | 0.59704 | 0.161339958 | 0.1 | 0.05 | 1 | 14 |
| hivep3b | 0.5972 | -0.10097336 | 0.075 | 0.15 | 1 | 14 |
| CU929070.1 | 0.59785 | 0.137048746 | 0.125 | 0.1 | 1 | 14 |
| mibp2 | 0.59789 | -0.15398155 | 0.65 | 0.7 | 1 | 14 |
| dynll2b | 0.59897 | 0.209188635 | 0.175 | 0.1 | 1 | 14 |
| dlc1 | 0.59899 | -0.17134305 | 0.05 | 0.1 | 1 | 14 |
| ccdc187 | 0.59921 | -0.16185561 | 0.1 | 0.15 | 1 | 14 |
| srgap3 | 0.59933 | 0.177604148 | 0.15 | 0.1 | 1 | 14 |
| cspg4 | 0.59935 | 0.246578363 | 0.2 | 0.1 | 1 | 14 |
| sema6e | 0.59948 | -0.10040213 | 0.075 | 0.15 | 1 | 14 |
| scaf11 | 0.59975 | 0.210367213 | 0.125 | 0.05 | 1 | 14 |
| adamts2 | 0.59985 | 0.213650335 | 0.2 | 0.1 | 1 | 14 |
| exosc6 | 0.59989 | -0.20479348 | 0.075 | 0.15 | 1 | 14 |
| ewsr1b | 0.6002 | -0.18885424 | 0.075 | 0.15 | 1 | 14 |
| si:ch211-106h4.12 | 0.60112 | -0.16212718 | 0.075 | 0.15 | 1 | 14 |
| hpgd | 0.60128 | 0.173156456 | 0.125 | 0.05 | 1 | 14 |
| afdna | 0.60173 | -0.13357003 | 0.075 | 0.15 | 1 | 14 |
| pkig | 0.60238 | 0.173688863 | 0.1 | 0.05 | 1 | 14 |
| naa50 | 0.60245 | -0.10403544 | 0.075 | 0.15 | 1 | 14 |
| tefb | 0.60258 | 0.176515496 | 0.15 | 0.1 | 1 | 14 |
| ier2a | 0.6029 | -0.18306454 | 0.4 | 0.45 | 1 | 14 |

| | | | | | | |
|---|---|---|---|---|---|---|
| nat9 | 0.60296 | 0.1550471 | 0.1 | 0.05 | 1 | 14 |
| irf2bpl | 0.60461 | -0.21155015 | 0.125 | 0.2 | 1 | 14 |
| pcdh7b | 0.6052 | -0.17410435 | 0.15 | 0.25 | 1 | 14 |
| tmem9b | 0.60555 | 0.141103633 | 0.075 | 0.05 | 1 | 14 |
| parp12b | 0.6057 | 0.163744716 | 0.125 | 0.05 | 1 | 14 |
| lmnb2 | 0.6064 | -0.16158923 | 0.15 | 0.2 | 1 | 14 |
| cdkn1a | 0.6069 | 0.203312417 | 0.225 | 0.15 | 1 | 14 |
| c1qtnf2 | 0.60812 | -0.10202212 | 0.125 | 0.2 | 1 | 14 |
| her6 | 0.60887 | 0.242581302 | 0.175 | 0.1 | 1 | 14 |
| slco2b1 | 0.60931 | 0.172685403 | 0.125 | 0.05 | 1 | 14 |
| crabp2b | 0.60949 | 0.227482355 | 0.525 | 0.4 | 1 | 14 |
| slc48a1b | 0.60994 | -0.19511338 | 0.075 | 0.15 | 1 | 14 |
| ap4s1 | 0.61034 | 0.148909323 | 0.1 | 0.05 | 1 | 14 |
| si:dkeyp-69b9.6 | 0.6111 | 0.173588043 | 0.125 | 0.05 | 1 | 14 |
| rpl8 | 0.61116 | 0.131369757 | 0.9 | 0.85 | 1 | 14 |
| tnfaip3 | 0.61121 | -0.18810912 | 0.075 | 0.15 | 1 | 14 |
| si:dkey-6n21.12 | 0.61144 | 0.18067744 | 0.15 | 0.1 | 1 | 14 |
| tfam | 0.61192 | -0.10636566 | 0.075 | 0.15 | 1 | 14 |
| sod2 | 0.61332 | 0.124829331 | 0.125 | 0.1 | 1 | 14 |
| ptpa | 0.61345 | 0.166143705 | 0.1 | 0.05 | 1 | 14 |
| yy1a | 0.61355 | 0.173701893 | 0.125 | 0.05 | 1 | 14 |
| st13 | 0.61388 | 0.166064771 | 0.125 | 0.05 | 1 | 14 |
| tor4aa | 0.6139 | -0.12871956 | 0.075 | 0.15 | 1 | 14 |
| gstm.2 | 0.61409 | 0.130342226 | 0.125 | 0.05 | 1 | 14 |
| sav1 | 0.61431 | 0.156498759 | 0.1 | 0.05 | 1 | 14 |
| CDK2AP1 | 0.61541 | 0.185099631 | 0.1 | 0.05 | 1 | 14 |
| dnajb11 | 0.61558 | 0.155313261 | 0.2 | 0.15 | 1 | 14 |
| snrnp25 | 0.61623 | -0.17683514 | 0.075 | 0.15 | 1 | 14 |
| tab1 | 0.61637 | 0.197053702 | 0.125 | 0.05 | 1 | 14 |
| polr3e | 0.61711 | 0.163044615 | 0.125 | 0.05 | 1 | 14 |
| tmem106ba | 0.61752 | -0.19468423 | 0.125 | 0.2 | 1 | 14 |
| fbxl6 | 0.61779 | -0.20999586 | 0.075 | 0.15 | 1 | 14 |
| eif4a3 | 0.61794 | 0.182802142 | 0.225 | 0.15 | 1 | 14 |
| tnrc6b | 0.61896 | -0.12292935 | 0.075 | 0.15 | 1 | 14 |
| psen1 | 0.61922 | -0.13406594 | 0.075 | 0.1 | 1 | 14 |
| acbd3 | 0.61947 | 0.142804688 | 0.1 | 0.05 | 1 | 14 |
| ivd | 0.61977 | 0.171977022 | 0.1 | 0.05 | 1 | 14 |
| dut | 0.62164 | -0.1735199 | 0.05 | 0.1 | 1 | 14 |
| si:dkey-67c22.2 | 0.62173 | -0.12532619 | 0.2 | 0.3 | 1 | 14 |
| cmtm7 | 0.62181 | -0.19018236 | 0.075 | 0.15 | 1 | 14 |
| tpm4a | 0.62202 | -0.18873947 | 0.65 | 0.75 | 1 | 14 |
| pros1 | 0.62218 | 0.17970497 | 0.125 | 0.05 | 1 | 14 |
| tsr2 | 0.62275 | 0.179557725 | 0.125 | 0.05 | 1 | 14 |
| ufm1 | 0.62291 | 0.145059596 | 0.125 | 0.05 | 1 | 14 |
| tspan4b | 0.62349 | -0.21084331 | 0.275 | 0.4 | 1 | 14 |
| tgfbr3 | 0.62362 | -0.21495529 | 0.275 | 0.4 | 1 | 14 |
| AL732488.1 | 0.62442 | 0.15812305 | 0.125 | 0.05 | 1 | 14 |
| nckap5l | 0.62454 | 0.20296601 | 0.1 | 0.05 | 1 | 14 |

| | | | | | | |
|---|---|---|---|---|---|---|
| camk2g1 | 0.62458 | -0.12420372 | 0.075 | 0.15 | 1 | 14 |
| rsl24d1 | 0.62492 | 0.190993486 | 0.525 | 0.45 | 1 | 14 |
| etf1b | 0.62515 | -0.21493625 | 0.25 | 0.35 | 1 | 14 |
| asip1 | 0.62533 | 0.258244657 | 0.425 | 0.3 | 1 | 14 |
| stom | 0.6258 | -0.21982989 | 0.2 | 0.3 | 1 | 14 |
| myl6 | 0.62582 | 0.169514361 | 0.525 | 0.4 | 1 | 14 |
| sdc2 | 0.62601 | 0.224208987 | 0.45 | 0.35 | 1 | 14 |
| zc3h13 | 0.62612 | 0.18824817 | 0.175 | 0.1 | 1 | 14 |
| zbtb37 | 0.62651 | 0.189651971 | 0.15 | 0.1 | 1 | 14 |
| ctnnb2 | 0.62667 | -0.11745595 | 0.25 | 0.35 | 1 | 14 |
| sucla2 | 0.62706 | 0.186882787 | 0.125 | 0.05 | 1 | 14 |
| rad21a | 0.62734 | -0.13609928 | 0.2 | 0.3 | 1 | 14 |
| hnrnpk | 0.62771 | -0.11792275 | 0.2 | 0.3 | 1 | 14 |
| syap1 | 0.62861 | -0.10166742 | 0.075 | 0.15 | 1 | 14 |
| cactin | 0.62866 | 0.134663178 | 0.075 | 0.05 | 1 | 14 |
| lsm14ab | 0.62891 | 0.14480957 | 0.125 | 0.05 | 1 | 14 |
| sdf4 | 0.62894 | 0.170748421 | 0.15 | 0.1 | 1 | 14 |
| atp2b1a | 0.6295 | 0.178211242 | 0.125 | 0.05 | 1 | 14 |
| si:ch211-231f6.6 | 0.63018 | 0.163247567 | 0.125 | 0.05 | 1 | 14 |
| foxp1b | 0.63056 | 0.172254502 | 0.125 | 0.05 | 1 | 14 |
| herc2 | 0.63103 | -0.13478749 | 0.075 | 0.15 | 1 | 14 |
| si:ch211-166a6.5 | 0.63159 | 0.156892551 | 0.125 | 0.05 | 1 | 14 |
| plxnb2a | 0.63163 | 0.156222167 | 0.125 | 0.05 | 1 | 14 |
| mrpl11 | 0.63178 | 0.144014488 | 0.125 | 0.05 | 1 | 14 |
| zgc:162730 | 0.63229 | -0.13202078 | 0.575 | 0.6 | 1 | 14 |
| arf2b | 0.63241 | -0.155515 | 0.15 | 0.25 | 1 | 14 |
| prrx1a | 0.63242 | 0.158684521 | 0.125 | 0.05 | 1 | 14 |
| sap30bp | 0.63254 | 0.122526274 | 0.125 | 0.05 | 1 | 14 |
| ankhd1 | 0.6327 | 0.124240242 | 0.075 | 0.05 | 1 | 14 |
| CU681842.1 | 0.63333 | -0.11155294 | 0.025 | 0.05 | 1 | 14 |
| LOC101885938 | 0.63333 | -0.13659977 | 0.025 | 0.05 | 1 | 14 |
| dnajc5ga | 0.63333 | -0.10811098 | 0.025 | 0.05 | 1 | 14 |
| pdxka | 0.63333 | -0.11064937 | 0.025 | 0.05 | 1 | 14 |
| serpinb14 | 0.63333 | -0.10907433 | 0.025 | 0.05 | 1 | 14 |
| si:ch211-195h23.3 | 0.63333 | -0.11945938 | 0.025 | 0.05 | 1 | 14 |
| slc35b1 | 0.63333 | -0.11227362 | 0.025 | 0.05 | 1 | 14 |
| vps26bl | 0.63333 | -0.11028039 | 0.025 | 0.05 | 1 | 14 |
| LOC110439512 | 0.63333 | -0.10621571 | 0.025 | 0.05 | 1 | 14 |
| ccdc142 | 0.63333 | -0.10520054 | 0.025 | 0.05 | 1 | 14 |
| coasy | 0.63333 | -0.10875934 | 0.025 | 0.05 | 1 | 14 |
| csrnp1b | 0.63333 | -0.12422681 | 0.025 | 0.05 | 1 | 14 |
| cyp27a1.4 | 0.63333 | -0.11920074 | 0.025 | 0.05 | 1 | 14 |
| ecm2 | 0.63333 | -0.13217771 | 0.025 | 0.05 | 1 | 14 |
| exosc4 | 0.63333 | -0.15400104 | 0.025 | 0.05 | 1 | 14 |
| gart | 0.63333 | -0.11006971 | 0.025 | 0.05 | 1 | 14 |
| gga1 | 0.63333 | -0.12308137 | 0.025 | 0.05 | 1 | 14 |
| klhl15 | 0.63333 | -0.11290593 | 0.025 | 0.05 | 1 | 14 |
| med8 | 0.63333 | -0.15146194 | 0.025 | 0.05 | 1 | 14 |

| | | | | | | |
|---|---|---|---|---|---|---|
| pigb | 0.63333 | -0.12295477 | 0.025 | 0.05 | 1 | 14 |
| thy1 | 0.63333 | -0.13129475 | 0.025 | 0.05 | 1 | 14 |
| tmem150aa | 0.63333 | -0.10767929 | 0.025 | 0.05 | 1 | 14 |
| ubp1 | 0.63333 | -0.12280781 | 0.025 | 0.05 | 1 | 14 |
| zgc:174904 | 0.63333 | -0.17021722 | 0.025 | 0.05 | 1 | 14 |
| BX569787.1 | 0.63333 | -0.1297745 | 0.025 | 0.05 | 1 | 14 |
| abi1a | 0.63333 | -0.1116977 | 0.025 | 0.05 | 1 | 14 |
| acad8 | 0.63333 | -0.10433434 | 0.025 | 0.05 | 1 | 14 |
| ankib1b | 0.63333 | -0.10444666 | 0.025 | 0.05 | 1 | 14 |
| arl3 | 0.63333 | -0.10672077 | 0.025 | 0.05 | 1 | 14 |
| atad2b | 0.63333 | -0.10760787 | 0.025 | 0.05 | 1 | 14 |
| chac2 | 0.63333 | -0.10317339 | 0.025 | 0.05 | 1 | 14 |
| efnb1 | 0.63333 | -0.14171408 | 0.025 | 0.05 | 1 | 14 |
| ephx1 | 0.63333 | -0.10800095 | 0.025 | 0.05 | 1 | 14 |
| itgb5 | 0.63333 | -0.13888708 | 0.025 | 0.05 | 1 | 14 |
| kcng3 | 0.63333 | -0.16236673 | 0.025 | 0.05 | 1 | 14 |
| maf1 | 0.63333 | -0.10679802 | 0.025 | 0.05 | 1 | 14 |
| mapk8a | 0.63333 | -0.1048238 | 0.025 | 0.05 | 1 | 14 |
| nphp3 | 0.63333 | -0.11170721 | 0.025 | 0.05 | 1 | 14 |
| pold3 | 0.63333 | -0.12351158 | 0.025 | 0.05 | 1 | 14 |
| scaf4b | 0.63333 | -0.10317146 | 0.025 | 0.05 | 1 | 14 |
| senp6a | 0.63333 | -0.10089249 | 0.025 | 0.05 | 1 | 14 |
| si:ch211-264f5.8 | 0.63333 | -0.11123323 | 0.025 | 0.05 | 1 | 14 |
| slit2 | 0.63333 | -0.1149322 | 0.025 | 0.05 | 1 | 14 |
| srrt | 0.63333 | -0.11476309 | 0.025 | 0.05 | 1 | 14 |
| ugt5b2 | 0.63333 | -0.1369599 | 0.025 | 0.05 | 1 | 14 |
| zgc:123010 | 0.63333 | -0.11015432 | 0.025 | 0.05 | 1 | 14 |
| XLOC-043124 | 0.63333 | -0.10417659 | 0.025 | 0.05 | 1 | 14 |
| cept1a | 0.63333 | -0.10498234 | 0.025 | 0.05 | 1 | 14 |
| stag1a | 0.63333 | -0.10392867 | 0.025 | 0.05 | 1 | 14 |
| cep131 | 0.63333 | -0.11293093 | 0.025 | 0.05 | 1 | 14 |
| fbxw2 | 0.63333 | -0.11717863 | 0.025 | 0.05 | 1 | 14 |
| rrp12 | 0.63333 | -0.1208721 | 0.025 | 0.05 | 1 | 14 |
| mtmr11 | 0.63335 | 0.176621409 | 0.1 | 0.05 | 1 | 14 |
| nuak1a | 0.63366 | -0.15370139 | 0.15 | 0.25 | 1 | 14 |
| tlr5b | 0.63394 | -0.13089282 | 0.075 | 0.1 | 1 | 14 |
| amotl2a | 0.63421 | -0.1297968 | 0.075 | 0.15 | 1 | 14 |
| cstf2 | 0.63433 | 0.167247501 | 0.125 | 0.05 | 1 | 14 |
| mrps16 | 0.63477 | 0.143973092 | 0.125 | 0.05 | 1 | 14 |
| zgc:92606 | 0.63565 | 0.15069059 | 0.125 | 0.05 | 1 | 14 |
| rps5 | 0.63587 | 0.10326713 | 0.95 | 0.9 | 1 | 14 |
| sult2st1 | 0.63591 | 0.134170748 | 0.1 | 0.05 | 1 | 14 |
| hvcn1 | 0.63646 | 0.136694485 | 0.125 | 0.1 | 1 | 14 |
| mcl1a | 0.63677 | 0.200842923 | 0.6 | 0.5 | 1 | 14 |
| egln1b | 0.6368 | 0.126947823 | 0.075 | 0.05 | 1 | 14 |
| nudcd2 | 0.63789 | 0.143134462 | 0.15 | 0.1 | 1 | 14 |
| hnrnpa0b | 0.63821 | 0.154389449 | 0.625 | 0.5 | 1 | 14 |
| chmp2a | 0.63833 | -0.19280446 | 0.15 | 0.25 | 1 | 14 |

| | | | | | | |
|---|---|---|---|---|---|---|
| selenof | 0.6385 | -0.16368715 | 0.575 | 0.7 | 1 | 14 |
| arhgap10 | 0.63916 | -0.12186405 | 0.075 | 0.15 | 1 | 14 |
| purab | 0.63988 | -0.10041996 | 0.125 | 0.2 | 1 | 14 |
| ube2e3 | 0.64 | 0.159106479 | 0.3 | 0.25 | 1 | 14 |
| srpx2 | 0.64018 | -0.17873409 | 0.15 | 0.25 | 1 | 14 |
| cct3 | 0.64022 | 0.201681758 | 0.35 | 0.25 | 1 | 14 |
| snx17 | 0.64081 | 0.166810629 | 0.2 | 0.15 | 1 | 14 |
| ndufb9 | 0.64138 | -0.19247515 | 0.3 | 0.4 | 1 | 14 |
| rrp9 | 0.6414 | 0.165069501 | 0.1 | 0.05 | 1 | 14 |
| lmcd1 | 0.64208 | -0.11906528 | 0.075 | 0.15 | 1 | 14 |
| nhsl2 | 0.64259 | -0.14609985 | 0.075 | 0.15 | 1 | 14 |
| XLOC-018082 | 0.64277 | 0.141425902 | 0.075 | 0.05 | 1 | 14 |
| psma3 | 0.64286 | 0.197724998 | 0.35 | 0.25 | 1 | 14 |
| ms4a17a.11 | 0.64394 | -0.20567067 | 0.15 | 0.25 | 1 | 14 |
| si:ch211-212k18.8 | 0.644 | -0.16068069 | 0.15 | 0.2 | 1 | 14 |
| si:ch73-308l14.2 | 0.64412 | 0.178789748 | 0.175 | 0.1 | 1 | 14 |
| hibadhb | 0.64419 | -0.16015962 | 0.075 | 0.15 | 1 | 14 |
| si:ch211-284f22.3 | 0.6445 | -0.19893487 | 0.075 | 0.15 | 1 | 14 |
| cox5aa | 0.64459 | 0.123765361 | 0.25 | 0.25 | 1 | 14 |
| si:ch211-196g2.4 | 0.6448 | 0.150834805 | 0.1 | 0.05 | 1 | 14 |
| nf2b | 0.64559 | -0.11745165 | 0.075 | 0.15 | 1 | 14 |
| lman1 | 0.64639 | -0.18336645 | 0.075 | 0.15 | 1 | 14 |
| atp6v1h | 0.64683 | -0.11543001 | 0.075 | 0.15 | 1 | 14 |
| eif3f | 0.64687 | 0.151357364 | 0.625 | 0.5 | 1 | 14 |
| smurf2 | 0.64712 | -0.18057929 | 0.075 | 0.15 | 1 | 14 |
| meis2b | 0.64719 | 0.126312889 | 0.425 | 0.4 | 1 | 14 |
| enah | 0.64827 | -0.10129524 | 0.075 | 0.15 | 1 | 14 |
| helz | 0.64884 | -0.20528675 | 0.075 | 0.15 | 1 | 14 |
| mkln1 | 0.64902 | 0.118847946 | 0.075 | 0.05 | 1 | 14 |
| selenom | 0.64902 | -0.14675725 | 0.2 | 0.3 | 1 | 14 |
| cltca | 0.64974 | -0.18843196 | 0.15 | 0.25 | 1 | 14 |
| hsp70.1 | 0.64993 | -0.16023836 | 0.075 | 0.15 | 1 | 14 |
| spop | 0.65028 | 0.169391779 | 0.1 | 0.05 | 1 | 14 |
| CABZ01068366.1 | 0.6503 | -0.19571201 | 0.15 | 0.25 | 1 | 14 |
| pbdc1 | 0.65039 | 0.187613465 | 0.25 | 0.15 | 1 | 14 |
| nono | 0.65047 | -0.16484965 | 0.15 | 0.25 | 1 | 14 |
| zgc:85777 | 0.65086 | -0.15431282 | 0.075 | 0.15 | 1 | 14 |
| pnn | 0.65103 | 0.118652653 | 0.175 | 0.1 | 1 | 14 |
| rnf8 | 0.6511 | 0.159700237 | 0.1 | 0.05 | 1 | 14 |
| ube2d3 | 0.65134 | 0.16499611 | 0.1 | 0.05 | 1 | 14 |
| atp5meb | 0.65146 | 0.168679442 | 0.475 | 0.35 | 1 | 14 |
| jpt2 | 0.65184 | -0.16127306 | 0.15 | 0.25 | 1 | 14 |
| tia1 | 0.65196 | -0.12077791 | 0.075 | 0.15 | 1 | 14 |
| mrps10 | 0.65208 | -0.16894482 | 0.15 | 0.25 | 1 | 14 |
| copa | 0.65216 | -0.17058447 | 0.15 | 0.25 | 1 | 14 |
| dab2ipb | 0.65254 | 0.14102861 | 0.075 | 0.05 | 1 | 14 |
| oaz1a | 0.65309 | 0.155698955 | 0.75 | 0.65 | 1 | 14 |
| msna | 0.65315 | -0.19301864 | 0.375 | 0.5 | 1 | 14 |

| | | | | | | |
|---|---|---|---|---|---|---|
| acvr1l | 0.65321 | -0.20235378 | 0.125 | 0.2 | 1 | 14 |
| rpl7 | 0.65383 | 0.129687356 | 0.8 | 0.75 | 1 | 14 |
| rgmd | 0.65445 | -0.15200574 | 0.075 | 0.15 | 1 | 14 |
| rab5c | 0.6545 | 0.171890834 | 0.25 | 0.15 | 1 | 14 |
| dctn5 | 0.65497 | -0.11818423 | 0.075 | 0.15 | 1 | 14 |
| cluap1 | 0.65563 | 0.119711195 | 0.075 | 0.05 | 1 | 14 |
| zgc:66388 | 0.65628 | 0.135334033 | 0.125 | 0.1 | 1 | 14 |
| fn1b | 0.65724 | -0.10720441 | 0.125 | 0.2 | 1 | 14 |
| twist3 | 0.65771 | -0.13527005 | 0.075 | 0.15 | 1 | 14 |
| tp53 | 0.65778 | 0.150399899 | 0.2 | 0.15 | 1 | 14 |
| znf385a | 0.65792 | -0.21770774 | 0.25 | 0.35 | 1 | 14 |
| fam32a | 0.65801 | 0.16765392 | 0.525 | 0.4 | 1 | 14 |
| pet100 | 0.65804 | 0.137300654 | 0.125 | 0.1 | 1 | 14 |
| cavin2b | 0.65823 | -0.19304578 | 0.425 | 0.55 | 1 | 14 |
| nhp2 | 0.65826 | 0.202050861 | 0.3 | 0.2 | 1 | 14 |
| psmd3 | 0.65829 | -0.20381264 | 0.125 | 0.2 | 1 | 14 |
| atrx | 0.65883 | 0.107526837 | 0.375 | 0.35 | 1 | 14 |
| pcbp2 | 0.6589 | -0.16051448 | 0.075 | 0.15 | 1 | 14 |
| map3k14a | 0.65972 | 0.163937371 | 0.1 | 0.05 | 1 | 14 |
| rhebl1 | 0.65981 | -0.17400995 | 0.075 | 0.15 | 1 | 14 |
| rca2.1 | 0.66105 | -0.17914932 | 0.075 | 0.15 | 1 | 14 |
| tmem126a | 0.66161 | 0.149512202 | 0.1 | 0.05 | 1 | 14 |
| aco2 | 0.6623 | -0.1770251 | 0.075 | 0.15 | 1 | 14 |
| atg13 | 0.66239 | 0.173539084 | 0.1 | 0.05 | 1 | 14 |
| pdzrn3b | 0.66314 | -0.14196852 | 0.075 | 0.15 | 1 | 14 |
| vstm4a | 0.66358 | -0.14707776 | 0.075 | 0.15 | 1 | 14 |
| elavl1a | 0.66361 | -0.14195179 | 0.2 | 0.3 | 1 | 14 |
| tardbpl | 0.66396 | 0.191762988 | 0.25 | 0.15 | 1 | 14 |
| cnot9 | 0.66408 | -0.12699568 | 0.075 | 0.15 | 1 | 14 |
| nxpe3 | 0.66421 | 0.122647262 | 0.075 | 0.05 | 1 | 14 |
| fstb | 0.66458 | 0.164270348 | 0.1 | 0.05 | 1 | 14 |
| mbd3a | 0.66503 | 0.164511082 | 0.15 | 0.1 | 1 | 14 |
| ldb1a | 0.66505 | -0.14778969 | 0.075 | 0.15 | 1 | 14 |
| mark3a | 0.66532 | 0.127975708 | 0.075 | 0.05 | 1 | 14 |
| apmap | 0.66636 | -0.12577415 | 0.075 | 0.15 | 1 | 14 |
| zc3h15 | 0.66643 | 0.12014428 | 0.1 | 0.05 | 1 | 14 |
| hirip3 | 0.66648 | 0.126301244 | 0.125 | 0.1 | 1 | 14 |
| rhogd | 0.66692 | -0.15529021 | 0.075 | 0.15 | 1 | 14 |
| hmgb2a | 0.66696 | 0.154888491 | 0.6 | 0.55 | 1 | 14 |
| tbc1d10aa | 0.66704 | -0.14691935 | 0.075 | 0.15 | 1 | 14 |
| p3h4 | 0.66707 | -0.19744287 | 0.075 | 0.15 | 1 | 14 |
| wfdc2 | 0.66737 | -0.1734436 | 0.65 | 0.75 | 1 | 14 |
| sept15 | 0.6675 | -0.14019921 | 0.075 | 0.15 | 1 | 14 |
| coq8a | 0.66791 | -0.10548245 | 0.075 | 0.15 | 1 | 14 |
| rrs1 | 0.66808 | -0.17699628 | 0.075 | 0.15 | 1 | 14 |
| hic1 | 0.66854 | 0.142622198 | 0.4 | 0.35 | 1 | 14 |
| hoxc9a | 0.6687 | -0.17833047 | 0.125 | 0.2 | 1 | 14 |
| nucb2a | 0.66877 | -0.19464326 | 0.2 | 0.3 | 1 | 14 |

| | | | | | | |
|---|---|---|---|---|---|---|
| zranb2 | 0.66923 | -0.13789022 | 0.075 | 0.15 | 1 | 14 |
| pes | 0.66972 | -0.14486406 | 0.075 | 0.15 | 1 | 14 |
| dohh | 0.66978 | 0.16079087 | 0.15 | 0.1 | 1 | 14 |
| creb1a | 0.67006 | -0.13819383 | 0.075 | 0.15 | 1 | 14 |
| aimp1 | 0.67043 | 0.153721447 | 0.2 | 0.15 | 1 | 14 |
| gnsa | 0.67044 | 0.151541179 | 0.15 | 0.1 | 1 | 14 |
| c1galt1b | 0.67066 | -0.15242448 | 0.075 | 0.15 | 1 | 14 |
| foxc1a | 0.67081 | 0.153114393 | 0.1 | 0.05 | 1 | 14 |
| nek7 | 0.67148 | -0.10803754 | 0.075 | 0.15 | 1 | 14 |
| rerea | 0.67155 | 0.151467141 | 0.175 | 0.1 | 1 | 14 |
| txndc15 | 0.67172 | -0.13070115 | 0.075 | 0.15 | 1 | 14 |
| mrpl17 | 0.67179 | -0.16528062 | 0.05 | 0.1 | 1 | 14 |
| asah1b | 0.67215 | -0.12422168 | 0.075 | 0.15 | 1 | 14 |
| tex264a | 0.67226 | 0.128908885 | 0.075 | 0.05 | 1 | 14 |
| nfyc | 0.67238 | -0.14964885 | 0.075 | 0.15 | 1 | 14 |
| thoc2 | 0.67245 | -0.11996466 | 0.075 | 0.15 | 1 | 14 |
| atp5f1e | 0.6725 | 0.100672612 | 0.6 | 0.6 | 1 | 14 |
| glyctk | 0.67265 | 0.122245602 | 0.075 | 0.05 | 1 | 14 |
| farsa | 0.67292 | -0.17126365 | 0.05 | 0.1 | 1 | 14 |
| cbfa2t3 | 0.6741 | -0.16989563 | 0.075 | 0.15 | 1 | 14 |
| tdp1 | 0.67417 | 0.124159632 | 0.075 | 0.05 | 1 | 14 |
| txndc17 | 0.67468 | -0.1110737 | 0.075 | 0.15 | 1 | 14 |
| tmtc1 | 0.67499 | -0.18436991 | 0.075 | 0.15 | 1 | 14 |
| ska2 | 0.67503 | -0.14455842 | 0.075 | 0.15 | 1 | 14 |
| sulf1 | 0.67508 | -0.16451303 | 0.075 | 0.15 | 1 | 14 |
| gigyf1b | 0.67528 | -0.14735266 | 0.075 | 0.15 | 1 | 14 |
| gtpbp4 | 0.67529 | -0.1438095 | 0.2 | 0.3 | 1 | 14 |
| sinhcafl | 0.67536 | -0.13136947 | 0.075 | 0.15 | 1 | 14 |
| cul4b | 0.6755 | -0.13959759 | 0.075 | 0.15 | 1 | 14 |
| stx4 | 0.67578 | -0.14724833 | 0.075 | 0.15 | 1 | 14 |
| rn7sk | 0.67591 | -0.16413977 | 0.075 | 0.15 | 1 | 14 |
| olfml1 | 0.6762 | -0.16911092 | 0.075 | 0.15 | 1 | 14 |
| gnptg | 0.6763 | -0.15833085 | 0.05 | 0.1 | 1 | 14 |
| eif4ebp1 | 0.67644 | -0.10577019 | 0.25 | 0.35 | 1 | 14 |
| elmsan1b | 0.67659 | -0.13183955 | 0.075 | 0.15 | 1 | 14 |
| ddr2a | 0.67666 | -0.17500024 | 0.075 | 0.15 | 1 | 14 |
| mrps21 | 0.67716 | 0.114036591 | 0.075 | 0.05 | 1 | 14 |
| zfand5b | 0.67725 | 0.136937623 | 0.125 | 0.1 | 1 | 14 |
| thbs1a | 0.67759 | 0.135724514 | 0.075 | 0.05 | 1 | 14 |
| gsta.1 | 0.67771 | -0.19212746 | 0.2 | 0.3 | 1 | 14 |
| slc3a2b | 0.67816 | 0.136816098 | 0.175 | 0.1 | 1 | 14 |
| arpin | 0.67897 | 0.16856344 | 0.1 | 0.05 | 1 | 14 |
| ammecr1 | 0.68044 | 0.121788047 | 0.175 | 0.15 | 1 | 14 |
| prkd2 | 0.6807 | 0.155678806 | 0.1 | 0.05 | 1 | 14 |
| rgl1 | 0.68163 | -0.12270431 | 0.075 | 0.1 | 1 | 14 |
| sap30l | 0.68215 | -0.14354747 | 0.2 | 0.3 | 1 | 14 |
| ciartb | 0.68274 | -0.17567614 | 0.05 | 0.1 | 1 | 14 |
| ckba | 0.68294 | -0.20267162 | 0.4 | 0.5 | 1 | 14 |

| | | | | | | |
|---|---|---|---|---|---|---|
| tcf20 | 0.68294 | 0.145820891 | 0.1 | 0.05 | 1 | 14 |
| snx14 | 0.68334 | -0.1244422 | 0.075 | 0.1 | 1 | 14 |
| kmt5aa | 0.68372 | 0.113865008 | 0.175 | 0.15 | 1 | 14 |
| glud1a | 0.68429 | 0.150837797 | 0.25 | 0.2 | 1 | 14 |
| cygb1 | 0.68488 | -0.1812816 | 0.2 | 0.3 | 1 | 14 |
| usp1 | 0.68615 | 0.141193066 | 0.1 | 0.05 | 1 | 14 |
| uqcrfs1 | 0.68697 | -0.17776299 | 0.175 | 0.25 | 1 | 14 |
| ppap2d | 0.68781 | -0.20022349 | 0.125 | 0.2 | 1 | 14 |
| lims1 | 0.68884 | 0.193290487 | 0.225 | 0.15 | 1 | 14 |
| aldh6a1 | 0.68943 | -0.15080704 | 0.125 | 0.2 | 1 | 14 |
| irf2bp2b | 0.68955 | 0.160116862 | 0.25 | 0.2 | 1 | 14 |
| arl6ip4 | 0.68974 | 0.11875346 | 0.075 | 0.05 | 1 | 14 |
| tmem119b | 0.69018 | 0.189984332 | 0.225 | 0.15 | 1 | 14 |
| mphosph10 | 0.69057 | 0.136367722 | 0.1 | 0.05 | 1 | 14 |
| rps27.2 | 0.69132 | 0.120181795 | 0.8 | 0.75 | 1 | 14 |
| aldoaa | 0.69174 | -0.20058447 | 0.2 | 0.3 | 1 | 14 |
| ptk7a | 0.69197 | 0.142630129 | 0.1 | 0.05 | 1 | 14 |
| NDUFB1 | 0.6921 | 0.137345978 | 0.4 | 0.35 | 1 | 14 |
| colec12 | 0.69259 | 0.151284851 | 0.3 | 0.25 | 1 | 14 |
| add1 | 0.69301 | 0.162173746 | 0.25 | 0.2 | 1 | 14 |
| ptprk | 0.69375 | -0.18726602 | 0.2 | 0.3 | 1 | 14 |
| appbp2 | 0.69417 | 0.174697506 | 0.1 | 0.05 | 1 | 14 |
| hipk2 | 0.69443 | 0.11892288 | 0.075 | 0.05 | 1 | 14 |
| tomm22 | 0.69499 | -0.16337352 | 0.125 | 0.2 | 1 | 14 |
| fap | 0.6952 | 0.15912679 | 0.1 | 0.05 | 1 | 14 |
| tcf12 | 0.69526 | -0.19070511 | 0.2 | 0.3 | 1 | 14 |
| MED9 | 0.69623 | 0.146934513 | 0.1 | 0.05 | 1 | 14 |
| zyx | 0.69628 | 0.200505243 | 0.35 | 0.25 | 1 | 14 |
| insig2 | 0.69629 | -0.15985669 | 0.05 | 0.1 | 1 | 14 |
| csnk2a1 | 0.69648 | -0.17490741 | 0.2 | 0.3 | 1 | 14 |
| cdc42ep4b | 0.69697 | 0.125674546 | 0.075 | 0.05 | 1 | 14 |
| ralab | 0.69707 | -0.15728886 | 0.05 | 0.1 | 1 | 14 |
| ccdc47 | 0.69731 | -0.14528381 | 0.05 | 0.1 | 1 | 14 |
| rmdn1 | 0.69917 | 0.151865586 | 0.175 | 0.1 | 1 | 14 |
| LO018340.1 | 0.70008 | -0.11074666 | 0.175 | 0.25 | 1 | 14 |
| wu:fb55g09 | 0.70091 | 0.187123113 | 0.3 | 0.2 | 1 | 14 |
| ifrd1 | 0.70118 | 0.144504672 | 0.15 | 0.1 | 1 | 14 |
| polr2f | 0.7012 | -0.10618333 | 0.125 | 0.2 | 1 | 14 |
| tspan14 | 0.70127 | 0.124168311 | 0.325 | 0.3 | 1 | 14 |
| sec61a1 | 0.70173 | 0.182147931 | 0.325 | 0.25 | 1 | 14 |
| fkbp1aa | 0.70241 | 0.117281663 | 0.75 | 0.7 | 1 | 14 |
| KIF2A | 0.70284 | -0.13920972 | 0.05 | 0.1 | 1 | 14 |
| nkap | 0.70325 | 0.109567088 | 0.125 | 0.1 | 1 | 14 |
| PDPR | 0.70331 | 0.128504152 | 0.175 | 0.1 | 1 | 14 |
| pnrc2 | 0.70387 | -0.12037344 | 0.7 | 0.8 | 1 | 14 |
| rab1aa | 0.70512 | -0.11407308 | 0.125 | 0.15 | 1 | 14 |
| bhlhe41 | 0.70513 | -0.16833008 | 0.1 | 0.15 | 1 | 14 |
| wdfy1 | 0.70583 | 0.132960498 | 0.075 | 0.05 | 1 | 14 |

| | | | | | | |
|---|---|---|---|---|---|---|
| CT025690.1 | 0.7062 | -0.16919111 | 0.05 | 0.1 | 1 | 14 |
| ctgfa | 0.70644 | 0.132096436 | 0.45 | 0.35 | 1 | 14 |
| wdr81 | 0.70662 | 0.143362245 | 0.1 | 0.05 | 1 | 14 |
| CABZ01113192.1 | 0.70663 | 0.164803922 | 0.1 | 0.05 | 1 | 14 |
| hadh | 0.70703 | 0.159516604 | 0.175 | 0.1 | 1 | 14 |
| gpat3 | 0.7071 | 0.124235355 | 0.075 | 0.05 | 1 | 14 |
| ccnd1 | 0.70809 | 0.107997503 | 0.175 | 0.15 | 1 | 14 |
| si:ch211-132b12.7 | 0.70858 | -0.17839338 | 0.125 | 0.2 | 1 | 14 |
| c1qbp | 0.70876 | 0.121723773 | 0.175 | 0.1 | 1 | 14 |
| hoxc8a | 0.70884 | -0.11392339 | 0.225 | 0.25 | 1 | 14 |
| si:dkeyp-75h12.5 | 0.70927 | 0.191380494 | 0.175 | 0.1 | 1 | 14 |
| kdm4b | 0.70968 | 0.139995885 | 0.1 | 0.05 | 1 | 14 |
| LEPROTL1 | 0.71085 | 0.131570855 | 0.1 | 0.05 | 1 | 14 |
| ccnt1 | 0.71118 | 0.143846506 | 0.1 | 0.05 | 1 | 14 |
| fxyd6l | 0.7113 | 0.16054449 | 0.35 | 0.25 | 1 | 14 |
| foxo1a | 0.71152 | -0.16381682 | 0.1 | 0.15 | 1 | 14 |
| si:ch211-216l23.2 | 0.7123 | 0.149205407 | 0.1 | 0.05 | 1 | 14 |
| fbxo25 | 0.71286 | 0.144928634 | 0.1 | 0.05 | 1 | 14 |
| cd164 | 0.71329 | -0.17722444 | 0.125 | 0.2 | 1 | 14 |
| zfp36l1b | 0.71333 | -0.19189554 | 0.225 | 0.3 | 1 | 14 |
| si:ch1073-224n8.1 | 0.71365 | 0.142157158 | 0.1 | 0.05 | 1 | 14 |
| tgfb2 | 0.71383 | -0.12169191 | 0.125 | 0.2 | 1 | 14 |
| myl12.2 | 0.7139 | 0.111224558 | 0.075 | 0.05 | 1 | 14 |
| rab30 | 0.71429 | -0.12875245 | 0.05 | 0.1 | 1 | 14 |
| cdh6 | 0.71518 | 0.126911732 | 0.125 | 0.1 | 1 | 14 |
| ciarta | 0.71535 | 0.161121995 | 0.25 | 0.2 | 1 | 14 |
| slc12a7b | 0.71844 | -0.13207608 | 0.05 | 0.1 | 1 | 14 |
| ufc1 | 0.71898 | 0.105688196 | 0.125 | 0.1 | 1 | 14 |
| fam204a | 0.71959 | 0.110165151 | 0.075 | 0.05 | 1 | 14 |
| rps27.1 | 0.72016 | 0.115119421 | 0.9 | 0.85 | 1 | 14 |
| cmip | 0.72037 | -0.11612512 | 0.075 | 0.1 | 1 | 14 |
| socs3a | 0.7205 | -0.17819489 | 0.525 | 0.6 | 1 | 14 |
| BX640522.2 | 0.72085 | -0.18749589 | 0.4 | 0.45 | 1 | 14 |
| usp22 | 0.72113 | 0.138063789 | 0.1 | 0.05 | 1 | 14 |
| rab5ab | 0.72119 | 0.16300916 | 0.15 | 0.1 | 1 | 14 |
| zgc:101858 | 0.72121 | 0.124959274 | 0.1 | 0.05 | 1 | 14 |
| fam84a | 0.7214 | 0.108436607 | 0.075 | 0.05 | 1 | 14 |
| slc30a1a | 0.72235 | -0.13980627 | 0.05 | 0.1 | 1 | 14 |
| col12a1b | 0.72256 | -0.12136145 | 0.425 | 0.45 | 1 | 14 |
| ctsk | 0.72304 | -0.18094046 | 0.25 | 0.35 | 1 | 14 |
| mycbp | 0.72328 | 0.170324516 | 0.175 | 0.1 | 1 | 14 |
| atrn | 0.72342 | -0.17331183 | 0.125 | 0.2 | 1 | 14 |
| ccnd2a | 0.72407 | 0.184355204 | 0.275 | 0.2 | 1 | 14 |
| tmem26a | 0.72439 | 0.134889419 | 0.1 | 0.05 | 1 | 14 |
| surf4l | 0.72441 | -0.11048196 | 0.175 | 0.25 | 1 | 14 |
| fxyd1 | 0.72454 | 0.185565412 | 0.175 | 0.1 | 1 | 14 |
| sc:d0202 | 0.7248 | 0.121798709 | 0.225 | 0.15 | 1 | 14 |
| zmiz1a | 0.72618 | 0.110803611 | 0.125 | 0.1 | 1 | 14 |

| | | | | | | |
|---|---|---|---|---|---|---|
| slc25a55a | 0.72636 | 0.158070359 | 0.1 | 0.05 | 1 | 14 |
| golgb1 | 0.72643 | 0.149216795 | 0.1 | 0.05 | 1 | 14 |
| commd1 | 0.72644 | 0.114387115 | 0.075 | 0.05 | 1 | 14 |
| zgc:100829 | 0.72708 | 0.117075277 | 0.1 | 0.05 | 1 | 14 |
| prpf40a | 0.72805 | 0.164543825 | 0.1 | 0.05 | 1 | 14 |
| shoc2 | 0.72854 | 0.110092502 | 0.175 | 0.15 | 1 | 14 |
| atf4a | 0.72956 | -0.10570415 | 0.125 | 0.2 | 1 | 14 |
| gstr | 0.72966 | -0.14910906 | 0.425 | 0.5 | 1 | 14 |
| zgc:158343 | 0.73028 | 0.188690488 | 0.35 | 0.25 | 1 | 14 |
| si:dkey-222f8.3 | 0.73061 | 0.132460115 | 0.175 | 0.1 | 1 | 14 |
| nop2 | 0.73086 | 0.132961882 | 0.15 | 0.1 | 1 | 14 |
| cst14b.1 | 0.73087 | -0.14811138 | 0.1 | 0.15 | 1 | 14 |
| gstm.3 | 0.73118 | -0.19850696 | 0.125 | 0.2 | 1 | 14 |
| ddx21 | 0.73125 | -0.12234218 | 0.125 | 0.2 | 1 | 14 |
| adgrl2a | 0.73168 | 0.152424594 | 0.15 | 0.1 | 1 | 14 |
| nucb1 | 0.7318 | 0.112669139 | 0.1 | 0.05 | 1 | 14 |
| os9 | 0.73207 | -0.13728624 | 0.05 | 0.1 | 1 | 14 |
| mycla | 0.73243 | 0.141889391 | 0.1 | 0.05 | 1 | 14 |
| prelid3b | 0.73263 | -0.1086338 | 0.125 | 0.15 | 1 | 14 |
| ccdc43 | 0.73282 | 0.114766248 | 0.1 | 0.05 | 1 | 14 |
| srpx | 0.73285 | 0.163574727 | 0.175 | 0.1 | 1 | 14 |
| mbnl1 | 0.73423 | 0.143301536 | 0.175 | 0.1 | 1 | 14 |
| wnt16 | 0.73424 | 0.162085353 | 0.175 | 0.1 | 1 | 14 |
| cyb5r1 | 0.73431 | 0.143991967 | 0.15 | 0.1 | 1 | 14 |
| rhogc | 0.73525 | 0.14625405 | 0.1 | 0.05 | 1 | 14 |
| sfrp2 | 0.73554 | -0.15224019 | 0.35 | 0.45 | 1 | 14 |
| aurkaip1 | 0.73606 | 0.113965701 | 0.1 | 0.05 | 1 | 14 |
| slc35d1a | 0.73645 | -0.1303396 | 0.05 | 0.1 | 1 | 14 |
| cpne1 | 0.73645 | -0.10283074 | 0.175 | 0.25 | 1 | 14 |
| psenen | 0.7366 | 0.168293614 | 0.225 | 0.15 | 1 | 14 |
| mpc2 | 0.73692 | 0.133894915 | 0.15 | 0.1 | 1 | 14 |
| bcl11aa | 0.73725 | 0.16043826 | 0.25 | 0.2 | 1 | 14 |
| si:ch211-74m13.3 | 0.73809 | -0.16021773 | 0.05 | 0.1 | 1 | 14 |
| bsdc1 | 0.73891 | -0.1067578 | 0.125 | 0.2 | 1 | 14 |
| polr1d | 0.73983 | 0.113908696 | 0.125 | 0.1 | 1 | 14 |
| arl6ip1 | 0.73985 | -0.14470163 | 0.1 | 0.15 | 1 | 14 |
| tsr1 | 0.73989 | -0.14227696 | 0.05 | 0.1 | 1 | 14 |
| fbxo11a | 0.74046 | 0.109836012 | 0.075 | 0.05 | 1 | 14 |
| kars | 0.74081 | 0.102969434 | 0.275 | 0.2 | 1 | 14 |
| smg7 | 0.74082 | -0.13126839 | 0.05 | 0.1 | 1 | 14 |
| mtus1b | 0.74095 | -0.12638213 | 0.05 | 0.1 | 1 | 14 |
| ndufb3 | 0.7413 | 0.166478765 | 0.375 | 0.3 | 1 | 14 |
| si:dkey-9i23.16 | 0.74223 | -0.13436201 | 0.125 | 0.2 | 1 | 14 |
| gapdhs | 0.74281 | -0.15453594 | 0.375 | 0.45 | 1 | 14 |
| zgc:103697 | 0.74341 | 0.101835235 | 0.075 | 0.05 | 1 | 14 |
| gng2 | 0.74356 | 0.142720392 | 0.1 | 0.05 | 1 | 14 |
| arhgap5 | 0.74389 | 0.122356321 | 0.1 | 0.05 | 1 | 14 |
| rel | 0.74396 | 0.119021604 | 0.1 | 0.05 | 1 | 14 |

| | | | | | | |
|---|---|---|---|---|---|---|
| dnaja2b | 0.74398 | 0.113669165 | 0.175 | 0.15 | 1 | 14 |
| naa25 | 0.74403 | 0.114949209 | 0.075 | 0.05 | 1 | 14 |
| cbx5 | 0.74406 | -0.15115048 | 0.3 | 0.4 | 1 | 14 |
| c1qtnf5 | 0.74482 | -0.15596933 | 0.125 | 0.2 | 1 | 14 |
| cdc42ep2 | 0.74485 | -0.10118793 | 0.05 | 0.1 | 1 | 14 |
| mphosph6 | 0.74509 | 0.114542661 | 0.1 | 0.05 | 1 | 14 |
| plekhg5a | 0.74574 | 0.100466829 | 0.075 | 0.05 | 1 | 14 |
| krt18b | 0.74583 | -0.12452303 | 0.6 | 0.7 | 1 | 14 |
| nck1a | 0.74584 | -0.15579528 | 0.1 | 0.15 | 1 | 14 |
| ssr2 | 0.74585 | -0.12629239 | 0.6 | 0.7 | 1 | 14 |
| prdx3 | 0.74591 | -0.14321526 | 0.125 | 0.2 | 1 | 14 |
| ebna1bp2 | 0.74608 | -0.10696658 | 0.125 | 0.2 | 1 | 14 |
| ldha | 0.74609 | -0.1377207 | 0.05 | 0.1 | 1 | 14 |
| kdelc2 | 0.74648 | 0.127098511 | 0.1 | 0.05 | 1 | 14 |
| acsl1b | 0.74659 | 0.109321956 | 0.075 | 0.05 | 1 | 14 |
| col15a1a | 0.74692 | -0.14475557 | 0.125 | 0.2 | 1 | 14 |
| mak16 | 0.74696 | 0.145234111 | 0.1 | 0.05 | 1 | 14 |
| badb | 0.74737 | -0.12232043 | 0.05 | 0.1 | 1 | 14 |
| brms1 | 0.74753 | -0.11696458 | 0.05 | 0.1 | 1 | 14 |
| calm3b | 0.74755 | 0.156657242 | 0.4 | 0.3 | 1 | 14 |
| megf8 | 0.74755 | 0.137199663 | 0.1 | 0.05 | 1 | 14 |
| rab6bb | 0.74758 | 0.115216261 | 0.075 | 0.05 | 1 | 14 |
| srgap2 | 0.7478 | -0.1313834 | 0.05 | 0.1 | 1 | 14 |
| adnp2b | 0.74821 | 0.116534668 | 0.075 | 0.05 | 1 | 14 |
| LOC103909972 | 0.74853 | -0.10766526 | 0.125 | 0.2 | 1 | 14 |
| tra2b | 0.74875 | 0.110476105 | 0.1 | 0.05 | 1 | 14 |
| ptbp1b | 0.74885 | -0.15464388 | 0.175 | 0.25 | 1 | 14 |
| cs | 0.74937 | -0.14407443 | 0.125 | 0.2 | 1 | 14 |
| krt94 | 0.74966 | -0.13514869 | 0.45 | 0.55 | 1 | 14 |
| si:rp71-1g18.1 | 0.74991 | 0.107078464 | 0.075 | 0.05 | 1 | 14 |
| ripor2 | 0.74993 | 0.137183648 | 0.1 | 0.05 | 1 | 14 |
| smco4 | 0.74999 | -0.13450387 | 0.125 | 0.2 | 1 | 14 |
| bcl2a | 0.75045 | -0.13855993 | 0.125 | 0.2 | 1 | 14 |
| wbp1 | 0.75066 | -0.14486765 | 0.05 | 0.1 | 1 | 14 |
| htatip2 | 0.75084 | 0.142023903 | 0.1 | 0.05 | 1 | 14 |
| nudcd3 | 0.75091 | 0.104675103 | 0.075 | 0.05 | 1 | 14 |
| cd82b | 0.75121 | -0.1527957 | 0.125 | 0.2 | 1 | 14 |
| atp5f1d | 0.75128 | 0.164237676 | 0.375 | 0.3 | 1 | 14 |
| prrc1 | 0.75136 | 0.113891231 | 0.075 | 0.05 | 1 | 14 |
| blmh | 0.75145 | -0.10143508 | 0.125 | 0.15 | 1 | 14 |
| phf3 | 0.75151 | -0.12976705 | 0.125 | 0.2 | 1 | 14 |
| rhoq | 0.75164 | -0.10541646 | 0.05 | 0.1 | 1 | 14 |
| atad5a | 0.75169 | 0.110393415 | 0.075 | 0.05 | 1 | 14 |
| si:dkey-57k2.7 | 0.75249 | -0.12900993 | 0.05 | 0.1 | 1 | 14 |
| ubfd1 | 0.75354 | 0.11672942 | 0.1 | 0.05 | 1 | 14 |
| zgc:153018 | 0.75372 | -0.14693969 | 0.125 | 0.2 | 1 | 14 |
| itsn1 | 0.75408 | 0.115075801 | 0.075 | 0.05 | 1 | 14 |
| zgc:163080 | 0.75433 | -0.12798413 | 0.125 | 0.2 | 1 | 14 |

| | | | | | | |
|---|---|---|---|---|---|---|
| sgce | 0.75435 | -0.14933786 | 0.125 | 0.2 | 1 | 14 |
| ankrd11 | 0.75442 | -0.14536707 | 0.125 | 0.2 | 1 | 14 |
| si:ch73-211e3.1 | 0.75462 | -0.1027781 | 0.05 | 0.1 | 1 | 14 |
| ptpn11a | 0.75523 | -0.14391451 | 0.05 | 0.1 | 1 | 14 |
| ext1a | 0.75547 | -0.14027888 | 0.05 | 0.1 | 1 | 14 |
| ctsa | 0.75564 | 0.166359197 | 0.225 | 0.15 | 1 | 14 |
| foxn3 | 0.75575 | 0.135185657 | 0.1 | 0.05 | 1 | 14 |
| gnav1 | 0.75608 | 0.104433567 | 0.075 | 0.05 | 1 | 14 |
| ubtf | 0.75654 | 0.147024078 | 0.15 | 0.1 | 1 | 14 |
| arhgap23a | 0.75669 | 0.126439537 | 0.1 | 0.05 | 1 | 14 |
| col5a3a | 0.75701 | -0.10569673 | 0.05 | 0.1 | 1 | 14 |
| cst3 | 0.75704 | -0.1481399 | 0.05 | 0.1 | 1 | 14 |
| ilk | 0.75719 | -0.11165698 | 0.175 | 0.25 | 1 | 14 |
| ufl1 | 0.75747 | -0.10528533 | 0.05 | 0.1 | 1 | 14 |
| BX927333.1 | 0.75759 | -0.1441547 | 0.05 | 0.1 | 1 | 14 |
| sfxn5b | 0.75796 | -0.1466073 | 0.05 | 0.1 | 1 | 14 |
| ube2v1 | 0.75816 | 0.121717097 | 0.5 | 0.45 | 1 | 14 |
| si:ch211-130m23.5 | 0.7585 | -0.14186954 | 0.05 | 0.1 | 1 | 14 |
| eif5a2 | 0.75866 | 0.105114521 | 0.575 | 0.55 | 1 | 14 |
| marcksa | 0.75932 | 0.117998882 | 0.225 | 0.2 | 1 | 14 |
| brd4 | 0.75982 | 0.109904184 | 0.125 | 0.1 | 1 | 14 |
| fbln2 | 0.76006 | 0.115048753 | 0.275 | 0.25 | 1 | 14 |
| ppp2r2d | 0.76026 | -0.12087905 | 0.05 | 0.1 | 1 | 14 |
| slc7a6os | 0.7616 | 0.143608061 | 0.1 | 0.05 | 1 | 14 |
| sntb2 | 0.76226 | 0.107429652 | 0.125 | 0.1 | 1 | 14 |
| tbxas1 | 0.76232 | -0.12926356 | 0.05 | 0.1 | 1 | 14 |
| btd | 0.76292 | 0.129745129 | 0.1 | 0.05 | 1 | 14 |
| sh3pxd2ab | 0.76338 | -0.13430007 | 0.05 | 0.1 | 1 | 14 |
| cdon | 0.76408 | 0.11867096 | 0.65 | 0.6 | 1 | 14 |
| si:ch211-197h24.9 | 0.76456 | 0.130914168 | 0.15 | 0.1 | 1 | 14 |
| col6a1 | 0.76466 | -0.14973978 | 0.525 | 0.6 | 1 | 14 |
| prpf38a | 0.76472 | 0.11874913 | 0.1 | 0.05 | 1 | 14 |
| psmd6 | 0.76478 | 0.141806131 | 0.15 | 0.1 | 1 | 14 |
| sp4 | 0.76522 | 0.148480934 | 0.1 | 0.05 | 1 | 14 |
| fkbp9 | 0.76555 | -0.16622027 | 0.325 | 0.4 | 1 | 14 |
| strn4 | 0.7656 | 0.119822959 | 0.075 | 0.05 | 1 | 14 |
| si:ch211-22i13.2 | 0.7656 | 0.129525741 | 0.15 | 0.1 | 1 | 14 |
| tead1b | 0.76579 | 0.136796708 | 0.225 | 0.15 | 1 | 14 |
| fbxw7 | 0.76631 | -0.13440592 | 0.05 | 0.1 | 1 | 14 |
| ttl | 0.76646 | -0.10107518 | 0.05 | 0.1 | 1 | 14 |
| si:ch211-57n23.1 | 0.76656 | -0.13156497 | 0.05 | 0.1 | 1 | 14 |
| fgfr1a | 0.76668 | -0.13350099 | 0.05 | 0.1 | 1 | 14 |
| akirin2 | 0.76669 | 0.108803631 | 0.1 | 0.05 | 1 | 14 |
| cyp2p7 | 0.76705 | -0.13068889 | 0.05 | 0.1 | 1 | 14 |
| dynll1 | 0.76707 | -0.12470155 | 0.55 | 0.6 | 1 | 14 |
| wrb | 0.76822 | 0.105620323 | 0.075 | 0.05 | 1 | 14 |
| rlim | 0.76828 | -0.11417165 | 0.05 | 0.1 | 1 | 14 |
| klf8 | 0.76844 | -0.10968655 | 0.05 | 0.1 | 1 | 14 |

| | | | | | | |
|---|---|---|---|---|---|---|
| si:dkey-238c7.16 | 0.76852 | 0.157349133 | 0.525 | 0.45 | 1 | 14 |
| zgc:154058 | 0.76878 | -0.10617009 | 0.05 | 0.1 | 1 | 14 |
| slc25a11 | 0.76883 | 0.108526833 | 0.1 | 0.05 | 1 | 14 |
| npr2 | 0.76894 | -0.13976523 | 0.05 | 0.1 | 1 | 14 |
| timm10 | 0.76895 | -0.13593834 | 0.1 | 0.15 | 1 | 14 |
| rnf121 | 0.76913 | -0.10363558 | 0.05 | 0.1 | 1 | 14 |
| si:ch211-37e10.2 | 0.76915 | -0.10166367 | 0.05 | 0.1 | 1 | 14 |
| cutc | 0.76924 | 0.13153024 | 0.1 | 0.05 | 1 | 14 |
| ddx39aa | 0.76931 | -0.14255936 | 0.175 | 0.25 | 1 | 14 |
| zgc:193541 | 0.76933 | 0.152244183 | 0.475 | 0.4 | 1 | 14 |
| fgfr2 | 0.76986 | -0.14569941 | 0.1 | 0.15 | 1 | 14 |
| drg1 | 0.77009 | 0.124065372 | 0.1 | 0.05 | 1 | 14 |
| wdr45 | 0.77018 | 0.103344483 | 0.125 | 0.1 | 1 | 14 |
| psmd11b | 0.77054 | 0.137724278 | 0.1 | 0.05 | 1 | 14 |
| npnt | 0.77064 | 0.120555202 | 0.1 | 0.05 | 1 | 14 |
| FO082781.1 | 0.77097 | -0.16036146 | 0.175 | 0.25 | 1 | 14 |
| gtf2f2a | 0.77181 | 0.128487815 | 0.1 | 0.05 | 1 | 14 |
| si:dkey-85k7.7 | 0.77223 | -0.14003506 | 0.05 | 0.1 | 1 | 14 |
| si:ch211-253b8.2 | 0.77236 | -0.13645512 | 0.05 | 0.1 | 1 | 14 |
| lrig1 | 0.77279 | -0.10881052 | 0.05 | 0.1 | 1 | 14 |
| ugdh | 0.77285 | -0.11658758 | 0.05 | 0.1 | 1 | 14 |
| fzd8a | 0.77287 | 0.115824509 | 0.075 | 0.05 | 1 | 14 |
| klf13 | 0.77385 | -0.11416887 | 0.075 | 0.1 | 1 | 14 |
| arglu1b | 0.77399 | -0.12280045 | 0.075 | 0.1 | 1 | 14 |
| loxa | 0.77399 | -0.17060624 | 0.275 | 0.35 | 1 | 14 |
| rdh14b | 0.77443 | -0.11106779 | 0.05 | 0.1 | 1 | 14 |
| acvr2aa | 0.77472 | -0.10286894 | 0.05 | 0.1 | 1 | 14 |
| cd248b | 0.77492 | -0.18749475 | 0.175 | 0.25 | 1 | 14 |
| dpp9 | 0.77516 | -0.10506359 | 0.05 | 0.1 | 1 | 14 |
| ttyh2 | 0.77518 | -0.12146604 | 0.05 | 0.1 | 1 | 14 |
| pum3 | 0.77529 | 0.136958108 | 0.1 | 0.05 | 1 | 14 |
| arcn1b | 0.77571 | -0.10297082 | 0.05 | 0.1 | 1 | 14 |
| lclat1 | 0.77596 | -0.12209512 | 0.05 | 0.1 | 1 | 14 |
| hoxa3a | 0.77601 | -0.10950909 | 0.05 | 0.1 | 1 | 14 |
| phldb2a | 0.77644 | 0.112703908 | 0.075 | 0.05 | 1 | 14 |
| slc36a4 | 0.77724 | -0.10911242 | 0.05 | 0.1 | 1 | 14 |
| per3 | 0.77729 | -0.15840699 | 0.175 | 0.25 | 1 | 14 |
| rab27a | 0.77742 | 0.106444932 | 0.1 | 0.05 | 1 | 14 |
| CU695232.1 | 0.77761 | -0.1255706 | 0.05 | 0.1 | 1 | 14 |
| arhgef4 | 0.77768 | -0.10981903 | 0.05 | 0.1 | 1 | 14 |
| arpc5lb | 0.77805 | -0.12244979 | 0.05 | 0.1 | 1 | 14 |
| tmem107 | 0.77837 | 0.11784525 | 0.1 | 0.05 | 1 | 14 |
| mtbl | 0.77842 | -0.10718999 | 0.05 | 0.1 | 1 | 14 |
| XLOC-004556 | 0.77852 | -0.13799001 | 0.15 | 0.2 | 1 | 14 |
| luc7l3 | 0.77857 | -0.10688661 | 0.05 | 0.1 | 1 | 14 |
| gabpa | 0.77869 | -0.12105118 | 0.05 | 0.1 | 1 | 14 |
| ece1 | 0.77908 | -0.11609472 | 0.05 | 0.1 | 1 | 14 |
| MDFI | 0.77933 | -0.13432214 | 0.05 | 0.1 | 1 | 14 |

| | | | | | | |
|---|---|---|---|---|---|---|
| col27a1b | 0.77972 | -0.13527063 | 0.05 | 0.1 | 1 | 14 |
| antxr1b | 0.77981 | -0.11598207 | 0.05 | 0.1 | 1 | 14 |
| trim8b | 0.78028 | -0.14296036 | 0.05 | 0.1 | 1 | 14 |
| rrp8 | 0.78034 | 0.101333252 | 0.075 | 0.05 | 1 | 14 |
| rgs4 | 0.78051 | -0.12549333 | 0.05 | 0.1 | 1 | 14 |
| tmem60 | 0.78051 | -0.13880779 | 0.05 | 0.1 | 1 | 14 |
| alg12 | 0.78054 | -0.11221526 | 0.05 | 0.1 | 1 | 14 |
| pex7 | 0.78058 | -0.11369355 | 0.05 | 0.1 | 1 | 14 |
| ncoa5 | 0.78067 | -0.10017004 | 0.05 | 0.1 | 1 | 14 |
| ar | 0.78069 | -0.10052069 | 0.05 | 0.1 | 1 | 14 |
| dusp6 | 0.78069 | -0.10415789 | 0.05 | 0.1 | 1 | 14 |
| zgc:113208 | 0.78098 | 0.113516457 | 0.1 | 0.05 | 1 | 14 |
| mef2d | 0.78157 | 0.132211093 | 0.1 | 0.05 | 1 | 14 |
| adh8b | 0.78161 | 0.150839216 | 0.275 | 0.2 | 1 | 14 |
| adamts1 | 0.78196 | 0.123203813 | 0.075 | 0.05 | 1 | 14 |
| pdcd10a | 0.7821 | 0.116262847 | 0.1 | 0.05 | 1 | 14 |
| mapre1b | 0.78258 | -0.13771447 | 0.15 | 0.2 | 1 | 14 |
| smarca2 | 0.78271 | 0.129369681 | 0.1 | 0.05 | 1 | 14 |
| ctsd | 0.78342 | -0.10619148 | 0.175 | 0.2 | 1 | 14 |
| ankrd52a | 0.78369 | 0.106534916 | 0.1 | 0.05 | 1 | 14 |
| nasp | 0.78376 | 0.132406953 | 0.15 | 0.1 | 1 | 14 |
| pdcd5 | 0.78464 | 0.139251897 | 0.15 | 0.1 | 1 | 14 |
| vapb | 0.78491 | 0.136542655 | 0.225 | 0.15 | 1 | 14 |
| tefa | 0.78508 | 0.12737623 | 0.225 | 0.15 | 1 | 14 |
| rtf1 | 0.78521 | 0.125959832 | 0.15 | 0.1 | 1 | 14 |
| foxp4 | 0.78534 | -0.16109459 | 0.225 | 0.3 | 1 | 14 |
| klf7b | 0.78568 | -0.15473485 | 0.175 | 0.25 | 1 | 14 |
| nxt2 | 0.78573 | 0.102661064 | 0.1 | 0.05 | 1 | 14 |
| nck1b | 0.78578 | 0.126288636 | 0.1 | 0.05 | 1 | 14 |
| dhrs4 | 0.78693 | 0.109731441 | 0.125 | 0.1 | 1 | 14 |
| zgc:175214 | 0.78693 | 0.134617574 | 0.1 | 0.05 | 1 | 14 |
| zgc:77086 | 0.78865 | -0.1021557 | 0.225 | 0.3 | 1 | 14 |
| sh3gl1b | 0.78902 | 0.122943999 | 0.1 | 0.05 | 1 | 14 |
| tcea1 | 0.78984 | 0.104647196 | 0.1 | 0.05 | 1 | 14 |
| si:dkeyp-55f12.3 | 0.79003 | 0.117717077 | 0.1 | 0.05 | 1 | 14 |
| hoxb3a | 0.79023 | 0.126272039 | 0.1 | 0.05 | 1 | 14 |
| akirin1 | 0.7904 | 0.102879397 | 0.1 | 0.05 | 1 | 14 |
| zgc:123105 | 0.79076 | 0.12139625 | 0.1 | 0.05 | 1 | 14 |
| prtfdc1 | 0.79153 | 0.127524145 | 0.1 | 0.05 | 1 | 14 |
| bod1 | 0.79184 | 0.104489863 | 0.1 | 0.05 | 1 | 14 |
| MYADM | 0.79328 | 0.135999843 | 0.1 | 0.05 | 1 | 14 |
| rgma | 0.79342 | 0.116556277 | 0.1 | 0.05 | 1 | 14 |
| snw1 | 0.79391 | 0.101265318 | 0.1 | 0.05 | 1 | 14 |
| phip | 0.79407 | 0.105107157 | 0.1 | 0.05 | 1 | 14 |
| cry1ab | 0.79441 | 0.105710079 | 0.1 | 0.05 | 1 | 14 |
| rabep1 | 0.79445 | 0.106651981 | 0.1 | 0.05 | 1 | 14 |
| flncb | 0.79452 | 0.110437954 | 0.1 | 0.05 | 1 | 14 |
| kirrel1a | 0.7953 | -0.14736522 | 0.175 | 0.25 | 1 | 14 |

| | | | | | | |
|---|---|---|---|---|---|---|
| si:dkey-42i9.4 | 0.7965 | -0.1262435 | 0.175 | 0.25 | 1 | 14 |
| agpat3 | 0.79677 | -0.11232261 | 0.075 | 0.1 | 1 | 14 |
| atp5mc3a | 0.79797 | 0.119127054 | 0.4 | 0.35 | 1 | 14 |
| polr2d | 0.79905 | 0.122602876 | 0.25 | 0.2 | 1 | 14 |
| rpn1 | 0.79963 | 0.103836586 | 0.125 | 0.1 | 1 | 14 |
| BX005309.1 | 0.80079 | 0.106151525 | 0.075 | 0.05 | 1 | 14 |
| chd4b | 0.80098 | -0.13801051 | 0.15 | 0.2 | 1 | 14 |
| AL954322.2 | 0.80221 | 0.132703757 | 0.15 | 0.1 | 1 | 14 |
| b2m | 0.8025 | -0.10634002 | 0.225 | 0.3 | 1 | 14 |
| parn | 0.80361 | 0.101035137 | 0.075 | 0.05 | 1 | 14 |
| eif2s1a | 0.80509 | 0.102292448 | 0.175 | 0.15 | 1 | 14 |
| mmp16a | 0.8055 | 0.139532888 | 0.25 | 0.2 | 1 | 14 |
| si:dkey-51e6.1 | 0.80563 | 0.116030544 | 0.15 | 0.1 | 1 | 14 |
| galnt2 | 0.80659 | 0.1205079 | 0.075 | 0.05 | 1 | 14 |
| si:ch211-246m6.4 | 0.80713 | 0.12578683 | 0.2 | 0.15 | 1 | 14 |
| ctdspl2b | 0.80719 | 0.108728957 | 0.075 | 0.05 | 1 | 14 |
| krt15 | 0.80756 | 0.1434427 | 0.075 | 0.05 | 1 | 14 |
| cope | 0.80773 | -0.10112852 | 0.125 | 0.15 | 1 | 14 |
| kank2 | 0.80793 | 0.123189825 | 0.275 | 0.2 | 1 | 14 |
| hdac4 | 0.808 | 0.101856468 | 0.075 | 0.05 | 1 | 14 |
| sptssa | 0.80925 | 0.122121443 | 0.15 | 0.1 | 1 | 14 |
| abl1 | 0.81214 | 0.132012289 | 0.15 | 0.1 | 1 | 14 |
| rsrc2 | 0.81225 | -0.1025834 | 0.075 | 0.1 | 1 | 14 |
| wnt11r | 0.81492 | -0.15024491 | 0.3 | 0.35 | 1 | 14 |
| atp10a | 0.8162 | -0.1248656 | 0.1 | 0.15 | 1 | 14 |
| utrn | 0.81647 | -0.13429705 | 0.15 | 0.2 | 1 | 14 |
| srrm1 | 0.81665 | -0.1476312 | 0.225 | 0.3 | 1 | 14 |
| pdgfrb | 0.81717 | -0.15398258 | 0.225 | 0.3 | 1 | 14 |
| fzd7b | 0.81838 | -0.13294207 | 0.225 | 0.3 | 1 | 14 |
| zgc:162698 | 0.81902 | -0.10437116 | 0.075 | 0.1 | 1 | 14 |
| psip1b | 0.82078 | 0.135636436 | 0.15 | 0.1 | 1 | 14 |
| cebpd | 0.82267 | 0.153244769 | 0.425 | 0.35 | 1 | 14 |
| ube2l3a | 0.82288 | 0.123726309 | 0.15 | 0.1 | 1 | 14 |
| rpe | 0.82313 | -0.11997846 | 0.1 | 0.15 | 1 | 14 |
| si:dkey-78l4.14 | 0.8233 | -0.11980787 | 0.1 | 0.15 | 1 | 14 |
| raraa | 0.82363 | 0.112617322 | 0.15 | 0.1 | 1 | 14 |
| gpc6a | 0.82388 | -0.12876905 | 0.15 | 0.2 | 1 | 14 |
| ndufb8 | 0.8277 | 0.126521167 | 0.2 | 0.15 | 1 | 14 |
| txnrd3 | 0.8288 | -0.12591134 | 0.1 | 0.15 | 1 | 14 |
| eif3jb | 0.83054 | 0.102316092 | 0.225 | 0.2 | 1 | 14 |
| dnmt3ab | 0.83261 | 0.127870617 | 0.35 | 0.3 | 1 | 14 |
| mbnl2 | 0.83316 | 0.112788803 | 0.125 | 0.1 | 1 | 14 |
| daam1b | 0.8334 | 0.13254136 | 0.15 | 0.1 | 1 | 14 |
| fcho2 | 0.83367 | 0.108196411 | 0.125 | 0.1 | 1 | 14 |
| ctdspl2a | 0.83423 | -0.10585571 | 0.1 | 0.15 | 1 | 14 |
| ctnnd1 | 0.83444 | 0.12689443 | 0.15 | 0.1 | 1 | 14 |
| gpm6ab | 0.83538 | -0.14011812 | 0.1 | 0.15 | 1 | 14 |
| hcfc1a | 0.83818 | 0.130021763 | 0.2 | 0.15 | 1 | 14 |

| | | | | | | |
|---|---|---|---|---|---|---|
| MMP23B | 0.83836 | -0.12558763 | 0.1 | 0.15 | 1 | 14 |
| csde1 | 0.83852 | 0.124691468 | 0.25 | 0.2 | 1 | 14 |
| ndufb7 | 0.83879 | -0.11881264 | 0.275 | 0.35 | 1 | 14 |
| pag1 | 0.83904 | 0.113466514 | 0.15 | 0.1 | 1 | 14 |
| reep5 | 0.84386 | 0.111594079 | 0.375 | 0.3 | 1 | 14 |
| qkia | 0.84465 | -0.11939156 | 0.475 | 0.55 | 1 | 14 |
| hif1al | 0.84561 | 0.123673585 | 0.25 | 0.2 | 1 | 14 |
| api5 | 0.84612 | -0.11565149 | 0.15 | 0.2 | 1 | 14 |
| smim15 | 0.84788 | 0.122046642 | 0.25 | 0.2 | 1 | 14 |
| ube2e1 | 0.85018 | 0.105332925 | 0.15 | 0.1 | 1 | 14 |
| zgc:174863 | 0.85063 | -0.12268299 | 0.1 | 0.15 | 1 | 14 |
| wsb1 | 0.85167 | 0.121437756 | 0.15 | 0.1 | 1 | 14 |
| knop1 | 0.85285 | 0.118038549 | 0.15 | 0.1 | 1 | 14 |
| cemip | 0.85335 | -0.11414755 | 0.175 | 0.2 | 1 | 14 |
| cx43.4 | 0.85337 | -0.11010254 | 0.1 | 0.15 | 1 | 14 |
| arpc2 | 0.85422 | -0.11273663 | 0.375 | 0.45 | 1 | 14 |
| slc43a2a | 0.8544 | 0.106223398 | 0.075 | 0.05 | 1 | 14 |
| hsf2 | 0.85488 | 0.113218699 | 0.15 | 0.1 | 1 | 14 |
| eif1ad | 0.85531 | -0.11135288 | 0.1 | 0.15 | 1 | 14 |
| hcfc1b | 0.85633 | 0.121169084 | 0.15 | 0.1 | 1 | 14 |
| tcf4 | 0.85639 | -0.10775211 | 0.1 | 0.15 | 1 | 14 |
| luzp1 | 0.85641 | -0.10644511 | 0.1 | 0.15 | 1 | 14 |
| bmper | 0.85937 | -0.10171627 | 0.075 | 0.1 | 1 | 14 |
| lsm7 | 0.85995 | 0.113598161 | 0.15 | 0.1 | 1 | 14 |
| zgc:77056 | 0.86056 | 0.11140778 | 0.15 | 0.1 | 1 | 14 |
| snai1a | 0.86073 | 0.106498837 | 0.15 | 0.1 | 1 | 14 |
| brk1 | 0.86174 | 0.113968645 | 0.2 | 0.15 | 1 | 14 |
| dkc1 | 0.86174 | 0.118345779 | 0.2 | 0.15 | 1 | 14 |
| sirt1 | 0.86332 | 0.100680344 | 0.075 | 0.05 | 1 | 14 |
| ccdc102a | 0.86429 | 0.103567558 | 0.075 | 0.05 | 1 | 14 |
| tuba8l4 | 0.86742 | 0.111915862 | 0.55 | 0.5 | 1 | 14 |
| usp10 | 0.86807 | -0.10657161 | 0.15 | 0.2 | 1 | 14 |
| sdha | 0.87275 | 0.116335573 | 0.25 | 0.2 | 1 | 14 |
| LOC103910438 | 0.87277 | 0.123734719 | 0.25 | 0.2 | 1 | 14 |
| cdc42l | 0.87376 | -0.11264 | 0.25 | 0.3 | 1 | 14 |
| snrnp70 | 0.87466 | 0.116217416 | 0.2 | 0.15 | 1 | 14 |
| ppp3r1b | 0.87716 | -0.1044863 | 0.15 | 0.2 | 1 | 14 |
| vim | 0.87776 | -0.11265612 | 0.15 | 0.2 | 1 | 14 |
| gstm.1 | 0.88316 | 0.103607276 | 0.45 | 0.4 | 1 | 14 |
| vasnb | 0.88401 | 0.109100092 | 0.2 | 0.15 | 1 | 14 |
| gpx4b | 0.88647 | 0.105062057 | 0.3 | 0.25 | 1 | 14 |
| p3h1 | 0.88957 | -0.10105689 | 0.15 | 0.2 | 1 | 14 |
| pawr | 0.89147 | 0.101164068 | 0.2 | 0.15 | 1 | 14 |
| sub1a | 0.90528 | 0.104159673 | 0.35 | 0.3 | 1 | 14 |
| si:ch73-335l21.4 | 0.90973 | 0.103379854 | 0.25 | 0.2 | 1 | 14 |
| fabp11a | 0.91829 | 0.103194527 | 0.075 | 0.05 | 1 | 14 |
| BX957297.2 | 1 | 0.106082632 | 0.05 | 0.05 | 1 | 14 |
| ccdc50 | 1 | 0.131938296 | 0.05 | 0.05 | 1 | 14 |

| | | | | | | |
|---|---|---|---|---|---|---|
| mt-cyb | 1.611383 | -1.41382767 | 0.194 | 0.957 | 3.25451217 | 15 |
| mt-nd3 | 2.407202 | -1.13113378 | 0.139 | 0.826 | 0.0048618 | 15 |
| mt-atp6 | 1.545915 | -1.02730027 | 0.472 | 0.957 | 0.0312229 | 15 |
| zgc:92744 | 4.984870 | 0.818309784 | 0.889 | 0.565 | 0.1006794 | 15 |
| spink2.1 | 1.819829 | -0.61184742 | 0 | 0.391 | 0.367551 | 15 |
| glula | 2.217700 | 0.986530625 | 0.667 | 0.087 | 0.447909 | 15 |
| hsp90ab1 | 2.710477 | 0.306599303 | 0.972 | 0.957 | 0.5474351 | 15 |
| psmb2 | 0.0001 | 0.376214899 | 0.5 | 0.13 | 1 | 15 |
| rpl26 | 0.00015 | 0.289252616 | 1 | 0.87 | 1 | 15 |
| edem1 | 0.00018 | 0.626018304 | 0.361 | 0 | 1 | 15 |
| rpl10 | 0.0002 | 0.380158619 | 0.917 | 0.783 | 1 | 15 |
| rpia | 0.00021 | 0.700146414 | 0.611 | 0.13 | 1 | 15 |
| fabp11a | 0.00023 | -0.80045777 | 0 | 0.304 | 1 | 15 |
| plxnb1a | 0.00023 | -0.58081851 | 0 | 0.304 | 1 | 15 |
| igf2b | 0.00029 | -0.23381208 | 0.194 | 0.174 | 1 | 15 |
| rps27.2 | 0.00031 | 0.316837221 | 0.944 | 0.87 | 1 | 15 |
| col12a1b | 0.00034 | -0.68554322 | 0.028 | 0.435 | 1 | 15 |
| psmg2 | 0.00036 | 0.501379959 | 0.333 | 0 | 1 | 15 |
| tspan7 | 0.00036 | 0.589796519 | 0.333 | 0 | 1 | 15 |
| vat1 | 0.0005 | -0.85384865 | 0.056 | 0.478 | 1 | 15 |
| rpl7a | 0.0005 | 0.309261741 | 0.972 | 0.87 | 1 | 15 |
| mt-co2 | 0.00055 | -0.53314655 | 0.75 | 1 | 1 | 15 |
| entpd1 | 0.00062 | -0.78099053 | 0.083 | 0.435 | 1 | 15 |
| XLOC-018328 | 0.00072 | 0.684606431 | 0.306 | 0 | 1 | 15 |
| MZT1 | 0.00072 | 0.461400925 | 0.306 | 0 | 1 | 15 |
| cyp17a1 | 0.00074 | 0.941453124 | 0.694 | 0.217 | 1 | 15 |
| rpl13a | 0.00076 | 0.283358152 | 1 | 0.913 | 1 | 15 |
| actn1 | 0.00077 | -0.36813253 | 0 | 0.261 | 1 | 15 |
| uqcrb | 0.00077 | -0.42471256 | 0 | 0.261 | 1 | 15 |
| cybrd1 | 0.00077 | -0.47584448 | 0 | 0.261 | 1 | 15 |
| adh8b | 0.00077 | -0.44771692 | 0 | 0.261 | 1 | 15 |
| dkk3b | 0.00077 | -0.43496589 | 0 | 0.261 | 1 | 15 |
| ms4a17a.8 | 0.00077 | -0.41810804 | 0 | 0.261 | 1 | 15 |
| ptprk | 0.00077 | -0.54867149 | 0 | 0.261 | 1 | 15 |
| ecm1b | 0.00086 | 0.555127279 | 0.944 | 0.696 | 1 | 15 |
| rps17 | 0.00089 | 0.270077771 | 0.917 | 0.913 | 1 | 15 |
| rps6 | 0.0009 | 0.273420672 | 1 | 0.913 | 1 | 15 |
| rps14 | 0.00102 | 0.282356401 | 0.972 | 0.913 | 1 | 15 |
| plxna4 | 0.00103 | 0.651092034 | 0.417 | 0.043 | 1 | 15 |
| mt-co1 | 0.00109 | -0.22826402 | 1 | 0.957 | 1 | 15 |
| fosab | 0.00128 | -0.47107494 | 0.778 | 1 | 1 | 15 |
| taf10 | 0.00141 | 0.438272187 | 0.278 | 0 | 1 | 15 |
| phb2b | 0.00141 | 0.452033894 | 0.278 | 0 | 1 | 15 |
| psmd11a | 0.00141 | 0.423146746 | 0.278 | 0 | 1 | 15 |
| rpl34 | 0.00143 | 0.200982501 | 1 | 0.957 | 1 | 15 |
| rps12 | 0.0015 | 0.372810414 | 1 | 0.783 | 1 | 15 |
| rpl32 | 0.00151 | 0.273884331 | 0.972 | 0.87 | 1 | 15 |
| sec61a1l | 0.00156 | 0.105358012 | 0.25 | 0.087 | 1 | 15 |

| | | | | | | |
|---|---|---|---|---|---|---|
| ackr4b | 0.00168 | 0.739810605 | 0.722 | 0.261 | 1 | 15 |
| rps13 | 0.00194 | 0.252940374 | 1 | 0.913 | 1 | 15 |
| mt-nd1 | 0.00211 | -0.66722276 | 0.5 | 0.87 | 1 | 15 |
| XLOC-010899 | 0.00251 | -0.4348809 | 0 | 0.217 | 1 | 15 |
| cyp3c1 | 0.00251 | -0.45869317 | 0 | 0.217 | 1 | 15 |
| cygb1 | 0.00251 | -0.49523638 | 0 | 0.217 | 1 | 15 |
| BX908782.2 | 0.00251 | -0.62127887 | 0 | 0.217 | 1 | 15 |
| tmem106a | 0.00251 | -0.40192618 | 0 | 0.217 | 1 | 15 |
| hspe1 | 0.00259 | 0.309169065 | 0.972 | 0.87 | 1 | 15 |
| rps23 | 0.00272 | 0.310965404 | 0.944 | 0.87 | 1 | 15 |
| bambia | 0.00272 | 0.819677904 | 0.583 | 0.174 | 1 | 15 |
| drd2a | 0.00274 | 0.495421586 | 0.25 | 0 | 1 | 15 |
| hsd17b1 | 0.00274 | 0.435296328 | 0.25 | 0 | 1 | 15 |
| si:ch211-210c8.7 | 0.00274 | 0.416668827 | 0.25 | 0 | 1 | 15 |
| rps18 | 0.00298 | 0.302702419 | 1 | 0.87 | 1 | 15 |
| rps2 | 0.00337 | 0.22674051 | 0.972 | 0.957 | 1 | 15 |
| cx34.5 | 0.00364 | 0.739067659 | 0.722 | 0.304 | 1 | 15 |
| dnmt3bb.1 | 0.00381 | 0.364812848 | 0.361 | 0.087 | 1 | 15 |
| aldh6a1 | 0.00399 | 0.379508463 | 0.361 | 0.043 | 1 | 15 |
| junba | 0.00403 | -0.36799373 | 0.861 | 0.957 | 1 | 15 |
| ppp2r1ba | 0.00414 | 0.530014799 | 0.444 | 0.13 | 1 | 15 |
| scarb2a | 0.00445 | -0.65086404 | 0.056 | 0.391 | 1 | 15 |
| rps5 | 0.00456 | 0.311192777 | 1 | 0.913 | 1 | 15 |
| rps9 | 0.00482 | 0.231025489 | 0.972 | 0.913 | 1 | 15 |
| rpl7 | 0.00484 | 0.295881321 | 0.889 | 0.826 | 1 | 15 |
| inka1a | 0.00524 | 0.391254988 | 0.222 | 0 | 1 | 15 |
| rxfp3.2a | 0.00524 | 0.431906219 | 0.222 | 0 | 1 | 15 |
| MED9 | 0.00524 | 0.355498719 | 0.222 | 0 | 1 | 15 |
| ap2a1 | 0.00524 | 0.38078071 | 0.222 | 0 | 1 | 15 |
| brd4 | 0.00524 | 0.380851661 | 0.222 | 0 | 1 | 15 |
| cops7a | 0.00524 | 0.367200003 | 0.222 | 0 | 1 | 15 |
| nop16 | 0.00524 | 0.342662367 | 0.222 | 0 | 1 | 15 |
| rap1ab | 0.00524 | 0.348359402 | 0.222 | 0 | 1 | 15 |
| gdpd4a | 0.00525 | -0.54514298 | 0.056 | 0.391 | 1 | 15 |
| gstm.3 | 0.0053 | -0.35064177 | 0.861 | 0.87 | 1 | 15 |
| bnc2 | 0.00549 | -0.61665408 | 0.139 | 0.478 | 1 | 15 |
| BX284696.1 | 0.00559 | -0.60815282 | 0.139 | 0.478 | 1 | 15 |
| ccl25b | 0.00559 | -0.59893198 | 0.028 | 0.217 | 1 | 15 |
| mmp16a | 0.0056 | -0.7002942 | 0.083 | 0.435 | 1 | 15 |
| tmem14ca | 0.00573 | 0.411048423 | 0.361 | 0.043 | 1 | 15 |
| krt8 | 0.00584 | -0.49164499 | 0.806 | 0.957 | 1 | 15 |
| st3gal3b | 0.00586 | -0.51900705 | 0.028 | 0.304 | 1 | 15 |
| tuba8l3 | 0.00591 | 0.653189832 | 0.722 | 0.304 | 1 | 15 |
| rpl3 | 0.00599 | 0.21225257 | 0.944 | 0.913 | 1 | 15 |
| eppk1 | 0.00627 | -0.64802974 | 0.111 | 0.478 | 1 | 15 |
| rps7 | 0.00638 | 0.186390154 | 0.972 | 0.957 | 1 | 15 |
| hsd3b7 | 0.00649 | 0.68393857 | 0.611 | 0.261 | 1 | 15 |
| rps28 | 0.00676 | 0.167039929 | 1 | 1 | 1 | 15 |

| | | | | | | |
|---|---|---|---|---|---|---|
| itga4 | 0.00685 | 0.454538961 | 0.333 | 0.043 | 1 | 15 |
| XLOC-043861 | 0.00688 | -0.48913853 | 0.028 | 0.304 | 1 | 15 |
| rasd1 | 0.00699 | 0.474393521 | 0.583 | 0.217 | 1 | 15 |
| ncam1a | 0.00727 | -0.57140269 | 0.028 | 0.304 | 1 | 15 |
| rpl19 | 0.00733 | 0.255140099 | 0.944 | 0.913 | 1 | 15 |
| rpl27a | 0.00751 | -0.33116555 | 0.833 | 0.913 | 1 | 15 |
| LOC101886015 | 0.00797 | -0.34516444 | 0 | 0.174 | 1 | 15 |
| irf1b | 0.00797 | -0.35185726 | 0 | 0.174 | 1 | 15 |
| cntfr | 0.00797 | -0.35853862 | 0 | 0.174 | 1 | 15 |
| arl6ip6 | 0.00797 | -0.33320983 | 0 | 0.174 | 1 | 15 |
| iqsec1b | 0.00797 | -0.35501944 | 0 | 0.174 | 1 | 15 |
| kank2 | 0.00797 | -0.3548252 | 0 | 0.174 | 1 | 15 |
| mboat7 | 0.00797 | -0.32053182 | 0 | 0.174 | 1 | 15 |
| mief2 | 0.00797 | -0.27888391 | 0 | 0.174 | 1 | 15 |
| ppp1cbl | 0.00797 | -0.24690714 | 0 | 0.174 | 1 | 15 |
| slc2a12 | 0.00797 | -0.36096543 | 0 | 0.174 | 1 | 15 |
| zbtb16b | 0.00797 | -0.31753123 | 0 | 0.174 | 1 | 15 |
| fbln5 | 0.00797 | -0.39922993 | 0 | 0.174 | 1 | 15 |
| fgfrl1b | 0.00797 | -0.34102611 | 0 | 0.174 | 1 | 15 |
| si:ch211-214p16.2 | 0.00797 | -0.25762365 | 0 | 0.174 | 1 | 15 |
| rasgef1ba | 0.00797 | -0.3077246 | 0 | 0.174 | 1 | 15 |
| rflnb | 0.00797 | -0.38411161 | 0 | 0.174 | 1 | 15 |
| actr2a | 0.00797 | -0.28472723 | 0 | 0.174 | 1 | 15 |
| dachd | 0.00797 | -0.31940515 | 0 | 0.174 | 1 | 15 |
| nt5c3a | 0.00797 | -0.2669329 | 0 | 0.174 | 1 | 15 |
| CR855311.1 | 0.00797 | -0.46697272 | 0 | 0.174 | 1 | 15 |
| rpl24 | 0.00835 | 0.200784724 | 0.972 | 0.957 | 1 | 15 |
| hmgb2a | 0.00837 | -0.46020891 | 0.556 | 0.609 | 1 | 15 |
| igflr1 | 0.00841 | 0.620598304 | 0.417 | 0.087 | 1 | 15 |
| rpl28 | 0.00859 | 0.182074279 | 1 | 0.957 | 1 | 15 |
| slc29a4 | 0.00866 | -0.48836029 | 0.028 | 0.304 | 1 | 15 |
| sppl3 | 0.00868 | 0.545511297 | 0.444 | 0.087 | 1 | 15 |
| XLOC-037780 | 0.00873 | 0.233562078 | 0.25 | 0.043 | 1 | 15 |
| slc43a3b | 0.00875 | -0.53198348 | 0.028 | 0.304 | 1 | 15 |
| metrnla | 0.009 | 0.555766648 | 0.361 | 0.043 | 1 | 15 |
| hspb1 | 0.00903 | -0.62876392 | 0.111 | 0.435 | 1 | 15 |
| map1lc3a | 0.00905 | 0.377754837 | 0.5 | 0.174 | 1 | 15 |
| myl12.1 | 0.00912 | -0.14369512 | 0.806 | 0.696 | 1 | 15 |
| jam2b | 0.00929 | -0.67130499 | 0.028 | 0.304 | 1 | 15 |
| fn1b | 0.0093 | -0.7230267 | 0.333 | 0.652 | 1 | 15 |
| usp5 | 0.00948 | -0.49741523 | 0.028 | 0.304 | 1 | 15 |
| chac1 | 0.0096 | 0.224425153 | 0.722 | 0.391 | 1 | 15 |
| tmem98 | 0.00965 | -0.47290341 | 0.028 | 0.304 | 1 | 15 |
| elovl1b | 0.00967 | -0.44313075 | 0.028 | 0.304 | 1 | 15 |
| mrpl54 | 0.0097 | 0.410853477 | 0.361 | 0.043 | 1 | 15 |
| si:ch1073-170o4.1 | 0.00991 | 0.289811364 | 0.194 | 0 | 1 | 15 |
| slc30a6 | 0.00991 | 0.356699805 | 0.194 | 0 | 1 | 15 |
| tmem231 | 0.00991 | 0.335817267 | 0.194 | 0 | 1 | 15 |

| | | | | | | |
|---|---|---|---|---|---|---|
| atp6ap1lb | 0.00991 | 0.30951974 | 0.194 | 0 | 1 | 15 |
| polr1e | 0.00991 | 0.340481239 | 0.194 | 0 | 1 | 15 |
| rpl7l1 | 0.00991 | 0.301320491 | 0.194 | 0 | 1 | 15 |
| rrp8 | 0.00991 | 0.316445205 | 0.194 | 0 | 1 | 15 |
| mex3b | 0.00991 | 0.339416412 | 0.194 | 0 | 1 | 15 |
| txnipb | 0.00991 | 0.29640014 | 0.194 | 0 | 1 | 15 |
| zgc:56106 | 0.00991 | 0.301913043 | 0.194 | 0 | 1 | 15 |
| chd9 | 0.00991 | 0.348042225 | 0.194 | 0 | 1 | 15 |
| chmp1a | 0.00991 | 0.363296806 | 0.194 | 0 | 1 | 15 |
| cwc27 | 0.00991 | 0.339529258 | 0.194 | 0 | 1 | 15 |
| erlec1 | 0.00991 | 0.341923861 | 0.194 | 0 | 1 | 15 |
| ppp1cb | 0.00991 | 0.374615543 | 0.194 | 0 | 1 | 15 |
| rfxank | 0.00991 | 0.312788061 | 0.194 | 0 | 1 | 15 |
| tapbpl | 0.00991 | 0.350531075 | 0.194 | 0 | 1 | 15 |
| rab1aa | 0.00991 | 0.298370009 | 0.194 | 0 | 1 | 15 |
| si:dkeyp-55f12.3 | 0.00991 | 0.316278605 | 0.194 | 0 | 1 | 15 |
| rplp2 | 0.01001 | 0.244376098 | 1 | 0.783 | 1 | 15 |
| rps15a | 0.01004 | 0.291746915 | 1 | 0.87 | 1 | 15 |
| rps16 | 0.01013 | 0.208077464 | 1 | 0.957 | 1 | 15 |
| rpl8 | 0.01045 | 0.290345053 | 0.944 | 0.826 | 1 | 15 |
| akap12b | 0.0107 | -0.48425669 | 0.5 | 0.87 | 1 | 15 |
| aebp1 | 0.01097 | -0.55124107 | 0.167 | 0.478 | 1 | 15 |
| rpl27 | 0.01104 | 0.18623096 | 0.944 | 0.913 | 1 | 15 |
| LOC103910903 | 0.0113 | 0.445232292 | 0.5 | 0.391 | 1 | 15 |
| rpl35a | 0.01147 | 0.222850734 | 0.972 | 0.913 | 1 | 15 |
| man2c1 | 0.01169 | 0.670604545 | 0.611 | 0.261 | 1 | 15 |
| rpl10a | 0.01173 | 0.264255935 | 0.972 | 0.87 | 1 | 15 |
| rps27a | 0.01195 | 0.303999049 | 0.972 | 0.783 | 1 | 15 |
| tnfsf10 | 0.01206 | -0.61627912 | 0.167 | 0.522 | 1 | 15 |
| eef1b2 | 0.01268 | 0.284317883 | 0.972 | 0.913 | 1 | 15 |
| egfra | 0.01317 | 0.44665535 | 0.333 | 0.043 | 1 | 15 |
| eif3f | 0.01345 | 0.467041784 | 0.861 | 0.609 | 1 | 15 |
| rpl23 | 0.01356 | 0.213938002 | 1 | 0.913 | 1 | 15 |
| rpl18a | 0.01366 | 0.240317832 | 1 | 0.913 | 1 | 15 |
| ap2m1a | 0.0137 | 0.433605467 | 0.917 | 0.652 | 1 | 15 |
| ldhba | 0.01371 | 0.528869625 | 0.444 | 0.13 | 1 | 15 |
| pdia6 | 0.01378 | -0.40158429 | 0.444 | 0.609 | 1 | 15 |
| junbb | 0.0139 | -0.55932822 | 0.139 | 0.478 | 1 | 15 |
| loxl2b | 0.01412 | -0.41989106 | 0.028 | 0.261 | 1 | 15 |
| fstl3 | 0.01425 | -0.44187934 | 0.056 | 0.348 | 1 | 15 |
| sostdc1a | 0.01448 | -0.69278255 | 0.056 | 0.348 | 1 | 15 |
| cldn5a | 0.01475 | -0.6160399 | 0.083 | 0.391 | 1 | 15 |
| cd81a | 0.01501 | -0.48085521 | 0.611 | 0.826 | 1 | 15 |
| spred2b | 0.01516 | -0.50661413 | 0.167 | 0.522 | 1 | 15 |
| jam2a | 0.01536 | 0.405108604 | 0.917 | 0.609 | 1 | 15 |
| ptprga | 0.01543 | -0.2707454 | 0.694 | 0.739 | 1 | 15 |
| col4a6 | 0.01554 | 0.597089198 | 0.417 | 0.087 | 1 | 15 |
| ddx42 | 0.01577 | 0.442092628 | 0.306 | 0.043 | 1 | 15 |

| | | | | | | |
|---|---|---|---|---|---|---|
| socs3b | 0.01587 | -0.60134544 | 0.167 | 0.522 | 1 | 15 |
| rps3 | 0.01588 | 0.353248123 | 0.972 | 0.783 | 1 | 15 |
| id1 | 0.01592 | -0.46463589 | 0.333 | 0.435 | 1 | 15 |
| rpl38 | 0.01595 | 0.271314864 | 1 | 0.87 | 1 | 15 |
| h3f3d | 0.016 | 0.241111585 | 1 | 0.87 | 1 | 15 |
| slco2b1 | 0.01611 | -0.64466977 | 0.083 | 0.391 | 1 | 15 |
| tjp2b | 0.01635 | -0.51922388 | 0.194 | 0.435 | 1 | 15 |
| srsf11 | 0.01659 | 0.122672714 | 0.417 | 0.217 | 1 | 15 |
| med28 | 0.0166 | 0.305900701 | 0.389 | 0.13 | 1 | 15 |
| rpl22 | 0.01661 | 0.262460331 | 0.944 | 0.87 | 1 | 15 |
| pleca | 0.01663 | -0.41210061 | 0.028 | 0.261 | 1 | 15 |
| zgc:100829 | 0.01685 | 0.540367311 | 0.5 | 0.217 | 1 | 15 |
| aga | 0.017 | 0.290644832 | 0.25 | 0.043 | 1 | 15 |
| si:ch211-199g17.2 | 0.017 | 0.486688092 | 0.528 | 0.174 | 1 | 15 |
| rpl35 | 0.01707 | 0.211079443 | 0.972 | 0.957 | 1 | 15 |
| eif3ja | 0.01712 | -0.42184196 | 0.389 | 0.565 | 1 | 15 |
| asb11 | 0.01718 | 0.613214076 | 0.528 | 0.174 | 1 | 15 |
| surf4l | 0.01721 | -0.56967271 | 0.083 | 0.391 | 1 | 15 |
| znf395b | 0.01731 | 0.41639043 | 0.333 | 0.043 | 1 | 15 |
| zgc:162730 | 0.01743 | -0.71628819 | 0.222 | 0.565 | 1 | 15 |
| XLOC-000431 | 0.01791 | -0.56680114 | 0.111 | 0.304 | 1 | 15 |
| cldn11a | 0.01844 | 0.235284439 | 0.972 | 0.957 | 1 | 15 |
| cmtm6 | 0.0185 | -0.53483394 | 0.222 | 0.565 | 1 | 15 |
| BX511129.1 | 0.01856 | 0.289999604 | 0.167 | 0 | 1 | 15 |
| atp2b1a | 0.01856 | 0.289887788 | 0.167 | 0 | 1 | 15 |
| c1qtnf5 | 0.01856 | 0.420945577 | 0.167 | 0 | 1 | 15 |
| ccz1 | 0.01856 | 0.34358289 | 0.167 | 0 | 1 | 15 |
| ciao2b | 0.01856 | 0.300108254 | 0.167 | 0 | 1 | 15 |
| cisd2 | 0.01856 | 0.264332956 | 0.167 | 0 | 1 | 15 |
| ednrab | 0.01856 | 0.288861678 | 0.167 | 0 | 1 | 15 |
| emc8 | 0.01856 | 0.270611952 | 0.167 | 0 | 1 | 15 |
| fhit | 0.01856 | 0.235062474 | 0.167 | 0 | 1 | 15 |
| guk1a | 0.01856 | 0.293212035 | 0.167 | 0 | 1 | 15 |
| im:7138535 | 0.01856 | 0.318484822 | 0.167 | 0 | 1 | 15 |
| immp2l | 0.01856 | 0.299014539 | 0.167 | 0 | 1 | 15 |
| letm2 | 0.01856 | 0.27503748 | 0.167 | 0 | 1 | 15 |
| mrpl38 | 0.01856 | 0.316646767 | 0.167 | 0 | 1 | 15 |
| nup153 | 0.01856 | 0.241597882 | 0.167 | 0 | 1 | 15 |
| pafah1b1b | 0.01856 | 0.351647285 | 0.167 | 0 | 1 | 15 |
| pdcl | 0.01856 | 0.278076466 | 0.167 | 0 | 1 | 15 |
| ppt2 | 0.01856 | 0.312551565 | 0.167 | 0 | 1 | 15 |
| psma6l | 0.01856 | 0.315383837 | 0.167 | 0 | 1 | 15 |
| riox2 | 0.01856 | 0.257644829 | 0.167 | 0 | 1 | 15 |
| rtf2 | 0.01856 | 0.287166831 | 0.167 | 0 | 1 | 15 |
| sardh | 0.01856 | 0.259470935 | 0.167 | 0 | 1 | 15 |
| tmem106bb | 0.01856 | 0.338258628 | 0.167 | 0 | 1 | 15 |
| trappc13 | 0.01856 | 0.281185294 | 0.167 | 0 | 1 | 15 |
| triap1 | 0.01856 | 0.275166666 | 0.167 | 0 | 1 | 15 |

| | | | | | | |
|---|---|---|---|---|---|---|
| ypel1 | 0.01856 | 0.335144247 | 0.167 | 0 | 1 | 15 |
| znf609a | 0.01856 | 0.279123962 | 0.167 | 0 | 1 | 15 |
| actr10 | 0.01856 | 0.285861555 | 0.167 | 0 | 1 | 15 |
| atrn | 0.01856 | 0.268600742 | 0.167 | 0 | 1 | 15 |
| cdh4 | 0.01856 | 0.39430634 | 0.167 | 0 | 1 | 15 |
| creb3l2 | 0.01856 | 0.352983738 | 0.167 | 0 | 1 | 15 |
| dhx16 | 0.01856 | 0.3191718 | 0.167 | 0 | 1 | 15 |
| eif4ba | 0.01856 | 0.322131644 | 0.167 | 0 | 1 | 15 |
| gps2 | 0.01856 | 0.297571244 | 0.167 | 0 | 1 | 15 |
| med20 | 0.01856 | 0.319226129 | 0.167 | 0 | 1 | 15 |
| oxa1l | 0.01856 | 0.332164485 | 0.167 | 0 | 1 | 15 |
| pgm1 | 0.01856 | 0.17125431 | 0.167 | 0 | 1 | 15 |
| ppm1la | 0.01856 | 0.261271898 | 0.167 | 0 | 1 | 15 |
| smarcc1a | 0.01856 | 0.279071579 | 0.167 | 0 | 1 | 15 |
| tns3.2 | 0.01856 | 0.335678071 | 0.167 | 0 | 1 | 15 |
| utp18 | 0.01856 | 0.263200382 | 0.167 | 0 | 1 | 15 |
| ccdc9 | 0.01856 | 0.283626867 | 0.167 | 0 | 1 | 15 |
| lap3 | 0.01856 | 0.289198373 | 0.167 | 0 | 1 | 15 |
| zgc:153151 | 0.01856 | 0.286086768 | 0.167 | 0 | 1 | 15 |
| cd74b | 0.01857 | -0.24932228 | 0.139 | 0.391 | 1 | 15 |
| rpl23a | 0.01888 | 0.250912604 | 1 | 0.87 | 1 | 15 |
| nr1d2a | 0.01909 | -0.44215961 | 0.139 | 0.391 | 1 | 15 |
| dusp14 | 0.01931 | 0.2379133 | 0.306 | 0.304 | 1 | 15 |
| jun | 0.01969 | -0.41872627 | 0.639 | 0.783 | 1 | 15 |
| UBL4A | 0.01974 | -0.23418621 | 0.028 | 0.217 | 1 | 15 |
| rpl13 | 0.02011 | 0.210998585 | 1 | 0.913 | 1 | 15 |
| ybx1 | 0.02062 | 0.222078366 | 0.972 | 0.957 | 1 | 15 |
| cfl1 | 0.02072 | -0.29150243 | 0.861 | 0.957 | 1 | 15 |
| mrpl30 | 0.02159 | 0.450305252 | 0.306 | 0.043 | 1 | 15 |
| rps15 | 0.02287 | 0.248437705 | 1 | 0.826 | 1 | 15 |
| ppa1b | 0.02296 | 0.399530009 | 0.25 | 0.043 | 1 | 15 |
| zmat2 | 0.02303 | -0.4822369 | 0.194 | 0.522 | 1 | 15 |
| foxl2a | 0.02316 | -0.5177551 | 0.167 | 0.478 | 1 | 15 |
| plk3 | 0.02321 | -0.44277849 | 0.028 | 0.261 | 1 | 15 |
| eif4a2 | 0.02353 | 0.430701526 | 0.306 | 0.043 | 1 | 15 |
| anxa1a | 0.02391 | -0.4266624 | 0.028 | 0.261 | 1 | 15 |
| islr2 | 0.02393 | 0.610780473 | 0.444 | 0.13 | 1 | 15 |
| tia1l | 0.02425 | 0.216960076 | 0.222 | 0.043 | 1 | 15 |
| rps6ka4 | 0.02439 | -0.43887809 | 0.028 | 0.261 | 1 | 15 |
| me3 | 0.02443 | -0.43274883 | 0.028 | 0.261 | 1 | 15 |
| c6 | 0.02454 | -0.46567386 | 0.583 | 0.826 | 1 | 15 |
| uqcrc1 | 0.02472 | 0.272256189 | 0.611 | 0.304 | 1 | 15 |
| flnb | 0.02495 | -0.24512913 | 0 | 0.13 | 1 | 15 |
| pkdc | 0.02495 | -0.20999263 | 0 | 0.13 | 1 | 15 |
| usp14 | 0.02495 | -0.21189322 | 0 | 0.13 | 1 | 15 |
| zgc:174904 | 0.02495 | -0.22793126 | 0 | 0.13 | 1 | 15 |
| LOC100534805 | 0.02495 | -0.2169766 | 0 | 0.13 | 1 | 15 |
| aftpha | 0.02495 | -0.23799865 | 0 | 0.13 | 1 | 15 |

| | | | | | | |
|---|---|---|---|---|---|---|
| arhgap42a | 0.02495 | -0.2314491 | 0 | 0.13 | 1 | 15 |
| casd1 | 0.02495 | -0.22668264 | 0 | 0.13 | 1 | 15 |
| ccnd2b | 0.02495 | -0.2839633 | 0 | 0.13 | 1 | 15 |
| ccnk | 0.02495 | -0.23117526 | 0 | 0.13 | 1 | 15 |
| cd302 | 0.02495 | -0.20720988 | 0 | 0.13 | 1 | 15 |
| col6a2 | 0.02495 | -0.25852316 | 0 | 0.13 | 1 | 15 |
| epha4a | 0.02495 | -0.23717549 | 0 | 0.13 | 1 | 15 |
| fam198b | 0.02495 | -0.1960956 | 0 | 0.13 | 1 | 15 |
| fat2 | 0.02495 | -0.27495138 | 0 | 0.13 | 1 | 15 |
| fscn1a | 0.02495 | -0.30925012 | 0 | 0.13 | 1 | 15 |
| gabra5 | 0.02495 | -0.29645031 | 0 | 0.13 | 1 | 15 |
| kdelr3 | 0.02495 | -0.24764987 | 0 | 0.13 | 1 | 15 |
| nmt2 | 0.02495 | -0.19105209 | 0 | 0.13 | 1 | 15 |
| pdgfrl | 0.02495 | -0.24562509 | 0 | 0.13 | 1 | 15 |
| per1a | 0.02495 | -0.27334453 | 0 | 0.13 | 1 | 15 |
| prkd2 | 0.02495 | -0.22048084 | 0 | 0.13 | 1 | 15 |
| prss23 | 0.02495 | -0.21042748 | 0 | 0.13 | 1 | 15 |
| rgs7a | 0.02495 | -0.221589 | 0 | 0.13 | 1 | 15 |
| si:ch211-194c3.5 | 0.02495 | -0.3353708 | 0 | 0.13 | 1 | 15 |
| si:dkey-16l2.20 | 0.02495 | -0.20322157 | 0 | 0.13 | 1 | 15 |
| si:dkey-240h12.3 | 0.02495 | -0.23014729 | 0 | 0.13 | 1 | 15 |
| slc35a3a | 0.02495 | -0.24739926 | 0 | 0.13 | 1 | 15 |
| sost | 0.02495 | -0.26992805 | 0 | 0.13 | 1 | 15 |
| tpm2 | 0.02495 | -0.2691507 | 0 | 0.13 | 1 | 15 |
| tspo | 0.02495 | -0.2020274 | 0 | 0.13 | 1 | 15 |
| txndc16 | 0.02495 | -0.25744474 | 0 | 0.13 | 1 | 15 |
| zgc:171459 | 0.02495 | -0.17824178 | 0 | 0.13 | 1 | 15 |
| zgc:171679 | 0.02495 | -0.2101673 | 0 | 0.13 | 1 | 15 |
| CABZ01032488.1 | 0.02495 | -0.18743409 | 0 | 0.13 | 1 | 15 |
| GNAZ | 0.02495 | -0.21382859 | 0 | 0.13 | 1 | 15 |
| HACD4 | 0.02495 | -0.23935139 | 0 | 0.13 | 1 | 15 |
| LO018154.1 | 0.02495 | -0.21022243 | 0 | 0.13 | 1 | 15 |
| LOC101886030 | 0.02495 | -0.25203252 | 0 | 0.13 | 1 | 15 |
| LOC103910015 | 0.02495 | -0.23377108 | 0 | 0.13 | 1 | 15 |
| NMNAT3 | 0.02495 | -0.17041286 | 0 | 0.13 | 1 | 15 |
| XLOC-032154 | 0.02495 | -0.26560859 | 0 | 0.13 | 1 | 15 |
| alkbh5 | 0.02495 | -0.18473058 | 0 | 0.13 | 1 | 15 |
| amigo1 | 0.02495 | -0.32987748 | 0 | 0.13 | 1 | 15 |
| ap1ar | 0.02495 | -0.27486913 | 0 | 0.13 | 1 | 15 |
| aven | 0.02495 | -0.2234003 | 0 | 0.13 | 1 | 15 |
| b4galt3 | 0.02495 | -0.22004109 | 0 | 0.13 | 1 | 15 |
| caln2 | 0.02495 | -0.38784693 | 0 | 0.13 | 1 | 15 |
| copb1 | 0.02495 | -0.23024975 | 0 | 0.13 | 1 | 15 |
| dzip1 | 0.02495 | -0.2473399 | 0 | 0.13 | 1 | 15 |
| emilin1a | 0.02495 | -0.31039489 | 0 | 0.13 | 1 | 15 |
| epb41l4a | 0.02495 | -0.19998396 | 0 | 0.13 | 1 | 15 |
| hic2 | 0.02495 | -0.25416653 | 0 | 0.13 | 1 | 15 |
| ino80db | 0.02495 | -0.24408396 | 0 | 0.13 | 1 | 15 |

| | | | | | | |
|---|---|---|---|---|---|---|
| katnbl1 | 0.02495 | -0.21490584 | 0 | 0.13 | 1 | 15 |
| klhl18 | 0.02495 | -0.24327231 | 0 | 0.13 | 1 | 15 |
| lmtk2 | 0.02495 | -0.24050183 | 0 | 0.13 | 1 | 15 |
| mgat4a | 0.02495 | -0.34326517 | 0 | 0.13 | 1 | 15 |
| mkln1 | 0.02495 | -0.19897009 | 0 | 0.13 | 1 | 15 |
| nkx3.3 | 0.02495 | -0.27154361 | 0 | 0.13 | 1 | 15 |
| pcdh10b | 0.02495 | -0.25106046 | 0 | 0.13 | 1 | 15 |
| pecr | 0.02495 | -0.19540498 | 0 | 0.13 | 1 | 15 |
| plekho2 | 0.02495 | -0.2498789 | 0 | 0.13 | 1 | 15 |
| plppr2b | 0.02495 | -0.2119915 | 0 | 0.13 | 1 | 15 |
| pls3 | 0.02495 | -0.21541357 | 0 | 0.13 | 1 | 15 |
| plxdc2 | 0.02495 | -0.21740179 | 0 | 0.13 | 1 | 15 |
| prtfdc1 | 0.02495 | -0.27168638 | 0 | 0.13 | 1 | 15 |
| reck | 0.02495 | -0.2809122 | 0 | 0.13 | 1 | 15 |
| rgs12b | 0.02495 | -0.26031107 | 0 | 0.13 | 1 | 15 |
| setdb1a | 0.02495 | -0.17343961 | 0 | 0.13 | 1 | 15 |
| si:ch211-199m9.3 | 0.02495 | -0.22856685 | 0 | 0.13 | 1 | 15 |
| si:ch73-127m5.2 | 0.02495 | -0.17678895 | 0 | 0.13 | 1 | 15 |
| si:ch73-141c7.1 | 0.02495 | -0.20575722 | 0 | 0.13 | 1 | 15 |
| si:dkey-28b4.8 | 0.02495 | -0.34575528 | 0 | 0.13 | 1 | 15 |
| si:dkey-30c15.10 | 0.02495 | -0.20746806 | 0 | 0.13 | 1 | 15 |
| si:dkey-6i22.5 | 0.02495 | -0.19613751 | 0 | 0.13 | 1 | 15 |
| spred2a | 0.02495 | -0.30641376 | 0 | 0.13 | 1 | 15 |
| trafd1 | 0.02495 | -0.17411003 | 0 | 0.13 | 1 | 15 |
| trmt2a | 0.02495 | -0.18471828 | 0 | 0.13 | 1 | 15 |
| vcam1b | 0.02495 | -0.20419363 | 0 | 0.13 | 1 | 15 |
| tmsb4x | 0.02502 | -0.1760652 | 1 | 1 | 1 | 15 |
| ywhabl | 0.02526 | 0.439475222 | 0.778 | 0.435 | 1 | 15 |
| htatsf1 | 0.02543 | 0.256853142 | 0.222 | 0.043 | 1 | 15 |
| serinc1 | 0.02544 | 0.454258813 | 0.444 | 0.13 | 1 | 15 |
| pdgfc | 0.02591 | -0.38669204 | 0.028 | 0.261 | 1 | 15 |
| mast2 | 0.02593 | -0.38743293 | 0.028 | 0.261 | 1 | 15 |
| esr2b | 0.02596 | -0.51125954 | 0.083 | 0.348 | 1 | 15 |
| sigmar1 | 0.02639 | 0.440856521 | 0.389 | 0.13 | 1 | 15 |
| si:ch73-22o12.1 | 0.02669 | 0.482037503 | 0.361 | 0.087 | 1 | 15 |
| mrps6 | 0.02675 | 0.433153677 | 0.389 | 0.087 | 1 | 15 |
| col1a2 | 0.0272 | -0.62651618 | 0.194 | 0.478 | 1 | 15 |
| prdx4 | 0.02747 | 0.141982768 | 0.583 | 0.304 | 1 | 15 |
| cebpb | 0.02749 | -0.64219598 | 0.194 | 0.522 | 1 | 15 |
| c1qbp | 0.02754 | 0.394607387 | 0.278 | 0.043 | 1 | 15 |
| knop1 | 0.02771 | 0.361837985 | 0.333 | 0.087 | 1 | 15 |
| nos1apa | 0.02799 | 0.490290017 | 0.444 | 0.13 | 1 | 15 |
| boc | 0.02809 | -0.5412038 | 0.167 | 0.435 | 1 | 15 |
| ndufa7 | 0.02834 | 0.149710268 | 0.25 | 0.087 | 1 | 15 |
| gtf2b | 0.02856 | 0.440338355 | 0.361 | 0.087 | 1 | 15 |
| sf3a2 | 0.02884 | 0.426348261 | 0.306 | 0.087 | 1 | 15 |
| eef1g | 0.02885 | 0.293792091 | 0.944 | 0.87 | 1 | 15 |
| appa | 0.02899 | 0.336605039 | 0.833 | 0.522 | 1 | 15 |

| | | | | | | |
|---|---|---|---|---|---|---|
| rnaseh1 | 0.02946 | 0.264368086 | 0.25 | 0.043 | 1 | 15 |
| rps11 | 0.02946 | 0.199949288 | 1 | 0.826 | 1 | 15 |
| phc2a | 0.02953 | -0.27463014 | 0.556 | 0.609 | 1 | 15 |
| schip1 | 0.02974 | -0.37240543 | 0.083 | 0.348 | 1 | 15 |
| ppifb | 0.03015 | 0.366602304 | 0.333 | 0.087 | 1 | 15 |
| naca | 0.03031 | 0.263655421 | 0.972 | 0.913 | 1 | 15 |
| nacc2 | 0.03083 | 0.426386204 | 0.306 | 0.043 | 1 | 15 |
| dbi | 0.03085 | -0.44940184 | 0.306 | 0.522 | 1 | 15 |
| gsdf | 0.03108 | 0.506749612 | 0.806 | 0.478 | 1 | 15 |
| cox7a1 | 0.03109 | 0.310548861 | 0.528 | 0.217 | 1 | 15 |
| cd276 | 0.0317 | -0.44073674 | 0.111 | 0.304 | 1 | 15 |
| psmb4 | 0.0317 | 0.429578426 | 0.556 | 0.217 | 1 | 15 |
| pmp22a | 0.03175 | -0.55799329 | 0.222 | 0.478 | 1 | 15 |
| arf5 | 0.03203 | -0.23318213 | 0.444 | 0.478 | 1 | 15 |
| psmd14 | 0.03211 | 0.430823449 | 0.444 | 0.13 | 1 | 15 |
| eif2s1b | 0.03233 | 0.453074261 | 0.639 | 0.304 | 1 | 15 |
| zswim5 | 0.03241 | 0.322867327 | 0.25 | 0.043 | 1 | 15 |
| col6a3 | 0.03292 | -0.55856317 | 0.167 | 0.478 | 1 | 15 |
| mphosph8 | 0.03407 | 0.408245751 | 0.667 | 0.478 | 1 | 15 |
| odc1 | 0.03415 | 0.560978973 | 0.639 | 0.304 | 1 | 15 |
| tubb4b | 0.0345 | -0.25513634 | 0.667 | 0.652 | 1 | 15 |
| BX901962.2 | 0.03454 | 0.221702784 | 0.139 | 0 | 1 | 15 |
| CU207221.1 | 0.03454 | 0.238701844 | 0.139 | 0 | 1 | 15 |
| CU929418.1 | 0.03454 | 0.238306077 | 0.139 | 0 | 1 | 15 |
| arpin | 0.03454 | 0.238994701 | 0.139 | 0 | 1 | 15 |
| cpeb4a | 0.03454 | 0.247300421 | 0.139 | 0 | 1 | 15 |
| fbxl7 | 0.03454 | 0.252023116 | 0.139 | 0 | 1 | 15 |
| lcorl | 0.03454 | 0.311128368 | 0.139 | 0 | 1 | 15 |
| mterf2 | 0.03454 | 0.284523362 | 0.139 | 0 | 1 | 15 |
| ncoa4 | 0.03454 | 0.177753164 | 0.139 | 0 | 1 | 15 |
| npm1b | 0.03454 | 0.210826172 | 0.139 | 0 | 1 | 15 |
| si:dkey-234h16.7 | 0.03454 | 0.233785441 | 0.139 | 0 | 1 | 15 |
| si:dkey-23k10.5 | 0.03454 | 0.235266339 | 0.139 | 0 | 1 | 15 |
| spred1 | 0.03454 | 0.250206767 | 0.139 | 0 | 1 | 15 |
| stxbp3 | 0.03454 | 0.233998313 | 0.139 | 0 | 1 | 15 |
| sympk | 0.03454 | 0.23188778 | 0.139 | 0 | 1 | 15 |
| tbx18 | 0.03454 | 0.299975274 | 0.139 | 0 | 1 | 15 |
| tfr1b | 0.03454 | 0.23405445 | 0.139 | 0 | 1 | 15 |
| zgc:171566 | 0.03454 | 0.270944654 | 0.139 | 0 | 1 | 15 |
| aph1b | 0.03454 | 0.231613764 | 0.139 | 0 | 1 | 15 |
| cds2 | 0.03454 | 0.231692109 | 0.139 | 0 | 1 | 15 |
| cltca | 0.03454 | 0.252364036 | 0.139 | 0 | 1 | 15 |
| coa1 | 0.03454 | 0.221105403 | 0.139 | 0 | 1 | 15 |
| coq8a | 0.03454 | 0.229813515 | 0.139 | 0 | 1 | 15 |
| gart | 0.03454 | 0.268649545 | 0.139 | 0 | 1 | 15 |
| gga1 | 0.03454 | 0.249699288 | 0.139 | 0 | 1 | 15 |
| hes6 | 0.03454 | 0.218221756 | 0.139 | 0 | 1 | 15 |
| hnrnpua | 0.03454 | 0.264281191 | 0.139 | 0 | 1 | 15 |

| | | | | | | |
|---|---|---|---|---|---|---|
| mbnl2 | 0.03454 | 0.253197196 | 0.139 | 0 | 1 | 15 |
| med6 | 0.03454 | 0.257798202 | 0.139 | 0 | 1 | 15 |
| minpp1b | 0.03454 | 0.266901823 | 0.139 | 0 | 1 | 15 |
| nagpa | 0.03454 | 0.276692242 | 0.139 | 0 | 1 | 15 |
| nfxl1 | 0.03454 | 0.284640881 | 0.139 | 0 | 1 | 15 |
| ppp1r3b | 0.03454 | 0.17078396 | 0.139 | 0 | 1 | 15 |
| prrg4 | 0.03454 | 0.255827686 | 0.139 | 0 | 1 | 15 |
| rab27a | 0.03454 | 0.248088272 | 0.139 | 0 | 1 | 15 |
| rpa1 | 0.03454 | 0.242435435 | 0.139 | 0 | 1 | 15 |
| sdk2b | 0.03454 | 0.273302759 | 0.139 | 0 | 1 | 15 |
| si:dkeyp-113d7.10 | 0.03454 | 0.254301163 | 0.139 | 0 | 1 | 15 |
| tex261 | 0.03454 | 0.216126311 | 0.139 | 0 | 1 | 15 |
| tma16 | 0.03454 | 0.220042521 | 0.139 | 0 | 1 | 15 |
| tmem183a | 0.03454 | 0.259165456 | 0.139 | 0 | 1 | 15 |
| tmem237b | 0.03454 | 0.272170498 | 0.139 | 0 | 1 | 15 |
| tnksa | 0.03454 | 0.255644794 | 0.139 | 0 | 1 | 15 |
| umps | 0.03454 | 0.252850773 | 0.139 | 0 | 1 | 15 |
| usb1 | 0.03454 | 0.279410554 | 0.139 | 0 | 1 | 15 |
| wnt4a | 0.03454 | 0.279594713 | 0.139 | 0 | 1 | 15 |
| zgc:112962 | 0.03454 | 0.252220156 | 0.139 | 0 | 1 | 15 |
| zgc:91944 | 0.03454 | 0.25154908 | 0.139 | 0 | 1 | 15 |
| edem2 | 0.03454 | 0.322574125 | 0.139 | 0 | 1 | 15 |
| kdm6ba | 0.03454 | 0.251716119 | 0.139 | 0 | 1 | 15 |
| mllt10 | 0.03454 | 0.234426581 | 0.139 | 0 | 1 | 15 |
| slc6a11a | 0.03454 | 0.245519702 | 0.139 | 0 | 1 | 15 |
| CABZ01113192.1 | 0.03454 | 0.225279286 | 0.139 | 0 | 1 | 15 |
| RASSF5 | 0.03454 | 0.245180317 | 0.139 | 0 | 1 | 15 |
| ddx54 | 0.03454 | 0.258572835 | 0.139 | 0 | 1 | 15 |
| dpm1 | 0.03454 | 0.221998798 | 0.139 | 0 | 1 | 15 |
| lrig2 | 0.03454 | 0.249663715 | 0.139 | 0 | 1 | 15 |
| ngrn | 0.03454 | 0.205118827 | 0.139 | 0 | 1 | 15 |
| plgrkt | 0.03454 | 0.206425889 | 0.139 | 0 | 1 | 15 |
| praf2 | 0.03454 | 0.239203067 | 0.139 | 0 | 1 | 15 |
| rars | 0.03454 | 0.257911924 | 0.139 | 0 | 1 | 15 |
| sbds | 0.03454 | 0.266444229 | 0.139 | 0 | 1 | 15 |
| smg6 | 0.03454 | 0.220119792 | 0.139 | 0 | 1 | 15 |
| taz | 0.03454 | 0.183410427 | 0.139 | 0 | 1 | 15 |
| tmem42a | 0.03454 | 0.245811622 | 0.139 | 0 | 1 | 15 |
| ttc4 | 0.03454 | 0.242177962 | 0.139 | 0 | 1 | 15 |
| yars | 0.03454 | 0.23334698 | 0.139 | 0 | 1 | 15 |
| zgc:136439 | 0.03454 | 0.220960702 | 0.139 | 0 | 1 | 15 |
| CU459056.1 | 0.03462 | -0.45351275 | 0.111 | 0.348 | 1 | 15 |
| sf3b2 | 0.03465 | 0.414102996 | 0.306 | 0.087 | 1 | 15 |
| cct3 | 0.0348 | 0.470240521 | 0.639 | 0.304 | 1 | 15 |
| aes | 0.03483 | -0.40927608 | 0.028 | 0.217 | 1 | 15 |
| mrpl32 | 0.03591 | 0.471749025 | 0.361 | 0.087 | 1 | 15 |
| rn7sk | 0.03605 | 0.437903245 | 0.833 | 0.522 | 1 | 15 |
| eef1db | 0.03695 | 0.348168106 | 0.806 | 0.652 | 1 | 15 |

| | | | | | | |
|---|---|---|---|---|---|---|
| CU074419.2 | 0.03751 | -0.36206981 | 0.028 | 0.217 | 1 | 15 |
| ndufb8 | 0.03779 | 0.32019143 | 0.611 | 0.304 | 1 | 15 |
| igf3 | 0.03786 | 0.604115579 | 0.667 | 0.348 | 1 | 15 |
| zc3h7ba | 0.03799 | 0.410149448 | 0.25 | 0.043 | 1 | 15 |
| ginm1 | 0.03807 | -0.25484831 | 0.278 | 0.348 | 1 | 15 |
| rbm19 | 0.03826 | 0.337711774 | 0.278 | 0.043 | 1 | 15 |
| kmt2cb | 0.03843 | -0.42297329 | 0.25 | 0.435 | 1 | 15 |
| col4a5 | 0.03954 | 0.426707814 | 0.361 | 0.087 | 1 | 15 |
| oaz1a | 0.03982 | 0.28744641 | 0.917 | 0.783 | 1 | 15 |
| fxyd6l | 0.03992 | -0.48343295 | 0.111 | 0.348 | 1 | 15 |
| sdf2l1 | 0.04001 | -0.37893017 | 0.111 | 0.261 | 1 | 15 |
| alpl | 0.04042 | 0.268199397 | 0.222 | 0.043 | 1 | 15 |
| col5a2a | 0.04046 | -0.50556384 | 0.083 | 0.348 | 1 | 15 |
| zgc:85777 | 0.04091 | -0.26837235 | 0.083 | 0.304 | 1 | 15 |
| hexim1 | 0.041 | 0.396985749 | 0.306 | 0.087 | 1 | 15 |
| ccnd2a | 0.04166 | 0.277201011 | 0.306 | 0.087 | 1 | 15 |
| ivns1abpa | 0.04188 | 0.369270727 | 0.278 | 0.043 | 1 | 15 |
| cd248a | 0.04252 | -0.5285521 | 0.111 | 0.391 | 1 | 15 |
| foxo3b | 0.04281 | 0.353483497 | 0.25 | 0.043 | 1 | 15 |
| krt18a.1 | 0.04289 | -0.30665406 | 0.833 | 0.87 | 1 | 15 |
| mcl1a | 0.043 | -0.35408746 | 0.778 | 0.913 | 1 | 15 |
| prdx2 | 0.04302 | -0.39173737 | 0.583 | 0.87 | 1 | 15 |
| ube2ia | 0.04343 | 0.436514914 | 0.389 | 0.174 | 1 | 15 |
| srsf4 | 0.04345 | 0.351636775 | 0.278 | 0.13 | 1 | 15 |
| proza | 0.04349 | -0.34678268 | 0.083 | 0.217 | 1 | 15 |
| tm2d1 | 0.04415 | 0.452659301 | 0.278 | 0.043 | 1 | 15 |
| s1pr2 | 0.04418 | -0.48140437 | 0.139 | 0.391 | 1 | 15 |
| pex5 | 0.0445 | -0.38989685 | 0.028 | 0.217 | 1 | 15 |
| zgc:114188 | 0.04479 | 0.211927483 | 1 | 0.913 | 1 | 15 |
| rad23b | 0.04536 | 0.465005019 | 0.444 | 0.174 | 1 | 15 |
| socs3a | 0.04617 | -0.26910216 | 0.889 | 0.913 | 1 | 15 |
| zgc:56095 | 0.04661 | -0.13882036 | 0.306 | 0.261 | 1 | 15 |
| rpl12 | 0.04663 | 0.194925104 | 0.972 | 0.913 | 1 | 15 |
| si:dkey-97a13.6 | 0.04689 | 0.136593174 | 0.167 | 0.043 | 1 | 15 |
| wbp2nl | 0.04804 | -0.40734213 | 0.083 | 0.304 | 1 | 15 |
| tspan18b | 0.04836 | -0.46471769 | 0.028 | 0.217 | 1 | 15 |
| fez1 | 0.04868 | 0.407411751 | 0.278 | 0.043 | 1 | 15 |
| vamp3 | 0.04904 | 0.235915828 | 0.5 | 0.261 | 1 | 15 |
| ptmaa | 0.04917 | -0.11076546 | 0.861 | 0.739 | 1 | 15 |
| atp2a2b | 0.04953 | 0.262486026 | 0.222 | 0.043 | 1 | 15 |
| ier2a | 0.04963 | -0.28500251 | 0.694 | 0.739 | 1 | 15 |
| mt-nd4 | 0.0498 | -0.33042126 | 0.778 | 0.87 | 1 | 15 |
| mtus1b | 0.04982 | 0.46165559 | 0.333 | 0.087 | 1 | 15 |
| CABZ01076667.1 | 0.05061 | 0.399024834 | 0.278 | 0.043 | 1 | 15 |
| rpl6 | 0.05069 | 0.213714301 | 0.972 | 0.913 | 1 | 15 |
| ndufa1 | 0.05093 | -0.23189615 | 0.444 | 0.435 | 1 | 15 |
| iscub | 0.05104 | 0.439591003 | 0.583 | 0.304 | 1 | 15 |
| pdcd5 | 0.05148 | 0.428564127 | 0.333 | 0.087 | 1 | 15 |

| | | | | | | |
|---|---|---|---|---|---|---|
| pip5k1ab | 0.05162 | -0.30671835 | 0.028 | 0.217 | 1 | 15 |
| npl | 0.05166 | 0.171205633 | 0.25 | 0.087 | 1 | 15 |
| hmg20a | 0.05233 | 0.387109142 | 0.25 | 0.043 | 1 | 15 |
| per1b | 0.05235 | -0.39559159 | 0.056 | 0.261 | 1 | 15 |
| eif6 | 0.05247 | 0.425720596 | 0.417 | 0.13 | 1 | 15 |
| glrx5 | 0.05277 | -0.34440892 | 0.361 | 0.478 | 1 | 15 |
| mrc2 | 0.05333 | -0.36457806 | 0.028 | 0.217 | 1 | 15 |
| tpp1 | 0.05364 | 0.175803104 | 0.25 | 0.087 | 1 | 15 |
| mrgbp | 0.05391 | 0.366199815 | 0.25 | 0.043 | 1 | 15 |
| CABZ01077217.1 | 0.05391 | -0.26589597 | 0.028 | 0.217 | 1 | 15 |
| CU694197.1 | 0.05413 | 0.308827565 | 0.389 | 0.261 | 1 | 15 |
| chst1 | 0.05474 | -0.25942218 | 0.028 | 0.174 | 1 | 15 |
| rbp4 | 0.05491 | -0.51505989 | 0.222 | 0.391 | 1 | 15 |
| rpl36 | 0.05551 | 0.154760402 | 0.972 | 0.913 | 1 | 15 |
| kdm6bb | 0.05601 | 0.311067917 | 0.222 | 0.043 | 1 | 15 |
| ccnl1b | 0.0562 | 0.355592867 | 0.306 | 0.087 | 1 | 15 |
| lnpep | 0.05631 | -0.38663488 | 0.028 | 0.217 | 1 | 15 |
| ckap5 | 0.05631 | -0.19457246 | 0.056 | 0.217 | 1 | 15 |
| bmp4 | 0.05634 | -0.34301825 | 0.083 | 0.304 | 1 | 15 |
| col1a1a | 0.05669 | -0.53881398 | 0.111 | 0.348 | 1 | 15 |
| arhgap35a | 0.05692 | -0.33030768 | 0.028 | 0.217 | 1 | 15 |
| cox6a2 | 0.05694 | -0.4637945 | 0.167 | 0.391 | 1 | 15 |
| cmbl | 0.05811 | 0.204670191 | 0.111 | 0.087 | 1 | 15 |
| prpf8 | 0.05873 | -0.32781493 | 0.028 | 0.217 | 1 | 15 |
| si:ch211-191j22.3 | 0.05886 | 0.271621774 | 0.5 | 0.391 | 1 | 15 |
| LOC110438024 | 0.05887 | 0.369683459 | 0.222 | 0.087 | 1 | 15 |
| rpl36a | 0.0592 | 0.122384813 | 1 | 1 | 1 | 15 |
| CR381686.3 | 0.05929 | 0.196838873 | 0.194 | 0.043 | 1 | 15 |
| fbn2b | 0.06031 | -0.37308977 | 0.194 | 0.391 | 1 | 15 |
| wu:fb16f03 | 0.06032 | -0.40742705 | 0.222 | 0.522 | 1 | 15 |
| fam49a | 0.06043 | -0.30777029 | 0.028 | 0.217 | 1 | 15 |
| ace | 0.06065 | -0.48789971 | 0.139 | 0.391 | 1 | 15 |
| zgc:195173 | 0.06077 | -0.37087318 | 0.639 | 0.826 | 1 | 15 |
| fem1c | 0.06116 | -0.27025296 | 0.028 | 0.217 | 1 | 15 |
| marcksb | 0.0612 | -0.45055572 | 0.139 | 0.391 | 1 | 15 |
| nr2f2 | 0.0614 | -0.42094426 | 0.222 | 0.522 | 1 | 15 |
| rnf146 | 0.0616 | -0.19075989 | 0.056 | 0.217 | 1 | 15 |
| cxcl12a | 0.06313 | -0.49108764 | 0.028 | 0.217 | 1 | 15 |
| fam120c | 0.06321 | 0.452239579 | 0.444 | 0.174 | 1 | 15 |
| btf3l4 | 0.06332 | 0.279638313 | 0.472 | 0.217 | 1 | 15 |
| tnfrsfa | 0.06341 | 0.419867611 | 0.444 | 0.217 | 1 | 15 |
| phactr4b | 0.06375 | -0.32887036 | 0.028 | 0.217 | 1 | 15 |
| actb1 | 0.06382 | -0.28118393 | 0.639 | 0.652 | 1 | 15 |
| smndc1 | 0.06383 | 0.428591528 | 0.417 | 0.174 | 1 | 15 |
| olfml3b | 0.06395 | -0.35236627 | 0.028 | 0.217 | 1 | 15 |
| abca2 | 0.06408 | 0.192861427 | 0.111 | 0 | 1 | 15 |
| LOC101883704 | 0.06408 | 0.199722079 | 0.111 | 0 | 1 | 15 |
| XLOC-023492 | 0.06408 | 0.165336864 | 0.111 | 0 | 1 | 15 |

| | | | | | | |
|---|---|---|---|---|---|---|
| abtb1 | 0.06408 | 0.199689035 | 0.111 | 0 | 1 | 15 |
| ache | 0.06408 | 0.198178717 | 0.111 | 0 | 1 | 15 |
| ccdc174 | 0.06408 | 0.195723895 | 0.111 | 0 | 1 | 15 |
| cwc25 | 0.06408 | 0.199020549 | 0.111 | 0 | 1 | 15 |
| dhrs12 | 0.06408 | 0.187445805 | 0.111 | 0 | 1 | 15 |
| dusp23b | 0.06408 | 0.177764378 | 0.111 | 0 | 1 | 15 |
| fam181b | 0.06408 | 0.213188947 | 0.111 | 0 | 1 | 15 |
| fcf1 | 0.06408 | 0.171663435 | 0.111 | 0 | 1 | 15 |
| hgh1 | 0.06408 | 0.183763908 | 0.111 | 0 | 1 | 15 |
| jade3 | 0.06408 | 0.205843303 | 0.111 | 0 | 1 | 15 |
| otud5b | 0.06408 | 0.212881786 | 0.111 | 0 | 1 | 15 |
| pex14 | 0.06408 | 0.178533012 | 0.111 | 0 | 1 | 15 |
| sephs2 | 0.06408 | 0.171852884 | 0.111 | 0 | 1 | 15 |
| si:ch211-15d5.12 | 0.06408 | 0.194311955 | 0.111 | 0 | 1 | 15 |
| tab2 | 0.06408 | 0.207637396 | 0.111 | 0 | 1 | 15 |
| timeless | 0.06408 | 0.18911519 | 0.111 | 0 | 1 | 15 |
| tprg1l | 0.06408 | 0.168772893 | 0.111 | 0 | 1 | 15 |
| ube3b | 0.06408 | 0.173317867 | 0.111 | 0 | 1 | 15 |
| zdhhc8b | 0.06408 | 0.221589757 | 0.111 | 0 | 1 | 15 |
| CABZ01072885.1 | 0.06408 | 0.210211659 | 0.111 | 0 | 1 | 15 |
| CR293507.1 | 0.06408 | 0.216078959 | 0.111 | 0 | 1 | 15 |
| XLOC-001102 | 0.06408 | 0.192699512 | 0.111 | 0 | 1 | 15 |
| aasdhppt | 0.06408 | 0.223199822 | 0.111 | 0 | 1 | 15 |
| arid4a | 0.06408 | 0.20343318 | 0.111 | 0 | 1 | 15 |
| bop1 | 0.06408 | 0.203987944 | 0.111 | 0 | 1 | 15 |
| cacul1 | 0.06408 | 0.210108848 | 0.111 | 0 | 1 | 15 |
| cdkn1a | 0.06408 | 0.176300265 | 0.111 | 0 | 1 | 15 |
| celf1 | 0.06408 | 0.163042261 | 0.111 | 0 | 1 | 15 |
| chpt1 | 0.06408 | 0.190137178 | 0.111 | 0 | 1 | 15 |
| cnppd1 | 0.06408 | 0.211581661 | 0.111 | 0 | 1 | 15 |
| dcun1d3 | 0.06408 | 0.172220129 | 0.111 | 0 | 1 | 15 |
| ddx10 | 0.06408 | 0.166145337 | 0.111 | 0 | 1 | 15 |
| ets1 | 0.06408 | 0.203899769 | 0.111 | 0 | 1 | 15 |
| gsk3aa | 0.06408 | 0.233232572 | 0.111 | 0 | 1 | 15 |
| hipk3a | 0.06408 | 0.17977248 | 0.111 | 0 | 1 | 15 |
| isca2 | 0.06408 | 0.111058329 | 0.111 | 0 | 1 | 15 |
| lats2 | 0.06408 | 0.198105658 | 0.111 | 0 | 1 | 15 |
| mtmr14 | 0.06408 | 0.238483875 | 0.111 | 0 | 1 | 15 |
| nipsnap3a | 0.06408 | 0.168437089 | 0.111 | 0 | 1 | 15 |
| rabggtb | 0.06408 | 0.187234767 | 0.111 | 0 | 1 | 15 |
| rnaseh2b | 0.06408 | 0.198754711 | 0.111 | 0 | 1 | 15 |
| sc:d189 | 0.06408 | 0.207191505 | 0.111 | 0 | 1 | 15 |
| si:ch73-248e21.1 | 0.06408 | 0.203603928 | 0.111 | 0 | 1 | 15 |
| si:dkey-23a13.22 | 0.06408 | 0.154555959 | 0.111 | 0 | 1 | 15 |
| si:dkeyp-115e12.7 | 0.06408 | 0.189393827 | 0.111 | 0 | 1 | 15 |
| ypel3 | 0.06408 | 0.189659496 | 0.111 | 0 | 1 | 15 |
| ywhae2 | 0.06408 | 0.185917984 | 0.111 | 0 | 1 | 15 |
| hbs1l | 0.06408 | 0.168356439 | 0.111 | 0 | 1 | 15 |

| | | | | | | |
|---|---|---|---|---|---|---|
| ptpmt1 | 0.06408 | 0.200192039 | 0.111 | 0 | 1 | 15 |
| si:dkeyp-68b7.5 | 0.06408 | 0.281112711 | 0.111 | 0 | 1 | 15 |
| uprt | 0.06408 | 0.18168796 | 0.111 | 0 | 1 | 15 |
| urm1 | 0.06408 | 0.181412707 | 0.111 | 0 | 1 | 15 |
| AK6 | 0.06408 | 0.192272212 | 0.111 | 0 | 1 | 15 |
| BX465227.1 | 0.06408 | 0.189329203 | 0.111 | 0 | 1 | 15 |
| CU693502.1 | 0.06408 | 0.195461684 | 0.111 | 0 | 1 | 15 |
| FBXO48 | 0.06408 | 0.105611549 | 0.111 | 0 | 1 | 15 |
| FP101887.1 | 0.06408 | 0.243589447 | 0.111 | 0 | 1 | 15 |
| LOC108190702 | 0.06408 | 0.214289769 | 0.111 | 0 | 1 | 15 |
| LOC110440070 | 0.06408 | 0.223641281 | 0.111 | 0 | 1 | 15 |
| XLOC-035413 | 0.06408 | 0.205003571 | 0.111 | 0 | 1 | 15 |
| abat | 0.06408 | 0.2378926 | 0.111 | 0 | 1 | 15 |
| agpat2 | 0.06408 | 0.179184798 | 0.111 | 0 | 1 | 15 |
| b3gnt2l | 0.06408 | 0.203927697 | 0.111 | 0 | 1 | 15 |
| bcar1 | 0.06408 | 0.20697322 | 0.111 | 0 | 1 | 15 |
| bcl11aa | 0.06408 | 0.209751447 | 0.111 | 0 | 1 | 15 |
| bcl9l | 0.06408 | 0.197514322 | 0.111 | 0 | 1 | 15 |
| c1galt1c1 | 0.06408 | 0.183305533 | 0.111 | 0 | 1 | 15 |
| cars2 | 0.06408 | 0.192631573 | 0.111 | 0 | 1 | 15 |
| cbx6b | 0.06408 | 0.221166217 | 0.111 | 0 | 1 | 15 |
| ccdc6a | 0.06408 | 0.204866015 | 0.111 | 0 | 1 | 15 |
| chmp2bb | 0.06408 | 0.204575632 | 0.111 | 0 | 1 | 15 |
| cisd1 | 0.06408 | 0.198532849 | 0.111 | 0 | 1 | 15 |
| coro2ba | 0.06408 | 0.173748821 | 0.111 | 0 | 1 | 15 |
| cpsf1 | 0.06408 | 0.210078735 | 0.111 | 0 | 1 | 15 |
| dhps | 0.06408 | 0.231524526 | 0.111 | 0 | 1 | 15 |
| dr1 | 0.06408 | 0.207910397 | 0.111 | 0 | 1 | 15 |
| dut | 0.06408 | 0.210113225 | 0.111 | 0 | 1 | 15 |
| efnb1 | 0.06408 | 0.166888628 | 0.111 | 0 | 1 | 15 |
| elavl1b | 0.06408 | 0.240403739 | 0.111 | 0 | 1 | 15 |
| erap2 | 0.06408 | 0.182870464 | 0.111 | 0 | 1 | 15 |
| fam173b | 0.06408 | 0.199487806 | 0.111 | 0 | 1 | 15 |
| fgd6 | 0.06408 | 0.182618497 | 0.111 | 0 | 1 | 15 |
| fuca1.1 | 0.06408 | 0.20930353 | 0.111 | 0 | 1 | 15 |
| gabpb2a | 0.06408 | 0.197617073 | 0.111 | 0 | 1 | 15 |
| galr2b | 0.06408 | 0.215777791 | 0.111 | 0 | 1 | 15 |
| gtf2e1 | 0.06408 | 0.211213104 | 0.111 | 0 | 1 | 15 |
| hcn2b | 0.06408 | 0.222215501 | 0.111 | 0 | 1 | 15 |
| igfbp7 | 0.06408 | 0.257951677 | 0.111 | 0 | 1 | 15 |
| ints14 | 0.06408 | 0.203695341 | 0.111 | 0 | 1 | 15 |
| ints8 | 0.06408 | 0.191734129 | 0.111 | 0 | 1 | 15 |
| jak2a | 0.06408 | 0.199918172 | 0.111 | 0 | 1 | 15 |
| klf11a | 0.06408 | 0.239683417 | 0.111 | 0 | 1 | 15 |
| klhl21 | 0.06408 | 0.194632682 | 0.111 | 0 | 1 | 15 |
| lamtor1 | 0.06408 | 0.185059001 | 0.111 | 0 | 1 | 15 |
| lrig1 | 0.06408 | 0.21484897 | 0.111 | 0 | 1 | 15 |
| lto1 | 0.06408 | 0.219998785 | 0.111 | 0 | 1 | 15 |

| | | | | | | |
|---|---|---|---|---|---|---|
| med17 | 0.06408 | 0.196954274 | 0.111 | 0 | 1 | 15 |
| med21 | 0.06408 | 0.170936309 | 0.111 | 0 | 1 | 15 |
| med23 | 0.06408 | 0.186596408 | 0.111 | 0 | 1 | 15 |
| mertka | 0.06408 | 0.203387997 | 0.111 | 0 | 1 | 15 |
| mpp5a | 0.06408 | 0.225584022 | 0.111 | 0 | 1 | 15 |
| mrps15 | 0.06408 | 0.170103323 | 0.111 | 0 | 1 | 15 |
| mut | 0.06408 | 0.20004609 | 0.111 | 0 | 1 | 15 |
| nelfb | 0.06408 | 0.231835191 | 0.111 | 0 | 1 | 15 |
| nlk1 | 0.06408 | 0.203646267 | 0.111 | 0 | 1 | 15 |
| noc2l | 0.06408 | 0.207605585 | 0.111 | 0 | 1 | 15 |
| nudt9 | 0.06408 | 0.24603959 | 0.111 | 0 | 1 | 15 |
| nup210 | 0.06408 | 0.205346143 | 0.111 | 0 | 1 | 15 |
| osgep | 0.06408 | 0.175945291 | 0.111 | 0 | 1 | 15 |
| pgp | 0.06408 | 0.16937245 | 0.111 | 0 | 1 | 15 |
| plekhb2 | 0.06408 | 0.196765951 | 0.111 | 0 | 1 | 15 |
| polr1a | 0.06408 | 0.218363587 | 0.111 | 0 | 1 | 15 |
| polr1d | 0.06408 | 0.178721033 | 0.111 | 0 | 1 | 15 |
| prkrip1 | 0.06408 | 0.180280608 | 0.111 | 0 | 1 | 15 |
| prox1a | 0.06408 | 0.264031189 | 0.111 | 0 | 1 | 15 |
| rab35b | 0.06408 | 0.202655518 | 0.111 | 0 | 1 | 15 |
| rabl6b | 0.06408 | 0.189412983 | 0.111 | 0 | 1 | 15 |
| rarab | 0.06408 | 0.189402875 | 0.111 | 0 | 1 | 15 |
| rassf8a | 0.06408 | 0.209302746 | 0.111 | 0 | 1 | 15 |
| rdh8a | 0.06408 | 0.174330901 | 0.111 | 0 | 1 | 15 |
| rgs2 | 0.06408 | 0.301123184 | 0.111 | 0 | 1 | 15 |
| rmnd1 | 0.06408 | 0.183021915 | 0.111 | 0 | 1 | 15 |
| rnaseh2a | 0.06408 | 0.176506535 | 0.111 | 0 | 1 | 15 |
| rnf111 | 0.06408 | 0.225132048 | 0.111 | 0 | 1 | 15 |
| rnf20 | 0.06408 | 0.198567261 | 0.111 | 0 | 1 | 15 |
| rspry1 | 0.06408 | 0.15730292 | 0.111 | 0 | 1 | 15 |
| samd4b | 0.06408 | 0.138381138 | 0.111 | 0 | 1 | 15 |
| sgk1 | 0.06408 | 0.215446592 | 0.111 | 0 | 1 | 15 |
| sh2b1 | 0.06408 | 0.21197857 | 0.111 | 0 | 1 | 15 |
| shmt2 | 0.06408 | 0.12365119 | 0.111 | 0 | 1 | 15 |
| si:ch211-266o15.1 | 0.06408 | 0.183321659 | 0.111 | 0 | 1 | 15 |
| si:ch211-89o9.6 | 0.06408 | 0.20524161 | 0.111 | 0 | 1 | 15 |
| si:dkey-122a22.2 | 0.06408 | 0.182047301 | 0.111 | 0 | 1 | 15 |
| si:dkey-24l11.2 | 0.06408 | 0.187641604 | 0.111 | 0 | 1 | 15 |
| si:dkey-7l6.3 | 0.06408 | 0.249961632 | 0.111 | 0 | 1 | 15 |
| si:dkey-90a24.1 | 0.06408 | 0.268103364 | 0.111 | 0 | 1 | 15 |
| si:rp71-1k13.6 | 0.06408 | 0.202722924 | 0.111 | 0 | 1 | 15 |
| slc39a9 | 0.06408 | 0.180162732 | 0.111 | 0 | 1 | 15 |
| slc7a10a | 0.06408 | 0.164089298 | 0.111 | 0 | 1 | 15 |
| slc9a3r1a | 0.06408 | 0.195569847 | 0.111 | 0 | 1 | 15 |
| smc5 | 0.06408 | 0.18403078 | 0.111 | 0 | 1 | 15 |
| tead1b | 0.06408 | 0.209099634 | 0.111 | 0 | 1 | 15 |
| tent5ba | 0.06408 | 0.218691355 | 0.111 | 0 | 1 | 15 |
| tmem187 | 0.06408 | 0.18512574 | 0.111 | 0 | 1 | 15 |

| | | | | | | |
|---|---|---|---|---|---|---|
| ttyh3a | 0.06408 | 0.179420564 | 0.111 | 0 | 1 | 15 |
| ubap2a | 0.06408 | 0.218864013 | 0.111 | 0 | 1 | 15 |
| xpot | 0.06408 | 0.164640559 | 0.111 | 0 | 1 | 15 |
| yipf1 | 0.06408 | 0.222901592 | 0.111 | 0 | 1 | 15 |
| zgc:101716 | 0.06408 | 0.147140525 | 0.111 | 0 | 1 | 15 |
| znf644a | 0.06408 | 0.191280898 | 0.111 | 0 | 1 | 15 |
| nrd1a | 0.0641 | -0.32690437 | 0.028 | 0.217 | 1 | 15 |
| hmgb1a | 0.06423 | 0.273333102 | 0.833 | 0.565 | 1 | 15 |
| gnai3 | 0.06426 | -0.30938825 | 0.028 | 0.217 | 1 | 15 |
| ncor2 | 0.06464 | -0.12985362 | 0.222 | 0.217 | 1 | 15 |
| errfi1a | 0.06472 | -0.40016569 | 0.028 | 0.217 | 1 | 15 |
| zgc:66433 | 0.06488 | -0.44595936 | 0.194 | 0.391 | 1 | 15 |
| pdgfrb | 0.06511 | -0.33437117 | 0.028 | 0.217 | 1 | 15 |
| stk19 | 0.06511 | -0.3310965 | 0.028 | 0.217 | 1 | 15 |
| zbtb46 | 0.06544 | 0.148465255 | 0.222 | 0.087 | 1 | 15 |
| rps4x | 0.06649 | 0.218982531 | 1 | 0.913 | 1 | 15 |
| hsf2 | 0.06738 | 0.398578138 | 0.306 | 0.087 | 1 | 15 |
| csnk1da | 0.06833 | 0.428255826 | 0.389 | 0.13 | 1 | 15 |
| hsp90b1 | 0.06888 | -0.2393785 | 0.306 | 0.348 | 1 | 15 |
| sema3ab | 0.06899 | 0.26755718 | 0.25 | 0.087 | 1 | 15 |
| cdh2 | 0.0693 | -0.33642519 | 0.417 | 0.565 | 1 | 15 |
| eif3ha | 0.06965 | 0.381290458 | 0.694 | 0.391 | 1 | 15 |
| ganab | 0.06999 | 0.155003627 | 0.222 | 0.087 | 1 | 15 |
| nlrc3l1 | 0.07026 | -0.21509247 | 0.028 | 0.174 | 1 | 15 |
| sypl2b | 0.07048 | 0.105562156 | 0.139 | 0.043 | 1 | 15 |
| tram2 | 0.07086 | -0.42638553 | 0.167 | 0.435 | 1 | 15 |
| rps29 | 0.07088 | 0.131275339 | 1 | 0.87 | 1 | 15 |
| tnrc6b | 0.07175 | -0.43932144 | 0.139 | 0.391 | 1 | 15 |
| stk17b | 0.07187 | 0.321392096 | 0.25 | 0.043 | 1 | 15 |
| vps29 | 0.07205 | -0.39437203 | 0.194 | 0.478 | 1 | 15 |
| rab1ab | 0.0722 | 0.400768981 | 0.556 | 0.261 | 1 | 15 |
| cd82b | 0.07259 | -0.38045801 | 0.056 | 0.261 | 1 | 15 |
| rps24 | 0.07293 | 0.162642588 | 0.972 | 0.913 | 1 | 15 |
| uqcrc2a | 0.07327 | 0.267899052 | 0.222 | 0.087 | 1 | 15 |
| ctnna1 | 0.07343 | 0.393199672 | 0.333 | 0.087 | 1 | 15 |
| rps8a | 0.0738 | 0.188421589 | 0.972 | 0.913 | 1 | 15 |
| cd164 | 0.07481 | -0.27759313 | 0.306 | 0.391 | 1 | 15 |
| sema4ba | 0.07591 | -0.3534327 | 0.056 | 0.261 | 1 | 15 |
| acvr1ba | 0.07629 | -0.32622225 | 0.056 | 0.261 | 1 | 15 |
| hspa5 | 0.0764 | -0.37312395 | 0.361 | 0.478 | 1 | 15 |
| s100a10b | 0.07674 | -0.39908506 | 0.5 | 0.609 | 1 | 15 |
| rpl37 | 0.07676 | 0.171369994 | 1 | 0.87 | 1 | 15 |
| mpzl2b | 0.07694 | 0.468791628 | 0.556 | 0.261 | 1 | 15 |
| sfswap | 0.07729 | -0.11939983 | 0.139 | 0.13 | 1 | 15 |
| ggcta | 0.07736 | 0.330669305 | 0.25 | 0.043 | 1 | 15 |
| st5 | 0.07741 | 0.207620321 | 0.194 | 0.043 | 1 | 15 |
| polr2a | 0.07776 | 0.377837891 | 0.25 | 0.043 | 1 | 15 |
| si:dkey-222f8.3 | 0.07784 | -0.29000082 | 0.056 | 0.261 | 1 | 15 |

| | | | | | | |
|---|---|---|---|---|---|---|
| LOC103910871 | 0.07802 | -0.17809081 | 0 | 0.087 | 1 | 15 |
| caska | 0.07802 | -0.20708766 | 0 | 0.087 | 1 | 15 |
| gnptg | 0.07802 | -0.13539909 | 0 | 0.087 | 1 | 15 |
| kif26ab | 0.07802 | -0.20878725 | 0 | 0.087 | 1 | 15 |
| si:ch211-108c6.2 | 0.07802 | -0.19945788 | 0 | 0.087 | 1 | 15 |
| tubgcp2 | 0.07802 | -0.13596964 | 0 | 0.087 | 1 | 15 |
| ucmaa | 0.07802 | -0.13557727 | 0 | 0.087 | 1 | 15 |
| CU550719.2 | 0.07802 | -0.14481858 | 0 | 0.087 | 1 | 15 |
| CU571074.1 | 0.07802 | -0.16945542 | 0 | 0.087 | 1 | 15 |
| CU984579.1 | 0.07802 | -0.14040607 | 0 | 0.087 | 1 | 15 |
| aak1b | 0.07802 | -0.163964 | 0 | 0.087 | 1 | 15 |
| arhgap32a | 0.07802 | -0.1931505 | 0 | 0.087 | 1 | 15 |
| card19 | 0.07802 | -0.13831599 | 0 | 0.087 | 1 | 15 |
| cd2ap | 0.07802 | -0.19701036 | 0 | 0.087 | 1 | 15 |
| efna1a | 0.07802 | -0.16572067 | 0 | 0.087 | 1 | 15 |
| maml1 | 0.07802 | -0.13517604 | 0 | 0.087 | 1 | 15 |
| p3h2 | 0.07802 | -0.14281036 | 0 | 0.087 | 1 | 15 |
| pdlim5b | 0.07802 | -0.16016869 | 0 | 0.087 | 1 | 15 |
| prr11 | 0.07802 | -0.18602684 | 0 | 0.087 | 1 | 15 |
| rab3db | 0.07802 | -0.15708164 | 0 | 0.087 | 1 | 15 |
| sb:cb81 | 0.07802 | -0.1715502 | 0 | 0.087 | 1 | 15 |
| slc25a40 | 0.07802 | -0.17369751 | 0 | 0.087 | 1 | 15 |
| BX901883.3 | 0.07802 | -0.14693718 | 0 | 0.087 | 1 | 15 |
| BX957297.2 | 0.07802 | -0.22410132 | 0 | 0.087 | 1 | 15 |
| CABZ01069022.2 | 0.07802 | -0.14640317 | 0 | 0.087 | 1 | 15 |
| CT583672.1 | 0.07802 | -0.14553913 | 0 | 0.087 | 1 | 15 |
| LOC101885935 | 0.07802 | -0.19860905 | 0 | 0.087 | 1 | 15 |
| LOC103910470 | 0.07802 | -0.15829284 | 0 | 0.087 | 1 | 15 |
| LOC796314 | 0.07802 | -0.15340534 | 0 | 0.087 | 1 | 15 |
| MFSD3 | 0.07802 | -0.14412978 | 0 | 0.087 | 1 | 15 |
| aldh4a1 | 0.07802 | -0.18443098 | 0 | 0.087 | 1 | 15 |
| arhgap29a | 0.07802 | -0.16434343 | 0 | 0.087 | 1 | 15 |
| bub1 | 0.07802 | -0.15143362 | 0 | 0.087 | 1 | 15 |
| cbx2 | 0.07802 | -0.18139208 | 0 | 0.087 | 1 | 15 |
| cdc20 | 0.07802 | -0.24403944 | 0 | 0.087 | 1 | 15 |
| cks1b | 0.07802 | -0.17671069 | 0 | 0.087 | 1 | 15 |
| dennd5b | 0.07802 | -0.13645914 | 0 | 0.087 | 1 | 15 |
| dlgap5 | 0.07802 | -0.17812084 | 0 | 0.087 | 1 | 15 |
| eif2ak3 | 0.07802 | -0.16778741 | 0 | 0.087 | 1 | 15 |
| kif20a | 0.07802 | -0.18161673 | 0 | 0.087 | 1 | 15 |
| kif23 | 0.07802 | -0.18154326 | 0 | 0.087 | 1 | 15 |
| mindy2 | 0.07802 | -0.22487372 | 0 | 0.087 | 1 | 15 |
| mki67 | 0.07802 | -0.17958801 | 0 | 0.087 | 1 | 15 |
| mtrr | 0.07802 | -0.15511625 | 0 | 0.087 | 1 | 15 |
| noc4l | 0.07802 | -0.16902317 | 0 | 0.087 | 1 | 15 |
| nusap1 | 0.07802 | -0.20857697 | 0 | 0.087 | 1 | 15 |
| pak1ip1 | 0.07802 | -0.18545629 | 0 | 0.087 | 1 | 15 |
| pccb | 0.07802 | -0.1408946 | 0 | 0.087 | 1 | 15 |

| | | | | | | |
|---|---|---|---|---|---|---|
| pcnt | 0.07802 | -0.1813928 | 0 | 0.087 | 1 | 15 |
| plekha7b | 0.07802 | -0.26952023 | 0 | 0.087 | 1 | 15 |
| sall1a | 0.07802 | -0.13752021 | 0 | 0.087 | 1 | 15 |
| sdr42e1 | 0.07802 | -0.17171914 | 0 | 0.087 | 1 | 15 |
| sema4bb | 0.07802 | -0.17281924 | 0 | 0.087 | 1 | 15 |
| sept9a | 0.07802 | -0.18372773 | 0 | 0.087 | 1 | 15 |
| si:ch211-266i6.3 | 0.07802 | -0.17929164 | 0 | 0.087 | 1 | 15 |
| slc50a1 | 0.07802 | -0.18143456 | 0 | 0.087 | 1 | 15 |
| ube2c | 0.07802 | -0.27963588 | 0 | 0.087 | 1 | 15 |
| ugt5b1 | 0.07802 | -0.12489088 | 0 | 0.087 | 1 | 15 |
| zbtb3 | 0.07802 | -0.2131528 | 0 | 0.087 | 1 | 15 |
| zgc:153184 | 0.07802 | -0.13532384 | 0 | 0.087 | 1 | 15 |
| zgc:194627 | 0.07802 | -0.37417872 | 0 | 0.087 | 1 | 15 |
| BX005183.3 | 0.07802 | -0.25448825 | 0 | 0.087 | 1 | 15 |
| CABZ01087948.1 | 0.07802 | -0.21254531 | 0 | 0.087 | 1 | 15 |
| CR936442.1 | 0.07802 | -0.2037627 | 0 | 0.087 | 1 | 15 |
| LOC101884506 | 0.07802 | -0.21925483 | 0 | 0.087 | 1 | 15 |
| RFESD (1 of many) | 0.07802 | -0.1747717 | 0 | 0.087 | 1 | 15 |
| XLOC-043151 | 0.07802 | -0.17137926 | 0 | 0.087 | 1 | 15 |
| abcc6a | 0.07802 | -0.2194738 | 0 | 0.087 | 1 | 15 |
| ajuba | 0.07802 | -0.12653133 | 0 | 0.087 | 1 | 15 |
| bicdl2 | 0.07802 | -0.14485768 | 0 | 0.087 | 1 | 15 |
| cdh24b | 0.07802 | -0.15967102 | 0 | 0.087 | 1 | 15 |
| clptm1l | 0.07802 | -0.21074934 | 0 | 0.087 | 1 | 15 |
| ephb4b | 0.07802 | -0.21291284 | 0 | 0.087 | 1 | 15 |
| gba | 0.07802 | -0.16759014 | 0 | 0.087 | 1 | 15 |
| kpna6 | 0.07802 | -0.22681943 | 0 | 0.087 | 1 | 15 |
| olfml2ba | 0.07802 | -0.25292806 | 0 | 0.087 | 1 | 15 |
| phf20a | 0.07802 | -0.15026634 | 0 | 0.087 | 1 | 15 |
| si:dkey-118j18.2 | 0.07802 | -0.17135939 | 0 | 0.087 | 1 | 15 |
| smad6a | 0.07802 | -0.17291188 | 0 | 0.087 | 1 | 15 |
| tbck | 0.07802 | -0.16741752 | 0 | 0.087 | 1 | 15 |
| tnfrsf19 | 0.07802 | -0.218526 | 0 | 0.087 | 1 | 15 |
| ublcp1 | 0.07802 | -0.23138757 | 0 | 0.087 | 1 | 15 |
| zgc:101783 | 0.07802 | -0.22376283 | 0 | 0.087 | 1 | 15 |
| enpp6 | 0.07802 | -0.18064542 | 0 | 0.087 | 1 | 15 |
| lrp2a | 0.07802 | -0.14969165 | 0 | 0.087 | 1 | 15 |
| nfatc3a | 0.07802 | -0.15075249 | 0 | 0.087 | 1 | 15 |
| CR753886.1 | 0.07802 | -0.12933555 | 0 | 0.087 | 1 | 15 |
| CU633745.1 | 0.07802 | -0.22697883 | 0 | 0.087 | 1 | 15 |
| LOC101885938 | 0.07802 | -0.12358079 | 0 | 0.087 | 1 | 15 |
| NCEH1 (1 of many) | 0.07802 | -0.23093219 | 0 | 0.087 | 1 | 15 |
| ankib1b | 0.07802 | -0.16325335 | 0 | 0.087 | 1 | 15 |
| dera | 0.07802 | -0.18385775 | 0 | 0.087 | 1 | 15 |
| drosha | 0.07802 | -0.22782255 | 0 | 0.087 | 1 | 15 |
| dusp2 | 0.07802 | -0.15869319 | 0 | 0.087 | 1 | 15 |
| fam173a | 0.07802 | -0.15170608 | 0 | 0.087 | 1 | 15 |
| fam199x | 0.07802 | -0.12810331 | 0 | 0.087 | 1 | 15 |

| | | | | | | |
|---|---|---|---|---|---|---|
| fkbp10a | 0.07802 | -0.19070203 | 0 | 0.087 | 1 | 15 |
| ftr51 | 0.07802 | -0.12037075 | 0 | 0.087 | 1 | 15 |
| or127-1 | 0.07802 | -0.13173302 | 0 | 0.087 | 1 | 15 |
| plch1 | 0.07802 | -0.14893697 | 0 | 0.087 | 1 | 15 |
| si:dkey-147f3.8 | 0.07802 | -0.25299706 | 0 | 0.087 | 1 | 15 |
| si:dkey-79d12.5 | 0.07802 | -0.16401806 | 0 | 0.087 | 1 | 15 |
| slc40a1 | 0.07802 | -0.11344517 | 0 | 0.087 | 1 | 15 |
| ttll1 | 0.07802 | -0.20326876 | 0 | 0.087 | 1 | 15 |
| zbtb12.2 | 0.07802 | -0.12817417 | 0 | 0.087 | 1 | 15 |
| zgc:158398 | 0.07802 | -0.1620732 | 0 | 0.087 | 1 | 15 |
| LOC101882475 | 0.07802 | -0.20001337 | 0 | 0.087 | 1 | 15 |
| LOC108182798 | 0.07802 | -0.12817884 | 0 | 0.087 | 1 | 15 |
| alp3 | 0.07802 | -0.11282236 | 0 | 0.087 | 1 | 15 |
| cyp26a1 | 0.07802 | -0.30348368 | 0 | 0.087 | 1 | 15 |
| dazap1 | 0.07802 | -0.11476476 | 0 | 0.087 | 1 | 15 |
| ftr93 | 0.07802 | -0.10835252 | 0 | 0.087 | 1 | 15 |
| gnpnat1 | 0.07802 | -0.12252292 | 0 | 0.087 | 1 | 15 |
| grb10b | 0.07802 | -0.21056287 | 0 | 0.087 | 1 | 15 |
| kcng2 | 0.07802 | -0.23469316 | 0 | 0.087 | 1 | 15 |
| mocs2 | 0.07802 | -0.20776231 | 0 | 0.087 | 1 | 15 |
| pkdcca | 0.07802 | -0.19400355 | 0 | 0.087 | 1 | 15 |
| pld1a | 0.07802 | -0.12606561 | 0 | 0.087 | 1 | 15 |
| ppp1r15a | 0.07802 | -0.19317766 | 0 | 0.087 | 1 | 15 |
| rxraa | 0.07802 | -0.13104285 | 0 | 0.087 | 1 | 15 |
| si:dkey-82o10.4 | 0.07802 | -0.1095679 | 0 | 0.087 | 1 | 15 |
| skila | 0.07802 | -0.15665889 | 0 | 0.087 | 1 | 15 |
| tnfrsf9a | 0.07802 | -0.2802522 | 0 | 0.087 | 1 | 15 |
| zgc:113531 | 0.07802 | -0.20954447 | 0 | 0.087 | 1 | 15 |
| znf1081 | 0.07802 | -0.12849486 | 0 | 0.087 | 1 | 15 |
| znf1173 | 0.07802 | -0.10884556 | 0 | 0.087 | 1 | 15 |
| AL935174.2 | 0.07802 | -0.1530048 | 0 | 0.087 | 1 | 15 |
| CABZ01038697.1 | 0.07802 | -0.15500795 | 0 | 0.087 | 1 | 15 |
| LOC108180286 | 0.07802 | -0.17543165 | 0 | 0.087 | 1 | 15 |
| LOC110439967 | 0.07802 | -0.17062055 | 0 | 0.087 | 1 | 15 |
| XLOC-024261 | 0.07802 | -0.15353248 | 0 | 0.087 | 1 | 15 |
| adamts10 | 0.07802 | -0.1748182 | 0 | 0.087 | 1 | 15 |
| bcl10 | 0.07802 | -0.17714469 | 0 | 0.087 | 1 | 15 |
| bet1 | 0.07802 | -0.16704229 | 0 | 0.087 | 1 | 15 |
| btbd6a | 0.07802 | -0.20484365 | 0 | 0.087 | 1 | 15 |
| c7a | 0.07802 | -0.18343466 | 0 | 0.087 | 1 | 15 |
| ca5a | 0.07802 | -0.19875885 | 0 | 0.087 | 1 | 15 |
| cdk17 | 0.07802 | -0.21148076 | 0 | 0.087 | 1 | 15 |
| cdkn1d | 0.07802 | -0.19353917 | 0 | 0.087 | 1 | 15 |
| ch25h | 0.07802 | -0.25152518 | 0 | 0.087 | 1 | 15 |
| col21a1 | 0.07802 | -0.2027858 | 0 | 0.087 | 1 | 15 |
| cry1ba | 0.07802 | -0.17023109 | 0 | 0.087 | 1 | 15 |
| ctu2 | 0.07802 | -0.13220052 | 0 | 0.087 | 1 | 15 |
| cyp2ad2 | 0.07802 | -0.13540307 | 0 | 0.087 | 1 | 15 |

| | | | | | | |
|---|---|---|---|---|---|---|
| dlg3 | 0.07802 | -0.18259405 | 0 | 0.087 | 1 | 15 |
| dtnbp1b | 0.07802 | -0.19012988 | 0 | 0.087 | 1 | 15 |
| ecpas | 0.07802 | -0.2087743 | 0 | 0.087 | 1 | 15 |
| edc4 | 0.07802 | -0.22048789 | 0 | 0.087 | 1 | 15 |
| ednraa | 0.07802 | -0.17669407 | 0 | 0.087 | 1 | 15 |
| efnb3b | 0.07802 | -0.24713721 | 0 | 0.087 | 1 | 15 |
| fbxo22 | 0.07802 | -0.15424963 | 0 | 0.087 | 1 | 15 |
| frmd4ba | 0.07802 | -0.21631301 | 0 | 0.087 | 1 | 15 |
| gpm6ab | 0.07802 | -0.17299423 | 0 | 0.087 | 1 | 15 |
| gxylt2 | 0.07802 | -0.192661 | 0 | 0.087 | 1 | 15 |
| igfbp1a | 0.07802 | -0.22157862 | 0 | 0.087 | 1 | 15 |
| ipmkb | 0.07802 | -0.18006696 | 0 | 0.087 | 1 | 15 |
| katnal1 | 0.07802 | -0.1239128 | 0 | 0.087 | 1 | 15 |
| magi2a | 0.07802 | -0.1730738 | 0 | 0.087 | 1 | 15 |
| mcoln1a | 0.07802 | -0.17121388 | 0 | 0.087 | 1 | 15 |
| mt-atp8 | 0.07802 | -0.17310042 | 0 | 0.087 | 1 | 15 |
| mycla | 0.07802 | -0.15334567 | 0 | 0.087 | 1 | 15 |
| narf | 0.07802 | -0.13435951 | 0 | 0.087 | 1 | 15 |
| ncdn | 0.07802 | -0.15825363 | 0 | 0.087 | 1 | 15 |
| nme6 | 0.07802 | -0.19693759 | 0 | 0.087 | 1 | 15 |
| nup93 | 0.07802 | -0.21098235 | 0 | 0.087 | 1 | 15 |
| oaz2a | 0.07802 | -0.13433706 | 0 | 0.087 | 1 | 15 |
| paip1 | 0.07802 | -0.14178203 | 0 | 0.087 | 1 | 15 |
| pane1 | 0.07802 | -0.1210422 | 0 | 0.087 | 1 | 15 |
| pemt | 0.07802 | -0.17387446 | 0 | 0.087 | 1 | 15 |
| plekhh1 | 0.07802 | -0.22831842 | 0 | 0.087 | 1 | 15 |
| ppm1db | 0.07802 | -0.14659821 | 0 | 0.087 | 1 | 15 |
| prpf18 | 0.07802 | -0.16280533 | 0 | 0.087 | 1 | 15 |
| prune | 0.07802 | -0.18153955 | 0 | 0.087 | 1 | 15 |
| rbm6 | 0.07802 | -0.17828249 | 0 | 0.087 | 1 | 15 |
| rgcc | 0.07802 | -0.16174931 | 0 | 0.087 | 1 | 15 |
| rgn | 0.07802 | -0.11847267 | 0 | 0.087 | 1 | 15 |
| rin3 | 0.07802 | -0.18032922 | 0 | 0.087 | 1 | 15 |
| rit1 | 0.07802 | -0.16106727 | 0 | 0.087 | 1 | 15 |
| rrm2 | 0.07802 | -0.11614851 | 0 | 0.087 | 1 | 15 |
| sft2d3 | 0.07802 | -0.16156511 | 0 | 0.087 | 1 | 15 |
| si:ch1073-159d7.13 | 0.07802 | -0.1647586 | 0 | 0.087 | 1 | 15 |
| si:ch211-130m23.5 | 0.07802 | -0.15101078 | 0 | 0.087 | 1 | 15 |
| si:ch73-25f10.6 | 0.07802 | -0.15327563 | 0 | 0.087 | 1 | 15 |
| si:dkey-187k19.2 | 0.07802 | -0.21430673 | 0 | 0.087 | 1 | 15 |
| si:dkey-20i20.2 | 0.07802 | -0.16361283 | 0 | 0.087 | 1 | 15 |
| si:dkey-75a21.2 | 0.07802 | -0.18416251 | 0 | 0.087 | 1 | 15 |
| slc25a36b | 0.07802 | -0.13549252 | 0 | 0.087 | 1 | 15 |
| snx30 | 0.07802 | -0.20960142 | 0 | 0.087 | 1 | 15 |
| snx33 | 0.07802 | -0.14726075 | 0 | 0.087 | 1 | 15 |
| srpx | 0.07802 | -0.20569558 | 0 | 0.087 | 1 | 15 |
| syt9b | 0.07802 | -0.19044803 | 0 | 0.087 | 1 | 15 |
| terb2 | 0.07802 | -0.1677977 | 0 | 0.087 | 1 | 15 |

| | | | | | | |
|---|---|---|---|---|---|---|
| tmbim1a | 0.07802 | -0.19741273 | 0 | 0.087 | 1 | 15 |
| tmem229b | 0.07802 | -0.12980032 | 0 | 0.087 | 1 | 15 |
| tmem259 | 0.07802 | -0.16264151 | 0 | 0.087 | 1 | 15 |
| tnfaip3 | 0.07802 | -0.1629863 | 0 | 0.087 | 1 | 15 |
| toe1 | 0.07802 | -0.13431559 | 0 | 0.087 | 1 | 15 |
| trdmt1 | 0.07802 | -0.16441216 | 0 | 0.087 | 1 | 15 |
| zdhhc18a | 0.07802 | -0.1450424 | 0 | 0.087 | 1 | 15 |
| zdhhc20b | 0.07802 | -0.16753991 | 0 | 0.087 | 1 | 15 |
| LOC405768 | 0.07802 | -0.20905602 | 0 | 0.087 | 1 | 15 |
| cyp46a1.2 | 0.07802 | -0.12171934 | 0 | 0.087 | 1 | 15 |
| inpp5d | 0.07802 | -0.12819982 | 0 | 0.087 | 1 | 15 |
| pmepa1 | 0.07802 | -0.25781217 | 0 | 0.087 | 1 | 15 |
| si:ch211-153a8.4 | 0.07802 | -0.13215294 | 0 | 0.087 | 1 | 15 |
| srek1ip1 | 0.07802 | -0.14430471 | 0 | 0.087 | 1 | 15 |
| CR387933.1 | 0.07802 | -0.1498154 | 0 | 0.087 | 1 | 15 |
| LOC101886337 | 0.07802 | -0.14732923 | 0 | 0.087 | 1 | 15 |
| bnip1b | 0.07802 | -0.12765925 | 0 | 0.087 | 1 | 15 |
| c1qtnf6a | 0.07802 | -0.16714693 | 0 | 0.087 | 1 | 15 |
| cahz | 0.07802 | -0.14482525 | 0 | 0.087 | 1 | 15 |
| cblc | 0.07802 | -0.15672201 | 0 | 0.087 | 1 | 15 |
| cdc42l2 | 0.07802 | -0.10883993 | 0 | 0.087 | 1 | 15 |
| col4a4 | 0.07802 | -0.15785789 | 0 | 0.087 | 1 | 15 |
| dchs1a | 0.07802 | -0.1369221 | 0 | 0.087 | 1 | 15 |
| e2f5 | 0.07802 | -0.10669593 | 0 | 0.087 | 1 | 15 |
| fnbp4 | 0.07802 | -0.14952644 | 0 | 0.087 | 1 | 15 |
| m6pr | 0.07802 | -0.11674401 | 0 | 0.087 | 1 | 15 |
| palld | 0.07802 | -0.13812227 | 0 | 0.087 | 1 | 15 |
| pard6a | 0.07802 | -0.13041324 | 0 | 0.087 | 1 | 15 |
| rab12 | 0.07802 | -0.13989806 | 0 | 0.087 | 1 | 15 |
| si:dkey-108k21.13 | 0.07802 | -0.17408083 | 0 | 0.087 | 1 | 15 |
| timp2b | 0.07802 | -0.12022239 | 0 | 0.087 | 1 | 15 |
| unc119.1 | 0.07802 | -0.13896347 | 0 | 0.087 | 1 | 15 |
| wdr18 | 0.07802 | -0.11993334 | 0 | 0.087 | 1 | 15 |
| wdr83 | 0.07802 | -0.15226376 | 0 | 0.087 | 1 | 15 |
| zgc:153292 | 0.07802 | -0.14746815 | 0 | 0.087 | 1 | 15 |
| ints12 | 0.07802 | -0.19117165 | 0 | 0.087 | 1 | 15 |
| kat6a | 0.07802 | -0.19414369 | 0 | 0.087 | 1 | 15 |
| rgs19 | 0.07802 | -0.16737651 | 0 | 0.087 | 1 | 15 |
| sptlc2b | 0.07802 | -0.20932949 | 0 | 0.087 | 1 | 15 |
| tmem169b | 0.07802 | -0.18599775 | 0 | 0.087 | 1 | 15 |
| LOC798120 | 0.07802 | -0.13776611 | 0 | 0.087 | 1 | 15 |
| agap1 | 0.07802 | -0.12043888 | 0 | 0.087 | 1 | 15 |
| atpaf1 | 0.07802 | -0.1250192 | 0 | 0.087 | 1 | 15 |
| brd1a | 0.07802 | -0.12266173 | 0 | 0.087 | 1 | 15 |
| dennd4c | 0.07802 | -0.18037517 | 0 | 0.087 | 1 | 15 |
| dock8 | 0.07802 | -0.15728329 | 0 | 0.087 | 1 | 15 |
| dusp28 | 0.07802 | -0.12089987 | 0 | 0.087 | 1 | 15 |
| fxn | 0.07802 | -0.15861569 | 0 | 0.087 | 1 | 15 |

| | | | | | | |
|---|---|---|---|---|---|---|
| ksr1b | 0.07802 | -0.12087973 | 0 | 0.087 | 1 | 15 |
| lpl | 0.07802 | -0.12811709 | 0 | 0.087 | 1 | 15 |
| march7 | 0.07802 | -0.13420529 | 0 | 0.087 | 1 | 15 |
| myrf | 0.07802 | -0.18378089 | 0 | 0.087 | 1 | 15 |
| nbeal2 | 0.07802 | -0.18115769 | 0 | 0.087 | 1 | 15 |
| nek1 | 0.07802 | -0.13882716 | 0 | 0.087 | 1 | 15 |
| nuak1a | 0.07802 | -0.15644844 | 0 | 0.087 | 1 | 15 |
| numb | 0.07802 | -0.15019336 | 0 | 0.087 | 1 | 15 |
| pald1a | 0.07802 | -0.16135928 | 0 | 0.087 | 1 | 15 |
| prkab1a | 0.07802 | -0.14430647 | 0 | 0.087 | 1 | 15 |
| rdh12 | 0.07802 | -0.15636257 | 0 | 0.087 | 1 | 15 |
| sec22ba | 0.07802 | -0.13167234 | 0 | 0.087 | 1 | 15 |
| si:cabz01021426.2 | 0.07802 | -0.14345247 | 0 | 0.087 | 1 | 15 |
| st3gal5 | 0.07802 | -0.14689387 | 0 | 0.087 | 1 | 15 |
| st6galnac5b | 0.07802 | -0.15495272 | 0 | 0.087 | 1 | 15 |
| tfb2m | 0.07802 | -0.14793598 | 0 | 0.087 | 1 | 15 |
| tmem88a | 0.07802 | -0.14982758 | 0 | 0.087 | 1 | 15 |
| vps9d1 | 0.07802 | -0.13272171 | 0 | 0.087 | 1 | 15 |
| wnt5b | 0.07802 | -0.15790721 | 0 | 0.087 | 1 | 15 |
| zgc:113436 | 0.07802 | -0.11999307 | 0 | 0.087 | 1 | 15 |
| zgc:153665 | 0.07802 | -0.15021766 | 0 | 0.087 | 1 | 15 |
| zgc:162858 | 0.07802 | -0.12638449 | 0 | 0.087 | 1 | 15 |
| zgc:77880 | 0.07802 | -0.18292628 | 0 | 0.087 | 1 | 15 |
| znf770 | 0.07802 | -0.12745834 | 0 | 0.087 | 1 | 15 |
| cdh1 | 0.07837 | -0.48249735 | 0.25 | 0.478 | 1 | 15 |
| btf3 | 0.07851 | 0.238383297 | 0.917 | 0.826 | 1 | 15 |
| b4galt6 | 0.07907 | 0.343432786 | 0.25 | 0.043 | 1 | 15 |
| snrpb2 | 0.07968 | 0.378799411 | 0.25 | 0.043 | 1 | 15 |
| znfx1 | 0.08027 | 0.398108472 | 0.25 | 0.043 | 1 | 15 |
| atg5 | 0.08109 | -0.37926114 | 0.056 | 0.261 | 1 | 15 |
| nup43 | 0.08111 | 0.132928253 | 0.167 | 0.043 | 1 | 15 |
| wdr82 | 0.08128 | 0.350289394 | 0.25 | 0.043 | 1 | 15 |
| ccng2 | 0.0814 | 0.325759765 | 0.306 | 0.087 | 1 | 15 |
| prkar2aa | 0.08143 | 0.34270172 | 0.639 | 0.348 | 1 | 15 |
| rps19 | 0.08171 | 0.179103241 | 0.972 | 0.913 | 1 | 15 |
| eif4g2b | 0.08196 | 0.346713251 | 0.25 | 0.043 | 1 | 15 |
| tmem9b | 0.08237 | 0.199670403 | 0.25 | 0.087 | 1 | 15 |
| kdm3b | 0.0831 | 0.21432303 | 0.194 | 0.043 | 1 | 15 |
| arxa | 0.08328 | -0.33822753 | 0.056 | 0.261 | 1 | 15 |
| eif4e1c | 0.08333 | 0.206951393 | 0.194 | 0.043 | 1 | 15 |
| smarcb1b | 0.08348 | -0.41404877 | 0.056 | 0.261 | 1 | 15 |
| rpl18 | 0.08359 | 0.202533916 | 0.972 | 0.913 | 1 | 15 |
| atp1b3b | 0.08396 | 0.438632264 | 0.528 | 0.304 | 1 | 15 |
| hdlbpa | 0.08399 | -0.22455316 | 0.028 | 0.174 | 1 | 15 |
| XLOC-005605 | 0.08445 | 0.274282874 | 0.861 | 0.609 | 1 | 15 |
| si:ch211-108d22.2 | 0.08454 | -0.32778419 | 0.056 | 0.261 | 1 | 15 |
| col5a1 | 0.08462 | -0.39401398 | 0.083 | 0.304 | 1 | 15 |
| p4hb | 0.08472 | -0.43962056 | 0.222 | 0.391 | 1 | 15 |

| | | | | | | |
|---|---|---|---|---|---|---|
| mkrn1 | 0.08478 | -0.34528158 | 0.111 | 0.348 | 1 | 15 |
| r3hdml | 0.08492 | 0.225719116 | 0.194 | 0.13 | 1 | 15 |
| tp53inp1 | 0.08534 | 0.441562669 | 0.5 | 0.217 | 1 | 15 |
| rps19bp1 | 0.08537 | 0.317856537 | 0.25 | 0.043 | 1 | 15 |
| robo2 | 0.08557 | 0.18083754 | 0.5 | 0.304 | 1 | 15 |
| adamts2 | 0.08581 | -0.42820441 | 0.139 | 0.348 | 1 | 15 |
| sept6 | 0.08605 | -0.39037773 | 0.139 | 0.391 | 1 | 15 |
| faua | 0.08641 | 0.179831071 | 1 | 0.913 | 1 | 15 |
| ndufab1b | 0.08643 | 0.376795741 | 0.444 | 0.217 | 1 | 15 |
| eva1ba | 0.0871 | -0.38740946 | 0.083 | 0.304 | 1 | 15 |
| bhlhe40 | 0.08713 | 0.113245491 | 0.472 | 0.304 | 1 | 15 |
| sf3b6 | 0.08769 | -0.22913526 | 0.417 | 0.478 | 1 | 15 |
| si:ch73-335l21.4 | 0.08822 | -0.52244944 | 0.25 | 0.478 | 1 | 15 |
| sf3b1 | 0.08823 | 0.223644674 | 0.194 | 0.043 | 1 | 15 |
| brd7 | 0.08829 | -0.36366434 | 0.139 | 0.391 | 1 | 15 |
| rwdd1 | 0.0889 | 0.288920516 | 0.472 | 0.217 | 1 | 15 |
| tomm5 | 0.08898 | 0.357382709 | 0.639 | 0.348 | 1 | 15 |
| si:ch211-166a6.5 | 0.08974 | -0.37951224 | 0.083 | 0.304 | 1 | 15 |
| rpl11 | 0.08995 | 0.17892231 | 1 | 0.913 | 1 | 15 |
| grhl1 | 0.09179 | -0.40279742 | 0.111 | 0.348 | 1 | 15 |
| eif2s1a | 0.09192 | 0.349193018 | 0.333 | 0.174 | 1 | 15 |
| zgc:162944 | 0.09229 | 0.359291145 | 0.778 | 0.522 | 1 | 15 |
| dpy30 | 0.09232 | 0.10269566 | 0.333 | 0.391 | 1 | 15 |
| cxcl18b | 0.09241 | -0.48784271 | 0.111 | 0.348 | 1 | 15 |
| sat1a.2 | 0.09259 | -0.39251449 | 0.472 | 0.739 | 1 | 15 |
| gstk1 | 0.09274 | -0.32967936 | 0.083 | 0.304 | 1 | 15 |
| snrpd1 | 0.09323 | 0.413483758 | 0.528 | 0.304 | 1 | 15 |
| eif1b | 0.09359 | 0.341348319 | 0.806 | 0.565 | 1 | 15 |
| rpl17 | 0.09362 | 0.154864707 | 0.944 | 0.913 | 1 | 15 |
| oaz1b | 0.09366 | 0.206898383 | 0.861 | 0.826 | 1 | 15 |
| rpl15 | 0.09403 | 0.217421709 | 0.917 | 0.826 | 1 | 15 |
| dhrs13l1 | 0.09414 | -0.34144698 | 0.083 | 0.261 | 1 | 15 |
| sltm | 0.09425 | -0.33849589 | 0.194 | 0.348 | 1 | 15 |
| actb2 | 0.09426 | -0.26153854 | 0.861 | 0.913 | 1 | 15 |
| ap1m1 | 0.09453 | 0.365189296 | 0.306 | 0.13 | 1 | 15 |
| decr1 | 0.09491 | 0.361537654 | 0.25 | 0.087 | 1 | 15 |
| cd63 | 0.09509 | -0.25069602 | 0.667 | 0.696 | 1 | 15 |
| marcksl1a | 0.09522 | -0.35235985 | 0.611 | 0.826 | 1 | 15 |
| pla2g15 | 0.09533 | -0.10534068 | 0.139 | 0.13 | 1 | 15 |
| fkbp2 | 0.09555 | 0.400355095 | 0.583 | 0.304 | 1 | 15 |
| h2afvb | 0.09578 | -0.36471766 | 0.639 | 0.783 | 1 | 15 |
| slc16a10 | 0.09597 | 0.278108657 | 0.361 | 0.13 | 1 | 15 |
| si:dkey-44g23.5 | 0.09653 | 0.269638481 | 0.222 | 0.043 | 1 | 15 |
| tbca | 0.09781 | -0.40459714 | 0.333 | 0.609 | 1 | 15 |
| pfn2 | 0.09881 | -0.3134895 | 0.528 | 0.652 | 1 | 15 |
| psma4 | 0.09923 | 0.19829992 | 0.472 | 0.261 | 1 | 15 |
| adipor2 | 0.09995 | 0.290550529 | 0.333 | 0.217 | 1 | 15 |
| ercc6 | 0.09998 | -0.33999947 | 0.028 | 0.174 | 1 | 15 |

| | | | | | | |
|---|---|---|---|---|---|---|
| hacd2 | 0.10097 | 0.341685206 | 0.306 | 0.087 | 1 | 15 |
| zgc:110626 | 0.10108 | -0.19592857 | 0.028 | 0.174 | 1 | 15 |
| anp32a | 0.10152 | 0.378760095 | 0.583 | 0.304 | 1 | 15 |
| atp5md | 0.1016 | -0.10634394 | 0.611 | 0.522 | 1 | 15 |
| sema4c | 0.10218 | -0.21418804 | 0.028 | 0.174 | 1 | 15 |
| col18a1a | 0.10247 | -0.41543669 | 0.25 | 0.522 | 1 | 15 |
| ifrd1 | 0.10317 | 0.300686609 | 0.306 | 0.087 | 1 | 15 |
| BOLA2 | 0.10362 | -0.32378435 | 0.083 | 0.217 | 1 | 15 |
| sema3b | 0.10367 | 0.26734993 | 0.278 | 0.087 | 1 | 15 |
| cx43 | 0.10377 | 0.473041777 | 0.667 | 0.435 | 1 | 15 |
| si:rp71-36a1.3 | 0.10407 | -0.34046691 | 0.083 | 0.261 | 1 | 15 |
| lama5 | 0.10435 | -0.38889571 | 0.25 | 0.522 | 1 | 15 |
| dnajc8 | 0.10444 | 0.305469381 | 0.306 | 0.087 | 1 | 15 |
| twf1a | 0.10515 | 0.244049684 | 0.278 | 0.087 | 1 | 15 |
| ahcy | 0.10538 | -0.10070695 | 0.222 | 0.174 | 1 | 15 |
| pnn | 0.10581 | 0.32879328 | 0.306 | 0.087 | 1 | 15 |
| snrpd2 | 0.10602 | 0.36259067 | 0.583 | 0.304 | 1 | 15 |
| eif4a1a | 0.10627 | 0.314160151 | 0.75 | 0.478 | 1 | 15 |
| rpl31 | 0.10647 | 0.172959061 | 0.944 | 0.913 | 1 | 15 |
| slc2a1b | 0.10647 | -0.31474995 | 0.028 | 0.13 | 1 | 15 |
| mrps23 | 0.10657 | 0.317333864 | 0.222 | 0.043 | 1 | 15 |
| si:ch211-225b11.1 | 0.10675 | 0.186026481 | 0.5 | 0.304 | 1 | 15 |
| snrnp27 | 0.10676 | 0.357084041 | 0.333 | 0.13 | 1 | 15 |
| ddx19 | 0.1072 | 0.288444616 | 0.222 | 0.087 | 1 | 15 |
| khdrbs1b | 0.1072 | 0.196501566 | 0.361 | 0.174 | 1 | 15 |
| chd2 | 0.10739 | 0.327034284 | 0.306 | 0.13 | 1 | 15 |
| snx1a | 0.10766 | -0.33810438 | 0.194 | 0.348 | 1 | 15 |
| nid1a | 0.10814 | -0.35100292 | 0.028 | 0.174 | 1 | 15 |
| nt5c2b | 0.10895 | 0.358871869 | 0.306 | 0.087 | 1 | 15 |
| eif5a2 | 0.10928 | 0.192196017 | 0.667 | 0.435 | 1 | 15 |
| slka | 0.10963 | 0.311776941 | 0.222 | 0.043 | 1 | 15 |
| nxn | 0.11006 | 0.342895952 | 0.306 | 0.087 | 1 | 15 |
| tia1 | 0.11039 | 0.338252037 | 0.361 | 0.13 | 1 | 15 |
| ralba | 0.11046 | -0.40631163 | 0.056 | 0.217 | 1 | 15 |
| rps27l | 0.11079 | 0.329850122 | 0.5 | 0.391 | 1 | 15 |
| hprt1 | 0.11087 | 0.324019871 | 0.306 | 0.087 | 1 | 15 |
| lgmn | 0.11137 | -0.34849545 | 0.167 | 0.348 | 1 | 15 |
| nfkbiab | 0.11147 | 0.213446553 | 0.583 | 0.348 | 1 | 15 |
| rad21a | 0.11151 | -0.39262979 | 0.333 | 0.609 | 1 | 15 |
| cct6a | 0.11163 | 0.333945154 | 0.528 | 0.261 | 1 | 15 |
| ubxn1 | 0.11187 | 0.184274744 | 0.222 | 0.174 | 1 | 15 |
| eif3d | 0.11234 | 0.335262192 | 0.778 | 0.565 | 1 | 15 |
| ilf3b | 0.11253 | 0.111212033 | 0.306 | 0.174 | 1 | 15 |
| usf1l | 0.11257 | -0.24281206 | 0.111 | 0.304 | 1 | 15 |
| camk2g1 | 0.1129 | 0.241401927 | 0.194 | 0.043 | 1 | 15 |
| tpm4a | 0.11305 | -0.25030899 | 0.389 | 0.478 | 1 | 15 |
| si:dkey-283j8.1 | 0.11332 | 0.39104923 | 0.222 | 0.043 | 1 | 15 |
| epha3 | 0.11377 | 0.10623596 | 0.139 | 0.043 | 1 | 15 |

| | | | | | | |
|---|---|---|---|---|---|---|
| synrg | 0.11387 | -0.22505598 | 0.028 | 0.174 | 1 | 15 |
| grb2b | 0.11392 | 0.364637332 | 0.361 | 0.13 | 1 | 15 |
| ppp3r1b | 0.11411 | 0.190811053 | 0.306 | 0.13 | 1 | 15 |
| ndufc2 | 0.11463 | 0.331115578 | 0.667 | 0.391 | 1 | 15 |
| musk | 0.11519 | 0.376074288 | 0.306 | 0.087 | 1 | 15 |
| ccdc124 | 0.11524 | -0.36791911 | 0.194 | 0.435 | 1 | 15 |
| tshz3b | 0.11529 | -0.13525665 | 0.056 | 0.174 | 1 | 15 |
| yaf2 | 0.11541 | -0.34167599 | 0.056 | 0.217 | 1 | 15 |
| si:dkey-89b17.4 | 0.11649 | 0.23418441 | 0.194 | 0.043 | 1 | 15 |
| inhbaa | 0.11662 | 0.392436204 | 0.528 | 0.261 | 1 | 15 |
| ndufb4 | 0.11679 | 0.382400325 | 0.611 | 0.348 | 1 | 15 |
| si:ch211-191i18.2 | 0.11738 | -0.27212575 | 0.083 | 0.261 | 1 | 15 |
| akap8l | 0.11819 | -0.32868619 | 0.083 | 0.261 | 1 | 15 |
| zgc:55943 | 0.11821 | 0.155298399 | 0.222 | 0.087 | 1 | 15 |
| tmed10 | 0.11847 | -0.38152719 | 0.444 | 0.609 | 1 | 15 |
| celf2 | 0.11868 | -0.39052283 | 0.222 | 0.391 | 1 | 15 |
| csnk2b | 0.11899 | 0.365082386 | 0.611 | 0.391 | 1 | 15 |
| scn4bb | 0.11908 | 0.286051865 | 0.278 | 0.087 | 1 | 15 |
| CABZ01072614.1 | 0.1191 | -0.36438802 | 0.361 | 0.435 | 1 | 15 |
| agfg1b | 0.11917 | 0.159273007 | 0.083 | 0 | 1 | 15 |
| golga1 | 0.11917 | 0.171580886 | 0.083 | 0 | 1 | 15 |
| hmgcrb | 0.11917 | 0.164346742 | 0.083 | 0 | 1 | 15 |
| ispd | 0.11917 | 0.186409808 | 0.083 | 0 | 1 | 15 |
| msl1a | 0.11917 | 0.157479956 | 0.083 | 0 | 1 | 15 |
| pex12 | 0.11917 | 0.140540883 | 0.083 | 0 | 1 | 15 |
| polrmt | 0.11917 | 0.153136115 | 0.083 | 0 | 1 | 15 |
| shroom4 | 0.11917 | 0.152322835 | 0.083 | 0 | 1 | 15 |
| usp22 | 0.11917 | 0.173109811 | 0.083 | 0 | 1 | 15 |
| zgc:92594 | 0.11917 | 0.204838971 | 0.083 | 0 | 1 | 15 |
| BX537358.1 | 0.11917 | 0.122909883 | 0.083 | 0 | 1 | 15 |
| celsr1a | 0.11917 | 0.168571871 | 0.083 | 0 | 1 | 15 |
| chchd3a | 0.11917 | 0.131103042 | 0.083 | 0 | 1 | 15 |
| crata | 0.11917 | 0.150091065 | 0.083 | 0 | 1 | 15 |
| cyb5b | 0.11917 | 0.121323359 | 0.083 | 0 | 1 | 15 |
| gdpd1 | 0.11917 | 0.147618756 | 0.083 | 0 | 1 | 15 |
| larsa | 0.11917 | 0.133413606 | 0.083 | 0 | 1 | 15 |
| ldlrad4a | 0.11917 | 0.134880647 | 0.083 | 0 | 1 | 15 |
| lgi1a | 0.11917 | 0.142117765 | 0.083 | 0 | 1 | 15 |
| mtif3 | 0.11917 | 0.178474642 | 0.083 | 0 | 1 | 15 |
| nagk | 0.11917 | 0.13006574 | 0.083 | 0 | 1 | 15 |
| pmm2 | 0.11917 | 0.117170923 | 0.083 | 0 | 1 | 15 |
| ppp1r2 | 0.11917 | 0.11979304 | 0.083 | 0 | 1 | 15 |
| rac1b | 0.11917 | 0.137296289 | 0.083 | 0 | 1 | 15 |
| si:ch1073-174d20.2 | 0.11917 | 0.142522782 | 0.083 | 0 | 1 | 15 |
| stox2a | 0.11917 | 0.183402234 | 0.083 | 0 | 1 | 15 |
| tox2 | 0.11917 | 0.14299379 | 0.083 | 0 | 1 | 15 |
| wapla | 0.11917 | 0.128591142 | 0.083 | 0 | 1 | 15 |
| wdr37 | 0.11917 | 0.179530191 | 0.083 | 0 | 1 | 15 |

| | | | | | | |
|---|---|---|---|---|---|---|
| CABZ01039304.1 | 0.11917 | 0.143566277 | 0.083 | 0 | 1 | 15 |
| CT956002.2 | 0.11917 | 0.205150183 | 0.083 | 0 | 1 | 15 |
| abraxas1 | 0.11917 | 0.12604448 | 0.083 | 0 | 1 | 15 |
| ankmy2a | 0.11917 | 0.133945139 | 0.083 | 0 | 1 | 15 |
| b3gat3 | 0.11917 | 0.139042235 | 0.083 | 0 | 1 | 15 |
| b4galt4 | 0.11917 | 0.149891337 | 0.083 | 0 | 1 | 15 |
| coq7 | 0.11917 | 0.153644944 | 0.083 | 0 | 1 | 15 |
| cpsf3 | 0.11917 | 0.161322621 | 0.083 | 0 | 1 | 15 |
| dhx37 | 0.11917 | 0.13558776 | 0.083 | 0 | 1 | 15 |
| eloa | 0.11917 | 0.14712603 | 0.083 | 0 | 1 | 15 |
| etfdh | 0.11917 | 0.149984876 | 0.083 | 0 | 1 | 15 |
| fastkd2 | 0.11917 | 0.132578782 | 0.083 | 0 | 1 | 15 |
| fbxo11b | 0.11917 | 0.131840595 | 0.083 | 0 | 1 | 15 |
| glmna | 0.11917 | 0.126048982 | 0.083 | 0 | 1 | 15 |
| gpn1 | 0.11917 | 0.133564448 | 0.083 | 0 | 1 | 15 |
| ift52 | 0.11917 | 0.181618169 | 0.083 | 0 | 1 | 15 |
| igf2bp3 | 0.11917 | 0.168878559 | 0.083 | 0 | 1 | 15 |
| itfg2 | 0.11917 | 0.168574243 | 0.083 | 0 | 1 | 15 |
| jade1 | 0.11917 | 0.129691241 | 0.083 | 0 | 1 | 15 |
| lama1 | 0.11917 | 0.157423498 | 0.083 | 0 | 1 | 15 |
| mrrf | 0.11917 | 0.129099074 | 0.083 | 0 | 1 | 15 |
| nfrkb | 0.11917 | 0.156256955 | 0.083 | 0 | 1 | 15 |
| nhsl1b | 0.11917 | 0.138402442 | 0.083 | 0 | 1 | 15 |
| nkain1 | 0.11917 | 0.136705077 | 0.083 | 0 | 1 | 15 |
| nom1 | 0.11917 | 0.146646655 | 0.083 | 0 | 1 | 15 |
| nsun2 | 0.11917 | 0.152230758 | 0.083 | 0 | 1 | 15 |
| numbl | 0.11917 | 0.159321282 | 0.083 | 0 | 1 | 15 |
| osbpl2a | 0.11917 | 0.156530549 | 0.083 | 0 | 1 | 15 |
| pag1 | 0.11917 | 0.14254919 | 0.083 | 0 | 1 | 15 |
| plekhg4 | 0.11917 | 0.166130152 | 0.083 | 0 | 1 | 15 |
| ppan | 0.11917 | 0.140976638 | 0.083 | 0 | 1 | 15 |
| ppp1r26 | 0.11917 | 0.138012115 | 0.083 | 0 | 1 | 15 |
| ptpn11b | 0.11917 | 0.191066645 | 0.083 | 0 | 1 | 15 |
| r3hdm1 | 0.11917 | 0.165980411 | 0.083 | 0 | 1 | 15 |
| rabl6a | 0.11917 | 0.160041467 | 0.083 | 0 | 1 | 15 |
| rims2b | 0.11917 | 0.151518505 | 0.083 | 0 | 1 | 15 |
| rnf141 | 0.11917 | 0.174291155 | 0.083 | 0 | 1 | 15 |
| rnf169 | 0.11917 | 0.125908056 | 0.083 | 0 | 1 | 15 |
| scamp1 | 0.11917 | 0.167170548 | 0.083 | 0 | 1 | 15 |
| scnm1 | 0.11917 | 0.132760321 | 0.083 | 0 | 1 | 15 |
| si:ch211-230g15.5 | 0.11917 | 0.130553596 | 0.083 | 0 | 1 | 15 |
| si:dkey-286h2.3 | 0.11917 | 0.133077406 | 0.083 | 0 | 1 | 15 |
| tpte | 0.11917 | 0.201543116 | 0.083 | 0 | 1 | 15 |
| usp30 | 0.11917 | 0.133678782 | 0.083 | 0 | 1 | 15 |
| zgc:153759 | 0.11917 | 0.105884303 | 0.083 | 0 | 1 | 15 |
| zgc:56235 | 0.11917 | 0.135369709 | 0.083 | 0 | 1 | 15 |
| zgc:65997 | 0.11917 | 0.1584596 | 0.083 | 0 | 1 | 15 |
| grtp1a | 0.11917 | 0.128772331 | 0.083 | 0 | 1 | 15 |

| | | | | | | |
|---|---|---|---|---|---|---|
| si:ch211-127i16.2 | 0.11917 | 0.158849316 | 0.083 | 0 | 1 | 15 |
| si:rp71-36a1.1 | 0.11917 | 0.179949944 | 0.083 | 0 | 1 | 15 |
| tex2 | 0.11917 | 0.162437306 | 0.083 | 0 | 1 | 15 |
| txlna | 0.11917 | 0.163129763 | 0.083 | 0 | 1 | 15 |
| zgc:153372 | 0.11917 | 0.137398511 | 0.083 | 0 | 1 | 15 |
| AL954696.1 | 0.11917 | 0.169714316 | 0.083 | 0 | 1 | 15 |
| XLOC-008119 | 0.11917 | 0.134612353 | 0.083 | 0 | 1 | 15 |
| bmf2 | 0.11917 | 0.137885528 | 0.083 | 0 | 1 | 15 |
| bmpr1aa | 0.11917 | 0.152291447 | 0.083 | 0 | 1 | 15 |
| cables1 | 0.11917 | 0.140215551 | 0.083 | 0 | 1 | 15 |
| calm1b | 0.11917 | 0.111373125 | 0.083 | 0 | 1 | 15 |
| chaf1a | 0.11917 | 0.154561475 | 0.083 | 0 | 1 | 15 |
| chic1 | 0.11917 | 0.174707722 | 0.083 | 0 | 1 | 15 |
| cipcb | 0.11917 | 0.151049615 | 0.083 | 0 | 1 | 15 |
| cox11 | 0.11917 | 0.153287462 | 0.083 | 0 | 1 | 15 |
| dhx58 | 0.11917 | 0.189844783 | 0.083 | 0 | 1 | 15 |
| dlgap4a | 0.11917 | 0.151987835 | 0.083 | 0 | 1 | 15 |
| efcab7 | 0.11917 | 0.145568023 | 0.083 | 0 | 1 | 15 |
| elp6 | 0.11917 | 0.131360589 | 0.083 | 0 | 1 | 15 |
| fbxl4 | 0.11917 | 0.171805695 | 0.083 | 0 | 1 | 15 |
| gltpa | 0.11917 | 0.142135577 | 0.083 | 0 | 1 | 15 |
| gtf3aa | 0.11917 | 0.109861549 | 0.083 | 0 | 1 | 15 |
| hddc2 | 0.11917 | 0.145323754 | 0.083 | 0 | 1 | 15 |
| hibch | 0.11917 | 0.140843038 | 0.083 | 0 | 1 | 15 |
| il10rb | 0.11917 | 0.137637441 | 0.083 | 0 | 1 | 15 |
| lhx9 | 0.11917 | 0.138654719 | 0.083 | 0 | 1 | 15 |
| lnpa | 0.11917 | 0.167338349 | 0.083 | 0 | 1 | 15 |
| lrrc8db | 0.11917 | 0.1503733 | 0.083 | 0 | 1 | 15 |
| map2k5 | 0.11917 | 0.14945876 | 0.083 | 0 | 1 | 15 |
| mbip | 0.11917 | 0.141958246 | 0.083 | 0 | 1 | 15 |
| mettl27 | 0.11917 | 0.167083383 | 0.083 | 0 | 1 | 15 |
| mgll | 0.11917 | 0.131475414 | 0.083 | 0 | 1 | 15 |
| nhsl1a | 0.11917 | 0.170270915 | 0.083 | 0 | 1 | 15 |
| nrp1a | 0.11917 | 0.157498569 | 0.083 | 0 | 1 | 15 |
| pars2 | 0.11917 | 0.201337494 | 0.083 | 0 | 1 | 15 |
| pcyt2 | 0.11917 | 0.145329156 | 0.083 | 0 | 1 | 15 |
| pgam1b | 0.11917 | 0.148148193 | 0.083 | 0 | 1 | 15 |
| prpf3 | 0.11917 | 0.147570317 | 0.083 | 0 | 1 | 15 |
| ric8b | 0.11917 | 0.161673249 | 0.083 | 0 | 1 | 15 |
| sept9b | 0.11917 | 0.15634935 | 0.083 | 0 | 1 | 15 |
| serhl | 0.11917 | 0.161322138 | 0.083 | 0 | 1 | 15 |
| si:ch211-128m15.3 | 0.11917 | 0.135731546 | 0.083 | 0 | 1 | 15 |
| si:ch211-239j9.1 | 0.11917 | 0.134261103 | 0.083 | 0 | 1 | 15 |
| si:dkey-34m19.3 | 0.11917 | 0.190434224 | 0.083 | 0 | 1 | 15 |
| slc25a44b | 0.11917 | 0.15048458 | 0.083 | 0 | 1 | 15 |
| slc27a1a | 0.11917 | 0.14184998 | 0.083 | 0 | 1 | 15 |
| slc35f2 | 0.11917 | 0.119055354 | 0.083 | 0 | 1 | 15 |
| soul3 | 0.11917 | 0.141063472 | 0.083 | 0 | 1 | 15 |

| | | | | | | |
|---|---|---|---|---|---|---|
| stx4 | 0.11917 | 0.134014521 | 0.083 | 0 | 1 | 15 |
| tmem79b | 0.11917 | 0.164972771 | 0.083 | 0 | 1 | 15 |
| usp1 | 0.11917 | 0.137179148 | 0.083 | 0 | 1 | 15 |
| BX682556.1 | 0.11917 | 0.145523109 | 0.083 | 0 | 1 | 15 |
| CU459056.2 | 0.11917 | 0.146665604 | 0.083 | 0 | 1 | 15 |
| LOC101885376 | 0.11917 | 0.164141585 | 0.083 | 0 | 1 | 15 |
| bach1a | 0.11917 | 0.149394219 | 0.083 | 0 | 1 | 15 |
| cd2bp2 | 0.11917 | 0.136339669 | 0.083 | 0 | 1 | 15 |
| efhd1 | 0.11917 | 0.146399674 | 0.083 | 0 | 1 | 15 |
| glra4a | 0.11917 | 0.195573593 | 0.083 | 0 | 1 | 15 |
| larp1 | 0.11917 | 0.173659977 | 0.083 | 0 | 1 | 15 |
| mblac1 | 0.11917 | 0.129582669 | 0.083 | 0 | 1 | 15 |
| mea1 | 0.11917 | 0.138570885 | 0.083 | 0 | 1 | 15 |
| mri1 | 0.11917 | 0.133038638 | 0.083 | 0 | 1 | 15 |
| mrps28 | 0.11917 | 0.125459703 | 0.083 | 0 | 1 | 15 |
| msh2 | 0.11917 | 0.168439716 | 0.083 | 0 | 1 | 15 |
| ncoa6 | 0.11917 | 0.146933789 | 0.083 | 0 | 1 | 15 |
| pcdh1b | 0.11917 | 0.198664262 | 0.083 | 0 | 1 | 15 |
| pigc | 0.11917 | 0.161752986 | 0.083 | 0 | 1 | 15 |
| pole4 | 0.11917 | 0.135613276 | 0.083 | 0 | 1 | 15 |
| polr3d | 0.11917 | 0.150110565 | 0.083 | 0 | 1 | 15 |
| pus1 | 0.11917 | 0.144298148 | 0.083 | 0 | 1 | 15 |
| rusc2 | 0.11917 | 0.162201814 | 0.083 | 0 | 1 | 15 |
| setd7 | 0.11917 | 0.164141585 | 0.083 | 0 | 1 | 15 |
| setdb1b | 0.11917 | 0.164625582 | 0.083 | 0 | 1 | 15 |
| si:ch211-207k7.4 | 0.11917 | 0.179531845 | 0.083 | 0 | 1 | 15 |
| tceanc2 | 0.11917 | 0.190903308 | 0.083 | 0 | 1 | 15 |
| tmem120a | 0.11917 | 0.16412548 | 0.083 | 0 | 1 | 15 |
| tsen54 | 0.11917 | 0.146198996 | 0.083 | 0 | 1 | 15 |
| ubxn8 | 0.11917 | 0.192657531 | 0.083 | 0 | 1 | 15 |
| uck2b | 0.11917 | 0.164542692 | 0.083 | 0 | 1 | 15 |
| utp15 | 0.11917 | 0.130045611 | 0.083 | 0 | 1 | 15 |
| waslb | 0.11917 | 0.16559508 | 0.083 | 0 | 1 | 15 |
| zfand1 | 0.11917 | 0.162484642 | 0.083 | 0 | 1 | 15 |
| zgc:110843 | 0.11917 | 0.115639721 | 0.083 | 0 | 1 | 15 |
| zgc:162972 | 0.11917 | 0.154798464 | 0.083 | 0 | 1 | 15 |
| afap1l1b | 0.11917 | 0.156133777 | 0.083 | 0 | 1 | 15 |
| alg3 | 0.11917 | 0.146600212 | 0.083 | 0 | 1 | 15 |
| eps8a | 0.11917 | 0.148327496 | 0.083 | 0 | 1 | 15 |
| gatad1 | 0.11917 | 0.128805948 | 0.083 | 0 | 1 | 15 |
| kirrel3l | 0.11917 | 0.179565887 | 0.083 | 0 | 1 | 15 |
| man2b2 | 0.11917 | 0.135557221 | 0.083 | 0 | 1 | 15 |
| mapk14a | 0.11917 | 0.1305099 | 0.083 | 0 | 1 | 15 |
| n6amt1 | 0.11917 | 0.127303254 | 0.083 | 0 | 1 | 15 |
| ptpa | 0.11917 | 0.141753363 | 0.083 | 0 | 1 | 15 |
| si:ch211-79k12.2 | 0.11917 | 0.151021317 | 0.083 | 0 | 1 | 15 |
| trim35-7 | 0.11917 | 0.112300272 | 0.083 | 0 | 1 | 15 |
| CU856539.1 | 0.11917 | 0.139133896 | 0.083 | 0 | 1 | 15 |

| | | | | | | |
|---|---|---|---|---|---|---|
| ascc3 | 0.11917 | 0.160050138 | 0.083 | 0 | 1 | 15 |
| atraid | 0.11917 | 0.10589225 | 0.083 | 0 | 1 | 15 |
| bloc1s4 | 0.11917 | 0.173104977 | 0.083 | 0 | 1 | 15 |
| btbd10a | 0.11917 | 0.159334432 | 0.083 | 0 | 1 | 15 |
| ccdc126 | 0.11917 | 0.142013196 | 0.083 | 0 | 1 | 15 |
| cdc27 | 0.11917 | 0.137383313 | 0.083 | 0 | 1 | 15 |
| cntn5 | 0.11917 | 0.175741376 | 0.083 | 0 | 1 | 15 |
| coq3 | 0.11917 | 0.164796975 | 0.083 | 0 | 1 | 15 |
| dbn1 | 0.11917 | 0.140894238 | 0.083 | 0 | 1 | 15 |
| ddx51 | 0.11917 | 0.164321625 | 0.083 | 0 | 1 | 15 |
| dglucy | 0.11917 | 0.149518094 | 0.083 | 0 | 1 | 15 |
| dhx29 | 0.11917 | 0.15009922 | 0.083 | 0 | 1 | 15 |
| eif2a | 0.11917 | 0.174837146 | 0.083 | 0 | 1 | 15 |
| fam214a | 0.11917 | 0.138167036 | 0.083 | 0 | 1 | 15 |
| frs2a | 0.11917 | 0.178988166 | 0.083 | 0 | 1 | 15 |
| fuca2 | 0.11917 | 0.144282462 | 0.083 | 0 | 1 | 15 |
| fyco1b | 0.11917 | 0.167136945 | 0.083 | 0 | 1 | 15 |
| inpp5b | 0.11917 | 0.138795348 | 0.083 | 0 | 1 | 15 |
| intu | 0.11917 | 0.145916244 | 0.083 | 0 | 1 | 15 |
| mast3a | 0.11917 | 0.184174191 | 0.083 | 0 | 1 | 15 |
| mtss1 | 0.11917 | 0.157048706 | 0.083 | 0 | 1 | 15 |
| myorg | 0.11917 | 0.124941234 | 0.083 | 0 | 1 | 15 |
| nlrc3l | 0.11917 | 0.151930046 | 0.083 | 0 | 1 | 15 |
| nrip1a | 0.11917 | 0.137341288 | 0.083 | 0 | 1 | 15 |
| ntf3 | 0.11917 | 0.161600212 | 0.083 | 0 | 1 | 15 |
| nxpe3 | 0.11917 | 0.145988881 | 0.083 | 0 | 1 | 15 |
| parga | 0.11917 | 0.160718506 | 0.083 | 0 | 1 | 15 |
| rab3gap2 | 0.11917 | 0.138179854 | 0.083 | 0 | 1 | 15 |
| rangap1b | 0.11917 | 0.139924538 | 0.083 | 0 | 1 | 15 |
| rfc5 | 0.11917 | 0.139365819 | 0.083 | 0 | 1 | 15 |
| rnf175 | 0.11917 | 0.145468652 | 0.083 | 0 | 1 | 15 |
| rp9 | 0.11917 | 0.185314052 | 0.083 | 0 | 1 | 15 |
| scyl1 | 0.11917 | 0.15501267 | 0.083 | 0 | 1 | 15 |
| setd3 | 0.11917 | 0.188482178 | 0.083 | 0 | 1 | 15 |
| si:ch211-113p18.3 | 0.11917 | 0.156582182 | 0.083 | 0 | 1 | 15 |
| si:dkey-16l2.16 | 0.11917 | 0.172793785 | 0.083 | 0 | 1 | 15 |
| si:dkey-230p4.1 | 0.11917 | 0.173202473 | 0.083 | 0 | 1 | 15 |
| si:dkey-260j18.2 | 0.11917 | 0.147131824 | 0.083 | 0 | 1 | 15 |
| si:dkey-51d8.6 | 0.11917 | 0.145922077 | 0.083 | 0 | 1 | 15 |
| si:dkey-94l16.4 | 0.11917 | 0.139165178 | 0.083 | 0 | 1 | 15 |
| tcerg1a | 0.11917 | 0.152112329 | 0.083 | 0 | 1 | 15 |
| tial1 | 0.11917 | 0.173674441 | 0.083 | 0 | 1 | 15 |
| tmem170a | 0.11917 | 0.155421991 | 0.083 | 0 | 1 | 15 |
| vps26b | 0.11917 | 0.195019314 | 0.083 | 0 | 1 | 15 |
| yme1l1a | 0.11917 | 0.145084871 | 0.083 | 0 | 1 | 15 |
| zcchc7 | 0.11917 | 0.180423637 | 0.083 | 0 | 1 | 15 |
| zgc:101851 | 0.11917 | 0.127206006 | 0.083 | 0 | 1 | 15 |
| zgc:103692 | 0.11917 | 0.152835563 | 0.083 | 0 | 1 | 15 |

| | | | | | | |
|---|---|---|---|---|---|---|
| zgc:113030 | 0.11917 | 0.139344027 | 0.083 | 0 | 1 | 15 |
| zgc:153521 | 0.11917 | 0.129140728 | 0.083 | 0 | 1 | 15 |
| zgc:162472 | 0.11917 | 0.175008336 | 0.083 | 0 | 1 | 15 |
| znf1183 | 0.11917 | 0.174743368 | 0.083 | 0 | 1 | 15 |
| znf148 | 0.11917 | 0.161382995 | 0.083 | 0 | 1 | 15 |
| CABZ01028771.1 | 0.11917 | 0.195211413 | 0.083 | 0 | 1 | 15 |
| CABZ01081752.3 | 0.11917 | 0.14247257 | 0.083 | 0 | 1 | 15 |
| CR388143.1 | 0.11917 | 0.156361279 | 0.083 | 0 | 1 | 15 |
| LOC101885018 | 0.11917 | 0.211952418 | 0.083 | 0 | 1 | 15 |
| LOC110437926 | 0.11917 | 0.19819031 | 0.083 | 0 | 1 | 15 |
| NINJ1 | 0.11917 | 0.153870629 | 0.083 | 0 | 1 | 15 |
| XLOC-004335 | 0.11917 | 0.194260791 | 0.083 | 0 | 1 | 15 |
| XLOC-034437 | 0.11917 | 0.129719625 | 0.083 | 0 | 1 | 15 |
| ZNF512B | 0.11917 | 0.12737416 | 0.083 | 0 | 1 | 15 |
| aatf | 0.11917 | 0.137437777 | 0.083 | 0 | 1 | 15 |
| abhd13 | 0.11917 | 0.140838289 | 0.083 | 0 | 1 | 15 |
| adcy6b | 0.11917 | 0.143026032 | 0.083 | 0 | 1 | 15 |
| adkb | 0.11917 | 0.136711452 | 0.083 | 0 | 1 | 15 |
| afg3l2 | 0.11917 | 0.156879381 | 0.083 | 0 | 1 | 15 |
| akt2l | 0.11917 | 0.142951226 | 0.083 | 0 | 1 | 15 |
| alas1 | 0.11917 | 0.158084703 | 0.083 | 0 | 1 | 15 |
| ankrd54 | 0.11917 | 0.142174781 | 0.083 | 0 | 1 | 15 |
| arhgef18a | 0.11917 | 0.153447307 | 0.083 | 0 | 1 | 15 |
| arl13b | 0.11917 | 0.1441298 | 0.083 | 0 | 1 | 15 |
| asap3 | 0.11917 | 0.175479555 | 0.083 | 0 | 1 | 15 |
| atg101 | 0.11917 | 0.131187035 | 0.083 | 0 | 1 | 15 |
| bcat2 | 0.11917 | 0.152352013 | 0.083 | 0 | 1 | 15 |
| bcl11ab | 0.11917 | 0.165131191 | 0.083 | 0 | 1 | 15 |
| bmper | 0.11917 | 0.181146271 | 0.083 | 0 | 1 | 15 |
| btg3 | 0.11917 | 0.181877435 | 0.083 | 0 | 1 | 15 |
| ccar1 | 0.11917 | 0.134362353 | 0.083 | 0 | 1 | 15 |
| ccdc127b | 0.11917 | 0.132526083 | 0.083 | 0 | 1 | 15 |
| cept1a | 0.11917 | 0.151350649 | 0.083 | 0 | 1 | 15 |
| clptm1 | 0.11917 | 0.136214772 | 0.083 | 0 | 1 | 15 |
| cyp27a1.4 | 0.11917 | 0.133792861 | 0.083 | 0 | 1 | 15 |
| dnaja1 | 0.11917 | 0.140609108 | 0.083 | 0 | 1 | 15 |
| dnm2a | 0.11917 | 0.145556982 | 0.083 | 0 | 1 | 15 |
| dync1i2b | 0.11917 | 0.158185177 | 0.083 | 0 | 1 | 15 |
| epc1b | 0.11917 | 0.122729251 | 0.083 | 0 | 1 | 15 |
| etv5a | 0.11917 | 0.139979781 | 0.083 | 0 | 1 | 15 |
| exd2 | 0.11917 | 0.156591263 | 0.083 | 0 | 1 | 15 |
| fam122b | 0.11917 | 0.19623909 | 0.083 | 0 | 1 | 15 |
| fam222bb | 0.11917 | 0.135583029 | 0.083 | 0 | 1 | 15 |
| fgfr1op | 0.11917 | 0.125320739 | 0.083 | 0 | 1 | 15 |
| fhdc3 | 0.11917 | 0.15645111 | 0.083 | 0 | 1 | 15 |
| foxj2 | 0.11917 | 0.181230649 | 0.083 | 0 | 1 | 15 |
| fzd10 | 0.11917 | 0.15238453 | 0.083 | 0 | 1 | 15 |
| gas2l1 | 0.11917 | 0.168341702 | 0.083 | 0 | 1 | 15 |

| | | | | | | |
|---|---|---|---|---|---|---|
| gramd1c | 0.11917 | 0.128854458 | 0.083 | 0 | 1 | 15 |
| grsf1 | 0.11917 | 0.134993222 | 0.083 | 0 | 1 | 15 |
| gsk3b | 0.11917 | 0.158188548 | 0.083 | 0 | 1 | 15 |
| ical1 | 0.11917 | 0.124939261 | 0.083 | 0 | 1 | 15 |
| itga2.3 | 0.11917 | 0.169175914 | 0.083 | 0 | 1 | 15 |
| lamp1b | 0.11917 | 0.139252472 | 0.083 | 0 | 1 | 15 |
| lsg1 | 0.11917 | 0.12881359 | 0.083 | 0 | 1 | 15 |
| mapk7 | 0.11917 | 0.169851071 | 0.083 | 0 | 1 | 15 |
| mdm1 | 0.11917 | 0.147651479 | 0.083 | 0 | 1 | 15 |
| med27 | 0.11917 | 0.156981361 | 0.083 | 0 | 1 | 15 |
| mier3b | 0.11917 | 0.155622165 | 0.083 | 0 | 1 | 15 |
| mipep | 0.11917 | 0.152546998 | 0.083 | 0 | 1 | 15 |
| nlgn2a | 0.11917 | 0.142011468 | 0.083 | 0 | 1 | 15 |
| noc3l | 0.11917 | 0.114642326 | 0.083 | 0 | 1 | 15 |
| nup98 | 0.11917 | 0.139506042 | 0.083 | 0 | 1 | 15 |
| pargl | 0.11917 | 0.145416167 | 0.083 | 0 | 1 | 15 |
| patl1 | 0.11917 | 0.154792907 | 0.083 | 0 | 1 | 15 |
| pebp1 | 0.11917 | 0.131234571 | 0.083 | 0 | 1 | 15 |
| pigh | 0.11917 | 0.155475076 | 0.083 | 0 | 1 | 15 |
| pla2g12a | 0.11917 | 0.167346075 | 0.083 | 0 | 1 | 15 |
| ppp2r2aa | 0.11917 | 0.180856591 | 0.083 | 0 | 1 | 15 |
| ppp2r5cb | 0.11917 | 0.161622629 | 0.083 | 0 | 1 | 15 |
| prkcdb | 0.11917 | 0.203168331 | 0.083 | 0 | 1 | 15 |
| rcc1l | 0.11917 | 0.15240362 | 0.083 | 0 | 1 | 15 |
| rsf1a | 0.11917 | 0.137217104 | 0.083 | 0 | 1 | 15 |
| sar1aa | 0.11917 | 0.133233066 | 0.083 | 0 | 1 | 15 |
| sec23ip | 0.11917 | 0.128962017 | 0.083 | 0 | 1 | 15 |
| setdb2 | 0.11917 | 0.11207329 | 0.083 | 0 | 1 | 15 |
| si:dkey-165a24.10 | 0.11917 | 0.149245137 | 0.083 | 0 | 1 | 15 |
| si:dkey-238i5.2 | 0.11917 | 0.14201379 | 0.083 | 0 | 1 | 15 |
| si:dkeyp-69c1.6 | 0.11917 | 0.181269801 | 0.083 | 0 | 1 | 15 |
| slc25a36a | 0.11917 | 0.126869738 | 0.083 | 0 | 1 | 15 |
| srgap3 | 0.11917 | 0.178418998 | 0.083 | 0 | 1 | 15 |
| stra6 | 0.11917 | 0.131802973 | 0.083 | 0 | 1 | 15 |
| sumf1 | 0.11917 | 0.161671592 | 0.083 | 0 | 1 | 15 |
| suz12b | 0.11917 | 0.154743271 | 0.083 | 0 | 1 | 15 |
| taf4a | 0.11917 | 0.166434906 | 0.083 | 0 | 1 | 15 |
| tgfb1a | 0.11917 | 0.162745808 | 0.083 | 0 | 1 | 15 |
| tmem86a | 0.11917 | 0.157251578 | 0.083 | 0 | 1 | 15 |
| trap1 | 0.11917 | 0.165611653 | 0.083 | 0 | 1 | 15 |
| trub1 | 0.11917 | 0.129352192 | 0.083 | 0 | 1 | 15 |
| tsr1 | 0.11917 | 0.150630318 | 0.083 | 0 | 1 | 15 |
| tubg1 | 0.11917 | 0.133628134 | 0.083 | 0 | 1 | 15 |
| utrn | 0.11917 | 0.142847222 | 0.083 | 0 | 1 | 15 |
| vamp2 | 0.11917 | 0.139863602 | 0.083 | 0 | 1 | 15 |
| zgc:110821 | 0.11917 | 0.160212927 | 0.083 | 0 | 1 | 15 |
| zgc:113149 | 0.11917 | 0.163930701 | 0.083 | 0 | 1 | 15 |
| zgc:174877 | 0.11917 | 0.17164519 | 0.083 | 0 | 1 | 15 |

| | | | | | | |
|---|---|---|---|---|---|---|
| znf1181 | 0.11917 | 0.150240979 | 0.083 | 0 | 1 | 15 |
| atpv0e2 | 0.11923 | 0.224789495 | 0.194 | 0.043 | 1 | 15 |
| sf3a3 | 0.1198 | 0.307772473 | 0.222 | 0.043 | 1 | 15 |
| cnn2 | 0.11997 | -0.39823871 | 0.167 | 0.391 | 1 | 15 |
| foxl2b | 0.12061 | 0.309983654 | 0.528 | 0.435 | 1 | 15 |
| dnmt3ab | 0.12065 | 0.378550971 | 0.361 | 0.13 | 1 | 15 |
| dnajc7 | 0.12109 | -0.33449579 | 0.028 | 0.174 | 1 | 15 |
| tubb5 | 0.12191 | -0.33652918 | 0.167 | 0.391 | 1 | 15 |
| lmf2b | 0.12195 | -0.25904001 | 0.028 | 0.174 | 1 | 15 |
| tmed9 | 0.12204 | 0.349131401 | 0.528 | 0.261 | 1 | 15 |
| stard3nl | 0.12215 | -0.21930107 | 0.028 | 0.174 | 1 | 15 |
| cndp2 | 0.12221 | -0.24869113 | 0.111 | 0.304 | 1 | 15 |
| gab2 | 0.12315 | -0.37851325 | 0.056 | 0.174 | 1 | 15 |
| matr3l1.1 | 0.12394 | 0.358619702 | 0.417 | 0.217 | 1 | 15 |
| srsf1b | 0.12401 | 0.278070492 | 0.278 | 0.174 | 1 | 15 |
| uchl5 | 0.12416 | 0.26545799 | 0.194 | 0.043 | 1 | 15 |
| vclb | 0.12446 | -0.30718937 | 0.028 | 0.174 | 1 | 15 |
| gtpbp4 | 0.1245 | 0.269838575 | 0.556 | 0.304 | 1 | 15 |
| odf2a | 0.12488 | -0.28352652 | 0.028 | 0.174 | 1 | 15 |
| rtn4a | 0.12535 | 0.324801455 | 0.528 | 0.348 | 1 | 15 |
| lactb | 0.12571 | -0.13016138 | 0.028 | 0.13 | 1 | 15 |
| ndufb5 | 0.12573 | -0.36544448 | 0.194 | 0.391 | 1 | 15 |
| psma3 | 0.12617 | -0.23808061 | 0.417 | 0.478 | 1 | 15 |
| hpf1 | 0.12631 | 0.310204789 | 0.222 | 0.043 | 1 | 15 |
| relb | 0.12692 | -0.2883137 | 0.028 | 0.174 | 1 | 15 |
| rps20 | 0.12694 | 0.126655043 | 0.944 | 0.913 | 1 | 15 |
| itpkcb | 0.12791 | 0.240852771 | 0.194 | 0.043 | 1 | 15 |
| ccnl1a | 0.1287 | -0.22666882 | 0.028 | 0.174 | 1 | 15 |
| ppp2r5eb | 0.12919 | 0.342992954 | 0.222 | 0.043 | 1 | 15 |
| si:ch211-235e9.6 | 0.12944 | 0.3378268 | 0.472 | 0.217 | 1 | 15 |
| ctsla | 0.12954 | -0.20604402 | 0.917 | 0.957 | 1 | 15 |
| adar | 0.12997 | -0.29144136 | 0.028 | 0.174 | 1 | 15 |
| si:dkey-29d8.3 | 0.13006 | -0.24755219 | 0.028 | 0.174 | 1 | 15 |
| BX548015.2 | 0.13018 | -0.2865687 | 0.028 | 0.174 | 1 | 15 |
| pik3r3b | 0.13078 | 0.340040658 | 0.194 | 0.043 | 1 | 15 |
| eif4a1b | 0.13107 | 0.113224084 | 0.722 | 0.522 | 1 | 15 |
| itm2bb | 0.13108 | 0.147265192 | 1 | 0.957 | 1 | 15 |
| nkrf | 0.1312 | 0.272551959 | 0.222 | 0.043 | 1 | 15 |
| dido1 | 0.13121 | 0.370553899 | 0.472 | 0.217 | 1 | 15 |
| si:ch73-42p12.2 | 0.13131 | 0.272321121 | 0.389 | 0.174 | 1 | 15 |
| egr1 | 0.13166 | -0.40729412 | 0.194 | 0.435 | 1 | 15 |
| ctsa | 0.1317 | 0.268596649 | 0.333 | 0.13 | 1 | 15 |
| ddx5 | 0.13189 | -0.13616269 | 0.667 | 0.652 | 1 | 15 |
| cacnb1 | 0.13192 | -0.27083307 | 0.028 | 0.174 | 1 | 15 |
| mrps14 | 0.13232 | -0.29427895 | 0.111 | 0.304 | 1 | 15 |
| trmt112 | 0.1325 | 0.329796052 | 0.361 | 0.13 | 1 | 15 |
| cdc42 | 0.13275 | -0.13183009 | 0.583 | 0.565 | 1 | 15 |
| gnl2 | 0.1328 | -0.32430431 | 0.056 | 0.217 | 1 | 15 |

| | | | | | | |
|---|---|---|---|---|---|---|
| cox6b2 | 0.13283 | 0.265580119 | 0.889 | 0.783 | 1 | 15 |
| gstt1b | 0.13293 | -0.37591408 | 0.361 | 0.609 | 1 | 15 |
| rps3a | 0.13355 | 0.179595135 | 0.972 | 0.957 | 1 | 15 |
| oip5-as1 | 0.13358 | 0.253411137 | 0.194 | 0.043 | 1 | 15 |
| CABZ01080702.1 | 0.13367 | -0.38814913 | 0.333 | 0.565 | 1 | 15 |
| fam13b | 0.13413 | 0.301394325 | 0.222 | 0.043 | 1 | 15 |
| asna1 | 0.13415 | -0.15231921 | 0.111 | 0.13 | 1 | 15 |
| gskip | 0.13419 | 0.326413361 | 0.417 | 0.174 | 1 | 15 |
| slc43a1a | 0.13422 | -0.3263835 | 0.028 | 0.174 | 1 | 15 |
| pde4ba | 0.13451 | 0.359925624 | 0.722 | 0.522 | 1 | 15 |
| ptp4a2b | 0.13475 | 0.301048437 | 0.556 | 0.304 | 1 | 15 |
| vdac3 | 0.13478 | 0.349723535 | 0.611 | 0.348 | 1 | 15 |
| ost4 | 0.13481 | -0.29084307 | 0.417 | 0.565 | 1 | 15 |
| gna12a | 0.13531 | 0.291448029 | 0.222 | 0.043 | 1 | 15 |
| mrpl18 | 0.13533 | 0.323118506 | 0.222 | 0.043 | 1 | 15 |
| pip4k2ca | 0.13572 | 0.343104032 | 0.222 | 0.043 | 1 | 15 |
| rbm42 | 0.13592 | -0.19564628 | 0.028 | 0.174 | 1 | 15 |
| zfyve21 | 0.13676 | 0.281387792 | 0.222 | 0.043 | 1 | 15 |
| fosb | 0.13686 | -0.45956437 | 0.139 | 0.304 | 1 | 15 |
| si:ch73-211e3.1 | 0.13689 | 0.291700527 | 0.222 | 0.043 | 1 | 15 |
| zgc:77650 | 0.13721 | 0.315598892 | 0.222 | 0.043 | 1 | 15 |
| rps27.1 | 0.1375 | 0.156710482 | 1 | 0.913 | 1 | 15 |
| si:ch211-214j24.10 | 0.13758 | 0.337738359 | 0.278 | 0.087 | 1 | 15 |
| mrpl52 | 0.13759 | 0.273234569 | 0.222 | 0.043 | 1 | 15 |
| cul4b | 0.13767 | 0.269191569 | 0.194 | 0.043 | 1 | 15 |
| adsl | 0.13788 | 0.263115813 | 0.222 | 0.043 | 1 | 15 |
| mat2aa | 0.13799 | -0.33209671 | 0.083 | 0.217 | 1 | 15 |
| b3gnt5a | 0.13815 | -0.30644252 | 0.083 | 0.217 | 1 | 15 |
| smap1 | 0.13834 | 0.315331227 | 0.194 | 0.043 | 1 | 15 |
| jagn1b | 0.13839 | -0.25223136 | 0.194 | 0.261 | 1 | 15 |
| nipblb | 0.13846 | 0.154481289 | 0.222 | 0.087 | 1 | 15 |
| serf2 | 0.13873 | -0.16820982 | 0.611 | 0.609 | 1 | 15 |
| pdhb | 0.13902 | 0.221362694 | 0.25 | 0.087 | 1 | 15 |
| hbp1 | 0.1393 | 0.334945888 | 0.417 | 0.261 | 1 | 15 |
| mcee | 0.13953 | -0.22653865 | 0.056 | 0.217 | 1 | 15 |
| sept2 | 0.13959 | -0.37108095 | 0.194 | 0.435 | 1 | 15 |
| elmo3 | 0.13979 | -0.32630458 | 0.028 | 0.174 | 1 | 15 |
| metap1 | 0.13981 | 0.227148852 | 0.194 | 0.043 | 1 | 15 |
| ndfip1 | 0.13983 | 0.337628529 | 0.611 | 0.348 | 1 | 15 |
| si:dkeyp-82a1.6 | 0.14027 | -0.30785417 | 0.028 | 0.174 | 1 | 15 |
| klf6a | 0.14045 | -0.30204354 | 0.056 | 0.174 | 1 | 15 |
| tpbgb | 0.14192 | -0.33418012 | 0.056 | 0.217 | 1 | 15 |
| mbd1b | 0.1422 | -0.27461839 | 0.028 | 0.174 | 1 | 15 |
| emg1 | 0.14235 | 0.206793243 | 0.194 | 0.043 | 1 | 15 |
| os9 | 0.14243 | -0.23988612 | 0.028 | 0.174 | 1 | 15 |
| zgc:56525 | 0.14248 | -0.25678308 | 0.028 | 0.174 | 1 | 15 |
| syngr2a | 0.14299 | -0.18310776 | 0.028 | 0.174 | 1 | 15 |
| hmgn6 | 0.14317 | -0.26098755 | 0.75 | 0.913 | 1 | 15 |

| | | | | | | |
|---|---|---|---|---|---|---|
| grinab | 0.14319 | -0.14216507 | 0.472 | 0.478 | 1 | 15 |
| cul3b | 0.14325 | -0.31918797 | 0.056 | 0.174 | 1 | 15 |
| thoc1 | 0.14335 | -0.22974141 | 0.028 | 0.174 | 1 | 15 |
| wsb1 | 0.14385 | 0.100401951 | 0.306 | 0.174 | 1 | 15 |
| rpl30 | 0.14415 | 0.243694944 | 0.944 | 0.783 | 1 | 15 |
| zgc:158343 | 0.14421 | -0.43006855 | 0.083 | 0.261 | 1 | 15 |
| fbxo34 | 0.14468 | -0.29940564 | 0.028 | 0.174 | 1 | 15 |
| znf217 | 0.145 | -0.12073457 | 0.028 | 0.13 | 1 | 15 |
| zgc:171480 | 0.14503 | 0.18760816 | 0.25 | 0.087 | 1 | 15 |
| aebp2 | 0.14528 | -0.36555621 | 0.222 | 0.435 | 1 | 15 |
| rps10 | 0.1455 | 0.152586709 | 1 | 0.913 | 1 | 15 |
| ctgfa | 0.14555 | -0.23608572 | 0.028 | 0.174 | 1 | 15 |
| pdcd4a | 0.14589 | 0.295943771 | 0.25 | 0.087 | 1 | 15 |
| CU539058.1 | 0.14612 | -0.22195648 | 0.028 | 0.174 | 1 | 15 |
| si:ch73-281n10.2 | 0.14619 | 0.317829651 | 0.833 | 0.609 | 1 | 15 |
| EML4 | 0.14631 | -0.30877637 | 0.028 | 0.174 | 1 | 15 |
| slc26a5 | 0.14635 | 0.277822122 | 0.167 | 0.043 | 1 | 15 |
| ss18l2 | 0.14731 | -0.23807805 | 0.056 | 0.217 | 1 | 15 |
| csnk1g2b | 0.14737 | -0.20360178 | 0.028 | 0.174 | 1 | 15 |
| mff | 0.14744 | 0.263553021 | 0.167 | 0.043 | 1 | 15 |
| vbp1 | 0.14745 | -0.20625675 | 0.139 | 0.174 | 1 | 15 |
| zgc:174906 | 0.14792 | -0.28901393 | 0.056 | 0.217 | 1 | 15 |
| golga4 | 0.14795 | 0.16425086 | 0.222 | 0.087 | 1 | 15 |
| nfkbib | 0.14796 | 0.273224546 | 0.167 | 0.043 | 1 | 15 |
| slc30a7 | 0.14818 | 0.285039041 | 0.194 | 0.043 | 1 | 15 |
| PRRC2B | 0.14839 | -0.24780328 | 0.028 | 0.174 | 1 | 15 |
| snu13b | 0.1486 | 0.33048664 | 0.472 | 0.304 | 1 | 15 |
| atp5mc1 | 0.14879 | 0.225373903 | 0.722 | 0.478 | 1 | 15 |
| tnrc6c2 | 0.14887 | 0.229290952 | 0.25 | 0.087 | 1 | 15 |
| rnf38 | 0.14953 | -0.23626001 | 0.028 | 0.174 | 1 | 15 |
| ttc1 | 0.14955 | -0.23072591 | 0.028 | 0.174 | 1 | 15 |
| csf1b | 0.1499 | -0.13433541 | 0.028 | 0.13 | 1 | 15 |
| ifngr1l | 0.14995 | -0.22810816 | 0.028 | 0.13 | 1 | 15 |
| spoplb | 0.14997 | -0.32884334 | 0.028 | 0.174 | 1 | 15 |
| nup50 | 0.15032 | 0.284183272 | 0.306 | 0.13 | 1 | 15 |
| si:zfos-1192g2.3 | 0.15069 | 0.184360472 | 0.333 | 0.261 | 1 | 15 |
| isg20 | 0.1507 | -0.2490285 | 0.056 | 0.217 | 1 | 15 |
| rnf128a | 0.15094 | 0.291929423 | 0.25 | 0.13 | 1 | 15 |
| nfya | 0.15143 | -0.23115634 | 0.028 | 0.174 | 1 | 15 |
| esf1 | 0.15159 | 0.272643906 | 0.167 | 0.043 | 1 | 15 |
| pds5a | 0.15162 | -0.21775654 | 0.028 | 0.174 | 1 | 15 |
| bdh1 | 0.15169 | -0.2398504 | 0.028 | 0.174 | 1 | 15 |
| FRMD1 | 0.1517 | -0.24892613 | 0.028 | 0.174 | 1 | 15 |
| lrba | 0.15174 | -0.26772815 | 0.028 | 0.174 | 1 | 15 |
| ttc9c | 0.15178 | -0.30612351 | 0.083 | 0.217 | 1 | 15 |
| abca3b | 0.15207 | -0.23128154 | 0.028 | 0.174 | 1 | 15 |
| psmc5 | 0.15209 | 0.321571317 | 0.361 | 0.217 | 1 | 15 |
| spsb4b | 0.1522 | -0.19615747 | 0.028 | 0.174 | 1 | 15 |

| | | | | | | |
|---|---|---|---|---|---|---|
| isoc1 | 0.1523 | -0.25834591 | 0.028 | 0.174 | 1 | 15 |
| cltb | 0.15232 | -0.26120742 | 0.028 | 0.174 | 1 | 15 |
| cript | 0.15234 | -0.23468999 | 0.028 | 0.174 | 1 | 15 |
| sec24c | 0.15241 | -0.21555625 | 0.028 | 0.174 | 1 | 15 |
| si:ch211-286b5.5 | 0.15372 | 0.229223514 | 0.556 | 0.478 | 1 | 15 |
| maip1 | 0.15384 | 0.212125871 | 0.194 | 0.043 | 1 | 15 |
| tomm20a | 0.15408 | 0.158808186 | 0.528 | 0.522 | 1 | 15 |
| si:dkeyp-113d7.1 | 0.15432 | -0.35754739 | 0.167 | 0.391 | 1 | 15 |
| si:ch73-37h15.2 | 0.15464 | 0.167142828 | 0.222 | 0.087 | 1 | 15 |
| CU929070.1 | 0.15519 | 0.200352414 | 0.194 | 0.043 | 1 | 15 |
| CABZ01021532.1 | 0.15526 | -0.43011293 | 0.139 | 0.348 | 1 | 15 |
| plekhf2 | 0.15584 | -0.35053807 | 0.056 | 0.217 | 1 | 15 |
| fam168b | 0.15628 | 0.176683274 | 0.222 | 0.174 | 1 | 15 |
| ola1 | 0.15676 | 0.295589861 | 0.639 | 0.391 | 1 | 15 |
| snrnp70 | 0.15698 | 0.352755055 | 0.333 | 0.13 | 1 | 15 |
| znf609b | 0.157 | -0.35920505 | 0.167 | 0.348 | 1 | 15 |
| krt15 | 0.15708 | -0.48753628 | 0.139 | 0.348 | 1 | 15 |
| zfand5b | 0.15727 | 0.258244215 | 0.25 | 0.087 | 1 | 15 |
| tars | 0.15735 | 0.291539507 | 0.194 | 0.043 | 1 | 15 |
| pfdn4 | 0.15757 | -0.23938574 | 0.306 | 0.391 | 1 | 15 |
| spag7 | 0.15801 | 0.180474059 | 0.167 | 0.043 | 1 | 15 |
| pgm2l1 | 0.15825 | 0.269542173 | 0.167 | 0.043 | 1 | 15 |
| wdr1 | 0.15931 | -0.35754568 | 0.139 | 0.348 | 1 | 15 |
| ctnnd1 | 0.15935 | -0.34776836 | 0.056 | 0.217 | 1 | 15 |
| mibp2 | 0.15994 | 0.266004084 | 0.833 | 0.609 | 1 | 15 |
| FP245456.1 | 0.15999 | 0.232427382 | 0.139 | 0.043 | 1 | 15 |
| wfikkn2a | 0.16005 | -0.31198539 | 0.056 | 0.217 | 1 | 15 |
| tmem39a | 0.1609 | 0.338590389 | 0.25 | 0.087 | 1 | 15 |
| tent4b | 0.16122 | 0.254773335 | 0.194 | 0.043 | 1 | 15 |
| ndufa4l | 0.16153 | 0.127843476 | 0.889 | 0.913 | 1 | 15 |
| arf1 | 0.16192 | 0.270186436 | 0.333 | 0.13 | 1 | 15 |
| wwtr1 | 0.16214 | -0.3248197 | 0.139 | 0.348 | 1 | 15 |
| anxa2a | 0.16278 | -0.46411004 | 0.361 | 0.609 | 1 | 15 |
| chmp7 | 0.16314 | 0.3179483 | 0.194 | 0.043 | 1 | 15 |
| nat8l | 0.16376 | -0.33473728 | 0.028 | 0.13 | 1 | 15 |
| arf6b | 0.16384 | 0.329193708 | 0.333 | 0.13 | 1 | 15 |
| cox7b | 0.16389 | 0.165304615 | 0.75 | 0.522 | 1 | 15 |
| rpl21 | 0.1641 | 0.190376267 | 0.944 | 0.87 | 1 | 15 |
| derl2 | 0.16483 | -0.12736917 | 0.167 | 0.174 | 1 | 15 |
| hsbp1a | 0.16485 | -0.31358555 | 0.083 | 0.217 | 1 | 15 |
| chd7 | 0.1653 | 0.339978287 | 0.361 | 0.174 | 1 | 15 |
| abhd15a | 0.16537 | -0.3216499 | 0.111 | 0.304 | 1 | 15 |
| sorbs1 | 0.16539 | -0.28399643 | 0.056 | 0.217 | 1 | 15 |
| ppp1r11 | 0.16543 | -0.31284758 | 0.056 | 0.174 | 1 | 15 |
| sub1a | 0.16549 | 0.298645235 | 0.722 | 0.478 | 1 | 15 |
| fosl1a | 0.16597 | -0.28398777 | 0.056 | 0.217 | 1 | 15 |
| arfgap2 | 0.16603 | -0.27545492 | 0.056 | 0.217 | 1 | 15 |
| grcc10 | 0.16606 | 0.294031568 | 0.278 | 0.13 | 1 | 15 |

| | | | | | | |
|---|---|---|---|---|---|---|
| ube2na | 0.1662 | 0.201161755 | 0.306 | 0.261 | 1 | 15 |
| CYTH2 | 0.16702 | -0.17846647 | 0.083 | 0.13 | 1 | 15 |
| calcoco1a | 0.16765 | 0.227793933 | 0.222 | 0.13 | 1 | 15 |
| zcrb1 | 0.1678 | 0.231915972 | 0.194 | 0.043 | 1 | 15 |
| porb | 0.16791 | -0.29862226 | 0.083 | 0.217 | 1 | 15 |
| anapc16 | 0.16835 | 0.233569501 | 0.417 | 0.217 | 1 | 15 |
| ppfibp1a | 0.16839 | -0.37178766 | 0.083 | 0.261 | 1 | 15 |
| ppiab | 0.16843 | 0.217659096 | 0.944 | 0.783 | 1 | 15 |
| tmem165 | 0.16852 | -0.30243219 | 0.167 | 0.304 | 1 | 15 |
| slc6a4a | 0.16885 | -0.15307778 | 0.111 | 0.13 | 1 | 15 |
| zgc:162509 | 0.169 | -0.30352313 | 0.139 | 0.348 | 1 | 15 |
| zgc:153867 | 0.16934 | -0.17540924 | 0.75 | 0.783 | 1 | 15 |
| si:ch211-212k18.9 | 0.1694 | -0.17214553 | 0.111 | 0.261 | 1 | 15 |
| fblim1 | 0.16962 | -0.2632318 | 0.083 | 0.261 | 1 | 15 |
| llph | 0.16966 | 0.333063494 | 0.278 | 0.087 | 1 | 15 |
| srpk1a | 0.16973 | 0.243131053 | 0.167 | 0.043 | 1 | 15 |
| dhdhl | 0.17011 | -0.18072384 | 0.083 | 0.13 | 1 | 15 |
| chmp2a | 0.17042 | -0.32203464 | 0.361 | 0.609 | 1 | 15 |
| tmem258 | 0.17084 | -0.13279914 | 0.556 | 0.522 | 1 | 15 |
| ywhah | 0.17088 | -0.32164862 | 0.139 | 0.348 | 1 | 15 |
| pabpc1b | 0.17105 | -0.30575321 | 0.194 | 0.348 | 1 | 15 |
| lsm6 | 0.17233 | 0.364753141 | 0.5 | 0.261 | 1 | 15 |
| krt18b | 0.17266 | -0.40693051 | 0.194 | 0.391 | 1 | 15 |
| acaa2 | 0.17305 | -0.2883866 | 0.139 | 0.348 | 1 | 15 |
| pdlim1 | 0.174 | -0.32852361 | 0.139 | 0.348 | 1 | 15 |
| ccng1 | 0.17423 | 0.183634936 | 0.611 | 0.391 | 1 | 15 |
| rbm4.2 | 0.17486 | 0.216998367 | 0.194 | 0.13 | 1 | 15 |
| atp5mf | 0.17497 | -0.31302016 | 0.361 | 0.609 | 1 | 15 |
| mapkapk2a | 0.17517 | -0.31093439 | 0.139 | 0.348 | 1 | 15 |
| pbx1a | 0.17535 | -0.25930143 | 0.056 | 0.217 | 1 | 15 |
| gadd45gip1 | 0.17539 | -0.29984027 | 0.139 | 0.348 | 1 | 15 |
| hnrnpa0l | 0.17548 | 0.161891153 | 0.944 | 0.783 | 1 | 15 |
| eif3i | 0.17558 | 0.310619118 | 0.639 | 0.391 | 1 | 15 |
| fntb | 0.17635 | 0.288557782 | 0.194 | 0.043 | 1 | 15 |
| krr1 | 0.17645 | 0.112251735 | 0.139 | 0.043 | 1 | 15 |
| zfhx3 | 0.17707 | -0.27752593 | 0.111 | 0.217 | 1 | 15 |
| nme2b.1 | 0.17707 | 0.110757404 | 0.944 | 0.957 | 1 | 15 |
| tmx4 | 0.1772 | 0.112402134 | 0.139 | 0.043 | 1 | 15 |
| fryl | 0.17749 | -0.34568274 | 0.028 | 0.13 | 1 | 15 |
| si:dkey-17m8.1 | 0.17753 | -0.34768576 | 0.222 | 0.435 | 1 | 15 |
| tsku | 0.17773 | -0.32483387 | 0.083 | 0.261 | 1 | 15 |
| slc29a1a | 0.17857 | -0.25305002 | 0.222 | 0.304 | 1 | 15 |
| txndc12 | 0.17879 | -0.14108509 | 0.194 | 0.217 | 1 | 15 |
| gtf2e2 | 0.17891 | 0.318078198 | 0.278 | 0.087 | 1 | 15 |
| pagr1 | 0.1791 | 0.293520524 | 0.278 | 0.087 | 1 | 15 |
| ccnd1 | 0.1794 | -0.28456003 | 0.111 | 0.217 | 1 | 15 |
| arl1 | 0.17942 | 0.251657113 | 0.167 | 0.043 | 1 | 15 |
| tbl1xr1b | 0.17965 | -0.29145946 | 0.056 | 0.217 | 1 | 15 |

| | | | | | | |
|---|---|---|---|---|---|---|
| u2surp | 0.17977 | -0.35451862 | 0.222 | 0.435 | 1 | 15 |
| rmdn3 | 0.17986 | -0.29328831 | 0.056 | 0.174 | 1 | 15 |
| ywhaqa | 0.18007 | 0.292939723 | 0.389 | 0.174 | 1 | 15 |
| abcd3a | 0.18077 | -0.2301367 | 0.056 | 0.217 | 1 | 15 |
| necap2 | 0.18085 | -0.25277533 | 0.056 | 0.217 | 1 | 15 |
| hddc3 | 0.18093 | 0.201983715 | 0.139 | 0.087 | 1 | 15 |
| ip6k2b | 0.18129 | 0.299204404 | 0.25 | 0.13 | 1 | 15 |
| adnpa | 0.18167 | -0.2810087 | 0.056 | 0.217 | 1 | 15 |
| FQ976913.1 | 0.18196 | -0.11385118 | 0.056 | 0.174 | 1 | 15 |
| si:dkeyp-75h12.5 | 0.18196 | -0.27250988 | 0.056 | 0.217 | 1 | 15 |
| cavin2a | 0.18205 | -0.26817152 | 0.056 | 0.217 | 1 | 15 |
| sh3gl1b | 0.18219 | -0.28219382 | 0.056 | 0.217 | 1 | 15 |
| strn4 | 0.1822 | -0.11032252 | 0.139 | 0.13 | 1 | 15 |
| rasa1a | 0.18231 | -0.26896845 | 0.056 | 0.174 | 1 | 15 |
| dspa | 0.18232 | -0.3348677 | 0.083 | 0.261 | 1 | 15 |
| zgc:110591 | 0.18243 | -0.27571995 | 0.083 | 0.261 | 1 | 15 |
| si:dkey-19e4.5 | 0.18251 | -0.31033167 | 0.111 | 0.304 | 1 | 15 |
| nup107 | 0.1836 | -0.28296009 | 0.056 | 0.174 | 1 | 15 |
| oga | 0.18386 | -0.32309214 | 0.167 | 0.304 | 1 | 15 |
| ctbp2a | 0.18395 | -0.29048867 | 0.111 | 0.304 | 1 | 15 |
| nop10 | 0.1841 | 0.313696487 | 0.556 | 0.391 | 1 | 15 |
| slc22a18 | 0.18446 | -0.19794207 | 0.083 | 0.13 | 1 | 15 |
| nfic | 0.18463 | -0.28123522 | 0.306 | 0.435 | 1 | 15 |
| anapc11 | 0.18467 | -0.30354247 | 0.083 | 0.261 | 1 | 15 |
| smurf2 | 0.1852 | -0.34786897 | 0.111 | 0.304 | 1 | 15 |
| tmem59 | 0.18532 | 0.33695514 | 0.444 | 0.217 | 1 | 15 |
| carhsp1 | 0.18556 | -0.30333502 | 0.111 | 0.304 | 1 | 15 |
| PYURF | 0.18573 | -0.32342928 | 0.167 | 0.304 | 1 | 15 |
| isg15 | 0.18602 | 0.117049557 | 0.139 | 0.13 | 1 | 15 |
| tnfrsf21 | 0.18603 | -0.10848217 | 0.083 | 0.087 | 1 | 15 |
| dars | 0.1862 | -0.27804363 | 0.083 | 0.261 | 1 | 15 |
| b2ml | 0.18622 | 0.214928419 | 0.889 | 0.696 | 1 | 15 |
| stmn1a | 0.18626 | -0.33840695 | 0.111 | 0.304 | 1 | 15 |
| hmox1a | 0.18654 | -0.21458087 | 0.028 | 0.13 | 1 | 15 |
| ube2d4 | 0.18663 | 0.178014485 | 0.278 | 0.13 | 1 | 15 |
| max | 0.18709 | 0.303820875 | 0.333 | 0.174 | 1 | 15 |
| trim25 | 0.18718 | 0.273504971 | 0.194 | 0.043 | 1 | 15 |
| mbnl1 | 0.18724 | -0.18747266 | 0.139 | 0.174 | 1 | 15 |
| ubr7 | 0.18754 | 0.238885791 | 0.167 | 0.043 | 1 | 15 |
| mrps30 | 0.18773 | 0.16305865 | 0.167 | 0.043 | 1 | 15 |
| rab5aa | 0.18794 | 0.296346374 | 0.194 | 0.043 | 1 | 15 |
| ndufb6 | 0.18813 | 0.308659356 | 0.361 | 0.217 | 1 | 15 |
| spon1b | 0.18822 | -0.2502083 | 0.083 | 0.261 | 1 | 15 |
| naa10 | 0.18831 | 0.303042865 | 0.389 | 0.217 | 1 | 15 |
| tufm | 0.18837 | 0.346609918 | 0.333 | 0.13 | 1 | 15 |
| tmem198b | 0.18846 | -0.28619812 | 0.056 | 0.174 | 1 | 15 |
| usp43b | 0.18906 | -0.29345867 | 0.028 | 0.13 | 1 | 15 |
| atg13 | 0.18935 | 0.241684986 | 0.194 | 0.043 | 1 | 15 |

| | | | | | | |
|---|---|---|---|---|---|---|
| inha | 0.18939 | 0.35392153 | 0.389 | 0.174 | 1 | 15 |
| DDX17 | 0.18941 | -0.14416078 | 0.056 | 0.174 | 1 | 15 |
| glulb | 0.18969 | 0.370921196 | 0.667 | 0.435 | 1 | 15 |
| zgc:110239 | 0.18972 | -0.29935827 | 0.083 | 0.261 | 1 | 15 |
| hmgb2b | 0.18986 | -0.16922833 | 0.833 | 0.826 | 1 | 15 |
| si:ch211-237c6.4 | 0.19013 | -0.26520214 | 0.083 | 0.261 | 1 | 15 |
| atp6v1e1a | 0.19018 | 0.240700542 | 0.222 | 0.13 | 1 | 15 |
| serbp1a | 0.19018 | 0.203214378 | 0.917 | 0.739 | 1 | 15 |
| mdka | 0.1902 | -0.35401531 | 0.083 | 0.261 | 1 | 15 |
| krt94 | 0.19024 | 0.197320928 | 0.917 | 0.739 | 1 | 15 |
| sec11a | 0.19048 | -0.15013943 | 0.528 | 0.522 | 1 | 15 |
| znf1104 | 0.19059 | 0.377152238 | 0.333 | 0.13 | 1 | 15 |
| spopla | 0.19082 | -0.25982423 | 0.028 | 0.13 | 1 | 15 |
| slirp | 0.19124 | 0.259450737 | 0.389 | 0.174 | 1 | 15 |
| cd3eap | 0.19182 | 0.226846735 | 0.167 | 0.043 | 1 | 15 |
| si:ch211-117c9.5 | 0.19214 | 0.114490693 | 0.306 | 0.174 | 1 | 15 |
| atp5if1b | 0.19255 | 0.296405765 | 0.417 | 0.261 | 1 | 15 |
| u2af1 | 0.19272 | 0.207270047 | 0.389 | 0.304 | 1 | 15 |
| gdi2 | 0.19279 | 0.288843356 | 0.444 | 0.217 | 1 | 15 |
| fam102ab | 0.19324 | 0.376648421 | 0.444 | 0.217 | 1 | 15 |
| zgc:77086 | 0.19326 | 0.16989143 | 0.389 | 0.217 | 1 | 15 |
| timm17a | 0.19403 | 0.271651532 | 0.194 | 0.043 | 1 | 15 |
| copz1 | 0.19458 | 0.302439887 | 0.444 | 0.217 | 1 | 15 |
| prodha | 0.19493 | -0.11033182 | 0.139 | 0.13 | 1 | 15 |
| chtopa | 0.19504 | 0.348289477 | 0.639 | 0.435 | 1 | 15 |
| vdac2 | 0.19583 | 0.326636594 | 0.667 | 0.435 | 1 | 15 |
| wu:fb55g09 | 0.19647 | -0.29794923 | 0.611 | 0.783 | 1 | 15 |
| dda1 | 0.19656 | 0.120936751 | 0.194 | 0.087 | 1 | 15 |
| exosc4 | 0.1966 | 0.267752307 | 0.194 | 0.043 | 1 | 15 |
| ppiaa | 0.19666 | 0.124457828 | 1 | 0.913 | 1 | 15 |
| znf706 | 0.19671 | 0.211358144 | 0.472 | 0.261 | 1 | 15 |
| rbm25a | 0.19747 | 0.273829178 | 0.194 | 0.043 | 1 | 15 |
| fam168a | 0.19781 | -0.34276462 | 0.222 | 0.435 | 1 | 15 |
| dynll2b | 0.19832 | 0.245130329 | 0.472 | 0.261 | 1 | 15 |
| wbp1 | 0.19888 | 0.222812688 | 0.25 | 0.087 | 1 | 15 |
| tgfbr3 | 0.19942 | -0.2734557 | 0.056 | 0.174 | 1 | 15 |
| rab41 | 0.19957 | 0.208330512 | 0.167 | 0.043 | 1 | 15 |
| ypel5 | 0.19976 | 0.24488387 | 0.167 | 0.043 | 1 | 15 |
| col4a3bpb | 0.1998 | -0.22330245 | 0.028 | 0.13 | 1 | 15 |
| eif3s6ip | 0.20003 | 0.34532366 | 0.472 | 0.261 | 1 | 15 |
| si:dkey-28b4.7 | 0.20025 | -0.20669045 | 0.444 | 0.522 | 1 | 15 |
| ewsr1a | 0.20099 | 0.24588901 | 0.278 | 0.174 | 1 | 15 |
| psme1 | 0.20123 | 0.295255636 | 0.583 | 0.348 | 1 | 15 |
| ddx27 | 0.20157 | 0.22166456 | 0.25 | 0.087 | 1 | 15 |
| hmgb1b | 0.20167 | 0.138308171 | 0.833 | 0.652 | 1 | 15 |
| top2b | 0.20168 | -0.2730302 | 0.194 | 0.304 | 1 | 15 |
| igfbp5b | 0.20179 | 0.412018794 | 0.361 | 0.174 | 1 | 15 |
| gigyf1b | 0.20263 | 0.227357434 | 0.25 | 0.087 | 1 | 15 |

| | | | | | | |
|---|---|---|---|---|---|---|
| mphosph6 | 0.20289 | 0.169447515 | 0.167 | 0.043 | 1 | 15 |
| aldoaa | 0.20311 | 0.309565945 | 0.556 | 0.391 | 1 | 15 |
| suds3 | 0.2037 | 0.252559714 | 0.194 | 0.043 | 1 | 15 |
| cmc1 | 0.20432 | 0.103350099 | 0.139 | 0.043 | 1 | 15 |
| calhm2 | 0.20437 | -0.27307167 | 0.056 | 0.13 | 1 | 15 |
| ezrb | 0.2047 | 0.261757463 | 0.194 | 0.043 | 1 | 15 |
| fas | 0.20479 | -0.10893133 | 0.083 | 0.087 | 1 | 15 |
| nup58 | 0.2053 | 0.107549196 | 0.139 | 0.043 | 1 | 15 |
| zgc:112255 | 0.20554 | 0.227253261 | 0.167 | 0.087 | 1 | 15 |
| si:ch211-13k12.2 | 0.20561 | 0.173617475 | 0.167 | 0.043 | 1 | 15 |
| yipf4 | 0.2057 | 0.120400197 | 0.306 | 0.174 | 1 | 15 |
| mogs | 0.2057 | 0.252881644 | 0.139 | 0.043 | 1 | 15 |
| npepps | 0.20572 | 0.201429001 | 0.167 | 0.043 | 1 | 15 |
| rnf4 | 0.20573 | 0.266799131 | 0.194 | 0.043 | 1 | 15 |
| rbm8a | 0.20583 | 0.268587905 | 0.694 | 0.565 | 1 | 15 |
| nfe2l3 | 0.20634 | -0.27917825 | 0.111 | 0.261 | 1 | 15 |
| rer1 | 0.20681 | -0.36816063 | 0.278 | 0.478 | 1 | 15 |
| csnk1db | 0.207 | 0.114030146 | 0.25 | 0.13 | 1 | 15 |
| mrpl34 | 0.2071 | 0.209595732 | 0.306 | 0.13 | 1 | 15 |
| lgals17 | 0.20718 | 0.200069027 | 0.417 | 0.391 | 1 | 15 |
| papola | 0.20753 | 0.24076692 | 0.194 | 0.043 | 1 | 15 |
| vps4a | 0.20799 | 0.126209658 | 0.139 | 0.043 | 1 | 15 |
| ak2 | 0.20824 | -0.10987191 | 0.389 | 0.348 | 1 | 15 |
| tomm22 | 0.20864 | 0.221212703 | 0.25 | 0.087 | 1 | 15 |
| bmpr1bb | 0.20869 | -0.26399089 | 0.028 | 0.13 | 1 | 15 |
| helz | 0.20871 | -0.2180862 | 0.056 | 0.13 | 1 | 15 |
| atp5if1a | 0.20879 | 0.31356412 | 0.583 | 0.435 | 1 | 15 |
| glod5 | 0.20919 | -0.29536477 | 0.111 | 0.261 | 1 | 15 |
| pabpc1a | 0.20955 | -0.10871113 | 0.889 | 0.87 | 1 | 15 |
| zfr | 0.20991 | 0.259570278 | 0.194 | 0.043 | 1 | 15 |
| si:ch1073-322p19.1 | 0.21029 | 0.232769775 | 0.194 | 0.043 | 1 | 15 |
| CABZ01039096.1 | 0.21058 | -0.16724459 | 0.028 | 0.13 | 1 | 15 |
| rps21 | 0.21064 | 0.160547783 | 0.972 | 0.87 | 1 | 15 |
| capzb | 0.21132 | 0.13510013 | 0.417 | 0.261 | 1 | 15 |
| fam110b | 0.21145 | 0.250685313 | 0.194 | 0.043 | 1 | 15 |
| mtfmt | 0.21146 | -0.29540651 | 0.028 | 0.13 | 1 | 15 |
| ptbp3 | 0.21154 | 0.259697703 | 0.306 | 0.13 | 1 | 15 |
| thrab | 0.21156 | 0.256613539 | 0.194 | 0.043 | 1 | 15 |
| pdk2b | 0.21204 | -0.15713333 | 0.083 | 0.217 | 1 | 15 |
| slc25a10 | 0.2122 | -0.12053357 | 0.028 | 0.13 | 1 | 15 |
| josd1 | 0.21226 | 0.285268902 | 0.167 | 0.043 | 1 | 15 |
| nfil3-5 | 0.21257 | -0.15972453 | 0.028 | 0.13 | 1 | 15 |
| napab | 0.21275 | 0.231771803 | 0.167 | 0.043 | 1 | 15 |
| hspg2 | 0.21313 | 0.139381057 | 0.361 | 0.217 | 1 | 15 |
| txndc15 | 0.21346 | -0.33040354 | 0.222 | 0.435 | 1 | 15 |
| zgc:56676 | 0.21353 | -0.23185076 | 0.222 | 0.304 | 1 | 15 |
| si:ch211-132g1.3 | 0.2137 | -0.16251268 | 0.056 | 0.174 | 1 | 15 |
| clstn1 | 0.21395 | 0.308874122 | 0.194 | 0.043 | 1 | 15 |

| | | | | | | |
|---|---|---|---|---|---|---|
| ktn1 | 0.21406 | 0.24101064 | 0.194 | 0.043 | 1 | 15 |
| zcchc17 | 0.21437 | 0.246713716 | 0.194 | 0.043 | 1 | 15 |
| dynll1 | 0.21474 | -0.23984381 | 0.5 | 0.565 | 1 | 15 |
| glrx | 0.21543 | -0.26445517 | 0.5 | 0.609 | 1 | 15 |
| hdac4 | 0.21554 | -0.27916985 | 0.194 | 0.304 | 1 | 15 |
| fdxr | 0.21651 | 0.26504353 | 0.361 | 0.174 | 1 | 15 |
| hmg20b | 0.21659 | 0.235409619 | 0.167 | 0.043 | 1 | 15 |
| zgc:153916 | 0.21665 | -0.156658 | 0.028 | 0.13 | 1 | 15 |
| ddost | 0.2176 | -0.28217324 | 0.111 | 0.217 | 1 | 15 |
| tpi1b | 0.2182 | -0.25891079 | 0.111 | 0.217 | 1 | 15 |
| zgc:195175 | 0.21828 | 0.227457804 | 0.25 | 0.174 | 1 | 15 |
| CR318588.1 | 0.21835 | 0.203099714 | 0.917 | 0.87 | 1 | 15 |
| lasp1 | 0.21877 | 0.202432211 | 0.333 | 0.261 | 1 | 15 |
| phactr4a | 0.21882 | -0.12320487 | 0.389 | 0.391 | 1 | 15 |
| top1l | 0.21887 | 0.219792014 | 0.472 | 0.391 | 1 | 15 |
| mob4 | 0.21912 | 0.226587972 | 0.278 | 0.217 | 1 | 15 |
| si:ch73-1a9.3 | 0.21958 | 0.165066674 | 0.972 | 0.957 | 1 | 15 |
| ponzr1 | 0.22031 | 0.178690596 | 0.861 | 0.826 | 1 | 15 |
| vma21 | 0.22133 | -0.32760082 | 0.278 | 0.478 | 1 | 15 |
| mif | 0.22163 | 0.320965429 | 0.611 | 0.391 | 1 | 15 |
| daam1a | 0.22268 | -0.16027983 | 0.028 | 0.13 | 1 | 15 |
| nabp1a | 0.22363 | 0.163311143 | 0.333 | 0.174 | 1 | 15 |
| rpz5 | 0.22376 | 0.155918128 | 0.694 | 0.696 | 1 | 15 |
| zgc:114181 | 0.22391 | -0.26924264 | 0.028 | 0.13 | 1 | 15 |
| coa5 | 0.22434 | 0.142543559 | 0.222 | 0.087 | 1 | 15 |
| AL732567.1 | 0.22453 | 0.114924013 | 0.056 | 0 | 1 | 15 |
| BX323824.1 | 0.22453 | 0.103292745 | 0.056 | 0 | 1 | 15 |
| BX530018.2 | 0.22453 | 0.107402825 | 0.056 | 0 | 1 | 15 |
| BX537342.1 | 0.22453 | 0.109802708 | 0.056 | 0 | 1 | 15 |
| BX957346.1 | 0.22453 | 0.109842243 | 0.056 | 0 | 1 | 15 |
| CABZ01063556.1 | 0.22453 | 0.133613502 | 0.056 | 0 | 1 | 15 |
| CABZ01078055.1 | 0.22453 | 0.111747819 | 0.056 | 0 | 1 | 15 |
| CABZ01086293.1 | 0.22453 | 0.10836135 | 0.056 | 0 | 1 | 15 |
| CR376783.1 | 0.22453 | 0.116488965 | 0.056 | 0 | 1 | 15 |
| CR450780.2 | 0.22453 | 0.123037404 | 0.056 | 0 | 1 | 15 |
| CR847984.1 | 0.22453 | 0.10672016 | 0.056 | 0 | 1 | 15 |
| CR855865.1 | 0.22453 | 0.124684743 | 0.056 | 0 | 1 | 15 |
| CT573342.1 | 0.22453 | 0.103115022 | 0.056 | 0 | 1 | 15 |
| CT737123.1 | 0.22453 | 0.10162466 | 0.056 | 0 | 1 | 15 |
| CU570894.2 | 0.22453 | 0.132611011 | 0.056 | 0 | 1 | 15 |
| CU856539.2 | 0.22453 | 0.117028257 | 0.056 | 0 | 1 | 15 |
| FO681288.1 | 0.22453 | 0.14842692 | 0.056 | 0 | 1 | 15 |
| FO704779.1 | 0.22453 | 0.104423208 | 0.056 | 0 | 1 | 15 |
| LOC100000365 | 0.22453 | 0.118340309 | 0.056 | 0 | 1 | 15 |
| LOC100536359 | 0.22453 | 0.124231677 | 0.056 | 0 | 1 | 15 |
| LOC101882111 | 0.22453 | 0.108184433 | 0.056 | 0 | 1 | 15 |
| LOC101883815 | 0.22453 | 0.104453434 | 0.056 | 0 | 1 | 15 |
| LOC101884686 | 0.22453 | 0.113949855 | 0.056 | 0 | 1 | 15 |

| | | | | | | |
|---|---|---|---|---|---|---|
| LOC101885478 | 0.22453 | 0.104618794 | 0.056 | 0 | 1 | 15 |
| LOC103911361 | 0.22453 | 0.139933837 | 0.056 | 0 | 1 | 15 |
| LOC108179575 | 0.22453 | 0.121363141 | 0.056 | 0 | 1 | 15 |
| LOC108180226 | 0.22453 | 0.115099024 | 0.056 | 0 | 1 | 15 |
| LOC108183500 | 0.22453 | 0.11868087 | 0.056 | 0 | 1 | 15 |
| LOC108190731 | 0.22453 | 0.122636622 | 0.056 | 0 | 1 | 15 |
| RANBP2 | 0.22453 | 0.11551394 | 0.056 | 0 | 1 | 15 |
| RF01268 | 0.22453 | 0.123442078 | 0.056 | 0 | 1 | 15 |
| SBNO2 | 0.22453 | 0.107248602 | 0.056 | 0 | 1 | 15 |
| SHKBP1 | 0.22453 | 0.110131118 | 0.056 | 0 | 1 | 15 |
| UBE2O | 0.22453 | 0.105462166 | 0.056 | 0 | 1 | 15 |
| XLOC-004796 | 0.22453 | 0.133241567 | 0.056 | 0 | 1 | 15 |
| XLOC-021534 | 0.22453 | 0.130434473 | 0.056 | 0 | 1 | 15 |
| XLOC-022032 | 0.22453 | 0.111716322 | 0.056 | 0 | 1 | 15 |
| XLOC-026761 | 0.22453 | 0.100418044 | 0.056 | 0 | 1 | 15 |
| XLOC-042512 | 0.22453 | 0.103115022 | 0.056 | 0 | 1 | 15 |
| abcb7 | 0.22453 | 0.100462534 | 0.056 | 0 | 1 | 15 |
| abhd4 | 0.22453 | 0.100557643 | 0.056 | 0 | 1 | 15 |
| adamtsl5 | 0.22453 | 0.124735118 | 0.056 | 0 | 1 | 15 |
| adgra2 | 0.22453 | 0.122324658 | 0.056 | 0 | 1 | 15 |
| adhfe1 | 0.22453 | 0.114090154 | 0.056 | 0 | 1 | 15 |
| adoa | 0.22453 | 0.100090525 | 0.056 | 0 | 1 | 15 |
| adprh | 0.22453 | 0.109802708 | 0.056 | 0 | 1 | 15 |
| ahi1 | 0.22453 | 0.112626578 | 0.056 | 0 | 1 | 15 |
| akap10 | 0.22453 | 0.122580388 | 0.056 | 0 | 1 | 15 |
| akap11 | 0.22453 | 0.1125981 | 0.056 | 0 | 1 | 15 |
| aldh16a1 | 0.22453 | 0.105057131 | 0.056 | 0 | 1 | 15 |
| alg1 | 0.22453 | 0.104423208 | 0.056 | 0 | 1 | 15 |
| ambra1a | 0.22453 | 0.119182974 | 0.056 | 0 | 1 | 15 |
| ankrd40 | 0.22453 | 0.103424968 | 0.056 | 0 | 1 | 15 |
| ankrd46b | 0.22453 | 0.100153441 | 0.056 | 0 | 1 | 15 |
| anks3 | 0.22453 | 0.149336476 | 0.056 | 0 | 1 | 15 |
| ap3m2 | 0.22453 | 0.110546469 | 0.056 | 0 | 1 | 15 |
| ap4e1 | 0.22453 | 0.104653243 | 0.056 | 0 | 1 | 15 |
| arhgef25b | 0.22453 | 0.138556549 | 0.056 | 0 | 1 | 15 |
| arl6ip5a | 0.22453 | 0.103746812 | 0.056 | 0 | 1 | 15 |
| arv1 | 0.22453 | 0.101781255 | 0.056 | 0 | 1 | 15 |
| atg7 | 0.22453 | 0.106267073 | 0.056 | 0 | 1 | 15 |
| atp6v1d | 0.22453 | 0.126154929 | 0.056 | 0 | 1 | 15 |
| auts2b | 0.22453 | 0.102005892 | 0.056 | 0 | 1 | 15 |
| bach2b | 0.22453 | 0.109212454 | 0.056 | 0 | 1 | 15 |
| baxa | 0.22453 | 0.10058427 | 0.056 | 0 | 1 | 15 |
| bbs5 | 0.22453 | 0.113189433 | 0.056 | 0 | 1 | 15 |
| bicc1b | 0.22453 | 0.106027923 | 0.056 | 0 | 1 | 15 |
| borcs8 | 0.22453 | 0.130022348 | 0.056 | 0 | 1 | 15 |
| bpgm | 0.22453 | 0.112996947 | 0.056 | 0 | 1 | 15 |
| ca8 | 0.22453 | 0.106242452 | 0.056 | 0 | 1 | 15 |
| cars | 0.22453 | 0.107205431 | 0.056 | 0 | 1 | 15 |

| | | | | | | |
|---|---|---|---|---|---|---|
| ccar2 | 0.22453 | 0.125697552 | 0.056 | 0 | 1 | 15 |
| ccdc3a | 0.22453 | 0.164441714 | 0.056 | 0 | 1 | 15 |
| ccl19a.1 | 0.22453 | 0.175714339 | 0.056 | 0 | 1 | 15 |
| cdr2l | 0.22453 | 0.120439131 | 0.056 | 0 | 1 | 15 |
| cebpz | 0.22453 | 0.128765758 | 0.056 | 0 | 1 | 15 |
| cep19 | 0.22453 | 0.10794079 | 0.056 | 0 | 1 | 15 |
| chrdl2 | 0.22453 | 0.116419173 | 0.056 | 0 | 1 | 15 |
| chst11 | 0.22453 | 0.147304421 | 0.056 | 0 | 1 | 15 |
| clint1b | 0.22453 | 0.105177361 | 0.056 | 0 | 1 | 15 |
| col4a2 | 0.22453 | 0.110361482 | 0.056 | 0 | 1 | 15 |
| coro1cb | 0.22453 | 0.105543559 | 0.056 | 0 | 1 | 15 |
| cplane2 | 0.22453 | 0.105062095 | 0.056 | 0 | 1 | 15 |
| ctdp1 | 0.22453 | 0.10072058 | 0.056 | 0 | 1 | 15 |
| ctso | 0.22453 | 0.121084784 | 0.056 | 0 | 1 | 15 |
| cttnbp2nlb | 0.22453 | 0.11386757 | 0.056 | 0 | 1 | 15 |
| cuedc1b | 0.22453 | 0.105063106 | 0.056 | 0 | 1 | 15 |
| cuedc2 | 0.22453 | 0.105725918 | 0.056 | 0 | 1 | 15 |
| cyp8b3 | 0.22453 | 0.134777662 | 0.056 | 0 | 1 | 15 |
| dact1 | 0.22453 | 0.10141586 | 0.056 | 0 | 1 | 15 |
| dapk1 | 0.22453 | 0.105722833 | 0.056 | 0 | 1 | 15 |
| dctd | 0.22453 | 0.104679946 | 0.056 | 0 | 1 | 15 |
| dctpp1 | 0.22453 | 0.140024845 | 0.056 | 0 | 1 | 15 |
| ddx31 | 0.22453 | 0.106554165 | 0.056 | 0 | 1 | 15 |
| dph7 | 0.22453 | 0.102775883 | 0.056 | 0 | 1 | 15 |
| dusp16 | 0.22453 | 0.133912614 | 0.056 | 0 | 1 | 15 |
| dusp26 | 0.22453 | 0.102771608 | 0.056 | 0 | 1 | 15 |
| eif2d | 0.22453 | 0.10672526 | 0.056 | 0 | 1 | 15 |
| elac1 | 0.22453 | 0.10427271 | 0.056 | 0 | 1 | 15 |
| ell | 0.22453 | 0.110908846 | 0.056 | 0 | 1 | 15 |
| elp2 | 0.22453 | 0.109050314 | 0.056 | 0 | 1 | 15 |
| enosf1 | 0.22453 | 0.110274651 | 0.056 | 0 | 1 | 15 |
| epas1b | 0.22453 | 0.100283238 | 0.056 | 0 | 1 | 15 |
| epn3a | 0.22453 | 0.111099342 | 0.056 | 0 | 1 | 15 |
| exoc6b | 0.22453 | 0.109681019 | 0.056 | 0 | 1 | 15 |
| f7 | 0.22453 | 0.118828754 | 0.056 | 0 | 1 | 15 |
| fam160b1 | 0.22453 | 0.102377055 | 0.056 | 0 | 1 | 15 |
| fam49bb | 0.22453 | 0.118011689 | 0.056 | 0 | 1 | 15 |
| fbxo3 | 0.22453 | 0.104148154 | 0.056 | 0 | 1 | 15 |
| fbxo33 | 0.22453 | 0.122426831 | 0.056 | 0 | 1 | 15 |
| fkbp10b | 0.22453 | 0.101391303 | 0.056 | 0 | 1 | 15 |
| fsta | 0.22453 | 0.114013019 | 0.056 | 0 | 1 | 15 |
| gapvd1 | 0.22453 | 0.102191935 | 0.056 | 0 | 1 | 15 |
| gbe1b | 0.22453 | 0.115027467 | 0.056 | 0 | 1 | 15 |
| ggh | 0.22453 | 0.160106804 | 0.056 | 0 | 1 | 15 |
| gkup | 0.22453 | 0.122179173 | 0.056 | 0 | 1 | 15 |
| gpr107 | 0.22453 | 0.110531372 | 0.056 | 0 | 1 | 15 |
| grpel1 | 0.22453 | 0.107877642 | 0.056 | 0 | 1 | 15 |
| hbegfa | 0.22453 | 0.117398226 | 0.056 | 0 | 1 | 15 |

| | | | | | | |
|---|---|---|---|---|---|---|
| hdac3 | 0.22453 | 0.12307739 | 0.056 | 0 | 1 | 15 |
| hdac8 | 0.22453 | 0.125832802 | 0.056 | 0 | 1 | 15 |
| heatr5a | 0.22453 | 0.112487004 | 0.056 | 0 | 1 | 15 |
| hspbap1 | 0.22453 | 0.10172225 | 0.056 | 0 | 1 | 15 |
| il11a | 0.22453 | 0.124450619 | 0.056 | 0 | 1 | 15 |
| impdh2 | 0.22453 | 0.112441572 | 0.056 | 0 | 1 | 15 |
| inpp5l | 0.22453 | 0.162860309 | 0.056 | 0 | 1 | 15 |
| itsn2b | 0.22453 | 0.149641155 | 0.056 | 0 | 1 | 15 |
| jazf1a | 0.22453 | 0.102034778 | 0.056 | 0 | 1 | 15 |
| kat8 | 0.22453 | 0.10091173 | 0.056 | 0 | 1 | 15 |
| kcmf1 | 0.22453 | 0.109072401 | 0.056 | 0 | 1 | 15 |
| kcnh7 | 0.22453 | 0.128178668 | 0.056 | 0 | 1 | 15 |
| kcnn1a | 0.22453 | 0.129434125 | 0.056 | 0 | 1 | 15 |
| kif3a | 0.22453 | 0.119140139 | 0.056 | 0 | 1 | 15 |
| kitlga | 0.22453 | 0.121606474 | 0.056 | 0 | 1 | 15 |
| lamb2l | 0.22453 | 0.140672541 | 0.056 | 0 | 1 | 15 |
| larp4b | 0.22453 | 0.108937584 | 0.056 | 0 | 1 | 15 |
| lmo4a | 0.22453 | 0.111414915 | 0.056 | 0 | 1 | 15 |
| lpcat4 | 0.22453 | 0.1041948 | 0.056 | 0 | 1 | 15 |
| madd | 0.22453 | 0.113328003 | 0.056 | 0 | 1 | 15 |
| map6a | 0.22453 | 0.114720896 | 0.056 | 0 | 1 | 15 |
| mat2ab | 0.22453 | 0.138183469 | 0.056 | 0 | 1 | 15 |
| minpp1a | 0.22453 | 0.127436857 | 0.056 | 0 | 1 | 15 |
| mms19 | 0.22453 | 0.106245908 | 0.056 | 0 | 1 | 15 |
| mon2 | 0.22453 | 0.10796607 | 0.056 | 0 | 1 | 15 |
| mre11a | 0.22453 | 0.112529736 | 0.056 | 0 | 1 | 15 |
| mrtfaa | 0.22453 | 0.113912985 | 0.056 | 0 | 1 | 15 |
| msmo1 | 0.22453 | 0.104967355 | 0.056 | 0 | 1 | 15 |
| mtpap | 0.22453 | 0.112921844 | 0.056 | 0 | 1 | 15 |
| mynn | 0.22453 | 0.111404744 | 0.056 | 0 | 1 | 15 |
| naa35 | 0.22453 | 0.104004394 | 0.056 | 0 | 1 | 15 |
| nbas | 0.22453 | 0.122430174 | 0.056 | 0 | 1 | 15 |
| nbr1a | 0.22453 | 0.111841771 | 0.056 | 0 | 1 | 15 |
| nemp2 | 0.22453 | 0.119297585 | 0.056 | 0 | 1 | 15 |
| nol12 | 0.22453 | 0.100049681 | 0.056 | 0 | 1 | 15 |
| nol9 | 0.22453 | 0.112186214 | 0.056 | 0 | 1 | 15 |
| odr4 | 0.22453 | 0.107801716 | 0.056 | 0 | 1 | 15 |
| osbpl10b | 0.22453 | 0.102296521 | 0.056 | 0 | 1 | 15 |
| p2rx4b | 0.22453 | 0.108491327 | 0.056 | 0 | 1 | 15 |
| patz1 | 0.22453 | 0.103289874 | 0.056 | 0 | 1 | 15 |
| pcolcea | 0.22453 | 0.113718937 | 0.056 | 0 | 1 | 15 |
| pdxdc1 | 0.22453 | 0.153047254 | 0.056 | 0 | 1 | 15 |
| phlda1 | 0.22453 | 0.108070208 | 0.056 | 0 | 1 | 15 |
| pigo | 0.22453 | 0.10590332 | 0.056 | 0 | 1 | 15 |
| pigu | 0.22453 | 0.115419934 | 0.056 | 0 | 1 | 15 |
| plxnb3 | 0.22453 | 0.117631998 | 0.056 | 0 | 1 | 15 |
| polr2h | 0.22453 | 0.115458055 | 0.056 | 0 | 1 | 15 |
| pomgnt1 | 0.22453 | 0.121084784 | 0.056 | 0 | 1 | 15 |

| | | | | | | |
|---|---|---|---|---|---|---|
| ppfia1 | 0.22453 | 0.103193925 | 0.056 | 0 | 1 | 15 |
| prepl | 0.22453 | 0.116771411 | 0.056 | 0 | 1 | 15 |
| psme4b | 0.22453 | 0.109652807 | 0.056 | 0 | 1 | 15 |
| ptcd1 | 0.22453 | 0.105302169 | 0.056 | 0 | 1 | 15 |
| ptgr2 | 0.22453 | 0.12453395 | 0.056 | 0 | 1 | 15 |
| qtrt1 | 0.22453 | 0.107613027 | 0.056 | 0 | 1 | 15 |
| rasa1b | 0.22453 | 0.102122295 | 0.056 | 0 | 1 | 15 |
| rbm18 | 0.22453 | 0.104160796 | 0.056 | 0 | 1 | 15 |
| rel | 0.22453 | 0.110406387 | 0.056 | 0 | 1 | 15 |
| retreg2 | 0.22453 | 0.14779723 | 0.056 | 0 | 1 | 15 |
| rnf103 | 0.22453 | 0.101113585 | 0.056 | 0 | 1 | 15 |
| rnf150a | 0.22453 | 0.110787356 | 0.056 | 0 | 1 | 15 |
| rnf181 | 0.22453 | 0.103594563 | 0.056 | 0 | 1 | 15 |
| rorab | 0.22453 | 0.112384611 | 0.056 | 0 | 1 | 15 |
| rpp25l | 0.22453 | 0.125009472 | 0.056 | 0 | 1 | 15 |
| rrp36 | 0.22453 | 0.103183419 | 0.056 | 0 | 1 | 15 |
| sbno1 | 0.22453 | 0.114117752 | 0.056 | 0 | 1 | 15 |
| scly | 0.22453 | 0.124814921 | 0.056 | 0 | 1 | 15 |
| si:busm1-57f23.1 | 0.22453 | 0.129917492 | 0.056 | 0 | 1 | 15 |
| si:ch211-110e21.4 | 0.22453 | 0.104618794 | 0.056 | 0 | 1 | 15 |
| si:ch211-113e8.11 | 0.22453 | 0.101075163 | 0.056 | 0 | 1 | 15 |
| si:ch211-203b8.6 | 0.22453 | 0.121507859 | 0.056 | 0 | 1 | 15 |
| si:ch211-209l18.2 | 0.22453 | 0.1293593 | 0.056 | 0 | 1 | 15 |
| si:ch211-266d19.4 | 0.22453 | 0.103071918 | 0.056 | 0 | 1 | 15 |
| si:ch211-281g13.5 | 0.22453 | 0.134223819 | 0.056 | 0 | 1 | 15 |
| si:ch211-40k21.9 | 0.22453 | 0.101432545 | 0.056 | 0 | 1 | 15 |
| si:ch73-144d13.5 | 0.22453 | 0.122430174 | 0.056 | 0 | 1 | 15 |
| si:ch73-181d5.4 | 0.22453 | 0.12431701 | 0.056 | 0 | 1 | 15 |
| si:ch73-191k20.3 | 0.22453 | 0.101128829 | 0.056 | 0 | 1 | 15 |
| si:ch73-193c12.2 | 0.22453 | 0.12055832 | 0.056 | 0 | 1 | 15 |
| si:ch73-199g24.2 | 0.22453 | 0.104260737 | 0.056 | 0 | 1 | 15 |
| si:ch73-252i11.1 | 0.22453 | 0.109625785 | 0.056 | 0 | 1 | 15 |
| si:ch73-278m9.1 | 0.22453 | 0.105063106 | 0.056 | 0 | 1 | 15 |
| si:ch73-368j24.8 | 0.22453 | 0.107634944 | 0.056 | 0 | 1 | 15 |
| si:ch73-390p7.2 | 0.22453 | 0.102957922 | 0.056 | 0 | 1 | 15 |
| si:dkey-172k15.11 | 0.22453 | 0.102445215 | 0.056 | 0 | 1 | 15 |
| si:dkey-182g1.10 | 0.22453 | 0.121637344 | 0.056 | 0 | 1 | 15 |
| si:dkey-182i3.10 | 0.22453 | 0.103753191 | 0.056 | 0 | 1 | 15 |
| si:dkey-23a13.5 | 0.22453 | 0.142892102 | 0.056 | 0 | 1 | 15 |
| si:dkey-247i3.6 | 0.22453 | 0.105725918 | 0.056 | 0 | 1 | 15 |
| si:dkey-29h14.10 | 0.22453 | 0.101316455 | 0.056 | 0 | 1 | 15 |
| si:dkey-85k7.7 | 0.22453 | 0.103341436 | 0.056 | 0 | 1 | 15 |
| slc26a11 | 0.22453 | 0.115890523 | 0.056 | 0 | 1 | 15 |
| slc2a11a | 0.22453 | 0.108521651 | 0.056 | 0 | 1 | 15 |
| slc31a2 | 0.22453 | 0.142401874 | 0.056 | 0 | 1 | 15 |
| slc35e3 | 0.22453 | 0.10427003 | 0.056 | 0 | 1 | 15 |
| slc37a4b | 0.22453 | 0.119486159 | 0.056 | 0 | 1 | 15 |
| smco4 | 0.22453 | 0.122057349 | 0.056 | 0 | 1 | 15 |

| | | | | | | |
|---|---|---|---|---|---|---|
| snupn | 0.22453 | 0.10156111 | 0.056 | 0 | 1 | 15 |
| snx14 | 0.22453 | 0.12933414 | 0.056 | 0 | 1 | 15 |
| sorcs2 | 0.22453 | 0.100322197 | 0.056 | 0 | 1 | 15 |
| sos2 | 0.22453 | 0.13313842 | 0.056 | 0 | 1 | 15 |
| spry4 | 0.22453 | 0.111387434 | 0.056 | 0 | 1 | 15 |
| spsb3a | 0.22453 | 0.100418044 | 0.056 | 0 | 1 | 15 |
| srp68 | 0.22453 | 0.113988102 | 0.056 | 0 | 1 | 15 |
| srsf6a | 0.22453 | 0.123994252 | 0.056 | 0 | 1 | 15 |
| st3gal3a | 0.22453 | 0.132597613 | 0.056 | 0 | 1 | 15 |
| syt2a | 0.22453 | 0.105991737 | 0.056 | 0 | 1 | 15 |
| tbk1 | 0.22453 | 0.100261932 | 0.056 | 0 | 1 | 15 |
| tdrkh | 0.22453 | 0.105004952 | 0.056 | 0 | 1 | 15 |
| tfip11 | 0.22453 | 0.132474703 | 0.056 | 0 | 1 | 15 |
| thbs1b | 0.22453 | 0.103418081 | 0.056 | 0 | 1 | 15 |
| tlr7 | 0.22453 | 0.104141429 | 0.056 | 0 | 1 | 15 |
| tmem131l | 0.22453 | 0.101026596 | 0.056 | 0 | 1 | 15 |
| tmem135 | 0.22453 | 0.124576327 | 0.056 | 0 | 1 | 15 |
| tmem260 | 0.22453 | 0.113810318 | 0.056 | 0 | 1 | 15 |
| tp53bp1 | 0.22453 | 0.124760801 | 0.056 | 0 | 1 | 15 |
| tpcn3 | 0.22453 | 0.161620804 | 0.056 | 0 | 1 | 15 |
| tpst1l | 0.22453 | 0.116641538 | 0.056 | 0 | 1 | 15 |
| traf3ip1 | 0.22453 | 0.130090388 | 0.056 | 0 | 1 | 15 |
| trappc9 | 0.22453 | 0.129434125 | 0.056 | 0 | 1 | 15 |
| trib2 | 0.22453 | 0.116371038 | 0.056 | 0 | 1 | 15 |
| trim35-1 | 0.22453 | 0.128013662 | 0.056 | 0 | 1 | 15 |
| tsc1a | 0.22453 | 0.106369757 | 0.056 | 0 | 1 | 15 |
| ttc8 | 0.22453 | 0.108869381 | 0.056 | 0 | 1 | 15 |
| ttl | 0.22453 | 0.100084951 | 0.056 | 0 | 1 | 15 |
| ube4b | 0.22453 | 0.110048643 | 0.056 | 0 | 1 | 15 |
| ufsp1 | 0.22453 | 0.116479193 | 0.056 | 0 | 1 | 15 |
| uggt2 | 0.22453 | 0.100803809 | 0.056 | 0 | 1 | 15 |
| unga | 0.22453 | 0.107330899 | 0.056 | 0 | 1 | 15 |
| vegfaa | 0.22453 | 0.125458095 | 0.056 | 0 | 1 | 15 |
| vps16 | 0.22453 | 0.104410999 | 0.056 | 0 | 1 | 15 |
| vps33b | 0.22453 | 0.104028894 | 0.056 | 0 | 1 | 15 |
| washc5 | 0.22453 | 0.103099188 | 0.056 | 0 | 1 | 15 |
| wrnip1 | 0.22453 | 0.139059774 | 0.056 | 0 | 1 | 15 |
| xpc | 0.22453 | 0.121974268 | 0.056 | 0 | 1 | 15 |
| zbtb37 | 0.22453 | 0.106880651 | 0.056 | 0 | 1 | 15 |
| zbtb43 | 0.22453 | 0.111496483 | 0.056 | 0 | 1 | 15 |
| zdhhc16b | 0.22453 | 0.11661533 | 0.056 | 0 | 1 | 15 |
| zgc:113102 | 0.22453 | 0.100239967 | 0.056 | 0 | 1 | 15 |
| zgc:113518 | 0.22453 | 0.108655509 | 0.056 | 0 | 1 | 15 |
| zgc:113921 | 0.22453 | 0.126260541 | 0.056 | 0 | 1 | 15 |
| zgc:136308 | 0.22453 | 0.105725918 | 0.056 | 0 | 1 | 15 |
| zgc:152830 | 0.22453 | 0.140172747 | 0.056 | 0 | 1 | 15 |
| zgc:153031 | 0.22453 | 0.106649975 | 0.056 | 0 | 1 | 15 |
| zgc:153215 | 0.22453 | 0.109221139 | 0.056 | 0 | 1 | 15 |

| | | | | | | |
|---|---|---|---|---|---|---|
| zgc:162898 | 0.22453 | 0.102309955 | 0.056 | 0 | 1 | 15 |
| zgc:174702 | 0.22453 | 0.103115022 | 0.056 | 0 | 1 | 15 |
| zgc:92313 | 0.22453 | 0.110349392 | 0.056 | 0 | 1 | 15 |
| znhit3 | 0.22453 | 0.107097664 | 0.056 | 0 | 1 | 15 |
| acp6 | 0.22453 | 0.147692856 | 0.056 | 0 | 1 | 15 |
| cnot4b | 0.22453 | 0.115662641 | 0.056 | 0 | 1 | 15 |
| cnot7 | 0.22453 | 0.130091051 | 0.056 | 0 | 1 | 15 |
| eps15l1a | 0.22453 | 0.102071589 | 0.056 | 0 | 1 | 15 |
| frmd8 | 0.22453 | 0.111889379 | 0.056 | 0 | 1 | 15 |
| gclc | 0.22453 | 0.10523556 | 0.056 | 0 | 1 | 15 |
| gpbp1l1 | 0.22453 | 0.102043067 | 0.056 | 0 | 1 | 15 |
| map4k3b | 0.22453 | 0.108281517 | 0.056 | 0 | 1 | 15 |
| mical3a | 0.22453 | 0.114349215 | 0.056 | 0 | 1 | 15 |
| phospho2 | 0.22453 | 0.145933812 | 0.056 | 0 | 1 | 15 |
| ppp1r3cb | 0.22453 | 0.109158447 | 0.056 | 0 | 1 | 15 |
| rgmb | 0.22453 | 0.108281517 | 0.056 | 0 | 1 | 15 |
| rock1 | 0.22453 | 0.130040617 | 0.056 | 0 | 1 | 15 |
| rrp9 | 0.22453 | 0.113154217 | 0.056 | 0 | 1 | 15 |
| shdb | 0.22453 | 0.102048618 | 0.056 | 0 | 1 | 15 |
| si:ch73-281f12.4 | 0.22453 | 0.110960636 | 0.056 | 0 | 1 | 15 |
| si:dkeyp-33b5.4 | 0.22453 | 0.111646874 | 0.056 | 0 | 1 | 15 |
| slc8a1b | 0.22453 | 0.142944805 | 0.056 | 0 | 1 | 15 |
| stx10 | 0.22453 | 0.11216494 | 0.056 | 0 | 1 | 15 |
| thap1 | 0.22453 | 0.10674946 | 0.056 | 0 | 1 | 15 |
| tmcc3 | 0.22453 | 0.115830012 | 0.056 | 0 | 1 | 15 |
| tsc1b | 0.22453 | 0.10488926 | 0.056 | 0 | 1 | 15 |
| ttll7 | 0.22453 | 0.111488067 | 0.056 | 0 | 1 | 15 |
| ulk1a | 0.22453 | 0.108045576 | 0.056 | 0 | 1 | 15 |
| unk | 0.22453 | 0.124867973 | 0.056 | 0 | 1 | 15 |
| wdr73 | 0.22453 | 0.1008654 | 0.056 | 0 | 1 | 15 |
| wipi2 | 0.22453 | 0.113470595 | 0.056 | 0 | 1 | 15 |
| xpo6 | 0.22453 | 0.108045576 | 0.056 | 0 | 1 | 15 |
| yif1b | 0.22453 | 0.119647503 | 0.056 | 0 | 1 | 15 |
| CABZ01067348.1 | 0.22453 | 0.115596147 | 0.056 | 0 | 1 | 15 |
| CR751602.1 | 0.22453 | 0.105287255 | 0.056 | 0 | 1 | 15 |
| CT737127.1 | 0.22453 | 0.111655113 | 0.056 | 0 | 1 | 15 |
| ERC1 | 0.22453 | 0.115596147 | 0.056 | 0 | 1 | 15 |
| apol1 | 0.22453 | 0.134213301 | 0.056 | 0 | 1 | 15 |
| borcs5 | 0.22453 | 0.130020679 | 0.056 | 0 | 1 | 15 |
| btbd7 | 0.22453 | 0.141825678 | 0.056 | 0 | 1 | 15 |
| ccndbp1 | 0.22453 | 0.12165835 | 0.056 | 0 | 1 | 15 |
| chsy1 | 0.22453 | 0.120178214 | 0.056 | 0 | 1 | 15 |
| dachb | 0.22453 | 0.104557521 | 0.056 | 0 | 1 | 15 |
| dgcr2 | 0.22453 | 0.148859663 | 0.056 | 0 | 1 | 15 |
| enc2 | 0.22453 | 0.113359988 | 0.056 | 0 | 1 | 15 |
| fastkd5 | 0.22453 | 0.124206881 | 0.056 | 0 | 1 | 15 |
| gb:ai877918 | 0.22453 | 0.102675565 | 0.056 | 0 | 1 | 15 |
| jmjd7 | 0.22453 | 0.104118182 | 0.056 | 0 | 1 | 15 |

| | | | | | | |
|---|---|---|---|---|---|---|
| nrip1b | 0.22453 | 0.129619587 | 0.056 | 0 | 1 | 15 |
| nudt3b | 0.22453 | 0.110500731 | 0.056 | 0 | 1 | 15 |
| pla2g7 | 0.22453 | 0.111781834 | 0.056 | 0 | 1 | 15 |
| pomt2 | 0.22453 | 0.133000005 | 0.056 | 0 | 1 | 15 |
| prdm4 | 0.22453 | 0.125075577 | 0.056 | 0 | 1 | 15 |
| rassf1 | 0.22453 | 0.126742052 | 0.056 | 0 | 1 | 15 |
| rbm27 | 0.22453 | 0.113907553 | 0.056 | 0 | 1 | 15 |
| ripply1 | 0.22453 | 0.136883239 | 0.056 | 0 | 1 | 15 |
| sfxn4 | 0.22453 | 0.163390094 | 0.056 | 0 | 1 | 15 |
| si:ch211-201o1.1 | 0.22453 | 0.142838533 | 0.056 | 0 | 1 | 15 |
| si:ch211-236l14.4 | 0.22453 | 0.102675565 | 0.056 | 0 | 1 | 15 |
| slc44a1a | 0.22453 | 0.102675565 | 0.056 | 0 | 1 | 15 |
| suox | 0.22453 | 0.124302263 | 0.056 | 0 | 1 | 15 |
| tcaim | 0.22453 | 0.116131135 | 0.056 | 0 | 1 | 15 |
| tsta3 | 0.22453 | 0.107055658 | 0.056 | 0 | 1 | 15 |
| zgc:171727 | 0.22453 | 0.111947968 | 0.056 | 0 | 1 | 15 |
| znf1164 | 0.22453 | 0.123883965 | 0.056 | 0 | 1 | 15 |
| znf207a | 0.22453 | 0.102554583 | 0.056 | 0 | 1 | 15 |
| XLOC-013197 | 0.22453 | 0.101527662 | 0.056 | 0 | 1 | 15 |
| elmod2 | 0.22453 | 0.131687028 | 0.056 | 0 | 1 | 15 |
| igdcc4 | 0.22453 | 0.10762285 | 0.056 | 0 | 1 | 15 |
| nedd9 | 0.22453 | 0.103411115 | 0.056 | 0 | 1 | 15 |
| zgc:77058 | 0.22453 | 0.105604724 | 0.056 | 0 | 1 | 15 |
| LOC110437744.1 | 0.22453 | 0.107714995 | 0.056 | 0 | 1 | 15 |
| XLOC-016198 | 0.22453 | 0.155152427 | 0.056 | 0 | 1 | 15 |
| fam206a | 0.22453 | 0.108780034 | 0.056 | 0 | 1 | 15 |
| mark1 | 0.22453 | 0.119951441 | 0.056 | 0 | 1 | 15 |
| ptpn3 | 0.22453 | 0.101297128 | 0.056 | 0 | 1 | 15 |
| taf1 | 0.22453 | 0.109440094 | 0.056 | 0 | 1 | 15 |
| utp4 | 0.22453 | 0.105671101 | 0.056 | 0 | 1 | 15 |
| XLOC-041269 | 0.22453 | 0.107386301 | 0.056 | 0 | 1 | 15 |
| dus4l | 0.22453 | 0.106694287 | 0.056 | 0 | 1 | 15 |
| cipca | 0.22453 | 0.117947421 | 0.056 | 0 | 1 | 15 |
| acsbg2 | 0.22453 | 0.104002597 | 0.056 | 0 | 1 | 15 |
| hipk3b | 0.22453 | 0.151539012 | 0.056 | 0 | 1 | 15 |
| hnrnph1 | 0.22453 | 0.107806549 | 0.056 | 0 | 1 | 15 |
| mark3b | 0.22453 | 0.105294926 | 0.056 | 0 | 1 | 15 |
| pde8a | 0.22453 | 0.119377825 | 0.056 | 0 | 1 | 15 |
| plppr5a | 0.22453 | 0.11546493 | 0.056 | 0 | 1 | 15 |
| polr1c | 0.22453 | 0.103096529 | 0.056 | 0 | 1 | 15 |
| slc22a15 | 0.22453 | 0.103235223 | 0.056 | 0 | 1 | 15 |
| slc30a5 | 0.22453 | 0.124293387 | 0.056 | 0 | 1 | 15 |
| tmem178 | 0.22453 | 0.105927327 | 0.056 | 0 | 1 | 15 |
| urb1 | 0.22453 | 0.119449432 | 0.056 | 0 | 1 | 15 |
| alyref | 0.22527 | 0.115340655 | 0.278 | 0.261 | 1 | 15 |
| nfe2l2a | 0.2258 | -0.18828546 | 0.083 | 0.13 | 1 | 15 |
| tegt | 0.2264 | 0.138745149 | 0.833 | 0.652 | 1 | 15 |
| mapk1 | 0.22664 | 0.178737654 | 0.167 | 0.043 | 1 | 15 |

| | | | | | | |
|---|---|---|---|---|---|---|
| agrn | 0.22669 | -0.21122848 | 0.389 | 0.435 | 1 | 15 |
| gde1 | 0.22708 | -0.24021103 | 0.028 | 0.13 | 1 | 15 |
| csgalnact1a | 0.22745 | -0.32467926 | 0.139 | 0.304 | 1 | 15 |
| NC-002333.17 | 0.22779 | -0.20854712 | 0.083 | 0.13 | 1 | 15 |
| atg3 | 0.22783 | 0.226630707 | 0.167 | 0.043 | 1 | 15 |
| amh | 0.22829 | -0.14176537 | 0.111 | 0.13 | 1 | 15 |
| tmem203 | 0.22843 | -0.15552081 | 0.056 | 0.174 | 1 | 15 |
| ssuh2.2 | 0.22844 | -0.17111711 | 0.028 | 0.13 | 1 | 15 |
| sap30bp | 0.22915 | -0.32299527 | 0.111 | 0.261 | 1 | 15 |
| cpn1 | 0.22937 | 0.264122655 | 0.778 | 0.565 | 1 | 15 |
| gstm.1 | 0.22997 | -0.23296309 | 0.75 | 0.913 | 1 | 15 |
| cct4 | 0.23157 | 0.285749067 | 0.639 | 0.478 | 1 | 15 |
| calm2b | 0.23162 | -0.30259157 | 0.528 | 0.739 | 1 | 15 |
| mrpl33 | 0.23192 | -0.15339295 | 0.222 | 0.261 | 1 | 15 |
| tmbim4 | 0.23208 | 0.20081453 | 0.139 | 0.043 | 1 | 15 |
| timp2a | 0.2322 | -0.11806093 | 0.306 | 0.304 | 1 | 15 |
| zfp36l1b | 0.23221 | -0.2953011 | 0.528 | 0.739 | 1 | 15 |
| prelid1a | 0.2331 | -0.27660858 | 0.139 | 0.304 | 1 | 15 |
| flna | 0.23315 | -0.17483508 | 0.194 | 0.217 | 1 | 15 |
| tnfaip6 | 0.23338 | 0.295753905 | 0.472 | 0.261 | 1 | 15 |
| tuba1b | 0.23378 | -0.25136368 | 0.028 | 0.13 | 1 | 15 |
| rbm5 | 0.23394 | 0.266175611 | 0.25 | 0.087 | 1 | 15 |
| ttll12 | 0.2344 | 0.351798179 | 0.194 | 0.087 | 1 | 15 |
| birc6 | 0.23461 | -0.13930338 | 0.028 | 0.13 | 1 | 15 |
| ccdc187 | 0.23482 | 0.290609544 | 0.472 | 0.261 | 1 | 15 |
| mir21-1 | 0.23492 | -0.34703522 | 0.222 | 0.391 | 1 | 15 |
| hsd3b1 | 0.23535 | 0.152535221 | 0.194 | 0.087 | 1 | 15 |
| si:ch211-155e24.3 | 0.23568 | -0.25371297 | 0.028 | 0.13 | 1 | 15 |
| sri | 0.23578 | -0.32515787 | 0.25 | 0.435 | 1 | 15 |
| si:ch211-198c19.3 | 0.23597 | -0.32616549 | 0.028 | 0.13 | 1 | 15 |
| cux1a | 0.23601 | -0.16064059 | 0.306 | 0.348 | 1 | 15 |
| tmed7 | 0.23653 | -0.14034903 | 0.583 | 0.609 | 1 | 15 |
| si:ch211-264f5.6 | 0.23661 | 0.100962986 | 0.194 | 0.087 | 1 | 15 |
| si:ch73-376l24.3 | 0.23667 | 0.242561058 | 0.556 | 0.348 | 1 | 15 |
| impad1 | 0.23688 | -0.25960202 | 0.028 | 0.13 | 1 | 15 |
| ier5l | 0.23697 | 0.146970947 | 0.111 | 0.087 | 1 | 15 |
| eif4ebp2 | 0.23699 | 0.214204068 | 0.5 | 0.304 | 1 | 15 |
| creg2 | 0.23748 | -0.2265608 | 0.056 | 0.13 | 1 | 15 |
| sephs1 | 0.23759 | 0.250967654 | 0.222 | 0.087 | 1 | 15 |
| rrp1 | 0.23788 | 0.236178576 | 0.25 | 0.087 | 1 | 15 |
| gnb1b | 0.23811 | -0.20510841 | 0.611 | 0.696 | 1 | 15 |
| slc25a3b | 0.23812 | 0.246942916 | 0.861 | 0.696 | 1 | 15 |
| gnai2a | 0.23907 | -0.23954797 | 0.028 | 0.13 | 1 | 15 |
| crk | 0.24106 | 0.240456264 | 0.25 | 0.087 | 1 | 15 |
| huwe1 | 0.24108 | -0.30120266 | 0.083 | 0.217 | 1 | 15 |
| skp1 | 0.24183 | 0.278075705 | 0.694 | 0.478 | 1 | 15 |
| rpl4 | 0.24186 | 0.202629266 | 0.944 | 0.826 | 1 | 15 |
| zgc:92664 | 0.24217 | 0.20964325 | 0.167 | 0.043 | 1 | 15 |

| | | | | | | |
|---|---|---|---|---|---|---|
| ptgr1 | 0.2423 | 0.255363506 | 0.306 | 0.13 | 1 | 15 |
| ntmt1 | 0.24268 | -0.26536377 | 0.056 | 0.174 | 1 | 15 |
| tyw3 | 0.2428 | 0.129597412 | 0.139 | 0.043 | 1 | 15 |
| tspan4a | 0.24303 | -0.21727705 | 0.111 | 0.174 | 1 | 15 |
| id3 | 0.24342 | -0.27008833 | 0.056 | 0.174 | 1 | 15 |
| ctsf | 0.24387 | 0.307793398 | 0.306 | 0.13 | 1 | 15 |
| timm23a | 0.24408 | 0.298046775 | 0.278 | 0.13 | 1 | 15 |
| metrn | 0.24451 | -0.3323834 | 0.083 | 0.217 | 1 | 15 |
| msra | 0.24462 | 0.25912842 | 0.278 | 0.13 | 1 | 15 |
| denr | 0.24469 | 0.299238821 | 0.444 | 0.261 | 1 | 15 |
| ndufa4 | 0.24502 | -0.26171015 | 0.028 | 0.13 | 1 | 15 |
| si:ch211-276i12.11 | 0.24518 | 0.204120287 | 0.222 | 0.13 | 1 | 15 |
| nudc | 0.24561 | -0.33551072 | 0.194 | 0.348 | 1 | 15 |
| pno1 | 0.24576 | -0.30325538 | 0.25 | 0.391 | 1 | 15 |
| si:dkey-188i13.9 | 0.24581 | 0.307694433 | 0.361 | 0.217 | 1 | 15 |
| si:dkey-22o12.2 | 0.24629 | 0.153468326 | 0.306 | 0.174 | 1 | 15 |
| kdelr2b | 0.24633 | -0.14371308 | 0.278 | 0.304 | 1 | 15 |
| ncl | 0.24792 | 0.312706514 | 0.639 | 0.478 | 1 | 15 |
| cst3 | 0.24854 | -0.22388716 | 0.028 | 0.13 | 1 | 15 |
| CU929259.1 | 0.24864 | 0.217515697 | 0.167 | 0.043 | 1 | 15 |
| phf6 | 0.24873 | -0.21815635 | 0.028 | 0.13 | 1 | 15 |
| polr2d | 0.24887 | 0.301555959 | 0.444 | 0.261 | 1 | 15 |
| zgc:63882 | 0.24897 | 0.251306557 | 0.167 | 0.043 | 1 | 15 |
| rack1 | 0.24923 | 0.169017579 | 0.944 | 0.87 | 1 | 15 |
| col6a1 | 0.24932 | -0.24785563 | 0.056 | 0.174 | 1 | 15 |
| lrp12 | 0.24947 | -0.24674411 | 0.028 | 0.13 | 1 | 15 |
| si:ch211-260e23.9 | 0.24969 | -0.26019357 | 0.056 | 0.174 | 1 | 15 |
| raraa | 0.24982 | -0.24941817 | 0.028 | 0.13 | 1 | 15 |
| abhd16a | 0.25012 | -0.18451355 | 0.083 | 0.217 | 1 | 15 |
| lrpap1 | 0.25058 | -0.2649124 | 0.194 | 0.391 | 1 | 15 |
| si:zfos-80g12.1 | 0.25069 | -0.24461281 | 0.056 | 0.174 | 1 | 15 |
| cnpy4 | 0.25079 | 0.187882644 | 0.167 | 0.043 | 1 | 15 |
| nr2f1a | 0.25118 | 0.187836155 | 0.833 | 0.652 | 1 | 15 |
| slc25a55a | 0.2514 | -0.30064475 | 0.167 | 0.348 | 1 | 15 |
| ppp2cb | 0.25168 | 0.280995912 | 0.611 | 0.391 | 1 | 15 |
| rerea | 0.25198 | -0.13733117 | 0.167 | 0.304 | 1 | 15 |
| ndufb3 | 0.252 | 0.271878114 | 0.611 | 0.391 | 1 | 15 |
| rnf170 | 0.25221 | -0.21596016 | 0.028 | 0.13 | 1 | 15 |
| cyp20a1 | 0.25374 | -0.22844105 | 0.056 | 0.174 | 1 | 15 |
| trim35-33 | 0.25381 | -0.22082981 | 0.028 | 0.13 | 1 | 15 |
| pgrmc1 | 0.254 | 0.25067179 | 0.861 | 0.696 | 1 | 15 |
| hnrnpr | 0.25443 | 0.204659039 | 0.278 | 0.13 | 1 | 15 |
| ctsc | 0.25445 | -0.25796833 | 0.028 | 0.13 | 1 | 15 |
| zgc:110366 | 0.25451 | -0.15467273 | 0.028 | 0.13 | 1 | 15 |
| ndufa2 | 0.25502 | 0.298015479 | 0.417 | 0.217 | 1 | 15 |
| sik2b | 0.25524 | -0.35084877 | 0.167 | 0.304 | 1 | 15 |
| ndufaf1 | 0.25535 | -0.25936795 | 0.056 | 0.174 | 1 | 15 |
| rrbp1a | 0.25622 | -0.26161537 | 0.194 | 0.304 | 1 | 15 |

| | | | | | | |
|---|---|---|---|---|---|---|
| cxxc5a | 0.25657 | 0.209952941 | 0.278 | 0.217 | 1 | 15 |
| med7 | 0.25667 | 0.161308554 | 0.139 | 0.087 | 1 | 15 |
| etfa | 0.25722 | -0.28182783 | 0.222 | 0.391 | 1 | 15 |
| ptgesl | 0.2573 | 0.108073592 | 0.139 | 0.13 | 1 | 15 |
| nfkb2 | 0.25735 | -0.22682212 | 0.028 | 0.13 | 1 | 15 |
| npm3 | 0.25778 | 0.270303051 | 0.361 | 0.174 | 1 | 15 |
| cd99 | 0.25837 | -0.31016721 | 0.139 | 0.304 | 1 | 15 |
| fkbp5 | 0.25848 | 0.28854925 | 0.472 | 0.261 | 1 | 15 |
| aip | 0.25853 | 0.116213805 | 0.25 | 0.13 | 1 | 15 |
| pdzd11 | 0.25864 | -0.11277761 | 0.139 | 0.261 | 1 | 15 |
| FO082781.1 | 0.25868 | -0.23259898 | 0.278 | 0.348 | 1 | 15 |
| tagln2 | 0.2587 | -0.28336212 | 0.111 | 0.261 | 1 | 15 |
| sh3glb1b | 0.25873 | 0.280644799 | 0.278 | 0.13 | 1 | 15 |
| mlf2 | 0.25881 | 0.283177882 | 0.222 | 0.087 | 1 | 15 |
| zgc:55262 | 0.25974 | 0.205299989 | 0.139 | 0.043 | 1 | 15 |
| setb | 0.26009 | 0.284977469 | 0.694 | 0.522 | 1 | 15 |
| sf3b4 | 0.2603 | 0.246655379 | 0.25 | 0.087 | 1 | 15 |
| slc44a2 | 0.26067 | -0.29078421 | 0.139 | 0.304 | 1 | 15 |
| polb | 0.26076 | 0.231463577 | 0.167 | 0.043 | 1 | 15 |
| uba52 | 0.26106 | 0.157383817 | 0.944 | 0.913 | 1 | 15 |
| cdk16 | 0.26135 | 0.209400075 | 0.167 | 0.043 | 1 | 15 |
| c4b | 0.2614 | -0.1134951 | 0.139 | 0.13 | 1 | 15 |
| cadm4 | 0.26155 | 0.293899168 | 0.25 | 0.087 | 1 | 15 |
| myclb | 0.26159 | 0.297978971 | 0.528 | 0.391 | 1 | 15 |
| txnipa | 0.26179 | 0.249005191 | 0.833 | 0.739 | 1 | 15 |
| commd9 | 0.26181 | -0.18333967 | 0.083 | 0.217 | 1 | 15 |
| ogdha | 0.26197 | 0.311926947 | 0.417 | 0.217 | 1 | 15 |
| ccdc25 | 0.26198 | 0.228039699 | 0.361 | 0.174 | 1 | 15 |
| arhgdig | 0.26242 | -0.21034944 | 0.222 | 0.261 | 1 | 15 |
| atp6v0d1 | 0.26269 | -0.1422896 | 0.167 | 0.174 | 1 | 15 |
| fzd7b | 0.26294 | -0.20183379 | 0.278 | 0.348 | 1 | 15 |
| fundc1 | 0.26359 | 0.243740153 | 0.333 | 0.217 | 1 | 15 |
| si:dkey-188i13.11 | 0.26367 | 0.173261712 | 0.472 | 0.435 | 1 | 15 |
| zc4h2 | 0.26474 | -0.30726042 | 0.194 | 0.348 | 1 | 15 |
| magt1 | 0.26504 | -0.2766431 | 0.278 | 0.478 | 1 | 15 |
| itgav | 0.2658 | -0.2891484 | 0.194 | 0.304 | 1 | 15 |
| tle2b | 0.2663 | -0.29938373 | 0.111 | 0.261 | 1 | 15 |
| lhfpl6 | 0.26634 | -0.16519765 | 0.028 | 0.13 | 1 | 15 |
| plp2 | 0.26638 | -0.18256933 | 0.083 | 0.13 | 1 | 15 |
| fut8a | 0.26646 | -0.24368705 | 0.028 | 0.13 | 1 | 15 |
| sult1st1 | 0.26654 | -0.13800441 | 0.028 | 0.13 | 1 | 15 |
| rfxap | 0.26705 | 0.275807355 | 0.361 | 0.174 | 1 | 15 |
| mrps26 | 0.26801 | 0.194694674 | 0.111 | 0.043 | 1 | 15 |
| h2afy2 | 0.26807 | -0.13801612 | 0.444 | 0.478 | 1 | 15 |
| htt | 0.2681 | -0.22207947 | 0.028 | 0.13 | 1 | 15 |
| zbtb33 | 0.26876 | 0.254814199 | 0.167 | 0.043 | 1 | 15 |
| hectd2 | 0.2688 | -0.10952994 | 0.083 | 0.087 | 1 | 15 |
| rhoj | 0.26881 | -0.30482244 | 0.139 | 0.304 | 1 | 15 |

| | | | | | | |
|---|---|---|---|---|---|---|
| tbcc | 0.269 | -0.18719296 | 0.028 | 0.13 | 1 | 15 |
| ccdc3b | 0.26943 | -0.1933743 | 0.028 | 0.13 | 1 | 15 |
| slc9a6a | 0.26996 | 0.206447611 | 0.111 | 0.043 | 1 | 15 |
| clns1a | 0.27045 | 0.133795707 | 0.111 | 0.087 | 1 | 15 |
| scfd1 | 0.27087 | 0.187919196 | 0.167 | 0.043 | 1 | 15 |
| coq10b | 0.27262 | -0.22478724 | 0.083 | 0.174 | 1 | 15 |
| vrk1 | 0.273 | -0.12823662 | 0.028 | 0.13 | 1 | 15 |
| edf1 | 0.27321 | 0.220007084 | 0.722 | 0.609 | 1 | 15 |
| u2af2a | 0.27383 | -0.20901792 | 0.111 | 0.174 | 1 | 15 |
| dbnlb | 0.27393 | 0.164202995 | 0.167 | 0.043 | 1 | 15 |
| eif4g2a | 0.27419 | -0.30803029 | 0.222 | 0.391 | 1 | 15 |
| flot1b | 0.27445 | -0.27572446 | 0.389 | 0.565 | 1 | 15 |
| plxnb2a | 0.27462 | -0.23221796 | 0.111 | 0.261 | 1 | 15 |
| fthl27 | 0.27492 | 0.131504561 | 0.917 | 0.913 | 1 | 15 |
| sart1 | 0.27503 | 0.199243005 | 0.194 | 0.13 | 1 | 15 |
| hadhaa | 0.27509 | 0.128952635 | 0.194 | 0.087 | 1 | 15 |
| lpar1 | 0.27509 | -0.20536937 | 0.333 | 0.391 | 1 | 15 |
| sdhb | 0.27539 | 0.154609326 | 0.278 | 0.261 | 1 | 15 |
| pfdn6 | 0.27565 | 0.206269947 | 0.306 | 0.217 | 1 | 15 |
| gadd45bb | 0.27598 | -0.19461142 | 0.028 | 0.13 | 1 | 15 |
| adipor1a | 0.27611 | -0.14968241 | 0.139 | 0.174 | 1 | 15 |
| si:ch211-194b1.1 | 0.27622 | 0.216850354 | 0.139 | 0.043 | 1 | 15 |
| capns1b | 0.27624 | 0.294891981 | 0.417 | 0.217 | 1 | 15 |
| si:ch211-253b8.2 | 0.27624 | -0.25083211 | 0.028 | 0.13 | 1 | 15 |
| rsu1 | 0.27637 | -0.19561032 | 0.083 | 0.13 | 1 | 15 |
| kif5ba | 0.27638 | -0.26065306 | 0.139 | 0.261 | 1 | 15 |
| nutf2l | 0.27658 | 0.258898214 | 0.306 | 0.174 | 1 | 15 |
| nudt3a | 0.27685 | -0.24145512 | 0.056 | 0.174 | 1 | 15 |
| parp8 | 0.27693 | -0.259535 | 0.056 | 0.174 | 1 | 15 |
| si:ch211-15b10.6 | 0.27731 | 0.272200777 | 0.361 | 0.217 | 1 | 15 |
| pacsin3 | 0.27735 | -0.26175056 | 0.028 | 0.13 | 1 | 15 |
| ube2d3 | 0.27796 | 0.208334854 | 0.139 | 0.043 | 1 | 15 |
| hmgb3a | 0.27818 | 0.176482046 | 0.75 | 0.565 | 1 | 15 |
| vgll4b | 0.27821 | -0.22289201 | 0.056 | 0.174 | 1 | 15 |
| mrps18c | 0.27823 | 0.222253437 | 0.139 | 0.043 | 1 | 15 |
| agpat5 | 0.27837 | 0.189773329 | 0.167 | 0.043 | 1 | 15 |
| cav1 | 0.2784 | -0.21475585 | 0.028 | 0.13 | 1 | 15 |
| mrpl58 | 0.2785 | 0.110136113 | 0.139 | 0.043 | 1 | 15 |
| rplp1 | 0.27872 | 0.116434078 | 1 | 0.957 | 1 | 15 |
| gng5 | 0.27919 | 0.128114564 | 0.556 | 0.391 | 1 | 15 |
| sypl1 | 0.27928 | -0.23868216 | 0.056 | 0.174 | 1 | 15 |
| pycr1b | 0.27969 | -0.14194166 | 0.056 | 0.087 | 1 | 15 |
| si:ch211-282b22.1 | 0.28007 | 0.242166963 | 0.222 | 0.087 | 1 | 15 |
| acvrl1 | 0.28054 | -0.2157423 | 0.028 | 0.13 | 1 | 15 |
| cavin2b | 0.28055 | -0.31589588 | 0.167 | 0.348 | 1 | 15 |
| dusp6 | 0.28098 | -0.25754576 | 0.111 | 0.217 | 1 | 15 |
| pkig | 0.28106 | -0.2517538 | 0.278 | 0.478 | 1 | 15 |
| atp1a3b | 0.28136 | -0.32583342 | 0.361 | 0.565 | 1 | 15 |

| | | | | | | |
|---|---|---|---|---|---|---|
| f3b | 0.2819 | -0.18623754 | 0.083 | 0.13 | 1 | 15 |
| nol4lb | 0.28194 | 0.297574049 | 0.222 | 0.087 | 1 | 15 |
| tcp11l2 | 0.2822 | -0.20817774 | 0.028 | 0.13 | 1 | 15 |
| znf318 | 0.28241 | 0.253820821 | 0.194 | 0.087 | 1 | 15 |
| s100v2 | 0.28258 | -0.25489295 | 0.306 | 0.435 | 1 | 15 |
| ufc1 | 0.28298 | -0.2199535 | 0.056 | 0.174 | 1 | 15 |
| crkl | 0.28306 | -0.24360175 | 0.083 | 0.217 | 1 | 15 |
| osbpl9 | 0.28351 | -0.23974254 | 0.028 | 0.13 | 1 | 15 |
| mmp15b | 0.28358 | -0.2525417 | 0.028 | 0.13 | 1 | 15 |
| pfkfb4b | 0.28364 | 0.248727097 | 0.611 | 0.522 | 1 | 15 |
| yes1 | 0.28374 | 0.222688755 | 0.167 | 0.043 | 1 | 15 |
| inip | 0.28376 | -0.27249386 | 0.083 | 0.217 | 1 | 15 |
| sec61a1 | 0.28499 | -0.22362377 | 0.083 | 0.217 | 1 | 15 |
| dnaja2a | 0.28504 | 0.288246178 | 0.556 | 0.348 | 1 | 15 |
| chmp6b | 0.28595 | -0.27237935 | 0.083 | 0.217 | 1 | 15 |
| arhgef4 | 0.2867 | -0.13843427 | 0.167 | 0.174 | 1 | 15 |
| tkfc | 0.28749 | -0.23165296 | 0.028 | 0.13 | 1 | 15 |
| pdgfra | 0.28751 | -0.26300877 | 0.056 | 0.174 | 1 | 15 |
| lrrc17 | 0.28783 | -0.30432529 | 0.222 | 0.348 | 1 | 15 |
| si:ch73-311h14.2 | 0.28818 | -0.22491983 | 0.028 | 0.13 | 1 | 15 |
| rbm4.3 | 0.28835 | 0.286002819 | 0.667 | 0.478 | 1 | 15 |
| tpt1 | 0.28847 | 0.205844225 | 0.917 | 0.783 | 1 | 15 |
| arl4d | 0.28895 | 0.187162428 | 0.333 | 0.261 | 1 | 15 |
| STMP1 | 0.28895 | 0.201265551 | 0.389 | 0.217 | 1 | 15 |
| pgap3 | 0.28913 | -0.14396932 | 0.056 | 0.087 | 1 | 15 |
| ncor1 | 0.28939 | -0.21884906 | 0.056 | 0.174 | 1 | 15 |
| znf281a | 0.28998 | -0.17106652 | 0.028 | 0.13 | 1 | 15 |
| serinc5 | 0.2902 | -0.348869 | 0.278 | 0.478 | 1 | 15 |
| timmdc1 | 0.2904 | -0.18163755 | 0.028 | 0.13 | 1 | 15 |
| si:ch211-39i22.1 | 0.29074 | -0.30155789 | 0.25 | 0.435 | 1 | 15 |
| josd2 | 0.29075 | -0.20893376 | 0.028 | 0.13 | 1 | 15 |
| ndufb10 | 0.29124 | -0.27941767 | 0.361 | 0.565 | 1 | 15 |
| fkbp14 | 0.29127 | -0.26230634 | 0.111 | 0.261 | 1 | 15 |
| serpinb1l1 | 0.29132 | -0.19205217 | 0.028 | 0.13 | 1 | 15 |
| naprt | 0.29133 | 0.237967798 | 0.167 | 0.043 | 1 | 15 |
| srp72 | 0.2915 | 0.237007863 | 0.139 | 0.043 | 1 | 15 |
| lypla2 | 0.29152 | -0.2750991 | 0.083 | 0.217 | 1 | 15 |
| poldip2 | 0.29182 | 0.114779586 | 0.194 | 0.087 | 1 | 15 |
| srsf1a | 0.29191 | 0.276036672 | 0.222 | 0.087 | 1 | 15 |
| ddx56 | 0.29205 | 0.120998335 | 0.139 | 0.043 | 1 | 15 |
| nob1 | 0.29223 | 0.199026609 | 0.167 | 0.043 | 1 | 15 |
| higd2a | 0.29252 | 0.250567891 | 0.167 | 0.043 | 1 | 15 |
| mtx2 | 0.29274 | 0.195961708 | 0.139 | 0.043 | 1 | 15 |
| mvb12a | 0.29303 | -0.14459343 | 0.028 | 0.13 | 1 | 15 |
| histh1l | 0.29364 | -0.28373579 | 0.083 | 0.217 | 1 | 15 |
| XLOC-005199 | 0.29416 | -0.16708831 | 0.028 | 0.13 | 1 | 15 |
| chmp5a | 0.29435 | -0.21335236 | 0.028 | 0.13 | 1 | 15 |
| dchs1b | 0.29542 | -0.25996485 | 0.028 | 0.13 | 1 | 15 |

| | | | | | | |
|---|---|---|---|---|---|---|
| arf6a | 0.29683 | 0.171065156 | 0.222 | 0.087 | 1 | 15 |
| mych | 0.29711 | -0.25859106 | 0.139 | 0.261 | 1 | 15 |
| marcksa | 0.29719 | 0.237023453 | 0.194 | 0.087 | 1 | 15 |
| pabpn1 | 0.29719 | -0.16646579 | 0.389 | 0.565 | 1 | 15 |
| mrps21 | 0.2972 | 0.218945686 | 0.167 | 0.043 | 1 | 15 |
| sfpq | 0.2973 | -0.31636056 | 0.222 | 0.391 | 1 | 15 |
| mfap2 | 0.29789 | -0.31621916 | 0.167 | 0.304 | 1 | 15 |
| si:dkey-58f10.11 | 0.29845 | -0.16352276 | 0.028 | 0.13 | 1 | 15 |
| dffb | 0.29872 | 0.231550881 | 0.167 | 0.043 | 1 | 15 |
| rbm4.1 | 0.29883 | 0.272869919 | 0.389 | 0.217 | 1 | 15 |
| XLOC-032256 | 0.2989 | -0.14081396 | 0.028 | 0.13 | 1 | 15 |
| fam129bb | 0.29915 | -0.22112609 | 0.028 | 0.13 | 1 | 15 |
| foxn3 | 0.30019 | 0.254558955 | 0.278 | 0.13 | 1 | 15 |
| cbfa2t3 | 0.30022 | -0.19726222 | 0.028 | 0.13 | 1 | 15 |
| LOC103910921 | 0.30043 | -0.18206848 | 0.028 | 0.13 | 1 | 15 |
| CABZ01087560.1 | 0.3005 | -0.14197851 | 0.167 | 0.304 | 1 | 15 |
| kpna4 | 0.30067 | 0.125518002 | 0.139 | 0.043 | 1 | 15 |
| il6st | 0.30067 | -0.26900308 | 0.139 | 0.304 | 1 | 15 |
| ngdn | 0.30107 | -0.24242722 | 0.167 | 0.261 | 1 | 15 |
| park7 | 0.30148 | -0.18953048 | 0.111 | 0.174 | 1 | 15 |
| smarcc2 | 0.30168 | 0.2420234 | 0.167 | 0.043 | 1 | 15 |
| col6a4a | 0.30232 | -0.23956388 | 0.028 | 0.13 | 1 | 15 |
| si:ch211-275j6.5 | 0.30235 | 0.198594258 | 0.139 | 0.043 | 1 | 15 |
| si:ch211-106h4.12 | 0.30244 | -0.28579592 | 0.028 | 0.13 | 1 | 15 |
| adgre5b.3 | 0.30282 | -0.20602703 | 0.028 | 0.13 | 1 | 15 |
| zbtb4 | 0.30301 | -0.17493959 | 0.083 | 0.217 | 1 | 15 |
| naxd | 0.30316 | 0.222254013 | 0.167 | 0.043 | 1 | 15 |
| appb | 0.3033 | -0.27789331 | 0.139 | 0.261 | 1 | 15 |
| dgkza | 0.30347 | -0.18909828 | 0.028 | 0.13 | 1 | 15 |
| grnb | 0.30349 | -0.25123468 | 0.056 | 0.13 | 1 | 15 |
| pkp3b | 0.30356 | -0.24182366 | 0.139 | 0.217 | 1 | 15 |
| nap1l1 | 0.30398 | 0.251866268 | 0.639 | 0.435 | 1 | 15 |
| sms | 0.30399 | 0.233146269 | 0.139 | 0.043 | 1 | 15 |
| fam76b | 0.30411 | 0.195277007 | 0.222 | 0.087 | 1 | 15 |
| eif4ebp1 | 0.30414 | 0.244831442 | 0.778 | 0.609 | 1 | 15 |
| tfdp2 | 0.30428 | 0.118468715 | 0.194 | 0.087 | 1 | 15 |
| ppme1 | 0.30485 | 0.222847279 | 0.167 | 0.043 | 1 | 15 |
| dock7 | 0.30505 | -0.15788057 | 0.083 | 0.13 | 1 | 15 |
| fam92a1 | 0.30522 | -0.13624885 | 0.028 | 0.13 | 1 | 15 |
| PAXBP1 | 0.30556 | -0.19992188 | 0.028 | 0.13 | 1 | 15 |
| LOC103910438 | 0.30563 | -0.1642246 | 0.361 | 0.391 | 1 | 15 |
| ptx3a | 0.30584 | -0.16434372 | 0.028 | 0.13 | 1 | 15 |
| kpnb3 | 0.30664 | 0.218557488 | 0.167 | 0.043 | 1 | 15 |
| zgc:112982 | 0.3067 | -0.17104986 | 0.028 | 0.13 | 1 | 15 |
| zgc:101679 | 0.30696 | -0.18644811 | 0.028 | 0.13 | 1 | 15 |
| CABZ01026681.1 | 0.30725 | -0.23331802 | 0.167 | 0.261 | 1 | 15 |
| coa7 | 0.30731 | -0.19940982 | 0.028 | 0.13 | 1 | 15 |
| copb2 | 0.30733 | 0.231628172 | 0.222 | 0.087 | 1 | 15 |

| | | | | | | |
|---|---|---|---|---|---|---|
| cnp | 0.30735 | -0.23190768 | 0.25 | 0.435 | 1 | 15 |
| rgl1 | 0.30763 | -0.22102629 | 0.056 | 0.174 | 1 | 15 |
| wipf1b | 0.30765 | -0.21263553 | 0.028 | 0.13 | 1 | 15 |
| ms4a17a.9 | 0.30774 | 0.187011998 | 0.167 | 0.087 | 1 | 15 |
| sumf2 | 0.30793 | -0.11475976 | 0.111 | 0.217 | 1 | 15 |
| zgc:172106 | 0.30809 | -0.25629553 | 0.028 | 0.13 | 1 | 15 |
| atp6v0cb | 0.30819 | 0.124558162 | 0.139 | 0.043 | 1 | 15 |
| ehmt1a | 0.30847 | -0.17556401 | 0.028 | 0.13 | 1 | 15 |
| slc16a9b | 0.30864 | 0.233274054 | 0.167 | 0.043 | 1 | 15 |
| rnf7 | 0.30881 | 0.276968151 | 0.361 | 0.217 | 1 | 15 |
| capza1b | 0.30883 | 0.229195069 | 0.361 | 0.261 | 1 | 15 |
| mfap5 | 0.30891 | -0.22122767 | 0.028 | 0.13 | 1 | 15 |
| sod2 | 0.30908 | 0.100006181 | 0.444 | 0.304 | 1 | 15 |
| ahnak | 0.30913 | -0.29161347 | 0.111 | 0.261 | 1 | 15 |
| foxp1a | 0.30936 | -0.16500916 | 0.028 | 0.13 | 1 | 15 |
| ddx21 | 0.30952 | 0.268904304 | 0.444 | 0.261 | 1 | 15 |
| kdm2bb | 0.30975 | -0.2057005 | 0.028 | 0.13 | 1 | 15 |
| pbx2 | 0.30991 | 0.201311196 | 0.139 | 0.043 | 1 | 15 |
| ids | 0.31032 | -0.20406761 | 0.028 | 0.13 | 1 | 15 |
| tpm1 | 0.31056 | 0.242249847 | 0.222 | 0.087 | 1 | 15 |
| bmp3 | 0.31066 | -0.17741061 | 0.028 | 0.13 | 1 | 15 |
| bloc1s1 | 0.31079 | 0.139712253 | 0.139 | 0.043 | 1 | 15 |
| hccsa.1 | 0.31083 | 0.206379502 | 0.139 | 0.043 | 1 | 15 |
| tox4a | 0.31097 | -0.15678057 | 0.028 | 0.13 | 1 | 15 |
| fmr1 | 0.31129 | 0.145314943 | 0.25 | 0.217 | 1 | 15 |
| csrnp1b | 0.31136 | -0.20455157 | 0.028 | 0.13 | 1 | 15 |
| dnal4b | 0.31141 | 0.21712087 | 0.167 | 0.043 | 1 | 15 |
| rps6ka3a | 0.31142 | -0.17220776 | 0.111 | 0.174 | 1 | 15 |
| prr14 | 0.31172 | -0.21949823 | 0.028 | 0.13 | 1 | 15 |
| trabd | 0.31203 | -0.16506889 | 0.028 | 0.13 | 1 | 15 |
| cavin1b | 0.31212 | -0.13844005 | 0.056 | 0.087 | 1 | 15 |
| nid2a | 0.31237 | -0.31207865 | 0.139 | 0.304 | 1 | 15 |
| dstyk | 0.31239 | 0.192620843 | 0.139 | 0.043 | 1 | 15 |
| gabpa | 0.31258 | 0.190606179 | 0.167 | 0.043 | 1 | 15 |
| slc4a7 | 0.31267 | -0.15760085 | 0.028 | 0.13 | 1 | 15 |
| neu1 | 0.31278 | 0.193717639 | 0.167 | 0.043 | 1 | 15 |
| lrrc20 | 0.3128 | 0.19126415 | 0.167 | 0.043 | 1 | 15 |
| CABZ01029938.1 | 0.3131 | 0.193191879 | 0.139 | 0.043 | 1 | 15 |
| tmem181 | 0.31314 | -0.22093503 | 0.028 | 0.13 | 1 | 15 |
| lmnb1 | 0.31335 | 0.227603206 | 0.167 | 0.043 | 1 | 15 |
| hirip3 | 0.31371 | -0.19571604 | 0.028 | 0.13 | 1 | 15 |
| gyg1a | 0.31411 | 0.102062429 | 0.139 | 0.043 | 1 | 15 |
| psmd3 | 0.31438 | 0.120395238 | 0.444 | 0.304 | 1 | 15 |
| pdxka | 0.3144 | -0.24359449 | 0.139 | 0.304 | 1 | 15 |
| mapre1b | 0.31468 | -0.25745776 | 0.139 | 0.304 | 1 | 15 |
| dsc2l | 0.31474 | -0.15726491 | 0.028 | 0.13 | 1 | 15 |
| rprmb | 0.31487 | -0.16415876 | 0.056 | 0.174 | 1 | 15 |
| sdf2 | 0.31504 | 0.208387781 | 0.167 | 0.043 | 1 | 15 |

| | | | | | | |
|---|---|---|---|---|---|---|
| fam172a | 0.31638 | -0.18043111 | 0.111 | 0.174 | 1 | 15 |
| fshr | 0.31644 | -0.27914697 | 0.056 | 0.174 | 1 | 15 |
| coil | 0.3166 | 0.175242753 | 0.167 | 0.043 | 1 | 15 |
| msi2a | 0.31664 | 0.293892279 | 0.444 | 0.261 | 1 | 15 |
| si:dkey-4e7.3 | 0.31714 | 0.237956096 | 0.167 | 0.043 | 1 | 15 |
| mrps16 | 0.31778 | 0.120374496 | 0.194 | 0.087 | 1 | 15 |
| acy1 | 0.31786 | -0.15963113 | 0.028 | 0.13 | 1 | 15 |
| BX569787.1 | 0.31836 | 0.182465051 | 0.361 | 0.217 | 1 | 15 |
| tnrc18 | 0.3185 | -0.24949955 | 0.111 | 0.261 | 1 | 15 |
| grb10a | 0.31855 | 0.233199938 | 0.167 | 0.043 | 1 | 15 |
| XLOC-018082 | 0.31855 | -0.15509812 | 0.028 | 0.13 | 1 | 15 |
| socs1a | 0.31861 | -0.18757396 | 0.056 | 0.174 | 1 | 15 |
| ube2g1a | 0.31865 | 0.207317024 | 0.167 | 0.043 | 1 | 15 |
| farp2 | 0.31866 | -0.21066841 | 0.028 | 0.13 | 1 | 15 |
| tcea2 | 0.31917 | -0.20313411 | 0.028 | 0.13 | 1 | 15 |
| lamtor5 | 0.31937 | -0.22367382 | 0.083 | 0.217 | 1 | 15 |
| CABZ01068362.1 | 0.31944 | -0.16298551 | 0.028 | 0.13 | 1 | 15 |
| virma | 0.31958 | -0.18173457 | 0.028 | 0.13 | 1 | 15 |
| slc38a10 | 0.31974 | 0.210297266 | 0.167 | 0.043 | 1 | 15 |
| tfam | 0.31987 | 0.128681301 | 0.25 | 0.13 | 1 | 15 |
| ndufs5 | 0.32016 | 0.228229592 | 0.5 | 0.304 | 1 | 15 |
| zeb2b | 0.32033 | -0.23663537 | 0.111 | 0.261 | 1 | 15 |
| ada | 0.32039 | -0.1813874 | 0.056 | 0.174 | 1 | 15 |
| bxdc2 | 0.32064 | 0.193478312 | 0.167 | 0.043 | 1 | 15 |
| mesd | 0.3208 | -0.2765853 | 0.194 | 0.348 | 1 | 15 |
| rnaset2 | 0.32081 | -0.1782257 | 0.083 | 0.217 | 1 | 15 |
| dnase1l1 | 0.32088 | -0.21512406 | 0.083 | 0.217 | 1 | 15 |
| wdr45b | 0.32095 | -0.14837876 | 0.028 | 0.13 | 1 | 15 |
| arfip1 | 0.32104 | 0.210185907 | 0.167 | 0.043 | 1 | 15 |
| riok2 | 0.32125 | 0.192969849 | 0.167 | 0.043 | 1 | 15 |
| smim20 | 0.32127 | 0.196740796 | 0.167 | 0.043 | 1 | 15 |
| sod1 | 0.32132 | 0.178868436 | 0.667 | 0.478 | 1 | 15 |
| si:ch211-262e15.1 | 0.32144 | -0.23230333 | 0.056 | 0.13 | 1 | 15 |
| csk | 0.32161 | -0.14436878 | 0.028 | 0.13 | 1 | 15 |
| snrnp48 | 0.32166 | -0.17350971 | 0.056 | 0.174 | 1 | 15 |
| PCMTD2 (1 of many) | 0.32187 | -0.16481924 | 0.028 | 0.13 | 1 | 15 |
| lsm1 | 0.3219 | 0.185856927 | 0.167 | 0.043 | 1 | 15 |
| piezo2a.2 | 0.322 | -0.22715294 | 0.028 | 0.13 | 1 | 15 |
| dffa | 0.32212 | 0.200116459 | 0.167 | 0.043 | 1 | 15 |
| si:ch211-162e15.3 | 0.32219 | 0.243725725 | 0.278 | 0.13 | 1 | 15 |
| vgll4a | 0.32223 | -0.23644598 | 0.056 | 0.174 | 1 | 15 |
| thbs3b | 0.32229 | 0.264898469 | 0.167 | 0.043 | 1 | 15 |
| fgfr2 | 0.32265 | -0.23111457 | 0.028 | 0.13 | 1 | 15 |
| si:ch211-106k21.5 | 0.32289 | -0.23829248 | 0.056 | 0.174 | 1 | 15 |
| dhrs13a.1 | 0.32294 | -0.13818741 | 0.056 | 0.087 | 1 | 15 |
| ugdh | 0.32296 | -0.19944028 | 0.028 | 0.13 | 1 | 15 |
| srcap | 0.32318 | -0.17103269 | 0.028 | 0.13 | 1 | 15 |
| fam3c | 0.32349 | 0.234346076 | 0.222 | 0.087 | 1 | 15 |

| | | | | | | |
|---|---|---|---|---|---|---|
| zgc:194981 | 0.3237 | 0.217170166 | 0.194 | 0.087 | 1 | 15 |
| stat1b | 0.32378 | -0.24333329 | 0.111 | 0.261 | 1 | 15 |
| nip7 | 0.32431 | -0.22844639 | 0.111 | 0.261 | 1 | 15 |
| RPL37A | 0.32443 | -0.14792971 | 0.028 | 0.13 | 1 | 15 |
| lix1l | 0.32479 | -0.13680506 | 0.028 | 0.13 | 1 | 15 |
| si:ch211-262i1.5 | 0.32504 | -0.16435601 | 0.028 | 0.13 | 1 | 15 |
| ipo4 | 0.32508 | -0.14318075 | 0.028 | 0.13 | 1 | 15 |
| tmem150aa | 0.3253 | -0.15957668 | 0.028 | 0.13 | 1 | 15 |
| inhbb | 0.32548 | 0.200548691 | 0.139 | 0.043 | 1 | 15 |
| wdr45 | 0.32549 | -0.1983115 | 0.028 | 0.13 | 1 | 15 |
| exosc5 | 0.32564 | -0.16406303 | 0.028 | 0.13 | 1 | 15 |
| adgra3 | 0.32568 | -0.20862138 | 0.028 | 0.13 | 1 | 15 |
| crcp | 0.3257 | -0.17199219 | 0.028 | 0.13 | 1 | 15 |
| si:ch211-178n15.1 | 0.32577 | -0.22989138 | 0.028 | 0.13 | 1 | 15 |
| nolc1 | 0.32595 | 0.231683142 | 0.222 | 0.087 | 1 | 15 |
| irf2bpl | 0.32596 | -0.16621571 | 0.028 | 0.13 | 1 | 15 |
| elmo1 | 0.3262 | -0.16692607 | 0.028 | 0.13 | 1 | 15 |
| pip5k1aa | 0.32621 | -0.17625404 | 0.028 | 0.13 | 1 | 15 |
| mxra8b | 0.32648 | -0.2152742 | 0.056 | 0.174 | 1 | 15 |
| si:ch211-282j22.3 | 0.3267 | -0.16999989 | 0.028 | 0.13 | 1 | 15 |
| entpd6 | 0.32672 | -0.17423142 | 0.028 | 0.13 | 1 | 15 |
| tefb | 0.32673 | -0.1927761 | 0.028 | 0.13 | 1 | 15 |
| asb12a | 0.32673 | -0.14470928 | 0.028 | 0.13 | 1 | 15 |
| pex3 | 0.32677 | -0.17236707 | 0.028 | 0.13 | 1 | 15 |
| smdt1b | 0.32677 | -0.10948742 | 0.417 | 0.565 | 1 | 15 |
| cpvl | 0.32678 | 0.221677734 | 0.194 | 0.087 | 1 | 15 |
| stard3 | 0.32679 | -0.21831214 | 0.028 | 0.13 | 1 | 15 |
| yipf3 | 0.3268 | -0.24637768 | 0.083 | 0.217 | 1 | 15 |
| gorab | 0.32681 | -0.17648178 | 0.028 | 0.13 | 1 | 15 |
| tmpob | 0.32688 | -0.21442282 | 0.028 | 0.13 | 1 | 15 |
| anapc15 | 0.32699 | -0.16360332 | 0.028 | 0.13 | 1 | 15 |
| znf532 | 0.327 | -0.22901178 | 0.028 | 0.13 | 1 | 15 |
| nf2b | 0.32701 | -0.15724404 | 0.028 | 0.13 | 1 | 15 |
| cfap36 | 0.32721 | 0.129713945 | 0.194 | 0.087 | 1 | 15 |
| BX548028.1 | 0.32722 | -0.29717521 | 0.167 | 0.304 | 1 | 15 |
| ctnnbip1 | 0.32812 | 0.134793165 | 0.194 | 0.087 | 1 | 15 |
| pttg1ipb | 0.32826 | -0.22056186 | 0.083 | 0.217 | 1 | 15 |
| mapk3 | 0.32881 | 0.170280164 | 0.194 | 0.13 | 1 | 15 |
| rab1ba | 0.329 | -0.23484359 | 0.056 | 0.174 | 1 | 15 |
| nomo | 0.32901 | -0.21547704 | 0.111 | 0.261 | 1 | 15 |
| ptena | 0.3293 | 0.181596043 | 0.222 | 0.174 | 1 | 15 |
| LOC100007100 | 0.32959 | -0.17460842 | 0.028 | 0.087 | 1 | 15 |
| dnajc3b | 0.32959 | -0.16084918 | 0.028 | 0.087 | 1 | 15 |
| rfx7b | 0.32959 | -0.11199697 | 0.028 | 0.087 | 1 | 15 |
| efna1b | 0.32959 | -0.15868128 | 0.028 | 0.087 | 1 | 15 |
| galnt2 | 0.32959 | -0.135369 | 0.028 | 0.087 | 1 | 15 |
| hmox2b | 0.32959 | -0.12588265 | 0.028 | 0.087 | 1 | 15 |
| pcsk5b | 0.32959 | -0.13372029 | 0.028 | 0.087 | 1 | 15 |

| | | | | | | |
|---|---|---|---|---|---|---|
| cul1b | 0.32959 | -0.10664886 | 0.028 | 0.087 | 1 | 15 |
| dsg2.1 | 0.32959 | -0.10156516 | 0.028 | 0.087 | 1 | 15 |
| sdhaf2 | 0.32959 | -0.16121945 | 0.028 | 0.087 | 1 | 15 |
| zmp:0000000735 | 0.32959 | -0.109266 | 0.028 | 0.087 | 1 | 15 |
| CR753876.1 | 0.32959 | -0.19720298 | 0.028 | 0.087 | 1 | 15 |
| NAV1 (1 of many) | 0.32959 | -0.1323668 | 0.028 | 0.087 | 1 | 15 |
| mpv17 | 0.32959 | -0.10500122 | 0.028 | 0.087 | 1 | 15 |
| nin | 0.32959 | -0.11106693 | 0.028 | 0.087 | 1 | 15 |
| XLOC-040582 | 0.32959 | -0.17836445 | 0.028 | 0.087 | 1 | 15 |
| acot20 | 0.32959 | -0.1161895 | 0.028 | 0.087 | 1 | 15 |
| adnp2b | 0.32959 | -0.13306884 | 0.028 | 0.087 | 1 | 15 |
| dus3l | 0.32959 | -0.13706341 | 0.028 | 0.087 | 1 | 15 |
| hpse | 0.32959 | -0.10475017 | 0.028 | 0.087 | 1 | 15 |
| LOC110438241 | 0.32959 | -0.20562436 | 0.028 | 0.087 | 1 | 15 |
| MTERF4 | 0.32959 | -0.12332842 | 0.028 | 0.087 | 1 | 15 |
| bach1b | 0.32959 | -0.11497025 | 0.028 | 0.087 | 1 | 15 |
| cdkn2c | 0.32959 | -0.14313751 | 0.028 | 0.087 | 1 | 15 |
| cers5 | 0.32959 | -0.10050518 | 0.028 | 0.087 | 1 | 15 |
| creb1b | 0.32959 | -0.10314981 | 0.028 | 0.087 | 1 | 15 |
| dhtkd1 | 0.32959 | -0.14177565 | 0.028 | 0.087 | 1 | 15 |
| epha6 | 0.32959 | -0.10772374 | 0.028 | 0.087 | 1 | 15 |
| evla | 0.32959 | -0.11918379 | 0.028 | 0.087 | 1 | 15 |
| fxr2 | 0.32959 | -0.1171327 | 0.028 | 0.087 | 1 | 15 |
| gtf2a1l | 0.32959 | -0.12276849 | 0.028 | 0.087 | 1 | 15 |
| herpud1 | 0.32959 | -0.13693079 | 0.028 | 0.087 | 1 | 15 |
| hsf1 | 0.32959 | -0.17569313 | 0.028 | 0.087 | 1 | 15 |
| lig3 | 0.32959 | -0.12701746 | 0.028 | 0.087 | 1 | 15 |
| mllt1b | 0.32959 | -0.13994025 | 0.028 | 0.087 | 1 | 15 |
| neo1b | 0.32959 | -0.10100816 | 0.028 | 0.087 | 1 | 15 |
| pbrm1l | 0.32959 | -0.11632686 | 0.028 | 0.087 | 1 | 15 |
| pde4d | 0.32959 | -0.11943068 | 0.028 | 0.087 | 1 | 15 |
| phf8 | 0.32959 | -0.18535394 | 0.028 | 0.087 | 1 | 15 |
| plce1 | 0.32959 | -0.14004326 | 0.028 | 0.087 | 1 | 15 |
| plxna2 | 0.32959 | -0.12492742 | 0.028 | 0.087 | 1 | 15 |
| pogzb | 0.32959 | -0.18287341 | 0.028 | 0.087 | 1 | 15 |
| rhbdf1a | 0.32959 | -0.13412633 | 0.028 | 0.087 | 1 | 15 |
| rhov | 0.32959 | -0.1518138 | 0.028 | 0.087 | 1 | 15 |
| rnf144b | 0.32959 | -0.11688996 | 0.028 | 0.087 | 1 | 15 |
| rnps1 | 0.32959 | -0.10463243 | 0.028 | 0.087 | 1 | 15 |
| saraf | 0.32959 | -0.17637878 | 0.028 | 0.087 | 1 | 15 |
| sgk3 | 0.32959 | -0.11272858 | 0.028 | 0.087 | 1 | 15 |
| si:ch73-92e7.4 | 0.32959 | -0.11150052 | 0.028 | 0.087 | 1 | 15 |
| si:dkey-172m14.1 | 0.32959 | -0.11878038 | 0.028 | 0.087 | 1 | 15 |
| slc35b3 | 0.32959 | -0.1227794 | 0.028 | 0.087 | 1 | 15 |
| slc39a3 | 0.32959 | -0.1667305 | 0.028 | 0.087 | 1 | 15 |
| srsf5b | 0.32959 | -0.14674765 | 0.028 | 0.087 | 1 | 15 |
| stag1b | 0.32959 | -0.11893471 | 0.028 | 0.087 | 1 | 15 |
| stx1b | 0.32959 | -0.10995199 | 0.028 | 0.087 | 1 | 15 |

| | | | | | | |
|---|---|---|---|---|---|---|
| tmem206 | 0.32959 | -0.10459789 | 0.028 | 0.087 | 1 | 15 |
| tmem222a | 0.32959 | -0.12699887 | 0.028 | 0.087 | 1 | 15 |
| trim13 | 0.32959 | -0.10130052 | 0.028 | 0.087 | 1 | 15 |
| wu:fb74b10 | 0.32959 | -0.13137197 | 0.028 | 0.087 | 1 | 15 |
| xirp2a | 0.32959 | -0.12966022 | 0.028 | 0.087 | 1 | 15 |
| zgc:110319 | 0.32959 | -0.11399174 | 0.028 | 0.087 | 1 | 15 |
| zgc:112977 | 0.32959 | -0.12570286 | 0.028 | 0.087 | 1 | 15 |
| zgc:91976 | 0.32959 | -0.19128003 | 0.028 | 0.087 | 1 | 15 |
| CABZ01078261.1 | 0.32959 | -0.23844698 | 0.028 | 0.087 | 1 | 15 |
| CR388029.1 | 0.32959 | -0.10309704 | 0.028 | 0.087 | 1 | 15 |
| FAR2 | 0.32959 | -0.16326146 | 0.028 | 0.087 | 1 | 15 |
| XLOC-024446 | 0.32959 | -0.14684552 | 0.028 | 0.087 | 1 | 15 |
| abhd10b | 0.32959 | -0.10246225 | 0.028 | 0.087 | 1 | 15 |
| acad8 | 0.32959 | -0.10490649 | 0.028 | 0.087 | 1 | 15 |
| ackr4a | 0.32959 | -0.14441815 | 0.028 | 0.087 | 1 | 15 |
| amacr | 0.32959 | -0.10886905 | 0.028 | 0.087 | 1 | 15 |
| amd1 | 0.32959 | -0.20869887 | 0.028 | 0.087 | 1 | 15 |
| arl8 | 0.32959 | -0.14288911 | 0.028 | 0.087 | 1 | 15 |
| asb3 | 0.32959 | -0.13998478 | 0.028 | 0.087 | 1 | 15 |
| dhdds | 0.32959 | -0.11084185 | 0.028 | 0.087 | 1 | 15 |
| dnajc3a | 0.32959 | -0.13601019 | 0.028 | 0.087 | 1 | 15 |
| ehbp1 | 0.32959 | -0.11051547 | 0.028 | 0.087 | 1 | 15 |
| exoc5 | 0.32959 | -0.12811829 | 0.028 | 0.087 | 1 | 15 |
| fopnl | 0.32959 | -0.10945171 | 0.028 | 0.087 | 1 | 15 |
| fto | 0.32959 | -0.21200557 | 0.028 | 0.087 | 1 | 15 |
| grk3 | 0.32959 | -0.13560255 | 0.028 | 0.087 | 1 | 15 |
| malt1 | 0.32959 | -0.11335303 | 0.028 | 0.087 | 1 | 15 |
| mbd4 | 0.32959 | -0.12769479 | 0.028 | 0.087 | 1 | 15 |
| mfn1b | 0.32959 | -0.112371 | 0.028 | 0.087 | 1 | 15 |
| nacc1b | 0.32959 | -0.12399162 | 0.028 | 0.087 | 1 | 15 |
| nrbp2a | 0.32959 | -0.11825045 | 0.028 | 0.087 | 1 | 15 |
| pard3ab | 0.32959 | -0.13596536 | 0.028 | 0.087 | 1 | 15 |
| pard6gb | 0.32959 | -0.16639113 | 0.028 | 0.087 | 1 | 15 |
| pmp22b | 0.32959 | -0.14709466 | 0.028 | 0.087 | 1 | 15 |
| polr1b | 0.32959 | -0.14883521 | 0.028 | 0.087 | 1 | 15 |
| qsox1 | 0.32959 | -0.1312117 | 0.028 | 0.087 | 1 | 15 |
| sept15 | 0.32959 | -0.11585261 | 0.028 | 0.087 | 1 | 15 |
| si:ch211-59d15.9 | 0.32959 | -0.10447575 | 0.028 | 0.087 | 1 | 15 |
| si:dkey-51a16.9 | 0.32959 | -0.1127723 | 0.028 | 0.087 | 1 | 15 |
| smarca5 | 0.32959 | -0.12064676 | 0.028 | 0.087 | 1 | 15 |
| syne2b | 0.32959 | -0.11606393 | 0.028 | 0.087 | 1 | 15 |
| tctn1 | 0.32959 | -0.11279953 | 0.028 | 0.087 | 1 | 15 |
| tgfbrap1 | 0.32959 | -0.19665922 | 0.028 | 0.087 | 1 | 15 |
| thrb | 0.32959 | -0.10968261 | 0.028 | 0.087 | 1 | 15 |
| tlr3 | 0.32959 | -0.11355657 | 0.028 | 0.087 | 1 | 15 |
| tmem205 | 0.32959 | -0.13705889 | 0.028 | 0.087 | 1 | 15 |
| tmem43 | 0.32959 | -0.11959552 | 0.028 | 0.087 | 1 | 15 |
| tnfsf10l | 0.32959 | -0.10497667 | 0.028 | 0.087 | 1 | 15 |

| | | | | | | |
|---|---|---|---|---|---|---|
| tpbga | 0.32959 | -0.19122132 | 0.028 | 0.087 | 1 | 15 |
| trim8a | 0.32959 | -0.10384932 | 0.028 | 0.087 | 1 | 15 |
| trip11 | 0.32959 | -0.10544664 | 0.028 | 0.087 | 1 | 15 |
| tubgcp3 | 0.32959 | -0.1099775 | 0.028 | 0.087 | 1 | 15 |
| ugp2b | 0.32959 | -0.1635816 | 0.028 | 0.087 | 1 | 15 |
| vars | 0.32959 | -0.1514817 | 0.028 | 0.087 | 1 | 15 |
| yme1l1b | 0.32959 | -0.17833133 | 0.028 | 0.087 | 1 | 15 |
| zbtb17 | 0.32959 | -0.1503241 | 0.028 | 0.087 | 1 | 15 |
| zfpm2b | 0.32959 | -0.10743453 | 0.028 | 0.087 | 1 | 15 |
| zfx | 0.32959 | -0.12115956 | 0.028 | 0.087 | 1 | 15 |
| zgc:77739 | 0.32959 | -0.10840272 | 0.028 | 0.087 | 1 | 15 |
| znf1159 | 0.32959 | -0.11941987 | 0.028 | 0.087 | 1 | 15 |
| CABZ01074397.1 | 0.32959 | -0.13587329 | 0.028 | 0.087 | 1 | 15 |
| CABZ01077220.2 | 0.32959 | -0.10231642 | 0.028 | 0.087 | 1 | 15 |
| LOC103911445 | 0.32959 | -0.11504571 | 0.028 | 0.087 | 1 | 15 |
| XLOC-004556 | 0.32959 | -0.15045749 | 0.028 | 0.087 | 1 | 15 |
| adcy7 | 0.32959 | -0.17191161 | 0.028 | 0.087 | 1 | 15 |
| alg12 | 0.32959 | -0.12410544 | 0.028 | 0.087 | 1 | 15 |
| ankib1a | 0.32959 | -0.10592032 | 0.028 | 0.087 | 1 | 15 |
| arhgap17b | 0.32959 | -0.11644279 | 0.028 | 0.087 | 1 | 15 |
| atp11a | 0.32959 | -0.22896612 | 0.028 | 0.087 | 1 | 15 |
| cyfip2 | 0.32959 | -0.13976524 | 0.028 | 0.087 | 1 | 15 |
| elovl1a | 0.32959 | -0.19444602 | 0.028 | 0.087 | 1 | 15 |
| hdac6 | 0.32959 | -0.10334502 | 0.028 | 0.087 | 1 | 15 |
| ift46 | 0.32959 | -0.1195264 | 0.028 | 0.087 | 1 | 15 |
| ikzf4 | 0.32959 | -0.10480189 | 0.028 | 0.087 | 1 | 15 |
| mthfd1l | 0.32959 | -0.15703118 | 0.028 | 0.087 | 1 | 15 |
| mtrex | 0.32959 | -0.12920433 | 0.028 | 0.087 | 1 | 15 |
| nfyal | 0.32959 | -0.15123223 | 0.028 | 0.087 | 1 | 15 |
| ribc1 | 0.32959 | -0.15796352 | 0.028 | 0.087 | 1 | 15 |
| senp6a | 0.32959 | -0.15737697 | 0.028 | 0.087 | 1 | 15 |
| sept8a | 0.32959 | -0.12232836 | 0.028 | 0.087 | 1 | 15 |
| si:dkeyp-38g8.5 | 0.32959 | -0.17669363 | 0.028 | 0.087 | 1 | 15 |
| slc35e1 | 0.32959 | -0.12882523 | 0.028 | 0.087 | 1 | 15 |
| sprydd7b | 0.32959 | -0.13010107 | 0.028 | 0.087 | 1 | 15 |
| srebf2 | 0.32959 | -0.13228796 | 0.028 | 0.087 | 1 | 15 |
| tbc1d23 | 0.32959 | -0.13590245 | 0.028 | 0.087 | 1 | 15 |
| tmem184ba | 0.32959 | -0.10309152 | 0.028 | 0.087 | 1 | 15 |
| tmem238a | 0.32959 | -0.16977204 | 0.028 | 0.087 | 1 | 15 |
| tmem9 | 0.32959 | -0.18453453 | 0.028 | 0.087 | 1 | 15 |
| tnfaip1 | 0.32959 | -0.15584127 | 0.028 | 0.087 | 1 | 15 |
| tp53inp2 | 0.32959 | -0.17705544 | 0.028 | 0.087 | 1 | 15 |
| vkorc1 | 0.32959 | -0.1246884 | 0.028 | 0.087 | 1 | 15 |
| zfyve9a | 0.32959 | -0.130509 | 0.028 | 0.087 | 1 | 15 |
| zgc:63587 | 0.32959 | -0.18929199 | 0.028 | 0.087 | 1 | 15 |
| zgc:77158 | 0.32959 | -0.11952925 | 0.028 | 0.087 | 1 | 15 |
| zmat5 | 0.32959 | -0.19087223 | 0.028 | 0.087 | 1 | 15 |
| bcorl1 | 0.32959 | -0.11604045 | 0.028 | 0.087 | 1 | 15 |

| | | | | | | |
|---|---|---|---|---|---|---|
| coasy | 0.32959 | -0.15711226 | 0.028 | 0.087 | 1 | 15 |
| hsdl1 | 0.32959 | -0.15464649 | 0.028 | 0.087 | 1 | 15 |
| plpp6 | 0.32959 | -0.15361534 | 0.028 | 0.087 | 1 | 15 |
| si:ch211-210c8.6 | 0.32959 | -0.10787146 | 0.028 | 0.087 | 1 | 15 |
| them6 | 0.32959 | -0.10791483 | 0.028 | 0.087 | 1 | 15 |
| CR847953.1 | 0.32959 | -0.14870529 | 0.028 | 0.087 | 1 | 15 |
| acvr2ba | 0.32959 | -0.11330936 | 0.028 | 0.087 | 1 | 15 |
| ankrd10b | 0.32959 | -0.12561638 | 0.028 | 0.087 | 1 | 15 |
| arid3b | 0.32959 | -0.12924086 | 0.028 | 0.087 | 1 | 15 |
| cachd1 | 0.32959 | -0.15834217 | 0.028 | 0.087 | 1 | 15 |
| coq6 | 0.32959 | -0.10238471 | 0.028 | 0.087 | 1 | 15 |
| dync1li1 | 0.32959 | -0.12019524 | 0.028 | 0.087 | 1 | 15 |
| evi5b | 0.32959 | -0.13523436 | 0.028 | 0.087 | 1 | 15 |
| ice1 | 0.32959 | -0.12640576 | 0.028 | 0.087 | 1 | 15 |
| ints2 | 0.32959 | -0.18026641 | 0.028 | 0.087 | 1 | 15 |
| lurap1 | 0.32959 | -0.15102323 | 0.028 | 0.087 | 1 | 15 |
| nckipsd | 0.32959 | -0.10950706 | 0.028 | 0.087 | 1 | 15 |
| prkd3 | 0.32959 | -0.1391165 | 0.028 | 0.087 | 1 | 15 |
| rnd3b | 0.32959 | -0.17625242 | 0.028 | 0.087 | 1 | 15 |
| s1pr5a | 0.32959 | -0.14688427 | 0.028 | 0.087 | 1 | 15 |
| sfxn2 | 0.32959 | -0.11445486 | 0.028 | 0.087 | 1 | 15 |
| si:dkey-102f14.5 | 0.32959 | -0.12150248 | 0.028 | 0.087 | 1 | 15 |
| si:dkey-66i24.9 | 0.32959 | -0.12012335 | 0.028 | 0.087 | 1 | 15 |
| slc7a6 | 0.32959 | -0.11334496 | 0.028 | 0.087 | 1 | 15 |
| snrnp25 | 0.32959 | -0.10229554 | 0.028 | 0.087 | 1 | 15 |
| stard8 | 0.32959 | -0.12750672 | 0.028 | 0.087 | 1 | 15 |
| tmem17 | 0.32959 | -0.1183877 | 0.028 | 0.087 | 1 | 15 |
| tmem63a | 0.32959 | -0.10586729 | 0.028 | 0.087 | 1 | 15 |
| tyms | 0.32959 | -0.10020306 | 0.028 | 0.087 | 1 | 15 |
| wdr26b | 0.32959 | -0.14798114 | 0.028 | 0.087 | 1 | 15 |
| AL935174.5 | 0.32959 | -0.13068159 | 0.028 | 0.087 | 1 | 15 |
| LOC103909810 | 0.32959 | -0.13044682 | 0.028 | 0.087 | 1 | 15 |
| elk1 | 0.32959 | -0.15132753 | 0.028 | 0.087 | 1 | 15 |
| gcat | 0.32959 | -0.13542673 | 0.028 | 0.087 | 1 | 15 |
| igf1ra | 0.32959 | -0.17110608 | 0.028 | 0.087 | 1 | 15 |
| lamb2 | 0.32959 | -0.14901291 | 0.028 | 0.087 | 1 | 15 |
| mark2a | 0.32959 | -0.10308126 | 0.028 | 0.087 | 1 | 15 |
| mrtfbb | 0.32959 | -0.13960878 | 0.028 | 0.087 | 1 | 15 |
| nadkb | 0.32959 | -0.13371484 | 0.028 | 0.087 | 1 | 15 |
| phlpp1 | 0.32959 | -0.13077577 | 0.028 | 0.087 | 1 | 15 |
| pwwp2a | 0.32959 | -0.15229042 | 0.028 | 0.087 | 1 | 15 |
| rdh12l | 0.32959 | -0.18370897 | 0.028 | 0.087 | 1 | 15 |
| rexo1 | 0.32959 | -0.13873202 | 0.028 | 0.087 | 1 | 15 |
| rprd2b | 0.32959 | -0.11194287 | 0.028 | 0.087 | 1 | 15 |
| si:ch211-218g4.2 | 0.32959 | -0.12514046 | 0.028 | 0.087 | 1 | 15 |
| si:dkey-16j16.4 | 0.32959 | -0.11320519 | 0.028 | 0.087 | 1 | 15 |
| hs3st1l2 | 0.32969 | -0.24441502 | 0.056 | 0.174 | 1 | 15 |
| tfap4 | 0.3305 | -0.21349681 | 0.083 | 0.217 | 1 | 15 |

| | | | | | | |
|---|---|---|---|---|---|---|
| serpina10a | 0.33095 | -0.23949411 | 0.056 | 0.174 | 1 | 15 |
| cct5 | 0.33148 | 0.265899109 | 0.639 | 0.478 | 1 | 15 |
| copg2 | 0.33266 | -0.27220666 | 0.194 | 0.348 | 1 | 15 |
| fam107b | 0.33271 | 0.29216429 | 0.278 | 0.13 | 1 | 15 |
| itga9 | 0.33281 | -0.12886948 | 0.25 | 0.261 | 1 | 15 |
| rab13 | 0.33298 | 0.121742324 | 0.194 | 0.087 | 1 | 15 |
| ap2s1 | 0.3337 | 0.237962704 | 0.278 | 0.13 | 1 | 15 |
| pum2 | 0.33405 | -0.22689792 | 0.056 | 0.13 | 1 | 15 |
| greb1 | 0.33456 | 0.249937759 | 0.222 | 0.087 | 1 | 15 |
| utp3 | 0.33483 | 0.13216329 | 0.194 | 0.087 | 1 | 15 |
| sec61b | 0.33577 | -0.23098493 | 0.611 | 0.783 | 1 | 15 |
| dvl3a | 0.3358 | -0.21357609 | 0.056 | 0.174 | 1 | 15 |
| mlec | 0.33581 | 0.181964132 | 0.139 | 0.043 | 1 | 15 |
| seh1l | 0.33589 | -0.24855154 | 0.111 | 0.261 | 1 | 15 |
| chd6 | 0.33593 | 0.222725977 | 0.333 | 0.174 | 1 | 15 |
| ppm1aa | 0.33593 | -0.25477594 | 0.083 | 0.217 | 1 | 15 |
| cd151 | 0.33611 | -0.30491913 | 0.222 | 0.348 | 1 | 15 |
| si:ch73-111k22.2 | 0.33623 | -0.21753474 | 0.083 | 0.217 | 1 | 15 |
| selenof | 0.33637 | 0.156817326 | 0.889 | 0.739 | 1 | 15 |
| emd | 0.33681 | -0.24434563 | 0.25 | 0.435 | 1 | 15 |
| gli2a | 0.33691 | -0.23480898 | 0.056 | 0.174 | 1 | 15 |
| jcada | 0.33699 | -0.20724852 | 0.194 | 0.261 | 1 | 15 |
| arhgap5 | 0.33707 | 0.266568813 | 0.278 | 0.13 | 1 | 15 |
| rpl5a | 0.33746 | 0.160494497 | 0.889 | 0.739 | 1 | 15 |
| sar1b | 0.33754 | -0.26529758 | 0.25 | 0.435 | 1 | 15 |
| gpx1a | 0.33754 | -0.2093942 | 0.111 | 0.261 | 1 | 15 |
| rnf19a | 0.33801 | 0.192495064 | 0.111 | 0.043 | 1 | 15 |
| kmt2ca | 0.33834 | -0.19902577 | 0.056 | 0.13 | 1 | 15 |
| gemin2 | 0.3384 | 0.226389876 | 0.139 | 0.043 | 1 | 15 |
| si:ch211-222k6.3 | 0.33851 | -0.19877525 | 0.056 | 0.174 | 1 | 15 |
| znf654 | 0.33857 | -0.17527 | 0.056 | 0.13 | 1 | 15 |
| gabarapl2 | 0.3386 | 0.182260387 | 0.25 | 0.174 | 1 | 15 |
| zmp:0000000711 | 0.33944 | 0.227485031 | 0.194 | 0.087 | 1 | 15 |
| hs2st1b | 0.33962 | -0.18689643 | 0.056 | 0.174 | 1 | 15 |
| atp6v1e1b | 0.33989 | -0.23719688 | 0.139 | 0.261 | 1 | 15 |
| gria2a | 0.34018 | 0.176172767 | 0.111 | 0.043 | 1 | 15 |
| map3k7 | 0.34035 | -0.13933595 | 0.056 | 0.087 | 1 | 15 |
| nectin1b | 0.34051 | 0.103645316 | 0.333 | 0.217 | 1 | 15 |
| myo1b | 0.34056 | 0.206290639 | 0.139 | 0.043 | 1 | 15 |
| cald1a | 0.34065 | -0.26532012 | 0.528 | 0.696 | 1 | 15 |
| uqcc2 | 0.34085 | -0.23642676 | 0.167 | 0.304 | 1 | 15 |
| ccni | 0.34089 | 0.131703272 | 0.5 | 0.348 | 1 | 15 |
| aplp2 | 0.34089 | -0.2237182 | 0.222 | 0.391 | 1 | 15 |
| znf827 | 0.34212 | -0.2244888 | 0.056 | 0.174 | 1 | 15 |
| znf622 | 0.34231 | 0.13141436 | 0.139 | 0.043 | 1 | 15 |
| faub | 0.34324 | 0.122997709 | 0.194 | 0.174 | 1 | 15 |
| ythdf2 | 0.34343 | -0.21433709 | 0.139 | 0.217 | 1 | 15 |
| lrp5.1 | 0.34347 | -0.23812893 | 0.056 | 0.174 | 1 | 15 |

| | | | | | | |
|---|---|---|---|---|---|---|
| pola1 | 0.34359 | 0.172798746 | 0.111 | 0.043 | 1 | 15 |
| tusc3 | 0.34399 | 0.154683902 | 0.25 | 0.13 | 1 | 15 |
| brd1b | 0.34404 | 0.190095343 | 0.139 | 0.043 | 1 | 15 |
| commd8 | 0.34424 | -0.22214903 | 0.111 | 0.217 | 1 | 15 |
| ppp1r37 | 0.34471 | 0.182246878 | 0.111 | 0.043 | 1 | 15 |
| zgc:92335 | 0.34471 | 0.157676843 | 0.139 | 0.087 | 1 | 15 |
| ubiad1 | 0.34505 | -0.14403896 | 0.056 | 0.174 | 1 | 15 |
| suclg2 | 0.34536 | -0.1893604 | 0.056 | 0.174 | 1 | 15 |
| znrf3 | 0.34543 | -0.17154956 | 0.056 | 0.174 | 1 | 15 |
| nckap1 | 0.34576 | 0.196456564 | 0.167 | 0.087 | 1 | 15 |
| eif5 | 0.34583 | 0.268250192 | 0.444 | 0.304 | 1 | 15 |
| ier3ip1 | 0.34592 | 0.262860263 | 0.556 | 0.391 | 1 | 15 |
| acp1 | 0.34596 | 0.211852916 | 0.222 | 0.087 | 1 | 15 |
| chd3 | 0.34644 | -0.27595525 | 0.389 | 0.565 | 1 | 15 |
| krtcap2 | 0.34661 | -0.22099845 | 0.528 | 0.609 | 1 | 15 |
| magi1b | 0.34684 | 0.143820721 | 0.194 | 0.087 | 1 | 15 |
| dedd | 0.34719 | -0.1743177 | 0.111 | 0.174 | 1 | 15 |
| znf207b | 0.34825 | 0.224327163 | 0.222 | 0.087 | 1 | 15 |
| lin37 | 0.34843 | -0.14752782 | 0.056 | 0.087 | 1 | 15 |
| mplkip | 0.34852 | 0.204806696 | 0.194 | 0.087 | 1 | 15 |
| exoc6 | 0.34876 | -0.24010909 | 0.083 | 0.217 | 1 | 15 |
| cflara | 0.34901 | -0.23516328 | 0.083 | 0.217 | 1 | 15 |
| si:ch211-1a19.3 | 0.34924 | -0.22573848 | 0.056 | 0.174 | 1 | 15 |
| phlda2 | 0.34985 | -0.23121562 | 0.139 | 0.217 | 1 | 15 |
| gnptab | 0.34994 | -0.25814112 | 0.056 | 0.174 | 1 | 15 |
| jupa | 0.35 | -0.1431748 | 0.139 | 0.261 | 1 | 15 |
| waplb | 0.3505 | 0.160646112 | 0.139 | 0.043 | 1 | 15 |
| nupr1a | 0.3516 | 0.209545235 | 0.222 | 0.087 | 1 | 15 |
| psmc6 | 0.35191 | -0.25833353 | 0.222 | 0.391 | 1 | 15 |
| ubc | 0.35229 | 0.157316995 | 0.917 | 0.783 | 1 | 15 |
| esr2a | 0.35234 | 0.165596156 | 0.139 | 0.043 | 1 | 15 |
| stk40 | 0.35261 | -0.17580277 | 0.056 | 0.174 | 1 | 15 |
| scube1 | 0.35292 | -0.24990227 | 0.083 | 0.217 | 1 | 15 |
| eif4eb | 0.35292 | 0.203748837 | 0.139 | 0.043 | 1 | 15 |
| ran | 0.35294 | 0.134606076 | 0.917 | 0.783 | 1 | 15 |
| ywhaz | 0.35341 | 0.146305189 | 0.917 | 0.783 | 1 | 15 |
| si:ch211-14a17.11 | 0.35357 | -0.19613721 | 0.139 | 0.217 | 1 | 15 |
| galnt11 | 0.35388 | -0.1997336 | 0.056 | 0.13 | 1 | 15 |
| ar | 0.3539 | 0.241024621 | 0.667 | 0.478 | 1 | 15 |
| rbpms2a | 0.35439 | -0.22248724 | 0.056 | 0.174 | 1 | 15 |
| emid1 | 0.35459 | -0.23811679 | 0.056 | 0.174 | 1 | 15 |
| sgce | 0.3547 | -0.10680872 | 0.167 | 0.174 | 1 | 15 |
| KIF2A | 0.35488 | -0.17441471 | 0.056 | 0.174 | 1 | 15 |
| kif1bp | 0.35497 | -0.22030484 | 0.056 | 0.174 | 1 | 15 |
| si:ch211-246m6.4 | 0.35521 | -0.25759453 | 0.278 | 0.391 | 1 | 15 |
| net1 | 0.35542 | -0.22378573 | 0.056 | 0.174 | 1 | 15 |
| unm-hu7910 | 0.35566 | -0.22050342 | 0.139 | 0.217 | 1 | 15 |
| bcas3 | 0.3563 | 0.140885129 | 0.139 | 0.043 | 1 | 15 |

| | | | | | | |
|---|---|---|---|---|---|---|
| nsrp1 | 0.35645 | -0.18216197 | 0.056 | 0.13 | 1 | 15 |
| rab6a | 0.35689 | 0.2326311 | 0.222 | 0.087 | 1 | 15 |
| tuba8l2 | 0.35692 | -0.16519442 | 0.083 | 0.13 | 1 | 15 |
| tfec | 0.357 | -0.11164502 | 0.083 | 0.087 | 1 | 15 |
| thop1 | 0.3571 | 0.125289163 | 0.111 | 0.087 | 1 | 15 |
| aamp | 0.35792 | -0.11832999 | 0.139 | 0.174 | 1 | 15 |
| znf593 | 0.35865 | 0.205411214 | 0.25 | 0.13 | 1 | 15 |
| gcn1 | 0.35877 | 0.203433034 | 0.139 | 0.043 | 1 | 15 |
| pard3bb | 0.35878 | 0.158047634 | 0.25 | 0.13 | 1 | 15 |
| ccnt1 | 0.35947 | -0.23998195 | 0.083 | 0.174 | 1 | 15 |
| hspa8 | 0.35976 | -0.1203303 | 0.944 | 1 | 1 | 15 |
| cry1aa | 0.36 | 0.227640339 | 0.139 | 0.043 | 1 | 15 |
| arf2a | 0.36021 | 0.269583334 | 0.333 | 0.174 | 1 | 15 |
| PDPR | 0.36063 | 0.182128538 | 0.139 | 0.043 | 1 | 15 |
| fdx1 | 0.36115 | 0.193363595 | 0.222 | 0.087 | 1 | 15 |
| nkain4 | 0.36116 | -0.18088943 | 0.056 | 0.13 | 1 | 15 |
| rnasekb | 0.36148 | 0.166463893 | 0.111 | 0.043 | 1 | 15 |
| sav1 | 0.36208 | 0.150700224 | 0.139 | 0.043 | 1 | 15 |
| nipsnap2 | 0.36217 | 0.131333683 | 0.25 | 0.13 | 1 | 15 |
| mycbp2 | 0.36333 | 0.222959746 | 0.222 | 0.087 | 1 | 15 |
| znf1124 | 0.36343 | 0.233630906 | 0.222 | 0.087 | 1 | 15 |
| axin2 | 0.36364 | -0.12862153 | 0.139 | 0.174 | 1 | 15 |
| fbxl14b | 0.36373 | 0.141924519 | 0.139 | 0.087 | 1 | 15 |
| rpp14 | 0.36378 | 0.135907876 | 0.139 | 0.043 | 1 | 15 |
| lygl1 | 0.36381 | -0.12100381 | 0.194 | 0.217 | 1 | 15 |
| sgta | 0.3643 | 0.118635214 | 0.194 | 0.087 | 1 | 15 |
| elf2b | 0.36472 | -0.10211002 | 0.083 | 0.174 | 1 | 15 |
| clic4 | 0.36521 | -0.1432302 | 0.222 | 0.261 | 1 | 15 |
| ddt | 0.36589 | 0.249872409 | 0.278 | 0.13 | 1 | 15 |
| eif4e3 | 0.36594 | 0.231924818 | 0.194 | 0.087 | 1 | 15 |
| ppp1caa | 0.366 | -0.12536814 | 0.306 | 0.435 | 1 | 15 |
| pdia4 | 0.36633 | -0.25717747 | 0.111 | 0.217 | 1 | 15 |
| gstr | 0.36663 | 0.218962847 | 0.278 | 0.13 | 1 | 15 |
| cib1 | 0.36731 | 0.137880805 | 0.139 | 0.043 | 1 | 15 |
| tdh | 0.36734 | 0.12780822 | 0.139 | 0.043 | 1 | 15 |
| CABZ01064859.1 | 0.36736 | 0.192246834 | 0.139 | 0.043 | 1 | 15 |
| arpp19a | 0.36805 | 0.238149836 | 0.222 | 0.087 | 1 | 15 |
| ubxn4 | 0.36808 | 0.225015894 | 0.222 | 0.087 | 1 | 15 |
| ptbp1a | 0.36819 | -0.27797642 | 0.222 | 0.391 | 1 | 15 |
| st13 | 0.36829 | 0.10111941 | 0.333 | 0.217 | 1 | 15 |
| sec13 | 0.36891 | 0.190617041 | 0.139 | 0.043 | 1 | 15 |
| cdk9 | 0.36893 | 0.254357204 | 0.278 | 0.13 | 1 | 15 |
| si:ch211-87m7.2 | 0.36928 | -0.15486336 | 0.056 | 0.087 | 1 | 15 |
| gas6 | 0.36955 | 0.232877507 | 0.333 | 0.174 | 1 | 15 |
| eef1e1 | 0.36979 | 0.165576585 | 0.139 | 0.043 | 1 | 15 |
| rsl24d1 | 0.37003 | 0.191108096 | 0.833 | 0.739 | 1 | 15 |
| usp10 | 0.37016 | -0.24647435 | 0.083 | 0.174 | 1 | 15 |
| prkab1b | 0.37062 | 0.104684654 | 0.472 | 0.348 | 1 | 15 |

| | | | | | | |
|---|---|---|---|---|---|---|
| CABZ01084501.3 | 0.37114 | 0.298675305 | 0.139 | 0.043 | 1 | 15 |
| cbwd | 0.37173 | 0.159806017 | 0.139 | 0.043 | 1 | 15 |
| smc1al | 0.3721 | 0.208514149 | 0.361 | 0.261 | 1 | 15 |
| spsb4a | 0.37241 | -0.13871253 | 0.139 | 0.174 | 1 | 15 |
| ciapin1 | 0.37271 | 0.205171787 | 0.194 | 0.087 | 1 | 15 |
| rap1aa | 0.37293 | -0.13635091 | 0.222 | 0.261 | 1 | 15 |
| hikeshi | 0.37301 | 0.223386085 | 0.333 | 0.174 | 1 | 15 |
| rpe | 0.37302 | 0.217978455 | 0.222 | 0.087 | 1 | 15 |
| lysmd2 | 0.3731 | 0.221042957 | 0.222 | 0.087 | 1 | 15 |
| SMIM10 | 0.3734 | 0.189829333 | 0.222 | 0.087 | 1 | 15 |
| si:ch73-91k6.2 | 0.37342 | 0.236247629 | 0.194 | 0.087 | 1 | 15 |
| zc3h15 | 0.37421 | 0.201957262 | 0.222 | 0.087 | 1 | 15 |
| pgd | 0.37452 | 0.224993663 | 0.389 | 0.217 | 1 | 15 |
| CABZ01090373.1 | 0.37564 | 0.133518423 | 0.194 | 0.087 | 1 | 15 |
| tsr2 | 0.37576 | 0.221829021 | 0.194 | 0.087 | 1 | 15 |
| nop2 | 0.37585 | 0.229127794 | 0.222 | 0.087 | 1 | 15 |
| clic1 | 0.37603 | -0.21653348 | 0.139 | 0.217 | 1 | 15 |
| lamp1a | 0.37628 | 0.200818767 | 0.222 | 0.087 | 1 | 15 |
| psip1b | 0.37682 | 0.279782884 | 0.333 | 0.174 | 1 | 15 |
| si:dkey-56m19.5 | 0.377 | -0.12611537 | 0.333 | 0.348 | 1 | 15 |
| psap | 0.37704 | -0.16611199 | 0.444 | 0.478 | 1 | 15 |
| pnp5a | 0.3775 | 0.256037262 | 0.278 | 0.13 | 1 | 15 |
| med30 | 0.37759 | -0.17466943 | 0.083 | 0.13 | 1 | 15 |
| dkc1 | 0.37768 | 0.207721803 | 0.278 | 0.13 | 1 | 15 |
| snx17 | 0.37792 | 0.283209784 | 0.389 | 0.217 | 1 | 15 |
| hspa4b | 0.37889 | -0.1186398 | 0.111 | 0.217 | 1 | 15 |
| zc3h18 | 0.37926 | 0.142817758 | 0.139 | 0.043 | 1 | 15 |
| wdr12 | 0.37956 | 0.190308073 | 0.194 | 0.087 | 1 | 15 |
| ptges3a | 0.37987 | 0.188809786 | 0.167 | 0.087 | 1 | 15 |
| CR391940.1 | 0.37987 | -0.1267135 | 0.056 | 0.087 | 1 | 15 |
| tinagl1 | 0.38036 | -0.17124625 | 0.056 | 0.087 | 1 | 15 |
| ugt5b2 | 0.38043 | -0.22572347 | 0.056 | 0.13 | 1 | 15 |
| tmem150ab | 0.38095 | 0.188684489 | 0.139 | 0.043 | 1 | 15 |
| atp6v1g1 | 0.38266 | 0.219799443 | 0.333 | 0.174 | 1 | 15 |
| plod2 | 0.38312 | -0.28071012 | 0.194 | 0.348 | 1 | 15 |
| furina | 0.38348 | -0.22013682 | 0.083 | 0.174 | 1 | 15 |
| ier5 | 0.38369 | -0.13477277 | 0.417 | 0.565 | 1 | 15 |
| abracl | 0.38398 | -0.12827042 | 0.111 | 0.13 | 1 | 15 |
| snrpc | 0.38493 | -0.24656234 | 0.167 | 0.304 | 1 | 15 |
| tcea1 | 0.38639 | 0.216676864 | 0.333 | 0.174 | 1 | 15 |
| nktr | 0.3864 | 0.243790974 | 0.333 | 0.174 | 1 | 15 |
| hint1 | 0.38651 | 0.239650653 | 0.556 | 0.435 | 1 | 15 |
| atp6v0b | 0.38705 | -0.15118376 | 0.306 | 0.348 | 1 | 15 |
| sdhaf3 | 0.38738 | 0.155674621 | 0.139 | 0.087 | 1 | 15 |
| sdha | 0.38739 | 0.237080887 | 0.333 | 0.174 | 1 | 15 |
| ndufb7 | 0.38757 | 0.210885342 | 0.528 | 0.348 | 1 | 15 |
| cbr1 | 0.38764 | -0.15245622 | 0.056 | 0.087 | 1 | 15 |
| her9 | 0.38783 | -0.10638858 | 0.083 | 0.087 | 1 | 15 |

| | | | | | | |
|---|---|---|---|---|---|---|
| irf9 | 0.38914 | 0.208220488 | 0.139 | 0.043 | 1 | 15 |
| ddx24 | 0.38938 | 0.253293915 | 0.333 | 0.174 | 1 | 15 |
| clcn7 | 0.39003 | 0.194356405 | 0.139 | 0.043 | 1 | 15 |
| si:dkey-23i12.7 | 0.39073 | -0.22438114 | 0.056 | 0.13 | 1 | 15 |
| eci1 | 0.39145 | 0.233735484 | 0.333 | 0.174 | 1 | 15 |
| cfl1l | 0.39195 | -0.16841256 | 0.056 | 0.13 | 1 | 15 |
| gpia | 0.39234 | 0.188437088 | 0.194 | 0.087 | 1 | 15 |
| snx27b | 0.39326 | 0.184988789 | 0.139 | 0.043 | 1 | 15 |
| mcrip1 | 0.3936 | 0.207522646 | 0.278 | 0.13 | 1 | 15 |
| gosr2 | 0.39378 | 0.240149136 | 0.278 | 0.13 | 1 | 15 |
| cs | 0.39407 | -0.2049754 | 0.167 | 0.261 | 1 | 15 |
| slc38a2 | 0.39526 | 0.241206772 | 0.25 | 0.13 | 1 | 15 |
| stt3b | 0.39535 | 0.205158579 | 0.25 | 0.13 | 1 | 15 |
| chic2 | 0.39563 | -0.22819593 | 0.111 | 0.217 | 1 | 15 |
| copa | 0.39598 | -0.12715799 | 0.194 | 0.217 | 1 | 15 |
| dad1 | 0.39679 | -0.17340244 | 0.778 | 0.87 | 1 | 15 |
| mmp2 | 0.3971 | 0.205184882 | 0.75 | 0.652 | 1 | 15 |
| MRPS12 | 0.39768 | -0.2502627 | 0.167 | 0.304 | 1 | 15 |
| slc7a3a | 0.39769 | 0.127842817 | 0.667 | 0.522 | 1 | 15 |
| abhd17c | 0.39788 | 0.168710487 | 0.139 | 0.043 | 1 | 15 |
| serpinh1b | 0.39818 | -0.16473669 | 0.056 | 0.087 | 1 | 15 |
| si:ch211-74m13.3 | 0.3983 | -0.19439195 | 0.167 | 0.304 | 1 | 15 |
| cox7a2a | 0.3986 | 0.165785301 | 0.611 | 0.435 | 1 | 15 |
| prelid3b | 0.39872 | -0.1870574 | 0.278 | 0.348 | 1 | 15 |
| si:dkey-19a16.7 | 0.39909 | -0.27017174 | 0.167 | 0.304 | 1 | 15 |
| mtdha | 0.40009 | -0.16412671 | 0.389 | 0.435 | 1 | 15 |
| prr12a | 0.40059 | -0.12818179 | 0.056 | 0.087 | 1 | 15 |
| cnpy2 | 0.4007 | 0.167000585 | 0.139 | 0.043 | 1 | 15 |
| pigb | 0.40128 | -0.13204877 | 0.056 | 0.087 | 1 | 15 |
| podxl | 0.40175 | -0.13745421 | 0.806 | 0.826 | 1 | 15 |
| syap1 | 0.40217 | -0.18499114 | 0.222 | 0.304 | 1 | 15 |
| ubqln4 | 0.4025 | -0.12463503 | 0.111 | 0.217 | 1 | 15 |
| ranbp1 | 0.40269 | 0.236518675 | 0.5 | 0.348 | 1 | 15 |
| mbd3b | 0.4027 | 0.20442473 | 0.139 | 0.043 | 1 | 15 |
| nucks1a | 0.4027 | 0.199561603 | 0.611 | 0.435 | 1 | 15 |
| rilpl2 | 0.40319 | 0.199883918 | 0.111 | 0.043 | 1 | 15 |
| polr3f | 0.40336 | 0.190903867 | 0.139 | 0.043 | 1 | 15 |
| puraa | 0.40364 | 0.149570317 | 0.25 | 0.13 | 1 | 15 |
| acads | 0.40449 | 0.165261284 | 0.139 | 0.043 | 1 | 15 |
| picalmb | 0.40449 | 0.176525578 | 0.139 | 0.043 | 1 | 15 |
| rpl29 | 0.40507 | 0.119296797 | 0.972 | 0.913 | 1 | 15 |
| cyc1 | 0.40547 | -0.23834166 | 0.361 | 0.522 | 1 | 15 |
| ppp3r1a | 0.40572 | 0.19186315 | 0.139 | 0.043 | 1 | 15 |
| ap1s2 | 0.40637 | -0.22816956 | 0.361 | 0.478 | 1 | 15 |
| atp5f1c | 0.40682 | 0.146422344 | 0.639 | 0.478 | 1 | 15 |
| wt1a | 0.407 | -0.27674335 | 0.333 | 0.478 | 1 | 15 |
| eif4ea | 0.40729 | 0.184524263 | 0.139 | 0.043 | 1 | 15 |
| ube2z | 0.40742 | 0.179087004 | 0.139 | 0.043 | 1 | 15 |

| | | | | | | |
|---|---|---|---|---|---|---|
| pggt1b | 0.40742 | 0.150253862 | 0.111 | 0.043 | 1 | 15 |
| zgc:92140 | 0.40756 | 0.11817532 | 0.361 | 0.348 | 1 | 15 |
| ebna1bp2 | 0.40785 | -0.10947645 | 0.111 | 0.217 | 1 | 15 |
| trappc3 | 0.40793 | 0.155702206 | 0.194 | 0.087 | 1 | 15 |
| atp5meb | 0.40806 | 0.178662711 | 0.694 | 0.522 | 1 | 15 |
| nsfl1c | 0.40823 | 0.2655307 | 0.361 | 0.217 | 1 | 15 |
| oaz2b | 0.40828 | 0.196632348 | 0.139 | 0.043 | 1 | 15 |
| timm8a | 0.40829 | 0.128259311 | 0.167 | 0.13 | 1 | 15 |
| hs6st2 | 0.40855 | 0.163448462 | 0.139 | 0.043 | 1 | 15 |
| plxna1a | 0.40919 | 0.195176814 | 0.194 | 0.087 | 1 | 15 |
| si:ch211-196l7.4 | 0.40965 | -0.21950657 | 0.639 | 0.783 | 1 | 15 |
| cdc73 | 0.40975 | 0.167176945 | 0.111 | 0.043 | 1 | 15 |
| snrpe | 0.40988 | 0.24333471 | 0.333 | 0.217 | 1 | 15 |
| stx6 | 0.41013 | 0.178457267 | 0.139 | 0.043 | 1 | 15 |
| phf11 | 0.41017 | 0.151762381 | 0.139 | 0.043 | 1 | 15 |
| rps25 | 0.41078 | 0.126320267 | 0.972 | 0.913 | 1 | 15 |
| sdc2 | 0.41119 | -0.19680871 | 0.056 | 0.13 | 1 | 15 |
| hmgn2 | 0.41129 | -0.34153517 | 0.472 | 0.609 | 1 | 15 |
| ddx39ab | 0.41142 | 0.114551779 | 0.389 | 0.261 | 1 | 15 |
| si:ch1073-325m22.2 | 0.41156 | 0.212972891 | 0.611 | 0.435 | 1 | 15 |
| atp1b1a | 0.41163 | -0.15137888 | 0.833 | 0.87 | 1 | 15 |
| smarce1 | 0.41179 | 0.222971793 | 0.556 | 0.435 | 1 | 15 |
| sntb2 | 0.41189 | -0.18196595 | 0.056 | 0.13 | 1 | 15 |
| parp1 | 0.41208 | 0.157580234 | 0.167 | 0.13 | 1 | 15 |
| gng10 | 0.41211 | -0.19053735 | 0.056 | 0.13 | 1 | 15 |
| alg5 | 0.4122 | -0.2307335 | 0.083 | 0.174 | 1 | 15 |
| adh5 | 0.41228 | -0.13019611 | 0.583 | 0.739 | 1 | 15 |
| si:dkey-33c12.4 | 0.41277 | 0.237646493 | 0.306 | 0.174 | 1 | 15 |
| per3 | 0.41277 | -0.22694588 | 0.111 | 0.217 | 1 | 15 |
| zgc:66472 | 0.41319 | 0.183957377 | 0.139 | 0.043 | 1 | 15 |
| ndrg1a | 0.41348 | -0.18400875 | 0.083 | 0.174 | 1 | 15 |
| gchfr | 0.41381 | 0.20672331 | 0.222 | 0.174 | 1 | 15 |
| fbxl14a | 0.41469 | 0.148790691 | 0.139 | 0.043 | 1 | 15 |
| pi15a | 0.41472 | 0.180560681 | 0.806 | 0.739 | 1 | 15 |
| actn4 | 0.41484 | -0.15324133 | 0.139 | 0.261 | 1 | 15 |
| uqcr10 | 0.4151 | 0.224521384 | 0.611 | 0.435 | 1 | 15 |
| ch25hl3 | 0.41596 | -0.14066977 | 0.222 | 0.348 | 1 | 15 |
| lypd6 | 0.41616 | 0.1069685 | 0.111 | 0.087 | 1 | 15 |
| fam192a | 0.41705 | -0.19047208 | 0.056 | 0.13 | 1 | 15 |
| calr3a | 0.4175 | -0.20116297 | 0.361 | 0.435 | 1 | 15 |
| phax | 0.41759 | 0.18554205 | 0.111 | 0.043 | 1 | 15 |
| yy1b | 0.41859 | 0.151850031 | 0.194 | 0.087 | 1 | 15 |
| fibpa | 0.41912 | 0.189363923 | 0.139 | 0.043 | 1 | 15 |
| npm1a | 0.41993 | 0.161230566 | 0.556 | 0.391 | 1 | 15 |
| arrdc3a | 0.42017 | 0.263155556 | 0.556 | 0.391 | 1 | 15 |
| ugt1b5 | 0.42193 | -0.25240224 | 0.194 | 0.348 | 1 | 15 |
| rras | 0.42225 | -0.18756979 | 0.306 | 0.391 | 1 | 15 |
| bmpr2a | 0.42259 | 0.166661561 | 0.25 | 0.13 | 1 | 15 |

| | | | | | | |
|---|---|---|---|---|---|---|
| nap1l4b | 0.42424 | 0.170590471 | 0.139 | 0.043 | 1 | 15 |
| cx43.4 | 0.42483 | -0.26643702 | 0.167 | 0.304 | 1 | 15 |
| susd6 | 0.42501 | 0.164208805 | 0.194 | 0.13 | 1 | 15 |
| zgc:92606 | 0.42512 | 0.180388772 | 0.194 | 0.087 | 1 | 15 |
| gng7 | 0.42523 | -0.21658557 | 0.194 | 0.304 | 1 | 15 |
| ylpm1 | 0.42594 | -0.22339653 | 0.139 | 0.217 | 1 | 15 |
| tspan14 | 0.42601 | -0.23083852 | 0.194 | 0.348 | 1 | 15 |
| FP016056.1 | 0.42761 | 0.110537506 | 0.167 | 0.087 | 1 | 15 |
| rab18b | 0.42787 | -0.19603231 | 0.139 | 0.217 | 1 | 15 |
| si:rp71-45k5.4 | 0.42796 | 0.182361904 | 0.361 | 0.217 | 1 | 15 |
| wbp1lb | 0.4285 | -0.21665661 | 0.111 | 0.217 | 1 | 15 |
| tbl2 | 0.42983 | -0.17193337 | 0.083 | 0.13 | 1 | 15 |
| tefa | 0.43028 | -0.25531605 | 0.222 | 0.348 | 1 | 15 |
| zgc:110158 | 0.43039 | -0.11440418 | 0.083 | 0.174 | 1 | 15 |
| raver2 | 0.43073 | -0.12934292 | 0.083 | 0.174 | 1 | 15 |
| pum3 | 0.43128 | 0.162254085 | 0.139 | 0.043 | 1 | 15 |
| kmt2bb | 0.43133 | 0.232648825 | 0.472 | 0.304 | 1 | 15 |
| cox15 | 0.43214 | 0.177696052 | 0.111 | 0.043 | 1 | 15 |
| gadd45ba | 0.43216 | -0.30485883 | 0.278 | 0.435 | 1 | 15 |
| psmg1 | 0.43229 | 0.175695107 | 0.278 | 0.217 | 1 | 15 |
| ifi35 | 0.43245 | 0.210966472 | 0.25 | 0.13 | 1 | 15 |
| tlk1b | 0.43287 | 0.21519835 | 0.25 | 0.13 | 1 | 15 |
| baz2a | 0.4329 | 0.140718094 | 0.139 | 0.043 | 1 | 15 |
| zbtb10 | 0.4329 | 0.149459059 | 0.111 | 0.043 | 1 | 15 |
| calm3a | 0.43304 | 0.183590639 | 0.139 | 0.043 | 1 | 15 |
| cox6c | 0.4332 | -0.21490829 | 0.556 | 0.652 | 1 | 15 |
| pbx1b | 0.43354 | -0.15476784 | 0.167 | 0.217 | 1 | 15 |
| rtf1 | 0.43356 | 0.183606253 | 0.306 | 0.174 | 1 | 15 |
| rnf213a | 0.43362 | -0.24507653 | 0.333 | 0.478 | 1 | 15 |
| pes | 0.43396 | 0.162006135 | 0.139 | 0.043 | 1 | 15 |
| lysmd4 | 0.43408 | 0.182601894 | 0.139 | 0.043 | 1 | 15 |
| mmp14b | 0.43415 | 0.168136615 | 0.139 | 0.043 | 1 | 15 |
| nxt2 | 0.43469 | -0.19494121 | 0.083 | 0.174 | 1 | 15 |
| gcsha | 0.43485 | -0.19789303 | 0.083 | 0.174 | 1 | 15 |
| adi1 | 0.43498 | 0.103050303 | 0.194 | 0.174 | 1 | 15 |
| fam53b | 0.43618 | -0.16463585 | 0.056 | 0.13 | 1 | 15 |
| mpc2 | 0.43645 | 0.214455152 | 0.306 | 0.174 | 1 | 15 |
| ankrd52a | 0.43674 | -0.14387968 | 0.139 | 0.174 | 1 | 15 |
| gnl3 | 0.43736 | 0.144275391 | 0.139 | 0.043 | 1 | 15 |
| cox14 | 0.43775 | 0.183012416 | 0.306 | 0.217 | 1 | 15 |
| hmbox1a | 0.43856 | 0.222784371 | 0.222 | 0.13 | 1 | 15 |
| zgc:66388 | 0.43887 | -0.20785806 | 0.111 | 0.217 | 1 | 15 |
| CABZ01113812.1 | 0.43899 | 0.184590573 | 0.111 | 0.043 | 1 | 15 |
| hkdc1 | 0.43919 | 0.16901654 | 0.139 | 0.043 | 1 | 15 |
| ifi30 | 0.4395 | -0.21325557 | 0.694 | 0.826 | 1 | 15 |
| psmd9 | 0.43951 | 0.153045664 | 0.111 | 0.043 | 1 | 15 |
| surf2 | 0.4396 | 0.170225326 | 0.167 | 0.087 | 1 | 15 |
| si:dkey-46a10.3 | 0.43991 | -0.14448232 | 0.167 | 0.217 | 1 | 15 |

| | | | | | | |
|---|---|---|---|---|---|---|
| pgs1 | 0.44132 | -0.15453977 | 0.083 | 0.13 | 1 | 15 |
| mgst1.1 | 0.44136 | 0.142605272 | 0.222 | 0.174 | 1 | 15 |
| dtd1 | 0.44152 | 0.234447386 | 0.194 | 0.087 | 1 | 15 |
| lrrc59 | 0.44156 | 0.121570884 | 0.139 | 0.043 | 1 | 15 |
| FO744854.1 | 0.44176 | -0.19865089 | 0.056 | 0.13 | 1 | 15 |
| mrpl4 | 0.44183 | 0.134532058 | 0.167 | 0.13 | 1 | 15 |
| rrbp1b | 0.44239 | -0.18629918 | 0.083 | 0.13 | 1 | 15 |
| atp11c | 0.44246 | 0.177958992 | 0.139 | 0.043 | 1 | 15 |
| usp9 | 0.4425 | 0.15384412 | 0.25 | 0.13 | 1 | 15 |
| lsm14ab | 0.44252 | 0.144145961 | 0.139 | 0.043 | 1 | 15 |
| nosip | 0.44265 | -0.19179631 | 0.083 | 0.174 | 1 | 15 |
| ube2e3 | 0.44276 | 0.118579464 | 0.611 | 0.478 | 1 | 15 |
| si:dkey-51e6.1 | 0.44351 | 0.218413351 | 0.25 | 0.13 | 1 | 15 |
| glmp | 0.4439 | 0.19572074 | 0.194 | 0.087 | 1 | 15 |
| foxp1b | 0.44409 | -0.20132909 | 0.056 | 0.13 | 1 | 15 |
| mkrn4 | 0.44422 | -0.1986769 | 0.083 | 0.174 | 1 | 15 |
| erp44 | 0.44678 | 0.234930085 | 0.361 | 0.217 | 1 | 15 |
| calub | 0.44679 | 0.182491695 | 0.139 | 0.043 | 1 | 15 |
| nansa | 0.44704 | 0.155378902 | 0.111 | 0.043 | 1 | 15 |
| chac2 | 0.44732 | 0.213837834 | 0.111 | 0.043 | 1 | 15 |
| trappc1 | 0.44735 | -0.22481925 | 0.361 | 0.522 | 1 | 15 |
| bace2 | 0.44748 | 0.168538201 | 0.139 | 0.043 | 1 | 15 |
| zc3h13 | 0.448 | -0.20610198 | 0.139 | 0.217 | 1 | 15 |
| fkbpl | 0.44924 | 0.158014403 | 0.111 | 0.043 | 1 | 15 |
| lancl1 | 0.44984 | 0.159072513 | 0.139 | 0.043 | 1 | 15 |
| anxa11b | 0.44986 | 0.119847411 | 0.472 | 0.348 | 1 | 15 |
| tmx1 | 0.45021 | 0.131532108 | 0.139 | 0.043 | 1 | 15 |
| eif3s10 | 0.45029 | -0.12492187 | 0.417 | 0.435 | 1 | 15 |
| naa38 | 0.45044 | 0.175917278 | 0.139 | 0.043 | 1 | 15 |
| slx1b | 0.45059 | -0.16931887 | 0.111 | 0.174 | 1 | 15 |
| slc43a2a | 0.45092 | -0.15071285 | 0.417 | 0.435 | 1 | 15 |
| brd8 | 0.45156 | 0.139779516 | 0.139 | 0.043 | 1 | 15 |
| mtap | 0.45162 | 0.186650245 | 0.25 | 0.13 | 1 | 15 |
| trmt61a | 0.45164 | 0.18236889 | 0.139 | 0.043 | 1 | 15 |
| col1a1b | 0.45203 | -0.15922081 | 0.278 | 0.304 | 1 | 15 |
| tcirg1b | 0.45206 | -0.20370844 | 0.083 | 0.174 | 1 | 15 |
| micall1a | 0.45209 | 0.141175827 | 0.139 | 0.043 | 1 | 15 |
| gtf2f2a | 0.45266 | -0.15790236 | 0.25 | 0.304 | 1 | 15 |
| otulinb | 0.45282 | 0.156842443 | 0.111 | 0.043 | 1 | 15 |
| tjp1a | 0.45286 | -0.21301013 | 0.167 | 0.261 | 1 | 15 |
| fbxl5 | 0.45313 | 0.133518775 | 0.139 | 0.043 | 1 | 15 |
| cxxc1a | 0.45314 | 0.174930766 | 0.111 | 0.043 | 1 | 15 |
| asph | 0.45342 | -0.25888303 | 0.167 | 0.304 | 1 | 15 |
| btr09 | 0.4536 | -0.1113388 | 0.083 | 0.174 | 1 | 15 |
| acbd6 | 0.45366 | 0.170764718 | 0.111 | 0.043 | 1 | 15 |
| nol11 | 0.45438 | 0.146707314 | 0.111 | 0.043 | 1 | 15 |
| zgc:91910 | 0.4547 | -0.17388204 | 0.583 | 0.652 | 1 | 15 |
| taf3 | 0.4548 | 0.204314511 | 0.194 | 0.087 | 1 | 15 |

| | | | | | | |
|---|---|---|---|---|---|---|
| zgc:123010 | 0.45584 | -0.20128253 | 0.111 | 0.174 | 1 | 15 |
| rmdn1 | 0.45613 | -0.22425303 | 0.25 | 0.348 | 1 | 15 |
| arhgdia | 0.45618 | -0.21860081 | 0.278 | 0.435 | 1 | 15 |
| pinx1 | 0.45621 | -0.15869462 | 0.139 | 0.217 | 1 | 15 |
| polr3h | 0.45637 | 0.175767917 | 0.194 | 0.087 | 1 | 15 |
| hp1bp3 | 0.45659 | -0.22065583 | 0.444 | 0.609 | 1 | 15 |
| crsp7 | 0.45688 | -0.19956826 | 0.056 | 0.13 | 1 | 15 |
| mgst3b | 0.4569 | -0.19835628 | 0.444 | 0.522 | 1 | 15 |
| sesn3 | 0.45691 | 0.149652206 | 0.111 | 0.043 | 1 | 15 |
| pim3 | 0.45762 | 0.184370323 | 0.139 | 0.043 | 1 | 15 |
| pgam1a | 0.45771 | -0.17256121 | 0.111 | 0.174 | 1 | 15 |
| aplp1 | 0.45782 | 0.160278482 | 0.139 | 0.043 | 1 | 15 |
| phf20l1 | 0.45826 | 0.168710461 | 0.139 | 0.043 | 1 | 15 |
| sash1a | 0.45834 | -0.12001506 | 0.056 | 0.087 | 1 | 15 |
| aimp1 | 0.45854 | -0.16272357 | 0.194 | 0.261 | 1 | 15 |
| klf11b | 0.45858 | 0.194516019 | 0.361 | 0.217 | 1 | 15 |
| pwwp2b | 0.45862 | 0.150434707 | 0.194 | 0.087 | 1 | 15 |
| ranbp10 | 0.45882 | 0.159494731 | 0.111 | 0.043 | 1 | 15 |
| gpalpp1 | 0.4591 | 0.160321332 | 0.139 | 0.043 | 1 | 15 |
| gpatch4 | 0.45955 | -0.14941632 | 0.139 | 0.174 | 1 | 15 |
| sft2d1 | 0.45958 | 0.149230966 | 0.139 | 0.043 | 1 | 15 |
| dock9b | 0.45958 | 0.15635117 | 0.139 | 0.043 | 1 | 15 |
| arcn1a | 0.45978 | 0.193964317 | 0.194 | 0.087 | 1 | 15 |
| sarnp | 0.45981 | -0.22386452 | 0.167 | 0.304 | 1 | 15 |
| fbrs | 0.45991 | -0.20274304 | 0.056 | 0.13 | 1 | 15 |
| phip | 0.45993 | 0.175857104 | 0.139 | 0.043 | 1 | 15 |
| ankrd12 | 0.46018 | 0.188681919 | 0.139 | 0.043 | 1 | 15 |
| CU695223.1 | 0.46019 | -0.20450966 | 0.056 | 0.13 | 1 | 15 |
| serinc2 | 0.46071 | -0.2323921 | 0.333 | 0.478 | 1 | 15 |
| otulina | 0.46155 | 0.127748014 | 0.139 | 0.043 | 1 | 15 |
| surf4 | 0.46162 | 0.192738373 | 0.139 | 0.043 | 1 | 15 |
| nmt1a | 0.46163 | -0.12298791 | 0.194 | 0.217 | 1 | 15 |
| mlst8 | 0.46166 | 0.147170071 | 0.139 | 0.043 | 1 | 15 |
| eif2b1 | 0.46175 | 0.143099724 | 0.111 | 0.043 | 1 | 15 |
| utp23 | 0.46183 | 0.18277228 | 0.139 | 0.043 | 1 | 15 |
| tubb2b | 0.46191 | -0.17470322 | 0.417 | 0.478 | 1 | 15 |
| si:dkey-112e7.2 | 0.46192 | 0.174134316 | 0.333 | 0.261 | 1 | 15 |
| mef2d | 0.46202 | -0.13179618 | 0.111 | 0.217 | 1 | 15 |
| ruvbl1 | 0.46205 | 0.137256006 | 0.139 | 0.043 | 1 | 15 |
| serpinb1l3 | 0.46256 | 0.162584635 | 0.139 | 0.043 | 1 | 15 |
| isoc2 | 0.46263 | 0.138298564 | 0.139 | 0.043 | 1 | 15 |
| anapc10 | 0.46291 | 0.155510542 | 0.167 | 0.087 | 1 | 15 |
| eif4a3 | 0.46298 | 0.187737764 | 0.361 | 0.217 | 1 | 15 |
| fam177a1 | 0.46303 | -0.18814982 | 0.111 | 0.217 | 1 | 15 |
| cc2d1b | 0.46362 | 0.142964172 | 0.111 | 0.043 | 1 | 15 |
| mrto4 | 0.46401 | 0.144165584 | 0.139 | 0.043 | 1 | 15 |
| ndnf | 0.46402 | -0.20377041 | 0.194 | 0.261 | 1 | 15 |
| chchd2 | 0.46469 | -0.12193113 | 0.139 | 0.174 | 1 | 15 |

| | | | | | | |
|---|---|---|---|---|---|---|
| sept7b | 0.46481 | 0.139257213 | 0.194 | 0.087 | 1 | 15 |
| xaf1 | 0.46503 | 0.171458179 | 0.167 | 0.087 | 1 | 15 |
| psme3 | 0.46546 | 0.152784131 | 0.139 | 0.043 | 1 | 15 |
| si:dkey-188i13.10 | 0.46567 | 0.216569498 | 0.639 | 0.478 | 1 | 15 |
| rab28 | 0.46574 | 0.158946541 | 0.139 | 0.043 | 1 | 15 |
| yif1a | 0.46619 | 0.209797851 | 0.139 | 0.043 | 1 | 15 |
| nop53 | 0.4662 | 0.199880819 | 0.556 | 0.391 | 1 | 15 |
| mrps18a | 0.46626 | 0.135800614 | 0.111 | 0.043 | 1 | 15 |
| sc5d | 0.4663 | 0.162338351 | 0.139 | 0.043 | 1 | 15 |
| btd | 0.4665 | 0.100847886 | 0.167 | 0.087 | 1 | 15 |
| apoob | 0.46651 | -0.1008826 | 0.083 | 0.174 | 1 | 15 |
| tmem141 | 0.46657 | 0.143797445 | 0.111 | 0.043 | 1 | 15 |
| rnf26 | 0.46673 | 0.205667453 | 0.139 | 0.043 | 1 | 15 |
| fam222a | 0.46677 | 0.101954091 | 0.111 | 0.087 | 1 | 15 |
| si:ch211-67e16.4 | 0.46682 | 0.160830626 | 0.139 | 0.043 | 1 | 15 |
| trrap | 0.46684 | 0.198220471 | 0.139 | 0.043 | 1 | 15 |
| pbxip1a | 0.46688 | 0.153327129 | 0.139 | 0.043 | 1 | 15 |
| CABZ01088149.1 | 0.4669 | 0.158278685 | 0.139 | 0.043 | 1 | 15 |
| neil1 | 0.46691 | 0.154084403 | 0.139 | 0.043 | 1 | 15 |
| BX957317.2 | 0.46715 | -0.10449471 | 0.056 | 0.13 | 1 | 15 |
| tm9sf2 | 0.46737 | -0.21814695 | 0.139 | 0.261 | 1 | 15 |
| psmc2 | 0.4675 | -0.21862911 | 0.194 | 0.304 | 1 | 15 |
| ssr3 | 0.46753 | -0.24264622 | 0.472 | 0.565 | 1 | 15 |
| nsa2 | 0.46764 | 0.152846147 | 0.833 | 0.696 | 1 | 15 |
| vkorc1l1 | 0.46776 | 0.112498431 | 0.167 | 0.13 | 1 | 15 |
| tmem106ba | 0.46782 | -0.20222316 | 0.167 | 0.304 | 1 | 15 |
| jam3b | 0.46833 | 0.230199354 | 0.417 | 0.261 | 1 | 15 |
| chchd10 | 0.46856 | -0.15153865 | 0.111 | 0.174 | 1 | 15 |
| trappc2l | 0.46865 | -0.14561906 | 0.25 | 0.304 | 1 | 15 |
| atp5pb | 0.46866 | 0.193410796 | 0.778 | 0.652 | 1 | 15 |
| prdx3 | 0.46909 | 0.193588767 | 0.5 | 0.391 | 1 | 15 |
| tmem176 | 0.46926 | -0.21102773 | 0.167 | 0.304 | 1 | 15 |
| gas1a | 0.46932 | -0.21107603 | 0.167 | 0.304 | 1 | 15 |
| bicd2 | 0.47009 | -0.20777555 | 0.056 | 0.13 | 1 | 15 |
| fnta | 0.47028 | 0.196923134 | 0.194 | 0.087 | 1 | 15 |
| lhfpl2a | 0.47042 | -0.22696467 | 0.083 | 0.174 | 1 | 15 |
| creb3l3l | 0.47146 | -0.2150697 | 0.111 | 0.217 | 1 | 15 |
| nhp2 | 0.47196 | 0.220096775 | 0.528 | 0.391 | 1 | 15 |
| tln1 | 0.47218 | -0.23641961 | 0.194 | 0.304 | 1 | 15 |
| glod4 | 0.47268 | 0.147829522 | 0.111 | 0.043 | 1 | 15 |
| XLOC-009815 | 0.47275 | -0.14639609 | 0.472 | 0.522 | 1 | 15 |
| prkcbp1l | 0.47292 | -0.11950028 | 0.111 | 0.13 | 1 | 15 |
| zgc:158482 | 0.47387 | 0.149351221 | 0.139 | 0.087 | 1 | 15 |
| ccdc47 | 0.47467 | -0.11289598 | 0.083 | 0.174 | 1 | 15 |
| acot13 | 0.47518 | -0.18514458 | 0.083 | 0.174 | 1 | 15 |
| LO018148.1 | 0.47602 | -0.16447859 | 0.056 | 0.13 | 1 | 15 |
| wee1 | 0.47652 | -0.12071225 | 0.056 | 0.087 | 1 | 15 |
| htatip2 | 0.47664 | -0.20867104 | 0.111 | 0.217 | 1 | 15 |

| | | | | | | |
|---|---|---|---|---|---|---|
| scamp2 | 0.47672 | 0.10480428 | 0.194 | 0.174 | 1 | 15 |
| rxrba | 0.47705 | -0.18911801 | 0.056 | 0.13 | 1 | 15 |
| hnrnpm | 0.47723 | -0.13407697 | 0.194 | 0.217 | 1 | 15 |
| elocb | 0.47724 | 0.195744892 | 0.639 | 0.478 | 1 | 15 |
| zgc:86598 | 0.47767 | 0.115588668 | 0.472 | 0.348 | 1 | 15 |
| phf10 | 0.47833 | -0.10145402 | 0.083 | 0.174 | 1 | 15 |
| creb1a | 0.47881 | 0.181718983 | 0.167 | 0.087 | 1 | 15 |
| ube2e1 | 0.4792 | -0.11942399 | 0.306 | 0.435 | 1 | 15 |
| apex1 | 0.47945 | 0.184814559 | 0.222 | 0.13 | 1 | 15 |
| svbp | 0.47946 | 0.158988864 | 0.111 | 0.043 | 1 | 15 |
| clk2a | 0.47996 | -0.1972075 | 0.083 | 0.174 | 1 | 15 |
| fkbp7 | 0.48017 | 0.103974896 | 0.222 | 0.13 | 1 | 15 |
| puf60b | 0.48074 | -0.18322319 | 0.194 | 0.261 | 1 | 15 |
| tpm3 | 0.48101 | -0.18372034 | 0.444 | 0.522 | 1 | 15 |
| crybg1a | 0.48102 | 0.138667724 | 0.222 | 0.174 | 1 | 15 |
| dldh | 0.48126 | 0.215037413 | 0.361 | 0.217 | 1 | 15 |
| id2b | 0.48163 | 0.238948702 | 0.306 | 0.174 | 1 | 15 |
| si:ch211-196c10.15 | 0.48168 | 0.156901625 | 0.111 | 0.043 | 1 | 15 |
| clgn | 0.48178 | -0.15976526 | 0.056 | 0.13 | 1 | 15 |
| abi2b | 0.4822 | -0.12711317 | 0.111 | 0.217 | 1 | 15 |
| rnf213b | 0.48229 | 0.134571875 | 0.222 | 0.13 | 1 | 15 |
| sptlc2a | 0.48355 | -0.16656357 | 0.111 | 0.174 | 1 | 15 |
| snrpa | 0.48392 | -0.22853101 | 0.111 | 0.217 | 1 | 15 |
| TST | 0.48402 | -0.10672398 | 0.056 | 0.13 | 1 | 15 |
| txn2 | 0.4843 | 0.201721049 | 0.306 | 0.174 | 1 | 15 |
| march2 | 0.48595 | 0.166544619 | 0.111 | 0.043 | 1 | 15 |
| rhoac | 0.48625 | 0.187206725 | 0.389 | 0.304 | 1 | 15 |
| smim19 | 0.48647 | -0.18376197 | 0.056 | 0.13 | 1 | 15 |
| sssca1 | 0.48648 | -0.1195615 | 0.083 | 0.174 | 1 | 15 |
| ube2d2 | 0.48683 | 0.194664184 | 0.361 | 0.217 | 1 | 15 |
| cyp11a2 | 0.48731 | 0.120901103 | 0.083 | 0.043 | 1 | 15 |
| fam171a2a | 0.4881 | -0.18570421 | 0.111 | 0.217 | 1 | 15 |
| rapgef2 | 0.48848 | -0.11939298 | 0.056 | 0.087 | 1 | 15 |
| trib3 | 0.48876 | -0.10841177 | 0.333 | 0.348 | 1 | 15 |
| nop58 | 0.48889 | 0.232529792 | 0.361 | 0.217 | 1 | 15 |
| si:ch73-109d9.1 | 0.48921 | -0.16155946 | 0.056 | 0.13 | 1 | 15 |
| srd5a3 | 0.48959 | 0.121413445 | 0.083 | 0.043 | 1 | 15 |
| emc2 | 0.49005 | 0.115052895 | 0.333 | 0.217 | 1 | 15 |
| si:ch211-66k16.27 | 0.49088 | -0.13278693 | 0.25 | 0.304 | 1 | 15 |
| fam89b | 0.49139 | 0.196915736 | 0.194 | 0.087 | 1 | 15 |
| ndufa12 | 0.49148 | 0.203473157 | 0.361 | 0.217 | 1 | 15 |
| fzr1a | 0.49183 | -0.16102414 | 0.111 | 0.174 | 1 | 15 |
| morf4l1 | 0.49309 | 0.206329722 | 0.361 | 0.217 | 1 | 15 |
| dnajc19 | 0.49339 | 0.187190757 | 0.361 | 0.217 | 1 | 15 |
| znf576.2 | 0.49364 | -0.17243395 | 0.139 | 0.261 | 1 | 15 |
| hnrnpc | 0.49377 | 0.218067071 | 0.361 | 0.217 | 1 | 15 |
| mpnd | 0.49384 | 0.135521588 | 0.111 | 0.043 | 1 | 15 |
| tshz2 | 0.49541 | -0.17625749 | 0.056 | 0.13 | 1 | 15 |

| | | | | | | |
|---|---|---|---|---|---|---|
| ilf3a | 0.49542 | 0.131119051 | 0.111 | 0.043 | 1 | 15 |
| psma6a | 0.49595 | 0.204914282 | 0.417 | 0.304 | 1 | 15 |
| si:ch1073-459j12.1 | 0.49602 | -0.20960115 | 0.139 | 0.261 | 1 | 15 |
| snrpg | 0.49627 | -0.12830357 | 0.444 | 0.478 | 1 | 15 |
| lin52 | 0.49633 | -0.10628158 | 0.083 | 0.174 | 1 | 15 |
| vapal | 0.49645 | 0.150050983 | 0.194 | 0.13 | 1 | 15 |
| rhoq | 0.4968 | 0.126124435 | 0.278 | 0.174 | 1 | 15 |
| sc:d217 | 0.49701 | 0.180210033 | 0.167 | 0.087 | 1 | 15 |
| sccpdhb | 0.49735 | 0.152976374 | 0.111 | 0.043 | 1 | 15 |
| si:ch211-173p18.3 | 0.49738 | 0.153796001 | 0.194 | 0.087 | 1 | 15 |
| brd2a | 0.49738 | -0.19849662 | 0.333 | 0.435 | 1 | 15 |
| zgc:162396 | 0.49743 | 0.131685412 | 0.111 | 0.043 | 1 | 15 |
| supt6h | 0.49753 | 0.139630513 | 0.139 | 0.087 | 1 | 15 |
| ccnh | 0.4978 | -0.1653184 | 0.056 | 0.13 | 1 | 15 |
| hnrpkl | 0.49812 | -0.12502148 | 0.361 | 0.391 | 1 | 15 |
| hist2h2l | 0.49858 | 0.185992144 | 0.639 | 0.565 | 1 | 15 |
| eif3k | 0.49873 | 0.13899942 | 0.556 | 0.522 | 1 | 15 |
| ecrg4a | 0.49878 | 0.18128482 | 0.833 | 0.783 | 1 | 15 |
| enc1 | 0.49916 | -0.24115656 | 0.361 | 0.478 | 1 | 15 |
| ptpn21 | 0.49917 | -0.12990761 | 0.083 | 0.174 | 1 | 15 |
| nars | 0.49955 | 0.18920872 | 0.278 | 0.174 | 1 | 15 |
| uqcc1 | 0.50006 | 0.167574012 | 0.111 | 0.043 | 1 | 15 |
| pnrc2 | 0.50008 | -0.11608235 | 0.917 | 0.957 | 1 | 15 |
| vdrb | 0.50018 | -0.22064755 | 0.222 | 0.348 | 1 | 15 |
| pik3ip1 | 0.5004 | 0.122717508 | 0.611 | 0.478 | 1 | 15 |
| gm2a | 0.50054 | -0.18777198 | 0.083 | 0.174 | 1 | 15 |
| lrp1ab | 0.50077 | -0.15772056 | 0.167 | 0.217 | 1 | 15 |
| LO016987.2 | 0.50083 | 0.190725119 | 0.111 | 0.043 | 1 | 15 |
| ift20 | 0.502 | 0.138270818 | 0.111 | 0.043 | 1 | 15 |
| kbtbd11 | 0.50252 | -0.11585657 | 0.056 | 0.087 | 1 | 15 |
| xrcc6 | 0.50253 | -0.13681088 | 0.083 | 0.13 | 1 | 15 |
| ergic2 | 0.50259 | 0.182302919 | 0.194 | 0.087 | 1 | 15 |
| si:dkey-250d21.1 | 0.50267 | 0.190618177 | 0.306 | 0.174 | 1 | 15 |
| psme2 | 0.50279 | -0.15554045 | 0.417 | 0.478 | 1 | 15 |
| sik1 | 0.50309 | -0.2197858 | 0.194 | 0.304 | 1 | 15 |
| brms1la | 0.50339 | -0.16760116 | 0.139 | 0.217 | 1 | 15 |
| slu7 | 0.50342 | -0.18274883 | 0.056 | 0.13 | 1 | 15 |
| golph3 | 0.50389 | 0.195482217 | 0.306 | 0.174 | 1 | 15 |
| ctsh | 0.50453 | -0.19526632 | 0.139 | 0.261 | 1 | 15 |
| trpt1 | 0.5047 | -0.18200055 | 0.056 | 0.13 | 1 | 15 |
| pbdc1 | 0.50494 | 0.117968988 | 0.472 | 0.348 | 1 | 15 |
| glud1a | 0.50495 | -0.13604812 | 0.111 | 0.217 | 1 | 15 |
| agpat3 | 0.5052 | -0.12000808 | 0.139 | 0.174 | 1 | 15 |
| rap1b | 0.50569 | 0.177174735 | 0.25 | 0.13 | 1 | 15 |
| pspc1 | 0.50591 | 0.130190938 | 0.139 | 0.087 | 1 | 15 |
| pex19 | 0.50593 | 0.159477501 | 0.111 | 0.043 | 1 | 15 |
| med19b | 0.50676 | -0.10960957 | 0.083 | 0.174 | 1 | 15 |
| tarbp2 | 0.50677 | 0.153459187 | 0.111 | 0.043 | 1 | 15 |

| | | | | | | |
|---|---|---|---|---|---|---|
| nfe2l1b | 0.5068 | -0.15895706 | 0.111 | 0.174 | 1 | 15 |
| bcl9 | 0.50685 | 0.152231604 | 0.111 | 0.043 | 1 | 15 |
| si:ch211-212k18.4 | 0.50739 | 0.174713179 | 0.194 | 0.087 | 1 | 15 |
| TMEM208 | 0.5077 | 0.147550646 | 0.194 | 0.087 | 1 | 15 |
| srrm1 | 0.50827 | 0.20201675 | 0.306 | 0.174 | 1 | 15 |
| FAT4 | 0.50885 | -0.17808407 | 0.083 | 0.174 | 1 | 15 |
| wrb | 0.50906 | -0.20330833 | 0.139 | 0.261 | 1 | 15 |
| akt1s1 | 0.50918 | 0.108043978 | 0.278 | 0.174 | 1 | 15 |
| zgc:171497 | 0.50933 | -0.16240519 | 0.111 | 0.174 | 1 | 15 |
| hsbp1b | 0.50943 | 0.193092993 | 0.5 | 0.348 | 1 | 15 |
| taf12 | 0.50967 | 0.208198185 | 0.333 | 0.217 | 1 | 15 |
| ptp4a2a | 0.50971 | -0.14336805 | 0.222 | 0.348 | 1 | 15 |
| crema | 0.50986 | -0.10706483 | 0.25 | 0.261 | 1 | 15 |
| slc1a3a | 0.51007 | 0.128315365 | 0.222 | 0.217 | 1 | 15 |
| tbc1d22b | 0.51021 | 0.121525582 | 0.083 | 0.043 | 1 | 15 |
| yy1a | 0.51024 | 0.180305015 | 0.25 | 0.174 | 1 | 15 |
| adma | 0.51029 | -0.20117603 | 0.083 | 0.174 | 1 | 15 |
| twistnb | 0.51047 | 0.130099836 | 0.111 | 0.043 | 1 | 15 |
| CABZ01034698.2 | 0.5107 | 0.181485268 | 0.194 | 0.087 | 1 | 15 |
| sptan1 | 0.51082 | -0.19391664 | 0.222 | 0.304 | 1 | 15 |
| phyh | 0.51101 | 0.17657922 | 0.194 | 0.087 | 1 | 15 |
| lxn | 0.51115 | 0.220431455 | 0.306 | 0.174 | 1 | 15 |
| api5 | 0.51141 | 0.169632147 | 0.25 | 0.13 | 1 | 15 |
| si:dkey-21a6.5 | 0.51151 | 0.170803947 | 0.25 | 0.13 | 1 | 15 |
| pskh1 | 0.51157 | -0.15317291 | 0.083 | 0.13 | 1 | 15 |
| eny2 | 0.51189 | -0.21101086 | 0.25 | 0.391 | 1 | 15 |
| large2 | 0.5122 | 0.171684567 | 0.194 | 0.087 | 1 | 15 |
| dyrk1ab | 0.51257 | -0.16743475 | 0.056 | 0.13 | 1 | 15 |
| naxe | 0.51298 | -0.1430404 | 0.056 | 0.13 | 1 | 15 |
| ciarta | 0.51306 | 0.127024528 | 0.278 | 0.174 | 1 | 15 |
| cnbpa | 0.51399 | 0.183383038 | 0.472 | 0.391 | 1 | 15 |
| oprd1a | 0.51416 | 0.168883017 | 0.194 | 0.087 | 1 | 15 |
| mrpl57 | 0.51421 | 0.196714467 | 0.306 | 0.174 | 1 | 15 |
| sp3b | 0.51422 | 0.137203109 | 0.083 | 0.043 | 1 | 15 |
| taf6 | 0.51424 | 0.131369736 | 0.111 | 0.043 | 1 | 15 |
| si:dkey-85k7.12 | 0.51431 | 0.151172229 | 0.111 | 0.043 | 1 | 15 |
| zc3h14 | 0.51451 | 0.172743166 | 0.194 | 0.087 | 1 | 15 |
| tmem243b | 0.51459 | 0.166960465 | 0.194 | 0.087 | 1 | 15 |
| myd88 | 0.51462 | 0.17323594 | 0.194 | 0.087 | 1 | 15 |
| zgc:77056 | 0.51487 | 0.175016395 | 0.194 | 0.087 | 1 | 15 |
| eif2b2 | 0.51514 | 0.161441594 | 0.194 | 0.087 | 1 | 15 |
| vps26a | 0.51535 | -0.18967705 | 0.083 | 0.174 | 1 | 15 |
| ube2q2 | 0.51557 | 0.212694652 | 0.306 | 0.174 | 1 | 15 |
| ttpal | 0.51633 | 0.167868263 | 0.167 | 0.087 | 1 | 15 |
| LOC108180076 | 0.51639 | -0.13478613 | 0.056 | 0.087 | 1 | 15 |
| si:ch1073-287p18.1 | 0.51668 | 0.137563028 | 0.111 | 0.043 | 1 | 15 |
| cherp | 0.51741 | 0.182578806 | 0.111 | 0.043 | 1 | 15 |
| rpn1 | 0.51761 | 0.139695898 | 0.528 | 0.391 | 1 | 15 |

| | | | | | | |
|---|---|---|---|---|---|---|
| cnot3a | 0.51762 | -0.16590615 | 0.056 | 0.13 | 1 | 15 |
| sdhc | 0.51787 | 0.104436082 | 0.417 | 0.304 | 1 | 15 |
| fbxo28 | 0.51787 | -0.1136755 | 0.194 | 0.217 | 1 | 15 |
| sirt7 | 0.51798 | 0.167360211 | 0.25 | 0.13 | 1 | 15 |
| slc31a1 | 0.51874 | -0.12986665 | 0.056 | 0.087 | 1 | 15 |
| cox6a1 | 0.51883 | 0.186646238 | 0.778 | 0.652 | 1 | 15 |
| atp5pd | 0.51886 | 0.209991439 | 0.611 | 0.478 | 1 | 15 |
| CABZ01086029.1 | 0.51892 | -0.16376769 | 0.111 | 0.174 | 1 | 15 |
| sel1l | 0.51893 | 0.145802035 | 0.111 | 0.043 | 1 | 15 |
| paqr3a | 0.5197 | -0.1944126 | 0.083 | 0.174 | 1 | 15 |
| si:ch73-71c20.5 | 0.51989 | 0.189224918 | 0.167 | 0.087 | 1 | 15 |
| dhrs7b | 0.52 | -0.15505308 | 0.056 | 0.13 | 1 | 15 |
| ncam2 | 0.52002 | 0.109737356 | 0.111 | 0.043 | 1 | 15 |
| rnf40 | 0.5215 | -0.15976607 | 0.056 | 0.13 | 1 | 15 |
| wbp1la | 0.52161 | 0.212027486 | 0.25 | 0.13 | 1 | 15 |
| ncalda | 0.52192 | -0.12907288 | 0.222 | 0.261 | 1 | 15 |
| nudt1 | 0.52192 | 0.139286833 | 0.111 | 0.043 | 1 | 15 |
| med4 | 0.52223 | -0.16567727 | 0.111 | 0.217 | 1 | 15 |
| dynlrb1 | 0.52249 | -0.16073804 | 0.417 | 0.565 | 1 | 15 |
| ndufab1a | 0.52251 | -0.17636271 | 0.278 | 0.348 | 1 | 15 |
| nrbf2b | 0.52268 | -0.12965245 | 0.111 | 0.217 | 1 | 15 |
| slc38a4 | 0.52309 | 0.176308577 | 0.25 | 0.13 | 1 | 15 |
| dcbld2 | 0.52314 | -0.18951957 | 0.139 | 0.217 | 1 | 15 |
| fbxo25 | 0.52339 | -0.14751016 | 0.111 | 0.217 | 1 | 15 |
| si:dkey-73n10.1 | 0.52344 | -0.14453721 | 0.222 | 0.348 | 1 | 15 |
| si:ch73-367f21.4 | 0.52344 | -0.10335278 | 0.056 | 0.13 | 1 | 15 |
| mylka | 0.52353 | 0.138802469 | 0.139 | 0.087 | 1 | 15 |
| atp5f1b | 0.5241 | 0.140107082 | 0.889 | 0.783 | 1 | 15 |
| si:ch211-284b7.3 | 0.52435 | 0.169879678 | 0.25 | 0.13 | 1 | 15 |
| CABZ01115113.1 | 0.52437 | 0.190356677 | 0.306 | 0.217 | 1 | 15 |
| pudp | 0.52447 | 0.117489212 | 0.111 | 0.043 | 1 | 15 |
| cstf3 | 0.5252 | 0.105024093 | 0.167 | 0.13 | 1 | 15 |
| cnpy1 | 0.52593 | -0.14827719 | 0.111 | 0.217 | 1 | 15 |
| pef1 | 0.52617 | 0.115523481 | 0.222 | 0.13 | 1 | 15 |
| cercam | 0.52671 | -0.11053093 | 0.056 | 0.13 | 1 | 15 |
| ldb1a | 0.527 | -0.10635042 | 0.167 | 0.261 | 1 | 15 |
| dap3 | 0.52735 | -0.14921228 | 0.056 | 0.13 | 1 | 15 |
| ddb2 | 0.52756 | -0.10894607 | 0.056 | 0.13 | 1 | 15 |
| zgc:153115 | 0.52814 | -0.20004978 | 0.083 | 0.174 | 1 | 15 |
| dpf2 | 0.52919 | 0.147087419 | 0.111 | 0.043 | 1 | 15 |
| ccdc85b | 0.52924 | -0.16793642 | 0.111 | 0.217 | 1 | 15 |
| ddah2 | 0.52952 | -0.10591822 | 0.194 | 0.217 | 1 | 15 |
| mob3c | 0.52976 | -0.17080539 | 0.056 | 0.13 | 1 | 15 |
| rhoua | 0.52983 | -0.16287674 | 0.056 | 0.13 | 1 | 15 |
| apool | 0.53092 | 0.113651556 | 0.083 | 0.043 | 1 | 15 |
| stxbp6 | 0.53118 | -0.17743817 | 0.111 | 0.217 | 1 | 15 |
| EARS2 | 0.53127 | 0.104233928 | 0.111 | 0.043 | 1 | 15 |
| smo | 0.53149 | -0.17688977 | 0.111 | 0.217 | 1 | 15 |

| | | | | | | |
|---|---|---|---|---|---|---|
| ttc26 | 0.53185 | 0.100773416 | 0.111 | 0.043 | 1 | 15 |
| kdm4b | 0.532 | -0.18088808 | 0.111 | 0.217 | 1 | 15 |
| myo7aa | 0.53207 | 0.118735442 | 0.083 | 0.043 | 1 | 15 |
| supt5h | 0.53262 | -0.19041148 | 0.083 | 0.174 | 1 | 15 |
| anxa13l | 0.53267 | 0.202425379 | 0.389 | 0.304 | 1 | 15 |
| FAM53C | 0.53294 | 0.158266518 | 0.111 | 0.043 | 1 | 15 |
| ndrg3b | 0.53309 | -0.1606787 | 0.083 | 0.174 | 1 | 15 |
| f10 | 0.53435 | 0.200714788 | 0.583 | 0.435 | 1 | 15 |
| btr31 | 0.5346 | -0.13036034 | 0.083 | 0.174 | 1 | 15 |
| sf3b3 | 0.5348 | 0.154628813 | 0.111 | 0.043 | 1 | 15 |
| abca5 | 0.53498 | -0.18839845 | 0.111 | 0.217 | 1 | 15 |
| ier2b | 0.5353 | -0.23966288 | 0.389 | 0.522 | 1 | 15 |
| mrpl9 | 0.53592 | -0.13835176 | 0.083 | 0.13 | 1 | 15 |
| tardbp | 0.53612 | 0.143987905 | 0.333 | 0.217 | 1 | 15 |
| mak16 | 0.5362 | -0.11296922 | 0.083 | 0.174 | 1 | 15 |
| zgc:66427 | 0.5369 | 0.129475325 | 0.111 | 0.043 | 1 | 15 |
| ino80e | 0.53692 | -0.16540979 | 0.056 | 0.13 | 1 | 15 |
| zgc:174268 | 0.537 | -0.12826884 | 0.083 | 0.174 | 1 | 15 |
| sae1 | 0.5373 | 0.131901006 | 0.111 | 0.043 | 1 | 15 |
| txnrd3 | 0.53736 | 0.139436655 | 0.111 | 0.043 | 1 | 15 |
| b2m | 0.53747 | 0.173478313 | 0.833 | 0.739 | 1 | 15 |
| chp1 | 0.53761 | -0.13381992 | 0.056 | 0.13 | 1 | 15 |
| lamc1 | 0.53776 | -0.20862912 | 0.306 | 0.391 | 1 | 15 |
| cdc34a | 0.53802 | -0.13494599 | 0.056 | 0.087 | 1 | 15 |
| stat2 | 0.53823 | -0.19821901 | 0.278 | 0.391 | 1 | 15 |
| zgc:112496 | 0.53881 | -0.10024313 | 0.056 | 0.13 | 1 | 15 |
| cybc1 | 0.53903 | 0.151768594 | 0.111 | 0.043 | 1 | 15 |
| pbx4 | 0.53972 | 0.165920734 | 0.333 | 0.261 | 1 | 15 |
| nelfe | 0.54051 | -0.10886559 | 0.083 | 0.174 | 1 | 15 |
| mov10b.1 | 0.54087 | -0.11617199 | 0.194 | 0.304 | 1 | 15 |
| tmub1 | 0.54128 | -0.18599206 | 0.111 | 0.217 | 1 | 15 |
| ifrd2 | 0.54173 | 0.172729607 | 0.167 | 0.087 | 1 | 15 |
| vegfab | 0.54234 | -0.20263321 | 0.083 | 0.174 | 1 | 15 |
| plagx | 0.54235 | -0.10418848 | 0.056 | 0.13 | 1 | 15 |
| gigyf2 | 0.54268 | -0.15127142 | 0.056 | 0.13 | 1 | 15 |
| phf23a | 0.54297 | -0.1771107 | 0.056 | 0.13 | 1 | 15 |
| ncoa5 | 0.543 | -0.14239851 | 0.111 | 0.217 | 1 | 15 |
| aco1 | 0.54355 | 0.137531617 | 0.111 | 0.043 | 1 | 15 |
| arpc1a | 0.54393 | 0.151473508 | 0.278 | 0.217 | 1 | 15 |
| mbd2 | 0.54407 | -0.11589195 | 0.056 | 0.087 | 1 | 15 |
| pgrmc2 | 0.54542 | -0.16417503 | 0.222 | 0.348 | 1 | 15 |
| fnbp1l | 0.54557 | 0.123796828 | 0.167 | 0.13 | 1 | 15 |
| plxna3 | 0.54557 | -0.1676698 | 0.056 | 0.13 | 1 | 15 |
| tbc1d4 | 0.54573 | -0.14952783 | 0.111 | 0.174 | 1 | 15 |
| rgs7bpb | 0.54674 | -0.18707338 | 0.111 | 0.217 | 1 | 15 |
| si:ch73-340n8.4 | 0.54719 | -0.10463121 | 0.056 | 0.13 | 1 | 15 |
| rnf13 | 0.54734 | -0.18551211 | 0.111 | 0.217 | 1 | 15 |
| ece2b | 0.54764 | 0.107785621 | 0.361 | 0.261 | 1 | 15 |

| | | | | | | |
|---|---|---|---|---|---|---|
| gpat3 | 0.54779 | -0.18429748 | 0.056 | 0.13 | 1 | 15 |
| vmp1 | 0.54786 | -0.18570415 | 0.111 | 0.217 | 1 | 15 |
| bcam | 0.54837 | 0.131788082 | 0.694 | 0.565 | 1 | 15 |
| usp42 | 0.54914 | -0.11914148 | 0.139 | 0.174 | 1 | 15 |
| serpinf1 | 0.54953 | -0.17288843 | 0.083 | 0.174 | 1 | 15 |
| erbin | 0.54962 | 0.130322278 | 0.278 | 0.174 | 1 | 15 |
| emc3 | 0.54994 | -0.15215937 | 0.083 | 0.174 | 1 | 15 |
| bud31 | 0.5507 | -0.108805 | 0.528 | 0.565 | 1 | 15 |
| prkag2a | 0.55088 | 0.148936477 | 0.139 | 0.087 | 1 | 15 |
| cct8 | 0.55155 | 0.180480558 | 0.444 | 0.304 | 1 | 15 |
| si:dkeyp-44a8.2 | 0.55213 | -0.16025855 | 0.139 | 0.217 | 1 | 15 |
| tomm70a | 0.5524 | -0.16368057 | 0.222 | 0.348 | 1 | 15 |
| kdm5c | 0.55299 | -0.15818976 | 0.111 | 0.174 | 1 | 15 |
| dpf3 | 0.55313 | -0.16627527 | 0.056 | 0.13 | 1 | 15 |
| tor1 | 0.5532 | -0.16688834 | 0.222 | 0.348 | 1 | 15 |
| fbxo44 | 0.55354 | 0.119976519 | 0.111 | 0.043 | 1 | 15 |
| laptm4b | 0.55433 | -0.15426566 | 0.056 | 0.13 | 1 | 15 |
| mrpl20 | 0.55448 | -0.11798175 | 0.222 | 0.261 | 1 | 15 |
| reep5 | 0.55509 | -0.13088381 | 0.361 | 0.391 | 1 | 15 |
| mt-nd5 | 0.55543 | 0.16372776 | 0.444 | 0.304 | 1 | 15 |
| htr1aa | 0.55551 | -0.15946368 | 0.083 | 0.174 | 1 | 15 |
| wdfy1 | 0.55592 | -0.16756839 | 0.167 | 0.261 | 1 | 15 |
| spop | 0.55672 | 0.102862809 | 0.111 | 0.043 | 1 | 15 |
| zfr2 | 0.55714 | -0.13344736 | 0.083 | 0.13 | 1 | 15 |
| psma8 | 0.55717 | -0.17625877 | 0.444 | 0.522 | 1 | 15 |
| tmem209 | 0.55736 | -0.18781033 | 0.083 | 0.174 | 1 | 15 |
| cnot9 | 0.55823 | 0.113476291 | 0.111 | 0.043 | 1 | 15 |
| fabp3 | 0.55824 | 0.124327923 | 0.111 | 0.087 | 1 | 15 |
| cep131 | 0.55901 | 0.109148673 | 0.111 | 0.043 | 1 | 15 |
| cst14b.1 | 0.55921 | 0.129156568 | 0.111 | 0.043 | 1 | 15 |
| trip12 | 0.55937 | -0.14808508 | 0.111 | 0.174 | 1 | 15 |
| ei24 | 0.55938 | 0.110503001 | 0.083 | 0.043 | 1 | 15 |
| sco2 | 0.55973 | -0.1489752 | 0.056 | 0.13 | 1 | 15 |
| pitpnbl | 0.55973 | -0.15788521 | 0.083 | 0.174 | 1 | 15 |
| extl2 | 0.5598 | -0.1609573 | 0.056 | 0.13 | 1 | 15 |
| ncln | 0.56014 | 0.145084993 | 0.167 | 0.087 | 1 | 15 |
| ctdsplb | 0.56037 | -0.10291645 | 0.167 | 0.261 | 1 | 15 |
| znf1001 | 0.56044 | 0.154960477 | 0.167 | 0.087 | 1 | 15 |
| pcsk7 | 0.56069 | -0.10901122 | 0.056 | 0.13 | 1 | 15 |
| gstm.2 | 0.56101 | -0.214207 | 0.194 | 0.304 | 1 | 15 |
| bcap31 | 0.56136 | -0.17499226 | 0.222 | 0.348 | 1 | 15 |
| ubr4 | 0.56217 | -0.14321175 | 0.167 | 0.217 | 1 | 15 |
| nde1 | 0.56218 | 0.106517419 | 0.083 | 0.043 | 1 | 15 |
| fbxl3a | 0.56304 | -0.15167232 | 0.083 | 0.174 | 1 | 15 |
| CABZ01086041.1 | 0.56307 | 0.152830072 | 0.111 | 0.043 | 1 | 15 |
| nae1 | 0.56356 | -0.14957177 | 0.083 | 0.174 | 1 | 15 |
| bccip | 0.56395 | 0.15225273 | 0.306 | 0.261 | 1 | 15 |
| mcm3ap | 0.56397 | 0.139787018 | 0.111 | 0.043 | 1 | 15 |

| | | | | | | |
|---|---|---|---|---|---|---|
| mrpl11 | 0.56461 | 0.171710921 | 0.306 | 0.217 | 1 | 15 |
| rabl2 | 0.56483 | 0.122327719 | 0.083 | 0.043 | 1 | 15 |
| bcl2l11 | 0.56485 | -0.11860929 | 0.056 | 0.13 | 1 | 15 |
| ndufs8a | 0.56518 | 0.169601657 | 0.222 | 0.13 | 1 | 15 |
| dele1 | 0.56533 | 0.101940323 | 0.111 | 0.043 | 1 | 15 |
| arih1 | 0.56551 | -0.17916956 | 0.083 | 0.174 | 1 | 15 |
| btr04 | 0.56621 | -0.13389149 | 0.083 | 0.13 | 1 | 15 |
| upf1 | 0.56677 | -0.12819963 | 0.056 | 0.13 | 1 | 15 |
| wdr83os | 0.56733 | -0.19200673 | 0.361 | 0.478 | 1 | 15 |
| pop5 | 0.56805 | -0.14695867 | 0.056 | 0.13 | 1 | 15 |
| hax1 | 0.56845 | -0.13442133 | 0.083 | 0.174 | 1 | 15 |
| ing2 | 0.56862 | -0.13346873 | 0.167 | 0.217 | 1 | 15 |
| si:ch211-281l24.3 | 0.56868 | 0.134829653 | 0.111 | 0.043 | 1 | 15 |
| dync1li2 | 0.56887 | -0.13432529 | 0.083 | 0.174 | 1 | 15 |
| phrf1 | 0.56897 | 0.125375396 | 0.083 | 0.043 | 1 | 15 |
| slc3a2b | 0.56917 | 0.134548272 | 0.472 | 0.348 | 1 | 15 |
| csde1 | 0.56943 | -0.19016684 | 0.306 | 0.435 | 1 | 15 |
| wu:fc75a09 | 0.5699 | -0.20481385 | 0.222 | 0.348 | 1 | 15 |
| hypk | 0.56993 | 0.202934521 | 0.528 | 0.391 | 1 | 15 |
| kdm5bb | 0.56996 | -0.15973938 | 0.056 | 0.13 | 1 | 15 |
| spen | 0.57001 | -0.12542863 | 0.056 | 0.087 | 1 | 15 |
| prmt5 | 0.57071 | 0.150207928 | 0.111 | 0.043 | 1 | 15 |
| LOC101885027 | 0.57166 | -0.13842741 | 0.056 | 0.13 | 1 | 15 |
| mcts1 | 0.57177 | -0.17441164 | 0.222 | 0.348 | 1 | 15 |
| rfc3 | 0.57224 | -0.14576004 | 0.056 | 0.13 | 1 | 15 |
| pdpk1b | 0.57237 | 0.147274277 | 0.111 | 0.043 | 1 | 15 |
| rac1a | 0.57303 | -0.17212592 | 0.222 | 0.348 | 1 | 15 |
| si:dkey-222b8.1 | 0.57329 | 0.122793542 | 0.139 | 0.087 | 1 | 15 |
| fuca1.2 | 0.57352 | -0.1295956 | 0.083 | 0.174 | 1 | 15 |
| wwp2 | 0.57363 | 0.150530611 | 0.111 | 0.043 | 1 | 15 |
| sipa1l2 | 0.57377 | 0.155738448 | 0.111 | 0.043 | 1 | 15 |
| ndfip2 | 0.57401 | -0.19334498 | 0.083 | 0.174 | 1 | 15 |
| hint2 | 0.57418 | -0.10343687 | 0.056 | 0.13 | 1 | 15 |
| polr2b | 0.57444 | -0.17487968 | 0.139 | 0.217 | 1 | 15 |
| vapb | 0.5745 | 0.166829564 | 0.389 | 0.261 | 1 | 15 |
| dcun1d5 | 0.57486 | 0.144969063 | 0.167 | 0.087 | 1 | 15 |
| reep3b | 0.57508 | 0.168798708 | 0.389 | 0.261 | 1 | 15 |
| pomp | 0.57518 | -0.14426556 | 0.556 | 0.609 | 1 | 15 |
| mrps5 | 0.57531 | -0.10979789 | 0.056 | 0.087 | 1 | 15 |
| vti1b | 0.57578 | 0.126693346 | 0.139 | 0.087 | 1 | 15 |
| naa30 | 0.57665 | -0.12277298 | 0.056 | 0.087 | 1 | 15 |
| mau2 | 0.57801 | -0.15368502 | 0.083 | 0.174 | 1 | 15 |
| ostc | 0.57808 | 0.175193527 | 0.556 | 0.478 | 1 | 15 |
| zgc:162816 | 0.57834 | 0.103614846 | 0.167 | 0.13 | 1 | 15 |
| eif2s3 | 0.57853 | 0.16704731 | 0.528 | 0.391 | 1 | 15 |
| cnnm2a | 0.57872 | 0.122931001 | 0.083 | 0.043 | 1 | 15 |
| pkp4 | 0.5789 | 0.115413608 | 0.083 | 0.043 | 1 | 15 |
| ndnl2 | 0.57891 | -0.1317998 | 0.083 | 0.174 | 1 | 15 |

| | | | | | | |
|---|---|---|---|---|---|---|
| pdcd6ip | 0.57915 | -0.11488038 | 0.056 | 0.13 | 1 | 15 |
| man2a2 | 0.57918 | -0.16234327 | 0.111 | 0.174 | 1 | 15 |
| map4k5 | 0.5795 | -0.1058927 | 0.056 | 0.087 | 1 | 15 |
| crebrf | 0.57981 | -0.13121711 | 0.083 | 0.13 | 1 | 15 |
| luc7l3 | 0.58026 | -0.16090128 | 0.056 | 0.13 | 1 | 15 |
| gmps | 0.58039 | -0.16526625 | 0.056 | 0.13 | 1 | 15 |
| laynb | 0.58041 | 0.139569055 | 0.111 | 0.043 | 1 | 15 |
| trappc4 | 0.58053 | -0.16078048 | 0.083 | 0.174 | 1 | 15 |
| hspb8 | 0.58179 | -0.14368742 | 0.083 | 0.174 | 1 | 15 |
| ece1 | 0.58214 | 0.105742231 | 0.083 | 0.043 | 1 | 15 |
| pyroxd2 | 0.58237 | 0.132408377 | 0.111 | 0.043 | 1 | 15 |
| dap1b | 0.58238 | 0.187474929 | 0.611 | 0.478 | 1 | 15 |
| zgc:101858 | 0.58271 | -0.15529762 | 0.083 | 0.174 | 1 | 15 |
| aldh2.2 | 0.58284 | -0.15770365 | 0.389 | 0.522 | 1 | 15 |
| hspd1 | 0.58297 | 0.17249404 | 0.694 | 0.609 | 1 | 15 |
| acad9 | 0.58342 | -0.1243489 | 0.056 | 0.13 | 1 | 15 |
| CU639469.1 | 0.58347 | 0.115968887 | 0.083 | 0.043 | 1 | 15 |
| znf516 | 0.5838 | -0.16527388 | 0.083 | 0.174 | 1 | 15 |
| cyhr1 | 0.58397 | 0.148871127 | 0.111 | 0.043 | 1 | 15 |
| ndufa11 | 0.58443 | -0.17816211 | 0.389 | 0.522 | 1 | 15 |
| riox1 | 0.58451 | -0.1682822 | 0.083 | 0.174 | 1 | 15 |
| ap1m2 | 0.58451 | 0.102684089 | 0.083 | 0.043 | 1 | 15 |
| eif2s2 | 0.58458 | -0.1200094 | 0.667 | 0.696 | 1 | 15 |
| pak4 | 0.5852 | -0.10529276 | 0.056 | 0.13 | 1 | 15 |
| gstz1 | 0.58536 | 0.107689206 | 0.083 | 0.043 | 1 | 15 |
| tax1bp3 | 0.58542 | -0.18108007 | 0.194 | 0.304 | 1 | 15 |
| kank4 | 0.58568 | 0.119957701 | 0.083 | 0.043 | 1 | 15 |
| ptprfb | 0.58638 | 0.139042286 | 0.278 | 0.174 | 1 | 15 |
| ubtd2 | 0.58642 | -0.11524169 | 0.056 | 0.13 | 1 | 15 |
| map7d1b | 0.58649 | -0.1310391 | 0.083 | 0.174 | 1 | 15 |
| cyb5r1 | 0.58667 | -0.12465489 | 0.083 | 0.174 | 1 | 15 |
| lhfpl2b | 0.5867 | -0.14660138 | 0.083 | 0.174 | 1 | 15 |
| fam32a | 0.58691 | 0.188855828 | 0.5 | 0.391 | 1 | 15 |
| kansl1b | 0.58692 | -0.1189559 | 0.139 | 0.174 | 1 | 15 |
| ndufb2 | 0.58692 | 0.116324819 | 0.417 | 0.304 | 1 | 15 |
| arl6ip4 | 0.58716 | -0.16203599 | 0.083 | 0.174 | 1 | 15 |
| bptf | 0.58731 | -0.15556616 | 0.083 | 0.174 | 1 | 15 |
| si:ch73-234b20.5 | 0.58735 | -0.13289519 | 0.083 | 0.13 | 1 | 15 |
| acadm | 0.58761 | 0.182178393 | 0.417 | 0.304 | 1 | 15 |
| nenf | 0.58765 | -0.15295818 | 0.083 | 0.174 | 1 | 15 |
| acot15 | 0.58784 | -0.17267238 | 0.083 | 0.174 | 1 | 15 |
| wt1b | 0.58793 | -0.17585756 | 0.083 | 0.174 | 1 | 15 |
| XLOC-011916 | 0.58812 | -0.17372455 | 0.083 | 0.174 | 1 | 15 |
| bbip1 | 0.58812 | -0.15617927 | 0.083 | 0.174 | 1 | 15 |
| tdp1 | 0.58815 | -0.14507412 | 0.083 | 0.174 | 1 | 15 |
| pak2b | 0.58816 | -0.14381448 | 0.083 | 0.174 | 1 | 15 |
| tra2a | 0.58828 | 0.162399778 | 0.306 | 0.217 | 1 | 15 |
| psmf1 | 0.5884 | 0.112620041 | 0.222 | 0.13 | 1 | 15 |

| | | | | | | |
|---|---|---|---|---|---|---|
| uqcrc2b | 0.58855 | -0.12984779 | 0.306 | 0.348 | 1 | 15 |
| drap1 | 0.58906 | 0.181067925 | 0.389 | 0.261 | 1 | 15 |
| march5l | 0.58936 | 0.15073362 | 0.167 | 0.087 | 1 | 15 |
| kdm2aa | 0.5894 | 0.12112724 | 0.083 | 0.043 | 1 | 15 |
| spg21 | 0.58946 | 0.126900744 | 0.194 | 0.13 | 1 | 15 |
| dcun1d4 | 0.58983 | 0.141801216 | 0.111 | 0.043 | 1 | 15 |
| u2af2b | 0.59019 | -0.11669867 | 0.278 | 0.391 | 1 | 15 |
| ankle2 | 0.59026 | -0.13752313 | 0.056 | 0.13 | 1 | 15 |
| gnai1 | 0.59043 | -0.10014524 | 0.194 | 0.217 | 1 | 15 |
| dgcr8 | 0.59048 | 0.141815122 | 0.111 | 0.043 | 1 | 15 |
| chmp4ba | 0.59052 | 0.153650992 | 0.167 | 0.087 | 1 | 15 |
| atp5mc3a | 0.59062 | 0.18847654 | 0.444 | 0.348 | 1 | 15 |
| rab8a | 0.59062 | -0.15576526 | 0.056 | 0.13 | 1 | 15 |
| sptssa | 0.59103 | 0.189625625 | 0.278 | 0.174 | 1 | 15 |
| pias4a | 0.59104 | 0.118025034 | 0.083 | 0.043 | 1 | 15 |
| ppp4r2b | 0.59143 | 0.14345299 | 0.111 | 0.043 | 1 | 15 |
| rbm28 | 0.59154 | -0.16340254 | 0.111 | 0.174 | 1 | 15 |
| aktip | 0.59179 | 0.149444208 | 0.111 | 0.043 | 1 | 15 |
| hmox2a | 0.59249 | 0.127393676 | 0.111 | 0.043 | 1 | 15 |
| wwc1 | 0.5928 | -0.1408452 | 0.056 | 0.13 | 1 | 15 |
| nucks1b | 0.59295 | 0.18154471 | 0.389 | 0.261 | 1 | 15 |
| ISCU (1 of many) | 0.59302 | 0.144293004 | 0.417 | 0.348 | 1 | 15 |
| sec24a | 0.59313 | 0.142498162 | 0.111 | 0.043 | 1 | 15 |
| psmc3 | 0.59391 | 0.130434833 | 0.222 | 0.174 | 1 | 15 |
| selenos | 0.59419 | -0.14710941 | 0.111 | 0.174 | 1 | 15 |
| tmem176l.3a | 0.59433 | -0.16528834 | 0.139 | 0.217 | 1 | 15 |
| pvrl2l | 0.59442 | 0.134288297 | 0.139 | 0.087 | 1 | 15 |
| zgc:174356 | 0.59472 | -0.16742109 | 0.056 | 0.13 | 1 | 15 |
| zfpm1 | 0.59516 | 0.175157752 | 0.306 | 0.217 | 1 | 15 |
| mbtps1 | 0.59557 | -0.12946411 | 0.056 | 0.087 | 1 | 15 |
| desi2 | 0.59598 | -0.14452812 | 0.056 | 0.13 | 1 | 15 |
| arrdc2 | 0.59598 | -0.16982702 | 0.167 | 0.261 | 1 | 15 |
| rbfa | 0.59605 | 0.141824462 | 0.111 | 0.043 | 1 | 15 |
| ldlrap1a | 0.59639 | -0.17415053 | 0.25 | 0.348 | 1 | 15 |
| col4a3bpa | 0.59672 | 0.126037692 | 0.111 | 0.043 | 1 | 15 |
| pkma | 0.59679 | 0.138614517 | 0.278 | 0.174 | 1 | 15 |
| fam234a | 0.59713 | 0.191880292 | 0.278 | 0.174 | 1 | 15 |
| smim12 | 0.59721 | 0.151578547 | 0.167 | 0.087 | 1 | 15 |
| hmgn7 | 0.59732 | 0.137398045 | 0.806 | 0.739 | 1 | 15 |
| myh10 | 0.59743 | 0.138579738 | 0.222 | 0.13 | 1 | 15 |
| il4r.1 | 0.59777 | -0.13285039 | 0.056 | 0.13 | 1 | 15 |
| trim69 | 0.59804 | -0.1430734 | 0.056 | 0.13 | 1 | 15 |
| cyp2n13 | 0.59884 | 0.163695376 | 0.167 | 0.087 | 1 | 15 |
| colec11 | 0.59894 | 0.134089632 | 0.111 | 0.043 | 1 | 15 |
| lsm7 | 0.59897 | -0.13251706 | 0.278 | 0.391 | 1 | 15 |
| atrnl1b | 0.59945 | 0.123332489 | 0.111 | 0.043 | 1 | 15 |
| si:dkey-23i12.5 | 0.60001 | -0.1439909 | 0.194 | 0.261 | 1 | 15 |
| poglut1 | 0.60077 | -0.13761112 | 0.056 | 0.13 | 1 | 15 |

| | | | | | | |
|---|---|---|---|---|---|---|
| aif1l | 0.60096 | -0.18421462 | 0.222 | 0.304 | 1 | 15 |
| adcy9 | 0.60126 | 0.10455355 | 0.083 | 0.043 | 1 | 15 |
| rnf2 | 0.60163 | -0.18047015 | 0.194 | 0.304 | 1 | 15 |
| pam | 0.60217 | 0.135927967 | 0.111 | 0.043 | 1 | 15 |
| nutf2 | 0.60223 | 0.148717062 | 0.167 | 0.087 | 1 | 15 |
| sh3pxd2aa | 0.60232 | -0.13759368 | 0.083 | 0.13 | 1 | 15 |
| vps11 | 0.6028 | 0.112632779 | 0.111 | 0.043 | 1 | 15 |
| CR352226.5 | 0.60316 | -0.10982138 | 0.056 | 0.13 | 1 | 15 |
| zyx | 0.60362 | -0.15674891 | 0.222 | 0.304 | 1 | 15 |
| ppp4r2a | 0.60376 | 0.143051473 | 0.111 | 0.043 | 1 | 15 |
| caprin1a | 0.60379 | -0.14122178 | 0.056 | 0.13 | 1 | 15 |
| pdcd6 | 0.60406 | -0.11354344 | 0.056 | 0.13 | 1 | 15 |
| abhd10a | 0.60421 | -0.13000872 | 0.056 | 0.13 | 1 | 15 |
| cox20 | 0.60443 | -0.11315376 | 0.056 | 0.13 | 1 | 15 |
| kita | 0.60459 | 0.145884422 | 0.111 | 0.043 | 1 | 15 |
| mdm4 | 0.60531 | -0.11571509 | 0.056 | 0.13 | 1 | 15 |
| smim15 | 0.60566 | 0.144715433 | 0.333 | 0.261 | 1 | 15 |
| slc25a26 | 0.60657 | 0.145522077 | 0.139 | 0.087 | 1 | 15 |
| paf1 | 0.60659 | -0.12924856 | 0.056 | 0.13 | 1 | 15 |
| leng1 | 0.60673 | -0.12878764 | 0.056 | 0.13 | 1 | 15 |
| psmd13 | 0.60728 | 0.133626408 | 0.194 | 0.13 | 1 | 15 |
| tpra | 0.60768 | -0.12195352 | 0.056 | 0.13 | 1 | 15 |
| homezb | 0.60863 | -0.12918227 | 0.056 | 0.13 | 1 | 15 |
| mob2a | 0.60875 | -0.12263263 | 0.056 | 0.13 | 1 | 15 |
| eef2k | 0.60887 | -0.11694865 | 0.056 | 0.13 | 1 | 15 |
| prkcz | 0.60904 | 0.136512542 | 0.111 | 0.043 | 1 | 15 |
| ube2l3a | 0.60936 | -0.17939469 | 0.194 | 0.304 | 1 | 15 |
| utp25 | 0.60949 | -0.11258513 | 0.056 | 0.13 | 1 | 15 |
| zgc:175088 | 0.60972 | 0.13413827 | 0.278 | 0.174 | 1 | 15 |
| mrps9 | 0.60988 | -0.15681343 | 0.139 | 0.217 | 1 | 15 |
| hnrnpabb | 0.60999 | 0.143273379 | 0.778 | 0.696 | 1 | 15 |
| nup160 | 0.61009 | 0.104088769 | 0.083 | 0.043 | 1 | 15 |
| ociad1 | 0.61019 | -0.12449291 | 0.056 | 0.13 | 1 | 15 |
| cmtr1 | 0.61025 | -0.11799977 | 0.083 | 0.13 | 1 | 15 |
| eif2b3 | 0.61038 | -0.11657159 | 0.056 | 0.13 | 1 | 15 |
| rbms3 | 0.6105 | -0.1002447 | 0.056 | 0.13 | 1 | 15 |
| pgk1 | 0.61079 | -0.15231356 | 0.167 | 0.261 | 1 | 15 |
| taok1b | 0.61113 | 0.143972754 | 0.111 | 0.043 | 1 | 15 |
| supt4h1 | 0.61149 | 0.156567589 | 0.222 | 0.13 | 1 | 15 |
| palm1b | 0.61151 | -0.13556758 | 0.056 | 0.13 | 1 | 15 |
| fahd2a | 0.61155 | 0.100980441 | 0.167 | 0.087 | 1 | 15 |
| rplp0 | 0.61175 | 0.148700996 | 0.861 | 0.783 | 1 | 15 |
| prkaa1 | 0.61184 | 0.141968145 | 0.111 | 0.043 | 1 | 15 |
| pithd1 | 0.612 | 0.103443175 | 0.139 | 0.087 | 1 | 15 |
| aqr | 0.61217 | 0.130219758 | 0.111 | 0.043 | 1 | 15 |
| znf362b | 0.61234 | 0.122787602 | 0.111 | 0.043 | 1 | 15 |
| ptpn1 | 0.61249 | 0.135145437 | 0.111 | 0.043 | 1 | 15 |
| XLOC-024248 | 0.61257 | -0.12631954 | 0.056 | 0.13 | 1 | 15 |

| | | | | | | |
|---|---|---|---|---|---|---|
| kdm5ba | 0.61259 | -0.15225822 | 0.056 | 0.13 | 1 | 15 |
| txndc9 | 0.61266 | 0.117925589 | 0.472 | 0.435 | 1 | 15 |
| fam210aa | 0.61286 | -0.11192218 | 0.056 | 0.087 | 1 | 15 |
| rtcb | 0.61293 | -0.13255201 | 0.056 | 0.13 | 1 | 15 |
| col27a1b | 0.61293 | -0.16775815 | 0.194 | 0.304 | 1 | 15 |
| mcm7 | 0.61321 | 0.113239206 | 0.111 | 0.043 | 1 | 15 |
| bud23 | 0.61333 | -0.1005416 | 0.056 | 0.13 | 1 | 15 |
| ramp2 | 0.61333 | -0.12468225 | 0.056 | 0.13 | 1 | 15 |
| flii | 0.61337 | -0.16431206 | 0.056 | 0.13 | 1 | 15 |
| si:ch211-150o23.2 | 0.61353 | -0.14257263 | 0.056 | 0.13 | 1 | 15 |
| paqr6 | 0.61359 | -0.14540514 | 0.056 | 0.13 | 1 | 15 |
| CABZ01074994.1 | 0.61361 | 0.131385025 | 0.111 | 0.043 | 1 | 15 |
| si:ch211-71n6.4 | 0.61413 | -0.11494362 | 0.056 | 0.13 | 1 | 15 |
| pdcd2 | 0.61418 | -0.13195747 | 0.056 | 0.13 | 1 | 15 |
| cdc34b | 0.6143 | 0.167047181 | 0.278 | 0.174 | 1 | 15 |
| atg16l1 | 0.61467 | -0.10477134 | 0.056 | 0.087 | 1 | 15 |
| e2f4 | 0.61486 | -0.13757982 | 0.056 | 0.13 | 1 | 15 |
| mrpl47 | 0.61503 | 0.113075714 | 0.222 | 0.13 | 1 | 15 |
| hiat1a | 0.61507 | -0.12685382 | 0.056 | 0.13 | 1 | 15 |
| tsn | 0.6151 | 0.109200629 | 0.083 | 0.043 | 1 | 15 |
| si:ch211-250c4.4 | 0.61519 | -0.15670806 | 0.056 | 0.13 | 1 | 15 |
| ciao2a | 0.61525 | -0.11451153 | 0.056 | 0.13 | 1 | 15 |
| si:ch1073-157b13.1 | 0.61541 | -0.11182863 | 0.056 | 0.13 | 1 | 15 |
| gata5 | 0.61555 | -0.1498398 | 0.056 | 0.13 | 1 | 15 |
| foxo4 | 0.61561 | -0.16076982 | 0.056 | 0.13 | 1 | 15 |
| pank2 | 0.61561 | -0.14015386 | 0.056 | 0.13 | 1 | 15 |
| dusp1 | 0.61565 | -0.12853735 | 0.056 | 0.13 | 1 | 15 |
| got1 | 0.61569 | 0.133581909 | 0.111 | 0.043 | 1 | 15 |
| borcs7 | 0.61589 | -0.10884674 | 0.056 | 0.087 | 1 | 15 |
| nck2a | 0.61591 | -0.11230416 | 0.056 | 0.13 | 1 | 15 |
| zgc:173607 | 0.61606 | -0.13903735 | 0.056 | 0.13 | 1 | 15 |
| si:ch211-261d7.3 | 0.61609 | -0.10395521 | 0.056 | 0.13 | 1 | 15 |
| dcn | 0.61613 | -0.14734382 | 0.056 | 0.13 | 1 | 15 |
| frem2a | 0.61616 | -0.13510555 | 0.056 | 0.13 | 1 | 15 |
| akr1a1b | 0.61617 | -0.12187554 | 0.056 | 0.13 | 1 | 15 |
| bmi1b | 0.61617 | -0.11429955 | 0.056 | 0.13 | 1 | 15 |
| kdm2ba | 0.61627 | 0.181184964 | 0.278 | 0.174 | 1 | 15 |
| fam114a2 | 0.61768 | 0.117371312 | 0.111 | 0.043 | 1 | 15 |
| zgc:153993 | 0.61793 | 0.128809956 | 0.111 | 0.043 | 1 | 15 |
| csrp1a | 0.61809 | -0.10513213 | 0.139 | 0.174 | 1 | 15 |
| fxr1 | 0.61818 | 0.112017849 | 0.111 | 0.087 | 1 | 15 |
| sgf29 | 0.61861 | 0.114355456 | 0.111 | 0.043 | 1 | 15 |
| asf1bb | 0.61905 | 0.1368847 | 0.111 | 0.043 | 1 | 15 |
| lsr | 0.61915 | -0.16229007 | 0.222 | 0.304 | 1 | 15 |
| blmh | 0.61986 | 0.152841932 | 0.278 | 0.174 | 1 | 15 |
| psmd5 | 0.62034 | 0.105375701 | 0.083 | 0.043 | 1 | 15 |
| srgap2 | 0.62073 | 0.135065718 | 0.111 | 0.043 | 1 | 15 |
| tdrd3 | 0.62077 | 0.105574093 | 0.111 | 0.043 | 1 | 15 |

| | | | | | | |
|---|---|---|---|---|---|---|
| trioa | 0.62125 | -0.11481314 | 0.056 | 0.087 | 1 | 15 |
| fam210b | 0.62133 | 0.102060648 | 0.111 | 0.043 | 1 | 15 |
| caskb | 0.62168 | 0.156753536 | 0.167 | 0.087 | 1 | 15 |
| cope | 0.62174 | 0.115405925 | 0.25 | 0.217 | 1 | 15 |
| tnpo1 | 0.62283 | -0.15013886 | 0.056 | 0.087 | 1 | 15 |
| psen1 | 0.62302 | 0.173886165 | 0.222 | 0.13 | 1 | 15 |
| hgsnat | 0.62329 | 0.136923783 | 0.083 | 0.043 | 1 | 15 |
| cpsf2 | 0.62338 | 0.12619331 | 0.111 | 0.043 | 1 | 15 |
| mgaa | 0.62453 | 0.142084053 | 0.167 | 0.087 | 1 | 15 |
| cbx1a | 0.62453 | 0.170046992 | 0.333 | 0.217 | 1 | 15 |
| alcamb | 0.62481 | 0.195121055 | 0.333 | 0.217 | 1 | 15 |
| eif5a | 0.62486 | 0.106453026 | 0.806 | 0.696 | 1 | 15 |
| kdm6al | 0.62507 | 0.133456673 | 0.111 | 0.043 | 1 | 15 |
| ercc1 | 0.62623 | 0.114676407 | 0.111 | 0.043 | 1 | 15 |
| socs2 | 0.6268 | 0.106434214 | 0.111 | 0.043 | 1 | 15 |
| si:ch73-138n13.1 | 0.62716 | -0.10602982 | 0.194 | 0.217 | 1 | 15 |
| dnajc5ga | 0.6272 | 0.107734798 | 0.111 | 0.043 | 1 | 15 |
| ssbp1 | 0.62762 | -0.13159964 | 0.083 | 0.13 | 1 | 15 |
| znhit1 | 0.62778 | -0.10660079 | 0.139 | 0.174 | 1 | 15 |
| clpp | 0.62782 | 0.132848783 | 0.278 | 0.174 | 1 | 15 |
| snx5 | 0.62817 | 0.103356966 | 0.167 | 0.087 | 1 | 15 |
| si:ch211-288g17.3 | 0.62842 | -0.15216389 | 0.389 | 0.435 | 1 | 15 |
| anxa4 | 0.62853 | -0.15353397 | 0.194 | 0.304 | 1 | 15 |
| yipf6 | 0.62857 | -0.16692551 | 0.167 | 0.261 | 1 | 15 |
| mrpl46 | 0.62863 | 0.163598274 | 0.222 | 0.13 | 1 | 15 |
| rnf185 | 0.62893 | 0.110020385 | 0.167 | 0.087 | 1 | 15 |
| arid1ab | 0.62895 | -0.107503 | 0.056 | 0.087 | 1 | 15 |
| usp8 | 0.62918 | 0.13298526 | 0.222 | 0.13 | 1 | 15 |
| vasna | 0.62975 | 0.128493186 | 0.111 | 0.043 | 1 | 15 |
| lamb1a | 0.63012 | -0.12304138 | 0.667 | 0.696 | 1 | 15 |
| si:zfos-169g10.3 | 0.63076 | 0.155464192 | 0.333 | 0.261 | 1 | 15 |
| map2k6 | 0.63088 | -0.17253025 | 0.194 | 0.304 | 1 | 15 |
| CABZ01008730.1 | 0.63089 | 0.110307284 | 0.111 | 0.043 | 1 | 15 |
| ube2e2 | 0.63115 | 0.139081661 | 0.222 | 0.13 | 1 | 15 |
| mrpl48 | 0.6316 | 0.157688978 | 0.111 | 0.043 | 1 | 15 |
| si:ch211-214p13.9 | 0.63161 | 0.127469501 | 0.111 | 0.043 | 1 | 15 |
| map2k1 | 0.63187 | 0.111016736 | 0.167 | 0.087 | 1 | 15 |
| rab7a | 0.63235 | 0.156529478 | 0.278 | 0.174 | 1 | 15 |
| pop7 | 0.6324 | 0.103135676 | 0.111 | 0.043 | 1 | 15 |
| amfra | 0.63283 | 0.139078487 | 0.111 | 0.043 | 1 | 15 |
| rbfox2 | 0.63286 | 0.125093416 | 0.222 | 0.13 | 1 | 15 |
| cdk5rap1 | 0.633 | 0.109110492 | 0.111 | 0.043 | 1 | 15 |
| grwd1 | 0.6334 | 0.125321592 | 0.111 | 0.043 | 1 | 15 |
| hnrnph1l | 0.6335 | 0.115751132 | 0.361 | 0.261 | 1 | 15 |
| cutc | 0.63377 | 0.107096895 | 0.111 | 0.043 | 1 | 15 |
| rnf41 | 0.63379 | 0.101630888 | 0.083 | 0.043 | 1 | 15 |
| pros1 | 0.6339 | 0.128617687 | 0.222 | 0.13 | 1 | 15 |
| CABZ01113954.1 | 0.63391 | 0.109518145 | 0.111 | 0.043 | 1 | 15 |

| | | | | | | |
|---|---|---|---|---|---|---|
| pitpnb | 0.63409 | 0.14813033 | 0.167 | 0.087 | 1 | 15 |
| pcmtd1 | 0.63412 | 0.100369551 | 0.139 | 0.087 | 1 | 15 |
| gzf1 | 0.6342 | 0.113138859 | 0.167 | 0.087 | 1 | 15 |
| tasp1 | 0.63439 | 0.123868485 | 0.111 | 0.043 | 1 | 15 |
| ppp5c | 0.6345 | 0.121382952 | 0.111 | 0.043 | 1 | 15 |
| psen2 | 0.63465 | 0.144008819 | 0.111 | 0.043 | 1 | 15 |
| hcfc1b | 0.63465 | -0.12749217 | 0.056 | 0.087 | 1 | 15 |
| arl2 | 0.63524 | 0.168581435 | 0.222 | 0.13 | 1 | 15 |
| znf330 | 0.63556 | -0.11509714 | 0.167 | 0.261 | 1 | 15 |
| zgc:110182 | 0.63564 | -0.12007922 | 0.361 | 0.391 | 1 | 15 |
| nr2c1 | 0.63587 | 0.122839177 | 0.111 | 0.043 | 1 | 15 |
| tmem107 | 0.63602 | 0.102566733 | 0.111 | 0.043 | 1 | 15 |
| upf3b | 0.63603 | 0.150812272 | 0.167 | 0.087 | 1 | 15 |
| kirrel1a | 0.63608 | 0.103527837 | 0.083 | 0.043 | 1 | 15 |
| ccnc | 0.63654 | 0.121414971 | 0.111 | 0.043 | 1 | 15 |
| acsl1a | 0.63694 | 0.121891538 | 0.111 | 0.043 | 1 | 15 |
| tle2a | 0.63711 | 0.130998282 | 0.139 | 0.087 | 1 | 15 |
| syncrip | 0.63718 | -0.14246902 | 0.528 | 0.652 | 1 | 15 |
| atp5f1d | 0.63858 | -0.12507857 | 0.5 | 0.522 | 1 | 15 |
| sypl2a | 0.63864 | 0.102788688 | 0.167 | 0.13 | 1 | 15 |
| ywhaqb | 0.63881 | 0.175826502 | 0.472 | 0.348 | 1 | 15 |
| sdad1 | 0.63884 | 0.104864522 | 0.111 | 0.043 | 1 | 15 |
| arhgef1b | 0.63911 | -0.20236503 | 0.278 | 0.391 | 1 | 15 |
| rab40b | 0.63913 | 0.115215941 | 0.111 | 0.043 | 1 | 15 |
| dynlt3 | 0.63914 | 0.127867331 | 0.278 | 0.217 | 1 | 15 |
| atic | 0.63916 | 0.12719087 | 0.111 | 0.043 | 1 | 15 |
| insig2 | 0.63953 | 0.153949516 | 0.111 | 0.043 | 1 | 15 |
| vcp | 0.63966 | 0.135767765 | 0.361 | 0.304 | 1 | 15 |
| XLOC-043857 | 0.63986 | -0.1228364 | 0.167 | 0.217 | 1 | 15 |
| sema7a | 0.64044 | 0.185868897 | 0.639 | 0.522 | 1 | 15 |
| si:ch211-125a15.1 | 0.64047 | 0.101340297 | 0.111 | 0.043 | 1 | 15 |
| selenop | 0.64056 | 0.115261708 | 0.111 | 0.043 | 1 | 15 |
| si:ch211-276i12.9 | 0.64067 | 0.101590409 | 0.111 | 0.043 | 1 | 15 |
| march5 | 0.64075 | 0.119654338 | 0.111 | 0.043 | 1 | 15 |
| larp4ab | 0.64105 | -0.17414924 | 0.167 | 0.261 | 1 | 15 |
| RASA2 | 0.6412 | 0.123287385 | 0.111 | 0.043 | 1 | 15 |
| eed | 0.64129 | 0.100454799 | 0.111 | 0.043 | 1 | 15 |
| arhgap35b | 0.64159 | 0.118255386 | 0.111 | 0.043 | 1 | 15 |
| tacc1 | 0.64207 | 0.101093401 | 0.111 | 0.043 | 1 | 15 |
| parp12a | 0.6423 | 0.122605484 | 0.361 | 0.261 | 1 | 15 |
| ppm1k | 0.64246 | 0.122540489 | 0.111 | 0.043 | 1 | 15 |
| msna | 0.64267 | -0.20329449 | 0.25 | 0.348 | 1 | 15 |
| pcca | 0.6429 | 0.11074332 | 0.111 | 0.043 | 1 | 15 |
| si:ch73-386h18.1 | 0.64294 | 0.110722384 | 0.083 | 0.043 | 1 | 15 |
| oxsm | 0.64314 | 0.110003046 | 0.111 | 0.043 | 1 | 15 |
| tmpoa | 0.64331 | 0.112902566 | 0.167 | 0.13 | 1 | 15 |
| psmd10 | 0.64331 | 0.111570765 | 0.111 | 0.043 | 1 | 15 |
| hccsb | 0.64333 | 0.116908455 | 0.111 | 0.043 | 1 | 15 |

| | | | | | | |
|---|---|---|---|---|---|---|
| mov10a | 0.64342 | 0.112975296 | 0.111 | 0.043 | 1 | 15 |
| naa40 | 0.64353 | 0.136539682 | 0.278 | 0.174 | 1 | 15 |
| tra2b | 0.64379 | 0.111957803 | 0.111 | 0.043 | 1 | 15 |
| clk4a | 0.64386 | 0.183376253 | 0.278 | 0.174 | 1 | 15 |
| eef2a.1 | 0.64388 | 0.115230775 | 0.111 | 0.043 | 1 | 15 |
| ell2 | 0.6441 | 0.113833462 | 0.111 | 0.043 | 1 | 15 |
| znf703 | 0.64443 | 0.119982612 | 0.111 | 0.043 | 1 | 15 |
| ubald1a | 0.64448 | 0.107917682 | 0.111 | 0.043 | 1 | 15 |
| si:dkey-85k15.4 | 0.64449 | 0.108065661 | 0.111 | 0.043 | 1 | 15 |
| ptenb | 0.6445 | -0.16562887 | 0.139 | 0.217 | 1 | 15 |
| cog4 | 0.6445 | 0.112848058 | 0.111 | 0.043 | 1 | 15 |
| hsd17b12b | 0.64456 | 0.110744809 | 0.111 | 0.043 | 1 | 15 |
| gid8a | 0.64457 | 0.101558231 | 0.111 | 0.043 | 1 | 15 |
| cops3 | 0.64457 | 0.104304168 | 0.111 | 0.043 | 1 | 15 |
| synm | 0.64458 | 0.123990471 | 0.111 | 0.043 | 1 | 15 |
| srsf3a | 0.64476 | 0.118674287 | 0.139 | 0.087 | 1 | 15 |
| wu:fj29h11 | 0.64559 | -0.10596914 | 0.139 | 0.217 | 1 | 15 |
| aff2 | 0.64559 | 0.143984686 | 0.222 | 0.13 | 1 | 15 |
| dctn2 | 0.64608 | 0.106015485 | 0.167 | 0.087 | 1 | 15 |
| mrpl27 | 0.64649 | 0.105915407 | 0.167 | 0.13 | 1 | 15 |
| irf2bp2a | 0.64663 | -0.10981878 | 0.278 | 0.304 | 1 | 15 |
| rrp7a | 0.64739 | 0.111625966 | 0.167 | 0.087 | 1 | 15 |
| fra10ac1 | 0.64792 | 0.111765402 | 0.083 | 0.043 | 1 | 15 |
| hcfc1a | 0.6486 | 0.168163045 | 0.278 | 0.174 | 1 | 15 |
| ppp2r2d | 0.6492 | -0.16345019 | 0.139 | 0.217 | 1 | 15 |
| MCMDC2 | 0.64923 | -0.13377564 | 0.111 | 0.174 | 1 | 15 |
| litaf | 0.64926 | -0.11462291 | 0.167 | 0.261 | 1 | 15 |
| myl9b | 0.6502 | 0.175823936 | 0.278 | 0.174 | 1 | 15 |
| adgrl3.1 | 0.65024 | 0.181177128 | 0.278 | 0.174 | 1 | 15 |
| ets2 | 0.65025 | 0.116803856 | 0.167 | 0.087 | 1 | 15 |
| gpn2 | 0.65027 | -0.10059574 | 0.056 | 0.087 | 1 | 15 |
| rbm22 | 0.65082 | 0.14029119 | 0.194 | 0.13 | 1 | 15 |
| pi4kb | 0.65185 | -0.14506016 | 0.167 | 0.261 | 1 | 15 |
| gmfb | 0.65246 | 0.163396191 | 0.389 | 0.304 | 1 | 15 |
| naa50 | 0.65246 | 0.135543202 | 0.194 | 0.13 | 1 | 15 |
| plcl1 | 0.6525 | 0.156661913 | 0.194 | 0.13 | 1 | 15 |
| eif3m | 0.653 | 0.144248241 | 0.75 | 0.652 | 1 | 15 |
| si:ch73-269m14.2 | 0.65302 | -0.12639221 | 0.167 | 0.261 | 1 | 15 |
| asah1b | 0.65311 | -0.11148495 | 0.278 | 0.304 | 1 | 15 |
| col16a1 | 0.65352 | 0.132491022 | 0.222 | 0.13 | 1 | 15 |
| arhgap1 | 0.65382 | -0.10442209 | 0.056 | 0.087 | 1 | 15 |
| crnkl1 | 0.65407 | 0.110860719 | 0.083 | 0.043 | 1 | 15 |
| CABZ01068367.1 | 0.65428 | 0.125735943 | 0.5 | 0.391 | 1 | 15 |
| manf | 0.65457 | 0.162868056 | 0.222 | 0.13 | 1 | 15 |
| ubr5 | 0.65478 | 0.128236133 | 0.222 | 0.13 | 1 | 15 |
| calm2a | 0.65484 | -0.12983894 | 0.167 | 0.217 | 1 | 15 |
| mia3 | 0.65493 | -0.12671039 | 0.083 | 0.13 | 1 | 15 |
| mccc2 | 0.65546 | -0.1105124 | 0.139 | 0.174 | 1 | 15 |

| | | | | | | |
|---|---|---|---|---|---|---|
| crebzf | 0.65594 | -0.10910735 | 0.333 | 0.435 | 1 | 15 |
| ctsba | 0.65624 | 0.16785524 | 0.444 | 0.348 | 1 | 15 |
| pmaip1 | 0.65631 | 0.118524084 | 0.167 | 0.087 | 1 | 15 |
| fbxo11a | 0.65636 | 0.118869485 | 0.167 | 0.087 | 1 | 15 |
| dpm3 | 0.65668 | 0.119452074 | 0.139 | 0.087 | 1 | 15 |
| postnb | 0.65673 | -0.2193676 | 0.278 | 0.391 | 1 | 15 |
| ikbke | 0.65681 | 0.117775637 | 0.139 | 0.087 | 1 | 15 |
| tdp2a | 0.65758 | 0.11000298 | 0.083 | 0.043 | 1 | 15 |
| hm13 | 0.65775 | -0.13000738 | 0.111 | 0.174 | 1 | 15 |
| mafba | 0.65778 | -0.1755236 | 0.444 | 0.565 | 1 | 15 |
| rcan1a | 0.65793 | -0.13657391 | 0.167 | 0.217 | 1 | 15 |
| ino80c | 0.65818 | 0.117400704 | 0.139 | 0.087 | 1 | 15 |
| ckba | 0.65924 | 0.167137942 | 0.278 | 0.217 | 1 | 15 |
| CU929037.3 | 0.66041 | 0.10540905 | 0.083 | 0.043 | 1 | 15 |
| nrp2a | 0.66102 | -0.1197637 | 0.139 | 0.217 | 1 | 15 |
| fosl2 | 0.66218 | -0.10102894 | 0.056 | 0.087 | 1 | 15 |
| pdk2a | 0.66251 | 0.175544744 | 0.222 | 0.13 | 1 | 15 |
| spcs1 | 0.66277 | -0.12186182 | 0.528 | 0.565 | 1 | 15 |
| si:ch211-256e16.11 | 0.664 | 0.107822674 | 0.083 | 0.043 | 1 | 15 |
| cpsf6 | 0.66404 | -0.15480721 | 0.278 | 0.391 | 1 | 15 |
| zfp36l1a | 0.66409 | -0.15706897 | 0.278 | 0.391 | 1 | 15 |
| aldh3a2a | 0.66458 | 0.127864539 | 0.167 | 0.087 | 1 | 15 |
| cnot6a | 0.66481 | 0.1530448 | 0.222 | 0.13 | 1 | 15 |
| zgc:77849 | 0.66527 | 0.150495833 | 0.306 | 0.217 | 1 | 15 |
| galnt1 | 0.66547 | 0.120734213 | 0.167 | 0.087 | 1 | 15 |
| psmc1a | 0.66578 | -0.15695388 | 0.306 | 0.391 | 1 | 15 |
| si:dkey-180p18.9 | 0.66613 | 0.11862336 | 0.167 | 0.087 | 1 | 15 |
| snapc1b | 0.6669 | 0.154443364 | 0.167 | 0.087 | 1 | 15 |
| mcfd2 | 0.66697 | 0.142092287 | 0.222 | 0.13 | 1 | 15 |
| swap70b | 0.66698 | 0.142051144 | 0.167 | 0.087 | 1 | 15 |
| pphln1 | 0.66722 | 0.116520838 | 0.167 | 0.087 | 1 | 15 |
| rchy1 | 0.66763 | 0.11267161 | 0.139 | 0.087 | 1 | 15 |
| pid1 | 0.66782 | 0.145977989 | 0.167 | 0.087 | 1 | 15 |
| dynll2a | 0.66856 | 0.154386722 | 0.222 | 0.13 | 1 | 15 |
| wu:fu71h07 | 0.66866 | 0.119097346 | 0.167 | 0.087 | 1 | 15 |
| irf2bp2b | 0.67004 | 0.152106506 | 0.639 | 0.522 | 1 | 15 |
| mob1a | 0.67012 | 0.102226462 | 0.306 | 0.217 | 1 | 15 |
| irf2bp1 | 0.67035 | -0.10693954 | 0.056 | 0.087 | 1 | 15 |
| selenok | 0.67036 | 0.115884581 | 0.5 | 0.391 | 1 | 15 |
| dlst | 0.67095 | 0.167466042 | 0.417 | 0.304 | 1 | 15 |
| mrps34 | 0.6714 | 0.117484557 | 0.139 | 0.087 | 1 | 15 |
| hspa4a | 0.67144 | 0.117279086 | 0.167 | 0.087 | 1 | 15 |
| tm2d2 | 0.67167 | 0.143132608 | 0.167 | 0.087 | 1 | 15 |
| sec62 | 0.67172 | -0.16873289 | 0.25 | 0.348 | 1 | 15 |
| amer1 | 0.672 | 0.142904558 | 0.167 | 0.087 | 1 | 15 |
| thtpa | 0.67244 | 0.139547879 | 0.167 | 0.087 | 1 | 15 |
| CR749775.2 | 0.6726 | -0.101589 | 0.056 | 0.087 | 1 | 15 |
| ube2a | 0.67261 | 0.119799575 | 0.167 | 0.087 | 1 | 15 |

| | | | | | | |
|---|---|---|---|---|---|---|
| yipf5 | 0.67266 | 0.14542791 | 0.167 | 0.087 | 1 | 15 |
| psma6b | 0.67274 | 0.137028189 | 0.278 | 0.217 | 1 | 15 |
| si:ch211-136a13.1 | 0.67301 | -0.20429313 | 0.167 | 0.261 | 1 | 15 |
| acin1b | 0.67345 | 0.117113646 | 0.139 | 0.087 | 1 | 15 |
| wdr5 | 0.67358 | -0.11874674 | 0.083 | 0.13 | 1 | 15 |
| arfrp1 | 0.67396 | 0.142069194 | 0.25 | 0.174 | 1 | 15 |
| cdkn1bb | 0.67494 | -0.15471746 | 0.167 | 0.261 | 1 | 15 |
| cmtm7 | 0.67518 | -0.11939595 | 0.083 | 0.13 | 1 | 15 |
| snrnp200 | 0.67522 | -0.13628995 | 0.111 | 0.174 | 1 | 15 |
| med11 | 0.6753 | 0.139260986 | 0.194 | 0.13 | 1 | 15 |
| csnk1a1 | 0.67549 | -0.14089463 | 0.5 | 0.565 | 1 | 15 |
| syf2 | 0.67656 | -0.12839794 | 0.333 | 0.391 | 1 | 15 |
| dhrs3a | 0.67728 | -0.17604189 | 0.167 | 0.261 | 1 | 15 |
| XLOC-015291 | 0.67736 | 0.163269345 | 0.306 | 0.217 | 1 | 15 |
| cops2 | 0.67772 | 0.1299685 | 0.194 | 0.13 | 1 | 15 |
| snrpa1 | 0.67799 | -0.14246826 | 0.194 | 0.261 | 1 | 15 |
| alcama | 0.67814 | 0.166620933 | 0.417 | 0.304 | 1 | 15 |
| psenen | 0.68127 | 0.165827692 | 0.417 | 0.304 | 1 | 15 |
| ndufa6 | 0.68152 | -0.16122797 | 0.417 | 0.522 | 1 | 15 |
| khsrp | 0.68419 | -0.12663794 | 0.167 | 0.261 | 1 | 15 |
| txnl4a | 0.68447 | 0.104866477 | 0.306 | 0.261 | 1 | 15 |
| npc2 | 0.68532 | -0.13390041 | 0.306 | 0.348 | 1 | 15 |
| faxdc2 | 0.68559 | 0.112779129 | 0.111 | 0.087 | 1 | 15 |
| gli3 | 0.686 | 0.138613887 | 0.194 | 0.13 | 1 | 15 |
| atp5l | 0.6863 | 0.10164838 | 0.833 | 0.739 | 1 | 15 |
| gng12a | 0.68663 | -0.15276443 | 0.167 | 0.261 | 1 | 15 |
| ogt.1 | 0.68664 | -0.14918416 | 0.167 | 0.261 | 1 | 15 |
| smim4 | 0.68733 | 0.128409979 | 0.083 | 0.043 | 1 | 15 |
| lrrc8da | 0.68795 | 0.118010158 | 0.083 | 0.043 | 1 | 15 |
| si:dkey-42i9.7 | 0.68877 | -0.145507 | 0.389 | 0.435 | 1 | 15 |
| serp1 | 0.68968 | 0.151343598 | 0.306 | 0.217 | 1 | 15 |
| glo1 | 0.69 | -0.13411647 | 0.139 | 0.217 | 1 | 15 |
| eci2 | 0.69215 | 0.139012339 | 0.25 | 0.174 | 1 | 15 |
| si:ch73-1a9.4 | 0.69235 | -0.10925744 | 0.056 | 0.087 | 1 | 15 |
| rhpn2 | 0.69272 | 0.121677291 | 0.139 | 0.087 | 1 | 15 |
| polr2eb | 0.69286 | 0.116393881 | 0.361 | 0.261 | 1 | 15 |
| si:dkey-54n8.2 | 0.69372 | 0.114672868 | 0.083 | 0.043 | 1 | 15 |
| eipr1 | 0.69412 | 0.103526992 | 0.083 | 0.043 | 1 | 15 |
| tiparp | 0.69585 | -0.13287852 | 0.083 | 0.13 | 1 | 15 |
| si:dkey-112a7.4 | 0.69604 | -0.14722534 | 0.194 | 0.261 | 1 | 15 |
| hnrnpul1l | 0.69714 | -0.10544131 | 0.139 | 0.217 | 1 | 15 |
| srpra | 0.69749 | -0.1288929 | 0.167 | 0.217 | 1 | 15 |
| cry1ab | 0.69803 | -0.15622581 | 0.111 | 0.174 | 1 | 15 |
| ssbp4 | 0.69826 | 0.100332119 | 0.083 | 0.043 | 1 | 15 |
| si:ch73-368j24.10.1 | 0.69863 | -0.1203902 | 0.083 | 0.13 | 1 | 15 |
| wu:fe05a04 | 0.69944 | -0.11844977 | 0.083 | 0.13 | 1 | 15 |
| ptbp1b | 0.6997 | 0.113996642 | 0.139 | 0.087 | 1 | 15 |
| dnajc10 | 0.70138 | 0.101577308 | 0.111 | 0.087 | 1 | 15 |

| | | | | | | |
|---|---|---|---|---|---|---|
| pafah1b1a | 0.70156 | -0.15609354 | 0.139 | 0.217 | 1 | 15 |
| hpn | 0.70162 | 0.172625987 | 0.194 | 0.13 | 1 | 15 |
| ctbp1 | 0.7018 | 0.137936007 | 0.361 | 0.304 | 1 | 15 |
| bsg | 0.70283 | -0.12106952 | 0.5 | 0.609 | 1 | 15 |
| cdip1 | 0.70449 | 0.121966075 | 0.139 | 0.087 | 1 | 15 |
| higd1a | 0.70522 | 0.120868352 | 0.444 | 0.391 | 1 | 15 |
| rsad2 | 0.70543 | 0.161178787 | 0.083 | 0.043 | 1 | 15 |
| rnf167 | 0.70646 | 0.11049168 | 0.139 | 0.087 | 1 | 15 |
| ubald2 | 0.70664 | -0.12596937 | 0.111 | 0.174 | 1 | 15 |
| msrb1b | 0.70701 | 0.10578546 | 0.139 | 0.087 | 1 | 15 |
| tcf3b | 0.70791 | -0.16167558 | 0.111 | 0.174 | 1 | 15 |
| si:dkey-286j15.3 | 0.70946 | -0.11866719 | 0.111 | 0.174 | 1 | 15 |
| acoxl | 0.71051 | 0.100586311 | 0.083 | 0.043 | 1 | 15 |
| ip6k2a | 0.71115 | -0.14562808 | 0.139 | 0.217 | 1 | 15 |
| mt-nd2 | 0.71222 | -0.11240904 | 0.639 | 0.739 | 1 | 15 |
| ube2v1 | 0.71237 | -0.13449632 | 0.5 | 0.609 | 1 | 15 |
| zgc:112163 | 0.7132 | 0.104050156 | 0.083 | 0.043 | 1 | 15 |
| emc10 | 0.71361 | -0.15629143 | 0.333 | 0.435 | 1 | 15 |
| ghra | 0.7139 | 0.10390736 | 0.167 | 0.13 | 1 | 15 |
| atp13a1 | 0.71532 | -0.10338918 | 0.083 | 0.13 | 1 | 15 |
| ezh2 | 0.7159 | -0.11012003 | 0.139 | 0.174 | 1 | 15 |
| timm8b | 0.71611 | -0.14146795 | 0.222 | 0.304 | 1 | 15 |
| ak1 | 0.71742 | -0.13492099 | 0.194 | 0.261 | 1 | 15 |
| atp2b2 | 0.7181 | 0.148466503 | 0.361 | 0.261 | 1 | 15 |
| gnb1a | 0.71948 | 0.146858236 | 0.361 | 0.261 | 1 | 15 |
| trappc5 | 0.7196 | 0.124528179 | 0.278 | 0.217 | 1 | 15 |
| perp | 0.72008 | -0.13547556 | 0.139 | 0.217 | 1 | 15 |
| arpc4l | 0.72098 | 0.144098604 | 0.361 | 0.261 | 1 | 15 |
| ndufs7 | 0.72137 | 0.13909073 | 0.472 | 0.391 | 1 | 15 |
| lsm5 | 0.72182 | 0.135201722 | 0.417 | 0.348 | 1 | 15 |
| pfn2l | 0.72265 | 0.131706198 | 0.639 | 0.565 | 1 | 15 |
| fam133b | 0.72266 | 0.132941378 | 0.278 | 0.217 | 1 | 15 |
| ppp3ccb | 0.72303 | -0.13458633 | 0.25 | 0.348 | 1 | 15 |
| srgap1b | 0.72307 | 0.119953524 | 0.139 | 0.087 | 1 | 15 |
| otud5a | 0.72329 | -0.14266442 | 0.111 | 0.174 | 1 | 15 |
| ssr1 | 0.72341 | -0.14849941 | 0.5 | 0.565 | 1 | 15 |
| marcksl1b | 0.72479 | -0.14024337 | 0.417 | 0.522 | 1 | 15 |
| LEPROTL1 | 0.72503 | 0.159426623 | 0.389 | 0.304 | 1 | 15 |
| pa2g4b | 0.72654 | 0.134907153 | 0.194 | 0.13 | 1 | 15 |
| bsdc1 | 0.72731 | -0.13171787 | 0.194 | 0.261 | 1 | 15 |
| CABZ01068358.1 | 0.72778 | -0.12559327 | 0.556 | 0.609 | 1 | 15 |
| klf13 | 0.72801 | -0.14150817 | 0.139 | 0.217 | 1 | 15 |
| ctsd | 0.72825 | 0.147489869 | 0.444 | 0.348 | 1 | 15 |
| cmtm3 | 0.72879 | 0.131468738 | 0.25 | 0.174 | 1 | 15 |
| rbm25b | 0.72905 | -0.12749824 | 0.111 | 0.174 | 1 | 15 |
| pdha1a | 0.72922 | 0.125959089 | 0.306 | 0.261 | 1 | 15 |
| namptb | 0.72978 | -0.11953186 | 0.083 | 0.13 | 1 | 15 |
| mcm6 | 0.73143 | -0.10265261 | 0.111 | 0.174 | 1 | 15 |

| | | | | | | |
|---|---|---|---|---|---|---|
| wls | 0.7315 | -0.13510717 | 0.417 | 0.522 | 1 | 15 |
| mcl1b | 0.73222 | -0.14603852 | 0.417 | 0.522 | 1 | 15 |
| zbtb2b | 0.73239 | 0.10624654 | 0.083 | 0.043 | 1 | 15 |
| psmd4b | 0.73366 | 0.107966066 | 0.222 | 0.174 | 1 | 15 |
| ifngr2 | 0.73402 | -0.11169513 | 0.167 | 0.217 | 1 | 15 |
| ddb1 | 0.73447 | -0.1145703 | 0.083 | 0.13 | 1 | 15 |
| lamtor2 | 0.73537 | -0.14091345 | 0.139 | 0.217 | 1 | 15 |
| ywhae1 | 0.73602 | 0.1185139 | 0.667 | 0.565 | 1 | 15 |
| cox5ab | 0.73638 | -0.13528922 | 0.472 | 0.565 | 1 | 15 |
| XLOC-034863 | 0.7371 | 0.119295718 | 0.194 | 0.13 | 1 | 15 |
| rhogd | 0.73786 | 0.122786795 | 0.194 | 0.13 | 1 | 15 |
| chd4b | 0.73805 | 0.150192039 | 0.306 | 0.217 | 1 | 15 |
| atf6 | 0.7389 | -0.12142889 | 0.222 | 0.261 | 1 | 15 |
| cox8b | 0.73925 | 0.130030475 | 0.444 | 0.348 | 1 | 15 |
| pcna | 0.73952 | 0.143557229 | 0.194 | 0.13 | 1 | 15 |
| atg12 | 0.74026 | 0.10058829 | 0.139 | 0.087 | 1 | 15 |
| sh3gl3b | 0.74028 | -0.12459717 | 0.139 | 0.217 | 1 | 15 |
| kremen1 | 0.74042 | -0.16146932 | 0.139 | 0.217 | 1 | 15 |
| btg2 | 0.74048 | -0.11481928 | 0.611 | 0.652 | 1 | 15 |
| kras | 0.74102 | -0.12629034 | 0.139 | 0.217 | 1 | 15 |
| otub1b | 0.74107 | -0.11351168 | 0.139 | 0.217 | 1 | 15 |
| CZQB01141835.1 | 0.74355 | 0.105016654 | 0.167 | 0.13 | 1 | 15 |
| cdh6 | 0.7444 | -0.10346135 | 0.083 | 0.13 | 1 | 15 |
| sf3b5 | 0.74448 | 0.120781479 | 0.444 | 0.348 | 1 | 15 |
| zfyve27 | 0.74471 | 0.112510544 | 0.194 | 0.13 | 1 | 15 |
| cab39l1 | 0.74495 | -0.1106139 | 0.083 | 0.13 | 1 | 15 |
| rnf10 | 0.74542 | 0.115260402 | 0.444 | 0.348 | 1 | 15 |
| cemip2 | 0.74642 | -0.16549908 | 0.111 | 0.174 | 1 | 15 |
| dpp7 | 0.7471 | -0.11871545 | 0.194 | 0.261 | 1 | 15 |
| pho | 0.74884 | -0.13207367 | 0.167 | 0.217 | 1 | 15 |
| zgc:123258 | 0.74924 | -0.10368366 | 0.056 | 0.087 | 1 | 15 |
| ap2b1 | 0.74959 | 0.134488386 | 0.139 | 0.087 | 1 | 15 |
| PDCL3 | 0.74968 | -0.10194056 | 0.139 | 0.174 | 1 | 15 |
| notch2 | 0.74979 | -0.13894184 | 0.25 | 0.304 | 1 | 15 |
| zgc:100832 | 0.75 | 0.108336619 | 0.083 | 0.043 | 1 | 15 |
| uxs1 | 0.75079 | 0.110816775 | 0.306 | 0.261 | 1 | 15 |
| rpf1 | 0.75128 | -0.1284008 | 0.167 | 0.217 | 1 | 15 |
| mtm1 | 0.75168 | -0.12341367 | 0.083 | 0.13 | 1 | 15 |
| cry-dash | 0.75272 | 0.101123234 | 0.139 | 0.087 | 1 | 15 |
| XLOC-016094 | 0.75369 | -0.10370638 | 0.222 | 0.304 | 1 | 15 |
| si:ch1073-429i10.3.1 | 0.75382 | -0.10694481 | 0.528 | 0.565 | 1 | 15 |
| chmp5b | 0.75385 | 0.123962735 | 0.444 | 0.391 | 1 | 15 |
| sdc4 | 0.75393 | -0.10595037 | 0.556 | 0.652 | 1 | 15 |
| mbnl3 | 0.75445 | 0.122673263 | 0.444 | 0.348 | 1 | 15 |
| plscr3b | 0.75716 | -0.11595931 | 0.111 | 0.174 | 1 | 15 |
| psmg3 | 0.75789 | -0.12373036 | 0.25 | 0.304 | 1 | 15 |
| hdac1 | 0.75811 | 0.143610143 | 0.444 | 0.348 | 1 | 15 |
| ppp4cb | 0.75865 | 0.131943804 | 0.25 | 0.174 | 1 | 15 |

| | | | | | | |
|---|---|---|---|---|---|---|
| abce1 | 0.76051 | 0.106882188 | 0.139 | 0.087 | 1 | 15 |
| anapc13 | 0.7615 | 0.117128295 | 0.139 | 0.087 | 1 | 15 |
| si:dkey-42i9.4 | 0.76171 | -0.10470191 | 0.528 | 0.565 | 1 | 15 |
| rca2.1 | 0.76178 | -0.13681321 | 0.222 | 0.304 | 1 | 15 |
| dram2b | 0.76415 | -0.1264236 | 0.111 | 0.174 | 1 | 15 |
| nfia | 0.76488 | -0.14867548 | 0.306 | 0.391 | 1 | 15 |
| si:ch73-109d9.2 | 0.76587 | 0.102492428 | 0.139 | 0.087 | 1 | 15 |
| tssc4 | 0.76719 | 0.121927392 | 0.25 | 0.174 | 1 | 15 |
| cited4a | 0.76924 | 0.110584584 | 0.139 | 0.087 | 1 | 15 |
| si:dkey-187e18.1 | 0.76961 | 0.138233339 | 0.472 | 0.391 | 1 | 15 |
| kat5b | 0.76972 | 0.141638453 | 0.194 | 0.13 | 1 | 15 |
| tmem256 | 0.7701 | 0.101667324 | 0.25 | 0.174 | 1 | 15 |
| tp53 | 0.77024 | 0.134503388 | 0.333 | 0.261 | 1 | 15 |
| si:ch211-194k22.8 | 0.77181 | -0.10754963 | 0.222 | 0.261 | 1 | 15 |
| tmem30ab | 0.77198 | -0.13071809 | 0.194 | 0.261 | 1 | 15 |
| ptp4a1 | 0.77555 | 0.13625641 | 0.389 | 0.304 | 1 | 15 |
| tmf1 | 0.77577 | -0.10987287 | 0.083 | 0.13 | 1 | 15 |
| adam17b | 0.77695 | -0.11467147 | 0.111 | 0.174 | 1 | 15 |
| ube2d2l | 0.777 | -0.10576009 | 0.083 | 0.13 | 1 | 15 |
| arhgef5 | 0.77798 | -0.13192588 | 0.083 | 0.13 | 1 | 15 |
| kctd5a | 0.77818 | 0.103059621 | 0.389 | 0.304 | 1 | 15 |
| uba1 | 0.77892 | -0.12260092 | 0.222 | 0.304 | 1 | 15 |
| cops9 | 0.77934 | -0.11265267 | 0.222 | 0.304 | 1 | 15 |
| trim47 | 0.7797 | 0.132464953 | 0.333 | 0.261 | 1 | 15 |
| rad54l2 | 0.78016 | -0.11497836 | 0.083 | 0.13 | 1 | 15 |
| foxo1a | 0.78039 | -0.1194478 | 0.194 | 0.217 | 1 | 15 |
| srrm2 | 0.7805 | -0.10715376 | 0.222 | 0.304 | 1 | 15 |
| prkar1ab | 0.78057 | -0.11804396 | 0.222 | 0.304 | 1 | 15 |
| si:dkey-67c22.2 | 0.78148 | -0.13132605 | 0.222 | 0.304 | 1 | 15 |
| zc3h11a | 0.78186 | -0.12754292 | 0.222 | 0.304 | 1 | 15 |
| arpp19b | 0.78339 | 0.119595845 | 0.25 | 0.174 | 1 | 15 |
| ilk | 0.78405 | -0.10505243 | 0.083 | 0.13 | 1 | 15 |
| ndufb9 | 0.78458 | 0.120685558 | 0.583 | 0.522 | 1 | 15 |
| pcyox1 | 0.78485 | -0.11656889 | 0.306 | 0.391 | 1 | 15 |
| taf7 | 0.78694 | -0.116483 | 0.111 | 0.174 | 1 | 15 |
| atp8a1 | 0.78787 | -0.11019216 | 0.111 | 0.174 | 1 | 15 |
| scaf1 | 0.78981 | 0.100912968 | 0.139 | 0.087 | 1 | 15 |
| LO017872.1 | 0.79143 | -0.11423268 | 0.111 | 0.174 | 1 | 15 |
| atrx | 0.79163 | 0.105938028 | 0.611 | 0.522 | 1 | 15 |
| kdm1a | 0.79164 | -0.10699673 | 0.306 | 0.391 | 1 | 15 |
| parn | 0.79278 | -0.10096634 | 0.111 | 0.174 | 1 | 15 |
| eif1axa | 0.79308 | -0.11693809 | 0.25 | 0.304 | 1 | 15 |
| mknk2a | 0.7933 | 0.103735842 | 0.139 | 0.087 | 1 | 15 |
| pdia5 | 0.79357 | -0.13980735 | 0.194 | 0.261 | 1 | 15 |
| rtca | 0.79466 | -0.11639188 | 0.111 | 0.174 | 1 | 15 |
| kdelr2a | 0.79476 | -0.12701511 | 0.194 | 0.261 | 1 | 15 |
| romo1 | 0.79512 | 0.101931432 | 0.611 | 0.522 | 1 | 15 |
| NDUFB1 | 0.79599 | -0.11675179 | 0.306 | 0.391 | 1 | 15 |

| | | | | | | |
|---|---|---|---|---|---|---|
| gabarapb | 0.79659 | 0.115366675 | 0.667 | 0.609 | 1 | 15 |
| skib | 0.79659 | 0.126106779 | 0.194 | 0.13 | 1 | 15 |
| nudt21 | 0.79671 | 0.105268754 | 0.361 | 0.304 | 1 | 15 |
| taf9 | 0.79693 | -0.1174932 | 0.194 | 0.261 | 1 | 15 |
| gb:bc139872 | 0.79834 | 0.116142062 | 0.139 | 0.087 | 1 | 15 |
| fth1a | 0.79857 | -0.1410172 | 0.444 | 0.522 | 1 | 15 |
| peli1b | 0.79899 | -0.11336921 | 0.083 | 0.13 | 1 | 15 |
| COLEC10 | 0.7994 | 0.137094857 | 0.278 | 0.217 | 1 | 15 |
| LOC553316 | 0.80032 | 0.137516719 | 0.389 | 0.304 | 1 | 15 |
| abca1a | 0.80191 | -0.12735009 | 0.167 | 0.217 | 1 | 15 |
| dnttip2 | 0.80288 | -0.10052133 | 0.083 | 0.13 | 1 | 15 |
| kpnb1 | 0.80349 | 0.11803182 | 0.194 | 0.13 | 1 | 15 |
| osr2 | 0.80523 | 0.111484283 | 0.306 | 0.261 | 1 | 15 |
| echs1 | 0.80595 | 0.1279692 | 0.333 | 0.261 | 1 | 15 |
| si:ch211-132p1.3 | 0.80631 | 0.140192154 | 0.333 | 0.261 | 1 | 15 |
| rarga | 0.8068 | -0.12075187 | 0.194 | 0.261 | 1 | 15 |
| si:dkey-220k22.1 | 0.80715 | 0.108826786 | 0.194 | 0.13 | 1 | 15 |
| fam208b | 0.80716 | -0.10108072 | 0.139 | 0.174 | 1 | 15 |
| limk2 | 0.80916 | 0.10401349 | 0.194 | 0.13 | 1 | 15 |
| ube2v2 | 0.80935 | 0.111242032 | 0.528 | 0.478 | 1 | 15 |
| fbxw11b | 0.80948 | 0.106991327 | 0.194 | 0.13 | 1 | 15 |
| aldh9a1a.1 | 0.81031 | -0.1195917 | 0.5 | 0.565 | 1 | 15 |
| dhrs3b | 0.81057 | -0.10005968 | 0.139 | 0.174 | 1 | 15 |
| shc1 | 0.81131 | 0.127369457 | 0.194 | 0.13 | 1 | 15 |
| mrpl17 | 0.81158 | 0.101205891 | 0.194 | 0.13 | 1 | 15 |
| dennd4a | 0.81164 | -0.10249113 | 0.056 | 0.087 | 1 | 15 |
| senp2 | 0.81164 | 0.113368254 | 0.194 | 0.13 | 1 | 15 |
| ophn1 | 0.81528 | 0.103182507 | 0.139 | 0.087 | 1 | 15 |
| ankhd1 | 0.81585 | 0.108498205 | 0.139 | 0.087 | 1 | 15 |
| zfand3 | 0.81641 | -0.11029994 | 0.278 | 0.348 | 1 | 15 |
| rbm14a | 0.81863 | 0.104334223 | 0.167 | 0.13 | 1 | 15 |
| tpp2 | 0.81898 | 0.106409775 | 0.083 | 0.043 | 1 | 15 |
| rnf24 | 0.81907 | -0.10771179 | 0.167 | 0.217 | 1 | 15 |
| sem1 | 0.82048 | -0.10084308 | 0.556 | 0.609 | 1 | 15 |
| ctdsp2 | 0.8205 | -0.11405956 | 0.361 | 0.435 | 1 | 15 |
| cyp2v1 | 0.8219 | -0.10501139 | 0.083 | 0.13 | 1 | 15 |
| laptm4a | 0.82349 | -0.11624062 | 0.361 | 0.435 | 1 | 15 |
| tob1a | 0.82426 | 0.112538132 | 0.472 | 0.391 | 1 | 15 |
| esd | 0.82709 | 0.1026918 | 0.472 | 0.391 | 1 | 15 |
| man1a2 | 0.82985 | 0.105448158 | 0.139 | 0.087 | 1 | 15 |
| dmrt1 | 0.83074 | 0.112673856 | 0.139 | 0.087 | 1 | 15 |
| eif1ad | 0.83216 | 0.108451243 | 0.333 | 0.261 | 1 | 15 |
| met | 0.83494 | 0.112269642 | 0.361 | 0.304 | 1 | 15 |
| ammecr1 | 0.83635 | -0.10592557 | 0.194 | 0.261 | 1 | 15 |
| slc4a4a | 0.83663 | -0.10533037 | 0.194 | 0.261 | 1 | 15 |
| ckap4 | 0.83683 | -0.10087631 | 0.194 | 0.261 | 1 | 15 |
| slc39a11 | 0.84258 | -0.1004861 | 0.083 | 0.13 | 1 | 15 |
| cnbpb | 0.84358 | 0.105627332 | 0.556 | 0.478 | 1 | 15 |

| | | | | | | |
|---|---|---|---|---|---|---|
| soga1 | 0.84425 | -0.11096426 | 0.083 | 0.13 | 1 | 15 |
| atf7ip | 0.84914 | -0.10532037 | 0.278 | 0.348 | 1 | 15 |
| hnrnpa1b | 0.85129 | -0.11480146 | 0.333 | 0.391 | 1 | 15 |
| nfyba | 0.85247 | 0.10152002 | 0.417 | 0.348 | 1 | 15 |
| add3a | 0.85274 | -0.10429944 | 0.361 | 0.435 | 1 | 15 |
| psmd1 | 0.85928 | -0.10160947 | 0.333 | 0.391 | 1 | 15 |
| fn1a | 0.86905 | 0.102385203 | 0.417 | 0.348 | 1 | 15 |
| lrp1aa | 0.87264 | 0.110050106 | 0.361 | 0.304 | 1 | 15 |
| tspan31 | 0.89185 | 0.102179358 | 0.222 | 0.174 | 1 | 15 |
| mt-cyb | 1.478125 | -1.35490293 | 0.171 | 1 | 2.98537009 | 16 |
| mt-nd3 | 3.662616 | -1.15695415 | 0.029 | 0.727 | 0.0007397 | 16 |
| mt-atp6 | 1.404470 | -1.10259249 | 0.371 | 0.955 | 0.0283661 | 16 |
| mt-co2 | 3.298009 | -0.89134744 | 0.4 | 0.955 | 0.666099 | 16 |
| rpl27a | 7.447372 | -0.43776599 | 0.857 | 1 | 1 | 16 |
| emx2 | 8.200032 | -0.68462102 | 0.257 | 0.818 | 1 | 16 |
| chac1 | 0.00018 | -0.73572752 | 0.486 | 0.909 | 1 | 16 |
| cox6b1 | 0.0002 | -0.43048295 | 0 | 0.318 | 1 | 16 |
| eif2s3 | 0.0003 | 0.701245036 | 0.714 | 0.182 | 1 | 16 |
| csrp1a | 0.00034 | -0.53536238 | 0.057 | 0.455 | 1 | 16 |
| ssr1 | 0.00064 | -0.66129186 | 0.486 | 0.773 | 1 | 16 |
| cahz | 0.0007 | -0.39268341 | 0 | 0.273 | 1 | 16 |
| crk | 0.00076 | 0.592159733 | 0.314 | 0 | 1 | 16 |
| tspan36 | 0.0008 | -0.59565142 | 0.143 | 0.455 | 1 | 16 |
| hspe1 | 0.0011 | 0.151084458 | 0.914 | 1 | 1 | 16 |
| abce1 | 0.00141 | -0.67344086 | 0.086 | 0.5 | 1 | 16 |
| appa | 0.0015 | 0.4732752 | 0.857 | 0.727 | 1 | 16 |
| mt-nd1 | 0.00179 | -0.733995 | 0.429 | 0.818 | 1 | 16 |
| aup1 | 0.0019 | -0.53200769 | 0.029 | 0.364 | 1 | 16 |
| rpl14 | 0.00201 | 0.124026547 | 1 | 0.773 | 1 | 16 |
| rsad2 | 0.00206 | -0.61871023 | 0.029 | 0.364 | 1 | 16 |
| suclg2 | 0.00228 | -0.49855706 | 0.029 | 0.364 | 1 | 16 |
| cx43 | 0.0023 | 0.488745118 | 0.971 | 0.636 | 1 | 16 |
| CABZ01078055.1 | 0.00233 | -0.43450711 | 0 | 0.227 | 1 | 16 |
| abhd17b | 0.00233 | -0.38022751 | 0 | 0.227 | 1 | 16 |
| dexi | 0.00233 | -0.41667877 | 0 | 0.227 | 1 | 16 |
| mtrr | 0.00233 | -0.37410475 | 0 | 0.227 | 1 | 16 |
| nkrf | 0.00233 | -0.44378282 | 0 | 0.227 | 1 | 16 |
| rae1 | 0.00233 | -0.31757549 | 0 | 0.227 | 1 | 16 |
| ripor1.1 | 0.00233 | -0.37920704 | 0 | 0.227 | 1 | 16 |
| swt1 | 0.00233 | -0.41921141 | 0 | 0.227 | 1 | 16 |
| tcf4 | 0.00233 | -0.40474562 | 0 | 0.227 | 1 | 16 |
| zgc:153215 | 0.00233 | -0.35192662 | 0 | 0.227 | 1 | 16 |
| CABZ01073795.1 | 0.00233 | -0.51009631 | 0 | 0.227 | 1 | 16 |
| serpinh1b | 0.00237 | -0.33452702 | 0.143 | 0.227 | 1 | 16 |
| txndc9 | 0.0025 | 0.4459825 | 0.514 | 0.136 | 1 | 16 |
| isg15 | 0.00252 | -0.70451361 | 0.229 | 0.682 | 1 | 16 |
| CR318588.1 | 0.00257 | -0.38037917 | 0.886 | 1 | 1 | 16 |
| apoob | 0.00287 | 0.454921001 | 0.257 | 0 | 1 | 16 |

| | | | | | | |
|---|---|---|---|---|---|---|
| arrdc3b | 0.00287 | 0.422780892 | 0.257 | 0 | 1 | 16 |
| tbx18 | 0.00287 | 0.593019794 | 0.257 | 0 | 1 | 16 |
| mt-nd4 | 0.00291 | -0.52213632 | 0.657 | 0.864 | 1 | 16 |
| ftr56 | 0.00372 | -0.42244465 | 0.029 | 0.318 | 1 | 16 |
| cfl1l | 0.00373 | -0.27633758 | 0.086 | 0.455 | 1 | 16 |
| twistnb | 0.00382 | -0.32769709 | 0.057 | 0.318 | 1 | 16 |
| fstl3 | 0.00404 | -0.62396133 | 0.086 | 0.455 | 1 | 16 |
| pdk2b | 0.00449 | 0.304353395 | 0.286 | 0.045 | 1 | 16 |
| surf4l | 0.00466 | -0.60081595 | 0.057 | 0.409 | 1 | 16 |
| tjp1a | 0.00488 | 0.623856558 | 0.571 | 0.182 | 1 | 16 |
| pim1 | 0.00503 | -0.63015881 | 0.114 | 0.5 | 1 | 16 |
| p4hb | 0.00509 | -0.60014008 | 0.371 | 0.636 | 1 | 16 |
| CABZ01021532.1 | 0.00521 | -0.72848398 | 0.257 | 0.636 | 1 | 16 |
| mrpl58 | 0.00548 | 0.370453464 | 0.229 | 0 | 1 | 16 |
| hmgcrb | 0.00548 | 0.420140612 | 0.229 | 0 | 1 | 16 |
| itcha | 0.00548 | 0.381712882 | 0.229 | 0 | 1 | 16 |
| ralab | 0.00548 | 0.40210692 | 0.229 | 0 | 1 | 16 |
| xrn2 | 0.00548 | 0.402238319 | 0.229 | 0 | 1 | 16 |
| thoc5 | 0.00548 | 0.349821287 | 0.229 | 0 | 1 | 16 |
| rnf213a | 0.00685 | -0.60050869 | 0.286 | 0.682 | 1 | 16 |
| eef1db | 0.00718 | -0.34001251 | 0.543 | 0.909 | 1 | 16 |
| si:dkey-42i9.4 | 0.0073 | 0.533251477 | 0.514 | 0.136 | 1 | 16 |
| ada2a | 0.00755 | -0.33250801 | 0 | 0.182 | 1 | 16 |
| bmt2 | 0.00755 | -0.2813014 | 0 | 0.182 | 1 | 16 |
| si:ch1073-170o4.1 | 0.00755 | -0.35124859 | 0 | 0.182 | 1 | 16 |
| tmem246 | 0.00755 | -0.27841834 | 0 | 0.182 | 1 | 16 |
| bin3 | 0.00755 | -0.33546207 | 0 | 0.182 | 1 | 16 |
| dnajc9 | 0.00755 | -0.29121509 | 0 | 0.182 | 1 | 16 |
| ppil2 | 0.00755 | -0.28573077 | 0 | 0.182 | 1 | 16 |
| ms4a17a.5 | 0.00755 | -0.29797287 | 0 | 0.182 | 1 | 16 |
| acss3 | 0.00755 | -0.33290784 | 0 | 0.182 | 1 | 16 |
| dkk1b | 0.00755 | -0.50077502 | 0 | 0.182 | 1 | 16 |
| gnptab | 0.00755 | -0.34753782 | 0 | 0.182 | 1 | 16 |
| lypd6 | 0.00755 | -0.29946432 | 0 | 0.182 | 1 | 16 |
| tsr3 | 0.00755 | -0.32248966 | 0 | 0.182 | 1 | 16 |
| igfbp1a | 0.00755 | -0.34562374 | 0 | 0.182 | 1 | 16 |
| nedd4l | 0.00755 | -0.28493142 | 0 | 0.182 | 1 | 16 |
| nup155 | 0.00755 | -0.33486324 | 0 | 0.182 | 1 | 16 |
| polr1e | 0.00755 | -0.3089026 | 0 | 0.182 | 1 | 16 |
| fut8a | 0.00755 | -0.33482589 | 0 | 0.182 | 1 | 16 |
| cx34.5 | 0.00765 | 0.755995385 | 0.543 | 0.182 | 1 | 16 |
| fkbp5 | 0.00781 | 0.444934859 | 0.4 | 0.091 | 1 | 16 |
| gstm.3 | 0.00793 | 0.415094291 | 0.943 | 0.727 | 1 | 16 |
| CU984600.2 | 0.00803 | -0.52221478 | 0.143 | 0.455 | 1 | 16 |
| igf1ra | 0.00872 | -0.46323448 | 0.029 | 0.318 | 1 | 16 |
| sf3b2 | 0.0096 | 0.319251838 | 0.286 | 0.045 | 1 | 16 |
| psmb6 | 0.00995 | 0.389639289 | 0.6 | 0.409 | 1 | 16 |
| foxo3b | 0.0101 | 0.532136096 | 0.371 | 0.045 | 1 | 16 |

| | | | | | | |
|---|---|---|---|---|---|---|
| ift20 | 0.01015 | -0.27742564 | 0.029 | 0.227 | 1 | 16 |
| nfil3-6 | 0.01032 | 0.391994499 | 0.2 | 0 | 1 | 16 |
| tango2 | 0.01032 | 0.316472742 | 0.2 | 0 | 1 | 16 |
| CU633762.1 | 0.01032 | 0.340113542 | 0.2 | 0 | 1 | 16 |
| akr1a1b | 0.01032 | 0.349779442 | 0.2 | 0 | 1 | 16 |
| cpne3 | 0.01032 | 0.362182465 | 0.2 | 0 | 1 | 16 |
| hddc3 | 0.01032 | 0.404651863 | 0.2 | 0 | 1 | 16 |
| malt1 | 0.01032 | 0.355126129 | 0.2 | 0 | 1 | 16 |
| arid1b | 0.01068 | -0.48855067 | 0.057 | 0.364 | 1 | 16 |
| arf2a | 0.01076 | 0.543351173 | 0.486 | 0.182 | 1 | 16 |
| ola1 | 0.01103 | 0.509341141 | 0.571 | 0.182 | 1 | 16 |
| tubb4b | 0.01136 | -0.29943787 | 0.743 | 1 | 1 | 16 |
| ctbp2a | 0.0116 | -0.4554131 | 0.114 | 0.455 | 1 | 16 |
| pfkfb4b | 0.01177 | 0.617274292 | 0.486 | 0.182 | 1 | 16 |
| LOC103910903 | 0.01186 | 0.362351506 | 0.857 | 0.773 | 1 | 16 |
| decr1 | 0.01239 | -0.13977031 | 0.514 | 0.5 | 1 | 16 |
| agrn | 0.01312 | -0.30816819 | 0.714 | 0.773 | 1 | 16 |
| si:ch211-1a19.3 | 0.01316 | -0.6786326 | 0.2 | 0.545 | 1 | 16 |
| mxa | 0.01316 | -0.50327402 | 0.086 | 0.409 | 1 | 16 |
| ssuh2rs1 | 0.0132 | -0.52388497 | 0.114 | 0.455 | 1 | 16 |
| qsox1 | 0.0138 | -0.32233815 | 0.029 | 0.227 | 1 | 16 |
| zgc:152791 | 0.0147 | -0.53354142 | 0.343 | 0.727 | 1 | 16 |
| mtdha | 0.01488 | 0.395006897 | 0.629 | 0.273 | 1 | 16 |
| arl8a | 0.01499 | -0.48052712 | 0.086 | 0.409 | 1 | 16 |
| xaf1 | 0.01502 | -0.51689497 | 0.114 | 0.455 | 1 | 16 |
| si:ch73-269m14.2 | 0.01509 | -0.50430402 | 0.086 | 0.409 | 1 | 16 |
| BX284696.1 | 0.01517 | -0.45650854 | 0.257 | 0.636 | 1 | 16 |
| tln1 | 0.01521 | 0.39265453 | 0.457 | 0.136 | 1 | 16 |
| nop10 | 0.01551 | -0.49257345 | 0.257 | 0.636 | 1 | 16 |
| zgc:100829 | 0.01595 | 0.615035169 | 0.514 | 0.182 | 1 | 16 |
| etnk1 | 0.01616 | 0.53701253 | 0.343 | 0.045 | 1 | 16 |
| ap2m1a | 0.01617 | 0.348425355 | 0.857 | 0.727 | 1 | 16 |
| snrpb2 | 0.01621 | 0.490002944 | 0.314 | 0.045 | 1 | 16 |
| ttpal | 0.01685 | 0.475751717 | 0.343 | 0.045 | 1 | 16 |
| lhfpl2a | 0.01694 | -0.53929643 | 0.257 | 0.636 | 1 | 16 |
| ccnd1 | 0.01734 | -0.53879088 | 0.257 | 0.545 | 1 | 16 |
| txnipa | 0.01753 | 0.46258746 | 0.857 | 0.636 | 1 | 16 |
| rpl32 | 0.01788 | 0.102705028 | 0.943 | 0.955 | 1 | 16 |
| mt-nd2 | 0.01864 | -0.52083325 | 0.543 | 0.864 | 1 | 16 |
| chpfa | 0.01883 | -0.3039971 | 0.029 | 0.227 | 1 | 16 |
| phc2a | 0.01883 | 0.351793846 | 0.543 | 0.455 | 1 | 16 |
| h1f0 | 0.01885 | -0.44811667 | 0.286 | 0.636 | 1 | 16 |
| glula | 0.0192 | 0.56533738 | 0.371 | 0.091 | 1 | 16 |
| ephb3 | 0.01925 | 0.309859751 | 0.171 | 0 | 1 | 16 |
| rab8a | 0.01925 | 0.285210454 | 0.171 | 0 | 1 | 16 |
| tnrc18 | 0.01925 | 0.335963385 | 0.171 | 0 | 1 | 16 |
| acat1 | 0.01925 | 0.284514232 | 0.171 | 0 | 1 | 16 |
| bag4 | 0.01925 | 0.334789687 | 0.171 | 0 | 1 | 16 |

| | | | | | | |
|---|---|---|---|---|---|---|
| hipk3b | 0.01925 | 0.272070855 | 0.171 | 0 | 1 | 16 |
| si:ch211-87m7.2 | 0.01925 | 0.311477432 | 0.171 | 0 | 1 | 16 |
| si:ch211-206k20.5 | 0.01925 | 0.305788235 | 0.171 | 0 | 1 | 16 |
| tprkb | 0.01925 | 0.251189231 | 0.171 | 0 | 1 | 16 |
| zfyve9a | 0.01925 | 0.307813273 | 0.171 | 0 | 1 | 16 |
| zgc:92242 | 0.01925 | 0.321893662 | 0.171 | 0 | 1 | 16 |
| cmasa | 0.01925 | 0.283212477 | 0.171 | 0 | 1 | 16 |
| cnot2 | 0.01925 | 0.294966624 | 0.171 | 0 | 1 | 16 |
| musk | 0.01925 | 0.380535802 | 0.171 | 0 | 1 | 16 |
| plekha1a | 0.01925 | 0.266910915 | 0.171 | 0 | 1 | 16 |
| ptpn11b | 0.01925 | 0.313733273 | 0.171 | 0 | 1 | 16 |
| xpo6 | 0.01925 | 0.30285269 | 0.171 | 0 | 1 | 16 |
| rnf220a | 0.01925 | 0.283466235 | 0.171 | 0 | 1 | 16 |
| serhl | 0.01925 | 0.297048539 | 0.171 | 0 | 1 | 16 |
| chrdl2 | 0.01976 | 0.412163341 | 0.314 | 0.045 | 1 | 16 |
| vta1 | 0.0201 | 0.449525578 | 0.314 | 0.045 | 1 | 16 |
| otub1b | 0.02044 | -0.27429359 | 0.086 | 0.318 | 1 | 16 |
| ece1 | 0.02102 | -0.40885984 | 0.029 | 0.273 | 1 | 16 |
| rin2 | 0.02169 | -0.41239313 | 0.029 | 0.273 | 1 | 16 |
| smoc2 | 0.02182 | -0.66943687 | 0.171 | 0.5 | 1 | 16 |
| cnp | 0.02225 | -0.46125544 | 0.229 | 0.591 | 1 | 16 |
| ndufa8 | 0.02249 | 0.33611995 | 0.571 | 0.409 | 1 | 16 |
| ankrd22 | 0.02261 | -0.419065 | 0.029 | 0.273 | 1 | 16 |
| her9 | 0.02307 | -0.53648422 | 0.057 | 0.227 | 1 | 16 |
| si:ch73-1a9.3 | 0.02317 | 0.280511104 | 1 | 0.955 | 1 | 16 |
| neu1 | 0.02342 | -0.32320555 | 0.029 | 0.227 | 1 | 16 |
| dcbld2 | 0.02351 | -0.38935522 | 0.029 | 0.273 | 1 | 16 |
| si:ch73-27e22.2 | 0.02353 | -0.43608305 | 0.029 | 0.273 | 1 | 16 |
| srprb | 0.02368 | -0.36839866 | 0.286 | 0.455 | 1 | 16 |
| tefb | 0.02393 | -0.35786728 | 0.029 | 0.273 | 1 | 16 |
| ecm1b | 0.02397 | 0.593905338 | 0.543 | 0.273 | 1 | 16 |
| glis2b | 0.02399 | -0.27879223 | 0 | 0.136 | 1 | 16 |
| ms4a17a.8 | 0.02399 | -0.28619638 | 0 | 0.136 | 1 | 16 |
| pdxp | 0.02399 | -0.23351525 | 0 | 0.136 | 1 | 16 |
| RANBP2 | 0.02399 | -0.22048287 | 0 | 0.136 | 1 | 16 |
| brox | 0.02399 | -0.23836019 | 0 | 0.136 | 1 | 16 |
| cables2b | 0.02399 | -0.29189588 | 0 | 0.136 | 1 | 16 |
| clk4a | 0.02399 | -0.27125463 | 0 | 0.136 | 1 | 16 |
| clptm1l | 0.02399 | -0.21957078 | 0 | 0.136 | 1 | 16 |
| cpeb4a | 0.02399 | -0.22761925 | 0 | 0.136 | 1 | 16 |
| dnajc18 | 0.02399 | -0.25851188 | 0 | 0.136 | 1 | 16 |
| edem3 | 0.02399 | -0.23784156 | 0 | 0.136 | 1 | 16 |
| eef2kmt | 0.02399 | -0.27481757 | 0 | 0.136 | 1 | 16 |
| emid1 | 0.02399 | -0.24249963 | 0 | 0.136 | 1 | 16 |
| fbxo7 | 0.02399 | -0.22003415 | 0 | 0.136 | 1 | 16 |
| fosb | 0.02399 | -0.35677923 | 0 | 0.136 | 1 | 16 |
| ftr93 | 0.02399 | -0.25062487 | 0 | 0.136 | 1 | 16 |
| gatad2ab | 0.02399 | -0.23526147 | 0 | 0.136 | 1 | 16 |

| | | | | | | |
|---|---|---|---|---|---|---|
| herpud2 | 0.02399 | -0.24037031 | 0 | 0.136 | 1 | 16 |
| hs6st1b | 0.02399 | -0.23480349 | 0 | 0.136 | 1 | 16 |
| intu | 0.02399 | -0.26340473 | 0 | 0.136 | 1 | 16 |
| mhc1zba | 0.02399 | -0.24798755 | 0 | 0.136 | 1 | 16 |
| mmd2a | 0.02399 | -0.27197703 | 0 | 0.136 | 1 | 16 |
| nup58 | 0.02399 | -0.21822064 | 0 | 0.136 | 1 | 16 |
| polr1c | 0.02399 | -0.2336893 | 0 | 0.136 | 1 | 16 |
| prmt6 | 0.02399 | -0.2800987 | 0 | 0.136 | 1 | 16 |
| prodha | 0.02399 | -0.21908917 | 0 | 0.136 | 1 | 16 |
| prpf3 | 0.02399 | -0.26346703 | 0 | 0.136 | 1 | 16 |
| r3hcc1l | 0.02399 | -0.2638796 | 0 | 0.136 | 1 | 16 |
| rbm45 | 0.02399 | -0.23543212 | 0 | 0.136 | 1 | 16 |
| rgmd | 0.02399 | -0.22472409 | 0 | 0.136 | 1 | 16 |
| scyl3 | 0.02399 | -0.22012539 | 0 | 0.136 | 1 | 16 |
| sh3bp5a | 0.02399 | -0.25293483 | 0 | 0.136 | 1 | 16 |
| si:ch73-238c9.1 | 0.02399 | -0.24298955 | 0 | 0.136 | 1 | 16 |
| snx4 | 0.02399 | -0.20763021 | 0 | 0.136 | 1 | 16 |
| trub2 | 0.02399 | -0.25738698 | 0 | 0.136 | 1 | 16 |
| tsen54 | 0.02399 | -0.24586537 | 0 | 0.136 | 1 | 16 |
| wdr18 | 0.02399 | -0.28286302 | 0 | 0.136 | 1 | 16 |
| wdr21 | 0.02399 | -0.26734186 | 0 | 0.136 | 1 | 16 |
| zgc:153372 | 0.02399 | -0.24601298 | 0 | 0.136 | 1 | 16 |
| cd83 | 0.02399 | -0.26631344 | 0 | 0.136 | 1 | 16 |
| fzd2 | 0.02399 | -0.27285844 | 0 | 0.136 | 1 | 16 |
| nfrkb | 0.02399 | -0.2183583 | 0 | 0.136 | 1 | 16 |
| nup98 | 0.02399 | -0.21399739 | 0 | 0.136 | 1 | 16 |
| tmc6b | 0.02399 | -0.20821893 | 0 | 0.136 | 1 | 16 |
| vsig10l | 0.02399 | -0.23073767 | 0 | 0.136 | 1 | 16 |
| CABZ01080371.2 | 0.02399 | -0.25620157 | 0 | 0.136 | 1 | 16 |
| CU856539.1 | 0.02399 | -0.23435374 | 0 | 0.136 | 1 | 16 |
| LOC101883097 | 0.02399 | -0.21863413 | 0 | 0.136 | 1 | 16 |
| aqp11 | 0.02399 | -0.2068081 | 0 | 0.136 | 1 | 16 |
| arntl1b | 0.02399 | -0.25144407 | 0 | 0.136 | 1 | 16 |
| ccl25b | 0.02399 | -0.16541969 | 0 | 0.136 | 1 | 16 |
| fam222a | 0.02399 | -0.24620535 | 0 | 0.136 | 1 | 16 |
| ftr91 | 0.02399 | -0.23577969 | 0 | 0.136 | 1 | 16 |
| relb | 0.02399 | -0.23293718 | 0 | 0.136 | 1 | 16 |
| si:ch211-10d23.5 | 0.02399 | -0.310573 | 0 | 0.136 | 1 | 16 |
| si:dkey-56d12.4 | 0.02399 | -0.21985256 | 0 | 0.136 | 1 | 16 |
| st3gal3b | 0.02399 | -0.28939156 | 0 | 0.136 | 1 | 16 |
| tmem135 | 0.02399 | -0.21109211 | 0 | 0.136 | 1 | 16 |
| traf2a | 0.02399 | -0.21890296 | 0 | 0.136 | 1 | 16 |
| wbp2 | 0.02399 | -0.2249061 | 0 | 0.136 | 1 | 16 |
| clptm1 | 0.02399 | -0.31563911 | 0 | 0.136 | 1 | 16 |
| irgq1 | 0.02399 | -0.26176042 | 0 | 0.136 | 1 | 16 |
| plpp7 | 0.02399 | -0.24682444 | 0 | 0.136 | 1 | 16 |
| LOC101884549 | 0.02399 | -0.23001049 | 0 | 0.136 | 1 | 16 |
| LOC108182857 | 0.02399 | -0.19388783 | 0 | 0.136 | 1 | 16 |

| | | | | | | |
|---|---|---|---|---|---|---|
| atrn | 0.02399 | -0.2478616 | 0 | 0.136 | 1 | 16 |
| gpx1a | 0.02399 | -0.10314289 | 0 | 0.136 | 1 | 16 |
| pcgf1 | 0.02399 | -0.23039758 | 0 | 0.136 | 1 | 16 |
| pitrm1 | 0.02399 | -0.20384182 | 0 | 0.136 | 1 | 16 |
| si:ch211-244b2.3 | 0.02399 | -0.22523909 | 0 | 0.136 | 1 | 16 |
| socs1b | 0.02399 | -0.2550784 | 0 | 0.136 | 1 | 16 |
| tbc1d10aa | 0.02399 | -0.21552714 | 0 | 0.136 | 1 | 16 |
| txn | 0.02399 | -0.2606888 | 0 | 0.136 | 1 | 16 |
| zgc:171497 | 0.02399 | -0.2129001 | 0 | 0.136 | 1 | 16 |
| si:ch73-181d5.4 | 0.02399 | -0.21406179 | 0 | 0.136 | 1 | 16 |
| si:dkey-15h8.16 | 0.02399 | -0.22858228 | 0 | 0.136 | 1 | 16 |
| tfa | 0.02399 | -0.21269995 | 0 | 0.136 | 1 | 16 |
| thoc6 | 0.02399 | -0.23902645 | 0 | 0.136 | 1 | 16 |
| tmem70 | 0.02399 | -0.21967456 | 0 | 0.136 | 1 | 16 |
| CR354588.1 | 0.02399 | -0.20981407 | 0 | 0.136 | 1 | 16 |
| LOC103910306 | 0.02399 | -0.23531376 | 0 | 0.136 | 1 | 16 |
| PLBD2 | 0.02399 | -0.29767406 | 0 | 0.136 | 1 | 16 |
| appl2 | 0.02399 | -0.24885926 | 0 | 0.136 | 1 | 16 |
| dpysl2b | 0.02399 | -0.22782164 | 0 | 0.136 | 1 | 16 |
| kalrna | 0.02399 | -0.28050563 | 0 | 0.136 | 1 | 16 |
| mbnl2 | 0.02399 | -0.25890361 | 0 | 0.136 | 1 | 16 |
| nadkb | 0.02399 | -0.25584697 | 0 | 0.136 | 1 | 16 |
| rab3gap2 | 0.02399 | -0.25572824 | 0 | 0.136 | 1 | 16 |
| reck | 0.02399 | -0.2414602 | 0 | 0.136 | 1 | 16 |
| rgma | 0.02399 | -0.27233886 | 0 | 0.136 | 1 | 16 |
| sash1a | 0.02399 | -0.25768099 | 0 | 0.136 | 1 | 16 |
| tbpl1 | 0.02399 | -0.20924084 | 0 | 0.136 | 1 | 16 |
| tep1 | 0.02399 | -0.25080115 | 0 | 0.136 | 1 | 16 |
| ube3a | 0.02399 | -0.23276037 | 0 | 0.136 | 1 | 16 |
| ubxn6 | 0.02399 | -0.22416892 | 0 | 0.136 | 1 | 16 |
| zgc:101783 | 0.02399 | -0.23886148 | 0 | 0.136 | 1 | 16 |
| zmiz2 | 0.02399 | -0.22554973 | 0 | 0.136 | 1 | 16 |
| znf1156 | 0.02399 | -0.21819298 | 0 | 0.136 | 1 | 16 |
| rbx1 | 0.02415 | 0.210306246 | 0.857 | 0.864 | 1 | 16 |
| slc7a3a | 0.02443 | 0.44188188 | 0.857 | 0.591 | 1 | 16 |
| cux1a | 0.02451 | -0.43430184 | 0.229 | 0.455 | 1 | 16 |
| si:ch211-222l21.1 | 0.02493 | -0.24032544 | 0.914 | 0.909 | 1 | 16 |
| ppa1b | 0.02532 | 0.142562638 | 0.314 | 0.136 | 1 | 16 |
| pfn2 | 0.02608 | 0.281710152 | 0.857 | 0.818 | 1 | 16 |
| tmem33 | 0.0264 | -0.50044267 | 0.2 | 0.545 | 1 | 16 |
| stx5a | 0.02785 | -0.1794872 | 0.029 | 0.182 | 1 | 16 |
| si:ch211-276k2.1 | 0.02805 | -0.57301699 | 0.143 | 0.455 | 1 | 16 |
| CABZ01113812.1 | 0.02826 | -0.41427822 | 0.086 | 0.364 | 1 | 16 |
| ogdha | 0.02859 | 0.497281965 | 0.4 | 0.091 | 1 | 16 |
| gig2e | 0.02869 | -0.45534697 | 0.171 | 0.5 | 1 | 16 |
| rpl22l1 | 0.029 | -0.24782893 | 0.686 | 0.955 | 1 | 16 |
| si:dkey-42i9.6 | 0.02947 | -0.49566911 | 0.2 | 0.455 | 1 | 16 |
| tob1a | 0.02959 | 0.371448676 | 0.6 | 0.5 | 1 | 16 |

| | | | | | | |
|---|---|---|---|---|---|---|
| psmd4a | 0.0299 | -0.41472366 | 0.086 | 0.364 | 1 | 16 |
| FAT4 | 0.02997 | 0.401588891 | 0.371 | 0.091 | 1 | 16 |
| gpia | 0.03022 | -0.38871637 | 0.286 | 0.636 | 1 | 16 |
| jun | 0.03066 | -0.23300441 | 0.514 | 0.5 | 1 | 16 |
| tia1 | 0.03156 | 0.453595518 | 0.314 | 0.045 | 1 | 16 |
| cdh1 | 0.03196 | -0.52224786 | 0.229 | 0.5 | 1 | 16 |
| pgam5 | 0.032 | -0.34230025 | 0.229 | 0.545 | 1 | 16 |
| eif1axb | 0.03247 | -0.26651347 | 0.6 | 0.682 | 1 | 16 |
| pdrg1 | 0.03287 | -0.27229661 | 0.029 | 0.227 | 1 | 16 |
| slc2a1b | 0.03355 | 0.526370484 | 0.429 | 0.136 | 1 | 16 |
| cbr1l | 0.03412 | -0.49451829 | 0.114 | 0.409 | 1 | 16 |
| bnip3lb | 0.03426 | 0.166756746 | 0.2 | 0.045 | 1 | 16 |
| h2afva | 0.03431 | 0.394148146 | 0.629 | 0.409 | 1 | 16 |
| tspan14 | 0.03529 | -0.41841651 | 0.086 | 0.364 | 1 | 16 |
| serpinh1a | 0.03549 | -0.42285486 | 0.086 | 0.364 | 1 | 16 |
| krt15 | 0.03565 | 0.287811407 | 0.143 | 0 | 1 | 16 |
| mpped2 | 0.03565 | 0.249225086 | 0.143 | 0 | 1 | 16 |
| rbfox1 | 0.03565 | 0.29670586 | 0.143 | 0 | 1 | 16 |
| si:ch73-305o9.3 | 0.03565 | 0.283838297 | 0.143 | 0 | 1 | 16 |
| zpr1 | 0.03565 | 0.218284335 | 0.143 | 0 | 1 | 16 |
| LOC103909810 | 0.03565 | 0.296571246 | 0.143 | 0 | 1 | 16 |
| bco2a | 0.03565 | 0.276927799 | 0.143 | 0 | 1 | 16 |
| camk1b | 0.03565 | 0.292685692 | 0.143 | 0 | 1 | 16 |
| ccar1 | 0.03565 | 0.249823814 | 0.143 | 0 | 1 | 16 |
| chchd6a | 0.03565 | 0.247617898 | 0.143 | 0 | 1 | 16 |
| clcn5a | 0.03565 | 0.271101193 | 0.143 | 0 | 1 | 16 |
| cpt1ab | 0.03565 | 0.254979899 | 0.143 | 0 | 1 | 16 |
| eif2a | 0.03565 | 0.301999481 | 0.143 | 0 | 1 | 16 |
| hsd17b1 | 0.03565 | 0.306225572 | 0.143 | 0 | 1 | 16 |
| jade1 | 0.03565 | 0.242374031 | 0.143 | 0 | 1 | 16 |
| kcnh4a | 0.03565 | 0.287318196 | 0.143 | 0 | 1 | 16 |
| cyp2p6 | 0.03565 | 0.264388343 | 0.143 | 0 | 1 | 16 |
| march7 | 0.03565 | 0.276478287 | 0.143 | 0 | 1 | 16 |
| mdm4 | 0.03565 | 0.213300989 | 0.143 | 0 | 1 | 16 |
| ndnf | 0.03565 | 0.254857323 | 0.143 | 0 | 1 | 16 |
| scaf4a | 0.03565 | 0.236432748 | 0.143 | 0 | 1 | 16 |
| XLOC-038073 | 0.03565 | 0.293764619 | 0.143 | 0 | 1 | 16 |
| aldh3a2a | 0.03565 | 0.256391275 | 0.143 | 0 | 1 | 16 |
| ehmt1b | 0.03565 | 0.296084062 | 0.143 | 0 | 1 | 16 |
| gdpd5b | 0.03565 | 0.257374563 | 0.143 | 0 | 1 | 16 |
| got2a | 0.03565 | 0.23053735 | 0.143 | 0 | 1 | 16 |
| larp4b | 0.03565 | 0.292303421 | 0.143 | 0 | 1 | 16 |
| lpcat3 | 0.03565 | 0.243735453 | 0.143 | 0 | 1 | 16 |
| pbxip1b | 0.03565 | 0.231210619 | 0.143 | 0 | 1 | 16 |
| pde8a | 0.03565 | 0.264930866 | 0.143 | 0 | 1 | 16 |
| polr2a | 0.03565 | 0.291953185 | 0.143 | 0 | 1 | 16 |
| sephs2 | 0.03565 | 0.212905807 | 0.143 | 0 | 1 | 16 |
| skia | 0.03565 | 0.269324065 | 0.143 | 0 | 1 | 16 |

| | | | | | | |
|---|---|---|---|---|---|---|
| slc1a3a | 0.03565 | 0.233540206 | 0.143 | 0 | 1 | 16 |
| snrnp48 | 0.03565 | 0.298476978 | 0.143 | 0 | 1 | 16 |
| srd5a2a | 0.03565 | 0.22100245 | 0.143 | 0 | 1 | 16 |
| tpd52l2a | 0.03565 | 0.29833596 | 0.143 | 0 | 1 | 16 |
| vamp4 | 0.03565 | 0.238261953 | 0.143 | 0 | 1 | 16 |
| zgc:162945 | 0.03565 | 0.245431262 | 0.143 | 0 | 1 | 16 |
| dpf2l | 0.03565 | 0.285792336 | 0.143 | 0 | 1 | 16 |
| nckipsd | 0.03565 | 0.240039414 | 0.143 | 0 | 1 | 16 |
| ppm1bb | 0.03565 | 0.280151982 | 0.143 | 0 | 1 | 16 |
| zgc:158517 | 0.03565 | 0.228519654 | 0.143 | 0 | 1 | 16 |
| ptpn11a | 0.03565 | 0.279216856 | 0.143 | 0 | 1 | 16 |
| CABZ01074203.1 | 0.03565 | 0.339440273 | 0.143 | 0 | 1 | 16 |
| gmppab | 0.03565 | 0.232619965 | 0.143 | 0 | 1 | 16 |
| ilk | 0.03565 | 0.208220516 | 0.143 | 0 | 1 | 16 |
| DDX17 | 0.03566 | -0.43211816 | 0.086 | 0.364 | 1 | 16 |
| CABZ01088149.1 | 0.03566 | -0.39571193 | 0.114 | 0.409 | 1 | 16 |
| ssr4 | 0.03592 | -0.10086593 | 0.886 | 0.773 | 1 | 16 |
| krt8 | 0.03611 | 0.368817495 | 0.857 | 0.727 | 1 | 16 |
| grb10b | 0.03614 | -0.29725086 | 0.029 | 0.227 | 1 | 16 |
| sirt7 | 0.03651 | -0.39687507 | 0.086 | 0.364 | 1 | 16 |
| sec13 | 0.03674 | -0.4152119 | 0.114 | 0.409 | 1 | 16 |
| btbd6b | 0.03751 | -0.42867808 | 0.086 | 0.364 | 1 | 16 |
| hspa5 | 0.03753 | -0.44910005 | 0.571 | 0.818 | 1 | 16 |
| baz1a | 0.03755 | -0.43491425 | 0.086 | 0.364 | 1 | 16 |
| ndufv2 | 0.03756 | -0.38128668 | 0.086 | 0.364 | 1 | 16 |
| rbm39a | 0.03795 | 0.507388863 | 0.486 | 0.182 | 1 | 16 |
| sec61a1 | 0.03805 | -0.40173739 | 0.114 | 0.409 | 1 | 16 |
| dspa | 0.03806 | -0.41771735 | 0.114 | 0.409 | 1 | 16 |
| prelid1a | 0.03918 | 0.234646926 | 0.286 | 0.091 | 1 | 16 |
| hprt1 | 0.03949 | 0.232409698 | 0.286 | 0.091 | 1 | 16 |
| ehd3 | 0.03957 | 0.530063486 | 0.371 | 0.091 | 1 | 16 |
| npm1a | 0.041 | -0.29692234 | 0.457 | 0.5 | 1 | 16 |
| mmp16a | 0.041 | -0.38156059 | 0.029 | 0.227 | 1 | 16 |
| rab1ab | 0.04106 | -0.41469553 | 0.257 | 0.591 | 1 | 16 |
| fblim1 | 0.04106 | 0.421805888 | 0.286 | 0.045 | 1 | 16 |
| ctsla | 0.04115 | -0.30915198 | 0.8 | 0.909 | 1 | 16 |
| aes | 0.04118 | -0.21571199 | 0.114 | 0.318 | 1 | 16 |
| ssr3 | 0.04138 | -0.48165073 | 0.543 | 0.727 | 1 | 16 |
| mt-co1 | 0.04211 | -0.37783088 | 0.743 | 0.909 | 1 | 16 |
| rbbp4 | 0.04224 | -0.38344444 | 0.171 | 0.318 | 1 | 16 |
| sod2 | 0.04228 | 0.359203326 | 0.543 | 0.227 | 1 | 16 |
| sept2 | 0.04332 | -0.46124409 | 0.257 | 0.545 | 1 | 16 |
| bhlhe40 | 0.04378 | 0.462921587 | 0.257 | 0.045 | 1 | 16 |
| ccnl1b | 0.04447 | -0.21267796 | 0.171 | 0.227 | 1 | 16 |
| timm9 | 0.04541 | -0.41984978 | 0.114 | 0.364 | 1 | 16 |
| mrpl45 | 0.04547 | -0.37510194 | 0.029 | 0.227 | 1 | 16 |
| plxna4 | 0.04561 | 0.306262441 | 0.486 | 0.227 | 1 | 16 |
| glyr1 | 0.04577 | -0.38003736 | 0.057 | 0.273 | 1 | 16 |

| | | | | | | |
|---|---|---|---|---|---|---|
| ndufb7 | 0.04606 | 0.352588977 | 0.686 | 0.5 | 1 | 16 |
| il6st | 0.04644 | -0.34104655 | 0.143 | 0.409 | 1 | 16 |
| zgc:113921 | 0.04719 | -0.41184593 | 0.029 | 0.227 | 1 | 16 |
| cebpd | 0.04763 | 0.372967927 | 0.629 | 0.5 | 1 | 16 |
| fosab | 0.04799 | -0.59802338 | 0.314 | 0.545 | 1 | 16 |
| amh | 0.04805 | -0.42407793 | 0.114 | 0.318 | 1 | 16 |
| ralba | 0.04809 | 0.41048479 | 0.286 | 0.045 | 1 | 16 |
| lpl | 0.0494 | -0.19185474 | 0.029 | 0.136 | 1 | 16 |
| slc43a1a | 0.04943 | -0.21282085 | 0.029 | 0.182 | 1 | 16 |
| pole4 | 0.05066 | -0.34301877 | 0.029 | 0.227 | 1 | 16 |
| mknk2b | 0.05077 | 0.317021761 | 0.429 | 0.364 | 1 | 16 |
| fez1 | 0.05089 | 0.438781983 | 0.686 | 0.455 | 1 | 16 |
| ube2al | 0.05105 | 0.415042048 | 0.286 | 0.045 | 1 | 16 |
| afap1l1a | 0.05111 | -0.3354059 | 0.114 | 0.227 | 1 | 16 |
| slc25a51b | 0.05212 | -0.38343027 | 0.029 | 0.227 | 1 | 16 |
| taf3 | 0.05213 | 0.419338963 | 0.286 | 0.045 | 1 | 16 |
| stip1 | 0.05249 | 0.103021396 | 0.257 | 0.273 | 1 | 16 |
| dlat | 0.05254 | -0.3253123 | 0.029 | 0.227 | 1 | 16 |
| polr1d | 0.05286 | -0.29571758 | 0.057 | 0.273 | 1 | 16 |
| si:ch211-114c12.2 | 0.0532 | -0.26867211 | 0.171 | 0.409 | 1 | 16 |
| nectin1b | 0.05355 | -0.40718046 | 0.514 | 0.773 | 1 | 16 |
| calua | 0.05375 | -0.42391072 | 0.114 | 0.364 | 1 | 16 |
| hmgn2 | 0.05455 | -0.18442346 | 0.543 | 0.818 | 1 | 16 |
| polr2b | 0.05554 | -0.38059349 | 0.057 | 0.273 | 1 | 16 |
| nme2b.1 | 0.05602 | 0.113751535 | 0.971 | 1 | 1 | 16 |
| slc35b4 | 0.05646 | -0.34556465 | 0.029 | 0.227 | 1 | 16 |
| zranb2 | 0.05649 | 0.425650425 | 0.457 | 0.182 | 1 | 16 |
| flot2a | 0.05658 | -0.19787097 | 0.314 | 0.364 | 1 | 16 |
| rsl1d1 | 0.05663 | 0.342586191 | 0.4 | 0.136 | 1 | 16 |
| pi4kb | 0.05694 | -0.44038075 | 0.057 | 0.273 | 1 | 16 |
| rhoaa | 0.05704 | -0.1686424 | 0.457 | 0.682 | 1 | 16 |
| LOC101883815 | 0.05726 | -0.32284471 | 0.029 | 0.227 | 1 | 16 |
| crnkl1 | 0.05727 | -0.32088978 | 0.029 | 0.227 | 1 | 16 |
| irf2bp1 | 0.05752 | -0.31042073 | 0.029 | 0.227 | 1 | 16 |
| ubb | 0.05766 | -0.15468689 | 0.629 | 0.864 | 1 | 16 |
| usp9 | 0.0584 | -0.4062504 | 0.086 | 0.318 | 1 | 16 |
| notch2 | 0.05913 | -0.21336375 | 0.171 | 0.364 | 1 | 16 |
| ubr4 | 0.05929 | -0.32484171 | 0.029 | 0.227 | 1 | 16 |
| qars | 0.05952 | -0.33049836 | 0.029 | 0.227 | 1 | 16 |
| hist2h2l | 0.05988 | 0.239549477 | 0.771 | 0.5 | 1 | 16 |
| coa5 | 0.06001 | 0.354913133 | 0.257 | 0.045 | 1 | 16 |
| arl5a | 0.06013 | -0.31962794 | 0.029 | 0.227 | 1 | 16 |
| namptb | 0.06049 | -0.36148594 | 0.029 | 0.227 | 1 | 16 |
| mfap3l | 0.06058 | -0.31533606 | 0.029 | 0.227 | 1 | 16 |
| XLOC-022032 | 0.06075 | -0.32072506 | 0.029 | 0.227 | 1 | 16 |
| cox6b2 | 0.06095 | 0.239586492 | 0.914 | 0.864 | 1 | 16 |
| mxb | 0.06109 | -0.34035782 | 0.029 | 0.227 | 1 | 16 |
| dlc1 | 0.06132 | -0.38704382 | 0.029 | 0.227 | 1 | 16 |

| | | | | | | |
|---|---|---|---|---|---|---|
| huwe1 | 0.06133 | -0.30472981 | 0.029 | 0.227 | 1 | 16 |
| rpl3 | 0.06258 | 0.201537376 | 1 | 0.864 | 1 | 16 |
| fam120c | 0.06267 | -0.20038069 | 0.229 | 0.455 | 1 | 16 |
| si:ch211-197h24.9 | 0.06355 | -0.47281481 | 0.343 | 0.636 | 1 | 16 |
| kctd5a | 0.06407 | -0.32656653 | 0.4 | 0.545 | 1 | 16 |
| ifi45 | 0.06407 | -0.30401875 | 0.8 | 0.864 | 1 | 16 |
| epsti1 | 0.06407 | -0.28562345 | 0.114 | 0.227 | 1 | 16 |
| ak1 | 0.0646 | -0.18473037 | 0.086 | 0.273 | 1 | 16 |
| si:ch211-213a13.1 | 0.06466 | -0.25834503 | 0.343 | 0.409 | 1 | 16 |
| emilin1a | 0.06473 | -0.38054772 | 0.429 | 0.591 | 1 | 16 |
| exoc1 | 0.06493 | -0.35059272 | 0.057 | 0.273 | 1 | 16 |
| mob1ba | 0.06516 | 0.372180349 | 0.229 | 0.045 | 1 | 16 |
| chmp5b | 0.06523 | 0.380132212 | 0.4 | 0.136 | 1 | 16 |
| xpo1a | 0.06553 | -0.16637643 | 0.029 | 0.136 | 1 | 16 |
| apeh | 0.06579 | 0.178679588 | 0.114 | 0 | 1 | 16 |
| atad2b | 0.06579 | 0.262621496 | 0.114 | 0 | 1 | 16 |
| cyp2n13 | 0.06579 | 0.226778119 | 0.114 | 0 | 1 | 16 |
| daam1a | 0.06579 | 0.224290589 | 0.114 | 0 | 1 | 16 |
| elavl1b | 0.06579 | 0.220158899 | 0.114 | 0 | 1 | 16 |
| fam173a | 0.06579 | 0.19716006 | 0.114 | 0 | 1 | 16 |
| mmadhc | 0.06579 | 0.195759096 | 0.114 | 0 | 1 | 16 |
| srebf2 | 0.06579 | 0.226917157 | 0.114 | 0 | 1 | 16 |
| tmem230a | 0.06579 | 0.20971815 | 0.114 | 0 | 1 | 16 |
| zgc:194469 | 0.06579 | 0.212523236 | 0.114 | 0 | 1 | 16 |
| LOC110439840 | 0.06579 | 0.191694962 | 0.114 | 0 | 1 | 16 |
| klhdc4 | 0.06579 | 0.176919435 | 0.114 | 0 | 1 | 16 |
| rab27a | 0.06579 | 0.181536587 | 0.114 | 0 | 1 | 16 |
| slc35e1 | 0.06579 | 0.206782373 | 0.114 | 0 | 1 | 16 |
| tmem129 | 0.06579 | 0.20893518 | 0.114 | 0 | 1 | 16 |
| tmem186 | 0.06579 | 0.208352078 | 0.114 | 0 | 1 | 16 |
| wacb | 0.06579 | 0.227653999 | 0.114 | 0 | 1 | 16 |
| CR450780.2 | 0.06579 | 0.211714232 | 0.114 | 0 | 1 | 16 |
| angel2 | 0.06579 | 0.217867159 | 0.114 | 0 | 1 | 16 |
| atp6v1c1a | 0.06579 | 0.196191631 | 0.114 | 0 | 1 | 16 |
| atxn3 | 0.06579 | 0.19290025 | 0.114 | 0 | 1 | 16 |
| cmtr2 | 0.06579 | 0.230341133 | 0.114 | 0 | 1 | 16 |
| cnnm2a | 0.06579 | 0.233901654 | 0.114 | 0 | 1 | 16 |
| coa1 | 0.06579 | 0.190116133 | 0.114 | 0 | 1 | 16 |
| cox16 | 0.06579 | 0.235891598 | 0.114 | 0 | 1 | 16 |
| entpd4 | 0.06579 | 0.207089401 | 0.114 | 0 | 1 | 16 |
| fam92a1 | 0.06579 | 0.200312324 | 0.114 | 0 | 1 | 16 |
| fndc3ba | 0.06579 | 0.190807644 | 0.114 | 0 | 1 | 16 |
| gatd3a | 0.06579 | 0.171977262 | 0.114 | 0 | 1 | 16 |
| gmppaa | 0.06579 | 0.190280732 | 0.114 | 0 | 1 | 16 |
| hat1 | 0.06579 | 0.194969556 | 0.114 | 0 | 1 | 16 |
| itpkcb | 0.06579 | 0.247677059 | 0.114 | 0 | 1 | 16 |
| larp6a | 0.06579 | 0.20692548 | 0.114 | 0 | 1 | 16 |
| lnx2a | 0.06579 | 0.248523956 | 0.114 | 0 | 1 | 16 |

| | | | | | | |
|---|---|---|---|---|---|---|
| mtss1la | 0.06579 | 0.206076871 | 0.114 | 0 | 1 | 16 |
| nck2b | 0.06579 | 0.191483183 | 0.114 | 0 | 1 | 16 |
| nudt7 | 0.06579 | 0.222304279 | 0.114 | 0 | 1 | 16 |
| pde6d | 0.06579 | 0.19392747 | 0.114 | 0 | 1 | 16 |
| pebp1 | 0.06579 | 0.184374563 | 0.114 | 0 | 1 | 16 |
| pik3r1 | 0.06579 | 0.22041839 | 0.114 | 0 | 1 | 16 |
| ppt2 | 0.06579 | 0.197530737 | 0.114 | 0 | 1 | 16 |
| ptprsa | 0.06579 | 0.220230809 | 0.114 | 0 | 1 | 16 |
| rbm39b | 0.06579 | 0.189696305 | 0.114 | 0 | 1 | 16 |
| sco1 | 0.06579 | 0.222956214 | 0.114 | 0 | 1 | 16 |
| scp2b | 0.06579 | 0.191635555 | 0.114 | 0 | 1 | 16 |
| sestd1 | 0.06579 | 0.27169146 | 0.114 | 0 | 1 | 16 |
| si:ch211-117c9.2 | 0.06579 | 0.231100139 | 0.114 | 0 | 1 | 16 |
| si:dkey-90a24.1 | 0.06579 | 0.302355882 | 0.114 | 0 | 1 | 16 |
| stxbp3 | 0.06579 | 0.232245633 | 0.114 | 0 | 1 | 16 |
| tbcb | 0.06579 | 0.204554409 | 0.114 | 0 | 1 | 16 |
| tulp4a | 0.06579 | 0.263681747 | 0.114 | 0 | 1 | 16 |
| uqcc3 | 0.06579 | 0.177790242 | 0.114 | 0 | 1 | 16 |
| zdhhc18a | 0.06579 | 0.198248261 | 0.114 | 0 | 1 | 16 |
| zgc:162472 | 0.06579 | 0.253153073 | 0.114 | 0 | 1 | 16 |
| ap3m1 | 0.06579 | 0.218538049 | 0.114 | 0 | 1 | 16 |
| atl2 | 0.06579 | 0.179092231 | 0.114 | 0 | 1 | 16 |
| baiap2b | 0.06579 | 0.230827175 | 0.114 | 0 | 1 | 16 |
| elovl1a | 0.06579 | 0.176551783 | 0.114 | 0 | 1 | 16 |
| gyg1a | 0.06579 | 0.19306481 | 0.114 | 0 | 1 | 16 |
| maf1 | 0.06579 | 0.176702326 | 0.114 | 0 | 1 | 16 |
| supt16h | 0.06579 | 0.204790673 | 0.114 | 0 | 1 | 16 |
| tpmt.1 | 0.06579 | 0.210826757 | 0.114 | 0 | 1 | 16 |
| uck2a | 0.06579 | 0.225051927 | 0.114 | 0 | 1 | 16 |
| BX465227.1 | 0.06579 | 0.22215642 | 0.114 | 0 | 1 | 16 |
| atp6v1f | 0.06579 | 0.197714948 | 0.114 | 0 | 1 | 16 |
| cutc | 0.06579 | 0.235759386 | 0.114 | 0 | 1 | 16 |
| kin | 0.06579 | 0.201123997 | 0.114 | 0 | 1 | 16 |
| nkd2a | 0.06579 | 0.218093212 | 0.114 | 0 | 1 | 16 |
| prkcz | 0.06579 | 0.227990263 | 0.114 | 0 | 1 | 16 |
| stoml2 | 0.06579 | 0.168869337 | 0.114 | 0 | 1 | 16 |
| sesn2 | 0.06579 | 0.186802726 | 0.114 | 0 | 1 | 16 |
| abi2b | 0.06579 | 0.190969815 | 0.114 | 0 | 1 | 16 |
| bloc1s5 | 0.06579 | 0.177106276 | 0.114 | 0 | 1 | 16 |
| cpsf4 | 0.06579 | 0.181215867 | 0.114 | 0 | 1 | 16 |
| hephl1a | 0.06588 | -0.33937549 | 0.057 | 0.273 | 1 | 16 |
| degs1 | 0.06614 | -0.37012418 | 0.229 | 0.409 | 1 | 16 |
| rrp8 | 0.06636 | -0.34444586 | 0.057 | 0.273 | 1 | 16 |
| eif2b1 | 0.06751 | -0.39588978 | 0.143 | 0.409 | 1 | 16 |
| atp5if1a | 0.06792 | -0.21346194 | 0.6 | 0.636 | 1 | 16 |
| arpc2 | 0.06843 | 0.120165439 | 0.543 | 0.364 | 1 | 16 |
| srsf1a | 0.06857 | 0.147487137 | 0.229 | 0.091 | 1 | 16 |
| csnk1a1 | 0.06905 | -0.40902768 | 0.457 | 0.727 | 1 | 16 |

| | | | | | | |
|---|---|---|---|---|---|---|
| si:dkey-239h2.3 | 0.0693 | -0.36041565 | 0.143 | 0.409 | 1 | 16 |
| dpp3 | 0.06935 | 0.490078624 | 0.543 | 0.273 | 1 | 16 |
| ddx21 | 0.0695 | 0.264892133 | 0.371 | 0.136 | 1 | 16 |
| lman1 | 0.06955 | -0.40564493 | 0.171 | 0.409 | 1 | 16 |
| timm23a | 0.07049 | 0.359581863 | 0.457 | 0.273 | 1 | 16 |
| capzb | 0.07067 | 0.424855122 | 0.629 | 0.364 | 1 | 16 |
| histh1l | 0.07075 | -0.33970729 | 0.086 | 0.227 | 1 | 16 |
| exosc4 | 0.07123 | -0.15639561 | 0.114 | 0.273 | 1 | 16 |
| dicer1 | 0.07123 | -0.21006579 | 0.057 | 0.227 | 1 | 16 |
| glud1b | 0.07163 | -0.15859712 | 0.143 | 0.182 | 1 | 16 |
| si:dkey-178e17.3 | 0.07174 | -0.40407744 | 0.171 | 0.455 | 1 | 16 |
| tjp2b | 0.07176 | -0.4043268 | 0.171 | 0.455 | 1 | 16 |
| mob4 | 0.07226 | 0.318585172 | 0.371 | 0.227 | 1 | 16 |
| sik1 | 0.07245 | -0.40326055 | 0.286 | 0.591 | 1 | 16 |
| pou2f1b | 0.07277 | -0.2387324 | 0.114 | 0.182 | 1 | 16 |
| arf6a | 0.07293 | -0.39261677 | 0.171 | 0.364 | 1 | 16 |
| ilf2 | 0.07311 | 0.38536806 | 0.714 | 0.5 | 1 | 16 |
| tfcp2 | 0.07315 | 0.301258543 | 0.229 | 0.045 | 1 | 16 |
| gabarapb | 0.07344 | 0.325121218 | 0.886 | 0.636 | 1 | 16 |
| si:ch211-191j22.3 | 0.07409 | -0.40719004 | 0.314 | 0.545 | 1 | 16 |
| adipor1b | 0.07409 | 0.198604771 | 0.2 | 0.045 | 1 | 16 |
| ptp4a2a | 0.07498 | 0.246728682 | 0.4 | 0.182 | 1 | 16 |
| slka | 0.07507 | 0.42937657 | 0.314 | 0.091 | 1 | 16 |
| zgc:162944 | 0.07507 | 0.304686103 | 0.886 | 0.682 | 1 | 16 |
| nfkbiab | 0.07509 | -0.42557305 | 0.343 | 0.636 | 1 | 16 |
| mrpl32 | 0.07526 | -0.20199908 | 0.2 | 0.409 | 1 | 16 |
| BX908782.2 | 0.0761 | -0.14886476 | 0 | 0.091 | 1 | 16 |
| LOC110440142 | 0.0761 | -0.15067583 | 0 | 0.091 | 1 | 16 |
| XLOC-022034 | 0.0761 | -0.1422386 | 0 | 0.091 | 1 | 16 |
| atxn10 | 0.0761 | -0.1492429 | 0 | 0.091 | 1 | 16 |
| borcs8 | 0.0761 | -0.12884346 | 0 | 0.091 | 1 | 16 |
| bpgm | 0.0761 | -0.15858323 | 0 | 0.091 | 1 | 16 |
| btr29 | 0.0761 | -0.14380313 | 0 | 0.091 | 1 | 16 |
| ccdc73 | 0.0761 | -0.14359497 | 0 | 0.091 | 1 | 16 |
| cd40 | 0.0761 | -0.17196472 | 0 | 0.091 | 1 | 16 |
| cdc42se1 | 0.0761 | -0.14926396 | 0 | 0.091 | 1 | 16 |
| chpt1 | 0.0761 | -0.14259591 | 0 | 0.091 | 1 | 16 |
| ddx55 | 0.0761 | -0.14289949 | 0 | 0.091 | 1 | 16 |
| dhx8 | 0.0761 | -0.18979067 | 0 | 0.091 | 1 | 16 |
| dpf2 | 0.0761 | -0.14659139 | 0 | 0.091 | 1 | 16 |
| dph6 | 0.0761 | -0.15259075 | 0 | 0.091 | 1 | 16 |
| dusp2 | 0.0761 | -0.11798868 | 0 | 0.091 | 1 | 16 |
| eef1a1a | 0.0761 | -0.16863596 | 0 | 0.091 | 1 | 16 |
| epha2a | 0.0761 | -0.14000133 | 0 | 0.091 | 1 | 16 |
| fbrs | 0.0761 | -0.13843786 | 0 | 0.091 | 1 | 16 |
| foxk1 | 0.0761 | -0.1499571 | 0 | 0.091 | 1 | 16 |
| hdac8 | 0.0761 | -0.14333302 | 0 | 0.091 | 1 | 16 |
| inhbab | 0.0761 | -0.14602128 | 0 | 0.091 | 1 | 16 |

| | | | | | | |
|---|---|---|---|---|---|---|
| ipo13 | 0.0761 | -0.14704182 | 0 | 0.091 | 1 | 16 |
| mical3b | 0.0761 | -0.13739728 | 0 | 0.091 | 1 | 16 |
| mipol1 | 0.0761 | -0.13792609 | 0 | 0.091 | 1 | 16 |
| mllt1a | 0.0761 | -0.14863948 | 0 | 0.091 | 1 | 16 |
| mnat1 | 0.0761 | -0.12423346 | 0 | 0.091 | 1 | 16 |
| mtbl | 0.0761 | -0.13569726 | 0 | 0.091 | 1 | 16 |
| nmt2 | 0.0761 | -0.13914092 | 0 | 0.091 | 1 | 16 |
| nsmaf | 0.0761 | -0.12753753 | 0 | 0.091 | 1 | 16 |
| nup35 | 0.0761 | -0.14025108 | 0 | 0.091 | 1 | 16 |
| rab11fip3 | 0.0761 | -0.18664638 | 0 | 0.091 | 1 | 16 |
| rfx7b | 0.0761 | -0.14071314 | 0 | 0.091 | 1 | 16 |
| rnf144ab | 0.0761 | -0.21974909 | 0 | 0.091 | 1 | 16 |
| si:ch211-155k24.9 | 0.0761 | -0.15462611 | 0 | 0.091 | 1 | 16 |
| si:ch211-40k21.5 | 0.0761 | -0.14339836 | 0 | 0.091 | 1 | 16 |
| si:ch211-71m22.1 | 0.0761 | -0.18800433 | 0 | 0.091 | 1 | 16 |
| si:ch73-190m4.1 | 0.0761 | -0.16162956 | 0 | 0.091 | 1 | 16 |
| si:dkey-181m9.8 | 0.0761 | -0.12517945 | 0 | 0.091 | 1 | 16 |
| si:dkey-58f10.14 | 0.0761 | -0.15066856 | 0 | 0.091 | 1 | 16 |
| taf5 | 0.0761 | -0.13062013 | 0 | 0.091 | 1 | 16 |
| tmeff2a | 0.0761 | -0.1858637 | 0 | 0.091 | 1 | 16 |
| tnfsf10l | 0.0761 | -0.20015015 | 0 | 0.091 | 1 | 16 |
| xpo4 | 0.0761 | -0.14399082 | 0 | 0.091 | 1 | 16 |
| zgc:162964 | 0.0761 | -0.13805954 | 0 | 0.091 | 1 | 16 |
| zgc:194655 | 0.0761 | -0.17800967 | 0 | 0.091 | 1 | 16 |
| ARHGEF17 | 0.0761 | -0.14588986 | 0 | 0.091 | 1 | 16 |
| LO016987.2 | 0.0761 | -0.19472203 | 0 | 0.091 | 1 | 16 |
| XLOC-007007 | 0.0761 | -0.15507843 | 0 | 0.091 | 1 | 16 |
| alkbh7 | 0.0761 | -0.15801471 | 0 | 0.091 | 1 | 16 |
| coq4 | 0.0761 | -0.1762248 | 0 | 0.091 | 1 | 16 |
| fbxw8 | 0.0761 | -0.15334246 | 0 | 0.091 | 1 | 16 |
| ftr06 | 0.0761 | -0.14254645 | 0 | 0.091 | 1 | 16 |
| miip | 0.0761 | -0.19582069 | 0 | 0.091 | 1 | 16 |
| p4ha1a | 0.0761 | -0.13708768 | 0 | 0.091 | 1 | 16 |
| pigw | 0.0761 | -0.15222818 | 0 | 0.091 | 1 | 16 |
| pknox1.2 | 0.0761 | -0.15954536 | 0 | 0.091 | 1 | 16 |
| rtn4b | 0.0761 | -0.19256304 | 0 | 0.091 | 1 | 16 |
| samm50l | 0.0761 | -0.14532128 | 0 | 0.091 | 1 | 16 |
| tm7sf3 | 0.0761 | -0.14799884 | 0 | 0.091 | 1 | 16 |
| znf576.1 | 0.0761 | -0.16056131 | 0 | 0.091 | 1 | 16 |
| rarres3l | 0.0761 | -0.14690313 | 0 | 0.091 | 1 | 16 |
| sema4ba | 0.0761 | -0.14204867 | 0 | 0.091 | 1 | 16 |
| LOC101885938 | 0.0761 | -0.17340616 | 0 | 0.091 | 1 | 16 |
| NDUFAF8 | 0.0761 | -0.21846223 | 0 | 0.091 | 1 | 16 |
| USP53 (1 of many) | 0.0761 | -0.15969967 | 0 | 0.091 | 1 | 16 |
| armh1 | 0.0761 | -0.14799715 | 0 | 0.091 | 1 | 16 |
| bicra | 0.0761 | -0.22025075 | 0 | 0.091 | 1 | 16 |
| ccdc93 | 0.0761 | -0.15597115 | 0 | 0.091 | 1 | 16 |
| col8a1a | 0.0761 | -0.18483179 | 0 | 0.091 | 1 | 16 |

| | | | | | | |
|---|---|---|---|---|---|---|
| cpne4b | 0.0761 | -0.16796439 | 0 | 0.091 | 1 | 16 |
| inpp5ka | 0.0761 | -0.17851337 | 0 | 0.091 | 1 | 16 |
| mad2l1 | 0.0761 | -0.14955574 | 0 | 0.091 | 1 | 16 |
| med13b | 0.0761 | -0.14767173 | 0 | 0.091 | 1 | 16 |
| notch3 | 0.0761 | -0.16138274 | 0 | 0.091 | 1 | 16 |
| nsd1a | 0.0761 | -0.15792954 | 0 | 0.091 | 1 | 16 |
| rab5b | 0.0761 | -0.15886872 | 0 | 0.091 | 1 | 16 |
| rdh8a | 0.0761 | -0.21910597 | 0 | 0.091 | 1 | 16 |
| si:dkey-40g16.5 | 0.0761 | -0.18704891 | 0 | 0.091 | 1 | 16 |
| si:dkey-71b5.7 | 0.0761 | -0.15518311 | 0 | 0.091 | 1 | 16 |
| si:dkey-82o10.4 | 0.0761 | -0.14236474 | 0 | 0.091 | 1 | 16 |
| sox18 | 0.0761 | -0.18024242 | 0 | 0.091 | 1 | 16 |
| tmem160 | 0.0761 | -0.17865611 | 0 | 0.091 | 1 | 16 |
| uap1 | 0.0761 | -0.15656225 | 0 | 0.091 | 1 | 16 |
| zgc:171572 | 0.0761 | -0.17051611 | 0 | 0.091 | 1 | 16 |
| grk6 | 0.0761 | -0.17543229 | 0 | 0.091 | 1 | 16 |
| il10rb | 0.0761 | -0.16723284 | 0 | 0.091 | 1 | 16 |
| piezo1 | 0.0761 | -0.16495187 | 0 | 0.091 | 1 | 16 |
| ppp4r2b | 0.0761 | -0.1761585 | 0 | 0.091 | 1 | 16 |
| si:ch211-107n13.1 | 0.0761 | -0.17212054 | 0 | 0.091 | 1 | 16 |
| tmem220 | 0.0761 | -0.17225554 | 0 | 0.091 | 1 | 16 |
| abhd2a | 0.0761 | -0.16898359 | 0 | 0.091 | 1 | 16 |
| arnt | 0.0761 | -0.16690649 | 0 | 0.091 | 1 | 16 |
| ccnc | 0.0761 | -0.20095236 | 0 | 0.091 | 1 | 16 |
| clp1 | 0.0761 | -0.1459476 | 0 | 0.091 | 1 | 16 |
| cyp2ae1 | 0.0761 | -0.15124788 | 0 | 0.091 | 1 | 16 |
| cyp2x7 | 0.0761 | -0.15257279 | 0 | 0.091 | 1 | 16 |
| dnajc21 | 0.0761 | -0.21357432 | 0 | 0.091 | 1 | 16 |
| dsg2.1 | 0.0761 | -0.15630922 | 0 | 0.091 | 1 | 16 |
| gcdha | 0.0761 | -0.2584677 | 0 | 0.091 | 1 | 16 |
| gdap2 | 0.0761 | -0.14346176 | 0 | 0.091 | 1 | 16 |
| ilvbl | 0.0761 | -0.20065332 | 0 | 0.091 | 1 | 16 |
| lrig3 | 0.0761 | -0.22049678 | 0 | 0.091 | 1 | 16 |
| mhc1zca | 0.0761 | -0.15216783 | 0 | 0.091 | 1 | 16 |
| mir181b-3 | 0.0761 | -0.21302353 | 0 | 0.091 | 1 | 16 |
| ndufaf4 | 0.0761 | -0.19685289 | 0 | 0.091 | 1 | 16 |
| pdcd2l | 0.0761 | -0.16499073 | 0 | 0.091 | 1 | 16 |
| pogzb | 0.0761 | -0.1506337 | 0 | 0.091 | 1 | 16 |
| rfk | 0.0761 | -0.19816107 | 0 | 0.091 | 1 | 16 |
| rh50 | 0.0761 | -0.20603159 | 0 | 0.091 | 1 | 16 |
| rnpepl1 | 0.0761 | -0.19595215 | 0 | 0.091 | 1 | 16 |
| sacm1lb | 0.0761 | -0.15778272 | 0 | 0.091 | 1 | 16 |
| si:ch211-175m2.5 | 0.0761 | -0.18623105 | 0 | 0.091 | 1 | 16 |
| si:ch211-212d10.2 | 0.0761 | -0.20286471 | 0 | 0.091 | 1 | 16 |
| si:dkey-29p10.4 | 0.0761 | -0.23423133 | 0 | 0.091 | 1 | 16 |
| slc26a5 | 0.0761 | -0.23287343 | 0 | 0.091 | 1 | 16 |
| toe1 | 0.0761 | -0.1494136 | 0 | 0.091 | 1 | 16 |
| tprg1 | 0.0761 | -0.19520957 | 0 | 0.091 | 1 | 16 |

| | | | | | | |
|---|---|---|---|---|---|---|
| wdr55 | 0.0761 | -0.15358104 | 0 | 0.091 | 1 | 16 |
| zgc:112977 | 0.0761 | -0.14941741 | 0 | 0.091 | 1 | 16 |
| znf1010 | 0.0761 | -0.20321469 | 0 | 0.091 | 1 | 16 |
| znf628 | 0.0761 | -0.1708198 | 0 | 0.091 | 1 | 16 |
| AL935153.1 | 0.0761 | -0.1586279 | 0 | 0.091 | 1 | 16 |
| CABZ01028771.1 | 0.0761 | -0.17154785 | 0 | 0.091 | 1 | 16 |
| CU693502.1 | 0.0761 | -0.14851585 | 0 | 0.091 | 1 | 16 |
| CU929391.1 | 0.0761 | -0.16436581 | 0 | 0.091 | 1 | 16 |
| FP085398.1 | 0.0761 | -0.19408324 | 0 | 0.091 | 1 | 16 |
| FQ378018.1 | 0.0761 | -0.19536402 | 0 | 0.091 | 1 | 16 |
| IZUMO1R | 0.0761 | -0.23275265 | 0 | 0.091 | 1 | 16 |
| XLOC-030237 | 0.0761 | -0.2027906 | 0 | 0.091 | 1 | 16 |
| abi1a | 0.0761 | -0.16756323 | 0 | 0.091 | 1 | 16 |
| abi2a | 0.0761 | -0.14517679 | 0 | 0.091 | 1 | 16 |
| aftpha | 0.0761 | -0.14231955 | 0 | 0.091 | 1 | 16 |
| birc6 | 0.0761 | -0.19741944 | 0 | 0.091 | 1 | 16 |
| cav2 | 0.0761 | -0.16502792 | 0 | 0.091 | 1 | 16 |
| clmn | 0.0761 | -0.18295185 | 0 | 0.091 | 1 | 16 |
| cop1 | 0.0761 | -0.13389858 | 0 | 0.091 | 1 | 16 |
| dnal4b | 0.0761 | -0.18145005 | 0 | 0.091 | 1 | 16 |
| dnmt1 | 0.0761 | -0.16698624 | 0 | 0.091 | 1 | 16 |
| dock6 | 0.0761 | -0.18248696 | 0 | 0.091 | 1 | 16 |
| exosc2 | 0.0761 | -0.172329 | 0 | 0.091 | 1 | 16 |
| fndc3a | 0.0761 | -0.25203961 | 0 | 0.091 | 1 | 16 |
| fuk | 0.0761 | -0.14583009 | 0 | 0.091 | 1 | 16 |
| gabpb2b | 0.0761 | -0.17605343 | 0 | 0.091 | 1 | 16 |
| grpel1 | 0.0761 | -0.13862261 | 0 | 0.091 | 1 | 16 |
| gxylt2 | 0.0761 | -0.16938314 | 0 | 0.091 | 1 | 16 |
| itgb1b.1 | 0.0761 | -0.14407807 | 0 | 0.091 | 1 | 16 |
| kaznb | 0.0761 | -0.18031746 | 0 | 0.091 | 1 | 16 |
| kctd7 | 0.0761 | -0.14558833 | 0 | 0.091 | 1 | 16 |
| ldb1b | 0.0761 | -0.1445656 | 0 | 0.091 | 1 | 16 |
| lrrfip1a | 0.0761 | -0.1324701 | 0 | 0.091 | 1 | 16 |
| mfn1b | 0.0761 | -0.19782808 | 0 | 0.091 | 1 | 16 |
| mical3a | 0.0761 | -0.23574983 | 0 | 0.091 | 1 | 16 |
| ndel1b | 0.0761 | -0.1954223 | 0 | 0.091 | 1 | 16 |
| nek6 | 0.0761 | -0.20418612 | 0 | 0.091 | 1 | 16 |
| nog1 | 0.0761 | -0.14525875 | 0 | 0.091 | 1 | 16 |
| nrg2a | 0.0761 | -0.18367786 | 0 | 0.091 | 1 | 16 |
| ormdl3 | 0.0761 | -0.13739561 | 0 | 0.091 | 1 | 16 |
| phf20a | 0.0761 | -0.20299709 | 0 | 0.091 | 1 | 16 |
| plaa | 0.0761 | -0.16939695 | 0 | 0.091 | 1 | 16 |
| ptprk | 0.0761 | -0.17677068 | 0 | 0.091 | 1 | 16 |
| ptprua | 0.0761 | -0.18291562 | 0 | 0.091 | 1 | 16 |
| rapgef1a | 0.0761 | -0.15180033 | 0 | 0.091 | 1 | 16 |
| rbpms2a | 0.0761 | -0.21624227 | 0 | 0.091 | 1 | 16 |
| rcbtb1 | 0.0761 | -0.22887112 | 0 | 0.091 | 1 | 16 |
| ribc1 | 0.0761 | -0.20951377 | 0 | 0.091 | 1 | 16 |

| | | | | | | |
|---|---|---|---|---|---|---|
| rpp38 | 0.0761 | -0.16268721 | 0 | 0.091 | 1 | 16 |
| sb:cb81 | 0.0761 | -0.17104697 | 0 | 0.091 | 1 | 16 |
| sdad1 | 0.0761 | -0.1607658 | 0 | 0.091 | 1 | 16 |
| sec22a | 0.0761 | -0.15149886 | 0 | 0.091 | 1 | 16 |
| sh2b3 | 0.0761 | -0.14049865 | 0 | 0.091 | 1 | 16 |
| si:busm1-194e12.11 | 0.0761 | -0.10866942 | 0 | 0.091 | 1 | 16 |
| si:ch211-261n11.5 | 0.0761 | -0.1682575 | 0 | 0.091 | 1 | 16 |
| si:dkey-68o6.6 | 0.0761 | -0.15395071 | 0 | 0.091 | 1 | 16 |
| si:dkeyp-120h9.1 | 0.0761 | -0.16454477 | 0 | 0.091 | 1 | 16 |
| slc35d1b | 0.0761 | -0.18205174 | 0 | 0.091 | 1 | 16 |
| slc39a11 | 0.0761 | -0.19929446 | 0 | 0.091 | 1 | 16 |
| slc48a1a | 0.0761 | -0.19410863 | 0 | 0.091 | 1 | 16 |
| strip1 | 0.0761 | -0.18079386 | 0 | 0.091 | 1 | 16 |
| tceanc2 | 0.0761 | -0.16767003 | 0 | 0.091 | 1 | 16 |
| tmem150c | 0.0761 | -0.20127328 | 0 | 0.091 | 1 | 16 |
| tpgs2 | 0.0761 | -0.19999036 | 0 | 0.091 | 1 | 16 |
| uxt | 0.0761 | -0.19462353 | 0 | 0.091 | 1 | 16 |
| vgll4b | 0.0761 | -0.16887675 | 0 | 0.091 | 1 | 16 |
| wwc3 | 0.0761 | -0.15542205 | 0 | 0.091 | 1 | 16 |
| zgc:113210 | 0.0761 | -0.17384893 | 0 | 0.091 | 1 | 16 |
| zgc:92873 | 0.0761 | -0.17633406 | 0 | 0.091 | 1 | 16 |
| BX950187.1 | 0.0761 | -0.20792747 | 0 | 0.091 | 1 | 16 |
| CABZ01076758.1 | 0.0761 | -0.1659899 | 0 | 0.091 | 1 | 16 |
| CABZ01086877.1 | 0.0761 | -0.17188485 | 0 | 0.091 | 1 | 16 |
| CR388132.2 | 0.0761 | -0.13808684 | 0 | 0.091 | 1 | 16 |
| XLOC-041870 | 0.0761 | -0.10321029 | 0 | 0.091 | 1 | 16 |
| adpgk2 | 0.0761 | -0.15471343 | 0 | 0.091 | 1 | 16 |
| afap1 | 0.0761 | -0.1624394 | 0 | 0.091 | 1 | 16 |
| alkbh5 | 0.0761 | -0.15255402 | 0 | 0.091 | 1 | 16 |
| atp11c | 0.0761 | -0.15853815 | 0 | 0.091 | 1 | 16 |
| atxn1a | 0.0761 | -0.16678407 | 0 | 0.091 | 1 | 16 |
| bach2b | 0.0761 | -0.15143956 | 0 | 0.091 | 1 | 16 |
| brwd1 | 0.0761 | -0.16572752 | 0 | 0.091 | 1 | 16 |
| casp9 | 0.0761 | -0.15251353 | 0 | 0.091 | 1 | 16 |
| ccnq | 0.0761 | -0.18466559 | 0 | 0.091 | 1 | 16 |
| cdca7a | 0.0761 | -0.14731874 | 0 | 0.091 | 1 | 16 |
| cog8 | 0.0761 | -0.15363071 | 0 | 0.091 | 1 | 16 |
| coq5 | 0.0761 | -0.16654696 | 0 | 0.091 | 1 | 16 |
| dhx57 | 0.0761 | -0.14857025 | 0 | 0.091 | 1 | 16 |
| dis3l2 | 0.0761 | -0.16090873 | 0 | 0.091 | 1 | 16 |
| efr3bb | 0.0761 | -0.1601748 | 0 | 0.091 | 1 | 16 |
| exosc1 | 0.0761 | -0.17579035 | 0 | 0.091 | 1 | 16 |
| fam8a1b | 0.0761 | -0.12943872 | 0 | 0.091 | 1 | 16 |
| fbxo22 | 0.0761 | -0.2049297 | 0 | 0.091 | 1 | 16 |
| fcho2 | 0.0761 | -0.14899946 | 0 | 0.091 | 1 | 16 |
| ftr22 | 0.0761 | -0.20538576 | 0 | 0.091 | 1 | 16 |
| gal3st1b | 0.0761 | -0.16030232 | 0 | 0.091 | 1 | 16 |
| gig2d | 0.0761 | -0.13125394 | 0 | 0.091 | 1 | 16 |

| | | | | | | |
|---|---|---|---|---|---|---|
| gnpda2 | 0.0761 | -0.16118013 | 0 | 0.091 | 1 | 16 |
| gtf3c6 | 0.0761 | -0.14816451 | 0 | 0.091 | 1 | 16 |
| hdhd3 | 0.0761 | -0.13776327 | 0 | 0.091 | 1 | 16 |
| hsd3b1 | 0.0761 | -0.16330354 | 0 | 0.091 | 1 | 16 |
| ikbkb | 0.0761 | -0.16432102 | 0 | 0.091 | 1 | 16 |
| ing2 | 0.0761 | -0.18174991 | 0 | 0.091 | 1 | 16 |
| ints1 | 0.0761 | -0.20791776 | 0 | 0.091 | 1 | 16 |
| ivns1abpb | 0.0761 | -0.12770955 | 0 | 0.091 | 1 | 16 |
| lima1a | 0.0761 | -0.18685126 | 0 | 0.091 | 1 | 16 |
| lin7c | 0.0761 | -0.2122312 | 0 | 0.091 | 1 | 16 |
| lmo7b | 0.0761 | -0.16109894 | 0 | 0.091 | 1 | 16 |
| mob2a | 0.0761 | -0.18275123 | 0 | 0.091 | 1 | 16 |
| mpp1 | 0.0761 | -0.15500307 | 0 | 0.091 | 1 | 16 |
| mtmr7a | 0.0761 | -0.13594676 | 0 | 0.091 | 1 | 16 |
| nkain4 | 0.0761 | -0.15538007 | 0 | 0.091 | 1 | 16 |
| nkiras2 | 0.0761 | -0.1738092 | 0 | 0.091 | 1 | 16 |
| noc3l | 0.0761 | -0.14679288 | 0 | 0.091 | 1 | 16 |
| osbpl2a | 0.0761 | -0.18173642 | 0 | 0.091 | 1 | 16 |
| osbpl9 | 0.0761 | -0.14272802 | 0 | 0.091 | 1 | 16 |
| pkp2 | 0.0761 | -0.18369323 | 0 | 0.091 | 1 | 16 |
| plekhh1 | 0.0761 | -0.16023481 | 0 | 0.091 | 1 | 16 |
| rbm26 | 0.0761 | -0.18381158 | 0 | 0.091 | 1 | 16 |
| rgs12b | 0.0761 | -0.15127351 | 0 | 0.091 | 1 | 16 |
| rnf144b | 0.0761 | -0.13297463 | 0 | 0.091 | 1 | 16 |
| rngtt | 0.0761 | -0.20675958 | 0 | 0.091 | 1 | 16 |
| rpn2 | 0.0761 | -0.15035097 | 0 | 0.091 | 1 | 16 |
| rtn1b | 0.0761 | -0.14392878 | 0 | 0.091 | 1 | 16 |
| rwdd3 | 0.0761 | -0.15760455 | 0 | 0.091 | 1 | 16 |
| sbno1 | 0.0761 | -0.14483462 | 0 | 0.091 | 1 | 16 |
| sec24d | 0.0761 | -0.12697197 | 0 | 0.091 | 1 | 16 |
| si:ch211-191d2.2 | 0.0761 | -0.17038594 | 0 | 0.091 | 1 | 16 |
| si:ch211-281g13.5 | 0.0761 | -0.17876085 | 0 | 0.091 | 1 | 16 |
| si:ch73-233k15.2 | 0.0761 | -0.24303093 | 0 | 0.091 | 1 | 16 |
| si:dkey-14d8.1 | 0.0761 | -0.16147859 | 0 | 0.091 | 1 | 16 |
| si:dkey-156n14.3 | 0.0761 | -0.15288586 | 0 | 0.091 | 1 | 16 |
| si:dkey-97o5.1 | 0.0761 | -0.15851516 | 0 | 0.091 | 1 | 16 |
| slc19a1 | 0.0761 | -0.1889643 | 0 | 0.091 | 1 | 16 |
| slc25a23a | 0.0761 | -0.20807567 | 0 | 0.091 | 1 | 16 |
| slc25a23b | 0.0761 | -0.16077921 | 0 | 0.091 | 1 | 16 |
| suz12a | 0.0761 | -0.20391333 | 0 | 0.091 | 1 | 16 |
| tefm | 0.0761 | -0.20228526 | 0 | 0.091 | 1 | 16 |
| ticam1 | 0.0761 | -0.16048461 | 0 | 0.091 | 1 | 16 |
| tigarb | 0.0761 | -0.17414465 | 0 | 0.091 | 1 | 16 |
| tmem106bb | 0.0761 | -0.16855265 | 0 | 0.091 | 1 | 16 |
| tmem138 | 0.0761 | -0.15215461 | 0 | 0.091 | 1 | 16 |
| tmem47 | 0.0761 | -0.13605023 | 0 | 0.091 | 1 | 16 |
| tnfaip2a | 0.0761 | -0.2565055 | 0 | 0.091 | 1 | 16 |
| trim35-3 | 0.0761 | -0.14204289 | 0 | 0.091 | 1 | 16 |

| | | | | | | |
|---|---|---|---|---|---|---|
| ttc38 | 0.0761 | -0.16638926 | 0 | 0.091 | 1 | 16 |
| txlng | 0.0761 | -0.15291575 | 0 | 0.091 | 1 | 16 |
| urod | 0.0761 | -0.1405865 | 0 | 0.091 | 1 | 16 |
| utp15 | 0.0761 | -0.2069382 | 0 | 0.091 | 1 | 16 |
| vac14 | 0.0761 | -0.15759576 | 0 | 0.091 | 1 | 16 |
| wdr36 | 0.0761 | -0.16121714 | 0 | 0.091 | 1 | 16 |
| wdr48b | 0.0761 | -0.15493055 | 0 | 0.091 | 1 | 16 |
| xrcc3 | 0.0761 | -0.14896704 | 0 | 0.091 | 1 | 16 |
| zgc:110782 | 0.0761 | -0.16518184 | 0 | 0.091 | 1 | 16 |
| zgc:113348 | 0.0761 | -0.20934555 | 0 | 0.091 | 1 | 16 |
| zgc:114174 | 0.0761 | -0.14015471 | 0 | 0.091 | 1 | 16 |
| zmp:0000000662 | 0.0761 | -0.13961625 | 0 | 0.091 | 1 | 16 |
| zmp:0000000912 | 0.0761 | -0.17973565 | 0 | 0.091 | 1 | 16 |
| znf644b | 0.0761 | -0.18764061 | 0 | 0.091 | 1 | 16 |
| znf654 | 0.0761 | -0.14368083 | 0 | 0.091 | 1 | 16 |
| BX544877.1 | 0.0761 | -0.156003 | 0 | 0.091 | 1 | 16 |
| CU074419.2 | 0.0761 | -0.1996567 | 0 | 0.091 | 1 | 16 |
| CU929418.1 | 0.0761 | -0.18023002 | 0 | 0.091 | 1 | 16 |
| FP102158.1 | 0.0761 | -0.18395033 | 0 | 0.091 | 1 | 16 |
| LOC793824 | 0.0761 | -0.16222301 | 0 | 0.091 | 1 | 16 |
| OSBPL8 | 0.0761 | -0.1920736 | 0 | 0.091 | 1 | 16 |
| XLOC-043151 | 0.0761 | -0.17266405 | 0 | 0.091 | 1 | 16 |
| agla | 0.0761 | -0.19193752 | 0 | 0.091 | 1 | 16 |
| alkbh3 | 0.0761 | -0.19103762 | 0 | 0.091 | 1 | 16 |
| arntl2 | 0.0761 | -0.19748266 | 0 | 0.091 | 1 | 16 |
| bmp7b | 0.0761 | -0.19302138 | 0 | 0.091 | 1 | 16 |
| brpf3a | 0.0761 | -0.17373861 | 0 | 0.091 | 1 | 16 |
| chtf18 | 0.0761 | -0.18673657 | 0 | 0.091 | 1 | 16 |
| clpxa | 0.0761 | -0.1934595 | 0 | 0.091 | 1 | 16 |
| cmpk2 | 0.0761 | -0.16285364 | 0 | 0.091 | 1 | 16 |
| cpt1aa | 0.0761 | -0.16329295 | 0 | 0.091 | 1 | 16 |
| creld1a | 0.0761 | -0.17554729 | 0 | 0.091 | 1 | 16 |
| cyb5b | 0.0761 | -0.18678688 | 0 | 0.091 | 1 | 16 |
| dnmt3ba | 0.0761 | -0.19029309 | 0 | 0.091 | 1 | 16 |
| ephb3a | 0.0761 | -0.17449764 | 0 | 0.091 | 1 | 16 |
| fen1 | 0.0761 | -0.15219693 | 0 | 0.091 | 1 | 16 |
| fip1l1a | 0.0761 | -0.16741306 | 0 | 0.091 | 1 | 16 |
| gbp2 | 0.0761 | -0.16819337 | 0 | 0.091 | 1 | 16 |
| hip1rb | 0.0761 | -0.18929644 | 0 | 0.091 | 1 | 16 |
| hmmr | 0.0761 | -0.14898588 | 0 | 0.091 | 1 | 16 |
| il34 | 0.0761 | -0.13515525 | 0 | 0.091 | 1 | 16 |
| junbb | 0.0761 | -0.19949387 | 0 | 0.091 | 1 | 16 |
| mcm5 | 0.0761 | -0.17937506 | 0 | 0.091 | 1 | 16 |
| mier1b | 0.0761 | -0.19193522 | 0 | 0.091 | 1 | 16 |
| mipep | 0.0761 | -0.16315715 | 0 | 0.091 | 1 | 16 |
| msh2 | 0.0761 | -0.198191 | 0 | 0.091 | 1 | 16 |
| qtrt2 | 0.0761 | -0.20608516 | 0 | 0.091 | 1 | 16 |
| rabgap1 | 0.0761 | -0.17507599 | 0 | 0.091 | 1 | 16 |

| | | | | | | |
|---|---|---|---|---|---|---|
| rad51 | 0.0761 | -0.18062918 | 0 | 0.091 | 1 | 16 |
| retreg2 | 0.0761 | -0.17107636 | 0 | 0.091 | 1 | 16 |
| sall1a | 0.0761 | -0.21060449 | 0 | 0.091 | 1 | 16 |
| si:dkey-110g7.8 | 0.0761 | -0.1692282 | 0 | 0.091 | 1 | 16 |
| slc25a25a | 0.0761 | -0.21282841 | 0 | 0.091 | 1 | 16 |
| tmco3 | 0.0761 | -0.17181353 | 0 | 0.091 | 1 | 16 |
| top1mt | 0.0761 | -0.15613206 | 0 | 0.091 | 1 | 16 |
| treh | 0.0761 | -0.18266931 | 0 | 0.091 | 1 | 16 |
| ube2t | 0.0761 | -0.18616092 | 0 | 0.091 | 1 | 16 |
| xpo7 | 0.0761 | -0.17953316 | 0 | 0.091 | 1 | 16 |
| xrcc4 | 0.0761 | -0.16350639 | 0 | 0.091 | 1 | 16 |
| zbtb12.2 | 0.0761 | -0.19823036 | 0 | 0.091 | 1 | 16 |
| zgc:158640 | 0.0761 | -0.12604336 | 0 | 0.091 | 1 | 16 |
| znf1005 | 0.0761 | -0.15026517 | 0 | 0.091 | 1 | 16 |
| dnajb11 | 0.07806 | -0.38119146 | 0.086 | 0.318 | 1 | 16 |
| si:rp71-36a1.3 | 0.07808 | -0.38724086 | 0.057 | 0.273 | 1 | 16 |
| marcksl1b | 0.07812 | 0.397344018 | 0.771 | 0.591 | 1 | 16 |
| si:dkey-78l4.14 | 0.07815 | -0.3457625 | 0.114 | 0.364 | 1 | 16 |
| tuba8l4 | 0.07815 | -0.34479109 | 0.429 | 0.727 | 1 | 16 |
| tspan18b | 0.07818 | -0.33458015 | 0.057 | 0.273 | 1 | 16 |
| rsrp1 | 0.07818 | -0.35324485 | 0.086 | 0.318 | 1 | 16 |
| hif1al | 0.07856 | -0.32597929 | 0.057 | 0.273 | 1 | 16 |
| mcl1b | 0.07902 | -0.43926814 | 0.257 | 0.545 | 1 | 16 |
| pwwp2b | 0.07937 | 0.323935937 | 0.257 | 0.045 | 1 | 16 |
| atp5fa1 | 0.07942 | 0.360148294 | 0.857 | 0.682 | 1 | 16 |
| txnrd3 | 0.0798 | -0.36299808 | 0.143 | 0.409 | 1 | 16 |
| gdpd4a | 0.07988 | 0.196729235 | 0.343 | 0.182 | 1 | 16 |
| kansl1b | 0.08044 | -0.35584552 | 0.086 | 0.273 | 1 | 16 |
| lxn | 0.08046 | -0.32481722 | 0.2 | 0.364 | 1 | 16 |
| si:dkeyp-27c8.2 | 0.0806 | -0.42780449 | 0.114 | 0.364 | 1 | 16 |
| efhd1 | 0.08079 | 0.305192011 | 0.314 | 0.182 | 1 | 16 |
| pkig | 0.0811 | 0.320908447 | 0.4 | 0.273 | 1 | 16 |
| zgc:153293 | 0.0813 | 0.332726292 | 0.229 | 0.045 | 1 | 16 |
| ewsr1b | 0.0817 | -0.31375598 | 0.429 | 0.727 | 1 | 16 |
| LOC103909972 | 0.0826 | -0.36793297 | 0.114 | 0.364 | 1 | 16 |
| xbp1 | 0.08269 | -0.33791685 | 0.657 | 0.773 | 1 | 16 |
| aldoaa | 0.08311 | 0.40703909 | 0.457 | 0.182 | 1 | 16 |
| wbp1la | 0.08323 | -0.38670535 | 0.114 | 0.364 | 1 | 16 |
| gata6 | 0.08352 | 0.28848535 | 0.543 | 0.455 | 1 | 16 |
| krt18a.1 | 0.08387 | 0.367947334 | 0.629 | 0.591 | 1 | 16 |
| wnt4a | 0.08415 | 0.382890226 | 0.257 | 0.045 | 1 | 16 |
| vangl2 | 0.08475 | -0.2769341 | 0.086 | 0.273 | 1 | 16 |
| rnf213b | 0.08561 | -0.37274528 | 0.086 | 0.318 | 1 | 16 |
| fbxo25 | 0.08567 | 0.337642585 | 0.2 | 0.045 | 1 | 16 |
| tbl2 | 0.08571 | -0.36765612 | 0.086 | 0.318 | 1 | 16 |
| gnaia | 0.08571 | -0.32848217 | 0.086 | 0.318 | 1 | 16 |
| lama5 | 0.08592 | -0.31542964 | 0.314 | 0.591 | 1 | 16 |
| parp12b | 0.08609 | -0.4319545 | 0.171 | 0.409 | 1 | 16 |

| | | | | | | |
|---|---|---|---|---|---|---|
| tmem258 | 0.08666 | -0.18374604 | 0.629 | 0.636 | 1 | 16 |
| socs3a | 0.08681 | -0.44973687 | 0.543 | 0.773 | 1 | 16 |
| kdm5bb | 0.08736 | 0.334153489 | 0.257 | 0.045 | 1 | 16 |
| clic2 | 0.08815 | -0.27907324 | 0.057 | 0.136 | 1 | 16 |
| ptenb | 0.08845 | -0.36488376 | 0.257 | 0.545 | 1 | 16 |
| irf10 | 0.08874 | -0.24605138 | 0.057 | 0.227 | 1 | 16 |
| cct5 | 0.0891 | 0.298063116 | 0.743 | 0.455 | 1 | 16 |
| cd63 | 0.08976 | -0.1596928 | 0.829 | 0.773 | 1 | 16 |
| eif3m | 0.09034 | 0.352544315 | 0.8 | 0.591 | 1 | 16 |
| cirbpb | 0.09058 | 0.198424402 | 1 | 0.909 | 1 | 16 |
| gpx4b | 0.09064 | 0.237396464 | 0.8 | 0.727 | 1 | 16 |
| psmd13 | 0.09097 | -0.3865692 | 0.343 | 0.591 | 1 | 16 |
| ap2s1 | 0.09112 | -0.14849541 | 0.343 | 0.364 | 1 | 16 |
| irf2 | 0.09176 | -0.34772805 | 0.2 | 0.364 | 1 | 16 |
| ptp4a1 | 0.09183 | -0.20904715 | 0.257 | 0.455 | 1 | 16 |
| igfbp5b | 0.09249 | -0.1444624 | 0.029 | 0.136 | 1 | 16 |
| reep5 | 0.09256 | 0.326938094 | 0.629 | 0.5 | 1 | 16 |
| si:dkey-73n10.1 | 0.0933 | -0.1000622 | 0.257 | 0.227 | 1 | 16 |
| vcp | 0.09333 | -0.41314888 | 0.257 | 0.455 | 1 | 16 |
| hbegfa | 0.09383 | -0.40246474 | 0.086 | 0.273 | 1 | 16 |
| zgc:110591 | 0.09397 | -0.36608585 | 0.114 | 0.318 | 1 | 16 |
| prkab1b | 0.09406 | -0.38246997 | 0.286 | 0.545 | 1 | 16 |
| socs3b | 0.09415 | -0.16073515 | 0.114 | 0.273 | 1 | 16 |
| hnrnpr | 0.09446 | 0.343845499 | 0.229 | 0.045 | 1 | 16 |
| gdi1 | 0.09544 | -0.33977786 | 0.2 | 0.364 | 1 | 16 |
| mfap2 | 0.09721 | -0.23329955 | 0.086 | 0.273 | 1 | 16 |
| eif4a1a | 0.09756 | -0.3350298 | 0.571 | 0.818 | 1 | 16 |
| adam17a | 0.09789 | -0.3473199 | 0.029 | 0.182 | 1 | 16 |
| rpa1 | 0.09822 | -0.31262715 | 0.114 | 0.318 | 1 | 16 |
| rpn1 | 0.09915 | -0.34092502 | 0.514 | 0.727 | 1 | 16 |
| hbp1 | 0.09941 | 0.432824984 | 0.314 | 0.091 | 1 | 16 |
| nol12 | 0.09967 | -0.38750671 | 0.086 | 0.273 | 1 | 16 |
| cbx7a | 0.09969 | -0.12390693 | 0.086 | 0.091 | 1 | 16 |
| capza1a | 0.0997 | 0.217242234 | 0.486 | 0.455 | 1 | 16 |
| gtf2f1 | 0.10069 | 0.312591372 | 0.314 | 0.091 | 1 | 16 |
| snrpe | 0.10088 | -0.24117436 | 0.457 | 0.727 | 1 | 16 |
| zc3h7ba | 0.1014 | -0.14734455 | 0.086 | 0.227 | 1 | 16 |
| jam2a | 0.10251 | 0.202005758 | 1 | 0.909 | 1 | 16 |
| nsd3 | 0.10309 | -0.37244215 | 0.029 | 0.182 | 1 | 16 |
| tssc4 | 0.10351 | 0.160021926 | 0.229 | 0.091 | 1 | 16 |
| cdkal1 | 0.10378 | -0.31403907 | 0.029 | 0.182 | 1 | 16 |
| ctsh | 0.10391 | -0.32560566 | 0.257 | 0.409 | 1 | 16 |
| zdhhc3a | 0.10416 | -0.22280872 | 0.029 | 0.182 | 1 | 16 |
| rcc1 | 0.10435 | -0.30422704 | 0.029 | 0.182 | 1 | 16 |
| hmgn7 | 0.10441 | 0.146430264 | 0.714 | 0.727 | 1 | 16 |
| nfe2l2a | 0.10462 | -0.40288704 | 0.114 | 0.318 | 1 | 16 |
| pin4 | 0.10466 | 0.400000878 | 0.371 | 0.136 | 1 | 16 |
| pbx1b | 0.10478 | -0.29614303 | 0.086 | 0.273 | 1 | 16 |

| | | | | | | |
|---|---|---|---|---|---|---|
| manbal | 0.1048 | 0.366594258 | 0.286 | 0.091 | 1 | 16 |
| clint1a | 0.10512 | 0.109998545 | 0.143 | 0.045 | 1 | 16 |
| rhoq | 0.10538 | 0.376125738 | 0.286 | 0.091 | 1 | 16 |
| erlin1 | 0.10546 | 0.387339924 | 0.286 | 0.091 | 1 | 16 |
| mcm7 | 0.10573 | -0.23836427 | 0.029 | 0.182 | 1 | 16 |
| trim35-14 | 0.10608 | -0.35183226 | 0.114 | 0.273 | 1 | 16 |
| rdh10a | 0.1064 | 0.294789689 | 0.8 | 0.545 | 1 | 16 |
| nanos1 | 0.10641 | -0.17634742 | 0.514 | 0.5 | 1 | 16 |
| g3bp1 | 0.1068 | -0.28968958 | 0.171 | 0.409 | 1 | 16 |
| mycbp | 0.10749 | 0.20739064 | 0.457 | 0.455 | 1 | 16 |
| snx3 | 0.10779 | 0.31913929 | 0.314 | 0.091 | 1 | 16 |
| mta3 | 0.10802 | -0.36010905 | 0.086 | 0.273 | 1 | 16 |
| psme1 | 0.10866 | 0.234391853 | 0.629 | 0.545 | 1 | 16 |
| kcng4a | 0.10893 | -0.27878542 | 0.029 | 0.182 | 1 | 16 |
| si:ch211-132p1.3 | 0.10919 | -0.43680715 | 0.314 | 0.545 | 1 | 16 |
| grinab | 0.10945 | -0.33280884 | 0.4 | 0.682 | 1 | 16 |
| phgdh | 0.11034 | -0.28448616 | 0.114 | 0.227 | 1 | 16 |
| axin2 | 0.11048 | -0.32222109 | 0.029 | 0.182 | 1 | 16 |
| echs1 | 0.11057 | 0.198811146 | 0.429 | 0.227 | 1 | 16 |
| dynlrb1 | 0.1112 | 0.397279361 | 0.571 | 0.318 | 1 | 16 |
| lypla2 | 0.11151 | -0.39262562 | 0.171 | 0.409 | 1 | 16 |
| mrps36 | 0.11291 | 0.371354267 | 0.457 | 0.227 | 1 | 16 |
| capn2a | 0.11312 | -0.14765554 | 0.029 | 0.136 | 1 | 16 |
| slc7a5 | 0.11327 | -0.26027139 | 0.029 | 0.182 | 1 | 16 |
| cmbl | 0.11346 | 0.320877966 | 0.229 | 0.045 | 1 | 16 |
| rbm5 | 0.11349 | 0.379063996 | 0.314 | 0.091 | 1 | 16 |
| snd1 | 0.11383 | -0.29484996 | 0.314 | 0.591 | 1 | 16 |
| si:ch211-244b2.4 | 0.1141 | -0.32617236 | 0.057 | 0.227 | 1 | 16 |
| junba | 0.11436 | -0.48561378 | 0.257 | 0.5 | 1 | 16 |
| bzw1b | 0.11477 | -0.13388946 | 0.514 | 0.5 | 1 | 16 |
| txndc15 | 0.11511 | -0.14631174 | 0.4 | 0.409 | 1 | 16 |
| srsf5a | 0.11555 | -0.25288477 | 0.686 | 0.773 | 1 | 16 |
| cpxm1a | 0.11591 | 0.392131317 | 0.314 | 0.091 | 1 | 16 |
| mrps22 | 0.11605 | -0.15074567 | 0.057 | 0.182 | 1 | 16 |
| hvcn1 | 0.11643 | 0.325158137 | 0.229 | 0.045 | 1 | 16 |
| nudt9 | 0.11665 | -0.28931816 | 0.029 | 0.182 | 1 | 16 |
| ncalda | 0.11672 | 0.210403568 | 0.257 | 0.091 | 1 | 16 |
| si:ch73-37h15.2 | 0.11738 | 0.406761121 | 0.286 | 0.091 | 1 | 16 |
| spcs3 | 0.11752 | 0.349647804 | 0.657 | 0.455 | 1 | 16 |
| tktb | 0.11784 | -0.36036057 | 0.2 | 0.455 | 1 | 16 |
| eif4g3b | 0.11822 | -0.24021974 | 0.057 | 0.227 | 1 | 16 |
| mia3 | 0.11946 | -0.25515381 | 0.029 | 0.182 | 1 | 16 |
| sprydr7a | 0.11971 | -0.2772785 | 0.029 | 0.182 | 1 | 16 |
| parp3 | 0.11997 | -0.25127489 | 0.114 | 0.318 | 1 | 16 |
| si:dkey-172h23.2 | 0.12073 | 0.362083082 | 0.286 | 0.136 | 1 | 16 |
| mob3c | 0.12102 | 0.319684812 | 0.229 | 0.045 | 1 | 16 |
| plxna1a | 0.12109 | -0.27932425 | 0.086 | 0.182 | 1 | 16 |
| dsc2l | 0.12115 | 0.382219431 | 0.229 | 0.045 | 1 | 16 |

| | | | | | | |
|---|---|---|---|---|---|---|
| AL954696.1 | 0.12167 | 0.134803142 | 0.086 | 0 | 1 | 16 |
| CU076099.1 | 0.12167 | 0.159254974 | 0.086 | 0 | 1 | 16 |
| LOC101883779 | 0.12167 | 0.151167189 | 0.086 | 0 | 1 | 16 |
| LOC101884106 | 0.12167 | 0.187297915 | 0.086 | 0 | 1 | 16 |
| LOC101886015 | 0.12167 | 0.141671354 | 0.086 | 0 | 1 | 16 |
| LOC108190726 | 0.12167 | 0.181221218 | 0.086 | 0 | 1 | 16 |
| XLOC-029426 | 0.12167 | 0.14011903 | 0.086 | 0 | 1 | 16 |
| akr7a3 | 0.12167 | 0.183218812 | 0.086 | 0 | 1 | 16 |
| arhgap33 | 0.12167 | 0.17025062 | 0.086 | 0 | 1 | 16 |
| ash2l | 0.12167 | 0.149371728 | 0.086 | 0 | 1 | 16 |
| atrnl1b | 0.12167 | 0.184680022 | 0.086 | 0 | 1 | 16 |
| cep131 | 0.12167 | 0.146026032 | 0.086 | 0 | 1 | 16 |
| coq7 | 0.12167 | 0.159746958 | 0.086 | 0 | 1 | 16 |
| crata | 0.12167 | 0.162251965 | 0.086 | 0 | 1 | 16 |
| csgalnact2 | 0.12167 | 0.152659103 | 0.086 | 0 | 1 | 16 |
| esama | 0.12167 | 0.145322684 | 0.086 | 0 | 1 | 16 |
| josd1 | 0.12167 | 0.143507593 | 0.086 | 0 | 1 | 16 |
| kcnq1 | 0.12167 | 0.171107979 | 0.086 | 0 | 1 | 16 |
| map2k5 | 0.12167 | 0.149970904 | 0.086 | 0 | 1 | 16 |
| nfixb | 0.12167 | 0.152911899 | 0.086 | 0 | 1 | 16 |
| pargl | 0.12167 | 0.159162912 | 0.086 | 0 | 1 | 16 |
| pgm2 | 0.12167 | 0.176108518 | 0.086 | 0 | 1 | 16 |
| phykpl | 0.12167 | 0.155210161 | 0.086 | 0 | 1 | 16 |
| pi4k2a | 0.12167 | 0.147507846 | 0.086 | 0 | 1 | 16 |
| prpf6 | 0.12167 | 0.155960923 | 0.086 | 0 | 1 | 16 |
| ptp4a3 | 0.12167 | 0.136882168 | 0.086 | 0 | 1 | 16 |
| raf1a | 0.12167 | 0.188275026 | 0.086 | 0 | 1 | 16 |
| si:ch211-133l5.8 | 0.12167 | 0.139250554 | 0.086 | 0 | 1 | 16 |
| si:ch73-347e22.4 | 0.12167 | 0.185513354 | 0.086 | 0 | 1 | 16 |
| slc22a15 | 0.12167 | 0.20415059 | 0.086 | 0 | 1 | 16 |
| slc25a43 | 0.12167 | 0.156492685 | 0.086 | 0 | 1 | 16 |
| suox | 0.12167 | 0.139855228 | 0.086 | 0 | 1 | 16 |
| tspan11 | 0.12167 | 0.170039661 | 0.086 | 0 | 1 | 16 |
| ubald2 | 0.12167 | 0.135948179 | 0.086 | 0 | 1 | 16 |
| washc5 | 0.12167 | 0.16524941 | 0.086 | 0 | 1 | 16 |
| zbtb33 | 0.12167 | 0.145872896 | 0.086 | 0 | 1 | 16 |
| zgc:110843 | 0.12167 | 0.133458706 | 0.086 | 0 | 1 | 16 |
| zgc:165481 | 0.12167 | 0.174303076 | 0.086 | 0 | 1 | 16 |
| zgc:172139 | 0.12167 | 0.16141634 | 0.086 | 0 | 1 | 16 |
| zmp:0000000652 | 0.12167 | 0.143078248 | 0.086 | 0 | 1 | 16 |
| AL935174.5 | 0.12167 | 0.178402072 | 0.086 | 0 | 1 | 16 |
| CT956002.2 | 0.12167 | 0.163977134 | 0.086 | 0 | 1 | 16 |
| CU694532.1 | 0.12167 | 0.181102391 | 0.086 | 0 | 1 | 16 |
| RASSF5 | 0.12167 | 0.152704085 | 0.086 | 0 | 1 | 16 |
| adcy2b | 0.12167 | 0.188705302 | 0.086 | 0 | 1 | 16 |
| adm2a | 0.12167 | 0.159322828 | 0.086 | 0 | 1 | 16 |
| agk | 0.12167 | 0.135579015 | 0.086 | 0 | 1 | 16 |
| btbd10a | 0.12167 | 0.149537451 | 0.086 | 0 | 1 | 16 |

| | | | | | | |
|---|---|---|---|---|---|---|
| camsap2b | 0.12167 | 0.14970412 | 0.086 | 0 | 1 | 16 |
| cks1b | 0.12167 | 0.108848618 | 0.086 | 0 | 1 | 16 |
| crispld1b | 0.12167 | 0.182369512 | 0.086 | 0 | 1 | 16 |
| dcaf5 | 0.12167 | 0.147908333 | 0.086 | 0 | 1 | 16 |
| elmsan1b | 0.12167 | 0.149420583 | 0.086 | 0 | 1 | 16 |
| fam219ab | 0.12167 | 0.155616029 | 0.086 | 0 | 1 | 16 |
| flvcr1 | 0.12167 | 0.181221409 | 0.086 | 0 | 1 | 16 |
| homezb | 0.12167 | 0.173919639 | 0.086 | 0 | 1 | 16 |
| immp1l | 0.12167 | 0.135299392 | 0.086 | 0 | 1 | 16 |
| ino80e | 0.12167 | 0.158834573 | 0.086 | 0 | 1 | 16 |
| nat15 | 0.12167 | 0.167549295 | 0.086 | 0 | 1 | 16 |
| nkd3 | 0.12167 | 0.14896938 | 0.086 | 0 | 1 | 16 |
| nxpe3 | 0.12167 | 0.183607227 | 0.086 | 0 | 1 | 16 |
| paf1 | 0.12167 | 0.140261989 | 0.086 | 0 | 1 | 16 |
| pard6gb | 0.12167 | 0.156000918 | 0.086 | 0 | 1 | 16 |
| picalmb | 0.12167 | 0.176568645 | 0.086 | 0 | 1 | 16 |
| pigc | 0.12167 | 0.156989988 | 0.086 | 0 | 1 | 16 |
| pnkd | 0.12167 | 0.204355884 | 0.086 | 0 | 1 | 16 |
| scap | 0.12167 | 0.152113665 | 0.086 | 0 | 1 | 16 |
| sdr16c5a | 0.12167 | 0.155218804 | 0.086 | 0 | 1 | 16 |
| si:ch211-269c21.2 | 0.12167 | 0.148739427 | 0.086 | 0 | 1 | 16 |
| si:dkey-262g12.10 | 0.12167 | 0.165000632 | 0.086 | 0 | 1 | 16 |
| stard4 | 0.12167 | 0.142133042 | 0.086 | 0 | 1 | 16 |
| tmem141 | 0.12167 | 0.17905922 | 0.086 | 0 | 1 | 16 |
| tmem56a | 0.12167 | 0.132323178 | 0.086 | 0 | 1 | 16 |
| vcam1b | 0.12167 | 0.200864287 | 0.086 | 0 | 1 | 16 |
| xpr1b | 0.12167 | 0.184501151 | 0.086 | 0 | 1 | 16 |
| CABZ01072952.1 | 0.12167 | 0.148715698 | 0.086 | 0 | 1 | 16 |
| CABZ01088274.1 | 0.12167 | 0.124370425 | 0.086 | 0 | 1 | 16 |
| XLOC-008119 | 0.12167 | 0.157880721 | 0.086 | 0 | 1 | 16 |
| XLOC-009787 | 0.12167 | 0.170706327 | 0.086 | 0 | 1 | 16 |
| XLOC-042228 | 0.12167 | 0.183306909 | 0.086 | 0 | 1 | 16 |
| aco1 | 0.12167 | 0.1568093 | 0.086 | 0 | 1 | 16 |
| alg8 | 0.12167 | 0.153781037 | 0.086 | 0 | 1 | 16 |
| asap1b | 0.12167 | 0.16402071 | 0.086 | 0 | 1 | 16 |
| cbx6a | 0.12167 | 0.15662287 | 0.086 | 0 | 1 | 16 |
| ccdc127b | 0.12167 | 0.150145921 | 0.086 | 0 | 1 | 16 |
| cnot4a | 0.12167 | 0.134194474 | 0.086 | 0 | 1 | 16 |
| coq8a | 0.12167 | 0.140292947 | 0.086 | 0 | 1 | 16 |
| dot1l | 0.12167 | 0.193925662 | 0.086 | 0 | 1 | 16 |
| dyrk1b | 0.12167 | 0.149048502 | 0.086 | 0 | 1 | 16 |
| ednrab | 0.12167 | 0.147205391 | 0.086 | 0 | 1 | 16 |
| elp5 | 0.12167 | 0.14810998 | 0.086 | 0 | 1 | 16 |
| fhdc3 | 0.12167 | 0.149338735 | 0.086 | 0 | 1 | 16 |
| gatd1 | 0.12167 | 0.134590986 | 0.086 | 0 | 1 | 16 |
| golga4 | 0.12167 | 0.178652233 | 0.086 | 0 | 1 | 16 |
| igf2r | 0.12167 | 0.16837502 | 0.086 | 0 | 1 | 16 |
| itga5 | 0.12167 | 0.171662945 | 0.086 | 0 | 1 | 16 |

| | | | | | | |
|---|---|---|---|---|---|---|
| mindy2 | 0.12167 | 0.206761328 | 0.086 | 0 | 1 | 16 |
| ncoa2 | 0.12167 | 0.154583619 | 0.086 | 0 | 1 | 16 |
| nf1b | 0.12167 | 0.193647097 | 0.086 | 0 | 1 | 16 |
| nipal3 | 0.12167 | 0.143371984 | 0.086 | 0 | 1 | 16 |
| npepps | 0.12167 | 0.175629009 | 0.086 | 0 | 1 | 16 |
| nsun3 | 0.12167 | 0.133802133 | 0.086 | 0 | 1 | 16 |
| nxn | 0.12167 | 0.17300191 | 0.086 | 0 | 1 | 16 |
| oprd1a | 0.12167 | 0.159666537 | 0.086 | 0 | 1 | 16 |
| rdh12l | 0.12167 | 0.182744425 | 0.086 | 0 | 1 | 16 |
| rgp1 | 0.12167 | 0.135211729 | 0.086 | 0 | 1 | 16 |
| rnf168 | 0.12167 | 0.132343665 | 0.086 | 0 | 1 | 16 |
| rxfp3.2a | 0.12167 | 0.241755664 | 0.086 | 0 | 1 | 16 |
| sap130a | 0.12167 | 0.208877394 | 0.086 | 0 | 1 | 16 |
| scml2 | 0.12167 | 0.146672304 | 0.086 | 0 | 1 | 16 |
| si:ch211-13c6.2 | 0.12167 | 0.185628861 | 0.086 | 0 | 1 | 16 |
| si:ch211-140b10.6 | 0.12167 | 0.150724815 | 0.086 | 0 | 1 | 16 |
| si:ch211-161m3.4 | 0.12167 | 0.171349044 | 0.086 | 0 | 1 | 16 |
| si:ch211-169p10.1 | 0.12167 | 0.174955418 | 0.086 | 0 | 1 | 16 |
| si:dkeyp-110g5.4 | 0.12167 | 0.142930711 | 0.086 | 0 | 1 | 16 |
| slc4a11 | 0.12167 | 0.238735026 | 0.086 | 0 | 1 | 16 |
| spryd3 | 0.12167 | 0.123041164 | 0.086 | 0 | 1 | 16 |
| tbc1d22b | 0.12167 | 0.13674384 | 0.086 | 0 | 1 | 16 |
| tom1l2a | 0.12167 | 0.157079074 | 0.086 | 0 | 1 | 16 |
| trmt61a | 0.12167 | 0.135095708 | 0.086 | 0 | 1 | 16 |
| vgll4a | 0.12167 | 0.162139794 | 0.086 | 0 | 1 | 16 |
| ypel3 | 0.12167 | 0.148762741 | 0.086 | 0 | 1 | 16 |
| zgc:112496 | 0.12167 | 0.151709657 | 0.086 | 0 | 1 | 16 |
| zgc:174890 | 0.12167 | 0.122826551 | 0.086 | 0 | 1 | 16 |
| znf993 | 0.12167 | 0.140137329 | 0.086 | 0 | 1 | 16 |
| HOOK3 | 0.12167 | 0.15531173 | 0.086 | 0 | 1 | 16 |
| ablim1b | 0.12167 | 0.164651995 | 0.086 | 0 | 1 | 16 |
| ahcy | 0.12167 | 0.129612502 | 0.086 | 0 | 1 | 16 |
| aven | 0.12167 | 0.13025555 | 0.086 | 0 | 1 | 16 |
| bcar1 | 0.12167 | 0.187169863 | 0.086 | 0 | 1 | 16 |
| cbx3b | 0.12167 | 0.126128085 | 0.086 | 0 | 1 | 16 |
| col5a1 | 0.12167 | 0.180044378 | 0.086 | 0 | 1 | 16 |
| crebrf | 0.12167 | 0.173062862 | 0.086 | 0 | 1 | 16 |
| dia1a | 0.12167 | 0.17375414 | 0.086 | 0 | 1 | 16 |
| dusp19b | 0.12167 | 0.194429821 | 0.086 | 0 | 1 | 16 |
| evi5b | 0.12167 | 0.14794524 | 0.086 | 0 | 1 | 16 |
| fryb | 0.12167 | 0.168358418 | 0.086 | 0 | 1 | 16 |
| ing5a | 0.12167 | 0.151722489 | 0.086 | 0 | 1 | 16 |
| inpp5b | 0.12167 | 0.206411376 | 0.086 | 0 | 1 | 16 |
| jak2b | 0.12167 | 0.163229338 | 0.086 | 0 | 1 | 16 |
| npl | 0.12167 | 0.127985634 | 0.086 | 0 | 1 | 16 |
| nr1d2a | 0.12167 | 0.147715786 | 0.086 | 0 | 1 | 16 |
| nrde2 | 0.12167 | 0.137857192 | 0.086 | 0 | 1 | 16 |
| oc90 | 0.12167 | 0.15773763 | 0.086 | 0 | 1 | 16 |

| | | | | | | |
|---|---|---|---|---|---|---|
| phf23b | 0.12167 | 0.168411388 | 0.086 | 0 | 1 | 16 |
| rpp25l | 0.12167 | 0.16463544 | 0.086 | 0 | 1 | 16 |
| shmt2 | 0.12167 | 0.164432985 | 0.086 | 0 | 1 | 16 |
| si:ch211-166a6.5 | 0.12167 | 0.153709145 | 0.086 | 0 | 1 | 16 |
| si:ch211-239j9.1 | 0.12167 | 0.17162243 | 0.086 | 0 | 1 | 16 |
| snapc4 | 0.12167 | 0.206307306 | 0.086 | 0 | 1 | 16 |
| sntb2 | 0.12167 | 0.199984465 | 0.086 | 0 | 1 | 16 |
| zer1 | 0.12167 | 0.199294875 | 0.086 | 0 | 1 | 16 |
| znf1015 | 0.12167 | 0.129870139 | 0.086 | 0 | 1 | 16 |
| adal | 0.12167 | 0.146786127 | 0.086 | 0 | 1 | 16 |
| adhfe1 | 0.12167 | 0.157115315 | 0.086 | 0 | 1 | 16 |
| agmo | 0.12167 | 0.14399081 | 0.086 | 0 | 1 | 16 |
| apaf1 | 0.12167 | 0.219537361 | 0.086 | 0 | 1 | 16 |
| atp6ap1lb | 0.12167 | 0.161309675 | 0.086 | 0 | 1 | 16 |
| bmpr2b | 0.12167 | 0.141945993 | 0.086 | 0 | 1 | 16 |
| cc2d1b | 0.12167 | 0.146852084 | 0.086 | 0 | 1 | 16 |
| ccdc160 | 0.12167 | 0.135130267 | 0.086 | 0 | 1 | 16 |
| cdcp1a | 0.12167 | 0.18404497 | 0.086 | 0 | 1 | 16 |
| chordc1b | 0.12167 | 0.14747796 | 0.086 | 0 | 1 | 16 |
| clasp2 | 0.12167 | 0.147444402 | 0.086 | 0 | 1 | 16 |
| clip2 | 0.12167 | 0.137561176 | 0.086 | 0 | 1 | 16 |
| cog2 | 0.12167 | 0.148058294 | 0.086 | 0 | 1 | 16 |
| cttnbp2 | 0.12167 | 0.139105064 | 0.086 | 0 | 1 | 16 |
| dstyk | 0.12167 | 0.160127701 | 0.086 | 0 | 1 | 16 |
| elp1 | 0.12167 | 0.157823942 | 0.086 | 0 | 1 | 16 |
| ext1c | 0.12167 | 0.133625867 | 0.086 | 0 | 1 | 16 |
| fshr | 0.12167 | 0.173284449 | 0.086 | 0 | 1 | 16 |
| glt8d1 | 0.12167 | 0.143109202 | 0.086 | 0 | 1 | 16 |
| igsf9a | 0.12167 | 0.216721577 | 0.086 | 0 | 1 | 16 |
| nup210 | 0.12167 | 0.146345496 | 0.086 | 0 | 1 | 16 |
| otud3 | 0.12167 | 0.124156437 | 0.086 | 0 | 1 | 16 |
| phf20l1 | 0.12167 | 0.191071561 | 0.086 | 0 | 1 | 16 |
| pnp5b | 0.12167 | 0.154059694 | 0.086 | 0 | 1 | 16 |
| ppp1r3b | 0.12167 | 0.108578901 | 0.086 | 0 | 1 | 16 |
| preb | 0.12167 | 0.159160465 | 0.086 | 0 | 1 | 16 |
| pts | 0.12167 | 0.167024992 | 0.086 | 0 | 1 | 16 |
| rbfox2 | 0.12167 | 0.154713339 | 0.086 | 0 | 1 | 16 |
| si:ch211-127m7.2 | 0.12167 | 0.16662029 | 0.086 | 0 | 1 | 16 |
| slc38a6 | 0.12167 | 0.171690888 | 0.086 | 0 | 1 | 16 |
| snx17 | 0.12167 | 0.193534564 | 0.086 | 0 | 1 | 16 |
| spa17 | 0.12167 | 0.148805134 | 0.086 | 0 | 1 | 16 |
| tdp1 | 0.12167 | 0.182075798 | 0.086 | 0 | 1 | 16 |
| thbs1a | 0.12167 | 0.223724226 | 0.086 | 0 | 1 | 16 |
| wnk1a | 0.12167 | 0.154530794 | 0.086 | 0 | 1 | 16 |
| zgc:101577 | 0.12167 | 0.190608152 | 0.086 | 0 | 1 | 16 |
| zgc:153031 | 0.12167 | 0.183570847 | 0.086 | 0 | 1 | 16 |
| znf1059 | 0.12167 | 0.153786401 | 0.086 | 0 | 1 | 16 |
| znhit1 | 0.12167 | 0.156427328 | 0.086 | 0 | 1 | 16 |

| | | | | | | |
|---|---|---|---|---|---|---|
| bach1b | 0.12167 | 0.13229601 | 0.086 | 0 | 1 | 16 |
| mex3b | 0.12167 | 0.133163783 | 0.086 | 0 | 1 | 16 |
| rpap1 | 0.12167 | 0.145163039 | 0.086 | 0 | 1 | 16 |
| LOC100149700 | 0.12167 | 0.178165734 | 0.086 | 0 | 1 | 16 |
| fan1 | 0.12167 | 0.139917542 | 0.086 | 0 | 1 | 16 |
| fitm2 | 0.12167 | 0.155257892 | 0.086 | 0 | 1 | 16 |
| nxf1 | 0.12167 | 0.172686991 | 0.086 | 0 | 1 | 16 |
| pkdcca | 0.12167 | 0.153655937 | 0.086 | 0 | 1 | 16 |
| spsb4b | 0.12167 | 0.152323965 | 0.086 | 0 | 1 | 16 |
| ttll7 | 0.12167 | 0.138936421 | 0.086 | 0 | 1 | 16 |
| zc3h10 | 0.12167 | 0.14405827 | 0.086 | 0 | 1 | 16 |
| FBXO48 | 0.12167 | 0.122825337 | 0.086 | 0 | 1 | 16 |
| acot15 | 0.12167 | 0.138747667 | 0.086 | 0 | 1 | 16 |
| bada | 0.12167 | 0.137321656 | 0.086 | 0 | 1 | 16 |
| mettl14 | 0.12167 | 0.181273281 | 0.086 | 0 | 1 | 16 |
| pmpca | 0.12167 | 0.171257865 | 0.086 | 0 | 1 | 16 |
| polr3d | 0.12167 | 0.139290152 | 0.086 | 0 | 1 | 16 |
| sirt3 | 0.12167 | 0.156026639 | 0.086 | 0 | 1 | 16 |
| twf1b | 0.12167 | 0.1475004 | 0.086 | 0 | 1 | 16 |
| xrcc5 | 0.12167 | 0.149067665 | 0.086 | 0 | 1 | 16 |
| zbtb11 | 0.12167 | 0.174531299 | 0.086 | 0 | 1 | 16 |
| znf1159 | 0.12167 | 0.136189183 | 0.086 | 0 | 1 | 16 |
| ADGRL3 | 0.12167 | 0.169161906 | 0.086 | 0 | 1 | 16 |
| BX323797.4 | 0.12167 | 0.145568831 | 0.086 | 0 | 1 | 16 |
| CABZ01008730.1 | 0.12167 | 0.161681723 | 0.086 | 0 | 1 | 16 |
| aptx | 0.12167 | 0.14981299 | 0.086 | 0 | 1 | 16 |
| c19h1orf109 | 0.12167 | 0.156538164 | 0.086 | 0 | 1 | 16 |
| ccdc115 | 0.12167 | 0.154888571 | 0.086 | 0 | 1 | 16 |
| ckap5 | 0.12167 | 0.202096938 | 0.086 | 0 | 1 | 16 |
| foxn2b | 0.12167 | 0.203800512 | 0.086 | 0 | 1 | 16 |
| g6pd | 0.12167 | 0.168589446 | 0.086 | 0 | 1 | 16 |
| gde1 | 0.12167 | 0.158349975 | 0.086 | 0 | 1 | 16 |
| gna11a | 0.12167 | 0.129179697 | 0.086 | 0 | 1 | 16 |
| hnrnph3 | 0.12167 | 0.190725294 | 0.086 | 0 | 1 | 16 |
| ice2 | 0.12167 | 0.176719869 | 0.086 | 0 | 1 | 16 |
| kmt2ca | 0.12167 | 0.145379458 | 0.086 | 0 | 1 | 16 |
| mef2aa | 0.12167 | 0.185359878 | 0.086 | 0 | 1 | 16 |
| mir22a | 0.12167 | 0.174368957 | 0.086 | 0 | 1 | 16 |
| mrpl15 | 0.12167 | 0.141639871 | 0.086 | 0 | 1 | 16 |
| ndufaf2 | 0.12167 | 0.136223573 | 0.086 | 0 | 1 | 16 |
| nts | 0.12167 | 0.270341773 | 0.086 | 0 | 1 | 16 |
| nudt18 | 0.12167 | 0.165777804 | 0.086 | 0 | 1 | 16 |
| pelp1 | 0.12167 | 0.144643776 | 0.086 | 0 | 1 | 16 |
| pisd | 0.12167 | 0.146042141 | 0.086 | 0 | 1 | 16 |
| raph1b | 0.12167 | 0.160024242 | 0.086 | 0 | 1 | 16 |
| scarb1 | 0.12167 | 0.154255082 | 0.086 | 0 | 1 | 16 |
| sfxn4 | 0.12167 | 0.15907469 | 0.086 | 0 | 1 | 16 |
| si:ch211-76m11.3 | 0.12167 | 0.13761657 | 0.086 | 0 | 1 | 16 |

| | | | | | | |
|---|---|---|---|---|---|---|
| si:ch73-340n8.4 | 0.12167 | 0.188880068 | 0.086 | 0 | 1 | 16 |
| si:dkey-184p9.7 | 0.12167 | 0.169369459 | 0.086 | 0 | 1 | 16 |
| si:rp71-19m20.1 | 0.12167 | 0.138748623 | 0.086 | 0 | 1 | 16 |
| slc2a12 | 0.12167 | 0.154701671 | 0.086 | 0 | 1 | 16 |
| slc34a2b | 0.12167 | 0.160647712 | 0.086 | 0 | 1 | 16 |
| slc7a10a | 0.12167 | 0.172367905 | 0.086 | 0 | 1 | 16 |
| snx19b | 0.12167 | 0.188327546 | 0.086 | 0 | 1 | 16 |
| srp14 | 0.12167 | 0.142384811 | 0.086 | 0 | 1 | 16 |
| suco | 0.12167 | 0.194008528 | 0.086 | 0 | 1 | 16 |
| syngr3a | 0.12167 | 0.17141632 | 0.086 | 0 | 1 | 16 |
| tcea3 | 0.12167 | 0.150398612 | 0.086 | 0 | 1 | 16 |
| timd4 | 0.12167 | 0.159939906 | 0.086 | 0 | 1 | 16 |
| vps36 | 0.12167 | 0.178540478 | 0.086 | 0 | 1 | 16 |
| wdsub1 | 0.12167 | 0.150645098 | 0.086 | 0 | 1 | 16 |
| znf1152 | 0.12167 | 0.175312696 | 0.086 | 0 | 1 | 16 |
| LO018197.1 | 0.12167 | 0.160066693 | 0.086 | 0 | 1 | 16 |
| rnf14 | 0.12167 | 0.158507719 | 0.086 | 0 | 1 | 16 |
| si:dkey-100n23.3 | 0.12167 | 0.134705055 | 0.086 | 0 | 1 | 16 |
| tmem206 | 0.12167 | 0.146710763 | 0.086 | 0 | 1 | 16 |
| zdhhc23b | 0.12167 | 0.133251623 | 0.086 | 0 | 1 | 16 |
| bcas2 | 0.1223 | 0.361167945 | 0.314 | 0.091 | 1 | 16 |
| ap1b1 | 0.12299 | -0.16211657 | 0.057 | 0.182 | 1 | 16 |
| crema | 0.1234 | 0.211481912 | 0.514 | 0.455 | 1 | 16 |
| tmem243b | 0.12394 | -0.25163976 | 0.057 | 0.227 | 1 | 16 |
| pmaip1 | 0.12422 | -0.30456291 | 0.029 | 0.182 | 1 | 16 |
| gli3 | 0.1249 | 0.373062317 | 0.2 | 0.045 | 1 | 16 |
| npm3 | 0.12521 | -0.34114587 | 0.257 | 0.409 | 1 | 16 |
| sdf2l1 | 0.12521 | -0.34479648 | 0.314 | 0.545 | 1 | 16 |
| clic1 | 0.12537 | 0.346985097 | 0.429 | 0.182 | 1 | 16 |
| bicd2 | 0.12543 | -0.24034293 | 0.029 | 0.182 | 1 | 16 |
| usp4 | 0.12558 | 0.270311753 | 0.2 | 0.091 | 1 | 16 |
| ptgr1 | 0.1257 | 0.411866748 | 0.457 | 0.227 | 1 | 16 |
| tsn | 0.12602 | -0.28802076 | 0.029 | 0.182 | 1 | 16 |
| rpf1 | 0.1265 | 0.357631809 | 0.229 | 0.045 | 1 | 16 |
| hsd3b7 | 0.12672 | 0.342168742 | 0.829 | 0.591 | 1 | 16 |
| psme3 | 0.12691 | 0.310677642 | 0.257 | 0.091 | 1 | 16 |
| golt1bb | 0.12691 | -0.35434705 | 0.2 | 0.455 | 1 | 16 |
| appb | 0.12705 | -0.2232266 | 0.286 | 0.5 | 1 | 16 |
| nol4lb | 0.1272 | 0.334177413 | 0.229 | 0.045 | 1 | 16 |
| rmdn1 | 0.12798 | -0.33718509 | 0.2 | 0.455 | 1 | 16 |
| fkbp9 | 0.12841 | -0.32654523 | 0.086 | 0.273 | 1 | 16 |
| setd5 | 0.12882 | -0.24590992 | 0.057 | 0.227 | 1 | 16 |
| caprin1a | 0.12903 | -0.31441868 | 0.086 | 0.182 | 1 | 16 |
| akap12b | 0.12916 | 0.416291483 | 0.371 | 0.136 | 1 | 16 |
| cyp17a1 | 0.13004 | 0.32473655 | 0.429 | 0.182 | 1 | 16 |
| iscub | 0.13044 | 0.33556697 | 0.486 | 0.273 | 1 | 16 |
| mycb | 0.13055 | -0.40481012 | 0.143 | 0.364 | 1 | 16 |
| cdkn1bb | 0.13087 | 0.363539026 | 0.343 | 0.136 | 1 | 16 |

| | | | | | | |
|---|---|---|---|---|---|---|
| m6pr | 0.13174 | -0.24189431 | 0.029 | 0.182 | 1 | 16 |
| mrpl38 | 0.13196 | -0.20496683 | 0.171 | 0.364 | 1 | 16 |
| sptbn2 | 0.13201 | -0.35782207 | 0.2 | 0.409 | 1 | 16 |
| rrbp1a | 0.13237 | -0.36018359 | 0.257 | 0.455 | 1 | 16 |
| yy1b | 0.13272 | -0.17970624 | 0.029 | 0.136 | 1 | 16 |
| rassf1 | 0.13356 | -0.27807377 | 0.029 | 0.182 | 1 | 16 |
| srpx | 0.13357 | -0.1630205 | 0.029 | 0.136 | 1 | 16 |
| spred2b | 0.13374 | -0.32722719 | 0.571 | 0.773 | 1 | 16 |
| hspa4a | 0.13419 | -0.32925522 | 0.057 | 0.227 | 1 | 16 |
| anxa11b | 0.13472 | -0.35357292 | 0.371 | 0.636 | 1 | 16 |
| si:ch211-197h24.6 | 0.13492 | -0.28030683 | 0.114 | 0.318 | 1 | 16 |
| cachd1 | 0.13526 | -0.17245547 | 0.029 | 0.136 | 1 | 16 |
| cpne1 | 0.1357 | 0.151306842 | 0.429 | 0.409 | 1 | 16 |
| si:ch211-246m6.4 | 0.13594 | -0.37164014 | 0.171 | 0.364 | 1 | 16 |
| ssb | 0.13626 | -0.32311588 | 0.143 | 0.364 | 1 | 16 |
| nek7 | 0.13667 | -0.10352999 | 0.114 | 0.227 | 1 | 16 |
| ctnnbip1 | 0.13699 | 0.293296764 | 0.229 | 0.045 | 1 | 16 |
| zgc:92481 | 0.13792 | -0.25209541 | 0.086 | 0.273 | 1 | 16 |
| med30 | 0.13831 | 0.310084207 | 0.229 | 0.045 | 1 | 16 |
| prdx3 | 0.13833 | 0.352993336 | 0.629 | 0.364 | 1 | 16 |
| CABZ01044053.1 | 0.13833 | 0.304812433 | 0.229 | 0.045 | 1 | 16 |
| XLOC-012680 | 0.13837 | -0.24602107 | 0.029 | 0.182 | 1 | 16 |
| psma6b | 0.1386 | -0.3700768 | 0.171 | 0.409 | 1 | 16 |
| naga | 0.13868 | -0.11899019 | 0.114 | 0.227 | 1 | 16 |
| ndc1 | 0.1391 | -0.26618931 | 0.029 | 0.182 | 1 | 16 |
| med6 | 0.13918 | -0.25860583 | 0.029 | 0.182 | 1 | 16 |
| cmc1 | 0.13938 | 0.319203467 | 0.286 | 0.091 | 1 | 16 |
| asf1bb | 0.13986 | -0.17505921 | 0.114 | 0.273 | 1 | 16 |
| ftr50 | 0.13991 | -0.21452704 | 0.029 | 0.182 | 1 | 16 |
| wnt9b | 0.13997 | -0.2051905 | 0.057 | 0.091 | 1 | 16 |
| slc20a2 | 0.14016 | 0.236120909 | 0.314 | 0.136 | 1 | 16 |
| cst14b.1 | 0.14021 | -0.22257185 | 0.029 | 0.182 | 1 | 16 |
| mlst8 | 0.14087 | 0.155884159 | 0.171 | 0.045 | 1 | 16 |
| rps4x | 0.14133 | 0.13582123 | 0.971 | 0.955 | 1 | 16 |
| rhogd | 0.14141 | -0.25470926 | 0.086 | 0.273 | 1 | 16 |
| CABZ01074307.1 | 0.14157 | -0.29469249 | 0.029 | 0.182 | 1 | 16 |
| ccdc85b | 0.14163 | 0.363242805 | 0.343 | 0.136 | 1 | 16 |
| hmgb1a | 0.14185 | 0.291695315 | 0.886 | 0.727 | 1 | 16 |
| XLOC-004335 | 0.14196 | 0.363479868 | 0.229 | 0.045 | 1 | 16 |
| atxn2l | 0.14201 | -0.27509902 | 0.029 | 0.182 | 1 | 16 |
| calub | 0.14223 | -0.36414485 | 0.143 | 0.318 | 1 | 16 |
| mapkapk2a | 0.14244 | 0.235017836 | 0.257 | 0.091 | 1 | 16 |
| pdcl | 0.14245 | 0.330746738 | 0.229 | 0.045 | 1 | 16 |
| ccni | 0.14267 | 0.360226323 | 0.686 | 0.455 | 1 | 16 |
| kcng1 | 0.14268 | -0.2793143 | 0.029 | 0.182 | 1 | 16 |
| tab1 | 0.14272 | 0.296151367 | 0.229 | 0.045 | 1 | 16 |
| rnf128a | 0.14279 | 0.326142581 | 0.229 | 0.045 | 1 | 16 |
| ap2a1 | 0.14281 | -0.22664454 | 0.171 | 0.364 | 1 | 16 |

| | | | | | | |
|---|---|---|---|---|---|---|
| myo6a | 0.14287 | -0.27236998 | 0.029 | 0.182 | 1 | 16 |
| commd5 | 0.14291 | 0.273452378 | 0.229 | 0.045 | 1 | 16 |
| tspan7 | 0.14308 | -0.25519512 | 0.029 | 0.182 | 1 | 16 |
| nfe2l1b | 0.14351 | -0.24376145 | 0.029 | 0.182 | 1 | 16 |
| psmb2 | 0.14376 | -0.21038439 | 0.4 | 0.455 | 1 | 16 |
| smfn | 0.14381 | -0.28043368 | 0.057 | 0.227 | 1 | 16 |
| siah1 | 0.14432 | -0.24360815 | 0.029 | 0.182 | 1 | 16 |
| letm2 | 0.14439 | -0.25141919 | 0.029 | 0.182 | 1 | 16 |
| sgk1 | 0.14445 | -0.25065397 | 0.029 | 0.182 | 1 | 16 |
| ier5 | 0.1445 | -0.3748104 | 0.143 | 0.364 | 1 | 16 |
| bola3 | 0.14455 | -0.2338814 | 0.029 | 0.182 | 1 | 16 |
| metrnla | 0.14473 | 0.307771591 | 0.571 | 0.455 | 1 | 16 |
| cybc1 | 0.14495 | -0.22467912 | 0.029 | 0.182 | 1 | 16 |
| AK6 | 0.14504 | -0.24017286 | 0.029 | 0.182 | 1 | 16 |
| bcl7bb | 0.1451 | 0.212147812 | 0.314 | 0.136 | 1 | 16 |
| rprd2a | 0.14522 | -0.25173485 | 0.029 | 0.182 | 1 | 16 |
| rnf115 | 0.14525 | -0.23920667 | 0.029 | 0.182 | 1 | 16 |
| etv5b | 0.14554 | -0.23928614 | 0.029 | 0.182 | 1 | 16 |
| stag1b | 0.14557 | -0.26152675 | 0.029 | 0.182 | 1 | 16 |
| chd4b | 0.14562 | -0.32253352 | 0.429 | 0.591 | 1 | 16 |
| ccnd2b | 0.14568 | -0.25797237 | 0.029 | 0.182 | 1 | 16 |
| XLOC-019096 | 0.14575 | -0.27864285 | 0.029 | 0.182 | 1 | 16 |
| cbx4 | 0.14575 | -0.24438594 | 0.029 | 0.182 | 1 | 16 |
| rangap1a | 0.14587 | -0.27319801 | 0.029 | 0.182 | 1 | 16 |
| hgs | 0.14587 | -0.30011146 | 0.114 | 0.318 | 1 | 16 |
| cyb5a | 0.14588 | -0.19452537 | 0.171 | 0.273 | 1 | 16 |
| cdk7 | 0.14592 | -0.24041545 | 0.029 | 0.182 | 1 | 16 |
| phf10 | 0.14593 | -0.24792399 | 0.029 | 0.182 | 1 | 16 |
| irf2a | 0.14625 | -0.15505329 | 0.086 | 0.227 | 1 | 16 |
| cnn3b | 0.14643 | -0.29570471 | 0.086 | 0.273 | 1 | 16 |
| limch1b | 0.14644 | -0.11112364 | 0.143 | 0.273 | 1 | 16 |
| eif4bb | 0.14653 | 0.347360529 | 0.343 | 0.136 | 1 | 16 |
| tmem43 | 0.1468 | -0.27590297 | 0.057 | 0.227 | 1 | 16 |
| kdelr2a | 0.14682 | -0.34445267 | 0.143 | 0.364 | 1 | 16 |
| zgc:112255 | 0.14718 | -0.16307632 | 0.114 | 0.136 | 1 | 16 |
| slc48a1b | 0.14764 | -0.15022904 | 0.057 | 0.182 | 1 | 16 |
| ppp1cab | 0.14772 | -0.30898759 | 0.286 | 0.545 | 1 | 16 |
| mdh2 | 0.14776 | 0.305730393 | 0.714 | 0.455 | 1 | 16 |
| usp10 | 0.14877 | -0.29571654 | 0.114 | 0.318 | 1 | 16 |
| ccdc126 | 0.14953 | -0.25351712 | 0.086 | 0.182 | 1 | 16 |
| arrb2b | 0.14976 | -0.33069987 | 0.086 | 0.273 | 1 | 16 |
| hs2st1b | 0.14982 | -0.16665387 | 0.086 | 0.227 | 1 | 16 |
| pdia6 | 0.14988 | -0.26368777 | 0.543 | 0.636 | 1 | 16 |
| pdcd10a | 0.14993 | -0.21454532 | 0.2 | 0.273 | 1 | 16 |
| pgr | 0.14997 | 0.134057922 | 0.2 | 0.182 | 1 | 16 |
| si:dkey-23i12.5 | 0.15021 | -0.34202904 | 0.143 | 0.364 | 1 | 16 |
| pin1 | 0.15028 | 0.200009953 | 0.571 | 0.5 | 1 | 16 |
| si:dkey-58f10.11 | 0.15053 | -0.34774656 | 0.057 | 0.227 | 1 | 16 |

| | | | | | | |
|---|---|---|---|---|---|---|
| ctbp1 | 0.15115 | 0.348997248 | 0.486 | 0.318 | 1 | 16 |
| si:dkey-42i9.7 | 0.15133 | -0.3941997 | 0.371 | 0.545 | 1 | 16 |
| ddx39ab | 0.15191 | -0.25863329 | 0.4 | 0.545 | 1 | 16 |
| ddx51 | 0.15208 | 0.124277378 | 0.143 | 0.045 | 1 | 16 |
| slc20a1b | 0.1522 | -0.305877 | 0.057 | 0.227 | 1 | 16 |
| cib1 | 0.15238 | -0.2576188 | 0.114 | 0.227 | 1 | 16 |
| tnpo2 | 0.15265 | -0.31820988 | 0.057 | 0.227 | 1 | 16 |
| sde2 | 0.15294 | 0.326307024 | 0.314 | 0.136 | 1 | 16 |
| pdia5 | 0.15299 | 0.289963175 | 0.286 | 0.091 | 1 | 16 |
| errfi1a | 0.15305 | 0.404127361 | 0.343 | 0.136 | 1 | 16 |
| ap1m1 | 0.15367 | -0.35315961 | 0.229 | 0.409 | 1 | 16 |
| rab8b | 0.15369 | -0.28254131 | 0.057 | 0.227 | 1 | 16 |
| apooa | 0.15389 | -0.19075348 | 0.057 | 0.136 | 1 | 16 |
| flna | 0.15406 | -0.3612572 | 0.086 | 0.273 | 1 | 16 |
| phf8 | 0.15448 | -0.26510632 | 0.086 | 0.273 | 1 | 16 |
| mccc2 | 0.15503 | -0.22959094 | 0.057 | 0.227 | 1 | 16 |
| CABZ01086029.1 | 0.15555 | -0.33087144 | 0.229 | 0.409 | 1 | 16 |
| podxl | 0.15582 | 0.415869548 | 0.657 | 0.409 | 1 | 16 |
| si:dkey-250d21.1 | 0.15613 | -0.23174763 | 0.171 | 0.364 | 1 | 16 |
| mrps9 | 0.15619 | -0.26322751 | 0.086 | 0.182 | 1 | 16 |
| abhd16a | 0.15633 | -0.16351714 | 0.029 | 0.136 | 1 | 16 |
| rab6a | 0.15656 | -0.33403962 | 0.171 | 0.364 | 1 | 16 |
| psmg2 | 0.15709 | -0.27875302 | 0.114 | 0.318 | 1 | 16 |
| mrpl33 | 0.15774 | 0.191799637 | 0.257 | 0.091 | 1 | 16 |
| sav1 | 0.15776 | -0.27439065 | 0.086 | 0.273 | 1 | 16 |
| ifi35 | 0.15785 | -0.26732767 | 0.257 | 0.364 | 1 | 16 |
| adam10a | 0.1579 | 0.31508042 | 0.8 | 0.591 | 1 | 16 |
| fkbp2 | 0.15798 | 0.13960576 | 0.629 | 0.636 | 1 | 16 |
| scpep1 | 0.15799 | -0.32442115 | 0.143 | 0.364 | 1 | 16 |
| rhoua | 0.15807 | -0.35065045 | 0.143 | 0.364 | 1 | 16 |
| cirbpa | 0.15835 | -0.18755535 | 0.943 | 0.955 | 1 | 16 |
| sec22bb | 0.15907 | -0.13703074 | 0.029 | 0.136 | 1 | 16 |
| mrpl57 | 0.15921 | 0.113690021 | 0.257 | 0.136 | 1 | 16 |
| gtf2e2 | 0.15977 | -0.30509454 | 0.2 | 0.318 | 1 | 16 |
| nap1l1 | 0.15999 | -0.10668869 | 0.457 | 0.636 | 1 | 16 |
| fas | 0.1601 | -0.32857856 | 0.143 | 0.364 | 1 | 16 |
| tpm2 | 0.16045 | -0.27672115 | 0.057 | 0.227 | 1 | 16 |
| adam19a | 0.16052 | -0.21082351 | 0.143 | 0.318 | 1 | 16 |
| tgfb2l | 0.16079 | -0.21743687 | 0.286 | 0.5 | 1 | 16 |
| birc2 | 0.16119 | -0.34800973 | 0.143 | 0.364 | 1 | 16 |
| ppiab | 0.1615 | 0.158593587 | 1 | 0.909 | 1 | 16 |
| clstn1 | 0.16167 | 0.319928519 | 0.2 | 0.045 | 1 | 16 |
| tmtc2a | 0.16219 | -0.33903534 | 0.086 | 0.273 | 1 | 16 |
| bzw1a | 0.16287 | -0.29209156 | 0.257 | 0.5 | 1 | 16 |
| fhl1b | 0.16296 | 0.348255326 | 0.257 | 0.091 | 1 | 16 |
| elovl1b | 0.16297 | -0.25434448 | 0.229 | 0.455 | 1 | 16 |
| eif3ba | 0.16321 | -0.28850947 | 0.6 | 0.773 | 1 | 16 |
| prdm11 | 0.16346 | -0.28863783 | 0.057 | 0.227 | 1 | 16 |

| | | | | | | |
|---|---|---|---|---|---|---|
| selenol | 0.16383 | -0.12103103 | 0.086 | 0.091 | 1 | 16 |
| si:ch73-335l21.4 | 0.16432 | -0.45010545 | 0.057 | 0.227 | 1 | 16 |
| zgc:55733 | 0.16439 | 0.290101023 | 0.286 | 0.091 | 1 | 16 |
| tmem115 | 0.16478 | 0.281546634 | 0.2 | 0.045 | 1 | 16 |
| plod1a | 0.16506 | -0.27268848 | 0.086 | 0.273 | 1 | 16 |
| arhgef2 | 0.16509 | -0.28597433 | 0.086 | 0.273 | 1 | 16 |
| atp1b1a | 0.16536 | -0.20150474 | 0.8 | 0.864 | 1 | 16 |
| lyrm4 | 0.16562 | -0.1553541 | 0.086 | 0.227 | 1 | 16 |
| znf827 | 0.16573 | -0.27517745 | 0.057 | 0.182 | 1 | 16 |
| matr3l1.1 | 0.1662 | -0.32653831 | 0.257 | 0.455 | 1 | 16 |
| rnf121 | 0.16632 | -0.35291302 | 0.114 | 0.318 | 1 | 16 |
| emc1 | 0.16633 | -0.30980746 | 0.086 | 0.273 | 1 | 16 |
| csde1 | 0.16639 | -0.23580501 | 0.343 | 0.455 | 1 | 16 |
| st6galnac5b | 0.16667 | 0.112927214 | 0.2 | 0.091 | 1 | 16 |
| zgc:162698 | 0.16696 | -0.291316 | 0.057 | 0.227 | 1 | 16 |
| col4a5 | 0.16751 | -0.30999383 | 0.343 | 0.5 | 1 | 16 |
| cdk2ap2 | 0.16928 | 0.325092936 | 0.4 | 0.182 | 1 | 16 |
| kpnb3 | 0.16939 | -0.2819326 | 0.057 | 0.227 | 1 | 16 |
| rp2 | 0.16995 | -0.24865035 | 0.057 | 0.227 | 1 | 16 |
| zfyve27 | 0.17015 | -0.11057169 | 0.171 | 0.182 | 1 | 16 |
| ppp4ca | 0.17021 | -0.26883129 | 0.057 | 0.227 | 1 | 16 |
| ptn | 0.17022 | -0.31495826 | 0.057 | 0.227 | 1 | 16 |
| chd2 | 0.17036 | -0.2932232 | 0.057 | 0.182 | 1 | 16 |
| sap30bp | 0.17077 | -0.31939366 | 0.114 | 0.318 | 1 | 16 |
| lamp1b | 0.17118 | -0.27074301 | 0.057 | 0.227 | 1 | 16 |
| dag1 | 0.17122 | 0.340965855 | 0.514 | 0.318 | 1 | 16 |
| abcf2a | 0.1717 | -0.26777544 | 0.086 | 0.273 | 1 | 16 |
| arrdc3a | 0.1719 | 0.297951895 | 0.486 | 0.409 | 1 | 16 |
| f10 | 0.1719 | 0.23894746 | 0.886 | 0.682 | 1 | 16 |
| stim1a | 0.172 | -0.28770517 | 0.086 | 0.273 | 1 | 16 |
| prkd2 | 0.17218 | 0.343036734 | 0.314 | 0.136 | 1 | 16 |
| tfe3a | 0.17224 | -0.28951582 | 0.057 | 0.227 | 1 | 16 |
| tdp2a | 0.17261 | -0.28829828 | 0.057 | 0.227 | 1 | 16 |
| cox7a1 | 0.1727 | 0.231842734 | 0.4 | 0.182 | 1 | 16 |
| LOC110438024 | 0.17299 | 0.393029881 | 0.6 | 0.364 | 1 | 16 |
| upf1 | 0.17302 | -0.29368195 | 0.086 | 0.227 | 1 | 16 |
| larp1b | 0.17304 | -0.25882777 | 0.057 | 0.227 | 1 | 16 |
| paqr3a | 0.17306 | -0.18901253 | 0.086 | 0.227 | 1 | 16 |
| anapc13 | 0.17306 | -0.29624449 | 0.057 | 0.227 | 1 | 16 |
| gstm.2 | 0.17326 | -0.33433745 | 0.229 | 0.455 | 1 | 16 |
| sdf4 | 0.17327 | -0.28975977 | 0.114 | 0.318 | 1 | 16 |
| cfap36 | 0.17354 | -0.33608713 | 0.114 | 0.318 | 1 | 16 |
| snrpd3l | 0.17386 | -0.1711959 | 0.371 | 0.409 | 1 | 16 |
| mrps34 | 0.17401 | 0.11506008 | 0.2 | 0.091 | 1 | 16 |
| nckap1 | 0.17458 | -0.30229856 | 0.086 | 0.273 | 1 | 16 |
| her6 | 0.17484 | -0.17429848 | 0.086 | 0.227 | 1 | 16 |
| si:dkey-6n6.1 | 0.17489 | 0.346100325 | 0.771 | 0.591 | 1 | 16 |
| tubb2b | 0.17493 | -0.31876858 | 0.343 | 0.591 | 1 | 16 |

| | | | | | | |
|---|---|---|---|---|---|---|
| fuca2 | 0.17595 | -0.29130361 | 0.086 | 0.227 | 1 | 16 |
| coa3a | 0.17623 | -0.35516705 | 0.143 | 0.318 | 1 | 16 |
| tnrc6c1 | 0.17696 | 0.239369958 | 0.571 | 0.455 | 1 | 16 |
| cry2 | 0.17731 | -0.15437073 | 0.057 | 0.182 | 1 | 16 |
| smarcd1 | 0.17781 | 0.23300064 | 0.2 | 0.045 | 1 | 16 |
| atp6v1ab | 0.17789 | -0.32500987 | 0.086 | 0.273 | 1 | 16 |
| ppp2r5eb | 0.17839 | -0.29080944 | 0.086 | 0.273 | 1 | 16 |
| si:ch1073-291c23.2 | 0.17848 | -0.16530457 | 0.029 | 0.136 | 1 | 16 |
| serpinb1l3 | 0.17849 | -0.19663612 | 0.2 | 0.273 | 1 | 16 |
| cdc5l | 0.17894 | -0.27509882 | 0.086 | 0.273 | 1 | 16 |
| aamp | 0.17901 | -0.29314651 | 0.086 | 0.273 | 1 | 16 |
| romo1 | 0.17912 | 0.18182992 | 0.771 | 0.773 | 1 | 16 |
| ccdc187 | 0.17986 | 0.243317842 | 0.629 | 0.409 | 1 | 16 |
| csnk2b | 0.18008 | 0.346564067 | 0.629 | 0.409 | 1 | 16 |
| ctdsp1 | 0.18013 | 0.275366805 | 0.286 | 0.091 | 1 | 16 |
| LOC108190056 | 0.18037 | -0.31494244 | 0.114 | 0.273 | 1 | 16 |
| nacc2 | 0.18062 | 0.295929454 | 0.286 | 0.091 | 1 | 16 |
| psmd1 | 0.18062 | -0.19972695 | 0.229 | 0.409 | 1 | 16 |
| rpl13 | 0.18086 | 0.113166952 | 1 | 0.955 | 1 | 16 |
| pcdh19 | 0.18128 | -0.26701131 | 0.029 | 0.136 | 1 | 16 |
| gstk1 | 0.18134 | 0.123734367 | 0.371 | 0.227 | 1 | 16 |
| btbd10b | 0.18155 | -0.23571525 | 0.171 | 0.364 | 1 | 16 |
| kdm6bb | 0.18201 | -0.15933798 | 0.2 | 0.364 | 1 | 16 |
| ppp6r3 | 0.18206 | 0.336817626 | 0.314 | 0.136 | 1 | 16 |
| ube2g1a | 0.18313 | -0.2184044 | 0.143 | 0.318 | 1 | 16 |
| spry4 | 0.184 | -0.23142248 | 0.143 | 0.318 | 1 | 16 |
| wu:fb55g09 | 0.1842 | 0.357438886 | 0.629 | 0.409 | 1 | 16 |
| cwf19l2 | 0.18435 | -0.10433968 | 0.114 | 0.227 | 1 | 16 |
| rtn4a | 0.1844 | 0.321776873 | 0.686 | 0.455 | 1 | 16 |
| vps28 | 0.18489 | -0.10839779 | 0.229 | 0.364 | 1 | 16 |
| alcamb | 0.18491 | 0.306175572 | 0.771 | 0.545 | 1 | 16 |
| ccnd2a | 0.18518 | -0.32275682 | 0.114 | 0.273 | 1 | 16 |
| praf2 | 0.18537 | 0.312787499 | 0.286 | 0.091 | 1 | 16 |
| CABZ01045617.1 | 0.18537 | 0.104160454 | 0.229 | 0.273 | 1 | 16 |
| nsfl1c | 0.18569 | -0.17747888 | 0.429 | 0.636 | 1 | 16 |
| si:ch211-236l14.4 | 0.18664 | 0.284147467 | 0.2 | 0.091 | 1 | 16 |
| pnrc2 | 0.18671 | 0.182090177 | 0.971 | 0.864 | 1 | 16 |
| hsbp1b | 0.18677 | 0.306853513 | 0.514 | 0.364 | 1 | 16 |
| sik2b | 0.18694 | -0.19076901 | 0.086 | 0.227 | 1 | 16 |
| ppdpfb | 0.1872 | 0.274772653 | 0.886 | 0.727 | 1 | 16 |
| wdr1 | 0.18746 | -0.25923307 | 0.143 | 0.273 | 1 | 16 |
| hypk | 0.18852 | 0.327633097 | 0.571 | 0.364 | 1 | 16 |
| zbtb8os | 0.18961 | 0.143748561 | 0.171 | 0.045 | 1 | 16 |
| pcna | 0.18964 | -0.23504874 | 0.143 | 0.227 | 1 | 16 |
| atf1 | 0.18972 | 0.319132705 | 0.257 | 0.091 | 1 | 16 |
| cnot9 | 0.19 | 0.250850934 | 0.2 | 0.045 | 1 | 16 |
| CABZ01115113.1 | 0.19005 | -0.34473795 | 0.2 | 0.409 | 1 | 16 |
| mta2 | 0.19028 | 0.274440088 | 0.2 | 0.045 | 1 | 16 |

| | | | | | | |
|---|---|---|---|---|---|---|
| map1lc3b | 0.19066 | 0.322956639 | 0.514 | 0.273 | 1 | 16 |
| ptges3b | 0.19091 | -0.27644317 | 0.486 | 0.727 | 1 | 16 |
| atp5mf | 0.19097 | -0.22060658 | 0.543 | 0.773 | 1 | 16 |
| hm13 | 0.1916 | -0.36039689 | 0.229 | 0.455 | 1 | 16 |
| cnpy1 | 0.1922 | -0.32926029 | 0.229 | 0.409 | 1 | 16 |
| FP102052.2 | 0.19245 | -0.16193311 | 0.057 | 0.182 | 1 | 16 |
| mphosph8 | 0.1926 | 0.290285582 | 0.771 | 0.591 | 1 | 16 |
| nutf2 | 0.19262 | 0.319057032 | 0.371 | 0.182 | 1 | 16 |
| eif2s1b | 0.19322 | 0.37089731 | 0.429 | 0.227 | 1 | 16 |
| tmem30aa | 0.19357 | -0.20259034 | 0.343 | 0.545 | 1 | 16 |
| sap18 | 0.19423 | -0.29676132 | 0.514 | 0.727 | 1 | 16 |
| ap2b1 | 0.19442 | 0.310415273 | 0.314 | 0.182 | 1 | 16 |
| jak2a | 0.19472 | 0.321455589 | 0.257 | 0.091 | 1 | 16 |
| znfx1 | 0.19485 | -0.35719939 | 0.286 | 0.5 | 1 | 16 |
| rbm4.1 | 0.19492 | 0.295795664 | 0.4 | 0.227 | 1 | 16 |
| phip | 0.19497 | 0.288469999 | 0.2 | 0.045 | 1 | 16 |
| kremen1 | 0.19506 | 0.291567472 | 0.2 | 0.045 | 1 | 16 |
| ubap2a | 0.19525 | -0.27408937 | 0.029 | 0.136 | 1 | 16 |
| poglut1 | 0.19527 | -0.23566024 | 0.057 | 0.182 | 1 | 16 |
| COX3 | 0.19598 | -0.22328478 | 0.857 | 0.955 | 1 | 16 |
| nhsl1a | 0.1961 | 0.306208553 | 0.2 | 0.045 | 1 | 16 |
| gbp1 | 0.19633 | -0.28402923 | 0.086 | 0.227 | 1 | 16 |
| nbas | 0.19637 | 0.279618261 | 0.2 | 0.045 | 1 | 16 |
| rps7 | 0.19705 | 0.123305902 | 1 | 0.909 | 1 | 16 |
| smad2 | 0.19803 | 0.264066879 | 0.2 | 0.045 | 1 | 16 |
| mpc2 | 0.19827 | -0.28819652 | 0.229 | 0.455 | 1 | 16 |
| fn1a | 0.2 | 0.316251682 | 0.657 | 0.455 | 1 | 16 |
| cnga1a | 0.20065 | 0.247815058 | 0.171 | 0.045 | 1 | 16 |
| smurf2 | 0.20118 | -0.17053419 | 0.171 | 0.318 | 1 | 16 |
| adgrl3.1 | 0.20137 | 0.144107229 | 0.429 | 0.409 | 1 | 16 |
| bcl6aa | 0.20146 | 0.245875163 | 0.143 | 0.045 | 1 | 16 |
| srrm2 | 0.20177 | 0.128152186 | 0.2 | 0.182 | 1 | 16 |
| pitpnb | 0.20181 | -0.22952271 | 0.143 | 0.318 | 1 | 16 |
| pag1 | 0.20201 | -0.21605553 | 0.143 | 0.318 | 1 | 16 |
| psmb5 | 0.20331 | -0.27658265 | 0.286 | 0.409 | 1 | 16 |
| babam2 | 0.20332 | -0.13073124 | 0.029 | 0.136 | 1 | 16 |
| cptp | 0.20334 | -0.27152401 | 0.057 | 0.182 | 1 | 16 |
| znf706 | 0.20451 | 0.293293873 | 0.457 | 0.227 | 1 | 16 |
| asph | 0.20458 | 0.250981382 | 0.486 | 0.273 | 1 | 16 |
| tnrc6a | 0.20551 | -0.22062192 | 0.257 | 0.455 | 1 | 16 |
| gnl3l | 0.2058 | 0.254667183 | 0.2 | 0.045 | 1 | 16 |
| snapc1b | 0.20702 | 0.260254644 | 0.171 | 0.045 | 1 | 16 |
| tsc22d2 | 0.20704 | -0.17391613 | 0.086 | 0.227 | 1 | 16 |
| cst14a.2 | 0.20716 | 0.291590885 | 0.6 | 0.364 | 1 | 16 |
| nfu1 | 0.20778 | 0.249794838 | 0.2 | 0.045 | 1 | 16 |
| fam177a1 | 0.20789 | 0.184685141 | 0.2 | 0.136 | 1 | 16 |
| psmb10 | 0.2081 | -0.27576175 | 0.257 | 0.409 | 1 | 16 |
| ccnl1a | 0.20821 | -0.24327744 | 0.029 | 0.136 | 1 | 16 |

| | | | | | | |
|---|---|---|---|---|---|---|
| igf2bp1 | 0.20845 | 0.25522436 | 0.2 | 0.045 | 1 | 16 |
| coa7 | 0.20884 | -0.22649127 | 0.029 | 0.136 | 1 | 16 |
| zgc:114181 | 0.2094 | -0.37817304 | 0.2 | 0.409 | 1 | 16 |
| dtd1 | 0.20952 | -0.15437692 | 0.057 | 0.182 | 1 | 16 |
| diaph2 | 0.2109 | 0.286252337 | 0.229 | 0.091 | 1 | 16 |
| mrpl54 | 0.21097 | -0.10034737 | 0.257 | 0.273 | 1 | 16 |
| timmdc1 | 0.21142 | 0.250453284 | 0.2 | 0.045 | 1 | 16 |
| calhm2 | 0.21149 | -0.17585468 | 0.057 | 0.182 | 1 | 16 |
| LOC553316 | 0.21152 | -0.3033392 | 0.514 | 0.727 | 1 | 16 |
| nucks1a | 0.2117 | -0.33111369 | 0.4 | 0.636 | 1 | 16 |
| meis3 | 0.21174 | -0.18084427 | 0.086 | 0.136 | 1 | 16 |
| slx1b | 0.21185 | -0.18451195 | 0.029 | 0.136 | 1 | 16 |
| ctnnb1 | 0.21192 | -0.29564018 | 0.457 | 0.636 | 1 | 16 |
| polr2c | 0.21322 | 0.221849962 | 0.2 | 0.045 | 1 | 16 |
| mmel1 | 0.21339 | 0.270963381 | 0.171 | 0.045 | 1 | 16 |
| cap1 | 0.21378 | -0.24685599 | 0.229 | 0.318 | 1 | 16 |
| stag2b | 0.21389 | -0.25893743 | 0.057 | 0.136 | 1 | 16 |
| farsa | 0.21436 | 0.110823825 | 0.2 | 0.091 | 1 | 16 |
| zgc:100832 | 0.21449 | -0.27202914 | 0.057 | 0.182 | 1 | 16 |
| mfge8a | 0.21453 | -0.21663914 | 0.057 | 0.136 | 1 | 16 |
| sdhb | 0.21512 | 0.333223967 | 0.429 | 0.227 | 1 | 16 |
| zbtb10 | 0.21516 | 0.287468929 | 0.229 | 0.091 | 1 | 16 |
| sigmar1 | 0.21546 | -0.3163893 | 0.257 | 0.455 | 1 | 16 |
| snx1a | 0.21574 | 0.324724591 | 0.4 | 0.227 | 1 | 16 |
| metrn | 0.2162 | 0.331434545 | 0.6 | 0.364 | 1 | 16 |
| ctps1a | 0.21686 | -0.12163118 | 0.143 | 0.273 | 1 | 16 |
| cox8a | 0.21696 | 0.148247593 | 0.829 | 0.818 | 1 | 16 |
| stat2 | 0.21745 | -0.30123157 | 0.2 | 0.409 | 1 | 16 |
| gak | 0.21765 | -0.23366013 | 0.029 | 0.136 | 1 | 16 |
| si:dkey-261h17.1 | 0.21787 | 0.302617818 | 0.2 | 0.045 | 1 | 16 |
| gb:ai877918 | 0.21836 | 0.238312519 | 0.2 | 0.045 | 1 | 16 |
| tnika | 0.2195 | 0.311331762 | 0.257 | 0.091 | 1 | 16 |
| tusc2b | 0.22027 | -0.18752009 | 0.086 | 0.227 | 1 | 16 |
| blzf1 | 0.22113 | -0.21172002 | 0.029 | 0.136 | 1 | 16 |
| si:busm1-266f07.2 | 0.22133 | -0.22588212 | 0.086 | 0.227 | 1 | 16 |
| bmp6 | 0.22221 | -0.37178317 | 0.286 | 0.5 | 1 | 16 |
| cox7a2a | 0.22308 | -0.13743217 | 0.571 | 0.545 | 1 | 16 |
| mrpl21 | 0.22314 | 0.179885387 | 0.171 | 0.045 | 1 | 16 |
| fam208b | 0.2233 | -0.27436176 | 0.086 | 0.227 | 1 | 16 |
| kif5bb | 0.22367 | -0.21540537 | 0.114 | 0.273 | 1 | 16 |
| tbca | 0.22389 | 0.2568455 | 0.829 | 0.636 | 1 | 16 |
| atrxl | 0.2239 | -0.29049976 | 0.086 | 0.227 | 1 | 16 |
| otub1a | 0.22431 | -0.21419859 | 0.143 | 0.318 | 1 | 16 |
| cldnc | 0.22444 | -0.3201058 | 0.314 | 0.545 | 1 | 16 |
| tiparp | 0.22456 | -0.21770025 | 0.057 | 0.136 | 1 | 16 |
| crybg1a | 0.22481 | 0.16306081 | 0.229 | 0.182 | 1 | 16 |
| cnot1 | 0.22545 | 0.192921872 | 0.143 | 0.091 | 1 | 16 |
| gsto2 | 0.22569 | 0.188145518 | 0.314 | 0.136 | 1 | 16 |

| | | | | | | |
|---|---|---|---|---|---|---|
| med19b | 0.22596 | 0.131191135 | 0.143 | 0.045 | 1 | 16 |
| hexb | 0.226 | -0.31449495 | 0.143 | 0.318 | 1 | 16 |
| psmd3 | 0.22628 | -0.14954713 | 0.286 | 0.455 | 1 | 16 |
| rps6ka3a | 0.22664 | 0.296299263 | 0.314 | 0.136 | 1 | 16 |
| sept10 | 0.22687 | -0.24907267 | 0.086 | 0.227 | 1 | 16 |
| tex261 | 0.22708 | -0.16140817 | 0.029 | 0.136 | 1 | 16 |
| gpr108 | 0.22738 | -0.28337927 | 0.114 | 0.273 | 1 | 16 |
| ajuba | 0.22782 | -0.16601882 | 0.029 | 0.136 | 1 | 16 |
| ANKRD50 | 0.22786 | 0.113683417 | 0.057 | 0 | 1 | 16 |
| CABZ01085201.1 | 0.22786 | 0.102336399 | 0.057 | 0 | 1 | 16 |
| alg9 | 0.22786 | 0.111426566 | 0.057 | 0 | 1 | 16 |
| dio1 | 0.22786 | 0.117567617 | 0.057 | 0 | 1 | 16 |
| fkbp10b | 0.22786 | 0.106599451 | 0.057 | 0 | 1 | 16 |
| phf23a | 0.22786 | 0.106542216 | 0.057 | 0 | 1 | 16 |
| senp7b | 0.22786 | 0.129945011 | 0.057 | 0 | 1 | 16 |
| si:dkey-16p19.5 | 0.22786 | 0.100570237 | 0.057 | 0 | 1 | 16 |
| tinagl1 | 0.22786 | 0.131470355 | 0.057 | 0 | 1 | 16 |
| tmem102 | 0.22786 | 0.109818128 | 0.057 | 0 | 1 | 16 |
| zgc:154075 | 0.22786 | 0.107339499 | 0.057 | 0 | 1 | 16 |
| FP102809.1 | 0.22786 | 0.114705131 | 0.057 | 0 | 1 | 16 |
| LOC110439310 | 0.22786 | 0.101852151 | 0.057 | 0 | 1 | 16 |
| auts2b | 0.22786 | 0.114336894 | 0.057 | 0 | 1 | 16 |
| cd74a | 0.22786 | 0.29424815 | 0.057 | 0 | 1 | 16 |
| celf5a | 0.22786 | 0.103485852 | 0.057 | 0 | 1 | 16 |
| commd6 | 0.22786 | 0.115915239 | 0.057 | 0 | 1 | 16 |
| exoc5 | 0.22786 | 0.103292705 | 0.057 | 0 | 1 | 16 |
| glcci1a | 0.22786 | 0.141111611 | 0.057 | 0 | 1 | 16 |
| kbtbd11 | 0.22786 | 0.115624809 | 0.057 | 0 | 1 | 16 |
| lix1l | 0.22786 | 0.11330326 | 0.057 | 0 | 1 | 16 |
| lnpk | 0.22786 | 0.104451275 | 0.057 | 0 | 1 | 16 |
| magi2a | 0.22786 | 0.118267889 | 0.057 | 0 | 1 | 16 |
| mettl17 | 0.22786 | 0.102920463 | 0.057 | 0 | 1 | 16 |
| nab2 | 0.22786 | 0.115131493 | 0.057 | 0 | 1 | 16 |
| pold2 | 0.22786 | 0.117800834 | 0.057 | 0 | 1 | 16 |
| ppp1r26 | 0.22786 | 0.115061541 | 0.057 | 0 | 1 | 16 |
| rdh14b | 0.22786 | 0.111490038 | 0.057 | 0 | 1 | 16 |
| si:ch211-195m9.3 | 0.22786 | 0.113548405 | 0.057 | 0 | 1 | 16 |
| slc6a16a | 0.22786 | 0.104605739 | 0.057 | 0 | 1 | 16 |
| ssh1a | 0.22786 | 0.11062716 | 0.057 | 0 | 1 | 16 |
| traf6 | 0.22786 | 0.115833898 | 0.057 | 0 | 1 | 16 |
| uevld | 0.22786 | 0.146260906 | 0.057 | 0 | 1 | 16 |
| vps72a | 0.22786 | 0.102650875 | 0.057 | 0 | 1 | 16 |
| zgc:103700 | 0.22786 | 0.157300425 | 0.057 | 0 | 1 | 16 |
| zgc:194627 | 0.22786 | 0.123809661 | 0.057 | 0 | 1 | 16 |
| BX001023.1 | 0.22786 | 0.100607584 | 0.057 | 0 | 1 | 16 |
| LOC569391 | 0.22786 | 0.142905656 | 0.057 | 0 | 1 | 16 |
| XLOC-004018 | 0.22786 | 0.109542969 | 0.057 | 0 | 1 | 16 |
| XLOC-031367 | 0.22786 | 0.128827477 | 0.057 | 0 | 1 | 16 |

| | | | | | | |
|---|---|---|---|---|---|---|
| XLOC-041832 | 0.22786 | 0.147714479 | 0.057 | 0 | 1 | 16 |
| XLOC-042565 | 0.22786 | 0.122510144 | 0.057 | 0 | 1 | 16 |
| atxn1b | 0.22786 | 0.11853817 | 0.057 | 0 | 1 | 16 |
| atxn7l2a | 0.22786 | 0.149006162 | 0.057 | 0 | 1 | 16 |
| casd1 | 0.22786 | 0.132308486 | 0.057 | 0 | 1 | 16 |
| caska | 0.22786 | 0.111568971 | 0.057 | 0 | 1 | 16 |
| clcn4 | 0.22786 | 0.100774213 | 0.057 | 0 | 1 | 16 |
| dram2b | 0.22786 | 0.132983609 | 0.057 | 0 | 1 | 16 |
| erbb3a | 0.22786 | 0.104418093 | 0.057 | 0 | 1 | 16 |
| fgd4a | 0.22786 | 0.103831921 | 0.057 | 0 | 1 | 16 |
| gria1b | 0.22786 | 0.132340595 | 0.057 | 0 | 1 | 16 |
| insra | 0.22786 | 0.105670124 | 0.057 | 0 | 1 | 16 |
| jmjd4 | 0.22786 | 0.127400616 | 0.057 | 0 | 1 | 16 |
| lama1 | 0.22786 | 0.101213052 | 0.057 | 0 | 1 | 16 |
| mapre2 | 0.22786 | 0.104346644 | 0.057 | 0 | 1 | 16 |
| pja2 | 0.22786 | 0.100947771 | 0.057 | 0 | 1 | 16 |
| pnpla6 | 0.22786 | 0.119037248 | 0.057 | 0 | 1 | 16 |
| rnf34a | 0.22786 | 0.119331778 | 0.057 | 0 | 1 | 16 |
| rrm2b | 0.22786 | 0.10731538 | 0.057 | 0 | 1 | 16 |
| si:ch1073-358c10.1 | 0.22786 | 0.111037134 | 0.057 | 0 | 1 | 16 |
| si:ch1073-363c19.3 | 0.22786 | 0.131716115 | 0.057 | 0 | 1 | 16 |
| si:ch211-134c9.2 | 0.22786 | 0.106841956 | 0.057 | 0 | 1 | 16 |
| si:ch211-212k18.8 | 0.22786 | 0.114991783 | 0.057 | 0 | 1 | 16 |
| si:ch211-246b8.5 | 0.22786 | 0.118627415 | 0.057 | 0 | 1 | 16 |
| si:dkey-154p10.3 | 0.22786 | 0.11111387 | 0.057 | 0 | 1 | 16 |
| smg1 | 0.22786 | 0.106216329 | 0.057 | 0 | 1 | 16 |
| stam | 0.22786 | 0.110608968 | 0.057 | 0 | 1 | 16 |
| taok1b | 0.22786 | 0.1178785 | 0.057 | 0 | 1 | 16 |
| tax1bp1a | 0.22786 | 0.109531393 | 0.057 | 0 | 1 | 16 |
| tfe3b | 0.22786 | 0.114834407 | 0.057 | 0 | 1 | 16 |
| trnt1 | 0.22786 | 0.134237701 | 0.057 | 0 | 1 | 16 |
| vps33b | 0.22786 | 0.127294523 | 0.057 | 0 | 1 | 16 |
| zgc:109744 | 0.22786 | 0.111812367 | 0.057 | 0 | 1 | 16 |
| zgc:113372 | 0.22786 | 0.102907613 | 0.057 | 0 | 1 | 16 |
| zgc:56556 | 0.22786 | 0.135900935 | 0.057 | 0 | 1 | 16 |
| zgc:66440 | 0.22786 | 0.128578185 | 0.057 | 0 | 1 | 16 |
| zgc:66443 | 0.22786 | 0.139006693 | 0.057 | 0 | 1 | 16 |
| zgc:77151 | 0.22786 | 0.10396387 | 0.057 | 0 | 1 | 16 |
| znf1184 | 0.22786 | 0.132340595 | 0.057 | 0 | 1 | 16 |
| znf532 | 0.22786 | 0.109991937 | 0.057 | 0 | 1 | 16 |
| BX511034.2 | 0.22786 | 0.10451574 | 0.057 | 0 | 1 | 16 |
| CR450808.4 | 0.22786 | 0.117843448 | 0.057 | 0 | 1 | 16 |
| abcc3 | 0.22786 | 0.11918555 | 0.057 | 0 | 1 | 16 |
| arhgef18b | 0.22786 | 0.117554232 | 0.057 | 0 | 1 | 16 |
| atraid | 0.22786 | 0.1177258 | 0.057 | 0 | 1 | 16 |
| brf1b | 0.22786 | 0.10451574 | 0.057 | 0 | 1 | 16 |
| ca9 | 0.22786 | 0.110467558 | 0.057 | 0 | 1 | 16 |
| cpeb2 | 0.22786 | 0.114605273 | 0.057 | 0 | 1 | 16 |

| | | | | | | |
|---|---|---|---|---|---|---|
| ctu1 | 0.22786 | 0.105675868 | 0.057 | 0 | 1 | 16 |
| fbxl4 | 0.22786 | 0.131921585 | 0.057 | 0 | 1 | 16 |
| ghra | 0.22786 | 0.135638249 | 0.057 | 0 | 1 | 16 |
| mettl2a | 0.22786 | 0.106049067 | 0.057 | 0 | 1 | 16 |
| mmp17a | 0.22786 | 0.139892345 | 0.057 | 0 | 1 | 16 |
| pm20d1.2 | 0.22786 | 0.130165305 | 0.057 | 0 | 1 | 16 |
| vars2 | 0.22786 | 0.119642463 | 0.057 | 0 | 1 | 16 |
| zgc:193711 | 0.22786 | 0.120875099 | 0.057 | 0 | 1 | 16 |
| zmp:0000000521 | 0.22786 | 0.120143729 | 0.057 | 0 | 1 | 16 |
| FBXO46 | 0.22786 | 0.110565722 | 0.057 | 0 | 1 | 16 |
| LOC103911404 | 0.22786 | 0.102330841 | 0.057 | 0 | 1 | 16 |
| adgra3 | 0.22786 | 0.116288905 | 0.057 | 0 | 1 | 16 |
| akap11 | 0.22786 | 0.112251033 | 0.057 | 0 | 1 | 16 |
| alg11 | 0.22786 | 0.115028647 | 0.057 | 0 | 1 | 16 |
| asxl2 | 0.22786 | 0.104130046 | 0.057 | 0 | 1 | 16 |
| b3gnt3.4 | 0.22786 | 0.112018787 | 0.057 | 0 | 1 | 16 |
| colgalt1 | 0.22786 | 0.106620801 | 0.057 | 0 | 1 | 16 |
| ehbp1l1a | 0.22786 | 0.101207146 | 0.057 | 0 | 1 | 16 |
| frmd6 | 0.22786 | 0.107392724 | 0.057 | 0 | 1 | 16 |
| gas1b | 0.22786 | 0.100302102 | 0.057 | 0 | 1 | 16 |
| ipo9 | 0.22786 | 0.104724974 | 0.057 | 0 | 1 | 16 |
| kat6b | 0.22786 | 0.102330841 | 0.057 | 0 | 1 | 16 |
| klhl13 | 0.22786 | 0.104769712 | 0.057 | 0 | 1 | 16 |
| krit1 | 0.22786 | 0.10740524 | 0.057 | 0 | 1 | 16 |
| ngrn | 0.22786 | 0.119874805 | 0.057 | 0 | 1 | 16 |
| nlk2 | 0.22786 | 0.129854691 | 0.057 | 0 | 1 | 16 |
| nucb1 | 0.22786 | 0.167745837 | 0.057 | 0 | 1 | 16 |
| nudcd1 | 0.22786 | 0.123531075 | 0.057 | 0 | 1 | 16 |
| osbpl2b | 0.22786 | 0.106814234 | 0.057 | 0 | 1 | 16 |
| patz1 | 0.22786 | 0.100598974 | 0.057 | 0 | 1 | 16 |
| pnpla8 | 0.22786 | 0.107051841 | 0.057 | 0 | 1 | 16 |
| sbf2 | 0.22786 | 0.110406128 | 0.057 | 0 | 1 | 16 |
| sc:d189 | 0.22786 | 0.124854752 | 0.057 | 0 | 1 | 16 |
| sgf29 | 0.22786 | 0.12946872 | 0.057 | 0 | 1 | 16 |
| si:ch211-195b15.8 | 0.22786 | 0.149558262 | 0.057 | 0 | 1 | 16 |
| si:ch211-197k17.3 | 0.22786 | 0.106748961 | 0.057 | 0 | 1 | 16 |
| si:ch73-266f23.3 | 0.22786 | 0.101834494 | 0.057 | 0 | 1 | 16 |
| si:dkey-100n23.5 | 0.22786 | 0.111961103 | 0.057 | 0 | 1 | 16 |
| si:dkey-247i3.6 | 0.22786 | 0.104210586 | 0.057 | 0 | 1 | 16 |
| si:dkey-264d12.4 | 0.22786 | 0.118802722 | 0.057 | 0 | 1 | 16 |
| sirt6 | 0.22786 | 0.104312458 | 0.057 | 0 | 1 | 16 |
| slc31a2 | 0.22786 | 0.141831371 | 0.057 | 0 | 1 | 16 |
| tgfbi | 0.22786 | 0.115507472 | 0.057 | 0 | 1 | 16 |
| tnfsf10l4 | 0.22786 | 0.100963141 | 0.057 | 0 | 1 | 16 |
| tyw5 | 0.22786 | 0.108588037 | 0.057 | 0 | 1 | 16 |
| zgc:77880 | 0.22786 | 0.100832613 | 0.057 | 0 | 1 | 16 |
| zmp:0000000529 | 0.22786 | 0.137589587 | 0.057 | 0 | 1 | 16 |
| CABZ01034698.1 | 0.22786 | 0.121203771 | 0.057 | 0 | 1 | 16 |

| | | | | | | |
|---|---|---|---|---|---|---|
| CABZ01077121.1 | 0.22786 | 0.100526928 | 0.057 | 0 | 1 | 16 |
| CU695232.1 | 0.22786 | 0.120550197 | 0.057 | 0 | 1 | 16 |
| LOC110438291 | 0.22786 | 0.106232417 | 0.057 | 0 | 1 | 16 |
| LOC110438482 | 0.22786 | 0.135867557 | 0.057 | 0 | 1 | 16 |
| LOC110439414 | 0.22786 | 0.129701804 | 0.057 | 0 | 1 | 16 |
| XLOC-002452 | 0.22786 | 0.131912245 | 0.057 | 0 | 1 | 16 |
| XLOC-005199 | 0.22786 | 0.172977875 | 0.057 | 0 | 1 | 16 |
| XLOC-038098 | 0.22786 | 0.106232417 | 0.057 | 0 | 1 | 16 |
| acbd4 | 0.22786 | 0.102171907 | 0.057 | 0 | 1 | 16 |
| adck1 | 0.22786 | 0.136247551 | 0.057 | 0 | 1 | 16 |
| afg3l2 | 0.22786 | 0.109907084 | 0.057 | 0 | 1 | 16 |
| akap10 | 0.22786 | 0.115474083 | 0.057 | 0 | 1 | 16 |
| cab39l1 | 0.22786 | 0.13269279 | 0.057 | 0 | 1 | 16 |
| cacna1c | 0.22786 | 0.109698351 | 0.057 | 0 | 1 | 16 |
| ccnt2a | 0.22786 | 0.104831825 | 0.057 | 0 | 1 | 16 |
| ctdspl2b | 0.22786 | 0.131790371 | 0.057 | 0 | 1 | 16 |
| dhodh | 0.22786 | 0.11053166 | 0.057 | 0 | 1 | 16 |
| dnm1l | 0.22786 | 0.102512249 | 0.057 | 0 | 1 | 16 |
| emsy | 0.22786 | 0.141483697 | 0.057 | 0 | 1 | 16 |
| exoc6b | 0.22786 | 0.100857585 | 0.057 | 0 | 1 | 16 |
| fam198a | 0.22786 | 0.10711982 | 0.057 | 0 | 1 | 16 |
| fbxo3 | 0.22786 | 0.120812704 | 0.057 | 0 | 1 | 16 |
| furinb | 0.22786 | 0.100688851 | 0.057 | 0 | 1 | 16 |
| iars | 0.22786 | 0.117660212 | 0.057 | 0 | 1 | 16 |
| im:7145024 | 0.22786 | 0.105896613 | 0.057 | 0 | 1 | 16 |
| ipcef1 | 0.22786 | 0.118338353 | 0.057 | 0 | 1 | 16 |
| lmna | 0.22786 | 0.131167586 | 0.057 | 0 | 1 | 16 |
| megf8 | 0.22786 | 0.125029427 | 0.057 | 0 | 1 | 16 |
| mertka | 0.22786 | 0.13108586 | 0.057 | 0 | 1 | 16 |
| mfap5 | 0.22786 | 0.10857591 | 0.057 | 0 | 1 | 16 |
| mtmr2 | 0.22786 | 0.124043447 | 0.057 | 0 | 1 | 16 |
| mut | 0.22786 | 0.120426216 | 0.057 | 0 | 1 | 16 |
| naif1 | 0.22786 | 0.136653102 | 0.057 | 0 | 1 | 16 |
| ncoa3 | 0.22786 | 0.132168961 | 0.057 | 0 | 1 | 16 |
| ndufaf1 | 0.22786 | 0.105618218 | 0.057 | 0 | 1 | 16 |
| nlgn2a | 0.22786 | 0.121203771 | 0.057 | 0 | 1 | 16 |
| oma1 | 0.22786 | 0.134764406 | 0.057 | 0 | 1 | 16 |
| ppargc1b | 0.22786 | 0.11523924 | 0.057 | 0 | 1 | 16 |
| ptk6a | 0.22786 | 0.104285247 | 0.057 | 0 | 1 | 16 |
| rarga | 0.22786 | 0.109575137 | 0.057 | 0 | 1 | 16 |
| rb1 | 0.22786 | 0.13285841 | 0.057 | 0 | 1 | 16 |
| rragd | 0.22786 | 0.132445184 | 0.057 | 0 | 1 | 16 |
| scamp2l | 0.22786 | 0.11741754 | 0.057 | 0 | 1 | 16 |
| sema6bb | 0.22786 | 0.144150781 | 0.057 | 0 | 1 | 16 |
| sgtb | 0.22786 | 0.100471102 | 0.057 | 0 | 1 | 16 |
| sh3bp4a | 0.22786 | 0.135867557 | 0.057 | 0 | 1 | 16 |
| si:ch211-130m23.5 | 0.22786 | 0.106296931 | 0.057 | 0 | 1 | 16 |
| si:ch211-93f2.1 | 0.22786 | 0.130721719 | 0.057 | 0 | 1 | 16 |

| | | | | | | |
|---|---|---|---|---|---|---|
| si:dkey-111e8.5 | 0.22786 | 0.120812704 | 0.057 | 0 | 1 | 16 |
| si:dkey-153k10.9 | 0.22786 | 0.106232417 | 0.057 | 0 | 1 | 16 |
| si:dkey-220k22.1 | 0.22786 | 0.110252649 | 0.057 | 0 | 1 | 16 |
| si:dkey-253i9.4 | 0.22786 | 0.105046237 | 0.057 | 0 | 1 | 16 |
| si:dkey-266m15.6 | 0.22786 | 0.135901782 | 0.057 | 0 | 1 | 16 |
| si:dkeyp-33b5.4 | 0.22786 | 0.110345353 | 0.057 | 0 | 1 | 16 |
| slc27a2a | 0.22786 | 0.111062167 | 0.057 | 0 | 1 | 16 |
| slit3 | 0.22786 | 0.117476936 | 0.057 | 0 | 1 | 16 |
| smad7 | 0.22786 | 0.104443431 | 0.057 | 0 | 1 | 16 |
| spata6 | 0.22786 | 0.110350833 | 0.057 | 0 | 1 | 16 |
| tbce | 0.22786 | 0.145757982 | 0.057 | 0 | 1 | 16 |
| thop1 | 0.22786 | 0.127726865 | 0.057 | 0 | 1 | 16 |
| thumpd2 | 0.22786 | 0.121203771 | 0.057 | 0 | 1 | 16 |
| tmem248 | 0.22786 | 0.111275831 | 0.057 | 0 | 1 | 16 |
| tsc1a | 0.22786 | 0.109874225 | 0.057 | 0 | 1 | 16 |
| tspearb | 0.22786 | 0.109698351 | 0.057 | 0 | 1 | 16 |
| BX470259.3 | 0.22786 | 0.13588342 | 0.057 | 0 | 1 | 16 |
| CU655961.2 | 0.22786 | 0.121675234 | 0.057 | 0 | 1 | 16 |
| FO904870.1 | 0.22786 | 0.120078188 | 0.057 | 0 | 1 | 16 |
| LOC101882402 | 0.22786 | 0.133119475 | 0.057 | 0 | 1 | 16 |
| LOC108190731 | 0.22786 | 0.117095582 | 0.057 | 0 | 1 | 16 |
| VASH1 | 0.22786 | 0.102609183 | 0.057 | 0 | 1 | 16 |
| XLOC-020730 | 0.22786 | 0.113601495 | 0.057 | 0 | 1 | 16 |
| XLOC-032154 | 0.22786 | 0.112540273 | 0.057 | 0 | 1 | 16 |
| acvr2ba | 0.22786 | 0.132593745 | 0.057 | 0 | 1 | 16 |
| atg4a | 0.22786 | 0.138172993 | 0.057 | 0 | 1 | 16 |
| baiap2a | 0.22786 | 0.14206703 | 0.057 | 0 | 1 | 16 |
| cenpo | 0.22786 | 0.110346498 | 0.057 | 0 | 1 | 16 |
| cplx2l | 0.22786 | 0.121844348 | 0.057 | 0 | 1 | 16 |
| cwc22 | 0.22786 | 0.118084089 | 0.057 | 0 | 1 | 16 |
| cyp26b1 | 0.22786 | 0.148169186 | 0.057 | 0 | 1 | 16 |
| dapk1 | 0.22786 | 0.108277078 | 0.057 | 0 | 1 | 16 |
| enpp5 | 0.22786 | 0.118496959 | 0.057 | 0 | 1 | 16 |
| f7 | 0.22786 | 0.122057307 | 0.057 | 0 | 1 | 16 |
| fam98a | 0.22786 | 0.138963129 | 0.057 | 0 | 1 | 16 |
| fbxo34 | 0.22786 | 0.119964281 | 0.057 | 0 | 1 | 16 |
| gdf7 | 0.22786 | 0.104923666 | 0.057 | 0 | 1 | 16 |
| ghrb | 0.22786 | 0.144611691 | 0.057 | 0 | 1 | 16 |
| glsa | 0.22786 | 0.108081405 | 0.057 | 0 | 1 | 16 |
| harbi1 | 0.22786 | 0.109058837 | 0.057 | 0 | 1 | 16 |
| il12rb2l | 0.22786 | 0.118961072 | 0.057 | 0 | 1 | 16 |
| itga2.3 | 0.22786 | 0.128444393 | 0.057 | 0 | 1 | 16 |
| kat7b | 0.22786 | 0.114461759 | 0.057 | 0 | 1 | 16 |
| kif22 | 0.22786 | 0.120770696 | 0.057 | 0 | 1 | 16 |
| klhl18 | 0.22786 | 0.125076743 | 0.057 | 0 | 1 | 16 |
| klhl21 | 0.22786 | 0.111792386 | 0.057 | 0 | 1 | 16 |
| klhl22 | 0.22786 | 0.111986854 | 0.057 | 0 | 1 | 16 |
| lancl1 | 0.22786 | 0.107056207 | 0.057 | 0 | 1 | 16 |

| | | | | | | |
|---|---|---|---|---|---|---|
| lrp2a | 0.22786 | 0.103495386 | 0.057 | 0 | 1 | 16 |
| lyrm2 | 0.22786 | 0.118602846 | 0.057 | 0 | 1 | 16 |
| macrod2 | 0.22786 | 0.121282631 | 0.057 | 0 | 1 | 16 |
| mest | 0.22786 | 0.127444168 | 0.057 | 0 | 1 | 16 |
| mettl24 | 0.22786 | 0.11026628 | 0.057 | 0 | 1 | 16 |
| mpp7a | 0.22786 | 0.112540273 | 0.057 | 0 | 1 | 16 |
| mta1 | 0.22786 | 0.119508391 | 0.057 | 0 | 1 | 16 |
| mtpap | 0.22786 | 0.110183155 | 0.057 | 0 | 1 | 16 |
| nin | 0.22786 | 0.122476247 | 0.057 | 0 | 1 | 16 |
| nrbp2b | 0.22786 | 0.116131008 | 0.057 | 0 | 1 | 16 |
| oaz2b | 0.22786 | 0.123653805 | 0.057 | 0 | 1 | 16 |
| pcmtd1 | 0.22786 | 0.111341856 | 0.057 | 0 | 1 | 16 |
| pdgfab | 0.22786 | 0.104389563 | 0.057 | 0 | 1 | 16 |
| phospho1 | 0.22786 | 0.113579858 | 0.057 | 0 | 1 | 16 |
| pkdc | 0.22786 | 0.119949171 | 0.057 | 0 | 1 | 16 |
| plppr5a | 0.22786 | 0.102419632 | 0.057 | 0 | 1 | 16 |
| ppm1ab | 0.22786 | 0.12928935 | 0.057 | 0 | 1 | 16 |
| ptdss2 | 0.22786 | 0.106378144 | 0.057 | 0 | 1 | 16 |
| qrich1 | 0.22786 | 0.117037832 | 0.057 | 0 | 1 | 16 |
| rangrf | 0.22786 | 0.10420455 | 0.057 | 0 | 1 | 16 |
| rasa1a | 0.22786 | 0.106737078 | 0.057 | 0 | 1 | 16 |
| rnps1 | 0.22786 | 0.144896219 | 0.057 | 0 | 1 | 16 |
| rpusd4 | 0.22786 | 0.128637711 | 0.057 | 0 | 1 | 16 |
| setd6 | 0.22786 | 0.101676605 | 0.057 | 0 | 1 | 16 |
| si:ch211-212c13.10 | 0.22786 | 0.10716352 | 0.057 | 0 | 1 | 16 |
| si:ch211-255f4.3 | 0.22786 | 0.126634823 | 0.057 | 0 | 1 | 16 |
| si:ch211-59d15.9 | 0.22786 | 0.106535395 | 0.057 | 0 | 1 | 16 |
| si:ch73-174h16.4 | 0.22786 | 0.10641758 | 0.057 | 0 | 1 | 16 |
| si:ch73-27e22.7 | 0.22786 | 0.130812355 | 0.057 | 0 | 1 | 16 |
| si:dkey-21a6.5 | 0.22786 | 0.106378144 | 0.057 | 0 | 1 | 16 |
| si:dkey-28a3.2 | 0.22786 | 0.112540273 | 0.057 | 0 | 1 | 16 |
| si:dkey-92i17.2 | 0.22786 | 0.118417111 | 0.057 | 0 | 1 | 16 |
| si:dkey-9i23.16 | 0.22786 | 0.108335212 | 0.057 | 0 | 1 | 16 |
| si:dkeyp-113d7.10 | 0.22786 | 0.130063536 | 0.057 | 0 | 1 | 16 |
| si:dkeyp-2e4.3 | 0.22786 | 0.101606826 | 0.057 | 0 | 1 | 16 |
| si:rp71-1g18.1 | 0.22786 | 0.110834675 | 0.057 | 0 | 1 | 16 |
| si:rp71-79p20.2 | 0.22786 | 0.100128733 | 0.057 | 0 | 1 | 16 |
| si:zfos-1011f11.2 | 0.22786 | 0.119856206 | 0.057 | 0 | 1 | 16 |
| si:zfos-1056e6.1 | 0.22786 | 0.120825845 | 0.057 | 0 | 1 | 16 |
| siah2l | 0.22786 | 0.106155799 | 0.057 | 0 | 1 | 16 |
| slc25a14 | 0.22786 | 0.139555483 | 0.057 | 0 | 1 | 16 |
| snx16 | 0.22786 | 0.107408958 | 0.057 | 0 | 1 | 16 |
| snx27a | 0.22786 | 0.118141795 | 0.057 | 0 | 1 | 16 |
| srgap1a | 0.22786 | 0.121415232 | 0.057 | 0 | 1 | 16 |
| stoml1 | 0.22786 | 0.107947325 | 0.057 | 0 | 1 | 16 |
| strada | 0.22786 | 0.105870076 | 0.057 | 0 | 1 | 16 |
| syt1a | 0.22786 | 0.149145565 | 0.057 | 0 | 1 | 16 |
| tbc1d14 | 0.22786 | 0.101502168 | 0.057 | 0 | 1 | 16 |

| | | | | | | |
|---|---|---|---|---|---|---|
| tha1 | 0.22786 | 0.124015829 | 0.057 | 0 | 1 | 16 |
| tmem150aa | 0.22786 | 0.100039183 | 0.057 | 0 | 1 | 16 |
| tnksb | 0.22786 | 0.1253409 | 0.057 | 0 | 1 | 16 |
| tp53bp1 | 0.22786 | 0.114311017 | 0.057 | 0 | 1 | 16 |
| trim3b | 0.22786 | 0.128444393 | 0.057 | 0 | 1 | 16 |
| ttyh2 | 0.22786 | 0.133276587 | 0.057 | 0 | 1 | 16 |
| usp6nl | 0.22786 | 0.110069454 | 0.057 | 0 | 1 | 16 |
| wu:fc21g02 | 0.22786 | 0.149145565 | 0.057 | 0 | 1 | 16 |
| zbtb41 | 0.22786 | 0.104596972 | 0.057 | 0 | 1 | 16 |
| zc3hc1 | 0.22786 | 0.14679145 | 0.057 | 0 | 1 | 16 |
| zgc:113149 | 0.22786 | 0.122734511 | 0.057 | 0 | 1 | 16 |
| zgc:113333 | 0.22786 | 0.104818185 | 0.057 | 0 | 1 | 16 |
| zgc:162898 | 0.22786 | 0.100084394 | 0.057 | 0 | 1 | 16 |
| zgc:174315 | 0.22786 | 0.104229423 | 0.057 | 0 | 1 | 16 |
| zgc:92594 | 0.22786 | 0.118215262 | 0.057 | 0 | 1 | 16 |
| znf1007 | 0.22786 | 0.111250909 | 0.057 | 0 | 1 | 16 |
| znf516 | 0.22786 | 0.107973991 | 0.057 | 0 | 1 | 16 |
| znf770 | 0.22786 | 0.115071382 | 0.057 | 0 | 1 | 16 |
| znrf3 | 0.22786 | 0.107990419 | 0.057 | 0 | 1 | 16 |
| XLOC-007665 | 0.22786 | 0.103170266 | 0.057 | 0 | 1 | 16 |
| fzr1b | 0.22786 | 0.112952055 | 0.057 | 0 | 1 | 16 |
| itprid2 | 0.22786 | 0.114639518 | 0.057 | 0 | 1 | 16 |
| khk | 0.22786 | 0.103689261 | 0.057 | 0 | 1 | 16 |
| oxsr1b | 0.22786 | 0.107541217 | 0.057 | 0 | 1 | 16 |
| polr3a | 0.22786 | 0.108990246 | 0.057 | 0 | 1 | 16 |
| tmem237b | 0.22786 | 0.10259724 | 0.057 | 0 | 1 | 16 |
| wdr44 | 0.22786 | 0.104434093 | 0.057 | 0 | 1 | 16 |
| wsb2 | 0.22786 | 0.101887059 | 0.057 | 0 | 1 | 16 |
| LOC101884172 | 0.22786 | 0.134597363 | 0.057 | 0 | 1 | 16 |
| LOC101886390 | 0.22786 | 0.113536953 | 0.057 | 0 | 1 | 16 |
| ints2 | 0.22786 | 0.103877821 | 0.057 | 0 | 1 | 16 |
| lrpprc | 0.22786 | 0.100224289 | 0.057 | 0 | 1 | 16 |
| parvab | 0.22786 | 0.103877821 | 0.057 | 0 | 1 | 16 |
| thyn1 | 0.22786 | 0.109618761 | 0.057 | 0 | 1 | 16 |
| tmem125 | 0.22786 | 0.113536953 | 0.057 | 0 | 1 | 16 |
| usp2a | 0.22786 | 0.104088036 | 0.057 | 0 | 1 | 16 |
| idi1 | 0.22786 | 0.107820977 | 0.057 | 0 | 1 | 16 |
| lrrc8db | 0.22786 | 0.135243353 | 0.057 | 0 | 1 | 16 |
| mgrn1b | 0.22786 | 0.124537648 | 0.057 | 0 | 1 | 16 |
| nars2 | 0.22786 | 0.117548731 | 0.057 | 0 | 1 | 16 |
| pds5b | 0.22786 | 0.112422368 | 0.057 | 0 | 1 | 16 |
| si:busm1-180g5.6 | 0.22786 | 0.115128247 | 0.057 | 0 | 1 | 16 |
| si:ch211-154o6.2 | 0.22786 | 0.111298055 | 0.057 | 0 | 1 | 16 |
| vwa8 | 0.22786 | 0.115074258 | 0.057 | 0 | 1 | 16 |
| zgc:174263 | 0.22786 | 0.115658522 | 0.057 | 0 | 1 | 16 |
| zgc:101583 | 0.22786 | 0.100101197 | 0.057 | 0 | 1 | 16 |
| zgc:152951 | 0.22786 | 0.103168707 | 0.057 | 0 | 1 | 16 |
| prpf4 | 0.22795 | -0.14747122 | 0.029 | 0.136 | 1 | 16 |

| | | | | | | |
|---|---|---|---|---|---|---|
| anxa13l | 0.22807 | 0.248447108 | 0.143 | 0.045 | 1 | 16 |
| wipf2b | 0.22919 | -0.25760899 | 0.057 | 0.182 | 1 | 16 |
| fstl1b | 0.2303 | -0.13600576 | 0.029 | 0.136 | 1 | 16 |
| zc4h2 | 0.23031 | -0.32050495 | 0.229 | 0.409 | 1 | 16 |
| psma6a | 0.23052 | 0.233652494 | 0.429 | 0.227 | 1 | 16 |
| CT573342.1 | 0.23061 | -0.19763525 | 0.029 | 0.136 | 1 | 16 |
| vamp2 | 0.23075 | 0.204375345 | 0.114 | 0.045 | 1 | 16 |
| esd | 0.23085 | 0.276124465 | 0.629 | 0.455 | 1 | 16 |
| LOC101885018 | 0.23091 | -0.1520379 | 0.029 | 0.136 | 1 | 16 |
| tomm5 | 0.23138 | -0.20009661 | 0.514 | 0.727 | 1 | 16 |
| yars | 0.23249 | -0.26597187 | 0.086 | 0.227 | 1 | 16 |
| asah1b | 0.23281 | -0.27671694 | 0.2 | 0.364 | 1 | 16 |
| pank1b | 0.23341 | 0.199101439 | 0.171 | 0.091 | 1 | 16 |
| tmeff1b | 0.23349 | -0.25276887 | 0.086 | 0.227 | 1 | 16 |
| lman2la | 0.23355 | -0.24839412 | 0.057 | 0.182 | 1 | 16 |
| eif4e2rs1 | 0.23358 | -0.2630995 | 0.057 | 0.182 | 1 | 16 |
| eif5 | 0.23412 | 0.319298117 | 0.6 | 0.409 | 1 | 16 |
| cstf2 | 0.23421 | -0.19500114 | 0.114 | 0.273 | 1 | 16 |
| zgc:165582 | 0.23439 | -0.25385193 | 0.029 | 0.136 | 1 | 16 |
| plekho2 | 0.23447 | -0.17403149 | 0.086 | 0.227 | 1 | 16 |
| cct3 | 0.23452 | 0.313545056 | 0.543 | 0.318 | 1 | 16 |
| eif5b | 0.23572 | -0.31985198 | 0.371 | 0.591 | 1 | 16 |
| uggt2 | 0.23663 | -0.24442439 | 0.057 | 0.182 | 1 | 16 |
| lgals17 | 0.23663 | -0.35709033 | 0.457 | 0.636 | 1 | 16 |
| slc43a2a | 0.23707 | 0.308345841 | 0.229 | 0.136 | 1 | 16 |
| tent4b | 0.23726 | -0.29451818 | 0.114 | 0.273 | 1 | 16 |
| rbsn | 0.23737 | -0.23035038 | 0.029 | 0.136 | 1 | 16 |
| si:ch211-212k18.4 | 0.23754 | -0.14153568 | 0.143 | 0.182 | 1 | 16 |
| bcl2a | 0.23776 | 0.276414593 | 0.171 | 0.045 | 1 | 16 |
| CU104709.1 | 0.23778 | -0.28510317 | 0.114 | 0.273 | 1 | 16 |
| pdcd6 | 0.23793 | -0.10519359 | 0.229 | 0.227 | 1 | 16 |
| atp1a3b | 0.23794 | -0.26056249 | 0.114 | 0.227 | 1 | 16 |
| tm9sf2 | 0.23806 | -0.30545827 | 0.2 | 0.409 | 1 | 16 |
| nrp2a | 0.23876 | -0.29910879 | 0.171 | 0.364 | 1 | 16 |
| jak1 | 0.23961 | 0.288340966 | 0.286 | 0.136 | 1 | 16 |
| atp5mc3b | 0.24018 | 0.134993297 | 0.829 | 0.864 | 1 | 16 |
| si:ch211-256e16.11 | 0.24067 | 0.117405685 | 0.143 | 0.136 | 1 | 16 |
| wu:fu71h07 | 0.24079 | 0.290875705 | 0.286 | 0.136 | 1 | 16 |
| ndufb9 | 0.24087 | 0.22579127 | 0.629 | 0.409 | 1 | 16 |
| tmem18 | 0.24093 | -0.28649638 | 0.171 | 0.364 | 1 | 16 |
| zc3h11a | 0.24103 | 0.124375738 | 0.2 | 0.091 | 1 | 16 |
| pabpn1 | 0.24137 | -0.28591229 | 0.457 | 0.682 | 1 | 16 |
| ctdsp2 | 0.24171 | -0.23581695 | 0.257 | 0.364 | 1 | 16 |
| iqgap1 | 0.24177 | 0.245046671 | 0.171 | 0.045 | 1 | 16 |
| mcrip1 | 0.24198 | -0.2115331 | 0.143 | 0.227 | 1 | 16 |
| si:ch73-234b20.5 | 0.24223 | 0.138155069 | 0.171 | 0.045 | 1 | 16 |
| ier2b | 0.24234 | -0.36124369 | 0.171 | 0.364 | 1 | 16 |
| calr | 0.24236 | -0.25789835 | 0.286 | 0.5 | 1 | 16 |

| | | | | | | |
|---|---|---|---|---|---|---|
| rbm25b | 0.24256 | -0.13419105 | 0.143 | 0.273 | 1 | 16 |
| CU462878.1 | 0.24269 | -0.15952278 | 0.029 | 0.136 | 1 | 16 |
| rab5ab | 0.24285 | 0.292874356 | 0.371 | 0.182 | 1 | 16 |
| usp14 | 0.24292 | -0.20476589 | 0.114 | 0.273 | 1 | 16 |
| slc16a10 | 0.24299 | 0.258309422 | 0.429 | 0.227 | 1 | 16 |
| blmh | 0.24301 | 0.174243925 | 0.4 | 0.318 | 1 | 16 |
| pip4k2ca | 0.24345 | -0.1843108 | 0.086 | 0.227 | 1 | 16 |
| psmb1 | 0.24346 | -0.23188533 | 0.686 | 0.864 | 1 | 16 |
| txlna | 0.24385 | -0.15318267 | 0.029 | 0.136 | 1 | 16 |
| mcm6 | 0.24521 | -0.26408819 | 0.114 | 0.227 | 1 | 16 |
| zgc:101716 | 0.24525 | -0.21886591 | 0.029 | 0.136 | 1 | 16 |
| pabpc1a | 0.24544 | 0.163717272 | 0.943 | 0.909 | 1 | 16 |
| rpl17 | 0.24559 | 0.133075666 | 1 | 0.955 | 1 | 16 |
| arglu1a | 0.24614 | 0.254884755 | 0.257 | 0.091 | 1 | 16 |
| mrpl9 | 0.24615 | -0.22092801 | 0.029 | 0.136 | 1 | 16 |
| dynll2a | 0.24621 | 0.196069282 | 0.257 | 0.182 | 1 | 16 |
| vps26c | 0.2463 | 0.226200846 | 0.257 | 0.136 | 1 | 16 |
| h1fx | 0.2464 | -0.36770309 | 0.2 | 0.364 | 1 | 16 |
| timp2a | 0.2465 | -0.25986287 | 0.171 | 0.364 | 1 | 16 |
| hyou1 | 0.24678 | -0.23304424 | 0.229 | 0.318 | 1 | 16 |
| tnfaip6 | 0.24684 | 0.317919007 | 0.257 | 0.091 | 1 | 16 |
| clic4 | 0.24723 | 0.13408523 | 0.2 | 0.091 | 1 | 16 |
| tomm22 | 0.24725 | -0.29665306 | 0.229 | 0.409 | 1 | 16 |
| smu1a | 0.24729 | 0.115330846 | 0.143 | 0.136 | 1 | 16 |
| cops5 | 0.24739 | 0.252800528 | 0.229 | 0.091 | 1 | 16 |
| CABZ01049847.1 | 0.24772 | 0.27764342 | 0.371 | 0.182 | 1 | 16 |
| pygo2 | 0.24793 | -0.2178274 | 0.029 | 0.136 | 1 | 16 |
| calr3b | 0.24799 | -0.1145964 | 0.514 | 0.5 | 1 | 16 |
| cd302 | 0.24801 | -0.16153294 | 0.029 | 0.136 | 1 | 16 |
| hadh | 0.24808 | 0.305093565 | 0.314 | 0.136 | 1 | 16 |
| psma8 | 0.24845 | 0.146466803 | 0.571 | 0.545 | 1 | 16 |
| seta | 0.24847 | -0.22219838 | 0.543 | 0.636 | 1 | 16 |
| kdm7ab | 0.24856 | 0.241400516 | 0.171 | 0.045 | 1 | 16 |
| arhgap17b | 0.24879 | -0.14900005 | 0.057 | 0.091 | 1 | 16 |
| siae | 0.24981 | -0.15741177 | 0.086 | 0.136 | 1 | 16 |
| ndufa5 | 0.25067 | 0.238333611 | 0.486 | 0.364 | 1 | 16 |
| phf5a | 0.25207 | -0.20550985 | 0.229 | 0.409 | 1 | 16 |
| brd9 | 0.2521 | 0.257961283 | 0.2 | 0.091 | 1 | 16 |
| chchd2 | 0.25301 | -0.1459949 | 0.257 | 0.409 | 1 | 16 |
| ak2 | 0.25312 | 0.194523882 | 0.629 | 0.409 | 1 | 16 |
| rhbdf1a | 0.25455 | -0.2276582 | 0.029 | 0.136 | 1 | 16 |
| ppp2r5d | 0.2548 | 0.191661383 | 0.257 | 0.182 | 1 | 16 |
| mhc1uba | 0.25515 | -0.2067401 | 0.057 | 0.182 | 1 | 16 |
| parp1 | 0.25542 | 0.270657144 | 0.171 | 0.045 | 1 | 16 |
| chd9 | 0.25552 | 0.321066537 | 0.314 | 0.136 | 1 | 16 |
| rbm28 | 0.25564 | 0.290088174 | 0.257 | 0.091 | 1 | 16 |
| si:dkey-222f8.3 | 0.25569 | 0.236863287 | 0.314 | 0.136 | 1 | 16 |
| strap | 0.25601 | 0.245627009 | 0.286 | 0.182 | 1 | 16 |

| | | | | | | |
|---|---|---|---|---|---|---|
| si:ch73-138n13.1 | 0.25606 | 0.246520226 | 0.286 | 0.182 | 1 | 16 |
| uchl5 | 0.25644 | -0.20752667 | 0.114 | 0.273 | 1 | 16 |
| calm2b | 0.25669 | 0.232030543 | 0.857 | 0.773 | 1 | 16 |
| extl2 | 0.25696 | -0.1723947 | 0.029 | 0.136 | 1 | 16 |
| si:dkey-240h12.3 | 0.25719 | -0.25224571 | 0.029 | 0.136 | 1 | 16 |
| zgc:171971 | 0.25769 | -0.22263844 | 0.029 | 0.136 | 1 | 16 |
| zbtb18 | 0.25776 | -0.21254696 | 0.029 | 0.136 | 1 | 16 |
| ddx18 | 0.2579 | 0.259922197 | 0.257 | 0.091 | 1 | 16 |
| prdx6 | 0.2583 | -0.21126614 | 0.429 | 0.636 | 1 | 16 |
| zgc:92606 | 0.25848 | 0.311309836 | 0.457 | 0.273 | 1 | 16 |
| ctsba | 0.25873 | -0.31350703 | 0.371 | 0.591 | 1 | 16 |
| tmsb | 0.25874 | -0.14936389 | 0.114 | 0.136 | 1 | 16 |
| dab2ipb | 0.25918 | -0.15370684 | 0.143 | 0.182 | 1 | 16 |
| gng2 | 0.25928 | -0.25381058 | 0.086 | 0.182 | 1 | 16 |
| sema3b | 0.25949 | 0.325304257 | 0.257 | 0.091 | 1 | 16 |
| slc22a31 | 0.25952 | -0.26852404 | 0.057 | 0.182 | 1 | 16 |
| atp11a | 0.2598 | 0.273698045 | 0.371 | 0.182 | 1 | 16 |
| ddt | 0.25987 | 0.224862361 | 0.429 | 0.318 | 1 | 16 |
| si:ch73-25f10.6 | 0.26009 | 0.171790512 | 0.171 | 0.045 | 1 | 16 |
| micu2 | 0.26051 | -0.2533423 | 0.057 | 0.182 | 1 | 16 |
| emd | 0.26056 | 0.208787066 | 0.514 | 0.318 | 1 | 16 |
| si:ch211-108d22.2 | 0.26094 | 0.218188634 | 0.143 | 0.045 | 1 | 16 |
| kpna4 | 0.26102 | 0.198588137 | 0.286 | 0.182 | 1 | 16 |
| dap1b | 0.26108 | -0.1335992 | 0.371 | 0.545 | 1 | 16 |
| pomp | 0.26114 | 0.117344964 | 0.857 | 0.864 | 1 | 16 |
| ca8 | 0.26146 | 0.185791201 | 0.229 | 0.091 | 1 | 16 |
| eif4g2b | 0.26146 | 0.271055877 | 0.171 | 0.045 | 1 | 16 |
| crtap | 0.26206 | -0.23137623 | 0.143 | 0.318 | 1 | 16 |
| gng5 | 0.26231 | 0.249999716 | 0.714 | 0.5 | 1 | 16 |
| cxxc1a | 0.26263 | -0.26822182 | 0.086 | 0.227 | 1 | 16 |
| sdhaf3 | 0.26297 | -0.15316955 | 0.029 | 0.136 | 1 | 16 |
| mmgt1 | 0.26317 | -0.23598113 | 0.4 | 0.5 | 1 | 16 |
| mrpl13 | 0.26341 | -0.23009277 | 0.143 | 0.318 | 1 | 16 |
| vclb | 0.26382 | -0.16879757 | 0.029 | 0.136 | 1 | 16 |
| si:ch211-160o17.4 | 0.26388 | -0.27754488 | 0.086 | 0.227 | 1 | 16 |
| slc39a7 | 0.26394 | 0.190016409 | 0.4 | 0.227 | 1 | 16 |
| tlk2 | 0.26401 | -0.18937786 | 0.029 | 0.136 | 1 | 16 |
| angpt1 | 0.26419 | -0.16622079 | 0.029 | 0.136 | 1 | 16 |
| gnal | 0.26447 | -0.29119241 | 0.114 | 0.273 | 1 | 16 |
| st7 | 0.2648 | -0.18520231 | 0.057 | 0.182 | 1 | 16 |
| clns1a | 0.26563 | 0.24223708 | 0.257 | 0.091 | 1 | 16 |
| si:ch211-156j16.1 | 0.26635 | 0.189643871 | 0.114 | 0.045 | 1 | 16 |
| slc25a3b | 0.26638 | 0.152412367 | 0.971 | 0.955 | 1 | 16 |
| cd74b | 0.267 | -0.39383952 | 0.286 | 0.5 | 1 | 16 |
| osgep | 0.2676 | -0.259833 | 0.143 | 0.318 | 1 | 16 |
| dnajc5ab | 0.26767 | -0.22749352 | 0.029 | 0.136 | 1 | 16 |
| fbxl14b | 0.26782 | 0.225920017 | 0.171 | 0.045 | 1 | 16 |
| wu:fj29h11 | 0.26842 | -0.27279192 | 0.171 | 0.364 | 1 | 16 |

| | | | | | | |
|---|---|---|---|---|---|---|
| mrpl53 | 0.26854 | 0.245611217 | 0.257 | 0.091 | 1 | 16 |
| rab3il1 | 0.26857 | -0.21992023 | 0.114 | 0.273 | 1 | 16 |
| usp1 | 0.26865 | -0.1970519 | 0.114 | 0.182 | 1 | 16 |
| prr13 | 0.26875 | -0.33045178 | 0.343 | 0.545 | 1 | 16 |
| anapc16 | 0.26896 | 0.260831309 | 0.343 | 0.182 | 1 | 16 |
| ralbp1 | 0.26911 | 0.275798518 | 0.429 | 0.273 | 1 | 16 |
| ppp4cb | 0.26961 | -0.30416518 | 0.143 | 0.318 | 1 | 16 |
| pcf11 | 0.2701 | 0.137651964 | 0.314 | 0.182 | 1 | 16 |
| tubb5 | 0.27031 | -0.3179837 | 0.143 | 0.318 | 1 | 16 |
| acaa2 | 0.27047 | -0.28237057 | 0.114 | 0.273 | 1 | 16 |
| thbs3b | 0.27071 | 0.290488087 | 0.257 | 0.091 | 1 | 16 |
| UTP14C | 0.27072 | -0.17312711 | 0.114 | 0.182 | 1 | 16 |
| gatad2b | 0.27105 | -0.31242325 | 0.229 | 0.409 | 1 | 16 |
| camk2g1 | 0.27144 | 0.235254163 | 0.171 | 0.045 | 1 | 16 |
| anp32e | 0.27205 | 0.24414418 | 0.314 | 0.136 | 1 | 16 |
| ppil1 | 0.27215 | 0.285462079 | 0.314 | 0.136 | 1 | 16 |
| islr2 | 0.27267 | 0.272514995 | 0.543 | 0.409 | 1 | 16 |
| ddx39aa | 0.27268 | -0.21156733 | 0.143 | 0.318 | 1 | 16 |
| rspry1 | 0.27298 | -0.2168844 | 0.029 | 0.136 | 1 | 16 |
| negr1 | 0.27337 | -0.21509389 | 0.029 | 0.136 | 1 | 16 |
| tfdp2 | 0.27369 | 0.276251173 | 0.257 | 0.091 | 1 | 16 |
| mrps14 | 0.27393 | 0.245319306 | 0.257 | 0.091 | 1 | 16 |
| optn | 0.2743 | -0.16249458 | 0.057 | 0.182 | 1 | 16 |
| ncor1 | 0.27436 | -0.30534348 | 0.143 | 0.318 | 1 | 16 |
| lsm8 | 0.27467 | -0.13750158 | 0.2 | 0.227 | 1 | 16 |
| arf2b | 0.27477 | 0.16168784 | 0.8 | 0.773 | 1 | 16 |
| slc35e3 | 0.27544 | -0.22619389 | 0.057 | 0.182 | 1 | 16 |
| CABZ01068367.1 | 0.27567 | 0.329524144 | 0.429 | 0.227 | 1 | 16 |
| tnfrsf1a | 0.27571 | -0.28402665 | 0.114 | 0.273 | 1 | 16 |
| fip1l1b | 0.27576 | 0.240049934 | 0.229 | 0.091 | 1 | 16 |
| STMP1 | 0.27577 | 0.280316944 | 0.6 | 0.409 | 1 | 16 |
| ildr1b | 0.27582 | -0.22255015 | 0.029 | 0.136 | 1 | 16 |
| aldh1a2 | 0.2759 | 0.212420399 | 0.943 | 0.864 | 1 | 16 |
| zc3h14 | 0.27613 | 0.20261797 | 0.171 | 0.045 | 1 | 16 |
| map1lc3a | 0.27614 | 0.244211301 | 0.371 | 0.182 | 1 | 16 |
| tmem189 | 0.27617 | 0.276952024 | 0.229 | 0.091 | 1 | 16 |
| ier2a | 0.277 | -0.37176167 | 0.257 | 0.409 | 1 | 16 |
| st5 | 0.27708 | -0.32042161 | 0.2 | 0.364 | 1 | 16 |
| pex7 | 0.2775 | -0.21335525 | 0.029 | 0.136 | 1 | 16 |
| plgrkt | 0.27776 | -0.21455583 | 0.029 | 0.136 | 1 | 16 |
| kars | 0.27803 | -0.29482415 | 0.314 | 0.5 | 1 | 16 |
| gtf2b | 0.27823 | 0.216325628 | 0.229 | 0.136 | 1 | 16 |
| slc44a5b | 0.27882 | -0.11514917 | 0.257 | 0.273 | 1 | 16 |
| pgam1a | 0.27893 | -0.16389766 | 0.086 | 0.136 | 1 | 16 |
| lamb2l | 0.27933 | 0.137234325 | 0.286 | 0.273 | 1 | 16 |
| XLOC-024446 | 0.27953 | -0.21889565 | 0.029 | 0.136 | 1 | 16 |
| stt3b | 0.27973 | -0.2754601 | 0.171 | 0.318 | 1 | 16 |
| drg2 | 0.28019 | 0.274165302 | 0.286 | 0.136 | 1 | 16 |

| | | | | | | |
|---|---|---|---|---|---|---|
| aldh2.1 | 0.28032 | -0.22993156 | 0.057 | 0.182 | 1 | 16 |
| dctn4 | 0.28039 | -0.20390071 | 0.114 | 0.273 | 1 | 16 |
| cfap20 | 0.28091 | -0.1939217 | 0.086 | 0.227 | 1 | 16 |
| isca2 | 0.28112 | -0.19722342 | 0.057 | 0.182 | 1 | 16 |
| psmd11a | 0.2816 | -0.1712608 | 0.057 | 0.182 | 1 | 16 |
| sptssa | 0.28165 | -0.27510698 | 0.171 | 0.318 | 1 | 16 |
| stau2 | 0.28174 | 0.326451426 | 0.429 | 0.227 | 1 | 16 |
| ell2 | 0.28234 | -0.16317972 | 0.029 | 0.136 | 1 | 16 |
| eef1da | 0.28236 | 0.205175124 | 0.743 | 0.727 | 1 | 16 |
| cbx1a | 0.28276 | 0.21599352 | 0.314 | 0.227 | 1 | 16 |
| si:rp71-36a1.5 | 0.28312 | -0.23904809 | 0.057 | 0.182 | 1 | 16 |
| fam3c | 0.28321 | -0.22835615 | 0.143 | 0.318 | 1 | 16 |
| zgc:114104 | 0.28329 | -0.20992845 | 0.029 | 0.136 | 1 | 16 |
| si:ch73-236c18.8 | 0.28336 | -0.17889473 | 0.029 | 0.136 | 1 | 16 |
| hnrnpabb | 0.284 | -0.20900824 | 0.714 | 0.818 | 1 | 16 |
| si:dkey-85k7.12 | 0.28439 | -0.24477647 | 0.057 | 0.182 | 1 | 16 |
| rhoub | 0.28461 | 0.241415468 | 0.143 | 0.045 | 1 | 16 |
| gstt1b | 0.28477 | 0.280493563 | 0.657 | 0.455 | 1 | 16 |
| eif3i | 0.28478 | 0.185156861 | 0.686 | 0.636 | 1 | 16 |
| atad1a | 0.28488 | -0.17698469 | 0.029 | 0.136 | 1 | 16 |
| ccdc90b | 0.28529 | -0.21749547 | 0.029 | 0.136 | 1 | 16 |
| slc4a1ap | 0.28549 | -0.15618257 | 0.029 | 0.136 | 1 | 16 |
| tmem65 | 0.28575 | -0.21132621 | 0.029 | 0.136 | 1 | 16 |
| mak16 | 0.28592 | -0.13635406 | 0.143 | 0.273 | 1 | 16 |
| gsk3b | 0.28691 | 0.196860663 | 0.229 | 0.091 | 1 | 16 |
| mpzl1l | 0.28692 | -0.19747427 | 0.029 | 0.136 | 1 | 16 |
| zgc:171480 | 0.28707 | -0.1470157 | 0.171 | 0.318 | 1 | 16 |
| rbm8a | 0.28709 | 0.22591947 | 0.829 | 0.682 | 1 | 16 |
| phb2b | 0.28712 | -0.16820854 | 0.057 | 0.182 | 1 | 16 |
| tmem145 | 0.28719 | -0.20139995 | 0.057 | 0.182 | 1 | 16 |
| mcts1 | 0.28738 | -0.24763551 | 0.257 | 0.455 | 1 | 16 |
| snx12 | 0.28783 | 0.279998499 | 0.4 | 0.227 | 1 | 16 |
| mbd3a | 0.28788 | -0.29398982 | 0.143 | 0.318 | 1 | 16 |
| man2b1 | 0.28795 | 0.239093905 | 0.171 | 0.045 | 1 | 16 |
| si:ch211-286b5.5 | 0.28798 | 0.234987252 | 0.657 | 0.545 | 1 | 16 |
| sympk | 0.28808 | 0.186457982 | 0.2 | 0.136 | 1 | 16 |
| LOC100536647 | 0.28823 | -0.25320992 | 0.057 | 0.182 | 1 | 16 |
| krt18b | 0.28838 | -0.12033975 | 0.143 | 0.273 | 1 | 16 |
| serp1 | 0.28853 | -0.14566809 | 0.286 | 0.318 | 1 | 16 |
| dock9b | 0.28867 | 0.25405345 | 0.171 | 0.045 | 1 | 16 |
| chac2 | 0.28904 | -0.17074073 | 0.029 | 0.136 | 1 | 16 |
| commd9 | 0.28907 | 0.250628139 | 0.229 | 0.091 | 1 | 16 |
| acy1 | 0.28918 | -0.21512864 | 0.057 | 0.182 | 1 | 16 |
| btr31 | 0.28924 | -0.16836002 | 0.057 | 0.182 | 1 | 16 |
| letm1 | 0.2893 | 0.216871187 | 0.171 | 0.045 | 1 | 16 |
| si:ch211-264f5.6 | 0.28933 | -0.24724042 | 0.086 | 0.227 | 1 | 16 |
| ldlrap1a | 0.28967 | 0.280892786 | 0.486 | 0.318 | 1 | 16 |
| gas6 | 0.28973 | 0.296212742 | 0.429 | 0.227 | 1 | 16 |

| | | | | | | |
|---|---|---|---|---|---|---|
| si:dkey-276j7.1 | 0.28997 | -0.21357589 | 0.171 | 0.273 | 1 | 16 |
| arpc5b | 0.29044 | 0.269176421 | 0.314 | 0.136 | 1 | 16 |
| si:ch211-276i12.11 | 0.29105 | -0.17684757 | 0.086 | 0.227 | 1 | 16 |
| srsf5b | 0.29113 | -0.21544731 | 0.086 | 0.227 | 1 | 16 |
| tob1b | 0.29119 | 0.306228389 | 0.314 | 0.136 | 1 | 16 |
| CR381686.3 | 0.29137 | -0.18975797 | 0.029 | 0.136 | 1 | 16 |
| mlec | 0.2916 | -0.24993845 | 0.143 | 0.318 | 1 | 16 |
| eif3g | 0.29166 | 0.205810775 | 0.657 | 0.455 | 1 | 16 |
| kxd1 | 0.29199 | -0.20467959 | 0.029 | 0.136 | 1 | 16 |
| larp4ab | 0.29205 | -0.3054261 | 0.171 | 0.318 | 1 | 16 |
| epb41l5 | 0.29225 | -0.24853911 | 0.114 | 0.227 | 1 | 16 |
| rcc2 | 0.29235 | 0.270229106 | 0.429 | 0.273 | 1 | 16 |
| si:ch211-156b7.4 | 0.29241 | -0.18558321 | 0.029 | 0.136 | 1 | 16 |
| pfdn4 | 0.29241 | 0.150990647 | 0.4 | 0.364 | 1 | 16 |
| nxt2 | 0.29258 | -0.27997309 | 0.143 | 0.318 | 1 | 16 |
| amn1 | 0.29268 | -0.15851752 | 0.029 | 0.136 | 1 | 16 |
| brd1b | 0.29272 | -0.23012957 | 0.029 | 0.136 | 1 | 16 |
| fgfr1op | 0.29297 | -0.21738823 | 0.057 | 0.182 | 1 | 16 |
| trmt10a | 0.2931 | -0.17826804 | 0.029 | 0.136 | 1 | 16 |
| gfer | 0.29363 | -0.17865905 | 0.029 | 0.136 | 1 | 16 |
| si:ch1073-340j19.1 | 0.29405 | 0.144746556 | 0.143 | 0.045 | 1 | 16 |
| sec61g | 0.29429 | -0.21360602 | 0.686 | 0.864 | 1 | 16 |
| myo10l3 | 0.29456 | -0.29178952 | 0.143 | 0.318 | 1 | 16 |
| uqcr10 | 0.29482 | 0.127418055 | 0.714 | 0.545 | 1 | 16 |
| CR457445.1 | 0.29484 | 0.227453714 | 0.171 | 0.045 | 1 | 16 |
| antxr2a | 0.29484 | -0.2722441 | 0.143 | 0.318 | 1 | 16 |
| coil | 0.29486 | -0.27074315 | 0.143 | 0.318 | 1 | 16 |
| bcl9 | 0.29512 | -0.1911573 | 0.029 | 0.136 | 1 | 16 |
| pdgfc | 0.29533 | -0.2188203 | 0.029 | 0.136 | 1 | 16 |
| pcdh10a | 0.29542 | 0.269765542 | 0.314 | 0.136 | 1 | 16 |
| ccdc12 | 0.29559 | -0.26364735 | 0.114 | 0.273 | 1 | 16 |
| pycr1b | 0.29589 | -0.17096548 | 0.057 | 0.182 | 1 | 16 |
| cndp2 | 0.296 | -0.13525 | 0.143 | 0.182 | 1 | 16 |
| ufm1 | 0.29627 | -0.2441318 | 0.143 | 0.318 | 1 | 16 |
| wdr33 | 0.29644 | -0.16080346 | 0.029 | 0.136 | 1 | 16 |
| ptgs2a | 0.29648 | -0.22118579 | 0.086 | 0.136 | 1 | 16 |
| si:ch211-11k18.4 | 0.29649 | -0.22449971 | 0.086 | 0.227 | 1 | 16 |
| npepl1 | 0.29672 | 0.137316273 | 0.143 | 0.045 | 1 | 16 |
| ube2kb | 0.29684 | 0.252234056 | 0.2 | 0.091 | 1 | 16 |
| gorasp2 | 0.29703 | -0.11256272 | 0.2 | 0.318 | 1 | 16 |
| cox6a1 | 0.2971 | 0.108326524 | 0.914 | 0.773 | 1 | 16 |
| homer3b | 0.29741 | 0.245647222 | 0.229 | 0.091 | 1 | 16 |
| psmc1b | 0.29756 | -0.17724073 | 0.114 | 0.182 | 1 | 16 |
| seh1l | 0.29802 | -0.27610105 | 0.114 | 0.273 | 1 | 16 |
| mkrn4 | 0.29816 | 0.23049181 | 0.171 | 0.045 | 1 | 16 |
| mkrn1 | 0.29869 | 0.177198165 | 0.171 | 0.045 | 1 | 16 |
| tmx2b | 0.29942 | -0.20733103 | 0.029 | 0.136 | 1 | 16 |
| cops4 | 0.29949 | -0.28006558 | 0.286 | 0.455 | 1 | 16 |

| | | | | | | |
|---|---|---|---|---|---|---|
| aebp2 | 0.2995 | 0.238360243 | 0.171 | 0.045 | 1 | 16 |
| timm17a | 0.29953 | 0.212232387 | 0.171 | 0.091 | 1 | 16 |
| cnppd1 | 0.2996 | -0.20200164 | 0.029 | 0.136 | 1 | 16 |
| cyp2ad3 | 0.29961 | 0.258032616 | 0.171 | 0.045 | 1 | 16 |
| tp53bp2a | 0.30006 | -0.24682913 | 0.057 | 0.182 | 1 | 16 |
| casc4 | 0.30011 | 0.127915775 | 0.143 | 0.045 | 1 | 16 |
| si:ch73-252i11.1 | 0.30056 | -0.18516483 | 0.057 | 0.182 | 1 | 16 |
| vamp5 | 0.30057 | -0.20449521 | 0.029 | 0.136 | 1 | 16 |
| pak4 | 0.30087 | -0.19097277 | 0.029 | 0.136 | 1 | 16 |
| adipor2 | 0.30094 | 0.22054337 | 0.229 | 0.091 | 1 | 16 |
| pigb | 0.30094 | -0.22673923 | 0.029 | 0.136 | 1 | 16 |
| wu:fc33b09 | 0.30095 | -0.16456148 | 0.029 | 0.136 | 1 | 16 |
| rab2a | 0.30103 | -0.22031253 | 0.429 | 0.545 | 1 | 16 |
| CU459094.2 | 0.30139 | -0.31850544 | 0.086 | 0.227 | 1 | 16 |
| fundc2 | 0.30142 | -0.16395489 | 0.371 | 0.545 | 1 | 16 |
| gng10 | 0.3015 | -0.11530416 | 0.114 | 0.136 | 1 | 16 |
| c6 | 0.30173 | -0.18966028 | 0.029 | 0.136 | 1 | 16 |
| itga3b | 0.30178 | 0.211221124 | 0.4 | 0.227 | 1 | 16 |
| sec24b | 0.30213 | -0.18383152 | 0.029 | 0.136 | 1 | 16 |
| atf7ip | 0.30247 | 0.20898585 | 0.314 | 0.227 | 1 | 16 |
| ube3c | 0.30258 | -0.21709845 | 0.029 | 0.136 | 1 | 16 |
| si:ch73-144d13.5 | 0.30315 | -0.18668747 | 0.029 | 0.136 | 1 | 16 |
| fam114a1 | 0.3032 | -0.1817984 | 0.029 | 0.136 | 1 | 16 |
| ube2g2 | 0.30376 | 0.189808317 | 0.171 | 0.045 | 1 | 16 |
| dnph1 | 0.30385 | -0.18845652 | 0.029 | 0.136 | 1 | 16 |
| dhrs3a | 0.30392 | -0.278783 | 0.229 | 0.409 | 1 | 16 |
| trmt13 | 0.30401 | -0.17023488 | 0.029 | 0.136 | 1 | 16 |
| jag1b | 0.30445 | -0.18703444 | 0.029 | 0.136 | 1 | 16 |
| aifm2 | 0.30449 | -0.2139387 | 0.057 | 0.182 | 1 | 16 |
| ivns1abpa | 0.30483 | -0.20048028 | 0.029 | 0.136 | 1 | 16 |
| thrap3b | 0.30486 | -0.25890366 | 0.086 | 0.227 | 1 | 16 |
| LOC103910015 | 0.30509 | -0.18086174 | 0.029 | 0.136 | 1 | 16 |
| si:dkey-97a13.6 | 0.30523 | -0.19351287 | 0.029 | 0.136 | 1 | 16 |
| CABZ01090373.1 | 0.30539 | -0.1894734 | 0.029 | 0.136 | 1 | 16 |
| lhx9 | 0.30564 | -0.23020229 | 0.029 | 0.136 | 1 | 16 |
| cldn11a | 0.30568 | 0.106429537 | 1 | 1 | 1 | 16 |
| si:ch73-368j24.10.1 | 0.30569 | -0.17584592 | 0.029 | 0.136 | 1 | 16 |
| polr3c | 0.30582 | -0.16631833 | 0.029 | 0.136 | 1 | 16 |
| syvn1 | 0.30594 | -0.15426794 | 0.029 | 0.136 | 1 | 16 |
| znf1124 | 0.30608 | 0.185315826 | 0.171 | 0.045 | 1 | 16 |
| amotl2a | 0.30622 | -0.18559143 | 0.029 | 0.136 | 1 | 16 |
| tctex1d2 | 0.30628 | -0.15815924 | 0.029 | 0.136 | 1 | 16 |
| ptprga | 0.30643 | -0.3085841 | 0.114 | 0.273 | 1 | 16 |
| mtr | 0.30651 | -0.16392762 | 0.029 | 0.136 | 1 | 16 |
| CABZ01073736.1 | 0.30681 | -0.21159142 | 0.029 | 0.136 | 1 | 16 |
| ugdh | 0.30714 | -0.19732812 | 0.057 | 0.182 | 1 | 16 |
| slirp | 0.30714 | -0.2662169 | 0.257 | 0.455 | 1 | 16 |
| ppme1 | 0.30714 | 0.137124658 | 0.257 | 0.136 | 1 | 16 |

| | | | | | | |
|---|---|---|---|---|---|---|
| lrrc17 | 0.30721 | -0.15149813 | 0.029 | 0.136 | 1 | 16 |
| mibp2 | 0.30734 | 0.271345583 | 0.8 | 0.682 | 1 | 16 |
| pias4a | 0.30734 | -0.23439893 | 0.086 | 0.227 | 1 | 16 |
| nifk | 0.30753 | -0.25190163 | 0.086 | 0.227 | 1 | 16 |
| tia1l | 0.30779 | -0.10160702 | 0.171 | 0.273 | 1 | 16 |
| zgc:56676 | 0.30787 | -0.16670053 | 0.2 | 0.364 | 1 | 16 |
| rnaseka | 0.30805 | -0.22487718 | 0.171 | 0.273 | 1 | 16 |
| kbtbd8 | 0.30806 | -0.20202688 | 0.029 | 0.136 | 1 | 16 |
| gabarapa | 0.3082 | 0.28104679 | 0.571 | 0.409 | 1 | 16 |
| nucks1b | 0.30926 | -0.22107497 | 0.229 | 0.409 | 1 | 16 |
| pou6f1 | 0.30926 | -0.19982929 | 0.029 | 0.136 | 1 | 16 |
| gnl3 | 0.30952 | -0.27878509 | 0.171 | 0.318 | 1 | 16 |
| cavin1b | 0.30954 | -0.17553074 | 0.029 | 0.136 | 1 | 16 |
| rchy1 | 0.30983 | -0.1672675 | 0.057 | 0.182 | 1 | 16 |
| nars | 0.31004 | -0.25723517 | 0.429 | 0.591 | 1 | 16 |
| ikbke | 0.31007 | -0.25917057 | 0.057 | 0.182 | 1 | 16 |
| si:dkey-219c3.2 | 0.31014 | -0.15331958 | 0.029 | 0.136 | 1 | 16 |
| arl3 | 0.31037 | -0.16324018 | 0.029 | 0.136 | 1 | 16 |
| paxx | 0.31037 | -0.21857322 | 0.057 | 0.182 | 1 | 16 |
| itsn1 | 0.31053 | 0.243980208 | 0.229 | 0.091 | 1 | 16 |
| mrps25 | 0.31097 | -0.21601808 | 0.171 | 0.273 | 1 | 16 |
| gng12a | 0.31104 | 0.121499683 | 0.2 | 0.091 | 1 | 16 |
| mapk9 | 0.31109 | -0.19860158 | 0.029 | 0.136 | 1 | 16 |
| pacsin2 | 0.31125 | -0.18634427 | 0.029 | 0.136 | 1 | 16 |
| hnrnpdl | 0.31132 | -0.19350535 | 0.029 | 0.136 | 1 | 16 |
| fibpa | 0.31158 | -0.19173805 | 0.029 | 0.136 | 1 | 16 |
| nfkbiaa | 0.31168 | -0.22316264 | 0.086 | 0.227 | 1 | 16 |
| src | 0.31259 | -0.19727355 | 0.029 | 0.136 | 1 | 16 |
| ndufb6 | 0.31263 | -0.17784504 | 0.514 | 0.591 | 1 | 16 |
| tmed10 | 0.31289 | -0.21464131 | 0.714 | 0.864 | 1 | 16 |
| LOC110438360 | 0.31292 | -0.17220407 | 0.029 | 0.136 | 1 | 16 |
| pus1 | 0.31312 | -0.2256101 | 0.114 | 0.273 | 1 | 16 |
| cnpy2 | 0.31319 | -0.16404918 | 0.029 | 0.136 | 1 | 16 |
| foxn3 | 0.31347 | 0.217620535 | 0.143 | 0.045 | 1 | 16 |
| gspt1 | 0.31353 | -0.18397817 | 0.029 | 0.136 | 1 | 16 |
| clta | 0.31359 | 0.240342809 | 0.4 | 0.273 | 1 | 16 |
| pom121 | 0.3136 | -0.15557228 | 0.029 | 0.136 | 1 | 16 |
| naa40 | 0.31384 | 0.230591216 | 0.229 | 0.091 | 1 | 16 |
| dnajb1a | 0.3145 | -0.18519423 | 0.057 | 0.136 | 1 | 16 |
| ift22 | 0.31468 | -0.16835959 | 0.029 | 0.136 | 1 | 16 |
| atp2b1a | 0.3147 | 0.222225255 | 0.2 | 0.091 | 1 | 16 |
| lin37 | 0.31483 | -0.16882484 | 0.029 | 0.136 | 1 | 16 |
| si:ch211-166g5.4 | 0.3155 | -0.19417103 | 0.029 | 0.136 | 1 | 16 |
| higd1a | 0.31571 | -0.19835253 | 0.486 | 0.682 | 1 | 16 |
| slc35a4 | 0.31588 | -0.16701821 | 0.029 | 0.136 | 1 | 16 |
| dus4l | 0.31588 | -0.16298992 | 0.029 | 0.136 | 1 | 16 |
| fbxl7 | 0.31592 | 0.215320145 | 0.171 | 0.045 | 1 | 16 |
| ankib1b | 0.31602 | -0.17459314 | 0.029 | 0.136 | 1 | 16 |

| | | | | | | |
|---|---|---|---|---|---|---|
| btr26 | 0.31602 | -0.1958014 | 0.029 | 0.136 | 1 | 16 |
| NFX1 | 0.31608 | -0.18771652 | 0.057 | 0.182 | 1 | 16 |
| gspt1l | 0.31619 | 0.230900973 | 0.2 | 0.091 | 1 | 16 |
| si:ch211-245h14.1 | 0.31627 | -0.22536565 | 0.086 | 0.227 | 1 | 16 |
| igf3 | 0.31628 | 0.200026982 | 0.914 | 0.773 | 1 | 16 |
| tcf7l1a | 0.3164 | -0.21133563 | 0.029 | 0.136 | 1 | 16 |
| timm8b | 0.3164 | 0.203736494 | 0.171 | 0.045 | 1 | 16 |
| hspg2 | 0.31671 | 0.256201726 | 0.314 | 0.182 | 1 | 16 |
| klc4 | 0.31677 | -0.15833489 | 0.029 | 0.136 | 1 | 16 |
| srsf1b | 0.31678 | -0.22954054 | 0.114 | 0.273 | 1 | 16 |
| wdr81 | 0.31678 | -0.16259125 | 0.029 | 0.136 | 1 | 16 |
| spast | 0.31679 | -0.18798357 | 0.029 | 0.136 | 1 | 16 |
| scaf1 | 0.31686 | -0.26948278 | 0.114 | 0.273 | 1 | 16 |
| stx4 | 0.31691 | -0.16344108 | 0.029 | 0.136 | 1 | 16 |
| atp6v1h | 0.3171 | -0.24714946 | 0.086 | 0.227 | 1 | 16 |
| arhgap20b | 0.3171 | 0.219791717 | 0.171 | 0.045 | 1 | 16 |
| LOC101886114 | 0.31718 | -0.17412767 | 0.029 | 0.136 | 1 | 16 |
| hyi | 0.31731 | -0.16410348 | 0.029 | 0.136 | 1 | 16 |
| tent5ba | 0.31734 | -0.1190098 | 0.314 | 0.455 | 1 | 16 |
| suz12b | 0.31758 | -0.18437708 | 0.029 | 0.136 | 1 | 16 |
| si:dkeyp-68b7.7 | 0.31764 | -0.18026296 | 0.029 | 0.136 | 1 | 16 |
| efcab2 | 0.31773 | -0.15521562 | 0.029 | 0.136 | 1 | 16 |
| rpp21 | 0.3178 | -0.14970598 | 0.029 | 0.136 | 1 | 16 |
| slc31a1 | 0.31784 | 0.224690404 | 0.171 | 0.045 | 1 | 16 |
| GDPGP1 | 0.31788 | -0.17443155 | 0.029 | 0.136 | 1 | 16 |
| LOC101883704 | 0.31788 | -0.19541947 | 0.029 | 0.136 | 1 | 16 |
| ucp2 | 0.31798 | 0.106876477 | 0.971 | 0.864 | 1 | 16 |
| rnf141 | 0.31798 | -0.19814904 | 0.057 | 0.182 | 1 | 16 |
| fmnl3 | 0.31798 | -0.17204628 | 0.029 | 0.136 | 1 | 16 |
| snap23.1 | 0.31798 | -0.19355062 | 0.029 | 0.136 | 1 | 16 |
| plekhb2 | 0.31802 | -0.16083426 | 0.029 | 0.136 | 1 | 16 |
| kri1 | 0.31841 | -0.1872624 | 0.057 | 0.182 | 1 | 16 |
| gusb | 0.31851 | 0.187176978 | 0.171 | 0.045 | 1 | 16 |
| tpp1 | 0.31917 | 0.131600868 | 0.286 | 0.273 | 1 | 16 |
| sltm | 0.31928 | 0.277652124 | 0.371 | 0.227 | 1 | 16 |
| ppan | 0.31949 | -0.21681416 | 0.057 | 0.182 | 1 | 16 |
| mesd | 0.3195 | -0.25416064 | 0.229 | 0.364 | 1 | 16 |
| nudt8 | 0.31957 | -0.21707807 | 0.057 | 0.182 | 1 | 16 |
| mrpl48 | 0.31962 | -0.22761619 | 0.114 | 0.273 | 1 | 16 |
| sar1b | 0.31984 | -0.24853208 | 0.343 | 0.545 | 1 | 16 |
| fam193a | 0.32004 | -0.19850067 | 0.057 | 0.136 | 1 | 16 |
| prpf39 | 0.32012 | -0.25171025 | 0.114 | 0.273 | 1 | 16 |
| kdm5ba | 0.32027 | 0.24743013 | 0.171 | 0.045 | 1 | 16 |
| zgc:86609 | 0.32035 | 0.180951117 | 0.171 | 0.045 | 1 | 16 |
| si:dkeyp-87e7.4 | 0.32063 | 0.220679717 | 0.143 | 0.045 | 1 | 16 |
| exosc5 | 0.32074 | -0.20086947 | 0.057 | 0.136 | 1 | 16 |
| fkbp3 | 0.3208 | 0.275718984 | 0.514 | 0.364 | 1 | 16 |
| sumf2 | 0.32083 | -0.24920572 | 0.114 | 0.273 | 1 | 16 |

| | | | | | | |
|---|---|---|---|---|---|---|
| ogg1 | 0.32102 | 0.208953198 | 0.171 | 0.045 | 1 | 16 |
| polr2gl | 0.32125 | -0.23308917 | 0.114 | 0.273 | 1 | 16 |
| gm2a | 0.32133 | -0.23927515 | 0.114 | 0.273 | 1 | 16 |
| daam1b | 0.32166 | 0.223219028 | 0.171 | 0.045 | 1 | 16 |
| nudt17 | 0.32217 | -0.10196059 | 0.029 | 0.091 | 1 | 16 |
| rbm17 | 0.32217 | -0.12908052 | 0.029 | 0.091 | 1 | 16 |
| CABZ01005379.1 | 0.32217 | -0.13114561 | 0.029 | 0.091 | 1 | 16 |
| CR382370.2 | 0.32217 | -0.13566784 | 0.029 | 0.091 | 1 | 16 |
| CU469420.3 | 0.32217 | -0.10899483 | 0.029 | 0.091 | 1 | 16 |
| DXO | 0.32217 | -0.10307378 | 0.029 | 0.091 | 1 | 16 |
| TMEM19 | 0.32217 | -0.10486313 | 0.029 | 0.091 | 1 | 16 |
| XLOC-000431 | 0.32217 | -0.10808124 | 0.029 | 0.091 | 1 | 16 |
| XLOC-037780 | 0.32217 | -0.13568691 | 0.029 | 0.091 | 1 | 16 |
| XLOC-040108 | 0.32217 | -0.12134729 | 0.029 | 0.091 | 1 | 16 |
| adprhl2 | 0.32217 | -0.11688696 | 0.029 | 0.091 | 1 | 16 |
| bop1 | 0.32217 | -0.10313415 | 0.029 | 0.091 | 1 | 16 |
| chchd7 | 0.32217 | -0.12074904 | 0.029 | 0.091 | 1 | 16 |
| chfr | 0.32217 | -0.15480165 | 0.029 | 0.091 | 1 | 16 |
| chst10 | 0.32217 | -0.11311497 | 0.029 | 0.091 | 1 | 16 |
| cntn2 | 0.32217 | -0.14078466 | 0.029 | 0.091 | 1 | 16 |
| col28a2b | 0.32217 | -0.17460524 | 0.029 | 0.091 | 1 | 16 |
| colec11 | 0.32217 | -0.13130463 | 0.029 | 0.091 | 1 | 16 |
| ctdspl3 | 0.32217 | -0.10254508 | 0.029 | 0.091 | 1 | 16 |
| dusp12 | 0.32217 | -0.10904694 | 0.029 | 0.091 | 1 | 16 |
| dusp7 | 0.32217 | -0.1135567 | 0.029 | 0.091 | 1 | 16 |
| dyrk3 | 0.32217 | -0.1416613 | 0.029 | 0.091 | 1 | 16 |
| eef2k | 0.32217 | -0.11378574 | 0.029 | 0.091 | 1 | 16 |
| egr1 | 0.32217 | -0.17288492 | 0.029 | 0.091 | 1 | 16 |
| fbxo8 | 0.32217 | -0.10935409 | 0.029 | 0.091 | 1 | 16 |
| gon4l | 0.32217 | -0.10693872 | 0.029 | 0.091 | 1 | 16 |
| gphna | 0.32217 | -0.11354287 | 0.029 | 0.091 | 1 | 16 |
| kdm2aa | 0.32217 | -0.13177623 | 0.029 | 0.091 | 1 | 16 |
| kdm5a | 0.32217 | -0.12097543 | 0.029 | 0.091 | 1 | 16 |
| lpcat4 | 0.32217 | -0.11266566 | 0.029 | 0.091 | 1 | 16 |
| ly75 | 0.32217 | -0.12184295 | 0.029 | 0.091 | 1 | 16 |
| macrod1 | 0.32217 | -0.11606778 | 0.029 | 0.091 | 1 | 16 |
| maml1 | 0.32217 | -0.12287306 | 0.029 | 0.091 | 1 | 16 |
| maml3 | 0.32217 | -0.13178342 | 0.029 | 0.091 | 1 | 16 |
| man2b2 | 0.32217 | -0.16144826 | 0.029 | 0.091 | 1 | 16 |
| mbip | 0.32217 | -0.19742482 | 0.029 | 0.091 | 1 | 16 |
| mettl16 | 0.32217 | -0.13405461 | 0.029 | 0.091 | 1 | 16 |
| mettl22 | 0.32217 | -0.10877905 | 0.029 | 0.091 | 1 | 16 |
| mier2 | 0.32217 | -0.11239814 | 0.029 | 0.091 | 1 | 16 |
| nudt22 | 0.32217 | -0.10106851 | 0.029 | 0.091 | 1 | 16 |
| olfml3a | 0.32217 | -0.11674395 | 0.029 | 0.091 | 1 | 16 |
| phf20b | 0.32217 | -0.12958389 | 0.029 | 0.091 | 1 | 16 |
| plxnb1a | 0.32217 | -0.10970425 | 0.029 | 0.091 | 1 | 16 |
| ppp1r1b | 0.32217 | -0.11039229 | 0.029 | 0.091 | 1 | 16 |

| | | | | | | |
|---|---|---|---|---|---|---|
| ppp3r1a | 0.32217 | -0.1041494 | 0.029 | 0.091 | 1 | 16 |
| pus10 | 0.32217 | -0.10682384 | 0.029 | 0.091 | 1 | 16 |
| rab3gap1 | 0.32217 | -0.12713118 | 0.029 | 0.091 | 1 | 16 |
| sfxn3 | 0.32217 | -0.12121211 | 0.029 | 0.091 | 1 | 16 |
| sh3yl1 | 0.32217 | -0.12598877 | 0.029 | 0.091 | 1 | 16 |
| si:dkey-237h12.3 | 0.32217 | -0.13031052 | 0.029 | 0.091 | 1 | 16 |
| si:dkey-78a14.4 | 0.32217 | -0.10347349 | 0.029 | 0.091 | 1 | 16 |
| slc27a1a | 0.32217 | -0.12134244 | 0.029 | 0.091 | 1 | 16 |
| slc33a1 | 0.32217 | -0.10803541 | 0.029 | 0.091 | 1 | 16 |
| stat5a | 0.32217 | -0.11086473 | 0.029 | 0.091 | 1 | 16 |
| syngr2a | 0.32217 | -0.12967749 | 0.029 | 0.091 | 1 | 16 |
| trappc10 | 0.32217 | -0.11785333 | 0.029 | 0.091 | 1 | 16 |
| trim35-7 | 0.32217 | -0.11220844 | 0.029 | 0.091 | 1 | 16 |
| tspy | 0.32217 | -0.1170417 | 0.029 | 0.091 | 1 | 16 |
| ubfd1 | 0.32217 | -0.12946678 | 0.029 | 0.091 | 1 | 16 |
| zbtb34 | 0.32217 | -0.15630613 | 0.029 | 0.091 | 1 | 16 |
| zfand2a | 0.32217 | -0.11463749 | 0.029 | 0.091 | 1 | 16 |
| zgc:112962 | 0.32217 | -0.14226569 | 0.029 | 0.091 | 1 | 16 |
| znf131 | 0.32217 | -0.12173523 | 0.029 | 0.091 | 1 | 16 |
| znf385c | 0.32217 | -0.10288365 | 0.029 | 0.091 | 1 | 16 |
| CR388143.1 | 0.32217 | -0.13143257 | 0.029 | 0.091 | 1 | 16 |
| CU469577.1 | 0.32217 | -0.12899492 | 0.029 | 0.091 | 1 | 16 |
| LO018148.1 | 0.32217 | -0.13089827 | 0.029 | 0.091 | 1 | 16 |
| abca1a | 0.32217 | -0.13125599 | 0.029 | 0.091 | 1 | 16 |
| ackr3b | 0.32217 | -0.1097087 | 0.029 | 0.091 | 1 | 16 |
| acoxl | 0.32217 | -0.10007569 | 0.029 | 0.091 | 1 | 16 |
| acsbg2 | 0.32217 | -0.16253908 | 0.029 | 0.091 | 1 | 16 |
| adamts10 | 0.32217 | -0.12563274 | 0.029 | 0.091 | 1 | 16 |
| ago3b | 0.32217 | -0.10834041 | 0.029 | 0.091 | 1 | 16 |
| bambib | 0.32217 | -0.15964409 | 0.029 | 0.091 | 1 | 16 |
| bcr | 0.32217 | -0.13871936 | 0.029 | 0.091 | 1 | 16 |
| cadm4 | 0.32217 | -0.15352691 | 0.029 | 0.091 | 1 | 16 |
| ccdc82 | 0.32217 | -0.10480195 | 0.029 | 0.091 | 1 | 16 |
| clcn3 | 0.32217 | -0.12582767 | 0.029 | 0.091 | 1 | 16 |
| cremb | 0.32217 | -0.10519953 | 0.029 | 0.091 | 1 | 16 |
| ddx49 | 0.32217 | -0.12385011 | 0.029 | 0.091 | 1 | 16 |
| ddx56 | 0.32217 | -0.11255478 | 0.029 | 0.091 | 1 | 16 |
| dek | 0.32217 | -0.14318639 | 0.029 | 0.091 | 1 | 16 |
| dhx16 | 0.32217 | -0.10973797 | 0.029 | 0.091 | 1 | 16 |
| dlg5a | 0.32217 | -0.11106917 | 0.029 | 0.091 | 1 | 16 |
| eif2b3 | 0.32217 | -0.12222671 | 0.029 | 0.091 | 1 | 16 |
| eif4e2 | 0.32217 | -0.16470697 | 0.029 | 0.091 | 1 | 16 |
| ep300b | 0.32217 | -0.10564651 | 0.029 | 0.091 | 1 | 16 |
| ephx5 | 0.32217 | -0.12945299 | 0.029 | 0.091 | 1 | 16 |
| fam117bb | 0.32217 | -0.13492907 | 0.029 | 0.091 | 1 | 16 |
| gopc | 0.32217 | -0.11363079 | 0.029 | 0.091 | 1 | 16 |
| gtf2e1 | 0.32217 | -0.12147419 | 0.029 | 0.091 | 1 | 16 |
| heatr5b | 0.32217 | -0.10175028 | 0.029 | 0.091 | 1 | 16 |

| | | | | | | |
|---|---|---|---|---|---|---|
| hells | 0.32217 | -0.19819549 | 0.029 | 0.091 | 1 | 16 |
| inppl1a | 0.32217 | -0.16170368 | 0.029 | 0.091 | 1 | 16 |
| katnal1 | 0.32217 | -0.12557239 | 0.029 | 0.091 | 1 | 16 |
| kctd2 | 0.32217 | -0.10672999 | 0.029 | 0.091 | 1 | 16 |
| kdelr3 | 0.32217 | -0.15720716 | 0.029 | 0.091 | 1 | 16 |
| ldlrad3 | 0.32217 | -0.1026103 | 0.029 | 0.091 | 1 | 16 |
| med24 | 0.32217 | -0.11197931 | 0.029 | 0.091 | 1 | 16 |
| nasp | 0.32217 | -0.10793546 | 0.029 | 0.091 | 1 | 16 |
| nfyc | 0.32217 | -0.10064847 | 0.029 | 0.091 | 1 | 16 |
| noc4l | 0.32217 | -0.10854128 | 0.029 | 0.091 | 1 | 16 |
| parp8 | 0.32217 | -0.11832074 | 0.029 | 0.091 | 1 | 16 |
| pcgf6 | 0.32217 | -0.13346752 | 0.029 | 0.091 | 1 | 16 |
| pcolce2b | 0.32217 | -0.121527 | 0.029 | 0.091 | 1 | 16 |
| peli1b | 0.32217 | -0.12148342 | 0.029 | 0.091 | 1 | 16 |
| phc2b | 0.32217 | -0.11020274 | 0.029 | 0.091 | 1 | 16 |
| pkz | 0.32217 | -0.12749698 | 0.029 | 0.091 | 1 | 16 |
| plekhg2 | 0.32217 | -0.11800929 | 0.029 | 0.091 | 1 | 16 |
| ppp1r13l | 0.32217 | -0.11000806 | 0.029 | 0.091 | 1 | 16 |
| prkacab | 0.32217 | -0.14281908 | 0.029 | 0.091 | 1 | 16 |
| pttg1ipb | 0.32217 | -0.11665346 | 0.029 | 0.091 | 1 | 16 |
| rab22a | 0.32217 | -0.11290393 | 0.029 | 0.091 | 1 | 16 |
| rell1 | 0.32217 | -0.14156654 | 0.029 | 0.091 | 1 | 16 |
| rnaseh2a | 0.32217 | -0.16168642 | 0.029 | 0.091 | 1 | 16 |
| rorb | 0.32217 | -0.11179103 | 0.029 | 0.091 | 1 | 16 |
| rps6kb1a | 0.32217 | -0.11729462 | 0.029 | 0.091 | 1 | 16 |
| si:ch211-196c10.15 | 0.32217 | -0.1174132 | 0.029 | 0.091 | 1 | 16 |
| si:ch211-216l23.2 | 0.32217 | -0.10272186 | 0.029 | 0.091 | 1 | 16 |
| si:ch211-243j20.2 | 0.32217 | -0.1028093 | 0.029 | 0.091 | 1 | 16 |
| si:ch211-255f4.5 | 0.32217 | -0.13739795 | 0.029 | 0.091 | 1 | 16 |
| si:ch211-66k16.28 | 0.32217 | -0.13435348 | 0.029 | 0.091 | 1 | 16 |
| si:ch211-67e16.4 | 0.32217 | -0.10166581 | 0.029 | 0.091 | 1 | 16 |
| si:ch73-147o17.1 | 0.32217 | -0.1053255 | 0.029 | 0.091 | 1 | 16 |
| si:ch73-308l14.2 | 0.32217 | -0.16823084 | 0.029 | 0.091 | 1 | 16 |
| si:dkey-237i9.1 | 0.32217 | -0.13841426 | 0.029 | 0.091 | 1 | 16 |
| si:dkey-32e6.3 | 0.32217 | -0.1391826 | 0.029 | 0.091 | 1 | 16 |
| si:dkey-77f5.3 | 0.32217 | -0.10753222 | 0.029 | 0.091 | 1 | 16 |
| slbp | 0.32217 | -0.11840314 | 0.029 | 0.091 | 1 | 16 |
| smc3 | 0.32217 | -0.10560207 | 0.029 | 0.091 | 1 | 16 |
| sult1st1 | 0.32217 | -0.14383716 | 0.029 | 0.091 | 1 | 16 |
| tap2t | 0.32217 | -0.1193356 | 0.029 | 0.091 | 1 | 16 |
| thap12a | 0.32217 | -0.10936437 | 0.029 | 0.091 | 1 | 16 |
| thy1 | 0.32217 | -0.16438392 | 0.029 | 0.091 | 1 | 16 |
| tmem241 | 0.32217 | -0.14581901 | 0.029 | 0.091 | 1 | 16 |
| trappc2 | 0.32217 | -0.10071053 | 0.029 | 0.091 | 1 | 16 |
| tsg101a | 0.32217 | -0.13926171 | 0.029 | 0.091 | 1 | 16 |
| vps37a | 0.32217 | -0.17753203 | 0.029 | 0.091 | 1 | 16 |
| zgc:91890 | 0.32217 | -0.10549105 | 0.029 | 0.091 | 1 | 16 |
| zrsr2 | 0.32217 | -0.11403112 | 0.029 | 0.091 | 1 | 16 |

| | | | | | | |
|---|---|---|---|---|---|---|
| CABZ01074307.2 | 0.32217 | -0.107131 | 0.029 | 0.091 | 1 | 16 |
| CR847984.1 | 0.32217 | -0.10581399 | 0.029 | 0.091 | 1 | 16 |
| CU467861.1 | 0.32217 | -0.10823661 | 0.029 | 0.091 | 1 | 16 |
| LOC110438394 | 0.32217 | -0.14845612 | 0.029 | 0.091 | 1 | 16 |
| SHKBP1 | 0.32217 | -0.10095799 | 0.029 | 0.091 | 1 | 16 |
| XLOC-028649 | 0.32217 | -0.1040565 | 0.029 | 0.091 | 1 | 16 |
| acvr2aa | 0.32217 | -0.12770582 | 0.029 | 0.091 | 1 | 16 |
| dedd | 0.32217 | -0.10913807 | 0.029 | 0.091 | 1 | 16 |
| fam219aa | 0.32217 | -0.12390792 | 0.029 | 0.091 | 1 | 16 |
| hmox2a | 0.32217 | -0.12941977 | 0.029 | 0.091 | 1 | 16 |
| ifih1 | 0.32217 | -0.10890184 | 0.029 | 0.091 | 1 | 16 |
| lats2 | 0.32217 | -0.13937942 | 0.029 | 0.091 | 1 | 16 |
| mprip | 0.32217 | -0.1552604 | 0.029 | 0.091 | 1 | 16 |
| pbrm1l | 0.32217 | -0.11202006 | 0.029 | 0.091 | 1 | 16 |
| pbxip1a | 0.32217 | -0.13043256 | 0.029 | 0.091 | 1 | 16 |
| ppwd1 | 0.32217 | -0.14384478 | 0.029 | 0.091 | 1 | 16 |
| rac3b | 0.32217 | -0.14989103 | 0.029 | 0.091 | 1 | 16 |
| rap1gap2b | 0.32217 | -0.11461419 | 0.029 | 0.091 | 1 | 16 |
| rgmb | 0.32217 | -0.17610405 | 0.029 | 0.091 | 1 | 16 |
| rsrc1 | 0.32217 | -0.12662361 | 0.029 | 0.091 | 1 | 16 |
| si:ch211-218g4.2 | 0.32217 | -0.14651999 | 0.029 | 0.091 | 1 | 16 |
| si:ch73-299h12.6 | 0.32217 | -0.10500937 | 0.029 | 0.091 | 1 | 16 |
| si:dkey-287g12.6 | 0.32217 | -0.12646963 | 0.029 | 0.091 | 1 | 16 |
| slc39a1 | 0.32217 | -0.14705841 | 0.029 | 0.091 | 1 | 16 |
| xkrx | 0.32217 | -0.11388108 | 0.029 | 0.091 | 1 | 16 |
| zcchc7 | 0.32217 | -0.12893981 | 0.029 | 0.091 | 1 | 16 |
| zfand1 | 0.32217 | -0.10682265 | 0.029 | 0.091 | 1 | 16 |
| zgc:153383 | 0.32217 | -0.12724602 | 0.029 | 0.091 | 1 | 16 |
| zgc:171679 | 0.32217 | -0.11647425 | 0.029 | 0.091 | 1 | 16 |
| XLOC-033403 | 0.32217 | -0.11455472 | 0.029 | 0.091 | 1 | 16 |
| hdac6 | 0.32217 | -0.16041444 | 0.029 | 0.091 | 1 | 16 |
| hif1an | 0.32217 | -0.11896111 | 0.029 | 0.091 | 1 | 16 |
| mbd5 | 0.32217 | -0.11158065 | 0.029 | 0.091 | 1 | 16 |
| rab32a | 0.32217 | -0.10795693 | 0.029 | 0.091 | 1 | 16 |
| rfx2 | 0.32217 | -0.15809145 | 0.029 | 0.091 | 1 | 16 |
| serpinb1 | 0.32217 | -0.10258977 | 0.029 | 0.091 | 1 | 16 |
| si:dkey-192g7.3 | 0.32217 | -0.1108845 | 0.029 | 0.091 | 1 | 16 |
| zgc:165555.3 | 0.32217 | -0.12488448 | 0.029 | 0.091 | 1 | 16 |
| zmp:0000000735 | 0.32217 | -0.17944667 | 0.029 | 0.091 | 1 | 16 |
| FP067440.1 | 0.32217 | -0.15120678 | 0.029 | 0.091 | 1 | 16 |
| LOC110437709 | 0.32217 | -0.14619011 | 0.029 | 0.091 | 1 | 16 |
| dctn5 | 0.32217 | -0.13208631 | 0.029 | 0.091 | 1 | 16 |
| ei24 | 0.32217 | -0.10144667 | 0.029 | 0.091 | 1 | 16 |
| fam83d | 0.32217 | -0.10543705 | 0.029 | 0.091 | 1 | 16 |
| mre11a | 0.32217 | -0.12973575 | 0.029 | 0.091 | 1 | 16 |
| si:ch211-284f22.3 | 0.32217 | -0.14252658 | 0.029 | 0.091 | 1 | 16 |
| trdn | 0.32217 | -0.1000491 | 0.029 | 0.091 | 1 | 16 |
| ube2r2 | 0.32217 | -0.17152885 | 0.029 | 0.091 | 1 | 16 |

| | | | | | | |
|---|---|---|---|---|---|---|
| SYDE2 | 0.32217 | -0.12856778 | 0.029 | 0.091 | 1 | 16 |
| TIMM21 | 0.32217 | -0.12844314 | 0.029 | 0.091 | 1 | 16 |
| akap1b | 0.32217 | -0.13578346 | 0.029 | 0.091 | 1 | 16 |
| asl | 0.32217 | -0.10523084 | 0.029 | 0.091 | 1 | 16 |
| cul1b | 0.32217 | -0.14198037 | 0.029 | 0.091 | 1 | 16 |
| dimt1l | 0.32217 | -0.10981021 | 0.029 | 0.091 | 1 | 16 |
| fadd | 0.32217 | -0.1441495 | 0.029 | 0.091 | 1 | 16 |
| gstt1a | 0.32217 | -0.11318061 | 0.029 | 0.091 | 1 | 16 |
| pecr | 0.32217 | -0.12725601 | 0.029 | 0.091 | 1 | 16 |
| prkab1a | 0.32217 | -0.10424954 | 0.029 | 0.091 | 1 | 16 |
| rab4b | 0.32217 | -0.11750462 | 0.029 | 0.091 | 1 | 16 |
| rras2 | 0.32217 | -0.13298229 | 0.029 | 0.091 | 1 | 16 |
| spag7 | 0.32217 | -0.10173555 | 0.029 | 0.091 | 1 | 16 |
| tmem176 | 0.32217 | -0.10444688 | 0.029 | 0.091 | 1 | 16 |
| tsg101b | 0.32217 | -0.10541991 | 0.029 | 0.091 | 1 | 16 |
| ubr3 | 0.32217 | -0.12517876 | 0.029 | 0.091 | 1 | 16 |
| ugt2a7 | 0.32217 | -0.11776444 | 0.029 | 0.091 | 1 | 16 |
| unc119.1 | 0.32217 | -0.12786883 | 0.029 | 0.091 | 1 | 16 |
| wdfy3 | 0.32217 | -0.1203946 | 0.029 | 0.091 | 1 | 16 |
| znf1014 | 0.32217 | -0.13410597 | 0.029 | 0.091 | 1 | 16 |
| CABZ01086293.1 | 0.32217 | -0.15808761 | 0.029 | 0.091 | 1 | 16 |
| CABZ01092746.1 | 0.32217 | -0.10131413 | 0.029 | 0.091 | 1 | 16 |
| F7 (1 of many) | 0.32217 | -0.13594336 | 0.029 | 0.091 | 1 | 16 |
| XLOC-034437 | 0.32217 | -0.13351792 | 0.029 | 0.091 | 1 | 16 |
| dennd5a | 0.32217 | -0.15983615 | 0.029 | 0.091 | 1 | 16 |
| fndc3bb | 0.32217 | -0.16591014 | 0.029 | 0.091 | 1 | 16 |
| gdpd5a | 0.32217 | -0.16601388 | 0.029 | 0.091 | 1 | 16 |
| glmna | 0.32217 | -0.10437977 | 0.029 | 0.091 | 1 | 16 |
| itpk1b | 0.32217 | -0.13023727 | 0.029 | 0.091 | 1 | 16 |
| mief2 | 0.32217 | -0.15620397 | 0.029 | 0.091 | 1 | 16 |
| nelfcd | 0.32217 | -0.10622753 | 0.029 | 0.091 | 1 | 16 |
| nlgn3a | 0.32217 | -0.1453662 | 0.029 | 0.091 | 1 | 16 |
| rab28 | 0.32217 | -0.15493075 | 0.029 | 0.091 | 1 | 16 |
| rasgef1bb | 0.32217 | -0.11658831 | 0.029 | 0.091 | 1 | 16 |
| rpap3 | 0.32217 | -0.12535958 | 0.029 | 0.091 | 1 | 16 |
| serinc2l | 0.32217 | -0.12825885 | 0.029 | 0.091 | 1 | 16 |
| sgk3 | 0.32217 | -0.11520106 | 0.029 | 0.091 | 1 | 16 |
| slc12a4 | 0.32217 | -0.10345901 | 0.029 | 0.091 | 1 | 16 |
| slc4a2a | 0.32217 | -0.12806169 | 0.029 | 0.091 | 1 | 16 |
| smad6a | 0.32217 | -0.12236575 | 0.029 | 0.091 | 1 | 16 |
| sptbn1 | 0.32217 | -0.15141306 | 0.029 | 0.091 | 1 | 16 |
| tcf7l2 | 0.32217 | -0.18246031 | 0.029 | 0.091 | 1 | 16 |
| tcp11l2 | 0.32217 | -0.10912015 | 0.029 | 0.091 | 1 | 16 |
| zgc:113436 | 0.32217 | -0.10769389 | 0.029 | 0.091 | 1 | 16 |
| zgc:136308 | 0.32217 | -0.15622918 | 0.029 | 0.091 | 1 | 16 |
| actr6 | 0.32217 | -0.11685464 | 0.029 | 0.091 | 1 | 16 |
| atg4b | 0.32217 | -0.15476675 | 0.029 | 0.091 | 1 | 16 |
| brf1a | 0.32217 | -0.11769686 | 0.029 | 0.091 | 1 | 16 |

| | | | | | | |
|---|---|---|---|---|---|---|
| plxnb1b | 0.32217 | -0.17492981 | 0.029 | 0.091 | 1 | 16 |
| qser1 | 0.32217 | -0.1307676 | 0.029 | 0.091 | 1 | 16 |
| sipa1l2.1 | 0.32217 | -0.14000822 | 0.029 | 0.091 | 1 | 16 |
| sv2 | 0.32217 | -0.1104547 | 0.029 | 0.091 | 1 | 16 |
| tmem106c | 0.32217 | -0.10517758 | 0.029 | 0.091 | 1 | 16 |
| wdfy2 | 0.32217 | -0.1448214 | 0.029 | 0.091 | 1 | 16 |
| wrap53 | 0.32217 | -0.14950424 | 0.029 | 0.091 | 1 | 16 |
| zgc:110366 | 0.32217 | -0.12953887 | 0.029 | 0.091 | 1 | 16 |
| gas1a | 0.32288 | 0.21693235 | 0.143 | 0.045 | 1 | 16 |
| fdps | 0.3229 | -0.2095768 | 0.086 | 0.182 | 1 | 16 |
| sept7b | 0.32392 | 0.210040227 | 0.457 | 0.273 | 1 | 16 |
| id1 | 0.32415 | -0.15571358 | 0.343 | 0.5 | 1 | 16 |
| med28 | 0.32423 | 0.222591181 | 0.4 | 0.227 | 1 | 16 |
| efnb1 | 0.32431 | -0.27971826 | 0.086 | 0.227 | 1 | 16 |
| stat1b | 0.32457 | -0.2709916 | 0.114 | 0.227 | 1 | 16 |
| top2b | 0.32523 | -0.23720575 | 0.086 | 0.227 | 1 | 16 |
| capns1b | 0.32534 | -0.25749313 | 0.486 | 0.682 | 1 | 16 |
| nup62l | 0.32535 | -0.21147594 | 0.086 | 0.227 | 1 | 16 |
| mrpl27 | 0.32559 | 0.198441561 | 0.171 | 0.045 | 1 | 16 |
| bmpr2a | 0.32563 | 0.142231038 | 0.457 | 0.455 | 1 | 16 |
| CT573476.1 | 0.32582 | -0.23367546 | 0.086 | 0.227 | 1 | 16 |
| pcyt2 | 0.32602 | 0.250777651 | 0.229 | 0.091 | 1 | 16 |
| kank3 | 0.32729 | 0.234369009 | 0.143 | 0.045 | 1 | 16 |
| cst3 | 0.32741 | -0.21899857 | 0.057 | 0.182 | 1 | 16 |
| smarcb1b | 0.3279 | 0.21551726 | 0.171 | 0.045 | 1 | 16 |
| rab18b | 0.32807 | 0.199695008 | 0.171 | 0.045 | 1 | 16 |
| pdk2a | 0.32827 | 0.187774991 | 0.229 | 0.091 | 1 | 16 |
| creb1a | 0.32897 | -0.18846778 | 0.057 | 0.136 | 1 | 16 |
| nop56 | 0.32906 | 0.17979168 | 0.257 | 0.182 | 1 | 16 |
| eppk1 | 0.32959 | 0.221234896 | 0.257 | 0.136 | 1 | 16 |
| rcn1 | 0.32974 | 0.198328252 | 0.171 | 0.045 | 1 | 16 |
| tmsb4x | 0.32975 | 0.107362706 | 0.943 | 0.955 | 1 | 16 |
| LOC110439817 | 0.3298 | -0.22463763 | 0.057 | 0.182 | 1 | 16 |
| fzd7b | 0.32981 | -0.23298297 | 0.057 | 0.182 | 1 | 16 |
| bxdc2 | 0.33022 | -0.21040676 | 0.143 | 0.273 | 1 | 16 |
| cbx5 | 0.33043 | 0.243856854 | 0.514 | 0.364 | 1 | 16 |
| ssbp3b | 0.33044 | -0.20692041 | 0.086 | 0.227 | 1 | 16 |
| sirt2 | 0.33086 | 0.192201222 | 0.171 | 0.045 | 1 | 16 |
| taok1a | 0.33088 | 0.21182731 | 0.171 | 0.045 | 1 | 16 |
| dcun1d4 | 0.33096 | 0.246279569 | 0.171 | 0.045 | 1 | 16 |
| gid8b | 0.3311 | 0.19186023 | 0.171 | 0.045 | 1 | 16 |
| elf1 | 0.33121 | -0.20957112 | 0.086 | 0.182 | 1 | 16 |
| yap1 | 0.33188 | 0.282453076 | 0.343 | 0.182 | 1 | 16 |
| rerea | 0.33218 | -0.2532168 | 0.086 | 0.227 | 1 | 16 |
| rab5if | 0.33221 | -0.13076602 | 0.286 | 0.318 | 1 | 16 |
| dars | 0.33243 | 0.177530323 | 0.286 | 0.136 | 1 | 16 |
| otud5a | 0.33309 | -0.18734574 | 0.057 | 0.182 | 1 | 16 |
| XLOC-035413 | 0.33399 | 0.11138154 | 0.257 | 0.227 | 1 | 16 |

| | | | | | | |
|---|---|---|---|---|---|---|
| dgcr2 | 0.33408 | -0.20470199 | 0.057 | 0.182 | 1 | 16 |
| prkacaa | 0.3347 | 0.23095926 | 0.229 | 0.091 | 1 | 16 |
| myef2 | 0.33473 | -0.25904319 | 0.086 | 0.227 | 1 | 16 |
| si:zfos-464b6.2 | 0.3348 | -0.17976908 | 0.171 | 0.318 | 1 | 16 |
| cicb | 0.33532 | 0.22558614 | 0.2 | 0.091 | 1 | 16 |
| plekhj1 | 0.33545 | -0.23025565 | 0.057 | 0.182 | 1 | 16 |
| rab13 | 0.33579 | -0.18104476 | 0.171 | 0.318 | 1 | 16 |
| tmem254 | 0.33748 | -0.20139878 | 0.086 | 0.227 | 1 | 16 |
| gtf2h5 | 0.33774 | 0.135984078 | 0.371 | 0.227 | 1 | 16 |
| pqlc3 | 0.3378 | -0.18637742 | 0.057 | 0.182 | 1 | 16 |
| arl1 | 0.33803 | -0.20874188 | 0.086 | 0.227 | 1 | 16 |
| mcm2 | 0.33819 | -0.2098939 | 0.086 | 0.227 | 1 | 16 |
| csgalnact1a | 0.33821 | 0.228807194 | 0.229 | 0.136 | 1 | 16 |
| ppig | 0.33847 | -0.13601819 | 0.114 | 0.227 | 1 | 16 |
| arl9 | 0.33848 | -0.19869377 | 0.057 | 0.182 | 1 | 16 |
| pla2g4aa | 0.33875 | -0.19875226 | 0.057 | 0.182 | 1 | 16 |
| rbck1 | 0.33905 | -0.23813746 | 0.086 | 0.227 | 1 | 16 |
| lman2lb | 0.33911 | -0.2295075 | 0.086 | 0.227 | 1 | 16 |
| adar | 0.33932 | -0.23108302 | 0.057 | 0.182 | 1 | 16 |
| afdna | 0.33934 | -0.25786221 | 0.057 | 0.182 | 1 | 16 |
| mrps33 | 0.33941 | -0.20269183 | 0.057 | 0.182 | 1 | 16 |
| pnpo | 0.33982 | -0.18360791 | 0.057 | 0.182 | 1 | 16 |
| fxr1 | 0.33985 | -0.21042442 | 0.057 | 0.182 | 1 | 16 |
| steap4 | 0.33985 | 0.265156722 | 0.229 | 0.136 | 1 | 16 |
| mgst1.2 | 0.34005 | -0.21584922 | 0.171 | 0.273 | 1 | 16 |
| fam133b | 0.34046 | 0.165785945 | 0.514 | 0.455 | 1 | 16 |
| p4ha2 | 0.34123 | -0.21264843 | 0.057 | 0.182 | 1 | 16 |
| tmem176l.3a | 0.34166 | 0.23966862 | 0.229 | 0.091 | 1 | 16 |
| cetn4 | 0.34182 | -0.20827567 | 0.057 | 0.182 | 1 | 16 |
| pdcd11 | 0.34209 | -0.1843406 | 0.057 | 0.182 | 1 | 16 |
| add3a | 0.34218 | -0.1582885 | 0.257 | 0.409 | 1 | 16 |
| ddit3 | 0.3423 | -0.21255463 | 0.057 | 0.182 | 1 | 16 |
| tmed5 | 0.34268 | 0.199015932 | 0.286 | 0.136 | 1 | 16 |
| lgals9l3 | 0.34305 | -0.25170639 | 0.486 | 0.682 | 1 | 16 |
| zdhhc6 | 0.34305 | -0.22192701 | 0.057 | 0.182 | 1 | 16 |
| faf2 | 0.34308 | -0.18454566 | 0.057 | 0.182 | 1 | 16 |
| ap3s1.1 | 0.34313 | -0.21365679 | 0.057 | 0.182 | 1 | 16 |
| dhx58 | 0.34321 | -0.19242262 | 0.057 | 0.182 | 1 | 16 |
| necap2 | 0.34323 | -0.20861773 | 0.057 | 0.182 | 1 | 16 |
| ccl39.7 | 0.34331 | -0.12724433 | 0.057 | 0.091 | 1 | 16 |
| ppie | 0.34343 | -0.20210427 | 0.057 | 0.182 | 1 | 16 |
| GGT7 | 0.34347 | -0.21040548 | 0.057 | 0.182 | 1 | 16 |
| selenot1a | 0.34366 | 0.154881955 | 0.371 | 0.227 | 1 | 16 |
| stk17b | 0.34509 | -0.16148614 | 0.171 | 0.318 | 1 | 16 |
| selenou1a | 0.34568 | -0.17731389 | 0.057 | 0.136 | 1 | 16 |
| CABZ01061591.1 | 0.34625 | 0.223483608 | 0.143 | 0.045 | 1 | 16 |
| ppm1aa | 0.34687 | 0.224312364 | 0.2 | 0.091 | 1 | 16 |
| psmb3 | 0.34689 | -0.24189142 | 0.571 | 0.727 | 1 | 16 |

| | | | | | | |
|---|---|---|---|---|---|---|
| klhdc3 | 0.34817 | 0.217752237 | 0.143 | 0.045 | 1 | 16 |
| tcf25 | 0.3482 | 0.198535885 | 0.229 | 0.091 | 1 | 16 |
| eif3ha | 0.34868 | 0.231018225 | 0.629 | 0.5 | 1 | 16 |
| trappc4 | 0.3491 | -0.190191 | 0.114 | 0.227 | 1 | 16 |
| naxe | 0.35 | 0.108907601 | 0.143 | 0.045 | 1 | 16 |
| ywhaqa | 0.35007 | 0.242630685 | 0.457 | 0.318 | 1 | 16 |
| tmem131 | 0.35018 | -0.12178852 | 0.086 | 0.182 | 1 | 16 |
| zgc:153713 | 0.35186 | 0.238220399 | 0.429 | 0.318 | 1 | 16 |
| cand1 | 0.35208 | 0.136576762 | 0.2 | 0.091 | 1 | 16 |
| si:dkey-44g23.5 | 0.35214 | 0.242589395 | 0.343 | 0.227 | 1 | 16 |
| sh3pxd2b | 0.35218 | -0.10823246 | 0.086 | 0.182 | 1 | 16 |
| arf1 | 0.3522 | -0.24449393 | 0.229 | 0.409 | 1 | 16 |
| gcshb | 0.35252 | 0.157982966 | 0.229 | 0.091 | 1 | 16 |
| si:ch73-109d9.2 | 0.3534 | -0.13521112 | 0.114 | 0.227 | 1 | 16 |
| si:zfos-1192g2.3 | 0.35352 | 0.156660385 | 0.286 | 0.227 | 1 | 16 |
| zgc:110339 | 0.35421 | 0.151536671 | 0.314 | 0.273 | 1 | 16 |
| pafah1b1a | 0.35536 | 0.234203603 | 0.229 | 0.091 | 1 | 16 |
| hkdc1 | 0.35584 | 0.231775592 | 0.286 | 0.182 | 1 | 16 |
| fam184b | 0.35603 | 0.176321342 | 0.257 | 0.182 | 1 | 16 |
| srgap3 | 0.3561 | -0.19774614 | 0.057 | 0.136 | 1 | 16 |
| si:dkey-180p18.9 | 0.35646 | -0.22519676 | 0.114 | 0.227 | 1 | 16 |
| c1galt1b | 0.35668 | -0.19907932 | 0.057 | 0.136 | 1 | 16 |
| chchd6b | 0.35679 | -0.20691374 | 0.2 | 0.364 | 1 | 16 |
| tead1b | 0.35782 | 0.232554335 | 0.2 | 0.091 | 1 | 16 |
| slc25a20 | 0.35865 | -0.20386878 | 0.171 | 0.273 | 1 | 16 |
| ube2l3b | 0.35885 | -0.22143309 | 0.4 | 0.591 | 1 | 16 |
| egln2 | 0.35903 | -0.13481521 | 0.057 | 0.091 | 1 | 16 |
| tor1 | 0.35932 | 0.167789577 | 0.371 | 0.227 | 1 | 16 |
| spoplb | 0.35968 | 0.158600814 | 0.143 | 0.091 | 1 | 16 |
| chd4a | 0.36068 | 0.19052785 | 0.543 | 0.455 | 1 | 16 |
| msl2b | 0.36086 | -0.20325663 | 0.057 | 0.136 | 1 | 16 |
| tnfrsf9a | 0.36209 | -0.15984154 | 0.143 | 0.273 | 1 | 16 |
| polr2d | 0.36214 | -0.17295359 | 0.257 | 0.318 | 1 | 16 |
| ccdc84 | 0.36291 | 0.138137355 | 0.143 | 0.045 | 1 | 16 |
| cdh4 | 0.36297 | -0.25401335 | 0.571 | 0.727 | 1 | 16 |
| arcn1b | 0.36332 | 0.137688805 | 0.229 | 0.182 | 1 | 16 |
| auh | 0.36453 | 0.219719948 | 0.286 | 0.136 | 1 | 16 |
| rps6ka4 | 0.36458 | 0.212670808 | 0.143 | 0.045 | 1 | 16 |
| CABZ01044746.1 | 0.36465 | -0.15770181 | 0.143 | 0.273 | 1 | 16 |
| jdp2b | 0.36482 | -0.13702143 | 0.114 | 0.136 | 1 | 16 |
| canx | 0.36509 | -0.154389 | 0.486 | 0.545 | 1 | 16 |
| cnpy3 | 0.36521 | -0.19957021 | 0.086 | 0.182 | 1 | 16 |
| si:dkey-188i13.9 | 0.36581 | -0.24946898 | 0.4 | 0.591 | 1 | 16 |
| pdlim1 | 0.36616 | 0.196879667 | 0.143 | 0.045 | 1 | 16 |
| pde7a | 0.36629 | 0.210894006 | 0.514 | 0.409 | 1 | 16 |
| actn4 | 0.36644 | 0.231446589 | 0.286 | 0.136 | 1 | 16 |
| ndrg3a | 0.36661 | -0.27178142 | 0.229 | 0.364 | 1 | 16 |
| gskip | 0.36679 | -0.26110983 | 0.2 | 0.364 | 1 | 16 |

| | | | | | | |
|---|---|---|---|---|---|---|
| si:dkey-187e18.1 | 0.36724 | 0.241554535 | 0.4 | 0.227 | 1 | 16 |
| znf292a | 0.3681 | 0.241283077 | 0.229 | 0.091 | 1 | 16 |
| ss18 | 0.36817 | 0.25512362 | 0.229 | 0.091 | 1 | 16 |
| naa10 | 0.36835 | -0.12349044 | 0.314 | 0.455 | 1 | 16 |
| rab5c | 0.36847 | -0.25012475 | 0.4 | 0.545 | 1 | 16 |
| slc38a2 | 0.36854 | 0.106190362 | 0.457 | 0.455 | 1 | 16 |
| bcat2 | 0.36867 | 0.231677161 | 0.229 | 0.091 | 1 | 16 |
| psap | 0.36877 | -0.13970758 | 0.429 | 0.591 | 1 | 16 |
| lgals3a | 0.36955 | -0.11269541 | 0.2 | 0.227 | 1 | 16 |
| htatsf1 | 0.36994 | 0.128574972 | 0.257 | 0.227 | 1 | 16 |
| cox14 | 0.36994 | 0.102967836 | 0.286 | 0.182 | 1 | 16 |
| znf362b | 0.37013 | 0.235908608 | 0.143 | 0.045 | 1 | 16 |
| si:zfos-169g10.3 | 0.37018 | -0.24912231 | 0.343 | 0.5 | 1 | 16 |
| rbmx | 0.37023 | -0.22816427 | 0.4 | 0.591 | 1 | 16 |
| bahcc1b | 0.37028 | 0.192839398 | 0.229 | 0.136 | 1 | 16 |
| rbms2a | 0.37042 | 0.260146573 | 0.229 | 0.091 | 1 | 16 |
| piezo2a.2 | 0.37066 | 0.246834825 | 0.229 | 0.091 | 1 | 16 |
| cdh2 | 0.37079 | 0.253668026 | 0.4 | 0.227 | 1 | 16 |
| si:dkey-51e6.1 | 0.37153 | -0.16004817 | 0.257 | 0.409 | 1 | 16 |
| btf3l4 | 0.37155 | 0.163584604 | 0.629 | 0.455 | 1 | 16 |
| si:dkey-33c12.4 | 0.37176 | -0.22650138 | 0.314 | 0.5 | 1 | 16 |
| si:ch211-194k22.8 | 0.37242 | -0.11689109 | 0.2 | 0.318 | 1 | 16 |
| hp1bp3 | 0.37262 | 0.12261473 | 0.743 | 0.591 | 1 | 16 |
| prps1a | 0.37292 | -0.25078736 | 0.2 | 0.364 | 1 | 16 |
| hnrnpk | 0.37293 | 0.23187695 | 0.543 | 0.409 | 1 | 16 |
| dnajc19 | 0.37307 | 0.251205651 | 0.343 | 0.182 | 1 | 16 |
| pi15a | 0.37348 | 0.157017993 | 0.943 | 0.864 | 1 | 16 |
| psmd12 | 0.37378 | -0.11255155 | 0.171 | 0.182 | 1 | 16 |
| fam168b | 0.37404 | 0.234423242 | 0.257 | 0.136 | 1 | 16 |
| psmd14 | 0.37409 | 0.22902936 | 0.429 | 0.318 | 1 | 16 |
| smarce1 | 0.3741 | -0.26254094 | 0.371 | 0.545 | 1 | 16 |
| tnksa | 0.37413 | 0.215219957 | 0.143 | 0.045 | 1 | 16 |
| si:ch211-217k17.7 | 0.37437 | 0.208823367 | 0.286 | 0.136 | 1 | 16 |
| LOC101884451 | 0.37437 | -0.28265619 | 0.314 | 0.5 | 1 | 16 |
| tufm | 0.37462 | 0.266101574 | 0.343 | 0.182 | 1 | 16 |
| tfg | 0.37494 | 0.205861934 | 0.286 | 0.136 | 1 | 16 |
| eipr1 | 0.37516 | 0.196913845 | 0.143 | 0.045 | 1 | 16 |
| brms1la | 0.37549 | 0.209944552 | 0.229 | 0.091 | 1 | 16 |
| zgc:113426 | 0.37604 | 0.227380332 | 0.143 | 0.045 | 1 | 16 |
| carm1 | 0.37605 | -0.12051145 | 0.086 | 0.182 | 1 | 16 |
| pum3 | 0.37632 | -0.19456135 | 0.086 | 0.182 | 1 | 16 |
| mapk14a | 0.37643 | -0.20761791 | 0.114 | 0.227 | 1 | 16 |
| igbp1 | 0.37658 | 0.142221622 | 0.143 | 0.045 | 1 | 16 |
| ppp1r37 | 0.3768 | 0.23584323 | 0.143 | 0.045 | 1 | 16 |
| tm2d3 | 0.37682 | -0.25629571 | 0.314 | 0.5 | 1 | 16 |
| bmp2b | 0.37736 | 0.251390266 | 0.143 | 0.045 | 1 | 16 |
| atp6v1e1a | 0.37779 | 0.218164231 | 0.343 | 0.182 | 1 | 16 |
| grb2b | 0.37834 | 0.227851235 | 0.229 | 0.091 | 1 | 16 |

| | | | | | | |
|---|---|---|---|---|---|---|
| ppp1r14bb | 0.37856 | 0.225375321 | 0.257 | 0.136 | 1 | 16 |
| XLOC-002129 | 0.37889 | -0.26709351 | 0.2 | 0.318 | 1 | 16 |
| rps11 | 0.37989 | 0.116478767 | 0.943 | 0.909 | 1 | 16 |
| nupr1a | 0.38039 | 0.244894761 | 0.257 | 0.136 | 1 | 16 |
| zc3h7bb | 0.38064 | -0.14899775 | 0.143 | 0.182 | 1 | 16 |
| ccl19a.1 | 0.38091 | -0.3139295 | 0.2 | 0.273 | 1 | 16 |
| pip4p1a | 0.38097 | 0.199674831 | 0.171 | 0.091 | 1 | 16 |
| CABZ01112656.1 | 0.38157 | 0.24522953 | 0.314 | 0.182 | 1 | 16 |
| NINJ1 | 0.38164 | 0.142863043 | 0.143 | 0.045 | 1 | 16 |
| brpf3b | 0.3819 | 0.210623317 | 0.143 | 0.045 | 1 | 16 |
| adipor1a | 0.3819 | -0.1425702 | 0.114 | 0.227 | 1 | 16 |
| dcaf7 | 0.38194 | -0.22580389 | 0.171 | 0.273 | 1 | 16 |
| nrg1 | 0.38237 | 0.206444593 | 0.114 | 0.045 | 1 | 16 |
| rnf19b | 0.38242 | -0.12116272 | 0.057 | 0.091 | 1 | 16 |
| pak2a | 0.38271 | 0.173472627 | 0.2 | 0.136 | 1 | 16 |
| rhpn2 | 0.38292 | 0.120098986 | 0.114 | 0.091 | 1 | 16 |
| bnc2 | 0.38329 | 0.245534286 | 0.229 | 0.091 | 1 | 16 |
| XLOC-043857 | 0.38341 | 0.234553036 | 0.229 | 0.091 | 1 | 16 |
| prkci | 0.38343 | -0.23483403 | 0.143 | 0.273 | 1 | 16 |
| tmub1 | 0.38353 | 0.227729943 | 0.257 | 0.136 | 1 | 16 |
| tnfrsfa | 0.38378 | 0.231031269 | 0.4 | 0.227 | 1 | 16 |
| usp36 | 0.38418 | -0.16808164 | 0.143 | 0.273 | 1 | 16 |
| rassf8a | 0.38483 | 0.110368526 | 0.143 | 0.136 | 1 | 16 |
| tmem9b | 0.38545 | 0.224626457 | 0.229 | 0.091 | 1 | 16 |
| rasl11a | 0.38551 | 0.24977862 | 0.229 | 0.091 | 1 | 16 |
| ndufa7 | 0.38561 | -0.13399732 | 0.429 | 0.455 | 1 | 16 |
| si:ch211-22i13.2 | 0.38642 | 0.206039204 | 0.229 | 0.091 | 1 | 16 |
| atp5f1e | 0.38717 | 0.20813965 | 0.857 | 0.727 | 1 | 16 |
| snrpa1 | 0.38721 | -0.25078225 | 0.171 | 0.318 | 1 | 16 |
| ube2v2 | 0.38725 | 0.201579118 | 0.8 | 0.636 | 1 | 16 |
| znf865 | 0.38756 | 0.184787278 | 0.114 | 0.045 | 1 | 16 |
| csrnp1b | 0.38763 | 0.260083643 | 0.229 | 0.091 | 1 | 16 |
| fntb | 0.38776 | 0.198026933 | 0.143 | 0.045 | 1 | 16 |
| cnn3a | 0.38779 | 0.281160681 | 0.229 | 0.091 | 1 | 16 |
| chmp6b | 0.38809 | 0.233968806 | 0.343 | 0.182 | 1 | 16 |
| creb3l3l | 0.3884 | 0.261779849 | 0.314 | 0.182 | 1 | 16 |
| gnb1b | 0.38892 | -0.20243315 | 0.657 | 0.818 | 1 | 16 |
| eif3jb | 0.39012 | 0.171986836 | 0.257 | 0.182 | 1 | 16 |
| dolpp1 | 0.39043 | -0.11621869 | 0.086 | 0.182 | 1 | 16 |
| phf11 | 0.39107 | -0.21734928 | 0.086 | 0.182 | 1 | 16 |
| cbx3a | 0.3923 | -0.17725231 | 0.6 | 0.773 | 1 | 16 |
| zgc:113208 | 0.39271 | 0.195530382 | 0.143 | 0.045 | 1 | 16 |
| si:ch211-235e9.6 | 0.39289 | -0.11677065 | 0.4 | 0.545 | 1 | 16 |
| abracl | 0.39344 | -0.2360298 | 0.2 | 0.364 | 1 | 16 |
| col16a1 | 0.39383 | 0.224477071 | 0.171 | 0.091 | 1 | 16 |
| ndufb10 | 0.39405 | -0.24300583 | 0.371 | 0.545 | 1 | 16 |
| pik3c2a | 0.39406 | 0.240658192 | 0.371 | 0.227 | 1 | 16 |
| churc1 | 0.39417 | 0.135886412 | 0.286 | 0.227 | 1 | 16 |

| | | | | | | |
|---|---|---|---|---|---|---|
| XLOC-016094 | 0.39516 | -0.24996668 | 0.143 | 0.273 | 1 | 16 |
| zgc:66448 | 0.39522 | -0.15359718 | 0.086 | 0.136 | 1 | 16 |
| pck2 | 0.39538 | -0.14010212 | 0.086 | 0.136 | 1 | 16 |
| ndufa10 | 0.39592 | -0.10838673 | 0.286 | 0.409 | 1 | 16 |
| spg21 | 0.39644 | -0.12671585 | 0.2 | 0.227 | 1 | 16 |
| ndufs6 | 0.39756 | -0.23748771 | 0.4 | 0.545 | 1 | 16 |
| prelid3b | 0.39766 | -0.22325893 | 0.2 | 0.364 | 1 | 16 |
| ndfip2 | 0.39771 | -0.17539127 | 0.143 | 0.273 | 1 | 16 |
| amd1 | 0.39908 | 0.249091667 | 0.343 | 0.182 | 1 | 16 |
| tgif1 | 0.39971 | 0.123784912 | 0.486 | 0.455 | 1 | 16 |
| msra | 0.39976 | 0.140615185 | 0.086 | 0.045 | 1 | 16 |
| flr | 0.39986 | 0.15658439 | 0.086 | 0.045 | 1 | 16 |
| gstm.1 | 0.40036 | 0.200272136 | 0.857 | 0.727 | 1 | 16 |
| fzr1a | 0.4005 | 0.210065196 | 0.2 | 0.091 | 1 | 16 |
| kdm4b | 0.40096 | 0.18760882 | 0.171 | 0.091 | 1 | 16 |
| mob3a | 0.40113 | 0.229953772 | 0.286 | 0.136 | 1 | 16 |
| selenow1 | 0.40162 | 0.211191256 | 0.8 | 0.682 | 1 | 16 |
| tmem198b | 0.4017 | -0.11356497 | 0.086 | 0.182 | 1 | 16 |
| znf1104 | 0.40218 | 0.265091063 | 0.286 | 0.136 | 1 | 16 |
| ggcta | 0.40331 | 0.185182003 | 0.143 | 0.045 | 1 | 16 |
| pdap1b | 0.40381 | 0.234490858 | 0.286 | 0.136 | 1 | 16 |
| atp6v1e1b | 0.40397 | 0.235748095 | 0.343 | 0.182 | 1 | 16 |
| ndfip1 | 0.40447 | 0.189416629 | 0.629 | 0.545 | 1 | 16 |
| nup107 | 0.4046 | 0.149110008 | 0.143 | 0.045 | 1 | 16 |
| txndc12 | 0.40469 | -0.10164362 | 0.4 | 0.409 | 1 | 16 |
| edf1 | 0.40549 | -0.17925363 | 0.657 | 0.818 | 1 | 16 |
| gpr155a | 0.40573 | 0.15268635 | 0.086 | 0.045 | 1 | 16 |
| ctdspla | 0.40588 | -0.14819332 | 0.086 | 0.136 | 1 | 16 |
| leng1 | 0.40613 | -0.2033313 | 0.086 | 0.182 | 1 | 16 |
| srsf9 | 0.40644 | 0.202180411 | 0.286 | 0.136 | 1 | 16 |
| mrpl36 | 0.40692 | 0.188533824 | 0.486 | 0.318 | 1 | 16 |
| vamp8 | 0.40706 | 0.221706468 | 0.286 | 0.136 | 1 | 16 |
| tmem223 | 0.40715 | 0.176326557 | 0.143 | 0.045 | 1 | 16 |
| naa15a | 0.4073 | 0.105066915 | 0.2 | 0.182 | 1 | 16 |
| si:ch211-210c8.7 | 0.40757 | 0.208058688 | 0.143 | 0.045 | 1 | 16 |
| arpc1a | 0.40812 | -0.16331044 | 0.343 | 0.5 | 1 | 16 |
| anxa2a | 0.40876 | 0.129099682 | 0.171 | 0.182 | 1 | 16 |
| rnft1 | 0.40886 | -0.16391451 | 0.057 | 0.136 | 1 | 16 |
| robo2 | 0.40927 | 0.254452679 | 0.429 | 0.273 | 1 | 16 |
| ncam2 | 0.40969 | 0.188876603 | 0.114 | 0.045 | 1 | 16 |
| zc3h6 | 0.40973 | 0.151590918 | 0.143 | 0.045 | 1 | 16 |
| cfdp1 | 0.40992 | -0.14679175 | 0.171 | 0.227 | 1 | 16 |
| gmfb | 0.41021 | 0.227050836 | 0.543 | 0.364 | 1 | 16 |
| mif | 0.41029 | 0.159255195 | 0.743 | 0.682 | 1 | 16 |
| fgl2a | 0.41044 | 0.270417635 | 0.257 | 0.136 | 1 | 16 |
| cdc37 | 0.41045 | 0.173694586 | 0.286 | 0.227 | 1 | 16 |
| stat1a | 0.41097 | -0.23686864 | 0.371 | 0.5 | 1 | 16 |
| si:ch211-219a4.3 | 0.41099 | -0.10041583 | 0.057 | 0.136 | 1 | 16 |

| | | | | | | |
|---|---|---|---|---|---|---|
| derl1 | 0.411 | 0.20172032 | 0.143 | 0.045 | 1 | 16 |
| utp23 | 0.41169 | 0.196262133 | 0.143 | 0.045 | 1 | 16 |
| nop53 | 0.41184 | 0.212109671 | 0.429 | 0.318 | 1 | 16 |
| trap1 | 0.41209 | 0.212038264 | 0.143 | 0.045 | 1 | 16 |
| BOLA2 | 0.41215 | -0.21614262 | 0.114 | 0.227 | 1 | 16 |
| atp6v1ba | 0.4122 | 0.106353514 | 0.114 | 0.091 | 1 | 16 |
| znf644a | 0.41272 | 0.168132356 | 0.114 | 0.045 | 1 | 16 |
| pdia4 | 0.41277 | -0.23377375 | 0.143 | 0.273 | 1 | 16 |
| tapbp.2 | 0.41295 | -0.2455191 | 0.114 | 0.227 | 1 | 16 |
| psmc6 | 0.41315 | -0.23873734 | 0.371 | 0.545 | 1 | 16 |
| zgc:77086 | 0.41347 | 0.187310841 | 0.429 | 0.273 | 1 | 16 |
| eif3s6ip | 0.41388 | 0.166247077 | 0.343 | 0.273 | 1 | 16 |
| si:ch211-225b11.1 | 0.41389 | 0.25078435 | 0.657 | 0.5 | 1 | 16 |
| nr0b1 | 0.41392 | 0.251823867 | 0.543 | 0.364 | 1 | 16 |
| sdhc | 0.41399 | 0.250272821 | 0.371 | 0.227 | 1 | 16 |
| tbc1d20 | 0.41405 | 0.196017301 | 0.143 | 0.045 | 1 | 16 |
| pbx3b | 0.41532 | 0.197216015 | 0.143 | 0.045 | 1 | 16 |
| pde4ba | 0.41548 | -0.239919 | 0.457 | 0.636 | 1 | 16 |
| dhrs1 | 0.41552 | -0.13162816 | 0.114 | 0.227 | 1 | 16 |
| dock7 | 0.41571 | 0.189027463 | 0.2 | 0.136 | 1 | 16 |
| scocb | 0.41582 | 0.1575995 | 0.143 | 0.045 | 1 | 16 |
| caprin1b | 0.41626 | -0.20310575 | 0.457 | 0.636 | 1 | 16 |
| atp5meb | 0.41685 | -0.17652011 | 0.8 | 0.864 | 1 | 16 |
| banf1 | 0.41707 | -0.19895594 | 0.286 | 0.455 | 1 | 16 |
| hikeshi | 0.41721 | -0.14537859 | 0.229 | 0.364 | 1 | 16 |
| tshz2 | 0.41769 | 0.209227475 | 0.2 | 0.091 | 1 | 16 |
| mafa | 0.41771 | -0.14941161 | 0.114 | 0.227 | 1 | 16 |
| MCUR1 | 0.41789 | 0.170668971 | 0.143 | 0.045 | 1 | 16 |
| rfxap | 0.41826 | 0.133282148 | 0.2 | 0.091 | 1 | 16 |
| wdr32 | 0.41835 | 0.188085446 | 0.143 | 0.045 | 1 | 16 |
| psmg3 | 0.41863 | -0.24068249 | 0.286 | 0.455 | 1 | 16 |
| u2af1 | 0.41865 | -0.19059208 | 0.371 | 0.545 | 1 | 16 |
| ube2f | 0.41879 | 0.104765436 | 0.2 | 0.182 | 1 | 16 |
| atf4a | 0.4189 | 0.102158733 | 0.286 | 0.182 | 1 | 16 |
| sdr16c5b | 0.41919 | 0.193906875 | 0.143 | 0.045 | 1 | 16 |
| urm1 | 0.41987 | -0.21262728 | 0.114 | 0.227 | 1 | 16 |
| syap1 | 0.42013 | 0.183721265 | 0.314 | 0.227 | 1 | 16 |
| stambpa | 0.42039 | -0.2024242 | 0.086 | 0.182 | 1 | 16 |
| si:ch211-85n16.3 | 0.42054 | -0.19855211 | 0.171 | 0.273 | 1 | 16 |
| ppih | 0.42058 | -0.17944957 | 0.171 | 0.318 | 1 | 16 |
| ppm1g | 0.42072 | 0.167035206 | 0.371 | 0.227 | 1 | 16 |
| kmt2bb | 0.42092 | 0.189727671 | 0.4 | 0.318 | 1 | 16 |
| nenf | 0.421 | -0.20229847 | 0.171 | 0.273 | 1 | 16 |
| zgc:77929 | 0.42137 | 0.200804545 | 0.257 | 0.136 | 1 | 16 |
| chchd4a | 0.42162 | 0.174476599 | 0.114 | 0.045 | 1 | 16 |
| CABZ01087560.1 | 0.42185 | 0.161405679 | 0.257 | 0.182 | 1 | 16 |
| pmm2 | 0.42186 | -0.168205 | 0.086 | 0.182 | 1 | 16 |
| CR318588.3 | 0.42193 | -0.10657406 | 0.657 | 0.636 | 1 | 16 |

| | | | | | | |
|---|---|---|---|---|---|---|
| arpc3 | 0.42201 | 0.211956436 | 0.714 | 0.545 | 1 | 16 |
| reep3a | 0.42219 | 0.19220112 | 0.143 | 0.045 | 1 | 16 |
| anp32a | 0.42277 | -0.24353088 | 0.371 | 0.545 | 1 | 16 |
| arhgap35a | 0.42288 | -0.2249918 | 0.143 | 0.273 | 1 | 16 |
| ube2e3 | 0.42302 | -0.15323986 | 0.543 | 0.591 | 1 | 16 |
| amfra | 0.4236 | -0.21123809 | 0.086 | 0.182 | 1 | 16 |
| impad1 | 0.42566 | -0.12230248 | 0.086 | 0.182 | 1 | 16 |
| btd | 0.42692 | 0.180865 | 0.143 | 0.045 | 1 | 16 |
| igf2b | 0.42738 | 0.278279404 | 0.371 | 0.227 | 1 | 16 |
| ppid | 0.42835 | -0.20402968 | 0.171 | 0.318 | 1 | 16 |
| pmp22b | 0.4285 | -0.22066431 | 0.114 | 0.227 | 1 | 16 |
| nip7 | 0.42972 | 0.14635873 | 0.171 | 0.136 | 1 | 16 |
| crip2 | 0.42979 | 0.184403437 | 0.143 | 0.045 | 1 | 16 |
| zyx | 0.43034 | -0.17999135 | 0.143 | 0.273 | 1 | 16 |
| si:ch1073-429i10.3.1 | 0.43058 | 0.119184349 | 0.657 | 0.636 | 1 | 16 |
| zgc:175088 | 0.43082 | 0.203971788 | 0.629 | 0.455 | 1 | 16 |
| eno1a | 0.43086 | 0.203487948 | 0.229 | 0.136 | 1 | 16 |
| srsf10b | 0.43091 | -0.19764038 | 0.114 | 0.227 | 1 | 16 |
| nup50 | 0.43118 | -0.20583419 | 0.114 | 0.227 | 1 | 16 |
| atp5pf | 0.43142 | 0.192040296 | 0.743 | 0.682 | 1 | 16 |
| tardbpl | 0.43152 | 0.173509242 | 0.371 | 0.227 | 1 | 16 |
| lrrc59 | 0.43193 | 0.223973405 | 0.286 | 0.182 | 1 | 16 |
| calm2a | 0.43237 | -0.18747037 | 0.286 | 0.364 | 1 | 16 |
| fsd1l | 0.43238 | -0.15822437 | 0.057 | 0.136 | 1 | 16 |
| zfand6 | 0.43277 | 0.187438954 | 0.143 | 0.045 | 1 | 16 |
| lrp1ab | 0.43281 | -0.18144966 | 0.114 | 0.182 | 1 | 16 |
| naa50 | 0.43283 | -0.23152678 | 0.114 | 0.227 | 1 | 16 |
| zcchc17 | 0.43303 | 0.171657308 | 0.143 | 0.045 | 1 | 16 |
| puf60a | 0.43319 | -0.22073363 | 0.286 | 0.455 | 1 | 16 |
| cul3b | 0.43334 | 0.196956605 | 0.143 | 0.045 | 1 | 16 |
| ncam1a | 0.43382 | -0.22868191 | 0.114 | 0.227 | 1 | 16 |
| ist1 | 0.43425 | -0.10670584 | 0.171 | 0.273 | 1 | 16 |
| sssca1 | 0.43425 | 0.209892592 | 0.257 | 0.136 | 1 | 16 |
| erbb2 | 0.43442 | -0.22439292 | 0.114 | 0.227 | 1 | 16 |
| rbpjb | 0.43443 | 0.167395979 | 0.143 | 0.091 | 1 | 16 |
| taf13 | 0.43455 | 0.232926298 | 0.571 | 0.409 | 1 | 16 |
| uqcrfs1 | 0.43526 | 0.170460002 | 0.571 | 0.5 | 1 | 16 |
| got2b | 0.43533 | 0.166290364 | 0.143 | 0.045 | 1 | 16 |
| zfhx3 | 0.43555 | 0.186460903 | 0.143 | 0.045 | 1 | 16 |
| smim19 | 0.43556 | 0.181790824 | 0.143 | 0.045 | 1 | 16 |
| cox4i1 | 0.43641 | 0.199582842 | 0.829 | 0.727 | 1 | 16 |
| aldh1l1 | 0.43691 | -0.11412322 | 0.057 | 0.091 | 1 | 16 |
| si:ch211-262i1.6 | 0.43704 | -0.20866499 | 0.171 | 0.273 | 1 | 16 |
| rab18a | 0.43706 | 0.19678666 | 0.143 | 0.045 | 1 | 16 |
| ccdc47 | 0.43714 | 0.120667821 | 0.171 | 0.136 | 1 | 16 |
| raf1b | 0.43753 | 0.139773318 | 0.143 | 0.045 | 1 | 16 |
| COX5B | 0.43779 | 0.129423594 | 0.771 | 0.773 | 1 | 16 |
| sulf1 | 0.43815 | -0.21987037 | 0.171 | 0.318 | 1 | 16 |

| | | | | | | |
|---|---|---|---|---|---|---|
| eci1 | 0.43824 | 0.237437932 | 0.257 | 0.136 | 1 | 16 |
| ywhae2 | 0.43865 | -0.23179472 | 0.171 | 0.318 | 1 | 16 |
| sept9b | 0.43904 | -0.12596769 | 0.057 | 0.091 | 1 | 16 |
| hcfc1a | 0.43909 | -0.14003849 | 0.171 | 0.227 | 1 | 16 |
| CR847566.1 | 0.43988 | 0.202370132 | 0.114 | 0.045 | 1 | 16 |
| pgrmc2 | 0.44029 | -0.22667525 | 0.171 | 0.318 | 1 | 16 |
| ip6k2a | 0.4403 | -0.17962106 | 0.171 | 0.227 | 1 | 16 |
| wdr5 | 0.44054 | -0.13251353 | 0.086 | 0.136 | 1 | 16 |
| arhgap1 | 0.44087 | 0.163047591 | 0.143 | 0.045 | 1 | 16 |
| b4galt1l | 0.44098 | 0.192501911 | 0.143 | 0.045 | 1 | 16 |
| rnf130 | 0.44142 | 0.103230359 | 0.486 | 0.364 | 1 | 16 |
| bmp1a | 0.44152 | -0.18148266 | 0.114 | 0.182 | 1 | 16 |
| pop7 | 0.44178 | -0.19820312 | 0.171 | 0.318 | 1 | 16 |
| ctcf | 0.44179 | 0.16481418 | 0.143 | 0.045 | 1 | 16 |
| tpm4a | 0.44205 | -0.24116962 | 0.514 | 0.682 | 1 | 16 |
| ndufa2 | 0.44221 | -0.22707878 | 0.314 | 0.455 | 1 | 16 |
| med4 | 0.44223 | -0.23785127 | 0.171 | 0.318 | 1 | 16 |
| slc29a1a | 0.44318 | -0.26263117 | 0.257 | 0.409 | 1 | 16 |
| agpat5 | 0.44336 | -0.15833356 | 0.057 | 0.136 | 1 | 16 |
| jtb | 0.44339 | -0.10286799 | 0.057 | 0.136 | 1 | 16 |
| eif4g2a | 0.44349 | 0.235187896 | 0.371 | 0.227 | 1 | 16 |
| dynll2b | 0.44359 | 0.212600662 | 0.629 | 0.5 | 1 | 16 |
| anapc15 | 0.44514 | 0.159892755 | 0.2 | 0.136 | 1 | 16 |
| ldhba | 0.44543 | -0.22453618 | 0.314 | 0.455 | 1 | 16 |
| rrs1 | 0.44546 | -0.2300075 | 0.143 | 0.273 | 1 | 16 |
| actr10 | 0.44554 | 0.177744251 | 0.143 | 0.045 | 1 | 16 |
| wdr82 | 0.44596 | -0.10710653 | 0.086 | 0.182 | 1 | 16 |
| grcc10 | 0.4464 | -0.2242314 | 0.171 | 0.318 | 1 | 16 |
| rbm7 | 0.44644 | 0.1923456 | 0.2 | 0.091 | 1 | 16 |
| dpm3 | 0.44651 | -0.20380487 | 0.171 | 0.318 | 1 | 16 |
| zdhhc5a | 0.4469 | 0.20515718 | 0.143 | 0.045 | 1 | 16 |
| stk35l | 0.44722 | 0.128717414 | 0.257 | 0.227 | 1 | 16 |
| higd2a | 0.44737 | 0.197970494 | 0.2 | 0.091 | 1 | 16 |
| gapdhs | 0.44778 | -0.13584367 | 0.571 | 0.591 | 1 | 16 |
| pdgfra | 0.4486 | 0.140056122 | 0.143 | 0.045 | 1 | 16 |
| sf3b6 | 0.44916 | -0.14093812 | 0.571 | 0.727 | 1 | 16 |
| arf6b | 0.44964 | -0.2251477 | 0.343 | 0.5 | 1 | 16 |
| s100v2 | 0.44991 | -0.2127369 | 0.543 | 0.682 | 1 | 16 |
| LOC103908925 | 0.45058 | 0.159780722 | 0.143 | 0.045 | 1 | 16 |
| klf3 | 0.45071 | 0.193372219 | 0.143 | 0.045 | 1 | 16 |
| col4a6 | 0.45171 | 0.127174515 | 0.486 | 0.455 | 1 | 16 |
| got1 | 0.45178 | -0.11893859 | 0.086 | 0.182 | 1 | 16 |
| cflara | 0.45181 | -0.19574903 | 0.171 | 0.273 | 1 | 16 |
| map2k2a | 0.45238 | -0.20527838 | 0.114 | 0.227 | 1 | 16 |
| btg2 | 0.45256 | 0.162949114 | 0.257 | 0.136 | 1 | 16 |
| cenpx | 0.45257 | -0.11337089 | 0.086 | 0.182 | 1 | 16 |
| snapc5 | 0.45308 | 0.14159862 | 0.143 | 0.045 | 1 | 16 |
| aldh9a1a.1 | 0.45311 | 0.204793815 | 0.6 | 0.545 | 1 | 16 |

| | | | | | | |
|---|---|---|---|---|---|---|
| pxmp2 | 0.45417 | -0.10954273 | 0.057 | 0.136 | 1 | 16 |
| usp8 | 0.45426 | -0.23198281 | 0.143 | 0.273 | 1 | 16 |
| cox6a2 | 0.45435 | 0.188620487 | 0.2 | 0.091 | 1 | 16 |
| zgc:153675 | 0.45439 | 0.168657942 | 0.143 | 0.045 | 1 | 16 |
| asb8 | 0.45447 | 0.16209311 | 0.143 | 0.045 | 1 | 16 |
| b3gnt2l | 0.45456 | 0.193974406 | 0.143 | 0.045 | 1 | 16 |
| si:ch211-196l7.4 | 0.45462 | 0.110551917 | 0.6 | 0.591 | 1 | 16 |
| sec61b | 0.45479 | -0.13572701 | 0.8 | 0.818 | 1 | 16 |
| pinx1 | 0.45488 | -0.20887865 | 0.286 | 0.409 | 1 | 16 |
| fkbp1ab | 0.4549 | 0.214461277 | 0.171 | 0.091 | 1 | 16 |
| fbn2b | 0.45511 | 0.240387395 | 0.343 | 0.227 | 1 | 16 |
| nktr | 0.45521 | 0.149884475 | 0.2 | 0.091 | 1 | 16 |
| ing3 | 0.45524 | 0.174851193 | 0.143 | 0.045 | 1 | 16 |
| zgc:194246 | 0.45532 | 0.162918252 | 0.171 | 0.091 | 1 | 16 |
| gnb2 | 0.45541 | 0.192141164 | 0.143 | 0.045 | 1 | 16 |
| triqk | 0.45556 | 0.179686877 | 0.2 | 0.136 | 1 | 16 |
| zgc:77262 | 0.45581 | -0.13017101 | 0.086 | 0.182 | 1 | 16 |
| hnrnpl | 0.45582 | -0.15155233 | 0.229 | 0.364 | 1 | 16 |
| lrrc1 | 0.45606 | 0.159874854 | 0.143 | 0.045 | 1 | 16 |
| zgc:112052 | 0.45651 | -0.14544313 | 0.057 | 0.136 | 1 | 16 |
| plod3 | 0.45668 | -0.1905493 | 0.143 | 0.273 | 1 | 16 |
| tmem205 | 0.45786 | -0.18517405 | 0.086 | 0.182 | 1 | 16 |
| macf1a | 0.45795 | 0.188977893 | 0.143 | 0.045 | 1 | 16 |
| ndufv3 | 0.45799 | 0.153314111 | 0.286 | 0.227 | 1 | 16 |
| zgc:171686 | 0.45848 | -0.17029494 | 0.114 | 0.227 | 1 | 16 |
| nop16 | 0.45884 | -0.17111145 | 0.114 | 0.227 | 1 | 16 |
| eif5a2 | 0.45898 | 0.123753405 | 0.886 | 0.864 | 1 | 16 |
| amdhd2 | 0.45904 | -0.18310053 | 0.114 | 0.227 | 1 | 16 |
| fubp1 | 0.45939 | 0.233945723 | 0.429 | 0.273 | 1 | 16 |
| XLOC-005605 | 0.46022 | -0.20154007 | 0.629 | 0.727 | 1 | 16 |
| farsb | 0.46031 | -0.16517888 | 0.114 | 0.182 | 1 | 16 |
| ddx3xb | 0.46044 | 0.118159617 | 0.257 | 0.227 | 1 | 16 |
| cntn5 | 0.46076 | 0.165845317 | 0.114 | 0.045 | 1 | 16 |
| sox9a | 0.46091 | 0.191140783 | 0.343 | 0.273 | 1 | 16 |
| pank2 | 0.46093 | 0.176952746 | 0.114 | 0.045 | 1 | 16 |
| cxxc5a | 0.46126 | 0.185636475 | 0.486 | 0.409 | 1 | 16 |
| tmem256 | 0.46127 | 0.176948899 | 0.371 | 0.227 | 1 | 16 |
| foxl2a | 0.46191 | -0.11227198 | 0.343 | 0.364 | 1 | 16 |
| zgc:162939 | 0.46195 | 0.151302737 | 0.143 | 0.045 | 1 | 16 |
| boc | 0.46197 | -0.17430997 | 0.114 | 0.227 | 1 | 16 |
| tram1 | 0.46222 | 0.190606202 | 0.314 | 0.227 | 1 | 16 |
| KIF2A | 0.46265 | 0.148022672 | 0.143 | 0.045 | 1 | 16 |
| pes | 0.46285 | -0.22075495 | 0.143 | 0.273 | 1 | 16 |
| sec22ba | 0.46293 | -0.16081696 | 0.2 | 0.273 | 1 | 16 |
| arl14ep | 0.46306 | 0.173846643 | 0.314 | 0.182 | 1 | 16 |
| kmt5b | 0.46319 | 0.170500665 | 0.143 | 0.045 | 1 | 16 |
| hpgd | 0.46326 | 0.148650304 | 0.143 | 0.045 | 1 | 16 |
| FAM53C | 0.46359 | 0.161638101 | 0.143 | 0.045 | 1 | 16 |

| | | | | | | |
|---|---|---|---|---|---|---|
| il15 | 0.46366 | 0.135339046 | 0.314 | 0.273 | 1 | 16 |
| ndufs5 | 0.46381 | -0.16051195 | 0.543 | 0.591 | 1 | 16 |
| sf3a1 | 0.46426 | 0.17522777 | 0.171 | 0.091 | 1 | 16 |
| cldn5a | 0.4649 | -0.11084055 | 0.057 | 0.136 | 1 | 16 |
| si:dkey-23k10.5 | 0.46497 | -0.11860171 | 0.057 | 0.136 | 1 | 16 |
| nol11 | 0.46497 | -0.10240214 | 0.057 | 0.136 | 1 | 16 |
| si:ch211-117c9.5 | 0.46522 | 0.224990954 | 0.629 | 0.5 | 1 | 16 |
| rsrc2 | 0.46523 | -0.13128025 | 0.2 | 0.318 | 1 | 16 |
| ptk7a | 0.46546 | -0.1325009 | 0.086 | 0.182 | 1 | 16 |
| zgc:92664 | 0.46562 | -0.15166373 | 0.114 | 0.227 | 1 | 16 |
| si:ch211-160o17.2 | 0.4658 | 0.190069986 | 0.143 | 0.045 | 1 | 16 |
| nosip | 0.46589 | 0.209339377 | 0.2 | 0.091 | 1 | 16 |
| vdac3 | 0.46589 | 0.216764226 | 0.571 | 0.409 | 1 | 16 |
| ago4 | 0.46717 | 0.180237602 | 0.143 | 0.045 | 1 | 16 |
| zgc:77158 | 0.46718 | 0.170254247 | 0.143 | 0.045 | 1 | 16 |
| spns1 | 0.46727 | 0.137283282 | 0.143 | 0.045 | 1 | 16 |
| cad | 0.46734 | 0.200669063 | 0.143 | 0.045 | 1 | 16 |
| exoc6 | 0.46735 | 0.117499049 | 0.114 | 0.091 | 1 | 16 |
| polr3glb | 0.46782 | 0.159431784 | 0.143 | 0.045 | 1 | 16 |
| pxk | 0.46813 | -0.11183969 | 0.057 | 0.136 | 1 | 16 |
| lims1 | 0.46829 | 0.172666793 | 0.143 | 0.045 | 1 | 16 |
| srsf6b | 0.46835 | -0.2095558 | 0.171 | 0.273 | 1 | 16 |
| chmp4bb | 0.46904 | 0.164961733 | 0.2 | 0.136 | 1 | 16 |
| si:ch211-162e15.3 | 0.46931 | 0.159213917 | 0.143 | 0.045 | 1 | 16 |
| limk2 | 0.46974 | 0.138469362 | 0.143 | 0.045 | 1 | 16 |
| ube2e1 | 0.46989 | -0.11822857 | 0.429 | 0.455 | 1 | 16 |
| ubtf | 0.47009 | 0.197363424 | 0.2 | 0.091 | 1 | 16 |
| zswim8 | 0.47025 | 0.184997127 | 0.2 | 0.091 | 1 | 16 |
| aida | 0.47025 | 0.140766852 | 0.143 | 0.045 | 1 | 16 |
| snrpb | 0.47073 | 0.19268039 | 0.771 | 0.636 | 1 | 16 |
| soga1 | 0.47084 | 0.186063668 | 0.2 | 0.091 | 1 | 16 |
| tgfb1a | 0.4711 | 0.180628513 | 0.114 | 0.045 | 1 | 16 |
| thap11 | 0.47118 | 0.163667241 | 0.143 | 0.045 | 1 | 16 |
| ythdf2 | 0.47135 | -0.17595064 | 0.057 | 0.136 | 1 | 16 |
| pam | 0.4715 | -0.20490034 | 0.2 | 0.273 | 1 | 16 |
| hdac4 | 0.47163 | 0.24206301 | 0.2 | 0.091 | 1 | 16 |
| srsf3b | 0.47168 | -0.19965642 | 0.514 | 0.636 | 1 | 16 |
| hmga2 | 0.47203 | 0.222238333 | 0.143 | 0.045 | 1 | 16 |
| setd2 | 0.47234 | 0.140610059 | 0.143 | 0.045 | 1 | 16 |
| stk38b | 0.4728 | 0.195366897 | 0.143 | 0.045 | 1 | 16 |
| med20 | 0.47312 | 0.150714669 | 0.143 | 0.045 | 1 | 16 |
| wdr13 | 0.47368 | 0.163106797 | 0.114 | 0.045 | 1 | 16 |
| rnf170 | 0.47381 | 0.155384148 | 0.143 | 0.045 | 1 | 16 |
| npm1b | 0.47393 | 0.164489422 | 0.171 | 0.091 | 1 | 16 |
| arcn1a | 0.47396 | 0.182211479 | 0.143 | 0.045 | 1 | 16 |
| cyfip1 | 0.47423 | 0.148671625 | 0.286 | 0.227 | 1 | 16 |
| idh3g | 0.47436 | -0.16598584 | 0.143 | 0.273 | 1 | 16 |
| rps27.2 | 0.4751 | -0.10313657 | 0.857 | 0.955 | 1 | 16 |

| | | | | | | |
|---|---|---|---|---|---|---|
| gpx1b | 0.4752 | 0.272329849 | 0.543 | 0.455 | 1 | 16 |
| ormdl2 | 0.47522 | 0.152883499 | 0.143 | 0.045 | 1 | 16 |
| zgc:63882 | 0.47526 | 0.159153697 | 0.257 | 0.136 | 1 | 16 |
| ccng2 | 0.47595 | 0.165209727 | 0.143 | 0.045 | 1 | 16 |
| FP074902.2 | 0.47608 | 0.166924321 | 0.143 | 0.045 | 1 | 16 |
| sccpdhb | 0.4761 | 0.151022114 | 0.143 | 0.045 | 1 | 16 |
| zgc:193541 | 0.47632 | -0.12084246 | 0.857 | 0.864 | 1 | 16 |
| echdc2 | 0.47634 | 0.133770069 | 0.143 | 0.045 | 1 | 16 |
| nras | 0.47663 | 0.16190889 | 0.143 | 0.045 | 1 | 16 |
| iah1 | 0.47665 | -0.16918911 | 0.057 | 0.136 | 1 | 16 |
| zgc:112982 | 0.47668 | 0.164864249 | 0.143 | 0.045 | 1 | 16 |
| axin1 | 0.4767 | -0.2168233 | 0.143 | 0.273 | 1 | 16 |
| robo3 | 0.47671 | 0.170564905 | 0.143 | 0.045 | 1 | 16 |
| zgc:175214 | 0.4772 | 0.171993272 | 0.114 | 0.045 | 1 | 16 |
| col4a1 | 0.47757 | -0.13478069 | 0.286 | 0.409 | 1 | 16 |
| vkorc1l1 | 0.47783 | -0.19434001 | 0.114 | 0.227 | 1 | 16 |
| sf3b4 | 0.47794 | -0.19092265 | 0.143 | 0.273 | 1 | 16 |
| trappc2l | 0.47818 | 0.226071463 | 0.257 | 0.136 | 1 | 16 |
| baxa | 0.47846 | -0.15968884 | 0.114 | 0.227 | 1 | 16 |
| arhgap17a | 0.47857 | 0.140983697 | 0.086 | 0.045 | 1 | 16 |
| araf | 0.47879 | 0.107139924 | 0.171 | 0.091 | 1 | 16 |
| pdk4 | 0.47919 | -0.11452417 | 0.057 | 0.091 | 1 | 16 |
| pitpnc1b | 0.4797 | 0.232021104 | 0.2 | 0.091 | 1 | 16 |
| ERC1 | 0.47994 | -0.10456641 | 0.057 | 0.136 | 1 | 16 |
| phyh | 0.48047 | 0.133749992 | 0.143 | 0.091 | 1 | 16 |
| cdk11b | 0.48075 | -0.17690488 | 0.086 | 0.182 | 1 | 16 |
| nr5a1b | 0.48085 | 0.207715527 | 0.2 | 0.091 | 1 | 16 |
| ahnak | 0.48085 | -0.22718752 | 0.343 | 0.5 | 1 | 16 |
| mphosph6 | 0.48134 | -0.10163318 | 0.171 | 0.273 | 1 | 16 |
| ebag9 | 0.48134 | 0.213100646 | 0.371 | 0.227 | 1 | 16 |
| si:ch211-74m13.3 | 0.48148 | -0.19362929 | 0.086 | 0.182 | 1 | 16 |
| cox19 | 0.48162 | -0.11902553 | 0.143 | 0.182 | 1 | 16 |
| ppp2r5ea | 0.48194 | -0.16212808 | 0.114 | 0.227 | 1 | 16 |
| riok1 | 0.4821 | -0.10757425 | 0.057 | 0.136 | 1 | 16 |
| jund | 0.48292 | -0.21078333 | 0.114 | 0.227 | 1 | 16 |
| sumo3b | 0.48308 | -0.1501765 | 0.086 | 0.136 | 1 | 16 |
| dchs1a | 0.4831 | -0.20594503 | 0.086 | 0.182 | 1 | 16 |
| elavl4 | 0.48412 | 0.199286398 | 0.171 | 0.091 | 1 | 16 |
| anxa13 | 0.48539 | -0.14877777 | 0.314 | 0.455 | 1 | 16 |
| ift172 | 0.48562 | -0.16830778 | 0.057 | 0.136 | 1 | 16 |
| lmbrd1 | 0.48608 | 0.187263769 | 0.2 | 0.091 | 1 | 16 |
| gadd45ba | 0.48608 | -0.13600355 | 0.2 | 0.227 | 1 | 16 |
| si:ch211-108c6.2 | 0.48623 | -0.1754103 | 0.057 | 0.136 | 1 | 16 |
| rcor1 | 0.48637 | -0.17928447 | 0.143 | 0.273 | 1 | 16 |
| fkbp7 | 0.48658 | -0.22114728 | 0.257 | 0.409 | 1 | 16 |
| hnrnph1l | 0.48662 | 0.193229345 | 0.371 | 0.227 | 1 | 16 |
| b4galt6 | 0.48688 | 0.179551924 | 0.343 | 0.273 | 1 | 16 |
| nedd8 | 0.48699 | -0.16102366 | 0.229 | 0.364 | 1 | 16 |

| | | | | | | |
|---|---|---|---|---|---|---|
| casc3 | 0.48761 | -0.14096854 | 0.086 | 0.182 | 1 | 16 |
| lsm1 | 0.4878 | -0.20707225 | 0.114 | 0.227 | 1 | 16 |
| ube2ia | 0.48785 | -0.23267291 | 0.314 | 0.455 | 1 | 16 |
| gna13b | 0.48792 | -0.11332644 | 0.057 | 0.136 | 1 | 16 |
| tm2d2 | 0.48802 | 0.21599357 | 0.257 | 0.136 | 1 | 16 |
| zgc:113411 | 0.48828 | -0.14325213 | 0.171 | 0.227 | 1 | 16 |
| pxnb | 0.48886 | 0.184004864 | 0.171 | 0.091 | 1 | 16 |
| zgc:92140 | 0.48913 | 0.231313319 | 0.457 | 0.318 | 1 | 16 |
| srrm1 | 0.48921 | 0.178045247 | 0.257 | 0.136 | 1 | 16 |
| slc35f6 | 0.48942 | 0.150725424 | 0.2 | 0.091 | 1 | 16 |
| cryl1 | 0.48977 | -0.13699286 | 0.086 | 0.136 | 1 | 16 |
| hspa8 | 0.48977 | -0.13172577 | 0.886 | 0.955 | 1 | 16 |
| zgc:153867 | 0.48998 | 0.146528328 | 0.857 | 0.727 | 1 | 16 |
| rbm34 | 0.49039 | -0.19093746 | 0.086 | 0.182 | 1 | 16 |
| tvp23b | 0.49049 | -0.20425663 | 0.143 | 0.273 | 1 | 16 |
| ctsa | 0.49051 | -0.14961049 | 0.314 | 0.455 | 1 | 16 |
| hspd1 | 0.49057 | 0.170113384 | 0.743 | 0.591 | 1 | 16 |
| ralaa | 0.49084 | -0.19358035 | 0.143 | 0.273 | 1 | 16 |
| smarca2 | 0.49085 | -0.22218667 | 0.143 | 0.273 | 1 | 16 |
| psmf1 | 0.49085 | -0.20685535 | 0.143 | 0.273 | 1 | 16 |
| asna1 | 0.49087 | -0.15306338 | 0.114 | 0.182 | 1 | 16 |
| prkar2aa | 0.49152 | -0.10010057 | 0.743 | 0.864 | 1 | 16 |
| mri1 | 0.4921 | -0.13408594 | 0.086 | 0.182 | 1 | 16 |
| prkar1ab | 0.49274 | 0.185052379 | 0.114 | 0.045 | 1 | 16 |
| zgc:174906 | 0.49334 | 0.192071864 | 0.2 | 0.091 | 1 | 16 |
| micall1a | 0.49345 | -0.15148806 | 0.057 | 0.136 | 1 | 16 |
| cox6c | 0.4937 | -0.18396551 | 0.6 | 0.682 | 1 | 16 |
| pcyox1 | 0.4939 | 0.172086833 | 0.457 | 0.364 | 1 | 16 |
| hnrnph1 | 0.49458 | -0.11472375 | 0.057 | 0.091 | 1 | 16 |
| tle2a | 0.4946 | -0.18977018 | 0.114 | 0.227 | 1 | 16 |
| manf | 0.49461 | -0.2207019 | 0.343 | 0.5 | 1 | 16 |
| rbm14b | 0.49479 | -0.13212336 | 0.086 | 0.136 | 1 | 16 |
| taf7 | 0.4948 | -0.10175699 | 0.171 | 0.273 | 1 | 16 |
| akt1s1 | 0.49492 | 0.18488305 | 0.257 | 0.136 | 1 | 16 |
| abhd10b | 0.49614 | -0.18112611 | 0.114 | 0.227 | 1 | 16 |
| utp3 | 0.49621 | 0.166955998 | 0.171 | 0.091 | 1 | 16 |
| dohh | 0.49649 | -0.13900999 | 0.114 | 0.227 | 1 | 16 |
| tial1 | 0.49657 | 0.158048121 | 0.171 | 0.091 | 1 | 16 |
| trmt112 | 0.49658 | -0.18429781 | 0.343 | 0.5 | 1 | 16 |
| commd3 | 0.49667 | 0.213597749 | 0.371 | 0.227 | 1 | 16 |
| march6 | 0.4973 | 0.207571142 | 0.2 | 0.091 | 1 | 16 |
| pfdn2 | 0.49737 | 0.210751348 | 0.314 | 0.182 | 1 | 16 |
| fgfr1op2 | 0.49762 | 0.178112863 | 0.2 | 0.091 | 1 | 16 |
| hmgb2a | 0.4978 | -0.19538525 | 0.629 | 0.773 | 1 | 16 |
| abhd13 | 0.49802 | 0.187800873 | 0.2 | 0.091 | 1 | 16 |
| slc3a2b | 0.49865 | -0.23871907 | 0.229 | 0.364 | 1 | 16 |
| itgav | 0.49885 | -0.15422272 | 0.114 | 0.182 | 1 | 16 |
| gpc4 | 0.49906 | 0.150530367 | 0.143 | 0.091 | 1 | 16 |

| | | | | | | |
|---|---|---|---|---|---|---|
| megf6a | 0.49944 | -0.13992792 | 0.086 | 0.182 | 1 | 16 |
| glg1b | 0.49956 | 0.149926511 | 0.114 | 0.045 | 1 | 16 |
| zgc:77056 | 0.49963 | 0.162596784 | 0.257 | 0.182 | 1 | 16 |
| eef2b | 0.49978 | 0.166162306 | 0.886 | 0.818 | 1 | 16 |
| vps25 | 0.49985 | -0.15941494 | 0.114 | 0.227 | 1 | 16 |
| khsrp | 0.49997 | -0.22173941 | 0.229 | 0.364 | 1 | 16 |
| sf3a2 | 0.50009 | -0.21510205 | 0.343 | 0.5 | 1 | 16 |
| prr12a | 0.50025 | -0.10195949 | 0.143 | 0.227 | 1 | 16 |
| emc10 | 0.50034 | -0.10535059 | 0.486 | 0.5 | 1 | 16 |
| dhrs4 | 0.50037 | -0.14688225 | 0.114 | 0.182 | 1 | 16 |
| eif3s10 | 0.5006 | -0.18668365 | 0.343 | 0.5 | 1 | 16 |
| zgc:162816 | 0.50065 | -0.17159817 | 0.114 | 0.227 | 1 | 16 |
| adssl | 0.5007 | -0.20268649 | 0.343 | 0.455 | 1 | 16 |
| cpsf2 | 0.50171 | -0.12893245 | 0.086 | 0.182 | 1 | 16 |
| coq10b | 0.50185 | 0.20558995 | 0.257 | 0.136 | 1 | 16 |
| prrc2c | 0.50236 | 0.183283423 | 0.171 | 0.091 | 1 | 16 |
| zmynd11 | 0.50242 | 0.194361134 | 0.2 | 0.091 | 1 | 16 |
| wu:fb16f03 | 0.50244 | -0.2198004 | 0.171 | 0.273 | 1 | 16 |
| zgc:163080 | 0.50264 | -0.18058668 | 0.229 | 0.318 | 1 | 16 |
| aimp2 | 0.50297 | 0.130956738 | 0.457 | 0.409 | 1 | 16 |
| cyb5r1 | 0.50401 | -0.18520773 | 0.143 | 0.227 | 1 | 16 |
| cplx2 | 0.50402 | -0.18485107 | 0.086 | 0.182 | 1 | 16 |
| lclat1 | 0.5049 | -0.11011783 | 0.057 | 0.136 | 1 | 16 |
| timm8a | 0.50516 | -0.11107943 | 0.143 | 0.182 | 1 | 16 |
| atp5pd | 0.50528 | -0.16910857 | 0.686 | 0.818 | 1 | 16 |
| sept6 | 0.50554 | -0.18261796 | 0.143 | 0.227 | 1 | 16 |
| hspb8 | 0.50637 | -0.11172413 | 0.114 | 0.136 | 1 | 16 |
| zgc:110783 | 0.50653 | -0.1293431 | 0.086 | 0.136 | 1 | 16 |
| zgc:158482 | 0.50686 | 0.195528189 | 0.257 | 0.136 | 1 | 16 |
| nap1l4a | 0.50695 | 0.161079476 | 0.257 | 0.136 | 1 | 16 |
| tspan4a | 0.5077 | -0.18097829 | 0.057 | 0.136 | 1 | 16 |
| casp8 | 0.50772 | -0.10992746 | 0.057 | 0.091 | 1 | 16 |
| rab14l | 0.50808 | 0.146180466 | 0.114 | 0.045 | 1 | 16 |
| cct8 | 0.50874 | 0.223756668 | 0.514 | 0.364 | 1 | 16 |
| col4a3bpb | 0.50898 | -0.10825811 | 0.057 | 0.091 | 1 | 16 |
| si:dkey-89b17.4 | 0.50911 | 0.191992478 | 0.2 | 0.091 | 1 | 16 |
| mrps7 | 0.50959 | 0.173307871 | 0.229 | 0.136 | 1 | 16 |
| vapal | 0.50969 | 0.219612495 | 0.314 | 0.182 | 1 | 16 |
| vat1 | 0.51007 | -0.21153672 | 0.486 | 0.636 | 1 | 16 |
| cbr1 | 0.51049 | 0.155866905 | 0.114 | 0.045 | 1 | 16 |
| irf2bp2a | 0.51057 | -0.2166018 | 0.286 | 0.409 | 1 | 16 |
| vrk1 | 0.51067 | -0.17414739 | 0.086 | 0.182 | 1 | 16 |
| rn7sk | 0.51139 | 0.296518568 | 0.457 | 0.318 | 1 | 16 |
| uxs1 | 0.51156 | 0.211505339 | 0.371 | 0.227 | 1 | 16 |
| utp18 | 0.51204 | -0.11956799 | 0.057 | 0.091 | 1 | 16 |
| kti12 | 0.5125 | 0.1461284 | 0.114 | 0.045 | 1 | 16 |
| UBL4A | 0.51254 | -0.18310745 | 0.114 | 0.227 | 1 | 16 |
| pspc1 | 0.51273 | 0.162967502 | 0.2 | 0.091 | 1 | 16 |

| | | | | | | |
|---|---|---|---|---|---|---|
| phf3 | 0.51311 | 0.134541594 | 0.086 | 0.045 | 1 | 16 |
| atrip | 0.51322 | -0.1761887 | 0.057 | 0.136 | 1 | 16 |
| si:ch211-194c3.5 | 0.51326 | -0.18863477 | 0.057 | 0.136 | 1 | 16 |
| si:ch211-241b2.1 | 0.51353 | -0.19258434 | 0.114 | 0.227 | 1 | 16 |
| zgc:158463 | 0.51377 | -0.20245388 | 0.4 | 0.455 | 1 | 16 |
| trmt6 | 0.51422 | -0.18216052 | 0.086 | 0.182 | 1 | 16 |
| hspa9 | 0.51511 | -0.2164316 | 0.2 | 0.318 | 1 | 16 |
| pgpep1 | 0.51533 | 0.173392961 | 0.2 | 0.091 | 1 | 16 |
| zgc:56525 | 0.51553 | -0.13249373 | 0.086 | 0.182 | 1 | 16 |
| sh3gl3b | 0.51566 | -0.1151211 | 0.057 | 0.136 | 1 | 16 |
| csnk1db | 0.516 | -0.16061443 | 0.2 | 0.273 | 1 | 16 |
| adamts9 | 0.51632 | -0.1053257 | 0.171 | 0.182 | 1 | 16 |
| tdrd3 | 0.51672 | -0.15998378 | 0.057 | 0.136 | 1 | 16 |
| atp5pb | 0.51702 | 0.173438033 | 0.829 | 0.727 | 1 | 16 |
| snx5 | 0.51702 | 0.167215969 | 0.257 | 0.136 | 1 | 16 |
| sema7a | 0.51715 | 0.201846071 | 0.257 | 0.182 | 1 | 16 |
| tomm40l | 0.51741 | 0.147438038 | 0.2 | 0.091 | 1 | 16 |
| slc25a16 | 0.51756 | 0.128164698 | 0.086 | 0.045 | 1 | 16 |
| CABZ01110881.1 | 0.51759 | 0.201705157 | 0.286 | 0.182 | 1 | 16 |
| oaz1b | 0.51769 | 0.126238545 | 0.943 | 0.909 | 1 | 16 |
| faub | 0.51795 | 0.190443176 | 0.2 | 0.091 | 1 | 16 |
| txndc5 | 0.51804 | -0.15454993 | 0.2 | 0.318 | 1 | 16 |
| CABZ01087514.1 | 0.5181 | 0.100431101 | 0.114 | 0.045 | 1 | 16 |
| mplkip | 0.51854 | -0.15707737 | 0.057 | 0.136 | 1 | 16 |
| nomo | 0.51889 | -0.15396753 | 0.057 | 0.136 | 1 | 16 |
| mycbp2 | 0.51918 | -0.16154419 | 0.114 | 0.227 | 1 | 16 |
| fbxo28 | 0.51926 | 0.179418605 | 0.171 | 0.091 | 1 | 16 |
| kif5ba | 0.51947 | -0.1386872 | 0.2 | 0.318 | 1 | 16 |
| tes | 0.51948 | 0.15310305 | 0.2 | 0.136 | 1 | 16 |
| prpf38a | 0.51966 | -0.16261275 | 0.114 | 0.227 | 1 | 16 |
| rpia | 0.51971 | 0.236533228 | 0.486 | 0.364 | 1 | 16 |
| gclm | 0.5198 | -0.16009505 | 0.114 | 0.182 | 1 | 16 |
| abhd4 | 0.51996 | 0.120058303 | 0.086 | 0.045 | 1 | 16 |
| calcoco1a | 0.52073 | 0.197017387 | 0.2 | 0.091 | 1 | 16 |
| lyar | 0.521 | -0.13666305 | 0.086 | 0.136 | 1 | 16 |
| zgc:92744 | 0.5212 | 0.15723369 | 0.771 | 0.727 | 1 | 16 |
| chmp4ba | 0.52127 | 0.148428533 | 0.171 | 0.091 | 1 | 16 |
| acbd3 | 0.52147 | 0.128951492 | 0.229 | 0.182 | 1 | 16 |
| slc25a36a | 0.52181 | 0.15650525 | 0.114 | 0.045 | 1 | 16 |
| ylpm1 | 0.52181 | 0.131966254 | 0.143 | 0.091 | 1 | 16 |
| wasla | 0.52186 | 0.17907044 | 0.2 | 0.091 | 1 | 16 |
| ndp | 0.52217 | -0.2092976 | 0.286 | 0.409 | 1 | 16 |
| ciapin1 | 0.52254 | -0.1616049 | 0.229 | 0.364 | 1 | 16 |
| polr2f | 0.52269 | -0.12398105 | 0.086 | 0.182 | 1 | 16 |
| nutf2l | 0.52281 | 0.148607468 | 0.2 | 0.091 | 1 | 16 |
| sirt1 | 0.52291 | 0.175266195 | 0.171 | 0.091 | 1 | 16 |
| oip5-as1 | 0.52295 | 0.195706907 | 0.2 | 0.091 | 1 | 16 |
| lnpep | 0.52328 | -0.16552367 | 0.086 | 0.136 | 1 | 16 |

| | | | | | | |
|---|---|---|---|---|---|---|
| arl6ip1 | 0.52372 | 0.1342608 | 0.686 | 0.545 | 1 | 16 |
| brms1 | 0.52382 | 0.100892597 | 0.114 | 0.045 | 1 | 16 |
| zgc:193812 | 0.52401 | 0.193366382 | 0.2 | 0.091 | 1 | 16 |
| tmco1 | 0.52405 | 0.178429904 | 0.2 | 0.091 | 1 | 16 |
| rbb4l | 0.52424 | -0.20178514 | 0.314 | 0.455 | 1 | 16 |
| park7 | 0.52426 | 0.178101755 | 0.171 | 0.091 | 1 | 16 |
| gtf2a2 | 0.52437 | 0.170237831 | 0.2 | 0.091 | 1 | 16 |
| glod5 | 0.52456 | -0.15168415 | 0.257 | 0.318 | 1 | 16 |
| ociad2 | 0.52463 | -0.18243409 | 0.114 | 0.227 | 1 | 16 |
| prpf40a | 0.52514 | 0.214661874 | 0.2 | 0.091 | 1 | 16 |
| tmtc3 | 0.52533 | -0.21046582 | 0.114 | 0.227 | 1 | 16 |
| slc39a6 | 0.52572 | 0.131411374 | 0.086 | 0.045 | 1 | 16 |
| pyroxd2 | 0.52576 | 0.146746884 | 0.114 | 0.045 | 1 | 16 |
| ppifb | 0.52591 | 0.134823678 | 0.2 | 0.091 | 1 | 16 |
| marcksa | 0.52594 | 0.219236896 | 0.2 | 0.091 | 1 | 16 |
| scamp3 | 0.52615 | 0.143025001 | 0.114 | 0.045 | 1 | 16 |
| stx12 | 0.52638 | -0.16891276 | 0.114 | 0.227 | 1 | 16 |
| mrto4 | 0.5265 | -0.17263054 | 0.114 | 0.227 | 1 | 16 |
| mrpl40 | 0.52651 | -0.15312672 | 0.086 | 0.182 | 1 | 16 |
| arpc5la | 0.52662 | -0.17157779 | 0.114 | 0.227 | 1 | 16 |
| cdkn2aipnl | 0.52666 | -0.11384961 | 0.171 | 0.273 | 1 | 16 |
| bzw2 | 0.52689 | -0.11461481 | 0.057 | 0.091 | 1 | 16 |
| rassf2a | 0.52706 | 0.164259818 | 0.171 | 0.091 | 1 | 16 |
| gemin2 | 0.52712 | -0.13068192 | 0.057 | 0.136 | 1 | 16 |
| bbip1 | 0.52734 | 0.188744229 | 0.286 | 0.182 | 1 | 16 |
| znf451 | 0.52762 | -0.15204955 | 0.086 | 0.182 | 1 | 16 |
| aff2 | 0.52778 | 0.174063433 | 0.2 | 0.091 | 1 | 16 |
| hmg20a | 0.52788 | 0.184953217 | 0.2 | 0.091 | 1 | 16 |
| rpz4 | 0.52895 | -0.1698525 | 0.143 | 0.227 | 1 | 16 |
| uqcrc1 | 0.52993 | 0.187994735 | 0.514 | 0.364 | 1 | 16 |
| pqbp1 | 0.53006 | -0.14912216 | 0.114 | 0.182 | 1 | 16 |
| josd2 | 0.53007 | -0.15586533 | 0.086 | 0.182 | 1 | 16 |
| aldh6a1 | 0.5301 | 0.160871201 | 0.257 | 0.136 | 1 | 16 |
| rest | 0.53136 | 0.142559576 | 0.114 | 0.045 | 1 | 16 |
| zgc:56106 | 0.53158 | -0.1063595 | 0.171 | 0.273 | 1 | 16 |
| mbd4 | 0.53174 | 0.154509649 | 0.114 | 0.045 | 1 | 16 |
| mao | 0.53191 | 0.122839275 | 0.086 | 0.045 | 1 | 16 |
| LEPROTL1 | 0.53197 | -0.11828931 | 0.371 | 0.409 | 1 | 16 |
| ppm1la | 0.53215 | 0.100767072 | 0.171 | 0.091 | 1 | 16 |
| mrpl34 | 0.53233 | 0.184942491 | 0.314 | 0.182 | 1 | 16 |
| tmed2 | 0.53243 | -0.17162099 | 0.629 | 0.727 | 1 | 16 |
| pgrmc1 | 0.53256 | 0.18180734 | 0.743 | 0.636 | 1 | 16 |
| syncripl | 0.53262 | -0.17132142 | 0.229 | 0.318 | 1 | 16 |
| tmem108 | 0.53281 | -0.10883147 | 0.057 | 0.136 | 1 | 16 |
| si:dkey-188i13.11 | 0.5334 | 0.198133613 | 0.657 | 0.545 | 1 | 16 |
| si:ch211-125a15.1 | 0.53345 | 0.113632133 | 0.086 | 0.045 | 1 | 16 |
| vps26a | 0.53412 | 0.192313212 | 0.229 | 0.136 | 1 | 16 |
| sp1 | 0.5343 | -0.16348623 | 0.057 | 0.136 | 1 | 16 |

| | | | | | | |
|---|---|---|---|---|---|---|
| si:dkey-199f5.8 | 0.53434 | -0.1814213 | 0.086 | 0.182 | 1 | 16 |
| ube2q2 | 0.53565 | 0.205237541 | 0.257 | 0.136 | 1 | 16 |
| ndufaf3 | 0.53585 | -0.15303715 | 0.057 | 0.136 | 1 | 16 |
| grtp1a | 0.53635 | -0.10705193 | 0.057 | 0.091 | 1 | 16 |
| anxa5b | 0.5367 | 0.201577981 | 0.429 | 0.318 | 1 | 16 |
| babam1 | 0.53676 | -0.16250453 | 0.057 | 0.136 | 1 | 16 |
| tm2d1 | 0.537 | -0.10450713 | 0.057 | 0.136 | 1 | 16 |
| si:ch73-61d6.3 | 0.53717 | 0.153857217 | 0.4 | 0.273 | 1 | 16 |
| wtap | 0.5372 | 0.182490435 | 0.257 | 0.136 | 1 | 16 |
| ndufs3 | 0.5385 | 0.149005008 | 0.343 | 0.273 | 1 | 16 |
| gale | 0.53857 | -0.10572827 | 0.057 | 0.136 | 1 | 16 |
| nipblb | 0.53857 | 0.173160043 | 0.171 | 0.091 | 1 | 16 |
| wsb1 | 0.53872 | 0.185017607 | 0.257 | 0.136 | 1 | 16 |
| tfam | 0.539 | 0.164486139 | 0.229 | 0.136 | 1 | 16 |
| fbxl6 | 0.53986 | 0.131213453 | 0.229 | 0.182 | 1 | 16 |
| wbp2nl | 0.5399 | 0.143084831 | 0.114 | 0.045 | 1 | 16 |
| tmem42a | 0.53996 | -0.11433931 | 0.057 | 0.091 | 1 | 16 |
| zgc:162339 | 0.54058 | 0.148188955 | 0.114 | 0.045 | 1 | 16 |
| sae1 | 0.54059 | -0.17887198 | 0.086 | 0.182 | 1 | 16 |
| hnrnpl2 | 0.54062 | 0.178679286 | 0.514 | 0.409 | 1 | 16 |
| CABZ01029938.1 | 0.54073 | -0.17092411 | 0.057 | 0.136 | 1 | 16 |
| rnf38 | 0.54096 | 0.158508245 | 0.114 | 0.045 | 1 | 16 |
| cd99 | 0.54158 | 0.157552808 | 0.114 | 0.045 | 1 | 16 |
| yes1 | 0.54232 | 0.145177962 | 0.086 | 0.045 | 1 | 16 |
| dazap2 | 0.54308 | -0.16281283 | 0.229 | 0.318 | 1 | 16 |
| mrpl14 | 0.54312 | -0.13396358 | 0.2 | 0.318 | 1 | 16 |
| tmpob | 0.54365 | 0.148072242 | 0.114 | 0.045 | 1 | 16 |
| irx3a | 0.54478 | -0.15078436 | 0.057 | 0.136 | 1 | 16 |
| nfybb | 0.54542 | 0.125857355 | 0.286 | 0.182 | 1 | 16 |
| cmc2 | 0.54575 | -0.12979551 | 0.057 | 0.136 | 1 | 16 |
| ttf1 | 0.54629 | -0.15075298 | 0.086 | 0.182 | 1 | 16 |
| dcp2 | 0.54631 | 0.185951418 | 0.229 | 0.136 | 1 | 16 |
| stimate | 0.54663 | 0.145462686 | 0.114 | 0.045 | 1 | 16 |
| borcs5 | 0.54669 | -0.15868373 | 0.057 | 0.136 | 1 | 16 |
| kank2 | 0.54693 | -0.10899489 | 0.057 | 0.136 | 1 | 16 |
| vdac1 | 0.54722 | -0.10687265 | 0.343 | 0.455 | 1 | 16 |
| ciao1 | 0.54756 | 0.169415093 | 0.171 | 0.091 | 1 | 16 |
| atp6ap1b | 0.54762 | -0.1467012 | 0.057 | 0.136 | 1 | 16 |
| faf1 | 0.54785 | -0.14881403 | 0.057 | 0.136 | 1 | 16 |
| wapla | 0.5481 | -0.11089009 | 0.057 | 0.136 | 1 | 16 |
| kmt5aa | 0.54837 | -0.18835474 | 0.229 | 0.364 | 1 | 16 |
| gar1 | 0.54849 | -0.12407516 | 0.171 | 0.273 | 1 | 16 |
| lfng | 0.54874 | -0.11325863 | 0.057 | 0.091 | 1 | 16 |
| meaf6 | 0.54922 | -0.17208416 | 0.171 | 0.273 | 1 | 16 |
| txndc17 | 0.54928 | -0.14897877 | 0.2 | 0.318 | 1 | 16 |
| zgc:63587 | 0.54928 | -0.13886985 | 0.086 | 0.182 | 1 | 16 |
| neo1a | 0.54983 | -0.15292156 | 0.057 | 0.136 | 1 | 16 |
| mbnl3 | 0.54985 | -0.15369495 | 0.657 | 0.727 | 1 | 16 |

| | | | | | | |
|---|---|---|---|---|---|---|
| hsd11b1la | 0.54997 | 0.148031215 | 0.2 | 0.136 | 1 | 16 |
| LOC101885027 | 0.5503 | -0.15548813 | 0.057 | 0.136 | 1 | 16 |
| syf2 | 0.55066 | -0.16999134 | 0.371 | 0.455 | 1 | 16 |
| ttll12 | 0.55094 | 0.139618045 | 0.286 | 0.182 | 1 | 16 |
| rxrba | 0.55184 | 0.107679591 | 0.171 | 0.091 | 1 | 16 |
| sdcbp2 | 0.55216 | 0.12101915 | 0.829 | 0.773 | 1 | 16 |
| purab | 0.55219 | 0.152499149 | 0.429 | 0.364 | 1 | 16 |
| smim14 | 0.55228 | -0.14200262 | 0.057 | 0.136 | 1 | 16 |
| rfc5 | 0.55305 | 0.127941391 | 0.114 | 0.045 | 1 | 16 |
| SBNO2 | 0.55316 | -0.18800116 | 0.086 | 0.182 | 1 | 16 |
| hmgb3a | 0.55318 | 0.16380319 | 0.771 | 0.682 | 1 | 16 |
| polr2k | 0.55325 | -0.20393294 | 0.2 | 0.318 | 1 | 16 |
| ppp1r10 | 0.55327 | -0.14715991 | 0.086 | 0.182 | 1 | 16 |
| prss35 | 0.55343 | 0.166041173 | 0.171 | 0.091 | 1 | 16 |
| egfra | 0.55348 | -0.16523607 | 0.086 | 0.182 | 1 | 16 |
| exosc8 | 0.55419 | -0.1133839 | 0.057 | 0.136 | 1 | 16 |
| SLC7A1 (1 of many) | 0.55514 | -0.10877379 | 0.057 | 0.136 | 1 | 16 |
| tarbp2 | 0.55536 | 0.149323034 | 0.114 | 0.045 | 1 | 16 |
| ppm1ba | 0.55559 | 0.114735911 | 0.171 | 0.091 | 1 | 16 |
| si:ch211-253h3.1 | 0.55618 | 0.143902052 | 0.086 | 0.045 | 1 | 16 |
| rap2b | 0.55639 | 0.125216234 | 0.143 | 0.091 | 1 | 16 |
| zgc:55781 | 0.55675 | -0.11948662 | 0.057 | 0.136 | 1 | 16 |
| ace | 0.55723 | 0.100677118 | 0.371 | 0.273 | 1 | 16 |
| trir | 0.55759 | -0.16058604 | 0.057 | 0.136 | 1 | 16 |
| golt1ba | 0.55768 | -0.10604939 | 0.057 | 0.136 | 1 | 16 |
| glulb | 0.55857 | 0.159592239 | 0.4 | 0.273 | 1 | 16 |
| smndc1 | 0.55879 | -0.13010821 | 0.286 | 0.409 | 1 | 16 |
| gps2 | 0.55889 | -0.1683912 | 0.086 | 0.182 | 1 | 16 |
| fgd5a | 0.55897 | 0.172393219 | 0.314 | 0.227 | 1 | 16 |
| tead3b | 0.55966 | -0.16937716 | 0.086 | 0.182 | 1 | 16 |
| gig2o | 0.55968 | -0.16016127 | 0.086 | 0.182 | 1 | 16 |
| eif1b | 0.56001 | -0.10007805 | 0.8 | 0.818 | 1 | 16 |
| ubxn7 | 0.56004 | -0.11778903 | 0.057 | 0.136 | 1 | 16 |
| LOC110437758 | 0.5601 | 0.149609685 | 0.114 | 0.045 | 1 | 16 |
| rabac1 | 0.5613 | -0.18066062 | 0.086 | 0.182 | 1 | 16 |
| ptbp3 | 0.56132 | -0.13646052 | 0.086 | 0.136 | 1 | 16 |
| cdc40 | 0.56139 | -0.12604585 | 0.086 | 0.182 | 1 | 16 |
| papola | 0.56171 | -0.12189231 | 0.143 | 0.182 | 1 | 16 |
| prrg4 | 0.56173 | -0.17769469 | 0.086 | 0.182 | 1 | 16 |
| zgc:85777 | 0.5623 | 0.146151285 | 0.114 | 0.045 | 1 | 16 |
| ZNF512B | 0.56235 | -0.12268829 | 0.057 | 0.136 | 1 | 16 |
| dph5 | 0.56277 | 0.121673932 | 0.086 | 0.045 | 1 | 16 |
| bida | 0.56316 | -0.12483145 | 0.086 | 0.182 | 1 | 16 |
| NAV1 (1 of many) | 0.56375 | -0.13331689 | 0.057 | 0.136 | 1 | 16 |
| aga | 0.56431 | -0.14014772 | 0.086 | 0.182 | 1 | 16 |
| tapbpl | 0.56438 | -0.18020511 | 0.086 | 0.182 | 1 | 16 |
| ttc9c | 0.5644 | -0.14422501 | 0.057 | 0.136 | 1 | 16 |
| slc30a1a | 0.56455 | -0.14914708 | 0.057 | 0.136 | 1 | 16 |

| | | | | | | |
|---|---|---|---|---|---|---|
| si:dkey-102m7.3 | 0.56531 | -0.18268271 | 0.086 | 0.182 | 1 | 16 |
| znf1181 | 0.5654 | 0.160955496 | 0.171 | 0.091 | 1 | 16 |
| ehbp1 | 0.56559 | -0.15128978 | 0.057 | 0.136 | 1 | 16 |
| srsf7a | 0.56575 | -0.17317767 | 0.4 | 0.5 | 1 | 16 |
| itga9 | 0.56601 | -0.14734493 | 0.143 | 0.182 | 1 | 16 |
| pdhb | 0.56636 | -0.18753031 | 0.086 | 0.182 | 1 | 16 |
| ppt1 | 0.5665 | 0.157593227 | 0.686 | 0.545 | 1 | 16 |
| dhdds | 0.56724 | -0.16426175 | 0.171 | 0.273 | 1 | 16 |
| serpinf1 | 0.56774 | 0.119374688 | 0.314 | 0.273 | 1 | 16 |
| mxra8a | 0.56805 | 0.194274903 | 0.657 | 0.545 | 1 | 16 |
| kif13ba | 0.56807 | -0.15911372 | 0.086 | 0.182 | 1 | 16 |
| ccdc58 | 0.56828 | -0.15034117 | 0.057 | 0.136 | 1 | 16 |
| CABZ01084501.3 | 0.56832 | 0.227183394 | 0.229 | 0.136 | 1 | 16 |
| elp6 | 0.56876 | -0.16184225 | 0.057 | 0.136 | 1 | 16 |
| BX005309.1 | 0.56907 | 0.134008736 | 0.229 | 0.182 | 1 | 16 |
| leo1 | 0.56935 | -0.14191386 | 0.086 | 0.182 | 1 | 16 |
| hnrnpd | 0.56983 | -0.16596399 | 0.086 | 0.182 | 1 | 16 |
| zgc:55262 | 0.56994 | 0.164983427 | 0.229 | 0.136 | 1 | 16 |
| atp6v0cb | 0.5701 | 0.188854926 | 0.343 | 0.227 | 1 | 16 |
| fra10ac1 | 0.57017 | -0.14265124 | 0.114 | 0.182 | 1 | 16 |
| si:ch211-132g1.3 | 0.5704 | 0.13428917 | 0.114 | 0.045 | 1 | 16 |
| cwc25 | 0.57046 | 0.134456401 | 0.114 | 0.045 | 1 | 16 |
| cttn | 0.57056 | -0.15373048 | 0.086 | 0.182 | 1 | 16 |
| tomm7 | 0.57081 | 0.125924042 | 0.543 | 0.409 | 1 | 16 |
| mcl1a | 0.57113 | -0.13666142 | 0.571 | 0.591 | 1 | 16 |
| unm-hu7910 | 0.57133 | 0.154162987 | 0.114 | 0.045 | 1 | 16 |
| gtf3c5 | 0.57138 | -0.16037833 | 0.086 | 0.182 | 1 | 16 |
| zgc:101744 | 0.57154 | -0.12307564 | 0.057 | 0.136 | 1 | 16 |
| appbp2 | 0.57165 | -0.14762542 | 0.086 | 0.182 | 1 | 16 |
| zcchc8 | 0.57176 | -0.15074165 | 0.086 | 0.182 | 1 | 16 |
| socs1a | 0.57182 | -0.17809161 | 0.086 | 0.182 | 1 | 16 |
| ddx47 | 0.5721 | -0.15022451 | 0.086 | 0.182 | 1 | 16 |
| c1galt1c1 | 0.57217 | -0.15183941 | 0.086 | 0.182 | 1 | 16 |
| srsf6a | 0.5722 | -0.12536129 | 0.171 | 0.273 | 1 | 16 |
| mrps18c | 0.57242 | -0.14889133 | 0.086 | 0.182 | 1 | 16 |
| drap1 | 0.57261 | 0.174847296 | 0.457 | 0.318 | 1 | 16 |
| pprc1 | 0.57263 | -0.18182114 | 0.086 | 0.182 | 1 | 16 |
| nipsnap3a | 0.57273 | -0.19078795 | 0.086 | 0.182 | 1 | 16 |
| glceb | 0.57274 | -0.16183584 | 0.086 | 0.182 | 1 | 16 |
| sil1 | 0.57283 | -0.15623062 | 0.086 | 0.182 | 1 | 16 |
| phrf1 | 0.57283 | -0.15935294 | 0.086 | 0.182 | 1 | 16 |
| polr3k | 0.57343 | -0.12642264 | 0.143 | 0.182 | 1 | 16 |
| ypel5 | 0.57345 | 0.128976022 | 0.2 | 0.136 | 1 | 16 |
| casp8ap2 | 0.57385 | -0.14428299 | 0.057 | 0.136 | 1 | 16 |
| CU459056.2 | 0.57393 | -0.16824633 | 0.057 | 0.136 | 1 | 16 |
| si:dkey-230p4.1 | 0.57404 | 0.106658475 | 0.114 | 0.045 | 1 | 16 |
| pcnp | 0.57583 | -0.11058069 | 0.057 | 0.136 | 1 | 16 |
| csrnp2 | 0.57609 | 0.123324961 | 0.229 | 0.182 | 1 | 16 |

| | | | | | | |
|---|---|---|---|---|---|---|
| rnf146 | 0.57635 | -0.11423703 | 0.057 | 0.091 | 1 | 16 |
| ciao2a | 0.5766 | -0.10965409 | 0.057 | 0.136 | 1 | 16 |
| lsm14ab | 0.57679 | 0.12292045 | 0.229 | 0.136 | 1 | 16 |
| atox1 | 0.57694 | 0.196382385 | 0.457 | 0.318 | 1 | 16 |
| zmat2 | 0.57703 | -0.13378932 | 0.314 | 0.364 | 1 | 16 |
| rhocb | 0.57724 | -0.1330645 | 0.257 | 0.318 | 1 | 16 |
| tp53rk | 0.57762 | 0.132377057 | 0.114 | 0.045 | 1 | 16 |
| ddx42 | 0.57798 | 0.1318057 | 0.143 | 0.091 | 1 | 16 |
| si:dkey-30g5.1 | 0.57822 | -0.12067156 | 0.057 | 0.136 | 1 | 16 |
| wls | 0.57852 | 0.105552187 | 0.571 | 0.455 | 1 | 16 |
| ndel1a | 0.57888 | -0.12598996 | 0.057 | 0.136 | 1 | 16 |
| gpm6bb | 0.57919 | 0.110508494 | 0.171 | 0.136 | 1 | 16 |
| ubqln4 | 0.57951 | -0.19501138 | 0.171 | 0.273 | 1 | 16 |
| rras | 0.57984 | -0.18545097 | 0.371 | 0.5 | 1 | 16 |
| pir | 0.58 | -0.14685506 | 0.057 | 0.136 | 1 | 16 |
| sat1a.1 | 0.58002 | 0.213010054 | 0.229 | 0.136 | 1 | 16 |
| mrpl24 | 0.58028 | -0.17264219 | 0.114 | 0.182 | 1 | 16 |
| drd2a | 0.58028 | 0.124422793 | 0.086 | 0.045 | 1 | 16 |
| zgc:153631 | 0.58033 | 0.151825938 | 0.114 | 0.045 | 1 | 16 |
| glipr2l | 0.58048 | -0.17675389 | 0.086 | 0.136 | 1 | 16 |
| cadm2a | 0.58086 | 0.189725258 | 0.657 | 0.545 | 1 | 16 |
| mrps16 | 0.5811 | -0.18953075 | 0.2 | 0.318 | 1 | 16 |
| myd88 | 0.58113 | 0.110001893 | 0.114 | 0.045 | 1 | 16 |
| arglu1b | 0.58125 | -0.13157011 | 0.171 | 0.273 | 1 | 16 |
| med11 | 0.58145 | -0.19582042 | 0.171 | 0.273 | 1 | 16 |
| si:ch211-51e12.7 | 0.58186 | 0.164754898 | 0.486 | 0.409 | 1 | 16 |
| mink1 | 0.58197 | 0.130873366 | 0.114 | 0.045 | 1 | 16 |
| ndufa12 | 0.58215 | -0.19380838 | 0.371 | 0.5 | 1 | 16 |
| dglucy | 0.58224 | 0.137766303 | 0.114 | 0.045 | 1 | 16 |
| lsm14aa | 0.58226 | -0.1309725 | 0.057 | 0.136 | 1 | 16 |
| adgrg1 | 0.58231 | 0.124957494 | 0.229 | 0.136 | 1 | 16 |
| prdm9 | 0.5826 | -0.1411211 | 0.057 | 0.136 | 1 | 16 |
| mtx1b | 0.58273 | -0.1203036 | 0.057 | 0.091 | 1 | 16 |
| fam129bb | 0.58312 | -0.16862007 | 0.057 | 0.136 | 1 | 16 |
| cmtm7 | 0.58363 | -0.11845808 | 0.057 | 0.136 | 1 | 16 |
| hsd11b2 | 0.58378 | -0.10415803 | 0.4 | 0.409 | 1 | 16 |
| pitpnbl | 0.58386 | -0.15522332 | 0.171 | 0.227 | 1 | 16 |
| aktip | 0.58476 | 0.165660977 | 0.171 | 0.091 | 1 | 16 |
| ccdc51 | 0.58562 | -0.11665781 | 0.057 | 0.136 | 1 | 16 |
| baz2a | 0.58563 | 0.144436066 | 0.114 | 0.045 | 1 | 16 |
| gabpa | 0.58594 | -0.15015697 | 0.2 | 0.318 | 1 | 16 |
| gtf2f2a | 0.58628 | 0.176465416 | 0.229 | 0.136 | 1 | 16 |
| kmt2cb | 0.5863 | -0.11597809 | 0.314 | 0.364 | 1 | 16 |
| nsrp1 | 0.58644 | 0.133382301 | 0.114 | 0.045 | 1 | 16 |
| mapre1a | 0.58654 | -0.10571998 | 0.057 | 0.136 | 1 | 16 |
| grnb | 0.58663 | 0.132021735 | 0.143 | 0.091 | 1 | 16 |
| acvr1l | 0.58665 | -0.11046499 | 0.057 | 0.136 | 1 | 16 |
| gpx8 | 0.58678 | -0.17241747 | 0.057 | 0.136 | 1 | 16 |

| | | | | | | |
|---|---|---|---|---|---|---|
| nr2f1a | 0.5868 | 0.161256706 | 0.229 | 0.136 | 1 | 16 |
| dkk3b | 0.58698 | 0.149033908 | 0.114 | 0.045 | 1 | 16 |
| si:ch211-51h9.7 | 0.58754 | -0.13804047 | 0.057 | 0.136 | 1 | 16 |
| sdhaf2 | 0.58815 | -0.1491578 | 0.114 | 0.182 | 1 | 16 |
| CDK2AP1 | 0.58842 | -0.1309658 | 0.057 | 0.136 | 1 | 16 |
| si:ch73-281f12.4 | 0.5887 | 0.152886025 | 0.114 | 0.045 | 1 | 16 |
| pwwp2a | 0.58873 | -0.1104651 | 0.057 | 0.136 | 1 | 16 |
| lpin2 | 0.5889 | -0.11718554 | 0.057 | 0.091 | 1 | 16 |
| rock2a | 0.58938 | 0.126413192 | 0.114 | 0.045 | 1 | 16 |
| pgap2 | 0.58969 | 0.171695329 | 0.171 | 0.091 | 1 | 16 |
| shroom4 | 0.58999 | -0.10918767 | 0.057 | 0.136 | 1 | 16 |
| baz1b | 0.59009 | -0.18586322 | 0.2 | 0.318 | 1 | 16 |
| borcs7 | 0.59018 | 0.142316883 | 0.171 | 0.091 | 1 | 16 |
| CU929037.3 | 0.59024 | -0.12976459 | 0.057 | 0.136 | 1 | 16 |
| man2c1 | 0.59036 | -0.13299174 | 0.057 | 0.136 | 1 | 16 |
| ubr7 | 0.59041 | -0.16611178 | 0.057 | 0.136 | 1 | 16 |
| pak1ip1 | 0.59049 | -0.14359855 | 0.057 | 0.136 | 1 | 16 |
| LOC110437926 | 0.5905 | 0.152213123 | 0.143 | 0.091 | 1 | 16 |
| atp6v0a1a | 0.59051 | -0.13911471 | 0.057 | 0.136 | 1 | 16 |
| cep350 | 0.59061 | -0.18338979 | 0.057 | 0.136 | 1 | 16 |
| ogt.1 | 0.5911 | -0.19339159 | 0.171 | 0.273 | 1 | 16 |
| gbp | 0.59127 | -0.17441146 | 0.229 | 0.318 | 1 | 16 |
| litaf | 0.59133 | -0.13620608 | 0.257 | 0.318 | 1 | 16 |
| mapk12a | 0.59144 | -0.10685713 | 0.057 | 0.136 | 1 | 16 |
| btbd7 | 0.5916 | -0.11223688 | 0.057 | 0.136 | 1 | 16 |
| smim7 | 0.59189 | 0.184258707 | 0.286 | 0.182 | 1 | 16 |
| ftr80 | 0.59206 | 0.156560446 | 0.171 | 0.091 | 1 | 16 |
| polr2j | 0.59211 | -0.13774996 | 0.057 | 0.136 | 1 | 16 |
| sh3kbp1 | 0.59216 | -0.12248956 | 0.057 | 0.136 | 1 | 16 |
| ldb1a | 0.59265 | -0.14014825 | 0.057 | 0.136 | 1 | 16 |
| ctnnbl1 | 0.59279 | 0.110205146 | 0.114 | 0.045 | 1 | 16 |
| fdxr | 0.59352 | 0.190441864 | 0.257 | 0.182 | 1 | 16 |
| tyw1 | 0.59363 | 0.124650443 | 0.114 | 0.045 | 1 | 16 |
| prps1b | 0.59405 | -0.13759597 | 0.057 | 0.136 | 1 | 16 |
| si:ch211-288g17.3 | 0.59415 | -0.2149325 | 0.457 | 0.591 | 1 | 16 |
| cygb1 | 0.59432 | 0.10588599 | 0.114 | 0.091 | 1 | 16 |
| ppp2r5cb | 0.59448 | -0.13134912 | 0.057 | 0.136 | 1 | 16 |
| smo | 0.59452 | -0.10305991 | 0.143 | 0.182 | 1 | 16 |
| kdm6al | 0.59454 | -0.12895867 | 0.057 | 0.136 | 1 | 16 |
| c1d | 0.59521 | 0.120790308 | 0.114 | 0.045 | 1 | 16 |
| csk | 0.59538 | -0.10731227 | 0.057 | 0.136 | 1 | 16 |
| tmem134 | 0.59541 | -0.16883734 | 0.143 | 0.227 | 1 | 16 |
| slc7a6 | 0.59556 | -0.12659031 | 0.057 | 0.136 | 1 | 16 |
| lsm12a | 0.59576 | 0.100035163 | 0.171 | 0.091 | 1 | 16 |
| mxe | 0.59582 | -0.13650511 | 0.057 | 0.136 | 1 | 16 |
| stard9 | 0.5961 | 0.168006867 | 0.171 | 0.091 | 1 | 16 |
| dync1li1 | 0.59615 | -0.12634625 | 0.057 | 0.136 | 1 | 16 |
| si:dkey-217d24.6 | 0.59617 | 0.145438722 | 0.114 | 0.045 | 1 | 16 |

| | | | | | | |
|---|---|---|---|---|---|---|
| rnf26 | 0.59623 | -0.14379916 | 0.057 | 0.136 | 1 | 16 |
| fbrsl1 | 0.59625 | -0.12377997 | 0.057 | 0.136 | 1 | 16 |
| cxcl18b | 0.5963 | -0.16460767 | 0.057 | 0.136 | 1 | 16 |
| cdc34b | 0.59636 | 0.152390951 | 0.114 | 0.045 | 1 | 16 |
| map2k4b | 0.59644 | 0.112295603 | 0.114 | 0.091 | 1 | 16 |
| tardbp | 0.59655 | 0.141700004 | 0.2 | 0.136 | 1 | 16 |
| uprt | 0.59689 | -0.11549924 | 0.086 | 0.136 | 1 | 16 |
| tmem101 | 0.59709 | -0.13914144 | 0.057 | 0.136 | 1 | 16 |
| ndufb2 | 0.5971 | 0.164721221 | 0.543 | 0.455 | 1 | 16 |
| nos1apa | 0.59732 | 0.199604797 | 0.543 | 0.409 | 1 | 16 |
| si:ch211-68a17.7 | 0.59804 | -0.12063037 | 0.171 | 0.273 | 1 | 16 |
| capza1b | 0.5981 | 0.170760283 | 0.514 | 0.409 | 1 | 16 |
| agpat4 | 0.59814 | -0.14639531 | 0.057 | 0.136 | 1 | 16 |
| krt4 | 0.59844 | -0.10900907 | 0.057 | 0.091 | 1 | 16 |
| chd1 | 0.59881 | -0.12929601 | 0.057 | 0.136 | 1 | 16 |
| hgh1 | 0.59882 | -0.1201798 | 0.057 | 0.136 | 1 | 16 |
| tradd | 0.59888 | -0.11329904 | 0.057 | 0.091 | 1 | 16 |
| brdt | 0.59922 | -0.14384475 | 0.057 | 0.136 | 1 | 16 |
| tnfrsf19 | 0.59924 | -0.17832911 | 0.2 | 0.318 | 1 | 16 |
| numbl | 0.59926 | -0.1293258 | 0.057 | 0.136 | 1 | 16 |
| mych | 0.59941 | -0.11075372 | 0.057 | 0.136 | 1 | 16 |
| mrpl20 | 0.59952 | -0.12200084 | 0.229 | 0.273 | 1 | 16 |
| gpm6ab | 0.59963 | -0.15364789 | 0.057 | 0.136 | 1 | 16 |
| dlgap1b | 0.60004 | -0.13134291 | 0.057 | 0.136 | 1 | 16 |
| mtor | 0.6005 | -0.13578419 | 0.057 | 0.136 | 1 | 16 |
| cct7 | 0.60078 | 0.145656095 | 0.543 | 0.409 | 1 | 16 |
| gna12a | 0.60099 | -0.13604696 | 0.057 | 0.136 | 1 | 16 |
| serpina10a | 0.60116 | 0.121391242 | 0.114 | 0.045 | 1 | 16 |
| si:ch211-150o23.3 | 0.60153 | -0.12603477 | 0.057 | 0.136 | 1 | 16 |
| ube2q1 | 0.60179 | -0.13125441 | 0.057 | 0.136 | 1 | 16 |
| sh2b1 | 0.60184 | 0.120075388 | 0.086 | 0.045 | 1 | 16 |
| plekha1b | 0.60191 | -0.12280639 | 0.057 | 0.136 | 1 | 16 |
| cs | 0.60206 | 0.172377324 | 0.229 | 0.136 | 1 | 16 |
| atp6v0d1 | 0.60213 | -0.14600634 | 0.057 | 0.136 | 1 | 16 |
| zgc:153935 | 0.60224 | -0.14814134 | 0.057 | 0.136 | 1 | 16 |
| zgc:110796 | 0.6023 | -0.14837122 | 0.057 | 0.136 | 1 | 16 |
| chchd4b | 0.60233 | 0.13971063 | 0.114 | 0.045 | 1 | 16 |
| rab7a | 0.60245 | -0.17594872 | 0.2 | 0.318 | 1 | 16 |
| nono | 0.6025 | -0.15791361 | 0.2 | 0.318 | 1 | 16 |
| arl4d | 0.60253 | 0.132656261 | 0.4 | 0.364 | 1 | 16 |
| oga | 0.6026 | -0.13156741 | 0.086 | 0.136 | 1 | 16 |
| zfp36l2 | 0.60263 | -0.13698622 | 0.057 | 0.136 | 1 | 16 |
| tax1bp3 | 0.60272 | 0.123820992 | 0.429 | 0.318 | 1 | 16 |
| pex26 | 0.60279 | -0.15721731 | 0.057 | 0.136 | 1 | 16 |
| nrd1a | 0.60281 | -0.12213353 | 0.057 | 0.136 | 1 | 16 |
| creb3l2 | 0.60281 | -0.15784703 | 0.057 | 0.136 | 1 | 16 |
| si:dkey-12j5.1 | 0.60288 | -0.14018964 | 0.057 | 0.136 | 1 | 16 |
| pros1 | 0.60295 | -0.17676852 | 0.057 | 0.136 | 1 | 16 |

| | | | | | | |
|---|---|---|---|---|---|---|
| cfl2 | 0.60302 | -0.13007258 | 0.057 | 0.136 | 1 | 16 |
| tbx2a | 0.60303 | 0.139849483 | 0.114 | 0.045 | 1 | 16 |
| ncl | 0.60305 | -0.12978102 | 0.6 | 0.727 | 1 | 16 |
| si:ch211-262h13.3 | 0.60324 | -0.12698579 | 0.057 | 0.136 | 1 | 16 |
| icmt | 0.60328 | -0.1419805 | 0.057 | 0.136 | 1 | 16 |
| agfg1b | 0.60331 | -0.12918575 | 0.057 | 0.136 | 1 | 16 |
| acss2 | 0.60332 | -0.12182117 | 0.057 | 0.136 | 1 | 16 |
| nsun2 | 0.60334 | -0.11788759 | 0.057 | 0.136 | 1 | 16 |
| rab23 | 0.60343 | -0.11913187 | 0.057 | 0.136 | 1 | 16 |
| si:ch211-140m22.7 | 0.60358 | -0.11951061 | 0.057 | 0.136 | 1 | 16 |
| ctdsplb | 0.60366 | -0.11722706 | 0.057 | 0.136 | 1 | 16 |
| kctd10 | 0.60368 | -0.153659 | 0.057 | 0.136 | 1 | 16 |
| acp1 | 0.60379 | -0.12240416 | 0.257 | 0.318 | 1 | 16 |
| pcmt | 0.60428 | 0.141763631 | 0.171 | 0.091 | 1 | 16 |
| dnaja2a | 0.60448 | 0.141380899 | 0.571 | 0.5 | 1 | 16 |
| tlk1a | 0.60471 | 0.1069535 | 0.086 | 0.045 | 1 | 16 |
| tollip | 0.60482 | 0.130092788 | 0.114 | 0.045 | 1 | 16 |
| elocb | 0.60539 | 0.101436061 | 0.571 | 0.545 | 1 | 16 |
| uri1 | 0.60556 | -0.19093251 | 0.2 | 0.318 | 1 | 16 |
| uqcrc2b | 0.60556 | -0.15450815 | 0.143 | 0.227 | 1 | 16 |
| coro1cb | 0.60608 | 0.132578039 | 0.114 | 0.045 | 1 | 16 |
| zgc:123010 | 0.60658 | -0.20640214 | 0.257 | 0.364 | 1 | 16 |
| kpnb1 | 0.60671 | -0.18479311 | 0.171 | 0.273 | 1 | 16 |
| ddx46 | 0.60693 | 0.119698505 | 0.114 | 0.045 | 1 | 16 |
| rnf4 | 0.60704 | 0.108833999 | 0.086 | 0.045 | 1 | 16 |
| dusp4 | 0.60708 | -0.20093285 | 0.2 | 0.318 | 1 | 16 |
| cited4b | 0.60724 | 0.11203792 | 0.114 | 0.045 | 1 | 16 |
| fam118b | 0.60733 | 0.166733298 | 0.171 | 0.091 | 1 | 16 |
| morc2 | 0.60847 | 0.11738085 | 0.086 | 0.045 | 1 | 16 |
| ada | 0.60873 | 0.111810196 | 0.114 | 0.045 | 1 | 16 |
| ncoa4 | 0.6091 | -0.13267219 | 0.143 | 0.182 | 1 | 16 |
| lrrfip2 | 0.60975 | 0.122220974 | 0.114 | 0.045 | 1 | 16 |
| ciao2b | 0.6109 | 0.106758885 | 0.114 | 0.045 | 1 | 16 |
| ascl1b | 0.61094 | 0.114655759 | 0.114 | 0.045 | 1 | 16 |
| supt4h1 | 0.61168 | -0.13177249 | 0.171 | 0.273 | 1 | 16 |
| pet100 | 0.61224 | 0.152227758 | 0.2 | 0.136 | 1 | 16 |
| zgc:123105 | 0.6125 | 0.169058643 | 0.229 | 0.136 | 1 | 16 |
| khdrbs1b | 0.61273 | -0.13564241 | 0.171 | 0.273 | 1 | 16 |
| nhp2 | 0.6132 | -0.14508966 | 0.286 | 0.409 | 1 | 16 |
| si:ch211-266d19.4 | 0.61361 | -0.11016483 | 0.057 | 0.091 | 1 | 16 |
| clpp | 0.61392 | -0.15516553 | 0.229 | 0.318 | 1 | 16 |
| fgfrl1a | 0.61415 | -0.18688205 | 0.343 | 0.409 | 1 | 16 |
| u2af2a | 0.6144 | -0.10439281 | 0.143 | 0.182 | 1 | 16 |
| mtap | 0.61451 | 0.127220571 | 0.229 | 0.136 | 1 | 16 |
| slmapa | 0.61473 | 0.12754104 | 0.114 | 0.045 | 1 | 16 |
| ube2d1a | 0.61489 | -0.12016849 | 0.114 | 0.136 | 1 | 16 |
| hsp90b1 | 0.61531 | -0.20053139 | 0.429 | 0.545 | 1 | 16 |
| rad21a | 0.61555 | -0.15231053 | 0.457 | 0.591 | 1 | 16 |

| | | | | | | |
|---|---|---|---|---|---|---|
| cisd1 | 0.61605 | 0.12980127 | 0.114 | 0.045 | 1 | 16 |
| hey2 | 0.61643 | 0.128614616 | 0.171 | 0.091 | 1 | 16 |
| pphln1 | 0.61647 | 0.10314014 | 0.171 | 0.091 | 1 | 16 |
| sem1 | 0.61659 | 0.128472589 | 0.743 | 0.682 | 1 | 16 |
| knop1 | 0.61695 | -0.13525011 | 0.171 | 0.227 | 1 | 16 |
| st3gal8 | 0.61706 | 0.103482317 | 0.086 | 0.045 | 1 | 16 |
| pcca | 0.6187 | 0.148416722 | 0.229 | 0.136 | 1 | 16 |
| gnav1 | 0.61961 | 0.182504537 | 0.171 | 0.091 | 1 | 16 |
| lasp1 | 0.62011 | 0.159181567 | 0.571 | 0.455 | 1 | 16 |
| tax1bp1b | 0.62048 | 0.161226716 | 0.286 | 0.182 | 1 | 16 |
| hnrnpaba | 0.62097 | 0.111830312 | 0.886 | 0.818 | 1 | 16 |
| myclb | 0.6211 | 0.107873949 | 0.171 | 0.136 | 1 | 16 |
| eif6 | 0.62139 | -0.15140649 | 0.286 | 0.409 | 1 | 16 |
| dzip1 | 0.62166 | 0.124413981 | 0.114 | 0.045 | 1 | 16 |
| csnk1da | 0.62202 | -0.13686058 | 0.257 | 0.318 | 1 | 16 |
| chmp7 | 0.62208 | 0.115348791 | 0.171 | 0.091 | 1 | 16 |
| arhgap5 | 0.62252 | -0.1763297 | 0.286 | 0.409 | 1 | 16 |
| usp22 | 0.62252 | 0.142220948 | 0.2 | 0.136 | 1 | 16 |
| XLOC-034394 | 0.62263 | 0.126629627 | 0.114 | 0.045 | 1 | 16 |
| aifm1 | 0.6228 | 0.128531777 | 0.143 | 0.091 | 1 | 16 |
| si:dkey-204f11.64 | 0.62293 | -0.12494617 | 0.171 | 0.273 | 1 | 16 |
| ppp6c | 0.62294 | -0.11239175 | 0.257 | 0.364 | 1 | 16 |
| lrp1aa | 0.62422 | 0.136664847 | 0.2 | 0.136 | 1 | 16 |
| srsf11 | 0.62488 | -0.11194619 | 0.457 | 0.5 | 1 | 16 |
| n4bp2l2 | 0.625 | 0.127580768 | 0.114 | 0.045 | 1 | 16 |
| rap1aa | 0.6255 | 0.12067864 | 0.371 | 0.318 | 1 | 16 |
| cox7b | 0.62573 | -0.14481395 | 0.743 | 0.818 | 1 | 16 |
| dda1 | 0.62576 | 0.10199639 | 0.257 | 0.227 | 1 | 16 |
| tmem203 | 0.62615 | 0.148972335 | 0.171 | 0.091 | 1 | 16 |
| smc1al | 0.62652 | -0.17242039 | 0.286 | 0.409 | 1 | 16 |
| nf2b | 0.62716 | 0.128061394 | 0.114 | 0.045 | 1 | 16 |
| txn2 | 0.62717 | -0.14130211 | 0.171 | 0.273 | 1 | 16 |
| ubap1 | 0.62718 | 0.118487004 | 0.114 | 0.045 | 1 | 16 |
| os9 | 0.62738 | 0.171846275 | 0.314 | 0.227 | 1 | 16 |
| spina | 0.62882 | 0.135289128 | 0.114 | 0.045 | 1 | 16 |
| phlda2 | 0.62937 | -0.18425869 | 0.286 | 0.409 | 1 | 16 |
| flj11011l | 0.62952 | -0.11382606 | 0.086 | 0.136 | 1 | 16 |
| puf60b | 0.6296 | 0.138199155 | 0.171 | 0.091 | 1 | 16 |
| ifngr2 | 0.62971 | 0.134876118 | 0.171 | 0.091 | 1 | 16 |
| dclk1a | 0.63027 | 0.134425209 | 0.114 | 0.045 | 1 | 16 |
| trub1 | 0.63086 | 0.126365041 | 0.114 | 0.045 | 1 | 16 |
| creld1b | 0.63106 | 0.137187538 | 0.114 | 0.045 | 1 | 16 |
| mrpl3 | 0.6316 | 0.126170776 | 0.114 | 0.045 | 1 | 16 |
| eif3c | 0.63203 | -0.15538397 | 0.371 | 0.5 | 1 | 16 |
| pcm1 | 0.63233 | 0.101797609 | 0.114 | 0.045 | 1 | 16 |
| brd2a | 0.63253 | -0.18093484 | 0.286 | 0.409 | 1 | 16 |
| rhoca | 0.63278 | 0.116514075 | 0.371 | 0.273 | 1 | 16 |
| ikzf2 | 0.6336 | 0.106497049 | 0.086 | 0.045 | 1 | 16 |

| | | | | | | |
|---|---|---|---|---|---|---|
| f11r.1 | 0.63397 | -0.17386263 | 0.371 | 0.5 | 1 | 16 |
| rab3db | 0.63414 | 0.107752925 | 0.114 | 0.045 | 1 | 16 |
| ptena | 0.63435 | 0.164837385 | 0.371 | 0.273 | 1 | 16 |
| ergic2 | 0.63496 | -0.13764294 | 0.171 | 0.273 | 1 | 16 |
| kpna6 | 0.63504 | 0.115476939 | 0.086 | 0.045 | 1 | 16 |
| parp2 | 0.63506 | 0.117521368 | 0.114 | 0.045 | 1 | 16 |
| lmnb2 | 0.63513 | -0.14922562 | 0.143 | 0.227 | 1 | 16 |
| znf385a | 0.63567 | 0.113268747 | 0.114 | 0.045 | 1 | 16 |
| ezrb | 0.63571 | 0.142521314 | 0.2 | 0.136 | 1 | 16 |
| med1 | 0.63584 | 0.121934634 | 0.086 | 0.045 | 1 | 16 |
| inha | 0.63585 | -0.16827271 | 0.171 | 0.227 | 1 | 16 |
| hmgcl | 0.63672 | 0.130228351 | 0.114 | 0.045 | 1 | 16 |
| si:ch73-42p12.2 | 0.63673 | 0.133860182 | 0.571 | 0.5 | 1 | 16 |
| slc36a1 | 0.63674 | 0.121134842 | 0.086 | 0.045 | 1 | 16 |
| trappc1 | 0.63718 | 0.174187243 | 0.514 | 0.409 | 1 | 16 |
| nudcd2 | 0.63721 | 0.174503696 | 0.343 | 0.227 | 1 | 16 |
| stard7 | 0.6373 | 0.125361035 | 0.114 | 0.045 | 1 | 16 |
| si:dkey-184a18.5 | 0.63739 | 0.117429309 | 0.114 | 0.045 | 1 | 16 |
| bmpr1bb | 0.63772 | 0.143409949 | 0.114 | 0.045 | 1 | 16 |
| si:dkey-56m19.5 | 0.63772 | -0.1890405 | 0.143 | 0.227 | 1 | 16 |
| morf4l1 | 0.63792 | -0.10814079 | 0.371 | 0.409 | 1 | 16 |
| cited4a | 0.63829 | 0.122389238 | 0.229 | 0.136 | 1 | 16 |
| ctdspl2a | 0.63831 | 0.135978786 | 0.114 | 0.045 | 1 | 16 |
| srp9 | 0.63907 | 0.114250339 | 0.171 | 0.136 | 1 | 16 |
| pcsk7 | 0.63942 | 0.126670536 | 0.114 | 0.045 | 1 | 16 |
| cdc26 | 0.6395 | -0.15214993 | 0.171 | 0.273 | 1 | 16 |
| lrpap1 | 0.63966 | -0.12178946 | 0.257 | 0.364 | 1 | 16 |
| snx27b | 0.63983 | 0.127377745 | 0.114 | 0.045 | 1 | 16 |
| ube2e2 | 0.63985 | 0.121573747 | 0.286 | 0.227 | 1 | 16 |
| slc4a4a | 0.64004 | -0.18317775 | 0.171 | 0.273 | 1 | 16 |
| vma21 | 0.64017 | 0.129324441 | 0.229 | 0.136 | 1 | 16 |
| txnl4b | 0.64023 | 0.125247344 | 0.114 | 0.045 | 1 | 16 |
| tk2 | 0.64066 | 0.12965076 | 0.114 | 0.045 | 1 | 16 |
| gzf1 | 0.64109 | 0.122917072 | 0.114 | 0.045 | 1 | 16 |
| sox5 | 0.64135 | -0.1312218 | 0.114 | 0.182 | 1 | 16 |
| ten1 | 0.64149 | 0.133422656 | 0.2 | 0.136 | 1 | 16 |
| sh3bgrl | 0.64169 | 0.155396903 | 0.229 | 0.136 | 1 | 16 |
| bloc1s1 | 0.64209 | 0.118656031 | 0.114 | 0.045 | 1 | 16 |
| rgs5b | 0.64248 | 0.11221084 | 0.171 | 0.091 | 1 | 16 |
| rab14 | 0.64271 | 0.16443194 | 0.371 | 0.273 | 1 | 16 |
| si:ch211-200p22.4 | 0.64271 | 0.10801313 | 0.114 | 0.045 | 1 | 16 |
| dnmt3bb.1 | 0.64278 | 0.132050737 | 0.371 | 0.318 | 1 | 16 |
| fam114a2 | 0.64295 | -0.11763749 | 0.086 | 0.136 | 1 | 16 |
| rybpa | 0.64317 | 0.103300126 | 0.114 | 0.045 | 1 | 16 |
| cers2a | 0.64328 | 0.122305814 | 0.114 | 0.045 | 1 | 16 |
| cica | 0.64349 | 0.122309489 | 0.114 | 0.045 | 1 | 16 |
| NC-002333.17 | 0.64359 | 0.136018213 | 0.171 | 0.091 | 1 | 16 |
| cers5 | 0.64376 | 0.106082145 | 0.114 | 0.045 | 1 | 16 |

| | | | | | | |
|---|---|---|---|---|---|---|
| tns3.2 | 0.64401 | 0.16246725 | 0.171 | 0.091 | 1 | 16 |
| irf2bpl | 0.64407 | 0.12511592 | 0.114 | 0.045 | 1 | 16 |
| ip6k2b | 0.64408 | 0.173717114 | 0.229 | 0.136 | 1 | 16 |
| kpna3 | 0.64409 | 0.110771645 | 0.171 | 0.091 | 1 | 16 |
| blcap | 0.6441 | 0.16031417 | 0.343 | 0.227 | 1 | 16 |
| abcd3a | 0.64421 | 0.128310778 | 0.114 | 0.045 | 1 | 16 |
| prkrip1 | 0.64463 | 0.13815438 | 0.114 | 0.045 | 1 | 16 |
| uso1 | 0.64503 | 0.125934505 | 0.114 | 0.045 | 1 | 16 |
| gid8a | 0.64503 | 0.126099277 | 0.114 | 0.045 | 1 | 16 |
| slc16a6b | 0.64522 | -0.10405205 | 0.057 | 0.091 | 1 | 16 |
| snw1 | 0.64528 | -0.14379418 | 0.2 | 0.273 | 1 | 16 |
| pa2g4b | 0.64533 | -0.11913971 | 0.257 | 0.364 | 1 | 16 |
| lcorl | 0.64595 | 0.11660169 | 0.229 | 0.182 | 1 | 16 |
| senp6a | 0.64637 | 0.123006837 | 0.114 | 0.045 | 1 | 16 |
| styx | 0.64645 | 0.125635826 | 0.114 | 0.045 | 1 | 16 |
| znf598 | 0.64649 | 0.118427543 | 0.114 | 0.045 | 1 | 16 |
| gdi2 | 0.64683 | 0.164183265 | 0.543 | 0.455 | 1 | 16 |
| si:ch211-199g17.2 | 0.64689 | -0.14252474 | 0.514 | 0.591 | 1 | 16 |
| sptlc2b | 0.64715 | -0.10842481 | 0.143 | 0.227 | 1 | 16 |
| hipk2 | 0.6472 | 0.104408563 | 0.086 | 0.045 | 1 | 16 |
| gosr2 | 0.64727 | -0.12143926 | 0.171 | 0.227 | 1 | 16 |
| psma4 | 0.64744 | -0.14701122 | 0.6 | 0.682 | 1 | 16 |
| mettl7a | 0.64795 | 0.127416175 | 0.171 | 0.091 | 1 | 16 |
| vrk2 | 0.64831 | 0.119961485 | 0.114 | 0.045 | 1 | 16 |
| ddx19 | 0.64857 | 0.107493557 | 0.171 | 0.136 | 1 | 16 |
| actr2a | 0.64935 | -0.15220647 | 0.2 | 0.273 | 1 | 16 |
| ndufs7 | 0.64936 | -0.12097299 | 0.429 | 0.5 | 1 | 16 |
| si:ch211-210c8.6 | 0.64939 | 0.126920162 | 0.143 | 0.091 | 1 | 16 |
| tuba8l | 0.64944 | -0.12388589 | 0.514 | 0.545 | 1 | 16 |
| pho | 0.64996 | -0.18135382 | 0.171 | 0.273 | 1 | 16 |
| zcrb1 | 0.65035 | 0.127032112 | 0.114 | 0.045 | 1 | 16 |
| mepce | 0.65039 | 0.120553071 | 0.086 | 0.045 | 1 | 16 |
| zhx2a | 0.65081 | 0.119530987 | 0.114 | 0.045 | 1 | 16 |
| amer1 | 0.65107 | 0.158072403 | 0.114 | 0.045 | 1 | 16 |
| atp6v0b | 0.65157 | 0.132536538 | 0.286 | 0.227 | 1 | 16 |
| osbp | 0.65185 | 0.114315128 | 0.143 | 0.091 | 1 | 16 |
| prkacbb | 0.65187 | 0.113119726 | 0.114 | 0.045 | 1 | 16 |
| pigk | 0.65202 | 0.123994894 | 0.114 | 0.045 | 1 | 16 |
| spen | 0.65241 | 0.115578626 | 0.114 | 0.045 | 1 | 16 |
| strn3 | 0.65263 | 0.115302216 | 0.114 | 0.045 | 1 | 16 |
| spint2 | 0.65283 | 0.113928889 | 0.143 | 0.091 | 1 | 16 |
| st13 | 0.65295 | -0.13569798 | 0.257 | 0.364 | 1 | 16 |
| bcl2l13 | 0.65302 | -0.1491016 | 0.114 | 0.182 | 1 | 16 |
| XLOC-032256 | 0.65305 | 0.113419007 | 0.114 | 0.045 | 1 | 16 |
| myl9a | 0.65319 | -0.10530269 | 0.143 | 0.227 | 1 | 16 |
| surf4 | 0.65321 | 0.109589932 | 0.114 | 0.045 | 1 | 16 |
| hbs1l | 0.65322 | 0.109221116 | 0.114 | 0.045 | 1 | 16 |
| dpyda.1 | 0.65336 | 0.116611933 | 0.114 | 0.045 | 1 | 16 |

| | | | | | | |
|---|---|---|---|---|---|---|
| ykt6 | 0.65338 | 0.144744277 | 0.114 | 0.045 | 1 | 16 |
| lrrc29 | 0.65341 | 0.111690927 | 0.114 | 0.045 | 1 | 16 |
| isca1 | 0.65341 | 0.103909887 | 0.114 | 0.045 | 1 | 16 |
| ubiad1 | 0.65366 | 0.109720403 | 0.114 | 0.045 | 1 | 16 |
| raver2 | 0.65378 | 0.12902873 | 0.114 | 0.045 | 1 | 16 |
| mapkapk5 | 0.65381 | 0.120901942 | 0.114 | 0.045 | 1 | 16 |
| emc9 | 0.65394 | 0.110600844 | 0.171 | 0.091 | 1 | 16 |
| srsf3a | 0.65397 | 0.103352356 | 0.114 | 0.045 | 1 | 16 |
| lrba | 0.6541 | 0.118614301 | 0.114 | 0.045 | 1 | 16 |
| fam49a | 0.65413 | 0.156317126 | 0.229 | 0.136 | 1 | 16 |
| znf839 | 0.65414 | 0.108104717 | 0.114 | 0.045 | 1 | 16 |
| bloc1s2 | 0.65423 | 0.121177206 | 0.114 | 0.045 | 1 | 16 |
| mrpl39 | 0.65429 | 0.117209459 | 0.114 | 0.045 | 1 | 16 |
| rab1ba | 0.65531 | 0.141633233 | 0.229 | 0.136 | 1 | 16 |
| plscr3b | 0.65606 | -0.17125542 | 0.171 | 0.273 | 1 | 16 |
| atp5mc3a | 0.65659 | -0.10051484 | 0.571 | 0.682 | 1 | 16 |
| bysl | 0.65672 | -0.15463393 | 0.171 | 0.273 | 1 | 16 |
| eef1akmt2 | 0.65696 | -0.10829399 | 0.086 | 0.136 | 1 | 16 |
| kctd9a | 0.65698 | 0.100093113 | 0.086 | 0.045 | 1 | 16 |
| copa | 0.65707 | 0.137613379 | 0.2 | 0.136 | 1 | 16 |
| serinc1 | 0.65782 | -0.1262665 | 0.314 | 0.364 | 1 | 16 |
| acad9 | 0.65798 | -0.10274008 | 0.057 | 0.091 | 1 | 16 |
| mfge8b | 0.6581 | -0.14820826 | 0.114 | 0.182 | 1 | 16 |
| calm3a | 0.65848 | 0.114434714 | 0.171 | 0.091 | 1 | 16 |
| timm44 | 0.65877 | 0.134951085 | 0.229 | 0.136 | 1 | 16 |
| mlf2 | 0.65892 | 0.143953787 | 0.229 | 0.136 | 1 | 16 |
| spry2 | 0.65955 | 0.168141833 | 0.571 | 0.455 | 1 | 16 |
| pfdn6 | 0.65984 | 0.135402551 | 0.343 | 0.273 | 1 | 16 |
| ube2d2l | 0.65991 | 0.144583033 | 0.286 | 0.227 | 1 | 16 |
| snrnp27 | 0.66015 | 0.144018769 | 0.2 | 0.136 | 1 | 16 |
| rpp30 | 0.66057 | 0.119725442 | 0.086 | 0.045 | 1 | 16 |
| si:ch211-284b7.3 | 0.66173 | 0.134350248 | 0.286 | 0.182 | 1 | 16 |
| cops2 | 0.66183 | 0.142222723 | 0.257 | 0.182 | 1 | 16 |
| polr2eb | 0.66185 | 0.106139671 | 0.314 | 0.273 | 1 | 16 |
| fam192a | 0.66205 | 0.123201975 | 0.143 | 0.091 | 1 | 16 |
| mapre1b | 0.66217 | 0.161805768 | 0.371 | 0.273 | 1 | 16 |
| si:dkey-6n21.12 | 0.66244 | 0.150617947 | 0.286 | 0.182 | 1 | 16 |
| zgc:162025 | 0.66268 | -0.10741836 | 0.143 | 0.227 | 1 | 16 |
| mogat2 | 0.66338 | -0.13007549 | 0.086 | 0.136 | 1 | 16 |
| ilf3b | 0.66339 | 0.104872406 | 0.371 | 0.273 | 1 | 16 |
| tagln2 | 0.66386 | 0.149721571 | 0.171 | 0.091 | 1 | 16 |
| etv5a | 0.66417 | 0.134524352 | 0.171 | 0.091 | 1 | 16 |
| tcf3b | 0.66446 | -0.14678414 | 0.171 | 0.273 | 1 | 16 |
| cdc14b | 0.6645 | -0.15414252 | 0.143 | 0.227 | 1 | 16 |
| si:ch211-198a12.6 | 0.66458 | -0.15263684 | 0.171 | 0.273 | 1 | 16 |
| pik3ip1 | 0.66458 | 0.124986353 | 0.171 | 0.091 | 1 | 16 |
| si:ch73-46j18.5 | 0.66466 | 0.12402833 | 0.8 | 0.727 | 1 | 16 |
| fkbp4 | 0.6651 | -0.14255561 | 0.143 | 0.227 | 1 | 16 |

| | | | | | | |
|---|---|---|---|---|---|---|
| atl3 | 0.66526 | -0.15805281 | 0.171 | 0.273 | 1 | 16 |
| arf4a | 0.66569 | 0.164606177 | 0.229 | 0.136 | 1 | 16 |
| pkp3b | 0.66584 | -0.13171567 | 0.257 | 0.364 | 1 | 16 |
| cdca4 | 0.666 | -0.14464876 | 0.229 | 0.318 | 1 | 16 |
| hnrnpa1b | 0.66661 | -0.17013848 | 0.4 | 0.5 | 1 | 16 |
| ndufa9a | 0.6669 | 0.140285553 | 0.343 | 0.273 | 1 | 16 |
| ddx39b | 0.66691 | 0.130795865 | 0.229 | 0.182 | 1 | 16 |
| wee1 | 0.66697 | -0.10532816 | 0.086 | 0.136 | 1 | 16 |
| arhgef18a | 0.66733 | 0.176109569 | 0.429 | 0.318 | 1 | 16 |
| zbtb20 | 0.66735 | 0.136617877 | 0.429 | 0.318 | 1 | 16 |
| chchd1 | 0.668 | -0.13210999 | 0.114 | 0.182 | 1 | 16 |
| ptges3a | 0.66808 | -0.12740143 | 0.229 | 0.318 | 1 | 16 |
| si:dkey-71l4.4 | 0.66826 | 0.117579231 | 0.086 | 0.045 | 1 | 16 |
| cebpg | 0.66845 | 0.123720272 | 0.171 | 0.091 | 1 | 16 |
| smap1 | 0.66895 | 0.144006182 | 0.2 | 0.136 | 1 | 16 |
| ube2v1 | 0.66905 | 0.14649099 | 0.686 | 0.591 | 1 | 16 |
| ccdc32 | 0.66916 | 0.146470015 | 0.171 | 0.091 | 1 | 16 |
| irf2bp2b | 0.66933 | 0.169825555 | 0.6 | 0.5 | 1 | 16 |
| glrx | 0.66947 | -0.12048578 | 0.429 | 0.545 | 1 | 16 |
| c1qbp | 0.66971 | -0.13440188 | 0.257 | 0.318 | 1 | 16 |
| tfr1a | 0.6698 | -0.17808304 | 0.086 | 0.136 | 1 | 16 |
| camlg | 0.67018 | 0.127927327 | 0.143 | 0.091 | 1 | 16 |
| fam207a | 0.67046 | -0.10785868 | 0.057 | 0.091 | 1 | 16 |
| lgi1a | 0.67119 | 0.130996822 | 0.171 | 0.091 | 1 | 16 |
| si:dkey-28b4.7 | 0.67155 | 0.129760331 | 0.429 | 0.318 | 1 | 16 |
| itpa | 0.67211 | 0.119752786 | 0.171 | 0.091 | 1 | 16 |
| psip1a | 0.67224 | 0.118391784 | 0.171 | 0.091 | 1 | 16 |
| LO018605.1 | 0.67306 | 0.116876313 | 0.086 | 0.045 | 1 | 16 |
| pno1 | 0.67318 | -0.12661171 | 0.257 | 0.364 | 1 | 16 |
| hmcn2 | 0.67482 | 0.146669269 | 0.229 | 0.136 | 1 | 16 |
| ap1g1 | 0.67523 | 0.104219688 | 0.086 | 0.045 | 1 | 16 |
| prpf31 | 0.6759 | -0.14681299 | 0.143 | 0.227 | 1 | 16 |
| ets2 | 0.67615 | 0.118886915 | 0.143 | 0.091 | 1 | 16 |
| dlg3 | 0.67636 | 0.126359205 | 0.086 | 0.045 | 1 | 16 |
| pcbd1 | 0.67669 | 0.102974031 | 0.171 | 0.091 | 1 | 16 |
| cox8b | 0.67683 | 0.125021902 | 0.229 | 0.136 | 1 | 16 |
| si:ch211-107o10.3 | 0.67769 | 0.114623568 | 0.171 | 0.091 | 1 | 16 |
| chtopa | 0.67796 | -0.13653702 | 0.429 | 0.545 | 1 | 16 |
| si:dkey-238i5.2 | 0.678 | 0.15572721 | 0.229 | 0.136 | 1 | 16 |
| glod4 | 0.67908 | 0.110975767 | 0.171 | 0.091 | 1 | 16 |
| zgc:195173 | 0.67909 | 0.182548461 | 0.429 | 0.318 | 1 | 16 |
| cavin2b | 0.68086 | -0.11404405 | 0.143 | 0.182 | 1 | 16 |
| taf11 | 0.68093 | 0.114670672 | 0.086 | 0.045 | 1 | 16 |
| sh3glb1b | 0.68094 | 0.133187673 | 0.171 | 0.091 | 1 | 16 |
| lamb1a | 0.68097 | -0.17640502 | 0.429 | 0.545 | 1 | 16 |
| add1 | 0.68105 | 0.143307827 | 0.229 | 0.136 | 1 | 16 |
| tanc2a | 0.6811 | 0.103780531 | 0.086 | 0.045 | 1 | 16 |
| mbnl1 | 0.68204 | 0.13882091 | 0.229 | 0.136 | 1 | 16 |

| | | | | | | |
|---|---|---|---|---|---|---|
| CR293507.1 | 0.68213 | 0.136277851 | 0.171 | 0.091 | 1 | 16 |
| MAP3K11 | 0.68237 | 0.112180261 | 0.086 | 0.045 | 1 | 16 |
| ctnna1 | 0.68269 | 0.102157872 | 0.314 | 0.227 | 1 | 16 |
| akirin2 | 0.68285 | 0.108694361 | 0.171 | 0.091 | 1 | 16 |
| me2 | 0.68289 | 0.109746185 | 0.086 | 0.045 | 1 | 16 |
| CABZ01026681.1 | 0.68303 | 0.151904558 | 0.229 | 0.136 | 1 | 16 |
| fam50a | 0.68327 | 0.14598021 | 0.229 | 0.136 | 1 | 16 |
| tpm3 | 0.68338 | -0.1068139 | 0.6 | 0.636 | 1 | 16 |
| irf7 | 0.68374 | -0.12145388 | 0.086 | 0.136 | 1 | 16 |
| paip2b | 0.6842 | -0.12118062 | 0.629 | 0.682 | 1 | 16 |
| katnb1 | 0.6843 | 0.127858737 | 0.171 | 0.091 | 1 | 16 |
| mrpl35 | 0.68457 | 0.135125055 | 0.171 | 0.091 | 1 | 16 |
| magi1b | 0.68474 | 0.135991324 | 0.171 | 0.091 | 1 | 16 |
| idh3a | 0.68492 | -0.1633253 | 0.143 | 0.227 | 1 | 16 |
| zgc:153993 | 0.68494 | 0.142525766 | 0.171 | 0.091 | 1 | 16 |
| CABZ01113192.1 | 0.68504 | 0.130851455 | 0.171 | 0.091 | 1 | 16 |
| mrps15 | 0.68507 | 0.122083343 | 0.171 | 0.091 | 1 | 16 |
| mrps10 | 0.68519 | 0.126953875 | 0.171 | 0.091 | 1 | 16 |
| MRPS12 | 0.68528 | 0.120858172 | 0.429 | 0.318 | 1 | 16 |
| tpcn2 | 0.6859 | -0.11585266 | 0.143 | 0.227 | 1 | 16 |
| twf1a | 0.68594 | -0.1278523 | 0.257 | 0.364 | 1 | 16 |
| gtpbp4 | 0.68619 | 0.106387716 | 0.4 | 0.364 | 1 | 16 |
| smim15 | 0.68744 | -0.14252724 | 0.429 | 0.545 | 1 | 16 |
| prrc1 | 0.68831 | 0.129994606 | 0.2 | 0.136 | 1 | 16 |
| mgll | 0.68837 | -0.108906 | 0.143 | 0.182 | 1 | 16 |
| znf318 | 0.68906 | -0.1112327 | 0.086 | 0.136 | 1 | 16 |
| scrib | 0.69071 | 0.100042506 | 0.086 | 0.045 | 1 | 16 |
| txnl4a | 0.69072 | -0.12955937 | 0.257 | 0.318 | 1 | 16 |
| kank4 | 0.69116 | 0.114705485 | 0.143 | 0.091 | 1 | 16 |
| atrx | 0.69258 | -0.12417067 | 0.571 | 0.636 | 1 | 16 |
| caap1 | 0.69301 | -0.12360681 | 0.143 | 0.227 | 1 | 16 |
| znf576.2 | 0.69318 | -0.12880021 | 0.2 | 0.273 | 1 | 16 |
| pof1b | 0.69396 | -0.10125242 | 0.057 | 0.091 | 1 | 16 |
| lsm4 | 0.69501 | -0.11257983 | 0.571 | 0.682 | 1 | 16 |
| vasnb | 0.69523 | -0.15351394 | 0.429 | 0.5 | 1 | 16 |
| dctn2 | 0.69592 | 0.106391203 | 0.143 | 0.091 | 1 | 16 |
| grb10a | 0.69607 | 0.102035419 | 0.086 | 0.045 | 1 | 16 |
| yipf6 | 0.69663 | 0.112283272 | 0.143 | 0.091 | 1 | 16 |
| cnbpb | 0.69734 | 0.140311184 | 0.514 | 0.455 | 1 | 16 |
| thumpd1 | 0.69828 | 0.103112105 | 0.143 | 0.091 | 1 | 16 |
| chka | 0.69857 | -0.10371796 | 0.4 | 0.5 | 1 | 16 |
| glo1 | 0.69878 | 0.156372772 | 0.429 | 0.318 | 1 | 16 |
| tprg1l | 0.69925 | 0.114432232 | 0.086 | 0.045 | 1 | 16 |
| denr | 0.69942 | -0.14286093 | 0.286 | 0.364 | 1 | 16 |
| eif4g1a | 0.69952 | 0.143575311 | 0.429 | 0.364 | 1 | 16 |
| cd151l | 0.70073 | -0.14029992 | 0.343 | 0.455 | 1 | 16 |
| hcfc1b | 0.70157 | -0.11737898 | 0.143 | 0.182 | 1 | 16 |
| ndufb3 | 0.70187 | 0.134747344 | 0.657 | 0.545 | 1 | 16 |

| | | | | | | |
|---|---|---|---|---|---|---|
| si:dkey-67c22.2 | 0.70203 | -0.11330142 | 0.229 | 0.273 | 1 | 16 |
| bckdk | 0.70439 | 0.147513685 | 0.343 | 0.273 | 1 | 16 |
| mdka | 0.70647 | -0.11790732 | 0.086 | 0.136 | 1 | 16 |
| cacul1 | 0.70697 | -0.1096155 | 0.086 | 0.136 | 1 | 16 |
| ccdc124 | 0.707 | -0.10121404 | 0.229 | 0.318 | 1 | 16 |
| crebbpb | 0.70722 | 0.108044858 | 0.086 | 0.045 | 1 | 16 |
| mdc1 | 0.7077 | 0.113693063 | 0.143 | 0.091 | 1 | 16 |
| inhbb | 0.70863 | 0.109913774 | 0.257 | 0.227 | 1 | 16 |
| bptf | 0.70889 | -0.14265952 | 0.143 | 0.227 | 1 | 16 |
| phlda1 | 0.70899 | 0.122132557 | 0.514 | 0.409 | 1 | 16 |
| pum1 | 0.70966 | 0.131539968 | 0.229 | 0.182 | 1 | 16 |
| dnaja1 | 0.7105 | -0.10062885 | 0.086 | 0.136 | 1 | 16 |
| polr2h | 0.71139 | -0.13611981 | 0.143 | 0.227 | 1 | 16 |
| net1 | 0.71173 | 0.103123499 | 0.086 | 0.045 | 1 | 16 |
| arpp19b | 0.7118 | -0.12778573 | 0.114 | 0.182 | 1 | 16 |
| zgc:86598 | 0.7121 | 0.127126664 | 0.371 | 0.273 | 1 | 16 |
| gmds | 0.71212 | -0.11756246 | 0.143 | 0.227 | 1 | 16 |
| hnrnpul1 | 0.71235 | -0.124348 | 0.143 | 0.227 | 1 | 16 |
| si:ch73-91k6.2 | 0.71266 | 0.118294012 | 0.2 | 0.136 | 1 | 16 |
| mrgbp | 0.71353 | -0.13704772 | 0.229 | 0.318 | 1 | 16 |
| bet1l | 0.71369 | -0.11250299 | 0.143 | 0.227 | 1 | 16 |
| apbb1 | 0.71446 | -0.10473739 | 0.057 | 0.091 | 1 | 16 |
| arhgef1b | 0.7147 | 0.117176769 | 0.314 | 0.273 | 1 | 16 |
| lbr | 0.71481 | -0.11009123 | 0.086 | 0.136 | 1 | 16 |
| eloca | 0.71506 | -0.14206946 | 0.143 | 0.227 | 1 | 16 |
| cops3 | 0.71605 | -0.1342599 | 0.143 | 0.227 | 1 | 16 |
| dus3l | 0.71637 | -0.12328216 | 0.143 | 0.227 | 1 | 16 |
| zgc:195175 | 0.71641 | 0.140159689 | 0.2 | 0.136 | 1 | 16 |
| snrnp200 | 0.71659 | -0.14732633 | 0.143 | 0.227 | 1 | 16 |
| fam3a | 0.71735 | -0.12058208 | 0.114 | 0.182 | 1 | 16 |
| usp7 | 0.71832 | -0.11747391 | 0.143 | 0.227 | 1 | 16 |
| arfrp1 | 0.71856 | 0.102465539 | 0.143 | 0.091 | 1 | 16 |
| stk35 | 0.71862 | 0.116007803 | 0.143 | 0.091 | 1 | 16 |
| si:dkeyp-113d7.1 | 0.71869 | -0.13954953 | 0.143 | 0.227 | 1 | 16 |
| fbxw11b | 0.71891 | -0.15162591 | 0.143 | 0.227 | 1 | 16 |
| rabif | 0.71958 | -0.12530654 | 0.143 | 0.227 | 1 | 16 |
| BX548028.1 | 0.7198 | -0.12553644 | 0.143 | 0.227 | 1 | 16 |
| tm9sf3 | 0.71997 | -0.13406597 | 0.143 | 0.227 | 1 | 16 |
| sec63 | 0.72001 | -0.13493378 | 0.143 | 0.227 | 1 | 16 |
| ifrd2 | 0.72032 | -0.13105521 | 0.143 | 0.227 | 1 | 16 |
| CABZ01058650.1 | 0.72207 | -0.11408194 | 0.143 | 0.227 | 1 | 16 |
| tmem209 | 0.72224 | 0.104538182 | 0.086 | 0.045 | 1 | 16 |
| zgc:162396 | 0.72229 | -0.12983439 | 0.143 | 0.227 | 1 | 16 |
| slc35a1 | 0.72299 | -0.12761153 | 0.143 | 0.227 | 1 | 16 |
| lamtor5 | 0.72313 | -0.12067582 | 0.143 | 0.227 | 1 | 16 |
| bud31 | 0.72315 | -0.13176146 | 0.457 | 0.545 | 1 | 16 |
| metap2b | 0.72346 | 0.121224406 | 0.514 | 0.455 | 1 | 16 |
| lyrm5a | 0.72355 | 0.1127729 | 0.143 | 0.091 | 1 | 16 |

| | | | | | | |
|---|---|---|---|---|---|---|
| prmt1 | 0.72386 | -0.13139031 | 0.4 | 0.455 | 1 | 16 |
| atp5po | 0.72426 | 0.147094526 | 0.571 | 0.5 | 1 | 16 |
| kdm5c | 0.72437 | -0.10647616 | 0.143 | 0.182 | 1 | 16 |
| rpf2 | 0.7244 | 0.107385646 | 0.286 | 0.227 | 1 | 16 |
| sec62 | 0.72451 | 0.131734038 | 0.286 | 0.227 | 1 | 16 |
| ddx61 | 0.72458 | -0.12972468 | 0.629 | 0.727 | 1 | 16 |
| si:ch211-195e19.1 | 0.72476 | 0.120458206 | 0.143 | 0.091 | 1 | 16 |
| serinc5 | 0.72668 | 0.153457869 | 0.257 | 0.182 | 1 | 16 |
| zfpm1 | 0.72681 | 0.101791673 | 0.086 | 0.045 | 1 | 16 |
| mzt2b | 0.727 | -0.11110349 | 0.086 | 0.136 | 1 | 16 |
| psmc5 | 0.72705 | -0.13907758 | 0.486 | 0.591 | 1 | 16 |
| dbt | 0.72706 | 0.107582958 | 0.143 | 0.091 | 1 | 16 |
| nfia | 0.72786 | 0.122427692 | 0.371 | 0.273 | 1 | 16 |
| tmem45a | 0.72804 | 0.139920154 | 0.2 | 0.136 | 1 | 16 |
| man1b1b | 0.72863 | 0.151276357 | 0.371 | 0.273 | 1 | 16 |
| smdt1b | 0.72986 | 0.138951694 | 0.543 | 0.455 | 1 | 16 |
| ttc3 | 0.73091 | -0.11115733 | 0.086 | 0.136 | 1 | 16 |
| zgc:113184 | 0.73094 | -0.109187 | 0.086 | 0.136 | 1 | 16 |
| tram2 | 0.73155 | -0.11878886 | 0.171 | 0.227 | 1 | 16 |
| pnp5a | 0.73161 | -0.14846894 | 0.229 | 0.318 | 1 | 16 |
| mecr | 0.73212 | -0.11471288 | 0.086 | 0.136 | 1 | 16 |
| mtx2 | 0.73213 | -0.13772304 | 0.2 | 0.273 | 1 | 16 |
| nfkbib | 0.73413 | 0.104351718 | 0.229 | 0.182 | 1 | 16 |
| kansl1a | 0.73452 | -0.12317132 | 0.057 | 0.091 | 1 | 16 |
| adrm1 | 0.73511 | 0.151270379 | 0.257 | 0.182 | 1 | 16 |
| mpdu1b | 0.73549 | -0.11899428 | 0.086 | 0.136 | 1 | 16 |
| zgc:194981 | 0.73549 | 0.109851746 | 0.143 | 0.091 | 1 | 16 |
| LOC103909115 | 0.73646 | 0.125140084 | 0.143 | 0.091 | 1 | 16 |
| zgc:77650 | 0.73689 | 0.14377916 | 0.314 | 0.227 | 1 | 16 |
| ganab | 0.73718 | -0.15231141 | 0.114 | 0.182 | 1 | 16 |
| ckba | 0.73771 | 0.125508859 | 0.286 | 0.227 | 1 | 16 |
| mogs | 0.73821 | -0.10069214 | 0.057 | 0.091 | 1 | 16 |
| dbi | 0.73833 | 0.143470069 | 0.686 | 0.591 | 1 | 16 |
| cdon | 0.73857 | 0.147566414 | 0.257 | 0.182 | 1 | 16 |
| tmed9 | 0.73888 | -0.12618519 | 0.6 | 0.682 | 1 | 16 |
| rrp12 | 0.73959 | -0.1230783 | 0.086 | 0.136 | 1 | 16 |
| slc25a24 | 0.74028 | -0.1136932 | 0.114 | 0.182 | 1 | 16 |
| nmt1a | 0.74065 | 0.133619878 | 0.371 | 0.273 | 1 | 16 |
| srp19 | 0.74131 | -0.11360614 | 0.314 | 0.409 | 1 | 16 |
| tenm3 | 0.74163 | 0.114655435 | 0.314 | 0.273 | 1 | 16 |
| cpda | 0.74203 | 0.101089197 | 0.086 | 0.045 | 1 | 16 |
| lgals3b | 0.74248 | 0.121856251 | 0.343 | 0.273 | 1 | 16 |
| chd7 | 0.7426 | -0.12278686 | 0.229 | 0.318 | 1 | 16 |
| mfap1 | 0.74332 | -0.11674347 | 0.229 | 0.318 | 1 | 16 |
| mrpl30 | 0.74411 | -0.13641355 | 0.314 | 0.409 | 1 | 16 |
| si:ch211-207d6.2 | 0.74615 | 0.17334925 | 0.543 | 0.455 | 1 | 16 |
| psmd10 | 0.74998 | -0.1306587 | 0.114 | 0.182 | 1 | 16 |
| foxp1a | 0.75076 | -0.12708645 | 0.114 | 0.182 | 1 | 16 |

| | | | | | | |
|---|---|---|---|---|---|---|
| si:dkeyp-44a8.2 | 0.75131 | 0.100086421 | 0.257 | 0.182 | 1 | 16 |
| aip | 0.75169 | 0.138920782 | 0.314 | 0.227 | 1 | 16 |
| psmd7 | 0.75393 | -0.11477876 | 0.314 | 0.409 | 1 | 16 |
| atp10a | 0.75443 | 0.121320047 | 0.314 | 0.227 | 1 | 16 |
| farp2 | 0.75619 | 0.119420425 | 0.2 | 0.136 | 1 | 16 |
| sdc2 | 0.75626 | 0.118723013 | 0.2 | 0.136 | 1 | 16 |
| arpc4l | 0.75692 | 0.136328231 | 0.4 | 0.318 | 1 | 16 |
| calm3b | 0.75716 | 0.109314995 | 0.771 | 0.682 | 1 | 16 |
| sdf2 | 0.75733 | 0.103219937 | 0.257 | 0.182 | 1 | 16 |
| lmf2b | 0.75756 | -0.1369814 | 0.2 | 0.273 | 1 | 16 |
| hsbp1a | 0.75818 | 0.122378026 | 0.286 | 0.227 | 1 | 16 |
| mt-nd5 | 0.75824 | -0.11204519 | 0.4 | 0.5 | 1 | 16 |
| rhoj | 0.75828 | 0.105414771 | 0.229 | 0.182 | 1 | 16 |
| frs2b | 0.75871 | -0.13001223 | 0.114 | 0.182 | 1 | 16 |
| fth1a | 0.75885 | -0.13469787 | 0.4 | 0.5 | 1 | 16 |
| nup93 | 0.7591 | -0.1119604 | 0.086 | 0.136 | 1 | 16 |
| nmi | 0.75913 | -0.12161678 | 0.229 | 0.318 | 1 | 16 |
| cald1a | 0.75926 | -0.1418248 | 0.229 | 0.318 | 1 | 16 |
| hnrpkl | 0.76008 | -0.12374485 | 0.2 | 0.273 | 1 | 16 |
| selenop | 0.76015 | 0.127035505 | 0.257 | 0.182 | 1 | 16 |
| si:ch211-276i12.9 | 0.76036 | -0.11185849 | 0.086 | 0.136 | 1 | 16 |
| tjap1 | 0.76037 | 0.101408698 | 0.143 | 0.091 | 1 | 16 |
| gmppb | 0.76106 | 0.102451425 | 0.143 | 0.091 | 1 | 16 |
| ppp6r2a | 0.76158 | -0.11180961 | 0.086 | 0.136 | 1 | 16 |
| ythdc1 | 0.76255 | -0.11739693 | 0.114 | 0.182 | 1 | 16 |
| stk11 | 0.76272 | -0.10506863 | 0.114 | 0.182 | 1 | 16 |
| fgfr2 | 0.76302 | -0.12223484 | 0.114 | 0.182 | 1 | 16 |
| rac1a | 0.76434 | 0.13323861 | 0.286 | 0.227 | 1 | 16 |
| nudt21 | 0.76441 | 0.131864731 | 0.4 | 0.318 | 1 | 16 |
| cdc42ep5 | 0.76455 | -0.11554986 | 0.114 | 0.182 | 1 | 16 |
| tmem245 | 0.76532 | -0.12074668 | 0.114 | 0.182 | 1 | 16 |
| card19 | 0.76535 | -0.12807412 | 0.114 | 0.182 | 1 | 16 |
| rnf2 | 0.76536 | 0.127214273 | 0.257 | 0.182 | 1 | 16 |
| ptpn9a | 0.76559 | -0.10248817 | 0.086 | 0.136 | 1 | 16 |
| chmp2a | 0.76633 | 0.135084711 | 0.4 | 0.318 | 1 | 16 |
| rps27l | 0.76639 | 0.128304103 | 0.543 | 0.455 | 1 | 16 |
| zc3h18 | 0.76679 | 0.115527645 | 0.2 | 0.136 | 1 | 16 |
| emc7 | 0.76707 | 0.116815421 | 0.314 | 0.227 | 1 | 16 |
| ndrg2 | 0.76708 | 0.137060986 | 0.457 | 0.364 | 1 | 16 |
| taf12 | 0.76728 | -0.12960733 | 0.314 | 0.409 | 1 | 16 |
| ubl7b | 0.7683 | -0.12677066 | 0.2 | 0.273 | 1 | 16 |
| znf593 | 0.76895 | 0.122482423 | 0.343 | 0.273 | 1 | 16 |
| abcf1 | 0.76939 | -0.12296378 | 0.429 | 0.5 | 1 | 16 |
| cyc1 | 0.76954 | -0.11894175 | 0.429 | 0.5 | 1 | 16 |
| gtf2a1 | 0.77126 | 0.129842615 | 0.2 | 0.136 | 1 | 16 |
| nsd1b | 0.77213 | 0.128265954 | 0.143 | 0.091 | 1 | 16 |
| ccdc25 | 0.77238 | 0.127956879 | 0.314 | 0.227 | 1 | 16 |
| yipf4 | 0.77243 | 0.127357744 | 0.314 | 0.227 | 1 | 16 |

| | | | | | | |
|---|---|---|---|---|---|---|
| cald1b | 0.77274 | -0.10934904 | 0.114 | 0.182 | 1 | 16 |
| sart1 | 0.77305 | 0.115158519 | 0.286 | 0.227 | 1 | 16 |
| tpst1 | 0.77425 | -0.11824484 | 0.114 | 0.182 | 1 | 16 |
| sat1a.2 | 0.7747 | 0.132167574 | 0.629 | 0.545 | 1 | 16 |
| cct4 | 0.77495 | -0.12005408 | 0.514 | 0.591 | 1 | 16 |
| adh5 | 0.77651 | -0.1186116 | 0.543 | 0.636 | 1 | 16 |
| aldh18a1 | 0.77657 | 0.110357274 | 0.086 | 0.045 | 1 | 16 |
| LOC108191614 | 0.77716 | -0.12004845 | 0.086 | 0.136 | 1 | 16 |
| srrt | 0.7773 | 0.126143415 | 0.143 | 0.091 | 1 | 16 |
| rnasekb | 0.77794 | -0.10265738 | 0.114 | 0.182 | 1 | 16 |
| smg5 | 0.77812 | -0.1057869 | 0.086 | 0.136 | 1 | 16 |
| gstr | 0.77839 | -0.10524426 | 0.4 | 0.455 | 1 | 16 |
| LOC103909284 | 0.77857 | -0.10571305 | 0.114 | 0.182 | 1 | 16 |
| as3mt | 0.7792 | -0.10600273 | 0.114 | 0.182 | 1 | 16 |
| hpf1 | 0.77975 | -0.11325471 | 0.114 | 0.182 | 1 | 16 |
| aplp2 | 0.78006 | 0.11096466 | 0.6 | 0.545 | 1 | 16 |
| eed | 0.78049 | -0.12009865 | 0.114 | 0.182 | 1 | 16 |
| slc38a3a | 0.78086 | 0.110741716 | 0.2 | 0.136 | 1 | 16 |
| mrpl44 | 0.78119 | -0.10214534 | 0.086 | 0.136 | 1 | 16 |
| snu13b | 0.78189 | -0.10973166 | 0.543 | 0.636 | 1 | 16 |
| znf609b | 0.78203 | -0.11060257 | 0.229 | 0.273 | 1 | 16 |
| nucb2a | 0.78397 | 0.128349526 | 0.457 | 0.364 | 1 | 16 |
| eno3 | 0.78414 | -0.10749048 | 0.371 | 0.409 | 1 | 16 |
| psenen | 0.78495 | 0.112402175 | 0.371 | 0.318 | 1 | 16 |
| ddost | 0.78555 | -0.13977767 | 0.371 | 0.455 | 1 | 16 |
| erlec1 | 0.78762 | 0.110771877 | 0.257 | 0.182 | 1 | 16 |
| hs3st3b1b | 0.78805 | 0.113200686 | 0.2 | 0.136 | 1 | 16 |
| cgnl1 | 0.78955 | 0.123402852 | 0.257 | 0.182 | 1 | 16 |
| naa20 | 0.78955 | 0.114933985 | 0.286 | 0.227 | 1 | 16 |
| rcan1a | 0.79006 | 0.108031708 | 0.257 | 0.182 | 1 | 16 |
| vps4b | 0.79073 | 0.11503493 | 0.257 | 0.182 | 1 | 16 |
| pcmtl | 0.79128 | -0.10318073 | 0.086 | 0.136 | 1 | 16 |
| me3 | 0.79134 | 0.103218054 | 0.143 | 0.091 | 1 | 16 |
| anxa11a | 0.79169 | -0.11035607 | 0.086 | 0.136 | 1 | 16 |
| srsf2a | 0.79451 | -0.1194768 | 0.457 | 0.545 | 1 | 16 |
| dnajc1 | 0.79469 | 0.107461986 | 0.286 | 0.227 | 1 | 16 |
| CU694197.1 | 0.7948 | 0.11643244 | 0.486 | 0.409 | 1 | 16 |
| pigf | 0.79582 | 0.118206374 | 0.2 | 0.136 | 1 | 16 |
| zgc:103697 | 0.79603 | 0.108816343 | 0.257 | 0.182 | 1 | 16 |
| phldb1a | 0.7964 | -0.10590624 | 0.086 | 0.136 | 1 | 16 |
| tmem167b | 0.79677 | 0.104105453 | 0.143 | 0.091 | 1 | 16 |
| cycsb | 0.798 | -0.10518513 | 0.486 | 0.545 | 1 | 16 |
| hmgb2b | 0.79994 | -0.12787584 | 0.686 | 0.727 | 1 | 16 |
| wdfy1 | 0.8007 | -0.11818878 | 0.2 | 0.273 | 1 | 16 |
| yipf3 | 0.80093 | 0.114000616 | 0.257 | 0.182 | 1 | 16 |
| ece2b | 0.80233 | 0.12419104 | 0.629 | 0.545 | 1 | 16 |
| rbbp6 | 0.80252 | -0.10399143 | 0.2 | 0.273 | 1 | 16 |
| zgc:158343 | 0.80433 | 0.114396569 | 0.2 | 0.136 | 1 | 16 |

| | | | | | | |
|---|---|---|---|---|---|---|
| imp3 | 0.80506 | 0.104512094 | 0.286 | 0.227 | 1 | 16 |
| XLOC-040516 | 0.8058 | 0.100216793 | 0.2 | 0.136 | 1 | 16 |
| rbm4.2 | 0.80648 | -0.10060093 | 0.229 | 0.273 | 1 | 16 |
| snrpd2 | 0.80747 | -0.11717152 | 0.457 | 0.545 | 1 | 16 |
| galnt11 | 0.80967 | -0.10056753 | 0.086 | 0.136 | 1 | 16 |
| thtpa | 0.81118 | -0.11611214 | 0.2 | 0.273 | 1 | 16 |
| pa2g4a | 0.8119 | -0.10222425 | 0.229 | 0.273 | 1 | 16 |
| aimp1 | 0.81392 | 0.112331501 | 0.286 | 0.227 | 1 | 16 |
| tmem30ab | 0.81403 | -0.10199653 | 0.371 | 0.455 | 1 | 16 |
| ppp2r2d | 0.81688 | -0.11792284 | 0.2 | 0.273 | 1 | 16 |
| prpsap1 | 0.81715 | 0.102571568 | 0.2 | 0.136 | 1 | 16 |
| mgst1.1 | 0.81797 | -0.10551023 | 0.2 | 0.273 | 1 | 16 |
| commd7 | 0.81797 | 0.103350922 | 0.229 | 0.182 | 1 | 16 |
| alg5 | 0.81809 | -0.10656037 | 0.2 | 0.273 | 1 | 16 |
| mgst3a | 0.81826 | -0.11450806 | 0.371 | 0.455 | 1 | 16 |
| zgc:162509 | 0.82162 | -0.11608892 | 0.286 | 0.364 | 1 | 16 |
| eci2 | 0.82232 | 0.100821679 | 0.2 | 0.136 | 1 | 16 |
| arhgef4 | 0.82363 | 0.109411655 | 0.2 | 0.136 | 1 | 16 |
| cemip2 | 0.82544 | 0.117993071 | 0.486 | 0.409 | 1 | 16 |
| dnase2 | 0.82606 | -0.10639373 | 0.086 | 0.136 | 1 | 16 |
| abl1 | 0.82644 | 0.100710058 | 0.143 | 0.091 | 1 | 16 |
| map2k1 | 0.82731 | 0.100431892 | 0.143 | 0.091 | 1 | 16 |
| mtpn | 0.82896 | 0.103770171 | 0.6 | 0.545 | 1 | 16 |
| ncoa5 | 0.83206 | 0.101170328 | 0.143 | 0.091 | 1 | 16 |
| ammecr1 | 0.83525 | -0.10201633 | 0.171 | 0.227 | 1 | 16 |
| tfap4 | 0.83653 | -0.10847765 | 0.086 | 0.136 | 1 | 16 |
| si:ch73-376l24.3 | 0.8368 | -0.10800869 | 0.257 | 0.318 | 1 | 16 |
| tsc22d3 | 0.83736 | -0.10041005 | 0.4 | 0.455 | 1 | 16 |
| taf9 | 0.84045 | 0.101683968 | 0.229 | 0.182 | 1 | 16 |
| cnbpa | 0.8481 | 0.103999586 | 0.486 | 0.409 | 1 | 16 |
| pgam1b | 0.84928 | 0.102973581 | 0.057 | 0.045 | 1 | 16 |
| mios | 0.84928 | 0.103355878 | 0.057 | 0.045 | 1 | 16 |
| pdgfaa | 0.85735 | -0.11623252 | 0.171 | 0.227 | 1 | 16 |
| ier3ip1 | 0.85931 | 0.104315461 | 0.571 | 0.5 | 1 | 16 |
| serinc2 | 0.86103 | -0.10092802 | 0.4 | 0.455 | 1 | 16 |
| bckdhb | 0.86149 | -0.10017203 | 0.171 | 0.227 | 1 | 16 |
| pik3r3a | 0.87784 | -0.10176671 | 0.257 | 0.318 | 1 | 16 |
| ndufa4l | 3.752670 | 0.722418478 | 1 | 0.39 | 0.7579268 | 17 |
| gstp1 | 0.00031 | 0.38897657 | 0.929 | 0.415 | 1 | 17 |
| selenow1 | 0.00037 | 0.973081599 | 0.786 | 0.195 | 1 | 17 |
| sycp3 | 0.00061 | 1.117013788 | 0.571 | 0.098 | 1 | 17 |
| uqcrc2b | 0.00073 | 0.934534416 | 0.571 | 0.073 | 1 | 17 |
| mrps18c | 0.00075 | 0.891078246 | 0.571 | 0.073 | 1 | 17 |
| polr2k | 0.00094 | 0.826542688 | 0.643 | 0.122 | 1 | 17 |
| CABZ01005379.1 | 0.00097 | 0.884503184 | 0.5 | 0.049 | 1 | 17 |
| mrpl51 | 0.00105 | 0.72695058 | 0.286 | 0 | 1 | 17 |
| rybpb | 0.00105 | 0.606869122 | 0.286 | 0 | 1 | 17 |
| si:ch211-277c7.7 | 0.00105 | 0.727742141 | 0.286 | 0 | 1 | 17 |

| | | | | | | |
|---|---|---|---|---|---|---|
| lhfpl2a | 0.00105 | 0.650677659 | 0.286 | 0 | 1 | 17 |
| ndufaf6 | 0.00105 | 0.606158432 | 0.286 | 0 | 1 | 17 |
| oxsr1a | 0.00105 | 0.636457994 | 0.286 | 0 | 1 | 17 |
| si:rp71-19m20.1 | 0.00105 | 0.68210583 | 0.286 | 0 | 1 | 17 |
| atp5f1c | 0.00143 | 0.858644781 | 0.857 | 0.317 | 1 | 17 |
| myl12.1 | 0.00145 | 0.834514965 | 0.786 | 0.244 | 1 | 17 |
| parp1 | 0.00209 | 0.778939573 | 0.643 | 0.146 | 1 | 17 |
| pcna | 0.00237 | 0.686630567 | 0.857 | 0.341 | 1 | 17 |
| psmb3 | 0.0031 | 0.723321765 | 0.714 | 0.22 | 1 | 17 |
| uqcrc1 | 0.00332 | 0.627583163 | 0.786 | 0.293 | 1 | 17 |
| dbi | 0.00359 | 0.771333969 | 0.5 | 0.073 | 1 | 17 |
| hmgb2a | 0.00378 | 0.113121733 | 1 | 0.707 | 1 | 17 |
| mcm2 | 0.0046 | 0.763895927 | 0.643 | 0.171 | 1 | 17 |
| cpsf3 | 0.00485 | 0.670491384 | 0.429 | 0.049 | 1 | 17 |
| dgcr6 | 0.00488 | 0.629775295 | 0.357 | 0.024 | 1 | 17 |
| bcas2 | 0.00508 | 0.786777009 | 0.429 | 0.049 | 1 | 17 |
| rtcb | 0.00513 | 0.807878153 | 0.429 | 0.049 | 1 | 17 |
| taf7 | 0.00523 | 0.902688975 | 0.429 | 0.049 | 1 | 17 |
| hsbp1b | 0.00531 | 0.723095482 | 0.429 | 0.049 | 1 | 17 |
| MAJIN | 0.00548 | 0.771054314 | 0.357 | 0.024 | 1 | 17 |
| nub1 | 0.00549 | 0.780426732 | 0.429 | 0.049 | 1 | 17 |
| chmp2a | 0.00551 | 0.763347287 | 0.429 | 0.049 | 1 | 17 |
| zgc:55943 | 0.00567 | 0.586793545 | 0.214 | 0 | 1 | 17 |
| hemgn | 0.00567 | 0.363128962 | 0.214 | 0 | 1 | 17 |
| myd88 | 0.00567 | 0.413683258 | 0.214 | 0 | 1 | 17 |
| smyd5 | 0.00567 | 0.483138962 | 0.214 | 0 | 1 | 17 |
| zgc:162898 | 0.00567 | 0.473194804 | 0.214 | 0 | 1 | 17 |
| zgc:174268 | 0.00567 | 0.45050312 | 0.214 | 0 | 1 | 17 |
| acat2 | 0.00567 | 0.455614126 | 0.214 | 0 | 1 | 17 |
| chd5 | 0.00567 | 0.421573488 | 0.214 | 0 | 1 | 17 |
| eaf1 | 0.00567 | 0.487602671 | 0.214 | 0 | 1 | 17 |
| ephx5 | 0.00567 | 0.428788397 | 0.214 | 0 | 1 | 17 |
| kctd9a | 0.00567 | 0.447260772 | 0.214 | 0 | 1 | 17 |
| kin | 0.00567 | 0.488404754 | 0.214 | 0 | 1 | 17 |
| smc6 | 0.00567 | 0.608508287 | 0.214 | 0 | 1 | 17 |
| CABZ01110881.1 | 0.00567 | 0.589798034 | 0.214 | 0 | 1 | 17 |
| bcl7bb | 0.00567 | 0.526631933 | 0.214 | 0 | 1 | 17 |
| lims1 | 0.00567 | 0.555071563 | 0.214 | 0 | 1 | 17 |
| pcmtl | 0.00567 | 0.545979512 | 0.214 | 0 | 1 | 17 |
| sqstm1 | 0.00567 | 0.495068846 | 0.214 | 0 | 1 | 17 |
| tfip11 | 0.00567 | 0.51446492 | 0.214 | 0 | 1 | 17 |
| ttc7b | 0.00567 | 0.557870215 | 0.214 | 0 | 1 | 17 |
| zc3h7bb | 0.00567 | 0.486582405 | 0.214 | 0 | 1 | 17 |
| CR812481.1 | 0.00567 | 0.404563482 | 0.214 | 0 | 1 | 17 |
| atad2b | 0.00567 | 0.461142761 | 0.214 | 0 | 1 | 17 |
| ccnl1a | 0.00567 | 0.434924024 | 0.214 | 0 | 1 | 17 |
| ddx11 | 0.00567 | 0.42493127 | 0.214 | 0 | 1 | 17 |
| golgb1 | 0.00567 | 0.49537674 | 0.214 | 0 | 1 | 17 |

| | | | | | | |
|---|---|---|---|---|---|---|
| hnrnpua | 0.00567 | 0.410063542 | 0.214 | 0 | 1 | 17 |
| kpnb3 | 0.00567 | 0.439632904 | 0.214 | 0 | 1 | 17 |
| mrps22 | 0.00567 | 0.397655014 | 0.214 | 0 | 1 | 17 |
| mycb | 0.00567 | 0.396014635 | 0.214 | 0 | 1 | 17 |
| oga | 0.00567 | 0.465982836 | 0.214 | 0 | 1 | 17 |
| rfx5 | 0.00567 | 0.453064666 | 0.214 | 0 | 1 | 17 |
| sgf29 | 0.00567 | 0.424225942 | 0.214 | 0 | 1 | 17 |
| tpm4b | 0.00567 | 0.465462761 | 0.214 | 0 | 1 | 17 |
| ttl | 0.00567 | 0.465731194 | 0.214 | 0 | 1 | 17 |
| zgc:112255 | 0.00567 | 0.43761614 | 0.214 | 0 | 1 | 17 |
| gmppab | 0.00567 | 0.463868576 | 0.214 | 0 | 1 | 17 |
| rbm5 | 0.00567 | 0.4511146 | 0.214 | 0 | 1 | 17 |
| wdhd1 | 0.00567 | 0.500319269 | 0.214 | 0 | 1 | 17 |
| brd8 | 0.00567 | 0.46987013 | 0.214 | 0 | 1 | 17 |
| elp5 | 0.00567 | 0.484062658 | 0.214 | 0 | 1 | 17 |
| fam102ba | 0.00567 | 0.474905282 | 0.214 | 0 | 1 | 17 |
| r3hdml | 0.00567 | 0.424208281 | 0.214 | 0 | 1 | 17 |
| si:ch211-199g17.2 | 0.00567 | 0.471474269 | 0.214 | 0 | 1 | 17 |
| spty2d1 | 0.00567 | 0.455752073 | 0.214 | 0 | 1 | 17 |
| tnni2b.1 | 0.00567 | 0.485815013 | 0.214 | 0 | 1 | 17 |
| xgb | 0.00567 | 0.449322243 | 0.214 | 0 | 1 | 17 |
| adal | 0.00567 | 0.545621499 | 0.214 | 0 | 1 | 17 |
| bloc1s5 | 0.00567 | 0.455533468 | 0.214 | 0 | 1 | 17 |
| dcaf15 | 0.00567 | 0.451913062 | 0.214 | 0 | 1 | 17 |
| kat7a | 0.00567 | 0.549050664 | 0.214 | 0 | 1 | 17 |
| lnpa | 0.00567 | 0.496394136 | 0.214 | 0 | 1 | 17 |
| sord | 0.00567 | 0.459655746 | 0.214 | 0 | 1 | 17 |
| zbtb7b | 0.00567 | 0.513672314 | 0.214 | 0 | 1 | 17 |
| si:ch211-214j24.10 | 0.00595 | 0.666337244 | 0.714 | 0.244 | 1 | 17 |
| serp1 | 0.00657 | 0.633095277 | 0.357 | 0.024 | 1 | 17 |
| snrpd2 | 0.00733 | 0.674805752 | 0.857 | 0.39 | 1 | 17 |
| lsm3 | 0.00773 | 0.432652901 | 0.786 | 0.366 | 1 | 17 |
| dnph1 | 0.00781 | 0.654622526 | 0.5 | 0.098 | 1 | 17 |
| cirbpb | 0.00814 | 0.185802693 | 0.929 | 0.61 | 1 | 17 |
| brk1 | 0.00842 | 0.708192628 | 0.786 | 0.317 | 1 | 17 |
| eif4h | 0.00849 | 0.654721015 | 0.5 | 0.098 | 1 | 17 |
| mrpl18 | 0.00874 | 0.697073666 | 0.5 | 0.098 | 1 | 17 |
| coq9 | 0.00904 | 0.717460655 | 0.5 | 0.098 | 1 | 17 |
| ndufa10 | 0.00922 | 0.779055712 | 0.5 | 0.098 | 1 | 17 |
| etfa | 0.00926 | 0.651860705 | 0.643 | 0.195 | 1 | 17 |
| hells | 0.00928 | 0.712022499 | 0.857 | 0.415 | 1 | 17 |
| sod1 | 0.00929 | 0.578719244 | 0.857 | 0.415 | 1 | 17 |
| ciapin1 | 0.00934 | 0.729076339 | 0.571 | 0.146 | 1 | 17 |
| cenpw | 0.01045 | 0.76281005 | 0.571 | 0.146 | 1 | 17 |
| sdcbp2 | 0.01062 | 0.51646569 | 0.571 | 0.171 | 1 | 17 |
| mt-co1 | 0.01063 | -0.77193186 | 0 | 0.268 | 1 | 17 |
| csde1 | 0.01154 | 0.606046118 | 0.429 | 0.073 | 1 | 17 |
| si:zfos-1192g2.3 | 0.01369 | 0.646596359 | 0.429 | 0.073 | 1 | 17 |

| | | | | | | |
|---|---|---|---|---|---|---|
| sf3a3 | 0.01516 | 0.72910565 | 0.429 | 0.073 | 1 | 17 |
| snw1 | 0.01535 | 0.657018501 | 0.5 | 0.122 | 1 | 17 |
| dcaf13 | 0.01602 | -0.65403076 | 0 | 0.244 | 1 | 17 |
| sdhdb | 0.01602 | -0.67630883 | 0 | 0.244 | 1 | 17 |
| cyb5r1 | 0.01638 | 0.493374097 | 0.357 | 0.049 | 1 | 17 |
| actb1 | 0.01733 | 0.226468879 | 0.786 | 0.439 | 1 | 17 |
| hmgb1b | 0.01783 | 0.384865038 | 1 | 0.683 | 1 | 17 |
| emd | 0.01876 | 0.664453198 | 0.5 | 0.122 | 1 | 17 |
| rmnd5b | 0.01879 | 0.542051611 | 0.357 | 0.049 | 1 | 17 |
| chtf18 | 0.01934 | 0.577355557 | 0.357 | 0.049 | 1 | 17 |
| mcm5 | 0.01948 | 0.700517862 | 0.5 | 0.122 | 1 | 17 |
| LOC101883959 | 0.01997 | 0.729291343 | 0.571 | 0.171 | 1 | 17 |
| wdr5 | 0.02017 | 0.535244807 | 0.357 | 0.049 | 1 | 17 |
| fopnl | 0.02045 | 0.668262525 | 0.357 | 0.049 | 1 | 17 |
| alas1 | 0.02078 | 0.674363321 | 0.357 | 0.049 | 1 | 17 |
| rps26 | 0.02084 | -0.1174861 | 0.857 | 0.707 | 1 | 17 |
| mtm1 | 0.02089 | 0.466351036 | 0.286 | 0.024 | 1 | 17 |
| trmt112 | 0.02121 | 0.466853037 | 0.857 | 0.463 | 1 | 17 |
| swt1 | 0.02141 | 0.720400537 | 0.357 | 0.049 | 1 | 17 |
| srsf6b | 0.02147 | 0.549727291 | 0.357 | 0.049 | 1 | 17 |
| npc2 | 0.02152 | 0.444821787 | 0.286 | 0.024 | 1 | 17 |
| zgc:193541 | 0.02154 | 0.665244176 | 0.786 | 0.366 | 1 | 17 |
| akirin1 | 0.02156 | 0.583520253 | 0.357 | 0.049 | 1 | 17 |
| GCA | 0.02165 | 0.62259499 | 0.357 | 0.049 | 1 | 17 |
| ilf2 | 0.02175 | 0.646384917 | 0.786 | 0.366 | 1 | 17 |
| cab39l | 0.02182 | 0.609461168 | 0.357 | 0.049 | 1 | 17 |
| mrpl54 | 0.02207 | 0.54330889 | 0.357 | 0.049 | 1 | 17 |
| trappc6b | 0.02224 | 0.647445547 | 0.357 | 0.049 | 1 | 17 |
| swi5 | 0.02229 | 0.574799859 | 0.357 | 0.049 | 1 | 17 |
| gabarapb | 0.02361 | 0.454682143 | 0.571 | 0.22 | 1 | 17 |
| wu:fb18f06 | 0.02379 | 0.503592374 | 0.286 | 0.024 | 1 | 17 |
| si:ch211-14a17.7 | 0.02397 | -0.60104252 | 0 | 0.22 | 1 | 17 |
| si:dkey-67c22.2 | 0.02397 | -0.62631003 | 0 | 0.22 | 1 | 17 |
| txndc9 | 0.02397 | -0.61777011 | 0 | 0.22 | 1 | 17 |
| coa5 | 0.02397 | -0.54411415 | 0 | 0.22 | 1 | 17 |
| rad9b | 0.02397 | -0.67719634 | 0 | 0.22 | 1 | 17 |
| wdr61 | 0.02537 | 0.539273054 | 0.286 | 0.024 | 1 | 17 |
| sptan1 | 0.02572 | 0.597325505 | 0.286 | 0.024 | 1 | 17 |
| phf19 | 0.02577 | 0.626924 | 0.286 | 0.024 | 1 | 17 |
| rpl7a | 0.02622 | 0.289571201 | 1 | 0.951 | 1 | 17 |
| zgc:91910 | 0.02623 | 0.583813188 | 0.643 | 0.244 | 1 | 17 |
| gins1 | 0.02632 | 0.677542895 | 0.286 | 0.024 | 1 | 17 |
| noc2l | 0.02643 | 0.66682065 | 0.286 | 0.024 | 1 | 17 |
| h2afy2 | 0.02658 | 0.598028537 | 0.643 | 0.244 | 1 | 17 |
| wdr34 | 0.02672 | 0.54897919 | 0.286 | 0.024 | 1 | 17 |
| prdx5 | 0.02675 | 0.513637806 | 0.286 | 0.024 | 1 | 17 |
| sltm | 0.02702 | 0.541622094 | 0.286 | 0.024 | 1 | 17 |
| hikeshi | 0.02721 | 0.559448917 | 0.286 | 0.024 | 1 | 17 |

| | | | | | | |
|---|---|---|---|---|---|---|
| snapc5 | 0.02722 | 0.646921874 | 0.286 | 0.024 | 1 | 17 |
| ube3a | 0.02734 | 0.626448269 | 0.286 | 0.024 | 1 | 17 |
| pin4 | 0.02737 | 0.547981445 | 0.286 | 0.024 | 1 | 17 |
| pole2 | 0.02742 | 0.553392377 | 0.286 | 0.024 | 1 | 17 |
| atf6 | 0.02748 | 0.510265151 | 0.286 | 0.024 | 1 | 17 |
| trmt1 | 0.02755 | 0.630550823 | 0.286 | 0.024 | 1 | 17 |
| cdc42l | 0.02769 | 0.543628638 | 0.429 | 0.098 | 1 | 17 |
| psmb2 | 0.02846 | 0.516178146 | 0.5 | 0.146 | 1 | 17 |
| DYNC2H1 | 0.02955 | 0.354241866 | 0.143 | 0 | 1 | 17 |
| TADA1 | 0.02955 | 0.343722218 | 0.143 | 0 | 1 | 17 |
| cbx3b | 0.02955 | 0.339970564 | 0.143 | 0 | 1 | 17 |
| klf11a | 0.02955 | 0.360252296 | 0.143 | 0 | 1 | 17 |
| tmem203 | 0.02955 | 0.286472804 | 0.143 | 0 | 1 | 17 |
| zgc:158254 | 0.02955 | 0.296999919 | 0.143 | 0 | 1 | 17 |
| znf1152 | 0.02955 | 0.366856952 | 0.143 | 0 | 1 | 17 |
| rpusd2 | 0.02955 | 0.391255872 | 0.143 | 0 | 1 | 17 |
| sytl1 | 0.02955 | 0.437111519 | 0.143 | 0 | 1 | 17 |
| thap12b | 0.02955 | 0.469498086 | 0.143 | 0 | 1 | 17 |
| wbp2nl | 0.02955 | 0.377508604 | 0.143 | 0 | 1 | 17 |
| CU927934.3 | 0.02955 | 0.337822494 | 0.143 | 0 | 1 | 17 |
| arhgef1b | 0.02955 | 0.295984369 | 0.143 | 0 | 1 | 17 |
| cebpz | 0.02955 | 0.297428946 | 0.143 | 0 | 1 | 17 |
| chd3 | 0.02955 | 0.335256637 | 0.143 | 0 | 1 | 17 |
| crata | 0.02955 | 0.341220842 | 0.143 | 0 | 1 | 17 |
| dbnlb | 0.02955 | 0.30889182 | 0.143 | 0 | 1 | 17 |
| dhx38 | 0.02955 | 0.355221426 | 0.143 | 0 | 1 | 17 |
| dnajc1 | 0.02955 | 0.331395147 | 0.143 | 0 | 1 | 17 |
| dpp7 | 0.02955 | 0.28895978 | 0.143 | 0 | 1 | 17 |
| fbxo18 | 0.02955 | 0.331194234 | 0.143 | 0 | 1 | 17 |
| hdhd3 | 0.02955 | 0.305460963 | 0.143 | 0 | 1 | 17 |
| irf2bp2b | 0.02955 | 0.31650509 | 0.143 | 0 | 1 | 17 |
| kpna3 | 0.02955 | 0.3116349 | 0.143 | 0 | 1 | 17 |
| ldb1b | 0.02955 | 0.346853517 | 0.143 | 0 | 1 | 17 |
| lmcd1 | 0.02955 | 0.387557518 | 0.143 | 0 | 1 | 17 |
| myh11a | 0.02955 | 0.379385911 | 0.143 | 0 | 1 | 17 |
| ntmt1 | 0.02955 | 0.368885712 | 0.143 | 0 | 1 | 17 |
| osbp | 0.02955 | 0.299264377 | 0.143 | 0 | 1 | 17 |
| prr14 | 0.02955 | 0.343884906 | 0.143 | 0 | 1 | 17 |
| rab8a | 0.02955 | 0.316176016 | 0.143 | 0 | 1 | 17 |
| rnf40 | 0.02955 | 0.333159269 | 0.143 | 0 | 1 | 17 |
| sec23ip | 0.02955 | 0.30290511 | 0.143 | 0 | 1 | 17 |
| sgk3 | 0.02955 | 0.327347741 | 0.143 | 0 | 1 | 17 |
| si:ch211-195o20.7 | 0.02955 | 0.249826406 | 0.143 | 0 | 1 | 17 |
| si:ch211-197k17.3 | 0.02955 | 0.299630623 | 0.143 | 0 | 1 | 17 |
| si:ch211-209a2.1 | 0.02955 | 0.30290511 | 0.143 | 0 | 1 | 17 |
| si:ch73-25f10.6 | 0.02955 | 0.296807883 | 0.143 | 0 | 1 | 17 |
| si:dkeyp-104h9.5 | 0.02955 | 0.344824241 | 0.143 | 0 | 1 | 17 |
| srbd1 | 0.02955 | 0.30290511 | 0.143 | 0 | 1 | 17 |

| | | | | | | |
|---|---|---|---|---|---|---|
| tmem60 | 0.02955 | 0.416831137 | 0.143 | 0 | 1 | 17 |
| tspan9b | 0.02955 | 0.408876462 | 0.143 | 0 | 1 | 17 |
| wapla | 0.02955 | 0.298274948 | 0.143 | 0 | 1 | 17 |
| wdr74 | 0.02955 | 0.302549629 | 0.143 | 0 | 1 | 17 |
| BX927282.2 | 0.02955 | 0.319439834 | 0.143 | 0 | 1 | 17 |
| ankrd29 | 0.02955 | 0.360164898 | 0.143 | 0 | 1 | 17 |
| ap1g2 | 0.02955 | 0.353777703 | 0.143 | 0 | 1 | 17 |
| asmt2 | 0.02955 | 0.268589515 | 0.143 | 0 | 1 | 17 |
| bin2b | 0.02955 | 0.340648398 | 0.143 | 0 | 1 | 17 |
| bnip4 | 0.02955 | 0.311738103 | 0.143 | 0 | 1 | 17 |
| btr09 | 0.02955 | 0.430000459 | 0.143 | 0 | 1 | 17 |
| caap1 | 0.02955 | 0.330422624 | 0.143 | 0 | 1 | 17 |
| capgb | 0.02955 | 0.365945051 | 0.143 | 0 | 1 | 17 |
| clul1 | 0.02955 | 0.397008556 | 0.143 | 0 | 1 | 17 |
| corin | 0.02955 | 0.242776356 | 0.143 | 0 | 1 | 17 |
| dctn1a | 0.02955 | 0.327437419 | 0.143 | 0 | 1 | 17 |
| fam207a | 0.02955 | 0.335467124 | 0.143 | 0 | 1 | 17 |
| fhl1b | 0.02955 | 0.397008556 | 0.143 | 0 | 1 | 17 |
| ftr98 | 0.02955 | 0.360164898 | 0.143 | 0 | 1 | 17 |
| hinfp | 0.02955 | 0.310758927 | 0.143 | 0 | 1 | 17 |
| ift20 | 0.02955 | 0.346316416 | 0.143 | 0 | 1 | 17 |
| inpp5ka | 0.02955 | 0.329451979 | 0.143 | 0 | 1 | 17 |
| kdm2aa | 0.02955 | 0.314892937 | 0.143 | 0 | 1 | 17 |
| kmt5aa | 0.02955 | 0.344749769 | 0.143 | 0 | 1 | 17 |
| nit1 | 0.02955 | 0.342732537 | 0.143 | 0 | 1 | 17 |
| nkd2a | 0.02955 | 0.346805333 | 0.143 | 0 | 1 | 17 |
| nucks1b | 0.02955 | 0.360901178 | 0.143 | 0 | 1 | 17 |
| pigv | 0.02955 | 0.394339235 | 0.143 | 0 | 1 | 17 |
| pja2 | 0.02955 | 0.391437665 | 0.143 | 0 | 1 | 17 |
| poli | 0.02955 | 0.341951387 | 0.143 | 0 | 1 | 17 |
| prmt3 | 0.02955 | 0.404866177 | 0.143 | 0 | 1 | 17 |
| ranbp9 | 0.02955 | 0.370692728 | 0.143 | 0 | 1 | 17 |
| rap1aa | 0.02955 | 0.44291765 | 0.143 | 0 | 1 | 17 |
| selenbp1 | 0.02955 | 0.365493258 | 0.143 | 0 | 1 | 17 |
| si:ch1073-228b5.2 | 0.02955 | 0.374972625 | 0.143 | 0 | 1 | 17 |
| si:dkeyp-51b7.3 | 0.02955 | 0.360164898 | 0.143 | 0 | 1 | 17 |
| snapc4 | 0.02955 | 0.431600637 | 0.143 | 0 | 1 | 17 |
| snd1 | 0.02955 | 0.303374591 | 0.143 | 0 | 1 | 17 |
| tmem120b | 0.02955 | 0.360164898 | 0.143 | 0 | 1 | 17 |
| trim35-40 | 0.02955 | 0.399235822 | 0.143 | 0 | 1 | 17 |
| ube2j2 | 0.02955 | 0.349129319 | 0.143 | 0 | 1 | 17 |
| uqcrb | 0.02955 | 0.395388649 | 0.143 | 0 | 1 | 17 |
| wdr66 | 0.02955 | 0.399235822 | 0.143 | 0 | 1 | 17 |
| wnt4a | 0.02955 | 0.34915759 | 0.143 | 0 | 1 | 17 |
| yme1l1a | 0.02955 | 0.349127467 | 0.143 | 0 | 1 | 17 |
| zgc:113054 | 0.02955 | 0.386466992 | 0.143 | 0 | 1 | 17 |
| znf367 | 0.02955 | 0.331922919 | 0.143 | 0 | 1 | 17 |
| zte38 | 0.02955 | 0.343878151 | 0.143 | 0 | 1 | 17 |

| | | | | | | |
|---|---|---|---|---|---|---|
| BX324164.3 | 0.02955 | 0.392522825 | 0.143 | 0 | 1 | 17 |
| BX465862.2 | 0.02955 | 0.352144871 | 0.143 | 0 | 1 | 17 |
| badb | 0.02955 | 0.311499595 | 0.143 | 0 | 1 | 17 |
| commd6 | 0.02955 | 0.317998676 | 0.143 | 0 | 1 | 17 |
| degs1 | 0.02955 | 0.349392673 | 0.143 | 0 | 1 | 17 |
| desi1a | 0.02955 | 0.211874589 | 0.143 | 0 | 1 | 17 |
| diablob | 0.02955 | 0.394105711 | 0.143 | 0 | 1 | 17 |
| dlgap4a | 0.02955 | 0.389758016 | 0.143 | 0 | 1 | 17 |
| drc1 | 0.02955 | 0.392522825 | 0.143 | 0 | 1 | 17 |
| dync1i1 | 0.02955 | 0.309867457 | 0.143 | 0 | 1 | 17 |
| elk1 | 0.02955 | 0.301307649 | 0.143 | 0 | 1 | 17 |
| epdr1 | 0.02955 | 0.309867457 | 0.143 | 0 | 1 | 17 |
| galnt7 | 0.02955 | 0.331646758 | 0.143 | 0 | 1 | 17 |
| hsd17b8 | 0.02955 | 0.287294762 | 0.143 | 0 | 1 | 17 |
| irf10 | 0.02955 | 0.380699644 | 0.143 | 0 | 1 | 17 |
| irf5 | 0.02955 | 0.34788058 | 0.143 | 0 | 1 | 17 |
| lrrc42 | 0.02955 | 0.387315543 | 0.143 | 0 | 1 | 17 |
| neil1 | 0.02955 | 0.349404177 | 0.143 | 0 | 1 | 17 |
| plekha5 | 0.02955 | 0.352144871 | 0.143 | 0 | 1 | 17 |
| polg | 0.02955 | 0.315058774 | 0.143 | 0 | 1 | 17 |
| ptbp2b | 0.02955 | 0.298562913 | 0.143 | 0 | 1 | 17 |
| scaper | 0.02955 | 0.32212027 | 0.143 | 0 | 1 | 17 |
| si:ch211-155e24.3 | 0.02955 | 0.283683761 | 0.143 | 0 | 1 | 17 |
| si:dkey-238f9.1 | 0.02955 | 0.389435573 | 0.143 | 0 | 1 | 17 |
| snx30 | 0.02955 | 0.338021709 | 0.143 | 0 | 1 | 17 |
| snx9b | 0.02955 | 0.357094246 | 0.143 | 0 | 1 | 17 |
| taf6 | 0.02955 | 0.318510747 | 0.143 | 0 | 1 | 17 |
| timmdc1 | 0.02955 | 0.333960166 | 0.143 | 0 | 1 | 17 |
| ube2ka | 0.02955 | 0.378906133 | 0.143 | 0 | 1 | 17 |
| wdr73 | 0.02955 | 0.352144871 | 0.143 | 0 | 1 | 17 |
| zdbf2 | 0.02955 | 0.347106123 | 0.143 | 0 | 1 | 17 |
| zgc:162239.1 | 0.02955 | 0.325227722 | 0.143 | 0 | 1 | 17 |
| acot14 | 0.02955 | 0.419123753 | 0.143 | 0 | 1 | 17 |
| ankrd11 | 0.02955 | 0.260695213 | 0.143 | 0 | 1 | 17 |
| arl8ba | 0.02955 | 0.356969372 | 0.143 | 0 | 1 | 17 |
| brip1 | 0.02955 | 0.284279041 | 0.143 | 0 | 1 | 17 |
| carm1l | 0.02955 | 0.259395678 | 0.143 | 0 | 1 | 17 |
| cutc | 0.02955 | 0.263164024 | 0.143 | 0 | 1 | 17 |
| cxxc1a | 0.02955 | 0.386016989 | 0.143 | 0 | 1 | 17 |
| cyth1b | 0.02955 | 0.354233756 | 0.143 | 0 | 1 | 17 |
| ddhd1a | 0.02955 | 0.288427808 | 0.143 | 0 | 1 | 17 |
| dlx6a | 0.02955 | 0.389191301 | 0.143 | 0 | 1 | 17 |
| dmap1 | 0.02955 | 0.282186849 | 0.143 | 0 | 1 | 17 |
| dnajc12 | 0.02955 | 0.259645424 | 0.143 | 0 | 1 | 17 |
| fam49a | 0.02955 | 0.258030373 | 0.143 | 0 | 1 | 17 |
| fstl5 | 0.02955 | 0.419123753 | 0.143 | 0 | 1 | 17 |
| golga7ba | 0.02955 | 0.265391391 | 0.143 | 0 | 1 | 17 |
| hps5 | 0.02955 | 0.310887604 | 0.143 | 0 | 1 | 17 |

| | | | | | | |
|---|---|---|---|---|---|---|
| ift52 | 0.02955 | 0.362594641 | 0.143 | 0 | 1 | 17 |
| ints12 | 0.02955 | 0.309022588 | 0.143 | 0 | 1 | 17 |
| kdm2ab | 0.02955 | 0.286384146 | 0.143 | 0 | 1 | 17 |
| mblac1 | 0.02955 | 0.261119511 | 0.143 | 0 | 1 | 17 |
| npl | 0.02955 | 0.269091319 | 0.143 | 0 | 1 | 17 |
| nr2c2ap | 0.02955 | 0.418323437 | 0.143 | 0 | 1 | 17 |
| oip5-as1 | 0.02955 | 0.404096106 | 0.143 | 0 | 1 | 17 |
| plekhj1 | 0.02955 | 0.276900547 | 0.143 | 0 | 1 | 17 |
| psme2 | 0.02955 | 0.252698192 | 0.143 | 0 | 1 | 17 |
| rbm11 | 0.02955 | 0.36703472 | 0.143 | 0 | 1 | 17 |
| rtn3 | 0.02955 | 0.413551309 | 0.143 | 0 | 1 | 17 |
| si:ch211-197h24.8 | 0.02955 | 0.262397748 | 0.143 | 0 | 1 | 17 |
| si:ch73-217b7.1 | 0.02955 | 0.246754189 | 0.143 | 0 | 1 | 17 |
| si:dkeyp-2e4.6 | 0.02955 | 0.36703472 | 0.143 | 0 | 1 | 17 |
| tab1 | 0.02955 | 0.400455134 | 0.143 | 0 | 1 | 17 |
| tmem248 | 0.02955 | 0.409619693 | 0.143 | 0 | 1 | 17 |
| tmem65 | 0.02955 | 0.259043163 | 0.143 | 0 | 1 | 17 |
| wasf2 | 0.02955 | 0.399885834 | 0.143 | 0 | 1 | 17 |
| wnt8b | 0.02955 | 0.288427808 | 0.143 | 0 | 1 | 17 |
| znf609b | 0.02955 | 0.265391391 | 0.143 | 0 | 1 | 17 |
| BX649250.1 | 0.02955 | 0.267519166 | 0.143 | 0 | 1 | 17 |
| CABZ01086877.1 | 0.02955 | 0.26260601 | 0.143 | 0 | 1 | 17 |
| CU929037.1 | 0.02955 | 0.326038903 | 0.143 | 0 | 1 | 17 |
| CU929391.1 | 0.02955 | 0.275884215 | 0.143 | 0 | 1 | 17 |
| LOC103909808 | 0.02955 | 0.263008923 | 0.143 | 0 | 1 | 17 |
| LOC110439840 | 0.02955 | 0.258221553 | 0.143 | 0 | 1 | 17 |
| ahctf1 | 0.02955 | 0.271661366 | 0.143 | 0 | 1 | 17 |
| asrgl1 | 0.02955 | 0.314244287 | 0.143 | 0 | 1 | 17 |
| atp2b1a | 0.02955 | 0.323541937 | 0.143 | 0 | 1 | 17 |
| atp6v1aa | 0.02955 | 0.312353538 | 0.143 | 0 | 1 | 17 |
| ccdc73 | 0.02955 | 0.275884215 | 0.143 | 0 | 1 | 17 |
| cfap298 | 0.02955 | 0.276861769 | 0.143 | 0 | 1 | 17 |
| clcn5b | 0.02955 | 0.240683345 | 0.143 | 0 | 1 | 17 |
| cldn5b | 0.02955 | 0.263297593 | 0.143 | 0 | 1 | 17 |
| cmbl | 0.02955 | 0.381278293 | 0.143 | 0 | 1 | 17 |
| commd9 | 0.02955 | 0.30093001 | 0.143 | 0 | 1 | 17 |
| crnkl1 | 0.02955 | 0.276695389 | 0.143 | 0 | 1 | 17 |
| ctc1 | 0.02955 | 0.377665371 | 0.143 | 0 | 1 | 17 |
| dennd2da | 0.02955 | 0.367161983 | 0.143 | 0 | 1 | 17 |
| dicer1 | 0.02955 | 0.319003291 | 0.143 | 0 | 1 | 17 |
| gcc1 | 0.02955 | 0.275534812 | 0.143 | 0 | 1 | 17 |
| gga3a | 0.02955 | 0.334204962 | 0.143 | 0 | 1 | 17 |
| gle1 | 0.02955 | 0.270000443 | 0.143 | 0 | 1 | 17 |
| gon4l | 0.02955 | 0.322979211 | 0.143 | 0 | 1 | 17 |
| hsd17b7 | 0.02955 | 0.345553902 | 0.143 | 0 | 1 | 17 |
| kif5bb | 0.02955 | 0.326038903 | 0.143 | 0 | 1 | 17 |
| ktn1 | 0.02955 | 0.365393429 | 0.143 | 0 | 1 | 17 |
| march6 | 0.02955 | 0.269742036 | 0.143 | 0 | 1 | 17 |

| | | | | | | |
|---|---|---|---|---|---|---|
| mier1a | 0.02955 | 0.347733205 | 0.143 | 0 | 1 | 17 |
| nipbla | 0.02955 | 0.258432289 | 0.143 | 0 | 1 | 17 |
| nnt | 0.02955 | 0.369620885 | 0.143 | 0 | 1 | 17 |
| nrarpa | 0.02955 | 0.39764851 | 0.143 | 0 | 1 | 17 |
| nsmfb | 0.02955 | 0.334204962 | 0.143 | 0 | 1 | 17 |
| ormdl1 | 0.02955 | 0.254409811 | 0.143 | 0 | 1 | 17 |
| paox1 | 0.02955 | 0.289239946 | 0.143 | 0 | 1 | 17 |
| pdk4 | 0.02955 | 0.259424795 | 0.143 | 0 | 1 | 17 |
| peli1b | 0.02955 | 0.272180445 | 0.143 | 0 | 1 | 17 |
| prcc | 0.02955 | 0.260994579 | 0.143 | 0 | 1 | 17 |
| prrc2a | 0.02955 | 0.269123969 | 0.143 | 0 | 1 | 17 |
| rcor1 | 0.02955 | 0.301952081 | 0.143 | 0 | 1 | 17 |
| rnf181 | 0.02955 | 0.268955079 | 0.143 | 0 | 1 | 17 |
| rp9 | 0.02955 | 0.36345385 | 0.143 | 0 | 1 | 17 |
| sgsh | 0.02955 | 0.322910575 | 0.143 | 0 | 1 | 17 |
| sh3bp5la | 0.02955 | 0.298221421 | 0.143 | 0 | 1 | 17 |
| si:ch211-152c8.4 | 0.02955 | 0.27142119 | 0.143 | 0 | 1 | 17 |
| si:ch73-299h12.9 | 0.02955 | 0.296168386 | 0.143 | 0 | 1 | 17 |
| si:ch73-347e22.4 | 0.02955 | 0.321055839 | 0.143 | 0 | 1 | 17 |
| si:dkey-288a3.2 | 0.02955 | 0.359157588 | 0.143 | 0 | 1 | 17 |
| slc20a1b | 0.02955 | 0.275489283 | 0.143 | 0 | 1 | 17 |
| slc7a10a | 0.02955 | 0.250023954 | 0.143 | 0 | 1 | 17 |
| tax1bp1b | 0.02955 | 0.268882208 | 0.143 | 0 | 1 | 17 |
| tbc1d14 | 0.02955 | 0.259570024 | 0.143 | 0 | 1 | 17 |
| trnau1apb | 0.02955 | 0.352466129 | 0.143 | 0 | 1 | 17 |
| usp47 | 0.02955 | 0.259521772 | 0.143 | 0 | 1 | 17 |
| usp7 | 0.02955 | 0.267238065 | 0.143 | 0 | 1 | 17 |
| vav2 | 0.02955 | 0.325825633 | 0.143 | 0 | 1 | 17 |
| vps41 | 0.02955 | 0.376328114 | 0.143 | 0 | 1 | 17 |
| xpo1a | 0.02955 | 0.32123398 | 0.143 | 0 | 1 | 17 |
| zgc:110319 | 0.02955 | 0.342458989 | 0.143 | 0 | 1 | 17 |
| znf384l | 0.02955 | 0.272797272 | 0.143 | 0 | 1 | 17 |
| birc5a | 0.02994 | 0.44573957 | 0.571 | 0.22 | 1 | 17 |
| pabpc1b | 0.03013 | 0.539211059 | 0.5 | 0.146 | 1 | 17 |
| clpp | 0.03086 | 0.584650974 | 0.429 | 0.098 | 1 | 17 |
| polr3f | 0.03152 | 0.699584105 | 0.429 | 0.098 | 1 | 17 |
| chd4a | 0.03203 | 0.570733735 | 0.5 | 0.146 | 1 | 17 |
| selenou1a | 0.03238 | 0.598483104 | 0.5 | 0.146 | 1 | 17 |
| atp5l | 0.03264 | 0.614502863 | 0.786 | 0.39 | 1 | 17 |
| naa40 | 0.0336 | 0.701380389 | 0.429 | 0.098 | 1 | 17 |
| h2afva | 0.0339 | 0.648318 | 0.714 | 0.317 | 1 | 17 |
| psma2 | 0.03403 | 0.615129797 | 0.5 | 0.146 | 1 | 17 |
| calm2b | 0.03415 | 0.201179979 | 1 | 0.756 | 1 | 17 |
| si:dkey-28b4.7 | 0.03428 | 0.429572759 | 0.357 | 0.073 | 1 | 17 |
| sub1b | 0.03442 | 0.406876335 | 0.929 | 0.585 | 1 | 17 |
| ap2m1a | 0.0346 | 0.626098279 | 0.5 | 0.146 | 1 | 17 |
| ube2d1b | 0.03499 | 0.262802011 | 0.571 | 0.268 | 1 | 17 |
| cct5 | 0.03515 | 0.363422163 | 0.857 | 0.512 | 1 | 17 |

| | | | | | | |
|---|---|---|---|---|---|---|
| cpsf2 | 0.03523 | 0.584021918 | 0.5 | 0.146 | 1 | 17 |
| eif4a1b | 0.03529 | 0.501728078 | 0.643 | 0.268 | 1 | 17 |
| CABZ01092156.1 | 0.03564 | -0.54343284 | 0 | 0.195 | 1 | 17 |
| mt-co2 | 0.03564 | -0.7934106 | 0 | 0.195 | 1 | 17 |
| CU929070.1 | 0.03564 | -0.54193592 | 0 | 0.195 | 1 | 17 |
| cnbpb | 0.03564 | -0.496596 | 0 | 0.195 | 1 | 17 |
| fkbp2 | 0.03564 | -0.51606985 | 0 | 0.195 | 1 | 17 |
| hnrnph1l | 0.03564 | -0.51892397 | 0 | 0.195 | 1 | 17 |
| si:ch211-102c2.7 | 0.03564 | -0.51208683 | 0 | 0.195 | 1 | 17 |
| zar1 | 0.03564 | -0.62648013 | 0 | 0.195 | 1 | 17 |
| ubtfl | 0.03564 | -0.53710855 | 0 | 0.195 | 1 | 17 |
| si:ch211-162e15.3 | 0.03626 | 0.451478426 | 0.357 | 0.073 | 1 | 17 |
| anp32e | 0.03745 | 0.521439209 | 0.357 | 0.073 | 1 | 17 |
| seta | 0.03905 | 0.550445584 | 0.857 | 0.488 | 1 | 17 |
| mphosph8 | 0.03925 | 0.411142 | 0.643 | 0.293 | 1 | 17 |
| eif3f | 0.03969 | 0.523554424 | 0.857 | 0.488 | 1 | 17 |
| eif2s2 | 0.04044 | 0.149305849 | 0.857 | 0.585 | 1 | 17 |
| ncl | 0.04112 | 0.364591231 | 0.643 | 0.317 | 1 | 17 |
| sap18 | 0.04134 | 0.672129882 | 0.714 | 0.341 | 1 | 17 |
| dpy30 | 0.04256 | 0.310909068 | 0.857 | 0.537 | 1 | 17 |
| nudc | 0.04334 | 0.264502764 | 0.714 | 0.415 | 1 | 17 |
| prdx3 | 0.04447 | 0.565925089 | 0.643 | 0.268 | 1 | 17 |
| h3f3c | 0.04506 | 0.525756103 | 0.714 | 0.341 | 1 | 17 |
| eif3c | 0.04616 | 0.516388275 | 0.429 | 0.122 | 1 | 17 |
| leng1 | 0.04706 | 0.551858077 | 0.357 | 0.073 | 1 | 17 |
| prmt5 | 0.04725 | 0.773607905 | 0.429 | 0.122 | 1 | 17 |
| eef1da | 0.04728 | 0.151877409 | 0.786 | 0.512 | 1 | 17 |
| mrpl44 | 0.04785 | 0.490636548 | 0.357 | 0.073 | 1 | 17 |
| idh3a | 0.04786 | 0.527142823 | 0.357 | 0.073 | 1 | 17 |
| sub1a | 0.04944 | 0.560453038 | 0.786 | 0.415 | 1 | 17 |
| tpd52l2a | 0.04958 | 0.642323036 | 0.357 | 0.073 | 1 | 17 |
| srsf5a | 0.05019 | 0.336520613 | 0.714 | 0.39 | 1 | 17 |
| cd3eap | 0.05078 | 0.631597086 | 0.357 | 0.073 | 1 | 17 |
| si:dkey-87o1.2 | 0.05088 | 0.375556621 | 0.286 | 0.049 | 1 | 17 |
| smc5 | 0.05149 | 0.567968967 | 0.357 | 0.073 | 1 | 17 |
| ddx61 | 0.05215 | 0.332659262 | 0.714 | 0.39 | 1 | 17 |
| psmd11b | 0.05228 | 0.645737211 | 0.357 | 0.073 | 1 | 17 |
| hspa4b | 0.05235 | 0.586413273 | 0.357 | 0.073 | 1 | 17 |
| tram2 | 0.05273 | -0.49780706 | 0 | 0.171 | 1 | 17 |
| ap2s1 | 0.05273 | -0.46329652 | 0 | 0.171 | 1 | 17 |
| nsun2 | 0.05273 | -0.47541239 | 0 | 0.171 | 1 | 17 |
| spink2.1 | 0.05273 | -0.44195075 | 0 | 0.171 | 1 | 17 |
| zmat5 | 0.05273 | -0.47241283 | 0 | 0.171 | 1 | 17 |
| hexim1 | 0.05273 | -0.44220974 | 0 | 0.171 | 1 | 17 |
| snap23.1 | 0.05273 | -0.43623322 | 0 | 0.171 | 1 | 17 |
| gstm.3 | 0.05273 | -0.47863011 | 0 | 0.171 | 1 | 17 |
| hnrnpa0b | 0.0528 | 0.119431098 | 0.786 | 0.537 | 1 | 17 |
| pfdn6 | 0.05297 | 0.47409493 | 0.643 | 0.293 | 1 | 17 |

| | | | | | | |
|---|---|---|---|---|---|---|
| odf3l2a | 0.05335 | 0.737013191 | 0.429 | 0.146 | 1 | 17 |
| lgals3a | 0.05355 | 0.539981934 | 0.357 | 0.073 | 1 | 17 |
| si:ch73-167f10.1 | 0.05362 | 0.52249957 | 0.786 | 0.61 | 1 | 17 |
| STMP1 | 0.05388 | 0.471466597 | 0.643 | 0.293 | 1 | 17 |
| pfn2l | 0.05457 | 0.581285567 | 0.571 | 0.22 | 1 | 17 |
| ost4 | 0.05642 | 0.5568439 | 0.571 | 0.22 | 1 | 17 |
| khdrbs1a | 0.05723 | 0.536828205 | 0.857 | 0.537 | 1 | 17 |
| mtap | 0.05927 | 0.550197986 | 0.5 | 0.171 | 1 | 17 |
| rpp21 | 0.05939 | 0.553467575 | 0.429 | 0.122 | 1 | 17 |
| apooa | 0.06093 | 0.543701119 | 0.429 | 0.122 | 1 | 17 |
| arl15a | 0.06136 | 0.618077441 | 0.429 | 0.122 | 1 | 17 |
| hsdl2 | 0.06136 | 0.548282986 | 0.429 | 0.122 | 1 | 17 |
| lyrm9 | 0.06188 | 0.502728746 | 0.429 | 0.122 | 1 | 17 |
| mtx2 | 0.06216 | 0.557327162 | 0.429 | 0.122 | 1 | 17 |
| mrps30 | 0.06264 | 0.406707736 | 0.286 | 0.049 | 1 | 17 |
| ndufb5 | 0.06296 | 0.475517942 | 0.643 | 0.293 | 1 | 17 |
| rcc2 | 0.06318 | 0.353155604 | 0.714 | 0.39 | 1 | 17 |
| tdrkh | 0.06408 | 0.588673067 | 0.5 | 0.171 | 1 | 17 |
| rad21a | 0.06409 | 0.655483385 | 0.429 | 0.122 | 1 | 17 |
| nme4 | 0.06418 | 0.563453761 | 0.5 | 0.171 | 1 | 17 |
| mrps25 | 0.06427 | 0.624592898 | 0.5 | 0.171 | 1 | 17 |
| gtf2h5 | 0.0644 | 0.591577217 | 0.429 | 0.122 | 1 | 17 |
| rbm39a | 0.06537 | 0.374231666 | 0.286 | 0.049 | 1 | 17 |
| si:ch211-126c2.4 | 0.06574 | 0.401457432 | 0.286 | 0.049 | 1 | 17 |
| psma5 | 0.0664 | 0.461919986 | 0.786 | 0.439 | 1 | 17 |
| nob1 | 0.06725 | 0.467184399 | 0.286 | 0.049 | 1 | 17 |
| rpl27 | 0.06748 | 0.317224891 | 1 | 0.78 | 1 | 17 |
| fen1 | 0.06791 | 0.543388217 | 0.643 | 0.293 | 1 | 17 |
| pin1 | 0.06809 | 0.41662001 | 0.5 | 0.195 | 1 | 17 |
| dynll2b | 0.06824 | 0.235519824 | 0.714 | 0.439 | 1 | 17 |
| tob1a | 0.06843 | 0.595912349 | 0.286 | 0.049 | 1 | 17 |
| ndufa6 | 0.06851 | 0.455844242 | 0.714 | 0.366 | 1 | 17 |
| coa3a | 0.06873 | 0.432505551 | 0.286 | 0.049 | 1 | 17 |
| si:ch211-226l4.6 | 0.0689 | 0.436442556 | 0.714 | 0.366 | 1 | 17 |
| chtf8 | 0.06919 | 0.698197431 | 0.286 | 0.049 | 1 | 17 |
| chchd2 | 0.06941 | 0.399022706 | 0.357 | 0.098 | 1 | 17 |
| senp3a | 0.06941 | 0.42888538 | 0.286 | 0.049 | 1 | 17 |
| ino80c | 0.06999 | 0.468333068 | 0.286 | 0.049 | 1 | 17 |
| fthl27 | 0.07006 | 0.259063265 | 0.786 | 0.488 | 1 | 17 |
| waslb | 0.07011 | 0.44417857 | 0.286 | 0.049 | 1 | 17 |
| si:ch211-176l24.4 | 0.07036 | 0.715836477 | 0.5 | 0.195 | 1 | 17 |
| as3mt | 0.07052 | 0.409619025 | 0.286 | 0.049 | 1 | 17 |
| selenoh | 0.07057 | 0.582116018 | 0.643 | 0.293 | 1 | 17 |
| senp2 | 0.07081 | 0.480737174 | 0.286 | 0.049 | 1 | 17 |
| morc3a | 0.0712 | 0.308280054 | 0.429 | 0.171 | 1 | 17 |
| rps23 | 0.07161 | 0.164101664 | 0.929 | 0.683 | 1 | 17 |
| rps26l | 0.07176 | 0.306183645 | 1 | 0.78 | 1 | 17 |
| rps4x | 0.07196 | 0.305274824 | 1 | 0.78 | 1 | 17 |

| | | | | | | |
|---|---|---|---|---|---|---|
| si:ch211-286b5.2 | 0.07254 | 0.500936124 | 0.286 | 0.049 | 1 | 17 |
| rmdn1 | 0.07264 | 0.383379848 | 0.286 | 0.049 | 1 | 17 |
| ndufa3 | 0.07264 | 0.399378548 | 0.643 | 0.317 | 1 | 17 |
| ccni | 0.07317 | 0.364629183 | 0.357 | 0.098 | 1 | 17 |
| sncb | 0.07343 | 0.419651689 | 0.286 | 0.049 | 1 | 17 |
| uba52 | 0.07346 | 0.391500945 | 0.929 | 0.634 | 1 | 17 |
| sfpq | 0.07352 | 0.384957242 | 0.357 | 0.098 | 1 | 17 |
| CR382383.2 | 0.07378 | 0.687277196 | 0.429 | 0.146 | 1 | 17 |
| cdkn1cb | 0.07474 | 0.428958305 | 0.286 | 0.049 | 1 | 17 |
| thoc1 | 0.07483 | 0.556036921 | 0.286 | 0.049 | 1 | 17 |
| pacsin3 | 0.07552 | 0.50572769 | 0.286 | 0.049 | 1 | 17 |
| nat9 | 0.07571 | 0.423902909 | 0.286 | 0.049 | 1 | 17 |
| exosc2 | 0.07594 | 0.482521129 | 0.286 | 0.049 | 1 | 17 |
| snu13b | 0.07602 | 0.389047935 | 0.857 | 0.537 | 1 | 17 |
| mrnip | 0.0761 | 0.537193353 | 0.286 | 0.049 | 1 | 17 |
| bnip2 | 0.07619 | 0.484804909 | 0.286 | 0.049 | 1 | 17 |
| gapdhs | 0.07642 | 0.12700461 | 0.786 | 0.561 | 1 | 17 |
| phc2a | 0.07645 | 0.602926615 | 0.286 | 0.049 | 1 | 17 |
| anapc13 | 0.07671 | 0.528556628 | 0.286 | 0.049 | 1 | 17 |
| pfkfb4b | 0.07691 | 0.482985811 | 0.286 | 0.049 | 1 | 17 |
| uqcrc2a | 0.07735 | 0.429308656 | 0.286 | 0.049 | 1 | 17 |
| coro1ca | 0.07741 | 0.480544689 | 0.286 | 0.049 | 1 | 17 |
| si:cabz01101003.1 | 0.07741 | 0.479083179 | 0.286 | 0.049 | 1 | 17 |
| gpd1a | 0.07744 | 0.429074659 | 0.286 | 0.049 | 1 | 17 |
| lrrc47 | 0.07747 | 0.578782167 | 0.286 | 0.049 | 1 | 17 |
| fnbp4 | 0.07754 | 0.509811801 | 0.286 | 0.049 | 1 | 17 |
| atf4b | 0.07774 | -0.38724618 | 0 | 0.146 | 1 | 17 |
| cox8b | 0.07774 | -0.44910792 | 0 | 0.146 | 1 | 17 |
| exosc1 | 0.07774 | -0.40797948 | 0 | 0.146 | 1 | 17 |
| gpd1b | 0.07774 | -0.33014026 | 0 | 0.146 | 1 | 17 |
| orc2 | 0.07774 | -0.39461923 | 0 | 0.146 | 1 | 17 |
| sdf2l1 | 0.07774 | -0.35681308 | 0 | 0.146 | 1 | 17 |
| slbp2 | 0.07774 | -0.42778665 | 0 | 0.146 | 1 | 17 |
| snip1 | 0.07774 | -0.39742811 | 0 | 0.146 | 1 | 17 |
| stx12 | 0.07774 | -0.43536468 | 0 | 0.146 | 1 | 17 |
| znf451 | 0.07774 | -0.42007502 | 0 | 0.146 | 1 | 17 |
| arf1 | 0.07774 | -0.36939162 | 0 | 0.146 | 1 | 17 |
| ctnnb2 | 0.07774 | -0.41763738 | 0 | 0.146 | 1 | 17 |
| dek | 0.07774 | -0.42714691 | 0 | 0.146 | 1 | 17 |
| fkbp1ab | 0.07774 | -0.45728368 | 0 | 0.146 | 1 | 17 |
| grcc10 | 0.07774 | -0.42376472 | 0 | 0.146 | 1 | 17 |
| nelfb | 0.07774 | -0.39829179 | 0 | 0.146 | 1 | 17 |
| nmt2 | 0.07774 | -0.42499616 | 0 | 0.146 | 1 | 17 |
| pcid2 | 0.07774 | -0.42622523 | 0 | 0.146 | 1 | 17 |
| ppm1la | 0.07774 | -0.42091366 | 0 | 0.146 | 1 | 17 |
| si:dkey-37o8.1 | 0.07774 | -0.45866164 | 0 | 0.146 | 1 | 17 |
| sri | 0.07774 | -0.43298474 | 0 | 0.146 | 1 | 17 |
| srprb | 0.07774 | -0.39659327 | 0 | 0.146 | 1 | 17 |

| | | | | | | |
|---|---|---|---|---|---|---|
| tars | 0.07774 | -0.47056051 | 0 | 0.146 | 1 | 17 |
| dcaf7 | 0.07774 | -0.38339904 | 0 | 0.146 | 1 | 17 |
| slc35f2 | 0.07774 | -0.35366177 | 0 | 0.146 | 1 | 17 |
| zswim7 | 0.07774 | -0.44959294 | 0 | 0.146 | 1 | 17 |
| gstk1 | 0.07775 | 0.472671681 | 0.286 | 0.049 | 1 | 17 |
| rpl24 | 0.07778 | 0.277073572 | 1 | 0.78 | 1 | 17 |
| calm2a | 0.07785 | 0.534329354 | 0.286 | 0.049 | 1 | 17 |
| smarcad1a | 0.07788 | 0.391427158 | 0.357 | 0.098 | 1 | 17 |
| gins2 | 0.07789 | 0.551402702 | 0.286 | 0.049 | 1 | 17 |
| ube2t | 0.07791 | 0.708213296 | 0.429 | 0.146 | 1 | 17 |
| rcor2 | 0.07802 | 0.522746549 | 0.286 | 0.049 | 1 | 17 |
| brwd1 | 0.07807 | 0.434916429 | 0.286 | 0.049 | 1 | 17 |
| cenpp | 0.0781 | 0.552849228 | 0.286 | 0.049 | 1 | 17 |
| ubc | 0.07815 | 0.264371226 | 1 | 0.78 | 1 | 17 |
| lsm7 | 0.07981 | 0.418907697 | 0.857 | 0.537 | 1 | 17 |
| ctsf | 0.08091 | 0.387120504 | 0.357 | 0.098 | 1 | 17 |
| si:dkey-40g16.5 | 0.08337 | 0.630286166 | 0.5 | 0.195 | 1 | 17 |
| eloca | 0.0834 | 0.472022215 | 0.5 | 0.195 | 1 | 17 |
| taf15 | 0.08347 | 0.434859057 | 0.643 | 0.317 | 1 | 17 |
| eif3m | 0.08374 | 0.494878686 | 0.571 | 0.244 | 1 | 17 |
| rab24 | 0.08382 | 0.32794253 | 0.214 | 0.024 | 1 | 17 |
| thumpd3 | 0.08388 | 0.603253337 | 0.214 | 0.024 | 1 | 17 |
| pnrc2 | 0.08392 | 0.307529916 | 0.643 | 0.341 | 1 | 17 |
| lsm12b | 0.08461 | 0.446308815 | 0.357 | 0.098 | 1 | 17 |
| psmb1 | 0.08471 | 0.196418195 | 0.571 | 0.317 | 1 | 17 |
| fam113 | 0.0848 | 0.342929452 | 0.214 | 0.024 | 1 | 17 |
| LOC100149647 | 0.08569 | 0.479113462 | 0.214 | 0.024 | 1 | 17 |
| cox7b | 0.08603 | 0.168088142 | 0.786 | 0.537 | 1 | 17 |
| nol12 | 0.08637 | 0.43188465 | 0.429 | 0.146 | 1 | 17 |
| rbm4.2 | 0.08701 | 0.446712918 | 0.357 | 0.098 | 1 | 17 |
| esco2 | 0.08787 | 0.586090131 | 0.571 | 0.244 | 1 | 17 |
| heatr3 | 0.08864 | 0.55965049 | 0.214 | 0.024 | 1 | 17 |
| edf1 | 0.08898 | 0.624480969 | 0.5 | 0.195 | 1 | 17 |
| glud1a | 0.08921 | 0.414085833 | 0.214 | 0.024 | 1 | 17 |
| rab11a | 0.08975 | 0.297994467 | 0.214 | 0.024 | 1 | 17 |
| rpl27a | 0.0901 | -0.59442885 | 0.357 | 0.585 | 1 | 17 |
| CABZ01113812.1 | 0.09025 | 0.393731395 | 0.214 | 0.024 | 1 | 17 |
| farsa | 0.09131 | 0.45826609 | 0.357 | 0.098 | 1 | 17 |
| usp4 | 0.09143 | 0.526847361 | 0.214 | 0.024 | 1 | 17 |
| idh3b | 0.09157 | 0.441120074 | 0.357 | 0.098 | 1 | 17 |
| caskb | 0.09164 | 0.52474974 | 0.214 | 0.024 | 1 | 17 |
| nsd2 | 0.09204 | 0.36171886 | 0.214 | 0.024 | 1 | 17 |
| rac1a | 0.09297 | 0.458691542 | 0.357 | 0.098 | 1 | 17 |
| ube2a | 0.09316 | 0.460682549 | 0.357 | 0.098 | 1 | 17 |
| nle1 | 0.09319 | 0.380470095 | 0.214 | 0.024 | 1 | 17 |
| zgc:113426 | 0.09319 | 0.417492616 | 0.214 | 0.024 | 1 | 17 |
| acot8 | 0.09333 | 0.440701765 | 0.214 | 0.024 | 1 | 17 |
| psmd5 | 0.09349 | 0.517869393 | 0.214 | 0.024 | 1 | 17 |

| | | | | | | |
|---|---|---|---|---|---|---|
| mrps33 | 0.09396 | 0.448240815 | 0.429 | 0.146 | 1 | 17 |
| srp54 | 0.09416 | 0.308699949 | 0.214 | 0.024 | 1 | 17 |
| ttc1 | 0.09419 | 0.318307057 | 0.214 | 0.024 | 1 | 17 |
| washc4 | 0.09458 | 0.394141112 | 0.214 | 0.024 | 1 | 17 |
| znrf2a | 0.09494 | 0.472835992 | 0.214 | 0.024 | 1 | 17 |
| psmb6 | 0.0955 | 0.391686252 | 0.571 | 0.268 | 1 | 17 |
| mmadhc | 0.09595 | 0.399949269 | 0.214 | 0.024 | 1 | 17 |
| prkci | 0.09596 | 0.467559445 | 0.214 | 0.024 | 1 | 17 |
| tmem160 | 0.09605 | 0.383428431 | 0.214 | 0.024 | 1 | 17 |
| ufc1 | 0.09633 | 0.340877565 | 0.5 | 0.22 | 1 | 17 |
| api5 | 0.09659 | 0.311516797 | 0.429 | 0.171 | 1 | 17 |
| nms | 0.09671 | 0.361366235 | 0.214 | 0.024 | 1 | 17 |
| kansl1b | 0.09675 | 0.312559597 | 0.214 | 0.024 | 1 | 17 |
| eci2 | 0.0969 | 0.37397689 | 0.214 | 0.024 | 1 | 17 |
| pcm1 | 0.09712 | 0.434967433 | 0.214 | 0.024 | 1 | 17 |
| tmem244 | 0.09728 | 0.374629177 | 0.214 | 0.024 | 1 | 17 |
| capns1b | 0.09729 | 0.522351707 | 0.5 | 0.195 | 1 | 17 |
| hdgfl2 | 0.09739 | 0.358272578 | 0.214 | 0.024 | 1 | 17 |
| rad51 | 0.09768 | 0.506022175 | 0.5 | 0.195 | 1 | 17 |
| XLOC-035884 | 0.09782 | 0.46497044 | 0.214 | 0.024 | 1 | 17 |
| nktr | 0.09811 | 0.452520823 | 0.214 | 0.024 | 1 | 17 |
| ndrg1a | 0.09832 | 0.343166247 | 0.214 | 0.024 | 1 | 17 |
| nt5c2a | 0.09842 | 0.538034403 | 0.214 | 0.024 | 1 | 17 |
| fbxo28 | 0.09844 | 0.367843245 | 0.214 | 0.024 | 1 | 17 |
| padi2 | 0.09853 | 0.427120296 | 0.214 | 0.024 | 1 | 17 |
| prrc1 | 0.0986 | 0.441067447 | 0.214 | 0.024 | 1 | 17 |
| CABZ01072614.1 | 0.09873 | 0.301535211 | 0.214 | 0.024 | 1 | 17 |
| noxo1b | 0.09877 | 0.499341207 | 0.357 | 0.098 | 1 | 17 |
| edc3 | 0.09885 | 0.387222477 | 0.214 | 0.024 | 1 | 17 |
| zgc:153293 | 0.09889 | 0.398965257 | 0.214 | 0.024 | 1 | 17 |
| si:ch211-125a15.1 | 0.09899 | 0.38151242 | 0.214 | 0.024 | 1 | 17 |
| arcn1b | 0.0991 | 0.444346358 | 0.214 | 0.024 | 1 | 17 |
| mgst3b | 0.09916 | 0.477732489 | 0.214 | 0.024 | 1 | 17 |
| mapkapk2a | 0.09925 | 0.398856591 | 0.214 | 0.024 | 1 | 17 |
| zgc:65873 | 0.09943 | 0.430403914 | 0.214 | 0.024 | 1 | 17 |
| ppp3r1b | 0.09947 | 0.369848281 | 0.214 | 0.024 | 1 | 17 |
| pdzk1ip1 | 0.09949 | 0.429762566 | 0.214 | 0.024 | 1 | 17 |
| eif1ad | 0.0995 | 0.48462299 | 0.5 | 0.195 | 1 | 17 |
| tpst1l | 0.09971 | 0.379001418 | 0.214 | 0.024 | 1 | 17 |
| cstf1 | 0.09995 | 0.473800856 | 0.214 | 0.024 | 1 | 17 |
| lta4h | 0.09995 | 0.344021037 | 0.214 | 0.024 | 1 | 17 |
| wdr45 | 0.09999 | 0.396815338 | 0.214 | 0.024 | 1 | 17 |
| znf106a | 0.10015 | 0.394329536 | 0.214 | 0.024 | 1 | 17 |
| telo2 | 0.10016 | 0.394329286 | 0.214 | 0.024 | 1 | 17 |
| eef1akmt1 | 0.10025 | 0.380505337 | 0.214 | 0.024 | 1 | 17 |
| rpp40 | 0.10058 | 0.512147563 | 0.214 | 0.024 | 1 | 17 |
| rbm33b | 0.10062 | 0.406677977 | 0.214 | 0.024 | 1 | 17 |
| srpk1a | 0.10075 | 0.313792522 | 0.286 | 0.073 | 1 | 17 |

| | | | | | | |
|---|---|---|---|---|---|---|
| mych | 0.10078 | 0.525468215 | 0.214 | 0.024 | 1 | 17 |
| mrpl22 | 0.10086 | 0.500462409 | 0.214 | 0.024 | 1 | 17 |
| cnep1r1 | 0.10089 | 0.459279461 | 0.214 | 0.024 | 1 | 17 |
| si:dkey-102m7.3 | 0.10089 | 0.492563083 | 0.429 | 0.146 | 1 | 17 |
| hsd17b10 | 0.10093 | 0.416795388 | 0.214 | 0.024 | 1 | 17 |
| anapc16 | 0.10094 | 0.439853608 | 0.214 | 0.024 | 1 | 17 |
| srfbp1 | 0.10095 | 0.431668153 | 0.214 | 0.024 | 1 | 17 |
| ccdc36 | 0.10095 | 0.476032827 | 0.214 | 0.024 | 1 | 17 |
| tprb | 0.10095 | 0.28090005 | 0.286 | 0.073 | 1 | 17 |
| zgc:153345 | 0.10096 | 0.416495457 | 0.214 | 0.024 | 1 | 17 |
| asl | 0.10098 | 0.439599958 | 0.214 | 0.024 | 1 | 17 |
| CR762420.1 | 0.101 | 0.435918037 | 0.214 | 0.024 | 1 | 17 |
| WDR70 | 0.10102 | 0.441712314 | 0.214 | 0.024 | 1 | 17 |
| umps | 0.10103 | 0.485140297 | 0.214 | 0.024 | 1 | 17 |
| usp44 | 0.10105 | 0.537449534 | 0.214 | 0.024 | 1 | 17 |
| eif2d | 0.10105 | 0.371348401 | 0.214 | 0.024 | 1 | 17 |
| dctn2 | 0.10106 | 0.366314917 | 0.214 | 0.024 | 1 | 17 |
| si:ch211-51h9.6 | 0.10107 | 0.455820858 | 0.214 | 0.024 | 1 | 17 |
| nedd1 | 0.10109 | 0.39990736 | 0.214 | 0.024 | 1 | 17 |
| tmem107l | 0.1011 | 0.379981101 | 0.214 | 0.024 | 1 | 17 |
| zgc:110224 | 0.1011 | 0.450394701 | 0.214 | 0.024 | 1 | 17 |
| ccnb3 | 0.10111 | 0.459552809 | 0.214 | 0.024 | 1 | 17 |
| cyb5r2 | 0.10111 | 0.472758733 | 0.214 | 0.024 | 1 | 17 |
| si:dkey-217d24.6 | 0.10112 | 0.476891754 | 0.214 | 0.024 | 1 | 17 |
| purba | 0.10112 | 0.459072518 | 0.214 | 0.024 | 1 | 17 |
| swsap1 | 0.10112 | 0.489785263 | 0.214 | 0.024 | 1 | 17 |
| cst14a.2 | 0.10125 | 0.539997507 | 0.357 | 0.098 | 1 | 17 |
| prpf38b | 0.1016 | 0.491859298 | 0.357 | 0.098 | 1 | 17 |
| sbds | 0.1025 | 0.561425078 | 0.5 | 0.195 | 1 | 17 |
| taf11 | 0.10359 | 0.519636977 | 0.357 | 0.098 | 1 | 17 |
| rad23b | 0.10365 | 0.360678148 | 0.643 | 0.341 | 1 | 17 |
| txnrd3 | 0.10411 | 0.552670964 | 0.357 | 0.098 | 1 | 17 |
| cops4 | 0.10517 | 0.481364283 | 0.429 | 0.146 | 1 | 17 |
| uba2 | 0.10676 | 0.519640571 | 0.429 | 0.146 | 1 | 17 |
| rpl3 | 0.1071 | 0.295405323 | 1 | 0.854 | 1 | 17 |
| naca | 0.10809 | 0.245744576 | 1 | 0.805 | 1 | 17 |
| snrpf | 0.10827 | 0.369549565 | 0.857 | 0.561 | 1 | 17 |
| ddx4 | 0.1087 | 0.310326665 | 1 | 0.854 | 1 | 17 |
| ndufs2 | 0.10924 | 0.559016299 | 0.429 | 0.146 | 1 | 17 |
| cnpy1 | 0.10971 | 0.304942208 | 0.286 | 0.073 | 1 | 17 |
| thap12a | 0.10975 | 0.299202853 | 0.286 | 0.073 | 1 | 17 |
| gnl3 | 0.11232 | 0.464830864 | 0.571 | 0.268 | 1 | 17 |
| rpl17 | 0.11277 | 0.120594851 | 0.929 | 0.732 | 1 | 17 |
| ap1s3b | 0.11442 | -0.30866191 | 0 | 0.122 | 1 | 17 |
| gdnfb | 0.11442 | -0.44693199 | 0 | 0.122 | 1 | 17 |
| gmps | 0.11442 | -0.35606484 | 0 | 0.122 | 1 | 17 |
| myclb | 0.11442 | -0.40086083 | 0 | 0.122 | 1 | 17 |
| si:ch73-189n23.1 | 0.11442 | -0.34561715 | 0 | 0.122 | 1 | 17 |

| | | | | | | |
|---|---|---|---|---|---|---|
| zgc:112265 | 0.11442 | -0.35978844 | 0 | 0.122 | 1 | 17 |
| depdc7b | 0.11442 | -0.29705648 | 0 | 0.122 | 1 | 17 |
| eno1b | 0.11442 | -0.3316705 | 0 | 0.122 | 1 | 17 |
| glipr2l | 0.11442 | -0.36917821 | 0 | 0.122 | 1 | 17 |
| slmapa | 0.11442 | -0.42267317 | 0 | 0.122 | 1 | 17 |
| ssr1 | 0.11442 | -0.25717889 | 0 | 0.122 | 1 | 17 |
| CR790368.1 | 0.11442 | -0.31113325 | 0 | 0.122 | 1 | 17 |
| MTERF4 | 0.11442 | -0.36089965 | 0 | 0.122 | 1 | 17 |
| elovl7b | 0.11442 | -0.36225965 | 0 | 0.122 | 1 | 17 |
| elp6 | 0.11442 | -0.37128772 | 0 | 0.122 | 1 | 17 |
| gemin8 | 0.11442 | -0.34439232 | 0 | 0.122 | 1 | 17 |
| gng5 | 0.11442 | -0.35214092 | 0 | 0.122 | 1 | 17 |
| gstz1 | 0.11442 | -0.28363921 | 0 | 0.122 | 1 | 17 |
| lias | 0.11442 | -0.36396035 | 0 | 0.122 | 1 | 17 |
| phyhd1 | 0.11442 | -0.34171961 | 0 | 0.122 | 1 | 17 |
| rab12 | 0.11442 | -0.34047641 | 0 | 0.122 | 1 | 17 |
| si:dkey-241l7.3 | 0.11442 | -0.32328919 | 0 | 0.122 | 1 | 17 |
| smarcb1b | 0.11442 | -0.31012196 | 0 | 0.122 | 1 | 17 |
| thg1l | 0.11442 | -0.31896937 | 0 | 0.122 | 1 | 17 |
| thrap3b | 0.11442 | -0.37438906 | 0 | 0.122 | 1 | 17 |
| timm8a | 0.11442 | -0.36785657 | 0 | 0.122 | 1 | 17 |
| tmub1 | 0.11442 | -0.33740867 | 0 | 0.122 | 1 | 17 |
| ttk | 0.11442 | -0.32557361 | 0 | 0.122 | 1 | 17 |
| ube2kb | 0.11442 | -0.36899262 | 0 | 0.122 | 1 | 17 |
| HOOK3 | 0.11442 | -0.34196235 | 0 | 0.122 | 1 | 17 |
| fabp11a | 0.11442 | -0.35443067 | 0 | 0.122 | 1 | 17 |
| pofut2 | 0.11442 | -0.3906383 | 0 | 0.122 | 1 | 17 |
| aldoaa | 0.11442 | -0.33322414 | 0 | 0.122 | 1 | 17 |
| eno1a | 0.11442 | -0.31997592 | 0 | 0.122 | 1 | 17 |
| kat8 | 0.11442 | -0.33637797 | 0 | 0.122 | 1 | 17 |
| kti12 | 0.11442 | -0.33066666 | 0 | 0.122 | 1 | 17 |
| metrnlb | 0.11442 | -0.28849302 | 0 | 0.122 | 1 | 17 |
| smarce1 | 0.11442 | -0.33313953 | 0 | 0.122 | 1 | 17 |
| tspan35 | 0.11442 | -0.42382983 | 0 | 0.122 | 1 | 17 |
| aldh3b1 | 0.11442 | -0.37916925 | 0 | 0.122 | 1 | 17 |
| bop1 | 0.11442 | -0.38990516 | 0 | 0.122 | 1 | 17 |
| zgc:171566 | 0.11442 | -0.37149354 | 0 | 0.122 | 1 | 17 |
| btbd10b | 0.11442 | -0.3504052 | 0 | 0.122 | 1 | 17 |
| ch25hl3 | 0.11442 | -0.39191295 | 0 | 0.122 | 1 | 17 |
| chuk | 0.11442 | -0.31835262 | 0 | 0.122 | 1 | 17 |
| cyb5a | 0.11442 | -0.33655034 | 0 | 0.122 | 1 | 17 |
| grsf1 | 0.11442 | -0.32408933 | 0 | 0.122 | 1 | 17 |
| nat10 | 0.11442 | -0.32907351 | 0 | 0.122 | 1 | 17 |
| nubp2 | 0.11442 | -0.33717407 | 0 | 0.122 | 1 | 17 |
| rrp15 | 0.11442 | -0.29585477 | 0 | 0.122 | 1 | 17 |
| ssna1 | 0.11442 | -0.38999097 | 0 | 0.122 | 1 | 17 |
| usp39 | 0.11442 | -0.34647731 | 0 | 0.122 | 1 | 17 |
| cx43.4 | 0.11442 | -0.39515388 | 0 | 0.122 | 1 | 17 |

| | | | | | | |
|---|---|---|---|---|---|---|
| si:ch211-161h7.4 | 0.11442 | -0.41068091 | 0 | 0.122 | 1 | 17 |
| hmgn2 | 0.11706 | 0.437625499 | 0.857 | 0.683 | 1 | 17 |
| anp32a | 0.11839 | 0.433291685 | 0.571 | 0.268 | 1 | 17 |
| ndufa11 | 0.1184 | 0.276758206 | 0.5 | 0.244 | 1 | 17 |
| atp5fa1 | 0.11954 | -0.37240997 | 0.571 | 0.659 | 1 | 17 |
| ppiab | 0.11969 | 0.489965691 | 0.786 | 0.488 | 1 | 17 |
| krtcap2 | 0.12198 | 0.245565668 | 0.571 | 0.317 | 1 | 17 |
| sf3b5 | 0.12299 | 0.320980556 | 0.571 | 0.293 | 1 | 17 |
| bccip | 0.1235 | 0.373731064 | 0.5 | 0.22 | 1 | 17 |
| ndel1a | 0.12635 | 0.340422724 | 0.286 | 0.073 | 1 | 17 |
| rabif | 0.12655 | 0.328446315 | 0.571 | 0.293 | 1 | 17 |
| eif4eb | 0.12682 | 0.554109246 | 0.571 | 0.268 | 1 | 17 |
| eno3 | 0.12866 | 0.287662703 | 0.643 | 0.366 | 1 | 17 |
| rplp0 | 0.12892 | 0.190337921 | 1 | 0.976 | 1 | 17 |
| BX649411.2 | 0.12901 | 0.228746285 | 0.143 | 0.024 | 1 | 17 |
| BX957362.1 | 0.12901 | 0.292066488 | 0.143 | 0.024 | 1 | 17 |
| CABZ01113954.1 | 0.12901 | 0.351510234 | 0.143 | 0.024 | 1 | 17 |
| CR786562.1 | 0.12901 | 0.201824263 | 0.143 | 0.024 | 1 | 17 |
| CU856358.1 | 0.12901 | 0.289080647 | 0.143 | 0.024 | 1 | 17 |
| LO018250.1 | 0.12901 | 0.189371075 | 0.143 | 0.024 | 1 | 17 |
| RUNDC1 | 0.12901 | 0.233051304 | 0.143 | 0.024 | 1 | 17 |
| abt1 | 0.12901 | 0.348510652 | 0.143 | 0.024 | 1 | 17 |
| acad9 | 0.12901 | 0.281576938 | 0.143 | 0.024 | 1 | 17 |
| antxr2a | 0.12901 | 0.18598546 | 0.143 | 0.024 | 1 | 17 |
| brd7 | 0.12901 | 0.22847789 | 0.143 | 0.024 | 1 | 17 |
| bud23 | 0.12901 | 0.287413546 | 0.143 | 0.024 | 1 | 17 |
| bzw2 | 0.12901 | 0.26636965 | 0.143 | 0.024 | 1 | 17 |
| camlg | 0.12901 | 0.260709562 | 0.143 | 0.024 | 1 | 17 |
| ccdc173 | 0.12901 | 0.317658116 | 0.143 | 0.024 | 1 | 17 |
| cdkal1 | 0.12901 | 0.311031009 | 0.143 | 0.024 | 1 | 17 |
| cluap1 | 0.12901 | 0.305316335 | 0.143 | 0.024 | 1 | 17 |
| coro1cb | 0.12901 | 0.319751498 | 0.143 | 0.024 | 1 | 17 |
| cpeb4b | 0.12901 | 0.219867457 | 0.143 | 0.024 | 1 | 17 |
| cry1bb | 0.12901 | 0.25191087 | 0.143 | 0.024 | 1 | 17 |
| cul3a | 0.12901 | 0.22840119 | 0.143 | 0.024 | 1 | 17 |
| dedd1 | 0.12901 | 0.210414735 | 0.143 | 0.024 | 1 | 17 |
| dmrt1 | 0.12901 | 0.195183413 | 0.143 | 0.024 | 1 | 17 |
| ebp | 0.12901 | 0.266397302 | 0.143 | 0.024 | 1 | 17 |
| eef1akmt2 | 0.12901 | 0.336078581 | 0.143 | 0.024 | 1 | 17 |
| emc10 | 0.12901 | 0.29037964 | 0.143 | 0.024 | 1 | 17 |
| emc6 | 0.12901 | 0.310519622 | 0.143 | 0.024 | 1 | 17 |
| fancl | 0.12901 | 0.355620695 | 0.143 | 0.024 | 1 | 17 |
| fntb | 0.12901 | 0.261859148 | 0.143 | 0.024 | 1 | 17 |
| fscn1a | 0.12901 | 0.22044019 | 0.143 | 0.024 | 1 | 17 |
| gramd1c | 0.12901 | 0.200656854 | 0.143 | 0.024 | 1 | 17 |
| hdac8 | 0.12901 | 0.280790463 | 0.143 | 0.024 | 1 | 17 |
| histh1l | 0.12901 | 0.142246351 | 0.143 | 0.024 | 1 | 17 |
| hnrnpm | 0.12901 | 0.164937524 | 0.143 | 0.024 | 1 | 17 |

| | | | | | | |
|---|---|---|---|---|---|---|
| ice1 | 0.12901 | 0.327084283 | 0.143 | 0.024 | 1 | 17 |
| inip | 0.12901 | 0.159292146 | 0.143 | 0.024 | 1 | 17 |
| jam2a | 0.12901 | 0.213352323 | 0.143 | 0.024 | 1 | 17 |
| kat2a | 0.12901 | 0.345711987 | 0.143 | 0.024 | 1 | 17 |
| las1l | 0.12901 | 0.199589561 | 0.143 | 0.024 | 1 | 17 |
| lmnb2 | 0.12901 | 0.331810067 | 0.143 | 0.024 | 1 | 17 |
| mbpb | 0.12901 | 0.305464903 | 0.143 | 0.024 | 1 | 17 |
| mcm10 | 0.12901 | 0.20137969 | 0.143 | 0.024 | 1 | 17 |
| mecr | 0.12901 | 0.17517955 | 0.143 | 0.024 | 1 | 17 |
| med18 | 0.12901 | 0.382966233 | 0.143 | 0.024 | 1 | 17 |
| mettl5 | 0.12901 | 0.275759887 | 0.143 | 0.024 | 1 | 17 |
| mlec | 0.12901 | 0.291790552 | 0.143 | 0.024 | 1 | 17 |
| naa35 | 0.12901 | 0.24942212 | 0.143 | 0.024 | 1 | 17 |
| ndufaf4 | 0.12901 | 0.216049223 | 0.143 | 0.024 | 1 | 17 |
| phlda3 | 0.12901 | 0.294626202 | 0.143 | 0.024 | 1 | 17 |
| polr3gla | 0.12901 | 0.269561243 | 0.143 | 0.024 | 1 | 17 |
| ptgr2 | 0.12901 | 0.224870361 | 0.143 | 0.024 | 1 | 17 |
| r3hdm1 | 0.12901 | 0.298888424 | 0.143 | 0.024 | 1 | 17 |
| rab41 | 0.12901 | 0.181774473 | 0.143 | 0.024 | 1 | 17 |
| rex1bd | 0.12901 | 0.257082082 | 0.143 | 0.024 | 1 | 17 |
| rrp1 | 0.12901 | 0.272190993 | 0.143 | 0.024 | 1 | 17 |
| samd13 | 0.12901 | 0.321807737 | 0.143 | 0.024 | 1 | 17 |
| si:dkey-152b24.6 | 0.12901 | 0.333798426 | 0.143 | 0.024 | 1 | 17 |
| si:dkey-188i13.9 | 0.12901 | 0.311710566 | 0.143 | 0.024 | 1 | 17 |
| si:dkey-238c7.16 | 0.12901 | 0.114592126 | 0.143 | 0.024 | 1 | 17 |
| si:dkey-274m17.3 | 0.12901 | 0.294564389 | 0.143 | 0.024 | 1 | 17 |
| si:dkey-286j15.3 | 0.12901 | 0.172942894 | 0.143 | 0.024 | 1 | 17 |
| smc4 | 0.12901 | 0.376144112 | 0.143 | 0.024 | 1 | 17 |
| spa17 | 0.12901 | 0.242682273 | 0.143 | 0.024 | 1 | 17 |
| spryd7b | 0.12901 | 0.289963427 | 0.143 | 0.024 | 1 | 17 |
| stag2b | 0.12901 | 0.223940055 | 0.143 | 0.024 | 1 | 17 |
| tbl1xr1a | 0.12901 | 0.220625805 | 0.143 | 0.024 | 1 | 17 |
| timm44 | 0.12901 | 0.25017324 | 0.143 | 0.024 | 1 | 17 |
| tmem17 | 0.12901 | 0.179083807 | 0.143 | 0.024 | 1 | 17 |
| toporsb | 0.12901 | 0.23333399 | 0.143 | 0.024 | 1 | 17 |
| tpm4a | 0.12901 | 0.128206261 | 0.143 | 0.024 | 1 | 17 |
| urm1 | 0.12901 | 0.264842701 | 0.143 | 0.024 | 1 | 17 |
| vamp3 | 0.12901 | 0.323769399 | 0.143 | 0.024 | 1 | 17 |
| vti1a | 0.12901 | 0.228475672 | 0.143 | 0.024 | 1 | 17 |
| xpo1b | 0.12901 | 0.313214964 | 0.143 | 0.024 | 1 | 17 |
| zcchc17 | 0.12901 | 0.178185739 | 0.143 | 0.024 | 1 | 17 |
| zgc:114119 | 0.12901 | 0.273561807 | 0.143 | 0.024 | 1 | 17 |
| zgc:123010 | 0.12901 | 0.169070124 | 0.143 | 0.024 | 1 | 17 |
| zgc:162962 | 0.12901 | 0.293622437 | 0.143 | 0.024 | 1 | 17 |
| zgc:92335 | 0.12901 | 0.208625534 | 0.143 | 0.024 | 1 | 17 |
| zswim5 | 0.12901 | 0.29427469 | 0.143 | 0.024 | 1 | 17 |
| BX248511.1 | 0.12901 | 0.281493652 | 0.143 | 0.024 | 1 | 17 |
| anapc10 | 0.12901 | 0.309372398 | 0.143 | 0.024 | 1 | 17 |

| | | | | | | |
|---|---|---|---|---|---|---|
| appa | 0.12901 | 0.253115666 | 0.143 | 0.024 | 1 | 17 |
| fam208ab | 0.12901 | 0.23249335 | 0.143 | 0.024 | 1 | 17 |
| miip | 0.12901 | 0.244923326 | 0.143 | 0.024 | 1 | 17 |
| ociad1 | 0.12901 | 0.290163683 | 0.143 | 0.024 | 1 | 17 |
| pds5a | 0.12901 | 0.310257146 | 0.143 | 0.024 | 1 | 17 |
| pgs1 | 0.12901 | 0.278738283 | 0.143 | 0.024 | 1 | 17 |
| ppp1r11 | 0.12901 | 0.239312426 | 0.143 | 0.024 | 1 | 17 |
| rgcc | 0.12901 | 0.205116561 | 0.143 | 0.024 | 1 | 17 |
| sdc4 | 0.12901 | 0.384012893 | 0.143 | 0.024 | 1 | 17 |
| ten1 | 0.12901 | 0.254812318 | 0.143 | 0.024 | 1 | 17 |
| tm2d3 | 0.12901 | 0.219414026 | 0.143 | 0.024 | 1 | 17 |
| tprg1l | 0.12901 | 0.366927224 | 0.143 | 0.024 | 1 | 17 |
| wasa | 0.12901 | 0.19799086 | 0.143 | 0.024 | 1 | 17 |
| e2f4 | 0.12901 | 0.246270381 | 0.143 | 0.024 | 1 | 17 |
| hcfc1a | 0.12901 | 0.24188796 | 0.143 | 0.024 | 1 | 17 |
| mark3a | 0.12901 | 0.272940199 | 0.143 | 0.024 | 1 | 17 |
| yipf2 | 0.12901 | 0.185045224 | 0.143 | 0.024 | 1 | 17 |
| cdc14b | 0.12901 | 0.228252 | 0.143 | 0.024 | 1 | 17 |
| cthrc1a | 0.12901 | 0.295612315 | 0.143 | 0.024 | 1 | 17 |
| dnajb1a | 0.12901 | 0.213751578 | 0.143 | 0.024 | 1 | 17 |
| si:dkey-69o16.5 | 0.12901 | 0.274353287 | 0.143 | 0.024 | 1 | 17 |
| arntl1a | 0.12901 | 0.239994676 | 0.143 | 0.024 | 1 | 17 |
| cacybp | 0.12901 | 0.19324792 | 0.143 | 0.024 | 1 | 17 |
| cenpq | 0.12901 | 0.21229029 | 0.143 | 0.024 | 1 | 17 |
| chd8 | 0.12901 | 0.286303131 | 0.143 | 0.024 | 1 | 17 |
| col12a1a | 0.12901 | 0.148330622 | 0.143 | 0.024 | 1 | 17 |
| crlf3 | 0.12901 | 0.186209614 | 0.143 | 0.024 | 1 | 17 |
| cul2 | 0.12901 | 0.25477493 | 0.143 | 0.024 | 1 | 17 |
| cxadr | 0.12901 | 0.115481355 | 0.143 | 0.024 | 1 | 17 |
| dhrs13l1 | 0.12901 | 0.272514841 | 0.143 | 0.024 | 1 | 17 |
| edrf1 | 0.12901 | 0.281789924 | 0.143 | 0.024 | 1 | 17 |
| ehhadh | 0.12901 | 0.187758228 | 0.143 | 0.024 | 1 | 17 |
| enpp2 | 0.12901 | 0.222214999 | 0.143 | 0.024 | 1 | 17 |
| fam49ba | 0.12901 | 0.132875843 | 0.143 | 0.024 | 1 | 17 |
| gmds | 0.12901 | 0.188280026 | 0.143 | 0.024 | 1 | 17 |
| grapb | 0.12901 | 0.201235664 | 0.143 | 0.024 | 1 | 17 |
| lpar2a | 0.12901 | 0.228495101 | 0.143 | 0.024 | 1 | 17 |
| mfge8a | 0.12901 | 0.303143261 | 0.143 | 0.024 | 1 | 17 |
| narfl | 0.12901 | 0.255135069 | 0.143 | 0.024 | 1 | 17 |
| nudcd1 | 0.12901 | 0.263192912 | 0.143 | 0.024 | 1 | 17 |
| pih1d1 | 0.12901 | 0.340620563 | 0.143 | 0.024 | 1 | 17 |
| pik3ca | 0.12901 | 0.196395513 | 0.143 | 0.024 | 1 | 17 |
| plekhb2 | 0.12901 | 0.294512213 | 0.143 | 0.024 | 1 | 17 |
| pola1 | 0.12901 | 0.289186416 | 0.143 | 0.024 | 1 | 17 |
| pot1 | 0.12901 | 0.225968104 | 0.143 | 0.024 | 1 | 17 |
| ppp1r12a | 0.12901 | 0.218470625 | 0.143 | 0.024 | 1 | 17 |
| prmt7 | 0.12901 | 0.250977666 | 0.143 | 0.024 | 1 | 17 |
| rab2a | 0.12901 | 0.155344038 | 0.143 | 0.024 | 1 | 17 |

| | | | | | | |
|---|---|---|---|---|---|---|
| rad54l2 | 0.12901 | 0.181367418 | 0.143 | 0.024 | 1 | 17 |
| rell1 | 0.12901 | 0.251491094 | 0.143 | 0.024 | 1 | 17 |
| sesn2 | 0.12901 | 0.216410628 | 0.143 | 0.024 | 1 | 17 |
| si:ch211-128m15.3 | 0.12901 | 0.192658534 | 0.143 | 0.024 | 1 | 17 |
| si:dkey-54n8.2 | 0.12901 | 0.23884168 | 0.143 | 0.024 | 1 | 17 |
| spast | 0.12901 | 0.28887325 | 0.143 | 0.024 | 1 | 17 |
| srp14 | 0.12901 | 0.302777046 | 0.143 | 0.024 | 1 | 17 |
| svbp | 0.12901 | 0.210598174 | 0.143 | 0.024 | 1 | 17 |
| triap1 | 0.12901 | 0.206346739 | 0.143 | 0.024 | 1 | 17 |
| usp36 | 0.12901 | 0.240740055 | 0.143 | 0.024 | 1 | 17 |
| cnpy3 | 0.12901 | 0.311142628 | 0.143 | 0.024 | 1 | 17 |
| dnajb6b | 0.12901 | 0.267148823 | 0.143 | 0.024 | 1 | 17 |
| fdps | 0.12901 | 0.159437599 | 0.143 | 0.024 | 1 | 17 |
| foxj3 | 0.12901 | 0.241240162 | 0.143 | 0.024 | 1 | 17 |
| gipc3 | 0.12901 | 0.292564656 | 0.143 | 0.024 | 1 | 17 |
| gpr63 | 0.12901 | 0.276011962 | 0.143 | 0.024 | 1 | 17 |
| hdac12 | 0.12901 | 0.246355822 | 0.143 | 0.024 | 1 | 17 |
| msmo1 | 0.12901 | 0.115224135 | 0.143 | 0.024 | 1 | 17 |
| mylpfb | 0.12901 | 0.211582228 | 0.143 | 0.024 | 1 | 17 |
| oser1 | 0.12901 | 0.280400802 | 0.143 | 0.024 | 1 | 17 |
| ovca2 | 0.12901 | 0.253899778 | 0.143 | 0.024 | 1 | 17 |
| pomt2 | 0.12901 | 0.223647958 | 0.143 | 0.024 | 1 | 17 |
| pwp1 | 0.12901 | 0.267434644 | 0.143 | 0.024 | 1 | 17 |
| rbm7 | 0.12901 | 0.210445653 | 0.143 | 0.024 | 1 | 17 |
| rpap1 | 0.12901 | 0.283137656 | 0.143 | 0.024 | 1 | 17 |
| si:ch1073-228h2.2 | 0.12901 | 0.256713136 | 0.143 | 0.024 | 1 | 17 |
| slc39a5 | 0.12901 | 0.240640428 | 0.143 | 0.024 | 1 | 17 |
| thumpd1 | 0.12901 | 0.281054438 | 0.143 | 0.024 | 1 | 17 |
| tut1 | 0.12901 | 0.227757622 | 0.143 | 0.024 | 1 | 17 |
| vps16 | 0.12901 | 0.254872523 | 0.143 | 0.024 | 1 | 17 |
| wrap73 | 0.12901 | 0.245940006 | 0.143 | 0.024 | 1 | 17 |
| zgc:113210 | 0.12901 | 0.282308496 | 0.143 | 0.024 | 1 | 17 |
| znf1035 | 0.12901 | 0.202198495 | 0.143 | 0.024 | 1 | 17 |
| brd4 | 0.12901 | 0.207771962 | 0.143 | 0.024 | 1 | 17 |
| chd4b | 0.12901 | 0.267681225 | 0.143 | 0.024 | 1 | 17 |
| clptm1 | 0.12901 | 0.282976083 | 0.143 | 0.024 | 1 | 17 |
| cpne2 | 0.12901 | 0.263909438 | 0.143 | 0.024 | 1 | 17 |
| dhcr7 | 0.12901 | 0.299242267 | 0.143 | 0.024 | 1 | 17 |
| dnal1 | 0.12901 | 0.266934824 | 0.143 | 0.024 | 1 | 17 |
| eif4g3a | 0.12901 | 0.273059808 | 0.143 | 0.024 | 1 | 17 |
| gatad1 | 0.12901 | 0.381055942 | 0.143 | 0.024 | 1 | 17 |
| lgi1a | 0.12901 | 0.281693185 | 0.143 | 0.024 | 1 | 17 |
| limk2 | 0.12901 | 0.124951175 | 0.143 | 0.024 | 1 | 17 |
| memo1 | 0.12901 | 0.267977496 | 0.143 | 0.024 | 1 | 17 |
| phf20b | 0.12901 | 0.210939485 | 0.143 | 0.024 | 1 | 17 |
| pik3r3b | 0.12901 | 0.275483767 | 0.143 | 0.024 | 1 | 17 |
| si:dkey-148f10.4 | 0.12901 | 0.290886739 | 0.143 | 0.024 | 1 | 17 |
| tango2 | 0.12901 | 0.278220068 | 0.143 | 0.024 | 1 | 17 |

| | | | | | | |
|---|---|---|---|---|---|---|
| tmem238a | 0.12901 | 0.223666513 | 0.143 | 0.024 | 1 | 17 |
| LOC100148648 | 0.12901 | 0.432105026 | 0.143 | 0.024 | 1 | 17 |
| adsl | 0.12901 | 0.323577894 | 0.143 | 0.024 | 1 | 17 |
| arrb2a | 0.12901 | 0.250521253 | 0.143 | 0.024 | 1 | 17 |
| baxa | 0.12901 | 0.251157318 | 0.143 | 0.024 | 1 | 17 |
| brf1a | 0.12901 | 0.220725153 | 0.143 | 0.024 | 1 | 17 |
| ccdc28a | 0.12901 | 0.23684649 | 0.143 | 0.024 | 1 | 17 |
| ccnk | 0.12901 | 0.271434418 | 0.143 | 0.024 | 1 | 17 |
| ccnt2b | 0.12901 | 0.226569973 | 0.143 | 0.024 | 1 | 17 |
| gipc2 | 0.12901 | 0.291186249 | 0.143 | 0.024 | 1 | 17 |
| hltf | 0.12901 | 0.217687951 | 0.143 | 0.024 | 1 | 17 |
| hmox2a | 0.12901 | 0.368908562 | 0.143 | 0.024 | 1 | 17 |
| ino80e | 0.12901 | 0.239471429 | 0.143 | 0.024 | 1 | 17 |
| iqck | 0.12901 | 0.339192386 | 0.143 | 0.024 | 1 | 17 |
| klhl20 | 0.12901 | 0.284222477 | 0.143 | 0.024 | 1 | 17 |
| ltv1 | 0.12901 | 0.220177635 | 0.143 | 0.024 | 1 | 17 |
| med19b | 0.12901 | 0.26251785 | 0.143 | 0.024 | 1 | 17 |
| mphosph10 | 0.12901 | 0.387340823 | 0.143 | 0.024 | 1 | 17 |
| mterf3 | 0.12901 | 0.312136176 | 0.143 | 0.024 | 1 | 17 |
| pold2 | 0.12901 | 0.295468745 | 0.143 | 0.024 | 1 | 17 |
| prpf3 | 0.12901 | 0.285018031 | 0.143 | 0.024 | 1 | 17 |
| rab5ab | 0.12901 | 0.241754747 | 0.143 | 0.024 | 1 | 17 |
| setd1a | 0.12901 | 0.343925777 | 0.143 | 0.024 | 1 | 17 |
| si:dkey-19e4.5 | 0.12901 | 0.286603797 | 0.143 | 0.024 | 1 | 17 |
| srpk1b | 0.12901 | 0.198676301 | 0.143 | 0.024 | 1 | 17 |
| zc3h7a | 0.12901 | 0.290501549 | 0.143 | 0.024 | 1 | 17 |
| zgc:113372 | 0.12901 | 0.264526301 | 0.143 | 0.024 | 1 | 17 |
| naa38 | 0.13034 | 0.338726 | 0.286 | 0.073 | 1 | 17 |
| psmf1 | 0.13106 | 0.24582723 | 0.357 | 0.146 | 1 | 17 |
| rbx1 | 0.13131 | -0.5417261 | 0.429 | 0.659 | 1 | 17 |
| tufm | 0.13181 | 0.401258863 | 0.286 | 0.073 | 1 | 17 |
| commd7 | 0.13304 | 0.374266526 | 0.429 | 0.171 | 1 | 17 |
| rpl28 | 0.13332 | 0.143652444 | 1 | 0.829 | 1 | 17 |
| rpl22l1 | 0.13398 | 0.188197196 | 0.714 | 0.463 | 1 | 17 |
| akr1a1b | 0.13468 | 0.411830717 | 0.286 | 0.073 | 1 | 17 |
| atp5pf | 0.13474 | 0.407741424 | 0.643 | 0.341 | 1 | 17 |
| dynlrb1 | 0.13541 | 0.260629597 | 0.643 | 0.39 | 1 | 17 |
| atp5mf | 0.13637 | 0.357321505 | 0.571 | 0.293 | 1 | 17 |
| pqbp1 | 0.13707 | 0.378804904 | 0.286 | 0.073 | 1 | 17 |
| cs | 0.13712 | 0.438297569 | 0.286 | 0.073 | 1 | 17 |
| alyref | 0.13736 | 0.43370031 | 0.286 | 0.073 | 1 | 17 |
| cox6b1 | 0.13848 | 0.589853568 | 0.429 | 0.171 | 1 | 17 |
| ptgr1 | 0.13876 | 0.324334064 | 0.571 | 0.293 | 1 | 17 |
| smc3 | 0.13888 | 0.396005136 | 0.286 | 0.073 | 1 | 17 |
| eef1e1 | 0.13974 | 0.355846984 | 0.286 | 0.073 | 1 | 17 |
| rpl23a | 0.14007 | 0.14372145 | 0.929 | 0.732 | 1 | 17 |
| dcun1d5 | 0.14029 | 0.169702369 | 0.571 | 0.341 | 1 | 17 |
| eif5a | 0.14101 | 0.492388003 | 0.714 | 0.415 | 1 | 17 |

| | | | | | | |
|---|---|---|---|---|---|---|
| rpl5b | 0.14147 | 0.306295305 | 0.929 | 0.683 | 1 | 17 |
| cdk11b | 0.14284 | 0.345728172 | 0.286 | 0.073 | 1 | 17 |
| tmed2 | 0.14341 | 0.276093841 | 0.429 | 0.195 | 1 | 17 |
| smndc1 | 0.14371 | 0.180533362 | 0.429 | 0.22 | 1 | 17 |
| hccsb | 0.14372 | 0.418671808 | 0.286 | 0.073 | 1 | 17 |
| gstr | 0.14397 | 0.358050913 | 0.571 | 0.293 | 1 | 17 |
| fam117ab | 0.14408 | 0.396889663 | 0.286 | 0.073 | 1 | 17 |
| zgc:63470 | 0.1447 | 0.628153151 | 0.286 | 0.073 | 1 | 17 |
| larp7 | 0.14559 | 0.458314273 | 0.5 | 0.22 | 1 | 17 |
| sart1 | 0.14584 | -0.48990344 | 0.286 | 0.39 | 1 | 17 |
| gpia | 0.1462 | 0.434154604 | 0.5 | 0.22 | 1 | 17 |
| ndufb10 | 0.14647 | 0.370181345 | 0.571 | 0.293 | 1 | 17 |
| ywhah | 0.14718 | 0.381155402 | 0.286 | 0.073 | 1 | 17 |
| hnrnph3 | 0.14838 | 0.362138275 | 0.286 | 0.073 | 1 | 17 |
| ctnnbl1 | 0.14872 | 0.400126284 | 0.286 | 0.073 | 1 | 17 |
| ndufa5 | 0.14978 | 0.429026073 | 0.429 | 0.171 | 1 | 17 |
| rps16 | 0.15021 | 0.221048591 | 1 | 0.829 | 1 | 17 |
| tomm34 | 0.15046 | 0.566527617 | 0.286 | 0.073 | 1 | 17 |
| si:ch211-241j8.2 | 0.15053 | 0.488369154 | 0.286 | 0.073 | 1 | 17 |
| dnajc9 | 0.15077 | 0.328239231 | 0.5 | 0.244 | 1 | 17 |
| mrps14 | 0.1508 | 0.440537236 | 0.429 | 0.171 | 1 | 17 |
| riox1 | 0.15114 | 0.387202753 | 0.286 | 0.073 | 1 | 17 |
| rab10 | 0.15195 | 0.486867803 | 0.5 | 0.22 | 1 | 17 |
| mrpl33 | 0.15234 | 0.47312614 | 0.5 | 0.22 | 1 | 17 |
| ube2e1 | 0.1524 | 0.40964623 | 0.429 | 0.171 | 1 | 17 |
| rbm46 | 0.1531 | 0.3933264 | 0.357 | 0.122 | 1 | 17 |
| hmgn6 | 0.15324 | 0.43985471 | 0.429 | 0.171 | 1 | 17 |
| fgfr1op | 0.15334 | 0.449955346 | 0.286 | 0.073 | 1 | 17 |
| dck | 0.15347 | 0.431583231 | 0.286 | 0.073 | 1 | 17 |
| sf1 | 0.15432 | 0.494666572 | 0.286 | 0.073 | 1 | 17 |
| AL929210.1 | 0.15584 | 0.178584158 | 0.071 | 0 | 1 | 17 |
| BX855614.1 | 0.15584 | 0.169209088 | 0.071 | 0 | 1 | 17 |
| CABZ01117348.1 | 0.15584 | 0.178584158 | 0.071 | 0 | 1 | 17 |
| CR759845.1 | 0.15584 | 0.178584158 | 0.071 | 0 | 1 | 17 |
| CR847971.1 | 0.15584 | 0.178584158 | 0.071 | 0 | 1 | 17 |
| FQ377903.3 | 0.15584 | 0.172990058 | 0.071 | 0 | 1 | 17 |
| GNAZ | 0.15584 | 0.216160118 | 0.071 | 0 | 1 | 17 |
| LOC100002706 | 0.15584 | 0.173417732 | 0.071 | 0 | 1 | 17 |
| LOC799914 | 0.15584 | 0.216960391 | 0.071 | 0 | 1 | 17 |
| XLOC-012772 | 0.15584 | 0.178584158 | 0.071 | 0 | 1 | 17 |
| acss1 | 0.15584 | 0.175071581 | 0.071 | 0 | 1 | 17 |
| adora2aa | 0.15584 | 0.17097557 | 0.071 | 0 | 1 | 17 |
| agfg1b | 0.15584 | 0.216960391 | 0.071 | 0 | 1 | 17 |
| agpat3 | 0.15584 | 0.176298519 | 0.071 | 0 | 1 | 17 |
| ankzf1 | 0.15584 | 0.175680815 | 0.071 | 0 | 1 | 17 |
| arrdc3b | 0.15584 | 0.162248422 | 0.071 | 0 | 1 | 17 |
| blvra | 0.15584 | 0.232251895 | 0.071 | 0 | 1 | 17 |
| ca16b | 0.15584 | 0.216960391 | 0.071 | 0 | 1 | 17 |

| | | | | | | |
|---|---|---|---|---|---|---|
| cacnb4b | 0.15584 | 0.216960391 | 0.071 | 0 | 1 | 17 |
| ccdc14 | 0.15584 | 0.171720189 | 0.071 | 0 | 1 | 17 |
| ccdc191 | 0.15584 | 0.175374291 | 0.071 | 0 | 1 | 17 |
| ccng2 | 0.15584 | 0.173245648 | 0.071 | 0 | 1 | 17 |
| cdkl5 | 0.15584 | 0.178584158 | 0.071 | 0 | 1 | 17 |
| cgas | 0.15584 | 0.178584158 | 0.071 | 0 | 1 | 17 |
| coro1b | 0.15584 | 0.178584158 | 0.071 | 0 | 1 | 17 |
| cst14b.1 | 0.15584 | 0.210390173 | 0.071 | 0 | 1 | 17 |
| ctcf | 0.15584 | 0.21550518 | 0.071 | 0 | 1 | 17 |
| ctdspl2a | 0.15584 | 0.176702433 | 0.071 | 0 | 1 | 17 |
| daxx | 0.15584 | 0.176459674 | 0.071 | 0 | 1 | 17 |
| dis3l | 0.15584 | 0.166705825 | 0.071 | 0 | 1 | 17 |
| dnmt3bb.1 | 0.15584 | 0.215642582 | 0.071 | 0 | 1 | 17 |
| eefsec | 0.15584 | 0.178584158 | 0.071 | 0 | 1 | 17 |
| ehd1a | 0.15584 | 0.214569439 | 0.071 | 0 | 1 | 17 |
| fam120b | 0.15584 | 0.216960391 | 0.071 | 0 | 1 | 17 |
| fryl | 0.15584 | 0.176360522 | 0.071 | 0 | 1 | 17 |
| gcdha | 0.15584 | 0.214832923 | 0.071 | 0 | 1 | 17 |
| ghrhra | 0.15584 | 0.216960391 | 0.071 | 0 | 1 | 17 |
| gorasp1b | 0.15584 | 0.178584158 | 0.071 | 0 | 1 | 17 |
| herpud1 | 0.15584 | 0.158355537 | 0.071 | 0 | 1 | 17 |
| il1b | 0.15584 | 0.207561515 | 0.071 | 0 | 1 | 17 |
| itpk1b | 0.15584 | 0.215966747 | 0.071 | 0 | 1 | 17 |
| lonrf1l | 0.15584 | 0.214447973 | 0.071 | 0 | 1 | 17 |
| lrrc39 | 0.15584 | 0.178584158 | 0.071 | 0 | 1 | 17 |
| maea | 0.15584 | 0.216325421 | 0.071 | 0 | 1 | 17 |
| mfsd2aa | 0.15584 | 0.170386104 | 0.071 | 0 | 1 | 17 |
| micall2a | 0.15584 | 0.178584158 | 0.071 | 0 | 1 | 17 |
| micu1 | 0.15584 | 0.178584158 | 0.071 | 0 | 1 | 17 |
| mosmob | 0.15584 | 0.178584158 | 0.071 | 0 | 1 | 17 |
| mrpl16 | 0.15584 | 0.174276174 | 0.071 | 0 | 1 | 17 |
| mynn | 0.15584 | 0.177273029 | 0.071 | 0 | 1 | 17 |
| naf1 | 0.15584 | 0.215875641 | 0.071 | 0 | 1 | 17 |
| nagpa | 0.15584 | 0.17755164 | 0.071 | 0 | 1 | 17 |
| nlrc3 | 0.15584 | 0.163775247 | 0.071 | 0 | 1 | 17 |
| olfml2ba | 0.15584 | 0.178584158 | 0.071 | 0 | 1 | 17 |
| opa3 | 0.15584 | 0.178584158 | 0.071 | 0 | 1 | 17 |
| osbpl9 | 0.15584 | 0.174185174 | 0.071 | 0 | 1 | 17 |
| pbld2 | 0.15584 | 0.178584158 | 0.071 | 0 | 1 | 17 |
| pcbp4 | 0.15584 | 0.178584158 | 0.071 | 0 | 1 | 17 |
| picalma | 0.15584 | 0.173786442 | 0.071 | 0 | 1 | 17 |
| pold3 | 0.15584 | 0.174246808 | 0.071 | 0 | 1 | 17 |
| ppm1da | 0.15584 | 0.216360426 | 0.071 | 0 | 1 | 17 |
| raraa | 0.15584 | 0.164542428 | 0.071 | 0 | 1 | 17 |
| rbm26 | 0.15584 | 0.178584158 | 0.071 | 0 | 1 | 17 |
| rel | 0.15584 | 0.174760664 | 0.071 | 0 | 1 | 17 |
| rimkla | 0.15584 | 0.215878138 | 0.071 | 0 | 1 | 17 |
| rlim | 0.15584 | 0.171937017 | 0.071 | 0 | 1 | 17 |

| | | | | | | |
|---|---|---|---|---|---|---|
| selp | 0.15584 | 0.210877233 | 0.071 | 0 | 1 | 17 |
| serinc1 | 0.15584 | 0.174802629 | 0.071 | 0 | 1 | 17 |
| si:ch211-162i8.5 | 0.15584 | 0.178584158 | 0.071 | 0 | 1 | 17 |
| si:ch211-194c3.5 | 0.15584 | 0.174044713 | 0.071 | 0 | 1 | 17 |
| si:ch211-222n22.1 | 0.15584 | 0.178584158 | 0.071 | 0 | 1 | 17 |
| si:ch211-244o22.2 | 0.15584 | 0.174738368 | 0.071 | 0 | 1 | 17 |
| si:ch211-85n16.3 | 0.15584 | 0.175690479 | 0.071 | 0 | 1 | 17 |
| si:dkey-30c15.2 | 0.15584 | 0.216960391 | 0.071 | 0 | 1 | 17 |
| si:dkeyp-118b1.2 | 0.15584 | 0.178584158 | 0.071 | 0 | 1 | 17 |
| slc25a40 | 0.15584 | 0.173963984 | 0.071 | 0 | 1 | 17 |
| slc25a42 | 0.15584 | 0.177872168 | 0.071 | 0 | 1 | 17 |
| slc9a7 | 0.15584 | 0.178584158 | 0.071 | 0 | 1 | 17 |
| smarcad1b | 0.15584 | 0.215068293 | 0.071 | 0 | 1 | 17 |
| stn1 | 0.15584 | 0.169851205 | 0.071 | 0 | 1 | 17 |
| syngr3b | 0.15584 | 0.171239035 | 0.071 | 0 | 1 | 17 |
| tmem120a | 0.15584 | 0.176347311 | 0.071 | 0 | 1 | 17 |
| tmem123 | 0.15584 | 0.178584158 | 0.071 | 0 | 1 | 17 |
| zgc:136864 | 0.15584 | 0.175116506 | 0.071 | 0 | 1 | 17 |
| zgc:153284 | 0.15584 | 0.212077804 | 0.071 | 0 | 1 | 17 |
| ARF5 (1 of many) | 0.15584 | 0.219129279 | 0.071 | 0 | 1 | 17 |
| BX511128.2 | 0.15584 | 0.219129279 | 0.071 | 0 | 1 | 17 |
| BX548046.1 | 0.15584 | 0.219129279 | 0.071 | 0 | 1 | 17 |
| CABZ01113192.1 | 0.15584 | 0.216889265 | 0.071 | 0 | 1 | 17 |
| CR392026.2 | 0.15584 | 0.208549247 | 0.071 | 0 | 1 | 17 |
| FAM124A | 0.15584 | 0.219129279 | 0.071 | 0 | 1 | 17 |
| GRM7 | 0.15584 | 0.219129279 | 0.071 | 0 | 1 | 17 |
| LOC103909742 | 0.15584 | 0.219129279 | 0.071 | 0 | 1 | 17 |
| LOC110438079 | 0.15584 | 0.219129279 | 0.071 | 0 | 1 | 17 |
| acot20 | 0.15584 | 0.248718121 | 0.071 | 0 | 1 | 17 |
| alg11 | 0.15584 | 0.217556076 | 0.071 | 0 | 1 | 17 |
| anapc7 | 0.15584 | 0.217052331 | 0.071 | 0 | 1 | 17 |
| bicra | 0.15584 | 0.252763154 | 0.071 | 0 | 1 | 17 |
| capn2a | 0.15584 | 0.212125775 | 0.071 | 0 | 1 | 17 |
| cog6 | 0.15584 | 0.219129279 | 0.071 | 0 | 1 | 17 |
| cuzd1.1 | 0.15584 | 0.219129279 | 0.071 | 0 | 1 | 17 |
| dcaf17 | 0.15584 | 0.252763154 | 0.071 | 0 | 1 | 17 |
| ddt | 0.15584 | 0.21462696 | 0.071 | 0 | 1 | 17 |
| dhx33 | 0.15584 | 0.219129279 | 0.071 | 0 | 1 | 17 |
| dkk1a | 0.15584 | 0.252763154 | 0.071 | 0 | 1 | 17 |
| drosha | 0.15584 | 0.219129279 | 0.071 | 0 | 1 | 17 |
| dtx4a | 0.15584 | 0.25173278 | 0.071 | 0 | 1 | 17 |
| eml1 | 0.15584 | 0.242002435 | 0.071 | 0 | 1 | 17 |
| ets2 | 0.15584 | 0.202184541 | 0.071 | 0 | 1 | 17 |
| f2rl1.2 | 0.15584 | 0.219129279 | 0.071 | 0 | 1 | 17 |
| foxi1 | 0.15584 | 0.252763154 | 0.071 | 0 | 1 | 17 |
| foxk1 | 0.15584 | 0.212219336 | 0.071 | 0 | 1 | 17 |
| galca | 0.15584 | 0.252763154 | 0.071 | 0 | 1 | 17 |
| gkup | 0.15584 | 0.251377194 | 0.071 | 0 | 1 | 17 |

| | | | | | | |
|---|---|---|---|---|---|---|
| gli3 | 0.15584 | 0.20988705 | 0.071 | 0 | 1 | 17 |
| gpat4 | 0.15584 | 0.252763154 | 0.071 | 0 | 1 | 17 |
| hbegfb | 0.15584 | 0.25041931 | 0.071 | 0 | 1 | 17 |
| hpca | 0.15584 | 0.219129279 | 0.071 | 0 | 1 | 17 |
| kcnc4 | 0.15584 | 0.252763154 | 0.071 | 0 | 1 | 17 |
| klc3 | 0.15584 | 0.252763154 | 0.071 | 0 | 1 | 17 |
| marcksl1a | 0.15584 | 0.203302945 | 0.071 | 0 | 1 | 17 |
| mgaa | 0.15584 | 0.252763154 | 0.071 | 0 | 1 | 17 |
| miga1 | 0.15584 | 0.216323576 | 0.071 | 0 | 1 | 17 |
| mipol1 | 0.15584 | 0.25144815 | 0.071 | 0 | 1 | 17 |
| mntb | 0.15584 | 0.219129279 | 0.071 | 0 | 1 | 17 |
| mybl2b | 0.15584 | 0.218718355 | 0.071 | 0 | 1 | 17 |
| mydgf | 0.15584 | 0.251986002 | 0.071 | 0 | 1 | 17 |
| nipa2 | 0.15584 | 0.25160012 | 0.071 | 0 | 1 | 17 |
| notch2 | 0.15584 | 0.215768219 | 0.071 | 0 | 1 | 17 |
| nprl2 | 0.15584 | 0.252763154 | 0.071 | 0 | 1 | 17 |
| pelp1 | 0.15584 | 0.218863622 | 0.071 | 0 | 1 | 17 |
| phactr2 | 0.15584 | 0.216169073 | 0.071 | 0 | 1 | 17 |
| pkp1b | 0.15584 | 0.219129279 | 0.071 | 0 | 1 | 17 |
| plekhm2 | 0.15584 | 0.219129279 | 0.071 | 0 | 1 | 17 |
| ppp1r26 | 0.15584 | 0.219129279 | 0.071 | 0 | 1 | 17 |
| rbl1 | 0.15584 | 0.252352877 | 0.071 | 0 | 1 | 17 |
| rhebl1 | 0.15584 | 0.215326851 | 0.071 | 0 | 1 | 17 |
| rps6ka3a | 0.15584 | 0.248157032 | 0.071 | 0 | 1 | 17 |
| sh3bgrl3 | 0.15584 | 0.25013712 | 0.071 | 0 | 1 | 17 |
| si:ch211-113g11.6 | 0.15584 | 0.252763154 | 0.071 | 0 | 1 | 17 |
| si:ch211-160o17.4 | 0.15584 | 0.217283401 | 0.071 | 0 | 1 | 17 |
| si:ch73-109d9.1 | 0.15584 | 0.219129279 | 0.071 | 0 | 1 | 17 |
| si:dkey-165n16.1 | 0.15584 | 0.251313757 | 0.071 | 0 | 1 | 17 |
| si:dkey-202e22.2 | 0.15584 | 0.252763154 | 0.071 | 0 | 1 | 17 |
| si:dkey-83m22.7 | 0.15584 | 0.219129279 | 0.071 | 0 | 1 | 17 |
| slc25a16 | 0.15584 | 0.251964703 | 0.071 | 0 | 1 | 17 |
| slc30a9 | 0.15584 | 0.252184099 | 0.071 | 0 | 1 | 17 |
| stap2b | 0.15584 | 0.252763154 | 0.071 | 0 | 1 | 17 |
| taz | 0.15584 | 0.249653627 | 0.071 | 0 | 1 | 17 |
| trappc12 | 0.15584 | 0.219129279 | 0.071 | 0 | 1 | 17 |
| tsen54 | 0.15584 | 0.252763154 | 0.071 | 0 | 1 | 17 |
| ubald1a | 0.15584 | 0.217774617 | 0.071 | 0 | 1 | 17 |
| ube2j1 | 0.15584 | 0.252763154 | 0.071 | 0 | 1 | 17 |
| usp42 | 0.15584 | 0.218521154 | 0.071 | 0 | 1 | 17 |
| wdr24 | 0.15584 | 0.219129279 | 0.071 | 0 | 1 | 17 |
| zbtb12.1 | 0.15584 | 0.213769675 | 0.071 | 0 | 1 | 17 |
| zfyve28 | 0.15584 | 0.219129279 | 0.071 | 0 | 1 | 17 |
| zgc:110269 | 0.15584 | 0.217799508 | 0.071 | 0 | 1 | 17 |
| zgc:165515 | 0.15584 | 0.251735917 | 0.071 | 0 | 1 | 17 |
| znf653 | 0.15584 | 0.215632856 | 0.071 | 0 | 1 | 17 |
| AL590151.2 | 0.15584 | 0.196138953 | 0.071 | 0 | 1 | 17 |
| AL772362.1 | 0.15584 | 0.158053911 | 0.071 | 0 | 1 | 17 |

| | | | | | | |
|---|---|---|---|---|---|---|
| AL928650.4 | 0.15584 | 0.165043554 | 0.071 | 0 | 1 | 17 |
| AL935174.2 | 0.15584 | 0.164016548 | 0.071 | 0 | 1 | 17 |
| AL953875.1 | 0.15584 | 0.170781964 | 0.071 | 0 | 1 | 17 |
| BX005389.4 | 0.15584 | 0.165043554 | 0.071 | 0 | 1 | 17 |
| BX284696.1 | 0.15584 | 0.169557508 | 0.071 | 0 | 1 | 17 |
| BX322555.3 | 0.15584 | 0.165043554 | 0.071 | 0 | 1 | 17 |
| BX323082.1 | 0.15584 | 0.153709502 | 0.071 | 0 | 1 | 17 |
| BX511166.1 | 0.15584 | 0.165043554 | 0.071 | 0 | 1 | 17 |
| BX571715.1 | 0.15584 | 0.170781964 | 0.071 | 0 | 1 | 17 |
| CABZ01026681.1 | 0.15584 | 0.22579364 | 0.071 | 0 | 1 | 17 |
| CABZ01038697.1 | 0.15584 | 0.149006961 | 0.071 | 0 | 1 | 17 |
| CABZ01055306.1 | 0.15584 | 0.153709502 | 0.071 | 0 | 1 | 17 |
| CABZ01058650.1 | 0.15584 | 0.15923292 | 0.071 | 0 | 1 | 17 |
| CABZ01084081.1 | 0.15584 | 0.156363071 | 0.071 | 0 | 1 | 17 |
| CABZ01084382.1 | 0.15584 | 0.277208742 | 0.071 | 0 | 1 | 17 |
| CABZ01085658.1 | 0.15584 | 0.102148633 | 0.071 | 0 | 1 | 17 |
| CABZ01090041.1 | 0.15584 | 0.170781964 | 0.071 | 0 | 1 | 17 |
| CABZ01090373.1 | 0.15584 | 0.148965844 | 0.071 | 0 | 1 | 17 |
| CABZ01112732.1 | 0.15584 | 0.165043554 | 0.071 | 0 | 1 | 17 |
| CR318601.1 | 0.15584 | 0.165043554 | 0.071 | 0 | 1 | 17 |
| CR381686.1 | 0.15584 | 0.170781964 | 0.071 | 0 | 1 | 17 |
| CR387933.1 | 0.15584 | 0.161653075 | 0.071 | 0 | 1 | 17 |
| CR753886.1 | 0.15584 | 0.147375547 | 0.071 | 0 | 1 | 17 |
| CR759918.1 | 0.15584 | 0.170781964 | 0.071 | 0 | 1 | 17 |
| CR954963.1 | 0.15584 | 0.153709502 | 0.071 | 0 | 1 | 17 |
| CT573429.2 | 0.15584 | 0.153709502 | 0.071 | 0 | 1 | 17 |
| CU179702.2 | 0.15584 | 0.206797871 | 0.071 | 0 | 1 | 17 |
| CU469420.4 | 0.15584 | 0.165043554 | 0.071 | 0 | 1 | 17 |
| CU639397.1 | 0.15584 | 0.182319933 | 0.071 | 0 | 1 | 17 |
| CU655961.4 | 0.15584 | 0.227921552 | 0.071 | 0 | 1 | 17 |
| FP102052.2 | 0.15584 | 0.162809956 | 0.071 | 0 | 1 | 17 |
| FQ377903.1 | 0.15584 | 0.153709502 | 0.071 | 0 | 1 | 17 |
| KHDRBS3 | 0.15584 | 0.196138953 | 0.071 | 0 | 1 | 17 |
| LO017820.1 | 0.15584 | 0.153709502 | 0.071 | 0 | 1 | 17 |
| LO018231.1 | 0.15584 | 0.153709502 | 0.071 | 0 | 1 | 17 |
| LOC100534832 | 0.15584 | 0.161515223 | 0.071 | 0 | 1 | 17 |
| LOC101882326 | 0.15584 | 0.165043554 | 0.071 | 0 | 1 | 17 |
| LOC101882596 | 0.15584 | 0.165043554 | 0.071 | 0 | 1 | 17 |
| LOC101882667 | 0.15584 | 0.165043554 | 0.071 | 0 | 1 | 17 |
| LOC101883779 | 0.15584 | 0.165043554 | 0.071 | 0 | 1 | 17 |
| LOC101885638 | 0.15584 | 0.165043554 | 0.071 | 0 | 1 | 17 |
| LOC103911938 | 0.15584 | 0.196138953 | 0.071 | 0 | 1 | 17 |
| LOC110437755 | 0.15584 | 0.277208742 | 0.071 | 0 | 1 | 17 |
| LOC110438072 | 0.15584 | 0.165043554 | 0.071 | 0 | 1 | 17 |
| LOC110438172 | 0.15584 | 0.165043554 | 0.071 | 0 | 1 | 17 |
| LOC560099 | 0.15584 | 0.165043554 | 0.071 | 0 | 1 | 17 |
| LRRC75A | 0.15584 | 0.18806628 | 0.071 | 0 | 1 | 17 |
| MDFI | 0.15584 | 0.165043554 | 0.071 | 0 | 1 | 17 |

| | | | | | | |
|---|---|---|---|---|---|---|
| NC-002333.17 | 0.15584 | 0.148155094 | 0.071 | 0 | 1 | 17 |
| NFX1 | 0.15584 | 0.163001282 | 0.071 | 0 | 1 | 17 |
| NHLRC3 | 0.15584 | 0.156412669 | 0.071 | 0 | 1 | 17 |
| PRRC2B | 0.15584 | 0.194822704 | 0.071 | 0 | 1 | 17 |
| RF00009 | 0.15584 | 0.157601532 | 0.071 | 0 | 1 | 17 |
| USP34 | 0.15584 | 0.215264094 | 0.071 | 0 | 1 | 17 |
| XLOC-002710 | 0.15584 | 0.165043554 | 0.071 | 0 | 1 | 17 |
| XLOC-003625 | 0.15584 | 0.165043554 | 0.071 | 0 | 1 | 17 |
| XLOC-004189 | 0.15584 | 0.170781964 | 0.071 | 0 | 1 | 17 |
| XLOC-014193 | 0.15584 | 0.196138953 | 0.071 | 0 | 1 | 17 |
| XLOC-014378 | 0.15584 | 0.165043554 | 0.071 | 0 | 1 | 17 |
| aars2 | 0.15584 | 0.165043554 | 0.071 | 0 | 1 | 17 |
| abat | 0.15584 | 0.154329664 | 0.071 | 0 | 1 | 17 |
| abcb10 | 0.15584 | 0.149597237 | 0.071 | 0 | 1 | 17 |
| abcc1 | 0.15584 | 0.165043554 | 0.071 | 0 | 1 | 17 |
| acaca | 0.15584 | 0.196138953 | 0.071 | 0 | 1 | 17 |
| acp2 | 0.15584 | 0.163800549 | 0.071 | 0 | 1 | 17 |
| acp6 | 0.15584 | 0.19517187 | 0.071 | 0 | 1 | 17 |
| adamtsl5 | 0.15584 | 0.165043554 | 0.071 | 0 | 1 | 17 |
| adcy7 | 0.15584 | 0.159451265 | 0.071 | 0 | 1 | 17 |
| adcyap1r1a | 0.15584 | 0.196138953 | 0.071 | 0 | 1 | 17 |
| adgre9 | 0.15584 | 0.277208742 | 0.071 | 0 | 1 | 17 |
| adgrv1 | 0.15584 | 0.170781964 | 0.071 | 0 | 1 | 17 |
| aebp2 | 0.15584 | 0.19290967 | 0.071 | 0 | 1 | 17 |
| aff4 | 0.15584 | 0.161260451 | 0.071 | 0 | 1 | 17 |
| aifm2 | 0.15584 | 0.144645423 | 0.071 | 0 | 1 | 17 |
| alas2 | 0.15584 | 0.20520242 | 0.071 | 0 | 1 | 17 |
| alg6 | 0.15584 | 0.1500334 | 0.071 | 0 | 1 | 17 |
| alms1 | 0.15584 | 0.160818565 | 0.071 | 0 | 1 | 17 |
| alpk2 | 0.15584 | 0.165043554 | 0.071 | 0 | 1 | 17 |
| als2a | 0.15584 | 0.151094935 | 0.071 | 0 | 1 | 17 |
| ampd3b | 0.15584 | 0.166863395 | 0.071 | 0 | 1 | 17 |
| ankmy2a | 0.15584 | 0.133251049 | 0.071 | 0 | 1 | 17 |
| anxa5a | 0.15584 | 0.107923408 | 0.071 | 0 | 1 | 17 |
| ap5m1 | 0.15584 | 0.147230938 | 0.071 | 0 | 1 | 17 |
| apba2b | 0.15584 | 0.165043554 | 0.071 | 0 | 1 | 17 |
| arhgap35a | 0.15584 | 0.146375146 | 0.071 | 0 | 1 | 17 |
| armc7 | 0.15584 | 0.158741268 | 0.071 | 0 | 1 | 17 |
| arntl2 | 0.15584 | 0.196138953 | 0.071 | 0 | 1 | 17 |
| arpp21 | 0.15584 | 0.146420414 | 0.071 | 0 | 1 | 17 |
| arv1 | 0.15584 | 0.153709502 | 0.071 | 0 | 1 | 17 |
| asah1a | 0.15584 | 0.10559198 | 0.071 | 0 | 1 | 17 |
| asb6 | 0.15584 | 0.165043554 | 0.071 | 0 | 1 | 17 |
| ascl1b | 0.15584 | 0.150015603 | 0.071 | 0 | 1 | 17 |
| asxl1 | 0.15584 | 0.163489297 | 0.071 | 0 | 1 | 17 |
| atg10 | 0.15584 | 0.149109464 | 0.071 | 0 | 1 | 17 |
| atg9b | 0.15584 | 0.153222659 | 0.071 | 0 | 1 | 17 |
| atp2a2a | 0.15584 | 0.227921552 | 0.071 | 0 | 1 | 17 |

| | | | | | | |
|---|---|---|---|---|---|---|
| atp6v0a1b | 0.15584 | 0.165043554 | 0.071 | 0 | 1 | 17 |
| atp6v1c1b | 0.15584 | 0.132979807 | 0.071 | 0 | 1 | 17 |
| bahd1 | 0.15584 | 0.149056866 | 0.071 | 0 | 1 | 17 |
| baiap2l2a | 0.15584 | 0.153709502 | 0.071 | 0 | 1 | 17 |
| bbs5 | 0.15584 | 0.227921552 | 0.071 | 0 | 1 | 17 |
| bmf2 | 0.15584 | 0.218899604 | 0.071 | 0 | 1 | 17 |
| braf | 0.15584 | 0.196138953 | 0.071 | 0 | 1 | 17 |
| brd1a | 0.15584 | 0.165043554 | 0.071 | 0 | 1 | 17 |
| btg1 | 0.15584 | 0.222496125 | 0.071 | 0 | 1 | 17 |
| c2cd2l | 0.15584 | 0.168409443 | 0.071 | 0 | 1 | 17 |
| c2cd3 | 0.15584 | 0.165043554 | 0.071 | 0 | 1 | 17 |
| c7b | 0.15584 | 0.165043554 | 0.071 | 0 | 1 | 17 |
| cachd1 | 0.15584 | 0.170781964 | 0.071 | 0 | 1 | 17 |
| carnmt1 | 0.15584 | 0.196138953 | 0.071 | 0 | 1 | 17 |
| casc1 | 0.15584 | 0.153709502 | 0.071 | 0 | 1 | 17 |
| caspa | 0.15584 | 0.139941805 | 0.071 | 0 | 1 | 17 |
| cd99 | 0.15584 | 0.192102488 | 0.071 | 0 | 1 | 17 |
| cdh7a | 0.15584 | 0.153709502 | 0.071 | 0 | 1 | 17 |
| cdk15 | 0.15584 | 0.153709502 | 0.071 | 0 | 1 | 17 |
| cenpn | 0.15584 | 0.194851835 | 0.071 | 0 | 1 | 17 |
| cep83 | 0.15584 | 0.165043554 | 0.071 | 0 | 1 | 17 |
| cers2b | 0.15584 | 0.193266069 | 0.071 | 0 | 1 | 17 |
| chd1l | 0.15584 | 0.153709502 | 0.071 | 0 | 1 | 17 |
| cipca | 0.15584 | 0.165043554 | 0.071 | 0 | 1 | 17 |
| clint1a | 0.15584 | 0.166679048 | 0.071 | 0 | 1 | 17 |
| clip2 | 0.15584 | 0.165043554 | 0.071 | 0 | 1 | 17 |
| clk4b | 0.15584 | 0.142271296 | 0.071 | 0 | 1 | 17 |
| clptm1l | 0.15584 | 0.152581279 | 0.071 | 0 | 1 | 17 |
| cmip | 0.15584 | 0.161472164 | 0.071 | 0 | 1 | 17 |
| cmpk2 | 0.15584 | 0.227921552 | 0.071 | 0 | 1 | 17 |
| cmtm6 | 0.15584 | 0.149100411 | 0.071 | 0 | 1 | 17 |
| cnga1b | 0.15584 | 0.218899604 | 0.071 | 0 | 1 | 17 |
| cnn1b | 0.15584 | 0.165043554 | 0.071 | 0 | 1 | 17 |
| cnot6a | 0.15584 | 0.196138953 | 0.071 | 0 | 1 | 17 |
| col11a1b | 0.15584 | 0.162425401 | 0.071 | 0 | 1 | 17 |
| commd8 | 0.15584 | 0.150472989 | 0.071 | 0 | 1 | 17 |
| coq3 | 0.15584 | 0.148593654 | 0.071 | 0 | 1 | 17 |
| coro1a | 0.15584 | 0.208045332 | 0.071 | 0 | 1 | 17 |
| cpeb2 | 0.15584 | 0.165043554 | 0.071 | 0 | 1 | 17 |
| cratb | 0.15584 | 0.151808297 | 0.071 | 0 | 1 | 17 |
| crebbpb | 0.15584 | 0.164190713 | 0.071 | 0 | 1 | 17 |
| cryz | 0.15584 | 0.223176562 | 0.071 | 0 | 1 | 17 |
| csnk1g1 | 0.15584 | 0.191791492 | 0.071 | 0 | 1 | 17 |
| ctdsp2 | 0.15584 | 0.144744673 | 0.071 | 0 | 1 | 17 |
| cxcr4a | 0.15584 | 0.190420924 | 0.071 | 0 | 1 | 17 |
| cyb561d2 | 0.15584 | 0.170781964 | 0.071 | 0 | 1 | 17 |
| cyp27a1.4 | 0.15584 | 0.227921552 | 0.071 | 0 | 1 | 17 |
| dap3 | 0.15584 | 0.151460149 | 0.071 | 0 | 1 | 17 |

| | | | | | | |
|---|---|---|---|---|---|---|
| dars2 | 0.15584 | 0.136879821 | 0.071 | 0 | 1 | 17 |
| dcps | 0.15584 | 0.226876128 | 0.071 | 0 | 1 | 17 |
| dennd1b | 0.15584 | 0.165043554 | 0.071 | 0 | 1 | 17 |
| dennd2db | 0.15584 | 0.165043554 | 0.071 | 0 | 1 | 17 |
| dgkza | 0.15584 | 0.163418338 | 0.071 | 0 | 1 | 17 |
| dip2bb | 0.15584 | 0.165043554 | 0.071 | 0 | 1 | 17 |
| dip2ca | 0.15584 | 0.161656854 | 0.071 | 0 | 1 | 17 |
| dmxl2 | 0.15584 | 0.13554963 | 0.071 | 0 | 1 | 17 |
| dnaaf4 | 0.15584 | 0.153709502 | 0.071 | 0 | 1 | 17 |
| dnajb1b | 0.15584 | 0.143907451 | 0.071 | 0 | 1 | 17 |
| dnajc17 | 0.15584 | 0.150737791 | 0.071 | 0 | 1 | 17 |
| dolpp1 | 0.15584 | 0.192327001 | 0.071 | 0 | 1 | 17 |
| draxin | 0.15584 | 0.227921552 | 0.071 | 0 | 1 | 17 |
| ece2a | 0.15584 | 0.275724923 | 0.071 | 0 | 1 | 17 |
| eif2ak1 | 0.15584 | 0.155823989 | 0.071 | 0 | 1 | 17 |
| elmod2 | 0.15584 | 0.165043554 | 0.071 | 0 | 1 | 17 |
| elovl5 | 0.15584 | 0.165043554 | 0.071 | 0 | 1 | 17 |
| emb | 0.15584 | 0.165043554 | 0.071 | 0 | 1 | 17 |
| enox2 | 0.15584 | 0.165043554 | 0.071 | 0 | 1 | 17 |
| eogt | 0.15584 | 0.153709502 | 0.071 | 0 | 1 | 17 |
| ep300b | 0.15584 | 0.16122031 | 0.071 | 0 | 1 | 17 |
| etv5a | 0.15584 | 0.227921552 | 0.071 | 0 | 1 | 17 |
| evi5b | 0.15584 | 0.168478942 | 0.071 | 0 | 1 | 17 |
| fam173a | 0.15584 | 0.149955454 | 0.071 | 0 | 1 | 17 |
| fam184b | 0.15584 | 0.138499744 | 0.071 | 0 | 1 | 17 |
| fam222bb | 0.15584 | 0.193191503 | 0.071 | 0 | 1 | 17 |
| fam3c | 0.15584 | 0.160879017 | 0.071 | 0 | 1 | 17 |
| fam43a | 0.15584 | 0.153709502 | 0.071 | 0 | 1 | 17 |
| fam45a | 0.15584 | 0.250644172 | 0.071 | 0 | 1 | 17 |
| fam83e | 0.15584 | 0.196138953 | 0.071 | 0 | 1 | 17 |
| fam83fb | 0.15584 | 0.153709502 | 0.071 | 0 | 1 | 17 |
| fam92a1 | 0.15584 | 0.196138953 | 0.071 | 0 | 1 | 17 |
| fam98a | 0.15584 | 0.225800855 | 0.071 | 0 | 1 | 17 |
| fam98b | 0.15584 | 0.153709502 | 0.071 | 0 | 1 | 17 |
| fastkd5 | 0.15584 | 0.277208742 | 0.071 | 0 | 1 | 17 |
| fbxl4 | 0.15584 | 0.166002109 | 0.071 | 0 | 1 | 17 |
| fbxo16 | 0.15584 | 0.153709502 | 0.071 | 0 | 1 | 17 |
| fbxo22 | 0.15584 | 0.158023253 | 0.071 | 0 | 1 | 17 |
| fbxo32 | 0.15584 | 0.153709502 | 0.071 | 0 | 1 | 17 |
| fbxo34 | 0.15584 | 0.151362541 | 0.071 | 0 | 1 | 17 |
| fbxo8 | 0.15584 | 0.192102793 | 0.071 | 0 | 1 | 17 |
| fer | 0.15584 | 0.162782596 | 0.071 | 0 | 1 | 17 |
| fgd1 | 0.15584 | 0.163757992 | 0.071 | 0 | 1 | 17 |
| fgf13a | 0.15584 | 0.153709502 | 0.071 | 0 | 1 | 17 |
| fgf8a | 0.15584 | 0.165043554 | 0.071 | 0 | 1 | 17 |
| fip1l1a | 0.15584 | 0.202951276 | 0.071 | 0 | 1 | 17 |
| fitm2 | 0.15584 | 0.282922337 | 0.071 | 0 | 1 | 17 |
| foxp3b | 0.15584 | 0.277208742 | 0.071 | 0 | 1 | 17 |

| | | | | | | |
|---|---|---|---|---|---|---|
| frem2b | 0.15584 | 0.165043554 | 0.071 | 0 | 1 | 17 |
| frem3 | 0.15584 | 0.227921552 | 0.071 | 0 | 1 | 17 |
| fzd3b | 0.15584 | 0.165043554 | 0.071 | 0 | 1 | 17 |
| fzd7b | 0.15584 | 0.161722807 | 0.071 | 0 | 1 | 17 |
| gabrd | 0.15584 | 0.170781964 | 0.071 | 0 | 1 | 17 |
| gatad2ab | 0.15584 | 0.226560116 | 0.071 | 0 | 1 | 17 |
| gga1 | 0.15584 | 0.158975346 | 0.071 | 0 | 1 | 17 |
| ghdc | 0.15584 | 0.165043554 | 0.071 | 0 | 1 | 17 |
| ginm1 | 0.15584 | 0.225249308 | 0.071 | 0 | 1 | 17 |
| git1 | 0.15584 | 0.163290621 | 0.071 | 0 | 1 | 17 |
| glg1a | 0.15584 | 0.163342725 | 0.071 | 0 | 1 | 17 |
| gnai2a | 0.15584 | 0.160842651 | 0.071 | 0 | 1 | 17 |
| gnal2 | 0.15584 | 0.166503817 | 0.071 | 0 | 1 | 17 |
| gnb2 | 0.15584 | 0.186102478 | 0.071 | 0 | 1 | 17 |
| got1 | 0.15584 | 0.143620986 | 0.071 | 0 | 1 | 17 |
| gpr141 | 0.15584 | 0.155753284 | 0.071 | 0 | 1 | 17 |
| gps1 | 0.15584 | 0.193790799 | 0.071 | 0 | 1 | 17 |
| grb10b | 0.15584 | 0.150044849 | 0.071 | 0 | 1 | 17 |
| grin2cb | 0.15584 | 0.165043554 | 0.071 | 0 | 1 | 17 |
| gsdmea | 0.15584 | 0.14377612 | 0.071 | 0 | 1 | 17 |
| gucy1b1 | 0.15584 | 0.153709502 | 0.071 | 0 | 1 | 17 |
| harbi1 | 0.15584 | 0.153709502 | 0.071 | 0 | 1 | 17 |
| hcfc1b | 0.15584 | 0.216673815 | 0.071 | 0 | 1 | 17 |
| hebp2 | 0.15584 | 0.145782145 | 0.071 | 0 | 1 | 17 |
| hectd2 | 0.15584 | 0.163089322 | 0.071 | 0 | 1 | 17 |
| hecw1b | 0.15584 | 0.170781964 | 0.071 | 0 | 1 | 17 |
| her9 | 0.15584 | 0.162357481 | 0.071 | 0 | 1 | 17 |
| hpcal1 | 0.15584 | 0.160571854 | 0.071 | 0 | 1 | 17 |
| hpse | 0.15584 | 0.227921552 | 0.071 | 0 | 1 | 17 |
| hsf2bp | 0.15584 | 0.153709502 | 0.071 | 0 | 1 | 17 |
| htt | 0.15584 | 0.167469571 | 0.071 | 0 | 1 | 17 |
| hykk.2 | 0.15584 | 0.165043554 | 0.071 | 0 | 1 | 17 |
| ica1 | 0.15584 | 0.20273242 | 0.071 | 0 | 1 | 17 |
| idh1 | 0.15584 | 0.159488741 | 0.071 | 0 | 1 | 17 |
| ifngr1l | 0.15584 | 0.124505253 | 0.071 | 0 | 1 | 17 |
| igfbp3 | 0.15584 | 0.162355177 | 0.071 | 0 | 1 | 17 |
| igsf9ba | 0.15584 | 0.170781964 | 0.071 | 0 | 1 | 17 |
| ikzf5 | 0.15584 | 0.191732191 | 0.071 | 0 | 1 | 17 |
| il15 | 0.15584 | 0.143794287 | 0.071 | 0 | 1 | 17 |
| il4r.1 | 0.15584 | 0.153021153 | 0.071 | 0 | 1 | 17 |
| ints9 | 0.15584 | 0.165043554 | 0.071 | 0 | 1 | 17 |
| irak1bp1 | 0.15584 | 0.196138953 | 0.071 | 0 | 1 | 17 |
| irf2 | 0.15584 | 0.154427782 | 0.071 | 0 | 1 | 17 |
| irx7 | 0.15584 | 0.165043554 | 0.071 | 0 | 1 | 17 |
| itgb1bp1 | 0.15584 | 0.194120661 | 0.071 | 0 | 1 | 17 |
| itk | 0.15584 | 0.165043554 | 0.071 | 0 | 1 | 17 |
| jmjd6 | 0.15584 | 0.196138953 | 0.071 | 0 | 1 | 17 |
| kazna | 0.15584 | 0.165043554 | 0.071 | 0 | 1 | 17 |

| | | | | | | |
|---|---|---|---|---|---|---|
| kbtbd4 | 0.15584 | 0.170781964 | 0.071 | 0 | 1 | 17 |
| kcnk6 | 0.15584 | 0.165043554 | 0.071 | 0 | 1 | 17 |
| kcnma1a | 0.15584 | 0.158149875 | 0.071 | 0 | 1 | 17 |
| kdf1a | 0.15584 | 0.196138953 | 0.071 | 0 | 1 | 17 |
| kdm2bb | 0.15584 | 0.146873461 | 0.071 | 0 | 1 | 17 |
| kdm6a | 0.15584 | 0.202456152 | 0.071 | 0 | 1 | 17 |
| kl | 0.15584 | 0.151584835 | 0.071 | 0 | 1 | 17 |
| laptm4b | 0.15584 | 0.134523665 | 0.071 | 0 | 1 | 17 |
| lcn15 | 0.15584 | 0.212100033 | 0.071 | 0 | 1 | 17 |
| lcorl | 0.15584 | 0.226934975 | 0.071 | 0 | 1 | 17 |
| lenep | 0.15584 | 0.196138953 | 0.071 | 0 | 1 | 17 |
| leng8 | 0.15584 | 0.227099805 | 0.071 | 0 | 1 | 17 |
| lepr | 0.15584 | 0.153709502 | 0.071 | 0 | 1 | 17 |
| lgals9l4 | 0.15584 | 0.18600264 | 0.071 | 0 | 1 | 17 |
| lhx8a | 0.15584 | 0.14289211 | 0.071 | 0 | 1 | 17 |
| limch1b | 0.15584 | 0.200051084 | 0.071 | 0 | 1 | 17 |
| lrch1 | 0.15584 | 0.165043554 | 0.071 | 0 | 1 | 17 |
| lrp10 | 0.15584 | 0.162874945 | 0.071 | 0 | 1 | 17 |
| lrwd1 | 0.15584 | 0.159930461 | 0.071 | 0 | 1 | 17 |
| map2k4b | 0.15584 | 0.148501108 | 0.071 | 0 | 1 | 17 |
| map3k21 | 0.15584 | 0.163293566 | 0.071 | 0 | 1 | 17 |
| map3k3 | 0.15584 | 0.165043554 | 0.071 | 0 | 1 | 17 |
| map4k2 | 0.15584 | 0.153709502 | 0.071 | 0 | 1 | 17 |
| map4l | 0.15584 | 0.165043554 | 0.071 | 0 | 1 | 17 |
| map9 | 0.15584 | 0.148305431 | 0.071 | 0 | 1 | 17 |
| mapk1 | 0.15584 | 0.164096894 | 0.071 | 0 | 1 | 17 |
| mapk4 | 0.15584 | 0.165043554 | 0.071 | 0 | 1 | 17 |
| marcksa | 0.15584 | 0.164340924 | 0.071 | 0 | 1 | 17 |
| mef2cb | 0.15584 | 0.141303424 | 0.071 | 0 | 1 | 17 |
| metrnla | 0.15584 | 0.119864929 | 0.071 | 0 | 1 | 17 |
| mfge8b | 0.15584 | 0.163564119 | 0.071 | 0 | 1 | 17 |
| mier3a | 0.15584 | 0.161780059 | 0.071 | 0 | 1 | 17 |
| mlf1 | 0.15584 | 0.227921552 | 0.071 | 0 | 1 | 17 |
| mmp16a | 0.15584 | 0.163986423 | 0.071 | 0 | 1 | 17 |
| mnat1 | 0.15584 | 0.146581794 | 0.071 | 0 | 1 | 17 |
| mplkip | 0.15584 | 0.168854668 | 0.071 | 0 | 1 | 17 |
| mrps2 | 0.15584 | 0.153561531 | 0.071 | 0 | 1 | 17 |
| mt-nd4l | 0.15584 | 0.155080172 | 0.071 | 0 | 1 | 17 |
| mtg1 | 0.15584 | 0.192237732 | 0.071 | 0 | 1 | 17 |
| mtmr9 | 0.15584 | 0.150670806 | 0.071 | 0 | 1 | 17 |
| mul1b | 0.15584 | 0.165043554 | 0.071 | 0 | 1 | 17 |
| mxra5b | 0.15584 | 0.165043554 | 0.071 | 0 | 1 | 17 |
| myhz1.2 | 0.15584 | 0.152658531 | 0.071 | 0 | 1 | 17 |
| myo18ab | 0.15584 | 0.156491165 | 0.071 | 0 | 1 | 17 |
| myo19 | 0.15584 | 0.153709502 | 0.071 | 0 | 1 | 17 |
| myof | 0.15584 | 0.165043554 | 0.071 | 0 | 1 | 17 |
| nadka | 0.15584 | 0.192081701 | 0.071 | 0 | 1 | 17 |
| nanos2 | 0.15584 | 0.196138953 | 0.071 | 0 | 1 | 17 |

| | | | | | | |
|---|---|---|---|---|---|---|
| ncam3 | 0.15584 | 0.165043554 | 0.071 | 0 | 1 | 17 |
| ncapg2 | 0.15584 | 0.15959704 | 0.071 | 0 | 1 | 17 |
| ncor1 | 0.15584 | 0.160208541 | 0.071 | 0 | 1 | 17 |
| nek8 | 0.15584 | 0.153709502 | 0.071 | 0 | 1 | 17 |
| neurl1ab | 0.15584 | 0.161451641 | 0.071 | 0 | 1 | 17 |
| nfe2 | 0.15584 | 0.152472965 | 0.071 | 0 | 1 | 17 |
| nfkbie | 0.15584 | 0.22598953 | 0.071 | 0 | 1 | 17 |
| nfxl1 | 0.15584 | 0.194977999 | 0.071 | 0 | 1 | 17 |
| nhsl1a | 0.15584 | 0.162760178 | 0.071 | 0 | 1 | 17 |
| nkx3-1 | 0.15584 | 0.15780012 | 0.071 | 0 | 1 | 17 |
| nlrc3l1 | 0.15584 | 0.165043554 | 0.071 | 0 | 1 | 17 |
| nme7 | 0.15584 | 0.190568546 | 0.071 | 0 | 1 | 17 |
| npb | 0.15584 | 0.165043554 | 0.071 | 0 | 1 | 17 |
| nppc | 0.15584 | 0.165043554 | 0.071 | 0 | 1 | 17 |
| nprl3 | 0.15584 | 0.196138953 | 0.071 | 0 | 1 | 17 |
| nsd1a | 0.15584 | 0.166813002 | 0.071 | 0 | 1 | 17 |
| nt5dc2 | 0.15584 | 0.166866487 | 0.071 | 0 | 1 | 17 |
| nup214 | 0.15584 | 0.163299143 | 0.071 | 0 | 1 | 17 |
| nxph2b | 0.15584 | 0.165043554 | 0.071 | 0 | 1 | 17 |
| oca2 | 0.15584 | 0.153709502 | 0.071 | 0 | 1 | 17 |
| ogdha | 0.15584 | 0.194062616 | 0.071 | 0 | 1 | 17 |
| opa1 | 0.15584 | 0.158756966 | 0.071 | 0 | 1 | 17 |
| osbpl2a | 0.15584 | 0.161574613 | 0.071 | 0 | 1 | 17 |
| oxr1b | 0.15584 | 0.226199698 | 0.071 | 0 | 1 | 17 |
| p3h2 | 0.15584 | 0.170781964 | 0.071 | 0 | 1 | 17 |
| paqr7b | 0.15584 | 0.187229064 | 0.071 | 0 | 1 | 17 |
| parp12a | 0.15584 | 0.161687192 | 0.071 | 0 | 1 | 17 |
| pcbp3 | 0.15584 | 0.165043554 | 0.071 | 0 | 1 | 17 |
| pcnt | 0.15584 | 0.214398691 | 0.071 | 0 | 1 | 17 |
| pdca | 0.15584 | 0.160122144 | 0.071 | 0 | 1 | 17 |
| pdcd2l | 0.15584 | 0.250652881 | 0.071 | 0 | 1 | 17 |
| per1b | 0.15584 | 0.162810202 | 0.071 | 0 | 1 | 17 |
| pex12 | 0.15584 | 0.154081895 | 0.071 | 0 | 1 | 17 |
| pex13 | 0.15584 | 0.149752332 | 0.071 | 0 | 1 | 17 |
| pgm3 | 0.15584 | 0.196138953 | 0.071 | 0 | 1 | 17 |
| phactr1 | 0.15584 | 0.165043554 | 0.071 | 0 | 1 | 17 |
| phc3 | 0.15584 | 0.153709502 | 0.071 | 0 | 1 | 17 |
| phf12a | 0.15584 | 0.165043554 | 0.071 | 0 | 1 | 17 |
| phkg1b | 0.15584 | 0.196138953 | 0.071 | 0 | 1 | 17 |
| pigo | 0.15584 | 0.165043554 | 0.071 | 0 | 1 | 17 |
| pik3ip1 | 0.15584 | 0.226592673 | 0.071 | 0 | 1 | 17 |
| pimr134 | 0.15584 | 0.165043554 | 0.071 | 0 | 1 | 17 |
| pla1a | 0.15584 | 0.153709502 | 0.071 | 0 | 1 | 17 |
| plecb | 0.15584 | 0.193910276 | 0.071 | 0 | 1 | 17 |
| plvapa | 0.15584 | 0.196138953 | 0.071 | 0 | 1 | 17 |
| pmepa1 | 0.15584 | 0.196138953 | 0.071 | 0 | 1 | 17 |
| ppp1r14ab | 0.15584 | 0.153709502 | 0.071 | 0 | 1 | 17 |
| ppp1r9ala | 0.15584 | 0.1596435 | 0.071 | 0 | 1 | 17 |

| | | | | | | |
|---|---|---|---|---|---|---|
| ppp2r2bb | 0.15584 | 0.165043554 | 0.071 | 0 | 1 | 17 |
| ppp5c | 0.15584 | 0.166588737 | 0.071 | 0 | 1 | 17 |
| prkcbp1l | 0.15584 | 0.194319097 | 0.071 | 0 | 1 | 17 |
| prune | 0.15584 | 0.165043554 | 0.071 | 0 | 1 | 17 |
| psd3l | 0.15584 | 0.166887128 | 0.071 | 0 | 1 | 17 |
| psma6l | 0.15584 | 0.205239945 | 0.071 | 0 | 1 | 17 |
| ptbp1b | 0.15584 | 0.16627875 | 0.071 | 0 | 1 | 17 |
| ptcd2 | 0.15584 | 0.193774971 | 0.071 | 0 | 1 | 17 |
| ptpmt1 | 0.15584 | 0.204587697 | 0.071 | 0 | 1 | 17 |
| ptprja | 0.15584 | 0.227921552 | 0.071 | 0 | 1 | 17 |
| pttg1ipb | 0.15584 | 0.122660437 | 0.071 | 0 | 1 | 17 |
| pvrl2l | 0.15584 | 0.164120035 | 0.071 | 0 | 1 | 17 |
| qpct | 0.15584 | 0.153709502 | 0.071 | 0 | 1 | 17 |
| qsox1 | 0.15584 | 0.142951473 | 0.071 | 0 | 1 | 17 |
| rab1ba | 0.15584 | 0.210307043 | 0.071 | 0 | 1 | 17 |
| rab23 | 0.15584 | 0.227258126 | 0.071 | 0 | 1 | 17 |
| rab33a | 0.15584 | 0.251362851 | 0.071 | 0 | 1 | 17 |
| rab34b | 0.15584 | 0.224612511 | 0.071 | 0 | 1 | 17 |
| rab40c | 0.15584 | 0.153709502 | 0.071 | 0 | 1 | 17 |
| rabep1 | 0.15584 | 0.161015542 | 0.071 | 0 | 1 | 17 |
| rabepk | 0.15584 | 0.196138953 | 0.071 | 0 | 1 | 17 |
| rasef | 0.15584 | 0.130420049 | 0.071 | 0 | 1 | 17 |
| rasgrf2b | 0.15584 | 0.170781964 | 0.071 | 0 | 1 | 17 |
| rbms3 | 0.15584 | 0.16270521 | 0.071 | 0 | 1 | 17 |
| rela | 0.15584 | 0.170781964 | 0.071 | 0 | 1 | 17 |
| reps1 | 0.15584 | 0.165043554 | 0.071 | 0 | 1 | 17 |
| rfk | 0.15584 | 0.205881103 | 0.071 | 0 | 1 | 17 |
| rgl1 | 0.15584 | 0.159680077 | 0.071 | 0 | 1 | 17 |
| rhbdf1a | 0.15584 | 0.15850642 | 0.071 | 0 | 1 | 17 |
| ribc2 | 0.15584 | 0.196138953 | 0.071 | 0 | 1 | 17 |
| rnf111 | 0.15584 | 0.153709502 | 0.071 | 0 | 1 | 17 |
| rnf126 | 0.15584 | 0.227921552 | 0.071 | 0 | 1 | 17 |
| rnf41 | 0.15584 | 0.14834823 | 0.071 | 0 | 1 | 17 |
| ropn1l | 0.15584 | 0.232506805 | 0.071 | 0 | 1 | 17 |
| rps6ka3b | 0.15584 | 0.162235061 | 0.071 | 0 | 1 | 17 |
| rragd | 0.15584 | 0.152378147 | 0.071 | 0 | 1 | 17 |
| rrbp1b | 0.15584 | 0.158914722 | 0.071 | 0 | 1 | 17 |
| rtn1b | 0.15584 | 0.131450426 | 0.071 | 0 | 1 | 17 |
| rxfp2a | 0.15584 | 0.143581483 | 0.071 | 0 | 1 | 17 |
| sars2 | 0.15584 | 0.196138953 | 0.071 | 0 | 1 | 17 |
| sc:d189 | 0.15584 | 0.153709502 | 0.071 | 0 | 1 | 17 |
| scamp1 | 0.15584 | 0.165144558 | 0.071 | 0 | 1 | 17 |
| scly | 0.15584 | 0.196138953 | 0.071 | 0 | 1 | 17 |
| scn1bb | 0.15584 | 0.165043554 | 0.071 | 0 | 1 | 17 |
| scyl3 | 0.15584 | 0.165043554 | 0.071 | 0 | 1 | 17 |
| selenon | 0.15584 | 0.135532679 | 0.071 | 0 | 1 | 17 |
| sema3b | 0.15584 | 0.196138953 | 0.071 | 0 | 1 | 17 |
| setdb1b | 0.15584 | 0.151995525 | 0.071 | 0 | 1 | 17 |

| | | | | | | |
|---|---|---|---|---|---|---|
| sfi1 | 0.15584 | 0.155620089 | 0.071 | 0 | 1 | 17 |
| si:cabz01007802.1 | 0.15584 | 0.165043554 | 0.071 | 0 | 1 | 17 |
| si:ch1073-70f20.1 | 0.15584 | 0.170781964 | 0.071 | 0 | 1 | 17 |
| si:ch211-106a19.1 | 0.15584 | 0.152171061 | 0.071 | 0 | 1 | 17 |
| si:ch211-106h4.9 | 0.15584 | 0.170781964 | 0.071 | 0 | 1 | 17 |
| si:ch211-108d22.2 | 0.15584 | 0.170781964 | 0.071 | 0 | 1 | 17 |
| si:ch211-133h13.1 | 0.15584 | 0.149323865 | 0.071 | 0 | 1 | 17 |
| si:ch211-137a8.2 | 0.15584 | 0.219752072 | 0.071 | 0 | 1 | 17 |
| si:ch211-137i24.12 | 0.15584 | 0.153709502 | 0.071 | 0 | 1 | 17 |
| si:ch211-15d5.12 | 0.15584 | 0.153200611 | 0.071 | 0 | 1 | 17 |
| si:ch211-163l21.7 | 0.15584 | 0.170781964 | 0.071 | 0 | 1 | 17 |
| si:ch211-173p18.3 | 0.15584 | 0.21772987 | 0.071 | 0 | 1 | 17 |
| si:ch211-195d17.2 | 0.15584 | 0.161300242 | 0.071 | 0 | 1 | 17 |
| si:ch211-197g15.7 | 0.15584 | 0.153709502 | 0.071 | 0 | 1 | 17 |
| si:ch211-197h24.9 | 0.15584 | 0.14861532 | 0.071 | 0 | 1 | 17 |
| si:ch211-207i20.2 | 0.15584 | 0.206797871 | 0.071 | 0 | 1 | 17 |
| si:ch211-218o21.4 | 0.15584 | 0.165043554 | 0.071 | 0 | 1 | 17 |
| si:ch211-239f4.1 | 0.15584 | 0.251402984 | 0.071 | 0 | 1 | 17 |
| si:ch211-241e1.5 | 0.15584 | 0.165043554 | 0.071 | 0 | 1 | 17 |
| si:ch211-272n13.3 | 0.15584 | 0.193856516 | 0.071 | 0 | 1 | 17 |
| si:ch211-284e13.5 | 0.15584 | 0.216706688 | 0.071 | 0 | 1 | 17 |
| si:ch211-285f17.1 | 0.15584 | 0.165043554 | 0.071 | 0 | 1 | 17 |
| si:ch211-33e4.3 | 0.15584 | 0.165043554 | 0.071 | 0 | 1 | 17 |
| si:ch211-66k16.27 | 0.15584 | 0.226330598 | 0.071 | 0 | 1 | 17 |
| si:ch73-132f6.5 | 0.15584 | 0.157167676 | 0.071 | 0 | 1 | 17 |
| si:ch73-1a9.4 | 0.15584 | 0.147760551 | 0.071 | 0 | 1 | 17 |
| si:ch73-376l24.6 | 0.15584 | 0.162912939 | 0.071 | 0 | 1 | 17 |
| si:ch73-92e7.4 | 0.15584 | 0.162962408 | 0.071 | 0 | 1 | 17 |
| si:dkey-172j4.3 | 0.15584 | 0.129550668 | 0.071 | 0 | 1 | 17 |
| si:dkey-183i3.6 | 0.15584 | 0.153709502 | 0.071 | 0 | 1 | 17 |
| si:dkey-188h10.3 | 0.15584 | 0.150235703 | 0.071 | 0 | 1 | 17 |
| si:dkey-197j19.6 | 0.15584 | 0.277208742 | 0.071 | 0 | 1 | 17 |
| si:dkey-19b23.10 | 0.15584 | 0.153709502 | 0.071 | 0 | 1 | 17 |
| si:dkey-22i16.9 | 0.15584 | 0.196138953 | 0.071 | 0 | 1 | 17 |
| si:dkey-237i9.1 | 0.15584 | 0.161310791 | 0.071 | 0 | 1 | 17 |
| si:dkey-237j10.2 | 0.15584 | 0.163573721 | 0.071 | 0 | 1 | 17 |
| si:dkey-250d21.1 | 0.15584 | 0.216799233 | 0.071 | 0 | 1 | 17 |
| si:dkey-262g12.12 | 0.15584 | 0.165043554 | 0.071 | 0 | 1 | 17 |
| si:dkey-262g12.7 | 0.15584 | 0.165043554 | 0.071 | 0 | 1 | 17 |
| si:dkey-26c10.5 | 0.15584 | 0.183811441 | 0.071 | 0 | 1 | 17 |
| si:dkey-32e23.4 | 0.15584 | 0.165043554 | 0.071 | 0 | 1 | 17 |
| si:dkey-66g10.2 | 0.15584 | 0.196138953 | 0.071 | 0 | 1 | 17 |
| si:dkey-6f10.3 | 0.15584 | 0.170781964 | 0.071 | 0 | 1 | 17 |
| si:dkey-6n21.12 | 0.15584 | 0.141640291 | 0.071 | 0 | 1 | 17 |
| si:dkey-73n8.3 | 0.15584 | 0.227921552 | 0.071 | 0 | 1 | 17 |
| si:dkey-77f5.10 | 0.15584 | 0.139206771 | 0.071 | 0 | 1 | 17 |
| si:dkey-77f5.8 | 0.15584 | 0.158404877 | 0.071 | 0 | 1 | 17 |
| si:dkey-79d12.4 | 0.15584 | 0.153709502 | 0.071 | 0 | 1 | 17 |

| | | | | | | |
|---|---|---|---|---|---|---|
| si:dkey-82i20.2 | 0.15584 | 0.16187512 | 0.071 | 0 | 1 | 17 |
| si:dkey-85a20.4 | 0.15584 | 0.196138953 | 0.071 | 0 | 1 | 17 |
| si:dkey-94e7.1 | 0.15584 | 0.165043554 | 0.071 | 0 | 1 | 17 |
| si:dkey-97o5.1 | 0.15584 | 0.147292359 | 0.071 | 0 | 1 | 17 |
| si:dkeyp-44a8.4 | 0.15584 | 0.18600264 | 0.071 | 0 | 1 | 17 |
| si:dkeyp-68b7.5 | 0.15584 | 0.165043554 | 0.071 | 0 | 1 | 17 |
| si:dkeyp-82a1.6 | 0.15584 | 0.193978872 | 0.071 | 0 | 1 | 17 |
| si:dkeyp-86d6.2 | 0.15584 | 0.165043554 | 0.071 | 0 | 1 | 17 |
| si:rp71-1d10.8 | 0.15584 | 0.165043554 | 0.071 | 0 | 1 | 17 |
| si:rp71-45g20.10 | 0.15584 | 0.170781964 | 0.071 | 0 | 1 | 17 |
| si:zfos-1056e6.1 | 0.15584 | 0.196138953 | 0.071 | 0 | 1 | 17 |
| sik2b | 0.15584 | 0.162836696 | 0.071 | 0 | 1 | 17 |
| sin3b | 0.15584 | 0.193358917 | 0.071 | 0 | 1 | 17 |
| ska3 | 0.15584 | 0.196138953 | 0.071 | 0 | 1 | 17 |
| slc15a4 | 0.15584 | 0.163168177 | 0.071 | 0 | 1 | 17 |
| slc16a1a | 0.15584 | 0.196138953 | 0.071 | 0 | 1 | 17 |
| slc17a9b | 0.15584 | 0.165043554 | 0.071 | 0 | 1 | 17 |
| slc38a5b | 0.15584 | 0.137253048 | 0.071 | 0 | 1 | 17 |
| slc39a7 | 0.15584 | 0.166908951 | 0.071 | 0 | 1 | 17 |
| slc44a1a | 0.15584 | 0.170781964 | 0.071 | 0 | 1 | 17 |
| slc6a14 | 0.15584 | 0.170781964 | 0.071 | 0 | 1 | 17 |
| slc6a6b | 0.15584 | 0.145897561 | 0.071 | 0 | 1 | 17 |
| slka | 0.15584 | 0.165043554 | 0.071 | 0 | 1 | 17 |
| smarca2 | 0.15584 | 0.163234917 | 0.071 | 0 | 1 | 17 |
| smc2 | 0.15584 | 0.169370947 | 0.071 | 0 | 1 | 17 |
| smchd1 | 0.15584 | 0.203487265 | 0.071 | 0 | 1 | 17 |
| smg5 | 0.15584 | 0.168379366 | 0.071 | 0 | 1 | 17 |
| snap25a | 0.15584 | 0.196138953 | 0.071 | 0 | 1 | 17 |
| sned1 | 0.15584 | 0.143970668 | 0.071 | 0 | 1 | 17 |
| snn | 0.15584 | 0.196138953 | 0.071 | 0 | 1 | 17 |
| snrka | 0.15584 | 0.160075097 | 0.071 | 0 | 1 | 17 |
| sntb1 | 0.15584 | 0.152291669 | 0.071 | 0 | 1 | 17 |
| snx16 | 0.15584 | 0.146370137 | 0.071 | 0 | 1 | 17 |
| snx25 | 0.15584 | 0.157315457 | 0.071 | 0 | 1 | 17 |
| snx27a | 0.15584 | 0.193605006 | 0.071 | 0 | 1 | 17 |
| sorl1 | 0.15584 | 0.162910438 | 0.071 | 0 | 1 | 17 |
| sox11a | 0.15584 | 0.143036391 | 0.071 | 0 | 1 | 17 |
| sox19a | 0.15584 | 0.138366852 | 0.071 | 0 | 1 | 17 |
| sox21a | 0.15584 | 0.151814678 | 0.071 | 0 | 1 | 17 |
| sp100.2 | 0.15584 | 0.196138953 | 0.071 | 0 | 1 | 17 |
| sp3a | 0.15584 | 0.16346394 | 0.071 | 0 | 1 | 17 |
| spaca9 | 0.15584 | 0.227921552 | 0.071 | 0 | 1 | 17 |
| sphk2 | 0.15584 | 0.134246837 | 0.071 | 0 | 1 | 17 |
| spi1b | 0.15584 | 0.156905533 | 0.071 | 0 | 1 | 17 |
| spout1 | 0.15584 | 0.156075485 | 0.071 | 0 | 1 | 17 |
| srek1 | 0.15584 | 0.150106963 | 0.071 | 0 | 1 | 17 |
| srgn | 0.15584 | 0.241338767 | 0.071 | 0 | 1 | 17 |
| st14a | 0.15584 | 0.165043554 | 0.071 | 0 | 1 | 17 |

| | | | | | | |
|---|---|---|---|---|---|---|
| st3gal1 | 0.15584 | 0.165043554 | 0.071 | 0 | 1 | 17 |
| st3gal8 | 0.15584 | 0.139808717 | 0.071 | 0 | 1 | 17 |
| stat3 | 0.15584 | 0.167946883 | 0.071 | 0 | 1 | 17 |
| stim2a | 0.15584 | 0.134150486 | 0.071 | 0 | 1 | 17 |
| stimate | 0.15584 | 0.162316178 | 0.071 | 0 | 1 | 17 |
| stk11ip | 0.15584 | 0.149349053 | 0.071 | 0 | 1 | 17 |
| stox2a | 0.15584 | 0.165043554 | 0.071 | 0 | 1 | 17 |
| stra6l | 0.15584 | 0.189868115 | 0.071 | 0 | 1 | 17 |
| stx16 | 0.15584 | 0.15845495 | 0.071 | 0 | 1 | 17 |
| suz12b | 0.15584 | 0.25081091 | 0.071 | 0 | 1 | 17 |
| syt1a | 0.15584 | 0.196138953 | 0.071 | 0 | 1 | 17 |
| sytl5 | 0.15584 | 0.165043554 | 0.071 | 0 | 1 | 17 |
| syvn1 | 0.15584 | 0.165345948 | 0.071 | 0 | 1 | 17 |
| tamm41 | 0.15584 | 0.165043554 | 0.071 | 0 | 1 | 17 |
| tbc1d10aa | 0.15584 | 0.167837941 | 0.071 | 0 | 1 | 17 |
| tbl1x | 0.15584 | 0.169245748 | 0.071 | 0 | 1 | 17 |
| tbpl1 | 0.15584 | 0.169660024 | 0.071 | 0 | 1 | 17 |
| tcp11l2 | 0.15584 | 0.160482246 | 0.071 | 0 | 1 | 17 |
| tctex1d1 | 0.15584 | 0.170781964 | 0.071 | 0 | 1 | 17 |
| tent4a | 0.15584 | 0.165043554 | 0.071 | 0 | 1 | 17 |
| tent5bb | 0.15584 | 0.159151615 | 0.071 | 0 | 1 | 17 |
| tfb2m | 0.15584 | 0.148744524 | 0.071 | 0 | 1 | 17 |
| ticrr | 0.15584 | 0.165043554 | 0.071 | 0 | 1 | 17 |
| timm29 | 0.15584 | 0.169350482 | 0.071 | 0 | 1 | 17 |
| tmem126a | 0.15584 | 0.249658019 | 0.071 | 0 | 1 | 17 |
| tmem220 | 0.15584 | 0.165720869 | 0.071 | 0 | 1 | 17 |
| tmem234 | 0.15584 | 0.142573557 | 0.071 | 0 | 1 | 17 |
| tmem30aa | 0.15584 | 0.225940151 | 0.071 | 0 | 1 | 17 |
| tmem30b | 0.15584 | 0.148118065 | 0.071 | 0 | 1 | 17 |
| tmem42a | 0.15584 | 0.1502509 | 0.071 | 0 | 1 | 17 |
| tmx2b | 0.15584 | 0.275626974 | 0.071 | 0 | 1 | 17 |
| tnfaip2a | 0.15584 | 0.153709502 | 0.071 | 0 | 1 | 17 |
| tnfaip6 | 0.15584 | 0.127178302 | 0.071 | 0 | 1 | 17 |
| tnksb | 0.15584 | 0.161813314 | 0.071 | 0 | 1 | 17 |
| top1 | 0.15584 | 0.152477387 | 0.071 | 0 | 1 | 17 |
| tp53bp1 | 0.15584 | 0.277208742 | 0.071 | 0 | 1 | 17 |
| tp53bp2a | 0.15584 | 0.162955803 | 0.071 | 0 | 1 | 17 |
| trim13 | 0.15584 | 0.170781964 | 0.071 | 0 | 1 | 17 |
| trim3a | 0.15584 | 0.153709502 | 0.071 | 0 | 1 | 17 |
| trip12 | 0.15584 | 0.249608212 | 0.071 | 0 | 1 | 17 |
| trpt1 | 0.15584 | 0.158879557 | 0.071 | 0 | 1 | 17 |
| tsfm | 0.15584 | 0.196138953 | 0.071 | 0 | 1 | 17 |
| tspan17 | 0.15584 | 0.151448901 | 0.071 | 0 | 1 | 17 |
| tspan9a | 0.15584 | 0.165043554 | 0.071 | 0 | 1 | 17 |
| ttc23 | 0.15584 | 0.165043554 | 0.071 | 0 | 1 | 17 |
| tubgcp2 | 0.15584 | 0.204683474 | 0.071 | 0 | 1 | 17 |
| txlnbb | 0.15584 | 0.170781964 | 0.071 | 0 | 1 | 17 |
| txndc11 | 0.15584 | 0.227921552 | 0.071 | 0 | 1 | 17 |

| | | | | | | |
|---|---|---|---|---|---|---|
| tyw3 | 0.15584 | 0.159764272 | 0.071 | 0 | 1 | 17 |
| ube3b | 0.15584 | 0.162040351 | 0.071 | 0 | 1 | 17 |
| ublcp1 | 0.15584 | 0.153709502 | 0.071 | 0 | 1 | 17 |
| uggt2 | 0.15584 | 0.150162696 | 0.071 | 0 | 1 | 17 |
| unkl | 0.15584 | 0.156354573 | 0.071 | 0 | 1 | 17 |
| vac14 | 0.15584 | 0.196138953 | 0.071 | 0 | 1 | 17 |
| vezf1b | 0.15584 | 0.168866883 | 0.071 | 0 | 1 | 17 |
| vgll2a | 0.15584 | 0.165043554 | 0.071 | 0 | 1 | 17 |
| vwa1 | 0.15584 | 0.160179546 | 0.071 | 0 | 1 | 17 |
| vwa8 | 0.15584 | 0.170781964 | 0.071 | 0 | 1 | 17 |
| wdr45b | 0.15584 | 0.163885522 | 0.071 | 0 | 1 | 17 |
| wu:fi36c02 | 0.15584 | 0.220193797 | 0.071 | 0 | 1 | 17 |
| wu:fu71h07 | 0.15584 | 0.143824068 | 0.071 | 0 | 1 | 17 |
| wwp2 | 0.15584 | 0.160068764 | 0.071 | 0 | 1 | 17 |
| xdh | 0.15584 | 0.153709502 | 0.071 | 0 | 1 | 17 |
| xkr8.2 | 0.15584 | 0.153709502 | 0.071 | 0 | 1 | 17 |
| yars2 | 0.15584 | 0.205893925 | 0.071 | 0 | 1 | 17 |
| yes1 | 0.15584 | 0.160927567 | 0.071 | 0 | 1 | 17 |
| zc2hc1a | 0.15584 | 0.194512181 | 0.071 | 0 | 1 | 17 |
| zc3h12a | 0.15584 | 0.165043554 | 0.071 | 0 | 1 | 17 |
| zgc:101016 | 0.15584 | 0.150594118 | 0.071 | 0 | 1 | 17 |
| zgc:110216 | 0.15584 | 0.226870909 | 0.071 | 0 | 1 | 17 |
| zgc:112356 | 0.15584 | 0.158285173 | 0.071 | 0 | 1 | 17 |
| zgc:113423 | 0.15584 | 0.153709502 | 0.071 | 0 | 1 | 17 |
| zgc:113436 | 0.15584 | 0.226233296 | 0.071 | 0 | 1 | 17 |
| zgc:113452 | 0.15584 | 0.170781964 | 0.071 | 0 | 1 | 17 |
| zgc:114041 | 0.15584 | 0.170781964 | 0.071 | 0 | 1 | 17 |
| zgc:114174 | 0.15584 | 0.159885234 | 0.071 | 0 | 1 | 17 |
| zgc:158463 | 0.15584 | 0.116089832 | 0.071 | 0 | 1 | 17 |
| zgc:162948 | 0.15584 | 0.153709502 | 0.071 | 0 | 1 | 17 |
| zgc:173705 | 0.15584 | 0.165043554 | 0.071 | 0 | 1 | 17 |
| zgc:193711 | 0.15584 | 0.163435982 | 0.071 | 0 | 1 | 17 |
| zgc:193790 | 0.15584 | 0.153709502 | 0.071 | 0 | 1 | 17 |
| zgc:193801 | 0.15584 | 0.153709502 | 0.071 | 0 | 1 | 17 |
| zgc:193807 | 0.15584 | 0.196138953 | 0.071 | 0 | 1 | 17 |
| zgc:91944 | 0.15584 | 0.163595397 | 0.071 | 0 | 1 | 17 |
| zmiz2 | 0.15584 | 0.163204431 | 0.071 | 0 | 1 | 17 |
| zmp:0000000624 | 0.15584 | 0.169074691 | 0.071 | 0 | 1 | 17 |
| zmp:0000000634 | 0.15584 | 0.165043554 | 0.071 | 0 | 1 | 17 |
| zmynd19 | 0.15584 | 0.15059569 | 0.071 | 0 | 1 | 17 |
| znf1010 | 0.15584 | 0.162302801 | 0.071 | 0 | 1 | 17 |
| znf1143 | 0.15584 | 0.170781964 | 0.071 | 0 | 1 | 17 |
| znf1154 | 0.15584 | 0.196138953 | 0.071 | 0 | 1 | 17 |
| znf280d | 0.15584 | 0.150311989 | 0.071 | 0 | 1 | 17 |
| znf281a | 0.15584 | 0.165043554 | 0.071 | 0 | 1 | 17 |
| znf536 | 0.15584 | 0.163447717 | 0.071 | 0 | 1 | 17 |
| znf692 | 0.15584 | 0.148103941 | 0.071 | 0 | 1 | 17 |
| znf839 | 0.15584 | 0.162579912 | 0.071 | 0 | 1 | 17 |

| | | | | | | |
|---|---|---|---|---|---|---|
| ACKR2 | 0.15584 | 0.173824046 | 0.071 | 0 | 1 | 17 |
| AL732488.1 | 0.15584 | 0.103326222 | 0.071 | 0 | 1 | 17 |
| AREL1 | 0.15584 | 0.111853761 | 0.071 | 0 | 1 | 17 |
| BX530077.1 | 0.15584 | 0.103161035 | 0.071 | 0 | 1 | 17 |
| BX569793.1 | 0.15584 | 0.123713846 | 0.071 | 0 | 1 | 17 |
| BX649423.1 | 0.15584 | 0.123713846 | 0.071 | 0 | 1 | 17 |
| BX957297.2 | 0.15584 | 0.120768256 | 0.071 | 0 | 1 | 17 |
| CABZ01033394.1 | 0.15584 | 0.123713846 | 0.071 | 0 | 1 | 17 |
| CABZ01052576.1 | 0.15584 | 0.115824774 | 0.071 | 0 | 1 | 17 |
| CABZ01078261.1 | 0.15584 | 0.118738164 | 0.071 | 0 | 1 | 17 |
| CR354547.1 | 0.15584 | 0.123713846 | 0.071 | 0 | 1 | 17 |
| CR388029.1 | 0.15584 | 0.123713846 | 0.071 | 0 | 1 | 17 |
| FP236542.1 | 0.15584 | 0.123713846 | 0.071 | 0 | 1 | 17 |
| LOC100332293 | 0.15584 | 0.123713846 | 0.071 | 0 | 1 | 17 |
| LOC101883704 | 0.15584 | 0.119226698 | 0.071 | 0 | 1 | 17 |
| LOC101883751 | 0.15584 | 0.173824046 | 0.071 | 0 | 1 | 17 |
| LOC101884756 | 0.15584 | 0.123713846 | 0.071 | 0 | 1 | 17 |
| LOC110439329 | 0.15584 | 0.123713846 | 0.071 | 0 | 1 | 17 |
| LOC570835 | 0.15584 | 0.171350415 | 0.071 | 0 | 1 | 17 |
| LOC796453 | 0.15584 | 0.167607021 | 0.071 | 0 | 1 | 17 |
| LTN1 | 0.15584 | 0.123713846 | 0.071 | 0 | 1 | 17 |
| MARF1 | 0.15584 | 0.115561949 | 0.071 | 0 | 1 | 17 |
| SHKBP1 | 0.15584 | 0.118035874 | 0.071 | 0 | 1 | 17 |
| XLOC-000946 | 0.15584 | 0.116102752 | 0.071 | 0 | 1 | 17 |
| XLOC-006367 | 0.15584 | 0.123713846 | 0.071 | 0 | 1 | 17 |
| XLOC-025815 | 0.15584 | 0.123713846 | 0.071 | 0 | 1 | 17 |
| XLOC-026761 | 0.15584 | 0.123713846 | 0.071 | 0 | 1 | 17 |
| XLOC-038013 | 0.15584 | 0.118365888 | 0.071 | 0 | 1 | 17 |
| XLOC-042514 | 0.15584 | 0.123713846 | 0.071 | 0 | 1 | 17 |
| abhd13 | 0.15584 | 0.113231711 | 0.071 | 0 | 1 | 17 |
| actr5 | 0.15584 | 0.123713846 | 0.071 | 0 | 1 | 17 |
| ago2 | 0.15584 | 0.121347509 | 0.071 | 0 | 1 | 17 |
| ankrd13a | 0.15584 | 0.172398707 | 0.071 | 0 | 1 | 17 |
| ankrd50l | 0.15584 | 0.173824046 | 0.071 | 0 | 1 | 17 |
| ap1s1 | 0.15584 | 0.116096545 | 0.071 | 0 | 1 | 17 |
| apeh | 0.15584 | 0.122638464 | 0.071 | 0 | 1 | 17 |
| areg | 0.15584 | 0.123713846 | 0.071 | 0 | 1 | 17 |
| arfip2a | 0.15584 | 0.101268872 | 0.071 | 0 | 1 | 17 |
| arhgap12a | 0.15584 | 0.116812143 | 0.071 | 0 | 1 | 17 |
| arhgap32b | 0.15584 | 0.123713846 | 0.071 | 0 | 1 | 17 |
| arhgef2 | 0.15584 | 0.12088272 | 0.071 | 0 | 1 | 17 |
| arl13b | 0.15584 | 0.114983659 | 0.071 | 0 | 1 | 17 |
| asphd2 | 0.15584 | 0.109147772 | 0.071 | 0 | 1 | 17 |
| atg4a | 0.15584 | 0.114407411 | 0.071 | 0 | 1 | 17 |
| atm | 0.15584 | 0.123713846 | 0.071 | 0 | 1 | 17 |
| b3galt6 | 0.15584 | 0.123713846 | 0.071 | 0 | 1 | 17 |
| b4galt2 | 0.15584 | 0.123713846 | 0.071 | 0 | 1 | 17 |
| bmp8a | 0.15584 | 0.115637601 | 0.071 | 0 | 1 | 17 |

| | | | | | | |
|---|---|---|---|---|---|---|
| bmpr2a | 0.15584 | 0.10628911 | 0.071 | 0 | 1 | 17 |
| brf1b | 0.15584 | 0.123713846 | 0.071 | 0 | 1 | 17 |
| c1galt1b | 0.15584 | 0.11901104 | 0.071 | 0 | 1 | 17 |
| cacna1ba | 0.15584 | 0.123713846 | 0.071 | 0 | 1 | 17 |
| cacna2d4b | 0.15584 | 0.123713846 | 0.071 | 0 | 1 | 17 |
| card19 | 0.15584 | 0.121425345 | 0.071 | 0 | 1 | 17 |
| ccdc167 | 0.15584 | 0.120012594 | 0.071 | 0 | 1 | 17 |
| ccdc18 | 0.15584 | 0.123713846 | 0.071 | 0 | 1 | 17 |
| cenpj | 0.15584 | 0.123713846 | 0.071 | 0 | 1 | 17 |
| cep131 | 0.15584 | 0.118968219 | 0.071 | 0 | 1 | 17 |
| cftr | 0.15584 | 0.123713846 | 0.071 | 0 | 1 | 17 |
| chmp5a | 0.15584 | 0.123713846 | 0.071 | 0 | 1 | 17 |
| col9a1a | 0.15584 | 0.156088724 | 0.071 | 0 | 1 | 17 |
| coq4 | 0.15584 | 0.118003228 | 0.071 | 0 | 1 | 17 |
| creb1b | 0.15584 | 0.121010023 | 0.071 | 0 | 1 | 17 |
| creld1a | 0.15584 | 0.123713846 | 0.071 | 0 | 1 | 17 |
| crtc3 | 0.15584 | 0.118338477 | 0.071 | 0 | 1 | 17 |
| csnk1g2a | 0.15584 | 0.121948877 | 0.071 | 0 | 1 | 17 |
| ctso | 0.15584 | 0.123713846 | 0.071 | 0 | 1 | 17 |
| cul4a | 0.15584 | 0.117114423 | 0.071 | 0 | 1 | 17 |
| cyth1a | 0.15584 | 0.113951809 | 0.071 | 0 | 1 | 17 |
| dclre1c | 0.15584 | 0.123713846 | 0.071 | 0 | 1 | 17 |
| dennd3a | 0.15584 | 0.123713846 | 0.071 | 0 | 1 | 17 |
| dna2 | 0.15584 | 0.118520749 | 0.071 | 0 | 1 | 17 |
| dnah1 | 0.15584 | 0.123713846 | 0.071 | 0 | 1 | 17 |
| dsg2.1 | 0.15584 | 0.111744941 | 0.071 | 0 | 1 | 17 |
| ebf2 | 0.15584 | 0.123713846 | 0.071 | 0 | 1 | 17 |
| elovl8b | 0.15584 | 0.112655258 | 0.071 | 0 | 1 | 17 |
| eml3 | 0.15584 | 0.115420403 | 0.071 | 0 | 1 | 17 |
| exoc8 | 0.15584 | 0.123713846 | 0.071 | 0 | 1 | 17 |
| fam102bb | 0.15584 | 0.114819043 | 0.071 | 0 | 1 | 17 |
| fam193b | 0.15584 | 0.123713846 | 0.071 | 0 | 1 | 17 |
| fam20b | 0.15584 | 0.123713846 | 0.071 | 0 | 1 | 17 |
| fbxl6 | 0.15584 | 0.121691834 | 0.071 | 0 | 1 | 17 |
| fcho1 | 0.15584 | 0.119081997 | 0.071 | 0 | 1 | 17 |
| fdxr | 0.15584 | 0.113950716 | 0.071 | 0 | 1 | 17 |
| fhod1 | 0.15584 | 0.123713846 | 0.071 | 0 | 1 | 17 |
| figla | 0.15584 | 0.123713846 | 0.071 | 0 | 1 | 17 |
| ftr67 | 0.15584 | 0.11327405 | 0.071 | 0 | 1 | 17 |
| fuk | 0.15584 | 0.113692809 | 0.071 | 0 | 1 | 17 |
| fzd6 | 0.15584 | 0.123713846 | 0.071 | 0 | 1 | 17 |
| fzr1b | 0.15584 | 0.111824901 | 0.071 | 0 | 1 | 17 |
| galnt12 | 0.15584 | 0.111589775 | 0.071 | 0 | 1 | 17 |
| gas1a | 0.15584 | 0.112701474 | 0.071 | 0 | 1 | 17 |
| gem | 0.15584 | 0.104737888 | 0.071 | 0 | 1 | 17 |
| golga4 | 0.15584 | 0.102697677 | 0.071 | 0 | 1 | 17 |
| gpatch1 | 0.15584 | 0.167959955 | 0.071 | 0 | 1 | 17 |
| gpbp1l1 | 0.15584 | 0.123713846 | 0.071 | 0 | 1 | 17 |

| | | | | | | |
|---|---|---|---|---|---|---|
| gpm6bb | 0.15584 | 0.109346485 | 0.071 | 0 | 1 | 17 |
| gtf2ird1 | 0.15584 | 0.123713846 | 0.071 | 0 | 1 | 17 |
| hacd2 | 0.15584 | 0.114994163 | 0.071 | 0 | 1 | 17 |
| hif1aa | 0.15584 | 0.123713846 | 0.071 | 0 | 1 | 17 |
| hivep2a | 0.15584 | 0.116615791 | 0.071 | 0 | 1 | 17 |
| huwe1 | 0.15584 | 0.172103343 | 0.071 | 0 | 1 | 17 |
| igdcc4 | 0.15584 | 0.112965122 | 0.071 | 0 | 1 | 17 |
| igsf8 | 0.15584 | 0.123713846 | 0.071 | 0 | 1 | 17 |
| insig2 | 0.15584 | 0.123713846 | 0.071 | 0 | 1 | 17 |
| ipo13 | 0.15584 | 0.122150297 | 0.071 | 0 | 1 | 17 |
| itga3a | 0.15584 | 0.123713846 | 0.071 | 0 | 1 | 17 |
| jkamp | 0.15584 | 0.121251087 | 0.071 | 0 | 1 | 17 |
| kdm5ba | 0.15584 | 0.110135748 | 0.071 | 0 | 1 | 17 |
| kdm6al | 0.15584 | 0.120110582 | 0.071 | 0 | 1 | 17 |
| kif1b | 0.15584 | 0.109817772 | 0.071 | 0 | 1 | 17 |
| lclat1 | 0.15584 | 0.116480505 | 0.071 | 0 | 1 | 17 |
| lmf2b | 0.15584 | 0.102892916 | 0.071 | 0 | 1 | 17 |
| lrrc8aa | 0.15584 | 0.120107708 | 0.071 | 0 | 1 | 17 |
| lsm11 | 0.15584 | 0.121736482 | 0.071 | 0 | 1 | 17 |
| mapk6 | 0.15584 | 0.1207727 | 0.071 | 0 | 1 | 17 |
| mapre2 | 0.15584 | 0.123713846 | 0.071 | 0 | 1 | 17 |
| mbd5 | 0.15584 | 0.120076162 | 0.071 | 0 | 1 | 17 |
| mcf2la | 0.15584 | 0.123713846 | 0.071 | 0 | 1 | 17 |
| mfap3l | 0.15584 | 0.118876065 | 0.071 | 0 | 1 | 17 |
| mfsd13a | 0.15584 | 0.123713846 | 0.071 | 0 | 1 | 17 |
| mis12 | 0.15584 | 0.119857126 | 0.071 | 0 | 1 | 17 |
| mmab | 0.15584 | 0.11436977 | 0.071 | 0 | 1 | 17 |
| mrpl58 | 0.15584 | 0.11955638 | 0.071 | 0 | 1 | 17 |
| mta1 | 0.15584 | 0.123713846 | 0.071 | 0 | 1 | 17 |
| mtor | 0.15584 | 0.122475388 | 0.071 | 0 | 1 | 17 |
| mtss1 | 0.15584 | 0.119314242 | 0.071 | 0 | 1 | 17 |
| mxd4 | 0.15584 | 0.115311563 | 0.071 | 0 | 1 | 17 |
| mycbp2 | 0.15584 | 0.119858489 | 0.071 | 0 | 1 | 17 |
| nbr1b | 0.15584 | 0.110247607 | 0.071 | 0 | 1 | 17 |
| nf2b | 0.15584 | 0.110830753 | 0.071 | 0 | 1 | 17 |
| ngfb | 0.15584 | 0.123713846 | 0.071 | 0 | 1 | 17 |
| ngly1 | 0.15584 | 0.12175679 | 0.071 | 0 | 1 | 17 |
| nipal3 | 0.15584 | 0.123713846 | 0.071 | 0 | 1 | 17 |
| noctb | 0.15584 | 0.123713846 | 0.071 | 0 | 1 | 17 |
| odf2b | 0.15584 | 0.115527952 | 0.071 | 0 | 1 | 17 |
| papolg | 0.15584 | 0.12136579 | 0.071 | 0 | 1 | 17 |
| parapinopsinb | 0.15584 | 0.123713846 | 0.071 | 0 | 1 | 17 |
| pccb | 0.15584 | 0.12250376 | 0.071 | 0 | 1 | 17 |
| pcdh1b | 0.15584 | 0.110234768 | 0.071 | 0 | 1 | 17 |
| pcif1 | 0.15584 | 0.123713846 | 0.071 | 0 | 1 | 17 |
| pdk3a | 0.15584 | 0.116929995 | 0.071 | 0 | 1 | 17 |
| pemt | 0.15584 | 0.114001974 | 0.071 | 0 | 1 | 17 |
| pgpep1 | 0.15584 | 0.119025852 | 0.071 | 0 | 1 | 17 |

| | | | | | | |
|---|---|---|---|---|---|---|
| pimr132 | 0.15584 | 0.123713846 | 0.071 | 0 | 1 | 17 |
| pink1 | 0.15584 | 0.173824046 | 0.071 | 0 | 1 | 17 |
| pla2g12a | 0.15584 | 0.123713846 | 0.071 | 0 | 1 | 17 |
| plcd4b | 0.15584 | 0.123713846 | 0.071 | 0 | 1 | 17 |
| plekhh1 | 0.15584 | 0.123713846 | 0.071 | 0 | 1 | 17 |
| plekho1b | 0.15584 | 0.116102321 | 0.071 | 0 | 1 | 17 |
| pnpla6 | 0.15584 | 0.121512196 | 0.071 | 0 | 1 | 17 |
| polr3c | 0.15584 | 0.107586936 | 0.071 | 0 | 1 | 17 |
| ppp1r3cb | 0.15584 | 0.110450397 | 0.071 | 0 | 1 | 17 |
| ppp2r3b | 0.15584 | 0.123713846 | 0.071 | 0 | 1 | 17 |
| prickle3 | 0.15584 | 0.123713846 | 0.071 | 0 | 1 | 17 |
| prkacaa | 0.15584 | 0.114221881 | 0.071 | 0 | 1 | 17 |
| prkar2ab | 0.15584 | 0.123713846 | 0.071 | 0 | 1 | 17 |
| ptena | 0.15584 | 0.117650883 | 0.071 | 0 | 1 | 17 |
| ptn | 0.15584 | 0.123713846 | 0.071 | 0 | 1 | 17 |
| ptpn11a | 0.15584 | 0.119311741 | 0.071 | 0 | 1 | 17 |
| rab27a | 0.15584 | 0.106614775 | 0.071 | 0 | 1 | 17 |
| ralgps2 | 0.15584 | 0.171422793 | 0.071 | 0 | 1 | 17 |
| rbpms | 0.15584 | 0.123713846 | 0.071 | 0 | 1 | 17 |
| rdh14b | 0.15584 | 0.11933353 | 0.071 | 0 | 1 | 17 |
| rec8a | 0.15584 | 0.123713846 | 0.071 | 0 | 1 | 17 |
| ripor1.1 | 0.15584 | 0.107863558 | 0.071 | 0 | 1 | 17 |
| rnaseka | 0.15584 | 0.164613265 | 0.071 | 0 | 1 | 17 |
| rnf11a | 0.15584 | 0.101735939 | 0.071 | 0 | 1 | 17 |
| rxfp1 | 0.15584 | 0.123713846 | 0.071 | 0 | 1 | 17 |
| ryr2b | 0.15584 | 0.123713846 | 0.071 | 0 | 1 | 17 |
| saal1 | 0.15584 | 0.118563107 | 0.071 | 0 | 1 | 17 |
| sap130a | 0.15584 | 0.12114796 | 0.071 | 0 | 1 | 17 |
| sar1aa | 0.15584 | 0.119791246 | 0.071 | 0 | 1 | 17 |
| sdf4 | 0.15584 | 0.112428186 | 0.071 | 0 | 1 | 17 |
| sema3e | 0.15584 | 0.123713846 | 0.071 | 0 | 1 | 17 |
| si:ch1073-296d18.1 | 0.15584 | 0.119683987 | 0.071 | 0 | 1 | 17 |
| si:ch211-191d2.2 | 0.15584 | 0.123713846 | 0.071 | 0 | 1 | 17 |
| si:ch211-66k16.28 | 0.15584 | 0.113200553 | 0.071 | 0 | 1 | 17 |
| si:ch211-71n6.4 | 0.15584 | 0.116843612 | 0.071 | 0 | 1 | 17 |
| si:ch73-144d13.7 | 0.15584 | 0.116325238 | 0.071 | 0 | 1 | 17 |
| si:ch73-213k20.5 | 0.15584 | 0.123713846 | 0.071 | 0 | 1 | 17 |
| si:ch73-221f6.3 | 0.15584 | 0.123713846 | 0.071 | 0 | 1 | 17 |
| si:dkey-111k8.3 | 0.15584 | 0.123713846 | 0.071 | 0 | 1 | 17 |
| si:dkey-37g12.1 | 0.15584 | 0.123713846 | 0.071 | 0 | 1 | 17 |
| si:dkey-37m8.3 | 0.15584 | 0.123713846 | 0.071 | 0 | 1 | 17 |
| si:dkey-73n10.1 | 0.15584 | 0.112949626 | 0.071 | 0 | 1 | 17 |
| sipa1l2.1 | 0.15584 | 0.123713846 | 0.071 | 0 | 1 | 17 |
| slc22a18 | 0.15584 | 0.117278263 | 0.071 | 0 | 1 | 17 |
| slc35f4 | 0.15584 | 0.123713846 | 0.071 | 0 | 1 | 17 |
| slc44a2 | 0.15584 | 0.120708472 | 0.071 | 0 | 1 | 17 |
| slc4a7 | 0.15584 | 0.123713846 | 0.071 | 0 | 1 | 17 |
| smg6 | 0.15584 | 0.120896889 | 0.071 | 0 | 1 | 17 |

| | | | | | | |
|---|---|---|---|---|---|---|
| snx27b | 0.15584 | 0.118129351 | 0.071 | 0 | 1 | 17 |
| spint1b | 0.15584 | 0.123713846 | 0.071 | 0 | 1 | 17 |
| srgap3 | 0.15584 | 0.118912017 | 0.071 | 0 | 1 | 17 |
| ssbp4 | 0.15584 | 0.123713846 | 0.071 | 0 | 1 | 17 |
| syngr2a | 0.15584 | 0.107125347 | 0.071 | 0 | 1 | 17 |
| taar19k | 0.15584 | 0.123713846 | 0.071 | 0 | 1 | 17 |
| tab3 | 0.15584 | 0.111515795 | 0.071 | 0 | 1 | 17 |
| tbc1d19 | 0.15584 | 0.123713846 | 0.071 | 0 | 1 | 17 |
| tbc1d31 | 0.15584 | 0.171594191 | 0.071 | 0 | 1 | 17 |
| tecpr1a | 0.15584 | 0.123713846 | 0.071 | 0 | 1 | 17 |
| tent5ba | 0.15584 | 0.102488768 | 0.071 | 0 | 1 | 17 |
| thumpd2 | 0.15584 | 0.173824046 | 0.071 | 0 | 1 | 17 |
| tln1 | 0.15584 | 0.117308587 | 0.071 | 0 | 1 | 17 |
| tmem106bb | 0.15584 | 0.123713846 | 0.071 | 0 | 1 | 17 |
| tmem145 | 0.15584 | 0.109442681 | 0.071 | 0 | 1 | 17 |
| tmem161a | 0.15584 | 0.12152272 | 0.071 | 0 | 1 | 17 |
| tmem186 | 0.15584 | 0.119741788 | 0.071 | 0 | 1 | 17 |
| triobpb | 0.15584 | 0.11597161 | 0.071 | 0 | 1 | 17 |
| tspan2a | 0.15584 | 0.119112695 | 0.071 | 0 | 1 | 17 |
| ttll9 | 0.15584 | 0.123713846 | 0.071 | 0 | 1 | 17 |
| tube1 | 0.15584 | 0.123713846 | 0.071 | 0 | 1 | 17 |
| tusc2b | 0.15584 | 0.116411999 | 0.071 | 0 | 1 | 17 |
| twsg1b | 0.15584 | 0.112468168 | 0.071 | 0 | 1 | 17 |
| ubn2b | 0.15584 | 0.119105088 | 0.071 | 0 | 1 | 17 |
| ulk1a | 0.15584 | 0.123713846 | 0.071 | 0 | 1 | 17 |
| unm-hu7910 | 0.15584 | 0.121108606 | 0.071 | 0 | 1 | 17 |
| wu:fj64h06 | 0.15584 | 0.123713846 | 0.071 | 0 | 1 | 17 |
| zbtb38 | 0.15584 | 0.123713846 | 0.071 | 0 | 1 | 17 |
| zbtb8os | 0.15584 | 0.114202545 | 0.071 | 0 | 1 | 17 |
| zc2hc1c | 0.15584 | 0.123713846 | 0.071 | 0 | 1 | 17 |
| zdhhc15b | 0.15584 | 0.11971439 | 0.071 | 0 | 1 | 17 |
| zgc:101731 | 0.15584 | 0.123713846 | 0.071 | 0 | 1 | 17 |
| zgc:153292 | 0.15584 | 0.104660367 | 0.071 | 0 | 1 | 17 |
| zgc:153901 | 0.15584 | 0.110875816 | 0.071 | 0 | 1 | 17 |
| zgc:154061 | 0.15584 | 0.117745286 | 0.071 | 0 | 1 | 17 |
| zgc:171704 | 0.15584 | 0.123713846 | 0.071 | 0 | 1 | 17 |
| zgc:56409 | 0.15584 | 0.123713846 | 0.071 | 0 | 1 | 17 |
| znf1153 | 0.15584 | 0.118654077 | 0.071 | 0 | 1 | 17 |
| znf318 | 0.15584 | 0.171107098 | 0.071 | 0 | 1 | 17 |
| AL929576.1 | 0.15584 | 0.202312984 | 0.071 | 0 | 1 | 17 |
| LO017872.1 | 0.15584 | 0.253755856 | 0.071 | 0 | 1 | 17 |
| LOC100534805 | 0.15584 | 0.156003571 | 0.071 | 0 | 1 | 17 |
| NDUFAF8 | 0.15584 | 0.198897321 | 0.071 | 0 | 1 | 17 |
| TCIM (1 of many) | 0.15584 | 0.144018569 | 0.071 | 0 | 1 | 17 |
| YTHDC2 | 0.15584 | 0.161428377 | 0.071 | 0 | 1 | 17 |
| abcg2c | 0.15584 | 0.149320487 | 0.071 | 0 | 1 | 17 |
| abraxas1 | 0.15584 | 0.159230092 | 0.071 | 0 | 1 | 17 |
| acox1 | 0.15584 | 0.151646907 | 0.071 | 0 | 1 | 17 |

| | | | | | | |
|---|---|---|---|---|---|---|
| akap7 | 0.15584 | 0.192598213 | 0.071 | 0 | 1 | 17 |
| akt3a | 0.15584 | 0.202312984 | 0.071 | 0 | 1 | 17 |
| anxa1b | 0.15584 | 0.153975877 | 0.071 | 0 | 1 | 17 |
| aptx | 0.15584 | 0.202312984 | 0.071 | 0 | 1 | 17 |
| b3glcta | 0.15584 | 0.201393547 | 0.071 | 0 | 1 | 17 |
| b4galt1l | 0.15584 | 0.254764204 | 0.071 | 0 | 1 | 17 |
| bach2b | 0.15584 | 0.158247747 | 0.071 | 0 | 1 | 17 |
| bbs2 | 0.15584 | 0.197464265 | 0.071 | 0 | 1 | 17 |
| btr25 | 0.15584 | 0.199979239 | 0.071 | 0 | 1 | 17 |
| cbx5 | 0.15584 | 0.156031177 | 0.071 | 0 | 1 | 17 |
| cc2d2a | 0.15584 | 0.202312984 | 0.071 | 0 | 1 | 17 |
| ccar2 | 0.15584 | 0.161428377 | 0.071 | 0 | 1 | 17 |
| ccdc186 | 0.15584 | 0.161428377 | 0.071 | 0 | 1 | 17 |
| cenpk | 0.15584 | 0.160374791 | 0.071 | 0 | 1 | 17 |
| cep192 | 0.15584 | 0.202312984 | 0.071 | 0 | 1 | 17 |
| cfap206 | 0.15584 | 0.161428377 | 0.071 | 0 | 1 | 17 |
| chp1 | 0.15584 | 0.157934798 | 0.071 | 0 | 1 | 17 |
| clint1b | 0.15584 | 0.202312984 | 0.071 | 0 | 1 | 17 |
| cpt1cb | 0.15584 | 0.161428377 | 0.071 | 0 | 1 | 17 |
| crema | 0.15584 | 0.200658158 | 0.071 | 0 | 1 | 17 |
| dapk3 | 0.15584 | 0.153256795 | 0.071 | 0 | 1 | 17 |
| dlg1l | 0.15584 | 0.123926256 | 0.071 | 0 | 1 | 17 |
| dusp2 | 0.15584 | 0.134032691 | 0.071 | 0 | 1 | 17 |
| e2f8 | 0.15584 | 0.202312984 | 0.071 | 0 | 1 | 17 |
| edem1 | 0.15584 | 0.145712544 | 0.071 | 0 | 1 | 17 |
| erbb2 | 0.15584 | 0.154616253 | 0.071 | 0 | 1 | 17 |
| etv4 | 0.15584 | 0.202312984 | 0.071 | 0 | 1 | 17 |
| exoc5 | 0.15584 | 0.159138266 | 0.071 | 0 | 1 | 17 |
| fbxo36a | 0.15584 | 0.180954858 | 0.071 | 0 | 1 | 17 |
| foxl2b | 0.15584 | 0.107977989 | 0.071 | 0 | 1 | 17 |
| foxq1b | 0.15584 | 0.130079163 | 0.071 | 0 | 1 | 17 |
| gak | 0.15584 | 0.218753995 | 0.071 | 0 | 1 | 17 |
| git2a | 0.15584 | 0.161428377 | 0.071 | 0 | 1 | 17 |
| gnpnat1 | 0.15584 | 0.196862156 | 0.071 | 0 | 1 | 17 |
| gpc2 | 0.15584 | 0.161428377 | 0.071 | 0 | 1 | 17 |
| grin2db | 0.15584 | 0.161428377 | 0.071 | 0 | 1 | 17 |
| hdac9b | 0.15584 | 0.157875171 | 0.071 | 0 | 1 | 17 |
| hexa | 0.15584 | 0.195076952 | 0.071 | 0 | 1 | 17 |
| hipk3a | 0.15584 | 0.251458557 | 0.071 | 0 | 1 | 17 |
| hkdc1 | 0.15584 | 0.202312984 | 0.071 | 0 | 1 | 17 |
| hmgb3a | 0.15584 | 0.193274736 | 0.071 | 0 | 1 | 17 |
| kif19 | 0.15584 | 0.161428377 | 0.071 | 0 | 1 | 17 |
| kif9 | 0.15584 | 0.161428377 | 0.071 | 0 | 1 | 17 |
| lgals3b | 0.15584 | 0.152968587 | 0.071 | 0 | 1 | 17 |
| lpcat1 | 0.15584 | 0.25382404 | 0.071 | 0 | 1 | 17 |
| maats1 | 0.15584 | 0.202312984 | 0.071 | 0 | 1 | 17 |
| man1a1 | 0.15584 | 0.252948258 | 0.071 | 0 | 1 | 17 |
| mast2 | 0.15584 | 0.251819656 | 0.071 | 0 | 1 | 17 |

| | | | | | | |
|---|---|---|---|---|---|---|
| mbnl1 | 0.15584 | 0.198418176 | 0.071 | 0 | 1 | 17 |
| mcrs1 | 0.15584 | 0.156826162 | 0.071 | 0 | 1 | 17 |
| mmachc | 0.15584 | 0.202312984 | 0.071 | 0 | 1 | 17 |
| mpi | 0.15584 | 0.158432142 | 0.071 | 0 | 1 | 17 |
| mrpl1 | 0.15584 | 0.200365854 | 0.071 | 0 | 1 | 17 |
| ms4a17a.6 | 0.15584 | 0.250525126 | 0.071 | 0 | 1 | 17 |
| mta2 | 0.15584 | 0.254127092 | 0.071 | 0 | 1 | 17 |
| mxb | 0.15584 | 0.161428377 | 0.071 | 0 | 1 | 17 |
| napepld | 0.15584 | 0.197127533 | 0.071 | 0 | 1 | 17 |
| ndufa4 | 0.15584 | 0.160181491 | 0.071 | 0 | 1 | 17 |
| ndufaf5 | 0.15584 | 0.159831095 | 0.071 | 0 | 1 | 17 |
| nek9 | 0.15584 | 0.159012809 | 0.071 | 0 | 1 | 17 |
| nfkbil1 | 0.15584 | 0.254764204 | 0.071 | 0 | 1 | 17 |
| nol4la | 0.15584 | 0.161428377 | 0.071 | 0 | 1 | 17 |
| nras | 0.15584 | 0.154828854 | 0.071 | 0 | 1 | 17 |
| nufip2 | 0.15584 | 0.253177417 | 0.071 | 0 | 1 | 17 |
| otulinb | 0.15584 | 0.156566141 | 0.071 | 0 | 1 | 17 |
| parvaa | 0.15584 | 0.19155116 | 0.071 | 0 | 1 | 17 |
| pkp3a | 0.15584 | 0.152748206 | 0.071 | 0 | 1 | 17 |
| pla2g6 | 0.15584 | 0.159255839 | 0.071 | 0 | 1 | 17 |
| plekhg4 | 0.15584 | 0.155773525 | 0.071 | 0 | 1 | 17 |
| ppm1k | 0.15584 | 0.161428377 | 0.071 | 0 | 1 | 17 |
| ppp1r13l | 0.15584 | 0.156781614 | 0.071 | 0 | 1 | 17 |
| ppp2r2aa | 0.15584 | 0.161428377 | 0.071 | 0 | 1 | 17 |
| ptpdc1a | 0.15584 | 0.202312984 | 0.071 | 0 | 1 | 17 |
| ptpn9a | 0.15584 | 0.157079084 | 0.071 | 0 | 1 | 17 |
| raf1a | 0.15584 | 0.159288519 | 0.071 | 0 | 1 | 17 |
| rassf8b | 0.15584 | 0.19701431 | 0.071 | 0 | 1 | 17 |
| rbks | 0.15584 | 0.160391581 | 0.071 | 0 | 1 | 17 |
| rfwd3 | 0.15584 | 0.161428377 | 0.071 | 0 | 1 | 17 |
| rilpl2 | 0.15584 | 0.157973274 | 0.071 | 0 | 1 | 17 |
| robo4 | 0.15584 | 0.161428377 | 0.071 | 0 | 1 | 17 |
| rpz5 | 0.15584 | 0.325578507 | 0.071 | 0 | 1 | 17 |
| rsf1a | 0.15584 | 0.202312984 | 0.071 | 0 | 1 | 17 |
| ryr3 | 0.15584 | 0.144168681 | 0.071 | 0 | 1 | 17 |
| sbf1 | 0.15584 | 0.200387733 | 0.071 | 0 | 1 | 17 |
| scap | 0.15584 | 0.161428377 | 0.071 | 0 | 1 | 17 |
| scarb2a | 0.15584 | 0.154745515 | 0.071 | 0 | 1 | 17 |
| sgms2a | 0.15584 | 0.250418206 | 0.071 | 0 | 1 | 17 |
| si:ch211-134m17.9 | 0.15584 | 0.196433301 | 0.071 | 0 | 1 | 17 |
| si:ch211-14a17.6 | 0.15584 | 0.194198323 | 0.071 | 0 | 1 | 17 |
| si:ch211-195h23.3 | 0.15584 | 0.198654627 | 0.071 | 0 | 1 | 17 |
| si:ch211-284e13.6 | 0.15584 | 0.310595619 | 0.071 | 0 | 1 | 17 |
| si:ch73-138n13.1 | 0.15584 | 0.253241169 | 0.071 | 0 | 1 | 17 |
| si:dkey-174n20.1 | 0.15584 | 0.161428377 | 0.071 | 0 | 1 | 17 |
| si:dkey-258f14.3 | 0.15584 | 0.253479637 | 0.071 | 0 | 1 | 17 |
| si:dkey-261m9.6 | 0.15584 | 0.202312984 | 0.071 | 0 | 1 | 17 |
| si:dkey-262g12.10 | 0.15584 | 0.161428377 | 0.071 | 0 | 1 | 17 |

| | | | | | | |
|---|---|---|---|---|---|---|
| si:dkey-29h14.10 | 0.15584 | 0.139749271 | 0.071 | 0 | 1 | 17 |
| si:dkeyp-110g5.4 | 0.15584 | 0.253247222 | 0.071 | 0 | 1 | 17 |
| si:dkeyp-69c1.6 | 0.15584 | 0.254764204 | 0.071 | 0 | 1 | 17 |
| slc26a5 | 0.15584 | 0.200706054 | 0.071 | 0 | 1 | 17 |
| slc38a8b | 0.15584 | 0.202312984 | 0.071 | 0 | 1 | 17 |
| snx1a | 0.15584 | 0.199863041 | 0.071 | 0 | 1 | 17 |
| spata5l1 | 0.15584 | 0.201043422 | 0.071 | 0 | 1 | 17 |
| stoml1 | 0.15584 | 0.158208381 | 0.071 | 0 | 1 | 17 |
| surf4 | 0.15584 | 0.200032335 | 0.071 | 0 | 1 | 17 |
| tbrg1 | 0.15584 | 0.252796351 | 0.071 | 0 | 1 | 17 |
| tbx2b | 0.15584 | 0.148243248 | 0.071 | 0 | 1 | 17 |
| tctn1 | 0.15584 | 0.157074467 | 0.071 | 0 | 1 | 17 |
| tmem187 | 0.15584 | 0.199426768 | 0.071 | 0 | 1 | 17 |
| tnpo3 | 0.15584 | 0.20001734 | 0.071 | 0 | 1 | 17 |
| tnrc6c2 | 0.15584 | 0.201456316 | 0.071 | 0 | 1 | 17 |
| tns2a | 0.15584 | 0.161428377 | 0.071 | 0 | 1 | 17 |
| tril | 0.15584 | 0.202312984 | 0.071 | 0 | 1 | 17 |
| trim16 | 0.15584 | 0.209181473 | 0.071 | 0 | 1 | 17 |
| trmo | 0.15584 | 0.159137389 | 0.071 | 0 | 1 | 17 |
| tspan12 | 0.15584 | 0.254764204 | 0.071 | 0 | 1 | 17 |
| uckl1b | 0.15584 | 0.201656232 | 0.071 | 0 | 1 | 17 |
| vps53 | 0.15584 | 0.156013149 | 0.071 | 0 | 1 | 17 |
| ywhae2 | 0.15584 | 0.200616727 | 0.071 | 0 | 1 | 17 |
| zc3hc1 | 0.15584 | 0.161428377 | 0.071 | 0 | 1 | 17 |
| zcrb1 | 0.15584 | 0.254281533 | 0.071 | 0 | 1 | 17 |
| zfyve21 | 0.15584 | 0.193116336 | 0.071 | 0 | 1 | 17 |
| zgc:101810 | 0.15584 | 0.196961504 | 0.071 | 0 | 1 | 17 |
| zgc:113274 | 0.15584 | 0.195361943 | 0.071 | 0 | 1 | 17 |
| zgc:152951 | 0.15584 | 0.144626964 | 0.071 | 0 | 1 | 17 |
| zgc:171220 | 0.15584 | 0.254764204 | 0.071 | 0 | 1 | 17 |
| zgc:195001 | 0.15584 | 0.254764204 | 0.071 | 0 | 1 | 17 |
| zgc:64051 | 0.15584 | 0.152872611 | 0.071 | 0 | 1 | 17 |
| zgc:92140 | 0.15584 | 0.195151777 | 0.071 | 0 | 1 | 17 |
| znf236 | 0.15584 | 0.158057559 | 0.071 | 0 | 1 | 17 |
| znf991 | 0.15584 | 0.19044418 | 0.071 | 0 | 1 | 17 |
| htatsf1 | 0.15607 | 0.529762795 | 0.286 | 0.073 | 1 | 17 |
| cox6b2 | 0.15668 | 0.244698198 | 0.714 | 0.463 | 1 | 17 |
| lsm5 | 0.15706 | 0.319529549 | 0.786 | 0.512 | 1 | 17 |
| gorasp2 | 0.15707 | 0.501009068 | 0.286 | 0.073 | 1 | 17 |
| fkbp8 | 0.15748 | 0.480274354 | 0.286 | 0.073 | 1 | 17 |
| nansa | 0.15765 | 0.473264185 | 0.286 | 0.073 | 1 | 17 |
| rpl23 | 0.15771 | 0.105932177 | 1 | 0.854 | 1 | 17 |
| zgc:173742 | 0.15818 | 0.373829169 | 0.571 | 0.293 | 1 | 17 |
| spata22 | 0.15828 | 0.421722144 | 0.286 | 0.073 | 1 | 17 |
| zgc:112496 | 0.15837 | 0.506457856 | 0.286 | 0.073 | 1 | 17 |
| fnta | 0.15891 | 0.454547997 | 0.286 | 0.073 | 1 | 17 |
| ip6k2a | 0.15899 | 0.486619384 | 0.286 | 0.073 | 1 | 17 |
| rbm28 | 0.15926 | 0.518425733 | 0.286 | 0.073 | 1 | 17 |

| | | | | | | |
|---|---|---|---|---|---|---|
| zgc:162297 | 0.15935 | 0.460958749 | 0.286 | 0.073 | 1 | 17 |
| mief2 | 0.15939 | 0.437742294 | 0.286 | 0.073 | 1 | 17 |
| elf2b | 0.15939 | 0.402099103 | 0.286 | 0.073 | 1 | 17 |
| gtf2e2 | 0.16056 | 0.280190057 | 0.429 | 0.195 | 1 | 17 |
| srsf2a | 0.16057 | 0.394187339 | 0.429 | 0.171 | 1 | 17 |
| si:dkey-92i17.2 | 0.16064 | 0.118321333 | 0.357 | 0.171 | 1 | 17 |
| med11 | 0.16192 | 0.150114131 | 0.357 | 0.171 | 1 | 17 |
| mrpl53 | 0.16194 | 0.433026051 | 0.357 | 0.122 | 1 | 17 |
| psmc1a | 0.16213 | -0.62916616 | 0.143 | 0.39 | 1 | 17 |
| mcm4 | 0.16247 | 0.427698426 | 0.429 | 0.171 | 1 | 17 |
| capzb | 0.16342 | 0.415097961 | 0.357 | 0.122 | 1 | 17 |
| arpc3 | 0.16412 | 0.358633356 | 0.643 | 0.366 | 1 | 17 |
| bbip1 | 0.16484 | 0.29774984 | 0.429 | 0.195 | 1 | 17 |
| b2m | 0.16565 | 0.461642651 | 0.357 | 0.122 | 1 | 17 |
| bloc1s3 | 0.16576 | 0.22393287 | 0.214 | 0.049 | 1 | 17 |
| mt-nd4 | 0.16621 | -0.66900489 | 0.071 | 0.293 | 1 | 17 |
| hnrnpub | 0.1673 | 0.234516397 | 0.357 | 0.146 | 1 | 17 |
| tdp1 | 0.1686 | -0.31359022 | 0 | 0.098 | 1 | 17 |
| farsb | 0.1686 | -0.304315 | 0 | 0.098 | 1 | 17 |
| wtap | 0.1686 | -0.31948029 | 0 | 0.098 | 1 | 17 |
| bcl2l13 | 0.1686 | -0.28440463 | 0 | 0.098 | 1 | 17 |
| bzw1a | 0.1686 | -0.33412925 | 0 | 0.098 | 1 | 17 |
| eif2b1 | 0.1686 | -0.30884611 | 0 | 0.098 | 1 | 17 |
| hectd1 | 0.1686 | -0.28433718 | 0 | 0.098 | 1 | 17 |
| isca2 | 0.1686 | -0.22461595 | 0 | 0.098 | 1 | 17 |
| mcfd2 | 0.1686 | -0.26558169 | 0 | 0.098 | 1 | 17 |
| mis18bp1 | 0.1686 | -0.31946426 | 0 | 0.098 | 1 | 17 |
| rbm4.1 | 0.1686 | -0.27924961 | 0 | 0.098 | 1 | 17 |
| sept10 | 0.1686 | -0.3113733 | 0 | 0.098 | 1 | 17 |
| SMIM10 | 0.1686 | -0.31330129 | 0 | 0.098 | 1 | 17 |
| abhd10a | 0.1686 | -0.29197302 | 0 | 0.098 | 1 | 17 |
| ctnnd1 | 0.1686 | -0.2927394 | 0 | 0.098 | 1 | 17 |
| dhrs4 | 0.1686 | -0.3324559 | 0 | 0.098 | 1 | 17 |
| gcat | 0.1686 | -0.31303514 | 0 | 0.098 | 1 | 17 |
| gpkow | 0.1686 | -0.26344477 | 0 | 0.098 | 1 | 17 |
| hnrpkl | 0.1686 | -0.26880308 | 0 | 0.098 | 1 | 17 |
| si:dkey-85k15.4 | 0.1686 | -0.29582514 | 0 | 0.098 | 1 | 17 |
| trmt5 | 0.1686 | -0.28399719 | 0 | 0.098 | 1 | 17 |
| ttc8 | 0.1686 | -0.26011688 | 0 | 0.098 | 1 | 17 |
| yif1a | 0.1686 | -0.35784974 | 0 | 0.098 | 1 | 17 |
| atp6ap2 | 0.1686 | -0.25304283 | 0 | 0.098 | 1 | 17 |
| clasrp | 0.1686 | -0.27325581 | 0 | 0.098 | 1 | 17 |
| srrt | 0.1686 | -0.30671149 | 0 | 0.098 | 1 | 17 |
| tra2b | 0.1686 | -0.25715284 | 0 | 0.098 | 1 | 17 |
| zp3a.2 | 0.1686 | -0.33087328 | 0 | 0.098 | 1 | 17 |
| CR925817.1 | 0.1686 | -0.33084996 | 0 | 0.098 | 1 | 17 |
| LOC101883356 | 0.1686 | -0.27075758 | 0 | 0.098 | 1 | 17 |
| ap1m1 | 0.1686 | -0.30763914 | 0 | 0.098 | 1 | 17 |

| | | | | | | |
|---|---|---|---|---|---|---|
| capns1a | 0.1686 | -0.28866315 | 0 | 0.098 | 1 | 17 |
| ccdc83 | 0.1686 | -0.32798346 | 0 | 0.098 | 1 | 17 |
| coq6 | 0.1686 | -0.29720886 | 0 | 0.098 | 1 | 17 |
| ftsj1 | 0.1686 | -0.30234917 | 0 | 0.098 | 1 | 17 |
| ggctb | 0.1686 | -0.33945313 | 0 | 0.098 | 1 | 17 |
| mettl21a | 0.1686 | -0.24797371 | 0 | 0.098 | 1 | 17 |
| nup37 | 0.1686 | -0.29455378 | 0 | 0.098 | 1 | 17 |
| tmem242 | 0.1686 | -0.26585894 | 0 | 0.098 | 1 | 17 |
| zgc:92630 | 0.1686 | -0.13966273 | 0 | 0.098 | 1 | 17 |
| cherp | 0.1686 | -0.27550945 | 0 | 0.098 | 1 | 17 |
| kif22 | 0.1686 | -0.31650797 | 0 | 0.098 | 1 | 17 |
| kif2c | 0.1686 | -0.29987606 | 0 | 0.098 | 1 | 17 |
| lrp13 | 0.1686 | -0.32621577 | 0 | 0.098 | 1 | 17 |
| si:dkey-90l23.1 | 0.1686 | -0.27406841 | 0 | 0.098 | 1 | 17 |
| si:dkeyp-98a7.9 | 0.1686 | -0.28740856 | 0 | 0.098 | 1 | 17 |
| spo11 | 0.1686 | -0.32648393 | 0 | 0.098 | 1 | 17 |
| tpx2 | 0.1686 | -0.29414114 | 0 | 0.098 | 1 | 17 |
| zp2.2 | 0.1686 | -0.27199631 | 0 | 0.098 | 1 | 17 |
| CABZ01038709.1 | 0.1686 | -0.26135635 | 0 | 0.098 | 1 | 17 |
| CABZ01072952.1 | 0.1686 | -0.28677063 | 0 | 0.098 | 1 | 17 |
| adar | 0.1686 | -0.34082285 | 0 | 0.098 | 1 | 17 |
| adh8b | 0.1686 | -0.21718366 | 0 | 0.098 | 1 | 17 |
| ahsa1b | 0.1686 | -0.26246503 | 0 | 0.098 | 1 | 17 |
| aldocb | 0.1686 | -0.25469018 | 0 | 0.098 | 1 | 17 |
| arpc5lb | 0.1686 | -0.27328033 | 0 | 0.098 | 1 | 17 |
| btg3 | 0.1686 | -0.26259393 | 0 | 0.098 | 1 | 17 |
| btr18 | 0.1686 | -0.24222249 | 0 | 0.098 | 1 | 17 |
| chordc1b | 0.1686 | -0.35331735 | 0 | 0.098 | 1 | 17 |
| cpm | 0.1686 | -0.28484532 | 0 | 0.098 | 1 | 17 |
| crebzf | 0.1686 | -0.3454346 | 0 | 0.098 | 1 | 17 |
| dlg3 | 0.1686 | -0.31260842 | 0 | 0.098 | 1 | 17 |
| dtd2 | 0.1686 | -0.31575737 | 0 | 0.098 | 1 | 17 |
| ecsit | 0.1686 | -0.26349568 | 0 | 0.098 | 1 | 17 |
| elac1 | 0.1686 | -0.29846098 | 0 | 0.098 | 1 | 17 |
| eloal | 0.1686 | -0.22183292 | 0 | 0.098 | 1 | 17 |
| erich2 | 0.1686 | -0.22601244 | 0 | 0.098 | 1 | 17 |
| fam84b | 0.1686 | -0.25038002 | 0 | 0.098 | 1 | 17 |
| ggnbp2 | 0.1686 | -0.26379414 | 0 | 0.098 | 1 | 17 |
| kri1 | 0.1686 | -0.32102552 | 0 | 0.098 | 1 | 17 |
| lbr | 0.1686 | -0.32680285 | 0 | 0.098 | 1 | 17 |
| lrrc59 | 0.1686 | -0.32550769 | 0 | 0.098 | 1 | 17 |
| mmgt1 | 0.1686 | -0.30271306 | 0 | 0.098 | 1 | 17 |
| mrps17 | 0.1686 | -0.26250173 | 0 | 0.098 | 1 | 17 |
| mus81 | 0.1686 | -0.27763843 | 0 | 0.098 | 1 | 17 |
| mzt2b | 0.1686 | -0.22546695 | 0 | 0.098 | 1 | 17 |
| nfatc2ip | 0.1686 | -0.26573025 | 0 | 0.098 | 1 | 17 |
| nol11 | 0.1686 | -0.24255017 | 0 | 0.098 | 1 | 17 |
| nup35 | 0.1686 | -0.26372539 | 0 | 0.098 | 1 | 17 |

| | | | | | | |
|---|---|---|---|---|---|---|
| ppardb | 0.1686 | -0.31533231 | 0 | 0.098 | 1 | 17 |
| rars | 0.1686 | -0.35877556 | 0 | 0.098 | 1 | 17 |
| rnpep | 0.1686 | -0.31944215 | 0 | 0.098 | 1 | 17 |
| sap30bp | 0.1686 | -0.35164848 | 0 | 0.098 | 1 | 17 |
| scrn3 | 0.1686 | -0.24497228 | 0 | 0.098 | 1 | 17 |
| si:ch211-12h2.8 | 0.1686 | -0.29658211 | 0 | 0.098 | 1 | 17 |
| si:ch211-246m6.4 | 0.1686 | -0.32637677 | 0 | 0.098 | 1 | 17 |
| si:ch73-340n8.4 | 0.1686 | -0.28334417 | 0 | 0.098 | 1 | 17 |
| si:dkey-241l7.2 | 0.1686 | -0.25760001 | 0 | 0.098 | 1 | 17 |
| terb2 | 0.1686 | -0.26325268 | 0 | 0.098 | 1 | 17 |
| trmt1l | 0.1686 | -0.33579495 | 0 | 0.098 | 1 | 17 |
| uap1 | 0.1686 | -0.24438063 | 0 | 0.098 | 1 | 17 |
| ubl7b | 0.1686 | -0.29893962 | 0 | 0.098 | 1 | 17 |
| zbtb40 | 0.1686 | -0.28800178 | 0 | 0.098 | 1 | 17 |
| zp3d.2 | 0.1686 | -0.32306338 | 0 | 0.098 | 1 | 17 |
| abhd17c | 0.1686 | -0.27280273 | 0 | 0.098 | 1 | 17 |
| aurka | 0.1686 | -0.30115851 | 0 | 0.098 | 1 | 17 |
| ccna2 | 0.1686 | -0.32620313 | 0 | 0.098 | 1 | 17 |
| dnajc2 | 0.1686 | -0.25184135 | 0 | 0.098 | 1 | 17 |
| usp3 | 0.1686 | -0.28907524 | 0 | 0.098 | 1 | 17 |
| CABZ01045617.1 | 0.1686 | -0.25315567 | 0 | 0.098 | 1 | 17 |
| actr8 | 0.1686 | -0.27590309 | 0 | 0.098 | 1 | 17 |
| atg13 | 0.1686 | -0.34965025 | 0 | 0.098 | 1 | 17 |
| cand1 | 0.1686 | -0.23454025 | 0 | 0.098 | 1 | 17 |
| cd151l | 0.1686 | -0.26599275 | 0 | 0.098 | 1 | 17 |
| cdc34b | 0.1686 | -0.27158784 | 0 | 0.098 | 1 | 17 |
| dnlz | 0.1686 | -0.27619706 | 0 | 0.098 | 1 | 17 |
| dtwd1 | 0.1686 | -0.28865665 | 0 | 0.098 | 1 | 17 |
| echdc2 | 0.1686 | -0.27009161 | 0 | 0.098 | 1 | 17 |
| eif4g1a | 0.1686 | -0.29110568 | 0 | 0.098 | 1 | 17 |
| fam53b | 0.1686 | -0.29861114 | 0 | 0.098 | 1 | 17 |
| mboat7 | 0.1686 | -0.2888495 | 0 | 0.098 | 1 | 17 |
| med15 | 0.1686 | -0.28180944 | 0 | 0.098 | 1 | 17 |
| ormdl2 | 0.1686 | -0.22140164 | 0 | 0.098 | 1 | 17 |
| ppil2 | 0.1686 | -0.26256036 | 0 | 0.098 | 1 | 17 |
| rab35b | 0.1686 | -0.3021395 | 0 | 0.098 | 1 | 17 |
| si:ch73-15b2.5 | 0.1686 | -0.26807676 | 0 | 0.098 | 1 | 17 |
| slc30a7 | 0.1686 | -0.33567553 | 0 | 0.098 | 1 | 17 |
| tmem254 | 0.1686 | -0.25309198 | 0 | 0.098 | 1 | 17 |
| txn | 0.1686 | -0.29595104 | 0 | 0.098 | 1 | 17 |
| vps45 | 0.1686 | -0.29636751 | 0 | 0.098 | 1 | 17 |
| qdprb1 | 0.17012 | 0.482661537 | 0.429 | 0.171 | 1 | 17 |
| hmga1a | 0.17023 | 0.522945091 | 0.429 | 0.171 | 1 | 17 |
| cox7a2l | 0.17024 | 0.561350326 | 0.357 | 0.122 | 1 | 17 |
| arpc2 | 0.17046 | 0.495467484 | 0.429 | 0.171 | 1 | 17 |
| eif3ba | 0.17052 | 0.491952662 | 0.429 | 0.171 | 1 | 17 |
| zgc:111986 | 0.17056 | 0.479016637 | 0.429 | 0.171 | 1 | 17 |
| atp5if1a | 0.1715 | -0.10082589 | 0.571 | 0.488 | 1 | 17 |

| | | | | | | |
|---|---|---|---|---|---|---|
| slc3a2b | 0.17275 | 0.255112698 | 0.214 | 0.049 | 1 | 17 |
| hadhaa | 0.17288 | 0.502934143 | 0.357 | 0.122 | 1 | 17 |
| polr1d | 0.17391 | 0.442066819 | 0.357 | 0.122 | 1 | 17 |
| srrm1 | 0.17424 | 0.492348641 | 0.357 | 0.122 | 1 | 17 |
| cct2 | 0.17426 | 0.115645215 | 0.714 | 0.512 | 1 | 17 |
| ndufs7 | 0.1752 | 0.422473197 | 0.571 | 0.293 | 1 | 17 |
| tk2 | 0.17525 | 0.456160924 | 0.357 | 0.122 | 1 | 17 |
| chchd6b | 0.1753 | 0.477375672 | 0.357 | 0.122 | 1 | 17 |
| dohh | 0.17553 | 0.446069427 | 0.357 | 0.122 | 1 | 17 |
| hnrnpk | 0.17568 | 0.485290873 | 0.357 | 0.122 | 1 | 17 |
| sf3b2 | 0.17586 | 0.167436647 | 0.357 | 0.171 | 1 | 17 |
| mt-cyb | 0.17591 | -0.72582091 | 0.071 | 0.268 | 1 | 17 |
| arpp19b | 0.17807 | -0.54688008 | 0.071 | 0.293 | 1 | 17 |
| rps18 | 0.17844 | 0.314460298 | 0.929 | 0.707 | 1 | 17 |
| nxt2 | 0.17888 | 0.31160894 | 0.429 | 0.195 | 1 | 17 |
| zgc:56304 | 0.17928 | -0.58881727 | 0.143 | 0.366 | 1 | 17 |
| zgc:103482 | 0.18074 | 0.429883003 | 0.571 | 0.293 | 1 | 17 |
| si:dkey-23i12.5 | 0.18121 | 0.175568332 | 0.357 | 0.171 | 1 | 17 |
| gstm.1 | 0.18142 | 0.362760142 | 0.786 | 0.512 | 1 | 17 |
| rps19bp1 | 0.18192 | 0.321214095 | 0.429 | 0.195 | 1 | 17 |
| dut | 0.18248 | 0.373314781 | 0.786 | 0.512 | 1 | 17 |
| rps15a | 0.18249 | 0.201503021 | 1 | 0.951 | 1 | 17 |
| rpl32 | 0.18265 | 0.227331234 | 1 | 0.878 | 1 | 17 |
| uqcrh | 0.18281 | 0.479872831 | 0.571 | 0.293 | 1 | 17 |
| ywhaqb | 0.18427 | -0.52455422 | 0.071 | 0.293 | 1 | 17 |
| CABZ01083945.1 | 0.18499 | 0.517501634 | 0.5 | 0.244 | 1 | 17 |
| psma4 | 0.18548 | 0.281217438 | 0.5 | 0.268 | 1 | 17 |
| rpsa | 0.18571 | 0.230818692 | 1 | 0.902 | 1 | 17 |
| hmgb2b | 0.18617 | 0.126658567 | 1 | 0.854 | 1 | 17 |
| hypk | 0.18674 | 0.343257181 | 0.5 | 0.244 | 1 | 17 |
| dad1 | 0.18749 | 0.173687886 | 0.5 | 0.293 | 1 | 17 |
| cdkn1bb | 0.18758 | 0.263782652 | 0.214 | 0.049 | 1 | 17 |
| zgc:114188 | 0.18758 | 0.292022138 | 0.929 | 0.707 | 1 | 17 |
| srsf1b | 0.18796 | 0.158628324 | 0.286 | 0.122 | 1 | 17 |
| ube2al | 0.18899 | 0.250615157 | 0.214 | 0.049 | 1 | 17 |
| nsmce2 | 0.18985 | 0.232072542 | 0.429 | 0.22 | 1 | 17 |
| nasp | 0.19105 | 0.486724361 | 0.643 | 0.366 | 1 | 17 |
| pld6 | 0.19144 | 0.291003725 | 0.571 | 0.317 | 1 | 17 |
| atpaf2 | 0.19222 | 0.261033046 | 0.214 | 0.049 | 1 | 17 |
| rps20 | 0.19259 | 0.127972014 | 1 | 0.854 | 1 | 17 |
| polr2l | 0.19433 | 0.278577242 | 0.714 | 0.463 | 1 | 17 |
| plgrkt | 0.19501 | 0.342166033 | 0.214 | 0.049 | 1 | 17 |
| insb | 0.19516 | 0.2291276 | 0.214 | 0.049 | 1 | 17 |
| iscub | 0.19542 | 0.277109287 | 0.357 | 0.146 | 1 | 17 |
| syce2 | 0.19543 | 0.562051666 | 0.5 | 0.268 | 1 | 17 |
| atp5meb | 0.19682 | 0.408084112 | 0.714 | 0.439 | 1 | 17 |
| tpi1a | 0.19715 | 0.265776432 | 0.214 | 0.049 | 1 | 17 |
| rbbp4 | 0.19735 | -0.54622901 | 0.214 | 0.366 | 1 | 17 |

| | | | | | | |
|---|---|---|---|---|---|---|
| eif1b | 0.1996 | 0.518422588 | 0.5 | 0.244 | 1 | 17 |
| rps24 | 0.2008 | 0.222847292 | 1 | 0.878 | 1 | 17 |
| si:zfos-169g10.3 | 0.20102 | 0.395409119 | 0.5 | 0.244 | 1 | 17 |
| tsr2 | 0.20169 | 0.353672578 | 0.429 | 0.195 | 1 | 17 |
| ssu72 | 0.20223 | 0.292698583 | 0.357 | 0.146 | 1 | 17 |
| msi2b | 0.20295 | 0.323935274 | 0.214 | 0.049 | 1 | 17 |
| ppp2cb | 0.20302 | 0.333223615 | 0.429 | 0.195 | 1 | 17 |
| rtraf | 0.20396 | 0.298280338 | 0.357 | 0.146 | 1 | 17 |
| phf5a | 0.20433 | 0.269281721 | 0.5 | 0.268 | 1 | 17 |
| timm10 | 0.20489 | 0.344374049 | 0.429 | 0.195 | 1 | 17 |
| ier3ip1 | 0.20499 | 0.324418961 | 0.286 | 0.098 | 1 | 17 |
| psmd4a | 0.2057 | 0.126363084 | 0.429 | 0.244 | 1 | 17 |
| mrps35 | 0.20594 | 0.283342425 | 0.214 | 0.049 | 1 | 17 |
| eri1 | 0.20598 | 0.309535794 | 0.214 | 0.049 | 1 | 17 |
| edc4 | 0.20761 | 0.27888282 | 0.286 | 0.098 | 1 | 17 |
| csnk2b | 0.20792 | 0.103031624 | 0.643 | 0.463 | 1 | 17 |
| bsg | 0.20797 | 0.388085434 | 0.429 | 0.195 | 1 | 17 |
| cyfip1 | 0.20833 | 0.348338762 | 0.214 | 0.049 | 1 | 17 |
| ndufab1a | 0.20862 | 0.248722637 | 0.571 | 0.341 | 1 | 17 |
| npm2b | 0.20866 | -0.62111564 | 0.143 | 0.366 | 1 | 17 |
| tmem230a | 0.20881 | 0.282462598 | 0.214 | 0.049 | 1 | 17 |
| eif4ebp1 | 0.20892 | 0.356786108 | 0.357 | 0.146 | 1 | 17 |
| asz1 | 0.20932 | -0.54741353 | 0.071 | 0.268 | 1 | 17 |
| BX511129.1 | 0.20954 | 0.278748065 | 0.214 | 0.049 | 1 | 17 |
| ube2v2 | 0.21023 | 0.222093058 | 0.643 | 0.415 | 1 | 17 |
| mrps24 | 0.21114 | 0.273527922 | 0.214 | 0.049 | 1 | 17 |
| trub2 | 0.21136 | 0.350274587 | 0.214 | 0.049 | 1 | 17 |
| bcl7a | 0.21165 | 0.279901507 | 0.214 | 0.049 | 1 | 17 |
| atp5mc1 | 0.21269 | 0.19385427 | 0.786 | 0.561 | 1 | 17 |
| bcat1 | 0.21286 | 0.448384969 | 0.214 | 0.049 | 1 | 17 |
| emc9 | 0.21334 | 0.27664979 | 0.286 | 0.098 | 1 | 17 |
| mrpl14 | 0.21339 | 0.324432424 | 0.429 | 0.195 | 1 | 17 |
| atp1b1a | 0.21383 | 0.286198079 | 0.214 | 0.049 | 1 | 17 |
| arl6ip1 | 0.21475 | -0.56260823 | 0.071 | 0.244 | 1 | 17 |
| gtf2h4 | 0.2148 | 0.293682474 | 0.214 | 0.049 | 1 | 17 |
| wdr4 | 0.21523 | 0.302582052 | 0.214 | 0.049 | 1 | 17 |
| aspscr1 | 0.21589 | 0.344260361 | 0.214 | 0.049 | 1 | 17 |
| atp5mc3a | 0.21617 | 0.142745658 | 0.571 | 0.39 | 1 | 17 |
| kcne4 | 0.21633 | 0.3232504 | 0.214 | 0.049 | 1 | 17 |
| smc1al | 0.2164 | 0.29228011 | 0.286 | 0.098 | 1 | 17 |
| vapb | 0.21729 | 0.259873497 | 0.286 | 0.098 | 1 | 17 |
| mrrf | 0.21733 | 0.295859277 | 0.214 | 0.049 | 1 | 17 |
| rab5c | 0.21832 | 0.323000836 | 0.214 | 0.049 | 1 | 17 |
| ppp1r7 | 0.21867 | 0.35302994 | 0.214 | 0.049 | 1 | 17 |
| ik | 0.21953 | 0.321046917 | 0.357 | 0.146 | 1 | 17 |
| glsb | 0.21968 | 0.383075421 | 0.214 | 0.049 | 1 | 17 |
| khdrbs1b | 0.21986 | 0.389003267 | 0.214 | 0.049 | 1 | 17 |
| sec61b | 0.22054 | 0.287932344 | 0.643 | 0.39 | 1 | 17 |

| | | | | | | |
|---|---|---|---|---|---|---|
| CABZ01080371.2 | 0.22067 | 0.394119994 | 0.214 | 0.049 | 1 | 17 |
| aggf1 | 0.22072 | 0.264276114 | 0.214 | 0.049 | 1 | 17 |
| znf207a | 0.22076 | 0.268927389 | 0.214 | 0.049 | 1 | 17 |
| tomm70a | 0.22088 | 0.287367733 | 0.286 | 0.098 | 1 | 17 |
| lamtor3 | 0.22176 | 0.265217978 | 0.286 | 0.098 | 1 | 17 |
| med8 | 0.22227 | 0.334588835 | 0.286 | 0.098 | 1 | 17 |
| AL954696.1 | 0.22234 | 0.376624322 | 0.214 | 0.049 | 1 | 17 |
| vrk1 | 0.22243 | 0.376386059 | 0.429 | 0.195 | 1 | 17 |
| tmem50a | 0.22318 | 0.352681006 | 0.214 | 0.049 | 1 | 17 |
| jpt1b | 0.2233 | -0.26623366 | 0.286 | 0.268 | 1 | 17 |
| snrpa | 0.22357 | 0.329183577 | 0.357 | 0.146 | 1 | 17 |
| bloc1s2 | 0.22421 | 0.412204332 | 0.214 | 0.049 | 1 | 17 |
| riox2 | 0.2249 | 0.348447736 | 0.214 | 0.049 | 1 | 17 |
| cdca7b | 0.22499 | 0.338180129 | 0.214 | 0.049 | 1 | 17 |
| msh2 | 0.22506 | 0.319263021 | 0.214 | 0.049 | 1 | 17 |
| golt1ba | 0.22518 | 0.42984787 | 0.214 | 0.049 | 1 | 17 |
| gnl1 | 0.22527 | 0.345175883 | 0.214 | 0.049 | 1 | 17 |
| orc6 | 0.22528 | 0.483474242 | 0.214 | 0.049 | 1 | 17 |
| nsmce1 | 0.22578 | 0.242054371 | 0.429 | 0.22 | 1 | 17 |
| ypel5 | 0.22586 | 0.289990721 | 0.286 | 0.098 | 1 | 17 |
| rpl19 | 0.226 | 0.197943165 | 1 | 0.927 | 1 | 17 |
| ciarta | 0.22608 | 0.38662844 | 0.214 | 0.049 | 1 | 17 |
| higd2a | 0.22615 | 0.378411763 | 0.214 | 0.049 | 1 | 17 |
| tyms | 0.22632 | 0.163058941 | 0.429 | 0.244 | 1 | 17 |
| cmasa | 0.22641 | 0.452235241 | 0.214 | 0.049 | 1 | 17 |
| rps17 | 0.22658 | 0.290352757 | 0.929 | 0.732 | 1 | 17 |
| phb2a | 0.22667 | 0.315036427 | 0.214 | 0.049 | 1 | 17 |
| si:dkeyp-55f12.3 | 0.22671 | 0.312514316 | 0.214 | 0.049 | 1 | 17 |
| dazap2 | 0.22709 | 0.320276253 | 0.214 | 0.049 | 1 | 17 |
| arl2bp | 0.2277 | 0.298909077 | 0.357 | 0.146 | 1 | 17 |
| rpp14 | 0.22794 | 0.34078929 | 0.286 | 0.098 | 1 | 17 |
| zgc:85789 | 0.22801 | 0.396002493 | 0.214 | 0.049 | 1 | 17 |
| efcab1 | 0.22802 | 0.371576971 | 0.214 | 0.049 | 1 | 17 |
| gpatch4 | 0.22882 | 0.449942516 | 0.214 | 0.049 | 1 | 17 |
| babam2 | 0.22883 | 0.382439789 | 0.214 | 0.049 | 1 | 17 |
| det1 | 0.22891 | 0.403637399 | 0.214 | 0.049 | 1 | 17 |
| map2k2a | 0.22899 | 0.399640914 | 0.214 | 0.049 | 1 | 17 |
| imp4 | 0.22912 | 0.461086445 | 0.214 | 0.049 | 1 | 17 |
| si:dkey-119f1.1 | 0.22912 | 0.398946811 | 0.214 | 0.049 | 1 | 17 |
| lyrm7 | 0.22919 | 0.363785017 | 0.214 | 0.049 | 1 | 17 |
| CABZ01118004.1 | 0.22978 | 0.346027743 | 0.214 | 0.049 | 1 | 17 |
| ctnnbip1 | 0.22995 | 0.35337843 | 0.214 | 0.049 | 1 | 17 |
| tuba1b | 0.23015 | 0.286310398 | 0.214 | 0.049 | 1 | 17 |
| ing5b | 0.23075 | 0.428843415 | 0.214 | 0.049 | 1 | 17 |
| actr6 | 0.23075 | 0.286647803 | 0.214 | 0.049 | 1 | 17 |
| hmces | 0.23106 | 0.36383275 | 0.214 | 0.049 | 1 | 17 |
| ap1s2 | 0.23106 | 0.371263607 | 0.214 | 0.049 | 1 | 17 |
| si:ch211-262e15.1 | 0.23117 | 0.339284984 | 0.214 | 0.049 | 1 | 17 |

| | | | | | | |
|---|---|---|---|---|---|---|
| COX7A2 (1 of many) | 0.23118 | 0.160926818 | 0.786 | 0.585 | 1 | 17 |
| adkb | 0.23137 | 0.411235494 | 0.214 | 0.049 | 1 | 17 |
| XLOC-014398 | 0.2314 | 0.363396865 | 0.214 | 0.049 | 1 | 17 |
| spc25 | 0.23142 | 0.362830346 | 0.214 | 0.049 | 1 | 17 |
| sept7a | 0.23147 | 0.368686021 | 0.214 | 0.049 | 1 | 17 |
| klhdc4 | 0.23149 | 0.371777576 | 0.214 | 0.049 | 1 | 17 |
| nap1l1 | 0.23238 | 0.164442868 | 0.857 | 0.659 | 1 | 17 |
| eif5a2 | 0.23298 | -0.12334704 | 0.643 | 0.585 | 1 | 17 |
| asf1ba | 0.23386 | 0.486622657 | 0.286 | 0.098 | 1 | 17 |
| pabpn1l | 0.23455 | -0.52102585 | 0.143 | 0.366 | 1 | 17 |
| rpl4 | 0.23511 | 0.154205668 | 0.857 | 0.659 | 1 | 17 |
| adad1 | 0.23564 | 0.305929255 | 0.5 | 0.268 | 1 | 17 |
| NC-002333.4 | 0.23566 | 0.182842784 | 0.286 | 0.317 | 1 | 17 |
| baz1a | 0.23597 | 0.221664072 | 0.429 | 0.22 | 1 | 17 |
| zgc:161973 | 0.23819 | 0.355021767 | 0.429 | 0.195 | 1 | 17 |
| prpf19 | 0.23898 | 0.378353378 | 0.286 | 0.098 | 1 | 17 |
| zgc:56493 | 0.24023 | -0.22869215 | 0.714 | 0.732 | 1 | 17 |
| nutf2l | 0.24025 | 0.336106469 | 0.786 | 0.537 | 1 | 17 |
| acaa1 | 0.24066 | 0.288058506 | 0.286 | 0.098 | 1 | 17 |
| ewsr1a | 0.24089 | 0.139112343 | 0.214 | 0.073 | 1 | 17 |
| syf2 | 0.24111 | 0.412705951 | 0.429 | 0.195 | 1 | 17 |
| si:ch211-235e9.6 | 0.2413 | 0.43189097 | 0.429 | 0.195 | 1 | 17 |
| ran | 0.24135 | 0.261813802 | 0.857 | 0.634 | 1 | 17 |
| cstf2 | 0.24154 | 0.413834965 | 0.429 | 0.195 | 1 | 17 |
| dctpp1 | 0.24189 | 0.318086538 | 0.5 | 0.268 | 1 | 17 |
| glrx | 0.24208 | -0.45103401 | 0.071 | 0.268 | 1 | 17 |
| COX3 | 0.2422 | -0.63798067 | 0.071 | 0.268 | 1 | 17 |
| rpl11 | 0.24314 | 0.204459536 | 1 | 0.902 | 1 | 17 |
| rab1ab | 0.24379 | 0.319994024 | 0.286 | 0.098 | 1 | 17 |
| pgam1a | 0.2447 | 0.344529309 | 0.357 | 0.146 | 1 | 17 |
| ywhabb | 0.24558 | 0.346705602 | 0.643 | 0.39 | 1 | 17 |
| hp1bp3 | 0.24675 | 0.408204355 | 0.571 | 0.317 | 1 | 17 |
| psmc6 | 0.24702 | 0.423874237 | 0.429 | 0.195 | 1 | 17 |
| minos1 | 0.24719 | 0.44838608 | 0.643 | 0.415 | 1 | 17 |
| cyc1 | 0.24731 | 0.36818178 | 0.357 | 0.146 | 1 | 17 |
| oaz1b | 0.24737 | 0.438453786 | 0.571 | 0.317 | 1 | 17 |
| dhx32a | 0.24976 | -0.2104411 | 0 | 0.073 | 1 | 17 |
| gspt1l | 0.24976 | -0.22404255 | 0 | 0.073 | 1 | 17 |
| CZQB01141835.1 | 0.24976 | -0.18709179 | 0 | 0.073 | 1 | 17 |
| alcama | 0.24976 | -0.27975815 | 0 | 0.073 | 1 | 17 |
| arl4aa | 0.24976 | -0.22634956 | 0 | 0.073 | 1 | 17 |
| armc4 | 0.24976 | -0.17383674 | 0 | 0.073 | 1 | 17 |
| asb12a | 0.24976 | -0.21122334 | 0 | 0.073 | 1 | 17 |
| ddb1 | 0.24976 | -0.14966705 | 0 | 0.073 | 1 | 17 |
| dpm3 | 0.24976 | -0.2297851 | 0 | 0.073 | 1 | 17 |
| exosc10 | 0.24976 | -0.1937582 | 0 | 0.073 | 1 | 17 |
| exosc7 | 0.24976 | -0.24518498 | 0 | 0.073 | 1 | 17 |
| extl2 | 0.24976 | -0.23850818 | 0 | 0.073 | 1 | 17 |

| | | | | | | |
|---|---|---|---|---|---|---|
| faf2 | 0.24976 | -0.17744557 | 0 | 0.073 | 1 | 17 |
| fam149a | 0.24976 | -0.21536991 | 0 | 0.073 | 1 | 17 |
| fuca2 | 0.24976 | -0.20828778 | 0 | 0.073 | 1 | 17 |
| gtf3ab | 0.24976 | -0.20989406 | 0 | 0.073 | 1 | 17 |
| hspa4l | 0.24976 | -0.20878216 | 0 | 0.073 | 1 | 17 |
| iars2 | 0.24976 | -0.17731772 | 0 | 0.073 | 1 | 17 |
| krt94 | 0.24976 | -0.25042353 | 0 | 0.073 | 1 | 17 |
| lgals9l3 | 0.24976 | -0.24995079 | 0 | 0.073 | 1 | 17 |
| mt-nd3 | 0.24976 | -0.22970316 | 0 | 0.073 | 1 | 17 |
| naxd | 0.24976 | -0.17114509 | 0 | 0.073 | 1 | 17 |
| nipsnap3a | 0.24976 | -0.19408966 | 0 | 0.073 | 1 | 17 |
| paqr3a | 0.24976 | -0.22225714 | 0 | 0.073 | 1 | 17 |
| sephs1 | 0.24976 | -0.18795141 | 0 | 0.073 | 1 | 17 |
| si:ch73-59c19.1 | 0.24976 | -0.2339002 | 0 | 0.073 | 1 | 17 |
| tmem70 | 0.24976 | -0.17241687 | 0 | 0.073 | 1 | 17 |
| vps26a | 0.24976 | -0.29282922 | 0 | 0.073 | 1 | 17 |
| CR318588.1 | 0.24976 | -0.23264869 | 0 | 0.073 | 1 | 17 |
| CR854989.1 | 0.24976 | -0.25750653 | 0 | 0.073 | 1 | 17 |
| LOC108181449 | 0.24976 | -0.19816333 | 0 | 0.073 | 1 | 17 |
| aclya | 0.24976 | -0.14422652 | 0 | 0.073 | 1 | 17 |
| aftpha | 0.24976 | -0.19876601 | 0 | 0.073 | 1 | 17 |
| ankrd54 | 0.24976 | -0.26563499 | 0 | 0.073 | 1 | 17 |
| asb8 | 0.24976 | -0.151202 | 0 | 0.073 | 1 | 17 |
| atp1a3a | 0.24976 | -0.21257676 | 0 | 0.073 | 1 | 17 |
| babam1 | 0.24976 | -0.2093899 | 0 | 0.073 | 1 | 17 |
| batf3 | 0.24976 | -0.18016906 | 0 | 0.073 | 1 | 17 |
| bbc3 | 0.24976 | -0.168096 | 0 | 0.073 | 1 | 17 |
| ccdc171 | 0.24976 | -0.19779229 | 0 | 0.073 | 1 | 17 |
| cdk20 | 0.24976 | -0.15545729 | 0 | 0.073 | 1 | 17 |
| cept1a | 0.24976 | -0.21719087 | 0 | 0.073 | 1 | 17 |
| chic2 | 0.24976 | -0.19686574 | 0 | 0.073 | 1 | 17 |
| chmp6b | 0.24976 | -0.18760821 | 0 | 0.073 | 1 | 17 |
| cmtr2 | 0.24976 | -0.260048 | 0 | 0.073 | 1 | 17 |
| cspp1b | 0.24976 | -0.22650817 | 0 | 0.073 | 1 | 17 |
| dclre1a | 0.24976 | -0.24316265 | 0 | 0.073 | 1 | 17 |
| dcp1b | 0.24976 | -0.26135771 | 0 | 0.073 | 1 | 17 |
| diabloa | 0.24976 | -0.23144335 | 0 | 0.073 | 1 | 17 |
| ech1 | 0.24976 | -0.18567768 | 0 | 0.073 | 1 | 17 |
| ell2 | 0.24976 | -0.26601484 | 0 | 0.073 | 1 | 17 |
| galnt18b | 0.24976 | -0.22419489 | 0 | 0.073 | 1 | 17 |
| gpatch8 | 0.24976 | -0.26919428 | 0 | 0.073 | 1 | 17 |
| gskip | 0.24976 | -0.21454919 | 0 | 0.073 | 1 | 17 |
| gsna | 0.24976 | -0.26631146 | 0 | 0.073 | 1 | 17 |
| hk2 | 0.24976 | -0.25321518 | 0 | 0.073 | 1 | 17 |
| ift80 | 0.24976 | -0.22049716 | 0 | 0.073 | 1 | 17 |
| ildr1a | 0.24976 | -0.24786547 | 0 | 0.073 | 1 | 17 |
| kbtbd8 | 0.24976 | -0.27849711 | 0 | 0.073 | 1 | 17 |
| kyat3 | 0.24976 | -0.19481183 | 0 | 0.073 | 1 | 17 |

| | | | | | | |
|---|---|---|---|---|---|---|
| lrrfip1a | 0.24976 | -0.24843292 | 0 | 0.073 | 1 | 17 |
| lyn | 0.24976 | -0.19866773 | 0 | 0.073 | 1 | 17 |
| march5l | 0.24976 | -0.1883878 | 0 | 0.073 | 1 | 17 |
| mdka | 0.24976 | -0.24234381 | 0 | 0.073 | 1 | 17 |
| mdm4 | 0.24976 | -0.23430071 | 0 | 0.073 | 1 | 17 |
| mknk2b | 0.24976 | -0.25252082 | 0 | 0.073 | 1 | 17 |
| mlst8 | 0.24976 | -0.21685773 | 0 | 0.073 | 1 | 17 |
| mrps15 | 0.24976 | -0.23954897 | 0 | 0.073 | 1 | 17 |
| n6amt1 | 0.24976 | -0.24939308 | 0 | 0.073 | 1 | 17 |
| ncoa5 | 0.24976 | -0.20262208 | 0 | 0.073 | 1 | 17 |
| nfil3 | 0.24976 | -0.25179534 | 0 | 0.073 | 1 | 17 |
| nfkb2 | 0.24976 | -0.17148895 | 0 | 0.073 | 1 | 17 |
| noc3l | 0.24976 | -0.17554287 | 0 | 0.073 | 1 | 17 |
| p4ha2 | 0.24976 | -0.19489195 | 0 | 0.073 | 1 | 17 |
| pars2 | 0.24976 | -0.25043687 | 0 | 0.073 | 1 | 17 |
| pcbd1 | 0.24976 | -0.19482419 | 0 | 0.073 | 1 | 17 |
| pomt1 | 0.24976 | -0.23745993 | 0 | 0.073 | 1 | 17 |
| pphln1 | 0.24976 | -0.22562679 | 0 | 0.073 | 1 | 17 |
| prkaa1 | 0.24976 | -0.17102932 | 0 | 0.073 | 1 | 17 |
| reep3b | 0.24976 | -0.19629219 | 0 | 0.073 | 1 | 17 |
| rogdi | 0.24976 | -0.17761106 | 0 | 0.073 | 1 | 17 |
| rras | 0.24976 | -0.21889388 | 0 | 0.073 | 1 | 17 |
| s100a10b | 0.24976 | -0.23058332 | 0 | 0.073 | 1 | 17 |
| sdhaf2 | 0.24976 | -0.21510759 | 0 | 0.073 | 1 | 17 |
| sfr1 | 0.24976 | -0.19848832 | 0 | 0.073 | 1 | 17 |
| si:ch211-194k22.8 | 0.24976 | -0.21166606 | 0 | 0.073 | 1 | 17 |
| si:ch211-197g15.6 | 0.24976 | -0.20425003 | 0 | 0.073 | 1 | 17 |
| si:ch211-199b20.3 | 0.24976 | -0.1825756 | 0 | 0.073 | 1 | 17 |
| si:ch211-40k21.5 | 0.24976 | -0.23654296 | 0 | 0.073 | 1 | 17 |
| si:dkey-147f3.8 | 0.24976 | -0.28877184 | 0 | 0.073 | 1 | 17 |
| si:dkeyp-98a7.8 | 0.24976 | -0.21799009 | 0 | 0.073 | 1 | 17 |
| ska1 | 0.24976 | -0.19025032 | 0 | 0.073 | 1 | 17 |
| snapc1a | 0.24976 | -0.18056558 | 0 | 0.073 | 1 | 17 |
| sp1 | 0.24976 | -0.16255116 | 0 | 0.073 | 1 | 17 |
| spop | 0.24976 | -0.20733348 | 0 | 0.073 | 1 | 17 |
| stau2 | 0.24976 | -0.18648376 | 0 | 0.073 | 1 | 17 |
| tbl2 | 0.24976 | -0.19983195 | 0 | 0.073 | 1 | 17 |
| tcta | 0.24976 | -0.23429289 | 0 | 0.073 | 1 | 17 |
| tdrd7a | 0.24976 | -0.1503474 | 0 | 0.073 | 1 | 17 |
| tlk1a | 0.24976 | -0.27057338 | 0 | 0.073 | 1 | 17 |
| tmem144b | 0.24976 | -0.17656885 | 0 | 0.073 | 1 | 17 |
| tmem222a | 0.24976 | -0.17033551 | 0 | 0.073 | 1 | 17 |
| trap1 | 0.24976 | -0.18385563 | 0 | 0.073 | 1 | 17 |
| ube2g1b | 0.24976 | -0.22487821 | 0 | 0.073 | 1 | 17 |
| upf3b | 0.24976 | -0.20137102 | 0 | 0.073 | 1 | 17 |
| wipi2 | 0.24976 | -0.25036642 | 0 | 0.073 | 1 | 17 |
| yif1b | 0.24976 | -0.17854204 | 0 | 0.073 | 1 | 17 |
| zgc:153499 | 0.24976 | -0.17412661 | 0 | 0.073 | 1 | 17 |

| | | | | | | |
|---|---|---|---|---|---|---|
| zgc:162344 | 0.24976 | -0.26471489 | 0 | 0.073 | 1 | 17 |
| zgc:92107 | 0.24976 | -0.20562701 | 0 | 0.073 | 1 | 17 |
| cd82b | 0.24976 | -0.19606172 | 0 | 0.073 | 1 | 17 |
| pip5k1ca | 0.24976 | -0.22163745 | 0 | 0.073 | 1 | 17 |
| ubqln4 | 0.24976 | -0.20637798 | 0 | 0.073 | 1 | 17 |
| BX510996.1 | 0.24976 | -0.24938646 | 0 | 0.073 | 1 | 17 |
| BX649337.1 | 0.24976 | -0.23564711 | 0 | 0.073 | 1 | 17 |
| LOC100331743 | 0.24976 | -0.18448027 | 0 | 0.073 | 1 | 17 |
| LOC101886766 | 0.24976 | -0.24383147 | 0 | 0.073 | 1 | 17 |
| TSTD1 | 0.24976 | -0.21271755 | 0 | 0.073 | 1 | 17 |
| aasdhppt | 0.24976 | -0.19194849 | 0 | 0.073 | 1 | 17 |
| acad11 | 0.24976 | -0.20100108 | 0 | 0.073 | 1 | 17 |
| acsbg2 | 0.24976 | -0.23561831 | 0 | 0.073 | 1 | 17 |
| acsl5 | 0.24976 | -0.20208045 | 0 | 0.073 | 1 | 17 |
| aldob | 0.24976 | -0.19731351 | 0 | 0.073 | 1 | 17 |
| anxa13l | 0.24976 | -0.2501139 | 0 | 0.073 | 1 | 17 |
| apex2 | 0.24976 | -0.23031494 | 0 | 0.073 | 1 | 17 |
| apmap | 0.24976 | -0.24894736 | 0 | 0.073 | 1 | 17 |
| apodb | 0.24976 | -0.21736794 | 0 | 0.073 | 1 | 17 |
| atad5a | 0.24976 | -0.27278761 | 0 | 0.073 | 1 | 17 |
| brap | 0.24976 | -0.26484586 | 0 | 0.073 | 1 | 17 |
| cd109 | 0.24976 | -0.29055987 | 0 | 0.073 | 1 | 17 |
| cdk5rap3 | 0.24976 | -0.25893785 | 0 | 0.073 | 1 | 17 |
| chsy1 | 0.24976 | -0.2877113 | 0 | 0.073 | 1 | 17 |
| cnga4 | 0.24976 | -0.21362569 | 0 | 0.073 | 1 | 17 |
| cox10 | 0.24976 | -0.18710605 | 0 | 0.073 | 1 | 17 |
| cry1ab | 0.24976 | -0.21134192 | 0 | 0.073 | 1 | 17 |
| crybg1a | 0.24976 | -0.2310527 | 0 | 0.073 | 1 | 17 |
| ctu2 | 0.24976 | -0.23192307 | 0 | 0.073 | 1 | 17 |
| dctn3 | 0.24976 | -0.22548928 | 0 | 0.073 | 1 | 17 |
| eapp | 0.24976 | -0.21864479 | 0 | 0.073 | 1 | 17 |
| efcab11 | 0.24976 | -0.18480583 | 0 | 0.073 | 1 | 17 |
| exoc2 | 0.24976 | -0.28579489 | 0 | 0.073 | 1 | 17 |
| flot2a | 0.24976 | -0.20329517 | 0 | 0.073 | 1 | 17 |
| g2e3 | 0.24976 | -0.20267154 | 0 | 0.073 | 1 | 17 |
| gtf2h2 | 0.24976 | -0.22061325 | 0 | 0.073 | 1 | 17 |
| hfm1 | 0.24976 | -0.26385092 | 0 | 0.073 | 1 | 17 |
| id2b | 0.24976 | -0.21621483 | 0 | 0.073 | 1 | 17 |
| igf2bp3 | 0.24976 | -0.20876933 | 0 | 0.073 | 1 | 17 |
| junba | 0.24976 | -0.20811883 | 0 | 0.073 | 1 | 17 |
| lbh | 0.24976 | -0.23062505 | 0 | 0.073 | 1 | 17 |
| mbd1b | 0.24976 | -0.22430974 | 0 | 0.073 | 1 | 17 |
| med7 | 0.24976 | -0.21925033 | 0 | 0.073 | 1 | 17 |
| oxr1a | 0.24976 | -0.26092678 | 0 | 0.073 | 1 | 17 |
| pigs | 0.24976 | -0.27303931 | 0 | 0.073 | 1 | 17 |
| pmm2 | 0.24976 | -0.20682428 | 0 | 0.073 | 1 | 17 |
| psme4b | 0.24976 | -0.24780666 | 0 | 0.073 | 1 | 17 |
| pxmp2 | 0.24976 | -0.25854946 | 0 | 0.073 | 1 | 17 |

| | | | | | | |
|---|---|---|---|---|---|---|
| rabggtb | 0.24976 | -0.22603626 | 0 | 0.073 | 1 | 17 |
| retsatl | 0.24976 | -0.28854899 | 0 | 0.073 | 1 | 17 |
| sdhaf4 | 0.24976 | -0.21297923 | 0 | 0.073 | 1 | 17 |
| sec61a1l | 0.24976 | -0.20007918 | 0 | 0.073 | 1 | 17 |
| si:ch211-196p9.1 | 0.24976 | -0.27292037 | 0 | 0.073 | 1 | 17 |
| si:ch73-174h16.4 | 0.24976 | -0.24861647 | 0 | 0.073 | 1 | 17 |
| si:ch73-376l24.2 | 0.24976 | -0.26535838 | 0 | 0.073 | 1 | 17 |
| slc25a33 | 0.24976 | -0.21894544 | 0 | 0.073 | 1 | 17 |
| ssbp3b | 0.24976 | -0.21367279 | 0 | 0.073 | 1 | 17 |
| st8sia7.1 | 0.24976 | -0.23305977 | 0 | 0.073 | 1 | 17 |
| tha1 | 0.24976 | -0.23045935 | 0 | 0.073 | 1 | 17 |
| tm9sf4 | 0.24976 | -0.21260516 | 0 | 0.073 | 1 | 17 |
| tmem201 | 0.24976 | -0.20476818 | 0 | 0.073 | 1 | 17 |
| tob1b | 0.24976 | -0.22943907 | 0 | 0.073 | 1 | 17 |
| tp53rk | 0.24976 | -0.23302826 | 0 | 0.073 | 1 | 17 |
| zgc:158316 | 0.24976 | -0.27921006 | 0 | 0.073 | 1 | 17 |
| zgc:65997 | 0.24976 | -0.15694097 | 0 | 0.073 | 1 | 17 |
| TMEM19 | 0.24976 | -0.21954079 | 0 | 0.073 | 1 | 17 |
| acp5a | 0.24976 | -0.20201074 | 0 | 0.073 | 1 | 17 |
| aifm4 | 0.24976 | -0.17520455 | 0 | 0.073 | 1 | 17 |
| chchd3a | 0.24976 | -0.18244621 | 0 | 0.073 | 1 | 17 |
| gnmt | 0.24976 | -0.15375279 | 0 | 0.073 | 1 | 17 |
| gsc | 0.24976 | -0.26097812 | 0 | 0.073 | 1 | 17 |
| hsd17b12a | 0.24976 | -0.20479742 | 0 | 0.073 | 1 | 17 |
| hyi | 0.24976 | -0.30556499 | 0 | 0.073 | 1 | 17 |
| ift46 | 0.24976 | -0.17906399 | 0 | 0.073 | 1 | 17 |
| ing3 | 0.24976 | -0.18238682 | 0 | 0.073 | 1 | 17 |
| itfg2 | 0.24976 | -0.17238549 | 0 | 0.073 | 1 | 17 |
| lamp2 | 0.24976 | -0.22691843 | 0 | 0.073 | 1 | 17 |
| larp4b | 0.24976 | -0.21835215 | 0 | 0.073 | 1 | 17 |
| magt1 | 0.24976 | -0.21326256 | 0 | 0.073 | 1 | 17 |
| mbd3a | 0.24976 | -0.2360106 | 0 | 0.073 | 1 | 17 |
| n4bp1 | 0.24976 | -0.2479913 | 0 | 0.073 | 1 | 17 |
| phpt1 | 0.24976 | -0.20583846 | 0 | 0.073 | 1 | 17 |
| plekhf2 | 0.24976 | -0.1563863 | 0 | 0.073 | 1 | 17 |
| plscr3b | 0.24976 | -0.22473036 | 0 | 0.073 | 1 | 17 |
| qpctla | 0.24976 | -0.18655954 | 0 | 0.073 | 1 | 17 |
| scamp2l | 0.24976 | -0.20038502 | 0 | 0.073 | 1 | 17 |
| si:ch211-114c12.2 | 0.24976 | -0.21799147 | 0 | 0.073 | 1 | 17 |
| si:dkey-21a6.5 | 0.24976 | -0.24502159 | 0 | 0.073 | 1 | 17 |
| spint1a | 0.24976 | -0.22010071 | 0 | 0.073 | 1 | 17 |
| tex261 | 0.24976 | -0.25344485 | 0 | 0.073 | 1 | 17 |
| tubgcp3 | 0.24976 | -0.21897446 | 0 | 0.073 | 1 | 17 |
| zgc:136254 | 0.24976 | -0.23867528 | 0 | 0.073 | 1 | 17 |
| atp6v1e1b | 0.24976 | -0.20431342 | 0 | 0.073 | 1 | 17 |
| si:dkey-93h22.8 | 0.24976 | -0.27585508 | 0 | 0.073 | 1 | 17 |
| bckdhb | 0.24976 | -0.2502291 | 0 | 0.073 | 1 | 17 |
| gtf3c6 | 0.24976 | -0.21340384 | 0 | 0.073 | 1 | 17 |

| | | | | | | |
|---|---|---|---|---|---|---|
| inpp5b | 0.24976 | -0.21263588 | 0 | 0.073 | 1 | 17 |
| larsb | 0.24976 | -0.19148618 | 0 | 0.073 | 1 | 17 |
| mkrn1 | 0.24976 | -0.26028753 | 0 | 0.073 | 1 | 17 |
| pde6h | 0.24976 | -0.28286374 | 0 | 0.073 | 1 | 17 |
| rdh10b | 0.24976 | -0.20705054 | 0 | 0.073 | 1 | 17 |
| rgs17 | 0.24976 | -0.25984976 | 0 | 0.073 | 1 | 17 |
| sec62 | 0.24976 | -0.24080782 | 0 | 0.073 | 1 | 17 |
| sypb | 0.24976 | -0.23002552 | 0 | 0.073 | 1 | 17 |
| tmem98 | 0.24976 | -0.27383116 | 0 | 0.073 | 1 | 17 |
| ube2w | 0.24976 | -0.22871396 | 0 | 0.073 | 1 | 17 |
| zc4h2 | 0.24976 | -0.22942542 | 0 | 0.073 | 1 | 17 |
| CR847984.1 | 0.24976 | -0.27463899 | 0 | 0.073 | 1 | 17 |
| CU467110.1 | 0.24976 | -0.22082098 | 0 | 0.073 | 1 | 17 |
| atp23 | 0.24976 | -0.24361545 | 0 | 0.073 | 1 | 17 |
| aup1 | 0.24976 | -0.23254416 | 0 | 0.073 | 1 | 17 |
| btbd2a | 0.24976 | -0.24566874 | 0 | 0.073 | 1 | 17 |
| ccdc47 | 0.24976 | -0.24447231 | 0 | 0.073 | 1 | 17 |
| copb2 | 0.24976 | -0.26701086 | 0 | 0.073 | 1 | 17 |
| coq10b | 0.24976 | -0.22699737 | 0 | 0.073 | 1 | 17 |
| crym | 0.24976 | -0.22447378 | 0 | 0.073 | 1 | 17 |
| exo1 | 0.24976 | -0.22861462 | 0 | 0.073 | 1 | 17 |
| faah2a | 0.24976 | -0.28304371 | 0 | 0.073 | 1 | 17 |
| gldc | 0.24976 | -0.24860109 | 0 | 0.073 | 1 | 17 |
| glmna | 0.24976 | -0.20436446 | 0 | 0.073 | 1 | 17 |
| homer2 | 0.24976 | -0.21388484 | 0 | 0.073 | 1 | 17 |
| lcmt1 | 0.24976 | -0.2320393 | 0 | 0.073 | 1 | 17 |
| metap1d | 0.24976 | -0.20120008 | 0 | 0.073 | 1 | 17 |
| mettl7a | 0.24976 | -0.26140409 | 0 | 0.073 | 1 | 17 |
| mier2 | 0.24976 | -0.24351767 | 0 | 0.073 | 1 | 17 |
| mkln1 | 0.24976 | -0.24222783 | 0 | 0.073 | 1 | 17 |
| morc3b | 0.24976 | -0.21047811 | 0 | 0.073 | 1 | 17 |
| mpped2 | 0.24976 | -0.21656362 | 0 | 0.073 | 1 | 17 |
| nf2a | 0.24976 | -0.21616615 | 0 | 0.073 | 1 | 17 |
| olah | 0.24976 | -0.19792598 | 0 | 0.073 | 1 | 17 |
| osgn1 | 0.24976 | -0.25359734 | 0 | 0.073 | 1 | 17 |
| pdcd11 | 0.24976 | -0.27515028 | 0 | 0.073 | 1 | 17 |
| pfn1 | 0.24976 | -0.163137 | 0 | 0.073 | 1 | 17 |
| rpusd4 | 0.24976 | -0.24232177 | 0 | 0.073 | 1 | 17 |
| selenow2a | 0.24976 | -0.2059035 | 0 | 0.073 | 1 | 17 |
| si:ch211-262h13.5 | 0.24976 | -0.22614731 | 0 | 0.073 | 1 | 17 |
| si:rp71-1g18.1 | 0.24976 | -0.26504514 | 0 | 0.073 | 1 | 17 |
| snrnp48 | 0.24976 | -0.26115604 | 0 | 0.073 | 1 | 17 |
| suv39h1b | 0.24976 | -0.23680025 | 0 | 0.073 | 1 | 17 |
| tacc3 | 0.24976 | -0.26576168 | 0 | 0.073 | 1 | 17 |
| tmem106c | 0.24976 | -0.26192645 | 0 | 0.073 | 1 | 17 |
| tmsb1 | 0.24976 | -0.23578739 | 0 | 0.073 | 1 | 17 |
| tollip | 0.24976 | -0.24367689 | 0 | 0.073 | 1 | 17 |
| yipf1 | 0.24976 | -0.25014909 | 0 | 0.073 | 1 | 17 |

| | | | | | | |
|---|---|---|---|---|---|---|
| zc3h13 | 0.24976 | -0.26756028 | 0 | 0.073 | 1 | 17 |
| zgc:193742 | 0.24976 | -0.23018043 | 0 | 0.073 | 1 | 17 |
| zgc:92481 | 0.24976 | -0.21442108 | 0 | 0.073 | 1 | 17 |
| ankrd28b | 0.24976 | -0.20910738 | 0 | 0.073 | 1 | 17 |
| rabac1 | 0.24976 | -0.22775936 | 0 | 0.073 | 1 | 17 |
| si:ch211-103n10.5 | 0.24976 | -0.24839567 | 0 | 0.073 | 1 | 17 |
| wdr92 | 0.24976 | -0.22663713 | 0 | 0.073 | 1 | 17 |
| zgc:86839 | 0.24976 | -0.24645887 | 0 | 0.073 | 1 | 17 |
| g6pd | 0.24976 | -0.27516026 | 0 | 0.073 | 1 | 17 |
| sem1 | 0.25004 | 0.199963114 | 0.571 | 0.366 | 1 | 17 |
| rcc1 | 0.25051 | 0.362313018 | 0.286 | 0.098 | 1 | 17 |
| gatd3a | 0.25236 | 0.36716121 | 0.286 | 0.098 | 1 | 17 |
| cops9 | 0.25247 | 0.348646456 | 0.286 | 0.098 | 1 | 17 |
| marcksl1b | 0.25249 | 0.356067635 | 0.286 | 0.098 | 1 | 17 |
| lamtor5 | 0.25287 | 0.347700812 | 0.357 | 0.146 | 1 | 17 |
| mrpl52 | 0.25314 | 0.2079611 | 0.286 | 0.122 | 1 | 17 |
| hormad1 | 0.25345 | 0.474029097 | 0.571 | 0.341 | 1 | 17 |
| psmb7 | 0.2541 | 0.428628804 | 0.643 | 0.39 | 1 | 17 |
| rpl10a | 0.256 | 0.186860727 | 1 | 0.878 | 1 | 17 |
| eef2b | 0.25662 | 0.392428521 | 0.714 | 0.463 | 1 | 17 |
| zgc:158803 | 0.25685 | -0.5268902 | 0.071 | 0.22 | 1 | 17 |
| myef2 | 0.25832 | 0.17114243 | 0.214 | 0.073 | 1 | 17 |
| ndufb4 | 0.25849 | 0.17760836 | 0.429 | 0.244 | 1 | 17 |
| lmnl3 | 0.25954 | 0.340604408 | 0.286 | 0.098 | 1 | 17 |
| idh3g | 0.25959 | 0.349416737 | 0.5 | 0.268 | 1 | 17 |
| cd2bp2 | 0.2599 | 0.38054332 | 0.357 | 0.146 | 1 | 17 |
| prelid1a | 0.26032 | 0.357549605 | 0.357 | 0.146 | 1 | 17 |
| rpl8 | 0.26138 | 0.188062536 | 1 | 0.878 | 1 | 17 |
| snrpd1 | 0.26161 | 0.38413421 | 0.786 | 0.561 | 1 | 17 |
| rpl26 | 0.26178 | 0.18094797 | 1 | 0.878 | 1 | 17 |
| yeats4 | 0.2623 | 0.375723953 | 0.357 | 0.146 | 1 | 17 |
| nelfcd | 0.2632 | 0.34398296 | 0.286 | 0.098 | 1 | 17 |
| prdx6 | 0.2634 | 0.333062507 | 0.286 | 0.098 | 1 | 17 |
| top2a | 0.2636 | 0.385338426 | 0.357 | 0.146 | 1 | 17 |
| hmgb1a | 0.26432 | 0.482733284 | 0.643 | 0.415 | 1 | 17 |
| fam204a | 0.2647 | 0.368063474 | 0.286 | 0.098 | 1 | 17 |
| cdca7a | 0.26479 | 0.460874083 | 0.357 | 0.146 | 1 | 17 |
| xrcc6 | 0.26643 | 0.412987254 | 0.357 | 0.146 | 1 | 17 |
| mt-nd2 | 0.26645 | -0.36415397 | 0.071 | 0.244 | 1 | 17 |
| naa50 | 0.26659 | 0.366829684 | 0.286 | 0.098 | 1 | 17 |
| nupr1a | 0.26665 | 0.454782419 | 0.357 | 0.146 | 1 | 17 |
| rfc4 | 0.26672 | 0.456531293 | 0.357 | 0.146 | 1 | 17 |
| spoplb | 0.26684 | 0.380314873 | 0.286 | 0.098 | 1 | 17 |
| hmgn3 | 0.26768 | 0.404096316 | 0.286 | 0.098 | 1 | 17 |
| ube2f | 0.26807 | 0.41149689 | 0.286 | 0.098 | 1 | 17 |
| stk31 | 0.26807 | 0.376707806 | 0.286 | 0.098 | 1 | 17 |
| vta1 | 0.26814 | 0.444001513 | 0.286 | 0.098 | 1 | 17 |
| tsn | 0.26818 | 0.435576476 | 0.286 | 0.098 | 1 | 17 |

| | | | | | | |
|---|---|---|---|---|---|---|
| calr3a | 0.26919 | 0.128553597 | 0.214 | 0.073 | 1 | 17 |
| ndufs5 | 0.2696 | 0.360884928 | 0.5 | 0.268 | 1 | 17 |
| got2b | 0.27064 | -0.50775398 | 0.071 | 0.244 | 1 | 17 |
| polr2d | 0.2729 | 0.349771814 | 0.5 | 0.268 | 1 | 17 |
| rtf2 | 0.273 | 0.279953642 | 0.429 | 0.22 | 1 | 17 |
| tuba8l4 | 0.2737 | 0.179559156 | 0.786 | 0.585 | 1 | 17 |
| rpl37 | 0.27379 | 0.170508815 | 1 | 0.878 | 1 | 17 |
| timm13 | 0.27438 | 0.293689066 | 0.429 | 0.22 | 1 | 17 |
| gar1 | 0.27457 | 0.229041046 | 0.286 | 0.122 | 1 | 17 |
| ldhba | 0.27462 | -0.51357829 | 0.143 | 0.293 | 1 | 17 |
| LOC101883594 | 0.27609 | 0.168036865 | 0.286 | 0.122 | 1 | 17 |
| cirbpa | 0.27623 | 0.163107104 | 0.929 | 0.756 | 1 | 17 |
| pnp5a | 0.27635 | 0.166981994 | 0.214 | 0.073 | 1 | 17 |
| sumo3a | 0.27881 | 0.138346777 | 0.643 | 0.463 | 1 | 17 |
| si:ch211-196l7.4 | 0.27939 | 0.149870349 | 0.357 | 0.195 | 1 | 17 |
| cct8 | 0.27951 | 0.177164743 | 0.643 | 0.439 | 1 | 17 |
| tomm6 | 0.28206 | -0.29888019 | 0.357 | 0.39 | 1 | 17 |
| pdia6 | 0.28213 | 0.145929162 | 0.214 | 0.073 | 1 | 17 |
| ube2ia | 0.28238 | 0.374870674 | 0.5 | 0.268 | 1 | 17 |
| si:dkey-151g10.6 | 0.28341 | 0.162593593 | 1 | 0.878 | 1 | 17 |
| snrpa1 | 0.28343 | 0.285817491 | 0.643 | 0.415 | 1 | 17 |
| uqcr10 | 0.28461 | -0.29040317 | 0.357 | 0.366 | 1 | 17 |
| dnd1 | 0.28502 | -0.47894249 | 0.214 | 0.341 | 1 | 17 |
| rps10 | 0.28621 | 0.24962742 | 0.929 | 0.805 | 1 | 17 |
| pole4 | 0.2865 | 0.127036337 | 0.357 | 0.195 | 1 | 17 |
| ccdc25 | 0.28716 | 0.300265248 | 0.429 | 0.22 | 1 | 17 |
| paip2b | 0.28745 | -0.16065332 | 0.571 | 0.537 | 1 | 17 |
| ube2l3b | 0.29024 | -0.4829849 | 0.071 | 0.244 | 1 | 17 |
| rps27.2 | 0.29043 | 0.263550453 | 0.714 | 0.488 | 1 | 17 |
| si:dkey-112a7.4 | 0.2915 | 0.167104141 | 0.214 | 0.073 | 1 | 17 |
| ewsr1b | 0.29225 | 0.283147096 | 0.429 | 0.22 | 1 | 17 |
| dnajc19 | 0.29593 | 0.149559975 | 0.214 | 0.073 | 1 | 17 |
| eif3s10 | 0.29612 | -0.47577233 | 0.071 | 0.244 | 1 | 17 |
| ppid | 0.29815 | 0.20799899 | 0.643 | 0.439 | 1 | 17 |
| tmed9 | 0.29891 | -0.53724248 | 0.071 | 0.22 | 1 | 17 |
| idh2 | 0.29955 | 0.247774149 | 0.5 | 0.293 | 1 | 17 |
| psmd8 | 0.29978 | 0.459690724 | 0.429 | 0.22 | 1 | 17 |
| spcs1 | 0.3026 | 0.320339762 | 0.429 | 0.22 | 1 | 17 |
| rpl31 | 0.30385 | 0.249045487 | 0.929 | 0.756 | 1 | 17 |
| ndufb3 | 0.30408 | 0.248872395 | 0.5 | 0.293 | 1 | 17 |
| chaf1a | 0.30628 | 0.115882402 | 0.5 | 0.341 | 1 | 17 |
| sumo2b | 0.30757 | 0.178871323 | 0.5 | 0.317 | 1 | 17 |
| rpa2 | 0.30978 | 0.34285491 | 0.571 | 0.341 | 1 | 17 |
| psma1 | 0.31316 | 0.480704642 | 0.429 | 0.22 | 1 | 17 |
| rbmx | 0.3137 | 0.317140339 | 0.429 | 0.22 | 1 | 17 |
| mrps9 | 0.31534 | -0.50915068 | 0.071 | 0.22 | 1 | 17 |
| slc25a3b | 0.31804 | 0.302537856 | 0.714 | 0.488 | 1 | 17 |
| eif1axa | 0.31831 | -0.41210974 | 0.071 | 0.244 | 1 | 17 |

| | | | | | | |
|---|---|---|---|---|---|---|
| srrm2 | 0.31856 | -0.43878352 | 0.286 | 0.415 | 1 | 17 |
| atp5mea | 0.3191 | 0.255393496 | 0.571 | 0.366 | 1 | 17 |
| rplp2 | 0.31977 | 0.302238109 | 0.857 | 0.683 | 1 | 17 |
| PAXBP1 | 0.32 | 0.206747213 | 0.214 | 0.073 | 1 | 17 |
| fbl | 0.32061 | -0.45185009 | 0.214 | 0.341 | 1 | 17 |
| uqcrq | 0.32305 | 0.376462524 | 0.571 | 0.341 | 1 | 17 |
| LOC110438491 | 0.32311 | 0.18826694 | 0.286 | 0.122 | 1 | 17 |
| rpf1 | 0.32319 | 0.386775911 | 0.571 | 0.341 | 1 | 17 |
| rer1 | 0.32341 | -0.47922372 | 0.071 | 0.244 | 1 | 17 |
| cnot9 | 0.32558 | -0.44084482 | 0.071 | 0.244 | 1 | 17 |
| vps4b | 0.32559 | -0.42331501 | 0.071 | 0.244 | 1 | 17 |
| gpatch11 | 0.32615 | 0.217195434 | 0.214 | 0.073 | 1 | 17 |
| rpl7 | 0.32627 | -0.17826847 | 0.786 | 0.805 | 1 | 17 |
| slc25a25a | 0.3266 | -0.39014813 | 0.143 | 0.195 | 1 | 17 |
| glud1b | 0.32667 | 0.251626518 | 0.357 | 0.171 | 1 | 17 |
| denr | 0.32759 | 0.341591158 | 0.429 | 0.22 | 1 | 17 |
| mrpl12 | 0.329 | 0.288129716 | 0.357 | 0.171 | 1 | 17 |
| CABZ01044053.1 | 0.32908 | 0.271015866 | 0.357 | 0.171 | 1 | 17 |
| stag3 | 0.3292 | 0.453165083 | 0.357 | 0.171 | 1 | 17 |
| si:dkey-180p18.9 | 0.32985 | 0.424352242 | 0.429 | 0.22 | 1 | 17 |
| pde6d | 0.33085 | 0.215368522 | 0.214 | 0.073 | 1 | 17 |
| ubb | 0.33193 | -0.27911705 | 0.357 | 0.39 | 1 | 17 |
| eif5b | 0.33216 | 0.228759494 | 0.429 | 0.244 | 1 | 17 |
| riok2 | 0.33282 | 0.31680403 | 0.357 | 0.171 | 1 | 17 |
| zw10 | 0.33325 | 0.227948833 | 0.214 | 0.073 | 1 | 17 |
| pdcd10a | 0.33546 | 0.233935044 | 0.214 | 0.073 | 1 | 17 |
| msto1 | 0.3371 | 0.164850493 | 0.214 | 0.073 | 1 | 17 |
| psma3 | 0.33788 | 0.417759521 | 0.429 | 0.22 | 1 | 17 |
| dmc1 | 0.33949 | 0.357144347 | 0.214 | 0.073 | 1 | 17 |
| thoc3 | 0.34022 | 0.186434891 | 0.214 | 0.073 | 1 | 17 |
| c1qbp | 0.34188 | -0.48039066 | 0.143 | 0.293 | 1 | 17 |
| suclg1 | 0.34216 | 0.206711024 | 0.286 | 0.122 | 1 | 17 |
| hibadhb | 0.34374 | 0.210258069 | 0.214 | 0.073 | 1 | 17 |
| FQ976913.1 | 0.34386 | 0.274899546 | 0.286 | 0.122 | 1 | 17 |
| gmfb | 0.34409 | 0.201654956 | 0.214 | 0.073 | 1 | 17 |
| tax1bp3 | 0.34466 | 0.106451995 | 0.286 | 0.146 | 1 | 17 |
| zgc:86598 | 0.34521 | -0.45281609 | 0.214 | 0.415 | 1 | 17 |
| crp2 | 0.34534 | -0.47659756 | 0.071 | 0.22 | 1 | 17 |
| setb | 0.34608 | 0.160804089 | 0.857 | 0.683 | 1 | 17 |
| rnf7 | 0.34739 | 0.223564249 | 0.214 | 0.073 | 1 | 17 |
| sucla2 | 0.34764 | 0.225231488 | 0.286 | 0.122 | 1 | 17 |
| dync1i2a | 0.34806 | 0.200983995 | 0.214 | 0.073 | 1 | 17 |
| vkorc1l1 | 0.34912 | 0.290106682 | 0.214 | 0.073 | 1 | 17 |
| ifi45 | 0.35016 | 0.178301827 | 0.357 | 0.195 | 1 | 17 |
| pcbp2 | 0.35028 | -0.44828924 | 0.071 | 0.22 | 1 | 17 |
| eif3d | 0.3504 | 0.317011581 | 0.5 | 0.293 | 1 | 17 |
| wdr12 | 0.35114 | -0.43942167 | 0.143 | 0.268 | 1 | 17 |
| xpc | 0.35126 | 0.216964886 | 0.214 | 0.073 | 1 | 17 |

| | | | | | | |
|---|---|---|---|---|---|---|
| tcp1 | 0.35159 | 0.221148931 | 0.714 | 0.512 | 1 | 17 |
| zgc:101851 | 0.35287 | -0.10949168 | 0.286 | 0.22 | 1 | 17 |
| adi1 | 0.35307 | 0.258613369 | 0.286 | 0.122 | 1 | 17 |
| ndufa1 | 0.35457 | 0.195769615 | 0.5 | 0.317 | 1 | 17 |
| arf2b | 0.355 | 0.427610827 | 0.286 | 0.122 | 1 | 17 |
| chchd7 | 0.35554 | 0.286446329 | 0.214 | 0.073 | 1 | 17 |
| dctn5 | 0.35571 | 0.22815801 | 0.214 | 0.073 | 1 | 17 |
| actb2 | 0.35584 | -0.44533727 | 0.286 | 0.488 | 1 | 17 |
| sae1 | 0.35606 | 0.155933196 | 0.286 | 0.146 | 1 | 17 |
| mt-atp6 | 0.35651 | -0.41071044 | 0.071 | 0.22 | 1 | 17 |
| nars | 0.35697 | -0.41399571 | 0.214 | 0.317 | 1 | 17 |
| xrcc5 | 0.35755 | 0.257075256 | 0.286 | 0.122 | 1 | 17 |
| pgk1 | 0.35846 | 0.317103525 | 0.357 | 0.171 | 1 | 17 |
| prpf39 | 0.35854 | 0.319555707 | 0.357 | 0.171 | 1 | 17 |
| mcm6 | 0.35888 | 0.176814716 | 0.357 | 0.195 | 1 | 17 |
| sec61g | 0.3592 | -0.18170502 | 0.429 | 0.366 | 1 | 17 |
| prdx4 | 0.35946 | 0.259640218 | 0.286 | 0.122 | 1 | 17 |
| pgghg | 0.36021 | 0.228226655 | 0.214 | 0.073 | 1 | 17 |
| cct3 | 0.36025 | 0.108628334 | 0.643 | 0.488 | 1 | 17 |
| dhx15 | 0.36038 | 0.258132966 | 0.286 | 0.122 | 1 | 17 |
| afmid | 0.36253 | 0.265617802 | 0.214 | 0.073 | 1 | 17 |
| nolc1 | 0.36265 | -0.31164778 | 0.214 | 0.244 | 1 | 17 |
| ranbp3b | 0.36366 | 0.238184862 | 0.214 | 0.073 | 1 | 17 |
| npepl1 | 0.36387 | 0.317723376 | 0.357 | 0.171 | 1 | 17 |
| rhoac | 0.36434 | 0.213734824 | 0.214 | 0.073 | 1 | 17 |
| rnaseh1 | 0.36435 | 0.280394035 | 0.286 | 0.122 | 1 | 17 |
| rps12 | 0.36453 | 0.143999245 | 1 | 0.902 | 1 | 17 |
| nup93 | 0.36484 | 0.147872881 | 0.286 | 0.146 | 1 | 17 |
| dynll2a | 0.36508 | -0.18679142 | 0.071 | 0.195 | 1 | 17 |
| spdya | 0.36546 | 0.405234534 | 0.286 | 0.122 | 1 | 17 |
| si:dkey-184a18.5 | 0.36613 | 0.248761506 | 0.214 | 0.073 | 1 | 17 |
| kdm7aa | 0.36674 | 0.245757342 | 0.214 | 0.073 | 1 | 17 |
| pwp2h | 0.36769 | 0.441056124 | 0.214 | 0.073 | 1 | 17 |
| psma6a | 0.36786 | 0.141763645 | 0.429 | 0.268 | 1 | 17 |
| smfn | 0.36853 | 0.210224444 | 0.5 | 0.317 | 1 | 17 |
| hnrnpabb | 0.37038 | -0.24992771 | 0.643 | 0.707 | 1 | 17 |
| mcts1 | 0.37042 | 0.400715686 | 0.357 | 0.171 | 1 | 17 |
| pfn2 | 0.3715 | 0.234690924 | 0.214 | 0.073 | 1 | 17 |
| ankrd10a | 0.37155 | 0.389688168 | 0.357 | 0.171 | 1 | 17 |
| ssr3 | 0.37155 | -0.24539085 | 0.143 | 0.098 | 1 | 17 |
| copz1 | 0.37183 | 0.375658379 | 0.214 | 0.073 | 1 | 17 |
| h1fx | 0.37231 | 0.183907479 | 0.214 | 0.073 | 1 | 17 |
| prdx1 | 0.37308 | 0.341609885 | 0.5 | 0.293 | 1 | 17 |
| si:ch211-212d10.2 | 0.37319 | 0.415827522 | 0.357 | 0.171 | 1 | 17 |
| psmc3 | 0.37321 | 0.102236878 | 0.357 | 0.22 | 1 | 17 |
| si:ch1073-325m22.2 | 0.37452 | 0.126737853 | 0.5 | 0.341 | 1 | 17 |
| LOC101886261 | 0.37527 | 0.356239318 | 0.214 | 0.073 | 1 | 17 |
| cnot2 | 0.37682 | 0.225204793 | 0.214 | 0.073 | 1 | 17 |

| | | | | | | |
|---|---|---|---|---|---|---|
| tspan3a | 0.37694 | 0.326963169 | 0.214 | 0.073 | 1 | 17 |
| surf1 | 0.37699 | 0.230631264 | 0.214 | 0.073 | 1 | 17 |
| AL935174.5 | 0.3774 | 0.272743306 | 0.214 | 0.073 | 1 | 17 |
| nosip | 0.37861 | 0.260176244 | 0.214 | 0.073 | 1 | 17 |
| nsmce4a | 0.37882 | 0.268701541 | 0.214 | 0.073 | 1 | 17 |
| LOC103909115 | 0.37883 | 0.271161645 | 0.214 | 0.073 | 1 | 17 |
| srsf7a | 0.37897 | 0.321229803 | 0.286 | 0.122 | 1 | 17 |
| tcea2 | 0.37924 | 0.430036331 | 0.214 | 0.073 | 1 | 17 |
| stard14 | 0.37937 | 0.213693445 | 0.214 | 0.073 | 1 | 17 |
| cox8a | 0.3803 | 0.171234884 | 0.429 | 0.268 | 1 | 17 |
| zgc:92594 | 0.38083 | 0.254763878 | 0.214 | 0.073 | 1 | 17 |
| synrg | 0.38094 | 0.276516581 | 0.214 | 0.073 | 1 | 17 |
| si:ch211-51e12.7 | 0.38145 | 0.256085103 | 0.643 | 0.439 | 1 | 17 |
| nsa2 | 0.38176 | 0.351408963 | 0.5 | 0.293 | 1 | 17 |
| mapk3 | 0.38228 | 0.252466914 | 0.214 | 0.073 | 1 | 17 |
| ppp1cab | 0.38232 | 0.284534509 | 0.214 | 0.073 | 1 | 17 |
| hnrnpc | 0.38254 | 0.289613006 | 0.286 | 0.122 | 1 | 17 |
| CT573476.1 | 0.38256 | 0.246283189 | 0.214 | 0.073 | 1 | 17 |
| ddx5 | 0.38264 | -0.35155576 | 0.286 | 0.366 | 1 | 17 |
| cwc27 | 0.38318 | 0.299106431 | 0.214 | 0.073 | 1 | 17 |
| anp32b | 0.3835 | -0.42994954 | 0.214 | 0.415 | 1 | 17 |
| rnf17 | 0.3836 | 0.260391432 | 0.214 | 0.073 | 1 | 17 |
| arl6ip4 | 0.3851 | 0.321458087 | 0.286 | 0.122 | 1 | 17 |
| aco2 | 0.38549 | 0.316768284 | 0.214 | 0.073 | 1 | 17 |
| rpl21 | 0.38556 | 0.121388277 | 1 | 0.902 | 1 | 17 |
| cnp | 0.38562 | 0.276847465 | 0.429 | 0.244 | 1 | 17 |
| h2afvb | 0.38566 | 0.218333034 | 0.929 | 0.78 | 1 | 17 |
| smn1 | 0.38627 | 0.306913161 | 0.286 | 0.122 | 1 | 17 |
| ptrh1 | 0.3865 | 0.25481657 | 0.214 | 0.073 | 1 | 17 |
| si:ch73-1a9.3 | 0.38653 | 0.240985741 | 0.5 | 0.317 | 1 | 17 |
| gstm.2 | 0.38709 | 0.275681877 | 0.214 | 0.073 | 1 | 17 |
| rpl18a | 0.38761 | 0.111369381 | 1 | 0.902 | 1 | 17 |
| CABZ01074453.1 | 0.38767 | 0.229124643 | 0.214 | 0.073 | 1 | 17 |
| aurkaip1 | 0.38777 | 0.36491457 | 0.214 | 0.073 | 1 | 17 |
| ddx27 | 0.38819 | 0.382870491 | 0.286 | 0.122 | 1 | 17 |
| gnaia | 0.3883 | 0.278502577 | 0.214 | 0.073 | 1 | 17 |
| tfdp1a | 0.38863 | 0.315225045 | 0.214 | 0.073 | 1 | 17 |
| srsf10b | 0.38943 | 0.346423916 | 0.214 | 0.073 | 1 | 17 |
| polr2gl | 0.3897 | 0.291046909 | 0.214 | 0.073 | 1 | 17 |
| zgc:101858 | 0.39017 | 0.33639714 | 0.214 | 0.073 | 1 | 17 |
| cdr2l | 0.39038 | 0.319656771 | 0.214 | 0.073 | 1 | 17 |
| msrb1a | 0.39039 | 0.252441673 | 0.214 | 0.073 | 1 | 17 |
| eef1a1a | 0.39093 | 0.335038528 | 0.214 | 0.073 | 1 | 17 |
| sephs2 | 0.3915 | 0.319741313 | 0.214 | 0.073 | 1 | 17 |
| zranb2 | 0.39195 | 0.330945821 | 0.286 | 0.122 | 1 | 17 |
| alad | 0.39198 | 0.31204075 | 0.214 | 0.073 | 1 | 17 |
| LOC103908877 | 0.39208 | 0.374223801 | 0.286 | 0.122 | 1 | 17 |
| cdc73 | 0.39215 | 0.333981038 | 0.214 | 0.073 | 1 | 17 |

| | | | | | | |
|---|---|---|---|---|---|---|
| prpf40a | 0.39224 | -0.13237704 | 0.214 | 0.171 | 1 | 17 |
| hes6 | 0.39231 | 0.303294049 | 0.214 | 0.073 | 1 | 17 |
| reep5 | 0.39237 | 0.339547496 | 0.214 | 0.073 | 1 | 17 |
| cdk2 | 0.39259 | 0.190839394 | 0.286 | 0.146 | 1 | 17 |
| ehd1b | 0.39259 | 0.353885041 | 0.214 | 0.073 | 1 | 17 |
| wdr3 | 0.39264 | 0.316229005 | 0.214 | 0.073 | 1 | 17 |
| tfpt | 0.39267 | 0.289790096 | 0.214 | 0.073 | 1 | 17 |
| tpd52l2b | 0.39273 | 0.309502128 | 0.214 | 0.073 | 1 | 17 |
| exosc5 | 0.39277 | 0.358781676 | 0.214 | 0.073 | 1 | 17 |
| mrpl38 | 0.3928 | 0.293267484 | 0.214 | 0.073 | 1 | 17 |
| arpc4 | 0.3928 | 0.329877453 | 0.214 | 0.073 | 1 | 17 |
| rsl1d1 | 0.39291 | 0.267681797 | 0.643 | 0.439 | 1 | 17 |
| park7 | 0.39325 | 0.399883638 | 0.286 | 0.122 | 1 | 17 |
| ccdc130 | 0.39357 | 0.383912702 | 0.286 | 0.122 | 1 | 17 |
| nifk | 0.3938 | 0.331949187 | 0.286 | 0.122 | 1 | 17 |
| sigmar1 | 0.39429 | -0.43742347 | 0.071 | 0.22 | 1 | 17 |
| acot13 | 0.39458 | 0.374003581 | 0.286 | 0.122 | 1 | 17 |
| st13 | 0.39462 | 0.339586679 | 0.286 | 0.122 | 1 | 17 |
| atf7ip2 | 0.39516 | 0.360178724 | 0.286 | 0.122 | 1 | 17 |
| fam162a | 0.39516 | 0.328074098 | 0.286 | 0.122 | 1 | 17 |
| ndufa8 | 0.39565 | 0.202560909 | 0.571 | 0.39 | 1 | 17 |
| pfdn4 | 0.39567 | -0.16624618 | 0.357 | 0.317 | 1 | 17 |
| arl1 | 0.39598 | 0.348891132 | 0.286 | 0.122 | 1 | 17 |
| tbcb | 0.39601 | 0.352350013 | 0.286 | 0.122 | 1 | 17 |
| cenps | 0.3961 | 0.33055177 | 0.286 | 0.122 | 1 | 17 |
| sh3glb2b | 0.39635 | 0.328654768 | 0.286 | 0.122 | 1 | 17 |
| spcs2 | 0.39674 | -0.1430971 | 0.214 | 0.171 | 1 | 17 |
| elocb | 0.39711 | 0.159669479 | 0.5 | 0.341 | 1 | 17 |
| rtca | 0.39879 | -0.37105599 | 0.286 | 0.39 | 1 | 17 |
| mad2l1 | 0.40149 | -0.44322955 | 0.143 | 0.268 | 1 | 17 |
| parp3 | 0.4016 | 0.185484876 | 0.429 | 0.268 | 1 | 17 |
| rps25 | 0.40413 | 0.222269751 | 0.929 | 0.805 | 1 | 17 |
| si:dkey-33c12.4 | 0.40571 | 0.180580709 | 0.429 | 0.268 | 1 | 17 |
| rpl14 | 0.40889 | 0.310688499 | 0.786 | 0.61 | 1 | 17 |
| emg1 | 0.40971 | -0.34891464 | 0.071 | 0.22 | 1 | 17 |
| mob4 | 0.41043 | -0.40906768 | 0.071 | 0.22 | 1 | 17 |
| mbd3b | 0.41099 | 0.209821319 | 0.357 | 0.195 | 1 | 17 |
| lsm6 | 0.41157 | 0.188942331 | 0.786 | 0.61 | 1 | 17 |
| dldh | 0.41181 | -0.39934682 | 0.071 | 0.22 | 1 | 17 |
| epcam | 0.41205 | -0.43276585 | 0.071 | 0.195 | 1 | 17 |
| ubxn1 | 0.414 | -0.3671047 | 0.071 | 0.22 | 1 | 17 |
| znf593 | 0.41456 | -0.37742402 | 0.071 | 0.22 | 1 | 17 |
| gid8b | 0.41457 | 0.158733941 | 0.286 | 0.146 | 1 | 17 |
| xbp1 | 0.41473 | -0.40432059 | 0.071 | 0.171 | 1 | 17 |
| gtf2a2 | 0.41481 | -0.37599637 | 0.071 | 0.22 | 1 | 17 |
| pdap1b | 0.41556 | 0.116123054 | 0.143 | 0.049 | 1 | 17 |
| cd63 | 0.41667 | -0.26138031 | 0.143 | 0.122 | 1 | 17 |
| nop53 | 0.41674 | 0.101979211 | 0.357 | 0.22 | 1 | 17 |

| | | | | | | |
|---|---|---|---|---|---|---|
| zgc:194246 | 0.41859 | 0.294269 | 0.429 | 0.244 | 1 | 17 |
| stmn1b | 0.42004 | 0.233780558 | 0.357 | 0.195 | 1 | 17 |
| mrpl41 | 0.42115 | 0.147180135 | 0.214 | 0.098 | 1 | 17 |
| utp23 | 0.42395 | 0.11363965 | 0.357 | 0.22 | 1 | 17 |
| rplp2l | 0.42545 | 0.150989303 | 1 | 0.927 | 1 | 17 |
| tmem176l.1 | 0.42917 | 0.117784784 | 0.143 | 0.049 | 1 | 17 |
| mcm7 | 0.43199 | -0.16965529 | 0.357 | 0.341 | 1 | 17 |
| cse1l | 0.43246 | -0.42034771 | 0.143 | 0.293 | 1 | 17 |
| ccdc6a | 0.43342 | 0.120030478 | 0.214 | 0.098 | 1 | 17 |
| lsm12a | 0.43421 | 0.360873677 | 0.429 | 0.244 | 1 | 17 |
| lin9 | 0.43499 | 0.114057187 | 0.143 | 0.049 | 1 | 17 |
| uqcrfs1 | 0.43524 | -0.22073128 | 0.357 | 0.366 | 1 | 17 |
| psme1 | 0.4353 | 0.121298932 | 0.214 | 0.098 | 1 | 17 |
| cops5 | 0.4355 | 0.313765442 | 0.429 | 0.244 | 1 | 17 |
| si:dkeyp-75h12.5 | 0.4356 | 0.11378488 | 0.143 | 0.049 | 1 | 17 |
| atp5if1b | 0.43574 | 0.241807067 | 0.357 | 0.195 | 1 | 17 |
| eif2s3 | 0.43672 | 0.222826364 | 0.357 | 0.195 | 1 | 17 |
| aurkb | 0.43961 | 0.244122565 | 0.357 | 0.195 | 1 | 17 |
| pfdn1 | 0.43986 | 0.253446324 | 0.5 | 0.317 | 1 | 17 |
| rnf10 | 0.44178 | 0.110892105 | 0.143 | 0.049 | 1 | 17 |
| ripply2 | 0.44202 | 0.32896259 | 0.143 | 0.049 | 1 | 17 |
| ddx18 | 0.44295 | 0.15988429 | 0.5 | 0.341 | 1 | 17 |
| caprin1a | 0.44367 | 0.163915909 | 0.286 | 0.146 | 1 | 17 |
| mrps26 | 0.4469 | 0.172857746 | 0.214 | 0.098 | 1 | 17 |
| hnrnpl2 | 0.44843 | -0.35372044 | 0.214 | 0.293 | 1 | 17 |
| fkbp3 | 0.44996 | 0.245851315 | 0.571 | 0.39 | 1 | 17 |
| spc24 | 0.45098 | 0.18856245 | 0.286 | 0.146 | 1 | 17 |
| mrps18b | 0.45105 | 0.164637235 | 0.214 | 0.098 | 1 | 17 |
| txlna | 0.45107 | 0.217356563 | 0.286 | 0.146 | 1 | 17 |
| pbdc1 | 0.45176 | 0.181607198 | 0.429 | 0.268 | 1 | 17 |
| si:ch73-95l15.5 | 0.4538 | 0.177067311 | 0.214 | 0.098 | 1 | 17 |
| zgc:101679 | 0.4545 | 0.110352449 | 0.143 | 0.049 | 1 | 17 |
| chek2 | 0.45464 | 0.141016972 | 0.214 | 0.098 | 1 | 17 |
| CU694197.1 | 0.45544 | 0.125778974 | 0.214 | 0.098 | 1 | 17 |
| prpf31 | 0.45637 | -0.39335247 | 0.143 | 0.244 | 1 | 17 |
| bcap31 | 0.45656 | 0.132067118 | 0.214 | 0.098 | 1 | 17 |
| pam16 | 0.45702 | 0.175006127 | 0.286 | 0.146 | 1 | 17 |
| znhit3 | 0.45931 | 0.17005252 | 0.214 | 0.098 | 1 | 17 |
| BX571701.1 | 0.46041 | 0.168010845 | 0.214 | 0.098 | 1 | 17 |
| zgc:194224 | 0.46094 | 0.104200866 | 0.143 | 0.049 | 1 | 17 |
| treh | 0.46142 | 0.353223243 | 0.143 | 0.049 | 1 | 17 |
| bxdc2 | 0.46174 | 0.209945968 | 0.286 | 0.146 | 1 | 17 |
| mesd | 0.46213 | 0.351991778 | 0.143 | 0.049 | 1 | 17 |
| rtn4a | 0.46214 | 0.178640727 | 0.286 | 0.146 | 1 | 17 |
| mrto4 | 0.46227 | 0.265356241 | 0.5 | 0.317 | 1 | 17 |
| supt3h | 0.46322 | 0.273552237 | 0.357 | 0.195 | 1 | 17 |
| ndufs3 | 0.46407 | 0.238810657 | 0.357 | 0.195 | 1 | 17 |
| BX005392.2 | 0.46455 | 0.183369946 | 0.071 | 0.024 | 1 | 17 |

| | | | | | | |
|---|---|---|---|---|---|---|
| BX465862.1 | 0.46455 | 0.122485955 | 0.071 | 0.024 | 1 | 17 |
| CABZ01072848.1 | 0.46455 | 0.170047787 | 0.071 | 0.024 | 1 | 17 |
| CABZ01078607.1 | 0.46455 | 0.118354465 | 0.071 | 0.024 | 1 | 17 |
| CR749162.1 | 0.46455 | 0.191202843 | 0.071 | 0.024 | 1 | 17 |
| CR749763.5 | 0.46455 | 0.132685599 | 0.071 | 0.024 | 1 | 17 |
| FP085398.1 | 0.46455 | 0.149094258 | 0.071 | 0.024 | 1 | 17 |
| LO018197.1 | 0.46455 | 0.120266892 | 0.071 | 0.024 | 1 | 17 |
| LOC101885027 | 0.46455 | 0.115535127 | 0.071 | 0.024 | 1 | 17 |
| LOC110439835 | 0.46455 | 0.129474046 | 0.071 | 0.024 | 1 | 17 |
| LOC797168 | 0.46455 | 0.107464336 | 0.071 | 0.024 | 1 | 17 |
| MCUR1 | 0.46455 | 0.213178136 | 0.071 | 0.024 | 1 | 17 |
| XLOC-007541 | 0.46455 | 0.122826738 | 0.071 | 0.024 | 1 | 17 |
| abcf2b | 0.46455 | 0.123113883 | 0.071 | 0.024 | 1 | 17 |
| acad8 | 0.46455 | 0.129258647 | 0.071 | 0.024 | 1 | 17 |
| acot9.1 | 0.46455 | 0.100995655 | 0.071 | 0.024 | 1 | 17 |
| amt | 0.46455 | 0.121727301 | 0.071 | 0.024 | 1 | 17 |
| angel2 | 0.46455 | 0.156870991 | 0.071 | 0.024 | 1 | 17 |
| arglu1b | 0.46455 | 0.101422914 | 0.071 | 0.024 | 1 | 17 |
| arhgap4b | 0.46455 | 0.125684455 | 0.071 | 0.024 | 1 | 17 |
| arhgef16 | 0.46455 | 0.13245342 | 0.071 | 0.024 | 1 | 17 |
| arih1l | 0.46455 | 0.148024121 | 0.071 | 0.024 | 1 | 17 |
| atg4b | 0.46455 | 0.205550086 | 0.071 | 0.024 | 1 | 17 |
| atp2b2 | 0.46455 | 0.12131046 | 0.071 | 0.024 | 1 | 17 |
| bada | 0.46455 | 0.126305227 | 0.071 | 0.024 | 1 | 17 |
| bcl2l1 | 0.46455 | 0.114892249 | 0.071 | 0.024 | 1 | 17 |
| bub1ba | 0.46455 | 0.148918765 | 0.071 | 0.024 | 1 | 17 |
| carm1 | 0.46455 | 0.114053441 | 0.071 | 0.024 | 1 | 17 |
| ccdc90b | 0.46455 | 0.127790559 | 0.071 | 0.024 | 1 | 17 |
| cdk5rap2 | 0.46455 | 0.102884067 | 0.071 | 0.024 | 1 | 17 |
| cep63 | 0.46455 | 0.20230532 | 0.071 | 0.024 | 1 | 17 |
| chic1 | 0.46455 | 0.143404401 | 0.071 | 0.024 | 1 | 17 |
| chid1 | 0.46455 | 0.170441675 | 0.071 | 0.024 | 1 | 17 |
| cicb | 0.46455 | 0.111772218 | 0.071 | 0.024 | 1 | 17 |
| cipcb | 0.46455 | 0.166138882 | 0.071 | 0.024 | 1 | 17 |
| cisd2 | 0.46455 | 0.207208908 | 0.071 | 0.024 | 1 | 17 |
| cry1aa | 0.46455 | 0.114674343 | 0.071 | 0.024 | 1 | 17 |
| cul3b | 0.46455 | 0.132773978 | 0.071 | 0.024 | 1 | 17 |
| dcaf11 | 0.46455 | 0.150208664 | 0.071 | 0.024 | 1 | 17 |
| ddr1 | 0.46455 | 0.118094933 | 0.071 | 0.024 | 1 | 17 |
| dnajc21 | 0.46455 | 0.182294937 | 0.071 | 0.024 | 1 | 17 |
| dnajc4 | 0.46455 | 0.126568826 | 0.071 | 0.024 | 1 | 17 |
| dync1li1 | 0.46455 | 0.169029732 | 0.071 | 0.024 | 1 | 17 |
| elp3 | 0.46455 | 0.16763381 | 0.071 | 0.024 | 1 | 17 |
| fgf12a | 0.46455 | 0.123951372 | 0.071 | 0.024 | 1 | 17 |
| golga2 | 0.46455 | 0.103329408 | 0.071 | 0.024 | 1 | 17 |
| gpr107 | 0.46455 | 0.121695866 | 0.071 | 0.024 | 1 | 17 |
| gtf3c4 | 0.46455 | 0.122606474 | 0.071 | 0.024 | 1 | 17 |
| htatip2 | 0.46455 | 0.117875944 | 0.071 | 0.024 | 1 | 17 |

| | | | | | | |
|---|---|---|---|---|---|---|
| impdh2 | 0.46455 | 0.117836019 | 0.071 | 0.024 | 1 | 17 |
| ints2 | 0.46455 | 0.11849156 | 0.071 | 0.024 | 1 | 17 |
| jade3 | 0.46455 | 0.123870362 | 0.071 | 0.024 | 1 | 17 |
| katna1 | 0.46455 | 0.108573302 | 0.071 | 0.024 | 1 | 17 |
| kif18a | 0.46455 | 0.120676329 | 0.071 | 0.024 | 1 | 17 |
| kif20ba | 0.46455 | 0.142625234 | 0.071 | 0.024 | 1 | 17 |
| kpna1 | 0.46455 | 0.100691884 | 0.071 | 0.024 | 1 | 17 |
| l3mbtl2 | 0.46455 | 0.117273901 | 0.071 | 0.024 | 1 | 17 |
| lrrfip2 | 0.46455 | 0.105563105 | 0.071 | 0.024 | 1 | 17 |
| lto1 | 0.46455 | 0.150069915 | 0.071 | 0.024 | 1 | 17 |
| lyrm2 | 0.46455 | 0.100200276 | 0.071 | 0.024 | 1 | 17 |
| mdga2a | 0.46455 | 0.119632462 | 0.071 | 0.024 | 1 | 17 |
| med17 | 0.46455 | 0.118784737 | 0.071 | 0.024 | 1 | 17 |
| mettl9 | 0.46455 | 0.190391291 | 0.071 | 0.024 | 1 | 17 |
| mgat4b | 0.46455 | 0.102800612 | 0.071 | 0.024 | 1 | 17 |
| mms22l | 0.46455 | 0.161277124 | 0.071 | 0.024 | 1 | 17 |
| mppe1 | 0.46455 | 0.182349225 | 0.071 | 0.024 | 1 | 17 |
| mrpl10 | 0.46455 | 0.142190217 | 0.071 | 0.024 | 1 | 17 |
| mtf2 | 0.46455 | 0.1187491 | 0.071 | 0.024 | 1 | 17 |
| mycla | 0.46455 | 0.139239027 | 0.071 | 0.024 | 1 | 17 |
| nadsyn1.1 | 0.46455 | 0.154210072 | 0.071 | 0.024 | 1 | 17 |
| naprt | 0.46455 | 0.147008937 | 0.071 | 0.024 | 1 | 17 |
| necap2 | 0.46455 | 0.173545263 | 0.071 | 0.024 | 1 | 17 |
| nelfe | 0.46455 | 0.140992489 | 0.071 | 0.024 | 1 | 17 |
| nfya | 0.46455 | 0.1778522 | 0.071 | 0.024 | 1 | 17 |
| nitr13 | 0.46455 | 0.131338918 | 0.071 | 0.024 | 1 | 17 |
| nop14 | 0.46455 | 0.179922003 | 0.071 | 0.024 | 1 | 17 |
| nsun3 | 0.46455 | 0.114371159 | 0.071 | 0.024 | 1 | 17 |
| parga | 0.46455 | 0.129269393 | 0.071 | 0.024 | 1 | 17 |
| pdf | 0.46455 | 0.153734511 | 0.071 | 0.024 | 1 | 17 |
| pih1d3 | 0.46455 | 0.11648763 | 0.071 | 0.024 | 1 | 17 |
| plcxd1 | 0.46455 | 0.120450926 | 0.071 | 0.024 | 1 | 17 |
| plekha3 | 0.46455 | 0.107085459 | 0.071 | 0.024 | 1 | 17 |
| plpp2b | 0.46455 | 0.143492863 | 0.071 | 0.024 | 1 | 17 |
| ppfia1 | 0.46455 | 0.120679015 | 0.071 | 0.024 | 1 | 17 |
| ppifa | 0.46455 | 0.125725651 | 0.071 | 0.024 | 1 | 17 |
| prr13 | 0.46455 | 0.177180775 | 0.071 | 0.024 | 1 | 17 |
| pter | 0.46455 | 0.107644466 | 0.071 | 0.024 | 1 | 17 |
| rasa3 | 0.46455 | 0.104243116 | 0.071 | 0.024 | 1 | 17 |
| rgs7bpb | 0.46455 | 0.149926526 | 0.071 | 0.024 | 1 | 17 |
| sccpdhb | 0.46455 | 0.14973642 | 0.071 | 0.024 | 1 | 17 |
| scoca | 0.46455 | 0.103514746 | 0.071 | 0.024 | 1 | 17 |
| selenot2 | 0.46455 | 0.172266747 | 0.071 | 0.024 | 1 | 17 |
| si:ch1073-143l10.2 | 0.46455 | 0.132971 | 0.071 | 0.024 | 1 | 17 |
| si:ch1073-357b18.4 | 0.46455 | 0.102437182 | 0.071 | 0.024 | 1 | 17 |
| si:ch211-139d20.3 | 0.46455 | 0.149584146 | 0.071 | 0.024 | 1 | 17 |
| si:ch211-188c16.1 | 0.46455 | 0.136263466 | 0.071 | 0.024 | 1 | 17 |
| si:ch211-214c7.5 | 0.46455 | 0.121762087 | 0.071 | 0.024 | 1 | 17 |

| | | | | | | |
|---|---|---|---|---|---|---|
| si:ch211-225g23.1 | 0.46455 | 0.152818519 | 0.071 | 0.024 | 1 | 17 |
| si:ch73-308l14.2 | 0.46455 | 0.195958404 | 0.071 | 0.024 | 1 | 17 |
| si:ch73-46j18.5 | 0.46455 | 0.108742118 | 0.071 | 0.024 | 1 | 17 |
| si:dkey-177p2.6 | 0.46455 | 0.127764876 | 0.071 | 0.024 | 1 | 17 |
| si:dkey-239h2.3 | 0.46455 | 0.121835811 | 0.071 | 0.024 | 1 | 17 |
| si:dkey-266m15.6 | 0.46455 | 0.116691659 | 0.071 | 0.024 | 1 | 17 |
| si:dkey-30k6.5 | 0.46455 | 0.105765993 | 0.071 | 0.024 | 1 | 17 |
| slc22a15 | 0.46455 | 0.148755687 | 0.071 | 0.024 | 1 | 17 |
| slc25a21 | 0.46455 | 0.169893235 | 0.071 | 0.024 | 1 | 17 |
| slkb | 0.46455 | 0.205087543 | 0.071 | 0.024 | 1 | 17 |
| smg9 | 0.46455 | 0.133211498 | 0.071 | 0.024 | 1 | 17 |
| st7l | 0.46455 | 0.130267901 | 0.071 | 0.024 | 1 | 17 |
| surf2 | 0.46455 | 0.154921143 | 0.071 | 0.024 | 1 | 17 |
| svild | 0.46455 | 0.152751416 | 0.071 | 0.024 | 1 | 17 |
| tmem231 | 0.46455 | 0.146336283 | 0.071 | 0.024 | 1 | 17 |
| trappc6bl | 0.46455 | 0.139984209 | 0.071 | 0.024 | 1 | 17 |
| tubg1 | 0.46455 | 0.11729749 | 0.071 | 0.024 | 1 | 17 |
| uri1 | 0.46455 | 0.124449903 | 0.071 | 0.024 | 1 | 17 |
| uros | 0.46455 | 0.167777764 | 0.071 | 0.024 | 1 | 17 |
| usp19 | 0.46455 | 0.14173741 | 0.071 | 0.024 | 1 | 17 |
| wdr55 | 0.46455 | 0.133850111 | 0.071 | 0.024 | 1 | 17 |
| wu:fj29h11 | 0.46455 | 0.142853797 | 0.071 | 0.024 | 1 | 17 |
| zgc:112962 | 0.46455 | 0.109395875 | 0.071 | 0.024 | 1 | 17 |
| zgc:113265 | 0.46455 | 0.133887254 | 0.071 | 0.024 | 1 | 17 |
| zgc:113886 | 0.46455 | 0.130249919 | 0.071 | 0.024 | 1 | 17 |
| zgc:114130 | 0.46455 | 0.100549773 | 0.071 | 0.024 | 1 | 17 |
| zgc:55733 | 0.46455 | 0.175579713 | 0.071 | 0.024 | 1 | 17 |
| zgc:76872 | 0.46455 | 0.125904527 | 0.071 | 0.024 | 1 | 17 |
| zgc:77262 | 0.46455 | 0.116549882 | 0.071 | 0.024 | 1 | 17 |
| zhx2a | 0.46455 | 0.118565484 | 0.071 | 0.024 | 1 | 17 |
| zmynd8 | 0.46455 | 0.140723804 | 0.071 | 0.024 | 1 | 17 |
| FP074902.2 | 0.46455 | 0.177011682 | 0.071 | 0.024 | 1 | 17 |
| LOC571912 | 0.46455 | 0.104378828 | 0.071 | 0.024 | 1 | 17 |
| adcy8 | 0.46455 | 0.113336527 | 0.071 | 0.024 | 1 | 17 |
| aldh2.1 | 0.46455 | 0.109543368 | 0.071 | 0.024 | 1 | 17 |
| amdhd2 | 0.46455 | 0.101532205 | 0.071 | 0.024 | 1 | 17 |
| armt1 | 0.46455 | 0.183676868 | 0.071 | 0.024 | 1 | 17 |
| baiap2a | 0.46455 | 0.149785544 | 0.071 | 0.024 | 1 | 17 |
| btr26 | 0.46455 | 0.106635012 | 0.071 | 0.024 | 1 | 17 |
| ccdc32 | 0.46455 | 0.176121422 | 0.071 | 0.024 | 1 | 17 |
| cfl2 | 0.46455 | 0.10586686 | 0.071 | 0.024 | 1 | 17 |
| clpb | 0.46455 | 0.174454248 | 0.071 | 0.024 | 1 | 17 |
| dachb | 0.46455 | 0.139540467 | 0.071 | 0.024 | 1 | 17 |
| dbnla | 0.46455 | 0.15548382 | 0.071 | 0.024 | 1 | 17 |
| dnai1.1 | 0.46455 | 0.170243785 | 0.071 | 0.024 | 1 | 17 |
| egln3 | 0.46455 | 0.112230818 | 0.071 | 0.024 | 1 | 17 |
| ergic2 | 0.46455 | 0.113614295 | 0.071 | 0.024 | 1 | 17 |
| fmr1 | 0.46455 | 0.208568796 | 0.071 | 0.024 | 1 | 17 |

| | | | | | | |
|---|---|---|---|---|---|---|
| gigyf2 | 0.46455 | 0.209678974 | 0.071 | 0.024 | 1 | 17 |
| ift74 | 0.46455 | 0.111940955 | 0.071 | 0.024 | 1 | 17 |
| ints8 | 0.46455 | 0.168649122 | 0.071 | 0.024 | 1 | 17 |
| kif23 | 0.46455 | 0.210589327 | 0.071 | 0.024 | 1 | 17 |
| lipia | 0.46455 | 0.105689859 | 0.071 | 0.024 | 1 | 17 |
| map7d1a | 0.46455 | 0.11292302 | 0.071 | 0.024 | 1 | 17 |
| me3 | 0.46455 | 0.108725651 | 0.071 | 0.024 | 1 | 17 |
| mrpl19 | 0.46455 | 0.114077696 | 0.071 | 0.024 | 1 | 17 |
| mtif3 | 0.46455 | 0.185772846 | 0.071 | 0.024 | 1 | 17 |
| oc90 | 0.46455 | 0.169591995 | 0.071 | 0.024 | 1 | 17 |
| phf6 | 0.46455 | 0.111791246 | 0.071 | 0.024 | 1 | 17 |
| pir | 0.46455 | 0.111342678 | 0.071 | 0.024 | 1 | 17 |
| rab7a | 0.46455 | 0.129734708 | 0.071 | 0.024 | 1 | 17 |
| rad23aa | 0.46455 | 0.115633822 | 0.071 | 0.024 | 1 | 17 |
| shroom4 | 0.46455 | 0.168985818 | 0.071 | 0.024 | 1 | 17 |
| si:ch211-121a2.2 | 0.46455 | 0.106142021 | 0.071 | 0.024 | 1 | 17 |
| si:ch211-207i20.3 | 0.46455 | 0.10202672 | 0.071 | 0.024 | 1 | 17 |
| si:dkey-66i24.9 | 0.46455 | 0.108040746 | 0.071 | 0.024 | 1 | 17 |
| snupn | 0.46455 | 0.210907425 | 0.071 | 0.024 | 1 | 17 |
| tmco3 | 0.46455 | 0.105463388 | 0.071 | 0.024 | 1 | 17 |
| tmem147 | 0.46455 | 0.212736493 | 0.071 | 0.024 | 1 | 17 |
| ubxn6 | 0.46455 | 0.142891889 | 0.071 | 0.024 | 1 | 17 |
| vps37b | 0.46455 | 0.103157695 | 0.071 | 0.024 | 1 | 17 |
| zfand6 | 0.46455 | 0.16888323 | 0.071 | 0.024 | 1 | 17 |
| zgc:152830 | 0.46455 | 0.179633348 | 0.071 | 0.024 | 1 | 17 |
| zgc:153352 | 0.46455 | 0.115143209 | 0.071 | 0.024 | 1 | 17 |
| zgc:153441 | 0.46455 | 0.111038609 | 0.071 | 0.024 | 1 | 17 |
| CABZ01076667.1 | 0.46455 | 0.127513762 | 0.071 | 0.024 | 1 | 17 |
| EARS2 | 0.46455 | 0.195387281 | 0.071 | 0.024 | 1 | 17 |
| LO018176.1 | 0.46455 | 0.138990425 | 0.071 | 0.024 | 1 | 17 |
| aagab | 0.46455 | 0.10065473 | 0.071 | 0.024 | 1 | 17 |
| aatf | 0.46455 | 0.147496507 | 0.071 | 0.024 | 1 | 17 |
| cd82a | 0.46455 | 0.121306686 | 0.071 | 0.024 | 1 | 17 |
| elovl4a | 0.46455 | 0.180586622 | 0.071 | 0.024 | 1 | 17 |
| ercc3 | 0.46455 | 0.154864209 | 0.071 | 0.024 | 1 | 17 |
| fbxo38 | 0.46455 | 0.111727626 | 0.071 | 0.024 | 1 | 17 |
| gdpd5a | 0.46455 | 0.130569126 | 0.071 | 0.024 | 1 | 17 |
| h2afx | 0.46455 | 0.13943726 | 0.071 | 0.024 | 1 | 17 |
| hmgxb4a | 0.46455 | 0.101930304 | 0.071 | 0.024 | 1 | 17 |
| hsbp1a | 0.46455 | 0.100050019 | 0.071 | 0.024 | 1 | 17 |
| ift22 | 0.46455 | 0.230772605 | 0.071 | 0.024 | 1 | 17 |
| impdh1b | 0.46455 | 0.180009686 | 0.071 | 0.024 | 1 | 17 |
| lrpap1 | 0.46455 | 0.157050681 | 0.071 | 0.024 | 1 | 17 |
| nol9 | 0.46455 | 0.101824897 | 0.071 | 0.024 | 1 | 17 |
| nup133 | 0.46455 | 0.102420654 | 0.071 | 0.024 | 1 | 17 |
| ppp2r3c | 0.46455 | 0.109198191 | 0.071 | 0.024 | 1 | 17 |
| si:ch211-198a12.6 | 0.46455 | 0.103769457 | 0.071 | 0.024 | 1 | 17 |
| slc25a28 | 0.46455 | 0.151769769 | 0.071 | 0.024 | 1 | 17 |

| | | | | | | |
|---|---|---|---|---|---|---|
| swap70a | 0.46455 | 0.162333744 | 0.071 | 0.024 | 1 | 17 |
| trmt61b | 0.46455 | 0.102741151 | 0.071 | 0.024 | 1 | 17 |
| zcchc4 | 0.46455 | 0.234494984 | 0.071 | 0.024 | 1 | 17 |
| znf511 | 0.46455 | 0.236195131 | 0.071 | 0.024 | 1 | 17 |
| CABZ01077555.1 | 0.46455 | 0.133070667 | 0.071 | 0.024 | 1 | 17 |
| cars | 0.46455 | 0.121993064 | 0.071 | 0.024 | 1 | 17 |
| hoxc11a | 0.46455 | 0.142935783 | 0.071 | 0.024 | 1 | 17 |
| nitr5 | 0.46455 | 0.134265557 | 0.071 | 0.024 | 1 | 17 |
| zc3h6 | 0.46455 | 0.135298565 | 0.071 | 0.024 | 1 | 17 |
| znf800a | 0.46455 | 0.132787371 | 0.071 | 0.024 | 1 | 17 |
| nudt17 | 0.46455 | 0.151889525 | 0.071 | 0.024 | 1 | 17 |
| bag1 | 0.46455 | 0.125529766 | 0.071 | 0.024 | 1 | 17 |
| gas6 | 0.46455 | 0.128658265 | 0.071 | 0.024 | 1 | 17 |
| klhl13 | 0.46455 | 0.130268182 | 0.071 | 0.024 | 1 | 17 |
| myo5ab | 0.46455 | 0.142329302 | 0.071 | 0.024 | 1 | 17 |
| pimr68 | 0.46455 | 0.130174663 | 0.071 | 0.024 | 1 | 17 |
| ppfibp1b | 0.46455 | 0.106252763 | 0.071 | 0.024 | 1 | 17 |
| ppl | 0.46455 | 0.130162866 | 0.071 | 0.024 | 1 | 17 |
| rc3h1b | 0.46455 | 0.179949336 | 0.071 | 0.024 | 1 | 17 |
| sardh | 0.46455 | 0.112723777 | 0.071 | 0.024 | 1 | 17 |
| si:ch211-160d20.3 | 0.46455 | 0.115607143 | 0.071 | 0.024 | 1 | 17 |
| si:dkey-42i9.8 | 0.46455 | 0.136753199 | 0.071 | 0.024 | 1 | 17 |
| slc35a1 | 0.46455 | 0.118917884 | 0.071 | 0.024 | 1 | 17 |
| syt9a | 0.46455 | 0.130174663 | 0.071 | 0.024 | 1 | 17 |
| tdrd12 | 0.46455 | 0.186332171 | 0.071 | 0.024 | 1 | 17 |
| tmem134 | 0.46455 | 0.151119835 | 0.071 | 0.024 | 1 | 17 |
| tom1l2b | 0.46455 | 0.102240413 | 0.071 | 0.024 | 1 | 17 |
| vamp1 | 0.46455 | 0.130853114 | 0.071 | 0.024 | 1 | 17 |
| vps33a | 0.46455 | 0.18752325 | 0.071 | 0.024 | 1 | 17 |
| zbtb16a | 0.46455 | 0.160741071 | 0.071 | 0.024 | 1 | 17 |
| zer1 | 0.46455 | 0.123098068 | 0.071 | 0.024 | 1 | 17 |
| zfp64 | 0.46455 | 0.165987593 | 0.071 | 0.024 | 1 | 17 |
| znf576.1 | 0.46455 | 0.11154134 | 0.071 | 0.024 | 1 | 17 |
| LOC101885675 | 0.46455 | 0.100592171 | 0.071 | 0.024 | 1 | 17 |
| LOC799656 | 0.46455 | 0.11963741 | 0.071 | 0.024 | 1 | 17 |
| RF00560 | 0.46455 | 0.112126926 | 0.071 | 0.024 | 1 | 17 |
| XLOC-043861 | 0.46455 | 0.131870559 | 0.071 | 0.024 | 1 | 17 |
| btbd9 | 0.46455 | 0.132843524 | 0.071 | 0.024 | 1 | 17 |
| cdk16 | 0.46455 | 0.102798488 | 0.071 | 0.024 | 1 | 17 |
| chkb | 0.46455 | 0.112244111 | 0.071 | 0.024 | 1 | 17 |
| cnot8 | 0.46455 | 0.140646596 | 0.071 | 0.024 | 1 | 17 |
| copb1 | 0.46455 | 0.111715916 | 0.071 | 0.024 | 1 | 17 |
| crlf1b | 0.46455 | 0.121325696 | 0.071 | 0.024 | 1 | 17 |
| cwf19l2 | 0.46455 | 0.114255419 | 0.071 | 0.024 | 1 | 17 |
| exoc3 | 0.46455 | 0.111356898 | 0.071 | 0.024 | 1 | 17 |
| fastkd2 | 0.46455 | 0.136782282 | 0.071 | 0.024 | 1 | 17 |
| gemin5 | 0.46455 | 0.136994036 | 0.071 | 0.024 | 1 | 17 |
| hectd3 | 0.46455 | 0.143383558 | 0.071 | 0.024 | 1 | 17 |

| | | | | | | |
|---|---|---|---|---|---|---|
| ino80db | 0.46455 | 0.107800383 | 0.071 | 0.024 | 1 | 17 |
| mettl13 | 0.46455 | 0.103280004 | 0.071 | 0.024 | 1 | 17 |
| mthfd2l | 0.46455 | 0.176607175 | 0.071 | 0.024 | 1 | 17 |
| naa30 | 0.46455 | 0.129643847 | 0.071 | 0.024 | 1 | 17 |
| nxpe3 | 0.46455 | 0.128136027 | 0.071 | 0.024 | 1 | 17 |
| onecut1 | 0.46455 | 0.151571784 | 0.071 | 0.024 | 1 | 17 |
| p3h3 | 0.46455 | 0.187344158 | 0.071 | 0.024 | 1 | 17 |
| pgm2l1 | 0.46455 | 0.144697926 | 0.071 | 0.024 | 1 | 17 |
| phf11 | 0.46455 | 0.163582662 | 0.071 | 0.024 | 1 | 17 |
| qtrt2 | 0.46455 | 0.119114871 | 0.071 | 0.024 | 1 | 17 |
| rab6ba | 0.46455 | 0.112431545 | 0.071 | 0.024 | 1 | 17 |
| rabl6a | 0.46455 | 0.118393928 | 0.071 | 0.024 | 1 | 17 |
| rassf1 | 0.46455 | 0.164261254 | 0.071 | 0.024 | 1 | 17 |
| rbms2b | 0.46455 | 0.104075105 | 0.071 | 0.024 | 1 | 17 |
| rufy3 | 0.46455 | 0.142907934 | 0.071 | 0.024 | 1 | 17 |
| sav1 | 0.46455 | 0.102700918 | 0.071 | 0.024 | 1 | 17 |
| sdc2 | 0.46455 | 0.102519497 | 0.071 | 0.024 | 1 | 17 |
| si:ch211-62a1.3 | 0.46455 | 0.177696535 | 0.071 | 0.024 | 1 | 17 |
| si:dkey-16l2.16 | 0.46455 | 0.105784016 | 0.071 | 0.024 | 1 | 17 |
| si:dkey-21c1.1 | 0.46455 | 0.115565461 | 0.071 | 0.024 | 1 | 17 |
| stk25b | 0.46455 | 0.115410362 | 0.071 | 0.024 | 1 | 17 |
| timeless | 0.46455 | 0.135415673 | 0.071 | 0.024 | 1 | 17 |
| tmcc1b | 0.46455 | 0.127372802 | 0.071 | 0.024 | 1 | 17 |
| zgc:112982 | 0.46455 | 0.130767538 | 0.071 | 0.024 | 1 | 17 |
| znf326 | 0.46455 | 0.107019628 | 0.071 | 0.024 | 1 | 17 |
| BX537342.1 | 0.46455 | 0.134035966 | 0.071 | 0.024 | 1 | 17 |
| chpf2 | 0.46455 | 0.132169101 | 0.071 | 0.024 | 1 | 17 |
| cryl1 | 0.46455 | 0.173973966 | 0.071 | 0.024 | 1 | 17 |
| flcn | 0.46455 | 0.15965735 | 0.071 | 0.024 | 1 | 17 |
| jam3a | 0.46455 | 0.175384614 | 0.071 | 0.024 | 1 | 17 |
| kat7b | 0.46455 | 0.112538716 | 0.071 | 0.024 | 1 | 17 |
| lpin2 | 0.46455 | 0.125725329 | 0.071 | 0.024 | 1 | 17 |
| minpp1b | 0.46455 | 0.15164608 | 0.071 | 0.024 | 1 | 17 |
| nt5c3a | 0.46455 | 0.165653984 | 0.071 | 0.024 | 1 | 17 |
| rb1cc1 | 0.46455 | 0.165762589 | 0.071 | 0.024 | 1 | 17 |
| rexo1 | 0.46455 | 0.15989314 | 0.071 | 0.024 | 1 | 17 |
| scpep1 | 0.46455 | 0.136550928 | 0.071 | 0.024 | 1 | 17 |
| scyl1 | 0.46455 | 0.135044431 | 0.071 | 0.024 | 1 | 17 |
| sfswap | 0.46455 | 0.105776732 | 0.071 | 0.024 | 1 | 17 |
| si:ch73-100l22.3 | 0.46455 | 0.186776171 | 0.071 | 0.024 | 1 | 17 |
| si:ch73-248e21.1 | 0.46455 | 0.181590657 | 0.071 | 0.024 | 1 | 17 |
| si:ch73-30l9.1 | 0.46455 | 0.188675368 | 0.071 | 0.024 | 1 | 17 |
| tmem177 | 0.46455 | 0.108746133 | 0.071 | 0.024 | 1 | 17 |
| tmem9b | 0.46455 | 0.130052595 | 0.071 | 0.024 | 1 | 17 |
| trabd | 0.46455 | 0.124670658 | 0.071 | 0.024 | 1 | 17 |
| twnk | 0.46455 | 0.131101409 | 0.071 | 0.024 | 1 | 17 |
| uprt | 0.46455 | 0.136473822 | 0.071 | 0.024 | 1 | 17 |
| uxs1 | 0.46455 | 0.174459328 | 0.071 | 0.024 | 1 | 17 |

| | | | | | | |
|---|---|---|---|---|---|---|
| zgc:56628 | 0.46455 | 0.116961405 | 0.071 | 0.024 | 1 | 17 |
| acss2 | 0.46455 | 0.144326722 | 0.071 | 0.024 | 1 | 17 |
| bckdhbl | 0.46455 | 0.123342707 | 0.071 | 0.024 | 1 | 17 |
| gnao1a | 0.46455 | 0.102906397 | 0.071 | 0.024 | 1 | 17 |
| lrrc29 | 0.46455 | 0.132896439 | 0.071 | 0.024 | 1 | 17 |
| pbrm1 | 0.46455 | 0.10440682 | 0.071 | 0.024 | 1 | 17 |
| ppm1bb | 0.46455 | 0.136870857 | 0.071 | 0.024 | 1 | 17 |
| prkacbb | 0.46455 | 0.134630976 | 0.071 | 0.024 | 1 | 17 |
| si:ch211-260e23.7 | 0.46455 | 0.146061136 | 0.071 | 0.024 | 1 | 17 |
| ttll12 | 0.46455 | 0.106441358 | 0.071 | 0.024 | 1 | 17 |
| marcksb | 0.46572 | 0.129708795 | 0.143 | 0.049 | 1 | 17 |
| fth1a | 0.46636 | -0.37633187 | 0.357 | 0.512 | 1 | 17 |
| cdca8 | 0.46645 | -0.40427869 | 0.143 | 0.268 | 1 | 17 |
| ppme1 | 0.46662 | 0.277321994 | 0.357 | 0.195 | 1 | 17 |
| ptmaa | 0.46789 | 0.199799681 | 0.286 | 0.146 | 1 | 17 |
| drg2 | 0.46951 | 0.191432705 | 0.429 | 0.268 | 1 | 17 |
| ywhaqa | 0.47004 | 0.289289293 | 0.5 | 0.317 | 1 | 17 |
| ranbp1 | 0.47033 | 0.193156834 | 0.5 | 0.341 | 1 | 17 |
| l2hgdh | 0.47057 | 0.1242551 | 0.143 | 0.049 | 1 | 17 |
| rpn1 | 0.47082 | -0.19744512 | 0.214 | 0.195 | 1 | 17 |
| ndufa7 | 0.47224 | 0.225666798 | 0.286 | 0.146 | 1 | 17 |
| chmp5b | 0.47284 | 0.231647333 | 0.286 | 0.146 | 1 | 17 |
| RSPH1 | 0.47384 | 0.115906631 | 0.143 | 0.049 | 1 | 17 |
| vdac1 | 0.47429 | 0.124176683 | 0.429 | 0.293 | 1 | 17 |
| ak5l | 0.47655 | 0.101982307 | 0.143 | 0.049 | 1 | 17 |
| zgc:66472 | 0.47759 | 0.205525831 | 0.286 | 0.146 | 1 | 17 |
| si:dkey-210j14.4 | 0.47785 | 0.117186694 | 0.143 | 0.049 | 1 | 17 |
| pold1 | 0.47838 | -0.37005317 | 0.071 | 0.171 | 1 | 17 |
| itm2ba | 0.47854 | -0.19240712 | 0.214 | 0.195 | 1 | 17 |
| mob1bb | 0.47977 | -0.40043518 | 0.143 | 0.293 | 1 | 17 |
| ccnd3 | 0.48153 | 0.148451028 | 0.357 | 0.22 | 1 | 17 |
| cdk2ap2 | 0.48162 | -0.33385864 | 0.143 | 0.195 | 1 | 17 |
| naxe | 0.4817 | 0.13943184 | 0.143 | 0.049 | 1 | 17 |
| ptmab | 0.48184 | -0.36839583 | 0.357 | 0.488 | 1 | 17 |
| LO018102.1 | 0.48239 | 0.139064747 | 0.214 | 0.098 | 1 | 17 |
| cbx1a | 0.4825 | 0.14488268 | 0.214 | 0.098 | 1 | 17 |
| snrpb | 0.48475 | 0.241429784 | 0.643 | 0.463 | 1 | 17 |
| cct6a | 0.4852 | -0.11859473 | 0.5 | 0.463 | 1 | 17 |
| arhgdia | 0.48578 | -0.28210275 | 0.143 | 0.293 | 1 | 17 |
| ptbp1a | 0.48849 | 0.310134732 | 0.357 | 0.195 | 1 | 17 |
| tmem30ab | 0.48909 | 0.179949715 | 0.143 | 0.049 | 1 | 17 |
| fam133b | 0.48931 | 0.340751308 | 0.357 | 0.195 | 1 | 17 |
| rps3a | 0.48936 | 0.12236284 | 0.929 | 0.805 | 1 | 17 |
| si:ch211-276i12.11 | 0.48995 | 0.320624959 | 0.357 | 0.195 | 1 | 17 |
| eif6 | 0.49002 | 0.291464638 | 0.357 | 0.195 | 1 | 17 |
| prkrip1 | 0.49138 | 0.127114856 | 0.143 | 0.049 | 1 | 17 |
| pes | 0.49212 | -0.364038 | 0.071 | 0.195 | 1 | 17 |
| spata6l | 0.49373 | 0.36085532 | 0.286 | 0.171 | 1 | 17 |

| | | | | | | |
|---|---|---|---|---|---|---|
| LOC103910438 | 0.49494 | 0.132023543 | 0.143 | 0.049 | 1 | 17 |
| h1f0 | 0.49522 | 0.220581769 | 0.286 | 0.146 | 1 | 17 |
| vdac2 | 0.49527 | 0.229362081 | 0.714 | 0.537 | 1 | 17 |
| chtopb | 0.49543 | 0.299718731 | 0.571 | 0.39 | 1 | 17 |
| ctdspl3 | 0.49572 | 0.104899913 | 0.143 | 0.049 | 1 | 17 |
| naa10 | 0.49574 | 0.274465391 | 0.571 | 0.39 | 1 | 17 |
| cnot3a | 0.49964 | 0.117648565 | 0.143 | 0.049 | 1 | 17 |
| ddx3xa | 0.50004 | 0.165775982 | 0.143 | 0.049 | 1 | 17 |
| ndufb7 | 0.50085 | 0.291295713 | 0.571 | 0.39 | 1 | 17 |
| cnppd1 | 0.50087 | 0.161904919 | 0.143 | 0.049 | 1 | 17 |
| nup62l | 0.5012 | -0.38283797 | 0.071 | 0.171 | 1 | 17 |
| terb1 | 0.50132 | 0.151865358 | 0.214 | 0.098 | 1 | 17 |
| si:dkey-31e10.5 | 0.50287 | 0.300934613 | 0.143 | 0.049 | 1 | 17 |
| nsrp1 | 0.50327 | 0.329674553 | 0.143 | 0.049 | 1 | 17 |
| ctdsplb | 0.50357 | 0.136410124 | 0.143 | 0.049 | 1 | 17 |
| gsta.1 | 0.50357 | 0.100037012 | 0.143 | 0.049 | 1 | 17 |
| ska2 | 0.50402 | 0.116574149 | 0.143 | 0.049 | 1 | 17 |
| isg20 | 0.50445 | -0.35644353 | 0.071 | 0.195 | 1 | 17 |
| tmem243b | 0.50492 | 0.131185291 | 0.143 | 0.049 | 1 | 17 |
| nedd8l | 0.50538 | 0.139563027 | 0.214 | 0.098 | 1 | 17 |
| tfg | 0.50557 | 0.146828464 | 0.143 | 0.049 | 1 | 17 |
| cox7a2a | 0.50629 | 0.176155275 | 0.571 | 0.415 | 1 | 17 |
| tubb4b | 0.5065 | 0.108075096 | 0.857 | 0.732 | 1 | 17 |
| dnajb12a | 0.50655 | 0.164307072 | 0.143 | 0.049 | 1 | 17 |
| armc1l | 0.50754 | 0.14917692 | 0.143 | 0.049 | 1 | 17 |
| zcchc2 | 0.5076 | 0.135407659 | 0.214 | 0.098 | 1 | 17 |
| polr3glb | 0.50773 | 0.126313978 | 0.143 | 0.049 | 1 | 17 |
| si:dkey-4e7.3 | 0.50803 | 0.152465843 | 0.143 | 0.049 | 1 | 17 |
| polr2h | 0.50949 | -0.35223999 | 0.143 | 0.293 | 1 | 17 |
| psma8 | 0.50963 | -0.25734051 | 0.5 | 0.585 | 1 | 17 |
| mrgbp | 0.5099 | 0.185024684 | 0.143 | 0.049 | 1 | 17 |
| cpsf1 | 0.51026 | 0.136484259 | 0.143 | 0.049 | 1 | 17 |
| si:ch211-207k7.4 | 0.51031 | 0.114582269 | 0.143 | 0.049 | 1 | 17 |
| utp15 | 0.5111 | -0.26756574 | 0.071 | 0.195 | 1 | 17 |
| psmd11a | 0.51187 | -0.3793394 | 0.071 | 0.171 | 1 | 17 |
| si:ch211-22i13.2 | 0.51241 | -0.28590362 | 0.143 | 0.171 | 1 | 17 |
| puf60b | 0.51309 | -0.3483479 | 0.071 | 0.171 | 1 | 17 |
| msl1a | 0.51467 | 0.112658808 | 0.143 | 0.049 | 1 | 17 |
| pias4a | 0.5152 | 0.193427501 | 0.143 | 0.049 | 1 | 17 |
| ube2d2 | 0.51564 | -0.22765922 | 0.357 | 0.39 | 1 | 17 |
| actl6a | 0.51644 | -0.28979195 | 0.071 | 0.195 | 1 | 17 |
| actr10 | 0.51731 | 0.117233362 | 0.143 | 0.049 | 1 | 17 |
| bsdc1 | 0.51737 | -0.33877193 | 0.071 | 0.195 | 1 | 17 |
| stag1b | 0.51786 | 0.125089703 | 0.143 | 0.049 | 1 | 17 |
| pank2 | 0.51833 | 0.192707954 | 0.143 | 0.049 | 1 | 17 |
| trappc1 | 0.5189 | 0.146594531 | 0.429 | 0.293 | 1 | 17 |
| igbp1 | 0.51919 | 0.299634761 | 0.286 | 0.146 | 1 | 17 |
| ybx1 | 0.51945 | 0.14266296 | 0.571 | 0.415 | 1 | 17 |

| | | | | | | |
|---|---|---|---|---|---|---|
| ppp1r10 | 0.51971 | -0.37107315 | 0.071 | 0.171 | 1 | 17 |
| prrg2 | 0.51973 | 0.1694842 | 0.143 | 0.049 | 1 | 17 |
| rasal2 | 0.51996 | 0.145012763 | 0.143 | 0.049 | 1 | 17 |
| cox7a1 | 0.52008 | 0.156114721 | 0.143 | 0.049 | 1 | 17 |
| yipf3 | 0.52057 | 0.193675618 | 0.143 | 0.049 | 1 | 17 |
| borcs7 | 0.52144 | 0.17294117 | 0.143 | 0.049 | 1 | 17 |
| esf1 | 0.52146 | 0.22077943 | 0.214 | 0.098 | 1 | 17 |
| bola1 | 0.52153 | -0.17232203 | 0.143 | 0.122 | 1 | 17 |
| psmd9 | 0.52179 | 0.196118616 | 0.143 | 0.049 | 1 | 17 |
| pimr109 | 0.52213 | 0.188026956 | 0.143 | 0.049 | 1 | 17 |
| e2f5 | 0.52233 | 0.190392972 | 0.357 | 0.22 | 1 | 17 |
| zgc:66474 | 0.52234 | 0.157800728 | 0.143 | 0.049 | 1 | 17 |
| knop1 | 0.52235 | 0.229073247 | 0.143 | 0.049 | 1 | 17 |
| mpv17 | 0.52283 | 0.138080318 | 0.143 | 0.049 | 1 | 17 |
| selenom | 0.52299 | 0.338369813 | 0.286 | 0.146 | 1 | 17 |
| sulf2a | 0.52301 | 0.318518436 | 0.143 | 0.049 | 1 | 17 |
| cops7a | 0.52385 | 0.278666799 | 0.286 | 0.146 | 1 | 17 |
| hspa9 | 0.52387 | 0.242319023 | 0.429 | 0.268 | 1 | 17 |
| rnf185 | 0.52418 | 0.190545796 | 0.214 | 0.098 | 1 | 17 |
| mibp | 0.52419 | 0.291134438 | 0.286 | 0.146 | 1 | 17 |
| hint2 | 0.52428 | 0.152869518 | 0.143 | 0.049 | 1 | 17 |
| taf12 | 0.52468 | -0.37986026 | 0.143 | 0.244 | 1 | 17 |
| etf1b | 0.52525 | 0.26559854 | 0.286 | 0.146 | 1 | 17 |
| zgc:110130 | 0.52633 | 0.162143481 | 0.214 | 0.098 | 1 | 17 |
| ankrd31 | 0.52637 | 0.14397667 | 0.143 | 0.049 | 1 | 17 |
| spata4 | 0.52656 | 0.161500944 | 0.214 | 0.098 | 1 | 17 |
| msra | 0.52671 | 0.171526894 | 0.143 | 0.049 | 1 | 17 |
| ccdc127b | 0.52685 | 0.171784858 | 0.143 | 0.049 | 1 | 17 |
| mcmbp | 0.52742 | 0.294074899 | 0.286 | 0.146 | 1 | 17 |
| pdrg1 | 0.5276 | 0.246322977 | 0.286 | 0.146 | 1 | 17 |
| ckap5 | 0.52775 | 0.122325999 | 0.143 | 0.049 | 1 | 17 |
| anapc11 | 0.52805 | 0.202151914 | 0.214 | 0.098 | 1 | 17 |
| kpna2 | 0.52818 | 0.217231025 | 0.5 | 0.341 | 1 | 17 |
| coa7 | 0.52874 | 0.123166737 | 0.143 | 0.049 | 1 | 17 |
| ifi30 | 0.52891 | 0.185204739 | 0.214 | 0.098 | 1 | 17 |
| ptp4a1 | 0.52965 | 0.234022575 | 0.429 | 0.268 | 1 | 17 |
| irak4 | 0.52991 | 0.105809076 | 0.143 | 0.049 | 1 | 17 |
| ift27 | 0.53014 | 0.142001582 | 0.143 | 0.049 | 1 | 17 |
| znf576.2 | 0.53016 | 0.202146455 | 0.143 | 0.049 | 1 | 17 |
| tomm20a | 0.53078 | 0.100681073 | 0.786 | 0.659 | 1 | 17 |
| uchl5 | 0.53078 | 0.191227776 | 0.143 | 0.049 | 1 | 17 |
| mfap1 | 0.53079 | 0.172570396 | 0.357 | 0.22 | 1 | 17 |
| cfap20 | 0.53133 | 0.165924725 | 0.143 | 0.049 | 1 | 17 |
| si:dkey-31g6.4 | 0.53162 | 0.28123738 | 0.143 | 0.049 | 1 | 17 |
| lyrm4 | 0.53186 | 0.274403033 | 0.286 | 0.146 | 1 | 17 |
| rnf2 | 0.53195 | 0.24785704 | 0.214 | 0.098 | 1 | 17 |
| tmem258 | 0.53231 | -0.31491748 | 0.286 | 0.366 | 1 | 17 |
| orc3 | 0.53252 | -0.32826283 | 0.071 | 0.171 | 1 | 17 |

| | | | | | | |
|---|---|---|---|---|---|---|
| bloc1s4 | 0.53253 | 0.277304294 | 0.143 | 0.049 | 1 | 17 |
| si:dkeyp-46h3.1 | 0.5326 | 0.258330039 | 0.143 | 0.049 | 1 | 17 |
| gnai1 | 0.5335 | 0.158468533 | 0.143 | 0.049 | 1 | 17 |
| zc3h4 | 0.53375 | 0.167408709 | 0.143 | 0.049 | 1 | 17 |
| tex10 | 0.53389 | 0.185607208 | 0.143 | 0.049 | 1 | 17 |
| rps6kb1b | 0.53455 | 0.182813791 | 0.143 | 0.049 | 1 | 17 |
| eif3ea | 0.53461 | 0.202229046 | 0.5 | 0.341 | 1 | 17 |
| acadm | 0.53471 | 0.180225936 | 0.357 | 0.22 | 1 | 17 |
| ptp4a2b | 0.53502 | 0.129479168 | 0.143 | 0.049 | 1 | 17 |
| katnal1 | 0.53505 | 0.147896698 | 0.143 | 0.049 | 1 | 17 |
| stip1 | 0.53509 | 0.154375475 | 0.429 | 0.293 | 1 | 17 |
| tmem11 | 0.53514 | 0.163693765 | 0.143 | 0.049 | 1 | 17 |
| dhdds | 0.53539 | 0.129243606 | 0.143 | 0.049 | 1 | 17 |
| rsf1b.1 | 0.53573 | 0.238761784 | 0.143 | 0.049 | 1 | 17 |
| smim13 | 0.5359 | 0.132374948 | 0.143 | 0.049 | 1 | 17 |
| hsd17b4 | 0.53594 | 0.144331743 | 0.143 | 0.049 | 1 | 17 |
| csnk1g2b | 0.53658 | 0.325858738 | 0.214 | 0.098 | 1 | 17 |
| irf7 | 0.53675 | 0.207803525 | 0.143 | 0.049 | 1 | 17 |
| selenof | 0.53683 | 0.346961512 | 0.357 | 0.22 | 1 | 17 |
| psmc5 | 0.53786 | 0.33567948 | 0.429 | 0.268 | 1 | 17 |
| sumf2 | 0.53804 | 0.279068273 | 0.143 | 0.049 | 1 | 17 |
| acads | 0.53841 | 0.264002757 | 0.143 | 0.049 | 1 | 17 |
| crip3 | 0.53886 | 0.101180159 | 0.143 | 0.049 | 1 | 17 |
| rps13 | 0.53918 | 0.213019486 | 0.857 | 0.732 | 1 | 17 |
| akap17a | 0.53941 | 0.191748382 | 0.143 | 0.049 | 1 | 17 |
| rnf212b | 0.53941 | 0.21357121 | 0.143 | 0.049 | 1 | 17 |
| smap1 | 0.54041 | 0.157411711 | 0.143 | 0.049 | 1 | 17 |
| ndufaf7 | 0.54131 | 0.313554701 | 0.143 | 0.049 | 1 | 17 |
| fam135a | 0.54133 | 0.13874215 | 0.143 | 0.049 | 1 | 17 |
| MCMDC2 | 0.54134 | 0.242394339 | 0.143 | 0.049 | 1 | 17 |
| ints11 | 0.54152 | 0.189909237 | 0.143 | 0.049 | 1 | 17 |
| mrpl2 | 0.54168 | 0.215180979 | 0.143 | 0.049 | 1 | 17 |
| stau1 | 0.54219 | 0.116197584 | 0.286 | 0.171 | 1 | 17 |
| dr1 | 0.54228 | 0.165017874 | 0.143 | 0.049 | 1 | 17 |
| ptgesl | 0.5425 | 0.309826814 | 0.214 | 0.098 | 1 | 17 |
| cpeb1b | 0.54302 | 0.192584537 | 0.214 | 0.098 | 1 | 17 |
| dynlt3 | 0.54315 | 0.284881118 | 0.429 | 0.268 | 1 | 17 |
| nabp1a | 0.54319 | 0.205416204 | 0.214 | 0.098 | 1 | 17 |
| wdr82 | 0.54336 | 0.276954196 | 0.429 | 0.268 | 1 | 17 |
| tpi1b | 0.54339 | 0.267392251 | 0.429 | 0.268 | 1 | 17 |
| ak1 | 0.54346 | 0.163790615 | 0.143 | 0.049 | 1 | 17 |
| nup50 | 0.54365 | 0.138849054 | 0.429 | 0.293 | 1 | 17 |
| mknk1 | 0.54384 | 0.202129815 | 0.143 | 0.049 | 1 | 17 |
| ascc3 | 0.54403 | 0.271687173 | 0.143 | 0.049 | 1 | 17 |
| smdt1b | 0.54412 | 0.185395853 | 0.143 | 0.049 | 1 | 17 |
| tada2a | 0.5442 | 0.208570935 | 0.143 | 0.049 | 1 | 17 |
| nfu1 | 0.5445 | 0.313684544 | 0.143 | 0.049 | 1 | 17 |
| mettl3 | 0.54462 | 0.235384562 | 0.143 | 0.049 | 1 | 17 |

| | | | | | | |
|---|---|---|---|---|---|---|
| si:dkey-6d5.1 | 0.54471 | 0.324245739 | 0.214 | 0.098 | 1 | 17 |
| emc8 | 0.54499 | 0.272218676 | 0.143 | 0.049 | 1 | 17 |
| gtpbp4 | 0.54501 | 0.307808574 | 0.429 | 0.268 | 1 | 17 |
| gars | 0.54537 | 0.202032897 | 0.214 | 0.098 | 1 | 17 |
| si:ch73-173h19.3 | 0.54544 | 0.15639424 | 0.143 | 0.049 | 1 | 17 |
| rgp1 | 0.54567 | -0.16808152 | 0.143 | 0.122 | 1 | 17 |
| ppp2r2d | 0.54583 | 0.174951981 | 0.143 | 0.049 | 1 | 17 |
| eif4e1c | 0.54606 | 0.306894377 | 0.429 | 0.268 | 1 | 17 |
| ndc1 | 0.54647 | -0.37489137 | 0.143 | 0.268 | 1 | 17 |
| tfdp1b | 0.54705 | 0.221658884 | 0.214 | 0.098 | 1 | 17 |
| khsrp | 0.54726 | 0.181338592 | 0.143 | 0.049 | 1 | 17 |
| jtb | 0.54732 | 0.178969229 | 0.143 | 0.049 | 1 | 17 |
| wu:fi37e09 | 0.54747 | 0.171995286 | 0.143 | 0.049 | 1 | 17 |
| nupl2 | 0.54764 | 0.280642757 | 0.143 | 0.049 | 1 | 17 |
| clic1 | 0.54774 | 0.182217246 | 0.143 | 0.049 | 1 | 17 |
| ccnf | 0.54826 | 0.25761593 | 0.143 | 0.049 | 1 | 17 |
| nsl1 | 0.54828 | 0.221622519 | 0.214 | 0.098 | 1 | 17 |
| tspo | 0.54837 | 0.219811218 | 0.143 | 0.049 | 1 | 17 |
| lipt1 | 0.54863 | 0.201014797 | 0.143 | 0.049 | 1 | 17 |
| noc4l | 0.54878 | 0.215696295 | 0.143 | 0.049 | 1 | 17 |
| smdt1a | 0.54885 | 0.199550122 | 0.143 | 0.049 | 1 | 17 |
| rab18b | 0.54899 | 0.166453345 | 0.143 | 0.049 | 1 | 17 |
| ankrd40 | 0.54901 | 0.235532529 | 0.214 | 0.098 | 1 | 17 |
| ube2q2 | 0.54914 | 0.229680959 | 0.143 | 0.049 | 1 | 17 |
| rab6bb | 0.54938 | 0.17423579 | 0.143 | 0.049 | 1 | 17 |
| cox6a1 | 0.54951 | -0.29681067 | 0.429 | 0.537 | 1 | 17 |
| rad50 | 0.54961 | 0.196383091 | 0.143 | 0.049 | 1 | 17 |
| ppp1r14aa | 0.54972 | 0.21901742 | 0.214 | 0.098 | 1 | 17 |
| acbd3 | 0.54974 | 0.168564771 | 0.143 | 0.049 | 1 | 17 |
| atg5 | 0.54974 | 0.202097946 | 0.143 | 0.049 | 1 | 17 |
| smim20 | 0.54977 | 0.192849602 | 0.214 | 0.098 | 1 | 17 |
| wasf3a | 0.54996 | 0.229831576 | 0.214 | 0.098 | 1 | 17 |
| naa15a | 0.5502 | 0.259478676 | 0.143 | 0.049 | 1 | 17 |
| rab11fip1b | 0.55038 | 0.163255862 | 0.143 | 0.049 | 1 | 17 |
| pias4b | 0.55044 | 0.221943349 | 0.143 | 0.049 | 1 | 17 |
| si:ch211-166a6.5 | 0.55049 | 0.163193 | 0.143 | 0.049 | 1 | 17 |
| commd4 | 0.55055 | 0.215345064 | 0.143 | 0.049 | 1 | 17 |
| brd9 | 0.55059 | 0.1975058 | 0.214 | 0.098 | 1 | 17 |
| smco4 | 0.55065 | 0.203501446 | 0.143 | 0.049 | 1 | 17 |
| cenpo | 0.55071 | 0.280184378 | 0.143 | 0.049 | 1 | 17 |
| pef1 | 0.55079 | 0.251952263 | 0.143 | 0.049 | 1 | 17 |
| scaf4a | 0.5508 | 0.19982225 | 0.214 | 0.098 | 1 | 17 |
| trmt61a | 0.55096 | 0.247928941 | 0.143 | 0.049 | 1 | 17 |
| rexo4 | 0.551 | -0.36421088 | 0.143 | 0.244 | 1 | 17 |
| smad2 | 0.55107 | 0.18260053 | 0.143 | 0.049 | 1 | 17 |
| CU469420.3 | 0.55142 | 0.29636828 | 0.143 | 0.049 | 1 | 17 |
| frem2a | 0.55142 | 0.154789435 | 0.143 | 0.049 | 1 | 17 |
| ndufs8b | 0.55168 | 0.299602604 | 0.214 | 0.098 | 1 | 17 |

| | | | | | | |
|---|---|---|---|---|---|---|
| kdm3b | 0.55177 | 0.18621676 | 0.143 | 0.049 | 1 | 17 |
| ankrd12 | 0.5519 | 0.156735553 | 0.143 | 0.049 | 1 | 17 |
| cptp | 0.552 | 0.188042717 | 0.143 | 0.049 | 1 | 17 |
| mpp2b | 0.55217 | 0.19402751 | 0.143 | 0.049 | 1 | 17 |
| kdm8 | 0.55223 | 0.21298319 | 0.143 | 0.049 | 1 | 17 |
| atg3 | 0.55257 | 0.229641564 | 0.143 | 0.049 | 1 | 17 |
| znf106b | 0.55264 | 0.268418721 | 0.143 | 0.049 | 1 | 17 |
| cat | 0.55269 | 0.243794972 | 0.143 | 0.049 | 1 | 17 |
| psmd14 | 0.55269 | -0.27586504 | 0.286 | 0.341 | 1 | 17 |
| acin1a | 0.55283 | 0.125700471 | 0.286 | 0.171 | 1 | 17 |
| lsg1 | 0.55286 | 0.180879373 | 0.143 | 0.049 | 1 | 17 |
| si:ch73-367f21.4 | 0.55293 | 0.215617202 | 0.143 | 0.049 | 1 | 17 |
| sf3b1 | 0.55296 | 0.25509119 | 0.143 | 0.049 | 1 | 17 |
| utp4 | 0.55299 | 0.230490972 | 0.143 | 0.049 | 1 | 17 |
| cndp2 | 0.55338 | 0.244482611 | 0.143 | 0.049 | 1 | 17 |
| cdk8 | 0.55341 | 0.215294838 | 0.143 | 0.049 | 1 | 17 |
| tac3b | 0.55363 | 0.181421258 | 0.143 | 0.049 | 1 | 17 |
| crsp7 | 0.55372 | 0.16494204 | 0.143 | 0.049 | 1 | 17 |
| pdgfd | 0.55375 | 0.194878616 | 0.143 | 0.049 | 1 | 17 |
| gtpbp1 | 0.55378 | 0.199463425 | 0.143 | 0.049 | 1 | 17 |
| rab8b | 0.55387 | 0.194996166 | 0.143 | 0.049 | 1 | 17 |
| hars | 0.554 | 0.193884786 | 0.143 | 0.049 | 1 | 17 |
| si:ch211-108c17.2 | 0.55414 | 0.205681393 | 0.143 | 0.049 | 1 | 17 |
| atl1 | 0.5543 | 0.205731511 | 0.143 | 0.049 | 1 | 17 |
| ctbp1 | 0.55431 | 0.259298528 | 0.214 | 0.098 | 1 | 17 |
| gpn2 | 0.55451 | 0.245409632 | 0.143 | 0.049 | 1 | 17 |
| setd4 | 0.55456 | 0.283497744 | 0.143 | 0.049 | 1 | 17 |
| ppp2r1ba | 0.55469 | 0.239111554 | 0.143 | 0.049 | 1 | 17 |
| cdk12 | 0.55475 | 0.204344643 | 0.143 | 0.049 | 1 | 17 |
| psmd7 | 0.55505 | -0.22333295 | 0.357 | 0.39 | 1 | 17 |
| eif2a | 0.55507 | 0.222265081 | 0.214 | 0.098 | 1 | 17 |
| atp6v1e1a | 0.55513 | 0.257161634 | 0.143 | 0.049 | 1 | 17 |
| eif4e2rs1 | 0.55518 | 0.234739563 | 0.143 | 0.049 | 1 | 17 |
| si:ch211-199g17.9 | 0.55518 | 0.205306679 | 0.143 | 0.049 | 1 | 17 |
| tfam | 0.55533 | 0.251199362 | 0.143 | 0.049 | 1 | 17 |
| mob1ba | 0.55547 | 0.242138198 | 0.143 | 0.049 | 1 | 17 |
| isoc2 | 0.55548 | 0.20093221 | 0.143 | 0.049 | 1 | 17 |
| mical2b | 0.55549 | 0.233263765 | 0.143 | 0.049 | 1 | 17 |
| ttc25 | 0.55557 | 0.208333657 | 0.143 | 0.049 | 1 | 17 |
| snx3 | 0.55565 | 0.196128604 | 0.143 | 0.049 | 1 | 17 |
| nvl | 0.55568 | 0.275660655 | 0.143 | 0.049 | 1 | 17 |
| trnau1apa | 0.55571 | 0.232969135 | 0.143 | 0.049 | 1 | 17 |
| zgc:123105 | 0.55571 | 0.291805532 | 0.214 | 0.098 | 1 | 17 |
| trib3 | 0.55571 | 0.192825711 | 0.143 | 0.049 | 1 | 17 |
| pigbos1 | 0.55586 | 0.217739415 | 0.143 | 0.049 | 1 | 17 |
| mgme1 | 0.55588 | 0.203033092 | 0.143 | 0.049 | 1 | 17 |
| cdkn1ca | 0.55588 | 0.227911103 | 0.143 | 0.049 | 1 | 17 |
| clk2b | 0.5559 | 0.213324664 | 0.143 | 0.049 | 1 | 17 |

| | | | | | | |
|---|---|---|---|---|---|---|
| coq7 | 0.5559 | 0.19263253 | 0.143 | 0.049 | 1 | 17 |
| oard1 | 0.5559 | 0.247317011 | 0.143 | 0.049 | 1 | 17 |
| si:dkey-12e7.4 | 0.5559 | 0.253962058 | 0.143 | 0.049 | 1 | 17 |
| blcap | 0.5559 | 0.241465838 | 0.143 | 0.049 | 1 | 17 |
| meaf6 | 0.55597 | -0.34941775 | 0.071 | 0.171 | 1 | 17 |
| prmt1 | 0.55618 | 0.235402569 | 0.5 | 0.341 | 1 | 17 |
| slirp | 0.55665 | 0.144327492 | 0.429 | 0.293 | 1 | 17 |
| myl12.2 | 0.55673 | -0.36806586 | 0.143 | 0.244 | 1 | 17 |
| rcc1l | 0.55676 | 0.224468532 | 0.214 | 0.098 | 1 | 17 |
| u2af2a | 0.55692 | 0.22729701 | 0.214 | 0.098 | 1 | 17 |
| psmd12 | 0.55751 | 0.280587849 | 0.214 | 0.098 | 1 | 17 |
| pinx1 | 0.55796 | 0.217690972 | 0.214 | 0.098 | 1 | 17 |
| cct7 | 0.55806 | -0.11462729 | 0.643 | 0.634 | 1 | 17 |
| abcf2a | 0.55846 | -0.2835503 | 0.214 | 0.268 | 1 | 17 |
| akap8l | 0.55852 | 0.206276961 | 0.214 | 0.098 | 1 | 17 |
| serbp1a | 0.5604 | 0.243431648 | 0.5 | 0.341 | 1 | 17 |
| arf6a | 0.56041 | 0.281943002 | 0.214 | 0.098 | 1 | 17 |
| ccdc137 | 0.56084 | 0.249477448 | 0.214 | 0.098 | 1 | 17 |
| lsm1 | 0.56085 | -0.35564902 | 0.214 | 0.341 | 1 | 17 |
| glod5 | 0.56138 | 0.250365669 | 0.214 | 0.098 | 1 | 17 |
| safb | 0.5614 | 0.266743281 | 0.214 | 0.098 | 1 | 17 |
| zgc:92606 | 0.56149 | 0.267611045 | 0.214 | 0.098 | 1 | 17 |
| cdc40 | 0.56159 | 0.29141003 | 0.214 | 0.098 | 1 | 17 |
| dhfr | 0.56165 | 0.285296194 | 0.214 | 0.098 | 1 | 17 |
| tbpl2 | 0.56185 | 0.234101519 | 0.214 | 0.098 | 1 | 17 |
| tprkb | 0.56187 | 0.257224537 | 0.214 | 0.098 | 1 | 17 |
| mrpl13 | 0.56191 | 0.242097556 | 0.214 | 0.098 | 1 | 17 |
| odc1 | 0.56254 | 0.205769583 | 0.357 | 0.22 | 1 | 17 |
| ppp6c | 0.56366 | -0.15571534 | 0.071 | 0.171 | 1 | 17 |
| zgc:56106 | 0.56526 | 0.111136491 | 0.357 | 0.244 | 1 | 17 |
| psap | 0.56727 | -0.31393513 | 0.071 | 0.122 | 1 | 17 |
| rps9 | 0.56887 | 0.100016684 | 1 | 0.951 | 1 | 17 |
| hspa5 | 0.56988 | 0.274935744 | 0.143 | 0.122 | 1 | 17 |
| polr2f | 0.57249 | -0.2348679 | 0.429 | 0.488 | 1 | 17 |
| mt-nd1 | 0.57312 | -0.22934568 | 0.071 | 0.171 | 1 | 17 |
| fkbp6 | 0.57626 | -0.33108365 | 0.143 | 0.268 | 1 | 17 |
| tmod2 | 0.5778 | -0.30405203 | 0.071 | 0.171 | 1 | 17 |
| tdrd9 | 0.57856 | 0.306469955 | 0.357 | 0.22 | 1 | 17 |
| nipsnap2 | 0.58035 | -0.29255488 | 0.143 | 0.195 | 1 | 17 |
| ndufb9 | 0.58039 | 0.265079073 | 0.5 | 0.341 | 1 | 17 |
| unga | 0.58201 | 0.154035021 | 0.286 | 0.171 | 1 | 17 |
| g3bp2 | 0.58269 | -0.3187089 | 0.071 | 0.171 | 1 | 17 |
| elavl2 | 0.58269 | 0.201280201 | 0.571 | 0.415 | 1 | 17 |
| vcp | 0.58506 | -0.27596615 | 0.286 | 0.341 | 1 | 17 |
| chrac1 | 0.58764 | 0.228913457 | 0.357 | 0.22 | 1 | 17 |
| prps1a | 0.58971 | -0.31311742 | 0.071 | 0.171 | 1 | 17 |
| zgc:162944 | 0.58986 | 0.319783937 | 0.429 | 0.293 | 1 | 17 |
| ssrp1a | 0.5909 | -0.11492415 | 0.357 | 0.341 | 1 | 17 |

| | | | | | | |
|---|---|---|---|---|---|---|
| rap2c | 0.59328 | -0.30659531 | 0.071 | 0.171 | 1 | 17 |
| aprt | 0.59371 | -0.32991106 | 0.143 | 0.268 | 1 | 17 |
| rpl36a | 0.59494 | 0.157435019 | 0.929 | 0.878 | 1 | 17 |
| hddc3 | 0.59518 | 0.173316901 | 0.429 | 0.293 | 1 | 17 |
| sh3glb1b | 0.59578 | -0.22093094 | 0.143 | 0.146 | 1 | 17 |
| nop56 | 0.59619 | -0.13503264 | 0.286 | 0.268 | 1 | 17 |
| pet100 | 0.59726 | -0.25119149 | 0.143 | 0.171 | 1 | 17 |
| ponzr1 | 0.59771 | -0.12102007 | 0.071 | 0.146 | 1 | 17 |
| tktb | 0.59791 | -0.29107932 | 0.286 | 0.366 | 1 | 17 |
| gnl2 | 0.59897 | -0.30280136 | 0.071 | 0.146 | 1 | 17 |
| aldh7a1 | 0.59935 | -0.18635293 | 0.286 | 0.293 | 1 | 17 |
| zgc:103697 | 0.60011 | -0.33582216 | 0.071 | 0.146 | 1 | 17 |
| eif3ja | 0.6011 | -0.21248604 | 0.286 | 0.317 | 1 | 17 |
| pim3 | 0.60164 | -0.29860809 | 0.071 | 0.171 | 1 | 17 |
| vdac3 | 0.60281 | -0.21702707 | 0.143 | 0.146 | 1 | 17 |
| tia1l | 0.60308 | -0.14686736 | 0.143 | 0.244 | 1 | 17 |
| rn7sk | 0.60674 | -0.2814653 | 0.071 | 0.171 | 1 | 17 |
| dele1 | 0.60758 | -0.33058841 | 0.071 | 0.171 | 1 | 17 |
| si:ch1073-429i10.3.1 | 0.6077 | -0.18664343 | 0.286 | 0.293 | 1 | 17 |
| btf3l4 | 0.6078 | 0.265396698 | 0.357 | 0.22 | 1 | 17 |
| rpia | 0.60783 | -0.3328163 | 0.214 | 0.341 | 1 | 17 |
| si:ch211-288g17.3 | 0.60812 | 0.300790428 | 0.357 | 0.22 | 1 | 17 |
| s100a1 | 0.60846 | -0.29919115 | 0.143 | 0.268 | 1 | 17 |
| adrm1 | 0.60848 | 0.176291898 | 0.286 | 0.171 | 1 | 17 |
| sms | 0.61143 | 0.195517931 | 0.429 | 0.293 | 1 | 17 |
| rpl12 | 0.61331 | -0.10228698 | 0.857 | 0.854 | 1 | 17 |
| zgc:136564 | 0.61384 | -0.24307206 | 0.071 | 0.171 | 1 | 17 |
| eif5 | 0.61454 | 0.119989295 | 0.357 | 0.244 | 1 | 17 |
| eif4bb | 0.61597 | -0.18058038 | 0.214 | 0.22 | 1 | 17 |
| zmat2 | 0.61694 | 0.167758733 | 0.286 | 0.171 | 1 | 17 |
| selenok | 0.61759 | -0.26950065 | 0.214 | 0.268 | 1 | 17 |
| cript | 0.61782 | 0.162842337 | 0.286 | 0.171 | 1 | 17 |
| rhocb | 0.61799 | -0.12641399 | 0.214 | 0.195 | 1 | 17 |
| cfl1 | 0.61971 | 0.174976549 | 0.5 | 0.366 | 1 | 17 |
| tomm40 | 0.62129 | 0.111084153 | 0.214 | 0.122 | 1 | 17 |
| ndufv2 | 0.62331 | -0.30473479 | 0.214 | 0.317 | 1 | 17 |
| rrp9 | 0.62511 | -0.24488132 | 0.071 | 0.171 | 1 | 17 |
| fubp1 | 0.62523 | -0.29964246 | 0.071 | 0.171 | 1 | 17 |
| rchy1 | 0.62756 | -0.26776713 | 0.071 | 0.171 | 1 | 17 |
| atp5mc3b | 0.62921 | 0.112181398 | 0.929 | 0.829 | 1 | 17 |
| hint1 | 0.62949 | 0.225533689 | 0.714 | 0.61 | 1 | 17 |
| mif | 0.63099 | -0.10518588 | 0.429 | 0.415 | 1 | 17 |
| thoc7 | 0.63144 | 0.17320286 | 0.286 | 0.171 | 1 | 17 |
| pabpc1a | 0.63146 | -0.13590715 | 0.643 | 0.659 | 1 | 17 |
| AK6 | 0.63166 | -0.28347731 | 0.071 | 0.171 | 1 | 17 |
| nap1l4a | 0.63225 | 0.108077156 | 0.214 | 0.122 | 1 | 17 |
| atp5f1d | 0.63471 | -0.15760296 | 0.429 | 0.439 | 1 | 17 |
| cfl1l | 0.63491 | -0.14922056 | 0.214 | 0.195 | 1 | 17 |

| | | | | | | |
|---|---|---|---|---|---|---|
| litaf | 0.63599 | -0.27799857 | 0.071 | 0.146 | 1 | 17 |
| taf13 | 0.63667 | 0.152372688 | 0.571 | 0.439 | 1 | 17 |
| ing2 | 0.63673 | 0.200845603 | 0.286 | 0.171 | 1 | 17 |
| sar1b | 0.63873 | -0.25654698 | 0.071 | 0.098 | 1 | 17 |
| alox5b.3 | 0.639 | -0.28683463 | 0.071 | 0.122 | 1 | 17 |
| lnpk | 0.63962 | 0.132519533 | 0.214 | 0.122 | 1 | 17 |
| eif2s1a | 0.64126 | 0.278567672 | 0.286 | 0.171 | 1 | 17 |
| lyar | 0.64183 | -0.11770048 | 0.214 | 0.195 | 1 | 17 |
| ppp4r2b | 0.64237 | 0.199061859 | 0.286 | 0.171 | 1 | 17 |
| adh5 | 0.64402 | 0.17861332 | 0.5 | 0.366 | 1 | 17 |
| rbm22 | 0.64883 | 0.219776761 | 0.286 | 0.171 | 1 | 17 |
| serbp1b | 0.65014 | -0.27304335 | 0.071 | 0.122 | 1 | 17 |
| dazl | 0.65076 | 0.136437987 | 0.429 | 0.317 | 1 | 17 |
| magoh | 0.6539 | 0.238104626 | 0.429 | 0.293 | 1 | 17 |
| fkbp4 | 0.65418 | 0.141736702 | 0.214 | 0.122 | 1 | 17 |
| ppdpfb | 0.65589 | 0.135943167 | 0.357 | 0.244 | 1 | 17 |
| ngdn | 0.65731 | 0.112167204 | 0.214 | 0.122 | 1 | 17 |
| spcs3 | 0.65751 | -0.29279472 | 0.071 | 0.146 | 1 | 17 |
| trappc3 | 0.65875 | 0.205147522 | 0.286 | 0.171 | 1 | 17 |
| atrx | 0.6591 | 0.145708712 | 0.214 | 0.122 | 1 | 17 |
| sycp1 | 0.65947 | -0.12409876 | 0.214 | 0.317 | 1 | 17 |
| tsen34 | 0.66108 | 0.136223333 | 0.214 | 0.122 | 1 | 17 |
| isy1 | 0.66134 | 0.278675914 | 0.143 | 0.073 | 1 | 17 |
| gtf2f2a | 0.66192 | 0.195225834 | 0.5 | 0.366 | 1 | 17 |
| acaa2 | 0.66271 | -0.28447887 | 0.071 | 0.146 | 1 | 17 |
| cwc25 | 0.66279 | 0.26246906 | 0.286 | 0.171 | 1 | 17 |
| hpf1 | 0.663 | 0.103332604 | 0.5 | 0.39 | 1 | 17 |
| mrpl47 | 0.66396 | 0.154273106 | 0.357 | 0.244 | 1 | 17 |
| elavl1a | 0.66463 | -0.28352888 | 0.071 | 0.146 | 1 | 17 |
| wdr83os | 0.66483 | 0.224569803 | 0.286 | 0.171 | 1 | 17 |
| si:dkey-185e18.7 | 0.6651 | 0.136215184 | 0.214 | 0.122 | 1 | 17 |
| cdv3 | 0.66623 | -0.31091907 | 0.143 | 0.244 | 1 | 17 |
| cdc6 | 0.66654 | 0.245204703 | 0.286 | 0.171 | 1 | 17 |
| mgmt | 0.66654 | 0.190976962 | 0.5 | 0.366 | 1 | 17 |
| ddx47 | 0.66657 | 0.249866088 | 0.286 | 0.171 | 1 | 17 |
| c1d | 0.66697 | 0.249913622 | 0.286 | 0.171 | 1 | 17 |
| srp9 | 0.66798 | -0.26817716 | 0.071 | 0.122 | 1 | 17 |
| xrcc1 | 0.66818 | 0.128174691 | 0.214 | 0.122 | 1 | 17 |
| manf | 0.66824 | 0.286676041 | 0.214 | 0.122 | 1 | 17 |
| pho | 0.6685 | -0.2041397 | 0.143 | 0.244 | 1 | 17 |
| tmem33 | 0.66868 | 0.160133952 | 0.214 | 0.122 | 1 | 17 |
| pkma | 0.67203 | 0.135489362 | 0.214 | 0.122 | 1 | 17 |
| ptrh2 | 0.67212 | -0.27792865 | 0.071 | 0.146 | 1 | 17 |
| zgc:114104 | 0.67288 | -0.26528165 | 0.071 | 0.122 | 1 | 17 |
| timm23a | 0.6742 | -0.16792612 | 0.5 | 0.537 | 1 | 17 |
| nuf2 | 0.67476 | 0.12808297 | 0.214 | 0.122 | 1 | 17 |
| usp1 | 0.67499 | 0.102699436 | 0.286 | 0.195 | 1 | 17 |
| uhrf1 | 0.67599 | -0.27306038 | 0.071 | 0.122 | 1 | 17 |

| | | | | | | |
|---|---|---|---|---|---|---|
| echs1 | 0.6786 | 0.151779312 | 0.214 | 0.122 | 1 | 17 |
| tmem14ca | 0.68068 | -0.26294229 | 0.143 | 0.195 | 1 | 17 |
| rpl9 | 0.68117 | -0.23958158 | 0.286 | 0.415 | 1 | 17 |
| ostc | 0.68161 | -0.17103555 | 0.214 | 0.22 | 1 | 17 |
| cmss1 | 0.68173 | 0.163017094 | 0.357 | 0.244 | 1 | 17 |
| LOC110440179 | 0.68178 | 0.133816382 | 0.214 | 0.122 | 1 | 17 |
| pabpn1 | 0.68228 | -0.22421519 | 0.286 | 0.415 | 1 | 17 |
| rrm2 | 0.68269 | 0.195817354 | 0.571 | 0.439 | 1 | 17 |
| eif4e1b | 0.68326 | 0.229088196 | 0.286 | 0.244 | 1 | 17 |
| eif4a1a | 0.68391 | 0.170886559 | 0.357 | 0.244 | 1 | 17 |
| ppa2 | 0.68478 | -0.25934283 | 0.143 | 0.195 | 1 | 17 |
| sept7b | 0.68654 | 0.157023638 | 0.214 | 0.122 | 1 | 17 |
| zgc:77086 | 0.68737 | -0.26969132 | 0.071 | 0.146 | 1 | 17 |
| gnb1b | 0.6894 | 0.140664263 | 0.214 | 0.122 | 1 | 17 |
| CABZ01115113.1 | 0.69032 | 0.272209618 | 0.143 | 0.073 | 1 | 17 |
| phax | 0.69082 | 0.178168896 | 0.357 | 0.244 | 1 | 17 |
| jpt2 | 0.69094 | 0.264152081 | 0.214 | 0.122 | 1 | 17 |
| cbx1b | 0.69219 | 0.155174921 | 0.214 | 0.122 | 1 | 17 |
| smc1b | 0.69233 | 0.256559867 | 0.357 | 0.244 | 1 | 17 |
| pomp | 0.69244 | 0.2144054 | 0.571 | 0.439 | 1 | 17 |
| nfyba | 0.69409 | 0.154932615 | 0.214 | 0.122 | 1 | 17 |
| hnrnpa0l | 0.69509 | 0.107523339 | 0.857 | 0.756 | 1 | 17 |
| gtf2h1 | 0.69632 | 0.242333708 | 0.143 | 0.073 | 1 | 17 |
| ddx42 | 0.69701 | -0.23237778 | 0.071 | 0.098 | 1 | 17 |
| cetn3 | 0.69753 | 0.17231346 | 0.214 | 0.122 | 1 | 17 |
| ttc5 | 0.69851 | -0.2541163 | 0.071 | 0.146 | 1 | 17 |
| gtf2h3 | 0.70117 | 0.15615369 | 0.214 | 0.122 | 1 | 17 |
| zgc:193690 | 0.70305 | 0.23919766 | 0.5 | 0.39 | 1 | 17 |
| bysl | 0.70316 | 0.117980301 | 0.286 | 0.195 | 1 | 17 |
| si:dkey-241l7.4 | 0.70462 | -0.25626002 | 0.071 | 0.146 | 1 | 17 |
| dnajc8 | 0.70515 | -0.15215932 | 0.214 | 0.22 | 1 | 17 |
| maip1 | 0.70538 | 0.254712906 | 0.143 | 0.073 | 1 | 17 |
| brms1la | 0.70603 | -0.27468762 | 0.071 | 0.146 | 1 | 17 |
| prpsap2 | 0.70643 | -0.25422227 | 0.071 | 0.146 | 1 | 17 |
| ctps1a | 0.70718 | 0.237268257 | 0.214 | 0.122 | 1 | 17 |
| uxt | 0.7072 | -0.24769288 | 0.071 | 0.146 | 1 | 17 |
| gpx4b | 0.70819 | 0.189113489 | 0.214 | 0.122 | 1 | 17 |
| rrp8 | 0.70857 | -0.27444947 | 0.071 | 0.146 | 1 | 17 |
| daam1a | 0.70912 | 0.101557629 | 0.143 | 0.073 | 1 | 17 |
| utp18 | 0.70927 | -0.26125406 | 0.071 | 0.122 | 1 | 17 |
| hax1 | 0.7094 | 0.187734887 | 0.214 | 0.122 | 1 | 17 |
| srsf9 | 0.70952 | -0.25630115 | 0.214 | 0.293 | 1 | 17 |
| polr1c | 0.70956 | -0.23977155 | 0.071 | 0.146 | 1 | 17 |
| ccdc58 | 0.70962 | 0.180703226 | 0.214 | 0.122 | 1 | 17 |
| alkbh4 | 0.70973 | 0.177295241 | 0.214 | 0.122 | 1 | 17 |
| ppan | 0.71037 | 0.105700404 | 0.143 | 0.073 | 1 | 17 |
| snapc2 | 0.71114 | 0.111468123 | 0.143 | 0.073 | 1 | 17 |
| mrpl57 | 0.71168 | -0.16665822 | 0.214 | 0.317 | 1 | 17 |

| | | | | | | |
|---|---|---|---|---|---|---|
| ndufv3 | 0.71199 | 0.108631193 | 0.143 | 0.073 | 1 | 17 |
| pnpo | 0.71215 | 0.105664694 | 0.143 | 0.073 | 1 | 17 |
| pfdn5 | 0.71311 | 0.170572338 | 0.214 | 0.122 | 1 | 17 |
| vps29 | 0.71336 | 0.167066537 | 0.214 | 0.122 | 1 | 17 |
| mars | 0.71358 | 0.119448444 | 0.143 | 0.073 | 1 | 17 |
| polr3h | 0.71426 | 0.230684151 | 0.214 | 0.122 | 1 | 17 |
| cmpk | 0.71434 | -0.26394085 | 0.143 | 0.22 | 1 | 17 |
| tuba8l2 | 0.71447 | -0.19716748 | 0.143 | 0.244 | 1 | 17 |
| ube2na | 0.71472 | 0.227479608 | 0.214 | 0.122 | 1 | 17 |
| hmgb3b | 0.71524 | 0.199333179 | 0.357 | 0.244 | 1 | 17 |
| nedd8 | 0.71585 | -0.18008115 | 0.357 | 0.39 | 1 | 17 |
| ubr7 | 0.7161 | -0.16253318 | 0.143 | 0.146 | 1 | 17 |
| rpl38 | 0.71616 | -0.11655989 | 0.857 | 0.902 | 1 | 17 |
| polr2eb | 0.71652 | -0.10826222 | 0.214 | 0.195 | 1 | 17 |
| kdm5c | 0.71668 | 0.184710782 | 0.214 | 0.122 | 1 | 17 |
| chmp1a | 0.71711 | -0.23701139 | 0.071 | 0.146 | 1 | 17 |
| baz1b | 0.71712 | -0.2395065 | 0.071 | 0.146 | 1 | 17 |
| ap1m2 | 0.71713 | 0.205962888 | 0.214 | 0.122 | 1 | 17 |
| sympk | 0.71749 | 0.239041057 | 0.214 | 0.122 | 1 | 17 |
| hnrnpl | 0.71756 | 0.21765798 | 0.214 | 0.122 | 1 | 17 |
| adssl | 0.71782 | 0.235114365 | 0.214 | 0.122 | 1 | 17 |
| xpot | 0.71798 | 0.217447734 | 0.214 | 0.122 | 1 | 17 |
| mrps23 | 0.718 | 0.180815469 | 0.214 | 0.122 | 1 | 17 |
| hdac1 | 0.71851 | -0.14215439 | 0.357 | 0.366 | 1 | 17 |
| slbp | 0.71894 | -0.24434219 | 0.286 | 0.366 | 1 | 17 |
| si:ch211-156b7.4 | 0.72028 | -0.155567 | 0.143 | 0.146 | 1 | 17 |
| si:dkey-188i13.11 | 0.72033 | 0.172151577 | 0.429 | 0.317 | 1 | 17 |
| yaf2 | 0.72093 | 0.107938974 | 0.143 | 0.073 | 1 | 17 |
| gkap1 | 0.72244 | 0.205457491 | 0.357 | 0.244 | 1 | 17 |
| smarca5 | 0.72324 | 0.105279022 | 0.143 | 0.073 | 1 | 17 |
| rpp30 | 0.72337 | 0.230554721 | 0.143 | 0.073 | 1 | 17 |
| nutf2 | 0.7247 | 0.111836251 | 0.143 | 0.073 | 1 | 17 |
| psmd4b | 0.72484 | -0.10206111 | 0.214 | 0.195 | 1 | 17 |
| mff | 0.72587 | 0.112664733 | 0.143 | 0.073 | 1 | 17 |
| pgp | 0.72699 | 0.117829347 | 0.143 | 0.073 | 1 | 17 |
| tomm5 | 0.7278 | -0.1508807 | 0.214 | 0.22 | 1 | 17 |
| qkia | 0.72799 | -0.24833247 | 0.071 | 0.146 | 1 | 17 |
| golph3 | 0.72821 | 0.226259452 | 0.143 | 0.073 | 1 | 17 |
| si:ch211-1o7.3 | 0.72831 | 0.102884245 | 0.143 | 0.073 | 1 | 17 |
| josd2 | 0.72885 | 0.119274837 | 0.143 | 0.073 | 1 | 17 |
| gdpd1 | 0.72939 | 0.100763911 | 0.143 | 0.073 | 1 | 17 |
| yipf5 | 0.72972 | -0.21183306 | 0.143 | 0.171 | 1 | 17 |
| prpf4 | 0.73029 | -0.10489545 | 0.143 | 0.122 | 1 | 17 |
| smu1a | 0.73032 | -0.25448983 | 0.214 | 0.293 | 1 | 17 |
| ddb2 | 0.73037 | -0.2381863 | 0.071 | 0.122 | 1 | 17 |
| top1mt | 0.7308 | -0.24365366 | 0.071 | 0.146 | 1 | 17 |
| cbr1l | 0.7317 | 0.113864544 | 0.143 | 0.073 | 1 | 17 |
| med22 | 0.73178 | 0.11213999 | 0.143 | 0.073 | 1 | 17 |

| | | | | | | |
|---|---|---|---|---|---|---|
| mrpl40 | 0.73198 | 0.151681096 | 0.143 | 0.073 | 1 | 17 |
| stk11 | 0.73322 | 0.128989341 | 0.143 | 0.073 | 1 | 17 |
| drg1 | 0.73335 | -0.24298617 | 0.071 | 0.146 | 1 | 17 |
| gb:bc139872 | 0.73346 | -0.24569237 | 0.071 | 0.146 | 1 | 17 |
| morc2 | 0.73378 | 0.23452777 | 0.143 | 0.073 | 1 | 17 |
| cdc123 | 0.73468 | 0.231834866 | 0.143 | 0.073 | 1 | 17 |
| pane1 | 0.73489 | -0.21135664 | 0.143 | 0.171 | 1 | 17 |
| sf3b3 | 0.73507 | -0.21698562 | 0.071 | 0.146 | 1 | 17 |
| nup58 | 0.7356 | -0.24837661 | 0.071 | 0.146 | 1 | 17 |
| ube2d3 | 0.73646 | 0.141439338 | 0.143 | 0.073 | 1 | 17 |
| trmt6 | 0.73677 | -0.21297323 | 0.071 | 0.146 | 1 | 17 |
| prpf38a | 0.73734 | -0.16230295 | 0.071 | 0.146 | 1 | 17 |
| zpr1 | 0.73838 | 0.145117401 | 0.143 | 0.073 | 1 | 17 |
| CDPF1 | 0.73852 | 0.211345889 | 0.143 | 0.073 | 1 | 17 |
| derl2 | 0.73887 | 0.13019431 | 0.143 | 0.073 | 1 | 17 |
| rab6a | 0.73978 | 0.101408466 | 0.143 | 0.073 | 1 | 17 |
| zgc:101744 | 0.74064 | -0.14934962 | 0.071 | 0.146 | 1 | 17 |
| osgep | 0.74065 | 0.156255117 | 0.143 | 0.073 | 1 | 17 |
| MED9 | 0.74072 | 0.207901269 | 0.143 | 0.073 | 1 | 17 |
| srsf2b | 0.74099 | 0.244978504 | 0.286 | 0.195 | 1 | 17 |
| vapal | 0.74124 | 0.147287926 | 0.143 | 0.073 | 1 | 17 |
| glo1 | 0.74197 | 0.131977981 | 0.286 | 0.195 | 1 | 17 |
| mad2l2 | 0.74202 | -0.18783239 | 0.214 | 0.244 | 1 | 17 |
| twistnb | 0.74209 | -0.16046502 | 0.071 | 0.146 | 1 | 17 |
| ndufs8a | 0.74215 | -0.10512921 | 0.143 | 0.122 | 1 | 17 |
| mat2aa | 0.74232 | 0.143960323 | 0.143 | 0.073 | 1 | 17 |
| asf1bb | 0.74244 | -0.24852121 | 0.143 | 0.22 | 1 | 17 |
| arid4a | 0.74269 | 0.11384566 | 0.143 | 0.073 | 1 | 17 |
| eif4ebp2 | 0.74314 | -0.21493884 | 0.071 | 0.146 | 1 | 17 |
| clns1a | 0.74354 | -0.22959766 | 0.214 | 0.268 | 1 | 17 |
| atp13a1 | 0.74383 | -0.21558087 | 0.071 | 0.146 | 1 | 17 |
| zgc:77929 | 0.74416 | -0.21646113 | 0.071 | 0.146 | 1 | 17 |
| si:dkeyp-50f7.2 | 0.74434 | -0.18854933 | 0.071 | 0.146 | 1 | 17 |
| rnaseh2b | 0.74538 | 0.250573241 | 0.286 | 0.195 | 1 | 17 |
| pdhx | 0.7462 | 0.107425235 | 0.143 | 0.073 | 1 | 17 |
| supt5h | 0.74624 | 0.128324146 | 0.143 | 0.073 | 1 | 17 |
| cdc7 | 0.74628 | 0.15931106 | 0.143 | 0.073 | 1 | 17 |
| commd3 | 0.74677 | 0.150142661 | 0.143 | 0.073 | 1 | 17 |
| znf598 | 0.747 | 0.112449298 | 0.143 | 0.073 | 1 | 17 |
| ttc4 | 0.7473 | 0.118097014 | 0.143 | 0.073 | 1 | 17 |
| bnip3la | 0.74756 | -0.20699261 | 0.071 | 0.146 | 1 | 17 |
| alcamb | 0.74794 | 0.149148037 | 0.143 | 0.073 | 1 | 17 |
| si:ch211-113p18.3 | 0.74798 | 0.141948101 | 0.143 | 0.073 | 1 | 17 |
| aamp | 0.74818 | 0.121584722 | 0.143 | 0.073 | 1 | 17 |
| dnajc18 | 0.7483 | 0.127145069 | 0.286 | 0.195 | 1 | 17 |
| csnk2a1 | 0.74882 | 0.218779024 | 0.429 | 0.317 | 1 | 17 |
| ash2l | 0.7492 | -0.21540181 | 0.071 | 0.146 | 1 | 17 |
| wu:fc33b09 | 0.7495 | -0.19685781 | 0.071 | 0.146 | 1 | 17 |

| | | | | | | |
|---|---|---|---|---|---|---|
| gabpa | 0.74951 | 0.142337929 | 0.143 | 0.073 | 1 | 17 |
| rnmt | 0.74951 | 0.13586464 | 0.143 | 0.073 | 1 | 17 |
| rab13 | 0.74989 | 0.117478737 | 0.143 | 0.073 | 1 | 17 |
| mrpl43 | 0.75009 | 0.148180375 | 0.143 | 0.073 | 1 | 17 |
| cdc20 | 0.75023 | -0.19755822 | 0.071 | 0.146 | 1 | 17 |
| ddx54 | 0.75026 | 0.141497413 | 0.143 | 0.073 | 1 | 17 |
| si:dkey-74k8.4 | 0.75039 | 0.128167349 | 0.143 | 0.073 | 1 | 17 |
| arpc4l | 0.75083 | -0.23813882 | 0.071 | 0.122 | 1 | 17 |
| mrps10 | 0.75099 | -0.23140202 | 0.071 | 0.146 | 1 | 17 |
| wdr46 | 0.75122 | -0.22534954 | 0.071 | 0.146 | 1 | 17 |
| zgc:92313 | 0.75124 | -0.18586998 | 0.071 | 0.146 | 1 | 17 |
| dars | 0.75145 | 0.133725429 | 0.143 | 0.073 | 1 | 17 |
| med21 | 0.75146 | -0.19843294 | 0.143 | 0.171 | 1 | 17 |
| yrdc | 0.75154 | 0.19159058 | 0.143 | 0.073 | 1 | 17 |
| zgc:110782 | 0.75278 | 0.119617683 | 0.143 | 0.073 | 1 | 17 |
| bag4 | 0.75287 | 0.190976972 | 0.143 | 0.073 | 1 | 17 |
| mrpl36 | 0.7531 | 0.221647399 | 0.143 | 0.073 | 1 | 17 |
| sept6 | 0.75336 | 0.194892524 | 0.143 | 0.073 | 1 | 17 |
| ddx43 | 0.7534 | 0.193500375 | 0.143 | 0.073 | 1 | 17 |
| atg12 | 0.75386 | 0.138820117 | 0.143 | 0.073 | 1 | 17 |
| trappc5 | 0.75408 | -0.24279407 | 0.071 | 0.122 | 1 | 17 |
| tuba1c | 0.75487 | 0.206682841 | 0.143 | 0.073 | 1 | 17 |
| mpped2a | 0.75488 | 0.184085213 | 0.143 | 0.073 | 1 | 17 |
| ube2ib | 0.7553 | -0.14947031 | 0.214 | 0.22 | 1 | 17 |
| arl14ep | 0.75554 | 0.179601632 | 0.143 | 0.073 | 1 | 17 |
| ppa1b | 0.75565 | 0.201858641 | 0.429 | 0.317 | 1 | 17 |
| vps36 | 0.75569 | 0.146721011 | 0.143 | 0.073 | 1 | 17 |
| med29 | 0.75603 | 0.166827578 | 0.143 | 0.073 | 1 | 17 |
| vps35 | 0.7562 | 0.171019357 | 0.143 | 0.073 | 1 | 17 |
| chmp4bb | 0.75626 | 0.13451212 | 0.143 | 0.073 | 1 | 17 |
| hgs | 0.75659 | 0.143344496 | 0.143 | 0.073 | 1 | 17 |
| tia1 | 0.75715 | 0.145632624 | 0.143 | 0.073 | 1 | 17 |
| scp2b | 0.7574 | 0.188046223 | 0.143 | 0.073 | 1 | 17 |
| arf6b | 0.7579 | 0.174208264 | 0.143 | 0.073 | 1 | 17 |
| tspan13a | 0.75791 | 0.103803808 | 0.143 | 0.073 | 1 | 17 |
| zgc:66160 | 0.75792 | 0.152000661 | 0.143 | 0.073 | 1 | 17 |
| egln1b | 0.7583 | 0.171579764 | 0.143 | 0.073 | 1 | 17 |
| si:ch211-262i1.6 | 0.75833 | 0.182401889 | 0.143 | 0.073 | 1 | 17 |
| tmco1 | 0.75834 | 0.18453166 | 0.143 | 0.073 | 1 | 17 |
| sumo2a | 0.75842 | 0.17014059 | 0.143 | 0.073 | 1 | 17 |
| si:ch73-7i4.1 | 0.75846 | 0.167638576 | 0.143 | 0.073 | 1 | 17 |
| mut | 0.75849 | 0.167362926 | 0.143 | 0.073 | 1 | 17 |
| LOC108180269 | 0.75865 | 0.177626378 | 0.143 | 0.073 | 1 | 17 |
| wu:fb59d01 | 0.75874 | 0.170931833 | 0.143 | 0.073 | 1 | 17 |
| dusp22b | 0.75886 | 0.175853595 | 0.143 | 0.073 | 1 | 17 |
| gtf2a1 | 0.7589 | 0.178251973 | 0.143 | 0.073 | 1 | 17 |
| nudcd3 | 0.7589 | 0.167703428 | 0.143 | 0.073 | 1 | 17 |
| txn2 | 0.7589 | 0.107025126 | 0.357 | 0.268 | 1 | 17 |

| | | | | | | |
|---|---|---|---|---|---|---|
| elac2 | 0.75895 | 0.158263476 | 0.143 | 0.073 | 1 | 17 |
| ndufs4 | 0.75896 | -0.14624961 | 0.214 | 0.22 | 1 | 17 |
| papola | 0.75897 | 0.149801131 | 0.143 | 0.073 | 1 | 17 |
| si:ch211-266i6.3 | 0.75898 | 0.153648272 | 0.143 | 0.073 | 1 | 17 |
| fam151b | 0.75901 | 0.151907005 | 0.143 | 0.073 | 1 | 17 |
| cdc26 | 0.75902 | 0.163803889 | 0.143 | 0.073 | 1 | 17 |
| snrnp40 | 0.75906 | 0.189979592 | 0.143 | 0.073 | 1 | 17 |
| taf10 | 0.75906 | 0.184138711 | 0.143 | 0.073 | 1 | 17 |
| pkp3b | 0.75907 | 0.19637043 | 0.143 | 0.073 | 1 | 17 |
| ddx51 | 0.75907 | 0.15657538 | 0.143 | 0.073 | 1 | 17 |
| zgc:86609 | 0.75959 | 0.142181165 | 0.071 | 0.049 | 1 | 17 |
| aldh16a1 | 0.75959 | 0.102929769 | 0.071 | 0.049 | 1 | 17 |
| arpc1b | 0.75959 | 0.103560844 | 0.071 | 0.049 | 1 | 17 |
| cdc27 | 0.75959 | 0.102073085 | 0.071 | 0.049 | 1 | 17 |
| cryba1l1 | 0.75959 | 0.126120522 | 0.071 | 0.049 | 1 | 17 |
| kcnip3b | 0.75959 | 0.102593365 | 0.071 | 0.049 | 1 | 17 |
| mettl17 | 0.75959 | 0.10242562 | 0.071 | 0.049 | 1 | 17 |
| pebp1 | 0.75959 | 0.121536231 | 0.071 | 0.049 | 1 | 17 |
| pelo | 0.75959 | 0.152655243 | 0.071 | 0.049 | 1 | 17 |
| poldip3 | 0.75959 | 0.136504761 | 0.071 | 0.049 | 1 | 17 |
| smarca4a | 0.75959 | 0.123134182 | 0.071 | 0.049 | 1 | 17 |
| trove2 | 0.75959 | 0.116485114 | 0.071 | 0.049 | 1 | 17 |
| vps8 | 0.75959 | 0.151018153 | 0.071 | 0.049 | 1 | 17 |
| zdhhc16a | 0.75959 | 0.104751438 | 0.071 | 0.049 | 1 | 17 |
| atp1b3a | 0.75959 | 0.10964514 | 0.071 | 0.049 | 1 | 17 |
| daw1 | 0.75959 | 0.100228719 | 0.071 | 0.049 | 1 | 17 |
| enoph1 | 0.75959 | 0.137527366 | 0.071 | 0.049 | 1 | 17 |
| lyrm1 | 0.75959 | 0.139465702 | 0.071 | 0.049 | 1 | 17 |
| nufip1 | 0.75959 | 0.109082472 | 0.071 | 0.049 | 1 | 17 |
| polh | 0.75959 | 0.102957274 | 0.071 | 0.049 | 1 | 17 |
| ptgis | 0.75959 | 0.107643054 | 0.071 | 0.049 | 1 | 17 |
| si:ch211-40k21.9 | 0.75959 | 0.137830904 | 0.071 | 0.049 | 1 | 17 |
| tmem214 | 0.75959 | 0.130126549 | 0.071 | 0.049 | 1 | 17 |
| celf2 | 0.75959 | 0.154958773 | 0.071 | 0.049 | 1 | 17 |
| rtn4ip1 | 0.75959 | 0.123849985 | 0.071 | 0.049 | 1 | 17 |
| snx10a | 0.75959 | 0.134890131 | 0.071 | 0.049 | 1 | 17 |
| tmem192 | 0.75959 | 0.130208784 | 0.071 | 0.049 | 1 | 17 |
| trim37 | 0.75959 | -0.14433563 | 0.071 | 0.049 | 1 | 17 |
| wipf2b | 0.75959 | 0.12162735 | 0.071 | 0.049 | 1 | 17 |
| etfdh | 0.75959 | 0.107995153 | 0.071 | 0.049 | 1 | 17 |
| isca1 | 0.75959 | 0.110999377 | 0.071 | 0.049 | 1 | 17 |
| msl1b | 0.75959 | 0.163644781 | 0.071 | 0.049 | 1 | 17 |
| srsf1a | 0.75959 | 0.113419197 | 0.071 | 0.049 | 1 | 17 |
| ctr9 | 0.75959 | 0.117932625 | 0.071 | 0.049 | 1 | 17 |
| nsun5 | 0.75959 | 0.155457078 | 0.071 | 0.049 | 1 | 17 |
| cbr4 | 0.75959 | 0.128294057 | 0.071 | 0.049 | 1 | 17 |
| haus1 | 0.75959 | 0.151899378 | 0.071 | 0.049 | 1 | 17 |
| dtd1 | 0.75959 | 0.105744659 | 0.071 | 0.049 | 1 | 17 |

| | | | | | | |
|---|---|---|---|---|---|---|
| mpnd | 0.75959 | 0.110966219 | 0.071 | 0.049 | 1 | 17 |
| ndel1b | 0.75959 | 0.103893613 | 0.071 | 0.049 | 1 | 17 |
| ntan1 | 0.75959 | 0.130935968 | 0.071 | 0.049 | 1 | 17 |
| metap2b | 0.76027 | -0.13244271 | 0.357 | 0.366 | 1 | 17 |
| bloc1s1 | 0.76046 | -0.1081574 | 0.143 | 0.122 | 1 | 17 |
| alg13 | 0.76217 | -0.23963048 | 0.071 | 0.122 | 1 | 17 |
| smim7 | 0.76323 | -0.23330679 | 0.071 | 0.122 | 1 | 17 |
| ppie | 0.7643 | 0.248893089 | 0.214 | 0.146 | 1 | 17 |
| napab | 0.76457 | -0.22692364 | 0.071 | 0.122 | 1 | 17 |
| hspe1 | 0.765 | 0.212704509 | 0.5 | 0.415 | 1 | 17 |
| hadhb | 0.76597 | 0.232144147 | 0.214 | 0.146 | 1 | 17 |
| smek1 | 0.7696 | -0.23110984 | 0.071 | 0.122 | 1 | 17 |
| pafah1b1a | 0.7703 | -0.2263551 | 0.071 | 0.122 | 1 | 17 |
| rfc5 | 0.77041 | -0.24757784 | 0.143 | 0.22 | 1 | 17 |
| s100u | 0.77162 | -0.16246708 | 0.071 | 0.073 | 1 | 17 |
| zgc:77739 | 0.77177 | -0.22592055 | 0.071 | 0.122 | 1 | 17 |
| csnk1a1 | 0.77187 | -0.22853815 | 0.286 | 0.366 | 1 | 17 |
| max | 0.77288 | -0.15126832 | 0.143 | 0.146 | 1 | 17 |
| map7d3 | 0.774 | -0.2250061 | 0.071 | 0.122 | 1 | 17 |
| hspa8 | 0.77433 | -0.10668263 | 0.857 | 0.902 | 1 | 17 |
| pafah1b1b | 0.77498 | -0.20331081 | 0.071 | 0.098 | 1 | 17 |
| nop58 | 0.77608 | -0.18656889 | 0.286 | 0.39 | 1 | 17 |
| atp5pd | 0.77635 | -0.10406017 | 0.5 | 0.512 | 1 | 17 |
| shmt1 | 0.77698 | -0.22365611 | 0.143 | 0.195 | 1 | 17 |
| ola1 | 0.77787 | 0.121856108 | 0.357 | 0.268 | 1 | 17 |
| gmppaa | 0.77817 | -0.2227155 | 0.071 | 0.122 | 1 | 17 |
| trir | 0.77857 | -0.1997963 | 0.071 | 0.098 | 1 | 17 |
| si:ch211-127m7.2 | 0.77868 | -0.22521748 | 0.071 | 0.122 | 1 | 17 |
| ccdc124 | 0.7787 | -0.15132666 | 0.143 | 0.146 | 1 | 17 |
| luc7l | 0.77919 | -0.20250968 | 0.071 | 0.098 | 1 | 17 |
| si:rp71-45k5.4 | 0.77929 | 0.1040058 | 0.357 | 0.268 | 1 | 17 |
| prkcsh | 0.78035 | -0.22295395 | 0.071 | 0.122 | 1 | 17 |
| chd7 | 0.78203 | -0.20101032 | 0.071 | 0.098 | 1 | 17 |
| zgc:110821 | 0.78318 | -0.2031991 | 0.071 | 0.098 | 1 | 17 |
| auh | 0.78324 | 0.170963377 | 0.286 | 0.195 | 1 | 17 |
| fdx1 | 0.7833 | -0.22157562 | 0.143 | 0.22 | 1 | 17 |
| eci1 | 0.78331 | -0.21383489 | 0.071 | 0.122 | 1 | 17 |
| smarca1 | 0.78497 | -0.23339167 | 0.071 | 0.122 | 1 | 17 |
| arf3a | 0.78508 | 0.221945073 | 0.143 | 0.098 | 1 | 17 |
| calr3b | 0.78643 | -0.19253775 | 0.071 | 0.122 | 1 | 17 |
| gcshb | 0.787 | 0.17217482 | 0.286 | 0.195 | 1 | 17 |
| slc25a12 | 0.78793 | -0.21805335 | 0.071 | 0.122 | 1 | 17 |
| dis3 | 0.78845 | -0.21468271 | 0.071 | 0.122 | 1 | 17 |
| pfdn2 | 0.78922 | -0.10689439 | 0.214 | 0.293 | 1 | 17 |
| gyg1a | 0.79022 | 0.118767485 | 0.357 | 0.268 | 1 | 17 |
| taf1a | 0.79231 | -0.19826536 | 0.071 | 0.098 | 1 | 17 |
| ubfd1 | 0.79378 | -0.21384704 | 0.071 | 0.122 | 1 | 17 |
| llph | 0.79417 | -0.1255678 | 0.286 | 0.293 | 1 | 17 |

| | | | | | | |
|---|---|---|---|---|---|---|
| si:dkey-78l4.14 | 0.79428 | -0.20353675 | 0.071 | 0.122 | 1 | 17 |
| kat5b | 0.79493 | -0.14128755 | 0.143 | 0.146 | 1 | 17 |
| adhfe1 | 0.79572 | -0.2003852 | 0.071 | 0.098 | 1 | 17 |
| yipf6 | 0.79649 | -0.14812604 | 0.071 | 0.073 | 1 | 17 |
| uchl3 | 0.79668 | 0.206961398 | 0.214 | 0.171 | 1 | 17 |
| dpp3 | 0.79757 | -0.20789743 | 0.071 | 0.122 | 1 | 17 |
| osbpl7 | 0.79933 | -0.1927337 | 0.071 | 0.098 | 1 | 17 |
| cenpx | 0.80017 | -0.21493819 | 0.143 | 0.22 | 1 | 17 |
| mdh1aa | 0.80097 | -0.15137937 | 0.357 | 0.39 | 1 | 17 |
| cdk13 | 0.8019 | -0.19219015 | 0.071 | 0.098 | 1 | 17 |
| cdh1 | 0.80243 | -0.20687341 | 0.071 | 0.122 | 1 | 17 |
| si:ch211-226h8.14 | 0.80343 | -0.20088527 | 0.071 | 0.122 | 1 | 17 |
| rnf168 | 0.8047 | -0.20786907 | 0.071 | 0.122 | 1 | 17 |
| atp5po | 0.80814 | 0.109787946 | 0.429 | 0.341 | 1 | 17 |
| lamtor2 | 0.80837 | -0.17708317 | 0.143 | 0.171 | 1 | 17 |
| hnrnpa3 | 0.80842 | -0.1822926 | 0.143 | 0.22 | 1 | 17 |
| calm1b | 0.80866 | -0.19461237 | 0.071 | 0.122 | 1 | 17 |
| ube2g1a | 0.80984 | -0.14670308 | 0.143 | 0.22 | 1 | 17 |
| zglp1 | 0.80993 | -0.15513717 | 0.143 | 0.146 | 1 | 17 |
| mibp2 | 0.80994 | -0.1610943 | 0.286 | 0.317 | 1 | 17 |
| r3hcc1 | 0.81041 | -0.13972294 | 0.071 | 0.073 | 1 | 17 |
| tspan18a | 0.81259 | -0.18300607 | 0.071 | 0.098 | 1 | 17 |
| rhoab | 0.81319 | 0.109526721 | 0.214 | 0.146 | 1 | 17 |
| phb | 0.81457 | -0.17506542 | 0.143 | 0.171 | 1 | 17 |
| ccdc12 | 0.81526 | -0.21146315 | 0.143 | 0.195 | 1 | 17 |
| wu:fa19b12 | 0.8153 | -0.19627241 | 0.071 | 0.122 | 1 | 17 |
| top1l | 0.81535 | -0.21060725 | 0.143 | 0.195 | 1 | 17 |
| strap | 0.81546 | -0.11952253 | 0.357 | 0.366 | 1 | 17 |
| hat1 | 0.81659 | -0.18251846 | 0.143 | 0.171 | 1 | 17 |
| pdcd6ip | 0.8169 | -0.19279461 | 0.071 | 0.122 | 1 | 17 |
| capza1b | 0.81713 | -0.16809879 | 0.214 | 0.244 | 1 | 17 |
| srsf6a | 0.81725 | -0.18975492 | 0.071 | 0.098 | 1 | 17 |
| rae1 | 0.81916 | 0.106749111 | 0.214 | 0.146 | 1 | 17 |
| zgc:153681 | 0.81925 | -0.19401128 | 0.071 | 0.122 | 1 | 17 |
| rpf2 | 0.8203 | -0.20329793 | 0.071 | 0.122 | 1 | 17 |
| aldh3a2a | 0.82147 | -0.18666646 | 0.071 | 0.122 | 1 | 17 |
| rcbtb1 | 0.82155 | -0.19484019 | 0.071 | 0.122 | 1 | 17 |
| eif1axb | 0.82282 | -0.19702576 | 0.286 | 0.366 | 1 | 17 |
| uchl1 | 0.82369 | 0.170047945 | 0.357 | 0.268 | 1 | 17 |
| arcn1a | 0.8247 | -0.18102859 | 0.071 | 0.098 | 1 | 17 |
| p4ha1b | 0.82483 | -0.19178853 | 0.071 | 0.122 | 1 | 17 |
| faub | 0.82658 | 0.115382265 | 0.429 | 0.341 | 1 | 17 |
| cdca9 | 0.82794 | 0.100919026 | 0.214 | 0.146 | 1 | 17 |
| vaspb | 0.82807 | -0.18157899 | 0.071 | 0.098 | 1 | 17 |
| zgc:92591 | 0.8281 | 0.182603796 | 0.143 | 0.098 | 1 | 17 |
| si:ch73-37h15.2 | 0.8285 | -0.18391743 | 0.071 | 0.098 | 1 | 17 |
| rps3 | 0.82875 | -0.10683313 | 0.786 | 0.854 | 1 | 17 |
| mrpl11 | 0.82904 | 0.10621305 | 0.214 | 0.146 | 1 | 17 |

| | | | | | | |
|---|---|---|---|---|---|---|
| maco1a | 0.8297 | -0.12384875 | 0.143 | 0.146 | 1 | 17 |
| eif2s1b | 0.83002 | -0.15657819 | 0.214 | 0.244 | 1 | 17 |
| cdkn3 | 0.83013 | -0.16924915 | 0.071 | 0.098 | 1 | 17 |
| timm9 | 0.83065 | -0.18234383 | 0.071 | 0.122 | 1 | 17 |
| ilf3b | 0.83243 | 0.118417172 | 0.214 | 0.146 | 1 | 17 |
| zgc:114123 | 0.83247 | 0.124674788 | 0.071 | 0.073 | 1 | 17 |
| serf2 | 0.833 | 0.184285833 | 0.357 | 0.293 | 1 | 17 |
| mrps21 | 0.83346 | -0.20038866 | 0.143 | 0.195 | 1 | 17 |
| castor2 | 0.834 | -0.13712441 | 0.071 | 0.073 | 1 | 17 |
| mrpl30 | 0.83447 | -0.16283642 | 0.143 | 0.171 | 1 | 17 |
| nop2 | 0.83447 | 0.181117 | 0.214 | 0.146 | 1 | 17 |
| med6 | 0.83548 | 0.110939955 | 0.214 | 0.146 | 1 | 17 |
| wee2 | 0.83636 | 0.122571936 | 0.214 | 0.146 | 1 | 17 |
| insig1 | 0.83696 | -0.1317684 | 0.071 | 0.073 | 1 | 17 |
| psmc1b | 0.83708 | 0.120324627 | 0.214 | 0.146 | 1 | 17 |
| aif1l | 0.83717 | 0.128576018 | 0.214 | 0.146 | 1 | 17 |
| wdr83 | 0.83721 | -0.17525474 | 0.071 | 0.122 | 1 | 17 |
| brf2 | 0.83822 | -0.1200652 | 0.143 | 0.146 | 1 | 17 |
| rsl24d1 | 0.83826 | 0.137287362 | 0.429 | 0.341 | 1 | 17 |
| hddc2 | 0.83842 | -0.14336779 | 0.214 | 0.293 | 1 | 17 |
| slc5a1 | 0.83863 | -0.12964821 | 0.071 | 0.073 | 1 | 17 |
| metap1 | 0.83883 | -0.17600484 | 0.071 | 0.122 | 1 | 17 |
| sec11a | 0.83936 | -0.13438726 | 0.143 | 0.146 | 1 | 17 |
| CABZ01084963.1 | 0.84037 | 0.193173616 | 0.143 | 0.098 | 1 | 17 |
| slc25a11 | 0.84053 | 0.176639866 | 0.214 | 0.146 | 1 | 17 |
| nudt21 | 0.8411 | -0.18685072 | 0.214 | 0.293 | 1 | 17 |
| gcsha | 0.84117 | -0.13907687 | 0.214 | 0.293 | 1 | 17 |
| hm13 | 0.84124 | 0.12640326 | 0.214 | 0.146 | 1 | 17 |
| os9 | 0.84126 | -0.18132208 | 0.071 | 0.122 | 1 | 17 |
| acat1 | 0.84131 | -0.15828362 | 0.071 | 0.122 | 1 | 17 |
| psmc3ip | 0.8416 | 0.144125725 | 0.214 | 0.146 | 1 | 17 |
| clgn | 0.84165 | -0.12714751 | 0.071 | 0.073 | 1 | 17 |
| ap2m1b | 0.84189 | 0.169844335 | 0.214 | 0.146 | 1 | 17 |
| mtfr1l | 0.84246 | -0.13187874 | 0.071 | 0.073 | 1 | 17 |
| stra6 | 0.84284 | -0.13964912 | 0.071 | 0.073 | 1 | 17 |
| wdr18 | 0.84296 | 0.185271117 | 0.143 | 0.098 | 1 | 17 |
| atic | 0.84347 | -0.16895408 | 0.071 | 0.122 | 1 | 17 |
| gpn1 | 0.84379 | -0.11879173 | 0.143 | 0.146 | 1 | 17 |
| ctnnb1 | 0.84405 | 0.203232904 | 0.143 | 0.098 | 1 | 17 |
| si:zfos-80g12.1 | 0.84407 | -0.12751827 | 0.071 | 0.073 | 1 | 17 |
| gins3 | 0.84423 | 0.161644339 | 0.214 | 0.146 | 1 | 17 |
| itpa | 0.84512 | -0.16375338 | 0.143 | 0.171 | 1 | 17 |
| grhpra | 0.84516 | -0.16115922 | 0.071 | 0.122 | 1 | 17 |
| rpa3 | 0.84526 | 0.157935737 | 0.429 | 0.341 | 1 | 17 |
| lsm14b | 0.84527 | -0.14165451 | 0.071 | 0.122 | 1 | 17 |
| UBL4A | 0.84546 | -0.1202466 | 0.143 | 0.146 | 1 | 17 |
| mob3a | 0.84563 | 0.14891994 | 0.214 | 0.146 | 1 | 17 |
| gpat2 | 0.84566 | 0.161090909 | 0.214 | 0.146 | 1 | 17 |

| | | | | | | |
|---|---|---|---|---|---|---|
| csnk1db | 0.84581 | -0.16388758 | 0.071 | 0.098 | 1 | 17 |
| znf76 | 0.84585 | -0.12787984 | 0.071 | 0.073 | 1 | 17 |
| trip13 | 0.84592 | 0.151817398 | 0.214 | 0.146 | 1 | 17 |
| sec13 | 0.84599 | 0.135512723 | 0.214 | 0.146 | 1 | 17 |
| pcgf6 | 0.84606 | -0.12048258 | 0.071 | 0.122 | 1 | 17 |
| gins4 | 0.84618 | -0.16846724 | 0.071 | 0.122 | 1 | 17 |
| ISCU (1 of many) | 0.84631 | 0.153273519 | 0.214 | 0.146 | 1 | 17 |
| ebna1bp2 | 0.84651 | -0.18557007 | 0.214 | 0.293 | 1 | 17 |
| ttf1 | 0.84661 | -0.16355731 | 0.143 | 0.171 | 1 | 17 |
| bzw1b | 0.84675 | -0.1774209 | 0.214 | 0.293 | 1 | 17 |
| lsm4 | 0.84675 | 0.143888786 | 0.643 | 0.561 | 1 | 17 |
| ssb | 0.84704 | -0.17938213 | 0.071 | 0.122 | 1 | 17 |
| UTP14C | 0.8472 | -0.10326899 | 0.071 | 0.122 | 1 | 17 |
| ifi46 | 0.84745 | -0.16808297 | 0.071 | 0.098 | 1 | 17 |
| ccnh | 0.84774 | -0.15427852 | 0.071 | 0.122 | 1 | 17 |
| polr1a | 0.84869 | -0.13033023 | 0.071 | 0.073 | 1 | 17 |
| ppp3ccb | 0.84927 | -0.12813048 | 0.071 | 0.073 | 1 | 17 |
| pask | 0.84989 | -0.17121332 | 0.071 | 0.098 | 1 | 17 |
| fam120c | 0.85023 | -0.12626031 | 0.071 | 0.073 | 1 | 17 |
| selenoj | 0.85038 | -0.16852101 | 0.071 | 0.122 | 1 | 17 |
| otub1a | 0.85098 | -0.1275443 | 0.071 | 0.073 | 1 | 17 |
| prim2 | 0.85117 | -0.11061984 | 0.071 | 0.122 | 1 | 17 |
| krcp | 0.85123 | -0.14920236 | 0.071 | 0.122 | 1 | 17 |
| arglu1a | 0.85154 | -0.1268512 | 0.071 | 0.073 | 1 | 17 |
| acadvl | 0.85182 | -0.16872307 | 0.071 | 0.122 | 1 | 17 |
| tmsb5 | 0.85212 | -0.15523751 | 0.071 | 0.122 | 1 | 17 |
| snrnp70 | 0.85289 | -0.1728724 | 0.071 | 0.122 | 1 | 17 |
| gemin6 | 0.85303 | -0.14930803 | 0.071 | 0.122 | 1 | 17 |
| cul1a | 0.85315 | -0.16655561 | 0.071 | 0.098 | 1 | 17 |
| phgdh | 0.85344 | -0.1566466 | 0.071 | 0.098 | 1 | 17 |
| si:ch211-191j22.3 | 0.85374 | -0.17281544 | 0.071 | 0.098 | 1 | 17 |
| nmt1a | 0.85433 | -0.1496531 | 0.071 | 0.122 | 1 | 17 |
| paics | 0.85527 | -0.16131601 | 0.071 | 0.122 | 1 | 17 |
| atp6v0ca | 0.85543 | -0.1597998 | 0.143 | 0.171 | 1 | 17 |
| zgc:66433 | 0.8562 | -0.10154408 | 0.071 | 0.122 | 1 | 17 |
| ndnl2 | 0.8565 | -0.12061627 | 0.071 | 0.122 | 1 | 17 |
| znf330 | 0.85683 | -0.11769009 | 0.071 | 0.122 | 1 | 17 |
| apip | 0.85746 | -0.11629644 | 0.071 | 0.122 | 1 | 17 |
| ckba | 0.85749 | -0.16107562 | 0.071 | 0.098 | 1 | 17 |
| sh3glb1a | 0.85874 | -0.12403062 | 0.071 | 0.073 | 1 | 17 |
| bms1 | 0.85916 | -0.15392639 | 0.071 | 0.122 | 1 | 17 |
| dnm1l | 0.85943 | -0.17203824 | 0.071 | 0.122 | 1 | 17 |
| rrp7a | 0.86036 | 0.180637124 | 0.143 | 0.098 | 1 | 17 |
| akap1a | 0.86055 | -0.18177963 | 0.071 | 0.122 | 1 | 17 |
| zgc:92818 | 0.86065 | -0.16658809 | 0.071 | 0.098 | 1 | 17 |
| snu13a | 0.86114 | -0.15199263 | 0.071 | 0.122 | 1 | 17 |
| nkrf | 0.86154 | -0.16768788 | 0.071 | 0.098 | 1 | 17 |
| si:dkey-30c15.10 | 0.86218 | -0.16375954 | 0.071 | 0.122 | 1 | 17 |

| | | | | | | |
|---|---|---|---|---|---|---|
| golga7 | 0.86312 | -0.16013025 | 0.071 | 0.098 | 1 | 17 |
| MRPS12 | 0.86339 | -0.1160442 | 0.071 | 0.122 | 1 | 17 |
| thap4 | 0.86401 | -0.14993945 | 0.071 | 0.122 | 1 | 17 |
| polr3k | 0.86423 | -0.14182544 | 0.071 | 0.122 | 1 | 17 |
| grpel2 | 0.86452 | 0.108068833 | 0.286 | 0.22 | 1 | 17 |
| si:ch1073-406l10.2 | 0.86464 | -0.12054576 | 0.071 | 0.073 | 1 | 17 |
| MZT1 | 0.86465 | -0.11231912 | 0.071 | 0.122 | 1 | 17 |
| si:ch211-10a23.2 | 0.86467 | -0.13121892 | 0.071 | 0.122 | 1 | 17 |
| waca | 0.86467 | -0.14611431 | 0.071 | 0.122 | 1 | 17 |
| lmnb1 | 0.86493 | -0.14517829 | 0.071 | 0.122 | 1 | 17 |
| btc | 0.86498 | -0.11458951 | 0.071 | 0.122 | 1 | 17 |
| sdha | 0.86501 | -0.11430544 | 0.071 | 0.122 | 1 | 17 |
| pdcd4b | 0.86529 | -0.13152245 | 0.071 | 0.122 | 1 | 17 |
| ppil1 | 0.86532 | -0.13379136 | 0.071 | 0.122 | 1 | 17 |
| anxa11b | 0.86535 | -0.12374627 | 0.071 | 0.122 | 1 | 17 |
| pgls | 0.86537 | -0.13711426 | 0.071 | 0.122 | 1 | 17 |
| ddx39aa | 0.86731 | 0.107332829 | 0.286 | 0.22 | 1 | 17 |
| cds2 | 0.86742 | -0.12452739 | 0.071 | 0.073 | 1 | 17 |
| otud5a | 0.86935 | -0.11973871 | 0.071 | 0.073 | 1 | 17 |
| sept4a | 0.86951 | 0.107355656 | 0.071 | 0.073 | 1 | 17 |
| prc1a | 0.87054 | -0.15027719 | 0.071 | 0.098 | 1 | 17 |
| psma6b | 0.87098 | -0.11523949 | 0.143 | 0.146 | 1 | 17 |
| ndufa9a | 0.87186 | 0.105603929 | 0.286 | 0.22 | 1 | 17 |
| dph3 | 0.87206 | -0.14501594 | 0.071 | 0.098 | 1 | 17 |
| med27 | 0.87229 | -0.11720672 | 0.071 | 0.073 | 1 | 17 |
| qars | 0.87237 | -0.15555886 | 0.071 | 0.098 | 1 | 17 |
| cyr61 | 0.87248 | -0.11377126 | 0.071 | 0.073 | 1 | 17 |
| ubap2l | 0.8743 | -0.11306867 | 0.071 | 0.073 | 1 | 17 |
| coil | 0.87471 | -0.14293113 | 0.214 | 0.244 | 1 | 17 |
| eif3s6ip | 0.87495 | 0.105379273 | 0.286 | 0.22 | 1 | 17 |
| mei4 | 0.87541 | -0.15186546 | 0.071 | 0.098 | 1 | 17 |
| paip1 | 0.87587 | -0.15086128 | 0.071 | 0.098 | 1 | 17 |
| nup43 | 0.87629 | -0.14624893 | 0.071 | 0.098 | 1 | 17 |
| nup107 | 0.87744 | 0.164826704 | 0.143 | 0.098 | 1 | 17 |
| srpra | 0.87887 | -0.10812915 | 0.143 | 0.146 | 1 | 17 |
| alkbh3 | 0.87909 | -0.16394805 | 0.143 | 0.195 | 1 | 17 |
| srp19 | 0.87939 | -0.15457803 | 0.143 | 0.195 | 1 | 17 |
| otub1b | 0.87962 | 0.142375189 | 0.143 | 0.122 | 1 | 17 |
| wrb | 0.87989 | -0.14428574 | 0.214 | 0.244 | 1 | 17 |
| mcm3 | 0.88042 | 0.156981007 | 0.286 | 0.244 | 1 | 17 |
| leo1 | 0.8807 | -0.11119005 | 0.071 | 0.073 | 1 | 17 |
| alg5 | 0.88077 | -0.15035597 | 0.071 | 0.098 | 1 | 17 |
| dkc1 | 0.88078 | 0.158101104 | 0.286 | 0.244 | 1 | 17 |
| ss18l2 | 0.8811 | 0.118776577 | 0.286 | 0.22 | 1 | 17 |
| dnajb11 | 0.88129 | -0.14009888 | 0.071 | 0.098 | 1 | 17 |
| ddx52 | 0.88193 | -0.16083455 | 0.143 | 0.195 | 1 | 17 |
| kdelr2b | 0.88253 | -0.15159356 | 0.071 | 0.098 | 1 | 17 |
| ndrg3a | 0.88273 | -0.14872079 | 0.071 | 0.098 | 1 | 17 |

| | | | | | | |
|---|---|---|---|---|---|---|
| stk35l | 0.88284 | -0.14321651 | 0.143 | 0.171 | 1 | 17 |
| sf3b4 | 0.88306 | -0.17017813 | 0.143 | 0.195 | 1 | 17 |
| hibadha | 0.88308 | 0.144549961 | 0.143 | 0.098 | 1 | 17 |
| nop16 | 0.88344 | 0.126829329 | 0.286 | 0.22 | 1 | 17 |
| hmgn7 | 0.88352 | 0.136238768 | 0.286 | 0.22 | 1 | 17 |
| pum3 | 0.88479 | -0.11118775 | 0.071 | 0.073 | 1 | 17 |
| chchd10 | 0.88502 | -0.18054862 | 0.143 | 0.195 | 1 | 17 |
| pip4k2aa | 0.88543 | 0.145383908 | 0.143 | 0.098 | 1 | 17 |
| polr1e | 0.88612 | 0.152080174 | 0.143 | 0.098 | 1 | 17 |
| elof1 | 0.88679 | -0.13398378 | 0.214 | 0.244 | 1 | 17 |
| saraf | 0.88734 | -0.10812202 | 0.071 | 0.073 | 1 | 17 |
| spg7 | 0.88818 | 0.156502476 | 0.143 | 0.098 | 1 | 17 |
| fam50a | 0.88824 | 0.141501376 | 0.143 | 0.098 | 1 | 17 |
| hnrnpa1a | 0.88894 | 0.166174682 | 0.143 | 0.098 | 1 | 17 |
| smim19 | 0.88917 | 0.151224594 | 0.143 | 0.098 | 1 | 17 |
| smarcc1a | 0.89007 | -0.14101134 | 0.071 | 0.098 | 1 | 17 |
| hsp90b1 | 0.89037 | -0.14137465 | 0.071 | 0.098 | 1 | 17 |
| kctd5a | 0.89047 | -0.14049509 | 0.071 | 0.098 | 1 | 17 |
| fance | 0.89077 | -0.13494726 | 0.071 | 0.098 | 1 | 17 |
| pa2g4b | 0.89151 | -0.14514089 | 0.143 | 0.171 | 1 | 17 |
| ubtf | 0.89186 | -0.14181511 | 0.214 | 0.244 | 1 | 17 |
| zgc:66024 | 0.89436 | -0.10531653 | 0.071 | 0.073 | 1 | 17 |
| itm2cb | 0.89457 | -0.11067821 | 0.071 | 0.073 | 1 | 17 |
| mrps31 | 0.89536 | 0.150839987 | 0.143 | 0.098 | 1 | 17 |
| LOC110440049 | 0.89593 | 0.128682325 | 0.143 | 0.098 | 1 | 17 |
| fbxw11b | 0.89598 | -0.14940868 | 0.071 | 0.098 | 1 | 17 |
| lrrc20 | 0.89622 | -0.14132554 | 0.071 | 0.098 | 1 | 17 |
| rrbp1a | 0.89684 | -0.10171673 | 0.071 | 0.073 | 1 | 17 |
| ncoa4 | 0.89694 | -0.14407551 | 0.071 | 0.098 | 1 | 17 |
| lsm8 | 0.89725 | 0.130561212 | 0.143 | 0.098 | 1 | 17 |
| pgam5 | 0.89738 | -0.13990525 | 0.357 | 0.415 | 1 | 17 |
| eif4ebp3l | 0.89747 | -0.10344257 | 0.071 | 0.073 | 1 | 17 |
| micu2 | 0.8975 | -0.14090339 | 0.071 | 0.098 | 1 | 17 |
| pno1 | 0.89804 | 0.108043164 | 0.357 | 0.293 | 1 | 17 |
| sars | 0.89816 | 0.125206622 | 0.143 | 0.098 | 1 | 17 |
| rab18a | 0.89858 | 0.102295093 | 0.143 | 0.098 | 1 | 17 |
| dnajb6a | 0.89898 | 0.126574126 | 0.143 | 0.098 | 1 | 17 |
| cib1 | 0.89911 | 0.129229203 | 0.143 | 0.098 | 1 | 17 |
| timm8b | 0.89912 | -0.13312283 | 0.214 | 0.244 | 1 | 17 |
| vbp1 | 0.89915 | -0.13511881 | 0.071 | 0.098 | 1 | 17 |
| si:ch211-218c6.8 | 0.89967 | -0.13663834 | 0.143 | 0.195 | 1 | 17 |
| lman1 | 0.89975 | -0.1076499 | 0.071 | 0.073 | 1 | 17 |
| CCNB1IP1 | 0.90023 | 0.1257326 | 0.143 | 0.098 | 1 | 17 |
| rfc2 | 0.90037 | 0.129814426 | 0.143 | 0.098 | 1 | 17 |
| ppp2r1bb | 0.90044 | 0.117637521 | 0.143 | 0.098 | 1 | 17 |
| ezh2 | 0.90048 | 0.111651446 | 0.143 | 0.098 | 1 | 17 |
| vars | 0.90077 | 0.121424738 | 0.143 | 0.098 | 1 | 17 |
| tspan4b | 0.90126 | 0.102714889 | 0.143 | 0.098 | 1 | 17 |

| | | | | | | |
|---|---|---|---|---|---|---|
| carhsp1 | 0.90127 | 0.102706235 | 0.143 | 0.098 | 1 | 17 |
| BOLA2 | 0.90134 | -0.13741804 | 0.071 | 0.098 | 1 | 17 |
| ddx39ab | 0.90142 | 0.106226466 | 0.143 | 0.098 | 1 | 17 |
| si:ch211-147d7.5 | 0.90155 | 0.105566032 | 0.143 | 0.098 | 1 | 17 |
| g3bp1 | 0.90156 | 0.113746621 | 0.143 | 0.098 | 1 | 17 |
| tardbpl | 0.90159 | 0.121734174 | 0.143 | 0.098 | 1 | 17 |
| CDK2AP1 | 0.90173 | -0.10082453 | 0.071 | 0.073 | 1 | 17 |
| pcgf1 | 0.9018 | 0.103249281 | 0.143 | 0.098 | 1 | 17 |
| cdc23 | 0.90202 | 0.119210561 | 0.143 | 0.098 | 1 | 17 |
| btr04 | 0.90216 | 0.138536977 | 0.143 | 0.098 | 1 | 17 |
| pole3 | 0.90305 | -0.13660827 | 0.143 | 0.195 | 1 | 17 |
| wars | 0.90319 | -0.1079359 | 0.071 | 0.073 | 1 | 17 |
| tp53 | 0.90495 | -0.13226278 | 0.143 | 0.195 | 1 | 17 |
| copa | 0.90557 | -0.131377 | 0.143 | 0.195 | 1 | 17 |
| fxr1 | 0.90707 | -0.13092482 | 0.071 | 0.098 | 1 | 17 |
| selenot1a | 0.90719 | -0.10389635 | 0.071 | 0.073 | 1 | 17 |
| xrn2 | 0.90834 | -0.13652646 | 0.071 | 0.098 | 1 | 17 |
| syap1 | 0.90837 | 0.139785554 | 0.214 | 0.171 | 1 | 17 |
| nucks1a | 0.90975 | -0.13782437 | 0.071 | 0.098 | 1 | 17 |
| mrpl32 | 0.91058 | 0.125144975 | 0.143 | 0.122 | 1 | 17 |
| ccdc106b | 0.91071 | -0.13569535 | 0.143 | 0.171 | 1 | 17 |
| si:dkeyp-87e7.4 | 0.91146 | -0.12855402 | 0.071 | 0.098 | 1 | 17 |
| dhx9 | 0.91147 | -0.11810289 | 0.071 | 0.098 | 1 | 17 |
| mrps18a | 0.91207 | -0.12992617 | 0.143 | 0.171 | 1 | 17 |
| ube4b | 0.91226 | -0.13039349 | 0.071 | 0.098 | 1 | 17 |
| sox19b | 0.91316 | -0.1314967 | 0.071 | 0.098 | 1 | 17 |
| pgrmc1 | 0.91419 | -0.13439686 | 0.071 | 0.098 | 1 | 17 |
| naa20 | 0.91463 | -0.12785643 | 0.143 | 0.171 | 1 | 17 |
| pprc1 | 0.91471 | -0.13179822 | 0.071 | 0.098 | 1 | 17 |
| dtnbp1a | 0.91472 | -0.12975072 | 0.071 | 0.098 | 1 | 17 |
| snx6 | 0.91587 | -0.1336892 | 0.071 | 0.098 | 1 | 17 |
| acyp1 | 0.91718 | -0.12063048 | 0.071 | 0.098 | 1 | 17 |
| tdrd1 | 0.91745 | 0.134305915 | 0.286 | 0.244 | 1 | 17 |
| kctd7 | 0.91802 | -0.12669415 | 0.071 | 0.098 | 1 | 17 |
| aes | 0.91839 | -0.12005988 | 0.071 | 0.098 | 1 | 17 |
| supt6h | 0.91941 | -0.12141483 | 0.071 | 0.098 | 1 | 17 |
| cplane2 | 0.91942 | -0.12696495 | 0.071 | 0.098 | 1 | 17 |
| adprhl2 | 0.91955 | -0.12445637 | 0.071 | 0.098 | 1 | 17 |
| hnrnpd | 0.92117 | -0.12814926 | 0.143 | 0.171 | 1 | 17 |
| hspa4a | 0.92217 | -0.11815755 | 0.214 | 0.244 | 1 | 17 |
| hist2h2l | 0.92369 | -0.12640973 | 0.071 | 0.098 | 1 | 17 |
| ccng1 | 0.92428 | -0.10675644 | 0.5 | 0.561 | 1 | 17 |
| wdr48b | 0.92714 | -0.11463059 | 0.071 | 0.098 | 1 | 17 |
| xaf1 | 0.92733 | -0.11950206 | 0.071 | 0.098 | 1 | 17 |
| pabpc1l | 0.92766 | -0.12885694 | 0.143 | 0.171 | 1 | 17 |
| zgc:112185 | 0.92814 | -0.11606061 | 0.071 | 0.098 | 1 | 17 |
| lin52 | 0.92907 | 0.122710715 | 0.286 | 0.244 | 1 | 17 |
| rnf4 | 0.93033 | -0.11538213 | 0.143 | 0.171 | 1 | 17 |

| | | | | | | |
|---|---|---|---|---|---|---|
| chmp2bb | 0.93056 | -0.11369811 | 0.071 | 0.098 | 1 | 17 |
| gale | 0.93183 | -0.11549912 | 0.071 | 0.098 | 1 | 17 |
| snrnp27 | 0.93292 | -0.10280101 | 0.429 | 0.463 | 1 | 17 |
| si:ch211-210c8.7 | 0.9337 | 0.116011906 | 0.214 | 0.171 | 1 | 17 |
| sumo3b | 0.93467 | -0.12096377 | 0.071 | 0.098 | 1 | 17 |
| ess2 | 0.9348 | -0.10732296 | 0.071 | 0.098 | 1 | 17 |
| ndufv1 | 0.93564 | -0.10557835 | 0.071 | 0.098 | 1 | 17 |
| psmg2 | 0.93719 | -0.11950532 | 0.071 | 0.098 | 1 | 17 |
| psmd6 | 0.93729 | 0.10234479 | 0.214 | 0.171 | 1 | 17 |
| adnpa | 0.93799 | -0.10622711 | 0.071 | 0.098 | 1 | 17 |
| fcf1 | 0.93828 | -0.11115006 | 0.071 | 0.098 | 1 | 17 |
| zgc:103692 | 0.93879 | -0.10346264 | 0.071 | 0.098 | 1 | 17 |
| cdkn1a | 0.9424 | -0.11110392 | 0.071 | 0.098 | 1 | 17 |
| cnbpa | 0.94491 | -0.10575925 | 0.286 | 0.317 | 1 | 17 |
| CABZ01086029.1 | 0.94596 | -0.10002896 | 0.071 | 0.098 | 1 | 17 |
| usp37 | 0.94635 | -0.10779232 | 0.071 | 0.098 | 1 | 17 |
| rrp36 | 0.94744 | 0.101490594 | 0.143 | 0.122 | 1 | 17 |
| supt16h | 0.95429 | -0.10253865 | 0.143 | 0.171 | 1 | 17 |
| taf9 | 0.95477 | -0.10310159 | 0.143 | 0.171 | 1 | 17 |
| mt-cyb | 3.705260 | -0.82924204 | 0.182 | 0.769 | 0.7483515 | 18 |
| mt-co2 | 9.245011 | -0.58983551 | 0.318 | 0.846 | 1 | 18 |
| ptx3a | 0.0002 | -0.79371617 | 0.182 | 0.731 | 1 | 18 |
| uqcrq | 0.00022 | -0.7622638 | 0.136 | 0.692 | 1 | 18 |
| si:ch1073-291c23.2 | 0.00043 | -0.82983543 | 0.091 | 0.615 | 1 | 18 |
| rtraf | 0.00061 | -0.47304027 | 0 | 0.346 | 1 | 18 |
| zgc:158343 | 0.00089 | -0.22010657 | 0.273 | 0.615 | 1 | 18 |
| si:ch211-218c6.8 | 0.00144 | -0.38154167 | 0 | 0.308 | 1 | 18 |
| srgn | 0.00186 | -0.35414411 | 0.091 | 0.462 | 1 | 18 |
| prr12b | 0.0022 | 0.525038305 | 0.273 | 0 | 1 | 18 |
| tob1b | 0.0022 | 0.513859547 | 0.273 | 0 | 1 | 18 |
| sarnp | 0.00236 | -0.62565854 | 0.091 | 0.538 | 1 | 18 |
| mt-atp6 | 0.00291 | -0.652145 | 0.5 | 0.923 | 1 | 18 |
| mmp14b | 0.00299 | -0.50659747 | 0.136 | 0.538 | 1 | 18 |
| si:ch211-51e12.7 | 0.00311 | 0.160240735 | 0.409 | 0.538 | 1 | 18 |
| mrpl23 | 0.0033 | -0.34917243 | 0 | 0.269 | 1 | 18 |
| ralbp1 | 0.0033 | -0.38564353 | 0 | 0.269 | 1 | 18 |
| wt1b | 0.0033 | -0.53945274 | 0 | 0.269 | 1 | 18 |
| ccl39.7 | 0.0033 | -0.4355177 | 0 | 0.269 | 1 | 18 |
| gpx4a | 0.0033 | -0.2333395 | 0 | 0.269 | 1 | 18 |
| mbd4 | 0.0033 | -0.43150269 | 0 | 0.269 | 1 | 18 |
| mmp11b | 0.0033 | -0.54449326 | 0 | 0.269 | 1 | 18 |
| nccrp1 | 0.0033 | -0.44943056 | 0 | 0.269 | 1 | 18 |
| purba | 0.0033 | -0.4140534 | 0 | 0.269 | 1 | 18 |
| sel1l | 0.0033 | -0.41087432 | 0 | 0.269 | 1 | 18 |
| tsg101b | 0.0033 | -0.4630746 | 0 | 0.269 | 1 | 18 |
| zgc:77650 | 0.0033 | -0.46753472 | 0 | 0.269 | 1 | 18 |
| zgc:66388 | 0.0033 | -0.39859261 | 0 | 0.269 | 1 | 18 |
| mesd | 0.00409 | -0.58020822 | 0.045 | 0.423 | 1 | 18 |

| | | | | | | |
|---|---|---|---|---|---|---|
| si:ch211-222l21.1 | 0.00435 | 0.503554903 | 0.773 | 0.692 | 1 | 18 |
| mrc2 | 0.00523 | -0.64531113 | 0.091 | 0.5 | 1 | 18 |
| zgc:193541 | 0.00547 | -0.51552343 | 0.409 | 0.846 | 1 | 18 |
| fam120c | 0.00549 | -0.4379676 | 0.091 | 0.423 | 1 | 18 |
| dnajc9 | 0.00554 | 0.507994379 | 0.273 | 0.077 | 1 | 18 |
| CABZ01072614.1 | 0.00596 | -0.73022833 | 0.136 | 0.538 | 1 | 18 |
| kctd10 | 0.00602 | 0.370283683 | 0.227 | 0 | 1 | 18 |
| ccbe1 | 0.00602 | 0.484155056 | 0.227 | 0 | 1 | 18 |
| lsm1 | 0.00602 | 0.395943015 | 0.227 | 0 | 1 | 18 |
| chrac1 | 0.00602 | 0.356887606 | 0.227 | 0 | 1 | 18 |
| nfkbiaa | 0.00602 | 0.418917386 | 0.227 | 0 | 1 | 18 |
| plrg1 | 0.00602 | 0.456314197 | 0.227 | 0 | 1 | 18 |
| gemin2 | 0.00602 | 0.472275271 | 0.227 | 0 | 1 | 18 |
| si:ch211-76m11.3 | 0.00602 | 0.382756132 | 0.227 | 0 | 1 | 18 |
| fbxo34 | 0.00661 | -0.34177964 | 0.045 | 0.308 | 1 | 18 |
| MCMDC2 | 0.0074 | -0.35565421 | 0 | 0.231 | 1 | 18 |
| cdk13 | 0.0074 | -0.33177129 | 0 | 0.231 | 1 | 18 |
| clic2 | 0.0074 | -0.3634808 | 0 | 0.231 | 1 | 18 |
| si:ch211-195o20.7 | 0.0074 | -0.41491018 | 0 | 0.231 | 1 | 18 |
| spon2b | 0.0074 | -0.66705887 | 0 | 0.231 | 1 | 18 |
| chmp1a | 0.0074 | -0.35466909 | 0 | 0.231 | 1 | 18 |
| egfra | 0.0074 | -0.3468259 | 0 | 0.231 | 1 | 18 |
| helz | 0.0074 | -0.40958055 | 0 | 0.231 | 1 | 18 |
| lin7c | 0.0074 | -0.42991677 | 0 | 0.231 | 1 | 18 |
| lman1 | 0.0074 | -0.34466394 | 0 | 0.231 | 1 | 18 |
| ms4a17a.8 | 0.0074 | -0.36075879 | 0 | 0.231 | 1 | 18 |
| plod1a | 0.0074 | -0.39725088 | 0 | 0.231 | 1 | 18 |
| rpf1 | 0.0074 | -0.36158632 | 0 | 0.231 | 1 | 18 |
| si:ch211-213a13.1 | 0.0074 | -0.36160359 | 0 | 0.231 | 1 | 18 |
| si:dkeyp-44a8.2 | 0.0074 | -0.38645108 | 0 | 0.231 | 1 | 18 |
| triqk | 0.0074 | -0.41389517 | 0 | 0.231 | 1 | 18 |
| vps35 | 0.0074 | -0.39566694 | 0 | 0.231 | 1 | 18 |
| wu:fu71h07 | 0.0074 | -0.39452698 | 0 | 0.231 | 1 | 18 |
| zeb1b | 0.0074 | -0.43866172 | 0 | 0.231 | 1 | 18 |
| zgc:56525 | 0.0074 | -0.40824525 | 0 | 0.231 | 1 | 18 |
| hspa12b | 0.00748 | -0.55609673 | 0.136 | 0.538 | 1 | 18 |
| anapc16 | 0.00816 | -0.64438249 | 0.136 | 0.538 | 1 | 18 |
| BX663503.3 | 0.00841 | -0.71944578 | 0.182 | 0.577 | 1 | 18 |
| tagln2 | 0.00878 | -0.63720875 | 0.409 | 0.769 | 1 | 18 |
| mt-nd4 | 0.00883 | -0.63049027 | 0.455 | 0.846 | 1 | 18 |
| gstt1b | 0.01055 | -0.60963143 | 0.227 | 0.577 | 1 | 18 |
| txnipa | 0.01131 | 0.329326231 | 0.545 | 0.192 | 1 | 18 |
| ldlrad3 | 0.01132 | 0.558387485 | 0.364 | 0.038 | 1 | 18 |
| chchd6b | 0.01134 | 0.461854502 | 0.364 | 0.038 | 1 | 18 |
| sned1 | 0.01249 | -0.54763319 | 0.273 | 0.692 | 1 | 18 |
| polr2eb | 0.01249 | -0.40288568 | 0.045 | 0.346 | 1 | 18 |
| lgals3b | 0.01363 | -0.49403056 | 0.091 | 0.462 | 1 | 18 |
| psma6b | 0.01363 | -0.31963936 | 0.045 | 0.308 | 1 | 18 |

| | | | | | | |
|---|---|---|---|---|---|---|
| zgc:195173 | 0.01368 | -0.47520115 | 0.364 | 0.769 | 1 | 18 |
| ier2a | 0.01465 | 0.639448566 | 0.818 | 0.462 | 1 | 18 |
| cct8 | 0.01473 | -0.11030812 | 0.409 | 0.654 | 1 | 18 |
| sub1b | 0.01554 | 0.567073641 | 0.364 | 0.077 | 1 | 18 |
| htra1a | 0.01598 | -0.62695835 | 0.364 | 0.692 | 1 | 18 |
| wscd2 | 0.01612 | 0.302457396 | 0.182 | 0 | 1 | 18 |
| traf7 | 0.01612 | 0.346598314 | 0.182 | 0 | 1 | 18 |
| CR384075.1 | 0.01612 | 0.403992001 | 0.182 | 0 | 1 | 18 |
| XLOC-042565 | 0.01612 | 0.430031062 | 0.182 | 0 | 1 | 18 |
| arrdc2 | 0.01612 | 0.276677543 | 0.182 | 0 | 1 | 18 |
| cbfb | 0.01612 | 0.299634764 | 0.182 | 0 | 1 | 18 |
| klf11a | 0.01612 | 0.372401849 | 0.182 | 0 | 1 | 18 |
| mgp | 0.01612 | 0.381946506 | 0.182 | 0 | 1 | 18 |
| msrb1b | 0.01612 | 0.332735418 | 0.182 | 0 | 1 | 18 |
| nup107 | 0.01612 | 0.345610258 | 0.182 | 0 | 1 | 18 |
| pard6gb | 0.01612 | 0.347734008 | 0.182 | 0 | 1 | 18 |
| smad5 | 0.01612 | 0.332849696 | 0.182 | 0 | 1 | 18 |
| stox2b | 0.01612 | 0.277484759 | 0.182 | 0 | 1 | 18 |
| supt16h | 0.01612 | 0.368822531 | 0.182 | 0 | 1 | 18 |
| creg1 | 0.01631 | -0.32293212 | 0 | 0.192 | 1 | 18 |
| fhl1b | 0.01631 | -0.35184138 | 0 | 0.192 | 1 | 18 |
| pmpcb | 0.01631 | -0.29810131 | 0 | 0.192 | 1 | 18 |
| si:ch211-282b22.1 | 0.01631 | -0.28574062 | 0 | 0.192 | 1 | 18 |
| smarcd2 | 0.01631 | -0.30795535 | 0 | 0.192 | 1 | 18 |
| srgap2 | 0.01631 | -0.33517747 | 0 | 0.192 | 1 | 18 |
| wdr12 | 0.01631 | -0.31205382 | 0 | 0.192 | 1 | 18 |
| htatip2 | 0.01631 | -0.29426199 | 0 | 0.192 | 1 | 18 |
| znf362a | 0.01631 | -0.4191139 | 0 | 0.192 | 1 | 18 |
| LOC108190702 | 0.01631 | -0.35286099 | 0 | 0.192 | 1 | 18 |
| abhd17c | 0.01631 | -0.28397059 | 0 | 0.192 | 1 | 18 |
| arhgap21a | 0.01631 | -0.33332156 | 0 | 0.192 | 1 | 18 |
| cfap45 | 0.01631 | -0.39959405 | 0 | 0.192 | 1 | 18 |
| cyp1c1 | 0.01631 | -0.39220488 | 0 | 0.192 | 1 | 18 |
| kat5b | 0.01631 | -0.34278607 | 0 | 0.192 | 1 | 18 |
| prss12 | 0.01631 | -0.39393431 | 0 | 0.192 | 1 | 18 |
| si:dkey-239n17.4 | 0.01631 | -0.30227556 | 0 | 0.192 | 1 | 18 |
| st7 | 0.01631 | -0.35010942 | 0 | 0.192 | 1 | 18 |
| tpbgb | 0.01631 | -0.36823956 | 0 | 0.192 | 1 | 18 |
| tra2b | 0.01631 | -0.27272639 | 0 | 0.192 | 1 | 18 |
| zgc:110783 | 0.01631 | -0.35705706 | 0 | 0.192 | 1 | 18 |
| LOC103910363 | 0.01631 | -0.35232688 | 0 | 0.192 | 1 | 18 |
| XLOC-032112 | 0.01631 | -0.34407476 | 0 | 0.192 | 1 | 18 |
| atp1b3a | 0.01631 | -0.31426119 | 0 | 0.192 | 1 | 18 |
| bcap29 | 0.01631 | -0.33999337 | 0 | 0.192 | 1 | 18 |
| bxdc2 | 0.01631 | -0.33462986 | 0 | 0.192 | 1 | 18 |
| cand1 | 0.01631 | -0.23598128 | 0 | 0.192 | 1 | 18 |
| dact3a | 0.01631 | -0.31734944 | 0 | 0.192 | 1 | 18 |
| dusp6 | 0.01631 | -0.33527686 | 0 | 0.192 | 1 | 18 |

| | | | | | | |
|---|---|---|---|---|---|---|
| eif2b1 | 0.01631 | -0.18006007 | 0 | 0.192 | 1 | 18 |
| fcho2 | 0.01631 | -0.30028294 | 0 | 0.192 | 1 | 18 |
| kdm5c | 0.01631 | -0.2722906 | 0 | 0.192 | 1 | 18 |
| plekho2 | 0.01631 | -0.36159427 | 0 | 0.192 | 1 | 18 |
| spen | 0.01631 | -0.40848873 | 0 | 0.192 | 1 | 18 |
| tes | 0.01631 | -0.28055377 | 0 | 0.192 | 1 | 18 |
| tmed1b | 0.01631 | -0.3201407 | 0 | 0.192 | 1 | 18 |
| nfkbiab | 0.01751 | 0.566452211 | 0.455 | 0.231 | 1 | 18 |
| fosab | 0.01779 | 0.234452009 | 0.909 | 0.923 | 1 | 18 |
| hmga1a | 0.01852 | 0.459690267 | 0.682 | 0.577 | 1 | 18 |
| alyref | 0.01908 | 0.61198012 | 0.409 | 0.115 | 1 | 18 |
| bicc1b | 0.01955 | -0.29612997 | 0.273 | 0.577 | 1 | 18 |
| ier2b | 0.02172 | 0.617833796 | 0.682 | 0.346 | 1 | 18 |
| cct5 | 0.02185 | -0.35161003 | 0.409 | 0.769 | 1 | 18 |
| hibadhb | 0.02263 | -0.44707576 | 0.045 | 0.346 | 1 | 18 |
| crip1 | 0.02301 | -0.70141657 | 0.5 | 0.731 | 1 | 18 |
| plod3 | 0.02333 | 0.354503323 | 0.273 | 0.038 | 1 | 18 |
| ctgfa | 0.02373 | 0.548100701 | 0.682 | 0.308 | 1 | 18 |
| rarga | 0.02385 | 0.270523439 | 0.273 | 0.231 | 1 | 18 |
| rab7a | 0.02393 | -0.45856096 | 0.045 | 0.346 | 1 | 18 |
| itm2ba | 0.02445 | -0.42243108 | 0.318 | 0.692 | 1 | 18 |
| mgst1.1 | 0.02467 | -0.24489808 | 0.273 | 0.538 | 1 | 18 |
| tia1l | 0.02481 | -0.44599783 | 0.045 | 0.346 | 1 | 18 |
| epha4l | 0.02547 | -0.44629443 | 0.045 | 0.346 | 1 | 18 |
| zgc:55943 | 0.02555 | -0.35714642 | 0.091 | 0.385 | 1 | 18 |
| npm3 | 0.02625 | -0.32360132 | 0.136 | 0.423 | 1 | 18 |
| hoxc8a | 0.02666 | -0.36804873 | 0.182 | 0.5 | 1 | 18 |
| dnmt3ab | 0.02687 | -0.44077164 | 0.182 | 0.538 | 1 | 18 |
| ifrd1 | 0.02815 | 0.41649524 | 0.318 | 0.038 | 1 | 18 |
| sult1st1 | 0.02829 | -0.53588242 | 0.091 | 0.385 | 1 | 18 |
| snrpe | 0.02873 | 0.520095813 | 0.682 | 0.346 | 1 | 18 |
| pdia6 | 0.02937 | -0.25542082 | 0.455 | 0.462 | 1 | 18 |
| tubb5 | 0.02954 | 0.610770027 | 0.5 | 0.154 | 1 | 18 |
| tmeff1b | 0.03059 | -0.4987077 | 0.182 | 0.538 | 1 | 18 |
| pora | 0.03132 | -0.51097725 | 0.136 | 0.462 | 1 | 18 |
| gemin8 | 0.0322 | 0.456304499 | 0.318 | 0.077 | 1 | 18 |
| krt18a.1 | 0.03386 | -0.35816373 | 0.045 | 0.308 | 1 | 18 |
| junbb | 0.03547 | -0.29385699 | 0 | 0.154 | 1 | 18 |
| larp4b | 0.03547 | -0.22356225 | 0 | 0.154 | 1 | 18 |
| si:ch211-260e23.9 | 0.03547 | -0.26276759 | 0 | 0.154 | 1 | 18 |
| ssbp1 | 0.03547 | -0.17717529 | 0 | 0.154 | 1 | 18 |
| CABZ01090890.1 | 0.03547 | -0.21656401 | 0 | 0.154 | 1 | 18 |
| appb | 0.03547 | -0.3630306 | 0 | 0.154 | 1 | 18 |
| arcn1b | 0.03547 | -0.24731869 | 0 | 0.154 | 1 | 18 |
| atp8a2 | 0.03547 | -0.2794159 | 0 | 0.154 | 1 | 18 |
| bnip3lb | 0.03547 | -0.27979052 | 0 | 0.154 | 1 | 18 |
| chst2b | 0.03547 | -0.27981296 | 0 | 0.154 | 1 | 18 |
| crip3 | 0.03547 | -0.28414309 | 0 | 0.154 | 1 | 18 |

| | | | | | | |
|---|---|---|---|---|---|---|
| frrs1b | 0.03547 | -0.2393873 | 0 | 0.154 | 1 | 18 |
| fryl | 0.03547 | -0.27913238 | 0 | 0.154 | 1 | 18 |
| gid8b | 0.03547 | -0.20897412 | 0 | 0.154 | 1 | 18 |
| mrpl41 | 0.03547 | -0.26796284 | 0 | 0.154 | 1 | 18 |
| nt5c2b | 0.03547 | -0.32369632 | 0 | 0.154 | 1 | 18 |
| otub1b | 0.03547 | -0.23098161 | 0 | 0.154 | 1 | 18 |
| paics | 0.03547 | -0.29351449 | 0 | 0.154 | 1 | 18 |
| ppp2r5ea | 0.03547 | -0.2570014 | 0 | 0.154 | 1 | 18 |
| sirt2 | 0.03547 | -0.25580897 | 0 | 0.154 | 1 | 18 |
| suclg2 | 0.03547 | -0.22154984 | 0 | 0.154 | 1 | 18 |
| sumf2 | 0.03547 | -0.26621489 | 0 | 0.154 | 1 | 18 |
| tmem198b | 0.03547 | -0.26592324 | 0 | 0.154 | 1 | 18 |
| traf3ip1 | 0.03547 | -0.23505296 | 0 | 0.154 | 1 | 18 |
| ubap2a | 0.03547 | -0.28787628 | 0 | 0.154 | 1 | 18 |
| uchl5 | 0.03547 | -0.19002488 | 0 | 0.154 | 1 | 18 |
| vkorc1l1 | 0.03547 | -0.2196455 | 0 | 0.154 | 1 | 18 |
| znf511 | 0.03547 | -0.23730292 | 0 | 0.154 | 1 | 18 |
| btbd10a | 0.03547 | -0.21737939 | 0 | 0.154 | 1 | 18 |
| dync1i2b | 0.03547 | -0.24266035 | 0 | 0.154 | 1 | 18 |
| kdm5bb | 0.03547 | -0.2481078 | 0 | 0.154 | 1 | 18 |
| naga | 0.03547 | -0.3082494 | 0 | 0.154 | 1 | 18 |
| CABZ01078261.1 | 0.03547 | -0.31012072 | 0 | 0.154 | 1 | 18 |
| FP101882.1 | 0.03547 | -0.33210615 | 0 | 0.154 | 1 | 18 |
| LOC571123 | 0.03547 | -0.30295816 | 0 | 0.154 | 1 | 18 |
| acaa1 | 0.03547 | -0.21653514 | 0 | 0.154 | 1 | 18 |
| ahdc1 | 0.03547 | -0.3096186 | 0 | 0.154 | 1 | 18 |
| appbp2 | 0.03547 | -0.22813019 | 0 | 0.154 | 1 | 18 |
| arhgef4 | 0.03547 | -0.30344961 | 0 | 0.154 | 1 | 18 |
| b9d1 | 0.03547 | -0.26901533 | 0 | 0.154 | 1 | 18 |
| cldn5a | 0.03547 | -0.33625057 | 0 | 0.154 | 1 | 18 |
| dclre1b | 0.03547 | -0.23068334 | 0 | 0.154 | 1 | 18 |
| dusp1 | 0.03547 | -0.24516918 | 0 | 0.154 | 1 | 18 |
| fam177a1 | 0.03547 | -0.21816239 | 0 | 0.154 | 1 | 18 |
| fdx2 | 0.03547 | -0.22232102 | 0 | 0.154 | 1 | 18 |
| foxj3 | 0.03547 | -0.22135898 | 0 | 0.154 | 1 | 18 |
| foxk1 | 0.03547 | -0.31847288 | 0 | 0.154 | 1 | 18 |
| foxk2 | 0.03547 | -0.32006018 | 0 | 0.154 | 1 | 18 |
| gabpa | 0.03547 | -0.28933899 | 0 | 0.154 | 1 | 18 |
| gpc1b | 0.03547 | -0.29213229 | 0 | 0.154 | 1 | 18 |
| mef2cb | 0.03547 | -0.19403954 | 0 | 0.154 | 1 | 18 |
| mrpl19 | 0.03547 | -0.23987194 | 0 | 0.154 | 1 | 18 |
| psph | 0.03547 | -0.13272032 | 0 | 0.154 | 1 | 18 |
| ptpra | 0.03547 | -0.29947289 | 0 | 0.154 | 1 | 18 |
| rab22a | 0.03547 | -0.26319502 | 0 | 0.154 | 1 | 18 |
| sept5a | 0.03547 | -0.22767877 | 0 | 0.154 | 1 | 18 |
| si:ch211-15b10.6 | 0.03547 | -0.26090721 | 0 | 0.154 | 1 | 18 |
| si:ch73-376l24.6 | 0.03547 | -0.32387605 | 0 | 0.154 | 1 | 18 |
| si:dkey-56m19.5 | 0.03547 | -0.30672143 | 0 | 0.154 | 1 | 18 |

| | | | | | | |
|---|---|---|---|---|---|---|
| slc12a4 | 0.03547 | -0.31049296 | 0 | 0.154 | 1 | 18 |
| smco4 | 0.03547 | -0.25983903 | 0 | 0.154 | 1 | 18 |
| socs6b | 0.03547 | -0.25503355 | 0 | 0.154 | 1 | 18 |
| tbc1d5 | 0.03547 | -0.2283098 | 0 | 0.154 | 1 | 18 |
| timmdc1 | 0.03547 | -0.2725141 | 0 | 0.154 | 1 | 18 |
| uba5 | 0.03547 | -0.22946372 | 0 | 0.154 | 1 | 18 |
| ubxn4 | 0.03547 | -0.3244554 | 0 | 0.154 | 1 | 18 |
| zgc:193742 | 0.03547 | -0.21841809 | 0 | 0.154 | 1 | 18 |
| AL954191.1 | 0.03547 | -0.28728726 | 0 | 0.154 | 1 | 18 |
| ATP11B | 0.03547 | -0.28636011 | 0 | 0.154 | 1 | 18 |
| BX470259.2 | 0.03547 | -0.27387938 | 0 | 0.154 | 1 | 18 |
| CU638870.1 | 0.03547 | -0.27481057 | 0 | 0.154 | 1 | 18 |
| LOC108180204 | 0.03547 | -0.22642679 | 0 | 0.154 | 1 | 18 |
| LOC110439840 | 0.03547 | -0.1649552 | 0 | 0.154 | 1 | 18 |
| acvr1l | 0.03547 | -0.22383304 | 0 | 0.154 | 1 | 18 |
| arid1b | 0.03547 | -0.24569264 | 0 | 0.154 | 1 | 18 |
| atrnl1b | 0.03547 | -0.294103 | 0 | 0.154 | 1 | 18 |
| cachd1 | 0.03547 | -0.2858243 | 0 | 0.154 | 1 | 18 |
| cbwd | 0.03547 | -0.19344373 | 0 | 0.154 | 1 | 18 |
| cep350 | 0.03547 | -0.30042776 | 0 | 0.154 | 1 | 18 |
| egfl7 | 0.03547 | -0.26182151 | 0 | 0.154 | 1 | 18 |
| eloca | 0.03547 | -0.24192812 | 0 | 0.154 | 1 | 18 |
| fam172a | 0.03547 | -0.27276314 | 0 | 0.154 | 1 | 18 |
| foxn3 | 0.03547 | -0.31621378 | 0 | 0.154 | 1 | 18 |
| gbp3 | 0.03547 | -0.22211642 | 0 | 0.154 | 1 | 18 |
| htr5ab | 0.03547 | -0.30725305 | 0 | 0.154 | 1 | 18 |
| lamp1a | 0.03547 | -0.36009817 | 0 | 0.154 | 1 | 18 |
| lyrm5a | 0.03547 | -0.24664979 | 0 | 0.154 | 1 | 18 |
| micu2 | 0.03547 | -0.21690843 | 0 | 0.154 | 1 | 18 |
| mrpl40 | 0.03547 | -0.24831132 | 0 | 0.154 | 1 | 18 |
| mvb12a | 0.03547 | -0.22373756 | 0 | 0.154 | 1 | 18 |
| pmp22b | 0.03547 | -0.32222423 | 0 | 0.154 | 1 | 18 |
| retsat | 0.03547 | -0.22016718 | 0 | 0.154 | 1 | 18 |
| rfx2 | 0.03547 | -0.23894099 | 0 | 0.154 | 1 | 18 |
| rnf8 | 0.03547 | -0.23070671 | 0 | 0.154 | 1 | 18 |
| si:ch211-166a6.5 | 0.03547 | -0.31526295 | 0 | 0.154 | 1 | 18 |
| si:dkey-264d12.4 | 0.03547 | -0.29693744 | 0 | 0.154 | 1 | 18 |
| stub1 | 0.03547 | -0.26375264 | 0 | 0.154 | 1 | 18 |
| top1 | 0.03547 | -0.27706462 | 0 | 0.154 | 1 | 18 |
| ttc1 | 0.03547 | -0.24929717 | 0 | 0.154 | 1 | 18 |
| twist3 | 0.03547 | -0.27909435 | 0 | 0.154 | 1 | 18 |
| wu:fc33b09 | 0.03547 | -0.221209 | 0 | 0.154 | 1 | 18 |
| xpot | 0.03547 | -0.25057755 | 0 | 0.154 | 1 | 18 |
| zgc:113372 | 0.03547 | -0.29821388 | 0 | 0.154 | 1 | 18 |
| sgk1 | 0.0358 | -0.51016193 | 0.091 | 0.346 | 1 | 18 |
| ssbp3b | 0.03667 | -0.33320339 | 0.091 | 0.346 | 1 | 18 |
| si:ch73-376l24.3 | 0.0379 | -0.32139804 | 0.182 | 0.462 | 1 | 18 |
| u2surp | 0.03886 | -0.51295323 | 0.136 | 0.462 | 1 | 18 |

| | | | | | | |
|---|---|---|---|---|---|---|
| mt-co1 | 0.03898 | -0.30363137 | 0.455 | 0.769 | 1 | 18 |
| psmd7 | 0.03949 | 0.308606426 | 0.318 | 0.231 | 1 | 18 |
| cavin2b | 0.03971 | -0.46593417 | 0.136 | 0.462 | 1 | 18 |
| cct2 | 0.03983 | -0.41134493 | 0.5 | 0.692 | 1 | 18 |
| ddit3 | 0.04036 | 0.608941504 | 0.636 | 0.346 | 1 | 18 |
| cdk2ap2 | 0.04043 | -0.23457155 | 0.045 | 0.269 | 1 | 18 |
| mob4 | 0.04124 | -0.14504161 | 0.091 | 0.269 | 1 | 18 |
| selenot2 | 0.04164 | -0.41922239 | 0.045 | 0.308 | 1 | 18 |
| tcf3b | 0.04231 | -0.12336932 | 0.045 | 0.192 | 1 | 18 |
| CU929259.1 | 0.04254 | 0.249412877 | 0.136 | 0 | 1 | 18 |
| acy3.1 | 0.04254 | 0.301666978 | 0.136 | 0 | 1 | 18 |
| cmss1 | 0.04254 | 0.274189008 | 0.136 | 0 | 1 | 18 |
| commd9 | 0.04254 | 0.267141472 | 0.136 | 0 | 1 | 18 |
| dnmbp | 0.04254 | 0.252814809 | 0.136 | 0 | 1 | 18 |
| gclc | 0.04254 | 0.258876621 | 0.136 | 0 | 1 | 18 |
| gpx8 | 0.04254 | 0.256689573 | 0.136 | 0 | 1 | 18 |
| nap1l4a | 0.04254 | 0.240566366 | 0.136 | 0 | 1 | 18 |
| pam16 | 0.04254 | 0.255455769 | 0.136 | 0 | 1 | 18 |
| poldip2 | 0.04254 | 0.20582842 | 0.136 | 0 | 1 | 18 |
| sh3glb1a | 0.04254 | 0.238233601 | 0.136 | 0 | 1 | 18 |
| si:ch211-214c7.4 | 0.04254 | 0.266228405 | 0.136 | 0 | 1 | 18 |
| smg9 | 0.04254 | 0.291341835 | 0.136 | 0 | 1 | 18 |
| wdr60 | 0.04254 | 0.281769386 | 0.136 | 0 | 1 | 18 |
| zgc:162472 | 0.04254 | 0.252486476 | 0.136 | 0 | 1 | 18 |
| akap17a | 0.04254 | 0.304942815 | 0.136 | 0 | 1 | 18 |
| ankrd52a | 0.04254 | 0.266948518 | 0.136 | 0 | 1 | 18 |
| casc3 | 0.04254 | 0.228728168 | 0.136 | 0 | 1 | 18 |
| chtf8 | 0.04254 | 0.249525407 | 0.136 | 0 | 1 | 18 |
| dlg3 | 0.04254 | 0.273075398 | 0.136 | 0 | 1 | 18 |
| dpy19l1l | 0.04254 | 0.29913216 | 0.136 | 0 | 1 | 18 |
| efhd2 | 0.04254 | 0.237258449 | 0.136 | 0 | 1 | 18 |
| efna1a | 0.04254 | 0.314891034 | 0.136 | 0 | 1 | 18 |
| epn2 | 0.04254 | 0.273831823 | 0.136 | 0 | 1 | 18 |
| fignl1 | 0.04254 | 0.234530371 | 0.136 | 0 | 1 | 18 |
| ing2 | 0.04254 | 0.220284973 | 0.136 | 0 | 1 | 18 |
| mapre1a | 0.04254 | 0.157599435 | 0.136 | 0 | 1 | 18 |
| myclb | 0.04254 | 0.265286905 | 0.136 | 0 | 1 | 18 |
| phactr1 | 0.04254 | 0.279624876 | 0.136 | 0 | 1 | 18 |
| prmt5 | 0.04254 | 0.253414871 | 0.136 | 0 | 1 | 18 |
| rc3h1b | 0.04254 | 0.279646404 | 0.136 | 0 | 1 | 18 |
| rfc4 | 0.04254 | 0.329670499 | 0.136 | 0 | 1 | 18 |
| scrn3 | 0.04254 | 0.191405492 | 0.136 | 0 | 1 | 18 |
| shmt1 | 0.04254 | 0.158614392 | 0.136 | 0 | 1 | 18 |
| si:zfos-80g12.1 | 0.04254 | 0.218446473 | 0.136 | 0 | 1 | 18 |
| trim8b | 0.04254 | 0.212663865 | 0.136 | 0 | 1 | 18 |
| zfr | 0.04254 | 0.262648616 | 0.136 | 0 | 1 | 18 |
| CABZ01088274.1 | 0.04254 | 0.218083363 | 0.136 | 0 | 1 | 18 |
| CU929418.1 | 0.04254 | 0.241326022 | 0.136 | 0 | 1 | 18 |

| | | | | | | |
|---|---|---|---|---|---|---|
| abtb1 | 0.04254 | 0.254983297 | 0.136 | 0 | 1 | 18 |
| cdc123 | 0.04254 | 0.257184642 | 0.136 | 0 | 1 | 18 |
| chico | 0.04254 | 0.243402747 | 0.136 | 0 | 1 | 18 |
| cops5 | 0.04254 | 0.214842123 | 0.136 | 0 | 1 | 18 |
| cox8b | 0.04254 | 0.233129508 | 0.136 | 0 | 1 | 18 |
| ddx39b | 0.04254 | 0.207157486 | 0.136 | 0 | 1 | 18 |
| fndc3bb | 0.04254 | 0.283002028 | 0.136 | 0 | 1 | 18 |
| fxyd1 | 0.04254 | 0.262089675 | 0.136 | 0 | 1 | 18 |
| gabpb2a | 0.04254 | 0.313830431 | 0.136 | 0 | 1 | 18 |
| glrx5 | 0.04254 | 0.206284711 | 0.136 | 0 | 1 | 18 |
| gmnn | 0.04254 | 0.218794285 | 0.136 | 0 | 1 | 18 |
| hsp70.1 | 0.04254 | 0.388609518 | 0.136 | 0 | 1 | 18 |
| isy1 | 0.04254 | 0.232287468 | 0.136 | 0 | 1 | 18 |
| lrrc1 | 0.04254 | 0.231777566 | 0.136 | 0 | 1 | 18 |
| mapre3b | 0.04254 | 0.247969537 | 0.136 | 0 | 1 | 18 |
| mcamb | 0.04254 | 0.272337041 | 0.136 | 0 | 1 | 18 |
| mrps23 | 0.04254 | 0.284091421 | 0.136 | 0 | 1 | 18 |
| mterf3 | 0.04254 | 0.270350661 | 0.136 | 0 | 1 | 18 |
| nudt8 | 0.04254 | 0.2788683 | 0.136 | 0 | 1 | 18 |
| pdzd11 | 0.04254 | 0.271921055 | 0.136 | 0 | 1 | 18 |
| prpsap1 | 0.04254 | 0.275908398 | 0.136 | 0 | 1 | 18 |
| rnaseh2c | 0.04254 | 0.263983141 | 0.136 | 0 | 1 | 18 |
| rnf126 | 0.04254 | 0.273970617 | 0.136 | 0 | 1 | 18 |
| sgtb | 0.04254 | 0.253296406 | 0.136 | 0 | 1 | 18 |
| si:dkey-118j18.2 | 0.04254 | 0.255692493 | 0.136 | 0 | 1 | 18 |
| sp3b | 0.04254 | 0.310432794 | 0.136 | 0 | 1 | 18 |
| tnika | 0.04254 | 0.229812658 | 0.136 | 0 | 1 | 18 |
| tp53inp2 | 0.04254 | 0.343097324 | 0.136 | 0 | 1 | 18 |
| ubl7b | 0.04254 | 0.307787879 | 0.136 | 0 | 1 | 18 |
| zgc:66472 | 0.04254 | 0.241930633 | 0.136 | 0 | 1 | 18 |
| ccna2 | 0.04254 | 0.263922088 | 0.136 | 0 | 1 | 18 |
| cdca8 | 0.04254 | 0.233507236 | 0.136 | 0 | 1 | 18 |
| cox10 | 0.04254 | 0.199172825 | 0.136 | 0 | 1 | 18 |
| dlgap5 | 0.04254 | 0.215148801 | 0.136 | 0 | 1 | 18 |
| ep300a | 0.04254 | 0.309555511 | 0.136 | 0 | 1 | 18 |
| hoxb2a | 0.04254 | 0.285239939 | 0.136 | 0 | 1 | 18 |
| lef1 | 0.04254 | 0.250314541 | 0.136 | 0 | 1 | 18 |
| nfil3-5 | 0.04254 | 0.224974978 | 0.136 | 0 | 1 | 18 |
| rab27a | 0.04254 | 0.218995838 | 0.136 | 0 | 1 | 18 |
| tfdp1a | 0.04254 | 0.235679301 | 0.136 | 0 | 1 | 18 |
| ppil1 | 0.04254 | 0.303343268 | 0.136 | 0 | 1 | 18 |
| si:dkey-10p5.10 | 0.04254 | 0.31711755 | 0.136 | 0 | 1 | 18 |
| efemp2a | 0.0428 | -0.51515294 | 0.136 | 0.423 | 1 | 18 |
| hnrnpa1a | 0.04371 | -0.4344074 | 0.227 | 0.577 | 1 | 18 |
| col1a1a | 0.04379 | -0.24424811 | 0.955 | 0.962 | 1 | 18 |
| pcyox1 | 0.04396 | -0.33790703 | 0.045 | 0.308 | 1 | 18 |
| cdv3 | 0.04494 | -0.39535258 | 0.227 | 0.385 | 1 | 18 |
| ddt | 0.04514 | -0.34209612 | 0.136 | 0.423 | 1 | 18 |

| | | | | | | |
|---|---|---|---|---|---|---|
| eif4a3 | 0.04556 | -0.45131257 | 0.227 | 0.577 | 1 | 18 |
| ift46 | 0.04572 | -0.30882399 | 0.045 | 0.269 | 1 | 18 |
| perp | 0.04608 | -0.43797852 | 0.045 | 0.308 | 1 | 18 |
| kif26ba | 0.0466 | -0.46061354 | 0.045 | 0.308 | 1 | 18 |
| sept6 | 0.04662 | -0.20510416 | 0.045 | 0.231 | 1 | 18 |
| pi15b | 0.0475 | -0.42292315 | 0.045 | 0.269 | 1 | 18 |
| AL732488.1 | 0.04761 | -0.4490359 | 0.045 | 0.308 | 1 | 18 |
| tmem30aa | 0.04807 | -0.3925523 | 0.045 | 0.308 | 1 | 18 |
| h2afvb | 0.04908 | -0.11963999 | 0.682 | 0.923 | 1 | 18 |
| cntfr | 0.04958 | -0.53888325 | 0.045 | 0.308 | 1 | 18 |
| zgc:56304 | 0.04973 | -0.15465354 | 0.182 | 0.385 | 1 | 18 |
| pkp3b | 0.05004 | -0.25649676 | 0.136 | 0.385 | 1 | 18 |
| smo | 0.05005 | 0.272993066 | 0.318 | 0.269 | 1 | 18 |
| h2afva | 0.05018 | 0.366590546 | 0.545 | 0.462 | 1 | 18 |
| banf1 | 0.05058 | 0.441812577 | 0.455 | 0.231 | 1 | 18 |
| fkbp9 | 0.05088 | -0.30636353 | 0.318 | 0.615 | 1 | 18 |
| hsp90b1 | 0.05099 | -0.4627843 | 0.273 | 0.423 | 1 | 18 |
| bsdc1 | 0.05176 | -0.42059624 | 0.091 | 0.385 | 1 | 18 |
| sdhc | 0.05181 | 0.203066744 | 0.182 | 0.192 | 1 | 18 |
| znf1035 | 0.05195 | -0.32002227 | 0.045 | 0.269 | 1 | 18 |
| tmsb2 | 0.05196 | -0.23099972 | 0.091 | 0.308 | 1 | 18 |
| zmat2 | 0.05226 | -0.27062579 | 0.273 | 0.577 | 1 | 18 |
| adamts15b | 0.05252 | -0.24311027 | 0.045 | 0.192 | 1 | 18 |
| pabpc1a | 0.05269 | -0.32981292 | 0.773 | 0.923 | 1 | 18 |
| rbx1 | 0.05283 | 0.257750234 | 0.455 | 0.462 | 1 | 18 |
| ywhaz | 0.05289 | 0.263528878 | 0.364 | 0.346 | 1 | 18 |
| mir21-1 | 0.05309 | 0.437635734 | 0.273 | 0.038 | 1 | 18 |
| mmp15a | 0.05356 | -0.31099321 | 0.045 | 0.269 | 1 | 18 |
| ctnnb1 | 0.05407 | -0.37255786 | 0.364 | 0.692 | 1 | 18 |
| spry2 | 0.05429 | 0.508999999 | 0.318 | 0.077 | 1 | 18 |
| yap1 | 0.05431 | 0.293079934 | 0.318 | 0.231 | 1 | 18 |
| si:dkeyp-113d7.1 | 0.0557 | -0.26056264 | 0.227 | 0.5 | 1 | 18 |
| atp5mf | 0.05609 | -0.37012306 | 0.273 | 0.615 | 1 | 18 |
| coa6 | 0.05812 | 0.317384401 | 0.273 | 0.038 | 1 | 18 |
| slit3 | 0.05825 | -0.51126502 | 0.136 | 0.423 | 1 | 18 |
| yipf5 | 0.0583 | -0.12296962 | 0.136 | 0.308 | 1 | 18 |
| maff | 0.05853 | -0.31047108 | 0.091 | 0.346 | 1 | 18 |
| sgf29 | 0.05914 | 0.436563667 | 0.227 | 0.038 | 1 | 18 |
| amer1 | 0.06034 | -0.28782716 | 0.045 | 0.269 | 1 | 18 |
| march2 | 0.06122 | -0.26054375 | 0.045 | 0.231 | 1 | 18 |
| ndufs5 | 0.06291 | -0.43451038 | 0.182 | 0.462 | 1 | 18 |
| ndufa1 | 0.06313 | -0.36076627 | 0.182 | 0.5 | 1 | 18 |
| sntb2 | 0.06366 | -0.21464264 | 0.136 | 0.346 | 1 | 18 |
| si:ch211-286b5.5 | 0.06447 | -0.38558695 | 0.182 | 0.5 | 1 | 18 |
| pdcd5 | 0.06554 | -0.31017491 | 0.091 | 0.346 | 1 | 18 |
| cst14b.1 | 0.06561 | -0.25047622 | 0.091 | 0.308 | 1 | 18 |
| fth1a | 0.06606 | -0.3000561 | 0.5 | 0.808 | 1 | 18 |
| sufu | 0.06618 | -0.31636947 | 0.045 | 0.269 | 1 | 18 |

| | | | | | | |
|---|---|---|---|---|---|---|
| parvaa | 0.06625 | -0.38006114 | 0.136 | 0.423 | 1 | 18 |
| krt18b | 0.06658 | -0.40890062 | 0.545 | 0.692 | 1 | 18 |
| nkd1 | 0.06717 | -0.24892136 | 0.091 | 0.308 | 1 | 18 |
| atp5f1e | 0.06743 | 0.427788846 | 0.773 | 0.462 | 1 | 18 |
| snrpd3l | 0.06756 | 0.41793599 | 0.318 | 0.115 | 1 | 18 |
| rwdd1 | 0.0677 | 0.317401393 | 0.364 | 0.269 | 1 | 18 |
| ackr3b | 0.06779 | 0.50987514 | 0.545 | 0.231 | 1 | 18 |
| rpl38 | 0.06852 | -0.16776906 | 0.955 | 0.962 | 1 | 18 |
| dnmt1 | 0.06863 | 0.168389688 | 0.182 | 0.038 | 1 | 18 |
| wwtr1 | 0.06874 | -0.44902233 | 0.045 | 0.269 | 1 | 18 |
| nap1l1 | 0.06925 | -0.32745727 | 0.5 | 0.808 | 1 | 18 |
| srsf7a | 0.06962 | -0.4050782 | 0.136 | 0.346 | 1 | 18 |
| arpc4l | 0.06976 | -0.42232466 | 0.136 | 0.423 | 1 | 18 |
| tp53 | 0.07009 | -0.40152856 | 0.091 | 0.346 | 1 | 18 |
| ubc | 0.07098 | 0.279874326 | 0.909 | 0.846 | 1 | 18 |
| pa2g4b | 0.07106 | 0.308446747 | 0.455 | 0.346 | 1 | 18 |
| si:ch211-212k18.8 | 0.07114 | -0.45178847 | 0.136 | 0.423 | 1 | 18 |
| dpysl2b | 0.07117 | -0.45557238 | 0.091 | 0.346 | 1 | 18 |
| ucp2 | 0.07136 | -0.33524646 | 0.318 | 0.423 | 1 | 18 |
| rpl24 | 0.07142 | 0.175309943 | 1 | 0.962 | 1 | 18 |
| mat2aa | 0.07147 | -0.20423235 | 0.091 | 0.269 | 1 | 18 |
| nova2 | 0.0721 | -0.42738051 | 0.136 | 0.423 | 1 | 18 |
| krtcap2 | 0.0727 | -0.13529113 | 0.455 | 0.692 | 1 | 18 |
| mrpl47 | 0.0743 | 0.150323864 | 0.091 | 0.154 | 1 | 18 |
| arglu1b | 0.07436 | -0.43637487 | 0.045 | 0.269 | 1 | 18 |
| cnbpa | 0.07548 | 0.103023718 | 0.409 | 0.5 | 1 | 18 |
| cdc42l | 0.07553 | 0.35157041 | 0.455 | 0.308 | 1 | 18 |
| si:ch211-212k18.9 | 0.07627 | -0.46821405 | 0.182 | 0.423 | 1 | 18 |
| CMKLR1 (1 of many) | 0.07669 | -0.19089576 | 0 | 0.115 | 1 | 18 |
| LOC110437774 | 0.07669 | -0.18875558 | 0 | 0.115 | 1 | 18 |
| adora2b | 0.07669 | -0.23073743 | 0 | 0.115 | 1 | 18 |
| b4galt1l | 0.07669 | -0.19245942 | 0 | 0.115 | 1 | 18 |
| cetn4 | 0.07669 | -0.21745639 | 0 | 0.115 | 1 | 18 |
| crebbpb | 0.07669 | -0.16087275 | 0 | 0.115 | 1 | 18 |
| csrnp2 | 0.07669 | -0.23761267 | 0 | 0.115 | 1 | 18 |
| gnptg | 0.07669 | -0.19959614 | 0 | 0.115 | 1 | 18 |
| nrarpb | 0.07669 | -0.21386118 | 0 | 0.115 | 1 | 18 |
| pgap3 | 0.07669 | -0.16526736 | 0 | 0.115 | 1 | 18 |
| prkd3 | 0.07669 | -0.25174525 | 0 | 0.115 | 1 | 18 |
| prpf31 | 0.07669 | -0.20808752 | 0 | 0.115 | 1 | 18 |
| sestd1 | 0.07669 | -0.17590326 | 0 | 0.115 | 1 | 18 |
| sfrp5 | 0.07669 | -0.27361187 | 0 | 0.115 | 1 | 18 |
| si:ch211-170d8.8 | 0.07669 | -0.21414766 | 0 | 0.115 | 1 | 18 |
| sort1a | 0.07669 | -0.25208577 | 0 | 0.115 | 1 | 18 |
| sun1 | 0.07669 | -0.21369176 | 0 | 0.115 | 1 | 18 |
| tcf3a | 0.07669 | -0.14526025 | 0 | 0.115 | 1 | 18 |
| thumpd3 | 0.07669 | -0.16086265 | 0 | 0.115 | 1 | 18 |
| zdhhc4 | 0.07669 | -0.16398109 | 0 | 0.115 | 1 | 18 |

| | | | | | | |
|---|---|---|---|---|---|---|
| ANKRD50 | 0.07669 | -0.19727399 | 0 | 0.115 | 1 | 18 |
| DXO | 0.07669 | -0.16325344 | 0 | 0.115 | 1 | 18 |
| GPATCH2 | 0.07669 | -0.21929526 | 0 | 0.115 | 1 | 18 |
| LOC100535208 | 0.07669 | -0.16405789 | 0 | 0.115 | 1 | 18 |
| LOC101884753 | 0.07669 | -0.20985203 | 0 | 0.115 | 1 | 18 |
| LOC108190731 | 0.07669 | -0.17394764 | 0 | 0.115 | 1 | 18 |
| TENM2 | 0.07669 | -0.18346023 | 0 | 0.115 | 1 | 18 |
| TTC28 | 0.07669 | -0.24539976 | 0 | 0.115 | 1 | 18 |
| XLOC-018981 | 0.07669 | -0.24209043 | 0 | 0.115 | 1 | 18 |
| acacb | 0.07669 | -0.19186364 | 0 | 0.115 | 1 | 18 |
| actr1 | 0.07669 | -0.2161749 | 0 | 0.115 | 1 | 18 |
| adhfe1 | 0.07669 | -0.16946346 | 0 | 0.115 | 1 | 18 |
| alkal2a | 0.07669 | -0.2517488 | 0 | 0.115 | 1 | 18 |
| alx4a | 0.07669 | -0.27271391 | 0 | 0.115 | 1 | 18 |
| ankhd1 | 0.07669 | -0.17749559 | 0 | 0.115 | 1 | 18 |
| aplp2 | 0.07669 | -0.16328779 | 0 | 0.115 | 1 | 18 |
| arhgap10 | 0.07669 | -0.19900781 | 0 | 0.115 | 1 | 18 |
| asl | 0.07669 | -0.14217586 | 0 | 0.115 | 1 | 18 |
| atf6 | 0.07669 | -0.1662033 | 0 | 0.115 | 1 | 18 |
| atf7b | 0.07669 | -0.22356008 | 0 | 0.115 | 1 | 18 |
| aup1 | 0.07669 | -0.16128746 | 0 | 0.115 | 1 | 18 |
| c1galt1b | 0.07669 | -0.15865999 | 0 | 0.115 | 1 | 18 |
| cab39l1 | 0.07669 | -0.15204137 | 0 | 0.115 | 1 | 18 |
| cars | 0.07669 | -0.19906377 | 0 | 0.115 | 1 | 18 |
| castor2 | 0.07669 | -0.18171644 | 0 | 0.115 | 1 | 18 |
| ccdc22 | 0.07669 | -0.1573902 | 0 | 0.115 | 1 | 18 |
| ccl44 | 0.07669 | -0.18902238 | 0 | 0.115 | 1 | 18 |
| ccng2 | 0.07669 | -0.22927518 | 0 | 0.115 | 1 | 18 |
| cemip2 | 0.07669 | -0.18636677 | 0 | 0.115 | 1 | 18 |
| chmp3 | 0.07669 | -0.1634057 | 0 | 0.115 | 1 | 18 |
| cmtr1 | 0.07669 | -0.25349883 | 0 | 0.115 | 1 | 18 |
| coa7 | 0.07669 | -0.2053769 | 0 | 0.115 | 1 | 18 |
| crot | 0.07669 | -0.16723513 | 0 | 0.115 | 1 | 18 |
| crsp7 | 0.07669 | -0.18939205 | 0 | 0.115 | 1 | 18 |
| dcaf6 | 0.07669 | -0.17122484 | 0 | 0.115 | 1 | 18 |
| dcbld2 | 0.07669 | -0.17575656 | 0 | 0.115 | 1 | 18 |
| dcun1d3 | 0.07669 | -0.2076444 | 0 | 0.115 | 1 | 18 |
| ddr2a | 0.07669 | -0.22681753 | 0 | 0.115 | 1 | 18 |
| ddx18 | 0.07669 | -0.17748143 | 0 | 0.115 | 1 | 18 |
| ddx56 | 0.07669 | -0.17143771 | 0 | 0.115 | 1 | 18 |
| dgcr6 | 0.07669 | -0.19816076 | 0 | 0.115 | 1 | 18 |
| dglucy | 0.07669 | -0.17340414 | 0 | 0.115 | 1 | 18 |
| dhrs12 | 0.07669 | -0.17847173 | 0 | 0.115 | 1 | 18 |
| dip2ca | 0.07669 | -0.1890263 | 0 | 0.115 | 1 | 18 |
| dnajb1a | 0.07669 | -0.1510695 | 0 | 0.115 | 1 | 18 |
| dock1 | 0.07669 | -0.27058476 | 0 | 0.115 | 1 | 18 |
| dyrk1ab | 0.07669 | -0.19859368 | 0 | 0.115 | 1 | 18 |
| dyrk1b | 0.07669 | -0.20714906 | 0 | 0.115 | 1 | 18 |

| | | | | | | |
|---|---|---|---|---|---|---|
| eef1akmt2 | 0.07669 | -0.14863648 | 0 | 0.115 | 1 | 18 |
| eif3hb | 0.07669 | -0.2151671 | 0 | 0.115 | 1 | 18 |
| eml1 | 0.07669 | -0.15547511 | 0 | 0.115 | 1 | 18 |
| ephx5 | 0.07669 | -0.17576102 | 0 | 0.115 | 1 | 18 |
| faf1 | 0.07669 | -0.18186418 | 0 | 0.115 | 1 | 18 |
| fbxw7 | 0.07669 | -0.21024287 | 0 | 0.115 | 1 | 18 |
| filip1b | 0.07669 | -0.27034949 | 0 | 0.115 | 1 | 18 |
| fuca2 | 0.07669 | -0.15543207 | 0 | 0.115 | 1 | 18 |
| ggcta | 0.07669 | -0.1642875 | 0 | 0.115 | 1 | 18 |
| gigyf1b | 0.07669 | -0.21951179 | 0 | 0.115 | 1 | 18 |
| gnpat | 0.07669 | -0.25299953 | 0 | 0.115 | 1 | 18 |
| gpatch3 | 0.07669 | -0.20243956 | 0 | 0.115 | 1 | 18 |
| hoxb1b | 0.07669 | -0.22088802 | 0 | 0.115 | 1 | 18 |
| hoxc4a | 0.07669 | -0.18351679 | 0 | 0.115 | 1 | 18 |
| htra3b | 0.07669 | -0.20437035 | 0 | 0.115 | 1 | 18 |
| ift57 | 0.07669 | -0.18276436 | 0 | 0.115 | 1 | 18 |
| igsf8 | 0.07669 | -0.19110986 | 0 | 0.115 | 1 | 18 |
| ints2 | 0.07669 | -0.22546743 | 0 | 0.115 | 1 | 18 |
| kdm6al | 0.07669 | -0.1779342 | 0 | 0.115 | 1 | 18 |
| kidins220b | 0.07669 | -0.17565634 | 0 | 0.115 | 1 | 18 |
| kif1bp | 0.07669 | -0.17284729 | 0 | 0.115 | 1 | 18 |
| kpna1 | 0.07669 | -0.21034863 | 0 | 0.115 | 1 | 18 |
| krr1 | 0.07669 | -0.11178267 | 0 | 0.115 | 1 | 18 |
| litaf | 0.07669 | -0.19385061 | 0 | 0.115 | 1 | 18 |
| lrba | 0.07669 | -0.19277714 | 0 | 0.115 | 1 | 18 |
| man2c1 | 0.07669 | -0.18418223 | 0 | 0.115 | 1 | 18 |
| mcee | 0.07669 | -0.18595546 | 0 | 0.115 | 1 | 18 |
| mcm3ap | 0.07669 | -0.19140743 | 0 | 0.115 | 1 | 18 |
| me3 | 0.07669 | -0.17200636 | 0 | 0.115 | 1 | 18 |
| med15 | 0.07669 | -0.17956536 | 0 | 0.115 | 1 | 18 |
| mogs | 0.07669 | -0.22249788 | 0 | 0.115 | 1 | 18 |
| mre11a | 0.07669 | -0.11446311 | 0 | 0.115 | 1 | 18 |
| mrpl15 | 0.07669 | -0.1123376 | 0 | 0.115 | 1 | 18 |
| mrps2 | 0.07669 | -0.170824 | 0 | 0.115 | 1 | 18 |
| mrps22 | 0.07669 | -0.14562023 | 0 | 0.115 | 1 | 18 |
| mrps24 | 0.07669 | -0.19081728 | 0 | 0.115 | 1 | 18 |
| mrps5 | 0.07669 | -0.15549248 | 0 | 0.115 | 1 | 18 |
| mrtfba | 0.07669 | -0.25409756 | 0 | 0.115 | 1 | 18 |
| mta3 | 0.07669 | -0.16299697 | 0 | 0.115 | 1 | 18 |
| ndrg3b | 0.07669 | -0.16718584 | 0 | 0.115 | 1 | 18 |
| ndufs8a | 0.07669 | -0.16502732 | 0 | 0.115 | 1 | 18 |
| nectin3b | 0.07669 | -0.20443471 | 0 | 0.115 | 1 | 18 |
| neurl1b | 0.07669 | -0.19922197 | 0 | 0.115 | 1 | 18 |
| nr1d2b | 0.07669 | -0.23684079 | 0 | 0.115 | 1 | 18 |
| nras | 0.07669 | -0.16527685 | 0 | 0.115 | 1 | 18 |
| otulina | 0.07669 | -0.23913414 | 0 | 0.115 | 1 | 18 |
| parp12b | 0.07669 | -0.1741238 | 0 | 0.115 | 1 | 18 |
| parp2 | 0.07669 | -0.18068534 | 0 | 0.115 | 1 | 18 |

| | | | | | | |
|---|---|---|---|---|---|---|
| pbrm1l | 0.07669 | -0.16504097 | 0 | 0.115 | 1 | 18 |
| peli1a | 0.07669 | -0.28158959 | 0 | 0.115 | 1 | 18 |
| pgls | 0.07669 | -0.14180849 | 0 | 0.115 | 1 | 18 |
| pgr | 0.07669 | -0.16296228 | 0 | 0.115 | 1 | 18 |
| pla2g4ab | 0.07669 | -0.17097696 | 0 | 0.115 | 1 | 18 |
| pla2g7 | 0.07669 | -0.15303402 | 0 | 0.115 | 1 | 18 |
| polr2b | 0.07669 | -0.15904327 | 0 | 0.115 | 1 | 18 |
| ppm1ab | 0.07669 | -0.27544948 | 0 | 0.115 | 1 | 18 |
| ppp4r2a | 0.07669 | -0.16823586 | 0 | 0.115 | 1 | 18 |
| ptprda | 0.07669 | -0.2887541 | 0 | 0.115 | 1 | 18 |
| ptprk | 0.07669 | -0.18775571 | 0 | 0.115 | 1 | 18 |
| rabgef1 | 0.07669 | -0.1727166 | 0 | 0.115 | 1 | 18 |
| raph1a | 0.07669 | -0.2244343 | 0 | 0.115 | 1 | 18 |
| rbm19 | 0.07669 | -0.19948241 | 0 | 0.115 | 1 | 18 |
| rhogb | 0.07669 | -0.19946251 | 0 | 0.115 | 1 | 18 |
| rin2 | 0.07669 | -0.22434932 | 0 | 0.115 | 1 | 18 |
| rnmt | 0.07669 | -0.1861953 | 0 | 0.115 | 1 | 18 |
| rpap1 | 0.07669 | -0.2180456 | 0 | 0.115 | 1 | 18 |
| rpap3 | 0.07669 | -0.18008467 | 0 | 0.115 | 1 | 18 |
| rtn3 | 0.07669 | -0.15620615 | 0 | 0.115 | 1 | 18 |
| s100w | 0.07669 | -0.15033461 | 0 | 0.115 | 1 | 18 |
| sacm1lb | 0.07669 | -0.21956891 | 0 | 0.115 | 1 | 18 |
| sec24c | 0.07669 | -0.15989399 | 0 | 0.115 | 1 | 18 |
| setbp1 | 0.07669 | -0.17803602 | 0 | 0.115 | 1 | 18 |
| setd1a | 0.07669 | -0.20477934 | 0 | 0.115 | 1 | 18 |
| sgsh | 0.07669 | -0.19485 | 0 | 0.115 | 1 | 18 |
| si:ch211-193c2.2 | 0.07669 | -0.1613616 | 0 | 0.115 | 1 | 18 |
| si:dkey-19e4.5 | 0.07669 | -0.17513721 | 0 | 0.115 | 1 | 18 |
| si:dkey-72l14.4 | 0.07669 | -0.15958424 | 0 | 0.115 | 1 | 18 |
| sik3 | 0.07669 | -0.17751165 | 0 | 0.115 | 1 | 18 |
| slc22a15 | 0.07669 | -0.20696217 | 0 | 0.115 | 1 | 18 |
| slc25a43 | 0.07669 | -0.17215323 | 0 | 0.115 | 1 | 18 |
| slc37a4a | 0.07669 | -0.12025949 | 0 | 0.115 | 1 | 18 |
| slka | 0.07669 | -0.18563937 | 0 | 0.115 | 1 | 18 |
| smfn | 0.07669 | -0.25522894 | 0 | 0.115 | 1 | 18 |
| smim8 | 0.07669 | -0.20561403 | 0 | 0.115 | 1 | 18 |
| smurf2 | 0.07669 | -0.19628495 | 0 | 0.115 | 1 | 18 |
| snapc3 | 0.07669 | -0.22196896 | 0 | 0.115 | 1 | 18 |
| snrkb | 0.07669 | -0.16735766 | 0 | 0.115 | 1 | 18 |
| sssca1 | 0.07669 | -0.18467854 | 0 | 0.115 | 1 | 18 |
| stard3nl | 0.07669 | -0.17749782 | 0 | 0.115 | 1 | 18 |
| stat2 | 0.07669 | -0.20012205 | 0 | 0.115 | 1 | 18 |
| stk38b | 0.07669 | -0.18712546 | 0 | 0.115 | 1 | 18 |
| strada | 0.07669 | -0.12714614 | 0 | 0.115 | 1 | 18 |
| syt11a | 0.07669 | -0.21092852 | 0 | 0.115 | 1 | 18 |
| tapbp.2 | 0.07669 | -0.16371353 | 0 | 0.115 | 1 | 18 |
| tbc1d4 | 0.07669 | -0.19872759 | 0 | 0.115 | 1 | 18 |
| tead3a | 0.07669 | -0.1663385 | 0 | 0.115 | 1 | 18 |

| | | | | | | |
|---|---|---|---|---|---|---|
| tmem135 | 0.07669 | -0.19701405 | 0 | 0.115 | 1 | 18 |
| tmem181 | 0.07669 | -0.1794567 | 0 | 0.115 | 1 | 18 |
| tmem256 | 0.07669 | -0.15134644 | 0 | 0.115 | 1 | 18 |
| tnfaip3 | 0.07669 | -0.18433817 | 0 | 0.115 | 1 | 18 |
| tpp2 | 0.07669 | -0.21726036 | 0 | 0.115 | 1 | 18 |
| tspo | 0.07669 | -0.16293665 | 0 | 0.115 | 1 | 18 |
| txndc16 | 0.07669 | -0.2066299 | 0 | 0.115 | 1 | 18 |
| ube2w | 0.07669 | -0.17345496 | 0 | 0.115 | 1 | 18 |
| ulk1a | 0.07669 | -0.23026705 | 0 | 0.115 | 1 | 18 |
| vapal | 0.07669 | -0.22977681 | 0 | 0.115 | 1 | 18 |
| vldlr | 0.07669 | -0.19279334 | 0 | 0.115 | 1 | 18 |
| wdr45 | 0.07669 | -0.18061717 | 0 | 0.115 | 1 | 18 |
| wdr45b | 0.07669 | -0.18984798 | 0 | 0.115 | 1 | 18 |
| wdr73 | 0.07669 | -0.18730984 | 0 | 0.115 | 1 | 18 |
| wnt11r | 0.07669 | -0.20258243 | 0 | 0.115 | 1 | 18 |
| wsb1 | 0.07669 | -0.19122645 | 0 | 0.115 | 1 | 18 |
| zgc:110843 | 0.07669 | -0.22311693 | 0 | 0.115 | 1 | 18 |
| zgc:112496 | 0.07669 | -0.20152053 | 0 | 0.115 | 1 | 18 |
| zgc:113436 | 0.07669 | -0.15323258 | 0 | 0.115 | 1 | 18 |
| zgc:158517 | 0.07669 | -0.18872387 | 0 | 0.115 | 1 | 18 |
| znf1124 | 0.07669 | -0.17617594 | 0 | 0.115 | 1 | 18 |
| znf362b | 0.07669 | -0.22334677 | 0 | 0.115 | 1 | 18 |
| znf609b | 0.07669 | -0.21385127 | 0 | 0.115 | 1 | 18 |
| lmbrd1 | 0.07669 | -0.19079721 | 0 | 0.115 | 1 | 18 |
| BX005254.1 | 0.07669 | -0.19613487 | 0 | 0.115 | 1 | 18 |
| LOC110440195 | 0.07669 | -0.20198465 | 0 | 0.115 | 1 | 18 |
| TCIM (1 of many) | 0.07669 | -0.38324844 | 0 | 0.115 | 1 | 18 |
| acvr1ba | 0.07669 | -0.19583817 | 0 | 0.115 | 1 | 18 |
| aifm2 | 0.07669 | -0.19264907 | 0 | 0.115 | 1 | 18 |
| camk2g2 | 0.07669 | -0.22200116 | 0 | 0.115 | 1 | 18 |
| ccdc186 | 0.07669 | -0.19885487 | 0 | 0.115 | 1 | 18 |
| cntn3a.1 | 0.07669 | -0.21754933 | 0 | 0.115 | 1 | 18 |
| ctif | 0.07669 | -0.19973395 | 0 | 0.115 | 1 | 18 |
| ctr9 | 0.07669 | -0.25178226 | 0 | 0.115 | 1 | 18 |
| cttn | 0.07669 | -0.22038066 | 0 | 0.115 | 1 | 18 |
| cyp3c1 | 0.07669 | -0.24318129 | 0 | 0.115 | 1 | 18 |
| ddx47 | 0.07669 | -0.12982322 | 0 | 0.115 | 1 | 18 |
| elovl1b | 0.07669 | -0.22689466 | 0 | 0.115 | 1 | 18 |
| emc1 | 0.07669 | -0.20522677 | 0 | 0.115 | 1 | 18 |
| eya3 | 0.07669 | -0.19541707 | 0 | 0.115 | 1 | 18 |
| gpr146 | 0.07669 | -0.22446284 | 0 | 0.115 | 1 | 18 |
| has2 | 0.07669 | -0.24236221 | 0 | 0.115 | 1 | 18 |
| herc2 | 0.07669 | -0.20209723 | 0 | 0.115 | 1 | 18 |
| ice1 | 0.07669 | -0.14294062 | 0 | 0.115 | 1 | 18 |
| ints11 | 0.07669 | -0.22248461 | 0 | 0.115 | 1 | 18 |
| man2a2 | 0.07669 | -0.20071581 | 0 | 0.115 | 1 | 18 |
| mgme1 | 0.07669 | -0.19858923 | 0 | 0.115 | 1 | 18 |
| mgrn1b | 0.07669 | -0.20041149 | 0 | 0.115 | 1 | 18 |

| | | | | | | |
|---|---|---|---|---|---|---|
| nmt2 | 0.07669 | -0.20973747 | 0 | 0.115 | 1 | 18 |
| phldb2a | 0.07669 | -0.20091092 | 0 | 0.115 | 1 | 18 |
| ppardb | 0.07669 | -0.14990382 | 0 | 0.115 | 1 | 18 |
| psd3l | 0.07669 | -0.19851673 | 0 | 0.115 | 1 | 18 |
| rab20 | 0.07669 | -0.11273627 | 0 | 0.115 | 1 | 18 |
| ric8b | 0.07669 | -0.24206468 | 0 | 0.115 | 1 | 18 |
| si:dkey-29d8.3 | 0.07669 | -0.21207581 | 0 | 0.115 | 1 | 18 |
| si:dkey-51e6.1 | 0.07669 | -0.19681545 | 0 | 0.115 | 1 | 18 |
| tbc1d22a | 0.07669 | -0.25903172 | 0 | 0.115 | 1 | 18 |
| tm9sf3 | 0.07669 | -0.19426793 | 0 | 0.115 | 1 | 18 |
| tnfrsf1a | 0.07669 | -0.2084021 | 0 | 0.115 | 1 | 18 |
| CABZ01068251.1 | 0.07669 | -0.17064639 | 0 | 0.115 | 1 | 18 |
| CABZ01086877.1 | 0.07669 | -0.25396791 | 0 | 0.115 | 1 | 18 |
| NC-002333.17 | 0.07669 | -0.20196148 | 0 | 0.115 | 1 | 18 |
| calm3a | 0.07669 | -0.21147295 | 0 | 0.115 | 1 | 18 |
| coro2ba | 0.07669 | -0.18960708 | 0 | 0.115 | 1 | 18 |
| cstf3 | 0.07669 | -0.22518253 | 0 | 0.115 | 1 | 18 |
| igf1rb | 0.07669 | -0.19115738 | 0 | 0.115 | 1 | 18 |
| si:ch211-151g22.1 | 0.07669 | -0.22179109 | 0 | 0.115 | 1 | 18 |
| thbs3a | 0.07669 | -0.23570309 | 0 | 0.115 | 1 | 18 |
| yipf2 | 0.07669 | -0.23116841 | 0 | 0.115 | 1 | 18 |
| zdhhc15b | 0.07669 | -0.23840372 | 0 | 0.115 | 1 | 18 |
| si:ch73-238c9.1 | 0.07681 | -0.15527774 | 0.091 | 0.269 | 1 | 18 |
| dkk3b | 0.07689 | -0.38621655 | 0.5 | 0.808 | 1 | 18 |
| si:ch73-42p12.2 | 0.07711 | -0.39207792 | 0.136 | 0.423 | 1 | 18 |
| ywhae2 | 0.07818 | -0.40562898 | 0.045 | 0.269 | 1 | 18 |
| zcchc24 | 0.07849 | -0.50678473 | 0.136 | 0.423 | 1 | 18 |
| bzw1b | 0.07884 | -0.35615201 | 0.227 | 0.538 | 1 | 18 |
| rad23b | 0.07961 | -0.15686634 | 0.318 | 0.538 | 1 | 18 |
| tsku | 0.07966 | -0.49475773 | 0.227 | 0.538 | 1 | 18 |
| si:dkey-222b8.1 | 0.08034 | 0.4663205 | 0.318 | 0.077 | 1 | 18 |
| jund | 0.0807 | -0.47854332 | 0.136 | 0.423 | 1 | 18 |
| cercam | 0.08112 | -0.40210729 | 0.045 | 0.269 | 1 | 18 |
| tmem50a | 0.08153 | -0.24074489 | 0.182 | 0.423 | 1 | 18 |
| rcn1 | 0.08167 | 0.402618674 | 0.318 | 0.154 | 1 | 18 |
| atp2b2 | 0.08229 | -0.43616717 | 0.136 | 0.423 | 1 | 18 |
| galnt1 | 0.08268 | -0.39887791 | 0.045 | 0.269 | 1 | 18 |
| lsm12b | 0.08309 | -0.12010309 | 0.273 | 0.462 | 1 | 18 |
| faub | 0.08366 | -0.28056572 | 0.045 | 0.269 | 1 | 18 |
| stau1 | 0.08371 | -0.42250494 | 0.227 | 0.538 | 1 | 18 |
| psmd14 | 0.08378 | 0.461340714 | 0.318 | 0.077 | 1 | 18 |
| inhbaa | 0.08443 | -0.13535183 | 0.045 | 0.154 | 1 | 18 |
| apbb1 | 0.0851 | -0.39150314 | 0.045 | 0.269 | 1 | 18 |
| hif1ab | 0.08515 | 0.187772131 | 0.318 | 0.308 | 1 | 18 |
| hmg20b | 0.08568 | -0.40128865 | 0.045 | 0.269 | 1 | 18 |
| eif4g2a | 0.08572 | -0.29650049 | 0.045 | 0.269 | 1 | 18 |
| npc1 | 0.08624 | 0.440939065 | 0.227 | 0.038 | 1 | 18 |
| rpl7l1 | 0.08627 | -0.35447601 | 0.045 | 0.269 | 1 | 18 |

| | | | | | | |
|---|---|---|---|---|---|---|
| sh3gl1b | 0.08652 | -0.35278239 | 0.045 | 0.269 | 1 | 18 |
| gpx1a | 0.08673 | -0.26251612 | 0.045 | 0.269 | 1 | 18 |
| rgcc | 0.08735 | -0.26458443 | 0.045 | 0.231 | 1 | 18 |
| f11r.1 | 0.0875 | 0.121225939 | 0.227 | 0.269 | 1 | 18 |
| qkia | 0.08847 | 0.252698216 | 0.727 | 0.654 | 1 | 18 |
| aars | 0.08866 | -0.35125325 | 0.045 | 0.269 | 1 | 18 |
| CT030188.1 | 0.08875 | -0.40161005 | 0.045 | 0.269 | 1 | 18 |
| CR847566.1 | 0.0891 | 0.333503897 | 0.864 | 0.731 | 1 | 18 |
| lpar2a | 0.08937 | -0.33873209 | 0.091 | 0.346 | 1 | 18 |
| etf1b | 0.08972 | -0.31116884 | 0.227 | 0.5 | 1 | 18 |
| ubtf | 0.08987 | -0.12097352 | 0.091 | 0.231 | 1 | 18 |
| mrps16 | 0.08996 | 0.21046444 | 0.136 | 0.115 | 1 | 18 |
| ube2l3a | 0.0902 | -0.17172817 | 0.136 | 0.346 | 1 | 18 |
| serbp1b | 0.09086 | -0.34647136 | 0.045 | 0.269 | 1 | 18 |
| ssu72 | 0.09086 | -0.27071198 | 0.045 | 0.269 | 1 | 18 |
| timm13 | 0.09086 | -0.22699589 | 0.045 | 0.269 | 1 | 18 |
| gmppb | 0.09102 | -0.32689998 | 0.045 | 0.269 | 1 | 18 |
| pigx | 0.09159 | -0.33670903 | 0.045 | 0.269 | 1 | 18 |
| prelid1a | 0.09166 | 0.347821877 | 0.318 | 0.192 | 1 | 18 |
| reep3a | 0.09174 | 0.399042057 | 0.318 | 0.077 | 1 | 18 |
| sdha | 0.092 | -0.31707168 | 0.045 | 0.269 | 1 | 18 |
| fthl27 | 0.09201 | -0.24921773 | 0.864 | 0.923 | 1 | 18 |
| zgc:77058 | 0.09296 | -0.35431734 | 0.045 | 0.269 | 1 | 18 |
| uqcrh | 0.09305 | -0.31347421 | 0.409 | 0.5 | 1 | 18 |
| zfand5a | 0.09325 | -0.33700311 | 0.045 | 0.269 | 1 | 18 |
| tspan18b | 0.09332 | -0.32006385 | 0.045 | 0.269 | 1 | 18 |
| tspan3a | 0.09339 | -0.36702754 | 0.045 | 0.269 | 1 | 18 |
| zgc:114119 | 0.0934 | 0.247752736 | 0.182 | 0.115 | 1 | 18 |
| eif1ad | 0.09366 | -0.36627525 | 0.091 | 0.346 | 1 | 18 |
| manbal | 0.09421 | -0.27453041 | 0.091 | 0.308 | 1 | 18 |
| hirip3 | 0.09487 | 0.225496588 | 0.182 | 0.192 | 1 | 18 |
| ppp6c | 0.09513 | -0.3914213 | 0.091 | 0.346 | 1 | 18 |
| zgc:110158 | 0.09536 | -0.40189307 | 0.091 | 0.346 | 1 | 18 |
| emc2 | 0.09552 | -0.36146599 | 0.091 | 0.346 | 1 | 18 |
| nr1d2a | 0.09566 | -0.23132362 | 0.091 | 0.269 | 1 | 18 |
| fkbp14 | 0.09595 | -0.35162254 | 0.091 | 0.346 | 1 | 18 |
| cdh2 | 0.09598 | -0.37142283 | 0.182 | 0.462 | 1 | 18 |
| ak2 | 0.0961 | -0.35478573 | 0.273 | 0.577 | 1 | 18 |
| eif4e1c | 0.09624 | 0.380127642 | 0.227 | 0.038 | 1 | 18 |
| neo1a | 0.09631 | -0.29580636 | 0.091 | 0.308 | 1 | 18 |
| epha4b | 0.09667 | 0.400000358 | 0.318 | 0.077 | 1 | 18 |
| dcun1d5 | 0.09693 | 0.293229783 | 0.227 | 0.115 | 1 | 18 |
| si:dkey-6n21.12 | 0.0973 | -0.28043487 | 0.091 | 0.308 | 1 | 18 |
| cnot3a | 0.09768 | 0.353074492 | 0.227 | 0.038 | 1 | 18 |
| hypk | 0.09875 | 0.273930235 | 0.364 | 0.308 | 1 | 18 |
| hspe1 | 0.09918 | 0.435147239 | 0.364 | 0.115 | 1 | 18 |
| si:dkey-266m15.6 | 0.09946 | 0.409668807 | 0.227 | 0.038 | 1 | 18 |
| kdelr2a | 0.10098 | -0.10209901 | 0.091 | 0.231 | 1 | 18 |

| | | | | | | |
|---|---|---|---|---|---|---|
| spon1b | 0.10101 | -0.25448636 | 0.773 | 0.962 | 1 | 18 |
| gabarapa | 0.10154 | -0.41590389 | 0.273 | 0.577 | 1 | 18 |
| aip | 0.10204 | 0.18742441 | 0.455 | 0.423 | 1 | 18 |
| egr1 | 0.10318 | 0.270938023 | 0.364 | 0.154 | 1 | 18 |
| spcs1 | 0.10393 | -0.26406464 | 0.409 | 0.692 | 1 | 18 |
| zgc:162944 | 0.104 | -0.36966669 | 0.318 | 0.5 | 1 | 18 |
| ier5 | 0.10475 | 0.31185333 | 0.545 | 0.423 | 1 | 18 |
| txndc15 | 0.10571 | -0.11851101 | 0.136 | 0.308 | 1 | 18 |
| ssr3 | 0.10597 | -0.43815045 | 0.364 | 0.615 | 1 | 18 |
| tuba8l | 0.1066 | -0.10567684 | 0.364 | 0.577 | 1 | 18 |
| mmp14a | 0.10664 | -0.17729214 | 0.227 | 0.423 | 1 | 18 |
| mpc1 | 0.10668 | -0.14056807 | 0.091 | 0.269 | 1 | 18 |
| eif3ja | 0.10676 | -0.14354045 | 0.318 | 0.538 | 1 | 18 |
| gpatch8 | 0.10706 | -0.39161538 | 0.091 | 0.308 | 1 | 18 |
| lsm6 | 0.10814 | 0.383000865 | 0.273 | 0.077 | 1 | 18 |
| lgals1l1 | 0.10839 | -0.18403102 | 0.045 | 0.192 | 1 | 18 |
| dynlt3 | 0.10849 | -0.33992345 | 0.318 | 0.615 | 1 | 18 |
| cap1 | 0.10883 | -0.40329998 | 0.136 | 0.346 | 1 | 18 |
| anxa11b | 0.10911 | -0.41199184 | 0.318 | 0.615 | 1 | 18 |
| dab2ipb | 0.10957 | 0.362078027 | 0.227 | 0.038 | 1 | 18 |
| nxn | 0.11006 | -0.173416 | 0.091 | 0.269 | 1 | 18 |
| ehd1b | 0.11024 | 0.361945364 | 0.227 | 0.038 | 1 | 18 |
| CU694197.1 | 0.11135 | -0.40363298 | 0.182 | 0.462 | 1 | 18 |
| ctnna1 | 0.11149 | -0.2503095 | 0.045 | 0.231 | 1 | 18 |
| brd4 | 0.1117 | -0.39869602 | 0.091 | 0.308 | 1 | 18 |
| rpl14 | 0.11201 | 0.204059471 | 1 | 0.923 | 1 | 18 |
| ckba | 0.11202 | 0.369178957 | 0.364 | 0.115 | 1 | 18 |
| BX539336.1 | 0.11217 | 0.212689982 | 0.091 | 0 | 1 | 18 |
| CABZ01061591.1 | 0.11217 | 0.150916848 | 0.091 | 0 | 1 | 18 |
| CABZ01064670.1 | 0.11217 | 0.153567607 | 0.091 | 0 | 1 | 18 |
| CABZ01078055.1 | 0.11217 | 0.137079457 | 0.091 | 0 | 1 | 18 |
| CABZ01088428.1 | 0.11217 | 0.158264448 | 0.091 | 0 | 1 | 18 |
| CR847984.1 | 0.11217 | 0.133210798 | 0.091 | 0 | 1 | 18 |
| FP101887.1 | 0.11217 | 0.174878839 | 0.091 | 0 | 1 | 18 |
| FP102158.1 | 0.11217 | 0.143009697 | 0.091 | 0 | 1 | 18 |
| LOC101884757 | 0.11217 | 0.17321956 | 0.091 | 0 | 1 | 18 |
| XLOC-007541 | 0.11217 | 0.153567607 | 0.091 | 0 | 1 | 18 |
| acads | 0.11217 | 0.20064742 | 0.091 | 0 | 1 | 18 |
| acsbg2 | 0.11217 | 0.141205364 | 0.091 | 0 | 1 | 18 |
| adamts10 | 0.11217 | 0.247546789 | 0.091 | 0 | 1 | 18 |
| adob | 0.11217 | 0.146142564 | 0.091 | 0 | 1 | 18 |
| agtr2 | 0.11217 | 0.16520088 | 0.091 | 0 | 1 | 18 |
| alms1 | 0.11217 | 0.178080272 | 0.091 | 0 | 1 | 18 |
| ankrd10a | 0.11217 | 0.151760422 | 0.091 | 0 | 1 | 18 |
| ap3s2 | 0.11217 | 0.176019972 | 0.091 | 0 | 1 | 18 |
| apopt1 | 0.11217 | 0.147104527 | 0.091 | 0 | 1 | 18 |
| arhgef2 | 0.11217 | 0.189341982 | 0.091 | 0 | 1 | 18 |
| arrdc3b | 0.11217 | 0.112350237 | 0.091 | 0 | 1 | 18 |

| | | | | | | |
|---|---|---|---|---|---|---|
| ascc1 | 0.11217 | 0.121563793 | 0.091 | 0 | 1 | 18 |
| asnsd1 | 0.11217 | 0.128891033 | 0.091 | 0 | 1 | 18 |
| aspm | 0.11217 | 0.200159304 | 0.091 | 0 | 1 | 18 |
| atp6v0cb | 0.11217 | 0.140923476 | 0.091 | 0 | 1 | 18 |
| atrn | 0.11217 | 0.171385442 | 0.091 | 0 | 1 | 18 |
| boka | 0.11217 | 0.211191014 | 0.091 | 0 | 1 | 18 |
| bub1ba | 0.11217 | 0.123622863 | 0.091 | 0 | 1 | 18 |
| ccdc93 | 0.11217 | 0.163696519 | 0.091 | 0 | 1 | 18 |
| cdk8 | 0.11217 | 0.153182443 | 0.091 | 0 | 1 | 18 |
| cetn3 | 0.11217 | 0.16352642 | 0.091 | 0 | 1 | 18 |
| chmp2bb | 0.11217 | 0.168349926 | 0.091 | 0 | 1 | 18 |
| clasp2 | 0.11217 | 0.198320108 | 0.091 | 0 | 1 | 18 |
| clk4b | 0.11217 | 0.180096432 | 0.091 | 0 | 1 | 18 |
| cntn5 | 0.11217 | 0.260373272 | 0.091 | 0 | 1 | 18 |
| coq8a | 0.11217 | 0.202716931 | 0.091 | 0 | 1 | 18 |
| cox18 | 0.11217 | 0.143274903 | 0.091 | 0 | 1 | 18 |
| cptp | 0.11217 | 0.147672788 | 0.091 | 0 | 1 | 18 |
| daam1b | 0.11217 | 0.317281903 | 0.091 | 0 | 1 | 18 |
| dcps | 0.11217 | 0.179794025 | 0.091 | 0 | 1 | 18 |
| ddr1 | 0.11217 | 0.276794164 | 0.091 | 0 | 1 | 18 |
| depdc1a | 0.11217 | 0.134063092 | 0.091 | 0 | 1 | 18 |
| dip2ba | 0.11217 | 0.222816216 | 0.091 | 0 | 1 | 18 |
| dnajb14 | 0.11217 | 0.211098885 | 0.091 | 0 | 1 | 18 |
| echdc2 | 0.11217 | 0.117348333 | 0.091 | 0 | 1 | 18 |
| edem2 | 0.11217 | 0.196461371 | 0.091 | 0 | 1 | 18 |
| edil3a | 0.11217 | 0.170877869 | 0.091 | 0 | 1 | 18 |
| eftud2 | 0.11217 | 0.156790685 | 0.091 | 0 | 1 | 18 |
| elmod2 | 0.11217 | 0.214964833 | 0.091 | 0 | 1 | 18 |
| etaa1 | 0.11217 | 0.150096131 | 0.091 | 0 | 1 | 18 |
| f3a | 0.11217 | 0.270874317 | 0.091 | 0 | 1 | 18 |
| fam207a | 0.11217 | 0.191033934 | 0.091 | 0 | 1 | 18 |
| fbrs | 0.11217 | 0.172499912 | 0.091 | 0 | 1 | 18 |
| fbxl14b | 0.11217 | 0.161218405 | 0.091 | 0 | 1 | 18 |
| frmd6 | 0.11217 | 0.134560189 | 0.091 | 0 | 1 | 18 |
| fuk | 0.11217 | 0.22493366 | 0.091 | 0 | 1 | 18 |
| gadd45ab | 0.11217 | 0.200942982 | 0.091 | 0 | 1 | 18 |
| gga1 | 0.11217 | 0.138509839 | 0.091 | 0 | 1 | 18 |
| gpkow | 0.11217 | 0.187350909 | 0.091 | 0 | 1 | 18 |
| hcfc1b | 0.11217 | 0.176054725 | 0.091 | 0 | 1 | 18 |
| hdac6 | 0.11217 | 0.215379674 | 0.091 | 0 | 1 | 18 |
| hpf1 | 0.11217 | 0.148198346 | 0.091 | 0 | 1 | 18 |
| ipo4 | 0.11217 | 0.174802443 | 0.091 | 0 | 1 | 18 |
| itpkcb | 0.11217 | 0.163788087 | 0.091 | 0 | 1 | 18 |
| jupa | 0.11217 | 0.214480632 | 0.091 | 0 | 1 | 18 |
| kif11 | 0.11217 | 0.189597236 | 0.091 | 0 | 1 | 18 |
| kif20a | 0.11217 | 0.243020807 | 0.091 | 0 | 1 | 18 |
| lin37 | 0.11217 | 0.110946959 | 0.091 | 0 | 1 | 18 |
| lrrc42 | 0.11217 | 0.141846815 | 0.091 | 0 | 1 | 18 |

| | | | | | | |
|---|---|---|---|---|---|---|
| mcat | 0.11217 | 0.195632762 | 0.091 | 0 | 1 | 18 |
| metrn | 0.11217 | 0.188531244 | 0.091 | 0 | 1 | 18 |
| mettl7a | 0.11217 | 0.145621815 | 0.091 | 0 | 1 | 18 |
| mob1ba | 0.11217 | 0.177892157 | 0.091 | 0 | 1 | 18 |
| mob1bb | 0.11217 | 0.216870597 | 0.091 | 0 | 1 | 18 |
| mon2 | 0.11217 | 0.160731305 | 0.091 | 0 | 1 | 18 |
| mrpl21 | 0.11217 | 0.14495937 | 0.091 | 0 | 1 | 18 |
| mus81 | 0.11217 | 0.127943435 | 0.091 | 0 | 1 | 18 |
| ncam1a | 0.11217 | 0.183561353 | 0.091 | 0 | 1 | 18 |
| ncapd2 | 0.11217 | 0.123348959 | 0.091 | 0 | 1 | 18 |
| ncaph2 | 0.11217 | 0.142182559 | 0.091 | 0 | 1 | 18 |
| ndufaf4 | 0.11217 | 0.154837007 | 0.091 | 0 | 1 | 18 |
| nle1 | 0.11217 | 0.168180447 | 0.091 | 0 | 1 | 18 |
| npb | 0.11217 | 0.233029095 | 0.091 | 0 | 1 | 18 |
| nsun2 | 0.11217 | 0.175183908 | 0.091 | 0 | 1 | 18 |
| ntn1a | 0.11217 | 0.222005994 | 0.091 | 0 | 1 | 18 |
| nup160 | 0.11217 | 0.210776781 | 0.091 | 0 | 1 | 18 |
| nusap1 | 0.11217 | 0.181472664 | 0.091 | 0 | 1 | 18 |
| odc1 | 0.11217 | 0.136009443 | 0.091 | 0 | 1 | 18 |
| odf2a | 0.11217 | 0.12179357 | 0.091 | 0 | 1 | 18 |
| ogfod1 | 0.11217 | 0.173102244 | 0.091 | 0 | 1 | 18 |
| paf1 | 0.11217 | 0.127833915 | 0.091 | 0 | 1 | 18 |
| pappab | 0.11217 | 0.157037113 | 0.091 | 0 | 1 | 18 |
| patz1 | 0.11217 | 0.198919027 | 0.091 | 0 | 1 | 18 |
| pbx2 | 0.11217 | 0.224212506 | 0.091 | 0 | 1 | 18 |
| pcolce2b | 0.11217 | 0.201663421 | 0.091 | 0 | 1 | 18 |
| pdcd6ip | 0.11217 | 0.22010519 | 0.091 | 0 | 1 | 18 |
| pde4ba | 0.11217 | 0.163313652 | 0.091 | 0 | 1 | 18 |
| pdk4 | 0.11217 | 0.150295561 | 0.091 | 0 | 1 | 18 |
| pid1 | 0.11217 | 0.163235821 | 0.091 | 0 | 1 | 18 |
| plk1 | 0.11217 | 0.130791734 | 0.091 | 0 | 1 | 18 |
| ppfia1 | 0.11217 | 0.22971939 | 0.091 | 0 | 1 | 18 |
| ppie | 0.11217 | 0.148268702 | 0.091 | 0 | 1 | 18 |
| preb | 0.11217 | 0.167596294 | 0.091 | 0 | 1 | 18 |
| pttg1 | 0.11217 | 0.164505065 | 0.091 | 0 | 1 | 18 |
| racgap1 | 0.11217 | 0.158379974 | 0.091 | 0 | 1 | 18 |
| rad54l | 0.11217 | 0.10862703 | 0.091 | 0 | 1 | 18 |
| rbmx2 | 0.11217 | 0.14417489 | 0.091 | 0 | 1 | 18 |
| rmdn3 | 0.11217 | 0.150096131 | 0.091 | 0 | 1 | 18 |
| rps6kal | 0.11217 | 0.210721379 | 0.091 | 0 | 1 | 18 |
| rspry1 | 0.11217 | 0.144548083 | 0.091 | 0 | 1 | 18 |
| sbds | 0.11217 | 0.133896614 | 0.091 | 0 | 1 | 18 |
| scaf4b | 0.11217 | 0.175338632 | 0.091 | 0 | 1 | 18 |
| sec22bb | 0.11217 | 0.193603977 | 0.091 | 0 | 1 | 18 |
| shisal1a | 0.11217 | 0.198723708 | 0.091 | 0 | 1 | 18 |
| si:cabz01071911.3 | 0.11217 | 0.149951338 | 0.091 | 0 | 1 | 18 |
| si:ch211-209f23.3 | 0.11217 | 0.245068191 | 0.091 | 0 | 1 | 18 |
| si:ch211-261n11.5 | 0.11217 | 0.143439972 | 0.091 | 0 | 1 | 18 |

| | | | | | | |
|---|---|---|---|---|---|---|
| si:ch211-275j6.5 | 0.11217 | 0.160446339 | 0.091 | 0 | 1 | 18 |
| si:ch211-63o20.7 | 0.11217 | 0.142182559 | 0.091 | 0 | 1 | 18 |
| si:ch211-68a17.7 | 0.11217 | 0.136209972 | 0.091 | 0 | 1 | 18 |
| si:dkey-121j17.6 | 0.11217 | 0.176358974 | 0.091 | 0 | 1 | 18 |
| si:dkey-14d8.1 | 0.11217 | 0.1351148 | 0.091 | 0 | 1 | 18 |
| si:dkey-163m14.2 | 0.11217 | 0.147387023 | 0.091 | 0 | 1 | 18 |
| si:dkey-262g12.10 | 0.11217 | 0.159674773 | 0.091 | 0 | 1 | 18 |
| si:dkey-66i24.9 | 0.11217 | 0.118603209 | 0.091 | 0 | 1 | 18 |
| si:dkey-6n21.13 | 0.11217 | 0.16826063 | 0.091 | 0 | 1 | 18 |
| si:dkey-7l6.3 | 0.11217 | 0.238188585 | 0.091 | 0 | 1 | 18 |
| slc12a7b | 0.11217 | 0.198994508 | 0.091 | 0 | 1 | 18 |
| slc16a5a | 0.11217 | 0.176358974 | 0.091 | 0 | 1 | 18 |
| slc25a36b | 0.11217 | 0.167411128 | 0.091 | 0 | 1 | 18 |
| snrnp35 | 0.11217 | 0.213440489 | 0.091 | 0 | 1 | 18 |
| snx18b | 0.11217 | 0.176358974 | 0.091 | 0 | 1 | 18 |
| spc25 | 0.11217 | 0.142182559 | 0.091 | 0 | 1 | 18 |
| sprtn | 0.11217 | 0.15358158 | 0.091 | 0 | 1 | 18 |
| srpk3 | 0.11217 | 0.220320775 | 0.091 | 0 | 1 | 18 |
| srrd | 0.11217 | 0.205086417 | 0.091 | 0 | 1 | 18 |
| srsf2b | 0.11217 | 0.142777114 | 0.091 | 0 | 1 | 18 |
| stam2 | 0.11217 | 0.162803264 | 0.091 | 0 | 1 | 18 |
| surf2 | 0.11217 | 0.18459041 | 0.091 | 0 | 1 | 18 |
| taf1a | 0.11217 | 0.159654213 | 0.091 | 0 | 1 | 18 |
| tbc1d23 | 0.11217 | 0.218031983 | 0.091 | 0 | 1 | 18 |
| tbcc | 0.11217 | 0.154735544 | 0.091 | 0 | 1 | 18 |
| tefb | 0.11217 | 0.217244435 | 0.091 | 0 | 1 | 18 |
| tfdp1b | 0.11217 | 0.17016106 | 0.091 | 0 | 1 | 18 |
| thoc3 | 0.11217 | 0.148305557 | 0.091 | 0 | 1 | 18 |
| tjp2b | 0.11217 | 0.213667042 | 0.091 | 0 | 1 | 18 |
| tmem145 | 0.11217 | 0.159671287 | 0.091 | 0 | 1 | 18 |
| tmem167b | 0.11217 | 0.173690227 | 0.091 | 0 | 1 | 18 |
| tmem67 | 0.11217 | 0.159674773 | 0.091 | 0 | 1 | 18 |
| tmtc1 | 0.11217 | 0.171476886 | 0.091 | 0 | 1 | 18 |
| tnpo1 | 0.11217 | 0.175370809 | 0.091 | 0 | 1 | 18 |
| tomm40l | 0.11217 | 0.12161473 | 0.091 | 0 | 1 | 18 |
| top2a | 0.11217 | 0.153557203 | 0.091 | 0 | 1 | 18 |
| tsn | 0.11217 | 0.134140775 | 0.091 | 0 | 1 | 18 |
| ttc7b | 0.11217 | 0.161600561 | 0.091 | 0 | 1 | 18 |
| tubg1 | 0.11217 | 0.137256155 | 0.091 | 0 | 1 | 18 |
| twist1b | 0.11217 | 0.152084816 | 0.091 | 0 | 1 | 18 |
| tyms | 0.11217 | 0.157013742 | 0.091 | 0 | 1 | 18 |
| ube2c | 0.11217 | 0.168521014 | 0.091 | 0 | 1 | 18 |
| uchl3 | 0.11217 | 0.174110926 | 0.091 | 0 | 1 | 18 |
| utrn | 0.11217 | 0.152766002 | 0.091 | 0 | 1 | 18 |
| vps25 | 0.11217 | 0.133692993 | 0.091 | 0 | 1 | 18 |
| vps4a | 0.11217 | 0.160446339 | 0.091 | 0 | 1 | 18 |
| wdfy3 | 0.11217 | 0.148232357 | 0.091 | 0 | 1 | 18 |
| wu:fc17b08 | 0.11217 | 0.125401073 | 0.091 | 0 | 1 | 18 |

| | | | | | | |
|---|---|---|---|---|---|---|
| xrcc1 | 0.11217 | 0.145131166 | 0.091 | 0 | 1 | 18 |
| yipf6 | 0.11217 | 0.164101706 | 0.091 | 0 | 1 | 18 |
| zgc:165409 | 0.11217 | 0.158457245 | 0.091 | 0 | 1 | 18 |
| zgc:175284 | 0.11217 | 0.151479578 | 0.091 | 0 | 1 | 18 |
| zgc:194627 | 0.11217 | 0.135743702 | 0.091 | 0 | 1 | 18 |
| zgc:55262 | 0.11217 | 0.170423921 | 0.091 | 0 | 1 | 18 |
| zgc:66475 | 0.11217 | 0.166984011 | 0.091 | 0 | 1 | 18 |
| zgc:77739 | 0.11217 | 0.143614329 | 0.091 | 0 | 1 | 18 |
| zgc:86839 | 0.11217 | 0.125908598 | 0.091 | 0 | 1 | 18 |
| zmynd11 | 0.11217 | 0.162752486 | 0.091 | 0 | 1 | 18 |
| zmynd19 | 0.11217 | 0.132649739 | 0.091 | 0 | 1 | 18 |
| znf131 | 0.11217 | 0.199171488 | 0.091 | 0 | 1 | 18 |
| znf142 | 0.11217 | 0.15366127 | 0.091 | 0 | 1 | 18 |
| znf770 | 0.11217 | 0.142564258 | 0.091 | 0 | 1 | 18 |
| CABZ01069162.1 | 0.11217 | 0.230626052 | 0.091 | 0 | 1 | 18 |
| FAR2 | 0.11217 | 0.182235247 | 0.091 | 0 | 1 | 18 |
| LOC101882640 | 0.11217 | 0.197606022 | 0.091 | 0 | 1 | 18 |
| LOC108182329 | 0.11217 | 0.179606365 | 0.091 | 0 | 1 | 18 |
| abcd3a | 0.11217 | 0.162323906 | 0.091 | 0 | 1 | 18 |
| abi2a | 0.11217 | 0.16144964 | 0.091 | 0 | 1 | 18 |
| ap1g1 | 0.11217 | 0.149243585 | 0.091 | 0 | 1 | 18 |
| arid2 | 0.11217 | 0.176598415 | 0.091 | 0 | 1 | 18 |
| atg12 | 0.11217 | 0.125286842 | 0.091 | 0 | 1 | 18 |
| bmt2 | 0.11217 | 0.195502249 | 0.091 | 0 | 1 | 18 |
| calm1b | 0.11217 | 0.183821195 | 0.091 | 0 | 1 | 18 |
| cbx6b | 0.11217 | 0.169632716 | 0.091 | 0 | 1 | 18 |
| cdkal1 | 0.11217 | 0.177786472 | 0.091 | 0 | 1 | 18 |
| cenpe | 0.11217 | 0.13005329 | 0.091 | 0 | 1 | 18 |
| cry-dash | 0.11217 | 0.211799845 | 0.091 | 0 | 1 | 18 |
| cul2 | 0.11217 | 0.158005215 | 0.091 | 0 | 1 | 18 |
| dcaf8 | 0.11217 | 0.198241029 | 0.091 | 0 | 1 | 18 |
| dck | 0.11217 | 0.149235696 | 0.091 | 0 | 1 | 18 |
| dctd | 0.11217 | 0.130711726 | 0.091 | 0 | 1 | 18 |
| ddx52 | 0.11217 | 0.115091381 | 0.091 | 0 | 1 | 18 |
| dpf2 | 0.11217 | 0.151838509 | 0.091 | 0 | 1 | 18 |
| elp3 | 0.11217 | 0.167860124 | 0.091 | 0 | 1 | 18 |
| epyc | 0.11217 | 0.236413734 | 0.091 | 0 | 1 | 18 |
| fam110b | 0.11217 | 0.216255673 | 0.091 | 0 | 1 | 18 |
| far1 | 0.11217 | 0.129342836 | 0.091 | 0 | 1 | 18 |
| golga2 | 0.11217 | 0.299678612 | 0.091 | 0 | 1 | 18 |
| gsk3aa | 0.11217 | 0.20566284 | 0.091 | 0 | 1 | 18 |
| hdac8 | 0.11217 | 0.179983806 | 0.091 | 0 | 1 | 18 |
| hoxc6a | 0.11217 | 0.159964648 | 0.091 | 0 | 1 | 18 |
| hs3st1l2 | 0.11217 | 0.127570875 | 0.091 | 0 | 1 | 18 |
| hsf1 | 0.11217 | 0.210293915 | 0.091 | 0 | 1 | 18 |
| irf2 | 0.11217 | 0.172172716 | 0.091 | 0 | 1 | 18 |
| itpkb | 0.11217 | 0.160883353 | 0.091 | 0 | 1 | 18 |
| jade1 | 0.11217 | 0.186432454 | 0.091 | 0 | 1 | 18 |

| | | | | | | |
|---|---|---|---|---|---|---|
| kalrna | 0.11217 | 0.182579883 | 0.091 | 0 | 1 | 18 |
| kcmf1 | 0.11217 | 0.202062251 | 0.091 | 0 | 1 | 18 |
| kdm2ab | 0.11217 | 0.205084912 | 0.091 | 0 | 1 | 18 |
| kif7 | 0.11217 | 0.163567646 | 0.091 | 0 | 1 | 18 |
| lto1 | 0.11217 | 0.184902672 | 0.091 | 0 | 1 | 18 |
| maf1 | 0.11217 | 0.159844426 | 0.091 | 0 | 1 | 18 |
| mcm9 | 0.11217 | 0.127976707 | 0.091 | 0 | 1 | 18 |
| med27 | 0.11217 | 0.131399244 | 0.091 | 0 | 1 | 18 |
| mettl5 | 0.11217 | 0.122579741 | 0.091 | 0 | 1 | 18 |
| mfn2 | 0.11217 | 0.180123104 | 0.091 | 0 | 1 | 18 |
| noc3l | 0.11217 | 0.151674364 | 0.091 | 0 | 1 | 18 |
| nt5dc2 | 0.11217 | 0.150713503 | 0.091 | 0 | 1 | 18 |
| nvl | 0.11217 | 0.175847663 | 0.091 | 0 | 1 | 18 |
| orc4 | 0.11217 | 0.148802186 | 0.091 | 0 | 1 | 18 |
| phf20l1 | 0.11217 | 0.137138003 | 0.091 | 0 | 1 | 18 |
| phf23a | 0.11217 | 0.160516371 | 0.091 | 0 | 1 | 18 |
| plekha3 | 0.11217 | 0.171675244 | 0.091 | 0 | 1 | 18 |
| pomgnt1 | 0.11217 | 0.197606022 | 0.091 | 0 | 1 | 18 |
| ppic | 0.11217 | 0.154213614 | 0.091 | 0 | 1 | 18 |
| prss23 | 0.11217 | 0.21503414 | 0.091 | 0 | 1 | 18 |
| ptcd3 | 0.11217 | 0.227693269 | 0.091 | 0 | 1 | 18 |
| r3hdml | 0.11217 | 0.181959755 | 0.091 | 0 | 1 | 18 |
| rrp9 | 0.11217 | 0.158762767 | 0.091 | 0 | 1 | 18 |
| samd11 | 0.11217 | 0.242180515 | 0.091 | 0 | 1 | 18 |
| sart3 | 0.11217 | 0.147397174 | 0.091 | 0 | 1 | 18 |
| sc:d217 | 0.11217 | 0.208021431 | 0.091 | 0 | 1 | 18 |
| si:ch211-113e8.10 | 0.11217 | 0.184902672 | 0.091 | 0 | 1 | 18 |
| si:ch211-175m2.5 | 0.11217 | 0.266247079 | 0.091 | 0 | 1 | 18 |
| si:ch211-220f16.2 | 0.11217 | 0.156294237 | 0.091 | 0 | 1 | 18 |
| si:ch211-255f4.9 | 0.11217 | 0.131913453 | 0.091 | 0 | 1 | 18 |
| si:ch73-127m5.2 | 0.11217 | 0.126119341 | 0.091 | 0 | 1 | 18 |
| sin3aa | 0.11217 | 0.20836933 | 0.091 | 0 | 1 | 18 |
| slc25a28 | 0.11217 | 0.197648779 | 0.091 | 0 | 1 | 18 |
| sptbn2 | 0.11217 | 0.155270622 | 0.091 | 0 | 1 | 18 |
| stk35l | 0.11217 | 0.185472686 | 0.091 | 0 | 1 | 18 |
| sumf1 | 0.11217 | 0.16237642 | 0.091 | 0 | 1 | 18 |
| taf1b | 0.11217 | 0.199203097 | 0.091 | 0 | 1 | 18 |
| tbc1d15 | 0.11217 | 0.142862136 | 0.091 | 0 | 1 | 18 |
| tbcelb | 0.11217 | 0.176604712 | 0.091 | 0 | 1 | 18 |
| tead3b | 0.11217 | 0.142322855 | 0.091 | 0 | 1 | 18 |
| thap12a | 0.11217 | 0.141076233 | 0.091 | 0 | 1 | 18 |
| tmem198a | 0.11217 | 0.147793281 | 0.091 | 0 | 1 | 18 |
| tom1l2a | 0.11217 | 0.179692874 | 0.091 | 0 | 1 | 18 |
| tsr3 | 0.11217 | 0.142341428 | 0.091 | 0 | 1 | 18 |
| wwc1 | 0.11217 | 0.139800005 | 0.091 | 0 | 1 | 18 |
| zgc:112185 | 0.11217 | 0.229349311 | 0.091 | 0 | 1 | 18 |
| zgc:92140 | 0.11217 | 0.17724431 | 0.091 | 0 | 1 | 18 |
| LOC108179023 | 0.11217 | 0.158561662 | 0.091 | 0 | 1 | 18 |

| | | | | | | |
|---|---|---|---|---|---|---|
| cdca7b | 0.11217 | 0.173703246 | 0.091 | 0 | 1 | 18 |
| cep57l1 | 0.11217 | 0.181215246 | 0.091 | 0 | 1 | 18 |
| cgnl1 | 0.11217 | 0.19500864 | 0.091 | 0 | 1 | 18 |
| ddx49 | 0.11217 | 0.165011389 | 0.091 | 0 | 1 | 18 |
| dnajc11a | 0.11217 | 0.165188176 | 0.091 | 0 | 1 | 18 |
| dus1l | 0.11217 | 0.167810009 | 0.091 | 0 | 1 | 18 |
| elk1 | 0.11217 | 0.140312383 | 0.091 | 0 | 1 | 18 |
| inppl1b | 0.11217 | 0.196864599 | 0.091 | 0 | 1 | 18 |
| msi2a | 0.11217 | 0.149349671 | 0.091 | 0 | 1 | 18 |
| myo18ab | 0.11217 | 0.18933039 | 0.091 | 0 | 1 | 18 |
| narfl | 0.11217 | 0.160191615 | 0.091 | 0 | 1 | 18 |
| pacsin2 | 0.11217 | 0.197668455 | 0.091 | 0 | 1 | 18 |
| patj | 0.11217 | 0.196689428 | 0.091 | 0 | 1 | 18 |
| pdcd7 | 0.11217 | 0.168020853 | 0.091 | 0 | 1 | 18 |
| pdk2b | 0.11217 | 0.19123319 | 0.091 | 0 | 1 | 18 |
| plekhb2 | 0.11217 | 0.19062029 | 0.091 | 0 | 1 | 18 |
| prdx1 | 0.11217 | 0.171648217 | 0.091 | 0 | 1 | 18 |
| prpf38a | 0.11217 | 0.184741963 | 0.091 | 0 | 1 | 18 |
| rab21 | 0.11217 | 0.17174673 | 0.091 | 0 | 1 | 18 |
| rcc1 | 0.11217 | 0.173340611 | 0.091 | 0 | 1 | 18 |
| reps1 | 0.11217 | 0.174454301 | 0.091 | 0 | 1 | 18 |
| rnf111 | 0.11217 | 0.180181735 | 0.091 | 0 | 1 | 18 |
| tmf1 | 0.11217 | 0.172273075 | 0.091 | 0 | 1 | 18 |
| vclb | 0.11217 | 0.163864311 | 0.091 | 0 | 1 | 18 |
| CU633745.1 | 0.11217 | 0.232927913 | 0.091 | 0 | 1 | 18 |
| apmap | 0.11217 | 0.141860171 | 0.091 | 0 | 1 | 18 |
| galt | 0.11217 | 0.179111902 | 0.091 | 0 | 1 | 18 |
| inpp5b | 0.11217 | 0.201539517 | 0.091 | 0 | 1 | 18 |
| prkd2 | 0.11217 | 0.234773065 | 0.091 | 0 | 1 | 18 |
| rnpepl1 | 0.11217 | 0.206536403 | 0.091 | 0 | 1 | 18 |
| zgc:91976 | 0.11217 | 0.136693885 | 0.091 | 0 | 1 | 18 |
| arl6ip6 | 0.11217 | 0.22842867 | 0.091 | 0 | 1 | 18 |
| otud7b | 0.11217 | 0.162554632 | 0.091 | 0 | 1 | 18 |
| tmem39b | 0.11217 | 0.207557632 | 0.091 | 0 | 1 | 18 |
| COX3 | 0.11298 | -0.37410349 | 0.727 | 0.923 | 1 | 18 |
| slc38a5b | 0.11341 | -0.40090998 | 0.182 | 0.462 | 1 | 18 |
| eif3k | 0.11351 | -0.40213369 | 0.318 | 0.577 | 1 | 18 |
| vipr2 | 0.11375 | -0.35843455 | 0.045 | 0.231 | 1 | 18 |
| cope | 0.11389 | -0.36437483 | 0.182 | 0.462 | 1 | 18 |
| osr1 | 0.11405 | 0.356777673 | 0.773 | 0.615 | 1 | 18 |
| pgam5 | 0.11426 | 0.386033198 | 0.318 | 0.154 | 1 | 18 |
| rab5ab | 0.11556 | 0.381149189 | 0.364 | 0.115 | 1 | 18 |
| angptl4 | 0.1169 | 0.396739616 | 0.409 | 0.269 | 1 | 18 |
| rbp4 | 0.1171 | -0.56721432 | 0.091 | 0.269 | 1 | 18 |
| tpm2 | 0.11806 | -0.17504173 | 0.045 | 0.192 | 1 | 18 |
| lima1a | 0.11833 | -0.40459739 | 0.045 | 0.231 | 1 | 18 |
| ndfip1 | 0.11862 | -0.22036982 | 0.136 | 0.346 | 1 | 18 |
| zgc:163030 | 0.11931 | 0.123461883 | 0.136 | 0.154 | 1 | 18 |

| | | | | | | |
|---|---|---|---|---|---|---|
| pfdn4 | 0.11945 | -0.230385 | 0.273 | 0.538 | 1 | 18 |
| psmd6 | 0.11962 | -0.19350186 | 0.182 | 0.385 | 1 | 18 |
| eif2s3 | 0.12003 | -0.26320429 | 0.545 | 0.808 | 1 | 18 |
| prpf19 | 0.12039 | 0.284148063 | 0.227 | 0.038 | 1 | 18 |
| srsf3b | 0.12057 | 0.292026093 | 0.364 | 0.308 | 1 | 18 |
| exosc6 | 0.12066 | 0.327428718 | 0.227 | 0.038 | 1 | 18 |
| zgc:111986 | 0.1207 | -0.44184546 | 0.227 | 0.5 | 1 | 18 |
| pabpc1b | 0.12078 | -0.42230684 | 0.182 | 0.423 | 1 | 18 |
| serpinf1 | 0.12093 | 0.301259544 | 0.273 | 0.077 | 1 | 18 |
| zgc:194469 | 0.12105 | 0.313175961 | 0.227 | 0.038 | 1 | 18 |
| fgfrl1a | 0.12126 | 0.349493442 | 0.227 | 0.038 | 1 | 18 |
| cd248a | 0.12147 | -0.36128342 | 0.227 | 0.5 | 1 | 18 |
| nipsnap2 | 0.12188 | 0.37115699 | 0.364 | 0.115 | 1 | 18 |
| cd81a | 0.12242 | -0.1307146 | 0.909 | 0.885 | 1 | 18 |
| si:dkey-239h2.3 | 0.12568 | 0.434138489 | 0.227 | 0.077 | 1 | 18 |
| u2af2b | 0.12724 | 0.331930671 | 0.5 | 0.346 | 1 | 18 |
| ndufb8 | 0.12738 | -0.19568216 | 0.364 | 0.615 | 1 | 18 |
| ugt5g1 | 0.12927 | -0.18998685 | 0.045 | 0.192 | 1 | 18 |
| atp5pf | 0.12969 | -0.2094641 | 0.273 | 0.538 | 1 | 18 |
| efnb1 | 0.12969 | -0.21377733 | 0.273 | 0.5 | 1 | 18 |
| aldh9a1a.1 | 0.12994 | 0.382688135 | 0.636 | 0.346 | 1 | 18 |
| arpp19b | 0.13051 | 0.32670266 | 0.364 | 0.269 | 1 | 18 |
| rhoab | 0.13057 | -0.19162906 | 0.455 | 0.692 | 1 | 18 |
| wu:fi46h04 | 0.13088 | -0.2542435 | 0.045 | 0.231 | 1 | 18 |
| CABZ01112656.1 | 0.13122 | -0.32481152 | 0.045 | 0.231 | 1 | 18 |
| fam107b | 0.13127 | -0.36895751 | 0.045 | 0.231 | 1 | 18 |
| ube2ib | 0.13158 | 0.107563023 | 0.545 | 0.577 | 1 | 18 |
| snrpd2 | 0.13224 | 0.315451735 | 0.5 | 0.385 | 1 | 18 |
| tprb | 0.13251 | 0.320896931 | 0.182 | 0.038 | 1 | 18 |
| txndc17 | 0.13276 | -0.32416355 | 0.045 | 0.231 | 1 | 18 |
| ctsd | 0.13291 | -0.30068252 | 0.136 | 0.385 | 1 | 18 |
| pprc1 | 0.13302 | 0.357863654 | 0.227 | 0.038 | 1 | 18 |
| rbms2b | 0.13313 | 0.399366743 | 0.273 | 0.077 | 1 | 18 |
| ywhabb | 0.13337 | 0.190343638 | 0.727 | 0.692 | 1 | 18 |
| inip | 0.13427 | 0.28451731 | 0.227 | 0.154 | 1 | 18 |
| polr2k | 0.13464 | -0.23343261 | 0.136 | 0.346 | 1 | 18 |
| nid1b | 0.13467 | -0.13502829 | 0.091 | 0.231 | 1 | 18 |
| marcksl1a | 0.13476 | -0.19695167 | 0.909 | 1 | 1 | 18 |
| nme3 | 0.13502 | -0.26739012 | 0.364 | 0.615 | 1 | 18 |
| aimp1 | 0.13503 | 0.257467193 | 0.318 | 0.192 | 1 | 18 |
| cebpd | 0.13559 | 0.499470072 | 0.5 | 0.231 | 1 | 18 |
| dcun1d1 | 0.13561 | 0.135613853 | 0.136 | 0.154 | 1 | 18 |
| cdc42bpb | 0.13574 | -0.2000694 | 0.045 | 0.192 | 1 | 18 |
| zgc:162730 | 0.13701 | 0.494335856 | 0.636 | 0.385 | 1 | 18 |
| ehmt1b | 0.13868 | 0.199874118 | 0.227 | 0.192 | 1 | 18 |
| znf292a | 0.13868 | -0.35996407 | 0.045 | 0.231 | 1 | 18 |
| si:ch73-335l21.4 | 0.13882 | 0.400890177 | 0.773 | 0.5 | 1 | 18 |
| vat1 | 0.13886 | -0.14262985 | 0.182 | 0.346 | 1 | 18 |

| | | | | | | |
|---|---|---|---|---|---|---|
| ponzr1 | 0.13943 | 0.247448582 | 0.227 | 0.231 | 1 | 18 |
| dbi | 0.13976 | -0.38505078 | 0.364 | 0.538 | 1 | 18 |
| si:dkey-26m3.3 | 0.14008 | -0.40562591 | 0.273 | 0.5 | 1 | 18 |
| ppp3ccb | 0.14054 | -0.1828737 | 0.045 | 0.192 | 1 | 18 |
| tmem258 | 0.14114 | -0.32050621 | 0.545 | 0.808 | 1 | 18 |
| jun | 0.14136 | 0.234727128 | 0.818 | 0.808 | 1 | 18 |
| smdt1b | 0.14153 | -0.27260136 | 0.182 | 0.423 | 1 | 18 |
| ddx3xa | 0.14171 | -0.30754101 | 0.045 | 0.231 | 1 | 18 |
| hivep3b | 0.14173 | 0.215578572 | 0.182 | 0.038 | 1 | 18 |
| lpar1 | 0.14201 | -0.3698198 | 0.136 | 0.385 | 1 | 18 |
| fkbp1ab | 0.14227 | -0.42105482 | 0.136 | 0.385 | 1 | 18 |
| tia1 | 0.14345 | -0.2905735 | 0.045 | 0.231 | 1 | 18 |
| ctsk | 0.14348 | -0.37698499 | 0.136 | 0.385 | 1 | 18 |
| puf60b | 0.1436 | -0.31578782 | 0.091 | 0.308 | 1 | 18 |
| cox6a1 | 0.14361 | -0.34749809 | 0.455 | 0.731 | 1 | 18 |
| sh3bgrl | 0.14361 | -0.36693586 | 0.136 | 0.385 | 1 | 18 |
| map1lc3a | 0.14365 | -0.36201241 | 0.045 | 0.231 | 1 | 18 |
| NDUFB1 | 0.1437 | 0.40739196 | 0.5 | 0.269 | 1 | 18 |
| eif4g2b | 0.14398 | -0.27739454 | 0.091 | 0.308 | 1 | 18 |
| nfyba | 0.14559 | -0.20388743 | 0.318 | 0.538 | 1 | 18 |
| si:dkey-33c12.4 | 0.14644 | 0.200719408 | 0.318 | 0.269 | 1 | 18 |
| atrx | 0.14683 | 0.254587736 | 0.727 | 0.615 | 1 | 18 |
| smu1a | 0.14745 | 0.389366811 | 0.273 | 0.077 | 1 | 18 |
| ulk2 | 0.14774 | 0.383861679 | 0.227 | 0.077 | 1 | 18 |
| ube2l3b | 0.14801 | 0.186154023 | 0.409 | 0.385 | 1 | 18 |
| zmiz1a | 0.1487 | -0.34620776 | 0.045 | 0.231 | 1 | 18 |
| mt-nd2 | 0.14889 | -0.33620987 | 0.5 | 0.769 | 1 | 18 |
| ndufa12 | 0.149 | -0.11152464 | 0.182 | 0.346 | 1 | 18 |
| ppfibp1b | 0.14904 | -0.26881285 | 0.045 | 0.231 | 1 | 18 |
| jag1b | 0.14968 | -0.21849744 | 0.227 | 0.423 | 1 | 18 |
| med19b | 0.15185 | 0.279682659 | 0.182 | 0.038 | 1 | 18 |
| psip1b | 0.15221 | -0.22415892 | 0.182 | 0.385 | 1 | 18 |
| june | 0.15286 | -0.2991958 | 0.045 | 0.231 | 1 | 18 |
| vta1 | 0.153 | -0.29972523 | 0.045 | 0.231 | 1 | 18 |
| setb | 0.15374 | 0.190216125 | 0.5 | 0.538 | 1 | 18 |
| eif3f | 0.15389 | 0.286185114 | 0.818 | 0.731 | 1 | 18 |
| uqcrc2b | 0.15396 | 0.184098691 | 0.182 | 0.154 | 1 | 18 |
| serinc5 | 0.15472 | 0.307801562 | 0.318 | 0.231 | 1 | 18 |
| chaf1a | 0.15472 | 0.35403055 | 0.227 | 0.115 | 1 | 18 |
| socs3b | 0.15486 | 0.265395778 | 0.273 | 0.192 | 1 | 18 |
| tmem100a | 0.15529 | -0.2545832 | 0.136 | 0.346 | 1 | 18 |
| zgc:77849 | 0.15597 | -0.32208459 | 0.273 | 0.538 | 1 | 18 |
| ndufa11 | 0.15644 | -0.37005496 | 0.136 | 0.346 | 1 | 18 |
| slx1b | 0.15647 | -0.25543658 | 0.045 | 0.231 | 1 | 18 |
| mpzl1l | 0.15651 | -0.32166954 | 0.045 | 0.231 | 1 | 18 |
| npm1b | 0.15765 | -0.1458322 | 0.091 | 0.231 | 1 | 18 |
| sart1 | 0.15813 | -0.23839568 | 0.136 | 0.346 | 1 | 18 |
| il6st | 0.15859 | -0.29296045 | 0.045 | 0.231 | 1 | 18 |

| | | | | | | |
|---|---|---|---|---|---|---|
| nsd3 | 0.15866 | -0.3259523 | 0.045 | 0.231 | 1 | 18 |
| pcolcea | 0.15882 | 0.23027329 | 0.909 | 0.692 | 1 | 18 |
| atp5mea | 0.15904 | -0.11632699 | 0.227 | 0.423 | 1 | 18 |
| acin1a | 0.15946 | -0.37159123 | 0.5 | 0.692 | 1 | 18 |
| atp6v1d | 0.15965 | -0.28086571 | 0.045 | 0.231 | 1 | 18 |
| fmr1 | 0.1601 | -0.32423119 | 0.091 | 0.308 | 1 | 18 |
| spry4 | 0.16054 | 0.309380497 | 0.273 | 0.077 | 1 | 18 |
| anapc13 | 0.1611 | -0.28465842 | 0.045 | 0.231 | 1 | 18 |
| fbxo28 | 0.16115 | -0.25907449 | 0.045 | 0.231 | 1 | 18 |
| ten1 | 0.16152 | -0.28912191 | 0.045 | 0.231 | 1 | 18 |
| faf2 | 0.16209 | -0.29551887 | 0.091 | 0.308 | 1 | 18 |
| brk1 | 0.16218 | 0.227750204 | 0.409 | 0.346 | 1 | 18 |
| serpine1 | 0.16234 | -0.39123776 | 0.045 | 0.231 | 1 | 18 |
| si:ch211-105c13.3 | 0.16253 | -0.32323218 | 0.227 | 0.462 | 1 | 18 |
| ptbp3 | 0.16259 | -0.31663073 | 0.045 | 0.231 | 1 | 18 |
| ablim1b | 0.16261 | -0.15051093 | 0.091 | 0.231 | 1 | 18 |
| hnrnpul1 | 0.16283 | -0.29506202 | 0.045 | 0.231 | 1 | 18 |
| akap8l | 0.16311 | -0.34381291 | 0.091 | 0.308 | 1 | 18 |
| fbxo9 | 0.16339 | -0.10533318 | 0.091 | 0.231 | 1 | 18 |
| rchy1 | 0.16356 | -0.2335396 | 0.045 | 0.231 | 1 | 18 |
| ndufb2 | 0.16383 | -0.22951494 | 0.227 | 0.462 | 1 | 18 |
| grk4 | 0.16394 | 0.190035422 | 0.136 | 0.115 | 1 | 18 |
| tnrc6b | 0.16428 | -0.3588518 | 0.045 | 0.231 | 1 | 18 |
| pno1 | 0.16433 | 0.121767349 | 0.182 | 0.192 | 1 | 18 |
| brd3b | 0.16475 | -0.31187842 | 0.091 | 0.308 | 1 | 18 |
| rab5c | 0.16485 | -0.30687582 | 0.091 | 0.308 | 1 | 18 |
| ptpn2b | 0.16493 | -0.24626191 | 0.045 | 0.231 | 1 | 18 |
| ifngr2 | 0.16507 | -0.33322496 | 0.091 | 0.308 | 1 | 18 |
| srgap3 | 0.16517 | -0.34162424 | 0.091 | 0.308 | 1 | 18 |
| twsg1a | 0.16536 | -0.31299016 | 0.045 | 0.231 | 1 | 18 |
| commd5 | 0.16539 | 0.309997038 | 0.182 | 0.038 | 1 | 18 |
| LOC110439623 | 0.16557 | -0.33979587 | 0.091 | 0.308 | 1 | 18 |
| angptl1a | 0.16566 | -0.4458713 | 0.091 | 0.308 | 1 | 18 |
| fis1 | 0.1657 | -0.29548201 | 0.091 | 0.308 | 1 | 18 |
| krt15 | 0.1657 | -0.40347931 | 0.136 | 0.308 | 1 | 18 |
| foxc1b | 0.16586 | -0.28890874 | 0.182 | 0.308 | 1 | 18 |
| tgfb1a | 0.16615 | -0.32444003 | 0.045 | 0.231 | 1 | 18 |
| scpep1 | 0.16632 | -0.28836686 | 0.045 | 0.231 | 1 | 18 |
| nktr | 0.16638 | -0.30907347 | 0.045 | 0.231 | 1 | 18 |
| akirin1 | 0.1664 | 0.103717197 | 0.273 | 0.308 | 1 | 18 |
| glrx | 0.16648 | 0.309088088 | 0.545 | 0.423 | 1 | 18 |
| ccnd1 | 0.16652 | -0.32924547 | 0.182 | 0.423 | 1 | 18 |
| si:ch211-195m9.3 | 0.16664 | -0.33119091 | 0.045 | 0.231 | 1 | 18 |
| nudt5 | 0.16665 | -0.29408062 | 0.045 | 0.231 | 1 | 18 |
| CABZ01066926.1 | 0.16684 | -0.137501 | 0 | 0.077 | 1 | 18 |
| CABZ01117348.1 | 0.16684 | -0.13706603 | 0 | 0.077 | 1 | 18 |
| CU695223.1 | 0.16684 | -0.15309597 | 0 | 0.077 | 1 | 18 |
| IYD | 0.16684 | -0.15536676 | 0 | 0.077 | 1 | 18 |

| | | | | | | |
|---|---|---|---|---|---|---|
| LO018340.1 | 0.16684 | -0.16424967 | 0 | 0.077 | 1 | 18 |
| LOC100332329 | 0.16684 | -0.10502536 | 0 | 0.077 | 1 | 18 |
| LOC100332847 | 0.16684 | -0.1230874 | 0 | 0.077 | 1 | 18 |
| LOC101882596 | 0.16684 | -0.11126417 | 0 | 0.077 | 1 | 18 |
| LOC108179036 | 0.16684 | -0.15097137 | 0 | 0.077 | 1 | 18 |
| LOC108181080 | 0.16684 | -0.12093142 | 0 | 0.077 | 1 | 18 |
| XLOC-001115 | 0.16684 | -0.13253394 | 0 | 0.077 | 1 | 18 |
| XLOC-004556 | 0.16684 | -0.19808498 | 0 | 0.077 | 1 | 18 |
| XLOC-034235 | 0.16684 | -0.12057253 | 0 | 0.077 | 1 | 18 |
| XLOC-042228 | 0.16684 | -0.13959169 | 0 | 0.077 | 1 | 18 |
| abhd10b | 0.16684 | -0.13596441 | 0 | 0.077 | 1 | 18 |
| agpat4 | 0.16684 | -0.10968891 | 0 | 0.077 | 1 | 18 |
| akt3b | 0.16684 | -0.16574811 | 0 | 0.077 | 1 | 18 |
| alg1 | 0.16684 | -0.10495743 | 0 | 0.077 | 1 | 18 |
| amacr | 0.16684 | -0.12722508 | 0 | 0.077 | 1 | 18 |
| ankrd54 | 0.16684 | -0.10263872 | 0 | 0.077 | 1 | 18 |
| arhgap42b | 0.16684 | -0.17272275 | 0 | 0.077 | 1 | 18 |
| atp13a2 | 0.16684 | -0.11610979 | 0 | 0.077 | 1 | 18 |
| atp8a1 | 0.16684 | -0.1427401 | 0 | 0.077 | 1 | 18 |
| bach2b | 0.16684 | -0.11183465 | 0 | 0.077 | 1 | 18 |
| baiap2a | 0.16684 | -0.179743 | 0 | 0.077 | 1 | 18 |
| bcl2l11 | 0.16684 | -0.14023878 | 0 | 0.077 | 1 | 18 |
| brinp2 | 0.16684 | -0.11328601 | 0 | 0.077 | 1 | 18 |
| cactin | 0.16684 | -0.13036441 | 0 | 0.077 | 1 | 18 |
| cbx8b | 0.16684 | -0.16620547 | 0 | 0.077 | 1 | 18 |
| ccdc142 | 0.16684 | -0.10588104 | 0 | 0.077 | 1 | 18 |
| ccdc174 | 0.16684 | -0.10070217 | 0 | 0.077 | 1 | 18 |
| ccnt1 | 0.16684 | -0.13319611 | 0 | 0.077 | 1 | 18 |
| cdk20 | 0.16684 | -0.12616501 | 0 | 0.077 | 1 | 18 |
| cerkl | 0.16684 | -0.16292286 | 0 | 0.077 | 1 | 18 |
| cers2a | 0.16684 | -0.1179961 | 0 | 0.077 | 1 | 18 |
| chad | 0.16684 | -0.1631369 | 0 | 0.077 | 1 | 18 |
| chrd | 0.16684 | -0.12393608 | 0 | 0.077 | 1 | 18 |
| cica | 0.16684 | -0.14946322 | 0 | 0.077 | 1 | 18 |
| cldnd1a | 0.16684 | -0.11072523 | 0 | 0.077 | 1 | 18 |
| cmtm7 | 0.16684 | -0.16937422 | 0 | 0.077 | 1 | 18 |
| cox15 | 0.16684 | -0.11494939 | 0 | 0.077 | 1 | 18 |
| creld1b | 0.16684 | -0.13880389 | 0 | 0.077 | 1 | 18 |
| cremb | 0.16684 | -0.12666754 | 0 | 0.077 | 1 | 18 |
| ctnnbl1 | 0.16684 | -0.13571121 | 0 | 0.077 | 1 | 18 |
| cybc1 | 0.16684 | -0.14860796 | 0 | 0.077 | 1 | 18 |
| dbt | 0.16684 | -0.12624383 | 0 | 0.077 | 1 | 18 |
| dctn5 | 0.16684 | -0.10741344 | 0 | 0.077 | 1 | 18 |
| dera | 0.16684 | -0.12607689 | 0 | 0.077 | 1 | 18 |
| dixdc1a | 0.16684 | -0.10743186 | 0 | 0.077 | 1 | 18 |
| dph5 | 0.16684 | -0.12436009 | 0 | 0.077 | 1 | 18 |
| dsg2l | 0.16684 | -0.11832668 | 0 | 0.077 | 1 | 18 |
| dub | 0.16684 | -0.12270157 | 0 | 0.077 | 1 | 18 |

| | | | | | | |
|---|---|---|---|---|---|---|
| egln1b | 0.16684 | -0.11728269 | 0 | 0.077 | 1 | 18 |
| eif4g3b | 0.16684 | -0.13134087 | 0 | 0.077 | 1 | 18 |
| elp4 | 0.16684 | -0.10684081 | 0 | 0.077 | 1 | 18 |
| enox1 | 0.16684 | -0.13775944 | 0 | 0.077 | 1 | 18 |
| epha2a | 0.16684 | -0.18680734 | 0 | 0.077 | 1 | 18 |
| erbb2 | 0.16684 | -0.17968442 | 0 | 0.077 | 1 | 18 |
| ercc5 | 0.16684 | -0.12436877 | 0 | 0.077 | 1 | 18 |
| exoc5 | 0.16684 | -0.14412341 | 0 | 0.077 | 1 | 18 |
| fam57a | 0.16684 | -0.12440065 | 0 | 0.077 | 1 | 18 |
| fbxo18 | 0.16684 | -0.12253813 | 0 | 0.077 | 1 | 18 |
| fgf18b | 0.16684 | -0.11303814 | 0 | 0.077 | 1 | 18 |
| fmo5 | 0.16684 | -0.14750313 | 0 | 0.077 | 1 | 18 |
| fus | 0.16684 | -0.10816905 | 0 | 0.077 | 1 | 18 |
| galm | 0.16684 | -0.13457816 | 0 | 0.077 | 1 | 18 |
| galns | 0.16684 | -0.14000798 | 0 | 0.077 | 1 | 18 |
| gcc1 | 0.16684 | -0.12066671 | 0 | 0.077 | 1 | 18 |
| gdpd4b | 0.16684 | -0.11446508 | 0 | 0.077 | 1 | 18 |
| gm2a | 0.16684 | -0.11304483 | 0 | 0.077 | 1 | 18 |
| gng13b | 0.16684 | -0.1955984 | 0 | 0.077 | 1 | 18 |
| gpn3 | 0.16684 | -0.15327749 | 0 | 0.077 | 1 | 18 |
| gsna | 0.16684 | -0.10882212 | 0 | 0.077 | 1 | 18 |
| gstcd | 0.16684 | -0.14825167 | 0 | 0.077 | 1 | 18 |
| hhipl2 | 0.16684 | -0.15285283 | 0 | 0.077 | 1 | 18 |
| hspg2 | 0.16684 | -0.14328199 | 0 | 0.077 | 1 | 18 |
| igf2a | 0.16684 | -0.21560686 | 0 | 0.077 | 1 | 18 |
| il16 | 0.16684 | -0.18584274 | 0 | 0.077 | 1 | 18 |
| inpp4aa | 0.16684 | -0.14353897 | 0 | 0.077 | 1 | 18 |
| insig1 | 0.16684 | -0.10160842 | 0 | 0.077 | 1 | 18 |
| invs | 0.16684 | -0.11100869 | 0 | 0.077 | 1 | 18 |
| irak1 | 0.16684 | -0.15231673 | 0 | 0.077 | 1 | 18 |
| itprip | 0.16684 | -0.13725491 | 0 | 0.077 | 1 | 18 |
| itpripl2 | 0.16684 | -0.1281952 | 0 | 0.077 | 1 | 18 |
| jtb | 0.16684 | -0.1336288 | 0 | 0.077 | 1 | 18 |
| khdc4 | 0.16684 | -0.14136824 | 0 | 0.077 | 1 | 18 |
| klf12b | 0.16684 | -0.1219524 | 0 | 0.077 | 1 | 18 |
| klhl24a | 0.16684 | -0.13763743 | 0 | 0.077 | 1 | 18 |
| klhl24b | 0.16684 | -0.13286951 | 0 | 0.077 | 1 | 18 |
| lmna | 0.16684 | -0.11037722 | 0 | 0.077 | 1 | 18 |
| lrch1 | 0.16684 | -0.11688809 | 0 | 0.077 | 1 | 18 |
| lyrm9 | 0.16684 | -0.10905242 | 0 | 0.077 | 1 | 18 |
| lzts2a | 0.16684 | -0.1136602 | 0 | 0.077 | 1 | 18 |
| m6pr | 0.16684 | -0.15475391 | 0 | 0.077 | 1 | 18 |
| map3k1 | 0.16684 | -0.17718692 | 0 | 0.077 | 1 | 18 |
| mat2al | 0.16684 | -0.12019646 | 0 | 0.077 | 1 | 18 |
| med8 | 0.16684 | -0.16412016 | 0 | 0.077 | 1 | 18 |
| mettl9 | 0.16684 | -0.12004779 | 0 | 0.077 | 1 | 18 |
| mid1ip1l | 0.16684 | -0.10263898 | 0 | 0.077 | 1 | 18 |
| mllt1b | 0.16684 | -0.11107916 | 0 | 0.077 | 1 | 18 |

| | | | | | | |
|---|---|---|---|---|---|---|
| mn1b | 0.16684 | -0.12304607 | 0 | 0.077 | 1 | 18 |
| mob2a | 0.16684 | -0.1747798 | 0 | 0.077 | 1 | 18 |
| mthfr | 0.16684 | -0.10453618 | 0 | 0.077 | 1 | 18 |
| mthfs | 0.16684 | -0.11790642 | 0 | 0.077 | 1 | 18 |
| mtrf1l | 0.16684 | -0.12076958 | 0 | 0.077 | 1 | 18 |
| mustn1a | 0.16684 | -0.13941053 | 0 | 0.077 | 1 | 18 |
| myo1ea | 0.16684 | -0.10511137 | 0 | 0.077 | 1 | 18 |
| nacad | 0.16684 | -0.12304607 | 0 | 0.077 | 1 | 18 |
| nat14 | 0.16684 | -0.15957451 | 0 | 0.077 | 1 | 18 |
| nckipsd | 0.16684 | -0.11750502 | 0 | 0.077 | 1 | 18 |
| nif3l1 | 0.16684 | -0.10436477 | 0 | 0.077 | 1 | 18 |
| nkiras2 | 0.16684 | -0.1065012 | 0 | 0.077 | 1 | 18 |
| npas2 | 0.16684 | -0.12513468 | 0 | 0.077 | 1 | 18 |
| nr2f6b | 0.16684 | -0.16713816 | 0 | 0.077 | 1 | 18 |
| nradd | 0.16684 | -0.18017837 | 0 | 0.077 | 1 | 18 |
| nthl1 | 0.16684 | -0.11442427 | 0 | 0.077 | 1 | 18 |
| ociad2 | 0.16684 | -0.15054154 | 0 | 0.077 | 1 | 18 |
| osbpl5 | 0.16684 | -0.14003882 | 0 | 0.077 | 1 | 18 |
| pak1 | 0.16684 | -0.11469735 | 0 | 0.077 | 1 | 18 |
| pcyt1aa | 0.16684 | -0.13261077 | 0 | 0.077 | 1 | 18 |
| pgfb | 0.16684 | -0.13757014 | 0 | 0.077 | 1 | 18 |
| phf11 | 0.16684 | -0.1404389 | 0 | 0.077 | 1 | 18 |
| pik3ca | 0.16684 | -0.11915806 | 0 | 0.077 | 1 | 18 |
| plagx | 0.16684 | -0.11093437 | 0 | 0.077 | 1 | 18 |
| plekhf2 | 0.16684 | -0.13508701 | 0 | 0.077 | 1 | 18 |
| plekhh1 | 0.16684 | -0.19328829 | 0 | 0.077 | 1 | 18 |
| ppp2r1bb | 0.16684 | -0.10277192 | 0 | 0.077 | 1 | 18 |
| ppp2r2ab | 0.16684 | -0.12367225 | 0 | 0.077 | 1 | 18 |
| psen1 | 0.16684 | -0.12271742 | 0 | 0.077 | 1 | 18 |
| qser1 | 0.16684 | -0.17129339 | 0 | 0.077 | 1 | 18 |
| rab3gap2 | 0.16684 | -0.10651261 | 0 | 0.077 | 1 | 18 |
| rab5b | 0.16684 | -0.11099815 | 0 | 0.077 | 1 | 18 |
| rabepk | 0.16684 | -0.11576744 | 0 | 0.077 | 1 | 18 |
| raf1b | 0.16684 | -0.10777872 | 0 | 0.077 | 1 | 18 |
| rbms1a | 0.16684 | -0.12007567 | 0 | 0.077 | 1 | 18 |
| rfx1b | 0.16684 | -0.16654656 | 0 | 0.077 | 1 | 18 |
| rgma | 0.16684 | -0.11009475 | 0 | 0.077 | 1 | 18 |
| rilp | 0.16684 | -0.13693909 | 0 | 0.077 | 1 | 18 |
| rnaset2 | 0.16684 | -0.11094933 | 0 | 0.077 | 1 | 18 |
| rnf14 | 0.16684 | -0.10495331 | 0 | 0.077 | 1 | 18 |
| rnf146 | 0.16684 | -0.11309312 | 0 | 0.077 | 1 | 18 |
| rnft1 | 0.16684 | -0.11135282 | 0 | 0.077 | 1 | 18 |
| rpz | 0.16684 | -0.14193036 | 0 | 0.077 | 1 | 18 |
| rpz4 | 0.16684 | -0.16963973 | 0 | 0.077 | 1 | 18 |
| rufy2 | 0.16684 | -0.16010884 | 0 | 0.077 | 1 | 18 |
| scml4 | 0.16684 | -0.11909469 | 0 | 0.077 | 1 | 18 |
| shisal1b | 0.16684 | -0.13702479 | 0 | 0.077 | 1 | 18 |
| si:ch1073-170o4.1 | 0.16684 | -0.1726233 | 0 | 0.077 | 1 | 18 |

| | | | | | | |
|---|---|---|---|---|---|---|
| si:ch1073-287p18.1 | 0.16684 | -0.17605586 | 0 | 0.077 | 1 | 18 |
| si:ch1073-357b18.4 | 0.16684 | -0.103285 | 0 | 0.077 | 1 | 18 |
| si:ch211-110e21.3 | 0.16684 | -0.12166576 | 0 | 0.077 | 1 | 18 |
| si:ch211-133l5.8 | 0.16684 | -0.15056683 | 0 | 0.077 | 1 | 18 |
| si:ch211-150o23.2 | 0.16684 | -0.10468997 | 0 | 0.077 | 1 | 18 |
| si:ch211-161h7.4 | 0.16684 | -0.17149791 | 0 | 0.077 | 1 | 18 |
| si:ch211-162i8.5 | 0.16684 | -0.14616933 | 0 | 0.077 | 1 | 18 |
| si:ch211-195b15.8 | 0.16684 | -0.16181751 | 0 | 0.077 | 1 | 18 |
| si:ch211-1a19.3 | 0.16684 | -0.1368728 | 0 | 0.077 | 1 | 18 |
| si:ch211-230g15.5 | 0.16684 | -0.15185866 | 0 | 0.077 | 1 | 18 |
| si:ch211-256e16.11 | 0.16684 | -0.11730388 | 0 | 0.077 | 1 | 18 |
| si:ch211-261d7.3 | 0.16684 | -0.13937159 | 0 | 0.077 | 1 | 18 |
| si:ch211-288d18.1 | 0.16684 | -0.12165268 | 0 | 0.077 | 1 | 18 |
| si:ch211-51h9.7 | 0.16684 | -0.13650352 | 0 | 0.077 | 1 | 18 |
| si:ch211-79k12.2 | 0.16684 | -0.11489695 | 0 | 0.077 | 1 | 18 |
| si:ch211-79l17.1 | 0.16684 | -0.14655167 | 0 | 0.077 | 1 | 18 |
| si:ch73-109d9.3 | 0.16684 | -0.13624247 | 0 | 0.077 | 1 | 18 |
| si:ch73-211l13.2 | 0.16684 | -0.11828915 | 0 | 0.077 | 1 | 18 |
| si:dkey-106g10.7 | 0.16684 | -0.11470177 | 0 | 0.077 | 1 | 18 |
| si:dkey-156n14.3 | 0.16684 | -0.12176189 | 0 | 0.077 | 1 | 18 |
| si:dkey-193c22.1 | 0.16684 | -0.10813819 | 0 | 0.077 | 1 | 18 |
| si:dkey-21c1.8 | 0.16684 | -0.19605239 | 0 | 0.077 | 1 | 18 |
| si:dkey-21c19.3 | 0.16684 | -0.16191113 | 0 | 0.077 | 1 | 18 |
| si:dkey-264d12.5 | 0.16684 | -0.1143874 | 0 | 0.077 | 1 | 18 |
| si:dkey-49n23.1 | 0.16684 | -0.11633897 | 0 | 0.077 | 1 | 18 |
| si:dkeyp-120h9.1 | 0.16684 | -0.10694153 | 0 | 0.077 | 1 | 18 |
| si:dkeyp-84f3.5 | 0.16684 | -0.17244658 | 0 | 0.077 | 1 | 18 |
| sik1 | 0.16684 | -0.15573558 | 0 | 0.077 | 1 | 18 |
| six4b | 0.16684 | -0.11328601 | 0 | 0.077 | 1 | 18 |
| slc25a55a | 0.16684 | -0.13309001 | 0 | 0.077 | 1 | 18 |
| slc2a8 | 0.16684 | -0.1122437 | 0 | 0.077 | 1 | 18 |
| slc40a1 | 0.16684 | -0.11488643 | 0 | 0.077 | 1 | 18 |
| slc45a4 | 0.16684 | -0.1102454 | 0 | 0.077 | 1 | 18 |
| smim19 | 0.16684 | -0.17457337 | 0 | 0.077 | 1 | 18 |
| sobpa | 0.16684 | -0.11046455 | 0 | 0.077 | 1 | 18 |
| sp2 | 0.16684 | -0.16229654 | 0 | 0.077 | 1 | 18 |
| sp4 | 0.16684 | -0.1109758 | 0 | 0.077 | 1 | 18 |
| spata7 | 0.16684 | -0.15137531 | 0 | 0.077 | 1 | 18 |
| spred2a | 0.16684 | -0.18171875 | 0 | 0.077 | 1 | 18 |
| spsb3a | 0.16684 | -0.17621779 | 0 | 0.077 | 1 | 18 |
| srxn1 | 0.16684 | -0.14835041 | 0 | 0.077 | 1 | 18 |
| stom | 0.16684 | -0.138294 | 0 | 0.077 | 1 | 18 |
| ston2 | 0.16684 | -0.14950218 | 0 | 0.077 | 1 | 18 |
| suco | 0.16684 | -0.11852458 | 0 | 0.077 | 1 | 18 |
| sugp1 | 0.16684 | -0.10956517 | 0 | 0.077 | 1 | 18 |
| sult1st6 | 0.16684 | -0.17618824 | 0 | 0.077 | 1 | 18 |
| supt20 | 0.16684 | -0.10381528 | 0 | 0.077 | 1 | 18 |
| supt7l | 0.16684 | -0.10435906 | 0 | 0.077 | 1 | 18 |

| | | | | | | |
|---|---|---|---|---|---|---|
| tbc1d20 | 0.16684 | -0.15455884 | 0 | 0.077 | 1 | 18 |
| tcaim | 0.16684 | -0.17405563 | 0 | 0.077 | 1 | 18 |
| tecpr2 | 0.16684 | -0.20008774 | 0 | 0.077 | 1 | 18 |
| tep1 | 0.16684 | -0.13348578 | 0 | 0.077 | 1 | 18 |
| tet3 | 0.16684 | -0.17497057 | 0 | 0.077 | 1 | 18 |
| tgfbi | 0.16684 | -0.20293061 | 0 | 0.077 | 1 | 18 |
| tha1 | 0.16684 | -0.10522126 | 0 | 0.077 | 1 | 18 |
| thap1 | 0.16684 | -0.11482906 | 0 | 0.077 | 1 | 18 |
| thoc6 | 0.16684 | -0.13593085 | 0 | 0.077 | 1 | 18 |
| thumpd1 | 0.16684 | -0.10921271 | 0 | 0.077 | 1 | 18 |
| tmem131l | 0.16684 | -0.12403433 | 0 | 0.077 | 1 | 18 |
| tmem161a | 0.16684 | -0.13094746 | 0 | 0.077 | 1 | 18 |
| tmem220 | 0.16684 | -0.10830854 | 0 | 0.077 | 1 | 18 |
| tpd52l2a | 0.16684 | -0.11738115 | 0 | 0.077 | 1 | 18 |
| tpmt.2 | 0.16684 | -0.14058585 | 0 | 0.077 | 1 | 18 |
| trafd1 | 0.16684 | -0.10062622 | 0 | 0.077 | 1 | 18 |
| trim24 | 0.16684 | -0.15234922 | 0 | 0.077 | 1 | 18 |
| trmt2b | 0.16684 | -0.15394973 | 0 | 0.077 | 1 | 18 |
| tspan17 | 0.16684 | -0.19587913 | 0 | 0.077 | 1 | 18 |
| ubn2a | 0.16684 | -0.13832174 | 0 | 0.077 | 1 | 18 |
| usp5 | 0.16684 | -0.15851589 | 0 | 0.077 | 1 | 18 |
| vkorc1 | 0.16684 | -0.16701846 | 0 | 0.077 | 1 | 18 |
| vps11 | 0.16684 | -0.14490088 | 0 | 0.077 | 1 | 18 |
| vps39 | 0.16684 | -0.15091521 | 0 | 0.077 | 1 | 18 |
| vps53 | 0.16684 | -0.10458164 | 0 | 0.077 | 1 | 18 |
| vstm4b | 0.16684 | -0.12223509 | 0 | 0.077 | 1 | 18 |
| wdr32 | 0.16684 | -0.14611807 | 0 | 0.077 | 1 | 18 |
| wwp1 | 0.16684 | -0.13844224 | 0 | 0.077 | 1 | 18 |
| yju2 | 0.16684 | -0.13157585 | 0 | 0.077 | 1 | 18 |
| ypel5 | 0.16684 | -0.12870283 | 0 | 0.077 | 1 | 18 |
| zdhhc5a | 0.16684 | -0.13713074 | 0 | 0.077 | 1 | 18 |
| zgc:153219 | 0.16684 | -0.10988697 | 0 | 0.077 | 1 | 18 |
| zgc:153292 | 0.16684 | -0.10892342 | 0 | 0.077 | 1 | 18 |
| zgc:174315 | 0.16684 | -0.13794073 | 0 | 0.077 | 1 | 18 |
| zgc:174888 | 0.16684 | -0.1506627 | 0 | 0.077 | 1 | 18 |
| zgc:194281 | 0.16684 | -0.1116873 | 0 | 0.077 | 1 | 18 |
| zgc:77838 | 0.16684 | -0.10084383 | 0 | 0.077 | 1 | 18 |
| zmp:0000000711 | 0.16684 | -0.12261663 | 0 | 0.077 | 1 | 18 |
| zmp:0000001082 | 0.16684 | -0.13230307 | 0 | 0.077 | 1 | 18 |
| zmp:0000001228 | 0.16684 | -0.10418033 | 0 | 0.077 | 1 | 18 |
| znf1158 | 0.16684 | -0.11231089 | 0 | 0.077 | 1 | 18 |
| znhit1 | 0.16684 | -0.10893917 | 0 | 0.077 | 1 | 18 |
| BX537342.1 | 0.16684 | -0.13302781 | 0 | 0.077 | 1 | 18 |
| CR352226.5 | 0.16684 | -0.11818248 | 0 | 0.077 | 1 | 18 |
| chd1 | 0.16684 | -0.1363649 | 0 | 0.077 | 1 | 18 |
| egln2 | 0.16684 | -0.12213719 | 0 | 0.077 | 1 | 18 |
| eml2 | 0.16684 | -0.14752499 | 0 | 0.077 | 1 | 18 |
| hoxb6b | 0.16684 | -0.14540549 | 0 | 0.077 | 1 | 18 |

| | | | | | | |
|---|---|---|---|---|---|---|
| ildr2 | 0.16684 | -0.11990621 | 0 | 0.077 | 1 | 18 |
| itpka | 0.16684 | -0.11123335 | 0 | 0.077 | 1 | 18 |
| mdn1 | 0.16684 | -0.1511182 | 0 | 0.077 | 1 | 18 |
| si:ch211-197h24.6 | 0.16684 | -0.11287011 | 0 | 0.077 | 1 | 18 |
| tcf15 | 0.16684 | -0.13744866 | 0 | 0.077 | 1 | 18 |
| ttbk2b | 0.16684 | -0.12876001 | 0 | 0.077 | 1 | 18 |
| LOC108190785 | 0.16684 | -0.16663724 | 0 | 0.077 | 1 | 18 |
| cav2 | 0.16684 | -0.12608422 | 0 | 0.077 | 1 | 18 |
| crebrf | 0.16684 | -0.11140476 | 0 | 0.077 | 1 | 18 |
| cyp2v1 | 0.16684 | -0.10740086 | 0 | 0.077 | 1 | 18 |
| hgd | 0.16684 | -0.10102846 | 0 | 0.077 | 1 | 18 |
| mrps15 | 0.16684 | -0.11100789 | 0 | 0.077 | 1 | 18 |
| nf1b | 0.16684 | -0.17148914 | 0 | 0.077 | 1 | 18 |
| piezo1 | 0.16684 | -0.11712795 | 0 | 0.077 | 1 | 18 |
| rgs7bpa | 0.16684 | -0.12851569 | 0 | 0.077 | 1 | 18 |
| setd5 | 0.16684 | -0.15414112 | 0 | 0.077 | 1 | 18 |
| si:dkey-234h16.7 | 0.16684 | -0.10641663 | 0 | 0.077 | 1 | 18 |
| trim25 | 0.16684 | -0.10019037 | 0 | 0.077 | 1 | 18 |
| xpr1b | 0.16684 | -0.16614557 | 0 | 0.077 | 1 | 18 |
| znf971 | 0.16684 | -0.11440309 | 0 | 0.077 | 1 | 18 |
| BX511061.1 | 0.16684 | -0.11134563 | 0 | 0.077 | 1 | 18 |
| BX572619.1 | 0.16684 | -0.14598943 | 0 | 0.077 | 1 | 18 |
| BX842699.1 | 0.16684 | -0.11981325 | 0 | 0.077 | 1 | 18 |
| CABZ01068366.1 | 0.16684 | -0.12353956 | 0 | 0.077 | 1 | 18 |
| CABZ01068600.1 | 0.16684 | -0.16224177 | 0 | 0.077 | 1 | 18 |
| CABZ01086041.1 | 0.16684 | -0.1468439 | 0 | 0.077 | 1 | 18 |
| CABZ01118004.1 | 0.16684 | -0.14296754 | 0 | 0.077 | 1 | 18 |
| CR376737.1 | 0.16684 | -0.11924992 | 0 | 0.077 | 1 | 18 |
| CR847953.1 | 0.16684 | -0.11757582 | 0 | 0.077 | 1 | 18 |
| CT737123.1 | 0.16684 | -0.14792756 | 0 | 0.077 | 1 | 18 |
| CU570894.2 | 0.16684 | -0.12331948 | 0 | 0.077 | 1 | 18 |
| CU660013.4 | 0.16684 | -0.12941742 | 0 | 0.077 | 1 | 18 |
| LOC101884549 | 0.16684 | -0.13490267 | 0 | 0.077 | 1 | 18 |
| PLEKHH3 | 0.16684 | -0.141851 | 0 | 0.077 | 1 | 18 |
| XLOC-006268 | 0.16684 | -0.15457905 | 0 | 0.077 | 1 | 18 |
| XLOC-028948 | 0.16684 | -0.11384676 | 0 | 0.077 | 1 | 18 |
| XLOC-039817 | 0.16684 | -0.10421537 | 0 | 0.077 | 1 | 18 |
| XLOC-042595 | 0.16684 | -0.16782335 | 0 | 0.077 | 1 | 18 |
| XLOC-044204 | 0.16684 | -0.13854994 | 0 | 0.077 | 1 | 18 |
| ZNF512B | 0.16684 | -0.16972977 | 0 | 0.077 | 1 | 18 |
| acap3a | 0.16684 | -0.18158861 | 0 | 0.077 | 1 | 18 |
| acsm3 | 0.16684 | -0.14208758 | 0 | 0.077 | 1 | 18 |
| ankib1a | 0.16684 | -0.13747284 | 0 | 0.077 | 1 | 18 |
| apoob | 0.16684 | -0.16685899 | 0 | 0.077 | 1 | 18 |
| arhgef28 | 0.16684 | -0.19556004 | 0 | 0.077 | 1 | 18 |
| arl16 | 0.16684 | -0.16029382 | 0 | 0.077 | 1 | 18 |
| atad1b | 0.16684 | -0.11579512 | 0 | 0.077 | 1 | 18 |
| atp23 | 0.16684 | -0.13761147 | 0 | 0.077 | 1 | 18 |

| | | | | | | |
|---|---|---|---|---|---|---|
| atp6v0a1a | 0.16684 | -0.15037691 | 0 | 0.077 | 1 | 18 |
| atpv0e2 | 0.16684 | -0.12069647 | 0 | 0.077 | 1 | 18 |
| becn1 | 0.16684 | -0.11657364 | 0 | 0.077 | 1 | 18 |
| blm | 0.16684 | -0.14196291 | 0 | 0.077 | 1 | 18 |
| btr31 | 0.16684 | -0.13865977 | 0 | 0.077 | 1 | 18 |
| cabin1 | 0.16684 | -0.1128079 | 0 | 0.077 | 1 | 18 |
| cacnb1 | 0.16684 | -0.10450616 | 0 | 0.077 | 1 | 18 |
| caska | 0.16684 | -0.16676745 | 0 | 0.077 | 1 | 18 |
| cbx8a | 0.16684 | -0.13429543 | 0 | 0.077 | 1 | 18 |
| ccnq | 0.16684 | -0.13565397 | 0 | 0.077 | 1 | 18 |
| chst10 | 0.16684 | -0.14507059 | 0 | 0.077 | 1 | 18 |
| ciartb | 0.16684 | -0.14283841 | 0 | 0.077 | 1 | 18 |
| cited2 | 0.16684 | -0.1205099 | 0 | 0.077 | 1 | 18 |
| clcn7 | 0.16684 | -0.14458121 | 0 | 0.077 | 1 | 18 |
| clic1 | 0.16684 | -0.14396719 | 0 | 0.077 | 1 | 18 |
| cltb | 0.16684 | -0.14398636 | 0 | 0.077 | 1 | 18 |
| cnrip1a | 0.16684 | -0.13193747 | 0 | 0.077 | 1 | 18 |
| colgalt1 | 0.16684 | -0.12428933 | 0 | 0.077 | 1 | 18 |
| cpt2 | 0.16684 | -0.14918701 | 0 | 0.077 | 1 | 18 |
| cpxm1b | 0.16684 | -0.12667167 | 0 | 0.077 | 1 | 18 |
| cx34.4 | 0.16684 | -0.12284792 | 0 | 0.077 | 1 | 18 |
| cyp2n13 | 0.16684 | -0.16757663 | 0 | 0.077 | 1 | 18 |
| cyp2p6 | 0.16684 | -0.13359964 | 0 | 0.077 | 1 | 18 |
| dag1 | 0.16684 | -0.13335263 | 0 | 0.077 | 1 | 18 |
| dcaf17 | 0.16684 | -0.183752 | 0 | 0.077 | 1 | 18 |
| dcdc2b | 0.16684 | -0.1237734 | 0 | 0.077 | 1 | 18 |
| dcp1b | 0.16684 | -0.12452618 | 0 | 0.077 | 1 | 18 |
| dhrs7b | 0.16684 | -0.14130883 | 0 | 0.077 | 1 | 18 |
| dmd | 0.16684 | -0.14519581 | 0 | 0.077 | 1 | 18 |
| dtnba | 0.16684 | -0.11252965 | 0 | 0.077 | 1 | 18 |
| dync1li1 | 0.16684 | -0.11662505 | 0 | 0.077 | 1 | 18 |
| efs | 0.16684 | -0.11515649 | 0 | 0.077 | 1 | 18 |
| erg | 0.16684 | -0.16734252 | 0 | 0.077 | 1 | 18 |
| ezh2 | 0.16684 | -0.10066566 | 0 | 0.077 | 1 | 18 |
| faim2a | 0.16684 | -0.20278176 | 0 | 0.077 | 1 | 18 |
| fam102bb | 0.16684 | -0.14356656 | 0 | 0.077 | 1 | 18 |
| fam20b | 0.16684 | -0.12902088 | 0 | 0.077 | 1 | 18 |
| fam8a1b | 0.16684 | -0.10566863 | 0 | 0.077 | 1 | 18 |
| fam98b | 0.16684 | -0.10858477 | 0 | 0.077 | 1 | 18 |
| fgfr1op2 | 0.16684 | -0.13933365 | 0 | 0.077 | 1 | 18 |
| fli1a | 0.16684 | -0.13269182 | 0 | 0.077 | 1 | 18 |
| fut8a | 0.16684 | -0.13665363 | 0 | 0.077 | 1 | 18 |
| gas2l3 | 0.16684 | -0.11426166 | 0 | 0.077 | 1 | 18 |
| glrx2 | 0.16684 | -0.14622896 | 0 | 0.077 | 1 | 18 |
| gne | 0.16684 | -0.19033226 | 0 | 0.077 | 1 | 18 |
| hccsb | 0.16684 | -0.11643635 | 0 | 0.077 | 1 | 18 |
| hipk3b | 0.16684 | -0.14596314 | 0 | 0.077 | 1 | 18 |
| hmox1a | 0.16684 | -0.116374 | 0 | 0.077 | 1 | 18 |

| | | | | | | |
|---|---|---|---|---|---|---|
| hoxb7a | 0.16684 | -0.16255811 | 0 | 0.077 | 1 | 18 |
| hsbp1l1 | 0.16684 | -0.17273969 | 0 | 0.077 | 1 | 18 |
| htra1b | 0.16684 | -0.12048344 | 0 | 0.077 | 1 | 18 |
| iars | 0.16684 | -0.10128293 | 0 | 0.077 | 1 | 18 |
| ibtk | 0.16684 | -0.11550242 | 0 | 0.077 | 1 | 18 |
| idh1 | 0.16684 | -0.13730386 | 0 | 0.077 | 1 | 18 |
| irak1bp1 | 0.16684 | -0.13234624 | 0 | 0.077 | 1 | 18 |
| irak4 | 0.16684 | -0.12361743 | 0 | 0.077 | 1 | 18 |
| irx2a | 0.16684 | -0.12614585 | 0 | 0.077 | 1 | 18 |
| jkamp | 0.16684 | -0.12807084 | 0 | 0.077 | 1 | 18 |
| jmjd7 | 0.16684 | -0.10857709 | 0 | 0.077 | 1 | 18 |
| kansl3 | 0.16684 | -0.12881887 | 0 | 0.077 | 1 | 18 |
| kat6a | 0.16684 | -0.13104605 | 0 | 0.077 | 1 | 18 |
| katna1 | 0.16684 | -0.13568554 | 0 | 0.077 | 1 | 18 |
| kcnk5b | 0.16684 | -0.12961546 | 0 | 0.077 | 1 | 18 |
| kdm4ab | 0.16684 | -0.13274415 | 0 | 0.077 | 1 | 18 |
| kif3a | 0.16684 | -0.12078743 | 0 | 0.077 | 1 | 18 |
| kitlga | 0.16684 | -0.11265458 | 0 | 0.077 | 1 | 18 |
| klf15 | 0.16684 | -0.10457623 | 0 | 0.077 | 1 | 18 |
| klf7a | 0.16684 | -0.13566984 | 0 | 0.077 | 1 | 18 |
| klhl8 | 0.16684 | -0.14089031 | 0 | 0.077 | 1 | 18 |
| ldb2b | 0.16684 | -0.1106543 | 0 | 0.077 | 1 | 18 |
| lman2la | 0.16684 | -0.14730446 | 0 | 0.077 | 1 | 18 |
| loxl2b | 0.16684 | -0.19264345 | 0 | 0.077 | 1 | 18 |
| lrch3 | 0.16684 | -0.21075283 | 0 | 0.077 | 1 | 18 |
| lrp1ba | 0.16684 | -0.14360065 | 0 | 0.077 | 1 | 18 |
| lrrc15 | 0.16684 | -0.11389233 | 0 | 0.077 | 1 | 18 |
| lrrc4ba | 0.16684 | -0.13534516 | 0 | 0.077 | 1 | 18 |
| lygl1 | 0.16684 | -0.13629013 | 0 | 0.077 | 1 | 18 |
| lyrm1 | 0.16684 | -0.10527828 | 0 | 0.077 | 1 | 18 |
| maco1b | 0.16684 | -0.14314381 | 0 | 0.077 | 1 | 18 |
| map2k4b | 0.16684 | -0.14555078 | 0 | 0.077 | 1 | 18 |
| mapk6 | 0.16684 | -0.16488535 | 0 | 0.077 | 1 | 18 |
| mapre2 | 0.16684 | -0.10197719 | 0 | 0.077 | 1 | 18 |
| mark2a | 0.16684 | -0.15290843 | 0 | 0.077 | 1 | 18 |
| mbtps2 | 0.16684 | -0.14686164 | 0 | 0.077 | 1 | 18 |
| mfsd8 | 0.16684 | -0.14333807 | 0 | 0.077 | 1 | 18 |
| mki67 | 0.16684 | -0.18968411 | 0 | 0.077 | 1 | 18 |
| mroh1 | 0.16684 | -0.20308957 | 0 | 0.077 | 1 | 18 |
| mrpl45 | 0.16684 | -0.14290986 | 0 | 0.077 | 1 | 18 |
| mtor | 0.16684 | -0.16502598 | 0 | 0.077 | 1 | 18 |
| mtrf1 | 0.16684 | -0.16006571 | 0 | 0.077 | 1 | 18 |
| myh9b | 0.16684 | -0.13382959 | 0 | 0.077 | 1 | 18 |
| myo1b | 0.16684 | -0.11388398 | 0 | 0.077 | 1 | 18 |
| nacc1a | 0.16684 | -0.19986 | 0 | 0.077 | 1 | 18 |
| nat8l | 0.16684 | -0.16197073 | 0 | 0.077 | 1 | 18 |
| ncaph | 0.16684 | -0.16043355 | 0 | 0.077 | 1 | 18 |
| nek9 | 0.16684 | -0.13671148 | 0 | 0.077 | 1 | 18 |

| | | | | | | |
|---|---|---|---|---|---|---|
| ngfb | 0.16684 | -0.10978161 | 0 | 0.077 | 1 | 18 |
| nog2 | 0.16684 | -0.121265 | 0 | 0.077 | 1 | 18 |
| ntrk2a | 0.16684 | -0.12804792 | 0 | 0.077 | 1 | 18 |
| nudcd1 | 0.16684 | -0.11517264 | 0 | 0.077 | 1 | 18 |
| nudt19 | 0.16684 | -0.12689006 | 0 | 0.077 | 1 | 18 |
| osbp | 0.16684 | -0.14133558 | 0 | 0.077 | 1 | 18 |
| p3h3 | 0.16684 | -0.14519307 | 0 | 0.077 | 1 | 18 |
| pcbd1 | 0.16684 | -0.128736 | 0 | 0.077 | 1 | 18 |
| pcm1 | 0.16684 | -0.15122836 | 0 | 0.077 | 1 | 18 |
| pcnt | 0.16684 | -0.16265758 | 0 | 0.077 | 1 | 18 |
| pex26 | 0.16684 | -0.11228953 | 0 | 0.077 | 1 | 18 |
| pi4k2a | 0.16684 | -0.12098466 | 0 | 0.077 | 1 | 18 |
| pi4kb | 0.16684 | -0.13677116 | 0 | 0.077 | 1 | 18 |
| pip4k2aa | 0.16684 | -0.15440691 | 0 | 0.077 | 1 | 18 |
| plk3 | 0.16684 | -0.1043533 | 0 | 0.077 | 1 | 18 |
| pold1 | 0.16684 | -0.15759837 | 0 | 0.077 | 1 | 18 |
| ppap2d | 0.16684 | -0.14370178 | 0 | 0.077 | 1 | 18 |
| ppp2r3c | 0.16684 | -0.10350934 | 0 | 0.077 | 1 | 18 |
| ppp4ca | 0.16684 | -0.15137993 | 0 | 0.077 | 1 | 18 |
| prkx | 0.16684 | -0.13438971 | 0 | 0.077 | 1 | 18 |
| prmt6 | 0.16684 | -0.10894394 | 0 | 0.077 | 1 | 18 |
| ptbp2a | 0.16684 | -0.12912406 | 0 | 0.077 | 1 | 18 |
| ptgs1 | 0.16684 | -0.13588131 | 0 | 0.077 | 1 | 18 |
| rbm14b | 0.16684 | -0.16643867 | 0 | 0.077 | 1 | 18 |
| rbm18 | 0.16684 | -0.11154916 | 0 | 0.077 | 1 | 18 |
| rcor1 | 0.16684 | -0.12315626 | 0 | 0.077 | 1 | 18 |
| rfx1a | 0.16684 | -0.14426921 | 0 | 0.077 | 1 | 18 |
| rhbdd1 | 0.16684 | -0.13927965 | 0 | 0.077 | 1 | 18 |
| rock2a | 0.16684 | -0.13417473 | 0 | 0.077 | 1 | 18 |
| rusc2 | 0.16684 | -0.12321333 | 0 | 0.077 | 1 | 18 |
| sap130a | 0.16684 | -0.14983384 | 0 | 0.077 | 1 | 18 |
| sapcd2 | 0.16684 | -0.16357663 | 0 | 0.077 | 1 | 18 |
| scarb1 | 0.16684 | -0.10790255 | 0 | 0.077 | 1 | 18 |
| sec31b | 0.16684 | -0.10450723 | 0 | 0.077 | 1 | 18 |
| secisbp2 | 0.16684 | -0.134556 | 0 | 0.077 | 1 | 18 |
| selenou1a | 0.16684 | -0.12553706 | 0 | 0.077 | 1 | 18 |
| sema3d | 0.16684 | -0.13055798 | 0 | 0.077 | 1 | 18 |
| sept9b | 0.16684 | -0.11066762 | 0 | 0.077 | 1 | 18 |
| serpinh2 | 0.16684 | -0.14118679 | 0 | 0.077 | 1 | 18 |
| sh3rf1 | 0.16684 | -0.17384513 | 0 | 0.077 | 1 | 18 |
| si:ch1073-392o20.2 | 0.16684 | -0.13839833 | 0 | 0.077 | 1 | 18 |
| si:ch211-113a14.27 | 0.16684 | -0.15364897 | 0 | 0.077 | 1 | 18 |
| si:ch211-11n16.2 | 0.16684 | -0.14088945 | 0 | 0.077 | 1 | 18 |
| si:ch211-136a13.1 | 0.16684 | -0.15654557 | 0 | 0.077 | 1 | 18 |
| si:ch211-199m9.3 | 0.16684 | -0.11666773 | 0 | 0.077 | 1 | 18 |
| si:ch211-250c4.4 | 0.16684 | -0.14974156 | 0 | 0.077 | 1 | 18 |
| si:ch73-347e22.4 | 0.16684 | -0.13216956 | 0 | 0.077 | 1 | 18 |
| si:dkey-117m1.4 | 0.16684 | -0.10475989 | 0 | 0.077 | 1 | 18 |

| | | | | | | |
|---|---|---|---|---|---|---|
| si:dkey-165n16.1 | 0.16684 | -0.1277655 | 0 | 0.077 | 1 | 18 |
| si:dkey-197c15.6 | 0.16684 | -0.11903154 | 0 | 0.077 | 1 | 18 |
| si:dkey-20i10.7 | 0.16684 | -0.11482571 | 0 | 0.077 | 1 | 18 |
| si:dkey-287g12.6 | 0.16684 | -0.11139663 | 0 | 0.077 | 1 | 18 |
| si:dkey-3h3.3 | 0.16684 | -0.13723897 | 0 | 0.077 | 1 | 18 |
| si:dkey-7j14.5 | 0.16684 | -0.18071707 | 0 | 0.077 | 1 | 18 |
| si:dkey-97o5.1 | 0.16684 | -0.13411296 | 0 | 0.077 | 1 | 18 |
| si:zfos-943e10.1 | 0.16684 | -0.13660008 | 0 | 0.077 | 1 | 18 |
| slc10a7 | 0.16684 | -0.10867862 | 0 | 0.077 | 1 | 18 |
| slc31a1 | 0.16684 | -0.14717769 | 0 | 0.077 | 1 | 18 |
| slc35a3a | 0.16684 | -0.11292756 | 0 | 0.077 | 1 | 18 |
| slc35b2 | 0.16684 | -0.11866349 | 0 | 0.077 | 1 | 18 |
| slc35e1 | 0.16684 | -0.15816665 | 0 | 0.077 | 1 | 18 |
| slco2b1 | 0.16684 | -0.13484793 | 0 | 0.077 | 1 | 18 |
| smarcc1b | 0.16684 | -0.10987961 | 0 | 0.077 | 1 | 18 |
| smim29 | 0.16684 | -0.10747654 | 0 | 0.077 | 1 | 18 |
| snapc2 | 0.16684 | -0.1037467 | 0 | 0.077 | 1 | 18 |
| snapc5 | 0.16684 | -0.13166438 | 0 | 0.077 | 1 | 18 |
| snx15 | 0.16684 | -0.1514784 | 0 | 0.077 | 1 | 18 |
| snx24 | 0.16684 | -0.13982587 | 0 | 0.077 | 1 | 18 |
| spata2 | 0.16684 | -0.13882031 | 0 | 0.077 | 1 | 18 |
| spop | 0.16684 | -0.11041256 | 0 | 0.077 | 1 | 18 |
| spsb1 | 0.16684 | -0.18990218 | 0 | 0.077 | 1 | 18 |
| sptbn1 | 0.16684 | -0.14571611 | 0 | 0.077 | 1 | 18 |
| st3gal3b | 0.16684 | -0.16794374 | 0 | 0.077 | 1 | 18 |
| st5 | 0.16684 | -0.17754875 | 0 | 0.077 | 1 | 18 |
| stag1b | 0.16684 | -0.11189995 | 0 | 0.077 | 1 | 18 |
| synj1 | 0.16684 | -0.14668795 | 0 | 0.077 | 1 | 18 |
| tcf7l1b | 0.16684 | -0.11362445 | 0 | 0.077 | 1 | 18 |
| tiparp | 0.16684 | -0.11399651 | 0 | 0.077 | 1 | 18 |
| tle3a | 0.16684 | -0.13311051 | 0 | 0.077 | 1 | 18 |
| tmem238a | 0.16684 | -0.12446734 | 0 | 0.077 | 1 | 18 |
| tmem53 | 0.16684 | -0.10776734 | 0 | 0.077 | 1 | 18 |
| tmem65 | 0.16684 | -0.14992267 | 0 | 0.077 | 1 | 18 |
| tmem86a | 0.16684 | -0.16178723 | 0 | 0.077 | 1 | 18 |
| tp53i11a | 0.16684 | -0.13517593 | 0 | 0.077 | 1 | 18 |
| tppp3 | 0.16684 | -0.2170057 | 0 | 0.077 | 1 | 18 |
| trnt1 | 0.16684 | -0.10855575 | 0 | 0.077 | 1 | 18 |
| tspan35 | 0.16684 | -0.16258406 | 0 | 0.077 | 1 | 18 |
| ttc37 | 0.16684 | -0.14019822 | 0 | 0.077 | 1 | 18 |
| vgll4b | 0.16684 | -0.15866499 | 0 | 0.077 | 1 | 18 |
| vps37a | 0.16684 | -0.15715152 | 0 | 0.077 | 1 | 18 |
| vps8 | 0.16684 | -0.11724507 | 0 | 0.077 | 1 | 18 |
| wasf3b | 0.16684 | -0.11606247 | 0 | 0.077 | 1 | 18 |
| xpc | 0.16684 | -0.178361 | 0 | 0.077 | 1 | 18 |
| ythdf3 | 0.16684 | -0.12903655 | 0 | 0.077 | 1 | 18 |
| zbtb39 | 0.16684 | -0.12680034 | 0 | 0.077 | 1 | 18 |
| zbtb46 | 0.16684 | -0.14790016 | 0 | 0.077 | 1 | 18 |

| | | | | | | |
|---|---|---|---|---|---|---|
| zgc:123305 | 0.16684 | -0.14377531 | 0 | 0.077 | 1 | 18 |
| zgc:153405 | 0.16684 | -0.16464138 | 0 | 0.077 | 1 | 18 |
| zgc:153935 | 0.16684 | -0.10898251 | 0 | 0.077 | 1 | 18 |
| zgc:162780 | 0.16684 | -0.11116069 | 0 | 0.077 | 1 | 18 |
| zgc:171497 | 0.16684 | -0.19217497 | 0 | 0.077 | 1 | 18 |
| zgc:77151 | 0.16684 | -0.14994145 | 0 | 0.077 | 1 | 18 |
| zmym2 | 0.16684 | -0.13440904 | 0 | 0.077 | 1 | 18 |
| znf1152 | 0.16684 | -0.11508615 | 0 | 0.077 | 1 | 18 |
| znf395a | 0.16684 | -0.13678381 | 0 | 0.077 | 1 | 18 |
| znf598 | 0.16684 | -0.13431208 | 0 | 0.077 | 1 | 18 |
| znf800b | 0.16684 | -0.16747821 | 0 | 0.077 | 1 | 18 |
| znf991 | 0.16684 | -0.12629859 | 0 | 0.077 | 1 | 18 |
| CABZ01055347.1 | 0.16684 | -0.12816553 | 0 | 0.077 | 1 | 18 |
| LOC556707 | 0.16684 | -0.19001521 | 0 | 0.077 | 1 | 18 |
| XLOC-029463 | 0.16684 | -0.12363646 | 0 | 0.077 | 1 | 18 |
| XLOC-031367 | 0.16684 | -0.17075059 | 0 | 0.077 | 1 | 18 |
| arhgap21b | 0.16684 | -0.22839954 | 0 | 0.077 | 1 | 18 |
| aurkaip1 | 0.16684 | -0.16843868 | 0 | 0.077 | 1 | 18 |
| dffb | 0.16684 | -0.1670453 | 0 | 0.077 | 1 | 18 |
| dhx8 | 0.16684 | -0.15069121 | 0 | 0.077 | 1 | 18 |
| dnajb4 | 0.16684 | -0.13295186 | 0 | 0.077 | 1 | 18 |
| dsc2l | 0.16684 | -0.18995312 | 0 | 0.077 | 1 | 18 |
| ech1 | 0.16684 | -0.10861449 | 0 | 0.077 | 1 | 18 |
| ehhadh | 0.16684 | -0.13124459 | 0 | 0.077 | 1 | 18 |
| fam57ba | 0.16684 | -0.14205249 | 0 | 0.077 | 1 | 18 |
| faxdc2 | 0.16684 | -0.15455005 | 0 | 0.077 | 1 | 18 |
| fbrsl1 | 0.16684 | -0.12119429 | 0 | 0.077 | 1 | 18 |
| fbxo3 | 0.16684 | -0.14897923 | 0 | 0.077 | 1 | 18 |
| idh3a | 0.16684 | -0.17505889 | 0 | 0.077 | 1 | 18 |
| lancl1 | 0.16684 | -0.11615224 | 0 | 0.077 | 1 | 18 |
| mgaa | 0.16684 | -0.15029766 | 0 | 0.077 | 1 | 18 |
| mhc1laa | 0.16684 | -0.1367561 | 0 | 0.077 | 1 | 18 |
| mtmr8 | 0.16684 | -0.11456965 | 0 | 0.077 | 1 | 18 |
| mybbp1a | 0.16684 | -0.14780785 | 0 | 0.077 | 1 | 18 |
| napbb | 0.16684 | -0.18736762 | 0 | 0.077 | 1 | 18 |
| nedd4l | 0.16684 | -0.11745888 | 0 | 0.077 | 1 | 18 |
| nfs1 | 0.16684 | -0.19433895 | 0 | 0.077 | 1 | 18 |
| nova1 | 0.16684 | -0.21976613 | 0 | 0.077 | 1 | 18 |
| nphs2 | 0.16684 | -0.18310354 | 0 | 0.077 | 1 | 18 |
| nrde2 | 0.16684 | -0.12597652 | 0 | 0.077 | 1 | 18 |
| pnpla7a | 0.16684 | -0.11529379 | 0 | 0.077 | 1 | 18 |
| polr3c | 0.16684 | -0.12297866 | 0 | 0.077 | 1 | 18 |
| ppp2r3a | 0.16684 | -0.12198961 | 0 | 0.077 | 1 | 18 |
| relb | 0.16684 | -0.12176071 | 0 | 0.077 | 1 | 18 |
| rictora | 0.16684 | -0.1309133 | 0 | 0.077 | 1 | 18 |
| sept4 | 0.16684 | -0.12454375 | 0 | 0.077 | 1 | 18 |
| si:ch211-132g1.3 | 0.16684 | -0.1951248 | 0 | 0.077 | 1 | 18 |
| si:ch211-154o6.2 | 0.16684 | -0.12023365 | 0 | 0.077 | 1 | 18 |

| | | | | | | |
|---|---|---|---|---|---|---|
| si:ch211-210c8.6 | 0.16684 | -0.1350108 | 0 | 0.077 | 1 | 18 |
| skilb | 0.16684 | -0.13068957 | 0 | 0.077 | 1 | 18 |
| slc25a34 | 0.16684 | -0.14132334 | 0 | 0.077 | 1 | 18 |
| slf1 | 0.16684 | -0.15478065 | 0 | 0.077 | 1 | 18 |
| tbpl1 | 0.16684 | -0.11425493 | 0 | 0.077 | 1 | 18 |
| tead1a | 0.16684 | -0.16842332 | 0 | 0.077 | 1 | 18 |
| tet1 | 0.16684 | -0.11774256 | 0 | 0.077 | 1 | 18 |
| tmem204 | 0.16684 | -0.18588731 | 0 | 0.077 | 1 | 18 |
| tmem88a | 0.16684 | -0.15072253 | 0 | 0.077 | 1 | 18 |
| tmtc4 | 0.16684 | -0.15372345 | 0 | 0.077 | 1 | 18 |
| tp53bp1 | 0.16684 | -0.1933685 | 0 | 0.077 | 1 | 18 |
| trmt61b | 0.16684 | -0.18091098 | 0 | 0.077 | 1 | 18 |
| zfand6 | 0.16684 | -0.11526344 | 0 | 0.077 | 1 | 18 |
| zgc:153039 | 0.16684 | -0.13828789 | 0 | 0.077 | 1 | 18 |
| znf1162 | 0.16684 | -0.15516986 | 0 | 0.077 | 1 | 18 |
| znf280d | 0.16684 | -0.11545398 | 0 | 0.077 | 1 | 18 |
| znf512 | 0.16684 | -0.13514209 | 0 | 0.077 | 1 | 18 |
| znf644a | 0.16684 | -0.20670846 | 0 | 0.077 | 1 | 18 |
| znf710b | 0.16684 | -0.12992778 | 0 | 0.077 | 1 | 18 |
| znf973 | 0.16684 | -0.14951841 | 0 | 0.077 | 1 | 18 |
| dpy30 | 0.16689 | 0.320553285 | 0.318 | 0.231 | 1 | 18 |
| eif5a2 | 0.16746 | -0.11725173 | 0.455 | 0.654 | 1 | 18 |
| tcf20 | 0.16794 | 0.368546849 | 0.227 | 0.077 | 1 | 18 |
| zfyve21 | 0.16805 | -0.16599943 | 0.136 | 0.308 | 1 | 18 |
| mccc1 | 0.16831 | 0.237109337 | 0.182 | 0.038 | 1 | 18 |
| XLOC-041269 | 0.16884 | -0.41306996 | 0.273 | 0.538 | 1 | 18 |
| hspa4b | 0.16908 | -0.177771 | 0.227 | 0.423 | 1 | 18 |
| spred2b | 0.16943 | -0.15297453 | 0.364 | 0.538 | 1 | 18 |
| sept8a | 0.16958 | 0.181016009 | 0.227 | 0.077 | 1 | 18 |
| MZT1 | 0.17014 | 0.352224428 | 0.227 | 0.077 | 1 | 18 |
| hnrnpa0a | 0.17033 | -0.10405639 | 0.364 | 0.538 | 1 | 18 |
| si:zfos-169g10.3 | 0.17065 | 0.12040298 | 0.318 | 0.346 | 1 | 18 |
| agrn | 0.17135 | -0.35730232 | 0.091 | 0.269 | 1 | 18 |
| ppa1b | 0.17262 | 0.283434572 | 0.273 | 0.077 | 1 | 18 |
| pdha1a | 0.17325 | 0.277938163 | 0.318 | 0.231 | 1 | 18 |
| metrnla | 0.17447 | 0.281029722 | 0.182 | 0.115 | 1 | 18 |
| rbm4.1 | 0.17471 | -0.33552083 | 0.182 | 0.423 | 1 | 18 |
| kdm4b | 0.17484 | -0.20671885 | 0.045 | 0.192 | 1 | 18 |
| zc3h11a | 0.17504 | 0.139358199 | 0.182 | 0.192 | 1 | 18 |
| zgc:63882 | 0.17647 | -0.10587413 | 0.045 | 0.154 | 1 | 18 |
| lrrc59 | 0.17729 | 0.228507963 | 0.273 | 0.077 | 1 | 18 |
| LOC103910438 | 0.17917 | -0.37300847 | 0.182 | 0.423 | 1 | 18 |
| pi15a | 0.17925 | -0.3224023 | 0.045 | 0.192 | 1 | 18 |
| fubp1 | 0.1795 | 0.339309163 | 0.364 | 0.192 | 1 | 18 |
| arf5 | 0.18011 | 0.199496822 | 0.455 | 0.423 | 1 | 18 |
| cst14a.2 | 0.18028 | 0.116638576 | 0.409 | 0.423 | 1 | 18 |
| cycsb | 0.18151 | 0.21041988 | 0.409 | 0.385 | 1 | 18 |
| manea | 0.18188 | -0.11224623 | 0.045 | 0.154 | 1 | 18 |

| | | | | | | |
|---|---|---|---|---|---|---|
| creb5b | 0.18221 | -0.19888103 | 0.136 | 0.154 | 1 | 18 |
| twist1a | 0.18364 | -0.33753157 | 0.182 | 0.423 | 1 | 18 |
| si:dkey-250d21.1 | 0.18544 | -0.3572656 | 0.136 | 0.346 | 1 | 18 |
| mrgbp | 0.18628 | -0.23305573 | 0.091 | 0.269 | 1 | 18 |
| barx1 | 0.18659 | 0.166828163 | 0.318 | 0.308 | 1 | 18 |
| hmgb3b | 0.18691 | 0.300138551 | 0.227 | 0.077 | 1 | 18 |
| mcm2 | 0.18721 | 0.446816605 | 0.273 | 0.077 | 1 | 18 |
| srsf3a | 0.18749 | 0.304892193 | 0.273 | 0.077 | 1 | 18 |
| mfge8a | 0.18784 | 0.300232307 | 0.182 | 0.077 | 1 | 18 |
| nkap | 0.18813 | -0.33707513 | 0.091 | 0.231 | 1 | 18 |
| acadm | 0.18848 | -0.31484649 | 0.182 | 0.423 | 1 | 18 |
| ppid | 0.18858 | 0.228554036 | 0.273 | 0.077 | 1 | 18 |
| nap1l4b | 0.18982 | -0.30694538 | 0.091 | 0.269 | 1 | 18 |
| aldh6a1 | 0.19035 | -0.18064183 | 0.182 | 0.192 | 1 | 18 |
| cd2bp2 | 0.19045 | 0.2175503 | 0.182 | 0.154 | 1 | 18 |
| rrm2 | 0.19064 | 0.104478542 | 0.136 | 0.038 | 1 | 18 |
| ccdc85b | 0.19139 | -0.23840799 | 0.091 | 0.269 | 1 | 18 |
| tnrc6c1 | 0.19201 | 0.24756635 | 0.409 | 0.346 | 1 | 18 |
| etfb | 0.19271 | 0.309638721 | 0.364 | 0.192 | 1 | 18 |
| adamts15a | 0.19282 | -0.39029074 | 0.091 | 0.269 | 1 | 18 |
| cdc42 | 0.19303 | -0.20820914 | 0.455 | 0.692 | 1 | 18 |
| uqcc1 | 0.19444 | 0.319990672 | 0.182 | 0.038 | 1 | 18 |
| uqcrc1 | 0.19466 | -0.20571092 | 0.091 | 0.269 | 1 | 18 |
| pepd | 0.19501 | -0.22553175 | 0.045 | 0.154 | 1 | 18 |
| pkig | 0.19545 | 0.223552818 | 0.364 | 0.308 | 1 | 18 |
| p4hb | 0.19575 | -0.40933035 | 0.273 | 0.5 | 1 | 18 |
| btf3l4 | 0.19619 | -0.16558855 | 0.273 | 0.462 | 1 | 18 |
| snrnp27 | 0.19653 | -0.19638599 | 0.091 | 0.269 | 1 | 18 |
| ddx6 | 0.19693 | 0.380184386 | 0.273 | 0.115 | 1 | 18 |
| tenm3 | 0.19806 | -0.37589449 | 0.409 | 0.615 | 1 | 18 |
| gmfb | 0.199 | 0.353139112 | 0.318 | 0.115 | 1 | 18 |
| tshz3b | 0.19908 | -0.23273508 | 0.045 | 0.192 | 1 | 18 |
| tril | 0.19971 | -0.34219776 | 0.045 | 0.192 | 1 | 18 |
| ndufa10 | 0.2002 | 0.156010987 | 0.273 | 0.269 | 1 | 18 |
| klf8 | 0.20028 | -0.29946334 | 0.045 | 0.192 | 1 | 18 |
| irf2bp2a | 0.20203 | 0.306893727 | 0.5 | 0.385 | 1 | 18 |
| slc25a11 | 0.20288 | 0.308695728 | 0.273 | 0.115 | 1 | 18 |
| tmed2 | 0.20429 | -0.35052558 | 0.364 | 0.577 | 1 | 18 |
| dedd | 0.20431 | -0.13768683 | 0.091 | 0.231 | 1 | 18 |
| tpt1 | 0.20443 | 0.147540063 | 1 | 0.885 | 1 | 18 |
| myef2 | 0.2047 | -0.2565502 | 0.136 | 0.346 | 1 | 18 |
| spryd4 | 0.20488 | 0.292647002 | 0.182 | 0.038 | 1 | 18 |
| picalmb | 0.20545 | -0.11196287 | 0.045 | 0.154 | 1 | 18 |
| nop56 | 0.20568 | 0.257769032 | 0.455 | 0.385 | 1 | 18 |
| snap23.1 | 0.20602 | -0.33993925 | 0.136 | 0.346 | 1 | 18 |
| ms4a17a.9 | 0.20661 | 0.338026096 | 0.318 | 0.115 | 1 | 18 |
| kctd1 | 0.20701 | 0.211831372 | 0.182 | 0.038 | 1 | 18 |
| hsp90ab1 | 0.20723 | 0.164775282 | 1 | 0.962 | 1 | 18 |

| | | | | | | |
|---|---|---|---|---|---|---|
| nme2b.1 | 0.20788 | 0.139434337 | 1 | 0.885 | 1 | 18 |
| uprt | 0.20803 | 0.303537479 | 0.182 | 0.038 | 1 | 18 |
| mrps35 | 0.20823 | 0.290910218 | 0.182 | 0.038 | 1 | 18 |
| arpc2 | 0.20823 | -0.10614049 | 0.364 | 0.538 | 1 | 18 |
| cpne1 | 0.20867 | -0.29878624 | 0.136 | 0.346 | 1 | 18 |
| mphosph8 | 0.20883 | 0.204027301 | 0.273 | 0.231 | 1 | 18 |
| mir214a | 0.20968 | -0.30506709 | 0.136 | 0.346 | 1 | 18 |
| dlgap4a | 0.20996 | 0.327240685 | 0.182 | 0.038 | 1 | 18 |
| rpz2 | 0.21003 | -0.19753718 | 0.045 | 0.192 | 1 | 18 |
| BX284696.1 | 0.21109 | -0.28352891 | 0.136 | 0.346 | 1 | 18 |
| grpel2 | 0.21185 | -0.16227041 | 0.136 | 0.308 | 1 | 18 |
| dlg5b.1 | 0.2119 | -0.22855645 | 0.045 | 0.192 | 1 | 18 |
| ube2d1b | 0.21214 | 0.219338164 | 0.318 | 0.269 | 1 | 18 |
| sec24a | 0.21449 | -0.11961984 | 0.045 | 0.154 | 1 | 18 |
| col5a1 | 0.21471 | -0.26485423 | 0.682 | 0.885 | 1 | 18 |
| ppme1 | 0.21478 | -0.23070735 | 0.091 | 0.269 | 1 | 18 |
| histh1l | 0.21524 | -0.12722451 | 0.136 | 0.115 | 1 | 18 |
| zgc:110591 | 0.21542 | -0.28588315 | 0.045 | 0.192 | 1 | 18 |
| atp5if1b | 0.21569 | -0.29341371 | 0.136 | 0.308 | 1 | 18 |
| pdgfrl | 0.21638 | -0.30510507 | 0.136 | 0.346 | 1 | 18 |
| colec12 | 0.21761 | -0.39634709 | 0.227 | 0.462 | 1 | 18 |
| ywhaba | 0.21805 | -0.12827311 | 0.364 | 0.538 | 1 | 18 |
| c1qtnf5 | 0.21816 | 0.161247824 | 0.773 | 0.577 | 1 | 18 |
| id2a | 0.21833 | 0.170586529 | 0.227 | 0.231 | 1 | 18 |
| ist1 | 0.21844 | 0.289845753 | 0.182 | 0.038 | 1 | 18 |
| rassf8a | 0.21923 | 0.357732578 | 0.318 | 0.115 | 1 | 18 |
| copa | 0.21935 | -0.2183269 | 0.136 | 0.308 | 1 | 18 |
| magoh | 0.21993 | 0.165058329 | 0.273 | 0.115 | 1 | 18 |
| sik2b | 0.22007 | -0.30371082 | 0.045 | 0.192 | 1 | 18 |
| meis1a | 0.22043 | -0.37440512 | 0.136 | 0.269 | 1 | 18 |
| zgc:171566 | 0.22059 | 0.19128191 | 0.182 | 0.154 | 1 | 18 |
| nedd8l | 0.22117 | -0.28233159 | 0.136 | 0.346 | 1 | 18 |
| lsm7 | 0.22145 | 0.170902973 | 0.409 | 0.423 | 1 | 18 |
| sap30l | 0.22162 | 0.11516609 | 0.364 | 0.385 | 1 | 18 |
| ddx27 | 0.22214 | 0.302659041 | 0.273 | 0.115 | 1 | 18 |
| laptm4a | 0.2222 | -0.2546909 | 0.091 | 0.269 | 1 | 18 |
| gtpbp4 | 0.22224 | -0.18431939 | 0.364 | 0.577 | 1 | 18 |
| asah1b | 0.22228 | 0.330025074 | 0.227 | 0.077 | 1 | 18 |
| zfhx4 | 0.22262 | 0.273192237 | 0.182 | 0.038 | 1 | 18 |
| rpf2 | 0.22279 | 0.281021228 | 0.227 | 0.115 | 1 | 18 |
| rps18 | 0.22313 | 0.132414914 | 1 | 0.923 | 1 | 18 |
| sox11a | 0.22315 | 0.311855256 | 0.318 | 0.115 | 1 | 18 |
| tcima | 0.22437 | 0.482544082 | 0.364 | 0.154 | 1 | 18 |
| sephs2 | 0.22465 | -0.1582033 | 0.136 | 0.154 | 1 | 18 |
| si:dkey-7j14.6 | 0.22523 | -0.32559774 | 0.409 | 0.654 | 1 | 18 |
| vapb | 0.22568 | -0.1561834 | 0.182 | 0.346 | 1 | 18 |
| tspan9b | 0.22569 | 0.132068716 | 0.136 | 0.038 | 1 | 18 |
| denr | 0.22601 | 0.104624249 | 0.318 | 0.346 | 1 | 18 |

| | | | | | | |
|---|---|---|---|---|---|---|
| mafba | 0.22603 | 0.21608801 | 0.318 | 0.269 | 1 | 18 |
| fam69aa | 0.22636 | -0.2894466 | 0.045 | 0.192 | 1 | 18 |
| pcdh18b | 0.22637 | 0.338602421 | 0.364 | 0.231 | 1 | 18 |
| ndnf | 0.22684 | -0.14138343 | 0.045 | 0.154 | 1 | 18 |
| adgrl2a | 0.22692 | 0.268167406 | 0.182 | 0.038 | 1 | 18 |
| MDFI | 0.22729 | -0.2540124 | 0.091 | 0.269 | 1 | 18 |
| MMP23B | 0.22735 | 0.174782044 | 0.227 | 0.192 | 1 | 18 |
| fabp3 | 0.22736 | -0.16437322 | 0.045 | 0.192 | 1 | 18 |
| rps6ka3b | 0.22755 | 0.328150612 | 0.182 | 0.038 | 1 | 18 |
| lhfpl2a | 0.22787 | -0.35730783 | 0.227 | 0.462 | 1 | 18 |
| reep5 | 0.22794 | 0.295480957 | 0.5 | 0.346 | 1 | 18 |
| strn3 | 0.22836 | -0.15684126 | 0.091 | 0.231 | 1 | 18 |
| smarcc2 | 0.22839 | 0.119674191 | 0.182 | 0.192 | 1 | 18 |
| cd63 | 0.22878 | -0.1914574 | 0.591 | 0.808 | 1 | 18 |
| ssr1 | 0.22886 | -0.32592415 | 0.636 | 0.808 | 1 | 18 |
| fbp1a | 0.22896 | 0.174270716 | 0.136 | 0.115 | 1 | 18 |
| ptgs2a | 0.22919 | -0.1484599 | 0.045 | 0.154 | 1 | 18 |
| rps23 | 0.2298 | 0.138682882 | 1 | 0.962 | 1 | 18 |
| nabp1a | 0.2298 | -0.24431672 | 0.091 | 0.269 | 1 | 18 |
| aebp1 | 0.23007 | -0.35346008 | 0.227 | 0.462 | 1 | 18 |
| h2afx1 | 0.23078 | -0.22465038 | 0.5 | 0.731 | 1 | 18 |
| wnt5b | 0.23109 | -0.33743657 | 0.136 | 0.346 | 1 | 18 |
| commd1 | 0.23128 | -0.31318688 | 0.045 | 0.192 | 1 | 18 |
| smn1 | 0.23129 | 0.194074237 | 0.182 | 0.038 | 1 | 18 |
| wdr83os | 0.23152 | 0.250034892 | 0.409 | 0.308 | 1 | 18 |
| sept9a | 0.23152 | -0.26860582 | 0.273 | 0.5 | 1 | 18 |
| gnav1 | 0.23227 | 0.314133217 | 0.227 | 0.077 | 1 | 18 |
| rpl13 | 0.2328 | 0.130164775 | 1 | 0.962 | 1 | 18 |
| mrpl34 | 0.23282 | 0.254491801 | 0.227 | 0.115 | 1 | 18 |
| atoh8 | 0.23301 | 0.392775422 | 0.591 | 0.385 | 1 | 18 |
| idh2 | 0.23346 | 0.292658448 | 0.5 | 0.423 | 1 | 18 |
| nfic | 0.23356 | 0.330242327 | 0.364 | 0.192 | 1 | 18 |
| si:ch211-222k6.3 | 0.23373 | -0.31155615 | 0.045 | 0.192 | 1 | 18 |
| sptan1 | 0.23402 | -0.32155058 | 0.136 | 0.346 | 1 | 18 |
| cd151l | 0.23432 | 0.143936972 | 0.273 | 0.269 | 1 | 18 |
| cox8a | 0.23449 | 0.32908235 | 0.682 | 0.5 | 1 | 18 |
| cops4 | 0.23457 | 0.193404825 | 0.273 | 0.231 | 1 | 18 |
| sult6b1 | 0.23512 | -0.14851206 | 0.273 | 0.269 | 1 | 18 |
| gata5 | 0.23537 | 0.364663983 | 0.364 | 0.154 | 1 | 18 |
| serpinh1b | 0.23543 | 0.341629475 | 0.409 | 0.308 | 1 | 18 |
| rpl7 | 0.23543 | -0.13700425 | 0.818 | 0.962 | 1 | 18 |
| TPGS1 | 0.23681 | 0.305278601 | 0.227 | 0.115 | 1 | 18 |
| spcs2 | 0.23683 | -0.13633355 | 0.318 | 0.5 | 1 | 18 |
| hipk2 | 0.23727 | 0.340383623 | 0.182 | 0.038 | 1 | 18 |
| psmc6 | 0.23739 | 0.345435049 | 0.364 | 0.154 | 1 | 18 |
| ptpn11a | 0.23813 | 0.226195283 | 0.182 | 0.038 | 1 | 18 |
| gpc6a | 0.23858 | 0.283240082 | 0.227 | 0.115 | 1 | 18 |
| larp1 | 0.23898 | 0.253753107 | 0.182 | 0.038 | 1 | 18 |

| | | | | | | |
|---|---|---|---|---|---|---|
| phyhiplb | 0.23904 | 0.30851824 | 0.364 | 0.154 | 1 | 18 |
| copz1 | 0.23912 | -0.24734058 | 0.273 | 0.5 | 1 | 18 |
| pet100 | 0.23941 | 0.324782 | 0.182 | 0.038 | 1 | 18 |
| rbm4.2 | 0.23981 | -0.27908559 | 0.045 | 0.192 | 1 | 18 |
| ash1l | 0.24028 | 0.358339711 | 0.364 | 0.154 | 1 | 18 |
| asip2b | 0.24094 | 0.460867649 | 0.409 | 0.231 | 1 | 18 |
| psma8 | 0.2411 | 0.322737265 | 0.5 | 0.308 | 1 | 18 |
| cul1a | 0.24123 | 0.251703284 | 0.227 | 0.077 | 1 | 18 |
| dcp2 | 0.24128 | 0.12056573 | 0.227 | 0.231 | 1 | 18 |
| pwwp2a | 0.24174 | 0.112580927 | 0.136 | 0.038 | 1 | 18 |
| crybg1a | 0.2421 | -0.20317142 | 0.045 | 0.192 | 1 | 18 |
| katnb1 | 0.24215 | -0.24941732 | 0.045 | 0.192 | 1 | 18 |
| psma5 | 0.24329 | -0.14397576 | 0.273 | 0.462 | 1 | 18 |
| CABZ01044053.1 | 0.24349 | 0.288308586 | 0.136 | 0.038 | 1 | 18 |
| psmb5 | 0.24403 | 0.277131473 | 0.409 | 0.308 | 1 | 18 |
| CABZ01115113.1 | 0.24466 | 0.247226788 | 0.182 | 0.038 | 1 | 18 |
| ndufs7 | 0.24467 | -0.14194859 | 0.182 | 0.346 | 1 | 18 |
| prkacab | 0.24505 | 0.268807065 | 0.182 | 0.038 | 1 | 18 |
| mt-nd1 | 0.24578 | -0.32265513 | 0.455 | 0.692 | 1 | 18 |
| tuba1b | 0.2459 | 0.370904551 | 0.364 | 0.192 | 1 | 18 |
| sf1 | 0.24659 | 0.247148559 | 0.364 | 0.269 | 1 | 18 |
| rsrc2 | 0.24685 | 0.323319821 | 0.318 | 0.154 | 1 | 18 |
| selenow2a | 0.24696 | 0.23594224 | 0.182 | 0.038 | 1 | 18 |
| pbx3b | 0.24702 | -0.29657901 | 0.182 | 0.385 | 1 | 18 |
| gnb1a | 0.24726 | 0.311591202 | 0.409 | 0.269 | 1 | 18 |
| mxd4 | 0.24774 | -0.19832162 | 0.045 | 0.192 | 1 | 18 |
| sumo3a | 0.24791 | -0.23899712 | 0.545 | 0.769 | 1 | 18 |
| plxna3 | 0.24861 | -0.20280374 | 0.045 | 0.192 | 1 | 18 |
| nutf2 | 0.24884 | 0.297250314 | 0.273 | 0.115 | 1 | 18 |
| mbtps1 | 0.24923 | -0.13297188 | 0.045 | 0.154 | 1 | 18 |
| atg101 | 0.25036 | 0.24509966 | 0.182 | 0.038 | 1 | 18 |
| rhebl1 | 0.25065 | -0.25168051 | 0.045 | 0.192 | 1 | 18 |
| ubb | 0.25147 | 0.153792168 | 0.591 | 0.577 | 1 | 18 |
| wdr61 | 0.25246 | 0.260245914 | 0.227 | 0.115 | 1 | 18 |
| cbx1a | 0.2525 | -0.1355878 | 0.227 | 0.385 | 1 | 18 |
| fam3a | 0.25288 | 0.232975616 | 0.182 | 0.038 | 1 | 18 |
| tbl1xr1a | 0.25298 | -0.27335077 | 0.091 | 0.269 | 1 | 18 |
| ube2e2 | 0.25321 | 0.229378842 | 0.136 | 0.077 | 1 | 18 |
| pnn | 0.25324 | -0.21498734 | 0.227 | 0.423 | 1 | 18 |
| suclg1 | 0.25338 | -0.16282145 | 0.273 | 0.269 | 1 | 18 |
| mrpl43 | 0.2536 | -0.18617296 | 0.045 | 0.192 | 1 | 18 |
| prdx6 | 0.25428 | -0.221803 | 0.182 | 0.385 | 1 | 18 |
| hist2h2l | 0.25429 | -0.30433023 | 0.091 | 0.269 | 1 | 18 |
| ntmt1 | 0.25447 | -0.219252 | 0.045 | 0.192 | 1 | 18 |
| crim1 | 0.25476 | -0.2307145 | 0.045 | 0.192 | 1 | 18 |
| tpm3 | 0.25487 | -0.2359746 | 0.591 | 0.808 | 1 | 18 |
| xrcc6 | 0.25495 | 0.202815236 | 0.227 | 0.154 | 1 | 18 |
| si:ch211-11k18.4 | 0.25515 | 0.144206673 | 0.182 | 0.038 | 1 | 18 |

| | | | | | | |
|---|---|---|---|---|---|---|
| cox7a2a | 0.25561 | 0.337659852 | 0.5 | 0.308 | 1 | 18 |
| clip2 | 0.25581 | 0.243794332 | 0.182 | 0.038 | 1 | 18 |
| hnrnpc | 0.25641 | 0.241322839 | 0.318 | 0.192 | 1 | 18 |
| tbx2a | 0.25645 | 0.263767328 | 0.182 | 0.038 | 1 | 18 |
| brms1la | 0.25647 | -0.28230659 | 0.045 | 0.192 | 1 | 18 |
| coch | 0.25651 | -0.2867978 | 0.727 | 0.808 | 1 | 18 |
| fbxl14a | 0.25672 | -0.19066076 | 0.045 | 0.192 | 1 | 18 |
| fam168a | 0.25695 | -0.33387399 | 0.364 | 0.577 | 1 | 18 |
| calhm2 | 0.25697 | -0.2164616 | 0.045 | 0.192 | 1 | 18 |
| rnd3a | 0.25706 | -0.35440197 | 0.182 | 0.346 | 1 | 18 |
| ppp1r14bb | 0.25714 | -0.15650017 | 0.091 | 0.231 | 1 | 18 |
| ttc26 | 0.25714 | -0.29256707 | 0.045 | 0.192 | 1 | 18 |
| nudt3a | 0.25744 | -0.25359867 | 0.045 | 0.192 | 1 | 18 |
| srsf10b | 0.2575 | 0.285750564 | 0.273 | 0.115 | 1 | 18 |
| supt4h1 | 0.25818 | -0.30006883 | 0.091 | 0.269 | 1 | 18 |
| tfe3a | 0.25827 | 0.266312046 | 0.182 | 0.038 | 1 | 18 |
| lysmd2 | 0.25866 | 0.245894443 | 0.182 | 0.038 | 1 | 18 |
| hoxb8a | 0.25888 | 0.113434374 | 0.136 | 0.154 | 1 | 18 |
| lsp1 | 0.25909 | -0.32652832 | 0.091 | 0.269 | 1 | 18 |
| gli3 | 0.25932 | -0.27646482 | 0.045 | 0.192 | 1 | 18 |
| szrd1 | 0.25948 | -0.22634044 | 0.045 | 0.192 | 1 | 18 |
| fbl | 0.26001 | 0.169441928 | 0.273 | 0.269 | 1 | 18 |
| itgb1bp1 | 0.26042 | -0.29227048 | 0.045 | 0.192 | 1 | 18 |
| ctdsp1 | 0.26073 | -0.25880795 | 0.045 | 0.192 | 1 | 18 |
| CU459056.1 | 0.26106 | -0.30958608 | 0.045 | 0.192 | 1 | 18 |
| f2rl2 | 0.26141 | -0.12472025 | 0.045 | 0.154 | 1 | 18 |
| FP102052.2 | 0.26157 | -0.28237911 | 0.091 | 0.269 | 1 | 18 |
| camk2g1 | 0.26181 | -0.27829022 | 0.045 | 0.192 | 1 | 18 |
| cox14 | 0.26207 | -0.24151849 | 0.091 | 0.269 | 1 | 18 |
| cenpw | 0.26229 | 0.187087774 | 0.182 | 0.038 | 1 | 18 |
| zfp36l2 | 0.26229 | -0.2572119 | 0.045 | 0.192 | 1 | 18 |
| hint2 | 0.26231 | -0.2502144 | 0.091 | 0.269 | 1 | 18 |
| slc25a51b | 0.26237 | 0.225135759 | 0.182 | 0.038 | 1 | 18 |
| tsr1 | 0.26257 | 0.2350177 | 0.182 | 0.038 | 1 | 18 |
| letm2 | 0.26262 | 0.234139568 | 0.182 | 0.038 | 1 | 18 |
| hmgn7 | 0.26263 | 0.199339479 | 0.955 | 0.846 | 1 | 18 |
| zdhhc20a | 0.26265 | -0.20810139 | 0.045 | 0.192 | 1 | 18 |
| rab41 | 0.26274 | 0.153400029 | 0.136 | 0.115 | 1 | 18 |
| snx9b | 0.26292 | -0.27884766 | 0.045 | 0.192 | 1 | 18 |
| nsfl1c | 0.26317 | -0.13521671 | 0.182 | 0.346 | 1 | 18 |
| aldoaa | 0.26333 | -0.1158958 | 0.136 | 0.269 | 1 | 18 |
| pdlim1 | 0.26357 | -0.31237198 | 0.182 | 0.385 | 1 | 18 |
| hddc3 | 0.26388 | 0.114106662 | 0.091 | 0.115 | 1 | 18 |
| nono | 0.26396 | -0.2469919 | 0.136 | 0.192 | 1 | 18 |
| eci1 | 0.26406 | -0.25302079 | 0.091 | 0.269 | 1 | 18 |
| hnrnpa0b | 0.26445 | 0.217347601 | 0.773 | 0.731 | 1 | 18 |
| mir199-3 | 0.26493 | -0.31665649 | 0.091 | 0.231 | 1 | 18 |
| imp4 | 0.26533 | -0.20228124 | 0.045 | 0.192 | 1 | 18 |

| | | | | | | |
|---|---|---|---|---|---|---|
| zgc:66160 | 0.2658 | -0.25797715 | 0.045 | 0.192 | 1 | 18 |
| prpf6 | 0.26597 | 0.259052569 | 0.136 | 0.038 | 1 | 18 |
| rac1a | 0.26611 | -0.25956134 | 0.182 | 0.385 | 1 | 18 |
| dnajc15 | 0.26633 | -0.22563848 | 0.045 | 0.192 | 1 | 18 |
| higd1a | 0.26637 | -0.14526566 | 0.182 | 0.346 | 1 | 18 |
| hvcn1 | 0.26655 | -0.26605169 | 0.045 | 0.192 | 1 | 18 |
| calr3b | 0.26677 | -0.22153492 | 0.273 | 0.462 | 1 | 18 |
| bnip4 | 0.26709 | -0.21342725 | 0.045 | 0.192 | 1 | 18 |
| rps9 | 0.26723 | 0.124912048 | 1 | 0.962 | 1 | 18 |
| ndufaf3 | 0.2674 | -0.23699455 | 0.091 | 0.269 | 1 | 18 |
| si:dkeyp-55f12.3 | 0.26847 | -0.26618909 | 0.091 | 0.269 | 1 | 18 |
| tmem107 | 0.26849 | -0.2515404 | 0.091 | 0.269 | 1 | 18 |
| zgc:158463 | 0.26851 | -0.29926167 | 0.091 | 0.269 | 1 | 18 |
| ythdc1 | 0.26854 | -0.29096612 | 0.045 | 0.154 | 1 | 18 |
| sf3a3 | 0.26935 | 0.253670875 | 0.136 | 0.038 | 1 | 18 |
| notch2 | 0.26951 | -0.12474704 | 0.273 | 0.423 | 1 | 18 |
| gstr | 0.2696 | 0.24417438 | 0.182 | 0.115 | 1 | 18 |
| kpna4 | 0.26984 | 0.232376238 | 0.182 | 0.115 | 1 | 18 |
| pycr1b | 0.27012 | -0.19530646 | 0.045 | 0.192 | 1 | 18 |
| tm9sf2 | 0.27053 | -0.20198237 | 0.045 | 0.192 | 1 | 18 |
| sf3b2 | 0.27111 | -0.24920208 | 0.091 | 0.269 | 1 | 18 |
| dctn3 | 0.27132 | -0.27414493 | 0.091 | 0.269 | 1 | 18 |
| rap2c | 0.27144 | -0.30007746 | 0.182 | 0.385 | 1 | 18 |
| unc50 | 0.27171 | -0.17923089 | 0.045 | 0.192 | 1 | 18 |
| mrpl57 | 0.27178 | -0.18647503 | 0.045 | 0.192 | 1 | 18 |
| snrpf | 0.27243 | 0.134756762 | 0.545 | 0.577 | 1 | 18 |
| si:ch211-262e15.1 | 0.2729 | -0.17157523 | 0.091 | 0.231 | 1 | 18 |
| dnajc3b | 0.27302 | -0.20683316 | 0.045 | 0.192 | 1 | 18 |
| ankrd40 | 0.27305 | 0.215642951 | 0.136 | 0.038 | 1 | 18 |
| akirin2 | 0.27354 | -0.23482617 | 0.045 | 0.192 | 1 | 18 |
| mcfd2 | 0.27354 | -0.28081393 | 0.045 | 0.192 | 1 | 18 |
| zgc:112356 | 0.27398 | 0.171545426 | 0.182 | 0.154 | 1 | 18 |
| arpc5a | 0.27402 | -0.21127741 | 0.045 | 0.192 | 1 | 18 |
| eif3d | 0.27423 | -0.10773042 | 0.636 | 0.615 | 1 | 18 |
| atp10a | 0.27502 | -0.28835344 | 0.045 | 0.192 | 1 | 18 |
| stxbp6 | 0.27528 | -0.24923308 | 0.045 | 0.192 | 1 | 18 |
| psat1 | 0.27572 | -0.18640777 | 0.045 | 0.192 | 1 | 18 |
| mrpl52 | 0.27573 | -0.20020489 | 0.045 | 0.192 | 1 | 18 |
| commd3 | 0.27611 | -0.19258119 | 0.045 | 0.192 | 1 | 18 |
| pitx2 | 0.27611 | 0.355433094 | 0.636 | 0.423 | 1 | 18 |
| rps16 | 0.27684 | 0.135602557 | 1 | 0.962 | 1 | 18 |
| atp2a2b | 0.27705 | -0.241118 | 0.045 | 0.192 | 1 | 18 |
| ormdl1 | 0.27726 | 0.260136135 | 0.227 | 0.077 | 1 | 18 |
| svbp | 0.27729 | -0.10309908 | 0.045 | 0.154 | 1 | 18 |
| arid5b | 0.27763 | 0.318561014 | 0.227 | 0.115 | 1 | 18 |
| taf13 | 0.27773 | 0.233920212 | 0.591 | 0.5 | 1 | 18 |
| slirp | 0.27773 | 0.344043987 | 0.364 | 0.192 | 1 | 18 |
| CR293507.1 | 0.27794 | -0.10324115 | 0.045 | 0.154 | 1 | 18 |

| | | | | | | |
|---|---|---|---|---|---|---|
| glis2a | 0.27797 | -0.33146736 | 0.045 | 0.192 | 1 | 18 |
| sh3glb1b | 0.27816 | -0.21532393 | 0.045 | 0.192 | 1 | 18 |
| junba | 0.27855 | 0.233065189 | 0.818 | 0.731 | 1 | 18 |
| fuca1.1 | 0.27879 | -0.22174955 | 0.045 | 0.192 | 1 | 18 |
| gskip | 0.27891 | 0.111133924 | 0.227 | 0.231 | 1 | 18 |
| ppp1cab | 0.27918 | -0.1988011 | 0.045 | 0.192 | 1 | 18 |
| axin2 | 0.27924 | 0.277114981 | 0.227 | 0.077 | 1 | 18 |
| prickle1b | 0.2796 | -0.24222189 | 0.045 | 0.192 | 1 | 18 |
| si:ch73-234b20.5 | 0.27972 | -0.13038928 | 0.045 | 0.154 | 1 | 18 |
| rpl18a | 0.28018 | 0.130923622 | 1 | 0.923 | 1 | 18 |
| si:dkey-276j7.1 | 0.28096 | -0.19167127 | 0.045 | 0.192 | 1 | 18 |
| iscub | 0.28097 | 0.257376841 | 0.227 | 0.077 | 1 | 18 |
| coro1cb | 0.2812 | 0.294059265 | 0.227 | 0.077 | 1 | 18 |
| naa20 | 0.28122 | 0.255390878 | 0.364 | 0.269 | 1 | 18 |
| si:ch211-204a13.2 | 0.28142 | 0.191554459 | 0.273 | 0.231 | 1 | 18 |
| eif3jb | 0.28159 | -0.25403136 | 0.045 | 0.192 | 1 | 18 |
| cebpb | 0.28174 | -0.17242521 | 0.045 | 0.192 | 1 | 18 |
| ube2d2l | 0.28174 | 0.245229319 | 0.227 | 0.115 | 1 | 18 |
| pde7a | 0.28218 | -0.25667482 | 0.045 | 0.192 | 1 | 18 |
| elnb | 0.28225 | -0.16705845 | 0.136 | 0.269 | 1 | 18 |
| smad3a | 0.28237 | -0.2883382 | 0.045 | 0.192 | 1 | 18 |
| calcoco1a | 0.28245 | -0.22863581 | 0.045 | 0.192 | 1 | 18 |
| sertad2b | 0.28245 | -0.24742332 | 0.045 | 0.192 | 1 | 18 |
| cnpy2 | 0.2825 | -0.22964305 | 0.045 | 0.192 | 1 | 18 |
| pbx4 | 0.28269 | -0.31124272 | 0.182 | 0.385 | 1 | 18 |
| zgc:153867 | 0.28285 | 0.170052634 | 0.955 | 0.808 | 1 | 18 |
| lcorl | 0.28302 | -0.25358641 | 0.045 | 0.192 | 1 | 18 |
| ms4a17a.11 | 0.28308 | -0.22927593 | 0.045 | 0.192 | 1 | 18 |
| arpc5b | 0.28309 | -0.21369231 | 0.045 | 0.192 | 1 | 18 |
| atp5meb | 0.28326 | 0.258292033 | 0.545 | 0.462 | 1 | 18 |
| smad2 | 0.28329 | -0.27690424 | 0.045 | 0.192 | 1 | 18 |
| atp5po | 0.28339 | -0.23374569 | 0.364 | 0.462 | 1 | 18 |
| march7 | 0.2834 | -0.26323774 | 0.045 | 0.192 | 1 | 18 |
| cd164 | 0.28343 | -0.22359176 | 0.045 | 0.192 | 1 | 18 |
| pfdn6 | 0.28344 | -0.21525907 | 0.045 | 0.192 | 1 | 18 |
| ugp2b | 0.28345 | -0.19940552 | 0.045 | 0.192 | 1 | 18 |
| endouc | 0.28348 | -0.2202436 | 0.045 | 0.192 | 1 | 18 |
| thrab | 0.28349 | -0.25494749 | 0.045 | 0.192 | 1 | 18 |
| dstyk | 0.28353 | -0.2193404 | 0.045 | 0.192 | 1 | 18 |
| ptp4a2a | 0.28423 | -0.24887091 | 0.182 | 0.385 | 1 | 18 |
| luc7l3 | 0.28541 | -0.17251019 | 0.091 | 0.231 | 1 | 18 |
| ank3b | 0.28549 | -0.25650343 | 0.045 | 0.154 | 1 | 18 |
| rab5if | 0.2861 | 0.28272051 | 0.227 | 0.115 | 1 | 18 |
| si:ch73-46j18.5 | 0.28645 | 0.183947653 | 0.818 | 0.769 | 1 | 18 |
| fam32a | 0.28665 | -0.1667854 | 0.455 | 0.654 | 1 | 18 |
| polr1c | 0.28693 | 0.282755942 | 0.136 | 0.038 | 1 | 18 |
| rpsa | 0.28784 | -0.10166192 | 1 | 1 | 1 | 18 |
| znf395b | 0.28806 | -0.21917724 | 0.136 | 0.308 | 1 | 18 |

| | | | | | | |
|---|---|---|---|---|---|---|
| sf3a2 | 0.28887 | -0.15479685 | 0.136 | 0.308 | 1 | 18 |
| ppp2cb | 0.28888 | 0.121134012 | 0.455 | 0.462 | 1 | 18 |
| si:dkey-42i9.4 | 0.28917 | 0.305065196 | 0.455 | 0.308 | 1 | 18 |
| ppp1cbl | 0.28933 | -0.12551159 | 0.045 | 0.154 | 1 | 18 |
| gpc4 | 0.28983 | 0.124163951 | 0.182 | 0.077 | 1 | 18 |
| cnn2 | 0.28984 | -0.37520173 | 0.318 | 0.5 | 1 | 18 |
| hmgn2 | 0.28994 | 0.214311553 | 0.818 | 0.885 | 1 | 18 |
| tpi1b | 0.29018 | -0.21057876 | 0.182 | 0.385 | 1 | 18 |
| atp5f1d | 0.2904 | -0.27851706 | 0.182 | 0.385 | 1 | 18 |
| flot1b | 0.29099 | -0.23587529 | 0.227 | 0.423 | 1 | 18 |
| ldhba | 0.29116 | 0.147438263 | 0.455 | 0.462 | 1 | 18 |
| mrps21 | 0.29159 | 0.158205043 | 0.091 | 0.077 | 1 | 18 |
| zgc:101851 | 0.29169 | -0.1077743 | 0.045 | 0.154 | 1 | 18 |
| mmp15b | 0.29191 | -0.32820036 | 0.227 | 0.346 | 1 | 18 |
| CABZ01076667.1 | 0.29191 | 0.262413392 | 0.182 | 0.077 | 1 | 18 |
| fhl1a | 0.29223 | -0.3862566 | 0.182 | 0.385 | 1 | 18 |
| csnk1a1 | 0.2936 | 0.303174754 | 0.636 | 0.423 | 1 | 18 |
| ppp1caa | 0.29407 | -0.20519533 | 0.136 | 0.308 | 1 | 18 |
| tspan14 | 0.29462 | -0.32153346 | 0.182 | 0.385 | 1 | 18 |
| dpysl3 | 0.29493 | -0.19077477 | 0.182 | 0.346 | 1 | 18 |
| phf2 | 0.29528 | 0.231672868 | 0.227 | 0.154 | 1 | 18 |
| tshz2 | 0.29544 | 0.349295726 | 0.273 | 0.115 | 1 | 18 |
| ier3ip1 | 0.29575 | 0.246472082 | 0.455 | 0.346 | 1 | 18 |
| XLOC-035413 | 0.29598 | -0.2876247 | 0.182 | 0.385 | 1 | 18 |
| lsm5 | 0.29635 | 0.284784934 | 0.364 | 0.231 | 1 | 18 |
| ifi30 | 0.2966 | -0.28967991 | 0.182 | 0.385 | 1 | 18 |
| klf12a | 0.29747 | -0.15801457 | 0.045 | 0.154 | 1 | 18 |
| ciapin1 | 0.2975 | 0.260853666 | 0.136 | 0.038 | 1 | 18 |
| rras | 0.29793 | 0.131710444 | 0.318 | 0.308 | 1 | 18 |
| sdc2 | 0.29834 | 0.115910283 | 0.136 | 0.154 | 1 | 18 |
| gb:bc139872 | 0.29934 | 0.225131331 | 0.273 | 0.192 | 1 | 18 |
| cxcl18b | 0.30027 | 0.278550104 | 0.136 | 0.038 | 1 | 18 |
| efna5b | 0.30073 | -0.17625283 | 0.091 | 0.231 | 1 | 18 |
| tm9sf4 | 0.30118 | -0.14639267 | 0.045 | 0.154 | 1 | 18 |
| ubfd1 | 0.3014 | 0.258931254 | 0.227 | 0.077 | 1 | 18 |
| rassf1 | 0.3032 | -0.28782073 | 0.045 | 0.154 | 1 | 18 |
| wu:fj13e08 | 0.30351 | 0.242228785 | 0.136 | 0.038 | 1 | 18 |
| mrps36 | 0.30355 | -0.13698407 | 0.182 | 0.346 | 1 | 18 |
| rab1ab | 0.304 | -0.29494508 | 0.318 | 0.538 | 1 | 18 |
| anp32b | 0.30437 | 0.195215598 | 0.455 | 0.423 | 1 | 18 |
| znf385a | 0.30475 | -0.26709875 | 0.227 | 0.423 | 1 | 18 |
| cacybp | 0.30529 | 0.197383437 | 0.227 | 0.192 | 1 | 18 |
| her9 | 0.30539 | -0.35891641 | 0.136 | 0.308 | 1 | 18 |
| oaz1b | 0.30545 | -0.27488955 | 0.545 | 0.692 | 1 | 18 |
| txndc12 | 0.30689 | -0.26205191 | 0.091 | 0.231 | 1 | 18 |
| ddx51 | 0.30692 | -0.14278082 | 0.045 | 0.154 | 1 | 18 |
| tnfsf10 | 0.30741 | -0.1615621 | 0.545 | 0.731 | 1 | 18 |
| lamtor4 | 0.30783 | -0.20034347 | 0.136 | 0.308 | 1 | 18 |

| | | | | | | |
|---|---|---|---|---|---|---|
| tor1 | 0.30844 | 0.244615779 | 0.273 | 0.154 | 1 | 18 |
| zgc:110366 | 0.3092 | 0.243622341 | 0.182 | 0.077 | 1 | 18 |
| mrpl54 | 0.30944 | 0.165191513 | 0.273 | 0.115 | 1 | 18 |
| zgc:110339 | 0.30949 | 0.291888776 | 0.364 | 0.231 | 1 | 18 |
| nup62l | 0.30962 | 0.215731995 | 0.227 | 0.077 | 1 | 18 |
| si:dkey-253d23.4 | 0.30966 | 0.211087458 | 0.136 | 0.077 | 1 | 18 |
| snrnp40 | 0.31016 | 0.283542497 | 0.227 | 0.077 | 1 | 18 |
| mrps10 | 0.31026 | 0.266512176 | 0.318 | 0.154 | 1 | 18 |
| prrx1a | 0.31034 | -0.16105663 | 0.045 | 0.154 | 1 | 18 |
| triap1 | 0.31081 | 0.224258049 | 0.136 | 0.038 | 1 | 18 |
| mccc2 | 0.31095 | -0.15035674 | 0.091 | 0.231 | 1 | 18 |
| wu:fb55g09 | 0.31207 | 0.320924063 | 0.591 | 0.385 | 1 | 18 |
| ndufs4 | 0.31249 | 0.262960783 | 0.364 | 0.231 | 1 | 18 |
| arrdc3a | 0.31377 | 0.281270034 | 0.273 | 0.154 | 1 | 18 |
| si:ch73-71c20.5 | 0.31383 | 0.265907314 | 0.227 | 0.077 | 1 | 18 |
| sept7a | 0.3144 | -0.12764504 | 0.136 | 0.269 | 1 | 18 |
| net1 | 0.31514 | 0.186977625 | 0.591 | 0.538 | 1 | 18 |
| trib3 | 0.31527 | 0.203452288 | 0.273 | 0.115 | 1 | 18 |
| eif1axb | 0.31581 | 0.183397726 | 0.455 | 0.423 | 1 | 18 |
| rps6kb1b | 0.31631 | 0.179764395 | 0.227 | 0.192 | 1 | 18 |
| sumo2b | 0.31649 | -0.27470514 | 0.591 | 0.769 | 1 | 18 |
| tomm7 | 0.31689 | 0.30057631 | 0.455 | 0.269 | 1 | 18 |
| si:ch1073-314i13.4 | 0.31732 | -0.11415383 | 0.045 | 0.154 | 1 | 18 |
| mast2 | 0.31787 | -0.14150998 | 0.045 | 0.154 | 1 | 18 |
| BX465227.1 | 0.3183 | -0.15615079 | 0.045 | 0.154 | 1 | 18 |
| cdc42ep5 | 0.31872 | 0.307866487 | 0.273 | 0.115 | 1 | 18 |
| ndufs3 | 0.31918 | 0.261149505 | 0.273 | 0.115 | 1 | 18 |
| rpn1 | 0.31995 | -0.17284831 | 0.318 | 0.5 | 1 | 18 |
| anxa11a | 0.32025 | -0.25574362 | 0.136 | 0.269 | 1 | 18 |
| id3 | 0.32028 | 0.343137633 | 0.591 | 0.385 | 1 | 18 |
| prdx3 | 0.32079 | 0.136726734 | 0.273 | 0.269 | 1 | 18 |
| polr3h | 0.32252 | 0.262323438 | 0.227 | 0.077 | 1 | 18 |
| pdgfra | 0.32352 | 0.17622455 | 0.455 | 0.423 | 1 | 18 |
| actr10 | 0.32355 | -0.1318053 | 0.045 | 0.154 | 1 | 18 |
| tagln | 0.32367 | -0.18396717 | 0.545 | 0.731 | 1 | 18 |
| antxr2a | 0.32396 | -0.287074 | 0.545 | 0.731 | 1 | 18 |
| emg1 | 0.32423 | -0.14075032 | 0.091 | 0.231 | 1 | 18 |
| mcrip1 | 0.32423 | -0.16111908 | 0.182 | 0.346 | 1 | 18 |
| slc39a11 | 0.32463 | 0.245187893 | 0.136 | 0.038 | 1 | 18 |
| ckap5 | 0.32722 | 0.276976924 | 0.227 | 0.077 | 1 | 18 |
| psmc3 | 0.32817 | 0.196540015 | 0.227 | 0.154 | 1 | 18 |
| zgc:153115 | 0.32847 | -0.26858035 | 0.045 | 0.154 | 1 | 18 |
| dhrs3a | 0.32977 | -0.25166579 | 0.091 | 0.231 | 1 | 18 |
| nr3c1 | 0.33052 | -0.25241087 | 0.136 | 0.308 | 1 | 18 |
| ost4 | 0.33053 | -0.29822904 | 0.364 | 0.577 | 1 | 18 |
| med19a | 0.33084 | 0.231321156 | 0.136 | 0.038 | 1 | 18 |
| si:ch211-191j22.3 | 0.33122 | -0.26240312 | 0.091 | 0.231 | 1 | 18 |
| wbp1 | 0.33129 | 0.228781275 | 0.227 | 0.077 | 1 | 18 |

| | | | | | | |
|---|---|---|---|---|---|---|
| rpl27a | 0.33202 | -0.14920663 | 0.909 | 0.962 | 1 | 18 |
| tpm4b | 0.33319 | 0.309024034 | 0.364 | 0.231 | 1 | 18 |
| ccng1 | 0.33345 | -0.25867714 | 0.364 | 0.577 | 1 | 18 |
| dynlrb1 | 0.33368 | 0.150431789 | 0.545 | 0.538 | 1 | 18 |
| elac1 | 0.33423 | 0.235903113 | 0.182 | 0.077 | 1 | 18 |
| lin52 | 0.3346 | 0.248928912 | 0.227 | 0.077 | 1 | 18 |
| gstp1 | 0.33491 | 0.221412402 | 0.364 | 0.308 | 1 | 18 |
| zgc:163080 | 0.33492 | -0.19714106 | 0.136 | 0.308 | 1 | 18 |
| ran | 0.33527 | 0.102060259 | 0.818 | 0.846 | 1 | 18 |
| golgb1 | 0.33576 | 0.271875109 | 0.273 | 0.115 | 1 | 18 |
| hivep1 | 0.33589 | 0.23305304 | 0.136 | 0.077 | 1 | 18 |
| cald1b | 0.33633 | -0.15345315 | 0.636 | 0.808 | 1 | 18 |
| b2ml | 0.33683 | -0.17069918 | 0.545 | 0.731 | 1 | 18 |
| ifrd2 | 0.33687 | -0.1043389 | 0.045 | 0.154 | 1 | 18 |
| rasgef1bb | 0.33699 | 0.325644487 | 0.318 | 0.154 | 1 | 18 |
| pias2 | 0.33715 | 0.264208223 | 0.227 | 0.077 | 1 | 18 |
| fblim1 | 0.33743 | 0.10733987 | 0.409 | 0.423 | 1 | 18 |
| ppp4cb | 0.33791 | 0.219651931 | 0.227 | 0.077 | 1 | 18 |
| cnih1 | 0.33795 | 0.251969305 | 0.227 | 0.077 | 1 | 18 |
| lama4 | 0.33909 | -0.24929202 | 0.045 | 0.154 | 1 | 18 |
| rasl12 | 0.33948 | 0.247256162 | 0.545 | 0.346 | 1 | 18 |
| tma7 | 0.34032 | 0.116389171 | 0.682 | 0.731 | 1 | 18 |
| emc6 | 0.34034 | 0.288517906 | 0.273 | 0.115 | 1 | 18 |
| mif | 0.34067 | -0.21648108 | 0.409 | 0.615 | 1 | 18 |
| slc4a10a | 0.34084 | 0.303872888 | 0.136 | 0.038 | 1 | 18 |
| peak1 | 0.34109 | -0.16474839 | 0.045 | 0.154 | 1 | 18 |
| cnn3a | 0.34155 | 0.26790613 | 0.591 | 0.462 | 1 | 18 |
| tead1b | 0.34351 | -0.17310257 | 0.227 | 0.385 | 1 | 18 |
| calm1a | 0.34493 | -0.30526457 | 0.227 | 0.385 | 1 | 18 |
| txnrd3 | 0.34548 | -0.12896744 | 0.045 | 0.154 | 1 | 18 |
| mfap5 | 0.3456 | -0.25314556 | 0.682 | 0.808 | 1 | 18 |
| cdk12 | 0.34607 | 0.245015449 | 0.136 | 0.038 | 1 | 18 |
| adam17b | 0.34614 | -0.24337777 | 0.045 | 0.154 | 1 | 18 |
| btg1 | 0.34623 | -0.24063323 | 0.409 | 0.615 | 1 | 18 |
| id2b | 0.34667 | 0.272107995 | 0.273 | 0.192 | 1 | 18 |
| ncoa6 | 0.34669 | 0.238197311 | 0.136 | 0.038 | 1 | 18 |
| bzw1a | 0.34686 | -0.1444419 | 0.136 | 0.269 | 1 | 18 |
| cebpg | 0.34737 | -0.12600791 | 0.182 | 0.308 | 1 | 18 |
| xbp1 | 0.34829 | 0.150211359 | 0.591 | 0.615 | 1 | 18 |
| atox1 | 0.34833 | 0.243819926 | 0.364 | 0.269 | 1 | 18 |
| tspan4a | 0.34835 | -0.2840159 | 0.136 | 0.308 | 1 | 18 |
| syncripl | 0.34898 | -0.25177238 | 0.182 | 0.269 | 1 | 18 |
| tln1 | 0.34953 | 0.20338056 | 0.318 | 0.269 | 1 | 18 |
| ddah2 | 0.34964 | 0.148105014 | 0.182 | 0.154 | 1 | 18 |
| lrp1aa | 0.34996 | -0.25586883 | 0.227 | 0.423 | 1 | 18 |
| eda | 0.35006 | -0.25559962 | 0.045 | 0.154 | 1 | 18 |
| cwc22 | 0.35044 | -0.23790762 | 0.045 | 0.154 | 1 | 18 |
| wls | 0.35178 | -0.28863475 | 0.227 | 0.423 | 1 | 18 |

| | | | | | | |
|---|---|---|---|---|---|---|
| mrpl14 | 0.35283 | 0.241327258 | 0.136 | 0.038 | 1 | 18 |
| zgc:56493 | 0.35319 | -0.21713089 | 0.727 | 0.885 | 1 | 18 |
| aph1b | 0.35377 | 0.243000862 | 0.136 | 0.038 | 1 | 18 |
| si:dkey-67c22.2 | 0.35389 | 0.238287563 | 0.227 | 0.115 | 1 | 18 |
| metap1 | 0.35484 | -0.17715066 | 0.091 | 0.231 | 1 | 18 |
| srsf1a | 0.35493 | 0.281372619 | 0.273 | 0.115 | 1 | 18 |
| si:dkey-9i23.16 | 0.35502 | -0.1474267 | 0.045 | 0.154 | 1 | 18 |
| psmd12 | 0.35535 | -0.23319096 | 0.136 | 0.308 | 1 | 18 |
| tmem18 | 0.35615 | -0.21449385 | 0.136 | 0.308 | 1 | 18 |
| cnih2 | 0.35617 | -0.12825494 | 0.045 | 0.154 | 1 | 18 |
| tsen34 | 0.3565 | -0.11497257 | 0.045 | 0.154 | 1 | 18 |
| prkab1b | 0.3569 | -0.24733868 | 0.409 | 0.615 | 1 | 18 |
| prelid3b | 0.3579 | 0.213415353 | 0.182 | 0.077 | 1 | 18 |
| gabarapl2 | 0.35814 | -0.18650531 | 0.182 | 0.346 | 1 | 18 |
| hdlbpa | 0.35818 | -0.27328859 | 0.455 | 0.654 | 1 | 18 |
| sav1 | 0.35849 | 0.116310499 | 0.273 | 0.269 | 1 | 18 |
| smim13 | 0.35914 | -0.18727797 | 0.091 | 0.231 | 1 | 18 |
| zgc:56106 | 0.35936 | 0.145962247 | 0.136 | 0.115 | 1 | 18 |
| cnbpb | 0.35944 | 0.259584167 | 0.591 | 0.462 | 1 | 18 |
| tpm4a | 0.35959 | 0.101415241 | 0.727 | 0.577 | 1 | 18 |
| caprin1a | 0.36001 | -0.24017287 | 0.136 | 0.308 | 1 | 18 |
| gnai1 | 0.36019 | 0.217338901 | 0.182 | 0.077 | 1 | 18 |
| mapkapk2a | 0.36025 | 0.171066375 | 0.273 | 0.231 | 1 | 18 |
| taf12 | 0.36063 | 0.283818818 | 0.455 | 0.269 | 1 | 18 |
| ciao2b | 0.36064 | -0.25426785 | 0.136 | 0.308 | 1 | 18 |
| capzb | 0.36093 | -0.27953661 | 0.273 | 0.462 | 1 | 18 |
| rhobtb4 | 0.36136 | -0.16916559 | 0.045 | 0.154 | 1 | 18 |
| tax1bp1b | 0.36158 | -0.17455728 | 0.091 | 0.231 | 1 | 18 |
| ubap2b | 0.36339 | 0.200407856 | 0.136 | 0.038 | 1 | 18 |
| ubtd2 | 0.36398 | -0.13301243 | 0.136 | 0.269 | 1 | 18 |
| ptmaa | 0.36399 | 0.221291746 | 0.864 | 0.769 | 1 | 18 |
| ninl | 0.36401 | 0.264766396 | 0.182 | 0.077 | 1 | 18 |
| ptp4a2b | 0.36407 | -0.25930037 | 0.091 | 0.231 | 1 | 18 |
| zgc:91910 | 0.36414 | 0.194207339 | 0.455 | 0.385 | 1 | 18 |
| uxs1 | 0.36425 | 0.226047489 | 0.136 | 0.038 | 1 | 18 |
| ctsa | 0.36429 | 0.25254543 | 0.273 | 0.115 | 1 | 18 |
| hint1 | 0.36446 | -0.23654149 | 0.455 | 0.654 | 1 | 18 |
| serf2 | 0.36466 | -0.2104309 | 0.682 | 0.769 | 1 | 18 |
| timm23a | 0.36512 | 0.227905646 | 0.318 | 0.231 | 1 | 18 |
| rplp0 | 0.3656 | 0.110616575 | 1 | 0.962 | 1 | 18 |
| rab11a | 0.366 | -0.1800478 | 0.182 | 0.346 | 1 | 18 |
| f3b | 0.36607 | 0.298957786 | 0.591 | 0.423 | 1 | 18 |
| eef1db | 0.36668 | -0.10858571 | 0.455 | 0.615 | 1 | 18 |
| tpm1 | 0.36672 | -0.35298765 | 0.273 | 0.462 | 1 | 18 |
| armc1 | 0.36686 | -0.26403006 | 0.045 | 0.154 | 1 | 18 |
| si:ch211-155e24.3 | 0.3669 | -0.2262137 | 0.091 | 0.231 | 1 | 18 |
| si:dkey-102m7.3 | 0.36717 | 0.262827331 | 0.318 | 0.192 | 1 | 18 |
| rpl11 | 0.36734 | 0.103384979 | 1 | 0.923 | 1 | 18 |

| | | | | | | |
|---|---|---|---|---|---|---|
| selenoh | 0.36757 | 0.147163279 | 0.227 | 0.231 | 1 | 18 |
| rabif | 0.36773 | 0.188406217 | 0.136 | 0.077 | 1 | 18 |
| hikeshi | 0.36806 | 0.259933714 | 0.318 | 0.154 | 1 | 18 |
| LEPROTL1 | 0.36837 | 0.165178163 | 0.227 | 0.192 | 1 | 18 |
| ubqln4 | 0.36922 | -0.13249491 | 0.045 | 0.154 | 1 | 18 |
| proza | 0.36942 | 0.243596972 | 0.273 | 0.154 | 1 | 18 |
| LOC110437782 | 0.36987 | 0.255243959 | 0.182 | 0.077 | 1 | 18 |
| hm13 | 0.37 | -0.13961905 | 0.136 | 0.269 | 1 | 18 |
| prdx5 | 0.37022 | 0.126976825 | 0.364 | 0.346 | 1 | 18 |
| pcnp | 0.37067 | 0.24456682 | 0.136 | 0.038 | 1 | 18 |
| sdc4 | 0.37098 | -0.19203432 | 0.5 | 0.692 | 1 | 18 |
| farsa | 0.37149 | 0.190748873 | 0.136 | 0.038 | 1 | 18 |
| si:ch211-39i22.1 | 0.37154 | -0.16607507 | 0.409 | 0.577 | 1 | 18 |
| anxa1a | 0.37164 | -0.1509319 | 0.273 | 0.423 | 1 | 18 |
| tpst1l | 0.37172 | -0.2532838 | 0.045 | 0.154 | 1 | 18 |
| smc1al | 0.37249 | 0.297657437 | 0.364 | 0.231 | 1 | 18 |
| sh3bgrl2 | 0.37295 | 0.28214394 | 0.318 | 0.192 | 1 | 18 |
| bcl2l1 | 0.37377 | -0.18333991 | 0.091 | 0.231 | 1 | 18 |
| npepps | 0.37378 | 0.211861574 | 0.136 | 0.038 | 1 | 18 |
| hmgb2a | 0.37396 | -0.17798505 | 0.591 | 0.769 | 1 | 18 |
| sptssa | 0.37434 | 0.23392305 | 0.182 | 0.115 | 1 | 18 |
| polr2l | 0.3745 | -0.11368448 | 0.273 | 0.423 | 1 | 18 |
| myofl | 0.37481 | -0.1615938 | 0.091 | 0.231 | 1 | 18 |
| h1fx | 0.37493 | -0.25440508 | 0.045 | 0.154 | 1 | 18 |
| cox4i1 | 0.37514 | 0.221567594 | 0.864 | 0.731 | 1 | 18 |
| c7a | 0.37517 | 0.219289003 | 0.136 | 0.038 | 1 | 18 |
| akr1a1b | 0.37533 | 0.216249461 | 0.136 | 0.038 | 1 | 18 |
| zgc:112437 | 0.37549 | -0.24187796 | 0.091 | 0.231 | 1 | 18 |
| fzd2 | 0.3755 | -0.25367067 | 0.273 | 0.462 | 1 | 18 |
| TADA1 | 0.37577 | 0.132588967 | 0.091 | 0.077 | 1 | 18 |
| gbp | 0.37607 | 0.291984102 | 0.409 | 0.231 | 1 | 18 |
| sap30bp | 0.37673 | -0.15669143 | 0.045 | 0.154 | 1 | 18 |
| ndufv1 | 0.37737 | -0.16296322 | 0.091 | 0.231 | 1 | 18 |
| her6 | 0.37773 | 0.144456318 | 0.136 | 0.038 | 1 | 18 |
| eif4eb | 0.37787 | 0.193620234 | 0.136 | 0.038 | 1 | 18 |
| cdkn1bb | 0.37818 | 0.284744613 | 0.318 | 0.154 | 1 | 18 |
| pigf | 0.37883 | -0.12340168 | 0.045 | 0.154 | 1 | 18 |
| stk25b | 0.37914 | -0.21520057 | 0.045 | 0.154 | 1 | 18 |
| smad1 | 0.37929 | 0.249218388 | 0.182 | 0.077 | 1 | 18 |
| mapre1b | 0.37979 | 0.243551772 | 0.273 | 0.115 | 1 | 18 |
| jak1 | 0.37991 | 0.272296228 | 0.273 | 0.115 | 1 | 18 |
| drg2 | 0.37996 | -0.1360746 | 0.045 | 0.154 | 1 | 18 |
| plxnb2a | 0.38003 | 0.255543482 | 0.273 | 0.115 | 1 | 18 |
| mrpl12 | 0.38077 | 0.22542313 | 0.227 | 0.115 | 1 | 18 |
| tardbpl | 0.38123 | -0.17651957 | 0.227 | 0.385 | 1 | 18 |
| clu | 0.38221 | 0.362492086 | 0.136 | 0.038 | 1 | 18 |
| stk17b | 0.38239 | 0.188554087 | 0.136 | 0.038 | 1 | 18 |
| arhgef12a | 0.3829 | 0.224987123 | 0.136 | 0.038 | 1 | 18 |

| | | | | | | |
|---|---|---|---|---|---|---|
| zc3h13 | 0.38299 | 0.135436147 | 0.273 | 0.154 | 1 | 18 |
| mt-nd3 | 0.38334 | -0.15159329 | 0.273 | 0.423 | 1 | 18 |
| dhrs13a.3 | 0.38372 | -0.13698217 | 0.045 | 0.154 | 1 | 18 |
| fam76b | 0.38416 | 0.220702122 | 0.136 | 0.038 | 1 | 18 |
| nup50 | 0.38428 | 0.210584122 | 0.136 | 0.038 | 1 | 18 |
| rsl1d1 | 0.38525 | -0.17015174 | 0.091 | 0.231 | 1 | 18 |
| btbd10b | 0.38557 | 0.147588724 | 0.182 | 0.154 | 1 | 18 |
| hcfc1a | 0.38585 | -0.27359611 | 0.136 | 0.269 | 1 | 18 |
| vdac2 | 0.38606 | -0.25569372 | 0.455 | 0.615 | 1 | 18 |
| cyr61 | 0.38618 | 0.240987815 | 0.409 | 0.231 | 1 | 18 |
| XLOC-011107 | 0.38643 | 0.227974833 | 0.136 | 0.077 | 1 | 18 |
| elmsan1b | 0.38698 | 0.237418287 | 0.136 | 0.038 | 1 | 18 |
| crebzf | 0.38704 | -0.11305234 | 0.182 | 0.308 | 1 | 18 |
| ube2g1a | 0.38737 | -0.23439795 | 0.091 | 0.231 | 1 | 18 |
| rnf7 | 0.38776 | 0.315349827 | 0.364 | 0.192 | 1 | 18 |
| cdc37 | 0.38792 | -0.10336065 | 0.091 | 0.192 | 1 | 18 |
| glud1a | 0.38824 | -0.20487427 | 0.182 | 0.346 | 1 | 18 |
| eif1b | 0.38854 | -0.2317128 | 0.636 | 0.808 | 1 | 18 |
| foxf1 | 0.38878 | 0.229961435 | 0.136 | 0.038 | 1 | 18 |
| fam210aa | 0.38987 | 0.187952137 | 0.136 | 0.038 | 1 | 18 |
| sar1b | 0.38999 | -0.27352223 | 0.318 | 0.462 | 1 | 18 |
| rbms3 | 0.39123 | -0.16446661 | 0.136 | 0.269 | 1 | 18 |
| si:ch211-204c21.1 | 0.3921 | -0.22263513 | 0.182 | 0.346 | 1 | 18 |
| lamtor3 | 0.39374 | -0.19565844 | 0.091 | 0.231 | 1 | 18 |
| antxr1c | 0.39403 | -0.23235214 | 0.091 | 0.231 | 1 | 18 |
| eif5 | 0.39423 | -0.20605381 | 0.273 | 0.462 | 1 | 18 |
| rpp14 | 0.39481 | 0.201233833 | 0.136 | 0.038 | 1 | 18 |
| zgc:101577 | 0.39486 | 0.205708694 | 0.136 | 0.038 | 1 | 18 |
| pggt1b | 0.39499 | 0.195397889 | 0.136 | 0.038 | 1 | 18 |
| porb | 0.39519 | 0.18644435 | 0.136 | 0.038 | 1 | 18 |
| sucla2 | 0.39522 | -0.16897704 | 0.091 | 0.231 | 1 | 18 |
| gnl2 | 0.39541 | -0.13721522 | 0.045 | 0.154 | 1 | 18 |
| uri1 | 0.39577 | -0.12820468 | 0.136 | 0.269 | 1 | 18 |
| fsta | 0.3959 | -0.19169586 | 0.045 | 0.154 | 1 | 18 |
| klf11b | 0.39594 | 0.249542742 | 0.136 | 0.038 | 1 | 18 |
| fundc2 | 0.39614 | -0.21217287 | 0.182 | 0.346 | 1 | 18 |
| morf4l1 | 0.39651 | 0.284873736 | 0.455 | 0.308 | 1 | 18 |
| btd | 0.39655 | -0.21264428 | 0.045 | 0.154 | 1 | 18 |
| epha3 | 0.39692 | 0.290458397 | 0.136 | 0.038 | 1 | 18 |
| canx | 0.39788 | -0.3160544 | 0.227 | 0.385 | 1 | 18 |
| pafah1b1b | 0.3979 | 0.252566349 | 0.318 | 0.154 | 1 | 18 |
| kctd12.2 | 0.39831 | -0.25889327 | 0.273 | 0.462 | 1 | 18 |
| tmem167a | 0.39872 | -0.26880978 | 0.273 | 0.462 | 1 | 18 |
| cdca7a | 0.39882 | 0.134512641 | 0.091 | 0.077 | 1 | 18 |
| FQ323156.1 | 0.39924 | -0.17694824 | 0.091 | 0.231 | 1 | 18 |
| pitpnbl | 0.39937 | -0.21755243 | 0.091 | 0.231 | 1 | 18 |
| snrpd1 | 0.39959 | 0.29008595 | 0.545 | 0.385 | 1 | 18 |
| rnaseh1 | 0.40019 | 0.141633098 | 0.136 | 0.115 | 1 | 18 |

| | | | | | | |
|---|---|---|---|---|---|---|
| cnp | 0.40064 | -0.26530308 | 0.273 | 0.423 | 1 | 18 |
| plgrkt | 0.40074 | -0.12128266 | 0.045 | 0.154 | 1 | 18 |
| tomm22 | 0.40119 | 0.263343265 | 0.273 | 0.154 | 1 | 18 |
| krt4 | 0.40128 | -0.1757809 | 0.045 | 0.154 | 1 | 18 |
| phf14 | 0.40155 | -0.22817214 | 0.045 | 0.115 | 1 | 18 |
| larp4ab | 0.40282 | -0.14645631 | 0.045 | 0.154 | 1 | 18 |
| cbx1b | 0.40306 | -0.25631079 | 0.091 | 0.231 | 1 | 18 |
| CU929237.1 | 0.40308 | -0.11898249 | 0.364 | 0.5 | 1 | 18 |
| skp1 | 0.40333 | 0.16924457 | 0.591 | 0.538 | 1 | 18 |
| cops7a | 0.40356 | 0.126584732 | 0.136 | 0.038 | 1 | 18 |
| adgra3 | 0.40379 | -0.11473068 | 0.091 | 0.192 | 1 | 18 |
| si:ch211-284b7.3 | 0.40401 | -0.16877772 | 0.091 | 0.231 | 1 | 18 |
| tns2a | 0.40413 | -0.24341556 | 0.091 | 0.231 | 1 | 18 |
| tab2 | 0.40438 | -0.21490307 | 0.045 | 0.154 | 1 | 18 |
| dus3l | 0.40443 | -0.15626696 | 0.045 | 0.154 | 1 | 18 |
| cdc5l | 0.40456 | 0.228519065 | 0.227 | 0.115 | 1 | 18 |
| per1b | 0.40482 | -0.24531262 | 0.091 | 0.231 | 1 | 18 |
| pdlim3a | 0.405 | -0.26505198 | 0.091 | 0.231 | 1 | 18 |
| znf609a | 0.40501 | 0.251035696 | 0.227 | 0.115 | 1 | 18 |
| cnot8 | 0.40506 | -0.21185283 | 0.091 | 0.231 | 1 | 18 |
| rap1b | 0.40525 | -0.19155082 | 0.318 | 0.385 | 1 | 18 |
| snrpg | 0.40525 | 0.283939785 | 0.455 | 0.269 | 1 | 18 |
| zgc:152830 | 0.40532 | -0.13405299 | 0.045 | 0.154 | 1 | 18 |
| nelfe | 0.40548 | -0.19234392 | 0.091 | 0.231 | 1 | 18 |
| alg6 | 0.40562 | -0.19994091 | 0.045 | 0.154 | 1 | 18 |
| naa40 | 0.40566 | 0.188764007 | 0.182 | 0.115 | 1 | 18 |
| capza1a | 0.40595 | -0.19949028 | 0.318 | 0.385 | 1 | 18 |
| dpf3 | 0.40599 | 0.197312526 | 0.136 | 0.038 | 1 | 18 |
| gng7 | 0.40772 | -0.26162873 | 0.182 | 0.308 | 1 | 18 |
| srfbp1 | 0.40788 | 0.157222397 | 0.136 | 0.038 | 1 | 18 |
| rpl12 | 0.40793 | 0.1012486 | 1 | 0.962 | 1 | 18 |
| si:dkey-40c11.2 | 0.40819 | -0.11167486 | 0.091 | 0.192 | 1 | 18 |
| ldb1b | 0.4088 | 0.137268388 | 0.136 | 0.038 | 1 | 18 |
| erp44 | 0.40909 | -0.25899659 | 0.182 | 0.346 | 1 | 18 |
| ece1 | 0.40945 | 0.141611073 | 0.591 | 0.577 | 1 | 18 |
| bckdhb | 0.41028 | -0.20490455 | 0.045 | 0.154 | 1 | 18 |
| si:dkey-159f12.2 | 0.41029 | 0.202904578 | 0.136 | 0.038 | 1 | 18 |
| nob1 | 0.41056 | 0.211687585 | 0.136 | 0.038 | 1 | 18 |
| tcf7l1a | 0.41067 | -0.22691925 | 0.045 | 0.154 | 1 | 18 |
| ddx3xb | 0.4109 | -0.19158465 | 0.045 | 0.154 | 1 | 18 |
| rab18b | 0.41184 | 0.194402683 | 0.136 | 0.038 | 1 | 18 |
| ptk7a | 0.41198 | -0.20668027 | 0.091 | 0.231 | 1 | 18 |
| calm3b | 0.41209 | 0.139212972 | 0.682 | 0.692 | 1 | 18 |
| esf1 | 0.41258 | -0.20086451 | 0.091 | 0.231 | 1 | 18 |
| CABZ01034698.2 | 0.41344 | -0.14865119 | 0.045 | 0.154 | 1 | 18 |
| rrp8 | 0.41407 | 0.133219504 | 0.136 | 0.038 | 1 | 18 |
| relt | 0.41491 | -0.23156105 | 0.091 | 0.231 | 1 | 18 |
| cacnb2a | 0.41527 | -0.21335832 | 0.091 | 0.231 | 1 | 18 |

| | | | | | | |
|---|---|---|---|---|---|---|
| nr2f2 | 0.41544 | 0.298949627 | 0.318 | 0.192 | 1 | 18 |
| hars | 0.41583 | -0.2015382 | 0.045 | 0.154 | 1 | 18 |
| si:ch211-108c6.2 | 0.41615 | -0.20366079 | 0.045 | 0.154 | 1 | 18 |
| thoc7 | 0.41698 | -0.18218823 | 0.091 | 0.231 | 1 | 18 |
| si:dkey-238c7.16 | 0.41699 | 0.226201833 | 0.273 | 0.192 | 1 | 18 |
| ap2m1b | 0.417 | -0.18793979 | 0.182 | 0.346 | 1 | 18 |
| dpm2 | 0.41732 | -0.21933165 | 0.091 | 0.231 | 1 | 18 |
| rpp38 | 0.4175 | -0.14108052 | 0.045 | 0.154 | 1 | 18 |
| psenen | 0.41769 | -0.20807967 | 0.091 | 0.231 | 1 | 18 |
| actl6a | 0.41779 | -0.18565374 | 0.091 | 0.231 | 1 | 18 |
| zfr2 | 0.41807 | 0.180607204 | 0.136 | 0.038 | 1 | 18 |
| entr1 | 0.41874 | -0.19717784 | 0.091 | 0.231 | 1 | 18 |
| tuba8l3 | 0.41876 | 0.158098258 | 0.455 | 0.423 | 1 | 18 |
| sh3pxd2b | 0.41877 | -0.22180876 | 0.091 | 0.231 | 1 | 18 |
| impad1 | 0.41911 | 0.205954586 | 0.136 | 0.038 | 1 | 18 |
| mknk2b | 0.41998 | 0.131650142 | 0.227 | 0.115 | 1 | 18 |
| kdelr3 | 0.42014 | -0.14691447 | 0.045 | 0.154 | 1 | 18 |
| dap1b | 0.42078 | -0.19485269 | 0.364 | 0.538 | 1 | 18 |
| phf10 | 0.42082 | 0.203669217 | 0.136 | 0.038 | 1 | 18 |
| rit1 | 0.42114 | -0.20953999 | 0.045 | 0.154 | 1 | 18 |
| gtf2a1l | 0.42141 | -0.10986622 | 0.091 | 0.192 | 1 | 18 |
| si:ch73-340n8.4 | 0.42143 | -0.20519603 | 0.045 | 0.154 | 1 | 18 |
| ppm1bb | 0.42148 | -0.17878361 | 0.045 | 0.154 | 1 | 18 |
| ptgs2b | 0.4219 | -0.20437773 | 0.045 | 0.154 | 1 | 18 |
| sec62 | 0.42206 | -0.20949725 | 0.182 | 0.346 | 1 | 18 |
| hdac1 | 0.42212 | 0.123571925 | 0.545 | 0.538 | 1 | 18 |
| nfya | 0.42229 | -0.2213162 | 0.045 | 0.154 | 1 | 18 |
| tars | 0.42311 | -0.20788747 | 0.045 | 0.154 | 1 | 18 |
| PAXBP1 | 0.42314 | 0.136841345 | 0.136 | 0.115 | 1 | 18 |
| bloc1s6 | 0.42328 | 0.174911603 | 0.136 | 0.038 | 1 | 18 |
| cry1ab | 0.42359 | 0.133489247 | 0.091 | 0.077 | 1 | 18 |
| vcp | 0.42376 | -0.25845324 | 0.182 | 0.346 | 1 | 18 |
| ncor2 | 0.42383 | -0.27702831 | 0.273 | 0.423 | 1 | 18 |
| si:dkey-180p18.9 | 0.42409 | 0.232374396 | 0.227 | 0.115 | 1 | 18 |
| rab10 | 0.4252 | -0.24006906 | 0.409 | 0.538 | 1 | 18 |
| si:ch73-109d9.2 | 0.42527 | -0.21387643 | 0.045 | 0.154 | 1 | 18 |
| ext2 | 0.42528 | -0.21766156 | 0.045 | 0.154 | 1 | 18 |
| igfbp2b | 0.42619 | -0.16727428 | 0.273 | 0.423 | 1 | 18 |
| bnip2 | 0.42633 | 0.188327009 | 0.136 | 0.038 | 1 | 18 |
| ak1 | 0.42683 | 0.23530196 | 0.273 | 0.154 | 1 | 18 |
| nup98 | 0.42711 | 0.149117298 | 0.182 | 0.077 | 1 | 18 |
| tada3l | 0.42764 | -0.11596305 | 0.045 | 0.154 | 1 | 18 |
| zgc:101569 | 0.4278 | 0.182004103 | 0.136 | 0.038 | 1 | 18 |
| ntd5 | 0.42787 | -0.16166648 | 0.045 | 0.154 | 1 | 18 |
| si:ch211-102c2.7 | 0.428 | 0.216174107 | 0.182 | 0.077 | 1 | 18 |
| mt-nd5 | 0.42852 | -0.28940625 | 0.227 | 0.385 | 1 | 18 |
| stxbp3 | 0.42916 | 0.124191448 | 0.136 | 0.038 | 1 | 18 |
| phf5a | 0.42919 | -0.25939108 | 0.273 | 0.385 | 1 | 18 |

| | | | | | | |
|---|---|---|---|---|---|---|
| dhrs1 | 0.42929 | -0.13853809 | 0.045 | 0.154 | 1 | 18 |
| mplkip | 0.42935 | -0.18314346 | 0.045 | 0.154 | 1 | 18 |
| mkrn4 | 0.42975 | 0.180559393 | 0.136 | 0.038 | 1 | 18 |
| slc39a6 | 0.43014 | 0.158556896 | 0.136 | 0.038 | 1 | 18 |
| eif4e2 | 0.43052 | -0.18108248 | 0.045 | 0.154 | 1 | 18 |
| pcbp2 | 0.43094 | -0.24266776 | 0.182 | 0.346 | 1 | 18 |
| ppp1r12a | 0.43117 | -0.13973826 | 0.045 | 0.154 | 1 | 18 |
| mxra8b | 0.43163 | -0.1307478 | 0.318 | 0.462 | 1 | 18 |
| si:ch211-214j24.10 | 0.43206 | -0.20279334 | 0.045 | 0.154 | 1 | 18 |
| wfdc1 | 0.43236 | 0.147703517 | 0.227 | 0.115 | 1 | 18 |
| prpf40a | 0.43248 | 0.192296579 | 0.136 | 0.038 | 1 | 18 |
| b2m | 0.43258 | -0.21890162 | 0.182 | 0.346 | 1 | 18 |
| BX511129.1 | 0.43261 | -0.14647799 | 0.045 | 0.154 | 1 | 18 |
| fut11 | 0.43268 | 0.189711025 | 0.136 | 0.038 | 1 | 18 |
| med10 | 0.43353 | -0.25569396 | 0.227 | 0.385 | 1 | 18 |
| eif3ea | 0.43367 | -0.23910437 | 0.545 | 0.692 | 1 | 18 |
| ciarta | 0.43465 | -0.23573949 | 0.182 | 0.346 | 1 | 18 |
| usf1 | 0.43469 | -0.14917351 | 0.045 | 0.154 | 1 | 18 |
| id1 | 0.43499 | 0.11832098 | 0.909 | 0.923 | 1 | 18 |
| tmed9 | 0.43572 | 0.128857999 | 0.409 | 0.269 | 1 | 18 |
| cops9 | 0.43577 | -0.25092168 | 0.182 | 0.346 | 1 | 18 |
| cog4 | 0.43579 | 0.178177825 | 0.136 | 0.038 | 1 | 18 |
| ephb4a | 0.43588 | -0.18380919 | 0.045 | 0.154 | 1 | 18 |
| bpnt1 | 0.43612 | -0.19778241 | 0.045 | 0.154 | 1 | 18 |
| prkar1aa | 0.43643 | -0.22345559 | 0.045 | 0.154 | 1 | 18 |
| csnk1g2a | 0.43711 | -0.19652683 | 0.045 | 0.154 | 1 | 18 |
| mrpl24 | 0.43717 | -0.14193308 | 0.045 | 0.154 | 1 | 18 |
| ednraa | 0.43719 | 0.192673412 | 0.182 | 0.115 | 1 | 18 |
| fam162a | 0.4375 | 0.172042371 | 0.136 | 0.038 | 1 | 18 |
| map3k7 | 0.43756 | 0.15319092 | 0.136 | 0.038 | 1 | 18 |
| arl6ip5b | 0.43848 | -0.24427165 | 0.318 | 0.5 | 1 | 18 |
| slc25a24 | 0.43864 | -0.21838737 | 0.045 | 0.154 | 1 | 18 |
| fahd2a | 0.4387 | 0.140605016 | 0.136 | 0.038 | 1 | 18 |
| papola | 0.43875 | 0.205339168 | 0.136 | 0.038 | 1 | 18 |
| rrp15 | 0.43923 | -0.19809972 | 0.045 | 0.154 | 1 | 18 |
| foxp4 | 0.43945 | -0.20520434 | 0.727 | 0.846 | 1 | 18 |
| sympk | 0.43971 | 0.159257055 | 0.136 | 0.038 | 1 | 18 |
| f11r.2 | 0.43984 | -0.20257522 | 0.045 | 0.154 | 1 | 18 |
| nfkbie | 0.44013 | -0.15959775 | 0.045 | 0.154 | 1 | 18 |
| ssbp4 | 0.44039 | -0.16333906 | 0.045 | 0.154 | 1 | 18 |
| gnas | 0.44043 | 0.216885276 | 0.136 | 0.038 | 1 | 18 |
| si:ch211-39k3.2 | 0.4405 | 0.138732259 | 0.136 | 0.038 | 1 | 18 |
| pelp1 | 0.44098 | 0.139089894 | 0.136 | 0.038 | 1 | 18 |
| cicb | 0.44146 | -0.18728654 | 0.045 | 0.154 | 1 | 18 |
| pde6d | 0.44169 | 0.226786363 | 0.136 | 0.038 | 1 | 18 |
| paqr3a | 0.44171 | -0.14181252 | 0.045 | 0.154 | 1 | 18 |
| dazap2 | 0.44173 | -0.15415884 | 0.045 | 0.154 | 1 | 18 |
| cox5ab | 0.44179 | -0.22766121 | 0.364 | 0.538 | 1 | 18 |

| | | | | | | |
|---|---|---|---|---|---|---|
| prkcsh | 0.44181 | -0.13843439 | 0.136 | 0.269 | 1 | 18 |
| ghrb | 0.44223 | 0.20884933 | 0.136 | 0.038 | 1 | 18 |
| abce1 | 0.44251 | 0.129789269 | 0.136 | 0.038 | 1 | 18 |
| dnajc1 | 0.44347 | 0.134935314 | 0.136 | 0.038 | 1 | 18 |
| si:ch211-71n6.4 | 0.44404 | -0.19038188 | 0.045 | 0.154 | 1 | 18 |
| fbxw11b | 0.44448 | -0.19914801 | 0.045 | 0.154 | 1 | 18 |
| gnsa | 0.44472 | -0.16460585 | 0.045 | 0.154 | 1 | 18 |
| desi2 | 0.44481 | -0.15960148 | 0.045 | 0.154 | 1 | 18 |
| usp9 | 0.44544 | -0.14129728 | 0.045 | 0.154 | 1 | 18 |
| nars | 0.44565 | 0.234583432 | 0.318 | 0.192 | 1 | 18 |
| qars | 0.44635 | -0.19989654 | 0.045 | 0.154 | 1 | 18 |
| cpvl | 0.44652 | 0.149850862 | 0.136 | 0.038 | 1 | 18 |
| bccip | 0.44656 | -0.1279223 | 0.273 | 0.423 | 1 | 18 |
| u2af2a | 0.44657 | -0.12527423 | 0.045 | 0.154 | 1 | 18 |
| zgc:136864 | 0.44667 | -0.16280258 | 0.045 | 0.154 | 1 | 18 |
| si:ch211-207i20.3 | 0.44669 | -0.14287457 | 0.045 | 0.154 | 1 | 18 |
| acbd3 | 0.44684 | -0.18218009 | 0.045 | 0.154 | 1 | 18 |
| olfml3b | 0.44699 | 0.152796976 | 0.455 | 0.308 | 1 | 18 |
| mlf2 | 0.44703 | -0.10157446 | 0.091 | 0.192 | 1 | 18 |
| porcnl | 0.44711 | -0.18586671 | 0.045 | 0.154 | 1 | 18 |
| si:ch211-173p18.3 | 0.44715 | -0.11638592 | 0.091 | 0.192 | 1 | 18 |
| sdhaf4 | 0.44723 | 0.177216689 | 0.136 | 0.038 | 1 | 18 |
| carhsp1 | 0.44723 | -0.19235089 | 0.045 | 0.154 | 1 | 18 |
| maco1a | 0.44729 | -0.18984401 | 0.045 | 0.154 | 1 | 18 |
| ptpn13 | 0.44737 | -0.2209247 | 0.045 | 0.115 | 1 | 18 |
| pak2a | 0.44788 | -0.1653502 | 0.045 | 0.154 | 1 | 18 |
| krt8 | 0.44834 | -0.22942073 | 0.727 | 0.846 | 1 | 18 |
| si:ch211-140m22.7 | 0.44851 | 0.100138151 | 0.182 | 0.192 | 1 | 18 |
| smg5 | 0.44903 | 0.172110138 | 0.136 | 0.038 | 1 | 18 |
| dicer1 | 0.4491 | -0.17250097 | 0.045 | 0.154 | 1 | 18 |
| ssb | 0.44941 | 0.135206947 | 0.136 | 0.115 | 1 | 18 |
| mcm3 | 0.44952 | 0.168931183 | 0.136 | 0.038 | 1 | 18 |
| pdcd6 | 0.44964 | -0.26361004 | 0.273 | 0.423 | 1 | 18 |
| trappc6b | 0.44974 | -0.17145719 | 0.045 | 0.154 | 1 | 18 |
| abi2b | 0.4498 | -0.16341781 | 0.045 | 0.154 | 1 | 18 |
| cox19 | 0.45013 | -0.18045998 | 0.045 | 0.154 | 1 | 18 |
| ddx19 | 0.45018 | -0.14549057 | 0.045 | 0.154 | 1 | 18 |
| ppp1r13l | 0.45027 | 0.147772646 | 0.182 | 0.077 | 1 | 18 |
| ddx21 | 0.45083 | -0.21437714 | 0.136 | 0.269 | 1 | 18 |
| agpat3 | 0.45111 | -0.17904712 | 0.045 | 0.154 | 1 | 18 |
| rnf2 | 0.45117 | 0.12455791 | 0.182 | 0.154 | 1 | 18 |
| si:dkey-78l4.14 | 0.45138 | 0.122494085 | 0.182 | 0.154 | 1 | 18 |
| mef2aa | 0.45157 | -0.20150458 | 0.045 | 0.154 | 1 | 18 |
| timm17a | 0.45158 | 0.174593773 | 0.136 | 0.038 | 1 | 18 |
| tp53inp1 | 0.45187 | -0.1571952 | 0.045 | 0.154 | 1 | 18 |
| serinc2l | 0.45204 | -0.17814178 | 0.045 | 0.154 | 1 | 18 |
| rfng | 0.45215 | -0.1616123 | 0.045 | 0.154 | 1 | 18 |
| creb1a | 0.45229 | -0.15618517 | 0.045 | 0.154 | 1 | 18 |

| | | | | | | |
|---|---|---|---|---|---|---|
| si:dkey-172m14.1 | 0.45238 | -0.19589061 | 0.045 | 0.154 | 1 | 18 |
| eaf2 | 0.45245 | 0.165763938 | 0.136 | 0.038 | 1 | 18 |
| usp1 | 0.45247 | 0.175371229 | 0.136 | 0.038 | 1 | 18 |
| znf593 | 0.45254 | -0.1278664 | 0.045 | 0.154 | 1 | 18 |
| yes1 | 0.45254 | 0.189867777 | 0.136 | 0.038 | 1 | 18 |
| nenf | 0.45269 | -0.17111674 | 0.045 | 0.154 | 1 | 18 |
| lrpap1 | 0.4527 | -0.14104639 | 0.182 | 0.308 | 1 | 18 |
| timm10 | 0.45272 | -0.16038292 | 0.045 | 0.154 | 1 | 18 |
| mtpn | 0.45307 | -0.24999431 | 0.227 | 0.385 | 1 | 18 |
| ehd2b | 0.45343 | 0.155237234 | 0.182 | 0.154 | 1 | 18 |
| pcid2 | 0.45349 | -0.16616683 | 0.045 | 0.154 | 1 | 18 |
| parp3 | 0.45385 | -0.15031067 | 0.045 | 0.154 | 1 | 18 |
| si:ch211-85n16.3 | 0.45395 | -0.13912367 | 0.045 | 0.154 | 1 | 18 |
| cisd2 | 0.45397 | 0.160121055 | 0.136 | 0.038 | 1 | 18 |
| yy1b | 0.45399 | -0.14591649 | 0.045 | 0.154 | 1 | 18 |
| cmc1 | 0.45404 | -0.18504482 | 0.045 | 0.154 | 1 | 18 |
| atg13 | 0.45416 | -0.18328228 | 0.045 | 0.154 | 1 | 18 |
| apex1 | 0.45424 | -0.14857057 | 0.045 | 0.154 | 1 | 18 |
| pdcd4a | 0.45441 | -0.1471672 | 0.045 | 0.154 | 1 | 18 |
| irf2bp2b | 0.45441 | -0.20989134 | 0.364 | 0.538 | 1 | 18 |
| ppp4r3b | 0.45446 | -0.172435 | 0.045 | 0.154 | 1 | 18 |
| hecw2b | 0.45453 | -0.16664827 | 0.045 | 0.154 | 1 | 18 |
| gtf3c6 | 0.45456 | -0.17258394 | 0.045 | 0.154 | 1 | 18 |
| tmem17 | 0.45458 | -0.16566677 | 0.045 | 0.154 | 1 | 18 |
| parp16 | 0.45464 | -0.16270076 | 0.045 | 0.154 | 1 | 18 |
| rpp25l | 0.45466 | -0.15137848 | 0.045 | 0.154 | 1 | 18 |
| map9 | 0.45468 | -0.15661651 | 0.045 | 0.154 | 1 | 18 |
| elavl1b | 0.45468 | -0.18212022 | 0.045 | 0.154 | 1 | 18 |
| rgl1 | 0.45469 | 0.202328298 | 0.182 | 0.115 | 1 | 18 |
| znf576.2 | 0.45473 | -0.16657691 | 0.045 | 0.154 | 1 | 18 |
| rbpms2a | 0.45473 | -0.17208175 | 0.045 | 0.154 | 1 | 18 |
| hsdl2 | 0.45476 | 0.154169769 | 0.136 | 0.038 | 1 | 18 |
| pgam1a | 0.45478 | -0.12304609 | 0.045 | 0.154 | 1 | 18 |
| pafah1b2 | 0.4548 | -0.21130638 | 0.045 | 0.154 | 1 | 18 |
| mrps31 | 0.4548 | -0.18301766 | 0.045 | 0.154 | 1 | 18 |
| jmjd1cb | 0.45481 | -0.18967554 | 0.045 | 0.154 | 1 | 18 |
| usp7 | 0.45481 | -0.20599189 | 0.045 | 0.154 | 1 | 18 |
| tdg.1 | 0.45481 | -0.16337193 | 0.045 | 0.154 | 1 | 18 |
| angpt2b | 0.45513 | -0.27621386 | 0.227 | 0.346 | 1 | 18 |
| raraa | 0.45514 | -0.10249332 | 0.136 | 0.231 | 1 | 18 |
| tjp1a | 0.4557 | -0.13708085 | 0.091 | 0.192 | 1 | 18 |
| emc10 | 0.45584 | 0.210773091 | 0.227 | 0.115 | 1 | 18 |
| si:dkey-46a10.3 | 0.45651 | 0.132363537 | 0.182 | 0.154 | 1 | 18 |
| pias4a | 0.45681 | 0.193967631 | 0.136 | 0.038 | 1 | 18 |
| scara3 | 0.45689 | 0.132804815 | 0.318 | 0.192 | 1 | 18 |
| trappc2l | 0.45719 | -0.14448761 | 0.136 | 0.269 | 1 | 18 |
| cmtm3 | 0.45741 | 0.15519601 | 0.273 | 0.231 | 1 | 18 |
| rxraa | 0.45751 | 0.118883746 | 0.091 | 0.077 | 1 | 18 |

| | | | | | | |
|---|---|---|---|---|---|---|
| srrt | 0.45765 | 0.215265096 | 0.136 | 0.038 | 1 | 18 |
| glg1a | 0.45814 | 0.18521954 | 0.136 | 0.038 | 1 | 18 |
| vasna | 0.45862 | -0.27522614 | 0.364 | 0.538 | 1 | 18 |
| lbr | 0.45931 | 0.135976124 | 0.136 | 0.038 | 1 | 18 |
| scml2 | 0.46014 | 0.164865494 | 0.136 | 0.038 | 1 | 18 |
| bud31 | 0.46036 | 0.111027622 | 0.364 | 0.346 | 1 | 18 |
| plcg1 | 0.46037 | 0.219436298 | 0.182 | 0.077 | 1 | 18 |
| ftsj3 | 0.46041 | 0.1587694 | 0.136 | 0.038 | 1 | 18 |
| fam171a2a | 0.46048 | 0.20048082 | 0.227 | 0.154 | 1 | 18 |
| drg1 | 0.46055 | 0.169459456 | 0.136 | 0.077 | 1 | 18 |
| med11 | 0.46066 | -0.16521122 | 0.227 | 0.385 | 1 | 18 |
| stk35 | 0.46148 | 0.232750097 | 0.227 | 0.115 | 1 | 18 |
| spc24 | 0.46153 | 0.149651874 | 0.136 | 0.038 | 1 | 18 |
| selenon | 0.46159 | 0.135892179 | 0.182 | 0.154 | 1 | 18 |
| epb41l5 | 0.46208 | 0.185898992 | 0.136 | 0.038 | 1 | 18 |
| traf4a | 0.46217 | 0.183534858 | 0.136 | 0.038 | 1 | 18 |
| ada | 0.46232 | 0.126755009 | 0.091 | 0.038 | 1 | 18 |
| clcn5a | 0.46232 | 0.12164649 | 0.091 | 0.038 | 1 | 18 |
| ctnnd1 | 0.46232 | 0.159074585 | 0.091 | 0.038 | 1 | 18 |
| cyfip1 | 0.46232 | 0.160724728 | 0.091 | 0.038 | 1 | 18 |
| foxo1a | 0.46232 | 0.138419435 | 0.091 | 0.038 | 1 | 18 |
| ift52 | 0.46232 | 0.143656694 | 0.091 | 0.038 | 1 | 18 |
| med7 | 0.46232 | 0.137581733 | 0.091 | 0.038 | 1 | 18 |
| pdcd2 | 0.46232 | 0.164186976 | 0.091 | 0.038 | 1 | 18 |
| phgdh | 0.46232 | 0.105452357 | 0.091 | 0.038 | 1 | 18 |
| ptrhd1 | 0.46232 | 0.105243607 | 0.091 | 0.038 | 1 | 18 |
| sfxn5b | 0.46232 | 0.131868663 | 0.091 | 0.038 | 1 | 18 |
| si:dkey-192l18.9 | 0.46232 | 0.158625035 | 0.091 | 0.038 | 1 | 18 |
| si:dkey-260j18.2 | 0.46232 | 0.105738871 | 0.091 | 0.038 | 1 | 18 |
| wu:fa19b12 | 0.46232 | 0.15590892 | 0.091 | 0.038 | 1 | 18 |
| CABZ01078600.1 | 0.46232 | 0.153970107 | 0.091 | 0.038 | 1 | 18 |
| LOC110438211 | 0.46232 | 0.153439629 | 0.091 | 0.038 | 1 | 18 |
| XLOC-043861 | 0.46232 | 0.128179865 | 0.091 | 0.038 | 1 | 18 |
| arhgap35b | 0.46232 | 0.114939571 | 0.091 | 0.038 | 1 | 18 |
| atp1a1b | 0.46232 | 0.173801881 | 0.091 | 0.038 | 1 | 18 |
| ca5a | 0.46232 | 0.109758311 | 0.091 | 0.038 | 1 | 18 |
| ccdc32 | 0.46232 | 0.153682346 | 0.091 | 0.038 | 1 | 18 |
| cdc73 | 0.46232 | 0.11291035 | 0.091 | 0.038 | 1 | 18 |
| ciao2a | 0.46232 | 0.142371383 | 0.091 | 0.038 | 1 | 18 |
| cks1b | 0.46232 | 0.136931334 | 0.091 | 0.038 | 1 | 18 |
| clockb | 0.46232 | 0.108769108 | 0.091 | 0.038 | 1 | 18 |
| dap3 | 0.46232 | 0.128229403 | 0.091 | 0.038 | 1 | 18 |
| dtwd1 | 0.46232 | 0.151501365 | 0.091 | 0.038 | 1 | 18 |
| dtx4a | 0.46232 | 0.124591513 | 0.091 | 0.038 | 1 | 18 |
| edc3 | 0.46232 | 0.155675994 | 0.091 | 0.038 | 1 | 18 |
| eml3 | 0.46232 | 0.118665354 | 0.091 | 0.038 | 1 | 18 |
| enah | 0.46232 | 0.188775852 | 0.091 | 0.038 | 1 | 18 |
| esr2b | 0.46232 | 0.194583909 | 0.091 | 0.038 | 1 | 18 |

| | | | | | | |
|---|---|---|---|---|---|---|
| fam208b | 0.46232 | 0.116244049 | 0.091 | 0.038 | 1 | 18 |
| fam3c | 0.46232 | 0.124080621 | 0.091 | 0.038 | 1 | 18 |
| fbxw11a | 0.46232 | 0.132971828 | 0.091 | 0.038 | 1 | 18 |
| flr | 0.46232 | 0.109885815 | 0.091 | 0.038 | 1 | 18 |
| fnta | 0.46232 | 0.142562374 | 0.091 | 0.038 | 1 | 18 |
| fundc1 | 0.46232 | 0.111384593 | 0.091 | 0.038 | 1 | 18 |
| gemin7 | 0.46232 | 0.154328897 | 0.091 | 0.038 | 1 | 18 |
| ghra | 0.46232 | 0.129398271 | 0.091 | 0.038 | 1 | 18 |
| gigyf1a | 0.46232 | 0.100966597 | 0.091 | 0.038 | 1 | 18 |
| gng10 | 0.46232 | 0.137612379 | 0.091 | 0.038 | 1 | 18 |
| golga1 | 0.46232 | 0.166432025 | 0.091 | 0.038 | 1 | 18 |
| gstz1 | 0.46232 | 0.120204795 | 0.091 | 0.038 | 1 | 18 |
| harbi1 | 0.46232 | 0.127795015 | 0.091 | 0.038 | 1 | 18 |
| idh3b | 0.46232 | 0.188564608 | 0.091 | 0.038 | 1 | 18 |
| irf9 | 0.46232 | 0.120737432 | 0.091 | 0.038 | 1 | 18 |
| itpk1b | 0.46232 | 0.21811234 | 0.091 | 0.038 | 1 | 18 |
| klhdc3 | 0.46232 | 0.16153733 | 0.091 | 0.038 | 1 | 18 |
| kmt2e | 0.46232 | 0.137802098 | 0.091 | 0.038 | 1 | 18 |
| lrrc3ca | 0.46232 | 0.107650845 | 0.091 | 0.038 | 1 | 18 |
| mapk1 | 0.46232 | 0.187273009 | 0.091 | 0.038 | 1 | 18 |
| mat2ab | 0.46232 | 0.134462211 | 0.091 | 0.038 | 1 | 18 |
| mbd1b | 0.46232 | 0.106949274 | 0.091 | 0.038 | 1 | 18 |
| med1 | 0.46232 | 0.136983246 | 0.091 | 0.038 | 1 | 18 |
| med22 | 0.46232 | 0.121278813 | 0.091 | 0.038 | 1 | 18 |
| mrpl46 | 0.46232 | 0.163794049 | 0.091 | 0.038 | 1 | 18 |
| mrpl53 | 0.46232 | 0.112485691 | 0.091 | 0.038 | 1 | 18 |
| nasp | 0.46232 | 0.170958422 | 0.091 | 0.038 | 1 | 18 |
| ncln | 0.46232 | 0.104382597 | 0.091 | 0.038 | 1 | 18 |
| ndc80 | 0.46232 | 0.130923261 | 0.091 | 0.038 | 1 | 18 |
| nfe2l3 | 0.46232 | 0.107220063 | 0.091 | 0.038 | 1 | 18 |
| nrip1b | 0.46232 | 0.143035912 | 0.091 | 0.038 | 1 | 18 |
| pdhb | 0.46232 | 0.157617592 | 0.091 | 0.038 | 1 | 18 |
| polr3glb | 0.46232 | 0.230334909 | 0.091 | 0.038 | 1 | 18 |
| psmg1 | 0.46232 | 0.127056834 | 0.091 | 0.038 | 1 | 18 |
| rab3gap1 | 0.46232 | 0.106290209 | 0.091 | 0.038 | 1 | 18 |
| rab8b | 0.46232 | 0.113487312 | 0.091 | 0.038 | 1 | 18 |
| rabl2 | 0.46232 | 0.116469054 | 0.091 | 0.038 | 1 | 18 |
| ralbb | 0.46232 | 0.102287978 | 0.091 | 0.038 | 1 | 18 |
| rex1bd | 0.46232 | 0.106167394 | 0.091 | 0.038 | 1 | 18 |
| rfx7b | 0.46232 | 0.144813174 | 0.091 | 0.038 | 1 | 18 |
| rnf40 | 0.46232 | 0.118819628 | 0.091 | 0.038 | 1 | 18 |
| rprd1a | 0.46232 | 0.190637013 | 0.091 | 0.038 | 1 | 18 |
| schip1 | 0.46232 | 0.15286861 | 0.091 | 0.038 | 1 | 18 |
| serpinh1a | 0.46232 | 0.173217188 | 0.091 | 0.038 | 1 | 18 |
| shisa4 | 0.46232 | 0.100521174 | 0.091 | 0.038 | 1 | 18 |
| si:ch211-106k21.5 | 0.46232 | 0.174210461 | 0.091 | 0.038 | 1 | 18 |
| si:ch211-196c10.15 | 0.46232 | 0.157590453 | 0.091 | 0.038 | 1 | 18 |
| si:dkey-21a6.5 | 0.46232 | 0.109734099 | 0.091 | 0.038 | 1 | 18 |

| | | | | | | |
|---|---|---|---|---|---|---|
| slc25a32a | 0.46232 | 0.170536618 | 0.091 | 0.038 | 1 | 18 |
| slc46a1 | 0.46232 | 0.176801769 | 0.091 | 0.038 | 1 | 18 |
| sorl1 | 0.46232 | 0.119719533 | 0.091 | 0.038 | 1 | 18 |
| specc1 | 0.46232 | 0.1375133 | 0.091 | 0.038 | 1 | 18 |
| syt3 | 0.46232 | 0.157910743 | 0.091 | 0.038 | 1 | 18 |
| tcf25 | 0.46232 | 0.179585222 | 0.091 | 0.038 | 1 | 18 |
| tfap4 | 0.46232 | 0.145651491 | 0.091 | 0.038 | 1 | 18 |
| tgfb2 | 0.46232 | 0.156241491 | 0.091 | 0.038 | 1 | 18 |
| tnrc6c2 | 0.46232 | 0.101646135 | 0.091 | 0.038 | 1 | 18 |
| trim71 | 0.46232 | 0.112580911 | 0.091 | 0.038 | 1 | 18 |
| ube4b | 0.46232 | 0.158759761 | 0.091 | 0.038 | 1 | 18 |
| vps36 | 0.46232 | 0.128301698 | 0.091 | 0.038 | 1 | 18 |
| wipi1 | 0.46232 | 0.101465985 | 0.091 | 0.038 | 1 | 18 |
| zbtb17 | 0.46232 | 0.107492917 | 0.091 | 0.038 | 1 | 18 |
| zgc:101566 | 0.46232 | 0.119040411 | 0.091 | 0.038 | 1 | 18 |
| zgc:193812 | 0.46232 | 0.110293513 | 0.091 | 0.038 | 1 | 18 |
| zgc:56585 | 0.46232 | 0.228585168 | 0.091 | 0.038 | 1 | 18 |
| zgc:66448 | 0.46232 | 0.167587182 | 0.091 | 0.038 | 1 | 18 |
| zgc:92907 | 0.46232 | 0.129830903 | 0.091 | 0.038 | 1 | 18 |
| abi1b | 0.46232 | 0.124257612 | 0.091 | 0.038 | 1 | 18 |
| bcl11aa | 0.46232 | 0.136290439 | 0.091 | 0.038 | 1 | 18 |
| bcor | 0.46232 | 0.174537637 | 0.091 | 0.038 | 1 | 18 |
| brd8 | 0.46232 | 0.11260291 | 0.091 | 0.038 | 1 | 18 |
| camk1b | 0.46232 | 0.1210316 | 0.091 | 0.038 | 1 | 18 |
| cdc20 | 0.46232 | 0.257575932 | 0.091 | 0.038 | 1 | 18 |
| cog6 | 0.46232 | 0.109084946 | 0.091 | 0.038 | 1 | 18 |
| crls1 | 0.46232 | 0.103340556 | 0.091 | 0.038 | 1 | 18 |
| ddx54 | 0.46232 | 0.126911671 | 0.091 | 0.038 | 1 | 18 |
| e2f4 | 0.46232 | 0.114159929 | 0.091 | 0.038 | 1 | 18 |
| exoc6 | 0.46232 | 0.112713931 | 0.091 | 0.038 | 1 | 18 |
| gpalpp1 | 0.46232 | 0.104676249 | 0.091 | 0.038 | 1 | 18 |
| im:7147486 | 0.46232 | 0.147370142 | 0.091 | 0.038 | 1 | 18 |
| ing3 | 0.46232 | 0.147914763 | 0.091 | 0.038 | 1 | 18 |
| intu | 0.46232 | 0.145310223 | 0.091 | 0.038 | 1 | 18 |
| kmt2a | 0.46232 | 0.146882236 | 0.091 | 0.038 | 1 | 18 |
| lig3 | 0.46232 | 0.129402414 | 0.091 | 0.038 | 1 | 18 |
| micu1 | 0.46232 | 0.105514473 | 0.091 | 0.038 | 1 | 18 |
| mmadhc | 0.46232 | 0.131732648 | 0.091 | 0.038 | 1 | 18 |
| msl2b | 0.46232 | 0.145183103 | 0.091 | 0.038 | 1 | 18 |
| myo9aa | 0.46232 | 0.108004793 | 0.091 | 0.038 | 1 | 18 |
| psmf1 | 0.46232 | 0.128960493 | 0.091 | 0.038 | 1 | 18 |
| rhogd | 0.46232 | 0.110409139 | 0.091 | 0.038 | 1 | 18 |
| ruvbl2 | 0.46232 | 0.149198191 | 0.091 | 0.038 | 1 | 18 |
| sez6l | 0.46232 | 0.151776236 | 0.091 | 0.038 | 1 | 18 |
| si:dkey-56d12.4 | 0.46232 | 0.105092214 | 0.091 | 0.038 | 1 | 18 |
| si:dkeyp-33b5.4 | 0.46232 | 0.116590856 | 0.091 | 0.038 | 1 | 18 |
| snphb | 0.46232 | 0.127661774 | 0.091 | 0.038 | 1 | 18 |
| stambpa | 0.46232 | 0.12777131 | 0.091 | 0.038 | 1 | 18 |

| | | | | | | |
|---|---|---|---|---|---|---|
| stox2a | 0.46232 | 0.106844211 | 0.091 | 0.038 | 1 | 18 |
| tcerg1a | 0.46232 | 0.121013821 | 0.091 | 0.038 | 1 | 18 |
| tfb2m | 0.46232 | 0.105713681 | 0.091 | 0.038 | 1 | 18 |
| ube2d3 | 0.46232 | 0.13997487 | 0.091 | 0.038 | 1 | 18 |
| usp4 | 0.46232 | 0.100504075 | 0.091 | 0.038 | 1 | 18 |
| arhgap42a | 0.46232 | 0.106487868 | 0.091 | 0.038 | 1 | 18 |
| cxxc5b | 0.46232 | 0.105517288 | 0.091 | 0.038 | 1 | 18 |
| gatad2ab | 0.46232 | 0.111990864 | 0.091 | 0.038 | 1 | 18 |
| smad6a | 0.46232 | 0.119957963 | 0.091 | 0.038 | 1 | 18 |
| XLOC-016094 | 0.46232 | 0.195865033 | 0.091 | 0.038 | 1 | 18 |
| aplnra | 0.46232 | 0.13396476 | 0.091 | 0.038 | 1 | 18 |
| c19h1orf109 | 0.46232 | 0.142769269 | 0.091 | 0.038 | 1 | 18 |
| cmtm4 | 0.46232 | 0.109594839 | 0.091 | 0.038 | 1 | 18 |
| cnppd1 | 0.46232 | 0.12938089 | 0.091 | 0.038 | 1 | 18 |
| eea1 | 0.46232 | 0.125319822 | 0.091 | 0.038 | 1 | 18 |
| gps2 | 0.46232 | 0.127637149 | 0.091 | 0.038 | 1 | 18 |
| hoxb3a | 0.46232 | 0.152096667 | 0.091 | 0.038 | 1 | 18 |
| impa1 | 0.46232 | 0.173779955 | 0.091 | 0.038 | 1 | 18 |
| klf7b | 0.46232 | 0.130100397 | 0.091 | 0.038 | 1 | 18 |
| mff | 0.46232 | 0.127039539 | 0.091 | 0.038 | 1 | 18 |
| mtx2 | 0.46232 | 0.117077822 | 0.091 | 0.038 | 1 | 18 |
| r3hcc1l | 0.46232 | 0.101624576 | 0.091 | 0.038 | 1 | 18 |
| rilpl2 | 0.46232 | 0.108996812 | 0.091 | 0.038 | 1 | 18 |
| si:ch211-271b14.1 | 0.46232 | 0.112855803 | 0.091 | 0.038 | 1 | 18 |
| si:dkey-109l4.3 | 0.46232 | 0.132256458 | 0.091 | 0.038 | 1 | 18 |
| si:dkey-187e18.1 | 0.46232 | 0.129315622 | 0.091 | 0.038 | 1 | 18 |
| stk40 | 0.46232 | 0.132761563 | 0.091 | 0.038 | 1 | 18 |
| fap | 0.46232 | 0.105201707 | 0.091 | 0.038 | 1 | 18 |
| ppp4r2b | 0.46232 | 0.118421194 | 0.091 | 0.038 | 1 | 18 |
| slkb | 0.46232 | 0.142756379 | 0.091 | 0.038 | 1 | 18 |
| zfpl1 | 0.46232 | 0.123579295 | 0.091 | 0.038 | 1 | 18 |
| shc1 | 0.46234 | 0.159800013 | 0.136 | 0.038 | 1 | 18 |
| pnrc2 | 0.4629 | -0.14759941 | 0.864 | 0.923 | 1 | 18 |
| dpt | 0.46314 | 0.242296683 | 0.136 | 0.038 | 1 | 18 |
| wdr36 | 0.46358 | 0.193638125 | 0.136 | 0.038 | 1 | 18 |
| mcmbp | 0.46431 | 0.157777506 | 0.136 | 0.038 | 1 | 18 |
| tmpoa | 0.46452 | 0.175831208 | 0.136 | 0.038 | 1 | 18 |
| sema3bl | 0.46501 | 0.231614263 | 0.136 | 0.038 | 1 | 18 |
| psmg2 | 0.46636 | 0.156109511 | 0.136 | 0.038 | 1 | 18 |
| aif1l | 0.46651 | 0.165982681 | 0.136 | 0.038 | 1 | 18 |
| ddx39ab | 0.46676 | -0.24205898 | 0.273 | 0.423 | 1 | 18 |
| dnajb6a | 0.46694 | 0.129103295 | 0.182 | 0.077 | 1 | 18 |
| cbx5 | 0.46696 | -0.13493009 | 0.364 | 0.5 | 1 | 18 |
| rpa2 | 0.46742 | 0.208855017 | 0.136 | 0.038 | 1 | 18 |
| rbm25b | 0.46744 | -0.13333806 | 0.182 | 0.308 | 1 | 18 |
| mta2 | 0.4675 | 0.16107202 | 0.136 | 0.038 | 1 | 18 |
| gna11b | 0.46824 | 0.215871404 | 0.182 | 0.077 | 1 | 18 |
| irf2a | 0.46891 | 0.19632512 | 0.136 | 0.038 | 1 | 18 |

| | | | | | | |
|---|---|---|---|---|---|---|
| kremen1 | 0.46895 | 0.169432839 | 0.136 | 0.038 | 1 | 18 |
| plpp5 | 0.46929 | -0.22309829 | 0.136 | 0.269 | 1 | 18 |
| surf4l | 0.4696 | -0.21279368 | 0.136 | 0.269 | 1 | 18 |
| tcp11l2 | 0.46971 | 0.148722802 | 0.136 | 0.038 | 1 | 18 |
| pds5b | 0.46985 | 0.12603142 | 0.091 | 0.077 | 1 | 18 |
| polr2c | 0.46988 | 0.200002262 | 0.182 | 0.115 | 1 | 18 |
| cdk9 | 0.46989 | 0.211476394 | 0.182 | 0.077 | 1 | 18 |
| galnt11 | 0.47031 | 0.171323927 | 0.136 | 0.038 | 1 | 18 |
| anapc10 | 0.4704 | 0.155492406 | 0.136 | 0.038 | 1 | 18 |
| marcksl1b | 0.47057 | 0.246726883 | 0.545 | 0.462 | 1 | 18 |
| map1sa | 0.47137 | 0.166340431 | 0.136 | 0.038 | 1 | 18 |
| fen1 | 0.47234 | 0.202774488 | 0.136 | 0.038 | 1 | 18 |
| keap1b | 0.4724 | 0.138453429 | 0.136 | 0.038 | 1 | 18 |
| ddx42 | 0.47248 | 0.206339505 | 0.227 | 0.115 | 1 | 18 |
| picalma | 0.47249 | 0.195420494 | 0.136 | 0.038 | 1 | 18 |
| clpxa | 0.4727 | 0.189381207 | 0.136 | 0.038 | 1 | 18 |
| atp6ap2 | 0.47279 | 0.144636325 | 0.136 | 0.038 | 1 | 18 |
| sem1 | 0.47299 | -0.16409845 | 0.591 | 0.654 | 1 | 18 |
| mrpl18 | 0.47359 | 0.139093179 | 0.136 | 0.038 | 1 | 18 |
| tspan4b | 0.47378 | -0.23317436 | 0.364 | 0.538 | 1 | 18 |
| stat6 | 0.47384 | 0.151616496 | 0.136 | 0.038 | 1 | 18 |
| loxl1 | 0.47386 | -0.11587388 | 0.5 | 0.5 | 1 | 18 |
| CABZ01039096.1 | 0.47409 | -0.23301402 | 0.364 | 0.538 | 1 | 18 |
| zgc:77929 | 0.4741 | 0.15197236 | 0.136 | 0.038 | 1 | 18 |
| ryk | 0.47435 | -0.20874956 | 0.227 | 0.385 | 1 | 18 |
| slc30a6 | 0.47447 | 0.204282878 | 0.136 | 0.038 | 1 | 18 |
| oser1 | 0.47488 | 0.194757059 | 0.136 | 0.038 | 1 | 18 |
| nfil3 | 0.47498 | 0.13285594 | 0.136 | 0.038 | 1 | 18 |
| cdc14ab | 0.47513 | 0.159766903 | 0.136 | 0.038 | 1 | 18 |
| cplx2l | 0.47515 | 0.208215143 | 0.136 | 0.038 | 1 | 18 |
| hs6st2 | 0.47516 | 0.179838178 | 0.136 | 0.038 | 1 | 18 |
| zgc:92744 | 0.47526 | 0.21269155 | 0.136 | 0.038 | 1 | 18 |
| lfng | 0.47533 | 0.229519573 | 0.182 | 0.077 | 1 | 18 |
| si:ch73-1a9.3 | 0.47556 | 0.171992893 | 0.909 | 0.846 | 1 | 18 |
| med28 | 0.47572 | 0.200283686 | 0.182 | 0.115 | 1 | 18 |
| cpn1 | 0.47702 | -0.22497742 | 0.227 | 0.385 | 1 | 18 |
| rbm4.3 | 0.47815 | 0.257447395 | 0.591 | 0.423 | 1 | 18 |
| mcm7 | 0.47893 | 0.120677494 | 0.136 | 0.115 | 1 | 18 |
| pfn2 | 0.47894 | -0.13801735 | 0.682 | 0.731 | 1 | 18 |
| ppm1la | 0.47975 | 0.185617632 | 0.182 | 0.115 | 1 | 18 |
| serp1 | 0.47997 | 0.117416207 | 0.364 | 0.231 | 1 | 18 |
| chchd2 | 0.48026 | 0.132737896 | 0.182 | 0.077 | 1 | 18 |
| anp32e | 0.48142 | -0.12166709 | 0.182 | 0.308 | 1 | 18 |
| rab11ba | 0.48198 | -0.24025515 | 0.227 | 0.346 | 1 | 18 |
| hmgb2b | 0.48284 | 0.133472665 | 0.682 | 0.692 | 1 | 18 |
| ube2ia | 0.48289 | 0.151341343 | 0.409 | 0.346 | 1 | 18 |
| rbm27 | 0.48307 | 0.151178523 | 0.182 | 0.077 | 1 | 18 |
| cul3b | 0.48386 | -0.21313 | 0.136 | 0.269 | 1 | 18 |

| | | | | | | |
|---|---|---|---|---|---|---|
| ephb3 | 0.48386 | -0.26109013 | 0.273 | 0.423 | 1 | 18 |
| cd276 | 0.4843 | -0.14656363 | 0.182 | 0.308 | 1 | 18 |
| anxa2a | 0.48491 | -0.23478238 | 0.409 | 0.462 | 1 | 18 |
| cops2 | 0.48619 | -0.10727388 | 0.091 | 0.192 | 1 | 18 |
| tusc2b | 0.48734 | -0.1075769 | 0.091 | 0.192 | 1 | 18 |
| nfixb | 0.48771 | -0.11236715 | 0.909 | 0.923 | 1 | 18 |
| adrm1 | 0.48795 | -0.16443407 | 0.136 | 0.269 | 1 | 18 |
| vdac3 | 0.48838 | -0.23529226 | 0.409 | 0.577 | 1 | 18 |
| bnc2 | 0.48857 | 0.228885812 | 0.318 | 0.231 | 1 | 18 |
| itga9 | 0.48908 | 0.245165035 | 0.182 | 0.077 | 1 | 18 |
| gpatch4 | 0.48971 | 0.126110641 | 0.136 | 0.115 | 1 | 18 |
| slc7a3a | 0.49068 | 0.201380987 | 0.182 | 0.115 | 1 | 18 |
| g3bp1 | 0.49077 | 0.178276218 | 0.273 | 0.192 | 1 | 18 |
| hmgb1a | 0.49094 | 0.213267791 | 0.818 | 0.692 | 1 | 18 |
| chac1 | 0.4911 | -0.11690673 | 0.227 | 0.385 | 1 | 18 |
| napab | 0.49224 | 0.235191824 | 0.273 | 0.154 | 1 | 18 |
| bmper | 0.49289 | -0.24616665 | 0.409 | 0.538 | 1 | 18 |
| btg2 | 0.49295 | 0.189947341 | 0.591 | 0.423 | 1 | 18 |
| gnl1 | 0.49337 | -0.2364514 | 0.091 | 0.192 | 1 | 18 |
| txndc5 | 0.49377 | -0.26899429 | 0.227 | 0.385 | 1 | 18 |
| lix1l | 0.49432 | 0.224587585 | 0.182 | 0.077 | 1 | 18 |
| rcan2 | 0.49546 | -0.22942112 | 0.227 | 0.385 | 1 | 18 |
| calub | 0.4955 | -0.20273257 | 0.136 | 0.231 | 1 | 18 |
| postna | 0.49658 | 0.116487809 | 0.091 | 0.077 | 1 | 18 |
| hnrpkl | 0.49738 | -0.15186583 | 0.182 | 0.308 | 1 | 18 |
| prpf18 | 0.49882 | -0.12279949 | 0.091 | 0.192 | 1 | 18 |
| srp54 | 0.49907 | 0.189734444 | 0.182 | 0.115 | 1 | 18 |
| s100a10b | 0.49919 | -0.16861597 | 0.864 | 0.923 | 1 | 18 |
| glmp | 0.49941 | 0.127697749 | 0.182 | 0.077 | 1 | 18 |
| hoxb5b | 0.49987 | 0.185062955 | 0.5 | 0.346 | 1 | 18 |
| actn1 | 0.50011 | -0.1060144 | 0.136 | 0.231 | 1 | 18 |
| arhgdig | 0.5016 | -0.13735126 | 0.318 | 0.462 | 1 | 18 |
| snx6 | 0.50167 | 0.164810367 | 0.182 | 0.115 | 1 | 18 |
| tuba1a | 0.50241 | -0.2948783 | 0.182 | 0.308 | 1 | 18 |
| ndufb9 | 0.50304 | -0.18081703 | 0.545 | 0.615 | 1 | 18 |
| taf7 | 0.50307 | 0.189454975 | 0.273 | 0.192 | 1 | 18 |
| kpnb3 | 0.5033 | -0.19735925 | 0.091 | 0.154 | 1 | 18 |
| ndufab1a | 0.50407 | 0.169591995 | 0.409 | 0.346 | 1 | 18 |
| mdh2 | 0.50436 | 0.165558247 | 0.545 | 0.385 | 1 | 18 |
| phip | 0.5044 | 0.193481212 | 0.182 | 0.077 | 1 | 18 |
| tfdp2 | 0.50462 | 0.173770557 | 0.182 | 0.077 | 1 | 18 |
| psmb2 | 0.50533 | -0.22170414 | 0.318 | 0.462 | 1 | 18 |
| ccnd2a | 0.50576 | -0.23195716 | 0.227 | 0.346 | 1 | 18 |
| oaz1a | 0.50752 | 0.132153418 | 0.864 | 0.731 | 1 | 18 |
| arl4d | 0.50759 | 0.189211921 | 0.182 | 0.077 | 1 | 18 |
| cd3eap | 0.50795 | -0.17389189 | 0.136 | 0.269 | 1 | 18 |
| ssr4 | 0.50863 | -0.18260611 | 0.773 | 0.846 | 1 | 18 |
| yaf2 | 0.5087 | -0.17703475 | 0.136 | 0.269 | 1 | 18 |

| | | | | | | |
|---|---|---|---|---|---|---|
| nfia | 0.50943 | -0.19152727 | 0.591 | 0.692 | 1 | 18 |
| cenpx | 0.51058 | -0.1745368 | 0.045 | 0.115 | 1 | 18 |
| dohh | 0.51059 | -0.10032209 | 0.091 | 0.192 | 1 | 18 |
| ip6k2b | 0.51068 | 0.16341745 | 0.273 | 0.231 | 1 | 18 |
| lmbr1l | 0.51071 | 0.171443635 | 0.136 | 0.077 | 1 | 18 |
| rexo4 | 0.51077 | 0.192048674 | 0.182 | 0.077 | 1 | 18 |
| cops8 | 0.51185 | -0.11223436 | 0.091 | 0.192 | 1 | 18 |
| kdm7ab | 0.51289 | 0.159764935 | 0.318 | 0.192 | 1 | 18 |
| erlec1 | 0.51308 | -0.17943459 | 0.227 | 0.308 | 1 | 18 |
| azin1b | 0.51334 | 0.175119751 | 0.182 | 0.115 | 1 | 18 |
| grb10a | 0.51366 | 0.226789184 | 0.273 | 0.154 | 1 | 18 |
| ptpa | 0.51368 | 0.191725196 | 0.136 | 0.077 | 1 | 18 |
| cpne3 | 0.51413 | -0.21372831 | 0.136 | 0.269 | 1 | 18 |
| papolg | 0.51425 | -0.14636235 | 0.091 | 0.192 | 1 | 18 |
| p4ha1b | 0.51473 | 0.196978461 | 0.182 | 0.077 | 1 | 18 |
| gdf10a | 0.51476 | 0.2271014 | 0.136 | 0.077 | 1 | 18 |
| sec61g | 0.51481 | 0.136801628 | 0.909 | 0.885 | 1 | 18 |
| loxa | 0.51592 | -0.25637045 | 0.364 | 0.5 | 1 | 18 |
| ddx39aa | 0.5161 | -0.11579721 | 0.364 | 0.5 | 1 | 18 |
| xpo4 | 0.51665 | 0.17430867 | 0.136 | 0.077 | 1 | 18 |
| wtap | 0.51675 | -0.21095915 | 0.136 | 0.231 | 1 | 18 |
| ccnc | 0.51721 | 0.160405326 | 0.182 | 0.077 | 1 | 18 |
| kpnb1 | 0.51725 | -0.20910755 | 0.136 | 0.269 | 1 | 18 |
| smim4 | 0.51773 | -0.19163364 | 0.091 | 0.192 | 1 | 18 |
| frs2b | 0.51783 | -0.21295974 | 0.136 | 0.269 | 1 | 18 |
| nr2f1a | 0.51811 | -0.12387682 | 0.409 | 0.538 | 1 | 18 |
| arl1 | 0.51848 | 0.214125175 | 0.318 | 0.192 | 1 | 18 |
| hectd1 | 0.51893 | 0.229843449 | 0.227 | 0.115 | 1 | 18 |
| cldn11a | 0.51913 | 0.229914254 | 0.636 | 0.538 | 1 | 18 |
| lnx1 | 0.51975 | -0.19509335 | 0.045 | 0.115 | 1 | 18 |
| bace2 | 0.52023 | -0.18490045 | 0.136 | 0.269 | 1 | 18 |
| COA4 | 0.52071 | -0.12505391 | 0.091 | 0.192 | 1 | 18 |
| bloc1s1 | 0.52089 | 0.122022381 | 0.091 | 0.077 | 1 | 18 |
| copb2 | 0.52105 | -0.18332251 | 0.136 | 0.269 | 1 | 18 |
| chd6 | 0.52108 | -0.21942166 | 0.136 | 0.269 | 1 | 18 |
| ier5l | 0.52124 | -0.22310591 | 0.136 | 0.269 | 1 | 18 |
| chd9 | 0.52151 | 0.11841867 | 0.091 | 0.077 | 1 | 18 |
| thoc5 | 0.52164 | 0.164694545 | 0.182 | 0.115 | 1 | 18 |
| mrpl32 | 0.5219 | 0.102204234 | 0.273 | 0.269 | 1 | 18 |
| XLOC-019096 | 0.52225 | -0.17382387 | 0.045 | 0.115 | 1 | 18 |
| cstf1 | 0.52273 | 0.108533639 | 0.091 | 0.077 | 1 | 18 |
| dnaja1 | 0.52285 | -0.16681074 | 0.045 | 0.115 | 1 | 18 |
| elocb | 0.52382 | 0.184730472 | 0.5 | 0.423 | 1 | 18 |
| pmaip1 | 0.52385 | 0.117634186 | 0.091 | 0.077 | 1 | 18 |
| twist2 | 0.52431 | 0.144580131 | 0.182 | 0.077 | 1 | 18 |
| atf5b | 0.52437 | -0.22196599 | 0.091 | 0.192 | 1 | 18 |
| cep290 | 0.52439 | 0.198575274 | 0.182 | 0.077 | 1 | 18 |
| mettl24 | 0.52458 | -0.10648796 | 0.045 | 0.115 | 1 | 18 |

| | | | | | | |
|---|---|---|---|---|---|---|
| glipr2l | 0.52461 | 0.192276043 | 0.364 | 0.308 | 1 | 18 |
| znf704 | 0.52547 | 0.200393401 | 0.227 | 0.115 | 1 | 18 |
| anp32a | 0.52615 | -0.10701361 | 0.409 | 0.538 | 1 | 18 |
| arl8ba | 0.52653 | -0.11410684 | 0.091 | 0.192 | 1 | 18 |
| yeats4 | 0.52666 | 0.201794821 | 0.273 | 0.192 | 1 | 18 |
| ppp6r3 | 0.52732 | 0.18331135 | 0.182 | 0.115 | 1 | 18 |
| map2k6 | 0.52843 | 0.223795123 | 0.273 | 0.154 | 1 | 18 |
| coa3a | 0.52998 | 0.151024423 | 0.227 | 0.192 | 1 | 18 |
| nifk | 0.53016 | 0.200989743 | 0.182 | 0.115 | 1 | 18 |
| ankrd12 | 0.53088 | -0.21222662 | 0.136 | 0.231 | 1 | 18 |
| itgav | 0.53107 | -0.13171885 | 0.091 | 0.192 | 1 | 18 |
| psmc5 | 0.53158 | -0.16837323 | 0.273 | 0.423 | 1 | 18 |
| fbxw4 | 0.53173 | 0.112663997 | 0.091 | 0.077 | 1 | 18 |
| ubxn7 | 0.5318 | 0.131967884 | 0.182 | 0.077 | 1 | 18 |
| mbd3b | 0.53254 | -0.12021051 | 0.227 | 0.346 | 1 | 18 |
| caap1 | 0.53266 | 0.186687708 | 0.182 | 0.077 | 1 | 18 |
| atp6v0ca | 0.53332 | -0.2326988 | 0.182 | 0.269 | 1 | 18 |
| gas1a | 0.53366 | -0.10838748 | 0.455 | 0.577 | 1 | 18 |
| si:ch73-37h15.2 | 0.53377 | 0.143468466 | 0.136 | 0.115 | 1 | 18 |
| srsf11 | 0.53625 | 0.204008782 | 0.409 | 0.308 | 1 | 18 |
| ahi1 | 0.53626 | -0.1228777 | 0.091 | 0.192 | 1 | 18 |
| socs3a | 0.5364 | 0.216085925 | 0.818 | 0.692 | 1 | 18 |
| CU929230.1 | 0.53653 | -0.11021199 | 0.045 | 0.115 | 1 | 18 |
| irf2bpl | 0.53727 | 0.168868228 | 0.227 | 0.192 | 1 | 18 |
| kdm6ba | 0.53733 | -0.19796359 | 0.045 | 0.115 | 1 | 18 |
| boc | 0.53758 | 0.178715762 | 0.727 | 0.654 | 1 | 18 |
| mex3b | 0.53835 | 0.118880679 | 0.682 | 0.654 | 1 | 18 |
| nsa2 | 0.53892 | -0.19208071 | 0.682 | 0.808 | 1 | 18 |
| alpl | 0.53914 | 0.110484976 | 0.091 | 0.077 | 1 | 18 |
| CU138547.1 | 0.54012 | 0.112425172 | 0.091 | 0.077 | 1 | 18 |
| mosmob | 0.54047 | 0.11170687 | 0.091 | 0.077 | 1 | 18 |
| eif2s1a | 0.5405 | 0.165445708 | 0.182 | 0.115 | 1 | 18 |
| sdf4 | 0.54069 | -0.18412564 | 0.091 | 0.192 | 1 | 18 |
| gatad2b | 0.54133 | -0.17217254 | 0.318 | 0.462 | 1 | 18 |
| ctdsp2 | 0.54174 | -0.19663191 | 0.273 | 0.423 | 1 | 18 |
| rnd1b | 0.54258 | 0.172883596 | 0.136 | 0.077 | 1 | 18 |
| flrt3 | 0.54281 | -0.14212736 | 0.091 | 0.192 | 1 | 18 |
| calr3a | 0.54282 | 0.1328223 | 0.455 | 0.423 | 1 | 18 |
| ormdl2 | 0.54341 | 0.110715293 | 0.091 | 0.077 | 1 | 18 |
| rcan1a | 0.54345 | 0.123726616 | 0.091 | 0.077 | 1 | 18 |
| ss18l2 | 0.54347 | 0.159642432 | 0.318 | 0.269 | 1 | 18 |
| flt4 | 0.54358 | -0.18878577 | 0.045 | 0.115 | 1 | 18 |
| sae1 | 0.54414 | 0.136315511 | 0.273 | 0.154 | 1 | 18 |
| FAT4 | 0.5442 | 0.172066241 | 0.227 | 0.115 | 1 | 18 |
| bmp1a | 0.5443 | 0.120605935 | 0.136 | 0.115 | 1 | 18 |
| rergla | 0.54445 | 0.275200526 | 0.227 | 0.115 | 1 | 18 |
| psmd1 | 0.54463 | 0.165577314 | 0.318 | 0.192 | 1 | 18 |
| rnf220a | 0.54466 | 0.215402795 | 0.227 | 0.115 | 1 | 18 |

| | | | | | | |
|---|---|---|---|---|---|---|
| mrpl4 | 0.54488 | 0.176574048 | 0.182 | 0.077 | 1 | 18 |
| celf2 | 0.5461 | -0.11863794 | 0.409 | 0.538 | 1 | 18 |
| mxra8a | 0.54634 | -0.23992492 | 0.318 | 0.462 | 1 | 18 |
| psmb7 | 0.5471 | 0.198129428 | 0.636 | 0.538 | 1 | 18 |
| sms | 0.54723 | 0.14912346 | 0.182 | 0.077 | 1 | 18 |
| rsu1 | 0.5476 | 0.166298947 | 0.182 | 0.077 | 1 | 18 |
| tomm40 | 0.5482 | 0.170701508 | 0.182 | 0.077 | 1 | 18 |
| mcl1a | 0.54823 | 0.214198604 | 0.273 | 0.192 | 1 | 18 |
| srsf4 | 0.54873 | -0.11398459 | 0.091 | 0.192 | 1 | 18 |
| mtmr14 | 0.54893 | 0.168097654 | 0.136 | 0.077 | 1 | 18 |
| slc7a6os | 0.5491 | 0.191960695 | 0.182 | 0.077 | 1 | 18 |
| gpm6bb | 0.54921 | 0.192890371 | 0.182 | 0.077 | 1 | 18 |
| phf8 | 0.54938 | 0.163521379 | 0.182 | 0.077 | 1 | 18 |
| commd7 | 0.54942 | 0.169405707 | 0.182 | 0.077 | 1 | 18 |
| ndufa4l | 0.54992 | -0.1874295 | 0.727 | 0.808 | 1 | 18 |
| cdipt | 0.54994 | 0.152766571 | 0.182 | 0.077 | 1 | 18 |
| fgf14 | 0.55008 | 0.186766745 | 0.182 | 0.077 | 1 | 18 |
| tmem254 | 0.55017 | 0.1578831 | 0.182 | 0.077 | 1 | 18 |
| sra1 | 0.55072 | 0.150707958 | 0.182 | 0.077 | 1 | 18 |
| znf703 | 0.55091 | 0.167966993 | 0.182 | 0.077 | 1 | 18 |
| cherp | 0.55106 | 0.212290512 | 0.182 | 0.077 | 1 | 18 |
| uck2b | 0.55108 | 0.167686404 | 0.182 | 0.077 | 1 | 18 |
| adh5 | 0.55117 | -0.19492197 | 0.545 | 0.654 | 1 | 18 |
| tmem214 | 0.5515 | 0.167479765 | 0.182 | 0.077 | 1 | 18 |
| sesn3 | 0.55172 | 0.187723501 | 0.182 | 0.077 | 1 | 18 |
| XLOC-042917 | 0.55173 | 0.206110784 | 0.182 | 0.077 | 1 | 18 |
| lrrc20 | 0.55173 | 0.160689571 | 0.182 | 0.077 | 1 | 18 |
| aoc2 | 0.55177 | -0.14955176 | 0.045 | 0.115 | 1 | 18 |
| decr1 | 0.55184 | 0.153500344 | 0.227 | 0.115 | 1 | 18 |
| hsbp1b | 0.55271 | -0.15035786 | 0.318 | 0.462 | 1 | 18 |
| si:ch73-390p7.2 | 0.55276 | 0.179966444 | 0.136 | 0.077 | 1 | 18 |
| tmem176l.1 | 0.55283 | -0.18550541 | 0.5 | 0.654 | 1 | 18 |
| anxa5b | 0.55289 | -0.15562817 | 0.318 | 0.462 | 1 | 18 |
| ddah1 | 0.55296 | 0.194632523 | 0.136 | 0.077 | 1 | 18 |
| ss18 | 0.55467 | 0.220109453 | 0.227 | 0.115 | 1 | 18 |
| sgcb | 0.55516 | -0.16201069 | 0.045 | 0.115 | 1 | 18 |
| mta1 | 0.55555 | -0.21759918 | 0.091 | 0.192 | 1 | 18 |
| eny2 | 0.55578 | 0.209655598 | 0.318 | 0.192 | 1 | 18 |
| tmem165 | 0.55601 | 0.191130565 | 0.227 | 0.115 | 1 | 18 |
| c1qtnf2 | 0.55656 | 0.212453126 | 0.227 | 0.115 | 1 | 18 |
| svild | 0.55689 | -0.23063719 | 0.136 | 0.231 | 1 | 18 |
| nell2a | 0.5569 | -0.25390701 | 0.091 | 0.192 | 1 | 18 |
| pdcd10a | 0.55701 | 0.168617142 | 0.136 | 0.077 | 1 | 18 |
| FO082781.1 | 0.55721 | -0.16447749 | 0.091 | 0.192 | 1 | 18 |
| krt94 | 0.5576 | -0.20947169 | 0.682 | 0.808 | 1 | 18 |
| hdgfl2 | 0.55769 | 0.192130268 | 0.227 | 0.115 | 1 | 18 |
| ccdc25 | 0.55886 | -0.1344858 | 0.091 | 0.192 | 1 | 18 |
| stt3b | 0.55898 | -0.16485451 | 0.091 | 0.154 | 1 | 18 |

| | | | | | | |
|---|---|---|---|---|---|---|
| thbs2b | 0.55904 | -0.24946466 | 0.091 | 0.192 | 1 | 18 |
| mtdhb | 0.56001 | -0.11537999 | 0.091 | 0.192 | 1 | 18 |
| eif1axa | 0.56213 | -0.18848806 | 0.091 | 0.192 | 1 | 18 |
| emc7 | 0.56232 | -0.17758604 | 0.091 | 0.192 | 1 | 18 |
| mmp16a | 0.5631 | 0.193352909 | 0.682 | 0.577 | 1 | 18 |
| nat9 | 0.56324 | 0.167621132 | 0.182 | 0.115 | 1 | 18 |
| cyb5a | 0.56324 | 0.20384712 | 0.273 | 0.192 | 1 | 18 |
| tmem263 | 0.56431 | -0.11792873 | 0.227 | 0.346 | 1 | 18 |
| acbd6 | 0.56443 | -0.17592876 | 0.045 | 0.115 | 1 | 18 |
| pik3r3b | 0.56464 | 0.230853883 | 0.227 | 0.115 | 1 | 18 |
| tmem218 | 0.5648 | -0.15404442 | 0.045 | 0.115 | 1 | 18 |
| naa50 | 0.5655 | 0.14858546 | 0.227 | 0.115 | 1 | 18 |
| ndufv2 | 0.56666 | -0.13033995 | 0.182 | 0.308 | 1 | 18 |
| ubap2l | 0.56711 | -0.11994952 | 0.136 | 0.231 | 1 | 18 |
| cdon | 0.56745 | -0.14785247 | 0.364 | 0.5 | 1 | 18 |
| si:ch211-133n4.4 | 0.56794 | -0.21424613 | 0.364 | 0.462 | 1 | 18 |
| FP102018.1 | 0.56856 | -0.21720192 | 0.182 | 0.308 | 1 | 18 |
| ap2s1 | 0.5701 | -0.2080754 | 0.182 | 0.308 | 1 | 18 |
| csnk2a1 | 0.57018 | 0.106941434 | 0.318 | 0.308 | 1 | 18 |
| h3f3a | 0.57078 | 0.188917572 | 0.682 | 0.615 | 1 | 18 |
| coq3 | 0.57089 | 0.165228941 | 0.182 | 0.115 | 1 | 18 |
| cdc14b | 0.57129 | 0.187177124 | 0.273 | 0.192 | 1 | 18 |
| tax1bp3 | 0.57182 | -0.17814972 | 0.318 | 0.462 | 1 | 18 |
| scp2b | 0.57196 | 0.122723059 | 0.136 | 0.115 | 1 | 18 |
| tmem147 | 0.57233 | -0.19215817 | 0.136 | 0.231 | 1 | 18 |
| zgc:112023 | 0.57239 | -0.13670864 | 0.091 | 0.192 | 1 | 18 |
| mrpl37 | 0.57251 | -0.16923175 | 0.045 | 0.115 | 1 | 18 |
| crtap | 0.57253 | -0.20915191 | 0.318 | 0.423 | 1 | 18 |
| zgc:103697 | 0.57279 | 0.134173121 | 0.136 | 0.077 | 1 | 18 |
| ctps1a | 0.57279 | 0.101072339 | 0.091 | 0.077 | 1 | 18 |
| kcnq5b | 0.57286 | 0.164662354 | 0.227 | 0.115 | 1 | 18 |
| zyx | 0.57339 | 0.107077325 | 0.318 | 0.308 | 1 | 18 |
| clpp | 0.5736 | -0.10834641 | 0.091 | 0.192 | 1 | 18 |
| gstm.1 | 0.57406 | 0.123184693 | 0.682 | 0.538 | 1 | 18 |
| etfa | 0.57436 | -0.18360098 | 0.182 | 0.308 | 1 | 18 |
| ptk2ab | 0.575 | -0.18008344 | 0.045 | 0.115 | 1 | 18 |
| cadm4 | 0.57501 | -0.22235316 | 0.227 | 0.346 | 1 | 18 |
| si:ch1073-429i10.3.1 | 0.57553 | 0.227051773 | 0.364 | 0.231 | 1 | 18 |
| mphosph10 | 0.57619 | 0.173269699 | 0.227 | 0.115 | 1 | 18 |
| mrps28 | 0.57634 | -0.17605757 | 0.091 | 0.154 | 1 | 18 |
| shoc2 | 0.57638 | -0.18325199 | 0.091 | 0.192 | 1 | 18 |
| fosb | 0.57659 | 0.249793145 | 0.409 | 0.269 | 1 | 18 |
| dnajc8 | 0.57722 | -0.19089278 | 0.091 | 0.192 | 1 | 18 |
| tfam | 0.57816 | -0.16654736 | 0.091 | 0.192 | 1 | 18 |
| acadvl | 0.57817 | 0.171206191 | 0.273 | 0.192 | 1 | 18 |
| mgst3a | 0.57892 | 0.198076782 | 0.227 | 0.115 | 1 | 18 |
| ubap1 | 0.58011 | -0.16756797 | 0.045 | 0.115 | 1 | 18 |
| wasf2 | 0.5811 | -0.13583755 | 0.091 | 0.192 | 1 | 18 |

| | | | | | | |
|---|---|---|---|---|---|---|
| lgals3a | 0.58115 | -0.18357378 | 0.091 | 0.192 | 1 | 18 |
| smarca2 | 0.58151 | -0.18610031 | 0.227 | 0.308 | 1 | 18 |
| wdr1 | 0.582 | -0.18937956 | 0.182 | 0.308 | 1 | 18 |
| mib1 | 0.58217 | -0.13677579 | 0.045 | 0.115 | 1 | 18 |
| rtn4a | 0.58219 | -0.20669883 | 0.318 | 0.423 | 1 | 18 |
| chtopa | 0.58268 | 0.15756086 | 0.318 | 0.192 | 1 | 18 |
| col1a1b | 0.58278 | -0.16367465 | 0.727 | 0.846 | 1 | 18 |
| ptges3a | 0.58319 | -0.11894363 | 0.091 | 0.192 | 1 | 18 |
| pnpla6 | 0.58356 | -0.12113911 | 0.091 | 0.115 | 1 | 18 |
| txndc9 | 0.58374 | -0.20375021 | 0.227 | 0.346 | 1 | 18 |
| prkar1ab | 0.58375 | -0.15182303 | 0.091 | 0.192 | 1 | 18 |
| eif4ebp3l | 0.58425 | -0.15356013 | 0.182 | 0.308 | 1 | 18 |
| sulf2b | 0.5843 | -0.16724009 | 0.045 | 0.115 | 1 | 18 |
| eef1da | 0.58475 | 0.171954333 | 0.773 | 0.692 | 1 | 18 |
| copg2 | 0.58495 | 0.177860099 | 0.227 | 0.115 | 1 | 18 |
| atf1 | 0.58558 | 0.158786593 | 0.227 | 0.115 | 1 | 18 |
| puf60a | 0.58585 | -0.1356931 | 0.227 | 0.346 | 1 | 18 |
| fam168b | 0.58598 | 0.177665475 | 0.364 | 0.231 | 1 | 18 |
| sec61a1 | 0.58609 | -0.22028627 | 0.318 | 0.423 | 1 | 18 |
| psmd10 | 0.58624 | -0.1278439 | 0.091 | 0.192 | 1 | 18 |
| si:ch211-237c6.4 | 0.58646 | 0.175081209 | 0.182 | 0.115 | 1 | 18 |
| LOC110438079 | 0.58675 | 0.129062241 | 0.136 | 0.115 | 1 | 18 |
| tbx2b | 0.58685 | 0.171767846 | 0.227 | 0.115 | 1 | 18 |
| zgc:92664 | 0.58724 | -0.21498817 | 0.273 | 0.385 | 1 | 18 |
| zgc:153675 | 0.58782 | 0.214683925 | 0.409 | 0.269 | 1 | 18 |
| rbfox1 | 0.58794 | 0.148421956 | 0.273 | 0.231 | 1 | 18 |
| clint1a | 0.58797 | 0.176618247 | 0.136 | 0.077 | 1 | 18 |
| fkbp1aa | 0.58847 | -0.16929523 | 0.682 | 0.808 | 1 | 18 |
| zgc:162025 | 0.58932 | -0.15112462 | 0.091 | 0.192 | 1 | 18 |
| eaf1 | 0.58954 | -0.16826779 | 0.091 | 0.154 | 1 | 18 |
| bmpr1ab | 0.58958 | 0.164538475 | 0.136 | 0.077 | 1 | 18 |
| cnot9 | 0.5902 | -0.14576309 | 0.091 | 0.192 | 1 | 18 |
| zgc:162509 | 0.59105 | -0.17329409 | 0.045 | 0.115 | 1 | 18 |
| ndufb3 | 0.59121 | -0.1058386 | 0.409 | 0.538 | 1 | 18 |
| cdh11 | 0.5914 | -0.23556803 | 0.227 | 0.346 | 1 | 18 |
| tubb2b | 0.59231 | 0.220219654 | 0.409 | 0.269 | 1 | 18 |
| si:dkey-172h23.2 | 0.59232 | -0.18069787 | 0.091 | 0.192 | 1 | 18 |
| BX640522.2 | 0.59249 | -0.11557982 | 0.727 | 0.846 | 1 | 18 |
| si:ch211-198c19.3 | 0.59264 | -0.16892957 | 0.091 | 0.192 | 1 | 18 |
| bcl2l10 | 0.59265 | -0.14013493 | 0.091 | 0.192 | 1 | 18 |
| pbx1a | 0.59291 | 0.207463322 | 0.636 | 0.5 | 1 | 18 |
| gng5 | 0.59293 | 0.15193296 | 0.682 | 0.538 | 1 | 18 |
| aldh1l1 | 0.59349 | -0.13213215 | 0.045 | 0.115 | 1 | 18 |
| dnajb11 | 0.59363 | -0.18129036 | 0.091 | 0.192 | 1 | 18 |
| nudt21 | 0.59373 | 0.156075537 | 0.318 | 0.192 | 1 | 18 |
| calua | 0.59427 | 0.206563926 | 0.364 | 0.269 | 1 | 18 |
| cdc34a | 0.59455 | 0.114548613 | 0.091 | 0.077 | 1 | 18 |
| dpm3 | 0.59494 | -0.18354012 | 0.091 | 0.192 | 1 | 18 |

| | | | | | | |
|---|---|---|---|---|---|---|
| chmp2a | 0.59526 | -0.17459287 | 0.182 | 0.308 | 1 | 18 |
| atf7ip | 0.59553 | 0.108901262 | 0.136 | 0.115 | 1 | 18 |
| lysmd4 | 0.59587 | -0.13894198 | 0.091 | 0.192 | 1 | 18 |
| tmem176l.3a | 0.5959 | -0.16595798 | 0.091 | 0.192 | 1 | 18 |
| kctd3 | 0.59591 | -0.13486703 | 0.091 | 0.192 | 1 | 18 |
| gsta.1 | 0.59612 | -0.21172003 | 0.182 | 0.308 | 1 | 18 |
| ctdspla | 0.59616 | 0.114559303 | 0.091 | 0.077 | 1 | 18 |
| gna12a | 0.59678 | -0.16398858 | 0.091 | 0.192 | 1 | 18 |
| phb2a | 0.59689 | 0.139274779 | 0.136 | 0.077 | 1 | 18 |
| rab40c | 0.59771 | -0.11010215 | 0.045 | 0.115 | 1 | 18 |
| pqlc3 | 0.59917 | -0.15259254 | 0.091 | 0.192 | 1 | 18 |
| chst2a | 0.5992 | -0.17119843 | 0.091 | 0.192 | 1 | 18 |
| rpz5 | 0.59936 | -0.21213302 | 0.182 | 0.308 | 1 | 18 |
| mrpl17 | 0.59938 | -0.15394631 | 0.091 | 0.192 | 1 | 18 |
| pon3.2 | 0.59955 | -0.14753442 | 0.091 | 0.192 | 1 | 18 |
| ppp2r5eb | 0.59958 | -0.16289475 | 0.182 | 0.308 | 1 | 18 |
| smg6 | 0.59983 | -0.18269634 | 0.091 | 0.192 | 1 | 18 |
| os9 | 0.59994 | -0.14838469 | 0.091 | 0.192 | 1 | 18 |
| fam198a | 0.60015 | -0.17641166 | 0.182 | 0.308 | 1 | 18 |
| mycb | 0.60032 | 0.220102268 | 0.318 | 0.192 | 1 | 18 |
| gtf2f2a | 0.60041 | -0.17002903 | 0.182 | 0.308 | 1 | 18 |
| rtf1 | 0.60051 | 0.191637458 | 0.364 | 0.231 | 1 | 18 |
| mapk3 | 0.60079 | -0.15113143 | 0.091 | 0.192 | 1 | 18 |
| rbpjb | 0.60083 | 0.19523524 | 0.273 | 0.154 | 1 | 18 |
| nck1b | 0.60102 | -0.17513863 | 0.091 | 0.192 | 1 | 18 |
| h1f0 | 0.60155 | -0.25713609 | 0.364 | 0.5 | 1 | 18 |
| npc2 | 0.60165 | 0.183476284 | 0.364 | 0.231 | 1 | 18 |
| psmc2 | 0.60199 | 0.179399176 | 0.273 | 0.154 | 1 | 18 |
| sh3kbp1 | 0.60222 | -0.14205426 | 0.091 | 0.192 | 1 | 18 |
| ralab | 0.60229 | -0.14125839 | 0.091 | 0.192 | 1 | 18 |
| hmbox1a | 0.60242 | -0.11382149 | 0.136 | 0.231 | 1 | 18 |
| lrrc17 | 0.60244 | -0.15869852 | 0.636 | 0.769 | 1 | 18 |
| cfap20 | 0.60247 | -0.14825381 | 0.091 | 0.192 | 1 | 18 |
| coro1ca | 0.60259 | 0.189032682 | 0.318 | 0.192 | 1 | 18 |
| map4k4 | 0.60363 | -0.16390289 | 0.091 | 0.192 | 1 | 18 |
| smad9 | 0.60399 | -0.18090939 | 0.091 | 0.192 | 1 | 18 |
| tmco1 | 0.60401 | -0.12817335 | 0.091 | 0.192 | 1 | 18 |
| ppig | 0.60411 | -0.15897859 | 0.091 | 0.192 | 1 | 18 |
| nanos1 | 0.60412 | 0.16034726 | 0.227 | 0.192 | 1 | 18 |
| pou2f1b | 0.60413 | -0.11890256 | 0.091 | 0.115 | 1 | 18 |
| stimate | 0.60418 | -0.13792574 | 0.091 | 0.192 | 1 | 18 |
| stx1b | 0.60429 | -0.16005002 | 0.091 | 0.192 | 1 | 18 |
| ndufc2 | 0.60447 | -0.1077769 | 0.273 | 0.385 | 1 | 18 |
| sept15 | 0.60451 | -0.15991648 | 0.091 | 0.192 | 1 | 18 |
| ewsr1a | 0.60461 | 0.190291148 | 0.318 | 0.192 | 1 | 18 |
| brd7 | 0.60489 | 0.202570487 | 0.318 | 0.192 | 1 | 18 |
| sertad4 | 0.6052 | -0.14845574 | 0.227 | 0.346 | 1 | 18 |
| ndufa7 | 0.60522 | 0.122611993 | 0.227 | 0.192 | 1 | 18 |

| | | | | | | |
|---|---|---|---|---|---|---|
| tmem126a | 0.60539 | -0.19520992 | 0.136 | 0.231 | 1 | 18 |
| si:dkey-88p24.11 | 0.60587 | 0.11292447 | 0.136 | 0.115 | 1 | 18 |
| cdca5 | 0.60611 | -0.17015615 | 0.045 | 0.115 | 1 | 18 |
| tmem176 | 0.60791 | 0.203500839 | 0.364 | 0.269 | 1 | 18 |
| cmbl | 0.60895 | -0.13334559 | 0.045 | 0.115 | 1 | 18 |
| immp2l | 0.60903 | -0.14625055 | 0.045 | 0.115 | 1 | 18 |
| uba1 | 0.60944 | 0.151136182 | 0.227 | 0.192 | 1 | 18 |
| hnrnpub | 0.61433 | -0.22069662 | 0.136 | 0.231 | 1 | 18 |
| kdm2ba | 0.61687 | 0.16276771 | 0.136 | 0.077 | 1 | 18 |
| aamp | 0.61833 | -0.15886095 | 0.045 | 0.115 | 1 | 18 |
| rprd1b | 0.61837 | 0.160169275 | 0.136 | 0.077 | 1 | 18 |
| stau2 | 0.61944 | 0.200154253 | 0.5 | 0.385 | 1 | 18 |
| crk | 0.61952 | -0.12323582 | 0.136 | 0.231 | 1 | 18 |
| sat1b | 0.6196 | -0.12982721 | 0.136 | 0.231 | 1 | 18 |
| marcksa | 0.62136 | 0.111714461 | 0.182 | 0.154 | 1 | 18 |
| znf740b | 0.62157 | -0.16447555 | 0.045 | 0.115 | 1 | 18 |
| babam1 | 0.62175 | -0.1298707 | 0.045 | 0.115 | 1 | 18 |
| myl12.2 | 0.62201 | 0.115542708 | 0.182 | 0.154 | 1 | 18 |
| cdkn1a | 0.62217 | -0.21580432 | 0.136 | 0.231 | 1 | 18 |
| atp5if1a | 0.62227 | -0.11700561 | 0.409 | 0.538 | 1 | 18 |
| psmd4b | 0.62235 | 0.134456342 | 0.273 | 0.231 | 1 | 18 |
| eif5b | 0.6229 | -0.20613926 | 0.227 | 0.346 | 1 | 18 |
| sept2 | 0.62302 | -0.17004822 | 0.5 | 0.577 | 1 | 18 |
| palld | 0.62334 | -0.14651883 | 0.182 | 0.231 | 1 | 18 |
| sdhda | 0.62376 | -0.14348086 | 0.045 | 0.115 | 1 | 18 |
| ranbp1 | 0.62393 | -0.10862299 | 0.455 | 0.577 | 1 | 18 |
| adka | 0.62475 | -0.1332607 | 0.045 | 0.115 | 1 | 18 |
| si:dkey-253i9.4 | 0.62622 | 0.101867386 | 0.091 | 0.077 | 1 | 18 |
| CABZ01088109.1 | 0.62703 | -0.14675586 | 0.045 | 0.115 | 1 | 18 |
| igf2bp2b | 0.62745 | -0.15630524 | 0.091 | 0.154 | 1 | 18 |
| ncbp2 | 0.62756 | 0.188505747 | 0.364 | 0.269 | 1 | 18 |
| rmnd5b | 0.62779 | 0.116419829 | 0.136 | 0.115 | 1 | 18 |
| lamb1a | 0.62793 | 0.175907792 | 0.273 | 0.192 | 1 | 18 |
| pttg1ipb | 0.6289 | -0.1015066 | 0.136 | 0.231 | 1 | 18 |
| si:ch211-197h24.9 | 0.62931 | 0.15124901 | 0.136 | 0.077 | 1 | 18 |
| si:dkey-11f4.14 | 0.63007 | 0.157876771 | 0.182 | 0.115 | 1 | 18 |
| crabp2b | 0.63013 | -0.19126314 | 0.182 | 0.269 | 1 | 18 |
| mrps27 | 0.6302 | 0.101335045 | 0.091 | 0.077 | 1 | 18 |
| ube2g1b | 0.63099 | 0.109165596 | 0.182 | 0.154 | 1 | 18 |
| pvrl2l | 0.63129 | 0.103037552 | 0.091 | 0.077 | 1 | 18 |
| fstb | 0.63146 | -0.22706424 | 0.045 | 0.115 | 1 | 18 |
| zgc:113102 | 0.63231 | -0.14279437 | 0.045 | 0.115 | 1 | 18 |
| slc22a18 | 0.63269 | -0.16894573 | 0.045 | 0.115 | 1 | 18 |
| myl6 | 0.63275 | 0.184847139 | 0.727 | 0.615 | 1 | 18 |
| si:ch211-276c2.4 | 0.63287 | 0.166894951 | 0.136 | 0.077 | 1 | 18 |
| six2a | 0.63288 | -0.11064888 | 0.045 | 0.115 | 1 | 18 |
| txnl4a | 0.63322 | 0.161243157 | 0.227 | 0.154 | 1 | 18 |
| zgc:175264 | 0.63397 | -0.15288804 | 0.045 | 0.115 | 1 | 18 |

| | | | | | | |
|---|---|---|---|---|---|---|
| ppfibp1a | 0.63421 | 0.131400878 | 0.227 | 0.192 | 1 | 18 |
| ttyh2 | 0.63448 | -0.10979469 | 0.045 | 0.115 | 1 | 18 |
| psmb6 | 0.63471 | -0.12658705 | 0.409 | 0.538 | 1 | 18 |
| rgmb | 0.63492 | -0.16997741 | 0.045 | 0.115 | 1 | 18 |
| polr2i | 0.63533 | -0.12339237 | 0.045 | 0.115 | 1 | 18 |
| lpp | 0.63562 | -0.13999139 | 0.045 | 0.115 | 1 | 18 |
| klf2a | 0.63592 | -0.1103297 | 0.455 | 0.462 | 1 | 18 |
| srsf2a | 0.63643 | -0.16659479 | 0.364 | 0.5 | 1 | 18 |
| cct4 | 0.63648 | -0.18606245 | 0.409 | 0.538 | 1 | 18 |
| ppp1r26 | 0.63707 | 0.164465783 | 0.136 | 0.077 | 1 | 18 |
| aes | 0.63712 | 0.103818405 | 0.091 | 0.077 | 1 | 18 |
| kif5ba | 0.63728 | -0.1538752 | 0.091 | 0.154 | 1 | 18 |
| snw1 | 0.63922 | -0.19607003 | 0.227 | 0.346 | 1 | 18 |
| fat1a | 0.6396 | -0.11666887 | 0.045 | 0.115 | 1 | 18 |
| etv4 | 0.63961 | 0.18988425 | 0.273 | 0.192 | 1 | 18 |
| pear1 | 0.64029 | -0.15606933 | 0.045 | 0.115 | 1 | 18 |
| eif2b4 | 0.6408 | -0.12964182 | 0.045 | 0.115 | 1 | 18 |
| spsb4a | 0.64156 | -0.1347298 | 0.045 | 0.115 | 1 | 18 |
| rbbp4 | 0.64208 | 0.178482877 | 0.364 | 0.308 | 1 | 18 |
| dpp7 | 0.64229 | -0.15581669 | 0.045 | 0.115 | 1 | 18 |
| prpf3 | 0.64229 | -0.13131068 | 0.045 | 0.115 | 1 | 18 |
| tex264a | 0.64289 | -0.1039374 | 0.045 | 0.115 | 1 | 18 |
| lmnb1 | 0.64304 | -0.11467928 | 0.136 | 0.231 | 1 | 18 |
| slit2 | 0.64316 | -0.10266863 | 0.045 | 0.115 | 1 | 18 |
| tm9sf1 | 0.64339 | -0.10192566 | 0.045 | 0.115 | 1 | 18 |
| timp2a | 0.64377 | 0.148158343 | 0.318 | 0.269 | 1 | 18 |
| zgc:56095 | 0.64395 | -0.13633362 | 0.227 | 0.269 | 1 | 18 |
| zgc:162698 | 0.644 | -0.1364618 | 0.045 | 0.115 | 1 | 18 |
| sirt7 | 0.64443 | -0.14870917 | 0.136 | 0.192 | 1 | 18 |
| taok1b | 0.64573 | -0.11753092 | 0.045 | 0.115 | 1 | 18 |
| gas6 | 0.64596 | 0.158822039 | 0.136 | 0.077 | 1 | 18 |
| mob3c | 0.64644 | 0.149653094 | 0.136 | 0.077 | 1 | 18 |
| tsen15 | 0.64686 | -0.1082323 | 0.045 | 0.115 | 1 | 18 |
| ptenb | 0.64724 | -0.10483008 | 0.318 | 0.423 | 1 | 18 |
| si:ch211-236l14.4 | 0.6474 | -0.11292745 | 0.045 | 0.115 | 1 | 18 |
| gadd45ba | 0.64741 | 0.207215583 | 0.545 | 0.423 | 1 | 18 |
| tpst1 | 0.64755 | -0.17183907 | 0.182 | 0.269 | 1 | 18 |
| pik3r2 | 0.64788 | 0.124160995 | 0.182 | 0.154 | 1 | 18 |
| klhl11 | 0.64856 | -0.10311781 | 0.045 | 0.115 | 1 | 18 |
| chsy1 | 0.64866 | -0.13125574 | 0.045 | 0.115 | 1 | 18 |
| yipf4 | 0.64866 | 0.111661387 | 0.182 | 0.154 | 1 | 18 |
| elovl6 | 0.64975 | -0.17925694 | 0.136 | 0.231 | 1 | 18 |
| ranbp9 | 0.65016 | -0.16951908 | 0.136 | 0.231 | 1 | 18 |
| eif4a1b | 0.65036 | 0.125003136 | 0.773 | 0.654 | 1 | 18 |
| MYO1D | 0.65038 | -0.11854285 | 0.045 | 0.115 | 1 | 18 |
| mrps33 | 0.65073 | 0.123095766 | 0.136 | 0.077 | 1 | 18 |
| slc43a1a | 0.65105 | 0.171264623 | 0.182 | 0.115 | 1 | 18 |
| tnrc6a | 0.65148 | -0.12115576 | 0.136 | 0.231 | 1 | 18 |

| | | | | | | |
|---|---|---|---|---|---|---|
| oaz2a | 0.65157 | -0.13831067 | 0.045 | 0.115 | 1 | 18 |
| atp5f1b | 0.65159 | -0.15561498 | 0.682 | 0.731 | 1 | 18 |
| zgc:163098 | 0.65192 | -0.11651902 | 0.045 | 0.115 | 1 | 18 |
| pdcl | 0.65194 | -0.11510038 | 0.136 | 0.231 | 1 | 18 |
| bmi1a | 0.65199 | -0.1162727 | 0.045 | 0.115 | 1 | 18 |
| syf2 | 0.65296 | -0.18205186 | 0.409 | 0.538 | 1 | 18 |
| mbd1a | 0.65296 | -0.13053127 | 0.045 | 0.115 | 1 | 18 |
| cib1 | 0.65347 | 0.164327816 | 0.182 | 0.115 | 1 | 18 |
| zgc:110626 | 0.65355 | -0.13223992 | 0.045 | 0.115 | 1 | 18 |
| rem1 | 0.65357 | -0.16221211 | 0.136 | 0.192 | 1 | 18 |
| pum3 | 0.65444 | -0.13633278 | 0.045 | 0.115 | 1 | 18 |
| rfc3 | 0.65484 | 0.105834894 | 0.091 | 0.077 | 1 | 18 |
| col27a1b | 0.65489 | -0.12017634 | 0.045 | 0.115 | 1 | 18 |
| gtf2f1 | 0.65615 | -0.12304753 | 0.136 | 0.231 | 1 | 18 |
| bet1l | 0.65621 | -0.13104503 | 0.045 | 0.115 | 1 | 18 |
| METTL18 | 0.65652 | -0.16600072 | 0.045 | 0.077 | 1 | 18 |
| ccdc97 | 0.65652 | -0.10941479 | 0.045 | 0.077 | 1 | 18 |
| ctso | 0.65652 | -0.11254487 | 0.045 | 0.077 | 1 | 18 |
| ppp1r21 | 0.65652 | -0.12350313 | 0.045 | 0.077 | 1 | 18 |
| prcc | 0.65652 | -0.11023426 | 0.045 | 0.077 | 1 | 18 |
| si:dkey-12l12.1 | 0.65652 | -0.11482706 | 0.045 | 0.077 | 1 | 18 |
| slc1a3a | 0.65652 | -0.14260359 | 0.045 | 0.077 | 1 | 18 |
| JMY | 0.65652 | -0.14246363 | 0.045 | 0.077 | 1 | 18 |
| adamts8a | 0.65652 | -0.16053196 | 0.045 | 0.077 | 1 | 18 |
| cnpy3 | 0.65652 | -0.10744334 | 0.045 | 0.077 | 1 | 18 |
| g2e3 | 0.65652 | -0.11591813 | 0.045 | 0.077 | 1 | 18 |
| nop16 | 0.65652 | -0.10156919 | 0.045 | 0.077 | 1 | 18 |
| pla2g15 | 0.65652 | -0.11374882 | 0.045 | 0.077 | 1 | 18 |
| sc:d0202 | 0.65652 | -0.12561236 | 0.045 | 0.077 | 1 | 18 |
| sec23a | 0.65652 | -0.11101253 | 0.045 | 0.077 | 1 | 18 |
| slc35f2 | 0.65652 | -0.12655508 | 0.045 | 0.077 | 1 | 18 |
| BX901883.3 | 0.65652 | -0.12096424 | 0.045 | 0.077 | 1 | 18 |
| hist1h4l | 0.65652 | -0.11913917 | 0.045 | 0.077 | 1 | 18 |
| mbd6 | 0.65652 | -0.1156164 | 0.045 | 0.077 | 1 | 18 |
| rap2aa | 0.65652 | -0.11826504 | 0.045 | 0.077 | 1 | 18 |
| rbm33a | 0.65652 | -0.10427 | 0.045 | 0.077 | 1 | 18 |
| rnf11b | 0.65652 | -0.11074779 | 0.045 | 0.077 | 1 | 18 |
| si:ch211-232b12.5 | 0.65652 | -0.16980324 | 0.045 | 0.077 | 1 | 18 |
| si:ch73-25f10.6 | 0.65652 | -0.10596278 | 0.045 | 0.077 | 1 | 18 |
| taf2 | 0.65652 | -0.12347153 | 0.045 | 0.077 | 1 | 18 |
| LOC110437917 | 0.65652 | -0.1046462 | 0.045 | 0.077 | 1 | 18 |
| cwf19l2 | 0.65652 | -0.12656964 | 0.045 | 0.077 | 1 | 18 |
| palm1a | 0.65652 | -0.10454931 | 0.045 | 0.077 | 1 | 18 |
| proser1 | 0.65652 | -0.10905705 | 0.045 | 0.077 | 1 | 18 |
| rflnb | 0.65652 | -0.13503469 | 0.045 | 0.077 | 1 | 18 |
| si:ch211-86h15.1 | 0.65652 | -0.10654565 | 0.045 | 0.077 | 1 | 18 |
| tmem35 | 0.65652 | -0.15622165 | 0.045 | 0.077 | 1 | 18 |
| ubxn6 | 0.65652 | -0.1185159 | 0.045 | 0.077 | 1 | 18 |

| | | | | | | |
|---|---|---|---|---|---|---|
| ada2a | 0.65652 | -0.13064018 | 0.045 | 0.077 | 1 | 18 |
| prr12a | 0.65652 | -0.11027496 | 0.045 | 0.077 | 1 | 18 |
| ubr5 | 0.65652 | -0.10073433 | 0.045 | 0.077 | 1 | 18 |
| dgcr8 | 0.65652 | -0.11024917 | 0.045 | 0.077 | 1 | 18 |
| CABZ01058650.1 | 0.6568 | -0.12089094 | 0.136 | 0.231 | 1 | 18 |
| znf410 | 0.65745 | 0.164287937 | 0.136 | 0.077 | 1 | 18 |
| baz1b | 0.65787 | 0.14018854 | 0.136 | 0.077 | 1 | 18 |
| mrps11 | 0.65791 | -0.13034619 | 0.045 | 0.115 | 1 | 18 |
| wbp2nl | 0.65832 | -0.17270299 | 0.182 | 0.269 | 1 | 18 |
| zfyve27 | 0.65842 | -0.13652473 | 0.045 | 0.115 | 1 | 18 |
| arf2a | 0.65888 | -0.17850443 | 0.091 | 0.154 | 1 | 18 |
| purab | 0.65909 | 0.106068113 | 0.136 | 0.115 | 1 | 18 |
| sorbs1 | 0.65918 | -0.10687645 | 0.045 | 0.115 | 1 | 18 |
| calr | 0.65968 | -0.19758709 | 0.318 | 0.423 | 1 | 18 |
| rxrga | 0.66019 | 0.130440643 | 0.136 | 0.077 | 1 | 18 |
| lgi1a | 0.66117 | -0.11454278 | 0.045 | 0.115 | 1 | 18 |
| hadhab | 0.66138 | -0.10958427 | 0.045 | 0.115 | 1 | 18 |
| fbn2b | 0.66162 | 0.143515844 | 0.773 | 0.654 | 1 | 18 |
| per3 | 0.66175 | -0.15217006 | 0.045 | 0.115 | 1 | 18 |
| atf4a | 0.66182 | -0.15406473 | 0.227 | 0.346 | 1 | 18 |
| dedd1 | 0.66209 | 0.182264298 | 0.545 | 0.423 | 1 | 18 |
| baz2a | 0.66248 | -0.15487913 | 0.045 | 0.115 | 1 | 18 |
| tab1 | 0.66249 | -0.10528908 | 0.045 | 0.115 | 1 | 18 |
| hdac4 | 0.66264 | -0.14883112 | 0.045 | 0.115 | 1 | 18 |
| nmt1a | 0.66297 | -0.11209968 | 0.045 | 0.115 | 1 | 18 |
| tmem189 | 0.66326 | 0.167941398 | 0.182 | 0.115 | 1 | 18 |
| glrx3 | 0.66372 | 0.141524691 | 0.182 | 0.115 | 1 | 18 |
| CABZ01072952.1 | 0.6639 | 0.103758132 | 0.091 | 0.077 | 1 | 18 |
| si:dkey-222f8.3 | 0.66424 | -0.1245771 | 0.045 | 0.115 | 1 | 18 |
| nudt17 | 0.66455 | 0.162275946 | 0.136 | 0.077 | 1 | 18 |
| atg4c | 0.66505 | -0.10986862 | 0.045 | 0.115 | 1 | 18 |
| bace1 | 0.66514 | -0.12886892 | 0.045 | 0.115 | 1 | 18 |
| cst3 | 0.66516 | 0.178960614 | 0.182 | 0.115 | 1 | 18 |
| creg2 | 0.66565 | 0.18062553 | 0.182 | 0.115 | 1 | 18 |
| kirrel1a | 0.66602 | -0.12103389 | 0.273 | 0.308 | 1 | 18 |
| macf1a | 0.66693 | -0.10474729 | 0.136 | 0.154 | 1 | 18 |
| kmt2bb | 0.66732 | -0.17891827 | 0.318 | 0.423 | 1 | 18 |
| extl2 | 0.66739 | -0.12522068 | 0.045 | 0.115 | 1 | 18 |
| scamp2 | 0.66748 | -0.10766559 | 0.045 | 0.115 | 1 | 18 |
| ptpn2a | 0.66762 | -0.13251334 | 0.045 | 0.115 | 1 | 18 |
| necap2 | 0.66822 | -0.15134342 | 0.045 | 0.115 | 1 | 18 |
| chd4b | 0.66865 | 0.156773329 | 0.227 | 0.154 | 1 | 18 |
| si:dkey-19a16.7 | 0.66876 | 0.152290983 | 0.136 | 0.077 | 1 | 18 |
| xrcc3 | 0.66885 | -0.11755619 | 0.045 | 0.115 | 1 | 18 |
| lamc1 | 0.66896 | 0.15492274 | 0.136 | 0.077 | 1 | 18 |
| angptl2a | 0.66898 | -0.14250833 | 0.045 | 0.115 | 1 | 18 |
| h3f3d | 0.66945 | 0.121188614 | 0.909 | 0.846 | 1 | 18 |
| sod1 | 0.66964 | 0.165687587 | 0.727 | 0.615 | 1 | 18 |

| | | | | | | |
|---|---|---|---|---|---|---|
| ndufaf5 | 0.66967 | 0.126306111 | 0.136 | 0.077 | 1 | 18 |
| rev3l | 0.66968 | -0.10720428 | 0.045 | 0.115 | 1 | 18 |
| igflr1 | 0.66972 | -0.17863895 | 0.409 | 0.538 | 1 | 18 |
| ell | 0.67022 | -0.11187489 | 0.045 | 0.115 | 1 | 18 |
| zgc:153018 | 0.67047 | -0.15474851 | 0.091 | 0.154 | 1 | 18 |
| fbln1 | 0.67048 | 0.16488872 | 0.182 | 0.115 | 1 | 18 |
| bbx | 0.67076 | -0.12132492 | 0.045 | 0.115 | 1 | 18 |
| med20 | 0.67094 | -0.11904538 | 0.045 | 0.115 | 1 | 18 |
| cry2 | 0.67097 | -0.10285084 | 0.045 | 0.115 | 1 | 18 |
| golga7 | 0.671 | 0.118165142 | 0.318 | 0.269 | 1 | 18 |
| ldb1a | 0.67118 | -0.10578524 | 0.045 | 0.115 | 1 | 18 |
| saraf | 0.67189 | -0.10847172 | 0.045 | 0.115 | 1 | 18 |
| bcl9 | 0.67194 | -0.15061453 | 0.136 | 0.231 | 1 | 18 |
| ccdc106a | 0.67202 | -0.11994006 | 0.045 | 0.115 | 1 | 18 |
| ep300b | 0.67204 | -0.10525836 | 0.045 | 0.115 | 1 | 18 |
| pld1a | 0.67259 | -0.13617672 | 0.045 | 0.115 | 1 | 18 |
| uchl1 | 0.67263 | -0.1076378 | 0.045 | 0.115 | 1 | 18 |
| rab1ba | 0.67302 | -0.11006948 | 0.045 | 0.115 | 1 | 18 |
| thap11 | 0.67302 | -0.13233021 | 0.045 | 0.115 | 1 | 18 |
| arl14ep | 0.6733 | -0.16221066 | 0.091 | 0.154 | 1 | 18 |
| gusb | 0.67347 | -0.10535509 | 0.045 | 0.115 | 1 | 18 |
| decr2 | 0.67352 | -0.1008923 | 0.045 | 0.115 | 1 | 18 |
| selenof | 0.6738 | 0.173425637 | 0.682 | 0.577 | 1 | 18 |
| dync1li2 | 0.67397 | -0.10000589 | 0.045 | 0.115 | 1 | 18 |
| snx27b | 0.674 | 0.154989766 | 0.136 | 0.077 | 1 | 18 |
| ccm2 | 0.67403 | -0.13144629 | 0.045 | 0.115 | 1 | 18 |
| zgc:101858 | 0.67429 | -0.12652434 | 0.045 | 0.115 | 1 | 18 |
| ptk2aa | 0.6744 | -0.11640315 | 0.045 | 0.115 | 1 | 18 |
| stx4 | 0.67453 | -0.1319242 | 0.045 | 0.115 | 1 | 18 |
| tmem107l | 0.67457 | -0.1208823 | 0.045 | 0.115 | 1 | 18 |
| zfpm1 | 0.67467 | -0.11284287 | 0.045 | 0.115 | 1 | 18 |
| afdna | 0.67538 | -0.13058101 | 0.045 | 0.115 | 1 | 18 |
| tango2 | 0.67579 | -0.11014335 | 0.045 | 0.115 | 1 | 18 |
| brd1b | 0.67581 | -0.11391456 | 0.045 | 0.115 | 1 | 18 |
| CU466287.2 | 0.67626 | 0.14118862 | 0.136 | 0.077 | 1 | 18 |
| aar2 | 0.67627 | -0.10581173 | 0.045 | 0.115 | 1 | 18 |
| LO018102.1 | 0.67639 | -0.10509371 | 0.045 | 0.115 | 1 | 18 |
| tox | 0.6764 | -0.12618846 | 0.045 | 0.115 | 1 | 18 |
| zgc:55558 | 0.67646 | -0.10417909 | 0.045 | 0.115 | 1 | 18 |
| XLOC-017111 | 0.67654 | -0.11239648 | 0.045 | 0.115 | 1 | 18 |
| bcl2a | 0.6766 | -0.12870092 | 0.045 | 0.115 | 1 | 18 |
| tjp1b | 0.67675 | -0.15317144 | 0.045 | 0.115 | 1 | 18 |
| arhgef12b | 0.67684 | -0.11970774 | 0.045 | 0.115 | 1 | 18 |
| ing1 | 0.67693 | -0.10345185 | 0.045 | 0.115 | 1 | 18 |
| dnal4b | 0.67694 | -0.12181271 | 0.045 | 0.115 | 1 | 18 |
| vaspb | 0.67706 | -0.10291893 | 0.045 | 0.115 | 1 | 18 |
| chd2 | 0.67708 | -0.13066205 | 0.045 | 0.115 | 1 | 18 |
| pskh1 | 0.67709 | -0.10752876 | 0.045 | 0.115 | 1 | 18 |

| | | | | | | |
|---|---|---|---|---|---|---|
| txnl4b | 0.67711 | -0.10138242 | 0.045 | 0.115 | 1 | 18 |
| fndc4b | 0.67717 | -0.14838506 | 0.045 | 0.115 | 1 | 18 |
| pex7 | 0.67724 | -0.12321586 | 0.045 | 0.115 | 1 | 18 |
| rbm12b | 0.67734 | -0.10552834 | 0.045 | 0.115 | 1 | 18 |
| flncb | 0.67736 | -0.13359025 | 0.045 | 0.115 | 1 | 18 |
| adamts12 | 0.67737 | -0.15034738 | 0.045 | 0.115 | 1 | 18 |
| amotl2a | 0.67744 | -0.13199163 | 0.045 | 0.115 | 1 | 18 |
| appa | 0.67796 | 0.107173853 | 0.136 | 0.115 | 1 | 18 |
| filip1l | 0.67798 | -0.20386852 | 0.273 | 0.385 | 1 | 18 |
| dse | 0.67828 | 0.157640631 | 0.182 | 0.115 | 1 | 18 |
| cast | 0.67853 | -0.13099225 | 0.136 | 0.231 | 1 | 18 |
| mdh1aa | 0.67862 | 0.156228058 | 0.591 | 0.5 | 1 | 18 |
| icn | 0.67862 | 0.19248332 | 0.727 | 0.615 | 1 | 18 |
| zranb2 | 0.6796 | -0.13436096 | 0.273 | 0.385 | 1 | 18 |
| cndp2 | 0.68228 | 0.109108156 | 0.182 | 0.154 | 1 | 18 |
| pak2b | 0.68237 | 0.120349223 | 0.227 | 0.192 | 1 | 18 |
| hmcn1 | 0.68289 | -0.15464954 | 0.136 | 0.231 | 1 | 18 |
| rnasekb | 0.68351 | -0.14305158 | 0.136 | 0.192 | 1 | 18 |
| vstm4a | 0.68363 | 0.163681804 | 0.182 | 0.115 | 1 | 18 |
| fstl1b | 0.68413 | -0.10538286 | 0.864 | 0.923 | 1 | 18 |
| nop58 | 0.6848 | -0.15840513 | 0.136 | 0.231 | 1 | 18 |
| isl2b | 0.68551 | -0.15211119 | 0.273 | 0.385 | 1 | 18 |
| lyar | 0.68638 | 0.118213865 | 0.136 | 0.077 | 1 | 18 |
| tusc3 | 0.6877 | -0.12201998 | 0.136 | 0.231 | 1 | 18 |
| ogdha | 0.68945 | -0.14318632 | 0.136 | 0.231 | 1 | 18 |
| chic1 | 0.69012 | 0.154646578 | 0.136 | 0.077 | 1 | 18 |
| ufc1 | 0.69066 | 0.134575581 | 0.136 | 0.077 | 1 | 18 |
| dhx15 | 0.6907 | -0.12672932 | 0.136 | 0.231 | 1 | 18 |
| dynll1 | 0.69108 | 0.166921459 | 0.682 | 0.577 | 1 | 18 |
| zc3h18 | 0.69196 | -0.14249723 | 0.136 | 0.231 | 1 | 18 |
| dldh | 0.69415 | -0.13071238 | 0.136 | 0.231 | 1 | 18 |
| sfpq | 0.69636 | -0.12874085 | 0.136 | 0.192 | 1 | 18 |
| mrps14 | 0.69643 | -0.13097057 | 0.136 | 0.231 | 1 | 18 |
| hnrnpm | 0.69666 | -0.14638253 | 0.136 | 0.231 | 1 | 18 |
| sccpdhb | 0.69686 | -0.13293254 | 0.136 | 0.231 | 1 | 18 |
| plpp3 | 0.69705 | -0.17202451 | 0.273 | 0.385 | 1 | 18 |
| gnl3 | 0.69736 | 0.16932933 | 0.182 | 0.115 | 1 | 18 |
| nkain1 | 0.69744 | -0.14785873 | 0.136 | 0.231 | 1 | 18 |
| ola1 | 0.6993 | 0.154421499 | 0.545 | 0.423 | 1 | 18 |
| eif3ha | 0.69942 | 0.140960424 | 0.455 | 0.385 | 1 | 18 |
| thoc2 | 0.69966 | -0.15739699 | 0.136 | 0.231 | 1 | 18 |
| si:ch73-111k22.2 | 0.69988 | 0.152203563 | 0.136 | 0.077 | 1 | 18 |
| actn4 | 0.70077 | 0.139062303 | 0.182 | 0.115 | 1 | 18 |
| baxa | 0.7008 | 0.140810584 | 0.227 | 0.154 | 1 | 18 |
| arf6a | 0.70112 | -0.13636943 | 0.136 | 0.231 | 1 | 18 |
| mmrn2a | 0.70159 | 0.120554289 | 0.273 | 0.231 | 1 | 18 |
| crlf3 | 0.7026 | -0.13924142 | 0.136 | 0.231 | 1 | 18 |
| arfrp1 | 0.70333 | 0.127972985 | 0.136 | 0.077 | 1 | 18 |

| | | | | | | |
|---|---|---|---|---|---|---|
| znf423 | 0.70378 | 0.150907927 | 0.182 | 0.115 | 1 | 18 |
| tmem251 | 0.70434 | 0.125738285 | 0.182 | 0.115 | 1 | 18 |
| XLOC-000431 | 0.70525 | -0.13738971 | 0.182 | 0.192 | 1 | 18 |
| actb1 | 0.70543 | 0.127043386 | 0.773 | 0.731 | 1 | 18 |
| arl2 | 0.70584 | 0.150012613 | 0.273 | 0.192 | 1 | 18 |
| gpat3 | 0.70618 | 0.100972221 | 0.136 | 0.115 | 1 | 18 |
| msna | 0.70661 | 0.138654011 | 0.636 | 0.577 | 1 | 18 |
| csnk1da | 0.70667 | 0.106625416 | 0.182 | 0.154 | 1 | 18 |
| krt91 | 0.70703 | 0.183905789 | 0.182 | 0.154 | 1 | 18 |
| bbip1 | 0.70709 | -0.1487805 | 0.091 | 0.154 | 1 | 18 |
| chmp1b | 0.70757 | 0.107748132 | 0.227 | 0.192 | 1 | 18 |
| polr2h | 0.70884 | 0.108884852 | 0.136 | 0.077 | 1 | 18 |
| cox6c | 0.70932 | -0.13221497 | 0.455 | 0.5 | 1 | 18 |
| rimkla | 0.70939 | 0.100931614 | 0.136 | 0.115 | 1 | 18 |
| daam1a | 0.7111 | 0.107490793 | 0.182 | 0.154 | 1 | 18 |
| brd3a | 0.71222 | 0.101818674 | 0.136 | 0.115 | 1 | 18 |
| si:dkey-17m8.1 | 0.71293 | 0.140617399 | 0.136 | 0.077 | 1 | 18 |
| ptprga | 0.7133 | -0.15829752 | 0.273 | 0.385 | 1 | 18 |
| ube2d2 | 0.71358 | 0.171577144 | 0.5 | 0.385 | 1 | 18 |
| selenot1b | 0.71359 | -0.13732151 | 0.227 | 0.308 | 1 | 18 |
| nfyc | 0.71453 | -0.13248961 | 0.136 | 0.192 | 1 | 18 |
| rab28 | 0.71555 | 0.136170973 | 0.136 | 0.077 | 1 | 18 |
| tcea1 | 0.71591 | 0.1876165 | 0.273 | 0.192 | 1 | 18 |
| cnot2 | 0.71596 | 0.15690411 | 0.136 | 0.077 | 1 | 18 |
| lemd3 | 0.7162 | 0.131422809 | 0.136 | 0.077 | 1 | 18 |
| col16a1 | 0.71657 | -0.12758594 | 0.227 | 0.269 | 1 | 18 |
| ccdc47 | 0.71703 | 0.118442581 | 0.136 | 0.077 | 1 | 18 |
| tfpt | 0.718 | 0.11855252 | 0.136 | 0.077 | 1 | 18 |
| flna | 0.71806 | 0.164872321 | 0.409 | 0.346 | 1 | 18 |
| yipf3 | 0.71897 | 0.143347978 | 0.136 | 0.077 | 1 | 18 |
| cpsf3 | 0.71928 | -0.13591003 | 0.091 | 0.154 | 1 | 18 |
| ctbp1 | 0.72009 | 0.140530202 | 0.455 | 0.385 | 1 | 18 |
| gars | 0.72093 | 0.125339351 | 0.182 | 0.115 | 1 | 18 |
| tbca | 0.72185 | -0.14748122 | 0.5 | 0.615 | 1 | 18 |
| ccni | 0.72222 | -0.1398631 | 0.455 | 0.538 | 1 | 18 |
| zc3h7bb | 0.72263 | 0.120971374 | 0.136 | 0.077 | 1 | 18 |
| phb | 0.72393 | 0.101269203 | 0.136 | 0.115 | 1 | 18 |
| tns3.2 | 0.7246 | 0.148819042 | 0.136 | 0.077 | 1 | 18 |
| hells | 0.7247 | 0.157211362 | 0.182 | 0.115 | 1 | 18 |
| csrnp1b | 0.72486 | 0.101907368 | 0.136 | 0.115 | 1 | 18 |
| gtf2h5 | 0.72581 | 0.103013172 | 0.136 | 0.077 | 1 | 18 |
| gtf2a1 | 0.72678 | -0.13662129 | 0.182 | 0.231 | 1 | 18 |
| meis2b | 0.72702 | -0.16055644 | 0.318 | 0.423 | 1 | 18 |
| LOC100536647 | 0.72747 | 0.12580723 | 0.182 | 0.115 | 1 | 18 |
| chmp4ba | 0.728 | 0.127910418 | 0.136 | 0.077 | 1 | 18 |
| mpc2 | 0.72942 | 0.136742137 | 0.182 | 0.115 | 1 | 18 |
| fam204a | 0.7297 | 0.128379093 | 0.182 | 0.115 | 1 | 18 |
| rsf1b.1 | 0.73032 | 0.100255744 | 0.182 | 0.154 | 1 | 18 |

| | | | | | | |
|---|---|---|---|---|---|---|
| ift74 | 0.73132 | 0.142976389 | 0.136 | 0.077 | 1 | 18 |
| srsf6b | 0.73226 | 0.168754165 | 0.273 | 0.192 | 1 | 18 |
| tomm20b | 0.73236 | -0.16116269 | 0.409 | 0.5 | 1 | 18 |
| slain2 | 0.73237 | 0.125870722 | 0.136 | 0.077 | 1 | 18 |
| romo1 | 0.73298 | 0.142867911 | 0.727 | 0.654 | 1 | 18 |
| urm1 | 0.73309 | 0.127105215 | 0.136 | 0.077 | 1 | 18 |
| nr2f6a | 0.73391 | -0.14363915 | 0.091 | 0.154 | 1 | 18 |
| col15a1a | 0.73393 | 0.126299663 | 0.318 | 0.269 | 1 | 18 |
| tfg | 0.73412 | -0.13261605 | 0.182 | 0.231 | 1 | 18 |
| si:ch211-276i12.9 | 0.73485 | 0.147909172 | 0.182 | 0.115 | 1 | 18 |
| snu13b | 0.7352 | 0.104954559 | 0.364 | 0.346 | 1 | 18 |
| adam19a | 0.73535 | -0.15422621 | 0.182 | 0.269 | 1 | 18 |
| hsbp1a | 0.73602 | 0.117962532 | 0.136 | 0.077 | 1 | 18 |
| chmp5b | 0.73753 | -0.10917787 | 0.227 | 0.269 | 1 | 18 |
| sgcd | 0.73851 | 0.116976854 | 0.136 | 0.077 | 1 | 18 |
| btg3 | 0.73897 | -0.15860082 | 0.182 | 0.269 | 1 | 18 |
| csnk1db | 0.73911 | -0.11853576 | 0.091 | 0.154 | 1 | 18 |
| uck2a | 0.73915 | -0.11733186 | 0.091 | 0.154 | 1 | 18 |
| itpr1a | 0.74027 | 0.17054494 | 0.136 | 0.077 | 1 | 18 |
| mydgf | 0.74035 | -0.11462224 | 0.182 | 0.269 | 1 | 18 |
| hgh1 | 0.74135 | -0.12986018 | 0.136 | 0.192 | 1 | 18 |
| pqbp1 | 0.74241 | 0.113496809 | 0.136 | 0.077 | 1 | 18 |
| dbpb | 0.74244 | 0.158188172 | 0.182 | 0.115 | 1 | 18 |
| kdelr2b | 0.74284 | -0.12069419 | 0.318 | 0.423 | 1 | 18 |
| top1l | 0.7443 | 0.142116902 | 0.364 | 0.308 | 1 | 18 |
| gdi2 | 0.74514 | -0.16738684 | 0.364 | 0.462 | 1 | 18 |
| ruvbl1 | 0.74518 | -0.11859366 | 0.091 | 0.154 | 1 | 18 |
| zgc:174863 | 0.74683 | 0.100104049 | 0.182 | 0.154 | 1 | 18 |
| phrf1 | 0.74799 | 0.147018124 | 0.136 | 0.077 | 1 | 18 |
| rpe | 0.74892 | 0.101875882 | 0.136 | 0.077 | 1 | 18 |
| XLOC-015238 | 0.74936 | -0.12275712 | 0.136 | 0.192 | 1 | 18 |
| eif3ba | 0.7494 | -0.12831238 | 0.636 | 0.692 | 1 | 18 |
| reck | 0.74971 | 0.13920477 | 0.136 | 0.077 | 1 | 18 |
| capza1b | 0.75043 | -0.15412401 | 0.318 | 0.423 | 1 | 18 |
| clk4a | 0.7506 | -0.12715773 | 0.091 | 0.154 | 1 | 18 |
| ndufaf1 | 0.75173 | 0.123628976 | 0.182 | 0.115 | 1 | 18 |
| elof1 | 0.7518 | -0.1428825 | 0.318 | 0.423 | 1 | 18 |
| aldh2.2 | 0.75224 | -0.13359072 | 0.091 | 0.154 | 1 | 18 |
| ccdc82 | 0.75226 | 0.14494659 | 0.227 | 0.154 | 1 | 18 |
| sf3b1 | 0.75279 | -0.12488732 | 0.182 | 0.269 | 1 | 18 |
| nucks1b | 0.75447 | -0.14668324 | 0.318 | 0.423 | 1 | 18 |
| col6a4a | 0.75463 | 0.136708102 | 0.591 | 0.538 | 1 | 18 |
| pfkfb4b | 0.75475 | -0.13950066 | 0.318 | 0.423 | 1 | 18 |
| exosc1 | 0.75531 | 0.126458676 | 0.136 | 0.077 | 1 | 18 |
| meis3 | 0.75599 | 0.150365964 | 0.227 | 0.154 | 1 | 18 |
| gng2 | 0.75707 | -0.13690943 | 0.182 | 0.231 | 1 | 18 |
| si:dkey-112e7.2 | 0.75724 | -0.11003212 | 0.182 | 0.269 | 1 | 18 |
| igf2b | 0.75775 | 0.16157307 | 0.591 | 0.5 | 1 | 18 |

| | | | | | | |
|---|---|---|---|---|---|---|
| glod4 | 0.75825 | 0.105788502 | 0.136 | 0.077 | 1 | 18 |
| abi3bpb | 0.75845 | -0.16891283 | 0.409 | 0.5 | 1 | 18 |
| polb | 0.75861 | 0.115933848 | 0.182 | 0.115 | 1 | 18 |
| aldh7a1 | 0.75914 | 0.102861042 | 0.136 | 0.077 | 1 | 18 |
| si:ch211-267e7.3 | 0.7606 | 0.14481071 | 0.182 | 0.115 | 1 | 18 |
| smtnl | 0.76079 | 0.12051472 | 0.136 | 0.077 | 1 | 18 |
| zgc:77056 | 0.76122 | -0.11671823 | 0.091 | 0.154 | 1 | 18 |
| cyb5r3 | 0.76246 | -0.12841355 | 0.091 | 0.154 | 1 | 18 |
| jpt1b | 0.76255 | 0.111499732 | 0.818 | 0.731 | 1 | 18 |
| prdx2 | 0.76271 | -0.12741095 | 0.682 | 0.769 | 1 | 18 |
| pdia3 | 0.76343 | -0.12881023 | 0.636 | 0.692 | 1 | 18 |
| fnbp1l | 0.76374 | 0.129565395 | 0.136 | 0.077 | 1 | 18 |
| srsf5a | 0.7639 | -0.1226374 | 0.591 | 0.692 | 1 | 18 |
| mia3 | 0.7642 | 0.11742599 | 0.136 | 0.077 | 1 | 18 |
| clns1a | 0.76457 | -0.12299381 | 0.182 | 0.269 | 1 | 18 |
| col6a1 | 0.76464 | 0.113673094 | 0.5 | 0.462 | 1 | 18 |
| plecb | 0.76472 | -0.13070934 | 0.091 | 0.154 | 1 | 18 |
| cacna1g | 0.76527 | -0.14856603 | 0.182 | 0.269 | 1 | 18 |
| stag2b | 0.76606 | -0.12587215 | 0.136 | 0.192 | 1 | 18 |
| znf503 | 0.76625 | 0.169740242 | 0.364 | 0.269 | 1 | 18 |
| akap12b | 0.76689 | -0.12656892 | 0.364 | 0.462 | 1 | 18 |
| ptp4a3 | 0.76775 | 0.132811946 | 0.182 | 0.115 | 1 | 18 |
| timm9 | 0.76805 | -0.12464552 | 0.136 | 0.192 | 1 | 18 |
| pcf11 | 0.7686 | 0.150643674 | 0.273 | 0.192 | 1 | 18 |
| ube2e1 | 0.76885 | -0.11949403 | 0.182 | 0.269 | 1 | 18 |
| phc2b | 0.76941 | 0.136491272 | 0.136 | 0.077 | 1 | 18 |
| med6 | 0.77 | 0.122409393 | 0.182 | 0.115 | 1 | 18 |
| fbln5 | 0.77092 | 0.128546325 | 0.136 | 0.077 | 1 | 18 |
| bckdhbl | 0.77093 | 0.11273644 | 0.318 | 0.269 | 1 | 18 |
| rnf44 | 0.77104 | -0.11069182 | 0.091 | 0.115 | 1 | 18 |
| ube2na | 0.77106 | -0.13873522 | 0.364 | 0.462 | 1 | 18 |
| tnfsf12 | 0.77117 | -0.1391953 | 0.182 | 0.269 | 1 | 18 |
| rab6a | 0.77139 | -0.12869578 | 0.182 | 0.269 | 1 | 18 |
| igf2bp3 | 0.77164 | 0.140616627 | 0.182 | 0.115 | 1 | 18 |
| meaf6 | 0.77173 | 0.134494496 | 0.455 | 0.385 | 1 | 18 |
| carm1 | 0.77215 | 0.136798458 | 0.182 | 0.115 | 1 | 18 |
| prdx4 | 0.77263 | -0.12454493 | 0.455 | 0.5 | 1 | 18 |
| bgna | 0.77272 | -0.10421363 | 0.227 | 0.308 | 1 | 18 |
| pim3 | 0.77352 | 0.138674752 | 0.455 | 0.385 | 1 | 18 |
| fbxo11a | 0.77356 | -0.13727722 | 0.091 | 0.154 | 1 | 18 |
| atp5fa1 | 0.77359 | -0.11920854 | 0.636 | 0.731 | 1 | 18 |
| tent5c | 0.7745 | -0.14906183 | 0.091 | 0.154 | 1 | 18 |
| zfp36l1b | 0.77694 | 0.113700333 | 0.273 | 0.231 | 1 | 18 |
| hspb1 | 0.77734 | -0.15340438 | 0.364 | 0.462 | 1 | 18 |
| naa15a | 0.77843 | 0.119439568 | 0.136 | 0.077 | 1 | 18 |
| ms4a17a.14 | 0.77867 | 0.115046347 | 0.136 | 0.077 | 1 | 18 |
| cox7c | 0.77923 | 0.111217468 | 0.818 | 0.769 | 1 | 18 |
| rnf185 | 0.77933 | -0.12631271 | 0.136 | 0.192 | 1 | 18 |

| | | | | | | |
|---|---|---|---|---|---|---|
| rpa3 | 0.77934 | 0.120126242 | 0.182 | 0.115 | 1 | 18 |
| ap2a1 | 0.77942 | 0.11608426 | 0.136 | 0.077 | 1 | 18 |
| GCA | 0.78068 | -0.11586135 | 0.364 | 0.462 | 1 | 18 |
| tsc22d3 | 0.78068 | -0.14115484 | 0.364 | 0.462 | 1 | 18 |
| rap1aa | 0.78078 | 0.132443195 | 0.318 | 0.231 | 1 | 18 |
| si:dkey-195m11.8 | 0.78137 | 0.124255407 | 0.136 | 0.077 | 1 | 18 |
| smim15 | 0.78147 | -0.10803694 | 0.364 | 0.462 | 1 | 18 |
| BX322530.1 | 0.78155 | -0.1268023 | 0.136 | 0.192 | 1 | 18 |
| zgc:92242 | 0.78162 | 0.120969842 | 0.136 | 0.077 | 1 | 18 |
| tuba8l4 | 0.78271 | 0.106942906 | 0.591 | 0.577 | 1 | 18 |
| arglu1a | 0.78294 | -0.13168986 | 0.182 | 0.231 | 1 | 18 |
| klf6a | 0.78372 | 0.160339736 | 0.5 | 0.423 | 1 | 18 |
| cfap97 | 0.78374 | 0.12437697 | 0.136 | 0.077 | 1 | 18 |
| pcna | 0.7847 | 0.129941828 | 0.318 | 0.231 | 1 | 18 |
| si:ch211-212k18.4 | 0.78515 | 0.142987671 | 0.136 | 0.077 | 1 | 18 |
| rbm7 | 0.78522 | 0.101983424 | 0.136 | 0.077 | 1 | 18 |
| hgs | 0.78604 | 0.11364564 | 0.136 | 0.077 | 1 | 18 |
| ube2h | 0.78633 | 0.121729772 | 0.182 | 0.115 | 1 | 18 |
| DDX17 | 0.78689 | 0.104241331 | 0.182 | 0.154 | 1 | 18 |
| gorasp2 | 0.78717 | -0.12708321 | 0.136 | 0.192 | 1 | 18 |
| UTP14C | 0.78767 | 0.13350758 | 0.136 | 0.077 | 1 | 18 |
| adgrl3.1 | 0.7878 | -0.10172712 | 0.227 | 0.308 | 1 | 18 |
| si:ch211-286o17.1 | 0.78862 | -0.14593551 | 0.364 | 0.462 | 1 | 18 |
| atf4b | 0.78877 | 0.130563089 | 0.636 | 0.538 | 1 | 18 |
| ccdc80 | 0.78883 | -0.12614873 | 0.182 | 0.231 | 1 | 18 |
| si:ch1073-325m22.2 | 0.78935 | -0.10910761 | 0.5 | 0.538 | 1 | 18 |
| nupr1a | 0.78969 | 0.125357207 | 0.227 | 0.154 | 1 | 18 |
| ddx61 | 0.7897 | -0.155355 | 0.409 | 0.5 | 1 | 18 |
| atp5mc3b | 0.79017 | 0.114317938 | 0.773 | 0.731 | 1 | 18 |
| mmp2 | 0.79038 | -0.14268753 | 0.364 | 0.462 | 1 | 18 |
| lrrc47 | 0.79047 | 0.112628932 | 0.136 | 0.077 | 1 | 18 |
| marveld1 | 0.79063 | 0.125327459 | 0.136 | 0.077 | 1 | 18 |
| wif1 | 0.79088 | 0.10702297 | 0.182 | 0.115 | 1 | 18 |
| nck1a | 0.79112 | 0.123959973 | 0.227 | 0.154 | 1 | 18 |
| snrpa1 | 0.79136 | 0.124801339 | 0.136 | 0.077 | 1 | 18 |
| kank2 | 0.79164 | 0.116325191 | 0.273 | 0.192 | 1 | 18 |
| blcap | 0.7921 | -0.1077643 | 0.091 | 0.154 | 1 | 18 |
| dnaja2b | 0.79254 | -0.11417535 | 0.182 | 0.231 | 1 | 18 |
| rpp21 | 0.79455 | -0.11568277 | 0.091 | 0.154 | 1 | 18 |
| ash2l | 0.79509 | -0.13157715 | 0.091 | 0.154 | 1 | 18 |
| sh3glb2b | 0.79542 | 0.106838622 | 0.136 | 0.077 | 1 | 18 |
| ccdc28a | 0.79575 | 0.10183774 | 0.136 | 0.077 | 1 | 18 |
| spg21 | 0.7963 | -0.11921296 | 0.091 | 0.154 | 1 | 18 |
| znf207a | 0.79631 | -0.10108858 | 0.091 | 0.154 | 1 | 18 |
| usp14 | 0.7965 | -0.10419576 | 0.227 | 0.269 | 1 | 18 |
| rybpb | 0.7965 | -0.10990387 | 0.091 | 0.154 | 1 | 18 |
| cnn3b | 0.79655 | -0.10046397 | 0.091 | 0.154 | 1 | 18 |
| zgc:113208 | 0.79662 | -0.14397151 | 0.091 | 0.154 | 1 | 18 |

| | | | | | | |
|---|---|---|---|---|---|---|
| pudp | 0.79698 | 0.121343337 | 0.182 | 0.115 | 1 | 18 |
| si:ch211-162e15.3 | 0.79723 | -0.1106453 | 0.091 | 0.154 | 1 | 18 |
| actr6 | 0.79758 | 0.120947421 | 0.136 | 0.077 | 1 | 18 |
| anapc11 | 0.79858 | 0.11592922 | 0.136 | 0.077 | 1 | 18 |
| pcdh11 | 0.79932 | 0.100428471 | 0.136 | 0.077 | 1 | 18 |
| arnt2 | 0.79945 | 0.106844101 | 0.136 | 0.077 | 1 | 18 |
| fgfr1a | 0.79955 | 0.121310116 | 0.136 | 0.077 | 1 | 18 |
| dnase1l1 | 0.79964 | 0.112424326 | 0.318 | 0.269 | 1 | 18 |
| knop1 | 0.79995 | -0.11856749 | 0.136 | 0.192 | 1 | 18 |
| gar1 | 0.79999 | 0.11875165 | 0.182 | 0.115 | 1 | 18 |
| stmn1a | 0.80097 | 0.164550605 | 0.136 | 0.077 | 1 | 18 |
| ythdf1 | 0.80109 | -0.10428527 | 0.091 | 0.154 | 1 | 18 |
| CABZ01039304.1 | 0.8011 | 0.118106199 | 0.136 | 0.077 | 1 | 18 |
| fzd4 | 0.8012 | -0.13750814 | 0.273 | 0.346 | 1 | 18 |
| gli2a | 0.80129 | 0.128506397 | 0.136 | 0.077 | 1 | 18 |
| LOC108191614 | 0.80143 | 0.107360803 | 0.136 | 0.077 | 1 | 18 |
| pcdh19 | 0.80199 | -0.10367506 | 0.091 | 0.154 | 1 | 18 |
| ufm1 | 0.80243 | 0.117009362 | 0.273 | 0.192 | 1 | 18 |
| hnrnpabb | 0.80264 | 0.126005893 | 0.636 | 0.577 | 1 | 18 |
| man2b1 | 0.80377 | -0.11837956 | 0.091 | 0.154 | 1 | 18 |
| cox7a2l | 0.8038 | 0.120383819 | 0.273 | 0.192 | 1 | 18 |
| npm1a | 0.80382 | 0.14550478 | 0.364 | 0.308 | 1 | 18 |
| fermt2 | 0.80408 | -0.12329326 | 0.091 | 0.154 | 1 | 18 |
| dhrs4 | 0.80448 | -0.10482074 | 0.091 | 0.154 | 1 | 18 |
| si:dkey-17o15.2 | 0.8045 | -0.11410649 | 0.091 | 0.154 | 1 | 18 |
| sema6e | 0.80451 | -0.1147813 | 0.091 | 0.154 | 1 | 18 |
| efna2b | 0.8048 | 0.140713792 | 0.227 | 0.154 | 1 | 18 |
| sat1a.2 | 0.80595 | -0.12709633 | 0.364 | 0.423 | 1 | 18 |
| vdrb | 0.80702 | 0.134079463 | 0.227 | 0.154 | 1 | 18 |
| tob1a | 0.80729 | -0.13251317 | 0.227 | 0.308 | 1 | 18 |
| si:ch73-138n13.1 | 0.80808 | 0.113255009 | 0.227 | 0.154 | 1 | 18 |
| srd5a3 | 0.80897 | 0.113316253 | 0.227 | 0.154 | 1 | 18 |
| gtf2e2 | 0.80918 | 0.106805964 | 0.182 | 0.115 | 1 | 18 |
| rbm39b | 0.80918 | 0.130059323 | 0.182 | 0.115 | 1 | 18 |
| ywhah | 0.81063 | -0.1337233 | 0.227 | 0.308 | 1 | 18 |
| psmd4a | 0.8108 | 0.10777922 | 0.182 | 0.115 | 1 | 18 |
| tomm6 | 0.8111 | -0.10018416 | 0.409 | 0.5 | 1 | 18 |
| bod1 | 0.81112 | 0.105503445 | 0.182 | 0.115 | 1 | 18 |
| serinc1 | 0.81145 | 0.111965506 | 0.227 | 0.154 | 1 | 18 |
| gstk1 | 0.81152 | 0.113313216 | 0.227 | 0.154 | 1 | 18 |
| mrps25 | 0.81153 | 0.111689035 | 0.182 | 0.115 | 1 | 18 |
| ddx5 | 0.81391 | -0.13591957 | 0.409 | 0.5 | 1 | 18 |
| si:ch73-269m14.2 | 0.81649 | -0.11227375 | 0.136 | 0.192 | 1 | 18 |
| abcc5 | 0.81776 | 0.103149231 | 0.182 | 0.154 | 1 | 18 |
| minos1 | 0.82005 | 0.100462717 | 0.227 | 0.192 | 1 | 18 |
| wt1a | 0.8214 | -0.13322139 | 0.136 | 0.192 | 1 | 18 |
| eif4e3 | 0.82145 | -0.10216948 | 0.182 | 0.231 | 1 | 18 |
| map1lc3b | 0.82153 | 0.126623701 | 0.409 | 0.346 | 1 | 18 |

| | | | | | | |
|---|---|---|---|---|---|---|
| anxa4 | 0.82356 | -0.11461537 | 0.545 | 0.615 | 1 | 18 |
| snai2 | 0.82379 | -0.1142926 | 0.136 | 0.192 | 1 | 18 |
| jagn1b | 0.82522 | -0.10020171 | 0.182 | 0.231 | 1 | 18 |
| kctd5a | 0.82751 | -0.12107612 | 0.364 | 0.423 | 1 | 18 |
| brd2a | 0.82904 | 0.10658012 | 0.273 | 0.231 | 1 | 18 |
| fgf24 | 0.82973 | 0.135713758 | 0.5 | 0.423 | 1 | 18 |
| myo1ca | 0.83013 | -0.11542963 | 0.136 | 0.192 | 1 | 18 |
| srsf9 | 0.83576 | 0.109400924 | 0.364 | 0.308 | 1 | 18 |
| fzd7b | 0.8372 | 0.133477419 | 0.455 | 0.385 | 1 | 18 |
| sox4a | 0.8387 | -0.12562792 | 0.273 | 0.346 | 1 | 18 |
| meis1b | 0.8445 | -0.10817058 | 0.455 | 0.538 | 1 | 18 |
| ahnak | 0.84482 | -0.10862336 | 0.455 | 0.538 | 1 | 18 |
| cstf2 | 0.84516 | -0.12389456 | 0.136 | 0.192 | 1 | 18 |
| tns1b | 0.84697 | -0.13290812 | 0.136 | 0.192 | 1 | 18 |
| rbm28 | 0.84832 | -0.11900686 | 0.136 | 0.192 | 1 | 18 |
| ppt1 | 0.84833 | 0.118780553 | 0.409 | 0.346 | 1 | 18 |
| kmt2cb | 0.84942 | -0.12490445 | 0.182 | 0.231 | 1 | 18 |
| spina | 0.84992 | -0.12115631 | 0.136 | 0.192 | 1 | 18 |
| zgc:153713 | 0.85284 | -0.1147168 | 0.273 | 0.346 | 1 | 18 |
| XLOC-013964 | 0.85629 | -0.10684821 | 0.091 | 0.115 | 1 | 18 |
| eif4a1a | 0.85641 | 0.113381992 | 0.5 | 0.423 | 1 | 18 |
| cyc1 | 0.85721 | -0.10232643 | 0.273 | 0.346 | 1 | 18 |
| ginm1 | 0.85968 | -0.11074905 | 0.182 | 0.231 | 1 | 18 |
| tpd52l2b | 0.85978 | 0.100863108 | 0.273 | 0.231 | 1 | 18 |
| pim1 | 0.85984 | -0.10339425 | 0.227 | 0.269 | 1 | 18 |
| ctdspl2b | 0.86586 | -0.11091284 | 0.136 | 0.192 | 1 | 18 |
| ptpn21 | 0.86681 | -0.11145642 | 0.136 | 0.192 | 1 | 18 |
| zfhx3 | 0.86769 | -0.11643766 | 0.318 | 0.385 | 1 | 18 |
| si:ch73-91k6.2 | 0.87214 | -0.10359201 | 0.136 | 0.192 | 1 | 18 |
| fibinb | 0.87459 | -0.10061883 | 0.455 | 0.5 | 1 | 18 |
| ccl25b | 0.88509 | 0.112466543 | 0.682 | 0.615 | 1 | 18 |
| zbtb20 | 0.88567 | -0.1065773 | 0.364 | 0.423 | 1 | 18 |
| arhgef1b | 0.90236 | 0.103516927 | 0.318 | 0.269 | 1 | 18 |
| mt-atp6 | 4.734967 | -1.15812265 | 0.129 | 0.824 | 0.0956321 | 19 |
| mt-co2 | 0.00052 | -1.08983485 | 0.258 | 0.824 | 1 | 19 |
| mt-cyb | 0.00112 | -1.02659736 | 0.226 | 0.765 | 1 | 19 |
| clasp2 | 0.00127 | -0.56669487 | 0 | 0.294 | 1 | 19 |
| arpc5b | 0.00273 | 0.875119231 | 0.516 | 0.059 | 1 | 19 |
| pomp | 0.00295 | -0.86335856 | 0.258 | 0.706 | 1 | 19 |
| sumo2b | 0.00392 | 0.793906012 | 0.484 | 0.059 | 1 | 19 |
| mt-nd2 | 0.00457 | -0.78142003 | 0.097 | 0.529 | 1 | 19 |
| brk1 | 0.00468 | -0.7603584 | 0.065 | 0.471 | 1 | 19 |
| atp2a2b | 0.00477 | -0.43429459 | 0 | 0.235 | 1 | 19 |
| cnot3b | 0.00477 | -0.46044851 | 0 | 0.235 | 1 | 19 |
| mrpl32 | 0.00477 | -0.41398986 | 0 | 0.235 | 1 | 19 |
| ncoa5 | 0.00477 | -0.41980017 | 0 | 0.235 | 1 | 19 |
| nono | 0.00477 | -0.58871888 | 0 | 0.235 | 1 | 19 |
| rsrc2 | 0.00477 | -0.52861998 | 0 | 0.235 | 1 | 19 |

| | | | | | | |
|---|---|---|---|---|---|---|
| si:ch73-226l13.2 | 0.00477 | -0.56991903 | 0 | 0.235 | 1 | 19 |
| tollip | 0.00477 | -0.43294493 | 0 | 0.235 | 1 | 19 |
| vat1 | 0.00477 | -0.47676084 | 0 | 0.235 | 1 | 19 |
| ssr1 | 0.00753 | -0.77715422 | 0.097 | 0.471 | 1 | 19 |
| COX3 | 0.00842 | -0.70327367 | 0.452 | 0.882 | 1 | 19 |
| mt-nd1 | 0.01078 | -0.74919655 | 0.129 | 0.529 | 1 | 19 |
| aimp2 | 0.01081 | -0.66508084 | 0.032 | 0.353 | 1 | 19 |
| LOC103911745 | 0.01487 | 0.576806369 | 0.226 | 0 | 1 | 19 |
| mycbp | 0.01487 | 0.502466759 | 0.226 | 0 | 1 | 19 |
| CR847936.10 | 0.01487 | 0.532844738 | 0.226 | 0 | 1 | 19 |
| snapc5 | 0.01487 | 0.449187448 | 0.226 | 0 | 1 | 19 |
| taf10 | 0.01487 | 0.562728133 | 0.226 | 0 | 1 | 19 |
| tra2a | 0.01487 | 0.435270957 | 0.226 | 0 | 1 | 19 |
| zmat2 | 0.01487 | 0.482011184 | 0.226 | 0 | 1 | 19 |
| pinx1 | 0.01512 | -0.54999843 | 0.065 | 0.412 | 1 | 19 |
| cnot9 | 0.0173 | -0.23979765 | 0 | 0.176 | 1 | 19 |
| gb:bc139872 | 0.0173 | -0.37070509 | 0 | 0.176 | 1 | 19 |
| hmg20b | 0.0173 | -0.49161836 | 0 | 0.176 | 1 | 19 |
| hnrnpub | 0.0173 | -0.32866043 | 0 | 0.176 | 1 | 19 |
| oat | 0.0173 | -0.28920708 | 0 | 0.176 | 1 | 19 |
| pik3cg | 0.0173 | -0.40883051 | 0 | 0.176 | 1 | 19 |
| rtraf | 0.0173 | -0.37235281 | 0 | 0.176 | 1 | 19 |
| si:dkey-147f3.8 | 0.0173 | -0.33713199 | 0 | 0.176 | 1 | 19 |
| si:dkey-253d23.4 | 0.0173 | -0.36974252 | 0 | 0.176 | 1 | 19 |
| si:dkey-78l4.1 | 0.0173 | -0.78784267 | 0 | 0.176 | 1 | 19 |
| slc29a2 | 0.0173 | -0.35167667 | 0 | 0.176 | 1 | 19 |
| ssna1 | 0.0173 | -0.33533478 | 0 | 0.176 | 1 | 19 |
| tank | 0.0173 | -0.35853527 | 0 | 0.176 | 1 | 19 |
| zgc:92161 | 0.0173 | -0.42075564 | 0 | 0.176 | 1 | 19 |
| crebbpa | 0.0173 | -0.27490142 | 0 | 0.176 | 1 | 19 |
| elmsan1a | 0.0173 | -0.37962179 | 0 | 0.176 | 1 | 19 |
| zdhhc21 | 0.0173 | -0.45356719 | 0 | 0.176 | 1 | 19 |
| gstm.2 | 0.0173 | -0.38146412 | 0 | 0.176 | 1 | 19 |
| lnpep | 0.0173 | -0.38692497 | 0 | 0.176 | 1 | 19 |
| ogdha | 0.0173 | -0.45526013 | 0 | 0.176 | 1 | 19 |
| si:ch1073-296d18.1 | 0.0173 | -0.3454943 | 0 | 0.176 | 1 | 19 |
| wdr36 | 0.0173 | -0.38394186 | 0 | 0.176 | 1 | 19 |
| ccnl1a | 0.0173 | -0.34834762 | 0 | 0.176 | 1 | 19 |
| dnajc3b | 0.0173 | -0.32325565 | 0 | 0.176 | 1 | 19 |
| gab2 | 0.0173 | -0.36128477 | 0 | 0.176 | 1 | 19 |
| gtf2b | 0.0173 | -0.26709725 | 0 | 0.176 | 1 | 19 |
| hexb | 0.0173 | -0.32330231 | 0 | 0.176 | 1 | 19 |
| lgals3a | 0.0173 | -0.45457807 | 0 | 0.176 | 1 | 19 |
| map7d3 | 0.0173 | -0.3381549 | 0 | 0.176 | 1 | 19 |
| prdx2 | 0.02533 | -0.65038853 | 0.226 | 0.588 | 1 | 19 |
| elmsan1b | 0.02649 | 0.442885054 | 0.194 | 0 | 1 | 19 |
| ndufb10 | 0.02649 | 0.473489356 | 0.194 | 0 | 1 | 19 |
| snrpc | 0.02649 | 0.356793417 | 0.194 | 0 | 1 | 19 |

| | | | | | | |
|---|---|---|---|---|---|---|
| u2af2b | 0.02649 | 0.400070883 | 0.194 | 0 | 1 | 19 |
| CABZ01117603.1 | 0.02649 | 0.428667654 | 0.194 | 0 | 1 | 19 |
| hint2 | 0.02649 | 0.350548843 | 0.194 | 0 | 1 | 19 |
| rab27a | 0.02649 | 0.391672248 | 0.194 | 0 | 1 | 19 |
| ndufc2 | 0.02649 | 0.409105223 | 0.194 | 0 | 1 | 19 |
| plp2 | 0.02649 | 0.429357474 | 0.194 | 0 | 1 | 19 |
| efhd2 | 0.03606 | -0.64383065 | 0.129 | 0.471 | 1 | 19 |
| smarce1 | 0.03621 | -0.46284915 | 0.032 | 0.294 | 1 | 19 |
| lcp1 | 0.03646 | -0.52721206 | 0.097 | 0.412 | 1 | 19 |
| erh | 0.03762 | -0.41319022 | 0.194 | 0.529 | 1 | 19 |
| ost4 | 0.03921 | -0.6488021 | 0.194 | 0.412 | 1 | 19 |
| srsf6b | 0.04046 | -0.43920939 | 0.065 | 0.353 | 1 | 19 |
| tm9sf4 | 0.04312 | -0.54290967 | 0.065 | 0.353 | 1 | 19 |
| hsp90b1 | 0.04324 | -0.57551446 | 0.065 | 0.353 | 1 | 19 |
| chd9 | 0.04681 | 0.326416485 | 0.161 | 0 | 1 | 19 |
| tmpob | 0.04681 | 0.288231082 | 0.161 | 0 | 1 | 19 |
| chd4b | 0.04681 | 0.349718626 | 0.161 | 0 | 1 | 19 |
| ndufs4 | 0.04681 | 0.292164479 | 0.161 | 0 | 1 | 19 |
| sort1b | 0.04681 | 0.344260116 | 0.161 | 0 | 1 | 19 |
| srgap3 | 0.04681 | 0.301007046 | 0.161 | 0 | 1 | 19 |
| tfr1b | 0.04681 | 0.352796239 | 0.161 | 0 | 1 | 19 |
| tspan33a | 0.04681 | 0.354194606 | 0.161 | 0 | 1 | 19 |
| CU467110.1 | 0.04681 | 0.39733305 | 0.161 | 0 | 1 | 19 |
| atp6ap2 | 0.04681 | 0.35135023 | 0.161 | 0 | 1 | 19 |
| dedd1 | 0.04681 | 0.345219994 | 0.161 | 0 | 1 | 19 |
| rhocb | 0.04681 | 0.301114658 | 0.161 | 0 | 1 | 19 |
| si:dkey-185m8.2 | 0.04681 | 0.347782721 | 0.161 | 0 | 1 | 19 |
| szrd1 | 0.04681 | 0.397157431 | 0.161 | 0 | 1 | 19 |
| zgc:165573 | 0.04681 | 0.324670559 | 0.161 | 0 | 1 | 19 |
| ccl35.1 | 0.04681 | 0.503005875 | 0.161 | 0 | 1 | 19 |
| chchd6b | 0.04681 | 0.252689393 | 0.161 | 0 | 1 | 19 |
| cyth4a | 0.04681 | 0.336560983 | 0.161 | 0 | 1 | 19 |
| ddx47 | 0.04681 | 0.316574999 | 0.161 | 0 | 1 | 19 |
| pam16 | 0.04681 | 0.323483426 | 0.161 | 0 | 1 | 19 |
| rpf2 | 0.04681 | 0.367557053 | 0.161 | 0 | 1 | 19 |
| si:dkey-78l4.7 | 0.04681 | 0.62682206 | 0.161 | 0 | 1 | 19 |
| hck | 0.04851 | 0.594295799 | 0.355 | 0.059 | 1 | 19 |
| eif5a2 | 0.0593 | -0.56293688 | 0.323 | 0.647 | 1 | 19 |
| BX470189.1 | 0.06172 | -0.23289863 | 0 | 0.118 | 1 | 19 |
| CABZ01046088.1 | 0.06172 | -0.22204056 | 0 | 0.118 | 1 | 19 |
| CABZ01081490.1 | 0.06172 | -0.28180684 | 0 | 0.118 | 1 | 19 |
| CR388166.2 | 0.06172 | -0.31861527 | 0 | 0.118 | 1 | 19 |
| LO018154.1 | 0.06172 | -0.16494194 | 0 | 0.118 | 1 | 19 |
| LOC100332329 | 0.06172 | -0.30046006 | 0 | 0.118 | 1 | 19 |
| LOC101884921 | 0.06172 | -0.27191741 | 0 | 0.118 | 1 | 19 |
| LOC101886015 | 0.06172 | -0.23933371 | 0 | 0.118 | 1 | 19 |
| acadl | 0.06172 | -0.23586109 | 0 | 0.118 | 1 | 19 |
| acp6 | 0.06172 | -0.26837114 | 0 | 0.118 | 1 | 19 |

| | | | | | | |
|---|---|---|---|---|---|---|
| adob | 0.06172 | -0.30808812 | 0 | 0.118 | 1 | 19 |
| akr7a3 | 0.06172 | -0.16920294 | 0 | 0.118 | 1 | 19 |
| anapc13 | 0.06172 | -0.18173977 | 0 | 0.118 | 1 | 19 |
| ascc1 | 0.06172 | -0.15872725 | 0 | 0.118 | 1 | 19 |
| atad5a | 0.06172 | -0.14070978 | 0 | 0.118 | 1 | 19 |
| atp13a1 | 0.06172 | -0.19477681 | 0 | 0.118 | 1 | 19 |
| atp6ap1b | 0.06172 | -0.20100541 | 0 | 0.118 | 1 | 19 |
| bag4 | 0.06172 | -0.30891475 | 0 | 0.118 | 1 | 19 |
| bbs1 | 0.06172 | -0.21704166 | 0 | 0.118 | 1 | 19 |
| bms1 | 0.06172 | -0.16322881 | 0 | 0.118 | 1 | 19 |
| bod1 | 0.06172 | -0.23580762 | 0 | 0.118 | 1 | 19 |
| c2cd2 | 0.06172 | -0.28671359 | 0 | 0.118 | 1 | 19 |
| capn7 | 0.06172 | -0.23838967 | 0 | 0.118 | 1 | 19 |
| ccl34b.1 | 0.06172 | -0.60911662 | 0 | 0.118 | 1 | 19 |
| ccl39.1 | 0.06172 | -0.36960564 | 0 | 0.118 | 1 | 19 |
| cdc40 | 0.06172 | -0.26857541 | 0 | 0.118 | 1 | 19 |
| cdc6 | 0.06172 | -0.14988487 | 0 | 0.118 | 1 | 19 |
| cdkal1 | 0.06172 | -0.33870362 | 0 | 0.118 | 1 | 19 |
| cecr2 | 0.06172 | -0.27506417 | 0 | 0.118 | 1 | 19 |
| cenpw | 0.06172 | -0.11634711 | 0 | 0.118 | 1 | 19 |
| cherp | 0.06172 | -0.30900902 | 0 | 0.118 | 1 | 19 |
| chst14 | 0.06172 | -0.38718421 | 0 | 0.118 | 1 | 19 |
| chtf18 | 0.06172 | -0.18984099 | 0 | 0.118 | 1 | 19 |
| ciao1 | 0.06172 | -0.15008413 | 0 | 0.118 | 1 | 19 |
| clns1a | 0.06172 | -0.35275746 | 0 | 0.118 | 1 | 19 |
| cox15 | 0.06172 | -0.30265289 | 0 | 0.118 | 1 | 19 |
| crk | 0.06172 | -0.2775785 | 0 | 0.118 | 1 | 19 |
| cstf1 | 0.06172 | -0.35979146 | 0 | 0.118 | 1 | 19 |
| ctcf | 0.06172 | -0.26176378 | 0 | 0.118 | 1 | 19 |
| cux1b | 0.06172 | -0.21669165 | 0 | 0.118 | 1 | 19 |
| dars | 0.06172 | -0.23733353 | 0 | 0.118 | 1 | 19 |
| dcps | 0.06172 | -0.16970972 | 0 | 0.118 | 1 | 19 |
| dhx9 | 0.06172 | -0.29482112 | 0 | 0.118 | 1 | 19 |
| diablob | 0.06172 | -0.22588739 | 0 | 0.118 | 1 | 19 |
| dnajc2 | 0.06172 | -0.32812033 | 0 | 0.118 | 1 | 19 |
| dnajc25 | 0.06172 | -0.34998294 | 0 | 0.118 | 1 | 19 |
| dnmt1 | 0.06172 | -0.16850155 | 0 | 0.118 | 1 | 19 |
| elmo3 | 0.06172 | -0.40035699 | 0 | 0.118 | 1 | 19 |
| emc8 | 0.06172 | -0.26517791 | 0 | 0.118 | 1 | 19 |
| fam53b | 0.06172 | -0.19427995 | 0 | 0.118 | 1 | 19 |
| gcat | 0.06172 | -0.20700428 | 0 | 0.118 | 1 | 19 |
| gins2 | 0.06172 | -0.2258828 | 0 | 0.118 | 1 | 19 |
| glud1a | 0.06172 | -0.26392506 | 0 | 0.118 | 1 | 19 |
| golgb1 | 0.06172 | -0.23524889 | 0 | 0.118 | 1 | 19 |
| gramd1a | 0.06172 | -0.21566704 | 0 | 0.118 | 1 | 19 |
| hat1 | 0.06172 | -0.22837068 | 0 | 0.118 | 1 | 19 |
| hdac3 | 0.06172 | -0.16699766 | 0 | 0.118 | 1 | 19 |
| hipk3a | 0.06172 | -0.15466564 | 0 | 0.118 | 1 | 19 |

| | | | | | | |
|---|---|---|---|---|---|---|
| hmg20a | 0.06172 | -0.2222921 | 0 | 0.118 | 1 | 19 |
| hmox2b | 0.06172 | -0.17105465 | 0 | 0.118 | 1 | 19 |
| hps3 | 0.06172 | -0.21249893 | 0 | 0.118 | 1 | 19 |
| hspa14 | 0.06172 | -0.26665749 | 0 | 0.118 | 1 | 19 |
| im:7140055 | 0.06172 | -0.16871961 | 0 | 0.118 | 1 | 19 |
| ipcef1 | 0.06172 | -0.38973736 | 0 | 0.118 | 1 | 19 |
| kdm1a | 0.06172 | -0.14181269 | 0 | 0.118 | 1 | 19 |
| kdm6bb | 0.06172 | -0.28705437 | 0 | 0.118 | 1 | 19 |
| lamtor4 | 0.06172 | -0.23435436 | 0 | 0.118 | 1 | 19 |
| lig1 | 0.06172 | -0.16285529 | 0 | 0.118 | 1 | 19 |
| lman1 | 0.06172 | -0.30566342 | 0 | 0.118 | 1 | 19 |
| lmo7b | 0.06172 | -0.2584342 | 0 | 0.118 | 1 | 19 |
| mau2 | 0.06172 | -0.29248878 | 0 | 0.118 | 1 | 19 |
| mboat2a | 0.06172 | -0.28903896 | 0 | 0.118 | 1 | 19 |
| mcm3 | 0.06172 | -0.19867163 | 0 | 0.118 | 1 | 19 |
| mcm7 | 0.06172 | -0.19702198 | 0 | 0.118 | 1 | 19 |
| med15 | 0.06172 | -0.24189762 | 0 | 0.118 | 1 | 19 |
| med20 | 0.06172 | -0.30333294 | 0 | 0.118 | 1 | 19 |
| mfap1 | 0.06172 | -0.12241007 | 0 | 0.118 | 1 | 19 |
| micu1 | 0.06172 | -0.25228982 | 0 | 0.118 | 1 | 19 |
| mis12 | 0.06172 | -0.18695234 | 0 | 0.118 | 1 | 19 |
| mlx | 0.06172 | -0.31655382 | 0 | 0.118 | 1 | 19 |
| mpped2a | 0.06172 | -0.27596555 | 0 | 0.118 | 1 | 19 |
| mrpl10 | 0.06172 | -0.24664475 | 0 | 0.118 | 1 | 19 |
| mybl2b | 0.06172 | -0.16956508 | 0 | 0.118 | 1 | 19 |
| myg1 | 0.06172 | -0.25301972 | 0 | 0.118 | 1 | 19 |
| nat10 | 0.06172 | -0.24052525 | 0 | 0.118 | 1 | 19 |
| naxd | 0.06172 | -0.16759058 | 0 | 0.118 | 1 | 19 |
| nbr1b | 0.06172 | -0.2356406 | 0 | 0.118 | 1 | 19 |
| neu1 | 0.06172 | -0.19509438 | 0 | 0.118 | 1 | 19 |
| ogfod1 | 0.06172 | -0.27524628 | 0 | 0.118 | 1 | 19 |
| parp2 | 0.06172 | -0.14440477 | 0 | 0.118 | 1 | 19 |
| pdcd2l | 0.06172 | -0.3465034 | 0 | 0.118 | 1 | 19 |
| pde7a | 0.06172 | -0.1537111 | 0 | 0.118 | 1 | 19 |
| pik3r5 | 0.06172 | -0.27226796 | 0 | 0.118 | 1 | 19 |
| polr1b | 0.06172 | -0.29445986 | 0 | 0.118 | 1 | 19 |
| ppm1g | 0.06172 | -0.40823885 | 0 | 0.118 | 1 | 19 |
| prpf6 | 0.06172 | -0.10161663 | 0 | 0.118 | 1 | 19 |
| prps1b | 0.06172 | -0.2112899 | 0 | 0.118 | 1 | 19 |
| pspc1 | 0.06172 | -0.15561628 | 0 | 0.118 | 1 | 19 |
| ptcd3 | 0.06172 | -0.26129911 | 0 | 0.118 | 1 | 19 |
| ptgis | 0.06172 | -0.24725134 | 0 | 0.118 | 1 | 19 |
| rab8b | 0.06172 | -0.29302063 | 0 | 0.118 | 1 | 19 |
| rbm22 | 0.06172 | -0.15639525 | 0 | 0.118 | 1 | 19 |
| rexo1 | 0.06172 | -0.28999494 | 0 | 0.118 | 1 | 19 |
| rnpc3 | 0.06172 | -0.14611305 | 0 | 0.118 | 1 | 19 |
| rrm1 | 0.06172 | -0.2021006 | 0 | 0.118 | 1 | 19 |
| rrm2 | 0.06172 | -0.27953931 | 0 | 0.118 | 1 | 19 |

| | | | | | | |
|---|---|---|---|---|---|---|
| sdhaf4 | 0.06172 | -0.24369131 | 0 | 0.118 | 1 | 19 |
| selenoj | 0.06172 | -0.11284202 | 0 | 0.118 | 1 | 19 |
| si:ch211-156b7.4 | 0.06172 | -0.26780036 | 0 | 0.118 | 1 | 19 |
| si:ch211-161m3.4 | 0.06172 | -0.15826569 | 0 | 0.118 | 1 | 19 |
| si:ch73-40a2.1 | 0.06172 | -0.29886972 | 0 | 0.118 | 1 | 19 |
| si:dkey-188i13.10 | 0.06172 | -0.39291746 | 0 | 0.118 | 1 | 19 |
| si:dkey-217f16.1 | 0.06172 | -0.24478437 | 0 | 0.118 | 1 | 19 |
| si:dkey-234h16.7 | 0.06172 | -0.26796844 | 0 | 0.118 | 1 | 19 |
| si:dkey-29h14.10 | 0.06172 | -0.29488858 | 0 | 0.118 | 1 | 19 |
| slbp | 0.06172 | -0.27753241 | 0 | 0.118 | 1 | 19 |
| slc16a6b | 0.06172 | -0.39084576 | 0 | 0.118 | 1 | 19 |
| slc29a1a | 0.06172 | -0.14650284 | 0 | 0.118 | 1 | 19 |
| slc35e3 | 0.06172 | -0.26675469 | 0 | 0.118 | 1 | 19 |
| smfn | 0.06172 | -0.1926527 | 0 | 0.118 | 1 | 19 |
| snx20 | 0.06172 | -0.17166312 | 0 | 0.118 | 1 | 19 |
| sp1 | 0.06172 | -0.36770347 | 0 | 0.118 | 1 | 19 |
| srp14 | 0.06172 | -0.31493588 | 0 | 0.118 | 1 | 19 |
| srsf1b | 0.06172 | -0.32309685 | 0 | 0.118 | 1 | 19 |
| st6gal2a | 0.06172 | -0.315223 | 0 | 0.118 | 1 | 19 |
| tarbp2 | 0.06172 | -0.27829927 | 0 | 0.118 | 1 | 19 |
| thg1l | 0.06172 | -0.16687352 | 0 | 0.118 | 1 | 19 |
| tk1 | 0.06172 | -0.20660007 | 0 | 0.118 | 1 | 19 |
| tor1 | 0.06172 | -0.23974743 | 0 | 0.118 | 1 | 19 |
| trioa | 0.06172 | -0.27458988 | 0 | 0.118 | 1 | 19 |
| trub1 | 0.06172 | -0.26732029 | 0 | 0.118 | 1 | 19 |
| tuba8l2 | 0.06172 | -0.17188432 | 0 | 0.118 | 1 | 19 |
| ubxn1 | 0.06172 | -0.25502539 | 0 | 0.118 | 1 | 19 |
| wdr46 | 0.06172 | -0.3666109 | 0 | 0.118 | 1 | 19 |
| xrcc6 | 0.06172 | -0.13926625 | 0 | 0.118 | 1 | 19 |
| zgc:101858 | 0.06172 | -0.30697989 | 0 | 0.118 | 1 | 19 |
| zgc:110540 | 0.06172 | -0.19717345 | 0 | 0.118 | 1 | 19 |
| zgc:112496 | 0.06172 | -0.1204 | 0 | 0.118 | 1 | 19 |
| zgc:193742 | 0.06172 | -0.16243464 | 0 | 0.118 | 1 | 19 |
| zgc:194981 | 0.06172 | -0.31061419 | 0 | 0.118 | 1 | 19 |
| zgc:92664 | 0.06172 | -0.26043066 | 0 | 0.118 | 1 | 19 |
| znf622 | 0.06172 | -0.20912138 | 0 | 0.118 | 1 | 19 |
| CABZ01029938.1 | 0.06172 | -0.3120696 | 0 | 0.118 | 1 | 19 |
| aff4 | 0.06172 | -0.32194837 | 0 | 0.118 | 1 | 19 |
| ap1ar | 0.06172 | -0.34233841 | 0 | 0.118 | 1 | 19 |
| foxo4 | 0.06172 | -0.34395886 | 0 | 0.118 | 1 | 19 |
| gbp1 | 0.06172 | -0.34627929 | 0 | 0.118 | 1 | 19 |
| ggps1 | 0.06172 | -0.32348393 | 0 | 0.118 | 1 | 19 |
| gmfg | 0.06172 | -0.34415627 | 0 | 0.118 | 1 | 19 |
| herpud2 | 0.06172 | -0.3184868 | 0 | 0.118 | 1 | 19 |
| ints2 | 0.06172 | -0.33926681 | 0 | 0.118 | 1 | 19 |
| rxrbb | 0.06172 | -0.35909547 | 0 | 0.118 | 1 | 19 |
| si:dkey-57k2.7 | 0.06172 | -0.34359008 | 0 | 0.118 | 1 | 19 |
| sp3a | 0.06172 | -0.31515099 | 0 | 0.118 | 1 | 19 |

| | | | | | | |
|---|---|---|---|---|---|---|
| supt6h | 0.06172 | -0.31185545 | 0 | 0.118 | 1 | 19 |
| vbp1 | 0.06172 | -0.28951624 | 0 | 0.118 | 1 | 19 |
| zgc:113411 | 0.06172 | -0.32076648 | 0 | 0.118 | 1 | 19 |
| CABZ01044053.1 | 0.06172 | -0.30184276 | 0 | 0.118 | 1 | 19 |
| CABZ01058650.1 | 0.06172 | -0.3068156 | 0 | 0.118 | 1 | 19 |
| COA4 | 0.06172 | -0.22133394 | 0 | 0.118 | 1 | 19 |
| CR786582.2 | 0.06172 | -0.24278395 | 0 | 0.118 | 1 | 19 |
| CU019662.1 | 0.06172 | -0.27994965 | 0 | 0.118 | 1 | 19 |
| acat1 | 0.06172 | -0.2257382 | 0 | 0.118 | 1 | 19 |
| aebp2 | 0.06172 | -0.18006816 | 0 | 0.118 | 1 | 19 |
| ddb1 | 0.06172 | -0.16931846 | 0 | 0.118 | 1 | 19 |
| dhdhl | 0.06172 | -0.23047291 | 0 | 0.118 | 1 | 19 |
| eif1axa | 0.06172 | -0.19294261 | 0 | 0.118 | 1 | 19 |
| fam114a2 | 0.06172 | -0.22455342 | 0 | 0.118 | 1 | 19 |
| fam204a | 0.06172 | -0.18814408 | 0 | 0.118 | 1 | 19 |
| fam3c | 0.06172 | -0.33765706 | 0 | 0.118 | 1 | 19 |
| fbxw11b | 0.06172 | -0.27478729 | 0 | 0.118 | 1 | 19 |
| flot2a | 0.06172 | -0.18549978 | 0 | 0.118 | 1 | 19 |
| hnrnpl | 0.06172 | -0.19784814 | 0 | 0.118 | 1 | 19 |
| itgav | 0.06172 | -0.27240887 | 0 | 0.118 | 1 | 19 |
| kansl1a | 0.06172 | -0.31373201 | 0 | 0.118 | 1 | 19 |
| kpna4 | 0.06172 | -0.27029977 | 0 | 0.118 | 1 | 19 |
| lmf2b | 0.06172 | -0.20368448 | 0 | 0.118 | 1 | 19 |
| lrrc20 | 0.06172 | -0.22707724 | 0 | 0.118 | 1 | 19 |
| lsg1 | 0.06172 | -0.14294517 | 0 | 0.118 | 1 | 19 |
| mfsd1 | 0.06172 | -0.2396832 | 0 | 0.118 | 1 | 19 |
| mrps23 | 0.06172 | -0.17274594 | 0 | 0.118 | 1 | 19 |
| mrps35 | 0.06172 | -0.19266914 | 0 | 0.118 | 1 | 19 |
| mtg1 | 0.06172 | -0.23456119 | 0 | 0.118 | 1 | 19 |
| nasp | 0.06172 | -0.20429769 | 0 | 0.118 | 1 | 19 |
| nudt3a | 0.06172 | -0.33261651 | 0 | 0.118 | 1 | 19 |
| nup43 | 0.06172 | -0.23768384 | 0 | 0.118 | 1 | 19 |
| pcf11 | 0.06172 | -0.31784806 | 0 | 0.118 | 1 | 19 |
| si:ch211-212d10.2 | 0.06172 | -0.18918051 | 0 | 0.118 | 1 | 19 |
| si:ch73-223f5.1 | 0.06172 | -0.20167507 | 0 | 0.118 | 1 | 19 |
| si:dkey-103e21.5 | 0.06172 | -0.28941353 | 0 | 0.118 | 1 | 19 |
| si:dkey-192g7.3 | 0.06172 | -0.20151061 | 0 | 0.118 | 1 | 19 |
| si:dkey-22o12.2 | 0.06172 | -0.26194982 | 0 | 0.118 | 1 | 19 |
| slc1a5 | 0.06172 | -0.24733666 | 0 | 0.118 | 1 | 19 |
| slc35a4 | 0.06172 | -0.27328666 | 0 | 0.118 | 1 | 19 |
| sod2 | 0.06172 | -0.13986403 | 0 | 0.118 | 1 | 19 |
| stk11 | 0.06172 | -0.17360259 | 0 | 0.118 | 1 | 19 |
| them4 | 0.06172 | -0.24581621 | 0 | 0.118 | 1 | 19 |
| tial1 | 0.06172 | -0.1841191 | 0 | 0.118 | 1 | 19 |
| trim33 | 0.06172 | -0.23570028 | 0 | 0.118 | 1 | 19 |
| trmt11 | 0.06172 | -0.26749982 | 0 | 0.118 | 1 | 19 |
| tshz2 | 0.06172 | -0.26047183 | 0 | 0.118 | 1 | 19 |
| ttc3 | 0.06172 | -0.34184007 | 0 | 0.118 | 1 | 19 |

| | | | | | | |
|---|---|---|---|---|---|---|
| tyrobp | 0.06172 | -0.20577235 | 0 | 0.118 | 1 | 19 |
| ubap1 | 0.06172 | -0.19876474 | 0 | 0.118 | 1 | 19 |
| ube2h | 0.06172 | -0.276327 | 0 | 0.118 | 1 | 19 |
| uri1 | 0.06172 | -0.18511238 | 0 | 0.118 | 1 | 19 |
| vdac3 | 0.06172 | -0.32353052 | 0 | 0.118 | 1 | 19 |
| vps37c | 0.06172 | -0.30787141 | 0 | 0.118 | 1 | 19 |
| yju2 | 0.06172 | -0.16225648 | 0 | 0.118 | 1 | 19 |
| bud23 | 0.06172 | -0.26559001 | 0 | 0.118 | 1 | 19 |
| rab41 | 0.06172 | -0.27680165 | 0 | 0.118 | 1 | 19 |
| tagln2 | 0.06309 | -0.33303853 | 0.29 | 0.588 | 1 | 19 |
| si:dkey-9i23.4 | 0.06475 | 0.400354778 | 0.29 | 0.294 | 1 | 19 |
| fth1a | 0.06839 | -0.59756914 | 0.161 | 0.471 | 1 | 19 |
| rpl12 | 0.07234 | 0.121728498 | 0.903 | 0.941 | 1 | 19 |
| mt-co1 | 0.07525 | -0.42462665 | 0.677 | 0.765 | 1 | 19 |
| h2afx1 | 0.0779 | 0.402803893 | 0.29 | 0.059 | 1 | 19 |
| sec61g | 0.08126 | -0.25725738 | 0.806 | 1 | 1 | 19 |
| irf2bpl | 0.08229 | 0.23838728 | 0.129 | 0 | 1 | 19 |
| tomm20a | 0.08229 | 0.318160196 | 0.129 | 0 | 1 | 19 |
| atox1 | 0.08229 | 0.22216453 | 0.129 | 0 | 1 | 19 |
| atp2b1a | 0.08229 | 0.370951922 | 0.129 | 0 | 1 | 19 |
| pcnx3 | 0.08229 | 0.234179413 | 0.129 | 0 | 1 | 19 |
| supt4h1 | 0.08229 | 0.214493799 | 0.129 | 0 | 1 | 19 |
| tp53inp1 | 0.08229 | 0.220696422 | 0.129 | 0 | 1 | 19 |
| wu:fb16f03 | 0.08229 | 0.284886407 | 0.129 | 0 | 1 | 19 |
| abi1a | 0.08229 | 0.317221389 | 0.129 | 0 | 1 | 19 |
| adgre13 | 0.08229 | 0.265255424 | 0.129 | 0 | 1 | 19 |
| atp6v1ba | 0.08229 | 0.286522128 | 0.129 | 0 | 1 | 19 |
| ciartb | 0.08229 | 0.258802994 | 0.129 | 0 | 1 | 19 |
| commd3 | 0.08229 | 0.272165903 | 0.129 | 0 | 1 | 19 |
| erbin | 0.08229 | 0.256742968 | 0.129 | 0 | 1 | 19 |
| gm2a | 0.08229 | 0.252803667 | 0.129 | 0 | 1 | 19 |
| mrps16 | 0.08229 | 0.381893839 | 0.129 | 0 | 1 | 19 |
| mta2 | 0.08229 | 0.293436665 | 0.129 | 0 | 1 | 19 |
| nup58 | 0.08229 | 0.299352538 | 0.129 | 0 | 1 | 19 |
| pdlim5b | 0.08229 | 0.245877935 | 0.129 | 0 | 1 | 19 |
| pitpnaa | 0.08229 | 0.313764476 | 0.129 | 0 | 1 | 19 |
| prdx6 | 0.08229 | 0.262826863 | 0.129 | 0 | 1 | 19 |
| psmd4a | 0.08229 | 0.240133114 | 0.129 | 0 | 1 | 19 |
| rab36 | 0.08229 | 0.295540956 | 0.129 | 0 | 1 | 19 |
| rassf2b | 0.08229 | 0.284515015 | 0.129 | 0 | 1 | 19 |
| sema7a | 0.08229 | 0.236072371 | 0.129 | 0 | 1 | 19 |
| si:ch211-112c15.8 | 0.08229 | 0.247332761 | 0.129 | 0 | 1 | 19 |
| si:ch211-246m6.4 | 0.08229 | 0.166444611 | 0.129 | 0 | 1 | 19 |
| si:dkey-118k5.3 | 0.08229 | 0.302186602 | 0.129 | 0 | 1 | 19 |
| si:dkey-180p18.9 | 0.08229 | 0.324654231 | 0.129 | 0 | 1 | 19 |
| sptlc1 | 0.08229 | 0.307285958 | 0.129 | 0 | 1 | 19 |
| sra1 | 0.08229 | 0.24936069 | 0.129 | 0 | 1 | 19 |
| tnrc6b | 0.08229 | 0.308446463 | 0.129 | 0 | 1 | 19 |

| | | | | | | |
|---|---|---|---|---|---|---|
| zfp36l2 | 0.08229 | 0.261168596 | 0.129 | 0 | 1 | 19 |
| aif1l | 0.08229 | 0.39904234 | 0.129 | 0 | 1 | 19 |
| amfra | 0.08229 | 0.264512842 | 0.129 | 0 | 1 | 19 |
| asb12a | 0.08229 | 0.237498979 | 0.129 | 0 | 1 | 19 |
| clint1a | 0.08229 | 0.229953093 | 0.129 | 0 | 1 | 19 |
| cox7a2l | 0.08229 | 0.265895386 | 0.129 | 0 | 1 | 19 |
| dhx16 | 0.08229 | 0.258759324 | 0.129 | 0 | 1 | 19 |
| dnaja2b | 0.08229 | 0.283930147 | 0.129 | 0 | 1 | 19 |
| eif4a2 | 0.08229 | 0.22132276 | 0.129 | 0 | 1 | 19 |
| fcer1g | 0.08229 | 0.325031215 | 0.129 | 0 | 1 | 19 |
| lmnb1 | 0.08229 | 0.249567256 | 0.129 | 0 | 1 | 19 |
| nkl.3 | 0.08229 | 0.453637399 | 0.129 | 0 | 1 | 19 |
| pdcd10b | 0.08229 | 0.227966295 | 0.129 | 0 | 1 | 19 |
| rspry1 | 0.08229 | 0.237943665 | 0.129 | 0 | 1 | 19 |
| safb | 0.08229 | 0.252564698 | 0.129 | 0 | 1 | 19 |
| scdb | 0.08229 | 0.282252007 | 0.129 | 0 | 1 | 19 |
| si:cabz01074946.1 | 0.08229 | 0.300074007 | 0.129 | 0 | 1 | 19 |
| si:dkey-222f8.3 | 0.08229 | 0.235283023 | 0.129 | 0 | 1 | 19 |
| si:dkey-78l4.5 | 0.08229 | 0.3924089 | 0.129 | 0 | 1 | 19 |
| smim7 | 0.08229 | 0.318345436 | 0.129 | 0 | 1 | 19 |
| sqstm1 | 0.08229 | 0.281166881 | 0.129 | 0 | 1 | 19 |
| triobpa | 0.08229 | 0.267815835 | 0.129 | 0 | 1 | 19 |
| tspan36 | 0.08229 | 0.245769869 | 0.129 | 0 | 1 | 19 |
| wdfy2 | 0.08229 | 0.250593771 | 0.129 | 0 | 1 | 19 |
| wdr61 | 0.08229 | 0.290606036 | 0.129 | 0 | 1 | 19 |
| zgc:77739 | 0.08229 | 0.243764735 | 0.129 | 0 | 1 | 19 |
| zgc:77929 | 0.08229 | 0.34242271 | 0.129 | 0 | 1 | 19 |
| eif4g2b | 0.08229 | 0.318159321 | 0.129 | 0 | 1 | 19 |
| gsto2 | 0.08229 | 0.248219324 | 0.129 | 0 | 1 | 19 |
| gtf2h5 | 0.08229 | 0.303908469 | 0.129 | 0 | 1 | 19 |
| hcfc1a | 0.08229 | 0.259217802 | 0.129 | 0 | 1 | 19 |
| hnrnpm | 0.08229 | 0.3012396 | 0.129 | 0 | 1 | 19 |
| jak2a | 0.08229 | 0.211059732 | 0.129 | 0 | 1 | 19 |
| si:busm1-194e12.11 | 0.08229 | 0.250340622 | 0.129 | 0 | 1 | 19 |
| si:ch211-239m17.1 | 0.08229 | 0.253577549 | 0.129 | 0 | 1 | 19 |
| slc25a20 | 0.08229 | 0.253423502 | 0.129 | 0 | 1 | 19 |
| atr | 0.08302 | -0.4835406 | 0.032 | 0.235 | 1 | 19 |
| smu1a | 0.0833 | -0.51706328 | 0.032 | 0.235 | 1 | 19 |
| ddx3xa | 0.09082 | -0.51652867 | 0.065 | 0.294 | 1 | 19 |
| gfi1ab | 0.09178 | -0.34640456 | 0.032 | 0.235 | 1 | 19 |
| ero1b | 0.09204 | -0.44888568 | 0.032 | 0.235 | 1 | 19 |
| naa40 | 0.0921 | -0.41770258 | 0.032 | 0.235 | 1 | 19 |
| adssl | 0.09237 | -0.4115246 | 0.032 | 0.235 | 1 | 19 |
| rac2 | 0.0926 | 0.500955822 | 0.71 | 0.471 | 1 | 19 |
| pdia4 | 0.09285 | -0.56412246 | 0.032 | 0.235 | 1 | 19 |
| cox7c | 0.09422 | -0.51270541 | 0.484 | 0.706 | 1 | 19 |
| sec61a1 | 0.09518 | -0.54380773 | 0.194 | 0.471 | 1 | 19 |
| junbb | 0.09569 | -0.45778035 | 0.032 | 0.235 | 1 | 19 |

| | | | | | | |
|---|---|---|---|---|---|---|
| tmx4 | 0.09671 | -0.50029043 | 0.032 | 0.235 | 1 | 19 |
| srp19 | 0.09721 | -0.51764162 | 0.065 | 0.294 | 1 | 19 |
| eif3i | 0.09771 | -0.40642212 | 0.226 | 0.529 | 1 | 19 |
| slc25a25a | 0.09805 | -0.39261392 | 0.032 | 0.235 | 1 | 19 |
| si:dkey-7j14.6 | 0.09813 | -0.37497721 | 0.032 | 0.235 | 1 | 19 |
| sf3b5 | 0.09844 | -0.52623642 | 0.097 | 0.353 | 1 | 19 |
| XLOC-043043 | 0.09927 | 0.520388353 | 0.774 | 0.471 | 1 | 19 |
| cyc1 | 0.1 | -0.50101351 | 0.065 | 0.294 | 1 | 19 |
| si:ch211-212k18.7 | 0.10025 | -0.36344888 | 0.032 | 0.235 | 1 | 19 |
| c1d | 0.10031 | -0.38379495 | 0.032 | 0.235 | 1 | 19 |
| dnajc1 | 0.10046 | -0.36891445 | 0.065 | 0.294 | 1 | 19 |
| rplp0 | 0.10058 | 0.100049221 | 0.935 | 0.765 | 1 | 19 |
| dock10 | 0.10123 | -0.45127229 | 0.032 | 0.235 | 1 | 19 |
| rpl27a | 0.10139 | -0.42110169 | 0.645 | 0.824 | 1 | 19 |
| rsu1 | 0.10435 | -0.46875411 | 0.065 | 0.294 | 1 | 19 |
| si:dkey-33i11.9 | 0.10657 | -0.50941892 | 0.355 | 0.529 | 1 | 19 |
| cd247l | 0.10682 | -0.52764451 | 0.194 | 0.471 | 1 | 19 |
| BX571675.1 | 0.10717 | -0.49981503 | 0.065 | 0.294 | 1 | 19 |
| ppp6c | 0.10752 | -0.47510025 | 0.065 | 0.294 | 1 | 19 |
| XLOC-042595 | 0.10784 | 0.383344434 | 0.355 | 0.118 | 1 | 19 |
| prkcda | 0.10872 | -0.39986947 | 0.065 | 0.294 | 1 | 19 |
| zgc:101583 | 0.109 | -0.4590058 | 0.065 | 0.294 | 1 | 19 |
| capgb | 0.10902 | -0.29587652 | 0.484 | 0.765 | 1 | 19 |
| slirp | 0.11032 | -0.3823472 | 0.065 | 0.294 | 1 | 19 |
| ezrb | 0.11082 | -0.46305245 | 0.065 | 0.294 | 1 | 19 |
| prelid3b | 0.1169 | -0.55119448 | 0.161 | 0.412 | 1 | 19 |
| cct2 | 0.11894 | -0.53953741 | 0.194 | 0.471 | 1 | 19 |
| ndufa1 | 0.1205 | -0.52762697 | 0.194 | 0.471 | 1 | 19 |
| si:dkey-11f4.14 | 0.12126 | 0.504232832 | 0.387 | 0.118 | 1 | 19 |
| cnn2 | 0.12458 | 0.437221525 | 0.29 | 0.059 | 1 | 19 |
| ywhaz | 0.12573 | 0.441918543 | 0.29 | 0.059 | 1 | 19 |
| faslg | 0.12595 | 0.330779597 | 0.419 | 0.176 | 1 | 19 |
| dusp5 | 0.12812 | -0.57465024 | 0.194 | 0.471 | 1 | 19 |
| arpc2 | 0.12862 | 0.407223005 | 0.71 | 0.529 | 1 | 19 |
| ndfip2 | 0.12888 | 0.391272151 | 0.29 | 0.059 | 1 | 19 |
| nfkbiaa | 0.12987 | -0.45740594 | 0.258 | 0.353 | 1 | 19 |
| rpl24 | 0.13002 | 0.121231636 | 0.935 | 0.765 | 1 | 19 |
| cebpb | 0.13198 | -0.26093777 | 0.548 | 0.529 | 1 | 19 |
| mt-nd3 | 0.13623 | -0.59370191 | 0.129 | 0.353 | 1 | 19 |
| tomm5 | 0.13763 | -0.33291723 | 0.161 | 0.412 | 1 | 19 |
| cd81a | 0.13789 | 0.273185899 | 0.258 | 0.059 | 1 | 19 |
| nfkbiab | 0.13795 | -0.46273512 | 0.194 | 0.471 | 1 | 19 |
| ahcy | 0.13843 | -0.11188589 | 0.032 | 0.176 | 1 | 19 |
| hmgb2a | 0.14039 | -0.20520624 | 0.742 | 0.706 | 1 | 19 |
| atp6v1e1a | 0.14475 | 0.211068235 | 0.097 | 0 | 1 | 19 |
| dbpb | 0.14475 | 0.202473211 | 0.097 | 0 | 1 | 19 |
| sh3glb1a | 0.14475 | 0.226325001 | 0.097 | 0 | 1 | 19 |
| slc25a24 | 0.14475 | 0.172848865 | 0.097 | 0 | 1 | 19 |

| | | | | | | |
|---|---|---|---|---|---|---|
| spink4 | 0.14475 | 0.275742725 | 0.097 | 0 | 1 | 19 |
| zgc:194224 | 0.14475 | 0.299957028 | 0.097 | 0 | 1 | 19 |
| CABZ01034698.2 | 0.14475 | 0.192687562 | 0.097 | 0 | 1 | 19 |
| LOC101885990 | 0.14475 | 0.241493212 | 0.097 | 0 | 1 | 19 |
| NC-002333.17 | 0.14475 | 0.162683459 | 0.097 | 0 | 1 | 19 |
| SCYL2 | 0.14475 | 0.196149608 | 0.097 | 0 | 1 | 19 |
| aip | 0.14475 | 0.187651982 | 0.097 | 0 | 1 | 19 |
| ap2m1b | 0.14475 | 0.215985545 | 0.097 | 0 | 1 | 19 |
| atf3 | 0.14475 | 0.231879612 | 0.097 | 0 | 1 | 19 |
| bloc1s1 | 0.14475 | 0.173054354 | 0.097 | 0 | 1 | 19 |
| cbr1l | 0.14475 | 0.191376996 | 0.097 | 0 | 1 | 19 |
| cops3 | 0.14475 | 0.291678688 | 0.097 | 0 | 1 | 19 |
| eif4bb | 0.14475 | 0.218102563 | 0.097 | 0 | 1 | 19 |
| fra10ac1 | 0.14475 | 0.150182937 | 0.097 | 0 | 1 | 19 |
| gnsb | 0.14475 | 0.197448558 | 0.097 | 0 | 1 | 19 |
| hcst | 0.14475 | 0.239994988 | 0.097 | 0 | 1 | 19 |
| irs1 | 0.14475 | 0.170696549 | 0.097 | 0 | 1 | 19 |
| kdm3b | 0.14475 | 0.157184653 | 0.097 | 0 | 1 | 19 |
| msl1a | 0.14475 | 0.258247946 | 0.097 | 0 | 1 | 19 |
| naa15a | 0.14475 | 0.185756441 | 0.097 | 0 | 1 | 19 |
| nfu1 | 0.14475 | 0.193760115 | 0.097 | 0 | 1 | 19 |
| nkl.2 | 0.14475 | 0.502571749 | 0.097 | 0 | 1 | 19 |
| nom1 | 0.14475 | 0.148382397 | 0.097 | 0 | 1 | 19 |
| pak1 | 0.14475 | 0.24472135 | 0.097 | 0 | 1 | 19 |
| pim1 | 0.14475 | 0.184610467 | 0.097 | 0 | 1 | 19 |
| pkn1a | 0.14475 | 0.251310441 | 0.097 | 0 | 1 | 19 |
| plcg1 | 0.14475 | 0.29095307 | 0.097 | 0 | 1 | 19 |
| ptpmt1 | 0.14475 | 0.137429321 | 0.097 | 0 | 1 | 19 |
| rcor1 | 0.14475 | 0.200011633 | 0.097 | 0 | 1 | 19 |
| sdc4 | 0.14475 | 0.224292986 | 0.097 | 0 | 1 | 19 |
| sgta | 0.14475 | 0.194452468 | 0.097 | 0 | 1 | 19 |
| si:ch211-195b13.1 | 0.14475 | 0.229549911 | 0.097 | 0 | 1 | 19 |
| si:dkey-159f12.2 | 0.14475 | 0.241103636 | 0.097 | 0 | 1 | 19 |
| si:dkey-276j7.1 | 0.14475 | 0.203311415 | 0.097 | 0 | 1 | 19 |
| si:dkeyp-28d2.4 | 0.14475 | 0.195983826 | 0.097 | 0 | 1 | 19 |
| snap23.1 | 0.14475 | 0.238873546 | 0.097 | 0 | 1 | 19 |
| srp9 | 0.14475 | 0.232497517 | 0.097 | 0 | 1 | 19 |
| tab1 | 0.14475 | 0.182072804 | 0.097 | 0 | 1 | 19 |
| timmdc1 | 0.14475 | 0.158006582 | 0.097 | 0 | 1 | 19 |
| tmcc3 | 0.14475 | 0.14046666 | 0.097 | 0 | 1 | 19 |
| tpcn2 | 0.14475 | 0.254931112 | 0.097 | 0 | 1 | 19 |
| tra2b | 0.14475 | 0.349834148 | 0.097 | 0 | 1 | 19 |
| traf1 | 0.14475 | 0.155761126 | 0.097 | 0 | 1 | 19 |
| trpc4apa | 0.14475 | 0.173885521 | 0.097 | 0 | 1 | 19 |
| waplb | 0.14475 | 0.264934695 | 0.097 | 0 | 1 | 19 |
| zgc:101577 | 0.14475 | 0.233618418 | 0.097 | 0 | 1 | 19 |
| zgc:110158 | 0.14475 | 0.210615302 | 0.097 | 0 | 1 | 19 |
| zgc:55262 | 0.14475 | 0.272706011 | 0.097 | 0 | 1 | 19 |

| | | | | | | |
|---|---|---|---|---|---|---|
| zgc:92744 | 0.14475 | 0.249545054 | 0.097 | 0 | 1 | 19 |
| amy2a | 0.14475 | 0.176615517 | 0.097 | 0 | 1 | 19 |
| rabgap1l | 0.14475 | 0.23387074 | 0.097 | 0 | 1 | 19 |
| tmem214 | 0.14475 | 0.222161678 | 0.097 | 0 | 1 | 19 |
| tpst1l | 0.14475 | 0.230055067 | 0.097 | 0 | 1 | 19 |
| tspan3a | 0.14475 | 0.259596646 | 0.097 | 0 | 1 | 19 |
| csgalnact1a | 0.14475 | 0.205519893 | 0.097 | 0 | 1 | 19 |
| castor1 | 0.14475 | 0.206938988 | 0.097 | 0 | 1 | 19 |
| cyfip2 | 0.14475 | 0.248771704 | 0.097 | 0 | 1 | 19 |
| kdm2ba | 0.14475 | 0.30030387 | 0.097 | 0 | 1 | 19 |
| lpcat3 | 0.14475 | 0.180584155 | 0.097 | 0 | 1 | 19 |
| metrnla | 0.14475 | 0.234846646 | 0.097 | 0 | 1 | 19 |
| orai1a | 0.14475 | 0.176024076 | 0.097 | 0 | 1 | 19 |
| pagr1 | 0.14475 | 0.243211543 | 0.097 | 0 | 1 | 19 |
| rabggtb | 0.14475 | 0.233504223 | 0.097 | 0 | 1 | 19 |
| si:ch211-243a20.4 | 0.14475 | 0.25753166 | 0.097 | 0 | 1 | 19 |
| tmub2 | 0.14475 | 0.245917438 | 0.097 | 0 | 1 | 19 |
| tsg101b | 0.14475 | 0.265587595 | 0.097 | 0 | 1 | 19 |
| zgc:194627 | 0.14475 | 0.235947515 | 0.097 | 0 | 1 | 19 |
| BX908750.1 | 0.14475 | 0.249161089 | 0.097 | 0 | 1 | 19 |
| CR513782.1 | 0.14475 | 0.190930434 | 0.097 | 0 | 1 | 19 |
| CU570889.1 | 0.14475 | 0.181252186 | 0.097 | 0 | 1 | 19 |
| LOC101884267 | 0.14475 | 0.294157168 | 0.097 | 0 | 1 | 19 |
| LOC108190761 | 0.14475 | 0.219714281 | 0.097 | 0 | 1 | 19 |
| XLOC-041870 | 0.14475 | 0.258021416 | 0.097 | 0 | 1 | 19 |
| aco2 | 0.14475 | 0.187757063 | 0.097 | 0 | 1 | 19 |
| adam10a | 0.14475 | 0.20214726 | 0.097 | 0 | 1 | 19 |
| anks1b | 0.14475 | 0.222881459 | 0.097 | 0 | 1 | 19 |
| apbb1ip | 0.14475 | 0.278734042 | 0.097 | 0 | 1 | 19 |
| atp6v1c1a | 0.14475 | 0.223679414 | 0.097 | 0 | 1 | 19 |
| baxa | 0.14475 | 0.242143032 | 0.097 | 0 | 1 | 19 |
| bcap29 | 0.14475 | 0.232086287 | 0.097 | 0 | 1 | 19 |
| ccdc171 | 0.14475 | 0.211294435 | 0.097 | 0 | 1 | 19 |
| ccdc88b | 0.14475 | 0.223203065 | 0.097 | 0 | 1 | 19 |
| cdc26 | 0.14475 | 0.204896263 | 0.097 | 0 | 1 | 19 |
| cflara | 0.14475 | 0.166401743 | 0.097 | 0 | 1 | 19 |
| crcp | 0.14475 | 0.245365231 | 0.097 | 0 | 1 | 19 |
| ddx23 | 0.14475 | 0.155612702 | 0.097 | 0 | 1 | 19 |
| dera | 0.14475 | 0.188957708 | 0.097 | 0 | 1 | 19 |
| dhx38 | 0.14475 | 0.192540746 | 0.097 | 0 | 1 | 19 |
| fam173a | 0.14475 | 0.187900134 | 0.097 | 0 | 1 | 19 |
| fam210b | 0.14475 | 0.227578635 | 0.097 | 0 | 1 | 19 |
| fbxl4 | 0.14475 | 0.169034681 | 0.097 | 0 | 1 | 19 |
| fbxo44 | 0.14475 | 0.166334079 | 0.097 | 0 | 1 | 19 |
| fpr1 | 0.14475 | 0.183517304 | 0.097 | 0 | 1 | 19 |
| galnt7 | 0.14475 | 0.177665751 | 0.097 | 0 | 1 | 19 |
| gigyf1a | 0.14475 | 0.177934572 | 0.097 | 0 | 1 | 19 |
| gusb | 0.14475 | 0.173775582 | 0.097 | 0 | 1 | 19 |

| | | | | | | |
|---|---|---|---|---|---|---|
| icn | 0.14475 | 0.319880814 | 0.097 | 0 | 1 | 19 |
| itpkb | 0.14475 | 0.236762747 | 0.097 | 0 | 1 | 19 |
| lrrc58b | 0.14475 | 0.155986579 | 0.097 | 0 | 1 | 19 |
| mavs | 0.14475 | 0.181871652 | 0.097 | 0 | 1 | 19 |
| mccc2 | 0.14475 | 0.167884753 | 0.097 | 0 | 1 | 19 |
| med21 | 0.14475 | 0.200659648 | 0.097 | 0 | 1 | 19 |
| mgmt | 0.14475 | 0.276408816 | 0.097 | 0 | 1 | 19 |
| mthfd2 | 0.14475 | 0.206023621 | 0.097 | 0 | 1 | 19 |
| mych | 0.14475 | 0.202279305 | 0.097 | 0 | 1 | 19 |
| naga | 0.14475 | 0.179528426 | 0.097 | 0 | 1 | 19 |
| napab | 0.14475 | 0.218447257 | 0.097 | 0 | 1 | 19 |
| ndufv1 | 0.14475 | 0.261699849 | 0.097 | 0 | 1 | 19 |
| phf23a | 0.14475 | 0.173255495 | 0.097 | 0 | 1 | 19 |
| qsox1 | 0.14475 | 0.256988989 | 0.097 | 0 | 1 | 19 |
| rab6ba | 0.14475 | 0.165360513 | 0.097 | 0 | 1 | 19 |
| rad50 | 0.14475 | 0.267843994 | 0.097 | 0 | 1 | 19 |
| rps6ka3b | 0.14475 | 0.275466447 | 0.097 | 0 | 1 | 19 |
| sh2d3ca | 0.14475 | 0.209669059 | 0.097 | 0 | 1 | 19 |
| shc1 | 0.14475 | 0.312321202 | 0.097 | 0 | 1 | 19 |
| si:dkey-222f2.7 | 0.14475 | 0.20562936 | 0.097 | 0 | 1 | 19 |
| si:dkey-260g12.1 | 0.14475 | 0.190350173 | 0.097 | 0 | 1 | 19 |
| si:dkey-27i16.2 | 0.14475 | 0.303913553 | 0.097 | 0 | 1 | 19 |
| si:dkey-78l4.8 | 0.14475 | 0.356572216 | 0.097 | 0 | 1 | 19 |
| sik2b | 0.14475 | 0.160743113 | 0.097 | 0 | 1 | 19 |
| slc30a7 | 0.14475 | 0.179573649 | 0.097 | 0 | 1 | 19 |
| snapin | 0.14475 | 0.206959085 | 0.097 | 0 | 1 | 19 |
| stard3nl | 0.14475 | 0.229441787 | 0.097 | 0 | 1 | 19 |
| taok3a | 0.14475 | 0.197839771 | 0.097 | 0 | 1 | 19 |
| tap2t | 0.14475 | 0.237274145 | 0.097 | 0 | 1 | 19 |
| tmtc1 | 0.14475 | 0.188443967 | 0.097 | 0 | 1 | 19 |
| tnrc6a | 0.14475 | 0.151325254 | 0.097 | 0 | 1 | 19 |
| tyk2 | 0.14475 | 0.247228776 | 0.097 | 0 | 1 | 19 |
| unc50 | 0.14475 | 0.249941707 | 0.097 | 0 | 1 | 19 |
| usp42 | 0.14475 | 0.21248512 | 0.097 | 0 | 1 | 19 |
| vil1 | 0.14475 | 0.183517304 | 0.097 | 0 | 1 | 19 |
| zgc:136864 | 0.14475 | 0.169014611 | 0.097 | 0 | 1 | 19 |
| zgc:92907 | 0.14475 | 0.247175447 | 0.097 | 0 | 1 | 19 |
| znf330 | 0.14475 | 0.201314674 | 0.097 | 0 | 1 | 19 |
| zyx | 0.14475 | 0.187767513 | 0.097 | 0 | 1 | 19 |
| aldh7a1 | 0.14475 | 0.224105301 | 0.097 | 0 | 1 | 19 |
| arap3 | 0.14475 | 0.154093229 | 0.097 | 0 | 1 | 19 |
| coq3 | 0.14475 | 0.163831625 | 0.097 | 0 | 1 | 19 |
| gatc | 0.14475 | 0.237813244 | 0.097 | 0 | 1 | 19 |
| gps1 | 0.14475 | 0.147815252 | 0.097 | 0 | 1 | 19 |
| ist1 | 0.14475 | 0.243043291 | 0.097 | 0 | 1 | 19 |
| lonrf1l | 0.14475 | 0.264851562 | 0.097 | 0 | 1 | 19 |
| march5l | 0.14475 | 0.217391593 | 0.097 | 0 | 1 | 19 |
| march7 | 0.14475 | 0.248820315 | 0.097 | 0 | 1 | 19 |

| | | | | | | |
|---|---|---|---|---|---|---|
| nitr5 | 0.14475 | 0.305647401 | 0.097 | 0 | 1 | 19 |
| parp16 | 0.14475 | 0.20134329 | 0.097 | 0 | 1 | 19 |
| plekhj1 | 0.14475 | 0.172145896 | 0.097 | 0 | 1 | 19 |
| ppp1r2 | 0.14475 | 0.156566708 | 0.097 | 0 | 1 | 19 |
| rnf14 | 0.14475 | 0.232379394 | 0.097 | 0 | 1 | 19 |
| si:ch211-284b7.3 | 0.14475 | 0.220881437 | 0.097 | 0 | 1 | 19 |
| tet3 | 0.14475 | 0.226649533 | 0.097 | 0 | 1 | 19 |
| vaspb | 0.14475 | 0.283522215 | 0.097 | 0 | 1 | 19 |
| dync1li2 | 0.14475 | 0.194008942 | 0.097 | 0 | 1 | 19 |
| foxp1b | 0.14475 | 0.181329708 | 0.097 | 0 | 1 | 19 |
| irf2 | 0.14475 | 0.278605776 | 0.097 | 0 | 1 | 19 |
| rbm5 | 0.14475 | 0.225209678 | 0.097 | 0 | 1 | 19 |
| selenos | 0.14475 | 0.17521555 | 0.097 | 0 | 1 | 19 |
| slc25a28 | 0.14475 | 0.19279844 | 0.097 | 0 | 1 | 19 |
| zgc:63882 | 0.14475 | 0.178038819 | 0.097 | 0 | 1 | 19 |
| PDPR | 0.14475 | 0.230202044 | 0.097 | 0 | 1 | 19 |
| ikzf2 | 0.14475 | 0.259339186 | 0.097 | 0 | 1 | 19 |
| si:dkey-239h2.3 | 0.14475 | 0.238798671 | 0.097 | 0 | 1 | 19 |
| pdxdc1 | 0.14475 | 0.194217745 | 0.097 | 0 | 1 | 19 |
| sbf1 | 0.14475 | 0.202117349 | 0.097 | 0 | 1 | 19 |
| sod1 | 0.14979 | 0.375167479 | 0.645 | 0.353 | 1 | 19 |
| atp1a1a.1 | 0.15322 | -0.55426305 | 0.065 | 0.235 | 1 | 19 |
| tmed2 | 0.15481 | -0.2952632 | 0.323 | 0.588 | 1 | 19 |
| ldha | 0.15836 | -0.41219412 | 0.161 | 0.412 | 1 | 19 |
| mdh2 | 0.15845 | 0.290863901 | 0.258 | 0.059 | 1 | 19 |
| arhgdia | 0.1585 | -0.40604354 | 0.419 | 0.706 | 1 | 19 |
| serp1 | 0.16161 | -0.42281551 | 0.226 | 0.294 | 1 | 19 |
| selenof | 0.16531 | 0.111759898 | 0.419 | 0.235 | 1 | 19 |
| chac1 | 0.16533 | -0.51579362 | 0.355 | 0.588 | 1 | 19 |
| hspa5 | 0.16591 | -0.42595979 | 0.161 | 0.412 | 1 | 19 |
| dnajb9b | 0.1682 | -0.50921382 | 0.097 | 0.294 | 1 | 19 |
| khdrbs1a | 0.1688 | -0.40986903 | 0.516 | 0.765 | 1 | 19 |
| ybx1 | 0.17227 | -0.36301724 | 0.71 | 0.882 | 1 | 19 |
| ranbp3b | 0.17401 | -0.45413293 | 0.032 | 0.176 | 1 | 19 |
| cirbpa | 0.17691 | -0.42318366 | 0.548 | 0.765 | 1 | 19 |
| si:zfos-2330d3.1 | 0.17699 | 0.541437901 | 0.226 | 0.059 | 1 | 19 |
| tob1a | 0.17719 | -0.44460444 | 0.129 | 0.353 | 1 | 19 |
| rtca | 0.17951 | -0.18909591 | 0.032 | 0.176 | 1 | 19 |
| clta | 0.17996 | -0.47063665 | 0.097 | 0.294 | 1 | 19 |
| ptpn6 | 0.18211 | -0.47622861 | 0.387 | 0.588 | 1 | 19 |
| tma7 | 0.1826 | -0.44478904 | 0.387 | 0.647 | 1 | 19 |
| tspan14 | 0.18342 | -0.48697907 | 0.097 | 0.294 | 1 | 19 |
| BX322787.1 | 0.18361 | -0.23019472 | 0.387 | 0.353 | 1 | 19 |
| bhlhe40 | 0.18438 | 0.452806224 | 0.677 | 0.412 | 1 | 19 |
| psmf1 | 0.18469 | -0.46783885 | 0.032 | 0.176 | 1 | 19 |
| hnrnpa0b | 0.18599 | -0.45566925 | 0.355 | 0.588 | 1 | 19 |
| btf3l4 | 0.18622 | -0.46358123 | 0.065 | 0.235 | 1 | 19 |
| si:dkey-78l4.6 | 0.1867 | 0.112607821 | 0.355 | 0.412 | 1 | 19 |

| | | | | | | |
|---|---|---|---|---|---|---|
| park7 | 0.18983 | -0.48346167 | 0.129 | 0.353 | 1 | 19 |
| sub1a | 0.19147 | -0.37239979 | 0.355 | 0.471 | 1 | 19 |
| dnaja2a | 0.19422 | 0.27819826 | 0.323 | 0.118 | 1 | 19 |
| fkbp5 | 0.19575 | -0.36415622 | 0.129 | 0.353 | 1 | 19 |
| snw1 | 0.19575 | -0.24230955 | 0.032 | 0.176 | 1 | 19 |
| zmp:0000001082 | 0.19892 | -0.52008228 | 0.065 | 0.235 | 1 | 19 |
| si:dkey-53k12.2 | 0.20016 | -0.4220575 | 0.129 | 0.353 | 1 | 19 |
| atf4b | 0.20193 | -0.447694 | 0.129 | 0.353 | 1 | 19 |
| dido1 | 0.20225 | -0.43583163 | 0.129 | 0.353 | 1 | 19 |
| zc3h15 | 0.20331 | -0.31223344 | 0.065 | 0.235 | 1 | 19 |
| bcl2l10 | 0.20452 | 0.299896466 | 0.387 | 0.176 | 1 | 19 |
| mt-nd4 | 0.20475 | -0.39896943 | 0.323 | 0.588 | 1 | 19 |
| plrg1 | 0.20753 | -0.4057005 | 0.032 | 0.176 | 1 | 19 |
| dldh | 0.21066 | -0.44553908 | 0.065 | 0.235 | 1 | 19 |
| grpel2 | 0.21193 | -0.2726691 | 0.097 | 0.294 | 1 | 19 |
| def6a | 0.21455 | -0.45201888 | 0.097 | 0.294 | 1 | 19 |
| mtap | 0.2148 | -0.3625626 | 0.032 | 0.176 | 1 | 19 |
| FBXO48 | 0.215 | -0.44421455 | 0.065 | 0.235 | 1 | 19 |
| rps26l | 0.21594 | -0.16684897 | 0.871 | 1 | 1 | 19 |
| arl1 | 0.21669 | -0.43083495 | 0.065 | 0.235 | 1 | 19 |
| acp5b | 0.21741 | 0.246543373 | 0.226 | 0.059 | 1 | 19 |
| ube2g2 | 0.21744 | -0.24297878 | 0.032 | 0.176 | 1 | 19 |
| eif2b2 | 0.21752 | -0.23277182 | 0.032 | 0.176 | 1 | 19 |
| p4hb | 0.21799 | -0.3488261 | 0.29 | 0.353 | 1 | 19 |
| bcl2a | 0.21813 | -0.24309762 | 0.032 | 0.176 | 1 | 19 |
| cass4 | 0.21997 | -0.42106607 | 0.065 | 0.235 | 1 | 19 |
| srsf9 | 0.22038 | -0.29598974 | 0.032 | 0.176 | 1 | 19 |
| pcyox1 | 0.22064 | -0.31229211 | 0.097 | 0.294 | 1 | 19 |
| ptp4a1 | 0.22143 | -0.47854168 | 0.161 | 0.353 | 1 | 19 |
| lsm4 | 0.2217 | -0.353627 | 0.097 | 0.294 | 1 | 19 |
| spcs2 | 0.22286 | -0.45139091 | 0.29 | 0.529 | 1 | 19 |
| yrk | 0.22321 | -0.11921968 | 0.29 | 0.235 | 1 | 19 |
| arpc4 | 0.22382 | 0.369912133 | 0.613 | 0.353 | 1 | 19 |
| cd44b | 0.22407 | -0.21079605 | 0.065 | 0.235 | 1 | 19 |
| AL929028.1 | 0.22544 | -0.14202205 | 0 | 0.059 | 1 | 19 |
| AREL1 | 0.22544 | -0.12895911 | 0 | 0.059 | 1 | 19 |
| CABZ01074397.1 | 0.22544 | -0.13793204 | 0 | 0.059 | 1 | 19 |
| LOC101882921 | 0.22544 | -0.14202205 | 0 | 0.059 | 1 | 19 |
| LOC110439849 | 0.22544 | -0.13564854 | 0 | 0.059 | 1 | 19 |
| acss1 | 0.22544 | -0.13592206 | 0 | 0.059 | 1 | 19 |
| asb13b | 0.22544 | -0.14202205 | 0 | 0.059 | 1 | 19 |
| atl2 | 0.22544 | -0.13307469 | 0 | 0.059 | 1 | 19 |
| atxn10 | 0.22544 | -0.139382 | 0 | 0.059 | 1 | 19 |
| bahcc1b | 0.22544 | -0.14056927 | 0 | 0.059 | 1 | 19 |
| capn15 | 0.22544 | -0.13745997 | 0 | 0.059 | 1 | 19 |
| ccdc43 | 0.22544 | -0.13547025 | 0 | 0.059 | 1 | 19 |
| ccna2 | 0.22544 | -0.12712997 | 0 | 0.059 | 1 | 19 |
| cnpy2 | 0.22544 | -0.12521573 | 0 | 0.059 | 1 | 19 |

| | | | | | | |
|---|---|---|---|---|---|---|
| ctdspl2a | 0.22544 | -0.18651577 | 0 | 0.059 | 1 | 19 |
| diabloa | 0.22544 | -0.11928128 | 0 | 0.059 | 1 | 19 |
| dpm3 | 0.22544 | -0.13404515 | 0 | 0.059 | 1 | 19 |
| e2f7 | 0.22544 | -0.12629672 | 0 | 0.059 | 1 | 19 |
| engase | 0.22544 | -0.13617921 | 0 | 0.059 | 1 | 19 |
| foxj3 | 0.22544 | -0.13987732 | 0 | 0.059 | 1 | 19 |
| gclm | 0.22544 | -0.13744577 | 0 | 0.059 | 1 | 19 |
| hadhab | 0.22544 | -0.13871418 | 0 | 0.059 | 1 | 19 |
| hgs | 0.22544 | -0.13943479 | 0 | 0.059 | 1 | 19 |
| hk1 | 0.22544 | -0.13947405 | 0 | 0.059 | 1 | 19 |
| hnrnpul1l | 0.22544 | -0.13606346 | 0 | 0.059 | 1 | 19 |
| ibtk | 0.22544 | -0.11628795 | 0 | 0.059 | 1 | 19 |
| il6st | 0.22544 | -0.13754095 | 0 | 0.059 | 1 | 19 |
| inip | 0.22544 | -0.13634628 | 0 | 0.059 | 1 | 19 |
| ints12 | 0.22544 | -0.13510431 | 0 | 0.059 | 1 | 19 |
| itprid2 | 0.22544 | -0.13252466 | 0 | 0.059 | 1 | 19 |
| kcnq5b | 0.22544 | -0.13988825 | 0 | 0.059 | 1 | 19 |
| mitd1 | 0.22544 | -0.13360512 | 0 | 0.059 | 1 | 19 |
| mtfp1 | 0.22544 | -0.13706507 | 0 | 0.059 | 1 | 19 |
| mtr | 0.22544 | -0.13058746 | 0 | 0.059 | 1 | 19 |
| mybbp1a | 0.22544 | -0.13523148 | 0 | 0.059 | 1 | 19 |
| naxe | 0.22544 | -0.12732895 | 0 | 0.059 | 1 | 19 |
| nomo | 0.22544 | -0.13835769 | 0 | 0.059 | 1 | 19 |
| nup107 | 0.22544 | -0.13503404 | 0 | 0.059 | 1 | 19 |
| palm1a | 0.22544 | -0.12918422 | 0 | 0.059 | 1 | 19 |
| papss2b | 0.22544 | -0.13927603 | 0 | 0.059 | 1 | 19 |
| parn | 0.22544 | -0.13290756 | 0 | 0.059 | 1 | 19 |
| pds5b | 0.22544 | -0.14018762 | 0 | 0.059 | 1 | 19 |
| pdss1 | 0.22544 | -0.14008312 | 0 | 0.059 | 1 | 19 |
| pdzd8 | 0.22544 | -0.14157503 | 0 | 0.059 | 1 | 19 |
| peak1 | 0.22544 | -0.14041128 | 0 | 0.059 | 1 | 19 |
| pgap1 | 0.22544 | -0.13656608 | 0 | 0.059 | 1 | 19 |
| plekhm3 | 0.22544 | -0.14202205 | 0 | 0.059 | 1 | 19 |
| plod3 | 0.22544 | -0.13145562 | 0 | 0.059 | 1 | 19 |
| pola2 | 0.22544 | -0.13587329 | 0 | 0.059 | 1 | 19 |
| polb | 0.22544 | -0.17844927 | 0 | 0.059 | 1 | 19 |
| pom121 | 0.22544 | -0.14141252 | 0 | 0.059 | 1 | 19 |
| rangap1a | 0.22544 | -0.13820748 | 0 | 0.059 | 1 | 19 |
| rictorb | 0.22544 | -0.14202205 | 0 | 0.059 | 1 | 19 |
| rrbp1b | 0.22544 | -0.13191028 | 0 | 0.059 | 1 | 19 |
| scyl1 | 0.22544 | -0.13696366 | 0 | 0.059 | 1 | 19 |
| si:ch211-261n11.5 | 0.22544 | -0.13504949 | 0 | 0.059 | 1 | 19 |
| si:ch73-1a9.4 | 0.22544 | -0.12364357 | 0 | 0.059 | 1 | 19 |
| si:dkey-204f11.64 | 0.22544 | -0.13523918 | 0 | 0.059 | 1 | 19 |
| si:dkey-6i22.5 | 0.22544 | -0.10110176 | 0 | 0.059 | 1 | 19 |
| slc25a18 | 0.22544 | -0.13497678 | 0 | 0.059 | 1 | 19 |
| slc5a11 | 0.22544 | -0.14051323 | 0 | 0.059 | 1 | 19 |
| snx2 | 0.22544 | -0.14058036 | 0 | 0.059 | 1 | 19 |

| | | | | | | |
|---|---|---|---|---|---|---|
| srm | 0.22544 | -0.1381378 | 0 | 0.059 | 1 | 19 |
| srsf6a | 0.22544 | -0.13741635 | 0 | 0.059 | 1 | 19 |
| sssca1 | 0.22544 | -0.12880622 | 0 | 0.059 | 1 | 19 |
| stimate | 0.22544 | -0.137279 | 0 | 0.059 | 1 | 19 |
| telo2 | 0.22544 | -0.13756714 | 0 | 0.059 | 1 | 19 |
| timm50 | 0.22544 | -0.13077249 | 0 | 0.059 | 1 | 19 |
| traf3ip2l | 0.22544 | -0.13067995 | 0 | 0.059 | 1 | 19 |
| twist3 | 0.22544 | -0.13969731 | 0 | 0.059 | 1 | 19 |
| ubash3bb | 0.22544 | -0.14202205 | 0 | 0.059 | 1 | 19 |
| ube2q1 | 0.22544 | -0.13478461 | 0 | 0.059 | 1 | 19 |
| ubqln4 | 0.22544 | -0.13744097 | 0 | 0.059 | 1 | 19 |
| usf2 | 0.22544 | -0.13740954 | 0 | 0.059 | 1 | 19 |
| wdr18 | 0.22544 | -0.13486098 | 0 | 0.059 | 1 | 19 |
| wdr33 | 0.22544 | -0.13662253 | 0 | 0.059 | 1 | 19 |
| znf16l | 0.22544 | -0.13866994 | 0 | 0.059 | 1 | 19 |
| BX000363.1 | 0.22544 | -0.19907565 | 0 | 0.059 | 1 | 19 |
| XLOC-005199 | 0.22544 | -0.19983535 | 0 | 0.059 | 1 | 19 |
| ankdd1a | 0.22544 | -0.19060778 | 0 | 0.059 | 1 | 19 |
| ccdc126 | 0.22544 | -0.19778411 | 0 | 0.059 | 1 | 19 |
| gramd2aa | 0.22544 | -0.19917051 | 0 | 0.059 | 1 | 19 |
| jmjd7 | 0.22544 | -0.19807058 | 0 | 0.059 | 1 | 19 |
| man1a1 | 0.22544 | -0.19901113 | 0 | 0.059 | 1 | 19 |
| nudcd2 | 0.22544 | -0.19476373 | 0 | 0.059 | 1 | 19 |
| pgm2 | 0.22544 | -0.1996541 | 0 | 0.059 | 1 | 19 |
| rrbp1a | 0.22544 | -0.19039131 | 0 | 0.059 | 1 | 19 |
| si:ch211-243a20.3 | 0.22544 | -0.19256543 | 0 | 0.059 | 1 | 19 |
| tango2 | 0.22544 | -0.19824504 | 0 | 0.059 | 1 | 19 |
| tmem150ab | 0.22544 | -0.19211394 | 0 | 0.059 | 1 | 19 |
| tnpo3 | 0.22544 | -0.19839096 | 0 | 0.059 | 1 | 19 |
| trib1 | 0.22544 | -0.20003262 | 0 | 0.059 | 1 | 19 |
| vps33b | 0.22544 | -0.1988792 | 0 | 0.059 | 1 | 19 |
| zgc:112083 | 0.22544 | -0.16263888 | 0 | 0.059 | 1 | 19 |
| BX000438.1 | 0.22544 | -0.22820359 | 0 | 0.059 | 1 | 19 |
| BX005421.3 | 0.22544 | -0.12736938 | 0 | 0.059 | 1 | 19 |
| BX322574.1 | 0.22544 | -0.12736938 | 0 | 0.059 | 1 | 19 |
| BX511129.1 | 0.22544 | -0.15215113 | 0 | 0.059 | 1 | 19 |
| BX544877.1 | 0.22544 | -0.16789267 | 0 | 0.059 | 1 | 19 |
| BX950187.1 | 0.22544 | -0.12253385 | 0 | 0.059 | 1 | 19 |
| BX957297.2 | 0.22544 | -0.19909057 | 0 | 0.059 | 1 | 19 |
| CABZ01030107.1 | 0.22544 | -0.19016408 | 0 | 0.059 | 1 | 19 |
| CABZ01054965.1 | 0.22544 | -0.12582176 | 0 | 0.059 | 1 | 19 |
| CABZ01058333.1 | 0.22544 | -0.19887833 | 0 | 0.059 | 1 | 19 |
| CABZ01072487.1 | 0.22544 | -0.20941797 | 0 | 0.059 | 1 | 19 |
| CABZ01078261.1 | 0.22544 | -0.11555311 | 0 | 0.059 | 1 | 19 |
| CABZ01080375.1 | 0.22544 | -0.16390839 | 0 | 0.059 | 1 | 19 |
| CABZ01085201.1 | 0.22544 | -0.1189198 | 0 | 0.059 | 1 | 19 |
| CABZ01088109.1 | 0.22544 | -0.16633734 | 0 | 0.059 | 1 | 19 |
| CABZ01088149.1 | 0.22544 | -0.19615972 | 0 | 0.059 | 1 | 19 |

| | | | | | | |
|---|---|---|---|---|---|---|
| CABZ01090361.1 | 0.22544 | -0.1185145 | 0 | 0.059 | 1 | 19 |
| CABZ01118738.1 | 0.22544 | -0.16504797 | 0 | 0.059 | 1 | 19 |
| CR293507.1 | 0.22544 | -0.16479426 | 0 | 0.059 | 1 | 19 |
| CR753886.1 | 0.22544 | -0.17102669 | 0 | 0.059 | 1 | 19 |
| CR759826.1 | 0.22544 | -0.12225285 | 0 | 0.059 | 1 | 19 |
| CU062501.1 | 0.22544 | -0.20429665 | 0 | 0.059 | 1 | 19 |
| CU469577.1 | 0.22544 | -0.16985436 | 0 | 0.059 | 1 | 19 |
| DDX17 | 0.22544 | -0.16139221 | 0 | 0.059 | 1 | 19 |
| EIF2AK4 | 0.22544 | -0.11647407 | 0 | 0.059 | 1 | 19 |
| FO082877.1 | 0.22544 | -0.11409606 | 0 | 0.059 | 1 | 19 |
| FO904870.1 | 0.22544 | -0.15479827 | 0 | 0.059 | 1 | 19 |
| FP016248.1 | 0.22544 | -0.16268575 | 0 | 0.059 | 1 | 19 |
| GFOD1 | 0.22544 | -0.12386094 | 0 | 0.059 | 1 | 19 |
| GK3P | 0.22544 | -0.2182189 | 0 | 0.059 | 1 | 19 |
| GPR180 | 0.22544 | -0.15479827 | 0 | 0.059 | 1 | 19 |
| LO017648.1 | 0.22544 | -0.1901231 | 0 | 0.059 | 1 | 19 |
| LO017938.1 | 0.22544 | -0.12736938 | 0 | 0.059 | 1 | 19 |
| LOC100536946 | 0.22544 | -0.22706202 | 0 | 0.059 | 1 | 19 |
| LOC101882159 | 0.22544 | -0.1629668 | 0 | 0.059 | 1 | 19 |
| LOC101883704 | 0.22544 | -0.10922228 | 0 | 0.059 | 1 | 19 |
| LOC101883948 | 0.22544 | -0.21788127 | 0 | 0.059 | 1 | 19 |
| LOC101884756 | 0.22544 | -0.21058728 | 0 | 0.059 | 1 | 19 |
| LOC101886002 | 0.22544 | -0.11393567 | 0 | 0.059 | 1 | 19 |
| LOC101886078 | 0.22544 | -0.19954535 | 0 | 0.059 | 1 | 19 |
| LOC103909899 | 0.22544 | -0.197525 | 0 | 0.059 | 1 | 19 |
| LOC103911715 | 0.22544 | -0.16390839 | 0 | 0.059 | 1 | 19 |
| LOC108179162 | 0.22544 | -0.19545493 | 0 | 0.059 | 1 | 19 |
| LOC108191465 | 0.22544 | -0.19495446 | 0 | 0.059 | 1 | 19 |
| LOC108191614 | 0.22544 | -0.16701897 | 0 | 0.059 | 1 | 19 |
| LOC110438518 | 0.22544 | -0.16985195 | 0 | 0.059 | 1 | 19 |
| LOC110439163 | 0.22544 | -0.19954535 | 0 | 0.059 | 1 | 19 |
| LOC110439310 | 0.22544 | -0.25688 | 0 | 0.059 | 1 | 19 |
| LOC110439320 | 0.22544 | -0.16952586 | 0 | 0.059 | 1 | 19 |
| LOC110439377 | 0.22544 | -0.12307472 | 0 | 0.059 | 1 | 19 |
| LOC570723 | 0.22544 | -0.19896747 | 0 | 0.059 | 1 | 19 |
| MAP3K11 | 0.22544 | -0.21036666 | 0 | 0.059 | 1 | 19 |
| PAQR9 | 0.22544 | -0.16298623 | 0 | 0.059 | 1 | 19 |
| RALGDS | 0.22544 | -0.19761421 | 0 | 0.059 | 1 | 19 |
| RF00560 | 0.22544 | -0.1905444 | 0 | 0.059 | 1 | 19 |
| RNF14 (1 of many) | 0.22544 | -0.10071153 | 0 | 0.059 | 1 | 19 |
| SBNO2 | 0.22544 | -0.1620476 | 0 | 0.059 | 1 | 19 |
| SFMBT1 | 0.22544 | -0.11756736 | 0 | 0.059 | 1 | 19 |
| SHKBP1 | 0.22544 | -0.19786095 | 0 | 0.059 | 1 | 19 |
| XLOC-000941 | 0.22544 | -0.16390839 | 0 | 0.059 | 1 | 19 |
| XLOC-001102 | 0.22544 | -0.12132979 | 0 | 0.059 | 1 | 19 |
| XLOC-004016 | 0.22544 | -0.1211566 | 0 | 0.059 | 1 | 19 |
| XLOC-006330 | 0.22544 | -0.11393567 | 0 | 0.059 | 1 | 19 |
| XLOC-011085 | 0.22544 | -0.12637926 | 0 | 0.059 | 1 | 19 |

| | | | | | | |
|---|---|---|---|---|---|---|
| XLOC-011107 | 0.22544 | -0.12477205 | 0 | 0.059 | 1 | 19 |
| XLOC-012680 | 0.22544 | -0.11874652 | 0 | 0.059 | 1 | 19 |
| XLOC-016094 | 0.22544 | -0.12479484 | 0 | 0.059 | 1 | 19 |
| XLOC-025819 | 0.22544 | -0.12225285 | 0 | 0.059 | 1 | 19 |
| XLOC-027844 | 0.22544 | -0.15479827 | 0 | 0.059 | 1 | 19 |
| XLOC-039817 | 0.22544 | -0.10983352 | 0 | 0.059 | 1 | 19 |
| XLOC-040756 | 0.22544 | -0.12225285 | 0 | 0.059 | 1 | 19 |
| XLOC-043834 | 0.22544 | -0.14032897 | 0 | 0.059 | 1 | 19 |
| XLOC-043861 | 0.22544 | -0.19849565 | 0 | 0.059 | 1 | 19 |
| abca5 | 0.22544 | -0.10948953 | 0 | 0.059 | 1 | 19 |
| abhd15a | 0.22544 | -0.19417149 | 0 | 0.059 | 1 | 19 |
| abhd16a | 0.22544 | -0.11600252 | 0 | 0.059 | 1 | 19 |
| abhd17b | 0.22544 | -0.16980252 | 0 | 0.059 | 1 | 19 |
| abhd2a | 0.22544 | -0.16299556 | 0 | 0.059 | 1 | 19 |
| abhd3 | 0.22544 | -0.16937692 | 0 | 0.059 | 1 | 19 |
| abi2b | 0.22544 | -0.11846852 | 0 | 0.059 | 1 | 19 |
| abraxas2 | 0.22544 | -0.11882253 | 0 | 0.059 | 1 | 19 |
| abt1 | 0.22544 | -0.10908187 | 0 | 0.059 | 1 | 19 |
| accs | 0.22544 | -0.19363404 | 0 | 0.059 | 1 | 19 |
| acp1 | 0.22544 | -0.16403947 | 0 | 0.059 | 1 | 19 |
| add1 | 0.22544 | -0.12448282 | 0 | 0.059 | 1 | 19 |
| adkb | 0.22544 | -0.16928351 | 0 | 0.059 | 1 | 19 |
| afg1lb | 0.22544 | -0.1507794 | 0 | 0.059 | 1 | 19 |
| ahcyl1 | 0.22544 | -0.12621514 | 0 | 0.059 | 1 | 19 |
| akap1b | 0.22544 | -0.16511237 | 0 | 0.059 | 1 | 19 |
| akap7 | 0.22544 | -0.14396724 | 0 | 0.059 | 1 | 19 |
| aldoab | 0.22544 | -0.12133635 | 0 | 0.059 | 1 | 19 |
| anapc5 | 0.22544 | -0.11570772 | 0 | 0.059 | 1 | 19 |
| ankrd52a | 0.22544 | -0.16138892 | 0 | 0.059 | 1 | 19 |
| ankrd54 | 0.22544 | -0.13889714 | 0 | 0.059 | 1 | 19 |
| aoc2 | 0.22544 | -0.11820389 | 0 | 0.059 | 1 | 19 |
| arfgap1 | 0.22544 | -0.15436185 | 0 | 0.059 | 1 | 19 |
| arfgef1 | 0.22544 | -0.17182102 | 0 | 0.059 | 1 | 19 |
| arglu1b | 0.22544 | -0.19830555 | 0 | 0.059 | 1 | 19 |
| arid1aa | 0.22544 | -0.12508051 | 0 | 0.059 | 1 | 19 |
| arl2 | 0.22544 | -0.21251469 | 0 | 0.059 | 1 | 19 |
| arpp19a | 0.22544 | -0.16044317 | 0 | 0.059 | 1 | 19 |
| arrdc2 | 0.22544 | -0.11913182 | 0 | 0.059 | 1 | 19 |
| asb8 | 0.22544 | -0.19032305 | 0 | 0.059 | 1 | 19 |
| ascc3 | 0.22544 | -0.16390839 | 0 | 0.059 | 1 | 19 |
| asnsd1 | 0.22544 | -0.16383247 | 0 | 0.059 | 1 | 19 |
| aspdh | 0.22544 | -0.16789267 | 0 | 0.059 | 1 | 19 |
| atxn1b | 0.22544 | -0.16089378 | 0 | 0.059 | 1 | 19 |
| atxn2l | 0.22544 | -0.16156995 | 0 | 0.059 | 1 | 19 |
| axin1 | 0.22544 | -0.17023326 | 0 | 0.059 | 1 | 19 |
| b4galt3 | 0.22544 | -0.16678792 | 0 | 0.059 | 1 | 19 |
| b4galt7 | 0.22544 | -0.19805446 | 0 | 0.059 | 1 | 19 |
| bmpr2a | 0.22544 | -0.15807997 | 0 | 0.059 | 1 | 19 |

| | | | | | | |
|---|---|---|---|---|---|---|
| borcs8 | 0.22544 | -0.1172348 | 0 | 0.059 | 1 | 19 |
| brcc3 | 0.22544 | -0.16854737 | 0 | 0.059 | 1 | 19 |
| brf2 | 0.22544 | -0.15013415 | 0 | 0.059 | 1 | 19 |
| brip1 | 0.22544 | -0.16670766 | 0 | 0.059 | 1 | 19 |
| brpf3b | 0.22544 | -0.10457014 | 0 | 0.059 | 1 | 19 |
| brwd1 | 0.22544 | -0.12356578 | 0 | 0.059 | 1 | 19 |
| c1galt1a | 0.22544 | -0.10540049 | 0 | 0.059 | 1 | 19 |
| cab39l | 0.22544 | -0.16493853 | 0 | 0.059 | 1 | 19 |
| calcoco2 | 0.22544 | -0.16806244 | 0 | 0.059 | 1 | 19 |
| calm3a | 0.22544 | -0.11969641 | 0 | 0.059 | 1 | 19 |
| calub | 0.22544 | -0.1187917 | 0 | 0.059 | 1 | 19 |
| camkk1a | 0.22544 | -0.19890732 | 0 | 0.059 | 1 | 19 |
| cant1a | 0.22544 | -0.12133051 | 0 | 0.059 | 1 | 19 |
| casc3 | 0.22544 | -0.11620003 | 0 | 0.059 | 1 | 19 |
| casp7 | 0.22544 | -0.19113765 | 0 | 0.059 | 1 | 19 |
| cblc | 0.22544 | -0.19954535 | 0 | 0.059 | 1 | 19 |
| cbx7b | 0.22544 | -0.17041933 | 0 | 0.059 | 1 | 19 |
| ccdc187 | 0.22544 | -0.15484013 | 0 | 0.059 | 1 | 19 |
| ccdc30 | 0.22544 | -0.15873392 | 0 | 0.059 | 1 | 19 |
| ccdc57 | 0.22544 | -0.11111584 | 0 | 0.059 | 1 | 19 |
| ccnq | 0.22544 | -0.14546743 | 0 | 0.059 | 1 | 19 |
| cd40lg | 0.22544 | -0.12693251 | 0 | 0.059 | 1 | 19 |
| cd81b | 0.22544 | -0.15479827 | 0 | 0.059 | 1 | 19 |
| cdc123 | 0.22544 | -0.15937916 | 0 | 0.059 | 1 | 19 |
| cdc27 | 0.22544 | -0.21761164 | 0 | 0.059 | 1 | 19 |
| cdkn1ba | 0.22544 | -0.16309832 | 0 | 0.059 | 1 | 19 |
| cds1 | 0.22544 | -0.17127293 | 0 | 0.059 | 1 | 19 |
| celf4 | 0.22544 | -0.11915943 | 0 | 0.059 | 1 | 19 |
| cfd | 0.22544 | -0.186452 | 0 | 0.059 | 1 | 19 |
| chaf1b | 0.22544 | -0.11766384 | 0 | 0.059 | 1 | 19 |
| chchd1 | 0.22544 | -0.21083562 | 0 | 0.059 | 1 | 19 |
| chd7 | 0.22544 | -0.11997052 | 0 | 0.059 | 1 | 19 |
| chordc1a | 0.22544 | -0.19864028 | 0 | 0.059 | 1 | 19 |
| clic2 | 0.22544 | -0.16155589 | 0 | 0.059 | 1 | 19 |
| clip1a | 0.22544 | -0.12533413 | 0 | 0.059 | 1 | 19 |
| cndp2 | 0.22544 | -0.12453395 | 0 | 0.059 | 1 | 19 |
| cnep1r1 | 0.22544 | -0.11341021 | 0 | 0.059 | 1 | 19 |
| col4a1 | 0.22544 | -0.12564566 | 0 | 0.059 | 1 | 19 |
| copz2 | 0.22544 | -0.16296863 | 0 | 0.059 | 1 | 19 |
| coq2 | 0.22544 | -0.11806009 | 0 | 0.059 | 1 | 19 |
| cox19 | 0.22544 | -0.12157136 | 0 | 0.059 | 1 | 19 |
| creb3l2 | 0.22544 | -0.1626981 | 0 | 0.059 | 1 | 19 |
| creld2 | 0.22544 | -0.16785152 | 0 | 0.059 | 1 | 19 |
| crlf1b | 0.22544 | -0.11732273 | 0 | 0.059 | 1 | 19 |
| cst14b.2 | 0.22544 | -0.11147866 | 0 | 0.059 | 1 | 19 |
| ctdsp2 | 0.22544 | -0.11528903 | 0 | 0.059 | 1 | 19 |
| ctnnbl1 | 0.22544 | -0.11579218 | 0 | 0.059 | 1 | 19 |
| ctns | 0.22544 | -0.16833655 | 0 | 0.059 | 1 | 19 |

| | | | | | | |
|---|---|---|---|---|---|---|
| cwf19l1 | 0.22544 | -0.12097467 | 0 | 0.059 | 1 | 19 |
| cxcr2 | 0.22544 | -0.12225285 | 0 | 0.059 | 1 | 19 |
| cyb5b | 0.22544 | -0.16422496 | 0 | 0.059 | 1 | 19 |
| cyth1a | 0.22544 | -0.21850819 | 0 | 0.059 | 1 | 19 |
| dag1 | 0.22544 | -0.11290621 | 0 | 0.059 | 1 | 19 |
| dennd6b | 0.22544 | -0.21891588 | 0 | 0.059 | 1 | 19 |
| dgcr8 | 0.22544 | -0.16513012 | 0 | 0.059 | 1 | 19 |
| dhrs1 | 0.22544 | -0.15706853 | 0 | 0.059 | 1 | 19 |
| dnah9l | 0.22544 | -0.16789267 | 0 | 0.059 | 1 | 19 |
| dnajb1a | 0.22544 | -0.16514195 | 0 | 0.059 | 1 | 19 |
| dnajc7 | 0.22544 | -0.19840325 | 0 | 0.059 | 1 | 19 |
| dnase1l1 | 0.22544 | -0.1655539 | 0 | 0.059 | 1 | 19 |
| dph2 | 0.22544 | -0.15081811 | 0 | 0.059 | 1 | 19 |
| drosha | 0.22544 | -0.12635418 | 0 | 0.059 | 1 | 19 |
| dtx4b | 0.22544 | -0.11250158 | 0 | 0.059 | 1 | 19 |
| dyrk1b | 0.22544 | -0.15744055 | 0 | 0.059 | 1 | 19 |
| eif4h | 0.22544 | -0.19171705 | 0 | 0.059 | 1 | 19 |
| eloa | 0.22544 | -0.12244333 | 0 | 0.059 | 1 | 19 |
| emc3 | 0.22544 | -0.16462767 | 0 | 0.059 | 1 | 19 |
| emid1 | 0.22544 | -0.17720202 | 0 | 0.059 | 1 | 19 |
| ep400 | 0.22544 | -0.16194446 | 0 | 0.059 | 1 | 19 |
| ergic1 | 0.22544 | -0.19830658 | 0 | 0.059 | 1 | 19 |
| esco1 | 0.22544 | -0.16275848 | 0 | 0.059 | 1 | 19 |
| esm1 | 0.22544 | -0.14437496 | 0 | 0.059 | 1 | 19 |
| ethe1 | 0.22544 | -0.11658778 | 0 | 0.059 | 1 | 19 |
| etv2 | 0.22544 | -0.22110389 | 0 | 0.059 | 1 | 19 |
| evlb | 0.22544 | -0.12587759 | 0 | 0.059 | 1 | 19 |
| exoc1 | 0.22544 | -0.15929538 | 0 | 0.059 | 1 | 19 |
| exoc3l2a | 0.22544 | -0.12821918 | 0 | 0.059 | 1 | 19 |
| ext2 | 0.22544 | -0.19565614 | 0 | 0.059 | 1 | 19 |
| ezh2 | 0.22544 | -0.11121564 | 0 | 0.059 | 1 | 19 |
| fabp11a | 0.22544 | -0.3844545 | 0 | 0.059 | 1 | 19 |
| fam129bb | 0.22544 | -0.16127935 | 0 | 0.059 | 1 | 19 |
| fam135a | 0.22544 | -0.22722801 | 0 | 0.059 | 1 | 19 |
| fam160b1 | 0.22544 | -0.15215607 | 0 | 0.059 | 1 | 19 |
| fam219b | 0.22544 | -0.16881397 | 0 | 0.059 | 1 | 19 |
| fam69ab | 0.22544 | -0.16390839 | 0 | 0.059 | 1 | 19 |
| fam91a1 | 0.22544 | -0.12190719 | 0 | 0.059 | 1 | 19 |
| fbxl14a | 0.22544 | -0.19091102 | 0 | 0.059 | 1 | 19 |
| fbxl3a | 0.22544 | -0.12403074 | 0 | 0.059 | 1 | 19 |
| fbxo25 | 0.22544 | -0.19261656 | 0 | 0.059 | 1 | 19 |
| fbxw5 | 0.22544 | -0.16767051 | 0 | 0.059 | 1 | 19 |
| fbxw8 | 0.22544 | -0.18457538 | 0 | 0.059 | 1 | 19 |
| fig4a | 0.22544 | -0.12117387 | 0 | 0.059 | 1 | 19 |
| fnip1 | 0.22544 | -0.17174979 | 0 | 0.059 | 1 | 19 |
| foxn2b | 0.22544 | -0.11792188 | 0 | 0.059 | 1 | 19 |
| ftr66 | 0.22544 | -0.21891588 | 0 | 0.059 | 1 | 19 |
| ftsj3 | 0.22544 | -0.11565359 | 0 | 0.059 | 1 | 19 |

| | | | | | | |
|---|---|---|---|---|---|---|
| fxyd1 | 0.22544 | -0.14972114 | 0 | 0.059 | 1 | 19 |
| gabpa | 0.22544 | -0.15993417 | 0 | 0.059 | 1 | 19 |
| gabpb2a | 0.22544 | -0.16610392 | 0 | 0.059 | 1 | 19 |
| galnt12 | 0.22544 | -0.12649965 | 0 | 0.059 | 1 | 19 |
| gata2a | 0.22544 | -0.17059673 | 0 | 0.059 | 1 | 19 |
| gba | 0.22544 | -0.19507059 | 0 | 0.059 | 1 | 19 |
| gbp4 | 0.22544 | -0.15479827 | 0 | 0.059 | 1 | 19 |
| ggact.2 | 0.22544 | -0.12060766 | 0 | 0.059 | 1 | 19 |
| glud1b | 0.22544 | -0.16138887 | 0 | 0.059 | 1 | 19 |
| gne | 0.22544 | -0.16068934 | 0 | 0.059 | 1 | 19 |
| gpalpp1 | 0.22544 | -0.21834698 | 0 | 0.059 | 1 | 19 |
| gpbp1l1 | 0.22544 | -0.1179314 | 0 | 0.059 | 1 | 19 |
| gpcpd1 | 0.22544 | -0.11277853 | 0 | 0.059 | 1 | 19 |
| gphna | 0.22544 | -0.12225285 | 0 | 0.059 | 1 | 19 |
| gpr174 | 0.22544 | -0.12145053 | 0 | 0.059 | 1 | 19 |
| gramd4b | 0.22544 | -0.17136593 | 0 | 0.059 | 1 | 19 |
| grk5l | 0.22544 | -0.12369096 | 0 | 0.059 | 1 | 19 |
| grn1 | 0.22544 | -0.1597521 | 0 | 0.059 | 1 | 19 |
| gtf3aa | 0.22544 | -0.15787846 | 0 | 0.059 | 1 | 19 |
| gtpbp10 | 0.22544 | -0.19020596 | 0 | 0.059 | 1 | 19 |
| gzf1 | 0.22544 | -0.19386822 | 0 | 0.059 | 1 | 19 |
| habp2 | 0.22544 | -0.12088874 | 0 | 0.059 | 1 | 19 |
| hacd2 | 0.22544 | -0.1962194 | 0 | 0.059 | 1 | 19 |
| haus7 | 0.22544 | -0.11475749 | 0 | 0.059 | 1 | 19 |
| hax1 | 0.22544 | -0.15496013 | 0 | 0.059 | 1 | 19 |
| hbegfa | 0.22544 | -0.15900261 | 0 | 0.059 | 1 | 19 |
| herc2 | 0.22544 | -0.21812097 | 0 | 0.059 | 1 | 19 |
| hopx | 0.22544 | -0.15871595 | 0 | 0.059 | 1 | 19 |
| hs3st3b1a | 0.22544 | -0.10220465 | 0 | 0.059 | 1 | 19 |
| hsd17b4 | 0.22544 | -0.12027596 | 0 | 0.059 | 1 | 19 |
| hsdl1 | 0.22544 | -0.12225285 | 0 | 0.059 | 1 | 19 |
| hsph1 | 0.22544 | -0.1209814 | 0 | 0.059 | 1 | 19 |
| ice1 | 0.22544 | -0.11273387 | 0 | 0.059 | 1 | 19 |
| il2rb | 0.22544 | -0.12685039 | 0 | 0.059 | 1 | 19 |
| il2rga | 0.22544 | -0.12118627 | 0 | 0.059 | 1 | 19 |
| im:7147486 | 0.22544 | -0.15695096 | 0 | 0.059 | 1 | 19 |
| ino80 | 0.22544 | -0.16714902 | 0 | 0.059 | 1 | 19 |
| inpp5b | 0.22544 | -0.11904831 | 0 | 0.059 | 1 | 19 |
| ip6k1 | 0.22544 | -0.14572609 | 0 | 0.059 | 1 | 19 |
| itgb1a | 0.22544 | -0.19091539 | 0 | 0.059 | 1 | 19 |
| jakmip1 | 0.22544 | -0.16227544 | 0 | 0.059 | 1 | 19 |
| kat14 | 0.22544 | -0.14529494 | 0 | 0.059 | 1 | 19 |
| kat7b | 0.22544 | -0.16575666 | 0 | 0.059 | 1 | 19 |
| katnbl1 | 0.22544 | -0.11802572 | 0 | 0.059 | 1 | 19 |
| kbtbd8 | 0.22544 | -0.18403096 | 0 | 0.059 | 1 | 19 |
| kcmf1 | 0.22544 | -0.11680383 | 0 | 0.059 | 1 | 19 |
| kdrl | 0.22544 | -0.20549721 | 0 | 0.059 | 1 | 19 |
| kif13ba | 0.22544 | -0.1687465 | 0 | 0.059 | 1 | 19 |

| | | | | | | |
|---|---|---|---|---|---|---|
| kif1ab | 0.22544 | -0.20941797 | 0 | 0.059 | 1 | 19 |
| klhdc4 | 0.22544 | -0.11664927 | 0 | 0.059 | 1 | 19 |
| klhl22 | 0.22544 | -0.19669354 | 0 | 0.059 | 1 | 19 |
| kras | 0.22544 | -0.16706263 | 0 | 0.059 | 1 | 19 |
| ksr1a | 0.22544 | -0.16390839 | 0 | 0.059 | 1 | 19 |
| kti12 | 0.22544 | -0.24008102 | 0 | 0.059 | 1 | 19 |
| lancl2 | 0.22544 | -0.16439521 | 0 | 0.059 | 1 | 19 |
| larp4aa | 0.22544 | -0.12030864 | 0 | 0.059 | 1 | 19 |
| ldb2a | 0.22544 | -0.13220928 | 0 | 0.059 | 1 | 19 |
| ldb3a | 0.22544 | -0.15479827 | 0 | 0.059 | 1 | 19 |
| lepr | 0.22544 | -0.11998481 | 0 | 0.059 | 1 | 19 |
| letm1 | 0.22544 | -0.21805858 | 0 | 0.059 | 1 | 19 |
| lgmn | 0.22544 | -0.16113211 | 0 | 0.059 | 1 | 19 |
| lig3 | 0.22544 | -0.1605813 | 0 | 0.059 | 1 | 19 |
| lpl | 0.22544 | -0.20006252 | 0 | 0.059 | 1 | 19 |
| lrch3 | 0.22544 | -0.11967399 | 0 | 0.059 | 1 | 19 |
| lrmp | 0.22544 | -0.14888083 | 0 | 0.059 | 1 | 19 |
| lrrc29 | 0.22544 | -0.11035019 | 0 | 0.059 | 1 | 19 |
| man2a2 | 0.22544 | -0.21832576 | 0 | 0.059 | 1 | 19 |
| mao | 0.22544 | -0.1617385 | 0 | 0.059 | 1 | 19 |
| map2k5 | 0.22544 | -0.16697542 | 0 | 0.059 | 1 | 19 |
| map4l | 0.22544 | -0.12225285 | 0 | 0.059 | 1 | 19 |
| mbd6 | 0.22544 | -0.15429767 | 0 | 0.059 | 1 | 19 |
| mcm6 | 0.22544 | -0.15093495 | 0 | 0.059 | 1 | 19 |
| mctp1b | 0.22544 | -0.17166162 | 0 | 0.059 | 1 | 19 |
| mdm1 | 0.22544 | -0.12085183 | 0 | 0.059 | 1 | 19 |
| mepce | 0.22544 | -0.12299123 | 0 | 0.059 | 1 | 19 |
| mettl15 | 0.22544 | -0.10158298 | 0 | 0.059 | 1 | 19 |
| mgst1.1 | 0.22544 | -0.16095677 | 0 | 0.059 | 1 | 19 |
| mia3 | 0.22544 | -0.11677434 | 0 | 0.059 | 1 | 19 |
| mlh3 | 0.22544 | -0.19803373 | 0 | 0.059 | 1 | 19 |
| mmrn2a | 0.22544 | -0.15479827 | 0 | 0.059 | 1 | 19 |
| mocs3 | 0.22544 | -0.17536953 | 0 | 0.059 | 1 | 19 |
| morc2 | 0.22544 | -0.21756297 | 0 | 0.059 | 1 | 19 |
| mrc1b | 0.22544 | -0.16587113 | 0 | 0.059 | 1 | 19 |
| mrpl37 | 0.22544 | -0.1150115 | 0 | 0.059 | 1 | 19 |
| mrpl38 | 0.22544 | -0.19140078 | 0 | 0.059 | 1 | 19 |
| mrpl44 | 0.22544 | -0.11330118 | 0 | 0.059 | 1 | 19 |
| mtbl | 0.22544 | -0.19381404 | 0 | 0.059 | 1 | 19 |
| mtmr8 | 0.22544 | -0.16158624 | 0 | 0.059 | 1 | 19 |
| mut | 0.22544 | -0.16409894 | 0 | 0.059 | 1 | 19 |
| myef2 | 0.22544 | -0.16850345 | 0 | 0.059 | 1 | 19 |
| naa30 | 0.22544 | -0.1557817 | 0 | 0.059 | 1 | 19 |
| nabp1a | 0.22544 | -0.10710245 | 0 | 0.059 | 1 | 19 |
| nae1 | 0.22544 | -0.19102717 | 0 | 0.059 | 1 | 19 |
| naprt | 0.22544 | -0.1069515 | 0 | 0.059 | 1 | 19 |
| nat14 | 0.22544 | -0.1627787 | 0 | 0.059 | 1 | 19 |
| nbr1a | 0.22544 | -0.21354129 | 0 | 0.059 | 1 | 19 |

| | | | | | | |
|---|---|---|---|---|---|---|
| ncapd2 | 0.22544 | -0.12403705 | 0 | 0.059 | 1 | 19 |
| ncor2 | 0.22544 | -0.19685724 | 0 | 0.059 | 1 | 19 |
| ndufaf6 | 0.22544 | -0.16770107 | 0 | 0.059 | 1 | 19 |
| nedd1 | 0.22544 | -0.13539674 | 0 | 0.059 | 1 | 19 |
| nelfe | 0.22544 | -0.10209615 | 0 | 0.059 | 1 | 19 |
| nfatc2ip | 0.22544 | -0.10032566 | 0 | 0.059 | 1 | 19 |
| nfya | 0.22544 | -0.16706299 | 0 | 0.059 | 1 | 19 |
| nlrc3l1 | 0.22544 | -0.12499838 | 0 | 0.059 | 1 | 19 |
| nr1d1 | 0.22544 | -0.19829086 | 0 | 0.059 | 1 | 19 |
| nsun3 | 0.22544 | -0.21718954 | 0 | 0.059 | 1 | 19 |
| nudt18 | 0.22544 | -0.12204404 | 0 | 0.059 | 1 | 19 |
| nuggc.1 | 0.22544 | -0.17158714 | 0 | 0.059 | 1 | 19 |
| nup210 | 0.22544 | -0.11952911 | 0 | 0.059 | 1 | 19 |
| nup98 | 0.22544 | -0.16250367 | 0 | 0.059 | 1 | 19 |
| nxf1 | 0.22544 | -0.19207502 | 0 | 0.059 | 1 | 19 |
| orai1b | 0.22544 | -0.19862471 | 0 | 0.059 | 1 | 19 |
| osbpl9 | 0.22544 | -0.17145375 | 0 | 0.059 | 1 | 19 |
| oser1 | 0.22544 | -0.17070759 | 0 | 0.059 | 1 | 19 |
| oxct1b | 0.22544 | -0.16390839 | 0 | 0.059 | 1 | 19 |
| pak4 | 0.22544 | -0.12177359 | 0 | 0.059 | 1 | 19 |
| paqr3a | 0.22544 | -0.15280742 | 0 | 0.059 | 1 | 19 |
| parla | 0.22544 | -0.19649673 | 0 | 0.059 | 1 | 19 |
| parp4 | 0.22544 | -0.12736938 | 0 | 0.059 | 1 | 19 |
| pbx2 | 0.22544 | -0.11692223 | 0 | 0.059 | 1 | 19 |
| pccb | 0.22544 | -0.18981111 | 0 | 0.059 | 1 | 19 |
| pcmtd1 | 0.22544 | -0.1692652 | 0 | 0.059 | 1 | 19 |
| pde12 | 0.22544 | -0.11584734 | 0 | 0.059 | 1 | 19 |
| pdk1 | 0.22544 | -0.16342078 | 0 | 0.059 | 1 | 19 |
| pdss2 | 0.22544 | -0.15501427 | 0 | 0.059 | 1 | 19 |
| pelp1 | 0.22544 | -0.21717775 | 0 | 0.059 | 1 | 19 |
| pfkpb | 0.22544 | -0.16148419 | 0 | 0.059 | 1 | 19 |
| pgap2 | 0.22544 | -0.16506292 | 0 | 0.059 | 1 | 19 |
| phax | 0.22544 | -0.11916122 | 0 | 0.059 | 1 | 19 |
| phf12b | 0.22544 | -0.12128422 | 0 | 0.059 | 1 | 19 |
| phf23b | 0.22544 | -0.18956288 | 0 | 0.059 | 1 | 19 |
| pik3r2 | 0.22544 | -0.11869689 | 0 | 0.059 | 1 | 19 |
| pikfyve | 0.22544 | -0.19878592 | 0 | 0.059 | 1 | 19 |
| plagl2 | 0.22544 | -0.21891588 | 0 | 0.059 | 1 | 19 |
| plekhf2 | 0.22544 | -0.11978552 | 0 | 0.059 | 1 | 19 |
| plpbp | 0.22544 | -0.19623839 | 0 | 0.059 | 1 | 19 |
| plscr3b | 0.22544 | -0.16156346 | 0 | 0.059 | 1 | 19 |
| plvapa | 0.22544 | -0.15479827 | 0 | 0.059 | 1 | 19 |
| plvapb | 0.22544 | -0.19680094 | 0 | 0.059 | 1 | 19 |
| plxnb3 | 0.22544 | -0.14437496 | 0 | 0.059 | 1 | 19 |
| pms2 | 0.22544 | -0.14100944 | 0 | 0.059 | 1 | 19 |
| pnp5b | 0.22544 | -0.16703789 | 0 | 0.059 | 1 | 19 |
| pnpla8 | 0.22544 | -0.1608019 | 0 | 0.059 | 1 | 19 |
| pofut2 | 0.22544 | -0.15633947 | 0 | 0.059 | 1 | 19 |

| | | | | | | |
|---|---|---|---|---|---|---|
| ppil1 | 0.22544 | -0.16302222 | 0 | 0.059 | 1 | 19 |
| ppp1cbl | 0.22544 | -0.19097247 | 0 | 0.059 | 1 | 19 |
| ppp1r3da | 0.22544 | -0.12559908 | 0 | 0.059 | 1 | 19 |
| ppp2r2aa | 0.22544 | -0.16082379 | 0 | 0.059 | 1 | 19 |
| ppp4r2b | 0.22544 | -0.16671384 | 0 | 0.059 | 1 | 19 |
| ppt2 | 0.22544 | -0.16345133 | 0 | 0.059 | 1 | 19 |
| prcc | 0.22544 | -0.16121013 | 0 | 0.059 | 1 | 19 |
| prdm1a | 0.22544 | -0.12670703 | 0 | 0.059 | 1 | 19 |
| preb | 0.22544 | -0.11952442 | 0 | 0.059 | 1 | 19 |
| pros1 | 0.22544 | -0.11977479 | 0 | 0.059 | 1 | 19 |
| prpf18 | 0.22544 | -0.12067569 | 0 | 0.059 | 1 | 19 |
| prpf40a | 0.22544 | -0.19752916 | 0 | 0.059 | 1 | 19 |
| ptk6b | 0.22544 | -0.22834736 | 0 | 0.059 | 1 | 19 |
| pwwp2a | 0.22544 | -0.16288418 | 0 | 0.059 | 1 | 19 |
| pxylp1 | 0.22544 | -0.1917629 | 0 | 0.059 | 1 | 19 |
| rab20 | 0.22544 | -0.11690867 | 0 | 0.059 | 1 | 19 |
| rab4a | 0.22544 | -0.16771273 | 0 | 0.059 | 1 | 19 |
| rad51 | 0.22544 | -0.11778564 | 0 | 0.059 | 1 | 19 |
| rad54l2 | 0.22544 | -0.11835236 | 0 | 0.059 | 1 | 19 |
| rap2aa | 0.22544 | -0.12446341 | 0 | 0.059 | 1 | 19 |
| rasa1a | 0.22544 | -0.12142834 | 0 | 0.059 | 1 | 19 |
| rbm17 | 0.22544 | -0.19463251 | 0 | 0.059 | 1 | 19 |
| rbm39a | 0.22544 | -0.11725208 | 0 | 0.059 | 1 | 19 |
| rbm6 | 0.22544 | -0.11717042 | 0 | 0.059 | 1 | 19 |
| rbp2a | 0.22544 | -0.34536156 | 0 | 0.059 | 1 | 19 |
| rbp7b | 0.22544 | -0.22164345 | 0 | 0.059 | 1 | 19 |
| rbpja | 0.22544 | -0.17066457 | 0 | 0.059 | 1 | 19 |
| rcl1 | 0.22544 | -0.10913221 | 0 | 0.059 | 1 | 19 |
| rcor3 | 0.22544 | -0.15237723 | 0 | 0.059 | 1 | 19 |
| rela | 0.22544 | -0.18913255 | 0 | 0.059 | 1 | 19 |
| rerea | 0.22544 | -0.1165397 | 0 | 0.059 | 1 | 19 |
| retsat | 0.22544 | -0.19735509 | 0 | 0.059 | 1 | 19 |
| rfxap | 0.22544 | -0.16675982 | 0 | 0.059 | 1 | 19 |
| rgcc | 0.22544 | -0.20787357 | 0 | 0.059 | 1 | 19 |
| rimkla | 0.22544 | -0.17105126 | 0 | 0.059 | 1 | 19 |
| riox2 | 0.22544 | -0.10946886 | 0 | 0.059 | 1 | 19 |
| rlim | 0.22544 | -0.11839099 | 0 | 0.059 | 1 | 19 |
| rnf126 | 0.22544 | -0.16082662 | 0 | 0.059 | 1 | 19 |
| rnf8 | 0.22544 | -0.21435801 | 0 | 0.059 | 1 | 19 |
| rps6kal | 0.22544 | -0.16417899 | 0 | 0.059 | 1 | 19 |
| rps6kb1b | 0.22544 | -0.11971516 | 0 | 0.059 | 1 | 19 |
| rpz4 | 0.22544 | -0.16683294 | 0 | 0.059 | 1 | 19 |
| rras | 0.22544 | -0.19126409 | 0 | 0.059 | 1 | 19 |
| rreb1b | 0.22544 | -0.11953875 | 0 | 0.059 | 1 | 19 |
| rtel1 | 0.22544 | -0.19734445 | 0 | 0.059 | 1 | 19 |
| rubcn | 0.22544 | -0.19771599 | 0 | 0.059 | 1 | 19 |
| rusc1 | 0.22544 | -0.1670549 | 0 | 0.059 | 1 | 19 |
| samd11 | 0.22544 | -0.12080059 | 0 | 0.059 | 1 | 19 |

| | | | | | | |
|---|---|---|---|---|---|---|
| scin | 0.22544 | -0.12638763 | 0 | 0.059 | 1 | 19 |
| scn8ab | 0.22544 | -0.11393567 | 0 | 0.059 | 1 | 19 |
| sdad1 | 0.22544 | -0.10965602 | 0 | 0.059 | 1 | 19 |
| sdha | 0.22544 | -0.10861981 | 0 | 0.059 | 1 | 19 |
| sec23ip | 0.22544 | -0.12076467 | 0 | 0.059 | 1 | 19 |
| sema3ga | 0.22544 | -0.16390839 | 0 | 0.059 | 1 | 19 |
| sharpin | 0.22544 | -0.16240792 | 0 | 0.059 | 1 | 19 |
| si:ch1073-340j19.1 | 0.22544 | -0.16241374 | 0 | 0.059 | 1 | 19 |
| si:ch1073-394i4.1 | 0.22544 | -0.16985195 | 0 | 0.059 | 1 | 19 |
| si:ch211-113e8.10 | 0.22544 | -0.16568976 | 0 | 0.059 | 1 | 19 |
| si:ch211-114l13.9 | 0.22544 | -0.125392 | 0 | 0.059 | 1 | 19 |
| si:ch211-195e19.1 | 0.22544 | -0.15905163 | 0 | 0.059 | 1 | 19 |
| si:ch211-198c19.3 | 0.22544 | -0.11366808 | 0 | 0.059 | 1 | 19 |
| si:ch211-203d1.3 | 0.22544 | -0.12666948 | 0 | 0.059 | 1 | 19 |
| si:ch211-204c21.1 | 0.22544 | -0.20968391 | 0 | 0.059 | 1 | 19 |
| si:ch211-210b2.1 | 0.22544 | -0.1400167 | 0 | 0.059 | 1 | 19 |
| si:ch211-213a13.2 | 0.22544 | -0.11962105 | 0 | 0.059 | 1 | 19 |
| si:ch211-215p11.1 | 0.22544 | -0.19954535 | 0 | 0.059 | 1 | 19 |
| si:ch211-229b6.1 | 0.22544 | -0.16293505 | 0 | 0.059 | 1 | 19 |
| si:ch211-253h3.1 | 0.22544 | -0.19584259 | 0 | 0.059 | 1 | 19 |
| si:ch211-59h6.1 | 0.22544 | -0.12225285 | 0 | 0.059 | 1 | 19 |
| si:ch211-66k16.2 | 0.22544 | -0.12225285 | 0 | 0.059 | 1 | 19 |
| si:ch211-74f19.2 | 0.22544 | -0.1985625 | 0 | 0.059 | 1 | 19 |
| si:ch73-109d9.3 | 0.22544 | -0.12035744 | 0 | 0.059 | 1 | 19 |
| si:ch73-173p19.1 | 0.22544 | -0.1901231 | 0 | 0.059 | 1 | 19 |
| si:ch73-191k20.3 | 0.22544 | -0.22850618 | 0 | 0.059 | 1 | 19 |
| si:ch73-223p23.2 | 0.22544 | -0.12697619 | 0 | 0.059 | 1 | 19 |
| si:ch73-27e22.7 | 0.22544 | -0.22855327 | 0 | 0.059 | 1 | 19 |
| si:dkey-117m1.4 | 0.22544 | -0.16209239 | 0 | 0.059 | 1 | 19 |
| si:dkey-13n15.11 | 0.22544 | -0.12605651 | 0 | 0.059 | 1 | 19 |
| si:dkey-14o6.2 | 0.22544 | -0.16201929 | 0 | 0.059 | 1 | 19 |
| si:dkey-183p4.10 | 0.22544 | -0.12654367 | 0 | 0.059 | 1 | 19 |
| si:dkey-222b8.1 | 0.22544 | -0.11809562 | 0 | 0.059 | 1 | 19 |
| si:dkey-238d18.5 | 0.22544 | -0.19954535 | 0 | 0.059 | 1 | 19 |
| si:dkey-249d8.1 | 0.22544 | -0.19713875 | 0 | 0.059 | 1 | 19 |
| si:dkey-251i10.1 | 0.22544 | -0.17164994 | 0 | 0.059 | 1 | 19 |
| si:dkey-33c14.7 | 0.22544 | -0.11104056 | 0 | 0.059 | 1 | 19 |
| si:dkey-66a8.7 | 0.22544 | -0.16889539 | 0 | 0.059 | 1 | 19 |
| si:dkey-69o16.5 | 0.22544 | -0.11741074 | 0 | 0.059 | 1 | 19 |
| si:dkey-76k16.6 | 0.22544 | -0.12242106 | 0 | 0.059 | 1 | 19 |
| si:dkey-7i4.8 | 0.22544 | -0.22872696 | 0 | 0.059 | 1 | 19 |
| si:dkey-93h22.8 | 0.22544 | -0.16452549 | 0 | 0.059 | 1 | 19 |
| si:rp71-79p20.2 | 0.22544 | -0.16109902 | 0 | 0.059 | 1 | 19 |
| ska3 | 0.22544 | -0.11485299 | 0 | 0.059 | 1 | 19 |
| slc11a2 | 0.22544 | -0.16706892 | 0 | 0.059 | 1 | 19 |
| slc15a4 | 0.22544 | -0.12511056 | 0 | 0.059 | 1 | 19 |
| slc16a13 | 0.22544 | -0.16062039 | 0 | 0.059 | 1 | 19 |
| slc25a16 | 0.22544 | -0.16195582 | 0 | 0.059 | 1 | 19 |

| | | | | | | |
|---|---|---|---|---|---|---|
| slc26a5 | 0.22544 | -0.21495834 | 0 | 0.059 | 1 | 19 |
| smc5 | 0.22544 | -0.13885135 | 0 | 0.059 | 1 | 19 |
| smpd3 | 0.22544 | -0.1721414 | 0 | 0.059 | 1 | 19 |
| snrnp48 | 0.22544 | -0.21711811 | 0 | 0.059 | 1 | 19 |
| snx27b | 0.22544 | -0.12154282 | 0 | 0.059 | 1 | 19 |
| soul3 | 0.22544 | -0.20983787 | 0 | 0.059 | 1 | 19 |
| spdl1 | 0.22544 | -0.11716612 | 0 | 0.059 | 1 | 19 |
| spoplb | 0.22544 | -0.19430987 | 0 | 0.059 | 1 | 19 |
| sprn2 | 0.22544 | -0.12736938 | 0 | 0.059 | 1 | 19 |
| st3gal4 | 0.22544 | -0.12705081 | 0 | 0.059 | 1 | 19 |
| stag2a | 0.22544 | -0.16814519 | 0 | 0.059 | 1 | 19 |
| stam2 | 0.22544 | -0.12004306 | 0 | 0.059 | 1 | 19 |
| stmn1a | 0.22544 | -0.24179535 | 0 | 0.059 | 1 | 19 |
| stx6 | 0.22544 | -0.19551713 | 0 | 0.059 | 1 | 19 |
| swi5 | 0.22544 | -0.11686053 | 0 | 0.059 | 1 | 19 |
| syvn1 | 0.22544 | -0.16429817 | 0 | 0.059 | 1 | 19 |
| taf1a | 0.22544 | -0.1643826 | 0 | 0.059 | 1 | 19 |
| tbce | 0.22544 | -0.19476643 | 0 | 0.059 | 1 | 19 |
| tceb2 | 0.22544 | -0.12065778 | 0 | 0.059 | 1 | 19 |
| tcf7 | 0.22544 | -0.17175478 | 0 | 0.059 | 1 | 19 |
| terf2ip | 0.22544 | -0.18105446 | 0 | 0.059 | 1 | 19 |
| tfa | 0.22544 | -0.32156643 | 0 | 0.059 | 1 | 19 |
| tgfbrap1 | 0.22544 | -0.12555398 | 0 | 0.059 | 1 | 19 |
| tgm5l | 0.22544 | -0.16535783 | 0 | 0.059 | 1 | 19 |
| thap11 | 0.22544 | -0.1495517 | 0 | 0.059 | 1 | 19 |
| thap12a | 0.22544 | -0.19610835 | 0 | 0.059 | 1 | 19 |
| thoc7 | 0.22544 | -0.11645432 | 0 | 0.059 | 1 | 19 |
| thy1 | 0.22544 | -0.15937 | 0 | 0.059 | 1 | 19 |
| timm10b | 0.22544 | -0.11925128 | 0 | 0.059 | 1 | 19 |
| tlr22 | 0.22544 | -0.19919796 | 0 | 0.059 | 1 | 19 |
| tm9sf2 | 0.22544 | -0.1956708 | 0 | 0.059 | 1 | 19 |
| tmc6a | 0.22544 | -0.16349101 | 0 | 0.059 | 1 | 19 |
| tmco4 | 0.22544 | -0.19219807 | 0 | 0.059 | 1 | 19 |
| tmco6 | 0.22544 | -0.13717153 | 0 | 0.059 | 1 | 19 |
| tmed5 | 0.22544 | -0.12083083 | 0 | 0.059 | 1 | 19 |
| tmem160 | 0.22544 | -0.11980923 | 0 | 0.059 | 1 | 19 |
| tmem184a | 0.22544 | -0.19096 | 0 | 0.059 | 1 | 19 |
| tmem50a | 0.22544 | -0.16116744 | 0 | 0.059 | 1 | 19 |
| tmub1 | 0.22544 | -0.11412648 | 0 | 0.059 | 1 | 19 |
| tnfaip1 | 0.22544 | -0.10110498 | 0 | 0.059 | 1 | 19 |
| tox | 0.22544 | -0.21026808 | 0 | 0.059 | 1 | 19 |
| tpd52l2a | 0.22544 | -0.10919622 | 0 | 0.059 | 1 | 19 |
| tpm4a | 0.22544 | -0.14850711 | 0 | 0.059 | 1 | 19 |
| tpmt.2 | 0.22544 | -0.19752864 | 0 | 0.059 | 1 | 19 |
| tpra | 0.22544 | -0.15857842 | 0 | 0.059 | 1 | 19 |
| tprb | 0.22544 | -0.21767136 | 0 | 0.059 | 1 | 19 |
| trap1 | 0.22544 | -0.11881707 | 0 | 0.059 | 1 | 19 |
| trim3a | 0.22544 | -0.15846059 | 0 | 0.059 | 1 | 19 |

| | | | | | | |
|---|---|---|---|---|---|---|
| trmt5 | 0.22544 | -0.19154192 | 0 | 0.059 | 1 | 19 |
| ttl | 0.22544 | -0.19556867 | 0 | 0.059 | 1 | 19 |
| twf1b | 0.22544 | -0.13220928 | 0 | 0.059 | 1 | 19 |
| txndc12 | 0.22544 | -0.1172756 | 0 | 0.059 | 1 | 19 |
| txndc5 | 0.22544 | -0.19572725 | 0 | 0.059 | 1 | 19 |
| txnl1 | 0.22544 | -0.21634264 | 0 | 0.059 | 1 | 19 |
| ubald2 | 0.22544 | -0.12524561 | 0 | 0.059 | 1 | 19 |
| ube2j1 | 0.22544 | -0.15503087 | 0 | 0.059 | 1 | 19 |
| ube2r2 | 0.22544 | -0.19661494 | 0 | 0.059 | 1 | 19 |
| ublcp1 | 0.22544 | -0.15224672 | 0 | 0.059 | 1 | 19 |
| ubn2b | 0.22544 | -0.16391278 | 0 | 0.059 | 1 | 19 |
| ubr4 | 0.22544 | -0.19827116 | 0 | 0.059 | 1 | 19 |
| ubtf | 0.22544 | -0.19126271 | 0 | 0.059 | 1 | 19 |
| ubxn7 | 0.22544 | -0.15540656 | 0 | 0.059 | 1 | 19 |
| uck1 | 0.22544 | -0.16656008 | 0 | 0.059 | 1 | 19 |
| uck2a | 0.22544 | -0.12060026 | 0 | 0.059 | 1 | 19 |
| ugdh | 0.22544 | -0.16066243 | 0 | 0.059 | 1 | 19 |
| uprt | 0.22544 | -0.16064995 | 0 | 0.059 | 1 | 19 |
| usp24 | 0.22544 | -0.1990834 | 0 | 0.059 | 1 | 19 |
| usp48 | 0.22544 | -0.11834492 | 0 | 0.059 | 1 | 19 |
| utp6 | 0.22544 | -0.10952211 | 0 | 0.059 | 1 | 19 |
| utrn | 0.22544 | -0.11790209 | 0 | 0.059 | 1 | 19 |
| vav3b | 0.22544 | -0.12567073 | 0 | 0.059 | 1 | 19 |
| vps26bl | 0.22544 | -0.170042 | 0 | 0.059 | 1 | 19 |
| vps53 | 0.22544 | -0.19635302 | 0 | 0.059 | 1 | 19 |
| vps54 | 0.22544 | -0.16945706 | 0 | 0.059 | 1 | 19 |
| vti1a | 0.22544 | -0.19437842 | 0 | 0.059 | 1 | 19 |
| wbp1 | 0.22544 | -0.1917658 | 0 | 0.059 | 1 | 19 |
| wdfy1 | 0.22544 | -0.15784176 | 0 | 0.059 | 1 | 19 |
| wdr21 | 0.22544 | -0.18926871 | 0 | 0.059 | 1 | 19 |
| wdr3 | 0.22544 | -0.11994407 | 0 | 0.059 | 1 | 19 |
| wdr45 | 0.22544 | -0.11857204 | 0 | 0.059 | 1 | 19 |
| wdr62 | 0.22544 | -0.11762331 | 0 | 0.059 | 1 | 19 |
| wsb1 | 0.22544 | -0.19833518 | 0 | 0.059 | 1 | 19 |
| wu:fc33b09 | 0.22544 | -0.11770254 | 0 | 0.059 | 1 | 19 |
| xpot | 0.22544 | -0.16622136 | 0 | 0.059 | 1 | 19 |
| yeats2 | 0.22544 | -0.16251477 | 0 | 0.059 | 1 | 19 |
| yme1l1a | 0.22544 | -0.11715597 | 0 | 0.059 | 1 | 19 |
| zbtb33 | 0.22544 | -0.15705723 | 0 | 0.059 | 1 | 19 |
| zdhhc15b | 0.22544 | -0.21383388 | 0 | 0.059 | 1 | 19 |
| zdhhc16b | 0.22544 | -0.130828 | 0 | 0.059 | 1 | 19 |
| zdhhc17 | 0.22544 | -0.16465343 | 0 | 0.059 | 1 | 19 |
| zdhhc3a | 0.22544 | -0.16960031 | 0 | 0.059 | 1 | 19 |
| zfat | 0.22544 | -0.22754041 | 0 | 0.059 | 1 | 19 |
| zfyve21 | 0.22544 | -0.11582865 | 0 | 0.059 | 1 | 19 |
| zgc:101559 | 0.22544 | -0.22733246 | 0 | 0.059 | 1 | 19 |
| zgc:101566 | 0.22544 | -0.16327038 | 0 | 0.059 | 1 | 19 |
| zgc:136564 | 0.22544 | -0.10762861 | 0 | 0.059 | 1 | 19 |

| | | | | | | |
|---|---|---|---|---|---|---|
| zgc:153292 | 0.22544 | -0.12875125 | 0 | 0.059 | 1 | 19 |
| zgc:153935 | 0.22544 | -0.16829805 | 0 | 0.059 | 1 | 19 |
| zgc:153990 | 0.22544 | -0.11755012 | 0 | 0.059 | 1 | 19 |
| zgc:161969 | 0.22544 | -0.16053533 | 0 | 0.059 | 1 | 19 |
| zgc:162509 | 0.22544 | -0.17797118 | 0 | 0.059 | 1 | 19 |
| zgc:76872 | 0.22544 | -0.20838216 | 0 | 0.059 | 1 | 19 |
| znf704 | 0.22544 | -0.17037063 | 0 | 0.059 | 1 | 19 |
| zwilch | 0.22544 | -0.12130474 | 0 | 0.059 | 1 | 19 |
| si:dkey-21e2.15 | 0.22618 | -0.12055067 | 0.29 | 0.412 | 1 | 19 |
| pgam5 | 0.22717 | -0.18351953 | 0.032 | 0.176 | 1 | 19 |
| ddx5 | 0.22721 | 0.437924991 | 0.516 | 0.294 | 1 | 19 |
| bloc1s6 | 0.22744 | -0.4235569 | 0.097 | 0.294 | 1 | 19 |
| fip1l1b | 0.22769 | -0.34948008 | 0.097 | 0.294 | 1 | 19 |
| psma2 | 0.22773 | 0.387569142 | 0.226 | 0.059 | 1 | 19 |
| mlec | 0.22819 | -0.37103777 | 0.097 | 0.294 | 1 | 19 |
| rps14 | 0.22929 | 0.18736786 | 0.935 | 0.765 | 1 | 19 |
| wtap | 0.22964 | -0.42029585 | 0.065 | 0.235 | 1 | 19 |
| syne1b | 0.23009 | -0.23384962 | 0.032 | 0.176 | 1 | 19 |
| dnajc8 | 0.23019 | -0.15582276 | 0.032 | 0.176 | 1 | 19 |
| cog7 | 0.23058 | -0.36187502 | 0.032 | 0.176 | 1 | 19 |
| iqgap1 | 0.23102 | -0.36474109 | 0.097 | 0.294 | 1 | 19 |
| zgc:136605 | 0.2311 | -0.48057413 | 0.097 | 0.235 | 1 | 19 |
| lamp1a | 0.23143 | -0.39529529 | 0.097 | 0.294 | 1 | 19 |
| samsn1a | 0.23147 | -0.42436038 | 0.097 | 0.294 | 1 | 19 |
| zgc:56304 | 0.23234 | -0.30699378 | 0.097 | 0.294 | 1 | 19 |
| eif4ebp1 | 0.23274 | 0.30307996 | 0.323 | 0.118 | 1 | 19 |
| chchd7 | 0.2334 | -0.37803984 | 0.065 | 0.235 | 1 | 19 |
| uqcrfs1 | 0.2347 | 0.402809828 | 0.323 | 0.118 | 1 | 19 |
| trnau1apb | 0.23481 | -0.33284343 | 0.065 | 0.235 | 1 | 19 |
| vps35 | 0.23568 | -0.24070194 | 0.065 | 0.235 | 1 | 19 |
| ola1 | 0.23658 | 0.378401814 | 0.29 | 0.118 | 1 | 19 |
| entpd4 | 0.23688 | -0.36784864 | 0.032 | 0.176 | 1 | 19 |
| ppiaa | 0.23701 | 0.252084307 | 0.935 | 0.765 | 1 | 19 |
| ppdpfb | 0.23761 | -0.33194377 | 0.452 | 0.529 | 1 | 19 |
| mmgt1 | 0.23769 | -0.29868246 | 0.065 | 0.235 | 1 | 19 |
| BX927081.3 | 0.238 | -0.27924237 | 0.032 | 0.176 | 1 | 19 |
| ptger4b | 0.23811 | -0.4346424 | 0.065 | 0.235 | 1 | 19 |
| XLOC-018356 | 0.23824 | -0.51460311 | 0.097 | 0.235 | 1 | 19 |
| tia1l | 0.23841 | -0.28111685 | 0.065 | 0.235 | 1 | 19 |
| mrpl33 | 0.23848 | -0.27125622 | 0.065 | 0.235 | 1 | 19 |
| LEPROTL1 | 0.23883 | 0.462977925 | 0.226 | 0.059 | 1 | 19 |
| rpl39 | 0.23894 | -0.14371734 | 0.903 | 0.941 | 1 | 19 |
| ube2ib | 0.23971 | -0.36890467 | 0.032 | 0.176 | 1 | 19 |
| acvr2aa | 0.24101 | -0.3324017 | 0.032 | 0.176 | 1 | 19 |
| nfil3 | 0.24117 | -0.311036 | 0.032 | 0.176 | 1 | 19 |
| tmem176l.1 | 0.24132 | -0.28105281 | 0.032 | 0.176 | 1 | 19 |
| agpat5 | 0.2416 | -0.25619511 | 0.065 | 0.235 | 1 | 19 |
| arpc1a | 0.2426 | -0.3539208 | 0.032 | 0.176 | 1 | 19 |

| | | | | | | |
|---|---|---|---|---|---|---|
| gcsha | 0.24297 | -0.27843557 | 0.032 | 0.176 | 1 | 19 |
| ubc | 0.243 | 0.42113407 | 0.677 | 0.471 | 1 | 19 |
| pip4k2aa | 0.24305 | -0.31221948 | 0.032 | 0.176 | 1 | 19 |
| si:ch211-191j22.3 | 0.24337 | -0.28923059 | 0.032 | 0.176 | 1 | 19 |
| si:dkey-56i24.1 | 0.24372 | -0.31831731 | 0.032 | 0.176 | 1 | 19 |
| myo1f | 0.24417 | -0.43787857 | 0.065 | 0.235 | 1 | 19 |
| lyar | 0.24419 | -0.24786135 | 0.065 | 0.235 | 1 | 19 |
| med19a | 0.24425 | 0.462199731 | 0.29 | 0.118 | 1 | 19 |
| ifi30 | 0.24462 | -0.3591233 | 0.065 | 0.235 | 1 | 19 |
| bysl | 0.24476 | -0.3249989 | 0.065 | 0.235 | 1 | 19 |
| mapk3 | 0.24495 | -0.28250859 | 0.065 | 0.235 | 1 | 19 |
| chd8 | 0.24531 | -0.30982794 | 0.032 | 0.176 | 1 | 19 |
| prkar1aa | 0.24531 | -0.23754652 | 0.032 | 0.176 | 1 | 19 |
| csnk1db | 0.24589 | -0.32269944 | 0.032 | 0.176 | 1 | 19 |
| arf2a | 0.24611 | -0.32418114 | 0.065 | 0.235 | 1 | 19 |
| rrp12 | 0.24615 | -0.36251835 | 0.032 | 0.176 | 1 | 19 |
| hsd17b10 | 0.24724 | -0.16604802 | 0.032 | 0.176 | 1 | 19 |
| akr1a1b | 0.24766 | -0.24177111 | 0.032 | 0.176 | 1 | 19 |
| hhex | 0.24787 | -0.41621204 | 0.065 | 0.235 | 1 | 19 |
| RASA2 | 0.24789 | -0.36169113 | 0.032 | 0.176 | 1 | 19 |
| cpox | 0.24798 | -0.31103522 | 0.032 | 0.176 | 1 | 19 |
| cyb5r2 | 0.24846 | -0.38687101 | 0.065 | 0.235 | 1 | 19 |
| grpel1 | 0.24853 | -0.22743984 | 0.032 | 0.176 | 1 | 19 |
| hnrnpl2 | 0.24867 | -0.31715544 | 0.032 | 0.176 | 1 | 19 |
| rnf41 | 0.24876 | -0.32395024 | 0.032 | 0.176 | 1 | 19 |
| si:ch1073-220m6.1 | 0.24892 | -0.3120279 | 0.032 | 0.176 | 1 | 19 |
| tmed1b | 0.24938 | -0.26631441 | 0.032 | 0.176 | 1 | 19 |
| cfdp1 | 0.2497 | -0.23485485 | 0.032 | 0.176 | 1 | 19 |
| lyrm4 | 0.24983 | -0.25254862 | 0.032 | 0.176 | 1 | 19 |
| dync1i2a | 0.24984 | -0.40523971 | 0.032 | 0.176 | 1 | 19 |
| calm2a | 0.25042 | -0.40928265 | 0.194 | 0.412 | 1 | 19 |
| ctsc | 0.2505 | -0.40205759 | 0.065 | 0.235 | 1 | 19 |
| ubtd2 | 0.25058 | -0.25864122 | 0.065 | 0.235 | 1 | 19 |
| trib3 | 0.25075 | -0.34246124 | 0.065 | 0.235 | 1 | 19 |
| tmem126a | 0.2519 | -0.32289609 | 0.065 | 0.235 | 1 | 19 |
| LOC100151196 | 0.25196 | -0.32215804 | 0.065 | 0.235 | 1 | 19 |
| def6c | 0.25233 | -0.30668892 | 0.065 | 0.235 | 1 | 19 |
| slc25a55a | 0.25279 | -0.30061363 | 0.065 | 0.235 | 1 | 19 |
| dus3l | 0.25281 | -0.29775027 | 0.065 | 0.235 | 1 | 19 |
| cebpa | 0.25289 | -0.39383063 | 0.065 | 0.235 | 1 | 19 |
| CU207241.1 | 0.25544 | -0.1346842 | 0.226 | 0.176 | 1 | 19 |
| LOC103909452 | 0.25736 | 0.113505335 | 0.065 | 0 | 1 | 19 |
| actn4 | 0.25736 | 0.109778698 | 0.065 | 0 | 1 | 19 |
| adnpa | 0.25736 | 0.177954266 | 0.065 | 0 | 1 | 19 |
| afmid | 0.25736 | 0.224166931 | 0.065 | 0 | 1 | 19 |
| alg2 | 0.25736 | 0.116291718 | 0.065 | 0 | 1 | 19 |
| alkbh4 | 0.25736 | 0.132271574 | 0.065 | 0 | 1 | 19 |
| apeh | 0.25736 | 0.135037136 | 0.065 | 0 | 1 | 19 |

| | | | | | | |
|---|---|---|---|---|---|---|
| aqp3a | 0.25736 | 0.205658241 | 0.065 | 0 | 1 | 19 |
| atp13a2 | 0.25736 | 0.132358508 | 0.065 | 0 | 1 | 19 |
| atp6v1h | 0.25736 | 0.120391841 | 0.065 | 0 | 1 | 19 |
| bola3 | 0.25736 | 0.109234789 | 0.065 | 0 | 1 | 19 |
| cbarpb | 0.25736 | 0.163705491 | 0.065 | 0 | 1 | 19 |
| ccdc58 | 0.25736 | 0.128045691 | 0.065 | 0 | 1 | 19 |
| cdc37 | 0.25736 | 0.154767206 | 0.065 | 0 | 1 | 19 |
| cers5 | 0.25736 | 0.132102865 | 0.065 | 0 | 1 | 19 |
| col14a1b | 0.25736 | 0.136278885 | 0.065 | 0 | 1 | 19 |
| cpa5 | 0.25736 | 0.317833979 | 0.065 | 0 | 1 | 19 |
| crfb4 | 0.25736 | 0.119844421 | 0.065 | 0 | 1 | 19 |
| dele1 | 0.25736 | 0.111547321 | 0.065 | 0 | 1 | 19 |
| eci1 | 0.25736 | 0.117113143 | 0.065 | 0 | 1 | 19 |
| eef1e1 | 0.25736 | 0.112432321 | 0.065 | 0 | 1 | 19 |
| ergic2 | 0.25736 | 0.138203705 | 0.065 | 0 | 1 | 19 |
| fryl | 0.25736 | 0.159377469 | 0.065 | 0 | 1 | 19 |
| gng10 | 0.25736 | 0.114709424 | 0.065 | 0 | 1 | 19 |
| gnpnat1 | 0.25736 | 0.104355969 | 0.065 | 0 | 1 | 19 |
| golt1ba | 0.25736 | 0.120406615 | 0.065 | 0 | 1 | 19 |
| hdgfl2 | 0.25736 | 0.126436002 | 0.065 | 0 | 1 | 19 |
| heatr5a | 0.25736 | 0.160375471 | 0.065 | 0 | 1 | 19 |
| hnrnpr | 0.25736 | 0.116915884 | 0.065 | 0 | 1 | 19 |
| irf7 | 0.25736 | 0.114185621 | 0.065 | 0 | 1 | 19 |
| loxl3b | 0.25736 | 0.114002738 | 0.065 | 0 | 1 | 19 |
| nutf2 | 0.25736 | 0.124143623 | 0.065 | 0 | 1 | 19 |
| pafah1b1b | 0.25736 | 0.150056231 | 0.065 | 0 | 1 | 19 |
| polr1e | 0.25736 | 0.179721931 | 0.065 | 0 | 1 | 19 |
| ppp1r8b | 0.25736 | 0.17215764 | 0.065 | 0 | 1 | 19 |
| prrc2c | 0.25736 | 0.106664034 | 0.065 | 0 | 1 | 19 |
| ptcd1 | 0.25736 | 0.119409359 | 0.065 | 0 | 1 | 19 |
| rab5b | 0.25736 | 0.167543384 | 0.065 | 0 | 1 | 19 |
| rbm7 | 0.25736 | 0.114557882 | 0.065 | 0 | 1 | 19 |
| rbmx | 0.25736 | 0.176007123 | 0.065 | 0 | 1 | 19 |
| rhbdd1 | 0.25736 | 0.125830002 | 0.065 | 0 | 1 | 19 |
| rprd1a | 0.25736 | 0.107702686 | 0.065 | 0 | 1 | 19 |
| rxrga | 0.25736 | 0.143800184 | 0.065 | 0 | 1 | 19 |
| sh3gl2a | 0.25736 | 0.122270943 | 0.065 | 0 | 1 | 19 |
| sh3glb1b | 0.25736 | 0.127369613 | 0.065 | 0 | 1 | 19 |
| si:ch211-215e19.3 | 0.25736 | 0.186946135 | 0.065 | 0 | 1 | 19 |
| si:ch211-225b11.1 | 0.25736 | 0.10952 | 0.065 | 0 | 1 | 19 |
| si:dkey-102m7.3 | 0.25736 | 0.145624761 | 0.065 | 0 | 1 | 19 |
| si:dkey-19e4.5 | 0.25736 | 0.121256117 | 0.065 | 0 | 1 | 19 |
| si:dkeyp-38g8.5 | 0.25736 | 0.131190039 | 0.065 | 0 | 1 | 19 |
| slc35e1 | 0.25736 | 0.13432759 | 0.065 | 0 | 1 | 19 |
| slc38a10 | 0.25736 | 0.124480331 | 0.065 | 0 | 1 | 19 |
| slc44a2 | 0.25736 | 0.120436371 | 0.065 | 0 | 1 | 19 |
| slka | 0.25736 | 0.181343202 | 0.065 | 0 | 1 | 19 |
| smc1al | 0.25736 | 0.22470816 | 0.065 | 0 | 1 | 19 |

| | | | | | | |
|---|---|---|---|---|---|---|
| srrm4 | 0.25736 | 0.12075024 | 0.065 | 0 | 1 | 19 |
| tbcb | 0.25736 | 0.15309825 | 0.065 | 0 | 1 | 19 |
| tent5ab | 0.25736 | 0.14870674 | 0.065 | 0 | 1 | 19 |
| tmem241 | 0.25736 | 0.136235698 | 0.065 | 0 | 1 | 19 |
| traf3 | 0.25736 | 0.123504289 | 0.065 | 0 | 1 | 19 |
| trappc6bl | 0.25736 | 0.132883412 | 0.065 | 0 | 1 | 19 |
| triqk | 0.25736 | 0.171100024 | 0.065 | 0 | 1 | 19 |
| tsen54 | 0.25736 | 0.136278885 | 0.065 | 0 | 1 | 19 |
| ube2a | 0.25736 | 0.128133094 | 0.065 | 0 | 1 | 19 |
| utp25 | 0.25736 | 0.117882137 | 0.065 | 0 | 1 | 19 |
| wdfy4 | 0.25736 | 0.132900059 | 0.065 | 0 | 1 | 19 |
| zgc:152863 | 0.25736 | 0.126988692 | 0.065 | 0 | 1 | 19 |
| zgc:174863 | 0.25736 | 0.128704315 | 0.065 | 0 | 1 | 19 |
| zgc:174945 | 0.25736 | 0.143074318 | 0.065 | 0 | 1 | 19 |
| zmp:0000000650 | 0.25736 | 0.122835646 | 0.065 | 0 | 1 | 19 |
| znf207b | 0.25736 | 0.129417735 | 0.065 | 0 | 1 | 19 |
| znhit1 | 0.25736 | 0.147225485 | 0.065 | 0 | 1 | 19 |
| zw10 | 0.25736 | 0.124798314 | 0.065 | 0 | 1 | 19 |
| adgrg1 | 0.25736 | 0.168290495 | 0.065 | 0 | 1 | 19 |
| anapc11 | 0.25736 | 0.143457569 | 0.065 | 0 | 1 | 19 |
| arih2 | 0.25736 | 0.12994763 | 0.065 | 0 | 1 | 19 |
| arl4aa | 0.25736 | 0.247347352 | 0.065 | 0 | 1 | 19 |
| arpin | 0.25736 | 0.143047605 | 0.065 | 0 | 1 | 19 |
| decr1 | 0.25736 | 0.104254347 | 0.065 | 0 | 1 | 19 |
| dpy19l1l | 0.25736 | 0.154473585 | 0.065 | 0 | 1 | 19 |
| elp6 | 0.25736 | 0.108823417 | 0.065 | 0 | 1 | 19 |
| hnrnpua | 0.25736 | 0.171280209 | 0.065 | 0 | 1 | 19 |
| ikbkb | 0.25736 | 0.172729924 | 0.065 | 0 | 1 | 19 |
| jarid2b | 0.25736 | 0.146801245 | 0.065 | 0 | 1 | 19 |
| kansl1b | 0.25736 | 0.197183481 | 0.065 | 0 | 1 | 19 |
| kdm5a | 0.25736 | 0.154473585 | 0.065 | 0 | 1 | 19 |
| krr1 | 0.25736 | 0.127938892 | 0.065 | 0 | 1 | 19 |
| lyst | 0.25736 | 0.126828527 | 0.065 | 0 | 1 | 19 |
| mcrip1 | 0.25736 | 0.157606996 | 0.065 | 0 | 1 | 19 |
| mob4 | 0.25736 | 0.184064393 | 0.065 | 0 | 1 | 19 |
| nab1b | 0.25736 | 0.195169238 | 0.065 | 0 | 1 | 19 |
| nbeal2 | 0.25736 | 0.150165532 | 0.065 | 0 | 1 | 19 |
| pcca | 0.25736 | 0.171316679 | 0.065 | 0 | 1 | 19 |
| psmc1a | 0.25736 | 0.184667763 | 0.065 | 0 | 1 | 19 |
| rab11bb | 0.25736 | 0.17551691 | 0.065 | 0 | 1 | 19 |
| rap2c | 0.25736 | 0.137031081 | 0.065 | 0 | 1 | 19 |
| sec31b | 0.25736 | 0.174924723 | 0.065 | 0 | 1 | 19 |
| sft2d1 | 0.25736 | 0.151243162 | 0.065 | 0 | 1 | 19 |
| si:dkey-111k8.3 | 0.25736 | 0.133867031 | 0.065 | 0 | 1 | 19 |
| si:dkey-188i13.11 | 0.25736 | 0.178298312 | 0.065 | 0 | 1 | 19 |
| si:dkey-217l24.1 | 0.25736 | 0.193423116 | 0.065 | 0 | 1 | 19 |
| srd5a3 | 0.25736 | 0.162813352 | 0.065 | 0 | 1 | 19 |
| stk11ip | 0.25736 | 0.150176137 | 0.065 | 0 | 1 | 19 |

| | | | | | | |
|---|---|---|---|---|---|---|
| taf4a | 0.25736 | 0.177261049 | 0.065 | 0 | 1 | 19 |
| xpc | 0.25736 | 0.172882662 | 0.065 | 0 | 1 | 19 |
| zdhhc4 | 0.25736 | 0.137353601 | 0.065 | 0 | 1 | 19 |
| zgc:153018 | 0.25736 | 0.121325337 | 0.065 | 0 | 1 | 19 |
| zgc:92360 | 0.25736 | 0.15014925 | 0.065 | 0 | 1 | 19 |
| BX248082.1 | 0.25736 | 0.21741867 | 0.065 | 0 | 1 | 19 |
| BX470259.2 | 0.25736 | 0.132735181 | 0.065 | 0 | 1 | 19 |
| CABZ01074453.1 | 0.25736 | 0.124785412 | 0.065 | 0 | 1 | 19 |
| CABZ01075242.1 | 0.25736 | 0.142290477 | 0.065 | 0 | 1 | 19 |
| CR318588.3 | 0.25736 | 0.164984313 | 0.065 | 0 | 1 | 19 |
| CR847543.1 | 0.25736 | 0.210952072 | 0.065 | 0 | 1 | 19 |
| CR847994.1 | 0.25736 | 0.176407779 | 0.065 | 0 | 1 | 19 |
| CU856520.1 | 0.25736 | 0.11999966 | 0.065 | 0 | 1 | 19 |
| FBXO46 | 0.25736 | 0.12096217 | 0.065 | 0 | 1 | 19 |
| LOC100534659 | 0.25736 | 0.150712412 | 0.065 | 0 | 1 | 19 |
| LOC101884098 | 0.25736 | 0.16404141 | 0.065 | 0 | 1 | 19 |
| LOC101885083 | 0.25736 | 0.117941602 | 0.065 | 0 | 1 | 19 |
| LOC103909972 | 0.25736 | 0.103260098 | 0.065 | 0 | 1 | 19 |
| LOC108190702 | 0.25736 | 0.136502443 | 0.065 | 0 | 1 | 19 |
| MARK4 | 0.25736 | 0.166400669 | 0.065 | 0 | 1 | 19 |
| MED9 | 0.25736 | 0.156572955 | 0.065 | 0 | 1 | 19 |
| TMEM208 | 0.25736 | 0.111180194 | 0.065 | 0 | 1 | 19 |
| UTP14C | 0.25736 | 0.157369443 | 0.065 | 0 | 1 | 19 |
| XLOC-019150 | 0.25736 | 0.169173524 | 0.065 | 0 | 1 | 19 |
| XLOC-039619 | 0.25736 | 0.141723697 | 0.065 | 0 | 1 | 19 |
| acad9 | 0.25736 | 0.121429358 | 0.065 | 0 | 1 | 19 |
| acsf2 | 0.25736 | 0.127170825 | 0.065 | 0 | 1 | 19 |
| adi1 | 0.25736 | 0.115148303 | 0.065 | 0 | 1 | 19 |
| adsl | 0.25736 | 0.106234359 | 0.065 | 0 | 1 | 19 |
| afap1l2 | 0.25736 | 0.242726873 | 0.065 | 0 | 1 | 19 |
| agr2 | 0.25736 | 0.162960773 | 0.065 | 0 | 1 | 19 |
| amd1 | 0.25736 | 0.186349325 | 0.065 | 0 | 1 | 19 |
| ano10a | 0.25736 | 0.138126184 | 0.065 | 0 | 1 | 19 |
| arhgef1b | 0.25736 | 0.121276881 | 0.065 | 0 | 1 | 19 |
| arid1ab | 0.25736 | 0.172631224 | 0.065 | 0 | 1 | 19 |
| armt1 | 0.25736 | 0.143702186 | 0.065 | 0 | 1 | 19 |
| atp10a | 0.25736 | 0.141469323 | 0.065 | 0 | 1 | 19 |
| atp1b2a | 0.25736 | 0.148116611 | 0.065 | 0 | 1 | 19 |
| atp2c1 | 0.25736 | 0.123678737 | 0.065 | 0 | 1 | 19 |
| atp6v1d | 0.25736 | 0.155260526 | 0.065 | 0 | 1 | 19 |
| b3gnt7l | 0.25736 | 0.114742357 | 0.065 | 0 | 1 | 19 |
| bbip1 | 0.25736 | 0.103588536 | 0.065 | 0 | 1 | 19 |
| bbx | 0.25736 | 0.229901982 | 0.065 | 0 | 1 | 19 |
| bcat2 | 0.25736 | 0.196091748 | 0.065 | 0 | 1 | 19 |
| becn1 | 0.25736 | 0.114970883 | 0.065 | 0 | 1 | 19 |
| bmi1b | 0.25736 | 0.209898661 | 0.065 | 0 | 1 | 19 |
| camk1b | 0.25736 | 0.240420223 | 0.065 | 0 | 1 | 19 |
| casp8 | 0.25736 | 0.146975662 | 0.065 | 0 | 1 | 19 |

| | | | | | | |
|---|---|---|---|---|---|---|
| cbwd | 0.25736 | 0.106827065 | 0.065 | 0 | 1 | 19 |
| ccnl1b | 0.25736 | 0.201580799 | 0.065 | 0 | 1 | 19 |
| ccz1 | 0.25736 | 0.1135767 | 0.065 | 0 | 1 | 19 |
| cdk9 | 0.25736 | 0.104532309 | 0.065 | 0 | 1 | 19 |
| cdkn3 | 0.25736 | 0.178601935 | 0.065 | 0 | 1 | 19 |
| chmp1b | 0.25736 | 0.143715646 | 0.065 | 0 | 1 | 19 |
| chsy1 | 0.25736 | 0.187223327 | 0.065 | 0 | 1 | 19 |
| chtopa | 0.25736 | 0.149981716 | 0.065 | 0 | 1 | 19 |
| cln8 | 0.25736 | 0.138770831 | 0.065 | 0 | 1 | 19 |
| coil | 0.25736 | 0.119895427 | 0.065 | 0 | 1 | 19 |
| creb3l3l | 0.25736 | 0.18871741 | 0.065 | 0 | 1 | 19 |
| cul1b | 0.25736 | 0.150216034 | 0.065 | 0 | 1 | 19 |
| dtnba | 0.25736 | 0.175818502 | 0.065 | 0 | 1 | 19 |
| dusp23b | 0.25736 | 0.189787713 | 0.065 | 0 | 1 | 19 |
| e4f1 | 0.25736 | 0.156779141 | 0.065 | 0 | 1 | 19 |
| eef2k | 0.25736 | 0.120638052 | 0.065 | 0 | 1 | 19 |
| egr2a | 0.25736 | 0.17324588 | 0.065 | 0 | 1 | 19 |
| ehbp1l1a | 0.25736 | 0.131860475 | 0.065 | 0 | 1 | 19 |
| eif3hb | 0.25736 | 0.104299874 | 0.065 | 0 | 1 | 19 |
| eif4ebp3 | 0.25736 | 0.139449773 | 0.065 | 0 | 1 | 19 |
| ell | 0.25736 | 0.105596192 | 0.065 | 0 | 1 | 19 |
| erlec1 | 0.25736 | 0.109464043 | 0.065 | 0 | 1 | 19 |
| esrra | 0.25736 | 0.116527491 | 0.065 | 0 | 1 | 19 |
| exosc9 | 0.25736 | 0.120768514 | 0.065 | 0 | 1 | 19 |
| fam76b | 0.25736 | 0.117700851 | 0.065 | 0 | 1 | 19 |
| fes | 0.25736 | 0.131155098 | 0.065 | 0 | 1 | 19 |
| fibpa | 0.25736 | 0.153379227 | 0.065 | 0 | 1 | 19 |
| flot1b | 0.25736 | 0.129303083 | 0.065 | 0 | 1 | 19 |
| fntb | 0.25736 | 0.166409056 | 0.065 | 0 | 1 | 19 |
| fut8a | 0.25736 | 0.20718409 | 0.065 | 0 | 1 | 19 |
| gamt | 0.25736 | 0.122220733 | 0.065 | 0 | 1 | 19 |
| gigyf2 | 0.25736 | 0.160363443 | 0.065 | 0 | 1 | 19 |
| gnal | 0.25736 | 0.10007458 | 0.065 | 0 | 1 | 19 |
| gsdmea | 0.25736 | 0.109594737 | 0.065 | 0 | 1 | 19 |
| gsr | 0.25736 | 0.103746732 | 0.065 | 0 | 1 | 19 |
| guk1a | 0.25736 | 0.164981632 | 0.065 | 0 | 1 | 19 |
| hectd3 | 0.25736 | 0.145791467 | 0.065 | 0 | 1 | 19 |
| hexa | 0.25736 | 0.128148385 | 0.065 | 0 | 1 | 19 |
| ifi46 | 0.25736 | 0.110832977 | 0.065 | 0 | 1 | 19 |
| igf2r | 0.25736 | 0.126492755 | 0.065 | 0 | 1 | 19 |
| ikbke | 0.25736 | 0.129497763 | 0.065 | 0 | 1 | 19 |
| il12rb2 | 0.25736 | 0.109357713 | 0.065 | 0 | 1 | 19 |
| il12rb2l | 0.25736 | 0.132718656 | 0.065 | 0 | 1 | 19 |
| itln1 | 0.25736 | 0.13194704 | 0.065 | 0 | 1 | 19 |
| ivd | 0.25736 | 0.105217863 | 0.065 | 0 | 1 | 19 |
| kctd13 | 0.25736 | 0.103154193 | 0.065 | 0 | 1 | 19 |
| kdm5c | 0.25736 | 0.148065842 | 0.065 | 0 | 1 | 19 |
| klc1b | 0.25736 | 0.118759516 | 0.065 | 0 | 1 | 19 |

| | | | | | | |
|---|---|---|---|---|---|---|
| klf13 | 0.25736 | 0.142875847 | 0.065 | 0 | 1 | 19 |
| klf2a | 0.25736 | 0.110480307 | 0.065 | 0 | 1 | 19 |
| klf7b | 0.25736 | 0.172599155 | 0.065 | 0 | 1 | 19 |
| kpna6 | 0.25736 | 0.152784838 | 0.065 | 0 | 1 | 19 |
| lancl1 | 0.25736 | 0.126126744 | 0.065 | 0 | 1 | 19 |
| larp1 | 0.25736 | 0.104534944 | 0.065 | 0 | 1 | 19 |
| larp7 | 0.25736 | 0.164453855 | 0.065 | 0 | 1 | 19 |
| lbr | 0.25736 | 0.14202694 | 0.065 | 0 | 1 | 19 |
| lgals2a | 0.25736 | 0.159419127 | 0.065 | 0 | 1 | 19 |
| lipea | 0.25736 | 0.107939303 | 0.065 | 0 | 1 | 19 |
| lrp10 | 0.25736 | 0.181007677 | 0.065 | 0 | 1 | 19 |
| madd | 0.25736 | 0.106321888 | 0.065 | 0 | 1 | 19 |
| mak16 | 0.25736 | 0.139672226 | 0.065 | 0 | 1 | 19 |
| map4k6 | 0.25736 | 0.184586075 | 0.065 | 0 | 1 | 19 |
| matr3l1.1 | 0.25736 | 0.122192932 | 0.065 | 0 | 1 | 19 |
| mcl1b | 0.25736 | 0.156856605 | 0.065 | 0 | 1 | 19 |
| med7 | 0.25736 | 0.155854367 | 0.065 | 0 | 1 | 19 |
| metap1 | 0.25736 | 0.118861841 | 0.065 | 0 | 1 | 19 |
| mrps18a | 0.25736 | 0.165745853 | 0.065 | 0 | 1 | 19 |
| mrps18c | 0.25736 | 0.140957057 | 0.065 | 0 | 1 | 19 |
| mrps36 | 0.25736 | 0.142814433 | 0.065 | 0 | 1 | 19 |
| msl2b | 0.25736 | 0.194479443 | 0.065 | 0 | 1 | 19 |
| mtrf1l | 0.25736 | 0.122487363 | 0.065 | 0 | 1 | 19 |
| ndnl2 | 0.25736 | 0.129152565 | 0.065 | 0 | 1 | 19 |
| ndufv2 | 0.25736 | 0.101800074 | 0.065 | 0 | 1 | 19 |
| nek7 | 0.25736 | 0.228136278 | 0.065 | 0 | 1 | 19 |
| nfat5a | 0.25736 | 0.102016427 | 0.065 | 0 | 1 | 19 |
| nfs1 | 0.25736 | 0.101384934 | 0.065 | 0 | 1 | 19 |
| nsun5 | 0.25736 | 0.113947847 | 0.065 | 0 | 1 | 19 |
| nt5c3a | 0.25736 | 0.106469226 | 0.065 | 0 | 1 | 19 |
| nudt3b | 0.25736 | 0.133326778 | 0.065 | 0 | 1 | 19 |
| osbpl3b | 0.25736 | 0.157988583 | 0.065 | 0 | 1 | 19 |
| oxr1a | 0.25736 | 0.141385143 | 0.065 | 0 | 1 | 19 |
| oxsr1a | 0.25736 | 0.105622326 | 0.065 | 0 | 1 | 19 |
| parm1 | 0.25736 | 0.112823697 | 0.065 | 0 | 1 | 19 |
| pcgf5b | 0.25736 | 0.192787258 | 0.065 | 0 | 1 | 19 |
| per1a | 0.25736 | 0.122724266 | 0.065 | 0 | 1 | 19 |
| per2 | 0.25736 | 0.158609686 | 0.065 | 0 | 1 | 19 |
| pex10 | 0.25736 | 0.154665726 | 0.065 | 0 | 1 | 19 |
| pgap3 | 0.25736 | 0.122296712 | 0.065 | 0 | 1 | 19 |
| phf8 | 0.25736 | 0.116391491 | 0.065 | 0 | 1 | 19 |
| phkg2 | 0.25736 | 0.117684051 | 0.065 | 0 | 1 | 19 |
| pi4k2b | 0.25736 | 0.117405069 | 0.065 | 0 | 1 | 19 |
| pih1d1 | 0.25736 | 0.187878574 | 0.065 | 0 | 1 | 19 |
| pik3r3a | 0.25736 | 0.124949417 | 0.065 | 0 | 1 | 19 |
| pla2g15 | 0.25736 | 0.144483895 | 0.065 | 0 | 1 | 19 |
| pnisr | 0.25736 | 0.116379485 | 0.065 | 0 | 1 | 19 |
| polr2c | 0.25736 | 0.134596209 | 0.065 | 0 | 1 | 19 |

| | | | | | | |
|---|---|---|---|---|---|---|
| polr3f | 0.25736 | 0.164256679 | 0.065 | 0 | 1 | 19 |
| ponzr6 | 0.25736 | 0.136502443 | 0.065 | 0 | 1 | 19 |
| prr12a | 0.25736 | 0.130368136 | 0.065 | 0 | 1 | 19 |
| pstpip1a | 0.25736 | 0.122487363 | 0.065 | 0 | 1 | 19 |
| pstpip2 | 0.25736 | 0.194165341 | 0.065 | 0 | 1 | 19 |
| ptpa | 0.25736 | 0.158602514 | 0.065 | 0 | 1 | 19 |
| purbb | 0.25736 | 0.124227374 | 0.065 | 0 | 1 | 19 |
| qtrt2 | 0.25736 | 0.154535061 | 0.065 | 0 | 1 | 19 |
| rab11a | 0.25736 | 0.171836547 | 0.065 | 0 | 1 | 19 |
| rab1aa | 0.25736 | 0.180892835 | 0.065 | 0 | 1 | 19 |
| rars | 0.25736 | 0.17141019 | 0.065 | 0 | 1 | 19 |
| rassf1 | 0.25736 | 0.114386393 | 0.065 | 0 | 1 | 19 |
| rgs14a | 0.25736 | 0.18911035 | 0.065 | 0 | 1 | 19 |
| rgs18 | 0.25736 | 0.137655545 | 0.065 | 0 | 1 | 19 |
| rilp | 0.25736 | 0.188294679 | 0.065 | 0 | 1 | 19 |
| rmdn3 | 0.25736 | 0.158200323 | 0.065 | 0 | 1 | 19 |
| rmnd1 | 0.25736 | 0.118631414 | 0.065 | 0 | 1 | 19 |
| rnf220a | 0.25736 | 0.156221829 | 0.065 | 0 | 1 | 19 |
| rpap1 | 0.25736 | 0.175446191 | 0.065 | 0 | 1 | 19 |
| rpp21 | 0.25736 | 0.113547162 | 0.065 | 0 | 1 | 19 |
| rrp36 | 0.25736 | 0.123792143 | 0.065 | 0 | 1 | 19 |
| rsf1b.1 | 0.25736 | 0.181559346 | 0.065 | 0 | 1 | 19 |
| runx3 | 0.25736 | 0.174767559 | 0.065 | 0 | 1 | 19 |
| scarb2a | 0.25736 | 0.117042183 | 0.065 | 0 | 1 | 19 |
| sccpdhb | 0.25736 | 0.111429273 | 0.065 | 0 | 1 | 19 |
| sco1 | 0.25736 | 0.120330822 | 0.065 | 0 | 1 | 19 |
| setdb1b | 0.25736 | 0.163957318 | 0.065 | 0 | 1 | 19 |
| sfxn3 | 0.25736 | 0.149138504 | 0.065 | 0 | 1 | 19 |
| shrprbck1r | 0.25736 | 0.139378043 | 0.065 | 0 | 1 | 19 |
| si:ch1073-440b2.1 | 0.25736 | 0.14741678 | 0.065 | 0 | 1 | 19 |
| si:ch211-113e8.11 | 0.25736 | 0.111862308 | 0.065 | 0 | 1 | 19 |
| si:ch211-160o17.4 | 0.25736 | 0.184622188 | 0.065 | 0 | 1 | 19 |
| si:ch211-214k5.3 | 0.25736 | 0.185634581 | 0.065 | 0 | 1 | 19 |
| si:ch211-256e16.11 | 0.25736 | 0.190952241 | 0.065 | 0 | 1 | 19 |
| si:ch211-262h13.3 | 0.25736 | 0.190522674 | 0.065 | 0 | 1 | 19 |
| si:ch211-282b22.1 | 0.25736 | 0.127473147 | 0.065 | 0 | 1 | 19 |
| si:ch211-282j22.3 | 0.25736 | 0.125964925 | 0.065 | 0 | 1 | 19 |
| si:ch211-59d17.3 | 0.25736 | 0.131244785 | 0.065 | 0 | 1 | 19 |
| si:dkey-172o19.2 | 0.25736 | 0.191974679 | 0.065 | 0 | 1 | 19 |
| si:dkey-21e2.11 | 0.25736 | 0.132141162 | 0.065 | 0 | 1 | 19 |
| si:dkey-25o1.7 | 0.25736 | 0.138041752 | 0.065 | 0 | 1 | 19 |
| si:dkey-287g12.6 | 0.25736 | 0.179085835 | 0.065 | 0 | 1 | 19 |
| si:dkey-28b4.7 | 0.25736 | 0.170385685 | 0.065 | 0 | 1 | 19 |
| slc35b4 | 0.25736 | 0.147094707 | 0.065 | 0 | 1 | 19 |
| slc39a9 | 0.25736 | 0.102022173 | 0.065 | 0 | 1 | 19 |
| slc43a1a | 0.25736 | 0.197823562 | 0.065 | 0 | 1 | 19 |
| slc4a2a | 0.25736 | 0.170659754 | 0.065 | 0 | 1 | 19 |
| slc4a4b | 0.25736 | 0.177983999 | 0.065 | 0 | 1 | 19 |

| | | | | | | |
|---|---|---|---|---|---|---|
| slc6a6a | 0.25736 | 0.139378043 | 0.065 | 0 | 1 | 19 |
| slc7a3b | 0.25736 | 0.147041802 | 0.065 | 0 | 1 | 19 |
| smc3 | 0.25736 | 0.113381721 | 0.065 | 0 | 1 | 19 |
| spryd3 | 0.25736 | 0.197052653 | 0.065 | 0 | 1 | 19 |
| sptan1 | 0.25736 | 0.123045989 | 0.065 | 0 | 1 | 19 |
| srp54 | 0.25736 | 0.111970969 | 0.065 | 0 | 1 | 19 |
| stard7 | 0.25736 | 0.141506196 | 0.065 | 0 | 1 | 19 |
| stk25b | 0.25736 | 0.163543075 | 0.065 | 0 | 1 | 19 |
| stx3a | 0.25736 | 0.139955041 | 0.065 | 0 | 1 | 19 |
| tbl2 | 0.25736 | 0.105572651 | 0.065 | 0 | 1 | 19 |
| tent4b | 0.25736 | 0.114052265 | 0.065 | 0 | 1 | 19 |
| tex264a | 0.25736 | 0.132461294 | 0.065 | 0 | 1 | 19 |
| tfg | 0.25736 | 0.161285391 | 0.065 | 0 | 1 | 19 |
| tle3a | 0.25736 | 0.108996593 | 0.065 | 0 | 1 | 19 |
| tm2d3 | 0.25736 | 0.114852542 | 0.065 | 0 | 1 | 19 |
| tmem106ba | 0.25736 | 0.103840781 | 0.065 | 0 | 1 | 19 |
| tmem165 | 0.25736 | 0.175592962 | 0.065 | 0 | 1 | 19 |
| tmem41b | 0.25736 | 0.121826308 | 0.065 | 0 | 1 | 19 |
| tmem65 | 0.25736 | 0.128714359 | 0.065 | 0 | 1 | 19 |
| tnip2 | 0.25736 | 0.22967674 | 0.065 | 0 | 1 | 19 |
| traf2b | 0.25736 | 0.157988583 | 0.065 | 0 | 1 | 19 |
| ttc39c | 0.25736 | 0.174496041 | 0.065 | 0 | 1 | 19 |
| ttll12 | 0.25736 | 0.13049976 | 0.065 | 0 | 1 | 19 |
| ube2d1a | 0.25736 | 0.165185365 | 0.065 | 0 | 1 | 19 |
| ube2d3 | 0.25736 | 0.19177023 | 0.065 | 0 | 1 | 19 |
| usp3 | 0.25736 | 0.11862807 | 0.065 | 0 | 1 | 19 |
| uspl1 | 0.25736 | 0.165615823 | 0.065 | 0 | 1 | 19 |
| utp23 | 0.25736 | 0.146092325 | 0.065 | 0 | 1 | 19 |
| uxs1 | 0.25736 | 0.116835892 | 0.065 | 0 | 1 | 19 |
| vcpip1 | 0.25736 | 0.190673554 | 0.065 | 0 | 1 | 19 |
| vps37b | 0.25736 | 0.122947153 | 0.065 | 0 | 1 | 19 |
| wdr41 | 0.25736 | 0.104558688 | 0.065 | 0 | 1 | 19 |
| wdr89 | 0.25736 | 0.113912208 | 0.065 | 0 | 1 | 19 |
| wee2 | 0.25736 | 0.174353336 | 0.065 | 0 | 1 | 19 |
| wu:fb74b10 | 0.25736 | 0.115402347 | 0.065 | 0 | 1 | 19 |
| xaf1 | 0.25736 | 0.109993402 | 0.065 | 0 | 1 | 19 |
| yipf1 | 0.25736 | 0.154379898 | 0.065 | 0 | 1 | 19 |
| yipf5 | 0.25736 | 0.107863845 | 0.065 | 0 | 1 | 19 |
| zbed4 | 0.25736 | 0.166389136 | 0.065 | 0 | 1 | 19 |
| zbtb11 | 0.25736 | 0.126869216 | 0.065 | 0 | 1 | 19 |
| zbtb20 | 0.25736 | 0.177725162 | 0.065 | 0 | 1 | 19 |
| zbtb32 | 0.25736 | 0.146872059 | 0.065 | 0 | 1 | 19 |
| zdhhc18a | 0.25736 | 0.123691564 | 0.065 | 0 | 1 | 19 |
| zdhhc6 | 0.25736 | 0.136897489 | 0.065 | 0 | 1 | 19 |
| zgc:114119 | 0.25736 | 0.149484296 | 0.065 | 0 | 1 | 19 |
| zgc:123258 | 0.25736 | 0.232567654 | 0.065 | 0 | 1 | 19 |
| zgc:152891 | 0.25736 | 0.159499587 | 0.065 | 0 | 1 | 19 |
| zgc:171967 | 0.25736 | 0.164317541 | 0.065 | 0 | 1 | 19 |

| | | | | | | |
|---|---|---|---|---|---|---|
| zgc:63587 | 0.25736 | 0.132487122 | 0.065 | 0 | 1 | 19 |
| zgc:85789 | 0.25736 | 0.105125655 | 0.065 | 0 | 1 | 19 |
| znf438 | 0.25736 | 0.122487363 | 0.065 | 0 | 1 | 19 |
| ehbp1l1b | 0.25736 | 0.16836473 | 0.065 | 0 | 1 | 19 |
| mkrn4 | 0.25736 | 0.164251099 | 0.065 | 0 | 1 | 19 |
| pik3r4 | 0.25736 | 0.165932726 | 0.065 | 0 | 1 | 19 |
| tfpt | 0.25736 | 0.146265428 | 0.065 | 0 | 1 | 19 |
| CR847906.1 | 0.25736 | 0.133817915 | 0.065 | 0 | 1 | 19 |
| FO834831.1 | 0.25736 | 0.133817915 | 0.065 | 0 | 1 | 19 |
| USP34 | 0.25736 | 0.233572208 | 0.065 | 0 | 1 | 19 |
| XLOC-011160 | 0.25736 | 0.158678932 | 0.065 | 0 | 1 | 19 |
| acbd3 | 0.25736 | 0.206705866 | 0.065 | 0 | 1 | 19 |
| atg13 | 0.25736 | 0.165761187 | 0.065 | 0 | 1 | 19 |
| atg3 | 0.25736 | 0.161308542 | 0.065 | 0 | 1 | 19 |
| brdt | 0.25736 | 0.176821263 | 0.065 | 0 | 1 | 19 |
| ccdc32 | 0.25736 | 0.113631398 | 0.065 | 0 | 1 | 19 |
| ccdc84 | 0.25736 | 0.17391557 | 0.065 | 0 | 1 | 19 |
| cdk5rap2 | 0.25736 | 0.124748888 | 0.065 | 0 | 1 | 19 |
| clec16a | 0.25736 | 0.176263447 | 0.065 | 0 | 1 | 19 |
| crata | 0.25736 | 0.126877882 | 0.065 | 0 | 1 | 19 |
| ddx49 | 0.25736 | 0.173366023 | 0.065 | 0 | 1 | 19 |
| dhtkd1 | 0.25736 | 0.110843074 | 0.065 | 0 | 1 | 19 |
| dnmt3ab | 0.25736 | 0.122966805 | 0.065 | 0 | 1 | 19 |
| fam126a | 0.25736 | 0.135066845 | 0.065 | 0 | 1 | 19 |
| fam50a | 0.25736 | 0.108816972 | 0.065 | 0 | 1 | 19 |
| flna | 0.25736 | 0.264956134 | 0.065 | 0 | 1 | 19 |
| g3bp1 | 0.25736 | 0.177340043 | 0.065 | 0 | 1 | 19 |
| gfm1 | 0.25736 | 0.130543173 | 0.065 | 0 | 1 | 19 |
| hccsb | 0.25736 | 0.116625199 | 0.065 | 0 | 1 | 19 |
| hectd1 | 0.25736 | 0.169659288 | 0.065 | 0 | 1 | 19 |
| ift81 | 0.25736 | 0.162654943 | 0.065 | 0 | 1 | 19 |
| itpr1b | 0.25736 | 0.124689601 | 0.065 | 0 | 1 | 19 |
| map3k14a | 0.25736 | 0.119090999 | 0.065 | 0 | 1 | 19 |
| march2 | 0.25736 | 0.133132791 | 0.065 | 0 | 1 | 19 |
| mfap3l | 0.25736 | 0.131162624 | 0.065 | 0 | 1 | 19 |
| mier2 | 0.25736 | 0.229815658 | 0.065 | 0 | 1 | 19 |
| mrpl28 | 0.25736 | 0.171268213 | 0.065 | 0 | 1 | 19 |
| mrpl46 | 0.25736 | 0.114852315 | 0.065 | 0 | 1 | 19 |
| mtch2 | 0.25736 | 0.152278977 | 0.065 | 0 | 1 | 19 |
| nudt9 | 0.25736 | 0.119212777 | 0.065 | 0 | 1 | 19 |
| otub1a | 0.25736 | 0.163004855 | 0.065 | 0 | 1 | 19 |
| oxa1l | 0.25736 | 0.118113457 | 0.065 | 0 | 1 | 19 |
| pmpcb | 0.25736 | 0.123203736 | 0.065 | 0 | 1 | 19 |
| ppp1r12a | 0.25736 | 0.137304703 | 0.065 | 0 | 1 | 19 |
| prf1.7 | 0.25736 | 0.273706912 | 0.065 | 0 | 1 | 19 |
| prnprs3 | 0.25736 | 0.140350956 | 0.065 | 0 | 1 | 19 |
| prss16 | 0.25736 | 0.176774482 | 0.065 | 0 | 1 | 19 |
| psmg1 | 0.25736 | 0.146061144 | 0.065 | 0 | 1 | 19 |

| | | | | | | |
|---|---|---|---|---|---|---|
| ralab | 0.25736 | 0.12393359 | 0.065 | 0 | 1 | 19 |
| rbbp8l | 0.25736 | 0.20810781 | 0.065 | 0 | 1 | 19 |
| rdh14a | 0.25736 | 0.126265385 | 0.065 | 0 | 1 | 19 |
| rprd1b | 0.25736 | 0.158846153 | 0.065 | 0 | 1 | 19 |
| sfmbt2 | 0.25736 | 0.126780735 | 0.065 | 0 | 1 | 19 |
| si:ch211-148l7.4 | 0.25736 | 0.190303525 | 0.065 | 0 | 1 | 19 |
| si:ch211-196f2.6 | 0.25736 | 0.171700416 | 0.065 | 0 | 1 | 19 |
| si:ch73-308l14.2 | 0.25736 | 0.165716564 | 0.065 | 0 | 1 | 19 |
| slc5a3b | 0.25736 | 0.140493422 | 0.065 | 0 | 1 | 19 |
| snf8 | 0.25736 | 0.133817915 | 0.065 | 0 | 1 | 19 |
| tnfaip8l2b | 0.25736 | 0.182765843 | 0.065 | 0 | 1 | 19 |
| tomm34 | 0.25736 | 0.171770157 | 0.065 | 0 | 1 | 19 |
| tox4a | 0.25736 | 0.133681356 | 0.065 | 0 | 1 | 19 |
| tpp2 | 0.25736 | 0.239659494 | 0.065 | 0 | 1 | 19 |
| tspan2b | 0.25736 | 0.144718713 | 0.065 | 0 | 1 | 19 |
| ttc13 | 0.25736 | 0.134961967 | 0.065 | 0 | 1 | 19 |
| ttf1 | 0.25736 | 0.129656009 | 0.065 | 0 | 1 | 19 |
| uap1 | 0.25736 | 0.115438316 | 0.065 | 0 | 1 | 19 |
| uba3 | 0.25736 | 0.117975142 | 0.065 | 0 | 1 | 19 |
| ube4b | 0.25736 | 0.200023309 | 0.065 | 0 | 1 | 19 |
| uckl1b | 0.25736 | 0.20147227 | 0.065 | 0 | 1 | 19 |
| xrn1 | 0.25736 | 0.131813285 | 0.065 | 0 | 1 | 19 |
| zgc:162025 | 0.25736 | 0.157598407 | 0.065 | 0 | 1 | 19 |
| fbxw7 | 0.25736 | 0.148520236 | 0.065 | 0 | 1 | 19 |
| il21r.1 | 0.25736 | 0.171167907 | 0.065 | 0 | 1 | 19 |
| ppargc1b | 0.25736 | 0.14739281 | 0.065 | 0 | 1 | 19 |
| scocb | 0.25736 | 0.182274897 | 0.065 | 0 | 1 | 19 |
| smim29 | 0.25736 | 0.1533994 | 0.065 | 0 | 1 | 19 |
| usp8 | 0.25736 | 0.169550949 | 0.065 | 0 | 1 | 19 |
| AL954696.1 | 0.25736 | 0.107127717 | 0.065 | 0 | 1 | 19 |
| LOC101883999 | 0.25736 | 0.112942841 | 0.065 | 0 | 1 | 19 |
| dbt | 0.25736 | 0.100261684 | 0.065 | 0 | 1 | 19 |
| ddx39b | 0.25736 | 0.104617242 | 0.065 | 0 | 1 | 19 |
| gak | 0.25736 | 0.101174976 | 0.065 | 0 | 1 | 19 |
| gkap1 | 0.25736 | 0.125412642 | 0.065 | 0 | 1 | 19 |
| glrx3 | 0.25736 | 0.120444423 | 0.065 | 0 | 1 | 19 |
| lyrm9 | 0.25736 | 0.141683934 | 0.065 | 0 | 1 | 19 |
| megf8l | 0.25736 | 0.108848525 | 0.065 | 0 | 1 | 19 |
| nktr | 0.25736 | 0.123439681 | 0.065 | 0 | 1 | 19 |
| rc3h1a | 0.25736 | 0.140246101 | 0.065 | 0 | 1 | 19 |
| sdhb | 0.25736 | 0.101513721 | 0.065 | 0 | 1 | 19 |
| snx9b | 0.25736 | 0.107447761 | 0.065 | 0 | 1 | 19 |
| srebf2 | 0.25736 | 0.106029302 | 0.065 | 0 | 1 | 19 |
| zgc:100846 | 0.25736 | 0.108088752 | 0.065 | 0 | 1 | 19 |
| zgc:114130 | 0.25736 | 0.104677727 | 0.065 | 0 | 1 | 19 |
| syncripl | 0.2584 | 0.258384864 | 0.226 | 0.059 | 1 | 19 |
| junba | 0.26133 | 0.170005797 | 0.355 | 0.176 | 1 | 19 |
| eif4g1a | 0.2626 | -0.40593879 | 0.129 | 0.235 | 1 | 19 |

| | | | | | | |
|---|---|---|---|---|---|---|
| rbm8a | 0.26384 | 0.396156467 | 0.323 | 0.118 | 1 | 19 |
| CU550719.2 | 0.26503 | -0.12714448 | 0.032 | 0.118 | 1 | 19 |
| atp6v0a1a | 0.26503 | -0.14548518 | 0.032 | 0.118 | 1 | 19 |
| b4galt6 | 0.26503 | -0.25722944 | 0.032 | 0.118 | 1 | 19 |
| chmp7 | 0.26503 | -0.30735016 | 0.032 | 0.118 | 1 | 19 |
| creg1 | 0.26503 | -0.28885821 | 0.032 | 0.118 | 1 | 19 |
| cript | 0.26503 | -0.2722761 | 0.032 | 0.118 | 1 | 19 |
| cybc1 | 0.26503 | -0.21319045 | 0.032 | 0.118 | 1 | 19 |
| cyth1b | 0.26503 | -0.19927088 | 0.032 | 0.118 | 1 | 19 |
| dph6 | 0.26503 | -0.20098931 | 0.032 | 0.118 | 1 | 19 |
| eaf1 | 0.26503 | -0.10268603 | 0.032 | 0.118 | 1 | 19 |
| faf2 | 0.26503 | -0.17421454 | 0.032 | 0.118 | 1 | 19 |
| golga5 | 0.26503 | -0.15993389 | 0.032 | 0.118 | 1 | 19 |
| gss | 0.26503 | -0.11656417 | 0.032 | 0.118 | 1 | 19 |
| hnrnpc | 0.26503 | -0.26449716 | 0.032 | 0.118 | 1 | 19 |
| hspd1 | 0.26503 | -0.22770635 | 0.032 | 0.118 | 1 | 19 |
| isca1 | 0.26503 | -0.14831295 | 0.032 | 0.118 | 1 | 19 |
| lamtor3 | 0.26503 | -0.15837348 | 0.032 | 0.118 | 1 | 19 |
| lman2 | 0.26503 | -0.20692915 | 0.032 | 0.118 | 1 | 19 |
| mcm5 | 0.26503 | -0.1499369 | 0.032 | 0.118 | 1 | 19 |
| mif4gdb | 0.26503 | -0.18616584 | 0.032 | 0.118 | 1 | 19 |
| mrps14 | 0.26503 | -0.23913628 | 0.032 | 0.118 | 1 | 19 |
| mtmr12 | 0.26503 | -0.12194635 | 0.032 | 0.118 | 1 | 19 |
| mvp | 0.26503 | -0.16200417 | 0.032 | 0.118 | 1 | 19 |
| nfil3-6 | 0.26503 | -0.33496897 | 0.032 | 0.118 | 1 | 19 |
| nsmce2 | 0.26503 | -0.10577414 | 0.032 | 0.118 | 1 | 19 |
| nt5dc2 | 0.26503 | -0.14513623 | 0.032 | 0.118 | 1 | 19 |
| nxpe3 | 0.26503 | -0.23672087 | 0.032 | 0.118 | 1 | 19 |
| pdxka | 0.26503 | -0.13723757 | 0.032 | 0.118 | 1 | 19 |
| phactr2 | 0.26503 | -0.15284852 | 0.032 | 0.118 | 1 | 19 |
| pigx | 0.26503 | -0.2058949 | 0.032 | 0.118 | 1 | 19 |
| pip5k1ba | 0.26503 | -0.23464021 | 0.032 | 0.118 | 1 | 19 |
| ptbp3 | 0.26503 | -0.12929448 | 0.032 | 0.118 | 1 | 19 |
| rab35b | 0.26503 | -0.15041132 | 0.032 | 0.118 | 1 | 19 |
| ralbp1 | 0.26503 | -0.10327672 | 0.032 | 0.118 | 1 | 19 |
| rpa3 | 0.26503 | -0.13419055 | 0.032 | 0.118 | 1 | 19 |
| rwdd1 | 0.26503 | -0.15465257 | 0.032 | 0.118 | 1 | 19 |
| si:zfos-943e10.1 | 0.26503 | -0.11553809 | 0.032 | 0.118 | 1 | 19 |
| st7 | 0.26503 | -0.18579521 | 0.032 | 0.118 | 1 | 19 |
| strap | 0.26503 | -0.13758905 | 0.032 | 0.118 | 1 | 19 |
| usp14 | 0.26503 | -0.14174099 | 0.032 | 0.118 | 1 | 19 |
| washc2c | 0.26503 | -0.24073963 | 0.032 | 0.118 | 1 | 19 |
| zgc:153118 | 0.26503 | -0.17967329 | 0.032 | 0.118 | 1 | 19 |
| abca2 | 0.26503 | -0.10061386 | 0.032 | 0.118 | 1 | 19 |
| acadvl | 0.26503 | -0.14006119 | 0.032 | 0.118 | 1 | 19 |
| bahd1 | 0.26503 | -0.25225039 | 0.032 | 0.118 | 1 | 19 |
| tob1b | 0.26503 | -0.26271832 | 0.032 | 0.118 | 1 | 19 |
| XLOC-043760 | 0.26503 | -0.37707018 | 0.032 | 0.118 | 1 | 19 |

| | | | | | | |
|---|---|---|---|---|---|---|
| ampd3b | 0.26503 | -0.29538548 | 0.032 | 0.118 | 1 | 19 |
| araf | 0.26503 | -0.17477309 | 0.032 | 0.118 | 1 | 19 |
| ctnnb2 | 0.26503 | -0.20803419 | 0.032 | 0.118 | 1 | 19 |
| klf11a | 0.26503 | -0.34860351 | 0.032 | 0.118 | 1 | 19 |
| lsm14aa | 0.26503 | -0.22885755 | 0.032 | 0.118 | 1 | 19 |
| mrps17 | 0.26503 | -0.2135949 | 0.032 | 0.118 | 1 | 19 |
| rrp7a | 0.26503 | -0.19653119 | 0.032 | 0.118 | 1 | 19 |
| taf7 | 0.26503 | -0.30229621 | 0.032 | 0.118 | 1 | 19 |
| BX284696.1 | 0.26503 | -0.14103961 | 0.032 | 0.118 | 1 | 19 |
| CABZ01087560.1 | 0.26503 | -0.14475297 | 0.032 | 0.118 | 1 | 19 |
| XLOC-037780 | 0.26503 | -0.20951748 | 0.032 | 0.118 | 1 | 19 |
| adgre5a | 0.26503 | -0.14528413 | 0.032 | 0.118 | 1 | 19 |
| alkbh5 | 0.26503 | -0.1160578 | 0.032 | 0.118 | 1 | 19 |
| apooa | 0.26503 | -0.20083545 | 0.032 | 0.118 | 1 | 19 |
| arf4a | 0.26503 | -0.14525476 | 0.032 | 0.118 | 1 | 19 |
| asf1ba | 0.26503 | -0.16874326 | 0.032 | 0.118 | 1 | 19 |
| asf1bb | 0.26503 | -0.19925201 | 0.032 | 0.118 | 1 | 19 |
| atf1 | 0.26503 | -0.24707355 | 0.032 | 0.118 | 1 | 19 |
| atf7ip | 0.26503 | -0.28503252 | 0.032 | 0.118 | 1 | 19 |
| bcap31 | 0.26503 | -0.24160653 | 0.032 | 0.118 | 1 | 19 |
| bcl11ba | 0.26503 | -0.21904282 | 0.032 | 0.118 | 1 | 19 |
| blk | 0.26503 | -0.23604566 | 0.032 | 0.118 | 1 | 19 |
| chchd2 | 0.26503 | -0.20140092 | 0.032 | 0.118 | 1 | 19 |
| chd3 | 0.26503 | -0.24172534 | 0.032 | 0.118 | 1 | 19 |
| chmp2bb | 0.26503 | -0.12596836 | 0.032 | 0.118 | 1 | 19 |
| cryl1 | 0.26503 | -0.15146081 | 0.032 | 0.118 | 1 | 19 |
| ddx54 | 0.26503 | -0.13071867 | 0.032 | 0.118 | 1 | 19 |
| ddx56 | 0.26503 | -0.1952066 | 0.032 | 0.118 | 1 | 19 |
| egln3 | 0.26503 | -0.14743535 | 0.032 | 0.118 | 1 | 19 |
| eif4ba | 0.26503 | -0.22402298 | 0.032 | 0.118 | 1 | 19 |
| eprs | 0.26503 | -0.10005028 | 0.032 | 0.118 | 1 | 19 |
| gsdmeb | 0.26503 | -0.23712208 | 0.032 | 0.118 | 1 | 19 |
| hcfc1b | 0.26503 | -0.16212201 | 0.032 | 0.118 | 1 | 19 |
| helz | 0.26503 | -0.13283125 | 0.032 | 0.118 | 1 | 19 |
| hspa4a | 0.26503 | -0.21003628 | 0.032 | 0.118 | 1 | 19 |
| inppl1a | 0.26503 | -0.25040365 | 0.032 | 0.118 | 1 | 19 |
| itprip | 0.26503 | -0.11302792 | 0.032 | 0.118 | 1 | 19 |
| kdm5ba | 0.26503 | -0.10934564 | 0.032 | 0.118 | 1 | 19 |
| kdm6ba | 0.26503 | -0.16773616 | 0.032 | 0.118 | 1 | 19 |
| kmt2bb | 0.26503 | -0.15546731 | 0.032 | 0.118 | 1 | 19 |
| luc7l3 | 0.26503 | -0.20416144 | 0.032 | 0.118 | 1 | 19 |
| mhc2dab | 0.26503 | -0.13334016 | 0.032 | 0.118 | 1 | 19 |
| mrpl36 | 0.26503 | -0.24712385 | 0.032 | 0.118 | 1 | 19 |
| mycla | 0.26503 | -0.20868095 | 0.032 | 0.118 | 1 | 19 |
| nedd4l | 0.26503 | -0.22276239 | 0.032 | 0.118 | 1 | 19 |
| nkap | 0.26503 | -0.12756621 | 0.032 | 0.118 | 1 | 19 |
| nmt1a | 0.26503 | -0.22246153 | 0.032 | 0.118 | 1 | 19 |
| nr1d2b | 0.26503 | -0.25786961 | 0.032 | 0.118 | 1 | 19 |

| | | | | | | |
|---|---|---|---|---|---|---|
| nr3c1 | 0.26503 | -0.26876882 | 0.032 | 0.118 | 1 | 19 |
| nsd3 | 0.26503 | -0.13178326 | 0.032 | 0.118 | 1 | 19 |
| olfml2ba | 0.26503 | -0.19725241 | 0.032 | 0.118 | 1 | 19 |
| pcna | 0.26503 | -0.23652783 | 0.032 | 0.118 | 1 | 19 |
| peli1b | 0.26503 | -0.12303804 | 0.032 | 0.118 | 1 | 19 |
| per3 | 0.26503 | -0.23706711 | 0.032 | 0.118 | 1 | 19 |
| pigbos1 | 0.26503 | -0.24583635 | 0.032 | 0.118 | 1 | 19 |
| pigo | 0.26503 | -0.18075462 | 0.032 | 0.118 | 1 | 19 |
| pigt | 0.26503 | -0.13493616 | 0.032 | 0.118 | 1 | 19 |
| ppp1cb | 0.26503 | -0.19680884 | 0.032 | 0.118 | 1 | 19 |
| ppp2cab | 0.26503 | -0.14379778 | 0.032 | 0.118 | 1 | 19 |
| prodha | 0.26503 | -0.2504743 | 0.032 | 0.118 | 1 | 19 |
| prpf39 | 0.26503 | -0.13000786 | 0.032 | 0.118 | 1 | 19 |
| prrc2a | 0.26503 | -0.1236997 | 0.032 | 0.118 | 1 | 19 |
| qpctla | 0.26503 | -0.13080815 | 0.032 | 0.118 | 1 | 19 |
| rel | 0.26503 | -0.16346928 | 0.032 | 0.118 | 1 | 19 |
| rnf121 | 0.26503 | -0.22253291 | 0.032 | 0.118 | 1 | 19 |
| rtn3 | 0.26503 | -0.27357257 | 0.032 | 0.118 | 1 | 19 |
| ruvbl2 | 0.26503 | -0.12085172 | 0.032 | 0.118 | 1 | 19 |
| scfd1 | 0.26503 | -0.15181281 | 0.032 | 0.118 | 1 | 19 |
| scyl3 | 0.26503 | -0.10450541 | 0.032 | 0.118 | 1 | 19 |
| sdhdb | 0.26503 | -0.17965022 | 0.032 | 0.118 | 1 | 19 |
| sec13 | 0.26503 | -0.11840924 | 0.032 | 0.118 | 1 | 19 |
| si:ch211-121a2.2 | 0.26503 | -0.21137328 | 0.032 | 0.118 | 1 | 19 |
| si:ch211-197h24.9 | 0.26503 | -0.19232229 | 0.032 | 0.118 | 1 | 19 |
| si:dkey-6e2.2 | 0.26503 | -0.2108235 | 0.032 | 0.118 | 1 | 19 |
| si:dkeyp-113d7.1 | 0.26503 | -0.22048342 | 0.032 | 0.118 | 1 | 19 |
| si:dkeyp-113d7.10 | 0.26503 | -0.17674189 | 0.032 | 0.118 | 1 | 19 |
| si:dkeyp-120h9.1 | 0.26503 | -0.26695496 | 0.032 | 0.118 | 1 | 19 |
| spopla | 0.26503 | -0.13914552 | 0.032 | 0.118 | 1 | 19 |
| srp68 | 0.26503 | -0.28741251 | 0.032 | 0.118 | 1 | 19 |
| st6galnac5b | 0.26503 | -0.1819264 | 0.032 | 0.118 | 1 | 19 |
| tax1bp1a | 0.26503 | -0.33840309 | 0.032 | 0.118 | 1 | 19 |
| tbl1xr1b | 0.26503 | -0.20179513 | 0.032 | 0.118 | 1 | 19 |
| tfap4 | 0.26503 | -0.19444166 | 0.032 | 0.118 | 1 | 19 |
| tiparp | 0.26503 | -0.18389204 | 0.032 | 0.118 | 1 | 19 |
| tmbim1a | 0.26503 | -0.17384148 | 0.032 | 0.118 | 1 | 19 |
| tmem203 | 0.26503 | -0.22225624 | 0.032 | 0.118 | 1 | 19 |
| trappc4 | 0.26503 | -0.21769798 | 0.032 | 0.118 | 1 | 19 |
| viml | 0.26503 | -0.27754596 | 0.032 | 0.118 | 1 | 19 |
| wdr5 | 0.26503 | -0.23330917 | 0.032 | 0.118 | 1 | 19 |
| zc3h4 | 0.26503 | -0.32682187 | 0.032 | 0.118 | 1 | 19 |
| zgc:55558 | 0.26503 | -0.27442736 | 0.032 | 0.118 | 1 | 19 |
| igf2bp2a | 0.26503 | -0.19925882 | 0.032 | 0.118 | 1 | 19 |
| kdm4b | 0.26503 | -0.22075673 | 0.032 | 0.118 | 1 | 19 |
| rpusd4 | 0.26503 | -0.26562769 | 0.032 | 0.118 | 1 | 19 |
| vta1 | 0.26503 | -0.19754868 | 0.032 | 0.118 | 1 | 19 |
| mbnl3 | 0.26503 | -0.16691613 | 0.032 | 0.118 | 1 | 19 |

| | | | | | | |
|---|---|---|---|---|---|---|
| pdcd4a | 0.26503 | -0.17047327 | 0.032 | 0.118 | 1 | 19 |
| FO704610.1 | 0.26503 | -0.15449234 | 0.032 | 0.118 | 1 | 19 |
| ano5b | 0.26503 | -0.19422435 | 0.032 | 0.118 | 1 | 19 |
| dnajc9 | 0.26503 | -0.21614546 | 0.032 | 0.118 | 1 | 19 |
| gnas | 0.26503 | -0.20099808 | 0.032 | 0.118 | 1 | 19 |
| mrpl53 | 0.26503 | -0.21609015 | 0.032 | 0.118 | 1 | 19 |
| nxt2 | 0.26503 | -0.20809034 | 0.032 | 0.118 | 1 | 19 |
| parp8 | 0.26503 | -0.30864643 | 0.032 | 0.118 | 1 | 19 |
| ppp6r3 | 0.26503 | -0.221535 | 0.032 | 0.118 | 1 | 19 |
| rraga | 0.26503 | -0.14538692 | 0.032 | 0.118 | 1 | 19 |
| sephs2 | 0.26503 | -0.2263753 | 0.032 | 0.118 | 1 | 19 |
| si:ch1073-170o4.1 | 0.26503 | -0.13995831 | 0.032 | 0.118 | 1 | 19 |
| si:ch1073-392o20.2 | 0.26503 | -0.12144239 | 0.032 | 0.118 | 1 | 19 |
| slc12a9 | 0.26503 | -0.23677833 | 0.032 | 0.118 | 1 | 19 |
| smarca4a | 0.26503 | -0.19547174 | 0.032 | 0.118 | 1 | 19 |
| smarcd2 | 0.26503 | -0.1854275 | 0.032 | 0.118 | 1 | 19 |
| smco4 | 0.26503 | -0.26404119 | 0.032 | 0.118 | 1 | 19 |
| spidr | 0.26503 | -0.21534877 | 0.032 | 0.118 | 1 | 19 |
| srcap | 0.26503 | -0.17978785 | 0.032 | 0.118 | 1 | 19 |
| susd6 | 0.26503 | -0.18076461 | 0.032 | 0.118 | 1 | 19 |
| timeless | 0.26503 | -0.15765745 | 0.032 | 0.118 | 1 | 19 |
| uqcrb | 0.26503 | -0.15189217 | 0.032 | 0.118 | 1 | 19 |
| zfand6 | 0.26503 | -0.17544882 | 0.032 | 0.118 | 1 | 19 |
| chmp1a | 0.26503 | -0.21141169 | 0.032 | 0.118 | 1 | 19 |
| gtf2a2 | 0.26503 | -0.23296402 | 0.032 | 0.118 | 1 | 19 |
| fkbp1aa | 0.26679 | -0.35193433 | 0.29 | 0.529 | 1 | 19 |
| sept6 | 0.26693 | -0.38829791 | 0.29 | 0.529 | 1 | 19 |
| CDPF1 | 0.27047 | -0.38685531 | 0.129 | 0.235 | 1 | 19 |
| tpi1b | 0.27065 | 0.409378489 | 0.581 | 0.353 | 1 | 19 |
| rpn1 | 0.27299 | -0.33241508 | 0.194 | 0.412 | 1 | 19 |
| si:dkey-118j18.2 | 0.27939 | -0.40810837 | 0.129 | 0.294 | 1 | 19 |
| calm2b | 0.28031 | -0.25062809 | 0.161 | 0.353 | 1 | 19 |
| ube2v1 | 0.28196 | -0.2542561 | 0.161 | 0.353 | 1 | 19 |
| prdx4 | 0.28211 | -0.15583683 | 0.226 | 0.176 | 1 | 19 |
| si:dkey-21e2.14 | 0.28227 | -0.22433477 | 0.258 | 0.235 | 1 | 19 |
| inpp5d | 0.28645 | 0.348021169 | 0.226 | 0.059 | 1 | 19 |
| cdk2ap2 | 0.289 | 0.341642631 | 0.226 | 0.059 | 1 | 19 |
| ndufa6 | 0.29205 | -0.39578731 | 0.258 | 0.471 | 1 | 19 |
| rpl13a | 0.29707 | 0.127413618 | 0.806 | 0.824 | 1 | 19 |
| tmed9 | 0.29892 | -0.41264653 | 0.129 | 0.294 | 1 | 19 |
| atp5if1b | 0.29944 | 0.395229181 | 0.581 | 0.412 | 1 | 19 |
| dynlt3 | 0.3006 | 0.167992096 | 0.194 | 0.059 | 1 | 19 |
| sec11a | 0.30687 | 0.347546633 | 0.452 | 0.235 | 1 | 19 |
| ssr3 | 0.30794 | -0.36516943 | 0.484 | 0.647 | 1 | 19 |
| serpinb1l1 | 0.30906 | 0.382884364 | 0.194 | 0.059 | 1 | 19 |
| arrb2b | 0.3093 | 0.427541933 | 0.355 | 0.176 | 1 | 19 |
| si:ch211-197h24.6 | 0.30974 | 0.40407901 | 0.161 | 0.059 | 1 | 19 |
| pfdn4 | 0.31064 | -0.11750444 | 0.194 | 0.353 | 1 | 19 |

| | | | | | | |
|---|---|---|---|---|---|---|
| FO203432.1 | 0.31166 | 0.2453351 | 0.355 | 0.176 | 1 | 19 |
| map1lc3b | 0.31335 | -0.16431143 | 0.129 | 0.294 | 1 | 19 |
| egr3 | 0.3139 | 0.423841314 | 0.161 | 0.059 | 1 | 19 |
| naa10 | 0.31552 | -0.23961428 | 0.161 | 0.353 | 1 | 19 |
| anp32a | 0.31632 | 0.256825594 | 0.29 | 0.118 | 1 | 19 |
| si:dkey-21e2.16 | 0.31759 | 0.134344911 | 0.29 | 0.294 | 1 | 19 |
| coro1a | 0.31884 | -0.26621017 | 0.71 | 0.882 | 1 | 19 |
| si:ch73-359m17.2 | 0.32021 | -0.43089307 | 0.129 | 0.294 | 1 | 19 |
| hmgn7 | 0.32065 | -0.39749713 | 0.323 | 0.471 | 1 | 19 |
| LOC103909115 | 0.32208 | 0.382018292 | 0.452 | 0.235 | 1 | 19 |
| nop10 | 0.32401 | -0.40445328 | 0.258 | 0.412 | 1 | 19 |
| MFAP4 (1 of many) | 0.3255 | 0.462215904 | 0.29 | 0.176 | 1 | 19 |
| ptprc | 0.32558 | 0.357643404 | 0.516 | 0.294 | 1 | 19 |
| eif3f | 0.32729 | 0.136788727 | 0.71 | 0.529 | 1 | 19 |
| serbp1a | 0.32836 | 0.316397502 | 0.742 | 0.588 | 1 | 19 |
| cops4 | 0.33176 | 0.321045209 | 0.194 | 0.059 | 1 | 19 |
| eno1a | 0.33228 | -0.3236306 | 0.258 | 0.471 | 1 | 19 |
| unm-hu7910 | 0.33253 | 0.197837316 | 0.194 | 0.059 | 1 | 19 |
| atp5mf | 0.33272 | -0.34562372 | 0.161 | 0.353 | 1 | 19 |
| gadd45ba | 0.33314 | -0.2680384 | 0.161 | 0.353 | 1 | 19 |
| ube2d2 | 0.33351 | -0.31566599 | 0.161 | 0.353 | 1 | 19 |
| max | 0.3341 | 0.162701866 | 0.194 | 0.059 | 1 | 19 |
| si:dkey-21e2.10 | 0.33427 | 0.39787494 | 0.29 | 0.235 | 1 | 19 |
| LOC100536213 | 0.33441 | -0.36813077 | 0.258 | 0.471 | 1 | 19 |
| nr4a1 | 0.33448 | -0.36273778 | 0.258 | 0.471 | 1 | 19 |
| cfbl | 0.33508 | -0.26603698 | 0.452 | 0.647 | 1 | 19 |
| crlf3 | 0.33515 | 0.15545245 | 0.194 | 0.059 | 1 | 19 |
| tubb4b | 0.33594 | -0.22005425 | 0.548 | 0.529 | 1 | 19 |
| rpl15 | 0.34002 | 0.107943942 | 0.935 | 0.941 | 1 | 19 |
| hnrnpa0l | 0.34239 | 0.367812299 | 0.645 | 0.471 | 1 | 19 |
| marcksl1a | 0.34306 | 0.337827558 | 0.29 | 0.118 | 1 | 19 |
| ywhaba | 0.34882 | -0.37622812 | 0.355 | 0.529 | 1 | 19 |
| faub | 0.35207 | -0.13164634 | 0.097 | 0.235 | 1 | 19 |
| ctss2.1 | 0.35247 | 0.145358941 | 0.323 | 0.176 | 1 | 19 |
| rps28 | 0.35355 | -0.18750512 | 0.839 | 0.941 | 1 | 19 |
| copz1 | 0.35415 | 0.224240746 | 0.194 | 0.059 | 1 | 19 |
| hmgn2 | 0.35543 | -0.23724273 | 0.742 | 0.765 | 1 | 19 |
| acin1b | 0.35567 | 0.339680741 | 0.194 | 0.059 | 1 | 19 |
| si:ch1073-429i10.3.1 | 0.35631 | 0.386974006 | 0.419 | 0.235 | 1 | 19 |
| ca2 | 0.35802 | -0.39926251 | 0.387 | 0.588 | 1 | 19 |
| id2a | 0.3582 | -0.17059894 | 0.194 | 0.176 | 1 | 19 |
| tnfaip8l2a | 0.35952 | -0.14078229 | 0.065 | 0.176 | 1 | 19 |
| cox7b | 0.3622 | -0.3479639 | 0.484 | 0.647 | 1 | 19 |
| pnp5a | 0.3629 | -0.33248422 | 0.161 | 0.235 | 1 | 19 |
| si:ch211-288g17.3 | 0.36359 | 0.347739714 | 0.194 | 0.118 | 1 | 19 |
| zgc:153713 | 0.36435 | 0.3984437 | 0.387 | 0.235 | 1 | 19 |
| gabarapb | 0.36507 | -0.36180676 | 0.323 | 0.529 | 1 | 19 |
| rpl22l1 | 0.36516 | 0.367987574 | 0.548 | 0.412 | 1 | 19 |

| | | | | | | |
|---|---|---|---|---|---|---|
| slc9a3r1a | 0.36766 | 0.308919876 | 0.194 | 0.059 | 1 | 19 |
| znf706 | 0.36973 | -0.29547725 | 0.129 | 0.294 | 1 | 19 |
| atf4a | 0.37113 | 0.409142103 | 0.355 | 0.176 | 1 | 19 |
| psmb4 | 0.37222 | -0.31978269 | 0.129 | 0.294 | 1 | 19 |
| ndufa11 | 0.37532 | 0.161085667 | 0.258 | 0.118 | 1 | 19 |
| zfp36l1a | 0.37636 | 0.365401123 | 0.516 | 0.353 | 1 | 19 |
| ewsr1b | 0.37686 | -0.27372814 | 0.129 | 0.294 | 1 | 19 |
| rsl24d1 | 0.37821 | 0.268609421 | 0.613 | 0.412 | 1 | 19 |
| hif1al | 0.37893 | -0.1465615 | 0.097 | 0.235 | 1 | 19 |
| chchd10 | 0.38128 | 0.310584773 | 0.161 | 0.059 | 1 | 19 |
| arpc1b | 0.38147 | -0.19685823 | 0.806 | 0.941 | 1 | 19 |
| pin1 | 0.38324 | -0.26962773 | 0.129 | 0.294 | 1 | 19 |
| si:dkey-262k9.2 | 0.38334 | -0.42689214 | 0.065 | 0.176 | 1 | 19 |
| hsf2 | 0.38381 | 0.274693729 | 0.194 | 0.059 | 1 | 19 |
| mir223 | 0.38396 | 0.221598027 | 0.194 | 0.059 | 1 | 19 |
| csk | 0.38429 | -0.36370918 | 0.226 | 0.412 | 1 | 19 |
| zgc:158343 | 0.38624 | -0.28680347 | 0.129 | 0.294 | 1 | 19 |
| si:dkey-21e2.8 | 0.38754 | 0.653211683 | 0.258 | 0.118 | 1 | 19 |
| rab5c | 0.38777 | -0.18229074 | 0.097 | 0.235 | 1 | 19 |
| manf | 0.38858 | -0.29448736 | 0.129 | 0.294 | 1 | 19 |
| pho | 0.38866 | 0.244427822 | 0.194 | 0.059 | 1 | 19 |
| ddost | 0.38914 | -0.34353369 | 0.129 | 0.294 | 1 | 19 |
| pabpn1 | 0.39108 | -0.25082575 | 0.129 | 0.294 | 1 | 19 |
| lpar5a | 0.39122 | -0.31157693 | 0.129 | 0.294 | 1 | 19 |
| hist2h2l | 0.39129 | -0.32743737 | 0.129 | 0.294 | 1 | 19 |
| fam120c | 0.39154 | -0.25120217 | 0.129 | 0.294 | 1 | 19 |
| lasp1 | 0.39188 | -0.30806599 | 0.516 | 0.706 | 1 | 19 |
| h2afvb | 0.39451 | -0.38423447 | 0.355 | 0.529 | 1 | 19 |
| ponzr1 | 0.39469 | 0.272519421 | 0.161 | 0.176 | 1 | 19 |
| cox8a | 0.39608 | 0.244935694 | 0.516 | 0.471 | 1 | 19 |
| litaf | 0.39653 | -0.28851159 | 0.226 | 0.412 | 1 | 19 |
| blcap | 0.39667 | 0.247957361 | 0.194 | 0.059 | 1 | 19 |
| ccr9a | 0.39677 | -0.3027523 | 0.226 | 0.294 | 1 | 19 |
| rpl30 | 0.39692 | -0.1223788 | 0.935 | 0.941 | 1 | 19 |
| lsm12b | 0.39741 | 0.244235334 | 0.194 | 0.059 | 1 | 19 |
| wasa | 0.39971 | 0.377748931 | 0.484 | 0.294 | 1 | 19 |
| nap1l4a | 0.39994 | -0.33238514 | 0.097 | 0.176 | 1 | 19 |
| dbi | 0.40205 | 0.273179921 | 0.484 | 0.294 | 1 | 19 |
| metap2b | 0.40221 | -0.32214088 | 0.226 | 0.412 | 1 | 19 |
| naa20 | 0.40331 | 0.18569471 | 0.194 | 0.059 | 1 | 19 |
| top2b | 0.40337 | -0.11610671 | 0.065 | 0.176 | 1 | 19 |
| samsn1b | 0.40355 | -0.3571246 | 0.065 | 0.176 | 1 | 19 |
| atp1b1a | 0.40382 | 0.332403321 | 0.355 | 0.176 | 1 | 19 |
| CT027744.2 | 0.40446 | -0.18813071 | 0.097 | 0.235 | 1 | 19 |
| mtor | 0.40451 | -0.34323279 | 0.065 | 0.176 | 1 | 19 |
| hspa4b | 0.40455 | -0.3157179 | 0.097 | 0.235 | 1 | 19 |
| atp5pb | 0.40526 | -0.33541123 | 0.29 | 0.471 | 1 | 19 |
| btk | 0.40565 | -0.21365631 | 0.226 | 0.235 | 1 | 19 |

| | | | | | | |
|---|---|---|---|---|---|---|
| bida | 0.40596 | 0.224482034 | 0.194 | 0.059 | 1 | 19 |
| si:ch73-248e21.7 | 0.40656 | 0.297206546 | 0.194 | 0.059 | 1 | 19 |
| si:dkey-262k9.4 | 0.40777 | 0.348427541 | 0.161 | 0.059 | 1 | 19 |
| scamp2 | 0.40878 | -0.1679371 | 0.194 | 0.353 | 1 | 19 |
| ap2m1a | 0.4109 | 0.299312191 | 0.355 | 0.176 | 1 | 19 |
| ier5 | 0.41151 | 0.220974765 | 0.194 | 0.059 | 1 | 19 |
| elof1 | 0.41163 | 0.242554072 | 0.194 | 0.059 | 1 | 19 |
| spink2.1 | 0.41166 | 0.223126891 | 0.194 | 0.059 | 1 | 19 |
| sltm | 0.41199 | -0.32552845 | 0.065 | 0.176 | 1 | 19 |
| cltca | 0.412 | 0.228936789 | 0.194 | 0.059 | 1 | 19 |
| CU075735.1 | 0.4121 | 0.339626224 | 0.194 | 0.059 | 1 | 19 |
| ube2na | 0.41251 | 0.353162722 | 0.355 | 0.176 | 1 | 19 |
| cox5ab | 0.41353 | -0.36751123 | 0.258 | 0.412 | 1 | 19 |
| b2m | 0.41358 | -0.34128199 | 0.226 | 0.353 | 1 | 19 |
| tnikb | 0.41359 | 0.110823256 | 0.161 | 0.059 | 1 | 19 |
| fabp3 | 0.41398 | -0.3693001 | 0.194 | 0.353 | 1 | 19 |
| il11b | 0.41821 | -0.29122765 | 0.452 | 0.647 | 1 | 19 |
| zbtb46 | 0.41979 | 0.108902044 | 0.161 | 0.059 | 1 | 19 |
| gsk3b | 0.42046 | 0.116825869 | 0.161 | 0.059 | 1 | 19 |
| CABZ01111959.1 | 0.42128 | -0.15900906 | 0.065 | 0.176 | 1 | 19 |
| vps26c | 0.42154 | -0.20901059 | 0.097 | 0.235 | 1 | 19 |
| CABZ01041494.1 | 0.422 | -0.18227253 | 0.065 | 0.176 | 1 | 19 |
| atp5pf | 0.4222 | -0.13221095 | 0.516 | 0.471 | 1 | 19 |
| atp5if1a | 0.42257 | 0.351659439 | 0.419 | 0.235 | 1 | 19 |
| pum2 | 0.42263 | -0.12857021 | 0.065 | 0.176 | 1 | 19 |
| si:dkey-8e10.2 | 0.42374 | -0.25821389 | 0.097 | 0.235 | 1 | 19 |
| timm10 | 0.42528 | -0.31778279 | 0.065 | 0.176 | 1 | 19 |
| uqcrq | 0.42545 | 0.278561647 | 0.484 | 0.294 | 1 | 19 |
| ndufb2 | 0.42562 | 0.175386554 | 0.258 | 0.118 | 1 | 19 |
| rhogb | 0.42569 | -0.34345661 | 0.323 | 0.471 | 1 | 19 |
| prkar2aa | 0.42835 | -0.33331161 | 0.065 | 0.176 | 1 | 19 |
| eif3ha | 0.42982 | -0.32019523 | 0.194 | 0.353 | 1 | 19 |
| hic1 | 0.42999 | -0.24509126 | 0.097 | 0.235 | 1 | 19 |
| mapre1a | 0.4317 | 0.314535062 | 0.484 | 0.294 | 1 | 19 |
| hif1ab | 0.43209 | 0.317341928 | 0.419 | 0.235 | 1 | 19 |
| eif4a3 | 0.43211 | -0.23255043 | 0.097 | 0.235 | 1 | 19 |
| cnih1 | 0.43231 | -0.14166006 | 0.065 | 0.176 | 1 | 19 |
| tomm6 | 0.43297 | -0.31283449 | 0.065 | 0.176 | 1 | 19 |
| eif3jb | 0.43517 | -0.31664301 | 0.129 | 0.235 | 1 | 19 |
| actb1 | 0.43554 | -0.12450283 | 0.935 | 1 | 1 | 19 |
| lypla2 | 0.43865 | -0.30910707 | 0.161 | 0.235 | 1 | 19 |
| srsf10b | 0.43937 | -0.11580734 | 0.065 | 0.176 | 1 | 19 |
| BX120005.1 | 0.43976 | -0.35051484 | 0.097 | 0.235 | 1 | 19 |
| FQ323156.1 | 0.4422 | -0.19653241 | 0.097 | 0.235 | 1 | 19 |
| si:ch211-105j21.9 | 0.44244 | 0.319202702 | 0.161 | 0.059 | 1 | 19 |
| surf4l | 0.44305 | -0.3206832 | 0.065 | 0.176 | 1 | 19 |
| calr3a | 0.44381 | -0.33024751 | 0.194 | 0.294 | 1 | 19 |
| adrm1 | 0.44454 | -0.27611051 | 0.065 | 0.176 | 1 | 19 |

| | | | | | | |
|---|---|---|---|---|---|---|
| pcbp2 | 0.44501 | -0.25382235 | 0.097 | 0.235 | 1 | 19 |
| tmem167a | 0.44532 | -0.256731 | 0.097 | 0.235 | 1 | 19 |
| h3f3d | 0.44596 | 0.175806403 | 0.548 | 0.529 | 1 | 19 |
| kmt5aa | 0.44603 | 0.242171407 | 0.194 | 0.176 | 1 | 19 |
| tescb | 0.44721 | 0.132562834 | 0.161 | 0.059 | 1 | 19 |
| srsf11 | 0.44754 | -0.26317776 | 0.097 | 0.235 | 1 | 19 |
| hmgb2b | 0.44756 | -0.28218491 | 0.097 | 0.235 | 1 | 19 |
| mylk5 | 0.44804 | -0.29962785 | 0.097 | 0.235 | 1 | 19 |
| map4k5 | 0.44807 | -0.23331493 | 0.097 | 0.235 | 1 | 19 |
| coa6 | 0.44864 | -0.22789166 | 0.097 | 0.235 | 1 | 19 |
| spcs1 | 0.44882 | -0.32366994 | 0.355 | 0.529 | 1 | 19 |
| psme2 | 0.44912 | -0.29105342 | 0.097 | 0.235 | 1 | 19 |
| rps3 | 0.44912 | 0.172953292 | 0.903 | 0.765 | 1 | 19 |
| khsrp | 0.44914 | -0.24239652 | 0.097 | 0.235 | 1 | 19 |
| scinlb | 0.45137 | 0.340453887 | 0.355 | 0.235 | 1 | 19 |
| phb | 0.45334 | 0.331470751 | 0.226 | 0.118 | 1 | 19 |
| cyba | 0.45426 | -0.31260702 | 0.258 | 0.353 | 1 | 19 |
| pfdn5 | 0.45641 | 0.266412927 | 0.29 | 0.235 | 1 | 19 |
| cdv3 | 0.45651 | -0.34472891 | 0.161 | 0.294 | 1 | 19 |
| tmc6b | 0.45805 | -0.30867491 | 0.065 | 0.176 | 1 | 19 |
| taf13 | 0.45819 | -0.29768192 | 0.097 | 0.176 | 1 | 19 |
| chmp2a | 0.45915 | -0.13819008 | 0.161 | 0.294 | 1 | 19 |
| cd74a | 0.46039 | -0.26006152 | 0.419 | 0.471 | 1 | 19 |
| LOC100150882 | 0.46153 | -0.44102634 | 0.29 | 0.471 | 1 | 19 |
| si:dkey-100n23.5 | 0.46197 | -0.16728125 | 0.065 | 0.176 | 1 | 19 |
| nucks1a | 0.46215 | -0.24984209 | 0.065 | 0.176 | 1 | 19 |
| dennd2da | 0.46245 | -0.26359633 | 0.065 | 0.176 | 1 | 19 |
| ddt | 0.46284 | 0.272246367 | 0.161 | 0.059 | 1 | 19 |
| gemin8 | 0.46379 | 0.253060145 | 0.161 | 0.059 | 1 | 19 |
| b2ml | 0.46437 | -0.30314796 | 0.29 | 0.471 | 1 | 19 |
| psmb10 | 0.46456 | -0.27552057 | 0.065 | 0.176 | 1 | 19 |
| sla1 | 0.4648 | 0.112767414 | 0.226 | 0.118 | 1 | 19 |
| tmco1 | 0.46623 | -0.17024527 | 0.129 | 0.118 | 1 | 19 |
| gapdhs | 0.46648 | 0.234005656 | 0.806 | 0.706 | 1 | 19 |
| pet100 | 0.46651 | -0.26901507 | 0.065 | 0.176 | 1 | 19 |
| ap1g1 | 0.46653 | -0.29310515 | 0.065 | 0.176 | 1 | 19 |
| rrp1 | 0.46693 | 0.128046845 | 0.161 | 0.059 | 1 | 19 |
| dusp1 | 0.46713 | -0.34861387 | 0.161 | 0.294 | 1 | 19 |
| CU855821.1 | 0.46749 | -0.30277202 | 0.065 | 0.176 | 1 | 19 |
| hiat1b | 0.46778 | -0.29864725 | 0.065 | 0.176 | 1 | 19 |
| zgc:112356 | 0.46839 | 0.272507768 | 0.226 | 0.176 | 1 | 19 |
| sat1a.2 | 0.46947 | 0.282384317 | 0.645 | 0.471 | 1 | 19 |
| derl2 | 0.46976 | 0.198751537 | 0.387 | 0.235 | 1 | 19 |
| eef1db | 0.47058 | -0.23004837 | 0.065 | 0.176 | 1 | 19 |
| nop53 | 0.47066 | -0.24152214 | 0.194 | 0.353 | 1 | 19 |
| psmb1 | 0.47074 | -0.33696561 | 0.258 | 0.412 | 1 | 19 |
| fkbp2 | 0.47083 | -0.35572921 | 0.065 | 0.176 | 1 | 19 |
| mob1a | 0.47123 | -0.31220122 | 0.194 | 0.353 | 1 | 19 |

| | | | | | | |
|---|---|---|---|---|---|---|
| gbp | 0.47162 | 0.263136171 | 0.161 | 0.059 | 1 | 19 |
| wu:fi37e09 | 0.47202 | -0.24963092 | 0.065 | 0.176 | 1 | 19 |
| zgc:63831 | 0.4724 | 0.218917344 | 0.258 | 0.118 | 1 | 19 |
| nutf2l | 0.47278 | -0.32743873 | 0.161 | 0.294 | 1 | 19 |
| atp6v0ca | 0.47514 | 0.241622215 | 0.258 | 0.118 | 1 | 19 |
| ckbb | 0.47625 | 0.320796534 | 0.258 | 0.118 | 1 | 19 |
| znf1035 | 0.47728 | -0.13928976 | 0.065 | 0.176 | 1 | 19 |
| cox6b2 | 0.47752 | -0.21144438 | 0.742 | 0.824 | 1 | 19 |
| mrps21 | 0.47893 | -0.12226918 | 0.065 | 0.176 | 1 | 19 |
| socs3a | 0.47997 | -0.26871872 | 0.355 | 0.412 | 1 | 19 |
| ciapin1 | 0.48104 | 0.188998256 | 0.097 | 0.118 | 1 | 19 |
| psmd8 | 0.48169 | -0.16823498 | 0.065 | 0.176 | 1 | 19 |
| pbdc1 | 0.48186 | 0.293644483 | 0.258 | 0.118 | 1 | 19 |
| CR855389.1 | 0.48227 | -0.33967028 | 0.097 | 0.176 | 1 | 19 |
| ube2nb | 0.48281 | -0.21316997 | 0.065 | 0.176 | 1 | 19 |
| aldocb | 0.48371 | -0.28906014 | 0.194 | 0.353 | 1 | 19 |
| hnrnpa1b | 0.48387 | 0.225814915 | 0.258 | 0.118 | 1 | 19 |
| ube2e1 | 0.48392 | -0.23033188 | 0.194 | 0.353 | 1 | 19 |
| vamp8 | 0.48418 | -0.25457496 | 0.065 | 0.176 | 1 | 19 |
| hmgn3 | 0.48477 | -0.33735146 | 0.129 | 0.235 | 1 | 19 |
| cnot1 | 0.48488 | 0.156076091 | 0.161 | 0.059 | 1 | 19 |
| snrpg | 0.48495 | -0.22975636 | 0.194 | 0.353 | 1 | 19 |
| zgc:91910 | 0.48498 | -0.24850876 | 0.194 | 0.353 | 1 | 19 |
| knop1 | 0.4859 | -0.22870845 | 0.065 | 0.176 | 1 | 19 |
| csrp1a | 0.48682 | 0.162771386 | 0.161 | 0.059 | 1 | 19 |
| atg12 | 0.48685 | -0.25602717 | 0.065 | 0.176 | 1 | 19 |
| twistnb | 0.48689 | -0.22595812 | 0.065 | 0.176 | 1 | 19 |
| hacd3 | 0.48815 | -0.28657488 | 0.065 | 0.176 | 1 | 19 |
| xbp1 | 0.48846 | -0.21915887 | 0.677 | 0.765 | 1 | 19 |
| si:dkey-237j10.2 | 0.48862 | 0.181632338 | 0.161 | 0.059 | 1 | 19 |
| BX005340.1 | 0.48862 | 0.125317264 | 0.161 | 0.059 | 1 | 19 |
| sf3b6 | 0.48978 | 0.257996257 | 0.258 | 0.118 | 1 | 19 |
| nolc1 | 0.49059 | -0.28533166 | 0.065 | 0.176 | 1 | 19 |
| cox7a2a | 0.49082 | 0.328250022 | 0.516 | 0.353 | 1 | 19 |
| si:dkey-187e18.1 | 0.49098 | -0.20873176 | 0.065 | 0.176 | 1 | 19 |
| nup62l | 0.49201 | -0.13575452 | 0.065 | 0.176 | 1 | 19 |
| atp6v0b | 0.49206 | -0.21340187 | 0.065 | 0.176 | 1 | 19 |
| mvb12a | 0.49226 | -0.26656626 | 0.065 | 0.176 | 1 | 19 |
| si:ch73-208g10.1 | 0.49296 | 0.28383048 | 0.258 | 0.118 | 1 | 19 |
| cox6a2 | 0.49316 | -0.23763188 | 0.161 | 0.176 | 1 | 19 |
| dip2ba | 0.49452 | -0.29036922 | 0.065 | 0.176 | 1 | 19 |
| rarga | 0.49458 | 0.276517292 | 0.258 | 0.118 | 1 | 19 |
| h2afy2 | 0.49512 | -0.24145809 | 0.065 | 0.176 | 1 | 19 |
| rps24 | 0.49531 | 0.101285899 | 0.903 | 0.882 | 1 | 19 |
| pgam1b | 0.49617 | -0.22532998 | 0.065 | 0.176 | 1 | 19 |
| si:ch73-281n10.2 | 0.49645 | -0.17679614 | 0.065 | 0.176 | 1 | 19 |
| psmd3 | 0.49662 | -0.25297724 | 0.065 | 0.176 | 1 | 19 |
| il13 | 0.49675 | -0.40679825 | 0.29 | 0.412 | 1 | 19 |

| | | | | | | |
|---|---|---|---|---|---|---|
| psmd13 | 0.49679 | -0.16330891 | 0.065 | 0.176 | 1 | 19 |
| pdia3 | 0.49702 | -0.28723353 | 0.355 | 0.529 | 1 | 19 |
| ikbkg | 0.4971 | -0.18241925 | 0.065 | 0.176 | 1 | 19 |
| rpl36a | 0.49746 | 0.150756049 | 0.935 | 0.824 | 1 | 19 |
| rab5ab | 0.49783 | -0.23990632 | 0.065 | 0.176 | 1 | 19 |
| fuca2 | 0.49821 | -0.22024267 | 0.065 | 0.176 | 1 | 19 |
| prkag1 | 0.49878 | -0.23703427 | 0.065 | 0.176 | 1 | 19 |
| uqcrc2a | 0.4988 | -0.14325831 | 0.065 | 0.176 | 1 | 19 |
| mcts1 | 0.49962 | -0.19925733 | 0.065 | 0.176 | 1 | 19 |
| krt8 | 0.49978 | 0.275825213 | 0.226 | 0.176 | 1 | 19 |
| hdac1 | 0.4998 | 0.316397865 | 0.258 | 0.176 | 1 | 19 |
| atp5meb | 0.49998 | -0.27074926 | 0.355 | 0.529 | 1 | 19 |
| ndufa5 | 0.50004 | -0.16072578 | 0.065 | 0.176 | 1 | 19 |
| rack1 | 0.50095 | 0.126873372 | 0.935 | 0.824 | 1 | 19 |
| mir21-1 | 0.50096 | -0.34126394 | 0.129 | 0.235 | 1 | 19 |
| emc1 | 0.50097 | -0.24574591 | 0.065 | 0.176 | 1 | 19 |
| mrps15 | 0.50127 | -0.2133683 | 0.065 | 0.176 | 1 | 19 |
| ccnt2b | 0.50133 | -0.21507111 | 0.065 | 0.176 | 1 | 19 |
| lyrm2 | 0.50158 | -0.24528047 | 0.065 | 0.176 | 1 | 19 |
| csnk1da | 0.50164 | -0.21995354 | 0.065 | 0.176 | 1 | 19 |
| eif4e1c | 0.50168 | -0.16318354 | 0.065 | 0.176 | 1 | 19 |
| sla2 | 0.50174 | 0.109829772 | 0.29 | 0.176 | 1 | 19 |
| dnttip2 | 0.50183 | -0.25450276 | 0.065 | 0.176 | 1 | 19 |
| chmp4bb | 0.50199 | -0.25764046 | 0.065 | 0.176 | 1 | 19 |
| cpsf6 | 0.50202 | -0.18589788 | 0.065 | 0.176 | 1 | 19 |
| ncoa3 | 0.50205 | -0.23363838 | 0.065 | 0.176 | 1 | 19 |
| elf1 | 0.5041 | -0.2804673 | 0.355 | 0.529 | 1 | 19 |
| si:ch211-277c7.6 | 0.50411 | -0.3598809 | 0.194 | 0.294 | 1 | 19 |
| thoc5 | 0.50536 | -0.26244827 | 0.129 | 0.176 | 1 | 19 |
| si:dkey-21e2.13 | 0.50956 | 0.143334693 | 0.387 | 0.294 | 1 | 19 |
| taf3 | 0.50996 | 0.243576256 | 0.161 | 0.059 | 1 | 19 |
| rpl27 | 0.51041 | -0.14177542 | 0.871 | 0.941 | 1 | 19 |
| tuba8l4 | 0.51205 | -0.30335236 | 0.355 | 0.471 | 1 | 19 |
| ndufa4 | 0.51223 | -0.32161668 | 0.161 | 0.294 | 1 | 19 |
| gpia | 0.5131 | -0.2994978 | 0.129 | 0.235 | 1 | 19 |
| CR381673.1 | 0.51313 | 0.247261954 | 0.161 | 0.059 | 1 | 19 |
| tomm7 | 0.51471 | -0.29101376 | 0.355 | 0.471 | 1 | 19 |
| sfpq | 0.51482 | 0.287455224 | 0.129 | 0.059 | 1 | 19 |
| jun | 0.51755 | -0.1019614 | 0.097 | 0.059 | 1 | 19 |
| llph | 0.51881 | -0.14276544 | 0.161 | 0.294 | 1 | 19 |
| nfe2l2a | 0.51914 | 0.254742973 | 0.161 | 0.059 | 1 | 19 |
| laptm4a | 0.51936 | -0.32972529 | 0.129 | 0.235 | 1 | 19 |
| tln1 | 0.5216 | -0.30499396 | 0.323 | 0.471 | 1 | 19 |
| id3 | 0.52168 | -0.22906708 | 0.419 | 0.471 | 1 | 19 |
| vps29 | 0.5227 | 0.282059374 | 0.258 | 0.176 | 1 | 19 |
| pak2a | 0.52309 | 0.20358989 | 0.161 | 0.059 | 1 | 19 |
| tmem30ab | 0.52476 | 0.156658339 | 0.161 | 0.059 | 1 | 19 |
| lamtor5 | 0.52629 | -0.29972127 | 0.161 | 0.294 | 1 | 19 |

| | | | | | | |
|---|---|---|---|---|---|---|
| dynlrb1 | 0.5291 | -0.29994214 | 0.226 | 0.353 | 1 | 19 |
| selenot1b | 0.52982 | -0.21910468 | 0.097 | 0.118 | 1 | 19 |
| oaz1a | 0.53016 | -0.26776945 | 0.516 | 0.647 | 1 | 19 |
| ppp1r18 | 0.53051 | 0.263014068 | 0.29 | 0.235 | 1 | 19 |
| ufm1 | 0.53059 | -0.17552354 | 0.161 | 0.294 | 1 | 19 |
| pabpc1a | 0.53101 | 0.233721652 | 0.774 | 0.647 | 1 | 19 |
| rps11 | 0.53489 | 0.150757578 | 0.935 | 0.882 | 1 | 19 |
| commd1 | 0.53638 | 0.259612555 | 0.097 | 0.059 | 1 | 19 |
| eif3k | 0.5384 | -0.30024465 | 0.161 | 0.294 | 1 | 19 |
| fbl | 0.53843 | -0.27386143 | 0.161 | 0.235 | 1 | 19 |
| rhoac | 0.5385 | -0.14290154 | 0.129 | 0.235 | 1 | 19 |
| cap1 | 0.53964 | -0.27020128 | 0.484 | 0.588 | 1 | 19 |
| ccdc25 | 0.54183 | 0.238907279 | 0.161 | 0.118 | 1 | 19 |
| serf2 | 0.54186 | 0.109182914 | 0.548 | 0.412 | 1 | 19 |
| celf2 | 0.54217 | 0.30138329 | 0.323 | 0.235 | 1 | 19 |
| ssr2 | 0.54347 | -0.28233034 | 0.419 | 0.529 | 1 | 19 |
| rpl37 | 0.54527 | -0.10857819 | 0.903 | 0.941 | 1 | 19 |
| ywhabl | 0.54648 | -0.21333921 | 0.484 | 0.647 | 1 | 19 |
| ptprjb.1 | 0.54672 | -0.27335693 | 0.065 | 0.118 | 1 | 19 |
| pygb | 0.54756 | 0.147123338 | 0.226 | 0.118 | 1 | 19 |
| ms4a17a.6 | 0.54763 | -0.34198681 | 0.161 | 0.294 | 1 | 19 |
| eef2b | 0.54931 | -0.21926028 | 0.613 | 0.765 | 1 | 19 |
| map2k6 | 0.54936 | 0.242757411 | 0.161 | 0.059 | 1 | 19 |
| srsf5a | 0.55008 | -0.17310663 | 0.161 | 0.294 | 1 | 19 |
| gadd45bb | 0.55019 | 0.17855875 | 0.161 | 0.059 | 1 | 19 |
| lrrfip1a | 0.55188 | 0.135449839 | 0.161 | 0.176 | 1 | 19 |
| si:ch211-162e15.3 | 0.55244 | 0.230767805 | 0.161 | 0.059 | 1 | 19 |
| hdlbpa | 0.55283 | -0.21819093 | 0.097 | 0.118 | 1 | 19 |
| arl5c | 0.55356 | -0.26084611 | 0.226 | 0.294 | 1 | 19 |
| ebna1bp2 | 0.55376 | 0.20142506 | 0.161 | 0.059 | 1 | 19 |
| txnipa | 0.55385 | -0.27478077 | 0.258 | 0.412 | 1 | 19 |
| hmgn6 | 0.5544 | -0.28134302 | 0.387 | 0.529 | 1 | 19 |
| eif1axb | 0.5556 | 0.268390277 | 0.452 | 0.294 | 1 | 19 |
| eif3ba | 0.55582 | -0.1436228 | 0.226 | 0.353 | 1 | 19 |
| uba52 | 0.55604 | -0.13529468 | 0.839 | 0.941 | 1 | 19 |
| rpl23a | 0.55605 | 0.155348563 | 0.935 | 0.882 | 1 | 19 |
| srp72 | 0.55688 | 0.180604658 | 0.161 | 0.059 | 1 | 19 |
| mbnl1 | 0.55732 | 0.293415515 | 0.387 | 0.235 | 1 | 19 |
| sumo3a | 0.55824 | -0.28703769 | 0.226 | 0.353 | 1 | 19 |
| txn | 0.55862 | -0.31741686 | 0.161 | 0.294 | 1 | 19 |
| rab14l | 0.55935 | -0.20995481 | 0.097 | 0.118 | 1 | 19 |
| hpdb | 0.55946 | -0.27686151 | 0.065 | 0.118 | 1 | 19 |
| unc13d | 0.56149 | 0.215725608 | 0.161 | 0.059 | 1 | 19 |
| uqcrc1 | 0.56157 | 0.198887224 | 0.161 | 0.059 | 1 | 19 |
| med28 | 0.56175 | 0.203552259 | 0.161 | 0.059 | 1 | 19 |
| rab5if | 0.56233 | 0.203761918 | 0.161 | 0.059 | 1 | 19 |
| rpl17 | 0.56236 | -0.11094536 | 0.871 | 0.882 | 1 | 19 |
| polr2d | 0.56279 | -0.29140463 | 0.161 | 0.294 | 1 | 19 |

| | | | | | | |
|---|---|---|---|---|---|---|
| sept12 | 0.56341 | 0.246540769 | 0.226 | 0.176 | 1 | 19 |
| srprb | 0.56358 | -0.16328681 | 0.226 | 0.235 | 1 | 19 |
| ddx21 | 0.56379 | -0.25709115 | 0.161 | 0.294 | 1 | 19 |
| eef1b2 | 0.56398 | 0.178959888 | 0.839 | 0.706 | 1 | 19 |
| ctsh | 0.56403 | 0.259982108 | 0.161 | 0.059 | 1 | 19 |
| sugt1 | 0.56441 | 0.20456571 | 0.161 | 0.059 | 1 | 19 |
| uqcrh | 0.56519 | -0.27208658 | 0.258 | 0.353 | 1 | 19 |
| anxa11a | 0.56537 | -0.23910345 | 0.161 | 0.294 | 1 | 19 |
| fosl2 | 0.56594 | -0.2531378 | 0.161 | 0.294 | 1 | 19 |
| rpl3 | 0.56648 | -0.12370202 | 0.806 | 0.824 | 1 | 19 |
| rtn4a | 0.56798 | -0.22648706 | 0.161 | 0.294 | 1 | 19 |
| gemin7 | 0.56853 | -0.29145609 | 0.097 | 0.176 | 1 | 19 |
| nme2b.2 | 0.56897 | -0.27845906 | 0.097 | 0.176 | 1 | 19 |
| skap2 | 0.57011 | 0.202943131 | 0.323 | 0.294 | 1 | 19 |
| rps5 | 0.57071 | 0.170213977 | 0.903 | 0.824 | 1 | 19 |
| si:ch211-102c2.4 | 0.57202 | -0.13030603 | 0.516 | 0.647 | 1 | 19 |
| zgc:112255 | 0.57494 | 0.247829055 | 0.129 | 0.059 | 1 | 19 |
| ddit3 | 0.57507 | 0.267601934 | 0.129 | 0.059 | 1 | 19 |
| cib1 | 0.57518 | -0.12821071 | 0.129 | 0.235 | 1 | 19 |
| vaspa | 0.5763 | -0.21188664 | 0.323 | 0.471 | 1 | 19 |
| atp5fa1 | 0.57675 | -0.18577303 | 0.226 | 0.235 | 1 | 19 |
| prkar1ab | 0.57849 | -0.17699005 | 0.161 | 0.176 | 1 | 19 |
| cox17 | 0.57868 | 0.14024739 | 0.613 | 0.471 | 1 | 19 |
| snrpd2 | 0.58012 | 0.144667122 | 0.161 | 0.176 | 1 | 19 |
| ikzf1 | 0.58133 | -0.2955476 | 0.129 | 0.235 | 1 | 19 |
| dynll1 | 0.58879 | -0.26338828 | 0.29 | 0.353 | 1 | 19 |
| atp5po | 0.58938 | -0.27162845 | 0.129 | 0.235 | 1 | 19 |
| eif2s1a | 0.59179 | 0.252724334 | 0.129 | 0.059 | 1 | 19 |
| rpl7l1 | 0.59389 | 0.235839115 | 0.129 | 0.059 | 1 | 19 |
| eif4a1b | 0.59405 | 0.103692468 | 0.323 | 0.353 | 1 | 19 |
| zgc:114188 | 0.5943 | -0.13300592 | 0.871 | 0.941 | 1 | 19 |
| hnrnpa1a | 0.59464 | -0.29170605 | 0.226 | 0.353 | 1 | 19 |
| h1fx | 0.59491 | -0.12625251 | 0.516 | 0.647 | 1 | 19 |
| atp5f1e | 0.59701 | -0.20044242 | 0.581 | 0.647 | 1 | 19 |
| rps26 | 0.59817 | -0.13888836 | 0.871 | 0.941 | 1 | 19 |
| si:ch211-212d10.1 | 0.59829 | -0.35431512 | 0.323 | 0.471 | 1 | 19 |
| rap1b | 0.59917 | 0.272040155 | 0.323 | 0.235 | 1 | 19 |
| eif5a | 0.59994 | 0.106934685 | 0.484 | 0.353 | 1 | 19 |
| prelid1a | 0.60345 | 0.240524637 | 0.129 | 0.059 | 1 | 19 |
| laptm5 | 0.6047 | -0.16657397 | 0.774 | 0.882 | 1 | 19 |
| yipf4 | 0.60504 | 0.239001792 | 0.129 | 0.059 | 1 | 19 |
| XLOC-000946 | 0.60627 | -0.25065345 | 0.065 | 0.118 | 1 | 19 |
| atp6v1e1b | 0.60735 | -0.19591749 | 0.097 | 0.118 | 1 | 19 |
| rbm28 | 0.60832 | -0.26511924 | 0.097 | 0.176 | 1 | 19 |
| NC-002333.4 | 0.61164 | -0.15527171 | 0.29 | 0.412 | 1 | 19 |
| eif3d | 0.61181 | -0.25389794 | 0.258 | 0.353 | 1 | 19 |
| ddx27 | 0.61223 | -0.12338191 | 0.129 | 0.235 | 1 | 19 |
| uqcr10 | 0.61496 | 0.252040428 | 0.29 | 0.176 | 1 | 19 |

| | | | | | | |
|---|---|---|---|---|---|---|
| zcchc7 | 0.61526 | -0.24367242 | 0.065 | 0.118 | 1 | 19 |
| si:dkey-206d17.12 | 0.61532 | -0.25498933 | 0.097 | 0.176 | 1 | 19 |
| gstk1 | 0.61625 | -0.27401896 | 0.129 | 0.235 | 1 | 19 |
| smdt1b | 0.61735 | -0.14235274 | 0.129 | 0.235 | 1 | 19 |
| bri3bp | 0.62025 | -0.16471477 | 0.129 | 0.235 | 1 | 19 |
| tomm20b | 0.62076 | 0.250641558 | 0.387 | 0.294 | 1 | 19 |
| sh3bgrl3 | 0.62083 | -0.25475691 | 0.226 | 0.353 | 1 | 19 |
| XLOC-026836 | 0.62314 | -0.1486039 | 0.29 | 0.294 | 1 | 19 |
| ucp2 | 0.62466 | -0.23160502 | 0.452 | 0.529 | 1 | 19 |
| eif3ja | 0.62559 | 0.243373925 | 0.226 | 0.118 | 1 | 19 |
| tmc8 | 0.62618 | 0.234490297 | 0.194 | 0.118 | 1 | 19 |
| tagapa | 0.62711 | -0.2698987 | 0.323 | 0.412 | 1 | 19 |
| tomm40 | 0.62768 | 0.21846177 | 0.129 | 0.059 | 1 | 19 |
| XLOC-025423 | 0.62891 | 0.294819126 | 0.419 | 0.294 | 1 | 19 |
| parvg | 0.62953 | 0.203821096 | 0.226 | 0.118 | 1 | 19 |
| midn | 0.6299 | -0.18037056 | 0.161 | 0.176 | 1 | 19 |
| zgc:77849 | 0.63031 | 0.202764955 | 0.161 | 0.118 | 1 | 19 |
| selenom | 0.63205 | 0.226335841 | 0.097 | 0.059 | 1 | 19 |
| csf3a | 0.6321 | -0.14924012 | 0.355 | 0.353 | 1 | 19 |
| ubap2l | 0.63217 | -0.19280685 | 0.097 | 0.118 | 1 | 19 |
| ndufab1a | 0.63252 | 0.153160233 | 0.226 | 0.118 | 1 | 19 |
| psma5 | 0.633 | -0.2342668 | 0.129 | 0.235 | 1 | 19 |
| si:ch73-234b20.5 | 0.63322 | 0.232458642 | 0.226 | 0.118 | 1 | 19 |
| snd1 | 0.63345 | 0.167323568 | 0.226 | 0.118 | 1 | 19 |
| snrpe | 0.6335 | -0.28308453 | 0.194 | 0.294 | 1 | 19 |
| nsa2 | 0.63386 | -0.2297726 | 0.387 | 0.529 | 1 | 19 |
| zgc:56585 | 0.63506 | 0.218501361 | 0.129 | 0.059 | 1 | 19 |
| eif4ebp2 | 0.63539 | -0.21826619 | 0.226 | 0.353 | 1 | 19 |
| calr3b | 0.63661 | -0.17803088 | 0.161 | 0.176 | 1 | 19 |
| XLOC-005449 | 0.63732 | 0.237144394 | 0.194 | 0.118 | 1 | 19 |
| eif4e3 | 0.63806 | 0.215924281 | 0.097 | 0.059 | 1 | 19 |
| fthl27 | 0.63832 | -0.2554445 | 0.419 | 0.529 | 1 | 19 |
| n4bp1 | 0.63987 | 0.225716092 | 0.129 | 0.059 | 1 | 19 |
| seta | 0.64017 | 0.200522959 | 0.194 | 0.118 | 1 | 19 |
| si:ch211-132b12.7 | 0.6416 | 0.220315842 | 0.129 | 0.059 | 1 | 19 |
| si:ch211-235e9.6 | 0.64202 | -0.17162027 | 0.129 | 0.235 | 1 | 19 |
| tnip1 | 0.64244 | 0.158827766 | 0.355 | 0.235 | 1 | 19 |
| sdf2l1 | 0.64257 | -0.18830142 | 0.129 | 0.235 | 1 | 19 |
| hyou1 | 0.64276 | -0.2132464 | 0.226 | 0.353 | 1 | 19 |
| nipsnap2 | 0.64413 | 0.179476675 | 0.129 | 0.118 | 1 | 19 |
| usf1l | 0.64434 | 0.230580667 | 0.194 | 0.118 | 1 | 19 |
| tnrc6c1 | 0.64487 | -0.21570519 | 0.129 | 0.235 | 1 | 19 |
| itga4 | 0.64558 | -0.17157733 | 0.129 | 0.235 | 1 | 19 |
| nptna | 0.64578 | -0.20246119 | 0.129 | 0.235 | 1 | 19 |
| trappc2l | 0.64603 | 0.255537307 | 0.194 | 0.118 | 1 | 19 |
| pwp1 | 0.64809 | 0.206186624 | 0.129 | 0.059 | 1 | 19 |
| si:dkeyp-68b7.12 | 0.64826 | -0.2562941 | 0.194 | 0.294 | 1 | 19 |
| lgals3b | 0.64834 | -0.21383012 | 0.129 | 0.235 | 1 | 19 |

| | | | | | | |
|---|---|---|---|---|---|---|
| mrps34 | 0.64887 | 0.216336451 | 0.129 | 0.059 | 1 | 19 |
| znf217 | 0.64911 | 0.195792241 | 0.129 | 0.059 | 1 | 19 |
| eif2s2 | 0.65136 | -0.2221683 | 0.452 | 0.588 | 1 | 19 |
| nupr1a | 0.65179 | -0.23583705 | 0.129 | 0.176 | 1 | 19 |
| h3f3a | 0.65211 | -0.1922419 | 0.129 | 0.235 | 1 | 19 |
| emc10 | 0.65233 | -0.11374529 | 0.129 | 0.118 | 1 | 19 |
| mif | 0.65269 | -0.13363759 | 0.29 | 0.294 | 1 | 19 |
| npc2 | 0.65351 | -0.24885524 | 0.258 | 0.353 | 1 | 19 |
| ndufv3 | 0.65392 | -0.22655413 | 0.097 | 0.176 | 1 | 19 |
| si:ch211-87m7.2 | 0.65424 | 0.100478741 | 0.129 | 0.059 | 1 | 19 |
| ankrd44 | 0.65451 | -0.11232053 | 0.129 | 0.118 | 1 | 19 |
| mydgf | 0.65465 | 0.210961654 | 0.129 | 0.059 | 1 | 19 |
| ywhaqa | 0.65466 | 0.165388988 | 0.355 | 0.235 | 1 | 19 |
| chmp6b | 0.65477 | 0.214706822 | 0.097 | 0.059 | 1 | 19 |
| mdh1aa | 0.65638 | 0.201864172 | 0.29 | 0.176 | 1 | 19 |
| lpcat2 | 0.65883 | -0.25456118 | 0.097 | 0.176 | 1 | 19 |
| edaradd | 0.65899 | -0.22835057 | 0.065 | 0.118 | 1 | 19 |
| ccr2 | 0.66064 | -0.16641016 | 0.194 | 0.294 | 1 | 19 |
| tomm22 | 0.6612 | -0.21975869 | 0.097 | 0.176 | 1 | 19 |
| lsm3 | 0.66155 | 0.225012191 | 0.129 | 0.059 | 1 | 19 |
| fundc2 | 0.66191 | 0.198258521 | 0.129 | 0.059 | 1 | 19 |
| cbx5 | 0.66255 | 0.183413416 | 0.129 | 0.118 | 1 | 19 |
| u2surp | 0.66327 | 0.191875847 | 0.129 | 0.059 | 1 | 19 |
| slc38a5a | 0.66376 | -0.23309669 | 0.097 | 0.176 | 1 | 19 |
| jpt2 | 0.66481 | 0.213849234 | 0.29 | 0.176 | 1 | 19 |
| ppp1caa | 0.66504 | -0.23143158 | 0.258 | 0.353 | 1 | 19 |
| abce1 | 0.66526 | 0.225354767 | 0.129 | 0.059 | 1 | 19 |
| osmr | 0.66579 | -0.24442766 | 0.065 | 0.118 | 1 | 19 |
| si:ch211-217k17.7 | 0.66711 | -0.22952473 | 0.097 | 0.176 | 1 | 19 |
| si:dkey-112e7.2 | 0.66741 | -0.16412459 | 0.323 | 0.353 | 1 | 19 |
| aldh9a1a.1 | 0.66832 | 0.154194918 | 0.29 | 0.176 | 1 | 19 |
| mef2d | 0.66894 | -0.12024368 | 0.129 | 0.118 | 1 | 19 |
| sh3bgrl | 0.66971 | 0.235625805 | 0.419 | 0.294 | 1 | 19 |
| hsbp1b | 0.67055 | 0.190547916 | 0.29 | 0.176 | 1 | 19 |
| setb | 0.67177 | -0.19117505 | 0.258 | 0.294 | 1 | 19 |
| hm13 | 0.67199 | 0.199092463 | 0.226 | 0.176 | 1 | 19 |
| arpp19b | 0.67317 | -0.21738868 | 0.097 | 0.176 | 1 | 19 |
| krtcap2 | 0.67319 | 0.163433021 | 0.484 | 0.353 | 1 | 19 |
| dkc1 | 0.67331 | 0.197749201 | 0.161 | 0.118 | 1 | 19 |
| AL954142.1 | 0.67336 | -0.18638415 | 0.258 | 0.294 | 1 | 19 |
| srsf2a | 0.67565 | -0.23026366 | 0.194 | 0.294 | 1 | 19 |
| BX510996.1 | 0.67726 | -0.1427146 | 0.032 | 0.059 | 1 | 19 |
| CABZ01117840.1 | 0.67726 | -0.10978794 | 0.032 | 0.059 | 1 | 19 |
| CU499330.1 | 0.67726 | -0.15745915 | 0.032 | 0.059 | 1 | 19 |
| ahctf1 | 0.67726 | -0.12683996 | 0.032 | 0.059 | 1 | 19 |
| arhgef12a | 0.67726 | -0.10802786 | 0.032 | 0.059 | 1 | 19 |
| atp11c | 0.67726 | -0.13265436 | 0.032 | 0.059 | 1 | 19 |
| chordc1b | 0.67726 | -0.16093691 | 0.032 | 0.059 | 1 | 19 |

| | | | | | | |
|---|---|---|---|---|---|---|
| dr1 | 0.67726 | -0.10897275 | 0.032 | 0.059 | 1 | 19 |
| esco2 | 0.67726 | 0.103141683 | 0.032 | 0.059 | 1 | 19 |
| grapa | 0.67726 | -0.10432242 | 0.032 | 0.059 | 1 | 19 |
| klhl6 | 0.67726 | -0.12609089 | 0.032 | 0.059 | 1 | 19 |
| klhl8 | 0.67726 | -0.10449303 | 0.032 | 0.059 | 1 | 19 |
| mzt2b | 0.67726 | -0.1037533 | 0.032 | 0.059 | 1 | 19 |
| plecb | 0.67726 | -0.22978162 | 0.032 | 0.059 | 1 | 19 |
| ptenb | 0.67726 | -0.10871651 | 0.032 | 0.059 | 1 | 19 |
| si:zfos-464b6.2 | 0.67726 | -0.10715579 | 0.032 | 0.059 | 1 | 19 |
| socs3b | 0.67726 | -0.13764886 | 0.032 | 0.059 | 1 | 19 |
| urod | 0.67726 | -0.11104992 | 0.032 | 0.059 | 1 | 19 |
| zdhhc23a | 0.67726 | -0.13171216 | 0.032 | 0.059 | 1 | 19 |
| CABZ01076667.1 | 0.67726 | -0.13618271 | 0.032 | 0.059 | 1 | 19 |
| LOC110439500 | 0.67726 | -0.12418829 | 0.032 | 0.059 | 1 | 19 |
| OSBPL8 | 0.67726 | -0.12361927 | 0.032 | 0.059 | 1 | 19 |
| acvr1l | 0.67726 | -0.14780913 | 0.032 | 0.059 | 1 | 19 |
| acy3.1 | 0.67726 | -0.11826719 | 0.032 | 0.059 | 1 | 19 |
| adipor1a | 0.67726 | -0.1129781 | 0.032 | 0.059 | 1 | 19 |
| arhgap29b | 0.67726 | -0.14103555 | 0.032 | 0.059 | 1 | 19 |
| atp8a1 | 0.67726 | -0.15752585 | 0.032 | 0.059 | 1 | 19 |
| atpv0e2 | 0.67726 | -0.14098484 | 0.032 | 0.059 | 1 | 19 |
| b4galnt1a | 0.67726 | -0.10757429 | 0.032 | 0.059 | 1 | 19 |
| brap | 0.67726 | -0.15911363 | 0.032 | 0.059 | 1 | 19 |
| btbd9 | 0.67726 | -0.14249512 | 0.032 | 0.059 | 1 | 19 |
| cops5 | 0.67726 | -0.13192833 | 0.032 | 0.059 | 1 | 19 |
| dnajc24 | 0.67726 | -0.10820783 | 0.032 | 0.059 | 1 | 19 |
| dusp7 | 0.67726 | -0.15864103 | 0.032 | 0.059 | 1 | 19 |
| elavl1b | 0.67726 | -0.13279758 | 0.032 | 0.059 | 1 | 19 |
| epc1b | 0.67726 | -0.12529142 | 0.032 | 0.059 | 1 | 19 |
| fam207a | 0.67726 | -0.12258768 | 0.032 | 0.059 | 1 | 19 |
| fis1 | 0.67726 | -0.15025306 | 0.032 | 0.059 | 1 | 19 |
| fut9b | 0.67726 | -0.10142843 | 0.032 | 0.059 | 1 | 19 |
| gltpa | 0.67726 | -0.13929381 | 0.032 | 0.059 | 1 | 19 |
| gmppb | 0.67726 | -0.11353377 | 0.032 | 0.059 | 1 | 19 |
| gorasp2 | 0.67726 | -0.12935722 | 0.032 | 0.059 | 1 | 19 |
| gpr141 | 0.67726 | -0.11075085 | 0.032 | 0.059 | 1 | 19 |
| gps2 | 0.67726 | -0.11138104 | 0.032 | 0.059 | 1 | 19 |
| ing1 | 0.67726 | -0.10943967 | 0.032 | 0.059 | 1 | 19 |
| lsm14ab | 0.67726 | -0.15340792 | 0.032 | 0.059 | 1 | 19 |
| mettl9 | 0.67726 | -0.12580553 | 0.032 | 0.059 | 1 | 19 |
| nkrf | 0.67726 | -0.12214691 | 0.032 | 0.059 | 1 | 19 |
| nop2 | 0.67726 | -0.12663703 | 0.032 | 0.059 | 1 | 19 |
| nrbf2b | 0.67726 | -0.11578571 | 0.032 | 0.059 | 1 | 19 |
| pelo | 0.67726 | -0.15086434 | 0.032 | 0.059 | 1 | 19 |
| pmm2 | 0.67726 | -0.10931894 | 0.032 | 0.059 | 1 | 19 |
| polr3k | 0.67726 | -0.10004034 | 0.032 | 0.059 | 1 | 19 |
| ppm1aa | 0.67726 | -0.10918119 | 0.032 | 0.059 | 1 | 19 |
| prkceb | 0.67726 | -0.10261667 | 0.032 | 0.059 | 1 | 19 |

| | | | | | | |
|---|---|---|---|---|---|---|
| rnaseh1 | 0.67726 | -0.14176614 | 0.032 | 0.059 | 1 | 19 |
| rtcb | 0.67726 | -0.14009658 | 0.032 | 0.059 | 1 | 19 |
| saraf | 0.67726 | -0.11798598 | 0.032 | 0.059 | 1 | 19 |
| sc:d217 | 0.67726 | -0.11152147 | 0.032 | 0.059 | 1 | 19 |
| si:ch211-194k22.8 | 0.67726 | -0.11184793 | 0.032 | 0.059 | 1 | 19 |
| si:ch211-25d12.7 | 0.67726 | -0.11146318 | 0.032 | 0.059 | 1 | 19 |
| si:ch211-93g21.1 | 0.67726 | -0.14500327 | 0.032 | 0.059 | 1 | 19 |
| slc25a11 | 0.67726 | -0.14363179 | 0.032 | 0.059 | 1 | 19 |
| slc25a43 | 0.67726 | -0.1630197 | 0.032 | 0.059 | 1 | 19 |
| snx17 | 0.67726 | -0.16028093 | 0.032 | 0.059 | 1 | 19 |
| spink2.2 | 0.67726 | -0.20468769 | 0.032 | 0.059 | 1 | 19 |
| tgm1l1 | 0.67726 | -0.13659832 | 0.032 | 0.059 | 1 | 19 |
| timm29 | 0.67726 | -0.14711425 | 0.032 | 0.059 | 1 | 19 |
| tmem181 | 0.67726 | -0.14990101 | 0.032 | 0.059 | 1 | 19 |
| tmem209 | 0.67726 | -0.14292297 | 0.032 | 0.059 | 1 | 19 |
| tradd | 0.67726 | -0.14887564 | 0.032 | 0.059 | 1 | 19 |
| ube2f | 0.67726 | -0.10646166 | 0.032 | 0.059 | 1 | 19 |
| ufsp2 | 0.67726 | -0.13202805 | 0.032 | 0.059 | 1 | 19 |
| urm1 | 0.67726 | -0.1145871 | 0.032 | 0.059 | 1 | 19 |
| uxt | 0.67726 | -0.11160457 | 0.032 | 0.059 | 1 | 19 |
| wdr83 | 0.67726 | -0.14863455 | 0.032 | 0.059 | 1 | 19 |
| wipi2 | 0.67726 | -0.14102474 | 0.032 | 0.059 | 1 | 19 |
| zgc:112052 | 0.67726 | -0.12023931 | 0.032 | 0.059 | 1 | 19 |
| zgc:194246 | 0.67726 | -0.14909153 | 0.032 | 0.059 | 1 | 19 |
| zgc:63470 | 0.67726 | -0.10858416 | 0.032 | 0.059 | 1 | 19 |
| znf593 | 0.67726 | -0.10708636 | 0.032 | 0.059 | 1 | 19 |
| CABZ01076275.1 | 0.67726 | -0.13619164 | 0.032 | 0.059 | 1 | 19 |
| CR354588.1 | 0.67726 | -0.14032315 | 0.032 | 0.059 | 1 | 19 |
| CR788248.1 | 0.67726 | -0.13500599 | 0.032 | 0.059 | 1 | 19 |
| CU570894.1 | 0.67726 | -0.13538287 | 0.032 | 0.059 | 1 | 19 |
| LOC101884753 | 0.67726 | -0.10348354 | 0.032 | 0.059 | 1 | 19 |
| LOC103909796 | 0.67726 | -0.18060139 | 0.032 | 0.059 | 1 | 19 |
| LOC103911445 | 0.67726 | -0.10288503 | 0.032 | 0.059 | 1 | 19 |
| LOC110438144 | 0.67726 | -0.16234345 | 0.032 | 0.059 | 1 | 19 |
| acaa1 | 0.67726 | -0.10094117 | 0.032 | 0.059 | 1 | 19 |
| azin1b | 0.67726 | -0.14716862 | 0.032 | 0.059 | 1 | 19 |
| clasrp | 0.67726 | -0.10162557 | 0.032 | 0.059 | 1 | 19 |
| cnot6a | 0.67726 | -0.10628024 | 0.032 | 0.059 | 1 | 19 |
| ctnnb1 | 0.67726 | -0.14217924 | 0.032 | 0.059 | 1 | 19 |
| cul3b | 0.67726 | -0.14591547 | 0.032 | 0.059 | 1 | 19 |
| dcp1b | 0.67726 | -0.14539119 | 0.032 | 0.059 | 1 | 19 |
| dpp3 | 0.67726 | -0.11134894 | 0.032 | 0.059 | 1 | 19 |
| ednraa | 0.67726 | -0.10029123 | 0.032 | 0.059 | 1 | 19 |
| elk1 | 0.67726 | -0.14289165 | 0.032 | 0.059 | 1 | 19 |
| ell2 | 0.67726 | -0.11130833 | 0.032 | 0.059 | 1 | 19 |
| enpp1 | 0.67726 | -0.10238859 | 0.032 | 0.059 | 1 | 19 |
| ero1a | 0.67726 | -0.13859745 | 0.032 | 0.059 | 1 | 19 |
| exoc7 | 0.67726 | -0.12987575 | 0.032 | 0.059 | 1 | 19 |

| | | | | | | |
|---|---|---|---|---|---|---|
| fem1c | 0.67726 | -0.13482743 | 0.032 | 0.059 | 1 | 19 |
| ganab | 0.67726 | -0.1229664 | 0.032 | 0.059 | 1 | 19 |
| gfi1aa | 0.67726 | -0.18471677 | 0.032 | 0.059 | 1 | 19 |
| gigyf1b | 0.67726 | -0.16862627 | 0.032 | 0.059 | 1 | 19 |
| gse1 | 0.67726 | -0.12064099 | 0.032 | 0.059 | 1 | 19 |
| gstp1 | 0.67726 | -0.11945235 | 0.032 | 0.059 | 1 | 19 |
| hdac6 | 0.67726 | -0.14227363 | 0.032 | 0.059 | 1 | 19 |
| hivep2a | 0.67726 | -0.13590379 | 0.032 | 0.059 | 1 | 19 |
| il1b | 0.67726 | -0.1689387 | 0.032 | 0.059 | 1 | 19 |
| impdh2 | 0.67726 | -0.10890882 | 0.032 | 0.059 | 1 | 19 |
| kdm2aa | 0.67726 | -0.11723738 | 0.032 | 0.059 | 1 | 19 |
| kif5ba | 0.67726 | -0.10455243 | 0.032 | 0.059 | 1 | 19 |
| mrpl19 | 0.67726 | -0.10254232 | 0.032 | 0.059 | 1 | 19 |
| mrpl41 | 0.67726 | -0.14551703 | 0.032 | 0.059 | 1 | 19 |
| polr2k | 0.67726 | -0.11547109 | 0.032 | 0.059 | 1 | 19 |
| ptdss1a | 0.67726 | -0.1190626 | 0.032 | 0.059 | 1 | 19 |
| ptrh2 | 0.67726 | -0.13499471 | 0.032 | 0.059 | 1 | 19 |
| rasal2 | 0.67726 | -0.10440645 | 0.032 | 0.059 | 1 | 19 |
| sgsh | 0.67726 | -0.12401754 | 0.032 | 0.059 | 1 | 19 |
| si:ch73-238c9.1 | 0.67726 | -0.10338894 | 0.032 | 0.059 | 1 | 19 |
| si:ch73-248e21.1 | 0.67726 | -0.1622413 | 0.032 | 0.059 | 1 | 19 |
| si:dkey-86e18.1 | 0.67726 | -0.12278606 | 0.032 | 0.059 | 1 | 19 |
| slc41a1 | 0.67726 | -0.10436462 | 0.032 | 0.059 | 1 | 19 |
| soul4 | 0.67726 | -0.12811172 | 0.032 | 0.059 | 1 | 19 |
| ssh2b | 0.67726 | -0.12517405 | 0.032 | 0.059 | 1 | 19 |
| stim1a | 0.67726 | -0.12433948 | 0.032 | 0.059 | 1 | 19 |
| stk40 | 0.67726 | -0.10687802 | 0.032 | 0.059 | 1 | 19 |
| surf1 | 0.67726 | -0.11292989 | 0.032 | 0.059 | 1 | 19 |
| tbc1d14 | 0.67726 | -0.10035593 | 0.032 | 0.059 | 1 | 19 |
| tcf12 | 0.67726 | -0.10136131 | 0.032 | 0.059 | 1 | 19 |
| tcirg1b | 0.67726 | -0.15310433 | 0.032 | 0.059 | 1 | 19 |
| tfe3b | 0.67726 | -0.13376153 | 0.032 | 0.059 | 1 | 19 |
| tmem106a | 0.67726 | -0.10258312 | 0.032 | 0.059 | 1 | 19 |
| tmem56a | 0.67726 | -0.13879062 | 0.032 | 0.059 | 1 | 19 |
| tsr1 | 0.67726 | -0.13266506 | 0.032 | 0.059 | 1 | 19 |
| ttll4 | 0.67726 | -0.10507861 | 0.032 | 0.059 | 1 | 19 |
| uggt1 | 0.67726 | -0.11238833 | 0.032 | 0.059 | 1 | 19 |
| vps36 | 0.67726 | -0.10602851 | 0.032 | 0.059 | 1 | 19 |
| xylt1 | 0.67726 | -0.10863585 | 0.032 | 0.059 | 1 | 19 |
| zeb1b | 0.67726 | -0.15579796 | 0.032 | 0.059 | 1 | 19 |
| zfand3 | 0.67726 | -0.10294061 | 0.032 | 0.059 | 1 | 19 |
| zgc:113232 | 0.67726 | -0.10387573 | 0.032 | 0.059 | 1 | 19 |
| zgc:77056 | 0.67726 | -0.11147591 | 0.032 | 0.059 | 1 | 19 |
| arhgap27 | 0.67744 | -0.17737925 | 0.097 | 0.118 | 1 | 19 |
| eif1b | 0.67803 | -0.20178429 | 0.355 | 0.412 | 1 | 19 |
| il4 | 0.67893 | -0.31283272 | 0.355 | 0.471 | 1 | 19 |
| arhgef3l | 0.67911 | -0.22338871 | 0.129 | 0.176 | 1 | 19 |
| mtpn | 0.67956 | -0.2397334 | 0.258 | 0.353 | 1 | 19 |

| | | | | | | |
|---|---|---|---|---|---|---|
| LOC108190717 | 0.68275 | -0.122436 | 0.097 | 0.176 | 1 | 19 |
| top1l | 0.68366 | -0.23179967 | 0.097 | 0.176 | 1 | 19 |
| sart3 | 0.68397 | 0.191529692 | 0.097 | 0.059 | 1 | 19 |
| fosab | 0.68415 | 0.171575538 | 0.419 | 0.294 | 1 | 19 |
| glipr1a | 0.68568 | 0.240555924 | 0.323 | 0.235 | 1 | 19 |
| kctd12.2 | 0.68585 | 0.205319857 | 0.355 | 0.235 | 1 | 19 |
| isg20 | 0.68607 | 0.208759729 | 0.129 | 0.059 | 1 | 19 |
| cd83 | 0.6868 | -0.21798118 | 0.097 | 0.176 | 1 | 19 |
| ywhabb | 0.68785 | -0.17227321 | 0.258 | 0.294 | 1 | 19 |
| cmc1 | 0.6886 | 0.185269744 | 0.129 | 0.059 | 1 | 19 |
| anxa11b | 0.68894 | 0.126990348 | 0.355 | 0.353 | 1 | 19 |
| znf395b | 0.69032 | -0.22850432 | 0.097 | 0.176 | 1 | 19 |
| kars | 0.69049 | 0.21366363 | 0.194 | 0.118 | 1 | 19 |
| eef1da | 0.69091 | 0.172635697 | 0.71 | 0.647 | 1 | 19 |
| irf2bp2a | 0.69286 | 0.216488537 | 0.323 | 0.235 | 1 | 19 |
| si:ch211-262i1.6 | 0.69318 | -0.21327471 | 0.097 | 0.176 | 1 | 19 |
| eif2s1b | 0.69433 | -0.21500113 | 0.258 | 0.353 | 1 | 19 |
| dnajc5ab | 0.69434 | 0.169498613 | 0.129 | 0.059 | 1 | 19 |
| fam162a | 0.69462 | -0.2032575 | 0.065 | 0.118 | 1 | 19 |
| mphosph6 | 0.69468 | -0.20252934 | 0.097 | 0.176 | 1 | 19 |
| nfkbie | 0.69496 | 0.192310354 | 0.226 | 0.176 | 1 | 19 |
| ebag9 | 0.69549 | 0.180473907 | 0.129 | 0.059 | 1 | 19 |
| CR753876.1 | 0.69722 | -0.17436696 | 0.194 | 0.294 | 1 | 19 |
| rbm39b | 0.69764 | -0.24418025 | 0.097 | 0.176 | 1 | 19 |
| atp5md | 0.6977 | -0.15710248 | 0.387 | 0.412 | 1 | 19 |
| syncrip | 0.69784 | -0.18347547 | 0.29 | 0.412 | 1 | 19 |
| adcy7 | 0.69802 | -0.20646029 | 0.065 | 0.118 | 1 | 19 |
| si:ch211-191c10.1 | 0.6986 | -0.11764908 | 0.355 | 0.353 | 1 | 19 |
| UBL4A | 0.6989 | -0.16989207 | 0.097 | 0.118 | 1 | 19 |
| srd5a2a | 0.69899 | -0.20080017 | 0.129 | 0.176 | 1 | 19 |
| si:ch211-198c19.1 | 0.69964 | 0.204876966 | 0.129 | 0.059 | 1 | 19 |
| bin2b | 0.6999 | 0.210217917 | 0.194 | 0.118 | 1 | 19 |
| nop16 | 0.7001 | 0.102491809 | 0.129 | 0.059 | 1 | 19 |
| ppp3r1a | 0.70018 | 0.10969293 | 0.129 | 0.059 | 1 | 19 |
| zgc:162698 | 0.70037 | -0.18499601 | 0.097 | 0.176 | 1 | 19 |
| nat9 | 0.701 | 0.173078973 | 0.129 | 0.059 | 1 | 19 |
| slc39a7 | 0.70115 | -0.2081112 | 0.065 | 0.118 | 1 | 19 |
| ube2g1b | 0.70194 | -0.20087385 | 0.065 | 0.118 | 1 | 19 |
| zgc:193541 | 0.70219 | -0.21876243 | 0.355 | 0.471 | 1 | 19 |
| FERMT3 (1 of many) | 0.70219 | 0.200355873 | 0.129 | 0.059 | 1 | 19 |
| slc25a5 | 0.70309 | 0.137697955 | 0.871 | 0.824 | 1 | 19 |
| skp1 | 0.70317 | 0.232984005 | 0.258 | 0.176 | 1 | 19 |
| tardbpl | 0.70355 | 0.134794295 | 0.129 | 0.059 | 1 | 19 |
| kpna3 | 0.70357 | -0.20745535 | 0.065 | 0.118 | 1 | 19 |
| erf | 0.70463 | -0.22894305 | 0.065 | 0.118 | 1 | 19 |
| wdr83os | 0.70589 | 0.216370104 | 0.194 | 0.118 | 1 | 19 |
| stxbp2 | 0.70617 | -0.14784461 | 0.194 | 0.294 | 1 | 19 |
| msra | 0.70655 | 0.191719758 | 0.129 | 0.059 | 1 | 19 |

| | | | | | | |
|---|---|---|---|---|---|---|
| denr | 0.70832 | -0.1861973 | 0.097 | 0.176 | 1 | 19 |
| mat2aa | 0.70924 | 0.210500902 | 0.129 | 0.059 | 1 | 19 |
| pdcd10a | 0.70944 | -0.18029818 | 0.097 | 0.176 | 1 | 19 |
| capn2a | 0.70973 | 0.208735298 | 0.129 | 0.059 | 1 | 19 |
| gna13b | 0.7111 | -0.15891352 | 0.161 | 0.176 | 1 | 19 |
| psenen | 0.71147 | 0.186177321 | 0.129 | 0.059 | 1 | 19 |
| serinc2 | 0.7117 | -0.20756726 | 0.065 | 0.118 | 1 | 19 |
| gsna | 0.71197 | -0.1101781 | 0.097 | 0.176 | 1 | 19 |
| il10ra | 0.71203 | -0.20673852 | 0.161 | 0.235 | 1 | 19 |
| sult5a1 | 0.7123 | 0.259129293 | 0.258 | 0.176 | 1 | 19 |
| slc25a33 | 0.7134 | 0.195453462 | 0.097 | 0.059 | 1 | 19 |
| ldhba | 0.71362 | -0.20984796 | 0.194 | 0.294 | 1 | 19 |
| ddx18 | 0.71377 | 0.109168455 | 0.097 | 0.118 | 1 | 19 |
| caprin1b | 0.71377 | 0.216404388 | 0.258 | 0.176 | 1 | 19 |
| ip6k2b | 0.71407 | -0.2185129 | 0.194 | 0.294 | 1 | 19 |
| si:ch1073-110a20.3 | 0.71446 | 0.182813115 | 0.129 | 0.059 | 1 | 19 |
| CABZ01109481.1 | 0.71518 | 0.12632336 | 0.129 | 0.059 | 1 | 19 |
| tdh | 0.71536 | -0.16170869 | 0.097 | 0.176 | 1 | 19 |
| LOC558761 | 0.71555 | 0.159149084 | 0.129 | 0.059 | 1 | 19 |
| fkbp3 | 0.71604 | -0.17417981 | 0.097 | 0.176 | 1 | 19 |
| pxna | 0.71636 | -0.22296914 | 0.161 | 0.235 | 1 | 19 |
| nsfl1c | 0.71751 | 0.147943898 | 0.129 | 0.118 | 1 | 19 |
| snrpb | 0.71792 | 0.205521274 | 0.323 | 0.235 | 1 | 19 |
| cops9 | 0.71873 | -0.17017577 | 0.097 | 0.176 | 1 | 19 |
| pfkpa | 0.71965 | 0.155232579 | 0.129 | 0.059 | 1 | 19 |
| wipf1b | 0.71971 | -0.12100535 | 0.097 | 0.176 | 1 | 19 |
| golga7 | 0.72021 | -0.10402935 | 0.097 | 0.176 | 1 | 19 |
| pfdn6 | 0.7203 | 0.200025635 | 0.194 | 0.118 | 1 | 19 |
| tec | 0.72037 | -0.2179239 | 0.097 | 0.176 | 1 | 19 |
| si:zfos-741a10.3 | 0.72058 | 0.177601426 | 0.129 | 0.059 | 1 | 19 |
| ndufb7 | 0.72092 | -0.21048306 | 0.194 | 0.294 | 1 | 19 |
| itm2bb | 0.72129 | 0.204927944 | 0.581 | 0.471 | 1 | 19 |
| lsm7 | 0.7218 | -0.12753407 | 0.097 | 0.176 | 1 | 19 |
| nup50 | 0.72238 | 0.125925192 | 0.129 | 0.059 | 1 | 19 |
| snx1a | 0.72277 | -0.17108996 | 0.097 | 0.176 | 1 | 19 |
| zgc:171480 | 0.723 | 0.174469341 | 0.097 | 0.059 | 1 | 19 |
| si:dkey-42i9.4 | 0.723 | 0.192406973 | 0.097 | 0.059 | 1 | 19 |
| naa50 | 0.72355 | 0.102791383 | 0.129 | 0.059 | 1 | 19 |
| alyref | 0.72468 | -0.13580986 | 0.097 | 0.176 | 1 | 19 |
| eif5b | 0.7258 | -0.18387027 | 0.129 | 0.176 | 1 | 19 |
| cyb5r3 | 0.72672 | 0.171378918 | 0.129 | 0.059 | 1 | 19 |
| rpa2 | 0.72791 | -0.16398709 | 0.161 | 0.176 | 1 | 19 |
| rgl2 | 0.72829 | 0.141367447 | 0.129 | 0.059 | 1 | 19 |
| ctss1 | 0.7291 | -0.1099312 | 0.29 | 0.294 | 1 | 19 |
| pdap1b | 0.72931 | 0.172018614 | 0.129 | 0.059 | 1 | 19 |
| ndufa3 | 0.7295 | -0.15251302 | 0.065 | 0.118 | 1 | 19 |
| mrpl20 | 0.73001 | 0.126987154 | 0.129 | 0.059 | 1 | 19 |
| slc6a6b | 0.73005 | -0.20622845 | 0.097 | 0.176 | 1 | 19 |

| | | | | | | |
|---|---|---|---|---|---|---|
| psmc3 | 0.73023 | 0.154521399 | 0.129 | 0.059 | 1 | 19 |
| nucb1 | 0.73051 | -0.20229725 | 0.097 | 0.176 | 1 | 19 |
| cct5 | 0.73098 | -0.20309491 | 0.355 | 0.471 | 1 | 19 |
| birc2 | 0.73101 | -0.15340982 | 0.097 | 0.176 | 1 | 19 |
| elavl1a | 0.73111 | 0.116604157 | 0.129 | 0.059 | 1 | 19 |
| tnfrsf9a | 0.73144 | -0.24285574 | 0.323 | 0.412 | 1 | 19 |
| pa2g4b | 0.73184 | 0.146111824 | 0.129 | 0.059 | 1 | 19 |
| fam210aa | 0.7319 | -0.20283072 | 0.065 | 0.118 | 1 | 19 |
| lamtor2 | 0.73191 | 0.16793998 | 0.161 | 0.118 | 1 | 19 |
| m6pr | 0.73252 | 0.15921023 | 0.129 | 0.059 | 1 | 19 |
| ptpn22 | 0.73252 | 0.195603915 | 0.129 | 0.059 | 1 | 19 |
| arl6ip5b | 0.73259 | -0.15751965 | 0.097 | 0.176 | 1 | 19 |
| jagn1b | 0.73278 | 0.148794119 | 0.129 | 0.059 | 1 | 19 |
| zgc:77650 | 0.7328 | 0.158686577 | 0.129 | 0.059 | 1 | 19 |
| si:ch1073-325m22.2 | 0.73282 | -0.18508862 | 0.355 | 0.471 | 1 | 19 |
| uqcrc2b | 0.73293 | 0.150923967 | 0.129 | 0.059 | 1 | 19 |
| tmx1 | 0.73302 | -0.17056775 | 0.097 | 0.176 | 1 | 19 |
| ran | 0.73377 | 0.174100025 | 0.645 | 0.529 | 1 | 19 |
| CR392341.1 | 0.73411 | -0.20337175 | 0.194 | 0.294 | 1 | 19 |
| gata3 | 0.73456 | 0.213368838 | 0.387 | 0.294 | 1 | 19 |
| rap1aa | 0.73481 | -0.17093145 | 0.194 | 0.294 | 1 | 19 |
| grb2b | 0.73492 | 0.123130705 | 0.129 | 0.059 | 1 | 19 |
| grasp | 0.73496 | 0.162528784 | 0.129 | 0.059 | 1 | 19 |
| ndufs3 | 0.73503 | 0.127930262 | 0.129 | 0.059 | 1 | 19 |
| glipr2 | 0.73509 | 0.125404727 | 0.129 | 0.059 | 1 | 19 |
| surf2 | 0.73518 | 0.111592003 | 0.129 | 0.059 | 1 | 19 |
| lyn | 0.73519 | 0.163672068 | 0.129 | 0.059 | 1 | 19 |
| rrs1 | 0.73531 | 0.110966471 | 0.129 | 0.059 | 1 | 19 |
| gnai2a | 0.73542 | -0.18928991 | 0.194 | 0.294 | 1 | 19 |
| egr1 | 0.73612 | -0.13147924 | 0.097 | 0.176 | 1 | 19 |
| gatad2b | 0.7363 | 0.129685226 | 0.129 | 0.059 | 1 | 19 |
| ppp1r9ba | 0.73631 | 0.130765853 | 0.129 | 0.059 | 1 | 19 |
| prkcbb | 0.73634 | -0.19139026 | 0.097 | 0.176 | 1 | 19 |
| si:dkey-12f6.5 | 0.73638 | -0.21212414 | 0.097 | 0.176 | 1 | 19 |
| nifk | 0.73648 | -0.18087646 | 0.065 | 0.118 | 1 | 19 |
| cnpy1 | 0.73673 | -0.16953222 | 0.194 | 0.294 | 1 | 19 |
| coa3a | 0.73703 | -0.11254591 | 0.097 | 0.176 | 1 | 19 |
| rab1ab | 0.73707 | -0.11099097 | 0.097 | 0.176 | 1 | 19 |
| srrm1 | 0.73708 | -0.14593223 | 0.097 | 0.176 | 1 | 19 |
| pgk1 | 0.73722 | -0.13752549 | 0.097 | 0.176 | 1 | 19 |
| zc3h13 | 0.73744 | -0.14753583 | 0.097 | 0.176 | 1 | 19 |
| chd4a | 0.73798 | -0.1212451 | 0.097 | 0.176 | 1 | 19 |
| ncoa4 | 0.73824 | -0.15754501 | 0.097 | 0.176 | 1 | 19 |
| ndufa7 | 0.73834 | -0.12088937 | 0.097 | 0.176 | 1 | 19 |
| suclg1 | 0.73844 | -0.13099374 | 0.097 | 0.176 | 1 | 19 |
| mrpl14 | 0.73844 | -0.16335717 | 0.097 | 0.176 | 1 | 19 |
| ehmt1b | 0.73851 | -0.1498061 | 0.097 | 0.176 | 1 | 19 |
| grap2a | 0.73857 | -0.1529609 | 0.097 | 0.176 | 1 | 19 |

| | | | | | | |
|---|---|---|---|---|---|---|
| ptbp1a | 0.73858 | -0.16739746 | 0.097 | 0.176 | 1 | 19 |
| dazap2 | 0.73858 | -0.15392311 | 0.097 | 0.176 | 1 | 19 |
| prpf8 | 0.73879 | 0.207267003 | 0.194 | 0.118 | 1 | 19 |
| lbh | 0.74115 | 0.164313676 | 0.194 | 0.176 | 1 | 19 |
| gmip | 0.74169 | -0.18880922 | 0.065 | 0.118 | 1 | 19 |
| dub | 0.74416 | 0.21947071 | 0.258 | 0.176 | 1 | 19 |
| sept7b | 0.74483 | -0.1953698 | 0.29 | 0.353 | 1 | 19 |
| atp5mc3b | 0.74529 | 0.192617382 | 0.581 | 0.471 | 1 | 19 |
| sdf4 | 0.74545 | -0.19818853 | 0.065 | 0.118 | 1 | 19 |
| oaz1b | 0.74588 | 0.126783897 | 0.581 | 0.471 | 1 | 19 |
| ube2ia | 0.74604 | -0.18149754 | 0.065 | 0.118 | 1 | 19 |
| eny2 | 0.74649 | 0.188491831 | 0.194 | 0.118 | 1 | 19 |
| higd1a | 0.74693 | -0.1905273 | 0.419 | 0.529 | 1 | 19 |
| taf9 | 0.74845 | -0.15617579 | 0.097 | 0.118 | 1 | 19 |
| ndufs8a | 0.74864 | -0.1861219 | 0.065 | 0.118 | 1 | 19 |
| lsm1 | 0.75051 | 0.15299509 | 0.097 | 0.059 | 1 | 19 |
| tram1 | 0.75274 | 0.198998716 | 0.194 | 0.118 | 1 | 19 |
| cct6a | 0.75354 | -0.17971491 | 0.29 | 0.353 | 1 | 19 |
| psme1 | 0.75421 | 0.141356362 | 0.194 | 0.176 | 1 | 19 |
| arpc4l | 0.75489 | 0.172308812 | 0.581 | 0.471 | 1 | 19 |
| selenow1 | 0.75506 | -0.13373535 | 0.419 | 0.529 | 1 | 19 |
| lsm5 | 0.75609 | 0.192569514 | 0.258 | 0.176 | 1 | 19 |
| MYADM | 0.75612 | -0.19243815 | 0.129 | 0.176 | 1 | 19 |
| btg1 | 0.7578 | 0.18409391 | 0.548 | 0.471 | 1 | 19 |
| pik3cd | 0.75804 | 0.174307452 | 0.194 | 0.118 | 1 | 19 |
| psma3 | 0.75898 | 0.184234691 | 0.323 | 0.235 | 1 | 19 |
| cwf19l2 | 0.75964 | -0.19245407 | 0.065 | 0.118 | 1 | 19 |
| tnpo2 | 0.75974 | -0.1684578 | 0.065 | 0.118 | 1 | 19 |
| spi1b | 0.7598 | -0.18385163 | 0.129 | 0.176 | 1 | 19 |
| cst14a.2 | 0.76008 | 0.143231559 | 0.258 | 0.235 | 1 | 19 |
| caspa | 0.76041 | 0.174400768 | 0.161 | 0.118 | 1 | 19 |
| xpnpep1 | 0.76086 | 0.160151578 | 0.097 | 0.059 | 1 | 19 |
| CU694197.1 | 0.76087 | 0.143030417 | 0.194 | 0.176 | 1 | 19 |
| zgc:64051 | 0.76091 | 0.140094279 | 0.581 | 0.471 | 1 | 19 |
| hspe1 | 0.76181 | 0.177664378 | 0.194 | 0.118 | 1 | 19 |
| smim4 | 0.76219 | 0.176813914 | 0.097 | 0.059 | 1 | 19 |
| dtnbp1a | 0.76385 | -0.1327753 | 0.161 | 0.176 | 1 | 19 |
| mbpb | 0.76424 | -0.13613553 | 0.161 | 0.176 | 1 | 19 |
| ddx39ab | 0.76425 | 0.18207167 | 0.194 | 0.118 | 1 | 19 |
| arf5 | 0.76495 | -0.18201276 | 0.258 | 0.353 | 1 | 19 |
| ugcg | 0.76553 | 0.140600185 | 0.129 | 0.118 | 1 | 19 |
| ndufb9 | 0.7663 | -0.11295569 | 0.258 | 0.353 | 1 | 19 |
| capza1b | 0.76652 | 0.185030746 | 0.419 | 0.353 | 1 | 19 |
| jak1 | 0.7668 | -0.17795782 | 0.065 | 0.118 | 1 | 19 |
| sf3a3 | 0.76717 | -0.16017064 | 0.097 | 0.118 | 1 | 19 |
| dpy30 | 0.76752 | 0.199218608 | 0.161 | 0.118 | 1 | 19 |
| nap1l1 | 0.76889 | -0.16322477 | 0.194 | 0.235 | 1 | 19 |
| LOC566587 | 0.7695 | -0.16457544 | 0.065 | 0.118 | 1 | 19 |

| | | | | | | |
|---|---|---|---|---|---|---|
| magoh | 0.77013 | 0.146800562 | 0.194 | 0.176 | 1 | 19 |
| crip1 | 0.7702 | 0.10799187 | 0.742 | 0.647 | 1 | 19 |
| notch2 | 0.77032 | -0.17168299 | 0.129 | 0.176 | 1 | 19 |
| sdhc | 0.77234 | -0.14865638 | 0.097 | 0.118 | 1 | 19 |
| ap2b1 | 0.77316 | -0.1781271 | 0.065 | 0.118 | 1 | 19 |
| XLOC-033137 | 0.77355 | 0.167295967 | 0.097 | 0.059 | 1 | 19 |
| sept2 | 0.77388 | 0.103766219 | 0.258 | 0.176 | 1 | 19 |
| tegt | 0.77395 | -0.17065079 | 0.258 | 0.353 | 1 | 19 |
| cdipt | 0.77413 | 0.195860008 | 0.194 | 0.118 | 1 | 19 |
| cct7 | 0.77492 | 0.189748948 | 0.387 | 0.294 | 1 | 19 |
| tmem39a | 0.77537 | -0.14880819 | 0.065 | 0.118 | 1 | 19 |
| BX901920.1 | 0.77544 | 0.189447078 | 0.194 | 0.118 | 1 | 19 |
| psmc5 | 0.77557 | -0.16184058 | 0.065 | 0.118 | 1 | 19 |
| CR318588.1 | 0.77591 | -0.10467616 | 0.065 | 0.118 | 1 | 19 |
| cmip | 0.77596 | -0.18067171 | 0.129 | 0.176 | 1 | 19 |
| rab14 | 0.77731 | -0.16167774 | 0.065 | 0.118 | 1 | 19 |
| gpn3 | 0.77732 | 0.166422234 | 0.097 | 0.059 | 1 | 19 |
| MZT1 | 0.77735 | 0.166056238 | 0.097 | 0.059 | 1 | 19 |
| sh3bp2 | 0.77797 | -0.18506913 | 0.129 | 0.176 | 1 | 19 |
| fli1b | 0.77984 | 0.161728132 | 0.097 | 0.059 | 1 | 19 |
| rer1 | 0.77997 | 0.121706166 | 0.194 | 0.118 | 1 | 19 |
| reep5 | 0.78019 | -0.12578757 | 0.484 | 0.588 | 1 | 19 |
| esf1 | 0.78051 | -0.18175999 | 0.065 | 0.118 | 1 | 19 |
| snrpa1 | 0.78197 | 0.169520458 | 0.161 | 0.118 | 1 | 19 |
| nop58 | 0.7833 | 0.127891777 | 0.258 | 0.176 | 1 | 19 |
| tes | 0.78331 | -0.14635877 | 0.258 | 0.353 | 1 | 19 |
| rca2.1 | 0.78351 | -0.16984575 | 0.129 | 0.176 | 1 | 19 |
| eif3m | 0.78531 | 0.105922519 | 0.419 | 0.412 | 1 | 19 |
| rbx1 | 0.78557 | -0.12939061 | 0.258 | 0.353 | 1 | 19 |
| fmnl1a | 0.78561 | -0.17545616 | 0.065 | 0.118 | 1 | 19 |
| rab11ba | 0.78603 | 0.116365308 | 0.194 | 0.118 | 1 | 19 |
| ptk2ba | 0.78688 | 0.162407792 | 0.097 | 0.059 | 1 | 19 |
| nap1l4b | 0.78828 | -0.15144097 | 0.065 | 0.118 | 1 | 19 |
| ches1 | 0.78845 | -0.15471536 | 0.065 | 0.118 | 1 | 19 |
| alas1 | 0.78865 | 0.157407881 | 0.194 | 0.118 | 1 | 19 |
| LOC100332187 | 0.78879 | -0.1808667 | 0.065 | 0.118 | 1 | 19 |
| anxa4 | 0.78916 | -0.15760846 | 0.258 | 0.353 | 1 | 19 |
| tpm3 | 0.78917 | -0.1062227 | 0.516 | 0.529 | 1 | 19 |
| ndufb6 | 0.78981 | 0.111713153 | 0.194 | 0.118 | 1 | 19 |
| anapc16 | 0.78999 | 0.143423627 | 0.194 | 0.118 | 1 | 19 |
| CABZ01055347.1 | 0.79018 | 0.132875853 | 0.194 | 0.118 | 1 | 19 |
| ap1s1 | 0.79021 | 0.147467466 | 0.194 | 0.118 | 1 | 19 |
| dusp2 | 0.79062 | -0.20242073 | 0.258 | 0.353 | 1 | 19 |
| arl6ip1 | 0.79088 | 0.192410337 | 0.258 | 0.176 | 1 | 19 |
| alox5ap | 0.79105 | -0.11110383 | 0.161 | 0.235 | 1 | 19 |
| ipo7 | 0.79138 | -0.1509396 | 0.065 | 0.118 | 1 | 19 |
| aurkaip1 | 0.7915 | -0.12142137 | 0.065 | 0.118 | 1 | 19 |
| ddx31 | 0.79178 | -0.17585751 | 0.065 | 0.118 | 1 | 19 |

| | | | | | | |
|---|---|---|---|---|---|---|
| osgn1 | 0.79186 | -0.149829 | 0.065 | 0.118 | 1 | 19 |
| aldh6a1 | 0.79206 | -0.15137243 | 0.065 | 0.118 | 1 | 19 |
| ccl33.3 | 0.79294 | -0.17349402 | 0.161 | 0.235 | 1 | 19 |
| micu2 | 0.79381 | -0.1738371 | 0.065 | 0.118 | 1 | 19 |
| cltcb | 0.79387 | -0.1440791 | 0.065 | 0.118 | 1 | 19 |
| si:dkey-177p2.6 | 0.79564 | -0.14825799 | 0.097 | 0.118 | 1 | 19 |
| sf3b2 | 0.79609 | -0.15313228 | 0.065 | 0.118 | 1 | 19 |
| CABZ01113812.1 | 0.7962 | -0.15879246 | 0.065 | 0.118 | 1 | 19 |
| cmpk | 0.79815 | -0.12516131 | 0.161 | 0.176 | 1 | 19 |
| slc20a1b | 0.79848 | -0.13323201 | 0.065 | 0.118 | 1 | 19 |
| vcp | 0.79944 | -0.14616133 | 0.161 | 0.235 | 1 | 19 |
| arf6a | 0.79975 | 0.170167626 | 0.258 | 0.176 | 1 | 19 |
| sh3bp1 | 0.7999 | -0.14905024 | 0.065 | 0.118 | 1 | 19 |
| atp6v0d1 | 0.80003 | -0.16011473 | 0.065 | 0.118 | 1 | 19 |
| serpine2 | 0.80009 | -0.14427468 | 0.161 | 0.176 | 1 | 19 |
| mrpl42 | 0.80025 | -0.13255261 | 0.065 | 0.118 | 1 | 19 |
| selenok | 0.80141 | -0.17623744 | 0.226 | 0.294 | 1 | 19 |
| cnbpb | 0.80162 | 0.167823761 | 0.226 | 0.176 | 1 | 19 |
| si:dkey-286j15.3 | 0.80202 | -0.15360949 | 0.065 | 0.118 | 1 | 19 |
| si:rp71-36a1.3 | 0.80314 | -0.14525697 | 0.065 | 0.118 | 1 | 19 |
| prps1a | 0.80329 | -0.14154996 | 0.065 | 0.118 | 1 | 19 |
| CABZ01084575.1 | 0.80358 | 0.144605604 | 0.387 | 0.294 | 1 | 19 |
| tmed7 | 0.8036 | 0.172153126 | 0.323 | 0.235 | 1 | 19 |
| txn2 | 0.8039 | -0.14509685 | 0.097 | 0.118 | 1 | 19 |
| ccng1 | 0.80412 | 0.181183528 | 0.323 | 0.235 | 1 | 19 |
| CABZ01073834.1 | 0.80427 | -0.13477813 | 0.323 | 0.353 | 1 | 19 |
| nansa | 0.80433 | -0.17245565 | 0.129 | 0.176 | 1 | 19 |
| ccl34b.4 | 0.80506 | 0.221309919 | 0.258 | 0.176 | 1 | 19 |
| rhoaa | 0.80512 | 0.152550671 | 0.161 | 0.118 | 1 | 19 |
| ptp4a2b | 0.80515 | 0.127133778 | 0.258 | 0.176 | 1 | 19 |
| ISCU (1 of many) | 0.80587 | 0.148111217 | 0.258 | 0.176 | 1 | 19 |
| rcc2 | 0.80593 | 0.144755219 | 0.097 | 0.059 | 1 | 19 |
| mcl1a | 0.80612 | -0.1032584 | 0.677 | 0.765 | 1 | 19 |
| fam107b | 0.80616 | -0.16348839 | 0.065 | 0.118 | 1 | 19 |
| wnk1b | 0.8062 | -0.1664867 | 0.065 | 0.118 | 1 | 19 |
| itm2cb | 0.80644 | 0.155857982 | 0.097 | 0.059 | 1 | 19 |
| psmd11b | 0.8071 | -0.15330018 | 0.065 | 0.118 | 1 | 19 |
| arhgef2 | 0.80752 | -0.13313276 | 0.065 | 0.118 | 1 | 19 |
| scgn | 0.80827 | -0.10187895 | 0.161 | 0.235 | 1 | 19 |
| myh11a | 0.80877 | -0.1730002 | 0.161 | 0.235 | 1 | 19 |
| zmp:0000000521 | 0.80926 | 0.150724135 | 0.097 | 0.059 | 1 | 19 |
| hypk | 0.80926 | -0.17289497 | 0.161 | 0.235 | 1 | 19 |
| zgc:153867 | 0.80956 | -0.10039926 | 0.677 | 0.765 | 1 | 19 |
| rhoab | 0.80993 | 0.13690198 | 0.323 | 0.294 | 1 | 19 |
| si:rp71-36a1.1 | 0.81038 | 0.12496069 | 0.129 | 0.118 | 1 | 19 |
| rnf130 | 0.81049 | -0.15903372 | 0.194 | 0.235 | 1 | 19 |
| kmt2d | 0.81062 | -0.12705119 | 0.065 | 0.118 | 1 | 19 |
| arhgef6 | 0.81144 | 0.162559528 | 0.161 | 0.118 | 1 | 19 |

| | | | | | | |
|---|---|---|---|---|---|---|
| prmt1 | 0.81185 | 0.15786543 | 0.323 | 0.235 | 1 | 19 |
| eif3g | 0.81193 | -0.17744013 | 0.226 | 0.294 | 1 | 19 |
| ak1 | 0.81283 | -0.1495818 | 0.065 | 0.118 | 1 | 19 |
| xiap | 0.81415 | -0.12733084 | 0.065 | 0.118 | 1 | 19 |
| ubash3ba | 0.81449 | 0.153561958 | 0.097 | 0.059 | 1 | 19 |
| gnb1b | 0.81535 | -0.11946161 | 0.323 | 0.412 | 1 | 19 |
| pdap1a | 0.81536 | 0.142009739 | 0.097 | 0.059 | 1 | 19 |
| shoc2 | 0.81545 | 0.149689772 | 0.097 | 0.059 | 1 | 19 |
| jpt1b | 0.81584 | -0.17600786 | 0.065 | 0.118 | 1 | 19 |
| lamp2 | 0.81584 | -0.15200507 | 0.097 | 0.118 | 1 | 19 |
| hars | 0.81599 | 0.150474921 | 0.097 | 0.059 | 1 | 19 |
| elmo1 | 0.81626 | 0.120714349 | 0.129 | 0.118 | 1 | 19 |
| plpp5 | 0.81689 | -0.10541117 | 0.065 | 0.118 | 1 | 19 |
| baz2a | 0.81693 | -0.12227272 | 0.065 | 0.118 | 1 | 19 |
| ssu72 | 0.81696 | -0.13069847 | 0.065 | 0.118 | 1 | 19 |
| prkcsh | 0.81714 | -0.1570508 | 0.161 | 0.235 | 1 | 19 |
| ppm1j | 0.81782 | -0.14057353 | 0.065 | 0.118 | 1 | 19 |
| serbp1b | 0.81812 | -0.13457434 | 0.097 | 0.118 | 1 | 19 |
| XLOC-010899 | 0.81821 | -0.17590884 | 0.29 | 0.353 | 1 | 19 |
| si:dkey-250d21.1 | 0.81825 | -0.17390698 | 0.129 | 0.176 | 1 | 19 |
| fbxl6 | 0.81839 | -0.10366059 | 0.065 | 0.118 | 1 | 19 |
| ngdn | 0.81845 | -0.12807052 | 0.065 | 0.118 | 1 | 19 |
| syne2b | 0.81845 | -0.15429328 | 0.065 | 0.118 | 1 | 19 |
| si:dkey-27h10.2 | 0.81871 | -0.1712278 | 0.129 | 0.176 | 1 | 19 |
| si:ch211-67e16.3 | 0.81901 | -0.14553547 | 0.065 | 0.118 | 1 | 19 |
| sec24a | 0.82042 | -0.13051968 | 0.065 | 0.118 | 1 | 19 |
| vdac2 | 0.82068 | 0.164563999 | 0.29 | 0.235 | 1 | 19 |
| myca | 0.82117 | -0.15387683 | 0.129 | 0.176 | 1 | 19 |
| mrps10 | 0.82135 | -0.13587894 | 0.097 | 0.118 | 1 | 19 |
| rnaseka | 0.82135 | -0.12462215 | 0.161 | 0.235 | 1 | 19 |
| ncf4 | 0.82142 | -0.12335487 | 0.065 | 0.118 | 1 | 19 |
| ubr7 | 0.82145 | -0.12839281 | 0.065 | 0.118 | 1 | 19 |
| mrpl11 | 0.82206 | 0.142271842 | 0.097 | 0.059 | 1 | 19 |
| polr2i | 0.82213 | -0.11592732 | 0.065 | 0.118 | 1 | 19 |
| LOC100148254 | 0.82227 | -0.1648383 | 0.129 | 0.176 | 1 | 19 |
| derl1 | 0.82278 | -0.1298312 | 0.065 | 0.118 | 1 | 19 |
| polr2f | 0.82287 | -0.15352426 | 0.194 | 0.235 | 1 | 19 |
| mpc1 | 0.82288 | -0.17007897 | 0.129 | 0.176 | 1 | 19 |
| LOC110437917 | 0.82308 | 0.142867701 | 0.097 | 0.059 | 1 | 19 |
| ctps1a | 0.82319 | -0.1010694 | 0.065 | 0.118 | 1 | 19 |
| dpm1 | 0.82323 | 0.13868814 | 0.097 | 0.059 | 1 | 19 |
| stk10 | 0.82332 | -0.11418093 | 0.065 | 0.118 | 1 | 19 |
| cox6b1 | 0.82389 | -0.12528369 | 0.387 | 0.412 | 1 | 19 |
| ufc1 | 0.824 | -0.14705428 | 0.161 | 0.235 | 1 | 19 |
| itsn2b | 0.82423 | -0.12870292 | 0.065 | 0.118 | 1 | 19 |
| cbfb | 0.82429 | -0.11048508 | 0.065 | 0.118 | 1 | 19 |
| eed | 0.82445 | -0.12747899 | 0.065 | 0.118 | 1 | 19 |
| si:zfos-169g10.3 | 0.82465 | -0.10319617 | 0.065 | 0.118 | 1 | 19 |

| | | | | | | |
|---|---|---|---|---|---|---|
| baz2ba | 0.82466 | 0.134017698 | 0.097 | 0.059 | 1 | 19 |
| itm2ba | 0.82468 | -0.15384427 | 0.161 | 0.235 | 1 | 19 |
| XLOC-013964 | 0.82472 | -0.17264087 | 0.161 | 0.235 | 1 | 19 |
| gskip | 0.82472 | -0.12141281 | 0.161 | 0.235 | 1 | 19 |
| selplg | 0.82474 | -0.12867889 | 0.161 | 0.235 | 1 | 19 |
| atp5mc3a | 0.82504 | -0.15096371 | 0.323 | 0.412 | 1 | 19 |
| dse | 0.82533 | -0.12340363 | 0.065 | 0.118 | 1 | 19 |
| tm9sf1 | 0.82546 | -0.11352992 | 0.065 | 0.118 | 1 | 19 |
| cox8b | 0.82546 | 0.13624295 | 0.194 | 0.176 | 1 | 19 |
| atrx | 0.82572 | -0.12966504 | 0.161 | 0.235 | 1 | 19 |
| sec31a | 0.82577 | -0.13980952 | 0.065 | 0.118 | 1 | 19 |
| CU571074.1 | 0.82586 | -0.10059441 | 0.065 | 0.118 | 1 | 19 |
| dnase1l4.2 | 0.82593 | -0.12179277 | 0.065 | 0.118 | 1 | 19 |
| nucb2a | 0.82658 | -0.11234774 | 0.065 | 0.118 | 1 | 19 |
| ftr50 | 0.8266 | -0.11874228 | 0.065 | 0.118 | 1 | 19 |
| rab10 | 0.82665 | -0.11828016 | 0.065 | 0.118 | 1 | 19 |
| sptbn2 | 0.82665 | -0.10034158 | 0.065 | 0.118 | 1 | 19 |
| impa1 | 0.82669 | -0.12184882 | 0.065 | 0.118 | 1 | 19 |
| mknk2b | 0.82676 | -0.13369754 | 0.065 | 0.118 | 1 | 19 |
| ppp4r3b | 0.82682 | -0.1120446 | 0.065 | 0.118 | 1 | 19 |
| dnajc5gb | 0.82683 | -0.12495229 | 0.065 | 0.118 | 1 | 19 |
| anp32b | 0.8301 | -0.13371229 | 0.161 | 0.176 | 1 | 19 |
| rpa1 | 0.83091 | 0.168690816 | 0.097 | 0.059 | 1 | 19 |
| hint1 | 0.83115 | -0.1559667 | 0.194 | 0.235 | 1 | 19 |
| si:ch73-62l21.1 | 0.83328 | 0.133121128 | 0.097 | 0.059 | 1 | 19 |
| XLOC-043857 | 0.83378 | 0.125893905 | 0.097 | 0.059 | 1 | 19 |
| nccrp1 | 0.83456 | -0.12602826 | 0.097 | 0.118 | 1 | 19 |
| casp3a | 0.83476 | 0.135566821 | 0.097 | 0.059 | 1 | 19 |
| aars | 0.83482 | 0.149506287 | 0.161 | 0.118 | 1 | 19 |
| zfand5a | 0.83746 | -0.1289407 | 0.097 | 0.118 | 1 | 19 |
| stat1a | 0.8404 | 0.12721257 | 0.097 | 0.059 | 1 | 19 |
| psma8 | 0.84086 | 0.153184877 | 0.161 | 0.118 | 1 | 19 |
| ppib | 0.84093 | 0.145772947 | 0.452 | 0.412 | 1 | 19 |
| atp5f1b | 0.84141 | -0.13725276 | 0.484 | 0.529 | 1 | 19 |
| timm8a | 0.84318 | 0.126711236 | 0.097 | 0.059 | 1 | 19 |
| mrpl39 | 0.84328 | 0.115321657 | 0.097 | 0.059 | 1 | 19 |
| mapkapk2a | 0.84624 | 0.113535888 | 0.129 | 0.118 | 1 | 19 |
| emc2 | 0.84702 | 0.116645347 | 0.097 | 0.059 | 1 | 19 |
| fndc3a | 0.84752 | 0.13983617 | 0.097 | 0.059 | 1 | 19 |
| gtf2f1 | 0.84797 | 0.115012496 | 0.097 | 0.059 | 1 | 19 |
| nhp2 | 0.84824 | 0.140904276 | 0.29 | 0.235 | 1 | 19 |
| eif4g2a | 0.84836 | 0.120726972 | 0.097 | 0.059 | 1 | 19 |
| spcs3 | 0.85031 | -0.11629013 | 0.323 | 0.353 | 1 | 19 |
| il16 | 0.85038 | 0.1555174 | 0.226 | 0.176 | 1 | 19 |
| hnrnpa0a | 0.8529 | 0.112217804 | 0.129 | 0.118 | 1 | 19 |
| imp4 | 0.85308 | 0.143199034 | 0.097 | 0.059 | 1 | 19 |
| si:ch73-1a9.3 | 0.85337 | -0.16895331 | 0.226 | 0.294 | 1 | 19 |
| il34 | 0.85508 | 0.136579545 | 0.258 | 0.235 | 1 | 19 |

| | | | | | | |
|---|---|---|---|---|---|---|
| ap1m1 | 0.85516 | -0.12473165 | 0.097 | 0.118 | 1 | 19 |
| mylipb | 0.8579 | 0.109579976 | 0.097 | 0.059 | 1 | 19 |
| dbnlb | 0.8585 | 0.154540435 | 0.161 | 0.118 | 1 | 19 |
| tmem189 | 0.85872 | 0.114626952 | 0.097 | 0.059 | 1 | 19 |
| psmd1 | 0.85917 | 0.115706272 | 0.097 | 0.059 | 1 | 19 |
| wbp1lb | 0.86085 | 0.116443083 | 0.097 | 0.059 | 1 | 19 |
| fcer1gl | 0.86217 | 0.128911826 | 0.581 | 0.529 | 1 | 19 |
| nip7 | 0.86234 | 0.110132657 | 0.097 | 0.059 | 1 | 19 |
| smarca5 | 0.86329 | 0.112173239 | 0.097 | 0.059 | 1 | 19 |
| tp53rk | 0.86441 | 0.140695411 | 0.097 | 0.059 | 1 | 19 |
| canx | 0.86626 | -0.15175396 | 0.226 | 0.294 | 1 | 19 |
| ubr5 | 0.86699 | 0.111653086 | 0.097 | 0.059 | 1 | 19 |
| mhc2b | 0.86737 | -0.14619918 | 0.129 | 0.176 | 1 | 19 |
| spg21 | 0.86787 | -0.11479422 | 0.097 | 0.118 | 1 | 19 |
| trappc13 | 0.86849 | 0.119169447 | 0.097 | 0.059 | 1 | 19 |
| mrps33 | 0.86871 | 0.103461996 | 0.097 | 0.059 | 1 | 19 |
| msna | 0.871 | -0.1307941 | 0.516 | 0.588 | 1 | 19 |
| acap1 | 0.87181 | -0.11627825 | 0.097 | 0.118 | 1 | 19 |
| tmod2 | 0.87232 | 0.122556025 | 0.097 | 0.059 | 1 | 19 |
| cirbpb | 0.87259 | -0.11334578 | 0.71 | 0.765 | 1 | 19 |
| stk17al | 0.87274 | 0.131733523 | 0.097 | 0.059 | 1 | 19 |
| cdkn2aipnl | 0.87383 | 0.106198401 | 0.129 | 0.118 | 1 | 19 |
| mphosph8 | 0.87558 | 0.119548058 | 0.097 | 0.059 | 1 | 19 |
| tmem59 | 0.87763 | -0.11995209 | 0.161 | 0.176 | 1 | 19 |
| tspan37 | 0.87802 | -0.11930379 | 0.097 | 0.118 | 1 | 19 |
| taf15 | 0.87912 | -0.12029647 | 0.129 | 0.176 | 1 | 19 |
| rad21a | 0.87973 | -0.10602492 | 0.161 | 0.176 | 1 | 19 |
| cox5aa | 0.87981 | -0.11634421 | 0.548 | 0.588 | 1 | 19 |
| ndufb5 | 0.88026 | 0.108061755 | 0.194 | 0.176 | 1 | 19 |
| ak2 | 0.88074 | -0.12979635 | 0.355 | 0.412 | 1 | 19 |
| sap30l | 0.88078 | 0.108181656 | 0.258 | 0.235 | 1 | 19 |
| gmps | 0.88262 | -0.11327162 | 0.097 | 0.118 | 1 | 19 |
| arcn1a | 0.88284 | 0.1104928 | 0.097 | 0.059 | 1 | 19 |
| ywhah | 0.88316 | 0.104836535 | 0.097 | 0.059 | 1 | 19 |
| zgc:66427 | 0.88397 | 0.116073114 | 0.097 | 0.059 | 1 | 19 |
| rp2 | 0.88603 | -0.11307995 | 0.097 | 0.118 | 1 | 19 |
| eif2s3 | 0.88616 | 0.123370047 | 0.484 | 0.412 | 1 | 19 |
| csnk1g2b | 0.88733 | 0.115093983 | 0.161 | 0.118 | 1 | 19 |
| arhgap20b | 0.88945 | 0.103475592 | 0.097 | 0.059 | 1 | 19 |
| ndufa4l | 0.88974 | -0.11328329 | 0.613 | 0.647 | 1 | 19 |
| fam49ba | 0.88979 | -0.10992717 | 0.323 | 0.353 | 1 | 19 |
| gtpbp4 | 0.89217 | 0.120667059 | 0.29 | 0.235 | 1 | 19 |
| mef2b | 0.8923 | 0.103932304 | 0.097 | 0.059 | 1 | 19 |
| polr2eb | 0.89412 | -0.11062742 | 0.129 | 0.176 | 1 | 19 |
| si:ch211-196l7.4 | 0.89415 | 0.122158485 | 0.419 | 0.353 | 1 | 19 |
| psd3l | 0.89431 | 0.106922181 | 0.097 | 0.059 | 1 | 19 |
| taf12 | 0.89506 | -0.1161131 | 0.129 | 0.176 | 1 | 19 |
| timm17a | 0.89575 | 0.116675501 | 0.097 | 0.059 | 1 | 19 |

| | | | | | | |
|---|---|---|---|---|---|---|
| BOLA2 | 0.89577 | 0.132131192 | 0.161 | 0.118 | 1 | 19 |
| trim35-28 | 0.8998 | 0.101117182 | 0.097 | 0.059 | 1 | 19 |
| bcl2l1 | 0.9008 | 0.115415521 | 0.161 | 0.118 | 1 | 19 |
| ubb | 0.90224 | 0.124895067 | 0.226 | 0.176 | 1 | 19 |
| eif3c | 0.90235 | 0.107907799 | 0.355 | 0.294 | 1 | 19 |
| bccip | 0.90313 | 0.107624736 | 0.161 | 0.118 | 1 | 19 |
| si:ch1073-443f11.2 | 0.90527 | 0.111073096 | 0.161 | 0.118 | 1 | 19 |
| syf2 | 0.90728 | 0.112091245 | 0.226 | 0.176 | 1 | 19 |
| pdia6 | 0.9078 | -0.10311503 | 0.129 | 0.176 | 1 | 19 |
| slc2a3b | 0.90813 | -0.10708674 | 0.129 | 0.176 | 1 | 19 |
| rbm4.3 | 0.90816 | 0.104412567 | 0.258 | 0.235 | 1 | 19 |
| nop56 | 0.9086 | 0.1151931 | 0.355 | 0.294 | 1 | 19 |
| si:dkey-21e2.3 | 0.90893 | 0.100785948 | 0.29 | 0.235 | 1 | 19 |
| ndufb4 | 0.91117 | -0.10723728 | 0.323 | 0.353 | 1 | 19 |
| atp5f1c | 0.91227 | 0.114506523 | 0.29 | 0.235 | 1 | 19 |
| cxcr4b | 0.9183 | 0.137209502 | 0.226 | 0.176 | 1 | 19 |
| ciarta | 0.92019 | 0.102682653 | 0.226 | 0.176 | 1 | 19 |
| c1qbp | 0.92185 | -0.10308507 | 0.194 | 0.235 | 1 | 19 |
| mob3a | 0.92322 | -0.10028231 | 0.355 | 0.412 | 1 | 19 |
| sarnp | 0.93308 | -0.10472649 | 0.194 | 0.235 | 1 | 19 |
| plekho1b | 0.93898 | 0.100775966 | 0.065 | 0.059 | 1 | 19 |
| riok2 | 0.93898 | 0.119043121 | 0.065 | 0.059 | 1 | 19 |
| scaf4b | 0.93898 | 0.150429964 | 0.065 | 0.059 | 1 | 19 |
| ssrp1a | 0.93898 | -0.10547751 | 0.065 | 0.059 | 1 | 19 |
| atg5 | 0.93898 | 0.103532959 | 0.065 | 0.059 | 1 | 19 |
| ccl38.6 | 0.93898 | -0.11456534 | 0.065 | 0.059 | 1 | 19 |
| cpsf3 | 0.93898 | 0.127117009 | 0.065 | 0.059 | 1 | 19 |
| ctdsp1 | 0.93898 | 0.118069113 | 0.065 | 0.059 | 1 | 19 |
| dcaf7 | 0.93898 | 0.109723457 | 0.065 | 0.059 | 1 | 19 |
| iars | 0.93898 | 0.113289481 | 0.065 | 0.059 | 1 | 19 |
| prkacab | 0.93898 | 0.103588315 | 0.065 | 0.059 | 1 | 19 |
| prkcq | 0.93898 | 0.116034711 | 0.065 | 0.059 | 1 | 19 |
| rexo4 | 0.93898 | 0.128405169 | 0.065 | 0.059 | 1 | 19 |
| taf11 | 0.93898 | 0.146781523 | 0.065 | 0.059 | 1 | 19 |
| usp37 | 0.93898 | 0.136067202 | 0.065 | 0.059 | 1 | 19 |
| gid8b | 0.93898 | -0.10528478 | 0.065 | 0.059 | 1 | 19 |
| pno1 | 0.93898 | 0.10092976 | 0.065 | 0.059 | 1 | 19 |
| ube2l3b | 0.93898 | 0.102948941 | 0.065 | 0.059 | 1 | 19 |
| casp8ap2 | 0.93898 | 0.147432356 | 0.065 | 0.059 | 1 | 19 |
| etfdh | 0.93898 | 0.129622259 | 0.065 | 0.059 | 1 | 19 |
| mrps7 | 0.93898 | -0.1083319 | 0.065 | 0.059 | 1 | 19 |
| rnaset2 | 0.93898 | 0.10628243 | 0.065 | 0.059 | 1 | 19 |
| sec61a1l | 0.93898 | 0.10419595 | 0.065 | 0.059 | 1 | 19 |
| spink2.3 | 0.93898 | 0.1302825 | 0.065 | 0.059 | 1 | 19 |
| zdhhc13 | 0.93898 | -0.1226741 | 0.065 | 0.059 | 1 | 19 |
| brms1la | 0.93898 | 0.161373242 | 0.065 | 0.059 | 1 | 19 |
| mt-co2 | 0.00013 | -1.06247651 | 0.094 | 0.667 | 1 | 20 |
| mt-nd4 | 0.00014 | -0.99623847 | 0 | 0.4 | 1 | 20 |

| | | | | | | |
|---|---|---|---|---|---|---|
| COX3 | 0.00017 | -1.28628273 | 0.125 | 0.733 | 1 | 20 |
| rps2 | 0.0003 | 0.490775999 | 1 | 0.933 | 1 | 20 |
| RASSF5 | 0.00039 | -1.17178088 | 0.094 | 0.533 | 1 | 20 |
| bzw1a | 0.00044 | 0.734044585 | 0.438 | 0 | 1 | 20 |
| rpl18 | 0.00046 | 0.594208732 | 0.906 | 0.733 | 1 | 20 |
| uba52 | 0.00058 | 0.574136664 | 1 | 0.6 | 1 | 20 |
| naa38 | 0.00066 | -0.58462278 | 0 | 0.333 | 1 | 20 |
| nifk | 0.00082 | 0.499071793 | 0.406 | 0 | 1 | 20 |
| blvrb | 0.0009 | -1.0708147 | 0.219 | 0.533 | 1 | 20 |
| rpl11 | 0.00105 | 0.572992922 | 1 | 0.8 | 1 | 20 |
| rplp0 | 0.00119 | 0.572529732 | 0.969 | 0.8 | 1 | 20 |
| mibp2 | 0.00125 | 0.691833157 | 0.906 | 0.4 | 1 | 20 |
| cct8 | 0.00134 | 0.824153839 | 0.719 | 0.267 | 1 | 20 |
| anp32b | 0.00135 | 0.54738555 | 0.531 | 0.067 | 1 | 20 |
| si:ch211-288g17.3 | 0.00136 | 0.367480318 | 0.469 | 0.067 | 1 | 20 |
| rps9 | 0.0014 | 0.668874997 | 0.969 | 0.533 | 1 | 20 |
| mybbp1a | 0.00149 | 0.480148718 | 0.375 | 0 | 1 | 20 |
| wdr83os | 0.00149 | 0.494298915 | 0.375 | 0 | 1 | 20 |
| hspe1 | 0.00149 | 0.633477384 | 0.375 | 0 | 1 | 20 |
| rpl35a | 0.00153 | 0.462293902 | 0.938 | 0.8 | 1 | 20 |
| nme2b.1 | 0.00239 | 0.366814998 | 0.938 | 0.867 | 1 | 20 |
| rps27.1 | 0.00247 | 0.241690028 | 0.906 | 0.933 | 1 | 20 |
| rpl7 | 0.00256 | 0.522819888 | 0.906 | 0.733 | 1 | 20 |
| cdk2ap2 | 0.00266 | 0.489728911 | 0.344 | 0 | 1 | 20 |
| mpc2 | 0.00266 | 0.422705548 | 0.344 | 0 | 1 | 20 |
| pdap1a | 0.00266 | 0.518624571 | 0.344 | 0 | 1 | 20 |
| stip1 | 0.00266 | 0.428449628 | 0.344 | 0 | 1 | 20 |
| zgc:162025 | 0.00266 | 0.347976214 | 0.344 | 0 | 1 | 20 |
| psmb4 | 0.00269 | 0.714871641 | 0.531 | 0.067 | 1 | 20 |
| npm1a | 0.00269 | 0.84885292 | 0.469 | 0.2 | 1 | 20 |
| rps27a | 0.00278 | 0.536683408 | 0.938 | 0.733 | 1 | 20 |
| rpl7a | 0.00279 | 0.583027974 | 0.969 | 0.667 | 1 | 20 |
| cavin1b | 0.00288 | -0.44847253 | 0 | 0.267 | 1 | 20 |
| cpn1 | 0.00288 | -0.86596744 | 0 | 0.267 | 1 | 20 |
| hcls1 | 0.00288 | -0.72252366 | 0 | 0.267 | 1 | 20 |
| rps16 | 0.00304 | 0.458604971 | 0.969 | 0.733 | 1 | 20 |
| gpia | 0.00321 | 0.272678446 | 0.438 | 0.067 | 1 | 20 |
| rpl19 | 0.00359 | 0.535280057 | 0.969 | 0.733 | 1 | 20 |
| ran | 0.00362 | 0.731034834 | 0.875 | 0.4 | 1 | 20 |
| rpl30 | 0.00362 | 0.443765509 | 0.906 | 0.8 | 1 | 20 |
| eif3f | 0.0039 | 0.663145128 | 0.906 | 0.467 | 1 | 20 |
| pnn | 0.00399 | 0.303001073 | 0.406 | 0.067 | 1 | 20 |
| rpl27 | 0.00399 | 0.515076256 | 0.938 | 0.733 | 1 | 20 |
| rps20 | 0.0041 | 0.580063581 | 0.938 | 0.667 | 1 | 20 |
| gnai1 | 0.00416 | -0.63407402 | 0.062 | 0.467 | 1 | 20 |
| med6 | 0.00467 | 0.551722796 | 0.312 | 0 | 1 | 20 |
| mrpl41 | 0.00467 | 0.350918701 | 0.312 | 0 | 1 | 20 |
| ndufa3 | 0.00528 | -0.2726544 | 0.344 | 0.2 | 1 | 20 |

| | | | | | | |
|---|---|---|---|---|---|---|
| bsg | 0.00535 | 0.113728212 | 0.5 | 0.2 | 1 | 20 |
| ybx1 | 0.00538 | 0.647810512 | 0.906 | 0.467 | 1 | 20 |
| cct2 | 0.00547 | 0.779429037 | 0.719 | 0.267 | 1 | 20 |
| tmem176l.1 | 0.00553 | -0.74153986 | 0.031 | 0.4 | 1 | 20 |
| rps23 | 0.0056 | 0.559900852 | 0.969 | 0.667 | 1 | 20 |
| mt-nd2 | 0.00561 | -0.73314382 | 0.062 | 0.467 | 1 | 20 |
| eef1b2 | 0.00561 | 0.515467768 | 0.969 | 0.6 | 1 | 20 |
| rpl21 | 0.00588 | 0.477851656 | 0.938 | 0.8 | 1 | 20 |
| nop58 | 0.006 | 0.737879792 | 0.625 | 0.333 | 1 | 20 |
| tmed10 | 0.00611 | -0.4933297 | 0.281 | 0.267 | 1 | 20 |
| rpl18a | 0.0062 | 0.380318854 | 0.969 | 0.867 | 1 | 20 |
| rpl22l1 | 0.00624 | 0.579254419 | 0.844 | 0.4 | 1 | 20 |
| ndufa1 | 0.00649 | -0.80831394 | 0.25 | 0.6 | 1 | 20 |
| rps6 | 0.00654 | 0.325173533 | 0.938 | 0.933 | 1 | 20 |
| mcl1a | 0.00659 | -0.60148748 | 0.719 | 1 | 1 | 20 |
| naca | 0.0066 | 0.454581012 | 0.938 | 0.733 | 1 | 20 |
| cxcr3.2 | 0.00689 | 0.237549676 | 0.312 | 0.067 | 1 | 20 |
| rps12 | 0.00689 | 0.472355698 | 0.844 | 0.733 | 1 | 20 |
| dido1 | 0.0075 | 0.530793264 | 0.469 | 0.067 | 1 | 20 |
| rps19 | 0.00753 | 0.432570199 | 1 | 0.8 | 1 | 20 |
| gstm.2 | 0.00759 | 0.251988844 | 0.375 | 0.067 | 1 | 20 |
| CR383676.1 | 0.00771 | -0.11353399 | 0.938 | 0.867 | 1 | 20 |
| rps11 | 0.0079 | 0.405463272 | 0.906 | 0.8 | 1 | 20 |
| rpia | 0.00808 | 0.350122832 | 0.281 | 0 | 1 | 20 |
| aars | 0.00808 | 0.330599154 | 0.281 | 0 | 1 | 20 |
| adka | 0.00808 | 0.420274075 | 0.281 | 0 | 1 | 20 |
| atf1 | 0.00808 | 0.361145677 | 0.281 | 0 | 1 | 20 |
| ccdc88b | 0.00808 | 0.348190569 | 0.281 | 0 | 1 | 20 |
| hsdl2 | 0.00808 | 0.372717337 | 0.281 | 0 | 1 | 20 |
| mrps14 | 0.00808 | 0.371900464 | 0.281 | 0 | 1 | 20 |
| mrto4 | 0.00808 | 0.340696869 | 0.281 | 0 | 1 | 20 |
| nipsnap2 | 0.00808 | 0.440896267 | 0.281 | 0 | 1 | 20 |
| ogt.1 | 0.00808 | 0.381607791 | 0.281 | 0 | 1 | 20 |
| pdcd6 | 0.00808 | 0.508889371 | 0.281 | 0 | 1 | 20 |
| polr1d | 0.00808 | 0.414996237 | 0.281 | 0 | 1 | 20 |
| rpap3 | 0.00808 | 0.430608849 | 0.281 | 0 | 1 | 20 |
| sumo3b | 0.00808 | 0.548710224 | 0.281 | 0 | 1 | 20 |
| supt3h | 0.00808 | 0.360234015 | 0.281 | 0 | 1 | 20 |
| tufm | 0.00808 | 0.388135206 | 0.281 | 0 | 1 | 20 |
| zgc:153317 | 0.00808 | 0.342679033 | 0.281 | 0 | 1 | 20 |
| rpl3 | 0.00828 | 0.246365011 | 0.938 | 0.933 | 1 | 20 |
| rps15 | 0.00889 | 0.376305417 | 0.969 | 0.867 | 1 | 20 |
| h3f3d | 0.00931 | 0.626586787 | 0.844 | 0.4 | 1 | 20 |
| ilk | 0.0097 | -0.45884974 | 0.188 | 0.2 | 1 | 20 |
| ndufs8a | 0.00973 | 0.278665944 | 0.375 | 0.067 | 1 | 20 |
| vamp8 | 0.01 | -0.85729297 | 0.031 | 0.333 | 1 | 20 |
| ndufb8 | 0.01015 | 0.484436416 | 0.656 | 0.2 | 1 | 20 |
| cox6b2 | 0.01016 | 0.364055879 | 0.906 | 0.533 | 1 | 20 |

| | | | | | | |
|---|---|---|---|---|---|---|
| map2k6 | 0.01024 | 0.212053363 | 0.344 | 0.067 | 1 | 20 |
| atp5meb | 0.01024 | 0.532217418 | 0.656 | 0.2 | 1 | 20 |
| rpl10a | 0.01026 | 0.384221672 | 0.875 | 0.8 | 1 | 20 |
| psmd7 | 0.01044 | 0.49378227 | 0.469 | 0.067 | 1 | 20 |
| suclg1 | 0.01061 | 0.387318073 | 0.438 | 0.067 | 1 | 20 |
| rpl17 | 0.01071 | 0.410834358 | 0.938 | 0.8 | 1 | 20 |
| rpl10 | 0.01098 | 0.303262355 | 0.969 | 0.933 | 1 | 20 |
| rps24 | 0.01115 | 0.531369837 | 0.906 | 0.733 | 1 | 20 |
| hmgb2b | 0.01131 | 0.837762938 | 0.625 | 0.2 | 1 | 20 |
| si:ch73-1a9.3 | 0.01131 | 0.635625409 | 0.781 | 0.333 | 1 | 20 |
| tln1 | 0.01142 | -0.78805435 | 0.375 | 0.8 | 1 | 20 |
| spcs2 | 0.01187 | 0.168349433 | 0.406 | 0.133 | 1 | 20 |
| faua | 0.01191 | 0.471203479 | 0.938 | 0.733 | 1 | 20 |
| ctsk | 0.01198 | -0.59941883 | 0 | 0.2 | 1 | 20 |
| reps2 | 0.01198 | -0.51318538 | 0 | 0.2 | 1 | 20 |
| XLOC-024516 | 0.01198 | -0.63748486 | 0 | 0.2 | 1 | 20 |
| XLOC-041870 | 0.01198 | -0.65331572 | 0 | 0.2 | 1 | 20 |
| dnajc5ga | 0.01198 | -0.41188544 | 0 | 0.2 | 1 | 20 |
| mt-cyb | 0.01198 | -0.54418496 | 0 | 0.2 | 1 | 20 |
| rbm27 | 0.01198 | -0.34759655 | 0 | 0.2 | 1 | 20 |
| tp53inp1 | 0.01198 | -0.45628626 | 0 | 0.2 | 1 | 20 |
| LOC103911445 | 0.01198 | -0.13020775 | 0 | 0.2 | 1 | 20 |
| XLOC-043761 | 0.01198 | -0.55304931 | 0 | 0.2 | 1 | 20 |
| ano5b | 0.01198 | -0.5919139 | 0 | 0.2 | 1 | 20 |
| pgrmc1 | 0.01198 | -0.45973453 | 0 | 0.2 | 1 | 20 |
| sec31b | 0.01198 | -0.32444251 | 0 | 0.2 | 1 | 20 |
| vaspa | 0.01198 | -0.31490043 | 0 | 0.2 | 1 | 20 |
| si:dkey-222f2.7 | 0.01198 | -0.4113394 | 0 | 0.2 | 1 | 20 |
| ubox5 | 0.01198 | -0.25504817 | 0 | 0.2 | 1 | 20 |
| XLOC-032002 | 0.01198 | -0.6273928 | 0 | 0.2 | 1 | 20 |
| zgc:66447 | 0.01198 | -0.41696403 | 0 | 0.2 | 1 | 20 |
| nop56 | 0.01203 | 0.735106129 | 0.625 | 0.267 | 1 | 20 |
| plek | 0.01204 | -0.80923112 | 0.219 | 0.667 | 1 | 20 |
| spred2b | 0.01224 | -0.84936277 | 0.125 | 0.467 | 1 | 20 |
| ppiaa | 0.01269 | 0.466853312 | 0.969 | 0.667 | 1 | 20 |
| atp5po | 0.01293 | 0.68683883 | 0.625 | 0.267 | 1 | 20 |
| rpl9 | 0.01347 | 0.416679428 | 0.875 | 0.733 | 1 | 20 |
| myh11a | 0.01362 | -0.75571709 | 0.25 | 0.467 | 1 | 20 |
| ier2a | 0.01375 | -0.68141612 | 0.531 | 0.667 | 1 | 20 |
| CABZ01072614.1 | 0.01378 | 0.204567279 | 0.25 | 0 | 1 | 20 |
| abcf2a | 0.01378 | 0.316115665 | 0.25 | 0 | 1 | 20 |
| amd1 | 0.01378 | 0.398156355 | 0.25 | 0 | 1 | 20 |
| coq3 | 0.01378 | 0.289893399 | 0.25 | 0 | 1 | 20 |
| fam210aa | 0.01378 | 0.399142335 | 0.25 | 0 | 1 | 20 |
| frg1 | 0.01378 | 0.347565817 | 0.25 | 0 | 1 | 20 |
| ranbp3b | 0.01378 | 0.278856801 | 0.25 | 0 | 1 | 20 |
| tuba1b | 0.01378 | 0.315248403 | 0.25 | 0 | 1 | 20 |
| yipf4 | 0.01378 | 0.380660603 | 0.25 | 0 | 1 | 20 |

| | | | | | | |
|---|---|---|---|---|---|---|
| ascc1 | 0.01378 | 0.408139897 | 0.25 | 0 | 1 | 20 |
| chchd10 | 0.01378 | 0.322740585 | 0.25 | 0 | 1 | 20 |
| chchd6b | 0.01378 | 0.298384807 | 0.25 | 0 | 1 | 20 |
| eaf2 | 0.01378 | 0.408124019 | 0.25 | 0 | 1 | 20 |
| emc7 | 0.01378 | 0.353197936 | 0.25 | 0 | 1 | 20 |
| msi2b | 0.01378 | 0.317374068 | 0.25 | 0 | 1 | 20 |
| naa20 | 0.01378 | 0.324323871 | 0.25 | 0 | 1 | 20 |
| pgd | 0.01378 | 0.401343489 | 0.25 | 0 | 1 | 20 |
| ppa1b | 0.01378 | 0.326310772 | 0.25 | 0 | 1 | 20 |
| sdhdb | 0.01378 | 0.395354238 | 0.25 | 0 | 1 | 20 |
| sod2 | 0.01378 | 0.365381398 | 0.25 | 0 | 1 | 20 |
| ube2g1b | 0.01378 | 0.37637907 | 0.25 | 0 | 1 | 20 |
| zgc:112356 | 0.01378 | 0.404725305 | 0.25 | 0 | 1 | 20 |
| CABZ01045617.1 | 0.01378 | 0.291932273 | 0.25 | 0 | 1 | 20 |
| XLOC-028370 | 0.01378 | 0.50809306 | 0.25 | 0 | 1 | 20 |
| dync1i2a | 0.01378 | 0.33916386 | 0.25 | 0 | 1 | 20 |
| mrps17 | 0.01378 | 0.344730123 | 0.25 | 0 | 1 | 20 |
| nsa2 | 0.01385 | 0.380337082 | 0.781 | 0.4 | 1 | 20 |
| rps3a | 0.01391 | 0.443356104 | 0.938 | 0.8 | 1 | 20 |
| hmgb1b | 0.01481 | 0.538445786 | 0.562 | 0.133 | 1 | 20 |
| rpl26 | 0.01493 | 0.427307827 | 0.938 | 0.733 | 1 | 20 |
| gpatch8 | 0.01516 | -0.51684271 | 0.125 | 0.2 | 1 | 20 |
| si:dkey-51e6.1 | 0.01581 | 0.666969246 | 0.438 | 0.067 | 1 | 20 |
| arf5 | 0.016 | -0.69996302 | 0.219 | 0.4 | 1 | 20 |
| rpl4 | 0.01635 | 0.586230799 | 0.875 | 0.533 | 1 | 20 |
| ddx5 | 0.0171 | -0.76053352 | 0.344 | 0.667 | 1 | 20 |
| arpc4 | 0.01739 | -0.69378899 | 0.312 | 0.6 | 1 | 20 |
| pbx1b | 0.01746 | -0.71687377 | 0.031 | 0.333 | 1 | 20 |
| eif4bb | 0.01771 | 0.603729721 | 0.5 | 0.133 | 1 | 20 |
| h3f3c | 0.01801 | 0.126459292 | 0.844 | 0.533 | 1 | 20 |
| rpsa | 0.01806 | 0.369329301 | 0.969 | 0.667 | 1 | 20 |
| rpl22 | 0.01806 | 0.427060068 | 0.906 | 0.733 | 1 | 20 |
| XLOC-025423 | 0.01835 | -0.67477727 | 0.188 | 0.333 | 1 | 20 |
| rpl24 | 0.01838 | 0.445104484 | 0.969 | 0.667 | 1 | 20 |
| fermt3b | 0.01845 | -0.72174247 | 0.062 | 0.4 | 1 | 20 |
| rpl32 | 0.01852 | 0.38284347 | 0.938 | 0.8 | 1 | 20 |
| rps7 | 0.01889 | 0.386855114 | 0.906 | 0.8 | 1 | 20 |
| ndufa4l | 0.01894 | 0.317500377 | 0.844 | 0.467 | 1 | 20 |
| cox17 | 0.01929 | 0.570408369 | 0.375 | 0.067 | 1 | 20 |
| eef1da | 0.01931 | 0.579384444 | 0.812 | 0.4 | 1 | 20 |
| rpl23a | 0.01991 | 0.435992075 | 0.938 | 0.733 | 1 | 20 |
| ppp1cbl | 0.02021 | -0.69378066 | 0.031 | 0.333 | 1 | 20 |
| prdx2 | 0.02033 | 0.383525728 | 0.844 | 0.467 | 1 | 20 |
| pfdn4 | 0.02044 | 0.627003463 | 0.625 | 0.2 | 1 | 20 |
| mrpl57 | 0.02056 | 0.547202261 | 0.469 | 0.133 | 1 | 20 |
| b2ml | 0.02151 | 0.341843598 | 0.719 | 0.333 | 1 | 20 |
| rps26 | 0.02176 | 0.512618169 | 0.875 | 0.6 | 1 | 20 |
| rsl1d1 | 0.02182 | 0.651647305 | 0.531 | 0.2 | 1 | 20 |

| | | | | | | |
|---|---|---|---|---|---|---|
| hnrnpabb | 0.02196 | 0.41689675 | 0.656 | 0.267 | 1 | 20 |
| zgc:64051 | 0.02216 | -0.70661132 | 0.406 | 0.6 | 1 | 20 |
| twistnb | 0.02245 | 0.365724253 | 0.375 | 0.067 | 1 | 20 |
| ube2nb | 0.02251 | 0.137288004 | 0.281 | 0.067 | 1 | 20 |
| mt-co1 | 0.02278 | -0.7313553 | 0.312 | 0.733 | 1 | 20 |
| adkb | 0.02322 | 0.289339638 | 0.219 | 0 | 1 | 20 |
| cmip | 0.02322 | 0.3098236 | 0.219 | 0 | 1 | 20 |
| ctps1a | 0.02322 | 0.238684366 | 0.219 | 0 | 1 | 20 |
| gemin8 | 0.02322 | 0.350920048 | 0.219 | 0 | 1 | 20 |
| mrpl37 | 0.02322 | 0.328032378 | 0.219 | 0 | 1 | 20 |
| pelp1 | 0.02322 | 0.302880761 | 0.219 | 0 | 1 | 20 |
| rtca | 0.02322 | 0.341454562 | 0.219 | 0 | 1 | 20 |
| si:dkeyp-113d7.1 | 0.02322 | 0.398520834 | 0.219 | 0 | 1 | 20 |
| arglu1a | 0.02322 | 0.300059666 | 0.219 | 0 | 1 | 20 |
| armt1 | 0.02322 | 0.401857078 | 0.219 | 0 | 1 | 20 |
| ccdc127b | 0.02322 | 0.333119799 | 0.219 | 0 | 1 | 20 |
| ddx52 | 0.02322 | 0.320902445 | 0.219 | 0 | 1 | 20 |
| e2f4 | 0.02322 | 0.272311428 | 0.219 | 0 | 1 | 20 |
| fbxo11a | 0.02322 | 0.369084821 | 0.219 | 0 | 1 | 20 |
| fntb | 0.02322 | 0.324341294 | 0.219 | 0 | 1 | 20 |
| gkap1 | 0.02322 | 0.360672342 | 0.219 | 0 | 1 | 20 |
| hdgfl2 | 0.02322 | 0.278795171 | 0.219 | 0 | 1 | 20 |
| hikeshi | 0.02322 | 0.295015134 | 0.219 | 0 | 1 | 20 |
| hsd17b10 | 0.02322 | 0.214761643 | 0.219 | 0 | 1 | 20 |
| htatsf1 | 0.02322 | 0.36792364 | 0.219 | 0 | 1 | 20 |
| jam2a | 0.02322 | 0.265957378 | 0.219 | 0 | 1 | 20 |
| lyrm4 | 0.02322 | 0.269627231 | 0.219 | 0 | 1 | 20 |
| med21 | 0.02322 | 0.23083512 | 0.219 | 0 | 1 | 20 |
| ndc1 | 0.02322 | 0.255123699 | 0.219 | 0 | 1 | 20 |
| rbm4.2 | 0.02322 | 0.318521968 | 0.219 | 0 | 1 | 20 |
| slc35f2 | 0.02322 | 0.284778291 | 0.219 | 0 | 1 | 20 |
| zcrb1 | 0.02322 | 0.299142628 | 0.219 | 0 | 1 | 20 |
| zgc:56106 | 0.02322 | 0.281808701 | 0.219 | 0 | 1 | 20 |
| BX005340.1 | 0.02322 | 0.209826643 | 0.219 | 0 | 1 | 20 |
| cmc1 | 0.02322 | 0.263267593 | 0.219 | 0 | 1 | 20 |
| dmac2l | 0.02322 | 0.310053645 | 0.219 | 0 | 1 | 20 |
| imp4 | 0.02322 | 0.294273198 | 0.219 | 0 | 1 | 20 |
| med11 | 0.02322 | 0.280536655 | 0.219 | 0 | 1 | 20 |
| mhc2dab | 0.02322 | 0.23175397 | 0.219 | 0 | 1 | 20 |
| nfybb | 0.02322 | 0.431084009 | 0.219 | 0 | 1 | 20 |
| ppme1 | 0.02322 | 0.230315037 | 0.219 | 0 | 1 | 20 |
| sdad1 | 0.02322 | 0.385415907 | 0.219 | 0 | 1 | 20 |
| spi2 | 0.02322 | 0.426615918 | 0.219 | 0 | 1 | 20 |
| tssc4 | 0.02322 | 0.250196859 | 0.219 | 0 | 1 | 20 |
| ufsp2 | 0.02322 | 0.250527527 | 0.219 | 0 | 1 | 20 |
| FQ323156.1 | 0.02322 | 0.251458464 | 0.219 | 0 | 1 | 20 |
| aldh2.2 | 0.02322 | 0.228157059 | 0.219 | 0 | 1 | 20 |
| heatr3 | 0.02322 | 0.242830774 | 0.219 | 0 | 1 | 20 |

| | | | | | | |
|---|---|---|---|---|---|---|
| klhdc4 | 0.02322 | 0.251168837 | 0.219 | 0 | 1 | 20 |
| lrrc59 | 0.02322 | 0.265208237 | 0.219 | 0 | 1 | 20 |
| zgc:110339 | 0.02322 | 0.206845231 | 0.219 | 0 | 1 | 20 |
| znhit3 | 0.02322 | 0.248382199 | 0.219 | 0 | 1 | 20 |
| prkar1ab | 0.02322 | 0.282491786 | 0.219 | 0 | 1 | 20 |
| hmgb3b | 0.02327 | 0.559280893 | 0.531 | 0.2 | 1 | 20 |
| rps21 | 0.02344 | 0.277678465 | 0.969 | 0.933 | 1 | 20 |
| rplp1 | 0.02401 | 0.416678645 | 0.969 | 0.733 | 1 | 20 |
| adh5 | 0.02421 | 0.578175294 | 0.688 | 0.267 | 1 | 20 |
| npm3 | 0.02491 | 0.196233493 | 0.406 | 0.133 | 1 | 20 |
| rpl23 | 0.02498 | 0.419089615 | 0.938 | 0.8 | 1 | 20 |
| glo1 | 0.02512 | 0.384485862 | 0.5 | 0.133 | 1 | 20 |
| cox7a2a | 0.02518 | 0.551682054 | 0.531 | 0.133 | 1 | 20 |
| rpl28 | 0.02597 | 0.409724966 | 0.906 | 0.733 | 1 | 20 |
| vdac2 | 0.02628 | 0.592344706 | 0.781 | 0.4 | 1 | 20 |
| rps14 | 0.02704 | 0.257286755 | 0.906 | 0.867 | 1 | 20 |
| rfx5 | 0.02717 | -0.62911215 | 0.094 | 0.267 | 1 | 20 |
| rps10 | 0.02738 | 0.444756676 | 0.875 | 0.667 | 1 | 20 |
| ankrd11 | 0.02754 | -0.63826634 | 0.188 | 0.4 | 1 | 20 |
| itm2ba | 0.02758 | -0.64976914 | 0.406 | 0.667 | 1 | 20 |
| pwp1 | 0.02763 | 0.307298407 | 0.344 | 0.067 | 1 | 20 |
| thbs1b | 0.02841 | -0.9377227 | 0.125 | 0.4 | 1 | 20 |
| hp1bp3 | 0.02871 | 0.527661005 | 0.531 | 0.2 | 1 | 20 |
| clpp | 0.02893 | 0.120284328 | 0.281 | 0.067 | 1 | 20 |
| rps3 | 0.02902 | 0.33213267 | 0.844 | 0.733 | 1 | 20 |
| eif5a2 | 0.02906 | 0.618814554 | 0.75 | 0.4 | 1 | 20 |
| sept12 | 0.02938 | -0.56526852 | 0.438 | 0.533 | 1 | 20 |
| ptpn2b | 0.02956 | -0.35522948 | 0.188 | 0.2 | 1 | 20 |
| rplp2l | 0.02961 | 0.481065369 | 0.938 | 0.667 | 1 | 20 |
| arf6b | 0.0303 | -0.37929617 | 0.25 | 0.267 | 1 | 20 |
| kmt5aa | 0.03032 | -0.10239 | 0.438 | 0.267 | 1 | 20 |
| rhocb | 0.03039 | -0.65742709 | 0.25 | 0.6 | 1 | 20 |
| atp5mc3b | 0.03047 | 0.556301673 | 0.844 | 0.467 | 1 | 20 |
| sap30l | 0.03118 | -0.64091506 | 0.219 | 0.533 | 1 | 20 |
| CU639469.1 | 0.03161 | -0.62696291 | 0.031 | 0.267 | 1 | 20 |
| med10 | 0.03167 | 0.488419675 | 0.406 | 0.067 | 1 | 20 |
| atp5f1c | 0.03172 | 0.573857053 | 0.594 | 0.2 | 1 | 20 |
| hsp90ab1 | 0.03185 | 0.356144376 | 0.969 | 0.8 | 1 | 20 |
| si:dkey-185m8.2 | 0.03187 | -0.64784717 | 0.188 | 0.4 | 1 | 20 |
| tpm1 | 0.03198 | -0.44144866 | 0.125 | 0.467 | 1 | 20 |
| rps27l | 0.03204 | 0.166317459 | 0.562 | 0.267 | 1 | 20 |
| rps27.2 | 0.03212 | 0.228865669 | 0.875 | 0.6 | 1 | 20 |
| mt-nd1 | 0.0329 | -0.69365951 | 0.062 | 0.333 | 1 | 20 |
| cfl1 | 0.03317 | -0.15633993 | 0.906 | 0.8 | 1 | 20 |
| rps25 | 0.03358 | 0.377093202 | 0.906 | 0.733 | 1 | 20 |
| LOC110438237 | 0.03386 | 0.103512331 | 0.25 | 0.067 | 1 | 20 |
| gapdhs | 0.03387 | -0.13870436 | 0.75 | 0.6 | 1 | 20 |
| zeb2a | 0.0339 | -0.58384161 | 0.125 | 0.267 | 1 | 20 |

| | | | | | | |
|---|---|---|---|---|---|---|
| fosab | 0.0348 | -0.65440265 | 0.406 | 0.8 | 1 | 20 |
| fbl | 0.03491 | 0.501698935 | 0.406 | 0.2 | 1 | 20 |
| hprt1 | 0.03547 | 0.152260093 | 0.312 | 0.067 | 1 | 20 |
| XLOC-043760 | 0.03577 | -0.73777058 | 0.031 | 0.2 | 1 | 20 |
| bud31 | 0.03589 | 0.539138907 | 0.5 | 0.133 | 1 | 20 |
| rpl37 | 0.03634 | 0.299603953 | 1 | 0.8 | 1 | 20 |
| tmem106ba | 0.03695 | -0.46663661 | 0.125 | 0.2 | 1 | 20 |
| c1qbp | 0.03712 | 0.449660397 | 0.406 | 0.067 | 1 | 20 |
| ch25hl3 | 0.03714 | -0.59356132 | 0.031 | 0.267 | 1 | 20 |
| pik3r3a | 0.03724 | -0.28930204 | 0.156 | 0.133 | 1 | 20 |
| ube2d1b | 0.03735 | 0.213420647 | 0.312 | 0.067 | 1 | 20 |
| ssr2 | 0.03866 | 0.595821425 | 0.531 | 0.2 | 1 | 20 |
| TST | 0.03877 | 0.292795312 | 0.188 | 0 | 1 | 20 |
| dnph1 | 0.03877 | 0.249330029 | 0.188 | 0 | 1 | 20 |
| fdx2 | 0.03877 | 0.254011319 | 0.188 | 0 | 1 | 20 |
| foxo4 | 0.03877 | 0.255430837 | 0.188 | 0 | 1 | 20 |
| gart | 0.03877 | 0.272454023 | 0.188 | 0 | 1 | 20 |
| gcn1 | 0.03877 | 0.246992627 | 0.188 | 0 | 1 | 20 |
| gyg1a | 0.03877 | 0.240782176 | 0.188 | 0 | 1 | 20 |
| ino80 | 0.03877 | 0.268280392 | 0.188 | 0 | 1 | 20 |
| lamtor3 | 0.03877 | 0.364014578 | 0.188 | 0 | 1 | 20 |
| prpsap1 | 0.03877 | 0.246271511 | 0.188 | 0 | 1 | 20 |
| rpp25l | 0.03877 | 0.208144218 | 0.188 | 0 | 1 | 20 |
| zfr | 0.03877 | 0.305649948 | 0.188 | 0 | 1 | 20 |
| bcap31 | 0.03877 | 0.229977461 | 0.188 | 0 | 1 | 20 |
| cd83 | 0.03877 | 0.28787819 | 0.188 | 0 | 1 | 20 |
| dcaf13 | 0.03877 | 0.238850718 | 0.188 | 0 | 1 | 20 |
| dnlz | 0.03877 | 0.166358454 | 0.188 | 0 | 1 | 20 |
| elp6 | 0.03877 | 0.249908478 | 0.188 | 0 | 1 | 20 |
| eny2 | 0.03877 | 0.248958106 | 0.188 | 0 | 1 | 20 |
| fblim1 | 0.03877 | 0.307459352 | 0.188 | 0 | 1 | 20 |
| gnptg | 0.03877 | 0.282094359 | 0.188 | 0 | 1 | 20 |
| hnrnpua | 0.03877 | 0.219223763 | 0.188 | 0 | 1 | 20 |
| med8 | 0.03877 | 0.336030952 | 0.188 | 0 | 1 | 20 |
| mgmt | 0.03877 | 0.238912692 | 0.188 | 0 | 1 | 20 |
| ms4a17a.6 | 0.03877 | 0.28928299 | 0.188 | 0 | 1 | 20 |
| polr2h | 0.03877 | 0.238583195 | 0.188 | 0 | 1 | 20 |
| ppp1r11 | 0.03877 | 0.249937268 | 0.188 | 0 | 1 | 20 |
| psmc1b | 0.03877 | 0.237718363 | 0.188 | 0 | 1 | 20 |
| rrp12 | 0.03877 | 0.199414903 | 0.188 | 0 | 1 | 20 |
| rtraf | 0.03877 | 0.208828647 | 0.188 | 0 | 1 | 20 |
| si:ch73-376l24.2 | 0.03877 | 0.245242567 | 0.188 | 0 | 1 | 20 |
| sms | 0.03877 | 0.260197539 | 0.188 | 0 | 1 | 20 |
| snu13a | 0.03877 | 0.242276964 | 0.188 | 0 | 1 | 20 |
| spi1a | 0.03877 | 0.38631184 | 0.188 | 0 | 1 | 20 |
| srsf6a | 0.03877 | 0.255763431 | 0.188 | 0 | 1 | 20 |
| stt3b | 0.03877 | 0.219296897 | 0.188 | 0 | 1 | 20 |
| tmem167a | 0.03877 | 0.365331082 | 0.188 | 0 | 1 | 20 |

| | | | | | | |
|---|---|---|---|---|---|---|
| zgc:101783 | 0.03877 | 0.214727118 | 0.188 | 0 | 1 | 20 |
| znf330 | 0.03877 | 0.217029771 | 0.188 | 0 | 1 | 20 |
| exosc9 | 0.03877 | 0.192264745 | 0.188 | 0 | 1 | 20 |
| mrpl40 | 0.03877 | 0.263353201 | 0.188 | 0 | 1 | 20 |
| rbmx2 | 0.03877 | 0.262505711 | 0.188 | 0 | 1 | 20 |
| snrnp40 | 0.03877 | 0.236502585 | 0.188 | 0 | 1 | 20 |
| sub1b | 0.03877 | 0.281150499 | 0.188 | 0 | 1 | 20 |
| tnrc18 | 0.03877 | 0.205068475 | 0.188 | 0 | 1 | 20 |
| trappc6b | 0.03877 | 0.275329742 | 0.188 | 0 | 1 | 20 |
| urod | 0.03877 | 0.219886126 | 0.188 | 0 | 1 | 20 |
| cyfip2 | 0.03877 | 0.247115635 | 0.188 | 0 | 1 | 20 |
| cyth1b | 0.03877 | 0.26591994 | 0.188 | 0 | 1 | 20 |
| fbxl3a | 0.03877 | 0.33077915 | 0.188 | 0 | 1 | 20 |
| gorasp2 | 0.03877 | 0.181848382 | 0.188 | 0 | 1 | 20 |
| isoc2 | 0.03877 | 0.289021835 | 0.188 | 0 | 1 | 20 |
| mrpl43 | 0.03877 | 0.290208226 | 0.188 | 0 | 1 | 20 |
| mrpl47 | 0.03877 | 0.233638183 | 0.188 | 0 | 1 | 20 |
| mrps27 | 0.03877 | 0.196035826 | 0.188 | 0 | 1 | 20 |
| prkar2aa | 0.03877 | 0.346908596 | 0.188 | 0 | 1 | 20 |
| scaf4a | 0.03877 | 0.244335635 | 0.188 | 0 | 1 | 20 |
| sdha | 0.03877 | 0.205011968 | 0.188 | 0 | 1 | 20 |
| sf3a3 | 0.03877 | 0.197949741 | 0.188 | 0 | 1 | 20 |
| si:ch211-196c10.15 | 0.03877 | 0.292435755 | 0.188 | 0 | 1 | 20 |
| snrnp200 | 0.03877 | 0.252212323 | 0.188 | 0 | 1 | 20 |
| suclg2 | 0.03877 | 0.197610258 | 0.188 | 0 | 1 | 20 |
| sufu | 0.03877 | 0.195712125 | 0.188 | 0 | 1 | 20 |
| wdr12 | 0.03877 | 0.210916661 | 0.188 | 0 | 1 | 20 |
| zgc:195175 | 0.03877 | 0.282232063 | 0.188 | 0 | 1 | 20 |
| si:ch1073-429i10.3.1 | 0.03924 | 0.105099449 | 0.625 | 0.333 | 1 | 20 |
| cox6a1 | 0.03961 | 0.285853574 | 0.75 | 0.4 | 1 | 20 |
| si:ch73-281n10.2 | 0.03971 | 0.589250887 | 0.656 | 0.267 | 1 | 20 |
| cct7 | 0.04003 | 0.576359998 | 0.656 | 0.267 | 1 | 20 |
| celf2 | 0.04048 | 0.576154578 | 0.719 | 0.333 | 1 | 20 |
| eif3ha | 0.04066 | 0.559947113 | 0.656 | 0.267 | 1 | 20 |
| mdh2 | 0.04125 | 0.468116777 | 0.719 | 0.333 | 1 | 20 |
| erh | 0.04277 | 0.559603584 | 0.562 | 0.2 | 1 | 20 |
| ppifb | 0.04318 | 0.579958178 | 0.625 | 0.267 | 1 | 20 |
| st13 | 0.04348 | 0.448230391 | 0.438 | 0.133 | 1 | 20 |
| serbp1a | 0.04402 | 0.454140045 | 0.906 | 0.6 | 1 | 20 |
| ptprc | 0.04413 | -0.26785173 | 0.188 | 0.133 | 1 | 20 |
| btf3 | 0.04438 | 0.365650122 | 0.844 | 0.667 | 1 | 20 |
| krtcap2 | 0.04439 | 0.520638284 | 0.375 | 0.067 | 1 | 20 |
| slc25a3b | 0.04465 | 0.387563648 | 0.812 | 0.467 | 1 | 20 |
| cd74a | 0.04473 | 0.515787211 | 0.375 | 0.067 | 1 | 20 |
| si:ch73-308l14.2 | 0.04473 | 0.288078449 | 0.344 | 0.067 | 1 | 20 |
| nucks1b | 0.04474 | 0.479920532 | 0.344 | 0.067 | 1 | 20 |
| rpl34 | 0.04485 | 0.400710051 | 0.906 | 0.733 | 1 | 20 |
| eif3c | 0.04495 | 0.520121923 | 0.562 | 0.333 | 1 | 20 |

| | | | | | | |
|---|---|---|---|---|---|---|
| arpc1b | 0.04553 | -0.24746212 | 0.812 | 0.733 | 1 | 20 |
| atp5mc3a | 0.04627 | 0.510345548 | 0.469 | 0.2 | 1 | 20 |
| rtn4a | 0.04688 | 0.481718454 | 0.406 | 0.2 | 1 | 20 |
| XLOC-026836 | 0.04709 | -0.60416692 | 0.219 | 0.333 | 1 | 20 |
| zgc:92907 | 0.0472 | -0.53726734 | 0.031 | 0.267 | 1 | 20 |
| hdac1 | 0.04735 | 0.534969078 | 0.531 | 0.2 | 1 | 20 |
| tmed2 | 0.04755 | 0.185021891 | 0.5 | 0.2 | 1 | 20 |
| nkap | 0.04795 | 0.483300935 | 0.344 | 0.067 | 1 | 20 |
| rpl39 | 0.04814 | 0.373305621 | 0.969 | 0.733 | 1 | 20 |
| rpl36 | 0.04817 | 0.321822732 | 0.938 | 0.8 | 1 | 20 |
| CR381673.1 | 0.04853 | -0.26667617 | 0 | 0.133 | 1 | 20 |
| LOC101883382 | 0.04853 | -0.26918499 | 0 | 0.133 | 1 | 20 |
| alox5ap | 0.04853 | -0.24970237 | 0 | 0.133 | 1 | 20 |
| arl5c | 0.04853 | -0.28498073 | 0 | 0.133 | 1 | 20 |
| catip | 0.04853 | -0.4362829 | 0 | 0.133 | 1 | 20 |
| cdc42se1 | 0.04853 | -0.30102804 | 0 | 0.133 | 1 | 20 |
| cfbl | 0.04853 | -0.24664445 | 0 | 0.133 | 1 | 20 |
| clk4a | 0.04853 | -0.4588089 | 0 | 0.133 | 1 | 20 |
| inpp5l | 0.04853 | -0.31520725 | 0 | 0.133 | 1 | 20 |
| osbpl5 | 0.04853 | -0.2883195 | 0 | 0.133 | 1 | 20 |
| pgm1 | 0.04853 | -0.24272903 | 0 | 0.133 | 1 | 20 |
| rabgap1 | 0.04853 | -0.23545386 | 0 | 0.133 | 1 | 20 |
| rgs5b | 0.04853 | -0.35657692 | 0 | 0.133 | 1 | 20 |
| rilpl2 | 0.04853 | -0.29478182 | 0 | 0.133 | 1 | 20 |
| rin3 | 0.04853 | -0.24887117 | 0 | 0.133 | 1 | 20 |
| scinlb | 0.04853 | -0.346041 | 0 | 0.133 | 1 | 20 |
| si:dkey-9k7.3 | 0.04853 | -0.2870935 | 0 | 0.133 | 1 | 20 |
| si:zfos-80g12.1 | 0.04853 | -0.31143431 | 0 | 0.133 | 1 | 20 |
| smarca1 | 0.04853 | -0.26329157 | 0 | 0.133 | 1 | 20 |
| spice1 | 0.04853 | -0.25846352 | 0 | 0.133 | 1 | 20 |
| swt1 | 0.04853 | -0.45368377 | 0 | 0.133 | 1 | 20 |
| ubl3b | 0.04853 | -0.30680092 | 0 | 0.133 | 1 | 20 |
| xylt1 | 0.04853 | -0.44681018 | 0 | 0.133 | 1 | 20 |
| zgc:113411 | 0.04853 | -0.42293442 | 0 | 0.133 | 1 | 20 |
| CIR1 | 0.04853 | -0.28372011 | 0 | 0.133 | 1 | 20 |
| COLGALT1 (1 of many) | 0.04853 | -0.28328479 | 0 | 0.133 | 1 | 20 |
| CR354588.1 | 0.04853 | -0.15302765 | 0 | 0.133 | 1 | 20 |
| CU856344.1 | 0.04853 | -0.13382424 | 0 | 0.133 | 1 | 20 |
| akr1a1b | 0.04853 | -0.13596499 | 0 | 0.133 | 1 | 20 |
| anxa1a | 0.04853 | -0.16956013 | 0 | 0.133 | 1 | 20 |
| bhlhe23 | 0.04853 | -0.17515497 | 0 | 0.133 | 1 | 20 |
| calm3a | 0.04853 | -0.1306047 | 0 | 0.133 | 1 | 20 |
| card9 | 0.04853 | -0.11343525 | 0 | 0.133 | 1 | 20 |
| chordc1b | 0.04853 | -0.11507838 | 0 | 0.133 | 1 | 20 |
| commd6 | 0.04853 | -0.26690229 | 0 | 0.133 | 1 | 20 |
| dyrk3 | 0.04853 | -0.11456473 | 0 | 0.133 | 1 | 20 |
| foxj3 | 0.04853 | -0.32154549 | 0 | 0.133 | 1 | 20 |
| gpr108 | 0.04853 | -0.14282853 | 0 | 0.133 | 1 | 20 |

| | | | | | | |
|---|---|---|---|---|---|---|
| il16 | 0.04853 | -0.13590549 | 0 | 0.133 | 1 | 20 |
| irak4 | 0.04853 | -0.10119236 | 0 | 0.133 | 1 | 20 |
| irgf1 | 0.04853 | -0.30398367 | 0 | 0.133 | 1 | 20 |
| kif1bp | 0.04853 | -0.238046 | 0 | 0.133 | 1 | 20 |
| lims2 | 0.04853 | -0.11839772 | 0 | 0.133 | 1 | 20 |
| mgat5 | 0.04853 | -0.16517821 | 0 | 0.133 | 1 | 20 |
| mocs2 | 0.04853 | -0.4879605 | 0 | 0.133 | 1 | 20 |
| necap1 | 0.04853 | -0.15317711 | 0 | 0.133 | 1 | 20 |
| oaz2b | 0.04853 | -0.29867505 | 0 | 0.133 | 1 | 20 |
| oga | 0.04853 | -0.27656403 | 0 | 0.133 | 1 | 20 |
| pik3cb | 0.04853 | -0.24784821 | 0 | 0.133 | 1 | 20 |
| pik3r5 | 0.04853 | -0.10052585 | 0 | 0.133 | 1 | 20 |
| ppp1r3b | 0.04853 | -0.23723054 | 0 | 0.133 | 1 | 20 |
| purbb | 0.04853 | -0.2898826 | 0 | 0.133 | 1 | 20 |
| rab11bb | 0.04853 | -0.14185929 | 0 | 0.133 | 1 | 20 |
| rasa1a | 0.04853 | -0.12258135 | 0 | 0.133 | 1 | 20 |
| si:dkey-192k22.2 | 0.04853 | -0.28019 | 0 | 0.133 | 1 | 20 |
| sik3 | 0.04853 | -0.15887374 | 0 | 0.133 | 1 | 20 |
| slc14a2 | 0.04853 | -0.12065243 | 0 | 0.133 | 1 | 20 |
| slc3a2a | 0.04853 | -0.2145906 | 0 | 0.133 | 1 | 20 |
| snx27b | 0.04853 | -0.29326063 | 0 | 0.133 | 1 | 20 |
| tacc1 | 0.04853 | -0.10898701 | 0 | 0.133 | 1 | 20 |
| tnfsf10 | 0.04853 | -0.24392334 | 0 | 0.133 | 1 | 20 |
| ushbp1 | 0.04853 | -0.14503682 | 0 | 0.133 | 1 | 20 |
| zfyve9a | 0.04853 | -0.25026912 | 0 | 0.133 | 1 | 20 |
| zgc:113294 | 0.04853 | -0.1019963 | 0 | 0.133 | 1 | 20 |
| zgc:153169 | 0.04853 | -0.10613671 | 0 | 0.133 | 1 | 20 |
| zgc:77838 | 0.04853 | -0.22621935 | 0 | 0.133 | 1 | 20 |
| calhm6 | 0.04853 | -0.4495911 | 0 | 0.133 | 1 | 20 |
| clic2 | 0.04853 | -0.42103026 | 0 | 0.133 | 1 | 20 |
| cx32.2 | 0.04853 | -0.42331494 | 0 | 0.133 | 1 | 20 |
| rad54l2 | 0.04853 | -0.26844877 | 0 | 0.133 | 1 | 20 |
| rhoh | 0.04853 | -0.2158998 | 0 | 0.133 | 1 | 20 |
| rp2 | 0.04853 | -0.25088684 | 0 | 0.133 | 1 | 20 |
| FP101882.1 | 0.04853 | -0.18113272 | 0 | 0.133 | 1 | 20 |
| b3glcta | 0.04853 | -0.1486001 | 0 | 0.133 | 1 | 20 |
| chd8 | 0.04853 | -0.15117193 | 0 | 0.133 | 1 | 20 |
| gab2 | 0.04853 | -0.14630302 | 0 | 0.133 | 1 | 20 |
| guf1 | 0.04853 | -0.14996833 | 0 | 0.133 | 1 | 20 |
| hsd11b2 | 0.04853 | -0.16487895 | 0 | 0.133 | 1 | 20 |
| id1 | 0.04853 | -0.38901233 | 0 | 0.133 | 1 | 20 |
| ints11 | 0.04853 | -0.14785418 | 0 | 0.133 | 1 | 20 |
| kat14 | 0.04853 | -0.14494139 | 0 | 0.133 | 1 | 20 |
| pdk1 | 0.04853 | -0.14880978 | 0 | 0.133 | 1 | 20 |
| rpz | 0.04853 | -0.38748457 | 0 | 0.133 | 1 | 20 |
| si:dkey-92i15.4 | 0.04853 | -0.15187704 | 0 | 0.133 | 1 | 20 |
| si:dkeyp-75h12.5 | 0.04853 | -0.42702934 | 0 | 0.133 | 1 | 20 |
| slc35d1a | 0.04853 | -0.1509425 | 0 | 0.133 | 1 | 20 |

| | | | | | | |
|---|---|---|---|---|---|---|
| suz12b | 0.04853 | -0.14993114 | 0 | 0.133 | 1 | 20 |
| NC-002333.17 | 0.04853 | -0.3889466 | 0 | 0.133 | 1 | 20 |
| ap4e1 | 0.04853 | -0.36919758 | 0 | 0.133 | 1 | 20 |
| mid1ip1b | 0.04853 | -0.44479077 | 0 | 0.133 | 1 | 20 |
| LOC108190717 | 0.04853 | -0.45764319 | 0 | 0.133 | 1 | 20 |
| chd1 | 0.04853 | -0.29177157 | 0 | 0.133 | 1 | 20 |
| iqgap2 | 0.04853 | -0.43083882 | 0 | 0.133 | 1 | 20 |
| plpp1a | 0.04853 | -0.42851092 | 0 | 0.133 | 1 | 20 |
| rpgrb | 0.04853 | -0.17799233 | 0 | 0.133 | 1 | 20 |
| sap130a | 0.04853 | -0.30211087 | 0 | 0.133 | 1 | 20 |
| sertad2b | 0.04853 | -0.42958097 | 0 | 0.133 | 1 | 20 |
| si:ch211-66e2.3 | 0.04853 | -0.27382208 | 0 | 0.133 | 1 | 20 |
| taf1 | 0.04853 | -0.32887001 | 0 | 0.133 | 1 | 20 |
| rps4x | 0.04891 | 0.409217799 | 0.906 | 0.733 | 1 | 20 |
| gars | 0.04903 | 0.471525407 | 0.469 | 0.133 | 1 | 20 |
| calm2a | 0.04904 | -0.63095886 | 0.188 | 0.533 | 1 | 20 |
| si:ch211-132p1.3 | 0.04927 | 0.463637628 | 0.375 | 0.067 | 1 | 20 |
| pfdn5 | 0.0493 | 0.509013501 | 0.469 | 0.133 | 1 | 20 |
| vat1 | 0.04953 | -0.56291772 | 0.094 | 0.267 | 1 | 20 |
| si:dkey-151g10.6 | 0.04954 | 0.513193192 | 0.906 | 0.6 | 1 | 20 |
| ppid | 0.04954 | 0.502867581 | 0.5 | 0.2 | 1 | 20 |
| si:ch1073-314i13.4 | 0.04965 | -0.20532073 | 0.188 | 0.133 | 1 | 20 |
| rpl35 | 0.05007 | 0.215170277 | 0.938 | 0.933 | 1 | 20 |
| ier2b | 0.05068 | -0.69077788 | 0.375 | 0.667 | 1 | 20 |
| rpl13 | 0.05089 | 0.284725091 | 0.938 | 0.867 | 1 | 20 |
| zgc:114188 | 0.05114 | 0.33109817 | 0.844 | 0.733 | 1 | 20 |
| mospd1 | 0.05225 | -0.57079018 | 0.031 | 0.267 | 1 | 20 |
| ddx39aa | 0.05261 | 0.468355309 | 0.438 | 0.133 | 1 | 20 |
| nip7 | 0.05291 | 0.469969358 | 0.375 | 0.133 | 1 | 20 |
| mtdha | 0.05291 | 0.570802229 | 0.406 | 0.133 | 1 | 20 |
| tmsb4x | 0.05297 | -0.31575603 | 0.906 | 0.933 | 1 | 20 |
| cst14a.2 | 0.05376 | -0.20000293 | 0.344 | 0.267 | 1 | 20 |
| fthl27 | 0.05433 | -0.47709298 | 0.625 | 0.733 | 1 | 20 |
| brd2a | 0.05477 | -0.64090957 | 0.125 | 0.4 | 1 | 20 |
| cd3eap | 0.05484 | 0.407271042 | 0.375 | 0.067 | 1 | 20 |
| psme1 | 0.05548 | 0.148692861 | 0.469 | 0.2 | 1 | 20 |
| lsm7 | 0.05562 | 0.501984228 | 0.375 | 0.067 | 1 | 20 |
| elf1 | 0.05569 | -0.46221229 | 0.094 | 0.4 | 1 | 20 |
| ptges3b | 0.05575 | 0.389360171 | 0.469 | 0.133 | 1 | 20 |
| zgc:56493 | 0.05621 | 0.402925184 | 0.906 | 0.6 | 1 | 20 |
| fbxo9 | 0.05636 | -0.46581002 | 0.031 | 0.267 | 1 | 20 |
| sumo2b | 0.05658 | -0.33501951 | 0.594 | 0.6 | 1 | 20 |
| eif3s6ip | 0.05668 | 0.478177217 | 0.562 | 0.2 | 1 | 20 |
| si:ch211-262i1.6 | 0.05681 | 0.138947169 | 0.25 | 0.067 | 1 | 20 |
| rac3a | 0.05683 | -0.7621018 | 0.062 | 0.333 | 1 | 20 |
| lsm6 | 0.05699 | 0.381513649 | 0.375 | 0.067 | 1 | 20 |
| rps13 | 0.05734 | 0.366305783 | 0.938 | 0.733 | 1 | 20 |
| hmgn6 | 0.05804 | 0.425551279 | 0.812 | 0.467 | 1 | 20 |

| | | | | | | |
|---|---|---|---|---|---|---|
| khdrbs1a | 0.05827 | 0.432851755 | 0.812 | 0.467 | 1 | 20 |
| sh3bgrl | 0.05856 | -0.39464957 | 0.375 | 0.4 | 1 | 20 |
| mplkip | 0.05866 | -0.56981813 | 0.031 | 0.2 | 1 | 20 |
| pabpc1a | 0.05885 | 0.467954281 | 0.812 | 0.467 | 1 | 20 |
| mcm2 | 0.05903 | 0.446531264 | 0.375 | 0.067 | 1 | 20 |
| mrpl32 | 0.05914 | 0.446377975 | 0.375 | 0.067 | 1 | 20 |
| rpl7l1 | 0.06007 | 0.456528324 | 0.344 | 0.067 | 1 | 20 |
| hypk | 0.0601 | 0.529111031 | 0.469 | 0.133 | 1 | 20 |
| keap1b | 0.06043 | -0.37093376 | 0.031 | 0.267 | 1 | 20 |
| rps17 | 0.06122 | 0.38685804 | 0.938 | 0.667 | 1 | 20 |
| riox2 | 0.06132 | 0.220272583 | 0.312 | 0.067 | 1 | 20 |
| vdac1 | 0.06138 | 0.473107556 | 0.344 | 0.067 | 1 | 20 |
| hnrnpub | 0.06145 | 0.206602452 | 0.312 | 0.067 | 1 | 20 |
| jdp2b | 0.06156 | -0.61033226 | 0.219 | 0.267 | 1 | 20 |
| hmga1a | 0.06188 | 0.32725317 | 0.781 | 0.467 | 1 | 20 |
| snrpd2 | 0.06226 | 0.500519114 | 0.406 | 0.2 | 1 | 20 |
| rps15a | 0.06278 | 0.397678613 | 0.938 | 0.8 | 1 | 20 |
| rpl15 | 0.06315 | 0.284242453 | 0.906 | 0.867 | 1 | 20 |
| rps5 | 0.06342 | 0.3901756 | 0.906 | 0.733 | 1 | 20 |
| ssb | 0.06364 | 0.385517762 | 0.312 | 0.067 | 1 | 20 |
| actb1 | 0.06405 | -0.49198211 | 0.656 | 0.933 | 1 | 20 |
| TCIM (1 of many) | 0.06429 | 0.318368906 | 0.156 | 0 | 1 | 20 |
| acsbg2 | 0.06429 | 0.218311991 | 0.156 | 0 | 1 | 20 |
| dnajc5gb | 0.06429 | 0.223299328 | 0.156 | 0 | 1 | 20 |
| dnajc7 | 0.06429 | 0.216971326 | 0.156 | 0 | 1 | 20 |
| ndufaf2 | 0.06429 | 0.25702015 | 0.156 | 0 | 1 | 20 |
| ccdc90b | 0.06429 | 0.206226848 | 0.156 | 0 | 1 | 20 |
| cnppd1 | 0.06429 | 0.203476341 | 0.156 | 0 | 1 | 20 |
| dnajb12a | 0.06429 | 0.208864868 | 0.156 | 0 | 1 | 20 |
| eif4ba | 0.06429 | 0.234559945 | 0.156 | 0 | 1 | 20 |
| gigyf2 | 0.06429 | 0.145979046 | 0.156 | 0 | 1 | 20 |
| ing3 | 0.06429 | 0.212485606 | 0.156 | 0 | 1 | 20 |
| morc2 | 0.06429 | 0.188608091 | 0.156 | 0 | 1 | 20 |
| mrps30 | 0.06429 | 0.16528617 | 0.156 | 0 | 1 | 20 |
| nipbla | 0.06429 | 0.204517437 | 0.156 | 0 | 1 | 20 |
| si:ch211-68a17.7 | 0.06429 | 0.214840428 | 0.156 | 0 | 1 | 20 |
| tex10 | 0.06429 | 0.182582086 | 0.156 | 0 | 1 | 20 |
| tnpo2 | 0.06429 | 0.18799739 | 0.156 | 0 | 1 | 20 |
| zgc:101858 | 0.06429 | 0.2524396 | 0.156 | 0 | 1 | 20 |
| zmp:0000000624 | 0.06429 | 0.178456454 | 0.156 | 0 | 1 | 20 |
| FP236812.2 | 0.06429 | 0.256505299 | 0.156 | 0 | 1 | 20 |
| adsl | 0.06429 | 0.25624277 | 0.156 | 0 | 1 | 20 |
| agap2 | 0.06429 | 0.219110142 | 0.156 | 0 | 1 | 20 |
| cux1a | 0.06429 | 0.265062601 | 0.156 | 0 | 1 | 20 |
| gata2b | 0.06429 | 0.196193634 | 0.156 | 0 | 1 | 20 |
| gpn1 | 0.06429 | 0.193275965 | 0.156 | 0 | 1 | 20 |
| rhebl1 | 0.06429 | 0.13893818 | 0.156 | 0 | 1 | 20 |
| srp9 | 0.06429 | 0.17798147 | 0.156 | 0 | 1 | 20 |

| | | | | | | |
|---|---|---|---|---|---|---|
| ssu72 | 0.06429 | 0.232016722 | 0.156 | 0 | 1 | 20 |
| szrd1 | 0.06429 | 0.292020703 | 0.156 | 0 | 1 | 20 |
| upf3b | 0.06429 | 0.298506149 | 0.156 | 0 | 1 | 20 |
| LOC108190785 | 0.06429 | 0.253888017 | 0.156 | 0 | 1 | 20 |
| XLOC-012680 | 0.06429 | 0.267885628 | 0.156 | 0 | 1 | 20 |
| casp3a | 0.06429 | 0.263354328 | 0.156 | 0 | 1 | 20 |
| ccdc9 | 0.06429 | 0.159559636 | 0.156 | 0 | 1 | 20 |
| cdc73 | 0.06429 | 0.201939294 | 0.156 | 0 | 1 | 20 |
| cenpx | 0.06429 | 0.289181197 | 0.156 | 0 | 1 | 20 |
| cmss1 | 0.06429 | 0.179923088 | 0.156 | 0 | 1 | 20 |
| dusp1 | 0.06429 | 0.181399591 | 0.156 | 0 | 1 | 20 |
| fam102ba | 0.06429 | 0.173900561 | 0.156 | 0 | 1 | 20 |
| gtf2a1 | 0.06429 | 0.224950975 | 0.156 | 0 | 1 | 20 |
| gtf3aa | 0.06429 | 0.184781312 | 0.156 | 0 | 1 | 20 |
| hcfc1b | 0.06429 | 0.155843363 | 0.156 | 0 | 1 | 20 |
| lyrm7 | 0.06429 | 0.156126096 | 0.156 | 0 | 1 | 20 |
| mlec | 0.06429 | 0.262410614 | 0.156 | 0 | 1 | 20 |
| mrps9 | 0.06429 | 0.182762247 | 0.156 | 0 | 1 | 20 |
| nat9 | 0.06429 | 0.142777281 | 0.156 | 0 | 1 | 20 |
| nle1 | 0.06429 | 0.192454092 | 0.156 | 0 | 1 | 20 |
| npepps | 0.06429 | 0.155452566 | 0.156 | 0 | 1 | 20 |
| nt5c3a | 0.06429 | 0.183918845 | 0.156 | 0 | 1 | 20 |
| pdhb | 0.06429 | 0.298736759 | 0.156 | 0 | 1 | 20 |
| pgam1b | 0.06429 | 0.201416888 | 0.156 | 0 | 1 | 20 |
| polr1c | 0.06429 | 0.235284068 | 0.156 | 0 | 1 | 20 |
| polr3k | 0.06429 | 0.204709754 | 0.156 | 0 | 1 | 20 |
| prpf3 | 0.06429 | 0.205493292 | 0.156 | 0 | 1 | 20 |
| ptgr1 | 0.06429 | 0.16747983 | 0.156 | 0 | 1 | 20 |
| pttg1 | 0.06429 | 0.277466891 | 0.156 | 0 | 1 | 20 |
| tfam | 0.06429 | 0.219095675 | 0.156 | 0 | 1 | 20 |
| tfr1b | 0.06429 | 0.219645366 | 0.156 | 0 | 1 | 20 |
| timm29 | 0.06429 | 0.186401068 | 0.156 | 0 | 1 | 20 |
| trmt1 | 0.06429 | 0.164418374 | 0.156 | 0 | 1 | 20 |
| tyw3 | 0.06429 | 0.224832793 | 0.156 | 0 | 1 | 20 |
| usp22 | 0.06429 | 0.213232302 | 0.156 | 0 | 1 | 20 |
| wdr45 | 0.06429 | 0.255085921 | 0.156 | 0 | 1 | 20 |
| ythdf2 | 0.06429 | 0.287558909 | 0.156 | 0 | 1 | 20 |
| zgc:101851 | 0.06429 | 0.188782291 | 0.156 | 0 | 1 | 20 |
| zgc:55262 | 0.06429 | 0.281536659 | 0.156 | 0 | 1 | 20 |
| znf142 | 0.06429 | 0.266557192 | 0.156 | 0 | 1 | 20 |
| PDCL3 | 0.06429 | 0.162557854 | 0.156 | 0 | 1 | 20 |
| dcun1d5 | 0.06429 | 0.227014631 | 0.156 | 0 | 1 | 20 |
| ddx56 | 0.06429 | 0.181900459 | 0.156 | 0 | 1 | 20 |
| npepl1 | 0.06429 | 0.206795138 | 0.156 | 0 | 1 | 20 |
| phf3 | 0.06429 | 0.26911973 | 0.156 | 0 | 1 | 20 |
| rbm38 | 0.06429 | 0.197213758 | 0.156 | 0 | 1 | 20 |
| reep3b | 0.06429 | 0.195780818 | 0.156 | 0 | 1 | 20 |
| rprd2b | 0.06429 | 0.285992354 | 0.156 | 0 | 1 | 20 |

| | | | | | | |
|---|---|---|---|---|---|---|
| rpusd4 | 0.06429 | 0.163771096 | 0.156 | 0 | 1 | 20 |
| sephs1 | 0.06429 | 0.221558896 | 0.156 | 0 | 1 | 20 |
| si:ch211-147a11.3 | 0.06429 | 0.235171488 | 0.156 | 0 | 1 | 20 |
| snrpa1 | 0.06429 | 0.154024351 | 0.156 | 0 | 1 | 20 |
| sssca1 | 0.06429 | 0.197397471 | 0.156 | 0 | 1 | 20 |
| thop1 | 0.06429 | 0.157024548 | 0.156 | 0 | 1 | 20 |
| trnau1apa | 0.06429 | 0.176752644 | 0.156 | 0 | 1 | 20 |
| trub1 | 0.06429 | 0.21344571 | 0.156 | 0 | 1 | 20 |
| uba2 | 0.06429 | 0.147237206 | 0.156 | 0 | 1 | 20 |
| zgc:55733 | 0.06429 | 0.21875314 | 0.156 | 0 | 1 | 20 |
| BX005309.1 | 0.06429 | 0.206751691 | 0.156 | 0 | 1 | 20 |
| glrx5 | 0.06429 | 0.236151565 | 0.156 | 0 | 1 | 20 |
| med12 | 0.06429 | 0.231709562 | 0.156 | 0 | 1 | 20 |
| mrm3b | 0.06429 | 0.203616297 | 0.156 | 0 | 1 | 20 |
| pank2 | 0.06429 | 0.211116345 | 0.156 | 0 | 1 | 20 |
| si:dkey-12j5.1 | 0.06429 | 0.210150086 | 0.156 | 0 | 1 | 20 |
| slc25a20 | 0.06429 | 0.295897099 | 0.156 | 0 | 1 | 20 |
| sptlc2b | 0.06429 | 0.176420202 | 0.156 | 0 | 1 | 20 |
| tent4b | 0.06429 | 0.200359184 | 0.156 | 0 | 1 | 20 |
| CABZ01068251.1 | 0.06429 | 0.218236868 | 0.156 | 0 | 1 | 20 |
| ap1b1 | 0.06429 | 0.206018213 | 0.156 | 0 | 1 | 20 |
| arhgef2 | 0.06429 | 0.251548243 | 0.156 | 0 | 1 | 20 |
| cd2bp2 | 0.06429 | 0.182109377 | 0.156 | 0 | 1 | 20 |
| crkl | 0.06429 | 0.214914069 | 0.156 | 0 | 1 | 20 |
| imp3 | 0.06429 | 0.161606868 | 0.156 | 0 | 1 | 20 |
| mroh1 | 0.06429 | 0.214872779 | 0.156 | 0 | 1 | 20 |
| nt5c2a | 0.06429 | 0.195895119 | 0.156 | 0 | 1 | 20 |
| nvl | 0.06429 | 0.217139334 | 0.156 | 0 | 1 | 20 |
| phb2b | 0.06429 | 0.227969593 | 0.156 | 0 | 1 | 20 |
| rcor1 | 0.06429 | 0.298943744 | 0.156 | 0 | 1 | 20 |
| retsat | 0.06429 | 0.212705333 | 0.156 | 0 | 1 | 20 |
| rnaseh2b | 0.06429 | 0.184889507 | 0.156 | 0 | 1 | 20 |
| setd2 | 0.06429 | 0.217680508 | 0.156 | 0 | 1 | 20 |
| si:ch211-276i12.9 | 0.06429 | 0.201294656 | 0.156 | 0 | 1 | 20 |
| si:dkey-103e21.5 | 0.06429 | 0.193983391 | 0.156 | 0 | 1 | 20 |
| siva1 | 0.06429 | 0.213949901 | 0.156 | 0 | 1 | 20 |
| taf11 | 0.06429 | 0.212414102 | 0.156 | 0 | 1 | 20 |
| znf622 | 0.06429 | 0.207654513 | 0.156 | 0 | 1 | 20 |
| myh9a | 0.06445 | -0.55394893 | 0.031 | 0.267 | 1 | 20 |
| eef1db | 0.06447 | 0.614166164 | 0.438 | 0.133 | 1 | 20 |
| rps18 | 0.06451 | 0.284108378 | 0.906 | 0.8 | 1 | 20 |
| tcp1 | 0.06493 | 0.521918643 | 0.625 | 0.267 | 1 | 20 |
| cct4 | 0.06519 | 0.543464075 | 0.688 | 0.333 | 1 | 20 |
| rpl36a | 0.06545 | 0.376268979 | 0.938 | 0.733 | 1 | 20 |
| ndufb2 | 0.06575 | 0.457025441 | 0.344 | 0.067 | 1 | 20 |
| alox5a | 0.06624 | -0.67715393 | 0.031 | 0.267 | 1 | 20 |
| atp5md | 0.06631 | 0.444544507 | 0.438 | 0.267 | 1 | 20 |
| mdh1aa | 0.06641 | 0.501379332 | 0.625 | 0.267 | 1 | 20 |

| | | | | | | |
|---|---|---|---|---|---|---|
| ssrp1a | 0.06653 | 0.449304195 | 0.281 | 0.067 | 1 | 20 |
| CR753876.1 | 0.06722 | -0.75583779 | 0.156 | 0.4 | 1 | 20 |
| rbm4.3 | 0.06728 | 0.284874631 | 0.656 | 0.333 | 1 | 20 |
| eif5 | 0.06741 | 0.509674904 | 0.594 | 0.267 | 1 | 20 |
| rac2 | 0.06742 | -0.20810409 | 0.531 | 0.467 | 1 | 20 |
| mt-atp6 | 0.06753 | -0.44041449 | 0.094 | 0.333 | 1 | 20 |
| ndufb4 | 0.06778 | 0.535980366 | 0.406 | 0.133 | 1 | 20 |
| ssr4 | 0.06811 | 0.467205622 | 0.625 | 0.267 | 1 | 20 |
| cox7c | 0.06946 | 0.197475705 | 0.625 | 0.333 | 1 | 20 |
| hars | 0.06985 | 0.498234635 | 0.312 | 0.067 | 1 | 20 |
| eif3g | 0.07028 | 0.444307081 | 0.75 | 0.4 | 1 | 20 |
| h2afvb | 0.07074 | 0.552831995 | 0.688 | 0.333 | 1 | 20 |
| soga1 | 0.07106 | -0.40198297 | 0.219 | 0.267 | 1 | 20 |
| rab1ab | 0.07181 | -0.44490201 | 0.281 | 0.333 | 1 | 20 |
| eif3k | 0.07231 | 0.455719828 | 0.688 | 0.333 | 1 | 20 |
| mych | 0.07249 | -0.52107481 | 0.281 | 0.4 | 1 | 20 |
| eif4a1b | 0.0727 | 0.466778072 | 0.688 | 0.333 | 1 | 20 |
| rps26l | 0.07287 | 0.325715262 | 0.938 | 0.8 | 1 | 20 |
| cdc26 | 0.07321 | 0.130872387 | 0.281 | 0.067 | 1 | 20 |
| sf1 | 0.07464 | 0.408425677 | 0.344 | 0.067 | 1 | 20 |
| ptges3a | 0.07507 | 0.146165373 | 0.25 | 0.067 | 1 | 20 |
| gnai2a | 0.07574 | -0.58565905 | 0.25 | 0.533 | 1 | 20 |
| ndufs7 | 0.07659 | 0.539422092 | 0.312 | 0.067 | 1 | 20 |
| sod1 | 0.07695 | 0.215894962 | 0.688 | 0.4 | 1 | 20 |
| pnrc2 | 0.07701 | -0.34014517 | 0.844 | 0.933 | 1 | 20 |
| psmb2 | 0.07701 | 0.30062042 | 0.594 | 0.267 | 1 | 20 |
| lsm14ab | 0.07731 | -0.25752567 | 0.219 | 0.2 | 1 | 20 |
| rpl5b | 0.07806 | 0.470440975 | 0.844 | 0.533 | 1 | 20 |
| psma1 | 0.07817 | 0.507694651 | 0.656 | 0.333 | 1 | 20 |
| hmgn7 | 0.07831 | 0.528636558 | 0.719 | 0.4 | 1 | 20 |
| rpl8 | 0.07875 | 0.381749368 | 0.938 | 0.733 | 1 | 20 |
| aimp2 | 0.0796 | 0.325013254 | 0.344 | 0.067 | 1 | 20 |
| rbm28 | 0.08059 | 0.478122099 | 0.531 | 0.2 | 1 | 20 |
| hmgb3a | 0.08126 | 0.440708152 | 0.406 | 0.133 | 1 | 20 |
| u2af2b | 0.0818 | -0.19360748 | 0.25 | 0.2 | 1 | 20 |
| anp32a | 0.08272 | -0.22745175 | 0.469 | 0.4 | 1 | 20 |
| gbp | 0.08344 | -0.29295639 | 0.625 | 0.6 | 1 | 20 |
| stmn1a | 0.08439 | 0.296668086 | 0.469 | 0.2 | 1 | 20 |
| eef1g | 0.0849 | 0.279830722 | 0.938 | 0.8 | 1 | 20 |
| psmg2 | 0.08517 | 0.452958237 | 0.438 | 0.133 | 1 | 20 |
| ppp6c | 0.08604 | -0.56916957 | 0.219 | 0.4 | 1 | 20 |
| psmg3 | 0.08692 | -0.30511657 | 0.25 | 0.267 | 1 | 20 |
| ube2v2 | 0.08715 | 0.253963853 | 0.562 | 0.267 | 1 | 20 |
| cul1a | 0.08735 | -0.38014328 | 0.156 | 0.2 | 1 | 20 |
| srprb | 0.08781 | 0.460556179 | 0.406 | 0.133 | 1 | 20 |
| rpl29 | 0.08822 | 0.198284925 | 0.938 | 0.933 | 1 | 20 |
| romo1 | 0.08832 | 0.36466002 | 0.719 | 0.4 | 1 | 20 |
| metap2b | 0.08838 | 0.519373347 | 0.656 | 0.333 | 1 | 20 |

| | | | | | | |
|---|---|---|---|---|---|---|
| rps6kb1b | 0.08934 | 0.340211236 | 0.344 | 0.067 | 1 | 20 |
| eif2s2 | 0.08966 | 0.332378973 | 0.656 | 0.333 | 1 | 20 |
| nudcd2 | 0.09068 | 0.437001035 | 0.312 | 0.067 | 1 | 20 |
| si:dkey-112e7.2 | 0.09084 | 0.520824868 | 0.688 | 0.4 | 1 | 20 |
| snu13b | 0.09272 | 0.550405193 | 0.5 | 0.267 | 1 | 20 |
| acin1a | 0.09322 | 0.122089895 | 0.5 | 0.267 | 1 | 20 |
| dusp5 | 0.09327 | -0.17249919 | 0.188 | 0.133 | 1 | 20 |
| caprin1b | 0.09398 | 0.489645622 | 0.5 | 0.267 | 1 | 20 |
| rps29 | 0.09461 | 0.317380899 | 0.969 | 0.8 | 1 | 20 |
| mrpl36 | 0.09466 | 0.496778428 | 0.5 | 0.2 | 1 | 20 |
| ddx61 | 0.09489 | 0.312542568 | 0.5 | 0.2 | 1 | 20 |
| rpl27a | 0.09506 | 0.111879596 | 0.719 | 0.533 | 1 | 20 |
| rpl6 | 0.09646 | 0.342957144 | 0.938 | 0.733 | 1 | 20 |
| irf2 | 0.09736 | -0.54136208 | 0.125 | 0.333 | 1 | 20 |
| oaz1b | 0.09738 | -0.2606507 | 0.656 | 0.6 | 1 | 20 |
| mir223 | 0.09809 | 0.386915427 | 0.5 | 0.2 | 1 | 20 |
| tagapb | 0.09821 | 0.44182956 | 0.438 | 0.133 | 1 | 20 |
| coro1a | 0.09927 | -0.15175833 | 0.375 | 0.267 | 1 | 20 |
| CR936442.1 | 0.09931 | -0.52797857 | 0.188 | 0.467 | 1 | 20 |
| gspt1l | 0.09987 | 0.408840142 | 0.438 | 0.133 | 1 | 20 |
| dok2 | 0.10017 | -0.51296236 | 0.031 | 0.2 | 1 | 20 |
| tomm5 | 0.10018 | 0.511307596 | 0.5 | 0.2 | 1 | 20 |
| sdc4 | 0.10074 | -0.48752567 | 0.156 | 0.267 | 1 | 20 |
| eif3ea | 0.1017 | 0.390943224 | 0.719 | 0.4 | 1 | 20 |
| myl12.2 | 0.10178 | -0.10423845 | 0.219 | 0.133 | 1 | 20 |
| sav1 | 0.10179 | -0.44623364 | 0.062 | 0.133 | 1 | 20 |
| sfpq | 0.103 | -0.24043415 | 0.156 | 0.133 | 1 | 20 |
| rbm8a | 0.10314 | 0.401769252 | 0.594 | 0.267 | 1 | 20 |
| rab10 | 0.10366 | 0.488712275 | 0.562 | 0.267 | 1 | 20 |
| arl6ip1 | 0.10381 | -0.45362869 | 0.562 | 0.733 | 1 | 20 |
| polr2f | 0.10398 | 0.379944946 | 0.406 | 0.2 | 1 | 20 |
| dpy30 | 0.10488 | -0.48507212 | 0.25 | 0.4 | 1 | 20 |
| GARNL3 | 0.10561 | -0.56342971 | 0.031 | 0.2 | 1 | 20 |
| histh1l | 0.10591 | -0.28459213 | 0.312 | 0.267 | 1 | 20 |
| ppp1r12a | 0.10614 | -0.5162236 | 0.062 | 0.2 | 1 | 20 |
| aldocb | 0.10624 | 0.19376677 | 0.125 | 0 | 1 | 20 |
| gnas | 0.10624 | 0.195682446 | 0.125 | 0 | 1 | 20 |
| lrmp | 0.10624 | 0.19076232 | 0.125 | 0 | 1 | 20 |
| mcmbp | 0.10624 | 0.159197544 | 0.125 | 0 | 1 | 20 |
| pdcd2 | 0.10624 | 0.190419527 | 0.125 | 0 | 1 | 20 |
| pmaip1 | 0.10624 | 0.205956968 | 0.125 | 0 | 1 | 20 |
| slc30a5 | 0.10624 | 0.162238379 | 0.125 | 0 | 1 | 20 |
| utp11l | 0.10624 | 0.182942999 | 0.125 | 0 | 1 | 20 |
| XLOC-005448 | 0.10624 | 0.158733467 | 0.125 | 0 | 1 | 20 |
| atp2a2b | 0.10624 | 0.130543003 | 0.125 | 0 | 1 | 20 |
| cad | 0.10624 | 0.195416521 | 0.125 | 0 | 1 | 20 |
| msl1a | 0.10624 | 0.153497175 | 0.125 | 0 | 1 | 20 |
| ndufaf4 | 0.10624 | 0.242247401 | 0.125 | 0 | 1 | 20 |

| | | | | | | |
|---|---|---|---|---|---|---|
| nedd8l | 0.10624 | 0.238047807 | 0.125 | 0 | 1 | 20 |
| CABZ01077218.1 | 0.10624 | 0.158667572 | 0.125 | 0 | 1 | 20 |
| CU074419.2 | 0.10624 | 0.162933601 | 0.125 | 0 | 1 | 20 |
| XLOC-011085 | 0.10624 | 0.194913017 | 0.125 | 0 | 1 | 20 |
| acot13 | 0.10624 | 0.179801223 | 0.125 | 0 | 1 | 20 |
| ankrd54 | 0.10624 | 0.232868672 | 0.125 | 0 | 1 | 20 |
| arid2 | 0.10624 | 0.156387338 | 0.125 | 0 | 1 | 20 |
| arl6ip5a | 0.10624 | 0.29495187 | 0.125 | 0 | 1 | 20 |
| borcs7 | 0.10624 | 0.149637512 | 0.125 | 0 | 1 | 20 |
| cat | 0.10624 | 0.133356084 | 0.125 | 0 | 1 | 20 |
| cbsa | 0.10624 | 0.112567304 | 0.125 | 0 | 1 | 20 |
| ccdc32 | 0.10624 | 0.137079134 | 0.125 | 0 | 1 | 20 |
| copb2 | 0.10624 | 0.147138373 | 0.125 | 0 | 1 | 20 |
| ddt | 0.10624 | 0.118315909 | 0.125 | 0 | 1 | 20 |
| dph6 | 0.10624 | 0.233898531 | 0.125 | 0 | 1 | 20 |
| dus1l | 0.10624 | 0.169448981 | 0.125 | 0 | 1 | 20 |
| dync1li2 | 0.10624 | 0.159486191 | 0.125 | 0 | 1 | 20 |
| fam210ab | 0.10624 | 0.142162062 | 0.125 | 0 | 1 | 20 |
| fam45a | 0.10624 | 0.199252876 | 0.125 | 0 | 1 | 20 |
| fip1l1b | 0.10624 | 0.185298327 | 0.125 | 0 | 1 | 20 |
| gemin2 | 0.10624 | 0.137225514 | 0.125 | 0 | 1 | 20 |
| gnl2 | 0.10624 | 0.154111673 | 0.125 | 0 | 1 | 20 |
| gsk3ab | 0.10624 | 0.199874604 | 0.125 | 0 | 1 | 20 |
| hibadhb | 0.10624 | 0.139674053 | 0.125 | 0 | 1 | 20 |
| inpp5b | 0.10624 | 0.150599336 | 0.125 | 0 | 1 | 20 |
| irf2a | 0.10624 | 0.204052575 | 0.125 | 0 | 1 | 20 |
| isy1 | 0.10624 | 0.206759425 | 0.125 | 0 | 1 | 20 |
| itgb1bp1 | 0.10624 | 0.215424878 | 0.125 | 0 | 1 | 20 |
| kansl1b | 0.10624 | 0.159023068 | 0.125 | 0 | 1 | 20 |
| kdm4b | 0.10624 | 0.228311559 | 0.125 | 0 | 1 | 20 |
| kyat3 | 0.10624 | 0.171253553 | 0.125 | 0 | 1 | 20 |
| marcksb | 0.10624 | 0.160747127 | 0.125 | 0 | 1 | 20 |
| med20 | 0.10624 | 0.198192551 | 0.125 | 0 | 1 | 20 |
| mesd | 0.10624 | 0.163197481 | 0.125 | 0 | 1 | 20 |
| mrpl24 | 0.10624 | 0.176410886 | 0.125 | 0 | 1 | 20 |
| mtdhb | 0.10624 | 0.15366788 | 0.125 | 0 | 1 | 20 |
| mtg2 | 0.10624 | 0.118908681 | 0.125 | 0 | 1 | 20 |
| mthfd1b | 0.10624 | 0.179853862 | 0.125 | 0 | 1 | 20 |
| n6amt1 | 0.10624 | 0.153866964 | 0.125 | 0 | 1 | 20 |
| ndufaf5 | 0.10624 | 0.175190649 | 0.125 | 0 | 1 | 20 |
| nfe2l3 | 0.10624 | 0.152143684 | 0.125 | 0 | 1 | 20 |
| nfxl1 | 0.10624 | 0.182377835 | 0.125 | 0 | 1 | 20 |
| nol11 | 0.10624 | 0.200427883 | 0.125 | 0 | 1 | 20 |
| nsrp1 | 0.10624 | 0.195781687 | 0.125 | 0 | 1 | 20 |
| nupl2 | 0.10624 | 0.203081101 | 0.125 | 0 | 1 | 20 |
| pcmt | 0.10624 | 0.251836927 | 0.125 | 0 | 1 | 20 |
| phb2a | 0.10624 | 0.133479192 | 0.125 | 0 | 1 | 20 |
| pitpnb | 0.10624 | 0.187516436 | 0.125 | 0 | 1 | 20 |

| | | | | | | |
|---|---|---|---|---|---|---|
| pon3.2 | 0.10624 | 0.182475668 | 0.125 | 0 | 1 | 20 |
| prpf4 | 0.10624 | 0.136417661 | 0.125 | 0 | 1 | 20 |
| psmd12 | 0.10624 | 0.164326039 | 0.125 | 0 | 1 | 20 |
| qtrt2 | 0.10624 | 0.138962503 | 0.125 | 0 | 1 | 20 |
| rab20 | 0.10624 | 0.247753214 | 0.125 | 0 | 1 | 20 |
| rab6a | 0.10624 | 0.17883793 | 0.125 | 0 | 1 | 20 |
| rnf121 | 0.10624 | 0.158116849 | 0.125 | 0 | 1 | 20 |
| sgf29 | 0.10624 | 0.163391173 | 0.125 | 0 | 1 | 20 |
| si:ch211-255f4.5 | 0.10624 | 0.130421588 | 0.125 | 0 | 1 | 20 |
| si:ch211-276i12.11 | 0.10624 | 0.240406688 | 0.125 | 0 | 1 | 20 |
| si:dkey-261j15.2 | 0.10624 | 0.210445941 | 0.125 | 0 | 1 | 20 |
| si:dkeyp-87e7.4 | 0.10624 | 0.201998831 | 0.125 | 0 | 1 | 20 |
| slc7a3a | 0.10624 | 0.106125489 | 0.125 | 0 | 1 | 20 |
| slc7a6os | 0.10624 | 0.1538183 | 0.125 | 0 | 1 | 20 |
| smc3 | 0.10624 | 0.144375532 | 0.125 | 0 | 1 | 20 |
| smco4 | 0.10624 | 0.113259337 | 0.125 | 0 | 1 | 20 |
| snapc1b | 0.10624 | 0.193175534 | 0.125 | 0 | 1 | 20 |
| srm | 0.10624 | 0.13885154 | 0.125 | 0 | 1 | 20 |
| sts | 0.10624 | 0.20903136 | 0.125 | 0 | 1 | 20 |
| sucla2 | 0.10624 | 0.174855336 | 0.125 | 0 | 1 | 20 |
| tiprl | 0.10624 | 0.173778365 | 0.125 | 0 | 1 | 20 |
| tmem69 | 0.10624 | 0.11286784 | 0.125 | 0 | 1 | 20 |
| trnt1 | 0.10624 | 0.161440327 | 0.125 | 0 | 1 | 20 |
| ttc1 | 0.10624 | 0.128729784 | 0.125 | 0 | 1 | 20 |
| wdr5 | 0.10624 | 0.202170638 | 0.125 | 0 | 1 | 20 |
| zbtb37 | 0.10624 | 0.189091506 | 0.125 | 0 | 1 | 20 |
| zdhhc23a | 0.10624 | 0.154682824 | 0.125 | 0 | 1 | 20 |
| zgc:110269 | 0.10624 | 0.162214983 | 0.125 | 0 | 1 | 20 |
| zgc:113184 | 0.10624 | 0.1315688 | 0.125 | 0 | 1 | 20 |
| zgc:161969 | 0.10624 | 0.156230231 | 0.125 | 0 | 1 | 20 |
| CU467861.1 | 0.10624 | 0.132507088 | 0.125 | 0 | 1 | 20 |
| LOC110438491 | 0.10624 | 0.117288904 | 0.125 | 0 | 1 | 20 |
| jmjd7 | 0.10624 | 0.161432469 | 0.125 | 0 | 1 | 20 |
| pou2f1b | 0.10624 | 0.17143329 | 0.125 | 0 | 1 | 20 |
| scnm1 | 0.10624 | 0.232058867 | 0.125 | 0 | 1 | 20 |
| si:dkey-172h23.2 | 0.10624 | 0.230422921 | 0.125 | 0 | 1 | 20 |
| stxbp3 | 0.10624 | 0.213357427 | 0.125 | 0 | 1 | 20 |
| zcchc7 | 0.10624 | 0.132368379 | 0.125 | 0 | 1 | 20 |
| ATP11B | 0.10624 | 0.145107757 | 0.125 | 0 | 1 | 20 |
| FP340257.1 | 0.10624 | 0.130945903 | 0.125 | 0 | 1 | 20 |
| XLOC-043861 | 0.10624 | 0.130544076 | 0.125 | 0 | 1 | 20 |
| aktip | 0.10624 | 0.232220754 | 0.125 | 0 | 1 | 20 |
| alg3 | 0.10624 | 0.247435257 | 0.125 | 0 | 1 | 20 |
| chtopa | 0.10624 | 0.16736886 | 0.125 | 0 | 1 | 20 |
| cnot3b | 0.10624 | 0.143004136 | 0.125 | 0 | 1 | 20 |
| ddx54 | 0.10624 | 0.144574353 | 0.125 | 0 | 1 | 20 |
| dhdds | 0.10624 | 0.150870549 | 0.125 | 0 | 1 | 20 |
| eef2a.1 | 0.10624 | 0.128392845 | 0.125 | 0 | 1 | 20 |

| | | | | | | |
|---|---|---|---|---|---|---|
| exosc7 | 0.10624 | 0.152510393 | 0.125 | 0 | 1 | 20 |
| gb:bc139872 | 0.10624 | 0.127028978 | 0.125 | 0 | 1 | 20 |
| hk1 | 0.10624 | 0.1371268 | 0.125 | 0 | 1 | 20 |
| jak2a | 0.10624 | 0.269877064 | 0.125 | 0 | 1 | 20 |
| med7 | 0.10624 | 0.158525244 | 0.125 | 0 | 1 | 20 |
| metap1d | 0.10624 | 0.195359703 | 0.125 | 0 | 1 | 20 |
| mpped2 | 0.10624 | 0.204798129 | 0.125 | 0 | 1 | 20 |
| mpped2a | 0.10624 | 0.156338181 | 0.125 | 0 | 1 | 20 |
| mrps28 | 0.10624 | 0.140035346 | 0.125 | 0 | 1 | 20 |
| nipsnap3a | 0.10624 | 0.200650368 | 0.125 | 0 | 1 | 20 |
| notch2 | 0.10624 | 0.204304245 | 0.125 | 0 | 1 | 20 |
| nsun5 | 0.10624 | 0.133320991 | 0.125 | 0 | 1 | 20 |
| pcyox1 | 0.10624 | 0.159990792 | 0.125 | 0 | 1 | 20 |
| ppp5c | 0.10624 | 0.219653936 | 0.125 | 0 | 1 | 20 |
| sdf2l1 | 0.10624 | 0.106045684 | 0.125 | 0 | 1 | 20 |
| sdr42e1 | 0.10624 | 0.141660955 | 0.125 | 0 | 1 | 20 |
| si:dkey-19e4.5 | 0.10624 | 0.153657747 | 0.125 | 0 | 1 | 20 |
| sppl3 | 0.10624 | 0.201095005 | 0.125 | 0 | 1 | 20 |
| trmt61a | 0.10624 | 0.211815636 | 0.125 | 0 | 1 | 20 |
| xpnpep1 | 0.10624 | 0.172021278 | 0.125 | 0 | 1 | 20 |
| xrcc6 | 0.10624 | 0.235682116 | 0.125 | 0 | 1 | 20 |
| zgc:101016 | 0.10624 | 0.148595883 | 0.125 | 0 | 1 | 20 |
| znf800b | 0.10624 | 0.182957034 | 0.125 | 0 | 1 | 20 |
| cavin2b | 0.10624 | 0.261707088 | 0.125 | 0 | 1 | 20 |
| golph3 | 0.10624 | 0.145117424 | 0.125 | 0 | 1 | 20 |
| hgh1 | 0.10624 | 0.132014112 | 0.125 | 0 | 1 | 20 |
| maip1 | 0.10624 | 0.143503784 | 0.125 | 0 | 1 | 20 |
| mrpl15 | 0.10624 | 0.182385127 | 0.125 | 0 | 1 | 20 |
| prpf38a | 0.10624 | 0.131350393 | 0.125 | 0 | 1 | 20 |
| si:ch211-132b12.7 | 0.10624 | 0.142820801 | 0.125 | 0 | 1 | 20 |
| CU570894.1 | 0.10624 | 0.136750715 | 0.125 | 0 | 1 | 20 |
| RASA2 | 0.10624 | 0.173762409 | 0.125 | 0 | 1 | 20 |
| XLOC-011450 | 0.10624 | 0.169405429 | 0.125 | 0 | 1 | 20 |
| aagab | 0.10624 | 0.18236471 | 0.125 | 0 | 1 | 20 |
| acat1 | 0.10624 | 0.263684767 | 0.125 | 0 | 1 | 20 |
| alg6 | 0.10624 | 0.130647578 | 0.125 | 0 | 1 | 20 |
| alkbh5 | 0.10624 | 0.164815094 | 0.125 | 0 | 1 | 20 |
| arfip1 | 0.10624 | 0.182196758 | 0.125 | 0 | 1 | 20 |
| atpaf2 | 0.10624 | 0.156300403 | 0.125 | 0 | 1 | 20 |
| brd1b | 0.10624 | 0.211031369 | 0.125 | 0 | 1 | 20 |
| cdkn1a | 0.10624 | 0.264503388 | 0.125 | 0 | 1 | 20 |
| ciao1 | 0.10624 | 0.210022075 | 0.125 | 0 | 1 | 20 |
| cks2 | 0.10624 | 0.209994617 | 0.125 | 0 | 1 | 20 |
| cops8 | 0.10624 | 0.139571244 | 0.125 | 0 | 1 | 20 |
| ctdp1 | 0.10624 | 0.13610253 | 0.125 | 0 | 1 | 20 |
| dele1 | 0.10624 | 0.163330249 | 0.125 | 0 | 1 | 20 |
| dnajc15 | 0.10624 | 0.142553077 | 0.125 | 0 | 1 | 20 |
| eif2b1 | 0.10624 | 0.119420042 | 0.125 | 0 | 1 | 20 |

| | | | | | | |
|---|---|---|---|---|---|---|
| elmsan1b | 0.10624 | 0.1979179 | 0.125 | 0 | 1 | 20 |
| fis1 | 0.10624 | 0.272944809 | 0.125 | 0 | 1 | 20 |
| fli1b | 0.10624 | 0.164749678 | 0.125 | 0 | 1 | 20 |
| ggh | 0.10624 | 0.16911902 | 0.125 | 0 | 1 | 20 |
| got1 | 0.10624 | 0.151009684 | 0.125 | 0 | 1 | 20 |
| gpd1l | 0.10624 | 0.14942411 | 0.125 | 0 | 1 | 20 |
| gpkow | 0.10624 | 0.153487939 | 0.125 | 0 | 1 | 20 |
| hmg20b | 0.10624 | 0.152658378 | 0.125 | 0 | 1 | 20 |
| hsd17b12b | 0.10624 | 0.117674325 | 0.125 | 0 | 1 | 20 |
| irf10 | 0.10624 | 0.117024233 | 0.125 | 0 | 1 | 20 |
| larsb | 0.10624 | 0.158736941 | 0.125 | 0 | 1 | 20 |
| lonrf1 | 0.10624 | 0.144944476 | 0.125 | 0 | 1 | 20 |
| lyrm9 | 0.10624 | 0.170548783 | 0.125 | 0 | 1 | 20 |
| mgst1.1 | 0.10624 | 0.211459425 | 0.125 | 0 | 1 | 20 |
| mrpl58 | 0.10624 | 0.128292669 | 0.125 | 0 | 1 | 20 |
| nek7 | 0.10624 | 0.188985458 | 0.125 | 0 | 1 | 20 |
| nup188 | 0.10624 | 0.14395475 | 0.125 | 0 | 1 | 20 |
| pik3r2 | 0.10624 | 0.165854875 | 0.125 | 0 | 1 | 20 |
| rab3ip | 0.10624 | 0.161658762 | 0.125 | 0 | 1 | 20 |
| rnmt | 0.10624 | 0.150413524 | 0.125 | 0 | 1 | 20 |
| rpp21 | 0.10624 | 0.140943035 | 0.125 | 0 | 1 | 20 |
| rxrba | 0.10624 | 0.122392464 | 0.125 | 0 | 1 | 20 |
| si:ch211-197k17.3 | 0.10624 | 0.17067233 | 0.125 | 0 | 1 | 20 |
| si:ch73-269m14.2 | 0.10624 | 0.186971988 | 0.125 | 0 | 1 | 20 |
| si:ch73-37h15.2 | 0.10624 | 0.189640964 | 0.125 | 0 | 1 | 20 |
| si:dkey-184a18.5 | 0.10624 | 0.170863368 | 0.125 | 0 | 1 | 20 |
| si:dkey-27b3.2 | 0.10624 | 0.10356238 | 0.125 | 0 | 1 | 20 |
| si:dkey-73n10.1 | 0.10624 | 0.180751125 | 0.125 | 0 | 1 | 20 |
| si:dkeyp-85e10.3 | 0.10624 | 0.174400266 | 0.125 | 0 | 1 | 20 |
| styx | 0.10624 | 0.156136418 | 0.125 | 0 | 1 | 20 |
| suox | 0.10624 | 0.207019515 | 0.125 | 0 | 1 | 20 |
| supt6h | 0.10624 | 0.171043459 | 0.125 | 0 | 1 | 20 |
| tango2 | 0.10624 | 0.131674366 | 0.125 | 0 | 1 | 20 |
| thoc3 | 0.10624 | 0.186166859 | 0.125 | 0 | 1 | 20 |
| tm9sf3 | 0.10624 | 0.150656732 | 0.125 | 0 | 1 | 20 |
| tpra | 0.10624 | 0.157092387 | 0.125 | 0 | 1 | 20 |
| tprkb | 0.10624 | 0.241209281 | 0.125 | 0 | 1 | 20 |
| trappc10 | 0.10624 | 0.187331227 | 0.125 | 0 | 1 | 20 |
| ubr3 | 0.10624 | 0.159502752 | 0.125 | 0 | 1 | 20 |
| vapal | 0.10624 | 0.272446689 | 0.125 | 0 | 1 | 20 |
| xpc | 0.10624 | 0.145425897 | 0.125 | 0 | 1 | 20 |
| zgc:92335 | 0.10624 | 0.16503092 | 0.125 | 0 | 1 | 20 |
| zgc:92590 | 0.10624 | 0.270717494 | 0.125 | 0 | 1 | 20 |
| zmp:0000001228 | 0.10624 | 0.213388865 | 0.125 | 0 | 1 | 20 |
| znf609b | 0.10624 | 0.142424737 | 0.125 | 0 | 1 | 20 |
| smc1al | 0.10684 | 0.42832018 | 0.406 | 0.133 | 1 | 20 |
| eif1b | 0.10697 | -0.28580181 | 0.594 | 0.6 | 1 | 20 |
| irf2bp2a | 0.10712 | -0.43752316 | 0.344 | 0.467 | 1 | 20 |

| | | | | | | |
|---|---|---|---|---|---|---|
| polr2l | 0.10906 | 0.312092098 | 0.406 | 0.133 | 1 | 20 |
| tomm20a | 0.10913 | -0.11527509 | 0.375 | 0.267 | 1 | 20 |
| si:ch211-213a13.1 | 0.10953 | 0.240654075 | 0.281 | 0.067 | 1 | 20 |
| btk | 0.10973 | -0.48271992 | 0.094 | 0.2 | 1 | 20 |
| polr3e | 0.11139 | 0.348415555 | 0.312 | 0.067 | 1 | 20 |
| edf1 | 0.11146 | -0.18643621 | 0.531 | 0.467 | 1 | 20 |
| plk3 | 0.11246 | -0.64154611 | 0.125 | 0.4 | 1 | 20 |
| eef1a1l1 | 0.11261 | 0.225327033 | 0.938 | 0.867 | 1 | 20 |
| adra2a | 0.11289 | -0.56699154 | 0.031 | 0.2 | 1 | 20 |
| naa10 | 0.11331 | 0.490883689 | 0.5 | 0.267 | 1 | 20 |
| jun | 0.11334 | -0.6171502 | 0.125 | 0.267 | 1 | 20 |
| crip1 | 0.11336 | 0.427792545 | 0.719 | 0.4 | 1 | 20 |
| h3f3a | 0.11359 | 0.454170699 | 0.656 | 0.333 | 1 | 20 |
| selenok | 0.11402 | -0.50961857 | 0.188 | 0.467 | 1 | 20 |
| krt18b | 0.11408 | 0.360616473 | 0.312 | 0.067 | 1 | 20 |
| llph | 0.11452 | 0.36240858 | 0.406 | 0.133 | 1 | 20 |
| apln | 0.11457 | -0.57039038 | 0.094 | 0.333 | 1 | 20 |
| cox7b | 0.11502 | 0.433172008 | 0.719 | 0.4 | 1 | 20 |
| si:ch211-102c2.4 | 0.11514 | -0.68750739 | 0.125 | 0.4 | 1 | 20 |
| zgc:174863 | 0.11518 | 0.361419573 | 0.312 | 0.067 | 1 | 20 |
| srgn | 0.11632 | -0.74292715 | 0.219 | 0.467 | 1 | 20 |
| hnrnpa1b | 0.11684 | 0.476655886 | 0.5 | 0.2 | 1 | 20 |
| btf3l4 | 0.11717 | 0.390445014 | 0.406 | 0.133 | 1 | 20 |
| rps28 | 0.11729 | 0.312772271 | 0.906 | 0.8 | 1 | 20 |
| shmt1 | 0.11753 | 0.370934219 | 0.312 | 0.067 | 1 | 20 |
| pdha1a | 0.11758 | -0.54003578 | 0.188 | 0.4 | 1 | 20 |
| COX5B | 0.11803 | 0.281730441 | 0.688 | 0.4 | 1 | 20 |
| dkc1 | 0.1184 | 0.461806153 | 0.469 | 0.267 | 1 | 20 |
| larp4ab | 0.11946 | 0.394919147 | 0.312 | 0.067 | 1 | 20 |
| fam133b | 0.1198 | 0.321450345 | 0.312 | 0.067 | 1 | 20 |
| nfyba | 0.12078 | -0.34898907 | 0.469 | 0.533 | 1 | 20 |
| psmg1 | 0.12078 | 0.246444897 | 0.312 | 0.067 | 1 | 20 |
| UBL4A | 0.12107 | -0.5495477 | 0.031 | 0.2 | 1 | 20 |
| acap1 | 0.12168 | -0.14540096 | 0.188 | 0.133 | 1 | 20 |
| hnrnpk | 0.1223 | 0.390139058 | 0.375 | 0.133 | 1 | 20 |
| ponzr1 | 0.12275 | 0.347180197 | 0.406 | 0.133 | 1 | 20 |
| si:ch211-226m16.2 | 0.12312 | -0.52097676 | 0.406 | 0.6 | 1 | 20 |
| tsc22d3 | 0.12329 | -0.55706708 | 0.312 | 0.6 | 1 | 20 |
| bloc1s5 | 0.12367 | -0.54401469 | 0.062 | 0.267 | 1 | 20 |
| chmp1b | 0.12385 | -0.55153817 | 0.219 | 0.4 | 1 | 20 |
| prmt1 | 0.12413 | 0.381550482 | 0.5 | 0.2 | 1 | 20 |
| polr2i | 0.1246 | -0.14713568 | 0.188 | 0.133 | 1 | 20 |
| plscr3b | 0.12515 | -0.46683836 | 0.031 | 0.2 | 1 | 20 |
| fn1b | 0.12538 | -0.28574513 | 0.062 | 0.267 | 1 | 20 |
| dnajc1 | 0.12582 | 0.343783322 | 0.281 | 0.067 | 1 | 20 |
| dnmt3bb.1 | 0.12608 | 0.508561833 | 0.344 | 0.133 | 1 | 20 |
| laptm4a | 0.12628 | -0.36128734 | 0.219 | 0.267 | 1 | 20 |
| arih2 | 0.1264 | -0.46210864 | 0.031 | 0.2 | 1 | 20 |

| | | | | | | |
|---|---|---|---|---|---|---|
| lsm12b | 0.12662 | 0.107289719 | 0.5 | 0.267 | 1 | 20 |
| napab | 0.12682 | 0.282393631 | 0.281 | 0.067 | 1 | 20 |
| smdt1b | 0.12699 | 0.224694772 | 0.375 | 0.133 | 1 | 20 |
| msra | 0.12723 | 0.339796978 | 0.312 | 0.067 | 1 | 20 |
| sntb2 | 0.12775 | -0.20544708 | 0.156 | 0.133 | 1 | 20 |
| nhp2 | 0.12793 | 0.395665021 | 0.469 | 0.333 | 1 | 20 |
| hbegfb | 0.12801 | -0.69044504 | 0.156 | 0.333 | 1 | 20 |
| cdkn2aipnl | 0.12859 | 0.235061959 | 0.375 | 0.133 | 1 | 20 |
| serinc2l | 0.12867 | -0.43706134 | 0.031 | 0.2 | 1 | 20 |
| ptk2ba | 0.12883 | -0.33533319 | 0.094 | 0.133 | 1 | 20 |
| si:ch211-214j24.10 | 0.12883 | 0.385525442 | 0.406 | 0.133 | 1 | 20 |
| parp1 | 0.12932 | 0.358478343 | 0.312 | 0.067 | 1 | 20 |
| calm1b | 0.12958 | -0.47357659 | 0.062 | 0.267 | 1 | 20 |
| adgrg1 | 0.12998 | -0.60453803 | 0.062 | 0.267 | 1 | 20 |
| dyrk1b | 0.13094 | -0.43296052 | 0.031 | 0.2 | 1 | 20 |
| myg1 | 0.13097 | 0.423436489 | 0.406 | 0.133 | 1 | 20 |
| sat1a.2 | 0.13158 | -0.42630349 | 0.438 | 0.533 | 1 | 20 |
| atrx | 0.13181 | 0.220022088 | 0.469 | 0.2 | 1 | 20 |
| cbr1l | 0.13192 | 0.313041719 | 0.312 | 0.067 | 1 | 20 |
| snrpd1 | 0.13198 | 0.406760503 | 0.5 | 0.2 | 1 | 20 |
| ap2m1a | 0.13199 | 0.366657065 | 0.5 | 0.2 | 1 | 20 |
| ndufa2 | 0.13262 | 0.374719045 | 0.281 | 0.067 | 1 | 20 |
| mcts1 | 0.13322 | 0.383468614 | 0.344 | 0.133 | 1 | 20 |
| reep5 | 0.13325 | -0.31463853 | 0.469 | 0.467 | 1 | 20 |
| coa3a | 0.13342 | 0.260238546 | 0.312 | 0.067 | 1 | 20 |
| sgk1 | 0.13344 | -0.4262139 | 0.062 | 0.267 | 1 | 20 |
| si:zfos-1192g2.3 | 0.13359 | 0.399964146 | 0.281 | 0.067 | 1 | 20 |
| aldh6a1 | 0.1343 | 0.299884729 | 0.312 | 0.067 | 1 | 20 |
| rpz5 | 0.13458 | -0.5232732 | 0.156 | 0.267 | 1 | 20 |
| si:ch73-42p12.2 | 0.13521 | 0.30691689 | 0.312 | 0.067 | 1 | 20 |
| ndufa12 | 0.13576 | 0.385348876 | 0.375 | 0.133 | 1 | 20 |
| si:ch211-281l24.3 | 0.13612 | -0.32844578 | 0.094 | 0.133 | 1 | 20 |
| rpl31 | 0.13618 | 0.401822357 | 0.844 | 0.6 | 1 | 20 |
| atp2b1a | 0.13633 | -0.53997707 | 0.156 | 0.4 | 1 | 20 |
| phf20b | 0.13745 | -0.18287689 | 0.156 | 0.133 | 1 | 20 |
| gosr2 | 0.13759 | 0.314925091 | 0.281 | 0.067 | 1 | 20 |
| ahcy | 0.13906 | 0.495160221 | 0.531 | 0.333 | 1 | 20 |
| snrpg | 0.13996 | 0.117027286 | 0.469 | 0.267 | 1 | 20 |
| ndufa6 | 0.14049 | 0.396234643 | 0.406 | 0.133 | 1 | 20 |
| cd63 | 0.14065 | -0.44331653 | 0.469 | 0.6 | 1 | 20 |
| si:ch73-91k6.2 | 0.14126 | -0.46168238 | 0.062 | 0.267 | 1 | 20 |
| ciarta | 0.14176 | -0.55908433 | 0.188 | 0.4 | 1 | 20 |
| rrs1 | 0.14211 | 0.336459414 | 0.281 | 0.067 | 1 | 20 |
| si:ch211-196l7.4 | 0.14294 | -0.48052121 | 0.625 | 0.733 | 1 | 20 |
| fth1a | 0.14303 | -0.41256207 | 0.188 | 0.467 | 1 | 20 |
| pomp | 0.14317 | 0.268817946 | 0.625 | 0.333 | 1 | 20 |
| arpc2 | 0.1435 | -0.20728025 | 0.781 | 0.733 | 1 | 20 |
| pinx1 | 0.14357 | 0.448352697 | 0.531 | 0.267 | 1 | 20 |

| | | | | | | |
|---|---|---|---|---|---|---|
| tpm4a | 0.14382 | -0.66025122 | 0.188 | 0.467 | 1 | 20 |
| cirbpb | 0.14455 | 0.304406986 | 0.906 | 0.667 | 1 | 20 |
| nop16 | 0.14471 | 0.316734037 | 0.25 | 0.133 | 1 | 20 |
| sf3b5 | 0.14489 | 0.190178262 | 0.375 | 0.133 | 1 | 20 |
| slirp | 0.14504 | 0.355448126 | 0.438 | 0.2 | 1 | 20 |
| alyref | 0.14565 | 0.450959626 | 0.25 | 0.067 | 1 | 20 |
| lgals3a | 0.14608 | -0.33117967 | 0.094 | 0.333 | 1 | 20 |
| mrpl18 | 0.14732 | 0.36663754 | 0.281 | 0.067 | 1 | 20 |
| ebna1bp2 | 0.14844 | 0.355959024 | 0.281 | 0.067 | 1 | 20 |
| rrp1 | 0.14878 | 0.371347365 | 0.406 | 0.2 | 1 | 20 |
| srsf10b | 0.14888 | 0.369278289 | 0.281 | 0.067 | 1 | 20 |
| gpatch4 | 0.1492 | 0.394475025 | 0.281 | 0.067 | 1 | 20 |
| ncf2 | 0.14921 | 0.458540021 | 0.281 | 0.133 | 1 | 20 |
| atp5f1b | 0.14941 | 0.346128543 | 0.812 | 0.533 | 1 | 20 |
| vsir | 0.14987 | -0.39426769 | 0.219 | 0.333 | 1 | 20 |
| pcna | 0.15077 | 0.336997327 | 0.375 | 0.133 | 1 | 20 |
| rhoac | 0.15106 | -0.53883426 | 0.312 | 0.533 | 1 | 20 |
| prelid1a | 0.15123 | 0.366616667 | 0.281 | 0.133 | 1 | 20 |
| mak16 | 0.15129 | 0.373202945 | 0.281 | 0.067 | 1 | 20 |
| fkbp2 | 0.15134 | 0.107569551 | 0.344 | 0.133 | 1 | 20 |
| commd7 | 0.15145 | 0.226390972 | 0.281 | 0.067 | 1 | 20 |
| tefa | 0.15159 | -0.24377758 | 0.344 | 0.333 | 1 | 20 |
| tagln2 | 0.1518 | -0.52174159 | 0.5 | 0.733 | 1 | 20 |
| lcorl | 0.15203 | -0.4052965 | 0.062 | 0.267 | 1 | 20 |
| tle2c | 0.1528 | -0.37885222 | 0.062 | 0.267 | 1 | 20 |
| mtap | 0.15375 | -0.50173297 | 0.188 | 0.4 | 1 | 20 |
| nudt9 | 0.15416 | 0.37937473 | 0.281 | 0.067 | 1 | 20 |
| ISCU (1 of many) | 0.15529 | -0.43765454 | 0.125 | 0.333 | 1 | 20 |
| znf593 | 0.15537 | 0.378033898 | 0.406 | 0.2 | 1 | 20 |
| ccdc47 | 0.15591 | -0.11570429 | 0.125 | 0.067 | 1 | 20 |
| tbcb | 0.156 | -0.42211753 | 0.031 | 0.2 | 1 | 20 |
| usp10 | 0.15608 | 0.366829388 | 0.25 | 0.067 | 1 | 20 |
| ndufb3 | 0.15622 | 0.24565213 | 0.469 | 0.2 | 1 | 20 |
| nup58 | 0.15665 | 0.115184171 | 0.219 | 0.067 | 1 | 20 |
| saraf | 0.15678 | -0.34137245 | 0.031 | 0.2 | 1 | 20 |
| tkfc | 0.15693 | -0.3339571 | 0.031 | 0.2 | 1 | 20 |
| CT030188.1 | 0.15718 | -0.33246184 | 0.031 | 0.2 | 1 | 20 |
| yipf6 | 0.15745 | -0.39983919 | 0.031 | 0.2 | 1 | 20 |
| si:ch211-119e14.1 | 0.1575 | 0.121726057 | 0.219 | 0.067 | 1 | 20 |
| hells | 0.15766 | 0.425855456 | 0.406 | 0.2 | 1 | 20 |
| LOC101886015 | 0.15786 | -0.54392978 | 0.031 | 0.2 | 1 | 20 |
| f5 | 0.15787 | -0.63502023 | 0.031 | 0.2 | 1 | 20 |
| ypel1 | 0.15807 | -0.1370446 | 0.031 | 0.2 | 1 | 20 |
| si:ch211-71m22.1 | 0.15819 | -0.33536191 | 0.094 | 0.133 | 1 | 20 |
| rhag | 0.15839 | -0.48248066 | 0.062 | 0.267 | 1 | 20 |
| stk17b | 0.15862 | -0.43731683 | 0.094 | 0.267 | 1 | 20 |
| nfkbiaa | 0.15892 | 0.379413655 | 0.281 | 0.067 | 1 | 20 |
| mrpl33 | 0.15912 | 0.309134294 | 0.281 | 0.067 | 1 | 20 |

| | | | | | | |
|---|---|---|---|---|---|---|
| kdm6ba | 0.15914 | -0.25419904 | 0.125 | 0.133 | 1 | 20 |
| XLOC-019062 | 0.15948 | -0.48995358 | 0.062 | 0.2 | 1 | 20 |
| elocb | 0.16022 | 0.296931664 | 0.375 | 0.133 | 1 | 20 |
| smarca4a | 0.16035 | 0.375594782 | 0.344 | 0.133 | 1 | 20 |
| polr2c | 0.16083 | -0.11718884 | 0.125 | 0.067 | 1 | 20 |
| tfdp2 | 0.1612 | -0.31522634 | 0.031 | 0.2 | 1 | 20 |
| si:dkey-261j4.4 | 0.16189 | -0.34362438 | 0.031 | 0.2 | 1 | 20 |
| klf7b | 0.16254 | -0.37191632 | 0.031 | 0.2 | 1 | 20 |
| lrrc8aa | 0.16259 | -0.11142966 | 0.125 | 0.067 | 1 | 20 |
| psma3 | 0.16262 | 0.461805763 | 0.531 | 0.267 | 1 | 20 |
| zgc:77056 | 0.16336 | -0.31905053 | 0.062 | 0.267 | 1 | 20 |
| eif4ebp2 | 0.16345 | 0.456909755 | 0.469 | 0.2 | 1 | 20 |
| emc2 | 0.16378 | 0.389166673 | 0.281 | 0.067 | 1 | 20 |
| cox4i1 | 0.16389 | 0.213553069 | 0.719 | 0.467 | 1 | 20 |
| CU499330.1 | 0.16435 | 0.509450552 | 0.5 | 0.267 | 1 | 20 |
| mettl5 | 0.16455 | -0.324857 | 0.031 | 0.2 | 1 | 20 |
| tes | 0.16492 | -0.41831321 | 0.062 | 0.133 | 1 | 20 |
| cirbpa | 0.16495 | 0.388620868 | 0.812 | 0.6 | 1 | 20 |
| pfn1 | 0.16504 | -0.22920206 | 0.969 | 1 | 1 | 20 |
| il10ra | 0.16546 | -0.32083871 | 0.031 | 0.2 | 1 | 20 |
| carhsp1 | 0.16649 | -0.18388989 | 0.312 | 0.267 | 1 | 20 |
| taf13 | 0.16778 | -0.36607672 | 0.219 | 0.267 | 1 | 20 |
| rap1b | 0.16799 | -0.47125948 | 0.625 | 0.733 | 1 | 20 |
| eif5a | 0.16808 | 0.215839937 | 0.688 | 0.467 | 1 | 20 |
| ergic2 | 0.16847 | -0.11332468 | 0.125 | 0.067 | 1 | 20 |
| ptmaa | 0.16907 | -0.23124267 | 0.938 | 1 | 1 | 20 |
| ndufab1a | 0.17059 | 0.365653974 | 0.312 | 0.133 | 1 | 20 |
| nutf2 | 0.17068 | -0.22630198 | 0.219 | 0.2 | 1 | 20 |
| sf3b2 | 0.17086 | -0.44451986 | 0.125 | 0.333 | 1 | 20 |
| aamp | 0.17091 | 0.327760053 | 0.312 | 0.133 | 1 | 20 |
| atp5if1a | 0.17091 | 0.364374372 | 0.625 | 0.333 | 1 | 20 |
| cd81a | 0.17211 | -0.3423446 | 0.062 | 0.267 | 1 | 20 |
| ddx21 | 0.17222 | 0.38533043 | 0.625 | 0.333 | 1 | 20 |
| yrk | 0.17242 | -0.53377089 | 0.094 | 0.267 | 1 | 20 |
| abce1 | 0.17294 | 0.322810598 | 0.344 | 0.2 | 1 | 20 |
| edem1 | 0.17313 | -0.37337314 | 0.062 | 0.133 | 1 | 20 |
| nuak1a | 0.17327 | -0.57993882 | 0.031 | 0.2 | 1 | 20 |
| fhl2a | 0.17334 | -0.63607228 | 0.062 | 0.267 | 1 | 20 |
| rap1gap2a | 0.17354 | -0.5972285 | 0.062 | 0.267 | 1 | 20 |
| ipo7 | 0.17358 | 0.377354882 | 0.344 | 0.133 | 1 | 20 |
| btbd10b | 0.17403 | -0.15162371 | 0.062 | 0.267 | 1 | 20 |
| vma21 | 0.17408 | 0.371551583 | 0.375 | 0.133 | 1 | 20 |
| rps8a | 0.17412 | 0.299543331 | 0.875 | 0.733 | 1 | 20 |
| agpat3 | 0.17437 | -0.25041067 | 0.062 | 0.267 | 1 | 20 |
| tpm3 | 0.17444 | -0.33747391 | 0.594 | 0.6 | 1 | 20 |
| macf1a | 0.17456 | -0.38669968 | 0.062 | 0.267 | 1 | 20 |
| celf4 | 0.17461 | -0.56194856 | 0.062 | 0.267 | 1 | 20 |
| si:dkey-46a10.3 | 0.17467 | -0.34548668 | 0.062 | 0.267 | 1 | 20 |

| | | | | | | |
|---|---|---|---|---|---|---|
| zgc:162730 | 0.17474 | -0.59966019 | 0.188 | 0.4 | 1 | 20 |
| mob4 | 0.17508 | 0.139394287 | 0.219 | 0.067 | 1 | 20 |
| apbb1ip | 0.17519 | -0.33136712 | 0.031 | 0.2 | 1 | 20 |
| CR847984.1 | 0.17586 | 0.146720143 | 0.094 | 0 | 1 | 20 |
| aclya | 0.17586 | 0.107944447 | 0.094 | 0 | 1 | 20 |
| apip | 0.17586 | 0.186060161 | 0.094 | 0 | 1 | 20 |
| atg4c | 0.17586 | 0.12592329 | 0.094 | 0 | 1 | 20 |
| cdca4 | 0.17586 | 0.124748763 | 0.094 | 0 | 1 | 20 |
| cdk7 | 0.17586 | 0.15652311 | 0.094 | 0 | 1 | 20 |
| cnnm2a | 0.17586 | 0.102069279 | 0.094 | 0 | 1 | 20 |
| coq10b | 0.17586 | 0.103078019 | 0.094 | 0 | 1 | 20 |
| ctdnep1a | 0.17586 | 0.119078677 | 0.094 | 0 | 1 | 20 |
| dpf2 | 0.17586 | 0.117897488 | 0.094 | 0 | 1 | 20 |
| dtnba | 0.17586 | 0.145885358 | 0.094 | 0 | 1 | 20 |
| ecd | 0.17586 | 0.110487862 | 0.094 | 0 | 1 | 20 |
| edc3 | 0.17586 | 0.165338358 | 0.094 | 0 | 1 | 20 |
| emsy | 0.17586 | 0.124313635 | 0.094 | 0 | 1 | 20 |
| erap1a | 0.17586 | 0.161265297 | 0.094 | 0 | 1 | 20 |
| ftsj1 | 0.17586 | 0.12167316 | 0.094 | 0 | 1 | 20 |
| fut9b | 0.17586 | 0.149680464 | 0.094 | 0 | 1 | 20 |
| gpr132b | 0.17586 | 0.120272355 | 0.094 | 0 | 1 | 20 |
| herc2 | 0.17586 | 0.141455012 | 0.094 | 0 | 1 | 20 |
| ighv1-4 | 0.17586 | 0.184138785 | 0.094 | 0 | 1 | 20 |
| jade1 | 0.17586 | 0.10012771 | 0.094 | 0 | 1 | 20 |
| mark3b | 0.17586 | 0.157951568 | 0.094 | 0 | 1 | 20 |
| mat2b | 0.17586 | 0.178378489 | 0.094 | 0 | 1 | 20 |
| mecr | 0.17586 | 0.128162669 | 0.094 | 0 | 1 | 20 |
| med18 | 0.17586 | 0.101119567 | 0.094 | 0 | 1 | 20 |
| mgrn1a | 0.17586 | 0.158990792 | 0.094 | 0 | 1 | 20 |
| mrpl10 | 0.17586 | 0.145148393 | 0.094 | 0 | 1 | 20 |
| mrpl3 | 0.17586 | 0.15534701 | 0.094 | 0 | 1 | 20 |
| mrps5 | 0.17586 | 0.174467456 | 0.094 | 0 | 1 | 20 |
| mtif3 | 0.17586 | 0.104809939 | 0.094 | 0 | 1 | 20 |
| myd88 | 0.17586 | 0.225037293 | 0.094 | 0 | 1 | 20 |
| mynn | 0.17586 | 0.113670072 | 0.094 | 0 | 1 | 20 |
| nfkbib | 0.17586 | 0.132228346 | 0.094 | 0 | 1 | 20 |
| nup35 | 0.17586 | 0.209008575 | 0.094 | 0 | 1 | 20 |
| p4hb | 0.17586 | 0.178744074 | 0.094 | 0 | 1 | 20 |
| parp12b | 0.17586 | 0.129549904 | 0.094 | 0 | 1 | 20 |
| pdhx | 0.17586 | 0.111433806 | 0.094 | 0 | 1 | 20 |
| pgap2 | 0.17586 | 0.109313455 | 0.094 | 0 | 1 | 20 |
| ppil2 | 0.17586 | 0.103839175 | 0.094 | 0 | 1 | 20 |
| prpf18 | 0.17586 | 0.130998825 | 0.094 | 0 | 1 | 20 |
| psmd9 | 0.17586 | 0.114153006 | 0.094 | 0 | 1 | 20 |
| ptcd3 | 0.17586 | 0.158211615 | 0.094 | 0 | 1 | 20 |
| rdh10b | 0.17586 | 0.100690833 | 0.094 | 0 | 1 | 20 |
| riox1 | 0.17586 | 0.128777286 | 0.094 | 0 | 1 | 20 |
| ripk3 | 0.17586 | 0.102597722 | 0.094 | 0 | 1 | 20 |

| | | | | | | |
|---|---|---|---|---|---|---|
| senp6a | 0.17586 | 0.106330086 | 0.094 | 0 | 1 | 20 |
| sh3gl1b | 0.17586 | 0.169844971 | 0.094 | 0 | 1 | 20 |
| slc16a13 | 0.17586 | 0.122321392 | 0.094 | 0 | 1 | 20 |
| smim12 | 0.17586 | 0.112234465 | 0.094 | 0 | 1 | 20 |
| spag7 | 0.17586 | 0.149367237 | 0.094 | 0 | 1 | 20 |
| srfbp1 | 0.17586 | 0.114311659 | 0.094 | 0 | 1 | 20 |
| ssbp1 | 0.17586 | 0.101680692 | 0.094 | 0 | 1 | 20 |
| stx12 | 0.17586 | 0.157760858 | 0.094 | 0 | 1 | 20 |
| sulf2b | 0.17586 | 0.144666341 | 0.094 | 0 | 1 | 20 |
| tent5c | 0.17586 | 0.151205599 | 0.094 | 0 | 1 | 20 |
| thg1l | 0.17586 | 0.109217619 | 0.094 | 0 | 1 | 20 |
| tm2d2 | 0.17586 | 0.211054883 | 0.094 | 0 | 1 | 20 |
| tmem53 | 0.17586 | 0.11976633 | 0.094 | 0 | 1 | 20 |
| tnksb | 0.17586 | 0.210260061 | 0.094 | 0 | 1 | 20 |
| tp53i11a | 0.17586 | 0.136213944 | 0.094 | 0 | 1 | 20 |
| usb1 | 0.17586 | 0.184214754 | 0.094 | 0 | 1 | 20 |
| wbp11 | 0.17586 | 0.120658859 | 0.094 | 0 | 1 | 20 |
| zgc:91944 | 0.17586 | 0.13744825 | 0.094 | 0 | 1 | 20 |
| CABZ01046093.1 | 0.17586 | 0.111280062 | 0.094 | 0 | 1 | 20 |
| CHST13 | 0.17586 | 0.195789455 | 0.094 | 0 | 1 | 20 |
| LOC101883999 | 0.17586 | 0.152290042 | 0.094 | 0 | 1 | 20 |
| XLOC-005449 | 0.17586 | 0.17991773 | 0.094 | 0 | 1 | 20 |
| acsl4b | 0.17586 | 0.157510176 | 0.094 | 0 | 1 | 20 |
| apmap | 0.17586 | 0.106572688 | 0.094 | 0 | 1 | 20 |
| arf1 | 0.17586 | 0.202781183 | 0.094 | 0 | 1 | 20 |
| btaf1 | 0.17586 | 0.128306925 | 0.094 | 0 | 1 | 20 |
| c1galt1b | 0.17586 | 0.116211965 | 0.094 | 0 | 1 | 20 |
| cdc123 | 0.17586 | 0.173415289 | 0.094 | 0 | 1 | 20 |
| cdc27 | 0.17586 | 0.136230215 | 0.094 | 0 | 1 | 20 |
| clybl | 0.17586 | 0.130895748 | 0.094 | 0 | 1 | 20 |
| cpox | 0.17586 | 0.128715294 | 0.094 | 0 | 1 | 20 |
| csnk1g2a | 0.17586 | 0.132165298 | 0.094 | 0 | 1 | 20 |
| ctnnbl1 | 0.17586 | 0.105431489 | 0.094 | 0 | 1 | 20 |
| dnajc11a | 0.17586 | 0.145811765 | 0.094 | 0 | 1 | 20 |
| dtx4b | 0.17586 | 0.119443783 | 0.094 | 0 | 1 | 20 |
| fam76b | 0.17586 | 0.129772042 | 0.094 | 0 | 1 | 20 |
| fas | 0.17586 | 0.136505756 | 0.094 | 0 | 1 | 20 |
| fbxo45 | 0.17586 | 0.11094404 | 0.094 | 0 | 1 | 20 |
| frs2b | 0.17586 | 0.135694141 | 0.094 | 0 | 1 | 20 |
| hadhab | 0.17586 | 0.145344793 | 0.094 | 0 | 1 | 20 |
| ipo13 | 0.17586 | 0.136113716 | 0.094 | 0 | 1 | 20 |
| josd2 | 0.17586 | 0.187310713 | 0.094 | 0 | 1 | 20 |
| lrrc8db | 0.17586 | 0.147707759 | 0.094 | 0 | 1 | 20 |
| march5l | 0.17586 | 0.137450874 | 0.094 | 0 | 1 | 20 |
| mef2ca | 0.17586 | 0.108485618 | 0.094 | 0 | 1 | 20 |
| naa30 | 0.17586 | 0.128223113 | 0.094 | 0 | 1 | 20 |
| parp3 | 0.17586 | 0.142961157 | 0.094 | 0 | 1 | 20 |
| pcgf1 | 0.17586 | 0.26246611 | 0.094 | 0 | 1 | 20 |

| | | | | | | |
|---|---|---|---|---|---|---|
| pdcd11 | 0.17586 | 0.10677999 | 0.094 | 0 | 1 | 20 |
| plekho2 | 0.17586 | 0.219844726 | 0.094 | 0 | 1 | 20 |
| rab1ba | 0.17586 | 0.17883013 | 0.094 | 0 | 1 | 20 |
| rbm14a | 0.17586 | 0.155301565 | 0.094 | 0 | 1 | 20 |
| riok1 | 0.17586 | 0.120018943 | 0.094 | 0 | 1 | 20 |
| rnf25 | 0.17586 | 0.110498081 | 0.094 | 0 | 1 | 20 |
| sae1 | 0.17586 | 0.111454496 | 0.094 | 0 | 1 | 20 |
| sec24a | 0.17586 | 0.131674477 | 0.094 | 0 | 1 | 20 |
| si:ch211-113e8.10 | 0.17586 | 0.198084473 | 0.094 | 0 | 1 | 20 |
| si:ch73-100l22.3 | 0.17586 | 0.212285088 | 0.094 | 0 | 1 | 20 |
| si:dkey-100n23.3 | 0.17586 | 0.267679158 | 0.094 | 0 | 1 | 20 |
| slc35b3 | 0.17586 | 0.139176687 | 0.094 | 0 | 1 | 20 |
| slc6a6b | 0.17586 | 0.112517964 | 0.094 | 0 | 1 | 20 |
| smyd2a | 0.17586 | 0.124054886 | 0.094 | 0 | 1 | 20 |
| spata2 | 0.17586 | 0.121102254 | 0.094 | 0 | 1 | 20 |
| srcap | 0.17586 | 0.150795819 | 0.094 | 0 | 1 | 20 |
| stambpa | 0.17586 | 0.124101401 | 0.094 | 0 | 1 | 20 |
| taf6l | 0.17586 | 0.136239488 | 0.094 | 0 | 1 | 20 |
| tfdp1b | 0.17586 | 0.207795722 | 0.094 | 0 | 1 | 20 |
| tfe3a | 0.17586 | 0.22431055 | 0.094 | 0 | 1 | 20 |
| tmem33 | 0.17586 | 0.204792972 | 0.094 | 0 | 1 | 20 |
| ttc4 | 0.17586 | 0.136799241 | 0.094 | 0 | 1 | 20 |
| tubg1 | 0.17586 | 0.135205979 | 0.094 | 0 | 1 | 20 |
| tusc2b | 0.17586 | 0.106709809 | 0.094 | 0 | 1 | 20 |
| usp24 | 0.17586 | 0.186679225 | 0.094 | 0 | 1 | 20 |
| zgc:66160 | 0.17586 | 0.194921045 | 0.094 | 0 | 1 | 20 |
| znf1001 | 0.17586 | 0.122682674 | 0.094 | 0 | 1 | 20 |
| znf292a | 0.17586 | 0.139183579 | 0.094 | 0 | 1 | 20 |
| LOC100333266 | 0.17586 | 0.128414268 | 0.094 | 0 | 1 | 20 |
| acss2 | 0.17586 | 0.174164572 | 0.094 | 0 | 1 | 20 |
| as3mt | 0.17586 | 0.176235326 | 0.094 | 0 | 1 | 20 |
| cfdp1 | 0.17586 | 0.166793935 | 0.094 | 0 | 1 | 20 |
| dusp12 | 0.17586 | 0.109093794 | 0.094 | 0 | 1 | 20 |
| edc4 | 0.17586 | 0.181462299 | 0.094 | 0 | 1 | 20 |
| lnpep | 0.17586 | 0.195401546 | 0.094 | 0 | 1 | 20 |
| lztfl1 | 0.17586 | 0.100678078 | 0.094 | 0 | 1 | 20 |
| mydgf | 0.17586 | 0.135864701 | 0.094 | 0 | 1 | 20 |
| ndufaf3 | 0.17586 | 0.138760209 | 0.094 | 0 | 1 | 20 |
| nup160 | 0.17586 | 0.132030357 | 0.094 | 0 | 1 | 20 |
| orc3 | 0.17586 | 0.118855646 | 0.094 | 0 | 1 | 20 |
| phf20l1 | 0.17586 | 0.147019909 | 0.094 | 0 | 1 | 20 |
| serpinb1l1 | 0.17586 | 0.167383212 | 0.094 | 0 | 1 | 20 |
| trir | 0.17586 | 0.154160252 | 0.094 | 0 | 1 | 20 |
| txndc17 | 0.17586 | 0.106254154 | 0.094 | 0 | 1 | 20 |
| vps41 | 0.17586 | 0.170941108 | 0.094 | 0 | 1 | 20 |
| BX248521.2 | 0.17586 | 0.178551377 | 0.094 | 0 | 1 | 20 |
| CABZ01088274.1 | 0.17586 | 0.112164933 | 0.094 | 0 | 1 | 20 |
| CSKMT | 0.17586 | 0.126662598 | 0.094 | 0 | 1 | 20 |

| | | | | | | |
|---|---|---|---|---|---|---|
| LOC101884690 | 0.17586 | 0.106317459 | 0.094 | 0 | 1 | 20 |
| adipor1b | 0.17586 | 0.146350593 | 0.094 | 0 | 1 | 20 |
| afmid | 0.17586 | 0.165534266 | 0.094 | 0 | 1 | 20 |
| ankrd10a | 0.17586 | 0.103010304 | 0.094 | 0 | 1 | 20 |
| badb | 0.17586 | 0.203115901 | 0.094 | 0 | 1 | 20 |
| chd2 | 0.17586 | 0.167531811 | 0.094 | 0 | 1 | 20 |
| cinp | 0.17586 | 0.132745679 | 0.094 | 0 | 1 | 20 |
| dgcr8 | 0.17586 | 0.118878821 | 0.094 | 0 | 1 | 20 |
| dnajc9 | 0.17586 | 0.164328032 | 0.094 | 0 | 1 | 20 |
| dtwd1 | 0.17586 | 0.110872601 | 0.094 | 0 | 1 | 20 |
| epb41a | 0.17586 | 0.143637607 | 0.094 | 0 | 1 | 20 |
| fbxw8 | 0.17586 | 0.110240674 | 0.094 | 0 | 1 | 20 |
| fhl3a | 0.17586 | 0.112622836 | 0.094 | 0 | 1 | 20 |
| fpgt | 0.17586 | 0.103250639 | 0.094 | 0 | 1 | 20 |
| iba57 | 0.17586 | 0.114394581 | 0.094 | 0 | 1 | 20 |
| im:7140055 | 0.17586 | 0.114418971 | 0.094 | 0 | 1 | 20 |
| ints1 | 0.17586 | 0.116903271 | 0.094 | 0 | 1 | 20 |
| map3k1 | 0.17586 | 0.177302471 | 0.094 | 0 | 1 | 20 |
| mrps34 | 0.17586 | 0.174639048 | 0.094 | 0 | 1 | 20 |
| nfatc2ip | 0.17586 | 0.110292801 | 0.094 | 0 | 1 | 20 |
| nudt1 | 0.17586 | 0.105563041 | 0.094 | 0 | 1 | 20 |
| polr3d | 0.17586 | 0.116166076 | 0.094 | 0 | 1 | 20 |
| ppp2r2ab | 0.17586 | 0.130156278 | 0.094 | 0 | 1 | 20 |
| rnf168 | 0.17586 | 0.115013618 | 0.094 | 0 | 1 | 20 |
| si:ch211-255f4.3 | 0.17586 | 0.159214051 | 0.094 | 0 | 1 | 20 |
| si:ch73-147o17.1 | 0.17586 | 0.230599587 | 0.094 | 0 | 1 | 20 |
| si:dkey-58f10.10 | 0.17586 | 0.130457584 | 0.094 | 0 | 1 | 20 |
| srpk1b | 0.17586 | 0.103135478 | 0.094 | 0 | 1 | 20 |
| tmed1b | 0.17586 | 0.167914563 | 0.094 | 0 | 1 | 20 |
| tmem187 | 0.17586 | 0.104126142 | 0.094 | 0 | 1 | 20 |
| trap1 | 0.17586 | 0.11913278 | 0.094 | 0 | 1 | 20 |
| trip12 | 0.17586 | 0.145088176 | 0.094 | 0 | 1 | 20 |
| trmt12 | 0.17586 | 0.109477672 | 0.094 | 0 | 1 | 20 |
| ubp1 | 0.17586 | 0.106157835 | 0.094 | 0 | 1 | 20 |
| utrn | 0.17586 | 0.146341411 | 0.094 | 0 | 1 | 20 |
| wbp1 | 0.17586 | 0.132813355 | 0.094 | 0 | 1 | 20 |
| wdfy1 | 0.17586 | 0.152002259 | 0.094 | 0 | 1 | 20 |
| zgc:173587 | 0.17586 | 0.108781732 | 0.094 | 0 | 1 | 20 |
| CABZ01084963.1 | 0.17586 | 0.108108569 | 0.094 | 0 | 1 | 20 |
| LOC100535806 | 0.17586 | 0.12316688 | 0.094 | 0 | 1 | 20 |
| adnpb | 0.17586 | 0.144596793 | 0.094 | 0 | 1 | 20 |
| arcn1b | 0.17586 | 0.135476604 | 0.094 | 0 | 1 | 20 |
| atg16l1 | 0.17586 | 0.187536879 | 0.094 | 0 | 1 | 20 |
| baxa | 0.17586 | 0.132617373 | 0.094 | 0 | 1 | 20 |
| cab39l1 | 0.17586 | 0.139172022 | 0.094 | 0 | 1 | 20 |
| casp8ap2 | 0.17586 | 0.105921918 | 0.094 | 0 | 1 | 20 |
| cbwd | 0.17586 | 0.108405163 | 0.094 | 0 | 1 | 20 |
| ccdc167 | 0.17586 | 0.109996303 | 0.094 | 0 | 1 | 20 |

| | | | | | | |
|---|---|---|---|---|---|---|
| cd99 | 0.17586 | 0.102548878 | 0.094 | 0 | 1 | 20 |
| cldn12 | 0.17586 | 0.144127809 | 0.094 | 0 | 1 | 20 |
| cox15 | 0.17586 | 0.181182312 | 0.094 | 0 | 1 | 20 |
| cpsf2 | 0.17586 | 0.11248885 | 0.094 | 0 | 1 | 20 |
| cybc1 | 0.17586 | 0.146731 | 0.094 | 0 | 1 | 20 |
| derl1 | 0.17586 | 0.118717199 | 0.094 | 0 | 1 | 20 |
| dus2 | 0.17586 | 0.123098054 | 0.094 | 0 | 1 | 20 |
| eftud2 | 0.17586 | 0.100555064 | 0.094 | 0 | 1 | 20 |
| elk4 | 0.17586 | 0.102082271 | 0.094 | 0 | 1 | 20 |
| elovl4a | 0.17586 | 0.110897801 | 0.094 | 0 | 1 | 20 |
| enox2 | 0.17586 | 0.113018269 | 0.094 | 0 | 1 | 20 |
| erap2 | 0.17586 | 0.146286429 | 0.094 | 0 | 1 | 20 |
| etfdh | 0.17586 | 0.144911302 | 0.094 | 0 | 1 | 20 |
| fbxw7 | 0.17586 | 0.210376605 | 0.094 | 0 | 1 | 20 |
| foxp1b | 0.17586 | 0.117124061 | 0.094 | 0 | 1 | 20 |
| gatb | 0.17586 | 0.114616294 | 0.094 | 0 | 1 | 20 |
| gpr55a | 0.17586 | 0.139877487 | 0.094 | 0 | 1 | 20 |
| gtf2h4 | 0.17586 | 0.130686956 | 0.094 | 0 | 1 | 20 |
| herpud2 | 0.17586 | 0.123014831 | 0.094 | 0 | 1 | 20 |
| hmgcrb | 0.17586 | 0.160333295 | 0.094 | 0 | 1 | 20 |
| hmox2b | 0.17586 | 0.107298301 | 0.094 | 0 | 1 | 20 |
| hpdl | 0.17586 | 0.104233106 | 0.094 | 0 | 1 | 20 |
| idh3g | 0.17586 | 0.103915751 | 0.094 | 0 | 1 | 20 |
| itprid2 | 0.17586 | 0.252971206 | 0.094 | 0 | 1 | 20 |
| larp4b | 0.17586 | 0.111347616 | 0.094 | 0 | 1 | 20 |
| lsg1 | 0.17586 | 0.109167092 | 0.094 | 0 | 1 | 20 |
| mcfd2 | 0.17586 | 0.152616777 | 0.094 | 0 | 1 | 20 |
| mettl2a | 0.17586 | 0.118248572 | 0.094 | 0 | 1 | 20 |
| mmadhc | 0.17586 | 0.131380484 | 0.094 | 0 | 1 | 20 |
| mrm2 | 0.17586 | 0.103443726 | 0.094 | 0 | 1 | 20 |
| mterf3 | 0.17586 | 0.113490792 | 0.094 | 0 | 1 | 20 |
| ncoa5 | 0.17586 | 0.127528514 | 0.094 | 0 | 1 | 20 |
| nras | 0.17586 | 0.134599778 | 0.094 | 0 | 1 | 20 |
| nsun2 | 0.17586 | 0.128398773 | 0.094 | 0 | 1 | 20 |
| parn | 0.17586 | 0.119211388 | 0.094 | 0 | 1 | 20 |
| pigp | 0.17586 | 0.108341882 | 0.094 | 0 | 1 | 20 |
| pip5k1ba | 0.17586 | 0.183694949 | 0.094 | 0 | 1 | 20 |
| ppardb | 0.17586 | 0.107694005 | 0.094 | 0 | 1 | 20 |
| prim1 | 0.17586 | 0.110670061 | 0.094 | 0 | 1 | 20 |
| rangap1b | 0.17586 | 0.10174196 | 0.094 | 0 | 1 | 20 |
| rbm4.1 | 0.17586 | 0.144177469 | 0.094 | 0 | 1 | 20 |
| rest | 0.17586 | 0.180444965 | 0.094 | 0 | 1 | 20 |
| shmt2 | 0.17586 | 0.189952996 | 0.094 | 0 | 1 | 20 |
| si:ch211-114c12.2 | 0.17586 | 0.111681687 | 0.094 | 0 | 1 | 20 |
| si:ch211-195b11.3 | 0.17586 | 0.366105483 | 0.094 | 0 | 1 | 20 |
| si:ch211-222k6.3 | 0.17586 | 0.104450614 | 0.094 | 0 | 1 | 20 |
| si:ch211-276c2.4 | 0.17586 | 0.10454848 | 0.094 | 0 | 1 | 20 |
| si:ch211-39k3.2 | 0.17586 | 0.101570065 | 0.094 | 0 | 1 | 20 |

| | | | | | | |
|---|---|---|---|---|---|---|
| si:ch73-285p12.4 | 0.17586 | 0.136289379 | 0.094 | 0 | 1 | 20 |
| si:dkey-110g7.8 | 0.17586 | 0.130122975 | 0.094 | 0 | 1 | 20 |
| si:dkey-154b15.1 | 0.17586 | 0.122039406 | 0.094 | 0 | 1 | 20 |
| si:dkey-72l14.4 | 0.17586 | 0.108623145 | 0.094 | 0 | 1 | 20 |
| si:rp71-1g18.1 | 0.17586 | 0.154678404 | 0.094 | 0 | 1 | 20 |
| slc25a12 | 0.17586 | 0.109683497 | 0.094 | 0 | 1 | 20 |
| slc5a6b | 0.17586 | 0.114466702 | 0.094 | 0 | 1 | 20 |
| specc1la | 0.17586 | 0.147022836 | 0.094 | 0 | 1 | 20 |
| spoplb | 0.17586 | 0.13501722 | 0.094 | 0 | 1 | 20 |
| stx5a | 0.17586 | 0.110191443 | 0.094 | 0 | 1 | 20 |
| taok2a | 0.17586 | 0.1016844 | 0.094 | 0 | 1 | 20 |
| tmem198b | 0.17586 | 0.119485583 | 0.094 | 0 | 1 | 20 |
| tnip1 | 0.17586 | 0.169527537 | 0.094 | 0 | 1 | 20 |
| trmt6 | 0.17586 | 0.133595875 | 0.094 | 0 | 1 | 20 |
| tsen34 | 0.17586 | 0.125131162 | 0.094 | 0 | 1 | 20 |
| tsg101a | 0.17586 | 0.100243629 | 0.094 | 0 | 1 | 20 |
| tyw5 | 0.17586 | 0.119011844 | 0.094 | 0 | 1 | 20 |
| uap1 | 0.17586 | 0.110527993 | 0.094 | 0 | 1 | 20 |
| ubap2a | 0.17586 | 0.115976842 | 0.094 | 0 | 1 | 20 |
| usp4 | 0.17586 | 0.119994885 | 0.094 | 0 | 1 | 20 |
| utp6 | 0.17586 | 0.121634152 | 0.094 | 0 | 1 | 20 |
| wu:fc33b09 | 0.17586 | 0.210948418 | 0.094 | 0 | 1 | 20 |
| ypel3 | 0.17586 | 0.15920688 | 0.094 | 0 | 1 | 20 |
| zc3h15 | 0.17586 | 0.119598147 | 0.094 | 0 | 1 | 20 |
| zc3h4 | 0.17586 | 0.123381457 | 0.094 | 0 | 1 | 20 |
| zdhhc20a | 0.17586 | 0.118139679 | 0.094 | 0 | 1 | 20 |
| zgc:112962 | 0.17586 | 0.134825741 | 0.094 | 0 | 1 | 20 |
| zgc:113518 | 0.17586 | 0.131757899 | 0.094 | 0 | 1 | 20 |
| zgc:153115 | 0.17586 | 0.111885765 | 0.094 | 0 | 1 | 20 |
| zgc:174906 | 0.17586 | 0.10711382 | 0.094 | 0 | 1 | 20 |
| zgc:193812 | 0.17586 | 0.106832721 | 0.094 | 0 | 1 | 20 |
| zgc:194659 | 0.17586 | 0.247116386 | 0.094 | 0 | 1 | 20 |
| zgc:194981 | 0.17586 | 0.109144946 | 0.094 | 0 | 1 | 20 |
| zmat4b | 0.17586 | 0.202472209 | 0.094 | 0 | 1 | 20 |
| zmiz2 | 0.17586 | 0.113712195 | 0.094 | 0 | 1 | 20 |
| znf511 | 0.17586 | 0.113080605 | 0.094 | 0 | 1 | 20 |
| znfl2a | 0.17586 | 0.255841908 | 0.094 | 0 | 1 | 20 |
| fahd2a | 0.17586 | 0.152175531 | 0.094 | 0 | 1 | 20 |
| fbxo3 | 0.17586 | 0.110950589 | 0.094 | 0 | 1 | 20 |
| hephl1a | 0.17586 | 0.121945312 | 0.094 | 0 | 1 | 20 |
| maea | 0.17586 | 0.15075089 | 0.094 | 0 | 1 | 20 |
| nde1 | 0.17586 | 0.164444512 | 0.094 | 0 | 1 | 20 |
| nudt12 | 0.17586 | 0.100536926 | 0.094 | 0 | 1 | 20 |
| si:ch73-248e21.1 | 0.17586 | 0.14092544 | 0.094 | 0 | 1 | 20 |
| taok3a | 0.17586 | 0.146633817 | 0.094 | 0 | 1 | 20 |
| eefsec | 0.17586 | 0.145937359 | 0.094 | 0 | 1 | 20 |
| mos | 0.17586 | 0.1398858 | 0.094 | 0 | 1 | 20 |
| mycla | 0.17586 | 0.12931923 | 0.094 | 0 | 1 | 20 |

| | | | | | | |
|---|---|---|---|---|---|---|
| rpgra | 0.17586 | 0.294990664 | 0.094 | 0 | 1 | 20 |
| runx3 | 0.17586 | 0.194716937 | 0.094 | 0 | 1 | 20 |
| srgap3 | 0.17586 | 0.168688442 | 0.094 | 0 | 1 | 20 |
| tgfbr2a | 0.17586 | 0.127444312 | 0.094 | 0 | 1 | 20 |
| FP017215.1 | 0.17586 | 0.133520123 | 0.094 | 0 | 1 | 20 |
| klhl6 | 0.17586 | 0.148550018 | 0.094 | 0 | 1 | 20 |
| atxn3 | 0.17586 | 0.116458551 | 0.094 | 0 | 1 | 20 |
| brwd3 | 0.17586 | 0.109798064 | 0.094 | 0 | 1 | 20 |
| c19h1orf109 | 0.17586 | 0.138098676 | 0.094 | 0 | 1 | 20 |
| hbegfa | 0.17586 | 0.124154665 | 0.094 | 0 | 1 | 20 |
| heca | 0.17586 | 0.107499353 | 0.094 | 0 | 1 | 20 |
| lmbr1 | 0.17586 | 0.134444695 | 0.094 | 0 | 1 | 20 |
| vezf1b | 0.17586 | 0.147010469 | 0.094 | 0 | 1 | 20 |
| slc44a2 | 0.17586 | -0.24210542 | 0.125 | 0.133 | 1 | 20 |
| ehd1a | 0.17689 | -0.49667441 | 0.062 | 0.2 | 1 | 20 |
| atp6v0b | 0.17699 | -0.44353009 | 0.094 | 0.267 | 1 | 20 |
| pggt1b | 0.17746 | -0.33733848 | 0.156 | 0.2 | 1 | 20 |
| cflara | 0.17781 | -0.27691324 | 0.031 | 0.2 | 1 | 20 |
| zfp36l2 | 0.17842 | -0.12690299 | 0.125 | 0.067 | 1 | 20 |
| nars | 0.17958 | 0.365054008 | 0.375 | 0.133 | 1 | 20 |
| si:ch211-51e12.7 | 0.17962 | 0.140459243 | 0.344 | 0.133 | 1 | 20 |
| khsrp | 0.18003 | 0.374551506 | 0.375 | 0.133 | 1 | 20 |
| si:ch1073-322p19.1 | 0.18017 | -0.31480965 | 0.031 | 0.2 | 1 | 20 |
| rpl13a | 0.18065 | 0.24911782 | 0.906 | 0.8 | 1 | 20 |
| pfdn2 | 0.18072 | 0.36470053 | 0.344 | 0.133 | 1 | 20 |
| arhgef1b | 0.18118 | 0.321182344 | 0.25 | 0.067 | 1 | 20 |
| si:ch73-376l24.6 | 0.18156 | 0.276637646 | 0.25 | 0.067 | 1 | 20 |
| ugcg | 0.182 | -0.39364559 | 0.031 | 0.2 | 1 | 20 |
| grpel2 | 0.1822 | 0.273478897 | 0.281 | 0.067 | 1 | 20 |
| mapk3 | 0.18244 | -0.11509685 | 0.188 | 0.133 | 1 | 20 |
| pdk4 | 0.18257 | -0.21033243 | 0.031 | 0.2 | 1 | 20 |
| tomm6 | 0.18258 | 0.359643737 | 0.344 | 0.133 | 1 | 20 |
| cebpb | 0.18263 | -0.34642366 | 0.375 | 0.4 | 1 | 20 |
| ccr9a | 0.18275 | 0.396109501 | 0.25 | 0.067 | 1 | 20 |
| pfkfb4b | 0.18309 | -0.4172021 | 0.25 | 0.4 | 1 | 20 |
| ifrd2 | 0.18358 | 0.306283901 | 0.281 | 0.067 | 1 | 20 |
| FO082781.1 | 0.18363 | 0.317243765 | 0.281 | 0.067 | 1 | 20 |
| thrap3b | 0.1842 | -0.4192664 | 0.344 | 0.467 | 1 | 20 |
| rab7a | 0.18489 | -0.19530696 | 0.031 | 0.2 | 1 | 20 |
| gstm.1 | 0.18495 | 0.415666731 | 0.406 | 0.2 | 1 | 20 |
| angpt1 | 0.18546 | -0.19632123 | 0.031 | 0.2 | 1 | 20 |
| psmd6 | 0.18569 | 0.276784722 | 0.281 | 0.067 | 1 | 20 |
| ubfd1 | 0.18599 | 0.299935372 | 0.281 | 0.067 | 1 | 20 |
| myl12.1 | 0.18616 | -0.32911773 | 0.812 | 0.933 | 1 | 20 |
| cd7al | 0.1863 | 0.320228264 | 0.375 | 0.133 | 1 | 20 |
| mcl1b | 0.18667 | -0.36448473 | 0.25 | 0.267 | 1 | 20 |
| csrnp1b | 0.18675 | -0.43933582 | 0.031 | 0.2 | 1 | 20 |
| diablob | 0.18694 | -0.37529355 | 0.031 | 0.2 | 1 | 20 |

| | | | | | | |
|---|---|---|---|---|---|---|
| taf15 | 0.18715 | 0.14043305 | 0.344 | 0.133 | 1 | 20 |
| mtss1 | 0.18731 | -0.52498002 | 0.031 | 0.2 | 1 | 20 |
| rchy1 | 0.18734 | -0.1522402 | 0.031 | 0.2 | 1 | 20 |
| mgst3b | 0.1874 | -0.25168024 | 0.281 | 0.267 | 1 | 20 |
| dhrs13a.1 | 0.1875 | -0.13347054 | 0.031 | 0.2 | 1 | 20 |
| mt-nd3 | 0.18752 | -0.40541738 | 0.031 | 0.2 | 1 | 20 |
| aup1 | 0.18754 | -0.24131187 | 0.031 | 0.2 | 1 | 20 |
| bmp16 | 0.18767 | -0.38980094 | 0.094 | 0.267 | 1 | 20 |
| tma16 | 0.18805 | 0.310518009 | 0.25 | 0.067 | 1 | 20 |
| myca | 0.18837 | 0.483094222 | 0.406 | 0.2 | 1 | 20 |
| prdx6 | 0.18855 | 0.289397439 | 0.438 | 0.2 | 1 | 20 |
| mcm5 | 0.1887 | 0.276563671 | 0.281 | 0.067 | 1 | 20 |
| setb | 0.18924 | 0.352646518 | 0.469 | 0.4 | 1 | 20 |
| knop1 | 0.19003 | 0.323585522 | 0.25 | 0.067 | 1 | 20 |
| cdc34a | 0.19081 | -0.30385468 | 0.094 | 0.133 | 1 | 20 |
| ier5 | 0.191 | -0.23176549 | 0.281 | 0.267 | 1 | 20 |
| snrpc | 0.19216 | 0.276904729 | 0.219 | 0.067 | 1 | 20 |
| ddx18 | 0.19226 | 0.293309066 | 0.312 | 0.267 | 1 | 20 |
| copz1 | 0.19228 | 0.396099543 | 0.375 | 0.2 | 1 | 20 |
| selenow1 | 0.19261 | -0.48722053 | 0.5 | 0.733 | 1 | 20 |
| ube2l3b | 0.19274 | 0.352418874 | 0.469 | 0.2 | 1 | 20 |
| bop1 | 0.19274 | 0.288564274 | 0.25 | 0.067 | 1 | 20 |
| atp5f1d | 0.19294 | 0.393005343 | 0.469 | 0.2 | 1 | 20 |
| sec11a | 0.19305 | 0.34114208 | 0.281 | 0.2 | 1 | 20 |
| cops7a | 0.19315 | -0.40001251 | 0.094 | 0.267 | 1 | 20 |
| rsf1b.1 | 0.19439 | 0.27546893 | 0.281 | 0.067 | 1 | 20 |
| jagn1b | 0.19583 | 0.360802005 | 0.25 | 0.067 | 1 | 20 |
| kars | 0.19663 | 0.356814001 | 0.375 | 0.2 | 1 | 20 |
| srrm2 | 0.19696 | 0.196006264 | 0.25 | 0.067 | 1 | 20 |
| eif3jb | 0.19738 | -0.41054577 | 0.188 | 0.4 | 1 | 20 |
| mrpl42 | 0.19825 | 0.380307466 | 0.438 | 0.2 | 1 | 20 |
| scamp2 | 0.19846 | -0.39064934 | 0.188 | 0.333 | 1 | 20 |
| mef2aa | 0.19859 | -0.44206408 | 0.406 | 0.533 | 1 | 20 |
| si:ch211-161c3.6 | 0.19866 | 0.323220582 | 0.5 | 0.267 | 1 | 20 |
| syk | 0.19884 | 0.138369607 | 0.219 | 0.067 | 1 | 20 |
| mbd3a | 0.19893 | 0.236041227 | 0.281 | 0.067 | 1 | 20 |
| ppp3ccb | 0.19991 | -0.21192157 | 0.156 | 0.133 | 1 | 20 |
| BX323824.1 | 0.19997 | -0.22395135 | 0 | 0.067 | 1 | 20 |
| CABZ01044746.1 | 0.19997 | -0.20873735 | 0 | 0.067 | 1 | 20 |
| CABZ01068600.1 | 0.19997 | -0.25116527 | 0 | 0.067 | 1 | 20 |
| CR339041.3 | 0.19997 | -0.25306972 | 0 | 0.067 | 1 | 20 |
| CT573148.1 | 0.19997 | -0.10242312 | 0 | 0.067 | 1 | 20 |
| CT583708.1 | 0.19997 | -0.24336394 | 0 | 0.067 | 1 | 20 |
| CU681842.1 | 0.19997 | -0.23196859 | 0 | 0.067 | 1 | 20 |
| CU929237.1 | 0.19997 | -0.23178282 | 0 | 0.067 | 1 | 20 |
| LOC110439915 | 0.19997 | -0.22902257 | 0 | 0.067 | 1 | 20 |
| XLOC-041857 | 0.19997 | -0.23449314 | 0 | 0.067 | 1 | 20 |
| acvr2aa | 0.19997 | -0.21247792 | 0 | 0.067 | 1 | 20 |

| | | | | | | |
|---|---|---|---|---|---|---|
| adam9 | 0.19997 | -0.28706535 | 0 | 0.067 | 1 | 20 |
| admb | 0.19997 | -0.23109194 | 0 | 0.067 | 1 | 20 |
| aldh1a2 | 0.19997 | -0.19524718 | 0 | 0.067 | 1 | 20 |
| arhgap36 | 0.19997 | -0.22853496 | 0 | 0.067 | 1 | 20 |
| arrdc1b | 0.19997 | -0.22913071 | 0 | 0.067 | 1 | 20 |
| bcl11ba | 0.19997 | -0.11951155 | 0 | 0.067 | 1 | 20 |
| cav1 | 0.19997 | -0.10030968 | 0 | 0.067 | 1 | 20 |
| ccny | 0.19997 | -0.21206199 | 0 | 0.067 | 1 | 20 |
| chp1 | 0.19997 | -0.22460542 | 0 | 0.067 | 1 | 20 |
| cldn5a | 0.19997 | -0.10209204 | 0 | 0.067 | 1 | 20 |
| cldnc | 0.19997 | -0.15540439 | 0 | 0.067 | 1 | 20 |
| col1a2 | 0.19997 | -0.16063011 | 0 | 0.067 | 1 | 20 |
| cptp | 0.19997 | -0.21029528 | 0 | 0.067 | 1 | 20 |
| cramp1 | 0.19997 | -0.17964604 | 0 | 0.067 | 1 | 20 |
| ddx20 | 0.19997 | -0.10750929 | 0 | 0.067 | 1 | 20 |
| dnajb9b | 0.19997 | -0.22821484 | 0 | 0.067 | 1 | 20 |
| dop1b | 0.19997 | -0.25376334 | 0 | 0.067 | 1 | 20 |
| eef2l2 | 0.19997 | -0.19221745 | 0 | 0.067 | 1 | 20 |
| foxl1 | 0.19997 | -0.21316822 | 0 | 0.067 | 1 | 20 |
| gpr156 | 0.19997 | -0.24336394 | 0 | 0.067 | 1 | 20 |
| hcst | 0.19997 | -0.11996945 | 0 | 0.067 | 1 | 20 |
| il13ra1 | 0.19997 | -0.20900326 | 0 | 0.067 | 1 | 20 |
| itgb1a | 0.19997 | -0.20818594 | 0 | 0.067 | 1 | 20 |
| jcada | 0.19997 | -0.25210017 | 0 | 0.067 | 1 | 20 |
| jund | 0.19997 | -0.22713971 | 0 | 0.067 | 1 | 20 |
| kcnab1b | 0.19997 | -0.13738727 | 0 | 0.067 | 1 | 20 |
| kcne4 | 0.19997 | -0.12982371 | 0 | 0.067 | 1 | 20 |
| krt4 | 0.19997 | -0.11118799 | 0 | 0.067 | 1 | 20 |
| lats1 | 0.19997 | -0.17964604 | 0 | 0.067 | 1 | 20 |
| ltbp1 | 0.19997 | -0.17534953 | 0 | 0.067 | 1 | 20 |
| luzp1 | 0.19997 | -0.21082876 | 0 | 0.067 | 1 | 20 |
| mfap2 | 0.19997 | -0.12203005 | 0 | 0.067 | 1 | 20 |
| mob1bb | 0.19997 | -0.24158738 | 0 | 0.067 | 1 | 20 |
| mon2 | 0.19997 | -0.22762114 | 0 | 0.067 | 1 | 20 |
| mtmr14 | 0.19997 | -0.2274666 | 0 | 0.067 | 1 | 20 |
| myct1b | 0.19997 | -0.3370597 | 0 | 0.067 | 1 | 20 |
| nubp1 | 0.19997 | -0.21316822 | 0 | 0.067 | 1 | 20 |
| nusap1 | 0.19997 | -0.21647409 | 0 | 0.067 | 1 | 20 |
| peak1 | 0.19997 | -0.25344791 | 0 | 0.067 | 1 | 20 |
| pex2 | 0.19997 | -0.25040048 | 0 | 0.067 | 1 | 20 |
| pik3c2b | 0.19997 | -0.12161295 | 0 | 0.067 | 1 | 20 |
| pmepa1 | 0.19997 | -0.23376984 | 0 | 0.067 | 1 | 20 |
| ptprja | 0.19997 | -0.21910862 | 0 | 0.067 | 1 | 20 |
| rac1b | 0.19997 | -0.20485559 | 0 | 0.067 | 1 | 20 |
| ralbb | 0.19997 | -0.23249385 | 0 | 0.067 | 1 | 20 |
| ralgapa1 | 0.19997 | -0.21032544 | 0 | 0.067 | 1 | 20 |
| rdh10a | 0.19997 | -0.19260459 | 0 | 0.067 | 1 | 20 |
| rnf128a | 0.19997 | -0.22638694 | 0 | 0.067 | 1 | 20 |

| | | | | | | |
|---|---|---|---|---|---|---|
| shc1 | 0.19997 | -0.23336245 | 0 | 0.067 | 1 | 20 |
| si:ch211-133h13.1 | 0.19997 | -0.24336394 | 0 | 0.067 | 1 | 20 |
| si:ch211-218g4.2 | 0.19997 | -0.20272665 | 0 | 0.067 | 1 | 20 |
| si:ch211-272n13.3 | 0.19997 | -0.24176187 | 0 | 0.067 | 1 | 20 |
| si:ch211-40k21.5 | 0.19997 | -0.12004135 | 0 | 0.067 | 1 | 20 |
| si:ch211-9d9.8 | 0.19997 | -0.23414745 | 0 | 0.067 | 1 | 20 |
| si:ch73-250a16.5 | 0.19997 | -0.1765098 | 0 | 0.067 | 1 | 20 |
| si:dkey-188i13.10 | 0.19997 | -0.28076411 | 0 | 0.067 | 1 | 20 |
| si:dkey-1b17.5 | 0.19997 | -0.11951155 | 0 | 0.067 | 1 | 20 |
| si:dkey-23k10.5 | 0.19997 | -0.22769662 | 0 | 0.067 | 1 | 20 |
| si:dkey-42i9.6 | 0.19997 | -0.23368385 | 0 | 0.067 | 1 | 20 |
| si:dkeyp-121d4.3 | 0.19997 | -0.20615299 | 0 | 0.067 | 1 | 20 |
| slc16a5a | 0.19997 | -0.25271407 | 0 | 0.067 | 1 | 20 |
| slc24a3 | 0.19997 | -0.21175697 | 0 | 0.067 | 1 | 20 |
| smad3b | 0.19997 | -0.22662041 | 0 | 0.067 | 1 | 20 |
| snrkb | 0.19997 | -0.21706188 | 0 | 0.067 | 1 | 20 |
| snx27a | 0.19997 | -0.21162252 | 0 | 0.067 | 1 | 20 |
| socs2 | 0.19997 | -0.20581823 | 0 | 0.067 | 1 | 20 |
| sos1 | 0.19997 | -0.20876692 | 0 | 0.067 | 1 | 20 |
| sostdc1a | 0.19997 | -0.12507155 | 0 | 0.067 | 1 | 20 |
| spink2.2 | 0.19997 | -0.1316337 | 0 | 0.067 | 1 | 20 |
| tbl2 | 0.19997 | -0.25293324 | 0 | 0.067 | 1 | 20 |
| tdh | 0.19997 | -0.21885723 | 0 | 0.067 | 1 | 20 |
| tmem35 | 0.19997 | -0.15695534 | 0 | 0.067 | 1 | 20 |
| tmem63ba | 0.19997 | -0.20954388 | 0 | 0.067 | 1 | 20 |
| tnikb | 0.19997 | -0.25203514 | 0 | 0.067 | 1 | 20 |
| tprn | 0.19997 | -0.23449314 | 0 | 0.067 | 1 | 20 |
| traf6 | 0.19997 | -0.23381009 | 0 | 0.067 | 1 | 20 |
| tsc22d1 | 0.19997 | -0.28028742 | 0 | 0.067 | 1 | 20 |
| tspan5a | 0.19997 | -0.21385524 | 0 | 0.067 | 1 | 20 |
| ttc7b | 0.19997 | -0.25262451 | 0 | 0.067 | 1 | 20 |
| ubtd1a | 0.19997 | -0.23109194 | 0 | 0.067 | 1 | 20 |
| vps37c | 0.19997 | -0.22681678 | 0 | 0.067 | 1 | 20 |
| wdr48b | 0.19997 | -0.22642751 | 0 | 0.067 | 1 | 20 |
| wu:fj29h11 | 0.19997 | -0.11463934 | 0 | 0.067 | 1 | 20 |
| xk | 0.19997 | -0.22395135 | 0 | 0.067 | 1 | 20 |
| zbtb34 | 0.19997 | -0.23209302 | 0 | 0.067 | 1 | 20 |
| zdhhc14 | 0.19997 | -0.25233153 | 0 | 0.067 | 1 | 20 |
| zer1 | 0.19997 | -0.24832844 | 0 | 0.067 | 1 | 20 |
| zgc:152791 | 0.19997 | -0.23250988 | 0 | 0.067 | 1 | 20 |
| zgc:56235 | 0.19997 | -0.20706409 | 0 | 0.067 | 1 | 20 |
| zgc:77151 | 0.19997 | -0.23140327 | 0 | 0.067 | 1 | 20 |
| znf1009 | 0.19997 | -0.23382393 | 0 | 0.067 | 1 | 20 |
| ATG14 | 0.19997 | -0.24324924 | 0 | 0.067 | 1 | 20 |
| BX511034.1 | 0.19997 | -0.17009436 | 0 | 0.067 | 1 | 20 |
| FO203432.1 | 0.19997 | -0.26098346 | 0 | 0.067 | 1 | 20 |
| LOC100536726 | 0.19997 | -0.24808522 | 0 | 0.067 | 1 | 20 |
| LOC101886390 | 0.19997 | -0.17441738 | 0 | 0.067 | 1 | 20 |

| | | | | | | |
|---|---|---|---|---|---|---|
| LOC110438970 | 0.19997 | -0.17103989 | 0 | 0.067 | 1 | 20 |
| LOC110439849 | 0.19997 | -0.16182489 | 0 | 0.067 | 1 | 20 |
| MFAP4 (1 of many) | 0.19997 | -0.37612207 | 0 | 0.067 | 1 | 20 |
| blnk | 0.19997 | -0.22918009 | 0 | 0.067 | 1 | 20 |
| cd28l | 0.19997 | -0.1707341 | 0 | 0.067 | 1 | 20 |
| cdc42ep5 | 0.19997 | -0.22126007 | 0 | 0.067 | 1 | 20 |
| cep57l1 | 0.19997 | -0.17299138 | 0 | 0.067 | 1 | 20 |
| cfd | 0.19997 | -0.1604582 | 0 | 0.067 | 1 | 20 |
| chkb | 0.19997 | -0.16380179 | 0 | 0.067 | 1 | 20 |
| crebrf | 0.19997 | -0.24770856 | 0 | 0.067 | 1 | 20 |
| dennd4c | 0.19997 | -0.17422936 | 0 | 0.067 | 1 | 20 |
| dnajb1a | 0.19997 | -0.17166068 | 0 | 0.067 | 1 | 20 |
| dnal1 | 0.19997 | -0.21709566 | 0 | 0.067 | 1 | 20 |
| egr2b | 0.19997 | -0.16139097 | 0 | 0.067 | 1 | 20 |
| ehbp1l1a | 0.19997 | -0.24765884 | 0 | 0.067 | 1 | 20 |
| enpp1 | 0.19997 | -0.2477563 | 0 | 0.067 | 1 | 20 |
| exoc8 | 0.19997 | -0.23380493 | 0 | 0.067 | 1 | 20 |
| f2rl2 | 0.19997 | -0.24777308 | 0 | 0.067 | 1 | 20 |
| gnao1b | 0.19997 | -0.23751137 | 0 | 0.067 | 1 | 20 |
| gnb2 | 0.19997 | -0.17197521 | 0 | 0.067 | 1 | 20 |
| ip6k1 | 0.19997 | -0.16139097 | 0 | 0.067 | 1 | 20 |
| lgals9l3 | 0.19997 | -0.15371318 | 0 | 0.067 | 1 | 20 |
| lrp1ba | 0.19997 | -0.2299322 | 0 | 0.067 | 1 | 20 |
| mapk9 | 0.19997 | -0.17227793 | 0 | 0.067 | 1 | 20 |
| mllt1a | 0.19997 | -0.2299322 | 0 | 0.067 | 1 | 20 |
| nkiras2 | 0.19997 | -0.24628082 | 0 | 0.067 | 1 | 20 |
| nrip1b | 0.19997 | -0.21539333 | 0 | 0.067 | 1 | 20 |
| nuf2 | 0.19997 | -0.17051682 | 0 | 0.067 | 1 | 20 |
| oip5-as1 | 0.19997 | -0.22673937 | 0 | 0.067 | 1 | 20 |
| pak4 | 0.19997 | -0.24648922 | 0 | 0.067 | 1 | 20 |
| pdzrn3b | 0.19997 | -0.20580618 | 0 | 0.067 | 1 | 20 |
| pi4k2a | 0.19997 | -0.24528441 | 0 | 0.067 | 1 | 20 |
| plekhg2 | 0.19997 | -0.23423138 | 0 | 0.067 | 1 | 20 |
| ppip5k1a | 0.19997 | -0.28267142 | 0 | 0.067 | 1 | 20 |
| prkag3a | 0.19997 | -0.23423138 | 0 | 0.067 | 1 | 20 |
| proser3 | 0.19997 | -0.17191428 | 0 | 0.067 | 1 | 20 |
| ptbp2b | 0.19997 | -0.23341644 | 0 | 0.067 | 1 | 20 |
| ptpdc1a | 0.19997 | -0.16957983 | 0 | 0.067 | 1 | 20 |
| ramp2 | 0.19997 | -0.20586832 | 0 | 0.067 | 1 | 20 |
| rgp1 | 0.19997 | -0.22243368 | 0 | 0.067 | 1 | 20 |
| rnf175 | 0.19997 | -0.23191463 | 0 | 0.067 | 1 | 20 |
| robo4 | 0.19997 | -0.24783353 | 0 | 0.067 | 1 | 20 |
| si:ch1073-110a20.3 | 0.19997 | -0.31351686 | 0 | 0.067 | 1 | 20 |
| si:ch211-251p5.5 | 0.19997 | -0.16843601 | 0 | 0.067 | 1 | 20 |
| si:ch211-63o20.7 | 0.19997 | -0.17097172 | 0 | 0.067 | 1 | 20 |
| si:dkey-211g8.4 | 0.19997 | -0.17284383 | 0 | 0.067 | 1 | 20 |
| si:dkey-4e7.3 | 0.19997 | -0.17177505 | 0 | 0.067 | 1 | 20 |
| si:dkey-56d12.4 | 0.19997 | -0.22538081 | 0 | 0.067 | 1 | 20 |

| | | | | | | |
|---|---|---|---|---|---|---|
| si:dkey-86e18.1 | 0.19997 | -0.1567975 | 0 | 0.067 | 1 | 20 |
| si:zfos-2330d3.1 | 0.19997 | -0.25771438 | 0 | 0.067 | 1 | 20 |
| sid1 | 0.19997 | -0.28697321 | 0 | 0.067 | 1 | 20 |
| slc37a1 | 0.19997 | -0.17441738 | 0 | 0.067 | 1 | 20 |
| st6galnac5b | 0.19997 | -0.1535187 | 0 | 0.067 | 1 | 20 |
| sugp1 | 0.19997 | -0.17293964 | 0 | 0.067 | 1 | 20 |
| syne3 | 0.19997 | -0.17320272 | 0 | 0.067 | 1 | 20 |
| tbc1d14 | 0.19997 | -0.22840607 | 0 | 0.067 | 1 | 20 |
| timp4.2 | 0.19997 | -0.16645491 | 0 | 0.067 | 1 | 20 |
| timp4.3 | 0.19997 | -0.10949299 | 0 | 0.067 | 1 | 20 |
| tpcn2 | 0.19997 | -0.13834587 | 0 | 0.067 | 1 | 20 |
| tpk1 | 0.19997 | -0.16786147 | 0 | 0.067 | 1 | 20 |
| trabd2a | 0.19997 | -0.24808522 | 0 | 0.067 | 1 | 20 |
| trappc9 | 0.19997 | -0.24780142 | 0 | 0.067 | 1 | 20 |
| wdr34 | 0.19997 | -0.22831514 | 0 | 0.067 | 1 | 20 |
| zap70 | 0.19997 | -0.21943126 | 0 | 0.067 | 1 | 20 |
| zbtb32 | 0.19997 | -0.16139097 | 0 | 0.067 | 1 | 20 |
| zgc:113436 | 0.19997 | -0.22374184 | 0 | 0.067 | 1 | 20 |
| zgc:113921 | 0.19997 | -0.23346319 | 0 | 0.067 | 1 | 20 |
| zgc:158463 | 0.19997 | -0.11473915 | 0 | 0.067 | 1 | 20 |
| zgc:175088 | 0.19997 | -0.24468744 | 0 | 0.067 | 1 | 20 |
| zgc:194839 | 0.19997 | -0.22538081 | 0 | 0.067 | 1 | 20 |
| zranb3 | 0.19997 | -0.17147954 | 0 | 0.067 | 1 | 20 |
| BX248113.1 | 0.19997 | -0.20711672 | 0 | 0.067 | 1 | 20 |
| BX957297.2 | 0.19997 | -0.24708608 | 0 | 0.067 | 1 | 20 |
| CABZ01001434.1 | 0.19997 | -0.259209 | 0 | 0.067 | 1 | 20 |
| CABZ01009512.2 | 0.19997 | -0.24488941 | 0 | 0.067 | 1 | 20 |
| CABZ01038709.1 | 0.19997 | -0.2474127 | 0 | 0.067 | 1 | 20 |
| CABZ01048402.1 | 0.19997 | -0.24722462 | 0 | 0.067 | 1 | 20 |
| CABZ01058046.1 | 0.19997 | -0.20591995 | 0 | 0.067 | 1 | 20 |
| CABZ01068358.1 | 0.19997 | -0.20376428 | 0 | 0.067 | 1 | 20 |
| CABZ01088367.1 | 0.19997 | -0.22343043 | 0 | 0.067 | 1 | 20 |
| CT573231.1 | 0.19997 | -0.20622725 | 0 | 0.067 | 1 | 20 |
| CT573342.2 | 0.19997 | -0.20711672 | 0 | 0.067 | 1 | 20 |
| CU459094.2 | 0.19997 | -0.16922784 | 0 | 0.067 | 1 | 20 |
| LO017872.1 | 0.19997 | -0.20576163 | 0 | 0.067 | 1 | 20 |
| LO018250.1 | 0.19997 | -0.18474422 | 0 | 0.067 | 1 | 20 |
| LO018493.1 | 0.19997 | -0.20666264 | 0 | 0.067 | 1 | 20 |
| PAQR9 | 0.19997 | -0.24852456 | 0 | 0.067 | 1 | 20 |
| SCYL2 | 0.19997 | -0.24577094 | 0 | 0.067 | 1 | 20 |
| SH2B2 | 0.19997 | -0.24898189 | 0 | 0.067 | 1 | 20 |
| XLOC-008233 | 0.19997 | -0.24898189 | 0 | 0.067 | 1 | 20 |
| XLOC-010337 | 0.19997 | -0.20644044 | 0 | 0.067 | 1 | 20 |
| XLOC-018356 | 0.19997 | -0.2066357 | 0 | 0.067 | 1 | 20 |
| XLOC-019150 | 0.19997 | -0.20582966 | 0 | 0.067 | 1 | 20 |
| XLOC-038035 | 0.19997 | -0.20711672 | 0 | 0.067 | 1 | 20 |
| ankrd46b | 0.19997 | -0.18370913 | 0 | 0.067 | 1 | 20 |
| ap3m1 | 0.19997 | -0.22117219 | 0 | 0.067 | 1 | 20 |

| | | | | | | |
|---|---|---|---|---|---|---|
| araf | 0.19997 | -0.24622991 | 0 | 0.067 | 1 | 20 |
| arcn1a | 0.19997 | -0.22243725 | 0 | 0.067 | 1 | 20 |
| arhgap31 | 0.19997 | -0.24639364 | 0 | 0.067 | 1 | 20 |
| atp6v0a1a | 0.19997 | -0.24845576 | 0 | 0.067 | 1 | 20 |
| b4galnt1a | 0.19997 | -0.20711672 | 0 | 0.067 | 1 | 20 |
| b4galt4 | 0.19997 | -0.20139656 | 0 | 0.067 | 1 | 20 |
| baiap2a | 0.19997 | -0.12895242 | 0 | 0.067 | 1 | 20 |
| camk4 | 0.19997 | -0.20711672 | 0 | 0.067 | 1 | 20 |
| cdk17 | 0.19997 | -0.20630812 | 0 | 0.067 | 1 | 20 |
| chrna1 | 0.19997 | -0.19671658 | 0 | 0.067 | 1 | 20 |
| chuk | 0.19997 | -0.20411964 | 0 | 0.067 | 1 | 20 |
| clasp1a | 0.19997 | -0.24295098 | 0 | 0.067 | 1 | 20 |
| ctsla | 0.19997 | -0.22653359 | 0 | 0.067 | 1 | 20 |
| cxcr3.1 | 0.19997 | -0.20654533 | 0 | 0.067 | 1 | 20 |
| cybb | 0.19997 | -0.20674036 | 0 | 0.067 | 1 | 20 |
| dcaf8 | 0.19997 | -0.24357134 | 0 | 0.067 | 1 | 20 |
| dctn5 | 0.19997 | -0.22091234 | 0 | 0.067 | 1 | 20 |
| dhrs13l1 | 0.19997 | -0.24206594 | 0 | 0.067 | 1 | 20 |
| dusp16 | 0.19997 | -0.20340426 | 0 | 0.067 | 1 | 20 |
| dync1i1 | 0.19997 | -0.25566174 | 0 | 0.067 | 1 | 20 |
| ehd1b | 0.19997 | -0.17106465 | 0 | 0.067 | 1 | 20 |
| ets2 | 0.19997 | -0.27613117 | 0 | 0.067 | 1 | 20 |
| exoc5 | 0.19997 | -0.20613201 | 0 | 0.067 | 1 | 20 |
| fam57a | 0.19997 | -0.24846228 | 0 | 0.067 | 1 | 20 |
| fbxo33 | 0.19997 | -0.24716665 | 0 | 0.067 | 1 | 20 |
| fpr1 | 0.19997 | -0.25769654 | 0 | 0.067 | 1 | 20 |
| fut7 | 0.19997 | -0.20673703 | 0 | 0.067 | 1 | 20 |
| g6pd | 0.19997 | -0.2229982 | 0 | 0.067 | 1 | 20 |
| galnt1 | 0.19997 | -0.20259838 | 0 | 0.067 | 1 | 20 |
| gdi1 | 0.19997 | -0.24031583 | 0 | 0.067 | 1 | 20 |
| glb1l | 0.19997 | -0.20651418 | 0 | 0.067 | 1 | 20 |
| glipr2 | 0.19997 | -0.22438719 | 0 | 0.067 | 1 | 20 |
| gpsm2 | 0.19997 | -0.24689969 | 0 | 0.067 | 1 | 20 |
| grn1 | 0.19997 | -0.28689583 | 0 | 0.067 | 1 | 20 |
| hdac9b | 0.19997 | -0.22439795 | 0 | 0.067 | 1 | 20 |
| hmbsb | 0.19997 | -0.29796635 | 0 | 0.067 | 1 | 20 |
| hps3 | 0.19997 | -0.20281507 | 0 | 0.067 | 1 | 20 |
| hsd11b1la | 0.19997 | -0.20519938 | 0 | 0.067 | 1 | 20 |
| hsdl1 | 0.19997 | -0.20711672 | 0 | 0.067 | 1 | 20 |
| hspa12b | 0.19997 | -0.25920583 | 0 | 0.067 | 1 | 20 |
| hyal2b | 0.19997 | -0.20599075 | 0 | 0.067 | 1 | 20 |
| ifngr1 | 0.19997 | -0.20711672 | 0 | 0.067 | 1 | 20 |
| il34 | 0.19997 | -0.22417165 | 0 | 0.067 | 1 | 20 |
| il4 | 0.19997 | -0.17507288 | 0 | 0.067 | 1 | 20 |
| irak3 | 0.19997 | -0.20672271 | 0 | 0.067 | 1 | 20 |
| itgae.1 | 0.19997 | -0.25931616 | 0 | 0.067 | 1 | 20 |
| june | 0.19997 | -0.22439124 | 0 | 0.067 | 1 | 20 |
| kat6a | 0.19997 | -0.24720557 | 0 | 0.067 | 1 | 20 |

| | | | | | | |
|---|---|---|---|---|---|---|
| laptm4b | 0.19997 | -0.2236266 | 0 | 0.067 | 1 | 20 |
| lcp2b | 0.19997 | -0.20669218 | 0 | 0.067 | 1 | 20 |
| lpar6b | 0.19997 | -0.25942929 | 0 | 0.067 | 1 | 20 |
| lygl1 | 0.19997 | -0.32848007 | 0 | 0.067 | 1 | 20 |
| marveld2a | 0.19997 | -0.19168506 | 0 | 0.067 | 1 | 20 |
| mical2a | 0.19997 | -0.19760141 | 0 | 0.067 | 1 | 20 |
| mpx | 0.19997 | -0.38865904 | 0 | 0.067 | 1 | 20 |
| mtmr2 | 0.19997 | -0.20448684 | 0 | 0.067 | 1 | 20 |
| myclb | 0.19997 | -0.23949045 | 0 | 0.067 | 1 | 20 |
| myo1cb | 0.19997 | -0.2014064 | 0 | 0.067 | 1 | 20 |
| nme2a | 0.19997 | -0.20711672 | 0 | 0.067 | 1 | 20 |
| noctb | 0.19997 | -0.19051441 | 0 | 0.067 | 1 | 20 |
| nr1d2b | 0.19997 | -0.20246117 | 0 | 0.067 | 1 | 20 |
| nrbp2a | 0.19997 | -0.2051361 | 0 | 0.067 | 1 | 20 |
| ormdl1 | 0.19997 | -0.24236482 | 0 | 0.067 | 1 | 20 |
| paqr7a | 0.19997 | -0.22440977 | 0 | 0.067 | 1 | 20 |
| pcdh9 | 0.19997 | -0.24898189 | 0 | 0.067 | 1 | 20 |
| pfkpb | 0.19997 | -0.24620107 | 0 | 0.067 | 1 | 20 |
| phlpp1 | 0.19997 | -0.24431099 | 0 | 0.067 | 1 | 20 |
| pkig | 0.19997 | -0.28761106 | 0 | 0.067 | 1 | 20 |
| ppm1e | 0.19997 | -0.20711672 | 0 | 0.067 | 1 | 20 |
| ppp1r21 | 0.19997 | -0.24561845 | 0 | 0.067 | 1 | 20 |
| prkcaa | 0.19997 | -0.24749943 | 0 | 0.067 | 1 | 20 |
| ptprga | 0.19997 | -0.24390794 | 0 | 0.067 | 1 | 20 |
| pvrl2l | 0.19997 | -0.20656588 | 0 | 0.067 | 1 | 20 |
| rapgef1a | 0.19997 | -0.22375955 | 0 | 0.067 | 1 | 20 |
| rasl11a | 0.19997 | -0.24308796 | 0 | 0.067 | 1 | 20 |
| rbm18 | 0.19997 | -0.24807606 | 0 | 0.067 | 1 | 20 |
| rbm47 | 0.19997 | -0.20485022 | 0 | 0.067 | 1 | 20 |
| rgs19 | 0.19997 | -0.20611791 | 0 | 0.067 | 1 | 20 |
| rps6ka3a | 0.19997 | -0.20498686 | 0 | 0.067 | 1 | 20 |
| rxrgb | 0.19997 | -0.20448317 | 0 | 0.067 | 1 | 20 |
| saal1 | 0.19997 | -0.24719825 | 0 | 0.067 | 1 | 20 |
| sc:d156 | 0.19997 | -0.22452008 | 0 | 0.067 | 1 | 20 |
| selenop | 0.19997 | -0.20318197 | 0 | 0.067 | 1 | 20 |
| sema6bb | 0.19997 | -0.24752603 | 0 | 0.067 | 1 | 20 |
| shc2 | 0.19997 | -0.2246755 | 0 | 0.067 | 1 | 20 |
| si:ch211-130m23.5 | 0.19997 | -0.20269275 | 0 | 0.067 | 1 | 20 |
| si:ch211-153b23.7 | 0.19997 | -0.20605727 | 0 | 0.067 | 1 | 20 |
| si:ch211-198m17.1 | 0.19997 | -0.2047936 | 0 | 0.067 | 1 | 20 |
| si:ch73-335l21.4 | 0.19997 | -0.25360544 | 0 | 0.067 | 1 | 20 |
| si:ch73-380l3.4 | 0.19997 | -0.20655355 | 0 | 0.067 | 1 | 20 |
| si:dkey-102c8.3 | 0.19997 | -0.17065805 | 0 | 0.067 | 1 | 20 |
| si:dkey-103j14.5 | 0.19997 | -0.24752603 | 0 | 0.067 | 1 | 20 |
| si:dkey-163m14.7 | 0.19997 | -0.27720841 | 0 | 0.067 | 1 | 20 |
| si:dkey-19b23.14 | 0.19997 | -0.22130777 | 0 | 0.067 | 1 | 20 |
| si:dkey-1c7.2 | 0.19997 | -0.25968191 | 0 | 0.067 | 1 | 20 |
| si:dkey-222n6.2 | 0.19997 | -0.21417157 | 0 | 0.067 | 1 | 20 |

| | | | | | | |
|---|---|---|---|---|---|---|
| si:dkey-3k24.5 | 0.19997 | -0.20623091 | 0 | 0.067 | 1 | 20 |
| si:dkey-53k12.2 | 0.19997 | -0.25956754 | 0 | 0.067 | 1 | 20 |
| slc15a4 | 0.19997 | -0.20593834 | 0 | 0.067 | 1 | 20 |
| slc25a23a | 0.19997 | -0.23467226 | 0 | 0.067 | 1 | 20 |
| slc6a6a | 0.19997 | -0.2039117 | 0 | 0.067 | 1 | 20 |
| slc9a3r1a | 0.19997 | -0.20558714 | 0 | 0.067 | 1 | 20 |
| sorl1 | 0.19997 | -0.20568797 | 0 | 0.067 | 1 | 20 |
| sowahd | 0.19997 | -0.20654565 | 0 | 0.067 | 1 | 20 |
| stag2a | 0.19997 | -0.24804965 | 0 | 0.067 | 1 | 20 |
| tmem176l.2 | 0.19997 | -0.2246755 | 0 | 0.067 | 1 | 20 |
| tmem237b | 0.19997 | -0.20244478 | 0 | 0.067 | 1 | 20 |
| traf7 | 0.19997 | -0.20631476 | 0 | 0.067 | 1 | 20 |
| trim2a | 0.19997 | -0.24359154 | 0 | 0.067 | 1 | 20 |
| trpv1 | 0.19997 | -0.20681623 | 0 | 0.067 | 1 | 20 |
| tshz1 | 0.19997 | -0.17065805 | 0 | 0.067 | 1 | 20 |
| tyk2 | 0.19997 | -0.22358519 | 0 | 0.067 | 1 | 20 |
| uba3 | 0.19997 | -0.20423109 | 0 | 0.067 | 1 | 20 |
| ubiad1 | 0.19997 | -0.19131965 | 0 | 0.067 | 1 | 20 |
| usp40 | 0.19997 | -0.24898189 | 0 | 0.067 | 1 | 20 |
| vps13d | 0.19997 | -0.20122288 | 0 | 0.067 | 1 | 20 |
| wasla | 0.19997 | -0.2435791 | 0 | 0.067 | 1 | 20 |
| wrap73 | 0.19997 | -0.24071429 | 0 | 0.067 | 1 | 20 |
| ywhae2 | 0.19997 | -0.24780326 | 0 | 0.067 | 1 | 20 |
| zbtb17 | 0.19997 | -0.20141041 | 0 | 0.067 | 1 | 20 |
| zdhhc13 | 0.19997 | -0.24603884 | 0 | 0.067 | 1 | 20 |
| zfp36l1b | 0.19997 | -0.1974871 | 0 | 0.067 | 1 | 20 |
| zgc:103438 | 0.19997 | -0.24565603 | 0 | 0.067 | 1 | 20 |
| zgc:112980 | 0.19997 | -0.20551021 | 0 | 0.067 | 1 | 20 |
| zgc:113149 | 0.19997 | -0.24815218 | 0 | 0.067 | 1 | 20 |
| zgc:113340 | 0.19997 | -0.21159314 | 0 | 0.067 | 1 | 20 |
| zgc:153118 | 0.19997 | -0.24589281 | 0 | 0.067 | 1 | 20 |
| zgc:165582 | 0.19997 | -0.20590814 | 0 | 0.067 | 1 | 20 |
| zmp:0000000650 | 0.19997 | -0.20254993 | 0 | 0.067 | 1 | 20 |
| zmp:0000001323 | 0.19997 | -0.23487252 | 0 | 0.067 | 1 | 20 |
| znf516 | 0.19997 | -0.20711672 | 0 | 0.067 | 1 | 20 |
| si:ch211-214p16.2 | 0.20013 | 0.140762066 | 0.594 | 0.4 | 1 | 20 |
| fkbp3 | 0.2007 | 0.326883264 | 0.406 | 0.267 | 1 | 20 |
| pkma | 0.20142 | -0.10225631 | 0.344 | 0.267 | 1 | 20 |
| exosc2 | 0.20171 | 0.181715231 | 0.25 | 0.067 | 1 | 20 |
| rbms1b | 0.20196 | -0.25483101 | 0.125 | 0.133 | 1 | 20 |
| eif5b | 0.20297 | 0.139981102 | 0.406 | 0.2 | 1 | 20 |
| ppp3r1a | 0.20417 | -0.17203417 | 0.094 | 0.067 | 1 | 20 |
| nop10 | 0.2051 | 0.438092124 | 0.438 | 0.2 | 1 | 20 |
| cwc27 | 0.20537 | 0.336394323 | 0.219 | 0.067 | 1 | 20 |
| aldoaa | 0.20705 | 0.269711263 | 0.719 | 0.467 | 1 | 20 |
| gstk1 | 0.20745 | 0.29764001 | 0.25 | 0.067 | 1 | 20 |
| actr2a | 0.20906 | -0.41004469 | 0.25 | 0.333 | 1 | 20 |
| si:ch211-197h24.9 | 0.20925 | 0.403147908 | 0.344 | 0.133 | 1 | 20 |

| | | | | | | |
|---|---|---|---|---|---|---|
| surf4l | 0.20929 | -0.10653777 | 0.125 | 0.067 | 1 | 20 |
| dhx37 | 0.20939 | 0.135497466 | 0.219 | 0.067 | 1 | 20 |
| atp5mc1 | 0.20943 | 0.430163726 | 0.531 | 0.267 | 1 | 20 |
| mrpl38 | 0.20974 | 0.22082046 | 0.25 | 0.067 | 1 | 20 |
| copb1 | 0.21044 | -0.3818805 | 0.031 | 0.133 | 1 | 20 |
| mbtps1 | 0.21044 | -0.41170311 | 0.031 | 0.133 | 1 | 20 |
| nr2f6a | 0.21044 | -0.2942638 | 0.031 | 0.133 | 1 | 20 |
| plp2 | 0.21044 | -0.19493467 | 0.031 | 0.133 | 1 | 20 |
| rab40c | 0.21044 | -0.16552779 | 0.031 | 0.133 | 1 | 20 |
| rps6ka1 | 0.21044 | -0.12788676 | 0.031 | 0.133 | 1 | 20 |
| si:ch211-191c10.1 | 0.21044 | -0.12365757 | 0.031 | 0.133 | 1 | 20 |
| CR318588.1 | 0.21044 | -0.22759948 | 0.031 | 0.133 | 1 | 20 |
| CU019662.1 | 0.21044 | -0.28495464 | 0.031 | 0.133 | 1 | 20 |
| FP102809.1 | 0.21044 | -0.14659032 | 0.031 | 0.133 | 1 | 20 |
| adcy6a | 0.21044 | -0.32369706 | 0.031 | 0.133 | 1 | 20 |
| arl6ip5b | 0.21044 | -0.30919324 | 0.031 | 0.133 | 1 | 20 |
| atg101 | 0.21044 | -0.25656316 | 0.031 | 0.133 | 1 | 20 |
| bmp2k | 0.21044 | -0.30870267 | 0.031 | 0.133 | 1 | 20 |
| calcoco1a | 0.21044 | -0.14889368 | 0.031 | 0.133 | 1 | 20 |
| camk2g1 | 0.21044 | -0.2535597 | 0.031 | 0.133 | 1 | 20 |
| chmp4ba | 0.21044 | -0.38495413 | 0.031 | 0.133 | 1 | 20 |
| crema | 0.21044 | -0.3383449 | 0.031 | 0.133 | 1 | 20 |
| cry2 | 0.21044 | -0.12858503 | 0.031 | 0.133 | 1 | 20 |
| dalrd3 | 0.21044 | -0.12266088 | 0.031 | 0.133 | 1 | 20 |
| desi1b | 0.21044 | -0.2492859 | 0.031 | 0.133 | 1 | 20 |
| elmod2 | 0.21044 | -0.22885276 | 0.031 | 0.133 | 1 | 20 |
| ero1b | 0.21044 | -0.43825949 | 0.031 | 0.133 | 1 | 20 |
| faf1 | 0.21044 | -0.25020631 | 0.031 | 0.133 | 1 | 20 |
| fastkd1 | 0.21044 | -0.27487428 | 0.031 | 0.133 | 1 | 20 |
| gadd45bb | 0.21044 | -0.30565802 | 0.031 | 0.133 | 1 | 20 |
| gnl1 | 0.21044 | -0.21586008 | 0.031 | 0.133 | 1 | 20 |
| hsbp1a | 0.21044 | -0.25992822 | 0.031 | 0.133 | 1 | 20 |
| mitd1 | 0.21044 | -0.19175576 | 0.031 | 0.133 | 1 | 20 |
| mvb12ba | 0.21044 | -0.24527447 | 0.031 | 0.133 | 1 | 20 |
| nme2b.2 | 0.21044 | -0.15855534 | 0.031 | 0.133 | 1 | 20 |
| nme7 | 0.21044 | -0.12223474 | 0.031 | 0.133 | 1 | 20 |
| ntan1 | 0.21044 | -0.15276469 | 0.031 | 0.133 | 1 | 20 |
| osgn1 | 0.21044 | -0.22456608 | 0.031 | 0.133 | 1 | 20 |
| paxx | 0.21044 | -0.17846871 | 0.031 | 0.133 | 1 | 20 |
| pdlim1 | 0.21044 | -0.18701384 | 0.031 | 0.133 | 1 | 20 |
| sgms1 | 0.21044 | -0.22563228 | 0.031 | 0.133 | 1 | 20 |
| si:ch211-173p18.3 | 0.21044 | -0.17181555 | 0.031 | 0.133 | 1 | 20 |
| si:dkey-261j4.3 | 0.21044 | -0.12739008 | 0.031 | 0.133 | 1 | 20 |
| slc12a7b | 0.21044 | -0.18525515 | 0.031 | 0.133 | 1 | 20 |
| slc38a2 | 0.21044 | -0.4399165 | 0.031 | 0.133 | 1 | 20 |
| socs3a | 0.21044 | -0.30812236 | 0.031 | 0.133 | 1 | 20 |
| tmem42b | 0.21044 | -0.33124509 | 0.031 | 0.133 | 1 | 20 |
| tnfrsf9a | 0.21044 | -0.2601589 | 0.031 | 0.133 | 1 | 20 |

| | | | | | | |
|---|---|---|---|---|---|---|
| ubash3ba | 0.21044 | -0.23918196 | 0.031 | 0.133 | 1 | 20 |
| ube2a | 0.21044 | -0.22262967 | 0.031 | 0.133 | 1 | 20 |
| ube2al | 0.21044 | -0.23855286 | 0.031 | 0.133 | 1 | 20 |
| vash2 | 0.21044 | -0.20233992 | 0.031 | 0.133 | 1 | 20 |
| zfand6 | 0.21044 | -0.24645114 | 0.031 | 0.133 | 1 | 20 |
| zgc:153018 | 0.21044 | -0.31747801 | 0.031 | 0.133 | 1 | 20 |
| zgc:162396 | 0.21044 | -0.38198312 | 0.031 | 0.133 | 1 | 20 |
| zgc:65873 | 0.21044 | -0.25119138 | 0.031 | 0.133 | 1 | 20 |
| zgc:91860 | 0.21044 | -0.38000556 | 0.031 | 0.133 | 1 | 20 |
| zmp:0000001114 | 0.21044 | -0.25180437 | 0.031 | 0.133 | 1 | 20 |
| LOC110437731 | 0.21044 | -0.40774418 | 0.031 | 0.133 | 1 | 20 |
| akap10 | 0.21044 | -0.34794775 | 0.031 | 0.133 | 1 | 20 |
| caspa | 0.21044 | -0.11124819 | 0.031 | 0.133 | 1 | 20 |
| celf1 | 0.21044 | -0.12719684 | 0.031 | 0.133 | 1 | 20 |
| chic1 | 0.21044 | -0.12451123 | 0.031 | 0.133 | 1 | 20 |
| dhfr | 0.21044 | -0.36032726 | 0.031 | 0.133 | 1 | 20 |
| egr3 | 0.21044 | -0.36560427 | 0.031 | 0.133 | 1 | 20 |
| eif3hb | 0.21044 | -0.3464578 | 0.031 | 0.133 | 1 | 20 |
| fhod1 | 0.21044 | -0.27708905 | 0.031 | 0.133 | 1 | 20 |
| gpat3 | 0.21044 | -0.11764949 | 0.031 | 0.133 | 1 | 20 |
| map2k4a | 0.21044 | -0.25278891 | 0.031 | 0.133 | 1 | 20 |
| mybl2b | 0.21044 | -0.16142543 | 0.031 | 0.133 | 1 | 20 |
| ndfip1l | 0.21044 | -0.21711201 | 0.031 | 0.133 | 1 | 20 |
| osbpl9 | 0.21044 | -0.1127938 | 0.031 | 0.133 | 1 | 20 |
| pacs1a | 0.21044 | -0.32440796 | 0.031 | 0.133 | 1 | 20 |
| papolg | 0.21044 | -0.27569833 | 0.031 | 0.133 | 1 | 20 |
| rasal2 | 0.21044 | -0.26203819 | 0.031 | 0.133 | 1 | 20 |
| rev3l | 0.21044 | -0.26815998 | 0.031 | 0.133 | 1 | 20 |
| si:ch1073-443f11.2 | 0.21044 | -0.21120225 | 0.031 | 0.133 | 1 | 20 |
| si:ch211-128m15.3 | 0.21044 | -0.13005398 | 0.031 | 0.133 | 1 | 20 |
| si:dkey-93h22.8 | 0.21044 | -0.11991801 | 0.031 | 0.133 | 1 | 20 |
| sptlc1 | 0.21044 | -0.11762292 | 0.031 | 0.133 | 1 | 20 |
| supt5h | 0.21044 | -0.27521855 | 0.031 | 0.133 | 1 | 20 |
| tcp11l2 | 0.21044 | -0.27648676 | 0.031 | 0.133 | 1 | 20 |
| tmem131l | 0.21044 | -0.22773587 | 0.031 | 0.133 | 1 | 20 |
| ube2d4 | 0.21044 | -0.20812961 | 0.031 | 0.133 | 1 | 20 |
| ubtd2 | 0.21044 | -0.22942865 | 0.031 | 0.133 | 1 | 20 |
| zgc:175264 | 0.21044 | -0.24174361 | 0.031 | 0.133 | 1 | 20 |
| col4a1 | 0.21044 | -0.25424698 | 0.031 | 0.133 | 1 | 20 |
| tapt1b | 0.21044 | -0.35701185 | 0.031 | 0.133 | 1 | 20 |
| JMY | 0.21044 | -0.26571763 | 0.031 | 0.133 | 1 | 20 |
| bcas3 | 0.21044 | -0.24858356 | 0.031 | 0.133 | 1 | 20 |
| dck | 0.21044 | -0.32231221 | 0.031 | 0.133 | 1 | 20 |
| dnm2a | 0.21044 | -0.26655244 | 0.031 | 0.133 | 1 | 20 |
| e4f1 | 0.21044 | -0.36801035 | 0.031 | 0.133 | 1 | 20 |
| fibpa | 0.21044 | -0.23475972 | 0.031 | 0.133 | 1 | 20 |
| ginm1 | 0.21044 | -0.23256612 | 0.031 | 0.133 | 1 | 20 |
| gnsb | 0.21044 | -0.26284883 | 0.031 | 0.133 | 1 | 20 |

| | | | | | | |
|---|---|---|---|---|---|---|
| map2k4b | 0.21044 | -0.22526726 | 0.031 | 0.133 | 1 | 20 |
| mest | 0.21044 | -0.18043186 | 0.031 | 0.133 | 1 | 20 |
| ppp1r18 | 0.21044 | -0.1100184 | 0.031 | 0.133 | 1 | 20 |
| ptpn4a | 0.21044 | -0.20969458 | 0.031 | 0.133 | 1 | 20 |
| rhbdl2 | 0.21044 | -0.2313539 | 0.031 | 0.133 | 1 | 20 |
| ripor1.1 | 0.21044 | -0.24182886 | 0.031 | 0.133 | 1 | 20 |
| sept8a | 0.21044 | -0.26768305 | 0.031 | 0.133 | 1 | 20 |
| si:dkey-276j7.1 | 0.21044 | -0.1991192 | 0.031 | 0.133 | 1 | 20 |
| tm9sf4 | 0.21044 | -0.17890845 | 0.031 | 0.133 | 1 | 20 |
| zgc:110843 | 0.21044 | -0.25865056 | 0.031 | 0.133 | 1 | 20 |
| znf710b | 0.21044 | -0.23974893 | 0.031 | 0.133 | 1 | 20 |
| AL935174.5 | 0.21044 | -0.21706575 | 0.031 | 0.133 | 1 | 20 |
| CABZ01005379.1 | 0.21044 | -0.13352085 | 0.031 | 0.133 | 1 | 20 |
| abhd18 | 0.21044 | -0.16104296 | 0.031 | 0.133 | 1 | 20 |
| aif1l | 0.21044 | -0.40476886 | 0.031 | 0.133 | 1 | 20 |
| arhgef12a | 0.21044 | -0.42201778 | 0.031 | 0.133 | 1 | 20 |
| bida | 0.21044 | -0.29712243 | 0.031 | 0.133 | 1 | 20 |
| chaf1b | 0.21044 | -0.20099077 | 0.031 | 0.133 | 1 | 20 |
| coro1ca | 0.21044 | -0.25714691 | 0.031 | 0.133 | 1 | 20 |
| cpeb4b | 0.21044 | -0.3795105 | 0.031 | 0.133 | 1 | 20 |
| cul3a | 0.21044 | -0.23583832 | 0.031 | 0.133 | 1 | 20 |
| engase | 0.21044 | -0.19566345 | 0.031 | 0.133 | 1 | 20 |
| hipk3a | 0.21044 | -0.24435419 | 0.031 | 0.133 | 1 | 20 |
| myct1a | 0.21044 | -0.35170907 | 0.031 | 0.133 | 1 | 20 |
| nfil3-6 | 0.21044 | -0.13038695 | 0.031 | 0.133 | 1 | 20 |
| paqr3a | 0.21044 | -0.20744919 | 0.031 | 0.133 | 1 | 20 |
| poli | 0.21044 | -0.10936399 | 0.031 | 0.133 | 1 | 20 |
| ppp1r7 | 0.21044 | -0.36833647 | 0.031 | 0.133 | 1 | 20 |
| pum1 | 0.21044 | -0.20315077 | 0.031 | 0.133 | 1 | 20 |
| scrib | 0.21044 | -0.20807719 | 0.031 | 0.133 | 1 | 20 |
| sdf4 | 0.21044 | -0.28116233 | 0.031 | 0.133 | 1 | 20 |
| si:dkey-234h16.7 | 0.21044 | -0.21661875 | 0.031 | 0.133 | 1 | 20 |
| si:dkey-24l11.2 | 0.21044 | -0.17022028 | 0.031 | 0.133 | 1 | 20 |
| si:dkeyp-68b7.7 | 0.21044 | -0.17205339 | 0.031 | 0.133 | 1 | 20 |
| slc39a11 | 0.21044 | -0.26944668 | 0.031 | 0.133 | 1 | 20 |
| slka | 0.21044 | -0.23522512 | 0.031 | 0.133 | 1 | 20 |
| tdg.1 | 0.21044 | -0.16743787 | 0.031 | 0.133 | 1 | 20 |
| tmem107 | 0.21044 | -0.15568308 | 0.031 | 0.133 | 1 | 20 |
| trim44 | 0.21044 | -0.24725751 | 0.031 | 0.133 | 1 | 20 |
| zfpm1 | 0.21044 | -0.26135462 | 0.031 | 0.133 | 1 | 20 |
| bcas2 | 0.21075 | 0.322427307 | 0.344 | 0.2 | 1 | 20 |
| rfc3 | 0.21079 | 0.346432841 | 0.281 | 0.133 | 1 | 20 |
| phf5a | 0.21182 | 0.379085453 | 0.375 | 0.133 | 1 | 20 |
| sb:cb81 | 0.21197 | 0.100247926 | 0.5 | 0.333 | 1 | 20 |
| smim8 | 0.21225 | -0.1032435 | 0.125 | 0.067 | 1 | 20 |
| nucks1a | 0.21268 | 0.141900796 | 0.344 | 0.133 | 1 | 20 |
| timm9 | 0.21348 | 0.248848534 | 0.25 | 0.067 | 1 | 20 |
| jpt2 | 0.21447 | -0.42669772 | 0.281 | 0.533 | 1 | 20 |

| | | | | | | |
|---|---|---|---|---|---|---|
| cbx3a | 0.2152 | 0.439761333 | 0.438 | 0.2 | 1 | 20 |
| eif3s10 | 0.21609 | 0.352020136 | 0.531 | 0.267 | 1 | 20 |
| wdr3 | 0.21629 | 0.264685196 | 0.219 | 0.067 | 1 | 20 |
| ndfip1 | 0.21668 | -0.37899237 | 0.406 | 0.467 | 1 | 20 |
| dohh | 0.21878 | 0.285117777 | 0.25 | 0.067 | 1 | 20 |
| hint1 | 0.22032 | 0.386840311 | 0.531 | 0.267 | 1 | 20 |
| coq8a | 0.22072 | -0.22874949 | 0.125 | 0.133 | 1 | 20 |
| taf9 | 0.22072 | 0.330377651 | 0.344 | 0.133 | 1 | 20 |
| tab1 | 0.22123 | 0.404755547 | 0.344 | 0.133 | 1 | 20 |
| cox5aa | 0.22175 | -0.12570349 | 0.656 | 0.6 | 1 | 20 |
| rpf2 | 0.22211 | 0.303171675 | 0.281 | 0.133 | 1 | 20 |
| tmem106bb | 0.22212 | -0.15234796 | 0.094 | 0.067 | 1 | 20 |
| elmo1 | 0.22215 | 0.339791523 | 0.25 | 0.067 | 1 | 20 |
| id2a | 0.22325 | 0.438820119 | 0.531 | 0.267 | 1 | 20 |
| zgc:92140 | 0.2237 | 0.170045982 | 0.219 | 0.067 | 1 | 20 |
| cotl1 | 0.22405 | -0.32887108 | 0.531 | 0.6 | 1 | 20 |
| nutf2l | 0.22572 | 0.416272923 | 0.531 | 0.333 | 1 | 20 |
| pdia6 | 0.22581 | 0.219949958 | 0.25 | 0.067 | 1 | 20 |
| fgl2a | 0.22591 | -0.63100765 | 0.219 | 0.467 | 1 | 20 |
| dennd2c | 0.22663 | -0.41633575 | 0.094 | 0.2 | 1 | 20 |
| got2b | 0.22837 | 0.35042211 | 0.219 | 0.067 | 1 | 20 |
| psme2 | 0.22897 | 0.190031389 | 0.5 | 0.267 | 1 | 20 |
| arhgdia | 0.22958 | -0.40794596 | 0.406 | 0.533 | 1 | 20 |
| si:dkey-67c22.2 | 0.23066 | -0.32479488 | 0.156 | 0.2 | 1 | 20 |
| map1lc3b | 0.23076 | -0.46050453 | 0.344 | 0.533 | 1 | 20 |
| ophn1 | 0.23325 | -0.13710457 | 0.281 | 0.2 | 1 | 20 |
| myl9b | 0.23475 | -0.53137358 | 0.188 | 0.333 | 1 | 20 |
| idh1 | 0.2348 | -0.37716537 | 0.125 | 0.333 | 1 | 20 |
| cald1a | 0.23494 | -0.54022243 | 0.219 | 0.467 | 1 | 20 |
| brd4 | 0.23509 | -0.20318115 | 0.219 | 0.2 | 1 | 20 |
| psmb10 | 0.23546 | 0.228576772 | 0.25 | 0.067 | 1 | 20 |
| eef2b | 0.23581 | 0.234535101 | 0.875 | 0.667 | 1 | 20 |
| si:cabz01074946.1 | 0.23583 | -0.45210559 | 0.188 | 0.333 | 1 | 20 |
| psenen | 0.23586 | 0.339069455 | 0.344 | 0.133 | 1 | 20 |
| hmgb2a | 0.23594 | 0.277578328 | 0.719 | 0.467 | 1 | 20 |
| utp23 | 0.23682 | 0.320538192 | 0.25 | 0.067 | 1 | 20 |
| kpnb3 | 0.2371 | 0.319332552 | 0.312 | 0.133 | 1 | 20 |
| ube2l3a | 0.23768 | 0.184852508 | 0.344 | 0.133 | 1 | 20 |
| lpp | 0.23799 | -0.50616109 | 0.125 | 0.333 | 1 | 20 |
| cpsf6 | 0.23838 | 0.306656431 | 0.281 | 0.133 | 1 | 20 |
| asb12a | 0.23879 | -0.38419841 | 0.156 | 0.333 | 1 | 20 |
| gp1bb | 0.23884 | -0.38550962 | 0.25 | 0.467 | 1 | 20 |
| rdh8a | 0.23891 | -0.36986986 | 0.062 | 0.133 | 1 | 20 |
| slc25a55a | 0.23905 | -0.35555762 | 0.219 | 0.267 | 1 | 20 |
| srd5a2a | 0.23932 | 0.369910997 | 0.281 | 0.133 | 1 | 20 |
| eif4ebp3l | 0.23941 | 0.288754077 | 0.25 | 0.067 | 1 | 20 |
| si:dkey-159f12.2 | 0.24021 | 0.316390847 | 0.219 | 0.067 | 1 | 20 |
| lcp2a | 0.24208 | 0.272924654 | 0.25 | 0.067 | 1 | 20 |

| | | | | | | |
|---|---|---|---|---|---|---|
| seta | 0.24228 | 0.339958217 | 0.594 | 0.333 | 1 | 20 |
| CABZ01034698.2 | 0.24233 | -0.40752364 | 0.062 | 0.2 | 1 | 20 |
| rnaseka | 0.24372 | -0.33319578 | 0.156 | 0.2 | 1 | 20 |
| srsf9 | 0.2442 | -0.17742122 | 0.375 | 0.333 | 1 | 20 |
| mrps6 | 0.24422 | 0.273768874 | 0.219 | 0.067 | 1 | 20 |
| serp1 | 0.24432 | 0.280817174 | 0.188 | 0.067 | 1 | 20 |
| fabp3 | 0.24449 | 0.406136831 | 0.594 | 0.333 | 1 | 20 |
| pim1 | 0.24462 | 0.146299235 | 0.312 | 0.133 | 1 | 20 |
| znf706 | 0.24494 | 0.156834318 | 0.344 | 0.133 | 1 | 20 |
| uchl5 | 0.24567 | 0.291450535 | 0.25 | 0.067 | 1 | 20 |
| tfap4 | 0.24574 | 0.157232043 | 0.219 | 0.067 | 1 | 20 |
| mrps2 | 0.24597 | 0.323240655 | 0.25 | 0.067 | 1 | 20 |
| csnk2a1 | 0.2461 | 0.307803738 | 0.344 | 0.133 | 1 | 20 |
| si:dkey-239h2.3 | 0.2476 | 0.401649207 | 0.5 | 0.267 | 1 | 20 |
| b3gnt3.4 | 0.24778 | -0.4934506 | 0.094 | 0.267 | 1 | 20 |
| ppat | 0.24851 | 0.287582782 | 0.25 | 0.067 | 1 | 20 |
| tmx1 | 0.24851 | 0.276413642 | 0.25 | 0.067 | 1 | 20 |
| actb2 | 0.24869 | -0.14019977 | 1 | 1 | 1 | 20 |
| trim35-28 | 0.24878 | -0.38913539 | 0.094 | 0.2 | 1 | 20 |
| snrpd3l | 0.25015 | 0.298770127 | 0.281 | 0.133 | 1 | 20 |
| hmha1b | 0.25079 | 0.305374529 | 0.344 | 0.133 | 1 | 20 |
| esd | 0.25106 | 0.251893323 | 0.219 | 0.2 | 1 | 20 |
| tra2a | 0.25217 | 0.314115679 | 0.344 | 0.2 | 1 | 20 |
| mcm7 | 0.25246 | 0.200761971 | 0.406 | 0.2 | 1 | 20 |
| baz1b | 0.25264 | 0.333873311 | 0.281 | 0.133 | 1 | 20 |
| CR855337.1 | 0.25292 | -0.39176333 | 0.094 | 0.2 | 1 | 20 |
| ddx27 | 0.25381 | 0.174360626 | 0.406 | 0.2 | 1 | 20 |
| srsf3b | 0.25474 | 0.267675692 | 0.438 | 0.2 | 1 | 20 |
| tubb4b | 0.25478 | -0.12708663 | 0.844 | 0.8 | 1 | 20 |
| bysl | 0.25499 | 0.303462526 | 0.438 | 0.2 | 1 | 20 |
| acadm | 0.255 | -0.14062543 | 0.094 | 0.267 | 1 | 20 |
| u2af2a | 0.25556 | 0.253264609 | 0.25 | 0.067 | 1 | 20 |
| pfdn1 | 0.25582 | 0.105039734 | 0.375 | 0.2 | 1 | 20 |
| tia1 | 0.25639 | 0.277232692 | 0.25 | 0.067 | 1 | 20 |
| meaf6 | 0.2566 | 0.182745355 | 0.25 | 0.067 | 1 | 20 |
| lpar1 | 0.2583 | -0.50932449 | 0.125 | 0.333 | 1 | 20 |
| h2afva | 0.25853 | 0.21396491 | 0.5 | 0.267 | 1 | 20 |
| prrc2c | 0.25897 | 0.20753607 | 0.25 | 0.067 | 1 | 20 |
| btg2 | 0.2594 | -0.29184218 | 0.438 | 0.467 | 1 | 20 |
| sdhb | 0.25941 | -0.21204481 | 0.281 | 0.267 | 1 | 20 |
| exosc1 | 0.25942 | 0.322670026 | 0.312 | 0.133 | 1 | 20 |
| gskip | 0.25946 | 0.275036594 | 0.219 | 0.067 | 1 | 20 |
| chd4b | 0.2596 | 0.315650758 | 0.281 | 0.133 | 1 | 20 |
| anp32e | 0.25968 | 0.176790429 | 0.406 | 0.2 | 1 | 20 |
| psma6a | 0.25976 | 0.353429812 | 0.281 | 0.133 | 1 | 20 |
| prpf19 | 0.26023 | 0.301996829 | 0.25 | 0.067 | 1 | 20 |
| rack1 | 0.26031 | 0.203878489 | 0.906 | 0.867 | 1 | 20 |
| atp5pd | 0.26041 | 0.203905151 | 0.562 | 0.333 | 1 | 20 |

| | | | | | | |
|---|---|---|---|---|---|---|
| gabarapb | 0.26044 | -0.41682975 | 0.469 | 0.667 | 1 | 20 |
| ncaldb | 0.2613 | -0.4153377 | 0.156 | 0.333 | 1 | 20 |
| lrrfip1a | 0.26184 | -0.33581784 | 0.125 | 0.333 | 1 | 20 |
| wu:fi46h04 | 0.26347 | 0.243702599 | 0.219 | 0.067 | 1 | 20 |
| uqcrfs1 | 0.26498 | 0.3657454 | 0.406 | 0.267 | 1 | 20 |
| utp3 | 0.26517 | 0.295852252 | 0.281 | 0.133 | 1 | 20 |
| rhogb | 0.26549 | -0.10757339 | 0.281 | 0.2 | 1 | 20 |
| ptmab | 0.26587 | 0.117446572 | 0.781 | 0.6 | 1 | 20 |
| mrpl53 | 0.2659 | -0.34218276 | 0.094 | 0.267 | 1 | 20 |
| wu:fb55g09 | 0.26603 | -0.28145055 | 0.594 | 0.6 | 1 | 20 |
| si:ch211-217k17.7 | 0.26669 | 0.315532457 | 0.438 | 0.2 | 1 | 20 |
| dad1 | 0.26757 | 0.152463809 | 0.406 | 0.2 | 1 | 20 |
| xpot | 0.26787 | -0.14702011 | 0.094 | 0.067 | 1 | 20 |
| ppih | 0.26813 | 0.240354358 | 0.25 | 0.067 | 1 | 20 |
| pno1 | 0.26891 | 0.296677754 | 0.438 | 0.2 | 1 | 20 |
| tmem251 | 0.26945 | -0.14538434 | 0.094 | 0.067 | 1 | 20 |
| sh3kbp1 | 0.26959 | -0.39585208 | 0.125 | 0.2 | 1 | 20 |
| nob1 | 0.26959 | 0.263619893 | 0.25 | 0.067 | 1 | 20 |
| glrx | 0.27089 | -0.233628 | 0.531 | 0.533 | 1 | 20 |
| golgb1 | 0.27104 | 0.275301177 | 0.25 | 0.067 | 1 | 20 |
| alas1 | 0.27145 | 0.303564785 | 0.219 | 0.067 | 1 | 20 |
| RSBN1 | 0.27183 | -0.14358526 | 0.094 | 0.067 | 1 | 20 |
| rhoab | 0.27279 | -0.37848963 | 0.406 | 0.533 | 1 | 20 |
| rcc1l | 0.27335 | -0.1443069 | 0.094 | 0.067 | 1 | 20 |
| tmem39b | 0.27478 | -0.37771456 | 0.094 | 0.2 | 1 | 20 |
| zfr2 | 0.27495 | -0.29901165 | 0.094 | 0.133 | 1 | 20 |
| psmd1 | 0.27559 | 0.235734609 | 0.25 | 0.067 | 1 | 20 |
| txnipa | 0.27571 | -0.3469872 | 0.531 | 0.6 | 1 | 20 |
| noc2l | 0.27724 | 0.13427619 | 0.219 | 0.067 | 1 | 20 |
| fbxw11b | 0.2778 | -0.14116504 | 0.094 | 0.067 | 1 | 20 |
| dnajc19 | 0.278 | 0.250244778 | 0.25 | 0.2 | 1 | 20 |
| si:dkey-187e18.1 | 0.27803 | -0.34636862 | 0.125 | 0.2 | 1 | 20 |
| uck2a | 0.27805 | 0.120314261 | 0.156 | 0.2 | 1 | 20 |
| arpc1a | 0.2781 | -0.2218572 | 0.125 | 0.133 | 1 | 20 |
| si:ch211-191j22.3 | 0.27811 | 0.103906057 | 0.219 | 0.067 | 1 | 20 |
| pa2g4b | 0.2801 | 0.312807381 | 0.312 | 0.2 | 1 | 20 |
| casc3 | 0.28043 | -0.30094227 | 0.062 | 0.133 | 1 | 20 |
| rpl12 | 0.28047 | 0.251555012 | 0.844 | 0.733 | 1 | 20 |
| snrpe | 0.28089 | 0.360916008 | 0.531 | 0.333 | 1 | 20 |
| sec22ba | 0.28107 | -0.1459541 | 0.094 | 0.067 | 1 | 20 |
| dera | 0.28152 | 0.128864565 | 0.188 | 0.067 | 1 | 20 |
| tfg | 0.28156 | -0.34703837 | 0.125 | 0.2 | 1 | 20 |
| aprt | 0.28158 | -0.26315669 | 0.094 | 0.133 | 1 | 20 |
| ccdc6a | 0.28201 | -0.14228896 | 0.094 | 0.067 | 1 | 20 |
| pgpep1 | 0.28221 | -0.27649056 | 0.094 | 0.267 | 1 | 20 |
| cd82a | 0.28231 | -0.25233733 | 0.062 | 0.2 | 1 | 20 |
| mrps11 | 0.2832 | 0.218097938 | 0.25 | 0.067 | 1 | 20 |
| pphln1 | 0.28356 | -0.37028891 | 0.156 | 0.267 | 1 | 20 |

| | | | | | | |
|---|---|---|---|---|---|---|
| si:ch73-234b20.5 | 0.28358 | -0.45397188 | 0.281 | 0.467 | 1 | 20 |
| gadd45ab | 0.28449 | 0.301958949 | 0.188 | 0.133 | 1 | 20 |
| dtnbp1a | 0.28471 | -0.32985729 | 0.062 | 0.2 | 1 | 20 |
| atl2 | 0.28567 | -0.14480146 | 0.094 | 0.067 | 1 | 20 |
| ubr7 | 0.28613 | -0.21571345 | 0.25 | 0.267 | 1 | 20 |
| ankrd12 | 0.28651 | -0.2887132 | 0.156 | 0.2 | 1 | 20 |
| idh2 | 0.28698 | 0.110880485 | 0.594 | 0.4 | 1 | 20 |
| ldhba | 0.28726 | 0.141955504 | 0.656 | 0.467 | 1 | 20 |
| pam16 | 0.28729 | 0.136782966 | 0.219 | 0.067 | 1 | 20 |
| vps52 | 0.28836 | -0.28584466 | 0.062 | 0.2 | 1 | 20 |
| b2m | 0.28848 | 0.348157845 | 0.5 | 0.267 | 1 | 20 |
| sh3glb1b | 0.28866 | -0.33683807 | 0.125 | 0.2 | 1 | 20 |
| si:dkey-52l18.4 | 0.2888 | -0.30258619 | 0.125 | 0.133 | 1 | 20 |
| zgc:101583 | 0.28957 | -0.30266749 | 0.062 | 0.133 | 1 | 20 |
| cdc14b | 0.28976 | -0.35919686 | 0.094 | 0.2 | 1 | 20 |
| rrp9 | 0.28987 | 0.242347658 | 0.188 | 0.067 | 1 | 20 |
| ppan | 0.29022 | 0.344746515 | 0.344 | 0.133 | 1 | 20 |
| mrgbp | 0.29025 | 0.244966008 | 0.219 | 0.067 | 1 | 20 |
| si:dkey-33c12.4 | 0.29055 | 0.230509763 | 0.344 | 0.267 | 1 | 20 |
| prkrip1 | 0.29158 | -0.266636 | 0.094 | 0.133 | 1 | 20 |
| gmfb | 0.29206 | -0.32102093 | 0.281 | 0.333 | 1 | 20 |
| pdcd10a | 0.29216 | -0.37413382 | 0.094 | 0.267 | 1 | 20 |
| ctbp1 | 0.29283 | -0.29576294 | 0.062 | 0.2 | 1 | 20 |
| trmt112 | 0.29316 | 0.284125533 | 0.344 | 0.133 | 1 | 20 |
| psmc4 | 0.29319 | 0.268883864 | 0.344 | 0.133 | 1 | 20 |
| tp53 | 0.29377 | 0.266724998 | 0.344 | 0.133 | 1 | 20 |
| nupr1a | 0.29381 | -0.42543384 | 0.25 | 0.467 | 1 | 20 |
| BX511246.1 | 0.29436 | 0.117306405 | 0.062 | 0 | 1 | 20 |
| BX649411.2 | 0.29436 | 0.11093637 | 0.062 | 0 | 1 | 20 |
| CT025742.1 | 0.29436 | 0.14912043 | 0.062 | 0 | 1 | 20 |
| CT573344.1 | 0.29436 | 0.133916393 | 0.062 | 0 | 1 | 20 |
| FP101861.1 | 0.29436 | 0.113510796 | 0.062 | 0 | 1 | 20 |
| GGT7 | 0.29436 | 0.129770973 | 0.062 | 0 | 1 | 20 |
| LOC110439416 | 0.29436 | 0.101876958 | 0.062 | 0 | 1 | 20 |
| NHLRC3 | 0.29436 | 0.109032488 | 0.062 | 0 | 1 | 20 |
| XLOC-032112 | 0.29436 | 0.210465626 | 0.062 | 0 | 1 | 20 |
| XLOC-043788 | 0.29436 | 0.113060668 | 0.062 | 0 | 1 | 20 |
| abhd4 | 0.29436 | 0.12059024 | 0.062 | 0 | 1 | 20 |
| adnp2b | 0.29436 | 0.150111292 | 0.062 | 0 | 1 | 20 |
| aga | 0.29436 | 0.112212225 | 0.062 | 0 | 1 | 20 |
| ankra2 | 0.29436 | 0.101499128 | 0.062 | 0 | 1 | 20 |
| ankrd44 | 0.29436 | 0.173551788 | 0.062 | 0 | 1 | 20 |
| apoob | 0.29436 | 0.131726015 | 0.062 | 0 | 1 | 20 |
| arhgap27 | 0.29436 | 0.15104302 | 0.062 | 0 | 1 | 20 |
| arhgef9b | 0.29436 | 0.247700376 | 0.062 | 0 | 1 | 20 |
| arih1 | 0.29436 | 0.157255859 | 0.062 | 0 | 1 | 20 |
| arl6ip6 | 0.29436 | 0.125060535 | 0.062 | 0 | 1 | 20 |
| atp6ap1b | 0.29436 | 0.124080749 | 0.062 | 0 | 1 | 20 |

| | | | | | | |
|---|---|---|---|---|---|---|
| bap1 | 0.29436 | 0.109151759 | 0.062 | 0 | 1 | 20 |
| bcor | 0.29436 | 0.166293926 | 0.062 | 0 | 1 | 20 |
| bpnt1 | 0.29436 | 0.10003261 | 0.062 | 0 | 1 | 20 |
| c6ast4 | 0.29436 | 0.270430216 | 0.062 | 0 | 1 | 20 |
| castor2 | 0.29436 | 0.158089505 | 0.062 | 0 | 1 | 20 |
| cbx6b | 0.29436 | 0.131448037 | 0.062 | 0 | 1 | 20 |
| ccdc137 | 0.29436 | 0.101617027 | 0.062 | 0 | 1 | 20 |
| cdc42ep4a | 0.29436 | 0.135487727 | 0.062 | 0 | 1 | 20 |
| cnrip1a | 0.29436 | 0.104558718 | 0.062 | 0 | 1 | 20 |
| commd9 | 0.29436 | 0.119576537 | 0.062 | 0 | 1 | 20 |
| cox18 | 0.29436 | 0.128986602 | 0.062 | 0 | 1 | 20 |
| cpa5 | 0.29436 | 0.256444754 | 0.062 | 0 | 1 | 20 |
| cpb1 | 0.29436 | 0.171126177 | 0.062 | 0 | 1 | 20 |
| dpm3 | 0.29436 | 0.107016788 | 0.062 | 0 | 1 | 20 |
| fbxo7 | 0.29436 | 0.106405981 | 0.062 | 0 | 1 | 20 |
| flii | 0.29436 | 0.121545604 | 0.062 | 0 | 1 | 20 |
| foxn3 | 0.29436 | 0.120460596 | 0.062 | 0 | 1 | 20 |
| frmd8 | 0.29436 | 0.121290772 | 0.062 | 0 | 1 | 20 |
| fto | 0.29436 | 0.111257262 | 0.062 | 0 | 1 | 20 |
| gna13a | 0.29436 | 0.152714779 | 0.062 | 0 | 1 | 20 |
| grk6 | 0.29436 | 0.130202756 | 0.062 | 0 | 1 | 20 |
| gsap | 0.29436 | 0.131471487 | 0.062 | 0 | 1 | 20 |
| hddc2 | 0.29436 | 0.114992829 | 0.062 | 0 | 1 | 20 |
| hddc3 | 0.29436 | 0.109598668 | 0.062 | 0 | 1 | 20 |
| hectd1 | 0.29436 | 0.107021364 | 0.062 | 0 | 1 | 20 |
| her6 | 0.29436 | 0.106153644 | 0.062 | 0 | 1 | 20 |
| hmbox1a | 0.29436 | 0.114452434 | 0.062 | 0 | 1 | 20 |
| kdm5ba | 0.29436 | 0.119693313 | 0.062 | 0 | 1 | 20 |
| lpcat2 | 0.29436 | 0.141329412 | 0.062 | 0 | 1 | 20 |
| lrp10 | 0.29436 | 0.162509504 | 0.062 | 0 | 1 | 20 |
| ltb4r | 0.29436 | 0.124400998 | 0.062 | 0 | 1 | 20 |
| ly6m2 | 0.29436 | 0.1698934 | 0.062 | 0 | 1 | 20 |
| macrod1 | 0.29436 | 0.116358414 | 0.062 | 0 | 1 | 20 |
| manea | 0.29436 | 0.117947742 | 0.062 | 0 | 1 | 20 |
| map7d3 | 0.29436 | 0.116500654 | 0.062 | 0 | 1 | 20 |
| mhc1uba | 0.29436 | 0.101566925 | 0.062 | 0 | 1 | 20 |
| mtx1a | 0.29436 | 0.142972368 | 0.062 | 0 | 1 | 20 |
| mylka | 0.29436 | 0.14599608 | 0.062 | 0 | 1 | 20 |
| nab1a | 0.29436 | 0.191335308 | 0.062 | 0 | 1 | 20 |
| nsd3 | 0.29436 | 0.103315279 | 0.062 | 0 | 1 | 20 |
| nsmce1 | 0.29436 | 0.110457704 | 0.062 | 0 | 1 | 20 |
| oxr1a | 0.29436 | 0.192401997 | 0.062 | 0 | 1 | 20 |
| pex13 | 0.29436 | 0.157340843 | 0.062 | 0 | 1 | 20 |
| phldb1b | 0.29436 | 0.112584732 | 0.062 | 0 | 1 | 20 |
| pla2g1b | 0.29436 | 0.215111782 | 0.062 | 0 | 1 | 20 |
| pold2 | 0.29436 | 0.104615569 | 0.062 | 0 | 1 | 20 |
| polg2 | 0.29436 | 0.110759661 | 0.062 | 0 | 1 | 20 |
| pou6f1 | 0.29436 | 0.143327065 | 0.062 | 0 | 1 | 20 |

| | | | | | | |
|---|---|---|---|---|---|---|
| ppm1ba | 0.29436 | 0.177922647 | 0.062 | 0 | 1 | 20 |
| ppp4r3b | 0.29436 | 0.100409844 | 0.062 | 0 | 1 | 20 |
| preb | 0.29436 | 0.108149031 | 0.062 | 0 | 1 | 20 |
| prss59.1 | 0.29436 | 0.376229015 | 0.062 | 0 | 1 | 20 |
| rab11fip1b | 0.29436 | 0.279257639 | 0.062 | 0 | 1 | 20 |
| rab43 | 0.29436 | 0.130861546 | 0.062 | 0 | 1 | 20 |
| rbm41 | 0.29436 | 0.170157635 | 0.062 | 0 | 1 | 20 |
| rffl | 0.29436 | 0.210483189 | 0.062 | 0 | 1 | 20 |
| rnf38 | 0.29436 | 0.13231987 | 0.062 | 0 | 1 | 20 |
| rpain | 0.29436 | 0.102757463 | 0.062 | 0 | 1 | 20 |
| sdhaf2 | 0.29436 | 0.148700173 | 0.062 | 0 | 1 | 20 |
| sec13 | 0.29436 | 0.117563521 | 0.062 | 0 | 1 | 20 |
| sec24d | 0.29436 | 0.115254893 | 0.062 | 0 | 1 | 20 |
| sema4ab | 0.29436 | 0.110879736 | 0.062 | 0 | 1 | 20 |
| si:ch211-150o23.2 | 0.29436 | 0.128419962 | 0.062 | 0 | 1 | 20 |
| si:ch211-154o6.2 | 0.29436 | 0.156650771 | 0.062 | 0 | 1 | 20 |
| si:ch211-160f23.5 | 0.29436 | 0.178699537 | 0.062 | 0 | 1 | 20 |
| si:ch211-168f7.5 | 0.29436 | 0.11776632 | 0.062 | 0 | 1 | 20 |
| si:ch211-243j20.2 | 0.29436 | 0.113431018 | 0.062 | 0 | 1 | 20 |
| si:ch73-236c18.8 | 0.29436 | 0.125398676 | 0.062 | 0 | 1 | 20 |
| si:ch73-359m17.2 | 0.29436 | 0.113580127 | 0.062 | 0 | 1 | 20 |
| si:dkey-171c9.3 | 0.29436 | 0.15146538 | 0.062 | 0 | 1 | 20 |
| slc27a1a | 0.29436 | 0.161688269 | 0.062 | 0 | 1 | 20 |
| slc5a8l | 0.29436 | 0.122335402 | 0.062 | 0 | 1 | 20 |
| smtnb | 0.29436 | 0.132755046 | 0.062 | 0 | 1 | 20 |
| ssuh2rs1 | 0.29436 | 0.102689524 | 0.062 | 0 | 1 | 20 |
| stau1 | 0.29436 | 0.115323772 | 0.062 | 0 | 1 | 20 |
| stk19 | 0.29436 | 0.103241571 | 0.062 | 0 | 1 | 20 |
| stx4 | 0.29436 | 0.126463646 | 0.062 | 0 | 1 | 20 |
| stx8 | 0.29436 | 0.134626117 | 0.062 | 0 | 1 | 20 |
| tbcc | 0.29436 | 0.124100168 | 0.062 | 0 | 1 | 20 |
| tbl1x | 0.29436 | 0.131278312 | 0.062 | 0 | 1 | 20 |
| tekt3 | 0.29436 | 0.132638176 | 0.062 | 0 | 1 | 20 |
| tjp2a | 0.29436 | 0.105899534 | 0.062 | 0 | 1 | 20 |
| tmem43 | 0.29436 | 0.103071094 | 0.062 | 0 | 1 | 20 |
| toporsb | 0.29436 | 0.126044736 | 0.062 | 0 | 1 | 20 |
| tpd52l2b | 0.29436 | 0.123643414 | 0.062 | 0 | 1 | 20 |
| trafd1 | 0.29436 | 0.129921048 | 0.062 | 0 | 1 | 20 |
| trmo | 0.29436 | 0.105277531 | 0.062 | 0 | 1 | 20 |
| tspan31 | 0.29436 | 0.10089173 | 0.062 | 0 | 1 | 20 |
| ube2q1 | 0.29436 | 0.126597944 | 0.062 | 0 | 1 | 20 |
| ube3a | 0.29436 | 0.105337342 | 0.062 | 0 | 1 | 20 |
| unga | 0.29436 | 0.109125056 | 0.062 | 0 | 1 | 20 |
| usf2 | 0.29436 | 0.166309296 | 0.062 | 0 | 1 | 20 |
| usp25 | 0.29436 | 0.101566925 | 0.062 | 0 | 1 | 20 |
| usp3 | 0.29436 | 0.110704488 | 0.062 | 0 | 1 | 20 |
| uvrag | 0.29436 | 0.119434557 | 0.062 | 0 | 1 | 20 |
| wdr44 | 0.29436 | 0.153825913 | 0.062 | 0 | 1 | 20 |

| | | | | | | |
|---|---|---|---|---|---|---|
| zcchc2 | 0.29436 | 0.14581658 | 0.062 | 0 | 1 | 20 |
| zgc:114081 | 0.29436 | 0.10745486 | 0.062 | 0 | 1 | 20 |
| zgc:162816 | 0.29436 | 0.113192564 | 0.062 | 0 | 1 | 20 |
| zgc:163057 | 0.29436 | 0.162605399 | 0.062 | 0 | 1 | 20 |
| zgc:92744 | 0.29436 | 0.236534507 | 0.062 | 0 | 1 | 20 |
| znf1117 | 0.29436 | 0.113926809 | 0.062 | 0 | 1 | 20 |
| MAP3K13 | 0.29436 | 0.114271974 | 0.062 | 0 | 1 | 20 |
| ap3s2 | 0.29436 | 0.113381442 | 0.062 | 0 | 1 | 20 |
| arhgef12b | 0.29436 | 0.145987685 | 0.062 | 0 | 1 | 20 |
| asah1b | 0.29436 | 0.118273783 | 0.062 | 0 | 1 | 20 |
| asf1ba | 0.29436 | 0.173400524 | 0.062 | 0 | 1 | 20 |
| ccl33.3 | 0.29436 | 0.177513673 | 0.062 | 0 | 1 | 20 |
| cica | 0.29436 | 0.121716774 | 0.062 | 0 | 1 | 20 |
| cluap1 | 0.29436 | 0.107520706 | 0.062 | 0 | 1 | 20 |
| dffa | 0.29436 | 0.165999913 | 0.062 | 0 | 1 | 20 |
| dnajc18 | 0.29436 | 0.156717248 | 0.062 | 0 | 1 | 20 |
| entr1 | 0.29436 | 0.193054049 | 0.062 | 0 | 1 | 20 |
| flcn | 0.29436 | 0.103917794 | 0.062 | 0 | 1 | 20 |
| mapk6 | 0.29436 | 0.164628133 | 0.062 | 0 | 1 | 20 |
| mettl15 | 0.29436 | 0.104432714 | 0.062 | 0 | 1 | 20 |
| mettl4 | 0.29436 | 0.100837501 | 0.062 | 0 | 1 | 20 |
| mtrr | 0.29436 | 0.102310795 | 0.062 | 0 | 1 | 20 |
| pdpk1b | 0.29436 | 0.153550585 | 0.062 | 0 | 1 | 20 |
| pgap1 | 0.29436 | 0.183752044 | 0.062 | 0 | 1 | 20 |
| pias2 | 0.29436 | 0.118437596 | 0.062 | 0 | 1 | 20 |
| rhoca | 0.29436 | 0.17455491 | 0.062 | 0 | 1 | 20 |
| scn2b | 0.29436 | 0.261329473 | 0.062 | 0 | 1 | 20 |
| spout1 | 0.29436 | 0.106490793 | 0.062 | 0 | 1 | 20 |
| tmem203 | 0.29436 | 0.184555013 | 0.062 | 0 | 1 | 20 |
| BX649250.1 | 0.29436 | 0.11325816 | 0.062 | 0 | 1 | 20 |
| CU929259.1 | 0.29436 | 0.14133266 | 0.062 | 0 | 1 | 20 |
| FO704750.1 | 0.29436 | 0.132702391 | 0.062 | 0 | 1 | 20 |
| atp6v1d | 0.29436 | 0.136052296 | 0.062 | 0 | 1 | 20 |
| cdc7 | 0.29436 | 0.101802572 | 0.062 | 0 | 1 | 20 |
| chek2 | 0.29436 | 0.126647084 | 0.062 | 0 | 1 | 20 |
| cyp20a1 | 0.29436 | 0.119852217 | 0.062 | 0 | 1 | 20 |
| dcun1d3 | 0.29436 | 0.126216075 | 0.062 | 0 | 1 | 20 |
| esco2 | 0.29436 | 0.140582268 | 0.062 | 0 | 1 | 20 |
| fam91a1 | 0.29436 | 0.145527977 | 0.062 | 0 | 1 | 20 |
| gnl3l | 0.29436 | 0.110237734 | 0.062 | 0 | 1 | 20 |
| gsdmeb | 0.29436 | 0.150181271 | 0.062 | 0 | 1 | 20 |
| isoc1 | 0.29436 | 0.150346404 | 0.062 | 0 | 1 | 20 |
| mlh1 | 0.29436 | 0.149466381 | 0.062 | 0 | 1 | 20 |
| naprt | 0.29436 | 0.110840764 | 0.062 | 0 | 1 | 20 |
| nme6 | 0.29436 | 0.101657094 | 0.062 | 0 | 1 | 20 |
| ppp1r26 | 0.29436 | 0.170298624 | 0.062 | 0 | 1 | 20 |
| psen2 | 0.29436 | 0.103169629 | 0.062 | 0 | 1 | 20 |
| rb1 | 0.29436 | 0.133349924 | 0.062 | 0 | 1 | 20 |

| | | | | | | |
|---|---|---|---|---|---|---|
| sh3bp5la | 0.29436 | 0.148216849 | 0.062 | 0 | 1 | 20 |
| smad2 | 0.29436 | 0.113587623 | 0.062 | 0 | 1 | 20 |
| spinb | 0.29436 | 0.135485733 | 0.062 | 0 | 1 | 20 |
| taf1b | 0.29436 | 0.115983617 | 0.062 | 0 | 1 | 20 |
| tank | 0.29436 | 0.148148171 | 0.062 | 0 | 1 | 20 |
| tmem115 | 0.29436 | 0.166527536 | 0.062 | 0 | 1 | 20 |
| tmx4 | 0.29436 | 0.115766234 | 0.062 | 0 | 1 | 20 |
| tnfrsf11a | 0.29436 | 0.116998474 | 0.062 | 0 | 1 | 20 |
| ube2w | 0.29436 | 0.146196172 | 0.062 | 0 | 1 | 20 |
| wdfy3 | 0.29436 | 0.163264453 | 0.062 | 0 | 1 | 20 |
| wipi1 | 0.29436 | 0.139998392 | 0.062 | 0 | 1 | 20 |
| wwp2 | 0.29436 | 0.16940521 | 0.062 | 0 | 1 | 20 |
| zbtb16b | 0.29436 | 0.105026213 | 0.062 | 0 | 1 | 20 |
| zc3hc1 | 0.29436 | 0.151454038 | 0.062 | 0 | 1 | 20 |
| zgc:56585 | 0.29436 | 0.100948669 | 0.062 | 0 | 1 | 20 |
| NINJ1 | 0.29436 | 0.180015689 | 0.062 | 0 | 1 | 20 |
| btr09 | 0.29436 | 0.104524957 | 0.062 | 0 | 1 | 20 |
| ctrl | 0.29436 | 0.232602466 | 0.062 | 0 | 1 | 20 |
| gpr35.2 | 0.29436 | 0.233617635 | 0.062 | 0 | 1 | 20 |
| mtus1b | 0.29436 | 0.184094834 | 0.062 | 0 | 1 | 20 |
| phlda2 | 0.29436 | 0.14493327 | 0.062 | 0 | 1 | 20 |
| ptgs1 | 0.29436 | 0.107194678 | 0.062 | 0 | 1 | 20 |
| retreg2 | 0.29436 | 0.14589661 | 0.062 | 0 | 1 | 20 |
| tuba8l3 | 0.29436 | 0.254836481 | 0.062 | 0 | 1 | 20 |
| zc3h10 | 0.29436 | 0.177754505 | 0.062 | 0 | 1 | 20 |
| CU856539.2 | 0.29436 | 0.106817631 | 0.062 | 0 | 1 | 20 |
| GDPGP1 | 0.29436 | 0.104836044 | 0.062 | 0 | 1 | 20 |
| ighd | 0.29436 | 0.122071629 | 0.062 | 0 | 1 | 20 |
| kdm8 | 0.29436 | 0.116112578 | 0.062 | 0 | 1 | 20 |
| mier3b | 0.29436 | 0.117894009 | 0.062 | 0 | 1 | 20 |
| ptpn22 | 0.29436 | 0.109371386 | 0.062 | 0 | 1 | 20 |
| rab24 | 0.29436 | 0.113179037 | 0.062 | 0 | 1 | 20 |
| sh3d21 | 0.29436 | 0.132799013 | 0.062 | 0 | 1 | 20 |
| tbc1d22a | 0.29436 | 0.114855313 | 0.062 | 0 | 1 | 20 |
| tox4a | 0.29436 | 0.13524635 | 0.062 | 0 | 1 | 20 |
| zc3h12b | 0.29436 | 0.102308835 | 0.062 | 0 | 1 | 20 |
| CABZ01113599.1 | 0.29436 | 0.121684789 | 0.062 | 0 | 1 | 20 |
| LOC108179634 | 0.29436 | 0.143032697 | 0.062 | 0 | 1 | 20 |
| adcy9 | 0.29436 | 0.145908922 | 0.062 | 0 | 1 | 20 |
| atp2c1 | 0.29436 | 0.16456779 | 0.062 | 0 | 1 | 20 |
| babam2 | 0.29436 | 0.130469216 | 0.062 | 0 | 1 | 20 |
| carm1l | 0.29436 | 0.138340661 | 0.062 | 0 | 1 | 20 |
| cuedc2 | 0.29436 | 0.101750459 | 0.062 | 0 | 1 | 20 |
| glmp | 0.29436 | 0.112638452 | 0.062 | 0 | 1 | 20 |
| hmgxb4a | 0.29436 | 0.101750459 | 0.062 | 0 | 1 | 20 |
| mef2d | 0.29436 | 0.170438283 | 0.062 | 0 | 1 | 20 |
| orai1a | 0.29436 | 0.189939328 | 0.062 | 0 | 1 | 20 |
| plcd1b | 0.29436 | 0.218856667 | 0.062 | 0 | 1 | 20 |

| | | | | | | |
|---|---|---|---|---|---|---|
| rasgrp2 | 0.29436 | 0.118238226 | 0.062 | 0 | 1 | 20 |
| rbms2a | 0.29436 | 0.111573214 | 0.062 | 0 | 1 | 20 |
| rce1a | 0.29436 | 0.175130753 | 0.062 | 0 | 1 | 20 |
| sec63 | 0.29436 | 0.119679851 | 0.062 | 0 | 1 | 20 |
| si:ch73-237c6.1 | 0.29436 | 0.109502044 | 0.062 | 0 | 1 | 20 |
| si:dkeyp-104h9.5 | 0.29436 | 0.108685844 | 0.062 | 0 | 1 | 20 |
| smg6 | 0.29436 | 0.140088134 | 0.062 | 0 | 1 | 20 |
| tmem56b | 0.29436 | 0.117755888 | 0.062 | 0 | 1 | 20 |
| tmx2b | 0.29436 | 0.110932536 | 0.062 | 0 | 1 | 20 |
| zzz3 | 0.29436 | 0.100087599 | 0.062 | 0 | 1 | 20 |
| gatad2ab | 0.29436 | 0.118493604 | 0.062 | 0 | 1 | 20 |
| kif22 | 0.29436 | 0.144244695 | 0.062 | 0 | 1 | 20 |
| nt5c2l1 | 0.29436 | 0.183545355 | 0.062 | 0 | 1 | 20 |
| pan3 | 0.29436 | 0.129601945 | 0.062 | 0 | 1 | 20 |
| rassf2b | 0.29436 | 0.177522151 | 0.062 | 0 | 1 | 20 |
| tdp2b | 0.29436 | 0.125812099 | 0.062 | 0 | 1 | 20 |
| zdhhc5a | 0.29436 | 0.142557251 | 0.062 | 0 | 1 | 20 |
| snrpf | 0.29452 | 0.36776623 | 0.625 | 0.4 | 1 | 20 |
| arpc5b | 0.29552 | -0.36482611 | 0.094 | 0.2 | 1 | 20 |
| gnai2b | 0.29571 | 0.346460827 | 0.281 | 0.133 | 1 | 20 |
| coq4 | 0.29583 | -0.1350249 | 0.094 | 0.067 | 1 | 20 |
| rer1 | 0.29763 | 0.254093104 | 0.219 | 0.067 | 1 | 20 |
| ppib | 0.29809 | 0.208895464 | 0.562 | 0.333 | 1 | 20 |
| qkia | 0.29854 | -0.35772898 | 0.375 | 0.467 | 1 | 20 |
| top1l | 0.2995 | 0.294264719 | 0.219 | 0.067 | 1 | 20 |
| mob1a | 0.29952 | -0.42651725 | 0.344 | 0.533 | 1 | 20 |
| gdi2 | 0.29972 | 0.152733549 | 0.469 | 0.267 | 1 | 20 |
| eif1axb | 0.30005 | 0.31272685 | 0.375 | 0.267 | 1 | 20 |
| atp6v1ba | 0.30017 | -0.31581096 | 0.062 | 0.2 | 1 | 20 |
| gnl3 | 0.30068 | 0.299691265 | 0.406 | 0.267 | 1 | 20 |
| nudt21 | 0.3008 | 0.294109452 | 0.219 | 0.133 | 1 | 20 |
| mrpl54 | 0.30085 | 0.34156344 | 0.375 | 0.267 | 1 | 20 |
| phb | 0.30111 | 0.324626041 | 0.281 | 0.133 | 1 | 20 |
| phc2a | 0.30215 | -0.31603869 | 0.062 | 0.133 | 1 | 20 |
| gigyf1b | 0.30238 | 0.315030446 | 0.219 | 0.067 | 1 | 20 |
| chac1 | 0.30281 | -0.41243974 | 0.094 | 0.267 | 1 | 20 |
| si:dkey-22o12.2 | 0.30333 | -0.15761726 | 0.156 | 0.133 | 1 | 20 |
| LOC110437917 | 0.30351 | -0.30728041 | 0.062 | 0.2 | 1 | 20 |
| uprt | 0.30356 | -0.33050341 | 0.125 | 0.2 | 1 | 20 |
| rpl5a | 0.30446 | 0.311114576 | 0.812 | 0.6 | 1 | 20 |
| ubald1a | 0.30466 | -0.13704615 | 0.094 | 0.067 | 1 | 20 |
| pin1 | 0.30501 | 0.203769099 | 0.406 | 0.2 | 1 | 20 |
| sumo3a | 0.30554 | 0.209563973 | 0.562 | 0.333 | 1 | 20 |
| mapk14a | 0.30649 | 0.222772817 | 0.219 | 0.067 | 1 | 20 |
| si:dkey-23i12.5 | 0.30693 | 0.280262526 | 0.5 | 0.267 | 1 | 20 |
| klf13 | 0.307 | -0.20052996 | 0.094 | 0.267 | 1 | 20 |
| yy1b | 0.30703 | -0.13633973 | 0.25 | 0.2 | 1 | 20 |
| rpn1 | 0.30731 | 0.338274326 | 0.469 | 0.267 | 1 | 20 |

| | | | | | | |
|---|---|---|---|---|---|---|
| mif | 0.30734 | 0.347424948 | 0.562 | 0.333 | 1 | 20 |
| eif4g1a | 0.30771 | 0.358343417 | 0.312 | 0.133 | 1 | 20 |
| CABZ01087560.1 | 0.30809 | 0.291095649 | 0.219 | 0.067 | 1 | 20 |
| cdk9 | 0.3097 | -0.36485973 | 0.062 | 0.2 | 1 | 20 |
| hnrpkl | 0.30984 | 0.27852244 | 0.344 | 0.267 | 1 | 20 |
| pes | 0.31014 | 0.229329716 | 0.219 | 0.067 | 1 | 20 |
| rpl38 | 0.31123 | 0.22905146 | 0.906 | 0.8 | 1 | 20 |
| tomm40 | 0.31171 | 0.251143562 | 0.25 | 0.133 | 1 | 20 |
| XLOC-043857 | 0.31385 | -0.42114855 | 0.125 | 0.267 | 1 | 20 |
| si:ch211-284f22.3 | 0.31394 | -0.28241164 | 0.156 | 0.2 | 1 | 20 |
| si:ch211-105j21.9 | 0.31409 | -0.14055811 | 0.156 | 0.133 | 1 | 20 |
| fam208b | 0.31416 | -0.12736054 | 0.094 | 0.067 | 1 | 20 |
| dnase1l1 | 0.31452 | -0.39527343 | 0.094 | 0.267 | 1 | 20 |
| fosb | 0.31462 | -0.52227272 | 0.156 | 0.333 | 1 | 20 |
| pnisr | 0.31506 | -0.35460782 | 0.156 | 0.267 | 1 | 20 |
| cmtm3 | 0.31641 | -0.16433492 | 0.062 | 0.2 | 1 | 20 |
| tardbp | 0.31685 | 0.273536864 | 0.344 | 0.2 | 1 | 20 |
| nap1l1 | 0.31717 | 0.260754483 | 0.625 | 0.4 | 1 | 20 |
| cbx8a | 0.31829 | -0.12855772 | 0.094 | 0.067 | 1 | 20 |
| smn1 | 0.31947 | 0.255005621 | 0.219 | 0.067 | 1 | 20 |
| nop53 | 0.31981 | 0.37863725 | 0.562 | 0.333 | 1 | 20 |
| pus7 | 0.32021 | 0.300533228 | 0.281 | 0.133 | 1 | 20 |
| ywhaba | 0.32084 | 0.157298256 | 0.531 | 0.333 | 1 | 20 |
| pa2g4a | 0.3217 | 0.32386277 | 0.375 | 0.2 | 1 | 20 |
| add1 | 0.32247 | -0.30421372 | 0.156 | 0.2 | 1 | 20 |
| si:zfos-169g10.3 | 0.32277 | 0.15682101 | 0.469 | 0.267 | 1 | 20 |
| si:ch211-260e23.9 | 0.32324 | -0.46709323 | 0.156 | 0.333 | 1 | 20 |
| si:dkey-7j14.6 | 0.32357 | -0.23155128 | 0.094 | 0.267 | 1 | 20 |
| si:dkeyp-68b7.12 | 0.32398 | -0.12231244 | 0.094 | 0.067 | 1 | 20 |
| ywhabl | 0.32442 | -0.32347851 | 0.5 | 0.6 | 1 | 20 |
| wasa | 0.32539 | -0.12369103 | 0.469 | 0.4 | 1 | 20 |
| uqcrh | 0.32617 | 0.323914849 | 0.562 | 0.333 | 1 | 20 |
| sec62 | 0.32645 | -0.26244563 | 0.094 | 0.267 | 1 | 20 |
| zgc:55781 | 0.32658 | -0.12445619 | 0.094 | 0.067 | 1 | 20 |
| nr1d2a | 0.32742 | -0.32344784 | 0.094 | 0.267 | 1 | 20 |
| srsf2a | 0.32787 | 0.352906358 | 0.531 | 0.333 | 1 | 20 |
| inka1b | 0.32806 | -0.48318095 | 0.156 | 0.333 | 1 | 20 |
| fam3a | 0.32829 | -0.13215296 | 0.094 | 0.067 | 1 | 20 |
| rbm19 | 0.32872 | 0.255033168 | 0.219 | 0.067 | 1 | 20 |
| mrpl48 | 0.32909 | 0.11088815 | 0.219 | 0.067 | 1 | 20 |
| serf2 | 0.32934 | -0.11912633 | 0.75 | 0.667 | 1 | 20 |
| gcnt4a | 0.33017 | -0.41740505 | 0.219 | 0.333 | 1 | 20 |
| ak2 | 0.33019 | 0.257074283 | 0.625 | 0.4 | 1 | 20 |
| zgc:110239 | 0.33071 | -0.13052669 | 0.094 | 0.067 | 1 | 20 |
| eif4e1c | 0.33116 | 0.210872101 | 0.219 | 0.2 | 1 | 20 |
| cct5 | 0.33126 | 0.329413492 | 0.688 | 0.467 | 1 | 20 |
| calr | 0.33142 | 0.212900424 | 0.219 | 0.067 | 1 | 20 |
| hnrnpc | 0.33145 | 0.271475298 | 0.219 | 0.067 | 1 | 20 |

| | | | | | | |
|---|---|---|---|---|---|---|
| zgc:113208 | 0.33212 | 0.259598014 | 0.219 | 0.067 | 1 | 20 |
| isca2 | 0.33236 | -0.32968158 | 0.062 | 0.133 | 1 | 20 |
| ptp4a1 | 0.33271 | 0.206016139 | 0.312 | 0.133 | 1 | 20 |
| psma6l | 0.33352 | 0.251405279 | 0.312 | 0.133 | 1 | 20 |
| bmp6 | 0.33443 | -0.29647223 | 0.062 | 0.2 | 1 | 20 |
| prdx4 | 0.33456 | 0.168949592 | 0.312 | 0.133 | 1 | 20 |
| mrpl30 | 0.33479 | 0.28977828 | 0.312 | 0.133 | 1 | 20 |
| calm3b | 0.33487 | -0.34938775 | 0.469 | 0.6 | 1 | 20 |
| casp8 | 0.33605 | 0.331757605 | 0.188 | 0.067 | 1 | 20 |
| sept15 | 0.33645 | -0.27772893 | 0.062 | 0.133 | 1 | 20 |
| ppp2r5cb | 0.3368 | -0.1218945 | 0.094 | 0.067 | 1 | 20 |
| cox6b1 | 0.33681 | 0.179097023 | 0.469 | 0.267 | 1 | 20 |
| ccng2 | 0.33707 | -0.12520771 | 0.094 | 0.067 | 1 | 20 |
| hint2 | 0.33723 | 0.251318932 | 0.219 | 0.067 | 1 | 20 |
| tax1bp1b | 0.33783 | 0.259569623 | 0.219 | 0.067 | 1 | 20 |
| gemin7 | 0.339 | 0.148642809 | 0.219 | 0.267 | 1 | 20 |
| kdm2bb | 0.33907 | -0.12634194 | 0.094 | 0.067 | 1 | 20 |
| susd1 | 0.34007 | 0.195540112 | 0.188 | 0.067 | 1 | 20 |
| hacd3 | 0.34148 | -0.45279927 | 0.25 | 0.467 | 1 | 20 |
| fam207a | 0.34348 | 0.258008376 | 0.281 | 0.133 | 1 | 20 |
| crlf3 | 0.34394 | 0.238774754 | 0.219 | 0.133 | 1 | 20 |
| ssbp3b | 0.34398 | 0.296433377 | 0.219 | 0.067 | 1 | 20 |
| mob3a | 0.3441 | 0.361823201 | 0.312 | 0.133 | 1 | 20 |
| zgc:66388 | 0.34496 | 0.173641345 | 0.312 | 0.133 | 1 | 20 |
| si:dkey-42i9.4 | 0.34517 | -0.2468991 | 0.375 | 0.4 | 1 | 20 |
| itpka | 0.34606 | -0.27742637 | 0.062 | 0.2 | 1 | 20 |
| sik1 | 0.34641 | -0.24685509 | 0.062 | 0.2 | 1 | 20 |
| rap2c | 0.34646 | -0.35968515 | 0.125 | 0.2 | 1 | 20 |
| eif4h | 0.34664 | 0.278111161 | 0.281 | 0.133 | 1 | 20 |
| zgc:103700 | 0.3467 | 0.205618609 | 0.219 | 0.067 | 1 | 20 |
| nol12 | 0.34719 | 0.100143568 | 0.125 | 0.2 | 1 | 20 |
| sox4a | 0.34726 | 0.186147514 | 0.312 | 0.133 | 1 | 20 |
| tktb | 0.34761 | -0.2281102 | 0.375 | 0.4 | 1 | 20 |
| fam49ba | 0.34891 | -0.3286593 | 0.188 | 0.333 | 1 | 20 |
| dpp3 | 0.34912 | 0.208457044 | 0.219 | 0.067 | 1 | 20 |
| creb3l3l | 0.34955 | -0.36390423 | 0.125 | 0.2 | 1 | 20 |
| ewsr1b | 0.35001 | 0.209017671 | 0.406 | 0.2 | 1 | 20 |
| nansa | 0.35093 | 0.22706937 | 0.188 | 0.067 | 1 | 20 |
| ccnd2a | 0.35153 | -0.31192963 | 0.312 | 0.333 | 1 | 20 |
| vdac3 | 0.35154 | 0.200895161 | 0.125 | 0.133 | 1 | 20 |
| ralba | 0.35164 | -0.11890544 | 0.094 | 0.067 | 1 | 20 |
| sh3bgrl3 | 0.35252 | 0.205473706 | 0.312 | 0.133 | 1 | 20 |
| coq9 | 0.35312 | -0.12269823 | 0.094 | 0.067 | 1 | 20 |
| ube2ib | 0.35371 | 0.111072437 | 0.438 | 0.267 | 1 | 20 |
| antxr2a | 0.35387 | -0.27283861 | 0.062 | 0.2 | 1 | 20 |
| mmgt1 | 0.35411 | 0.245046527 | 0.312 | 0.133 | 1 | 20 |
| nmi | 0.35435 | 0.255638716 | 0.312 | 0.133 | 1 | 20 |
| cap1 | 0.35477 | -0.35094425 | 0.469 | 0.533 | 1 | 20 |

| | | | | | | |
|---|---|---|---|---|---|---|
| banf1 | 0.35518 | 0.355140484 | 0.438 | 0.267 | 1 | 20 |
| tram1 | 0.3553 | 0.258826245 | 0.188 | 0.067 | 1 | 20 |
| ube2v1 | 0.35547 | 0.192141803 | 0.469 | 0.267 | 1 | 20 |
| map2k2a | 0.35557 | -0.10884338 | 0.094 | 0.067 | 1 | 20 |
| ciao2b | 0.356 | -0.36370083 | 0.062 | 0.2 | 1 | 20 |
| ruvbl1 | 0.35606 | 0.248551906 | 0.25 | 0.133 | 1 | 20 |
| nudcd3 | 0.35609 | 0.23469836 | 0.219 | 0.067 | 1 | 20 |
| zgc:171686 | 0.35612 | -0.33558124 | 0.25 | 0.333 | 1 | 20 |
| dcps | 0.35623 | -0.3616459 | 0.125 | 0.2 | 1 | 20 |
| ptpn11b | 0.35646 | 0.281063255 | 0.219 | 0.067 | 1 | 20 |
| prpf31 | 0.35781 | -0.12151518 | 0.094 | 0.067 | 1 | 20 |
| atp1b1a | 0.3591 | 0.315698711 | 0.438 | 0.267 | 1 | 20 |
| chd3 | 0.35982 | 0.188076191 | 0.25 | 0.267 | 1 | 20 |
| chrac1 | 0.35999 | 0.264719475 | 0.281 | 0.133 | 1 | 20 |
| myocd | 0.36013 | -0.24791248 | 0.094 | 0.133 | 1 | 20 |
| hmgn3 | 0.36042 | 0.22101657 | 0.219 | 0.067 | 1 | 20 |
| klf6a | 0.36116 | -0.4321554 | 0.188 | 0.333 | 1 | 20 |
| COX7A2 (1 of many) | 0.36197 | 0.290118524 | 0.438 | 0.267 | 1 | 20 |
| actn1 | 0.36251 | -0.42600417 | 0.156 | 0.333 | 1 | 20 |
| hsbp1b | 0.36251 | -0.18580979 | 0.344 | 0.333 | 1 | 20 |
| vps26c | 0.36304 | 0.214388981 | 0.188 | 0.067 | 1 | 20 |
| si:ch211-235e9.6 | 0.36316 | 0.293999344 | 0.375 | 0.2 | 1 | 20 |
| zgc:158803 | 0.36393 | 0.264008106 | 0.188 | 0.067 | 1 | 20 |
| tmed7 | 0.36399 | 0.220622031 | 0.469 | 0.267 | 1 | 20 |
| ngdn | 0.36453 | 0.230365866 | 0.281 | 0.2 | 1 | 20 |
| smarce1 | 0.36546 | 0.354878684 | 0.375 | 0.2 | 1 | 20 |
| cstf2 | 0.36836 | -0.22483831 | 0.156 | 0.333 | 1 | 20 |
| eif2s1a | 0.36933 | -0.3227175 | 0.219 | 0.4 | 1 | 20 |
| ccdc58 | 0.36948 | 0.227274793 | 0.188 | 0.067 | 1 | 20 |
| cox6a2 | 0.36972 | 0.276177724 | 0.219 | 0.133 | 1 | 20 |
| prdx5 | 0.37018 | 0.36027126 | 0.531 | 0.333 | 1 | 20 |
| fam162a | 0.37073 | 0.233696208 | 0.281 | 0.2 | 1 | 20 |
| pus1 | 0.37101 | 0.209150851 | 0.188 | 0.067 | 1 | 20 |
| polr2k | 0.37171 | 0.233681872 | 0.312 | 0.133 | 1 | 20 |
| snip1 | 0.3721 | 0.189684359 | 0.188 | 0.067 | 1 | 20 |
| mrtfab | 0.37217 | -0.1805117 | 0.062 | 0.2 | 1 | 20 |
| XLOC-000431 | 0.37234 | -0.44604094 | 0.344 | 0.467 | 1 | 20 |
| pef1 | 0.37246 | -0.32531401 | 0.125 | 0.267 | 1 | 20 |
| gtf2b | 0.37261 | 0.287050462 | 0.156 | 0.067 | 1 | 20 |
| timm17a | 0.37406 | 0.203445029 | 0.219 | 0.067 | 1 | 20 |
| csnk1g2b | 0.37413 | -0.24141843 | 0.062 | 0.2 | 1 | 20 |
| mrpl46 | 0.37432 | 0.194485488 | 0.219 | 0.067 | 1 | 20 |
| hnrnpm | 0.37477 | 0.208869676 | 0.219 | 0.067 | 1 | 20 |
| zgc:56095 | 0.37498 | -0.21053018 | 0.125 | 0.133 | 1 | 20 |
| psmd8 | 0.37513 | 0.160071446 | 0.219 | 0.067 | 1 | 20 |
| rabggtb | 0.37516 | -0.26019752 | 0.062 | 0.2 | 1 | 20 |
| srsf4 | 0.37544 | 0.186001758 | 0.219 | 0.2 | 1 | 20 |
| ccdc43 | 0.37622 | 0.222352605 | 0.219 | 0.067 | 1 | 20 |

| | | | | | | |
|---|---|---|---|---|---|---|
| puraa | 0.3763 | -0.21681953 | 0.125 | 0.133 | 1 | 20 |
| arpc4l | 0.37649 | -0.29853778 | 0.5 | 0.533 | 1 | 20 |
| samd11 | 0.37665 | -0.3120554 | 0.125 | 0.267 | 1 | 20 |
| atg13 | 0.37672 | -0.10937162 | 0.094 | 0.067 | 1 | 20 |
| zgc:123258 | 0.37697 | -0.2429697 | 0.062 | 0.2 | 1 | 20 |
| flna | 0.3776 | -0.13361168 | 0.094 | 0.2 | 1 | 20 |
| impa1 | 0.37779 | -0.11183223 | 0.094 | 0.067 | 1 | 20 |
| si:ch211-212k18.4 | 0.37819 | 0.235027043 | 0.312 | 0.133 | 1 | 20 |
| lrrc47 | 0.37843 | 0.239176891 | 0.188 | 0.067 | 1 | 20 |
| rwdd1 | 0.37886 | 0.275360418 | 0.312 | 0.133 | 1 | 20 |
| gmps | 0.37902 | 0.172096726 | 0.219 | 0.067 | 1 | 20 |
| rap1ab | 0.37922 | -0.18587831 | 0.125 | 0.133 | 1 | 20 |
| ssr3 | 0.37928 | 0.277833324 | 0.188 | 0.133 | 1 | 20 |
| pum2 | 0.3803 | -0.11721047 | 0.094 | 0.067 | 1 | 20 |
| cct3 | 0.38061 | 0.182823911 | 0.656 | 0.467 | 1 | 20 |
| mcm4 | 0.38133 | 0.200445745 | 0.219 | 0.067 | 1 | 20 |
| srsf11 | 0.38141 | 0.146594893 | 0.281 | 0.133 | 1 | 20 |
| cited4a | 0.38156 | -0.26384818 | 0.062 | 0.2 | 1 | 20 |
| zgc:194312 | 0.3817 | -0.19093865 | 0.062 | 0.2 | 1 | 20 |
| arglu1b | 0.38181 | 0.20160914 | 0.219 | 0.067 | 1 | 20 |
| supt16h | 0.38201 | 0.189951344 | 0.219 | 0.067 | 1 | 20 |
| uqcc2 | 0.38338 | 0.18852022 | 0.219 | 0.067 | 1 | 20 |
| mvb12a | 0.38427 | -0.31396348 | 0.062 | 0.2 | 1 | 20 |
| calr3b | 0.38501 | 0.297448789 | 0.281 | 0.133 | 1 | 20 |
| crfb4 | 0.38516 | -0.22495223 | 0.188 | 0.2 | 1 | 20 |
| thoc1 | 0.38545 | -0.21206246 | 0.062 | 0.2 | 1 | 20 |
| pwp2h | 0.38582 | 0.194227462 | 0.312 | 0.133 | 1 | 20 |
| wdr36 | 0.38595 | 0.193760645 | 0.219 | 0.067 | 1 | 20 |
| STMP1 | 0.3861 | -0.18256757 | 0.312 | 0.267 | 1 | 20 |
| pmp22b | 0.38656 | -0.34418968 | 0.375 | 0.467 | 1 | 20 |
| canx | 0.38678 | 0.216589339 | 0.281 | 0.133 | 1 | 20 |
| ctdsp2 | 0.38745 | 0.271092187 | 0.312 | 0.267 | 1 | 20 |
| akap8l | 0.38833 | -0.1860447 | 0.062 | 0.2 | 1 | 20 |
| cib1 | 0.38862 | -0.11519683 | 0.094 | 0.067 | 1 | 20 |
| sdr16c5b | 0.38862 | 0.159348846 | 0.219 | 0.067 | 1 | 20 |
| ap1s2 | 0.38877 | 0.18314523 | 0.219 | 0.067 | 1 | 20 |
| cox6c | 0.38945 | 0.312337775 | 0.656 | 0.467 | 1 | 20 |
| tmsb1 | 0.38972 | 0.348893967 | 0.219 | 0.067 | 1 | 20 |
| mrpl4 | 0.39007 | 0.170136531 | 0.219 | 0.067 | 1 | 20 |
| DDX17 | 0.39062 | 0.228835959 | 0.219 | 0.067 | 1 | 20 |
| slc25a51b | 0.3912 | 0.229811408 | 0.219 | 0.067 | 1 | 20 |
| cyp4f3 | 0.39121 | 0.175003324 | 0.219 | 0.067 | 1 | 20 |
| lamtor2 | 0.39122 | 0.148497473 | 0.219 | 0.067 | 1 | 20 |
| pprc1 | 0.39125 | 0.189022837 | 0.219 | 0.067 | 1 | 20 |
| ndufaf1 | 0.3915 | -0.24552116 | 0.094 | 0.133 | 1 | 20 |
| rcc2 | 0.39165 | -0.30048118 | 0.156 | 0.333 | 1 | 20 |
| adi1 | 0.3919 | 0.243445145 | 0.188 | 0.067 | 1 | 20 |
| bzw1b | 0.39213 | 0.310863809 | 0.469 | 0.267 | 1 | 20 |

| | | | | | | |
|---|---|---|---|---|---|---|
| zranb2 | 0.39242 | 0.274922118 | 0.188 | 0.067 | 1 | 20 |
| nolc1 | 0.39248 | 0.248847324 | 0.281 | 0.133 | 1 | 20 |
| spi1b | 0.39248 | -0.21990428 | 0.25 | 0.267 | 1 | 20 |
| selenof | 0.3925 | 0.200459815 | 0.469 | 0.267 | 1 | 20 |
| ube2g1a | 0.3925 | -0.10671295 | 0.094 | 0.067 | 1 | 20 |
| naa40 | 0.39341 | 0.246553391 | 0.312 | 0.133 | 1 | 20 |
| ankrd13c | 0.39352 | -0.2510316 | 0.062 | 0.2 | 1 | 20 |
| slc25a26 | 0.39445 | -0.1050273 | 0.062 | 0.2 | 1 | 20 |
| immp2l | 0.39449 | 0.224655452 | 0.188 | 0.067 | 1 | 20 |
| prkcda | 0.39451 | -0.20524253 | 0.062 | 0.2 | 1 | 20 |
| mrpl14 | 0.39474 | 0.233308864 | 0.312 | 0.133 | 1 | 20 |
| thoc7 | 0.39478 | -0.16301852 | 0.062 | 0.2 | 1 | 20 |
| si:dkey-78l4.14 | 0.39509 | 0.261650709 | 0.188 | 0.067 | 1 | 20 |
| CR318588.3 | 0.39558 | -0.45703773 | 0.062 | 0.2 | 1 | 20 |
| CABZ01029938.1 | 0.39628 | -0.30673613 | 0.062 | 0.2 | 1 | 20 |
| nup88 | 0.39654 | -0.3086373 | 0.062 | 0.2 | 1 | 20 |
| DHDH | 0.39656 | -0.15851509 | 0.062 | 0.2 | 1 | 20 |
| ipcef1 | 0.39666 | -0.21777832 | 0.062 | 0.2 | 1 | 20 |
| trib3 | 0.39679 | -0.37089858 | 0.125 | 0.267 | 1 | 20 |
| ccnt2a | 0.39696 | -0.19796339 | 0.062 | 0.2 | 1 | 20 |
| ubl3a | 0.39697 | -0.31037453 | 0.062 | 0.2 | 1 | 20 |
| ubc | 0.39701 | -0.1154002 | 0.719 | 0.667 | 1 | 20 |
| rpz2 | 0.39839 | -0.1326569 | 0.156 | 0.133 | 1 | 20 |
| cnot8 | 0.39858 | -0.14453197 | 0.156 | 0.133 | 1 | 20 |
| echdc2 | 0.39925 | 0.254118317 | 0.156 | 0.067 | 1 | 20 |
| kdelr2b | 0.40098 | 0.263260967 | 0.188 | 0.133 | 1 | 20 |
| hnrnpul1l | 0.40148 | 0.214300288 | 0.188 | 0.067 | 1 | 20 |
| ndufb7 | 0.40253 | 0.276146839 | 0.469 | 0.267 | 1 | 20 |
| vps28 | 0.40274 | 0.171530609 | 0.25 | 0.267 | 1 | 20 |
| polr2d | 0.40294 | 0.254239718 | 0.469 | 0.267 | 1 | 20 |
| limd2 | 0.40318 | -0.30942251 | 0.281 | 0.333 | 1 | 20 |
| phf10 | 0.40324 | 0.244199158 | 0.188 | 0.133 | 1 | 20 |
| sltm | 0.40505 | -0.20789388 | 0.25 | 0.267 | 1 | 20 |
| shq1 | 0.40529 | -0.21751849 | 0.125 | 0.133 | 1 | 20 |
| psma2 | 0.40542 | 0.320205882 | 0.406 | 0.267 | 1 | 20 |
| ddx39ab | 0.40583 | 0.293757388 | 0.469 | 0.333 | 1 | 20 |
| btg3 | 0.40611 | 0.195964485 | 0.188 | 0.067 | 1 | 20 |
| bccip | 0.4064 | 0.251815572 | 0.469 | 0.267 | 1 | 20 |
| xbp1 | 0.407 | -0.25375844 | 0.188 | 0.2 | 1 | 20 |
| grinab | 0.40708 | 0.295743319 | 0.188 | 0.067 | 1 | 20 |
| gatad2b | 0.40711 | 0.253219235 | 0.469 | 0.267 | 1 | 20 |
| ip6k2a | 0.4078 | -0.32490323 | 0.062 | 0.133 | 1 | 20 |
| churc1 | 0.40816 | 0.258474485 | 0.281 | 0.133 | 1 | 20 |
| ndufs6 | 0.40817 | -0.23412364 | 0.281 | 0.333 | 1 | 20 |
| sem1 | 0.40862 | 0.119067541 | 0.438 | 0.267 | 1 | 20 |
| rcan1a | 0.40862 | -0.33557305 | 0.094 | 0.2 | 1 | 20 |
| spcs1 | 0.40905 | 0.115833638 | 0.5 | 0.333 | 1 | 20 |
| mrpl28 | 0.40946 | -0.11103269 | 0.094 | 0.067 | 1 | 20 |

| | | | | | | |
|---|---|---|---|---|---|---|
| brd9 | 0.40947 | -0.30138361 | 0.094 | 0.2 | 1 | 20 |
| ywhaqa | 0.40948 | 0.323098118 | 0.438 | 0.267 | 1 | 20 |
| slc25a5 | 0.41185 | 0.15803599 | 0.938 | 0.867 | 1 | 20 |
| UTP14C | 0.41206 | 0.11458056 | 0.219 | 0.267 | 1 | 20 |
| mir21-1 | 0.41215 | -0.14645753 | 0.156 | 0.133 | 1 | 20 |
| taf7 | 0.41231 | -0.23843342 | 0.094 | 0.133 | 1 | 20 |
| ncl | 0.41323 | 0.218395467 | 0.656 | 0.467 | 1 | 20 |
| tmem110l | 0.41325 | -0.20056574 | 0.125 | 0.133 | 1 | 20 |
| mff | 0.41448 | 0.212133432 | 0.188 | 0.067 | 1 | 20 |
| ilf2 | 0.41517 | -0.13972771 | 0.438 | 0.4 | 1 | 20 |
| pum3 | 0.41549 | 0.268254416 | 0.25 | 0.133 | 1 | 20 |
| gsto2 | 0.41591 | 0.314363406 | 0.281 | 0.133 | 1 | 20 |
| ndrg3b | 0.41629 | -0.28333711 | 0.062 | 0.133 | 1 | 20 |
| chchd1 | 0.41693 | 0.220930242 | 0.219 | 0.133 | 1 | 20 |
| ucp2 | 0.41779 | -0.22241857 | 0.656 | 0.667 | 1 | 20 |
| rbpjb | 0.41803 | 0.233462953 | 0.188 | 0.067 | 1 | 20 |
| bloc1s1 | 0.41839 | 0.199451557 | 0.156 | 0.133 | 1 | 20 |
| timm23a | 0.41896 | 0.186267977 | 0.281 | 0.267 | 1 | 20 |
| ncor2 | 0.41949 | 0.212379396 | 0.188 | 0.067 | 1 | 20 |
| exosc4 | 0.41954 | 0.219747685 | 0.188 | 0.067 | 1 | 20 |
| uqcrc2a | 0.42019 | 0.242978057 | 0.25 | 0.2 | 1 | 20 |
| itgb3b | 0.42152 | -0.4622027 | 0.188 | 0.333 | 1 | 20 |
| gtf2f1 | 0.42166 | 0.241846499 | 0.281 | 0.133 | 1 | 20 |
| hbs1l | 0.42197 | -0.33553554 | 0.125 | 0.267 | 1 | 20 |
| zgc:91910 | 0.42291 | 0.238436528 | 0.281 | 0.2 | 1 | 20 |
| dbi | 0.42339 | 0.193656867 | 0.531 | 0.333 | 1 | 20 |
| ftr50 | 0.42423 | 0.21532744 | 0.188 | 0.067 | 1 | 20 |
| ppp2r5eb | 0.42473 | -0.11970672 | 0.188 | 0.133 | 1 | 20 |
| wu:fi37e09 | 0.42496 | -0.2601709 | 0.156 | 0.2 | 1 | 20 |
| tmem11 | 0.425 | 0.2533832 | 0.219 | 0.133 | 1 | 20 |
| echs1 | 0.42614 | 0.11644409 | 0.281 | 0.133 | 1 | 20 |
| slc38a5b | 0.42764 | 0.222729214 | 0.25 | 0.133 | 1 | 20 |
| zgc:86598 | 0.42802 | 0.119138078 | 0.281 | 0.133 | 1 | 20 |
| ier3ip1 | 0.42855 | 0.268295098 | 0.375 | 0.267 | 1 | 20 |
| micu2 | 0.42911 | -0.10532784 | 0.094 | 0.067 | 1 | 20 |
| fscn1a | 0.42987 | 0.245182114 | 0.188 | 0.067 | 1 | 20 |
| chmp1a | 0.43113 | -0.10299932 | 0.094 | 0.067 | 1 | 20 |
| adrm1 | 0.43185 | -0.18492357 | 0.188 | 0.2 | 1 | 20 |
| ccnl1b | 0.43196 | -0.10646781 | 0.094 | 0.067 | 1 | 20 |
| id3 | 0.43342 | 0.336856571 | 0.25 | 0.133 | 1 | 20 |
| pdia3 | 0.43388 | 0.118177973 | 0.438 | 0.267 | 1 | 20 |
| rrm2 | 0.43527 | 0.234654874 | 0.188 | 0.067 | 1 | 20 |
| mrps18c | 0.43582 | 0.205543819 | 0.156 | 0.067 | 1 | 20 |
| sh2b3 | 0.4361 | -0.34527219 | 0.125 | 0.2 | 1 | 20 |
| per1b | 0.43701 | 0.255638577 | 0.188 | 0.067 | 1 | 20 |
| hm13 | 0.43702 | 0.262200251 | 0.188 | 0.067 | 1 | 20 |
| mcm3 | 0.43865 | 0.197781972 | 0.156 | 0.067 | 1 | 20 |
| arl1 | 0.43877 | -0.28110759 | 0.094 | 0.2 | 1 | 20 |

| | | | | | | |
|---|---|---|---|---|---|---|
| wasf2 | 0.43927 | 0.147112708 | 0.125 | 0.133 | 1 | 20 |
| psmb1 | 0.44062 | 0.230280036 | 0.531 | 0.333 | 1 | 20 |
| ost4 | 0.44144 | 0.151990605 | 0.281 | 0.133 | 1 | 20 |
| sept7b | 0.44159 | 0.220682048 | 0.219 | 0.2 | 1 | 20 |
| psmc2 | 0.44216 | 0.160252232 | 0.281 | 0.133 | 1 | 20 |
| gtpbp4 | 0.44225 | 0.300086606 | 0.656 | 0.467 | 1 | 20 |
| proser1 | 0.44254 | -0.10040943 | 0.094 | 0.067 | 1 | 20 |
| chtf8 | 0.44294 | 0.169812265 | 0.125 | 0.133 | 1 | 20 |
| tmem59 | 0.44364 | -0.33558396 | 0.219 | 0.333 | 1 | 20 |
| rtf1 | 0.44377 | 0.238504554 | 0.375 | 0.2 | 1 | 20 |
| rsl24d1 | 0.44457 | 0.206248884 | 0.875 | 0.733 | 1 | 20 |
| eif3m | 0.44611 | 0.225624353 | 0.719 | 0.533 | 1 | 20 |
| arpp19b | 0.44668 | 0.223158673 | 0.219 | 0.133 | 1 | 20 |
| vmp1 | 0.44674 | -0.28360357 | 0.062 | 0.133 | 1 | 20 |
| skap2 | 0.44684 | -0.38039075 | 0.125 | 0.267 | 1 | 20 |
| rmdn1 | 0.44743 | 0.210644161 | 0.281 | 0.133 | 1 | 20 |
| cmtm6 | 0.44773 | 0.314983197 | 0.188 | 0.067 | 1 | 20 |
| cs | 0.44798 | 0.163803266 | 0.188 | 0.067 | 1 | 20 |
| ubap2b | 0.44897 | 0.217985733 | 0.125 | 0.067 | 1 | 20 |
| itpa | 0.45014 | 0.197377445 | 0.188 | 0.133 | 1 | 20 |
| gna11b | 0.45022 | -0.15031169 | 0.156 | 0.133 | 1 | 20 |
| eif3i | 0.45056 | 0.240275038 | 0.594 | 0.4 | 1 | 20 |
| aurkaip1 | 0.45067 | 0.168175998 | 0.125 | 0.133 | 1 | 20 |
| atp5pb | 0.4511 | 0.260301814 | 0.719 | 0.533 | 1 | 20 |
| snrnp70 | 0.4512 | -0.13657745 | 0.219 | 0.2 | 1 | 20 |
| ptena | 0.45189 | 0.237938191 | 0.188 | 0.067 | 1 | 20 |
| abcf1 | 0.45299 | 0.274975668 | 0.406 | 0.267 | 1 | 20 |
| sept6 | 0.45335 | 0.122149227 | 0.5 | 0.333 | 1 | 20 |
| tomm7 | 0.45357 | 0.293059542 | 0.438 | 0.267 | 1 | 20 |
| atp5fa1 | 0.45414 | 0.252631159 | 0.719 | 0.533 | 1 | 20 |
| insra | 0.45478 | -0.10360871 | 0.094 | 0.067 | 1 | 20 |
| hpf1 | 0.45494 | 0.228946383 | 0.188 | 0.067 | 1 | 20 |
| si:rp71-36a1.1 | 0.45592 | 0.211536463 | 0.188 | 0.067 | 1 | 20 |
| ammecr1 | 0.45604 | 0.213523817 | 0.188 | 0.067 | 1 | 20 |
| rnaseh2a | 0.45668 | 0.228864087 | 0.188 | 0.067 | 1 | 20 |
| ftsj3 | 0.45673 | 0.192890485 | 0.312 | 0.267 | 1 | 20 |
| tmsb | 0.45874 | 0.214728772 | 0.188 | 0.067 | 1 | 20 |
| cry-dash | 0.45886 | 0.163810134 | 0.188 | 0.067 | 1 | 20 |
| atp6v1g1 | 0.45911 | -0.27319142 | 0.219 | 0.267 | 1 | 20 |
| psmb5 | 0.45921 | -0.1576948 | 0.125 | 0.133 | 1 | 20 |
| si:dkeyp-55f12.3 | 0.46131 | 0.20749152 | 0.188 | 0.133 | 1 | 20 |
| gnpda1 | 0.46139 | 0.188438635 | 0.094 | 0.067 | 1 | 20 |
| spsb4b | 0.46193 | -0.26654703 | 0.062 | 0.133 | 1 | 20 |
| idh3a | 0.4623 | 0.156815767 | 0.188 | 0.067 | 1 | 20 |
| si:ch1073-325m22.2 | 0.46282 | 0.238284695 | 0.594 | 0.4 | 1 | 20 |
| rpf1 | 0.46294 | 0.145740245 | 0.188 | 0.2 | 1 | 20 |
| ubtf | 0.463 | 0.238770762 | 0.219 | 0.133 | 1 | 20 |
| usp36 | 0.46458 | 0.242747637 | 0.375 | 0.2 | 1 | 20 |

| | | | | | | |
|---|---|---|---|---|---|---|
| nr2c2ap | 0.46516 | 0.158073759 | 0.188 | 0.067 | 1 | 20 |
| sdcbp2 | 0.46573 | 0.159786747 | 0.281 | 0.133 | 1 | 20 |
| psmc1a | 0.46638 | 0.196700155 | 0.188 | 0.067 | 1 | 20 |
| zgc:162509 | 0.46665 | -0.10727032 | 0.156 | 0.133 | 1 | 20 |
| wdr43 | 0.46836 | 0.243595045 | 0.312 | 0.2 | 1 | 20 |
| nudt5 | 0.46923 | 0.191899157 | 0.188 | 0.067 | 1 | 20 |
| eif3d | 0.47053 | 0.262504657 | 0.656 | 0.467 | 1 | 20 |
| cd74b | 0.47086 | 0.279616478 | 0.375 | 0.2 | 1 | 20 |
| si:dkey-180p18.9 | 0.47119 | -0.25989614 | 0.281 | 0.4 | 1 | 20 |
| ttf1 | 0.47311 | 0.189967129 | 0.188 | 0.067 | 1 | 20 |
| tprb | 0.47363 | -0.22994902 | 0.094 | 0.133 | 1 | 20 |
| sf3a1 | 0.47379 | 0.249319014 | 0.25 | 0.133 | 1 | 20 |
| paics | 0.47436 | 0.249238915 | 0.25 | 0.133 | 1 | 20 |
| rnf14 | 0.4756 | -0.22925235 | 0.125 | 0.267 | 1 | 20 |
| srsf1a | 0.47566 | 0.203500065 | 0.156 | 0.067 | 1 | 20 |
| tsr1 | 0.47644 | 0.178122709 | 0.188 | 0.067 | 1 | 20 |
| ppp2r2d | 0.47649 | 0.182482504 | 0.188 | 0.067 | 1 | 20 |
| pet100 | 0.47675 | 0.140300303 | 0.094 | 0.133 | 1 | 20 |
| ip6k2b | 0.47748 | -0.3034902 | 0.188 | 0.267 | 1 | 20 |
| usf1l | 0.47754 | -0.25929964 | 0.219 | 0.267 | 1 | 20 |
| ctgfa | 0.47757 | -0.51606733 | 0.125 | 0.267 | 1 | 20 |
| snrnp35 | 0.47774 | -0.26852977 | 0.062 | 0.133 | 1 | 20 |
| eif2a | 0.47789 | 0.202935535 | 0.188 | 0.067 | 1 | 20 |
| arrb2b | 0.47814 | -0.30417683 | 0.25 | 0.4 | 1 | 20 |
| nfkbie | 0.47856 | -0.27807201 | 0.125 | 0.267 | 1 | 20 |
| tfpia | 0.47879 | -0.33761833 | 0.094 | 0.2 | 1 | 20 |
| rbbp5 | 0.47925 | 0.173165402 | 0.188 | 0.067 | 1 | 20 |
| zfand3 | 0.47993 | -0.14292467 | 0.125 | 0.267 | 1 | 20 |
| rfx1b | 0.48109 | -0.29899283 | 0.094 | 0.2 | 1 | 20 |
| ypel5 | 0.48295 | -0.27588845 | 0.094 | 0.2 | 1 | 20 |
| zc4h2 | 0.48365 | 0.126211439 | 0.25 | 0.267 | 1 | 20 |
| cox8a | 0.48399 | 0.272956809 | 0.625 | 0.467 | 1 | 20 |
| atf4b | 0.4841 | -0.26641713 | 0.406 | 0.467 | 1 | 20 |
| si:dkey-112a7.4 | 0.48423 | -0.25530873 | 0.062 | 0.133 | 1 | 20 |
| magoh | 0.48433 | 0.181852939 | 0.188 | 0.067 | 1 | 20 |
| prmt5 | 0.48468 | 0.174805609 | 0.188 | 0.067 | 1 | 20 |
| aco2 | 0.48639 | -0.23423255 | 0.094 | 0.2 | 1 | 20 |
| fgfr1op2 | 0.48666 | -0.29668624 | 0.188 | 0.333 | 1 | 20 |
| sugt1 | 0.4871 | -0.23084121 | 0.125 | 0.267 | 1 | 20 |
| sap18 | 0.4873 | 0.251164935 | 0.438 | 0.267 | 1 | 20 |
| glrx3 | 0.4874 | 0.164591091 | 0.188 | 0.133 | 1 | 20 |
| smarcb1b | 0.48871 | 0.172462831 | 0.188 | 0.067 | 1 | 20 |
| lig1 | 0.48911 | 0.220275053 | 0.125 | 0.067 | 1 | 20 |
| litaf | 0.49015 | 0.159642085 | 0.188 | 0.067 | 1 | 20 |
| cbx5 | 0.49093 | 0.188051315 | 0.438 | 0.267 | 1 | 20 |
| foxo3b | 0.49223 | 0.154444883 | 0.219 | 0.2 | 1 | 20 |
| rcc1 | 0.49348 | 0.146288256 | 0.188 | 0.067 | 1 | 20 |
| pycard | 0.49358 | -0.2189203 | 0.25 | 0.267 | 1 | 20 |

| | | | | | | |
|---|---|---|---|---|---|---|
| glyr1 | 0.49393 | -0.12702734 | 0.156 | 0.133 | 1 | 20 |
| ap2m1b | 0.49406 | 0.166912792 | 0.125 | 0.133 | 1 | 20 |
| ndufa11 | 0.49441 | 0.230725902 | 0.438 | 0.267 | 1 | 20 |
| spg21 | 0.49531 | -0.12420122 | 0.125 | 0.267 | 1 | 20 |
| eif4a2 | 0.49567 | -0.17202048 | 0.125 | 0.133 | 1 | 20 |
| mphosph10 | 0.49603 | 0.115799106 | 0.188 | 0.067 | 1 | 20 |
| ptenb | 0.49603 | 0.170866679 | 0.188 | 0.067 | 1 | 20 |
| tuba8l | 0.49612 | 0.137190902 | 0.5 | 0.333 | 1 | 20 |
| urm1 | 0.49622 | 0.166686566 | 0.188 | 0.067 | 1 | 20 |
| tomm40l | 0.49622 | 0.183431945 | 0.188 | 0.067 | 1 | 20 |
| capza1b | 0.49649 | -0.29043069 | 0.531 | 0.667 | 1 | 20 |
| ctsh | 0.4966 | -0.25284039 | 0.188 | 0.267 | 1 | 20 |
| eif4g2b | 0.49687 | -0.16295843 | 0.125 | 0.267 | 1 | 20 |
| pik3ip1 | 0.49761 | -0.22950009 | 0.094 | 0.133 | 1 | 20 |
| cops4 | 0.4977 | 0.157186683 | 0.188 | 0.067 | 1 | 20 |
| si:dkey-188h10.3 | 0.49881 | 0.168813824 | 0.156 | 0.133 | 1 | 20 |
| atg12 | 0.49887 | -0.30802842 | 0.094 | 0.2 | 1 | 20 |
| ubr5 | 0.49897 | 0.22459498 | 0.156 | 0.067 | 1 | 20 |
| rpp30 | 0.49917 | 0.159129031 | 0.188 | 0.067 | 1 | 20 |
| mtpn | 0.49964 | -0.26521001 | 0.344 | 0.4 | 1 | 20 |
| XLOC-035413 | 0.50002 | 0.156090812 | 0.188 | 0.067 | 1 | 20 |
| chchd4a | 0.50035 | 0.163680628 | 0.188 | 0.067 | 1 | 20 |
| ormdl2 | 0.50103 | 0.170572849 | 0.188 | 0.067 | 1 | 20 |
| acp1 | 0.50131 | 0.180648315 | 0.188 | 0.067 | 1 | 20 |
| mrpl27 | 0.50146 | 0.183978132 | 0.188 | 0.067 | 1 | 20 |
| pgam1a | 0.50291 | 0.140876231 | 0.5 | 0.333 | 1 | 20 |
| brk1 | 0.50313 | 0.252076503 | 0.406 | 0.267 | 1 | 20 |
| egr1 | 0.50354 | -0.10674217 | 0.094 | 0.067 | 1 | 20 |
| mgst1.2 | 0.50471 | -0.20330642 | 0.406 | 0.4 | 1 | 20 |
| hectd3 | 0.50492 | -0.1869784 | 0.188 | 0.2 | 1 | 20 |
| zgc:77739 | 0.50496 | 0.24573655 | 0.219 | 0.133 | 1 | 20 |
| pias4a | 0.50598 | 0.173130238 | 0.188 | 0.067 | 1 | 20 |
| zgc:66433 | 0.50624 | -0.1014503 | 0.062 | 0.133 | 1 | 20 |
| ap3d1 | 0.50659 | 0.212307759 | 0.156 | 0.067 | 1 | 20 |
| cct6a | 0.50722 | 0.140167318 | 0.562 | 0.4 | 1 | 20 |
| cdkn1bb | 0.50766 | 0.228191238 | 0.188 | 0.067 | 1 | 20 |
| rad21a | 0.50776 | 0.110240777 | 0.344 | 0.2 | 1 | 20 |
| rel | 0.50797 | -0.25458479 | 0.062 | 0.133 | 1 | 20 |
| ska2 | 0.50865 | -0.25458203 | 0.062 | 0.133 | 1 | 20 |
| ywhah | 0.50884 | 0.225367347 | 0.219 | 0.133 | 1 | 20 |
| ddx23 | 0.50904 | 0.144939737 | 0.188 | 0.067 | 1 | 20 |
| AL954142.1 | 0.50943 | -0.10018081 | 0.219 | 0.2 | 1 | 20 |
| rcl1 | 0.51088 | 0.143949421 | 0.188 | 0.067 | 1 | 20 |
| dctd | 0.51202 | 0.133579318 | 0.125 | 0.133 | 1 | 20 |
| itm2bb | 0.51204 | -0.2169595 | 0.5 | 0.533 | 1 | 20 |
| tmem14ca | 0.51225 | 0.222949781 | 0.188 | 0.067 | 1 | 20 |
| pdcd4a | 0.51237 | 0.224591846 | 0.281 | 0.133 | 1 | 20 |
| rangap1a | 0.51251 | 0.193148888 | 0.156 | 0.067 | 1 | 20 |

| | | | | | | |
|---|---|---|---|---|---|---|
| cmtm7 | 0.5134 | -0.23698069 | 0.094 | 0.2 | 1 | 20 |
| csf1rb | 0.51374 | 0.215929015 | 0.281 | 0.133 | 1 | 20 |
| p2ry10 | 0.5141 | 0.228086684 | 0.188 | 0.067 | 1 | 20 |
| psmb3 | 0.51453 | 0.212063903 | 0.25 | 0.133 | 1 | 20 |
| cebpg | 0.51498 | -0.27510567 | 0.062 | 0.133 | 1 | 20 |
| cul4b | 0.51564 | 0.163294254 | 0.188 | 0.067 | 1 | 20 |
| rpe | 0.51579 | 0.131153007 | 0.188 | 0.067 | 1 | 20 |
| dctn3 | 0.51583 | -0.30809132 | 0.25 | 0.4 | 1 | 20 |
| atp5l | 0.51607 | 0.269825294 | 0.625 | 0.467 | 1 | 20 |
| senp2 | 0.51752 | 0.178533391 | 0.125 | 0.067 | 1 | 20 |
| emc10 | 0.51782 | -0.16478337 | 0.125 | 0.133 | 1 | 20 |
| h2afx1 | 0.51814 | 0.192681485 | 0.5 | 0.333 | 1 | 20 |
| rtf2 | 0.51829 | 0.210978537 | 0.188 | 0.067 | 1 | 20 |
| LOC103911430 | 0.51833 | -0.2914978 | 0.094 | 0.2 | 1 | 20 |
| kdm2aa | 0.51874 | -0.22970814 | 0.062 | 0.133 | 1 | 20 |
| rap1aa | 0.51881 | 0.118762053 | 0.344 | 0.2 | 1 | 20 |
| higd2a | 0.51892 | 0.155302542 | 0.188 | 0.067 | 1 | 20 |
| si:dkey-222f8.3 | 0.51976 | 0.129052033 | 0.188 | 0.067 | 1 | 20 |
| magt1 | 0.51991 | -0.23133636 | 0.219 | 0.333 | 1 | 20 |
| ctdspl2b | 0.51999 | 0.199513748 | 0.156 | 0.067 | 1 | 20 |
| prkab1b | 0.52022 | 0.223162162 | 0.438 | 0.267 | 1 | 20 |
| blvra | 0.52062 | -0.27961974 | 0.062 | 0.133 | 1 | 20 |
| hnrnpl | 0.52123 | 0.1288088 | 0.188 | 0.067 | 1 | 20 |
| polr2eb | 0.52168 | 0.174284491 | 0.188 | 0.067 | 1 | 20 |
| chd9 | 0.52177 | 0.135854324 | 0.188 | 0.067 | 1 | 20 |
| zgc:163080 | 0.52178 | 0.161475961 | 0.188 | 0.067 | 1 | 20 |
| nsfl1c | 0.52184 | 0.120036105 | 0.188 | 0.067 | 1 | 20 |
| ndufb6 | 0.52239 | 0.205680148 | 0.25 | 0.133 | 1 | 20 |
| tspan7 | 0.52244 | -0.35263516 | 0.156 | 0.267 | 1 | 20 |
| tomm22 | 0.52252 | 0.110550426 | 0.188 | 0.067 | 1 | 20 |
| hif1ab | 0.52252 | 0.135286764 | 0.188 | 0.067 | 1 | 20 |
| zgc:114119 | 0.52405 | 0.143299539 | 0.156 | 0.067 | 1 | 20 |
| tmem50a | 0.52563 | -0.20595598 | 0.094 | 0.133 | 1 | 20 |
| sub1a | 0.52581 | 0.139175828 | 0.625 | 0.467 | 1 | 20 |
| mrps35 | 0.52637 | 0.243760479 | 0.219 | 0.133 | 1 | 20 |
| si:ch73-248e21.5 | 0.52771 | 0.233755394 | 0.25 | 0.133 | 1 | 20 |
| selenot2 | 0.52785 | 0.181506425 | 0.156 | 0.133 | 1 | 20 |
| h1fx | 0.52808 | -0.20978861 | 0.25 | 0.4 | 1 | 20 |
| sv2 | 0.53044 | -0.23849472 | 0.062 | 0.133 | 1 | 20 |
| arl8ba | 0.53213 | -0.21737224 | 0.094 | 0.2 | 1 | 20 |
| tob1a | 0.53219 | -0.25787677 | 0.188 | 0.333 | 1 | 20 |
| ywhaqb | 0.5324 | -0.22543107 | 0.312 | 0.333 | 1 | 20 |
| tmem39a | 0.53328 | -0.17164838 | 0.125 | 0.133 | 1 | 20 |
| trip10a | 0.53446 | -0.30462869 | 0.062 | 0.133 | 1 | 20 |
| pho | 0.53451 | 0.137904553 | 0.344 | 0.2 | 1 | 20 |
| skap1 | 0.53468 | 0.172278609 | 0.094 | 0.067 | 1 | 20 |
| atox1 | 0.53522 | 0.163342455 | 0.156 | 0.133 | 1 | 20 |
| plrg1 | 0.53553 | 0.123184703 | 0.188 | 0.2 | 1 | 20 |

| | | | | | | |
|---|---|---|---|---|---|---|
| h2afy2 | 0.53698 | -0.26455878 | 0.219 | 0.267 | 1 | 20 |
| hmbsa | 0.53704 | -0.23166397 | 0.094 | 0.133 | 1 | 20 |
| pde7a | 0.53899 | -0.27584044 | 0.25 | 0.333 | 1 | 20 |
| lpar5a | 0.5395 | 0.17656159 | 0.156 | 0.133 | 1 | 20 |
| dnttip2 | 0.53963 | 0.166888475 | 0.219 | 0.133 | 1 | 20 |
| atp5mea | 0.54003 | 0.196308775 | 0.188 | 0.133 | 1 | 20 |
| zbtb8os | 0.54013 | 0.2174903 | 0.219 | 0.133 | 1 | 20 |
| dusp2 | 0.54039 | 0.254267801 | 0.156 | 0.067 | 1 | 20 |
| srrm1 | 0.541 | 0.205424928 | 0.281 | 0.2 | 1 | 20 |
| ptpn6 | 0.54112 | -0.22550706 | 0.25 | 0.267 | 1 | 20 |
| mrpl35 | 0.54171 | 0.173587566 | 0.156 | 0.067 | 1 | 20 |
| LOC103909972 | 0.54195 | -0.24751594 | 0.25 | 0.333 | 1 | 20 |
| nfe2 | 0.54198 | -0.1269687 | 0.281 | 0.267 | 1 | 20 |
| creb1a | 0.54254 | 0.206131097 | 0.156 | 0.067 | 1 | 20 |
| nosip | 0.54313 | -0.20958737 | 0.094 | 0.2 | 1 | 20 |
| b3galt2 | 0.54395 | 0.172301873 | 0.094 | 0.067 | 1 | 20 |
| ociad1 | 0.54433 | 0.233231386 | 0.156 | 0.067 | 1 | 20 |
| akirin2 | 0.5445 | 0.155845229 | 0.156 | 0.067 | 1 | 20 |
| c1d | 0.54541 | 0.202768382 | 0.156 | 0.067 | 1 | 20 |
| hnrnpa0a | 0.54546 | 0.254053583 | 0.156 | 0.067 | 1 | 20 |
| pabpn1 | 0.54593 | 0.178527554 | 0.5 | 0.333 | 1 | 20 |
| nol10 | 0.54622 | 0.167718874 | 0.156 | 0.067 | 1 | 20 |
| grasp | 0.54626 | -0.25768228 | 0.125 | 0.2 | 1 | 20 |
| aip | 0.54636 | 0.179135955 | 0.188 | 0.133 | 1 | 20 |
| spty2d1 | 0.54671 | 0.191395569 | 0.156 | 0.067 | 1 | 20 |
| gabarapa | 0.54836 | -0.29602702 | 0.25 | 0.4 | 1 | 20 |
| fosl2 | 0.54924 | -0.2041542 | 0.094 | 0.133 | 1 | 20 |
| etfb | 0.54996 | 0.203558537 | 0.25 | 0.133 | 1 | 20 |
| uqcrq | 0.55006 | 0.147099811 | 0.562 | 0.4 | 1 | 20 |
| btg1 | 0.55024 | -0.27525918 | 0.688 | 0.8 | 1 | 20 |
| sept2 | 0.55085 | -0.19176602 | 0.469 | 0.467 | 1 | 20 |
| aimp1 | 0.55124 | 0.107906675 | 0.344 | 0.2 | 1 | 20 |
| psme3 | 0.55192 | 0.18940463 | 0.156 | 0.067 | 1 | 20 |
| klhdc3 | 0.55297 | 0.211711009 | 0.125 | 0.067 | 1 | 20 |
| tomm70a | 0.5535 | 0.222277827 | 0.25 | 0.2 | 1 | 20 |
| ubb | 0.55481 | -0.21503626 | 0.469 | 0.467 | 1 | 20 |
| txndc5 | 0.55509 | -0.22252716 | 0.094 | 0.2 | 1 | 20 |
| psip1b | 0.5556 | 0.23780633 | 0.312 | 0.2 | 1 | 20 |
| sptan1 | 0.55561 | 0.15758557 | 0.156 | 0.067 | 1 | 20 |
| tomm20b | 0.55648 | 0.247818674 | 0.469 | 0.333 | 1 | 20 |
| twf1a | 0.55695 | -0.14662407 | 0.438 | 0.4 | 1 | 20 |
| clic1 | 0.55712 | 0.241056972 | 0.5 | 0.333 | 1 | 20 |
| fnbp4 | 0.55771 | 0.179015779 | 0.156 | 0.067 | 1 | 20 |
| zgc:110591 | 0.55789 | 0.183194005 | 0.25 | 0.2 | 1 | 20 |
| cnot9 | 0.55889 | 0.208325867 | 0.219 | 0.133 | 1 | 20 |
| ndufb9 | 0.55942 | 0.214259865 | 0.5 | 0.333 | 1 | 20 |
| capza1a | 0.56042 | 0.125676101 | 0.469 | 0.333 | 1 | 20 |
| mphosph6 | 0.56088 | 0.211670416 | 0.25 | 0.133 | 1 | 20 |

| | | | | | | |
|---|---|---|---|---|---|---|
| znf1035 | 0.56108 | 0.167969921 | 0.156 | 0.067 | 1 | 20 |
| mbpb | 0.56118 | 0.260630971 | 0.281 | 0.2 | 1 | 20 |
| CABZ01115113.1 | 0.56126 | 0.168414319 | 0.25 | 0.133 | 1 | 20 |
| aifm1 | 0.56186 | -0.21611843 | 0.062 | 0.133 | 1 | 20 |
| tsr2 | 0.56304 | 0.187865972 | 0.25 | 0.2 | 1 | 20 |
| tmem258 | 0.56707 | 0.165729372 | 0.562 | 0.4 | 1 | 20 |
| slc25a11 | 0.56753 | -0.10546378 | 0.156 | 0.133 | 1 | 20 |
| TMEM19 | 0.56764 | 0.202208275 | 0.156 | 0.067 | 1 | 20 |
| zgc:158564 | 0.56836 | 0.20915335 | 0.156 | 0.067 | 1 | 20 |
| gins2 | 0.56971 | 0.170342801 | 0.125 | 0.067 | 1 | 20 |
| ap2s1 | 0.56995 | 0.175504681 | 0.219 | 0.2 | 1 | 20 |
| ndrg1a | 0.57093 | -0.21546879 | 0.094 | 0.133 | 1 | 20 |
| nme3 | 0.57151 | 0.218973538 | 0.25 | 0.133 | 1 | 20 |
| rnf11a | 0.57221 | -0.18525444 | 0.094 | 0.133 | 1 | 20 |
| flot1b | 0.57238 | -0.26829577 | 0.25 | 0.333 | 1 | 20 |
| smek1 | 0.57353 | -0.28937739 | 0.094 | 0.2 | 1 | 20 |
| tmpob | 0.57443 | 0.161465748 | 0.156 | 0.067 | 1 | 20 |
| csnk1da | 0.57444 | -0.23160997 | 0.094 | 0.2 | 1 | 20 |
| ap4s1 | 0.57444 | 0.173203284 | 0.156 | 0.133 | 1 | 20 |
| degs1 | 0.57444 | -0.23259667 | 0.062 | 0.133 | 1 | 20 |
| si:ch211-140m22.7 | 0.57453 | 0.185860716 | 0.156 | 0.067 | 1 | 20 |
| trappc6bl | 0.57567 | 0.133930504 | 0.156 | 0.067 | 1 | 20 |
| cbfb | 0.57594 | 0.113794067 | 0.25 | 0.133 | 1 | 20 |
| runx1t1 | 0.57614 | 0.180289055 | 0.125 | 0.067 | 1 | 20 |
| slc25a34 | 0.57774 | 0.186986213 | 0.156 | 0.067 | 1 | 20 |
| ccdc25 | 0.57802 | 0.205693426 | 0.312 | 0.2 | 1 | 20 |
| hnrnpa0l | 0.57809 | -0.14030878 | 0.812 | 0.867 | 1 | 20 |
| si:ch211-14a17.11 | 0.57926 | 0.231949297 | 0.344 | 0.2 | 1 | 20 |
| anapc16 | 0.58007 | 0.219487857 | 0.25 | 0.133 | 1 | 20 |
| tma7 | 0.58035 | 0.209663302 | 0.562 | 0.4 | 1 | 20 |
| yipf5 | 0.5805 | -0.20097575 | 0.094 | 0.2 | 1 | 20 |
| cnot6a | 0.58058 | 0.200031863 | 0.156 | 0.133 | 1 | 20 |
| cd40 | 0.58075 | 0.179686524 | 0.156 | 0.067 | 1 | 20 |
| cdv3 | 0.58145 | -0.16527743 | 0.312 | 0.333 | 1 | 20 |
| fam177a1 | 0.58156 | -0.23936921 | 0.125 | 0.2 | 1 | 20 |
| ddx24 | 0.58243 | 0.188786794 | 0.219 | 0.133 | 1 | 20 |
| ola1 | 0.58257 | 0.223426993 | 0.562 | 0.4 | 1 | 20 |
| glcci1a | 0.58275 | -0.26922527 | 0.094 | 0.2 | 1 | 20 |
| ppp1cab | 0.58335 | -0.24339797 | 0.156 | 0.267 | 1 | 20 |
| BOLA2 | 0.58444 | 0.171914983 | 0.156 | 0.067 | 1 | 20 |
| parvb | 0.58469 | 0.108763844 | 0.219 | 0.133 | 1 | 20 |
| apex1 | 0.58521 | 0.177498642 | 0.281 | 0.2 | 1 | 20 |
| oxsr1a | 0.5853 | -0.20364963 | 0.094 | 0.2 | 1 | 20 |
| znf410 | 0.58538 | 0.15741735 | 0.125 | 0.067 | 1 | 20 |
| rhof | 0.58656 | -0.16533319 | 0.125 | 0.133 | 1 | 20 |
| cnbpb | 0.58689 | 0.129518469 | 0.406 | 0.267 | 1 | 20 |
| farsa | 0.58711 | 0.131731414 | 0.25 | 0.267 | 1 | 20 |
| pafah1b1a | 0.58736 | -0.26931117 | 0.125 | 0.2 | 1 | 20 |

| | | | | | | |
|---|---|---|---|---|---|---|
| cpsf3 | 0.58763 | 0.166157507 | 0.125 | 0.067 | 1 | 20 |
| gtf2a2 | 0.58763 | -0.20059793 | 0.094 | 0.2 | 1 | 20 |
| coa6 | 0.58823 | 0.155328053 | 0.156 | 0.067 | 1 | 20 |
| ppig | 0.58923 | 0.14732315 | 0.219 | 0.2 | 1 | 20 |
| pnpla7a | 0.59046 | -0.21427799 | 0.062 | 0.133 | 1 | 20 |
| NC-002333.4 | 0.5907 | 0.237659208 | 0.625 | 0.467 | 1 | 20 |
| kmt5b | 0.59117 | 0.19487389 | 0.094 | 0.067 | 1 | 20 |
| mhc2b | 0.59173 | 0.18942871 | 0.25 | 0.133 | 1 | 20 |
| rhoaa | 0.5918 | 0.195909332 | 0.344 | 0.2 | 1 | 20 |
| nr3c1 | 0.59209 | -0.183864 | 0.125 | 0.133 | 1 | 20 |
| emg1 | 0.59257 | -0.1737929 | 0.094 | 0.2 | 1 | 20 |
| LOC100536213 | 0.59344 | 0.217717429 | 0.125 | 0.067 | 1 | 20 |
| ruvbl2 | 0.59385 | -0.11344955 | 0.156 | 0.133 | 1 | 20 |
| ttc5 | 0.59419 | 0.156136511 | 0.094 | 0.067 | 1 | 20 |
| siah1 | 0.59497 | -0.29122402 | 0.125 | 0.2 | 1 | 20 |
| mccc2 | 0.59509 | 0.120670035 | 0.156 | 0.067 | 1 | 20 |
| nup93 | 0.59596 | 0.146691591 | 0.156 | 0.133 | 1 | 20 |
| gar1 | 0.59677 | 0.178103269 | 0.188 | 0.133 | 1 | 20 |
| cdca7a | 0.59684 | 0.140439752 | 0.156 | 0.067 | 1 | 20 |
| polr3h | 0.59693 | 0.142722037 | 0.156 | 0.133 | 1 | 20 |
| vps26b | 0.59702 | 0.203155536 | 0.156 | 0.067 | 1 | 20 |
| zgc:66475 | 0.59734 | -0.1886497 | 0.094 | 0.2 | 1 | 20 |
| cdc34b | 0.59748 | 0.151229241 | 0.25 | 0.133 | 1 | 20 |
| ppt1 | 0.59834 | -0.15373682 | 0.406 | 0.4 | 1 | 20 |
| chaf1a | 0.59843 | 0.179918125 | 0.156 | 0.067 | 1 | 20 |
| im:7136021 | 0.59938 | 0.12971213 | 0.156 | 0.067 | 1 | 20 |
| arl14ep | 0.59985 | -0.18299258 | 0.094 | 0.2 | 1 | 20 |
| tmem147 | 0.60051 | 0.158346962 | 0.125 | 0.067 | 1 | 20 |
| u2surp | 0.60084 | 0.174523177 | 0.25 | 0.2 | 1 | 20 |
| ANKRD50 | 0.60096 | -0.19963364 | 0.031 | 0.067 | 1 | 20 |
| BX321875.1 | 0.60096 | -0.17268439 | 0.031 | 0.067 | 1 | 20 |
| BX470189.1 | 0.60096 | -0.19217213 | 0.031 | 0.067 | 1 | 20 |
| BX908750.1 | 0.60096 | -0.18914433 | 0.031 | 0.067 | 1 | 20 |
| CABZ01084575.1 | 0.60096 | -0.13438562 | 0.031 | 0.067 | 1 | 20 |
| CR848749.2 | 0.60096 | -0.11366794 | 0.031 | 0.067 | 1 | 20 |
| CU467110.1 | 0.60096 | -0.17987778 | 0.031 | 0.067 | 1 | 20 |
| LOC101883913 | 0.60096 | -0.17636653 | 0.031 | 0.067 | 1 | 20 |
| LOC101885018 | 0.60096 | -0.19264696 | 0.031 | 0.067 | 1 | 20 |
| LOC101885027 | 0.60096 | -0.11405034 | 0.031 | 0.067 | 1 | 20 |
| LOC101885907 | 0.60096 | -0.1377129 | 0.031 | 0.067 | 1 | 20 |
| MARK4 | 0.60096 | -0.18168348 | 0.031 | 0.067 | 1 | 20 |
| XLOC-000946 | 0.60096 | -0.15027427 | 0.031 | 0.067 | 1 | 20 |
| XLOC-022032 | 0.60096 | -0.16643309 | 0.031 | 0.067 | 1 | 20 |
| XLOC-026010 | 0.60096 | -0.23535513 | 0.031 | 0.067 | 1 | 20 |
| adora2aa | 0.60096 | -0.15966911 | 0.031 | 0.067 | 1 | 20 |
| adprm | 0.60096 | -0.17001005 | 0.031 | 0.067 | 1 | 20 |
| ano6 | 0.60096 | -0.197134 | 0.031 | 0.067 | 1 | 20 |
| baz2ba | 0.60096 | -0.18189894 | 0.031 | 0.067 | 1 | 20 |

| | | | | | | |
|---|---|---|---|---|---|---|
| bcl9l | 0.60096 | -0.20878001 | 0.031 | 0.067 | 1 | 20 |
| bnip3lb | 0.60096 | -0.14805506 | 0.031 | 0.067 | 1 | 20 |
| btbd7 | 0.60096 | -0.15608086 | 0.031 | 0.067 | 1 | 20 |
| cbx3b | 0.60096 | -0.20917181 | 0.031 | 0.067 | 1 | 20 |
| ccdc160 | 0.60096 | -0.12102803 | 0.031 | 0.067 | 1 | 20 |
| cenpn | 0.60096 | -0.14046273 | 0.031 | 0.067 | 1 | 20 |
| coro1b | 0.60096 | -0.15779408 | 0.031 | 0.067 | 1 | 20 |
| ctdsplb | 0.60096 | -0.21542277 | 0.031 | 0.067 | 1 | 20 |
| dhrs12 | 0.60096 | -0.19314159 | 0.031 | 0.067 | 1 | 20 |
| dhrs7b | 0.60096 | -0.17521466 | 0.031 | 0.067 | 1 | 20 |
| enc2 | 0.60096 | -0.16696707 | 0.031 | 0.067 | 1 | 20 |
| entpd1 | 0.60096 | -0.13013728 | 0.031 | 0.067 | 1 | 20 |
| fam57ba | 0.60096 | -0.17010092 | 0.031 | 0.067 | 1 | 20 |
| fam89b | 0.60096 | -0.17031565 | 0.031 | 0.067 | 1 | 20 |
| fdxr | 0.60096 | -0.22026244 | 0.031 | 0.067 | 1 | 20 |
| g3bp2 | 0.60096 | -0.19926538 | 0.031 | 0.067 | 1 | 20 |
| ggps1 | 0.60096 | -0.18163324 | 0.031 | 0.067 | 1 | 20 |
| idh3b | 0.60096 | -0.20817214 | 0.031 | 0.067 | 1 | 20 |
| il6r | 0.60096 | -0.14455914 | 0.031 | 0.067 | 1 | 20 |
| incenp | 0.60096 | -0.10217773 | 0.031 | 0.067 | 1 | 20 |
| ing5b | 0.60096 | -0.13602634 | 0.031 | 0.067 | 1 | 20 |
| ints13 | 0.60096 | -0.16882191 | 0.031 | 0.067 | 1 | 20 |
| kat7a | 0.60096 | -0.10047265 | 0.031 | 0.067 | 1 | 20 |
| lemd3 | 0.60096 | -0.13539191 | 0.031 | 0.067 | 1 | 20 |
| lnpk | 0.60096 | -0.18532319 | 0.031 | 0.067 | 1 | 20 |
| mafba | 0.60096 | -0.19577575 | 0.031 | 0.067 | 1 | 20 |
| mlf1 | 0.60096 | -0.10122809 | 0.031 | 0.067 | 1 | 20 |
| mpc1 | 0.60096 | -0.181534 | 0.031 | 0.067 | 1 | 20 |
| nbeal2 | 0.60096 | -0.22852613 | 0.031 | 0.067 | 1 | 20 |
| nfyc | 0.60096 | -0.20975005 | 0.031 | 0.067 | 1 | 20 |
| nrbf2b | 0.60096 | -0.21054704 | 0.031 | 0.067 | 1 | 20 |
| pdxdc1 | 0.60096 | -0.17203219 | 0.031 | 0.067 | 1 | 20 |
| pex14 | 0.60096 | -0.17708061 | 0.031 | 0.067 | 1 | 20 |
| phip | 0.60096 | -0.17017761 | 0.031 | 0.067 | 1 | 20 |
| pigrl2.3 | 0.60096 | -0.19239534 | 0.031 | 0.067 | 1 | 20 |
| pkn1a | 0.60096 | -0.21886134 | 0.031 | 0.067 | 1 | 20 |
| prkcha | 0.60096 | -0.20692874 | 0.031 | 0.067 | 1 | 20 |
| prrc1 | 0.60096 | -0.20647926 | 0.031 | 0.067 | 1 | 20 |
| ptger2b | 0.60096 | -0.17308649 | 0.031 | 0.067 | 1 | 20 |
| r3hcc1l | 0.60096 | -0.13729544 | 0.031 | 0.067 | 1 | 20 |
| rad18 | 0.60096 | -0.19272846 | 0.031 | 0.067 | 1 | 20 |
| rgl2 | 0.60096 | -0.11321811 | 0.031 | 0.067 | 1 | 20 |
| sec22bb | 0.60096 | -0.20821776 | 0.031 | 0.067 | 1 | 20 |
| sele | 0.60096 | -0.16060449 | 0.031 | 0.067 | 1 | 20 |
| si:ch211-114l13.9 | 0.60096 | -0.17680731 | 0.031 | 0.067 | 1 | 20 |
| si:ch211-212o1.2 | 0.60096 | -0.21521376 | 0.031 | 0.067 | 1 | 20 |
| si:ch211-261d7.3 | 0.60096 | -0.22677396 | 0.031 | 0.067 | 1 | 20 |
| si:ch73-109d9.1 | 0.60096 | -0.11089052 | 0.031 | 0.067 | 1 | 20 |

| | | | | | | |
|---|---|---|---|---|---|---|
| si:ch73-109d9.2 | 0.60096 | -0.18711609 | 0.031 | 0.067 | 1 | 20 |
| si:dkey-16j16.4 | 0.60096 | -0.19999537 | 0.031 | 0.067 | 1 | 20 |
| slc25a44a | 0.60096 | -0.12492769 | 0.031 | 0.067 | 1 | 20 |
| slc25a44b | 0.60096 | -0.16368362 | 0.031 | 0.067 | 1 | 20 |
| slc35e3 | 0.60096 | -0.18750959 | 0.031 | 0.067 | 1 | 20 |
| slc4a1ap | 0.60096 | -0.19232762 | 0.031 | 0.067 | 1 | 20 |
| snx14 | 0.60096 | -0.20879965 | 0.031 | 0.067 | 1 | 20 |
| ssr1 | 0.60096 | 0.100607659 | 0.031 | 0.067 | 1 | 20 |
| susd6 | 0.60096 | -0.16575534 | 0.031 | 0.067 | 1 | 20 |
| tcerg1a | 0.60096 | -0.20475533 | 0.031 | 0.067 | 1 | 20 |
| thap11 | 0.60096 | -0.20885766 | 0.031 | 0.067 | 1 | 20 |
| ttc33 | 0.60096 | -0.16576261 | 0.031 | 0.067 | 1 | 20 |
| unk | 0.60096 | -0.20411618 | 0.031 | 0.067 | 1 | 20 |
| usp42 | 0.60096 | -0.19617772 | 0.031 | 0.067 | 1 | 20 |
| wbp1lb | 0.60096 | -0.1609882 | 0.031 | 0.067 | 1 | 20 |
| zfand5a | 0.60096 | -0.14306384 | 0.031 | 0.067 | 1 | 20 |
| zfyve19 | 0.60096 | -0.16271261 | 0.031 | 0.067 | 1 | 20 |
| zgc:113263 | 0.60096 | -0.17968643 | 0.031 | 0.067 | 1 | 20 |
| zgc:153990 | 0.60096 | -0.15739369 | 0.031 | 0.067 | 1 | 20 |
| zgc:162200 | 0.60096 | -0.13971223 | 0.031 | 0.067 | 1 | 20 |
| zgc:162780 | 0.60096 | -0.2151073 | 0.031 | 0.067 | 1 | 20 |
| zgc:162952 | 0.60096 | -0.18425289 | 0.031 | 0.067 | 1 | 20 |
| BX005183.3 | 0.60096 | -0.10897254 | 0.031 | 0.067 | 1 | 20 |
| CABZ01032488.1 | 0.60096 | -0.22475477 | 0.031 | 0.067 | 1 | 20 |
| CABZ01074307.2 | 0.60096 | -0.18720238 | 0.031 | 0.067 | 1 | 20 |
| CU633745.1 | 0.60096 | -0.1997964 | 0.031 | 0.067 | 1 | 20 |
| CU856520.1 | 0.60096 | -0.17628445 | 0.031 | 0.067 | 1 | 20 |
| LOC110440146 | 0.60096 | -0.13656772 | 0.031 | 0.067 | 1 | 20 |
| LOC799656 | 0.60096 | -0.15847827 | 0.031 | 0.067 | 1 | 20 |
| aatka | 0.60096 | -0.13422736 | 0.031 | 0.067 | 1 | 20 |
| akt2 | 0.60096 | -0.20060931 | 0.031 | 0.067 | 1 | 20 |
| amfra | 0.60096 | -0.23174873 | 0.031 | 0.067 | 1 | 20 |
| asic1b | 0.60096 | -0.10103424 | 0.031 | 0.067 | 1 | 20 |
| bach1b | 0.60096 | -0.14134643 | 0.031 | 0.067 | 1 | 20 |
| bod1l1 | 0.60096 | -0.20171461 | 0.031 | 0.067 | 1 | 20 |
| bri3bp | 0.60096 | -0.21654367 | 0.031 | 0.067 | 1 | 20 |
| brwd1 | 0.60096 | -0.11314349 | 0.031 | 0.067 | 1 | 20 |
| camk2b1 | 0.60096 | -0.17331809 | 0.031 | 0.067 | 1 | 20 |
| caprin1a | 0.60096 | -0.10869102 | 0.031 | 0.067 | 1 | 20 |
| ccndbp1 | 0.60096 | -0.21584299 | 0.031 | 0.067 | 1 | 20 |
| ccr6a | 0.60096 | -0.18840367 | 0.031 | 0.067 | 1 | 20 |
| clcn3 | 0.60096 | -0.20685832 | 0.031 | 0.067 | 1 | 20 |
| cnep1r1 | 0.60096 | -0.21817162 | 0.031 | 0.067 | 1 | 20 |
| csrnp1a | 0.60096 | -0.18133025 | 0.031 | 0.067 | 1 | 20 |
| cytip | 0.60096 | -0.14321008 | 0.031 | 0.067 | 1 | 20 |
| dapk3 | 0.60096 | -0.20875117 | 0.031 | 0.067 | 1 | 20 |
| dhrs13a.3 | 0.60096 | -0.17791269 | 0.031 | 0.067 | 1 | 20 |
| fbxo34 | 0.60096 | -0.20233717 | 0.031 | 0.067 | 1 | 20 |

| | | | | | | |
|---|---|---|---|---|---|---|
| irs2b | 0.60096 | -0.1747786 | 0.031 | 0.067 | 1 | 20 |
| jak2b | 0.60096 | -0.18389639 | 0.031 | 0.067 | 1 | 20 |
| junbb | 0.60096 | -0.25387923 | 0.031 | 0.067 | 1 | 20 |
| kif5bb | 0.60096 | -0.19891789 | 0.031 | 0.067 | 1 | 20 |
| mark2b | 0.60096 | -0.15114206 | 0.031 | 0.067 | 1 | 20 |
| nck1a | 0.60096 | -0.21984519 | 0.031 | 0.067 | 1 | 20 |
| nelfe | 0.60096 | -0.19085423 | 0.031 | 0.067 | 1 | 20 |
| pcyt1aa | 0.60096 | -0.21997486 | 0.031 | 0.067 | 1 | 20 |
| pgrmc2 | 0.60096 | -0.20255208 | 0.031 | 0.067 | 1 | 20 |
| phrf1 | 0.60096 | -0.10220362 | 0.031 | 0.067 | 1 | 20 |
| pik3c3 | 0.60096 | -0.1780286 | 0.031 | 0.067 | 1 | 20 |
| pik3ca | 0.60096 | -0.19936773 | 0.031 | 0.067 | 1 | 20 |
| plekhf2 | 0.60096 | -0.12551043 | 0.031 | 0.067 | 1 | 20 |
| ppp1r12c | 0.60096 | -0.20493091 | 0.031 | 0.067 | 1 | 20 |
| prkab1a | 0.60096 | -0.24018727 | 0.031 | 0.067 | 1 | 20 |
| prkag1 | 0.60096 | -0.20567378 | 0.031 | 0.067 | 1 | 20 |
| pter | 0.60096 | -0.17797552 | 0.031 | 0.067 | 1 | 20 |
| ptpn21 | 0.60096 | -0.17716174 | 0.031 | 0.067 | 1 | 20 |
| rhogc | 0.60096 | -0.14537948 | 0.031 | 0.067 | 1 | 20 |
| scocb | 0.60096 | -0.22041147 | 0.031 | 0.067 | 1 | 20 |
| selplg | 0.60096 | -0.18613514 | 0.031 | 0.067 | 1 | 20 |
| si:ch211-114c17.1 | 0.60096 | -0.1867062 | 0.031 | 0.067 | 1 | 20 |
| si:ch73-264p11.1 | 0.60096 | -0.11650541 | 0.031 | 0.067 | 1 | 20 |
| slc10a7 | 0.60096 | -0.12928993 | 0.031 | 0.067 | 1 | 20 |
| tmem183a | 0.60096 | -0.11053274 | 0.031 | 0.067 | 1 | 20 |
| trrap | 0.60096 | -0.18093794 | 0.031 | 0.067 | 1 | 20 |
| twist3 | 0.60096 | -0.18478997 | 0.031 | 0.067 | 1 | 20 |
| uap1l1 | 0.60096 | -0.10910351 | 0.031 | 0.067 | 1 | 20 |
| wdr83 | 0.60096 | -0.1455116 | 0.031 | 0.067 | 1 | 20 |
| znf346 | 0.60096 | -0.20035151 | 0.031 | 0.067 | 1 | 20 |
| DGKD | 0.60096 | -0.20463997 | 0.031 | 0.067 | 1 | 20 |
| LOC108180269 | 0.60096 | -0.1085011 | 0.031 | 0.067 | 1 | 20 |
| XLOC-023492 | 0.60096 | -0.17127777 | 0.031 | 0.067 | 1 | 20 |
| XLOC-043918 | 0.60096 | -0.15900595 | 0.031 | 0.067 | 1 | 20 |
| abcc3 | 0.60096 | -0.19085033 | 0.031 | 0.067 | 1 | 20 |
| aftpha | 0.60096 | -0.21514616 | 0.031 | 0.067 | 1 | 20 |
| aifm2 | 0.60096 | -0.13449525 | 0.031 | 0.067 | 1 | 20 |
| arl16 | 0.60096 | -0.17727004 | 0.031 | 0.067 | 1 | 20 |
| asphd2 | 0.60096 | -0.20020425 | 0.031 | 0.067 | 1 | 20 |
| bnip2 | 0.60096 | -0.20796487 | 0.031 | 0.067 | 1 | 20 |
| cdh30 | 0.60096 | -0.21383217 | 0.031 | 0.067 | 1 | 20 |
| chmp7 | 0.60096 | -0.16421897 | 0.031 | 0.067 | 1 | 20 |
| ctnnb2 | 0.60096 | -0.13621396 | 0.031 | 0.067 | 1 | 20 |
| ddah2 | 0.60096 | -0.15683288 | 0.031 | 0.067 | 1 | 20 |
| gripap1 | 0.60096 | -0.20000876 | 0.031 | 0.067 | 1 | 20 |
| gstp1 | 0.60096 | -0.25277352 | 0.031 | 0.067 | 1 | 20 |
| hvcn1 | 0.60096 | -0.15429012 | 0.031 | 0.067 | 1 | 20 |
| kctd13 | 0.60096 | -0.18335834 | 0.031 | 0.067 | 1 | 20 |

| | | | | | | |
|---|---|---|---|---|---|---|
| mpnd | 0.60096 | -0.16291447 | 0.031 | 0.067 | 1 | 20 |
| mtm1 | 0.60096 | -0.18337734 | 0.031 | 0.067 | 1 | 20 |
| mxd4 | 0.60096 | -0.14064033 | 0.031 | 0.067 | 1 | 20 |
| npsn | 0.60096 | -0.2220542 | 0.031 | 0.067 | 1 | 20 |
| numb | 0.60096 | -0.17506439 | 0.031 | 0.067 | 1 | 20 |
| pacsin3 | 0.60096 | -0.18780977 | 0.031 | 0.067 | 1 | 20 |
| pdpk1a | 0.60096 | -0.21346984 | 0.031 | 0.067 | 1 | 20 |
| ppp1r37 | 0.60096 | -0.21925376 | 0.031 | 0.067 | 1 | 20 |
| ppp2r3c | 0.60096 | -0.19774834 | 0.031 | 0.067 | 1 | 20 |
| prkar1aa | 0.60096 | -0.19984852 | 0.031 | 0.067 | 1 | 20 |
| rag1 | 0.60096 | -0.31664912 | 0.031 | 0.067 | 1 | 20 |
| si:dkey-16l2.16 | 0.60096 | -0.21947317 | 0.031 | 0.067 | 1 | 20 |
| si:dkey-7l6.3 | 0.60096 | -0.12051266 | 0.031 | 0.067 | 1 | 20 |
| trib2 | 0.60096 | -0.22416467 | 0.031 | 0.067 | 1 | 20 |
| usp39 | 0.60096 | -0.12247215 | 0.031 | 0.067 | 1 | 20 |
| vps26a | 0.60096 | -0.13676167 | 0.031 | 0.067 | 1 | 20 |
| zgc:91976 | 0.60096 | -0.19456365 | 0.031 | 0.067 | 1 | 20 |
| LOC103910015 | 0.60096 | -0.15607997 | 0.031 | 0.067 | 1 | 20 |
| TSTD1 | 0.60096 | -0.1879814 | 0.031 | 0.067 | 1 | 20 |
| fam3c | 0.60096 | -0.15663913 | 0.031 | 0.067 | 1 | 20 |
| jakmip1 | 0.60096 | -0.1870009 | 0.031 | 0.067 | 1 | 20 |
| lin7c | 0.60096 | -0.21733728 | 0.031 | 0.067 | 1 | 20 |
| mxi1 | 0.60096 | -0.19810923 | 0.031 | 0.067 | 1 | 20 |
| snrka | 0.60096 | -0.20824049 | 0.031 | 0.067 | 1 | 20 |
| src | 0.60096 | -0.13990684 | 0.031 | 0.067 | 1 | 20 |
| znf598 | 0.60096 | -0.21867257 | 0.031 | 0.067 | 1 | 20 |
| csrnp2 | 0.60096 | -0.18700103 | 0.031 | 0.067 | 1 | 20 |
| faub | 0.60096 | -0.19330534 | 0.031 | 0.067 | 1 | 20 |
| gng2 | 0.60096 | -0.2274131 | 0.031 | 0.067 | 1 | 20 |
| nfrkb | 0.60096 | -0.18265797 | 0.031 | 0.067 | 1 | 20 |
| sec61a1 | 0.60096 | -0.17040517 | 0.031 | 0.067 | 1 | 20 |
| tbc1d15 | 0.60096 | -0.19843383 | 0.031 | 0.067 | 1 | 20 |
| zfand2a | 0.60096 | -0.1967114 | 0.031 | 0.067 | 1 | 20 |
| MYADM | 0.60096 | -0.21088035 | 0.031 | 0.067 | 1 | 20 |
| aldh5a1 | 0.60096 | -0.18475871 | 0.031 | 0.067 | 1 | 20 |
| bms1 | 0.60154 | 0.107142891 | 0.25 | 0.133 | 1 | 20 |
| chchd6a | 0.60194 | 0.177116105 | 0.156 | 0.067 | 1 | 20 |
| unc50 | 0.60255 | -0.19021653 | 0.094 | 0.133 | 1 | 20 |
| rspry1 | 0.60291 | -0.2074723 | 0.062 | 0.133 | 1 | 20 |
| sypl2b | 0.60345 | 0.202869592 | 0.219 | 0.133 | 1 | 20 |
| nrf1 | 0.60356 | -0.20079157 | 0.062 | 0.133 | 1 | 20 |
| ch25hl2 | 0.60495 | 0.146871727 | 0.094 | 0.067 | 1 | 20 |
| ap3s1.1 | 0.60532 | 0.148864117 | 0.219 | 0.2 | 1 | 20 |
| LOC108179023 | 0.60546 | 0.179287795 | 0.156 | 0.067 | 1 | 20 |
| cnpy3 | 0.60648 | -0.2005131 | 0.094 | 0.133 | 1 | 20 |
| tnr | 0.60722 | 0.213399075 | 0.156 | 0.067 | 1 | 20 |
| gpr183a | 0.60741 | 0.165706892 | 0.219 | 0.133 | 1 | 20 |
| selenot1b | 0.60851 | 0.180072264 | 0.219 | 0.133 | 1 | 20 |

| | | | | | | |
|---|---|---|---|---|---|---|
| CABZ01113192.1 | 0.60862 | 0.166653271 | 0.156 | 0.067 | 1 | 20 |
| sfxn4 | 0.60897 | -0.20846226 | 0.062 | 0.133 | 1 | 20 |
| ndufa8 | 0.60932 | 0.168329227 | 0.25 | 0.2 | 1 | 20 |
| vamp3 | 0.60939 | -0.19316021 | 0.312 | 0.333 | 1 | 20 |
| ccdc12 | 0.60968 | 0.139321334 | 0.156 | 0.133 | 1 | 20 |
| fdps | 0.61161 | -0.16090967 | 0.094 | 0.2 | 1 | 20 |
| ogfrl2 | 0.61188 | 0.145874207 | 0.156 | 0.067 | 1 | 20 |
| BX957362.1 | 0.61191 | 0.148323469 | 0.156 | 0.067 | 1 | 20 |
| rpa2 | 0.61231 | 0.186426667 | 0.125 | 0.067 | 1 | 20 |
| ppp2r1ba | 0.61257 | -0.19288335 | 0.094 | 0.2 | 1 | 20 |
| ifi30 | 0.61303 | 0.160509727 | 0.156 | 0.133 | 1 | 20 |
| smndc1 | 0.61305 | -0.22798097 | 0.281 | 0.4 | 1 | 20 |
| h2afy | 0.61305 | 0.173010322 | 0.094 | 0.067 | 1 | 20 |
| card19 | 0.61312 | -0.26259571 | 0.062 | 0.133 | 1 | 20 |
| srp68 | 0.61338 | 0.182340706 | 0.156 | 0.067 | 1 | 20 |
| ino80c | 0.61402 | -0.11162521 | 0.094 | 0.2 | 1 | 20 |
| XLOC-013964 | 0.61402 | -0.11625434 | 0.344 | 0.467 | 1 | 20 |
| gtf2f2a | 0.61403 | -0.1871547 | 0.219 | 0.267 | 1 | 20 |
| si:dkey-48p11.3 | 0.61416 | 0.124227055 | 0.156 | 0.067 | 1 | 20 |
| cnbpa | 0.61421 | 0.233483878 | 0.594 | 0.467 | 1 | 20 |
| rap2b | 0.61423 | -0.19355615 | 0.094 | 0.2 | 1 | 20 |
| nmt1a | 0.61428 | 0.143556517 | 0.219 | 0.2 | 1 | 20 |
| atpaf1 | 0.61433 | 0.178386253 | 0.156 | 0.067 | 1 | 20 |
| mrps10 | 0.61471 | -0.11825053 | 0.281 | 0.267 | 1 | 20 |
| trmt5 | 0.61533 | 0.141721932 | 0.156 | 0.067 | 1 | 20 |
| srp19 | 0.61579 | -0.26147229 | 0.156 | 0.267 | 1 | 20 |
| calr3a | 0.61631 | 0.167380358 | 0.25 | 0.133 | 1 | 20 |
| cables2b | 0.61651 | -0.16920176 | 0.094 | 0.2 | 1 | 20 |
| cisd2 | 0.61652 | -0.21502805 | 0.062 | 0.133 | 1 | 20 |
| calm1a | 0.61665 | -0.1817089 | 0.375 | 0.4 | 1 | 20 |
| mta3 | 0.6167 | 0.20214655 | 0.125 | 0.067 | 1 | 20 |
| akirin1 | 0.61676 | -0.17820737 | 0.094 | 0.2 | 1 | 20 |
| si:ch211-214p13.9 | 0.61738 | -0.30827579 | 0.094 | 0.2 | 1 | 20 |
| chico | 0.61785 | -0.19299552 | 0.094 | 0.133 | 1 | 20 |
| ctdsp1 | 0.61787 | -0.21843383 | 0.094 | 0.2 | 1 | 20 |
| eci2 | 0.61791 | -0.14083727 | 0.094 | 0.2 | 1 | 20 |
| si:dkey-56i24.1 | 0.61794 | 0.170047831 | 0.156 | 0.067 | 1 | 20 |
| arpc5lb | 0.61886 | 0.151818944 | 0.188 | 0.133 | 1 | 20 |
| pudp | 0.61886 | 0.143104446 | 0.125 | 0.067 | 1 | 20 |
| nasp | 0.61937 | 0.124450122 | 0.25 | 0.133 | 1 | 20 |
| cyb5r1 | 0.61995 | 0.155947928 | 0.156 | 0.067 | 1 | 20 |
| patl1 | 0.62003 | 0.16994066 | 0.156 | 0.067 | 1 | 20 |
| hnrnpul1 | 0.62078 | 0.177683561 | 0.156 | 0.067 | 1 | 20 |
| syngr2a | 0.62142 | -0.19151936 | 0.188 | 0.2 | 1 | 20 |
| ddx51 | 0.62174 | 0.143207279 | 0.156 | 0.067 | 1 | 20 |
| ptbp1a | 0.62179 | 0.131704556 | 0.25 | 0.133 | 1 | 20 |
| smap1 | 0.62207 | 0.162848672 | 0.094 | 0.067 | 1 | 20 |
| ice2 | 0.62247 | 0.161977217 | 0.156 | 0.067 | 1 | 20 |

| | | | | | | |
|---|---|---|---|---|---|---|
| caap1 | 0.62348 | 0.179893999 | 0.125 | 0.067 | 1 | 20 |
| tbl1xr1a | 0.62529 | -0.17251374 | 0.062 | 0.133 | 1 | 20 |
| eif2b2 | 0.62538 | 0.16162984 | 0.156 | 0.067 | 1 | 20 |
| csde1 | 0.62679 | -0.12689077 | 0.281 | 0.267 | 1 | 20 |
| rrp15 | 0.62708 | 0.157508553 | 0.188 | 0.133 | 1 | 20 |
| serinc1 | 0.62723 | -0.19405591 | 0.062 | 0.133 | 1 | 20 |
| meis1b | 0.62745 | 0.116038872 | 0.188 | 0.2 | 1 | 20 |
| marcksl1a | 0.62778 | -0.19355249 | 0.219 | 0.2 | 1 | 20 |
| dnajb11 | 0.62809 | 0.136052884 | 0.125 | 0.067 | 1 | 20 |
| pfn2l | 0.62844 | 0.116818017 | 0.156 | 0.067 | 1 | 20 |
| slc16a10 | 0.62859 | -0.12246536 | 0.219 | 0.2 | 1 | 20 |
| kctd5a | 0.62983 | -0.1389742 | 0.406 | 0.4 | 1 | 20 |
| def6a | 0.62986 | 0.125076085 | 0.156 | 0.067 | 1 | 20 |
| arhgap4b | 0.63002 | 0.133852824 | 0.156 | 0.067 | 1 | 20 |
| snrnp25 | 0.63003 | 0.180449642 | 0.125 | 0.067 | 1 | 20 |
| cebpa | 0.63085 | 0.176655199 | 0.25 | 0.133 | 1 | 20 |
| hadhb | 0.6312 | 0.14958364 | 0.219 | 0.133 | 1 | 20 |
| gmip | 0.63151 | -0.18295783 | 0.062 | 0.133 | 1 | 20 |
| lsm5 | 0.63186 | 0.187382635 | 0.406 | 0.267 | 1 | 20 |
| usf1 | 0.6319 | -0.19768132 | 0.156 | 0.2 | 1 | 20 |
| zgc:77849 | 0.63203 | -0.20638449 | 0.281 | 0.333 | 1 | 20 |
| fcer1gl | 0.63307 | -0.24858318 | 0.156 | 0.267 | 1 | 20 |
| sf3b4 | 0.63331 | 0.143581265 | 0.156 | 0.067 | 1 | 20 |
| hspa5 | 0.6334 | 0.141853659 | 0.25 | 0.133 | 1 | 20 |
| cul3b | 0.63442 | 0.127350428 | 0.125 | 0.067 | 1 | 20 |
| epc2 | 0.63468 | -0.18298402 | 0.094 | 0.133 | 1 | 20 |
| tpte | 0.63485 | -0.29164755 | 0.062 | 0.133 | 1 | 20 |
| ash2l | 0.63491 | 0.128257926 | 0.125 | 0.067 | 1 | 20 |
| mrpl45 | 0.63492 | 0.179537151 | 0.188 | 0.133 | 1 | 20 |
| AL954696.1 | 0.63511 | 0.133018077 | 0.156 | 0.067 | 1 | 20 |
| uri1 | 0.63576 | 0.167433378 | 0.188 | 0.133 | 1 | 20 |
| tlk1b | 0.63599 | -0.20104061 | 0.062 | 0.133 | 1 | 20 |
| emp2 | 0.63604 | -0.20154434 | 0.062 | 0.133 | 1 | 20 |
| gnb1a | 0.6365 | 0.22550084 | 0.406 | 0.267 | 1 | 20 |
| rrp36 | 0.63699 | 0.125870092 | 0.219 | 0.2 | 1 | 20 |
| hsf2 | 0.63723 | 0.131623194 | 0.156 | 0.067 | 1 | 20 |
| CABZ01079663.1 | 0.63753 | 0.166547276 | 0.156 | 0.067 | 1 | 20 |
| ech1 | 0.63892 | -0.17960618 | 0.094 | 0.133 | 1 | 20 |
| gadd45ga | 0.63902 | 0.181255411 | 0.219 | 0.133 | 1 | 20 |
| prpf8 | 0.63944 | 0.166875837 | 0.125 | 0.067 | 1 | 20 |
| cfap20 | 0.63982 | 0.106265051 | 0.125 | 0.133 | 1 | 20 |
| cdc42 | 0.63988 | 0.193599768 | 0.406 | 0.267 | 1 | 20 |
| paip2b | 0.63992 | 0.204342793 | 0.406 | 0.267 | 1 | 20 |
| erlin1 | 0.64038 | 0.157880557 | 0.156 | 0.067 | 1 | 20 |
| surf6 | 0.64056 | 0.186887963 | 0.156 | 0.067 | 1 | 20 |
| dnajc24 | 0.64058 | 0.133299709 | 0.156 | 0.067 | 1 | 20 |
| aqr | 0.64136 | 0.142773507 | 0.125 | 0.067 | 1 | 20 |
| eif1axa | 0.64204 | 0.146928104 | 0.188 | 0.133 | 1 | 20 |

| | | | | | | |
|---|---|---|---|---|---|---|
| sarnp | 0.64256 | 0.168760075 | 0.406 | 0.267 | 1 | 20 |
| ccdc59 | 0.6429 | 0.149627933 | 0.156 | 0.133 | 1 | 20 |
| actn4 | 0.64336 | -0.16263876 | 0.188 | 0.2 | 1 | 20 |
| tnfaip8l2a | 0.64356 | -0.17310873 | 0.156 | 0.2 | 1 | 20 |
| acadl | 0.6439 | 0.174903847 | 0.125 | 0.067 | 1 | 20 |
| jpt1b | 0.64541 | -0.23174374 | 0.156 | 0.2 | 1 | 20 |
| utp15 | 0.64553 | 0.151934235 | 0.125 | 0.067 | 1 | 20 |
| zbtb18 | 0.64614 | -0.18946805 | 0.062 | 0.133 | 1 | 20 |
| sars | 0.64751 | 0.107734854 | 0.25 | 0.267 | 1 | 20 |
| si:rp71-68n21.9 | 0.64792 | -0.21007556 | 0.219 | 0.333 | 1 | 20 |
| rgs2 | 0.64803 | -0.21039708 | 0.156 | 0.2 | 1 | 20 |
| lamtor4 | 0.64876 | -0.19080139 | 0.156 | 0.267 | 1 | 20 |
| bod1 | 0.64924 | 0.150930305 | 0.188 | 0.133 | 1 | 20 |
| igf2bp1 | 0.64998 | -0.20709566 | 0.156 | 0.267 | 1 | 20 |
| ctr9 | 0.65059 | -0.20094521 | 0.125 | 0.2 | 1 | 20 |
| si:dkey-66i24.7 | 0.65182 | 0.137430724 | 0.125 | 0.067 | 1 | 20 |
| ythdc1 | 0.65236 | -0.20474166 | 0.062 | 0.133 | 1 | 20 |
| leo1 | 0.65253 | 0.146244031 | 0.156 | 0.067 | 1 | 20 |
| cyba | 0.65329 | 0.177951737 | 0.188 | 0.133 | 1 | 20 |
| apeh | 0.65357 | 0.122585081 | 0.125 | 0.133 | 1 | 20 |
| tmc8 | 0.65366 | -0.17526946 | 0.094 | 0.133 | 1 | 20 |
| FP101887.1 | 0.65439 | 0.141754941 | 0.125 | 0.067 | 1 | 20 |
| rrp7a | 0.65441 | 0.138877462 | 0.188 | 0.133 | 1 | 20 |
| ndufv1 | 0.65489 | 0.180444926 | 0.188 | 0.133 | 1 | 20 |
| CU694197.1 | 0.65537 | -0.13798994 | 0.219 | 0.2 | 1 | 20 |
| zgc:111986 | 0.6556 | 0.191818789 | 0.469 | 0.333 | 1 | 20 |
| sf3b1 | 0.65561 | -0.18777046 | 0.094 | 0.133 | 1 | 20 |
| nol7 | 0.65611 | 0.117567693 | 0.156 | 0.067 | 1 | 20 |
| lamp1a | 0.65619 | -0.21520321 | 0.156 | 0.267 | 1 | 20 |
| dpp9 | 0.65687 | 0.141872979 | 0.125 | 0.067 | 1 | 20 |
| stag2b | 0.65712 | 0.143654081 | 0.156 | 0.067 | 1 | 20 |
| si:busm1-194e12.11 | 0.65733 | 0.101669427 | 0.156 | 0.067 | 1 | 20 |
| men1 | 0.65756 | 0.136798972 | 0.156 | 0.067 | 1 | 20 |
| si:ch211-85n16.3 | 0.65895 | 0.123883305 | 0.125 | 0.067 | 1 | 20 |
| vkorc1l1 | 0.659 | -0.17270745 | 0.062 | 0.133 | 1 | 20 |
| LOC553316 | 0.65919 | -0.14284923 | 0.125 | 0.133 | 1 | 20 |
| tra2b | 0.65966 | 0.11438051 | 0.156 | 0.067 | 1 | 20 |
| fam50a | 0.66021 | 0.127051975 | 0.125 | 0.067 | 1 | 20 |
| atp1a1a.1 | 0.66156 | 0.119405232 | 0.156 | 0.067 | 1 | 20 |
| eif4g2a | 0.66244 | 0.112068958 | 0.156 | 0.067 | 1 | 20 |
| rbm6 | 0.66248 | 0.114627464 | 0.156 | 0.067 | 1 | 20 |
| LOC110439840 | 0.66267 | -0.17449326 | 0.094 | 0.133 | 1 | 20 |
| polr3gla | 0.6628 | 0.116529938 | 0.156 | 0.067 | 1 | 20 |
| mrpl12 | 0.66294 | 0.176497879 | 0.312 | 0.2 | 1 | 20 |
| dnajc2 | 0.6634 | 0.119322062 | 0.125 | 0.067 | 1 | 20 |
| tapbpl | 0.66348 | 0.157508457 | 0.125 | 0.067 | 1 | 20 |
| ell | 0.66355 | -0.17337896 | 0.062 | 0.133 | 1 | 20 |
| smox | 0.66468 | -0.19169981 | 0.062 | 0.133 | 1 | 20 |

| | | | | | | |
|---|---|---|---|---|---|---|
| cicb | 0.66475 | -0.17880417 | 0.094 | 0.133 | 1 | 20 |
| zgc:77929 | 0.66483 | 0.118500828 | 0.156 | 0.067 | 1 | 20 |
| mrpl17 | 0.66565 | 0.116176485 | 0.156 | 0.067 | 1 | 20 |
| ppp1r10 | 0.6664 | 0.102054504 | 0.156 | 0.067 | 1 | 20 |
| ralab | 0.66645 | -0.19860498 | 0.094 | 0.133 | 1 | 20 |
| sptlc2a | 0.66691 | 0.109771683 | 0.156 | 0.067 | 1 | 20 |
| isg20 | 0.6674 | 0.155234059 | 0.25 | 0.2 | 1 | 20 |
| phf20a | 0.66748 | 0.134564444 | 0.125 | 0.067 | 1 | 20 |
| tbk1 | 0.66764 | -0.19990697 | 0.062 | 0.133 | 1 | 20 |
| tmem189 | 0.66777 | 0.104601604 | 0.156 | 0.067 | 1 | 20 |
| slc30a4 | 0.66802 | 0.100700623 | 0.156 | 0.067 | 1 | 20 |
| s1pr5b | 0.66845 | 0.160315758 | 0.156 | 0.067 | 1 | 20 |
| hccsb | 0.66904 | -0.17473125 | 0.062 | 0.133 | 1 | 20 |
| pole4 | 0.66914 | 0.170784503 | 0.156 | 0.133 | 1 | 20 |
| txn2 | 0.66922 | 0.134897273 | 0.156 | 0.067 | 1 | 20 |
| cox7a2l | 0.66965 | 0.207826463 | 0.344 | 0.267 | 1 | 20 |
| ndufs4 | 0.66985 | 0.185595233 | 0.312 | 0.2 | 1 | 20 |
| tmem243b | 0.67034 | 0.153656444 | 0.125 | 0.067 | 1 | 20 |
| rbbp6 | 0.6707 | -0.17724213 | 0.156 | 0.267 | 1 | 20 |
| cab39l | 0.6707 | 0.122383196 | 0.125 | 0.067 | 1 | 20 |
| cnot2 | 0.67103 | -0.15964233 | 0.062 | 0.133 | 1 | 20 |
| ehd4 | 0.67106 | 0.108158176 | 0.156 | 0.067 | 1 | 20 |
| helz | 0.67121 | -0.18247969 | 0.188 | 0.267 | 1 | 20 |
| tmem106a | 0.67145 | -0.19498485 | 0.062 | 0.133 | 1 | 20 |
| sqstm1 | 0.67152 | -0.11013745 | 0.062 | 0.133 | 1 | 20 |
| sept9b | 0.67156 | -0.23076289 | 0.156 | 0.267 | 1 | 20 |
| si:ch211-194k22.8 | 0.67209 | 0.116027397 | 0.156 | 0.067 | 1 | 20 |
| purab | 0.67226 | -0.15995821 | 0.156 | 0.2 | 1 | 20 |
| dnmt1 | 0.67226 | 0.139128798 | 0.094 | 0.067 | 1 | 20 |
| tubb2b | 0.67333 | -0.18755042 | 0.438 | 0.467 | 1 | 20 |
| ndufa5 | 0.67383 | -0.16817209 | 0.25 | 0.267 | 1 | 20 |
| tspan2a | 0.67413 | 0.13224109 | 0.094 | 0.067 | 1 | 20 |
| stk35l | 0.6749 | -0.12392634 | 0.125 | 0.133 | 1 | 20 |
| pitpnaa | 0.67517 | -0.18709218 | 0.125 | 0.2 | 1 | 20 |
| sumo1 | 0.67544 | -0.12397539 | 0.281 | 0.267 | 1 | 20 |
| nfkbiab | 0.67575 | 0.160269467 | 0.469 | 0.333 | 1 | 20 |
| psma8 | 0.67658 | 0.16538174 | 0.469 | 0.333 | 1 | 20 |
| pak2a | 0.67669 | -0.18261518 | 0.062 | 0.133 | 1 | 20 |
| ctdspl2a | 0.67711 | 0.175568703 | 0.094 | 0.067 | 1 | 20 |
| ufl1 | 0.67711 | -0.15881935 | 0.062 | 0.133 | 1 | 20 |
| tbca | 0.67713 | 0.219388299 | 0.375 | 0.267 | 1 | 20 |
| pi4k2b | 0.6773 | -0.15170222 | 0.125 | 0.133 | 1 | 20 |
| si:dkey-261h17.1 | 0.67735 | 0.234604976 | 0.531 | 0.4 | 1 | 20 |
| decr1 | 0.67801 | 0.143296603 | 0.219 | 0.133 | 1 | 20 |
| atg10 | 0.67803 | 0.148754726 | 0.094 | 0.067 | 1 | 20 |
| ctsba | 0.67829 | -0.10917119 | 0.156 | 0.267 | 1 | 20 |
| igf2bp3 | 0.67894 | 0.175389701 | 0.219 | 0.133 | 1 | 20 |
| smc5 | 0.67929 | 0.149158075 | 0.094 | 0.067 | 1 | 20 |

| | | | | | | |
|---|---|---|---|---|---|---|
| ckbb | 0.67933 | -0.13987089 | 0.188 | 0.2 | 1 | 20 |
| dldh | 0.68041 | -0.10883485 | 0.281 | 0.267 | 1 | 20 |
| dynlrb1 | 0.68072 | -0.13218694 | 0.469 | 0.467 | 1 | 20 |
| si:ch73-238c9.1 | 0.68291 | 0.133101113 | 0.094 | 0.067 | 1 | 20 |
| wdr74 | 0.68402 | -0.16822418 | 0.094 | 0.133 | 1 | 20 |
| tmem126a | 0.68494 | 0.105187301 | 0.125 | 0.133 | 1 | 20 |
| copg2 | 0.68562 | 0.157253155 | 0.125 | 0.067 | 1 | 20 |
| snrpb2 | 0.68564 | 0.117713681 | 0.219 | 0.2 | 1 | 20 |
| nt5dc2 | 0.68587 | -0.13086662 | 0.125 | 0.133 | 1 | 20 |
| wtap | 0.68615 | 0.151558932 | 0.219 | 0.133 | 1 | 20 |
| mcm6 | 0.68655 | 0.157008963 | 0.312 | 0.2 | 1 | 20 |
| mid1ip1l | 0.68655 | -0.19441804 | 0.094 | 0.133 | 1 | 20 |
| p2rx7 | 0.68698 | -0.12079879 | 0.062 | 0.133 | 1 | 20 |
| chmp5b | 0.68845 | 0.215420998 | 0.25 | 0.2 | 1 | 20 |
| si:ch211-210c8.7 | 0.68863 | 0.115813181 | 0.125 | 0.067 | 1 | 20 |
| stat5a | 0.6888 | -0.17576606 | 0.062 | 0.133 | 1 | 20 |
| cyb5r2 | 0.68891 | 0.141448306 | 0.094 | 0.067 | 1 | 20 |
| prpf4bb | 0.6898 | 0.122099088 | 0.094 | 0.067 | 1 | 20 |
| rasal3 | 0.6912 | 0.119426606 | 0.125 | 0.133 | 1 | 20 |
| PYURF | 0.69157 | 0.140663921 | 0.219 | 0.133 | 1 | 20 |
| rsrp1 | 0.69196 | -0.20538836 | 0.094 | 0.133 | 1 | 20 |
| tmem30aa | 0.69353 | 0.154008809 | 0.156 | 0.133 | 1 | 20 |
| ehbp1l1b | 0.69472 | -0.16833623 | 0.062 | 0.133 | 1 | 20 |
| cops5 | 0.69484 | 0.150188284 | 0.125 | 0.067 | 1 | 20 |
| ak1 | 0.69501 | -0.14864602 | 0.188 | 0.2 | 1 | 20 |
| arhgap25 | 0.69635 | -0.14692599 | 0.094 | 0.133 | 1 | 20 |
| XLOC-014336 | 0.69656 | 0.164828325 | 0.312 | 0.2 | 1 | 20 |
| capzb | 0.69686 | -0.14852703 | 0.469 | 0.467 | 1 | 20 |
| kif5ba | 0.69734 | -0.23401717 | 0.125 | 0.2 | 1 | 20 |
| tipin | 0.69744 | -0.13137186 | 0.062 | 0.133 | 1 | 20 |
| ranbp1 | 0.6981 | 0.107690222 | 0.656 | 0.533 | 1 | 20 |
| gins1 | 0.69924 | -0.14611276 | 0.062 | 0.133 | 1 | 20 |
| clint1a | 0.69967 | -0.18005527 | 0.062 | 0.133 | 1 | 20 |
| tcf25 | 0.70035 | -0.16526886 | 0.094 | 0.133 | 1 | 20 |
| sde2 | 0.70065 | 0.180020885 | 0.125 | 0.067 | 1 | 20 |
| nrd1a | 0.7012 | -0.17319224 | 0.094 | 0.133 | 1 | 20 |
| timm13 | 0.7025 | 0.130153217 | 0.188 | 0.133 | 1 | 20 |
| pip4k2ca | 0.70317 | 0.141388724 | 0.156 | 0.133 | 1 | 20 |
| timm10 | 0.70326 | 0.12942438 | 0.25 | 0.2 | 1 | 20 |
| rraga | 0.70534 | 0.145406962 | 0.188 | 0.133 | 1 | 20 |
| ndufc2 | 0.70567 | 0.177407965 | 0.312 | 0.267 | 1 | 20 |
| ppp3r1b | 0.70592 | 0.14251831 | 0.188 | 0.133 | 1 | 20 |
| wdr82 | 0.70598 | -0.1260866 | 0.125 | 0.133 | 1 | 20 |
| slc3a2b | 0.70601 | 0.175078607 | 0.312 | 0.2 | 1 | 20 |
| sptlc3 | 0.70607 | 0.172813192 | 0.125 | 0.067 | 1 | 20 |
| mrpl11 | 0.70754 | 0.122174796 | 0.312 | 0.2 | 1 | 20 |
| arpc3 | 0.7076 | -0.20741281 | 0.531 | 0.6 | 1 | 20 |
| ifrd1 | 0.70765 | -0.10697289 | 0.156 | 0.133 | 1 | 20 |

| | | | | | | |
|---|---|---|---|---|---|---|
| hspa4b | 0.70774 | -0.1664669 | 0.281 | 0.333 | 1 | 20 |
| wipf1b | 0.70775 | 0.129446127 | 0.281 | 0.267 | 1 | 20 |
| hhex | 0.70799 | 0.165765664 | 0.25 | 0.2 | 1 | 20 |
| dnmt3ab | 0.7089 | 0.130914982 | 0.125 | 0.067 | 1 | 20 |
| cox14 | 0.70892 | 0.133728918 | 0.188 | 0.133 | 1 | 20 |
| pim3 | 0.71052 | 0.110067234 | 0.125 | 0.067 | 1 | 20 |
| mov10a | 0.7107 | 0.126322849 | 0.125 | 0.067 | 1 | 20 |
| slc25a25a | 0.71108 | -0.16774155 | 0.062 | 0.133 | 1 | 20 |
| CR293507.1 | 0.71162 | 0.125221426 | 0.094 | 0.067 | 1 | 20 |
| plac8l1 | 0.7117 | -0.25677534 | 0.125 | 0.2 | 1 | 20 |
| selenom | 0.71183 | 0.103647962 | 0.125 | 0.067 | 1 | 20 |
| lsm8 | 0.71246 | 0.166906899 | 0.312 | 0.2 | 1 | 20 |
| cry1aa | 0.71313 | -0.1522855 | 0.062 | 0.133 | 1 | 20 |
| selenow2a | 0.71355 | 0.120123192 | 0.156 | 0.133 | 1 | 20 |
| rnf41 | 0.7136 | -0.17175457 | 0.062 | 0.133 | 1 | 20 |
| si:dkey-286j15.3 | 0.71418 | 0.145142579 | 0.094 | 0.067 | 1 | 20 |
| tspan14 | 0.71423 | -0.17973724 | 0.094 | 0.133 | 1 | 20 |
| mrpl21 | 0.71452 | 0.112823243 | 0.094 | 0.067 | 1 | 20 |
| spry2 | 0.7151 | -0.18168945 | 0.062 | 0.133 | 1 | 20 |
| yeats4 | 0.71535 | 0.120624324 | 0.094 | 0.067 | 1 | 20 |
| emd | 0.71546 | 0.163587889 | 0.094 | 0.067 | 1 | 20 |
| hist2h2l | 0.71643 | 0.145138156 | 0.312 | 0.2 | 1 | 20 |
| tmem141 | 0.71669 | -0.1452165 | 0.062 | 0.133 | 1 | 20 |
| hmgn2 | 0.71672 | 0.150704463 | 0.906 | 0.867 | 1 | 20 |
| ngly1 | 0.71882 | -0.14416987 | 0.062 | 0.133 | 1 | 20 |
| atp6ap2 | 0.72019 | -0.12945137 | 0.188 | 0.2 | 1 | 20 |
| kmt2a | 0.72109 | -0.14136375 | 0.156 | 0.2 | 1 | 20 |
| ifi35 | 0.72164 | 0.145055509 | 0.156 | 0.133 | 1 | 20 |
| cand1 | 0.72165 | 0.12647407 | 0.094 | 0.067 | 1 | 20 |
| h2afx | 0.72165 | -0.16739311 | 0.094 | 0.133 | 1 | 20 |
| si:dkey-88p24.11 | 0.72207 | -0.16424368 | 0.125 | 0.2 | 1 | 20 |
| trappc4 | 0.72239 | -0.10656407 | 0.062 | 0.133 | 1 | 20 |
| derl2 | 0.72284 | -0.23803028 | 0.281 | 0.4 | 1 | 20 |
| FBXO48 | 0.72343 | 0.163951598 | 0.125 | 0.067 | 1 | 20 |
| ino80e | 0.72344 | -0.16676082 | 0.094 | 0.133 | 1 | 20 |
| syap1 | 0.72411 | -0.12173347 | 0.125 | 0.133 | 1 | 20 |
| pdcl | 0.72484 | 0.10769681 | 0.125 | 0.067 | 1 | 20 |
| gzf1 | 0.72508 | -0.20417751 | 0.125 | 0.2 | 1 | 20 |
| gtf2e2 | 0.7253 | 0.117194911 | 0.156 | 0.133 | 1 | 20 |
| trappc5 | 0.72613 | -0.17026931 | 0.094 | 0.133 | 1 | 20 |
| si:ch211-22i13.2 | 0.72713 | 0.177525542 | 0.188 | 0.133 | 1 | 20 |
| bloc1s6 | 0.72746 | -0.17564728 | 0.219 | 0.267 | 1 | 20 |
| rnf7 | 0.72767 | 0.115311453 | 0.125 | 0.067 | 1 | 20 |
| csrp1a | 0.72798 | -0.14713805 | 0.344 | 0.333 | 1 | 20 |
| pik3cd | 0.72868 | 0.1439152 | 0.125 | 0.067 | 1 | 20 |
| npm1b | 0.72871 | -0.18059684 | 0.219 | 0.267 | 1 | 20 |
| plcg2 | 0.72886 | -0.12313281 | 0.062 | 0.133 | 1 | 20 |
| dnajb6a | 0.73022 | 0.110905936 | 0.125 | 0.133 | 1 | 20 |

| | | | | | | |
|---|---|---|---|---|---|---|
| dgkh | 0.73031 | 0.145268948 | 0.125 | 0.067 | 1 | 20 |
| dut | 0.73068 | -0.19377633 | 0.188 | 0.267 | 1 | 20 |
| si:ch211-156b7.4 | 0.73081 | 0.115872505 | 0.094 | 0.067 | 1 | 20 |
| napga | 0.73253 | 0.118517075 | 0.094 | 0.067 | 1 | 20 |
| blf | 0.73293 | 0.228573671 | 0.188 | 0.133 | 1 | 20 |
| ube2g2 | 0.73354 | 0.133827132 | 0.125 | 0.133 | 1 | 20 |
| si:ch211-195h23.3 | 0.73467 | -0.11230655 | 0.062 | 0.133 | 1 | 20 |
| gosr1 | 0.73488 | -0.10314621 | 0.062 | 0.133 | 1 | 20 |
| pop4 | 0.73494 | -0.10489435 | 0.062 | 0.133 | 1 | 20 |
| hspa8 | 0.73557 | 0.128673872 | 0.875 | 0.8 | 1 | 20 |
| ap2b1 | 0.73562 | 0.131414117 | 0.094 | 0.067 | 1 | 20 |
| ap1g1 | 0.7357 | 0.115888703 | 0.094 | 0.067 | 1 | 20 |
| mrps18a | 0.73625 | -0.17281274 | 0.125 | 0.2 | 1 | 20 |
| thtpa | 0.73645 | 0.169903235 | 0.094 | 0.067 | 1 | 20 |
| gpx4b | 0.73648 | 0.200168475 | 0.344 | 0.267 | 1 | 20 |
| rbpms | 0.73667 | -0.20062208 | 0.062 | 0.133 | 1 | 20 |
| lrrc8da | 0.73696 | -0.21056226 | 0.156 | 0.2 | 1 | 20 |
| rbm7 | 0.73758 | 0.118314887 | 0.094 | 0.067 | 1 | 20 |
| asap1b | 0.73772 | -0.18508956 | 0.094 | 0.133 | 1 | 20 |
| ppm1aa | 0.73803 | 0.153765013 | 0.125 | 0.067 | 1 | 20 |
| nr4a1 | 0.73812 | -0.18764979 | 0.375 | 0.4 | 1 | 20 |
| lpxn | 0.73842 | 0.145956294 | 0.094 | 0.067 | 1 | 20 |
| krt8 | 0.73877 | -0.1995344 | 0.062 | 0.133 | 1 | 20 |
| mat2aa | 0.73894 | -0.16752943 | 0.156 | 0.2 | 1 | 20 |
| fhl1a | 0.73927 | -0.18699435 | 0.062 | 0.133 | 1 | 20 |
| cdk11b | 0.7393 | 0.12625536 | 0.125 | 0.067 | 1 | 20 |
| ralbp1 | 0.73944 | 0.107348348 | 0.125 | 0.133 | 1 | 20 |
| gmppaa | 0.73968 | -0.14010102 | 0.062 | 0.133 | 1 | 20 |
| klf2a | 0.73975 | -0.20160188 | 0.062 | 0.133 | 1 | 20 |
| ppp4ca | 0.74032 | 0.111988436 | 0.094 | 0.067 | 1 | 20 |
| kdm1a | 0.74047 | -0.12585877 | 0.188 | 0.2 | 1 | 20 |
| GNAZ | 0.74111 | 0.131351573 | 0.094 | 0.067 | 1 | 20 |
| tpd52l2a | 0.74233 | -0.16494155 | 0.094 | 0.133 | 1 | 20 |
| csnk1a1 | 0.74261 | -0.11014224 | 0.406 | 0.4 | 1 | 20 |
| rnpc3 | 0.74331 | 0.124906128 | 0.094 | 0.067 | 1 | 20 |
| bcl2l13 | 0.74347 | 0.130252583 | 0.125 | 0.067 | 1 | 20 |
| poldip2 | 0.7461 | -0.160569 | 0.094 | 0.133 | 1 | 20 |
| hexb | 0.74668 | -0.12767747 | 0.094 | 0.133 | 1 | 20 |
| ccdc124 | 0.74704 | 0.119329623 | 0.094 | 0.067 | 1 | 20 |
| rbm14b | 0.74798 | 0.125289817 | 0.125 | 0.067 | 1 | 20 |
| ddx42 | 0.74803 | -0.15180985 | 0.156 | 0.2 | 1 | 20 |
| ppp1caa | 0.75085 | -0.10679886 | 0.344 | 0.333 | 1 | 20 |
| spcs3 | 0.75147 | 0.136379165 | 0.375 | 0.267 | 1 | 20 |
| ten1 | 0.75155 | 0.10582061 | 0.094 | 0.067 | 1 | 20 |
| denr | 0.75163 | 0.11173216 | 0.375 | 0.267 | 1 | 20 |
| zc3h13 | 0.75167 | 0.124573766 | 0.219 | 0.133 | 1 | 20 |
| zgc:162698 | 0.75177 | 0.115407162 | 0.125 | 0.067 | 1 | 20 |
| ube2f | 0.7528 | 0.137412494 | 0.125 | 0.067 | 1 | 20 |

| | | | | | | |
|---|---|---|---|---|---|---|
| max | 0.75287 | 0.181611202 | 0.438 | 0.333 | 1 | 20 |
| ccnl1a | 0.75302 | -0.1540079 | 0.094 | 0.133 | 1 | 20 |
| ccdc174 | 0.75407 | 0.131694791 | 0.125 | 0.067 | 1 | 20 |
| kri1 | 0.75644 | 0.121913102 | 0.125 | 0.067 | 1 | 20 |
| zgc:194224 | 0.75647 | 0.116681881 | 0.125 | 0.067 | 1 | 20 |
| sept7a | 0.75664 | 0.121458831 | 0.219 | 0.133 | 1 | 20 |
| mrps24 | 0.75738 | 0.133262413 | 0.125 | 0.067 | 1 | 20 |
| coil | 0.75752 | 0.108791884 | 0.125 | 0.067 | 1 | 20 |
| si:ch211-212k18.7 | 0.7576 | -0.11005131 | 0.125 | 0.133 | 1 | 20 |
| serinc5 | 0.7578 | -0.14713361 | 0.094 | 0.133 | 1 | 20 |
| cnp | 0.75781 | -0.13678022 | 0.219 | 0.267 | 1 | 20 |
| serpinb1l3 | 0.75815 | -0.17591647 | 0.094 | 0.133 | 1 | 20 |
| ppm1da | 0.75827 | 0.120567969 | 0.125 | 0.067 | 1 | 20 |
| ltv1 | 0.75841 | 0.114778434 | 0.156 | 0.133 | 1 | 20 |
| rab35b | 0.75858 | 0.142384671 | 0.094 | 0.067 | 1 | 20 |
| matr3l1.1 | 0.75921 | -0.11950143 | 0.188 | 0.2 | 1 | 20 |
| sgta | 0.75959 | 0.115675421 | 0.219 | 0.133 | 1 | 20 |
| eif6 | 0.75966 | 0.129474673 | 0.375 | 0.267 | 1 | 20 |
| dpm1 | 0.75974 | -0.14346872 | 0.125 | 0.2 | 1 | 20 |
| usp9 | 0.76083 | 0.1004429 | 0.125 | 0.067 | 1 | 20 |
| utp18 | 0.76099 | 0.107970775 | 0.125 | 0.067 | 1 | 20 |
| znf207a | 0.76104 | 0.148706259 | 0.094 | 0.067 | 1 | 20 |
| rnf213a | 0.76187 | 0.107075003 | 0.219 | 0.2 | 1 | 20 |
| uqcc3 | 0.76221 | 0.119850427 | 0.094 | 0.067 | 1 | 20 |
| cmpk | 0.76271 | 0.128647513 | 0.188 | 0.133 | 1 | 20 |
| tmc6b | 0.76276 | 0.112401427 | 0.125 | 0.067 | 1 | 20 |
| commd3 | 0.76289 | 0.134824256 | 0.188 | 0.133 | 1 | 20 |
| rnf130 | 0.76378 | 0.152424863 | 0.094 | 0.067 | 1 | 20 |
| peak3 | 0.7641 | -0.13084723 | 0.125 | 0.2 | 1 | 20 |
| cox20 | 0.76418 | 0.116433797 | 0.125 | 0.067 | 1 | 20 |
| si:busm1-266f07.2 | 0.76461 | 0.196321826 | 0.281 | 0.2 | 1 | 20 |
| sart1 | 0.76497 | 0.110615719 | 0.219 | 0.2 | 1 | 20 |
| alg5 | 0.76499 | 0.114228983 | 0.156 | 0.133 | 1 | 20 |
| lsm4 | 0.76512 | -0.14120769 | 0.25 | 0.267 | 1 | 20 |
| thumpd1 | 0.76881 | 0.126045545 | 0.188 | 0.133 | 1 | 20 |
| raf1b | 0.76912 | -0.15491013 | 0.094 | 0.133 | 1 | 20 |
| sccpdhb | 0.76974 | -0.14632874 | 0.156 | 0.2 | 1 | 20 |
| rmnd5b | 0.77156 | -0.10623954 | 0.125 | 0.133 | 1 | 20 |
| iqgap1 | 0.77159 | -0.11589887 | 0.125 | 0.133 | 1 | 20 |
| cox10 | 0.77181 | 0.119723466 | 0.125 | 0.067 | 1 | 20 |
| bri3 | 0.77269 | 0.14671514 | 0.094 | 0.067 | 1 | 20 |
| chchd3a | 0.77287 | 0.137658577 | 0.094 | 0.067 | 1 | 20 |
| nccrp1 | 0.77318 | -0.18003571 | 0.125 | 0.2 | 1 | 20 |
| pdrg1 | 0.77352 | 0.110280146 | 0.094 | 0.067 | 1 | 20 |
| rab11ba | 0.77424 | 0.119240488 | 0.156 | 0.133 | 1 | 20 |
| dars | 0.77437 | 0.145732471 | 0.281 | 0.2 | 1 | 20 |
| bcl11aa | 0.77461 | 0.126337617 | 0.125 | 0.067 | 1 | 20 |
| zgc:92664 | 0.7749 | 0.108293282 | 0.219 | 0.133 | 1 | 20 |

| | | | | | | |
|---|---|---|---|---|---|---|
| chmp4bb | 0.77544 | 0.107666686 | 0.094 | 0.067 | 1 | 20 |
| surf2 | 0.77722 | -0.11584351 | 0.25 | 0.267 | 1 | 20 |
| msi2a | 0.77727 | 0.169498103 | 0.219 | 0.133 | 1 | 20 |
| sirt1 | 0.77738 | 0.109389238 | 0.125 | 0.067 | 1 | 20 |
| bud23 | 0.77762 | 0.114929493 | 0.094 | 0.067 | 1 | 20 |
| zgc:152774 | 0.77777 | 0.102462988 | 0.219 | 0.133 | 1 | 20 |
| naa15a | 0.77799 | 0.120339135 | 0.188 | 0.133 | 1 | 20 |
| kptn | 0.77803 | 0.112312789 | 0.125 | 0.067 | 1 | 20 |
| hspd1 | 0.77808 | 0.152855688 | 0.281 | 0.2 | 1 | 20 |
| metap1 | 0.7786 | 0.129380113 | 0.25 | 0.2 | 1 | 20 |
| krit1 | 0.77897 | 0.126974914 | 0.094 | 0.067 | 1 | 20 |
| mrps31 | 0.77927 | -0.1345818 | 0.094 | 0.133 | 1 | 20 |
| sec22a | 0.7803 | -0.13590132 | 0.094 | 0.133 | 1 | 20 |
| MRPS12 | 0.78034 | 0.144190651 | 0.344 | 0.267 | 1 | 20 |
| rhogd | 0.78089 | 0.118338869 | 0.125 | 0.067 | 1 | 20 |
| itga2b | 0.7812 | -0.23403029 | 0.188 | 0.267 | 1 | 20 |
| mrps25 | 0.78171 | 0.110140619 | 0.125 | 0.067 | 1 | 20 |
| tarbp2 | 0.78224 | 0.117174723 | 0.125 | 0.067 | 1 | 20 |
| ptp4a2b | 0.78257 | 0.107093453 | 0.125 | 0.067 | 1 | 20 |
| papola | 0.78326 | 0.114668435 | 0.188 | 0.133 | 1 | 20 |
| atp2a3 | 0.78348 | -0.13143166 | 0.125 | 0.2 | 1 | 20 |
| glud1b | 0.78461 | -0.16403611 | 0.188 | 0.267 | 1 | 20 |
| sar1b | 0.78559 | -0.13275333 | 0.156 | 0.2 | 1 | 20 |
| gatd3a | 0.78563 | 0.106179143 | 0.125 | 0.067 | 1 | 20 |
| rprd1b | 0.78585 | -0.11772604 | 0.094 | 0.133 | 1 | 20 |
| rrp8 | 0.78741 | 0.139247552 | 0.25 | 0.2 | 1 | 20 |
| wnt2 | 0.78752 | -0.17234365 | 0.094 | 0.133 | 1 | 20 |
| rab5aa | 0.78753 | 0.109845955 | 0.125 | 0.067 | 1 | 20 |
| prdx3 | 0.78754 | 0.129390404 | 0.219 | 0.2 | 1 | 20 |
| aldh7a1 | 0.78797 | -0.10769632 | 0.188 | 0.2 | 1 | 20 |
| NDUFB1 | 0.78817 | 0.144992513 | 0.438 | 0.333 | 1 | 20 |
| nenf | 0.78823 | 0.114232889 | 0.156 | 0.133 | 1 | 20 |
| herpud1 | 0.78929 | -0.10832121 | 0.125 | 0.133 | 1 | 20 |
| slc35a1 | 0.78992 | -0.15131464 | 0.094 | 0.133 | 1 | 20 |
| tbl3 | 0.78996 | 0.101985154 | 0.094 | 0.067 | 1 | 20 |
| kdm6bb | 0.79043 | 0.166227163 | 0.438 | 0.333 | 1 | 20 |
| dedd1 | 0.79267 | -0.13820125 | 0.188 | 0.267 | 1 | 20 |
| tmem18 | 0.79389 | -0.13279551 | 0.156 | 0.2 | 1 | 20 |
| cacybp | 0.79396 | -0.14534614 | 0.094 | 0.133 | 1 | 20 |
| hexim1 | 0.794 | -0.12995398 | 0.094 | 0.133 | 1 | 20 |
| immt | 0.795 | 0.111279383 | 0.125 | 0.067 | 1 | 20 |
| si:ch211-197h24.8 | 0.79537 | 0.115301518 | 0.125 | 0.067 | 1 | 20 |
| cyth4b | 0.79609 | -0.14329098 | 0.094 | 0.133 | 1 | 20 |
| znf576.2 | 0.79665 | -0.11071742 | 0.125 | 0.133 | 1 | 20 |
| rnf10 | 0.79684 | -0.14995561 | 0.156 | 0.2 | 1 | 20 |
| cript | 0.79836 | 0.102767644 | 0.094 | 0.067 | 1 | 20 |
| lyar | 0.79854 | 0.101059553 | 0.281 | 0.267 | 1 | 20 |
| xkrx | 0.8001 | 0.131492461 | 0.094 | 0.067 | 1 | 20 |

| | | | | | | |
|---|---|---|---|---|---|---|
| ogg1 | 0.8002 | 0.1001088 | 0.125 | 0.067 | 1 | 20 |
| mtmr8 | 0.80065 | -0.12136503 | 0.094 | 0.133 | 1 | 20 |
| brms1la | 0.80084 | -0.10732194 | 0.125 | 0.133 | 1 | 20 |
| rbbp4 | 0.80093 | 0.125439739 | 0.156 | 0.133 | 1 | 20 |
| zgc:153867 | 0.80116 | -0.11676874 | 0.781 | 0.8 | 1 | 20 |
| cmc2 | 0.80125 | -0.1362485 | 0.094 | 0.133 | 1 | 20 |
| svila | 0.80291 | -0.10046413 | 0.125 | 0.133 | 1 | 20 |
| MCMDC2 | 0.80355 | 0.136801163 | 0.375 | 0.333 | 1 | 20 |
| naga | 0.80422 | 0.105204909 | 0.094 | 0.067 | 1 | 20 |
| agpat5 | 0.80545 | -0.15984209 | 0.094 | 0.133 | 1 | 20 |
| huwe1 | 0.80556 | 0.100479792 | 0.125 | 0.067 | 1 | 20 |
| stxbp2 | 0.80558 | 0.118683798 | 0.188 | 0.133 | 1 | 20 |
| gfi1ab | 0.80564 | -0.15986544 | 0.125 | 0.2 | 1 | 20 |
| bbip1 | 0.80574 | -0.1142665 | 0.094 | 0.133 | 1 | 20 |
| syncrip | 0.8099 | 0.123270497 | 0.5 | 0.4 | 1 | 20 |
| zbtb46 | 0.81073 | 0.113593193 | 0.094 | 0.067 | 1 | 20 |
| rfc4 | 0.81076 | 0.111609737 | 0.094 | 0.067 | 1 | 20 |
| zgc:77086 | 0.81243 | 0.131295122 | 0.125 | 0.067 | 1 | 20 |
| rnf2 | 0.81291 | 0.10355622 | 0.094 | 0.067 | 1 | 20 |
| puf60a | 0.81534 | -0.16359415 | 0.188 | 0.267 | 1 | 20 |
| ufd1l | 0.81567 | 0.108331942 | 0.094 | 0.067 | 1 | 20 |
| zgc:110540 | 0.8162 | 0.157648416 | 0.125 | 0.067 | 1 | 20 |
| il6st | 0.81647 | -0.10352694 | 0.125 | 0.133 | 1 | 20 |
| setdb2 | 0.8173 | 0.123909432 | 0.188 | 0.133 | 1 | 20 |
| fam49a | 0.81738 | 0.113197562 | 0.281 | 0.2 | 1 | 20 |
| tradd | 0.8178 | 0.104806129 | 0.156 | 0.133 | 1 | 20 |
| si:dkeyp-97b10.3 | 0.81832 | -0.11999189 | 0.094 | 0.133 | 1 | 20 |
| prkacba | 0.81835 | -0.11223588 | 0.188 | 0.267 | 1 | 20 |
| si:ch211-203d1.3 | 0.81869 | 0.138970219 | 0.125 | 0.067 | 1 | 20 |
| nuggc.3 | 0.81965 | 0.104756128 | 0.094 | 0.067 | 1 | 20 |
| tubgcp3 | 0.81999 | 0.122789016 | 0.094 | 0.067 | 1 | 20 |
| znhit1 | 0.82028 | 0.108240924 | 0.125 | 0.067 | 1 | 20 |
| ube4b | 0.82154 | 0.109129941 | 0.094 | 0.067 | 1 | 20 |
| ube2ia | 0.82196 | -0.12578046 | 0.25 | 0.333 | 1 | 20 |
| dynll2b | 0.82253 | 0.103365736 | 0.125 | 0.067 | 1 | 20 |
| si:dkeyp-113d7.10 | 0.82277 | 0.10532159 | 0.094 | 0.067 | 1 | 20 |
| zbtb20 | 0.82297 | -0.10677958 | 0.188 | 0.267 | 1 | 20 |
| pin4 | 0.82494 | 0.134275682 | 0.188 | 0.133 | 1 | 20 |
| rbm39a | 0.8263 | -0.12426342 | 0.375 | 0.4 | 1 | 20 |
| mgaa | 0.82634 | -0.12562961 | 0.094 | 0.133 | 1 | 20 |
| rab5if | 0.827 | 0.101664802 | 0.281 | 0.2 | 1 | 20 |
| mtch2 | 0.82722 | 0.100065429 | 0.188 | 0.133 | 1 | 20 |
| morf4l1 | 0.82777 | 0.139503644 | 0.344 | 0.267 | 1 | 20 |
| taf12 | 0.82905 | 0.144621585 | 0.312 | 0.267 | 1 | 20 |
| si:ch211-286b5.5 | 0.83099 | -0.11378596 | 0.094 | 0.133 | 1 | 20 |
| rab8b | 0.83121 | -0.1554705 | 0.156 | 0.2 | 1 | 20 |
| LOC103910438 | 0.83317 | -0.15428215 | 0.219 | 0.267 | 1 | 20 |
| g3bp1 | 0.83388 | 0.115757536 | 0.281 | 0.2 | 1 | 20 |

| | | | | | | |
|---|---|---|---|---|---|---|
| rab5c | 0.83563 | -0.1331129 | 0.25 | 0.267 | 1 | 20 |
| chd6 | 0.83791 | 0.102402579 | 0.094 | 0.067 | 1 | 20 |
| snrpb | 0.83927 | 0.10076067 | 0.688 | 0.6 | 1 | 20 |
| strap | 0.84105 | 0.103141038 | 0.188 | 0.133 | 1 | 20 |
| tia1l | 0.84115 | -0.11675297 | 0.094 | 0.133 | 1 | 20 |
| vaspb | 0.84226 | -0.15988048 | 0.156 | 0.2 | 1 | 20 |
| med28 | 0.843 | 0.101564675 | 0.188 | 0.133 | 1 | 20 |
| bahcc1b | 0.84329 | 0.111425083 | 0.188 | 0.133 | 1 | 20 |
| mthfs | 0.84603 | -0.12900495 | 0.094 | 0.133 | 1 | 20 |
| mrpl20 | 0.8469 | 0.126082587 | 0.312 | 0.267 | 1 | 20 |
| zc3h11a | 0.84803 | 0.122870678 | 0.094 | 0.067 | 1 | 20 |
| pde4ba | 0.84957 | -0.13029538 | 0.094 | 0.133 | 1 | 20 |
| prps1a | 0.85053 | 0.120263923 | 0.375 | 0.333 | 1 | 20 |
| lzts2a | 0.85119 | -0.12896384 | 0.094 | 0.133 | 1 | 20 |
| ctsd | 0.85188 | -0.11974485 | 0.094 | 0.133 | 1 | 20 |
| desi2 | 0.85222 | 0.106046861 | 0.094 | 0.067 | 1 | 20 |
| psip1a | 0.85241 | -0.13044457 | 0.094 | 0.133 | 1 | 20 |
| fgfr1op | 0.85292 | 0.112563468 | 0.094 | 0.067 | 1 | 20 |
| psmd13 | 0.85548 | -0.12159681 | 0.219 | 0.267 | 1 | 20 |
| tp53bp1 | 0.85571 | 0.10721528 | 0.094 | 0.067 | 1 | 20 |
| fbp1a | 0.85729 | -0.1036946 | 0.25 | 0.267 | 1 | 20 |
| arid5b | 0.85777 | -0.11592908 | 0.094 | 0.133 | 1 | 20 |
| spns2 | 0.85821 | -0.15581268 | 0.094 | 0.133 | 1 | 20 |
| cbfa2t3 | 0.85886 | 0.122766283 | 0.312 | 0.267 | 1 | 20 |
| kmt2bb | 0.85912 | -0.13721855 | 0.219 | 0.267 | 1 | 20 |
| pbxip1b | 0.86063 | -0.12907795 | 0.094 | 0.133 | 1 | 20 |
| psma6b | 0.86136 | -0.11535985 | 0.219 | 0.267 | 1 | 20 |
| si:ch1073-157b13.1 | 0.86173 | 0.106582483 | 0.094 | 0.067 | 1 | 20 |
| pcbp2 | 0.86332 | -0.15140254 | 0.219 | 0.267 | 1 | 20 |
| baxb | 0.86432 | 0.133872911 | 0.094 | 0.067 | 1 | 20 |
| npc2 | 0.86439 | -0.10522526 | 0.188 | 0.2 | 1 | 20 |
| srsf6b | 0.86507 | 0.110492481 | 0.344 | 0.267 | 1 | 20 |
| brd3b | 0.86624 | -0.12061097 | 0.156 | 0.2 | 1 | 20 |
| zgc:162939 | 0.87393 | -0.12434918 | 0.094 | 0.133 | 1 | 20 |
| pak2b | 0.87494 | 0.101036623 | 0.156 | 0.133 | 1 | 20 |
| ipmkb | 0.87621 | -0.12531889 | 0.094 | 0.133 | 1 | 20 |
| rbx1 | 0.88069 | 0.116005625 | 0.469 | 0.4 | 1 | 20 |
| surf1 | 0.88173 | -0.1089994 | 0.094 | 0.133 | 1 | 20 |
| mphosph8 | 0.89508 | 0.118169059 | 0.312 | 0.267 | 1 | 20 |
| rab11a | 0.89636 | -0.13098671 | 0.219 | 0.267 | 1 | 20 |
| lrrfip2 | 0.90026 | -0.11690277 | 0.094 | 0.133 | 1 | 20 |
| fam120c | 0.9007 | -0.11675717 | 0.156 | 0.2 | 1 | 20 |
| cdkn2c | 0.90154 | -0.12961054 | 0.094 | 0.133 | 1 | 20 |
| BX284696.1 | 0.90656 | -0.10777215 | 0.281 | 0.333 | 1 | 20 |
| zgc:92606 | 0.91288 | -0.10400514 | 0.156 | 0.2 | 1 | 20 |
| bcl2l10 | 0.91606 | -0.13221777 | 0.344 | 0.4 | 1 | 20 |
| p2ry13 | 0.92321 | -0.1208458 | 0.094 | 0.133 | 1 | 20 |
| fli1a | 0.93184 | 0.106800845 | 0.312 | 0.267 | 1 | 20 |

| | | | | | | |
|---|---|---|---|---|---|---|
| cnn2 | 0.93712 | -0.10523786 | 0.219 | 0.267 | 1 | 20 |
| mras | 0.95768 | -0.11808537 | 0.062 | 0.067 | 1 | 20 |
| nub1 | 0.95768 | 0.118760962 | 0.062 | 0.067 | 1 | 20 |
| ston2 | 0.95768 | 0.12306333 | 0.062 | 0.067 | 1 | 20 |
| cog6 | 0.95768 | 0.119541609 | 0.062 | 0.067 | 1 | 20 |
| il11b | 0.95768 | -0.1539947 | 0.062 | 0.067 | 1 | 20 |
| kansl3 | 0.95768 | -0.19237784 | 0.062 | 0.067 | 1 | 20 |
| nup214 | 0.95768 | -0.18113878 | 0.062 | 0.067 | 1 | 20 |
| plekha3 | 0.95768 | -0.17257952 | 0.062 | 0.067 | 1 | 20 |
| sc:d217 | 0.95768 | -0.11035886 | 0.062 | 0.067 | 1 | 20 |
| si:dkeyp-110e4.6 | 0.95768 | -0.10850864 | 0.062 | 0.067 | 1 | 20 |
| tm2d1 | 0.95768 | -0.15251709 | 0.062 | 0.067 | 1 | 20 |
| tmem150ab | 0.95768 | -0.12323855 | 0.062 | 0.067 | 1 | 20 |
| LO018102.1 | 0.95768 | 0.108262562 | 0.062 | 0.067 | 1 | 20 |
| LOC103908877 | 0.95768 | -0.1170179 | 0.062 | 0.067 | 1 | 20 |
| arhgef1a | 0.95768 | -0.14564736 | 0.062 | 0.067 | 1 | 20 |
| arrdc3a | 0.95768 | -0.12844462 | 0.062 | 0.067 | 1 | 20 |
| bcap29 | 0.95768 | -0.16023362 | 0.062 | 0.067 | 1 | 20 |
| ccnt1 | 0.95768 | 0.103004279 | 0.062 | 0.067 | 1 | 20 |
| cdk2 | 0.95768 | -0.15790859 | 0.062 | 0.067 | 1 | 20 |
| cdkn1d | 0.95768 | -0.1851128 | 0.062 | 0.067 | 1 | 20 |
| fbrsl1 | 0.95768 | -0.12408648 | 0.062 | 0.067 | 1 | 20 |
| fxr2 | 0.95768 | -0.14861264 | 0.062 | 0.067 | 1 | 20 |
| gig2i | 0.95768 | -0.15037262 | 0.062 | 0.067 | 1 | 20 |
| ikzf5 | 0.95768 | -0.12731091 | 0.062 | 0.067 | 1 | 20 |
| ldb1b | 0.95768 | -0.11419029 | 0.062 | 0.067 | 1 | 20 |
| pitpnc1b | 0.95768 | -0.16231569 | 0.062 | 0.067 | 1 | 20 |
| pptc7b | 0.95768 | -0.16136707 | 0.062 | 0.067 | 1 | 20 |
| rarga | 0.95768 | -0.16239028 | 0.062 | 0.067 | 1 | 20 |
| si:zfos-741a10.3 | 0.95768 | -0.14413599 | 0.062 | 0.067 | 1 | 20 |
| sla1 | 0.95768 | -0.10585503 | 0.062 | 0.067 | 1 | 20 |
| spink2.1 | 0.95768 | 0.191810742 | 0.062 | 0.067 | 1 | 20 |
| stk24b | 0.95768 | -0.16391308 | 0.062 | 0.067 | 1 | 20 |
| tmem131 | 0.95768 | -0.19335328 | 0.062 | 0.067 | 1 | 20 |
| zgc:123105 | 0.95768 | 0.129870286 | 0.062 | 0.067 | 1 | 20 |
| zgc:63587 | 0.95768 | -0.11556093 | 0.062 | 0.067 | 1 | 20 |
| znf76 | 0.95768 | -0.16274337 | 0.062 | 0.067 | 1 | 20 |
| CABZ01110881.1 | 0.95768 | -0.13008822 | 0.062 | 0.067 | 1 | 20 |
| CR383662.8 | 0.95768 | -0.13032784 | 0.062 | 0.067 | 1 | 20 |
| XLOC-022621 | 0.95768 | -0.14406635 | 0.062 | 0.067 | 1 | 20 |
| abhd12 | 0.95768 | -0.1519491 | 0.062 | 0.067 | 1 | 20 |
| abhd16a | 0.95768 | -0.1311007 | 0.062 | 0.067 | 1 | 20 |
| ccdc15 | 0.95768 | -0.14297988 | 0.062 | 0.067 | 1 | 20 |
| chid1 | 0.95768 | 0.120654042 | 0.062 | 0.067 | 1 | 20 |
| cyfip1 | 0.95768 | -0.11835883 | 0.062 | 0.067 | 1 | 20 |
| elmo3 | 0.95768 | -0.15110122 | 0.062 | 0.067 | 1 | 20 |
| epc1b | 0.95768 | 0.105587971 | 0.062 | 0.067 | 1 | 20 |
| furina | 0.95768 | -0.21570235 | 0.062 | 0.067 | 1 | 20 |

| | | | | | | |
|---|---|---|---|---|---|---|
| gsna | 0.95768 | -0.12194926 | 0.062 | 0.067 | 1 | 20 |
| hdac8 | 0.95768 | -0.10275325 | 0.062 | 0.067 | 1 | 20 |
| jak1 | 0.95768 | -0.18547079 | 0.062 | 0.067 | 1 | 20 |
| kat5b | 0.95768 | 0.142433696 | 0.062 | 0.067 | 1 | 20 |
| lpcat1 | 0.95768 | -0.1172395 | 0.062 | 0.067 | 1 | 20 |
| march5 | 0.95768 | 0.109990171 | 0.062 | 0.067 | 1 | 20 |
| mysm1 | 0.95768 | -0.18567611 | 0.062 | 0.067 | 1 | 20 |
| pag1 | 0.95768 | -0.15652313 | 0.062 | 0.067 | 1 | 20 |
| pdcd4b | 0.95768 | 0.147844554 | 0.062 | 0.067 | 1 | 20 |
| ppm1k | 0.95768 | -0.16740753 | 0.062 | 0.067 | 1 | 20 |
| rab41 | 0.95768 | -0.1005858 | 0.062 | 0.067 | 1 | 20 |
| rell1 | 0.95768 | -0.15161998 | 0.062 | 0.067 | 1 | 20 |
| sh3glb1a | 0.95768 | -0.14409776 | 0.062 | 0.067 | 1 | 20 |
| slain2 | 0.95768 | -0.14800899 | 0.062 | 0.067 | 1 | 20 |
| slc7a3b | 0.95768 | -0.15743481 | 0.062 | 0.067 | 1 | 20 |
| snx17 | 0.95768 | -0.10826822 | 0.062 | 0.067 | 1 | 20 |
| tdrd3 | 0.95768 | -0.17554648 | 0.062 | 0.067 | 1 | 20 |
| tnpo3 | 0.95768 | -0.14446888 | 0.062 | 0.067 | 1 | 20 |
| vhl | 0.95768 | -0.12714107 | 0.062 | 0.067 | 1 | 20 |
| zc3h18 | 0.95768 | -0.1045737 | 0.062 | 0.067 | 1 | 20 |
| zgc:162339 | 0.95768 | -0.14335191 | 0.062 | 0.067 | 1 | 20 |
| znf296 | 0.95768 | -0.17397292 | 0.062 | 0.067 | 1 | 20 |
| bloc1s2 | 0.95768 | 0.114932216 | 0.062 | 0.067 | 1 | 20 |
| calcoco2 | 0.95768 | -0.1003994 | 0.062 | 0.067 | 1 | 20 |
| si:dkey-9i23.15 | 0.95768 | 0.178332268 | 0.062 | 0.067 | 1 | 20 |
| ubald2 | 0.95768 | 0.115186589 | 0.062 | 0.067 | 1 | 20 |
| anapc13 | 0.95768 | -0.11449753 | 0.062 | 0.067 | 1 | 20 |
| bhlhe40 | 0.95768 | -0.14816491 | 0.062 | 0.067 | 1 | 20 |
| csnk1g1 | 0.95768 | -0.12618817 | 0.062 | 0.067 | 1 | 20 |
| enoph1 | 0.95768 | -0.13836285 | 0.062 | 0.067 | 1 | 20 |
| fam173a | 0.95768 | -0.15655844 | 0.062 | 0.067 | 1 | 20 |
| itsn2b | 0.95768 | -0.17123477 | 0.062 | 0.067 | 1 | 20 |
| klf3 | 0.95768 | -0.13524424 | 0.062 | 0.067 | 1 | 20 |
| mrps7 | 0.95768 | -0.18305172 | 0.062 | 0.067 | 1 | 20 |
| mzt2b | 0.95768 | -0.16313972 | 0.062 | 0.067 | 1 | 20 |
| patj | 0.95768 | -0.18478087 | 0.062 | 0.067 | 1 | 20 |
| seh1l | 0.95768 | -0.14662807 | 0.062 | 0.067 | 1 | 20 |
| tbc1d10aa | 0.95768 | -0.13765427 | 0.062 | 0.067 | 1 | 20 |
| tmem263 | 0.95768 | -0.13320184 | 0.062 | 0.067 | 1 | 20 |
| vegfc | 0.95768 | -0.101939 | 0.062 | 0.067 | 1 | 20 |
| anapc1 | 0.95768 | -0.16932122 | 0.062 | 0.067 | 1 | 20 |
| asap3 | 0.95768 | -0.14248332 | 0.062 | 0.067 | 1 | 20 |
| bada | 0.95768 | -0.13033473 | 0.062 | 0.067 | 1 | 20 |
| chic2 | 0.95768 | -0.15613115 | 0.062 | 0.067 | 1 | 20 |
| chpt1 | 0.95768 | -0.2125625 | 0.062 | 0.067 | 1 | 20 |
| cwc22 | 0.95768 | -0.13524568 | 0.062 | 0.067 | 1 | 20 |
| dscc1 | 0.95768 | -0.10501902 | 0.062 | 0.067 | 1 | 20 |
| eef1akmt1 | 0.95768 | -0.13289695 | 0.062 | 0.067 | 1 | 20 |

| | | | | | | |
|---|---|---|---|---|---|---|
| emc1 | 0.95768 | -0.11999209 | 0.062 | 0.067 | 1 | 20 |
| htatip2 | 0.95768 | 0.128364377 | 0.062 | 0.067 | 1 | 20 |
| ilf3a | 0.95768 | -0.12994209 | 0.062 | 0.067 | 1 | 20 |
| ncaph2 | 0.95768 | -0.10547917 | 0.062 | 0.067 | 1 | 20 |
| necap2 | 0.95768 | -0.13683867 | 0.062 | 0.067 | 1 | 20 |
| pip5k1aa | 0.95768 | -0.15009794 | 0.062 | 0.067 | 1 | 20 |
| ppp4r2b | 0.95768 | -0.1730669 | 0.062 | 0.067 | 1 | 20 |
| pqbp1 | 0.95768 | -0.16335508 | 0.062 | 0.067 | 1 | 20 |
| rgs14a | 0.95768 | -0.17050453 | 0.062 | 0.067 | 1 | 20 |
| snap23.1 | 0.95768 | -0.16128718 | 0.062 | 0.067 | 1 | 20 |
| snx30 | 0.95768 | 0.11711738 | 0.062 | 0.067 | 1 | 20 |
| stx18 | 0.95768 | -0.15618366 | 0.062 | 0.067 | 1 | 20 |
| cmasa | 0.95768 | -0.15741509 | 0.062 | 0.067 | 1 | 20 |
| ctsz | 0.95768 | -0.14582866 | 0.062 | 0.067 | 1 | 20 |
| hgsnat | 0.95768 | 0.112783553 | 0.062 | 0.067 | 1 | 20 |
| mntb | 0.95768 | -0.16266049 | 0.062 | 0.067 | 1 | 20 |
| si:dkeyp-115e12.6 | 0.95768 | 0.1104764 | 0.062 | 0.067 | 1 | 20 |
| si:dkeyp-69c1.6 | 0.95768 | -0.12688893 | 0.062 | 0.067 | 1 | 20 |
| si:rp71-36a1.3 | 0.95768 | -0.11382525 | 0.062 | 0.067 | 1 | 20 |
| tmem30ab | 0.95768 | -0.15808076 | 0.062 | 0.067 | 1 | 20 |
| zgc:92744 | 9.101670 | 0.92908088 | 0.471 | 0 | 1 | 21 |
| COX3 | 0.00248 | -0.66158148 | 0.176 | 0.652 | 1 | 21 |
| limd2 | 0.00495 | -0.70692728 | 0 | 0.304 | 1 | 21 |
| zgc:152830 | 0.00495 | -0.64945069 | 0 | 0.304 | 1 | 21 |
| cnn2 | 0.00639 | -0.78893749 | 0.353 | 0.783 | 1 | 21 |
| u2af2b | 0.01059 | -0.58741406 | 0 | 0.261 | 1 | 21 |
| pfdn2 | 0.01068 | 0.511721766 | 0.235 | 0 | 1 | 21 |
| rab44 | 0.01068 | 0.689985316 | 0.235 | 0 | 1 | 21 |
| eif5 | 0.01068 | 0.551962671 | 0.235 | 0 | 1 | 21 |
| fam162a | 0.01068 | 0.467374254 | 0.235 | 0 | 1 | 21 |
| rab10 | 0.01068 | 0.464243043 | 0.235 | 0 | 1 | 21 |
| si:dkey-261h17.1 | 0.01068 | 0.638834156 | 0.235 | 0 | 1 | 21 |
| crlf3 | 0.01993 | 0.769920962 | 0.471 | 0.087 | 1 | 21 |
| mt-nd4 | 0.02096 | -0.40952138 | 0.118 | 0.478 | 1 | 21 |
| dpy30 | 0.02217 | -0.32533437 | 0 | 0.217 | 1 | 21 |
| selenot2 | 0.02217 | -0.55941605 | 0 | 0.217 | 1 | 21 |
| cacybp | 0.02217 | -0.56610119 | 0 | 0.217 | 1 | 21 |
| atp5mf | 0.02217 | -0.53044614 | 0 | 0.217 | 1 | 21 |
| dynlt3 | 0.02217 | -0.50108927 | 0 | 0.217 | 1 | 21 |
| plpp5 | 0.02217 | -0.56024508 | 0 | 0.217 | 1 | 21 |
| hmgb2b | 0.02217 | -0.17711333 | 0 | 0.217 | 1 | 21 |
| pgam1a | 0.02217 | -0.54867271 | 0 | 0.217 | 1 | 21 |
| sf3b6 | 0.02217 | -0.43810463 | 0 | 0.217 | 1 | 21 |
| sf3b5 | 0.02217 | -0.48827528 | 0 | 0.217 | 1 | 21 |
| vsir | 0.02217 | -0.52888362 | 0 | 0.217 | 1 | 21 |
| ndufb3 | 0.02217 | -0.47412364 | 0 | 0.217 | 1 | 21 |
| zyx | 0.02217 | -0.48019939 | 0 | 0.217 | 1 | 21 |
| atp5f1e | 0.02343 | -0.51474397 | 0.294 | 0.696 | 1 | 21 |

| | | | | | | |
|---|---|---|---|---|---|---|
| clic1 | 0.02937 | -0.6729958 | 0.235 | 0.652 | 1 | 21 |
| ak4 | 0.0317 | 0.377060212 | 0.176 | 0 | 1 | 21 |
| cmtm6 | 0.0317 | 0.417590689 | 0.176 | 0 | 1 | 21 |
| elmo1 | 0.0317 | 0.439751929 | 0.176 | 0 | 1 | 21 |
| prps1a | 0.0317 | 0.450676256 | 0.176 | 0 | 1 | 21 |
| si:ch73-71c20.5 | 0.0317 | 0.34570893 | 0.176 | 0 | 1 | 21 |
| cybc1 | 0.0317 | 0.512529552 | 0.176 | 0 | 1 | 21 |
| mycbp | 0.0317 | 0.422747336 | 0.176 | 0 | 1 | 21 |
| tpmt.1 | 0.0317 | 0.414669728 | 0.176 | 0 | 1 | 21 |
| calm3a | 0.0317 | 0.423819349 | 0.176 | 0 | 1 | 21 |
| ctsba | 0.0317 | 0.476568428 | 0.176 | 0 | 1 | 21 |
| lypla2 | 0.0317 | 0.435253161 | 0.176 | 0 | 1 | 21 |
| ndufs3 | 0.0317 | 0.402052549 | 0.176 | 0 | 1 | 21 |
| pkn1a | 0.0317 | 0.504644904 | 0.176 | 0 | 1 | 21 |
| sept7a | 0.0317 | 0.408670148 | 0.176 | 0 | 1 | 21 |
| uqcc2 | 0.0317 | 0.359331794 | 0.176 | 0 | 1 | 21 |
| etfa | 0.0317 | 0.441559988 | 0.176 | 0 | 1 | 21 |
| nap1l1 | 0.0317 | 0.390932143 | 0.176 | 0 | 1 | 21 |
| psmd13 | 0.0317 | 0.449709711 | 0.176 | 0 | 1 | 21 |
| si:ch211-51h9.7 | 0.0317 | 0.454298537 | 0.176 | 0 | 1 | 21 |
| sypl2b | 0.0317 | 0.549990771 | 0.176 | 0 | 1 | 21 |
| elac1 | 0.0317 | 0.414451553 | 0.176 | 0 | 1 | 21 |
| med22 | 0.0317 | 0.377440627 | 0.176 | 0 | 1 | 21 |
| rsl1d1 | 0.0317 | 0.415960848 | 0.176 | 0 | 1 | 21 |
| znf576.2 | 0.0317 | 0.356434549 | 0.176 | 0 | 1 | 21 |
| fubp1 | 0.0317 | 0.425279234 | 0.176 | 0 | 1 | 21 |
| gskip | 0.035 | 0.667364413 | 0.353 | 0.043 | 1 | 21 |
| atp5mc3b | 0.03812 | 0.569170357 | 0.765 | 0.565 | 1 | 21 |
| cdc42l | 0.03959 | -0.53838965 | 0.471 | 0.696 | 1 | 21 |
| dicp1.1 | 0.04216 | -0.7333382 | 0.118 | 0.478 | 1 | 21 |
| ranbp1 | 0.04563 | 0.649572072 | 0.294 | 0.043 | 1 | 21 |
| psmb7 | 0.0457 | -0.37055745 | 0 | 0.174 | 1 | 21 |
| CABZ01079663.1 | 0.0457 | -0.4729693 | 0 | 0.174 | 1 | 21 |
| LOC103911430 | 0.0457 | -0.51456399 | 0 | 0.174 | 1 | 21 |
| XLOC-043760 | 0.0457 | -0.45831556 | 0 | 0.174 | 1 | 21 |
| il16 | 0.0457 | -0.46721262 | 0 | 0.174 | 1 | 21 |
| mtdha | 0.0457 | -0.43950257 | 0 | 0.174 | 1 | 21 |
| nxt2 | 0.0457 | -0.40153614 | 0 | 0.174 | 1 | 21 |
| snrpb2 | 0.0457 | -0.36997345 | 0 | 0.174 | 1 | 21 |
| tmub1 | 0.0457 | -0.45067896 | 0 | 0.174 | 1 | 21 |
| zdhhc23a | 0.0457 | -0.47383804 | 0 | 0.174 | 1 | 21 |
| zgc:152791 | 0.0457 | -0.54983669 | 0 | 0.174 | 1 | 21 |
| ciarta | 0.0457 | -0.34943403 | 0 | 0.174 | 1 | 21 |
| denr | 0.0457 | -0.37752573 | 0 | 0.174 | 1 | 21 |
| mrps2 | 0.0457 | -0.32297118 | 0 | 0.174 | 1 | 21 |
| si:ch211-59d17.3 | 0.0457 | -0.40575438 | 0 | 0.174 | 1 | 21 |
| rsl24d1 | 0.04572 | -0.60608828 | 0.412 | 0.783 | 1 | 21 |
| npsn | 0.0461 | 0.720376747 | 0.353 | 0.087 | 1 | 21 |

| | | | | | | |
|---|---|---|---|---|---|---|
| btg1 | 0.05099 | -0.41255721 | 0.353 | 0.696 | 1 | 21 |
| atp5f1d | 0.0533 | 0.66706471 | 0.471 | 0.174 | 1 | 21 |
| ptpn6 | 0.05332 | -0.49937329 | 0.529 | 0.87 | 1 | 21 |
| glrx | 0.0534 | -0.66559731 | 0.294 | 0.652 | 1 | 21 |
| sh3kbp1 | 0.05477 | -0.57587623 | 0.059 | 0.348 | 1 | 21 |
| rpl7a | 0.05496 | -0.13990865 | 0.824 | 1 | 1 | 21 |
| cfbl | 0.0554 | 0.630361373 | 0.588 | 0.217 | 1 | 21 |
| ucp2 | 0.05715 | -0.34088163 | 0.412 | 0.739 | 1 | 21 |
| fhit | 0.06128 | -0.64374787 | 0.059 | 0.348 | 1 | 21 |
| rpl10a | 0.07033 | 0.237372168 | 1 | 0.783 | 1 | 21 |
| cnpy3 | 0.07163 | -0.59389811 | 0.059 | 0.348 | 1 | 21 |
| ppiab | 0.07255 | -0.27309285 | 0.706 | 0.957 | 1 | 21 |
| rps27.1 | 0.07409 | -0.28193087 | 0.706 | 0.957 | 1 | 21 |
| rpl35a | 0.07782 | -0.16739895 | 0.824 | 1 | 1 | 21 |
| prdx2 | 0.07847 | -0.58938516 | 0.235 | 0.565 | 1 | 21 |
| rpl8 | 0.07945 | -0.26073466 | 0.824 | 1 | 1 | 21 |
| lcp1 | 0.08539 | -0.61102204 | 0.294 | 0.609 | 1 | 21 |
| rpl29 | 0.08663 | -0.33128287 | 0.706 | 0.957 | 1 | 21 |
| zgc:175088 | 0.08667 | 0.527380676 | 0.294 | 0.043 | 1 | 21 |
| fkbp3 | 0.09095 | -0.61697902 | 0.059 | 0.304 | 1 | 21 |
| esd | 0.09284 | 0.295427221 | 0.118 | 0 | 1 | 21 |
| nktr | 0.09284 | 0.279632882 | 0.118 | 0 | 1 | 21 |
| nsd1a | 0.09284 | 0.279196111 | 0.118 | 0 | 1 | 21 |
| CT027744.2 | 0.09284 | 0.428311094 | 0.118 | 0 | 1 | 21 |
| abcd3a | 0.09284 | 0.234558889 | 0.118 | 0 | 1 | 21 |
| abhd3 | 0.09284 | 0.365146568 | 0.118 | 0 | 1 | 21 |
| acp1 | 0.09284 | 0.271281851 | 0.118 | 0 | 1 | 21 |
| adsl | 0.09284 | 0.231549413 | 0.118 | 0 | 1 | 21 |
| bin1b | 0.09284 | 0.391673315 | 0.118 | 0 | 1 | 21 |
| casp3a | 0.09284 | 0.32905875 | 0.118 | 0 | 1 | 21 |
| crfb4 | 0.09284 | 0.324758702 | 0.118 | 0 | 1 | 21 |
| ddx27 | 0.09284 | 0.269482139 | 0.118 | 0 | 1 | 21 |
| dusp1 | 0.09284 | 0.438810658 | 0.118 | 0 | 1 | 21 |
| eif4ba | 0.09284 | 0.437932405 | 0.118 | 0 | 1 | 21 |
| epg5 | 0.09284 | 0.315691096 | 0.118 | 0 | 1 | 21 |
| etf1b | 0.09284 | 0.297956017 | 0.118 | 0 | 1 | 21 |
| faf2 | 0.09284 | 0.231128361 | 0.118 | 0 | 1 | 21 |
| fam53b | 0.09284 | 0.368417436 | 0.118 | 0 | 1 | 21 |
| fbl | 0.09284 | 0.415121763 | 0.118 | 0 | 1 | 21 |
| fkbp2 | 0.09284 | 0.30758204 | 0.118 | 0 | 1 | 21 |
| fmnl1a | 0.09284 | 0.353027309 | 0.118 | 0 | 1 | 21 |
| gatd1 | 0.09284 | 0.271688471 | 0.118 | 0 | 1 | 21 |
| gatm | 0.09284 | 0.192795651 | 0.118 | 0 | 1 | 21 |
| gmppb | 0.09284 | 0.277037026 | 0.118 | 0 | 1 | 21 |
| hexim1 | 0.09284 | 0.291666423 | 0.118 | 0 | 1 | 21 |
| hikeshi | 0.09284 | 0.294293036 | 0.118 | 0 | 1 | 21 |
| insra | 0.09284 | 0.239577258 | 0.118 | 0 | 1 | 21 |
| lamtor3 | 0.09284 | 0.28815361 | 0.118 | 0 | 1 | 21 |

| | | | | | | |
|---|---|---|---|---|---|---|
| mgmt | 0.09284 | 0.274430868 | 0.118 | 0 | 1 | 21 |
| mxi1 | 0.09284 | 0.326133489 | 0.118 | 0 | 1 | 21 |
| opa1 | 0.09284 | 0.305316757 | 0.118 | 0 | 1 | 21 |
| pag1 | 0.09284 | 0.327600493 | 0.118 | 0 | 1 | 21 |
| phactr4a | 0.09284 | 0.283512121 | 0.118 | 0 | 1 | 21 |
| prdx3 | 0.09284 | 0.213486438 | 0.118 | 0 | 1 | 21 |
| psmd4b | 0.09284 | 0.21940887 | 0.118 | 0 | 1 | 21 |
| ptgs2b | 0.09284 | 0.392644801 | 0.118 | 0 | 1 | 21 |
| rnf185 | 0.09284 | 0.296240999 | 0.118 | 0 | 1 | 21 |
| samd1a | 0.09284 | 0.19117325 | 0.118 | 0 | 1 | 21 |
| scpep1 | 0.09284 | 0.287906262 | 0.118 | 0 | 1 | 21 |
| sgta | 0.09284 | 0.273885876 | 0.118 | 0 | 1 | 21 |
| sirt7 | 0.09284 | 0.292759959 | 0.118 | 0 | 1 | 21 |
| snx27a | 0.09284 | 0.308042021 | 0.118 | 0 | 1 | 21 |
| srrm1 | 0.09284 | 0.349567942 | 0.118 | 0 | 1 | 21 |
| ssbp1 | 0.09284 | 0.485419483 | 0.118 | 0 | 1 | 21 |
| taf7 | 0.09284 | 0.231969437 | 0.118 | 0 | 1 | 21 |
| tefb | 0.09284 | 0.324419402 | 0.118 | 0 | 1 | 21 |
| tspan3a | 0.09284 | 0.274514649 | 0.118 | 0 | 1 | 21 |
| uba3 | 0.09284 | 0.218414585 | 0.118 | 0 | 1 | 21 |
| ubxn4 | 0.09284 | 0.239997958 | 0.118 | 0 | 1 | 21 |
| utp18 | 0.09284 | 0.397433426 | 0.118 | 0 | 1 | 21 |
| vma21 | 0.09284 | 0.300885981 | 0.118 | 0 | 1 | 21 |
| zfand3 | 0.09284 | 0.345426947 | 0.118 | 0 | 1 | 21 |
| znf511 | 0.09284 | 0.3230227 | 0.118 | 0 | 1 | 21 |
| znrf1 | 0.09284 | 0.36364879 | 0.118 | 0 | 1 | 21 |
| dnajc1 | 0.09284 | 0.254283698 | 0.118 | 0 | 1 | 21 |
| kpnb1 | 0.09284 | 0.251902538 | 0.118 | 0 | 1 | 21 |
| nfil3-6 | 0.09284 | 0.343660171 | 0.118 | 0 | 1 | 21 |
| pdcd2 | 0.09284 | 0.241518802 | 0.118 | 0 | 1 | 21 |
| ppp1r7 | 0.09284 | 0.315584279 | 0.118 | 0 | 1 | 21 |
| txndc12 | 0.09284 | 0.314213472 | 0.118 | 0 | 1 | 21 |
| zmp:0000000991 | 0.09284 | 0.367958902 | 0.118 | 0 | 1 | 21 |
| LOC108180076 | 0.09284 | 0.348419768 | 0.118 | 0 | 1 | 21 |
| TSTD1 | 0.09284 | 0.258039141 | 0.118 | 0 | 1 | 21 |
| XLOC-043761 | 0.09284 | 0.257099899 | 0.118 | 0 | 1 | 21 |
| agpat3 | 0.09284 | 0.342199205 | 0.118 | 0 | 1 | 21 |
| ankrd54 | 0.09284 | 0.254970289 | 0.118 | 0 | 1 | 21 |
| arpc1a | 0.09284 | 0.255686793 | 0.118 | 0 | 1 | 21 |
| atp6v1d | 0.09284 | 0.260810079 | 0.118 | 0 | 1 | 21 |
| cdc73 | 0.09284 | 0.306286352 | 0.118 | 0 | 1 | 21 |
| dffa | 0.09284 | 0.348997342 | 0.118 | 0 | 1 | 21 |
| eif5b | 0.09284 | 0.356066315 | 0.118 | 0 | 1 | 21 |
| elf2a | 0.09284 | 0.427920248 | 0.118 | 0 | 1 | 21 |
| gadd45ab | 0.09284 | 0.24651292 | 0.118 | 0 | 1 | 21 |
| grn1 | 0.09284 | 0.314472423 | 0.118 | 0 | 1 | 21 |
| gtf2e2 | 0.09284 | 0.283034737 | 0.118 | 0 | 1 | 21 |
| kdm7ab | 0.09284 | 0.386157675 | 0.118 | 0 | 1 | 21 |

| | | | | | | |
|---|---|---|---|---|---|---|
| map2k6 | 0.09284 | 0.309863607 | 0.118 | 0 | 1 | 21 |
| mbnl3 | 0.09284 | 0.306875045 | 0.118 | 0 | 1 | 21 |
| mrps7 | 0.09284 | 0.234399762 | 0.118 | 0 | 1 | 21 |
| mrtfba | 0.09284 | 0.276792921 | 0.118 | 0 | 1 | 21 |
| mtmr12 | 0.09284 | 0.25636801 | 0.118 | 0 | 1 | 21 |
| nop16 | 0.09284 | 0.2417887 | 0.118 | 0 | 1 | 21 |
| pim2 | 0.09284 | 0.264954168 | 0.118 | 0 | 1 | 21 |
| polr2f | 0.09284 | 0.235762538 | 0.118 | 0 | 1 | 21 |
| rab1ba | 0.09284 | 0.262236123 | 0.118 | 0 | 1 | 21 |
| rab4b | 0.09284 | 0.332221884 | 0.118 | 0 | 1 | 21 |
| rrp1 | 0.09284 | 0.219271702 | 0.118 | 0 | 1 | 21 |
| scp2a | 0.09284 | 0.166262868 | 0.118 | 0 | 1 | 21 |
| si:dkey-177p2.6 | 0.09284 | 0.315884217 | 0.118 | 0 | 1 | 21 |
| srrt | 0.09284 | 0.273336077 | 0.118 | 0 | 1 | 21 |
| tcf25 | 0.09284 | 0.326893705 | 0.118 | 0 | 1 | 21 |
| ube2g2 | 0.09284 | 0.25896893 | 0.118 | 0 | 1 | 21 |
| zc3h18 | 0.09284 | 0.37838679 | 0.118 | 0 | 1 | 21 |
| zgc:112433 | 0.09284 | 0.350473317 | 0.118 | 0 | 1 | 21 |
| zgc:158343 | 0.09284 | 0.286705536 | 0.118 | 0 | 1 | 21 |
| zgc:66475 | 0.09284 | 0.257253439 | 0.118 | 0 | 1 | 21 |
| snx3 | 0.09284 | 0.285163256 | 0.118 | 0 | 1 | 21 |
| vps41 | 0.09284 | 0.330723772 | 0.118 | 0 | 1 | 21 |
| yeats4 | 0.09284 | 0.285018009 | 0.118 | 0 | 1 | 21 |
| cltca | 0.09284 | 0.394415819 | 0.118 | 0 | 1 | 21 |
| ints10 | 0.09284 | 0.268920205 | 0.118 | 0 | 1 | 21 |
| mrpl33 | 0.09284 | 0.282185763 | 0.118 | 0 | 1 | 21 |
| nosip | 0.09284 | 0.256979333 | 0.118 | 0 | 1 | 21 |
| oxct1a | 0.09284 | 0.317297915 | 0.118 | 0 | 1 | 21 |
| zgc:100918 | 0.09284 | 0.263054534 | 0.118 | 0 | 1 | 21 |
| zgc:85858 | 0.09284 | 0.310837788 | 0.118 | 0 | 1 | 21 |
| CABZ01115113.1 | 0.09284 | 0.268764924 | 0.118 | 0 | 1 | 21 |
| leng8 | 0.09284 | 0.279173796 | 0.118 | 0 | 1 | 21 |
| osgn1 | 0.09284 | 0.27921407 | 0.118 | 0 | 1 | 21 |
| zgc:153901 | 0.09284 | 0.273091084 | 0.118 | 0 | 1 | 21 |
| bxdc2 | 0.09284 | 0.243881759 | 0.118 | 0 | 1 | 21 |
| canx | 0.09284 | 0.362978443 | 0.118 | 0 | 1 | 21 |
| irs1 | 0.09284 | 0.424182607 | 0.118 | 0 | 1 | 21 |
| mt-nd5 | 0.09284 | 0.262670989 | 0.118 | 0 | 1 | 21 |
| prps1b | 0.09284 | 0.356178422 | 0.118 | 0 | 1 | 21 |
| slka | 0.09284 | 0.384027819 | 0.118 | 0 | 1 | 21 |
| spink2.1 | 0.09284 | 0.305665414 | 0.118 | 0 | 1 | 21 |
| thrap3b | 0.09284 | 0.285609386 | 0.118 | 0 | 1 | 21 |
| unc93b1 | 0.09284 | 0.424043058 | 0.118 | 0 | 1 | 21 |
| bpifcl | 0.09338 | -0.27797954 | 0 | 0.13 | 1 | 21 |
| pcca | 0.09338 | -0.28016868 | 0 | 0.13 | 1 | 21 |
| plcg2 | 0.09338 | -0.31893807 | 0 | 0.13 | 1 | 21 |
| vbp1 | 0.09338 | -0.26537899 | 0 | 0.13 | 1 | 21 |
| clk4b | 0.09338 | -0.30771716 | 0 | 0.13 | 1 | 21 |

| | | | | | | |
|---|---|---|---|---|---|---|
| naa38 | 0.09338 | -0.34334972 | 0 | 0.13 | 1 | 21 |
| nmi | 0.09338 | -0.33139877 | 0 | 0.13 | 1 | 21 |
| ppp6c | 0.09338 | -0.33235889 | 0 | 0.13 | 1 | 21 |
| psma6b | 0.09338 | -0.30102273 | 0 | 0.13 | 1 | 21 |
| apooa | 0.09338 | -0.31545567 | 0 | 0.13 | 1 | 21 |
| elof1 | 0.09338 | -0.27655442 | 0 | 0.13 | 1 | 21 |
| impa1 | 0.09338 | -0.33439968 | 0 | 0.13 | 1 | 21 |
| otub1b | 0.09338 | -0.28515432 | 0 | 0.13 | 1 | 21 |
| ube2g1a | 0.09338 | -0.32562307 | 0 | 0.13 | 1 | 21 |
| FBXO48 | 0.09338 | -0.29292125 | 0 | 0.13 | 1 | 21 |
| LOC110438870 | 0.09338 | -0.36270822 | 0 | 0.13 | 1 | 21 |
| agap2 | 0.09338 | -0.37612774 | 0 | 0.13 | 1 | 21 |
| ahcyl1 | 0.09338 | -0.3471675 | 0 | 0.13 | 1 | 21 |
| ap2m1b | 0.09338 | -0.32767704 | 0 | 0.13 | 1 | 21 |
| b3gnt2l | 0.09338 | -0.33285533 | 0 | 0.13 | 1 | 21 |
| b3gnt3.4 | 0.09338 | -0.36264272 | 0 | 0.13 | 1 | 21 |
| bbip1 | 0.09338 | -0.27311327 | 0 | 0.13 | 1 | 21 |
| cab39l1 | 0.09338 | -0.30938202 | 0 | 0.13 | 1 | 21 |
| cbx5 | 0.09338 | -0.28838079 | 0 | 0.13 | 1 | 21 |
| ccnd2a | 0.09338 | -0.29745901 | 0 | 0.13 | 1 | 21 |
| cdca4 | 0.09338 | -0.32556494 | 0 | 0.13 | 1 | 21 |
| chmp6b | 0.09338 | -0.32985145 | 0 | 0.13 | 1 | 21 |
| clns1a | 0.09338 | -0.27528841 | 0 | 0.13 | 1 | 21 |
| cmtm3 | 0.09338 | -0.33379395 | 0 | 0.13 | 1 | 21 |
| csrnp1b | 0.09338 | -0.37412599 | 0 | 0.13 | 1 | 21 |
| cyb5r3 | 0.09338 | -0.36977027 | 0 | 0.13 | 1 | 21 |
| dctn5 | 0.09338 | -0.23975697 | 0 | 0.13 | 1 | 21 |
| dpm2 | 0.09338 | -0.3662405 | 0 | 0.13 | 1 | 21 |
| dtx4a | 0.09338 | -0.2558015 | 0 | 0.13 | 1 | 21 |
| dub | 0.09338 | -0.37498529 | 0 | 0.13 | 1 | 21 |
| eif3ja | 0.09338 | -0.34937245 | 0 | 0.13 | 1 | 21 |
| eif4g1a | 0.09338 | -0.33457775 | 0 | 0.13 | 1 | 21 |
| ell | 0.09338 | -0.29458767 | 0 | 0.13 | 1 | 21 |
| fabp3 | 0.09338 | -0.23697203 | 0 | 0.13 | 1 | 21 |
| fdx1 | 0.09338 | -0.31900811 | 0 | 0.13 | 1 | 21 |
| fdx1b | 0.09338 | -0.30196614 | 0 | 0.13 | 1 | 21 |
| ftr14 | 0.09338 | -0.2747889 | 0 | 0.13 | 1 | 21 |
| gar1 | 0.09338 | -0.33126605 | 0 | 0.13 | 1 | 21 |
| gcn1 | 0.09338 | -0.33139113 | 0 | 0.13 | 1 | 21 |
| got1 | 0.09338 | -0.25865972 | 0 | 0.13 | 1 | 21 |
| grpel2 | 0.09338 | -0.39405411 | 0 | 0.13 | 1 | 21 |
| hsdl2 | 0.09338 | -0.2411743 | 0 | 0.13 | 1 | 21 |
| ilf3b | 0.09338 | -0.29709365 | 0 | 0.13 | 1 | 21 |
| lcp2a | 0.09338 | -0.3361375 | 0 | 0.13 | 1 | 21 |
| med31 | 0.09338 | -0.34466314 | 0 | 0.13 | 1 | 21 |
| mfsd6b | 0.09338 | -0.30537961 | 0 | 0.13 | 1 | 21 |
| mrpl20 | 0.09338 | -0.29483757 | 0 | 0.13 | 1 | 21 |
| mrpl23 | 0.09338 | -0.32414613 | 0 | 0.13 | 1 | 21 |

| | | | | | | |
|---|---|---|---|---|---|---|
| mrpl28 | 0.09338 | -0.28707844 | 0 | 0.13 | 1 | 21 |
| mrtfaa | 0.09338 | -0.34559899 | 0 | 0.13 | 1 | 21 |
| mycla | 0.09338 | -0.25848469 | 0 | 0.13 | 1 | 21 |
| naa20 | 0.09338 | -0.35734838 | 0 | 0.13 | 1 | 21 |
| nkd1 | 0.09338 | -0.32600754 | 0 | 0.13 | 1 | 21 |
| p4hb | 0.09338 | -0.36661016 | 0 | 0.13 | 1 | 21 |
| pias4a | 0.09338 | -0.26131981 | 0 | 0.13 | 1 | 21 |
| polr3h | 0.09338 | -0.27241409 | 0 | 0.13 | 1 | 21 |
| ppp2r1ba | 0.09338 | -0.30701496 | 0 | 0.13 | 1 | 21 |
| prpf4 | 0.09338 | -0.28436664 | 0 | 0.13 | 1 | 21 |
| pstpip2 | 0.09338 | -0.2455268 | 0 | 0.13 | 1 | 21 |
| rab5ab | 0.09338 | -0.32125041 | 0 | 0.13 | 1 | 21 |
| rcc2 | 0.09338 | -0.34087006 | 0 | 0.13 | 1 | 21 |
| rrp15 | 0.09338 | -0.33387181 | 0 | 0.13 | 1 | 21 |
| sap18 | 0.09338 | -0.33175932 | 0 | 0.13 | 1 | 21 |
| si:dkey-165a24.10 | 0.09338 | -0.25344879 | 0 | 0.13 | 1 | 21 |
| si:dkey-211g8.4 | 0.09338 | -0.31604895 | 0 | 0.13 | 1 | 21 |
| si:dkey-222f8.3 | 0.09338 | -0.30107813 | 0 | 0.13 | 1 | 21 |
| si:dkey-51e6.1 | 0.09338 | -0.39325594 | 0 | 0.13 | 1 | 21 |
| si:dkey-69o16.5 | 0.09338 | -0.2671996 | 0 | 0.13 | 1 | 21 |
| skia | 0.09338 | -0.35572905 | 0 | 0.13 | 1 | 21 |
| snx5 | 0.09338 | -0.35062582 | 0 | 0.13 | 1 | 21 |
| stk24b | 0.09338 | -0.3021998 | 0 | 0.13 | 1 | 21 |
| timm8a | 0.09338 | -0.28173073 | 0 | 0.13 | 1 | 21 |
| trnau1apb | 0.09338 | -0.30604977 | 0 | 0.13 | 1 | 21 |
| tufm | 0.09338 | -0.29318692 | 0 | 0.13 | 1 | 21 |
| uba1 | 0.09338 | -0.34950876 | 0 | 0.13 | 1 | 21 |
| uqcrb | 0.09338 | -0.30350591 | 0 | 0.13 | 1 | 21 |
| zc3h14 | 0.09338 | -0.36131996 | 0 | 0.13 | 1 | 21 |
| ckbb | 0.09725 | -0.22613517 | 0.824 | 1 | 1 | 21 |
| ywhaqa | 0.10032 | -0.43930392 | 0.059 | 0.304 | 1 | 21 |
| XLOC-026836 | 0.10612 | -0.58326547 | 0.235 | 0.565 | 1 | 21 |
| cycsb | 0.10998 | -0.55675499 | 0.059 | 0.304 | 1 | 21 |
| rpl18 | 0.11219 | 0.178320639 | 0.882 | 0.913 | 1 | 21 |
| naa10 | 0.12077 | 0.280459353 | 0.353 | 0.13 | 1 | 21 |
| eif5a | 0.1208 | 0.379115871 | 0.706 | 0.391 | 1 | 21 |
| stk17al | 0.12304 | -0.49345106 | 0.059 | 0.304 | 1 | 21 |
| arhgdia | 0.12439 | -0.39155946 | 0.471 | 0.609 | 1 | 21 |
| eif3i | 0.1261 | -0.52473584 | 0.176 | 0.478 | 1 | 21 |
| crip1 | 0.12675 | 0.423430256 | 0.706 | 0.522 | 1 | 21 |
| mdh2 | 0.12809 | 0.527356668 | 0.412 | 0.13 | 1 | 21 |
| nedd8 | 0.12989 | -0.22938462 | 0.059 | 0.217 | 1 | 21 |
| tra2a | 0.12993 | -0.47375539 | 0.059 | 0.304 | 1 | 21 |
| mt-cyb | 0.1301 | -0.59190053 | 0.118 | 0.391 | 1 | 21 |
| rpl5a | 0.13198 | -0.33287169 | 0.765 | 0.913 | 1 | 21 |
| si:ch211-220f21.3 | 0.13425 | 0.510662838 | 0.706 | 0.391 | 1 | 21 |
| eif3c | 0.13733 | -0.3383354 | 0.118 | 0.348 | 1 | 21 |
| vaspa | 0.14194 | -0.27194557 | 0.824 | 0.957 | 1 | 21 |

| | | | | | | |
|---|---|---|---|---|---|---|
| oaz1b | 0.14389 | -0.34118047 | 0.529 | 0.609 | 1 | 21 |
| rhoaa | 0.14412 | 0.340062075 | 0.235 | 0.043 | 1 | 21 |
| cirbpa | 0.14794 | -0.52091624 | 0.353 | 0.609 | 1 | 21 |
| selenow1 | 0.15066 | 0.431625371 | 0.529 | 0.391 | 1 | 21 |
| actb1 | 0.15347 | -0.10197189 | 1 | 1 | 1 | 21 |
| atp5md | 0.15504 | 0.367669252 | 0.588 | 0.478 | 1 | 21 |
| naca | 0.15572 | -0.16102191 | 0.765 | 0.957 | 1 | 21 |
| fth1a | 0.15574 | 0.402458838 | 0.471 | 0.391 | 1 | 21 |
| snrnp27 | 0.15942 | 0.350263649 | 0.235 | 0.043 | 1 | 21 |
| gstm.2 | 0.16192 | 0.373071223 | 0.235 | 0.043 | 1 | 21 |
| tomm20b | 0.16473 | 0.208650623 | 0.471 | 0.261 | 1 | 21 |
| CU207241.1 | 0.16554 | 0.370563412 | 0.235 | 0.043 | 1 | 21 |
| ywhabb | 0.16573 | 0.389214614 | 0.529 | 0.261 | 1 | 21 |
| gpx4b | 0.16579 | 0.514946848 | 0.353 | 0.13 | 1 | 21 |
| zgc:193541 | 0.16612 | -0.48225584 | 0.353 | 0.652 | 1 | 21 |
| xbp1 | 0.16877 | 0.337412536 | 0.235 | 0.043 | 1 | 21 |
| smap1 | 0.17125 | 0.429163718 | 0.235 | 0.043 | 1 | 21 |
| atp5fa1 | 0.17158 | -0.23616359 | 0.176 | 0.391 | 1 | 21 |
| ccng1 | 0.1743 | -0.52963177 | 0.294 | 0.565 | 1 | 21 |
| scamp2 | 0.17584 | 0.34176147 | 0.294 | 0.087 | 1 | 21 |
| hdlbpa | 0.17586 | 0.105247093 | 0.176 | 0.043 | 1 | 21 |
| cfl1 | 0.17713 | -0.1753604 | 0.941 | 1 | 1 | 21 |
| rps20 | 0.18025 | -0.21629664 | 0.765 | 0.957 | 1 | 21 |
| si:ch73-281n10.2 | 0.18109 | 0.483905092 | 0.294 | 0.087 | 1 | 21 |
| krt18b | 0.1841 | 0.404714204 | 0.235 | 0.043 | 1 | 21 |
| bcam | 0.18922 | 0.473102641 | 0.235 | 0.043 | 1 | 21 |
| mob1a | 0.19076 | 0.468841295 | 0.647 | 0.391 | 1 | 21 |
| arpc1b | 0.19083 | -0.16108528 | 0.941 | 1 | 1 | 21 |
| rps27l | 0.19113 | 0.47849199 | 0.235 | 0.043 | 1 | 21 |
| CU467110.1 | 0.19149 | -0.24167256 | 0 | 0.087 | 1 | 21 |
| anxa4 | 0.19149 | -0.24304376 | 0 | 0.087 | 1 | 21 |
| ddx3xb | 0.19149 | -0.24665264 | 0 | 0.087 | 1 | 21 |
| irak3 | 0.19149 | -0.24868107 | 0 | 0.087 | 1 | 21 |
| lyrm4 | 0.19149 | -0.23139763 | 0 | 0.087 | 1 | 21 |
| zgc:172090 | 0.19149 | -0.24840309 | 0 | 0.087 | 1 | 21 |
| BOLA2 | 0.19149 | -0.22420899 | 0 | 0.087 | 1 | 21 |
| CU855821.1 | 0.19149 | -0.24920219 | 0 | 0.087 | 1 | 21 |
| LOC110439924 | 0.19149 | -0.17561679 | 0 | 0.087 | 1 | 21 |
| arid4a | 0.19149 | -0.2956471 | 0 | 0.087 | 1 | 21 |
| cdk13 | 0.19149 | -0.23236214 | 0 | 0.087 | 1 | 21 |
| cndp2 | 0.19149 | -0.17770164 | 0 | 0.087 | 1 | 21 |
| cyfip2 | 0.19149 | -0.22229065 | 0 | 0.087 | 1 | 21 |
| dldh | 0.19149 | -0.16599578 | 0 | 0.087 | 1 | 21 |
| endog | 0.19149 | -0.27935445 | 0 | 0.087 | 1 | 21 |
| fgd4a | 0.19149 | -0.27162448 | 0 | 0.087 | 1 | 21 |
| mob1bb | 0.19149 | -0.22835151 | 0 | 0.087 | 1 | 21 |
| mphosph8 | 0.19149 | -0.24352071 | 0 | 0.087 | 1 | 21 |
| nfu1 | 0.19149 | -0.29725255 | 0 | 0.087 | 1 | 21 |

| | | | | | | |
|---|---|---|---|---|---|---|
| nt5dc2 | 0.19149 | -0.17246741 | 0 | 0.087 | 1 | 21 |
| pop4 | 0.19149 | -0.17229432 | 0 | 0.087 | 1 | 21 |
| prdx6 | 0.19149 | -0.23429466 | 0 | 0.087 | 1 | 21 |
| prpf4bb | 0.19149 | -0.24679761 | 0 | 0.087 | 1 | 21 |
| rcor1 | 0.19149 | -0.25148685 | 0 | 0.087 | 1 | 21 |
| rpa3 | 0.19149 | -0.22469208 | 0 | 0.087 | 1 | 21 |
| serinc2 | 0.19149 | -0.2085795 | 0 | 0.087 | 1 | 21 |
| si:dkey-188i13.10 | 0.19149 | -0.16355729 | 0 | 0.087 | 1 | 21 |
| si:dkeyp-44a8.2 | 0.19149 | -0.27515197 | 0 | 0.087 | 1 | 21 |
| si:rp71-45k5.4 | 0.19149 | -0.15972762 | 0 | 0.087 | 1 | 21 |
| slc25a55a | 0.19149 | -0.278195 | 0 | 0.087 | 1 | 21 |
| sod2 | 0.19149 | -0.29215842 | 0 | 0.087 | 1 | 21 |
| srsf9 | 0.19149 | -0.16045145 | 0 | 0.087 | 1 | 21 |
| tax1bp1b | 0.19149 | -0.20305501 | 0 | 0.087 | 1 | 21 |
| tbl3 | 0.19149 | -0.16589484 | 0 | 0.087 | 1 | 21 |
| ube2ka | 0.19149 | -0.25940187 | 0 | 0.087 | 1 | 21 |
| uqcrc2b | 0.19149 | -0.24707001 | 0 | 0.087 | 1 | 21 |
| zgc:113184 | 0.19149 | -0.26371303 | 0 | 0.087 | 1 | 21 |
| zgc:56106 | 0.19149 | -0.22441438 | 0 | 0.087 | 1 | 21 |
| BX908750.1 | 0.19149 | -0.21249789 | 0 | 0.087 | 1 | 21 |
| LOC103910015 | 0.19149 | -0.18482367 | 0 | 0.087 | 1 | 21 |
| LOC110437917 | 0.19149 | -0.20681532 | 0 | 0.087 | 1 | 21 |
| actr10 | 0.19149 | -0.18031717 | 0 | 0.087 | 1 | 21 |
| ap4s1 | 0.19149 | -0.23983004 | 0 | 0.087 | 1 | 21 |
| asb8 | 0.19149 | -0.24431257 | 0 | 0.087 | 1 | 21 |
| ccdc47 | 0.19149 | -0.20097349 | 0 | 0.087 | 1 | 21 |
| dohh | 0.19149 | -0.18783034 | 0 | 0.087 | 1 | 21 |
| emc8 | 0.19149 | -0.21938621 | 0 | 0.087 | 1 | 21 |
| erlin1 | 0.19149 | -0.20458347 | 0 | 0.087 | 1 | 21 |
| fahd2a | 0.19149 | -0.20757324 | 0 | 0.087 | 1 | 21 |
| fbxo28 | 0.19149 | -0.19708375 | 0 | 0.087 | 1 | 21 |
| flot2a | 0.19149 | -0.20505362 | 0 | 0.087 | 1 | 21 |
| fntb | 0.19149 | -0.20618231 | 0 | 0.087 | 1 | 21 |
| mff | 0.19149 | -0.2069572 | 0 | 0.087 | 1 | 21 |
| mycb | 0.19149 | -0.2204561 | 0 | 0.087 | 1 | 21 |
| nipsnap2 | 0.19149 | -0.19517403 | 0 | 0.087 | 1 | 21 |
| pdcd10a | 0.19149 | -0.19918888 | 0 | 0.087 | 1 | 21 |
| pi4k2b | 0.19149 | -0.20367685 | 0 | 0.087 | 1 | 21 |
| ppa1b | 0.19149 | -0.18703019 | 0 | 0.087 | 1 | 21 |
| pphln1 | 0.19149 | -0.21472051 | 0 | 0.087 | 1 | 21 |
| puf60b | 0.19149 | -0.19379508 | 0 | 0.087 | 1 | 21 |
| rcbtb1 | 0.19149 | -0.19890634 | 0 | 0.087 | 1 | 21 |
| sccpdhb | 0.19149 | -0.19738223 | 0 | 0.087 | 1 | 21 |
| sdhc | 0.19149 | -0.19239918 | 0 | 0.087 | 1 | 21 |
| selplg | 0.19149 | -0.28289699 | 0 | 0.087 | 1 | 21 |
| sh3gl1a | 0.19149 | -0.20909606 | 0 | 0.087 | 1 | 21 |
| si:ch211-162e15.3 | 0.19149 | -0.27664769 | 0 | 0.087 | 1 | 21 |
| si:dkey-222h21.2 | 0.19149 | -0.20897092 | 0 | 0.087 | 1 | 21 |

| | | | | | | |
|---|---|---|---|---|---|---|
| srgap3 | 0.19149 | -0.2130868 | 0 | 0.087 | 1 | 21 |
| tcea1 | 0.19149 | -0.19583934 | 0 | 0.087 | 1 | 21 |
| tia1 | 0.19149 | -0.20427364 | 0 | 0.087 | 1 | 21 |
| tsr2 | 0.19149 | -0.21885808 | 0 | 0.087 | 1 | 21 |
| txnl4b | 0.19149 | -0.24077952 | 0 | 0.087 | 1 | 21 |
| utp11l | 0.19149 | -0.16263412 | 0 | 0.087 | 1 | 21 |
| BX005245.1 | 0.19149 | -0.25474502 | 0 | 0.087 | 1 | 21 |
| CU570894.1 | 0.19149 | -0.25762601 | 0 | 0.087 | 1 | 21 |
| XLOC-032112 | 0.19149 | -0.21039443 | 0 | 0.087 | 1 | 21 |
| adora2aa | 0.19149 | -0.2524491 | 0 | 0.087 | 1 | 21 |
| btg2 | 0.19149 | -0.20066133 | 0 | 0.087 | 1 | 21 |
| c1qbp | 0.19149 | -0.30400349 | 0 | 0.087 | 1 | 21 |
| cdk17 | 0.19149 | -0.24261248 | 0 | 0.087 | 1 | 21 |
| ciartb | 0.19149 | -0.19933347 | 0 | 0.087 | 1 | 21 |
| cmtm7 | 0.19149 | -0.18320435 | 0 | 0.087 | 1 | 21 |
| csf2rb | 0.19149 | -0.24541004 | 0 | 0.087 | 1 | 21 |
| cwf19l2 | 0.19149 | -0.20944758 | 0 | 0.087 | 1 | 21 |
| dctn2 | 0.19149 | -0.22722926 | 0 | 0.087 | 1 | 21 |
| def6c | 0.19149 | -0.22644083 | 0 | 0.087 | 1 | 21 |
| dok1b | 0.19149 | -0.23373307 | 0 | 0.087 | 1 | 21 |
| dynlrb1 | 0.19149 | -0.17841274 | 0 | 0.087 | 1 | 21 |
| fbxo9 | 0.19149 | -0.28747284 | 0 | 0.087 | 1 | 21 |
| fuom | 0.19149 | -0.22546092 | 0 | 0.087 | 1 | 21 |
| gemin2 | 0.19149 | -0.16986584 | 0 | 0.087 | 1 | 21 |
| gemin8 | 0.19149 | -0.23749438 | 0 | 0.087 | 1 | 21 |
| gmfg | 0.19149 | -0.25665513 | 0 | 0.087 | 1 | 21 |
| hadh | 0.19149 | -0.14057909 | 0 | 0.087 | 1 | 21 |
| hadhb | 0.19149 | -0.1860829 | 0 | 0.087 | 1 | 21 |
| hspe1 | 0.19149 | -0.15947193 | 0 | 0.087 | 1 | 21 |
| idh3b | 0.19149 | -0.17967772 | 0 | 0.087 | 1 | 21 |
| kctd10 | 0.19149 | -0.26766838 | 0 | 0.087 | 1 | 21 |
| klf7b | 0.19149 | -0.24449141 | 0 | 0.087 | 1 | 21 |
| larp1b | 0.19149 | -0.20517297 | 0 | 0.087 | 1 | 21 |
| lpin1 | 0.19149 | -0.22992588 | 0 | 0.087 | 1 | 21 |
| mapk3 | 0.19149 | -0.22526654 | 0 | 0.087 | 1 | 21 |
| mrpl21 | 0.19149 | -0.15025802 | 0 | 0.087 | 1 | 21 |
| mta3 | 0.19149 | -0.25073598 | 0 | 0.087 | 1 | 21 |
| nagk | 0.19149 | -0.2332232 | 0 | 0.087 | 1 | 21 |
| nars | 0.19149 | -0.22037941 | 0 | 0.087 | 1 | 21 |
| nat6 | 0.19149 | -0.21471229 | 0 | 0.087 | 1 | 21 |
| nup62l | 0.19149 | -0.26419123 | 0 | 0.087 | 1 | 21 |
| ost4 | 0.19149 | -0.16830497 | 0 | 0.087 | 1 | 21 |
| pes | 0.19149 | -0.27095686 | 0 | 0.087 | 1 | 21 |
| phpt1 | 0.19149 | -0.24817544 | 0 | 0.087 | 1 | 21 |
| prkcda | 0.19149 | -0.18191239 | 0 | 0.087 | 1 | 21 |
| ralab | 0.19149 | -0.26329086 | 0 | 0.087 | 1 | 21 |
| rfx5 | 0.19149 | -0.18248331 | 0 | 0.087 | 1 | 21 |
| rxfp2a | 0.19149 | -0.14610432 | 0 | 0.087 | 1 | 21 |

| | | | | | | |
|---|---|---|---|---|---|---|
| sart3 | 0.19149 | -0.1935717 | 0 | 0.087 | 1 | 21 |
| scdb | 0.19149 | -0.23153269 | 0 | 0.087 | 1 | 21 |
| sec13 | 0.19149 | -0.14035705 | 0 | 0.087 | 1 | 21 |
| selenbp1 | 0.19149 | -0.22487763 | 0 | 0.087 | 1 | 21 |
| sept7b | 0.19149 | -0.27454341 | 0 | 0.087 | 1 | 21 |
| si:ch1073-392o20.2 | 0.19149 | -0.24981885 | 0 | 0.087 | 1 | 21 |
| si:ch211-14a17.11 | 0.19149 | -0.1897958 | 0 | 0.087 | 1 | 21 |
| si:ch211-203d1.3 | 0.19149 | -0.15688677 | 0 | 0.087 | 1 | 21 |
| si:dkey-234h16.7 | 0.19149 | -0.25891166 | 0 | 0.087 | 1 | 21 |
| si:dkey-52l18.4 | 0.19149 | -0.23321816 | 0 | 0.087 | 1 | 21 |
| slc38a10 | 0.19149 | -0.20651625 | 0 | 0.087 | 1 | 21 |
| snu13a | 0.19149 | -0.25389536 | 0 | 0.087 | 1 | 21 |
| snx4 | 0.19149 | -0.25145328 | 0 | 0.087 | 1 | 21 |
| srp68 | 0.19149 | -0.21408456 | 0 | 0.087 | 1 | 21 |
| ssbp3b | 0.19149 | -0.26379167 | 0 | 0.087 | 1 | 21 |
| stimate | 0.19149 | -0.19681019 | 0 | 0.087 | 1 | 21 |
| stx5a | 0.19149 | -0.17679408 | 0 | 0.087 | 1 | 21 |
| thoc1 | 0.19149 | -0.1868408 | 0 | 0.087 | 1 | 21 |
| timm10 | 0.19149 | -0.21190765 | 0 | 0.087 | 1 | 21 |
| timm17a | 0.19149 | -0.21521674 | 0 | 0.087 | 1 | 21 |
| tmf1 | 0.19149 | -0.13121744 | 0 | 0.087 | 1 | 21 |
| utp4 | 0.19149 | -0.26647599 | 0 | 0.087 | 1 | 21 |
| zgc:113149 | 0.19149 | -0.23274982 | 0 | 0.087 | 1 | 21 |
| zgc:153012 | 0.19149 | -0.22667597 | 0 | 0.087 | 1 | 21 |
| zgc:194312 | 0.19149 | -0.1847277 | 0 | 0.087 | 1 | 21 |
| zgc:66440 | 0.19149 | -0.15030608 | 0 | 0.087 | 1 | 21 |
| BX322618.1 | 0.19149 | -0.24059787 | 0 | 0.087 | 1 | 21 |
| CABZ01068251.1 | 0.19149 | -0.17038074 | 0 | 0.087 | 1 | 21 |
| MED9 | 0.19149 | -0.17224682 | 0 | 0.087 | 1 | 21 |
| PDCL3 | 0.19149 | -0.18338683 | 0 | 0.087 | 1 | 21 |
| XLOC-000431 | 0.19149 | -0.18950785 | 0 | 0.087 | 1 | 21 |
| acbd6 | 0.19149 | -0.2095376 | 0 | 0.087 | 1 | 21 |
| adap2 | 0.19149 | -0.18071949 | 0 | 0.087 | 1 | 21 |
| atad1b | 0.19149 | -0.22342209 | 0 | 0.087 | 1 | 21 |
| b4galnt1a | 0.19149 | -0.19835579 | 0 | 0.087 | 1 | 21 |
| birc2 | 0.19149 | -0.17211387 | 0 | 0.087 | 1 | 21 |
| cdc123 | 0.19149 | -0.19983769 | 0 | 0.087 | 1 | 21 |
| cept1a | 0.19149 | -0.23646774 | 0 | 0.087 | 1 | 21 |
| dcun1d5 | 0.19149 | -0.17160799 | 0 | 0.087 | 1 | 21 |
| dnm1l | 0.19149 | -0.20256996 | 0 | 0.087 | 1 | 21 |
| ell2 | 0.19149 | -0.25149772 | 0 | 0.087 | 1 | 21 |
| gorasp2 | 0.19149 | -0.25000845 | 0 | 0.087 | 1 | 21 |
| gstk1 | 0.19149 | -0.23428956 | 0 | 0.087 | 1 | 21 |
| hic1 | 0.19149 | -0.17496038 | 0 | 0.087 | 1 | 21 |
| ier2b | 0.19149 | -0.28710623 | 0 | 0.087 | 1 | 21 |
| ikbkg | 0.19149 | -0.22159774 | 0 | 0.087 | 1 | 21 |
| ino80c | 0.19149 | -0.30068392 | 0 | 0.087 | 1 | 21 |
| mcm7 | 0.19149 | -0.16858773 | 0 | 0.087 | 1 | 21 |

| | | | | | | |
|---|---|---|---|---|---|---|
| mgst1.2 | 0.19149 | -0.23911541 | 0 | 0.087 | 1 | 21 |
| ncalda | 0.19149 | -0.22068497 | 0 | 0.087 | 1 | 21 |
| nipa2 | 0.19149 | -0.20004532 | 0 | 0.087 | 1 | 21 |
| nutf2 | 0.19149 | -0.19327653 | 0 | 0.087 | 1 | 21 |
| pabpn1 | 0.19149 | -0.20085626 | 0 | 0.087 | 1 | 21 |
| parp1 | 0.19149 | -0.13364535 | 0 | 0.087 | 1 | 21 |
| pdcd5 | 0.19149 | -0.24358688 | 0 | 0.087 | 1 | 21 |
| pggt1b | 0.19149 | -0.20438967 | 0 | 0.087 | 1 | 21 |
| pgm2l1 | 0.19149 | -0.19389163 | 0 | 0.087 | 1 | 21 |
| pigf | 0.19149 | -0.1665856 | 0 | 0.087 | 1 | 21 |
| pik3ap1 | 0.19149 | -0.21916362 | 0 | 0.087 | 1 | 21 |
| plp2 | 0.19149 | -0.21328347 | 0 | 0.087 | 1 | 21 |
| ppp1r12a | 0.19149 | -0.21792524 | 0 | 0.087 | 1 | 21 |
| pqlc3 | 0.19149 | -0.179881 | 0 | 0.087 | 1 | 21 |
| psma2 | 0.19149 | -0.1792587 | 0 | 0.087 | 1 | 21 |
| qkia | 0.19149 | -0.16775205 | 0 | 0.087 | 1 | 21 |
| rab8a | 0.19149 | -0.24083663 | 0 | 0.087 | 1 | 21 |
| rarab | 0.19149 | -0.23777675 | 0 | 0.087 | 1 | 21 |
| rcan1a | 0.19149 | -0.23532046 | 0 | 0.087 | 1 | 21 |
| rchy1 | 0.19149 | -0.20862419 | 0 | 0.087 | 1 | 21 |
| rfc3 | 0.19149 | -0.25487432 | 0 | 0.087 | 1 | 21 |
| rhogd | 0.19149 | -0.17517995 | 0 | 0.087 | 1 | 21 |
| rxrba | 0.19149 | -0.17353694 | 0 | 0.087 | 1 | 21 |
| si:dkey-262k9.2 | 0.19149 | -0.2394407 | 0 | 0.087 | 1 | 21 |
| si:dkeyp-82a1.6 | 0.19149 | -0.24569818 | 0 | 0.087 | 1 | 21 |
| slc30a7 | 0.19149 | -0.20836831 | 0 | 0.087 | 1 | 21 |
| snrpa | 0.19149 | -0.16224157 | 0 | 0.087 | 1 | 21 |
| srsf4 | 0.19149 | -0.20408753 | 0 | 0.087 | 1 | 21 |
| stat6 | 0.19149 | -0.30930721 | 0 | 0.087 | 1 | 21 |
| stx18 | 0.19149 | -0.16840483 | 0 | 0.087 | 1 | 21 |
| taf13 | 0.19149 | -0.2056701 | 0 | 0.087 | 1 | 21 |
| taf6 | 0.19149 | -0.24228093 | 0 | 0.087 | 1 | 21 |
| tbc1d5 | 0.19149 | -0.21642314 | 0 | 0.087 | 1 | 21 |
| trappc4 | 0.19149 | -0.21997438 | 0 | 0.087 | 1 | 21 |
| tsg101b | 0.19149 | -0.18090798 | 0 | 0.087 | 1 | 21 |
| ube2a | 0.19149 | -0.20526664 | 0 | 0.087 | 1 | 21 |
| ube2e2 | 0.19149 | -0.26944366 | 0 | 0.087 | 1 | 21 |
| ube2ia | 0.19149 | -0.1965124 | 0 | 0.087 | 1 | 21 |
| unkl | 0.19149 | -0.18028836 | 0 | 0.087 | 1 | 21 |
| uqcc3 | 0.19149 | -0.17357004 | 0 | 0.087 | 1 | 21 |
| vps25 | 0.19149 | -0.17059461 | 0 | 0.087 | 1 | 21 |
| zgc:163080 | 0.19149 | -0.19003998 | 0 | 0.087 | 1 | 21 |
| zmp:0000001082 | 0.19149 | -0.17741215 | 0 | 0.087 | 1 | 21 |
| BX323557.1 | 0.19149 | -0.18175099 | 0 | 0.087 | 1 | 21 |
| BX571825.4 | 0.19149 | -0.21540293 | 0 | 0.087 | 1 | 21 |
| CABZ01074397.1 | 0.19149 | -0.21199339 | 0 | 0.087 | 1 | 21 |
| CABZ01087514.1 | 0.19149 | -0.21622871 | 0 | 0.087 | 1 | 21 |
| CR388166.2 | 0.19149 | -0.22928599 | 0 | 0.087 | 1 | 21 |

| | | | | | | |
|---|---|---|---|---|---|---|
| LOC108190717 | 0.19149 | -0.20963632 | 0 | 0.087 | 1 | 21 |
| LOC110440216 | 0.19149 | -0.20515111 | 0 | 0.087 | 1 | 21 |
| OSBPL8 | 0.19149 | -0.21931204 | 0 | 0.087 | 1 | 21 |
| XLOC-035413 | 0.19149 | -0.20306628 | 0 | 0.087 | 1 | 21 |
| abhd13 | 0.19149 | -0.1966323 | 0 | 0.087 | 1 | 21 |
| arhgap4a | 0.19149 | -0.21189718 | 0 | 0.087 | 1 | 21 |
| atp6v1h | 0.19149 | -0.24458888 | 0 | 0.087 | 1 | 21 |
| blzf1 | 0.19149 | -0.19433291 | 0 | 0.087 | 1 | 21 |
| btbd10b | 0.19149 | -0.16273578 | 0 | 0.087 | 1 | 21 |
| cnot7 | 0.19149 | -0.22960036 | 0 | 0.087 | 1 | 21 |
| cpsf6 | 0.19149 | -0.19574845 | 0 | 0.087 | 1 | 21 |
| efhd2 | 0.19149 | -0.27680966 | 0 | 0.087 | 1 | 21 |
| eif4a3 | 0.19149 | -0.22805227 | 0 | 0.087 | 1 | 21 |
| fcf1 | 0.19149 | -0.18765132 | 0 | 0.087 | 1 | 21 |
| gna13b | 0.19149 | -0.17066516 | 0 | 0.087 | 1 | 21 |
| gpn1 | 0.19149 | -0.19847705 | 0 | 0.087 | 1 | 21 |
| gpn3 | 0.19149 | -0.20968307 | 0 | 0.087 | 1 | 21 |
| grsf1 | 0.19149 | -0.2677293 | 0 | 0.087 | 1 | 21 |
| hmgb3a | 0.19149 | -0.17765089 | 0 | 0.087 | 1 | 21 |
| hps5 | 0.19149 | -0.2456634 | 0 | 0.087 | 1 | 21 |
| iah1 | 0.19149 | -0.19975412 | 0 | 0.087 | 1 | 21 |
| ier5 | 0.19149 | -0.28104425 | 0 | 0.087 | 1 | 21 |
| ifngr2 | 0.19149 | -0.21774239 | 0 | 0.087 | 1 | 21 |
| ilk | 0.19149 | -0.16573251 | 0 | 0.087 | 1 | 21 |
| irf5 | 0.19149 | -0.21010996 | 0 | 0.087 | 1 | 21 |
| junba | 0.19149 | -0.19355751 | 0 | 0.087 | 1 | 21 |
| kdm1a | 0.19149 | -0.20741396 | 0 | 0.087 | 1 | 21 |
| kpna3 | 0.19149 | -0.24742889 | 0 | 0.087 | 1 | 21 |
| lpxn | 0.19149 | -0.24683813 | 0 | 0.087 | 1 | 21 |
| lyrm2 | 0.19149 | -0.19697868 | 0 | 0.087 | 1 | 21 |
| manf | 0.19149 | -0.24442986 | 0 | 0.087 | 1 | 21 |
| med17 | 0.19149 | -0.24904918 | 0 | 0.087 | 1 | 21 |
| mkrn1 | 0.19149 | -0.27198845 | 0 | 0.087 | 1 | 21 |
| mob1ba | 0.19149 | -0.16365634 | 0 | 0.087 | 1 | 21 |
| mob2b | 0.19149 | -0.20161158 | 0 | 0.087 | 1 | 21 |
| mrpl27 | 0.19149 | -0.20307713 | 0 | 0.087 | 1 | 21 |
| ncbp2 | 0.19149 | -0.17623031 | 0 | 0.087 | 1 | 21 |
| ndufs8a | 0.19149 | -0.16106816 | 0 | 0.087 | 1 | 21 |
| nek6 | 0.19149 | -0.24396077 | 0 | 0.087 | 1 | 21 |
| nmt1a | 0.19149 | -0.16880414 | 0 | 0.087 | 1 | 21 |
| ociad1 | 0.19149 | -0.23253543 | 0 | 0.087 | 1 | 21 |
| otub1a | 0.19149 | -0.2054429 | 0 | 0.087 | 1 | 21 |
| pbdc1 | 0.19149 | -0.19955292 | 0 | 0.087 | 1 | 21 |
| pgrmc2 | 0.19149 | -0.16958246 | 0 | 0.087 | 1 | 21 |
| pigc | 0.19149 | -0.28577326 | 0 | 0.087 | 1 | 21 |
| ppm1bb | 0.19149 | -0.17690877 | 0 | 0.087 | 1 | 21 |
| ppp1r12c | 0.19149 | -0.27710131 | 0 | 0.087 | 1 | 21 |
| prkcbb | 0.19149 | -0.28586994 | 0 | 0.087 | 1 | 21 |

| | | | | | | |
|---|---|---|---|---|---|---|
| prpf19 | 0.19149 | -0.2104389 | 0 | 0.087 | 1 | 21 |
| rnf14 | 0.19149 | -0.21835596 | 0 | 0.087 | 1 | 21 |
| si:ch211-85n16.3 | 0.19149 | -0.18413159 | 0 | 0.087 | 1 | 21 |
| slc5a8l | 0.19149 | -0.19605597 | 0 | 0.087 | 1 | 21 |
| smim12 | 0.19149 | -0.20234525 | 0 | 0.087 | 1 | 21 |
| tbl1xr1a | 0.19149 | -0.21046158 | 0 | 0.087 | 1 | 21 |
| tomm70a | 0.19149 | -0.17330446 | 0 | 0.087 | 1 | 21 |
| yars | 0.19149 | -0.24370639 | 0 | 0.087 | 1 | 21 |
| yipf2 | 0.19149 | -0.24753405 | 0 | 0.087 | 1 | 21 |
| yipf5 | 0.19149 | -0.19403263 | 0 | 0.087 | 1 | 21 |
| znf207a | 0.19149 | -0.24766439 | 0 | 0.087 | 1 | 21 |
| bnip3lb | 0.19149 | -0.16135681 | 0 | 0.087 | 1 | 21 |
| ctdsp2 | 0.19149 | -0.2080846 | 0 | 0.087 | 1 | 21 |
| dnajc9 | 0.19149 | -0.15474958 | 0 | 0.087 | 1 | 21 |
| ewsr1a | 0.19149 | -0.14808408 | 0 | 0.087 | 1 | 21 |
| si:dkey-6n6.2 | 0.19149 | -0.16230052 | 0 | 0.087 | 1 | 21 |
| lsm8 | 0.1935 | 0.401641143 | 0.235 | 0.043 | 1 | 21 |
| rgs13 | 0.19459 | 0.507488852 | 0.588 | 0.304 | 1 | 21 |
| kctd12.2 | 0.19465 | -0.43571824 | 0.471 | 0.739 | 1 | 21 |
| rpl19 | 0.19628 | -0.24057699 | 0.824 | 0.913 | 1 | 21 |
| MRPS12 | 0.20118 | -0.41560716 | 0.059 | 0.261 | 1 | 21 |
| lasp1 | 0.20147 | 0.215027638 | 0.706 | 0.478 | 1 | 21 |
| ap3m1 | 0.20202 | 0.198984117 | 0.176 | 0.043 | 1 | 21 |
| arpc4 | 0.20241 | -0.35949309 | 0.647 | 0.87 | 1 | 21 |
| idh2 | 0.20269 | -0.42336672 | 0.529 | 0.739 | 1 | 21 |
| TSTA3 (1 of many) | 0.20345 | -0.4669322 | 0.059 | 0.261 | 1 | 21 |
| h1fx | 0.20495 | -0.21771035 | 0.647 | 0.652 | 1 | 21 |
| cox6b1 | 0.20503 | -0.53497755 | 0.118 | 0.348 | 1 | 21 |
| sept2 | 0.20671 | -0.49656269 | 0.176 | 0.348 | 1 | 21 |
| rpl14 | 0.20712 | -0.20809214 | 0.882 | 0.957 | 1 | 21 |
| ier3ip1 | 0.20988 | -0.39934553 | 0.059 | 0.261 | 1 | 21 |
| eif3k | 0.21134 | -0.44554195 | 0.059 | 0.261 | 1 | 21 |
| FO393424.2 | 0.21731 | -0.46881252 | 0.059 | 0.261 | 1 | 21 |
| si:rp71-68n21.9 | 0.21798 | -0.41811546 | 0.059 | 0.261 | 1 | 21 |
| mif | 0.21814 | -0.50829091 | 0.118 | 0.348 | 1 | 21 |
| cib1 | 0.21989 | -0.40016441 | 0.059 | 0.261 | 1 | 21 |
| mt-nd3 | 0.21993 | -0.43760278 | 0.118 | 0.348 | 1 | 21 |
| gnai2a | 0.22008 | -0.41863995 | 0.059 | 0.261 | 1 | 21 |
| cd81a | 0.22037 | -0.40491989 | 0.059 | 0.261 | 1 | 21 |
| arpc4l | 0.22584 | -0.28573203 | 0.706 | 0.913 | 1 | 21 |
| fam49ba | 0.22624 | -0.27377757 | 0.588 | 0.826 | 1 | 21 |
| copz1 | 0.23177 | 0.391721583 | 0.353 | 0.13 | 1 | 21 |
| psap | 0.23187 | -0.46267742 | 0.118 | 0.348 | 1 | 21 |
| fosab | 0.23422 | -0.29330821 | 0.118 | 0.304 | 1 | 21 |
| zgc:162730 | 0.2375 | -0.22807231 | 0.882 | 0.957 | 1 | 21 |
| hcls1 | 0.23769 | -0.35657391 | 0.529 | 0.783 | 1 | 21 |
| rpl21 | 0.23972 | -0.16019831 | 0.941 | 1 | 1 | 21 |
| eif3ha | 0.24063 | 0.378809889 | 0.353 | 0.13 | 1 | 21 |

| | | | | | | |
|---|---|---|---|---|---|---|
| emc10 | 0.24359 | -0.13580776 | 0.059 | 0.174 | 1 | 21 |
| psma1 | 0.24474 | 0.3839137 | 0.353 | 0.13 | 1 | 21 |
| cox7c | 0.24522 | -0.35124554 | 0.529 | 0.652 | 1 | 21 |
| ran | 0.24906 | -0.29401094 | 0.588 | 0.826 | 1 | 21 |
| rpl27a | 0.25004 | -0.32204564 | 0.588 | 0.826 | 1 | 21 |
| XLOC-032002 | 0.25012 | -0.29011707 | 0.059 | 0.217 | 1 | 21 |
| arhgdig | 0.25248 | -0.3206611 | 0.588 | 0.826 | 1 | 21 |
| CR383676.1 | 0.25329 | 0.736035939 | 0.176 | 0.087 | 1 | 21 |
| psme1 | 0.25787 | -0.28537806 | 0.118 | 0.304 | 1 | 21 |
| CU972454.4 | 0.26874 | 0.342661797 | 0.176 | 0.13 | 1 | 21 |
| uba52 | 0.26927 | -0.11596045 | 0.706 | 0.87 | 1 | 21 |
| abracl | 0.26935 | -0.22287831 | 0.412 | 0.609 | 1 | 21 |
| nop53 | 0.27031 | -0.3241669 | 0.176 | 0.391 | 1 | 21 |
| CR381673.1 | 0.27082 | -0.24209788 | 0.353 | 0.565 | 1 | 21 |
| calm2a | 0.27302 | 0.214286745 | 0.353 | 0.174 | 1 | 21 |
| fcer1g | 0.27493 | -0.41152905 | 0.353 | 0.609 | 1 | 21 |
| tmc6a | 0.27617 | -0.28623629 | 0.059 | 0.217 | 1 | 21 |
| rpl13 | 0.27664 | -0.1007254 | 0.824 | 0.957 | 1 | 21 |
| BX908782.3 | 0.27829 | 0.129176194 | 0.059 | 0 | 1 | 21 |
| CABZ01078055.1 | 0.27829 | 0.126650324 | 0.059 | 0 | 1 | 21 |
| CR293507.1 | 0.27829 | 0.133717334 | 0.059 | 0 | 1 | 21 |
| CR450785.1 | 0.27829 | 0.178213357 | 0.059 | 0 | 1 | 21 |
| CR450808.1 | 0.27829 | 0.102279354 | 0.059 | 0 | 1 | 21 |
| CT025690.1 | 0.27829 | 0.122246398 | 0.059 | 0 | 1 | 21 |
| CZQB01141835.1 | 0.27829 | 0.127466213 | 0.059 | 0 | 1 | 21 |
| GNAZ | 0.27829 | 0.121155577 | 0.059 | 0 | 1 | 21 |
| LO018102.1 | 0.27829 | 0.219341878 | 0.059 | 0 | 1 | 21 |
| LO018176.1 | 0.27829 | 0.1316214 | 0.059 | 0 | 1 | 21 |
| LOC100007100 | 0.27829 | 0.139441619 | 0.059 | 0 | 1 | 21 |
| LOC100333241 | 0.27829 | 0.213329179 | 0.059 | 0 | 1 | 21 |
| LOC101882159 | 0.27829 | 0.275526126 | 0.059 | 0 | 1 | 21 |
| LOC101884253 | 0.27829 | 0.139441619 | 0.059 | 0 | 1 | 21 |
| LOC110438970 | 0.27829 | 0.108135583 | 0.059 | 0 | 1 | 21 |
| LOC110439829 | 0.27829 | 0.139441619 | 0.059 | 0 | 1 | 21 |
| LOC110440146 | 0.27829 | 0.118955607 | 0.059 | 0 | 1 | 21 |
| LOC793824 | 0.27829 | 0.117884299 | 0.059 | 0 | 1 | 21 |
| MCMDC2 | 0.27829 | 0.12982888 | 0.059 | 0 | 1 | 21 |
| XLOC-000946 | 0.27829 | 0.137223952 | 0.059 | 0 | 1 | 21 |
| XLOC-008119 | 0.27829 | 0.171149442 | 0.059 | 0 | 1 | 21 |
| XLOC-013197 | 0.27829 | 0.275526126 | 0.059 | 0 | 1 | 21 |
| XLOC-013854 | 0.27829 | 0.275526126 | 0.059 | 0 | 1 | 21 |
| XLOC-041980 | 0.27829 | 0.136698203 | 0.059 | 0 | 1 | 21 |
| aamdc | 0.27829 | 0.118677461 | 0.059 | 0 | 1 | 21 |
| abca1a | 0.27829 | 0.217807007 | 0.059 | 0 | 1 | 21 |
| abcg1 | 0.27829 | 0.222356983 | 0.059 | 0 | 1 | 21 |
| abhd2a | 0.27829 | 0.115761704 | 0.059 | 0 | 1 | 21 |
| acadl | 0.27829 | 0.137059661 | 0.059 | 0 | 1 | 21 |
| acbd3 | 0.27829 | 0.178535686 | 0.059 | 0 | 1 | 21 |

| | | | | | | |
|---|---|---|---|---|---|---|
| acin1a | 0.27829 | 0.13254224 | 0.059 | 0 | 1 | 21 |
| acsbg2 | 0.27829 | 0.123750279 | 0.059 | 0 | 1 | 21 |
| acsl4a | 0.27829 | 0.129669334 | 0.059 | 0 | 1 | 21 |
| adam10a | 0.27829 | 0.274801609 | 0.059 | 0 | 1 | 21 |
| adipor1a | 0.27829 | 0.178535335 | 0.059 | 0 | 1 | 21 |
| adipor1b | 0.27829 | 0.120280841 | 0.059 | 0 | 1 | 21 |
| aes | 0.27829 | 0.219351687 | 0.059 | 0 | 1 | 21 |
| afg3l2 | 0.27829 | 0.182250218 | 0.059 | 0 | 1 | 21 |
| akt2l | 0.27829 | 0.108686451 | 0.059 | 0 | 1 | 21 |
| akt3b | 0.27829 | 0.275526126 | 0.059 | 0 | 1 | 21 |
| aldh6a1 | 0.27829 | 0.112472076 | 0.059 | 0 | 1 | 21 |
| alkbh4 | 0.27829 | 0.182495816 | 0.059 | 0 | 1 | 21 |
| alyref | 0.27829 | 0.117983993 | 0.059 | 0 | 1 | 21 |
| anapc5 | 0.27829 | 0.124270615 | 0.059 | 0 | 1 | 21 |
| ap1g1 | 0.27829 | 0.113595651 | 0.059 | 0 | 1 | 21 |
| ap3b1a | 0.27829 | 0.274539601 | 0.059 | 0 | 1 | 21 |
| aqr | 0.27829 | 0.137418257 | 0.059 | 0 | 1 | 21 |
| arhgap31 | 0.27829 | 0.213329179 | 0.059 | 0 | 1 | 21 |
| arhgef12a | 0.27829 | 0.180607872 | 0.059 | 0 | 1 | 21 |
| arid3b | 0.27829 | 0.122246398 | 0.059 | 0 | 1 | 21 |
| arl4aa | 0.27829 | 0.150325809 | 0.059 | 0 | 1 | 21 |
| atf7ip | 0.27829 | 0.115017081 | 0.059 | 0 | 1 | 21 |
| atg101 | 0.27829 | 0.27386962 | 0.059 | 0 | 1 | 21 |
| atp2a2b | 0.27829 | 0.135994231 | 0.059 | 0 | 1 | 21 |
| atxn2l | 0.27829 | 0.275178761 | 0.059 | 0 | 1 | 21 |
| bap1 | 0.27829 | 0.222356983 | 0.059 | 0 | 1 | 21 |
| bcat2 | 0.27829 | 0.120748125 | 0.059 | 0 | 1 | 21 |
| bcl7ba | 0.27829 | 0.176978663 | 0.059 | 0 | 1 | 21 |
| bicdl2 | 0.27829 | 0.275526126 | 0.059 | 0 | 1 | 21 |
| birc6 | 0.27829 | 0.166090008 | 0.059 | 0 | 1 | 21 |
| bmi1b | 0.27829 | 0.126211206 | 0.059 | 0 | 1 | 21 |
| borcs7 | 0.27829 | 0.178469689 | 0.059 | 0 | 1 | 21 |
| brd4 | 0.27829 | 0.138459535 | 0.059 | 0 | 1 | 21 |
| bri3 | 0.27829 | 0.116097506 | 0.059 | 0 | 1 | 21 |
| btf3l4 | 0.27829 | 0.104799576 | 0.059 | 0 | 1 | 21 |
| btr31 | 0.27829 | 0.139441619 | 0.059 | 0 | 1 | 21 |
| c6ast1 | 0.27829 | 0.139441619 | 0.059 | 0 | 1 | 21 |
| catip | 0.27829 | 0.124270615 | 0.059 | 0 | 1 | 21 |
| cbwd | 0.27829 | 0.173657152 | 0.059 | 0 | 1 | 21 |
| cdc14b | 0.27829 | 0.118099415 | 0.059 | 0 | 1 | 21 |
| cdc26 | 0.27829 | 0.175609423 | 0.059 | 0 | 1 | 21 |
| cdv3 | 0.27829 | 0.168397763 | 0.059 | 0 | 1 | 21 |
| cebpg | 0.27829 | 0.107478533 | 0.059 | 0 | 1 | 21 |
| chchd6b | 0.27829 | 0.11049876 | 0.059 | 0 | 1 | 21 |
| chm | 0.27829 | 0.176700377 | 0.059 | 0 | 1 | 21 |
| chmp5b | 0.27829 | 0.177781262 | 0.059 | 0 | 1 | 21 |
| chtopa | 0.27829 | 0.11356224 | 0.059 | 0 | 1 | 21 |
| clasrp | 0.27829 | 0.118098307 | 0.059 | 0 | 1 | 21 |

| | | | | | | |
|---|---|---|---|---|---|---|
| cldn12 | 0.27829 | 0.123393518 | 0.059 | 0 | 1 | 21 |
| clic5b | 0.27829 | 0.120138264 | 0.059 | 0 | 1 | 21 |
| clptm1l | 0.27829 | 0.125992923 | 0.059 | 0 | 1 | 21 |
| comta | 0.27829 | 0.182495816 | 0.059 | 0 | 1 | 21 |
| cpne3 | 0.27829 | 0.123342661 | 0.059 | 0 | 1 | 21 |
| csgalnact1a | 0.27829 | 0.274687929 | 0.059 | 0 | 1 | 21 |
| csnk1g2b | 0.27829 | 0.120836946 | 0.059 | 0 | 1 | 21 |
| csnk2a2a | 0.27829 | 0.120138264 | 0.059 | 0 | 1 | 21 |
| ctdp1 | 0.27829 | 0.267948224 | 0.059 | 0 | 1 | 21 |
| ctps1a | 0.27829 | 0.11134532 | 0.059 | 0 | 1 | 21 |
| ctsz | 0.27829 | 0.134751359 | 0.059 | 0 | 1 | 21 |
| cyth4a | 0.27829 | 0.165448812 | 0.059 | 0 | 1 | 21 |
| dbn1 | 0.27829 | 0.126686894 | 0.059 | 0 | 1 | 21 |
| dbpb | 0.27829 | 0.110732462 | 0.059 | 0 | 1 | 21 |
| dctn1a | 0.27829 | 0.137361929 | 0.059 | 0 | 1 | 21 |
| ddi2 | 0.27829 | 0.138081866 | 0.059 | 0 | 1 | 21 |
| dhps | 0.27829 | 0.11582183 | 0.059 | 0 | 1 | 21 |
| dhrs13a.1 | 0.27829 | 0.274297654 | 0.059 | 0 | 1 | 21 |
| dkk2 | 0.27829 | 0.275526126 | 0.059 | 0 | 1 | 21 |
| dnajb1b | 0.27829 | 0.202955141 | 0.059 | 0 | 1 | 21 |
| dnajc5ab | 0.27829 | 0.122246398 | 0.059 | 0 | 1 | 21 |
| dock11 | 0.27829 | 0.115761704 | 0.059 | 0 | 1 | 21 |
| dot1l | 0.27829 | 0.117531581 | 0.059 | 0 | 1 | 21 |
| dtwd1 | 0.27829 | 0.108605088 | 0.059 | 0 | 1 | 21 |
| dusp5 | 0.27829 | 0.135310214 | 0.059 | 0 | 1 | 21 |
| dut | 0.27829 | 0.165815769 | 0.059 | 0 | 1 | 21 |
| eaf2 | 0.27829 | 0.116559067 | 0.059 | 0 | 1 | 21 |
| eci1 | 0.27829 | 0.175458165 | 0.059 | 0 | 1 | 21 |
| eef2k | 0.27829 | 0.182844792 | 0.059 | 0 | 1 | 21 |
| eif4e2rs1 | 0.27829 | 0.133788316 | 0.059 | 0 | 1 | 21 |
| eipr1 | 0.27829 | 0.111293069 | 0.059 | 0 | 1 | 21 |
| enoph1 | 0.27829 | 0.137160301 | 0.059 | 0 | 1 | 21 |
| epsti1 | 0.27829 | 0.273826739 | 0.059 | 0 | 1 | 21 |
| exoc2 | 0.27829 | 0.130561734 | 0.059 | 0 | 1 | 21 |
| fam102bb | 0.27829 | 0.112120921 | 0.059 | 0 | 1 | 21 |
| fibpa | 0.27829 | 0.132254182 | 0.059 | 0 | 1 | 21 |
| flad1 | 0.27829 | 0.184885032 | 0.059 | 0 | 1 | 21 |
| fli1b | 0.27829 | 0.120611924 | 0.059 | 0 | 1 | 21 |
| foxo3b | 0.27829 | 0.126211592 | 0.059 | 0 | 1 | 21 |
| foxp3a | 0.27829 | 0.272689464 | 0.059 | 0 | 1 | 21 |
| gatd3a | 0.27829 | 0.103813625 | 0.059 | 0 | 1 | 21 |
| gemin7 | 0.27829 | 0.110621932 | 0.059 | 0 | 1 | 21 |
| ggnbp2 | 0.27829 | 0.165914411 | 0.059 | 0 | 1 | 21 |
| gmps | 0.27829 | 0.119288495 | 0.059 | 0 | 1 | 21 |
| gpalpp1 | 0.27829 | 0.116946158 | 0.059 | 0 | 1 | 21 |
| gpat3 | 0.27829 | 0.172152426 | 0.059 | 0 | 1 | 21 |
| gtf2e1 | 0.27829 | 0.223386839 | 0.059 | 0 | 1 | 21 |
| gtf2f1 | 0.27829 | 0.273806818 | 0.059 | 0 | 1 | 21 |

| | | | | | | |
|---|---|---|---|---|---|---|
| hirip3 | 0.27829 | 0.113336462 | 0.059 | 0 | 1 | 21 |
| hmgcl | 0.27829 | 0.219918534 | 0.059 | 0 | 1 | 21 |
| im:6904045 | 0.27829 | 0.115761704 | 0.059 | 0 | 1 | 21 |
| im:7136021 | 0.27829 | 0.122246398 | 0.059 | 0 | 1 | 21 |
| ing4 | 0.27829 | 0.113831535 | 0.059 | 0 | 1 | 21 |
| inpp4aa | 0.27829 | 0.122246398 | 0.059 | 0 | 1 | 21 |
| ints13 | 0.27829 | 0.171149442 | 0.059 | 0 | 1 | 21 |
| irf2 | 0.27829 | 0.111366004 | 0.059 | 0 | 1 | 21 |
| irf2a | 0.27829 | 0.11858508 | 0.059 | 0 | 1 | 21 |
| ivd | 0.27829 | 0.179496609 | 0.059 | 0 | 1 | 21 |
| josd1 | 0.27829 | 0.216916486 | 0.059 | 0 | 1 | 21 |
| kdm6ba | 0.27829 | 0.132858179 | 0.059 | 0 | 1 | 21 |
| kmt2bb | 0.27829 | 0.16813259 | 0.059 | 0 | 1 | 21 |
| kpna1 | 0.27829 | 0.273673471 | 0.059 | 0 | 1 | 21 |
| lbr | 0.27829 | 0.12806135 | 0.059 | 0 | 1 | 21 |
| letm2 | 0.27829 | 0.132555857 | 0.059 | 0 | 1 | 21 |
| lin37 | 0.27829 | 0.131964605 | 0.059 | 0 | 1 | 21 |
| lsm14aa | 0.27829 | 0.132803481 | 0.059 | 0 | 1 | 21 |
| ltv1 | 0.27829 | 0.212733459 | 0.059 | 0 | 1 | 21 |
| malt3 | 0.27829 | 0.16313589 | 0.059 | 0 | 1 | 21 |
| map3k7 | 0.27829 | 0.13567236 | 0.059 | 0 | 1 | 21 |
| matr3l1.1 | 0.27829 | 0.110198699 | 0.059 | 0 | 1 | 21 |
| mcl1b | 0.27829 | 0.168168684 | 0.059 | 0 | 1 | 21 |
| mcm5 | 0.27829 | 0.163274408 | 0.059 | 0 | 1 | 21 |
| med24 | 0.27829 | 0.12557055 | 0.059 | 0 | 1 | 21 |
| med27 | 0.27829 | 0.215244073 | 0.059 | 0 | 1 | 21 |
| mfap1 | 0.27829 | 0.114337651 | 0.059 | 0 | 1 | 21 |
| mmab | 0.27829 | 0.129465746 | 0.059 | 0 | 1 | 21 |
| mon1bb | 0.27829 | 0.139441619 | 0.059 | 0 | 1 | 21 |
| mpped2a | 0.27829 | 0.132233789 | 0.059 | 0 | 1 | 21 |
| mrps18a | 0.27829 | 0.131546817 | 0.059 | 0 | 1 | 21 |
| msra | 0.27829 | 0.218219348 | 0.059 | 0 | 1 | 21 |
| mtch2 | 0.27829 | 0.133692283 | 0.059 | 0 | 1 | 21 |
| mylipa | 0.27829 | 0.154141807 | 0.059 | 0 | 1 | 21 |
| myo19 | 0.27829 | 0.139441619 | 0.059 | 0 | 1 | 21 |
| naa15a | 0.27829 | 0.117611132 | 0.059 | 0 | 1 | 21 |
| naa30 | 0.27829 | 0.114840937 | 0.059 | 0 | 1 | 21 |
| nde1 | 0.27829 | 0.133133563 | 0.059 | 0 | 1 | 21 |
| ndufv2 | 0.27829 | 0.108970661 | 0.059 | 0 | 1 | 21 |
| nob1 | 0.27829 | 0.100837883 | 0.059 | 0 | 1 | 21 |
| nop2 | 0.27829 | 0.114452455 | 0.059 | 0 | 1 | 21 |
| nutf2l | 0.27829 | 0.126432014 | 0.059 | 0 | 1 | 21 |
| ogfrl1 | 0.27829 | 0.184885032 | 0.059 | 0 | 1 | 21 |
| parn | 0.27829 | 0.221655625 | 0.059 | 0 | 1 | 21 |
| pbxip1b | 0.27829 | 0.160267959 | 0.059 | 0 | 1 | 21 |
| pdcd4a | 0.27829 | 0.112458085 | 0.059 | 0 | 1 | 21 |
| pde7a | 0.27829 | 0.200289382 | 0.059 | 0 | 1 | 21 |
| pdlim5b | 0.27829 | 0.120284477 | 0.059 | 0 | 1 | 21 |

| | | | | | | |
|---|---|---|---|---|---|---|
| perp | 0.27829 | 0.131392597 | 0.059 | 0 | 1 | 21 |
| pex3 | 0.27829 | 0.118026187 | 0.059 | 0 | 1 | 21 |
| phf8 | 0.27829 | 0.116521101 | 0.059 | 0 | 1 | 21 |
| pigbos1 | 0.27829 | 0.102674597 | 0.059 | 0 | 1 | 21 |
| pigk | 0.27829 | 0.135927455 | 0.059 | 0 | 1 | 21 |
| pigu | 0.27829 | 0.122246398 | 0.059 | 0 | 1 | 21 |
| pik3cb | 0.27829 | 0.119313781 | 0.059 | 0 | 1 | 21 |
| plaa | 0.27829 | 0.114364801 | 0.059 | 0 | 1 | 21 |
| porb | 0.27829 | 0.137401554 | 0.059 | 0 | 1 | 21 |
| pou6f1 | 0.27829 | 0.128233871 | 0.059 | 0 | 1 | 21 |
| ppargc1b | 0.27829 | 0.119233822 | 0.059 | 0 | 1 | 21 |
| ppp2r5ea | 0.27829 | 0.123393288 | 0.059 | 0 | 1 | 21 |
| prdm2b | 0.27829 | 0.275526126 | 0.059 | 0 | 1 | 21 |
| prkar1aa | 0.27829 | 0.275130656 | 0.059 | 0 | 1 | 21 |
| proza | 0.27829 | 0.183796929 | 0.059 | 0 | 1 | 21 |
| prpf18 | 0.27829 | 0.12818401 | 0.059 | 0 | 1 | 21 |
| prpf3 | 0.27829 | 0.274540171 | 0.059 | 0 | 1 | 21 |
| prrc1 | 0.27829 | 0.121044768 | 0.059 | 0 | 1 | 21 |
| ptcd1 | 0.27829 | 0.107010011 | 0.059 | 0 | 1 | 21 |
| ptpn11a | 0.27829 | 0.135738595 | 0.059 | 0 | 1 | 21 |
| pxdc1b | 0.27829 | 0.116934164 | 0.059 | 0 | 1 | 21 |
| rab2a | 0.27829 | 0.105643571 | 0.059 | 0 | 1 | 21 |
| rap2b | 0.27829 | 0.275348445 | 0.059 | 0 | 1 | 21 |
| rbm19 | 0.27829 | 0.131220089 | 0.059 | 0 | 1 | 21 |
| rbm28 | 0.27829 | 0.112677768 | 0.059 | 0 | 1 | 21 |
| rnaseh2b | 0.27829 | 0.273674999 | 0.059 | 0 | 1 | 21 |
| rnf167 | 0.27829 | 0.120334284 | 0.059 | 0 | 1 | 21 |
| rpusd4 | 0.27829 | 0.122246398 | 0.059 | 0 | 1 | 21 |
| rrm2b | 0.27829 | 0.121861196 | 0.059 | 0 | 1 | 21 |
| safb | 0.27829 | 0.135201296 | 0.059 | 0 | 1 | 21 |
| sde2 | 0.27829 | 0.209092093 | 0.059 | 0 | 1 | 21 |
| sec23ip | 0.27829 | 0.139441619 | 0.059 | 0 | 1 | 21 |
| sf3b2 | 0.27829 | 0.11503136 | 0.059 | 0 | 1 | 21 |
| sf3b4 | 0.27829 | 0.218427256 | 0.059 | 0 | 1 | 21 |
| shq1 | 0.27829 | 0.120138264 | 0.059 | 0 | 1 | 21 |
| si:ch1073-296d18.1 | 0.27829 | 0.270640403 | 0.059 | 0 | 1 | 21 |
| si:ch211-195b11.3 | 0.27829 | 0.235173729 | 0.059 | 0 | 1 | 21 |
| si:ch211-212o1.2 | 0.27829 | 0.222356983 | 0.059 | 0 | 1 | 21 |
| si:ch211-269k10.2 | 0.27829 | 0.122246398 | 0.059 | 0 | 1 | 21 |
| si:ch211-276i12.11 | 0.27829 | 0.177122416 | 0.059 | 0 | 1 | 21 |
| si:ch211-66k16.27 | 0.27829 | 0.17885432 | 0.059 | 0 | 1 | 21 |
| si:ch73-25f10.6 | 0.27829 | 0.115761704 | 0.059 | 0 | 1 | 21 |
| si:ch73-91k6.2 | 0.27829 | 0.116193317 | 0.059 | 0 | 1 | 21 |
| si:dkey-106l3.7 | 0.27829 | 0.172864414 | 0.059 | 0 | 1 | 21 |
| si:dkey-111e8.5 | 0.27829 | 0.122246398 | 0.059 | 0 | 1 | 21 |
| si:dkey-118j18.2 | 0.27829 | 0.108446874 | 0.059 | 0 | 1 | 21 |
| si:dkey-159f12.2 | 0.27829 | 0.121479283 | 0.059 | 0 | 1 | 21 |
| si:dkey-169i5.4 | 0.27829 | 0.122246398 | 0.059 | 0 | 1 | 21 |

| | | | | | | |
|---|---|---|---|---|---|---|
| si:dkey-237j10.2 | 0.27829 | 0.182871978 | 0.059 | 0 | 1 | 21 |
| si:dkey-46a10.3 | 0.27829 | 0.115988156 | 0.059 | 0 | 1 | 21 |
| si:dkey-56e3.3 | 0.27829 | 0.176027808 | 0.059 | 0 | 1 | 21 |
| si:dkey-72l14.4 | 0.27829 | 0.122771422 | 0.059 | 0 | 1 | 21 |
| si:dkey-92f12.2 | 0.27829 | 0.275526126 | 0.059 | 0 | 1 | 21 |
| si:dkeyp-69e1.8 | 0.27829 | 0.123842627 | 0.059 | 0 | 1 | 21 |
| sik1 | 0.27829 | 0.182359356 | 0.059 | 0 | 1 | 21 |
| ska2 | 0.27829 | 0.274787801 | 0.059 | 0 | 1 | 21 |
| slain2 | 0.27829 | 0.119944509 | 0.059 | 0 | 1 | 21 |
| slc39a9 | 0.27829 | 0.127973102 | 0.059 | 0 | 1 | 21 |
| slc8a2a | 0.27829 | 0.222356983 | 0.059 | 0 | 1 | 21 |
| slc9a3r1a | 0.27829 | 0.152363964 | 0.059 | 0 | 1 | 21 |
| slco5a1b | 0.27829 | 0.122246398 | 0.059 | 0 | 1 | 21 |
| smad3b | 0.27829 | 0.179502471 | 0.059 | 0 | 1 | 21 |
| smarcb1b | 0.27829 | 0.274207346 | 0.059 | 0 | 1 | 21 |
| snip1 | 0.27829 | 0.113691941 | 0.059 | 0 | 1 | 21 |
| snn | 0.27829 | 0.275526126 | 0.059 | 0 | 1 | 21 |
| snrkb | 0.27829 | 0.163570186 | 0.059 | 0 | 1 | 21 |
| snx1a | 0.27829 | 0.117741064 | 0.059 | 0 | 1 | 21 |
| spop | 0.27829 | 0.181993658 | 0.059 | 0 | 1 | 21 |
| srp72 | 0.27829 | 0.129050116 | 0.059 | 0 | 1 | 21 |
| srsf10b | 0.27829 | 0.128748868 | 0.059 | 0 | 1 | 21 |
| stim2b | 0.27829 | 0.176342188 | 0.059 | 0 | 1 | 21 |
| sucla2 | 0.27829 | 0.177201997 | 0.059 | 0 | 1 | 21 |
| surf1 | 0.27829 | 0.181238004 | 0.059 | 0 | 1 | 21 |
| susd6 | 0.27829 | 0.174003178 | 0.059 | 0 | 1 | 21 |
| syt1a | 0.27829 | 0.182495816 | 0.059 | 0 | 1 | 21 |
| szrd1 | 0.27829 | 0.115513076 | 0.059 | 0 | 1 | 21 |
| tbc1d25 | 0.27829 | 0.132173631 | 0.059 | 0 | 1 | 21 |
| tfdp1a | 0.27829 | 0.197128812 | 0.059 | 0 | 1 | 21 |
| tfe3a | 0.27829 | 0.137280859 | 0.059 | 0 | 1 | 21 |
| timmdc1 | 0.27829 | 0.113527127 | 0.059 | 0 | 1 | 21 |
| tiparp | 0.27829 | 0.180791053 | 0.059 | 0 | 1 | 21 |
| tlr19 | 0.27829 | 0.139441619 | 0.059 | 0 | 1 | 21 |
| tmem150ab | 0.27829 | 0.178213357 | 0.059 | 0 | 1 | 21 |
| tmem176l.1 | 0.27829 | 0.205376966 | 0.059 | 0 | 1 | 21 |
| tmem263 | 0.27829 | 0.114453519 | 0.059 | 0 | 1 | 21 |
| tmem266 | 0.27829 | 0.178213357 | 0.059 | 0 | 1 | 21 |
| tmem35 | 0.27829 | 0.182495816 | 0.059 | 0 | 1 | 21 |
| tmem41b | 0.27829 | 0.12818401 | 0.059 | 0 | 1 | 21 |
| tmem56a | 0.27829 | 0.274216855 | 0.059 | 0 | 1 | 21 |
| tnrc6c1 | 0.27829 | 0.109304904 | 0.059 | 0 | 1 | 21 |
| tpd52l1 | 0.27829 | 0.182980508 | 0.059 | 0 | 1 | 21 |
| tpd52l2a | 0.27829 | 0.128496029 | 0.059 | 0 | 1 | 21 |
| traf1 | 0.27829 | 0.13486007 | 0.059 | 0 | 1 | 21 |
| trappc9 | 0.27829 | 0.12818401 | 0.059 | 0 | 1 | 21 |
| tspan31 | 0.27829 | 0.269280126 | 0.059 | 0 | 1 | 21 |
| twist1a | 0.27829 | 0.163570186 | 0.059 | 0 | 1 | 21 |

| | | | | | | |
|---|---|---|---|---|---|---|
| txn | 0.27829 | 0.272011575 | 0.059 | 0 | 1 | 21 |
| txndc17 | 0.27829 | 0.100870812 | 0.059 | 0 | 1 | 21 |
| u2surp | 0.27829 | 0.178045759 | 0.059 | 0 | 1 | 21 |
| ube2al | 0.27829 | 0.109089106 | 0.059 | 0 | 1 | 21 |
| ube2z | 0.27829 | 0.274846096 | 0.059 | 0 | 1 | 21 |
| ube3c | 0.27829 | 0.21798966 | 0.059 | 0 | 1 | 21 |
| ubtd1a | 0.27829 | 0.184885032 | 0.059 | 0 | 1 | 21 |
| unm-hu7910 | 0.27829 | 0.179599506 | 0.059 | 0 | 1 | 21 |
| usf2 | 0.27829 | 0.171393684 | 0.059 | 0 | 1 | 21 |
| usp42 | 0.27829 | 0.118944223 | 0.059 | 0 | 1 | 21 |
| vaspb | 0.27829 | 0.132238319 | 0.059 | 0 | 1 | 21 |
| vps26b | 0.27829 | 0.125322005 | 0.059 | 0 | 1 | 21 |
| wdr4 | 0.27829 | 0.139441619 | 0.059 | 0 | 1 | 21 |
| wdr62 | 0.27829 | 0.124270615 | 0.059 | 0 | 1 | 21 |
| wdr89 | 0.27829 | 0.11560839 | 0.059 | 0 | 1 | 21 |
| xiap | 0.27829 | 0.112232632 | 0.059 | 0 | 1 | 21 |
| xkrx | 0.27829 | 0.117840208 | 0.059 | 0 | 1 | 21 |
| yif1a | 0.27829 | 0.105406915 | 0.059 | 0 | 1 | 21 |
| zfand2a | 0.27829 | 0.18136556 | 0.059 | 0 | 1 | 21 |
| zgc:100832 | 0.27829 | 0.110503228 | 0.059 | 0 | 1 | 21 |
| zgc:101679 | 0.27829 | 0.126486906 | 0.059 | 0 | 1 | 21 |
| zgc:112496 | 0.27829 | 0.126013086 | 0.059 | 0 | 1 | 21 |
| zgc:152774 | 0.27829 | 0.137890002 | 0.059 | 0 | 1 | 21 |
| zgc:153993 | 0.27829 | 0.198478772 | 0.059 | 0 | 1 | 21 |
| zgc:162200 | 0.27829 | 0.12818401 | 0.059 | 0 | 1 | 21 |
| zgc:172106 | 0.27829 | 0.274920871 | 0.059 | 0 | 1 | 21 |
| zgc:174268 | 0.27829 | 0.118729167 | 0.059 | 0 | 1 | 21 |
| zgc:175264 | 0.27829 | 0.138608783 | 0.059 | 0 | 1 | 21 |
| zgc:55733 | 0.27829 | 0.217223824 | 0.059 | 0 | 1 | 21 |
| zgc:56095 | 0.27829 | 0.128835273 | 0.059 | 0 | 1 | 21 |
| zgc:56304 | 0.27829 | 0.172193047 | 0.059 | 0 | 1 | 21 |
| zgc:66388 | 0.27829 | 0.137380702 | 0.059 | 0 | 1 | 21 |
| zgc:92907 | 0.27829 | 0.274317136 | 0.059 | 0 | 1 | 21 |
| znf1178 | 0.27829 | 0.12818401 | 0.059 | 0 | 1 | 21 |
| znf296 | 0.27829 | 0.115337717 | 0.059 | 0 | 1 | 21 |
| zpr1 | 0.27829 | 0.151836371 | 0.059 | 0 | 1 | 21 |
| AL840638.1 | 0.27829 | 0.12958591 | 0.059 | 0 | 1 | 21 |
| BX571680.1 | 0.27829 | 0.203928488 | 0.059 | 0 | 1 | 21 |
| CABZ01039096.1 | 0.27829 | 0.148041387 | 0.059 | 0 | 1 | 21 |
| CABZ01078607.1 | 0.27829 | 0.167015163 | 0.059 | 0 | 1 | 21 |
| CABZ01083448.1 | 0.27829 | 0.101319041 | 0.059 | 0 | 1 | 21 |
| CABZ01088109.1 | 0.27829 | 0.129321947 | 0.059 | 0 | 1 | 21 |
| CABZ01088274.1 | 0.27829 | 0.149187066 | 0.059 | 0 | 1 | 21 |
| CABZ01092156.1 | 0.27829 | 0.248088348 | 0.059 | 0 | 1 | 21 |
| CABZ01102170.1 | 0.27829 | 0.164954976 | 0.059 | 0 | 1 | 21 |
| CABZ01112656.1 | 0.27829 | 0.134647855 | 0.059 | 0 | 1 | 21 |
| CDK2AP1 | 0.27829 | 0.177372848 | 0.059 | 0 | 1 | 21 |
| CR382370.2 | 0.27829 | 0.202533818 | 0.059 | 0 | 1 | 21 |

| | | | | | | |
|---|---|---|---|---|---|---|
| CR383662.3 | 0.27829 | 0.167015163 | 0.059 | 0 | 1 | 21 |
| CR387919.1 | 0.27829 | 0.253295085 | 0.059 | 0 | 1 | 21 |
| CR847844.1 | 0.27829 | 0.196512165 | 0.059 | 0 | 1 | 21 |
| CU659670.1 | 0.27829 | 0.161647544 | 0.059 | 0 | 1 | 21 |
| CU694481.1 | 0.27829 | 0.253295085 | 0.059 | 0 | 1 | 21 |
| HUS1B | 0.27829 | 0.162800982 | 0.059 | 0 | 1 | 21 |
| LOC101882640 | 0.27829 | 0.132596887 | 0.059 | 0 | 1 | 21 |
| LOC101882843 | 0.27829 | 0.147563014 | 0.059 | 0 | 1 | 21 |
| LOC101883913 | 0.27829 | 0.163466795 | 0.059 | 0 | 1 | 21 |
| LOC101884710 | 0.27829 | 0.152165215 | 0.059 | 0 | 1 | 21 |
| LOC103910438 | 0.27829 | 0.167961439 | 0.059 | 0 | 1 | 21 |
| LOC108179036 | 0.27829 | 0.252087767 | 0.059 | 0 | 1 | 21 |
| LOC110438236 | 0.27829 | 0.203928488 | 0.059 | 0 | 1 | 21 |
| MZT1 | 0.27829 | 0.117867756 | 0.059 | 0 | 1 | 21 |
| RALGDS | 0.27829 | 0.160293657 | 0.059 | 0 | 1 | 21 |
| RPL37A | 0.27829 | 0.127189979 | 0.059 | 0 | 1 | 21 |
| TMEM19 | 0.27829 | 0.161840524 | 0.059 | 0 | 1 | 21 |
| XLOC-005448 | 0.27829 | 0.165189769 | 0.059 | 0 | 1 | 21 |
| XLOC-006268 | 0.27829 | 0.201726351 | 0.059 | 0 | 1 | 21 |
| XLOC-011085 | 0.27829 | 0.2526997 | 0.059 | 0 | 1 | 21 |
| XLOC-013964 | 0.27829 | 0.299707388 | 0.059 | 0 | 1 | 21 |
| XLOC-028370 | 0.27829 | 0.164012613 | 0.059 | 0 | 1 | 21 |
| XLOC-032256 | 0.27829 | 0.253295085 | 0.059 | 0 | 1 | 21 |
| XLOC-032772 | 0.27829 | 0.132596887 | 0.059 | 0 | 1 | 21 |
| XLOC-033809 | 0.27829 | 0.249421143 | 0.059 | 0 | 1 | 21 |
| XLOC-042868 | 0.27829 | 0.203928488 | 0.059 | 0 | 1 | 21 |
| XLOC-043861 | 0.27829 | 0.251905864 | 0.059 | 0 | 1 | 21 |
| abcf1 | 0.27829 | 0.120737064 | 0.059 | 0 | 1 | 21 |
| aclya | 0.27829 | 0.130308178 | 0.059 | 0 | 1 | 21 |
| actr1 | 0.27829 | 0.252395472 | 0.059 | 0 | 1 | 21 |
| adar | 0.27829 | 0.129598546 | 0.059 | 0 | 1 | 21 |
| agpat5 | 0.27829 | 0.201891139 | 0.059 | 0 | 1 | 21 |
| aif1l | 0.27829 | 0.164668952 | 0.059 | 0 | 1 | 21 |
| ajuba | 0.27829 | 0.16754844 | 0.059 | 0 | 1 | 21 |
| akt2 | 0.27829 | 0.201234609 | 0.059 | 0 | 1 | 21 |
| aldh16a1 | 0.27829 | 0.11222792 | 0.059 | 0 | 1 | 21 |
| aldh7a1 | 0.27829 | 0.100362293 | 0.059 | 0 | 1 | 21 |
| ap2b1 | 0.27829 | 0.201713107 | 0.059 | 0 | 1 | 21 |
| ap3s2 | 0.27829 | 0.11261477 | 0.059 | 0 | 1 | 21 |
| arhgap20b | 0.27829 | 0.155950248 | 0.059 | 0 | 1 | 21 |
| arl11 | 0.27829 | 0.153429089 | 0.059 | 0 | 1 | 21 |
| arl2 | 0.27829 | 0.14869322 | 0.059 | 0 | 1 | 21 |
| armh3 | 0.27829 | 0.163331495 | 0.059 | 0 | 1 | 21 |
| asap1b | 0.27829 | 0.203494545 | 0.059 | 0 | 1 | 21 |
| asnsd1 | 0.27829 | 0.145881556 | 0.059 | 0 | 1 | 21 |
| atp1a1a.1 | 0.27829 | 0.170440337 | 0.059 | 0 | 1 | 21 |
| atp2c1 | 0.27829 | 0.208742345 | 0.059 | 0 | 1 | 21 |
| atp6v0a2b | 0.27829 | 0.253295085 | 0.059 | 0 | 1 | 21 |

| | | | | | | |
|---|---|---|---|---|---|---|
| atxn1l | 0.27829 | 0.203928488 | 0.059 | 0 | 1 | 21 |
| atxn7 | 0.27829 | 0.157589042 | 0.059 | 0 | 1 | 21 |
| b3glcta | 0.27829 | 0.164794727 | 0.059 | 0 | 1 | 21 |
| b3gnt5b | 0.27829 | 0.149165009 | 0.059 | 0 | 1 | 21 |
| bach1a | 0.27829 | 0.253295085 | 0.059 | 0 | 1 | 21 |
| baz2a | 0.27829 | 0.194887621 | 0.059 | 0 | 1 | 21 |
| bckdhb | 0.27829 | 0.20477104 | 0.059 | 0 | 1 | 21 |
| bhlhe40 | 0.27829 | 0.201661919 | 0.059 | 0 | 1 | 21 |
| blk | 0.27829 | 0.16632641 | 0.059 | 0 | 1 | 21 |
| bloc1s4 | 0.27829 | 0.205971948 | 0.059 | 0 | 1 | 21 |
| bop1 | 0.27829 | 0.154441933 | 0.059 | 0 | 1 | 21 |
| brwd1 | 0.27829 | 0.157589042 | 0.059 | 0 | 1 | 21 |
| bud23 | 0.27829 | 0.16177048 | 0.059 | 0 | 1 | 21 |
| c2cd2 | 0.27829 | 0.203928488 | 0.059 | 0 | 1 | 21 |
| calr3a | 0.27829 | 0.157417644 | 0.059 | 0 | 1 | 21 |
| casp8 | 0.27829 | 0.130756625 | 0.059 | 0 | 1 | 21 |
| casp9 | 0.27829 | 0.161303925 | 0.059 | 0 | 1 | 21 |
| caspa | 0.27829 | 0.163426998 | 0.059 | 0 | 1 | 21 |
| cbr1l | 0.27829 | 0.156385909 | 0.059 | 0 | 1 | 21 |
| ccdc9 | 0.27829 | 0.152909086 | 0.059 | 0 | 1 | 21 |
| ccnh | 0.27829 | 0.109328256 | 0.059 | 0 | 1 | 21 |
| cdadc1 | 0.27829 | 0.167015163 | 0.059 | 0 | 1 | 21 |
| cdc42ep4a | 0.27829 | 0.132596887 | 0.059 | 0 | 1 | 21 |
| cel.1 | 0.27829 | 0.177972495 | 0.059 | 0 | 1 | 21 |
| cers2b | 0.27829 | 0.149040526 | 0.059 | 0 | 1 | 21 |
| chmp5a | 0.27829 | 0.16754844 | 0.059 | 0 | 1 | 21 |
| cited4b | 0.27829 | 0.12856882 | 0.059 | 0 | 1 | 21 |
| clcn7 | 0.27829 | 0.196602902 | 0.059 | 0 | 1 | 21 |
| coa7 | 0.27829 | 0.138900753 | 0.059 | 0 | 1 | 21 |
| coasy | 0.27829 | 0.132596887 | 0.059 | 0 | 1 | 21 |
| col14a1b | 0.27829 | 0.16754844 | 0.059 | 0 | 1 | 21 |
| coq10b | 0.27829 | 0.151482341 | 0.059 | 0 | 1 | 21 |
| cox19 | 0.27829 | 0.202347045 | 0.059 | 0 | 1 | 21 |
| cpb1 | 0.27829 | 0.134565676 | 0.059 | 0 | 1 | 21 |
| cptp | 0.27829 | 0.201565962 | 0.059 | 0 | 1 | 21 |
| cryz | 0.27829 | 0.14220336 | 0.059 | 0 | 1 | 21 |
| cst14b.2 | 0.27829 | 0.133639372 | 0.059 | 0 | 1 | 21 |
| ctif | 0.27829 | 0.205579644 | 0.059 | 0 | 1 | 21 |
| cux1b | 0.27829 | 0.152158561 | 0.059 | 0 | 1 | 21 |
| cxcr3.3 | 0.27829 | 0.157589042 | 0.059 | 0 | 1 | 21 |
| cyhr1 | 0.27829 | 0.194786677 | 0.059 | 0 | 1 | 21 |
| cylda | 0.27829 | 0.251942223 | 0.059 | 0 | 1 | 21 |
| ddx24 | 0.27829 | 0.125371975 | 0.059 | 0 | 1 | 21 |
| ddx3xa | 0.27829 | 0.137705333 | 0.059 | 0 | 1 | 21 |
| dennd4c | 0.27829 | 0.154233281 | 0.059 | 0 | 1 | 21 |
| dera | 0.27829 | 0.111719177 | 0.059 | 0 | 1 | 21 |
| derl3 | 0.27829 | 0.136164904 | 0.059 | 0 | 1 | 21 |
| dgkza | 0.27829 | 0.251035685 | 0.059 | 0 | 1 | 21 |

| | | | | | | |
|---|---|---|---|---|---|---|
| dnaja3a | 0.27829 | 0.14741959 | 0.059 | 0 | 1 | 21 |
| dnajc5gb | 0.27829 | 0.129011968 | 0.059 | 0 | 1 | 21 |
| dnph1 | 0.27829 | 0.14756174 | 0.059 | 0 | 1 | 21 |
| dok4 | 0.27829 | 0.132596887 | 0.059 | 0 | 1 | 21 |
| dpp7 | 0.27829 | 0.205650437 | 0.059 | 0 | 1 | 21 |
| dse | 0.27829 | 0.153254228 | 0.059 | 0 | 1 | 21 |
| dsela | 0.27829 | 0.203928488 | 0.059 | 0 | 1 | 21 |
| dtd1 | 0.27829 | 0.20116149 | 0.059 | 0 | 1 | 21 |
| dus4l | 0.27829 | 0.191645078 | 0.059 | 0 | 1 | 21 |
| dync1li1 | 0.27829 | 0.199989349 | 0.059 | 0 | 1 | 21 |
| eci2 | 0.27829 | 0.114951583 | 0.059 | 0 | 1 | 21 |
| edrf1 | 0.27829 | 0.249747188 | 0.059 | 0 | 1 | 21 |
| eef1akmt1 | 0.27829 | 0.10432716 | 0.059 | 0 | 1 | 21 |
| egln2 | 0.27829 | 0.189443778 | 0.059 | 0 | 1 | 21 |
| eif3hb | 0.27829 | 0.147774816 | 0.059 | 0 | 1 | 21 |
| eif4a2 | 0.27829 | 0.134191267 | 0.059 | 0 | 1 | 21 |
| eif4h | 0.27829 | 0.19655545 | 0.059 | 0 | 1 | 21 |
| enosf1 | 0.27829 | 0.151297949 | 0.059 | 0 | 1 | 21 |
| epb41l5 | 0.27829 | 0.148444442 | 0.059 | 0 | 1 | 21 |
| epdl2 | 0.27829 | 0.16283601 | 0.059 | 0 | 1 | 21 |
| epn2 | 0.27829 | 0.131206433 | 0.059 | 0 | 1 | 21 |
| ercc5 | 0.27829 | 0.16754844 | 0.059 | 0 | 1 | 21 |
| fam13b | 0.27829 | 0.203001276 | 0.059 | 0 | 1 | 21 |
| fam207a | 0.27829 | 0.163729198 | 0.059 | 0 | 1 | 21 |
| fam3c | 0.27829 | 0.154857149 | 0.059 | 0 | 1 | 21 |
| fbxo34 | 0.27829 | 0.202430049 | 0.059 | 0 | 1 | 21 |
| fbxo7 | 0.27829 | 0.157589042 | 0.059 | 0 | 1 | 21 |
| fbxw11a | 0.27829 | 0.153429089 | 0.059 | 0 | 1 | 21 |
| fkbp8 | 0.27829 | 0.203261033 | 0.059 | 0 | 1 | 21 |
| foxk1 | 0.27829 | 0.148167408 | 0.059 | 0 | 1 | 21 |
| ftr06 | 0.27829 | 0.201234609 | 0.059 | 0 | 1 | 21 |
| fuca2 | 0.27829 | 0.129864227 | 0.059 | 0 | 1 | 21 |
| gadd45bb | 0.27829 | 0.201204706 | 0.059 | 0 | 1 | 21 |
| gadd45gip1 | 0.27829 | 0.155992957 | 0.059 | 0 | 1 | 21 |
| galnt1 | 0.27829 | 0.201217015 | 0.059 | 0 | 1 | 21 |
| galnt6 | 0.27829 | 0.203928488 | 0.059 | 0 | 1 | 21 |
| gata3 | 0.27829 | 0.129290237 | 0.059 | 0 | 1 | 21 |
| gatad2b | 0.27829 | 0.25256658 | 0.059 | 0 | 1 | 21 |
| get4 | 0.27829 | 0.125168919 | 0.059 | 0 | 1 | 21 |
| gkap1 | 0.27829 | 0.15893842 | 0.059 | 0 | 1 | 21 |
| glod4 | 0.27829 | 0.159095237 | 0.059 | 0 | 1 | 21 |
| gnai2b | 0.27829 | 0.151508915 | 0.059 | 0 | 1 | 21 |
| gne | 0.27829 | 0.252373415 | 0.059 | 0 | 1 | 21 |
| golga5 | 0.27829 | 0.200801575 | 0.059 | 0 | 1 | 21 |
| golt1ba | 0.27829 | 0.156265429 | 0.059 | 0 | 1 | 21 |
| gpr108 | 0.27829 | 0.124104434 | 0.059 | 0 | 1 | 21 |
| gpsm2l | 0.27829 | 0.245634397 | 0.059 | 0 | 1 | 21 |
| grk3 | 0.27829 | 0.201980384 | 0.059 | 0 | 1 | 21 |

| | | | | | | |
|---|---|---|---|---|---|---|
| gsr | 0.27829 | 0.129256504 | 0.059 | 0 | 1 | 21 |
| hagh | 0.27829 | 0.151396953 | 0.059 | 0 | 1 | 21 |
| hif1al | 0.27829 | 0.157259514 | 0.059 | 0 | 1 | 21 |
| hnrnpm | 0.27829 | 0.128109252 | 0.059 | 0 | 1 | 21 |
| hnrnpua | 0.27829 | 0.128903642 | 0.059 | 0 | 1 | 21 |
| hnrpkl | 0.27829 | 0.150138219 | 0.059 | 0 | 1 | 21 |
| huwe1 | 0.27829 | 0.127503489 | 0.059 | 0 | 1 | 21 |
| hvcn1 | 0.27829 | 0.186313033 | 0.059 | 0 | 1 | 21 |
| hyal2a | 0.27829 | 0.200148876 | 0.059 | 0 | 1 | 21 |
| hyi | 0.27829 | 0.141933925 | 0.059 | 0 | 1 | 21 |
| icn | 0.27829 | 0.182912349 | 0.059 | 0 | 1 | 21 |
| id1 | 0.27829 | 0.166424674 | 0.059 | 0 | 1 | 21 |
| ift20 | 0.27829 | 0.251125652 | 0.059 | 0 | 1 | 21 |
| ik | 0.27829 | 0.194264637 | 0.059 | 0 | 1 | 21 |
| il21r.1 | 0.27829 | 0.21398314 | 0.059 | 0 | 1 | 21 |
| impdh2 | 0.27829 | 0.162085806 | 0.059 | 0 | 1 | 21 |
| inpp5f | 0.27829 | 0.189581764 | 0.059 | 0 | 1 | 21 |
| inpp5l | 0.27829 | 0.252159493 | 0.059 | 0 | 1 | 21 |
| ints12 | 0.27829 | 0.165250808 | 0.059 | 0 | 1 | 21 |
| ip6k2b | 0.27829 | 0.197823006 | 0.059 | 0 | 1 | 21 |
| ivns1abpa | 0.27829 | 0.249484215 | 0.059 | 0 | 1 | 21 |
| ivns1abpb | 0.27829 | 0.150196562 | 0.059 | 0 | 1 | 21 |
| jazf1b | 0.27829 | 0.125330228 | 0.059 | 0 | 1 | 21 |
| khnyn | 0.27829 | 0.201044887 | 0.059 | 0 | 1 | 21 |
| klhl15 | 0.27829 | 0.164156543 | 0.059 | 0 | 1 | 21 |
| kmt2cb | 0.27829 | 0.20102622 | 0.059 | 0 | 1 | 21 |
| kti12 | 0.27829 | 0.143463605 | 0.059 | 0 | 1 | 21 |
| lbh | 0.27829 | 0.163382683 | 0.059 | 0 | 1 | 21 |
| lcorl | 0.27829 | 0.252095708 | 0.059 | 0 | 1 | 21 |
| lman1 | 0.27829 | 0.126872989 | 0.059 | 0 | 1 | 21 |
| lmtk2 | 0.27829 | 0.157589042 | 0.059 | 0 | 1 | 21 |
| lnpep | 0.27829 | 0.252855131 | 0.059 | 0 | 1 | 21 |
| lrp10 | 0.27829 | 0.202674307 | 0.059 | 0 | 1 | 21 |
| lrrc29 | 0.27829 | 0.200280715 | 0.059 | 0 | 1 | 21 |
| lrrc40 | 0.27829 | 0.19510382 | 0.059 | 0 | 1 | 21 |
| luc7l | 0.27829 | 0.142028576 | 0.059 | 0 | 1 | 21 |
| map3k4 | 0.27829 | 0.162243631 | 0.059 | 0 | 1 | 21 |
| map4k4 | 0.27829 | 0.149015734 | 0.059 | 0 | 1 | 21 |
| mapk6 | 0.27829 | 0.163862323 | 0.059 | 0 | 1 | 21 |
| mau2 | 0.27829 | 0.12820681 | 0.059 | 0 | 1 | 21 |
| mbnl2 | 0.27829 | 0.203017449 | 0.059 | 0 | 1 | 21 |
| mcrip1 | 0.27829 | 0.119702232 | 0.059 | 0 | 1 | 21 |
| mecp2 | 0.27829 | 0.251440909 | 0.059 | 0 | 1 | 21 |
| med20 | 0.27829 | 0.12479761 | 0.059 | 0 | 1 | 21 |
| memo1 | 0.27829 | 0.247663883 | 0.059 | 0 | 1 | 21 |
| mettl3 | 0.27829 | 0.162656368 | 0.059 | 0 | 1 | 21 |
| mfn1b | 0.27829 | 0.198862747 | 0.059 | 0 | 1 | 21 |
| mgaa | 0.27829 | 0.253295085 | 0.059 | 0 | 1 | 21 |

| | | | | | | |
|---|---|---|---|---|---|---|
| minpp1b | 0.27829 | 0.158324829 | 0.059 | 0 | 1 | 21 |
| mlf2 | 0.27829 | 0.158452099 | 0.059 | 0 | 1 | 21 |
| mrpl41 | 0.27829 | 0.145592009 | 0.059 | 0 | 1 | 21 |
| mrpl42 | 0.27829 | 0.145146869 | 0.059 | 0 | 1 | 21 |
| mrps9 | 0.27829 | 0.201043894 | 0.059 | 0 | 1 | 21 |
| msrb1b | 0.27829 | 0.174344555 | 0.059 | 0 | 1 | 21 |
| mta2 | 0.27829 | 0.127588961 | 0.059 | 0 | 1 | 21 |
| mtfr1l | 0.27829 | 0.139448704 | 0.059 | 0 | 1 | 21 |
| mycbp2 | 0.27829 | 0.202822082 | 0.059 | 0 | 1 | 21 |
| nadk2 | 0.27829 | 0.157589042 | 0.059 | 0 | 1 | 21 |
| nap1l4a | 0.27829 | 0.14733835 | 0.059 | 0 | 1 | 21 |
| napab | 0.27829 | 0.160548118 | 0.059 | 0 | 1 | 21 |
| ndel1a | 0.27829 | 0.241871542 | 0.059 | 0 | 1 | 21 |
| neil1 | 0.27829 | 0.158852126 | 0.059 | 0 | 1 | 21 |
| nelfe | 0.27829 | 0.143393972 | 0.059 | 0 | 1 | 21 |
| nfatc3b | 0.27829 | 0.162014313 | 0.059 | 0 | 1 | 21 |
| nfya | 0.27829 | 0.158852126 | 0.059 | 0 | 1 | 21 |
| nkrf | 0.27829 | 0.160632537 | 0.059 | 0 | 1 | 21 |
| nmt2 | 0.27829 | 0.116864231 | 0.059 | 0 | 1 | 21 |
| nol9 | 0.27829 | 0.206347213 | 0.059 | 0 | 1 | 21 |
| nr1d2b | 0.27829 | 0.160503732 | 0.059 | 0 | 1 | 21 |
| nr2c2ap | 0.27829 | 0.154020807 | 0.059 | 0 | 1 | 21 |
| nt5c3a | 0.27829 | 0.122686533 | 0.059 | 0 | 1 | 21 |
| nvl | 0.27829 | 0.153429089 | 0.059 | 0 | 1 | 21 |
| ocrl | 0.27829 | 0.163331495 | 0.059 | 0 | 1 | 21 |
| ogdha | 0.27829 | 0.129235892 | 0.059 | 0 | 1 | 21 |
| optn | 0.27829 | 0.124946711 | 0.059 | 0 | 1 | 21 |
| orai1a | 0.27829 | 0.148673095 | 0.059 | 0 | 1 | 21 |
| osbpl11 | 0.27829 | 0.157589042 | 0.059 | 0 | 1 | 21 |
| oser1 | 0.27829 | 0.202246287 | 0.059 | 0 | 1 | 21 |
| pagr1 | 0.27829 | 0.160664086 | 0.059 | 0 | 1 | 21 |
| pak1ip1 | 0.27829 | 0.159320027 | 0.059 | 0 | 1 | 21 |
| park7 | 0.27829 | 0.208509326 | 0.059 | 0 | 1 | 21 |
| pbx4 | 0.27829 | 0.115430522 | 0.059 | 0 | 1 | 21 |
| pcmt | 0.27829 | 0.160830001 | 0.059 | 0 | 1 | 21 |
| pdia6 | 0.27829 | 0.118178956 | 0.059 | 0 | 1 | 21 |
| pdp2 | 0.27829 | 0.152165215 | 0.059 | 0 | 1 | 21 |
| pds5a | 0.27829 | 0.252507602 | 0.059 | 0 | 1 | 21 |
| pepd | 0.27829 | 0.201576186 | 0.059 | 0 | 1 | 21 |
| pex11a | 0.27829 | 0.152165215 | 0.059 | 0 | 1 | 21 |
| pfkpb | 0.27829 | 0.201861493 | 0.059 | 0 | 1 | 21 |
| phf20a | 0.27829 | 0.132596887 | 0.059 | 0 | 1 | 21 |
| pik3cd | 0.27829 | 0.163906572 | 0.059 | 0 | 1 | 21 |
| pir | 0.27829 | 0.139018997 | 0.059 | 0 | 1 | 21 |
| pisd | 0.27829 | 0.142827743 | 0.059 | 0 | 1 | 21 |
| pitpnbl | 0.27829 | 0.117785548 | 0.059 | 0 | 1 | 21 |
| pkig | 0.27829 | 0.210103135 | 0.059 | 0 | 1 | 21 |
| plk1 | 0.27829 | 0.249994193 | 0.059 | 0 | 1 | 21 |

| | | | | | | |
|---|---|---|---|---|---|---|
| plpp1a | 0.27829 | 0.198780738 | 0.059 | 0 | 1 | 21 |
| plxnb1a | 0.27829 | 0.203928488 | 0.059 | 0 | 1 | 21 |
| pnisr | 0.27829 | 0.195818509 | 0.059 | 0 | 1 | 21 |
| pnpo | 0.27829 | 0.153552894 | 0.059 | 0 | 1 | 21 |
| polr1d | 0.27829 | 0.1641295 | 0.059 | 0 | 1 | 21 |
| polr2i | 0.27829 | 0.123098058 | 0.059 | 0 | 1 | 21 |
| pop5 | 0.27829 | 0.191153445 | 0.059 | 0 | 1 | 21 |
| ppip5k1b | 0.27829 | 0.129316717 | 0.059 | 0 | 1 | 21 |
| ppp1r11 | 0.27829 | 0.13551383 | 0.059 | 0 | 1 | 21 |
| ppp6r3 | 0.27829 | 0.200670841 | 0.059 | 0 | 1 | 21 |
| ppt2 | 0.27829 | 0.143669608 | 0.059 | 0 | 1 | 21 |
| pptc7b | 0.27829 | 0.203928488 | 0.059 | 0 | 1 | 21 |
| prf1.7 | 0.27829 | 0.163297268 | 0.059 | 0 | 1 | 21 |
| prkar1b | 0.27829 | 0.144316977 | 0.059 | 0 | 1 | 21 |
| psmd12 | 0.27829 | 0.119698667 | 0.059 | 0 | 1 | 21 |
| ptenb | 0.27829 | 0.126947019 | 0.059 | 0 | 1 | 21 |
| ptgdsb.1 | 0.27829 | 0.122311367 | 0.059 | 0 | 1 | 21 |
| ptprq | 0.27829 | 0.167015163 | 0.059 | 0 | 1 | 21 |
| pum2 | 0.27829 | 0.201489565 | 0.059 | 0 | 1 | 21 |
| purba | 0.27829 | 0.17032291 | 0.059 | 0 | 1 | 21 |
| rab13 | 0.27829 | 0.128580771 | 0.059 | 0 | 1 | 21 |
| rab6a | 0.27829 | 0.114849335 | 0.059 | 0 | 1 | 21 |
| rab6bb | 0.27829 | 0.12329392 | 0.059 | 0 | 1 | 21 |
| rabggta | 0.27829 | 0.125800733 | 0.059 | 0 | 1 | 21 |
| rabl3 | 0.27829 | 0.129875182 | 0.059 | 0 | 1 | 21 |
| ranbp9 | 0.27829 | 0.163416834 | 0.059 | 0 | 1 | 21 |
| rap1gap2b | 0.27829 | 0.167015163 | 0.059 | 0 | 1 | 21 |
| rassf2b | 0.27829 | 0.162935921 | 0.059 | 0 | 1 | 21 |
| rbm4.2 | 0.27829 | 0.203611632 | 0.059 | 0 | 1 | 21 |
| rbm7 | 0.27829 | 0.154104909 | 0.059 | 0 | 1 | 21 |
| rcn1 | 0.27829 | 0.148041387 | 0.059 | 0 | 1 | 21 |
| rel | 0.27829 | 0.154076427 | 0.059 | 0 | 1 | 21 |
| rfxank | 0.27829 | 0.150473032 | 0.059 | 0 | 1 | 21 |
| rgs8 | 0.27829 | 0.253295085 | 0.059 | 0 | 1 | 21 |
| rhbdl2 | 0.27829 | 0.130811522 | 0.059 | 0 | 1 | 21 |
| rnaseh1 | 0.27829 | 0.161073933 | 0.059 | 0 | 1 | 21 |
| rnaseh2a | 0.27829 | 0.15626588 | 0.059 | 0 | 1 | 21 |
| rnf213a | 0.27829 | 0.164666818 | 0.059 | 0 | 1 | 21 |
| rnf40 | 0.27829 | 0.200577881 | 0.059 | 0 | 1 | 21 |
| rnf44 | 0.27829 | 0.201899525 | 0.059 | 0 | 1 | 21 |
| rpl7l1 | 0.27829 | 0.107503268 | 0.059 | 0 | 1 | 21 |
| rps6kb1a | 0.27829 | 0.116579878 | 0.059 | 0 | 1 | 21 |
| rrp9 | 0.27829 | 0.25142566 | 0.059 | 0 | 1 | 21 |
| s100a10b | 0.27829 | 0.177272229 | 0.059 | 0 | 1 | 21 |
| s1pr1 | 0.27829 | 0.149015734 | 0.059 | 0 | 1 | 21 |
| sars | 0.27829 | 0.143519855 | 0.059 | 0 | 1 | 21 |
| sash3 | 0.27829 | 0.204391784 | 0.059 | 0 | 1 | 21 |
| scp2b | 0.27829 | 0.202688769 | 0.059 | 0 | 1 | 21 |

| | | | | | | |
|---|---|---|---|---|---|---|
| sdad1 | 0.27829 | 0.201497403 | 0.059 | 0 | 1 | 21 |
| sdr16c5b | 0.27829 | 0.143672889 | 0.059 | 0 | 1 | 21 |
| sec14l1 | 0.27829 | 0.148679813 | 0.059 | 0 | 1 | 21 |
| serp1 | 0.27829 | 0.113793178 | 0.059 | 0 | 1 | 21 |
| serpina1l | 0.27829 | 0.135853973 | 0.059 | 0 | 1 | 21 |
| serpinb14 | 0.27829 | 0.153429089 | 0.059 | 0 | 1 | 21 |
| setdb2 | 0.27829 | 0.156894603 | 0.059 | 0 | 1 | 21 |
| sgk3 | 0.27829 | 0.162202954 | 0.059 | 0 | 1 | 21 |
| sh3glb1a | 0.27829 | 0.154248673 | 0.059 | 0 | 1 | 21 |
| sh3glb2b | 0.27829 | 0.203610678 | 0.059 | 0 | 1 | 21 |
| si:cabz01071911.4 | 0.27829 | 0.167015163 | 0.059 | 0 | 1 | 21 |
| si:ch211-107o10.3 | 0.27829 | 0.156872164 | 0.059 | 0 | 1 | 21 |
| si:ch211-11k18.4 | 0.27829 | 0.126055542 | 0.059 | 0 | 1 | 21 |
| si:ch211-129c21.1 | 0.27829 | 0.203928488 | 0.059 | 0 | 1 | 21 |
| si:ch211-13c6.2 | 0.27829 | 0.148444442 | 0.059 | 0 | 1 | 21 |
| si:ch211-197h24.9 | 0.27829 | 0.192262019 | 0.059 | 0 | 1 | 21 |
| si:ch211-198a12.6 | 0.27829 | 0.148444442 | 0.059 | 0 | 1 | 21 |
| si:ch211-214j24.10 | 0.27829 | 0.153590368 | 0.059 | 0 | 1 | 21 |
| si:ch211-219a4.3 | 0.27829 | 0.202379572 | 0.059 | 0 | 1 | 21 |
| si:ch211-262i1.6 | 0.27829 | 0.127241716 | 0.059 | 0 | 1 | 21 |
| si:ch211-57i17.1 | 0.27829 | 0.199753158 | 0.059 | 0 | 1 | 21 |
| si:ch73-109d9.2 | 0.27829 | 0.149689276 | 0.059 | 0 | 1 | 21 |
| si:ch73-147o17.1 | 0.27829 | 0.120604216 | 0.059 | 0 | 1 | 21 |
| si:ch73-238c9.1 | 0.27829 | 0.145108412 | 0.059 | 0 | 1 | 21 |
| si:ch73-248e21.5 | 0.27829 | 0.155305158 | 0.059 | 0 | 1 | 21 |
| si:ch73-334d15.2 | 0.27829 | 0.165195603 | 0.059 | 0 | 1 | 21 |
| si:dkey-204f11.64 | 0.27829 | 0.194320954 | 0.059 | 0 | 1 | 21 |
| si:dkey-217f16.1 | 0.27829 | 0.203928488 | 0.059 | 0 | 1 | 21 |
| si:dkey-239n17.4 | 0.27829 | 0.163331495 | 0.059 | 0 | 1 | 21 |
| si:dkey-25g12.4 | 0.27829 | 0.203928488 | 0.059 | 0 | 1 | 21 |
| si:dkey-30g5.1 | 0.27829 | 0.150594138 | 0.059 | 0 | 1 | 21 |
| si:dkey-32e23.4 | 0.27829 | 0.132596887 | 0.059 | 0 | 1 | 21 |
| si:dkey-56d12.4 | 0.27829 | 0.16754844 | 0.059 | 0 | 1 | 21 |
| si:dkey-76b14.2 | 0.27829 | 0.167015163 | 0.059 | 0 | 1 | 21 |
| si:dkey-77g12.4 | 0.27829 | 0.121257655 | 0.059 | 0 | 1 | 21 |
| si:dkey-7j14.6 | 0.27829 | 0.110822793 | 0.059 | 0 | 1 | 21 |
| si:dkeyp-110g5.4 | 0.27829 | 0.126186728 | 0.059 | 0 | 1 | 21 |
| si:dkeyp-38g8.5 | 0.27829 | 0.203688832 | 0.059 | 0 | 1 | 21 |
| si:dkeyp-55f12.3 | 0.27829 | 0.191118316 | 0.059 | 0 | 1 | 21 |
| si:rp71-36a1.5 | 0.27829 | 0.131313 | 0.059 | 0 | 1 | 21 |
| sinhcafl | 0.27829 | 0.248878628 | 0.059 | 0 | 1 | 21 |
| slc12a2 | 0.27829 | 0.132596887 | 0.059 | 0 | 1 | 21 |
| slc12a9 | 0.27829 | 0.20788812 | 0.059 | 0 | 1 | 21 |
| slc1a5 | 0.27829 | 0.250798252 | 0.059 | 0 | 1 | 21 |
| slc22a18 | 0.27829 | 0.198692383 | 0.059 | 0 | 1 | 21 |
| slc25a1a | 0.27829 | 0.170000734 | 0.059 | 0 | 1 | 21 |
| slc25a36a | 0.27829 | 0.250608276 | 0.059 | 0 | 1 | 21 |
| slc43a1a | 0.27829 | 0.252402233 | 0.059 | 0 | 1 | 21 |

| | | | | | | |
|---|---|---|---|---|---|---|
| slc7a6os | 0.27829 | 0.14864428 | 0.059 | 0 | 1 | 21 |
| smad2 | 0.27829 | 0.129592128 | 0.059 | 0 | 1 | 21 |
| smad3a | 0.27829 | 0.160298064 | 0.059 | 0 | 1 | 21 |
| smarcb1a | 0.27829 | 0.251592425 | 0.059 | 0 | 1 | 21 |
| smchd1 | 0.27829 | 0.202343703 | 0.059 | 0 | 1 | 21 |
| snw1 | 0.27829 | 0.149614156 | 0.059 | 0 | 1 | 21 |
| snx12 | 0.27829 | 0.148786439 | 0.059 | 0 | 1 | 21 |
| snx20 | 0.27829 | 0.155145301 | 0.059 | 0 | 1 | 21 |
| socs1a | 0.27829 | 0.166172781 | 0.059 | 0 | 1 | 21 |
| sod3a | 0.27829 | 0.163331495 | 0.059 | 0 | 1 | 21 |
| sorl1 | 0.27829 | 0.200973068 | 0.059 | 0 | 1 | 21 |
| sort1b | 0.27829 | 0.20177049 | 0.059 | 0 | 1 | 21 |
| sp1 | 0.27829 | 0.145528049 | 0.059 | 0 | 1 | 21 |
| sp100.1 | 0.27829 | 0.167015163 | 0.059 | 0 | 1 | 21 |
| spg11 | 0.27829 | 0.144316977 | 0.059 | 0 | 1 | 21 |
| spink4 | 0.27829 | 0.118536395 | 0.059 | 0 | 1 | 21 |
| sppl3 | 0.27829 | 0.147544976 | 0.059 | 0 | 1 | 21 |
| spred2b | 0.27829 | 0.13778506 | 0.059 | 0 | 1 | 21 |
| spryd3 | 0.27829 | 0.129965846 | 0.059 | 0 | 1 | 21 |
| srsf5b | 0.27829 | 0.251201624 | 0.059 | 0 | 1 | 21 |
| sssca1 | 0.27829 | 0.154323261 | 0.059 | 0 | 1 | 21 |
| stag2a | 0.27829 | 0.203928488 | 0.059 | 0 | 1 | 21 |
| stat5b | 0.27829 | 0.128976447 | 0.059 | 0 | 1 | 21 |
| stk40 | 0.27829 | 0.145212925 | 0.059 | 0 | 1 | 21 |
| stx5al | 0.27829 | 0.197353667 | 0.059 | 0 | 1 | 21 |
| suds3 | 0.27829 | 0.20229976 | 0.059 | 0 | 1 | 21 |
| sv2 | 0.27829 | 0.148963536 | 0.059 | 0 | 1 | 21 |
| svbp | 0.27829 | 0.136906855 | 0.059 | 0 | 1 | 21 |
| taf3 | 0.27829 | 0.147780648 | 0.059 | 0 | 1 | 21 |
| tbcd | 0.27829 | 0.200739969 | 0.059 | 0 | 1 | 21 |
| tcerg1a | 0.27829 | 0.148444442 | 0.059 | 0 | 1 | 21 |
| tecpr2 | 0.27829 | 0.163331495 | 0.059 | 0 | 1 | 21 |
| terfa | 0.27829 | 0.200019402 | 0.059 | 0 | 1 | 21 |
| tfb1m | 0.27829 | 0.144316977 | 0.059 | 0 | 1 | 21 |
| thap4 | 0.27829 | 0.127857521 | 0.059 | 0 | 1 | 21 |
| thg1l | 0.27829 | 0.125911067 | 0.059 | 0 | 1 | 21 |
| tm9sf2 | 0.27829 | 0.132487997 | 0.059 | 0 | 1 | 21 |
| tmcc1b | 0.27829 | 0.154441933 | 0.059 | 0 | 1 | 21 |
| tmed5 | 0.27829 | 0.12993567 | 0.059 | 0 | 1 | 21 |
| tmem131 | 0.27829 | 0.202318864 | 0.059 | 0 | 1 | 21 |
| tmem251 | 0.27829 | 0.159772955 | 0.059 | 0 | 1 | 21 |
| tmem63a | 0.27829 | 0.193394183 | 0.059 | 0 | 1 | 21 |
| tmem9b | 0.27829 | 0.202567543 | 0.059 | 0 | 1 | 21 |
| tnfb | 0.27829 | 0.161277361 | 0.059 | 0 | 1 | 21 |
| tnrc6b | 0.27829 | 0.203959377 | 0.059 | 0 | 1 | 21 |
| tnrc6c2 | 0.27829 | 0.130507113 | 0.059 | 0 | 1 | 21 |
| tnxba | 0.27829 | 0.164829583 | 0.059 | 0 | 1 | 21 |
| toe1 | 0.27829 | 0.126728778 | 0.059 | 0 | 1 | 21 |

| | | | | | | |
|---|---|---|---|---|---|---|
| tomm34 | 0.27829 | 0.127031432 | 0.059 | 0 | 1 | 21 |
| tor4aa | 0.27829 | 0.152165215 | 0.059 | 0 | 1 | 21 |
| tox | 0.27829 | 0.148938632 | 0.059 | 0 | 1 | 21 |
| tra2b | 0.27829 | 0.199221025 | 0.059 | 0 | 1 | 21 |
| trafd1 | 0.27829 | 0.200462646 | 0.059 | 0 | 1 | 21 |
| trappc2l | 0.27829 | 0.200982312 | 0.059 | 0 | 1 | 21 |
| trioa | 0.27829 | 0.199662208 | 0.059 | 0 | 1 | 21 |
| trpc4apa | 0.27829 | 0.194786677 | 0.059 | 0 | 1 | 21 |
| tsc22d2 | 0.27829 | 0.252461084 | 0.059 | 0 | 1 | 21 |
| tspan37 | 0.27829 | 0.131502404 | 0.059 | 0 | 1 | 21 |
| tspy | 0.27829 | 0.149924272 | 0.059 | 0 | 1 | 21 |
| ttc3 | 0.27829 | 0.251536252 | 0.059 | 0 | 1 | 21 |
| ttc4 | 0.27829 | 0.248957744 | 0.059 | 0 | 1 | 21 |
| txnl1 | 0.27829 | 0.128080596 | 0.059 | 0 | 1 | 21 |
| uap1 | 0.27829 | 0.154838564 | 0.059 | 0 | 1 | 21 |
| uba5 | 0.27829 | 0.148520351 | 0.059 | 0 | 1 | 21 |
| ubap1 | 0.27829 | 0.148628184 | 0.059 | 0 | 1 | 21 |
| usp10 | 0.27829 | 0.252456241 | 0.059 | 0 | 1 | 21 |
| usp3 | 0.27829 | 0.129972717 | 0.059 | 0 | 1 | 21 |
| utp15 | 0.27829 | 0.137007493 | 0.059 | 0 | 1 | 21 |
| vhll | 0.27829 | 0.252561275 | 0.059 | 0 | 1 | 21 |
| vps26a | 0.27829 | 0.128651594 | 0.059 | 0 | 1 | 21 |
| wdr36 | 0.27829 | 0.165345079 | 0.059 | 0 | 1 | 21 |
| xrn2 | 0.27829 | 0.130540341 | 0.059 | 0 | 1 | 21 |
| zbtb17 | 0.27829 | 0.162438073 | 0.059 | 0 | 1 | 21 |
| zbtb7a | 0.27829 | 0.249660194 | 0.059 | 0 | 1 | 21 |
| zc3h4 | 0.27829 | 0.162254677 | 0.059 | 0 | 1 | 21 |
| zcrb1 | 0.27829 | 0.163591947 | 0.059 | 0 | 1 | 21 |
| zdhhc4 | 0.27829 | 0.137007493 | 0.059 | 0 | 1 | 21 |
| zeb2a | 0.27829 | 0.157362423 | 0.059 | 0 | 1 | 21 |
| zfand6 | 0.27829 | 0.144852331 | 0.059 | 0 | 1 | 21 |
| zfyve19 | 0.27829 | 0.129655529 | 0.059 | 0 | 1 | 21 |
| zgc:101569 | 0.27829 | 0.138136266 | 0.059 | 0 | 1 | 21 |
| zgc:101583 | 0.27829 | 0.12979529 | 0.059 | 0 | 1 | 21 |
| zgc:101783 | 0.27829 | 0.125819758 | 0.059 | 0 | 1 | 21 |
| zgc:110269 | 0.27829 | 0.163234124 | 0.059 | 0 | 1 | 21 |
| zgc:112160 | 0.27829 | 0.180574642 | 0.059 | 0 | 1 | 21 |
| zgc:153733 | 0.27829 | 0.150621177 | 0.059 | 0 | 1 | 21 |
| zgc:162339 | 0.27829 | 0.163294673 | 0.059 | 0 | 1 | 21 |
| zgc:162698 | 0.27829 | 0.202794781 | 0.059 | 0 | 1 | 21 |
| zgc:171967 | 0.27829 | 0.162288672 | 0.059 | 0 | 1 | 21 |
| zgc:173915 | 0.27829 | 0.201234609 | 0.059 | 0 | 1 | 21 |
| zgc:63470 | 0.27829 | 0.160969473 | 0.059 | 0 | 1 | 21 |
| zgc:65873 | 0.27829 | 0.121106274 | 0.059 | 0 | 1 | 21 |
| zgc:65997 | 0.27829 | 0.153552894 | 0.059 | 0 | 1 | 21 |
| zgc:66448 | 0.27829 | 0.154076427 | 0.059 | 0 | 1 | 21 |
| zgc:92590 | 0.27829 | 0.11587664 | 0.059 | 0 | 1 | 21 |
| zhx3 | 0.27829 | 0.200288756 | 0.059 | 0 | 1 | 21 |

| | | | | | | |
|---|---|---|---|---|---|---|
| znf1133 | 0.27829 | 0.203928488 | 0.059 | 0 | 1 | 21 |
| znf143a | 0.27829 | 0.20301251 | 0.059 | 0 | 1 | 21 |
| znf410 | 0.27829 | 0.139387539 | 0.059 | 0 | 1 | 21 |
| zranb2 | 0.27829 | 0.252546374 | 0.059 | 0 | 1 | 21 |
| COQ10A | 0.27829 | 0.148712852 | 0.059 | 0 | 1 | 21 |
| CR847906.1 | 0.27829 | 0.148712852 | 0.059 | 0 | 1 | 21 |
| FP102052.2 | 0.27829 | 0.144614039 | 0.059 | 0 | 1 | 21 |
| adck5 | 0.27829 | 0.148712852 | 0.059 | 0 | 1 | 21 |
| anapc15 | 0.27829 | 0.144236603 | 0.059 | 0 | 1 | 21 |
| anapc7 | 0.27829 | 0.140270948 | 0.059 | 0 | 1 | 21 |
| aprt | 0.27829 | 0.139822413 | 0.059 | 0 | 1 | 21 |
| ash2l | 0.27829 | 0.141915705 | 0.059 | 0 | 1 | 21 |
| chkb | 0.27829 | 0.10911736 | 0.059 | 0 | 1 | 21 |
| dnajb1a | 0.27829 | 0.1391645 | 0.059 | 0 | 1 | 21 |
| dnajc18 | 0.27829 | 0.140270948 | 0.059 | 0 | 1 | 21 |
| dnajc5ga | 0.27829 | 0.132648236 | 0.059 | 0 | 1 | 21 |
| eloa | 0.27829 | 0.148712852 | 0.059 | 0 | 1 | 21 |
| erbin | 0.27829 | 0.145806789 | 0.059 | 0 | 1 | 21 |
| fam234b | 0.27829 | 0.148712852 | 0.059 | 0 | 1 | 21 |
| fis1 | 0.27829 | 0.138402288 | 0.059 | 0 | 1 | 21 |
| glsa | 0.27829 | 0.144614039 | 0.059 | 0 | 1 | 21 |
| kat5b | 0.27829 | 0.143629319 | 0.059 | 0 | 1 | 21 |
| magi1b | 0.27829 | 0.148712852 | 0.059 | 0 | 1 | 21 |
| mapkapk3 | 0.27829 | 0.141034486 | 0.059 | 0 | 1 | 21 |
| march7 | 0.27829 | 0.144865449 | 0.059 | 0 | 1 | 21 |
| med28 | 0.27829 | 0.131200005 | 0.059 | 0 | 1 | 21 |
| mpp1 | 0.27829 | 0.142501368 | 0.059 | 0 | 1 | 21 |
| mrps16 | 0.27829 | 0.140526507 | 0.059 | 0 | 1 | 21 |
| mrps22 | 0.27829 | 0.119754461 | 0.059 | 0 | 1 | 21 |
| mrps30 | 0.27829 | 0.139122595 | 0.059 | 0 | 1 | 21 |
| mrps33 | 0.27829 | 0.142785469 | 0.059 | 0 | 1 | 21 |
| ms4a17a.11 | 0.27829 | 0.119569075 | 0.059 | 0 | 1 | 21 |
| ndufaf3 | 0.27829 | 0.131975985 | 0.059 | 0 | 1 | 21 |
| nlrc3 | 0.27829 | 0.142411881 | 0.059 | 0 | 1 | 21 |
| nol10 | 0.27829 | 0.141264319 | 0.059 | 0 | 1 | 21 |
| ormdl1 | 0.27829 | 0.135792235 | 0.059 | 0 | 1 | 21 |
| pdrg1 | 0.27829 | 0.138624463 | 0.059 | 0 | 1 | 21 |
| phc2a | 0.27829 | 0.13969013 | 0.059 | 0 | 1 | 21 |
| rgs17 | 0.27829 | 0.143614508 | 0.059 | 0 | 1 | 21 |
| rictorb | 0.27829 | 0.148712852 | 0.059 | 0 | 1 | 21 |
| rnf13 | 0.27829 | 0.144614039 | 0.059 | 0 | 1 | 21 |
| rnf20 | 0.27829 | 0.139009026 | 0.059 | 0 | 1 | 21 |
| si:dkey-202e17.1 | 0.27829 | 0.148712852 | 0.059 | 0 | 1 | 21 |
| spata2 | 0.27829 | 0.147116792 | 0.059 | 0 | 1 | 21 |
| ssrp1a | 0.27829 | 0.114605825 | 0.059 | 0 | 1 | 21 |
| tex30 | 0.27829 | 0.140270948 | 0.059 | 0 | 1 | 21 |
| tle2c | 0.27829 | 0.146672982 | 0.059 | 0 | 1 | 21 |
| tmem120b | 0.27829 | 0.148712852 | 0.059 | 0 | 1 | 21 |

| | | | | | | |
|---|---|---|---|---|---|---|
| tnfaip3 | 0.27829 | 0.148712852 | 0.059 | 0 | 1 | 21 |
| ube2w | 0.27829 | 0.147454661 | 0.059 | 0 | 1 | 21 |
| vegfd | 0.27829 | 0.148712852 | 0.059 | 0 | 1 | 21 |
| wdtc1 | 0.27829 | 0.135653444 | 0.059 | 0 | 1 | 21 |
| zfpl1 | 0.27829 | 0.147374526 | 0.059 | 0 | 1 | 21 |
| nfkbie | 0.28393 | 0.510328118 | 0.294 | 0.13 | 1 | 21 |
| mt-co1 | 0.29543 | -0.25408134 | 0.235 | 0.435 | 1 | 21 |
| vps29 | 0.29549 | 0.200416617 | 0.176 | 0.043 | 1 | 21 |
| psmg3 | 0.29579 | 0.254348606 | 0.235 | 0.087 | 1 | 21 |
| mob3a | 0.2968 | 0.341637771 | 0.588 | 0.348 | 1 | 21 |
| ndufb9 | 0.2984 | 0.243351441 | 0.412 | 0.217 | 1 | 21 |
| phf5a | 0.29917 | -0.32262275 | 0.059 | 0.217 | 1 | 21 |
| skap2 | 0.29964 | -0.40201906 | 0.353 | 0.565 | 1 | 21 |
| hint1 | 0.30135 | -0.45916661 | 0.059 | 0.217 | 1 | 21 |
| LOC108181220 | 0.30168 | -0.43388868 | 0.059 | 0.217 | 1 | 21 |
| srgn | 0.30511 | 0.420400859 | 0.647 | 0.435 | 1 | 21 |
| rpl39 | 0.3098 | -0.1102447 | 0.941 | 0.957 | 1 | 21 |
| ppp2r5eb | 0.30994 | -0.3906409 | 0.059 | 0.217 | 1 | 21 |
| selenot1b | 0.31018 | -0.38691096 | 0.059 | 0.217 | 1 | 21 |
| rpl38 | 0.31142 | -0.22316193 | 0.824 | 0.957 | 1 | 21 |
| rhocb | 0.31181 | 0.191422499 | 0.176 | 0.043 | 1 | 21 |
| mfng | 0.312 | 0.429931606 | 0.176 | 0.043 | 1 | 21 |
| plekho2 | 0.31788 | 0.476723535 | 0.176 | 0.043 | 1 | 21 |
| ndufs4 | 0.31949 | 0.404899698 | 0.235 | 0.087 | 1 | 21 |
| psmb2 | 0.31976 | -0.11707696 | 0.059 | 0.174 | 1 | 21 |
| atp5po | 0.32341 | -0.42716548 | 0.118 | 0.304 | 1 | 21 |
| ndufa4l | 0.32481 | 0.403267594 | 0.529 | 0.348 | 1 | 21 |
| clta | 0.32543 | -0.28741882 | 0.059 | 0.217 | 1 | 21 |
| mt-nd2 | 0.3261 | -0.28857391 | 0.059 | 0.217 | 1 | 21 |
| ndufs6 | 0.32614 | 0.344801857 | 0.176 | 0.043 | 1 | 21 |
| CABZ01090890.1 | 0.33631 | 0.35260998 | 0.176 | 0.043 | 1 | 21 |
| itga4 | 0.33685 | -0.41631322 | 0.059 | 0.217 | 1 | 21 |
| arpc5b | 0.33724 | 0.138598104 | 0.765 | 0.739 | 1 | 21 |
| rps29 | 0.33964 | -0.14969066 | 0.882 | 0.957 | 1 | 21 |
| si:ch211-214p16.1 | 0.34016 | -0.22852289 | 0.706 | 0.783 | 1 | 21 |
| pnrc2 | 0.34027 | 0.210828161 | 0.941 | 0.783 | 1 | 21 |
| msna | 0.3409 | -0.40796183 | 0.294 | 0.522 | 1 | 21 |
| cct4 | 0.34094 | -0.33939857 | 0.118 | 0.304 | 1 | 21 |
| cmpk | 0.34198 | 0.206894869 | 0.294 | 0.13 | 1 | 21 |
| cdc42 | 0.34374 | 0.329590579 | 0.529 | 0.304 | 1 | 21 |
| sub1b | 0.34501 | -0.23351845 | 0.059 | 0.217 | 1 | 21 |
| prkacab | 0.34541 | 0.243911484 | 0.235 | 0.087 | 1 | 21 |
| alas1 | 0.3456 | 0.404125494 | 0.176 | 0.043 | 1 | 21 |
| itga1 | 0.34582 | -0.17840492 | 0.353 | 0.522 | 1 | 21 |
| nop56 | 0.34649 | -0.34180692 | 0.059 | 0.217 | 1 | 21 |
| znf395b | 0.34678 | 0.251828607 | 0.176 | 0.043 | 1 | 21 |
| CU694197.1 | 0.34754 | -0.34791399 | 0.059 | 0.217 | 1 | 21 |
| celf2 | 0.34853 | 0.261876677 | 0.176 | 0.043 | 1 | 21 |

| | | | | | | |
|---|---|---|---|---|---|---|
| eif4a1a | 0.34985 | -0.39168279 | 0.059 | 0.217 | 1 | 21 |
| u2af1 | 0.35051 | 0.387538062 | 0.235 | 0.087 | 1 | 21 |
| spcs1 | 0.35069 | -0.37562882 | 0.059 | 0.217 | 1 | 21 |
| fgfr1op2 | 0.35195 | 0.384035193 | 0.176 | 0.043 | 1 | 21 |
| snrpe | 0.35215 | -0.12534703 | 0.059 | 0.174 | 1 | 21 |
| hnrnpk | 0.35217 | -0.13002067 | 0.059 | 0.174 | 1 | 21 |
| hmgn7 | 0.35236 | -0.33371958 | 0.353 | 0.478 | 1 | 21 |
| cdipt | 0.35312 | 0.2475915 | 0.176 | 0.043 | 1 | 21 |
| dnmt3ab | 0.35373 | -0.32899597 | 0.059 | 0.217 | 1 | 21 |
| psme2 | 0.35403 | -0.3144478 | 0.059 | 0.217 | 1 | 21 |
| zgc:152891 | 0.35412 | -0.34753183 | 0.059 | 0.217 | 1 | 21 |
| CABZ01046427.2 | 0.35417 | -0.37056394 | 0.059 | 0.217 | 1 | 21 |
| zgc:110591 | 0.35655 | 0.399267996 | 0.176 | 0.043 | 1 | 21 |
| hspa8 | 0.35859 | -0.12085377 | 0.824 | 0.957 | 1 | 21 |
| tomm5 | 0.35873 | -0.35545971 | 0.118 | 0.304 | 1 | 21 |
| arf2b | 0.36159 | 0.323803515 | 0.176 | 0.043 | 1 | 21 |
| pet100 | 0.362 | 0.325325446 | 0.176 | 0.043 | 1 | 21 |
| ubb | 0.36657 | 0.349436232 | 0.176 | 0.043 | 1 | 21 |
| rps19bp1 | 0.36988 | 0.294184449 | 0.176 | 0.043 | 1 | 21 |
| CR391910.2 | 0.37125 | 0.354824193 | 0.176 | 0.043 | 1 | 21 |
| si:ch1073-358c10.1 | 0.37536 | 0.288715553 | 0.176 | 0.043 | 1 | 21 |
| tagapb | 0.37603 | 0.451917069 | 0.235 | 0.087 | 1 | 21 |
| eif3ea | 0.37628 | 0.392431021 | 0.471 | 0.261 | 1 | 21 |
| si:ch211-136m16.8 | 0.37684 | 0.293442919 | 0.176 | 0.043 | 1 | 21 |
| rps21 | 0.37691 | -0.13879798 | 0.941 | 1 | 1 | 21 |
| rbm38 | 0.37822 | 0.339304632 | 0.176 | 0.043 | 1 | 21 |
| bcl10 | 0.37825 | 0.302841879 | 0.176 | 0.043 | 1 | 21 |
| me2 | 0.37828 | 0.338612746 | 0.176 | 0.043 | 1 | 21 |
| mylipb | 0.37879 | 0.286720636 | 0.176 | 0.043 | 1 | 21 |
| arf5 | 0.38041 | 0.2855273 | 0.176 | 0.043 | 1 | 21 |
| tob1a | 0.38201 | 0.270453769 | 0.176 | 0.043 | 1 | 21 |
| BX682234.1 | 0.38288 | 0.377404045 | 0.235 | 0.087 | 1 | 21 |
| bcas2 | 0.38311 | 0.269834244 | 0.176 | 0.043 | 1 | 21 |
| hsp90b1 | 0.38342 | 0.322394008 | 0.176 | 0.043 | 1 | 21 |
| SLA (1 of many) | 0.38589 | 0.351114123 | 0.176 | 0.043 | 1 | 21 |
| cox4i1 | 0.38603 | -0.29900312 | 0.529 | 0.739 | 1 | 21 |
| tmem14ca | 0.38674 | 0.300069463 | 0.176 | 0.043 | 1 | 21 |
| ap1s2 | 0.38701 | 0.272886551 | 0.176 | 0.043 | 1 | 21 |
| chd4b | 0.38703 | 0.413663523 | 0.176 | 0.043 | 1 | 21 |
| ube2v1 | 0.38806 | -0.19110752 | 0.118 | 0.261 | 1 | 21 |
| rpl17 | 0.38808 | -0.1713944 | 0.824 | 0.957 | 1 | 21 |
| mdh1aa | 0.38808 | 0.273487242 | 0.294 | 0.13 | 1 | 21 |
| CR318588.1 | 0.38814 | -0.46918608 | 0.412 | 0.565 | 1 | 21 |
| ostc | 0.38916 | 0.384741636 | 0.176 | 0.043 | 1 | 21 |
| taf9 | 0.38927 | 0.328536028 | 0.176 | 0.043 | 1 | 21 |
| tmx1 | 0.38975 | -0.15834462 | 0.059 | 0.174 | 1 | 21 |
| ralbp1 | 0.39002 | 0.32046426 | 0.176 | 0.043 | 1 | 21 |
| MFAP4 (1 of many) | 0.3904 | 0.179913485 | 0.118 | 0.043 | 1 | 21 |

| | | | | | | |
|---|---|---|---|---|---|---|
| arl8bb | 0.3904 | 0.163004287 | 0.118 | 0.043 | 1 | 21 |
| baxb | 0.3904 | 0.198099039 | 0.118 | 0.043 | 1 | 21 |
| ciapin1 | 0.3904 | 0.242897984 | 0.118 | 0.043 | 1 | 21 |
| cisd2 | 0.3904 | 0.174306986 | 0.118 | 0.043 | 1 | 21 |
| dnajb11 | 0.3904 | 0.142641019 | 0.118 | 0.043 | 1 | 21 |
| gabarapl2 | 0.3904 | 0.15971103 | 0.118 | 0.043 | 1 | 21 |
| hm13 | 0.3904 | 0.317403458 | 0.118 | 0.043 | 1 | 21 |
| hmha1b | 0.3904 | 0.132238381 | 0.118 | 0.043 | 1 | 21 |
| iscub | 0.3904 | 0.153738537 | 0.118 | 0.043 | 1 | 21 |
| mrps6 | 0.3904 | 0.143318104 | 0.118 | 0.043 | 1 | 21 |
| rin3 | 0.3904 | 0.225628274 | 0.118 | 0.043 | 1 | 21 |
| rit1 | 0.3904 | 0.196414212 | 0.118 | 0.043 | 1 | 21 |
| tmem165 | 0.3904 | 0.1763187 | 0.118 | 0.043 | 1 | 21 |
| wdr43 | 0.3904 | 0.278580472 | 0.118 | 0.043 | 1 | 21 |
| CU570889.1 | 0.3904 | 0.336150673 | 0.118 | 0.043 | 1 | 21 |
| LOC101886073 | 0.3904 | 0.195479814 | 0.118 | 0.043 | 1 | 21 |
| baxa | 0.3904 | 0.265079406 | 0.118 | 0.043 | 1 | 21 |
| dcaf7 | 0.3904 | 0.15419415 | 0.118 | 0.043 | 1 | 21 |
| grna | 0.3904 | 0.245692895 | 0.118 | 0.043 | 1 | 21 |
| mrpl11 | 0.3904 | 0.126305355 | 0.118 | 0.043 | 1 | 21 |
| mrps35 | 0.3904 | 0.1884013 | 0.118 | 0.043 | 1 | 21 |
| plrg1 | 0.3904 | 0.152185774 | 0.118 | 0.043 | 1 | 21 |
| psmd1 | 0.3904 | 0.142527439 | 0.118 | 0.043 | 1 | 21 |
| sema4ab | 0.3904 | 0.226811445 | 0.118 | 0.043 | 1 | 21 |
| snd1 | 0.3904 | 0.130463138 | 0.118 | 0.043 | 1 | 21 |
| timm13 | 0.3904 | 0.18474875 | 0.118 | 0.043 | 1 | 21 |
| tnfsf13b | 0.3904 | 0.179291787 | 0.118 | 0.043 | 1 | 21 |
| ugp2b | 0.3904 | 0.283665592 | 0.118 | 0.043 | 1 | 21 |
| znf740b | 0.3904 | 0.226826827 | 0.118 | 0.043 | 1 | 21 |
| akirin1 | 0.3904 | 0.227822841 | 0.118 | 0.043 | 1 | 21 |
| anapc11 | 0.3904 | 0.145330566 | 0.118 | 0.043 | 1 | 21 |
| cat | 0.3904 | 0.160041233 | 0.118 | 0.043 | 1 | 21 |
| chmp2bb | 0.3904 | 0.129228193 | 0.118 | 0.043 | 1 | 21 |
| cipcb | 0.3904 | 0.282328458 | 0.118 | 0.043 | 1 | 21 |
| cox16 | 0.3904 | 0.102186589 | 0.118 | 0.043 | 1 | 21 |
| dnajc19 | 0.3904 | 0.195460547 | 0.118 | 0.043 | 1 | 21 |
| eif4e3 | 0.3904 | 0.19227718 | 0.118 | 0.043 | 1 | 21 |
| fam50a | 0.3904 | 0.143167453 | 0.118 | 0.043 | 1 | 21 |
| gpr137ba | 0.3904 | 0.135686746 | 0.118 | 0.043 | 1 | 21 |
| hmgb3b | 0.3904 | 0.183674063 | 0.118 | 0.043 | 1 | 21 |
| hnrnpa1a | 0.3904 | 0.161106353 | 0.118 | 0.043 | 1 | 21 |
| hp1bp3 | 0.3904 | 0.235749631 | 0.118 | 0.043 | 1 | 21 |
| iqgap1 | 0.3904 | 0.322298663 | 0.118 | 0.043 | 1 | 21 |
| mbd2 | 0.3904 | 0.289371289 | 0.118 | 0.043 | 1 | 21 |
| mrps28 | 0.3904 | 0.225111875 | 0.118 | 0.043 | 1 | 21 |
| ndufa10 | 0.3904 | 0.226700792 | 0.118 | 0.043 | 1 | 21 |
| pno1 | 0.3904 | 0.109771495 | 0.118 | 0.043 | 1 | 21 |
| ppp3ccb | 0.3904 | 0.281506767 | 0.118 | 0.043 | 1 | 21 |

| | | | | | | |
|---|---|---|---|---|---|---|
| psmc4 | 0.3904 | 0.24761286 | 0.118 | 0.043 | 1 | 21 |
| rangap1a | 0.3904 | 0.246507597 | 0.118 | 0.043 | 1 | 21 |
| rer1 | 0.3904 | 0.173577089 | 0.118 | 0.043 | 1 | 21 |
| rnf130 | 0.3904 | 0.20750716 | 0.118 | 0.043 | 1 | 21 |
| runx3 | 0.3904 | 0.245695397 | 0.118 | 0.043 | 1 | 21 |
| srsf11 | 0.3904 | 0.164962207 | 0.118 | 0.043 | 1 | 21 |
| taf12 | 0.3904 | 0.156086809 | 0.118 | 0.043 | 1 | 21 |
| trappc1 | 0.3904 | 0.272732863 | 0.118 | 0.043 | 1 | 21 |
| txnipa | 0.3904 | 0.138739158 | 0.118 | 0.043 | 1 | 21 |
| wbp2nl | 0.3904 | 0.148163877 | 0.118 | 0.043 | 1 | 21 |
| wdr61 | 0.3904 | 0.141745162 | 0.118 | 0.043 | 1 | 21 |
| zgc:174863 | 0.3904 | 0.298512932 | 0.118 | 0.043 | 1 | 21 |
| aamp | 0.3904 | 0.104400143 | 0.118 | 0.043 | 1 | 21 |
| adssl | 0.3904 | 0.120025771 | 0.118 | 0.043 | 1 | 21 |
| csnk1db | 0.3904 | 0.298603662 | 0.118 | 0.043 | 1 | 21 |
| cst3 | 0.3904 | 0.174956771 | 0.118 | 0.043 | 1 | 21 |
| eif4e1c | 0.3904 | 0.161277296 | 0.118 | 0.043 | 1 | 21 |
| lamtor2 | 0.3904 | 0.215998157 | 0.118 | 0.043 | 1 | 21 |
| ppp3r1b | 0.3904 | 0.207781038 | 0.118 | 0.043 | 1 | 21 |
| psmc2 | 0.3904 | 0.203912578 | 0.118 | 0.043 | 1 | 21 |
| psmd14 | 0.3904 | 0.126460771 | 0.118 | 0.043 | 1 | 21 |
| ptbp1a | 0.3904 | 0.225510326 | 0.118 | 0.043 | 1 | 21 |
| tbca | 0.3904 | 0.121993476 | 0.118 | 0.043 | 1 | 21 |
| tktb | 0.3904 | 0.203811067 | 0.118 | 0.043 | 1 | 21 |
| tm2d3 | 0.3904 | 0.104816877 | 0.118 | 0.043 | 1 | 21 |
| tprkb | 0.3904 | 0.134326194 | 0.118 | 0.043 | 1 | 21 |
| txndc9 | 0.3904 | 0.199715512 | 0.118 | 0.043 | 1 | 21 |
| zfand5a | 0.3904 | 0.218405724 | 0.118 | 0.043 | 1 | 21 |
| cd164 | 0.3904 | 0.253688902 | 0.118 | 0.043 | 1 | 21 |
| cnbpa | 0.3904 | 0.20424567 | 0.118 | 0.043 | 1 | 21 |
| fbp1a | 0.3904 | 0.110850999 | 0.118 | 0.043 | 1 | 21 |
| gpd1c | 0.3904 | 0.284005196 | 0.118 | 0.043 | 1 | 21 |
| h2afy2 | 0.3904 | 0.167157214 | 0.118 | 0.043 | 1 | 21 |
| histh1l | 0.3904 | 0.226525047 | 0.118 | 0.043 | 1 | 21 |
| mpc2 | 0.3904 | 0.149872745 | 0.118 | 0.043 | 1 | 21 |
| p2ry10 | 0.3904 | 0.213721044 | 0.118 | 0.043 | 1 | 21 |
| pdcd6ip | 0.3904 | 0.212401569 | 0.118 | 0.043 | 1 | 21 |
| pdk2b | 0.3904 | 0.139128817 | 0.118 | 0.043 | 1 | 21 |
| polr2l | 0.3904 | 0.244325991 | 0.118 | 0.043 | 1 | 21 |
| prkacba | 0.3904 | 0.247160274 | 0.118 | 0.043 | 1 | 21 |
| rbmx | 0.3904 | 0.103777023 | 0.118 | 0.043 | 1 | 21 |
| selenoj | 0.3904 | 0.186412714 | 0.118 | 0.043 | 1 | 21 |
| si:ch211-226m16.2 | 0.3904 | 0.146121201 | 0.118 | 0.043 | 1 | 21 |
| slu7 | 0.3904 | 0.270247537 | 0.118 | 0.043 | 1 | 21 |
| smc1al | 0.3904 | 0.312254698 | 0.118 | 0.043 | 1 | 21 |
| spcs2 | 0.3904 | 0.282002151 | 0.118 | 0.043 | 1 | 21 |
| ssb | 0.3904 | 0.129775421 | 0.118 | 0.043 | 1 | 21 |
| tab1 | 0.3904 | 0.270042719 | 0.118 | 0.043 | 1 | 21 |

| | | | | | | |
|---|---|---|---|---|---|---|
| STMP1 | 0.3904 | 0.176938523 | 0.118 | 0.043 | 1 | 21 |
| cxxc1a | 0.3904 | 0.185482857 | 0.118 | 0.043 | 1 | 21 |
| emd | 0.3904 | 0.262200261 | 0.118 | 0.043 | 1 | 21 |
| hyal3 | 0.3904 | 0.14005688 | 0.118 | 0.043 | 1 | 21 |
| ikzf1 | 0.3904 | 0.275555359 | 0.118 | 0.043 | 1 | 21 |
| il10ra | 0.3904 | 0.237070624 | 0.118 | 0.043 | 1 | 21 |
| mief1 | 0.3904 | 0.208407859 | 0.118 | 0.043 | 1 | 21 |
| mpeg1.1 | 0.3904 | 0.337443317 | 0.118 | 0.043 | 1 | 21 |
| odc1 | 0.3904 | 0.327328042 | 0.118 | 0.043 | 1 | 21 |
| uqcrc1 | 0.3904 | 0.145900604 | 0.118 | 0.043 | 1 | 21 |
| alox12 | 0.3904 | 0.118604324 | 0.118 | 0.043 | 1 | 21 |
| pa2g4a | 0.39049 | 0.251306029 | 0.176 | 0.043 | 1 | 21 |
| dkc1 | 0.39062 | 0.327411057 | 0.176 | 0.043 | 1 | 21 |
| nup50 | 0.39075 | 0.293141613 | 0.176 | 0.043 | 1 | 21 |
| mir223 | 0.39101 | 0.404421208 | 0.176 | 0.043 | 1 | 21 |
| LOC100148254 | 0.39105 | 0.291319283 | 0.176 | 0.043 | 1 | 21 |
| arpc2 | 0.39363 | 0.137641504 | 0.941 | 0.913 | 1 | 21 |
| ube2ib | 0.39459 | -0.37788764 | 0.118 | 0.261 | 1 | 21 |
| sec61g | 0.3967 | -0.2611507 | 0.706 | 0.87 | 1 | 21 |
| tsc22d3 | 0.39887 | 0.375084288 | 0.235 | 0.087 | 1 | 21 |
| rac2 | 0.39896 | 0.167879562 | 0.824 | 0.783 | 1 | 21 |
| gosr2 | 0.40108 | 0.388945299 | 0.176 | 0.087 | 1 | 21 |
| tpm1 | 0.40451 | -0.39074102 | 0.059 | 0.174 | 1 | 21 |
| XLOC-018356 | 0.40574 | 0.447197338 | 0.294 | 0.13 | 1 | 21 |
| cope | 0.40671 | 0.354748337 | 0.235 | 0.087 | 1 | 21 |
| LOC108190565 | 0.40883 | -0.37924827 | 0.235 | 0.435 | 1 | 21 |
| sumo3a | 0.4106 | 0.380324836 | 0.353 | 0.174 | 1 | 21 |
| tpt1 | 0.41127 | 0.119260798 | 1 | 0.913 | 1 | 21 |
| eno1a | 0.41152 | 0.365068172 | 0.353 | 0.174 | 1 | 21 |
| tardbpl | 0.41374 | 0.276155954 | 0.118 | 0.087 | 1 | 21 |
| tpi1b | 0.41399 | 0.373298472 | 0.353 | 0.174 | 1 | 21 |
| paip2b | 0.41804 | 0.233587291 | 0.294 | 0.261 | 1 | 21 |
| cxcr4b | 0.41865 | 0.36575712 | 0.412 | 0.217 | 1 | 21 |
| csnk1a1 | 0.42038 | 0.34202745 | 0.235 | 0.087 | 1 | 21 |
| si:zfos-741a10.3 | 0.42242 | 0.379318402 | 0.235 | 0.087 | 1 | 21 |
| sugt1 | 0.42434 | 0.369507572 | 0.235 | 0.087 | 1 | 21 |
| commd3 | 0.42464 | 0.304242975 | 0.235 | 0.174 | 1 | 21 |
| rps2 | 0.42574 | 0.107731729 | 1 | 0.913 | 1 | 21 |
| ccni | 0.42746 | -0.32400363 | 0.471 | 0.652 | 1 | 21 |
| h2afvb | 0.42921 | -0.18701546 | 0.353 | 0.348 | 1 | 21 |
| elocb | 0.43251 | 0.365742054 | 0.353 | 0.174 | 1 | 21 |
| magt1 | 0.4327 | 0.354330695 | 0.235 | 0.087 | 1 | 21 |
| nfyba | 0.43337 | 0.289778475 | 0.235 | 0.087 | 1 | 21 |
| nfybb | 0.43351 | 0.348962949 | 0.235 | 0.087 | 1 | 21 |
| ldha | 0.43388 | 0.298696559 | 0.235 | 0.087 | 1 | 21 |
| ppp1cab | 0.43462 | 0.343930408 | 0.235 | 0.087 | 1 | 21 |
| cnbpb | 0.43511 | -0.1418193 | 0.235 | 0.217 | 1 | 21 |
| banf1 | 0.43616 | 0.285767985 | 0.235 | 0.087 | 1 | 21 |

| | | | | | | |
|---|---|---|---|---|---|---|
| si:ch73-380l3.2 | 0.4365 | 0.374709225 | 0.294 | 0.174 | 1 | 21 |
| DHDH | 0.4372 | 0.279012473 | 0.235 | 0.087 | 1 | 21 |
| atf4a | 0.43774 | 0.338758088 | 0.235 | 0.087 | 1 | 21 |
| si:dkey-151g10.6 | 0.4392 | -0.10226964 | 0.941 | 1 | 1 | 21 |
| anxa11a | 0.43991 | 0.349852842 | 0.294 | 0.13 | 1 | 21 |
| rpl4 | 0.44196 | 0.182779303 | 0.824 | 0.652 | 1 | 21 |
| rraga | 0.44431 | -0.36980879 | 0.059 | 0.174 | 1 | 21 |
| myo1f | 0.44467 | 0.368707802 | 0.235 | 0.13 | 1 | 21 |
| ccdc124 | 0.44479 | 0.351230377 | 0.235 | 0.13 | 1 | 21 |
| gdi2 | 0.44587 | -0.31047923 | 0.353 | 0.478 | 1 | 21 |
| cst14a.2 | 0.44667 | 0.329313362 | 0.294 | 0.13 | 1 | 21 |
| oaz1a | 0.45015 | -0.35022681 | 0.353 | 0.522 | 1 | 21 |
| si:dkey-9i23.4 | 0.45243 | -0.40967588 | 0.059 | 0.174 | 1 | 21 |
| rps19 | 0.45368 | -0.10287625 | 0.941 | 0.957 | 1 | 21 |
| cox8a | 0.4544 | 0.357876317 | 0.471 | 0.304 | 1 | 21 |
| scinlb | 0.45461 | -0.38832986 | 0.118 | 0.261 | 1 | 21 |
| hprt1 | 0.45547 | -0.31558341 | 0.059 | 0.174 | 1 | 21 |
| arpc3 | 0.46001 | -0.17463627 | 0.824 | 0.913 | 1 | 21 |
| rps15a | 0.46603 | -0.14678701 | 0.882 | 0.957 | 1 | 21 |
| tmem59 | 0.46692 | -0.35108824 | 0.118 | 0.261 | 1 | 21 |
| si:busm1-266f07.2 | 0.46723 | -0.31290492 | 0.412 | 0.609 | 1 | 21 |
| bysl | 0.47008 | 0.246444127 | 0.118 | 0.087 | 1 | 21 |
| eif4a1b | 0.47274 | -0.28437293 | 0.176 | 0.348 | 1 | 21 |
| tspan14 | 0.47568 | -0.34713983 | 0.059 | 0.174 | 1 | 21 |
| si:ch211-102c2.4 | 0.47616 | -0.16339377 | 0.882 | 0.957 | 1 | 21 |
| rpl22l1 | 0.4774 | -0.29409479 | 0.176 | 0.348 | 1 | 21 |
| gabarapb | 0.47946 | 0.27812362 | 0.706 | 0.522 | 1 | 21 |
| psmf1 | 0.47957 | -0.31744386 | 0.059 | 0.174 | 1 | 21 |
| mt-co2 | 0.48059 | -0.35519366 | 0.294 | 0.478 | 1 | 21 |
| grasp | 0.4815 | 0.370515059 | 0.353 | 0.217 | 1 | 21 |
| mef2aa | 0.48389 | -0.23310307 | 0.059 | 0.174 | 1 | 21 |
| sftpbb | 0.48839 | -0.10517183 | 0.353 | 0.478 | 1 | 21 |
| mtpn | 0.48841 | -0.28645094 | 0.294 | 0.478 | 1 | 21 |
| hnrnpa0l | 0.48903 | -0.30214994 | 0.529 | 0.696 | 1 | 21 |
| gnb1b | 0.49005 | -0.24794232 | 0.118 | 0.261 | 1 | 21 |
| cct7 | 0.4917 | 0.317190717 | 0.294 | 0.174 | 1 | 21 |
| eif3g | 0.49205 | 0.178676685 | 0.353 | 0.217 | 1 | 21 |
| eef2b | 0.49587 | 0.166303143 | 0.882 | 0.826 | 1 | 21 |
| jpt2 | 0.49662 | -0.32565418 | 0.118 | 0.261 | 1 | 21 |
| si:zfos-169g10.3 | 0.49811 | -0.27615732 | 0.059 | 0.174 | 1 | 21 |
| rhoab | 0.49836 | -0.35787101 | 0.235 | 0.391 | 1 | 21 |
| ssu72 | 0.50042 | -0.28956657 | 0.059 | 0.174 | 1 | 21 |
| fam49bb | 0.503 | -0.25590874 | 0.059 | 0.174 | 1 | 21 |
| si:ch73-27e22.7 | 0.50303 | -0.25422137 | 0.059 | 0.174 | 1 | 21 |
| prmt1 | 0.50675 | -0.29751551 | 0.118 | 0.261 | 1 | 21 |
| ifrd1 | 0.50842 | -0.2383479 | 0.059 | 0.174 | 1 | 21 |
| calm3b | 0.51143 | -0.23170576 | 0.471 | 0.652 | 1 | 21 |
| ndufb10 | 0.51172 | 0.286781213 | 0.176 | 0.087 | 1 | 21 |

| | | | | | | |
|---|---|---|---|---|---|---|
| rpl12 | 0.51222 | -0.153559 | 0.882 | 0.957 | 1 | 21 |
| dynll1 | 0.51356 | 0.360076915 | 0.353 | 0.217 | 1 | 21 |
| hypk | 0.51387 | -0.28122703 | 0.118 | 0.261 | 1 | 21 |
| btk | 0.51478 | -0.27376588 | 0.118 | 0.261 | 1 | 21 |
| cct6a | 0.51621 | -0.27609763 | 0.118 | 0.261 | 1 | 21 |
| mcu | 0.51629 | 0.349574982 | 0.176 | 0.087 | 1 | 21 |
| rplp0 | 0.51718 | -0.14085033 | 0.882 | 0.87 | 1 | 21 |
| wipf1a | 0.51751 | -0.27732999 | 0.471 | 0.652 | 1 | 21 |
| ppp2cb | 0.51773 | -0.25792498 | 0.059 | 0.174 | 1 | 21 |
| pcmtd1 | 0.51913 | -0.29956759 | 0.118 | 0.261 | 1 | 21 |
| si:ch211-93g21.1 | 0.51976 | -0.28965069 | 0.059 | 0.174 | 1 | 21 |
| ewsr1b | 0.52066 | -0.14614763 | 0.176 | 0.304 | 1 | 21 |
| prpsap2 | 0.52099 | -0.25772918 | 0.059 | 0.174 | 1 | 21 |
| cct3 | 0.52192 | -0.28481135 | 0.118 | 0.261 | 1 | 21 |
| rab7a | 0.52194 | -0.2912628 | 0.118 | 0.261 | 1 | 21 |
| rap1gap2a | 0.5222 | -0.30025801 | 0.059 | 0.174 | 1 | 21 |
| dad1 | 0.52257 | -0.30084844 | 0.235 | 0.391 | 1 | 21 |
| gtf2b | 0.52289 | -0.27141074 | 0.059 | 0.174 | 1 | 21 |
| ndufa8 | 0.52303 | -0.27922912 | 0.059 | 0.174 | 1 | 21 |
| mvb12a | 0.52317 | -0.26142725 | 0.059 | 0.174 | 1 | 21 |
| romo1 | 0.52366 | 0.295070148 | 0.588 | 0.435 | 1 | 21 |
| tpd52l2b | 0.52639 | -0.20055652 | 0.059 | 0.174 | 1 | 21 |
| atp6v0b | 0.5267 | -0.27657216 | 0.059 | 0.174 | 1 | 21 |
| rhogb | 0.52761 | -0.14877523 | 0.471 | 0.478 | 1 | 21 |
| adgre5b.3 | 0.52849 | 0.275657247 | 0.118 | 0.087 | 1 | 21 |
| tagapa | 0.52879 | -0.31043842 | 0.059 | 0.174 | 1 | 21 |
| eomesa | 0.529 | 0.304298615 | 0.588 | 0.435 | 1 | 21 |
| pxna | 0.53026 | 0.287662866 | 0.471 | 0.348 | 1 | 21 |
| pycard | 0.53056 | 0.333790443 | 0.353 | 0.217 | 1 | 21 |
| ube2d1b | 0.5326 | -0.23125331 | 0.059 | 0.174 | 1 | 21 |
| hnrnpa1b | 0.5328 | -0.24976755 | 0.059 | 0.174 | 1 | 21 |
| ubtd2 | 0.53397 | -0.29602145 | 0.059 | 0.174 | 1 | 21 |
| ssr4 | 0.53476 | -0.27804299 | 0.118 | 0.217 | 1 | 21 |
| pgk1 | 0.53509 | -0.2683283 | 0.059 | 0.174 | 1 | 21 |
| gnb1a | 0.5351 | -0.25902533 | 0.059 | 0.174 | 1 | 21 |
| arpp19b | 0.53528 | -0.24173791 | 0.059 | 0.174 | 1 | 21 |
| hck | 0.53543 | -0.24986527 | 0.059 | 0.174 | 1 | 21 |
| apbb1ip | 0.53554 | -0.26226232 | 0.059 | 0.174 | 1 | 21 |
| psma5 | 0.53558 | -0.3011867 | 0.176 | 0.304 | 1 | 21 |
| ccr2 | 0.53573 | -0.300834 | 0.059 | 0.174 | 1 | 21 |
| ufd1l | 0.53582 | -0.19626857 | 0.059 | 0.174 | 1 | 21 |
| lamp2 | 0.53582 | -0.24881428 | 0.059 | 0.174 | 1 | 21 |
| si:ch73-42p12.2 | 0.53599 | -0.22543128 | 0.059 | 0.174 | 1 | 21 |
| gtf2f2a | 0.53599 | -0.22386643 | 0.059 | 0.174 | 1 | 21 |
| mhc2dab | 0.53784 | -0.26336031 | 0.235 | 0.304 | 1 | 21 |
| fgl2a | 0.5394 | 0.125667887 | 0.176 | 0.087 | 1 | 21 |
| cct5 | 0.54106 | -0.27650109 | 0.294 | 0.391 | 1 | 21 |
| vamp3 | 0.54127 | -0.34201626 | 0.176 | 0.304 | 1 | 21 |

| | | | | | | |
|---|---|---|---|---|---|---|
| mbpb | 0.54533 | 0.253589085 | 0.235 | 0.174 | 1 | 21 |
| gpam | 0.547 | 0.243160193 | 0.118 | 0.087 | 1 | 21 |
| selenok | 0.54844 | 0.30670333 | 0.235 | 0.13 | 1 | 21 |
| si:dkey-262k9.4 | 0.54922 | -0.12980988 | 0.118 | 0.217 | 1 | 21 |
| ndufa6 | 0.55013 | 0.132975186 | 0.118 | 0.13 | 1 | 21 |
| zgc:91910 | 0.5527 | -0.2776096 | 0.176 | 0.261 | 1 | 21 |
| eif4ebp2 | 0.55326 | -0.23928838 | 0.353 | 0.522 | 1 | 21 |
| hmga1a | 0.55477 | -0.16056837 | 0.176 | 0.304 | 1 | 21 |
| si:ch1073-429i10.3.1 | 0.5582 | 0.32041763 | 0.471 | 0.304 | 1 | 21 |
| ywhae1 | 0.55874 | -0.24797332 | 0.235 | 0.391 | 1 | 21 |
| itgb2 | 0.55915 | 0.239690223 | 0.765 | 0.609 | 1 | 21 |
| wdr1 | 0.56154 | -0.14086785 | 0.471 | 0.609 | 1 | 21 |
| parvg | 0.56287 | -0.17879654 | 0.294 | 0.435 | 1 | 21 |
| cd63 | 0.56292 | -0.23083964 | 0.529 | 0.696 | 1 | 21 |
| zgc:153713 | 0.56337 | -0.31328053 | 0.235 | 0.391 | 1 | 21 |
| rps18 | 0.5641 | -0.11043895 | 0.941 | 0.957 | 1 | 21 |
| CR753876.1 | 0.56591 | -0.25870072 | 0.529 | 0.652 | 1 | 21 |
| ywhabl | 0.56689 | -0.31978171 | 0.294 | 0.435 | 1 | 21 |
| rap1aa | 0.56819 | -0.28375764 | 0.118 | 0.217 | 1 | 21 |
| cox5ab | 0.57232 | -0.27430191 | 0.353 | 0.478 | 1 | 21 |
| ppt1 | 0.57344 | -0.27980725 | 0.235 | 0.391 | 1 | 21 |
| LOC101884267 | 0.57559 | 0.277922706 | 0.176 | 0.13 | 1 | 21 |
| hmgn6 | 0.57652 | -0.27187381 | 0.235 | 0.391 | 1 | 21 |
| washc2c | 0.57652 | 0.153600652 | 0.529 | 0.391 | 1 | 21 |
| ppp2cab | 0.57838 | 0.161444676 | 0.235 | 0.13 | 1 | 21 |
| fundc1 | 0.57976 | 0.231730079 | 0.176 | 0.087 | 1 | 21 |
| ctss1 | 0.58022 | 0.151456832 | 0.176 | 0.174 | 1 | 21 |
| btf3 | 0.58023 | -0.19875464 | 0.706 | 0.826 | 1 | 21 |
| CABZ01109481.1 | 0.58178 | -0.28896591 | 0.059 | 0.13 | 1 | 21 |
| si:ch211-214p16.2 | 0.58331 | 0.231858707 | 0.471 | 0.391 | 1 | 21 |
| uqcrh | 0.58608 | -0.17785709 | 0.235 | 0.261 | 1 | 21 |
| grap2b | 0.58715 | -0.21694396 | 0.647 | 0.783 | 1 | 21 |
| sesn1 | 0.59088 | 0.312274098 | 0.235 | 0.13 | 1 | 21 |
| lcp2b | 0.59508 | 0.269816352 | 0.588 | 0.435 | 1 | 21 |
| cox7a2a | 0.59577 | 0.188896435 | 0.353 | 0.217 | 1 | 21 |
| si:ch1073-443f11.2 | 0.59671 | -0.29022595 | 0.118 | 0.217 | 1 | 21 |
| si:ch211-213a13.1 | 0.59804 | -0.16029751 | 0.118 | 0.217 | 1 | 21 |
| cab39l | 0.59877 | -0.25425631 | 0.059 | 0.13 | 1 | 21 |
| ndufb7 | 0.60284 | -0.23736961 | 0.118 | 0.174 | 1 | 21 |
| ccr9a | 0.60396 | 0.274635819 | 0.412 | 0.261 | 1 | 21 |
| csde1 | 0.60512 | -0.30230007 | 0.118 | 0.217 | 1 | 21 |
| zgc:136864 | 0.6092 | -0.26024512 | 0.059 | 0.13 | 1 | 21 |
| cd74a | 0.61 | 0.117651447 | 0.941 | 0.913 | 1 | 21 |
| yrk | 0.61356 | -0.2080753 | 0.294 | 0.435 | 1 | 21 |
| laptm4a | 0.6165 | 0.142157125 | 0.176 | 0.087 | 1 | 21 |
| kctd5a | 0.61754 | 0.314197088 | 0.176 | 0.087 | 1 | 21 |
| minos1 | 0.61808 | 0.250054199 | 0.176 | 0.087 | 1 | 21 |
| coro1a | 0.61844 | -0.11845743 | 0.882 | 0.957 | 1 | 21 |

| | | | | | | |
|---|---|---|---|---|---|---|
| rp2 | 0.61929 | 0.274665303 | 0.176 | 0.087 | 1 | 21 |
| psmb3 | 0.62364 | -0.26387836 | 0.118 | 0.217 | 1 | 21 |
| si:ch211-188c18.1 | 0.62517 | -0.26683628 | 0.059 | 0.13 | 1 | 21 |
| gm2a | 0.62638 | -0.25926338 | 0.412 | 0.565 | 1 | 21 |
| ctss2.1 | 0.62638 | 0.195006819 | 0.294 | 0.174 | 1 | 21 |
| ss18l2 | 0.62664 | 0.124630371 | 0.176 | 0.087 | 1 | 21 |
| nccrp1 | 0.62709 | 0.163690793 | 0.176 | 0.087 | 1 | 21 |
| si:dkey-187e18.1 | 0.62826 | 0.277099814 | 0.176 | 0.087 | 1 | 21 |
| rab11a | 0.62844 | -0.26108163 | 0.059 | 0.13 | 1 | 21 |
| anp32e | 0.62944 | 0.267863528 | 0.176 | 0.087 | 1 | 21 |
| eaf1 | 0.63322 | -0.25031003 | 0.059 | 0.13 | 1 | 21 |
| pik3r3a | 0.63325 | 0.277944514 | 0.176 | 0.087 | 1 | 21 |
| sh3bgrl3 | 0.63339 | 0.26787644 | 0.471 | 0.348 | 1 | 21 |
| ube2d2 | 0.63416 | -0.25619674 | 0.235 | 0.304 | 1 | 21 |
| rhoac | 0.63701 | 0.261451615 | 0.294 | 0.217 | 1 | 21 |
| hnrnpaba | 0.63712 | 0.254109235 | 0.412 | 0.304 | 1 | 21 |
| zgc:56676 | 0.63893 | -0.21837831 | 0.176 | 0.304 | 1 | 21 |
| rpl9 | 0.6409 | -0.13457984 | 0.706 | 0.826 | 1 | 21 |
| prkar2aa | 0.64122 | 0.130805804 | 0.176 | 0.087 | 1 | 21 |
| snrpf | 0.64202 | 0.155924518 | 0.294 | 0.174 | 1 | 21 |
| ube2l3b | 0.64297 | 0.239021739 | 0.294 | 0.217 | 1 | 21 |
| tbcb | 0.64334 | -0.24429621 | 0.059 | 0.13 | 1 | 21 |
| arf6b | 0.64393 | -0.22753169 | 0.176 | 0.304 | 1 | 21 |
| si:dkey-222f2.7 | 0.64533 | -0.30292673 | 0.176 | 0.304 | 1 | 21 |
| BX005340.1 | 0.64567 | 0.169874211 | 0.235 | 0.13 | 1 | 21 |
| tmed7 | 0.64602 | -0.218814 | 0.176 | 0.304 | 1 | 21 |
| plek | 0.64731 | -0.11775167 | 0.059 | 0.13 | 1 | 21 |
| vdac2 | 0.64844 | -0.24158031 | 0.353 | 0.435 | 1 | 21 |
| laptm5 | 0.64906 | -0.13320301 | 0.588 | 0.609 | 1 | 21 |
| eif3s10 | 0.6497 | 0.268255592 | 0.176 | 0.087 | 1 | 21 |
| csk | 0.65032 | -0.25280451 | 0.294 | 0.435 | 1 | 21 |
| bloc1s6 | 0.65114 | 0.259052262 | 0.176 | 0.087 | 1 | 21 |
| si:ch211-277c7.6 | 0.65125 | 0.182223472 | 0.176 | 0.087 | 1 | 21 |
| dazap2 | 0.65225 | 0.302745517 | 0.176 | 0.087 | 1 | 21 |
| slc25a3b | 0.65349 | -0.20108979 | 0.529 | 0.609 | 1 | 21 |
| eef1g | 0.65401 | 0.123160185 | 0.765 | 0.739 | 1 | 21 |
| mhc2b | 0.65435 | 0.253578215 | 0.294 | 0.174 | 1 | 21 |
| smfn | 0.65454 | -0.23727598 | 0.059 | 0.13 | 1 | 21 |
| serf2 | 0.65594 | -0.14725813 | 0.529 | 0.565 | 1 | 21 |
| pcna | 0.6563 | -0.22468933 | 0.059 | 0.13 | 1 | 21 |
| pin1 | 0.65676 | -0.1820514 | 0.118 | 0.217 | 1 | 21 |
| itm2bb | 0.65865 | 0.195485973 | 0.706 | 0.565 | 1 | 21 |
| suclg1 | 0.65895 | 0.251004682 | 0.235 | 0.13 | 1 | 21 |
| nr4a1 | 0.66091 | 0.204358934 | 0.118 | 0.087 | 1 | 21 |
| mdkb | 0.66171 | 0.28747379 | 0.294 | 0.174 | 1 | 21 |
| sarnp | 0.66206 | 0.156143668 | 0.176 | 0.087 | 1 | 21 |
| max | 0.6624 | 0.251550519 | 0.176 | 0.087 | 1 | 21 |
| sftpba | 0.66365 | -0.13942443 | 0.706 | 0.826 | 1 | 21 |

| | | | | | | |
|---|---|---|---|---|---|---|
| eif1b | 0.66554 | 0.132778215 | 0.412 | 0.391 | 1 | 21 |
| ctsh | 0.66583 | -0.12024288 | 0.059 | 0.13 | 1 | 21 |
| ndufa3 | 0.66867 | 0.122366212 | 0.235 | 0.13 | 1 | 21 |
| hnrnpabb | 0.66942 | 0.166739122 | 0.294 | 0.261 | 1 | 21 |
| smdt1b | 0.66964 | -0.13969079 | 0.059 | 0.13 | 1 | 21 |
| spcs3 | 0.67194 | 0.248966628 | 0.235 | 0.174 | 1 | 21 |
| pam16 | 0.67339 | 0.195882013 | 0.118 | 0.087 | 1 | 21 |
| rpl13a | 0.675 | -0.10236931 | 0.882 | 0.957 | 1 | 21 |
| seta | 0.67521 | 0.256366607 | 0.176 | 0.087 | 1 | 21 |
| dctn3 | 0.67792 | -0.10060935 | 0.059 | 0.13 | 1 | 21 |
| ndufab1b | 0.6801 | 0.214596208 | 0.235 | 0.13 | 1 | 21 |
| ipcef1 | 0.68018 | 0.229502037 | 0.235 | 0.13 | 1 | 21 |
| glo1 | 0.68023 | 0.139313603 | 0.412 | 0.391 | 1 | 21 |
| adh5 | 0.68032 | 0.170341106 | 0.235 | 0.13 | 1 | 21 |
| edf1 | 0.68148 | -0.27013227 | 0.118 | 0.217 | 1 | 21 |
| prkab1b | 0.68187 | -0.22970226 | 0.118 | 0.217 | 1 | 21 |
| ube2l3a | 0.683 | -0.2554629 | 0.118 | 0.217 | 1 | 21 |
| AL954142.1 | 0.68429 | -0.24393669 | 0.118 | 0.217 | 1 | 21 |
| zgc:111986 | 0.68498 | 0.224521767 | 0.176 | 0.087 | 1 | 21 |
| cwc25 | 0.68499 | -0.25180363 | 0.059 | 0.13 | 1 | 21 |
| atp5pf | 0.68511 | -0.25084474 | 0.353 | 0.478 | 1 | 21 |
| zgc:112356 | 0.68564 | -0.27689062 | 0.118 | 0.217 | 1 | 21 |
| LOC110438754 | 0.68598 | -0.14619064 | 0.118 | 0.13 | 1 | 21 |
| rps27.2 | 0.68619 | -0.20274465 | 0.647 | 0.739 | 1 | 21 |
| gtpbp4 | 0.68867 | -0.25180128 | 0.118 | 0.217 | 1 | 21 |
| eif3jb | 0.68894 | 0.233151038 | 0.235 | 0.13 | 1 | 21 |
| cox8b | 0.68971 | -0.19507443 | 0.176 | 0.217 | 1 | 21 |
| si:ch73-380l3.4 | 0.69002 | 0.108892801 | 0.353 | 0.261 | 1 | 21 |
| psmc6 | 0.69145 | 0.187130531 | 0.176 | 0.087 | 1 | 21 |
| elovl1b | 0.69245 | 0.227864143 | 0.235 | 0.13 | 1 | 21 |
| myh11a | 0.69319 | 0.224247871 | 0.176 | 0.087 | 1 | 21 |
| XLOC-023492 | 0.69437 | 0.226073695 | 0.176 | 0.13 | 1 | 21 |
| crema | 0.69534 | -0.24324583 | 0.059 | 0.13 | 1 | 21 |
| alox5a | 0.69577 | 0.246056745 | 0.176 | 0.087 | 1 | 21 |
| gpr141 | 0.69605 | -0.13669686 | 0.059 | 0.13 | 1 | 21 |
| ap1s1 | 0.69607 | -0.18581599 | 0.118 | 0.217 | 1 | 21 |
| eif4bb | 0.69637 | -0.20954707 | 0.059 | 0.13 | 1 | 21 |
| scgn | 0.69686 | 0.201671712 | 0.176 | 0.087 | 1 | 21 |
| wu:fb55g09 | 0.69703 | -0.20108937 | 0.059 | 0.13 | 1 | 21 |
| sh3glb1b | 0.69707 | 0.196721482 | 0.176 | 0.087 | 1 | 21 |
| hif1ab | 0.6975 | -0.22171276 | 0.118 | 0.217 | 1 | 21 |
| dnaja2b | 0.69771 | 0.205828258 | 0.118 | 0.087 | 1 | 21 |
| usf1l | 0.6982 | -0.20672626 | 0.059 | 0.13 | 1 | 21 |
| si:ch73-248e21.7 | 0.69878 | 0.213262152 | 0.176 | 0.087 | 1 | 21 |
| stk17b | 0.69908 | 0.248718988 | 0.471 | 0.348 | 1 | 21 |
| npm3 | 0.69928 | 0.156740368 | 0.176 | 0.087 | 1 | 21 |
| XLOC-043043 | 0.69941 | 0.186973072 | 0.176 | 0.087 | 1 | 21 |
| snrpd2 | 0.69963 | 0.177207803 | 0.176 | 0.087 | 1 | 21 |

| | | | | | | |
|---|---|---|---|---|---|---|
| faub | 0.70027 | 0.193676224 | 0.176 | 0.087 | 1 | 21 |
| serpinb1l3 | 0.70079 | 0.212567509 | 0.176 | 0.087 | 1 | 21 |
| lgals2a | 0.7008 | 0.251809865 | 0.176 | 0.087 | 1 | 21 |
| ndufc2 | 0.70092 | 0.252102663 | 0.176 | 0.087 | 1 | 21 |
| atp6ap2 | 0.70368 | -0.20357948 | 0.059 | 0.13 | 1 | 21 |
| ptpn1 | 0.70431 | 0.172724666 | 0.176 | 0.087 | 1 | 21 |
| ddx39aa | 0.70621 | -0.21952962 | 0.118 | 0.217 | 1 | 21 |
| CR855389.1 | 0.70623 | -0.22636922 | 0.118 | 0.217 | 1 | 21 |
| zgc:171480 | 0.70635 | 0.192341953 | 0.176 | 0.087 | 1 | 21 |
| slc25a33 | 0.70636 | 0.239200107 | 0.176 | 0.087 | 1 | 21 |
| CDPF1 | 0.70642 | 0.190856898 | 0.176 | 0.087 | 1 | 21 |
| slc25a20 | 0.70645 | 0.176650321 | 0.176 | 0.087 | 1 | 21 |
| ppp1caa | 0.70669 | -0.19096787 | 0.118 | 0.217 | 1 | 21 |
| nop10 | 0.70703 | -0.21305582 | 0.118 | 0.217 | 1 | 21 |
| tubb4b | 0.70851 | -0.20714282 | 0.529 | 0.652 | 1 | 21 |
| si:ch211-195h23.3 | 0.71064 | 0.193988873 | 0.118 | 0.087 | 1 | 21 |
| arl5c | 0.71272 | -0.12102659 | 0.059 | 0.13 | 1 | 21 |
| arl6ip1 | 0.7154 | -0.20462824 | 0.118 | 0.174 | 1 | 21 |
| il7r | 0.71647 | -0.15354918 | 0.059 | 0.13 | 1 | 21 |
| tes | 0.71953 | -0.20861784 | 0.059 | 0.13 | 1 | 21 |
| dnajc4 | 0.71972 | -0.18278904 | 0.059 | 0.13 | 1 | 21 |
| sept6 | 0.71997 | 0.235598315 | 0.353 | 0.261 | 1 | 21 |
| pdap1a | 0.72065 | -0.16612855 | 0.059 | 0.13 | 1 | 21 |
| rnf7 | 0.72205 | -0.13794102 | 0.059 | 0.13 | 1 | 21 |
| rwdd1 | 0.72374 | -0.21021861 | 0.118 | 0.174 | 1 | 21 |
| zmat2 | 0.72511 | -0.18422731 | 0.059 | 0.13 | 1 | 21 |
| arrb2b | 0.72593 | 0.138497105 | 0.235 | 0.217 | 1 | 21 |
| LOC103909115 | 0.72621 | 0.2274369 | 0.471 | 0.348 | 1 | 21 |
| si:ch211-246m6.4 | 0.7279 | -0.12564112 | 0.059 | 0.13 | 1 | 21 |
| si:ch211-153b23.7 | 0.72844 | -0.18841868 | 0.059 | 0.13 | 1 | 21 |
| znf451 | 0.72858 | -0.20577562 | 0.059 | 0.13 | 1 | 21 |
| CABZ01046432.1 | 0.73077 | 0.195843379 | 0.176 | 0.13 | 1 | 21 |
| lsm12b | 0.73225 | -0.19846555 | 0.059 | 0.13 | 1 | 21 |
| mrpl32 | 0.73227 | -0.17820808 | 0.059 | 0.13 | 1 | 21 |
| blcap | 0.7323 | 0.199196115 | 0.176 | 0.13 | 1 | 21 |
| cox6a1 | 0.73302 | -0.15287563 | 0.529 | 0.652 | 1 | 21 |
| cox7b | 0.73323 | 0.193736238 | 0.471 | 0.348 | 1 | 21 |
| cox17 | 0.73436 | -0.18771351 | 0.471 | 0.565 | 1 | 21 |
| rplp2l | 0.73442 | 0.106863916 | 0.941 | 0.87 | 1 | 21 |
| sec61b | 0.73503 | 0.198856789 | 0.471 | 0.348 | 1 | 21 |
| pik3r1 | 0.73518 | -0.10500168 | 0.176 | 0.261 | 1 | 21 |
| rasal3 | 0.73583 | -0.23877022 | 0.059 | 0.13 | 1 | 21 |
| mtap | 0.73584 | -0.17799272 | 0.059 | 0.13 | 1 | 21 |
| sap30l | 0.73597 | 0.216756579 | 0.235 | 0.174 | 1 | 21 |
| rtn4a | 0.736 | -0.18104376 | 0.059 | 0.13 | 1 | 21 |
| pou2f2a | 0.73634 | -0.14440101 | 0.059 | 0.13 | 1 | 21 |
| bin2b | 0.73657 | -0.18418074 | 0.059 | 0.13 | 1 | 21 |
| ddx21 | 0.73667 | -0.15948675 | 0.059 | 0.13 | 1 | 21 |

| | | | | | | |
|---|---|---|---|---|---|---|
| acadvl | 0.73793 | -0.11933555 | 0.059 | 0.13 | 1 | 21 |
| ndufa11 | 0.73862 | -0.21190094 | 0.059 | 0.13 | 1 | 21 |
| fam49al | 0.73926 | -0.21458348 | 0.059 | 0.13 | 1 | 21 |
| GCA | 0.74038 | -0.21053005 | 0.235 | 0.348 | 1 | 21 |
| bcl7bb | 0.74091 | -0.12321347 | 0.059 | 0.087 | 1 | 21 |
| pdcl | 0.74091 | -0.13685917 | 0.059 | 0.087 | 1 | 21 |
| tmc8 | 0.74091 | -0.13576929 | 0.059 | 0.087 | 1 | 21 |
| trim35-28 | 0.74091 | -0.10500309 | 0.059 | 0.087 | 1 | 21 |
| chac1 | 0.74091 | -0.18056982 | 0.059 | 0.087 | 1 | 21 |
| gsna | 0.74091 | -0.10298741 | 0.059 | 0.087 | 1 | 21 |
| pfkfb4b | 0.74091 | -0.15155341 | 0.059 | 0.087 | 1 | 21 |
| pik3r5 | 0.74091 | 0.120038502 | 0.059 | 0.087 | 1 | 21 |
| psmd7 | 0.74091 | -0.17918978 | 0.059 | 0.087 | 1 | 21 |
| rab27a | 0.74091 | -0.11080182 | 0.059 | 0.087 | 1 | 21 |
| u2af2a | 0.74091 | -0.11991597 | 0.059 | 0.087 | 1 | 21 |
| vcp | 0.74091 | -0.12724952 | 0.059 | 0.087 | 1 | 21 |
| aars | 0.74091 | -0.10227432 | 0.059 | 0.087 | 1 | 21 |
| anp32a | 0.74091 | -0.11108955 | 0.059 | 0.087 | 1 | 21 |
| cbx1a | 0.74091 | -0.1348085 | 0.059 | 0.087 | 1 | 21 |
| gbp1 | 0.74091 | -0.13243026 | 0.059 | 0.087 | 1 | 21 |
| gtf2a1 | 0.74091 | -0.10345551 | 0.059 | 0.087 | 1 | 21 |
| ifi46 | 0.74091 | -0.19790072 | 0.059 | 0.087 | 1 | 21 |
| irf2bp2a | 0.74091 | -0.16559673 | 0.059 | 0.087 | 1 | 21 |
| klhdc3 | 0.74091 | -0.11923965 | 0.059 | 0.087 | 1 | 21 |
| nkap | 0.74091 | -0.12164612 | 0.059 | 0.087 | 1 | 21 |
| ostf1 | 0.74091 | -0.12142499 | 0.059 | 0.087 | 1 | 21 |
| prkcsh | 0.74091 | -0.13793934 | 0.059 | 0.087 | 1 | 21 |
| si:dkey-23i12.5 | 0.74091 | -0.12438371 | 0.059 | 0.087 | 1 | 21 |
| tdh | 0.74091 | -0.10982481 | 0.059 | 0.087 | 1 | 21 |
| vps26c | 0.74091 | -0.12020371 | 0.059 | 0.087 | 1 | 21 |
| abcf2a | 0.74091 | -0.14003706 | 0.059 | 0.087 | 1 | 21 |
| cdk9 | 0.74091 | -0.1007796 | 0.059 | 0.087 | 1 | 21 |
| mcm2 | 0.74091 | -0.12550615 | 0.059 | 0.087 | 1 | 21 |
| ppih | 0.74091 | 0.108665283 | 0.059 | 0.087 | 1 | 21 |
| si:dkey-27b3.2 | 0.74091 | -0.1090414 | 0.059 | 0.087 | 1 | 21 |
| znf330 | 0.74091 | -0.12453411 | 0.059 | 0.087 | 1 | 21 |
| ciao2a | 0.74091 | -0.11572071 | 0.059 | 0.087 | 1 | 21 |
| ube2q2 | 0.74091 | -0.1765657 | 0.059 | 0.087 | 1 | 21 |
| aldh9a1a.1 | 0.74091 | -0.13950131 | 0.059 | 0.087 | 1 | 21 |
| dpp3 | 0.74091 | -0.11261245 | 0.059 | 0.087 | 1 | 21 |
| mark2a | 0.74091 | -0.11171081 | 0.059 | 0.087 | 1 | 21 |
| strap | 0.74091 | -0.1036255 | 0.059 | 0.087 | 1 | 21 |
| phlda2 | 0.74137 | 0.182585249 | 0.118 | 0.087 | 1 | 21 |
| aimp1 | 0.74167 | -0.1583176 | 0.059 | 0.13 | 1 | 21 |
| cbx3a | 0.74193 | -0.14222697 | 0.059 | 0.13 | 1 | 21 |
| cops9 | 0.74302 | -0.18051422 | 0.059 | 0.13 | 1 | 21 |
| chmp2a | 0.74447 | -0.16632056 | 0.059 | 0.13 | 1 | 21 |
| vps4b | 0.74538 | 0.175147554 | 0.118 | 0.087 | 1 | 21 |

| | | | | | | |
|---|---|---|---|---|---|---|
| gapdhs | 0.74556 | -0.16038328 | 0.588 | 0.652 | 1 | 21 |
| farsa | 0.74655 | -0.17157134 | 0.059 | 0.13 | 1 | 21 |
| metap2b | 0.74674 | -0.16036307 | 0.059 | 0.13 | 1 | 21 |
| atp5l | 0.74715 | -0.17869454 | 0.412 | 0.478 | 1 | 21 |
| rap2c | 0.7473 | -0.15276114 | 0.059 | 0.13 | 1 | 21 |
| psma4 | 0.74783 | -0.20910354 | 0.059 | 0.13 | 1 | 21 |
| calhm6 | 0.74801 | -0.1309311 | 0.059 | 0.13 | 1 | 21 |
| FQ323156.1 | 0.74859 | -0.19460946 | 0.059 | 0.13 | 1 | 21 |
| srsf2a | 0.74906 | -0.18616278 | 0.059 | 0.13 | 1 | 21 |
| lsm7 | 0.74919 | -0.15047737 | 0.059 | 0.13 | 1 | 21 |
| bud31 | 0.74953 | -0.12182115 | 0.059 | 0.13 | 1 | 21 |
| hnrnph1l | 0.74954 | -0.15911968 | 0.059 | 0.13 | 1 | 21 |
| psma8 | 0.7499 | -0.19338462 | 0.118 | 0.174 | 1 | 21 |
| rpa2 | 0.74994 | -0.13606822 | 0.059 | 0.13 | 1 | 21 |
| selenot1a | 0.75067 | -0.16788238 | 0.059 | 0.13 | 1 | 21 |
| rras | 0.75105 | -0.15287049 | 0.059 | 0.13 | 1 | 21 |
| med6 | 0.75108 | -0.14154122 | 0.059 | 0.13 | 1 | 21 |
| si:dkey-12f6.5 | 0.75116 | -0.18716252 | 0.059 | 0.13 | 1 | 21 |
| si:dkeyp-68b7.12 | 0.75118 | -0.15384074 | 0.059 | 0.13 | 1 | 21 |
| gpx1a | 0.75137 | -0.10795473 | 0.176 | 0.174 | 1 | 21 |
| capns1a | 0.75191 | -0.16265444 | 0.059 | 0.13 | 1 | 21 |
| pa2g4b | 0.75216 | -0.14198249 | 0.059 | 0.13 | 1 | 21 |
| irf2bp2b | 0.75216 | -0.16356259 | 0.059 | 0.13 | 1 | 21 |
| sdhb | 0.75218 | -0.17398241 | 0.059 | 0.13 | 1 | 21 |
| gabarapa | 0.75445 | -0.11229435 | 0.294 | 0.304 | 1 | 21 |
| b2m | 0.75518 | -0.15204178 | 0.353 | 0.391 | 1 | 21 |
| eif2s1b | 0.76115 | -0.16289051 | 0.176 | 0.217 | 1 | 21 |
| eif4ebp1 | 0.76586 | -0.1857038 | 0.471 | 0.565 | 1 | 21 |
| fam32a | 0.7705 | -0.18975802 | 0.176 | 0.261 | 1 | 21 |
| ndufs7 | 0.77356 | -0.16494953 | 0.176 | 0.217 | 1 | 21 |
| tuba8l | 0.77369 | 0.180761197 | 0.176 | 0.13 | 1 | 21 |
| sat1a.2 | 0.77523 | -0.16040287 | 0.412 | 0.522 | 1 | 21 |
| gstp1 | 0.77669 | 0.110015684 | 0.353 | 0.348 | 1 | 21 |
| mapre1a | 0.78155 | -0.14418902 | 0.588 | 0.696 | 1 | 21 |
| bzw1b | 0.78176 | -0.18689927 | 0.118 | 0.174 | 1 | 21 |
| eif2s2 | 0.78282 | 0.164572737 | 0.353 | 0.304 | 1 | 21 |
| ndufa2 | 0.78284 | 0.190921386 | 0.176 | 0.13 | 1 | 21 |
| XLOC-005449 | 0.78419 | 0.17861296 | 0.118 | 0.087 | 1 | 21 |
| tma7 | 0.79158 | -0.13962155 | 0.176 | 0.261 | 1 | 21 |
| nucb2a | 0.79529 | 0.160198354 | 0.118 | 0.087 | 1 | 21 |
| kmt5aa | 0.79631 | 0.167455476 | 0.118 | 0.087 | 1 | 21 |
| higd2a | 0.79738 | -0.18192064 | 0.118 | 0.174 | 1 | 21 |
| atp5f1c | 0.79769 | 0.12789597 | 0.353 | 0.261 | 1 | 21 |
| rpl5b | 0.79963 | -0.14307624 | 0.647 | 0.739 | 1 | 21 |
| eif2s3 | 0.8013 | -0.14350094 | 0.294 | 0.391 | 1 | 21 |
| itpkb | 0.80281 | 0.178136417 | 0.118 | 0.087 | 1 | 21 |
| ptges3b | 0.80286 | -0.18265525 | 0.118 | 0.174 | 1 | 21 |
| pkma | 0.805 | -0.14735463 | 0.294 | 0.391 | 1 | 21 |

| | | | | | | |
|---|---|---|---|---|---|---|
| ndufb8 | 0.80634 | 0.174192603 | 0.353 | 0.261 | 1 | 21 |
| nupr1a | 0.80655 | 0.158199931 | 0.176 | 0.13 | 1 | 21 |
| si:ch211-215e19.4 | 0.80823 | -0.19169059 | 0.176 | 0.261 | 1 | 21 |
| h2afx1 | 0.80887 | -0.20919617 | 0.118 | 0.174 | 1 | 21 |
| med10 | 0.81086 | 0.1693854 | 0.235 | 0.174 | 1 | 21 |
| ifi30 | 0.81311 | 0.146226352 | 0.294 | 0.261 | 1 | 21 |
| znf706 | 0.81385 | -0.12478334 | 0.118 | 0.13 | 1 | 21 |
| rabif | 0.81584 | 0.145796741 | 0.118 | 0.087 | 1 | 21 |
| zgc:63831 | 0.81704 | -0.16833618 | 0.176 | 0.261 | 1 | 21 |
| ywhaba | 0.8178 | 0.123241954 | 0.529 | 0.435 | 1 | 21 |
| npc2 | 0.81802 | -0.11727945 | 0.118 | 0.13 | 1 | 21 |
| inpp5d | 0.81852 | 0.161610255 | 0.118 | 0.087 | 1 | 21 |
| tegt | 0.81915 | -0.17891825 | 0.118 | 0.174 | 1 | 21 |
| brk1 | 0.81954 | -0.18126116 | 0.118 | 0.174 | 1 | 21 |
| fthl27 | 0.82024 | 0.156015562 | 0.353 | 0.261 | 1 | 21 |
| ilf2 | 0.82161 | 0.146012115 | 0.118 | 0.087 | 1 | 21 |
| ssr3 | 0.82258 | 0.140550863 | 0.353 | 0.261 | 1 | 21 |
| gstr | 0.82282 | 0.146111735 | 0.118 | 0.087 | 1 | 21 |
| degs1 | 0.8229 | 0.154084872 | 0.118 | 0.087 | 1 | 21 |
| rab5c | 0.82485 | 0.155162764 | 0.176 | 0.13 | 1 | 21 |
| si:ch73-1a9.3 | 0.82622 | 0.199328909 | 0.235 | 0.174 | 1 | 21 |
| h3f3d | 0.82632 | -0.11625626 | 0.588 | 0.609 | 1 | 21 |
| si:ch211-288g17.3 | 0.83232 | -0.18290535 | 0.118 | 0.174 | 1 | 21 |
| cbfb | 0.83258 | 0.147327328 | 0.059 | 0.043 | 1 | 21 |
| eloca | 0.83258 | 0.108699342 | 0.059 | 0.043 | 1 | 21 |
| plxnc1 | 0.83258 | 0.132419471 | 0.059 | 0.043 | 1 | 21 |
| ppp6r2a | 0.83258 | 0.159402041 | 0.059 | 0.043 | 1 | 21 |
| pspc1 | 0.83258 | 0.185133085 | 0.059 | 0.043 | 1 | 21 |
| si:dkey-16l2.16 | 0.83258 | 0.130865244 | 0.059 | 0.043 | 1 | 21 |
| ubald1a | 0.83258 | 0.134674139 | 0.059 | 0.043 | 1 | 21 |
| ago2 | 0.83258 | 0.159819979 | 0.059 | 0.043 | 1 | 21 |
| bpnt1 | 0.83258 | 0.102566552 | 0.059 | 0.043 | 1 | 21 |
| chchd7 | 0.83258 | 0.105648126 | 0.059 | 0.043 | 1 | 21 |
| chd3 | 0.83258 | 0.167968647 | 0.059 | 0.043 | 1 | 21 |
| cops5 | 0.83258 | 0.121025813 | 0.059 | 0.043 | 1 | 21 |
| ddx46 | 0.83258 | 0.101341252 | 0.059 | 0.043 | 1 | 21 |
| epc2 | 0.83258 | 0.155828137 | 0.059 | 0.043 | 1 | 21 |
| nedd9 | 0.83258 | 0.104041374 | 0.059 | 0.043 | 1 | 21 |
| pole4 | 0.83258 | 0.176290526 | 0.059 | 0.043 | 1 | 21 |
| ppp2r2d | 0.83258 | 0.122801556 | 0.059 | 0.043 | 1 | 21 |
| sco1 | 0.83258 | 0.151964227 | 0.059 | 0.043 | 1 | 21 |
| si:ch211-199g17.2 | 0.83258 | 0.1409081 | 0.059 | 0.043 | 1 | 21 |
| slc35a4 | 0.83258 | 0.119995036 | 0.059 | 0.043 | 1 | 21 |
| slc6a6b | 0.83258 | 0.117276193 | 0.059 | 0.043 | 1 | 21 |
| tmem131l | 0.83258 | 0.111646321 | 0.059 | 0.043 | 1 | 21 |
| znf593 | 0.83258 | 0.177856365 | 0.059 | 0.043 | 1 | 21 |
| znf827 | 0.83258 | 0.138640932 | 0.059 | 0.043 | 1 | 21 |
| calr3b | 0.83258 | 0.168804074 | 0.059 | 0.043 | 1 | 21 |

| | | | | | | |
|---|---|---|---|---|---|---|
| csnk1g2a | 0.83258 | 0.101739792 | 0.059 | 0.043 | 1 | 21 |
| ecm1b | 0.83258 | 0.208348484 | 0.059 | 0.043 | 1 | 21 |
| pitpnb | 0.83258 | 0.141102105 | 0.059 | 0.043 | 1 | 21 |
| sdhaf4 | 0.83258 | 0.149455032 | 0.059 | 0.043 | 1 | 21 |
| si:ch211-220f21.1 | 0.83258 | 0.145241557 | 0.059 | 0.043 | 1 | 21 |
| txndc15 | 0.83258 | 0.211911479 | 0.059 | 0.043 | 1 | 21 |
| zgc:194246 | 0.83258 | 0.191187269 | 0.059 | 0.043 | 1 | 21 |
| atf6 | 0.83258 | 0.133265124 | 0.059 | 0.043 | 1 | 21 |
| atox1 | 0.83258 | 0.175108883 | 0.059 | 0.043 | 1 | 21 |
| cox10 | 0.83258 | 0.102393897 | 0.059 | 0.043 | 1 | 21 |
| dcp1b | 0.83258 | 0.112616893 | 0.059 | 0.043 | 1 | 21 |
| mlec | 0.83258 | 0.172549883 | 0.059 | 0.043 | 1 | 21 |
| nucks1b | 0.83258 | 0.100219613 | 0.059 | 0.043 | 1 | 21 |
| pcmtl | 0.83258 | 0.110174982 | 0.059 | 0.043 | 1 | 21 |
| rex1bd | 0.83258 | 0.165018704 | 0.059 | 0.043 | 1 | 21 |
| sf1 | 0.83258 | 0.107372001 | 0.059 | 0.043 | 1 | 21 |
| spink2.4 | 0.83258 | 0.108486702 | 0.059 | 0.043 | 1 | 21 |
| sec11a | 0.83387 | 0.19936766 | 0.235 | 0.174 | 1 | 21 |
| sf3b1 | 0.83446 | 0.146917842 | 0.118 | 0.087 | 1 | 21 |
| ccdc28a | 0.83483 | 0.122760718 | 0.294 | 0.217 | 1 | 21 |
| sla1 | 0.83551 | 0.117204476 | 0.294 | 0.217 | 1 | 21 |
| atp5pd | 0.83556 | -0.10940923 | 0.294 | 0.304 | 1 | 21 |
| h3f3c | 0.83665 | -0.15979954 | 0.353 | 0.435 | 1 | 21 |
| RASSF5 | 0.84372 | -0.11875028 | 0.235 | 0.261 | 1 | 21 |
| psmb6 | 0.84596 | 0.17352896 | 0.294 | 0.217 | 1 | 21 |
| ak1 | 0.85159 | -0.14423699 | 0.294 | 0.348 | 1 | 21 |
| atp6v1g1 | 0.85422 | 0.14492312 | 0.353 | 0.304 | 1 | 21 |
| eif3ba | 0.85423 | 0.12392091 | 0.294 | 0.217 | 1 | 21 |
| ube2na | 0.85434 | -0.10328648 | 0.118 | 0.13 | 1 | 21 |
| sptssa | 0.85597 | -0.15687926 | 0.118 | 0.174 | 1 | 21 |
| gng5 | 0.85828 | -0.156336 | 0.353 | 0.435 | 1 | 21 |
| eif5a2 | 0.85939 | -0.15993612 | 0.235 | 0.304 | 1 | 21 |
| arf1 | 0.86223 | -0.15726816 | 0.118 | 0.174 | 1 | 21 |
| snrpb | 0.86465 | -0.10353194 | 0.118 | 0.13 | 1 | 21 |
| litaf | 0.86513 | -0.10223591 | 0.118 | 0.13 | 1 | 21 |
| FERMT3 (1 of many) | 0.86521 | 0.100393576 | 0.235 | 0.174 | 1 | 21 |
| actn4 | 0.86614 | -0.15742374 | 0.118 | 0.174 | 1 | 21 |
| atp5if1a | 0.86822 | -0.11463466 | 0.353 | 0.435 | 1 | 21 |
| csrp1a | 0.86905 | -0.14151689 | 0.118 | 0.174 | 1 | 21 |
| arl6ip5b | 0.86992 | -0.14959502 | 0.176 | 0.217 | 1 | 21 |
| capza1a | 0.87006 | 0.145257658 | 0.471 | 0.391 | 1 | 21 |
| cct8 | 0.87113 | -0.10508007 | 0.118 | 0.174 | 1 | 21 |
| vamp8 | 0.87333 | -0.10655923 | 0.235 | 0.261 | 1 | 21 |
| marcksl1a | 0.87344 | -0.16873416 | 0.118 | 0.174 | 1 | 21 |
| psmb1 | 0.87409 | -0.14719406 | 0.118 | 0.174 | 1 | 21 |
| mych | 0.87431 | -0.14233831 | 0.176 | 0.217 | 1 | 21 |
| eif1axa | 0.87457 | 0.148797768 | 0.118 | 0.087 | 1 | 21 |
| tpm3 | 0.87568 | -0.11740179 | 0.588 | 0.652 | 1 | 21 |

| | | | | | | |
|---|---|---|---|---|---|---|
| psmc3 | 0.87578 | -0.14334914 | 0.235 | 0.304 | 1 | 21 |
| rnf10 | 0.87646 | -0.14010658 | 0.118 | 0.174 | 1 | 21 |
| dtnbp1a | 0.87723 | -0.15972854 | 0.235 | 0.304 | 1 | 21 |
| eif6 | 0.87766 | -0.13135956 | 0.118 | 0.174 | 1 | 21 |
| ptprea | 0.8784 | -0.12918225 | 0.118 | 0.174 | 1 | 21 |
| eno3 | 0.87859 | -0.10307636 | 0.529 | 0.609 | 1 | 21 |
| atp5mc3a | 0.88243 | -0.12016527 | 0.118 | 0.174 | 1 | 21 |
| tomm7 | 0.88292 | -0.12982482 | 0.118 | 0.174 | 1 | 21 |
| dock8 | 0.88429 | 0.144185026 | 0.118 | 0.087 | 1 | 21 |
| mgst3b | 0.8844 | -0.12711752 | 0.118 | 0.174 | 1 | 21 |
| ipo7 | 0.88505 | -0.10239679 | 0.118 | 0.174 | 1 | 21 |
| LOC100535743 | 0.8864 | 0.118130221 | 0.235 | 0.174 | 1 | 21 |
| BX120005.1 | 0.88641 | 0.17016565 | 0.176 | 0.13 | 1 | 21 |
| cct2 | 0.88665 | -0.13920082 | 0.235 | 0.304 | 1 | 21 |
| uckl1b | 0.88677 | 0.119747594 | 0.118 | 0.087 | 1 | 21 |
| socs3a | 0.8876 | 0.154279305 | 0.176 | 0.13 | 1 | 21 |
| ephx2 | 0.88768 | 0.138323199 | 0.412 | 0.348 | 1 | 21 |
| lrrfip1a | 0.88858 | 0.104942448 | 0.235 | 0.217 | 1 | 21 |
| pomp | 0.88872 | -0.11200233 | 0.235 | 0.304 | 1 | 21 |
| tmem167a | 0.88903 | -0.13255663 | 0.176 | 0.217 | 1 | 21 |
| sem1 | 0.88947 | 0.113171545 | 0.294 | 0.261 | 1 | 21 |
| caprin1b | 0.88984 | 0.125140564 | 0.235 | 0.174 | 1 | 21 |
| ddx5 | 0.89252 | 0.125494093 | 0.235 | 0.174 | 1 | 21 |
| sod1 | 0.89349 | -0.13209748 | 0.412 | 0.478 | 1 | 21 |
| sumo2b | 0.89977 | 0.114357675 | 0.588 | 0.522 | 1 | 21 |
| higd1a | 0.90023 | 0.12077527 | 0.118 | 0.087 | 1 | 21 |
| hacd3 | 0.90068 | 0.12706043 | 0.118 | 0.087 | 1 | 21 |
| ppp1r18 | 0.90157 | -0.11337244 | 0.353 | 0.391 | 1 | 21 |
| pfdn4 | 0.9019 | -0.12093495 | 0.176 | 0.217 | 1 | 21 |
| scaf1 | 0.90313 | 0.118010121 | 0.118 | 0.087 | 1 | 21 |
| pfdn1 | 0.90601 | 0.105399051 | 0.118 | 0.087 | 1 | 21 |
| si:ch211-217k17.7 | 0.90843 | 0.109747558 | 0.118 | 0.087 | 1 | 21 |
| dusp22b | 0.90908 | 0.115630862 | 0.118 | 0.087 | 1 | 21 |
| reep5 | 0.9126 | 0.12339569 | 0.176 | 0.13 | 1 | 21 |
| churc1 | 0.91363 | -0.12508698 | 0.176 | 0.217 | 1 | 21 |
| ak2 | 0.91462 | -0.1029691 | 0.412 | 0.478 | 1 | 21 |
| prdx5 | 0.91607 | 0.109334981 | 0.176 | 0.13 | 1 | 21 |
| rhogc | 0.91976 | 0.111073486 | 0.118 | 0.087 | 1 | 21 |
| COX7A2 (1 of many) | 0.92211 | 0.103982584 | 0.176 | 0.13 | 1 | 21 |
| srsf3b | 0.93138 | -0.10735925 | 0.176 | 0.217 | 1 | 21 |
| lxn | 0.93274 | 0.104852204 | 0.118 | 0.087 | 1 | 21 |
| cyba | 0.94199 | -0.10374071 | 0.176 | 0.217 | 1 | 21 |
| eif1axb | 0.9434 | -0.10376665 | 0.176 | 0.217 | 1 | 21 |
| grb2b | 0.94362 | 0.101718455 | 0.118 | 0.087 | 1 | 21 |
| si:ch211-196l7.4 | 0.94653 | -0.10106894 | 0.176 | 0.217 | 1 | 21 |
| mt-co2 | 3.384045 | -1.36211132 | 0.15 | 0.9 | 0.0683476 | 22 |
| eno1a | 0.00053 | -0.82899149 | 0 | 0.4 | 1 | 22 |
| mt-cyb | 0.00064 | -0.92414153 | 0.15 | 0.7 | 1 | 22 |

| | | | | | | |
|---|---|---|---|---|---|---|
| dynlrb1 | 0.00143 | -0.61287493 | 0 | 0.35 | 1 | 22 |
| ppp2cab | 0.00143 | -0.61304696 | 0 | 0.35 | 1 | 22 |
| rtn4a | 0.00143 | 0.717385168 | 0.35 | 0 | 1 | 22 |
| si:dkeyp-19e1.3 | 0.00143 | -0.62901634 | 0 | 0.35 | 1 | 22 |
| khdrbs1a | 0.00264 | -0.65640186 | 0.4 | 0.9 | 1 | 22 |
| pabpc1a | 0.00302 | -0.73952901 | 0.5 | 0.9 | 1 | 22 |
| eef1a1l1 | 0.00319 | -0.3221115 | 0.95 | 1 | 1 | 22 |
| lcp2b | 0.00364 | -0.85506314 | 0.05 | 0.5 | 1 | 22 |
| LOC101886459 | 0.00374 | -0.54804167 | 0 | 0.3 | 1 | 22 |
| lman2 | 0.00374 | -0.57746258 | 0 | 0.3 | 1 | 22 |
| arl11 | 0.00374 | -0.70761156 | 0 | 0.3 | 1 | 22 |
| kdm4b | 0.00374 | -0.54210579 | 0 | 0.3 | 1 | 22 |
| tmed10 | 0.00374 | -0.4820044 | 0 | 0.3 | 1 | 22 |
| FO393424.2 | 0.00374 | -0.66164046 | 0 | 0.3 | 1 | 22 |
| ppdpfb | 0.00494 | -0.70788637 | 0.25 | 0.75 | 1 | 22 |
| si:dkey-262k9.4 | 0.00554 | -0.71765637 | 0.1 | 0.55 | 1 | 22 |
| rpl29 | 0.00827 | -0.25057228 | 0.95 | 0.95 | 1 | 22 |
| sept6 | 0.00893 | -0.62380847 | 0.2 | 0.65 | 1 | 22 |
| capzb | 0.00898 | -0.51451763 | 0.55 | 0.95 | 1 | 22 |
| ostc | 0.00946 | -0.41810921 | 0 | 0.25 | 1 | 22 |
| ptp4a1 | 0.00946 | -0.46660027 | 0 | 0.25 | 1 | 22 |
| capns1b | 0.00946 | -0.35592342 | 0 | 0.25 | 1 | 22 |
| pef1 | 0.00946 | -0.53130714 | 0 | 0.25 | 1 | 22 |
| usp36 | 0.00946 | -0.42987576 | 0 | 0.25 | 1 | 22 |
| anapc16 | 0.00946 | -0.50424894 | 0 | 0.25 | 1 | 22 |
| nek6 | 0.00946 | -0.55061869 | 0 | 0.25 | 1 | 22 |
| safb | 0.00946 | -0.51431248 | 0 | 0.25 | 1 | 22 |
| ppp4cb | 0.00946 | -0.46376029 | 0 | 0.25 | 1 | 22 |
| uqcrh | 0.00946 | -0.37307278 | 0 | 0.25 | 1 | 22 |
| nkap | 0.00946 | 0.640227443 | 0.25 | 0 | 1 | 22 |
| arf6b | 0.01428 | -0.83043461 | 0.1 | 0.5 | 1 | 22 |
| COX3 | 0.01722 | -0.55634887 | 0.5 | 0.9 | 1 | 22 |
| XLOC-043760 | 0.01794 | -0.64924705 | 0.05 | 0.4 | 1 | 22 |
| rpl27a | 0.01983 | -0.67286567 | 0.45 | 0.8 | 1 | 22 |
| CT027744.2 | 0.0201 | -0.45492879 | 0.1 | 0.45 | 1 | 22 |
| atp5pf | 0.02074 | -0.42253668 | 0.2 | 0.6 | 1 | 22 |
| cmip | 0.02097 | -0.62406187 | 0.05 | 0.4 | 1 | 22 |
| rpl39 | 0.02105 | -0.29037023 | 0.95 | 1 | 1 | 22 |
| mt-atp6 | 0.0211 | -0.62573244 | 0.2 | 0.6 | 1 | 22 |
| acin1a | 0.02166 | -0.68657702 | 0.05 | 0.4 | 1 | 22 |
| CABZ01069022.2 | 0.02332 | -0.40283244 | 0 | 0.2 | 1 | 22 |
| copb1 | 0.02332 | 0.506458067 | 0.2 | 0 | 1 | 22 |
| ndufa12 | 0.02332 | -0.33981001 | 0 | 0.2 | 1 | 22 |
| pfdn5 | 0.02332 | -0.36530019 | 0 | 0.2 | 1 | 22 |
| phf5a | 0.02332 | -0.43958531 | 0 | 0.2 | 1 | 22 |
| ppp1cbl | 0.02332 | -0.42745727 | 0 | 0.2 | 1 | 22 |
| si:ch211-106e7.2 | 0.02332 | 0.465425237 | 0.2 | 0 | 1 | 22 |
| ythdf1 | 0.02332 | -0.46202469 | 0 | 0.2 | 1 | 22 |

| | | | | | | |
|---|---|---|---|---|---|---|
| dok1b | 0.02332 | -0.3237813 | 0 | 0.2 | 1 | 22 |
| mbnl3 | 0.02332 | -0.35167022 | 0 | 0.2 | 1 | 22 |
| mfap1 | 0.02332 | -0.29823474 | 0 | 0.2 | 1 | 22 |
| rpa1 | 0.02332 | -0.36923122 | 0 | 0.2 | 1 | 22 |
| skilb | 0.02332 | -0.42682234 | 0 | 0.2 | 1 | 22 |
| aco2 | 0.02332 | -0.35403646 | 0 | 0.2 | 1 | 22 |
| ahcy | 0.02332 | -0.35573545 | 0 | 0.2 | 1 | 22 |
| bloc1s6 | 0.02332 | -0.42424807 | 0 | 0.2 | 1 | 22 |
| cdk2ap2 | 0.02332 | -0.33455511 | 0 | 0.2 | 1 | 22 |
| cgrrf1 | 0.02332 | -0.41626984 | 0 | 0.2 | 1 | 22 |
| ehbp1l1b | 0.02332 | 0.503964872 | 0.2 | 0 | 1 | 22 |
| eif4g2b | 0.02332 | -0.39639288 | 0 | 0.2 | 1 | 22 |
| ets2 | 0.02332 | -0.46470604 | 0 | 0.2 | 1 | 22 |
| gorasp2 | 0.02332 | 0.445710206 | 0.2 | 0 | 1 | 22 |
| lims1 | 0.02332 | -0.51191097 | 0 | 0.2 | 1 | 22 |
| lpin2 | 0.02332 | -0.26158345 | 0 | 0.2 | 1 | 22 |
| mgst1.2 | 0.02332 | -0.4327743 | 0 | 0.2 | 1 | 22 |
| nars | 0.02332 | -0.37272061 | 0 | 0.2 | 1 | 22 |
| npepl1 | 0.02332 | -0.44874175 | 0 | 0.2 | 1 | 22 |
| nutf2l | 0.02332 | -0.31931218 | 0 | 0.2 | 1 | 22 |
| pdcd10a | 0.02332 | -0.41815011 | 0 | 0.2 | 1 | 22 |
| pfkpb | 0.02332 | -0.43207187 | 0 | 0.2 | 1 | 22 |
| ppp2cb | 0.02332 | -0.37661815 | 0 | 0.2 | 1 | 22 |
| psmc6 | 0.02332 | -0.26360413 | 0 | 0.2 | 1 | 22 |
| psmd11a | 0.02332 | -0.30534605 | 0 | 0.2 | 1 | 22 |
| rbmx | 0.02332 | -0.49529266 | 0 | 0.2 | 1 | 22 |
| rpn1 | 0.02332 | -0.32523575 | 0 | 0.2 | 1 | 22 |
| si:ch73-158p21.3 | 0.02332 | -0.35941088 | 0 | 0.2 | 1 | 22 |
| si:ch73-208g10.1 | 0.02332 | -0.37475426 | 0 | 0.2 | 1 | 22 |
| si:dkey-42i9.4 | 0.02332 | 0.487531768 | 0.2 | 0 | 1 | 22 |
| stimate | 0.02332 | 0.475760334 | 0.2 | 0 | 1 | 22 |
| tirap | 0.02332 | -0.37965327 | 0 | 0.2 | 1 | 22 |
| tomm20a | 0.02332 | -0.46875432 | 0 | 0.2 | 1 | 22 |
| unc119b | 0.02332 | -0.42076666 | 0 | 0.2 | 1 | 22 |
| LOC100537499 | 0.02332 | -0.38515822 | 0 | 0.2 | 1 | 22 |
| arhgap10 | 0.02332 | -0.43318072 | 0 | 0.2 | 1 | 22 |
| prdx5 | 0.02332 | -0.39323714 | 0 | 0.2 | 1 | 22 |
| ccr9a | 0.02553 | -0.52355268 | 0.3 | 0.7 | 1 | 22 |
| rps21 | 0.02554 | -0.41098549 | 0.8 | 0.95 | 1 | 22 |
| setb | 0.02593 | -0.51081947 | 0.2 | 0.6 | 1 | 22 |
| XLOC-032002 | 0.02885 | -0.67035802 | 0.2 | 0.6 | 1 | 22 |
| limd2 | 0.03074 | -0.73430966 | 0.05 | 0.35 | 1 | 22 |
| rps11 | 0.03321 | -0.26786261 | 0.75 | 1 | 1 | 22 |
| mt-nd1 | 0.03361 | -0.58483718 | 0.1 | 0.45 | 1 | 22 |
| uba52 | 0.03402 | -0.4560858 | 0.65 | 0.95 | 1 | 22 |
| itm2bb | 0.03424 | -0.267261 | 0.75 | 1 | 1 | 22 |
| mych | 0.03578 | -0.617763 | 0.25 | 0.65 | 1 | 22 |
| eno3 | 0.03606 | -0.61157809 | 0.3 | 0.7 | 1 | 22 |

| | | | | | | |
|---|---|---|---|---|---|---|
| ptprc | 0.04404 | -0.3153213 | 0.65 | 0.95 | 1 | 22 |
| ccdc28a | 0.04457 | -0.50343418 | 0.05 | 0.35 | 1 | 22 |
| rhoab | 0.04489 | -0.10675401 | 0.4 | 0.65 | 1 | 22 |
| psmb2 | 0.04489 | -0.46586534 | 0.05 | 0.35 | 1 | 22 |
| npsn | 0.04747 | 0.728759145 | 0.35 | 0.05 | 1 | 22 |
| ndfip1 | 0.04773 | -0.55351337 | 0.05 | 0.35 | 1 | 22 |
| eif3ea | 0.05215 | -0.31394787 | 0.15 | 0.45 | 1 | 22 |
| rpl5b | 0.05221 | -0.48681468 | 0.6 | 0.9 | 1 | 22 |
| mt-nd4 | 0.0542 | -0.22987213 | 0.25 | 0.55 | 1 | 22 |
| fkbp5 | 0.05632 | -0.43370906 | 0.1 | 0.4 | 1 | 22 |
| kdm6a | 0.05649 | -0.30381815 | 0 | 0.15 | 1 | 22 |
| si:ch211-241f5.3 | 0.05649 | -0.28596688 | 0 | 0.15 | 1 | 22 |
| smndc1 | 0.05649 | -0.23486926 | 0 | 0.15 | 1 | 22 |
| LOC103909373 | 0.05649 | -0.34600963 | 0 | 0.15 | 1 | 22 |
| LOC110440142 | 0.05649 | -0.30481562 | 0 | 0.15 | 1 | 22 |
| SHKBP1 | 0.05649 | -0.30930449 | 0 | 0.15 | 1 | 22 |
| alas1 | 0.05649 | -0.33615929 | 0 | 0.15 | 1 | 22 |
| arid1ab | 0.05649 | 0.318057241 | 0.15 | 0 | 1 | 22 |
| atp6v1ba | 0.05649 | -0.26269864 | 0 | 0.15 | 1 | 22 |
| b3gnt5b | 0.05649 | 0.32212972 | 0.15 | 0 | 1 | 22 |
| ccdc167 | 0.05649 | -0.28907532 | 0 | 0.15 | 1 | 22 |
| cetn3 | 0.05649 | 0.314258699 | 0.15 | 0 | 1 | 22 |
| cmpk | 0.05649 | -0.25837962 | 0 | 0.15 | 1 | 22 |
| ctnnb1 | 0.05649 | -0.21861204 | 0 | 0.15 | 1 | 22 |
| ctss1 | 0.05649 | 0.372969736 | 0.15 | 0 | 1 | 22 |
| dkk2 | 0.05649 | -0.29459548 | 0 | 0.15 | 1 | 22 |
| dnajc1 | 0.05649 | -0.32623888 | 0 | 0.15 | 1 | 22 |
| drg2 | 0.05649 | -0.2862463 | 0 | 0.15 | 1 | 22 |
| galnt6 | 0.05649 | -0.300218 | 0 | 0.15 | 1 | 22 |
| golgb1 | 0.05649 | 0.323470365 | 0.15 | 0 | 1 | 22 |
| hiat1a | 0.05649 | -0.33285974 | 0 | 0.15 | 1 | 22 |
| ilf3a | 0.05649 | 0.334816642 | 0.15 | 0 | 1 | 22 |
| irak3 | 0.05649 | 0.367596961 | 0.15 | 0 | 1 | 22 |
| jak2a | 0.05649 | -0.34138196 | 0 | 0.15 | 1 | 22 |
| lrch3 | 0.05649 | -0.42743902 | 0 | 0.15 | 1 | 22 |
| lrrk1 | 0.05649 | -0.29459548 | 0 | 0.15 | 1 | 22 |
| maff | 0.05649 | -0.36704274 | 0 | 0.15 | 1 | 22 |
| ndufaf3 | 0.05649 | -0.23310139 | 0 | 0.15 | 1 | 22 |
| npm3 | 0.05649 | -0.2676291 | 0 | 0.15 | 1 | 22 |
| nudt5 | 0.05649 | 0.39681038 | 0.15 | 0 | 1 | 22 |
| nudt8 | 0.05649 | -0.2329468 | 0 | 0.15 | 1 | 22 |
| orai1a | 0.05649 | -0.23031873 | 0 | 0.15 | 1 | 22 |
| phf20b | 0.05649 | 0.437916899 | 0.15 | 0 | 1 | 22 |
| ptpn1 | 0.05649 | -0.26663964 | 0 | 0.15 | 1 | 22 |
| rabif | 0.05649 | -0.17727203 | 0 | 0.15 | 1 | 22 |
| ralbb | 0.05649 | -0.33812545 | 0 | 0.15 | 1 | 22 |
| rassf2a | 0.05649 | -0.33264241 | 0 | 0.15 | 1 | 22 |
| rbm14b | 0.05649 | -0.45852204 | 0 | 0.15 | 1 | 22 |

| | | | | | | |
|---|---|---|---|---|---|---|
| s1pr2 | 0.05649 | -0.46108269 | 0 | 0.15 | 1 | 22 |
| sc:d217 | 0.05649 | -0.31054443 | 0 | 0.15 | 1 | 22 |
| sec13 | 0.05649 | -0.20368842 | 0 | 0.15 | 1 | 22 |
| si:dkey-147f3.8 | 0.05649 | -0.37133505 | 0 | 0.15 | 1 | 22 |
| si:dkey-238d18.10 | 0.05649 | 0.355873028 | 0.15 | 0 | 1 | 22 |
| siah1 | 0.05649 | -0.3059347 | 0 | 0.15 | 1 | 22 |
| slc41a1 | 0.05649 | 0.332939898 | 0.15 | 0 | 1 | 22 |
| sptlc2a | 0.05649 | -0.37294737 | 0 | 0.15 | 1 | 22 |
| tex30 | 0.05649 | -0.32540608 | 0 | 0.15 | 1 | 22 |
| ube2ka | 0.05649 | -0.26450942 | 0 | 0.15 | 1 | 22 |
| uqcrc1 | 0.05649 | -0.33813939 | 0 | 0.15 | 1 | 22 |
| vma21 | 0.05649 | -0.2252103 | 0 | 0.15 | 1 | 22 |
| vps35 | 0.05649 | -0.28815252 | 0 | 0.15 | 1 | 22 |
| yrdc | 0.05649 | -0.18444077 | 0 | 0.15 | 1 | 22 |
| zc3h13 | 0.05649 | -0.32774669 | 0 | 0.15 | 1 | 22 |
| zc3h7bb | 0.05649 | -0.30181444 | 0 | 0.15 | 1 | 22 |
| CR396595.1 | 0.05649 | 0.401293935 | 0.15 | 0 | 1 | 22 |
| FP101887.1 | 0.05649 | -0.31838991 | 0 | 0.15 | 1 | 22 |
| LOC101885053 | 0.05649 | -0.31644232 | 0 | 0.15 | 1 | 22 |
| adora2aa | 0.05649 | 0.357673028 | 0.15 | 0 | 1 | 22 |
| cmtm6 | 0.05649 | -0.22389264 | 0 | 0.15 | 1 | 22 |
| cnot2 | 0.05649 | -0.35642707 | 0 | 0.15 | 1 | 22 |
| cops9 | 0.05649 | -0.28203902 | 0 | 0.15 | 1 | 22 |
| cremb | 0.05649 | -0.28477252 | 0 | 0.15 | 1 | 22 |
| cyb5r3 | 0.05649 | -0.2992307 | 0 | 0.15 | 1 | 22 |
| ddx3xb | 0.05649 | -0.29003512 | 0 | 0.15 | 1 | 22 |
| dlat | 0.05649 | -0.2411305 | 0 | 0.15 | 1 | 22 |
| dnajc8 | 0.05649 | -0.22269213 | 0 | 0.15 | 1 | 22 |
| dohh | 0.05649 | -0.33599892 | 0 | 0.15 | 1 | 22 |
| fam135a | 0.05649 | -0.24159412 | 0 | 0.15 | 1 | 22 |
| fbxo28 | 0.05649 | -0.30716225 | 0 | 0.15 | 1 | 22 |
| fosl2 | 0.05649 | -0.27652363 | 0 | 0.15 | 1 | 22 |
| il12rb2 | 0.05649 | 0.314101203 | 0.15 | 0 | 1 | 22 |
| ints13 | 0.05649 | 0.33354377 | 0.15 | 0 | 1 | 22 |
| itgav | 0.05649 | -0.36211321 | 0 | 0.15 | 1 | 22 |
| jpt1b | 0.05649 | -0.3272954 | 0 | 0.15 | 1 | 22 |
| kdm5ba | 0.05649 | -0.30453094 | 0 | 0.15 | 1 | 22 |
| kdm6ba | 0.05649 | -0.3398974 | 0 | 0.15 | 1 | 22 |
| matr3l1.1 | 0.05649 | -0.30028896 | 0 | 0.15 | 1 | 22 |
| naxe | 0.05649 | -0.21447322 | 0 | 0.15 | 1 | 22 |
| ophn1 | 0.05649 | 0.292883809 | 0.15 | 0 | 1 | 22 |
| ostf1 | 0.05649 | 0.40885586 | 0.15 | 0 | 1 | 22 |
| p2ry11 | 0.05649 | 0.327233341 | 0.15 | 0 | 1 | 22 |
| pkig | 0.05649 | 0.425297426 | 0.15 | 0 | 1 | 22 |
| prrc2a | 0.05649 | 0.322632651 | 0.15 | 0 | 1 | 22 |
| psen1 | 0.05649 | -0.37263714 | 0 | 0.15 | 1 | 22 |
| sarnp | 0.05649 | -0.25022 | 0 | 0.15 | 1 | 22 |
| serp1 | 0.05649 | -0.2357132 | 0 | 0.15 | 1 | 22 |

| | | | | | | |
|---|---|---|---|---|---|---|
| si:ch211-165f21.2 | 0.05649 | 0.408671512 | 0.15 | 0 | 1 | 22 |
| sidt2 | 0.05649 | -0.31993837 | 0 | 0.15 | 1 | 22 |
| slc6a7 | 0.05649 | 0.335191288 | 0.15 | 0 | 1 | 22 |
| snip1 | 0.05649 | -0.23951568 | 0 | 0.15 | 1 | 22 |
| st7 | 0.05649 | 0.348798759 | 0.15 | 0 | 1 | 22 |
| tbk1 | 0.05649 | 0.406553726 | 0.15 | 0 | 1 | 22 |
| tmem14ca | 0.05649 | 0.290267959 | 0.15 | 0 | 1 | 22 |
| tmem243b | 0.05649 | 0.403104344 | 0.15 | 0 | 1 | 22 |
| tp53rk | 0.05649 | -0.32366882 | 0 | 0.15 | 1 | 22 |
| wdr46 | 0.05649 | -0.3045886 | 0 | 0.15 | 1 | 22 |
| zgc:165555.3 | 0.05649 | 0.380303222 | 0.15 | 0 | 1 | 22 |
| znf217 | 0.05649 | -0.36083283 | 0 | 0.15 | 1 | 22 |
| RPL37A | 0.05649 | -0.26009834 | 0 | 0.15 | 1 | 22 |
| adka | 0.05649 | -0.23587744 | 0 | 0.15 | 1 | 22 |
| borcs8 | 0.05649 | -0.23589932 | 0 | 0.15 | 1 | 22 |
| cipcb | 0.05649 | -0.25128181 | 0 | 0.15 | 1 | 22 |
| cyb5a | 0.05649 | -0.27322512 | 0 | 0.15 | 1 | 22 |
| dbr1 | 0.05649 | -0.28133378 | 0 | 0.15 | 1 | 22 |
| ddt | 0.05649 | -0.28717347 | 0 | 0.15 | 1 | 22 |
| dnase2 | 0.05649 | -0.35229814 | 0 | 0.15 | 1 | 22 |
| ell | 0.05649 | 0.473796925 | 0.15 | 0 | 1 | 22 |
| gskip | 0.05649 | -0.34566328 | 0 | 0.15 | 1 | 22 |
| gtf2b | 0.05649 | -0.2143968 | 0 | 0.15 | 1 | 22 |
| hyal1 | 0.05649 | 0.316870664 | 0.15 | 0 | 1 | 22 |
| irf5 | 0.05649 | -0.29449937 | 0 | 0.15 | 1 | 22 |
| luc7l3 | 0.05649 | -0.27083479 | 0 | 0.15 | 1 | 22 |
| mcrip1 | 0.05649 | -0.26538568 | 0 | 0.15 | 1 | 22 |
| mycbp | 0.05649 | 0.318397812 | 0.15 | 0 | 1 | 22 |
| myef2 | 0.05649 | -0.22964958 | 0 | 0.15 | 1 | 22 |
| ndufa9a | 0.05649 | -0.25575886 | 0 | 0.15 | 1 | 22 |
| nr1d1 | 0.05649 | -0.3691423 | 0 | 0.15 | 1 | 22 |
| pdcl | 0.05649 | -0.29638524 | 0 | 0.15 | 1 | 22 |
| pla2g4ab | 0.05649 | -0.30905917 | 0 | 0.15 | 1 | 22 |
| prpf38b | 0.05649 | -0.25854132 | 0 | 0.15 | 1 | 22 |
| psmc1a | 0.05649 | -0.20052736 | 0 | 0.15 | 1 | 22 |
| pstpip2 | 0.05649 | -0.24309362 | 0 | 0.15 | 1 | 22 |
| rin3 | 0.05649 | -0.35842704 | 0 | 0.15 | 1 | 22 |
| srpk1a | 0.05649 | -0.32000792 | 0 | 0.15 | 1 | 22 |
| stx12 | 0.05649 | -0.26142681 | 0 | 0.15 | 1 | 22 |
| supt6h | 0.05649 | -0.31212849 | 0 | 0.15 | 1 | 22 |
| trnau1apb | 0.05649 | 0.365050958 | 0.15 | 0 | 1 | 22 |
| uprt | 0.05649 | 0.330656391 | 0.15 | 0 | 1 | 22 |
| vps26c | 0.05649 | -0.26951949 | 0 | 0.15 | 1 | 22 |
| wdr43 | 0.05649 | -0.27625648 | 0 | 0.15 | 1 | 22 |
| pex19 | 0.05649 | 0.39607894 | 0.15 | 0 | 1 | 22 |
| strn4 | 0.05649 | 0.424312551 | 0.15 | 0 | 1 | 22 |
| CR759918.1 | 0.05649 | 0.297418421 | 0.15 | 0 | 1 | 22 |
| add1 | 0.05649 | 0.358019922 | 0.15 | 0 | 1 | 22 |

| | | | | | | |
|---|---|---|---|---|---|---|
| chmp2bb | 0.05649 | -0.30936088 | 0 | 0.15 | 1 | 22 |
| ckap5 | 0.05649 | 0.33728268 | 0.15 | 0 | 1 | 22 |
| dop1b | 0.05649 | 0.30659758 | 0.15 | 0 | 1 | 22 |
| eif3hb | 0.05649 | -0.27907816 | 0 | 0.15 | 1 | 22 |
| ggps1 | 0.05649 | -0.35341829 | 0 | 0.15 | 1 | 22 |
| got1 | 0.05649 | -0.28884899 | 0 | 0.15 | 1 | 22 |
| klf11a | 0.05649 | -0.26603793 | 0 | 0.15 | 1 | 22 |
| kmt2a | 0.05649 | 0.359362954 | 0.15 | 0 | 1 | 22 |
| mtm1 | 0.05649 | -0.26592487 | 0 | 0.15 | 1 | 22 |
| nupr1a | 0.05649 | -0.26523852 | 0 | 0.15 | 1 | 22 |
| per3 | 0.05649 | 0.428046325 | 0.15 | 0 | 1 | 22 |
| plat | 0.05649 | -0.33779104 | 0 | 0.15 | 1 | 22 |
| rgs14a | 0.05649 | 0.358043467 | 0.15 | 0 | 1 | 22 |
| ss18l2 | 0.05649 | -0.25781275 | 0 | 0.15 | 1 | 22 |
| tra2a | 0.05649 | 0.317216359 | 0.15 | 0 | 1 | 22 |
| tufm | 0.05649 | -0.3123266 | 0 | 0.15 | 1 | 22 |
| vapal | 0.05649 | -0.23496425 | 0 | 0.15 | 1 | 22 |
| zgc:152774 | 0.05649 | 0.379547162 | 0.15 | 0 | 1 | 22 |
| XLOC-018356 | 0.05664 | 0.353123661 | 0.95 | 0.8 | 1 | 22 |
| si:ch211-196l7.4 | 0.05866 | 0.555783051 | 0.7 | 0.45 | 1 | 22 |
| cdc42l | 0.06036 | -0.24601022 | 0.5 | 0.8 | 1 | 22 |
| atp5f1c | 0.06201 | -0.55248262 | 0.1 | 0.4 | 1 | 22 |
| rps5 | 0.06243 | -0.41025266 | 0.7 | 0.95 | 1 | 22 |
| lcp1 | 0.06403 | -0.44081684 | 0.3 | 0.65 | 1 | 22 |
| cox6b1 | 0.06428 | -0.45879274 | 0.2 | 0.55 | 1 | 22 |
| rps13 | 0.06458 | -0.29018449 | 0.8 | 1 | 1 | 22 |
| eomesa | 0.0647 | -0.25880275 | 0.5 | 0.8 | 1 | 22 |
| arl6ip1 | 0.06515 | 0.581255355 | 0.4 | 0.2 | 1 | 22 |
| tpt1 | 0.06628 | -0.21084364 | 0.9 | 0.9 | 1 | 22 |
| hdac1 | 0.06664 | 0.613188147 | 0.4 | 0.1 | 1 | 22 |
| BX005183.3 | 0.06711 | -0.42840082 | 0.35 | 0.7 | 1 | 22 |
| spink2.1 | 0.06785 | -0.63765426 | 0.1 | 0.35 | 1 | 22 |
| csnk2b | 0.068 | -0.41582593 | 0.1 | 0.4 | 1 | 22 |
| sec61b | 0.06839 | -0.47708167 | 0.2 | 0.55 | 1 | 22 |
| faua | 0.06841 | -0.4600527 | 0.65 | 0.9 | 1 | 22 |
| si:dkey-262k9.2 | 0.06866 | -0.55151722 | 0.2 | 0.55 | 1 | 22 |
| erh | 0.07183 | 0.29699977 | 0.4 | 0.4 | 1 | 22 |
| si:dkey-11f4.14 | 0.07213 | 0.65457962 | 0.5 | 0.2 | 1 | 22 |
| si:ch211-66e2.3 | 0.07236 | -0.41008611 | 0.05 | 0.3 | 1 | 22 |
| rpl13a | 0.07268 | -0.39648625 | 0.7 | 0.95 | 1 | 22 |
| ola1 | 0.07303 | -0.54737626 | 0.1 | 0.4 | 1 | 22 |
| rab5ab | 0.07455 | 0.540861004 | 0.3 | 0.05 | 1 | 22 |
| si:ch211-212k18.7 | 0.07652 | -0.53922325 | 0.25 | 0.6 | 1 | 22 |
| cnp | 0.07713 | -0.180631 | 0.4 | 0.65 | 1 | 22 |
| XLOC-005449 | 0.07716 | 0.7089578 | 0.3 | 0.05 | 1 | 22 |
| ccl35.1 | 0.07732 | -0.37861668 | 0.75 | 0.95 | 1 | 22 |
| rps15a | 0.07741 | -0.26387822 | 0.85 | 1 | 1 | 22 |
| srsf11 | 0.07817 | -0.3576112 | 0.05 | 0.3 | 1 | 22 |

| | | | | | | |
|---|---|---|---|---|---|---|
| kctd12.2 | 0.07857 | -0.23591887 | 0.7 | 0.95 | 1 | 22 |
| nr1d2a | 0.08001 | -0.46541183 | 0.15 | 0.45 | 1 | 22 |
| rps28 | 0.08047 | -0.20291939 | 0.9 | 0.95 | 1 | 22 |
| gpx4b | 0.08145 | -0.51272121 | 0.1 | 0.4 | 1 | 22 |
| yeats4 | 0.08164 | -0.33987979 | 0.05 | 0.3 | 1 | 22 |
| uqcr10 | 0.08339 | 0.457826906 | 0.4 | 0.3 | 1 | 22 |
| XLOC-025423 | 0.08373 | 0.355748103 | 0.5 | 0.45 | 1 | 22 |
| tmx4 | 0.08428 | 0.648808917 | 0.3 | 0.05 | 1 | 22 |
| pparg | 0.08561 | -0.41455955 | 0.05 | 0.3 | 1 | 22 |
| cst14a.2 | 0.08569 | -0.3586779 | 0.05 | 0.3 | 1 | 22 |
| ran | 0.08602 | -0.25005249 | 0.5 | 0.8 | 1 | 22 |
| sf3b6 | 0.08788 | 0.593493194 | 0.35 | 0.1 | 1 | 22 |
| ndfip1l | 0.08801 | -0.45950325 | 0.05 | 0.3 | 1 | 22 |
| ptges3b | 0.08816 | -0.53866481 | 0.05 | 0.3 | 1 | 22 |
| rpl15 | 0.09255 | -0.34489704 | 0.75 | 0.95 | 1 | 22 |
| rpl18a | 0.09275 | -0.35635918 | 0.8 | 0.95 | 1 | 22 |
| si:dkey-151g10.6 | 0.09299 | -0.18825955 | 0.95 | 1 | 1 | 22 |
| shoc2 | 0.09349 | -0.47358504 | 0.05 | 0.3 | 1 | 22 |
| mt-nd2 | 0.0938 | -0.29052018 | 0.15 | 0.45 | 1 | 22 |
| mycb | 0.09492 | -0.4353384 | 0.05 | 0.3 | 1 | 22 |
| dock8 | 0.09506 | -0.56512326 | 0.05 | 0.3 | 1 | 22 |
| si:dkey-51e6.1 | 0.09623 | -0.42373433 | 0.05 | 0.3 | 1 | 22 |
| sap30l | 0.09655 | -0.16241455 | 0.1 | 0.3 | 1 | 22 |
| mtap | 0.09719 | -0.41759796 | 0.05 | 0.3 | 1 | 22 |
| ywhae1 | 0.09879 | -0.59495506 | 0.05 | 0.3 | 1 | 22 |
| tes | 0.09879 | -0.48980496 | 0.05 | 0.3 | 1 | 22 |
| npm1a | 0.09998 | -0.13598108 | 0.1 | 0.3 | 1 | 22 |
| ier5 | 0.10253 | -0.11437125 | 0.25 | 0.45 | 1 | 22 |
| pnrc2 | 0.10279 | -0.23303333 | 0.9 | 1 | 1 | 22 |
| itga1 | 0.1045 | -0.37034403 | 0.3 | 0.6 | 1 | 22 |
| CABZ01111959.1 | 0.1063 | -0.6167275 | 0.15 | 0.45 | 1 | 22 |
| ndufc2 | 0.10696 | -0.40281528 | 0.15 | 0.45 | 1 | 22 |
| dynlt3 | 0.10796 | -0.43424355 | 0.15 | 0.45 | 1 | 22 |
| si:ch211-214k5.3 | 0.10816 | -0.34767144 | 0.3 | 0.6 | 1 | 22 |
| sept2 | 0.10879 | -0.50887284 | 0.15 | 0.45 | 1 | 22 |
| b3gnt2l | 0.11009 | -0.49469763 | 0.15 | 0.45 | 1 | 22 |
| grasp | 0.11034 | -0.41043672 | 0.2 | 0.5 | 1 | 22 |
| si:ch73-27e22.7 | 0.11045 | -0.56712041 | 0.15 | 0.45 | 1 | 22 |
| atp5md | 0.11199 | -0.52597214 | 0.3 | 0.5 | 1 | 22 |
| rpl12 | 0.11243 | -0.33737884 | 0.75 | 0.95 | 1 | 22 |
| rps29 | 0.11389 | -0.18728171 | 0.95 | 1 | 1 | 22 |
| clic1 | 0.11411 | 0.53177818 | 0.6 | 0.35 | 1 | 22 |
| rad23b | 0.11716 | -0.29722633 | 0.1 | 0.35 | 1 | 22 |
| gm2a | 0.11765 | 0.271014782 | 0.75 | 0.7 | 1 | 22 |
| brk1 | 0.11784 | -0.29976329 | 0.05 | 0.25 | 1 | 22 |
| CU855821.1 | 0.11833 | -0.59492105 | 0.2 | 0.5 | 1 | 22 |
| si:dkey-9i23.4 | 0.12052 | -0.12440243 | 0.05 | 0.2 | 1 | 22 |
| hmga1a | 0.12096 | 0.455221225 | 0.35 | 0.25 | 1 | 22 |

| | | | | | | |
|---|---|---|---|---|---|---|
| ppp6c | 0.1213 | -0.35286509 | 0.1 | 0.35 | 1 | 22 |
| ubb | 0.12146 | 0.412843893 | 0.6 | 0.45 | 1 | 22 |
| atp5mc3b | 0.12186 | -0.39749347 | 0.65 | 0.8 | 1 | 22 |
| rab5c | 0.12254 | -0.52321202 | 0.1 | 0.35 | 1 | 22 |
| rpl10 | 0.12399 | -0.1044065 | 0.85 | 1 | 1 | 22 |
| actr2a | 0.12852 | -0.42497891 | 0.25 | 0.55 | 1 | 22 |
| glipr1a | 0.13062 | -0.4743137 | 0.2 | 0.5 | 1 | 22 |
| cox5ab | 0.13065 | -0.38642139 | 0.1 | 0.35 | 1 | 22 |
| CR383676.1 | 0.13157 | 0.144483255 | 1 | 0.95 | 1 | 22 |
| arhgdia | 0.1331 | -0.39232446 | 0.5 | 0.8 | 1 | 22 |
| elavl1a | 0.13318 | 0.510592302 | 0.25 | 0.05 | 1 | 22 |
| rbm8a | 0.13455 | 0.440107734 | 0.2 | 0.1 | 1 | 22 |
| hmgn6 | 0.13537 | -0.55592228 | 0.25 | 0.55 | 1 | 22 |
| BX000363.1 | 0.13645 | 0.329241918 | 0.1 | 0 | 1 | 22 |
| BX248410.1 | 0.13645 | -0.21891348 | 0 | 0.1 | 1 | 22 |
| BX571825.4 | 0.13645 | -0.27044895 | 0 | 0.1 | 1 | 22 |
| CABZ01085658.1 | 0.13645 | -0.21640183 | 0 | 0.1 | 1 | 22 |
| CABZ01087514.1 | 0.13645 | -0.17413569 | 0 | 0.1 | 1 | 22 |
| CABZ01113954.1 | 0.13645 | -0.18341463 | 0 | 0.1 | 1 | 22 |
| CT583625.5 | 0.13645 | 0.249331649 | 0.1 | 0 | 1 | 22 |
| CU681842.1 | 0.13645 | 0.328167567 | 0.1 | 0 | 1 | 22 |
| FAM83G | 0.13645 | -0.25566322 | 0 | 0.1 | 1 | 22 |
| FP074902.2 | 0.13645 | -0.18970315 | 0 | 0.1 | 1 | 22 |
| LOC100332221 | 0.13645 | -0.18457107 | 0 | 0.1 | 1 | 22 |
| LOC108180269 | 0.13645 | -0.15612527 | 0 | 0.1 | 1 | 22 |
| LOC110437917 | 0.13645 | -0.15976144 | 0 | 0.1 | 1 | 22 |
| LOC110439915 | 0.13645 | -0.20997749 | 0 | 0.1 | 1 | 22 |
| RANBP2 | 0.13645 | -0.16338284 | 0 | 0.1 | 1 | 22 |
| XLOC-022621 | 0.13645 | 0.210480319 | 0.1 | 0 | 1 | 22 |
| XLOC-032112 | 0.13645 | -0.19994199 | 0 | 0.1 | 1 | 22 |
| XLOC-043857 | 0.13645 | -0.15985183 | 0 | 0.1 | 1 | 22 |
| acsbg2 | 0.13645 | -0.19125094 | 0 | 0.1 | 1 | 22 |
| ada | 0.13645 | -0.1439229 | 0 | 0.1 | 1 | 22 |
| adnp2b | 0.13645 | -0.16058465 | 0 | 0.1 | 1 | 22 |
| adora1b | 0.13645 | -0.15557037 | 0 | 0.1 | 1 | 22 |
| aff4 | 0.13645 | -0.1498898 | 0 | 0.1 | 1 | 22 |
| akirin2 | 0.13645 | -0.18395794 | 0 | 0.1 | 1 | 22 |
| ankfy1 | 0.13645 | -0.24512905 | 0 | 0.1 | 1 | 22 |
| ankrd13a | 0.13645 | -0.19958046 | 0 | 0.1 | 1 | 22 |
| anks1b | 0.13645 | 0.327485565 | 0.1 | 0 | 1 | 22 |
| arhgef12b | 0.13645 | -0.29315769 | 0 | 0.1 | 1 | 22 |
| arl1 | 0.13645 | 0.196263002 | 0.1 | 0 | 1 | 22 |
| arl6ip5a | 0.13645 | -0.22278398 | 0 | 0.1 | 1 | 22 |
| arl8bb | 0.13645 | -0.24727942 | 0 | 0.1 | 1 | 22 |
| arpc1a | 0.13645 | -0.17564989 | 0 | 0.1 | 1 | 22 |
| atad1b | 0.13645 | -0.19882443 | 0 | 0.1 | 1 | 22 |
| atg12 | 0.13645 | -0.20718454 | 0 | 0.1 | 1 | 22 |
| axin1 | 0.13645 | -0.24008682 | 0 | 0.1 | 1 | 22 |

| | | | | | | |
|---|---|---|---|---|---|---|
| badb | 0.13645 | 0.259219814 | 0.1 | 0 | 1 | 22 |
| baz1b | 0.13645 | 0.264243111 | 0.1 | 0 | 1 | 22 |
| brd3a | 0.13645 | 0.266664742 | 0.1 | 0 | 1 | 22 |
| brd7 | 0.13645 | -0.15467442 | 0 | 0.1 | 1 | 22 |
| btf3l4 | 0.13645 | -0.17853338 | 0 | 0.1 | 1 | 22 |
| c1galt1b | 0.13645 | 0.233166968 | 0.1 | 0 | 1 | 22 |
| cdkn1a | 0.13645 | 0.239329432 | 0.1 | 0 | 1 | 22 |
| cept1b | 0.13645 | -0.27341915 | 0 | 0.1 | 1 | 22 |
| cflara | 0.13645 | -0.37910066 | 0 | 0.1 | 1 | 22 |
| chuk | 0.13645 | -0.15357164 | 0 | 0.1 | 1 | 22 |
| cip2a | 0.13645 | -0.30663461 | 0 | 0.1 | 1 | 22 |
| clk2a | 0.13645 | 0.222279717 | 0.1 | 0 | 1 | 22 |
| cmc2 | 0.13645 | -0.21446579 | 0 | 0.1 | 1 | 22 |
| cpsf3 | 0.13645 | -0.16052953 | 0 | 0.1 | 1 | 22 |
| cry-dash | 0.13645 | 0.224731446 | 0.1 | 0 | 1 | 22 |
| cry1aa | 0.13645 | -0.1979872 | 0 | 0.1 | 1 | 22 |
| dffa | 0.13645 | -0.1665661 | 0 | 0.1 | 1 | 22 |
| dlgap4a | 0.13645 | -0.23956038 | 0 | 0.1 | 1 | 22 |
| dpm2 | 0.13645 | -0.19035197 | 0 | 0.1 | 1 | 22 |
| dse | 0.13645 | -0.24712042 | 0 | 0.1 | 1 | 22 |
| dync1li2 | 0.13645 | -0.17193389 | 0 | 0.1 | 1 | 22 |
| ehmt2 | 0.13645 | -0.20301293 | 0 | 0.1 | 1 | 22 |
| eif3ja | 0.13645 | -0.13388375 | 0 | 0.1 | 1 | 22 |
| epn1 | 0.13645 | -0.16032092 | 0 | 0.1 | 1 | 22 |
| exoc1 | 0.13645 | -0.22321817 | 0 | 0.1 | 1 | 22 |
| exosc10 | 0.13645 | -0.21482212 | 0 | 0.1 | 1 | 22 |
| exosc4 | 0.13645 | -0.18946595 | 0 | 0.1 | 1 | 22 |
| ext2 | 0.13645 | -0.28167951 | 0 | 0.1 | 1 | 22 |
| ezh2 | 0.13645 | -0.19524663 | 0 | 0.1 | 1 | 22 |
| fam193b | 0.13645 | -0.25881486 | 0 | 0.1 | 1 | 22 |
| fam53b | 0.13645 | -0.17746637 | 0 | 0.1 | 1 | 22 |
| farsb | 0.13645 | 0.195584614 | 0.1 | 0 | 1 | 22 |
| fbxo5 | 0.13645 | 0.246284584 | 0.1 | 0 | 1 | 22 |
| fip1l1b | 0.13645 | -0.23349257 | 0 | 0.1 | 1 | 22 |
| ftr50 | 0.13645 | -0.22085985 | 0 | 0.1 | 1 | 22 |
| gadd45gip1 | 0.13645 | -0.17334845 | 0 | 0.1 | 1 | 22 |
| gemin7 | 0.13645 | -0.21410659 | 0 | 0.1 | 1 | 22 |
| gltpa | 0.13645 | -0.15370795 | 0 | 0.1 | 1 | 22 |
| glyr1 | 0.13645 | -0.19306396 | 0 | 0.1 | 1 | 22 |
| gnai2b | 0.13645 | -0.21141198 | 0 | 0.1 | 1 | 22 |
| gnl3 | 0.13645 | -0.14653052 | 0 | 0.1 | 1 | 22 |
| gpatch4 | 0.13645 | -0.30226516 | 0 | 0.1 | 1 | 22 |
| gpia | 0.13645 | -0.16916267 | 0 | 0.1 | 1 | 22 |
| gps2 | 0.13645 | -0.1884895 | 0 | 0.1 | 1 | 22 |
| gstk1 | 0.13645 | 0.278633031 | 0.1 | 0 | 1 | 22 |
| hdac3 | 0.13645 | -0.14103449 | 0 | 0.1 | 1 | 22 |
| hdgfl2 | 0.13645 | -0.13269855 | 0 | 0.1 | 1 | 22 |
| hint2 | 0.13645 | -0.15588668 | 0 | 0.1 | 1 | 22 |

| | | | | | | |
|---|---|---|---|---|---|---|
| hnrnpr | 0.13645 | -0.18105782 | 0 | 0.1 | 1 | 22 |
| hnrnpul1l | 0.13645 | -0.18937378 | 0 | 0.1 | 1 | 22 |
| hsd17b10 | 0.13645 | -0.18908369 | 0 | 0.1 | 1 | 22 |
| hyou1 | 0.13645 | -0.26140485 | 0 | 0.1 | 1 | 22 |
| iah1 | 0.13645 | 0.238159327 | 0.1 | 0 | 1 | 22 |
| icmt | 0.13645 | 0.233935666 | 0.1 | 0 | 1 | 22 |
| idh1 | 0.13645 | -0.16495588 | 0 | 0.1 | 1 | 22 |
| ift88 | 0.13645 | -0.15652652 | 0 | 0.1 | 1 | 22 |
| igf2bp2a | 0.13645 | 0.27064953 | 0.1 | 0 | 1 | 22 |
| il4r.1 | 0.13645 | -0.16839618 | 0 | 0.1 | 1 | 22 |
| inpp5b | 0.13645 | -0.30325632 | 0 | 0.1 | 1 | 22 |
| itpr3 | 0.13645 | -0.17208991 | 0 | 0.1 | 1 | 22 |
| kat8 | 0.13645 | 0.197582798 | 0.1 | 0 | 1 | 22 |
| kif21b | 0.13645 | -0.33017458 | 0 | 0.1 | 1 | 22 |
| knop1 | 0.13645 | -0.22353836 | 0 | 0.1 | 1 | 22 |
| kpna4 | 0.13645 | 0.291830013 | 0.1 | 0 | 1 | 22 |
| leo1 | 0.13645 | -0.28567252 | 0 | 0.1 | 1 | 22 |
| lifrb | 0.13645 | -0.31921571 | 0 | 0.1 | 1 | 22 |
| lin37 | 0.13645 | -0.17007387 | 0 | 0.1 | 1 | 22 |
| lmtk2 | 0.13645 | 0.289667032 | 0.1 | 0 | 1 | 22 |
| lrch4 | 0.13645 | -0.23702695 | 0 | 0.1 | 1 | 22 |
| lsm3 | 0.13645 | -0.15279851 | 0 | 0.1 | 1 | 22 |
| maf1 | 0.13645 | -0.17387997 | 0 | 0.1 | 1 | 22 |
| mdc1 | 0.13645 | -0.19489673 | 0 | 0.1 | 1 | 22 |
| med19a | 0.13645 | -0.25183325 | 0 | 0.1 | 1 | 22 |
| mgst1.1 | 0.13645 | -0.25448618 | 0 | 0.1 | 1 | 22 |
| mid1ip1l | 0.13645 | -0.26996053 | 0 | 0.1 | 1 | 22 |
| morc3a | 0.13645 | 0.2335612 | 0.1 | 0 | 1 | 22 |
| mri1 | 0.13645 | -0.24774508 | 0 | 0.1 | 1 | 22 |
| mrpl16 | 0.13645 | -0.18540254 | 0 | 0.1 | 1 | 22 |
| mrpl21 | 0.13645 | -0.11620771 | 0 | 0.1 | 1 | 22 |
| mrpl45 | 0.13645 | -0.21235085 | 0 | 0.1 | 1 | 22 |
| mrps25 | 0.13645 | -0.21670284 | 0 | 0.1 | 1 | 22 |
| mtfr1l | 0.13645 | -0.28028508 | 0 | 0.1 | 1 | 22 |
| mtmr4 | 0.13645 | -0.25968961 | 0 | 0.1 | 1 | 22 |
| mxi1 | 0.13645 | -0.19407712 | 0 | 0.1 | 1 | 22 |
| myo18ab | 0.13645 | -0.19038689 | 0 | 0.1 | 1 | 22 |
| mzt2b | 0.13645 | -0.2168293 | 0 | 0.1 | 1 | 22 |
| naa20 | 0.13645 | -0.17053181 | 0 | 0.1 | 1 | 22 |
| nasp | 0.13645 | -0.25242021 | 0 | 0.1 | 1 | 22 |
| ncaldb | 0.13645 | -0.19377536 | 0 | 0.1 | 1 | 22 |
| ncor2 | 0.13645 | 0.27745204 | 0.1 | 0 | 1 | 22 |
| ndufs2 | 0.13645 | 0.254002965 | 0.1 | 0 | 1 | 22 |
| nedd9 | 0.13645 | -0.26804047 | 0 | 0.1 | 1 | 22 |
| nek7 | 0.13645 | -0.16376525 | 0 | 0.1 | 1 | 22 |
| nfs1 | 0.13645 | -0.28410988 | 0 | 0.1 | 1 | 22 |
| nipa2 | 0.13645 | -0.21639767 | 0 | 0.1 | 1 | 22 |
| nol7 | 0.13645 | -0.21385208 | 0 | 0.1 | 1 | 22 |

| | | | | | | |
|---|---|---|---|---|---|---|
| nop16 | 0.13645 | -0.20302178 | 0 | 0.1 | 1 | 22 |
| nrbp2a | 0.13645 | -0.2204298 | 0 | 0.1 | 1 | 22 |
| nsmaf | 0.13645 | -0.12926717 | 0 | 0.1 | 1 | 22 |
| nudc | 0.13645 | -0.20512088 | 0 | 0.1 | 1 | 22 |
| os9 | 0.13645 | -0.24149394 | 0 | 0.1 | 1 | 22 |
| otud3 | 0.13645 | -0.21055625 | 0 | 0.1 | 1 | 22 |
| pa2g4a | 0.13645 | -0.24204605 | 0 | 0.1 | 1 | 22 |
| pag1 | 0.13645 | -0.31151778 | 0 | 0.1 | 1 | 22 |
| pam16 | 0.13645 | -0.20938015 | 0 | 0.1 | 1 | 22 |
| parp16 | 0.13645 | -0.22342653 | 0 | 0.1 | 1 | 22 |
| pcgf1 | 0.13645 | 0.27529509 | 0.1 | 0 | 1 | 22 |
| pdxkb | 0.13645 | -0.18620456 | 0 | 0.1 | 1 | 22 |
| pex7 | 0.13645 | -0.20351622 | 0 | 0.1 | 1 | 22 |
| pias4a | 0.13645 | -0.24294355 | 0 | 0.1 | 1 | 22 |
| pmm2 | 0.13645 | -0.14866293 | 0 | 0.1 | 1 | 22 |
| polr2h | 0.13645 | -0.1518843 | 0 | 0.1 | 1 | 22 |
| pptc7b | 0.13645 | -0.19964444 | 0 | 0.1 | 1 | 22 |
| prpf18 | 0.13645 | 0.295701169 | 0.1 | 0 | 1 | 22 |
| psme4b | 0.13645 | -0.148438 | 0 | 0.1 | 1 | 22 |
| psmg2 | 0.13645 | -0.20026222 | 0 | 0.1 | 1 | 22 |
| pwwp2b | 0.13645 | 0.242242846 | 0.1 | 0 | 1 | 22 |
| rab9b | 0.13645 | -0.21606458 | 0 | 0.1 | 1 | 22 |
| rangap1a | 0.13645 | -0.16794899 | 0 | 0.1 | 1 | 22 |
| rasal3 | 0.13645 | 0.203126533 | 0.1 | 0 | 1 | 22 |
| rfxank | 0.13645 | -0.23755963 | 0 | 0.1 | 1 | 22 |
| riok2 | 0.13645 | -0.1520318 | 0 | 0.1 | 1 | 22 |
| rnf20 | 0.13645 | -0.14825234 | 0 | 0.1 | 1 | 22 |
| rnf213b | 0.13645 | -0.20143228 | 0 | 0.1 | 1 | 22 |
| rnps1 | 0.13645 | -0.24722635 | 0 | 0.1 | 1 | 22 |
| rprd1b | 0.13645 | 0.207419619 | 0.1 | 0 | 1 | 22 |
| rrp15 | 0.13645 | -0.18442116 | 0 | 0.1 | 1 | 22 |
| sacm1lb | 0.13645 | -0.19329101 | 0 | 0.1 | 1 | 22 |
| scaf4a | 0.13645 | -0.28256631 | 0 | 0.1 | 1 | 22 |
| sdr42e1 | 0.13645 | -0.21652262 | 0 | 0.1 | 1 | 22 |
| sec24c | 0.13645 | -0.28439553 | 0 | 0.1 | 1 | 22 |
| selenoh | 0.13645 | 0.244581379 | 0.1 | 0 | 1 | 22 |
| sfxn3 | 0.13645 | -0.192579 | 0 | 0.1 | 1 | 22 |
| sgf29 | 0.13645 | 0.251605207 | 0.1 | 0 | 1 | 22 |
| si:ch1073-322p19.1 | 0.13645 | -0.18812326 | 0 | 0.1 | 1 | 22 |
| si:ch1073-392o20.2 | 0.13645 | -0.17542743 | 0 | 0.1 | 1 | 22 |
| si:ch211-130m23.5 | 0.13645 | -0.14892547 | 0 | 0.1 | 1 | 22 |
| si:ch211-162i8.5 | 0.13645 | -0.23536806 | 0 | 0.1 | 1 | 22 |
| si:ch211-276i12.11 | 0.13645 | 0.24800564 | 0.1 | 0 | 1 | 22 |
| si:ch73-238c9.1 | 0.13645 | -0.18633912 | 0 | 0.1 | 1 | 22 |
| si:ch73-71c20.5 | 0.13645 | -0.14567233 | 0 | 0.1 | 1 | 22 |
| si:dkey-100n23.3 | 0.13645 | 0.249781133 | 0.1 | 0 | 1 | 22 |
| si:dkey-111k8.3 | 0.13645 | -0.17376881 | 0 | 0.1 | 1 | 22 |
| si:dkey-16l2.16 | 0.13645 | -0.24074043 | 0 | 0.1 | 1 | 22 |

| | | | | | | |
|---|---|---|---|---|---|---|
| si:dkey-180p18.9 | 0.13645 | -0.25184398 | 0 | 0.1 | 1 | 22 |
| si:dkey-1c7.2 | 0.13645 | -0.20728513 | 0 | 0.1 | 1 | 22 |
| si:dkey-276j7.1 | 0.13645 | -0.33778296 | 0 | 0.1 | 1 | 22 |
| si:dkey-93h22.8 | 0.13645 | -0.21626663 | 0 | 0.1 | 1 | 22 |
| si:dkeyp-113d7.1 | 0.13645 | -0.16811629 | 0 | 0.1 | 1 | 22 |
| si:dkeyp-113d7.10 | 0.13645 | 0.304965089 | 0.1 | 0 | 1 | 22 |
| si:dkeyp-87e7.4 | 0.13645 | 0.291836375 | 0.1 | 0 | 1 | 22 |
| smad3a | 0.13645 | 0.258548701 | 0.1 | 0 | 1 | 22 |
| smu1a | 0.13645 | -0.13829264 | 0 | 0.1 | 1 | 22 |
| snapc3 | 0.13645 | -0.16624466 | 0 | 0.1 | 1 | 22 |
| snrnp25 | 0.13645 | -0.17427846 | 0 | 0.1 | 1 | 22 |
| snrnp70 | 0.13645 | -0.17575967 | 0 | 0.1 | 1 | 22 |
| snx14 | 0.13645 | -0.21392372 | 0 | 0.1 | 1 | 22 |
| snx30 | 0.13645 | -0.19658962 | 0 | 0.1 | 1 | 22 |
| spns3 | 0.13645 | -0.20321245 | 0 | 0.1 | 1 | 22 |
| srsf6a | 0.13645 | -0.23946984 | 0 | 0.1 | 1 | 22 |
| ssbp3a | 0.13645 | -0.20064434 | 0 | 0.1 | 1 | 22 |
| ssbp3b | 0.13645 | -0.28686042 | 0 | 0.1 | 1 | 22 |
| stam2 | 0.13645 | -0.2030882 | 0 | 0.1 | 1 | 22 |
| stat2 | 0.13645 | -0.23375137 | 0 | 0.1 | 1 | 22 |
| stim2b | 0.13645 | -0.15852644 | 0 | 0.1 | 1 | 22 |
| taf3 | 0.13645 | -0.16525972 | 0 | 0.1 | 1 | 22 |
| tagln2 | 0.13645 | -0.19101079 | 0 | 0.1 | 1 | 22 |
| thap12a | 0.13645 | -0.24853647 | 0 | 0.1 | 1 | 22 |
| tia1l | 0.13645 | -0.21918633 | 0 | 0.1 | 1 | 22 |
| tial1 | 0.13645 | 0.228884149 | 0.1 | 0 | 1 | 22 |
| timm23a | 0.13645 | -0.19940732 | 0 | 0.1 | 1 | 22 |
| timm9 | 0.13645 | -0.1860129 | 0 | 0.1 | 1 | 22 |
| tiparp | 0.13645 | -0.2753405 | 0 | 0.1 | 1 | 22 |
| tmem106c | 0.13645 | -0.29002339 | 0 | 0.1 | 1 | 22 |
| tmem168b | 0.13645 | -0.23165543 | 0 | 0.1 | 1 | 22 |
| tmem42b | 0.13645 | -0.16725553 | 0 | 0.1 | 1 | 22 |
| tomm40 | 0.13645 | -0.16209879 | 0 | 0.1 | 1 | 22 |
| tox4b | 0.13645 | -0.32765552 | 0 | 0.1 | 1 | 22 |
| tpd52l2a | 0.13645 | -0.2165617 | 0 | 0.1 | 1 | 22 |
| trappc4 | 0.13645 | -0.16681266 | 0 | 0.1 | 1 | 22 |
| trappc6bl | 0.13645 | -0.31089245 | 0 | 0.1 | 1 | 22 |
| trir | 0.13645 | -0.19412245 | 0 | 0.1 | 1 | 22 |
| ttc9c | 0.13645 | -0.16588178 | 0 | 0.1 | 1 | 22 |
| ube2ib | 0.13645 | -0.21172363 | 0 | 0.1 | 1 | 22 |
| ulk2 | 0.13645 | -0.29051587 | 0 | 0.1 | 1 | 22 |
| usp22 | 0.13645 | 0.252262243 | 0.1 | 0 | 1 | 22 |
| vapb | 0.13645 | -0.22799094 | 0 | 0.1 | 1 | 22 |
| vgll4b | 0.13645 | -0.23534366 | 0 | 0.1 | 1 | 22 |
| vps26a | 0.13645 | -0.14863821 | 0 | 0.1 | 1 | 22 |
| wdtc1 | 0.13645 | 0.209056193 | 0.1 | 0 | 1 | 22 |
| wu:fi46h04 | 0.13645 | -0.15366029 | 0 | 0.1 | 1 | 22 |
| wwp1 | 0.13645 | -0.18032471 | 0 | 0.1 | 1 | 22 |

| | | | | | | |
|---|---|---|---|---|---|---|
| zc3h15 | 0.13645 | -0.16364273 | 0 | 0.1 | 1 | 22 |
| zgc:111986 | 0.13645 | -0.22840336 | 0 | 0.1 | 1 | 22 |
| zgc:112962 | 0.13645 | -0.2222343 | 0 | 0.1 | 1 | 22 |
| zgc:113436 | 0.13645 | 0.234403933 | 0.1 | 0 | 1 | 22 |
| zgc:66474 | 0.13645 | -0.26116844 | 0 | 0.1 | 1 | 22 |
| zgc:92481 | 0.13645 | -0.21463065 | 0 | 0.1 | 1 | 22 |
| zhx3 | 0.13645 | 0.28766823 | 0.1 | 0 | 1 | 22 |
| znf236 | 0.13645 | -0.20307177 | 0 | 0.1 | 1 | 22 |
| znf318 | 0.13645 | 0.261368454 | 0.1 | 0 | 1 | 22 |
| znf609a | 0.13645 | -0.2442173 | 0 | 0.1 | 1 | 22 |
| znf704 | 0.13645 | -0.29582222 | 0 | 0.1 | 1 | 22 |
| znf710b | 0.13645 | 0.252904257 | 0.1 | 0 | 1 | 22 |
| znf800b | 0.13645 | -0.13200314 | 0 | 0.1 | 1 | 22 |
| LO018340.1 | 0.13645 | 0.243517007 | 0.1 | 0 | 1 | 22 |
| LOC103909972 | 0.13645 | -0.24751068 | 0 | 0.1 | 1 | 22 |
| XLOC-035413 | 0.13645 | -0.13239968 | 0 | 0.1 | 1 | 22 |
| actr1 | 0.13645 | -0.18532684 | 0 | 0.1 | 1 | 22 |
| ada2a | 0.13645 | -0.19436142 | 0 | 0.1 | 1 | 22 |
| adipor1a | 0.13645 | -0.20133114 | 0 | 0.1 | 1 | 22 |
| aimp2 | 0.13645 | -0.18385284 | 0 | 0.1 | 1 | 22 |
| akt2l | 0.13645 | -0.17163657 | 0 | 0.1 | 1 | 22 |
| arhgap36 | 0.13645 | 0.223470025 | 0.1 | 0 | 1 | 22 |
| ccr10 | 0.13645 | -0.20110257 | 0 | 0.1 | 1 | 22 |
| cd59 | 0.13645 | 0.256485124 | 0.1 | 0 | 1 | 22 |
| cmtm3 | 0.13645 | -0.18265234 | 0 | 0.1 | 1 | 22 |
| coa3a | 0.13645 | 0.202416764 | 0.1 | 0 | 1 | 22 |
| dcaf7 | 0.13645 | -0.25272087 | 0 | 0.1 | 1 | 22 |
| ddx10 | 0.13645 | 0.237432448 | 0.1 | 0 | 1 | 22 |
| ero1b | 0.13645 | 0.23131757 | 0.1 | 0 | 1 | 22 |
| flii | 0.13645 | -0.13811039 | 0 | 0.1 | 1 | 22 |
| glrx3 | 0.13645 | 0.210482734 | 0.1 | 0 | 1 | 22 |
| gtf2h5 | 0.13645 | 0.242068474 | 0.1 | 0 | 1 | 22 |
| hsp70.3 | 0.13645 | 0.198749638 | 0.1 | 0 | 1 | 22 |
| jagn1b | 0.13645 | 0.235556734 | 0.1 | 0 | 1 | 22 |
| katnb1 | 0.13645 | -0.11651093 | 0 | 0.1 | 1 | 22 |
| lgals9l1 | 0.13645 | -0.14108364 | 0 | 0.1 | 1 | 22 |
| maco1b | 0.13645 | -0.14113125 | 0 | 0.1 | 1 | 22 |
| mccc2 | 0.13645 | -0.12494557 | 0 | 0.1 | 1 | 22 |
| metap1 | 0.13645 | -0.11194075 | 0 | 0.1 | 1 | 22 |
| mrps35 | 0.13645 | -0.18431016 | 0 | 0.1 | 1 | 22 |
| nacc2 | 0.13645 | -0.18711649 | 0 | 0.1 | 1 | 22 |
| nol12 | 0.13645 | 0.241437338 | 0.1 | 0 | 1 | 22 |
| nucb2a | 0.13645 | -0.20617303 | 0 | 0.1 | 1 | 22 |
| ogdhb | 0.13645 | -0.29588644 | 0 | 0.1 | 1 | 22 |
| oxsr1a | 0.13645 | 0.243618512 | 0.1 | 0 | 1 | 22 |
| pdzd11 | 0.13645 | -0.17119376 | 0 | 0.1 | 1 | 22 |
| pnpla6 | 0.13645 | 0.232244908 | 0.1 | 0 | 1 | 22 |
| rnf38 | 0.13645 | -0.1930811 | 0 | 0.1 | 1 | 22 |

| | | | | | | |
|---|---|---|---|---|---|---|
| sco2 | 0.13645 | -0.16421942 | 0 | 0.1 | 1 | 22 |
| sec23ip | 0.13645 | -0.19004818 | 0 | 0.1 | 1 | 22 |
| serpinb1l3 | 0.13645 | -0.18361919 | 0 | 0.1 | 1 | 22 |
| si:ch211-150o23.2 | 0.13645 | -0.18737447 | 0 | 0.1 | 1 | 22 |
| si:ch211-157j23.3 | 0.13645 | -0.14678249 | 0 | 0.1 | 1 | 22 |
| si:ch211-157p22.10 | 0.13645 | -0.2560112 | 0 | 0.1 | 1 | 22 |
| smg5 | 0.13645 | 0.270501049 | 0.1 | 0 | 1 | 22 |
| smim7 | 0.13645 | 0.207751193 | 0.1 | 0 | 1 | 22 |
| tarbp2 | 0.13645 | -0.18346531 | 0 | 0.1 | 1 | 22 |
| tbc1d14 | 0.13645 | 0.251963301 | 0.1 | 0 | 1 | 22 |
| tbl3 | 0.13645 | -0.20088657 | 0 | 0.1 | 1 | 22 |
| tceanc2 | 0.13645 | -0.18125858 | 0 | 0.1 | 1 | 22 |
| tdh | 0.13645 | 0.194978943 | 0.1 | 0 | 1 | 22 |
| timm8b | 0.13645 | -0.21040033 | 0 | 0.1 | 1 | 22 |
| tomm40l | 0.13645 | -0.13343019 | 0 | 0.1 | 1 | 22 |
| ubl3a | 0.13645 | -0.19456846 | 0 | 0.1 | 1 | 22 |
| ufd1l | 0.13645 | -0.1841927 | 0 | 0.1 | 1 | 22 |
| yaf2 | 0.13645 | -0.1758563 | 0 | 0.1 | 1 | 22 |
| zgc:77838 | 0.13645 | -0.12136358 | 0 | 0.1 | 1 | 22 |
| znf740b | 0.13645 | -0.17744773 | 0 | 0.1 | 1 | 22 |
| arglu1b | 0.13645 | -0.22054035 | 0 | 0.1 | 1 | 22 |
| arih2 | 0.13645 | -0.25510817 | 0 | 0.1 | 1 | 22 |
| bola1 | 0.13645 | -0.19720706 | 0 | 0.1 | 1 | 22 |
| coq10b | 0.13645 | -0.25242571 | 0 | 0.1 | 1 | 22 |
| exoc2 | 0.13645 | -0.22727543 | 0 | 0.1 | 1 | 22 |
| gyg1a | 0.13645 | 0.234758023 | 0.1 | 0 | 1 | 22 |
| mat2aa | 0.13645 | -0.27171541 | 0 | 0.1 | 1 | 22 |
| mvb12ba | 0.13645 | -0.20860514 | 0 | 0.1 | 1 | 22 |
| osbpl5 | 0.13645 | 0.150659653 | 0.1 | 0 | 1 | 22 |
| papss1 | 0.13645 | -0.21137989 | 0 | 0.1 | 1 | 22 |
| pgrmc1 | 0.13645 | -0.2626282 | 0 | 0.1 | 1 | 22 |
| rnaset2 | 0.13645 | 0.29019616 | 0.1 | 0 | 1 | 22 |
| rnf11b | 0.13645 | -0.27162738 | 0 | 0.1 | 1 | 22 |
| senp3b | 0.13645 | -0.20664138 | 0 | 0.1 | 1 | 22 |
| twist3 | 0.13645 | 0.264373723 | 0.1 | 0 | 1 | 22 |
| ubald1b | 0.13645 | -0.26917049 | 0 | 0.1 | 1 | 22 |
| unkl | 0.13645 | 0.291742653 | 0.1 | 0 | 1 | 22 |
| alg12 | 0.13645 | 0.257943309 | 0.1 | 0 | 1 | 22 |
| ammecr1 | 0.13645 | 0.29505265 | 0.1 | 0 | 1 | 22 |
| c1d | 0.13645 | -0.13860825 | 0 | 0.1 | 1 | 22 |
| cab39l | 0.13645 | 0.261470681 | 0.1 | 0 | 1 | 22 |
| dnajb9b | 0.13645 | 0.291943162 | 0.1 | 0 | 1 | 22 |
| dyrk1ab | 0.13645 | 0.310234439 | 0.1 | 0 | 1 | 22 |
| etnk1 | 0.13645 | -0.2893276 | 0 | 0.1 | 1 | 22 |
| gmfg | 0.13645 | -0.19777582 | 0 | 0.1 | 1 | 22 |
| grsf1 | 0.13645 | 0.304408008 | 0.1 | 0 | 1 | 22 |
| pes | 0.13645 | 0.281224765 | 0.1 | 0 | 1 | 22 |
| rmnd5b | 0.13645 | -0.24055886 | 0 | 0.1 | 1 | 22 |

| | | | | | | |
|---|---|---|---|---|---|---|
| sin3ab | 0.13645 | -0.31747736 | 0 | 0.1 | 1 | 22 |
| sorl1 | 0.13645 | 0.313180829 | 0.1 | 0 | 1 | 22 |
| trim33l | 0.13645 | 0.300410268 | 0.1 | 0 | 1 | 22 |
| tsc22d1 | 0.13645 | 0.242091242 | 0.1 | 0 | 1 | 22 |
| wdr89 | 0.13645 | -0.23505492 | 0 | 0.1 | 1 | 22 |
| zdbf2 | 0.13645 | 0.24801667 | 0.1 | 0 | 1 | 22 |
| clpxb | 0.13645 | -0.23275841 | 0 | 0.1 | 1 | 22 |
| eif4e2rs1 | 0.13645 | -0.23831589 | 0 | 0.1 | 1 | 22 |
| psma6b | 0.13645 | -0.23968326 | 0 | 0.1 | 1 | 22 |
| rinl | 0.13645 | -0.27220075 | 0 | 0.1 | 1 | 22 |
| BX284666.1 | 0.13645 | 0.254666237 | 0.1 | 0 | 1 | 22 |
| CU467110.1 | 0.13645 | 0.295482593 | 0.1 | 0 | 1 | 22 |
| acsl1a | 0.13645 | 0.319197725 | 0.1 | 0 | 1 | 22 |
| dbf4 | 0.13645 | 0.296326318 | 0.1 | 0 | 1 | 22 |
| gpn3 | 0.13645 | 0.214596303 | 0.1 | 0 | 1 | 22 |
| scrib | 0.13645 | 0.262189592 | 0.1 | 0 | 1 | 22 |
| si:ch211-226m16.2 | 0.13645 | 0.255490238 | 0.1 | 0 | 1 | 22 |
| vmp1 | 0.13645 | 0.205651164 | 0.1 | 0 | 1 | 22 |
| BX005450.1 | 0.13645 | 0.290928126 | 0.1 | 0 | 1 | 22 |
| BX957362.1 | 0.13645 | -0.23493548 | 0 | 0.1 | 1 | 22 |
| CABZ01069162.1 | 0.13645 | 0.281784212 | 0.1 | 0 | 1 | 22 |
| CCDC127 | 0.13645 | 0.194344467 | 0.1 | 0 | 1 | 22 |
| CU019662.1 | 0.13645 | -0.2538165 | 0 | 0.1 | 1 | 22 |
| CU462878.2 | 0.13645 | 0.287638852 | 0.1 | 0 | 1 | 22 |
| CU694481.1 | 0.13645 | -0.20852355 | 0 | 0.1 | 1 | 22 |
| FO393424.4 | 0.13645 | -0.21588396 | 0 | 0.1 | 1 | 22 |
| FQ976913.1 | 0.13645 | 0.315022595 | 0.1 | 0 | 1 | 22 |
| LO017872.1 | 0.13645 | 0.187112833 | 0.1 | 0 | 1 | 22 |
| LOC100148254 | 0.13645 | -0.19353942 | 0 | 0.1 | 1 | 22 |
| LOC101883052 | 0.13645 | -0.19171963 | 0 | 0.1 | 1 | 22 |
| LOC103910871 | 0.13645 | -0.1619718 | 0 | 0.1 | 1 | 22 |
| MAP3K11 | 0.13645 | 0.198073392 | 0.1 | 0 | 1 | 22 |
| SIKE1 | 0.13645 | 0.241881183 | 0.1 | 0 | 1 | 22 |
| SIPA1 | 0.13645 | -0.26594832 | 0 | 0.1 | 1 | 22 |
| XLOC-001102 | 0.13645 | 0.218697881 | 0.1 | 0 | 1 | 22 |
| XLOC-014477 | 0.13645 | 0.251105818 | 0.1 | 0 | 1 | 22 |
| XLOC-043670 | 0.13645 | 0.271155098 | 0.1 | 0 | 1 | 22 |
| abhd15a | 0.13645 | -0.20242138 | 0 | 0.1 | 1 | 22 |
| abhd17b | 0.13645 | -0.18463155 | 0 | 0.1 | 1 | 22 |
| adap2 | 0.13645 | -0.29882726 | 0 | 0.1 | 1 | 22 |
| adtrp1 | 0.13645 | 0.226390746 | 0.1 | 0 | 1 | 22 |
| akr1a1b | 0.13645 | 0.247286187 | 0.1 | 0 | 1 | 22 |
| anapc11 | 0.13645 | 0.259502061 | 0.1 | 0 | 1 | 22 |
| anln | 0.13645 | 0.267908353 | 0.1 | 0 | 1 | 22 |
| ap2s1 | 0.13645 | -0.17145769 | 0 | 0.1 | 1 | 22 |
| arhgap42a | 0.13645 | -0.1787542 | 0 | 0.1 | 1 | 22 |
| arhgef18b | 0.13645 | -0.190473 | 0 | 0.1 | 1 | 22 |
| arih1 | 0.13645 | -0.17186209 | 0 | 0.1 | 1 | 22 |

| | | | | | | |
|---|---|---|---|---|---|---|
| aspm | 0.13645 | 0.268713061 | 0.1 | 0 | 1 | 22 |
| azin1b | 0.13645 | -0.20536451 | 0 | 0.1 | 1 | 22 |
| b4galt1l | 0.13645 | 0.289478983 | 0.1 | 0 | 1 | 22 |
| baz2ba | 0.13645 | -0.18062391 | 0 | 0.1 | 1 | 22 |
| bckdha | 0.13645 | -0.22574275 | 0 | 0.1 | 1 | 22 |
| bicra | 0.13645 | 0.227211408 | 0.1 | 0 | 1 | 22 |
| birc5a | 0.13645 | 0.243420329 | 0.1 | 0 | 1 | 22 |
| capn2a | 0.13645 | -0.23973014 | 0 | 0.1 | 1 | 22 |
| cbx8b | 0.13645 | -0.26651926 | 0 | 0.1 | 1 | 22 |
| cd209 | 0.13645 | -0.2182686 | 0 | 0.1 | 1 | 22 |
| cdipt | 0.13645 | -0.18333085 | 0 | 0.1 | 1 | 22 |
| cdk11b | 0.13645 | -0.23563216 | 0 | 0.1 | 1 | 22 |
| chd3 | 0.13645 | -0.24516232 | 0 | 0.1 | 1 | 22 |
| cited4b | 0.13645 | 0.205594279 | 0.1 | 0 | 1 | 22 |
| cks2 | 0.13645 | 0.255803293 | 0.1 | 0 | 1 | 22 |
| clasp2 | 0.13645 | 0.209063801 | 0.1 | 0 | 1 | 22 |
| cndp2 | 0.13645 | 0.224285829 | 0.1 | 0 | 1 | 22 |
| cnot1 | 0.13645 | -0.14432111 | 0 | 0.1 | 1 | 22 |
| cnot3b | 0.13645 | -0.24437133 | 0 | 0.1 | 1 | 22 |
| cnot6a | 0.13645 | 0.233034645 | 0.1 | 0 | 1 | 22 |
| cnot8 | 0.13645 | -0.2053544 | 0 | 0.1 | 1 | 22 |
| cnppd1 | 0.13645 | 0.301941909 | 0.1 | 0 | 1 | 22 |
| col4a3bpb | 0.13645 | -0.17700326 | 0 | 0.1 | 1 | 22 |
| cops7a | 0.13645 | -0.18242843 | 0 | 0.1 | 1 | 22 |
| coq5 | 0.13645 | 0.203810115 | 0.1 | 0 | 1 | 22 |
| cox20 | 0.13645 | -0.18654087 | 0 | 0.1 | 1 | 22 |
| csnk1db | 0.13645 | 0.25235639 | 0.1 | 0 | 1 | 22 |
| ctbp1 | 0.13645 | -0.24154549 | 0 | 0.1 | 1 | 22 |
| ctdspl3 | 0.13645 | 0.174128563 | 0.1 | 0 | 1 | 22 |
| cyp46a1.2 | 0.13645 | -0.18111211 | 0 | 0.1 | 1 | 22 |
| ddost | 0.13645 | -0.17335759 | 0 | 0.1 | 1 | 22 |
| derl1 | 0.13645 | -0.17426675 | 0 | 0.1 | 1 | 22 |
| dlgap5 | 0.13645 | 0.210926062 | 0.1 | 0 | 1 | 22 |
| eif2b4 | 0.13645 | 0.329518766 | 0.1 | 0 | 1 | 22 |
| eif4e1c | 0.13645 | -0.15082094 | 0 | 0.1 | 1 | 22 |
| epn2 | 0.13645 | 0.276592783 | 0.1 | 0 | 1 | 22 |
| exoc4 | 0.13645 | 0.241339663 | 0.1 | 0 | 1 | 22 |
| f11r.1 | 0.13645 | 0.322905057 | 0.1 | 0 | 1 | 22 |
| fahd2a | 0.13645 | -0.20245936 | 0 | 0.1 | 1 | 22 |
| fam126a | 0.13645 | -0.24404289 | 0 | 0.1 | 1 | 22 |
| fbxw11b | 0.13645 | -0.26029342 | 0 | 0.1 | 1 | 22 |
| foxo3b | 0.13645 | 0.19245432 | 0.1 | 0 | 1 | 22 |
| ftr24 | 0.13645 | 0.186434613 | 0.1 | 0 | 1 | 22 |
| furina | 0.13645 | 0.284728733 | 0.1 | 0 | 1 | 22 |
| gabarapl2 | 0.13645 | 0.269659843 | 0.1 | 0 | 1 | 22 |
| gak | 0.13645 | 0.279936017 | 0.1 | 0 | 1 | 22 |
| gnl3l | 0.13645 | 0.295099836 | 0.1 | 0 | 1 | 22 |
| gnptg | 0.13645 | -0.17954179 | 0 | 0.1 | 1 | 22 |

| | | | | | | |
|---|---|---|---|---|---|---|
| got2b | 0.13645 | -0.28221611 | 0 | 0.1 | 1 | 22 |
| gpr137c | 0.13645 | -0.16566197 | 0 | 0.1 | 1 | 22 |
| gsdmeb | 0.13645 | -0.21862559 | 0 | 0.1 | 1 | 22 |
| gtf2e1 | 0.13645 | -0.1610799 | 0 | 0.1 | 1 | 22 |
| gtf3c6 | 0.13645 | 0.233218936 | 0.1 | 0 | 1 | 22 |
| gtse1 | 0.13645 | 0.22007404 | 0.1 | 0 | 1 | 22 |
| hcfc1b | 0.13645 | -0.32717896 | 0 | 0.1 | 1 | 22 |
| hddc3 | 0.13645 | 0.233302844 | 0.1 | 0 | 1 | 22 |
| helz | 0.13645 | -0.21088045 | 0 | 0.1 | 1 | 22 |
| hif1an | 0.13645 | -0.16536349 | 0 | 0.1 | 1 | 22 |
| htr5ab | 0.13645 | 0.200069185 | 0.1 | 0 | 1 | 22 |
| ikbke | 0.13645 | -0.21486615 | 0 | 0.1 | 1 | 22 |
| il2rgb | 0.13645 | -0.19663044 | 0 | 0.1 | 1 | 22 |
| illr1 | 0.13645 | 0.329657382 | 0.1 | 0 | 1 | 22 |
| irak4 | 0.13645 | 0.208231151 | 0.1 | 0 | 1 | 22 |
| itpka | 0.13645 | 0.211534934 | 0.1 | 0 | 1 | 22 |
| kctd6b | 0.13645 | 0.285009483 | 0.1 | 0 | 1 | 22 |
| kif20bb | 0.13645 | 0.22007404 | 0.1 | 0 | 1 | 22 |
| kin | 0.13645 | 0.268374967 | 0.1 | 0 | 1 | 22 |
| klf12a | 0.13645 | 0.242340861 | 0.1 | 0 | 1 | 22 |
| kmt2ca | 0.13645 | 0.260168835 | 0.1 | 0 | 1 | 22 |
| ktn1 | 0.13645 | 0.330543838 | 0.1 | 0 | 1 | 22 |
| letm1 | 0.13645 | -0.23693136 | 0 | 0.1 | 1 | 22 |
| letm2 | 0.13645 | -0.24967775 | 0 | 0.1 | 1 | 22 |
| limk1a | 0.13645 | 0.226980785 | 0.1 | 0 | 1 | 22 |
| lpp | 0.13645 | -0.26977744 | 0 | 0.1 | 1 | 22 |
| lsm4 | 0.13645 | -0.15543312 | 0 | 0.1 | 1 | 22 |
| malt2 | 0.13645 | -0.21366815 | 0 | 0.1 | 1 | 22 |
| map3k12 | 0.13645 | 0.25355796 | 0.1 | 0 | 1 | 22 |
| mapk3 | 0.13645 | -0.17058591 | 0 | 0.1 | 1 | 22 |
| mapkapk5 | 0.13645 | -0.19850648 | 0 | 0.1 | 1 | 22 |
| mbtps2 | 0.13645 | -0.20424368 | 0 | 0.1 | 1 | 22 |
| mdm2 | 0.13645 | -0.23564326 | 0 | 0.1 | 1 | 22 |
| mif4gda | 0.13645 | 0.275096537 | 0.1 | 0 | 1 | 22 |
| mpc1 | 0.13645 | -0.27404744 | 0 | 0.1 | 1 | 22 |
| mpp6b | 0.13645 | 0.199748089 | 0.1 | 0 | 1 | 22 |
| mrpl4 | 0.13645 | -0.18730584 | 0 | 0.1 | 1 | 22 |
| mrps16 | 0.13645 | 0.207186861 | 0.1 | 0 | 1 | 22 |
| mrps17 | 0.13645 | -0.1810987 | 0 | 0.1 | 1 | 22 |
| mrps24 | 0.13645 | -0.18222191 | 0 | 0.1 | 1 | 22 |
| mrps6 | 0.13645 | -0.1486753 | 0 | 0.1 | 1 | 22 |
| mrtfbb | 0.13645 | -0.22247611 | 0 | 0.1 | 1 | 22 |
| myorg | 0.13645 | 0.21159331 | 0.1 | 0 | 1 | 22 |
| nat9 | 0.13645 | -0.17682791 | 0 | 0.1 | 1 | 22 |
| naxd | 0.13645 | -0.1926295 | 0 | 0.1 | 1 | 22 |
| ncbp2 | 0.13645 | 0.278109878 | 0.1 | 0 | 1 | 22 |
| ndnl2 | 0.13645 | -0.10195475 | 0 | 0.1 | 1 | 22 |
| ngdn | 0.13645 | -0.21803155 | 0 | 0.1 | 1 | 22 |

| | | | | | | |
|---|---|---|---|---|---|---|
| nkd1 | 0.13645 | -0.23148959 | 0 | 0.1 | 1 | 22 |
| nkl.3 | 0.13645 | 0.31801199 | 0.1 | 0 | 1 | 22 |
| nolc1 | 0.13645 | -0.17109018 | 0 | 0.1 | 1 | 22 |
| nsd1a | 0.13645 | -0.2301744 | 0 | 0.1 | 1 | 22 |
| nudcd3 | 0.13645 | -0.18528809 | 0 | 0.1 | 1 | 22 |
| nutf2 | 0.13645 | -0.17348278 | 0 | 0.1 | 1 | 22 |
| parp2 | 0.13645 | 0.205450808 | 0.1 | 0 | 1 | 22 |
| pbxip1b | 0.13645 | 0.169417778 | 0.1 | 0 | 1 | 22 |
| pde7a | 0.13645 | 0.224795867 | 0.1 | 0 | 1 | 22 |
| pik3ca | 0.13645 | 0.265921034 | 0.1 | 0 | 1 | 22 |
| pnn | 0.13645 | 0.388905653 | 0.1 | 0 | 1 | 22 |
| pole4 | 0.13645 | -0.27346903 | 0 | 0.1 | 1 | 22 |
| psmd3 | 0.13645 | 0.247211274 | 0.1 | 0 | 1 | 22 |
| ptpn2a | 0.13645 | 0.235737248 | 0.1 | 0 | 1 | 22 |
| ptpn2b | 0.13645 | -0.21014072 | 0 | 0.1 | 1 | 22 |
| pttg1 | 0.13645 | -0.1453286 | 0 | 0.1 | 1 | 22 |
| rab27a | 0.13645 | -0.28783593 | 0 | 0.1 | 1 | 22 |
| rabgap1 | 0.13645 | 0.278277809 | 0.1 | 0 | 1 | 22 |
| rap2b | 0.13645 | -0.16014352 | 0 | 0.1 | 1 | 22 |
| rap2c | 0.13645 | -0.13883119 | 0 | 0.1 | 1 | 22 |
| rdh14b | 0.13645 | 0.235189047 | 0.1 | 0 | 1 | 22 |
| rfx5 | 0.13645 | 0.244173708 | 0.1 | 0 | 1 | 22 |
| rnf145b | 0.13645 | 0.262100817 | 0.1 | 0 | 1 | 22 |
| rnf181 | 0.13645 | -0.24566891 | 0 | 0.1 | 1 | 22 |
| rnf4 | 0.13645 | 0.218291253 | 0.1 | 0 | 1 | 22 |
| rtraf | 0.13645 | -0.23980871 | 0 | 0.1 | 1 | 22 |
| samm50 | 0.13645 | -0.16393701 | 0 | 0.1 | 1 | 22 |
| sap30bp | 0.13645 | -0.20990716 | 0 | 0.1 | 1 | 22 |
| shq1 | 0.13645 | -0.28206733 | 0 | 0.1 | 1 | 22 |
| si:ch1073-357b18.4 | 0.13645 | -0.22780529 | 0 | 0.1 | 1 | 22 |
| si:ch211-160o17.4 | 0.13645 | 0.273096792 | 0.1 | 0 | 1 | 22 |
| si:ch211-194k22.8 | 0.13645 | 0.243766297 | 0.1 | 0 | 1 | 22 |
| si:ch211-209f23.6 | 0.13645 | 0.257001124 | 0.1 | 0 | 1 | 22 |
| si:ch211-210b2.4 | 0.13645 | 0.228800977 | 0.1 | 0 | 1 | 22 |
| si:ch211-244b2.4 | 0.13645 | -0.21188148 | 0 | 0.1 | 1 | 22 |
| si:ch211-259g3.4 | 0.13645 | 0.29374128 | 0.1 | 0 | 1 | 22 |
| si:ch211-276i12.9 | 0.13645 | 0.26177088 | 0.1 | 0 | 1 | 22 |
| si:ch211-8c17.3 | 0.13645 | -0.21180433 | 0 | 0.1 | 1 | 22 |
| si:dkey-171c9.3 | 0.13645 | 0.302497804 | 0.1 | 0 | 1 | 22 |
| si:dkey-204f11.64 | 0.13645 | 0.26420511 | 0.1 | 0 | 1 | 22 |
| si:dkey-239h2.3 | 0.13645 | 0.231250994 | 0.1 | 0 | 1 | 22 |
| si:dkey-26m3.3 | 0.13645 | -0.21600707 | 0 | 0.1 | 1 | 22 |
| si:dkey-27b3.2 | 0.13645 | 0.22176656 | 0.1 | 0 | 1 | 22 |
| si:dkeyp-69e1.8 | 0.13645 | -0.1416259 | 0 | 0.1 | 1 | 22 |
| ska2 | 0.13645 | 0.230098873 | 0.1 | 0 | 1 | 22 |
| slain2 | 0.13645 | -0.16184187 | 0 | 0.1 | 1 | 22 |
| slc35a4 | 0.13645 | -0.21049573 | 0 | 0.1 | 1 | 22 |
| slc35b2 | 0.13645 | -0.16246245 | 0 | 0.1 | 1 | 22 |

| | | | | | | |
|---|---|---|---|---|---|---|
| slc7a8a | 0.13645 | -0.16138084 | 0 | 0.1 | 1 | 22 |
| smad2 | 0.13645 | -0.18132309 | 0 | 0.1 | 1 | 22 |
| smarcd1 | 0.13645 | -0.15991928 | 0 | 0.1 | 1 | 22 |
| smfn | 0.13645 | -0.16915764 | 0 | 0.1 | 1 | 22 |
| spdl1 | 0.13645 | 0.250898713 | 0.1 | 0 | 1 | 22 |
| spsb4b | 0.13645 | -0.21210386 | 0 | 0.1 | 1 | 22 |
| sptbn2 | 0.13645 | -0.17776182 | 0 | 0.1 | 1 | 22 |
| sptlc2b | 0.13645 | -0.15948533 | 0 | 0.1 | 1 | 22 |
| srp19 | 0.13645 | 0.192584209 | 0.1 | 0 | 1 | 22 |
| srp72 | 0.13645 | -0.20490277 | 0 | 0.1 | 1 | 22 |
| srsf10b | 0.13645 | -0.122508 | 0 | 0.1 | 1 | 22 |
| st3gal3a | 0.13645 | 0.218418467 | 0.1 | 0 | 1 | 22 |
| stk40 | 0.13645 | -0.21270106 | 0 | 0.1 | 1 | 22 |
| syne1b | 0.13645 | -0.24235442 | 0 | 0.1 | 1 | 22 |
| taf7 | 0.13645 | -0.1666685 | 0 | 0.1 | 1 | 22 |
| tango2 | 0.13645 | -0.28750076 | 0 | 0.1 | 1 | 22 |
| taok1a | 0.13645 | -0.24967008 | 0 | 0.1 | 1 | 22 |
| tfr1b | 0.13645 | -0.21090886 | 0 | 0.1 | 1 | 22 |
| tkfc | 0.13645 | 0.11101071 | 0.1 | 0 | 1 | 22 |
| tmem11 | 0.13645 | -0.1681843 | 0 | 0.1 | 1 | 22 |
| tmem131 | 0.13645 | 0.296175334 | 0.1 | 0 | 1 | 22 |
| tmem231 | 0.13645 | 0.19635741 | 0.1 | 0 | 1 | 22 |
| tmem30ab | 0.13645 | 0.280770254 | 0.1 | 0 | 1 | 22 |
| tomm70a | 0.13645 | 0.209844912 | 0.1 | 0 | 1 | 22 |
| trpt1 | 0.13645 | 0.190055961 | 0.1 | 0 | 1 | 22 |
| ttc7b | 0.13645 | -0.15360353 | 0 | 0.1 | 1 | 22 |
| txlna | 0.13645 | 0.246596217 | 0.1 | 0 | 1 | 22 |
| ube2g2 | 0.13645 | -0.1624778 | 0 | 0.1 | 1 | 22 |
| ube2h | 0.13645 | -0.22784762 | 0 | 0.1 | 1 | 22 |
| ube2q1 | 0.13645 | 0.141010347 | 0.1 | 0 | 1 | 22 |
| ubqln4 | 0.13645 | 0.21096781 | 0.1 | 0 | 1 | 22 |
| ubxn1 | 0.13645 | -0.19275346 | 0 | 0.1 | 1 | 22 |
| unk | 0.13645 | -0.26772162 | 0 | 0.1 | 1 | 22 |
| uqcrc2b | 0.13645 | -0.15952613 | 0 | 0.1 | 1 | 22 |
| usp25 | 0.13645 | 0.228800977 | 0.1 | 0 | 1 | 22 |
| usp5 | 0.13645 | 0.271514729 | 0.1 | 0 | 1 | 22 |
| usp7 | 0.13645 | -0.24268184 | 0 | 0.1 | 1 | 22 |
| utp23 | 0.13645 | -0.27570733 | 0 | 0.1 | 1 | 22 |
| vbp1 | 0.13645 | -0.2298416 | 0 | 0.1 | 1 | 22 |
| wdr11 | 0.13645 | 0.134559113 | 0.1 | 0 | 1 | 22 |
| wu:fu71h07 | 0.13645 | 0.235799432 | 0.1 | 0 | 1 | 22 |
| xpo4 | 0.13645 | -0.1571226 | 0 | 0.1 | 1 | 22 |
| yy1a | 0.13645 | -0.29687592 | 0 | 0.1 | 1 | 22 |
| zc3h14 | 0.13645 | 0.194208852 | 0.1 | 0 | 1 | 22 |
| zfx | 0.13645 | 0.20962612 | 0.1 | 0 | 1 | 22 |
| zgc:110239 | 0.13645 | 0.263402745 | 0.1 | 0 | 1 | 22 |
| zgc:110843 | 0.13645 | -0.2323513 | 0 | 0.1 | 1 | 22 |
| zgc:112255 | 0.13645 | 0.304952033 | 0.1 | 0 | 1 | 22 |

| | | | | | | |
|---|---|---|---|---|---|---|
| zgc:77151 | 0.13645 | -0.20335946 | 0 | 0.1 | 1 | 22 |
| zgc:77262 | 0.13645 | -0.21290542 | 0 | 0.1 | 1 | 22 |
| zgc:92360 | 0.13645 | -0.21887391 | 0 | 0.1 | 1 | 22 |
| zgc:92907 | 0.13645 | -0.2075384 | 0 | 0.1 | 1 | 22 |
| zmat5 | 0.13645 | -0.17971735 | 0 | 0.1 | 1 | 22 |
| znf827 | 0.13645 | -0.20365816 | 0 | 0.1 | 1 | 22 |
| akirin1 | 0.13663 | -0.26225452 | 0.1 | 0.3 | 1 | 22 |
| si:ch211-277c7.6 | 0.13677 | -0.16391773 | 0.4 | 0.6 | 1 | 22 |
| rps25 | 0.13683 | -0.33776779 | 0.8 | 0.9 | 1 | 22 |
| cd74b | 0.13751 | 0.135881938 | 1 | 1 | 1 | 22 |
| NC-002333.4 | 0.13802 | -0.32727349 | 0.1 | 0.3 | 1 | 22 |
| XLOC-010899 | 0.13879 | 0.543667934 | 0.55 | 0.3 | 1 | 22 |
| rpl31 | 0.14052 | -0.13010388 | 0.95 | 0.9 | 1 | 22 |
| egr3 | 0.14125 | -0.16719887 | 0.2 | 0.4 | 1 | 22 |
| notch2 | 0.14146 | 0.339942308 | 0.35 | 0.3 | 1 | 22 |
| spcs2 | 0.14294 | 0.525518113 | 0.3 | 0.15 | 1 | 22 |
| rhogb | 0.14317 | -0.37851422 | 0.4 | 0.7 | 1 | 22 |
| h2afy2 | 0.14385 | -0.33836943 | 0.1 | 0.35 | 1 | 22 |
| rps2 | 0.14526 | -0.21946729 | 0.85 | 1 | 1 | 22 |
| si:ch211-119e14.1 | 0.14745 | -0.41533093 | 0.3 | 0.6 | 1 | 22 |
| prdx2 | 0.1488 | -0.33075798 | 0.45 | 0.75 | 1 | 22 |
| psap | 0.15009 | 0.392842616 | 0.5 | 0.4 | 1 | 22 |
| rps3a | 0.1503 | -0.28162104 | 0.65 | 0.9 | 1 | 22 |
| atp5po | 0.15387 | 0.490795593 | 0.5 | 0.3 | 1 | 22 |
| serbp1a | 0.1544 | -0.40540843 | 0.55 | 0.8 | 1 | 22 |
| psmb6 | 0.15481 | -0.41251864 | 0.1 | 0.35 | 1 | 22 |
| rbm4.3 | 0.15544 | -0.45585898 | 0.1 | 0.35 | 1 | 22 |
| atp5mf | 0.15556 | -0.43036647 | 0.1 | 0.35 | 1 | 22 |
| phlda2 | 0.1559 | -0.22894673 | 0.05 | 0.25 | 1 | 22 |
| eif3s10 | 0.15592 | -0.48950577 | 0.1 | 0.35 | 1 | 22 |
| cfbl | 0.15626 | -0.40004696 | 0.4 | 0.7 | 1 | 22 |
| rbm39a | 0.15975 | -0.35134512 | 0.05 | 0.25 | 1 | 22 |
| ndufb3 | 0.16176 | 0.499994186 | 0.45 | 0.25 | 1 | 22 |
| hint1 | 0.16503 | -0.28160371 | 0.05 | 0.25 | 1 | 22 |
| eif2s2 | 0.16519 | 0.2257835 | 0.35 | 0.4 | 1 | 22 |
| mknk2b | 0.16569 | -0.18732202 | 0.15 | 0.35 | 1 | 22 |
| bcam | 0.16787 | 0.488688121 | 0.25 | 0.05 | 1 | 22 |
| slc25a3b | 0.1689 | -0.20258623 | 0.45 | 0.7 | 1 | 22 |
| rpl30 | 0.16937 | -0.18558674 | 0.75 | 0.95 | 1 | 22 |
| mhc2dab | 0.1711 | -0.16011078 | 0.25 | 0.45 | 1 | 22 |
| rpl26 | 0.17375 | -0.19884181 | 0.9 | 1 | 1 | 22 |
| itga4 | 0.17428 | -0.28207004 | 0.3 | 0.55 | 1 | 22 |
| rps12 | 0.1756 | -0.39530395 | 0.6 | 0.85 | 1 | 22 |
| ddx61 | 0.17686 | 0.249237821 | 0.4 | 0.4 | 1 | 22 |
| zgc:114188 | 0.17714 | -0.28060314 | 0.75 | 0.95 | 1 | 22 |
| tox | 0.17971 | -0.31001542 | 0.05 | 0.25 | 1 | 22 |
| parvg | 0.18017 | 0.509390482 | 0.4 | 0.15 | 1 | 22 |
| arf5 | 0.18032 | -0.37390836 | 0.05 | 0.25 | 1 | 22 |

| | | | | | | |
|---|---|---|---|---|---|---|
| arhgap33 | 0.18068 | -0.30636137 | 0.05 | 0.25 | 1 | 22 |
| npc2 | 0.18106 | 0.465535813 | 0.25 | 0.05 | 1 | 22 |
| rpl11 | 0.18109 | -0.26217824 | 0.75 | 0.8 | 1 | 22 |
| gpr137ba | 0.18232 | -0.51351439 | 0.15 | 0.4 | 1 | 22 |
| rpl28 | 0.18236 | -0.1961692 | 0.9 | 0.95 | 1 | 22 |
| CR391910.2 | 0.18306 | -0.26256018 | 0.1 | 0.3 | 1 | 22 |
| si:ch211-51e12.7 | 0.18382 | -0.26072382 | 0.05 | 0.25 | 1 | 22 |
| ube2ia | 0.18422 | -0.29903226 | 0.05 | 0.25 | 1 | 22 |
| gstr | 0.18477 | -0.27453379 | 0.05 | 0.25 | 1 | 22 |
| tnrc6c1 | 0.18542 | -0.50688507 | 0.15 | 0.4 | 1 | 22 |
| rab11ba | 0.18559 | -0.35247461 | 0.05 | 0.25 | 1 | 22 |
| dbnlb | 0.18727 | -0.4595248 | 0.05 | 0.25 | 1 | 22 |
| eif6 | 0.18738 | -0.38954148 | 0.05 | 0.25 | 1 | 22 |
| wu:fi37e09 | 0.18789 | -0.34486291 | 0.2 | 0.45 | 1 | 22 |
| sik1 | 0.18789 | 0.538176796 | 0.4 | 0.15 | 1 | 22 |
| si:dkeyp-68b7.12 | 0.18816 | -0.41129571 | 0.05 | 0.25 | 1 | 22 |
| hic1 | 0.18827 | 0.488234804 | 0.25 | 0.05 | 1 | 22 |
| rps3 | 0.18928 | -0.37276459 | 0.65 | 0.85 | 1 | 22 |
| lmnb1 | 0.18968 | -0.34755256 | 0.05 | 0.25 | 1 | 22 |
| LOC101886073 | 0.19034 | -0.46769296 | 0.05 | 0.25 | 1 | 22 |
| fth1a | 0.19035 | -0.41420986 | 0.15 | 0.4 | 1 | 22 |
| mtmr12 | 0.19054 | 0.411486919 | 0.25 | 0.05 | 1 | 22 |
| bcl2l13 | 0.19062 | -0.36335848 | 0.05 | 0.25 | 1 | 22 |
| hspd1 | 0.19076 | -0.43826883 | 0.05 | 0.25 | 1 | 22 |
| slc25a20 | 0.1909 | -0.33340921 | 0.05 | 0.25 | 1 | 22 |
| cisd2 | 0.19112 | -0.37618211 | 0.05 | 0.25 | 1 | 22 |
| cycsb | 0.19225 | -0.32927795 | 0.2 | 0.45 | 1 | 22 |
| tma7 | 0.19257 | -0.15829898 | 0.25 | 0.45 | 1 | 22 |
| fybb | 0.19274 | -0.39338056 | 0.15 | 0.4 | 1 | 22 |
| nop10 | 0.19357 | 0.276657513 | 0.3 | 0.3 | 1 | 22 |
| il10ra | 0.1941 | -0.51028832 | 0.25 | 0.5 | 1 | 22 |
| cirbpb | 0.19514 | 0.366399897 | 0.85 | 0.65 | 1 | 22 |
| metap2b | 0.19708 | -0.39617529 | 0.15 | 0.4 | 1 | 22 |
| synrg | 0.19798 | 0.445463981 | 0.2 | 0.05 | 1 | 22 |
| rpl19 | 0.20084 | -0.29607261 | 0.75 | 0.9 | 1 | 22 |
| si:ch73-380l3.2 | 0.20089 | 0.124706495 | 0.3 | 0.35 | 1 | 22 |
| eif3k | 0.20153 | -0.42312704 | 0.15 | 0.4 | 1 | 22 |
| dtnbp1a | 0.20213 | -0.15872273 | 0.25 | 0.45 | 1 | 22 |
| sod2 | 0.20279 | 0.447420611 | 0.25 | 0.1 | 1 | 22 |
| bbip1 | 0.20313 | 0.44897879 | 0.25 | 0.1 | 1 | 22 |
| rpl38 | 0.20353 | -0.24869305 | 0.85 | 0.95 | 1 | 22 |
| pcmtd1 | 0.20372 | 0.493532714 | 0.3 | 0.1 | 1 | 22 |
| rps14 | 0.20489 | -0.23058913 | 0.8 | 0.85 | 1 | 22 |
| nap1l4b | 0.20681 | -0.23945412 | 0.1 | 0.3 | 1 | 22 |
| rgs13 | 0.20797 | 0.411841885 | 0.7 | 0.55 | 1 | 22 |
| fkbp3 | 0.20838 | -0.22248444 | 0.1 | 0.3 | 1 | 22 |
| bin2b | 0.2087 | -0.13834606 | 0.1 | 0.25 | 1 | 22 |
| rab10 | 0.20911 | -0.23601401 | 0.1 | 0.3 | 1 | 22 |

| | | | | | | |
|---|---|---|---|---|---|---|
| lypla2 | 0.20939 | -0.14857395 | 0.05 | 0.2 | 1 | 22 |
| mef2aa | 0.21201 | 0.489064245 | 0.3 | 0.1 | 1 | 22 |
| ssr3 | 0.21394 | -0.2635937 | 0.2 | 0.45 | 1 | 22 |
| prkar2aa | 0.21681 | -0.17500085 | 0.2 | 0.4 | 1 | 22 |
| rpl37 | 0.22144 | -0.20927857 | 0.9 | 1 | 1 | 22 |
| snrpe | 0.22193 | -0.10101991 | 0.1 | 0.25 | 1 | 22 |
| rap1aa | 0.22307 | 0.355944618 | 0.2 | 0.15 | 1 | 22 |
| fosab | 0.22357 | 0.380062287 | 0.75 | 0.6 | 1 | 22 |
| brd2a | 0.22414 | -0.1759524 | 0.1 | 0.25 | 1 | 22 |
| eif4a1b | 0.22425 | 0.298539861 | 0.45 | 0.4 | 1 | 22 |
| mibp2 | 0.22469 | -0.4217415 | 0.55 | 0.75 | 1 | 22 |
| tnrc6b | 0.2247 | 0.457804538 | 0.2 | 0.05 | 1 | 22 |
| eif5a | 0.22555 | 0.379238038 | 0.55 | 0.4 | 1 | 22 |
| mt-co1 | 0.22562 | -0.25935718 | 0.8 | 0.85 | 1 | 22 |
| CR753876.1 | 0.22618 | -0.27236185 | 0.8 | 0.95 | 1 | 22 |
| nsa2 | 0.22683 | -0.11282182 | 0.45 | 0.65 | 1 | 22 |
| serf2 | 0.23083 | -0.42612304 | 0.4 | 0.65 | 1 | 22 |
| ndufa3 | 0.23111 | 0.443301293 | 0.3 | 0.15 | 1 | 22 |
| ppp3ccb | 0.23151 | -0.31437713 | 0.1 | 0.3 | 1 | 22 |
| pho | 0.23186 | 0.255380493 | 0.25 | 0.25 | 1 | 22 |
| ndufa7 | 0.23229 | -0.2174925 | 0.05 | 0.2 | 1 | 22 |
| eef1g | 0.2323 | -0.31255633 | 0.55 | 0.8 | 1 | 22 |
| sept7b | 0.23426 | -0.29483359 | 0.1 | 0.3 | 1 | 22 |
| pfdn1 | 0.23574 | -0.16001424 | 0.05 | 0.2 | 1 | 22 |
| myo1f | 0.23602 | -0.36631461 | 0.25 | 0.5 | 1 | 22 |
| ndufab1b | 0.23626 | 0.171960678 | 0.15 | 0.2 | 1 | 22 |
| psmg3 | 0.23677 | 0.414215099 | 0.25 | 0.1 | 1 | 22 |
| ywhaqb | 0.239 | 0.464952187 | 0.3 | 0.1 | 1 | 22 |
| trim25 | 0.23935 | 0.449432612 | 0.2 | 0.05 | 1 | 22 |
| CABZ01046432.1 | 0.2424 | -0.30538448 | 0.1 | 0.3 | 1 | 22 |
| wbp2nl | 0.24266 | -0.12995944 | 0.1 | 0.25 | 1 | 22 |
| arpc4 | 0.24335 | -0.18455705 | 0.65 | 0.85 | 1 | 22 |
| rnf130 | 0.24399 | -0.47876501 | 0.1 | 0.3 | 1 | 22 |
| cox17 | 0.24406 | -0.35008336 | 0.25 | 0.5 | 1 | 22 |
| rpl7a | 0.24422 | -0.20436571 | 0.7 | 0.9 | 1 | 22 |
| tnfb | 0.24442 | -0.1482677 | 0.15 | 0.3 | 1 | 22 |
| ndufb9 | 0.24535 | -0.2915791 | 0.1 | 0.3 | 1 | 22 |
| rpl34 | 0.24604 | -0.25230539 | 0.8 | 0.95 | 1 | 22 |
| hmgn7 | 0.24674 | 0.198359305 | 0.45 | 0.45 | 1 | 22 |
| rps4x | 0.24681 | -0.30148713 | 0.7 | 0.9 | 1 | 22 |
| oaz1b | 0.24747 | -0.36734071 | 0.45 | 0.7 | 1 | 22 |
| dicp1.1 | 0.24847 | -0.21051237 | 0.3 | 0.5 | 1 | 22 |
| hsbp1b | 0.24908 | -0.35177393 | 0.1 | 0.3 | 1 | 22 |
| map4k5 | 0.25086 | -0.19662867 | 0.05 | 0.2 | 1 | 22 |
| BX901920.1 | 0.25355 | -0.27784365 | 0.15 | 0.35 | 1 | 22 |
| rpl23 | 0.25557 | -0.17957255 | 0.9 | 1 | 1 | 22 |
| tardbp | 0.2574 | -0.20270051 | 0.05 | 0.2 | 1 | 22 |
| atp2a3 | 0.25894 | 0.419552646 | 0.3 | 0.1 | 1 | 22 |

| | | | | | | |
|---|---|---|---|---|---|---|
| skap2 | 0.25989 | -0.30088051 | 0.35 | 0.6 | 1 | 22 |
| fcer1g | 0.25991 | -0.34475054 | 0.5 | 0.75 | 1 | 22 |
| sod1 | 0.26078 | -0.41894941 | 0.3 | 0.55 | 1 | 22 |
| psmb3 | 0.2608 | -0.25356553 | 0.05 | 0.2 | 1 | 22 |
| crema | 0.26089 | 0.418210118 | 0.6 | 0.4 | 1 | 22 |
| plcg2 | 0.26135 | -0.21917817 | 0.05 | 0.2 | 1 | 22 |
| srgn | 0.26213 | 0.272057534 | 0.9 | 0.7 | 1 | 22 |
| mdkb | 0.26469 | 0.399062495 | 0.3 | 0.1 | 1 | 22 |
| ndufb4 | 0.26582 | -0.2776778 | 0.1 | 0.3 | 1 | 22 |
| ctss2.1 | 0.26764 | -0.1855143 | 0.45 | 0.65 | 1 | 22 |
| wu:fb55g09 | 0.26795 | -0.32863658 | 0.35 | 0.6 | 1 | 22 |
| zgc:174945 | 0.26818 | -0.34594848 | 0.1 | 0.3 | 1 | 22 |
| herpud1 | 0.26849 | 0.408258214 | 0.25 | 0.15 | 1 | 22 |
| seta | 0.26877 | 0.457113468 | 0.35 | 0.15 | 1 | 22 |
| aldoab | 0.26931 | -0.38291006 | 0.1 | 0.3 | 1 | 22 |
| kmt5aa | 0.26969 | -0.35307833 | 0.1 | 0.3 | 1 | 22 |
| kctd5a | 0.26983 | 0.413310976 | 0.4 | 0.25 | 1 | 22 |
| selenok | 0.2718 | -0.17109452 | 0.05 | 0.2 | 1 | 22 |
| selenof | 0.27274 | 0.408201097 | 0.3 | 0.15 | 1 | 22 |
| ctsba | 0.27286 | -0.15447021 | 0.1 | 0.25 | 1 | 22 |
| rab8b | 0.27359 | -0.31012757 | 0.1 | 0.3 | 1 | 22 |
| tp53 | 0.27389 | -0.22255589 | 0.05 | 0.2 | 1 | 22 |
| aldh6a1 | 0.2748 | -0.42970299 | 0.05 | 0.2 | 1 | 22 |
| ndufa4l | 0.27666 | 0.257580479 | 0.5 | 0.45 | 1 | 22 |
| rad21a | 0.27674 | -0.34735149 | 0.1 | 0.3 | 1 | 22 |
| RASSF5 | 0.27678 | 0.280369378 | 0.4 | 0.35 | 1 | 22 |
| chac1 | 0.27707 | -0.31776895 | 0.4 | 0.65 | 1 | 22 |
| mob3a | 0.27741 | -0.39505104 | 0.35 | 0.6 | 1 | 22 |
| rhocb | 0.27996 | -0.22912783 | 0.05 | 0.2 | 1 | 22 |
| ncoa4 | 0.2807 | 0.357391031 | 0.2 | 0.05 | 1 | 22 |
| kdm2ba | 0.28224 | 0.324806978 | 0.2 | 0.15 | 1 | 22 |
| rpl8 | 0.28254 | -0.1064581 | 0.8 | 0.95 | 1 | 22 |
| ptprea | 0.28266 | 0.450272987 | 0.45 | 0.25 | 1 | 22 |
| il7r | 0.28385 | -0.28722676 | 0.15 | 0.35 | 1 | 22 |
| cbx3a | 0.28766 | -0.28927326 | 0.15 | 0.35 | 1 | 22 |
| eif4g2a | 0.28827 | -0.19726317 | 0.05 | 0.2 | 1 | 22 |
| BX682234.1 | 0.28879 | -0.41036934 | 0.15 | 0.35 | 1 | 22 |
| mob4 | 0.29077 | -0.177365 | 0.05 | 0.2 | 1 | 22 |
| gabarapa | 0.29105 | -0.18321163 | 0.3 | 0.5 | 1 | 22 |
| h1fx | 0.29136 | -0.19062232 | 0.55 | 0.75 | 1 | 22 |
| tmpob | 0.29207 | 0.377970949 | 0.2 | 0.05 | 1 | 22 |
| CABZ01109481.1 | 0.2948 | 0.42861792 | 0.35 | 0.2 | 1 | 22 |
| pdpk1a | 0.29548 | -0.26475026 | 0.05 | 0.2 | 1 | 22 |
| lrrfip1a | 0.29573 | -0.18266754 | 0.4 | 0.6 | 1 | 22 |
| slc25a51b | 0.29602 | 0.370934858 | 0.15 | 0.05 | 1 | 22 |
| eif3ha | 0.29612 | -0.29445265 | 0.15 | 0.35 | 1 | 22 |
| atp6v0b | 0.29619 | -0.13158503 | 0.1 | 0.25 | 1 | 22 |
| tob1a | 0.2966 | -0.43985871 | 0.25 | 0.45 | 1 | 22 |

| | | | | | | |
|---|---|---|---|---|---|---|
| ppp1r18 | 0.29955 | 0.332005744 | 0.35 | 0.15 | 1 | 22 |
| atf1 | 0.3021 | 0.33806627 | 0.2 | 0.05 | 1 | 22 |
| MYADM | 0.3025 | 0.34252094 | 0.2 | 0.05 | 1 | 22 |
| med29 | 0.30365 | 0.366213995 | 0.2 | 0.05 | 1 | 22 |
| mtch2 | 0.3041 | -0.17235398 | 0.05 | 0.2 | 1 | 22 |
| rps26l | 0.30498 | -0.22378626 | 0.8 | 0.95 | 1 | 22 |
| smdt1b | 0.30697 | -0.18145024 | 0.1 | 0.25 | 1 | 22 |
| selenot1a | 0.30717 | -0.28376742 | 0.15 | 0.35 | 1 | 22 |
| FERMT3 (1 of many) | 0.30753 | -0.20021092 | 0.1 | 0.25 | 1 | 22 |
| nptna | 0.30806 | 0.183065076 | 0.45 | 0.45 | 1 | 22 |
| zgc:158564 | 0.30815 | 0.268225153 | 0.2 | 0.05 | 1 | 22 |
| plekhj1 | 0.3084 | -0.23430023 | 0.05 | 0.2 | 1 | 22 |
| si:dkey-206d17.12 | 0.30899 | -0.14169163 | 0.2 | 0.35 | 1 | 22 |
| mvb12a | 0.30974 | 0.362891807 | 0.2 | 0.05 | 1 | 22 |
| rgs18 | 0.31035 | -0.42962977 | 0.15 | 0.35 | 1 | 22 |
| si:dkey-187e18.1 | 0.31037 | -0.22891607 | 0.05 | 0.2 | 1 | 22 |
| ywhaqa | 0.31085 | -0.15292366 | 0.1 | 0.25 | 1 | 22 |
| selplg | 0.3119 | -0.31673059 | 0.15 | 0.35 | 1 | 22 |
| tardbpl | 0.31353 | 0.391670926 | 0.2 | 0.05 | 1 | 22 |
| edf1 | 0.31369 | 0.399449469 | 0.35 | 0.15 | 1 | 22 |
| ppp2r5eb | 0.31397 | -0.22039673 | 0.05 | 0.2 | 1 | 22 |
| arhgdig | 0.31439 | -0.20287207 | 0.75 | 0.8 | 1 | 22 |
| rpl32 | 0.31623 | -0.21405751 | 0.8 | 0.95 | 1 | 22 |
| calm2a | 0.31691 | 0.400614823 | 0.3 | 0.15 | 1 | 22 |
| psme2 | 0.3176 | 0.389893921 | 0.25 | 0.1 | 1 | 22 |
| grap2b | 0.31781 | -0.20811394 | 0.55 | 0.75 | 1 | 22 |
| bloc1s2 | 0.31803 | -0.24914896 | 0.05 | 0.2 | 1 | 22 |
| si:zfos-741a10.3 | 0.31891 | 0.36398514 | 0.35 | 0.25 | 1 | 22 |
| fhit | 0.31895 | 0.423007243 | 0.2 | 0.05 | 1 | 22 |
| sumo3a | 0.31955 | -0.23334902 | 0.05 | 0.2 | 1 | 22 |
| rpl6 | 0.3205 | -0.21327333 | 0.8 | 0.95 | 1 | 22 |
| si:ch73-376l24.3 | 0.32065 | -0.20706475 | 0.05 | 0.2 | 1 | 22 |
| rps24 | 0.3208 | -0.21692114 | 0.8 | 0.95 | 1 | 22 |
| tmc8 | 0.32305 | -0.24862078 | 0.05 | 0.2 | 1 | 22 |
| ndufa8 | 0.32333 | -0.2564063 | 0.05 | 0.2 | 1 | 22 |
| eif4a3 | 0.32393 | -0.23625712 | 0.05 | 0.2 | 1 | 22 |
| wdr19 | 0.32394 | 0.37504324 | 0.25 | 0.15 | 1 | 22 |
| rplp0 | 0.3244 | -0.29023629 | 0.65 | 0.85 | 1 | 22 |
| tmed7 | 0.32452 | -0.15049549 | 0.15 | 0.3 | 1 | 22 |
| cxcr4b | 0.32453 | 0.382877697 | 0.7 | 0.5 | 1 | 22 |
| impa1 | 0.32455 | -0.20626366 | 0.05 | 0.2 | 1 | 22 |
| LOC110438754 | 0.32476 | -0.26661083 | 0.05 | 0.2 | 1 | 22 |
| atp5if1b | 0.32501 | 0.395953167 | 0.35 | 0.2 | 1 | 22 |
| commd7 | 0.32556 | -0.25095555 | 0.05 | 0.2 | 1 | 22 |
| tmem59 | 0.32571 | 0.23125212 | 0.25 | 0.25 | 1 | 22 |
| arpc5b | 0.32639 | -0.20463655 | 0.6 | 0.8 | 1 | 22 |
| tax1bp1b | 0.32675 | -0.27975562 | 0.05 | 0.2 | 1 | 22 |
| lamtor2 | 0.32752 | -0.22249189 | 0.05 | 0.2 | 1 | 22 |

| | | | | | | |
|---|---|---|---|---|---|---|
| pkn1a | 0.32774 | -0.31565314 | 0.05 | 0.2 | 1 | 22 |
| psma1 | 0.32784 | 0.399356392 | 0.4 | 0.25 | 1 | 22 |
| tspan3a | 0.32798 | -0.24408496 | 0.05 | 0.2 | 1 | 22 |
| ndufb10 | 0.32847 | -0.27678332 | 0.05 | 0.2 | 1 | 22 |
| si:ch211-102c2.4 | 0.32909 | 0.100855873 | 0.95 | 0.95 | 1 | 22 |
| atp6v1g1 | 0.32949 | 0.359427129 | 0.45 | 0.3 | 1 | 22 |
| wnk1a | 0.32969 | 0.356363509 | 0.2 | 0.05 | 1 | 22 |
| lrmp | 0.32989 | -0.21897484 | 0.05 | 0.2 | 1 | 22 |
| cct3 | 0.33077 | 0.407795299 | 0.3 | 0.15 | 1 | 22 |
| tagapb | 0.33082 | 0.312327787 | 0.8 | 0.6 | 1 | 22 |
| rpl3 | 0.33129 | -0.28269379 | 0.65 | 0.85 | 1 | 22 |
| slc11a2 | 0.33195 | -0.22007442 | 0.05 | 0.2 | 1 | 22 |
| plxnc1 | 0.33238 | -0.35010247 | 0.15 | 0.35 | 1 | 22 |
| crebzf | 0.33257 | -0.28407246 | 0.05 | 0.2 | 1 | 22 |
| ier3ip1 | 0.33308 | 0.300222607 | 0.2 | 0.05 | 1 | 22 |
| lgals2a | 0.33309 | 0.387852385 | 0.2 | 0.05 | 1 | 22 |
| adcy7 | 0.33348 | -0.4141642 | 0.05 | 0.2 | 1 | 22 |
| itm2ba | 0.33348 | 0.214124299 | 0.3 | 0.3 | 1 | 22 |
| smc1al | 0.33352 | -0.26627784 | 0.05 | 0.2 | 1 | 22 |
| wee2 | 0.33448 | -0.28746037 | 0.05 | 0.2 | 1 | 22 |
| zgc:91910 | 0.33459 | 0.383873222 | 0.25 | 0.1 | 1 | 22 |
| prmt1 | 0.33472 | 0.370532485 | 0.15 | 0.05 | 1 | 22 |
| si:dkeyp-80c12.4 | 0.33519 | -0.30110074 | 0.05 | 0.2 | 1 | 22 |
| eif4ebp2 | 0.33529 | -0.37080589 | 0.15 | 0.35 | 1 | 22 |
| fgd4a | 0.33534 | -0.36828209 | 0.15 | 0.35 | 1 | 22 |
| zgc:174863 | 0.33547 | -0.18545921 | 0.1 | 0.25 | 1 | 22 |
| ppargc1b | 0.33566 | -0.27900412 | 0.05 | 0.2 | 1 | 22 |
| fli1b | 0.33722 | 0.374314006 | 0.2 | 0.05 | 1 | 22 |
| nocta | 0.33771 | -0.38548607 | 0.15 | 0.35 | 1 | 22 |
| parp8 | 0.33799 | -0.2909342 | 0.05 | 0.2 | 1 | 22 |
| BX470116.1 | 0.3393 | 0.12107271 | 0.05 | 0 | 1 | 22 |
| CABZ01079302.1 | 0.3393 | 0.144720172 | 0.05 | 0 | 1 | 22 |
| CABZ01079427.1 | 0.3393 | 0.14203593 | 0.05 | 0 | 1 | 22 |
| CR387919.1 | 0.3393 | 0.164499116 | 0.05 | 0 | 1 | 22 |
| CR847994.1 | 0.3393 | 0.132797662 | 0.05 | 0 | 1 | 22 |
| CR855337.1 | 0.3393 | 0.132797662 | 0.05 | 0 | 1 | 22 |
| CT583708.1 | 0.3393 | 0.116949861 | 0.05 | 0 | 1 | 22 |
| CU856539.2 | 0.3393 | 0.129234235 | 0.05 | 0 | 1 | 22 |
| EML4 | 0.3393 | 0.184256764 | 0.05 | 0 | 1 | 22 |
| FP102783.1 | 0.3393 | 0.132797662 | 0.05 | 0 | 1 | 22 |
| LOC103909167 | 0.3393 | 0.132797662 | 0.05 | 0 | 1 | 22 |
| LOC103910363 | 0.3393 | 0.142878505 | 0.05 | 0 | 1 | 22 |
| LOC103911601 | 0.3393 | 0.144720172 | 0.05 | 0 | 1 | 22 |
| LOC108179036 | 0.3393 | 0.128861068 | 0.05 | 0 | 1 | 22 |
| LOC108182329 | 0.3393 | 0.122813994 | 0.05 | 0 | 1 | 22 |
| LOC110438330 | 0.3393 | 0.144720172 | 0.05 | 0 | 1 | 22 |
| XLOC-015481 | 0.3393 | 0.132797662 | 0.05 | 0 | 1 | 22 |
| XLOC-026010 | 0.3393 | 0.127734995 | 0.05 | 0 | 1 | 22 |

| | | | | | | |
|---|---|---|---|---|---|---|
| XLOC-042487 | 0.3393 | 0.132797662 | 0.05 | 0 | 1 | 22 |
| abcc3 | 0.3393 | 0.141197952 | 0.05 | 0 | 1 | 22 |
| abhd2a | 0.3393 | 0.141866653 | 0.05 | 0 | 1 | 22 |
| acp2 | 0.3393 | 0.142971503 | 0.05 | 0 | 1 | 22 |
| actr5 | 0.3393 | 0.144720172 | 0.05 | 0 | 1 | 22 |
| agps | 0.3393 | 0.139201425 | 0.05 | 0 | 1 | 22 |
| akap8l | 0.3393 | 0.12440346 | 0.05 | 0 | 1 | 22 |
| aldh7a1 | 0.3393 | 0.157849702 | 0.05 | 0 | 1 | 22 |
| arfgap2 | 0.3393 | 0.139139724 | 0.05 | 0 | 1 | 22 |
| arnt2 | 0.3393 | 0.127003156 | 0.05 | 0 | 1 | 22 |
| ascc3 | 0.3393 | 0.14086338 | 0.05 | 0 | 1 | 22 |
| atp13a2 | 0.3393 | 0.144720172 | 0.05 | 0 | 1 | 22 |
| atxn7l2a | 0.3393 | 0.132797662 | 0.05 | 0 | 1 | 22 |
| ccl39.7 | 0.3393 | 0.135694967 | 0.05 | 0 | 1 | 22 |
| chd2 | 0.3393 | 0.142214336 | 0.05 | 0 | 1 | 22 |
| ciao1 | 0.3393 | 0.140453072 | 0.05 | 0 | 1 | 22 |
| clptm1 | 0.3393 | 0.129852987 | 0.05 | 0 | 1 | 22 |
| clpxa | 0.3393 | 0.127937474 | 0.05 | 0 | 1 | 22 |
| commd1 | 0.3393 | 0.127689821 | 0.05 | 0 | 1 | 22 |
| cryz | 0.3393 | 0.124817026 | 0.05 | 0 | 1 | 22 |
| cul4b | 0.3393 | 0.141780972 | 0.05 | 0 | 1 | 22 |
| dcps | 0.3393 | 0.128553169 | 0.05 | 0 | 1 | 22 |
| dhdds | 0.3393 | 0.138299658 | 0.05 | 0 | 1 | 22 |
| dnajc10 | 0.3393 | 0.130783064 | 0.05 | 0 | 1 | 22 |
| dnajc18 | 0.3393 | 0.112227223 | 0.05 | 0 | 1 | 22 |
| dnal1 | 0.3393 | 0.164499116 | 0.05 | 0 | 1 | 22 |
| dus1l | 0.3393 | 0.140643252 | 0.05 | 0 | 1 | 22 |
| ezrb | 0.3393 | 0.129308519 | 0.05 | 0 | 1 | 22 |
| fam207a | 0.3393 | 0.143471487 | 0.05 | 0 | 1 | 22 |
| fis1 | 0.3393 | 0.139392729 | 0.05 | 0 | 1 | 22 |
| gnal | 0.3393 | 0.129996891 | 0.05 | 0 | 1 | 22 |
| gosr1 | 0.3393 | 0.128818142 | 0.05 | 0 | 1 | 22 |
| gucd1 | 0.3393 | 0.14221655 | 0.05 | 0 | 1 | 22 |
| hecw1b | 0.3393 | 0.132797662 | 0.05 | 0 | 1 | 22 |
| hlfa | 0.3393 | 0.121321034 | 0.05 | 0 | 1 | 22 |
| htatip2 | 0.3393 | 0.163183293 | 0.05 | 0 | 1 | 22 |
| immt | 0.3393 | 0.139840375 | 0.05 | 0 | 1 | 22 |
| itpkca | 0.3393 | 0.135943445 | 0.05 | 0 | 1 | 22 |
| itpr1a | 0.3393 | 0.132797662 | 0.05 | 0 | 1 | 22 |
| katnbl1 | 0.3393 | 0.143600768 | 0.05 | 0 | 1 | 22 |
| klf8 | 0.3393 | 0.162042905 | 0.05 | 0 | 1 | 22 |
| klhl12 | 0.3393 | 0.132797662 | 0.05 | 0 | 1 | 22 |
| klhl24a | 0.3393 | 0.125203244 | 0.05 | 0 | 1 | 22 |
| klhl32 | 0.3393 | 0.144720172 | 0.05 | 0 | 1 | 22 |
| ldb1a | 0.3393 | 0.115550186 | 0.05 | 0 | 1 | 22 |
| lrpap1 | 0.3393 | 0.127670241 | 0.05 | 0 | 1 | 22 |
| lrrc45 | 0.3393 | 0.164499116 | 0.05 | 0 | 1 | 22 |
| med18 | 0.3393 | 0.115802561 | 0.05 | 0 | 1 | 22 |

| | | | | | | |
|---|---|---|---|---|---|---|
| mef2cb | 0.3393 | 0.128875349 | 0.05 | 0 | 1 | 22 |
| mhc1uba | 0.3393 | 0.164499116 | 0.05 | 0 | 1 | 22 |
| mier3b | 0.3393 | 0.144720172 | 0.05 | 0 | 1 | 22 |
| mif4gdb | 0.3393 | 0.139743513 | 0.05 | 0 | 1 | 22 |
| mrpl27 | 0.3393 | 0.126960122 | 0.05 | 0 | 1 | 22 |
| mrpl39 | 0.3393 | 0.128441313 | 0.05 | 0 | 1 | 22 |
| mrps26 | 0.3393 | 0.115141382 | 0.05 | 0 | 1 | 22 |
| mrps27 | 0.3393 | 0.154447517 | 0.05 | 0 | 1 | 22 |
| mta3 | 0.3393 | 0.138768998 | 0.05 | 0 | 1 | 22 |
| mxd | 0.3393 | 0.140414719 | 0.05 | 0 | 1 | 22 |
| myg1 | 0.3393 | 0.123481905 | 0.05 | 0 | 1 | 22 |
| myo1cb | 0.3393 | 0.124525348 | 0.05 | 0 | 1 | 22 |
| naa15a | 0.3393 | 0.162988844 | 0.05 | 0 | 1 | 22 |
| nfil3-6 | 0.3393 | 0.128572776 | 0.05 | 0 | 1 | 22 |
| nme6 | 0.3393 | 0.144720172 | 0.05 | 0 | 1 | 22 |
| nol6 | 0.3393 | 0.115247616 | 0.05 | 0 | 1 | 22 |
| nucb1 | 0.3393 | 0.131512022 | 0.05 | 0 | 1 | 22 |
| nudt19 | 0.3393 | 0.144720172 | 0.05 | 0 | 1 | 22 |
| nup160 | 0.3393 | 0.121750055 | 0.05 | 0 | 1 | 22 |
| pdzd8 | 0.3393 | 0.132797662 | 0.05 | 0 | 1 | 22 |
| pgm1 | 0.3393 | 0.117827743 | 0.05 | 0 | 1 | 22 |
| pih1d1 | 0.3393 | 0.129494053 | 0.05 | 0 | 1 | 22 |
| pip4k2ca | 0.3393 | 0.162303409 | 0.05 | 0 | 1 | 22 |
| pmvk | 0.3393 | 0.129904416 | 0.05 | 0 | 1 | 22 |
| pou6f1 | 0.3393 | 0.159995392 | 0.05 | 0 | 1 | 22 |
| ppp4r2a | 0.3393 | 0.130372721 | 0.05 | 0 | 1 | 22 |
| praf2 | 0.3393 | 0.124453671 | 0.05 | 0 | 1 | 22 |
| psmd4b | 0.3393 | 0.141170576 | 0.05 | 0 | 1 | 22 |
| ptenb | 0.3393 | 0.129107463 | 0.05 | 0 | 1 | 22 |
| rbks | 0.3393 | 0.129079566 | 0.05 | 0 | 1 | 22 |
| rexo4 | 0.3393 | 0.182759522 | 0.05 | 0 | 1 | 22 |
| ripk2 | 0.3393 | 0.164499116 | 0.05 | 0 | 1 | 22 |
| rrn3 | 0.3393 | 0.143250391 | 0.05 | 0 | 1 | 22 |
| si:ch211-207k7.4 | 0.3393 | 0.138813296 | 0.05 | 0 | 1 | 22 |
| si:ch211-220f21.2 | 0.3393 | 0.148534956 | 0.05 | 0 | 1 | 22 |
| si:ch211-79k12.2 | 0.3393 | 0.158231576 | 0.05 | 0 | 1 | 22 |
| si:ch73-199g24.2 | 0.3393 | 0.132797662 | 0.05 | 0 | 1 | 22 |
| si:dkey-172j4.3 | 0.3393 | 0.143244723 | 0.05 | 0 | 1 | 22 |
| si:dkey-187k19.2 | 0.3393 | 0.132797662 | 0.05 | 0 | 1 | 22 |
| si:dkey-188i13.11 | 0.3393 | 0.130519337 | 0.05 | 0 | 1 | 22 |
| si:dkey-24l11.2 | 0.3393 | 0.135655638 | 0.05 | 0 | 1 | 22 |
| si:dkey-66a8.7 | 0.3393 | 0.128445176 | 0.05 | 0 | 1 | 22 |
| si:dkey-69o16.5 | 0.3393 | 0.127289815 | 0.05 | 0 | 1 | 22 |
| si:dkey-78p8.1 | 0.3393 | 0.132797662 | 0.05 | 0 | 1 | 22 |
| sinup | 0.3393 | 0.132797662 | 0.05 | 0 | 1 | 22 |
| slc10a7 | 0.3393 | 0.142281301 | 0.05 | 0 | 1 | 22 |
| slc30a7 | 0.3393 | 0.128560009 | 0.05 | 0 | 1 | 22 |
| slc31a1 | 0.3393 | 0.129130196 | 0.05 | 0 | 1 | 22 |

| | | | | | | |
|---|---|---|---|---|---|---|
| slc6a1l | 0.3393 | 0.144720172 | 0.05 | 0 | 1 | 22 |
| slc7a6os | 0.3393 | 0.141328431 | 0.05 | 0 | 1 | 22 |
| snrpb2 | 0.3393 | 0.142731027 | 0.05 | 0 | 1 | 22 |
| soul4 | 0.3393 | 0.127172206 | 0.05 | 0 | 1 | 22 |
| spoplb | 0.3393 | 0.131317031 | 0.05 | 0 | 1 | 22 |
| ssna1 | 0.3393 | 0.14051764 | 0.05 | 0 | 1 | 22 |
| supt3h | 0.3393 | 0.133100326 | 0.05 | 0 | 1 | 22 |
| tent2 | 0.3393 | 0.162222357 | 0.05 | 0 | 1 | 22 |
| tmed5 | 0.3393 | 0.131051139 | 0.05 | 0 | 1 | 22 |
| tmem120a | 0.3393 | 0.139455889 | 0.05 | 0 | 1 | 22 |
| tmem187 | 0.3393 | 0.124856312 | 0.05 | 0 | 1 | 22 |
| tmem201 | 0.3393 | 0.128542966 | 0.05 | 0 | 1 | 22 |
| tmem259 | 0.3393 | 0.127564552 | 0.05 | 0 | 1 | 22 |
| tnfrsf21 | 0.3393 | 0.144056961 | 0.05 | 0 | 1 | 22 |
| tomm34 | 0.3393 | 0.162378695 | 0.05 | 0 | 1 | 22 |
| ttpal | 0.3393 | 0.131587853 | 0.05 | 0 | 1 | 22 |
| ube2e2 | 0.3393 | 0.115426712 | 0.05 | 0 | 1 | 22 |
| ugt2a5 | 0.3393 | 0.159301007 | 0.05 | 0 | 1 | 22 |
| vangl2 | 0.3393 | 0.122572352 | 0.05 | 0 | 1 | 22 |
| wars2 | 0.3393 | 0.132797662 | 0.05 | 0 | 1 | 22 |
| wu:fb16f03 | 0.3393 | 0.127324288 | 0.05 | 0 | 1 | 22 |
| ythdf3 | 0.3393 | 0.130920092 | 0.05 | 0 | 1 | 22 |
| zdhhc4 | 0.3393 | 0.141959974 | 0.05 | 0 | 1 | 22 |
| zfhx3 | 0.3393 | 0.144720172 | 0.05 | 0 | 1 | 22 |
| zgc:153115 | 0.3393 | 0.143801863 | 0.05 | 0 | 1 | 22 |
| zgc:162339 | 0.3393 | 0.142709215 | 0.05 | 0 | 1 | 22 |
| zgc:86609 | 0.3393 | 0.152017633 | 0.05 | 0 | 1 | 22 |
| znf408 | 0.3393 | 0.164499116 | 0.05 | 0 | 1 | 22 |
| ASTE1 | 0.3393 | 0.175202231 | 0.05 | 0 | 1 | 22 |
| BX005005.2 | 0.3393 | -0.12126462 | 0 | 0.05 | 1 | 22 |
| BX248410.3 | 0.3393 | 0.207835778 | 0.05 | 0 | 1 | 22 |
| BX649300.1 | 0.3393 | 0.108026449 | 0.05 | 0 | 1 | 22 |
| BX908804.3 | 0.3393 | 0.146434687 | 0.05 | 0 | 1 | 22 |
| CABZ01029938.1 | 0.3393 | 0.105217342 | 0.05 | 0 | 1 | 22 |
| CABZ01058046.1 | 0.3393 | -0.12114246 | 0 | 0.05 | 1 | 22 |
| CABZ01061501.1 | 0.3393 | 0.17107891 | 0.05 | 0 | 1 | 22 |
| CABZ01072487.1 | 0.3393 | 0.148432793 | 0.05 | 0 | 1 | 22 |
| CABZ01080371.1 | 0.3393 | 0.108026449 | 0.05 | 0 | 1 | 22 |
| CABZ01084963.1 | 0.3393 | -0.10489753 | 0 | 0.05 | 1 | 22 |
| CR628365.1 | 0.3393 | -0.1151297 | 0 | 0.05 | 1 | 22 |
| CR848788.1 | 0.3393 | -0.23128805 | 0 | 0.05 | 1 | 22 |
| CT583652.1 | 0.3393 | -0.17616871 | 0 | 0.05 | 1 | 22 |
| CU929418.1 | 0.3393 | 0.144594151 | 0.05 | 0 | 1 | 22 |
| DOCK4 (1 of many) | 0.3393 | -0.12270515 | 0 | 0.05 | 1 | 22 |
| FP236454.1 | 0.3393 | 0.103198636 | 0.05 | 0 | 1 | 22 |
| HOOK3 | 0.3393 | -0.15491253 | 0 | 0.05 | 1 | 22 |
| LOC101883382 | 0.3393 | -0.12240242 | 0 | 0.05 | 1 | 22 |
| LOC101883913 | 0.3393 | 0.124614633 | 0.05 | 0 | 1 | 22 |

| | | | | | | |
|---|---|---|---|---|---|---|
| LOC101884045 | 0.3393 | 0.103198636 | 0.05 | 0 | 1 | 22 |
| LOC103909810 | 0.3393 | -0.12016261 | 0 | 0.05 | 1 | 22 |
| LOC108190423 | 0.3393 | 0.108026449 | 0.05 | 0 | 1 | 22 |
| LOC110437782 | 0.3393 | 0.108026449 | 0.05 | 0 | 1 | 22 |
| LOC110438172 | 0.3393 | 0.135020872 | 0.05 | 0 | 1 | 22 |
| LOC110440198 | 0.3393 | 0.103198636 | 0.05 | 0 | 1 | 22 |
| LOC793824 | 0.3393 | 0.146570376 | 0.05 | 0 | 1 | 22 |
| NMNAT3 | 0.3393 | -0.11711857 | 0 | 0.05 | 1 | 22 |
| PDCL3 | 0.3393 | -0.11256826 | 0 | 0.05 | 1 | 22 |
| XLOC-005448 | 0.3393 | 0.102389919 | 0.05 | 0 | 1 | 22 |
| XLOC-008119 | 0.3393 | -0.11923632 | 0 | 0.05 | 1 | 22 |
| XLOC-011272 | 0.3393 | -0.12209445 | 0 | 0.05 | 1 | 22 |
| XLOC-040582 | 0.3393 | 0.148432793 | 0.05 | 0 | 1 | 22 |
| XLOC-042565 | 0.3393 | 0.148432793 | 0.05 | 0 | 1 | 22 |
| XLOC-042868 | 0.3393 | 0.148432793 | 0.05 | 0 | 1 | 22 |
| XLOC-044090 | 0.3393 | 0.148432793 | 0.05 | 0 | 1 | 22 |
| XLOC-044204 | 0.3393 | -0.17202482 | 0 | 0.05 | 1 | 22 |
| abca3b | 0.3393 | 0.206333394 | 0.05 | 0 | 1 | 22 |
| abhd13 | 0.3393 | 0.248570833 | 0.05 | 0 | 1 | 22 |
| acads | 0.3393 | -0.18926285 | 0 | 0.05 | 1 | 22 |
| acvr2ab | 0.3393 | 0.169467141 | 0.05 | 0 | 1 | 22 |
| acvr2bb | 0.3393 | -0.12007184 | 0 | 0.05 | 1 | 22 |
| agfg1a | 0.3393 | -0.17573867 | 0 | 0.05 | 1 | 22 |
| alg11 | 0.3393 | 0.145969298 | 0.05 | 0 | 1 | 22 |
| alg3 | 0.3393 | 0.132792566 | 0.05 | 0 | 1 | 22 |
| alg8 | 0.3393 | 0.103198636 | 0.05 | 0 | 1 | 22 |
| ankrd37 | 0.3393 | 0.103198636 | 0.05 | 0 | 1 | 22 |
| ankrd46a | 0.3393 | -0.10899963 | 0 | 0.05 | 1 | 22 |
| ankrd52a | 0.3393 | -0.17494211 | 0 | 0.05 | 1 | 22 |
| ano6 | 0.3393 | 0.167371836 | 0.05 | 0 | 1 | 22 |
| arcn1b | 0.3393 | -0.18784437 | 0 | 0.05 | 1 | 22 |
| arhgap20b | 0.3393 | -0.12226721 | 0 | 0.05 | 1 | 22 |
| arhgap29b | 0.3393 | 0.105799149 | 0.05 | 0 | 1 | 22 |
| arid2 | 0.3393 | -0.10066049 | 0 | 0.05 | 1 | 22 |
| atf7b | 0.3393 | -0.11429214 | 0 | 0.05 | 1 | 22 |
| atg13 | 0.3393 | -0.18858366 | 0 | 0.05 | 1 | 22 |
| atp6v0a2b | 0.3393 | 0.108026449 | 0.05 | 0 | 1 | 22 |
| aven | 0.3393 | 0.101649322 | 0.05 | 0 | 1 | 22 |
| bag5 | 0.3393 | -0.18769096 | 0 | 0.05 | 1 | 22 |
| bcas3 | 0.3393 | -0.17719271 | 0 | 0.05 | 1 | 22 |
| bcorl1 | 0.3393 | 0.137316119 | 0.05 | 0 | 1 | 22 |
| blvra | 0.3393 | -0.15839047 | 0 | 0.05 | 1 | 22 |
| blzf1 | 0.3393 | -0.10928877 | 0 | 0.05 | 1 | 22 |
| c2cd2 | 0.3393 | 0.207835778 | 0.05 | 0 | 1 | 22 |
| cadm1b | 0.3393 | 0.103198636 | 0.05 | 0 | 1 | 22 |
| cars | 0.3393 | -0.10892327 | 0 | 0.05 | 1 | 22 |
| cbfb | 0.3393 | -0.11468357 | 0 | 0.05 | 1 | 22 |
| cbx6b | 0.3393 | -0.12032283 | 0 | 0.05 | 1 | 22 |

| | | | | | | |
|---|---|---|---|---|---|---|
| ccne1 | 0.3393 | 0.148432793 | 0.05 | 0 | 1 | 22 |
| cdc27 | 0.3393 | -0.18918383 | 0 | 0.05 | 1 | 22 |
| cdk12 | 0.3393 | 0.17107891 | 0.05 | 0 | 1 | 22 |
| cdk13 | 0.3393 | 0.168724035 | 0.05 | 0 | 1 | 22 |
| cdk6 | 0.3393 | 0.103198636 | 0.05 | 0 | 1 | 22 |
| cep131 | 0.3393 | 0.166003487 | 0.05 | 0 | 1 | 22 |
| cep85l | 0.3393 | -0.15926559 | 0 | 0.05 | 1 | 22 |
| cept1a | 0.3393 | 0.103954847 | 0.05 | 0 | 1 | 22 |
| clasrp | 0.3393 | -0.11727591 | 0 | 0.05 | 1 | 22 |
| cnot6l | 0.3393 | -0.17583376 | 0 | 0.05 | 1 | 22 |
| cnpy2 | 0.3393 | -0.10094533 | 0 | 0.05 | 1 | 22 |
| coro1b | 0.3393 | -0.15433834 | 0 | 0.05 | 1 | 22 |
| cox11 | 0.3393 | 0.100972461 | 0.05 | 0 | 1 | 22 |
| cpt2 | 0.3393 | 0.167535628 | 0.05 | 0 | 1 | 22 |
| crtc3 | 0.3393 | -0.17546011 | 0 | 0.05 | 1 | 22 |
| csnk2a2b | 0.3393 | 0.206860636 | 0.05 | 0 | 1 | 22 |
| ctsc | 0.3393 | 0.100814502 | 0.05 | 0 | 1 | 22 |
| cul3b | 0.3393 | -0.18783963 | 0 | 0.05 | 1 | 22 |
| cyp27a7 | 0.3393 | -0.18936254 | 0 | 0.05 | 1 | 22 |
| cyp2u1 | 0.3393 | -0.12270515 | 0 | 0.05 | 1 | 22 |
| dcaf12 | 0.3393 | 0.103198636 | 0.05 | 0 | 1 | 22 |
| ddi2 | 0.3393 | 0.101651622 | 0.05 | 0 | 1 | 22 |
| ddx23 | 0.3393 | -0.10290607 | 0 | 0.05 | 1 | 22 |
| ddx49 | 0.3393 | -0.12058725 | 0 | 0.05 | 1 | 22 |
| ddx51 | 0.3393 | 0.127324985 | 0.05 | 0 | 1 | 22 |
| dido1 | 0.3393 | 0.16844154 | 0.05 | 0 | 1 | 22 |
| dmtf1 | 0.3393 | 0.103198636 | 0.05 | 0 | 1 | 22 |
| dock3 | 0.3393 | -0.16597964 | 0 | 0.05 | 1 | 22 |
| dot1l | 0.3393 | -0.11412507 | 0 | 0.05 | 1 | 22 |
| dtx4b | 0.3393 | -0.10439198 | 0 | 0.05 | 1 | 22 |
| dusp23b | 0.3393 | 0.10455904 | 0.05 | 0 | 1 | 22 |
| dyrk1b | 0.3393 | 0.204945905 | 0.05 | 0 | 1 | 22 |
| ebpl | 0.3393 | -0.15716183 | 0 | 0.05 | 1 | 22 |
| egr1 | 0.3393 | 0.145671701 | 0.05 | 0 | 1 | 22 |
| ehbp1l1a | 0.3393 | -0.11450339 | 0 | 0.05 | 1 | 22 |
| eif2a | 0.3393 | 0.10680784 | 0.05 | 0 | 1 | 22 |
| elk4 | 0.3393 | -0.11956399 | 0 | 0.05 | 1 | 22 |
| elp2 | 0.3393 | 0.11884585 | 0.05 | 0 | 1 | 22 |
| elp5 | 0.3393 | 0.100206352 | 0.05 | 0 | 1 | 22 |
| enpp5 | 0.3393 | 0.128097017 | 0.05 | 0 | 1 | 22 |
| entpd4 | 0.3393 | 0.106621532 | 0.05 | 0 | 1 | 22 |
| epg5 | 0.3393 | 0.168156492 | 0.05 | 0 | 1 | 22 |
| ercc1 | 0.3393 | -0.10767118 | 0 | 0.05 | 1 | 22 |
| erlec1 | 0.3393 | -0.11775362 | 0 | 0.05 | 1 | 22 |
| etfbkmt | 0.3393 | 0.108026449 | 0.05 | 0 | 1 | 22 |
| ewsr1a | 0.3393 | 0.125046329 | 0.05 | 0 | 1 | 22 |
| exoc7 | 0.3393 | 0.105008958 | 0.05 | 0 | 1 | 22 |
| fam117ab | 0.3393 | -0.11463055 | 0 | 0.05 | 1 | 22 |

| | | | | | | |
|---|---|---|---|---|---|---|
| fam172a | 0.3393 | -0.17373858 | 0 | 0.05 | 1 | 22 |
| fam193a | 0.3393 | 0.114591487 | 0.05 | 0 | 1 | 22 |
| fam78ab | 0.3393 | -0.12194543 | 0 | 0.05 | 1 | 22 |
| fam84b | 0.3393 | -0.10725787 | 0 | 0.05 | 1 | 22 |
| fam98b | 0.3393 | 0.135020872 | 0.05 | 0 | 1 | 22 |
| fbxw11a | 0.3393 | 0.132832355 | 0.05 | 0 | 1 | 22 |
| foxj2 | 0.3393 | 0.10165029 | 0.05 | 0 | 1 | 22 |
| foxn3 | 0.3393 | 0.132646276 | 0.05 | 0 | 1 | 22 |
| fpgt | 0.3393 | 0.132597804 | 0.05 | 0 | 1 | 22 |
| fuca1.2 | 0.3393 | -0.15895891 | 0 | 0.05 | 1 | 22 |
| fut9b | 0.3393 | -0.11183365 | 0 | 0.05 | 1 | 22 |
| g3bp1 | 0.3393 | -0.11949792 | 0 | 0.05 | 1 | 22 |
| gatad2ab | 0.3393 | 0.131629079 | 0.05 | 0 | 1 | 22 |
| gbp1 | 0.3393 | 0.10657942 | 0.05 | 0 | 1 | 22 |
| gcat | 0.3393 | 0.1271077 | 0.05 | 0 | 1 | 22 |
| gclc | 0.3393 | 0.104453798 | 0.05 | 0 | 1 | 22 |
| gcsha | 0.3393 | 0.13886686 | 0.05 | 0 | 1 | 22 |
| gdi1 | 0.3393 | 0.148432793 | 0.05 | 0 | 1 | 22 |
| ggcx | 0.3393 | 0.17107891 | 0.05 | 0 | 1 | 22 |
| got2a | 0.3393 | 0.16762245 | 0.05 | 0 | 1 | 22 |
| gramd4b | 0.3393 | -0.15964042 | 0 | 0.05 | 1 | 22 |
| gsk3aa | 0.3393 | 0.103198636 | 0.05 | 0 | 1 | 22 |
| gtf2h4 | 0.3393 | -0.10462256 | 0 | 0.05 | 1 | 22 |
| gtf2ird1 | 0.3393 | 0.103198636 | 0.05 | 0 | 1 | 22 |
| hadhab | 0.3393 | -0.1750478 | 0 | 0.05 | 1 | 22 |
| hbegfa | 0.3393 | -0.11718696 | 0 | 0.05 | 1 | 22 |
| hdac9b | 0.3393 | 0.107536663 | 0.05 | 0 | 1 | 22 |
| hdlbpb | 0.3393 | 0.103198636 | 0.05 | 0 | 1 | 22 |
| hecw2b | 0.3393 | -0.15740917 | 0 | 0.05 | 1 | 22 |
| herpud2 | 0.3393 | -0.11927529 | 0 | 0.05 | 1 | 22 |
| hmgcrb | 0.3393 | -0.18893773 | 0 | 0.05 | 1 | 22 |
| hps3 | 0.3393 | -0.15705742 | 0 | 0.05 | 1 | 22 |
| ibtk | 0.3393 | -0.10300974 | 0 | 0.05 | 1 | 22 |
| ifi35 | 0.3393 | -0.1106344 | 0 | 0.05 | 1 | 22 |
| ifng1 | 0.3393 | -0.12270515 | 0 | 0.05 | 1 | 22 |
| igf2bp2b | 0.3393 | 0.147077043 | 0.05 | 0 | 1 | 22 |
| ik | 0.3393 | 0.144325208 | 0.05 | 0 | 1 | 22 |
| illr4 | 0.3393 | -0.11183521 | 0 | 0.05 | 1 | 22 |
| im:7138535 | 0.3393 | 0.105202823 | 0.05 | 0 | 1 | 22 |
| ino80e | 0.3393 | -0.18842123 | 0 | 0.05 | 1 | 22 |
| inpp4aa | 0.3393 | 0.103198636 | 0.05 | 0 | 1 | 22 |
| ints2 | 0.3393 | -0.17476289 | 0 | 0.05 | 1 | 22 |
| ippk | 0.3393 | -0.12049927 | 0 | 0.05 | 1 | 22 |
| isca1 | 0.3393 | 0.120283204 | 0.05 | 0 | 1 | 22 |
| itprip | 0.3393 | 0.100794476 | 0.05 | 0 | 1 | 22 |
| jak3 | 0.3393 | -0.11437719 | 0 | 0.05 | 1 | 22 |
| jakmip1 | 0.3393 | 0.205479312 | 0.05 | 0 | 1 | 22 |
| jkamp | 0.3393 | 0.133659759 | 0.05 | 0 | 1 | 22 |

| | | | | | | |
|---|---|---|---|---|---|---|
| junbb | 0.3393 | -0.11363736 | 0 | 0.05 | 1 | 22 |
| kansl1b | 0.3393 | -0.12168538 | 0 | 0.05 | 1 | 22 |
| kat2b | 0.3393 | 0.108026449 | 0.05 | 0 | 1 | 22 |
| kdelr2a | 0.3393 | 0.101327058 | 0.05 | 0 | 1 | 22 |
| kdm3b | 0.3393 | -0.17490667 | 0 | 0.05 | 1 | 22 |
| kdm6al | 0.3393 | -0.17516388 | 0 | 0.05 | 1 | 22 |
| kif3a | 0.3393 | -0.10899724 | 0 | 0.05 | 1 | 22 |
| kif5bb | 0.3393 | -0.11339936 | 0 | 0.05 | 1 | 22 |
| klhdc10 | 0.3393 | -0.10858202 | 0 | 0.05 | 1 | 22 |
| klhl11 | 0.3393 | -0.18973615 | 0 | 0.05 | 1 | 22 |
| kpna1 | 0.3393 | -0.1201087 | 0 | 0.05 | 1 | 22 |
| larp4b | 0.3393 | -0.1087422 | 0 | 0.05 | 1 | 22 |
| ldlrap1b | 0.3393 | 0.207835778 | 0.05 | 0 | 1 | 22 |
| lman2lb | 0.3393 | 0.207223131 | 0.05 | 0 | 1 | 22 |
| lmf1 | 0.3393 | 0.145410854 | 0.05 | 0 | 1 | 22 |
| lrp12 | 0.3393 | 0.207835778 | 0.05 | 0 | 1 | 22 |
| lrrc8db | 0.3393 | 0.108026449 | 0.05 | 0 | 1 | 22 |
| lsm14aa | 0.3393 | -0.17491945 | 0 | 0.05 | 1 | 22 |
| mak | 0.3393 | -0.10451267 | 0 | 0.05 | 1 | 22 |
| map2k4b | 0.3393 | 0.128425813 | 0.05 | 0 | 1 | 22 |
| mapk14a | 0.3393 | 0.170214054 | 0.05 | 0 | 1 | 22 |
| mapk14b | 0.3393 | -0.1703476 | 0 | 0.05 | 1 | 22 |
| mast3a | 0.3393 | 0.148432793 | 0.05 | 0 | 1 | 22 |
| mavs | 0.3393 | 0.108026449 | 0.05 | 0 | 1 | 22 |
| max | 0.3393 | -0.11155649 | 0 | 0.05 | 1 | 22 |
| mbd1b | 0.3393 | -0.1106837 | 0 | 0.05 | 1 | 22 |
| mbtps1 | 0.3393 | -0.11992441 | 0 | 0.05 | 1 | 22 |
| mcf2l2 | 0.3393 | -0.19004069 | 0 | 0.05 | 1 | 22 |
| med21 | 0.3393 | 0.16930018 | 0.05 | 0 | 1 | 22 |
| mgat2 | 0.3393 | 0.103198636 | 0.05 | 0 | 1 | 22 |
| mgat3b | 0.3393 | -0.11503576 | 0 | 0.05 | 1 | 22 |
| mgat4b | 0.3393 | -0.14862338 | 0 | 0.05 | 1 | 22 |
| mgmt | 0.3393 | 0.205464422 | 0.05 | 0 | 1 | 22 |
| mkrn4 | 0.3393 | 0.206579467 | 0.05 | 0 | 1 | 22 |
| msc | 0.3393 | 0.103198636 | 0.05 | 0 | 1 | 22 |
| msi2a | 0.3393 | -0.11756842 | 0 | 0.05 | 1 | 22 |
| mtg2 | 0.3393 | -0.18510145 | 0 | 0.05 | 1 | 22 |
| mtmr1a | 0.3393 | 0.103198636 | 0.05 | 0 | 1 | 22 |
| mycla | 0.3393 | 0.103198636 | 0.05 | 0 | 1 | 22 |
| naa30 | 0.3393 | -0.17227805 | 0 | 0.05 | 1 | 22 |
| nab1a | 0.3393 | 0.167626783 | 0.05 | 0 | 1 | 22 |
| nadkb | 0.3393 | 0.129636558 | 0.05 | 0 | 1 | 22 |
| nat15 | 0.3393 | -0.17334143 | 0 | 0.05 | 1 | 22 |
| ndel1b | 0.3393 | -0.11758534 | 0 | 0.05 | 1 | 22 |
| nepro | 0.3393 | 0.103198636 | 0.05 | 0 | 1 | 22 |
| nipal3 | 0.3393 | -0.12149167 | 0 | 0.05 | 1 | 22 |
| nisch | 0.3393 | -0.17518646 | 0 | 0.05 | 1 | 22 |
| nktr | 0.3393 | -0.11263392 | 0 | 0.05 | 1 | 22 |

| | | | | | | |
|---|---|---|---|---|---|---|
| noa1 | 0.3393 | 0.131710784 | 0.05 | 0 | 1 | 22 |
| nrbp1 | 0.3393 | 0.17107891 | 0.05 | 0 | 1 | 22 |
| nsfl1c | 0.3393 | -0.10358736 | 0 | 0.05 | 1 | 22 |
| nt5c1bb | 0.3393 | 0.103198636 | 0.05 | 0 | 1 | 22 |
| nubp1 | 0.3393 | -0.12270515 | 0 | 0.05 | 1 | 22 |
| nudcd2 | 0.3393 | -0.10432698 | 0 | 0.05 | 1 | 22 |
| nudt17 | 0.3393 | -0.16079674 | 0 | 0.05 | 1 | 22 |
| nufip1 | 0.3393 | 0.129176315 | 0.05 | 0 | 1 | 22 |
| nup50 | 0.3393 | 0.131477306 | 0.05 | 0 | 1 | 22 |
| oip5-as1 | 0.3393 | -0.11375672 | 0 | 0.05 | 1 | 22 |
| opcml | 0.3393 | -0.10652972 | 0 | 0.05 | 1 | 22 |
| otud4 | 0.3393 | -0.17289876 | 0 | 0.05 | 1 | 22 |
| otud7b | 0.3393 | 0.167217689 | 0.05 | 0 | 1 | 22 |
| pacs1a | 0.3393 | 0.207145549 | 0.05 | 0 | 1 | 22 |
| patz1 | 0.3393 | -0.12122167 | 0 | 0.05 | 1 | 22 |
| pbx2 | 0.3393 | -0.10770166 | 0 | 0.05 | 1 | 22 |
| pcid2 | 0.3393 | -0.10302958 | 0 | 0.05 | 1 | 22 |
| pcm1 | 0.3393 | -0.10080195 | 0 | 0.05 | 1 | 22 |
| pcmt | 0.3393 | -0.16455335 | 0 | 0.05 | 1 | 22 |
| pdcd4b | 0.3393 | -0.11312757 | 0 | 0.05 | 1 | 22 |
| pgap3 | 0.3393 | 0.170091354 | 0.05 | 0 | 1 | 22 |
| phactr4b | 0.3393 | -0.14732833 | 0 | 0.05 | 1 | 22 |
| phospho2 | 0.3393 | 0.152339668 | 0.05 | 0 | 1 | 22 |
| pick1 | 0.3393 | 0.121434547 | 0.05 | 0 | 1 | 22 |
| pigc | 0.3393 | 0.100314785 | 0.05 | 0 | 1 | 22 |
| pign | 0.3393 | 0.168033877 | 0.05 | 0 | 1 | 22 |
| pik3cd | 0.3393 | -0.11511556 | 0 | 0.05 | 1 | 22 |
| plekhf2 | 0.3393 | 0.17107891 | 0.05 | 0 | 1 | 22 |
| plrg1 | 0.3393 | -0.10480724 | 0 | 0.05 | 1 | 22 |
| ppa2 | 0.3393 | -0.10840227 | 0 | 0.05 | 1 | 22 |
| ppp1r3da | 0.3393 | -0.18956988 | 0 | 0.05 | 1 | 22 |
| ppp2r1ba | 0.3393 | -0.11239477 | 0 | 0.05 | 1 | 22 |
| ppp2r2d | 0.3393 | 0.106963484 | 0.05 | 0 | 1 | 22 |
| ppp4r1 | 0.3393 | 0.105501508 | 0.05 | 0 | 1 | 22 |
| prdm10 | 0.3393 | 0.202936558 | 0.05 | 0 | 1 | 22 |
| prickle3 | 0.3393 | -0.17249907 | 0 | 0.05 | 1 | 22 |
| prkacaa | 0.3393 | 0.144307107 | 0.05 | 0 | 1 | 22 |
| prune | 0.3393 | 0.103198636 | 0.05 | 0 | 1 | 22 |
| psmd1 | 0.3393 | -0.10993463 | 0 | 0.05 | 1 | 22 |
| ptgis | 0.3393 | -0.15937519 | 0 | 0.05 | 1 | 22 |
| ptpn9a | 0.3393 | -0.11427012 | 0 | 0.05 | 1 | 22 |
| purg | 0.3393 | -0.11966977 | 0 | 0.05 | 1 | 22 |
| rangrf | 0.3393 | -0.15640097 | 0 | 0.05 | 1 | 22 |
| rarab | 0.3393 | -0.12081769 | 0 | 0.05 | 1 | 22 |
| rbm12b | 0.3393 | -0.17301942 | 0 | 0.05 | 1 | 22 |
| rbms1b | 0.3393 | -0.17559611 | 0 | 0.05 | 1 | 22 |
| rcc1l | 0.3393 | -0.11244734 | 0 | 0.05 | 1 | 22 |
| rcor1 | 0.3393 | 0.105180818 | 0.05 | 0 | 1 | 22 |

| | | | | | | |
|---|---|---|---|---|---|---|
| rhobtb4 | 0.3393 | 0.139528196 | 0.05 | 0 | 1 | 22 |
| rnf115 | 0.3393 | -0.17510573 | 0 | 0.05 | 1 | 22 |
| rnf126 | 0.3393 | -0.11925089 | 0 | 0.05 | 1 | 22 |
| rnf145a | 0.3393 | 0.16606492 | 0.05 | 0 | 1 | 22 |
| rnf180 | 0.3393 | 0.107742544 | 0.05 | 0 | 1 | 22 |
| rnf6 | 0.3393 | -0.1141602 | 0 | 0.05 | 1 | 22 |
| rnft2 | 0.3393 | 0.207835778 | 0.05 | 0 | 1 | 22 |
| rock1 | 0.3393 | 0.133801306 | 0.05 | 0 | 1 | 22 |
| rreb1a | 0.3393 | 0.146021198 | 0.05 | 0 | 1 | 22 |
| rtca | 0.3393 | 0.130395581 | 0.05 | 0 | 1 | 22 |
| rtf2 | 0.3393 | 0.138352921 | 0.05 | 0 | 1 | 22 |
| rwdd | 0.3393 | -0.11689762 | 0 | 0.05 | 1 | 22 |
| sacm1la | 0.3393 | -0.12029287 | 0 | 0.05 | 1 | 22 |
| sdhaf4 | 0.3393 | 0.146506882 | 0.05 | 0 | 1 | 22 |
| sec31b | 0.3393 | 0.103198636 | 0.05 | 0 | 1 | 22 |
| senp7b | 0.3393 | 0.206542563 | 0.05 | 0 | 1 | 22 |
| sept9b | 0.3393 | 0.207335122 | 0.05 | 0 | 1 | 22 |
| sf3b2 | 0.3393 | 0.145381465 | 0.05 | 0 | 1 | 22 |
| sgpl1 | 0.3393 | 0.17107891 | 0.05 | 0 | 1 | 22 |
| sh3bp2 | 0.3393 | 0.106507 | 0.05 | 0 | 1 | 22 |
| sh3bp5la | 0.3393 | 0.104243138 | 0.05 | 0 | 1 | 22 |
| si:ch1073-224n8.1 | 0.3393 | 0.108026449 | 0.05 | 0 | 1 | 22 |
| si:ch211-114l13.12 | 0.3393 | 0.17107891 | 0.05 | 0 | 1 | 22 |
| si:ch211-117c9.5 | 0.3393 | 0.145089109 | 0.05 | 0 | 1 | 22 |
| si:ch211-121a2.2 | 0.3393 | 0.124050689 | 0.05 | 0 | 1 | 22 |
| si:ch211-132b12.8 | 0.3393 | -0.16938938 | 0 | 0.05 | 1 | 22 |
| si:ch211-132g1.3 | 0.3393 | -0.16015689 | 0 | 0.05 | 1 | 22 |
| si:ch211-154o6.2 | 0.3393 | 0.10329654 | 0.05 | 0 | 1 | 22 |
| si:ch211-161m3.6 | 0.3393 | 0.103198636 | 0.05 | 0 | 1 | 22 |
| si:ch211-196g2.4 | 0.3393 | -0.1546493 | 0 | 0.05 | 1 | 22 |
| si:ch211-197h24.6 | 0.3393 | -0.11549142 | 0 | 0.05 | 1 | 22 |
| si:ch211-212o1.2 | 0.3393 | 0.207835778 | 0.05 | 0 | 1 | 22 |
| si:ch211-214e3.5 | 0.3393 | 0.17107891 | 0.05 | 0 | 1 | 22 |
| si:ch211-215e19.3 | 0.3393 | 0.135020872 | 0.05 | 0 | 1 | 22 |
| si:ch211-223a21.3 | 0.3393 | -0.16015689 | 0 | 0.05 | 1 | 22 |
| si:ch211-227e10.3 | 0.3393 | -0.15890975 | 0 | 0.05 | 1 | 22 |
| si:ch211-22i13.2 | 0.3393 | 0.102240189 | 0.05 | 0 | 1 | 22 |
| si:ch211-238n5.4 | 0.3393 | 0.205803384 | 0.05 | 0 | 1 | 22 |
| si:ch211-269k10.5 | 0.3393 | 0.103198636 | 0.05 | 0 | 1 | 22 |
| si:ch211-281l24.3 | 0.3393 | -0.17429956 | 0 | 0.05 | 1 | 22 |
| si:ch211-79l17.1 | 0.3393 | 0.135020872 | 0.05 | 0 | 1 | 22 |
| si:ch73-193c12.2 | 0.3393 | -0.10819475 | 0 | 0.05 | 1 | 22 |
| si:ch73-248e21.1 | 0.3393 | -0.17463877 | 0 | 0.05 | 1 | 22 |
| si:ch73-375g18.1 | 0.3393 | -0.1122772 | 0 | 0.05 | 1 | 22 |
| si:dkey-117m1.4 | 0.3393 | 0.166980599 | 0.05 | 0 | 1 | 22 |
| si:dkey-230p4.1 | 0.3393 | -0.15845809 | 0 | 0.05 | 1 | 22 |
| si:dkey-239n17.4 | 0.3393 | -0.1200607 | 0 | 0.05 | 1 | 22 |
| si:dkey-260j18.2 | 0.3393 | -0.18931467 | 0 | 0.05 | 1 | 22 |

| | | | | | | |
|---|---|---|---|---|---|---|
| si:dkey-33c9.6 | 0.3393 | 0.148432793 | 0.05 | 0 | 1 | 22 |
| si:dkey-5n18.1 | 0.3393 | 0.127710849 | 0.05 | 0 | 1 | 22 |
| si:dkey-6b12.5 | 0.3393 | -0.14736105 | 0 | 0.05 | 1 | 22 |
| si:dkey-77f5.3 | 0.3393 | -0.17361898 | 0 | 0.05 | 1 | 22 |
| si:dkey-78l4.6 | 0.3393 | 0.176038073 | 0.05 | 0 | 1 | 22 |
| si:dkey-82o10.4 | 0.3393 | 0.103198636 | 0.05 | 0 | 1 | 22 |
| si:dkey-88p24.11 | 0.3393 | -0.18096587 | 0 | 0.05 | 1 | 22 |
| si:dkeyp-44a8.2 | 0.3393 | -0.12012953 | 0 | 0.05 | 1 | 22 |
| si:dkeyp-75h12.2 | 0.3393 | -0.15207516 | 0 | 0.05 | 1 | 22 |
| sin3aa | 0.3393 | 0.108026449 | 0.05 | 0 | 1 | 22 |
| sirt4 | 0.3393 | 0.138464809 | 0.05 | 0 | 1 | 22 |
| slc16a1b | 0.3393 | 0.132453442 | 0.05 | 0 | 1 | 22 |
| slc5a11 | 0.3393 | 0.17107891 | 0.05 | 0 | 1 | 22 |
| slkb | 0.3393 | 0.142498664 | 0.05 | 0 | 1 | 22 |
| smarcal1 | 0.3393 | -0.12826354 | 0 | 0.05 | 1 | 22 |
| smg6 | 0.3393 | 0.206965018 | 0.05 | 0 | 1 | 22 |
| smurf2 | 0.3393 | -0.10960753 | 0 | 0.05 | 1 | 22 |
| snrpa | 0.3393 | 0.141222818 | 0.05 | 0 | 1 | 22 |
| sntb2 | 0.3393 | -0.12069772 | 0 | 0.05 | 1 | 22 |
| snx2 | 0.3393 | 0.136375246 | 0.05 | 0 | 1 | 22 |
| snx29 | 0.3393 | -0.17536567 | 0 | 0.05 | 1 | 22 |
| snx6 | 0.3393 | -0.1711311 | 0 | 0.05 | 1 | 22 |
| spns1 | 0.3393 | 0.165806532 | 0.05 | 0 | 1 | 22 |
| sprtn | 0.3393 | 0.206119469 | 0.05 | 0 | 1 | 22 |
| spryd4 | 0.3393 | 0.126005738 | 0.05 | 0 | 1 | 22 |
| srm | 0.3393 | -0.15284896 | 0 | 0.05 | 1 | 22 |
| stag2a | 0.3393 | -0.15899305 | 0 | 0.05 | 1 | 22 |
| stx5al | 0.3393 | -0.17139326 | 0 | 0.05 | 1 | 22 |
| stx6 | 0.3393 | -0.11222832 | 0 | 0.05 | 1 | 22 |
| surf2 | 0.3393 | -0.11215947 | 0 | 0.05 | 1 | 22 |
| swi5 | 0.3393 | 0.101307724 | 0.05 | 0 | 1 | 22 |
| tbc1d24 | 0.3393 | 0.207835778 | 0.05 | 0 | 1 | 22 |
| tbx21 | 0.3393 | -0.16221861 | 0 | 0.05 | 1 | 22 |
| tbx2b | 0.3393 | -0.12948262 | 0 | 0.05 | 1 | 22 |
| tcerg1a | 0.3393 | 0.12721832 | 0.05 | 0 | 1 | 22 |
| tcf7l1a | 0.3393 | 0.100859142 | 0.05 | 0 | 1 | 22 |
| tefm | 0.3393 | 0.206871927 | 0.05 | 0 | 1 | 22 |
| tent5ab | 0.3393 | -0.15996431 | 0 | 0.05 | 1 | 22 |
| tfam | 0.3393 | 0.127133329 | 0.05 | 0 | 1 | 22 |
| timm29 | 0.3393 | 0.132341252 | 0.05 | 0 | 1 | 22 |
| tlr4ba | 0.3393 | 0.108026449 | 0.05 | 0 | 1 | 22 |
| tm9sf4 | 0.3393 | 0.101068069 | 0.05 | 0 | 1 | 22 |
| tmem115 | 0.3393 | -0.10997265 | 0 | 0.05 | 1 | 22 |
| tmem241 | 0.3393 | 0.135020872 | 0.05 | 0 | 1 | 22 |
| tnfaip2a | 0.3393 | 0.103198636 | 0.05 | 0 | 1 | 22 |
| tnikb | 0.3393 | 0.13257115 | 0.05 | 0 | 1 | 22 |
| tnrc18 | 0.3393 | -0.1158068 | 0 | 0.05 | 1 | 22 |
| triap1 | 0.3393 | 0.142168651 | 0.05 | 0 | 1 | 22 |

| | | | | | | |
|---|---|---|---|---|---|---|
| trim71 | 0.3393 | 0.108026449 | 0.05 | 0 | 1 | 22 |
| tsen15 | 0.3393 | 0.131467262 | 0.05 | 0 | 1 | 22 |
| tsen34 | 0.3393 | -0.17312801 | 0 | 0.05 | 1 | 22 |
| tsku | 0.3393 | 0.204251461 | 0.05 | 0 | 1 | 22 |
| ttc13 | 0.3393 | 0.206280932 | 0.05 | 0 | 1 | 22 |
| ttc39c | 0.3393 | 0.147849322 | 0.05 | 0 | 1 | 22 |
| uap1l1 | 0.3393 | -0.15579731 | 0 | 0.05 | 1 | 22 |
| ubap2l | 0.3393 | -0.17420604 | 0 | 0.05 | 1 | 22 |
| ube2w | 0.3393 | -0.17185766 | 0 | 0.05 | 1 | 22 |
| ubfd1 | 0.3393 | -0.1013654 | 0 | 0.05 | 1 | 22 |
| ufl1 | 0.3393 | -0.11464227 | 0 | 0.05 | 1 | 22 |
| ugt5b2 | 0.3393 | 0.207725901 | 0.05 | 0 | 1 | 22 |
| upf3b | 0.3393 | 0.130784149 | 0.05 | 0 | 1 | 22 |
| uri1 | 0.3393 | 0.145448907 | 0.05 | 0 | 1 | 22 |
| usb1 | 0.3393 | 0.102749736 | 0.05 | 0 | 1 | 22 |
| usp11 | 0.3393 | 0.17107891 | 0.05 | 0 | 1 | 22 |
| usp46 | 0.3393 | -0.18976355 | 0 | 0.05 | 1 | 22 |
| vps25 | 0.3393 | -0.12001211 | 0 | 0.05 | 1 | 22 |
| vwa8 | 0.3393 | -0.11305818 | 0 | 0.05 | 1 | 22 |
| wdr6 | 0.3393 | -0.15615597 | 0 | 0.05 | 1 | 22 |
| wdr61 | 0.3393 | 0.132237587 | 0.05 | 0 | 1 | 22 |
| wsb1 | 0.3393 | 0.144587066 | 0.05 | 0 | 1 | 22 |
| wsb2 | 0.3393 | -0.15807055 | 0 | 0.05 | 1 | 22 |
| xpo1b | 0.3393 | -0.11260918 | 0 | 0.05 | 1 | 22 |
| zbtb37 | 0.3393 | -0.12172591 | 0 | 0.05 | 1 | 22 |
| zbtb43 | 0.3393 | -0.12200359 | 0 | 0.05 | 1 | 22 |
| zbtb7b | 0.3393 | 0.147486815 | 0.05 | 0 | 1 | 22 |
| zc3h7a | 0.3393 | -0.11199388 | 0 | 0.05 | 1 | 22 |
| zdhhc13 | 0.3393 | -0.18801413 | 0 | 0.05 | 1 | 22 |
| zdhhc17 | 0.3393 | -0.10831595 | 0 | 0.05 | 1 | 22 |
| zdhhc5a | 0.3393 | 0.169616438 | 0.05 | 0 | 1 | 22 |
| zfand4 | 0.3393 | -0.17616871 | 0 | 0.05 | 1 | 22 |
| zfyve19 | 0.3393 | 0.100079975 | 0.05 | 0 | 1 | 22 |
| zgc:101016 | 0.3393 | -0.11480833 | 0 | 0.05 | 1 | 22 |
| zgc:152830 | 0.3393 | -0.11459252 | 0 | 0.05 | 1 | 22 |
| zgc:153142 | 0.3393 | 0.17107891 | 0.05 | 0 | 1 | 22 |
| zgc:153409 | 0.3393 | 0.17107891 | 0.05 | 0 | 1 | 22 |
| zgc:162025 | 0.3393 | -0.16991873 | 0 | 0.05 | 1 | 22 |
| zgc:171857 | 0.3393 | 0.17107891 | 0.05 | 0 | 1 | 22 |
| zgc:173737 | 0.3393 | -0.15207516 | 0 | 0.05 | 1 | 22 |
| zgc:194285 | 0.3393 | 0.100247432 | 0.05 | 0 | 1 | 22 |
| zgc:194981 | 0.3393 | 0.133963105 | 0.05 | 0 | 1 | 22 |
| zgc:195175 | 0.3393 | 0.135020872 | 0.05 | 0 | 1 | 22 |
| zgc:66160 | 0.3393 | 0.100933591 | 0.05 | 0 | 1 | 22 |
| zgc:85932 | 0.3393 | -0.11086509 | 0 | 0.05 | 1 | 22 |
| znf1008 | 0.3393 | 0.17107891 | 0.05 | 0 | 1 | 22 |
| znf1014 | 0.3393 | 0.102148018 | 0.05 | 0 | 1 | 22 |
| znf1059 | 0.3393 | -0.17433145 | 0 | 0.05 | 1 | 22 |

| | | | | | | |
|---|---|---|---|---|---|---|
| znf106a | 0.3393 | 0.208186006 | 0.05 | 0 | 1 | 22 |
| znf1183 | 0.3393 | 0.103198636 | 0.05 | 0 | 1 | 22 |
| znf292a | 0.3393 | 0.204129854 | 0.05 | 0 | 1 | 22 |
| znf395a | 0.3393 | -0.15525523 | 0 | 0.05 | 1 | 22 |
| znf622 | 0.3393 | -0.18901122 | 0 | 0.05 | 1 | 22 |
| zswim8 | 0.3393 | -0.1069278 | 0 | 0.05 | 1 | 22 |
| zzz3 | 0.3393 | 0.131147405 | 0.05 | 0 | 1 | 22 |
| AL732488.1 | 0.3393 | 0.164416634 | 0.05 | 0 | 1 | 22 |
| BOLA2 | 0.3393 | 0.144400773 | 0.05 | 0 | 1 | 22 |
| BX005340.1 | 0.3393 | -0.11746502 | 0 | 0.05 | 1 | 22 |
| BX322587.2 | 0.3393 | 0.13597632 | 0.05 | 0 | 1 | 22 |
| BX548015.2 | 0.3393 | -0.21596073 | 0 | 0.05 | 1 | 22 |
| BX649250.1 | 0.3393 | 0.136227215 | 0.05 | 0 | 1 | 22 |
| BX908804.2 | 0.3393 | 0.139575893 | 0.05 | 0 | 1 | 22 |
| CABZ01002690.1 | 0.3393 | 0.132006068 | 0.05 | 0 | 1 | 22 |
| CABZ01038697.1 | 0.3393 | -0.14903766 | 0 | 0.05 | 1 | 22 |
| CABZ01063556.1 | 0.3393 | 0.132972464 | 0.05 | 0 | 1 | 22 |
| CABZ01073736.1 | 0.3393 | 0.166493715 | 0.05 | 0 | 1 | 22 |
| CABZ01078600.1 | 0.3393 | 0.154985092 | 0.05 | 0 | 1 | 22 |
| CABZ01088271.1 | 0.3393 | 0.130266926 | 0.05 | 0 | 1 | 22 |
| CABZ01088367.1 | 0.3393 | 0.132468051 | 0.05 | 0 | 1 | 22 |
| CR388166.2 | 0.3393 | -0.11849905 | 0 | 0.05 | 1 | 22 |
| CR392341.1 | 0.3393 | -0.11845561 | 0 | 0.05 | 1 | 22 |
| CR392341.4 | 0.3393 | 0.115105188 | 0.05 | 0 | 1 | 22 |
| CR550307.1 | 0.3393 | 0.139575893 | 0.05 | 0 | 1 | 22 |
| CR788316.3 | 0.3393 | 0.119178432 | 0.05 | 0 | 1 | 22 |
| CR847906.1 | 0.3393 | -0.11833039 | 0 | 0.05 | 1 | 22 |
| CU694532.1 | 0.3393 | 0.114757657 | 0.05 | 0 | 1 | 22 |
| CU929230.1 | 0.3393 | -0.14869641 | 0 | 0.05 | 1 | 22 |
| FP101861.1 | 0.3393 | 0.118005184 | 0.05 | 0 | 1 | 22 |
| GNAZ | 0.3393 | -0.14914634 | 0 | 0.05 | 1 | 22 |
| LO018197.1 | 0.3393 | 0.17000603 | 0.05 | 0 | 1 | 22 |
| LOC100006926 | 0.3393 | -0.12054202 | 0 | 0.05 | 1 | 22 |
| LOC100332329 | 0.3393 | 0.146139584 | 0.05 | 0 | 1 | 22 |
| LOC101883185 | 0.3393 | 0.13597632 | 0.05 | 0 | 1 | 22 |
| LOC101883815 | 0.3393 | 0.132709338 | 0.05 | 0 | 1 | 22 |
| LOC101884404 | 0.3393 | 0.113359691 | 0.05 | 0 | 1 | 22 |
| LOC101884451 | 0.3393 | 0.139208216 | 0.05 | 0 | 1 | 22 |
| LOC101884921 | 0.3393 | 0.122611796 | 0.05 | 0 | 1 | 22 |
| LOC101884966 | 0.3393 | -0.14929361 | 0 | 0.05 | 1 | 22 |
| LOC101886111 | 0.3393 | -0.1413181 | 0 | 0.05 | 1 | 22 |
| LOC101886337 | 0.3393 | 0.115827051 | 0.05 | 0 | 1 | 22 |
| LOC103909796 | 0.3393 | -0.1906243 | 0 | 0.05 | 1 | 22 |
| LOC103911652 | 0.3393 | -0.10804838 | 0 | 0.05 | 1 | 22 |
| LOC110437938 | 0.3393 | 0.170926212 | 0.05 | 0 | 1 | 22 |
| LOC110438236 | 0.3393 | 0.170926212 | 0.05 | 0 | 1 | 22 |
| LOC110439500 | 0.3393 | 0.129152241 | 0.05 | 0 | 1 | 22 |
| LOC110439924 | 0.3393 | 0.124422477 | 0.05 | 0 | 1 | 22 |

| | | | | | | |
|---|---|---|---|---|---|---|
| LOC562049 | 0.3393 | 0.124422477 | 0.05 | 0 | 1 | 22 |
| MRPS12 | 0.3393 | 0.128244437 | 0.05 | 0 | 1 | 22 |
| NHLRC3 | 0.3393 | -0.14376955 | 0 | 0.05 | 1 | 22 |
| PDE3B | 0.3393 | 0.119051119 | 0.05 | 0 | 1 | 22 |
| RALGDS | 0.3393 | -0.1031117 | 0 | 0.05 | 1 | 22 |
| RUNDC1 | 0.3393 | -0.10730465 | 0 | 0.05 | 1 | 22 |
| SEMA4F | 0.3393 | 0.118005184 | 0.05 | 0 | 1 | 22 |
| SPAG9 | 0.3393 | 0.143546667 | 0.05 | 0 | 1 | 22 |
| TST | 0.3393 | 0.108182039 | 0.05 | 0 | 1 | 22 |
| USP34 | 0.3393 | -0.14535533 | 0 | 0.05 | 1 | 22 |
| XLOC-004796 | 0.3393 | 0.147876456 | 0.05 | 0 | 1 | 22 |
| XLOC-009640 | 0.3393 | 0.144161418 | 0.05 | 0 | 1 | 22 |
| XLOC-011160 | 0.3393 | -0.12054202 | 0 | 0.05 | 1 | 22 |
| XLOC-038013 | 0.3393 | 0.128352416 | 0.05 | 0 | 1 | 22 |
| XLOC-043545 | 0.3393 | 0.121740846 | 0.05 | 0 | 1 | 22 |
| ZNF462 | 0.3393 | 0.124422477 | 0.05 | 0 | 1 | 22 |
| ZNF821 | 0.3393 | 0.170926212 | 0.05 | 0 | 1 | 22 |
| abhd12 | 0.3393 | 0.169812176 | 0.05 | 0 | 1 | 22 |
| aclya | 0.3393 | 0.116007743 | 0.05 | 0 | 1 | 22 |
| acot8 | 0.3393 | 0.119857252 | 0.05 | 0 | 1 | 22 |
| acss2 | 0.3393 | 0.118480209 | 0.05 | 0 | 1 | 22 |
| acvr1l | 0.3393 | -0.1162051 | 0 | 0.05 | 1 | 22 |
| ada2b | 0.3393 | 0.141986026 | 0.05 | 0 | 1 | 22 |
| adam17a | 0.3393 | -0.11830174 | 0 | 0.05 | 1 | 22 |
| adprhl2 | 0.3393 | 0.163641018 | 0.05 | 0 | 1 | 22 |
| aga | 0.3393 | -0.10383463 | 0 | 0.05 | 1 | 22 |
| agla | 0.3393 | 0.141986026 | 0.05 | 0 | 1 | 22 |
| ago2 | 0.3393 | -0.11926225 | 0 | 0.05 | 1 | 22 |
| ahctf1 | 0.3393 | 0.13658352 | 0.05 | 0 | 1 | 22 |
| aip | 0.3393 | 0.134743356 | 0.05 | 0 | 1 | 22 |
| akt2 | 0.3393 | 0.123247502 | 0.05 | 0 | 1 | 22 |
| alg5 | 0.3393 | -0.11570522 | 0 | 0.05 | 1 | 22 |
| alg9 | 0.3393 | -0.11300965 | 0 | 0.05 | 1 | 22 |
| alox12 | 0.3393 | 0.120764316 | 0.05 | 0 | 1 | 22 |
| angel2 | 0.3393 | -0.21555924 | 0 | 0.05 | 1 | 22 |
| ankrd28b | 0.3393 | 0.135580844 | 0.05 | 0 | 1 | 22 |
| ano10b | 0.3393 | 0.114579284 | 0.05 | 0 | 1 | 22 |
| anxa2a | 0.3393 | 0.108488404 | 0.05 | 0 | 1 | 22 |
| ap3b1a | 0.3393 | 0.169009506 | 0.05 | 0 | 1 | 22 |
| ap3s1.1 | 0.3393 | 0.116600656 | 0.05 | 0 | 1 | 22 |
| api5 | 0.3393 | -0.14183233 | 0 | 0.05 | 1 | 22 |
| apoob | 0.3393 | 0.15113227 | 0.05 | 0 | 1 | 22 |
| apopt1 | 0.3393 | 0.113655516 | 0.05 | 0 | 1 | 22 |
| appbp2 | 0.3393 | 0.137997555 | 0.05 | 0 | 1 | 22 |
| arf4a | 0.3393 | 0.133048709 | 0.05 | 0 | 1 | 22 |
| arfgef1 | 0.3393 | 0.106787428 | 0.05 | 0 | 1 | 22 |
| arfgef2 | 0.3393 | -0.11706534 | 0 | 0.05 | 1 | 22 |
| arglu1a | 0.3393 | -0.11640316 | 0 | 0.05 | 1 | 22 |

| | | | | | | |
|---|---|---|---|---|---|---|
| arhgap17a | 0.3393 | 0.118005184 | 0.05 | 0 | 1 | 22 |
| arid1aa | 0.3393 | 0.112253519 | 0.05 | 0 | 1 | 22 |
| arid1b | 0.3393 | 0.113967406 | 0.05 | 0 | 1 | 22 |
| asah1b | 0.3393 | 0.1340289 | 0.05 | 0 | 1 | 22 |
| asb8 | 0.3393 | 0.160269239 | 0.05 | 0 | 1 | 22 |
| asf1bb | 0.3393 | 0.135484612 | 0.05 | 0 | 1 | 22 |
| atf6 | 0.3393 | 0.133641525 | 0.05 | 0 | 1 | 22 |
| atg101 | 0.3393 | 0.117664107 | 0.05 | 0 | 1 | 22 |
| atp11c | 0.3393 | 0.170167022 | 0.05 | 0 | 1 | 22 |
| atp13a1 | 0.3393 | 0.117505958 | 0.05 | 0 | 1 | 22 |
| atp2c1 | 0.3393 | 0.166493715 | 0.05 | 0 | 1 | 22 |
| atp6v1c1a | 0.3393 | -0.10699207 | 0 | 0.05 | 1 | 22 |
| atp6v1e1a | 0.3393 | 0.114429465 | 0.05 | 0 | 1 | 22 |
| atp7a | 0.3393 | -0.14887914 | 0 | 0.05 | 1 | 22 |
| atp9b | 0.3393 | 0.139575893 | 0.05 | 0 | 1 | 22 |
| atxn3 | 0.3393 | 0.134282294 | 0.05 | 0 | 1 | 22 |
| atxn7 | 0.3393 | 0.13597632 | 0.05 | 0 | 1 | 22 |
| babam2 | 0.3393 | 0.132495406 | 0.05 | 0 | 1 | 22 |
| batf3 | 0.3393 | 0.141986026 | 0.05 | 0 | 1 | 22 |
| baxb | 0.3393 | -0.11732362 | 0 | 0.05 | 1 | 22 |
| bbox1 | 0.3393 | 0.118005184 | 0.05 | 0 | 1 | 22 |
| bcat2 | 0.3393 | 0.116890324 | 0.05 | 0 | 1 | 22 |
| bida | 0.3393 | 0.164728762 | 0.05 | 0 | 1 | 22 |
| birc2 | 0.3393 | 0.112130251 | 0.05 | 0 | 1 | 22 |
| bnip3lb | 0.3393 | -0.14569275 | 0 | 0.05 | 1 | 22 |
| bola3 | 0.3393 | -0.13277432 | 0 | 0.05 | 1 | 22 |
| brca2 | 0.3393 | 0.139575893 | 0.05 | 0 | 1 | 22 |
| brms1la | 0.3393 | 0.162470844 | 0.05 | 0 | 1 | 22 |
| brpf3a | 0.3393 | -0.10540649 | 0 | 0.05 | 1 | 22 |
| btbd3a | 0.3393 | 0.13597632 | 0.05 | 0 | 1 | 22 |
| bxdc2 | 0.3393 | -0.11605625 | 0 | 0.05 | 1 | 22 |
| c1qtnf2 | 0.3393 | -0.10080222 | 0 | 0.05 | 1 | 22 |
| cacul1 | 0.3393 | -0.14758984 | 0 | 0.05 | 1 | 22 |
| caprin1a | 0.3393 | 0.17012641 | 0.05 | 0 | 1 | 22 |
| casp8 | 0.3393 | 0.127625182 | 0.05 | 0 | 1 | 22 |
| caspa | 0.3393 | 0.117816668 | 0.05 | 0 | 1 | 22 |
| ccdc174 | 0.3393 | 0.114667445 | 0.05 | 0 | 1 | 22 |
| ccdc93 | 0.3393 | -0.10520051 | 0 | 0.05 | 1 | 22 |
| ccl39.3 | 0.3393 | 0.133792854 | 0.05 | 0 | 1 | 22 |
| ccna2 | 0.3393 | 0.161971913 | 0.05 | 0 | 1 | 22 |
| ccnb2 | 0.3393 | 0.131598654 | 0.05 | 0 | 1 | 22 |
| ccne2 | 0.3393 | 0.139575893 | 0.05 | 0 | 1 | 22 |
| ccnf | 0.3393 | 0.129824629 | 0.05 | 0 | 1 | 22 |
| ccnt2a | 0.3393 | -0.11434971 | 0 | 0.05 | 1 | 22 |
| ccny | 0.3393 | -0.11894732 | 0 | 0.05 | 1 | 22 |
| ccser2a | 0.3393 | 0.119337053 | 0.05 | 0 | 1 | 22 |
| cd2bp2 | 0.3393 | 0.135221509 | 0.05 | 0 | 1 | 22 |
| cdk1 | 0.3393 | 0.125710743 | 0.05 | 0 | 1 | 22 |

| | | | | | | |
|---|---|---|---|---|---|---|
| cdk5rap3 | 0.3393 | 0.101749255 | 0.05 | 0 | 1 | 22 |
| cdk8 | 0.3393 | 0.159011212 | 0.05 | 0 | 1 | 22 |
| cdkn1d | 0.3393 | 0.143207905 | 0.05 | 0 | 1 | 22 |
| cdkn2aipnl | 0.3393 | -0.10293346 | 0 | 0.05 | 1 | 22 |
| cdkn2c | 0.3393 | 0.127537033 | 0.05 | 0 | 1 | 22 |
| cebpz | 0.3393 | 0.131069977 | 0.05 | 0 | 1 | 22 |
| cenpo | 0.3393 | 0.136948199 | 0.05 | 0 | 1 | 22 |
| cep57l1 | 0.3393 | 0.139575893 | 0.05 | 0 | 1 | 22 |
| cfap20 | 0.3393 | -0.1454998 | 0 | 0.05 | 1 | 22 |
| chchd2 | 0.3393 | 0.123445501 | 0.05 | 0 | 1 | 22 |
| chd1 | 0.3393 | 0.132619169 | 0.05 | 0 | 1 | 22 |
| chd1l | 0.3393 | 0.119857252 | 0.05 | 0 | 1 | 22 |
| ciartb | 0.3393 | -0.1165382 | 0 | 0.05 | 1 | 22 |
| clic5b | 0.3393 | -0.1384912 | 0 | 0.05 | 1 | 22 |
| cln8 | 0.3393 | -0.14885595 | 0 | 0.05 | 1 | 22 |
| clns1a | 0.3393 | 0.131548405 | 0.05 | 0 | 1 | 22 |
| clockb | 0.3393 | 0.132006137 | 0.05 | 0 | 1 | 22 |
| cnot4b | 0.3393 | 0.169960807 | 0.05 | 0 | 1 | 22 |
| cnsta | 0.3393 | 0.124422477 | 0.05 | 0 | 1 | 22 |
| cog2 | 0.3393 | 0.16752038 | 0.05 | 0 | 1 | 22 |
| commd4 | 0.3393 | 0.132225912 | 0.05 | 0 | 1 | 22 |
| cox16 | 0.3393 | -0.14786492 | 0 | 0.05 | 1 | 22 |
| cramp1 | 0.3393 | 0.105366508 | 0.05 | 0 | 1 | 22 |
| crebbpb | 0.3393 | -0.11776519 | 0 | 0.05 | 1 | 22 |
| crnkl1 | 0.3393 | -0.11686587 | 0 | 0.05 | 1 | 22 |
| crot | 0.3393 | 0.104660539 | 0.05 | 0 | 1 | 22 |
| crtc1b | 0.3393 | 0.141986026 | 0.05 | 0 | 1 | 22 |
| ctps1a | 0.3393 | 0.134138779 | 0.05 | 0 | 1 | 22 |
| ctps1b | 0.3393 | -0.10383942 | 0 | 0.05 | 1 | 22 |
| ctsd | 0.3393 | 0.117192265 | 0.05 | 0 | 1 | 22 |
| cuedc2 | 0.3393 | 0.139575893 | 0.05 | 0 | 1 | 22 |
| cyp3c1 | 0.3393 | -0.1036695 | 0 | 0.05 | 1 | 22 |
| cyp4f3 | 0.3393 | -0.1192185 | 0 | 0.05 | 1 | 22 |
| dbpa | 0.3393 | 0.13597632 | 0.05 | 0 | 1 | 22 |
| dbt | 0.3393 | -0.14358281 | 0 | 0.05 | 1 | 22 |
| dck | 0.3393 | 0.103957543 | 0.05 | 0 | 1 | 22 |
| dclre1b | 0.3393 | 0.13411824 | 0.05 | 0 | 1 | 22 |
| dcp1b | 0.3393 | 0.145233737 | 0.05 | 0 | 1 | 22 |
| dctpp1 | 0.3393 | 0.126336433 | 0.05 | 0 | 1 | 22 |
| ddb1 | 0.3393 | 0.108114501 | 0.05 | 0 | 1 | 22 |
| ddx24 | 0.3393 | -0.11449473 | 0 | 0.05 | 1 | 22 |
| dek | 0.3393 | 0.144779634 | 0.05 | 0 | 1 | 22 |
| dhrs12 | 0.3393 | -0.12806452 | 0 | 0.05 | 1 | 22 |
| dhrs7 | 0.3393 | 0.109058376 | 0.05 | 0 | 1 | 22 |
| dhx38 | 0.3393 | 0.114565695 | 0.05 | 0 | 1 | 22 |
| dhx8 | 0.3393 | 0.134863947 | 0.05 | 0 | 1 | 22 |
| dmac2l | 0.3393 | 0.114543772 | 0.05 | 0 | 1 | 22 |
| dna2 | 0.3393 | 0.11706451 | 0.05 | 0 | 1 | 22 |

| | | | | | | |
|---|---|---|---|---|---|---|
| dnaja2b | 0.3393 | 0.1338443 | 0.05 | 0 | 1 | 22 |
| dnajb12a | 0.3393 | 0.13694294 | 0.05 | 0 | 1 | 22 |
| dnajc21 | 0.3393 | 0.136623626 | 0.05 | 0 | 1 | 22 |
| dnajc5ab | 0.3393 | 0.118005184 | 0.05 | 0 | 1 | 22 |
| dnlz | 0.3393 | -0.14229169 | 0 | 0.05 | 1 | 22 |
| dnm1l | 0.3393 | 0.140697866 | 0.05 | 0 | 1 | 22 |
| dok6 | 0.3393 | -0.10686823 | 0 | 0.05 | 1 | 22 |
| drg1 | 0.3393 | -0.1117883 | 0 | 0.05 | 1 | 22 |
| drosha | 0.3393 | -0.10716431 | 0 | 0.05 | 1 | 22 |
| dtd1 | 0.3393 | 0.161247296 | 0.05 | 0 | 1 | 22 |
| dtx2 | 0.3393 | -0.11536665 | 0 | 0.05 | 1 | 22 |
| dtx4a | 0.3393 | 0.11055105 | 0.05 | 0 | 1 | 22 |
| dus4l | 0.3393 | 0.116872551 | 0.05 | 0 | 1 | 22 |
| e2f8 | 0.3393 | 0.139575893 | 0.05 | 0 | 1 | 22 |
| eea1 | 0.3393 | -0.11453626 | 0 | 0.05 | 1 | 22 |
| egr2a | 0.3393 | 0.114884089 | 0.05 | 0 | 1 | 22 |
| ehd1a | 0.3393 | 0.11570195 | 0.05 | 0 | 1 | 22 |
| ehmt1b | 0.3393 | 0.14523863 | 0.05 | 0 | 1 | 22 |
| eif1axa | 0.3393 | -0.11490435 | 0 | 0.05 | 1 | 22 |
| eif2b1 | 0.3393 | -0.10240874 | 0 | 0.05 | 1 | 22 |
| eif4enif1 | 0.3393 | 0.13597632 | 0.05 | 0 | 1 | 22 |
| eif4h | 0.3393 | -0.11289128 | 0 | 0.05 | 1 | 22 |
| elf2a | 0.3393 | 0.117329297 | 0.05 | 0 | 1 | 22 |
| elmod2 | 0.3393 | -0.14096936 | 0 | 0.05 | 1 | 22 |
| emc3 | 0.3393 | 0.11521635 | 0.05 | 0 | 1 | 22 |
| enc1 | 0.3393 | 0.131984119 | 0.05 | 0 | 1 | 22 |
| ep300b | 0.3393 | 0.135383092 | 0.05 | 0 | 1 | 22 |
| epc2 | 0.3393 | 0.132581867 | 0.05 | 0 | 1 | 22 |
| eprs | 0.3393 | -0.11608796 | 0 | 0.05 | 1 | 22 |
| ern1 | 0.3393 | -0.11431702 | 0 | 0.05 | 1 | 22 |
| esrra | 0.3393 | 0.122830501 | 0.05 | 0 | 1 | 22 |
| ess2 | 0.3393 | 0.124422477 | 0.05 | 0 | 1 | 22 |
| exoc6 | 0.3393 | 0.116649964 | 0.05 | 0 | 1 | 22 |
| exosc1 | 0.3393 | -0.11756974 | 0 | 0.05 | 1 | 22 |
| extl3 | 0.3393 | 0.147876456 | 0.05 | 0 | 1 | 22 |
| faah2a | 0.3393 | -0.21513427 | 0 | 0.05 | 1 | 22 |
| fam102bb | 0.3393 | 0.130830609 | 0.05 | 0 | 1 | 22 |
| fam107b | 0.3393 | 0.133341117 | 0.05 | 0 | 1 | 22 |
| fam208ab | 0.3393 | 0.124422477 | 0.05 | 0 | 1 | 22 |
| fam83d | 0.3393 | 0.176201111 | 0.05 | 0 | 1 | 22 |
| fastk | 0.3393 | -0.14808287 | 0 | 0.05 | 1 | 22 |
| fbrs | 0.3393 | 0.164798658 | 0.05 | 0 | 1 | 22 |
| fbxl5 | 0.3393 | 0.108283088 | 0.05 | 0 | 1 | 22 |
| fbxo42 | 0.3393 | 0.170926212 | 0.05 | 0 | 1 | 22 |
| fcf1 | 0.3393 | 0.163328873 | 0.05 | 0 | 1 | 22 |
| fgd4b | 0.3393 | 0.146866487 | 0.05 | 0 | 1 | 22 |
| fibpa | 0.3393 | 0.134985416 | 0.05 | 0 | 1 | 22 |
| foxp1a | 0.3393 | -0.10808405 | 0 | 0.05 | 1 | 22 |

| | | | | | | |
|---|---|---|---|---|---|---|
| ftr63 | 0.3393 | 0.13597632 | 0.05 | 0 | 1 | 22 |
| fxr2 | 0.3393 | 0.116165369 | 0.05 | 0 | 1 | 22 |
| g2e3 | 0.3393 | 0.129822189 | 0.05 | 0 | 1 | 22 |
| gapvd1 | 0.3393 | -0.14847225 | 0 | 0.05 | 1 | 22 |
| gar1 | 0.3393 | 0.134857122 | 0.05 | 0 | 1 | 22 |
| gb:bc139872 | 0.3393 | -0.10076156 | 0 | 0.05 | 1 | 22 |
| gcna | 0.3393 | 0.130573299 | 0.05 | 0 | 1 | 22 |
| gemin2 | 0.3393 | 0.138091096 | 0.05 | 0 | 1 | 22 |
| gpatch1 | 0.3393 | 0.168311128 | 0.05 | 0 | 1 | 22 |
| gpn1 | 0.3393 | 0.113174809 | 0.05 | 0 | 1 | 22 |
| gpr55a | 0.3393 | 0.170926212 | 0.05 | 0 | 1 | 22 |
| gps1 | 0.3393 | 0.117275613 | 0.05 | 0 | 1 | 22 |
| grapa | 0.3393 | 0.139972106 | 0.05 | 0 | 1 | 22 |
| grk4 | 0.3393 | 0.158999843 | 0.05 | 0 | 1 | 22 |
| grpel1 | 0.3393 | 0.146230174 | 0.05 | 0 | 1 | 22 |
| gtf3aa | 0.3393 | 0.128679653 | 0.05 | 0 | 1 | 22 |
| h2afx | 0.3393 | 0.126624393 | 0.05 | 0 | 1 | 22 |
| hadhb | 0.3393 | 0.129388103 | 0.05 | 0 | 1 | 22 |
| hbp1 | 0.3393 | 0.140502893 | 0.05 | 0 | 1 | 22 |
| heatr5b | 0.3393 | 0.116444626 | 0.05 | 0 | 1 | 22 |
| hic1l | 0.3393 | -0.10492045 | 0 | 0.05 | 1 | 22 |
| hmgxb4a | 0.3393 | 0.139575893 | 0.05 | 0 | 1 | 22 |
| hoxb5a | 0.3393 | 0.132033762 | 0.05 | 0 | 1 | 22 |
| hsd17b4 | 0.3393 | 0.13295359 | 0.05 | 0 | 1 | 22 |
| hsd17b8 | 0.3393 | -0.1436304 | 0 | 0.05 | 1 | 22 |
| hyal2a | 0.3393 | 0.112666401 | 0.05 | 0 | 1 | 22 |
| ifrd2 | 0.3393 | 0.143449309 | 0.05 | 0 | 1 | 22 |
| ift20 | 0.3393 | 0.135237022 | 0.05 | 0 | 1 | 22 |
| il13 | 0.3393 | 0.11066651 | 0.05 | 0 | 1 | 22 |
| il2rb | 0.3393 | -0.14906831 | 0 | 0.05 | 1 | 22 |
| il6st | 0.3393 | 0.141003235 | 0.05 | 0 | 1 | 22 |
| incenp | 0.3393 | 0.134413644 | 0.05 | 0 | 1 | 22 |
| ints8 | 0.3393 | 0.113444109 | 0.05 | 0 | 1 | 22 |
| ipmkb | 0.3393 | -0.10591079 | 0 | 0.05 | 1 | 22 |
| ipo11 | 0.3393 | 0.161916348 | 0.05 | 0 | 1 | 22 |
| ipo9 | 0.3393 | 0.135184607 | 0.05 | 0 | 1 | 22 |
| irf9 | 0.3393 | 0.169485431 | 0.05 | 0 | 1 | 22 |
| isy1 | 0.3393 | 0.120284021 | 0.05 | 0 | 1 | 22 |
| itgae.2 | 0.3393 | 0.141986026 | 0.05 | 0 | 1 | 22 |
| itpripl2 | 0.3393 | 0.132545726 | 0.05 | 0 | 1 | 22 |
| ivns1abpa | 0.3393 | -0.10338586 | 0 | 0.05 | 1 | 22 |
| kcnab2a | 0.3393 | 0.114476005 | 0.05 | 0 | 1 | 22 |
| kdelc1 | 0.3393 | -0.14612396 | 0 | 0.05 | 1 | 22 |
| kdm5bb | 0.3393 | 0.163046713 | 0.05 | 0 | 1 | 22 |
| keap1a | 0.3393 | -0.10677828 | 0 | 0.05 | 1 | 22 |
| kidins220b | 0.3393 | 0.124422477 | 0.05 | 0 | 1 | 22 |
| kif14 | 0.3393 | 0.133639904 | 0.05 | 0 | 1 | 22 |
| kif22 | 0.3393 | 0.133808448 | 0.05 | 0 | 1 | 22 |

| | | | | | | |
|---|---|---|---|---|---|---|
| klhl20 | 0.3393 | -0.14710713 | 0 | 0.05 | 1 | 22 |
| klhl21 | 0.3393 | 0.142582964 | 0.05 | 0 | 1 | 22 |
| klhl6 | 0.3393 | -0.1480027 | 0 | 0.05 | 1 | 22 |
| krcp | 0.3393 | 0.108850947 | 0.05 | 0 | 1 | 22 |
| krr1 | 0.3393 | 0.176140042 | 0.05 | 0 | 1 | 22 |
| ksr1a | 0.3393 | -0.2577573 | 0 | 0.05 | 1 | 22 |
| ksr1b | 0.3393 | -0.11245402 | 0 | 0.05 | 1 | 22 |
| lactb | 0.3393 | 0.166493715 | 0.05 | 0 | 1 | 22 |
| larp4aa | 0.3393 | 0.111571097 | 0.05 | 0 | 1 | 22 |
| las1l | 0.3393 | -0.11454833 | 0 | 0.05 | 1 | 22 |
| lcorl | 0.3393 | 0.168628497 | 0.05 | 0 | 1 | 22 |
| lman1 | 0.3393 | 0.113008618 | 0.05 | 0 | 1 | 22 |
| lpar5b | 0.3393 | 0.138986643 | 0.05 | 0 | 1 | 22 |
| lpcat4 | 0.3393 | 0.13919105 | 0.05 | 0 | 1 | 22 |
| lrch1 | 0.3393 | 0.119857252 | 0.05 | 0 | 1 | 22 |
| lrrc40 | 0.3393 | 0.126396537 | 0.05 | 0 | 1 | 22 |
| lrrfip2 | 0.3393 | 0.128271175 | 0.05 | 0 | 1 | 22 |
| lyn | 0.3393 | 0.12200299 | 0.05 | 0 | 1 | 22 |
| lztr1 | 0.3393 | -0.14879084 | 0 | 0.05 | 1 | 22 |
| m6pr | 0.3393 | 0.119400688 | 0.05 | 0 | 1 | 22 |
| mad1l1 | 0.3393 | 0.127317039 | 0.05 | 0 | 1 | 22 |
| mad2l1 | 0.3393 | 0.127642325 | 0.05 | 0 | 1 | 22 |
| maea | 0.3393 | 0.138588289 | 0.05 | 0 | 1 | 22 |
| maml3 | 0.3393 | -0.10681385 | 0 | 0.05 | 1 | 22 |
| map3k20 | 0.3393 | 0.124422477 | 0.05 | 0 | 1 | 22 |
| mat2b | 0.3393 | -0.10360234 | 0 | 0.05 | 1 | 22 |
| mbnl2 | 0.3393 | 0.122679537 | 0.05 | 0 | 1 | 22 |
| mcfd2 | 0.3393 | 0.144853214 | 0.05 | 0 | 1 | 22 |
| mcm7 | 0.3393 | 0.127988032 | 0.05 | 0 | 1 | 22 |
| mcmbp | 0.3393 | -0.14679065 | 0 | 0.05 | 1 | 22 |
| med4 | 0.3393 | 0.122387715 | 0.05 | 0 | 1 | 22 |
| med7 | 0.3393 | 0.164520803 | 0.05 | 0 | 1 | 22 |
| memo1 | 0.3393 | 0.132873773 | 0.05 | 0 | 1 | 22 |
| mettl9 | 0.3393 | -0.11380568 | 0 | 0.05 | 1 | 22 |
| mfap3l | 0.3393 | 0.166521743 | 0.05 | 0 | 1 | 22 |
| mfsd6a | 0.3393 | -0.21050389 | 0 | 0.05 | 1 | 22 |
| mgat5 | 0.3393 | 0.119857252 | 0.05 | 0 | 1 | 22 |
| mgrn1b | 0.3393 | -0.11977094 | 0 | 0.05 | 1 | 22 |
| mhc1zda | 0.3393 | 0.166493715 | 0.05 | 0 | 1 | 22 |
| mis18bp1 | 0.3393 | 0.139575893 | 0.05 | 0 | 1 | 22 |
| mlec | 0.3393 | 0.13750243 | 0.05 | 0 | 1 | 22 |
| mlf2 | 0.3393 | -0.14562001 | 0 | 0.05 | 1 | 22 |
| mrpl10 | 0.3393 | 0.137264878 | 0.05 | 0 | 1 | 22 |
| mrpl20 | 0.3393 | 0.165984225 | 0.05 | 0 | 1 | 22 |
| mrpl3 | 0.3393 | -0.14659183 | 0 | 0.05 | 1 | 22 |
| mrpl35 | 0.3393 | -0.10341067 | 0 | 0.05 | 1 | 22 |
| mrpl44 | 0.3393 | -0.11296678 | 0 | 0.05 | 1 | 22 |
| mrpl58 | 0.3393 | -0.14717087 | 0 | 0.05 | 1 | 22 |

| | | | | | | |
|---|---|---|---|---|---|---|
| msl2a | 0.3393 | 0.110216479 | 0.05 | 0 | 1 | 22 |
| msl2b | 0.3393 | -0.10396184 | 0 | 0.05 | 1 | 22 |
| msrb1b | 0.3393 | 0.119484201 | 0.05 | 0 | 1 | 22 |
| mtfp1 | 0.3393 | 0.134124906 | 0.05 | 0 | 1 | 22 |
| mtrex | 0.3393 | 0.109812261 | 0.05 | 0 | 1 | 22 |
| mxd3 | 0.3393 | 0.134025011 | 0.05 | 0 | 1 | 22 |
| myclb | 0.3393 | 0.159733154 | 0.05 | 0 | 1 | 22 |
| myh9a | 0.3393 | 0.146667527 | 0.05 | 0 | 1 | 22 |
| ncbp3 | 0.3393 | 0.122018765 | 0.05 | 0 | 1 | 22 |
| ncoa2 | 0.3393 | 0.115892489 | 0.05 | 0 | 1 | 22 |
| ncoa5 | 0.3393 | 0.122575529 | 0.05 | 0 | 1 | 22 |
| ndc1 | 0.3393 | 0.102490183 | 0.05 | 0 | 1 | 22 |
| nelfcd | 0.3393 | -0.10651974 | 0 | 0.05 | 1 | 22 |
| neu1 | 0.3393 | 0.1674254 | 0.05 | 0 | 1 | 22 |
| nf1a | 0.3393 | 0.139575893 | 0.05 | 0 | 1 | 22 |
| nfkbib | 0.3393 | -0.10334931 | 0 | 0.05 | 1 | 22 |
| nif3l1 | 0.3393 | 0.132840244 | 0.05 | 0 | 1 | 22 |
| nkiras2 | 0.3393 | 0.16571449 | 0.05 | 0 | 1 | 22 |
| nlk2 | 0.3393 | -0.21451367 | 0 | 0.05 | 1 | 22 |
| nmt1a | 0.3393 | -0.11856689 | 0 | 0.05 | 1 | 22 |
| nol8 | 0.3393 | 0.111841878 | 0.05 | 0 | 1 | 22 |
| nosip | 0.3393 | 0.145819942 | 0.05 | 0 | 1 | 22 |
| nr2f6a | 0.3393 | 0.14241021 | 0.05 | 0 | 1 | 22 |
| nrd1b | 0.3393 | -0.14724819 | 0 | 0.05 | 1 | 22 |
| nrf1 | 0.3393 | 0.13093172 | 0.05 | 0 | 1 | 22 |
| nsfb | 0.3393 | 0.138628125 | 0.05 | 0 | 1 | 22 |
| nsmce4a | 0.3393 | 0.131657497 | 0.05 | 0 | 1 | 22 |
| nub1 | 0.3393 | 0.11490479 | 0.05 | 0 | 1 | 22 |
| nuggc.2 | 0.3393 | 0.13597632 | 0.05 | 0 | 1 | 22 |
| nup153 | 0.3393 | -0.10493562 | 0 | 0.05 | 1 | 22 |
| nup43 | 0.3393 | 0.10435947 | 0.05 | 0 | 1 | 22 |
| ogfrl1 | 0.3393 | -0.21616311 | 0 | 0.05 | 1 | 22 |
| osbpl2b | 0.3393 | 0.139601322 | 0.05 | 0 | 1 | 22 |
| oxa1l | 0.3393 | 0.169223827 | 0.05 | 0 | 1 | 22 |
| paip1 | 0.3393 | 0.131694149 | 0.05 | 0 | 1 | 22 |
| papolg | 0.3393 | -0.10097953 | 0 | 0.05 | 1 | 22 |
| parp3 | 0.3393 | 0.115428248 | 0.05 | 0 | 1 | 22 |
| pbrm1l | 0.3393 | 0.126787258 | 0.05 | 0 | 1 | 22 |
| pcmtl | 0.3393 | 0.1222552 | 0.05 | 0 | 1 | 22 |
| pcnt | 0.3393 | -0.14880561 | 0 | 0.05 | 1 | 22 |
| pcnx1 | 0.3393 | 0.118005184 | 0.05 | 0 | 1 | 22 |
| pde12 | 0.3393 | 0.139575893 | 0.05 | 0 | 1 | 22 |
| pdk3a | 0.3393 | 0.133365777 | 0.05 | 0 | 1 | 22 |
| pdlim1 | 0.3393 | 0.115992683 | 0.05 | 0 | 1 | 22 |
| pex10 | 0.3393 | -0.147692 | 0 | 0.05 | 1 | 22 |
| pex14 | 0.3393 | 0.141927102 | 0.05 | 0 | 1 | 22 |
| pex26 | 0.3393 | 0.140309262 | 0.05 | 0 | 1 | 22 |
| pex3 | 0.3393 | -0.10606226 | 0 | 0.05 | 1 | 22 |

| | | | | | | |
|---|---|---|---|---|---|---|
| pgap1 | 0.3393 | 0.133147487 | 0.05 | 0 | 1 | 22 |
| phf14 | 0.3393 | 0.109757021 | 0.05 | 0 | 1 | 22 |
| phf3 | 0.3393 | 0.110184461 | 0.05 | 0 | 1 | 22 |
| phf6 | 0.3393 | -0.11624134 | 0 | 0.05 | 1 | 22 |
| pi4k2b | 0.3393 | 0.131235903 | 0.05 | 0 | 1 | 22 |
| pias2 | 0.3393 | 0.136065425 | 0.05 | 0 | 1 | 22 |
| pigb | 0.3393 | -0.14886151 | 0 | 0.05 | 1 | 22 |
| pigu | 0.3393 | 0.118005184 | 0.05 | 0 | 1 | 22 |
| pigx | 0.3393 | -0.14396469 | 0 | 0.05 | 1 | 22 |
| pknox1.2 | 0.3393 | 0.13597632 | 0.05 | 0 | 1 | 22 |
| pkp3b | 0.3393 | 0.111810286 | 0.05 | 0 | 1 | 22 |
| plaa | 0.3393 | 0.135612 | 0.05 | 0 | 1 | 22 |
| plcb3 | 0.3393 | 0.143729848 | 0.05 | 0 | 1 | 22 |
| plekha3 | 0.3393 | 0.115595912 | 0.05 | 0 | 1 | 22 |
| plekhm1 | 0.3393 | -0.1421108 | 0 | 0.05 | 1 | 22 |
| plin2 | 0.3393 | -0.14852099 | 0 | 0.05 | 1 | 22 |
| plod3 | 0.3393 | 0.128445625 | 0.05 | 0 | 1 | 22 |
| plxnb2a | 0.3393 | 0.144193512 | 0.05 | 0 | 1 | 22 |
| pola2 | 0.3393 | -0.1040998 | 0 | 0.05 | 1 | 22 |
| pold1 | 0.3393 | -0.10298801 | 0 | 0.05 | 1 | 22 |
| pole2 | 0.3393 | -0.1045318 | 0 | 0.05 | 1 | 22 |
| pole3 | 0.3393 | 0.131291847 | 0.05 | 0 | 1 | 22 |
| polr2c | 0.3393 | 0.108361612 | 0.05 | 0 | 1 | 22 |
| polr3c | 0.3393 | 0.111379294 | 0.05 | 0 | 1 | 22 |
| polr3h | 0.3393 | 0.10780572 | 0.05 | 0 | 1 | 22 |
| ppcdc | 0.3393 | 0.116206935 | 0.05 | 0 | 1 | 22 |
| pphln1 | 0.3393 | 0.138743315 | 0.05 | 0 | 1 | 22 |
| ppm1ba | 0.3393 | 0.138130207 | 0.05 | 0 | 1 | 22 |
| ppm1j | 0.3393 | 0.11613974 | 0.05 | 0 | 1 | 22 |
| ppm1k | 0.3393 | -0.10176861 | 0 | 0.05 | 1 | 22 |
| ppp1r12c | 0.3393 | 0.140031561 | 0.05 | 0 | 1 | 22 |
| ppp1r8b | 0.3393 | 0.134046023 | 0.05 | 0 | 1 | 22 |
| ppp1r9a | 0.3393 | 0.108520272 | 0.05 | 0 | 1 | 22 |
| ppp1r9ba | 0.3393 | 0.170569821 | 0.05 | 0 | 1 | 22 |
| ppp3r1b | 0.3393 | 0.111476045 | 0.05 | 0 | 1 | 22 |
| ppp4r3b | 0.3393 | 0.132470722 | 0.05 | 0 | 1 | 22 |
| prc1a | 0.3393 | 0.127924351 | 0.05 | 0 | 1 | 22 |
| primpol | 0.3393 | 0.133166131 | 0.05 | 0 | 1 | 22 |
| prkd3 | 0.3393 | -0.21208481 | 0 | 0.05 | 1 | 22 |
| prpf19 | 0.3393 | 0.134214232 | 0.05 | 0 | 1 | 22 |
| prpf4ba | 0.3393 | 0.141986026 | 0.05 | 0 | 1 | 22 |
| prps1b | 0.3393 | 0.134337041 | 0.05 | 0 | 1 | 22 |
| psma6l | 0.3393 | -0.11652169 | 0 | 0.05 | 1 | 22 |
| psmf1 | 0.3393 | 0.134626765 | 0.05 | 0 | 1 | 22 |
| ptgr2 | 0.3393 | 0.124422477 | 0.05 | 0 | 1 | 22 |
| ptprr | 0.3393 | -0.21638233 | 0 | 0.05 | 1 | 22 |
| ptrhd1 | 0.3393 | -0.14648044 | 0 | 0.05 | 1 | 22 |
| pts | 0.3393 | 0.124422477 | 0.05 | 0 | 1 | 22 |

| | | | | | | |
|---|---|---|---|---|---|---|
| pudp | 0.3393 | -0.14615049 | 0 | 0.05 | 1 | 22 |
| pum3 | 0.3393 | 0.135323193 | 0.05 | 0 | 1 | 22 |
| purab | 0.3393 | -0.10522422 | 0 | 0.05 | 1 | 22 |
| pygo2 | 0.3393 | 0.133573887 | 0.05 | 0 | 1 | 22 |
| qdprb1 | 0.3393 | -0.13448146 | 0 | 0.05 | 1 | 22 |
| rab4a | 0.3393 | 0.115081536 | 0.05 | 0 | 1 | 22 |
| rab6ba | 0.3393 | 0.13597632 | 0.05 | 0 | 1 | 22 |
| rabgef1l | 0.3393 | -0.17303045 | 0 | 0.05 | 1 | 22 |
| rabl3 | 0.3393 | 0.146498466 | 0.05 | 0 | 1 | 22 |
| rabl6b | 0.3393 | 0.118005184 | 0.05 | 0 | 1 | 22 |
| rad51ap1 | 0.3393 | 0.137176839 | 0.05 | 0 | 1 | 22 |
| rad9a | 0.3393 | -0.102735 | 0 | 0.05 | 1 | 22 |
| ralab | 0.3393 | -0.14638894 | 0 | 0.05 | 1 | 22 |
| ralba | 0.3393 | 0.118005184 | 0.05 | 0 | 1 | 22 |
| ralgapa2 | 0.3393 | 0.169056365 | 0.05 | 0 | 1 | 22 |
| ranbp3b | 0.3393 | 0.132923183 | 0.05 | 0 | 1 | 22 |
| raraa | 0.3393 | -0.10463962 | 0 | 0.05 | 1 | 22 |
| rbm28 | 0.3393 | -0.10280474 | 0 | 0.05 | 1 | 22 |
| rbm39b | 0.3393 | -0.21595133 | 0 | 0.05 | 1 | 22 |
| rbm42 | 0.3393 | -0.1125778 | 0 | 0.05 | 1 | 22 |
| rbmx2 | 0.3393 | 0.113412786 | 0.05 | 0 | 1 | 22 |
| rbpja | 0.3393 | 0.167225997 | 0.05 | 0 | 1 | 22 |
| rc3h2 | 0.3393 | 0.170926212 | 0.05 | 0 | 1 | 22 |
| rcn2 | 0.3393 | 0.141986026 | 0.05 | 0 | 1 | 22 |
| reck | 0.3393 | 0.103990773 | 0.05 | 0 | 1 | 22 |
| rell1 | 0.3393 | 0.144019681 | 0.05 | 0 | 1 | 22 |
| rev3l | 0.3393 | -0.14694462 | 0 | 0.05 | 1 | 22 |
| rfc1 | 0.3393 | 0.136594085 | 0.05 | 0 | 1 | 22 |
| rfc2 | 0.3393 | 0.162500838 | 0.05 | 0 | 1 | 22 |
| rgs17 | 0.3393 | 0.162511314 | 0.05 | 0 | 1 | 22 |
| rimkla | 0.3393 | -0.10637191 | 0 | 0.05 | 1 | 22 |
| rmi1 | 0.3393 | 0.129713343 | 0.05 | 0 | 1 | 22 |
| rnf169 | 0.3393 | 0.129471161 | 0.05 | 0 | 1 | 22 |
| rnf7 | 0.3393 | 0.125685877 | 0.05 | 0 | 1 | 22 |
| rnpc3 | 0.3393 | 0.124746801 | 0.05 | 0 | 1 | 22 |
| rpain | 0.3393 | 0.117976468 | 0.05 | 0 | 1 | 22 |
| rpgrb | 0.3393 | 0.111614643 | 0.05 | 0 | 1 | 22 |
| rps6ka4 | 0.3393 | 0.165400743 | 0.05 | 0 | 1 | 22 |
| rps6kb1b | 0.3393 | 0.133825487 | 0.05 | 0 | 1 | 22 |
| rrm2 | 0.3393 | 0.214476037 | 0.05 | 0 | 1 | 22 |
| rtn4ip1 | 0.3393 | 0.14445404 | 0.05 | 0 | 1 | 22 |
| rxrba | 0.3393 | -0.11641054 | 0 | 0.05 | 1 | 22 |
| samd13 | 0.3393 | 0.137193353 | 0.05 | 0 | 1 | 22 |
| samd4b | 0.3393 | -0.10460088 | 0 | 0.05 | 1 | 22 |
| sat1a.1 | 0.3393 | -0.14817524 | 0 | 0.05 | 1 | 22 |
| scarb2a | 0.3393 | 0.133286356 | 0.05 | 0 | 1 | 22 |
| scfd1 | 0.3393 | 0.100293571 | 0.05 | 0 | 1 | 22 |
| scml2 | 0.3393 | 0.141986026 | 0.05 | 0 | 1 | 22 |

| | | | | | | |
|---|---|---|---|---|---|---|
| scocb | 0.3393 | 0.166493715 | 0.05 | 0 | 1 | 22 |
| scpep1 | 0.3393 | -0.14813966 | 0 | 0.05 | 1 | 22 |
| sdc2 | 0.3393 | -0.21606642 | 0 | 0.05 | 1 | 22 |
| senp2 | 0.3393 | 0.163498714 | 0.05 | 0 | 1 | 22 |
| setd3 | 0.3393 | 0.136031918 | 0.05 | 0 | 1 | 22 |
| sfpq | 0.3393 | 0.128567841 | 0.05 | 0 | 1 | 22 |
| sh3gl3a | 0.3393 | -0.10736464 | 0 | 0.05 | 1 | 22 |
| si:ch1073-104i17.1 | 0.3393 | 0.124422477 | 0.05 | 0 | 1 | 22 |
| si:ch1073-170o4.1 | 0.3393 | 0.164102933 | 0.05 | 0 | 1 | 22 |
| si:ch211-107m4.1 | 0.3393 | 0.13597632 | 0.05 | 0 | 1 | 22 |
| si:ch211-113a14.18 | 0.3393 | 0.13597632 | 0.05 | 0 | 1 | 22 |
| si:ch211-11k18.4 | 0.3393 | 0.135692284 | 0.05 | 0 | 1 | 22 |
| si:ch211-156b7.4 | 0.3393 | -0.14246658 | 0 | 0.05 | 1 | 22 |
| si:ch211-160d20.3 | 0.3393 | 0.119884753 | 0.05 | 0 | 1 | 22 |
| si:ch211-161c3.6 | 0.3393 | -0.14629657 | 0 | 0.05 | 1 | 22 |
| si:ch211-171l17.4 | 0.3393 | 0.141986026 | 0.05 | 0 | 1 | 22 |
| si:ch211-191c10.1 | 0.3393 | 0.121978275 | 0.05 | 0 | 1 | 22 |
| si:ch211-191c10.3 | 0.3393 | 0.13597632 | 0.05 | 0 | 1 | 22 |
| si:ch211-193c2.2 | 0.3393 | 0.113373137 | 0.05 | 0 | 1 | 22 |
| si:ch211-195d17.2 | 0.3393 | -0.2161186 | 0 | 0.05 | 1 | 22 |
| si:ch211-207i20.2 | 0.3393 | -0.10085878 | 0 | 0.05 | 1 | 22 |
| si:ch211-210g13.5 | 0.3393 | 0.135012825 | 0.05 | 0 | 1 | 22 |
| si:ch211-212k18.4 | 0.3393 | 0.135747453 | 0.05 | 0 | 1 | 22 |
| si:ch211-216l23.2 | 0.3393 | 0.165209144 | 0.05 | 0 | 1 | 22 |
| si:ch211-227e10.6 | 0.3393 | 0.119857252 | 0.05 | 0 | 1 | 22 |
| si:ch211-227n13.3 | 0.3393 | 0.130948507 | 0.05 | 0 | 1 | 22 |
| si:ch211-230g15.5 | 0.3393 | 0.135133761 | 0.05 | 0 | 1 | 22 |
| si:ch211-261n11.7 | 0.3393 | -0.14943887 | 0 | 0.05 | 1 | 22 |
| si:ch211-262e15.1 | 0.3393 | 0.168515234 | 0.05 | 0 | 1 | 22 |
| si:ch211-271b14.1 | 0.3393 | 0.130350455 | 0.05 | 0 | 1 | 22 |
| si:ch211-272n13.3 | 0.3393 | -0.20948907 | 0 | 0.05 | 1 | 22 |
| si:ch211-275j6.5 | 0.3393 | -0.10558395 | 0 | 0.05 | 1 | 22 |
| si:ch211-282j22.3 | 0.3393 | -0.1064195 | 0 | 0.05 | 1 | 22 |
| si:ch211-57i17.1 | 0.3393 | 0.132796567 | 0.05 | 0 | 1 | 22 |
| si:ch211-59d17.3 | 0.3393 | 0.166493715 | 0.05 | 0 | 1 | 22 |
| si:ch211-66i15.4 | 0.3393 | 0.124422477 | 0.05 | 0 | 1 | 22 |
| si:ch211-67e16.4 | 0.3393 | 0.142859181 | 0.05 | 0 | 1 | 22 |
| si:ch211-69g19.2 | 0.3393 | 0.13240287 | 0.05 | 0 | 1 | 22 |
| si:ch211-71m22.1 | 0.3393 | 0.131560573 | 0.05 | 0 | 1 | 22 |
| si:ch73-109i22.2 | 0.3393 | -0.11898155 | 0 | 0.05 | 1 | 22 |
| si:ch73-147o17.1 | 0.3393 | -0.10163221 | 0 | 0.05 | 1 | 22 |
| si:ch73-233m11.2 | 0.3393 | 0.141986026 | 0.05 | 0 | 1 | 22 |
| si:ch73-252i11.1 | 0.3393 | -0.11510527 | 0 | 0.05 | 1 | 22 |
| si:ch73-368j24.10.1 | 0.3393 | -0.14873658 | 0 | 0.05 | 1 | 22 |
| si:ch73-91k6.2 | 0.3393 | 0.113519899 | 0.05 | 0 | 1 | 22 |
| si:dkey-105i14.1 | 0.3393 | 0.117085362 | 0.05 | 0 | 1 | 22 |
| si:dkey-108k21.19 | 0.3393 | 0.139575893 | 0.05 | 0 | 1 | 22 |
| si:dkey-109j17.5 | 0.3393 | 0.13597632 | 0.05 | 0 | 1 | 22 |

| | | | | | | |
|---|---|---|---|---|---|---|
| si:dkey-14d8.1 | 0.3393 | 0.119170704 | 0.05 | 0 | 1 | 22 |
| si:dkey-14o6.2 | 0.3393 | -0.11835124 | 0 | 0.05 | 1 | 22 |
| si:dkey-188i13.8 | 0.3393 | 0.188554623 | 0.05 | 0 | 1 | 22 |
| si:dkey-193c22.1 | 0.3393 | 0.116163223 | 0.05 | 0 | 1 | 22 |
| si:dkey-1c7.1 | 0.3393 | 0.156550666 | 0.05 | 0 | 1 | 22 |
| si:dkey-211g8.4 | 0.3393 | -0.120371 | 0 | 0.05 | 1 | 22 |
| si:dkey-217d24.6 | 0.3393 | 0.170926212 | 0.05 | 0 | 1 | 22 |
| si:dkey-229b18.3 | 0.3393 | 0.117243151 | 0.05 | 0 | 1 | 22 |
| si:dkey-24p1.1 | 0.3393 | 0.13597632 | 0.05 | 0 | 1 | 22 |
| si:dkey-253i9.4 | 0.3393 | 0.170926212 | 0.05 | 0 | 1 | 22 |
| si:dkey-261p22.1 | 0.3393 | 0.119857252 | 0.05 | 0 | 1 | 22 |
| si:dkey-29p9.3 | 0.3393 | 0.139575893 | 0.05 | 0 | 1 | 22 |
| si:dkey-31f5.8 | 0.3393 | 0.165994235 | 0.05 | 0 | 1 | 22 |
| si:dkey-40c23.2 | 0.3393 | 0.129842643 | 0.05 | 0 | 1 | 22 |
| si:dkey-48p11.3 | 0.3393 | -0.11158937 | 0 | 0.05 | 1 | 22 |
| si:dkey-53k12.1 | 0.3393 | 0.16547451 | 0.05 | 0 | 1 | 22 |
| si:dkey-6e2.2 | 0.3393 | 0.143268018 | 0.05 | 0 | 1 | 22 |
| si:dkey-7i4.1 | 0.3393 | 0.118005184 | 0.05 | 0 | 1 | 22 |
| si:dkey-92i17.2 | 0.3393 | 0.128116026 | 0.05 | 0 | 1 | 22 |
| si:dkey-93n13.2 | 0.3393 | 0.116876225 | 0.05 | 0 | 1 | 22 |
| si:dkeyp-104h9.5 | 0.3393 | 0.146730128 | 0.05 | 0 | 1 | 22 |
| si:dkeyp-84f3.5 | 0.3393 | 0.106606653 | 0.05 | 0 | 1 | 22 |
| si:rp71-46j2.7 | 0.3393 | 0.137621144 | 0.05 | 0 | 1 | 22 |
| si:zfos-452g4.1 | 0.3393 | 0.13597632 | 0.05 | 0 | 1 | 22 |
| si:zfos-943e10.1 | 0.3393 | 0.145171796 | 0.05 | 0 | 1 | 22 |
| skp2 | 0.3393 | -0.10402936 | 0 | 0.05 | 1 | 22 |
| slc12a9 | 0.3393 | -0.10750483 | 0 | 0.05 | 1 | 22 |
| slc16a5a | 0.3393 | -0.21582304 | 0 | 0.05 | 1 | 22 |
| slc25a12 | 0.3393 | -0.11736433 | 0 | 0.05 | 1 | 22 |
| slc25a24 | 0.3393 | 0.114885155 | 0.05 | 0 | 1 | 22 |
| slc25a36a | 0.3393 | 0.146025422 | 0.05 | 0 | 1 | 22 |
| slc29a2 | 0.3393 | 0.131194414 | 0.05 | 0 | 1 | 22 |
| slc2a1b | 0.3393 | -0.211274 | 0 | 0.05 | 1 | 22 |
| slc33a1 | 0.3393 | 0.139575893 | 0.05 | 0 | 1 | 22 |
| slc39a3 | 0.3393 | 0.137460478 | 0.05 | 0 | 1 | 22 |
| slc5a8l | 0.3393 | 0.112571517 | 0.05 | 0 | 1 | 22 |
| slc7a3b | 0.3393 | 0.170328384 | 0.05 | 0 | 1 | 22 |
| slc9a6a | 0.3393 | -0.14385454 | 0 | 0.05 | 1 | 22 |
| slco5a1b | 0.3393 | 0.119857252 | 0.05 | 0 | 1 | 22 |
| smc2 | 0.3393 | 0.124707827 | 0.05 | 0 | 1 | 22 |
| smc5 | 0.3393 | 0.137914991 | 0.05 | 0 | 1 | 22 |
| smc6 | 0.3393 | 0.133414643 | 0.05 | 0 | 1 | 22 |
| snap47 | 0.3393 | 0.115021495 | 0.05 | 0 | 1 | 22 |
| snapc4 | 0.3393 | -0.10249283 | 0 | 0.05 | 1 | 22 |
| snf8 | 0.3393 | -0.10581104 | 0 | 0.05 | 1 | 22 |
| snrpc | 0.3393 | 0.136626018 | 0.05 | 0 | 1 | 22 |
| snu13a | 0.3393 | -0.10033511 | 0 | 0.05 | 1 | 22 |
| snw1 | 0.3393 | 0.110694481 | 0.05 | 0 | 1 | 22 |

| | | | | | | |
|---|---|---|---|---|---|---|
| snx15 | 0.3393 | 0.170926212 | 0.05 | 0 | 1 | 22 |
| snx9a | 0.3393 | 0.163331588 | 0.05 | 0 | 1 | 22 |
| socs6a | 0.3393 | 0.146052215 | 0.05 | 0 | 1 | 22 |
| sos2 | 0.3393 | 0.129938795 | 0.05 | 0 | 1 | 22 |
| sp2 | 0.3393 | 0.111150585 | 0.05 | 0 | 1 | 22 |
| specc1la | 0.3393 | 0.100090756 | 0.05 | 0 | 1 | 22 |
| spop | 0.3393 | -0.14251033 | 0 | 0.05 | 1 | 22 |
| spryd3 | 0.3393 | 0.165352494 | 0.05 | 0 | 1 | 22 |
| spsb1 | 0.3393 | 0.133252421 | 0.05 | 0 | 1 | 22 |
| srebf2 | 0.3393 | 0.133744852 | 0.05 | 0 | 1 | 22 |
| srp14 | 0.3393 | 0.122354841 | 0.05 | 0 | 1 | 22 |
| ssrp1a | 0.3393 | -0.14108022 | 0 | 0.05 | 1 | 22 |
| st6galnac5b | 0.3393 | 0.114850669 | 0.05 | 0 | 1 | 22 |
| stau2 | 0.3393 | -0.14679275 | 0 | 0.05 | 1 | 22 |
| strap | 0.3393 | 0.165852278 | 0.05 | 0 | 1 | 22 |
| stub1 | 0.3393 | 0.167226272 | 0.05 | 0 | 1 | 22 |
| suclg2 | 0.3393 | 0.127521974 | 0.05 | 0 | 1 | 22 |
| surf1 | 0.3393 | -0.14786671 | 0 | 0.05 | 1 | 22 |
| svild | 0.3393 | 0.143144758 | 0.05 | 0 | 1 | 22 |
| swap70a | 0.3393 | -0.11684559 | 0 | 0.05 | 1 | 22 |
| sytl3 | 0.3393 | 0.170926212 | 0.05 | 0 | 1 | 22 |
| syvn1 | 0.3393 | -0.10268571 | 0 | 0.05 | 1 | 22 |
| tab2 | 0.3393 | -0.11850514 | 0 | 0.05 | 1 | 22 |
| tab3 | 0.3393 | 0.158921257 | 0.05 | 0 | 1 | 22 |
| tada2a | 0.3393 | -0.10218561 | 0 | 0.05 | 1 | 22 |
| taf6 | 0.3393 | 0.120809491 | 0.05 | 0 | 1 | 22 |
| tbc1d10b | 0.3393 | 0.144521493 | 0.05 | 0 | 1 | 22 |
| tbc1d23 | 0.3393 | 0.119857252 | 0.05 | 0 | 1 | 22 |
| tbcelb | 0.3393 | 0.121078295 | 0.05 | 0 | 1 | 22 |
| tcirg1b | 0.3393 | -0.11934329 | 0 | 0.05 | 1 | 22 |
| tdrkh | 0.3393 | 0.160434794 | 0.05 | 0 | 1 | 22 |
| tent5bb | 0.3393 | -0.1004311 | 0 | 0.05 | 1 | 22 |
| terf1 | 0.3393 | 0.162765842 | 0.05 | 0 | 1 | 22 |
| tex2 | 0.3393 | 0.113093119 | 0.05 | 0 | 1 | 22 |
| tex261 | 0.3393 | 0.163544074 | 0.05 | 0 | 1 | 22 |
| tfe3a | 0.3393 | 0.118057646 | 0.05 | 0 | 1 | 22 |
| thap1 | 0.3393 | -0.1059533 | 0 | 0.05 | 1 | 22 |
| thap7 | 0.3393 | 0.145355355 | 0.05 | 0 | 1 | 22 |
| tlk1a | 0.3393 | 0.117604294 | 0.05 | 0 | 1 | 22 |
| tmed4 | 0.3393 | 0.13694746 | 0.05 | 0 | 1 | 22 |
| tmem102 | 0.3393 | -0.10428092 | 0 | 0.05 | 1 | 22 |
| tmem106ba | 0.3393 | -0.11409459 | 0 | 0.05 | 1 | 22 |
| tmem126a | 0.3393 | -0.10171346 | 0 | 0.05 | 1 | 22 |
| tmem135 | 0.3393 | 0.141357016 | 0.05 | 0 | 1 | 22 |
| tmem183a | 0.3393 | -0.10010318 | 0 | 0.05 | 1 | 22 |
| tmem203 | 0.3393 | 0.124460337 | 0.05 | 0 | 1 | 22 |
| tmem209 | 0.3393 | -0.11866209 | 0 | 0.05 | 1 | 22 |
| tmem53 | 0.3393 | 0.110615864 | 0.05 | 0 | 1 | 22 |

| | | | | | | |
|---|---|---|---|---|---|---|
| tmem69 | 0.3393 | -0.10407123 | 0 | 0.05 | 1 | 22 |
| tmpoa | 0.3393 | 0.166458602 | 0.05 | 0 | 1 | 22 |
| tmub2 | 0.3393 | 0.13569588 | 0.05 | 0 | 1 | 22 |
| tnip2 | 0.3393 | -0.14706024 | 0 | 0.05 | 1 | 22 |
| top2a | 0.3393 | 0.127681889 | 0.05 | 0 | 1 | 22 |
| tpk1 | 0.3393 | -0.14640486 | 0 | 0.05 | 1 | 22 |
| tpp2 | 0.3393 | 0.112588156 | 0.05 | 0 | 1 | 22 |
| traf7 | 0.3393 | 0.133886964 | 0.05 | 0 | 1 | 22 |
| trappc10 | 0.3393 | 0.112795062 | 0.05 | 0 | 1 | 22 |
| trappc11 | 0.3393 | 0.145744322 | 0.05 | 0 | 1 | 22 |
| trioa | 0.3393 | 0.135559415 | 0.05 | 0 | 1 | 22 |
| trmt44 | 0.3393 | 0.164509561 | 0.05 | 0 | 1 | 22 |
| trpc4apa | 0.3393 | 0.113279011 | 0.05 | 0 | 1 | 22 |
| trub2 | 0.3393 | -0.10392237 | 0 | 0.05 | 1 | 22 |
| tsr1 | 0.3393 | 0.115052555 | 0.05 | 0 | 1 | 22 |
| ubac2 | 0.3393 | 0.139575893 | 0.05 | 0 | 1 | 22 |
| ube2d2l | 0.3393 | 0.120900143 | 0.05 | 0 | 1 | 22 |
| ube3a | 0.3393 | 0.134128975 | 0.05 | 0 | 1 | 22 |
| ubl3b | 0.3393 | 0.114068445 | 0.05 | 0 | 1 | 22 |
| ubr2 | 0.3393 | 0.115688476 | 0.05 | 0 | 1 | 22 |
| ubr3 | 0.3393 | 0.120758478 | 0.05 | 0 | 1 | 22 |
| ugp2a | 0.3393 | 0.128565354 | 0.05 | 0 | 1 | 22 |
| upp1 | 0.3393 | -0.10732169 | 0 | 0.05 | 1 | 22 |
| usf1 | 0.3393 | 0.143310559 | 0.05 | 0 | 1 | 22 |
| usf2 | 0.3393 | 0.118166685 | 0.05 | 0 | 1 | 22 |
| usp19 | 0.3393 | -0.21526226 | 0 | 0.05 | 1 | 22 |
| usp24 | 0.3393 | 0.137124351 | 0.05 | 0 | 1 | 22 |
| usp37 | 0.3393 | 0.154595955 | 0.05 | 0 | 1 | 22 |
| usp44 | 0.3393 | 0.134611885 | 0.05 | 0 | 1 | 22 |
| utp18 | 0.3393 | -0.11437423 | 0 | 0.05 | 1 | 22 |
| utp25 | 0.3393 | -0.10531106 | 0 | 0.05 | 1 | 22 |
| vps52 | 0.3393 | 0.13498001 | 0.05 | 0 | 1 | 22 |
| vps53 | 0.3393 | -0.11787441 | 0 | 0.05 | 1 | 22 |
| vrk3 | 0.3393 | 0.131432463 | 0.05 | 0 | 1 | 22 |
| vti1a | 0.3393 | 0.124422477 | 0.05 | 0 | 1 | 22 |
| wdhd1 | 0.3393 | 0.159891415 | 0.05 | 0 | 1 | 22 |
| wdr33 | 0.3393 | -0.14665376 | 0 | 0.05 | 1 | 22 |
| wdr45 | 0.3393 | 0.145031883 | 0.05 | 0 | 1 | 22 |
| wrb | 0.3393 | 0.131536345 | 0.05 | 0 | 1 | 22 |
| wscd2 | 0.3393 | 0.124422477 | 0.05 | 0 | 1 | 22 |
| wu:fa19b12 | 0.3393 | 0.111972686 | 0.05 | 0 | 1 | 22 |
| xpc | 0.3393 | 0.115149941 | 0.05 | 0 | 1 | 22 |
| xpr1b | 0.3393 | 0.117744764 | 0.05 | 0 | 1 | 22 |
| xrn1 | 0.3393 | 0.134713707 | 0.05 | 0 | 1 | 22 |
| yju2 | 0.3393 | -0.14095594 | 0 | 0.05 | 1 | 22 |
| yme1l1a | 0.3393 | 0.130090288 | 0.05 | 0 | 1 | 22 |
| ypel1 | 0.3393 | -0.14988166 | 0 | 0.05 | 1 | 22 |
| zbtb1 | 0.3393 | -0.2160486 | 0 | 0.05 | 1 | 22 |

| | | | | | | |
|---|---|---|---|---|---|---|
| zbtb8a | 0.3393 | 0.167804549 | 0.05 | 0 | 1 | 22 |
| zdhhc20b | 0.3393 | -0.13217559 | 0 | 0.05 | 1 | 22 |
| zfp36l2 | 0.3393 | 0.133240602 | 0.05 | 0 | 1 | 22 |
| zfr2 | 0.3393 | 0.137509221 | 0.05 | 0 | 1 | 22 |
| zgc:100846 | 0.3393 | -0.14930389 | 0 | 0.05 | 1 | 22 |
| zgc:101783 | 0.3393 | 0.13555426 | 0.05 | 0 | 1 | 22 |
| zgc:103759 | 0.3393 | 0.141986026 | 0.05 | 0 | 1 | 22 |
| zgc:112496 | 0.3393 | -0.10817783 | 0 | 0.05 | 1 | 22 |
| zgc:113149 | 0.3393 | 0.167566564 | 0.05 | 0 | 1 | 22 |
| zgc:152863 | 0.3393 | 0.114524257 | 0.05 | 0 | 1 | 22 |
| zgc:153631 | 0.3393 | 0.134175008 | 0.05 | 0 | 1 | 22 |
| zgc:153935 | 0.3393 | 0.120663585 | 0.05 | 0 | 1 | 22 |
| zgc:162971 | 0.3393 | 0.13597632 | 0.05 | 0 | 1 | 22 |
| zgc:163080 | 0.3393 | 0.148450038 | 0.05 | 0 | 1 | 22 |
| zgc:165508 | 0.3393 | 0.124970135 | 0.05 | 0 | 1 | 22 |
| zgc:165604 | 0.3393 | 0.139575893 | 0.05 | 0 | 1 | 22 |
| zgc:171599 | 0.3393 | -0.21517203 | 0 | 0.05 | 1 | 22 |
| zgc:174972 | 0.3393 | 0.138400995 | 0.05 | 0 | 1 | 22 |
| zgc:194312 | 0.3393 | 0.116782569 | 0.05 | 0 | 1 | 22 |
| zgc:55733 | 0.3393 | -0.11324835 | 0 | 0.05 | 1 | 22 |
| zgc:56556 | 0.3393 | 0.141986026 | 0.05 | 0 | 1 | 22 |
| zgc:76872 | 0.3393 | 0.112302469 | 0.05 | 0 | 1 | 22 |
| zgc:77086 | 0.3393 | -0.11598798 | 0 | 0.05 | 1 | 22 |
| zgc:77880 | 0.3393 | 0.169522796 | 0.05 | 0 | 1 | 22 |
| zgc:86764 | 0.3393 | 0.127605392 | 0.05 | 0 | 1 | 22 |
| zmiz1a | 0.3393 | -0.10433325 | 0 | 0.05 | 1 | 22 |
| znf1010 | 0.3393 | 0.120166238 | 0.05 | 0 | 1 | 22 |
| znf1156 | 0.3393 | 0.13597632 | 0.05 | 0 | 1 | 22 |
| znf16l | 0.3393 | 0.132782294 | 0.05 | 0 | 1 | 22 |
| znf362a | 0.3393 | 0.105913419 | 0.05 | 0 | 1 | 22 |
| znf367 | 0.3393 | 0.123668463 | 0.05 | 0 | 1 | 22 |
| znf770 | 0.3393 | -0.10300565 | 0 | 0.05 | 1 | 22 |
| znfl1l | 0.3393 | 0.137435207 | 0.05 | 0 | 1 | 22 |
| zranb2 | 0.3393 | 0.117000003 | 0.05 | 0 | 1 | 22 |
| AL954741.2 | 0.3393 | -0.12642333 | 0 | 0.05 | 1 | 22 |
| ATG14 | 0.3393 | -0.11259197 | 0 | 0.05 | 1 | 22 |
| BX927081.3 | 0.3393 | -0.13527833 | 0 | 0.05 | 1 | 22 |
| BX957297.2 | 0.3393 | -0.10956207 | 0 | 0.05 | 1 | 22 |
| CABZ01045617.1 | 0.3393 | 0.16371361 | 0.05 | 0 | 1 | 22 |
| CABZ01081752.1 | 0.3393 | -0.10875322 | 0 | 0.05 | 1 | 22 |
| CU929070.1 | 0.3393 | -0.12844189 | 0 | 0.05 | 1 | 22 |
| FO904898.3 | 0.3393 | -0.12941792 | 0 | 0.05 | 1 | 22 |
| FP236734.1 | 0.3393 | -0.13639368 | 0 | 0.05 | 1 | 22 |
| FQ311908.1 | 0.3393 | -0.1100848 | 0 | 0.05 | 1 | 22 |
| LOC101883948 | 0.3393 | -0.10551368 | 0 | 0.05 | 1 | 22 |
| LOC101885445 | 0.3393 | -0.13661303 | 0 | 0.05 | 1 | 22 |
| LOC103910921 | 0.3393 | -0.10090571 | 0 | 0.05 | 1 | 22 |
| LOC108182798 | 0.3393 | -0.12098929 | 0 | 0.05 | 1 | 22 |

| | | | | | | |
|---|---|---|---|---|---|---|
| LOC110440075 | 0.3393 | -0.12349732 | 0 | 0.05 | 1 | 22 |
| XLOC-006268 | 0.3393 | -0.12896285 | 0 | 0.05 | 1 | 22 |
| XLOC-017111 | 0.3393 | -0.10441833 | 0 | 0.05 | 1 | 22 |
| XLOC-024750 | 0.3393 | -0.12261112 | 0 | 0.05 | 1 | 22 |
| XLOC-038098 | 0.3393 | -0.12246705 | 0 | 0.05 | 1 | 22 |
| XLOC-043918 | 0.3393 | -0.13661303 | 0 | 0.05 | 1 | 22 |
| abhd17ab | 0.3393 | -0.1205665 | 0 | 0.05 | 1 | 22 |
| acbd3 | 0.3393 | -0.10570665 | 0 | 0.05 | 1 | 22 |
| adck5 | 0.3393 | -0.12884229 | 0 | 0.05 | 1 | 22 |
| adipor2 | 0.3393 | -0.13269992 | 0 | 0.05 | 1 | 22 |
| adnp2a | 0.3393 | -0.13322894 | 0 | 0.05 | 1 | 22 |
| agpat3 | 0.3393 | -0.10025243 | 0 | 0.05 | 1 | 22 |
| akap17a | 0.3393 | -0.13388615 | 0 | 0.05 | 1 | 22 |
| aldh3a2a | 0.3393 | -0.11577864 | 0 | 0.05 | 1 | 22 |
| alg10 | 0.3393 | 0.17117911 | 0.05 | 0 | 1 | 22 |
| ambra1b | 0.3393 | -0.10729096 | 0 | 0.05 | 1 | 22 |
| ap2b1 | 0.3393 | -0.10498458 | 0 | 0.05 | 1 | 22 |
| aprt | 0.3393 | -0.11765053 | 0 | 0.05 | 1 | 22 |
| arl5a | 0.3393 | -0.12287986 | 0 | 0.05 | 1 | 22 |
| ascl1b | 0.3393 | 0.163910111 | 0.05 | 0 | 1 | 22 |
| atg3 | 0.3393 | -0.10424274 | 0 | 0.05 | 1 | 22 |
| atraid | 0.3393 | -0.1192022 | 0 | 0.05 | 1 | 22 |
| atxn10 | 0.3393 | -0.12444901 | 0 | 0.05 | 1 | 22 |
| bin1b | 0.3393 | -0.10251029 | 0 | 0.05 | 1 | 22 |
| bloc1s1 | 0.3393 | -0.12313723 | 0 | 0.05 | 1 | 22 |
| bloc1s4 | 0.3393 | -0.12882835 | 0 | 0.05 | 1 | 22 |
| bmi1a | 0.3393 | -0.10482011 | 0 | 0.05 | 1 | 22 |
| bnip2 | 0.3393 | -0.12803245 | 0 | 0.05 | 1 | 22 |
| brcc3 | 0.3393 | -0.12298934 | 0 | 0.05 | 1 | 22 |
| brd1a | 0.3393 | -0.12556013 | 0 | 0.05 | 1 | 22 |
| brd8 | 0.3393 | -0.10260295 | 0 | 0.05 | 1 | 22 |
| btr09 | 0.3393 | -0.10906229 | 0 | 0.05 | 1 | 22 |
| camk2g2 | 0.3393 | -0.12378591 | 0 | 0.05 | 1 | 22 |
| carm1 | 0.3393 | -0.10363962 | 0 | 0.05 | 1 | 22 |
| cbx1a | 0.3393 | -0.12151998 | 0 | 0.05 | 1 | 22 |
| ccdc22 | 0.3393 | -0.12728234 | 0 | 0.05 | 1 | 22 |
| ccl38a.5 | 0.3393 | -0.10942982 | 0 | 0.05 | 1 | 22 |
| cd82a | 0.3393 | 0.152389917 | 0.05 | 0 | 1 | 22 |
| cep135 | 0.3393 | -0.12840975 | 0 | 0.05 | 1 | 22 |
| cep290 | 0.3393 | -0.12463313 | 0 | 0.05 | 1 | 22 |
| cers2a | 0.3393 | -0.13461801 | 0 | 0.05 | 1 | 22 |
| cers5 | 0.3393 | -0.12540407 | 0 | 0.05 | 1 | 22 |
| cfdp1 | 0.3393 | -0.10153944 | 0 | 0.05 | 1 | 22 |
| cflarb | 0.3393 | -0.12562379 | 0 | 0.05 | 1 | 22 |
| ch25hl2 | 0.3393 | -0.12285995 | 0 | 0.05 | 1 | 22 |
| chchd1 | 0.3393 | -0.11923209 | 0 | 0.05 | 1 | 22 |
| chic1 | 0.3393 | -0.12257604 | 0 | 0.05 | 1 | 22 |
| chst12a | 0.3393 | -0.10768353 | 0 | 0.05 | 1 | 22 |

| | | | | | | |
|---|---|---|---|---|---|---|
| cish | 0.3393 | -0.10696828 | 0 | 0.05 | 1 | 22 |
| clcn5b | 0.3393 | -0.12386808 | 0 | 0.05 | 1 | 22 |
| clip2 | 0.3393 | -0.11988253 | 0 | 0.05 | 1 | 22 |
| cog8 | 0.3393 | 0.165752861 | 0.05 | 0 | 1 | 22 |
| dap3 | 0.3393 | 0.166245073 | 0.05 | 0 | 1 | 22 |
| ddx18 | 0.3393 | -0.12233591 | 0 | 0.05 | 1 | 22 |
| ddx56 | 0.3393 | -0.12255763 | 0 | 0.05 | 1 | 22 |
| dedd | 0.3393 | -0.12305946 | 0 | 0.05 | 1 | 22 |
| dennd4a | 0.3393 | -0.12596225 | 0 | 0.05 | 1 | 22 |
| dhx37 | 0.3393 | -0.11670478 | 0 | 0.05 | 1 | 22 |
| dicer1 | 0.3393 | -0.1226735 | 0 | 0.05 | 1 | 22 |
| dnase1l4.2 | 0.3393 | -0.13203481 | 0 | 0.05 | 1 | 22 |
| dnttip2 | 0.3393 | -0.12374461 | 0 | 0.05 | 1 | 22 |
| dph5 | 0.3393 | -0.11764647 | 0 | 0.05 | 1 | 22 |
| e2f1 | 0.3393 | -0.12585917 | 0 | 0.05 | 1 | 22 |
| eepd1 | 0.3393 | -0.10696183 | 0 | 0.05 | 1 | 22 |
| eif4ea | 0.3393 | -0.12413314 | 0 | 0.05 | 1 | 22 |
| elmsan1b | 0.3393 | -0.10648735 | 0 | 0.05 | 1 | 22 |
| eps15l1a | 0.3393 | -0.12050141 | 0 | 0.05 | 1 | 22 |
| fam57a | 0.3393 | -0.10728824 | 0 | 0.05 | 1 | 22 |
| fbxl20 | 0.3393 | -0.12967203 | 0 | 0.05 | 1 | 22 |
| fbxo25 | 0.3393 | -0.11638888 | 0 | 0.05 | 1 | 22 |
| fbxo33 | 0.3393 | -0.1238722 | 0 | 0.05 | 1 | 22 |
| fcho2 | 0.3393 | -0.13440626 | 0 | 0.05 | 1 | 22 |
| fdps | 0.3393 | -0.12589017 | 0 | 0.05 | 1 | 22 |
| fgd1 | 0.3393 | -0.12135252 | 0 | 0.05 | 1 | 22 |
| foxk1 | 0.3393 | -0.12447896 | 0 | 0.05 | 1 | 22 |
| ftr51 | 0.3393 | -0.11320004 | 0 | 0.05 | 1 | 22 |
| fxr1 | 0.3393 | -0.12916274 | 0 | 0.05 | 1 | 22 |
| g6pd | 0.3393 | -0.13450127 | 0 | 0.05 | 1 | 22 |
| get4 | 0.3393 | 0.168308324 | 0.05 | 0 | 1 | 22 |
| gig2p | 0.3393 | -0.12027782 | 0 | 0.05 | 1 | 22 |
| gins2 | 0.3393 | -0.12133477 | 0 | 0.05 | 1 | 22 |
| glmp | 0.3393 | -0.13605324 | 0 | 0.05 | 1 | 22 |
| gnptab | 0.3393 | 0.169526 | 0.05 | 0 | 1 | 22 |
| golga4 | 0.3393 | -0.13661303 | 0 | 0.05 | 1 | 22 |
| golga5 | 0.3393 | -0.1268268 | 0 | 0.05 | 1 | 22 |
| gorab | 0.3393 | -0.11929293 | 0 | 0.05 | 1 | 22 |
| gpatch11 | 0.3393 | -0.10177815 | 0 | 0.05 | 1 | 22 |
| grna | 0.3393 | 0.169947389 | 0.05 | 0 | 1 | 22 |
| gsk3ab | 0.3393 | -0.10132398 | 0 | 0.05 | 1 | 22 |
| gtf3c4 | 0.3393 | -0.1222084 | 0 | 0.05 | 1 | 22 |
| hace1 | 0.3393 | -0.10875322 | 0 | 0.05 | 1 | 22 |
| hadhaa | 0.3393 | 0.169345089 | 0.05 | 0 | 1 | 22 |
| helb | 0.3393 | -0.10148911 | 0 | 0.05 | 1 | 22 |
| hemk1 | 0.3393 | -0.12341313 | 0 | 0.05 | 1 | 22 |
| herc1 | 0.3393 | -0.11554543 | 0 | 0.05 | 1 | 22 |
| hgh1 | 0.3393 | -0.11883953 | 0 | 0.05 | 1 | 22 |

| | | | | | | |
|---|---|---|---|---|---|---|
| hmbox1a | 0.3393 | -0.12059818 | 0 | 0.05 | 1 | 22 |
| hmg20a | 0.3393 | -0.11872201 | 0 | 0.05 | 1 | 22 |
| hnrnph3 | 0.3393 | -0.10395515 | 0 | 0.05 | 1 | 22 |
| igbp1 | 0.3393 | -0.12276228 | 0 | 0.05 | 1 | 22 |
| igf2r | 0.3393 | -0.10875322 | 0 | 0.05 | 1 | 22 |
| ing4 | 0.3393 | -0.12786292 | 0 | 0.05 | 1 | 22 |
| ino80da | 0.3393 | -0.1240283 | 0 | 0.05 | 1 | 22 |
| inpp5l | 0.3393 | -0.12426424 | 0 | 0.05 | 1 | 22 |
| insig1 | 0.3393 | -0.12174335 | 0 | 0.05 | 1 | 22 |
| itpkcb | 0.3393 | -0.11863435 | 0 | 0.05 | 1 | 22 |
| kcnh7 | 0.3393 | -0.12967203 | 0 | 0.05 | 1 | 22 |
| khnyn | 0.3393 | -0.1350948 | 0 | 0.05 | 1 | 22 |
| kri1 | 0.3393 | -0.10087108 | 0 | 0.05 | 1 | 22 |
| larp1b | 0.3393 | -0.12566034 | 0 | 0.05 | 1 | 22 |
| ldb1b | 0.3393 | -0.13343517 | 0 | 0.05 | 1 | 22 |
| lemd3 | 0.3393 | -0.12449138 | 0 | 0.05 | 1 | 22 |
| lmbrd1 | 0.3393 | -0.1213428 | 0 | 0.05 | 1 | 22 |
| lrrc58b | 0.3393 | -0.13568773 | 0 | 0.05 | 1 | 22 |
| lyrm1 | 0.3393 | -0.10130962 | 0 | 0.05 | 1 | 22 |
| lyst | 0.3393 | -0.12454516 | 0 | 0.05 | 1 | 22 |
| macf1a | 0.3393 | -0.12750572 | 0 | 0.05 | 1 | 22 |
| madd | 0.3393 | -0.10426444 | 0 | 0.05 | 1 | 22 |
| manf | 0.3393 | -0.1561754 | 0 | 0.05 | 1 | 22 |
| mapk8ip3 | 0.3393 | -0.12578569 | 0 | 0.05 | 1 | 22 |
| mars | 0.3393 | -0.12411839 | 0 | 0.05 | 1 | 22 |
| mgat1a | 0.3393 | -0.12219314 | 0 | 0.05 | 1 | 22 |
| mlst8 | 0.3393 | -0.10564912 | 0 | 0.05 | 1 | 22 |
| mpnd | 0.3393 | -0.10538639 | 0 | 0.05 | 1 | 22 |
| mrpl15 | 0.3393 | -0.12451117 | 0 | 0.05 | 1 | 22 |
| mrps11 | 0.3393 | -0.12037197 | 0 | 0.05 | 1 | 22 |
| mrps28 | 0.3393 | -0.10131144 | 0 | 0.05 | 1 | 22 |
| msi2b | 0.3393 | -0.10649388 | 0 | 0.05 | 1 | 22 |
| mtif2 | 0.3393 | -0.11349304 | 0 | 0.05 | 1 | 22 |
| n4bp2l2 | 0.3393 | -0.11956531 | 0 | 0.05 | 1 | 22 |
| ncapg | 0.3393 | -0.12708313 | 0 | 0.05 | 1 | 22 |
| ncoa6 | 0.3393 | -0.10624365 | 0 | 0.05 | 1 | 22 |
| neil1 | 0.3393 | -0.12270725 | 0 | 0.05 | 1 | 22 |
| nfrkb | 0.3393 | -0.10326346 | 0 | 0.05 | 1 | 22 |
| nrbf2b | 0.3393 | -0.12817708 | 0 | 0.05 | 1 | 22 |
| nt5c2a | 0.3393 | -0.12514353 | 0 | 0.05 | 1 | 22 |
| nxpe3 | 0.3393 | -0.13620431 | 0 | 0.05 | 1 | 22 |
| orc3 | 0.3393 | -0.12081732 | 0 | 0.05 | 1 | 22 |
| ormdl2 | 0.3393 | -0.11781496 | 0 | 0.05 | 1 | 22 |
| pak1ip1 | 0.3393 | -0.1127475 | 0 | 0.05 | 1 | 22 |
| paqr3b | 0.3393 | -0.12281868 | 0 | 0.05 | 1 | 22 |
| paqr4a | 0.3393 | -0.10618411 | 0 | 0.05 | 1 | 22 |
| parp4 | 0.3393 | -0.12642333 | 0 | 0.05 | 1 | 22 |
| paxx | 0.3393 | -0.10475315 | 0 | 0.05 | 1 | 22 |

| | | | | | | |
|---|---|---|---|---|---|---|
| pdhb | 0.3393 | -0.13452831 | 0 | 0.05 | 1 | 22 |
| phf2 | 0.3393 | -0.12527196 | 0 | 0.05 | 1 | 22 |
| picalma | 0.3393 | -0.12873961 | 0 | 0.05 | 1 | 22 |
| pigp | 0.3393 | -0.10566412 | 0 | 0.05 | 1 | 22 |
| plekhb2 | 0.3393 | -0.13267027 | 0 | 0.05 | 1 | 22 |
| plp2 | 0.3393 | -0.1093746 | 0 | 0.05 | 1 | 22 |
| polr2b | 0.3393 | -0.12478502 | 0 | 0.05 | 1 | 22 |
| ppip5k1a | 0.3393 | -0.10391694 | 0 | 0.05 | 1 | 22 |
| ppp1r7 | 0.3393 | -0.12107556 | 0 | 0.05 | 1 | 22 |
| ppp4r2b | 0.3393 | -0.12008972 | 0 | 0.05 | 1 | 22 |
| pqlc3 | 0.3393 | -0.1356409 | 0 | 0.05 | 1 | 22 |
| prkcdb | 0.3393 | -0.12600349 | 0 | 0.05 | 1 | 22 |
| prkn | 0.3393 | -0.11078151 | 0 | 0.05 | 1 | 22 |
| psda | 0.3393 | -0.12967203 | 0 | 0.05 | 1 | 22 |
| psmd2 | 0.3393 | -0.12335764 | 0 | 0.05 | 1 | 22 |
| ptgesl | 0.3393 | -0.13038295 | 0 | 0.05 | 1 | 22 |
| rbm7 | 0.3393 | -0.10416586 | 0 | 0.05 | 1 | 22 |
| rexo1 | 0.3393 | -0.12043836 | 0 | 0.05 | 1 | 22 |
| rfk | 0.3393 | -0.12494349 | 0 | 0.05 | 1 | 22 |
| rhot1a | 0.3393 | -0.12562885 | 0 | 0.05 | 1 | 22 |
| rps6ka3b | 0.3393 | -0.10178582 | 0 | 0.05 | 1 | 22 |
| rps6kc1 | 0.3393 | -0.12642333 | 0 | 0.05 | 1 | 22 |
| rragcb | 0.3393 | -0.12967203 | 0 | 0.05 | 1 | 22 |
| rrm1 | 0.3393 | -0.12209558 | 0 | 0.05 | 1 | 22 |
| rtcb | 0.3393 | -0.12948133 | 0 | 0.05 | 1 | 22 |
| s1pr1 | 0.3393 | -0.11699077 | 0 | 0.05 | 1 | 22 |
| serac1 | 0.3393 | -0.12336314 | 0 | 0.05 | 1 | 22 |
| sfmbt2 | 0.3393 | -0.12460254 | 0 | 0.05 | 1 | 22 |
| sfxn2 | 0.3393 | -0.10333479 | 0 | 0.05 | 1 | 22 |
| si:cabz01102082.1 | 0.3393 | -0.1358593 | 0 | 0.05 | 1 | 22 |
| si:ch1073-287p18.1 | 0.3393 | -0.11953674 | 0 | 0.05 | 1 | 22 |
| si:ch1073-358c10.1 | 0.3393 | 0.168301047 | 0.05 | 0 | 1 | 22 |
| si:ch211-13c6.2 | 0.3393 | 0.166939755 | 0.05 | 0 | 1 | 22 |
| si:ch211-200p22.4 | 0.3393 | -0.1228308 | 0 | 0.05 | 1 | 22 |
| si:ch211-91p5.3 | 0.3393 | -0.1038045 | 0 | 0.05 | 1 | 22 |
| si:ch73-92e7.4 | 0.3393 | -0.11893815 | 0 | 0.05 | 1 | 22 |
| si:dkey-192k22.2 | 0.3393 | -0.12642333 | 0 | 0.05 | 1 | 22 |
| si:dkey-42i9.7 | 0.3393 | -0.10950927 | 0 | 0.05 | 1 | 22 |
| si:dkey-46g23.1 | 0.3393 | -0.1353129 | 0 | 0.05 | 1 | 22 |
| si:dkey-71l1.1 | 0.3393 | -0.10920104 | 0 | 0.05 | 1 | 22 |
| si:dkey-83k24.5 | 0.3393 | -0.12744178 | 0 | 0.05 | 1 | 22 |
| si:dkey-86e18.1 | 0.3393 | -0.12088101 | 0 | 0.05 | 1 | 22 |
| si:dkey-90l23.1 | 0.3393 | 0.17117911 | 0.05 | 0 | 1 | 22 |
| si:dkey-94l16.4 | 0.3393 | -0.12642333 | 0 | 0.05 | 1 | 22 |
| si:dkeyp-53d3.5 | 0.3393 | -0.12967203 | 0 | 0.05 | 1 | 22 |
| slc37a2 | 0.3393 | -0.12525315 | 0 | 0.05 | 1 | 22 |
| slmapa | 0.3393 | -0.12002463 | 0 | 0.05 | 1 | 22 |
| snx13 | 0.3393 | -0.12295078 | 0 | 0.05 | 1 | 22 |

| | | | | | | |
|---|---|---|---|---|---|---|
| snx16 | 0.3393 | -0.12151003 | 0 | 0.05 | 1 | 22 |
| snx17 | 0.3393 | -0.13358706 | 0 | 0.05 | 1 | 22 |
| snx27b | 0.3393 | -0.12926657 | 0 | 0.05 | 1 | 22 |
| spa17 | 0.3393 | -0.12903922 | 0 | 0.05 | 1 | 22 |
| spen | 0.3393 | -0.10495202 | 0 | 0.05 | 1 | 22 |
| sptan1 | 0.3393 | -0.12087569 | 0 | 0.05 | 1 | 22 |
| sptlc3 | 0.3393 | -0.12490893 | 0 | 0.05 | 1 | 22 |
| sra1 | 0.3393 | -0.16284782 | 0 | 0.05 | 1 | 22 |
| ssh2a | 0.3393 | -0.13621059 | 0 | 0.05 | 1 | 22 |
| ssu72 | 0.3393 | 0.167799355 | 0.05 | 0 | 1 | 22 |
| stag1b | 0.3393 | -0.10497973 | 0 | 0.05 | 1 | 22 |
| stat3 | 0.3393 | -0.18967084 | 0 | 0.05 | 1 | 22 |
| stat4 | 0.3393 | -0.1081499 | 0 | 0.05 | 1 | 22 |
| stat5b | 0.3393 | -0.12609299 | 0 | 0.05 | 1 | 22 |
| stim1b | 0.3393 | -0.11869994 | 0 | 0.05 | 1 | 22 |
| stx7l | 0.3393 | -0.13245811 | 0 | 0.05 | 1 | 22 |
| stx8 | 0.3393 | -0.12593607 | 0 | 0.05 | 1 | 22 |
| sult2st1 | 0.3393 | -0.12162503 | 0 | 0.05 | 1 | 22 |
| sult2st3 | 0.3393 | -0.10551045 | 0 | 0.05 | 1 | 22 |
| supv3l1 | 0.3393 | -0.11419606 | 0 | 0.05 | 1 | 22 |
| suv39h1b | 0.3393 | -0.13240422 | 0 | 0.05 | 1 | 22 |
| taf1 | 0.3393 | -0.13313187 | 0 | 0.05 | 1 | 22 |
| taf8 | 0.3393 | -0.10556497 | 0 | 0.05 | 1 | 22 |
| taf9 | 0.3393 | -0.12120673 | 0 | 0.05 | 1 | 22 |
| taok3a | 0.3393 | -0.13566627 | 0 | 0.05 | 1 | 22 |
| tbc1d5 | 0.3393 | -0.12386728 | 0 | 0.05 | 1 | 22 |
| tbcd | 0.3393 | -0.12444801 | 0 | 0.05 | 1 | 22 |
| tbl1xr1b | 0.3393 | 0.169266526 | 0.05 | 0 | 1 | 22 |
| tcima | 0.3393 | -0.1310367 | 0 | 0.05 | 1 | 22 |
| tet2 | 0.3393 | -0.11487435 | 0 | 0.05 | 1 | 22 |
| tgfbrap1 | 0.3393 | -0.13540038 | 0 | 0.05 | 1 | 22 |
| tlr19 | 0.3393 | 0.17117911 | 0.05 | 0 | 1 | 22 |
| tlr8b | 0.3393 | -0.12967203 | 0 | 0.05 | 1 | 22 |
| tmem229b | 0.3393 | -0.12605584 | 0 | 0.05 | 1 | 22 |
| tmem245 | 0.3393 | -0.12158827 | 0 | 0.05 | 1 | 22 |
| tmem263 | 0.3393 | -0.10181985 | 0 | 0.05 | 1 | 22 |
| tmem33 | 0.3393 | -0.12039315 | 0 | 0.05 | 1 | 22 |
| tmem39a | 0.3393 | -0.12174302 | 0 | 0.05 | 1 | 22 |
| tmem68 | 0.3393 | 0.17117911 | 0.05 | 0 | 1 | 22 |
| tnfrsf9b | 0.3393 | -0.12563376 | 0 | 0.05 | 1 | 22 |
| top3b | 0.3393 | -0.10479611 | 0 | 0.05 | 1 | 22 |
| tpcn1 | 0.3393 | -0.16338989 | 0 | 0.05 | 1 | 22 |
| tprg1l | 0.3393 | -0.12393267 | 0 | 0.05 | 1 | 22 |
| trabd | 0.3393 | -0.12458223 | 0 | 0.05 | 1 | 22 |
| tradd | 0.3393 | -0.10717723 | 0 | 0.05 | 1 | 22 |
| traf5 | 0.3393 | -0.1219696 | 0 | 0.05 | 1 | 22 |
| trib2 | 0.3393 | -0.12518605 | 0 | 0.05 | 1 | 22 |
| trib3 | 0.3393 | -0.12371071 | 0 | 0.05 | 1 | 22 |

| | | | | | | |
|---|---|---|---|---|---|---|
| trim8a | 0.3393 | -0.12775139 | 0 | 0.05 | 1 | 22 |
| trnau1apa | 0.3393 | -0.12770183 | 0 | 0.05 | 1 | 22 |
| trrap | 0.3393 | -0.10580719 | 0 | 0.05 | 1 | 22 |
| tspan33a | 0.3393 | -0.12866729 | 0 | 0.05 | 1 | 22 |
| ttc3 | 0.3393 | -0.10188386 | 0 | 0.05 | 1 | 22 |
| ttl | 0.3393 | -0.12235335 | 0 | 0.05 | 1 | 22 |
| ube2d3 | 0.3393 | -0.1159794 | 0 | 0.05 | 1 | 22 |
| ube2d4 | 0.3393 | -0.10561483 | 0 | 0.05 | 1 | 22 |
| ubxn2a | 0.3393 | -0.1100848 | 0 | 0.05 | 1 | 22 |
| ugp2b | 0.3393 | -0.13529674 | 0 | 0.05 | 1 | 22 |
| ulk1b | 0.3393 | -0.10759247 | 0 | 0.05 | 1 | 22 |
| unc50 | 0.3393 | -0.12602602 | 0 | 0.05 | 1 | 22 |
| uso1 | 0.3393 | -0.12446781 | 0 | 0.05 | 1 | 22 |
| usp12a | 0.3393 | -0.1223158 | 0 | 0.05 | 1 | 22 |
| usp8 | 0.3393 | 0.169975371 | 0.05 | 0 | 1 | 22 |
| vars | 0.3393 | -0.12162581 | 0 | 0.05 | 1 | 22 |
| vclb | 0.3393 | -0.12846368 | 0 | 0.05 | 1 | 22 |
| vrk1 | 0.3393 | -0.12213005 | 0 | 0.05 | 1 | 22 |
| wacb | 0.3393 | -0.11654969 | 0 | 0.05 | 1 | 22 |
| waslb | 0.3393 | -0.11441218 | 0 | 0.05 | 1 | 22 |
| wbp1la | 0.3393 | -0.13304269 | 0 | 0.05 | 1 | 22 |
| wdr3 | 0.3393 | 0.167998651 | 0.05 | 0 | 1 | 22 |
| wdr91 | 0.3393 | 0.169809963 | 0.05 | 0 | 1 | 22 |
| wu:fb74b10 | 0.3393 | -0.1270157 | 0 | 0.05 | 1 | 22 |
| zbtb8os | 0.3393 | -0.12369259 | 0 | 0.05 | 1 | 22 |
| zdhhc3a | 0.3393 | -0.12687472 | 0 | 0.05 | 1 | 22 |
| zfpm1 | 0.3393 | -0.10974707 | 0 | 0.05 | 1 | 22 |
| zgc:153654 | 0.3393 | -0.12589671 | 0 | 0.05 | 1 | 22 |
| zgc:162193 | 0.3393 | -0.12601121 | 0 | 0.05 | 1 | 22 |
| zgc:172128 | 0.3393 | -0.12312538 | 0 | 0.05 | 1 | 22 |
| zgc:174904 | 0.3393 | -0.13572856 | 0 | 0.05 | 1 | 22 |
| zgc:195282 | 0.3393 | -0.12496379 | 0 | 0.05 | 1 | 22 |
| zgc:65895 | 0.3393 | -0.13544077 | 0 | 0.05 | 1 | 22 |
| zgc:66475 | 0.3393 | -0.12439818 | 0 | 0.05 | 1 | 22 |
| zgc:77287 | 0.3393 | -0.1212975 | 0 | 0.05 | 1 | 22 |
| zgc:77849 | 0.3393 | -0.13245048 | 0 | 0.05 | 1 | 22 |
| zmym4 | 0.3393 | -0.12689054 | 0 | 0.05 | 1 | 22 |
| znf1007 | 0.3393 | -0.13356802 | 0 | 0.05 | 1 | 22 |
| znf1104 | 0.3393 | -0.10932627 | 0 | 0.05 | 1 | 22 |
| znf438 | 0.3393 | -0.11266003 | 0 | 0.05 | 1 | 22 |
| znf511 | 0.3393 | -0.1183388 | 0 | 0.05 | 1 | 22 |
| mob1a | 0.3395 | -0.21008759 | 0.4 | 0.6 | 1 | 22 |
| rps27l | 0.3401 | 0.194359753 | 0.2 | 0.25 | 1 | 22 |
| eif4bb | 0.34117 | -0.23925663 | 0.05 | 0.2 | 1 | 22 |
| nccrp1 | 0.34137 | -0.35481523 | 0.05 | 0.2 | 1 | 22 |
| il21r.1 | 0.34215 | -0.36993802 | 0.05 | 0.2 | 1 | 22 |
| stxbp2 | 0.34229 | -0.33126181 | 0.05 | 0.2 | 1 | 22 |
| mrps14 | 0.34274 | -0.22744721 | 0.05 | 0.2 | 1 | 22 |

| | | | | | | |
|---|---|---|---|---|---|---|
| slc38a2 | 0.34303 | -0.26489066 | 0.05 | 0.2 | 1 | 22 |
| rpl27 | 0.34303 | -0.12659509 | 0.75 | 0.9 | 1 | 22 |
| hprt1 | 0.34307 | 0.301870684 | 0.2 | 0.05 | 1 | 22 |
| rps8a | 0.34359 | -0.22477687 | 0.75 | 0.85 | 1 | 22 |
| coro1a | 0.34378 | -0.13789438 | 0.95 | 1 | 1 | 22 |
| tmem18 | 0.3439 | -0.24251839 | 0.05 | 0.2 | 1 | 22 |
| spcs3 | 0.34398 | 0.105995391 | 0.3 | 0.35 | 1 | 22 |
| lcp2a | 0.34408 | -0.25614113 | 0.05 | 0.2 | 1 | 22 |
| crkl | 0.34408 | -0.28257251 | 0.05 | 0.2 | 1 | 22 |
| med10 | 0.34428 | -0.23337976 | 0.05 | 0.2 | 1 | 22 |
| capns1a | 0.34473 | -0.26162364 | 0.05 | 0.2 | 1 | 22 |
| heca | 0.34481 | 0.310480263 | 0.2 | 0.05 | 1 | 22 |
| ptbp3 | 0.34483 | -0.29298898 | 0.05 | 0.2 | 1 | 22 |
| rpl7l1 | 0.34483 | -0.34257463 | 0.05 | 0.2 | 1 | 22 |
| dock10 | 0.34487 | 0.29956985 | 0.2 | 0.05 | 1 | 22 |
| styx | 0.34629 | 0.341848288 | 0.15 | 0.05 | 1 | 22 |
| si:ch73-1a9.3 | 0.34691 | -0.41446506 | 0.2 | 0.35 | 1 | 22 |
| zmat2 | 0.34776 | -0.13535091 | 0.1 | 0.25 | 1 | 22 |
| eif1axb | 0.34912 | 0.139039422 | 0.15 | 0.2 | 1 | 22 |
| CU972454.4 | 0.34993 | -0.43231432 | 0.25 | 0.45 | 1 | 22 |
| themis2 | 0.35032 | 0.411851228 | 0.3 | 0.15 | 1 | 22 |
| smarce1 | 0.35262 | -0.17595513 | 0.1 | 0.25 | 1 | 22 |
| psme1 | 0.35596 | 0.211394129 | 0.25 | 0.25 | 1 | 22 |
| scinlb | 0.35596 | -0.28601207 | 0.2 | 0.4 | 1 | 22 |
| ikzf2 | 0.35813 | -0.31373983 | 0.2 | 0.4 | 1 | 22 |
| si:ch211-136m16.8 | 0.35883 | -0.25695281 | 0.1 | 0.25 | 1 | 22 |
| atp1b1a | 0.36006 | 0.323619062 | 0.4 | 0.2 | 1 | 22 |
| ash1l | 0.36014 | 0.369564903 | 0.15 | 0.05 | 1 | 22 |
| naa10 | 0.36278 | 0.105355335 | 0.25 | 0.3 | 1 | 22 |
| pgpep1 | 0.36324 | -0.11005076 | 0.05 | 0.15 | 1 | 22 |
| LOC110438870 | 0.36422 | -0.31719711 | 0.2 | 0.4 | 1 | 22 |
| tnfrsf9a | 0.36607 | -0.12676318 | 0.05 | 0.15 | 1 | 22 |
| snrpd1 | 0.3667 | 0.158198693 | 0.15 | 0.2 | 1 | 22 |
| arpp19b | 0.36686 | 0.353740654 | 0.15 | 0.05 | 1 | 22 |
| idh2 | 0.36741 | -0.32514752 | 0.5 | 0.65 | 1 | 22 |
| rplp2 | 0.36764 | -0.10531282 | 1 | 1 | 1 | 22 |
| clta | 0.36855 | 0.392406803 | 0.25 | 0.1 | 1 | 22 |
| cap1 | 0.36875 | -0.2598559 | 0.6 | 0.7 | 1 | 22 |
| rpl17 | 0.36883 | -0.24836478 | 0.75 | 0.85 | 1 | 22 |
| tagapa | 0.3735 | -0.2402835 | 0.45 | 0.65 | 1 | 22 |
| rbx1 | 0.37454 | 0.345249203 | 0.3 | 0.2 | 1 | 22 |
| si:busm1-194e12.11 | 0.37534 | 0.31490825 | 0.35 | 0.25 | 1 | 22 |
| nme2b.1 | 0.37584 | -0.20706649 | 0.85 | 0.95 | 1 | 22 |
| arpc4l | 0.37614 | -0.11503039 | 0.4 | 0.55 | 1 | 22 |
| ube2v2 | 0.37767 | 0.34429586 | 0.55 | 0.4 | 1 | 22 |
| si:dkey-9i23.5 | 0.37825 | 0.378725523 | 0.15 | 0.05 | 1 | 22 |
| spi1b | 0.38231 | -0.26387962 | 0.6 | 0.8 | 1 | 22 |
| atp5mc1 | 0.38253 | -0.307627 | 0.2 | 0.4 | 1 | 22 |

| | | | | | | |
|---|---|---|---|---|---|---|
| c1qa | 0.38302 | -0.38167599 | 0.1 | 0.25 | 1 | 22 |
| nav3 | 0.38343 | 0.301732751 | 0.15 | 0.05 | 1 | 22 |
| eif3i | 0.38359 | -0.2798797 | 0.25 | 0.45 | 1 | 22 |
| wasa | 0.38372 | 0.377127337 | 0.6 | 0.4 | 1 | 22 |
| ndufb8 | 0.38518 | -0.24152695 | 0.25 | 0.45 | 1 | 22 |
| tomm7 | 0.38595 | -0.13199229 | 0.2 | 0.35 | 1 | 22 |
| msna | 0.38679 | 0.185730585 | 0.95 | 0.85 | 1 | 22 |
| dnajb1b | 0.3869 | -0.23155443 | 0.1 | 0.25 | 1 | 22 |
| ntan1 | 0.38794 | 0.36896377 | 0.15 | 0.05 | 1 | 22 |
| rps23 | 0.39038 | -0.13415773 | 0.75 | 0.9 | 1 | 22 |
| lbr | 0.39186 | 0.295088815 | 0.2 | 0.15 | 1 | 22 |
| gpc1b | 0.39459 | 0.163310184 | 0.15 | 0.05 | 1 | 22 |
| vdrb | 0.39469 | 0.346275972 | 0.15 | 0.05 | 1 | 22 |
| mrpl52 | 0.39487 | 0.3231099 | 0.15 | 0.05 | 1 | 22 |
| vaspa | 0.3949 | -0.22929298 | 0.75 | 0.9 | 1 | 22 |
| hmha1b | 0.39662 | -0.22906568 | 0.1 | 0.25 | 1 | 22 |
| hif1ab | 0.40007 | 0.286833982 | 0.2 | 0.15 | 1 | 22 |
| pak2a | 0.40112 | -0.27886525 | 0.25 | 0.45 | 1 | 22 |
| rps15 | 0.40125 | -0.23960607 | 0.75 | 0.9 | 1 | 22 |
| lgals3a | 0.40153 | 0.347247756 | 0.15 | 0.05 | 1 | 22 |
| XLOC-041870 | 0.4036 | -0.28039999 | 0.3 | 0.5 | 1 | 22 |
| agap2 | 0.40419 | 0.360837425 | 0.25 | 0.1 | 1 | 22 |
| myh11a | 0.40645 | 0.23528182 | 0.45 | 0.4 | 1 | 22 |
| nhp2 | 0.40772 | 0.19823566 | 0.2 | 0.2 | 1 | 22 |
| atp5f1e | 0.40779 | -0.27163525 | 0.55 | 0.75 | 1 | 22 |
| unc93b1 | 0.40799 | -0.18269773 | 0.15 | 0.3 | 1 | 22 |
| ywhabl | 0.40859 | -0.34460048 | 0.35 | 0.55 | 1 | 22 |
| ppifb | 0.4103 | -0.1284287 | 0.25 | 0.4 | 1 | 22 |
| elmo1 | 0.41033 | -0.2297246 | 0.1 | 0.25 | 1 | 22 |
| glo1 | 0.41069 | 0.35341482 | 0.45 | 0.3 | 1 | 22 |
| mbnl1 | 0.41228 | -0.12771577 | 0.4 | 0.55 | 1 | 22 |
| rab11fip4b | 0.41248 | -0.35889258 | 0.1 | 0.25 | 1 | 22 |
| snrpb | 0.41394 | -0.17571145 | 0.1 | 0.25 | 1 | 22 |
| washc2c | 0.41395 | -0.25994795 | 0.35 | 0.55 | 1 | 22 |
| rpl22 | 0.41404 | -0.11579629 | 0.85 | 0.85 | 1 | 22 |
| rpl5a | 0.41517 | -0.2213992 | 0.75 | 0.9 | 1 | 22 |
| prpf39 | 0.41701 | 0.311013253 | 0.15 | 0.05 | 1 | 22 |
| rap1gap2a | 0.41721 | -0.34507823 | 0.1 | 0.25 | 1 | 22 |
| psmc3 | 0.4176 | 0.324396602 | 0.2 | 0.1 | 1 | 22 |
| srsf2a | 0.41778 | 0.35190786 | 0.4 | 0.25 | 1 | 22 |
| unm-hu7910 | 0.41979 | -0.23170214 | 0.1 | 0.25 | 1 | 22 |
| tuba8l4 | 0.41987 | 0.216135809 | 0.45 | 0.45 | 1 | 22 |
| ccdc88b | 0.42096 | 0.382679594 | 0.25 | 0.1 | 1 | 22 |
| uqcrq | 0.42222 | -0.2495017 | 0.35 | 0.55 | 1 | 22 |
| baz1a | 0.42237 | 0.338027166 | 0.2 | 0.1 | 1 | 22 |
| fam49ba | 0.4235 | -0.29222799 | 0.5 | 0.7 | 1 | 22 |
| tnpo3 | 0.42416 | -0.10175304 | 0.05 | 0.15 | 1 | 22 |
| dedd1 | 0.42571 | 0.366708078 | 0.25 | 0.1 | 1 | 22 |

| | | | | | | |
|---|---|---|---|---|---|---|
| pde4ba | 0.42605 | 0.277375834 | 0.25 | 0.1 | 1 | 22 |
| hp1bp3 | 0.42647 | -0.17638282 | 0.15 | 0.3 | 1 | 22 |
| chtopa | 0.42721 | 0.321181749 | 0.2 | 0.1 | 1 | 22 |
| mki67 | 0.42807 | 0.324312461 | 0.15 | 0.05 | 1 | 22 |
| adam19a | 0.4285 | -0.10347003 | 0.05 | 0.15 | 1 | 22 |
| cct4 | 0.42866 | 0.186001431 | 0.2 | 0.2 | 1 | 22 |
| myd88 | 0.42948 | 0.315897051 | 0.35 | 0.25 | 1 | 22 |
| si:zfos-169g10.3 | 0.42957 | 0.360090754 | 0.3 | 0.15 | 1 | 22 |
| sftpbb | 0.42973 | -0.26386307 | 0.5 | 0.7 | 1 | 22 |
| rab5aa | 0.4317 | -0.31138468 | 0.1 | 0.25 | 1 | 22 |
| tm2d2 | 0.4323 | -0.10701783 | 0.05 | 0.15 | 1 | 22 |
| magt1 | 0.43293 | -0.17574803 | 0.2 | 0.35 | 1 | 22 |
| btf3 | 0.43298 | -0.135831 | 0.75 | 0.9 | 1 | 22 |
| szrd1 | 0.43571 | -0.10433547 | 0.05 | 0.15 | 1 | 22 |
| irs1 | 0.43623 | 0.364069446 | 0.25 | 0.1 | 1 | 22 |
| stk35 | 0.43624 | -0.3482081 | 0.1 | 0.25 | 1 | 22 |
| znf1035 | 0.43728 | 0.183792578 | 0.15 | 0.15 | 1 | 22 |
| ndufs3 | 0.4378 | -0.3151111 | 0.1 | 0.25 | 1 | 22 |
| ankib1b | 0.43787 | -0.34634178 | 0.05 | 0.15 | 1 | 22 |
| psmc5 | 0.43896 | -0.10044254 | 0.05 | 0.15 | 1 | 22 |
| wdr1 | 0.44017 | -0.27265494 | 0.4 | 0.6 | 1 | 22 |
| rpl36a | 0.44036 | -0.18520256 | 0.8 | 0.85 | 1 | 22 |
| rsrp1 | 0.44145 | 0.324343621 | 0.55 | 0.35 | 1 | 22 |
| cnn2 | 0.44251 | -0.31710896 | 0.35 | 0.55 | 1 | 22 |
| XLOC-026836 | 0.44258 | -0.16063507 | 0.75 | 0.9 | 1 | 22 |
| tmem167a | 0.44266 | 0.322289523 | 0.25 | 0.15 | 1 | 22 |
| rps20 | 0.44294 | -0.21431423 | 0.8 | 0.9 | 1 | 22 |
| aimp1 | 0.44336 | 0.300672607 | 0.25 | 0.1 | 1 | 22 |
| tcp1 | 0.44386 | -0.16202804 | 0.2 | 0.35 | 1 | 22 |
| cops4 | 0.44489 | -0.10646027 | 0.05 | 0.15 | 1 | 22 |
| sh3bgrl | 0.44545 | -0.28285253 | 0.4 | 0.6 | 1 | 22 |
| aars | 0.44603 | -0.11982059 | 0.05 | 0.15 | 1 | 22 |
| slc25a33 | 0.44617 | -0.26795508 | 0.1 | 0.25 | 1 | 22 |
| NDUFB1 | 0.44639 | -0.25795193 | 0.1 | 0.25 | 1 | 22 |
| runx3 | 0.4464 | -0.39021996 | 0.15 | 0.3 | 1 | 22 |
| b2m | 0.44854 | 0.235742003 | 0.3 | 0.25 | 1 | 22 |
| dbi | 0.44858 | 0.184809773 | 0.3 | 0.3 | 1 | 22 |
| ewsr1b | 0.44862 | -0.37077269 | 0.15 | 0.3 | 1 | 22 |
| ifngr2 | 0.44985 | -0.26006086 | 0.1 | 0.25 | 1 | 22 |
| tox2 | 0.44996 | -0.31620602 | 0.1 | 0.25 | 1 | 22 |
| eif2s1b | 0.45001 | -0.23401622 | 0.1 | 0.25 | 1 | 22 |
| AL954142.1 | 0.45074 | -0.28963812 | 0.1 | 0.25 | 1 | 22 |
| znf395b | 0.45168 | 0.306591573 | 0.25 | 0.1 | 1 | 22 |
| arhgap4a | 0.45224 | -0.29090786 | 0.1 | 0.25 | 1 | 22 |
| pdk2b | 0.45269 | -0.36195392 | 0.1 | 0.25 | 1 | 22 |
| rpl23a | 0.4527 | -0.17400838 | 0.75 | 0.9 | 1 | 22 |
| adh5 | 0.4529 | -0.22113262 | 0.1 | 0.25 | 1 | 22 |
| eif3jb | 0.45347 | -0.23365867 | 0.1 | 0.25 | 1 | 22 |

| | | | | | | |
|---|---|---|---|---|---|---|
| suz12b | 0.45397 | 0.207229576 | 0.15 | 0.05 | 1 | 22 |
| rplp2l | 0.45638 | -0.18311866 | 0.85 | 0.9 | 1 | 22 |
| lpin1 | 0.45743 | -0.10940923 | 0.05 | 0.15 | 1 | 22 |
| eif3g | 0.45862 | -0.17521373 | 0.2 | 0.35 | 1 | 22 |
| mdh2 | 0.45869 | -0.19220746 | 0.15 | 0.3 | 1 | 22 |
| taf12 | 0.45941 | 0.184780804 | 0.15 | 0.15 | 1 | 22 |
| tec | 0.4607 | -0.33739905 | 0.05 | 0.15 | 1 | 22 |
| ube2v1 | 0.46077 | 0.185747973 | 0.15 | 0.15 | 1 | 22 |
| LOC101882805 | 0.46118 | 0.318125466 | 0.25 | 0.15 | 1 | 22 |
| ube2g1a | 0.46637 | 0.273654453 | 0.15 | 0.05 | 1 | 22 |
| gdi2 | 0.46662 | 0.158228763 | 0.45 | 0.45 | 1 | 22 |
| si:ch211-213a13.1 | 0.46683 | 0.346866058 | 0.3 | 0.15 | 1 | 22 |
| tuba8l | 0.46854 | 0.287512759 | 0.25 | 0.2 | 1 | 22 |
| vps28 | 0.46855 | 0.283460575 | 0.15 | 0.05 | 1 | 22 |
| top2b | 0.46913 | 0.294415106 | 0.15 | 0.05 | 1 | 22 |
| si:dkey-32n7.4 | 0.46962 | 0.364817712 | 0.15 | 0.05 | 1 | 22 |
| pxna | 0.47013 | 0.288903393 | 0.35 | 0.25 | 1 | 22 |
| lamtor4 | 0.47144 | -0.13348683 | 0.05 | 0.15 | 1 | 22 |
| gata3 | 0.47167 | -0.14357789 | 0.05 | 0.15 | 1 | 22 |
| ago4 | 0.47301 | -0.14192656 | 0.05 | 0.15 | 1 | 22 |
| ino80c | 0.47548 | 0.283095673 | 0.15 | 0.05 | 1 | 22 |
| gpr141 | 0.47592 | -0.13983791 | 0.05 | 0.15 | 1 | 22 |
| hnrnpub | 0.47618 | 0.184733536 | 0.15 | 0.15 | 1 | 22 |
| dad1 | 0.47824 | 0.293284322 | 0.4 | 0.3 | 1 | 22 |
| si:rp71-68n21.9 | 0.47854 | 0.224057341 | 0.35 | 0.3 | 1 | 22 |
| cyth4b | 0.47921 | 0.292268096 | 0.15 | 0.05 | 1 | 22 |
| gid8b | 0.4799 | -0.10158195 | 0.05 | 0.15 | 1 | 22 |
| b4galt3 | 0.48005 | -0.2767616 | 0.1 | 0.2 | 1 | 22 |
| CABZ01046427.2 | 0.48392 | -0.13747399 | 0.35 | 0.5 | 1 | 22 |
| si:dkey-112e7.2 | 0.48519 | -0.15601514 | 0.3 | 0.45 | 1 | 22 |
| mt-nd3 | 0.48657 | -0.1708638 | 0.15 | 0.3 | 1 | 22 |
| tnrc6a | 0.48921 | 0.284108075 | 0.15 | 0.1 | 1 | 22 |
| tmem106a | 0.49199 | 0.298909006 | 0.15 | 0.05 | 1 | 22 |
| si:dkey-23i12.5 | 0.49207 | -0.21241922 | 0.15 | 0.3 | 1 | 22 |
| mpeg1.1 | 0.49752 | -0.330633 | 0.25 | 0.4 | 1 | 22 |
| LOC108190565 | 0.49998 | -0.15519682 | 0.35 | 0.5 | 1 | 22 |
| rab7a | 0.50044 | -0.32186538 | 0.1 | 0.2 | 1 | 22 |
| ccdc124 | 0.5018 | -0.12452545 | 0.05 | 0.15 | 1 | 22 |
| paip2b | 0.50397 | -0.22218232 | 0.15 | 0.3 | 1 | 22 |
| adgre5b.3 | 0.50455 | 0.172257644 | 0.2 | 0.2 | 1 | 22 |
| tubb2b | 0.50603 | 0.324978724 | 0.3 | 0.15 | 1 | 22 |
| tsr2 | 0.50675 | 0.297328275 | 0.15 | 0.05 | 1 | 22 |
| mef2d | 0.50679 | 0.276811624 | 0.15 | 0.1 | 1 | 22 |
| hikeshi | 0.5069 | 0.25948723 | 0.15 | 0.05 | 1 | 22 |
| hps5 | 0.50696 | -0.23336532 | 0.15 | 0.3 | 1 | 22 |
| ccl35.2 | 0.50737 | -0.1266246 | 0.05 | 0.15 | 1 | 22 |
| igf2bp3 | 0.50766 | -0.16746654 | 0.05 | 0.15 | 1 | 22 |
| cox7c | 0.50772 | -0.18428269 | 0.65 | 0.7 | 1 | 22 |

| | | | | | | |
|---|---|---|---|---|---|---|
| fgfr1op2 | 0.5078 | -0.1310925 | 0.05 | 0.15 | 1 | 22 |
| si:dkey-177p2.6 | 0.50846 | -0.34846139 | 0.15 | 0.25 | 1 | 22 |
| mkrn1 | 0.50872 | 0.276553594 | 0.15 | 0.05 | 1 | 22 |
| lamtor3 | 0.50901 | -0.11198235 | 0.05 | 0.15 | 1 | 22 |
| ssr1 | 0.5092 | -0.18736528 | 0.05 | 0.15 | 1 | 22 |
| rpl35 | 0.511 | -0.1107025 | 0.9 | 0.9 | 1 | 22 |
| ppp1caa | 0.51106 | -0.25120943 | 0.15 | 0.3 | 1 | 22 |
| cnbpb | 0.51134 | 0.286681504 | 0.3 | 0.15 | 1 | 22 |
| inpp5d | 0.51159 | -0.33741828 | 0.05 | 0.15 | 1 | 22 |
| taf11 | 0.51217 | 0.254760623 | 0.15 | 0.05 | 1 | 22 |
| gadd45ba | 0.51234 | 0.312725764 | 0.3 | 0.2 | 1 | 22 |
| pet100 | 0.51559 | -0.1338135 | 0.05 | 0.15 | 1 | 22 |
| galnt7 | 0.51632 | -0.11990084 | 0.1 | 0.2 | 1 | 22 |
| rer1 | 0.51665 | 0.277106927 | 0.15 | 0.1 | 1 | 22 |
| oaz1a | 0.51684 | -0.28276879 | 0.45 | 0.6 | 1 | 22 |
| nr1d2b | 0.51793 | -0.17812005 | 0.05 | 0.15 | 1 | 22 |
| aif1l | 0.51848 | 0.247936372 | 0.15 | 0.1 | 1 | 22 |
| bcl7bb | 0.51979 | -0.12156288 | 0.05 | 0.15 | 1 | 22 |
| vamp3 | 0.52002 | -0.24238177 | 0.15 | 0.3 | 1 | 22 |
| si:ch211-197k17.3 | 0.52044 | 0.291456122 | 0.3 | 0.15 | 1 | 22 |
| syncripl | 0.52056 | -0.14402896 | 0.05 | 0.15 | 1 | 22 |
| st3gal5 | 0.521 | 0.295173202 | 0.25 | 0.15 | 1 | 22 |
| srrm2 | 0.52103 | -0.11740077 | 0.05 | 0.15 | 1 | 22 |
| churc1 | 0.52156 | -0.25586335 | 0.15 | 0.3 | 1 | 22 |
| grap2a | 0.52287 | 0.23688778 | 0.15 | 0.05 | 1 | 22 |
| rhogd | 0.52289 | -0.28404956 | 0.05 | 0.15 | 1 | 22 |
| fbl | 0.52313 | -0.27226771 | 0.05 | 0.15 | 1 | 22 |
| hmgn2 | 0.52365 | -0.29214188 | 0.65 | 0.8 | 1 | 22 |
| sulf2b | 0.52481 | -0.15081574 | 0.05 | 0.15 | 1 | 22 |
| arhgap15 | 0.52509 | -0.33663853 | 0.1 | 0.2 | 1 | 22 |
| cenpx | 0.5255 | 0.239583585 | 0.15 | 0.1 | 1 | 22 |
| tob1b | 0.52551 | 0.326847242 | 0.2 | 0.1 | 1 | 22 |
| lpar6b | 0.52585 | 0.163573725 | 0.15 | 0.05 | 1 | 22 |
| pa2g4b | 0.52591 | -0.15447933 | 0.05 | 0.15 | 1 | 22 |
| grpel2 | 0.52607 | 0.183664551 | 0.1 | 0.1 | 1 | 22 |
| kpnb1 | 0.52611 | -0.17170968 | 0.05 | 0.15 | 1 | 22 |
| cdkn1bb | 0.52663 | -0.1429121 | 0.5 | 0.65 | 1 | 22 |
| dusp22b | 0.52693 | 0.300844688 | 0.25 | 0.15 | 1 | 22 |
| fbxo34 | 0.52782 | -0.13566125 | 0.05 | 0.15 | 1 | 22 |
| unc13d | 0.52791 | -0.29211326 | 0.05 | 0.15 | 1 | 22 |
| rp2 | 0.52997 | -0.13488823 | 0.05 | 0.15 | 1 | 22 |
| mvp | 0.53007 | -0.12746735 | 0.05 | 0.15 | 1 | 22 |
| fes | 0.53025 | -0.25362297 | 0.05 | 0.15 | 1 | 22 |
| pank1b | 0.53103 | -0.1986432 | 0.05 | 0.15 | 1 | 22 |
| tln1 | 0.53104 | 0.106593777 | 0.6 | 0.6 | 1 | 22 |
| calm1a | 0.53125 | -0.24215103 | 0.05 | 0.15 | 1 | 22 |
| papola | 0.53234 | 0.277115411 | 0.15 | 0.05 | 1 | 22 |
| ccnl1a | 0.53244 | -0.26540292 | 0.05 | 0.15 | 1 | 22 |

| | | | | | | |
|---|---|---|---|---|---|---|
| rps27a | 0.53306 | -0.1482836 | 0.85 | 0.9 | 1 | 22 |
| myo5ab | 0.53315 | 0.27967294 | 0.15 | 0.05 | 1 | 22 |
| XLOC-043043 | 0.53336 | 0.222665001 | 0.4 | 0.25 | 1 | 22 |
| rpa2 | 0.53393 | -0.22368217 | 0.05 | 0.15 | 1 | 22 |
| zgc:64051 | 0.53548 | -0.16934268 | 0.65 | 0.8 | 1 | 22 |
| pdha1a | 0.53657 | 0.130161626 | 0.15 | 0.05 | 1 | 22 |
| si:dkey-85k7.12 | 0.53694 | 0.302018411 | 0.15 | 0.05 | 1 | 22 |
| slc25a25a | 0.53804 | -0.15551834 | 0.05 | 0.15 | 1 | 22 |
| si:busm1-266f07.2 | 0.53853 | 0.229206902 | 0.8 | 0.65 | 1 | 22 |
| si:ch211-197h24.9 | 0.53895 | -0.11478895 | 0.05 | 0.15 | 1 | 22 |
| amd1 | 0.53915 | 0.160693098 | 0.15 | 0.15 | 1 | 22 |
| sumo2b | 0.54009 | 0.264801399 | 0.4 | 0.3 | 1 | 22 |
| CU570889.1 | 0.54099 | 0.272952656 | 0.15 | 0.05 | 1 | 22 |
| eif3c | 0.54105 | -0.32187231 | 0.2 | 0.35 | 1 | 22 |
| mef2b | 0.54113 | -0.20938406 | 0.2 | 0.35 | 1 | 22 |
| CU571074.1 | 0.5412 | -0.1307692 | 0.05 | 0.15 | 1 | 22 |
| bri3 | 0.54148 | -0.15908362 | 0.05 | 0.15 | 1 | 22 |
| kctd10 | 0.54343 | -0.13776659 | 0.05 | 0.15 | 1 | 22 |
| iqgap1 | 0.54385 | 0.250240453 | 0.15 | 0.05 | 1 | 22 |
| ccdc32 | 0.54491 | 0.196288376 | 0.15 | 0.05 | 1 | 22 |
| atp2b1a | 0.54505 | 0.276992934 | 0.15 | 0.05 | 1 | 22 |
| idh3b | 0.54522 | 0.246776635 | 0.15 | 0.1 | 1 | 22 |
| rgs3a | 0.54542 | 0.169414082 | 0.1 | 0.1 | 1 | 22 |
| bag1 | 0.54561 | -0.22910418 | 0.05 | 0.15 | 1 | 22 |
| arhgap22 | 0.54587 | -0.28955486 | 0.05 | 0.15 | 1 | 22 |
| zdhhc18a | 0.54706 | -0.1878215 | 0.05 | 0.15 | 1 | 22 |
| ndufs7 | 0.5471 | 0.15836988 | 0.25 | 0.25 | 1 | 22 |
| zgc:171497 | 0.54711 | 0.255032642 | 0.15 | 0.05 | 1 | 22 |
| rps9 | 0.54719 | -0.16184172 | 0.65 | 0.8 | 1 | 22 |
| mphosph8 | 0.5472 | 0.27561167 | 0.15 | 0.05 | 1 | 22 |
| CABZ01076667.1 | 0.54776 | -0.23191731 | 0.05 | 0.15 | 1 | 22 |
| atp1a1a.1 | 0.54857 | -0.15825131 | 0.05 | 0.15 | 1 | 22 |
| ddx21 | 0.54945 | -0.21607362 | 0.05 | 0.15 | 1 | 22 |
| dgkza | 0.54983 | -0.18035439 | 0.05 | 0.15 | 1 | 22 |
| mrpl13 | 0.54987 | 0.250382862 | 0.15 | 0.05 | 1 | 22 |
| cyth4a | 0.5501 | -0.14483166 | 0.05 | 0.15 | 1 | 22 |
| trappc1 | 0.55042 | -0.15698124 | 0.05 | 0.15 | 1 | 22 |
| CABZ01112656.1 | 0.55058 | -0.2178104 | 0.05 | 0.15 | 1 | 22 |
| ndufa6 | 0.55061 | 0.276147597 | 0.35 | 0.2 | 1 | 22 |
| rsl1d1 | 0.55105 | -0.14019725 | 0.05 | 0.15 | 1 | 22 |
| si:ch211-195b13.1 | 0.5512 | -0.23861569 | 0.2 | 0.35 | 1 | 22 |
| cox7b | 0.55127 | -0.2668613 | 0.45 | 0.6 | 1 | 22 |
| cyfip2 | 0.5515 | 0.266990614 | 0.15 | 0.05 | 1 | 22 |
| si:ch211-87m7.2 | 0.55235 | -0.144451 | 0.05 | 0.15 | 1 | 22 |
| zgc:56585 | 0.55243 | 0.218379308 | 0.15 | 0.05 | 1 | 22 |
| bpifcl | 0.55271 | -0.10007026 | 0.15 | 0.25 | 1 | 22 |
| BX296551.1 | 0.5531 | 0.150217022 | 0.1 | 0.05 | 1 | 22 |
| CABZ01088428.1 | 0.5531 | 0.142232303 | 0.1 | 0.05 | 1 | 22 |

| | | | | | | |
|---|---|---|---|---|---|---|
| CABZ01110881.1 | 0.5531 | 0.104151848 | 0.1 | 0.05 | 1 | 22 |
| DDX17 | 0.5531 | 0.152438555 | 0.1 | 0.05 | 1 | 22 |
| FQ323156.1 | 0.5531 | 0.131801965 | 0.1 | 0.05 | 1 | 22 |
| LOC110437825 | 0.5531 | 0.13371657 | 0.1 | 0.05 | 1 | 22 |
| acin1b | 0.5531 | 0.132447346 | 0.1 | 0.05 | 1 | 22 |
| armc1l | 0.5531 | 0.152190822 | 0.1 | 0.05 | 1 | 22 |
| atp6v1d | 0.5531 | 0.179989422 | 0.1 | 0.05 | 1 | 22 |
| camk2g1 | 0.5531 | 0.198206641 | 0.1 | 0.05 | 1 | 22 |
| cdkn3 | 0.5531 | 0.131568944 | 0.1 | 0.05 | 1 | 22 |
| cep350 | 0.5531 | 0.174044046 | 0.1 | 0.05 | 1 | 22 |
| cica | 0.5531 | 0.156597937 | 0.1 | 0.05 | 1 | 22 |
| ddx47 | 0.5531 | 0.145376002 | 0.1 | 0.05 | 1 | 22 |
| dhx16 | 0.5531 | 0.145940765 | 0.1 | 0.05 | 1 | 22 |
| dusp5 | 0.5531 | 0.178965398 | 0.1 | 0.05 | 1 | 22 |
| ebna1bp2 | 0.5531 | 0.164120419 | 0.1 | 0.05 | 1 | 22 |
| eif3s6ip | 0.5531 | 0.142750179 | 0.1 | 0.05 | 1 | 22 |
| eml3 | 0.5531 | 0.193817874 | 0.1 | 0.05 | 1 | 22 |
| ergic3 | 0.5531 | 0.186540994 | 0.1 | 0.05 | 1 | 22 |
| fam162a | 0.5531 | 0.16832059 | 0.1 | 0.05 | 1 | 22 |
| fam210aa | 0.5531 | 0.201019052 | 0.1 | 0.05 | 1 | 22 |
| fam214b | 0.5531 | 0.129631265 | 0.1 | 0.05 | 1 | 22 |
| gmfb | 0.5531 | 0.142786501 | 0.1 | 0.05 | 1 | 22 |
| gnl1 | 0.5531 | 0.205546591 | 0.1 | 0.05 | 1 | 22 |
| hipk3a | 0.5531 | 0.208900961 | 0.1 | 0.05 | 1 | 22 |
| hnrnpl2 | 0.5531 | 0.227356316 | 0.1 | 0.05 | 1 | 22 |
| ilk | 0.5531 | 0.178186687 | 0.1 | 0.05 | 1 | 22 |
| ipo7 | 0.5531 | -0.13734491 | 0.05 | 0.1 | 1 | 22 |
| irak1 | 0.5531 | 0.115880281 | 0.1 | 0.05 | 1 | 22 |
| itpkb | 0.5531 | 0.146661913 | 0.1 | 0.05 | 1 | 22 |
| kars | 0.5531 | 0.138419743 | 0.1 | 0.05 | 1 | 22 |
| lpar5a | 0.5531 | 0.158771308 | 0.1 | 0.05 | 1 | 22 |
| march6 | 0.5531 | 0.13036163 | 0.1 | 0.05 | 1 | 22 |
| mcl1b | 0.5531 | 0.121331749 | 0.1 | 0.05 | 1 | 22 |
| ndc80 | 0.5531 | 0.217777443 | 0.1 | 0.05 | 1 | 22 |
| per1a | 0.5531 | 0.131064255 | 0.1 | 0.05 | 1 | 22 |
| pip4k2aa | 0.5531 | 0.161041377 | 0.1 | 0.05 | 1 | 22 |
| pmaip1 | 0.5531 | 0.200031703 | 0.1 | 0.05 | 1 | 22 |
| prdx3 | 0.5531 | 0.143368459 | 0.1 | 0.05 | 1 | 22 |
| ptk6b | 0.5531 | 0.107533566 | 0.1 | 0.05 | 1 | 22 |
| ralgapa1 | 0.5531 | 0.105185089 | 0.1 | 0.05 | 1 | 22 |
| rasal2 | 0.5531 | 0.131078632 | 0.1 | 0.05 | 1 | 22 |
| rbm5 | 0.5531 | 0.190485925 | 0.1 | 0.05 | 1 | 22 |
| si:ch73-248e21.5 | 0.5531 | 0.106187459 | 0.1 | 0.05 | 1 | 22 |
| si:dkey-102m7.3 | 0.5531 | -0.12257504 | 0.05 | 0.1 | 1 | 22 |
| si:dkey-165a24.10 | 0.5531 | 0.158671382 | 0.1 | 0.05 | 1 | 22 |
| slc6a6a | 0.5531 | 0.166340657 | 0.1 | 0.05 | 1 | 22 |
| stat1a | 0.5531 | 0.177200593 | 0.1 | 0.05 | 1 | 22 |
| tab1 | 0.5531 | 0.180924795 | 0.1 | 0.05 | 1 | 22 |

| | | | | | | |
|---|---|---|---|---|---|---|
| tspan14 | 0.5531 | 0.205774703 | 0.1 | 0.05 | 1 | 22 |
| txnl4a | 0.5531 | 0.188284776 | 0.1 | 0.05 | 1 | 22 |
| ube3c | 0.5531 | 0.13123386 | 0.1 | 0.05 | 1 | 22 |
| zgc:153012 | 0.5531 | 0.202070149 | 0.1 | 0.05 | 1 | 22 |
| zgc:63882 | 0.5531 | 0.138324883 | 0.1 | 0.05 | 1 | 22 |
| znrf1 | 0.5531 | 0.14731128 | 0.1 | 0.05 | 1 | 22 |
| BX908750.1 | 0.5531 | -0.20614356 | 0.05 | 0.1 | 1 | 22 |
| CHST13 | 0.5531 | 0.206074932 | 0.1 | 0.05 | 1 | 22 |
| CR388029.1 | 0.5531 | 0.175276877 | 0.1 | 0.05 | 1 | 22 |
| RASA2 | 0.5531 | 0.13433818 | 0.1 | 0.05 | 1 | 22 |
| XLOC-011085 | 0.5531 | 0.104534083 | 0.1 | 0.05 | 1 | 22 |
| XLOC-013964 | 0.5531 | 0.133806537 | 0.1 | 0.05 | 1 | 22 |
| abcg1 | 0.5531 | -0.10760477 | 0.05 | 0.1 | 1 | 22 |
| anapc10 | 0.5531 | -0.13609421 | 0.05 | 0.1 | 1 | 22 |
| atp2b1b | 0.5531 | 0.129651355 | 0.1 | 0.05 | 1 | 22 |
| baz2a | 0.5531 | 0.114406733 | 0.1 | 0.05 | 1 | 22 |
| birc6 | 0.5531 | 0.10774132 | 0.1 | 0.05 | 1 | 22 |
| bptf | 0.5531 | 0.135007253 | 0.1 | 0.05 | 1 | 22 |
| brd9 | 0.5531 | 0.172295469 | 0.1 | 0.05 | 1 | 22 |
| ccnd2b | 0.5531 | 0.124583479 | 0.1 | 0.05 | 1 | 22 |
| cdv3 | 0.5531 | 0.114095555 | 0.1 | 0.05 | 1 | 22 |
| cks1b | 0.5531 | 0.234842324 | 0.1 | 0.05 | 1 | 22 |
| cst3 | 0.5531 | 0.140604152 | 0.1 | 0.05 | 1 | 22 |
| fam76b | 0.5531 | 0.138077929 | 0.1 | 0.05 | 1 | 22 |
| gfi1aa | 0.5531 | -0.1009875 | 0.05 | 0.1 | 1 | 22 |
| gpatch8 | 0.5531 | 0.161032917 | 0.1 | 0.05 | 1 | 22 |
| hax1 | 0.5531 | 0.170217263 | 0.1 | 0.05 | 1 | 22 |
| lonrf1l | 0.5531 | -0.17666552 | 0.05 | 0.1 | 1 | 22 |
| map1lc3b | 0.5531 | -0.1080698 | 0.05 | 0.1 | 1 | 22 |
| mgst3a | 0.5531 | 0.124536846 | 0.1 | 0.05 | 1 | 22 |
| mpped2 | 0.5531 | 0.176719881 | 0.1 | 0.05 | 1 | 22 |
| mtor | 0.5531 | 0.15231388 | 0.1 | 0.05 | 1 | 22 |
| osbpl9 | 0.5531 | 0.159404763 | 0.1 | 0.05 | 1 | 22 |
| pak2b | 0.5531 | 0.105212357 | 0.1 | 0.05 | 1 | 22 |
| polr2k | 0.5531 | -0.10023559 | 0.05 | 0.1 | 1 | 22 |
| prpf4bb | 0.5531 | 0.109068896 | 0.1 | 0.05 | 1 | 22 |
| rit1 | 0.5531 | 0.115875423 | 0.1 | 0.05 | 1 | 22 |
| sbf1 | 0.5531 | 0.10141009 | 0.1 | 0.05 | 1 | 22 |
| sdf4 | 0.5531 | -0.17940243 | 0.05 | 0.1 | 1 | 22 |
| serinc2 | 0.5531 | 0.145940756 | 0.1 | 0.05 | 1 | 22 |
| serpine2 | 0.5531 | 0.12618895 | 0.1 | 0.05 | 1 | 22 |
| si:ch211-147a11.3 | 0.5531 | 0.150469166 | 0.1 | 0.05 | 1 | 22 |
| si:ch211-191j22.3 | 0.5531 | 0.100843324 | 0.1 | 0.05 | 1 | 22 |
| si:ch211-269k10.2 | 0.5531 | 0.149390642 | 0.1 | 0.05 | 1 | 22 |
| si:dkey-28k24.2 | 0.5531 | 0.132468293 | 0.1 | 0.05 | 1 | 22 |
| srgap2 | 0.5531 | -0.10413796 | 0.05 | 0.1 | 1 | 22 |
| srsf9 | 0.5531 | -0.11590573 | 0.05 | 0.1 | 1 | 22 |
| terfa | 0.5531 | -0.11913773 | 0.05 | 0.1 | 1 | 22 |

| | | | | | | |
|---|---|---|---|---|---|---|
| tmc6b | 0.5531 | 0.10770512 | 0.1 | 0.05 | 1 | 22 |
| wbp11 | 0.5531 | 0.160527089 | 0.1 | 0.05 | 1 | 22 |
| zgc:92664 | 0.5531 | -0.12106498 | 0.05 | 0.1 | 1 | 22 |
| CABZ01035279.1 | 0.5531 | 0.136382911 | 0.1 | 0.05 | 1 | 22 |
| CU639397.1 | 0.5531 | 0.264956668 | 0.1 | 0.05 | 1 | 22 |
| PDPR | 0.5531 | 0.196779505 | 0.1 | 0.05 | 1 | 22 |
| brms1 | 0.5531 | 0.180027138 | 0.1 | 0.05 | 1 | 22 |
| btbd7 | 0.5531 | 0.139401929 | 0.1 | 0.05 | 1 | 22 |
| calm3a | 0.5531 | 0.15296193 | 0.1 | 0.05 | 1 | 22 |
| daxx | 0.5531 | 0.208413174 | 0.1 | 0.05 | 1 | 22 |
| eef2k | 0.5531 | -0.11045328 | 0.05 | 0.1 | 1 | 22 |
| gnaia | 0.5531 | 0.194296238 | 0.1 | 0.05 | 1 | 22 |
| hat1 | 0.5531 | 0.181281786 | 0.1 | 0.05 | 1 | 22 |
| mcts1 | 0.5531 | 0.165215971 | 0.1 | 0.05 | 1 | 22 |
| micu1 | 0.5531 | -0.10445263 | 0.05 | 0.1 | 1 | 22 |
| rab8a | 0.5531 | 0.220415322 | 0.1 | 0.05 | 1 | 22 |
| rbm38 | 0.5531 | 0.172344384 | 0.1 | 0.05 | 1 | 22 |
| si:ch211-108d22.2 | 0.5531 | 0.131131733 | 0.1 | 0.05 | 1 | 22 |
| slc16a6b | 0.5531 | -0.19923935 | 0.05 | 0.1 | 1 | 22 |
| snx27a | 0.5531 | 0.16834951 | 0.1 | 0.05 | 1 | 22 |
| znf207b | 0.5531 | 0.20912147 | 0.1 | 0.05 | 1 | 22 |
| abcc5 | 0.5531 | 0.155545309 | 0.1 | 0.05 | 1 | 22 |
| abhd3 | 0.5531 | -0.1110824 | 0.05 | 0.1 | 1 | 22 |
| ap1m1 | 0.5531 | 0.187586046 | 0.1 | 0.05 | 1 | 22 |
| atpv0e2 | 0.5531 | 0.113605588 | 0.1 | 0.05 | 1 | 22 |
| brd4 | 0.5531 | 0.183453102 | 0.1 | 0.05 | 1 | 22 |
| carmil3 | 0.5531 | -0.15522326 | 0.05 | 0.1 | 1 | 22 |
| cenpw | 0.5531 | 0.105475609 | 0.1 | 0.05 | 1 | 22 |
| chmp6b | 0.5531 | -0.11000007 | 0.05 | 0.1 | 1 | 22 |
| cmtm7 | 0.5531 | -0.20457324 | 0.05 | 0.1 | 1 | 22 |
| coq4 | 0.5531 | 0.220882785 | 0.1 | 0.05 | 1 | 22 |
| csnk1da | 0.5531 | -0.15944798 | 0.05 | 0.1 | 1 | 22 |
| ddx39aa | 0.5531 | -0.12875822 | 0.05 | 0.1 | 1 | 22 |
| desi2 | 0.5531 | 0.181018205 | 0.1 | 0.05 | 1 | 22 |
| dock11 | 0.5531 | -0.18809467 | 0.05 | 0.1 | 1 | 22 |
| etfa | 0.5531 | 0.171166624 | 0.1 | 0.05 | 1 | 22 |
| gramd2aa | 0.5531 | 0.168278409 | 0.1 | 0.05 | 1 | 22 |
| grk3 | 0.5531 | 0.132108617 | 0.1 | 0.05 | 1 | 22 |
| gspt1l | 0.5531 | -0.1535785 | 0.05 | 0.1 | 1 | 22 |
| gtf2e2 | 0.5531 | 0.184047958 | 0.1 | 0.05 | 1 | 22 |
| gtf2f2a | 0.5531 | 0.225721968 | 0.1 | 0.05 | 1 | 22 |
| lysmd3 | 0.5531 | -0.13631161 | 0.05 | 0.1 | 1 | 22 |
| man1a2 | 0.5531 | 0.194836276 | 0.1 | 0.05 | 1 | 22 |
| man1b1b | 0.5531 | -0.11626678 | 0.05 | 0.1 | 1 | 22 |
| mbpb | 0.5531 | 0.257110474 | 0.1 | 0.05 | 1 | 22 |
| megf8 | 0.5531 | 0.159090823 | 0.1 | 0.05 | 1 | 22 |
| mesd | 0.5531 | -0.15683728 | 0.05 | 0.1 | 1 | 22 |
| mir21-1 | 0.5531 | 0.146236628 | 0.1 | 0.05 | 1 | 22 |

| | | | | | | |
|---|---|---|---|---|---|---|
| mrpl12 | 0.5531 | 0.114379128 | 0.1 | 0.05 | 1 | 22 |
| mycbp2 | 0.5531 | 0.150571638 | 0.1 | 0.05 | 1 | 22 |
| ndufv2 | 0.5531 | -0.11238503 | 0.05 | 0.1 | 1 | 22 |
| ppm1aa | 0.5531 | -0.10378809 | 0.05 | 0.1 | 1 | 22 |
| prkacab | 0.5531 | -0.10316614 | 0.05 | 0.1 | 1 | 22 |
| ptena | 0.5531 | -0.16907281 | 0.05 | 0.1 | 1 | 22 |
| rassf2b | 0.5531 | 0.215086772 | 0.1 | 0.05 | 1 | 22 |
| rps6ka3a | 0.5531 | -0.13587759 | 0.05 | 0.1 | 1 | 22 |
| rsrc2 | 0.5531 | 0.162536462 | 0.1 | 0.05 | 1 | 22 |
| rsu1 | 0.5531 | -0.12286225 | 0.05 | 0.1 | 1 | 22 |
| samsn1a | 0.5531 | 0.220077992 | 0.1 | 0.05 | 1 | 22 |
| sh3glb1b | 0.5531 | 0.144206872 | 0.1 | 0.05 | 1 | 22 |
| si:ch211-233a24.2 | 0.5531 | -0.10624794 | 0.05 | 0.1 | 1 | 22 |
| si:rp71-36a1.5 | 0.5531 | 0.109214993 | 0.1 | 0.05 | 1 | 22 |
| slka | 0.5531 | 0.13690883 | 0.1 | 0.05 | 1 | 22 |
| smg7 | 0.5531 | 0.143300606 | 0.1 | 0.05 | 1 | 22 |
| snx12 | 0.5531 | -0.1387884 | 0.05 | 0.1 | 1 | 22 |
| sox4b | 0.5531 | -0.12106301 | 0.05 | 0.1 | 1 | 22 |
| spryd7a | 0.5531 | -0.1698496 | 0.05 | 0.1 | 1 | 22 |
| tbkbp1 | 0.5531 | -0.16480335 | 0.05 | 0.1 | 1 | 22 |
| tmem189 | 0.5531 | 0.239495389 | 0.1 | 0.05 | 1 | 22 |
| tmtc1 | 0.5531 | -0.1111219 | 0.05 | 0.1 | 1 | 22 |
| tmub1 | 0.5531 | -0.10345307 | 0.05 | 0.1 | 1 | 22 |
| tnfsf13b | 0.5531 | -0.1078595 | 0.05 | 0.1 | 1 | 22 |
| trappc2l | 0.5531 | -0.1215359 | 0.05 | 0.1 | 1 | 22 |
| ube2al | 0.5531 | -0.1576406 | 0.05 | 0.1 | 1 | 22 |
| ube2r2 | 0.5531 | 0.229375239 | 0.1 | 0.05 | 1 | 22 |
| wdr83os | 0.5531 | 0.182202346 | 0.1 | 0.05 | 1 | 22 |
| zgc:112356 | 0.5531 | 0.133753736 | 0.1 | 0.05 | 1 | 22 |
| zgc:153215 | 0.5531 | -0.10347851 | 0.05 | 0.1 | 1 | 22 |
| zgc:56235 | 0.5531 | -0.14991542 | 0.05 | 0.1 | 1 | 22 |
| abca1a | 0.5531 | -0.10034067 | 0.05 | 0.1 | 1 | 22 |
| arrdc3b | 0.5531 | -0.16400043 | 0.05 | 0.1 | 1 | 22 |
| irf2 | 0.5531 | -0.27730533 | 0.05 | 0.1 | 1 | 22 |
| mon2 | 0.5531 | -0.10473801 | 0.05 | 0.1 | 1 | 22 |
| si:ch211-262i1.6 | 0.5531 | 0.167243874 | 0.1 | 0.05 | 1 | 22 |
| slc44a1b | 0.5531 | -0.14182828 | 0.05 | 0.1 | 1 | 22 |
| snrpg | 0.5531 | 0.283828153 | 0.1 | 0.05 | 1 | 22 |
| LOC101882159 | 0.5531 | -0.14862596 | 0.05 | 0.1 | 1 | 22 |
| b3gnt3.4 | 0.5531 | -0.11491661 | 0.05 | 0.1 | 1 | 22 |
| bpgm | 0.5531 | -0.11678386 | 0.05 | 0.1 | 1 | 22 |
| bzw1a | 0.5531 | -0.161516 | 0.05 | 0.1 | 1 | 22 |
| cpeb4b | 0.5531 | -0.22668323 | 0.05 | 0.1 | 1 | 22 |
| fbxo4 | 0.5531 | -0.10654933 | 0.05 | 0.1 | 1 | 22 |
| fryl | 0.5531 | -0.10355408 | 0.05 | 0.1 | 1 | 22 |
| itpk1b | 0.5531 | -0.10604018 | 0.05 | 0.1 | 1 | 22 |
| lrrc59 | 0.5531 | -0.13846694 | 0.05 | 0.1 | 1 | 22 |
| mcm6 | 0.5531 | -0.12320807 | 0.05 | 0.1 | 1 | 22 |

| | | | | | | |
|---|---|---|---|---|---|---|
| mtmr11 | 0.5531 | -0.1844223 | 0.05 | 0.1 | 1 | 22 |
| naa40 | 0.5531 | -0.11911832 | 0.05 | 0.1 | 1 | 22 |
| nfu1 | 0.5531 | -0.13187284 | 0.05 | 0.1 | 1 | 22 |
| pxdc1b | 0.5531 | -0.1137607 | 0.05 | 0.1 | 1 | 22 |
| rnf19b | 0.5531 | -0.1144049 | 0.05 | 0.1 | 1 | 22 |
| sh3bp1 | 0.5531 | -0.14365691 | 0.05 | 0.1 | 1 | 22 |
| si:dkey-42i9.6 | 0.5531 | -0.1920837 | 0.05 | 0.1 | 1 | 22 |
| slc6a9 | 0.5531 | -0.14124274 | 0.05 | 0.1 | 1 | 22 |
| smarcb1a | 0.5531 | 0.104257428 | 0.1 | 0.05 | 1 | 22 |
| snx9b | 0.5531 | -0.14344995 | 0.05 | 0.1 | 1 | 22 |
| stt3b | 0.5531 | -0.11928776 | 0.05 | 0.1 | 1 | 22 |
| stx11a | 0.5531 | -0.23427059 | 0.05 | 0.1 | 1 | 22 |
| tmed8 | 0.5531 | -0.16528892 | 0.05 | 0.1 | 1 | 22 |
| tmem256 | 0.5531 | -0.12760722 | 0.05 | 0.1 | 1 | 22 |
| tnip1 | 0.5531 | -0.10545632 | 0.05 | 0.1 | 1 | 22 |
| ubap2b | 0.5531 | -0.14656874 | 0.05 | 0.1 | 1 | 22 |
| uqcc3 | 0.5531 | -0.14970101 | 0.05 | 0.1 | 1 | 22 |
| wipi2 | 0.5531 | -0.20399906 | 0.05 | 0.1 | 1 | 22 |
| CDPF1 | 0.5531 | 0.176841414 | 0.1 | 0.05 | 1 | 22 |
| LOC103910015 | 0.5531 | 0.166892929 | 0.1 | 0.05 | 1 | 22 |
| ankrd44 | 0.5531 | 0.178141514 | 0.1 | 0.05 | 1 | 22 |
| cd3eap | 0.5531 | 0.164720766 | 0.1 | 0.05 | 1 | 22 |
| cdc37 | 0.5531 | 0.233543083 | 0.1 | 0.05 | 1 | 22 |
| csnk1a1 | 0.5531 | 0.172275183 | 0.1 | 0.05 | 1 | 22 |
| ctcf | 0.5531 | 0.19296255 | 0.1 | 0.05 | 1 | 22 |
| dnajc9 | 0.5531 | -0.11819658 | 0.05 | 0.1 | 1 | 22 |
| hnrnpa1a | 0.5531 | -0.10728102 | 0.05 | 0.1 | 1 | 22 |
| krt18b | 0.5531 | 0.250745133 | 0.1 | 0.05 | 1 | 22 |
| lnpep | 0.5531 | 0.183265479 | 0.1 | 0.05 | 1 | 22 |
| nufip2 | 0.5531 | 0.12228235 | 0.1 | 0.05 | 1 | 22 |
| pkn1b | 0.5531 | -0.14264286 | 0.05 | 0.1 | 1 | 22 |
| ppp1cab | 0.5531 | 0.179959776 | 0.1 | 0.05 | 1 | 22 |
| si:ch211-134c9.2 | 0.5531 | 0.185653384 | 0.1 | 0.05 | 1 | 22 |
| si:ch211-196c10.15 | 0.5531 | 0.193767543 | 0.1 | 0.05 | 1 | 22 |
| si:ch211-198c19.3 | 0.5531 | -0.12052037 | 0.05 | 0.1 | 1 | 22 |
| smim12 | 0.5531 | -0.13905915 | 0.05 | 0.1 | 1 | 22 |
| spag1a | 0.5531 | 0.169660292 | 0.1 | 0.05 | 1 | 22 |
| srsf1a | 0.5531 | 0.125187534 | 0.1 | 0.05 | 1 | 22 |
| zgc:103692 | 0.5531 | 0.165925871 | 0.1 | 0.05 | 1 | 22 |
| slc35c2 | 0.5531 | -0.189283 | 0.05 | 0.1 | 1 | 22 |
| si:dkey-24p1.7 | 0.55311 | 0.231391729 | 0.15 | 0.05 | 1 | 22 |
| COA4 | 0.55397 | 0.228589777 | 0.15 | 0.1 | 1 | 22 |
| ak2 | 0.55445 | -0.15525347 | 0.25 | 0.4 | 1 | 22 |
| ppp1r12a | 0.55459 | -0.19661801 | 0.05 | 0.15 | 1 | 22 |
| timm10 | 0.55466 | -0.17169998 | 0.05 | 0.15 | 1 | 22 |
| zgc:56676 | 0.55507 | -0.11319693 | 0.1 | 0.2 | 1 | 22 |
| zgc:100918 | 0.55513 | -0.15657439 | 0.05 | 0.15 | 1 | 22 |
| LOC110440146 | 0.55581 | -0.17328106 | 0.05 | 0.15 | 1 | 22 |

| | | | | | | |
|---|---|---|---|---|---|---|
| fam13b | 0.55617 | -0.24219895 | 0.05 | 0.15 | 1 | 22 |
| dldh | 0.55617 | -0.16794508 | 0.05 | 0.15 | 1 | 22 |
| si:ch73-109d9.3 | 0.55641 | 0.251302991 | 0.15 | 0.05 | 1 | 22 |
| epc1b | 0.55679 | -0.18373867 | 0.05 | 0.15 | 1 | 22 |
| supt4h1 | 0.55809 | 0.238046201 | 0.15 | 0.05 | 1 | 22 |
| ier2a | 0.55842 | 0.161666567 | 0.45 | 0.45 | 1 | 22 |
| sp4 | 0.55866 | -0.18901742 | 0.05 | 0.15 | 1 | 22 |
| tram1 | 0.55873 | 0.285341291 | 0.25 | 0.15 | 1 | 22 |
| rxfp2a | 0.55902 | 0.247015901 | 0.15 | 0.05 | 1 | 22 |
| xiap | 0.55945 | 0.23970256 | 0.15 | 0.05 | 1 | 22 |
| dpf2 | 0.55978 | -0.25620823 | 0.05 | 0.15 | 1 | 22 |
| hipk3b | 0.56003 | 0.247290037 | 0.15 | 0.05 | 1 | 22 |
| nr3c1 | 0.56004 | -0.2256759 | 0.05 | 0.15 | 1 | 22 |
| ppp1cb | 0.56024 | -0.17745997 | 0.05 | 0.15 | 1 | 22 |
| saraf | 0.56055 | -0.20219381 | 0.05 | 0.15 | 1 | 22 |
| uqcc2 | 0.56077 | -0.18655749 | 0.05 | 0.15 | 1 | 22 |
| ccdc12 | 0.56082 | -0.14476396 | 0.05 | 0.15 | 1 | 22 |
| jmjd1cb | 0.56083 | 0.257661378 | 0.15 | 0.05 | 1 | 22 |
| txndc15 | 0.56091 | -0.18059119 | 0.05 | 0.15 | 1 | 22 |
| sh3glb1a | 0.56118 | -0.19844377 | 0.05 | 0.15 | 1 | 22 |
| parp1 | 0.56152 | 0.18545334 | 0.15 | 0.15 | 1 | 22 |
| snx1a | 0.56204 | -0.16487731 | 0.05 | 0.15 | 1 | 22 |
| csrnp1b | 0.5624 | 0.272625531 | 0.35 | 0.25 | 1 | 22 |
| glud1a | 0.5624 | -0.21267433 | 0.05 | 0.15 | 1 | 22 |
| rbm4.2 | 0.56273 | 0.179657022 | 0.15 | 0.05 | 1 | 22 |
| LOC101883999 | 0.56361 | -0.22394648 | 0.05 | 0.15 | 1 | 22 |
| pik3r5 | 0.56366 | -0.23247107 | 0.05 | 0.15 | 1 | 22 |
| tmem106bb | 0.56367 | 0.252774123 | 0.15 | 0.05 | 1 | 22 |
| si:ch211-93g21.1 | 0.56426 | -0.19679033 | 0.05 | 0.15 | 1 | 22 |
| mid1ip1b | 0.56433 | -0.25287946 | 0.05 | 0.15 | 1 | 22 |
| zgc:110591 | 0.5649 | -0.19604442 | 0.05 | 0.15 | 1 | 22 |
| clk4a | 0.56505 | 0.216386941 | 0.15 | 0.05 | 1 | 22 |
| wwp2 | 0.56542 | 0.244155724 | 0.15 | 0.05 | 1 | 22 |
| trappc5 | 0.56554 | -0.18350622 | 0.05 | 0.15 | 1 | 22 |
| clptm1l | 0.56568 | -0.19971604 | 0.05 | 0.15 | 1 | 22 |
| c1qbp | 0.56576 | -0.18936743 | 0.05 | 0.15 | 1 | 22 |
| ptger4b | 0.56671 | -0.26406224 | 0.05 | 0.15 | 1 | 22 |
| adssl | 0.56691 | 0.226183379 | 0.15 | 0.05 | 1 | 22 |
| atf4a | 0.56743 | -0.27773037 | 0.2 | 0.35 | 1 | 22 |
| susd6 | 0.56768 | -0.22652486 | 0.05 | 0.15 | 1 | 22 |
| BX548028.1 | 0.56784 | -0.26585867 | 0.05 | 0.15 | 1 | 22 |
| ccdc25 | 0.56804 | -0.18171608 | 0.05 | 0.15 | 1 | 22 |
| psmd13 | 0.56815 | -0.21168616 | 0.05 | 0.15 | 1 | 22 |
| scgn | 0.56818 | -0.22773463 | 0.05 | 0.15 | 1 | 22 |
| ier2b | 0.56824 | 0.207626495 | 0.55 | 0.5 | 1 | 22 |
| itgb1bp1 | 0.56828 | 0.177476582 | 0.15 | 0.05 | 1 | 22 |
| meaf6 | 0.56832 | -0.17710821 | 0.05 | 0.15 | 1 | 22 |
| ppan | 0.56833 | 0.24975438 | 0.15 | 0.05 | 1 | 22 |

| | | | | | | |
|---|---|---|---|---|---|---|
| sumo1 | 0.56837 | -0.18006304 | 0.05 | 0.15 | 1 | 22 |
| snrpd3l | 0.56886 | -0.18975364 | 0.05 | 0.15 | 1 | 22 |
| eaf1 | 0.56901 | -0.16218887 | 0.05 | 0.15 | 1 | 22 |
| zgc:56304 | 0.56916 | -0.18534684 | 0.05 | 0.15 | 1 | 22 |
| banf1 | 0.56932 | 0.229112857 | 0.15 | 0.05 | 1 | 22 |
| txndc9 | 0.56934 | -0.22069441 | 0.05 | 0.15 | 1 | 22 |
| sowahd | 0.56945 | 0.201190646 | 0.15 | 0.05 | 1 | 22 |
| LOC103911430 | 0.56948 | 0.134019379 | 0.25 | 0.15 | 1 | 22 |
| vcp | 0.56956 | -0.24408435 | 0.05 | 0.15 | 1 | 22 |
| cass4 | 0.56983 | -0.21979653 | 0.05 | 0.15 | 1 | 22 |
| si:ch73-42p12.2 | 0.56989 | -0.15533554 | 0.05 | 0.15 | 1 | 22 |
| btbd10b | 0.56999 | 0.282697269 | 0.2 | 0.1 | 1 | 22 |
| mcm4 | 0.57004 | -0.21681708 | 0.05 | 0.15 | 1 | 22 |
| atp5f1d | 0.57008 | 0.28521623 | 0.4 | 0.25 | 1 | 22 |
| CU856520.1 | 0.57011 | -0.26863856 | 0.05 | 0.15 | 1 | 22 |
| smim4 | 0.57016 | 0.236837307 | 0.15 | 0.05 | 1 | 22 |
| LOC110440216 | 0.57052 | -0.25127936 | 0.05 | 0.15 | 1 | 22 |
| sort1b | 0.57052 | -0.24708111 | 0.05 | 0.15 | 1 | 22 |
| tpst1l | 0.57054 | -0.19420502 | 0.05 | 0.15 | 1 | 22 |
| abcf2a | 0.57057 | -0.23315033 | 0.05 | 0.15 | 1 | 22 |
| cdc26 | 0.5706 | 0.209016829 | 0.15 | 0.05 | 1 | 22 |
| jam3b | 0.5706 | 0.227552091 | 0.15 | 0.05 | 1 | 22 |
| ddx6 | 0.57061 | 0.199921238 | 0.15 | 0.05 | 1 | 22 |
| sptssa | 0.57063 | -0.23024407 | 0.05 | 0.15 | 1 | 22 |
| st3gal4 | 0.57064 | -0.17044877 | 0.05 | 0.15 | 1 | 22 |
| psmd6 | 0.57064 | 0.202687968 | 0.15 | 0.05 | 1 | 22 |
| ctdsp1 | 0.57066 | 0.212533997 | 0.15 | 0.05 | 1 | 22 |
| bsdc1 | 0.57066 | -0.19150636 | 0.05 | 0.15 | 1 | 22 |
| rnaset2l | 0.57137 | 0.300049503 | 0.45 | 0.3 | 1 | 22 |
| thrap3b | 0.5726 | 0.166703581 | 0.2 | 0.2 | 1 | 22 |
| rpl4 | 0.57308 | -0.11660685 | 0.85 | 0.85 | 1 | 22 |
| cs | 0.57451 | 0.249265475 | 0.2 | 0.1 | 1 | 22 |
| crip1 | 0.57533 | 0.26591102 | 0.65 | 0.5 | 1 | 22 |
| dapp1 | 0.57886 | 0.252986636 | 0.15 | 0.1 | 1 | 22 |
| CMKLR1 (1 of many) | 0.57948 | 0.230414533 | 0.2 | 0.15 | 1 | 22 |
| gapdhs | 0.57987 | 0.206828634 | 0.75 | 0.65 | 1 | 22 |
| gstp1 | 0.58155 | -0.23032816 | 0.25 | 0.4 | 1 | 22 |
| osgn1 | 0.58294 | 0.292527835 | 0.25 | 0.15 | 1 | 22 |
| glrx | 0.58329 | -0.19420813 | 0.45 | 0.5 | 1 | 22 |
| rps7 | 0.5834 | -0.16868678 | 0.8 | 0.9 | 1 | 22 |
| ak1 | 0.58423 | -0.303296 | 0.15 | 0.25 | 1 | 22 |
| eif1b | 0.58427 | 0.113653826 | 0.6 | 0.6 | 1 | 22 |
| arpc5a | 0.58493 | 0.101590941 | 0.55 | 0.55 | 1 | 22 |
| emc7 | 0.58498 | -0.11669868 | 0.1 | 0.2 | 1 | 22 |
| ddx39ab | 0.58759 | 0.229066891 | 0.15 | 0.1 | 1 | 22 |
| si:ch1073-325m22.2 | 0.58763 | -0.18336023 | 0.25 | 0.4 | 1 | 22 |
| anp32a | 0.58835 | -0.25734139 | 0.25 | 0.4 | 1 | 22 |
| nucks1a | 0.5897 | -0.10999255 | 0.1 | 0.2 | 1 | 22 |

| | | | | | | |
|---|---|---|---|---|---|---|
| si:ch211-215e19.4 | 0.59143 | -0.24052799 | 0.25 | 0.4 | 1 | 22 |
| higd2a | 0.59233 | -0.12821228 | 0.1 | 0.2 | 1 | 22 |
| mrtfaa | 0.59395 | -0.16521718 | 0.1 | 0.2 | 1 | 22 |
| hspa8 | 0.59425 | -0.16863682 | 0.85 | 0.9 | 1 | 22 |
| LOC101884267 | 0.59434 | -0.19683506 | 0.35 | 0.5 | 1 | 22 |
| cox6c | 0.59466 | -0.17789026 | 0.3 | 0.45 | 1 | 22 |
| srsf3b | 0.59606 | 0.143223025 | 0.25 | 0.25 | 1 | 22 |
| eif4a1a | 0.59736 | 0.26933569 | 0.3 | 0.2 | 1 | 22 |
| eif2s3 | 0.59772 | -0.23537242 | 0.25 | 0.4 | 1 | 22 |
| dctn2 | 0.60357 | -0.11698833 | 0.1 | 0.2 | 1 | 22 |
| mapre1a | 0.60378 | -0.16908214 | 0.55 | 0.7 | 1 | 22 |
| CR855389.1 | 0.60479 | -0.25035025 | 0.1 | 0.2 | 1 | 22 |
| LOC103909115 | 0.60551 | 0.252651918 | 0.6 | 0.45 | 1 | 22 |
| rfc5 | 0.60615 | 0.22605626 | 0.15 | 0.1 | 1 | 22 |
| romo1 | 0.60878 | 0.119166551 | 0.6 | 0.6 | 1 | 22 |
| hyal3 | 0.61143 | 0.174061928 | 0.2 | 0.1 | 1 | 22 |
| plk3 | 0.61154 | -0.26962402 | 0.15 | 0.25 | 1 | 22 |
| cnbpa | 0.61206 | 0.158716184 | 0.1 | 0.1 | 1 | 22 |
| rpl9 | 0.61331 | -0.11366284 | 0.85 | 0.85 | 1 | 22 |
| snrpd2 | 0.61413 | -0.12162724 | 0.1 | 0.2 | 1 | 22 |
| lasp1 | 0.61449 | -0.20676618 | 0.55 | 0.7 | 1 | 22 |
| dusp1 | 0.61533 | 0.276786193 | 0.5 | 0.35 | 1 | 22 |
| pstpip1b | 0.61543 | -0.25896209 | 0.1 | 0.2 | 1 | 22 |
| kmt2d | 0.61587 | 0.227687729 | 0.15 | 0.1 | 1 | 22 |
| slc43a3b | 0.61802 | -0.11632544 | 0.1 | 0.2 | 1 | 22 |
| si:ch73-380l3.4 | 0.61807 | 0.206706824 | 0.7 | 0.55 | 1 | 22 |
| btk | 0.6183 | -0.16090851 | 0.1 | 0.2 | 1 | 22 |
| stk17b | 0.61842 | -0.20050715 | 0.55 | 0.7 | 1 | 22 |
| calm2b | 0.61916 | 0.259976973 | 0.4 | 0.3 | 1 | 22 |
| inpp5f | 0.62074 | -0.16443317 | 0.1 | 0.2 | 1 | 22 |
| sri | 0.62263 | -0.11908535 | 0.1 | 0.2 | 1 | 22 |
| fam210b | 0.62362 | -0.24782844 | 0.2 | 0.3 | 1 | 22 |
| rnf44 | 0.62388 | 0.273821086 | 0.2 | 0.1 | 1 | 22 |
| atp5l | 0.62719 | 0.229743626 | 0.6 | 0.45 | 1 | 22 |
| ndfip2 | 0.62732 | -0.10493469 | 0.15 | 0.25 | 1 | 22 |
| TSTA3 (1 of many) | 0.62835 | -0.22738057 | 0.1 | 0.15 | 1 | 22 |
| hmgb1a | 0.62938 | -0.25487884 | 0.1 | 0.2 | 1 | 22 |
| hnrnpa0a | 0.62982 | 0.214017286 | 0.15 | 0.1 | 1 | 22 |
| hck | 0.63083 | -0.23087926 | 0.35 | 0.5 | 1 | 22 |
| GCA | 0.6311 | -0.20193716 | 0.4 | 0.55 | 1 | 22 |
| socs3a | 0.63237 | -0.23237761 | 0.4 | 0.55 | 1 | 22 |
| map2k2a | 0.63478 | 0.256592478 | 0.2 | 0.1 | 1 | 22 |
| arl5c | 0.63649 | 0.229052952 | 0.55 | 0.4 | 1 | 22 |
| yrk | 0.63687 | -0.22844562 | 0.4 | 0.55 | 1 | 22 |
| nudt21 | 0.63846 | -0.1076964 | 0.15 | 0.25 | 1 | 22 |
| eef2b | 0.6413 | -0.15809757 | 0.8 | 0.9 | 1 | 22 |
| hmgb2b | 0.64242 | 0.245965666 | 0.2 | 0.1 | 1 | 22 |
| si:dkey-185m8.2 | 0.64595 | -0.13741943 | 0.15 | 0.25 | 1 | 22 |

| | | | | | | |
|---|---|---|---|---|---|---|
| bcl2l10 | 0.64599 | -0.26019408 | 0.1 | 0.2 | 1 | 22 |
| jak1 | 0.64629 | 0.26597072 | 0.25 | 0.15 | 1 | 22 |
| gigyf1b | 0.64633 | 0.14597402 | 0.1 | 0.1 | 1 | 22 |
| atp5pb | 0.64711 | 0.159830691 | 0.4 | 0.4 | 1 | 22 |
| btg1 | 0.6475 | 0.175054902 | 0.85 | 0.75 | 1 | 22 |
| adam17b | 0.64872 | 0.229519143 | 0.15 | 0.1 | 1 | 22 |
| ndufb2 | 0.64882 | -0.2534693 | 0.15 | 0.25 | 1 | 22 |
| glipr2 | 0.65306 | -0.17224293 | 0.1 | 0.2 | 1 | 22 |
| csgalnact1a | 0.65366 | 0.135959624 | 0.15 | 0.15 | 1 | 22 |
| gabarapb | 0.65427 | 0.177522853 | 0.55 | 0.5 | 1 | 22 |
| commd3 | 0.65442 | 0.119309615 | 0.3 | 0.3 | 1 | 22 |
| ube2g1b | 0.65448 | -0.18045741 | 0.1 | 0.2 | 1 | 22 |
| cacybp | 0.65479 | -0.22748116 | 0.1 | 0.2 | 1 | 22 |
| srgap3 | 0.65507 | 0.230028698 | 0.2 | 0.15 | 1 | 22 |
| vsir | 0.65566 | 0.232020757 | 0.5 | 0.4 | 1 | 22 |
| ptk2ba | 0.65639 | -0.1925186 | 0.1 | 0.2 | 1 | 22 |
| ppib | 0.65692 | 0.11216312 | 0.35 | 0.35 | 1 | 22 |
| krtcap2 | 0.66113 | -0.22349537 | 0.1 | 0.2 | 1 | 22 |
| grb2b | 0.66161 | -0.26246135 | 0.15 | 0.25 | 1 | 22 |
| sat1a.2 | 0.66334 | -0.20285621 | 0.6 | 0.7 | 1 | 22 |
| psma2 | 0.66453 | 0.157425085 | 0.2 | 0.1 | 1 | 22 |
| pacsin2 | 0.66522 | -0.18115287 | 0.1 | 0.2 | 1 | 22 |
| CABZ01073834.1 | 0.66534 | 0.168347402 | 0.2 | 0.1 | 1 | 22 |
| parp12b | 0.667 | 0.225334278 | 0.2 | 0.1 | 1 | 22 |
| apbb1ip | 0.66832 | -0.21919746 | 0.1 | 0.2 | 1 | 22 |
| ube2l3a | 0.66859 | -0.21331199 | 0.1 | 0.2 | 1 | 22 |
| psma3 | 0.66862 | -0.27534063 | 0.1 | 0.2 | 1 | 22 |
| mrpl54 | 0.66887 | -0.15598677 | 0.1 | 0.2 | 1 | 22 |
| taf13 | 0.66939 | 0.173299552 | 0.2 | 0.1 | 1 | 22 |
| si:dkey-52l18.4 | 0.66945 | -0.13055521 | 0.15 | 0.25 | 1 | 22 |
| csf2rb | 0.66965 | 0.228844849 | 0.2 | 0.1 | 1 | 22 |
| srrm1 | 0.6698 | 0.249823368 | 0.25 | 0.15 | 1 | 22 |
| ubash3ba | 0.67011 | -0.22255472 | 0.1 | 0.2 | 1 | 22 |
| ssr2 | 0.6705 | 0.235624333 | 0.25 | 0.15 | 1 | 22 |
| prkab1b | 0.67059 | -0.18642965 | 0.1 | 0.2 | 1 | 22 |
| gnai1 | 0.67073 | -0.16865305 | 0.1 | 0.2 | 1 | 22 |
| ndufa2 | 0.67074 | 0.2300903 | 0.2 | 0.1 | 1 | 22 |
| pcf11 | 0.67094 | 0.229078484 | 0.2 | 0.1 | 1 | 22 |
| ss18 | 0.67119 | -0.15983014 | 0.1 | 0.2 | 1 | 22 |
| eif5 | 0.67129 | 0.17068975 | 0.2 | 0.1 | 1 | 22 |
| pstpip1a | 0.67136 | -0.12458031 | 0.15 | 0.25 | 1 | 22 |
| sdhc | 0.67136 | 0.196060754 | 0.2 | 0.1 | 1 | 22 |
| zmp:0000001082 | 0.67221 | -0.19564609 | 0.1 | 0.2 | 1 | 22 |
| dnaja2a | 0.67229 | 0.116923693 | 0.3 | 0.3 | 1 | 22 |
| stk17al | 0.67327 | -0.20421293 | 0.1 | 0.2 | 1 | 22 |
| dbn1 | 0.67343 | -0.18622426 | 0.1 | 0.2 | 1 | 22 |
| dpy30 | 0.67352 | -0.15950227 | 0.1 | 0.2 | 1 | 22 |
| cpsf6 | 0.6743 | 0.20214445 | 0.2 | 0.1 | 1 | 22 |

| | | | | | | |
|---|---|---|---|---|---|---|
| rac2 | 0.67462 | -0.13599152 | 0.7 | 0.75 | 1 | 22 |
| snu13b | 0.67471 | -0.17975078 | 0.1 | 0.2 | 1 | 22 |
| atrx | 0.67471 | 0.20260026 | 0.2 | 0.1 | 1 | 22 |
| coa6 | 0.67485 | 0.192263061 | 0.2 | 0.1 | 1 | 22 |
| midn | 0.67496 | -0.20505194 | 0.1 | 0.2 | 1 | 22 |
| malt3 | 0.67508 | 0.208819036 | 0.2 | 0.1 | 1 | 22 |
| prkar1ab | 0.67646 | 0.224653664 | 0.2 | 0.15 | 1 | 22 |
| thoc2 | 0.67657 | -0.13933145 | 0.15 | 0.25 | 1 | 22 |
| si:dkeyp-104b3.21 | 0.67669 | 0.22029343 | 0.15 | 0.1 | 1 | 22 |
| pdia3 | 0.67791 | -0.1200199 | 0.15 | 0.25 | 1 | 22 |
| pycard | 0.67995 | -0.18333145 | 0.3 | 0.35 | 1 | 22 |
| BX120005.1 | 0.68074 | 0.10455095 | 0.7 | 0.7 | 1 | 22 |
| plekho1b | 0.6811 | 0.264799413 | 0.25 | 0.15 | 1 | 22 |
| srsf5a | 0.68218 | 0.122030962 | 0.35 | 0.35 | 1 | 22 |
| rab44 | 0.68304 | 0.170713067 | 0.5 | 0.45 | 1 | 22 |
| sub1a | 0.68837 | 0.220834114 | 0.45 | 0.35 | 1 | 22 |
| ywhabb | 0.68863 | 0.227632393 | 0.4 | 0.3 | 1 | 22 |
| cox7a2l | 0.68916 | -0.1041945 | 0.25 | 0.35 | 1 | 22 |
| rplp1 | 0.69063 | -0.12803702 | 0.9 | 0.95 | 1 | 22 |
| XLOC-000431 | 0.69069 | 0.133982311 | 0.2 | 0.2 | 1 | 22 |
| atp6ap2 | 0.69069 | -0.1085959 | 0.15 | 0.25 | 1 | 22 |
| gna13b | 0.69118 | -0.24358521 | 0.15 | 0.25 | 1 | 22 |
| zgc:158343 | 0.69224 | 0.164014257 | 0.25 | 0.15 | 1 | 22 |
| anxa11b | 0.69472 | 0.127412972 | 0.15 | 0.15 | 1 | 22 |
| copz1 | 0.69604 | 0.210030232 | 0.25 | 0.2 | 1 | 22 |
| cblb | 0.69646 | -0.16919515 | 0.15 | 0.25 | 1 | 22 |
| prdx4 | 0.69868 | 0.198662 | 0.15 | 0.1 | 1 | 22 |
| si:ch73-234b20.5 | 0.69896 | 0.190664292 | 0.15 | 0.1 | 1 | 22 |
| kdm7ab | 0.70205 | 0.23240761 | 0.25 | 0.15 | 1 | 22 |
| ndufs5 | 0.70336 | -0.20176065 | 0.15 | 0.25 | 1 | 22 |
| pcca | 0.70354 | 0.127170513 | 0.1 | 0.1 | 1 | 22 |
| dctn3 | 0.70362 | 0.21023481 | 0.25 | 0.15 | 1 | 22 |
| med6 | 0.70494 | 0.122933359 | 0.15 | 0.15 | 1 | 22 |
| iscub | 0.70552 | 0.237090335 | 0.25 | 0.15 | 1 | 22 |
| uckl1b | 0.70756 | 0.216086677 | 0.15 | 0.1 | 1 | 22 |
| vipas39 | 0.71078 | 0.124009163 | 0.15 | 0.15 | 1 | 22 |
| ap2a1 | 0.71172 | 0.195012784 | 0.25 | 0.15 | 1 | 22 |
| naca | 0.71369 | 0.166024314 | 0.8 | 0.7 | 1 | 22 |
| si:ch211-203d1.3 | 0.714 | 0.114632399 | 0.15 | 0.15 | 1 | 22 |
| ndufa1 | 0.71442 | -0.10109353 | 0.2 | 0.3 | 1 | 22 |
| CABZ01046425.1 | 0.71553 | -0.18330976 | 0.15 | 0.25 | 1 | 22 |
| cct8 | 0.71556 | 0.118091064 | 0.3 | 0.2 | 1 | 22 |
| ldha | 0.71895 | -0.16408351 | 0.15 | 0.25 | 1 | 22 |
| zdhhc23a | 0.71929 | 0.201229291 | 0.25 | 0.2 | 1 | 22 |
| mir223 | 0.72086 | 0.110552603 | 0.5 | 0.5 | 1 | 22 |
| si:ch211-153b23.7 | 0.72105 | -0.23574686 | 0.15 | 0.25 | 1 | 22 |
| psma8 | 0.72115 | 0.191731832 | 0.2 | 0.15 | 1 | 22 |
| psmb4 | 0.72134 | 0.197121022 | 0.2 | 0.15 | 1 | 22 |

| | | | | | | |
|---|---|---|---|---|---|---|
| jun | 0.72141 | -0.17001837 | 0.15 | 0.25 | 1 | 22 |
| tefa | 0.72176 | -0.18359417 | 0.15 | 0.25 | 1 | 22 |
| gtpbp4 | 0.72181 | 0.203372489 | 0.2 | 0.15 | 1 | 22 |
| tcea1 | 0.72352 | 0.122896795 | 0.1 | 0.1 | 1 | 22 |
| fgl2a | 0.72491 | 0.190910479 | 0.3 | 0.25 | 1 | 22 |
| si:cabz01036006.1 | 0.72551 | 0.183579574 | 0.35 | 0.3 | 1 | 22 |
| tpm1 | 0.72612 | 0.166649117 | 0.25 | 0.15 | 1 | 22 |
| arhgef6 | 0.72613 | -0.1550223 | 0.15 | 0.25 | 1 | 22 |
| calm3b | 0.72684 | -0.20340816 | 0.4 | 0.5 | 1 | 22 |
| uqcrfs1 | 0.7273 | -0.16065192 | 0.15 | 0.25 | 1 | 22 |
| mmgt1 | 0.72819 | -0.16997955 | 0.15 | 0.25 | 1 | 22 |
| sec61g | 0.7295 | -0.14448204 | 0.75 | 0.8 | 1 | 22 |
| mrtfab | 0.72955 | -0.17015998 | 0.15 | 0.25 | 1 | 22 |
| ephx2 | 0.7301 | 0.161138189 | 0.5 | 0.45 | 1 | 22 |
| gnb1b | 0.73042 | -0.19330101 | 0.4 | 0.5 | 1 | 22 |
| hspa5 | 0.73154 | -0.23258659 | 0.15 | 0.25 | 1 | 22 |
| si:dkey-261h17.1 | 0.73158 | -0.14629433 | 0.15 | 0.25 | 1 | 22 |
| irf2bp2a | 0.73169 | -0.19822525 | 0.15 | 0.25 | 1 | 22 |
| mif | 0.73171 | -0.17518947 | 0.15 | 0.25 | 1 | 22 |
| elf1 | 0.73171 | 0.143548601 | 0.3 | 0.2 | 1 | 22 |
| irf2a | 0.73295 | 0.180395295 | 0.15 | 0.1 | 1 | 22 |
| tmem176l.1 | 0.7331 | 0.121549389 | 0.1 | 0.1 | 1 | 22 |
| gnb1a | 0.73462 | 0.223632874 | 0.35 | 0.25 | 1 | 22 |
| ap2m1a | 0.73503 | 0.198053738 | 0.15 | 0.1 | 1 | 22 |
| fubp1 | 0.73724 | 0.119915622 | 0.1 | 0.1 | 1 | 22 |
| slc46a2 | 0.7381 | 0.114045198 | 0.1 | 0.1 | 1 | 22 |
| gpx1a | 0.73835 | -0.20492313 | 0.25 | 0.35 | 1 | 22 |
| arfip1 | 0.73882 | 0.207404039 | 0.15 | 0.1 | 1 | 22 |
| csk | 0.74027 | -0.20708007 | 0.45 | 0.55 | 1 | 22 |
| pgam1a | 0.74346 | 0.10916908 | 0.2 | 0.2 | 1 | 22 |
| bud31 | 0.74541 | 0.179313903 | 0.15 | 0.1 | 1 | 22 |
| tpi1b | 0.74777 | 0.193299872 | 0.5 | 0.4 | 1 | 22 |
| rpl14 | 0.74793 | 0.111240265 | 0.95 | 0.9 | 1 | 22 |
| si:ch211-220f21.3 | 0.74826 | 0.114175371 | 0.5 | 0.4 | 1 | 22 |
| oxct1a | 0.74993 | 0.18803717 | 0.2 | 0.15 | 1 | 22 |
| sh3kbp1 | 0.75123 | -0.20679351 | 0.2 | 0.3 | 1 | 22 |
| pik3cg | 0.75219 | 0.117345455 | 0.1 | 0.1 | 1 | 22 |
| rps6 | 0.75225 | -0.11475102 | 0.7 | 0.8 | 1 | 22 |
| zc3h12a | 0.75292 | 0.114675329 | 0.1 | 0.1 | 1 | 22 |
| b2ml | 0.75304 | 0.211739077 | 0.4 | 0.3 | 1 | 22 |
| mbd2 | 0.75339 | 0.188050595 | 0.15 | 0.1 | 1 | 22 |
| elocb | 0.75371 | 0.174253278 | 0.2 | 0.15 | 1 | 22 |
| ptpn6 | 0.75391 | -0.11033706 | 0.7 | 0.8 | 1 | 22 |
| syap1 | 0.755 | 0.113572099 | 0.1 | 0.1 | 1 | 22 |
| mapkapk2a | 0.75549 | -0.14686287 | 0.2 | 0.3 | 1 | 22 |
| si:ch73-269m14.2 | 0.75566 | 0.173123176 | 0.3 | 0.2 | 1 | 22 |
| tnfrsf1a | 0.75569 | 0.184726245 | 0.2 | 0.15 | 1 | 22 |
| sypl2b | 0.75594 | 0.116990984 | 0.2 | 0.2 | 1 | 22 |

| | | | | | | |
|---|---|---|---|---|---|---|
| tipin | 0.75765 | 0.181356419 | 0.15 | 0.1 | 1 | 22 |
| XLOC-023492 | 0.75847 | 0.169641767 | 0.3 | 0.25 | 1 | 22 |
| jpt2 | 0.76079 | -0.1334443 | 0.25 | 0.35 | 1 | 22 |
| gse1 | 0.76283 | 0.114823402 | 0.1 | 0.1 | 1 | 22 |
| histh1l | 0.76439 | -0.18251067 | 0.1 | 0.15 | 1 | 22 |
| rbbp6 | 0.76494 | -0.18397583 | 0.2 | 0.3 | 1 | 22 |
| mt-nd5 | 0.76511 | 0.106089426 | 0.15 | 0.15 | 1 | 22 |
| gmps | 0.76515 | 0.113320016 | 0.1 | 0.1 | 1 | 22 |
| selenow1 | 0.76605 | 0.172907172 | 0.45 | 0.4 | 1 | 22 |
| fam49bb | 0.76611 | -0.1772295 | 0.2 | 0.3 | 1 | 22 |
| syf2 | 0.76619 | -0.1651378 | 0.2 | 0.3 | 1 | 22 |
| hmgb2a | 0.76619 | 0.125206792 | 0.8 | 0.7 | 1 | 22 |
| hnrpkl | 0.76808 | 0.109331308 | 0.1 | 0.1 | 1 | 22 |
| zgc:162944 | 0.77039 | 0.206339524 | 0.4 | 0.3 | 1 | 22 |
| si:ch1073-429i10.3.1 | 0.77235 | 0.198567029 | 0.45 | 0.35 | 1 | 22 |
| sema7a | 0.77343 | 0.105180687 | 0.15 | 0.15 | 1 | 22 |
| arf6a | 0.77416 | 0.136753466 | 0.4 | 0.3 | 1 | 22 |
| cox6b2 | 0.77535 | -0.10251116 | 0.65 | 0.75 | 1 | 22 |
| sem1 | 0.77596 | 0.182428904 | 0.45 | 0.35 | 1 | 22 |
| ywhaba | 0.77679 | -0.10808741 | 0.45 | 0.55 | 1 | 22 |
| atp6v1e1b | 0.77735 | 0.169422186 | 0.25 | 0.2 | 1 | 22 |
| si:rp71-45k5.4 | 0.77876 | 0.109763728 | 0.1 | 0.1 | 1 | 22 |
| twf1a | 0.78133 | 0.160076956 | 0.4 | 0.35 | 1 | 22 |
| u2af1 | 0.78161 | 0.175104052 | 0.2 | 0.15 | 1 | 22 |
| ing1 | 0.78242 | 0.107915128 | 0.1 | 0.1 | 1 | 22 |
| eif3d | 0.78303 | -0.14710736 | 0.25 | 0.35 | 1 | 22 |
| dynll1 | 0.78583 | -0.15330523 | 0.25 | 0.35 | 1 | 22 |
| copg2 | 0.78583 | 0.181106531 | 0.15 | 0.1 | 1 | 22 |
| ube2na | 0.78605 | -0.15971659 | 0.25 | 0.35 | 1 | 22 |
| ip6k2b | 0.78652 | 0.181431189 | 0.15 | 0.1 | 1 | 22 |
| irf7 | 0.78655 | 0.177636671 | 0.15 | 0.1 | 1 | 22 |
| pou2f2a | 0.78667 | 0.194731757 | 0.4 | 0.3 | 1 | 22 |
| larp4ab | 0.78683 | 0.171221331 | 0.2 | 0.15 | 1 | 22 |
| cox4i1 | 0.78821 | -0.11724136 | 0.65 | 0.75 | 1 | 22 |
| gng5 | 0.79151 | 0.179806823 | 0.55 | 0.45 | 1 | 22 |
| abi3a | 0.79181 | -0.12952284 | 0.35 | 0.45 | 1 | 22 |
| cox6a1 | 0.79183 | -0.11538064 | 0.5 | 0.6 | 1 | 22 |
| anxa11a | 0.79281 | 0.174043967 | 0.2 | 0.15 | 1 | 22 |
| ikzf1 | 0.79307 | 0.170192291 | 0.15 | 0.1 | 1 | 22 |
| slc3a2b | 0.79424 | 0.186253907 | 0.2 | 0.15 | 1 | 22 |
| ccng1 | 0.79501 | -0.1862394 | 0.35 | 0.45 | 1 | 22 |
| kpna3 | 0.79531 | -0.17974833 | 0.1 | 0.15 | 1 | 22 |
| pdcd10b | 0.79783 | 0.109301715 | 0.1 | 0.1 | 1 | 22 |
| sept7a | 0.79881 | 0.101430596 | 0.1 | 0.1 | 1 | 22 |
| efhd2 | 0.80084 | 0.170690295 | 0.4 | 0.3 | 1 | 22 |
| ifi46 | 0.80096 | 0.174987651 | 0.2 | 0.15 | 1 | 22 |
| bcas2 | 0.80115 | 0.163675265 | 0.15 | 0.1 | 1 | 22 |
| prelid3b | 0.80187 | 0.150395864 | 0.4 | 0.3 | 1 | 22 |

| | | | | | | |
|---|---|---|---|---|---|---|
| arrb2b | 0.80384 | -0.13641749 | 0.35 | 0.45 | 1 | 22 |
| mov10a | 0.80427 | 0.105942043 | 0.1 | 0.1 | 1 | 22 |
| nup62l | 0.80678 | -0.10103627 | 0.1 | 0.1 | 1 | 22 |
| cebpb | 0.81139 | -0.17840779 | 0.45 | 0.55 | 1 | 22 |
| ppt1 | 0.81144 | 0.172988108 | 0.45 | 0.35 | 1 | 22 |
| rap1b | 0.81149 | -0.14982243 | 0.35 | 0.45 | 1 | 22 |
| fabp3 | 0.81534 | 0.133821068 | 0.15 | 0.1 | 1 | 22 |
| psd3l | 0.81668 | 0.159693616 | 0.15 | 0.1 | 1 | 22 |
| anp32e | 0.82183 | 0.143475462 | 0.15 | 0.1 | 1 | 22 |
| dusp7 | 0.82258 | 0.147417392 | 0.15 | 0.1 | 1 | 22 |
| mrps21 | 0.8239 | 0.147078432 | 0.15 | 0.1 | 1 | 22 |
| ndufb5 | 0.82621 | 0.157713524 | 0.25 | 0.2 | 1 | 22 |
| il12rb2l | 0.82842 | 0.161235536 | 0.15 | 0.1 | 1 | 22 |
| si:ch211-199g17.2 | 0.83003 | -0.14690114 | 0.4 | 0.45 | 1 | 22 |
| pcna | 0.83127 | -0.17295439 | 0.1 | 0.15 | 1 | 22 |
| sucla2 | 0.83141 | 0.153954804 | 0.15 | 0.1 | 1 | 22 |
| stip1 | 0.83245 | -0.14921536 | 0.1 | 0.15 | 1 | 22 |
| eif4a2 | 0.83475 | 0.159089888 | 0.15 | 0.1 | 1 | 22 |
| cct5 | 0.83737 | 0.146162947 | 0.35 | 0.3 | 1 | 22 |
| ap1s1 | 0.83911 | 0.148926555 | 0.2 | 0.15 | 1 | 22 |
| hnrnpk | 0.84002 | 0.165465423 | 0.2 | 0.15 | 1 | 22 |
| vps29 | 0.84014 | 0.154300858 | 0.15 | 0.1 | 1 | 22 |
| cox15 | 0.84203 | 0.150954883 | 0.15 | 0.1 | 1 | 22 |
| zgc:66427 | 0.84489 | -0.1518091 | 0.1 | 0.15 | 1 | 22 |
| psenen | 0.8476 | 0.143472244 | 0.15 | 0.1 | 1 | 22 |
| ebag9 | 0.85003 | -0.1629747 | 0.15 | 0.2 | 1 | 22 |
| cybc1 | 0.853 | 0.152936465 | 0.15 | 0.1 | 1 | 22 |
| si:ch73-376l24.2 | 0.85498 | 0.145736667 | 0.15 | 0.1 | 1 | 22 |
| sept8a | 0.85651 | 0.156279506 | 0.15 | 0.1 | 1 | 22 |
| cope | 0.85679 | 0.142245687 | 0.15 | 0.1 | 1 | 22 |
| atp5f1b | 0.85748 | 0.140098143 | 0.4 | 0.35 | 1 | 22 |
| marcksl1a | 0.85863 | -0.1478909 | 0.15 | 0.2 | 1 | 22 |
| ndufab1a | 0.85983 | 0.131664749 | 0.15 | 0.1 | 1 | 22 |
| rhoaa | 0.86302 | -0.13111699 | 0.35 | 0.4 | 1 | 22 |
| atp6v1h | 0.86347 | 0.137010376 | 0.15 | 0.1 | 1 | 22 |
| alyref | 0.86898 | 0.137361091 | 0.2 | 0.15 | 1 | 22 |
| u2af2b | 0.86974 | 0.139213168 | 0.2 | 0.15 | 1 | 22 |
| zeb2a | 0.87347 | 0.152584925 | 0.15 | 0.1 | 1 | 22 |
| si:cabz01074946.1 | 0.8744 | -0.14883932 | 0.15 | 0.2 | 1 | 22 |
| si:dkey-33c12.4 | 0.87491 | -0.143767 | 0.2 | 0.25 | 1 | 22 |
| ndufs6 | 0.87632 | 0.133779982 | 0.15 | 0.1 | 1 | 22 |
| fbxl6 | 0.87704 | -0.14181358 | 0.1 | 0.15 | 1 | 22 |
| fam120c | 0.87722 | 0.137497613 | 0.15 | 0.1 | 1 | 22 |
| fthl27 | 0.87868 | 0.131803143 | 0.35 | 0.3 | 1 | 22 |
| fam57ba | 0.879 | 0.133902966 | 0.15 | 0.1 | 1 | 22 |
| arf1 | 0.87916 | -0.11758831 | 0.1 | 0.15 | 1 | 22 |
| acox1 | 0.87984 | -0.12458744 | 0.1 | 0.15 | 1 | 22 |
| mrpl36 | 0.87993 | 0.118188002 | 0.15 | 0.1 | 1 | 22 |

| | | | | | | |
|---|---|---|---|---|---|---|
| sltm | 0.88345 | 0.135294724 | 0.15 | 0.1 | 1 | 22 |
| LOC103910438 | 0.88453 | 0.136784628 | 0.2 | 0.15 | 1 | 22 |
| anp32b | 0.88482 | -0.14189662 | 0.2 | 0.25 | 1 | 22 |
| etf1b | 0.88601 | -0.12500994 | 0.1 | 0.15 | 1 | 22 |
| efna2a | 0.88611 | 0.124036676 | 0.15 | 0.1 | 1 | 22 |
| ccnl1b | 0.88666 | -0.11232324 | 0.1 | 0.15 | 1 | 22 |
| ndufs4 | 0.88792 | -0.13407682 | 0.15 | 0.2 | 1 | 22 |
| asap1b | 0.88797 | -0.12578404 | 0.1 | 0.15 | 1 | 22 |
| si:ch73-334d15.2 | 0.88955 | -0.10760936 | 0.1 | 0.15 | 1 | 22 |
| si:ch211-198a12.6 | 0.89006 | -0.12602124 | 0.15 | 0.2 | 1 | 22 |
| tktb | 0.89064 | -0.11160639 | 0.1 | 0.15 | 1 | 22 |
| txnipa | 0.89113 | 0.128604396 | 0.15 | 0.1 | 1 | 22 |
| selenos | 0.89149 | 0.113610343 | 0.15 | 0.1 | 1 | 22 |
| tmem56a | 0.89244 | -0.12230494 | 0.1 | 0.15 | 1 | 22 |
| arl6ip5b | 0.8925 | 0.100633665 | 0.15 | 0.1 | 1 | 22 |
| rgl2 | 0.89269 | -0.11058404 | 0.1 | 0.15 | 1 | 22 |
| cdca4 | 0.89288 | 0.124492843 | 0.15 | 0.1 | 1 | 22 |
| caprin1b | 0.89304 | -0.10677514 | 0.1 | 0.15 | 1 | 22 |
| ppp6r3 | 0.89304 | 0.115905066 | 0.15 | 0.1 | 1 | 22 |
| FP085398.1 | 0.89304 | -0.11484741 | 0.1 | 0.15 | 1 | 22 |
| tyk2 | 0.89306 | 0.114949935 | 0.15 | 0.1 | 1 | 22 |
| laptm4a | 0.89509 | -0.13308553 | 0.2 | 0.25 | 1 | 22 |
| zgc:152891 | 0.89567 | -0.13738435 | 0.2 | 0.25 | 1 | 22 |
| atp5mc3a | 0.8973 | 0.12954038 | 0.25 | 0.2 | 1 | 22 |
| ndufb6 | 0.90406 | 0.130116995 | 0.2 | 0.15 | 1 | 22 |
| slc25a55a | 0.90596 | 0.118201129 | 0.2 | 0.15 | 1 | 22 |
| pik3r3a | 0.90749 | -0.11490132 | 0.15 | 0.2 | 1 | 22 |
| fermt3b | 0.908 | 0.121911906 | 0.2 | 0.15 | 1 | 22 |
| rwdd1 | 0.90923 | 0.113488681 | 0.2 | 0.15 | 1 | 22 |
| jdp2b | 0.90965 | 0.12787155 | 0.35 | 0.3 | 1 | 22 |
| crlf3 | 0.91056 | 0.113694491 | 0.3 | 0.25 | 1 | 22 |
| ube2d2 | 0.91354 | -0.12404778 | 0.25 | 0.3 | 1 | 22 |
| si:ch211-243a20.4 | 0.91419 | 0.118830092 | 0.25 | 0.2 | 1 | 22 |
| tmem50a | 0.91529 | -0.1075476 | 0.15 | 0.2 | 1 | 22 |
| ist1 | 0.91586 | -0.10443212 | 0.15 | 0.2 | 1 | 22 |
| tomm5 | 0.91763 | -0.10621065 | 0.15 | 0.2 | 1 | 22 |
| znf706 | 0.91764 | -0.1018015 | 0.15 | 0.2 | 1 | 22 |
| si:ch73-264p11.1 | 0.91805 | -0.11157382 | 0.35 | 0.4 | 1 | 22 |
| rhoac | 0.91836 | 0.116649154 | 0.25 | 0.2 | 1 | 22 |
| pik3r1 | 0.91913 | -0.10910458 | 0.2 | 0.25 | 1 | 22 |
| nfkbiaa | 0.92326 | -0.12583054 | 0.25 | 0.3 | 1 | 22 |
| ccr2 | 0.92667 | 0.116505954 | 0.25 | 0.2 | 1 | 22 |
| cox7a2a | 0.92694 | 0.111057814 | 0.35 | 0.3 | 1 | 22 |
| reep5 | 0.92808 | -0.10737367 | 0.2 | 0.25 | 1 | 22 |
| ciarta | 0.93389 | -0.10750584 | 0.25 | 0.3 | 1 | 22 |
| knstrn | 1 | 0.109489806 | 0.05 | 0.05 | 1 | 22 |
| AK6 | 1 | 0.110622232 | 0.05 | 0.05 | 1 | 22 |
| itfg2 | 1 | 0.103705917 | 0.05 | 0.05 | 1 | 22 |

| | | | | | | |
|---|---|---|---|---|---|---|
| zgc:172106 | 1 | 0.110048862 | 0.05 | 0.05 | 1 | 22 |
| fam98a | 1 | 0.119519652 | 0.05 | 0.05 | 1 | 22 |
| asap2b | 1 | 0.136030857 | 0.05 | 0.05 | 1 | 22 |
| chmp7 | 1 | 0.106938044 | 0.05 | 0.05 | 1 | 22 |
| dph3 | 1 | -0.13240947 | 0.05 | 0.05 | 1 | 22 |
| edc4 | 1 | 0.15496426 | 0.05 | 0.05 | 1 | 22 |
| gmppaa | 1 | 0.116030694 | 0.05 | 0.05 | 1 | 22 |
| pccb | 1 | 0.117107452 | 0.05 | 0.05 | 1 | 22 |
| pdia6 | 1 | 0.108408639 | 0.05 | 0.05 | 1 | 22 |
| plpp1a | 1 | -0.12789003 | 0.05 | 0.05 | 1 | 22 |
| pspc1 | 1 | 0.104179495 | 0.05 | 0.05 | 1 | 22 |
| rcan1a | 1 | 0.141237118 | 0.05 | 0.05 | 1 | 22 |
| snapin | 1 | 0.10261502 | 0.05 | 0.05 | 1 | 22 |
| ssbp2 | 1 | 0.109700708 | 0.05 | 0.05 | 1 | 22 |
| ttc5 | 1 | 0.105981618 | 0.05 | 0.05 | 1 | 22 |
| usf1l | 1 | 0.150226033 | 0.05 | 0.05 | 1 | 22 |
| PAQR9 | 1 | -0.11603106 | 0.05 | 0.05 | 1 | 22 |
| XLOC-019150 | 1 | -0.11078149 | 0.05 | 0.05 | 1 | 22 |
| bcap31 | 1 | 0.14770063 | 0.05 | 0.05 | 1 | 22 |
| coa5 | 1 | 0.107015058 | 0.05 | 0.05 | 1 | 22 |
| mt-nd3 | 2.537017 | -1.23750499 | 0 | 0.625 | 0.5124013 | 23 |
| mt-atp6 | 2.943779 | -1.52830863 | 0.161 | 1 | 0.5945551 | 23 |
| psmb4 | 3.098350 | 0.294434758 | 0.71 | 0.25 | 0.6257738 | 23 |
| mt-co2 | 7.038129 | -1.45973859 | 0.194 | 1 | 1 | 23 |
| rpl28 | 9.678982 | 0.238556495 | 0.968 | 1 | 1 | 23 |
| mt-cyb | 0.00025 | -1.29469342 | 0.129 | 0.875 | 1 | 23 |
| LOC101884451 | 0.00026 | -0.91848343 | 0 | 0.5 | 1 | 23 |
| nrp2a | 0.00026 | -0.84551944 | 0 | 0.5 | 1 | 23 |
| strn3 | 0.00026 | -0.84783405 | 0 | 0.5 | 1 | 23 |
| cahz | 0.00026 | -0.85104618 | 0 | 0.5 | 1 | 23 |
| XLOC-015291 | 0.0003 | -0.90236371 | 0.129 | 0.875 | 1 | 23 |
| ppdpfb | 0.00047 | 0.552127569 | 0.968 | 0.875 | 1 | 23 |
| zgc:114188 | 0.00057 | 0.216057018 | 1 | 1 | 1 | 23 |
| tcima | 0.00058 | -1.1808746 | 0.032 | 0.5 | 1 | 23 |
| mt-nd4 | 0.0006 | -1.10869203 | 0.387 | 1 | 1 | 23 |
| rps7 | 0.00061 | 0.576712267 | 0.968 | 0.75 | 1 | 23 |
| cnbpa | 0.00066 | 0.877392135 | 0.613 | 0 | 1 | 23 |
| mt-co1 | 0.00074 | -1.04772528 | 0.613 | 1 | 1 | 23 |
| rpl19 | 0.00092 | 0.434432054 | 0.968 | 0.875 | 1 | 23 |
| nop53 | 0.00093 | 1.07938851 | 0.774 | 0.125 | 1 | 23 |
| mt-nd1 | 0.00105 | -0.88281423 | 0.323 | 1 | 1 | 23 |
| sesn1 | 0.0011 | 1.0712788 | 0.581 | 0 | 1 | 23 |
| si:dkey-112e7.2 | 0.00117 | 0.365638049 | 0.581 | 0.125 | 1 | 23 |
| bnc2 | 0.00118 | -0.86928044 | 0.355 | 0.875 | 1 | 23 |
| rps16 | 0.00124 | 0.190116261 | 0.968 | 1 | 1 | 23 |
| rpl26 | 0.0013 | 0.256231889 | 1 | 1 | 1 | 23 |
| rps4x | 0.00177 | 0.184249514 | 0.968 | 1 | 1 | 23 |
| atp1a1a.1 | 0.00185 | -0.8805452 | 0.29 | 0.875 | 1 | 23 |

| | | | | | | |
|---|---|---|---|---|---|---|
| atp10a | 0.00189 | -0.8583618 | 0.129 | 0.75 | 1 | 23 |
| wwp2 | 0.00226 | -0.72256759 | 0 | 0.375 | 1 | 23 |
| ccdc181 | 0.00226 | -0.72162461 | 0 | 0.375 | 1 | 23 |
| strip1 | 0.00226 | -0.6607895 | 0 | 0.375 | 1 | 23 |
| zmp:0000000912 | 0.00226 | -0.76055897 | 0 | 0.375 | 1 | 23 |
| skib | 0.00248 | -0.81468241 | 0.065 | 0.625 | 1 | 23 |
| faua | 0.0026 | 0.424710946 | 1 | 1 | 1 | 23 |
| rpl8 | 0.0027 | 0.458338703 | 0.968 | 0.75 | 1 | 23 |
| rpl7 | 0.0028 | 0.641043947 | 0.968 | 0.625 | 1 | 23 |
| rpl32 | 0.00281 | 0.510528316 | 0.968 | 0.75 | 1 | 23 |
| ndufa2 | 0.00285 | 0.602961081 | 0.516 | 0 | 1 | 23 |
| romo1 | 0.00288 | 0.646225824 | 0.871 | 0.25 | 1 | 23 |
| id2a | 0.00295 | -0.63038795 | 0.677 | 0.875 | 1 | 23 |
| rpl35a | 0.00354 | 0.24029233 | 1 | 1 | 1 | 23 |
| COX3 | 0.0037 | -0.94708483 | 0.581 | 1 | 1 | 23 |
| si:ch211-166a6.5 | 0.00371 | -0.78942527 | 0.065 | 0.625 | 1 | 23 |
| h3f3c | 0.00384 | 0.771350421 | 0.935 | 0.375 | 1 | 23 |
| dhrs3a | 0.00389 | -0.84712203 | 0.258 | 0.75 | 1 | 23 |
| dock1 | 0.00395 | -0.84578066 | 0.032 | 0.5 | 1 | 23 |
| akap1b | 0.00414 | -0.81924948 | 0.032 | 0.5 | 1 | 23 |
| nme2b.1 | 0.00451 | 0.168021776 | 0.968 | 1 | 1 | 23 |
| vps4b | 0.00459 | 0.183460378 | 0.387 | 0.125 | 1 | 23 |
| rpl30 | 0.00462 | 0.277040574 | 0.968 | 0.875 | 1 | 23 |
| rpl11 | 0.00481 | 0.187437619 | 1 | 1 | 1 | 23 |
| tpt1 | 0.00488 | 0.769700371 | 0.968 | 0.5 | 1 | 23 |
| rpl18a | 0.00502 | 0.300140042 | 0.968 | 0.875 | 1 | 23 |
| DDX17 | 0.00506 | -0.79429887 | 0.032 | 0.5 | 1 | 23 |
| rps23 | 0.00537 | 0.409987547 | 1 | 0.875 | 1 | 23 |
| snrpa | 0.00555 | -0.47010942 | 0.355 | 0.5 | 1 | 23 |
| fabp3 | 0.0061 | -0.70756297 | 0.129 | 0.625 | 1 | 23 |
| eif4bb | 0.00673 | 0.634486841 | 0.452 | 0 | 1 | 23 |
| serinc5 | 0.00673 | 0.717022813 | 0.452 | 0 | 1 | 23 |
| cope | 0.00673 | 0.643596476 | 0.452 | 0 | 1 | 23 |
| jpt2 | 0.00673 | 0.69197405 | 0.452 | 0 | 1 | 23 |
| rps20 | 0.00728 | 0.162613473 | 1 | 1 | 1 | 23 |
| rpl22 | 0.00733 | 0.68886656 | 0.968 | 0.5 | 1 | 23 |
| actn4 | 0.00775 | -0.7676185 | 0.097 | 0.625 | 1 | 23 |
| rps12 | 0.00784 | 0.465043011 | 0.968 | 0.75 | 1 | 23 |
| nat8l | 0.00785 | -0.86844843 | 0.032 | 0.5 | 1 | 23 |
| akt2 | 0.00789 | -0.69029766 | 0.032 | 0.5 | 1 | 23 |
| nectin1b | 0.00793 | -0.77308874 | 0.032 | 0.5 | 1 | 23 |
| ybx1 | 0.0081 | 0.307730081 | 0.968 | 0.875 | 1 | 23 |
| rpl3 | 0.00839 | 0.174261256 | 0.968 | 1 | 1 | 23 |
| rpl23a | 0.00855 | 0.376850192 | 0.968 | 0.875 | 1 | 23 |
| gtpbp4 | 0.00863 | 0.884459118 | 0.613 | 0.125 | 1 | 23 |
| cct7 | 0.00882 | 0.742542633 | 0.774 | 0.375 | 1 | 23 |
| rpl10 | 0.00883 | 0.461482356 | 0.968 | 0.75 | 1 | 23 |
| nop10 | 0.00886 | 0.106916022 | 0.516 | 0.25 | 1 | 23 |

| | | | | | | |
|---|---|---|---|---|---|---|
| fubp1 | 0.00889 | -0.78272904 | 0.097 | 0.625 | 1 | 23 |
| brd3a | 0.00947 | -0.57314475 | 0.258 | 0.5 | 1 | 23 |
| isg15 | 0.00958 | -0.59070422 | 0.129 | 0.625 | 1 | 23 |
| vat1 | 0.00982 | -0.31513947 | 0.387 | 0.375 | 1 | 23 |
| smad2 | 0.00998 | -0.67066798 | 0.097 | 0.625 | 1 | 23 |
| mt-nd5 | 0.01003 | -0.21333768 | 0.387 | 0.25 | 1 | 23 |
| eif3ea | 0.01005 | 0.777674927 | 0.839 | 0.375 | 1 | 23 |
| chmp5b | 0.01007 | 0.539383161 | 0.419 | 0 | 1 | 23 |
| ppid | 0.01007 | 0.640296629 | 0.419 | 0 | 1 | 23 |
| rab5c | 0.01007 | 0.630194137 | 0.419 | 0 | 1 | 23 |
| srprb | 0.01007 | 0.542486325 | 0.419 | 0 | 1 | 23 |
| rpl21 | 0.0103 | 0.233853145 | 1 | 1 | 1 | 23 |
| pfn2 | 0.0107 | -0.65979354 | 0.581 | 1 | 1 | 23 |
| ggnbp2 | 0.01077 | -0.71888501 | 0.065 | 0.5 | 1 | 23 |
| rplp1 | 0.01163 | 0.124152349 | 1 | 1 | 1 | 23 |
| mtbl | 0.0118 | -0.51342682 | 0.032 | 0.375 | 1 | 23 |
| rps8a | 0.01193 | 0.437266966 | 1 | 0.75 | 1 | 23 |
| stat1a | 0.01209 | -0.45288605 | 0.355 | 0.5 | 1 | 23 |
| magoh | 0.01222 | -0.40786199 | 0.258 | 0.375 | 1 | 23 |
| rpl10a | 0.01223 | 0.315156814 | 0.968 | 0.875 | 1 | 23 |
| ywhaz | 0.01229 | 0.747601114 | 0.806 | 0.25 | 1 | 23 |
| hnrnpl | 0.01271 | -0.76588807 | 0.194 | 0.75 | 1 | 23 |
| esr2a | 0.01275 | -0.76115141 | 0.194 | 0.75 | 1 | 23 |
| si:ch211-199g17.2 | 0.01303 | -0.59118555 | 0.323 | 0.625 | 1 | 23 |
| rplp0 | 0.01338 | 0.233381885 | 0.968 | 1 | 1 | 23 |
| CABZ01072614.1 | 0.01342 | -0.73957145 | 0.194 | 0.75 | 1 | 23 |
| rpl34 | 0.01418 | 0.308482266 | 1 | 0.875 | 1 | 23 |
| rps3 | 0.01427 | 0.425016265 | 0.968 | 0.75 | 1 | 23 |
| slc12a7b | 0.01475 | -0.77786232 | 0.065 | 0.5 | 1 | 23 |
| si:dkeyp-44a8.2 | 0.01479 | -0.74278848 | 0.129 | 0.5 | 1 | 23 |
| CU459056.1 | 0.01483 | 0.633775447 | 0.387 | 0 | 1 | 23 |
| adh8b | 0.01483 | 0.499467951 | 0.387 | 0 | 1 | 23 |
| ccdc25 | 0.01483 | 0.597362815 | 0.387 | 0 | 1 | 23 |
| ctdsp2 | 0.01483 | 0.651830069 | 0.387 | 0 | 1 | 23 |
| gsto2 | 0.01483 | 0.507115047 | 0.387 | 0 | 1 | 23 |
| hspa9 | 0.01483 | 0.609063131 | 0.387 | 0 | 1 | 23 |
| rbmx | 0.01483 | 0.645680876 | 0.387 | 0 | 1 | 23 |
| zgc:153372 | 0.01483 | 0.635804807 | 0.387 | 0 | 1 | 23 |
| ndufs5 | 0.01489 | -0.60065082 | 0.323 | 0.625 | 1 | 23 |
| mt-nd2 | 0.01521 | -0.9771631 | 0.323 | 0.75 | 1 | 23 |
| ppp2r1ba | 0.0163 | -0.31498938 | 0.258 | 0.25 | 1 | 23 |
| smek1 | 0.01653 | -0.13256031 | 0.194 | 0.125 | 1 | 23 |
| rps15a | 0.01694 | 0.177622169 | 1 | 1 | 1 | 23 |
| fuom | 0.01704 | -0.31299702 | 0 | 0.25 | 1 | 23 |
| igfbp1a | 0.01704 | -0.42993194 | 0 | 0.25 | 1 | 23 |
| plekhb2 | 0.01704 | -0.3207835 | 0 | 0.25 | 1 | 23 |
| ppap2d | 0.01704 | -0.35476159 | 0 | 0.25 | 1 | 23 |
| spoplb | 0.01704 | -0.33850187 | 0 | 0.25 | 1 | 23 |

| | | | | | | |
|---|---|---|---|---|---|---|
| csf1a | 0.01704 | -0.49728888 | 0 | 0.25 | 1 | 23 |
| fbln2 | 0.01704 | -0.42228671 | 0 | 0.25 | 1 | 23 |
| foxo1a | 0.01704 | -0.42340342 | 0 | 0.25 | 1 | 23 |
| pip5k1ab | 0.01704 | -0.42884256 | 0 | 0.25 | 1 | 23 |
| CU660013.4 | 0.01704 | -0.49095462 | 0 | 0.25 | 1 | 23 |
| bach1a | 0.01704 | -0.40126392 | 0 | 0.25 | 1 | 23 |
| cdr2a | 0.01704 | -0.49396399 | 0 | 0.25 | 1 | 23 |
| cog7 | 0.01704 | -0.47033183 | 0 | 0.25 | 1 | 23 |
| ipo11 | 0.01704 | -0.4060717 | 0 | 0.25 | 1 | 23 |
| nelfb | 0.01704 | -0.4057011 | 0 | 0.25 | 1 | 23 |
| phc2b | 0.01704 | -0.42034594 | 0 | 0.25 | 1 | 23 |
| sesn3 | 0.01704 | -0.40432993 | 0 | 0.25 | 1 | 23 |
| sh3gl1b | 0.01704 | -0.4067374 | 0 | 0.25 | 1 | 23 |
| tmed4 | 0.01704 | -0.41683479 | 0 | 0.25 | 1 | 23 |
| tprg1l | 0.01704 | -0.42074214 | 0 | 0.25 | 1 | 23 |
| traf3ip2l | 0.01704 | -0.54118934 | 0 | 0.25 | 1 | 23 |
| tspo | 0.01704 | -0.3415451 | 0 | 0.25 | 1 | 23 |
| urod | 0.01704 | -0.37229113 | 0 | 0.25 | 1 | 23 |
| CABZ01086041.1 | 0.01704 | -0.41641676 | 0 | 0.25 | 1 | 23 |
| CT027638.1 | 0.01704 | -0.26316738 | 0 | 0.25 | 1 | 23 |
| LOC108179575 | 0.01704 | -0.4980611 | 0 | 0.25 | 1 | 23 |
| ankrd28b | 0.01704 | -0.38108923 | 0 | 0.25 | 1 | 23 |
| birc6 | 0.01704 | -0.39576573 | 0 | 0.25 | 1 | 23 |
| bmp2k | 0.01704 | -0.49792394 | 0 | 0.25 | 1 | 23 |
| cbarpb | 0.01704 | -0.53585466 | 0 | 0.25 | 1 | 23 |
| cdkn1ba | 0.01704 | -0.40247779 | 0 | 0.25 | 1 | 23 |
| cfap298 | 0.01704 | -0.381996 | 0 | 0.25 | 1 | 23 |
| dcun1d4 | 0.01704 | -0.3843031 | 0 | 0.25 | 1 | 23 |
| dusp22b | 0.01704 | -0.45507509 | 0 | 0.25 | 1 | 23 |
| eef2a.1 | 0.01704 | -0.41494456 | 0 | 0.25 | 1 | 23 |
| epc1b | 0.01704 | -0.41882377 | 0 | 0.25 | 1 | 23 |
| gabpb2a | 0.01704 | -0.46138693 | 0 | 0.25 | 1 | 23 |
| lpgat1 | 0.01704 | -0.44993593 | 0 | 0.25 | 1 | 23 |
| mavs | 0.01704 | -0.46873676 | 0 | 0.25 | 1 | 23 |
| me3 | 0.01704 | -0.32657989 | 0 | 0.25 | 1 | 23 |
| nab2 | 0.01704 | -0.4822061 | 0 | 0.25 | 1 | 23 |
| pan3 | 0.01704 | -0.40317323 | 0 | 0.25 | 1 | 23 |
| pglyrp2 | 0.01704 | -0.42294898 | 0 | 0.25 | 1 | 23 |
| plaa | 0.01704 | -0.37361324 | 0 | 0.25 | 1 | 23 |
| retreg3 | 0.01704 | -0.41514078 | 0 | 0.25 | 1 | 23 |
| sh3yl1 | 0.01704 | -0.38793737 | 0 | 0.25 | 1 | 23 |
| slc6a6b | 0.01704 | -0.44303168 | 0 | 0.25 | 1 | 23 |
| sptbn1 | 0.01704 | -0.46569467 | 0 | 0.25 | 1 | 23 |
| tfb2m | 0.01704 | -0.39693143 | 0 | 0.25 | 1 | 23 |
| usp47 | 0.01704 | -0.45491591 | 0 | 0.25 | 1 | 23 |
| zbtb11 | 0.01704 | -0.44994708 | 0 | 0.25 | 1 | 23 |
| BX548015.2 | 0.01704 | -0.57110131 | 0 | 0.25 | 1 | 23 |
| LO018197.1 | 0.01704 | -0.55171087 | 0 | 0.25 | 1 | 23 |

| | | | | | | |
|---|---|---|---|---|---|---|
| atg101 | 0.01704 | -0.50401334 | 0 | 0.25 | 1 | 23 |
| ccdc114 | 0.01704 | -0.48906022 | 0 | 0.25 | 1 | 23 |
| cerk | 0.01704 | -0.46462855 | 0 | 0.25 | 1 | 23 |
| clip1a | 0.01704 | -0.35426263 | 0 | 0.25 | 1 | 23 |
| fam49ba | 0.01704 | -0.27617757 | 0 | 0.25 | 1 | 23 |
| fntb | 0.01704 | -0.48128739 | 0 | 0.25 | 1 | 23 |
| hint3 | 0.01704 | -0.57329422 | 0 | 0.25 | 1 | 23 |
| mibp | 0.01704 | -0.41413868 | 0 | 0.25 | 1 | 23 |
| mstnb | 0.01704 | -0.44390742 | 0 | 0.25 | 1 | 23 |
| ndufaf4 | 0.01704 | -0.53370805 | 0 | 0.25 | 1 | 23 |
| nucb1 | 0.01704 | -0.35235185 | 0 | 0.25 | 1 | 23 |
| pbxip1b | 0.01704 | -0.53337048 | 0 | 0.25 | 1 | 23 |
| rnpc3 | 0.01704 | -0.47852704 | 0 | 0.25 | 1 | 23 |
| rnps1 | 0.01704 | -0.3474825 | 0 | 0.25 | 1 | 23 |
| scaf11 | 0.01704 | -0.57087358 | 0 | 0.25 | 1 | 23 |
| slco5a1b | 0.01704 | -0.47573368 | 0 | 0.25 | 1 | 23 |
| timm29 | 0.01704 | -0.38954252 | 0 | 0.25 | 1 | 23 |
| zgc:165555.3 | 0.01704 | -0.44824143 | 0 | 0.25 | 1 | 23 |
| peli1b | 0.01704 | -0.45485765 | 0 | 0.25 | 1 | 23 |
| BX569793.1 | 0.01704 | -0.34291525 | 0 | 0.25 | 1 | 23 |
| phactr2 | 0.01704 | -0.58022418 | 0 | 0.25 | 1 | 23 |
| pnpla7a | 0.01704 | -0.43249551 | 0 | 0.25 | 1 | 23 |
| si:dkey-40c23.2 | 0.01704 | -0.43018695 | 0 | 0.25 | 1 | 23 |
| fam162a | 0.01741 | -0.76642258 | 0.065 | 0.5 | 1 | 23 |
| coa3a | 0.01764 | -0.67954692 | 0.161 | 0.5 | 1 | 23 |
| eif3m | 0.01768 | 0.614991489 | 0.935 | 0.5 | 1 | 23 |
| rnf10 | 0.01807 | -0.62880884 | 0.452 | 0.75 | 1 | 23 |
| stk35l | 0.01837 | -0.65883542 | 0.065 | 0.5 | 1 | 23 |
| tia1 | 0.01943 | -0.68146386 | 0.323 | 0.75 | 1 | 23 |
| sphk2 | 0.01952 | -0.58366883 | 0.032 | 0.375 | 1 | 23 |
| ackr3b | 0.01965 | -0.25089766 | 0.613 | 0.625 | 1 | 23 |
| rpl12 | 0.02079 | 0.197245545 | 1 | 1 | 1 | 23 |
| cnpy4 | 0.02155 | 0.528779633 | 0.355 | 0 | 1 | 23 |
| nos1apa | 0.02155 | 0.587778743 | 0.355 | 0 | 1 | 23 |
| rrp1 | 0.02155 | 0.535832584 | 0.355 | 0 | 1 | 23 |
| slc25a11 | 0.02155 | 0.498488761 | 0.355 | 0 | 1 | 23 |
| atox1 | 0.02155 | 0.525701333 | 0.355 | 0 | 1 | 23 |
| fzd2 | 0.02155 | 0.540915348 | 0.355 | 0 | 1 | 23 |
| gtf2f2a | 0.02155 | 0.578171322 | 0.355 | 0 | 1 | 23 |
| mrpl32 | 0.02155 | 0.486034812 | 0.355 | 0 | 1 | 23 |
| ndufv3 | 0.02155 | 0.477521031 | 0.355 | 0 | 1 | 23 |
| pdk2b | 0.02155 | 0.463263585 | 0.355 | 0 | 1 | 23 |
| eif4e1c | 0.02192 | -0.61269658 | 0.129 | 0.625 | 1 | 23 |
| serbp1a | 0.02247 | 0.22037782 | 0.935 | 1 | 1 | 23 |
| rps5 | 0.02257 | 0.353338899 | 1 | 1 | 1 | 23 |
| tyms | 0.02398 | -0.30251991 | 0.129 | 0.125 | 1 | 23 |
| cotl1 | 0.02415 | -0.54034735 | 0.226 | 0.75 | 1 | 23 |
| pik3r3b | 0.02417 | -0.7331787 | 0.226 | 0.75 | 1 | 23 |

| | | | | | | |
|---|---|---|---|---|---|---|
| rps24 | 0.02421 | 0.108767096 | 0.968 | 1 | 1 | 23 |
| rps19 | 0.02426 | 0.120993494 | 0.968 | 1 | 1 | 23 |
| xbp1 | 0.02463 | 0.203117848 | 0.581 | 0.75 | 1 | 23 |
| BX957297.2 | 0.02487 | -0.71848823 | 0.097 | 0.5 | 1 | 23 |
| pqbp1 | 0.02489 | -0.63851631 | 0.097 | 0.375 | 1 | 23 |
| hdlbpa | 0.02534 | 0.15725004 | 0.419 | 0.125 | 1 | 23 |
| psmd12 | 0.0255 | -0.33077671 | 0.194 | 0.25 | 1 | 23 |
| rpl23 | 0.02723 | 0.296706635 | 1 | 0.875 | 1 | 23 |
| rps27.1 | 0.0274 | 0.329276886 | 0.968 | 0.875 | 1 | 23 |
| tubb2b | 0.0275 | -0.4419964 | 0.613 | 0.625 | 1 | 23 |
| podxl | 0.02837 | -0.61511267 | 0.581 | 0.875 | 1 | 23 |
| wsb1 | 0.02905 | -0.72764257 | 0.097 | 0.5 | 1 | 23 |
| vps25 | 0.02909 | -0.67053749 | 0.097 | 0.5 | 1 | 23 |
| rps6 | 0.02917 | 0.314757861 | 1 | 0.875 | 1 | 23 |
| fkbp1aa | 0.02935 | 0.168304454 | 1 | 0.75 | 1 | 23 |
| btg1 | 0.02957 | -0.49935964 | 0.613 | 1 | 1 | 23 |
| rps17 | 0.02967 | 0.169882535 | 1 | 1 | 1 | 23 |
| mrpl53 | 0.03016 | -0.19228122 | 0.161 | 0.125 | 1 | 23 |
| spry4 | 0.03033 | -0.68379535 | 0.032 | 0.375 | 1 | 23 |
| dbi | 0.03041 | -0.71317184 | 0.419 | 0.75 | 1 | 23 |
| csnk1db | 0.03094 | 0.440578421 | 0.323 | 0 | 1 | 23 |
| hsf2 | 0.03094 | 0.493674082 | 0.323 | 0 | 1 | 23 |
| isg20 | 0.03094 | 0.45339913 | 0.323 | 0 | 1 | 23 |
| meis2b | 0.03094 | 0.506254301 | 0.323 | 0 | 1 | 23 |
| ndufb6 | 0.03094 | 0.395906688 | 0.323 | 0 | 1 | 23 |
| slc39a7 | 0.03094 | 0.461938623 | 0.323 | 0 | 1 | 23 |
| sypl2a | 0.03094 | 0.55047325 | 0.323 | 0 | 1 | 23 |
| tmem14ca | 0.03094 | 0.405904435 | 0.323 | 0 | 1 | 23 |
| wdr61 | 0.03094 | 0.495775472 | 0.323 | 0 | 1 | 23 |
| gstm.2 | 0.03094 | 0.523625512 | 0.323 | 0 | 1 | 23 |
| map1lc3a | 0.03094 | 0.490879361 | 0.323 | 0 | 1 | 23 |
| pbx1b | 0.03094 | 0.504983684 | 0.323 | 0 | 1 | 23 |
| PYURF | 0.03094 | 0.446303103 | 0.323 | 0 | 1 | 23 |
| emc2 | 0.03094 | 0.489437943 | 0.323 | 0 | 1 | 23 |
| ccdc12 | 0.03094 | 0.527758149 | 0.323 | 0 | 1 | 23 |
| med31 | 0.03094 | 0.454403279 | 0.323 | 0 | 1 | 23 |
| crebbpb | 0.03155 | -0.60366214 | 0.032 | 0.375 | 1 | 23 |
| lmnb2 | 0.03211 | -0.72336582 | 0.129 | 0.5 | 1 | 23 |
| nhp2 | 0.03297 | 0.733925647 | 0.613 | 0.125 | 1 | 23 |
| atp5mc3b | 0.03324 | 0.691920001 | 0.742 | 0.5 | 1 | 23 |
| spred2b | 0.0338 | -0.5022572 | 0.452 | 0.625 | 1 | 23 |
| srrt | 0.03394 | -0.58324346 | 0.032 | 0.375 | 1 | 23 |
| fam13b | 0.03413 | -0.70066182 | 0.065 | 0.375 | 1 | 23 |
| flna | 0.03433 | -0.53309507 | 0.387 | 0.625 | 1 | 23 |
| rpl5a | 0.03444 | 0.496127633 | 0.968 | 0.625 | 1 | 23 |
| eif3f | 0.03455 | 0.558206894 | 0.935 | 0.625 | 1 | 23 |
| map3k1 | 0.03509 | -0.59725273 | 0.032 | 0.375 | 1 | 23 |
| ndufs8a | 0.03527 | -0.39874289 | 0.258 | 0.375 | 1 | 23 |

| | | | | | | |
|---|---|---|---|---|---|---|
| os9 | 0.0354 | -0.60719636 | 0.065 | 0.375 | 1 | 23 |
| wscd2 | 0.03559 | -0.71098227 | 0.097 | 0.5 | 1 | 23 |
| scrib | 0.03559 | -0.57761956 | 0.032 | 0.375 | 1 | 23 |
| rbbp4 | 0.03587 | 0.301812583 | 0.452 | 0.125 | 1 | 23 |
| sec61g | 0.03646 | 0.584200594 | 0.935 | 0.625 | 1 | 23 |
| pkp3b | 0.03649 | -0.5133693 | 0.032 | 0.375 | 1 | 23 |
| ap2a1 | 0.03705 | -0.54100045 | 0.129 | 0.375 | 1 | 23 |
| cox6c | 0.03707 | -0.22589087 | 0.645 | 0.5 | 1 | 23 |
| alcamb | 0.03728 | -0.15817291 | 0.161 | 0.125 | 1 | 23 |
| elmsan1b | 0.03733 | -0.61398261 | 0.032 | 0.375 | 1 | 23 |
| spty2d1 | 0.03749 | -0.6058482 | 0.032 | 0.375 | 1 | 23 |
| dusp1 | 0.0376 | -0.60253691 | 0.065 | 0.375 | 1 | 23 |
| clasrp | 0.03776 | -0.52246263 | 0.032 | 0.375 | 1 | 23 |
| tomm5 | 0.03803 | 0.488495653 | 0.71 | 0.25 | 1 | 23 |
| pgr | 0.03862 | -0.61563308 | 0.032 | 0.375 | 1 | 23 |
| bmpr2a | 0.03886 | -0.63859887 | 0.032 | 0.375 | 1 | 23 |
| usp43b | 0.03984 | -0.59365173 | 0.258 | 0.75 | 1 | 23 |
| tdrd1 | 0.04036 | -0.51069443 | 0.032 | 0.375 | 1 | 23 |
| emx2 | 0.04075 | -0.36593652 | 0.484 | 0.625 | 1 | 23 |
| pin1 | 0.04087 | 0.467028889 | 0.516 | 0.125 | 1 | 23 |
| gpx1a | 0.0414 | -0.27889947 | 0.032 | 0.375 | 1 | 23 |
| rpl9 | 0.04194 | 0.148065226 | 1 | 1 | 1 | 23 |
| ppp1r12a | 0.04206 | -0.171773 | 0.161 | 0.125 | 1 | 23 |
| sfpq | 0.04213 | -0.55542694 | 0.323 | 0.5 | 1 | 23 |
| si:dkey-23k10.5 | 0.04217 | -0.68886004 | 0.032 | 0.375 | 1 | 23 |
| rbm8a | 0.04226 | 0.494227208 | 0.742 | 0.5 | 1 | 23 |
| si:rp71-36a1.5 | 0.04226 | -0.70829226 | 0.065 | 0.375 | 1 | 23 |
| rab6ba | 0.04242 | -0.53836722 | 0.032 | 0.375 | 1 | 23 |
| smu1a | 0.0426 | -0.49367002 | 0.032 | 0.375 | 1 | 23 |
| cpeb4b | 0.04262 | -0.56238957 | 0.032 | 0.375 | 1 | 23 |
| col1a1a | 0.04341 | 0.876025386 | 0.548 | 0.125 | 1 | 23 |
| ctnnb1 | 0.04347 | -0.44289082 | 0.484 | 0.625 | 1 | 23 |
| uqcrh | 0.04383 | -0.63009847 | 0.548 | 0.75 | 1 | 23 |
| cebpg | 0.04392 | -0.36461474 | 0.161 | 0.25 | 1 | 23 |
| commd5 | 0.04395 | 0.384499221 | 0.29 | 0 | 1 | 23 |
| eif1ad | 0.04395 | 0.469671934 | 0.29 | 0 | 1 | 23 |
| h1fx | 0.04395 | 0.443579127 | 0.29 | 0 | 1 | 23 |
| ipo7 | 0.04395 | 0.440101419 | 0.29 | 0 | 1 | 23 |
| lnx2b | 0.04395 | 0.434653853 | 0.29 | 0 | 1 | 23 |
| map7d1b | 0.04395 | 0.432920095 | 0.29 | 0 | 1 | 23 |
| prkd2 | 0.04395 | 0.539200969 | 0.29 | 0 | 1 | 23 |
| rab1ba | 0.04395 | 0.333377298 | 0.29 | 0 | 1 | 23 |
| rabac1 | 0.04395 | 0.444533614 | 0.29 | 0 | 1 | 23 |
| rap2c | 0.04395 | 0.464991317 | 0.29 | 0 | 1 | 23 |
| si:ch211-222k6.3 | 0.04395 | 0.414088853 | 0.29 | 0 | 1 | 23 |
| tmpob | 0.04395 | 0.498055739 | 0.29 | 0 | 1 | 23 |
| rraga | 0.04395 | 0.416267412 | 0.29 | 0 | 1 | 23 |
| dars | 0.04395 | 0.44043984 | 0.29 | 0 | 1 | 23 |

| | | | | | | |
|---|---|---|---|---|---|---|
| hmgb3b | 0.04395 | 0.518285973 | 0.29 | 0 | 1 | 23 |
| iscub | 0.04395 | 0.482807541 | 0.29 | 0 | 1 | 23 |
| mrps6 | 0.04395 | 0.476920441 | 0.29 | 0 | 1 | 23 |
| psap | 0.04395 | 0.412791542 | 0.29 | 0 | 1 | 23 |
| ptges3a | 0.04395 | 0.408132915 | 0.29 | 0 | 1 | 23 |
| zgc:123258 | 0.04395 | 0.493038619 | 0.29 | 0 | 1 | 23 |
| rpl13 | 0.04399 | 0.257406449 | 1 | 0.875 | 1 | 23 |
| rps2 | 0.04442 | 0.252409551 | 1 | 1 | 1 | 23 |
| slc11a2 | 0.04465 | -0.41184209 | 0.032 | 0.375 | 1 | 23 |
| snx27b | 0.0449 | -0.52318687 | 0.032 | 0.375 | 1 | 23 |
| rnf167 | 0.04549 | -0.45424747 | 0.032 | 0.375 | 1 | 23 |
| fndc3ba | 0.04549 | -0.63333446 | 0.032 | 0.375 | 1 | 23 |
| gstt1a | 0.04576 | -0.3509038 | 0.032 | 0.375 | 1 | 23 |
| fam208ab | 0.04576 | -0.46045186 | 0.032 | 0.375 | 1 | 23 |
| ifi45 | 0.04596 | -0.23482675 | 0.839 | 0.75 | 1 | 23 |
| si:dkey-85k7.12 | 0.04628 | -0.58805839 | 0.097 | 0.5 | 1 | 23 |
| fzd7a | 0.04637 | -0.49708197 | 0.161 | 0.375 | 1 | 23 |
| cd276 | 0.04642 | -0.63274977 | 0.161 | 0.5 | 1 | 23 |
| ywhabl | 0.04652 | -0.48330648 | 0.613 | 0.875 | 1 | 23 |
| cap1 | 0.04715 | -0.50466164 | 0.194 | 0.375 | 1 | 23 |
| zgc:158463 | 0.0472 | -0.65466704 | 0.161 | 0.375 | 1 | 23 |
| dsg2.1 | 0.04749 | -0.53188978 | 0.097 | 0.25 | 1 | 23 |
| ash1l | 0.04758 | -0.56584291 | 0.194 | 0.5 | 1 | 23 |
| pabpc1a | 0.04766 | 0.350801451 | 1 | 0.75 | 1 | 23 |
| zgc:162698 | 0.04799 | -0.60103951 | 0.161 | 0.5 | 1 | 23 |
| gata6 | 0.04838 | -0.2448052 | 0.226 | 0.25 | 1 | 23 |
| tmem107 | 0.05054 | -0.30233647 | 0.194 | 0.25 | 1 | 23 |
| mrpl52 | 0.05083 | -0.51507725 | 0.258 | 0.625 | 1 | 23 |
| ppih | 0.05136 | -0.47134069 | 0.097 | 0.25 | 1 | 23 |
| bmp6 | 0.05181 | 0.61390236 | 0.548 | 0.125 | 1 | 23 |
| zgc:100832 | 0.05184 | -0.61911897 | 0.065 | 0.375 | 1 | 23 |
| rps26l | 0.05239 | 0.18728473 | 0.968 | 1 | 1 | 23 |
| dhdhl | 0.05327 | -0.32251099 | 0.194 | 0.25 | 1 | 23 |
| rpl27 | 0.05337 | 0.318144871 | 1 | 0.875 | 1 | 23 |
| si:dkey-250d21.1 | 0.05425 | -0.51898924 | 0.452 | 0.875 | 1 | 23 |
| rpp21 | 0.05456 | -0.14285827 | 0.161 | 0.125 | 1 | 23 |
| tcf3b | 0.05481 | -0.5910822 | 0.161 | 0.5 | 1 | 23 |
| txndc9 | 0.05497 | 0.622908594 | 0.516 | 0.125 | 1 | 23 |
| epcam | 0.05547 | -0.35255018 | 0.226 | 0.25 | 1 | 23 |
| sumo3b | 0.05552 | -0.48068822 | 0.065 | 0.375 | 1 | 23 |
| tnika | 0.05563 | 0.136787434 | 0.323 | 0.125 | 1 | 23 |
| rps9 | 0.05564 | 0.31147436 | 1 | 0.875 | 1 | 23 |
| cct5 | 0.05619 | 0.59561535 | 0.742 | 0.375 | 1 | 23 |
| cdh2 | 0.0562 | -0.50245949 | 0.548 | 0.875 | 1 | 23 |
| fkbp2 | 0.05653 | -0.56861499 | 0.258 | 0.625 | 1 | 23 |
| serf2 | 0.05718 | 0.572988087 | 0.774 | 0.5 | 1 | 23 |
| rmdn1 | 0.05743 | -0.56119459 | 0.323 | 0.75 | 1 | 23 |
| cct4 | 0.05807 | 0.419148007 | 0.806 | 0.375 | 1 | 23 |

| | | | | | | |
|---|---|---|---|---|---|---|
| ssrp1a | 0.05833 | -0.59104847 | 0.129 | 0.5 | 1 | 23 |
| pcbd1 | 0.0584 | 0.117327461 | 0.387 | 0.125 | 1 | 23 |
| tnip1 | 0.05881 | -0.59825585 | 0.194 | 0.5 | 1 | 23 |
| rbpms2b | 0.05895 | -0.52640455 | 0.29 | 0.75 | 1 | 23 |
| si:ch211-108c6.2 | 0.05898 | -0.64220993 | 0.129 | 0.5 | 1 | 23 |
| tbca | 0.05901 | 0.622953384 | 0.645 | 0.25 | 1 | 23 |
| prps1a | 0.05935 | 0.320890438 | 0.452 | 0.125 | 1 | 23 |
| tomm20a | 0.05969 | 0.592918575 | 0.484 | 0.125 | 1 | 23 |
| si:ch73-375g18.1 | 0.06075 | -0.59284799 | 0.065 | 0.375 | 1 | 23 |
| ndufa6 | 0.0608 | -0.51389653 | 0.581 | 0.875 | 1 | 23 |
| trdn | 0.0618 | -0.56384242 | 0.097 | 0.375 | 1 | 23 |
| nkrf | 0.06186 | 0.378830019 | 0.258 | 0 | 1 | 23 |
| psmf1 | 0.06186 | 0.406584901 | 0.258 | 0 | 1 | 23 |
| CT573476.1 | 0.06186 | 0.424003575 | 0.258 | 0 | 1 | 23 |
| cd3eap | 0.06186 | 0.424911472 | 0.258 | 0 | 1 | 23 |
| chd9 | 0.06186 | 0.366827028 | 0.258 | 0 | 1 | 23 |
| cnot2 | 0.06186 | 0.331964477 | 0.258 | 0 | 1 | 23 |
| cnpy1 | 0.06186 | 0.40286613 | 0.258 | 0 | 1 | 23 |
| creb1a | 0.06186 | 0.442632994 | 0.258 | 0 | 1 | 23 |
| f10 | 0.06186 | 0.425476583 | 0.258 | 0 | 1 | 23 |
| fkbp7 | 0.06186 | 0.403469144 | 0.258 | 0 | 1 | 23 |
| gclm | 0.06186 | 0.385661227 | 0.258 | 0 | 1 | 23 |
| grwd1 | 0.06186 | 0.438138432 | 0.258 | 0 | 1 | 23 |
| hells | 0.06186 | 0.595140052 | 0.258 | 0 | 1 | 23 |
| itga9 | 0.06186 | 0.51279041 | 0.258 | 0 | 1 | 23 |
| knop1 | 0.06186 | 0.38704616 | 0.258 | 0 | 1 | 23 |
| lrrc59 | 0.06186 | 0.431153975 | 0.258 | 0 | 1 | 23 |
| lsm12a | 0.06186 | 0.406816195 | 0.258 | 0 | 1 | 23 |
| lyrm9 | 0.06186 | 0.394932111 | 0.258 | 0 | 1 | 23 |
| magi1b | 0.06186 | 0.401508834 | 0.258 | 0 | 1 | 23 |
| med6 | 0.06186 | 0.39787136 | 0.258 | 0 | 1 | 23 |
| mrps14 | 0.06186 | 0.364083669 | 0.258 | 0 | 1 | 23 |
| mrps17 | 0.06186 | 0.369877277 | 0.258 | 0 | 1 | 23 |
| nob1 | 0.06186 | 0.372248803 | 0.258 | 0 | 1 | 23 |
| nutf2l | 0.06186 | 0.325327445 | 0.258 | 0 | 1 | 23 |
| paics | 0.06186 | 0.369818263 | 0.258 | 0 | 1 | 23 |
| rrs1 | 0.06186 | 0.356432831 | 0.258 | 0 | 1 | 23 |
| sept7a | 0.06186 | 0.359726732 | 0.258 | 0 | 1 | 23 |
| si:ch1073-157b13.1 | 0.06186 | 0.412982871 | 0.258 | 0 | 1 | 23 |
| tcf25 | 0.06186 | 0.423062978 | 0.258 | 0 | 1 | 23 |
| tmx1 | 0.06186 | 0.395891669 | 0.258 | 0 | 1 | 23 |
| blcap | 0.06186 | 0.338092851 | 0.258 | 0 | 1 | 23 |
| desi2 | 0.06186 | 0.421717377 | 0.258 | 0 | 1 | 23 |
| gpatch8 | 0.06186 | 0.440582392 | 0.258 | 0 | 1 | 23 |
| hoxb5a | 0.06186 | 0.482579753 | 0.258 | 0 | 1 | 23 |
| nenf | 0.06186 | 0.410106486 | 0.258 | 0 | 1 | 23 |
| nfkbib | 0.06186 | 0.430048834 | 0.258 | 0 | 1 | 23 |
| nudcd2 | 0.06186 | 0.412041794 | 0.258 | 0 | 1 | 23 |

| | | | | | | |
|---|---|---|---|---|---|---|
| pdha1a | 0.06186 | 0.382285643 | 0.258 | 0 | 1 | 23 |
| rtf1 | 0.06186 | 0.417342734 | 0.258 | 0 | 1 | 23 |
| trir | 0.06186 | 0.46511386 | 0.258 | 0 | 1 | 23 |
| wdr1 | 0.06186 | 0.340658636 | 0.258 | 0 | 1 | 23 |
| wdr82 | 0.06186 | 0.339479113 | 0.258 | 0 | 1 | 23 |
| zc3h13 | 0.06186 | 0.41601266 | 0.258 | 0 | 1 | 23 |
| bahcc1b | 0.06186 | 0.399458286 | 0.258 | 0 | 1 | 23 |
| surf2 | 0.06186 | 0.436404051 | 0.258 | 0 | 1 | 23 |
| tars | 0.06186 | 0.4167174 | 0.258 | 0 | 1 | 23 |
| egr1 | 0.06362 | -0.35501341 | 0.581 | 0.625 | 1 | 23 |
| cox8a | 0.06437 | -0.11130766 | 0.774 | 0.625 | 1 | 23 |
| tjp1b | 0.06451 | -0.54205479 | 0.065 | 0.375 | 1 | 23 |
| polr2l | 0.06472 | -0.21934054 | 0.194 | 0.5 | 1 | 23 |
| pkma | 0.06499 | -0.59192385 | 0.129 | 0.5 | 1 | 23 |
| uchl1 | 0.06601 | -0.39982431 | 0.129 | 0.5 | 1 | 23 |
| lingo2a | 0.06805 | -0.13867758 | 0.161 | 0.125 | 1 | 23 |
| rbm25b | 0.06826 | -0.27856966 | 0.194 | 0.25 | 1 | 23 |
| snrnp70 | 0.06842 | -0.62169842 | 0.226 | 0.625 | 1 | 23 |
| ndrg1a | 0.06907 | -0.16421965 | 0.419 | 0.375 | 1 | 23 |
| klf11a | 0.06914 | -0.14315728 | 0.161 | 0.125 | 1 | 23 |
| smarcb1b | 0.06961 | -0.53414314 | 0.194 | 0.625 | 1 | 23 |
| agap3 | 0.07026 | -0.63191867 | 0.065 | 0.375 | 1 | 23 |
| zbtb4 | 0.07045 | -0.63411645 | 0.065 | 0.375 | 1 | 23 |
| ndufa11 | 0.07157 | 0.102709206 | 0.452 | 0.25 | 1 | 23 |
| antxr1c | 0.07178 | -0.59243458 | 0.097 | 0.375 | 1 | 23 |
| npm1b | 0.07324 | -0.34146908 | 0.161 | 0.25 | 1 | 23 |
| LOC103909284 | 0.07346 | -0.55974837 | 0.452 | 0.75 | 1 | 23 |
| sf3a1 | 0.07388 | -0.54732042 | 0.129 | 0.5 | 1 | 23 |
| efna1a | 0.07396 | -0.60046871 | 0.065 | 0.375 | 1 | 23 |
| eif3d | 0.07465 | 0.620352828 | 0.677 | 0.375 | 1 | 23 |
| BX323458.1 | 0.07466 | -0.43278537 | 0.032 | 0.25 | 1 | 23 |
| FAM83G | 0.07466 | -0.44251758 | 0.032 | 0.25 | 1 | 23 |
| FO704858.1 | 0.07466 | -0.48070928 | 0.032 | 0.25 | 1 | 23 |
| KIF2A | 0.07466 | -0.41353188 | 0.032 | 0.25 | 1 | 23 |
| LOC101882642 | 0.07466 | -0.57180201 | 0.032 | 0.25 | 1 | 23 |
| LOC103910363 | 0.07466 | -0.25080512 | 0.032 | 0.25 | 1 | 23 |
| LOC103911818 | 0.07466 | -0.44654495 | 0.032 | 0.25 | 1 | 23 |
| MED9 | 0.07466 | -0.28096088 | 0.032 | 0.25 | 1 | 23 |
| akirin2 | 0.07466 | -0.47889457 | 0.032 | 0.25 | 1 | 23 |
| amdhd2 | 0.07466 | -0.30468941 | 0.032 | 0.25 | 1 | 23 |
| arl13b | 0.07466 | -0.2787715 | 0.032 | 0.25 | 1 | 23 |
| arl6ip4 | 0.07466 | -0.26859467 | 0.032 | 0.25 | 1 | 23 |
| asap1b | 0.07466 | -0.47604999 | 0.032 | 0.25 | 1 | 23 |
| cactin | 0.07466 | -0.37693872 | 0.032 | 0.25 | 1 | 23 |
| cdc37 | 0.07466 | -0.29846857 | 0.032 | 0.25 | 1 | 23 |
| cdh1 | 0.07466 | -0.34320883 | 0.032 | 0.25 | 1 | 23 |
| cfap36 | 0.07466 | -0.22310695 | 0.032 | 0.25 | 1 | 23 |
| clgn | 0.07466 | -0.40088649 | 0.032 | 0.25 | 1 | 23 |

| | | | | | | |
|---|---|---|---|---|---|---|
| crnkl1 | 0.07466 | -0.36353257 | 0.032 | 0.25 | 1 | 23 |
| cttn | 0.07466 | -0.32807229 | 0.032 | 0.25 | 1 | 23 |
| cyp27a7 | 0.07466 | -0.37056715 | 0.032 | 0.25 | 1 | 23 |
| ddrgk1 | 0.07466 | -0.40619441 | 0.032 | 0.25 | 1 | 23 |
| dip2ba | 0.07466 | -0.40541057 | 0.032 | 0.25 | 1 | 23 |
| exosc7 | 0.07466 | -0.37891075 | 0.032 | 0.25 | 1 | 23 |
| fbxo3 | 0.07466 | -0.3905488 | 0.032 | 0.25 | 1 | 23 |
| fbxw5 | 0.07466 | -0.49719071 | 0.032 | 0.25 | 1 | 23 |
| fhdc3 | 0.07466 | -0.42900357 | 0.032 | 0.25 | 1 | 23 |
| ftr76 | 0.07466 | -0.41235269 | 0.032 | 0.25 | 1 | 23 |
| gclc | 0.07466 | -0.3241044 | 0.032 | 0.25 | 1 | 23 |
| glg1a | 0.07466 | -0.33684001 | 0.032 | 0.25 | 1 | 23 |
| gna11a | 0.07466 | -0.43801505 | 0.032 | 0.25 | 1 | 23 |
| gtf2h4 | 0.07466 | -0.40257659 | 0.032 | 0.25 | 1 | 23 |
| gtf3aa | 0.07466 | -0.45618281 | 0.032 | 0.25 | 1 | 23 |
| hic2 | 0.07466 | -0.345745 | 0.032 | 0.25 | 1 | 23 |
| hoxa9b | 0.07466 | -0.33688945 | 0.032 | 0.25 | 1 | 23 |
| hsp70.3 | 0.07466 | -0.40013461 | 0.032 | 0.25 | 1 | 23 |
| ints8 | 0.07466 | -0.43769416 | 0.032 | 0.25 | 1 | 23 |
| ipcef1 | 0.07466 | -0.39058912 | 0.032 | 0.25 | 1 | 23 |
| itpkcb | 0.07466 | -0.49812065 | 0.032 | 0.25 | 1 | 23 |
| kank1a | 0.07466 | -0.40189679 | 0.032 | 0.25 | 1 | 23 |
| limk2 | 0.07466 | -0.42930151 | 0.032 | 0.25 | 1 | 23 |
| lrp5.1 | 0.07466 | -0.50792531 | 0.032 | 0.25 | 1 | 23 |
| med21 | 0.07466 | -0.33634386 | 0.032 | 0.25 | 1 | 23 |
| med30 | 0.07466 | -0.30214932 | 0.032 | 0.25 | 1 | 23 |
| mettl7a | 0.07466 | -0.34116475 | 0.032 | 0.25 | 1 | 23 |
| mib1 | 0.07466 | -0.38121765 | 0.032 | 0.25 | 1 | 23 |
| mrpl46 | 0.07466 | -0.27438097 | 0.032 | 0.25 | 1 | 23 |
| nop2 | 0.07466 | -0.503851 | 0.032 | 0.25 | 1 | 23 |
| nup160 | 0.07466 | -0.45965082 | 0.032 | 0.25 | 1 | 23 |
| ormdl2 | 0.07466 | -0.33683572 | 0.032 | 0.25 | 1 | 23 |
| phrf1 | 0.07466 | -0.4042694 | 0.032 | 0.25 | 1 | 23 |
| pithd1 | 0.07466 | -0.25026972 | 0.032 | 0.25 | 1 | 23 |
| pls3 | 0.07466 | -0.44189341 | 0.032 | 0.25 | 1 | 23 |
| plxnb2a | 0.07466 | -0.33727538 | 0.032 | 0.25 | 1 | 23 |
| ppan | 0.07466 | -0.27459671 | 0.032 | 0.25 | 1 | 23 |
| prpf18 | 0.07466 | -0.51963204 | 0.032 | 0.25 | 1 | 23 |
| prrc2a | 0.07466 | -0.32294598 | 0.032 | 0.25 | 1 | 23 |
| rabgef1 | 0.07466 | -0.29398211 | 0.032 | 0.25 | 1 | 23 |
| rfx1b | 0.07466 | -0.23736244 | 0.032 | 0.25 | 1 | 23 |
| rimkla | 0.07466 | -0.32213071 | 0.032 | 0.25 | 1 | 23 |
| rit1 | 0.07466 | -0.37803626 | 0.032 | 0.25 | 1 | 23 |
| rnf19a | 0.07466 | -0.43154823 | 0.032 | 0.25 | 1 | 23 |
| sept15 | 0.07466 | -0.40060692 | 0.032 | 0.25 | 1 | 23 |
| sfxn5b | 0.07466 | -0.448575 | 0.032 | 0.25 | 1 | 23 |
| sh2b3 | 0.07466 | -0.30944062 | 0.032 | 0.25 | 1 | 23 |
| si:ch211-194e18.2 | 0.07466 | -0.45133298 | 0.032 | 0.25 | 1 | 23 |

| | | | | | | |
|---|---|---|---|---|---|---|
| si:ch211-74m13.3 | 0.07466 | -0.30984209 | 0.032 | 0.25 | 1 | 23 |
| si:ch73-103b11.2 | 0.07466 | -0.38301338 | 0.032 | 0.25 | 1 | 23 |
| si:dkey-12e7.1 | 0.07466 | -0.37917297 | 0.032 | 0.25 | 1 | 23 |
| si:dkey-13n15.11 | 0.07466 | -0.46216324 | 0.032 | 0.25 | 1 | 23 |
| sipa1l3 | 0.07466 | -0.34986606 | 0.032 | 0.25 | 1 | 23 |
| slc29a1a | 0.07466 | -0.47878426 | 0.032 | 0.25 | 1 | 23 |
| slc38a4 | 0.07466 | -0.35613599 | 0.032 | 0.25 | 1 | 23 |
| smg6 | 0.07466 | -0.34442869 | 0.032 | 0.25 | 1 | 23 |
| snrnp40 | 0.07466 | -0.31116266 | 0.032 | 0.25 | 1 | 23 |
| snx27a | 0.07466 | -0.49395658 | 0.032 | 0.25 | 1 | 23 |
| socs1a | 0.07466 | -0.24754212 | 0.032 | 0.25 | 1 | 23 |
| sp3b | 0.07466 | -0.33670554 | 0.032 | 0.25 | 1 | 23 |
| spock1 | 0.07466 | -0.40493999 | 0.032 | 0.25 | 1 | 23 |
| spsb4b | 0.07466 | -0.41537839 | 0.032 | 0.25 | 1 | 23 |
| srp9 | 0.07466 | -0.44242536 | 0.032 | 0.25 | 1 | 23 |
| stambpa | 0.07466 | -0.35447658 | 0.032 | 0.25 | 1 | 23 |
| sympk | 0.07466 | -0.40941011 | 0.032 | 0.25 | 1 | 23 |
| synm | 0.07466 | -0.44455643 | 0.032 | 0.25 | 1 | 23 |
| thoc7 | 0.07466 | -0.26099098 | 0.032 | 0.25 | 1 | 23 |
| tm9sf1 | 0.07466 | -0.31681137 | 0.032 | 0.25 | 1 | 23 |
| trappc11 | 0.07466 | -0.34169842 | 0.032 | 0.25 | 1 | 23 |
| trim35-28 | 0.07466 | -0.27204924 | 0.032 | 0.25 | 1 | 23 |
| tspan4b | 0.07466 | -0.25344324 | 0.032 | 0.25 | 1 | 23 |
| txn | 0.07466 | -0.41784967 | 0.032 | 0.25 | 1 | 23 |
| ube2d3 | 0.07466 | -0.32469845 | 0.032 | 0.25 | 1 | 23 |
| ube2f | 0.07466 | -0.27707701 | 0.032 | 0.25 | 1 | 23 |
| ugp2b | 0.07466 | -0.444407 | 0.032 | 0.25 | 1 | 23 |
| unc50 | 0.07466 | -0.28477534 | 0.032 | 0.25 | 1 | 23 |
| usp8 | 0.07466 | -0.49533037 | 0.032 | 0.25 | 1 | 23 |
| washc3 | 0.07466 | -0.30328032 | 0.032 | 0.25 | 1 | 23 |
| wwp1 | 0.07466 | -0.37960556 | 0.032 | 0.25 | 1 | 23 |
| zc3h12a | 0.07466 | -0.28592274 | 0.032 | 0.25 | 1 | 23 |
| zgc:158640 | 0.07466 | -0.29075405 | 0.032 | 0.25 | 1 | 23 |
| znf292b | 0.07466 | -0.28567469 | 0.032 | 0.25 | 1 | 23 |
| CU655842.1 | 0.07466 | -0.40409841 | 0.032 | 0.25 | 1 | 23 |
| hyal2b | 0.07466 | -0.41630291 | 0.032 | 0.25 | 1 | 23 |
| tsen2 | 0.07466 | -0.41056668 | 0.032 | 0.25 | 1 | 23 |
| alkbh3 | 0.07466 | -0.3220671 | 0.032 | 0.25 | 1 | 23 |
| leng8 | 0.07466 | -0.38838023 | 0.032 | 0.25 | 1 | 23 |
| lgalslb | 0.07466 | -0.30629421 | 0.032 | 0.25 | 1 | 23 |
| rnf166 | 0.07466 | -0.55124411 | 0.032 | 0.25 | 1 | 23 |
| slmapa | 0.07466 | -0.35161321 | 0.032 | 0.25 | 1 | 23 |
| tmeff2a | 0.07466 | -0.27746336 | 0.032 | 0.25 | 1 | 23 |
| zgc:162972 | 0.07466 | -0.26776832 | 0.032 | 0.25 | 1 | 23 |
| p4hb | 0.07572 | -0.70275202 | 0.161 | 0.5 | 1 | 23 |
| dut | 0.07645 | -0.31793536 | 0.194 | 0.25 | 1 | 23 |
| bahcc1a | 0.07688 | -0.59433869 | 0.065 | 0.375 | 1 | 23 |
| pdcd10b | 0.07688 | -0.35603248 | 0.065 | 0.375 | 1 | 23 |

| | | | | | | |
|---|---|---|---|---|---|---|
| rack1 | 0.0776 | 0.166136921 | 1 | 1 | 1 | 23 |
| b2m | 0.07763 | 0.399410296 | 0.871 | 0.75 | 1 | 23 |
| tuba8l4 | 0.07768 | 0.352348714 | 0.774 | 0.375 | 1 | 23 |
| eny2 | 0.07809 | -0.51323505 | 0.29 | 0.5 | 1 | 23 |
| cflara | 0.07832 | -0.61096472 | 0.129 | 0.5 | 1 | 23 |
| NC-002333.4 | 0.0786 | -0.48577484 | 0.871 | 1 | 1 | 23 |
| nme7 | 0.07882 | -0.51047668 | 0.065 | 0.375 | 1 | 23 |
| nfil3 | 0.07927 | -0.63089249 | 0.129 | 0.5 | 1 | 23 |
| dock7 | 0.07928 | -0.51806716 | 0.065 | 0.375 | 1 | 23 |
| psmb7 | 0.07939 | 0.569035574 | 0.774 | 0.375 | 1 | 23 |
| nanos1 | 0.08059 | 0.382949825 | 0.839 | 0.75 | 1 | 23 |
| nupr1a | 0.08098 | -0.28502931 | 0.194 | 0.25 | 1 | 23 |
| tbcb | 0.08251 | -0.51179613 | 0.226 | 0.5 | 1 | 23 |
| prpf38b | 0.08388 | -0.55880427 | 0.161 | 0.5 | 1 | 23 |
| rps27a | 0.08436 | 0.194547054 | 1 | 1 | 1 | 23 |
| mtdhb | 0.08538 | -0.54688744 | 0.065 | 0.375 | 1 | 23 |
| ralbb | 0.08548 | -0.22413868 | 0.226 | 0.25 | 1 | 23 |
| MRPS12 | 0.08642 | 0.304503941 | 0.226 | 0 | 1 | 23 |
| gas6 | 0.08642 | 0.354043422 | 0.226 | 0 | 1 | 23 |
| hacd2 | 0.08642 | 0.33430537 | 0.226 | 0 | 1 | 23 |
| larp4ab | 0.08642 | 0.338992778 | 0.226 | 0 | 1 | 23 |
| letm2 | 0.08642 | 0.339764675 | 0.226 | 0 | 1 | 23 |
| nfkbie | 0.08642 | 0.330512877 | 0.226 | 0 | 1 | 23 |
| rsu1 | 0.08642 | 0.341558919 | 0.226 | 0 | 1 | 23 |
| ube2d1a | 0.08642 | 0.342296011 | 0.226 | 0 | 1 | 23 |
| ufc1 | 0.08642 | 0.270022668 | 0.226 | 0 | 1 | 23 |
| zgc:92140 | 0.08642 | 0.373816544 | 0.226 | 0 | 1 | 23 |
| CU929070.1 | 0.08642 | 0.399008255 | 0.226 | 0 | 1 | 23 |
| FO082781.1 | 0.08642 | 0.364602611 | 0.226 | 0 | 1 | 23 |
| ap2b1 | 0.08642 | 0.447331351 | 0.226 | 0 | 1 | 23 |
| asf1bb | 0.08642 | 0.337931754 | 0.226 | 0 | 1 | 23 |
| cops4 | 0.08642 | 0.351397468 | 0.226 | 0 | 1 | 23 |
| entr1 | 0.08642 | 0.352374359 | 0.226 | 0 | 1 | 23 |
| ndufa7 | 0.08642 | 0.303115349 | 0.226 | 0 | 1 | 23 |
| rps19bp1 | 0.08642 | 0.347270436 | 0.226 | 0 | 1 | 23 |
| tmem176 | 0.08642 | 0.366036257 | 0.226 | 0 | 1 | 23 |
| zgc:110591 | 0.08642 | 0.36266428 | 0.226 | 0 | 1 | 23 |
| AL954322.2 | 0.08642 | 0.403545801 | 0.226 | 0 | 1 | 23 |
| anks1b | 0.08642 | 0.404874806 | 0.226 | 0 | 1 | 23 |
| cab39l | 0.08642 | 0.310779262 | 0.226 | 0 | 1 | 23 |
| ccdc59 | 0.08642 | 0.34205556 | 0.226 | 0 | 1 | 23 |
| csnk1g2b | 0.08642 | 0.382098176 | 0.226 | 0 | 1 | 23 |
| ctdsp1 | 0.08642 | 0.34410969 | 0.226 | 0 | 1 | 23 |
| ctsz | 0.08642 | 0.311521891 | 0.226 | 0 | 1 | 23 |
| dctn2 | 0.08642 | 0.33196616 | 0.226 | 0 | 1 | 23 |
| dynll2a | 0.08642 | 0.38193521 | 0.226 | 0 | 1 | 23 |
| eif4a2 | 0.08642 | 0.306643329 | 0.226 | 0 | 1 | 23 |
| elavl1b | 0.08642 | 0.389124276 | 0.226 | 0 | 1 | 23 |

| | | | | | | |
|---|---|---|---|---|---|---|
| fgl2a | 0.08642 | 0.447935337 | 0.226 | 0 | 1 | 23 |
| fibinb | 0.08642 | 0.385019 | 0.226 | 0 | 1 | 23 |
| fkbp14 | 0.08642 | 0.354851537 | 0.226 | 0 | 1 | 23 |
| idh3b | 0.08642 | 0.338430174 | 0.226 | 0 | 1 | 23 |
| il6st | 0.08642 | 0.397611383 | 0.226 | 0 | 1 | 23 |
| irf2 | 0.08642 | 0.35768627 | 0.226 | 0 | 1 | 23 |
| lamp1a | 0.08642 | 0.369859596 | 0.226 | 0 | 1 | 23 |
| lmcd1 | 0.08642 | 0.388692848 | 0.226 | 0 | 1 | 23 |
| matr3l1.1 | 0.08642 | 0.391899187 | 0.226 | 0 | 1 | 23 |
| ndnl2 | 0.08642 | 0.351583701 | 0.226 | 0 | 1 | 23 |
| nfybb | 0.08642 | 0.355783979 | 0.226 | 0 | 1 | 23 |
| pak4 | 0.08642 | 0.379173585 | 0.226 | 0 | 1 | 23 |
| pelo | 0.08642 | 0.336070717 | 0.226 | 0 | 1 | 23 |
| rab28 | 0.08642 | 0.387368998 | 0.226 | 0 | 1 | 23 |
| rbms2b | 0.08642 | 0.338993654 | 0.226 | 0 | 1 | 23 |
| sec62 | 0.08642 | 0.402554857 | 0.226 | 0 | 1 | 23 |
| szrd1 | 0.08642 | 0.301732355 | 0.226 | 0 | 1 | 23 |
| tmem33 | 0.08642 | 0.326997421 | 0.226 | 0 | 1 | 23 |
| uqcrc2b | 0.08642 | 0.356359912 | 0.226 | 0 | 1 | 23 |
| vapb | 0.08642 | 0.352994702 | 0.226 | 0 | 1 | 23 |
| wbp1lb | 0.08642 | 0.363122611 | 0.226 | 0 | 1 | 23 |
| ylpm1 | 0.08642 | 0.38171691 | 0.226 | 0 | 1 | 23 |
| zgc:110339 | 0.08642 | 0.326301418 | 0.226 | 0 | 1 | 23 |
| zgc:65873 | 0.08642 | 0.346818653 | 0.226 | 0 | 1 | 23 |
| ada2a | 0.08642 | 0.390882231 | 0.226 | 0 | 1 | 23 |
| aup1 | 0.08642 | 0.318807943 | 0.226 | 0 | 1 | 23 |
| cd164 | 0.08642 | 0.349188407 | 0.226 | 0 | 1 | 23 |
| commd3 | 0.08642 | 0.281386859 | 0.226 | 0 | 1 | 23 |
| ddx42 | 0.08642 | 0.37313952 | 0.226 | 0 | 1 | 23 |
| dnaja2b | 0.08642 | 0.361610627 | 0.226 | 0 | 1 | 23 |
| dtnba | 0.08642 | 0.290506462 | 0.226 | 0 | 1 | 23 |
| lyar | 0.08642 | 0.309312425 | 0.226 | 0 | 1 | 23 |
| med29 | 0.08642 | 0.358933387 | 0.226 | 0 | 1 | 23 |
| mplkip | 0.08642 | 0.344490892 | 0.226 | 0 | 1 | 23 |
| ms4a17a.9 | 0.08642 | 0.409547543 | 0.226 | 0 | 1 | 23 |
| nansa | 0.08642 | 0.336208606 | 0.226 | 0 | 1 | 23 |
| nat9 | 0.08642 | 0.355629314 | 0.226 | 0 | 1 | 23 |
| ndnf | 0.08642 | 0.418659774 | 0.226 | 0 | 1 | 23 |
| phb | 0.08642 | 0.364595071 | 0.226 | 0 | 1 | 23 |
| rab1bb | 0.08642 | 0.325201593 | 0.226 | 0 | 1 | 23 |
| rasl11a | 0.08642 | 0.361527921 | 0.226 | 0 | 1 | 23 |
| rbfox3a | 0.08642 | 0.411524491 | 0.226 | 0 | 1 | 23 |
| si:ch211-251p5.5 | 0.08642 | 0.407315976 | 0.226 | 0 | 1 | 23 |
| si:dkey-46a10.3 | 0.08642 | 0.347520063 | 0.226 | 0 | 1 | 23 |
| tmem183a | 0.08642 | 0.363754153 | 0.226 | 0 | 1 | 23 |
| top2b | 0.08642 | 0.361778683 | 0.226 | 0 | 1 | 23 |
| ubap2l | 0.08642 | 0.410665166 | 0.226 | 0 | 1 | 23 |
| wnt4a | 0.08642 | 0.436234939 | 0.226 | 0 | 1 | 23 |

| | | | | | | |
|---|---|---|---|---|---|---|
| zgc:92664 | 0.08642 | 0.327718022 | 0.226 | 0 | 1 | 23 |
| XLOC-001102 | 0.08642 | 0.365344081 | 0.226 | 0 | 1 | 23 |
| ddx47 | 0.08642 | 0.353564004 | 0.226 | 0 | 1 | 23 |
| sv2bb | 0.08642 | 0.409589732 | 0.226 | 0 | 1 | 23 |
| znf827 | 0.08642 | 0.38115395 | 0.226 | 0 | 1 | 23 |
| ar | 0.08694 | -0.58914279 | 0.323 | 0.75 | 1 | 23 |
| pgrmc1 | 0.08738 | 0.62763288 | 0.484 | 0.125 | 1 | 23 |
| ptprsa | 0.08769 | -0.57014501 | 0.226 | 0.625 | 1 | 23 |
| dbn1 | 0.0878 | -0.46844857 | 0.323 | 0.75 | 1 | 23 |
| rbms3 | 0.08794 | -0.18968039 | 0.129 | 0.125 | 1 | 23 |
| kdm4b | 0.08818 | -0.12164458 | 0.161 | 0.125 | 1 | 23 |
| rbm28 | 0.08822 | -0.12433074 | 0.29 | 0.25 | 1 | 23 |
| nfkbiab | 0.08932 | 0.466237044 | 0.645 | 0.25 | 1 | 23 |
| rab1ab | 0.08939 | -0.54717014 | 0.29 | 0.625 | 1 | 23 |
| clic4 | 0.09115 | -0.22033246 | 0.387 | 0.375 | 1 | 23 |
| rpl17 | 0.09118 | 0.258947574 | 1 | 1 | 1 | 23 |
| lsm12b | 0.09126 | -0.50273713 | 0.323 | 0.75 | 1 | 23 |
| snd1 | 0.0913 | -0.40293309 | 0.258 | 0.375 | 1 | 23 |
| rbm39b | 0.09141 | -0.11696569 | 0.161 | 0.125 | 1 | 23 |
| snrpd2 | 0.0922 | 0.562585014 | 0.677 | 0.25 | 1 | 23 |
| rnf44 | 0.09222 | -0.5269934 | 0.323 | 0.75 | 1 | 23 |
| kansl1a | 0.09238 | -0.28547534 | 0.097 | 0.125 | 1 | 23 |
| fer | 0.09244 | -0.52838776 | 0.065 | 0.375 | 1 | 23 |
| inka1a | 0.09267 | 0.530545453 | 0.323 | 0.125 | 1 | 23 |
| tram1 | 0.09281 | -0.53193359 | 0.129 | 0.5 | 1 | 23 |
| cox6b2 | 0.09295 | 0.355047367 | 0.774 | 0.75 | 1 | 23 |
| LOC103910438 | 0.09444 | -0.33129481 | 0.29 | 0.375 | 1 | 23 |
| si:ch73-335l21.4 | 0.09468 | -0.15343817 | 0.581 | 0.5 | 1 | 23 |
| sema3b | 0.09534 | -0.49607929 | 0.129 | 0.375 | 1 | 23 |
| c7a | 0.0954 | -0.48446933 | 0.065 | 0.375 | 1 | 23 |
| h3f3d | 0.09569 | 0.164938121 | 0.968 | 1 | 1 | 23 |
| eif3s6ip | 0.09582 | 0.610178894 | 0.516 | 0.125 | 1 | 23 |
| kdm1a | 0.09608 | -0.62447993 | 0.129 | 0.5 | 1 | 23 |
| cyc1 | 0.09687 | -0.17406194 | 0.29 | 0.25 | 1 | 23 |
| cdv3 | 0.09808 | -0.40520417 | 0.323 | 0.5 | 1 | 23 |
| brd7 | 0.09816 | 0.197578813 | 0.355 | 0.125 | 1 | 23 |
| ube2d2 | 0.09819 | -0.40274294 | 0.355 | 0.5 | 1 | 23 |
| parn | 0.09836 | -0.48212469 | 0.065 | 0.375 | 1 | 23 |
| itm2bb | 0.09844 | 0.34226524 | 0.742 | 0.375 | 1 | 23 |
| golgb1 | 0.09866 | -0.45187547 | 0.161 | 0.375 | 1 | 23 |
| med11 | 0.09887 | 0.204286198 | 0.355 | 0.125 | 1 | 23 |
| XLOC-043857 | 0.09917 | -0.3007641 | 0.194 | 0.25 | 1 | 23 |
| acin1a | 0.0992 | -0.37476495 | 0.581 | 0.75 | 1 | 23 |
| nsa2 | 0.10069 | 0.443945056 | 0.871 | 0.5 | 1 | 23 |
| smc3 | 0.10091 | -0.54143648 | 0.065 | 0.375 | 1 | 23 |
| antxr2a | 0.10115 | -0.5255514 | 0.355 | 0.625 | 1 | 23 |
| vangl2 | 0.10165 | -0.39203013 | 0.129 | 0.25 | 1 | 23 |
| fbxl6 | 0.10184 | 0.601492902 | 0.484 | 0.125 | 1 | 23 |

| | | | | | | |
|---|---|---|---|---|---|---|
| marcksb | 0.1029 | -0.49524349 | 0.387 | 0.625 | 1 | 23 |
| abcf2a | 0.10336 | -0.4295661 | 0.065 | 0.375 | 1 | 23 |
| afap1 | 0.10387 | -0.52606292 | 0.065 | 0.375 | 1 | 23 |
| psma4 | 0.1061 | 0.433567694 | 0.774 | 0.375 | 1 | 23 |
| zgc:100918 | 0.10716 | -0.46700429 | 0.129 | 0.375 | 1 | 23 |
| ptmaa | 0.10722 | 0.219118915 | 1 | 1 | 1 | 23 |
| tdp2a | 0.10745 | -0.42426579 | 0.065 | 0.375 | 1 | 23 |
| pik3c2a | 0.10764 | -0.44347618 | 0.065 | 0.375 | 1 | 23 |
| atp6ap2 | 0.10806 | 0.370615099 | 0.452 | 0.125 | 1 | 23 |
| timm23a | 0.10832 | 0.473855433 | 0.452 | 0.125 | 1 | 23 |
| hnrnpa0a | 0.10869 | -0.31343838 | 0.387 | 0.75 | 1 | 23 |
| ppme1 | 0.111 | -0.47186591 | 0.065 | 0.375 | 1 | 23 |
| ier2a | 0.1117 | 0.488698975 | 0.903 | 0.625 | 1 | 23 |
| gdi2 | 0.11196 | 0.433293302 | 0.484 | 0.125 | 1 | 23 |
| smad3a | 0.11209 | -0.47245011 | 0.065 | 0.375 | 1 | 23 |
| mdh2 | 0.11233 | 0.551617355 | 0.516 | 0.125 | 1 | 23 |
| dsc2l | 0.11273 | -0.53608559 | 0.065 | 0.375 | 1 | 23 |
| ppp2r1bb | 0.1129 | -0.39658121 | 0.065 | 0.375 | 1 | 23 |
| ywhae2 | 0.1129 | -0.41021924 | 0.065 | 0.375 | 1 | 23 |
| foxo4 | 0.11377 | -0.55172481 | 0.194 | 0.5 | 1 | 23 |
| lyrm4 | 0.114 | -0.26508166 | 0.194 | 0.25 | 1 | 23 |
| rad21a | 0.11433 | 0.265458793 | 0.71 | 0.375 | 1 | 23 |
| pbx3b | 0.1146 | -0.37666531 | 0.129 | 0.25 | 1 | 23 |
| eif5b | 0.11501 | -0.27379662 | 0.323 | 0.375 | 1 | 23 |
| zgc:193541 | 0.11529 | -0.42350901 | 0.774 | 0.875 | 1 | 23 |
| gng7 | 0.11568 | -0.11319579 | 0.161 | 0.125 | 1 | 23 |
| cst14a.2 | 0.1159 | 0.29579736 | 0.581 | 0.25 | 1 | 23 |
| paplna | 0.11634 | -0.47320061 | 0.129 | 0.375 | 1 | 23 |
| rpl39 | 0.11639 | 0.245767869 | 1 | 0.875 | 1 | 23 |
| ppp3ccb | 0.11689 | -0.14542322 | 0.29 | 0.25 | 1 | 23 |
| ppib | 0.11741 | 0.28686267 | 1 | 0.875 | 1 | 23 |
| meis3 | 0.11753 | 0.286820047 | 0.387 | 0.125 | 1 | 23 |
| chkb | 0.11768 | -0.39484874 | 0.097 | 0.25 | 1 | 23 |
| rpl24 | 0.11852 | 0.223565831 | 1 | 0.875 | 1 | 23 |
| si:ch211-196l7.4 | 0.11909 | 0.119405145 | 0.903 | 0.625 | 1 | 23 |
| CABZ01110881.1 | 0.12003 | 0.291589649 | 0.194 | 0 | 1 | 23 |
| TSTD1 | 0.12003 | 0.301768172 | 0.194 | 0 | 1 | 23 |
| ankrd10b | 0.12003 | 0.31796244 | 0.194 | 0 | 1 | 23 |
| ankrd54 | 0.12003 | 0.342010787 | 0.194 | 0 | 1 | 23 |
| ch25hl3 | 0.12003 | 0.318538964 | 0.194 | 0 | 1 | 23 |
| chd2 | 0.12003 | 0.295600937 | 0.194 | 0 | 1 | 23 |
| eif4ba | 0.12003 | 0.299756339 | 0.194 | 0 | 1 | 23 |
| ezh2 | 0.12003 | 0.315247924 | 0.194 | 0 | 1 | 23 |
| fam114a2 | 0.12003 | 0.323539908 | 0.194 | 0 | 1 | 23 |
| gng2 | 0.12003 | 0.301370639 | 0.194 | 0 | 1 | 23 |
| pdia4 | 0.12003 | 0.265411641 | 0.194 | 0 | 1 | 23 |
| rnf24 | 0.12003 | 0.296313038 | 0.194 | 0 | 1 | 23 |
| setbp1 | 0.12003 | 0.310302022 | 0.194 | 0 | 1 | 23 |

| | | | | | | |
|---|---|---|---|---|---|---|
| si:ch73-234b20.5 | 0.12003 | 0.230302569 | 0.194 | 0 | 1 | 23 |
| tpi1b | 0.12003 | 0.260524013 | 0.194 | 0 | 1 | 23 |
| ufm1 | 0.12003 | 0.268595266 | 0.194 | 0 | 1 | 23 |
| zfand3 | 0.12003 | 0.307051787 | 0.194 | 0 | 1 | 23 |
| ccz1 | 0.12003 | 0.300617838 | 0.194 | 0 | 1 | 23 |
| fkbp9 | 0.12003 | 0.321066533 | 0.194 | 0 | 1 | 23 |
| gcdhb | 0.12003 | 0.268868943 | 0.194 | 0 | 1 | 23 |
| nup98 | 0.12003 | 0.301116896 | 0.194 | 0 | 1 | 23 |
| PDCL3 | 0.12003 | 0.339561263 | 0.194 | 0 | 1 | 23 |
| XLOC-037780 | 0.12003 | 0.278375984 | 0.194 | 0 | 1 | 23 |
| ahi1 | 0.12003 | 0.353729636 | 0.194 | 0 | 1 | 23 |
| akt1s1 | 0.12003 | 0.293332701 | 0.194 | 0 | 1 | 23 |
| aldh7a1 | 0.12003 | 0.269991211 | 0.194 | 0 | 1 | 23 |
| arl9 | 0.12003 | 0.341744875 | 0.194 | 0 | 1 | 23 |
| caln2 | 0.12003 | 0.314613637 | 0.194 | 0 | 1 | 23 |
| camk2g1 | 0.12003 | 0.307540592 | 0.194 | 0 | 1 | 23 |
| cercam | 0.12003 | 0.299618469 | 0.194 | 0 | 1 | 23 |
| cnn3a | 0.12003 | 0.284782696 | 0.194 | 0 | 1 | 23 |
| commd2 | 0.12003 | 0.286149511 | 0.194 | 0 | 1 | 23 |
| cpsf2 | 0.12003 | 0.270216837 | 0.194 | 0 | 1 | 23 |
| dnajc2 | 0.12003 | 0.332072332 | 0.194 | 0 | 1 | 23 |
| fam3c | 0.12003 | 0.320730985 | 0.194 | 0 | 1 | 23 |
| jak1 | 0.12003 | 0.32770912 | 0.194 | 0 | 1 | 23 |
| mat2ab | 0.12003 | 0.273240688 | 0.194 | 0 | 1 | 23 |
| mrps18c | 0.12003 | 0.32121414 | 0.194 | 0 | 1 | 23 |
| mrto4 | 0.12003 | 0.270808356 | 0.194 | 0 | 1 | 23 |
| ms4a17a.11 | 0.12003 | 0.297196523 | 0.194 | 0 | 1 | 23 |
| mto1 | 0.12003 | 0.374442962 | 0.194 | 0 | 1 | 23 |
| nifk | 0.12003 | 0.306096887 | 0.194 | 0 | 1 | 23 |
| nol12 | 0.12003 | 0.310649956 | 0.194 | 0 | 1 | 23 |
| pds5a | 0.12003 | 0.299885165 | 0.194 | 0 | 1 | 23 |
| prpf8 | 0.12003 | 0.428666593 | 0.194 | 0 | 1 | 23 |
| rab7a | 0.12003 | 0.322426244 | 0.194 | 0 | 1 | 23 |
| rassf8b | 0.12003 | 0.306471853 | 0.194 | 0 | 1 | 23 |
| si:ch211-140m22.7 | 0.12003 | 0.268953603 | 0.194 | 0 | 1 | 23 |
| si:ch211-206k20.5 | 0.12003 | 0.310027716 | 0.194 | 0 | 1 | 23 |
| srsf1a | 0.12003 | 0.289403776 | 0.194 | 0 | 1 | 23 |
| supt3h | 0.12003 | 0.307720611 | 0.194 | 0 | 1 | 23 |
| tlk1b | 0.12003 | 0.318285456 | 0.194 | 0 | 1 | 23 |
| ube2g2 | 0.12003 | 0.291716482 | 0.194 | 0 | 1 | 23 |
| ythdc1 | 0.12003 | 0.309108118 | 0.194 | 0 | 1 | 23 |
| zgc:56106 | 0.12003 | 0.270848161 | 0.194 | 0 | 1 | 23 |
| znf330 | 0.12003 | 0.313190537 | 0.194 | 0 | 1 | 23 |
| znhit1 | 0.12003 | 0.273101826 | 0.194 | 0 | 1 | 23 |
| arfrp1 | 0.12003 | 0.274105009 | 0.194 | 0 | 1 | 23 |
| ddx3xb | 0.12003 | 0.400604934 | 0.194 | 0 | 1 | 23 |
| ephb4a | 0.12003 | 0.315070269 | 0.194 | 0 | 1 | 23 |
| gnai1 | 0.12003 | 0.281700588 | 0.194 | 0 | 1 | 23 |

| | | | | | | |
|---|---|---|---|---|---|---|
| imp3 | 0.12003 | 0.341910075 | 0.194 | 0 | 1 | 23 |
| map2k4a | 0.12003 | 0.270238045 | 0.194 | 0 | 1 | 23 |
| mbpb | 0.12003 | 0.289165981 | 0.194 | 0 | 1 | 23 |
| mrpl11 | 0.12003 | 0.306138182 | 0.194 | 0 | 1 | 23 |
| mrps24 | 0.12003 | 0.323849437 | 0.194 | 0 | 1 | 23 |
| mta3 | 0.12003 | 0.294190827 | 0.194 | 0 | 1 | 23 |
| prkacaa | 0.12003 | 0.287852398 | 0.194 | 0 | 1 | 23 |
| pttg1ipb | 0.12003 | 0.286866289 | 0.194 | 0 | 1 | 23 |
| smad5 | 0.12003 | 0.328300306 | 0.194 | 0 | 1 | 23 |
| zgc:161969 | 0.12003 | 0.351631014 | 0.194 | 0 | 1 | 23 |
| BX569787.1 | 0.12003 | 0.290103553 | 0.194 | 0 | 1 | 23 |
| CABZ01039304.1 | 0.12003 | 0.356883392 | 0.194 | 0 | 1 | 23 |
| acaa2 | 0.12003 | 0.302377702 | 0.194 | 0 | 1 | 23 |
| anapc11 | 0.12003 | 0.306820562 | 0.194 | 0 | 1 | 23 |
| aqr | 0.12003 | 0.302638462 | 0.194 | 0 | 1 | 23 |
| atg3 | 0.12003 | 0.328355609 | 0.194 | 0 | 1 | 23 |
| chic2 | 0.12003 | 0.313152204 | 0.194 | 0 | 1 | 23 |
| clpp | 0.12003 | 0.301305007 | 0.194 | 0 | 1 | 23 |
| cmbl | 0.12003 | 0.322592971 | 0.194 | 0 | 1 | 23 |
| creb3l3l | 0.12003 | 0.272546497 | 0.194 | 0 | 1 | 23 |
| ctdspla | 0.12003 | 0.310925975 | 0.194 | 0 | 1 | 23 |
| ctps1a | 0.12003 | 0.319933569 | 0.194 | 0 | 1 | 23 |
| dhx9 | 0.12003 | 0.303387128 | 0.194 | 0 | 1 | 23 |
| gemin7 | 0.12003 | 0.274416563 | 0.194 | 0 | 1 | 23 |
| lztfl1 | 0.12003 | 0.308951611 | 0.194 | 0 | 1 | 23 |
| mcfd2 | 0.12003 | 0.360752321 | 0.194 | 0 | 1 | 23 |
| mgmt | 0.12003 | 0.302395399 | 0.194 | 0 | 1 | 23 |
| mycn | 0.12003 | 0.372959173 | 0.194 | 0 | 1 | 23 |
| olfml3b | 0.12003 | 0.356720439 | 0.194 | 0 | 1 | 23 |
| phf2 | 0.12003 | 0.267790004 | 0.194 | 0 | 1 | 23 |
| prpf4bb | 0.12003 | 0.315007285 | 0.194 | 0 | 1 | 23 |
| rgl1 | 0.12003 | 0.309963926 | 0.194 | 0 | 1 | 23 |
| rnd3b | 0.12003 | 0.30779108 | 0.194 | 0 | 1 | 23 |
| sf3a3 | 0.12003 | 0.345847174 | 0.194 | 0 | 1 | 23 |
| stat1b | 0.12003 | 0.396752011 | 0.194 | 0 | 1 | 23 |
| sypl2b | 0.12003 | 0.281169312 | 0.194 | 0 | 1 | 23 |
| taf10 | 0.12003 | 0.274531069 | 0.194 | 0 | 1 | 23 |
| uqcc3 | 0.12003 | 0.262457587 | 0.194 | 0 | 1 | 23 |
| vma21 | 0.12003 | 0.280775986 | 0.194 | 0 | 1 | 23 |
| zc3h18 | 0.12003 | 0.344955073 | 0.194 | 0 | 1 | 23 |
| zgc:175214 | 0.12003 | 0.305199838 | 0.194 | 0 | 1 | 23 |
| rsl24d1 | 0.12056 | 0.446507808 | 0.903 | 0.625 | 1 | 23 |
| AL840638.1 | 0.12136 | -0.40629498 | 0 | 0.125 | 1 | 23 |
| CABZ01078737.1 | 0.12136 | -0.31782846 | 0 | 0.125 | 1 | 23 |
| CABZ01079550.1 | 0.12136 | -0.31912187 | 0 | 0.125 | 1 | 23 |
| CR749775.2 | 0.12136 | -0.31844117 | 0 | 0.125 | 1 | 23 |
| DNAH10 | 0.12136 | -0.31912187 | 0 | 0.125 | 1 | 23 |
| GAN | 0.12136 | -0.31741118 | 0 | 0.125 | 1 | 23 |

| | | | | | | |
|---|---|---|---|---|---|---|
| LOC101884469 | 0.12136 | -0.3011849 | 0 | 0.125 | 1 | 23 |
| LOC103911404 | 0.12136 | -0.4056701 | 0 | 0.125 | 1 | 23 |
| XLOC-035371 | 0.12136 | -0.31765077 | 0 | 0.125 | 1 | 23 |
| abhd17ab | 0.12136 | -0.31704304 | 0 | 0.125 | 1 | 23 |
| acvr2aa | 0.12136 | -0.31843292 | 0 | 0.125 | 1 | 23 |
| adamts10 | 0.12136 | -0.31884969 | 0 | 0.125 | 1 | 23 |
| adck5 | 0.12136 | -0.31813243 | 0 | 0.125 | 1 | 23 |
| afg1la | 0.12136 | -0.31680472 | 0 | 0.125 | 1 | 23 |
| btbd11a | 0.12136 | -0.31912187 | 0 | 0.125 | 1 | 23 |
| cdc25b | 0.12136 | -0.31172087 | 0 | 0.125 | 1 | 23 |
| cpt1aa | 0.12136 | -0.31829764 | 0 | 0.125 | 1 | 23 |
| cuedc1b | 0.12136 | -0.31845985 | 0 | 0.125 | 1 | 23 |
| cyth3a | 0.12136 | -0.29796883 | 0 | 0.125 | 1 | 23 |
| dhx29 | 0.12136 | -0.31731727 | 0 | 0.125 | 1 | 23 |
| dnajc25 | 0.12136 | -0.31684156 | 0 | 0.125 | 1 | 23 |
| e2f7 | 0.12136 | -0.3174663 | 0 | 0.125 | 1 | 23 |
| espl1 | 0.12136 | -0.31638744 | 0 | 0.125 | 1 | 23 |
| etaa1 | 0.12136 | -0.31849765 | 0 | 0.125 | 1 | 23 |
| fam129bb | 0.12136 | -0.31660691 | 0 | 0.125 | 1 | 23 |
| fancd2 | 0.12136 | -0.31331 | 0 | 0.125 | 1 | 23 |
| fgfr4 | 0.12136 | -0.31753037 | 0 | 0.125 | 1 | 23 |
| fthl28 | 0.12136 | -0.31862506 | 0 | 0.125 | 1 | 23 |
| gkup | 0.12136 | -0.31804581 | 0 | 0.125 | 1 | 23 |
| gpat4 | 0.12136 | -0.31505038 | 0 | 0.125 | 1 | 23 |
| her6 | 0.12136 | -0.31659354 | 0 | 0.125 | 1 | 23 |
| hist1h4l | 0.12136 | -0.31777108 | 0 | 0.125 | 1 | 23 |
| ints10 | 0.12136 | -0.3159389 | 0 | 0.125 | 1 | 23 |
| kif20ba | 0.12136 | -0.31760308 | 0 | 0.125 | 1 | 23 |
| lamtor1 | 0.12136 | -0.31418628 | 0 | 0.125 | 1 | 23 |
| lancl2 | 0.12136 | -0.31767112 | 0 | 0.125 | 1 | 23 |
| large1 | 0.12136 | -0.31796632 | 0 | 0.125 | 1 | 23 |
| lpl | 0.12136 | -0.312002 | 0 | 0.125 | 1 | 23 |
| maco1b | 0.12136 | -0.31821814 | 0 | 0.125 | 1 | 23 |
| map9 | 0.12136 | -0.31617975 | 0 | 0.125 | 1 | 23 |
| mcm3ap | 0.12136 | -0.3181314 | 0 | 0.125 | 1 | 23 |
| mgll | 0.12136 | -0.31695905 | 0 | 0.125 | 1 | 23 |
| mthfd2 | 0.12136 | -0.31521226 | 0 | 0.125 | 1 | 23 |
| myo1b | 0.12136 | -0.31807248 | 0 | 0.125 | 1 | 23 |
| nbn | 0.12136 | -0.31822702 | 0 | 0.125 | 1 | 23 |
| ncapd2 | 0.12136 | -0.31713847 | 0 | 0.125 | 1 | 23 |
| nr1h3 | 0.12136 | -0.31830587 | 0 | 0.125 | 1 | 23 |
| nrarpa | 0.12136 | -0.3184062 | 0 | 0.125 | 1 | 23 |
| pane1 | 0.12136 | -0.30365933 | 0 | 0.125 | 1 | 23 |
| pcnx1 | 0.12136 | -0.31743637 | 0 | 0.125 | 1 | 23 |
| phlda1 | 0.12136 | -0.31484627 | 0 | 0.125 | 1 | 23 |
| plxnb1a | 0.12136 | -0.31714547 | 0 | 0.125 | 1 | 23 |
| pola2 | 0.12136 | -0.31731384 | 0 | 0.125 | 1 | 23 |
| pole | 0.12136 | -0.31682376 | 0 | 0.125 | 1 | 23 |

| | | | | | | |
|---|---|---|---|---|---|---|
| ppp1r21 | 0.12136 | -0.31325831 | 0 | 0.125 | 1 | 23 |
| ptdss1a | 0.12136 | -0.31690054 | 0 | 0.125 | 1 | 23 |
| si:dkey-111e8.1 | 0.12136 | -0.31253421 | 0 | 0.125 | 1 | 23 |
| si:dkey-261j15.2 | 0.12136 | -0.31834527 | 0 | 0.125 | 1 | 23 |
| slc25a35 | 0.12136 | -0.31741338 | 0 | 0.125 | 1 | 23 |
| slc40a1 | 0.12136 | -0.30008956 | 0 | 0.125 | 1 | 23 |
| slc4a2a | 0.12136 | -0.31680391 | 0 | 0.125 | 1 | 23 |
| slc7a14a | 0.12136 | -0.31912187 | 0 | 0.125 | 1 | 23 |
| smad6b | 0.12136 | -0.3150949 | 0 | 0.125 | 1 | 23 |
| sntb1 | 0.12136 | -0.31637291 | 0 | 0.125 | 1 | 23 |
| styx | 0.12136 | -0.31550784 | 0 | 0.125 | 1 | 23 |
| tacc3 | 0.12136 | -0.31617268 | 0 | 0.125 | 1 | 23 |
| tent4a | 0.12136 | -0.31912187 | 0 | 0.125 | 1 | 23 |
| tnfsf13b | 0.12136 | -0.31592075 | 0 | 0.125 | 1 | 23 |
| top2a | 0.12136 | -0.4033976 | 0 | 0.125 | 1 | 23 |
| trappc10 | 0.12136 | -0.31726344 | 0 | 0.125 | 1 | 23 |
| trim59 | 0.12136 | -0.31824421 | 0 | 0.125 | 1 | 23 |
| ttf2 | 0.12136 | -0.3145221 | 0 | 0.125 | 1 | 23 |
| ube2t | 0.12136 | -0.30790006 | 0 | 0.125 | 1 | 23 |
| unc45a | 0.12136 | -0.31783312 | 0 | 0.125 | 1 | 23 |
| usp37 | 0.12136 | -0.31773285 | 0 | 0.125 | 1 | 23 |
| washc5 | 0.12136 | -0.31805767 | 0 | 0.125 | 1 | 23 |
| xrcc3 | 0.12136 | -0.31594605 | 0 | 0.125 | 1 | 23 |
| zgc:194285 | 0.12136 | -0.3146231 | 0 | 0.125 | 1 | 23 |
| zgc:86764 | 0.12136 | -0.31662118 | 0 | 0.125 | 1 | 23 |
| ANKRD50 | 0.12136 | -0.26396064 | 0 | 0.125 | 1 | 23 |
| BX470229.1 | 0.12136 | -0.26434869 | 0 | 0.125 | 1 | 23 |
| CABZ01068362.1 | 0.12136 | -0.19684109 | 0 | 0.125 | 1 | 23 |
| CABZ01069162.1 | 0.12136 | -0.36058137 | 0 | 0.125 | 1 | 23 |
| CABZ01074453.1 | 0.12136 | -0.18733981 | 0 | 0.125 | 1 | 23 |
| CABZ01078767.1 | 0.12136 | -0.27701263 | 0 | 0.125 | 1 | 23 |
| CABZ01087629.1 | 0.12136 | -0.19354509 | 0 | 0.125 | 1 | 23 |
| CR388369.1 | 0.12136 | -0.26817843 | 0 | 0.125 | 1 | 23 |
| CR848667.1 | 0.12136 | -0.26453631 | 0 | 0.125 | 1 | 23 |
| CR936442.1 | 0.12136 | -0.2250177 | 0 | 0.125 | 1 | 23 |
| CT030017.2 | 0.12136 | -0.26431107 | 0 | 0.125 | 1 | 23 |
| CU062501.1 | 0.12136 | -0.34804249 | 0 | 0.125 | 1 | 23 |
| CU694481.1 | 0.12136 | -0.26432957 | 0 | 0.125 | 1 | 23 |
| CU914487.1 | 0.12136 | -0.25598266 | 0 | 0.125 | 1 | 23 |
| FP236197.1 | 0.12136 | -0.18852519 | 0 | 0.125 | 1 | 23 |
| LO018183.1 | 0.12136 | -0.27197196 | 0 | 0.125 | 1 | 23 |
| LOC100537147 | 0.12136 | -0.26192981 | 0 | 0.125 | 1 | 23 |
| LOC101884919 | 0.12136 | -0.26711849 | 0 | 0.125 | 1 | 23 |
| LOC101885430 | 0.12136 | -0.26571271 | 0 | 0.125 | 1 | 23 |
| LOC101885938 | 0.12136 | -0.25660904 | 0 | 0.125 | 1 | 23 |
| LOC101886015 | 0.12136 | -0.26277258 | 0 | 0.125 | 1 | 23 |
| LOC107195253 | 0.12136 | -0.25675059 | 0 | 0.125 | 1 | 23 |
| LOC110438040 | 0.12136 | -0.2638994 | 0 | 0.125 | 1 | 23 |

| | | | | | | |
|---|---|---|---|---|---|---|
| LOC110439111 | 0.12136 | -0.2424177 | 0 | 0.125 | 1 | 23 |
| USP34 | 0.12136 | -0.26267492 | 0 | 0.125 | 1 | 23 |
| VASH1 | 0.12136 | -0.26571271 | 0 | 0.125 | 1 | 23 |
| XLOC-004843 | 0.12136 | -0.2610995 | 0 | 0.125 | 1 | 23 |
| XLOC-007665 | 0.12136 | -0.19057029 | 0 | 0.125 | 1 | 23 |
| XLOC-014378 | 0.12136 | -0.27701263 | 0 | 0.125 | 1 | 23 |
| XLOC-018061 | 0.12136 | -0.25038348 | 0 | 0.125 | 1 | 23 |
| XLOC-023916 | 0.12136 | -0.27701263 | 0 | 0.125 | 1 | 23 |
| XLOC-025565 | 0.12136 | -0.27701263 | 0 | 0.125 | 1 | 23 |
| XLOC-025573 | 0.12136 | -0.27202678 | 0 | 0.125 | 1 | 23 |
| XLOC-031182 | 0.12136 | -0.27701263 | 0 | 0.125 | 1 | 23 |
| XLOC-033953 | 0.12136 | -0.26571271 | 0 | 0.125 | 1 | 23 |
| XLOC-043124 | 0.12136 | -0.27701263 | 0 | 0.125 | 1 | 23 |
| XLOC-044048 | 0.12136 | -0.14819955 | 0 | 0.125 | 1 | 23 |
| acap1 | 0.12136 | -0.26882126 | 0 | 0.125 | 1 | 23 |
| alx1 | 0.12136 | -0.21731719 | 0 | 0.125 | 1 | 23 |
| ankmy1 | 0.12136 | -0.25038348 | 0 | 0.125 | 1 | 23 |
| arf4a | 0.12136 | -0.18952008 | 0 | 0.125 | 1 | 23 |
| arhgef5 | 0.12136 | -0.26003935 | 0 | 0.125 | 1 | 23 |
| arid3a | 0.12136 | -0.25827569 | 0 | 0.125 | 1 | 23 |
| armc9 | 0.12136 | -0.21731719 | 0 | 0.125 | 1 | 23 |
| arntl1b | 0.12136 | -0.19507246 | 0 | 0.125 | 1 | 23 |
| atat1 | 0.12136 | -0.27701263 | 0 | 0.125 | 1 | 23 |
| atm | 0.12136 | -0.26440994 | 0 | 0.125 | 1 | 23 |
| atp6ap1b | 0.12136 | -0.19165728 | 0 | 0.125 | 1 | 23 |
| atp6v0a2b | 0.12136 | -0.26052864 | 0 | 0.125 | 1 | 23 |
| atp6v0cb | 0.12136 | -0.17908531 | 0 | 0.125 | 1 | 23 |
| bcor | 0.12136 | -0.27214057 | 0 | 0.125 | 1 | 23 |
| btr04 | 0.12136 | -0.18441281 | 0 | 0.125 | 1 | 23 |
| bzw2 | 0.12136 | -0.1870544 | 0 | 0.125 | 1 | 23 |
| c1qtnf12 | 0.12136 | -0.27581838 | 0 | 0.125 | 1 | 23 |
| c2cd2 | 0.12136 | -0.26262769 | 0 | 0.125 | 1 | 23 |
| ca10b | 0.12136 | -0.27701263 | 0 | 0.125 | 1 | 23 |
| cacnb1 | 0.12136 | -0.26114504 | 0 | 0.125 | 1 | 23 |
| ccni2 | 0.12136 | -0.26571271 | 0 | 0.125 | 1 | 23 |
| cdc42ep2 | 0.12136 | -0.26120986 | 0 | 0.125 | 1 | 23 |
| cdk15 | 0.12136 | -0.23483429 | 0 | 0.125 | 1 | 23 |
| chordc1a | 0.12136 | -0.26359127 | 0 | 0.125 | 1 | 23 |
| clpxb | 0.12136 | -0.14126511 | 0 | 0.125 | 1 | 23 |
| crocc2 | 0.12136 | -0.27701263 | 0 | 0.125 | 1 | 23 |
| ctdp1 | 0.12136 | -0.27469101 | 0 | 0.125 | 1 | 23 |
| ctdspl3 | 0.12136 | -0.18693436 | 0 | 0.125 | 1 | 23 |
| dcaf15 | 0.12136 | -0.26219925 | 0 | 0.125 | 1 | 23 |
| ddr2l | 0.12136 | -0.27536606 | 0 | 0.125 | 1 | 23 |
| ddx39b | 0.12136 | -0.13357893 | 0 | 0.125 | 1 | 23 |
| def6c | 0.12136 | -0.26749187 | 0 | 0.125 | 1 | 23 |
| dhrs11b | 0.12136 | -0.12243507 | 0 | 0.125 | 1 | 23 |
| diabloa | 0.12136 | -0.18627559 | 0 | 0.125 | 1 | 23 |

| | | | | | | |
|---|---|---|---|---|---|---|
| disp1 | 0.12136 | -0.2636374 | 0 | 0.125 | 1 | 23 |
| dnah1 | 0.12136 | -0.26377731 | 0 | 0.125 | 1 | 23 |
| dpep2 | 0.12136 | -0.24671259 | 0 | 0.125 | 1 | 23 |
| dyrk3 | 0.12136 | -0.26321952 | 0 | 0.125 | 1 | 23 |
| egr3 | 0.12136 | -0.24946653 | 0 | 0.125 | 1 | 23 |
| ehmt2 | 0.12136 | -0.19125449 | 0 | 0.125 | 1 | 23 |
| etv5b | 0.12136 | -0.2622186 | 0 | 0.125 | 1 | 23 |
| exoc2 | 0.12136 | -0.17609887 | 0 | 0.125 | 1 | 23 |
| f11r.2 | 0.12136 | -0.19078688 | 0 | 0.125 | 1 | 23 |
| fam129ba | 0.12136 | -0.26929005 | 0 | 0.125 | 1 | 23 |
| foxred2 | 0.12136 | -0.26389392 | 0 | 0.125 | 1 | 23 |
| galca | 0.12136 | -0.26935594 | 0 | 0.125 | 1 | 23 |
| ggcx | 0.12136 | -0.26365725 | 0 | 0.125 | 1 | 23 |
| gmeb1 | 0.12136 | -0.19348639 | 0 | 0.125 | 1 | 23 |
| gpam | 0.12136 | -0.27007073 | 0 | 0.125 | 1 | 23 |
| gpc2 | 0.12136 | -0.26331042 | 0 | 0.125 | 1 | 23 |
| gpr146 | 0.12136 | -0.27533481 | 0 | 0.125 | 1 | 23 |
| hic1l | 0.12136 | -0.26161115 | 0 | 0.125 | 1 | 23 |
| hivep1 | 0.12136 | -0.3498205 | 0 | 0.125 | 1 | 23 |
| hivep2a | 0.12136 | -0.19351889 | 0 | 0.125 | 1 | 23 |
| hoxd3a | 0.12136 | -0.27701263 | 0 | 0.125 | 1 | 23 |
| hpn | 0.12136 | -0.19295927 | 0 | 0.125 | 1 | 23 |
| hsd11b1la | 0.12136 | -0.19500049 | 0 | 0.125 | 1 | 23 |
| ift80 | 0.12136 | -0.19266971 | 0 | 0.125 | 1 | 23 |
| ift81 | 0.12136 | -0.18883614 | 0 | 0.125 | 1 | 23 |
| ippk | 0.12136 | -0.35005686 | 0 | 0.125 | 1 | 23 |
| jcada | 0.12136 | -0.19350727 | 0 | 0.125 | 1 | 23 |
| kbtbd8 | 0.12136 | -0.26354998 | 0 | 0.125 | 1 | 23 |
| kcnq5a | 0.12136 | -0.26571271 | 0 | 0.125 | 1 | 23 |
| kmt2d | 0.12136 | -0.25854519 | 0 | 0.125 | 1 | 23 |
| l3mbtl1b | 0.12136 | -0.26279766 | 0 | 0.125 | 1 | 23 |
| lig4 | 0.12136 | -0.26066972 | 0 | 0.125 | 1 | 23 |
| lrrc45 | 0.12136 | -0.19199308 | 0 | 0.125 | 1 | 23 |
| lrrc6 | 0.12136 | -0.26241382 | 0 | 0.125 | 1 | 23 |
| maip1 | 0.12136 | -0.25940007 | 0 | 0.125 | 1 | 23 |
| map3k20 | 0.12136 | -0.26771516 | 0 | 0.125 | 1 | 23 |
| mark3b | 0.12136 | -0.26223693 | 0 | 0.125 | 1 | 23 |
| mblac2 | 0.12136 | -0.16989376 | 0 | 0.125 | 1 | 23 |
| mir301b | 0.12136 | -0.26571271 | 0 | 0.125 | 1 | 23 |
| mmp23bb | 0.12136 | -0.26219443 | 0 | 0.125 | 1 | 23 |
| mpp1 | 0.12136 | -0.18696671 | 0 | 0.125 | 1 | 23 |
| mpp5a | 0.12136 | -0.26502317 | 0 | 0.125 | 1 | 23 |
| mrtfbb | 0.12136 | -0.26410955 | 0 | 0.125 | 1 | 23 |
| msh3 | 0.12136 | -0.2737485 | 0 | 0.125 | 1 | 23 |
| mtpap | 0.12136 | -0.26337643 | 0 | 0.125 | 1 | 23 |
| mtrf1 | 0.12136 | -0.17933815 | 0 | 0.125 | 1 | 23 |
| mxb | 0.12136 | -0.25722797 | 0 | 0.125 | 1 | 23 |
| mylz3 | 0.12136 | -0.26866947 | 0 | 0.125 | 1 | 23 |

| | | | | | | |
|---|---|---|---|---|---|---|
| naa25 | 0.12136 | -0.26312179 | 0 | 0.125 | 1 | 23 |
| nbeal1 | 0.12136 | -0.26506179 | 0 | 0.125 | 1 | 23 |
| ncoa1 | 0.12136 | -0.26434869 | 0 | 0.125 | 1 | 23 |
| ncoa3 | 0.12136 | -0.26505604 | 0 | 0.125 | 1 | 23 |
| nell2a | 0.12136 | -0.27385827 | 0 | 0.125 | 1 | 23 |
| nf1a | 0.12136 | -0.27094119 | 0 | 0.125 | 1 | 23 |
| nlk2 | 0.12136 | -0.26274211 | 0 | 0.125 | 1 | 23 |
| nlrc3l1 | 0.12136 | -0.1950318 | 0 | 0.125 | 1 | 23 |
| npffr2b | 0.12136 | -0.27701263 | 0 | 0.125 | 1 | 23 |
| nrbp2a | 0.12136 | -0.26262043 | 0 | 0.125 | 1 | 23 |
| pacs2 | 0.12136 | -0.26262246 | 0 | 0.125 | 1 | 23 |
| palm2 | 0.12136 | -0.27260783 | 0 | 0.125 | 1 | 23 |
| pcnx3 | 0.12136 | -0.274037 | 0 | 0.125 | 1 | 23 |
| pcp4l1 | 0.12136 | -0.20592777 | 0 | 0.125 | 1 | 23 |
| pde12 | 0.12136 | -0.18656377 | 0 | 0.125 | 1 | 23 |
| pdk3a | 0.12136 | -0.23483429 | 0 | 0.125 | 1 | 23 |
| pex7 | 0.12136 | -0.26096917 | 0 | 0.125 | 1 | 23 |
| phlda3 | 0.12136 | -0.27479181 | 0 | 0.125 | 1 | 23 |
| pi4k2b | 0.12136 | -0.18105425 | 0 | 0.125 | 1 | 23 |
| pink1 | 0.12136 | -0.1952604 | 0 | 0.125 | 1 | 23 |
| pld6 | 0.12136 | -0.1142075 | 0 | 0.125 | 1 | 23 |
| pold1 | 0.12136 | -0.26057608 | 0 | 0.125 | 1 | 23 |
| ppm1ab | 0.12136 | -0.19017927 | 0 | 0.125 | 1 | 23 |
| ppp1r13l | 0.12136 | -0.27473475 | 0 | 0.125 | 1 | 23 |
| prcc | 0.12136 | -0.27490487 | 0 | 0.125 | 1 | 23 |
| ptgs2a | 0.12136 | -0.27602058 | 0 | 0.125 | 1 | 23 |
| pycr1a | 0.12136 | -0.27701263 | 0 | 0.125 | 1 | 23 |
| pyroxd1 | 0.12136 | -0.2602799 | 0 | 0.125 | 1 | 23 |
| rab12 | 0.12136 | -0.18099297 | 0 | 0.125 | 1 | 23 |
| raf1a | 0.12136 | -0.19071814 | 0 | 0.125 | 1 | 23 |
| rbbp8l | 0.12136 | -0.24867493 | 0 | 0.125 | 1 | 23 |
| rbck1 | 0.12136 | -0.19237132 | 0 | 0.125 | 1 | 23 |
| rgp1 | 0.12136 | -0.2615588 | 0 | 0.125 | 1 | 23 |
| rhot1a | 0.12136 | -0.19131745 | 0 | 0.125 | 1 | 23 |
| rnf165a | 0.12136 | -0.21731719 | 0 | 0.125 | 1 | 23 |
| rnf165b | 0.12136 | -0.2643185 | 0 | 0.125 | 1 | 23 |
| rps6ka4 | 0.12136 | -0.26399152 | 0 | 0.125 | 1 | 23 |
| rspo3 | 0.12136 | -0.19549702 | 0 | 0.125 | 1 | 23 |
| s100w | 0.12136 | -0.26649589 | 0 | 0.125 | 1 | 23 |
| samd4b | 0.12136 | -0.26810672 | 0 | 0.125 | 1 | 23 |
| sardh | 0.12136 | -0.25429479 | 0 | 0.125 | 1 | 23 |
| scaf4b | 0.12136 | -0.26891408 | 0 | 0.125 | 1 | 23 |
| scai | 0.12136 | -0.26490128 | 0 | 0.125 | 1 | 23 |
| scn4ba | 0.12136 | -0.18777112 | 0 | 0.125 | 1 | 23 |
| sdhda | 0.12136 | -0.18875201 | 0 | 0.125 | 1 | 23 |
| sema3d | 0.12136 | -0.36245079 | 0 | 0.125 | 1 | 23 |
| serpine3 | 0.12136 | -0.26516245 | 0 | 0.125 | 1 | 23 |
| sestd1 | 0.12136 | -0.26313023 | 0 | 0.125 | 1 | 23 |

| | | | | | | |
|---|---|---|---|---|---|---|
| sgip1a | 0.12136 | -0.26402328 | 0 | 0.125 | 1 | 23 |
| shmt2 | 0.12136 | -0.18638755 | 0 | 0.125 | 1 | 23 |
| shrprbck1r | 0.12136 | -0.26169568 | 0 | 0.125 | 1 | 23 |
| si:cabz01063885.1 | 0.12136 | -0.26462236 | 0 | 0.125 | 1 | 23 |
| si:ch211-105j21.9 | 0.12136 | -0.25825426 | 0 | 0.125 | 1 | 23 |
| si:ch211-148l7.4 | 0.12136 | -0.2628523 | 0 | 0.125 | 1 | 23 |
| si:ch211-181d7.3 | 0.12136 | -0.26442495 | 0 | 0.125 | 1 | 23 |
| si:ch211-188c16.1 | 0.12136 | -0.19304759 | 0 | 0.125 | 1 | 23 |
| si:ch211-1i11.3 | 0.12136 | -0.27701263 | 0 | 0.125 | 1 | 23 |
| si:ch211-227p7.1 | 0.12136 | -0.25869638 | 0 | 0.125 | 1 | 23 |
| si:ch211-262i1.3 | 0.12136 | -0.34285233 | 0 | 0.125 | 1 | 23 |
| si:ch211-264f5.6 | 0.12136 | -0.19163059 | 0 | 0.125 | 1 | 23 |
| si:ch211-66k16.28 | 0.12136 | -0.27406716 | 0 | 0.125 | 1 | 23 |
| si:ch73-233k15.2 | 0.12136 | -0.18857244 | 0 | 0.125 | 1 | 23 |
| si:dkey-111k8.3 | 0.12136 | -0.19246595 | 0 | 0.125 | 1 | 23 |
| si:dkey-14d8.1 | 0.12136 | -0.19067564 | 0 | 0.125 | 1 | 23 |
| si:dkey-174i8.1 | 0.12136 | -0.27701263 | 0 | 0.125 | 1 | 23 |
| si:dkey-184a18.5 | 0.12136 | -0.18849744 | 0 | 0.125 | 1 | 23 |
| si:dkey-204l11.1 | 0.12136 | -0.26388883 | 0 | 0.125 | 1 | 23 |
| si:dkey-261e22.4 | 0.12136 | -0.25051151 | 0 | 0.125 | 1 | 23 |
| si:dkey-29p23.1 | 0.12136 | -0.27701263 | 0 | 0.125 | 1 | 23 |
| si:dkey-66i24.8 | 0.12136 | -0.18859317 | 0 | 0.125 | 1 | 23 |
| si:dkey-85n7.6 | 0.12136 | -0.26525372 | 0 | 0.125 | 1 | 23 |
| si:dkey-94l16.4 | 0.12136 | -0.19684109 | 0 | 0.125 | 1 | 23 |
| si:dkeyp-53d3.5 | 0.12136 | -0.27701263 | 0 | 0.125 | 1 | 23 |
| si:dkeyp-9d4.3 | 0.12136 | -0.26844359 | 0 | 0.125 | 1 | 23 |
| si:zfos-1011f11.2 | 0.12136 | -0.27640368 | 0 | 0.125 | 1 | 23 |
| si:zfos-452g4.1 | 0.12136 | -0.2637975 | 0 | 0.125 | 1 | 23 |
| slc20a1b | 0.12136 | -0.26150904 | 0 | 0.125 | 1 | 23 |
| slc5a8l | 0.12136 | -0.26279872 | 0 | 0.125 | 1 | 23 |
| slc6a16b | 0.12136 | -0.26313667 | 0 | 0.125 | 1 | 23 |
| snu13a | 0.12136 | -0.25852171 | 0 | 0.125 | 1 | 23 |
| snx30 | 0.12136 | -0.18527447 | 0 | 0.125 | 1 | 23 |
| soul5 | 0.12136 | -0.27657085 | 0 | 0.125 | 1 | 23 |
| sox9a | 0.12136 | -0.19433902 | 0 | 0.125 | 1 | 23 |
| sox9b | 0.12136 | -0.26571271 | 0 | 0.125 | 1 | 23 |
| sp1 | 0.12136 | -0.26011469 | 0 | 0.125 | 1 | 23 |
| spata6 | 0.12136 | -0.26499772 | 0 | 0.125 | 1 | 23 |
| spidr | 0.12136 | -0.19082868 | 0 | 0.125 | 1 | 23 |
| spns2 | 0.12136 | -0.19684109 | 0 | 0.125 | 1 | 23 |
| st3gal3b | 0.12136 | -0.27413078 | 0 | 0.125 | 1 | 23 |
| stard3 | 0.12136 | -0.2702155 | 0 | 0.125 | 1 | 23 |
| stard9 | 0.12136 | -0.19563241 | 0 | 0.125 | 1 | 23 |
| supt7l | 0.12136 | -0.18157252 | 0 | 0.125 | 1 | 23 |
| syndig1l | 0.12136 | -0.26571271 | 0 | 0.125 | 1 | 23 |
| syne1b | 0.12136 | -0.25760893 | 0 | 0.125 | 1 | 23 |
| synj1 | 0.12136 | -0.19538707 | 0 | 0.125 | 1 | 23 |
| tada2b | 0.12136 | -0.19498333 | 0 | 0.125 | 1 | 23 |

| | | | | | | |
|---|---|---|---|---|---|---|
| tagapa | 0.12136 | -0.21327674 | 0 | 0.125 | 1 | 23 |
| tbc1d24 | 0.12136 | -0.26361539 | 0 | 0.125 | 1 | 23 |
| tbc1d2b | 0.12136 | -0.26355064 | 0 | 0.125 | 1 | 23 |
| tet1 | 0.12136 | -0.19381686 | 0 | 0.125 | 1 | 23 |
| tgfbr1a | 0.12136 | -0.2707259 | 0 | 0.125 | 1 | 23 |
| tgfbrap1 | 0.12136 | -0.2639172 | 0 | 0.125 | 1 | 23 |
| thraa | 0.12136 | -0.27185115 | 0 | 0.125 | 1 | 23 |
| tjap1 | 0.12136 | -0.19242089 | 0 | 0.125 | 1 | 23 |
| tlk2 | 0.12136 | -0.18911285 | 0 | 0.125 | 1 | 23 |
| tmem102 | 0.12136 | -0.25670113 | 0 | 0.125 | 1 | 23 |
| tmem60 | 0.12136 | -0.25818152 | 0 | 0.125 | 1 | 23 |
| tnfrsf9a | 0.12136 | -0.34163598 | 0 | 0.125 | 1 | 23 |
| tnks1bp1 | 0.12136 | -0.18101599 | 0 | 0.125 | 1 | 23 |
| tnksa | 0.12136 | -0.26724225 | 0 | 0.125 | 1 | 23 |
| tor2a | 0.12136 | -0.27224807 | 0 | 0.125 | 1 | 23 |
| tpx2 | 0.12136 | -0.26183308 | 0 | 0.125 | 1 | 23 |
| trpv4 | 0.12136 | -0.27434255 | 0 | 0.125 | 1 | 23 |
| ttc38 | 0.12136 | -0.27334661 | 0 | 0.125 | 1 | 23 |
| ttll9 | 0.12136 | -0.21731719 | 0 | 0.125 | 1 | 23 |
| tyro3 | 0.12136 | -0.26434869 | 0 | 0.125 | 1 | 23 |
| uacaa | 0.12136 | -0.27429908 | 0 | 0.125 | 1 | 23 |
| ube2j2 | 0.12136 | -0.27002544 | 0 | 0.125 | 1 | 23 |
| ugt5e1 | 0.12136 | -0.19077014 | 0 | 0.125 | 1 | 23 |
| urb1 | 0.12136 | -0.19488843 | 0 | 0.125 | 1 | 23 |
| usp54a | 0.12136 | -0.19033748 | 0 | 0.125 | 1 | 23 |
| uxs1 | 0.12136 | -0.19314989 | 0 | 0.125 | 1 | 23 |
| vgll4a | 0.12136 | -0.27096289 | 0 | 0.125 | 1 | 23 |
| washc2c | 0.12136 | -0.25812318 | 0 | 0.125 | 1 | 23 |
| wnt11r | 0.12136 | -0.27507057 | 0 | 0.125 | 1 | 23 |
| wrap73 | 0.12136 | -0.1873433 | 0 | 0.125 | 1 | 23 |
| zbtb24 | 0.12136 | -0.26571271 | 0 | 0.125 | 1 | 23 |
| zc3h3 | 0.12136 | -0.26403207 | 0 | 0.125 | 1 | 23 |
| zgc:113090 | 0.12136 | -0.27701263 | 0 | 0.125 | 1 | 23 |
| zgc:162344 | 0.12136 | -0.25060257 | 0 | 0.125 | 1 | 23 |
| zgc:173486 | 0.12136 | -0.19311957 | 0 | 0.125 | 1 | 23 |
| znf1174 | 0.12136 | -0.17352563 | 0 | 0.125 | 1 | 23 |
| znf362a | 0.12136 | -0.19581344 | 0 | 0.125 | 1 | 23 |
| znf513a | 0.12136 | -0.1941332 | 0 | 0.125 | 1 | 23 |
| zranb3 | 0.12136 | -0.26113189 | 0 | 0.125 | 1 | 23 |
| BX511061.1 | 0.12136 | -0.36074943 | 0 | 0.125 | 1 | 23 |
| BX530721.1 | 0.12136 | -0.15233076 | 0 | 0.125 | 1 | 23 |
| BX569798.1 | 0.12136 | -0.1578638 | 0 | 0.125 | 1 | 23 |
| BX649500.1 | 0.12136 | -0.35533198 | 0 | 0.125 | 1 | 23 |
| BX901883.3 | 0.12136 | -0.18705238 | 0 | 0.125 | 1 | 23 |
| BX901962.2 | 0.12136 | -0.17973008 | 0 | 0.125 | 1 | 23 |
| CABZ01057496.1 | 0.12136 | -0.18705238 | 0 | 0.125 | 1 | 23 |
| CABZ01064670.1 | 0.12136 | -0.15883309 | 0 | 0.125 | 1 | 23 |
| CABZ01066720.1 | 0.12136 | -0.36048338 | 0 | 0.125 | 1 | 23 |

| | | | | | | |
|---|---|---|---|---|---|---|
| CABZ01080371.2 | 0.12136 | -0.18281624 | 0 | 0.125 | 1 | 23 |
| CABZ01085201.1 | 0.12136 | -0.36090771 | 0 | 0.125 | 1 | 23 |
| CABZ01113812.1 | 0.12136 | -0.1552875 | 0 | 0.125 | 1 | 23 |
| CR382370.2 | 0.12136 | -0.18295907 | 0 | 0.125 | 1 | 23 |
| CR753876.1 | 0.12136 | -0.13269073 | 0 | 0.125 | 1 | 23 |
| CR847936.1 | 0.12136 | -0.14804502 | 0 | 0.125 | 1 | 23 |
| CT573793.2 | 0.12136 | -0.14899391 | 0 | 0.125 | 1 | 23 |
| FO704721.1 | 0.12136 | -0.15925638 | 0 | 0.125 | 1 | 23 |
| FO907087.1 | 0.12136 | -0.18410557 | 0 | 0.125 | 1 | 23 |
| FP243383.1 | 0.12136 | -0.18467721 | 0 | 0.125 | 1 | 23 |
| JMY | 0.12136 | -0.1641204 | 0 | 0.125 | 1 | 23 |
| LO018608.1 | 0.12136 | -0.16486193 | 0 | 0.125 | 1 | 23 |
| LOC101882607 | 0.12136 | -0.16486193 | 0 | 0.125 | 1 | 23 |
| LOC101884549 | 0.12136 | -0.36142513 | 0 | 0.125 | 1 | 23 |
| LOC101884606 | 0.12136 | -0.1576778 | 0 | 0.125 | 1 | 23 |
| LOC101884686 | 0.12136 | -0.16113133 | 0 | 0.125 | 1 | 23 |
| LOC101886860 | 0.12136 | -0.11616559 | 0 | 0.125 | 1 | 23 |
| LOC101886929 | 0.12136 | -0.18705238 | 0 | 0.125 | 1 | 23 |
| LOC101887070 | 0.12136 | -0.1812101 | 0 | 0.125 | 1 | 23 |
| LOC103908666 | 0.12136 | -0.16385728 | 0 | 0.125 | 1 | 23 |
| LOC103911563 | 0.12136 | -0.23464031 | 0 | 0.125 | 1 | 23 |
| LOC108180286 | 0.12136 | -0.18514748 | 0 | 0.125 | 1 | 23 |
| LOC110437877 | 0.12136 | -0.35676712 | 0 | 0.125 | 1 | 23 |
| XLOC-008367 | 0.12136 | -0.26604787 | 0 | 0.125 | 1 | 23 |
| XLOC-010289 | 0.12136 | -0.15754441 | 0 | 0.125 | 1 | 23 |
| XLOC-019150 | 0.12136 | -0.1423352 | 0 | 0.125 | 1 | 23 |
| XLOC-029426 | 0.12136 | -0.1841805 | 0 | 0.125 | 1 | 23 |
| ZNF512B | 0.12136 | -0.15867758 | 0 | 0.125 | 1 | 23 |
| abcc3 | 0.12136 | -0.3616707 | 0 | 0.125 | 1 | 23 |
| abhd18 | 0.12136 | -0.18196342 | 0 | 0.125 | 1 | 23 |
| adam19a | 0.12136 | -0.18384543 | 0 | 0.125 | 1 | 23 |
| adamts12 | 0.12136 | -0.16232542 | 0 | 0.125 | 1 | 23 |
| adh8a | 0.12136 | -0.11484615 | 0 | 0.125 | 1 | 23 |
| aggf1 | 0.12136 | -0.360891 | 0 | 0.125 | 1 | 23 |
| ap5m1 | 0.12136 | -0.15071424 | 0 | 0.125 | 1 | 23 |
| apaf1 | 0.12136 | -0.16188019 | 0 | 0.125 | 1 | 23 |
| apln | 0.12136 | -0.17341345 | 0 | 0.125 | 1 | 23 |
| apool | 0.12136 | -0.18171994 | 0 | 0.125 | 1 | 23 |
| arhgap21a | 0.12136 | -0.15580876 | 0 | 0.125 | 1 | 23 |
| arhgef10la | 0.12136 | -0.16043415 | 0 | 0.125 | 1 | 23 |
| arl6ip5a | 0.12136 | -0.18275645 | 0 | 0.125 | 1 | 23 |
| armc1 | 0.12136 | -0.15166488 | 0 | 0.125 | 1 | 23 |
| armc10 | 0.12136 | -0.1833191 | 0 | 0.125 | 1 | 23 |
| armc1l | 0.12136 | -0.16977938 | 0 | 0.125 | 1 | 23 |
| arrdc3b | 0.12136 | -0.17717284 | 0 | 0.125 | 1 | 23 |
| ash2l | 0.12136 | -0.15683988 | 0 | 0.125 | 1 | 23 |
| bahd1 | 0.12136 | -0.16345503 | 0 | 0.125 | 1 | 23 |
| bbs2 | 0.12136 | -0.18276985 | 0 | 0.125 | 1 | 23 |

| | | | | | | |
|---|---|---|---|---|---|---|
| bbs7 | 0.12136 | -0.36171066 | 0 | 0.125 | 1 | 23 |
| bcl2a | 0.12136 | -0.16206415 | 0 | 0.125 | 1 | 23 |
| bcl6aa | 0.12136 | -0.18075473 | 0 | 0.125 | 1 | 23 |
| c1galt1b | 0.12136 | -0.15916164 | 0 | 0.125 | 1 | 23 |
| ccdc186 | 0.12136 | -0.1583101 | 0 | 0.125 | 1 | 23 |
| ccdc65 | 0.12136 | -0.36164156 | 0 | 0.125 | 1 | 23 |
| cfl2 | 0.12136 | -0.17140697 | 0 | 0.125 | 1 | 23 |
| cmtr2 | 0.12136 | -0.18143973 | 0 | 0.125 | 1 | 23 |
| cnot10 | 0.12136 | -0.15695191 | 0 | 0.125 | 1 | 23 |
| coq7 | 0.12136 | -0.1474219 | 0 | 0.125 | 1 | 23 |
| cox4i2 | 0.12136 | -0.14430943 | 0 | 0.125 | 1 | 23 |
| crb3a | 0.12136 | -0.1613081 | 0 | 0.125 | 1 | 23 |
| creld1a | 0.12136 | -0.16486193 | 0 | 0.125 | 1 | 23 |
| cxadr | 0.12136 | -0.17617101 | 0 | 0.125 | 1 | 23 |
| cyb5r2 | 0.12136 | -0.14669331 | 0 | 0.125 | 1 | 23 |
| dcp1a | 0.12136 | -0.16293212 | 0 | 0.125 | 1 | 23 |
| deptor | 0.12136 | -0.15573053 | 0 | 0.125 | 1 | 23 |
| dguok | 0.12136 | -0.17040926 | 0 | 0.125 | 1 | 23 |
| dhodh | 0.12136 | -0.17999497 | 0 | 0.125 | 1 | 23 |
| dis3 | 0.12136 | -0.26321103 | 0 | 0.125 | 1 | 23 |
| dlg5b.1 | 0.12136 | -0.16020878 | 0 | 0.125 | 1 | 23 |
| dusp19b | 0.12136 | -0.15932197 | 0 | 0.125 | 1 | 23 |
| edc3 | 0.12136 | -0.15840819 | 0 | 0.125 | 1 | 23 |
| eef1a1a | 0.12136 | -0.15713662 | 0 | 0.125 | 1 | 23 |
| ei24 | 0.12136 | -0.18427889 | 0 | 0.125 | 1 | 23 |
| enc3 | 0.12136 | -0.15761409 | 0 | 0.125 | 1 | 23 |
| eral1 | 0.12136 | -0.1522059 | 0 | 0.125 | 1 | 23 |
| ercc8 | 0.12136 | -0.17939375 | 0 | 0.125 | 1 | 23 |
| extl2 | 0.12136 | -0.15927137 | 0 | 0.125 | 1 | 23 |
| fam174b | 0.12136 | -0.16858135 | 0 | 0.125 | 1 | 23 |
| fam8a1b | 0.12136 | -0.15248034 | 0 | 0.125 | 1 | 23 |
| fbxl4 | 0.12136 | -0.13625186 | 0 | 0.125 | 1 | 23 |
| galnt2 | 0.12136 | -0.16323982 | 0 | 0.125 | 1 | 23 |
| gatd1 | 0.12136 | -0.1596283 | 0 | 0.125 | 1 | 23 |
| ggact.2 | 0.12136 | -0.17075938 | 0 | 0.125 | 1 | 23 |
| ggps1 | 0.12136 | -0.16171107 | 0 | 0.125 | 1 | 23 |
| gna12a | 0.12136 | -0.26411438 | 0 | 0.125 | 1 | 23 |
| gnb4b | 0.12136 | -0.36082413 | 0 | 0.125 | 1 | 23 |
| gpbar1 | 0.12136 | -0.18364653 | 0 | 0.125 | 1 | 23 |
| gpd1l | 0.12136 | -0.15847314 | 0 | 0.125 | 1 | 23 |
| gpr137ba | 0.12136 | -0.18381231 | 0 | 0.125 | 1 | 23 |
| gpr161 | 0.12136 | -0.36253987 | 0 | 0.125 | 1 | 23 |
| gtf3c2 | 0.12136 | -0.17952416 | 0 | 0.125 | 1 | 23 |
| gtpbp1l | 0.12136 | -0.22673463 | 0 | 0.125 | 1 | 23 |
| gtpbp2b | 0.12136 | -0.18399385 | 0 | 0.125 | 1 | 23 |
| hagh | 0.12136 | -0.34888913 | 0 | 0.125 | 1 | 23 |
| hdac6 | 0.12136 | -0.17700965 | 0 | 0.125 | 1 | 23 |
| helz2 | 0.12136 | -0.16486193 | 0 | 0.125 | 1 | 23 |

| | | | | | | |
|---|---|---|---|---|---|---|
| hmbsa | 0.12136 | -0.23150653 | 0 | 0.125 | 1 | 23 |
| hoxc5a | 0.12136 | -0.35730439 | 0 | 0.125 | 1 | 23 |
| hoxd9a | 0.12136 | -0.31389867 | 0 | 0.125 | 1 | 23 |
| hrasb | 0.12136 | -0.15489575 | 0 | 0.125 | 1 | 23 |
| irak1bp1 | 0.12136 | -0.18073429 | 0 | 0.125 | 1 | 23 |
| itcha | 0.12136 | -0.18237191 | 0 | 0.125 | 1 | 23 |
| itgb1b | 0.12136 | -0.16486193 | 0 | 0.125 | 1 | 23 |
| itprip | 0.12136 | -0.18348695 | 0 | 0.125 | 1 | 23 |
| kank4 | 0.12136 | -0.15508721 | 0 | 0.125 | 1 | 23 |
| kansl3 | 0.12136 | -0.18397099 | 0 | 0.125 | 1 | 23 |
| katnbl1 | 0.12136 | -0.15767769 | 0 | 0.125 | 1 | 23 |
| klhl24a | 0.12136 | -0.15088773 | 0 | 0.125 | 1 | 23 |
| l2hgdh | 0.12136 | -0.18240795 | 0 | 0.125 | 1 | 23 |
| leng1 | 0.12136 | -0.15621795 | 0 | 0.125 | 1 | 23 |
| lhfpl6 | 0.12136 | -0.1768049 | 0 | 0.125 | 1 | 23 |
| lrrc3ca | 0.12136 | -0.18705238 | 0 | 0.125 | 1 | 23 |
| mad1l1 | 0.12136 | -0.17673422 | 0 | 0.125 | 1 | 23 |
| med24 | 0.12136 | -0.16071806 | 0 | 0.125 | 1 | 23 |
| mhc1zfa | 0.12136 | -0.18705238 | 0 | 0.125 | 1 | 23 |
| mindy2 | 0.12136 | -0.16486193 | 0 | 0.125 | 1 | 23 |
| mon1bb | 0.12136 | -0.18289919 | 0 | 0.125 | 1 | 23 |
| ms4a17a.8 | 0.12136 | -0.26428337 | 0 | 0.125 | 1 | 23 |
| msi1 | 0.12136 | -0.15215989 | 0 | 0.125 | 1 | 23 |
| msl1a | 0.12136 | -0.15530019 | 0 | 0.125 | 1 | 23 |
| mul1a | 0.12136 | -0.15455381 | 0 | 0.125 | 1 | 23 |
| myo18ab | 0.12136 | -0.18112713 | 0 | 0.125 | 1 | 23 |
| myorg | 0.12136 | -0.18331662 | 0 | 0.125 | 1 | 23 |
| naglu | 0.12136 | -0.16037603 | 0 | 0.125 | 1 | 23 |
| nckap5l | 0.12136 | -0.18479833 | 0 | 0.125 | 1 | 23 |
| ncoa2 | 0.12136 | -0.16157175 | 0 | 0.125 | 1 | 23 |
| nktr | 0.12136 | -0.18275135 | 0 | 0.125 | 1 | 23 |
| nme5 | 0.12136 | -0.15803024 | 0 | 0.125 | 1 | 23 |
| nme8 | 0.12136 | -0.16486193 | 0 | 0.125 | 1 | 23 |
| nufip1 | 0.12136 | -0.18216042 | 0 | 0.125 | 1 | 23 |
| odf2b | 0.12136 | -0.18574073 | 0 | 0.125 | 1 | 23 |
| paf1 | 0.12136 | -0.23591155 | 0 | 0.125 | 1 | 23 |
| pamr1 | 0.12136 | -0.16486193 | 0 | 0.125 | 1 | 23 |
| pank4 | 0.12136 | -0.17801016 | 0 | 0.125 | 1 | 23 |
| pappaa | 0.12136 | -0.18705238 | 0 | 0.125 | 1 | 23 |
| pde4d | 0.12136 | -0.23407873 | 0 | 0.125 | 1 | 23 |
| pgam1b | 0.12136 | -0.16094071 | 0 | 0.125 | 1 | 23 |
| pigl | 0.12136 | -0.18406111 | 0 | 0.125 | 1 | 23 |
| pign | 0.12136 | -0.14029944 | 0 | 0.125 | 1 | 23 |
| pitx2 | 0.12136 | -0.17867939 | 0 | 0.125 | 1 | 23 |
| pknox1.2 | 0.12136 | -0.16136096 | 0 | 0.125 | 1 | 23 |
| pld3 | 0.12136 | -0.18705238 | 0 | 0.125 | 1 | 23 |
| pnkp | 0.12136 | -0.1737058 | 0 | 0.125 | 1 | 23 |
| polr3c | 0.12136 | -0.1863868 | 0 | 0.125 | 1 | 23 |

| | | | | | | |
|---|---|---|---|---|---|---|
| polrmt | 0.12136 | -0.18274978 | 0 | 0.125 | 1 | 23 |
| pou6f1 | 0.12136 | -0.18705238 | 0 | 0.125 | 1 | 23 |
| prkcda | 0.12136 | -0.13974084 | 0 | 0.125 | 1 | 23 |
| ptger3 | 0.12136 | -0.29193239 | 0 | 0.125 | 1 | 23 |
| pygb | 0.12136 | -0.15871802 | 0 | 0.125 | 1 | 23 |
| qrich1 | 0.12136 | -0.18254397 | 0 | 0.125 | 1 | 23 |
| rabgap1 | 0.12136 | -0.16140605 | 0 | 0.125 | 1 | 23 |
| raf1b | 0.12136 | -0.1792937 | 0 | 0.125 | 1 | 23 |
| rasl11b | 0.12136 | -0.18568531 | 0 | 0.125 | 1 | 23 |
| rassf8a | 0.12136 | -0.15693327 | 0 | 0.125 | 1 | 23 |
| rbks | 0.12136 | -0.15426475 | 0 | 0.125 | 1 | 23 |
| rictora | 0.12136 | -0.17793785 | 0 | 0.125 | 1 | 23 |
| rnf111 | 0.12136 | -0.36037744 | 0 | 0.125 | 1 | 23 |
| rnf121 | 0.12136 | -0.15887445 | 0 | 0.125 | 1 | 23 |
| rorb | 0.12136 | -0.15672701 | 0 | 0.125 | 1 | 23 |
| rpain | 0.12136 | -0.17408141 | 0 | 0.125 | 1 | 23 |
| rpia | 0.12136 | -0.13765957 | 0 | 0.125 | 1 | 23 |
| sar1aa | 0.12136 | -0.16243835 | 0 | 0.125 | 1 | 23 |
| sat2b | 0.12136 | -0.15909183 | 0 | 0.125 | 1 | 23 |
| saxo2 | 0.12136 | -0.36253987 | 0 | 0.125 | 1 | 23 |
| sb:cb81 | 0.12136 | -0.14547036 | 0 | 0.125 | 1 | 23 |
| sbds | 0.12136 | -0.15991429 | 0 | 0.125 | 1 | 23 |
| scp2a | 0.12136 | -0.14626769 | 0 | 0.125 | 1 | 23 |
| scyl1 | 0.12136 | -0.15220692 | 0 | 0.125 | 1 | 23 |
| si:busm1-105l16.2 | 0.12136 | -0.15868713 | 0 | 0.125 | 1 | 23 |
| si:ch211-12e13.12 | 0.12136 | -0.15837458 | 0 | 0.125 | 1 | 23 |
| si:ch211-161m3.6 | 0.12136 | -0.18705238 | 0 | 0.125 | 1 | 23 |
| si:ch211-167b20.8 | 0.12136 | -0.17705545 | 0 | 0.125 | 1 | 23 |
| si:ch211-191d2.2 | 0.12136 | -0.16144396 | 0 | 0.125 | 1 | 23 |
| si:ch211-256e16.11 | 0.12136 | -0.36202422 | 0 | 0.125 | 1 | 23 |
| si:ch211-262i1.5 | 0.12136 | -0.15823971 | 0 | 0.125 | 1 | 23 |
| si:ch211-91p5.3 | 0.12136 | -0.16148899 | 0 | 0.125 | 1 | 23 |
| si:ch73-14h10.2 | 0.12136 | -0.18515321 | 0 | 0.125 | 1 | 23 |
| si:ch73-252i11.1 | 0.12136 | -0.1802179 | 0 | 0.125 | 1 | 23 |
| si:dkey-10h3.2 | 0.12136 | -0.13387705 | 0 | 0.125 | 1 | 23 |
| si:dkey-172k15.5 | 0.12136 | -0.17255103 | 0 | 0.125 | 1 | 23 |
| si:dkey-187k19.2 | 0.12136 | -0.1604459 | 0 | 0.125 | 1 | 23 |
| si:dkey-19f4.2 | 0.12136 | -0.16486193 | 0 | 0.125 | 1 | 23 |
| si:dkey-204f11.64 | 0.12136 | -0.15335937 | 0 | 0.125 | 1 | 23 |
| si:dkey-219c3.2 | 0.12136 | -0.17815055 | 0 | 0.125 | 1 | 23 |
| si:dkey-30k6.5 | 0.12136 | -0.26167102 | 0 | 0.125 | 1 | 23 |
| si:dkey-6e2.2 | 0.12136 | -0.16062517 | 0 | 0.125 | 1 | 23 |
| si:dkey-74k8.4 | 0.12136 | -0.13179271 | 0 | 0.125 | 1 | 23 |
| si:dkey-85a20.4 | 0.12136 | -0.18024265 | 0 | 0.125 | 1 | 23 |
| si:zfos-80g12.1 | 0.12136 | -0.15889054 | 0 | 0.125 | 1 | 23 |
| sirt3 | 0.12136 | -0.18304682 | 0 | 0.125 | 1 | 23 |
| slc17a6b | 0.12136 | -0.36253987 | 0 | 0.125 | 1 | 23 |
| slc25a17 | 0.12136 | -0.1635111 | 0 | 0.125 | 1 | 23 |

| | | | | | | |
|---|---|---|---|---|---|---|
| slc25a40 | 0.12136 | -0.15141896 | 0 | 0.125 | 1 | 23 |
| slc2a10 | 0.12136 | -0.18299525 | 0 | 0.125 | 1 | 23 |
| slc39a10 | 0.12136 | -0.15944928 | 0 | 0.125 | 1 | 23 |
| smad3b | 0.12136 | -0.17910639 | 0 | 0.125 | 1 | 23 |
| smg9 | 0.12136 | -0.15291279 | 0 | 0.125 | 1 | 23 |
| sned1 | 0.12136 | -0.16872718 | 0 | 0.125 | 1 | 23 |
| snx33 | 0.12136 | -0.14682621 | 0 | 0.125 | 1 | 23 |
| sowahd | 0.12136 | -0.12150035 | 0 | 0.125 | 1 | 23 |
| spag1a | 0.12136 | -0.15114216 | 0 | 0.125 | 1 | 23 |
| spryd3 | 0.12136 | -0.16183794 | 0 | 0.125 | 1 | 23 |
| stat5a | 0.12136 | -0.15474745 | 0 | 0.125 | 1 | 23 |
| stxbp5a | 0.12136 | -0.23757371 | 0 | 0.125 | 1 | 23 |
| sult1st5 | 0.12136 | -0.16063718 | 0 | 0.125 | 1 | 23 |
| svep1 | 0.12136 | -0.36133126 | 0 | 0.125 | 1 | 23 |
| swap70b | 0.12136 | -0.1721212 | 0 | 0.125 | 1 | 23 |
| taco1 | 0.12136 | -0.18060324 | 0 | 0.125 | 1 | 23 |
| tbc1d23 | 0.12136 | -0.18163002 | 0 | 0.125 | 1 | 23 |
| tbl2 | 0.12136 | -0.1809134 | 0 | 0.125 | 1 | 23 |
| tefm | 0.12136 | -0.17928487 | 0 | 0.125 | 1 | 23 |
| tent5c | 0.12136 | -0.17041532 | 0 | 0.125 | 1 | 23 |
| tgfbr2a | 0.12136 | -0.15764426 | 0 | 0.125 | 1 | 23 |
| tle3b | 0.12136 | -0.16330136 | 0 | 0.125 | 1 | 23 |
| tmem68 | 0.12136 | -0.16486193 | 0 | 0.125 | 1 | 23 |
| tmem97 | 0.12136 | -0.15858838 | 0 | 0.125 | 1 | 23 |
| tnikb | 0.12136 | -0.16096771 | 0 | 0.125 | 1 | 23 |
| tradd | 0.12136 | -0.1569611 | 0 | 0.125 | 1 | 23 |
| trap1 | 0.12136 | -0.18444635 | 0 | 0.125 | 1 | 23 |
| trim3b | 0.12136 | -0.36253987 | 0 | 0.125 | 1 | 23 |
| triob | 0.12136 | -0.16132212 | 0 | 0.125 | 1 | 23 |
| tubgcp4 | 0.12136 | -0.18057973 | 0 | 0.125 | 1 | 23 |
| twist3 | 0.12136 | -0.18183876 | 0 | 0.125 | 1 | 23 |
| uba7 | 0.12136 | -0.16086614 | 0 | 0.125 | 1 | 23 |
| ube2j1 | 0.12136 | -0.15850182 | 0 | 0.125 | 1 | 23 |
| ufsp1 | 0.12136 | -0.15576339 | 0 | 0.125 | 1 | 23 |
| usp48 | 0.12136 | -0.15957424 | 0 | 0.125 | 1 | 23 |
| uspl1 | 0.12136 | -0.17936492 | 0 | 0.125 | 1 | 23 |
| vangl1 | 0.12136 | -0.18705238 | 0 | 0.125 | 1 | 23 |
| vcpkmt | 0.12136 | -0.10945514 | 0 | 0.125 | 1 | 23 |
| vps13a | 0.12136 | -0.2651502 | 0 | 0.125 | 1 | 23 |
| vps39 | 0.12136 | -0.18563349 | 0 | 0.125 | 1 | 23 |
| wdr18 | 0.12136 | -0.15206762 | 0 | 0.125 | 1 | 23 |
| wdr37 | 0.12136 | -0.36127579 | 0 | 0.125 | 1 | 23 |
| wif1 | 0.12136 | -0.18705238 | 0 | 0.125 | 1 | 23 |
| xab2 | 0.12136 | -0.35960705 | 0 | 0.125 | 1 | 23 |
| xpo6 | 0.12136 | -0.36032153 | 0 | 0.125 | 1 | 23 |
| xrn1 | 0.12136 | -0.16098731 | 0 | 0.125 | 1 | 23 |
| yrdc | 0.12136 | -0.14358085 | 0 | 0.125 | 1 | 23 |
| zgc:101577 | 0.12136 | -0.1578886 | 0 | 0.125 | 1 | 23 |

| | | | | | | |
|---|---|---|---|---|---|---|
| zgc:112980 | 0.12136 | -0.17748891 | 0 | 0.125 | 1 | 23 |
| zgc:113210 | 0.12136 | -0.15456844 | 0 | 0.125 | 1 | 23 |
| zgc:114174 | 0.12136 | -0.15532769 | 0 | 0.125 | 1 | 23 |
| zgc:153169 | 0.12136 | -0.17934171 | 0 | 0.125 | 1 | 23 |
| zmiz2 | 0.12136 | -0.26299277 | 0 | 0.125 | 1 | 23 |
| znf1007 | 0.12136 | -0.18436425 | 0 | 0.125 | 1 | 23 |
| znf1113 | 0.12136 | -0.36253987 | 0 | 0.125 | 1 | 23 |
| znf1124 | 0.12136 | -0.16158019 | 0 | 0.125 | 1 | 23 |
| znf1130 | 0.12136 | -0.36253987 | 0 | 0.125 | 1 | 23 |
| znf1181 | 0.12136 | -0.15490034 | 0 | 0.125 | 1 | 23 |
| znhit6 | 0.12136 | -0.17957087 | 0 | 0.125 | 1 | 23 |
| ACSF3 | 0.12136 | -0.22403963 | 0 | 0.125 | 1 | 23 |
| BX005060.1 | 0.12136 | -0.21938756 | 0 | 0.125 | 1 | 23 |
| CABZ01072309.1 | 0.12136 | -0.22587884 | 0 | 0.125 | 1 | 23 |
| CR391986.1 | 0.12136 | -0.30018985 | 0 | 0.125 | 1 | 23 |
| LOC103910447 | 0.12136 | -0.21778512 | 0 | 0.125 | 1 | 23 |
| LOC108184012 | 0.12136 | -0.22587884 | 0 | 0.125 | 1 | 23 |
| TIMM21 | 0.12136 | -0.22415612 | 0 | 0.125 | 1 | 23 |
| XLOC-004482 | 0.12136 | -0.22587884 | 0 | 0.125 | 1 | 23 |
| XLOC-024377 | 0.12136 | -0.22478943 | 0 | 0.125 | 1 | 23 |
| XLOC-028935 | 0.12136 | -0.22587884 | 0 | 0.125 | 1 | 23 |
| XLOC-035493 | 0.12136 | -0.21648103 | 0 | 0.125 | 1 | 23 |
| adam11 | 0.12136 | -0.22587884 | 0 | 0.125 | 1 | 23 |
| ankfy1 | 0.12136 | -0.22460816 | 0 | 0.125 | 1 | 23 |
| atxn10 | 0.12136 | -0.22428536 | 0 | 0.125 | 1 | 23 |
| bach2b | 0.12136 | -0.22211005 | 0 | 0.125 | 1 | 23 |
| bicral | 0.12136 | -0.22241468 | 0 | 0.125 | 1 | 23 |
| capn1a | 0.12136 | -0.22295846 | 0 | 0.125 | 1 | 23 |
| ccbe1 | 0.12136 | -0.22377865 | 0 | 0.125 | 1 | 23 |
| ccdc13 | 0.12136 | -0.21868594 | 0 | 0.125 | 1 | 23 |
| cdc23 | 0.12136 | -0.22365737 | 0 | 0.125 | 1 | 23 |
| cdc42bpb | 0.12136 | -0.22399931 | 0 | 0.125 | 1 | 23 |
| cgna | 0.12136 | -0.21744296 | 0 | 0.125 | 1 | 23 |
| chst2a | 0.12136 | -0.22233135 | 0 | 0.125 | 1 | 23 |
| cse1l | 0.12136 | -0.3057878 | 0 | 0.125 | 1 | 23 |
| dcaf17 | 0.12136 | -0.22490772 | 0 | 0.125 | 1 | 23 |
| ddx11 | 0.12136 | -0.21973147 | 0 | 0.125 | 1 | 23 |
| ddx20 | 0.12136 | -0.30669489 | 0 | 0.125 | 1 | 23 |
| esyt2a | 0.12136 | -0.22334119 | 0 | 0.125 | 1 | 23 |
| fam160b1 | 0.12136 | -0.22316218 | 0 | 0.125 | 1 | 23 |
| fgf10b | 0.12136 | -0.22114222 | 0 | 0.125 | 1 | 23 |
| gdpd5a | 0.12136 | -0.22194387 | 0 | 0.125 | 1 | 23 |
| gigyf2 | 0.12136 | -0.22278282 | 0 | 0.125 | 1 | 23 |
| gmnn | 0.12136 | -0.21776305 | 0 | 0.125 | 1 | 23 |
| gsk3aa | 0.12136 | -0.22473689 | 0 | 0.125 | 1 | 23 |
| hdhd3 | 0.12136 | -0.21819194 | 0 | 0.125 | 1 | 23 |
| hykk.2 | 0.12136 | -0.21430997 | 0 | 0.125 | 1 | 23 |
| ints13 | 0.12136 | -0.22237616 | 0 | 0.125 | 1 | 23 |

| | | | | | | |
|---|---|---|---|---|---|---|
| lcmt1 | 0.12136 | -0.21671679 | 0 | 0.125 | 1 | 23 |
| lgr4 | 0.12136 | -0.21848641 | 0 | 0.125 | 1 | 23 |
| lipeb | 0.12136 | -0.22241708 | 0 | 0.125 | 1 | 23 |
| mrm3a | 0.12136 | -0.22469851 | 0 | 0.125 | 1 | 23 |
| mtss1la | 0.12136 | -0.22502903 | 0 | 0.125 | 1 | 23 |
| natd1 | 0.12136 | -0.20082383 | 0 | 0.125 | 1 | 23 |
| nrp2b | 0.12136 | -0.2247685 | 0 | 0.125 | 1 | 23 |
| otud4 | 0.12136 | -0.22313163 | 0 | 0.125 | 1 | 23 |
| plekha1b | 0.12136 | -0.22348721 | 0 | 0.125 | 1 | 23 |
| ppm1e | 0.12136 | -0.22587884 | 0 | 0.125 | 1 | 23 |
| ppp2r2aa | 0.12136 | -0.22094458 | 0 | 0.125 | 1 | 23 |
| psph | 0.12136 | -0.21300896 | 0 | 0.125 | 1 | 23 |
| scamp5a | 0.12136 | -0.20100941 | 0 | 0.125 | 1 | 23 |
| setx | 0.12136 | -0.22379682 | 0 | 0.125 | 1 | 23 |
| si:ch211-188f17.1 | 0.12136 | -0.22486802 | 0 | 0.125 | 1 | 23 |
| si:ch73-173p19.1 | 0.12136 | -0.22422534 | 0 | 0.125 | 1 | 23 |
| si:ch73-266f23.1 | 0.12136 | -0.22587884 | 0 | 0.125 | 1 | 23 |
| si:ch73-278m9.1 | 0.12136 | -0.22587884 | 0 | 0.125 | 1 | 23 |
| si:ch73-27e22.1 | 0.12136 | -0.22333536 | 0 | 0.125 | 1 | 23 |
| si:dkey-206f10.1 | 0.12136 | -0.21498397 | 0 | 0.125 | 1 | 23 |
| si:dkey-229e3.2 | 0.12136 | -0.22366066 | 0 | 0.125 | 1 | 23 |
| si:dkey-244a7.1 | 0.12136 | -0.22343875 | 0 | 0.125 | 1 | 23 |
| si:dkey-78a14.4 | 0.12136 | -0.2218829 | 0 | 0.125 | 1 | 23 |
| si:zfos-2326c3.2 | 0.12136 | -0.21783825 | 0 | 0.125 | 1 | 23 |
| slc25a38b | 0.12136 | -0.22239034 | 0 | 0.125 | 1 | 23 |
| snap47 | 0.12136 | -0.1898297 | 0 | 0.125 | 1 | 23 |
| st6galnac5a | 0.12136 | -0.22587884 | 0 | 0.125 | 1 | 23 |
| suco | 0.12136 | -0.22148428 | 0 | 0.125 | 1 | 23 |
| syne1a | 0.12136 | -0.22400683 | 0 | 0.125 | 1 | 23 |
| tbc1d12b | 0.12136 | -0.22330797 | 0 | 0.125 | 1 | 23 |
| tmem201 | 0.12136 | -0.2190385 | 0 | 0.125 | 1 | 23 |
| tpcn3 | 0.12136 | -0.22364201 | 0 | 0.125 | 1 | 23 |
| trim107 | 0.12136 | -0.22277529 | 0 | 0.125 | 1 | 23 |
| trip4 | 0.12136 | -0.22195601 | 0 | 0.125 | 1 | 23 |
| trit1 | 0.12136 | -0.2210711 | 0 | 0.125 | 1 | 23 |
| trub1 | 0.12136 | -0.21701663 | 0 | 0.125 | 1 | 23 |
| ttc9c | 0.12136 | -0.21731732 | 0 | 0.125 | 1 | 23 |
| txnl4b | 0.12136 | -0.22341519 | 0 | 0.125 | 1 | 23 |
| ulk4 | 0.12136 | -0.22417068 | 0 | 0.125 | 1 | 23 |
| virma | 0.12136 | -0.2217917 | 0 | 0.125 | 1 | 23 |
| vps45 | 0.12136 | -0.22046719 | 0 | 0.125 | 1 | 23 |
| vti1a | 0.12136 | -0.22498634 | 0 | 0.125 | 1 | 23 |
| wrap53 | 0.12136 | -0.21820293 | 0 | 0.125 | 1 | 23 |
| zgc:92594 | 0.12136 | -0.21025764 | 0 | 0.125 | 1 | 23 |
| zmym4 | 0.12136 | -0.22407785 | 0 | 0.125 | 1 | 23 |
| znf740a | 0.12136 | -0.22190254 | 0 | 0.125 | 1 | 23 |
| mcl1b | 0.12147 | -0.10285849 | 0.613 | 0.5 | 1 | 23 |
| prpf6 | 0.12156 | -0.55723689 | 0.097 | 0.375 | 1 | 23 |

| | | | | | | |
|---|---|---|---|---|---|---|
| med13a | 0.12168 | -0.48288285 | 0.129 | 0.375 | 1 | 23 |
| psip1b | 0.12187 | -0.40971711 | 0.355 | 0.5 | 1 | 23 |
| ndfip1l | 0.12217 | -0.51486234 | 0.129 | 0.375 | 1 | 23 |
| eppk1 | 0.12304 | -0.44505242 | 0.129 | 0.25 | 1 | 23 |
| letm1 | 0.12327 | -0.18897249 | 0.129 | 0.125 | 1 | 23 |
| si:ch1073-392o20.2 | 0.1234 | -0.44742414 | 0.065 | 0.25 | 1 | 23 |
| dag1 | 0.12372 | -0.58245554 | 0.097 | 0.375 | 1 | 23 |
| kdm6bb | 0.12373 | -0.45787781 | 0.194 | 0.375 | 1 | 23 |
| nfe2l1b | 0.12422 | -0.247672 | 0.097 | 0.125 | 1 | 23 |
| ldhba | 0.12439 | -0.19898086 | 0.516 | 0.5 | 1 | 23 |
| kctd12.2 | 0.12471 | -0.49562527 | 0.484 | 0.75 | 1 | 23 |
| tnrc18 | 0.12475 | -0.50992522 | 0.097 | 0.375 | 1 | 23 |
| rpl15 | 0.1248 | 0.139547234 | 0.968 | 1 | 1 | 23 |
| zgc:114181 | 0.12504 | -0.59197873 | 0.258 | 0.625 | 1 | 23 |
| actl6a | 0.12533 | -0.45765278 | 0.161 | 0.5 | 1 | 23 |
| gig2e | 0.12567 | -0.54553013 | 0.161 | 0.5 | 1 | 23 |
| flot2a | 0.12614 | 0.468204341 | 0.484 | 0.125 | 1 | 23 |
| pfn2l | 0.12656 | -0.42388522 | 0.613 | 0.875 | 1 | 23 |
| map4k4 | 0.12702 | -0.49388449 | 0.161 | 0.5 | 1 | 23 |
| stag2b | 0.1274 | -0.25462159 | 0.097 | 0.125 | 1 | 23 |
| ypel5 | 0.12799 | -0.44402658 | 0.097 | 0.375 | 1 | 23 |
| cpne3 | 0.1286 | 0.132512462 | 0.29 | 0.125 | 1 | 23 |
| si:dkey-188i13.10 | 0.12871 | -0.17504199 | 0.548 | 0.5 | 1 | 23 |
| pbx4 | 0.12916 | -0.45795752 | 0.355 | 0.625 | 1 | 23 |
| gnl3 | 0.12951 | -0.15280118 | 0.29 | 0.25 | 1 | 23 |
| faub | 0.12954 | -0.51995019 | 0.161 | 0.5 | 1 | 23 |
| aplp2 | 0.12956 | -0.43994916 | 0.258 | 0.5 | 1 | 23 |
| selenof | 0.13031 | -0.1444976 | 0.839 | 0.75 | 1 | 23 |
| rnf213a | 0.13103 | -0.64200747 | 0.258 | 0.625 | 1 | 23 |
| fosb | 0.13113 | -0.51466127 | 0.581 | 0.875 | 1 | 23 |
| itgb1bp1 | 0.13202 | -0.43484257 | 0.097 | 0.25 | 1 | 23 |
| per1b | 0.13229 | -0.25547193 | 0.097 | 0.125 | 1 | 23 |
| ndufs2 | 0.13282 | -0.35028894 | 0.258 | 0.375 | 1 | 23 |
| eif3s10 | 0.13286 | 0.157658355 | 0.484 | 0.25 | 1 | 23 |
| jupa | 0.13341 | -0.16971083 | 0.129 | 0.125 | 1 | 23 |
| lrrc17 | 0.1335 | 0.575950614 | 0.484 | 0.125 | 1 | 23 |
| eif2s3 | 0.13353 | 0.333196339 | 0.839 | 0.5 | 1 | 23 |
| si:ch211-13c6.2 | 0.13406 | -0.16993754 | 0.129 | 0.125 | 1 | 23 |
| hnrnpa1a | 0.13492 | -0.47859007 | 0.387 | 0.75 | 1 | 23 |
| atrx | 0.13502 | 0.408872807 | 0.677 | 0.5 | 1 | 23 |
| cox4i1 | 0.13558 | 0.455972273 | 0.839 | 0.5 | 1 | 23 |
| lamc1 | 0.13613 | -0.42191888 | 0.097 | 0.375 | 1 | 23 |
| tmem59 | 0.13664 | 0.452869406 | 0.419 | 0.125 | 1 | 23 |
| arpc4l | 0.13769 | 0.377125161 | 0.452 | 0.125 | 1 | 23 |
| zc3h7bb | 0.13771 | -0.52129973 | 0.097 | 0.375 | 1 | 23 |
| rpl35 | 0.13864 | 0.263454691 | 1 | 0.875 | 1 | 23 |
| rsf1b.1 | 0.13882 | -0.20058858 | 0.226 | 0.25 | 1 | 23 |
| mdka | 0.13906 | 0.153618821 | 0.774 | 0.875 | 1 | 23 |

| | | | | | | |
|---|---|---|---|---|---|---|
| tomm70a | 0.13921 | -0.35507343 | 0.226 | 0.375 | 1 | 23 |
| cox7a2a | 0.14021 | 0.416757347 | 0.484 | 0.125 | 1 | 23 |
| hsbp1b | 0.14116 | -0.14485904 | 0.419 | 0.375 | 1 | 23 |
| snrpa1 | 0.14127 | -0.50131028 | 0.194 | 0.5 | 1 | 23 |
| hnrnpaba | 0.14251 | 0.400831001 | 0.839 | 0.625 | 1 | 23 |
| h2afva | 0.14281 | -0.4137291 | 0.452 | 0.625 | 1 | 23 |
| xpo1b | 0.14303 | -0.43769179 | 0.097 | 0.25 | 1 | 23 |
| msna | 0.14337 | -0.4488714 | 0.516 | 0.875 | 1 | 23 |
| col4a6 | 0.14371 | -0.30346319 | 0.097 | 0.375 | 1 | 23 |
| dnmt3ab | 0.14388 | -0.34450791 | 0.258 | 0.375 | 1 | 23 |
| foxp4 | 0.14433 | -0.52277372 | 0.387 | 0.75 | 1 | 23 |
| zgc:153311 | 0.14448 | -0.49544244 | 0.097 | 0.25 | 1 | 23 |
| mtpn | 0.14451 | 0.500552473 | 0.419 | 0.125 | 1 | 23 |
| trappc1 | 0.14482 | -0.22927175 | 0.484 | 0.5 | 1 | 23 |
| fkbp3 | 0.14637 | 0.510859741 | 0.452 | 0.125 | 1 | 23 |
| syf2 | 0.14783 | 0.461649694 | 0.484 | 0.125 | 1 | 23 |
| prkab1b | 0.14812 | 0.463677632 | 0.516 | 0.25 | 1 | 23 |
| elocb | 0.14877 | 0.360143254 | 0.742 | 0.375 | 1 | 23 |
| anxa5b | 0.14901 | 0.534213008 | 0.484 | 0.125 | 1 | 23 |
| cetn3 | 0.14951 | -0.42627515 | 0.065 | 0.25 | 1 | 23 |
| cox8b | 0.14994 | -0.47807703 | 0.419 | 0.625 | 1 | 23 |
| si:ch211-214j24.10 | 0.15079 | 0.514435299 | 0.419 | 0.125 | 1 | 23 |
| mcts1 | 0.15118 | 0.391299607 | 0.452 | 0.125 | 1 | 23 |
| crebrf | 0.15164 | -0.23658052 | 0.097 | 0.125 | 1 | 23 |
| LOC110438360 | 0.15167 | -0.23042719 | 0.097 | 0.125 | 1 | 23 |
| slc38a5b | 0.1519 | -0.4860353 | 0.323 | 0.5 | 1 | 23 |
| csnk2b | 0.15221 | -0.37495754 | 0.581 | 0.75 | 1 | 23 |
| mast2 | 0.15266 | -0.43650709 | 0.065 | 0.25 | 1 | 23 |
| mphosph6 | 0.15316 | -0.5069753 | 0.129 | 0.375 | 1 | 23 |
| nudc | 0.1533 | 0.440615694 | 0.452 | 0.25 | 1 | 23 |
| yy1a | 0.15366 | -0.44190856 | 0.29 | 0.5 | 1 | 23 |
| prdx6 | 0.15449 | 0.373782392 | 0.613 | 0.25 | 1 | 23 |
| tatdn1 | 0.1546 | -0.4066882 | 0.065 | 0.25 | 1 | 23 |
| si:ch211-207d6.2 | 0.15467 | -0.35383584 | 0.581 | 0.625 | 1 | 23 |
| boc | 0.15585 | 0.469340714 | 0.452 | 0.125 | 1 | 23 |
| psmg2 | 0.15686 | -0.21056899 | 0.226 | 0.25 | 1 | 23 |
| zgc:194246 | 0.15759 | 0.305498727 | 0.226 | 0.125 | 1 | 23 |
| ctbp1 | 0.15773 | -0.34239492 | 0.29 | 0.375 | 1 | 23 |
| ndufs3 | 0.15785 | -0.38881691 | 0.29 | 0.5 | 1 | 23 |
| aclya | 0.15825 | -0.3848044 | 0.097 | 0.25 | 1 | 23 |
| mrpl58 | 0.15832 | -0.29140239 | 0.161 | 0.25 | 1 | 23 |
| si:ch211-22i13.2 | 0.15859 | 0.51462873 | 0.452 | 0.125 | 1 | 23 |
| zc3h15 | 0.16009 | -0.43598075 | 0.161 | 0.5 | 1 | 23 |
| psma5 | 0.16043 | 0.388264014 | 0.387 | 0.25 | 1 | 23 |
| asah1b | 0.16045 | -0.43437798 | 0.258 | 0.625 | 1 | 23 |
| wbp1 | 0.16055 | -0.57871498 | 0.097 | 0.375 | 1 | 23 |
| arhgdig | 0.16093 | 0.262801938 | 0.548 | 0.25 | 1 | 23 |
| cnbpb | 0.16177 | 0.421105845 | 0.613 | 0.25 | 1 | 23 |

| | | | | | | |
|---|---|---|---|---|---|---|
| psmb2 | 0.16212 | 0.175782172 | 0.645 | 0.375 | 1 | 23 |
| wdr43 | 0.16224 | -0.1622389 | 0.129 | 0.125 | 1 | 23 |
| glulb | 0.16249 | -0.47509916 | 0.161 | 0.5 | 1 | 23 |
| ost4 | 0.16275 | -0.51331829 | 0.355 | 0.625 | 1 | 23 |
| nt5c2b | 0.16366 | -0.50633841 | 0.161 | 0.5 | 1 | 23 |
| usp14 | 0.16398 | -0.31498414 | 0.161 | 0.25 | 1 | 23 |
| oga | 0.16413 | -0.54337722 | 0.097 | 0.375 | 1 | 23 |
| ppp1r10 | 0.16417 | -0.40465881 | 0.161 | 0.5 | 1 | 23 |
| snx3 | 0.1649 | 0.220402023 | 0.355 | 0.125 | 1 | 23 |
| iqca1 | 0.16607 | 0.294809515 | 0.161 | 0 | 1 | 23 |
| cdc42ep5 | 0.16607 | 0.282857602 | 0.161 | 0 | 1 | 23 |
| pik3ip1 | 0.16607 | 0.285069132 | 0.161 | 0 | 1 | 23 |
| ssbp4 | 0.16607 | 0.366960885 | 0.161 | 0 | 1 | 23 |
| ube2a | 0.16607 | 0.252626032 | 0.161 | 0 | 1 | 23 |
| AL935174.5 | 0.16607 | 0.267617081 | 0.161 | 0 | 1 | 23 |
| CU459056.2 | 0.16607 | 0.253276642 | 0.161 | 0 | 1 | 23 |
| CU929259.1 | 0.16607 | 0.253415954 | 0.161 | 0 | 1 | 23 |
| FP102052.2 | 0.16607 | 0.242195479 | 0.161 | 0 | 1 | 23 |
| LOC100004944 | 0.16607 | 0.284835856 | 0.161 | 0 | 1 | 23 |
| LOC110438518 | 0.16607 | 0.231441825 | 0.161 | 0 | 1 | 23 |
| MCUR1 | 0.16607 | 0.249152154 | 0.161 | 0 | 1 | 23 |
| actr6 | 0.16607 | 0.223200582 | 0.161 | 0 | 1 | 23 |
| ajuba | 0.16607 | 0.302079098 | 0.161 | 0 | 1 | 23 |
| ak4 | 0.16607 | 0.220871695 | 0.161 | 0 | 1 | 23 |
| aldh2.1 | 0.16607 | 0.254110855 | 0.161 | 0 | 1 | 23 |
| ap2m1b | 0.16607 | 0.280634632 | 0.161 | 0 | 1 | 23 |
| ap3s2 | 0.16607 | 0.23608017 | 0.161 | 0 | 1 | 23 |
| api5 | 0.16607 | 0.218703525 | 0.161 | 0 | 1 | 23 |
| arl14ep | 0.16607 | 0.245115448 | 0.161 | 0 | 1 | 23 |
| atf1 | 0.16607 | 0.24289777 | 0.161 | 0 | 1 | 23 |
| atg13 | 0.16607 | 0.26670696 | 0.161 | 0 | 1 | 23 |
| bida | 0.16607 | 0.237185556 | 0.161 | 0 | 1 | 23 |
| bnip4 | 0.16607 | 0.235885619 | 0.161 | 0 | 1 | 23 |
| borcs7 | 0.16607 | 0.264123 | 0.161 | 0 | 1 | 23 |
| cadm2a | 0.16607 | 0.282831605 | 0.161 | 0 | 1 | 23 |
| calhm2 | 0.16607 | 0.273734189 | 0.161 | 0 | 1 | 23 |
| calua | 0.16607 | 0.256440189 | 0.161 | 0 | 1 | 23 |
| carm1 | 0.16607 | 0.313360139 | 0.161 | 0 | 1 | 23 |
| cbx8a | 0.16607 | 0.265279705 | 0.161 | 0 | 1 | 23 |
| ccnyl1 | 0.16607 | 0.248291256 | 0.161 | 0 | 1 | 23 |
| cers5 | 0.16607 | 0.243553908 | 0.161 | 0 | 1 | 23 |
| chchd4a | 0.16607 | 0.242169591 | 0.161 | 0 | 1 | 23 |
| chmp2bb | 0.16607 | 0.228712958 | 0.161 | 0 | 1 | 23 |
| clk4b | 0.16607 | 0.193052617 | 0.161 | 0 | 1 | 23 |
| clstn1 | 0.16607 | 0.295972224 | 0.161 | 0 | 1 | 23 |
| crip3 | 0.16607 | 0.240186332 | 0.161 | 0 | 1 | 23 |
| crkl | 0.16607 | 0.252936294 | 0.161 | 0 | 1 | 23 |
| cry1ab | 0.16607 | 0.284661585 | 0.161 | 0 | 1 | 23 |

| | | | | | | |
|---|---|---|---|---|---|---|
| cwc27 | 0.16607 | 0.244079481 | 0.161 | 0 | 1 | 23 |
| cyr61 | 0.16607 | 0.440869214 | 0.161 | 0 | 1 | 23 |
| dcun1d2a | 0.16607 | 0.248557625 | 0.161 | 0 | 1 | 23 |
| ddx51 | 0.16607 | 0.301755888 | 0.161 | 0 | 1 | 23 |
| ddx52 | 0.16607 | 0.192660898 | 0.161 | 0 | 1 | 23 |
| dnajc1 | 0.16607 | 0.250529025 | 0.161 | 0 | 1 | 23 |
| dpf2l | 0.16607 | 0.275204528 | 0.161 | 0 | 1 | 23 |
| dtnbp1a | 0.16607 | 0.239549047 | 0.161 | 0 | 1 | 23 |
| dusp8a | 0.16607 | 0.283007444 | 0.161 | 0 | 1 | 23 |
| efna1b | 0.16607 | 0.296073571 | 0.161 | 0 | 1 | 23 |
| elf2a | 0.16607 | 0.219760142 | 0.161 | 0 | 1 | 23 |
| exosc6 | 0.16607 | 0.230152681 | 0.161 | 0 | 1 | 23 |
| ezrb | 0.16607 | 0.268084413 | 0.161 | 0 | 1 | 23 |
| fam193a | 0.16607 | 0.300436959 | 0.161 | 0 | 1 | 23 |
| fam50a | 0.16607 | 0.240596315 | 0.161 | 0 | 1 | 23 |
| fdx2 | 0.16607 | 0.20425928 | 0.161 | 0 | 1 | 23 |
| fkbp1b | 0.16607 | 0.293455097 | 0.161 | 0 | 1 | 23 |
| gnas | 0.16607 | 0.298676196 | 0.161 | 0 | 1 | 23 |
| gnl3l | 0.16607 | 0.236145292 | 0.161 | 0 | 1 | 23 |
| gpat3 | 0.16607 | 0.207747903 | 0.161 | 0 | 1 | 23 |
| gpatch3 | 0.16607 | 0.337649602 | 0.161 | 0 | 1 | 23 |
| grb2a | 0.16607 | 0.246411754 | 0.161 | 0 | 1 | 23 |
| grk4 | 0.16607 | 0.272053936 | 0.161 | 0 | 1 | 23 |
| hadh | 0.16607 | 0.215227176 | 0.161 | 0 | 1 | 23 |
| herpud2 | 0.16607 | 0.203630094 | 0.161 | 0 | 1 | 23 |
| hgs | 0.16607 | 0.300604053 | 0.161 | 0 | 1 | 23 |
| hnrnpul1 | 0.16607 | 0.239555065 | 0.161 | 0 | 1 | 23 |
| homeza | 0.16607 | 0.249271989 | 0.161 | 0 | 1 | 23 |
| hspa14 | 0.16607 | 0.231961885 | 0.161 | 0 | 1 | 23 |
| ift20 | 0.16607 | 0.272813757 | 0.161 | 0 | 1 | 23 |
| immp2l | 0.16607 | 0.250791225 | 0.161 | 0 | 1 | 23 |
| impdh1b | 0.16607 | 0.258533496 | 0.161 | 0 | 1 | 23 |
| ip6k2b | 0.16607 | 0.245062494 | 0.161 | 0 | 1 | 23 |
| irf10 | 0.16607 | 0.270993219 | 0.161 | 0 | 1 | 23 |
| kat7a | 0.16607 | 0.263536581 | 0.161 | 0 | 1 | 23 |
| kdm5bb | 0.16607 | 0.328111335 | 0.161 | 0 | 1 | 23 |
| kdsr | 0.16607 | 0.288400155 | 0.161 | 0 | 1 | 23 |
| lancl1 | 0.16607 | 0.261308171 | 0.161 | 0 | 1 | 23 |
| lipia | 0.16607 | 0.241053641 | 0.161 | 0 | 1 | 23 |
| lrrc47 | 0.16607 | 0.248447263 | 0.161 | 0 | 1 | 23 |
| ltbp1 | 0.16607 | 0.240988092 | 0.161 | 0 | 1 | 23 |
| maff | 0.16607 | 0.251112399 | 0.161 | 0 | 1 | 23 |
| mapk3 | 0.16607 | 0.279862851 | 0.161 | 0 | 1 | 23 |
| mecr | 0.16607 | 0.230710413 | 0.161 | 0 | 1 | 23 |
| mgst1.2 | 0.16607 | 0.192921126 | 0.161 | 0 | 1 | 23 |
| mpc1 | 0.16607 | 0.208832888 | 0.161 | 0 | 1 | 23 |
| mrpl2 | 0.16607 | 0.27833868 | 0.161 | 0 | 1 | 23 |
| mrps15 | 0.16607 | 0.234758702 | 0.161 | 0 | 1 | 23 |

| | | | | | | |
|---|---|---|---|---|---|---|
| mrps16 | 0.16607 | 0.275520829 | 0.161 | 0 | 1 | 23 |
| mrps18b | 0.16607 | 0.252789878 | 0.161 | 0 | 1 | 23 |
| mrps28 | 0.16607 | 0.236577534 | 0.161 | 0 | 1 | 23 |
| mxi1 | 0.16607 | 0.218967233 | 0.161 | 0 | 1 | 23 |
| myd88 | 0.16607 | 0.247662692 | 0.161 | 0 | 1 | 23 |
| naa38 | 0.16607 | 0.24784838 | 0.161 | 0 | 1 | 23 |
| naa50 | 0.16607 | 0.255650052 | 0.161 | 0 | 1 | 23 |
| nipa2 | 0.16607 | 0.30536329 | 0.161 | 0 | 1 | 23 |
| nipblb | 0.16607 | 0.239217934 | 0.161 | 0 | 1 | 23 |
| nt5dc2 | 0.16607 | 0.260327302 | 0.161 | 0 | 1 | 23 |
| nufip2 | 0.16607 | 0.246973739 | 0.161 | 0 | 1 | 23 |
| pafah1b3 | 0.16607 | 0.224793342 | 0.161 | 0 | 1 | 23 |
| parp3 | 0.16607 | 0.275505137 | 0.161 | 0 | 1 | 23 |
| pdcl | 0.16607 | 0.27576051 | 0.161 | 0 | 1 | 23 |
| pdpk1b | 0.16607 | 0.272814233 | 0.161 | 0 | 1 | 23 |
| pdzd11 | 0.16607 | 0.287137959 | 0.161 | 0 | 1 | 23 |
| pgam5 | 0.16607 | 0.226870143 | 0.161 | 0 | 1 | 23 |
| phax | 0.16607 | 0.22909437 | 0.161 | 0 | 1 | 23 |
| phf6 | 0.16607 | 0.245043175 | 0.161 | 0 | 1 | 23 |
| polr3glb | 0.16607 | 0.246634107 | 0.161 | 0 | 1 | 23 |
| polr3k | 0.16607 | 0.246989892 | 0.161 | 0 | 1 | 23 |
| prrg2 | 0.16607 | 0.248251525 | 0.161 | 0 | 1 | 23 |
| pter | 0.16607 | 0.248298579 | 0.161 | 0 | 1 | 23 |
| rab27a | 0.16607 | 0.246030037 | 0.161 | 0 | 1 | 23 |
| rab3gap1 | 0.16607 | 0.254724854 | 0.161 | 0 | 1 | 23 |
| ranbp3b | 0.16607 | 0.224495655 | 0.161 | 0 | 1 | 23 |
| rca2.1 | 0.16607 | 0.236978336 | 0.161 | 0 | 1 | 23 |
| rex1bd | 0.16607 | 0.296031161 | 0.161 | 0 | 1 | 23 |
| rnasekb | 0.16607 | 0.261854658 | 0.161 | 0 | 1 | 23 |
| rnf168 | 0.16607 | 0.27325046 | 0.161 | 0 | 1 | 23 |
| rsad2 | 0.16607 | 0.380920087 | 0.161 | 0 | 1 | 23 |
| sdad1 | 0.16607 | 0.247812181 | 0.161 | 0 | 1 | 23 |
| sdf2l1 | 0.16607 | 0.209246685 | 0.161 | 0 | 1 | 23 |
| sec13 | 0.16607 | 0.21087583 | 0.161 | 0 | 1 | 23 |
| sertad4 | 0.16607 | 0.247411371 | 0.161 | 0 | 1 | 23 |
| setdb2 | 0.16607 | 0.238450442 | 0.161 | 0 | 1 | 23 |
| si:ch1073-459j12.1 | 0.16607 | 0.33880739 | 0.161 | 0 | 1 | 23 |
| si:ch211-150o23.2 | 0.16607 | 0.254179227 | 0.161 | 0 | 1 | 23 |
| si:ch211-195m9.3 | 0.16607 | 0.399077811 | 0.161 | 0 | 1 | 23 |
| si:ch211-204a13.2 | 0.16607 | 0.284690233 | 0.161 | 0 | 1 | 23 |
| si:ch211-218c6.8 | 0.16607 | 0.23616503 | 0.161 | 0 | 1 | 23 |
| si:ch211-236l14.4 | 0.16607 | 0.304776043 | 0.161 | 0 | 1 | 23 |
| si:ch211-262i1.6 | 0.16607 | 0.244928909 | 0.161 | 0 | 1 | 23 |
| si:ch211-271b14.1 | 0.16607 | 0.236180649 | 0.161 | 0 | 1 | 23 |
| si:ch73-269m14.2 | 0.16607 | 0.277283284 | 0.161 | 0 | 1 | 23 |
| si:dkey-230p4.1 | 0.16607 | 0.317547675 | 0.161 | 0 | 1 | 23 |
| si:dkey-264d12.4 | 0.16607 | 0.289539082 | 0.161 | 0 | 1 | 23 |
| si:dkey-44g23.5 | 0.16607 | 0.237431127 | 0.161 | 0 | 1 | 23 |

| | | | | | | |
|---|---|---|---|---|---|---|
| si:dkey-57k2.7 | 0.16607 | 0.273875748 | 0.161 | 0 | 1 | 23 |
| si:dkeyp-69e1.8 | 0.16607 | 0.238070817 | 0.161 | 0 | 1 | 23 |
| sirt1 | 0.16607 | 0.276019333 | 0.161 | 0 | 1 | 23 |
| slc20a2 | 0.16607 | 0.278589294 | 0.161 | 0 | 1 | 23 |
| slc30a7 | 0.16607 | 0.272183488 | 0.161 | 0 | 1 | 23 |
| smap1 | 0.16607 | 0.254476873 | 0.161 | 0 | 1 | 23 |
| smim8 | 0.16607 | 0.221432409 | 0.161 | 0 | 1 | 23 |
| srsf4 | 0.16607 | 0.271365588 | 0.161 | 0 | 1 | 23 |
| ssbp3b | 0.16607 | 0.292211873 | 0.161 | 0 | 1 | 23 |
| ssu72 | 0.16607 | 0.261848254 | 0.161 | 0 | 1 | 23 |
| sugt1 | 0.16607 | 0.298187927 | 0.161 | 0 | 1 | 23 |
| tfpia | 0.16607 | 0.277341361 | 0.161 | 0 | 1 | 23 |
| timmdc1 | 0.16607 | 0.223643688 | 0.161 | 0 | 1 | 23 |
| tmed5 | 0.16607 | 0.235142241 | 0.161 | 0 | 1 | 23 |
| tmem263 | 0.16607 | 0.323679224 | 0.161 | 0 | 1 | 23 |
| tmub1 | 0.16607 | 0.238696156 | 0.161 | 0 | 1 | 23 |
| tob1b | 0.16607 | 0.298080401 | 0.161 | 0 | 1 | 23 |
| tomm34 | 0.16607 | 0.2828397 | 0.161 | 0 | 1 | 23 |
| trappc5 | 0.16607 | 0.285765818 | 0.161 | 0 | 1 | 23 |
| trub2 | 0.16607 | 0.244892602 | 0.161 | 0 | 1 | 23 |
| tshz2 | 0.16607 | 0.279343436 | 0.161 | 0 | 1 | 23 |
| uchl5 | 0.16607 | 0.257400146 | 0.161 | 0 | 1 | 23 |
| vps26a | 0.16607 | 0.253468925 | 0.161 | 0 | 1 | 23 |
| waca | 0.16607 | 0.239590825 | 0.161 | 0 | 1 | 23 |
| wu:fi04e12 | 0.16607 | 0.281014885 | 0.161 | 0 | 1 | 23 |
| xirp2a | 0.16607 | 0.28585275 | 0.161 | 0 | 1 | 23 |
| zcchc17 | 0.16607 | 0.321630506 | 0.161 | 0 | 1 | 23 |
| zcrb1 | 0.16607 | 0.254410924 | 0.161 | 0 | 1 | 23 |
| zdbf2 | 0.16607 | 0.26478881 | 0.161 | 0 | 1 | 23 |
| zfhx4 | 0.16607 | 0.244336562 | 0.161 | 0 | 1 | 23 |
| zfpm1 | 0.16607 | 0.290436914 | 0.161 | 0 | 1 | 23 |
| zgc:153215 | 0.16607 | 0.298096982 | 0.161 | 0 | 1 | 23 |
| zgc:153293 | 0.16607 | 0.298362111 | 0.161 | 0 | 1 | 23 |
| zgc:163098 | 0.16607 | 0.232636027 | 0.161 | 0 | 1 | 23 |
| zgc:85777 | 0.16607 | 0.238324581 | 0.161 | 0 | 1 | 23 |
| eif4e2rs1 | 0.16607 | 0.230597936 | 0.161 | 0 | 1 | 23 |
| mospd1 | 0.16607 | 0.230955771 | 0.161 | 0 | 1 | 23 |
| msra | 0.16607 | 0.263170933 | 0.161 | 0 | 1 | 23 |
| myca | 0.16607 | 0.321525372 | 0.161 | 0 | 1 | 23 |
| otud6b | 0.16607 | 0.245189351 | 0.161 | 0 | 1 | 23 |
| pcmtd1 | 0.16607 | 0.233636326 | 0.161 | 0 | 1 | 23 |
| slc43a2a | 0.16607 | 0.300813914 | 0.161 | 0 | 1 | 23 |
| smim20 | 0.16607 | 0.26555497 | 0.161 | 0 | 1 | 23 |
| sumf2 | 0.16607 | 0.277143976 | 0.161 | 0 | 1 | 23 |
| zgc:114045 | 0.16607 | 0.238734377 | 0.161 | 0 | 1 | 23 |
| lsm1 | 0.16607 | 0.241011607 | 0.161 | 0 | 1 | 23 |
| si:dkey-88p24.11 | 0.16607 | 0.235220544 | 0.161 | 0 | 1 | 23 |
| METTL18 | 0.16607 | 0.252341234 | 0.161 | 0 | 1 | 23 |

| | | | | | | |
|---|---|---|---|---|---|---|
| bag3 | 0.16607 | 0.329777426 | 0.161 | 0 | 1 | 23 |
| bbip1 | 0.16607 | 0.239612921 | 0.161 | 0 | 1 | 23 |
| cmpk | 0.16607 | 0.231891983 | 0.161 | 0 | 1 | 23 |
| crtap | 0.16607 | 0.304273971 | 0.161 | 0 | 1 | 23 |
| ddx3xa | 0.16607 | 0.279292171 | 0.161 | 0 | 1 | 23 |
| emp2 | 0.16607 | 0.257256573 | 0.161 | 0 | 1 | 23 |
| fam207a | 0.16607 | 0.285525059 | 0.161 | 0 | 1 | 23 |
| glrx3 | 0.16607 | 0.252153333 | 0.161 | 0 | 1 | 23 |
| hdac3 | 0.16607 | 0.293230936 | 0.161 | 0 | 1 | 23 |
| kansl1b | 0.16607 | 0.223069085 | 0.161 | 0 | 1 | 23 |
| nr2f6b | 0.16607 | 0.26866747 | 0.161 | 0 | 1 | 23 |
| pkmyt1 | 0.16607 | 0.272747492 | 0.161 | 0 | 1 | 23 |
| prpsap1 | 0.16607 | 0.26034382 | 0.161 | 0 | 1 | 23 |
| sgk1 | 0.16607 | 0.25536519 | 0.161 | 0 | 1 | 23 |
| si:ch211-216l23.2 | 0.16607 | 0.263657026 | 0.161 | 0 | 1 | 23 |
| si:ch73-109d9.2 | 0.16607 | 0.26248437 | 0.161 | 0 | 1 | 23 |
| si:ch73-376l24.6 | 0.16607 | 0.279730417 | 0.161 | 0 | 1 | 23 |
| si:dkey-70b23.2 | 0.16607 | 0.240530025 | 0.161 | 0 | 1 | 23 |
| si:dkeyp-55f12.3 | 0.16607 | 0.232396562 | 0.161 | 0 | 1 | 23 |
| slc22a18 | 0.16607 | 0.257582171 | 0.161 | 0 | 1 | 23 |
| srm | 0.16607 | 0.249087373 | 0.161 | 0 | 1 | 23 |
| tcea2 | 0.16607 | 0.242509242 | 0.161 | 0 | 1 | 23 |
| ypel1 | 0.16607 | 0.256507555 | 0.161 | 0 | 1 | 23 |
| bckdk | 0.16607 | 0.267482385 | 0.161 | 0 | 1 | 23 |
| hccsb | 0.16607 | 0.247868347 | 0.161 | 0 | 1 | 23 |
| cacybp | 0.16607 | 0.276695187 | 0.161 | 0 | 1 | 23 |
| itgb4 | 0.16607 | 0.303802314 | 0.161 | 0 | 1 | 23 |
| si:ch73-25f10.6 | 0.16607 | 0.250271359 | 0.161 | 0 | 1 | 23 |
| rpl36a | 0.16636 | 0.17075253 | 1 | 1 | 1 | 23 |
| si:dkeyp-121d4.3 | 0.16664 | -0.46343075 | 0.065 | 0.25 | 1 | 23 |
| uqcrq | 0.16669 | -0.50626266 | 0.387 | 0.625 | 1 | 23 |
| CABZ01068358.1 | 0.16722 | -0.3225012 | 0.097 | 0.375 | 1 | 23 |
| tax1bp3 | 0.16732 | 0.493606635 | 0.581 | 0.25 | 1 | 23 |
| si:dkey-4e7.3 | 0.16749 | -0.32970349 | 0.097 | 0.375 | 1 | 23 |
| pfdn5 | 0.16768 | -0.13535523 | 0.419 | 0.375 | 1 | 23 |
| pim1 | 0.16876 | -0.19488751 | 0.387 | 0.375 | 1 | 23 |
| tead3a | 0.1695 | 0.505291352 | 0.323 | 0.125 | 1 | 23 |
| cdkn1a | 0.16986 | 0.18258796 | 0.323 | 0.125 | 1 | 23 |
| h2afx1 | 0.1711 | -0.3595329 | 0.742 | 0.875 | 1 | 23 |
| CABZ01090890.1 | 0.17131 | -0.33388818 | 0.226 | 0.375 | 1 | 23 |
| amotl2a | 0.17208 | -0.50229145 | 0.097 | 0.375 | 1 | 23 |
| coro1ca | 0.17276 | -0.55961924 | 0.29 | 0.625 | 1 | 23 |
| gabarapa | 0.17288 | 0.309415258 | 0.419 | 0.125 | 1 | 23 |
| nprl2 | 0.17362 | -0.14741193 | 0.129 | 0.125 | 1 | 23 |
| actb1 | 0.17492 | -0.28144799 | 0.742 | 1 | 1 | 23 |
| tfe3a | 0.1751 | -0.48993348 | 0.065 | 0.25 | 1 | 23 |
| ISCU (1 of many) | 0.17578 | 0.314093763 | 0.387 | 0.125 | 1 | 23 |
| tuba1a | 0.1762 | -0.41812001 | 0.097 | 0.375 | 1 | 23 |

| | | | | | | |
|---|---|---|---|---|---|---|
| usp9 | 0.17673 | -0.40296552 | 0.097 | 0.375 | 1 | 23 |
| ddx5 | 0.17738 | -0.42887529 | 0.645 | 0.875 | 1 | 23 |
| rangap1a | 0.17752 | -0.36159568 | 0.097 | 0.375 | 1 | 23 |
| itm2ba | 0.17767 | -0.31201745 | 0.645 | 0.75 | 1 | 23 |
| adssl | 0.1778 | 0.428312018 | 0.29 | 0.125 | 1 | 23 |
| mcm7 | 0.17898 | -0.4737884 | 0.29 | 0.625 | 1 | 23 |
| tjp2a | 0.1805 | -0.1454085 | 0.129 | 0.125 | 1 | 23 |
| ctnnb2 | 0.18164 | 0.365823688 | 0.581 | 0.25 | 1 | 23 |
| ifngr2 | 0.18169 | -0.23806816 | 0.194 | 0.25 | 1 | 23 |
| s100v2 | 0.1817 | 0.358372125 | 0.71 | 0.625 | 1 | 23 |
| snrpd1 | 0.18171 | -0.22337491 | 0.581 | 0.875 | 1 | 23 |
| spint1a | 0.18185 | -0.34357521 | 0.194 | 0.375 | 1 | 23 |
| babam2 | 0.18288 | -0.32780888 | 0.097 | 0.375 | 1 | 23 |
| lama5 | 0.18306 | -0.46764686 | 0.419 | 0.75 | 1 | 23 |
| arpc1b | 0.1835 | -0.5591769 | 0.097 | 0.375 | 1 | 23 |
| pomp | 0.18365 | -0.31386961 | 0.806 | 1 | 1 | 23 |
| denr | 0.18369 | -0.22062475 | 0.323 | 0.375 | 1 | 23 |
| slu7 | 0.18389 | -0.1038962 | 0.161 | 0.125 | 1 | 23 |
| tango2 | 0.18417 | 0.112672467 | 0.29 | 0.125 | 1 | 23 |
| msh2 | 0.18428 | -0.3697527 | 0.097 | 0.25 | 1 | 23 |
| rpa2 | 0.18434 | -0.2298061 | 0.129 | 0.125 | 1 | 23 |
| eef2b | 0.18461 | -0.33784233 | 0.806 | 1 | 1 | 23 |
| vdac3 | 0.18534 | 0.462964474 | 0.581 | 0.25 | 1 | 23 |
| palm1b | 0.1856 | -0.45466441 | 0.065 | 0.25 | 1 | 23 |
| sri | 0.18566 | 0.406603673 | 0.484 | 0.25 | 1 | 23 |
| rbm22 | 0.18605 | -0.35022225 | 0.097 | 0.375 | 1 | 23 |
| CABZ01115113.1 | 0.18648 | 0.476323304 | 0.419 | 0.125 | 1 | 23 |
| ctdsplb | 0.18662 | -0.43126342 | 0.065 | 0.25 | 1 | 23 |
| luc7l3 | 0.18687 | -0.15809242 | 0.129 | 0.125 | 1 | 23 |
| ginm1 | 0.18708 | 0.424927361 | 0.419 | 0.125 | 1 | 23 |
| dnajc7 | 0.18718 | -0.14720339 | 0.258 | 0.25 | 1 | 23 |
| mau2 | 0.18723 | -0.43786414 | 0.129 | 0.375 | 1 | 23 |
| fgfr3 | 0.18725 | 0.207256482 | 0.129 | 0.125 | 1 | 23 |
| meis1b | 0.18758 | -0.44281175 | 0.419 | 0.625 | 1 | 23 |
| apc | 0.18772 | -0.46235585 | 0.065 | 0.25 | 1 | 23 |
| slc1a3a | 0.18801 | -0.30129821 | 0.097 | 0.375 | 1 | 23 |
| eef1b2 | 0.18962 | 0.303475913 | 0.968 | 0.75 | 1 | 23 |
| lrrc8aa | 0.19007 | -0.45913912 | 0.097 | 0.375 | 1 | 23 |
| jag1b | 0.1901 | -0.14190982 | 0.129 | 0.125 | 1 | 23 |
| purab | 0.19028 | -0.50549973 | 0.226 | 0.5 | 1 | 23 |
| ucp2 | 0.19039 | 0.463980592 | 0.71 | 0.5 | 1 | 23 |
| ergic3 | 0.19076 | -0.40765161 | 0.065 | 0.25 | 1 | 23 |
| alcama | 0.19098 | -0.36393034 | 0.29 | 0.625 | 1 | 23 |
| cox17 | 0.19154 | -0.34480141 | 0.097 | 0.375 | 1 | 23 |
| wwtr1 | 0.19178 | -0.38775764 | 0.226 | 0.375 | 1 | 23 |
| taf12 | 0.19196 | 0.333497066 | 0.419 | 0.125 | 1 | 23 |
| chd3 | 0.19357 | 0.45699787 | 0.71 | 0.375 | 1 | 23 |
| aip | 0.19373 | 0.504673077 | 0.452 | 0.125 | 1 | 23 |

| | | | | | | |
|---|---|---|---|---|---|---|
| bin3 | 0.19415 | -0.39503787 | 0.097 | 0.375 | 1 | 23 |
| pdia5 | 0.1944 | -0.28045916 | 0.161 | 0.25 | 1 | 23 |
| ehmt1b | 0.19457 | -0.37784073 | 0.355 | 0.5 | 1 | 23 |
| wt1a | 0.19514 | -0.47971253 | 0.452 | 0.75 | 1 | 23 |
| cox7c | 0.19679 | -0.30762162 | 0.774 | 1 | 1 | 23 |
| lrp10 | 0.1968 | -0.38870466 | 0.097 | 0.375 | 1 | 23 |
| tmod2 | 0.19712 | -0.39494123 | 0.097 | 0.375 | 1 | 23 |
| rpl18 | 0.19726 | 0.218953437 | 1 | 0.875 | 1 | 23 |
| eif4ebp1 | 0.19832 | -0.10589469 | 0.71 | 0.625 | 1 | 23 |
| m6pr | 0.19881 | -0.140847 | 0.129 | 0.125 | 1 | 23 |
| tegt | 0.19945 | -0.39243168 | 0.645 | 0.875 | 1 | 23 |
| hif1an | 0.20123 | -0.36221293 | 0.097 | 0.25 | 1 | 23 |
| yipf5 | 0.20135 | -0.38361371 | 0.097 | 0.375 | 1 | 23 |
| tprb | 0.20144 | -0.21838398 | 0.097 | 0.125 | 1 | 23 |
| tomm20b | 0.2016 | 0.333169448 | 0.71 | 0.375 | 1 | 23 |
| gadd45ba | 0.2018 | 0.417337812 | 0.871 | 0.625 | 1 | 23 |
| myo1ca | 0.20198 | -0.38460021 | 0.097 | 0.25 | 1 | 23 |
| rps26 | 0.20268 | 0.211877099 | 1 | 0.875 | 1 | 23 |
| clk4a | 0.20278 | -0.14219976 | 0.129 | 0.125 | 1 | 23 |
| foxk2 | 0.20324 | -0.38510639 | 0.097 | 0.25 | 1 | 23 |
| rab11a | 0.20381 | -0.23002446 | 0.452 | 0.5 | 1 | 23 |
| cyp26b1 | 0.20437 | -0.22385543 | 0.097 | 0.125 | 1 | 23 |
| ube2e3 | 0.20449 | 0.327555969 | 0.903 | 0.625 | 1 | 23 |
| snx2 | 0.20469 | -0.38032447 | 0.065 | 0.25 | 1 | 23 |
| cdk11b | 0.20491 | -0.5054204 | 0.129 | 0.375 | 1 | 23 |
| ptbp1a | 0.20523 | 0.480949146 | 0.387 | 0.125 | 1 | 23 |
| ttc33 | 0.20527 | -0.17723466 | 0.065 | 0.25 | 1 | 23 |
| klf6a | 0.20575 | -0.5004479 | 0.387 | 0.625 | 1 | 23 |
| tspan3a | 0.20666 | 0.320609849 | 0.548 | 0.25 | 1 | 23 |
| arrdc3a | 0.20742 | -0.40679906 | 0.097 | 0.375 | 1 | 23 |
| zgc:63882 | 0.20782 | -0.38912822 | 0.097 | 0.375 | 1 | 23 |
| gstk1 | 0.20811 | 0.119906413 | 0.29 | 0.125 | 1 | 23 |
| XLOC-016094 | 0.20823 | -0.20857694 | 0.097 | 0.125 | 1 | 23 |
| ddx23 | 0.20903 | -0.18754492 | 0.065 | 0.25 | 1 | 23 |
| rhoac | 0.21015 | 0.431735803 | 0.548 | 0.25 | 1 | 23 |
| dynlrb1 | 0.21017 | 0.305308841 | 0.548 | 0.25 | 1 | 23 |
| smo | 0.2102 | -0.22748515 | 0.194 | 0.25 | 1 | 23 |
| thumpd3 | 0.21021 | -0.39537138 | 0.097 | 0.25 | 1 | 23 |
| mars | 0.21052 | -0.44102711 | 0.129 | 0.375 | 1 | 23 |
| mcrip1 | 0.21157 | -0.25261143 | 0.129 | 0.375 | 1 | 23 |
| spina | 0.21196 | -0.49144068 | 0.129 | 0.375 | 1 | 23 |
| eif4a3 | 0.21311 | 0.456640594 | 0.419 | 0.125 | 1 | 23 |
| rab6a | 0.2133 | -0.16101564 | 0.226 | 0.25 | 1 | 23 |
| cpsf6 | 0.21375 | -0.44000263 | 0.29 | 0.625 | 1 | 23 |
| fgfr1a | 0.21382 | -0.37943392 | 0.097 | 0.25 | 1 | 23 |
| s100a10b | 0.21415 | 0.177269493 | 0.774 | 0.5 | 1 | 23 |
| hm13 | 0.21424 | 0.441734896 | 0.419 | 0.125 | 1 | 23 |
| LO018188.1 | 0.21506 | -0.46098202 | 0.194 | 0.375 | 1 | 23 |

| | | | | | | |
|---|---|---|---|---|---|---|
| rbm4.3 | 0.21506 | 0.428210468 | 0.774 | 0.5 | 1 | 23 |
| ralyl | 0.21512 | -0.10871377 | 0.161 | 0.125 | 1 | 23 |
| celf2 | 0.21528 | -0.30119981 | 0.645 | 0.75 | 1 | 23 |
| eno3 | 0.21587 | -0.36569325 | 0.29 | 0.375 | 1 | 23 |
| zgc:162730 | 0.2165 | -0.43028722 | 0.645 | 0.75 | 1 | 23 |
| rad23ab | 0.21668 | -0.42137036 | 0.065 | 0.25 | 1 | 23 |
| sem1 | 0.21842 | 0.342929831 | 0.806 | 0.625 | 1 | 23 |
| f11r.1 | 0.21969 | 0.236126446 | 0.645 | 0.625 | 1 | 23 |
| fxyd1 | 0.21991 | -0.12646521 | 0.161 | 0.125 | 1 | 23 |
| gdpd5b | 0.22039 | -0.21096808 | 0.097 | 0.125 | 1 | 23 |
| ddx24 | 0.22072 | -0.45136826 | 0.226 | 0.5 | 1 | 23 |
| eif3g | 0.22151 | 0.395647215 | 0.806 | 0.5 | 1 | 23 |
| DST | 0.22166 | -0.45169612 | 0.065 | 0.25 | 1 | 23 |
| scpep1 | 0.22173 | -0.14249651 | 0.129 | 0.125 | 1 | 23 |
| suclg2 | 0.22219 | -0.33139494 | 0.065 | 0.25 | 1 | 23 |
| gnai2a | 0.22246 | 0.465036468 | 0.419 | 0.125 | 1 | 23 |
| CR318588.3 | 0.22268 | -0.54574933 | 0.194 | 0.5 | 1 | 23 |
| dynll1 | 0.22337 | -0.41268109 | 0.645 | 0.875 | 1 | 23 |
| psmc1b | 0.2234 | 0.284598675 | 0.387 | 0.125 | 1 | 23 |
| cox7b | 0.22398 | -0.27574247 | 0.839 | 0.875 | 1 | 23 |
| naa10 | 0.22431 | 0.387161569 | 0.581 | 0.25 | 1 | 23 |
| si:dkey-6n21.12 | 0.22449 | 0.462088034 | 0.419 | 0.125 | 1 | 23 |
| nkain1 | 0.22474 | -0.13740121 | 0.129 | 0.125 | 1 | 23 |
| nnt | 0.2265 | -0.41539429 | 0.065 | 0.25 | 1 | 23 |
| rap1aa | 0.22655 | -0.38197823 | 0.29 | 0.5 | 1 | 23 |
| anxa4 | 0.22679 | 0.440189915 | 0.581 | 0.25 | 1 | 23 |
| dnajc9 | 0.22749 | 0.44945948 | 0.355 | 0.125 | 1 | 23 |
| pfdn1 | 0.22758 | 0.428670974 | 0.387 | 0.125 | 1 | 23 |
| mycb | 0.22813 | 0.510159434 | 0.581 | 0.25 | 1 | 23 |
| mbtps1 | 0.22859 | -0.20935841 | 0.097 | 0.125 | 1 | 23 |
| mrps36 | 0.22868 | -0.35195564 | 0.194 | 0.375 | 1 | 23 |
| cinp | 0.22953 | 0.186528369 | 0.129 | 0 | 1 | 23 |
| glb1l | 0.22953 | 0.19591415 | 0.129 | 0 | 1 | 23 |
| gstz1 | 0.22953 | 0.109300074 | 0.129 | 0 | 1 | 23 |
| rrbp1b | 0.22953 | 0.192867375 | 0.129 | 0 | 1 | 23 |
| si:ch211-225b7.5 | 0.22953 | 0.259296135 | 0.129 | 0 | 1 | 23 |
| tmem9 | 0.22953 | 0.199696599 | 0.129 | 0 | 1 | 23 |
| ttpa | 0.22953 | 0.215180903 | 0.129 | 0 | 1 | 23 |
| znf395b | 0.22953 | 0.220471505 | 0.129 | 0 | 1 | 23 |
| CR388143.1 | 0.22953 | 0.203763699 | 0.129 | 0 | 1 | 23 |
| EML4 | 0.22953 | 0.183251072 | 0.129 | 0 | 1 | 23 |
| MYADM | 0.22953 | 0.259970823 | 0.129 | 0 | 1 | 23 |
| XLOC-004016 | 0.22953 | 0.206457782 | 0.129 | 0 | 1 | 23 |
| aagab | 0.22953 | 0.212884592 | 0.129 | 0 | 1 | 23 |
| abhd15a | 0.22953 | 0.286414056 | 0.129 | 0 | 1 | 23 |
| ablim1b | 0.22953 | 0.206970892 | 0.129 | 0 | 1 | 23 |
| ahsa1b | 0.22953 | 0.19613863 | 0.129 | 0 | 1 | 23 |
| anapc15 | 0.22953 | 0.225388205 | 0.129 | 0 | 1 | 23 |

| | | | | | | |
|---|---|---|---|---|---|---|
| appbp2 | 0.22953 | 0.190247161 | 0.129 | 0 | 1 | 23 |
| arhgap35b | 0.22953 | 0.243455746 | 0.129 | 0 | 1 | 23 |
| bcar1 | 0.22953 | 0.21342903 | 0.129 | 0 | 1 | 23 |
| ca10a | 0.22953 | 0.253793413 | 0.129 | 0 | 1 | 23 |
| cav2 | 0.22953 | 0.245380259 | 0.129 | 0 | 1 | 23 |
| ccdc28a | 0.22953 | 0.247622925 | 0.129 | 0 | 1 | 23 |
| chchd7 | 0.22953 | 0.214150662 | 0.129 | 0 | 1 | 23 |
| chm | 0.22953 | 0.203543367 | 0.129 | 0 | 1 | 23 |
| cipcb | 0.22953 | 0.204219524 | 0.129 | 0 | 1 | 23 |
| cops5 | 0.22953 | 0.19303081 | 0.129 | 0 | 1 | 23 |
| cox10 | 0.22953 | 0.181568542 | 0.129 | 0 | 1 | 23 |
| dab2ipb | 0.22953 | 0.197903233 | 0.129 | 0 | 1 | 23 |
| dek | 0.22953 | 0.165882778 | 0.129 | 0 | 1 | 23 |
| dhrsx | 0.22953 | 0.187446697 | 0.129 | 0 | 1 | 23 |
| dnajb12a | 0.22953 | 0.189324163 | 0.129 | 0 | 1 | 23 |
| dnajc3b | 0.22953 | 0.216741868 | 0.129 | 0 | 1 | 23 |
| dpm1 | 0.22953 | 0.207893857 | 0.129 | 0 | 1 | 23 |
| dr1 | 0.22953 | 0.207924666 | 0.129 | 0 | 1 | 23 |
| dusp6 | 0.22953 | 0.183192961 | 0.129 | 0 | 1 | 23 |
| epb41a | 0.22953 | 0.195692978 | 0.129 | 0 | 1 | 23 |
| flii | 0.22953 | 0.181264488 | 0.129 | 0 | 1 | 23 |
| gcsha | 0.22953 | 0.202321896 | 0.129 | 0 | 1 | 23 |
| glyctk | 0.22953 | 0.17908233 | 0.129 | 0 | 1 | 23 |
| glyr1 | 0.22953 | 0.18618988 | 0.129 | 0 | 1 | 23 |
| gosr2 | 0.22953 | 0.221926077 | 0.129 | 0 | 1 | 23 |
| gtf2h3 | 0.22953 | 0.242711598 | 0.129 | 0 | 1 | 23 |
| hddc3 | 0.22953 | 0.195527612 | 0.129 | 0 | 1 | 23 |
| hspbp1 | 0.22953 | 0.199702947 | 0.129 | 0 | 1 | 23 |
| il10rb | 0.22953 | 0.220601782 | 0.129 | 0 | 1 | 23 |
| ino80c | 0.22953 | 0.204737739 | 0.129 | 0 | 1 | 23 |
| inpp5l | 0.22953 | 0.192919575 | 0.129 | 0 | 1 | 23 |
| insra | 0.22953 | 0.198188726 | 0.129 | 0 | 1 | 23 |
| ireb2 | 0.22953 | 0.196675521 | 0.129 | 0 | 1 | 23 |
| kdm3b | 0.22953 | 0.200388085 | 0.129 | 0 | 1 | 23 |
| kdm6ba | 0.22953 | 0.20708772 | 0.129 | 0 | 1 | 23 |
| klf7b | 0.22953 | 0.238017829 | 0.129 | 0 | 1 | 23 |
| klf9 | 0.22953 | 0.257782962 | 0.129 | 0 | 1 | 23 |
| kmt2ca | 0.22953 | 0.186199673 | 0.129 | 0 | 1 | 23 |
| ktn1 | 0.22953 | 0.196023863 | 0.129 | 0 | 1 | 23 |
| lamp1b | 0.22953 | 0.228227153 | 0.129 | 0 | 1 | 23 |
| lrrc20 | 0.22953 | 0.197636755 | 0.129 | 0 | 1 | 23 |
| magixa | 0.22953 | 0.189173548 | 0.129 | 0 | 1 | 23 |
| mapk4 | 0.22953 | 0.26897889 | 0.129 | 0 | 1 | 23 |
| mbd4 | 0.22953 | 0.224255802 | 0.129 | 0 | 1 | 23 |
| mcat | 0.22953 | 0.192084122 | 0.129 | 0 | 1 | 23 |
| med25 | 0.22953 | 0.241906884 | 0.129 | 0 | 1 | 23 |
| men1 | 0.22953 | 0.214609726 | 0.129 | 0 | 1 | 23 |
| mpped2 | 0.22953 | 0.198378907 | 0.129 | 0 | 1 | 23 |

| | | | | | | |
|---|---|---|---|---|---|---|
| mrpl37 | 0.22953 | 0.204074149 | 0.129 | 0 | 1 | 23 |
| naa40 | 0.22953 | 0.282305479 | 0.129 | 0 | 1 | 23 |
| nol10 | 0.22953 | 0.206128687 | 0.129 | 0 | 1 | 23 |
| nol4lb | 0.22953 | 0.257861629 | 0.129 | 0 | 1 | 23 |
| nr2f6a | 0.22953 | 0.191983318 | 0.129 | 0 | 1 | 23 |
| nsrp1 | 0.22953 | 0.20247646 | 0.129 | 0 | 1 | 23 |
| ociad1 | 0.22953 | 0.180548325 | 0.129 | 0 | 1 | 23 |
| pcid2 | 0.22953 | 0.19818696 | 0.129 | 0 | 1 | 23 |
| pcm1 | 0.22953 | 0.223851577 | 0.129 | 0 | 1 | 23 |
| pdk2a | 0.22953 | 0.18504365 | 0.129 | 0 | 1 | 23 |
| pgm1 | 0.22953 | 0.127022748 | 0.129 | 0 | 1 | 23 |
| plekha6 | 0.22953 | 0.20983297 | 0.129 | 0 | 1 | 23 |
| plpp1a | 0.22953 | 0.260696429 | 0.129 | 0 | 1 | 23 |
| prdm11 | 0.22953 | 0.246115726 | 0.129 | 0 | 1 | 23 |
| prdm2b | 0.22953 | 0.288956188 | 0.129 | 0 | 1 | 23 |
| prrc1 | 0.22953 | 0.20892872 | 0.129 | 0 | 1 | 23 |
| rab3db | 0.22953 | 0.191815756 | 0.129 | 0 | 1 | 23 |
| rbfox2 | 0.22953 | 0.231194134 | 0.129 | 0 | 1 | 23 |
| rbm48 | 0.22953 | 0.216491984 | 0.129 | 0 | 1 | 23 |
| rexo4 | 0.22953 | 0.191799111 | 0.129 | 0 | 1 | 23 |
| rfxap | 0.22953 | 0.181126236 | 0.129 | 0 | 1 | 23 |
| rgs7bpb | 0.22953 | 0.209377432 | 0.129 | 0 | 1 | 23 |
| rnaseh2c | 0.22953 | 0.202255537 | 0.129 | 0 | 1 | 23 |
| rnf141 | 0.22953 | 0.211551742 | 0.129 | 0 | 1 | 23 |
| rxrba | 0.22953 | 0.20063067 | 0.129 | 0 | 1 | 23 |
| shdb | 0.22953 | 0.173522486 | 0.129 | 0 | 1 | 23 |
| shmt1 | 0.22953 | 0.171149556 | 0.129 | 0 | 1 | 23 |
| si:ch211-275j6.5 | 0.22953 | 0.257339407 | 0.129 | 0 | 1 | 23 |
| si:ch73-1a9.4 | 0.22953 | 0.219532401 | 0.129 | 0 | 1 | 23 |
| si:dkey-16p19.5 | 0.22953 | 0.211170165 | 0.129 | 0 | 1 | 23 |
| sin3b | 0.22953 | 0.236053501 | 0.129 | 0 | 1 | 23 |
| slc25a1a | 0.22953 | 0.176356721 | 0.129 | 0 | 1 | 23 |
| slc38a10 | 0.22953 | 0.230592015 | 0.129 | 0 | 1 | 23 |
| smim29 | 0.22953 | 0.196777731 | 0.129 | 0 | 1 | 23 |
| spopla | 0.22953 | 0.187276049 | 0.129 | 0 | 1 | 23 |
| svild | 0.22953 | 0.191312131 | 0.129 | 0 | 1 | 23 |
| tarbp2 | 0.22953 | 0.179976027 | 0.129 | 0 | 1 | 23 |
| tgfb2 | 0.22953 | 0.280083497 | 0.129 | 0 | 1 | 23 |
| thrab | 0.22953 | 0.22587913 | 0.129 | 0 | 1 | 23 |
| tnfsf12 | 0.22953 | 0.233790471 | 0.129 | 0 | 1 | 23 |
| tram2 | 0.22953 | 0.214600361 | 0.129 | 0 | 1 | 23 |
| trim33 | 0.22953 | 0.198550311 | 0.129 | 0 | 1 | 23 |
| ube2kb | 0.22953 | 0.18728481 | 0.129 | 0 | 1 | 23 |
| usp10 | 0.22953 | 0.203383361 | 0.129 | 0 | 1 | 23 |
| wu:fc33b09 | 0.22953 | 0.248715637 | 0.129 | 0 | 1 | 23 |
| ythdf2 | 0.22953 | 0.244009257 | 0.129 | 0 | 1 | 23 |
| zgc:153675 | 0.22953 | 0.180856011 | 0.129 | 0 | 1 | 23 |
| znf451 | 0.22953 | 0.201514885 | 0.129 | 0 | 1 | 23 |

| | | | | | | |
|---|---|---|---|---|---|---|
| DHDH | 0.22953 | 0.163943575 | 0.129 | 0 | 1 | 23 |
| FP074902.2 | 0.22953 | 0.270601269 | 0.129 | 0 | 1 | 23 |
| FQ976913.1 | 0.22953 | 0.178758442 | 0.129 | 0 | 1 | 23 |
| LOC110437758 | 0.22953 | 0.240051951 | 0.129 | 0 | 1 | 23 |
| LOC110437917 | 0.22953 | 0.207468869 | 0.129 | 0 | 1 | 23 |
| abca2 | 0.22953 | 0.238948306 | 0.129 | 0 | 1 | 23 |
| adnp2b | 0.22953 | 0.201315301 | 0.129 | 0 | 1 | 23 |
| aldh1l1 | 0.22953 | 0.111098965 | 0.129 | 0 | 1 | 23 |
| ap3m2 | 0.22953 | 0.196039373 | 0.129 | 0 | 1 | 23 |
| apbb1 | 0.22953 | 0.200909509 | 0.129 | 0 | 1 | 23 |
| arl2 | 0.22953 | 0.190932099 | 0.129 | 0 | 1 | 23 |
| asnsd1 | 0.22953 | 0.182036067 | 0.129 | 0 | 1 | 23 |
| azin1b | 0.22953 | 0.196290598 | 0.129 | 0 | 1 | 23 |
| bod1 | 0.22953 | 0.216204845 | 0.129 | 0 | 1 | 23 |
| cbr4 | 0.22953 | 0.178844629 | 0.129 | 0 | 1 | 23 |
| cnot1 | 0.22953 | 0.193912926 | 0.129 | 0 | 1 | 23 |
| cops8 | 0.22953 | 0.187847901 | 0.129 | 0 | 1 | 23 |
| crebbpa | 0.22953 | 0.190328497 | 0.129 | 0 | 1 | 23 |
| ctdspl2a | 0.22953 | 0.271410731 | 0.129 | 0 | 1 | 23 |
| dclk1a | 0.22953 | 0.284667468 | 0.129 | 0 | 1 | 23 |
| decr2 | 0.22953 | 0.234525864 | 0.129 | 0 | 1 | 23 |
| derl2 | 0.22953 | 0.215966239 | 0.129 | 0 | 1 | 23 |
| dhfr | 0.22953 | 0.229598446 | 0.129 | 0 | 1 | 23 |
| eri1 | 0.22953 | 0.19772506 | 0.129 | 0 | 1 | 23 |
| fbxl5 | 0.22953 | 0.159910965 | 0.129 | 0 | 1 | 23 |
| gsk3b | 0.22953 | 0.209704168 | 0.129 | 0 | 1 | 23 |
| hadhaa | 0.22953 | 0.205537212 | 0.129 | 0 | 1 | 23 |
| imp4 | 0.22953 | 0.220082518 | 0.129 | 0 | 1 | 23 |
| ing1 | 0.22953 | 0.204702666 | 0.129 | 0 | 1 | 23 |
| ing3 | 0.22953 | 0.245603677 | 0.129 | 0 | 1 | 23 |
| itgb1a | 0.22953 | 0.245566183 | 0.129 | 0 | 1 | 23 |
| kcmf1 | 0.22953 | 0.197862328 | 0.129 | 0 | 1 | 23 |
| kctd6b | 0.22953 | 0.215738901 | 0.129 | 0 | 1 | 23 |
| leo1 | 0.22953 | 0.180799273 | 0.129 | 0 | 1 | 23 |
| lin52 | 0.22953 | 0.173049257 | 0.129 | 0 | 1 | 23 |
| lin7c | 0.22953 | 0.187367045 | 0.129 | 0 | 1 | 23 |
| lzts1 | 0.22953 | 0.174880209 | 0.129 | 0 | 1 | 23 |
| med15 | 0.22953 | 0.200611605 | 0.129 | 0 | 1 | 23 |
| ncalda | 0.22953 | 0.206126107 | 0.129 | 0 | 1 | 23 |
| nfatc3a | 0.22953 | 0.201506914 | 0.129 | 0 | 1 | 23 |
| nupl2 | 0.22953 | 0.207508157 | 0.129 | 0 | 1 | 23 |
| parp12a | 0.22953 | 0.231070001 | 0.129 | 0 | 1 | 23 |
| pdap1b | 0.22953 | 0.221463004 | 0.129 | 0 | 1 | 23 |
| pgp | 0.22953 | 0.177368472 | 0.129 | 0 | 1 | 23 |
| pitpnaa | 0.22953 | 0.19931981 | 0.129 | 0 | 1 | 23 |
| plpp5 | 0.22953 | 0.19522461 | 0.129 | 0 | 1 | 23 |
| polr3h | 0.22953 | 0.179850325 | 0.129 | 0 | 1 | 23 |
| pop7 | 0.22953 | 0.211019642 | 0.129 | 0 | 1 | 23 |

| | | | | | | |
|---|---|---|---|---|---|---|
| ppp3r1a | 0.22953 | 0.19863049 | 0.129 | 0 | 1 | 23 |
| qpctla | 0.22953 | 0.221027391 | 0.129 | 0 | 1 | 23 |
| rbm34 | 0.22953 | 0.189973522 | 0.129 | 0 | 1 | 23 |
| rbm42 | 0.22953 | 0.198728746 | 0.129 | 0 | 1 | 23 |
| rbms1b | 0.22953 | 0.224636231 | 0.129 | 0 | 1 | 23 |
| rcor1 | 0.22953 | 0.188323137 | 0.129 | 0 | 1 | 23 |
| rdh14b | 0.22953 | 0.234668105 | 0.129 | 0 | 1 | 23 |
| rnf8 | 0.22953 | 0.21050897 | 0.129 | 0 | 1 | 23 |
| rpp25l | 0.22953 | 0.189808993 | 0.129 | 0 | 1 | 23 |
| sap30bp | 0.22953 | 0.2060576 | 0.129 | 0 | 1 | 23 |
| sccpdhb | 0.22953 | 0.177158471 | 0.129 | 0 | 1 | 23 |
| sec23a | 0.22953 | 0.226424871 | 0.129 | 0 | 1 | 23 |
| selenot1a | 0.22953 | 0.17151636 | 0.129 | 0 | 1 | 23 |
| si:dkey-23i12.5 | 0.22953 | 0.175801918 | 0.129 | 0 | 1 | 23 |
| si:dkey-26c10.5 | 0.22953 | 0.404886837 | 0.129 | 0 | 1 | 23 |
| si:dkeyp-87e7.4 | 0.22953 | 0.225763757 | 0.129 | 0 | 1 | 23 |
| slc35b4 | 0.22953 | 0.209399518 | 0.129 | 0 | 1 | 23 |
| smfn | 0.22953 | 0.175019201 | 0.129 | 0 | 1 | 23 |
| sppl3 | 0.22953 | 0.202357626 | 0.129 | 0 | 1 | 23 |
| spryd4 | 0.22953 | 0.180627575 | 0.129 | 0 | 1 | 23 |
| strada | 0.22953 | 0.179331225 | 0.129 | 0 | 1 | 23 |
| tecrb | 0.22953 | 0.180757955 | 0.129 | 0 | 1 | 23 |
| tmem11 | 0.22953 | 0.16615133 | 0.129 | 0 | 1 | 23 |
| tyw1 | 0.22953 | 0.162846555 | 0.129 | 0 | 1 | 23 |
| ube2g1b | 0.22953 | 0.217821157 | 0.129 | 0 | 1 | 23 |
| uqcc1 | 0.22953 | 0.20609625 | 0.129 | 0 | 1 | 23 |
| utp3 | 0.22953 | 0.19480893 | 0.129 | 0 | 1 | 23 |
| zgc:103697 | 0.22953 | 0.19993037 | 0.129 | 0 | 1 | 23 |
| zgc:110269 | 0.22953 | 0.190420681 | 0.129 | 0 | 1 | 23 |
| zgc:162509 | 0.22953 | 0.186539708 | 0.129 | 0 | 1 | 23 |
| znf362b | 0.22953 | 0.219258618 | 0.129 | 0 | 1 | 23 |
| CABZ01086877.1 | 0.22953 | 0.250171424 | 0.129 | 0 | 1 | 23 |
| CABZ01088149.1 | 0.22953 | 0.189973878 | 0.129 | 0 | 1 | 23 |
| LO018340.1 | 0.22953 | 0.304027893 | 0.129 | 0 | 1 | 23 |
| actr2a | 0.22953 | 0.15995399 | 0.129 | 0 | 1 | 23 |
| adamts15a | 0.22953 | 0.291703781 | 0.129 | 0 | 1 | 23 |
| adamts2 | 0.22953 | 0.203896645 | 0.129 | 0 | 1 | 23 |
| arhgef12b | 0.22953 | 0.281515148 | 0.129 | 0 | 1 | 23 |
| arrb2b | 0.22953 | 0.20175605 | 0.129 | 0 | 1 | 23 |
| aspscr1 | 0.22953 | 0.211810533 | 0.129 | 0 | 1 | 23 |
| atp6v1d | 0.22953 | 0.177605839 | 0.129 | 0 | 1 | 23 |
| atpaf1 | 0.22953 | 0.200248394 | 0.129 | 0 | 1 | 23 |
| atxn3 | 0.22953 | 0.200555827 | 0.129 | 0 | 1 | 23 |
| btg3 | 0.22953 | 0.17825783 | 0.129 | 0 | 1 | 23 |
| cmss1 | 0.22953 | 0.203050063 | 0.129 | 0 | 1 | 23 |
| cox16 | 0.22953 | 0.173673318 | 0.129 | 0 | 1 | 23 |
| cyp20a1 | 0.22953 | 0.18691992 | 0.129 | 0 | 1 | 23 |
| ddx54 | 0.22953 | 0.20829213 | 0.129 | 0 | 1 | 23 |

| | | | | | | |
|---|---|---|---|---|---|---|
| dnajc4 | 0.22953 | 0.204382133 | 0.129 | 0 | 1 | 23 |
| dnajc5gb | 0.22953 | 0.179818671 | 0.129 | 0 | 1 | 23 |
| dvl3a | 0.22953 | 0.21740385 | 0.129 | 0 | 1 | 23 |
| epc2 | 0.22953 | 0.189628817 | 0.129 | 0 | 1 | 23 |
| exoc1 | 0.22953 | 0.233304014 | 0.129 | 0 | 1 | 23 |
| fam173a | 0.22953 | 0.2046289 | 0.129 | 0 | 1 | 23 |
| fam49a | 0.22953 | 0.190126743 | 0.129 | 0 | 1 | 23 |
| frs2b | 0.22953 | 0.202844087 | 0.129 | 0 | 1 | 23 |
| gnsb | 0.22953 | 0.192969964 | 0.129 | 0 | 1 | 23 |
| hat1 | 0.22953 | 0.18616517 | 0.129 | 0 | 1 | 23 |
| hcfc1b | 0.22953 | 0.270766277 | 0.129 | 0 | 1 | 23 |
| ilf3a | 0.22953 | 0.226285008 | 0.129 | 0 | 1 | 23 |
| kdm7aa | 0.22953 | 0.218806089 | 0.129 | 0 | 1 | 23 |
| lmtk2 | 0.22953 | 0.201349856 | 0.129 | 0 | 1 | 23 |
| lsm8 | 0.22953 | 0.14786894 | 0.129 | 0 | 1 | 23 |
| maea | 0.22953 | 0.218664456 | 0.129 | 0 | 1 | 23 |
| mif4gda | 0.22953 | 0.22774377 | 0.129 | 0 | 1 | 23 |
| mrps9 | 0.22953 | 0.199802538 | 0.129 | 0 | 1 | 23 |
| mthfd1l | 0.22953 | 0.228408516 | 0.129 | 0 | 1 | 23 |
| ndc1 | 0.22953 | 0.17884253 | 0.129 | 0 | 1 | 23 |
| ndufv1 | 0.22953 | 0.22920535 | 0.129 | 0 | 1 | 23 |
| npepps | 0.22953 | 0.213389679 | 0.129 | 0 | 1 | 23 |
| pdf | 0.22953 | 0.222519176 | 0.129 | 0 | 1 | 23 |
| pdk4 | 0.22953 | 0.248989465 | 0.129 | 0 | 1 | 23 |
| pgrmc2 | 0.22953 | 0.205274279 | 0.129 | 0 | 1 | 23 |
| phkg2 | 0.22953 | 0.161106834 | 0.129 | 0 | 1 | 23 |
| pou2f1b | 0.22953 | 0.212473215 | 0.129 | 0 | 1 | 23 |
| prkrip1 | 0.22953 | 0.243883955 | 0.129 | 0 | 1 | 23 |
| prmt6 | 0.22953 | 0.183321458 | 0.129 | 0 | 1 | 23 |
| psme3 | 0.22953 | 0.248641785 | 0.129 | 0 | 1 | 23 |
| ptrh2 | 0.22953 | 0.209859625 | 0.129 | 0 | 1 | 23 |
| qdprb1 | 0.22953 | 0.19956531 | 0.129 | 0 | 1 | 23 |
| rac3a | 0.22953 | 0.274425576 | 0.129 | 0 | 1 | 23 |
| rassf1 | 0.22953 | 0.260085927 | 0.129 | 0 | 1 | 23 |
| rbfox1 | 0.22953 | 0.186217078 | 0.129 | 0 | 1 | 23 |
| rbm12b | 0.22953 | 0.220372665 | 0.129 | 0 | 1 | 23 |
| rbm14a | 0.22953 | 0.196217844 | 0.129 | 0 | 1 | 23 |
| rbms2a | 0.22953 | 0.192293681 | 0.129 | 0 | 1 | 23 |
| rnaseh2a | 0.22953 | 0.239278909 | 0.129 | 0 | 1 | 23 |
| rnd1b | 0.22953 | 0.207842196 | 0.129 | 0 | 1 | 23 |
| rpp14 | 0.22953 | 0.214851548 | 0.129 | 0 | 1 | 23 |
| rpz4 | 0.22953 | 0.194919328 | 0.129 | 0 | 1 | 23 |
| selenoj | 0.22953 | 0.16033123 | 0.129 | 0 | 1 | 23 |
| selenou1a | 0.22953 | 0.178552084 | 0.129 | 0 | 1 | 23 |
| si:ch1073-340j19.1 | 0.22953 | 0.178605591 | 0.129 | 0 | 1 | 23 |
| si:dkey-253d23.3 | 0.22953 | 0.225444694 | 0.129 | 0 | 1 | 23 |
| slc12a2 | 0.22953 | 0.220104028 | 0.129 | 0 | 1 | 23 |
| slc25a51b | 0.22953 | 0.224878046 | 0.129 | 0 | 1 | 23 |

| | | | | | | |
|---|---|---|---|---|---|---|
| sox11a | 0.22953 | 0.175487962 | 0.129 | 0 | 1 | 23 |
| tead1b | 0.22953 | 0.231803859 | 0.129 | 0 | 1 | 23 |
| tfap4 | 0.22953 | 0.209600566 | 0.129 | 0 | 1 | 23 |
| tigarb | 0.22953 | 0.19045403 | 0.129 | 0 | 1 | 23 |
| tm9sf5 | 0.22953 | 0.230928603 | 0.129 | 0 | 1 | 23 |
| tmcc3 | 0.22953 | 0.204164595 | 0.129 | 0 | 1 | 23 |
| tmem108 | 0.22953 | 0.239182135 | 0.129 | 0 | 1 | 23 |
| tmem141 | 0.22953 | 0.207157034 | 0.129 | 0 | 1 | 23 |
| tp53i11a | 0.22953 | 0.210270798 | 0.129 | 0 | 1 | 23 |
| trappc4 | 0.22953 | 0.150821539 | 0.129 | 0 | 1 | 23 |
| ttyh2 | 0.22953 | 0.250092835 | 0.129 | 0 | 1 | 23 |
| ube2z | 0.22953 | 0.259497499 | 0.129 | 0 | 1 | 23 |
| usp25 | 0.22953 | 0.225996479 | 0.129 | 0 | 1 | 23 |
| vclb | 0.22953 | 0.229639078 | 0.129 | 0 | 1 | 23 |
| vkorc1 | 0.22953 | 0.219640171 | 0.129 | 0 | 1 | 23 |
| wrb | 0.22953 | 0.213051814 | 0.129 | 0 | 1 | 23 |
| zgc:158517 | 0.22953 | 0.199086428 | 0.129 | 0 | 1 | 23 |
| zgpat | 0.22953 | 0.185404375 | 0.129 | 0 | 1 | 23 |
| zrsr2 | 0.22953 | 0.246357645 | 0.129 | 0 | 1 | 23 |
| acot13 | 0.22953 | 0.20923856 | 0.129 | 0 | 1 | 23 |
| ehd1b | 0.22953 | 0.222057274 | 0.129 | 0 | 1 | 23 |
| minpp1b | 0.22953 | 0.221641957 | 0.129 | 0 | 1 | 23 |
| mycla | 0.22953 | 0.21676375 | 0.129 | 0 | 1 | 23 |
| prim2 | 0.22953 | 0.21137284 | 0.129 | 0 | 1 | 23 |
| si:ch211-248e11.2 | 0.22953 | 0.239397011 | 0.129 | 0 | 1 | 23 |
| si:ch73-211e3.1 | 0.22953 | 0.193139583 | 0.129 | 0 | 1 | 23 |
| tapbp.2 | 0.22953 | 0.207836871 | 0.129 | 0 | 1 | 23 |
| wipi2 | 0.22953 | 0.224756148 | 0.129 | 0 | 1 | 23 |
| CR759885.1 | 0.22953 | 0.256236135 | 0.129 | 0 | 1 | 23 |
| LOC108182329 | 0.22953 | 0.212718476 | 0.129 | 0 | 1 | 23 |
| arrb2a | 0.22953 | 0.220615642 | 0.129 | 0 | 1 | 23 |
| asna1 | 0.22953 | 0.260664717 | 0.129 | 0 | 1 | 23 |
| atad2b | 0.22953 | 0.188495445 | 0.129 | 0 | 1 | 23 |
| atf7b | 0.22953 | 0.194459658 | 0.129 | 0 | 1 | 23 |
| bcas3 | 0.22953 | 0.2219645 | 0.129 | 0 | 1 | 23 |
| bop1 | 0.22953 | 0.209946754 | 0.129 | 0 | 1 | 23 |
| capsla | 0.22953 | 0.204713708 | 0.129 | 0 | 1 | 23 |
| cops7a | 0.22953 | 0.20468993 | 0.129 | 0 | 1 | 23 |
| erap1a | 0.22953 | 0.187676849 | 0.129 | 0 | 1 | 23 |
| faf1 | 0.22953 | 0.195378238 | 0.129 | 0 | 1 | 23 |
| fgfrl1a | 0.22953 | 0.240824775 | 0.129 | 0 | 1 | 23 |
| gabpa | 0.22953 | 0.252465261 | 0.129 | 0 | 1 | 23 |
| kat5b | 0.22953 | 0.208362448 | 0.129 | 0 | 1 | 23 |
| krr1 | 0.22953 | 0.188714186 | 0.129 | 0 | 1 | 23 |
| lsp1 | 0.22953 | 0.22610556 | 0.129 | 0 | 1 | 23 |
| map3k7 | 0.22953 | 0.212882447 | 0.129 | 0 | 1 | 23 |
| mthfd2l | 0.22953 | 0.177203874 | 0.129 | 0 | 1 | 23 |
| pdhx | 0.22953 | 0.179902422 | 0.129 | 0 | 1 | 23 |

| | | | | | | |
|---|---|---|---|---|---|---|
| pdss1 | 0.22953 | 0.1944386 | 0.129 | 0 | 1 | 23 |
| plekha1a | 0.22953 | 0.211174906 | 0.129 | 0 | 1 | 23 |
| rab36 | 0.22953 | 0.240255886 | 0.129 | 0 | 1 | 23 |
| rnf130 | 0.22953 | 0.203292846 | 0.129 | 0 | 1 | 23 |
| sdr39u1 | 0.22953 | 0.221076299 | 0.129 | 0 | 1 | 23 |
| sel1l | 0.22953 | 0.227289628 | 0.129 | 0 | 1 | 23 |
| si:ch211-212k18.9 | 0.22953 | 0.213858535 | 0.129 | 0 | 1 | 23 |
| si:dkey-40c11.2 | 0.22953 | 0.194597249 | 0.129 | 0 | 1 | 23 |
| si:dkey-48p11.3 | 0.22953 | 0.196915129 | 0.129 | 0 | 1 | 23 |
| si:dkey-69o16.5 | 0.22953 | 0.151375121 | 0.129 | 0 | 1 | 23 |
| sirt7 | 0.22953 | 0.189198649 | 0.129 | 0 | 1 | 23 |
| smim19 | 0.22953 | 0.217107479 | 0.129 | 0 | 1 | 23 |
| srpk1a | 0.22953 | 0.233881635 | 0.129 | 0 | 1 | 23 |
| stx3a | 0.22953 | 0.186959741 | 0.129 | 0 | 1 | 23 |
| tmem251 | 0.22953 | 0.183973603 | 0.129 | 0 | 1 | 23 |
| tspy | 0.22953 | 0.244086505 | 0.129 | 0 | 1 | 23 |
| wnt7aa | 0.22953 | 0.202933595 | 0.129 | 0 | 1 | 23 |
| ankrd22 | 0.22953 | 0.209082464 | 0.129 | 0 | 1 | 23 |
| bloc1s2 | 0.22953 | 0.221204483 | 0.129 | 0 | 1 | 23 |
| cdkn1bb | 0.22953 | 0.242778337 | 0.129 | 0 | 1 | 23 |
| cwf19l2 | 0.22953 | 0.27386093 | 0.129 | 0 | 1 | 23 |
| exosc9 | 0.22953 | 0.222566528 | 0.129 | 0 | 1 | 23 |
| gdpd1 | 0.22953 | 0.213730004 | 0.129 | 0 | 1 | 23 |
| mrpl3 | 0.22953 | 0.203423166 | 0.129 | 0 | 1 | 23 |
| mtor | 0.22953 | 0.249282073 | 0.129 | 0 | 1 | 23 |
| sntb2 | 0.22953 | 0.265539636 | 0.129 | 0 | 1 | 23 |
| srp54 | 0.22953 | 0.197038758 | 0.129 | 0 | 1 | 23 |
| tmem231 | 0.22953 | 0.219942364 | 0.129 | 0 | 1 | 23 |
| zgc:172139 | 0.22953 | 0.239659422 | 0.129 | 0 | 1 | 23 |
| zgc:77929 | 0.22953 | 0.231204098 | 0.129 | 0 | 1 | 23 |
| rhbdf1a | 0.22972 | -0.35314828 | 0.097 | 0.25 | 1 | 23 |
| fstl1b | 0.22987 | 0.49005206 | 0.387 | 0.125 | 1 | 23 |
| copb2 | 0.23 | -0.19967041 | 0.226 | 0.25 | 1 | 23 |
| rps3a | 0.23027 | 0.260881797 | 0.968 | 0.875 | 1 | 23 |
| CR376783.1 | 0.23117 | -0.27804337 | 0.161 | 0.25 | 1 | 23 |
| add1 | 0.23231 | -0.250035 | 0.323 | 0.375 | 1 | 23 |
| cct2 | 0.23287 | 0.352778223 | 0.839 | 0.625 | 1 | 23 |
| tspan18b | 0.23404 | 0.433037264 | 0.355 | 0.125 | 1 | 23 |
| aldh2.2 | 0.23438 | 0.189455264 | 0.323 | 0.125 | 1 | 23 |
| rpl38 | 0.23453 | 0.198389027 | 1 | 0.875 | 1 | 23 |
| kmt2e | 0.23483 | -0.34662225 | 0.097 | 0.25 | 1 | 23 |
| coro1cb | 0.23483 | -0.43561784 | 0.065 | 0.25 | 1 | 23 |
| esf1 | 0.23497 | -0.40876242 | 0.161 | 0.375 | 1 | 23 |
| ncbp2 | 0.2353 | 0.140033618 | 0.29 | 0.125 | 1 | 23 |
| atp5mf | 0.23562 | -0.35249858 | 0.581 | 0.75 | 1 | 23 |
| metap2b | 0.23639 | -0.24791097 | 0.452 | 0.5 | 1 | 23 |
| dlat | 0.23728 | -0.39049998 | 0.097 | 0.25 | 1 | 23 |
| eif3ja | 0.23744 | -0.38123829 | 0.419 | 0.75 | 1 | 23 |

| | | | | | | |
|---|---|---|---|---|---|---|
| ube2d1b | 0.23768 | 0.268909203 | 0.387 | 0.125 | 1 | 23 |
| wu:fj29h11 | 0.23779 | -0.39760944 | 0.065 | 0.25 | 1 | 23 |
| si:ch211-198a12.6 | 0.23879 | -0.2249533 | 0.194 | 0.25 | 1 | 23 |
| pole3 | 0.23906 | -0.21611937 | 0.194 | 0.25 | 1 | 23 |
| GCA | 0.23929 | 0.345572512 | 0.419 | 0.25 | 1 | 23 |
| atf3 | 0.23995 | -0.14357964 | 0.129 | 0.125 | 1 | 23 |
| ngdn | 0.24002 | 0.445310006 | 0.581 | 0.25 | 1 | 23 |
| ddx19 | 0.24019 | -0.34620283 | 0.129 | 0.25 | 1 | 23 |
| rbbp6 | 0.24102 | -0.35233524 | 0.419 | 0.75 | 1 | 23 |
| uso1 | 0.24172 | -0.37961671 | 0.097 | 0.25 | 1 | 23 |
| CU694197.1 | 0.24234 | 0.299927906 | 0.548 | 0.25 | 1 | 23 |
| tufm | 0.243 | 0.142426312 | 0.29 | 0.125 | 1 | 23 |
| psma1 | 0.24361 | -0.31534166 | 0.71 | 0.875 | 1 | 23 |
| syncrip | 0.24426 | -0.35523221 | 0.581 | 0.875 | 1 | 23 |
| CABZ01028771.1 | 0.24475 | -0.3590898 | 0.065 | 0.25 | 1 | 23 |
| atp5f1e | 0.24496 | -0.30180148 | 0.677 | 0.75 | 1 | 23 |
| setd5 | 0.24558 | -0.20805711 | 0.097 | 0.125 | 1 | 23 |
| sirt5 | 0.2463 | -0.20252399 | 0.097 | 0.125 | 1 | 23 |
| arvcfb | 0.24656 | -0.14455347 | 0.258 | 0.25 | 1 | 23 |
| UTP14C | 0.24721 | -0.35549648 | 0.194 | 0.375 | 1 | 23 |
| cdc34a | 0.24828 | -0.18424953 | 0.065 | 0.25 | 1 | 23 |
| sar1b | 0.24901 | -0.37757011 | 0.194 | 0.375 | 1 | 23 |
| znf593 | 0.24959 | -0.17937549 | 0.226 | 0.25 | 1 | 23 |
| cwc22 | 0.24978 | -0.40796536 | 0.065 | 0.25 | 1 | 23 |
| prkacbb | 0.25008 | -0.29373129 | 0.161 | 0.25 | 1 | 23 |
| tle2a | 0.25149 | -0.38849637 | 0.194 | 0.375 | 1 | 23 |
| mif | 0.25153 | 0.293552723 | 0.677 | 0.375 | 1 | 23 |
| utrn | 0.25191 | -0.402369 | 0.065 | 0.25 | 1 | 23 |
| ccdc90b | 0.25236 | -0.42659668 | 0.226 | 0.5 | 1 | 23 |
| si:ch73-22o12.1 | 0.25326 | -0.20753564 | 0.226 | 0.25 | 1 | 23 |
| ndufab1b | 0.25487 | 0.434839802 | 0.419 | 0.125 | 1 | 23 |
| top1l | 0.25545 | -0.40718382 | 0.452 | 0.75 | 1 | 23 |
| si:zfos-169g10.3 | 0.25553 | 0.415603876 | 0.387 | 0.125 | 1 | 23 |
| thtpa | 0.25569 | -0.16718202 | 0.29 | 0.5 | 1 | 23 |
| capns1a | 0.25569 | 0.41901949 | 0.387 | 0.125 | 1 | 23 |
| acin1b | 0.25605 | -0.36824448 | 0.097 | 0.25 | 1 | 23 |
| tenm3 | 0.25727 | -0.21802885 | 0.194 | 0.25 | 1 | 23 |
| cfap20 | 0.2574 | 0.204742801 | 0.323 | 0.125 | 1 | 23 |
| f3b | 0.25797 | -0.13601007 | 0.516 | 0.5 | 1 | 23 |
| si:ch211-1a19.3 | 0.25831 | -0.46579133 | 0.323 | 0.625 | 1 | 23 |
| zfp36l1b | 0.25852 | -0.25360699 | 0.419 | 0.5 | 1 | 23 |
| sat1b | 0.25875 | -0.19637993 | 0.097 | 0.125 | 1 | 23 |
| gadd45ab | 0.26055 | -0.38309782 | 0.097 | 0.25 | 1 | 23 |
| eef1g | 0.26178 | 0.127536577 | 1 | 1 | 1 | 23 |
| lasp1 | 0.26197 | -0.17019417 | 0.516 | 0.5 | 1 | 23 |
| bcl2l10 | 0.26211 | 0.297207484 | 0.387 | 0.125 | 1 | 23 |
| tmem258 | 0.26213 | 0.179381217 | 0.742 | 0.5 | 1 | 23 |
| pa2g4b | 0.26301 | 0.437958002 | 0.387 | 0.125 | 1 | 23 |

| | | | | | | |
|---|---|---|---|---|---|---|
| jund | 0.26319 | 0.21924386 | 0.323 | 0.125 | 1 | 23 |
| capzb | 0.26375 | -0.11610437 | 0.581 | 0.5 | 1 | 23 |
| sf3b6 | 0.26387 | 0.365850589 | 0.774 | 0.5 | 1 | 23 |
| rps15 | 0.26401 | 0.140018295 | 1 | 1 | 1 | 23 |
| gsta.1 | 0.26405 | -0.18864893 | 0.065 | 0.25 | 1 | 23 |
| prrc2c | 0.26435 | -0.22191979 | 0.194 | 0.25 | 1 | 23 |
| lig1 | 0.26467 | -0.18808782 | 0.097 | 0.125 | 1 | 23 |
| pnrc2 | 0.2656 | -0.20192015 | 0.968 | 1 | 1 | 23 |
| ppp2r5d | 0.2657 | -0.12026392 | 0.097 | 0.25 | 1 | 23 |
| LOC103910871 | 0.26633 | -0.32490578 | 0.097 | 0.25 | 1 | 23 |
| si:dkey-6f10.3 | 0.26652 | -0.41672328 | 0.065 | 0.25 | 1 | 23 |
| nup153 | 0.2667 | -0.37988924 | 0.065 | 0.25 | 1 | 23 |
| tsr2 | 0.26692 | -0.30766008 | 0.065 | 0.25 | 1 | 23 |
| zmynd11 | 0.26758 | -0.15738778 | 0.226 | 0.25 | 1 | 23 |
| kpnb3 | 0.26788 | -0.1938311 | 0.194 | 0.25 | 1 | 23 |
| snx10a | 0.26793 | -0.48698187 | 0.129 | 0.375 | 1 | 23 |
| rpl7a | 0.27016 | 0.131773849 | 1 | 1 | 1 | 23 |
| nfic | 0.27053 | -0.4424641 | 0.387 | 0.625 | 1 | 23 |
| ncoa4 | 0.27081 | -0.28865152 | 0.129 | 0.25 | 1 | 23 |
| meaf6 | 0.27141 | 0.358943515 | 0.452 | 0.25 | 1 | 23 |
| zdhhc21 | 0.27153 | -0.13134286 | 0.129 | 0.125 | 1 | 23 |
| fbxo11a | 0.27195 | -0.1945528 | 0.065 | 0.25 | 1 | 23 |
| clta | 0.27206 | 0.344116991 | 0.355 | 0.125 | 1 | 23 |
| snrpf | 0.27229 | -0.30902072 | 0.71 | 0.875 | 1 | 23 |
| dab1a | 0.27256 | -0.22893835 | 0.065 | 0.25 | 1 | 23 |
| ahnak | 0.2735 | -0.31304099 | 0.355 | 0.5 | 1 | 23 |
| rcan1a | 0.27386 | -0.27079886 | 0.161 | 0.25 | 1 | 23 |
| hmg20a | 0.27413 | -0.40698554 | 0.129 | 0.375 | 1 | 23 |
| psma8 | 0.27448 | 0.4119707 | 0.677 | 0.375 | 1 | 23 |
| mir21-1 | 0.27483 | -0.12054881 | 0.419 | 0.375 | 1 | 23 |
| btbd6b | 0.27501 | -0.28657801 | 0.194 | 0.25 | 1 | 23 |
| hexb | 0.27531 | -0.33007907 | 0.097 | 0.25 | 1 | 23 |
| nmt1a | 0.27576 | -0.37551213 | 0.129 | 0.375 | 1 | 23 |
| rpsa | 0.27584 | 0.12889968 | 1 | 0.875 | 1 | 23 |
| appa | 0.27743 | -0.36455642 | 0.355 | 0.5 | 1 | 23 |
| ift22 | 0.27825 | -0.11983491 | 0.129 | 0.125 | 1 | 23 |
| plpp2b | 0.27849 | -0.36332009 | 0.097 | 0.25 | 1 | 23 |
| srsf11 | 0.27868 | -0.16074531 | 0.387 | 0.375 | 1 | 23 |
| eloca | 0.27883 | -0.24217077 | 0.129 | 0.375 | 1 | 23 |
| ahcyl2 | 0.27907 | -0.40903155 | 0.065 | 0.25 | 1 | 23 |
| rtca | 0.2796 | 0.349379213 | 0.258 | 0.125 | 1 | 23 |
| erlin1 | 0.28002 | -0.18408958 | 0.097 | 0.125 | 1 | 23 |
| hypk | 0.28205 | 0.213443634 | 0.645 | 0.375 | 1 | 23 |
| psmb6 | 0.28237 | 0.398703182 | 0.645 | 0.375 | 1 | 23 |
| sp4 | 0.2826 | -0.34169089 | 0.097 | 0.25 | 1 | 23 |
| namptb | 0.28261 | -0.44839155 | 0.065 | 0.25 | 1 | 23 |
| shc1 | 0.28422 | -0.3677992 | 0.161 | 0.375 | 1 | 23 |
| ccdc125 | 0.28444 | -0.36016416 | 0.065 | 0.25 | 1 | 23 |

| | | | | | | |
|---|---|---|---|---|---|---|
| uchl3 | 0.28483 | -0.13238575 | 0.129 | 0.125 | 1 | 23 |
| pam16 | 0.28506 | 0.306597728 | 0.29 | 0.125 | 1 | 23 |
| yif1a | 0.28546 | -0.36384475 | 0.065 | 0.25 | 1 | 23 |
| mrpl35 | 0.28569 | -0.21775612 | 0.065 | 0.25 | 1 | 23 |
| cs | 0.28587 | -0.41633357 | 0.129 | 0.375 | 1 | 23 |
| fam133b | 0.28637 | -0.22261997 | 0.677 | 0.75 | 1 | 23 |
| apex1 | 0.28644 | -0.18087445 | 0.484 | 0.5 | 1 | 23 |
| id1 | 0.2866 | 0.484222696 | 0.516 | 0.5 | 1 | 23 |
| cul4b | 0.28664 | -0.24987631 | 0.161 | 0.25 | 1 | 23 |
| lrriq1 | 0.28677 | -0.38857084 | 0.129 | 0.25 | 1 | 23 |
| zfp36l1a | 0.28722 | -0.12762534 | 0.548 | 0.5 | 1 | 23 |
| rsrp1 | 0.28778 | 0.348867675 | 0.323 | 0.125 | 1 | 23 |
| CR847566.1 | 0.28802 | -0.45538227 | 0.226 | 0.5 | 1 | 23 |
| lcorl | 0.28891 | -0.30895441 | 0.129 | 0.375 | 1 | 23 |
| laptm4a | 0.28898 | -0.30421527 | 0.355 | 0.5 | 1 | 23 |
| ptp4a2b | 0.28908 | -0.16919126 | 0.226 | 0.25 | 1 | 23 |
| glrx5 | 0.28974 | 0.115398658 | 0.29 | 0.125 | 1 | 23 |
| chordc1b | 0.29024 | -0.18588926 | 0.097 | 0.125 | 1 | 23 |
| gnl2 | 0.29067 | -0.18984001 | 0.097 | 0.125 | 1 | 23 |
| fundc1 | 0.29146 | -0.35472696 | 0.129 | 0.375 | 1 | 23 |
| map2k6 | 0.29156 | -0.11543701 | 0.194 | 0.375 | 1 | 23 |
| cyp26a1 | 0.29195 | -0.31151593 | 0.129 | 0.25 | 1 | 23 |
| ndufb3 | 0.29308 | 0.293529496 | 0.677 | 0.375 | 1 | 23 |
| med20 | 0.29396 | -0.36631048 | 0.065 | 0.25 | 1 | 23 |
| nudcd1 | 0.29432 | -0.34707714 | 0.065 | 0.25 | 1 | 23 |
| stau1 | 0.2948 | -0.38918633 | 0.129 | 0.375 | 1 | 23 |
| arl5a | 0.2957 | -0.19081748 | 0.097 | 0.125 | 1 | 23 |
| atp5f1d | 0.2959 | -0.31544915 | 0.548 | 0.625 | 1 | 23 |
| fbl | 0.2961 | -0.38911363 | 0.387 | 0.625 | 1 | 23 |
| nipsnap2 | 0.29689 | 0.374265312 | 0.387 | 0.125 | 1 | 23 |
| arpc2 | 0.29689 | 0.342961601 | 0.677 | 0.5 | 1 | 23 |
| ss18 | 0.29832 | -0.3965353 | 0.226 | 0.5 | 1 | 23 |
| slc29a4 | 0.29873 | -0.36389207 | 0.097 | 0.25 | 1 | 23 |
| rpl5b | 0.29923 | 0.129286834 | 1 | 0.875 | 1 | 23 |
| rpz | 0.29924 | -0.35445571 | 0.129 | 0.375 | 1 | 23 |
| si:dkeyp-38g8.5 | 0.29928 | -0.37387096 | 0.097 | 0.25 | 1 | 23 |
| ccdc85b | 0.29935 | -0.35409107 | 0.161 | 0.375 | 1 | 23 |
| plxna4 | 0.30018 | -0.35827305 | 0.065 | 0.25 | 1 | 23 |
| cetn4 | 0.30083 | -0.38376399 | 0.065 | 0.25 | 1 | 23 |
| zfr2 | 0.30088 | -0.2685561 | 0.161 | 0.25 | 1 | 23 |
| zgc:56095 | 0.30181 | -0.32911784 | 0.097 | 0.25 | 1 | 23 |
| lhfpl2b | 0.30234 | -0.2801658 | 0.129 | 0.375 | 1 | 23 |
| fen1 | 0.30274 | -0.12124387 | 0.129 | 0.125 | 1 | 23 |
| fth1a | 0.30287 | -0.41629496 | 0.452 | 0.75 | 1 | 23 |
| parp8 | 0.30313 | -0.37023425 | 0.065 | 0.25 | 1 | 23 |
| zgc:152830 | 0.30336 | -0.34467509 | 0.065 | 0.25 | 1 | 23 |
| XLOC-025423 | 0.30358 | -0.11938291 | 0.29 | 0.25 | 1 | 23 |
| atp5md | 0.30384 | -0.33824949 | 0.323 | 0.625 | 1 | 23 |

| | | | | | | |
|---|---|---|---|---|---|---|
| spen | 0.30425 | -0.18062978 | 0.097 | 0.125 | 1 | 23 |
| cdc42 | 0.30449 | 0.107672497 | 0.581 | 0.375 | 1 | 23 |
| gfer | 0.30471 | -0.35353876 | 0.065 | 0.25 | 1 | 23 |
| nr0b1 | 0.30486 | -0.41756288 | 0.129 | 0.375 | 1 | 23 |
| st14a | 0.30486 | -0.33324555 | 0.065 | 0.25 | 1 | 23 |
| psmc5 | 0.30489 | 0.375371322 | 0.645 | 0.375 | 1 | 23 |
| ccnjl | 0.30605 | -0.38392451 | 0.065 | 0.25 | 1 | 23 |
| arl8bb | 0.3062 | -0.32509719 | 0.097 | 0.25 | 1 | 23 |
| glrx | 0.30625 | 0.388296362 | 0.71 | 0.5 | 1 | 23 |
| pdia6 | 0.30651 | 0.326898718 | 0.387 | 0.125 | 1 | 23 |
| snrpd3l | 0.30653 | -0.18838541 | 0.452 | 0.5 | 1 | 23 |
| zfand5a | 0.30672 | -0.26502862 | 0.065 | 0.25 | 1 | 23 |
| acsl1a | 0.30696 | -0.35187399 | 0.065 | 0.25 | 1 | 23 |
| CABZ01045617.1 | 0.30804 | -0.43652121 | 0.194 | 0.375 | 1 | 23 |
| hnrnpa0l | 0.30827 | 0.148106242 | 0.935 | 0.75 | 1 | 23 |
| hif1al | 0.30834 | -0.30667824 | 0.065 | 0.25 | 1 | 23 |
| axin2 | 0.30863 | -0.38236602 | 0.065 | 0.25 | 1 | 23 |
| rhocb | 0.30887 | -0.29729486 | 0.29 | 0.375 | 1 | 23 |
| cdk5rap1 | 0.30889 | -0.26139512 | 0.065 | 0.25 | 1 | 23 |
| si:ch211-222l21.1 | 0.30895 | -0.11415778 | 1 | 1 | 1 | 23 |
| col5a1 | 0.30931 | 0.399453569 | 0.387 | 0.25 | 1 | 23 |
| stxbp6 | 0.30935 | 0.20182853 | 0.323 | 0.125 | 1 | 23 |
| dusp2 | 0.31097 | -0.34257859 | 0.161 | 0.25 | 1 | 23 |
| rpl6 | 0.3116 | 0.141363197 | 1 | 1 | 1 | 23 |
| efnb1 | 0.31196 | -0.34330046 | 0.323 | 0.5 | 1 | 23 |
| txndc17 | 0.31225 | -0.22983559 | 0.29 | 0.375 | 1 | 23 |
| zbtb20 | 0.31228 | -0.31001847 | 0.613 | 0.75 | 1 | 23 |
| tcf12 | 0.31243 | -0.38165438 | 0.129 | 0.375 | 1 | 23 |
| cltcb | 0.31259 | -0.26002317 | 0.161 | 0.25 | 1 | 23 |
| supt16h | 0.31353 | -0.42229953 | 0.194 | 0.375 | 1 | 23 |
| lmo1 | 0.31372 | -0.11093356 | 0.129 | 0.125 | 1 | 23 |
| stmn1a | 0.31438 | -0.53296261 | 0.258 | 0.5 | 1 | 23 |
| lzts2a | 0.3147 | -0.20057235 | 0.194 | 0.25 | 1 | 23 |
| rab11fip4b | 0.31614 | -0.16215463 | 0.097 | 0.125 | 1 | 23 |
| fam204a | 0.31646 | -0.31116925 | 0.129 | 0.375 | 1 | 23 |
| cbx1a | 0.31721 | 0.368640488 | 0.387 | 0.125 | 1 | 23 |
| h2afvb | 0.31723 | 0.215282895 | 0.968 | 1 | 1 | 23 |
| tpd52l2b | 0.31782 | 0.338573798 | 0.323 | 0.125 | 1 | 23 |
| AL954696.1 | 0.31817 | 0.153877986 | 0.097 | 0 | 1 | 23 |
| CABZ01008730.1 | 0.31817 | 0.16891556 | 0.097 | 0 | 1 | 23 |
| CABZ01061591.1 | 0.31817 | 0.15593057 | 0.097 | 0 | 1 | 23 |
| CABZ01068600.1 | 0.31817 | 0.146885301 | 0.097 | 0 | 1 | 23 |
| CABZ01077218.1 | 0.31817 | 0.163040082 | 0.097 | 0 | 1 | 23 |
| CABZ01081730.1 | 0.31817 | 0.178210602 | 0.097 | 0 | 1 | 23 |
| CABZ01113192.1 | 0.31817 | 0.16771966 | 0.097 | 0 | 1 | 23 |
| CABZ01113954.1 | 0.31817 | 0.178449494 | 0.097 | 0 | 1 | 23 |
| FO082877.1 | 0.31817 | 0.137982807 | 0.097 | 0 | 1 | 23 |
| LO017951.1 | 0.31817 | 0.185900209 | 0.097 | 0 | 1 | 23 |

| | | | | | | |
|---|---|---|---|---|---|---|
| LOC101883704 | 0.31817 | 0.160326056 | 0.097 | 0 | 1 | 23 |
| LOC110438237 | 0.31817 | 0.157537801 | 0.097 | 0 | 1 | 23 |
| MTERF4 | 0.31817 | 0.135038836 | 0.097 | 0 | 1 | 23 |
| SCARF2 | 0.31817 | 0.16742197 | 0.097 | 0 | 1 | 23 |
| WDR7 | 0.31817 | 0.145580044 | 0.097 | 0 | 1 | 23 |
| XLOC-038035 | 0.31817 | 0.195314577 | 0.097 | 0 | 1 | 23 |
| abhd17b | 0.31817 | 0.15494099 | 0.097 | 0 | 1 | 23 |
| acad8 | 0.31817 | 0.156248361 | 0.097 | 0 | 1 | 23 |
| acads | 0.31817 | 0.147078726 | 0.097 | 0 | 1 | 23 |
| ackr4b | 0.31817 | 0.207986963 | 0.097 | 0 | 1 | 23 |
| adob | 0.31817 | 0.144705817 | 0.097 | 0 | 1 | 23 |
| agpat2 | 0.31817 | 0.153944965 | 0.097 | 0 | 1 | 23 |
| aldh3a1 | 0.31817 | 0.182744827 | 0.097 | 0 | 1 | 23 |
| ap3m1 | 0.31817 | 0.149245545 | 0.097 | 0 | 1 | 23 |
| apoob | 0.31817 | 0.129254428 | 0.097 | 0 | 1 | 23 |
| arid1aa | 0.31817 | 0.184476947 | 0.097 | 0 | 1 | 23 |
| armh3 | 0.31817 | 0.144736547 | 0.097 | 0 | 1 | 23 |
| bcl2l1 | 0.31817 | 0.138688255 | 0.097 | 0 | 1 | 23 |
| bmp2b | 0.31817 | 0.134857592 | 0.097 | 0 | 1 | 23 |
| c1qtnf2 | 0.31817 | 0.171140109 | 0.097 | 0 | 1 | 23 |
| calm1b | 0.31817 | 0.151179754 | 0.097 | 0 | 1 | 23 |
| casc4 | 0.31817 | 0.174403035 | 0.097 | 0 | 1 | 23 |
| castor2 | 0.31817 | 0.150663207 | 0.097 | 0 | 1 | 23 |
| cbwd | 0.31817 | 0.156318514 | 0.097 | 0 | 1 | 23 |
| ccdc88c | 0.31817 | 0.155941531 | 0.097 | 0 | 1 | 23 |
| ccl39.7 | 0.31817 | 0.12808619 | 0.097 | 0 | 1 | 23 |
| cebpz | 0.31817 | 0.183821947 | 0.097 | 0 | 1 | 23 |
| cep170aa | 0.31817 | 0.163259343 | 0.097 | 0 | 1 | 23 |
| cherp | 0.31817 | 0.172151288 | 0.097 | 0 | 1 | 23 |
| cpxm1a | 0.31817 | 0.160459539 | 0.097 | 0 | 1 | 23 |
| crcp | 0.31817 | 0.14494221 | 0.097 | 0 | 1 | 23 |
| cul1a | 0.31817 | 0.148315146 | 0.097 | 0 | 1 | 23 |
| ddb2 | 0.31817 | 0.152785993 | 0.097 | 0 | 1 | 23 |
| ddx10 | 0.31817 | 0.157500717 | 0.097 | 0 | 1 | 23 |
| dnajc5ga | 0.31817 | 0.148901406 | 0.097 | 0 | 1 | 23 |
| dnm1l | 0.31817 | 0.157886873 | 0.097 | 0 | 1 | 23 |
| dph6 | 0.31817 | 0.130107884 | 0.097 | 0 | 1 | 23 |
| dph7 | 0.31817 | 0.154033012 | 0.097 | 0 | 1 | 23 |
| dtd1 | 0.31817 | 0.169759204 | 0.097 | 0 | 1 | 23 |
| dtymk | 0.31817 | 0.173714958 | 0.097 | 0 | 1 | 23 |
| dvl2 | 0.31817 | 0.157722602 | 0.097 | 0 | 1 | 23 |
| eef1akmt2 | 0.31817 | 0.159202146 | 0.097 | 0 | 1 | 23 |
| eif4g3a | 0.31817 | 0.167510911 | 0.097 | 0 | 1 | 23 |
| elf2b | 0.31817 | 0.177622122 | 0.097 | 0 | 1 | 23 |
| enc1 | 0.31817 | 0.204004929 | 0.097 | 0 | 1 | 23 |
| exoc6 | 0.31817 | 0.172668139 | 0.097 | 0 | 1 | 23 |
| fam102ab | 0.31817 | 0.157316698 | 0.097 | 0 | 1 | 23 |
| fam219b | 0.31817 | 0.142258014 | 0.097 | 0 | 1 | 23 |

| | | | | | | |
|---|---|---|---|---|---|---|
| frmd8 | 0.31817 | 0.158264953 | 0.097 | 0 | 1 | 23 |
| fuca1.1 | 0.31817 | 0.184803466 | 0.097 | 0 | 1 | 23 |
| fzd10 | 0.31817 | 0.228864394 | 0.097 | 0 | 1 | 23 |
| fzr1a | 0.31817 | 0.171258884 | 0.097 | 0 | 1 | 23 |
| galnt1 | 0.31817 | 0.181998829 | 0.097 | 0 | 1 | 23 |
| get4 | 0.31817 | 0.130153636 | 0.097 | 0 | 1 | 23 |
| gfod2 | 0.31817 | 0.152980021 | 0.097 | 0 | 1 | 23 |
| gins2 | 0.31817 | 0.139022393 | 0.097 | 0 | 1 | 23 |
| glis2b | 0.31817 | 0.171479812 | 0.097 | 0 | 1 | 23 |
| gphnb | 0.31817 | 0.152187405 | 0.097 | 0 | 1 | 23 |
| gpr37b | 0.31817 | 0.183549225 | 0.097 | 0 | 1 | 23 |
| gtpbp1 | 0.31817 | 0.197831423 | 0.097 | 0 | 1 | 23 |
| hpf1 | 0.31817 | 0.144705262 | 0.097 | 0 | 1 | 23 |
| im:7136021 | 0.31817 | 0.177502608 | 0.097 | 0 | 1 | 23 |
| impdh2 | 0.31817 | 0.116842101 | 0.097 | 0 | 1 | 23 |
| ipo8 | 0.31817 | 0.141383279 | 0.097 | 0 | 1 | 23 |
| irx3a | 0.31817 | 0.20843492 | 0.097 | 0 | 1 | 23 |
| kctd2 | 0.31817 | 0.163485701 | 0.097 | 0 | 1 | 23 |
| lipt1 | 0.31817 | 0.168087621 | 0.097 | 0 | 1 | 23 |
| lnpep | 0.31817 | 0.171603169 | 0.097 | 0 | 1 | 23 |
| lpar2a | 0.31817 | 0.149481327 | 0.097 | 0 | 1 | 23 |
| luzp1 | 0.31817 | 0.231115792 | 0.097 | 0 | 1 | 23 |
| man2b1 | 0.31817 | 0.203384335 | 0.097 | 0 | 1 | 23 |
| map4k5 | 0.31817 | 0.135727821 | 0.097 | 0 | 1 | 23 |
| mapk8b | 0.31817 | 0.165396732 | 0.097 | 0 | 1 | 23 |
| mbd2 | 0.31817 | 0.16170331 | 0.097 | 0 | 1 | 23 |
| mbnl2 | 0.31817 | 0.145104221 | 0.097 | 0 | 1 | 23 |
| metap1d | 0.31817 | 0.207606886 | 0.097 | 0 | 1 | 23 |
| mex3b | 0.31817 | 0.196774189 | 0.097 | 0 | 1 | 23 |
| micall1a | 0.31817 | 0.155626448 | 0.097 | 0 | 1 | 23 |
| mrps22 | 0.31817 | 0.137598557 | 0.097 | 0 | 1 | 23 |
| mrps31 | 0.31817 | 0.136237906 | 0.097 | 0 | 1 | 23 |
| mrps34 | 0.31817 | 0.15926795 | 0.097 | 0 | 1 | 23 |
| mrrf | 0.31817 | 0.149331283 | 0.097 | 0 | 1 | 23 |
| nacc1a | 0.31817 | 0.138391093 | 0.097 | 0 | 1 | 23 |
| nacc2 | 0.31817 | 0.17708105 | 0.097 | 0 | 1 | 23 |
| nadkb | 0.31817 | 0.135342098 | 0.097 | 0 | 1 | 23 |
| napga | 0.31817 | 0.177795804 | 0.097 | 0 | 1 | 23 |
| nr5a1b | 0.31817 | 0.177808862 | 0.097 | 0 | 1 | 23 |
| nrg2a | 0.31817 | 0.166087279 | 0.097 | 0 | 1 | 23 |
| nsun2 | 0.31817 | 0.148440892 | 0.097 | 0 | 1 | 23 |
| ntng2b | 0.31817 | 0.163259343 | 0.097 | 0 | 1 | 23 |
| nuak1b | 0.31817 | 0.128654134 | 0.097 | 0 | 1 | 23 |
| nup58 | 0.31817 | 0.141656337 | 0.097 | 0 | 1 | 23 |
| orc3 | 0.31817 | 0.153277048 | 0.097 | 0 | 1 | 23 |
| osr1 | 0.31817 | 0.153367462 | 0.097 | 0 | 1 | 23 |
| pbrm1 | 0.31817 | 0.142176263 | 0.097 | 0 | 1 | 23 |
| pbrm1l | 0.31817 | 0.15304949 | 0.097 | 0 | 1 | 23 |

| | | | | | | |
|---|---|---|---|---|---|---|
| pde6d | 0.31817 | 0.155484096 | 0.097 | 0 | 1 | 23 |
| pim3 | 0.31817 | 0.168562388 | 0.097 | 0 | 1 | 23 |
| pla2g6 | 0.31817 | 0.18225568 | 0.097 | 0 | 1 | 23 |
| pold3 | 0.31817 | 0.159919964 | 0.097 | 0 | 1 | 23 |
| prickle3 | 0.31817 | 0.202458554 | 0.097 | 0 | 1 | 23 |
| prss23 | 0.31817 | 0.192380605 | 0.097 | 0 | 1 | 23 |
| rab4a | 0.31817 | 0.142211148 | 0.097 | 0 | 1 | 23 |
| rabep1 | 0.31817 | 0.161960103 | 0.097 | 0 | 1 | 23 |
| ralaa | 0.31817 | 0.123853781 | 0.097 | 0 | 1 | 23 |
| ranbp10 | 0.31817 | 0.16296511 | 0.097 | 0 | 1 | 23 |
| rbm18 | 0.31817 | 0.143991603 | 0.097 | 0 | 1 | 23 |
| rgs3b | 0.31817 | 0.173716885 | 0.097 | 0 | 1 | 23 |
| rnf114 | 0.31817 | 0.180488835 | 0.097 | 0 | 1 | 23 |
| rnf11b | 0.31817 | 0.150794007 | 0.097 | 0 | 1 | 23 |
| rnf14 | 0.31817 | 0.13362242 | 0.097 | 0 | 1 | 23 |
| rybpb | 0.31817 | 0.196456791 | 0.097 | 0 | 1 | 23 |
| scamp2 | 0.31817 | 0.186716601 | 0.097 | 0 | 1 | 23 |
| si:ch211-196c10.15 | 0.31817 | 0.19806892 | 0.097 | 0 | 1 | 23 |
| si:ch211-212k18.4 | 0.31817 | 0.16997387 | 0.097 | 0 | 1 | 23 |
| si:ch211-214p13.9 | 0.31817 | 0.144161498 | 0.097 | 0 | 1 | 23 |
| si:ch211-225b11.1 | 0.31817 | 0.192623576 | 0.097 | 0 | 1 | 23 |
| si:ch211-226m16.2 | 0.31817 | 0.167278853 | 0.097 | 0 | 1 | 23 |
| si:ch211-233m11.1 | 0.31817 | 0.202193948 | 0.097 | 0 | 1 | 23 |
| si:ch73-144d13.4 | 0.31817 | 0.199884388 | 0.097 | 0 | 1 | 23 |
| si:ch73-340n8.4 | 0.31817 | 0.15096067 | 0.097 | 0 | 1 | 23 |
| si:ch73-367f21.4 | 0.31817 | 0.188669814 | 0.097 | 0 | 1 | 23 |
| si:dkey-237h12.3 | 0.31817 | 0.171198517 | 0.097 | 0 | 1 | 23 |
| si:dkey-74k8.3 | 0.31817 | 0.145235787 | 0.097 | 0 | 1 | 23 |
| si:dkey-9i23.15 | 0.31817 | 0.175874489 | 0.097 | 0 | 1 | 23 |
| slc25a33 | 0.31817 | 0.145982581 | 0.097 | 0 | 1 | 23 |
| slc37a2 | 0.31817 | 0.145576644 | 0.097 | 0 | 1 | 23 |
| slc7a6os | 0.31817 | 0.154778993 | 0.097 | 0 | 1 | 23 |
| sostdc1a | 0.31817 | 0.200244029 | 0.097 | 0 | 1 | 23 |
| sra1 | 0.31817 | 0.175326826 | 0.097 | 0 | 1 | 23 |
| stx4 | 0.31817 | 0.148864927 | 0.097 | 0 | 1 | 23 |
| taf5 | 0.31817 | 0.154148912 | 0.097 | 0 | 1 | 23 |
| tapbpl | 0.31817 | 0.133669755 | 0.097 | 0 | 1 | 23 |
| tasp1 | 0.31817 | 0.183315482 | 0.097 | 0 | 1 | 23 |
| tbc1d20 | 0.31817 | 0.158909369 | 0.097 | 0 | 1 | 23 |
| terfa | 0.31817 | 0.148465698 | 0.097 | 0 | 1 | 23 |
| tex264a | 0.31817 | 0.148967843 | 0.097 | 0 | 1 | 23 |
| tfdp1b | 0.31817 | 0.147054582 | 0.097 | 0 | 1 | 23 |
| tfip11 | 0.31817 | 0.139187382 | 0.097 | 0 | 1 | 23 |
| thap4 | 0.31817 | 0.145860427 | 0.097 | 0 | 1 | 23 |
| thy1 | 0.31817 | 0.153515525 | 0.097 | 0 | 1 | 23 |
| tkfc | 0.31817 | 0.167625001 | 0.097 | 0 | 1 | 23 |
| tmem135 | 0.31817 | 0.114956582 | 0.097 | 0 | 1 | 23 |
| tmem176l.3a | 0.31817 | 0.144935753 | 0.097 | 0 | 1 | 23 |

| | | | | | | |
|---|---|---|---|---|---|---|
| tnfsf10l4 | 0.31817 | 0.165806525 | 0.097 | 0 | 1 | 23 |
| toporsa | 0.31817 | 0.171533776 | 0.097 | 0 | 1 | 23 |
| tpst1l | 0.31817 | 0.176582082 | 0.097 | 0 | 1 | 23 |
| tsn | 0.31817 | 0.196401744 | 0.097 | 0 | 1 | 23 |
| ttyh3a | 0.31817 | 0.162125047 | 0.097 | 0 | 1 | 23 |
| tubg1 | 0.31817 | 0.143077955 | 0.097 | 0 | 1 | 23 |
| u2af2a | 0.31817 | 0.140058829 | 0.097 | 0 | 1 | 23 |
| uggt1 | 0.31817 | 0.158561436 | 0.097 | 0 | 1 | 23 |
| vhll | 0.31817 | 0.15154999 | 0.097 | 0 | 1 | 23 |
| vps36 | 0.31817 | 0.155982047 | 0.097 | 0 | 1 | 23 |
| vps52 | 0.31817 | 0.208098896 | 0.097 | 0 | 1 | 23 |
| vti1b | 0.31817 | 0.166946792 | 0.097 | 0 | 1 | 23 |
| vwa8 | 0.31817 | 0.151501747 | 0.097 | 0 | 1 | 23 |
| wu:fe05a04 | 0.31817 | 0.230699203 | 0.097 | 0 | 1 | 23 |
| wu:fi46h04 | 0.31817 | 0.157420405 | 0.097 | 0 | 1 | 23 |
| wwc3 | 0.31817 | 0.150848909 | 0.097 | 0 | 1 | 23 |
| xpr1b | 0.31817 | 0.19027265 | 0.097 | 0 | 1 | 23 |
| yipf1 | 0.31817 | 0.155065947 | 0.097 | 0 | 1 | 23 |
| zbtb17 | 0.31817 | 0.177095974 | 0.097 | 0 | 1 | 23 |
| zbtb33 | 0.31817 | 0.149297708 | 0.097 | 0 | 1 | 23 |
| zgc:158564 | 0.31817 | 0.155571476 | 0.097 | 0 | 1 | 23 |
| zgc:158803 | 0.31817 | 0.184384164 | 0.097 | 0 | 1 | 23 |
| zgc:193711 | 0.31817 | 0.13713259 | 0.097 | 0 | 1 | 23 |
| zgc:66427 | 0.31817 | 0.154441984 | 0.097 | 0 | 1 | 23 |
| zgc:66479 | 0.31817 | 0.139488242 | 0.097 | 0 | 1 | 23 |
| zswim5 | 0.31817 | 0.200493923 | 0.097 | 0 | 1 | 23 |
| BX465227.1 | 0.31817 | 0.169191058 | 0.097 | 0 | 1 | 23 |
| CABZ01005379.1 | 0.31817 | 0.167143557 | 0.097 | 0 | 1 | 23 |
| CABZ01055306.1 | 0.31817 | 0.188209772 | 0.097 | 0 | 1 | 23 |
| CABZ01078055.1 | 0.31817 | 0.148339883 | 0.097 | 0 | 1 | 23 |
| CABZ01081752.1 | 0.31817 | 0.143068805 | 0.097 | 0 | 1 | 23 |
| CR391991.4 | 0.31817 | 0.178791665 | 0.097 | 0 | 1 | 23 |
| CR855389.1 | 0.31817 | 0.109784531 | 0.097 | 0 | 1 | 23 |
| CU467856.3 | 0.31817 | 0.181656827 | 0.097 | 0 | 1 | 23 |
| CU571318.1 | 0.31817 | 0.144580909 | 0.097 | 0 | 1 | 23 |
| FBXO48 | 0.31817 | 0.120778543 | 0.097 | 0 | 1 | 23 |
| LOC101882334 | 0.31817 | 0.191832676 | 0.097 | 0 | 1 | 23 |
| LOC103909112 | 0.31817 | 0.156002956 | 0.097 | 0 | 1 | 23 |
| LOC110438079 | 0.31817 | 0.207133097 | 0.097 | 0 | 1 | 23 |
| SMIM10 | 0.31817 | 0.135508799 | 0.097 | 0 | 1 | 23 |
| XLOC-007541 | 0.31817 | 0.16703597 | 0.097 | 0 | 1 | 23 |
| XLOC-033403 | 0.31817 | 0.160482835 | 0.097 | 0 | 1 | 23 |
| aars | 0.31817 | 0.155710734 | 0.097 | 0 | 1 | 23 |
| acot15 | 0.31817 | 0.165937248 | 0.097 | 0 | 1 | 23 |
| adhfe1 | 0.31817 | 0.139953363 | 0.097 | 0 | 1 | 23 |
| afap1l2 | 0.31817 | 0.205224284 | 0.097 | 0 | 1 | 23 |
| agmo | 0.31817 | 0.132505505 | 0.097 | 0 | 1 | 23 |
| ago2 | 0.31817 | 0.19317608 | 0.097 | 0 | 1 | 23 |

| | | | | | | |
|---|---|---|---|---|---|---|
| agpat4 | 0.31817 | 0.154237564 | 0.097 | 0 | 1 | 23 |
| ak3 | 0.31817 | 0.118792131 | 0.097 | 0 | 1 | 23 |
| alpl | 0.31817 | 0.209247661 | 0.097 | 0 | 1 | 23 |
| anapc13 | 0.31817 | 0.142794912 | 0.097 | 0 | 1 | 23 |
| ankrd46a | 0.31817 | 0.134742623 | 0.097 | 0 | 1 | 23 |
| ap3b1a | 0.31817 | 0.148685749 | 0.097 | 0 | 1 | 23 |
| araf | 0.31817 | 0.152413848 | 0.097 | 0 | 1 | 23 |
| arhgap42a | 0.31817 | 0.14011451 | 0.097 | 0 | 1 | 23 |
| arpc5lb | 0.31817 | 0.158827466 | 0.097 | 0 | 1 | 23 |
| atg4a | 0.31817 | 0.162576423 | 0.097 | 0 | 1 | 23 |
| atp2c1 | 0.31817 | 0.186441534 | 0.097 | 0 | 1 | 23 |
| atxn2l | 0.31817 | 0.176061737 | 0.097 | 0 | 1 | 23 |
| b9d1 | 0.31817 | 0.143921443 | 0.097 | 0 | 1 | 23 |
| bag1 | 0.31817 | 0.141936783 | 0.097 | 0 | 1 | 23 |
| bag2 | 0.31817 | 0.161996443 | 0.097 | 0 | 1 | 23 |
| bag5 | 0.31817 | 0.161230354 | 0.097 | 0 | 1 | 23 |
| baiap2a | 0.31817 | 0.151813939 | 0.097 | 0 | 1 | 23 |
| bckdhbl | 0.31817 | 0.169900061 | 0.097 | 0 | 1 | 23 |
| bcorl1 | 0.31817 | 0.191970308 | 0.097 | 0 | 1 | 23 |
| bet1l | 0.31817 | 0.160822382 | 0.097 | 0 | 1 | 23 |
| brwd3 | 0.31817 | 0.131223705 | 0.097 | 0 | 1 | 23 |
| btc | 0.31817 | 0.137283965 | 0.097 | 0 | 1 | 23 |
| capn12 | 0.31817 | 0.183934463 | 0.097 | 0 | 1 | 23 |
| cars | 0.31817 | 0.193592069 | 0.097 | 0 | 1 | 23 |
| cars2 | 0.31817 | 0.164679109 | 0.097 | 0 | 1 | 23 |
| casp3a | 0.31817 | 0.164824701 | 0.097 | 0 | 1 | 23 |
| cav1 | 0.31817 | 0.138789852 | 0.097 | 0 | 1 | 23 |
| cbx3b | 0.31817 | 0.186514114 | 0.097 | 0 | 1 | 23 |
| cbx7a | 0.31817 | 0.190669679 | 0.097 | 0 | 1 | 23 |
| ccdc137 | 0.31817 | 0.14042615 | 0.097 | 0 | 1 | 23 |
| ccdc173 | 0.31817 | 0.207143789 | 0.097 | 0 | 1 | 23 |
| ccdc43 | 0.31817 | 0.153175753 | 0.097 | 0 | 1 | 23 |
| ccdc58 | 0.31817 | 0.160962172 | 0.097 | 0 | 1 | 23 |
| cdk13 | 0.31817 | 0.132089124 | 0.097 | 0 | 1 | 23 |
| celf1 | 0.31817 | 0.133946676 | 0.097 | 0 | 1 | 23 |
| celsr1a | 0.31817 | 0.158926341 | 0.097 | 0 | 1 | 23 |
| cenpw | 0.31817 | 0.121068173 | 0.097 | 0 | 1 | 23 |
| cep72 | 0.31817 | 0.11499341 | 0.097 | 0 | 1 | 23 |
| chp1 | 0.31817 | 0.155053064 | 0.097 | 0 | 1 | 23 |
| chst2b | 0.31817 | 0.142037907 | 0.097 | 0 | 1 | 23 |
| ciao2b | 0.31817 | 0.149063333 | 0.097 | 0 | 1 | 23 |
| cish | 0.31817 | 0.163621655 | 0.097 | 0 | 1 | 23 |
| clip2 | 0.31817 | 0.141153294 | 0.097 | 0 | 1 | 23 |
| cln6a | 0.31817 | 0.129352582 | 0.097 | 0 | 1 | 23 |
| cluap1 | 0.31817 | 0.114835969 | 0.097 | 0 | 1 | 23 |
| cmtm7 | 0.31817 | 0.133532754 | 0.097 | 0 | 1 | 23 |
| commd8 | 0.31817 | 0.13615852 | 0.097 | 0 | 1 | 23 |
| cpvl | 0.31817 | 0.150080376 | 0.097 | 0 | 1 | 23 |

| | | | | | | |
|---|---|---|---|---|---|---|
| crls1 | 0.31817 | 0.148574924 | 0.097 | 0 | 1 | 23 |
| cyfip1 | 0.31817 | 0.143686319 | 0.097 | 0 | 1 | 23 |
| dctd | 0.31817 | 0.19861816 | 0.097 | 0 | 1 | 23 |
| dedd | 0.31817 | 0.149416833 | 0.097 | 0 | 1 | 23 |
| dhcr7 | 0.31817 | 0.159512177 | 0.097 | 0 | 1 | 23 |
| dhrs12 | 0.31817 | 0.154122044 | 0.097 | 0 | 1 | 23 |
| dlgap1b | 0.31817 | 0.15976584 | 0.097 | 0 | 1 | 23 |
| dnajc3a | 0.31817 | 0.147738214 | 0.097 | 0 | 1 | 23 |
| dnal1 | 0.31817 | 0.163037194 | 0.097 | 0 | 1 | 23 |
| dnal4b | 0.31817 | 0.174813309 | 0.097 | 0 | 1 | 23 |
| dyrk2 | 0.31817 | 0.164732122 | 0.097 | 0 | 1 | 23 |
| e2f4 | 0.31817 | 0.159467017 | 0.097 | 0 | 1 | 23 |
| edem1 | 0.31817 | 0.123671162 | 0.097 | 0 | 1 | 23 |
| eif2b3 | 0.31817 | 0.172446232 | 0.097 | 0 | 1 | 23 |
| epha4l | 0.31817 | 0.126075202 | 0.097 | 0 | 1 | 23 |
| ephx2 | 0.31817 | 0.164230822 | 0.097 | 0 | 1 | 23 |
| fbln5 | 0.31817 | 0.125245008 | 0.097 | 0 | 1 | 23 |
| fbxw7 | 0.31817 | 0.159711814 | 0.097 | 0 | 1 | 23 |
| fhod3b | 0.31817 | 0.161891406 | 0.097 | 0 | 1 | 23 |
| foxk1 | 0.31817 | 0.128024577 | 0.097 | 0 | 1 | 23 |
| foxl2b | 0.31817 | 0.16455571 | 0.097 | 0 | 1 | 23 |
| foxp1a | 0.31817 | 0.12369958 | 0.097 | 0 | 1 | 23 |
| ftr91 | 0.31817 | 0.191364284 | 0.097 | 0 | 1 | 23 |
| gak | 0.31817 | 0.16466394 | 0.097 | 0 | 1 | 23 |
| gigyf1a | 0.31817 | 0.180534267 | 0.097 | 0 | 1 | 23 |
| golga4 | 0.31817 | 0.170144854 | 0.097 | 0 | 1 | 23 |
| gpatch1 | 0.31817 | 0.160061578 | 0.097 | 0 | 1 | 23 |
| hip1 | 0.31817 | 0.147742948 | 0.097 | 0 | 1 | 23 |
| hnrnpua | 0.31817 | 0.151729871 | 0.097 | 0 | 1 | 23 |
| hpca | 0.31817 | 0.152812986 | 0.097 | 0 | 1 | 23 |
| hsd17b10 | 0.31817 | 0.127633687 | 0.097 | 0 | 1 | 23 |
| ildr2 | 0.31817 | 0.141839444 | 0.097 | 0 | 1 | 23 |
| ino80e | 0.31817 | 0.134206785 | 0.097 | 0 | 1 | 23 |
| inpp4aa | 0.31817 | 0.153944859 | 0.097 | 0 | 1 | 23 |
| kdm6a | 0.31817 | 0.149430843 | 0.097 | 0 | 1 | 23 |
| klhl8 | 0.31817 | 0.206818313 | 0.097 | 0 | 1 | 23 |
| lama4 | 0.31817 | 0.147439457 | 0.097 | 0 | 1 | 23 |
| ldlrad3 | 0.31817 | 0.145497209 | 0.097 | 0 | 1 | 23 |
| lpar2b | 0.31817 | 0.165880257 | 0.097 | 0 | 1 | 23 |
| lrba | 0.31817 | 0.167045155 | 0.097 | 0 | 1 | 23 |
| lrp1aa | 0.31817 | 0.17090137 | 0.097 | 0 | 1 | 23 |
| lyrm7 | 0.31817 | 0.143991087 | 0.097 | 0 | 1 | 23 |
| mbd5 | 0.31817 | 0.223079524 | 0.097 | 0 | 1 | 23 |
| mboat2b | 0.31817 | 0.168842624 | 0.097 | 0 | 1 | 23 |
| mdc1 | 0.31817 | 0.145089107 | 0.097 | 0 | 1 | 23 |
| mea1 | 0.31817 | 0.199626149 | 0.097 | 0 | 1 | 23 |
| mepce | 0.31817 | 0.166328803 | 0.097 | 0 | 1 | 23 |
| mfap3l | 0.31817 | 0.14589826 | 0.097 | 0 | 1 | 23 |

| | | | | | | |
|---|---|---|---|---|---|---|
| mff | 0.31817 | 0.151703395 | 0.097 | 0 | 1 | 23 |
| mical3a | 0.31817 | 0.190797534 | 0.097 | 0 | 1 | 23 |
| mlst8 | 0.31817 | 0.148638555 | 0.097 | 0 | 1 | 23 |
| mob1bb | 0.31817 | 0.149376192 | 0.097 | 0 | 1 | 23 |
| mpp6b | 0.31817 | 0.173445894 | 0.097 | 0 | 1 | 23 |
| mrpl4 | 0.31817 | 0.141603312 | 0.097 | 0 | 1 | 23 |
| mrpl9 | 0.31817 | 0.135681517 | 0.097 | 0 | 1 | 23 |
| mrps2 | 0.31817 | 0.130454969 | 0.097 | 0 | 1 | 23 |
| mrps27 | 0.31817 | 0.126673033 | 0.097 | 0 | 1 | 23 |
| mta2 | 0.31817 | 0.198366851 | 0.097 | 0 | 1 | 23 |
| mtm1 | 0.31817 | 0.206260643 | 0.097 | 0 | 1 | 23 |
| mylipb | 0.31817 | 0.136292713 | 0.097 | 0 | 1 | 23 |
| mynn | 0.31817 | 0.144598484 | 0.097 | 0 | 1 | 23 |
| nap1l4a | 0.31817 | 0.121355864 | 0.097 | 0 | 1 | 23 |
| nat10 | 0.31817 | 0.146975762 | 0.097 | 0 | 1 | 23 |
| nav3 | 0.31817 | 0.153083855 | 0.097 | 0 | 1 | 23 |
| ndst2b | 0.31817 | 0.20234322 | 0.097 | 0 | 1 | 23 |
| ndufaf1 | 0.31817 | 0.115351523 | 0.097 | 0 | 1 | 23 |
| nek7 | 0.31817 | 0.135773036 | 0.097 | 0 | 1 | 23 |
| nelfe | 0.31817 | 0.131693703 | 0.097 | 0 | 1 | 23 |
| nfil3-5 | 0.31817 | 0.141207054 | 0.097 | 0 | 1 | 23 |
| nfil3-6 | 0.31817 | 0.178342369 | 0.097 | 0 | 1 | 23 |
| nfu1 | 0.31817 | 0.151721013 | 0.097 | 0 | 1 | 23 |
| nipsnap3a | 0.31817 | 0.132206695 | 0.097 | 0 | 1 | 23 |
| nom1 | 0.31817 | 0.136593998 | 0.097 | 0 | 1 | 23 |
| nr0b2a | 0.31817 | 0.139260043 | 0.097 | 0 | 1 | 23 |
| nr1d2b | 0.31817 | 0.168005055 | 0.097 | 0 | 1 | 23 |
| nsd2 | 0.31817 | 0.183522897 | 0.097 | 0 | 1 | 23 |
| nup37 | 0.31817 | 0.168658317 | 0.097 | 0 | 1 | 23 |
| nxpe3 | 0.31817 | 0.2019351 | 0.097 | 0 | 1 | 23 |
| ophn1 | 0.31817 | 0.142108502 | 0.097 | 0 | 1 | 23 |
| pak2a | 0.31817 | 0.171010146 | 0.097 | 0 | 1 | 23 |
| pard6b | 0.31817 | 0.129698121 | 0.097 | 0 | 1 | 23 |
| park7 | 0.31817 | 0.147658154 | 0.097 | 0 | 1 | 23 |
| pcnp | 0.31817 | 0.125988455 | 0.097 | 0 | 1 | 23 |
| pdcd2 | 0.31817 | 0.126040928 | 0.097 | 0 | 1 | 23 |
| pdlim1 | 0.31817 | 0.146991515 | 0.097 | 0 | 1 | 23 |
| pdxp | 0.31817 | 0.104940517 | 0.097 | 0 | 1 | 23 |
| peak1 | 0.31817 | 0.156157295 | 0.097 | 0 | 1 | 23 |
| pgam1a | 0.31817 | 0.146292983 | 0.097 | 0 | 1 | 23 |
| phf21ab | 0.31817 | 0.144754554 | 0.097 | 0 | 1 | 23 |
| pias1a | 0.31817 | 0.181529882 | 0.097 | 0 | 1 | 23 |
| pip4p1a | 0.31817 | 0.175153933 | 0.097 | 0 | 1 | 23 |
| plcg1 | 0.31817 | 0.149290908 | 0.097 | 0 | 1 | 23 |
| plp2 | 0.31817 | 0.145423475 | 0.097 | 0 | 1 | 23 |
| plpp3 | 0.31817 | 0.100050108 | 0.097 | 0 | 1 | 23 |
| polr2h | 0.31817 | 0.129363411 | 0.097 | 0 | 1 | 23 |
| pom121 | 0.31817 | 0.158422139 | 0.097 | 0 | 1 | 23 |

| | | | | | | |
|---|---|---|---|---|---|---|
| pomk | 0.31817 | 0.153362919 | 0.097 | 0 | 1 | 23 |
| pomt2 | 0.31817 | 0.142671929 | 0.097 | 0 | 1 | 23 |
| ppm1da | 0.31817 | 0.164971405 | 0.097 | 0 | 1 | 23 |
| ppp2r3c | 0.31817 | 0.118164881 | 0.097 | 0 | 1 | 23 |
| ppp3cca | 0.31817 | 0.182559378 | 0.097 | 0 | 1 | 23 |
| prex2 | 0.31817 | 0.209626257 | 0.097 | 0 | 1 | 23 |
| prmt3 | 0.31817 | 0.184477168 | 0.097 | 0 | 1 | 23 |
| prmt5 | 0.31817 | 0.143590697 | 0.097 | 0 | 1 | 23 |
| prpf38a | 0.31817 | 0.172019532 | 0.097 | 0 | 1 | 23 |
| pudp | 0.31817 | 0.139742328 | 0.097 | 0 | 1 | 23 |
| rabl6b | 0.31817 | 0.144076619 | 0.097 | 0 | 1 | 23 |
| rac1b | 0.31817 | 0.146387651 | 0.097 | 0 | 1 | 23 |
| rangap1b | 0.31817 | 0.14369224 | 0.097 | 0 | 1 | 23 |
| rbm19 | 0.31817 | 0.153001277 | 0.097 | 0 | 1 | 23 |
| rerea | 0.31817 | 0.176044588 | 0.097 | 0 | 1 | 23 |
| rgs7a | 0.31817 | 0.156015441 | 0.097 | 0 | 1 | 23 |
| rhogb | 0.31817 | 0.10276665 | 0.097 | 0 | 1 | 23 |
| rmnd5b | 0.31817 | 0.152024786 | 0.097 | 0 | 1 | 23 |
| rnft1 | 0.31817 | 0.138652082 | 0.097 | 0 | 1 | 23 |
| rps6kal | 0.31817 | 0.141724917 | 0.097 | 0 | 1 | 23 |
| samm50l | 0.31817 | 0.120488598 | 0.097 | 0 | 1 | 23 |
| sc:d217 | 0.31817 | 0.134613349 | 0.097 | 0 | 1 | 23 |
| sdr16c5b | 0.31817 | 0.131236367 | 0.097 | 0 | 1 | 23 |
| sema3fa | 0.31817 | 0.172895067 | 0.097 | 0 | 1 | 23 |
| serpina10a | 0.31817 | 0.166524924 | 0.097 | 0 | 1 | 23 |
| serpinh2 | 0.31817 | 0.152277936 | 0.097 | 0 | 1 | 23 |
| si:ch211-113p18.3 | 0.31817 | 0.169025014 | 0.097 | 0 | 1 | 23 |
| si:ch211-150o23.3 | 0.31817 | 0.148170381 | 0.097 | 0 | 1 | 23 |
| si:ch211-157b11.12 | 0.31817 | 0.173716288 | 0.097 | 0 | 1 | 23 |
| si:ch211-196g2.4 | 0.31817 | 0.14231804 | 0.097 | 0 | 1 | 23 |
| si:dkey-109l4.3 | 0.31817 | 0.140234573 | 0.097 | 0 | 1 | 23 |
| si:dkey-159f12.2 | 0.31817 | 0.149990723 | 0.097 | 0 | 1 | 23 |
| si:dkey-16l2.20 | 0.31817 | 0.168131672 | 0.097 | 0 | 1 | 23 |
| si:dkey-172k15.3 | 0.31817 | 0.125863528 | 0.097 | 0 | 1 | 23 |
| si:dkey-172m14.1 | 0.31817 | 0.157976136 | 0.097 | 0 | 1 | 23 |
| si:dkey-22h13.2 | 0.31817 | 0.160036482 | 0.097 | 0 | 1 | 23 |
| si:dkey-247i3.6 | 0.31817 | 0.152845666 | 0.097 | 0 | 1 | 23 |
| si:dkey-28b4.8 | 0.31817 | 0.1601809 | 0.097 | 0 | 1 | 23 |
| si:dkey-31c13.1 | 0.31817 | 0.141136792 | 0.097 | 0 | 1 | 23 |
| si:dkey-52l18.4 | 0.31817 | 0.154610798 | 0.097 | 0 | 1 | 23 |
| si:dkey-79d12.5 | 0.31817 | 0.115968775 | 0.097 | 0 | 1 | 23 |
| sik3 | 0.31817 | 0.176934496 | 0.097 | 0 | 1 | 23 |
| slc25a36a | 0.31817 | 0.158211521 | 0.097 | 0 | 1 | 23 |
| slc27a1a | 0.31817 | 0.141520137 | 0.097 | 0 | 1 | 23 |
| slc35a4 | 0.31817 | 0.159344013 | 0.097 | 0 | 1 | 23 |
| snrnp35 | 0.31817 | 0.166051761 | 0.097 | 0 | 1 | 23 |
| spag9a | 0.31817 | 0.164525645 | 0.097 | 0 | 1 | 23 |
| spata17 | 0.31817 | 0.164526716 | 0.097 | 0 | 1 | 23 |

| | | | | | | |
|---|---|---|---|---|---|---|
| spata5l1 | 0.31817 | 0.147475643 | 0.097 | 0 | 1 | 23 |
| spon1b | 0.31817 | 0.148860413 | 0.097 | 0 | 1 | 23 |
| srfbp1 | 0.31817 | 0.168093627 | 0.097 | 0 | 1 | 23 |
| srsf2b | 0.31817 | 0.203467418 | 0.097 | 0 | 1 | 23 |
| stag1a | 0.31817 | 0.143942423 | 0.097 | 0 | 1 | 23 |
| stat3 | 0.31817 | 0.132135998 | 0.097 | 0 | 1 | 23 |
| synrg | 0.31817 | 0.179568673 | 0.097 | 0 | 1 | 23 |
| tbl1xr1b | 0.31817 | 0.154219039 | 0.097 | 0 | 1 | 23 |
| tcf7l2 | 0.31817 | 0.1769458 | 0.097 | 0 | 1 | 23 |
| tead3b | 0.31817 | 0.161083959 | 0.097 | 0 | 1 | 23 |
| tex2 | 0.31817 | 0.139423206 | 0.097 | 0 | 1 | 23 |
| tfb1m | 0.31817 | 0.160796524 | 0.097 | 0 | 1 | 23 |
| thoc3 | 0.31817 | 0.189984136 | 0.097 | 0 | 1 | 23 |
| tlr3 | 0.31817 | 0.13468708 | 0.097 | 0 | 1 | 23 |
| tm9sf2 | 0.31817 | 0.16302748 | 0.097 | 0 | 1 | 23 |
| tmem126a | 0.31817 | 0.126125346 | 0.097 | 0 | 1 | 23 |
| tmem184ba | 0.31817 | 0.153113155 | 0.097 | 0 | 1 | 23 |
| tmem192 | 0.31817 | 0.171773488 | 0.097 | 0 | 1 | 23 |
| tmem203 | 0.31817 | 0.135360996 | 0.097 | 0 | 1 | 23 |
| tmsb1 | 0.31817 | 0.224693442 | 0.097 | 0 | 1 | 23 |
| tmtc2a | 0.31817 | 0.138541854 | 0.097 | 0 | 1 | 23 |
| tmtc3 | 0.31817 | 0.143761798 | 0.097 | 0 | 1 | 23 |
| tmub2 | 0.31817 | 0.163555117 | 0.097 | 0 | 1 | 23 |
| tp53bp1 | 0.31817 | 0.187431637 | 0.097 | 0 | 1 | 23 |
| tpp2 | 0.31817 | 0.11748804 | 0.097 | 0 | 1 | 23 |
| tprkb | 0.31817 | 0.123165734 | 0.097 | 0 | 1 | 23 |
| tsg101a | 0.31817 | 0.155498967 | 0.097 | 0 | 1 | 23 |
| txlna | 0.31817 | 0.149446389 | 0.097 | 0 | 1 | 23 |
| txnrd3 | 0.31817 | 0.102992103 | 0.097 | 0 | 1 | 23 |
| tyw3 | 0.31817 | 0.151044359 | 0.097 | 0 | 1 | 23 |
| uba3 | 0.31817 | 0.170953121 | 0.097 | 0 | 1 | 23 |
| ubald2 | 0.31817 | 0.169204001 | 0.097 | 0 | 1 | 23 |
| ube2q1 | 0.31817 | 0.139662462 | 0.097 | 0 | 1 | 23 |
| ube2ql1 | 0.31817 | 0.191380163 | 0.097 | 0 | 1 | 23 |
| ubtd2 | 0.31817 | 0.159453642 | 0.097 | 0 | 1 | 23 |
| uck2a | 0.31817 | 0.1736076 | 0.097 | 0 | 1 | 23 |
| uck2b | 0.31817 | 0.114499222 | 0.097 | 0 | 1 | 23 |
| unga | 0.31817 | 0.139723919 | 0.097 | 0 | 1 | 23 |
| usf1 | 0.31817 | 0.147040562 | 0.097 | 0 | 1 | 23 |
| usp1 | 0.31817 | 0.142275356 | 0.097 | 0 | 1 | 23 |
| utp15 | 0.31817 | 0.123545973 | 0.097 | 0 | 1 | 23 |
| utp4 | 0.31817 | 0.148093156 | 0.097 | 0 | 1 | 23 |
| uvrag | 0.31817 | 0.225555324 | 0.097 | 0 | 1 | 23 |
| vps26b | 0.31817 | 0.172044305 | 0.097 | 0 | 1 | 23 |
| wasf3b | 0.31817 | 0.143492029 | 0.097 | 0 | 1 | 23 |
| wfdc1 | 0.31817 | 0.157484274 | 0.097 | 0 | 1 | 23 |
| zcchc7 | 0.31817 | 0.156155173 | 0.097 | 0 | 1 | 23 |
| zdhhc6 | 0.31817 | 0.151895545 | 0.097 | 0 | 1 | 23 |

| | | | | | | |
|---|---|---|---|---|---|---|
| zer1 | 0.31817 | 0.157467673 | 0.097 | 0 | 1 | 23 |
| zgc:110366 | 0.31817 | 0.157856345 | 0.097 | 0 | 1 | 23 |
| zgc:112148 | 0.31817 | 0.176852044 | 0.097 | 0 | 1 | 23 |
| zgc:114119 | 0.31817 | 0.156364537 | 0.097 | 0 | 1 | 23 |
| zgc:153916 | 0.31817 | 0.157977377 | 0.097 | 0 | 1 | 23 |
| zgc:153990 | 0.31817 | 0.175625709 | 0.097 | 0 | 1 | 23 |
| zgc:66484 | 0.31817 | 0.139866785 | 0.097 | 0 | 1 | 23 |
| zmp:0000001316 | 0.31817 | 0.14623277 | 0.097 | 0 | 1 | 23 |
| zmpste24 | 0.31817 | 0.16224407 | 0.097 | 0 | 1 | 23 |
| znf1010 | 0.31817 | 0.164317069 | 0.097 | 0 | 1 | 23 |
| znf280d | 0.31817 | 0.139472777 | 0.097 | 0 | 1 | 23 |
| znf622 | 0.31817 | 0.141752475 | 0.097 | 0 | 1 | 23 |
| ankrd40 | 0.31817 | 0.176034957 | 0.097 | 0 | 1 | 23 |
| arid4a | 0.31817 | 0.176591863 | 0.097 | 0 | 1 | 23 |
| capn2a | 0.31817 | 0.172207489 | 0.097 | 0 | 1 | 23 |
| card19 | 0.31817 | 0.165878832 | 0.097 | 0 | 1 | 23 |
| ccdc126 | 0.31817 | 0.152689911 | 0.097 | 0 | 1 | 23 |
| cfap126 | 0.31817 | 0.194241067 | 0.097 | 0 | 1 | 23 |
| erlec1 | 0.31817 | 0.159382798 | 0.097 | 0 | 1 | 23 |
| fem1b | 0.31817 | 0.167279555 | 0.097 | 0 | 1 | 23 |
| fermt2 | 0.31817 | 0.129379246 | 0.097 | 0 | 1 | 23 |
| lrp12 | 0.31817 | 0.148003548 | 0.097 | 0 | 1 | 23 |
| mtrex | 0.31817 | 0.160043653 | 0.097 | 0 | 1 | 23 |
| pkig | 0.31817 | 0.165622981 | 0.097 | 0 | 1 | 23 |
| ppm1la | 0.31817 | 0.144950486 | 0.097 | 0 | 1 | 23 |
| rb1cc1 | 0.31817 | 0.183273323 | 0.097 | 0 | 1 | 23 |
| robo1 | 0.31817 | 0.172834166 | 0.097 | 0 | 1 | 23 |
| rprd2b | 0.31817 | 0.214113261 | 0.097 | 0 | 1 | 23 |
| sash1a | 0.31817 | 0.196252827 | 0.097 | 0 | 1 | 23 |
| si:dkey-31f5.8 | 0.31817 | 0.152588434 | 0.097 | 0 | 1 | 23 |
| smad6a | 0.31817 | 0.160750857 | 0.097 | 0 | 1 | 23 |
| snrnp200 | 0.31817 | 0.142991975 | 0.097 | 0 | 1 | 23 |
| stambpb | 0.31817 | 0.15908866 | 0.097 | 0 | 1 | 23 |
| timm13 | 0.31817 | 0.179189986 | 0.097 | 0 | 1 | 23 |
| tmx2b | 0.31817 | 0.16001453 | 0.097 | 0 | 1 | 23 |
| tppp3 | 0.31817 | 0.171385142 | 0.097 | 0 | 1 | 23 |
| ttll6 | 0.31817 | 0.197327624 | 0.097 | 0 | 1 | 23 |
| waslb | 0.31817 | 0.165098996 | 0.097 | 0 | 1 | 23 |
| zdhhc17 | 0.31817 | 0.16854876 | 0.097 | 0 | 1 | 23 |
| zgc:85858 | 0.31817 | 0.202999547 | 0.097 | 0 | 1 | 23 |
| znf1156 | 0.31817 | 0.19257926 | 0.097 | 0 | 1 | 23 |
| CABZ01033394.1 | 0.31817 | 0.224771086 | 0.097 | 0 | 1 | 23 |
| CABZ01068367.1 | 0.31817 | 0.176634042 | 0.097 | 0 | 1 | 23 |
| CU655961.2 | 0.31817 | 0.219452183 | 0.097 | 0 | 1 | 23 |
| CU681842.1 | 0.31817 | 0.145328431 | 0.097 | 0 | 1 | 23 |
| LOC110438491 | 0.31817 | 0.115659414 | 0.097 | 0 | 1 | 23 |
| XLOC-034538 | 0.31817 | 0.163012093 | 0.097 | 0 | 1 | 23 |
| abhd17c | 0.31817 | 0.148244856 | 0.097 | 0 | 1 | 23 |

| | | | | | | |
|---|---|---|---|---|---|---|
| acat1 | 0.31817 | 0.131024676 | 0.097 | 0 | 1 | 23 |
| acot18 | 0.31817 | 0.145389901 | 0.097 | 0 | 1 | 23 |
| acvr1ba | 0.31817 | 0.159075253 | 0.097 | 0 | 1 | 23 |
| ahr2 | 0.31817 | 0.206661461 | 0.097 | 0 | 1 | 23 |
| ap4s1 | 0.31817 | 0.165988278 | 0.097 | 0 | 1 | 23 |
| arhgap32a | 0.31817 | 0.152983693 | 0.097 | 0 | 1 | 23 |
| arid5b | 0.31817 | 0.167646183 | 0.097 | 0 | 1 | 23 |
| asb8 | 0.31817 | 0.149884209 | 0.097 | 0 | 1 | 23 |
| atp2a2b | 0.31817 | 0.203116608 | 0.097 | 0 | 1 | 23 |
| atp2b1a | 0.31817 | 0.194434566 | 0.097 | 0 | 1 | 23 |
| atpv0e2 | 0.31817 | 0.172666574 | 0.097 | 0 | 1 | 23 |
| atrxl | 0.31817 | 0.15495298 | 0.097 | 0 | 1 | 23 |
| atxn1a | 0.31817 | 0.172235996 | 0.097 | 0 | 1 | 23 |
| bmi1b | 0.31817 | 0.170444394 | 0.097 | 0 | 1 | 23 |
| bud23 | 0.31817 | 0.158104826 | 0.097 | 0 | 1 | 23 |
| caap1 | 0.31817 | 0.171728238 | 0.097 | 0 | 1 | 23 |
| ccdc22 | 0.31817 | 0.14783889 | 0.097 | 0 | 1 | 23 |
| ccdc85cb | 0.31817 | 0.169371881 | 0.097 | 0 | 1 | 23 |
| ch25h | 0.31817 | 0.213996425 | 0.097 | 0 | 1 | 23 |
| chad | 0.31817 | 0.203159222 | 0.097 | 0 | 1 | 23 |
| cmc4 | 0.31817 | 0.177566437 | 0.097 | 0 | 1 | 23 |
| cnot4b | 0.31817 | 0.170448173 | 0.097 | 0 | 1 | 23 |
| cryz | 0.31817 | 0.137533998 | 0.097 | 0 | 1 | 23 |
| dgkza | 0.31817 | 0.164504107 | 0.097 | 0 | 1 | 23 |
| dnaaf1 | 0.31817 | 0.196054802 | 0.097 | 0 | 1 | 23 |
| ech1 | 0.31817 | 0.169908875 | 0.097 | 0 | 1 | 23 |
| ehd1a | 0.31817 | 0.143924138 | 0.097 | 0 | 1 | 23 |
| erap2 | 0.31817 | 0.149301974 | 0.097 | 0 | 1 | 23 |
| f2rl1.2 | 0.31817 | 0.202921483 | 0.097 | 0 | 1 | 23 |
| fam199x | 0.31817 | 0.146122858 | 0.097 | 0 | 1 | 23 |
| frzb | 0.31817 | 0.153341449 | 0.097 | 0 | 1 | 23 |
| fzd9b | 0.31817 | 0.158554929 | 0.097 | 0 | 1 | 23 |
| gabpb2b | 0.31817 | 0.162196469 | 0.097 | 0 | 1 | 23 |
| gale | 0.31817 | 0.167521381 | 0.097 | 0 | 1 | 23 |
| gpr108 | 0.31817 | 0.15390179 | 0.097 | 0 | 1 | 23 |
| gtf2a1l | 0.31817 | 0.1587436 | 0.097 | 0 | 1 | 23 |
| gzf1 | 0.31817 | 0.217525025 | 0.097 | 0 | 1 | 23 |
| hsd17b12b | 0.31817 | 0.152917417 | 0.097 | 0 | 1 | 23 |
| htra3b | 0.31817 | 0.188128041 | 0.097 | 0 | 1 | 23 |
| icn | 0.31817 | 0.364869899 | 0.097 | 0 | 1 | 23 |
| invs | 0.31817 | 0.153002887 | 0.097 | 0 | 1 | 23 |
| itpr1a | 0.31817 | 0.156300747 | 0.097 | 0 | 1 | 23 |
| kdelr2a | 0.31817 | 0.145313891 | 0.097 | 0 | 1 | 23 |
| kif6 | 0.31817 | 0.162851971 | 0.097 | 0 | 1 | 23 |
| lman2la | 0.31817 | 0.143189805 | 0.097 | 0 | 1 | 23 |
| mad2l2 | 0.31817 | 0.155947559 | 0.097 | 0 | 1 | 23 |
| mark2b | 0.31817 | 0.17805127 | 0.097 | 0 | 1 | 23 |
| med19b | 0.31817 | 0.1765619 | 0.097 | 0 | 1 | 23 |

| | | | | | | |
|---|---|---|---|---|---|---|
| metrn | 0.31817 | 0.156203941 | 0.097 | 0 | 1 | 23 |
| mthfs | 0.31817 | 0.210048888 | 0.097 | 0 | 1 | 23 |
| mycbpap | 0.31817 | 0.21655027 | 0.097 | 0 | 1 | 23 |
| myo10 | 0.31817 | 0.15828077 | 0.097 | 0 | 1 | 23 |
| ndufs4 | 0.31817 | 0.119814749 | 0.097 | 0 | 1 | 23 |
| neil1 | 0.31817 | 0.163615567 | 0.097 | 0 | 1 | 23 |
| oxr1b | 0.31817 | 0.15966883 | 0.097 | 0 | 1 | 23 |
| paox1 | 0.31817 | 0.150545639 | 0.097 | 0 | 1 | 23 |
| pds5b | 0.31817 | 0.182121026 | 0.097 | 0 | 1 | 23 |
| pdzrn3b | 0.31817 | 0.167589161 | 0.097 | 0 | 1 | 23 |
| peli3 | 0.31817 | 0.139338342 | 0.097 | 0 | 1 | 23 |
| pla2g7 | 0.31817 | 0.128202878 | 0.097 | 0 | 1 | 23 |
| prkcbp1l | 0.31817 | 0.148501972 | 0.097 | 0 | 1 | 23 |
| prmt7 | 0.31817 | 0.131946449 | 0.097 | 0 | 1 | 23 |
| qtrt2 | 0.31817 | 0.145631422 | 0.097 | 0 | 1 | 23 |
| rasgef1ba | 0.31817 | 0.17777188 | 0.097 | 0 | 1 | 23 |
| rassf7b | 0.31817 | 0.178193646 | 0.097 | 0 | 1 | 23 |
| relch | 0.31817 | 0.149644323 | 0.097 | 0 | 1 | 23 |
| rtkn2b | 0.31817 | 0.163733819 | 0.097 | 0 | 1 | 23 |
| scamp4 | 0.31817 | 0.146720233 | 0.097 | 0 | 1 | 23 |
| sdr42e1 | 0.31817 | 0.219237091 | 0.097 | 0 | 1 | 23 |
| sec22ba | 0.31817 | 0.134667221 | 0.097 | 0 | 1 | 23 |
| sec23ip | 0.31817 | 0.163117512 | 0.097 | 0 | 1 | 23 |
| shroom4 | 0.31817 | 0.167529688 | 0.097 | 0 | 1 | 23 |
| si:ch211-207k7.4 | 0.31817 | 0.162987668 | 0.097 | 0 | 1 | 23 |
| si:ch211-89o9.6 | 0.31817 | 0.171883604 | 0.097 | 0 | 1 | 23 |
| si:ch73-386h18.1 | 0.31817 | 0.189610313 | 0.097 | 0 | 1 | 23 |
| si:dkey-38p12.3 | 0.31817 | 0.171183051 | 0.097 | 0 | 1 | 23 |
| slc12a4 | 0.31817 | 0.151614655 | 0.097 | 0 | 1 | 23 |
| slc25a28 | 0.31817 | 0.185182301 | 0.097 | 0 | 1 | 23 |
| spag6 | 0.31817 | 0.196054802 | 0.097 | 0 | 1 | 23 |
| stxbp3 | 0.31817 | 0.146686006 | 0.097 | 0 | 1 | 23 |
| swt1 | 0.31817 | 0.179353064 | 0.097 | 0 | 1 | 23 |
| tab2 | 0.31817 | 0.153351252 | 0.097 | 0 | 1 | 23 |
| taf6 | 0.31817 | 0.165924996 | 0.097 | 0 | 1 | 23 |
| tbc1d10b | 0.31817 | 0.123643775 | 0.097 | 0 | 1 | 23 |
| tex9 | 0.31817 | 0.149591013 | 0.097 | 0 | 1 | 23 |
| tgfbi | 0.31817 | 0.179217416 | 0.097 | 0 | 1 | 23 |
| tiam2a | 0.31817 | 0.16080799 | 0.097 | 0 | 1 | 23 |
| tmem63ba | 0.31817 | 0.19649346 | 0.097 | 0 | 1 | 23 |
| ubap1 | 0.31817 | 0.199120094 | 0.097 | 0 | 1 | 23 |
| ufsp2 | 0.31817 | 0.144840094 | 0.097 | 0 | 1 | 23 |
| vwa1 | 0.31817 | 0.19718168 | 0.097 | 0 | 1 | 23 |
| wdr32 | 0.31817 | 0.149858125 | 0.097 | 0 | 1 | 23 |
| wdr75 | 0.31817 | 0.150102991 | 0.097 | 0 | 1 | 23 |
| wu:fa19b12 | 0.31817 | 0.141642216 | 0.097 | 0 | 1 | 23 |
| wu:fc21g02 | 0.31817 | 0.159288433 | 0.097 | 0 | 1 | 23 |
| zeb1b | 0.31817 | 0.15125355 | 0.097 | 0 | 1 | 23 |

| | | | | | | |
|---|---|---|---|---|---|---|
| zgc:163030 | 0.31817 | 0.153210867 | 0.097 | 0 | 1 | 23 |
| zgc:171459 | 0.31817 | 0.149947714 | 0.097 | 0 | 1 | 23 |
| zgc:77838 | 0.31817 | 0.135976989 | 0.097 | 0 | 1 | 23 |
| zmat5 | 0.31817 | 0.123115171 | 0.097 | 0 | 1 | 23 |
| znf131 | 0.31817 | 0.168548127 | 0.097 | 0 | 1 | 23 |
| znf207b | 0.31817 | 0.162714607 | 0.097 | 0 | 1 | 23 |
| elof1 | 0.31817 | 0.24496285 | 0.645 | 0.375 | 1 | 23 |
| sept8a | 0.31954 | -0.39071758 | 0.129 | 0.375 | 1 | 23 |
| gatc | 0.3198 | 0.409847491 | 0.323 | 0.125 | 1 | 23 |
| zgc:195173 | 0.31981 | 0.388438299 | 0.645 | 0.375 | 1 | 23 |
| actr1 | 0.32014 | -0.11291174 | 0.129 | 0.125 | 1 | 23 |
| commd7 | 0.32042 | -0.35850618 | 0.097 | 0.25 | 1 | 23 |
| hnrnpub | 0.3207 | -0.38150019 | 0.258 | 0.5 | 1 | 23 |
| ncor1 | 0.32084 | -0.31327143 | 0.129 | 0.25 | 1 | 23 |
| esco1 | 0.32104 | -0.29422165 | 0.065 | 0.25 | 1 | 23 |
| gemin6 | 0.3211 | -0.36650449 | 0.065 | 0.25 | 1 | 23 |
| dctn3 | 0.3211 | 0.371800393 | 0.387 | 0.125 | 1 | 23 |
| slc25a24 | 0.32123 | -0.34884288 | 0.065 | 0.25 | 1 | 23 |
| stox1 | 0.32164 | -0.38056876 | 0.065 | 0.25 | 1 | 23 |
| ryk | 0.32173 | -0.32925558 | 0.065 | 0.25 | 1 | 23 |
| si:ch211-68a17.7 | 0.32174 | -0.3226237 | 0.129 | 0.25 | 1 | 23 |
| serpinh1b | 0.32191 | -0.39090755 | 0.129 | 0.375 | 1 | 23 |
| cdon | 0.32272 | 0.186558883 | 0.29 | 0.125 | 1 | 23 |
| cldn12 | 0.32295 | -0.39348282 | 0.226 | 0.5 | 1 | 23 |
| gas1b | 0.32295 | -0.15363409 | 0.097 | 0.25 | 1 | 23 |
| pi15a | 0.32314 | -0.19852649 | 0.516 | 0.75 | 1 | 23 |
| macf1a | 0.32325 | -0.38692637 | 0.129 | 0.375 | 1 | 23 |
| adar | 0.32328 | -0.34022719 | 0.065 | 0.25 | 1 | 23 |
| kars | 0.32341 | -0.27061867 | 0.29 | 0.375 | 1 | 23 |
| rnf185 | 0.32354 | -0.33368637 | 0.129 | 0.375 | 1 | 23 |
| kif5ba | 0.32363 | 0.145673496 | 0.29 | 0.125 | 1 | 23 |
| kdm5c | 0.32381 | -0.33200697 | 0.129 | 0.375 | 1 | 23 |
| foxo3b | 0.32382 | 0.450225228 | 0.387 | 0.125 | 1 | 23 |
| si:ch211-237c6.4 | 0.32412 | -0.34679586 | 0.129 | 0.375 | 1 | 23 |
| tpst1 | 0.32476 | -0.32932051 | 0.129 | 0.375 | 1 | 23 |
| ywhae1 | 0.32511 | 0.328578615 | 0.645 | 0.5 | 1 | 23 |
| relb | 0.3254 | -0.32828861 | 0.065 | 0.25 | 1 | 23 |
| hoxb8a | 0.32594 | 0.3824646 | 0.355 | 0.125 | 1 | 23 |
| naca | 0.32648 | 0.214647912 | 0.935 | 0.875 | 1 | 23 |
| cpne1 | 0.32683 | 0.36362214 | 0.387 | 0.125 | 1 | 23 |
| pcsk7 | 0.32698 | -0.18642631 | 0.097 | 0.125 | 1 | 23 |
| ifrd2 | 0.3271 | -0.2253494 | 0.065 | 0.25 | 1 | 23 |
| ptgr1 | 0.32825 | 0.338742992 | 0.387 | 0.125 | 1 | 23 |
| hoxc3a | 0.32873 | -0.31565545 | 0.129 | 0.25 | 1 | 23 |
| mrpl20 | 0.32873 | -0.24220221 | 0.355 | 0.625 | 1 | 23 |
| hdac1 | 0.32875 | -0.26909079 | 0.613 | 0.875 | 1 | 23 |
| ulk2 | 0.32885 | -0.20963149 | 0.323 | 0.375 | 1 | 23 |
| sod2 | 0.32894 | 0.342385303 | 0.323 | 0.125 | 1 | 23 |

| | | | | | | |
|---|---|---|---|---|---|---|
| higd1a | 0.32894 | 0.481483858 | 0.516 | 0.25 | 1 | 23 |
| mmp14b | 0.32897 | 0.373480969 | 0.323 | 0.125 | 1 | 23 |
| lgals17 | 0.32904 | -0.40480908 | 0.226 | 0.5 | 1 | 23 |
| ndufb9 | 0.32907 | -0.37482925 | 0.484 | 0.75 | 1 | 23 |
| cmtm3 | 0.33083 | 0.163647083 | 0.29 | 0.125 | 1 | 23 |
| ywhaba | 0.33112 | -0.24112389 | 0.581 | 0.625 | 1 | 23 |
| clic2 | 0.33152 | -0.31459749 | 0.065 | 0.25 | 1 | 23 |
| canx | 0.33269 | 0.408053991 | 0.387 | 0.125 | 1 | 23 |
| stk11 | 0.33284 | -0.34220354 | 0.065 | 0.25 | 1 | 23 |
| CABZ01092156.1 | 0.33289 | -0.2253477 | 0.161 | 0.375 | 1 | 23 |
| zgc:112356 | 0.33361 | -0.11722416 | 0.097 | 0.25 | 1 | 23 |
| eif4ebp2 | 0.33404 | 0.367916352 | 0.484 | 0.25 | 1 | 23 |
| klhdc4 | 0.33417 | -0.31878338 | 0.065 | 0.25 | 1 | 23 |
| tax1bp1b | 0.33458 | 0.379819173 | 0.355 | 0.125 | 1 | 23 |
| dnmt3bb.1 | 0.33511 | 0.103554812 | 0.258 | 0.125 | 1 | 23 |
| syncripl | 0.33515 | -0.39316459 | 0.226 | 0.5 | 1 | 23 |
| ier3ip1 | 0.33552 | 0.217683115 | 0.484 | 0.25 | 1 | 23 |
| tp53 | 0.33682 | 0.107711822 | 0.548 | 0.375 | 1 | 23 |
| csnk1a1 | 0.33756 | -0.39615313 | 0.355 | 0.625 | 1 | 23 |
| pfdn6 | 0.33782 | 0.339774162 | 0.323 | 0.125 | 1 | 23 |
| taf15 | 0.33862 | -0.29848061 | 0.452 | 0.625 | 1 | 23 |
| prpf19 | 0.33903 | -0.36748044 | 0.097 | 0.25 | 1 | 23 |
| tuba1c | 0.33976 | -0.12078741 | 0.129 | 0.125 | 1 | 23 |
| necap2 | 0.34044 | -0.302819 | 0.129 | 0.25 | 1 | 23 |
| mtdha | 0.34078 | 0.223490746 | 0.323 | 0.125 | 1 | 23 |
| si:ch211-210c8.7 | 0.3408 | -0.33902738 | 0.065 | 0.25 | 1 | 23 |
| znf207a | 0.34082 | -0.1707535 | 0.097 | 0.125 | 1 | 23 |
| mrps30 | 0.34095 | -0.11706464 | 0.129 | 0.125 | 1 | 23 |
| as3mt | 0.34106 | -0.35377971 | 0.065 | 0.25 | 1 | 23 |
| neurl1b | 0.34127 | -0.35485771 | 0.065 | 0.25 | 1 | 23 |
| cnn2 | 0.34163 | 0.410725751 | 0.484 | 0.25 | 1 | 23 |
| lamb2 | 0.34204 | 0.110886735 | 0.258 | 0.125 | 1 | 23 |
| magt1 | 0.34235 | 0.188600917 | 0.323 | 0.125 | 1 | 23 |
| prelid3b | 0.34269 | -0.21220988 | 0.226 | 0.25 | 1 | 23 |
| srrm1 | 0.34313 | -0.39839128 | 0.355 | 0.625 | 1 | 23 |
| zgc:153713 | 0.34397 | -0.3131426 | 0.258 | 0.375 | 1 | 23 |
| rn7sk | 0.34402 | 0.123076001 | 0.419 | 0.25 | 1 | 23 |
| u2af1 | 0.34441 | -0.26713568 | 0.516 | 0.625 | 1 | 23 |
| ube2v2 | 0.34468 | -0.22768927 | 0.677 | 0.75 | 1 | 23 |
| si:dkey-56d12.4 | 0.34483 | -0.24214722 | 0.065 | 0.25 | 1 | 23 |
| ola1 | 0.34506 | -0.34736655 | 0.452 | 0.625 | 1 | 23 |
| mboat7 | 0.34531 | -0.21218119 | 0.065 | 0.25 | 1 | 23 |
| josd1 | 0.34567 | -0.35201413 | 0.065 | 0.25 | 1 | 23 |
| arhgap29a | 0.34591 | -0.33328092 | 0.065 | 0.25 | 1 | 23 |
| zgc:153867 | 0.34616 | 0.334607309 | 0.839 | 0.625 | 1 | 23 |
| rab5if | 0.3467 | 0.33576121 | 0.355 | 0.125 | 1 | 23 |
| mboat2a | 0.34676 | -0.25924583 | 0.065 | 0.25 | 1 | 23 |
| mllt10 | 0.34688 | -0.32760739 | 0.065 | 0.25 | 1 | 23 |

| | | | | | | |
|---|---|---|---|---|---|---|
| eif4a1a | 0.34696 | -0.17136441 | 0.516 | 0.5 | 1 | 23 |
| LOC101883507 | 0.34728 | -0.34987829 | 0.097 | 0.25 | 1 | 23 |
| rilpl2 | 0.34759 | -0.33914047 | 0.097 | 0.25 | 1 | 23 |
| polr2f | 0.34794 | 0.378741482 | 0.516 | 0.25 | 1 | 23 |
| myl9b | 0.34813 | 0.201881256 | 0.452 | 0.25 | 1 | 23 |
| si:dkey-56m19.5 | 0.34844 | 0.309538814 | 0.226 | 0.125 | 1 | 23 |
| dnajb11 | 0.34878 | -0.11276525 | 0.129 | 0.125 | 1 | 23 |
| arid1ab | 0.34999 | -0.35680702 | 0.065 | 0.25 | 1 | 23 |
| baz1a | 0.35 | -0.37358092 | 0.258 | 0.5 | 1 | 23 |
| pcna | 0.35133 | -0.45085009 | 0.258 | 0.375 | 1 | 23 |
| ikzf2 | 0.35164 | -0.28962703 | 0.129 | 0.25 | 1 | 23 |
| idh3g | 0.35254 | -0.30356984 | 0.065 | 0.25 | 1 | 23 |
| ube2na | 0.35268 | 0.386656871 | 0.484 | 0.25 | 1 | 23 |
| pleca | 0.35428 | -0.23792712 | 0.452 | 0.5 | 1 | 23 |
| cct3 | 0.35506 | 0.169785232 | 0.742 | 0.5 | 1 | 23 |
| tnrc6c2 | 0.35507 | -0.11148502 | 0.129 | 0.125 | 1 | 23 |
| col8a1a | 0.35645 | -0.29830668 | 0.065 | 0.25 | 1 | 23 |
| prkd3 | 0.35802 | -0.2140599 | 0.032 | 0.125 | 1 | 23 |
| sec24b | 0.35802 | -0.21370752 | 0.032 | 0.125 | 1 | 23 |
| si:dkey-21c1.8 | 0.35802 | -0.21655777 | 0.032 | 0.125 | 1 | 23 |
| tusc3 | 0.35802 | -0.21211766 | 0.032 | 0.125 | 1 | 23 |
| FBXO46 | 0.35802 | -0.21955844 | 0.032 | 0.125 | 1 | 23 |
| amd1 | 0.35802 | -0.29690384 | 0.032 | 0.125 | 1 | 23 |
| calub | 0.35802 | -0.22399222 | 0.032 | 0.125 | 1 | 23 |
| cerkl | 0.35802 | -0.21835949 | 0.032 | 0.125 | 1 | 23 |
| gal3st1b | 0.35802 | -0.21602208 | 0.032 | 0.125 | 1 | 23 |
| il17rd | 0.35802 | -0.2205026 | 0.032 | 0.125 | 1 | 23 |
| mhc1zda | 0.35802 | -0.21703254 | 0.032 | 0.125 | 1 | 23 |
| mmp11a | 0.35802 | -0.15581127 | 0.032 | 0.125 | 1 | 23 |
| papss1 | 0.35802 | -0.20205688 | 0.032 | 0.125 | 1 | 23 |
| recql | 0.35802 | -0.22134227 | 0.032 | 0.125 | 1 | 23 |
| tmem198b | 0.35802 | -0.22838522 | 0.032 | 0.125 | 1 | 23 |
| zfyve9a | 0.35802 | -0.22302763 | 0.032 | 0.125 | 1 | 23 |
| ARHGEF17 | 0.35802 | -0.30124593 | 0.032 | 0.125 | 1 | 23 |
| CT737123.1 | 0.35802 | -0.22056701 | 0.032 | 0.125 | 1 | 23 |
| RALGDS | 0.35802 | -0.16034291 | 0.032 | 0.125 | 1 | 23 |
| adnpa | 0.35802 | -0.21872089 | 0.032 | 0.125 | 1 | 23 |
| ankib1b | 0.35802 | -0.22356079 | 0.032 | 0.125 | 1 | 23 |
| asxl2 | 0.35802 | -0.17097014 | 0.032 | 0.125 | 1 | 23 |
| bicc1b | 0.35802 | -0.21347009 | 0.032 | 0.125 | 1 | 23 |
| caska | 0.35802 | -0.22980239 | 0.032 | 0.125 | 1 | 23 |
| casp8 | 0.35802 | -0.22037382 | 0.032 | 0.125 | 1 | 23 |
| cbx6a | 0.35802 | -0.29090386 | 0.032 | 0.125 | 1 | 23 |
| ciz1b | 0.35802 | -0.18375028 | 0.032 | 0.125 | 1 | 23 |
| cmtr1 | 0.35802 | -0.23417755 | 0.032 | 0.125 | 1 | 23 |
| cwf19l1 | 0.35802 | -0.22168026 | 0.032 | 0.125 | 1 | 23 |
| cyp2ad3 | 0.35802 | -0.22720339 | 0.032 | 0.125 | 1 | 23 |
| cyp2p10 | 0.35802 | -0.20331129 | 0.032 | 0.125 | 1 | 23 |

| | | | | | | |
|---|---|---|---|---|---|---|
| ddx28 | 0.35802 | -0.21584171 | 0.032 | 0.125 | 1 | 23 |
| dennd4c | 0.35802 | -0.22825444 | 0.032 | 0.125 | 1 | 23 |
| fstb | 0.35802 | -0.21272997 | 0.032 | 0.125 | 1 | 23 |
| galnt11 | 0.35802 | -0.20480147 | 0.032 | 0.125 | 1 | 23 |
| hnrnph1 | 0.35802 | -0.20243674 | 0.032 | 0.125 | 1 | 23 |
| kremen1 | 0.35802 | -0.20765174 | 0.032 | 0.125 | 1 | 23 |
| lmf2a | 0.35802 | -0.21897252 | 0.032 | 0.125 | 1 | 23 |
| map1sa | 0.35802 | -0.35893089 | 0.032 | 0.125 | 1 | 23 |
| mef2ca | 0.35802 | -0.24442709 | 0.032 | 0.125 | 1 | 23 |
| mylipa | 0.35802 | -0.1115823 | 0.032 | 0.125 | 1 | 23 |
| nup205 | 0.35802 | -0.15413992 | 0.032 | 0.125 | 1 | 23 |
| ppm1ba | 0.35802 | -0.22499312 | 0.032 | 0.125 | 1 | 23 |
| qsox2 | 0.35802 | -0.19213606 | 0.032 | 0.125 | 1 | 23 |
| rab18b | 0.35802 | -0.21320927 | 0.032 | 0.125 | 1 | 23 |
| rbm14b | 0.35802 | -0.2241435 | 0.032 | 0.125 | 1 | 23 |
| scoca | 0.35802 | -0.21306765 | 0.032 | 0.125 | 1 | 23 |
| sertad2b | 0.35802 | -0.23389795 | 0.032 | 0.125 | 1 | 23 |
| si:dkey-234h16.7 | 0.35802 | -0.21202829 | 0.032 | 0.125 | 1 | 23 |
| si:dkeyp-113d7.10 | 0.35802 | -0.31821208 | 0.032 | 0.125 | 1 | 23 |
| tbc1d4 | 0.35802 | -0.17627295 | 0.032 | 0.125 | 1 | 23 |
| tcf20 | 0.35802 | -0.2336298 | 0.032 | 0.125 | 1 | 23 |
| tsc1b | 0.35802 | -0.1493092 | 0.032 | 0.125 | 1 | 23 |
| tti2 | 0.35802 | -0.20339116 | 0.032 | 0.125 | 1 | 23 |
| unk | 0.35802 | -0.20322439 | 0.032 | 0.125 | 1 | 23 |
| wdr26a | 0.35802 | -0.15029089 | 0.032 | 0.125 | 1 | 23 |
| xrra1 | 0.35802 | -0.21927403 | 0.032 | 0.125 | 1 | 23 |
| zcchc24 | 0.35802 | -0.22725084 | 0.032 | 0.125 | 1 | 23 |
| zgc:174877 | 0.35802 | -0.22678742 | 0.032 | 0.125 | 1 | 23 |
| CR786577.1 | 0.35802 | -0.24911115 | 0.032 | 0.125 | 1 | 23 |
| cgas | 0.35802 | -0.12394092 | 0.032 | 0.125 | 1 | 23 |
| crym | 0.35802 | -0.13418297 | 0.032 | 0.125 | 1 | 23 |
| dhrs1 | 0.35802 | -0.12689395 | 0.032 | 0.125 | 1 | 23 |
| dlgap4b | 0.35802 | -0.1220147 | 0.032 | 0.125 | 1 | 23 |
| eda | 0.35802 | -0.15929868 | 0.032 | 0.125 | 1 | 23 |
| fam151b | 0.35802 | -0.16036691 | 0.032 | 0.125 | 1 | 23 |
| ift74 | 0.35802 | -0.12488051 | 0.032 | 0.125 | 1 | 23 |
| myo1eb | 0.35802 | -0.17721959 | 0.032 | 0.125 | 1 | 23 |
| nudt4a | 0.35802 | -0.12126988 | 0.032 | 0.125 | 1 | 23 |
| rrp9 | 0.35802 | -0.15113154 | 0.032 | 0.125 | 1 | 23 |
| si:ch73-368j24.10.1 | 0.35802 | -0.29946217 | 0.032 | 0.125 | 1 | 23 |
| slc10a3 | 0.35802 | -0.13167965 | 0.032 | 0.125 | 1 | 23 |
| tox4b | 0.35802 | -0.17122227 | 0.032 | 0.125 | 1 | 23 |
| vamp4 | 0.35802 | -0.30804032 | 0.032 | 0.125 | 1 | 23 |
| zgc:173575 | 0.35802 | -0.17498 | 0.032 | 0.125 | 1 | 23 |
| ATP11B | 0.35802 | -0.26500865 | 0.032 | 0.125 | 1 | 23 |
| BX470189.1 | 0.35802 | -0.11607689 | 0.032 | 0.125 | 1 | 23 |
| CABZ01092982.1 | 0.35802 | -0.20577645 | 0.032 | 0.125 | 1 | 23 |
| CR751227.2 | 0.35802 | -0.11853775 | 0.032 | 0.125 | 1 | 23 |

| | | | | | | |
|---|---|---|---|---|---|---|
| LOC100333482 | 0.35802 | -0.26511897 | 0.032 | 0.125 | 1 | 23 |
| LOC101886876 | 0.35802 | -0.21489661 | 0.032 | 0.125 | 1 | 23 |
| LOC110437744.1 | 0.35802 | -0.11689151 | 0.032 | 0.125 | 1 | 23 |
| XLOC-004018 | 0.35802 | -0.11341886 | 0.032 | 0.125 | 1 | 23 |
| accs | 0.35802 | -0.21277718 | 0.032 | 0.125 | 1 | 23 |
| adsl | 0.35802 | -0.21744398 | 0.032 | 0.125 | 1 | 23 |
| akap7 | 0.35802 | -0.10603332 | 0.032 | 0.125 | 1 | 23 |
| aktip | 0.35802 | -0.10294195 | 0.032 | 0.125 | 1 | 23 |
| alkbh5 | 0.35802 | -0.18845076 | 0.032 | 0.125 | 1 | 23 |
| ap1ar | 0.35802 | -0.24950853 | 0.032 | 0.125 | 1 | 23 |
| asf1ba | 0.35802 | -0.28087584 | 0.032 | 0.125 | 1 | 23 |
| cad | 0.35802 | -0.2502393 | 0.032 | 0.125 | 1 | 23 |
| cat | 0.35802 | -0.12797771 | 0.032 | 0.125 | 1 | 23 |
| chpt1 | 0.35802 | -0.25670892 | 0.032 | 0.125 | 1 | 23 |
| cop1 | 0.35802 | -0.2571241 | 0.032 | 0.125 | 1 | 23 |
| coq5 | 0.35802 | -0.21718361 | 0.032 | 0.125 | 1 | 23 |
| crfb4 | 0.35802 | -0.21504319 | 0.032 | 0.125 | 1 | 23 |
| cry-dash | 0.35802 | -0.12918209 | 0.032 | 0.125 | 1 | 23 |
| ctdspl2b | 0.35802 | -0.24868249 | 0.032 | 0.125 | 1 | 23 |
| cyp2aa6 | 0.35802 | -0.10837319 | 0.032 | 0.125 | 1 | 23 |
| dennd4a | 0.35802 | -0.11284367 | 0.032 | 0.125 | 1 | 23 |
| dgcr8 | 0.35802 | -0.19031062 | 0.032 | 0.125 | 1 | 23 |
| dok1b | 0.35802 | -0.1392204 | 0.032 | 0.125 | 1 | 23 |
| dph5 | 0.35802 | -0.23157241 | 0.032 | 0.125 | 1 | 23 |
| dus2 | 0.35802 | -0.18685863 | 0.032 | 0.125 | 1 | 23 |
| entpd4 | 0.35802 | -0.21017369 | 0.032 | 0.125 | 1 | 23 |
| fam98a | 0.35802 | -0.1285811 | 0.032 | 0.125 | 1 | 23 |
| foxj3 | 0.35802 | -0.25080044 | 0.032 | 0.125 | 1 | 23 |
| galcb | 0.35802 | -0.23595877 | 0.032 | 0.125 | 1 | 23 |
| hcfc2 | 0.35802 | -0.11092487 | 0.032 | 0.125 | 1 | 23 |
| hebp2 | 0.35802 | -0.12196714 | 0.032 | 0.125 | 1 | 23 |
| il1rapl1a | 0.35802 | -0.10335906 | 0.032 | 0.125 | 1 | 23 |
| klhl3 | 0.35802 | -0.10676797 | 0.032 | 0.125 | 1 | 23 |
| lmo3 | 0.35802 | -0.24344175 | 0.032 | 0.125 | 1 | 23 |
| map7d3 | 0.35802 | -0.1164578 | 0.032 | 0.125 | 1 | 23 |
| mhc1zba | 0.35802 | -0.19815057 | 0.032 | 0.125 | 1 | 23 |
| mis12 | 0.35802 | -0.2782387 | 0.032 | 0.125 | 1 | 23 |
| mocs2 | 0.35802 | -0.21012073 | 0.032 | 0.125 | 1 | 23 |
| mrpl41 | 0.35802 | -0.11321711 | 0.032 | 0.125 | 1 | 23 |
| mthfr | 0.35802 | -0.12302623 | 0.032 | 0.125 | 1 | 23 |
| mtmr12 | 0.35802 | -0.22376961 | 0.032 | 0.125 | 1 | 23 |
| mycbp2 | 0.35802 | -0.10921246 | 0.032 | 0.125 | 1 | 23 |
| nedd1 | 0.35802 | -0.33831234 | 0.032 | 0.125 | 1 | 23 |
| piezo1 | 0.35802 | -0.30196806 | 0.032 | 0.125 | 1 | 23 |
| prkcbb | 0.35802 | -0.12119926 | 0.032 | 0.125 | 1 | 23 |
| rfx5 | 0.35802 | -0.11224109 | 0.032 | 0.125 | 1 | 23 |
| rrm1 | 0.35802 | -0.27849403 | 0.032 | 0.125 | 1 | 23 |
| sbno1 | 0.35802 | -0.11252764 | 0.032 | 0.125 | 1 | 23 |

| | | | | | | |
|---|---|---|---|---|---|---|
| si:ch1073-416d2.4 | 0.35802 | -0.11522832 | 0.032 | 0.125 | 1 | 23 |
| si:ch211-107o10.3 | 0.35802 | -0.11484846 | 0.032 | 0.125 | 1 | 23 |
| si:dkey-148f10.4 | 0.35802 | -0.20929985 | 0.032 | 0.125 | 1 | 23 |
| si:dkey-188h10.3 | 0.35802 | -0.11088999 | 0.032 | 0.125 | 1 | 23 |
| smarcd3b | 0.35802 | -0.11173576 | 0.032 | 0.125 | 1 | 23 |
| srebf2 | 0.35802 | -0.20835677 | 0.032 | 0.125 | 1 | 23 |
| timp2b | 0.35802 | -0.2158408 | 0.032 | 0.125 | 1 | 23 |
| tmem214 | 0.35802 | -0.10878744 | 0.032 | 0.125 | 1 | 23 |
| tmem35 | 0.35802 | -0.10817847 | 0.032 | 0.125 | 1 | 23 |
| trappc6b | 0.35802 | -0.11034471 | 0.032 | 0.125 | 1 | 23 |
| ubac1 | 0.35802 | -0.11537241 | 0.032 | 0.125 | 1 | 23 |
| ube3d | 0.35802 | -0.10078583 | 0.032 | 0.125 | 1 | 23 |
| ubxn7 | 0.35802 | -0.11655072 | 0.032 | 0.125 | 1 | 23 |
| ugt5c3 | 0.35802 | -0.21083088 | 0.032 | 0.125 | 1 | 23 |
| vps33a | 0.35802 | -0.24501477 | 0.032 | 0.125 | 1 | 23 |
| wdr44 | 0.35802 | -0.18860523 | 0.032 | 0.125 | 1 | 23 |
| wdr62 | 0.35802 | -0.21747871 | 0.032 | 0.125 | 1 | 23 |
| zgc:101679 | 0.35802 | -0.26868977 | 0.032 | 0.125 | 1 | 23 |
| zgc:113102 | 0.35802 | -0.2038682 | 0.032 | 0.125 | 1 | 23 |
| zgc:113265 | 0.35802 | -0.11222457 | 0.032 | 0.125 | 1 | 23 |
| zgc:173548 | 0.35802 | -0.11510666 | 0.032 | 0.125 | 1 | 23 |
| BX323557.1 | 0.35802 | -0.27743211 | 0.032 | 0.125 | 1 | 23 |
| BX470238.1 | 0.35802 | -0.18594287 | 0.032 | 0.125 | 1 | 23 |
| BX511034.2 | 0.35802 | -0.14393398 | 0.032 | 0.125 | 1 | 23 |
| BX842699.1 | 0.35802 | -0.11763801 | 0.032 | 0.125 | 1 | 23 |
| CABZ01072952.1 | 0.35802 | -0.16927236 | 0.032 | 0.125 | 1 | 23 |
| CABZ01074397.1 | 0.35802 | -0.14140372 | 0.032 | 0.125 | 1 | 23 |
| CABZ01076275.1 | 0.35802 | -0.22795528 | 0.032 | 0.125 | 1 | 23 |
| CABZ01081752.3 | 0.35802 | -0.15677753 | 0.032 | 0.125 | 1 | 23 |
| CABZ01083943.1 | 0.35802 | -0.15008419 | 0.032 | 0.125 | 1 | 23 |
| CU467861.1 | 0.35802 | -0.16896633 | 0.032 | 0.125 | 1 | 23 |
| CU984579.1 | 0.35802 | -0.14247544 | 0.032 | 0.125 | 1 | 23 |
| FP067440.1 | 0.35802 | -0.14490051 | 0.032 | 0.125 | 1 | 23 |
| FQ377903.1 | 0.35802 | -0.16950097 | 0.032 | 0.125 | 1 | 23 |
| SHKBP1 | 0.35802 | -0.25145362 | 0.032 | 0.125 | 1 | 23 |
| XLOC-014397 | 0.35802 | -0.16691747 | 0.032 | 0.125 | 1 | 23 |
| aass | 0.35802 | -0.16106878 | 0.032 | 0.125 | 1 | 23 |
| abcc4 | 0.35802 | -0.12188809 | 0.032 | 0.125 | 1 | 23 |
| acaca | 0.35802 | -0.16704067 | 0.032 | 0.125 | 1 | 23 |
| acsl1b | 0.35802 | -0.26673823 | 0.032 | 0.125 | 1 | 23 |
| adipor1b | 0.35802 | -0.12319047 | 0.032 | 0.125 | 1 | 23 |
| alg3 | 0.35802 | -0.10153651 | 0.032 | 0.125 | 1 | 23 |
| ambra1a | 0.35802 | -0.22365278 | 0.032 | 0.125 | 1 | 23 |
| apoeb | 0.35802 | -0.27070804 | 0.032 | 0.125 | 1 | 23 |
| appl2 | 0.35802 | -0.17604181 | 0.032 | 0.125 | 1 | 23 |
| arfgap2 | 0.35802 | -0.16818236 | 0.032 | 0.125 | 1 | 23 |
| arhgap42b | 0.35802 | -0.26291971 | 0.032 | 0.125 | 1 | 23 |
| asic1b | 0.35802 | -0.22265994 | 0.032 | 0.125 | 1 | 23 |

| | | | | | | |
|---|---|---|---|---|---|---|
| atf5a | 0.35802 | -0.17344937 | 0.032 | 0.125 | 1 | 23 |
| atg10 | 0.35802 | -0.17553964 | 0.032 | 0.125 | 1 | 23 |
| bicra | 0.35802 | -0.27603319 | 0.032 | 0.125 | 1 | 23 |
| brcc3 | 0.35802 | -0.16491863 | 0.032 | 0.125 | 1 | 23 |
| bri3bp | 0.35802 | -0.15243186 | 0.032 | 0.125 | 1 | 23 |
| brinp3b | 0.35802 | -0.17549113 | 0.032 | 0.125 | 1 | 23 |
| cachd1 | 0.35802 | -0.14939417 | 0.032 | 0.125 | 1 | 23 |
| camta1a | 0.35802 | -0.25370001 | 0.032 | 0.125 | 1 | 23 |
| cbr1 | 0.35802 | -0.27102923 | 0.032 | 0.125 | 1 | 23 |
| ccdc171 | 0.35802 | -0.21911542 | 0.032 | 0.125 | 1 | 23 |
| ccl19a.1 | 0.35802 | -0.25082663 | 0.032 | 0.125 | 1 | 23 |
| cd40 | 0.35802 | -0.15207678 | 0.032 | 0.125 | 1 | 23 |
| cdk5rap2 | 0.35802 | -0.26627005 | 0.032 | 0.125 | 1 | 23 |
| cep85l | 0.35802 | -0.11158463 | 0.032 | 0.125 | 1 | 23 |
| cfap300 | 0.35802 | -0.21707638 | 0.032 | 0.125 | 1 | 23 |
| cfap43 | 0.35802 | -0.14207397 | 0.032 | 0.125 | 1 | 23 |
| chtf18 | 0.35802 | -0.26933626 | 0.032 | 0.125 | 1 | 23 |
| ciartb | 0.35802 | -0.10545604 | 0.032 | 0.125 | 1 | 23 |
| cisd2 | 0.35802 | -0.1081481 | 0.032 | 0.125 | 1 | 23 |
| clasp1a | 0.35802 | -0.15254629 | 0.032 | 0.125 | 1 | 23 |
| cmip | 0.35802 | -0.13187376 | 0.032 | 0.125 | 1 | 23 |
| cnot3a | 0.35802 | -0.25324613 | 0.032 | 0.125 | 1 | 23 |
| cylda | 0.35802 | -0.27368747 | 0.032 | 0.125 | 1 | 23 |
| dcun1d2b | 0.35802 | -0.26386456 | 0.032 | 0.125 | 1 | 23 |
| ddx55 | 0.35802 | -0.17241958 | 0.032 | 0.125 | 1 | 23 |
| dglucy | 0.35802 | -0.21286544 | 0.032 | 0.125 | 1 | 23 |
| dok6 | 0.35802 | -0.16855188 | 0.032 | 0.125 | 1 | 23 |
| dpm2 | 0.35802 | -0.11888743 | 0.032 | 0.125 | 1 | 23 |
| dync1i2b | 0.35802 | -0.26836274 | 0.032 | 0.125 | 1 | 23 |
| eef1akmt1 | 0.35802 | -0.10557344 | 0.032 | 0.125 | 1 | 23 |
| ehd3 | 0.35802 | -0.11029283 | 0.032 | 0.125 | 1 | 23 |
| ergic1 | 0.35802 | -0.26347271 | 0.032 | 0.125 | 1 | 23 |
| exoc3 | 0.35802 | -0.16709921 | 0.032 | 0.125 | 1 | 23 |
| exosc10 | 0.35802 | -0.13575583 | 0.032 | 0.125 | 1 | 23 |
| fam118b | 0.35802 | -0.21931845 | 0.032 | 0.125 | 1 | 23 |
| fam171a1 | 0.35802 | -0.21562083 | 0.032 | 0.125 | 1 | 23 |
| fam49bb | 0.35802 | -0.14439856 | 0.032 | 0.125 | 1 | 23 |
| fam57a | 0.35802 | -0.1942569 | 0.032 | 0.125 | 1 | 23 |
| fat2 | 0.35802 | -0.16105995 | 0.032 | 0.125 | 1 | 23 |
| fndc3a | 0.35802 | -0.13439602 | 0.032 | 0.125 | 1 | 23 |
| frs2a | 0.35802 | -0.20814113 | 0.032 | 0.125 | 1 | 23 |
| glceb | 0.35802 | -0.13037713 | 0.032 | 0.125 | 1 | 23 |
| gpr155a | 0.35802 | -0.14653713 | 0.032 | 0.125 | 1 | 23 |
| gramd1a | 0.35802 | -0.16986642 | 0.032 | 0.125 | 1 | 23 |
| grip1 | 0.35802 | -0.130372 | 0.032 | 0.125 | 1 | 23 |
| gripap1 | 0.35802 | -0.10962529 | 0.032 | 0.125 | 1 | 23 |
| hmox2b | 0.35802 | -0.14793205 | 0.032 | 0.125 | 1 | 23 |
| hspa4a | 0.35802 | -0.21308527 | 0.032 | 0.125 | 1 | 23 |

| | | | | | | |
|---|---|---|---|---|---|---|
| ifih1 | 0.35802 | -0.15945125 | 0.032 | 0.125 | 1 | 23 |
| inppl1a | 0.35802 | -0.14543324 | 0.032 | 0.125 | 1 | 23 |
| ints12 | 0.35802 | -0.10901988 | 0.032 | 0.125 | 1 | 23 |
| irf1b | 0.35802 | -0.2400744 | 0.032 | 0.125 | 1 | 23 |
| irx5a | 0.35802 | -0.18139318 | 0.032 | 0.125 | 1 | 23 |
| itga3b | 0.35802 | -0.26521785 | 0.032 | 0.125 | 1 | 23 |
| jmjd6 | 0.35802 | -0.25396208 | 0.032 | 0.125 | 1 | 23 |
| kazna | 0.35802 | -0.26830468 | 0.032 | 0.125 | 1 | 23 |
| khdc4 | 0.35802 | -0.17582472 | 0.032 | 0.125 | 1 | 23 |
| klf12a | 0.35802 | -0.13776524 | 0.032 | 0.125 | 1 | 23 |
| klf13 | 0.35802 | -0.2149598 | 0.032 | 0.125 | 1 | 23 |
| kmt5b | 0.35802 | -0.17590226 | 0.032 | 0.125 | 1 | 23 |
| kpna2 | 0.35802 | -0.20383246 | 0.032 | 0.125 | 1 | 23 |
| larsb | 0.35802 | -0.15304905 | 0.032 | 0.125 | 1 | 23 |
| lats2 | 0.35802 | -0.10698321 | 0.032 | 0.125 | 1 | 23 |
| lin37 | 0.35802 | -0.2777865 | 0.032 | 0.125 | 1 | 23 |
| lonp1 | 0.35802 | -0.15266521 | 0.032 | 0.125 | 1 | 23 |
| magi3a | 0.35802 | -0.21782861 | 0.032 | 0.125 | 1 | 23 |
| mapk14a | 0.35802 | -0.11972303 | 0.032 | 0.125 | 1 | 23 |
| med7 | 0.35802 | -0.14609851 | 0.032 | 0.125 | 1 | 23 |
| mettl6 | 0.35802 | -0.1591987 | 0.032 | 0.125 | 1 | 23 |
| mfsd10 | 0.35802 | -0.19031552 | 0.032 | 0.125 | 1 | 23 |
| mier3a | 0.35802 | -0.26919043 | 0.032 | 0.125 | 1 | 23 |
| mrpl40 | 0.35802 | -0.1395057 | 0.032 | 0.125 | 1 | 23 |
| msl1b | 0.35802 | -0.16930473 | 0.032 | 0.125 | 1 | 23 |
| mtmr4 | 0.35802 | -0.16286097 | 0.032 | 0.125 | 1 | 23 |
| nfil3-2 | 0.35802 | -0.13138655 | 0.032 | 0.125 | 1 | 23 |
| nfxl1 | 0.35802 | -0.26805894 | 0.032 | 0.125 | 1 | 23 |
| nova2 | 0.35802 | -0.12051743 | 0.032 | 0.125 | 1 | 23 |
| nsun4 | 0.35802 | -0.14151071 | 0.032 | 0.125 | 1 | 23 |
| oma1 | 0.35802 | -0.27380955 | 0.032 | 0.125 | 1 | 23 |
| pdcd7 | 0.35802 | -0.15916524 | 0.032 | 0.125 | 1 | 23 |
| pdlim4 | 0.35802 | -0.11237108 | 0.032 | 0.125 | 1 | 23 |
| pgap1 | 0.35802 | -0.14150715 | 0.032 | 0.125 | 1 | 23 |
| phf8 | 0.35802 | -0.1722088 | 0.032 | 0.125 | 1 | 23 |
| pigh | 0.35802 | -0.10444541 | 0.032 | 0.125 | 1 | 23 |
| polr1c | 0.35802 | -0.16652309 | 0.032 | 0.125 | 1 | 23 |
| ppil4 | 0.35802 | -0.12626811 | 0.032 | 0.125 | 1 | 23 |
| ppm1k | 0.35802 | -0.26822685 | 0.032 | 0.125 | 1 | 23 |
| ppp1r9ba | 0.35802 | -0.13976956 | 0.032 | 0.125 | 1 | 23 |
| ppp2cab | 0.35802 | -0.22662219 | 0.032 | 0.125 | 1 | 23 |
| ppt2 | 0.35802 | -0.13939851 | 0.032 | 0.125 | 1 | 23 |
| prim1 | 0.35802 | -0.26881329 | 0.032 | 0.125 | 1 | 23 |
| proser1 | 0.35802 | -0.10550552 | 0.032 | 0.125 | 1 | 23 |
| prpf4 | 0.35802 | -0.17708565 | 0.032 | 0.125 | 1 | 23 |
| ptdss2 | 0.35802 | -0.12943255 | 0.032 | 0.125 | 1 | 23 |
| qtrt1 | 0.35802 | -0.14771012 | 0.032 | 0.125 | 1 | 23 |
| r3hdm1 | 0.35802 | -0.1215671 | 0.032 | 0.125 | 1 | 23 |

| | | | | | | |
|---|---|---|---|---|---|---|
| rab42b | 0.35802 | -0.15886723 | 0.032 | 0.125 | 1 | 23 |
| rab8a | 0.35802 | -0.17245983 | 0.032 | 0.125 | 1 | 23 |
| rad50 | 0.35802 | -0.26049364 | 0.032 | 0.125 | 1 | 23 |
| rell1 | 0.35802 | -0.17180241 | 0.032 | 0.125 | 1 | 23 |
| rgs12b | 0.35802 | -0.25337624 | 0.032 | 0.125 | 1 | 23 |
| rictorb | 0.35802 | -0.27706195 | 0.032 | 0.125 | 1 | 23 |
| rnf38 | 0.35802 | -0.10627835 | 0.032 | 0.125 | 1 | 23 |
| rnpep | 0.35802 | -0.16836981 | 0.032 | 0.125 | 1 | 23 |
| robo3 | 0.35802 | -0.165121 | 0.032 | 0.125 | 1 | 23 |
| rock2a | 0.35802 | -0.13047776 | 0.032 | 0.125 | 1 | 23 |
| rps6ka3a | 0.35802 | -0.1642402 | 0.032 | 0.125 | 1 | 23 |
| rps6ka3b | 0.35802 | -0.24413697 | 0.032 | 0.125 | 1 | 23 |
| rptor | 0.35802 | -0.26967597 | 0.032 | 0.125 | 1 | 23 |
| rrp12 | 0.35802 | -0.16045951 | 0.032 | 0.125 | 1 | 23 |
| saraf | 0.35802 | -0.13039486 | 0.032 | 0.125 | 1 | 23 |
| sart3 | 0.35802 | -0.24935194 | 0.032 | 0.125 | 1 | 23 |
| sc:d0202 | 0.35802 | -0.221903 | 0.032 | 0.125 | 1 | 23 |
| sema7a | 0.35802 | -0.10207922 | 0.032 | 0.125 | 1 | 23 |
| setd1ba | 0.35802 | -0.1788054 | 0.032 | 0.125 | 1 | 23 |
| sfxn3 | 0.35802 | -0.26408936 | 0.032 | 0.125 | 1 | 23 |
| sgsm3 | 0.35802 | -0.10090741 | 0.032 | 0.125 | 1 | 23 |
| si:ch211-162i8.5 | 0.35802 | -0.14957151 | 0.032 | 0.125 | 1 | 23 |
| si:ch211-194c3.5 | 0.35802 | -0.26755026 | 0.032 | 0.125 | 1 | 23 |
| si:ch211-219a4.3 | 0.35802 | -0.11460311 | 0.032 | 0.125 | 1 | 23 |
| si:ch211-225h24.2 | 0.35802 | -0.15770262 | 0.032 | 0.125 | 1 | 23 |
| si:ch211-230g15.5 | 0.35802 | -0.1142529 | 0.032 | 0.125 | 1 | 23 |
| si:ch211-232b12.5 | 0.35802 | -0.14530915 | 0.032 | 0.125 | 1 | 23 |
| si:ch211-261d7.3 | 0.35802 | -0.1446648 | 0.032 | 0.125 | 1 | 23 |
| si:ch211-71m22.1 | 0.35802 | -0.18080703 | 0.032 | 0.125 | 1 | 23 |
| si:ch73-103b9.2 | 0.35802 | -0.13847173 | 0.032 | 0.125 | 1 | 23 |
| si:ch73-223f5.1 | 0.35802 | -0.21783781 | 0.032 | 0.125 | 1 | 23 |
| si:ch73-358j7.2 | 0.35802 | -0.22213874 | 0.032 | 0.125 | 1 | 23 |
| si:dkey-188i13.8 | 0.35802 | -0.26021002 | 0.032 | 0.125 | 1 | 23 |
| si:dkey-253d23.11 | 0.35802 | -0.14094273 | 0.032 | 0.125 | 1 | 23 |
| si:dkey-82i20.2 | 0.35802 | -0.13610543 | 0.032 | 0.125 | 1 | 23 |
| si:dkeyp-104f11.6 | 0.35802 | -0.11805553 | 0.032 | 0.125 | 1 | 23 |
| si:zfos-464b6.2 | 0.35802 | -0.16299555 | 0.032 | 0.125 | 1 | 23 |
| slc10a7 | 0.35802 | -0.15200463 | 0.032 | 0.125 | 1 | 23 |
| slc38a3b | 0.35802 | -0.10508213 | 0.032 | 0.125 | 1 | 23 |
| slc4a1ap | 0.35802 | -0.14435184 | 0.032 | 0.125 | 1 | 23 |
| sp3a | 0.35802 | -0.12756749 | 0.032 | 0.125 | 1 | 23 |
| spata2 | 0.35802 | -0.27358597 | 0.032 | 0.125 | 1 | 23 |
| srpk1b | 0.35802 | -0.17114701 | 0.032 | 0.125 | 1 | 23 |
| sssca1 | 0.35802 | -0.22411582 | 0.032 | 0.125 | 1 | 23 |
| st7l | 0.35802 | -0.12121014 | 0.032 | 0.125 | 1 | 23 |
| stam2 | 0.35802 | -0.10423449 | 0.032 | 0.125 | 1 | 23 |
| sufu | 0.35802 | -0.15428595 | 0.032 | 0.125 | 1 | 23 |
| sugp1 | 0.35802 | -0.25389615 | 0.032 | 0.125 | 1 | 23 |

| | | | | | | |
|---|---|---|---|---|---|---|
| tfe3b | 0.35802 | -0.13053633 | 0.032 | 0.125 | 1 | 23 |
| thoc6 | 0.35802 | -0.27529081 | 0.032 | 0.125 | 1 | 23 |
| tmem131l | 0.35802 | -0.12981643 | 0.032 | 0.125 | 1 | 23 |
| tmem145 | 0.35802 | -0.22092787 | 0.032 | 0.125 | 1 | 23 |
| tmem220 | 0.35802 | -0.12389631 | 0.032 | 0.125 | 1 | 23 |
| tmem39b | 0.35802 | -0.17987529 | 0.032 | 0.125 | 1 | 23 |
| tmem70 | 0.35802 | -0.17370261 | 0.032 | 0.125 | 1 | 23 |
| tmprss4a | 0.35802 | -0.15126221 | 0.032 | 0.125 | 1 | 23 |
| tollip | 0.35802 | -0.25972025 | 0.032 | 0.125 | 1 | 23 |
| tom1 | 0.35802 | -0.13034991 | 0.032 | 0.125 | 1 | 23 |
| tp53i11b | 0.35802 | -0.26685833 | 0.032 | 0.125 | 1 | 23 |
| trim8a | 0.35802 | -0.14702275 | 0.032 | 0.125 | 1 | 23 |
| trrap | 0.35802 | -0.24566597 | 0.032 | 0.125 | 1 | 23 |
| ttc8 | 0.35802 | -0.26057785 | 0.032 | 0.125 | 1 | 23 |
| tuba1b | 0.35802 | -0.10560853 | 0.032 | 0.125 | 1 | 23 |
| ube2c | 0.35802 | -0.30356013 | 0.032 | 0.125 | 1 | 23 |
| usp18 | 0.35802 | -0.19931549 | 0.032 | 0.125 | 1 | 23 |
| usp19 | 0.35802 | -0.12782939 | 0.032 | 0.125 | 1 | 23 |
| utp6 | 0.35802 | -0.25103587 | 0.032 | 0.125 | 1 | 23 |
| vhl | 0.35802 | -0.24905748 | 0.032 | 0.125 | 1 | 23 |
| vps35l | 0.35802 | -0.1217225 | 0.032 | 0.125 | 1 | 23 |
| wars2 | 0.35802 | -0.26728199 | 0.032 | 0.125 | 1 | 23 |
| wdr66 | 0.35802 | -0.15738094 | 0.032 | 0.125 | 1 | 23 |
| xrcc5 | 0.35802 | -0.2738459 | 0.032 | 0.125 | 1 | 23 |
| zbtb2b | 0.35802 | -0.17312907 | 0.032 | 0.125 | 1 | 23 |
| zbtb43 | 0.35802 | -0.12656516 | 0.032 | 0.125 | 1 | 23 |
| zbtb44 | 0.35802 | -0.12631234 | 0.032 | 0.125 | 1 | 23 |
| zdhhc23b | 0.35802 | -0.10914363 | 0.032 | 0.125 | 1 | 23 |
| zdhhc24 | 0.35802 | -0.17087024 | 0.032 | 0.125 | 1 | 23 |
| zgc:101566 | 0.35802 | -0.18007547 | 0.032 | 0.125 | 1 | 23 |
| zgc:112185 | 0.35802 | -0.1470138 | 0.032 | 0.125 | 1 | 23 |
| zgc:112496 | 0.35802 | -0.14383638 | 0.032 | 0.125 | 1 | 23 |
| zgc:113518 | 0.35802 | -0.13133192 | 0.032 | 0.125 | 1 | 23 |
| zgc:162472 | 0.35802 | -0.10840594 | 0.032 | 0.125 | 1 | 23 |
| zgc:172106 | 0.35802 | -0.26815194 | 0.032 | 0.125 | 1 | 23 |
| zgc:193742 | 0.35802 | -0.26942493 | 0.032 | 0.125 | 1 | 23 |
| zmp:0000000521 | 0.35802 | -0.13211385 | 0.032 | 0.125 | 1 | 23 |
| znf1041 | 0.35802 | -0.14602499 | 0.032 | 0.125 | 1 | 23 |
| znf16l | 0.35802 | -0.22365373 | 0.032 | 0.125 | 1 | 23 |
| znf574 | 0.35802 | -0.27998666 | 0.032 | 0.125 | 1 | 23 |
| znf576.2 | 0.35802 | -0.17272139 | 0.032 | 0.125 | 1 | 23 |
| znf687a | 0.35802 | -0.15937945 | 0.032 | 0.125 | 1 | 23 |
| dydc2 | 0.35802 | -0.15488328 | 0.032 | 0.125 | 1 | 23 |
| hephl1a | 0.35802 | -0.1613409 | 0.032 | 0.125 | 1 | 23 |
| klhl7 | 0.35802 | -0.11525636 | 0.032 | 0.125 | 1 | 23 |
| psmd10 | 0.35802 | -0.11559291 | 0.032 | 0.125 | 1 | 23 |
| si:ch211-234c11.3 | 0.35802 | -0.25608906 | 0.032 | 0.125 | 1 | 23 |
| si:dkey-165a24.10 | 0.35802 | -0.15481168 | 0.032 | 0.125 | 1 | 23 |

| | | | | | | |
|---|---|---|---|---|---|---|
| si:dkey-287g12.6 | 0.35802 | -0.12153617 | 0.032 | 0.125 | 1 | 23 |
| GK3P | 0.35802 | -0.1409211 | 0.032 | 0.125 | 1 | 23 |
| ankra2 | 0.35802 | -0.11696933 | 0.032 | 0.125 | 1 | 23 |
| cd9a | 0.35802 | -0.17623466 | 0.032 | 0.125 | 1 | 23 |
| cept1a | 0.35802 | -0.12938641 | 0.032 | 0.125 | 1 | 23 |
| cyfip2 | 0.35802 | -0.15182831 | 0.032 | 0.125 | 1 | 23 |
| daxx | 0.35802 | -0.1263537 | 0.032 | 0.125 | 1 | 23 |
| foxl2a | 0.35802 | -0.13707233 | 0.032 | 0.125 | 1 | 23 |
| ift88 | 0.35802 | -0.12738394 | 0.032 | 0.125 | 1 | 23 |
| kyat3 | 0.35802 | -0.12558984 | 0.032 | 0.125 | 1 | 23 |
| marveld3 | 0.35802 | -0.12773912 | 0.032 | 0.125 | 1 | 23 |
| mfge8a | 0.35802 | -0.13091701 | 0.032 | 0.125 | 1 | 23 |
| ndufaf2 | 0.35802 | -0.13974455 | 0.032 | 0.125 | 1 | 23 |
| nhsl1b | 0.35802 | -0.13543773 | 0.032 | 0.125 | 1 | 23 |
| nup93 | 0.35802 | -0.11950799 | 0.032 | 0.125 | 1 | 23 |
| oxsm | 0.35802 | -0.14128623 | 0.032 | 0.125 | 1 | 23 |
| si:ch211-218g23.6 | 0.35802 | -0.12340622 | 0.032 | 0.125 | 1 | 23 |
| si:ch73-92e7.4 | 0.35802 | -0.11869016 | 0.032 | 0.125 | 1 | 23 |
| trim35-33 | 0.35802 | -0.14106834 | 0.032 | 0.125 | 1 | 23 |
| trmt10b | 0.35802 | -0.14542611 | 0.032 | 0.125 | 1 | 23 |
| tspan5a | 0.35802 | -0.14765768 | 0.032 | 0.125 | 1 | 23 |
| zgc:77058 | 0.35802 | -0.13451747 | 0.032 | 0.125 | 1 | 23 |
| AL732488.1 | 0.35802 | -0.19633452 | 0.032 | 0.125 | 1 | 23 |
| BX000701.1 | 0.35802 | -0.31138918 | 0.032 | 0.125 | 1 | 23 |
| CABZ01117348.1 | 0.35802 | -0.13448658 | 0.032 | 0.125 | 1 | 23 |
| CR381686.3 | 0.35802 | -0.31088396 | 0.032 | 0.125 | 1 | 23 |
| CR388132.2 | 0.35802 | -0.13165202 | 0.032 | 0.125 | 1 | 23 |
| CR786571.2 | 0.35802 | -0.20998695 | 0.032 | 0.125 | 1 | 23 |
| CR847994.1 | 0.35802 | -0.13486832 | 0.032 | 0.125 | 1 | 23 |
| CT990618.1 | 0.35802 | -0.12629453 | 0.032 | 0.125 | 1 | 23 |
| CU633745.1 | 0.35802 | -0.13449877 | 0.032 | 0.125 | 1 | 23 |
| CU929418.1 | 0.35802 | -0.24905448 | 0.032 | 0.125 | 1 | 23 |
| FP016056.1 | 0.35802 | -0.29406929 | 0.032 | 0.125 | 1 | 23 |
| LOC100332329 | 0.35802 | -0.21906524 | 0.032 | 0.125 | 1 | 23 |
| LOC100333241 | 0.35802 | -0.22026402 | 0.032 | 0.125 | 1 | 23 |
| LOC101883319 | 0.35802 | -0.19869724 | 0.032 | 0.125 | 1 | 23 |
| LOC101883815 | 0.35802 | -0.11802829 | 0.032 | 0.125 | 1 | 23 |
| LOC103909205 | 0.35802 | -0.31163387 | 0.032 | 0.125 | 1 | 23 |
| LOC103909928 | 0.35802 | -0.12939237 | 0.032 | 0.125 | 1 | 23 |
| LOC103911996 | 0.35802 | -0.13257003 | 0.032 | 0.125 | 1 | 23 |
| LOC566587 | 0.35802 | -0.22195476 | 0.032 | 0.125 | 1 | 23 |
| RSBN1 | 0.35802 | -0.19875332 | 0.032 | 0.125 | 1 | 23 |
| SBNO2 | 0.35802 | -0.13585515 | 0.032 | 0.125 | 1 | 23 |
| SPAG9 | 0.35802 | -0.2163558 | 0.032 | 0.125 | 1 | 23 |
| XLOC-006330 | 0.35802 | -0.14373898 | 0.032 | 0.125 | 1 | 23 |
| XLOC-013964 | 0.35802 | -0.25536474 | 0.032 | 0.125 | 1 | 23 |
| adka | 0.35802 | -0.11580357 | 0.032 | 0.125 | 1 | 23 |
| adnpb | 0.35802 | -0.13826036 | 0.032 | 0.125 | 1 | 23 |

| | | | | | | |
|---|---|---|---|---|---|---|
| alx4a | 0.35802 | -0.13045299 | 0.032 | 0.125 | 1 | 23 |
| atg4c | 0.35802 | -0.12421018 | 0.032 | 0.125 | 1 | 23 |
| b4galt7 | 0.35802 | -0.12303514 | 0.032 | 0.125 | 1 | 23 |
| bach1b | 0.35802 | -0.13567271 | 0.032 | 0.125 | 1 | 23 |
| bcl9 | 0.35802 | -0.19916776 | 0.032 | 0.125 | 1 | 23 |
| bri3 | 0.35802 | -0.11643842 | 0.032 | 0.125 | 1 | 23 |
| cables2b | 0.35802 | -0.11067309 | 0.032 | 0.125 | 1 | 23 |
| camk2g2 | 0.35802 | -0.32679378 | 0.032 | 0.125 | 1 | 23 |
| cand1 | 0.35802 | -0.14395997 | 0.032 | 0.125 | 1 | 23 |
| cbln2b | 0.35802 | -0.26884145 | 0.032 | 0.125 | 1 | 23 |
| ccdc40 | 0.35802 | -0.21122339 | 0.032 | 0.125 | 1 | 23 |
| ccdc6a | 0.35802 | -0.12846763 | 0.032 | 0.125 | 1 | 23 |
| cdk8 | 0.35802 | -0.11081722 | 0.032 | 0.125 | 1 | 23 |
| cep290 | 0.35802 | -0.10562172 | 0.032 | 0.125 | 1 | 23 |
| chmp2ba | 0.35802 | -0.12183431 | 0.032 | 0.125 | 1 | 23 |
| chst14 | 0.35802 | -0.11717022 | 0.032 | 0.125 | 1 | 23 |
| commd6 | 0.35802 | -0.13147081 | 0.032 | 0.125 | 1 | 23 |
| crsp7 | 0.35802 | -0.30156072 | 0.032 | 0.125 | 1 | 23 |
| cstf3 | 0.35802 | -0.21274366 | 0.032 | 0.125 | 1 | 23 |
| cul4a | 0.35802 | -0.11774212 | 0.032 | 0.125 | 1 | 23 |
| cyhr1 | 0.35802 | -0.12522868 | 0.032 | 0.125 | 1 | 23 |
| cyp2aa7 | 0.35802 | -0.30827788 | 0.032 | 0.125 | 1 | 23 |
| cyyr1 | 0.35802 | -0.11849399 | 0.032 | 0.125 | 1 | 23 |
| dennd3a | 0.35802 | -0.21193953 | 0.032 | 0.125 | 1 | 23 |
| dhrs3b | 0.35802 | -0.21730451 | 0.032 | 0.125 | 1 | 23 |
| dhx37 | 0.35802 | -0.20081551 | 0.032 | 0.125 | 1 | 23 |
| dtx4b | 0.35802 | -0.14000578 | 0.032 | 0.125 | 1 | 23 |
| edem3 | 0.35802 | -0.19760813 | 0.032 | 0.125 | 1 | 23 |
| efs | 0.35802 | -0.13949618 | 0.032 | 0.125 | 1 | 23 |
| eloa | 0.35802 | -0.1195163 | 0.032 | 0.125 | 1 | 23 |
| emc9 | 0.35802 | -0.20459184 | 0.032 | 0.125 | 1 | 23 |
| ephb3a | 0.35802 | -0.32546379 | 0.032 | 0.125 | 1 | 23 |
| fbxo21 | 0.35802 | -0.21086195 | 0.032 | 0.125 | 1 | 23 |
| fbxw11a | 0.35802 | -0.31200465 | 0.032 | 0.125 | 1 | 23 |
| fgfr1op | 0.35802 | -0.29844755 | 0.032 | 0.125 | 1 | 23 |
| fis1 | 0.35802 | -0.1287928 | 0.032 | 0.125 | 1 | 23 |
| fxr1 | 0.35802 | -0.19681416 | 0.032 | 0.125 | 1 | 23 |
| fzd1 | 0.35802 | -0.13390389 | 0.032 | 0.125 | 1 | 23 |
| gnai2b | 0.35802 | -0.12360573 | 0.032 | 0.125 | 1 | 23 |
| gnaq | 0.35802 | -0.10333063 | 0.032 | 0.125 | 1 | 23 |
| gne | 0.35802 | -0.21185706 | 0.032 | 0.125 | 1 | 23 |
| gpn2 | 0.35802 | -0.19444344 | 0.032 | 0.125 | 1 | 23 |
| hps3 | 0.35802 | -0.14721615 | 0.032 | 0.125 | 1 | 23 |
| hs2st1b | 0.35802 | -0.13879291 | 0.032 | 0.125 | 1 | 23 |
| hsf1 | 0.35802 | -0.37781198 | 0.032 | 0.125 | 1 | 23 |
| ikbkb | 0.35802 | -0.12372351 | 0.032 | 0.125 | 1 | 23 |
| irgq1 | 0.35802 | -0.28796612 | 0.032 | 0.125 | 1 | 23 |
| itga1 | 0.35802 | -0.13890954 | 0.032 | 0.125 | 1 | 23 |

| | | | | | | |
|---|---|---|---|---|---|---|
| jmjd7 | 0.35802 | -0.1239305 | 0.032 | 0.125 | 1 | 23 |
| kat7b | 0.35802 | -0.12880495 | 0.032 | 0.125 | 1 | 23 |
| kbtbd2 | 0.35802 | -0.13206408 | 0.032 | 0.125 | 1 | 23 |
| kdelc1 | 0.35802 | -0.13070844 | 0.032 | 0.125 | 1 | 23 |
| kdm6al | 0.35802 | -0.30998779 | 0.032 | 0.125 | 1 | 23 |
| larp4aa | 0.35802 | -0.22464763 | 0.032 | 0.125 | 1 | 23 |
| maf1 | 0.35802 | -0.22010064 | 0.032 | 0.125 | 1 | 23 |
| mark2a | 0.35802 | -0.21002161 | 0.032 | 0.125 | 1 | 23 |
| mbtps2 | 0.35802 | -0.21891697 | 0.032 | 0.125 | 1 | 23 |
| mgat2 | 0.35802 | -0.1396031 | 0.032 | 0.125 | 1 | 23 |
| mhc1uka | 0.35802 | -0.12034589 | 0.032 | 0.125 | 1 | 23 |
| mlxip | 0.35802 | -0.12787162 | 0.032 | 0.125 | 1 | 23 |
| mns1 | 0.35802 | -0.28452442 | 0.032 | 0.125 | 1 | 23 |
| mon2 | 0.35802 | -0.19682448 | 0.032 | 0.125 | 1 | 23 |
| mrm2 | 0.35802 | -0.21344531 | 0.032 | 0.125 | 1 | 23 |
| ms4a17a.5 | 0.35802 | -0.20034024 | 0.032 | 0.125 | 1 | 23 |
| mtmr6 | 0.35802 | -0.19987069 | 0.032 | 0.125 | 1 | 23 |
| nanp | 0.35802 | -0.11036286 | 0.032 | 0.125 | 1 | 23 |
| ncdn | 0.35802 | -0.10562242 | 0.032 | 0.125 | 1 | 23 |
| ndel1a | 0.35802 | -0.10828086 | 0.032 | 0.125 | 1 | 23 |
| ndst2a | 0.35802 | -0.12373814 | 0.032 | 0.125 | 1 | 23 |
| ndufaf6 | 0.35802 | -0.1383443 | 0.032 | 0.125 | 1 | 23 |
| nomo | 0.35802 | -0.20128716 | 0.032 | 0.125 | 1 | 23 |
| npc1 | 0.35802 | -0.20983588 | 0.032 | 0.125 | 1 | 23 |
| ogfrl2 | 0.35802 | -0.31530545 | 0.032 | 0.125 | 1 | 23 |
| orc5 | 0.35802 | -0.18158382 | 0.032 | 0.125 | 1 | 23 |
| pbx2 | 0.35802 | -0.30037487 | 0.032 | 0.125 | 1 | 23 |
| pdcd2l | 0.35802 | -0.14351588 | 0.032 | 0.125 | 1 | 23 |
| pecr | 0.35802 | -0.19982637 | 0.032 | 0.125 | 1 | 23 |
| pes | 0.35802 | -0.30993502 | 0.032 | 0.125 | 1 | 23 |
| pja2 | 0.35802 | -0.12795076 | 0.032 | 0.125 | 1 | 23 |
| pla2g15 | 0.35802 | -0.13392146 | 0.032 | 0.125 | 1 | 23 |
| plcb3 | 0.35802 | -0.20308243 | 0.032 | 0.125 | 1 | 23 |
| plcd4b | 0.35802 | -0.21280559 | 0.032 | 0.125 | 1 | 23 |
| plekhj1 | 0.35802 | -0.28608698 | 0.032 | 0.125 | 1 | 23 |
| polr1a | 0.35802 | -0.11236286 | 0.032 | 0.125 | 1 | 23 |
| ppp5c | 0.35802 | -0.11141092 | 0.032 | 0.125 | 1 | 23 |
| prrx1a | 0.35802 | -0.12976469 | 0.032 | 0.125 | 1 | 23 |
| psen1 | 0.35802 | -0.12371345 | 0.032 | 0.125 | 1 | 23 |
| ptpn13 | 0.35802 | -0.26643984 | 0.032 | 0.125 | 1 | 23 |
| pttg1 | 0.35802 | -0.31788774 | 0.032 | 0.125 | 1 | 23 |
| pum3 | 0.35802 | -0.1689819 | 0.032 | 0.125 | 1 | 23 |
| pwp2h | 0.35802 | -0.11540933 | 0.032 | 0.125 | 1 | 23 |
| qki2 | 0.35802 | -0.20095025 | 0.032 | 0.125 | 1 | 23 |
| rad51c | 0.35802 | -0.21478967 | 0.032 | 0.125 | 1 | 23 |
| rad54l2 | 0.35802 | -0.21219628 | 0.032 | 0.125 | 1 | 23 |
| rbm15 | 0.35802 | -0.3280201 | 0.032 | 0.125 | 1 | 23 |
| rbm27 | 0.35802 | -0.32197755 | 0.032 | 0.125 | 1 | 23 |

| | | | | | | |
|---|---|---|---|---|---|---|
| rfc1 | 0.35802 | -0.14791291 | 0.032 | 0.125 | 1 | 23 |
| rftn2 | 0.35802 | -0.29861815 | 0.032 | 0.125 | 1 | 23 |
| rint1 | 0.35802 | -0.14062901 | 0.032 | 0.125 | 1 | 23 |
| rnf169 | 0.35802 | -0.21823532 | 0.032 | 0.125 | 1 | 23 |
| rpp30 | 0.35802 | -0.11844198 | 0.032 | 0.125 | 1 | 23 |
| s1pr5a | 0.35802 | -0.26273119 | 0.032 | 0.125 | 1 | 23 |
| samd11 | 0.35802 | -0.29789448 | 0.032 | 0.125 | 1 | 23 |
| samd13 | 0.35802 | -0.21319121 | 0.032 | 0.125 | 1 | 23 |
| sdhaf4 | 0.35802 | -0.1005706 | 0.032 | 0.125 | 1 | 23 |
| si:ch1073-228b5.2 | 0.35802 | -0.10353139 | 0.032 | 0.125 | 1 | 23 |
| si:ch211-238n5.4 | 0.35802 | -0.19856572 | 0.032 | 0.125 | 1 | 23 |
| si:ch211-274p24.2 | 0.35802 | -0.21160747 | 0.032 | 0.125 | 1 | 23 |
| si:ch73-109d9.3 | 0.35802 | -0.14134492 | 0.032 | 0.125 | 1 | 23 |
| si:dkey-10c21.1 | 0.35802 | -0.13594501 | 0.032 | 0.125 | 1 | 23 |
| si:dkey-29h14.10 | 0.35802 | -0.12587687 | 0.032 | 0.125 | 1 | 23 |
| si:dkey-85k15.6 | 0.35802 | -0.1320201 | 0.032 | 0.125 | 1 | 23 |
| si:rp71-1g18.1 | 0.35802 | -0.12795487 | 0.032 | 0.125 | 1 | 23 |
| si:rp71-36a1.3 | 0.35802 | -0.14090873 | 0.032 | 0.125 | 1 | 23 |
| si:rp71-79p20.2 | 0.35802 | -0.13781377 | 0.032 | 0.125 | 1 | 23 |
| slc38a2 | 0.35802 | -0.12717991 | 0.032 | 0.125 | 1 | 23 |
| slc38a6 | 0.35802 | -0.13627259 | 0.032 | 0.125 | 1 | 23 |
| slit2 | 0.35802 | -0.12197318 | 0.032 | 0.125 | 1 | 23 |
| smchd1 | 0.35802 | -0.11860024 | 0.032 | 0.125 | 1 | 23 |
| snapc3 | 0.35802 | -0.13388428 | 0.032 | 0.125 | 1 | 23 |
| snrka | 0.35802 | -0.21520507 | 0.032 | 0.125 | 1 | 23 |
| snx9b | 0.35802 | -0.11070499 | 0.032 | 0.125 | 1 | 23 |
| sobpa | 0.35802 | -0.19840016 | 0.032 | 0.125 | 1 | 23 |
| stim1a | 0.35802 | -0.14203766 | 0.032 | 0.125 | 1 | 23 |
| tada3l | 0.35802 | -0.19771185 | 0.032 | 0.125 | 1 | 23 |
| tbcelb | 0.35802 | -0.21349696 | 0.032 | 0.125 | 1 | 23 |
| tbx18 | 0.35802 | -0.31737556 | 0.032 | 0.125 | 1 | 23 |
| terf1 | 0.35802 | -0.21593801 | 0.032 | 0.125 | 1 | 23 |
| tmem45a | 0.35802 | -0.12762555 | 0.032 | 0.125 | 1 | 23 |
| tom1l2b | 0.35802 | -0.24331817 | 0.032 | 0.125 | 1 | 23 |
| tpmt.1 | 0.35802 | -0.10629377 | 0.032 | 0.125 | 1 | 23 |
| trmt11 | 0.35802 | -0.21878933 | 0.032 | 0.125 | 1 | 23 |
| uap1 | 0.35802 | -0.21570641 | 0.032 | 0.125 | 1 | 23 |
| ube3a | 0.35802 | -0.12025458 | 0.032 | 0.125 | 1 | 23 |
| ubxn4 | 0.35802 | -0.12890664 | 0.032 | 0.125 | 1 | 23 |
| ufd1l | 0.35802 | -0.32168058 | 0.032 | 0.125 | 1 | 23 |
| wdr6 | 0.35802 | -0.31598832 | 0.032 | 0.125 | 1 | 23 |
| yod1 | 0.35802 | -0.3151094 | 0.032 | 0.125 | 1 | 23 |
| zbtb7c | 0.35802 | -0.21016191 | 0.032 | 0.125 | 1 | 23 |
| zc3h11a | 0.35802 | -0.20162864 | 0.032 | 0.125 | 1 | 23 |
| zgc:153409 | 0.35802 | -0.19906421 | 0.032 | 0.125 | 1 | 23 |
| zgc:162898 | 0.35802 | -0.1106846 | 0.032 | 0.125 | 1 | 23 |
| zgc:194281 | 0.35802 | -0.21379373 | 0.032 | 0.125 | 1 | 23 |
| zgc:76872 | 0.35802 | -0.2159398 | 0.032 | 0.125 | 1 | 23 |

| | | | | | | |
|---|---|---|---|---|---|---|
| zgc:77287 | 0.35802 | -0.13903935 | 0.032 | 0.125 | 1 | 23 |
| zgc:77439 | 0.35802 | -0.24728095 | 0.032 | 0.125 | 1 | 23 |
| zmp:0000000711 | 0.35802 | -0.12325784 | 0.032 | 0.125 | 1 | 23 |
| znf1081 | 0.35802 | -0.34271839 | 0.032 | 0.125 | 1 | 23 |
| znf1173 | 0.35802 | -0.13987134 | 0.032 | 0.125 | 1 | 23 |
| znf507 | 0.35802 | -0.13549177 | 0.032 | 0.125 | 1 | 23 |
| etf1b | 0.35807 | -0.22046396 | 0.548 | 0.625 | 1 | 23 |
| srsf5a | 0.35825 | -0.31493987 | 0.581 | 0.75 | 1 | 23 |
| frem2b | 0.35832 | -0.13973075 | 0.419 | 0.625 | 1 | 23 |
| csnk1da | 0.35857 | -0.17884826 | 0.097 | 0.125 | 1 | 23 |
| picalmb | 0.35932 | -0.35078186 | 0.065 | 0.25 | 1 | 23 |
| smndc1 | 0.35934 | 0.148333418 | 0.258 | 0.25 | 1 | 23 |
| gli2a | 0.35953 | -0.27401593 | 0.065 | 0.25 | 1 | 23 |
| RANBP2 | 0.35973 | -0.24058492 | 0.065 | 0.25 | 1 | 23 |
| ubap2b | 0.36057 | -0.25307018 | 0.161 | 0.25 | 1 | 23 |
| ap3s1.1 | 0.361 | -0.22996819 | 0.065 | 0.25 | 1 | 23 |
| gpd1c | 0.36202 | -0.2384443 | 0.065 | 0.25 | 1 | 23 |
| si:dkey-102m7.3 | 0.36253 | -0.3055171 | 0.129 | 0.25 | 1 | 23 |
| hiat1b | 0.363 | -0.21781165 | 0.065 | 0.25 | 1 | 23 |
| ugt1b5 | 0.36307 | 0.253731522 | 0.484 | 0.25 | 1 | 23 |
| atp2a3 | 0.36318 | -0.35038065 | 0.097 | 0.25 | 1 | 23 |
| si:ch211-262h13.3 | 0.36321 | -0.23806711 | 0.065 | 0.25 | 1 | 23 |
| e2f5 | 0.36335 | 0.137215535 | 0.129 | 0.125 | 1 | 23 |
| cyba | 0.36405 | -0.15777642 | 0.097 | 0.125 | 1 | 23 |
| sat1a.2 | 0.36414 | -0.31634014 | 0.226 | 0.375 | 1 | 23 |
| si:ch211-284b7.3 | 0.36417 | 0.309115696 | 0.323 | 0.125 | 1 | 23 |
| ranbp9 | 0.36425 | -0.26993782 | 0.161 | 0.375 | 1 | 23 |
| eif5a2 | 0.36501 | 0.20532513 | 0.742 | 0.5 | 1 | 23 |
| ormdl1 | 0.36525 | -0.30655157 | 0.065 | 0.25 | 1 | 23 |
| srgap2 | 0.36526 | -0.22743588 | 0.065 | 0.25 | 1 | 23 |
| hipk3b | 0.36536 | -0.16916726 | 0.097 | 0.125 | 1 | 23 |
| hipk2 | 0.36549 | -0.3461087 | 0.065 | 0.25 | 1 | 23 |
| pdik1l | 0.36577 | -0.33877429 | 0.065 | 0.25 | 1 | 23 |
| polr2c | 0.36594 | -0.21217212 | 0.065 | 0.25 | 1 | 23 |
| flot1b | 0.3661 | 0.384621518 | 0.452 | 0.25 | 1 | 23 |
| qsox1 | 0.36641 | -0.21558521 | 0.065 | 0.25 | 1 | 23 |
| si:dkey-71b5.6 | 0.36643 | -0.21465179 | 0.065 | 0.25 | 1 | 23 |
| cnot9 | 0.3667 | -0.31553478 | 0.065 | 0.25 | 1 | 23 |
| glrx2 | 0.36673 | -0.22629234 | 0.065 | 0.25 | 1 | 23 |
| LO017872.1 | 0.36699 | -0.294247 | 0.065 | 0.25 | 1 | 23 |
| zgc:92818 | 0.3678 | -0.35072283 | 0.194 | 0.375 | 1 | 23 |
| LO018176.1 | 0.36799 | -0.31773763 | 0.065 | 0.25 | 1 | 23 |
| fxr2 | 0.36842 | -0.33406755 | 0.065 | 0.25 | 1 | 23 |
| ndufaf3 | 0.36938 | -0.22896897 | 0.065 | 0.25 | 1 | 23 |
| acox1 | 0.36951 | -0.31494901 | 0.065 | 0.25 | 1 | 23 |
| luc7l | 0.36974 | -0.32351368 | 0.129 | 0.25 | 1 | 23 |
| smarcad1a | 0.37102 | -0.32366712 | 0.065 | 0.25 | 1 | 23 |
| mical3b | 0.37102 | -0.2666076 | 0.065 | 0.25 | 1 | 23 |

| | | | | | | |
|---|---|---|---|---|---|---|
| ier2b | 0.37123 | -0.13283863 | 0.839 | 1 | 1 | 23 |
| ppp1r2 | 0.37136 | -0.21220529 | 0.065 | 0.25 | 1 | 23 |
| sirt2 | 0.37147 | -0.22507112 | 0.065 | 0.25 | 1 | 23 |
| anxa2a | 0.37151 | -0.43764532 | 0.387 | 0.625 | 1 | 23 |
| myl12.2 | 0.37176 | 0.298537664 | 0.419 | 0.25 | 1 | 23 |
| ganab | 0.37179 | -0.312413 | 0.065 | 0.25 | 1 | 23 |
| ralgapa1 | 0.37206 | -0.33650304 | 0.065 | 0.25 | 1 | 23 |
| ndfip2 | 0.3723 | -0.30645285 | 0.065 | 0.25 | 1 | 23 |
| kctd3 | 0.37232 | -0.27719087 | 0.065 | 0.25 | 1 | 23 |
| ches1 | 0.37247 | -0.27179235 | 0.065 | 0.25 | 1 | 23 |
| hsbp1l1 | 0.37249 | -0.33998754 | 0.065 | 0.25 | 1 | 23 |
| lrch4 | 0.37357 | -0.32115734 | 0.065 | 0.25 | 1 | 23 |
| cbx5 | 0.37362 | 0.1490015 | 0.484 | 0.5 | 1 | 23 |
| ptbp1b | 0.37407 | -0.31888777 | 0.226 | 0.375 | 1 | 23 |
| tbk1 | 0.37418 | -0.27973675 | 0.065 | 0.25 | 1 | 23 |
| atp6v1ba | 0.37471 | -0.30868526 | 0.065 | 0.25 | 1 | 23 |
| spcs1 | 0.37482 | -0.3460251 | 0.516 | 0.75 | 1 | 23 |
| med18 | 0.37491 | -0.33490306 | 0.097 | 0.25 | 1 | 23 |
| ess2 | 0.37526 | -0.26783772 | 0.065 | 0.25 | 1 | 23 |
| lsm6 | 0.3755 | 0.289169787 | 0.387 | 0.25 | 1 | 23 |
| zgc:123010 | 0.3756 | -0.29321363 | 0.065 | 0.25 | 1 | 23 |
| larp7 | 0.37565 | -0.16150973 | 0.097 | 0.125 | 1 | 23 |
| cebpa | 0.37574 | -0.30450811 | 0.065 | 0.25 | 1 | 23 |
| st5 | 0.37576 | 0.124066086 | 0.258 | 0.125 | 1 | 23 |
| LOC110439840 | 0.37614 | -0.22653213 | 0.065 | 0.25 | 1 | 23 |
| cnih1 | 0.37644 | -0.12874712 | 0.258 | 0.25 | 1 | 23 |
| copa | 0.37644 | -0.28937709 | 0.129 | 0.25 | 1 | 23 |
| zgc:77086 | 0.37644 | 0.284109633 | 0.516 | 0.375 | 1 | 23 |
| tpp1 | 0.37724 | -0.23545447 | 0.065 | 0.25 | 1 | 23 |
| mrps21 | 0.37741 | -0.35153291 | 0.161 | 0.375 | 1 | 23 |
| vdrb | 0.37776 | -0.42268377 | 0.161 | 0.375 | 1 | 23 |
| fam122b | 0.37857 | -0.32158082 | 0.065 | 0.25 | 1 | 23 |
| arhgdia | 0.37867 | 0.277633104 | 0.452 | 0.375 | 1 | 23 |
| nxt2 | 0.37924 | 0.30094589 | 0.323 | 0.125 | 1 | 23 |
| zgc:113184 | 0.37985 | -0.22158307 | 0.161 | 0.375 | 1 | 23 |
| map3k21 | 0.37993 | -0.28988735 | 0.065 | 0.25 | 1 | 23 |
| dpp7 | 0.38002 | -0.27349462 | 0.065 | 0.25 | 1 | 23 |
| dad1 | 0.38022 | -0.2247115 | 0.71 | 0.75 | 1 | 23 |
| nit2 | 0.38025 | -0.21097875 | 0.065 | 0.25 | 1 | 23 |
| ece2b | 0.38034 | -0.10183462 | 0.129 | 0.125 | 1 | 23 |
| sfrp2 | 0.38037 | -0.22739933 | 0.065 | 0.25 | 1 | 23 |
| dhrs4 | 0.38046 | -0.28461692 | 0.065 | 0.25 | 1 | 23 |
| ncoa6 | 0.38117 | -0.28055301 | 0.065 | 0.25 | 1 | 23 |
| nabp1a | 0.38121 | -0.3275079 | 0.226 | 0.375 | 1 | 23 |
| tjp2b | 0.38132 | -0.34501614 | 0.355 | 0.625 | 1 | 23 |
| herc1 | 0.38133 | -0.28229289 | 0.065 | 0.25 | 1 | 23 |
| tpm1 | 0.38139 | -0.33287671 | 0.194 | 0.375 | 1 | 23 |
| mmp14a | 0.38191 | -0.33632763 | 0.065 | 0.25 | 1 | 23 |

| | | | | | | |
|---|---|---|---|---|---|---|
| rhebl1 | 0.38237 | -0.1053705 | 0.129 | 0.125 | 1 | 23 |
| efemp2a | 0.38245 | -0.30566342 | 0.065 | 0.25 | 1 | 23 |
| si:ch1073-127d16.1 | 0.38249 | -0.30391199 | 0.065 | 0.25 | 1 | 23 |
| PAXBP1 | 0.38271 | -0.28123664 | 0.065 | 0.25 | 1 | 23 |
| rasa3 | 0.38276 | -0.233764 | 0.065 | 0.25 | 1 | 23 |
| asb12a | 0.38281 | -0.22301379 | 0.065 | 0.25 | 1 | 23 |
| ebag9 | 0.38285 | -0.369506 | 0.161 | 0.375 | 1 | 23 |
| si:ch73-127m5.2 | 0.38291 | -0.27576067 | 0.065 | 0.25 | 1 | 23 |
| si:ch73-111k22.2 | 0.38312 | -0.29785669 | 0.065 | 0.25 | 1 | 23 |
| fus | 0.38319 | -0.30483681 | 0.065 | 0.25 | 1 | 23 |
| ppiab | 0.38325 | -0.17152742 | 0.839 | 1 | 1 | 23 |
| rab32a | 0.38376 | -0.27683602 | 0.065 | 0.25 | 1 | 23 |
| csnk2a1 | 0.38379 | 0.192430025 | 0.452 | 0.25 | 1 | 23 |
| nrbf2b | 0.38384 | -0.26847857 | 0.065 | 0.25 | 1 | 23 |
| tnfrsf1a | 0.38402 | -0.31874053 | 0.065 | 0.25 | 1 | 23 |
| cnot3b | 0.38419 | -0.28668922 | 0.065 | 0.25 | 1 | 23 |
| trim25 | 0.38422 | -0.28228727 | 0.065 | 0.25 | 1 | 23 |
| tk2 | 0.3844 | -0.27101425 | 0.065 | 0.25 | 1 | 23 |
| pigp | 0.38455 | -0.28295257 | 0.065 | 0.25 | 1 | 23 |
| rcn2 | 0.38458 | -0.26082611 | 0.065 | 0.25 | 1 | 23 |
| si:ch211-276c2.4 | 0.38467 | -0.23133218 | 0.065 | 0.25 | 1 | 23 |
| mtch2 | 0.38483 | -0.30582046 | 0.226 | 0.375 | 1 | 23 |
| si:ch211-137i24.10 | 0.38495 | -0.3550013 | 0.065 | 0.25 | 1 | 23 |
| slc25a43 | 0.38501 | -0.31067793 | 0.065 | 0.25 | 1 | 23 |
| mapk7 | 0.38502 | -0.28455356 | 0.065 | 0.25 | 1 | 23 |
| dpp9 | 0.38506 | -0.25720974 | 0.065 | 0.25 | 1 | 23 |
| rnf26 | 0.38506 | -0.3028486 | 0.065 | 0.25 | 1 | 23 |
| MDFI | 0.38525 | 0.344953559 | 0.419 | 0.25 | 1 | 23 |
| nap1l1 | 0.38534 | -0.32211597 | 0.484 | 0.75 | 1 | 23 |
| si:ch211-194b1.1 | 0.38559 | -0.38487382 | 0.161 | 0.375 | 1 | 23 |
| hnrnpr | 0.38565 | -0.26118678 | 0.258 | 0.375 | 1 | 23 |
| lsm3 | 0.38588 | -0.25971406 | 0.29 | 0.375 | 1 | 23 |
| hnrpkl | 0.3863 | -0.33993442 | 0.387 | 0.5 | 1 | 23 |
| ssr3 | 0.38638 | 0.162890306 | 0.323 | 0.375 | 1 | 23 |
| eif4g2b | 0.38647 | 0.338066266 | 0.323 | 0.125 | 1 | 23 |
| zbtb10 | 0.38836 | -0.34071693 | 0.097 | 0.25 | 1 | 23 |
| snrpb2 | 0.39147 | -0.20406341 | 0.226 | 0.25 | 1 | 23 |
| eif5 | 0.39252 | -0.30904111 | 0.548 | 0.75 | 1 | 23 |
| tbl1xr1a | 0.39354 | -0.4262186 | 0.161 | 0.375 | 1 | 23 |
| zgc:92606 | 0.39357 | 0.317829831 | 0.258 | 0.125 | 1 | 23 |
| ppp1cb | 0.39477 | -0.25839317 | 0.161 | 0.25 | 1 | 23 |
| eif3c | 0.39502 | -0.28313623 | 0.516 | 0.625 | 1 | 23 |
| irf8 | 0.39509 | -0.44335932 | 0.161 | 0.375 | 1 | 23 |
| etfa | 0.39517 | -0.14722794 | 0.226 | 0.25 | 1 | 23 |
| thrap3b | 0.3984 | 0.255020022 | 0.258 | 0.125 | 1 | 23 |
| si:dkey-180p18.9 | 0.39857 | 0.281642659 | 0.29 | 0.125 | 1 | 23 |
| col6a3 | 0.39935 | 0.319337606 | 0.29 | 0.125 | 1 | 23 |
| sec63 | 0.39945 | -0.11395335 | 0.258 | 0.25 | 1 | 23 |

| | | | | | | |
|---|---|---|---|---|---|---|
| ddit3 | 0.39956 | 0.34832448 | 0.323 | 0.125 | 1 | 23 |
| ccdc167 | 0.39979 | -0.16142551 | 0.097 | 0.125 | 1 | 23 |
| ube2al | 0.39999 | -0.31249563 | 0.194 | 0.375 | 1 | 23 |
| eif4g1a | 0.40039 | 0.173035688 | 0.29 | 0.125 | 1 | 23 |
| nme3 | 0.40073 | 0.343504353 | 0.581 | 0.375 | 1 | 23 |
| snrpg | 0.40165 | 0.319017912 | 0.742 | 0.5 | 1 | 23 |
| phf20b | 0.40171 | -0.10959432 | 0.129 | 0.125 | 1 | 23 |
| pde4ba | 0.40183 | -0.39938804 | 0.419 | 0.625 | 1 | 23 |
| tjp3 | 0.40203 | -0.14516919 | 0.097 | 0.25 | 1 | 23 |
| rnf7 | 0.40207 | 0.31969999 | 0.355 | 0.125 | 1 | 23 |
| nkd1 | 0.40223 | 0.204981445 | 0.29 | 0.125 | 1 | 23 |
| mrpl12 | 0.40267 | -0.14759008 | 0.226 | 0.25 | 1 | 23 |
| nop16 | 0.4039 | -0.31339556 | 0.226 | 0.375 | 1 | 23 |
| XLOC-040582 | 0.40474 | -0.13689316 | 0.097 | 0.25 | 1 | 23 |
| snrnp27 | 0.40481 | 0.111558675 | 0.355 | 0.375 | 1 | 23 |
| yipf6 | 0.4055 | -0.15319107 | 0.097 | 0.25 | 1 | 23 |
| wu:fu71h07 | 0.40555 | -0.16869069 | 0.097 | 0.125 | 1 | 23 |
| cebpd | 0.40625 | -0.13626938 | 0.645 | 0.625 | 1 | 23 |
| hmbox1a | 0.40652 | -0.3563201 | 0.161 | 0.375 | 1 | 23 |
| tomm7 | 0.40683 | 0.355276913 | 0.355 | 0.125 | 1 | 23 |
| si:ch211-108d22.2 | 0.40698 | -0.17491342 | 0.097 | 0.125 | 1 | 23 |
| postnb | 0.40712 | 0.535113717 | 0.355 | 0.125 | 1 | 23 |
| rab2a | 0.40716 | 0.329152282 | 0.452 | 0.25 | 1 | 23 |
| smc1al | 0.40737 | 0.199151789 | 0.323 | 0.25 | 1 | 23 |
| pde7a | 0.40834 | -0.35314325 | 0.258 | 0.5 | 1 | 23 |
| mrps18a | 0.4085 | -0.33420923 | 0.194 | 0.375 | 1 | 23 |
| myh10 | 0.40851 | 0.369069558 | 0.355 | 0.125 | 1 | 23 |
| lrrc51 | 0.4095 | -0.20565365 | 0.194 | 0.25 | 1 | 23 |
| ap2s1 | 0.40952 | -0.16234253 | 0.226 | 0.25 | 1 | 23 |
| CR318588.1 | 0.4098 | -0.28964351 | 0.806 | 0.875 | 1 | 23 |
| snrpe | 0.41032 | -0.20394954 | 0.645 | 0.875 | 1 | 23 |
| smarcc1a | 0.41058 | -0.30210807 | 0.129 | 0.25 | 1 | 23 |
| si:ch1073-291c23.2 | 0.41069 | 0.329882458 | 0.323 | 0.125 | 1 | 23 |
| mrpl18 | 0.41147 | -0.10533507 | 0.258 | 0.25 | 1 | 23 |
| serinc1 | 0.41165 | -0.38446447 | 0.258 | 0.5 | 1 | 23 |
| tpd52l2a | 0.4121 | 0.129225758 | 0.226 | 0.25 | 1 | 23 |
| hist2h2l | 0.41247 | 0.324471368 | 0.645 | 0.5 | 1 | 23 |
| rarab | 0.4126 | -0.15477653 | 0.097 | 0.125 | 1 | 23 |
| psmd1 | 0.41289 | 0.35073195 | 0.355 | 0.125 | 1 | 23 |
| ints1 | 0.41303 | -0.16464633 | 0.129 | 0.125 | 1 | 23 |
| ywhaqb | 0.41371 | -0.25113157 | 0.645 | 0.875 | 1 | 23 |
| zgc:55558 | 0.41473 | -0.3383715 | 0.097 | 0.25 | 1 | 23 |
| tuba8l | 0.41497 | -0.40636555 | 0.387 | 0.625 | 1 | 23 |
| pard3bb | 0.41518 | 0.102543633 | 0.387 | 0.25 | 1 | 23 |
| inhbaa | 0.41562 | 0.422723693 | 0.355 | 0.125 | 1 | 23 |
| socs3b | 0.41565 | 0.384764226 | 0.355 | 0.125 | 1 | 23 |
| zbtb37 | 0.41605 | -0.16022954 | 0.097 | 0.125 | 1 | 23 |
| grcc10 | 0.41622 | 0.192642521 | 0.323 | 0.25 | 1 | 23 |

| | | | | | | |
|---|---|---|---|---|---|---|
| wu:fc27b11 | 0.41694 | 0.308697795 | 0.419 | 0.25 | 1 | 23 |
| hmgb1a | 0.41748 | 0.182016749 | 0.839 | 0.625 | 1 | 23 |
| atp5meb | 0.41748 | -0.26996935 | 0.677 | 0.75 | 1 | 23 |
| gtf2a2 | 0.41808 | -0.3259943 | 0.194 | 0.375 | 1 | 23 |
| cfap45 | 0.41809 | -0.42558837 | 0.161 | 0.375 | 1 | 23 |
| arl16 | 0.41828 | -0.1572041 | 0.097 | 0.125 | 1 | 23 |
| gar1 | 0.41878 | -0.28122308 | 0.161 | 0.375 | 1 | 23 |
| CZQB01141835.1 | 0.41913 | -0.28339259 | 0.226 | 0.375 | 1 | 23 |
| rgs5b | 0.41975 | 0.259775946 | 0.29 | 0.25 | 1 | 23 |
| rbx1 | 0.42019 | 0.285653574 | 0.645 | 0.5 | 1 | 23 |
| psmd8 | 0.42055 | 0.36478562 | 0.484 | 0.25 | 1 | 23 |
| h2afx | 0.4209 | -0.11544873 | 0.129 | 0.125 | 1 | 23 |
| LOC103910015 | 0.42151 | -0.12507166 | 0.129 | 0.25 | 1 | 23 |
| sdha | 0.42194 | 0.114291144 | 0.258 | 0.125 | 1 | 23 |
| atp6v0ca | 0.42281 | -0.2376798 | 0.29 | 0.375 | 1 | 23 |
| mrpl34 | 0.42368 | 0.321315379 | 0.323 | 0.125 | 1 | 23 |
| zgc:175088 | 0.42424 | 0.215731144 | 0.194 | 0.125 | 1 | 23 |
| fam168a | 0.42453 | -0.23476211 | 0.516 | 0.625 | 1 | 23 |
| wdr3 | 0.42471 | -0.10944492 | 0.129 | 0.125 | 1 | 23 |
| dis3l2 | 0.42509 | -0.29628005 | 0.194 | 0.375 | 1 | 23 |
| chka | 0.4251 | -0.32513869 | 0.194 | 0.375 | 1 | 23 |
| hspa8 | 0.42549 | -0.17606055 | 0.903 | 1 | 1 | 23 |
| ncam1a | 0.4261 | -0.34389865 | 0.258 | 0.5 | 1 | 23 |
| nedd8 | 0.42707 | -0.16222473 | 0.355 | 0.375 | 1 | 23 |
| brd3b | 0.42712 | -0.21547923 | 0.323 | 0.375 | 1 | 23 |
| ptprga | 0.42718 | -0.36460495 | 0.258 | 0.5 | 1 | 23 |
| rpl4 | 0.42754 | 0.10888903 | 1 | 1 | 1 | 23 |
| phf11 | 0.42814 | -0.36022364 | 0.097 | 0.25 | 1 | 23 |
| ifnphi1 | 0.42887 | -0.17133256 | 0.097 | 0.125 | 1 | 23 |
| ftr82 | 0.4296 | 0.247139652 | 0.613 | 0.375 | 1 | 23 |
| rbp5 | 0.42962 | 0.161764687 | 0.29 | 0.125 | 1 | 23 |
| hspe1 | 0.42998 | -0.33213169 | 0.516 | 0.75 | 1 | 23 |
| sf3a2 | 0.4302 | -0.13224923 | 0.387 | 0.375 | 1 | 23 |
| tcp1 | 0.43088 | 0.27190351 | 0.806 | 0.625 | 1 | 23 |
| calm2a | 0.43098 | 0.321162869 | 0.258 | 0.125 | 1 | 23 |
| lgals3a | 0.43244 | -0.11180818 | 0.258 | 0.25 | 1 | 23 |
| nutf2 | 0.43327 | -0.3225509 | 0.226 | 0.375 | 1 | 23 |
| alad | 0.43343 | -0.25030125 | 0.129 | 0.25 | 1 | 23 |
| rac1a | 0.43403 | -0.33894272 | 0.29 | 0.5 | 1 | 23 |
| mrpl57 | 0.4344 | -0.1178096 | 0.323 | 0.5 | 1 | 23 |
| atg5 | 0.43473 | -0.29687142 | 0.097 | 0.25 | 1 | 23 |
| nr3c1 | 0.43494 | -0.32449292 | 0.194 | 0.375 | 1 | 23 |
| mafba | 0.43508 | 0.314208501 | 0.323 | 0.125 | 1 | 23 |
| stard13b | 0.43514 | -0.29721256 | 0.097 | 0.25 | 1 | 23 |
| BOLA2 | 0.4353 | 0.245095786 | 0.258 | 0.125 | 1 | 23 |
| cirbpa | 0.43542 | -0.1673404 | 0.968 | 1 | 1 | 23 |
| ccdc187 | 0.43616 | -0.31618299 | 0.194 | 0.375 | 1 | 23 |
| si:dkey-42i9.4 | 0.43728 | 0.360954512 | 0.613 | 0.375 | 1 | 23 |

| | | | | | | |
|---|---|---|---|---|---|---|
| vps26c | 0.43734 | -0.26509993 | 0.129 | 0.25 | 1 | 23 |
| eif3i | 0.43765 | 0.172739319 | 0.71 | 0.5 | 1 | 23 |
| arl4d | 0.43809 | 0.34197792 | 0.419 | 0.25 | 1 | 23 |
| sgta | 0.43834 | 0.291436903 | 0.323 | 0.125 | 1 | 23 |
| rhogd | 0.43879 | -0.26481758 | 0.226 | 0.375 | 1 | 23 |
| nf2b | 0.43941 | -0.1571542 | 0.097 | 0.25 | 1 | 23 |
| mxra8a | 0.4395 | 0.259688597 | 0.484 | 0.25 | 1 | 23 |
| suds3 | 0.44033 | -0.14476087 | 0.097 | 0.25 | 1 | 23 |
| eif6 | 0.44099 | 0.251699884 | 0.323 | 0.125 | 1 | 23 |
| sept10 | 0.44141 | 0.128358902 | 0.258 | 0.125 | 1 | 23 |
| zgc:194469 | 0.44227 | -0.30213124 | 0.097 | 0.25 | 1 | 23 |
| seh1l | 0.44245 | -0.34802572 | 0.29 | 0.5 | 1 | 23 |
| NC-002333.17 | 0.4425 | -0.24612772 | 0.129 | 0.25 | 1 | 23 |
| rad23b | 0.44354 | 0.338794191 | 0.613 | 0.375 | 1 | 23 |
| nrde2 | 0.444 | -0.15503625 | 0.097 | 0.125 | 1 | 23 |
| mcl1a | 0.44409 | -0.29651574 | 0.677 | 0.75 | 1 | 23 |
| actb2 | 0.44493 | 0.175736275 | 0.935 | 0.875 | 1 | 23 |
| XLOC-014336 | 0.44541 | 0.100945491 | 0.065 | 0 | 1 | 23 |
| coq9 | 0.44541 | 0.122262065 | 0.065 | 0 | 1 | 23 |
| cul1b | 0.44541 | 0.104674272 | 0.065 | 0 | 1 | 23 |
| dhrs7 | 0.44541 | 0.121528214 | 0.065 | 0 | 1 | 23 |
| pcyt2 | 0.44541 | 0.119844274 | 0.065 | 0 | 1 | 23 |
| sinhcaf | 0.44541 | 0.134135972 | 0.065 | 0 | 1 | 23 |
| tbc1d10aa | 0.44541 | 0.110797808 | 0.065 | 0 | 1 | 23 |
| tmem178b | 0.44541 | 0.105377189 | 0.065 | 0 | 1 | 23 |
| tnfaip6 | 0.44541 | 0.128894468 | 0.065 | 0 | 1 | 23 |
| ttc26 | 0.44541 | 0.123489315 | 0.065 | 0 | 1 | 23 |
| ttll7 | 0.44541 | 0.134135972 | 0.065 | 0 | 1 | 23 |
| tuft1a | 0.44541 | 0.108361576 | 0.065 | 0 | 1 | 23 |
| waplb | 0.44541 | 0.105747915 | 0.065 | 0 | 1 | 23 |
| zgc:66443 | 0.44541 | 0.131226346 | 0.065 | 0 | 1 | 23 |
| znf1152 | 0.44541 | 0.131226346 | 0.065 | 0 | 1 | 23 |
| BX322661.2 | 0.44541 | 0.13682642 | 0.065 | 0 | 1 | 23 |
| BX548046.2 | 0.44541 | 0.103696593 | 0.065 | 0 | 1 | 23 |
| BX901920.1 | 0.44541 | 0.110374389 | 0.065 | 0 | 1 | 23 |
| BX936308.1 | 0.44541 | 0.145946688 | 0.065 | 0 | 1 | 23 |
| CABZ01034698.1 | 0.44541 | 0.104813237 | 0.065 | 0 | 1 | 23 |
| CABZ01068248.1 | 0.44541 | 0.110727837 | 0.065 | 0 | 1 | 23 |
| CABZ01085139.1 | 0.44541 | 0.104813237 | 0.065 | 0 | 1 | 23 |
| CABZ01088271.1 | 0.44541 | 0.130796123 | 0.065 | 0 | 1 | 23 |
| CR318607.1 | 0.44541 | 0.118087066 | 0.065 | 0 | 1 | 23 |
| CR382300.1 | 0.44541 | 0.12774394 | 0.065 | 0 | 1 | 23 |
| CR383662.8 | 0.44541 | 0.160870212 | 0.065 | 0 | 1 | 23 |
| CU459094.2 | 0.44541 | 0.11805517 | 0.065 | 0 | 1 | 23 |
| CU929037.3 | 0.44541 | 0.129043175 | 0.065 | 0 | 1 | 23 |
| FQ311908.1 | 0.44541 | 0.152458605 | 0.065 | 0 | 1 | 23 |
| LOC100001832 | 0.44541 | 0.114553604 | 0.065 | 0 | 1 | 23 |
| LOC101884172 | 0.44541 | 0.11113496 | 0.065 | 0 | 1 | 23 |

| | | | | | | |
|---|---|---|---|---|---|---|
| LOC103909808 | 0.44541 | 0.113321868 | 0.065 | 0 | 1 | 23 |
| MAP3K11 | 0.44541 | 0.161174884 | 0.065 | 0 | 1 | 23 |
| RF01268 | 0.44541 | 0.119535021 | 0.065 | 0 | 1 | 23 |
| RNF14 (1 of many) | 0.44541 | 0.117827114 | 0.065 | 0 | 1 | 23 |
| SCYL2 | 0.44541 | 0.104173221 | 0.065 | 0 | 1 | 23 |
| SEMA4F | 0.44541 | 0.124967187 | 0.065 | 0 | 1 | 23 |
| TADA1 | 0.44541 | 0.107053061 | 0.065 | 0 | 1 | 23 |
| XLOC-005449 | 0.44541 | 0.16975157 | 0.065 | 0 | 1 | 23 |
| XLOC-024248 | 0.44541 | 0.108752916 | 0.065 | 0 | 1 | 23 |
| abi2a | 0.44541 | 0.105037492 | 0.065 | 0 | 1 | 23 |
| acbd3 | 0.44541 | 0.10844595 | 0.065 | 0 | 1 | 23 |
| ache | 0.44541 | 0.139810206 | 0.065 | 0 | 1 | 23 |
| acsm3 | 0.44541 | 0.115279795 | 0.065 | 0 | 1 | 23 |
| acvr1l | 0.44541 | 0.146238199 | 0.065 | 0 | 1 | 23 |
| adss | 0.44541 | 0.157338822 | 0.065 | 0 | 1 | 23 |
| agk | 0.44541 | 0.112074631 | 0.065 | 0 | 1 | 23 |
| ak9 | 0.44541 | 0.104017114 | 0.065 | 0 | 1 | 23 |
| amn1 | 0.44541 | 0.103556109 | 0.065 | 0 | 1 | 23 |
| anapc2 | 0.44541 | 0.100816295 | 0.065 | 0 | 1 | 23 |
| ankib1a | 0.44541 | 0.102827668 | 0.065 | 0 | 1 | 23 |
| ankrd52a | 0.44541 | 0.138393138 | 0.065 | 0 | 1 | 23 |
| ano5b | 0.44541 | 0.134977122 | 0.065 | 0 | 1 | 23 |
| arl6 | 0.44541 | 0.10889758 | 0.065 | 0 | 1 | 23 |
| arnt2 | 0.44541 | 0.141781228 | 0.065 | 0 | 1 | 23 |
| atg4b | 0.44541 | 0.116028068 | 0.065 | 0 | 1 | 23 |
| atp6v1f | 0.44541 | 0.104352656 | 0.065 | 0 | 1 | 23 |
| bco1 | 0.44541 | 0.112660577 | 0.065 | 0 | 1 | 23 |
| bmp1a | 0.44541 | 0.118828122 | 0.065 | 0 | 1 | 23 |
| btbd10a | 0.44541 | 0.110123199 | 0.065 | 0 | 1 | 23 |
| btbd6a | 0.44541 | 0.115279795 | 0.065 | 0 | 1 | 23 |
| c1galt1c1 | 0.44541 | 0.10801826 | 0.065 | 0 | 1 | 23 |
| cab39l1 | 0.44541 | 0.101023242 | 0.065 | 0 | 1 | 23 |
| cadm4 | 0.44541 | 0.13539459 | 0.065 | 0 | 1 | 23 |
| calcrlb | 0.44541 | 0.156437103 | 0.065 | 0 | 1 | 23 |
| cavin2a | 0.44541 | 0.100931774 | 0.065 | 0 | 1 | 23 |
| cbx2 | 0.44541 | 0.112588152 | 0.065 | 0 | 1 | 23 |
| ccdc191 | 0.44541 | 0.121992166 | 0.065 | 0 | 1 | 23 |
| ccdc30 | 0.44541 | 0.134274769 | 0.065 | 0 | 1 | 23 |
| ccdc80 | 0.44541 | 0.129644614 | 0.065 | 0 | 1 | 23 |
| ccnd2b | 0.44541 | 0.125065914 | 0.065 | 0 | 1 | 23 |
| ccndbp1 | 0.44541 | 0.109966436 | 0.065 | 0 | 1 | 23 |
| ccnt1 | 0.44541 | 0.138477462 | 0.065 | 0 | 1 | 23 |
| cd248b | 0.44541 | 0.116856341 | 0.065 | 0 | 1 | 23 |
| cdc42se1 | 0.44541 | 0.134172118 | 0.065 | 0 | 1 | 23 |
| cenpo | 0.44541 | 0.112097508 | 0.065 | 0 | 1 | 23 |
| cfap57 | 0.44541 | 0.112105857 | 0.065 | 0 | 1 | 23 |
| chchd3b | 0.44541 | 0.101528275 | 0.065 | 0 | 1 | 23 |
| chp2 | 0.44541 | 0.111347902 | 0.065 | 0 | 1 | 23 |

| | | | | | | |
|---|---|---|---|---|---|---|
| cldn7a | 0.44541 | 0.116131922 | 0.065 | 0 | 1 | 23 |
| cluha | 0.44541 | 0.149233329 | 0.065 | 0 | 1 | 23 |
| cnih4 | 0.44541 | 0.105681409 | 0.065 | 0 | 1 | 23 |
| col4a3bpa | 0.44541 | 0.112018592 | 0.065 | 0 | 1 | 23 |
| cpne4a | 0.44541 | 0.104580354 | 0.065 | 0 | 1 | 23 |
| cpt1ab | 0.44541 | 0.11031987 | 0.065 | 0 | 1 | 23 |
| cramp1 | 0.44541 | 0.143590883 | 0.065 | 0 | 1 | 23 |
| crema | 0.44541 | 0.120473354 | 0.065 | 0 | 1 | 23 |
| csgalnact1a | 0.44541 | 0.116631601 | 0.065 | 0 | 1 | 23 |
| csnk2a2b | 0.44541 | 0.109703128 | 0.065 | 0 | 1 | 23 |
| cspp1a | 0.44541 | 0.123417047 | 0.065 | 0 | 1 | 23 |
| ctsc | 0.44541 | 0.136415994 | 0.065 | 0 | 1 | 23 |
| cul3a | 0.44541 | 0.111998421 | 0.065 | 0 | 1 | 23 |
| cul5a | 0.44541 | 0.113453279 | 0.065 | 0 | 1 | 23 |
| cutc | 0.44541 | 0.110227486 | 0.065 | 0 | 1 | 23 |
| cyp2p7 | 0.44541 | 0.107250694 | 0.065 | 0 | 1 | 23 |
| dact1 | 0.44541 | 0.104181279 | 0.065 | 0 | 1 | 23 |
| dbr1 | 0.44541 | 0.107140114 | 0.065 | 0 | 1 | 23 |
| dclre1b | 0.44541 | 0.154373371 | 0.065 | 0 | 1 | 23 |
| dcun1d1 | 0.44541 | 0.10411196 | 0.065 | 0 | 1 | 23 |
| ddhd2 | 0.44541 | 0.104749626 | 0.065 | 0 | 1 | 23 |
| dennd6aa | 0.44541 | 0.145720353 | 0.065 | 0 | 1 | 23 |
| dexi | 0.44541 | 0.117466171 | 0.065 | 0 | 1 | 23 |
| dhx34 | 0.44541 | 0.103787391 | 0.065 | 0 | 1 | 23 |
| dhx57 | 0.44541 | 0.108357432 | 0.065 | 0 | 1 | 23 |
| dixdc1b | 0.44541 | 0.116533517 | 0.065 | 0 | 1 | 23 |
| dlg1 | 0.44541 | 0.114219523 | 0.065 | 0 | 1 | 23 |
| dlg1l | 0.44541 | 0.100718228 | 0.065 | 0 | 1 | 23 |
| dlgap4a | 0.44541 | 0.105883893 | 0.065 | 0 | 1 | 23 |
| dnajc10 | 0.44541 | 0.112093145 | 0.065 | 0 | 1 | 23 |
| dnali1 | 0.44541 | 0.110959729 | 0.065 | 0 | 1 | 23 |
| dolpp1 | 0.44541 | 0.11887806 | 0.065 | 0 | 1 | 23 |
| dpf3 | 0.44541 | 0.113985302 | 0.065 | 0 | 1 | 23 |
| dph2 | 0.44541 | 0.129187717 | 0.065 | 0 | 1 | 23 |
| dtx4a | 0.44541 | 0.116258318 | 0.065 | 0 | 1 | 23 |
| dusp22a | 0.44541 | 0.1107077 | 0.065 | 0 | 1 | 23 |
| dyrk1ab | 0.44541 | 0.107459467 | 0.065 | 0 | 1 | 23 |
| eaf1 | 0.44541 | 0.130822565 | 0.065 | 0 | 1 | 23 |
| ednraa | 0.44541 | 0.1202407 | 0.065 | 0 | 1 | 23 |
| efl1 | 0.44541 | 0.109600254 | 0.065 | 0 | 1 | 23 |
| efnb3b | 0.44541 | 0.108014314 | 0.065 | 0 | 1 | 23 |
| egfra | 0.44541 | 0.106244355 | 0.065 | 0 | 1 | 23 |
| eif4g3b | 0.44541 | 0.113613883 | 0.065 | 0 | 1 | 23 |
| elp5 | 0.44541 | 0.100689 | 0.065 | 0 | 1 | 23 |
| entpd6 | 0.44541 | 0.101538308 | 0.065 | 0 | 1 | 23 |
| exoc6b | 0.44541 | 0.115444186 | 0.065 | 0 | 1 | 23 |
| fam184b | 0.44541 | 0.116105246 | 0.065 | 0 | 1 | 23 |
| fam196ab | 0.44541 | 0.101622311 | 0.065 | 0 | 1 | 23 |

| | | | | | | |
|---|---|---|---|---|---|---|
| fap | 0.44541 | 0.109445325 | 0.065 | 0 | 1 | 23 |
| fbxo34 | 0.44541 | 0.101127352 | 0.065 | 0 | 1 | 23 |
| fbxw11b | 0.44541 | 0.100573498 | 0.065 | 0 | 1 | 23 |
| fbxw2 | 0.44541 | 0.147740067 | 0.065 | 0 | 1 | 23 |
| flot2b | 0.44541 | 0.117624286 | 0.065 | 0 | 1 | 23 |
| flr | 0.44541 | 0.113936982 | 0.065 | 0 | 1 | 23 |
| flrt3 | 0.44541 | 0.14983726 | 0.065 | 0 | 1 | 23 |
| fnbp1l | 0.44541 | 0.110166755 | 0.065 | 0 | 1 | 23 |
| foxn2b | 0.44541 | 0.104914895 | 0.065 | 0 | 1 | 23 |
| ftr98 | 0.44541 | 0.16383393 | 0.065 | 0 | 1 | 23 |
| fuca1.2 | 0.44541 | 0.127583235 | 0.065 | 0 | 1 | 23 |
| fut11 | 0.44541 | 0.12482589 | 0.065 | 0 | 1 | 23 |
| gab2 | 0.44541 | 0.129585726 | 0.065 | 0 | 1 | 23 |
| gadd45bb | 0.44541 | 0.127668751 | 0.065 | 0 | 1 | 23 |
| gart | 0.44541 | 0.112858829 | 0.065 | 0 | 1 | 23 |
| gatd3a | 0.44541 | 0.130145298 | 0.065 | 0 | 1 | 23 |
| gbe1b | 0.44541 | 0.107234377 | 0.065 | 0 | 1 | 23 |
| gcn1 | 0.44541 | 0.117575243 | 0.065 | 0 | 1 | 23 |
| gdpd4a | 0.44541 | 0.135732224 | 0.065 | 0 | 1 | 23 |
| ghra | 0.44541 | 0.124487168 | 0.065 | 0 | 1 | 23 |
| gng13b | 0.44541 | 0.106533716 | 0.065 | 0 | 1 | 23 |
| gpank1 | 0.44541 | 0.130873539 | 0.065 | 0 | 1 | 23 |
| gpr107 | 0.44541 | 0.113216121 | 0.065 | 0 | 1 | 23 |
| gpr27 | 0.44541 | 0.120561446 | 0.065 | 0 | 1 | 23 |
| gps1 | 0.44541 | 0.110035554 | 0.065 | 0 | 1 | 23 |
| gpx8 | 0.44541 | 0.108132752 | 0.065 | 0 | 1 | 23 |
| hapln1b | 0.44541 | 0.103772223 | 0.065 | 0 | 1 | 23 |
| hapln3 | 0.44541 | 0.132037516 | 0.065 | 0 | 1 | 23 |
| has2 | 0.44541 | 0.159967459 | 0.065 | 0 | 1 | 23 |
| hdac8 | 0.44541 | 0.130141668 | 0.065 | 0 | 1 | 23 |
| heatr1 | 0.44541 | 0.127737771 | 0.065 | 0 | 1 | 23 |
| heatr6 | 0.44541 | 0.145946688 | 0.065 | 0 | 1 | 23 |
| hs6st2 | 0.44541 | 0.124944348 | 0.065 | 0 | 1 | 23 |
| htra1a | 0.44541 | 0.105662504 | 0.065 | 0 | 1 | 23 |
| id3 | 0.44541 | 0.107926536 | 0.065 | 0 | 1 | 23 |
| igf2bp1 | 0.44541 | 0.107759049 | 0.065 | 0 | 1 | 23 |
| igfbp2b | 0.44541 | 0.108192721 | 0.065 | 0 | 1 | 23 |
| igflr1 | 0.44541 | 0.110858185 | 0.065 | 0 | 1 | 23 |
| ikzf5 | 0.44541 | 0.117114236 | 0.065 | 0 | 1 | 23 |
| im:7140055 | 0.44541 | 0.111739231 | 0.065 | 0 | 1 | 23 |
| inavab | 0.44541 | 0.101831747 | 0.065 | 0 | 1 | 23 |
| inhbb | 0.44541 | 0.111005495 | 0.065 | 0 | 1 | 23 |
| irge4 | 0.44541 | 0.111008053 | 0.065 | 0 | 1 | 23 |
| isoc1 | 0.44541 | 0.148367276 | 0.065 | 0 | 1 | 23 |
| itga2.3 | 0.44541 | 0.12047578 | 0.065 | 0 | 1 | 23 |
| itsn1 | 0.44541 | 0.124415585 | 0.065 | 0 | 1 | 23 |
| jmjd4 | 0.44541 | 0.135551193 | 0.065 | 0 | 1 | 23 |
| kank3 | 0.44541 | 0.111931437 | 0.065 | 0 | 1 | 23 |

| | | | | | | |
|---|---|---|---|---|---|---|
| kctd9b | 0.44541 | 0.100505814 | 0.065 | 0 | 1 | 23 |
| kif1bp | 0.44541 | 0.113101272 | 0.065 | 0 | 1 | 23 |
| kifc3 | 0.44541 | 0.100313309 | 0.065 | 0 | 1 | 23 |
| klf2a | 0.44541 | 0.109657483 | 0.065 | 0 | 1 | 23 |
| klhl13 | 0.44541 | 0.107895642 | 0.065 | 0 | 1 | 23 |
| lacc1 | 0.44541 | 0.115279795 | 0.065 | 0 | 1 | 23 |
| lactb | 0.44541 | 0.11977553 | 0.065 | 0 | 1 | 23 |
| larp4b | 0.44541 | 0.10621899 | 0.065 | 0 | 1 | 23 |
| lclat1 | 0.44541 | 0.103977505 | 0.065 | 0 | 1 | 23 |
| lgi1a | 0.44541 | 0.108243461 | 0.065 | 0 | 1 | 23 |
| loxl2b | 0.44541 | 0.152830704 | 0.065 | 0 | 1 | 23 |
| lrrc1 | 0.44541 | 0.122685996 | 0.065 | 0 | 1 | 23 |
| lrrc58b | 0.44541 | 0.104439719 | 0.065 | 0 | 1 | 23 |
| lrrtm1 | 0.44541 | 0.163376493 | 0.065 | 0 | 1 | 23 |
| lrsam1 | 0.44541 | 0.118221857 | 0.065 | 0 | 1 | 23 |
| lrtm2a | 0.44541 | 0.126354316 | 0.065 | 0 | 1 | 23 |
| lsg1 | 0.44541 | 0.104792753 | 0.065 | 0 | 1 | 23 |
| maco1a | 0.44541 | 0.121276171 | 0.065 | 0 | 1 | 23 |
| mapk1 | 0.44541 | 0.104278353 | 0.065 | 0 | 1 | 23 |
| mapre2 | 0.44541 | 0.129043175 | 0.065 | 0 | 1 | 23 |
| mcmbp | 0.44541 | 0.100474452 | 0.065 | 0 | 1 | 23 |
| mdm2 | 0.44541 | 0.168601248 | 0.065 | 0 | 1 | 23 |
| memo1 | 0.44541 | 0.103982996 | 0.065 | 0 | 1 | 23 |
| mettl17 | 0.44541 | 0.103126433 | 0.065 | 0 | 1 | 23 |
| mettl21a | 0.44541 | 0.110296606 | 0.065 | 0 | 1 | 23 |
| mettl24 | 0.44541 | 0.11068955 | 0.065 | 0 | 1 | 23 |
| mgme1 | 0.44541 | 0.133005621 | 0.065 | 0 | 1 | 23 |
| mief1 | 0.44541 | 0.103439623 | 0.065 | 0 | 1 | 23 |
| mmp11b | 0.44541 | 0.119226672 | 0.065 | 0 | 1 | 23 |
| mogat2 | 0.44541 | 0.135223013 | 0.065 | 0 | 1 | 23 |
| morn2 | 0.44541 | 0.110959729 | 0.065 | 0 | 1 | 23 |
| mrpl38 | 0.44541 | 0.11452774 | 0.065 | 0 | 1 | 23 |
| mrpl48 | 0.44541 | 0.122271092 | 0.065 | 0 | 1 | 23 |
| mt-atp8 | 0.44541 | 0.139006519 | 0.065 | 0 | 1 | 23 |
| mtfp1 | 0.44541 | 0.115739989 | 0.065 | 0 | 1 | 23 |
| mtfr1l | 0.44541 | 0.102209515 | 0.065 | 0 | 1 | 23 |
| mtif3 | 0.44541 | 0.11165756 | 0.065 | 0 | 1 | 23 |
| mtrf1l | 0.44541 | 0.100466132 | 0.065 | 0 | 1 | 23 |
| mxd4 | 0.44541 | 0.132498437 | 0.065 | 0 | 1 | 23 |
| myo9aa | 0.44541 | 0.141892353 | 0.065 | 0 | 1 | 23 |
| myrip | 0.44541 | 0.134275218 | 0.065 | 0 | 1 | 23 |
| nbas | 0.44541 | 0.122560836 | 0.065 | 0 | 1 | 23 |
| nefla | 0.44541 | 0.111683064 | 0.065 | 0 | 1 | 23 |
| nfe2l2a | 0.44541 | 0.121560302 | 0.065 | 0 | 1 | 23 |
| nfya | 0.44541 | 0.109526598 | 0.065 | 0 | 1 | 23 |
| nin | 0.44541 | 0.122017136 | 0.065 | 0 | 1 | 23 |
| nrip1a | 0.44541 | 0.1409635 | 0.065 | 0 | 1 | 23 |
| nudt8 | 0.44541 | 0.116767943 | 0.065 | 0 | 1 | 23 |

| | | | | | | |
|---|---|---|---|---|---|---|
| ogg1 | 0.44541 | 0.107362491 | 0.065 | 0 | 1 | 23 |
| orai1b | 0.44541 | 0.102649697 | 0.065 | 0 | 1 | 23 |
| osbpl2a | 0.44541 | 0.14977206 | 0.065 | 0 | 1 | 23 |
| p4ha2 | 0.44541 | 0.114229738 | 0.065 | 0 | 1 | 23 |
| pacsin2 | 0.44541 | 0.117637668 | 0.065 | 0 | 1 | 23 |
| padi2 | 0.44541 | 0.112587316 | 0.065 | 0 | 1 | 23 |
| pard6a | 0.44541 | 0.143641812 | 0.065 | 0 | 1 | 23 |
| pargl | 0.44541 | 0.105497945 | 0.065 | 0 | 1 | 23 |
| pcdh10a | 0.44541 | 0.150695194 | 0.065 | 0 | 1 | 23 |
| pcmt | 0.44541 | 0.111133812 | 0.065 | 0 | 1 | 23 |
| pdgfd | 0.44541 | 0.138158214 | 0.065 | 0 | 1 | 23 |
| pdgfrb | 0.44541 | 0.134639892 | 0.065 | 0 | 1 | 23 |
| pgghg | 0.44541 | 0.11001277 | 0.065 | 0 | 1 | 23 |
| phf23a | 0.44541 | 0.154898154 | 0.065 | 0 | 1 | 23 |
| phgdh | 0.44541 | 0.110884763 | 0.065 | 0 | 1 | 23 |
| pigw | 0.44541 | 0.126820212 | 0.065 | 0 | 1 | 23 |
| pip5k1aa | 0.44541 | 0.125815953 | 0.065 | 0 | 1 | 23 |
| plekha5 | 0.44541 | 0.116533971 | 0.065 | 0 | 1 | 23 |
| polr1d | 0.44541 | 0.14986143 | 0.065 | 0 | 1 | 23 |
| polr2j | 0.44541 | 0.12292663 | 0.065 | 0 | 1 | 23 |
| pora | 0.44541 | 0.116665255 | 0.065 | 0 | 1 | 23 |
| postna | 0.44541 | 0.116131922 | 0.065 | 0 | 1 | 23 |
| ppat | 0.44541 | 0.103251175 | 0.065 | 0 | 1 | 23 |
| ppp1r8b | 0.44541 | 0.112213208 | 0.065 | 0 | 1 | 23 |
| ppp2r3b | 0.44541 | 0.11113496 | 0.065 | 0 | 1 | 23 |
| proza | 0.44541 | 0.104390234 | 0.065 | 0 | 1 | 23 |
| pskh1 | 0.44541 | 0.108601829 | 0.065 | 0 | 1 | 23 |
| ptpn23a | 0.44541 | 0.108667732 | 0.065 | 0 | 1 | 23 |
| ptpn2b | 0.44541 | 0.122515978 | 0.065 | 0 | 1 | 23 |
| quo | 0.44541 | 0.153083172 | 0.065 | 0 | 1 | 23 |
| rab11bb | 0.44541 | 0.100313309 | 0.065 | 0 | 1 | 23 |
| rab24 | 0.44541 | 0.126743406 | 0.065 | 0 | 1 | 23 |
| raph1b | 0.44541 | 0.101490209 | 0.065 | 0 | 1 | 23 |
| rasgrp3 | 0.44541 | 0.11693928 | 0.065 | 0 | 1 | 23 |
| rb1 | 0.44541 | 0.106762829 | 0.065 | 0 | 1 | 23 |
| rfxank | 0.44541 | 0.11518178 | 0.065 | 0 | 1 | 23 |
| rgs12a | 0.44541 | 0.129116662 | 0.065 | 0 | 1 | 23 |
| rgs3a | 0.44541 | 0.108773709 | 0.065 | 0 | 1 | 23 |
| rhpn2 | 0.44541 | 0.100151218 | 0.065 | 0 | 1 | 23 |
| ribc2 | 0.44541 | 0.142472998 | 0.065 | 0 | 1 | 23 |
| rmnd1 | 0.44541 | 0.108518379 | 0.065 | 0 | 1 | 23 |
| rnf144b | 0.44541 | 0.116182148 | 0.065 | 0 | 1 | 23 |
| rtn4rl2a | 0.44541 | 0.112105857 | 0.065 | 0 | 1 | 23 |
| s100v1 | 0.44541 | 0.158719698 | 0.065 | 0 | 1 | 23 |
| sc5d | 0.44541 | 0.109475069 | 0.065 | 0 | 1 | 23 |
| sgk3 | 0.44541 | 0.119549514 | 0.065 | 0 | 1 | 23 |
| sh2d5 | 0.44541 | 0.115193815 | 0.065 | 0 | 1 | 23 |
| si:ch1073-358c10.1 | 0.44541 | 0.10871721 | 0.065 | 0 | 1 | 23 |

| | | | | | | |
|---|---|---|---|---|---|---|
| si:ch211-154o6.2 | 0.44541 | 0.107678179 | 0.065 | 0 | 1 | 23 |
| si:ch211-166e11.5 | 0.44541 | 0.146340071 | 0.065 | 0 | 1 | 23 |
| si:ch211-178n15.1 | 0.44541 | 0.105943021 | 0.065 | 0 | 1 | 23 |
| si:ch211-195b15.8 | 0.44541 | 0.10254116 | 0.065 | 0 | 1 | 23 |
| si:ch211-195e19.1 | 0.44541 | 0.103554343 | 0.065 | 0 | 1 | 23 |
| si:ch211-198c19.3 | 0.44541 | 0.11257235 | 0.065 | 0 | 1 | 23 |
| si:ch211-220f16.2 | 0.44541 | 0.126820212 | 0.065 | 0 | 1 | 23 |
| si:ch211-226h8.6 | 0.44541 | 0.100835186 | 0.065 | 0 | 1 | 23 |
| si:ch211-266o15.1 | 0.44541 | 0.123098082 | 0.065 | 0 | 1 | 23 |
| si:ch211-76m11.8 | 0.44541 | 0.112660577 | 0.065 | 0 | 1 | 23 |
| si:dkey-106g10.7 | 0.44541 | 0.108197768 | 0.065 | 0 | 1 | 23 |
| si:dkey-192l18.9 | 0.44541 | 0.130257279 | 0.065 | 0 | 1 | 23 |
| si:dkey-206d17.12 | 0.44541 | 0.101688931 | 0.065 | 0 | 1 | 23 |
| si:dkey-239n17.4 | 0.44541 | 0.115078368 | 0.065 | 0 | 1 | 23 |
| si:dkey-23a13.5 | 0.44541 | 0.12448193 | 0.065 | 0 | 1 | 23 |
| si:dkey-256h2.1 | 0.44541 | 0.107353813 | 0.065 | 0 | 1 | 23 |
| si:dkey-29d8.3 | 0.44541 | 0.110949557 | 0.065 | 0 | 1 | 23 |
| si:dkey-29p10.4 | 0.44541 | 0.135998277 | 0.065 | 0 | 1 | 23 |
| si:dkey-30g5.1 | 0.44541 | 0.15775141 | 0.065 | 0 | 1 | 23 |
| si:dkey-51a16.9 | 0.44541 | 0.108029044 | 0.065 | 0 | 1 | 23 |
| si:dkey-56e3.3 | 0.44541 | 0.110646577 | 0.065 | 0 | 1 | 23 |
| si:dkey-93h22.8 | 0.44541 | 0.101689769 | 0.065 | 0 | 1 | 23 |
| si:dkeyp-104h9.5 | 0.44541 | 0.111919221 | 0.065 | 0 | 1 | 23 |
| si:dkeyp-69c1.7 | 0.44541 | 0.145588997 | 0.065 | 0 | 1 | 23 |
| si:dkeyp-84f3.5 | 0.44541 | 0.106759232 | 0.065 | 0 | 1 | 23 |
| si:rp71-19m20.1 | 0.44541 | 0.106743304 | 0.065 | 0 | 1 | 23 |
| slc22a31 | 0.44541 | 0.112240297 | 0.065 | 0 | 1 | 23 |
| slc35a1 | 0.44541 | 0.114481979 | 0.065 | 0 | 1 | 23 |
| slc35e1 | 0.44541 | 0.142108529 | 0.065 | 0 | 1 | 23 |
| slc36a4 | 0.44541 | 0.115279795 | 0.065 | 0 | 1 | 23 |
| slc39a13 | 0.44541 | 0.108029382 | 0.065 | 0 | 1 | 23 |
| slc45a4 | 0.44541 | 0.104192437 | 0.065 | 0 | 1 | 23 |
| slc48a1b | 0.44541 | 0.108916419 | 0.065 | 0 | 1 | 23 |
| slc7a10a | 0.44541 | 0.104507583 | 0.065 | 0 | 1 | 23 |
| smg7 | 0.44541 | 0.101123221 | 0.065 | 0 | 1 | 23 |
| smpd4 | 0.44541 | 0.115450632 | 0.065 | 0 | 1 | 23 |
| snai1a | 0.44541 | 0.139566461 | 0.065 | 0 | 1 | 23 |
| snai1b | 0.44541 | 0.143844008 | 0.065 | 0 | 1 | 23 |
| snai2 | 0.44541 | 0.112527204 | 0.065 | 0 | 1 | 23 |
| snap23.2 | 0.44541 | 0.141578638 | 0.065 | 0 | 1 | 23 |
| snapc1b | 0.44541 | 0.107271708 | 0.065 | 0 | 1 | 23 |
| snapin | 0.44541 | 0.10875603 | 0.065 | 0 | 1 | 23 |
| socs6a | 0.44541 | 0.109618531 | 0.065 | 0 | 1 | 23 |
| soga3a | 0.44541 | 0.108029044 | 0.065 | 0 | 1 | 23 |
| sox4b | 0.44541 | 0.148555494 | 0.065 | 0 | 1 | 23 |
| spag7 | 0.44541 | 0.109560032 | 0.065 | 0 | 1 | 23 |
| spred2a | 0.44541 | 0.109217316 | 0.065 | 0 | 1 | 23 |
| sprtn | 0.44541 | 0.101327535 | 0.065 | 0 | 1 | 23 |

| | | | | | | |
|---|---|---|---|---|---|---|
| spsb4a | 0.44541 | 0.112866621 | 0.065 | 0 | 1 | 23 |
| srebf1 | 0.44541 | 0.114602266 | 0.065 | 0 | 1 | 23 |
| ssrp1b | 0.44541 | 0.109349072 | 0.065 | 0 | 1 | 23 |
| stag2a | 0.44541 | 0.112746708 | 0.065 | 0 | 1 | 23 |
| stub1 | 0.44541 | 0.100697937 | 0.065 | 0 | 1 | 23 |
| susd6 | 0.44541 | 0.112796812 | 0.065 | 0 | 1 | 23 |
| sv2 | 0.44541 | 0.125072451 | 0.065 | 0 | 1 | 23 |
| syt11a | 0.44541 | 0.134258288 | 0.065 | 0 | 1 | 23 |
| sytl5 | 0.44541 | 0.112639111 | 0.065 | 0 | 1 | 23 |
| tada2a | 0.44541 | 0.111854434 | 0.065 | 0 | 1 | 23 |
| taf1b | 0.44541 | 0.104813237 | 0.065 | 0 | 1 | 23 |
| tango6 | 0.44541 | 0.127478927 | 0.065 | 0 | 1 | 23 |
| tbcc | 0.44541 | 0.144142007 | 0.065 | 0 | 1 | 23 |
| tctex1d1 | 0.44541 | 0.140645631 | 0.065 | 0 | 1 | 23 |
| tctex1d2 | 0.44541 | 0.132644423 | 0.065 | 0 | 1 | 23 |
| tex10 | 0.44541 | 0.118624903 | 0.065 | 0 | 1 | 23 |
| thap1 | 0.44541 | 0.125731058 | 0.065 | 0 | 1 | 23 |
| thumpd1 | 0.44541 | 0.108570192 | 0.065 | 0 | 1 | 23 |
| ticam1 | 0.44541 | 0.115279795 | 0.065 | 0 | 1 | 23 |
| tle3a | 0.44541 | 0.107093723 | 0.065 | 0 | 1 | 23 |
| tmem160 | 0.44541 | 0.115730597 | 0.065 | 0 | 1 | 23 |
| tmem170a | 0.44541 | 0.111299884 | 0.065 | 0 | 1 | 23 |
| tmem218 | 0.44541 | 0.170544947 | 0.065 | 0 | 1 | 23 |
| tmem62 | 0.44541 | 0.110959729 | 0.065 | 0 | 1 | 23 |
| tmem9b | 0.44541 | 0.144456886 | 0.065 | 0 | 1 | 23 |
| tnfaip2a | 0.44541 | 0.100226432 | 0.065 | 0 | 1 | 23 |
| tnip2 | 0.44541 | 0.163149559 | 0.065 | 0 | 1 | 23 |
| tnpo2 | 0.44541 | 0.158202446 | 0.065 | 0 | 1 | 23 |
| tnpo3 | 0.44541 | 0.102569725 | 0.065 | 0 | 1 | 23 |
| tpbgb | 0.44541 | 0.136754297 | 0.065 | 0 | 1 | 23 |
| trim47 | 0.44541 | 0.10564224 | 0.065 | 0 | 1 | 23 |
| trioa | 0.44541 | 0.102288805 | 0.065 | 0 | 1 | 23 |
| trmt10c | 0.44541 | 0.108200109 | 0.065 | 0 | 1 | 23 |
| trmt1l | 0.44541 | 0.112810428 | 0.065 | 0 | 1 | 23 |
| trmt6 | 0.44541 | 0.111830489 | 0.065 | 0 | 1 | 23 |
| trove2 | 0.44541 | 0.110233594 | 0.065 | 0 | 1 | 23 |
| trpt1 | 0.44541 | 0.106395023 | 0.065 | 0 | 1 | 23 |
| tsen34 | 0.44541 | 0.117514593 | 0.065 | 0 | 1 | 23 |
| ttc1 | 0.44541 | 0.103943059 | 0.065 | 0 | 1 | 23 |
| ttc13 | 0.44541 | 0.15609597 | 0.065 | 0 | 1 | 23 |
| ttc4 | 0.44541 | 0.115129651 | 0.065 | 0 | 1 | 23 |
| ttpal | 0.44541 | 0.143153056 | 0.065 | 0 | 1 | 23 |
| ttyh2l | 0.44541 | 0.118472173 | 0.065 | 0 | 1 | 23 |
| txndc16 | 0.44541 | 0.123207701 | 0.065 | 0 | 1 | 23 |
| ubap2a | 0.44541 | 0.101079342 | 0.065 | 0 | 1 | 23 |
| ugcg | 0.44541 | 0.112832213 | 0.065 | 0 | 1 | 23 |
| uggt2 | 0.44541 | 0.105288384 | 0.065 | 0 | 1 | 23 |
| ugt2a5 | 0.44541 | 0.12078611 | 0.065 | 0 | 1 | 23 |

| | | | | | | |
|---|---|---|---|---|---|---|
| urm1 | 0.44541 | 0.101461365 | 0.065 | 0 | 1 | 23 |
| usp42 | 0.44541 | 0.107829297 | 0.065 | 0 | 1 | 23 |
| utp23 | 0.44541 | 0.104843356 | 0.065 | 0 | 1 | 23 |
| uvssa | 0.44541 | 0.130444013 | 0.065 | 0 | 1 | 23 |
| vegfab | 0.44541 | 0.118057122 | 0.065 | 0 | 1 | 23 |
| wipi1 | 0.44541 | 0.139595137 | 0.065 | 0 | 1 | 23 |
| zc2hc1a | 0.44541 | 0.109511964 | 0.065 | 0 | 1 | 23 |
| zc3h12b | 0.44541 | 0.134596333 | 0.065 | 0 | 1 | 23 |
| zdhhc4 | 0.44541 | 0.119477082 | 0.065 | 0 | 1 | 23 |
| zgc:101851 | 0.44541 | 0.105717722 | 0.065 | 0 | 1 | 23 |
| zgc:109986 | 0.44541 | 0.105082065 | 0.065 | 0 | 1 | 23 |
| zgc:110130 | 0.44541 | 0.121762047 | 0.065 | 0 | 1 | 23 |
| zgc:110329 | 0.44541 | 0.109101946 | 0.065 | 0 | 1 | 23 |
| zgc:112977 | 0.44541 | 0.108480906 | 0.065 | 0 | 1 | 23 |
| zgc:113149 | 0.44541 | 0.136972911 | 0.065 | 0 | 1 | 23 |
| zgc:113220 | 0.44541 | 0.138010004 | 0.065 | 0 | 1 | 23 |
| zgc:113333 | 0.44541 | 0.108435145 | 0.065 | 0 | 1 | 23 |
| zgc:113436 | 0.44541 | 0.111654474 | 0.065 | 0 | 1 | 23 |
| zgc:113921 | 0.44541 | 0.135276723 | 0.065 | 0 | 1 | 23 |
| zgc:136864 | 0.44541 | 0.118516858 | 0.065 | 0 | 1 | 23 |
| zgc:153184 | 0.44541 | 0.125625056 | 0.065 | 0 | 1 | 23 |
| zgc:154075 | 0.44541 | 0.102088147 | 0.065 | 0 | 1 | 23 |
| zgc:162952 | 0.44541 | 0.115688633 | 0.065 | 0 | 1 | 23 |
| zgc:165515 | 0.44541 | 0.104526337 | 0.065 | 0 | 1 | 23 |
| zgc:174680 | 0.44541 | 0.129803693 | 0.065 | 0 | 1 | 23 |
| zgc:175264 | 0.44541 | 0.120989188 | 0.065 | 0 | 1 | 23 |
| zgc:63470 | 0.44541 | 0.115644338 | 0.065 | 0 | 1 | 23 |
| zgc:66475 | 0.44541 | 0.108737034 | 0.065 | 0 | 1 | 23 |
| zgc:91849 | 0.44541 | 0.146496521 | 0.065 | 0 | 1 | 23 |
| zmp:0000000735 | 0.44541 | 0.133443218 | 0.065 | 0 | 1 | 23 |
| znf1114 | 0.44541 | 0.11529711 | 0.065 | 0 | 1 | 23 |
| znf277 | 0.44541 | 0.109681497 | 0.065 | 0 | 1 | 23 |
| znf385c | 0.44541 | 0.100226432 | 0.065 | 0 | 1 | 23 |
| znf971 | 0.44541 | 0.114975564 | 0.065 | 0 | 1 | 23 |
| KCNIP1 | 0.44541 | 0.115626171 | 0.065 | 0 | 1 | 23 |
| LOC570723 | 0.44541 | 0.110851091 | 0.065 | 0 | 1 | 23 |
| alkbh4 | 0.44541 | 0.114078783 | 0.065 | 0 | 1 | 23 |
| atg9a | 0.44541 | 0.13454493 | 0.065 | 0 | 1 | 23 |
| cabcoco1 | 0.44541 | 0.127622003 | 0.065 | 0 | 1 | 23 |
| casc1 | 0.44541 | 0.138289597 | 0.065 | 0 | 1 | 23 |
| chd8 | 0.44541 | 0.126587194 | 0.065 | 0 | 1 | 23 |
| chn2 | 0.44541 | 0.101749488 | 0.065 | 0 | 1 | 23 |
| coq8a | 0.44541 | 0.113217858 | 0.065 | 0 | 1 | 23 |
| dnajc17 | 0.44541 | 0.102494816 | 0.065 | 0 | 1 | 23 |
| exosc1 | 0.44541 | 0.118043977 | 0.065 | 0 | 1 | 23 |
| gb:ai877918 | 0.44541 | 0.13546088 | 0.065 | 0 | 1 | 23 |
| jag2a | 0.44541 | 0.10932855 | 0.065 | 0 | 1 | 23 |
| lingo1b | 0.44541 | 0.115760283 | 0.065 | 0 | 1 | 23 |

| | | | | | | |
|---|---|---|---|---|---|---|
| lztr1 | 0.44541 | 0.121923502 | 0.065 | 0 | 1 | 23 |
| map3k19 | 0.44541 | 0.128974063 | 0.065 | 0 | 1 | 23 |
| naa35 | 0.44541 | 0.121266117 | 0.065 | 0 | 1 | 23 |
| nek9 | 0.44541 | 0.148076207 | 0.065 | 0 | 1 | 23 |
| npl | 0.44541 | 0.102937597 | 0.065 | 0 | 1 | 23 |
| nt5e | 0.44541 | 0.125414045 | 0.065 | 0 | 1 | 23 |
| nub1 | 0.44541 | 0.126044327 | 0.065 | 0 | 1 | 23 |
| nudt18 | 0.44541 | 0.123857311 | 0.065 | 0 | 1 | 23 |
| nup50 | 0.44541 | 0.11716834 | 0.065 | 0 | 1 | 23 |
| pih1d2 | 0.44541 | 0.115991367 | 0.065 | 0 | 1 | 23 |
| rlf | 0.44541 | 0.128366607 | 0.065 | 0 | 1 | 23 |
| ropn1l | 0.44541 | 0.102804339 | 0.065 | 0 | 1 | 23 |
| sh3glb2b | 0.44541 | 0.121576464 | 0.065 | 0 | 1 | 23 |
| si:ch211-147d7.5 | 0.44541 | 0.107887698 | 0.065 | 0 | 1 | 23 |
| srcap | 0.44541 | 0.125907639 | 0.065 | 0 | 1 | 23 |
| tbr1a | 0.44541 | 0.12353032 | 0.065 | 0 | 1 | 23 |
| tmem138 | 0.44541 | 0.110865361 | 0.065 | 0 | 1 | 23 |
| tmem185 | 0.44541 | 0.111734733 | 0.065 | 0 | 1 | 23 |
| xpo1a | 0.44541 | 0.118125228 | 0.065 | 0 | 1 | 23 |
| CABZ01079427.1 | 0.44541 | 0.100556889 | 0.065 | 0 | 1 | 23 |
| LOC100536647 | 0.44541 | 0.109483285 | 0.065 | 0 | 1 | 23 |
| LOC108184061 | 0.44541 | 0.10896618 | 0.065 | 0 | 1 | 23 |
| USP53 (1 of many) | 0.44541 | 0.110763225 | 0.065 | 0 | 1 | 23 |
| XLOC-031367 | 0.44541 | 0.121399192 | 0.065 | 0 | 1 | 23 |
| acsf2 | 0.44541 | 0.11521821 | 0.065 | 0 | 1 | 23 |
| cux1b | 0.44541 | 0.116002034 | 0.065 | 0 | 1 | 23 |
| dusp7 | 0.44541 | 0.108612423 | 0.065 | 0 | 1 | 23 |
| hgh1 | 0.44541 | 0.100403418 | 0.065 | 0 | 1 | 23 |
| kdelr3 | 0.44541 | 0.112446797 | 0.065 | 0 | 1 | 23 |
| mtmr2 | 0.44541 | 0.105251106 | 0.065 | 0 | 1 | 23 |
| phc1 | 0.44541 | 0.105494406 | 0.065 | 0 | 1 | 23 |
| serinc2l | 0.44541 | 0.106028305 | 0.065 | 0 | 1 | 23 |
| si:ch73-199g24.2 | 0.44541 | 0.113844538 | 0.065 | 0 | 1 | 23 |
| si:dkey-182g1.10 | 0.44541 | 0.117436372 | 0.065 | 0 | 1 | 23 |
| zgc:113364 | 0.44541 | 0.101814866 | 0.065 | 0 | 1 | 23 |
| znf1059 | 0.44541 | 0.113758473 | 0.065 | 0 | 1 | 23 |
| AL954741.2 | 0.44541 | 0.128193818 | 0.065 | 0 | 1 | 23 |
| BX957346.1 | 0.44541 | 0.100707125 | 0.065 | 0 | 1 | 23 |
| CABZ01038697.1 | 0.44541 | 0.124185876 | 0.065 | 0 | 1 | 23 |
| CABZ01074307.1 | 0.44541 | 0.100445533 | 0.065 | 0 | 1 | 23 |
| CABZ01074307.2 | 0.44541 | 0.1617274 | 0.065 | 0 | 1 | 23 |
| CABZ01109081.1 | 0.44541 | 0.106195477 | 0.065 | 0 | 1 | 23 |
| EML5 | 0.44541 | 0.102531972 | 0.065 | 0 | 1 | 23 |
| LOC100333826 | 0.44541 | 0.102111719 | 0.065 | 0 | 1 | 23 |
| LOC100334612 | 0.44541 | 0.123417499 | 0.065 | 0 | 1 | 23 |
| LOC101884799 | 0.44541 | 0.113954369 | 0.065 | 0 | 1 | 23 |
| LOC101886822 | 0.44541 | 0.106563068 | 0.065 | 0 | 1 | 23 |
| RAMP1 | 0.44541 | 0.132310419 | 0.065 | 0 | 1 | 23 |

| | | | | | | |
|---|---|---|---|---|---|---|
| XLOC-004796 | 0.44541 | 0.106785324 | 0.065 | 0 | 1 | 23 |
| abhd2a | 0.44541 | 0.120654939 | 0.065 | 0 | 1 | 23 |
| acacb | 0.44541 | 0.115983727 | 0.065 | 0 | 1 | 23 |
| aff3 | 0.44541 | 0.114896368 | 0.065 | 0 | 1 | 23 |
| akap11 | 0.44541 | 0.112884037 | 0.065 | 0 | 1 | 23 |
| ankrd12 | 0.44541 | 0.107035092 | 0.065 | 0 | 1 | 23 |
| apbb2b | 0.44541 | 0.106892162 | 0.065 | 0 | 1 | 23 |
| aprt | 0.44541 | 0.109514592 | 0.065 | 0 | 1 | 23 |
| atg7 | 0.44541 | 0.1077942 | 0.065 | 0 | 1 | 23 |
| bcl6ab | 0.44541 | 0.172393482 | 0.065 | 0 | 1 | 23 |
| bdh1 | 0.44541 | 0.100598808 | 0.065 | 0 | 1 | 23 |
| brpf3a | 0.44541 | 0.131129088 | 0.065 | 0 | 1 | 23 |
| cbfa2t3 | 0.44541 | 0.107136393 | 0.065 | 0 | 1 | 23 |
| ccdc103 | 0.44541 | 0.108887991 | 0.065 | 0 | 1 | 23 |
| ccdc57 | 0.44541 | 0.106011564 | 0.065 | 0 | 1 | 23 |
| ccdc83 | 0.44541 | 0.111752781 | 0.065 | 0 | 1 | 23 |
| cdkn1ca | 0.44541 | 0.160249335 | 0.065 | 0 | 1 | 23 |
| cenps | 0.44541 | 0.167961539 | 0.065 | 0 | 1 | 23 |
| cep68 | 0.44541 | 0.117193636 | 0.065 | 0 | 1 | 23 |
| cep97 | 0.44541 | 0.112456313 | 0.065 | 0 | 1 | 23 |
| cfap70 | 0.44541 | 0.104040224 | 0.065 | 0 | 1 | 23 |
| chmp4ba | 0.44541 | 0.107339498 | 0.065 | 0 | 1 | 23 |
| cib2 | 0.44541 | 0.133813823 | 0.065 | 0 | 1 | 23 |
| cited2 | 0.44541 | 0.16086946 | 0.065 | 0 | 1 | 23 |
| clcn4 | 0.44541 | 0.138507878 | 0.065 | 0 | 1 | 23 |
| clmn | 0.44541 | 0.107712331 | 0.065 | 0 | 1 | 23 |
| cnep1r1 | 0.44541 | 0.120286292 | 0.065 | 0 | 1 | 23 |
| cnppd1 | 0.44541 | 0.102427405 | 0.065 | 0 | 1 | 23 |
| col12a1a | 0.44541 | 0.107666624 | 0.065 | 0 | 1 | 23 |
| col15a1a | 0.44541 | 0.108656319 | 0.065 | 0 | 1 | 23 |
| copb1 | 0.44541 | 0.100873051 | 0.065 | 0 | 1 | 23 |
| cpda | 0.44541 | 0.111386509 | 0.065 | 0 | 1 | 23 |
| dbpa | 0.44541 | 0.104637935 | 0.065 | 0 | 1 | 23 |
| ddr1 | 0.44541 | 0.112195533 | 0.065 | 0 | 1 | 23 |
| dnajc21 | 0.44541 | 0.102408871 | 0.065 | 0 | 1 | 23 |
| dpm3 | 0.44541 | 0.123946492 | 0.065 | 0 | 1 | 23 |
| dscc1 | 0.44541 | 0.107270666 | 0.065 | 0 | 1 | 23 |
| eif2ak3 | 0.44541 | 0.117264398 | 0.065 | 0 | 1 | 23 |
| elmo3 | 0.44541 | 0.115839892 | 0.065 | 0 | 1 | 23 |
| epg5 | 0.44541 | 0.11713138 | 0.065 | 0 | 1 | 23 |
| epn2 | 0.44541 | 0.100598492 | 0.065 | 0 | 1 | 23 |
| fam184a | 0.44541 | 0.11298113 | 0.065 | 0 | 1 | 23 |
| fam208aa | 0.44541 | 0.117130425 | 0.065 | 0 | 1 | 23 |
| fem1c | 0.44541 | 0.125523687 | 0.065 | 0 | 1 | 23 |
| fermt1 | 0.44541 | 0.117495241 | 0.065 | 0 | 1 | 23 |
| fgf18b | 0.44541 | 0.110832667 | 0.065 | 0 | 1 | 23 |
| fndc4b | 0.44541 | 0.127549186 | 0.065 | 0 | 1 | 23 |
| fosl2 | 0.44541 | 0.10430756 | 0.065 | 0 | 1 | 23 |

| | | | | | | |
|---|---|---|---|---|---|---|
| frrs1b | 0.44541 | 0.115804753 | 0.065 | 0 | 1 | 23 |
| gata5 | 0.44541 | 0.12862997 | 0.065 | 0 | 1 | 23 |
| glipr2l | 0.44541 | 0.111737938 | 0.065 | 0 | 1 | 23 |
| gpr63 | 0.44541 | 0.111521016 | 0.065 | 0 | 1 | 23 |
| grasp | 0.44541 | 0.164358054 | 0.065 | 0 | 1 | 23 |
| hax1 | 0.44541 | 0.144143559 | 0.065 | 0 | 1 | 23 |
| hexa | 0.44541 | 0.115874017 | 0.065 | 0 | 1 | 23 |
| hira | 0.44541 | 0.131139657 | 0.065 | 0 | 1 | 23 |
| hoxc6b | 0.44541 | 0.151255732 | 0.065 | 0 | 1 | 23 |
| hspb7 | 0.44541 | 0.117424491 | 0.065 | 0 | 1 | 23 |
| ift57 | 0.44541 | 0.158080365 | 0.065 | 0 | 1 | 23 |
| ints3 | 0.44541 | 0.116525992 | 0.065 | 0 | 1 | 23 |
| iqcd | 0.44541 | 0.104040224 | 0.065 | 0 | 1 | 23 |
| ivns1abpa | 0.44541 | 0.119967151 | 0.065 | 0 | 1 | 23 |
| kcnk6 | 0.44541 | 0.109834565 | 0.065 | 0 | 1 | 23 |
| laptm4b | 0.44541 | 0.122999976 | 0.065 | 0 | 1 | 23 |
| lin7a | 0.44541 | 0.125817053 | 0.065 | 0 | 1 | 23 |
| lin7b | 0.44541 | 0.108029501 | 0.065 | 0 | 1 | 23 |
| lonrf1 | 0.44541 | 0.13562106 | 0.065 | 0 | 1 | 23 |
| lrrc40 | 0.44541 | 0.124463244 | 0.065 | 0 | 1 | 23 |
| lrrc8db | 0.44541 | 0.129113358 | 0.065 | 0 | 1 | 23 |
| lyrm5a | 0.44541 | 0.174304994 | 0.065 | 0 | 1 | 23 |
| lzic | 0.44541 | 0.111363588 | 0.065 | 0 | 1 | 23 |
| manea | 0.44541 | 0.117291468 | 0.065 | 0 | 1 | 23 |
| map3k14a | 0.44541 | 0.163729023 | 0.065 | 0 | 1 | 23 |
| mccc1 | 0.44541 | 0.137517027 | 0.065 | 0 | 1 | 23 |
| mcrs1 | 0.44541 | 0.109782571 | 0.065 | 0 | 1 | 23 |
| mdm4 | 0.44541 | 0.104044532 | 0.065 | 0 | 1 | 23 |
| mettl23 | 0.44541 | 0.109400039 | 0.065 | 0 | 1 | 23 |
| mfsd6a | 0.44541 | 0.144062815 | 0.065 | 0 | 1 | 23 |
| mrs2 | 0.44541 | 0.110781066 | 0.065 | 0 | 1 | 23 |
| mxra8b | 0.44541 | 0.12551574 | 0.065 | 0 | 1 | 23 |
| myo10l3 | 0.44541 | 0.125484518 | 0.065 | 0 | 1 | 23 |
| nadsyn1.1 | 0.44541 | 0.124773503 | 0.065 | 0 | 1 | 23 |
| necap1 | 0.44541 | 0.114219978 | 0.065 | 0 | 1 | 23 |
| nfrkb | 0.44541 | 0.136805618 | 0.065 | 0 | 1 | 23 |
| nkx2.3 | 0.44541 | 0.150659031 | 0.065 | 0 | 1 | 23 |
| nptna | 0.44541 | 0.127578636 | 0.065 | 0 | 1 | 23 |
| nrf1 | 0.44541 | 0.121531481 | 0.065 | 0 | 1 | 23 |
| nsd1b | 0.44541 | 0.117147429 | 0.065 | 0 | 1 | 23 |
| nudt9 | 0.44541 | 0.108760225 | 0.065 | 0 | 1 | 23 |
| nup210 | 0.44541 | 0.162876962 | 0.065 | 0 | 1 | 23 |
| oxnad1 | 0.44541 | 0.10346624 | 0.065 | 0 | 1 | 23 |
| pcgf6 | 0.44541 | 0.113349429 | 0.065 | 0 | 1 | 23 |
| pdp2 | 0.44541 | 0.113290438 | 0.065 | 0 | 1 | 23 |
| pkn2 | 0.44541 | 0.109726057 | 0.065 | 0 | 1 | 23 |
| plagx | 0.44541 | 0.126841026 | 0.065 | 0 | 1 | 23 |
| pofut1 | 0.44541 | 0.110969302 | 0.065 | 0 | 1 | 23 |

| | | | | | | |
|---|---|---|---|---|---|---|
| ppfibp1b | 0.44541 | 0.126377448 | 0.065 | 0 | 1 | 23 |
| pprc1 | 0.44541 | 0.107644166 | 0.065 | 0 | 1 | 23 |
| prr7 | 0.44541 | 0.117034279 | 0.065 | 0 | 1 | 23 |
| rab23 | 0.44541 | 0.104344963 | 0.065 | 0 | 1 | 23 |
| rab35b | 0.44541 | 0.104563939 | 0.065 | 0 | 1 | 23 |
| rffl | 0.44541 | 0.109589403 | 0.065 | 0 | 1 | 23 |
| rhoj | 0.44541 | 0.10506118 | 0.065 | 0 | 1 | 23 |
| rrm2b | 0.44541 | 0.105341769 | 0.065 | 0 | 1 | 23 |
| scn4bb | 0.44541 | 0.114738625 | 0.065 | 0 | 1 | 23 |
| sgtb | 0.44541 | 0.100578014 | 0.065 | 0 | 1 | 23 |
| sh2b1 | 0.44541 | 0.118009157 | 0.065 | 0 | 1 | 23 |
| si:busm1-180g5.6 | 0.44541 | 0.114838671 | 0.065 | 0 | 1 | 23 |
| si:ch211-256e16.3 | 0.44541 | 0.113954369 | 0.065 | 0 | 1 | 23 |
| si:ch211-276i12.9 | 0.44541 | 0.123561521 | 0.065 | 0 | 1 | 23 |
| si:ch211-87m7.2 | 0.44541 | 0.100970066 | 0.065 | 0 | 1 | 23 |
| si:dkey-21o19.2 | 0.44541 | 0.102390501 | 0.065 | 0 | 1 | 23 |
| si:dkey-237i9.1 | 0.44541 | 0.123496405 | 0.065 | 0 | 1 | 23 |
| si:dkey-238f9.1 | 0.44541 | 0.149872569 | 0.065 | 0 | 1 | 23 |
| si:dkey-253d23.4 | 0.44541 | 0.106008443 | 0.065 | 0 | 1 | 23 |
| si:dkey-27b3.4 | 0.44541 | 0.111521016 | 0.065 | 0 | 1 | 23 |
| si:dkey-46g23.1 | 0.44541 | 0.115885055 | 0.065 | 0 | 1 | 23 |
| si:dkey-84k17.2 | 0.44541 | 0.145827318 | 0.065 | 0 | 1 | 23 |
| slc30a4 | 0.44541 | 0.156163429 | 0.065 | 0 | 1 | 23 |
| slc35b1 | 0.44541 | 0.131940929 | 0.065 | 0 | 1 | 23 |
| slka | 0.44541 | 0.115350651 | 0.065 | 0 | 1 | 23 |
| smim12 | 0.44541 | 0.106271436 | 0.065 | 0 | 1 | 23 |
| smyd2a | 0.44541 | 0.132627025 | 0.065 | 0 | 1 | 23 |
| snx25 | 0.44541 | 0.104610309 | 0.065 | 0 | 1 | 23 |
| spire1a | 0.44541 | 0.147521599 | 0.065 | 0 | 1 | 23 |
| sprydd7b | 0.44541 | 0.108029501 | 0.065 | 0 | 1 | 23 |
| surf4l | 0.44541 | 0.124023837 | 0.065 | 0 | 1 | 23 |
| tbc1d22a | 0.44541 | 0.12788769 | 0.065 | 0 | 1 | 23 |
| tceanc2 | 0.44541 | 0.101708691 | 0.065 | 0 | 1 | 23 |
| tcf7l1a | 0.44541 | 0.112296432 | 0.065 | 0 | 1 | 23 |
| tent2 | 0.44541 | 0.123333906 | 0.065 | 0 | 1 | 23 |
| tmem106bb | 0.44541 | 0.117264398 | 0.065 | 0 | 1 | 23 |
| tmem150ab | 0.44541 | 0.112304323 | 0.065 | 0 | 1 | 23 |
| tmem181 | 0.44541 | 0.106563068 | 0.065 | 0 | 1 | 23 |
| tmem184c | 0.44541 | 0.100262141 | 0.065 | 0 | 1 | 23 |
| tnfaip3 | 0.44541 | 0.127867184 | 0.065 | 0 | 1 | 23 |
| trdmt1 | 0.44541 | 0.10736437 | 0.065 | 0 | 1 | 23 |
| ubald1b | 0.44541 | 0.102209703 | 0.065 | 0 | 1 | 23 |
| ubn2b | 0.44541 | 0.104020914 | 0.065 | 0 | 1 | 23 |
| unc13ba | 0.44541 | 0.138507878 | 0.065 | 0 | 1 | 23 |
| wdr33 | 0.44541 | 0.14371461 | 0.065 | 0 | 1 | 23 |
| xrcc4 | 0.44541 | 0.111960142 | 0.065 | 0 | 1 | 23 |
| zfyve21 | 0.44541 | 0.114247072 | 0.065 | 0 | 1 | 23 |
| zgc:158320 | 0.44541 | 0.130394354 | 0.065 | 0 | 1 | 23 |

| | | | | | | |
|---|---|---|---|---|---|---|
| zgc:172145 | 0.44541 | 0.114838671 | 0.065 | 0 | 1 | 23 |
| zgc:173705 | 0.44541 | 0.128193818 | 0.065 | 0 | 1 | 23 |
| zgc:56699 | 0.44541 | 0.108982898 | 0.065 | 0 | 1 | 23 |
| zgc:66472 | 0.44541 | 0.110949943 | 0.065 | 0 | 1 | 23 |
| zgc:77056 | 0.44541 | 0.109089844 | 0.065 | 0 | 1 | 23 |
| zgc:77880 | 0.44541 | 0.113471358 | 0.065 | 0 | 1 | 23 |
| zmp:0000000624 | 0.44541 | 0.133036686 | 0.065 | 0 | 1 | 23 |
| znf1015 | 0.44541 | 0.108806077 | 0.065 | 0 | 1 | 23 |
| znf346 | 0.44541 | 0.124839783 | 0.065 | 0 | 1 | 23 |
| znf423 | 0.44541 | 0.115767215 | 0.065 | 0 | 1 | 23 |
| znf704 | 0.44541 | 0.122204955 | 0.065 | 0 | 1 | 23 |
| znhit2 | 0.44541 | 0.116361521 | 0.065 | 0 | 1 | 23 |
| znrf3 | 0.44541 | 0.106563068 | 0.065 | 0 | 1 | 23 |
| arl4cb | 0.44541 | 0.106443855 | 0.065 | 0 | 1 | 23 |
| atp6v0d1 | 0.44541 | 0.137506862 | 0.065 | 0 | 1 | 23 |
| axin1 | 0.44541 | 0.10030833 | 0.065 | 0 | 1 | 23 |
| ercc2 | 0.44541 | 0.107402759 | 0.065 | 0 | 1 | 23 |
| fam92a1 | 0.44541 | 0.108805394 | 0.065 | 0 | 1 | 23 |
| hdx | 0.44541 | 0.112470058 | 0.065 | 0 | 1 | 23 |
| iffo1b | 0.44541 | 0.103449293 | 0.065 | 0 | 1 | 23 |
| ispd | 0.44541 | 0.105262104 | 0.065 | 0 | 1 | 23 |
| serpinb14 | 0.44541 | 0.10562052 | 0.065 | 0 | 1 | 23 |
| si:ch73-236c18.8 | 0.44541 | 0.130707337 | 0.065 | 0 | 1 | 23 |
| slc35e3 | 0.44541 | 0.1022278 | 0.065 | 0 | 1 | 23 |
| tnksb | 0.44541 | 0.103906682 | 0.065 | 0 | 1 | 23 |
| CT583625.5 | 0.44541 | 0.122630383 | 0.065 | 0 | 1 | 23 |
| CU464124.1 | 0.44541 | 0.117698407 | 0.065 | 0 | 1 | 23 |
| alox12 | 0.44541 | 0.166251629 | 0.065 | 0 | 1 | 23 |
| ccdc169 | 0.44541 | 0.122630383 | 0.065 | 0 | 1 | 23 |
| cisd1 | 0.44541 | 0.107621283 | 0.065 | 0 | 1 | 23 |
| eaf2 | 0.44541 | 0.105322547 | 0.065 | 0 | 1 | 23 |
| fam177a1 | 0.44541 | 0.120702157 | 0.065 | 0 | 1 | 23 |
| fhl3a | 0.44541 | 0.118924705 | 0.065 | 0 | 1 | 23 |
| fitm2 | 0.44541 | 0.123943163 | 0.065 | 0 | 1 | 23 |
| gsk3ab | 0.44541 | 0.110341569 | 0.065 | 0 | 1 | 23 |
| kdm2aa | 0.44541 | 0.102239141 | 0.065 | 0 | 1 | 23 |
| lmna | 0.44541 | 0.106327554 | 0.065 | 0 | 1 | 23 |
| mest | 0.44541 | 0.10677979 | 0.065 | 0 | 1 | 23 |
| pgm3 | 0.44541 | 0.113104677 | 0.065 | 0 | 1 | 23 |
| ppie | 0.44541 | 0.114906378 | 0.065 | 0 | 1 | 23 |
| prkacab | 0.44541 | 0.108604868 | 0.065 | 0 | 1 | 23 |
| ptpn1 | 0.44541 | 0.104628357 | 0.065 | 0 | 1 | 23 |
| rnf128a | 0.44541 | 0.121551411 | 0.065 | 0 | 1 | 23 |
| roraa | 0.44541 | 0.140876801 | 0.065 | 0 | 1 | 23 |
| si:ch211-197g15.6 | 0.44541 | 0.109270202 | 0.065 | 0 | 1 | 23 |
| si:dkey-260j18.2 | 0.44541 | 0.11640068 | 0.065 | 0 | 1 | 23 |
| smtnl | 0.44541 | 0.106327554 | 0.065 | 0 | 1 | 23 |
| sorbs3 | 0.44541 | 0.109917799 | 0.065 | 0 | 1 | 23 |

| | | | | | | |
|---|---|---|---|---|---|---|
| spns1 | 0.44541 | 0.117094397 | 0.065 | 0 | 1 | 23 |
| tchp | 0.44541 | 0.123570478 | 0.065 | 0 | 1 | 23 |
| trpc4apa | 0.44541 | 0.114176692 | 0.065 | 0 | 1 | 23 |
| tsen15 | 0.44541 | 0.116014336 | 0.065 | 0 | 1 | 23 |
| zgc:171967 | 0.44541 | 0.109627973 | 0.065 | 0 | 1 | 23 |
| CABZ01054965.1 | 0.44541 | 0.113337535 | 0.065 | 0 | 1 | 23 |
| CR457445.1 | 0.44541 | 0.141939631 | 0.065 | 0 | 1 | 23 |
| CU929237.1 | 0.44541 | 0.130650509 | 0.065 | 0 | 1 | 23 |
| LO016987.2 | 0.44541 | 0.120279574 | 0.065 | 0 | 1 | 23 |
| XLOC-041870 | 0.44541 | 0.108642454 | 0.065 | 0 | 1 | 23 |
| arl8a | 0.44541 | 0.137236126 | 0.065 | 0 | 1 | 23 |
| bloc1s4 | 0.44541 | 0.137120052 | 0.065 | 0 | 1 | 23 |
| chdh | 0.44541 | 0.190129268 | 0.065 | 0 | 1 | 23 |
| fam210ab | 0.44541 | 0.111266221 | 0.065 | 0 | 1 | 23 |
| gosr1 | 0.44541 | 0.114383373 | 0.065 | 0 | 1 | 23 |
| grb10b | 0.44541 | 0.130908796 | 0.065 | 0 | 1 | 23 |
| mapkapk5 | 0.44541 | 0.111857441 | 0.065 | 0 | 1 | 23 |
| mrps5 | 0.44541 | 0.119898634 | 0.065 | 0 | 1 | 23 |
| nkx3.2 | 0.44541 | 0.120368337 | 0.065 | 0 | 1 | 23 |
| paqr3b | 0.44541 | 0.117327565 | 0.065 | 0 | 1 | 23 |
| ppwd1 | 0.44541 | 0.10575662 | 0.065 | 0 | 1 | 23 |
| rc3h1a | 0.44541 | 0.121758428 | 0.065 | 0 | 1 | 23 |
| senp3a | 0.44541 | 0.128068178 | 0.065 | 0 | 1 | 23 |
| si:ch211-152c8.4 | 0.44541 | 0.109795512 | 0.065 | 0 | 1 | 23 |
| si:ch73-211l13.2 | 0.44541 | 0.116202657 | 0.065 | 0 | 1 | 23 |
| si:ch73-221f6.4 | 0.44541 | 0.128593622 | 0.065 | 0 | 1 | 23 |
| slc43a1a | 0.44541 | 0.13831699 | 0.065 | 0 | 1 | 23 |
| stam | 0.44541 | 0.111872145 | 0.065 | 0 | 1 | 23 |
| thap11 | 0.44541 | 0.134650393 | 0.065 | 0 | 1 | 23 |
| tnc | 0.44541 | 0.149516459 | 0.065 | 0 | 1 | 23 |
| tns1b | 0.44541 | 0.121758428 | 0.065 | 0 | 1 | 23 |
| vezf1a | 0.44541 | 0.121758428 | 0.065 | 0 | 1 | 23 |
| LOC108191465 | 0.44541 | 0.135321749 | 0.065 | 0 | 1 | 23 |
| mapk12a | 0.44541 | 0.113297525 | 0.065 | 0 | 1 | 23 |
| met | 0.44541 | 0.127922017 | 0.065 | 0 | 1 | 23 |
| rab11fip1a | 0.44541 | 0.126344747 | 0.065 | 0 | 1 | 23 |
| si:ch211-160d20.3 | 0.44541 | 0.125973954 | 0.065 | 0 | 1 | 23 |
| snx7 | 0.44541 | 0.126600742 | 0.065 | 0 | 1 | 23 |
| vps41 | 0.44541 | 0.114776556 | 0.065 | 0 | 1 | 23 |
| ccdc130 | 0.44541 | 0.110593256 | 0.065 | 0 | 1 | 23 |
| cica | 0.44541 | 0.192627205 | 0.065 | 0 | 1 | 23 |
| rbsn | 0.44541 | 0.104051846 | 0.065 | 0 | 1 | 23 |
| spag8 | 0.44541 | 0.110593256 | 0.065 | 0 | 1 | 23 |
| taf4a | 0.44541 | 0.10543759 | 0.065 | 0 | 1 | 23 |
| znf576.1 | 0.44541 | 0.113817069 | 0.065 | 0 | 1 | 23 |
| CABZ01038494.1 | 0.44541 | 0.123987579 | 0.065 | 0 | 1 | 23 |
| CABZ01080375.1 | 0.44541 | 0.121280449 | 0.065 | 0 | 1 | 23 |
| tomm22 | 0.44566 | 0.30314299 | 0.452 | 0.25 | 1 | 23 |

| | | | | | | |
|---|---|---|---|---|---|---|
| uqcr10 | 0.44583 | 0.100953074 | 0.548 | 0.375 | 1 | 23 |
| ndufb5 | 0.44609 | -0.10105495 | 0.387 | 0.375 | 1 | 23 |
| sf3b2 | 0.44623 | -0.31369557 | 0.387 | 0.625 | 1 | 23 |
| ilf2 | 0.44641 | 0.368804619 | 0.484 | 0.25 | 1 | 23 |
| si:ch211-162e15.3 | 0.44642 | -0.25285131 | 0.161 | 0.375 | 1 | 23 |
| rgmd | 0.44669 | -0.1620959 | 0.097 | 0.125 | 1 | 23 |
| nudcd3 | 0.44692 | -0.28170671 | 0.161 | 0.375 | 1 | 23 |
| sf3b1 | 0.44732 | 0.239745992 | 0.323 | 0.125 | 1 | 23 |
| grhl1 | 0.44798 | 0.166526379 | 0.419 | 0.25 | 1 | 23 |
| mfap2 | 0.44834 | -0.12397221 | 0.516 | 0.5 | 1 | 23 |
| h3f3a | 0.44845 | -0.11769304 | 0.677 | 0.625 | 1 | 23 |
| si:dkey-78l4.14 | 0.44883 | 0.129074242 | 0.258 | 0.125 | 1 | 23 |
| mt-nd4l | 0.44989 | -0.24675894 | 0.097 | 0.25 | 1 | 23 |
| per1a | 0.45084 | -0.29676327 | 0.097 | 0.25 | 1 | 23 |
| rab5ab | 0.45091 | -0.15989741 | 0.226 | 0.25 | 1 | 23 |
| usp22 | 0.45104 | -0.28807369 | 0.161 | 0.375 | 1 | 23 |
| epb41l5 | 0.4525 | -0.3222942 | 0.161 | 0.375 | 1 | 23 |
| xrn2 | 0.45263 | -0.28931089 | 0.161 | 0.375 | 1 | 23 |
| psmd11a | 0.45272 | -0.29269295 | 0.161 | 0.375 | 1 | 23 |
| agrn | 0.45274 | -0.31069145 | 0.516 | 0.75 | 1 | 23 |
| nol11 | 0.45323 | -0.26989902 | 0.161 | 0.375 | 1 | 23 |
| fam84a | 0.45345 | -0.38797614 | 0.097 | 0.25 | 1 | 23 |
| fmo5 | 0.45394 | -0.14513058 | 0.097 | 0.125 | 1 | 23 |
| hmgb2a | 0.45425 | -0.29271801 | 0.677 | 0.875 | 1 | 23 |
| sik1 | 0.45454 | -0.32663171 | 0.194 | 0.375 | 1 | 23 |
| thbs1b | 0.45456 | -0.21822153 | 0.097 | 0.25 | 1 | 23 |
| tmem30aa | 0.45529 | 0.1268101 | 0.258 | 0.125 | 1 | 23 |
| blmh | 0.45562 | -0.26517888 | 0.129 | 0.25 | 1 | 23 |
| psmc6 | 0.45597 | 0.264740143 | 0.516 | 0.375 | 1 | 23 |
| dnaja2a | 0.4561 | 0.224334749 | 0.613 | 0.5 | 1 | 23 |
| rbm39a | 0.45656 | -0.21650629 | 0.548 | 0.625 | 1 | 23 |
| sh3bp5la | 0.45659 | -0.13868892 | 0.129 | 0.125 | 1 | 23 |
| paip2b | 0.45662 | 0.205402197 | 0.677 | 0.625 | 1 | 23 |
| rab10 | 0.45764 | 0.352739663 | 0.613 | 0.375 | 1 | 23 |
| mrpl22 | 0.45921 | -0.17413297 | 0.097 | 0.25 | 1 | 23 |
| scarb2a | 0.45979 | -0.31515273 | 0.097 | 0.25 | 1 | 23 |
| mut | 0.46026 | -0.15127411 | 0.097 | 0.125 | 1 | 23 |
| brd4 | 0.46046 | -0.33029001 | 0.194 | 0.375 | 1 | 23 |
| bsg | 0.46049 | 0.18123065 | 0.71 | 0.5 | 1 | 23 |
| mov10b.1 | 0.46061 | -0.18746128 | 0.194 | 0.25 | 1 | 23 |
| fn1b | 0.46087 | -0.18973343 | 0.71 | 0.75 | 1 | 23 |
| gbp | 0.46279 | -0.33695074 | 0.323 | 0.5 | 1 | 23 |
| srpra | 0.46321 | 0.110219731 | 0.129 | 0.125 | 1 | 23 |
| plscr3b | 0.4635 | 0.23151464 | 0.355 | 0.25 | 1 | 23 |
| sqstm1 | 0.4637 | 0.241580446 | 0.258 | 0.125 | 1 | 23 |
| malt1 | 0.46414 | -0.2913491 | 0.129 | 0.25 | 1 | 23 |
| ogdha | 0.46418 | -0.28352002 | 0.129 | 0.25 | 1 | 23 |
| LOC101882596 | 0.46466 | -0.15254151 | 0.097 | 0.125 | 1 | 23 |

| | | | | | | |
|---|---|---|---|---|---|---|
| dohh | 0.46537 | -0.31163723 | 0.194 | 0.375 | 1 | 23 |
| tbx2b | 0.46657 | 0.326622783 | 0.387 | 0.25 | 1 | 23 |
| lig3 | 0.4671 | -0.14189824 | 0.097 | 0.125 | 1 | 23 |
| wt1b | 0.46766 | 0.317470013 | 0.29 | 0.125 | 1 | 23 |
| ehd2b | 0.46787 | 0.148814983 | 0.258 | 0.25 | 1 | 23 |
| pabpc1b | 0.46816 | -0.20587519 | 0.097 | 0.25 | 1 | 23 |
| btf3l4 | 0.46848 | -0.14652592 | 0.484 | 0.5 | 1 | 23 |
| ctsf | 0.46914 | 0.324806857 | 0.29 | 0.125 | 1 | 23 |
| pvrl2l | 0.46939 | -0.29676131 | 0.097 | 0.25 | 1 | 23 |
| caprin1b | 0.46974 | 0.293687262 | 0.613 | 0.375 | 1 | 23 |
| gstm.1 | 0.46979 | 0.117914342 | 0.806 | 0.625 | 1 | 23 |
| ywhah | 0.4706 | -0.16232692 | 0.484 | 0.5 | 1 | 23 |
| hmg20b | 0.47097 | -0.28778654 | 0.194 | 0.375 | 1 | 23 |
| usp36 | 0.47114 | 0.326113149 | 0.29 | 0.125 | 1 | 23 |
| hnrnpk | 0.4733 | 0.305577679 | 0.516 | 0.375 | 1 | 23 |
| si:ch211-217k17.7 | 0.47341 | 0.162678779 | 0.29 | 0.125 | 1 | 23 |
| pho | 0.47377 | 0.190714363 | 0.29 | 0.125 | 1 | 23 |
| cct8 | 0.47478 | 0.260273233 | 0.613 | 0.375 | 1 | 23 |
| snx1a | 0.47517 | 0.275713955 | 0.323 | 0.125 | 1 | 23 |
| atp6v1e1a | 0.4753 | -0.24463474 | 0.194 | 0.25 | 1 | 23 |
| si:dkey-42i9.6 | 0.47546 | -0.19318589 | 0.097 | 0.25 | 1 | 23 |
| arf5 | 0.47573 | 0.305816177 | 0.419 | 0.25 | 1 | 23 |
| rer1 | 0.47583 | 0.308167237 | 0.323 | 0.125 | 1 | 23 |
| rhoua | 0.47782 | -0.29285621 | 0.129 | 0.25 | 1 | 23 |
| zgc:194981 | 0.47813 | 0.302999316 | 0.323 | 0.125 | 1 | 23 |
| psma2 | 0.47821 | 0.291367129 | 0.484 | 0.25 | 1 | 23 |
| zgc:77849 | 0.47943 | 0.306252086 | 0.484 | 0.25 | 1 | 23 |
| itgav | 0.47959 | 0.338315766 | 0.484 | 0.25 | 1 | 23 |
| tpm3 | 0.48089 | -0.24489419 | 0.677 | 0.875 | 1 | 23 |
| eif3ba | 0.48153 | 0.310804666 | 0.71 | 0.5 | 1 | 23 |
| jam2a | 0.48244 | 0.328591766 | 0.355 | 0.25 | 1 | 23 |
| sept6 | 0.48312 | -0.2902626 | 0.194 | 0.375 | 1 | 23 |
| lmo4a | 0.48456 | 0.102708146 | 0.129 | 0.125 | 1 | 23 |
| plk3 | 0.48487 | -0.23743521 | 0.194 | 0.25 | 1 | 23 |
| nop58 | 0.48573 | 0.304388183 | 0.613 | 0.375 | 1 | 23 |
| abce1 | 0.4859 | 0.275487798 | 0.29 | 0.125 | 1 | 23 |
| ttf1 | 0.48738 | -0.28656535 | 0.097 | 0.25 | 1 | 23 |
| cst14b.1 | 0.48746 | -0.15170541 | 0.097 | 0.125 | 1 | 23 |
| atp5pd | 0.48768 | 0.29599925 | 0.581 | 0.5 | 1 | 23 |
| dgcr6 | 0.48781 | -0.28225774 | 0.129 | 0.25 | 1 | 23 |
| sptlc2a | 0.488 | -0.26699197 | 0.097 | 0.25 | 1 | 23 |
| dhrs13a.3 | 0.4882 | 0.28189349 | 0.226 | 0.125 | 1 | 23 |
| brdt | 0.48862 | 0.20912579 | 0.29 | 0.125 | 1 | 23 |
| col4a1 | 0.48954 | 0.329241449 | 0.323 | 0.125 | 1 | 23 |
| nucks1b | 0.49078 | -0.19159765 | 0.548 | 0.625 | 1 | 23 |
| mak16 | 0.49114 | 0.142219667 | 0.258 | 0.125 | 1 | 23 |
| si:dkey-19e4.5 | 0.49213 | 0.242114277 | 0.226 | 0.125 | 1 | 23 |
| si:dkey-51e6.1 | 0.49221 | 0.284768505 | 0.323 | 0.125 | 1 | 23 |

| | | | | | | |
|---|---|---|---|---|---|---|
| emd | 0.49272 | 0.263610037 | 0.516 | 0.375 | 1 | 23 |
| ncl | 0.49277 | 0.263096707 | 0.742 | 0.625 | 1 | 23 |
| rbb4l | 0.49334 | -0.10273386 | 0.226 | 0.375 | 1 | 23 |
| fosab | 0.49338 | -0.12245139 | 1 | 1 | 1 | 23 |
| edil3a | 0.4934 | 0.187497636 | 0.161 | 0.125 | 1 | 23 |
| mrpl47 | 0.49457 | -0.13791298 | 0.097 | 0.125 | 1 | 23 |
| lsm5 | 0.49493 | 0.283854969 | 0.419 | 0.25 | 1 | 23 |
| ranbp1 | 0.49517 | -0.21590755 | 0.581 | 0.625 | 1 | 23 |
| mfap1 | 0.49525 | -0.24381046 | 0.161 | 0.25 | 1 | 23 |
| ndufa8 | 0.49573 | 0.101682233 | 0.355 | 0.375 | 1 | 23 |
| taf9 | 0.49578 | 0.236825061 | 0.355 | 0.25 | 1 | 23 |
| rab11fip1b | 0.49636 | -0.25498613 | 0.129 | 0.25 | 1 | 23 |
| cox5ab | 0.49685 | -0.17579999 | 0.516 | 0.5 | 1 | 23 |
| mtap | 0.49713 | 0.272628312 | 0.323 | 0.125 | 1 | 23 |
| spry2 | 0.49738 | 0.292708385 | 0.258 | 0.125 | 1 | 23 |
| pbdc1 | 0.49811 | 0.149508098 | 0.419 | 0.25 | 1 | 23 |
| rtn3 | 0.49826 | 0.275022417 | 0.323 | 0.125 | 1 | 23 |
| si:ch211-76m11.3 | 0.49831 | -0.21047014 | 0.161 | 0.25 | 1 | 23 |
| slx1b | 0.49862 | -0.28898822 | 0.194 | 0.375 | 1 | 23 |
| psme2 | 0.49887 | 0.293398057 | 0.516 | 0.375 | 1 | 23 |
| ankrd13c | 0.49911 | -0.28418637 | 0.097 | 0.25 | 1 | 23 |
| sdhc | 0.50009 | 0.294731886 | 0.323 | 0.125 | 1 | 23 |
| gins4 | 0.50058 | -0.14074926 | 0.097 | 0.125 | 1 | 23 |
| phc2a | 0.50127 | 0.30485263 | 0.387 | 0.25 | 1 | 23 |
| zgc:55781 | 0.50167 | -0.25165872 | 0.129 | 0.25 | 1 | 23 |
| zgc:110182 | 0.50264 | 0.1521177 | 0.806 | 0.625 | 1 | 23 |
| cfl1 | 0.50273 | -0.14755551 | 0.871 | 0.875 | 1 | 23 |
| si:dkeyp-113d7.1 | 0.50309 | 0.256053208 | 0.258 | 0.125 | 1 | 23 |
| krt15 | 0.50348 | 0.449793763 | 0.452 | 0.25 | 1 | 23 |
| wbp2nl | 0.50357 | -0.17209924 | 0.194 | 0.25 | 1 | 23 |
| ctsk | 0.50401 | 0.30998475 | 0.323 | 0.125 | 1 | 23 |
| ddx18 | 0.50401 | 0.193382179 | 0.29 | 0.125 | 1 | 23 |
| sf1 | 0.50447 | -0.21022105 | 0.161 | 0.25 | 1 | 23 |
| mapk14b | 0.50462 | -0.31256823 | 0.097 | 0.25 | 1 | 23 |
| cd82a | 0.50508 | -0.35430709 | 0.194 | 0.375 | 1 | 23 |
| edf1 | 0.50588 | -0.21523887 | 0.774 | 0.875 | 1 | 23 |
| pxna | 0.50613 | 0.247158854 | 0.29 | 0.125 | 1 | 23 |
| efnb3a | 0.50715 | -0.26767756 | 0.129 | 0.25 | 1 | 23 |
| rtraf | 0.50911 | -0.18217935 | 0.194 | 0.375 | 1 | 23 |
| zgc:113208 | 0.5096 | -0.17742905 | 0.097 | 0.25 | 1 | 23 |
| BX284638.1 | 0.51172 | -0.23187646 | 0.161 | 0.25 | 1 | 23 |
| eif3k | 0.51183 | 0.230470049 | 0.581 | 0.5 | 1 | 23 |
| tnfrsfa | 0.51211 | 0.279494608 | 0.29 | 0.125 | 1 | 23 |
| gstp1 | 0.51239 | 0.311620404 | 0.581 | 0.375 | 1 | 23 |
| irs1 | 0.51241 | -0.26595772 | 0.226 | 0.375 | 1 | 23 |
| clic1 | 0.51278 | -0.28356283 | 0.323 | 0.5 | 1 | 23 |
| bckdhb | 0.5134 | -0.26810034 | 0.226 | 0.375 | 1 | 23 |
| aldoaa | 0.51342 | 0.280425898 | 0.387 | 0.25 | 1 | 23 |

| | | | | | | |
|---|---|---|---|---|---|---|
| soga1 | 0.51413 | 0.104562837 | 0.129 | 0.125 | 1 | 23 |
| hoxa3a | 0.51476 | -0.26042046 | 0.097 | 0.25 | 1 | 23 |
| pdia3 | 0.51493 | 0.147621661 | 0.774 | 0.75 | 1 | 23 |
| psmd4a | 0.51562 | 0.210183739 | 0.226 | 0.125 | 1 | 23 |
| idh2 | 0.51582 | -0.18888373 | 0.613 | 0.625 | 1 | 23 |
| ndufs7 | 0.51602 | -0.26243096 | 0.29 | 0.5 | 1 | 23 |
| cstf1 | 0.51689 | 0.198277668 | 0.194 | 0.125 | 1 | 23 |
| ier5 | 0.51753 | 0.238590815 | 0.452 | 0.25 | 1 | 23 |
| khdrbs1b | 0.51766 | -0.30696002 | 0.323 | 0.5 | 1 | 23 |
| lgmn | 0.51786 | 0.219595721 | 0.29 | 0.125 | 1 | 23 |
| klf3 | 0.51807 | -0.18930437 | 0.097 | 0.25 | 1 | 23 |
| esr2b | 0.51924 | -0.3046164 | 0.419 | 0.625 | 1 | 23 |
| ldlrad4a | 0.52016 | -0.10333576 | 0.129 | 0.25 | 1 | 23 |
| tmsb4x | 0.52064 | -0.14903238 | 1 | 1 | 1 | 23 |
| sap30l | 0.52089 | 0.28869398 | 0.548 | 0.375 | 1 | 23 |
| arih2 | 0.52109 | -0.10113515 | 0.129 | 0.25 | 1 | 23 |
| copg2 | 0.52124 | -0.20457193 | 0.194 | 0.375 | 1 | 23 |
| acadl | 0.5223 | -0.20233983 | 0.194 | 0.375 | 1 | 23 |
| si:dkey-42p14.3 | 0.52244 | -0.13960452 | 0.097 | 0.125 | 1 | 23 |
| ppm1aa | 0.52256 | 0.198683604 | 0.323 | 0.25 | 1 | 23 |
| vasna | 0.52294 | -0.29666619 | 0.226 | 0.375 | 1 | 23 |
| akap12b | 0.52365 | 0.373216906 | 0.452 | 0.25 | 1 | 23 |
| mmp15b | 0.52373 | 0.130338627 | 0.387 | 0.25 | 1 | 23 |
| hsbp1a | 0.52389 | -0.22886548 | 0.548 | 0.625 | 1 | 23 |
| si:dkey-276j7.1 | 0.52493 | 0.237902465 | 0.226 | 0.125 | 1 | 23 |
| dlst | 0.52551 | 0.176779385 | 0.323 | 0.25 | 1 | 23 |
| arglu1b | 0.52624 | -0.30865317 | 0.194 | 0.375 | 1 | 23 |
| btbd10b | 0.52656 | 0.179708431 | 0.161 | 0.125 | 1 | 23 |
| chd4b | 0.52659 | -0.14480363 | 0.452 | 0.625 | 1 | 23 |
| fam107b | 0.52664 | 0.230320363 | 0.452 | 0.25 | 1 | 23 |
| thoc5 | 0.52749 | -0.25243924 | 0.097 | 0.25 | 1 | 23 |
| rai14 | 0.52749 | -0.2159522 | 0.161 | 0.25 | 1 | 23 |
| si:dkey-188i13.9 | 0.52776 | -0.31526954 | 0.129 | 0.25 | 1 | 23 |
| bysl | 0.52839 | 0.203827363 | 0.29 | 0.125 | 1 | 23 |
| glo1 | 0.52845 | 0.13553359 | 0.419 | 0.25 | 1 | 23 |
| ckap4 | 0.52905 | -0.30511479 | 0.29 | 0.5 | 1 | 23 |
| vps35 | 0.52935 | -0.16056814 | 0.194 | 0.25 | 1 | 23 |
| aurkaip1 | 0.52939 | 0.274301463 | 0.29 | 0.125 | 1 | 23 |
| tagln2 | 0.52962 | 0.219411234 | 0.452 | 0.25 | 1 | 23 |
| naxe | 0.52963 | -0.172047 | 0.097 | 0.25 | 1 | 23 |
| si:ch211-191j22.3 | 0.53108 | -0.26957824 | 0.355 | 0.5 | 1 | 23 |
| rcn1 | 0.53125 | -0.2600102 | 0.097 | 0.25 | 1 | 23 |
| irf9 | 0.53158 | -0.12747449 | 0.129 | 0.25 | 1 | 23 |
| nars | 0.53213 | 0.160232589 | 0.419 | 0.25 | 1 | 23 |
| hsdl2 | 0.53303 | -0.24316167 | 0.097 | 0.25 | 1 | 23 |
| zgc:91910 | 0.5336 | 0.257893497 | 0.71 | 0.5 | 1 | 23 |
| slc4a4a | 0.53425 | -0.25019476 | 0.29 | 0.5 | 1 | 23 |
| arxa | 0.53438 | -0.14670012 | 0.097 | 0.125 | 1 | 23 |

| | | | | | | |
|---|---|---|---|---|---|---|
| zgc:112437 | 0.53455 | 0.167238488 | 0.258 | 0.125 | 1 | 23 |
| naga | 0.53469 | -0.25988209 | 0.097 | 0.25 | 1 | 23 |
| ralab | 0.53472 | -0.17706687 | 0.194 | 0.25 | 1 | 23 |
| lmnb1 | 0.53525 | -0.17432158 | 0.097 | 0.25 | 1 | 23 |
| ctsd | 0.5359 | -0.2072106 | 0.419 | 0.5 | 1 | 23 |
| hmgb3a | 0.53636 | 0.17030014 | 0.677 | 0.5 | 1 | 23 |
| LOC553316 | 0.53697 | -0.15613807 | 0.452 | 0.5 | 1 | 23 |
| nmi | 0.53786 | -0.24846483 | 0.29 | 0.5 | 1 | 23 |
| cx43.4 | 0.53795 | 0.233978341 | 0.226 | 0.125 | 1 | 23 |
| hspd1 | 0.53813 | -0.15738204 | 0.226 | 0.25 | 1 | 23 |
| oip5-as1 | 0.53816 | -0.2524424 | 0.129 | 0.25 | 1 | 23 |
| timm8a | 0.53817 | -0.23917004 | 0.097 | 0.25 | 1 | 23 |
| col6a4a | 0.53856 | 0.288200249 | 0.258 | 0.125 | 1 | 23 |
| dync1i2a | 0.54001 | -0.24359666 | 0.097 | 0.25 | 1 | 23 |
| iah1 | 0.54084 | 0.137647584 | 0.161 | 0.125 | 1 | 23 |
| eprs | 0.54106 | -0.20141096 | 0.097 | 0.25 | 1 | 23 |
| pik3r1 | 0.54116 | -0.25127641 | 0.097 | 0.25 | 1 | 23 |
| si:ch211-105c13.3 | 0.54147 | -0.29367944 | 0.29 | 0.5 | 1 | 23 |
| mrpl17 | 0.54168 | -0.14101158 | 0.097 | 0.125 | 1 | 23 |
| gdi1 | 0.54171 | -0.23550518 | 0.097 | 0.25 | 1 | 23 |
| atp5po | 0.54181 | 0.209992155 | 0.323 | 0.25 | 1 | 23 |
| ssr1 | 0.54181 | 0.270570799 | 0.419 | 0.25 | 1 | 23 |
| tmpoa | 0.54349 | -0.23199048 | 0.097 | 0.25 | 1 | 23 |
| tmem30ab | 0.54421 | 0.253855226 | 0.258 | 0.125 | 1 | 23 |
| nedd4l | 0.54432 | -0.27409846 | 0.097 | 0.25 | 1 | 23 |
| ccdc82 | 0.54448 | -0.28386176 | 0.097 | 0.25 | 1 | 23 |
| ctnnd1 | 0.54478 | -0.2165018 | 0.194 | 0.375 | 1 | 23 |
| ing4 | 0.54487 | -0.25369381 | 0.129 | 0.25 | 1 | 23 |
| stx18 | 0.54515 | -0.17425758 | 0.097 | 0.25 | 1 | 23 |
| ifi35 | 0.54698 | -0.21069094 | 0.097 | 0.25 | 1 | 23 |
| mgaa | 0.54748 | -0.169622 | 0.097 | 0.125 | 1 | 23 |
| abcc5 | 0.54753 | -0.27850047 | 0.129 | 0.25 | 1 | 23 |
| ndufa10 | 0.54758 | 0.169782028 | 0.194 | 0.125 | 1 | 23 |
| lamp2 | 0.54767 | -0.27836522 | 0.29 | 0.5 | 1 | 23 |
| gabarapb | 0.54797 | 0.238038126 | 0.71 | 0.5 | 1 | 23 |
| praf2 | 0.54811 | 0.234522087 | 0.258 | 0.125 | 1 | 23 |
| mrpl27 | 0.54813 | -0.2476503 | 0.097 | 0.25 | 1 | 23 |
| cul2 | 0.54814 | -0.21738075 | 0.097 | 0.25 | 1 | 23 |
| si:ch211-154o6.3 | 0.54826 | -0.11349143 | 0.129 | 0.25 | 1 | 23 |
| cox6a1 | 0.55096 | 0.266134851 | 0.774 | 0.625 | 1 | 23 |
| tle2b | 0.5517 | -0.30220358 | 0.194 | 0.375 | 1 | 23 |
| zfr | 0.55259 | -0.22176614 | 0.097 | 0.25 | 1 | 23 |
| ctr9 | 0.55291 | -0.25120497 | 0.097 | 0.25 | 1 | 23 |
| pdgfra | 0.55299 | 0.129667529 | 0.129 | 0.125 | 1 | 23 |
| XLOC-000431 | 0.5531 | -0.22264007 | 0.129 | 0.25 | 1 | 23 |
| mettl5 | 0.5536 | -0.19963889 | 0.097 | 0.25 | 1 | 23 |
| napab | 0.55418 | -0.24454279 | 0.226 | 0.375 | 1 | 23 |
| hs3st3b1b | 0.55419 | -0.27281115 | 0.097 | 0.25 | 1 | 23 |

| | | | | | | |
|---|---|---|---|---|---|---|
| stau2 | 0.55453 | 0.236346239 | 0.29 | 0.125 | 1 | 23 |
| tmsb | 0.55561 | -0.26605007 | 0.387 | 0.5 | 1 | 23 |
| si:rp71-45k5.4 | 0.55652 | -0.25679712 | 0.452 | 0.625 | 1 | 23 |
| trip12 | 0.55675 | -0.2161407 | 0.161 | 0.25 | 1 | 23 |
| ptpn21 | 0.55676 | -0.26962452 | 0.097 | 0.25 | 1 | 23 |
| kdm2bb | 0.55696 | -0.22912293 | 0.097 | 0.25 | 1 | 23 |
| sh3bgrl | 0.55864 | -0.22088313 | 0.29 | 0.375 | 1 | 23 |
| huwe1 | 0.55887 | -0.30175378 | 0.194 | 0.375 | 1 | 23 |
| bada | 0.55912 | -0.22853581 | 0.097 | 0.25 | 1 | 23 |
| cfap100 | 0.55934 | -0.24743468 | 0.097 | 0.25 | 1 | 23 |
| ubtf | 0.55943 | 0.258096046 | 0.29 | 0.125 | 1 | 23 |
| LOC110437774 | 0.56016 | -0.25507197 | 0.097 | 0.25 | 1 | 23 |
| alg5 | 0.56027 | -0.20689163 | 0.097 | 0.25 | 1 | 23 |
| anxa11a | 0.5611 | 0.222800587 | 0.258 | 0.125 | 1 | 23 |
| tnfsf10 | 0.5622 | -0.2547899 | 0.161 | 0.25 | 1 | 23 |
| pih1d1 | 0.56223 | -0.19566988 | 0.097 | 0.25 | 1 | 23 |
| si:dkey-187e18.1 | 0.56225 | -0.23717538 | 0.097 | 0.25 | 1 | 23 |
| grm4 | 0.56226 | -0.13975589 | 0.129 | 0.25 | 1 | 23 |
| zswim8 | 0.56228 | -0.22640219 | 0.097 | 0.25 | 1 | 23 |
| hoxc8a | 0.56272 | 0.164718533 | 0.258 | 0.125 | 1 | 23 |
| hikeshi | 0.56323 | -0.1963654 | 0.194 | 0.375 | 1 | 23 |
| sox4a | 0.56336 | -0.2240553 | 0.548 | 0.625 | 1 | 23 |
| ip6k2a | 0.56359 | -0.29236272 | 0.097 | 0.25 | 1 | 23 |
| rbfa | 0.56384 | -0.21181666 | 0.097 | 0.25 | 1 | 23 |
| zgc:66160 | 0.56443 | -0.18192411 | 0.097 | 0.25 | 1 | 23 |
| epsti1 | 0.56479 | -0.25037989 | 0.097 | 0.25 | 1 | 23 |
| lmf2b | 0.5649 | -0.17858577 | 0.097 | 0.25 | 1 | 23 |
| aif1l | 0.56498 | -0.24301051 | 0.097 | 0.25 | 1 | 23 |
| lamtor5 | 0.5651 | -0.20440991 | 0.097 | 0.25 | 1 | 23 |
| dtwd1 | 0.56543 | -0.21960405 | 0.097 | 0.25 | 1 | 23 |
| lipf | 0.56585 | 0.210435423 | 0.226 | 0.125 | 1 | 23 |
| lysmd4 | 0.56616 | -0.21002983 | 0.097 | 0.25 | 1 | 23 |
| tm2d1 | 0.56626 | -0.19431907 | 0.097 | 0.25 | 1 | 23 |
| arfgap1 | 0.56673 | -0.18803448 | 0.097 | 0.25 | 1 | 23 |
| zbtb8os | 0.56674 | -0.21079861 | 0.097 | 0.25 | 1 | 23 |
| tspan7 | 0.56684 | -0.24781303 | 0.097 | 0.25 | 1 | 23 |
| psmd13 | 0.56689 | -0.12814809 | 0.484 | 0.5 | 1 | 23 |
| mpped2a | 0.56707 | -0.14560453 | 0.097 | 0.125 | 1 | 23 |
| fam32a | 0.56751 | 0.28699925 | 0.452 | 0.25 | 1 | 23 |
| med10 | 0.56801 | 0.255887108 | 0.452 | 0.25 | 1 | 23 |
| ran | 0.56848 | -0.13233416 | 0.968 | 1 | 1 | 23 |
| hyou1 | 0.56931 | -0.16196309 | 0.194 | 0.25 | 1 | 23 |
| krt18b | 0.56992 | -0.27915946 | 0.419 | 0.625 | 1 | 23 |
| irf2a | 0.57043 | 0.150933601 | 0.161 | 0.125 | 1 | 23 |
| UBL4A | 0.57077 | 0.127511298 | 0.258 | 0.125 | 1 | 23 |
| zfand5b | 0.57118 | 0.276597077 | 0.452 | 0.25 | 1 | 23 |
| zgc:162944 | 0.57191 | 0.230762779 | 0.613 | 0.5 | 1 | 23 |
| tsc22d2 | 0.57213 | -0.24695049 | 0.129 | 0.25 | 1 | 23 |

| | | | | | | |
|---|---|---|---|---|---|---|
| hmgn6 | 0.57223 | 0.145830093 | 0.968 | 0.875 | 1 | 23 |
| hoxb3a | 0.57228 | -0.27046361 | 0.194 | 0.375 | 1 | 23 |
| bzw1b | 0.57229 | -0.21154148 | 0.548 | 0.75 | 1 | 23 |
| arf1 | 0.57278 | 0.104202899 | 0.355 | 0.375 | 1 | 23 |
| si:ch211-51e12.7 | 0.57303 | 0.114328348 | 0.516 | 0.5 | 1 | 23 |
| snap23.1 | 0.57363 | -0.13094894 | 0.097 | 0.125 | 1 | 23 |
| usp5 | 0.57561 | 0.294764214 | 0.29 | 0.125 | 1 | 23 |
| clasp2 | 0.57674 | -0.14033132 | 0.097 | 0.125 | 1 | 23 |
| csrnp1b | 0.57683 | -0.27184938 | 0.194 | 0.375 | 1 | 23 |
| ciao1 | 0.57744 | -0.19154513 | 0.161 | 0.25 | 1 | 23 |
| tfg | 0.57812 | -0.12263998 | 0.129 | 0.25 | 1 | 23 |
| BNC1 | 0.5806 | -0.2480633 | 0.129 | 0.25 | 1 | 23 |
| cd151l | 0.58086 | -0.23187336 | 0.419 | 0.625 | 1 | 23 |
| carhsp1 | 0.58103 | -0.24940454 | 0.194 | 0.375 | 1 | 23 |
| sptbn2 | 0.58141 | 0.28226239 | 0.258 | 0.125 | 1 | 23 |
| faf2 | 0.58172 | -0.14847657 | 0.226 | 0.375 | 1 | 23 |
| ddost | 0.58206 | 0.175271061 | 0.419 | 0.25 | 1 | 23 |
| inpp5ka | 0.58207 | -0.13610684 | 0.097 | 0.125 | 1 | 23 |
| eif2s2 | 0.58217 | 0.16487693 | 0.774 | 0.75 | 1 | 23 |
| tor1 | 0.58233 | 0.139564842 | 0.161 | 0.125 | 1 | 23 |
| chtopa | 0.58405 | -0.24942776 | 0.419 | 0.625 | 1 | 23 |
| c1qbp | 0.58513 | 0.284884209 | 0.29 | 0.125 | 1 | 23 |
| decr1 | 0.58553 | -0.22097193 | 0.161 | 0.25 | 1 | 23 |
| lysmd2 | 0.58601 | -0.249705 | 0.129 | 0.25 | 1 | 23 |
| pa2g4a | 0.5871 | 0.216694781 | 0.226 | 0.125 | 1 | 23 |
| rtn4b | 0.58741 | -0.11656347 | 0.129 | 0.25 | 1 | 23 |
| mrps35 | 0.58784 | -0.23940243 | 0.129 | 0.25 | 1 | 23 |
| ptp4a1 | 0.5886 | -0.1821528 | 0.323 | 0.5 | 1 | 23 |
| hmga2 | 0.58969 | 0.306043273 | 0.29 | 0.125 | 1 | 23 |
| zgc:112023 | 0.59001 | -0.11428695 | 0.129 | 0.25 | 1 | 23 |
| ube2l3a | 0.59297 | 0.248991938 | 0.258 | 0.125 | 1 | 23 |
| rpz5 | 0.59437 | -0.13796819 | 0.871 | 0.875 | 1 | 23 |
| calm2b | 0.59472 | -0.20979796 | 0.806 | 0.875 | 1 | 23 |
| tmed2 | 0.59509 | 0.176970869 | 0.419 | 0.375 | 1 | 23 |
| emc10 | 0.59515 | 0.133590052 | 0.29 | 0.25 | 1 | 23 |
| snw1 | 0.59553 | 0.294562756 | 0.29 | 0.125 | 1 | 23 |
| angptl4 | 0.59629 | -0.1264753 | 0.516 | 0.5 | 1 | 23 |
| sept2 | 0.59645 | 0.280250379 | 0.548 | 0.375 | 1 | 23 |
| tmeff1b | 0.59686 | -0.27734886 | 0.581 | 0.75 | 1 | 23 |
| yeats4 | 0.59753 | -0.12551008 | 0.226 | 0.25 | 1 | 23 |
| gcga | 0.59754 | 0.17241275 | 0.258 | 0.125 | 1 | 23 |
| ywhaqa | 0.59763 | 0.264671122 | 0.677 | 0.5 | 1 | 23 |
| kctd5a | 0.59764 | 0.169451187 | 0.548 | 0.375 | 1 | 23 |
| smarcb1a | 0.59779 | 0.150267553 | 0.258 | 0.125 | 1 | 23 |
| BX005309.1 | 0.59802 | -0.20895251 | 0.065 | 0.125 | 1 | 23 |
| BX465862.1 | 0.59802 | -0.1603341 | 0.065 | 0.125 | 1 | 23 |
| arhgap35a | 0.59802 | -0.13180848 | 0.065 | 0.125 | 1 | 23 |
| diaph2 | 0.59802 | -0.16576978 | 0.065 | 0.125 | 1 | 23 |

| | | | | | | |
|---|---|---|---|---|---|---|
| fam131bb | 0.59802 | -0.17667709 | 0.065 | 0.125 | 1 | 23 |
| iqce | 0.59802 | -0.12481942 | 0.065 | 0.125 | 1 | 23 |
| oxsr1a | 0.59802 | -0.18537366 | 0.065 | 0.125 | 1 | 23 |
| pcdh7a | 0.59802 | -0.16088124 | 0.065 | 0.125 | 1 | 23 |
| rasgef1bb | 0.59802 | -0.1428244 | 0.065 | 0.125 | 1 | 23 |
| si:dkey-266m15.5 | 0.59802 | -0.17268619 | 0.065 | 0.125 | 1 | 23 |
| zgc:154058 | 0.59802 | -0.1704321 | 0.065 | 0.125 | 1 | 23 |
| CABZ01088109.1 | 0.59802 | -0.18685397 | 0.065 | 0.125 | 1 | 23 |
| CT956002.2 | 0.59802 | -0.26704741 | 0.065 | 0.125 | 1 | 23 |
| PLBD2 | 0.59802 | -0.19857509 | 0.065 | 0.125 | 1 | 23 |
| abhd17aa | 0.59802 | -0.30598483 | 0.065 | 0.125 | 1 | 23 |
| acot9.1 | 0.59802 | -0.21066517 | 0.065 | 0.125 | 1 | 23 |
| ap1s3b | 0.59802 | -0.14859404 | 0.065 | 0.125 | 1 | 23 |
| casc3 | 0.59802 | -0.1195937 | 0.065 | 0.125 | 1 | 23 |
| ccdc73 | 0.59802 | -0.15549142 | 0.065 | 0.125 | 1 | 23 |
| cd302 | 0.59802 | -0.15833082 | 0.065 | 0.125 | 1 | 23 |
| cdc14b | 0.59802 | -0.27248397 | 0.065 | 0.125 | 1 | 23 |
| cog2 | 0.59802 | -0.1154665 | 0.065 | 0.125 | 1 | 23 |
| ctif | 0.59802 | -0.18358722 | 0.065 | 0.125 | 1 | 23 |
| cttnbp2nlb | 0.59802 | -0.17921839 | 0.065 | 0.125 | 1 | 23 |
| dusp4 | 0.59802 | -0.20842719 | 0.065 | 0.125 | 1 | 23 |
| dync1li1 | 0.59802 | -0.14620391 | 0.065 | 0.125 | 1 | 23 |
| enc2 | 0.59802 | -0.16377464 | 0.065 | 0.125 | 1 | 23 |
| fam192a | 0.59802 | -0.15928157 | 0.065 | 0.125 | 1 | 23 |
| fam198a | 0.59802 | -0.17292879 | 0.065 | 0.125 | 1 | 23 |
| frem2a | 0.59802 | -0.18516575 | 0.065 | 0.125 | 1 | 23 |
| grb2b | 0.59802 | -0.1445948 | 0.065 | 0.125 | 1 | 23 |
| hoxb10a | 0.59802 | -0.15185599 | 0.065 | 0.125 | 1 | 23 |
| ildr1b | 0.59802 | -0.15820563 | 0.065 | 0.125 | 1 | 23 |
| ilk | 0.59802 | -0.13441422 | 0.065 | 0.125 | 1 | 23 |
| isoc2 | 0.59802 | -0.1122594 | 0.065 | 0.125 | 1 | 23 |
| jarid2b | 0.59802 | -0.1129223 | 0.065 | 0.125 | 1 | 23 |
| ldlrap1a | 0.59802 | -0.20342136 | 0.065 | 0.125 | 1 | 23 |
| msh6 | 0.59802 | -0.13253512 | 0.065 | 0.125 | 1 | 23 |
| n6amt1 | 0.59802 | -0.20529059 | 0.065 | 0.125 | 1 | 23 |
| naa15a | 0.59802 | -0.21198124 | 0.065 | 0.125 | 1 | 23 |
| nlrc3l | 0.59802 | -0.18105545 | 0.065 | 0.125 | 1 | 23 |
| ntan1 | 0.59802 | -0.16027592 | 0.065 | 0.125 | 1 | 23 |
| pard6gb | 0.59802 | -0.16842848 | 0.065 | 0.125 | 1 | 23 |
| pigt | 0.59802 | -0.15800645 | 0.065 | 0.125 | 1 | 23 |
| pptc7b | 0.59802 | -0.20695458 | 0.065 | 0.125 | 1 | 23 |
| rab3gap2 | 0.59802 | -0.20896234 | 0.065 | 0.125 | 1 | 23 |
| rab40c | 0.59802 | -0.15375667 | 0.065 | 0.125 | 1 | 23 |
| rab4b | 0.59802 | -0.12019352 | 0.065 | 0.125 | 1 | 23 |
| riok1 | 0.59802 | -0.17689022 | 0.065 | 0.125 | 1 | 23 |
| rnd3a | 0.59802 | -0.22767559 | 0.065 | 0.125 | 1 | 23 |
| s100u | 0.59802 | -0.14782446 | 0.065 | 0.125 | 1 | 23 |
| sars | 0.59802 | -0.2573565 | 0.065 | 0.125 | 1 | 23 |

| | | | | | | |
|---|---|---|---|---|---|---|
| selenot1b | 0.59802 | -0.15184706 | 0.065 | 0.125 | 1 | 23 |
| sept9a | 0.59802 | -0.16186937 | 0.065 | 0.125 | 1 | 23 |
| si:ch211-11n16.2 | 0.59802 | -0.17657168 | 0.065 | 0.125 | 1 | 23 |
| si:ch211-281l24.3 | 0.59802 | -0.134698 | 0.065 | 0.125 | 1 | 23 |
| slc4a4b | 0.59802 | -0.1432265 | 0.065 | 0.125 | 1 | 23 |
| slkb | 0.59802 | -0.19487183 | 0.065 | 0.125 | 1 | 23 |
| socs6b | 0.59802 | -0.16510269 | 0.065 | 0.125 | 1 | 23 |
| sorl1 | 0.59802 | -0.26059789 | 0.065 | 0.125 | 1 | 23 |
| stk40 | 0.59802 | -0.15560847 | 0.065 | 0.125 | 1 | 23 |
| tp53bp2a | 0.59802 | -0.18403769 | 0.065 | 0.125 | 1 | 23 |
| ttc32 | 0.59802 | -0.16901511 | 0.065 | 0.125 | 1 | 23 |
| vps37b | 0.59802 | -0.16037469 | 0.065 | 0.125 | 1 | 23 |
| vrk3 | 0.59802 | -0.1385166 | 0.065 | 0.125 | 1 | 23 |
| xpo4 | 0.59802 | -0.18546088 | 0.065 | 0.125 | 1 | 23 |
| zgc:113030 | 0.59802 | -0.17856874 | 0.065 | 0.125 | 1 | 23 |
| znf292a | 0.59802 | -0.16070241 | 0.065 | 0.125 | 1 | 23 |
| znf367 | 0.59802 | -0.1834635 | 0.065 | 0.125 | 1 | 23 |
| CABZ01077217.1 | 0.59802 | -0.19562935 | 0.065 | 0.125 | 1 | 23 |
| LOC101884921 | 0.59802 | -0.14832361 | 0.065 | 0.125 | 1 | 23 |
| XLOC-004556 | 0.59802 | -0.1284348 | 0.065 | 0.125 | 1 | 23 |
| XLOC-011450 | 0.59802 | -0.12829627 | 0.065 | 0.125 | 1 | 23 |
| XLOC-022032 | 0.59802 | -0.12308819 | 0.065 | 0.125 | 1 | 23 |
| aatf | 0.59802 | -0.15262367 | 0.065 | 0.125 | 1 | 23 |
| acvr2ab | 0.59802 | -0.1398521 | 0.065 | 0.125 | 1 | 23 |
| ankhb | 0.59802 | -0.25151882 | 0.065 | 0.125 | 1 | 23 |
| ankle2 | 0.59802 | -0.14995646 | 0.065 | 0.125 | 1 | 23 |
| apeh | 0.59802 | -0.22231329 | 0.065 | 0.125 | 1 | 23 |
| aplp1 | 0.59802 | -0.25126772 | 0.065 | 0.125 | 1 | 23 |
| arhgap29b | 0.59802 | -0.15926544 | 0.065 | 0.125 | 1 | 23 |
| arrdc1a | 0.59802 | -0.14232842 | 0.065 | 0.125 | 1 | 23 |
| ascc3 | 0.59802 | -0.11342279 | 0.065 | 0.125 | 1 | 23 |
| bambia | 0.59802 | -0.2243561 | 0.065 | 0.125 | 1 | 23 |
| borcs5 | 0.59802 | -0.13363935 | 0.065 | 0.125 | 1 | 23 |
| ccdc51 | 0.59802 | -0.12798054 | 0.065 | 0.125 | 1 | 23 |
| cdk12 | 0.59802 | -0.15044592 | 0.065 | 0.125 | 1 | 23 |
| chd5 | 0.59802 | -0.17926244 | 0.065 | 0.125 | 1 | 23 |
| chico | 0.59802 | -0.24484885 | 0.065 | 0.125 | 1 | 23 |
| chst10 | 0.59802 | -0.20518163 | 0.065 | 0.125 | 1 | 23 |
| cltb | 0.59802 | -0.14973914 | 0.065 | 0.125 | 1 | 23 |
| coa7 | 0.59802 | -0.1283021 | 0.065 | 0.125 | 1 | 23 |
| col11a1b | 0.59802 | -0.1478292 | 0.065 | 0.125 | 1 | 23 |
| col4a3bpb | 0.59802 | -0.13714476 | 0.065 | 0.125 | 1 | 23 |
| cyb5r3 | 0.59802 | -0.15901484 | 0.065 | 0.125 | 1 | 23 |
| dbpb | 0.59802 | -0.27885262 | 0.065 | 0.125 | 1 | 23 |
| dgcr2 | 0.59802 | -0.29999415 | 0.065 | 0.125 | 1 | 23 |
| dhps | 0.59802 | -0.1181486 | 0.065 | 0.125 | 1 | 23 |
| dhx16 | 0.59802 | -0.12115914 | 0.065 | 0.125 | 1 | 23 |
| dhx58 | 0.59802 | -0.21261769 | 0.065 | 0.125 | 1 | 23 |

| | | | | | | |
|---|---|---|---|---|---|---|
| dnaja3a | 0.59802 | -0.13925907 | 0.065 | 0.125 | 1 | 23 |
| dpf2 | 0.59802 | -0.25485219 | 0.065 | 0.125 | 1 | 23 |
| ep400 | 0.59802 | -0.15682316 | 0.065 | 0.125 | 1 | 23 |
| ephx5 | 0.59802 | -0.24766245 | 0.065 | 0.125 | 1 | 23 |
| etnk1 | 0.59802 | -0.15219545 | 0.065 | 0.125 | 1 | 23 |
| evi5b | 0.59802 | -0.1280079 | 0.065 | 0.125 | 1 | 23 |
| fdxr | 0.59802 | -0.25120646 | 0.065 | 0.125 | 1 | 23 |
| fut8a | 0.59802 | -0.10899928 | 0.065 | 0.125 | 1 | 23 |
| gatad2ab | 0.59802 | -0.25329749 | 0.065 | 0.125 | 1 | 23 |
| gbp1 | 0.59802 | -0.16831193 | 0.065 | 0.125 | 1 | 23 |
| glod4 | 0.59802 | -0.14459906 | 0.065 | 0.125 | 1 | 23 |
| gnpda2 | 0.59802 | -0.15061816 | 0.065 | 0.125 | 1 | 23 |
| gnptab | 0.59802 | -0.20237003 | 0.065 | 0.125 | 1 | 23 |
| golph3 | 0.59802 | -0.25404503 | 0.065 | 0.125 | 1 | 23 |
| gpr158a | 0.59802 | -0.20546731 | 0.065 | 0.125 | 1 | 23 |
| hibch | 0.59802 | -0.15029507 | 0.065 | 0.125 | 1 | 23 |
| im:7151449 | 0.59802 | -0.20847538 | 0.065 | 0.125 | 1 | 23 |
| isca2 | 0.59802 | -0.20071019 | 0.065 | 0.125 | 1 | 23 |
| ivns1abpb | 0.59802 | -0.11589742 | 0.065 | 0.125 | 1 | 23 |
| kat6a | 0.59802 | -0.21095557 | 0.065 | 0.125 | 1 | 23 |
| katnb1 | 0.59802 | -0.1604287 | 0.065 | 0.125 | 1 | 23 |
| klf8 | 0.59802 | -0.17651339 | 0.065 | 0.125 | 1 | 23 |
| kxd1 | 0.59802 | -0.23366746 | 0.065 | 0.125 | 1 | 23 |
| lrrfip2 | 0.59802 | -0.1642645 | 0.065 | 0.125 | 1 | 23 |
| ly75 | 0.59802 | -0.24918319 | 0.065 | 0.125 | 1 | 23 |
| mad2l1 | 0.59802 | -0.22988966 | 0.065 | 0.125 | 1 | 23 |
| man2a2 | 0.59802 | -0.12766793 | 0.065 | 0.125 | 1 | 23 |
| mapk9 | 0.59802 | -0.15273702 | 0.065 | 0.125 | 1 | 23 |
| mbtd1 | 0.59802 | -0.14725402 | 0.065 | 0.125 | 1 | 23 |
| med12 | 0.59802 | -0.18094323 | 0.065 | 0.125 | 1 | 23 |
| mettl1 | 0.59802 | -0.11954971 | 0.065 | 0.125 | 1 | 23 |
| mkln1 | 0.59802 | -0.1258662 | 0.065 | 0.125 | 1 | 23 |
| mmp15a | 0.59802 | -0.17525284 | 0.065 | 0.125 | 1 | 23 |
| mrpl16 | 0.59802 | -0.20057427 | 0.065 | 0.125 | 1 | 23 |
| mxe | 0.59802 | -0.16190549 | 0.065 | 0.125 | 1 | 23 |
| mybbp1a | 0.59802 | -0.19889514 | 0.065 | 0.125 | 1 | 23 |
| nab1b | 0.59802 | -0.18613193 | 0.065 | 0.125 | 1 | 23 |
| nf2a | 0.59802 | -0.12883655 | 0.065 | 0.125 | 1 | 23 |
| nol7 | 0.59802 | -0.17432564 | 0.065 | 0.125 | 1 | 23 |
| nsd1a | 0.59802 | -0.15053026 | 0.065 | 0.125 | 1 | 23 |
| ntmt1 | 0.59802 | -0.27432057 | 0.065 | 0.125 | 1 | 23 |
| nup43 | 0.59802 | -0.13433806 | 0.065 | 0.125 | 1 | 23 |
| osgep | 0.59802 | -0.10304105 | 0.065 | 0.125 | 1 | 23 |
| oxa1l | 0.59802 | -0.12042975 | 0.065 | 0.125 | 1 | 23 |
| pdgfrl | 0.59802 | -0.168733 | 0.065 | 0.125 | 1 | 23 |
| per3 | 0.59802 | -0.16123809 | 0.065 | 0.125 | 1 | 23 |
| pgk1 | 0.59802 | -0.16473243 | 0.065 | 0.125 | 1 | 23 |
| phb2b | 0.59802 | -0.12214022 | 0.065 | 0.125 | 1 | 23 |

| | | | | | | |
|---|---|---|---|---|---|---|
| pkdccb | 0.59802 | -0.16938048 | 0.065 | 0.125 | 1 | 23 |
| polr2a | 0.59802 | -0.24773006 | 0.065 | 0.125 | 1 | 23 |
| ppm1bb | 0.59802 | -0.14127563 | 0.065 | 0.125 | 1 | 23 |
| ppp2r5ea | 0.59802 | -0.15368243 | 0.065 | 0.125 | 1 | 23 |
| purbb | 0.59802 | -0.15645948 | 0.065 | 0.125 | 1 | 23 |
| rasal2 | 0.59802 | -0.27249217 | 0.065 | 0.125 | 1 | 23 |
| rc3h1b | 0.59802 | -0.14071474 | 0.065 | 0.125 | 1 | 23 |
| rdh10b | 0.59802 | -0.12149435 | 0.065 | 0.125 | 1 | 23 |
| rest | 0.59802 | -0.15838856 | 0.065 | 0.125 | 1 | 23 |
| rfc5 | 0.59802 | -0.17863005 | 0.065 | 0.125 | 1 | 23 |
| rrm2 | 0.59802 | -0.36937869 | 0.065 | 0.125 | 1 | 23 |
| sco1 | 0.59802 | -0.12743415 | 0.065 | 0.125 | 1 | 23 |
| selenol | 0.59802 | -0.11054347 | 0.065 | 0.125 | 1 | 23 |
| sfxn4 | 0.59802 | -0.13114218 | 0.065 | 0.125 | 1 | 23 |
| sh3bgrl2 | 0.59802 | -0.19988231 | 0.065 | 0.125 | 1 | 23 |
| si:cabz01101003.1 | 0.59802 | -0.1512727 | 0.065 | 0.125 | 1 | 23 |
| si:ch211-161h7.4 | 0.59802 | -0.16838978 | 0.065 | 0.125 | 1 | 23 |
| si:ch211-71n6.4 | 0.59802 | -0.14963081 | 0.065 | 0.125 | 1 | 23 |
| si:ch73-237c6.1 | 0.59802 | -0.17013283 | 0.065 | 0.125 | 1 | 23 |
| si:ch73-257c13.2 | 0.59802 | -0.10315471 | 0.065 | 0.125 | 1 | 23 |
| si:ch73-311h14.2 | 0.59802 | -0.21051365 | 0.065 | 0.125 | 1 | 23 |
| si:dkey-101k6.5 | 0.59802 | -0.17734028 | 0.065 | 0.125 | 1 | 23 |
| si:dkey-154b15.1 | 0.59802 | -0.1112834 | 0.065 | 0.125 | 1 | 23 |
| slc25a25a | 0.59802 | -0.14957584 | 0.065 | 0.125 | 1 | 23 |
| smurf2 | 0.59802 | -0.24769469 | 0.065 | 0.125 | 1 | 23 |
| snx13 | 0.59802 | -0.14754419 | 0.065 | 0.125 | 1 | 23 |
| sppl2 | 0.59802 | -0.13344941 | 0.065 | 0.125 | 1 | 23 |
| sptlc1 | 0.59802 | -0.10818548 | 0.065 | 0.125 | 1 | 23 |
| sptlc3 | 0.59802 | -0.123201 | 0.065 | 0.125 | 1 | 23 |
| stk17b | 0.59802 | -0.29697987 | 0.065 | 0.125 | 1 | 23 |
| stox2a | 0.59802 | -0.10680687 | 0.065 | 0.125 | 1 | 23 |
| stx2a | 0.59802 | -0.1562061 | 0.065 | 0.125 | 1 | 23 |
| sult1st6 | 0.59802 | -0.13226046 | 0.065 | 0.125 | 1 | 23 |
| taf2 | 0.59802 | -0.12810103 | 0.065 | 0.125 | 1 | 23 |
| tcerg1a | 0.59802 | -0.11757575 | 0.065 | 0.125 | 1 | 23 |
| tcirg1b | 0.59802 | -0.17178099 | 0.065 | 0.125 | 1 | 23 |
| tgfbr3 | 0.59802 | -0.35550254 | 0.065 | 0.125 | 1 | 23 |
| thap12a | 0.59802 | -0.12790322 | 0.065 | 0.125 | 1 | 23 |
| tmem147 | 0.59802 | -0.26591141 | 0.065 | 0.125 | 1 | 23 |
| tmem222a | 0.59802 | -0.17402867 | 0.065 | 0.125 | 1 | 23 |
| tnrc6b | 0.59802 | -0.20533705 | 0.065 | 0.125 | 1 | 23 |
| trpm6 | 0.59802 | -0.16888792 | 0.065 | 0.125 | 1 | 23 |
| tspan36 | 0.59802 | -0.11658862 | 0.065 | 0.125 | 1 | 23 |
| vkorc1l1 | 0.59802 | -0.17559273 | 0.065 | 0.125 | 1 | 23 |
| wu:fi36c02 | 0.59802 | -0.18904543 | 0.065 | 0.125 | 1 | 23 |
| xiap | 0.59802 | -0.15027788 | 0.065 | 0.125 | 1 | 23 |
| yes1 | 0.59802 | -0.16522447 | 0.065 | 0.125 | 1 | 23 |
| ythdf3 | 0.59802 | -0.21606457 | 0.065 | 0.125 | 1 | 23 |

| | | | | | | |
|---|---|---|---|---|---|---|
| zbtb21 | 0.59802 | -0.11680686 | 0.065 | 0.125 | 1 | 23 |
| zfand4 | 0.59802 | -0.12651688 | 0.065 | 0.125 | 1 | 23 |
| zgc:101130 | 0.59802 | -0.14645778 | 0.065 | 0.125 | 1 | 23 |
| zgc:113411 | 0.59802 | -0.12205654 | 0.065 | 0.125 | 1 | 23 |
| zgc:77151 | 0.59802 | -0.1504629 | 0.065 | 0.125 | 1 | 23 |
| zgc:85936 | 0.59802 | -0.13965369 | 0.065 | 0.125 | 1 | 23 |
| zhx3 | 0.59802 | -0.22789453 | 0.065 | 0.125 | 1 | 23 |
| znf1094 | 0.59802 | -0.12119177 | 0.065 | 0.125 | 1 | 23 |
| znf326 | 0.59802 | -0.21656203 | 0.065 | 0.125 | 1 | 23 |
| znf609a | 0.59802 | -0.18290604 | 0.065 | 0.125 | 1 | 23 |
| znf770 | 0.59802 | -0.11293228 | 0.065 | 0.125 | 1 | 23 |
| CABZ01083448.1 | 0.59802 | -0.14034341 | 0.065 | 0.125 | 1 | 23 |
| FP102158.1 | 0.59802 | -0.16957401 | 0.065 | 0.125 | 1 | 23 |
| LOC100330186 | 0.59802 | -0.18037598 | 0.065 | 0.125 | 1 | 23 |
| LOC101886337 | 0.59802 | -0.25738481 | 0.065 | 0.125 | 1 | 23 |
| LOC110437782 | 0.59802 | -0.15529844 | 0.065 | 0.125 | 1 | 23 |
| LOC110439998 | 0.59802 | -0.13275263 | 0.065 | 0.125 | 1 | 23 |
| XLOC-043861 | 0.59802 | -0.19581593 | 0.065 | 0.125 | 1 | 23 |
| XLOC-044204 | 0.59802 | -0.30853621 | 0.065 | 0.125 | 1 | 23 |
| abhd14a | 0.59802 | -0.14796074 | 0.065 | 0.125 | 1 | 23 |
| acbd6 | 0.59802 | -0.15482466 | 0.065 | 0.125 | 1 | 23 |
| bend5 | 0.59802 | -0.23972284 | 0.065 | 0.125 | 1 | 23 |
| bloc1s1 | 0.59802 | -0.21561465 | 0.065 | 0.125 | 1 | 23 |
| cntfr | 0.59802 | -0.10393684 | 0.065 | 0.125 | 1 | 23 |
| cpox | 0.59802 | -0.18147652 | 0.065 | 0.125 | 1 | 23 |
| crk | 0.59802 | -0.17343137 | 0.065 | 0.125 | 1 | 23 |
| cry1ba | 0.59802 | -0.14456603 | 0.065 | 0.125 | 1 | 23 |
| dcbld2 | 0.59802 | -0.13527181 | 0.065 | 0.125 | 1 | 23 |
| dcps | 0.59802 | -0.18643496 | 0.065 | 0.125 | 1 | 23 |
| dnmt3ba | 0.59802 | -0.23116034 | 0.065 | 0.125 | 1 | 23 |
| dnph1 | 0.59802 | -0.21034725 | 0.065 | 0.125 | 1 | 23 |
| ece1 | 0.59802 | -0.2025105 | 0.065 | 0.125 | 1 | 23 |
| enoph1 | 0.59802 | -0.13032761 | 0.065 | 0.125 | 1 | 23 |
| fbrsl1 | 0.59802 | -0.21475077 | 0.065 | 0.125 | 1 | 23 |
| gmps | 0.59802 | -0.22116089 | 0.065 | 0.125 | 1 | 23 |
| hbs1l | 0.59802 | -0.21069211 | 0.065 | 0.125 | 1 | 23 |
| igdcc3 | 0.59802 | -0.16228123 | 0.065 | 0.125 | 1 | 23 |
| mafa | 0.59802 | -0.22045349 | 0.065 | 0.125 | 1 | 23 |
| mettl14 | 0.59802 | -0.17268769 | 0.065 | 0.125 | 1 | 23 |
| msto1 | 0.59802 | -0.11346527 | 0.065 | 0.125 | 1 | 23 |
| nfkb2 | 0.59802 | -0.12964316 | 0.065 | 0.125 | 1 | 23 |
| osbpl10b | 0.59802 | -0.14485064 | 0.065 | 0.125 | 1 | 23 |
| pcdh19 | 0.59802 | -0.19485226 | 0.065 | 0.125 | 1 | 23 |
| pdgfab | 0.59802 | -0.10392526 | 0.065 | 0.125 | 1 | 23 |
| ppil1 | 0.59802 | -0.17153181 | 0.065 | 0.125 | 1 | 23 |
| prr12b | 0.59802 | -0.18201313 | 0.065 | 0.125 | 1 | 23 |
| rnf11a | 0.59802 | -0.12677673 | 0.065 | 0.125 | 1 | 23 |
| rnf213b | 0.59802 | -0.10100782 | 0.065 | 0.125 | 1 | 23 |

| | | | | | | |
|---|---|---|---|---|---|---|
| rps6ka5 | 0.59802 | -0.14987471 | 0.065 | 0.125 | 1 | 23 |
| setdb1a | 0.59802 | -0.14104365 | 0.065 | 0.125 | 1 | 23 |
| sfrp1a | 0.59802 | -0.26612103 | 0.065 | 0.125 | 1 | 23 |
| si:ch211-197h24.6 | 0.59802 | -0.20849708 | 0.065 | 0.125 | 1 | 23 |
| si:dkey-10p5.10 | 0.59802 | -0.15490539 | 0.065 | 0.125 | 1 | 23 |
| slbp | 0.59802 | -0.3164423 | 0.065 | 0.125 | 1 | 23 |
| slc30a6 | 0.59802 | -0.12867388 | 0.065 | 0.125 | 1 | 23 |
| slco5a1a | 0.59802 | -0.15670932 | 0.065 | 0.125 | 1 | 23 |
| stard8 | 0.59802 | -0.11186038 | 0.065 | 0.125 | 1 | 23 |
| strn4 | 0.59802 | -0.12608373 | 0.065 | 0.125 | 1 | 23 |
| tnfrsf19 | 0.59802 | -0.20887603 | 0.065 | 0.125 | 1 | 23 |
| tpm4b | 0.59802 | -0.17274749 | 0.065 | 0.125 | 1 | 23 |
| trappc3 | 0.59802 | -0.22582717 | 0.065 | 0.125 | 1 | 23 |
| trim8b | 0.59802 | -0.16541294 | 0.065 | 0.125 | 1 | 23 |
| BX005254.1 | 0.59802 | -0.13091456 | 0.065 | 0.125 | 1 | 23 |
| XLOC-040069 | 0.59802 | -0.21763271 | 0.065 | 0.125 | 1 | 23 |
| cep63 | 0.59802 | -0.1765223 | 0.065 | 0.125 | 1 | 23 |
| fam84b | 0.59802 | -0.15559179 | 0.065 | 0.125 | 1 | 23 |
| gatb | 0.59802 | -0.16731103 | 0.065 | 0.125 | 1 | 23 |
| grb10a | 0.59802 | -0.17655374 | 0.065 | 0.125 | 1 | 23 |
| iars | 0.59802 | -0.19011036 | 0.065 | 0.125 | 1 | 23 |
| ide | 0.59802 | -0.16903147 | 0.065 | 0.125 | 1 | 23 |
| lrrcc1 | 0.59802 | -0.12785173 | 0.065 | 0.125 | 1 | 23 |
| myof | 0.59802 | -0.11535596 | 0.065 | 0.125 | 1 | 23 |
| si:dkey-16p21.8 | 0.59802 | -0.11801922 | 0.065 | 0.125 | 1 | 23 |
| slain2 | 0.59802 | -0.18348376 | 0.065 | 0.125 | 1 | 23 |
| znf384l | 0.59802 | -0.14900547 | 0.065 | 0.125 | 1 | 23 |
| gorab | 0.59802 | -0.21238884 | 0.065 | 0.125 | 1 | 23 |
| cox14 | 0.59824 | 0.165031262 | 0.419 | 0.25 | 1 | 23 |
| fmr1 | 0.59876 | 0.218154529 | 0.258 | 0.125 | 1 | 23 |
| shoc2 | 0.59904 | 0.222107441 | 0.226 | 0.125 | 1 | 23 |
| kdelr2b | 0.59956 | 0.230521013 | 0.29 | 0.125 | 1 | 23 |
| rbpjb | 0.59991 | -0.12832118 | 0.097 | 0.125 | 1 | 23 |
| exoc5 | 0.60048 | -0.13688744 | 0.097 | 0.125 | 1 | 23 |
| cops9 | 0.60116 | 0.236644166 | 0.29 | 0.125 | 1 | 23 |
| myclb | 0.60165 | 0.277025454 | 0.29 | 0.125 | 1 | 23 |
| zgc:92335 | 0.60171 | -0.12594415 | 0.097 | 0.125 | 1 | 23 |
| vdac1 | 0.60278 | 0.187234652 | 0.452 | 0.375 | 1 | 23 |
| acvr2ba | 0.60327 | -0.12225916 | 0.097 | 0.125 | 1 | 23 |
| cd151 | 0.60374 | 0.162117087 | 0.194 | 0.125 | 1 | 23 |
| sec22bb | 0.60381 | -0.12176972 | 0.097 | 0.125 | 1 | 23 |
| lbr | 0.60413 | -0.13340659 | 0.097 | 0.125 | 1 | 23 |
| zgc:163080 | 0.60485 | 0.192889442 | 0.29 | 0.125 | 1 | 23 |
| nfkbiaa | 0.60541 | 0.300200273 | 0.29 | 0.125 | 1 | 23 |
| mkrn1 | 0.60637 | -0.26088328 | 0.258 | 0.375 | 1 | 23 |
| LOC103909115 | 0.6064 | -0.22563861 | 0.129 | 0.25 | 1 | 23 |
| dcn | 0.60657 | 0.353906315 | 0.29 | 0.125 | 1 | 23 |
| mrpl42 | 0.60672 | 0.214480529 | 0.29 | 0.125 | 1 | 23 |

| | | | | | | |
|---|---|---|---|---|---|---|
| mrpl54 | 0.60673 | 0.206883555 | 0.29 | 0.125 | 1 | 23 |
| zgc:92313 | 0.60736 | 0.221070243 | 0.226 | 0.125 | 1 | 23 |
| mrps7 | 0.60803 | 0.228035289 | 0.258 | 0.125 | 1 | 23 |
| gng12a | 0.60895 | 0.208119245 | 0.258 | 0.125 | 1 | 23 |
| thoc1 | 0.61017 | 0.206372033 | 0.226 | 0.125 | 1 | 23 |
| col18a1a | 0.61127 | -0.12949694 | 0.097 | 0.125 | 1 | 23 |
| cox7a2l | 0.61288 | 0.179742915 | 0.419 | 0.25 | 1 | 23 |
| fryb | 0.61297 | -0.22342995 | 0.161 | 0.25 | 1 | 23 |
| rwdd1 | 0.613 | 0.17791345 | 0.419 | 0.25 | 1 | 23 |
| tprg1 | 0.61371 | -0.11406762 | 0.097 | 0.125 | 1 | 23 |
| cct6a | 0.61426 | 0.23161789 | 0.645 | 0.5 | 1 | 23 |
| tardbp | 0.61432 | -0.22995184 | 0.355 | 0.5 | 1 | 23 |
| phf10 | 0.61573 | -0.11909398 | 0.097 | 0.125 | 1 | 23 |
| eif4g2a | 0.61606 | -0.23169406 | 0.258 | 0.375 | 1 | 23 |
| kmt2cb | 0.61647 | -0.20823084 | 0.29 | 0.375 | 1 | 23 |
| timm8b | 0.61658 | 0.223901746 | 0.258 | 0.125 | 1 | 23 |
| srsf3b | 0.61695 | -0.14042661 | 0.742 | 0.75 | 1 | 23 |
| tagln | 0.61836 | -0.12415837 | 0.129 | 0.25 | 1 | 23 |
| ndufc2 | 0.61868 | 0.255946319 | 0.548 | 0.375 | 1 | 23 |
| tvp23b | 0.61941 | 0.158585877 | 0.194 | 0.125 | 1 | 23 |
| elavl1a | 0.61949 | -0.179445 | 0.323 | 0.5 | 1 | 23 |
| khdrbs1a | 0.61958 | -0.13033173 | 0.935 | 1 | 1 | 23 |
| ggcta | 0.62015 | -0.18879279 | 0.161 | 0.25 | 1 | 23 |
| prmt1 | 0.62073 | -0.16537386 | 0.452 | 0.5 | 1 | 23 |
| sucla2 | 0.6214 | 0.136903932 | 0.161 | 0.125 | 1 | 23 |
| zgc:158343 | 0.62144 | 0.119312335 | 0.387 | 0.25 | 1 | 23 |
| dcaf7 | 0.62179 | -0.15876275 | 0.323 | 0.375 | 1 | 23 |
| si:ch211-39k3.2 | 0.62215 | 0.185293677 | 0.226 | 0.125 | 1 | 23 |
| tpm4a | 0.62242 | 0.141733349 | 0.774 | 0.75 | 1 | 23 |
| marcksl1b | 0.62245 | -0.27191949 | 0.323 | 0.5 | 1 | 23 |
| eif1b | 0.62303 | -0.13779632 | 0.839 | 0.875 | 1 | 23 |
| raver2 | 0.62326 | -0.1195862 | 0.097 | 0.125 | 1 | 23 |
| churc1 | 0.62378 | 0.121562825 | 0.387 | 0.25 | 1 | 23 |
| ndufa3 | 0.62385 | -0.19890634 | 0.548 | 0.625 | 1 | 23 |
| znf1001 | 0.62424 | 0.145886512 | 0.161 | 0.125 | 1 | 23 |
| tra2a | 0.62561 | -0.245682 | 0.387 | 0.5 | 1 | 23 |
| nck1b | 0.62667 | 0.119694593 | 0.29 | 0.25 | 1 | 23 |
| march6 | 0.62687 | 0.231266784 | 0.226 | 0.125 | 1 | 23 |
| tomm6 | 0.62688 | 0.160250996 | 0.419 | 0.375 | 1 | 23 |
| CABZ01112656.1 | 0.6271 | 0.154760946 | 0.194 | 0.125 | 1 | 23 |
| MCMDC2 | 0.62809 | 0.199504524 | 0.226 | 0.125 | 1 | 23 |
| sarnp | 0.62825 | -0.25429285 | 0.226 | 0.375 | 1 | 23 |
| znfx1 | 0.62923 | -0.12192925 | 0.226 | 0.25 | 1 | 23 |
| tmem167a | 0.62935 | 0.188983425 | 0.226 | 0.125 | 1 | 23 |
| smarcd1 | 0.63007 | -0.24875077 | 0.323 | 0.5 | 1 | 23 |
| inip | 0.63053 | 0.185561241 | 0.226 | 0.125 | 1 | 23 |
| bud31 | 0.63086 | 0.17835978 | 0.419 | 0.25 | 1 | 23 |
| kdm7ab | 0.63087 | 0.148801527 | 0.161 | 0.125 | 1 | 23 |

| | | | | | | |
|---|---|---|---|---|---|---|
| atp5f1c | 0.63126 | 0.248483622 | 0.645 | 0.5 | 1 | 23 |
| atp5mea | 0.6327 | -0.2571029 | 0.355 | 0.5 | 1 | 23 |
| ppm1g | 0.63341 | -0.14017408 | 0.226 | 0.375 | 1 | 23 |
| spcs3 | 0.63351 | 0.216228461 | 0.484 | 0.375 | 1 | 23 |
| rpn1 | 0.63362 | -0.24339579 | 0.355 | 0.5 | 1 | 23 |
| ext2 | 0.6345 | 0.176331289 | 0.258 | 0.125 | 1 | 23 |
| ftsj3 | 0.63584 | -0.18388962 | 0.161 | 0.25 | 1 | 23 |
| fzd4 | 0.63617 | -0.19105518 | 0.194 | 0.25 | 1 | 23 |
| hoxb6a | 0.63634 | 0.218622604 | 0.226 | 0.125 | 1 | 23 |
| ciapin1 | 0.6367 | 0.224508761 | 0.258 | 0.125 | 1 | 23 |
| etfb | 0.63725 | 0.19141107 | 0.355 | 0.25 | 1 | 23 |
| mgst3b | 0.63725 | -0.15940108 | 0.548 | 0.625 | 1 | 23 |
| slc25a55a | 0.63753 | -0.24617524 | 0.323 | 0.5 | 1 | 23 |
| anapc16 | 0.63827 | -0.11716886 | 0.129 | 0.25 | 1 | 23 |
| irf2bp2b | 0.6384 | 0.198979077 | 0.419 | 0.25 | 1 | 23 |
| exosc4 | 0.6402 | 0.179047993 | 0.226 | 0.125 | 1 | 23 |
| si:ch211-133n4.4 | 0.64025 | 0.23252264 | 0.258 | 0.125 | 1 | 23 |
| tgfbr2b | 0.6404 | -0.12308016 | 0.097 | 0.125 | 1 | 23 |
| tia1l | 0.64053 | -0.26261012 | 0.129 | 0.25 | 1 | 23 |
| cd248a | 0.64314 | -0.14615683 | 0.129 | 0.25 | 1 | 23 |
| rpe | 0.64405 | 0.158725812 | 0.194 | 0.125 | 1 | 23 |
| tubb5 | 0.64438 | -0.1047422 | 0.258 | 0.375 | 1 | 23 |
| si:rp71-36a1.1 | 0.64496 | -0.23985891 | 0.161 | 0.25 | 1 | 23 |
| commd9 | 0.6452 | 0.123419027 | 0.161 | 0.125 | 1 | 23 |
| wdr45 | 0.64558 | -0.24901896 | 0.226 | 0.375 | 1 | 23 |
| ftr14 | 0.64609 | 0.189346631 | 0.226 | 0.125 | 1 | 23 |
| crim1 | 0.64721 | -0.24255897 | 0.129 | 0.25 | 1 | 23 |
| chmp2a | 0.64723 | 0.227803847 | 0.258 | 0.125 | 1 | 23 |
| zgc:55733 | 0.64865 | 0.179215987 | 0.194 | 0.125 | 1 | 23 |
| pros1 | 0.64977 | -0.20464379 | 0.161 | 0.25 | 1 | 23 |
| tmed10 | 0.65003 | 0.171779998 | 0.452 | 0.375 | 1 | 23 |
| pax2a | 0.65006 | 0.183428112 | 0.258 | 0.125 | 1 | 23 |
| ap2m1a | 0.65055 | 0.182661278 | 0.677 | 0.5 | 1 | 23 |
| tardbpl | 0.65065 | -0.23551253 | 0.452 | 0.625 | 1 | 23 |
| xaf1 | 0.65087 | 0.11658783 | 0.258 | 0.25 | 1 | 23 |
| tmem256 | 0.65151 | 0.218492064 | 0.258 | 0.125 | 1 | 23 |
| jam2b | 0.65188 | -0.24259185 | 0.323 | 0.5 | 1 | 23 |
| lxn | 0.65209 | 0.120929934 | 0.226 | 0.125 | 1 | 23 |
| igf2bp2b | 0.65256 | -0.225414 | 0.226 | 0.375 | 1 | 23 |
| cdk9 | 0.65276 | 0.181258216 | 0.226 | 0.125 | 1 | 23 |
| dnase1l1 | 0.65285 | -0.18581668 | 0.129 | 0.25 | 1 | 23 |
| mych | 0.65292 | 0.130531183 | 0.29 | 0.25 | 1 | 23 |
| ndufs6 | 0.65315 | 0.230616467 | 0.516 | 0.375 | 1 | 23 |
| uprt | 0.65337 | 0.153600743 | 0.194 | 0.125 | 1 | 23 |
| rpl36 | 0.65431 | 0.111156577 | 0.968 | 0.875 | 1 | 23 |
| pik3ca | 0.65432 | -0.11987257 | 0.097 | 0.125 | 1 | 23 |
| ehmt1a | 0.65441 | -0.21323288 | 0.161 | 0.25 | 1 | 23 |
| tmem238a | 0.65451 | 0.212252081 | 0.419 | 0.25 | 1 | 23 |

| | | | | | | |
|---|---|---|---|---|---|---|
| rap1b | 0.65501 | 0.243762693 | 0.516 | 0.375 | 1 | 23 |
| NPAS3 | 0.65504 | -0.15034636 | 0.129 | 0.25 | 1 | 23 |
| cebpb | 0.65521 | -0.22565418 | 0.581 | 0.625 | 1 | 23 |
| tmed9 | 0.65607 | 0.196280587 | 0.226 | 0.125 | 1 | 23 |
| hmcn2 | 0.65667 | 0.190178098 | 0.194 | 0.125 | 1 | 23 |
| u2af2b | 0.65669 | 0.167833557 | 0.258 | 0.125 | 1 | 23 |
| rsrc2 | 0.65824 | -0.21998319 | 0.226 | 0.375 | 1 | 23 |
| zgc:110158 | 0.65836 | -0.11885364 | 0.097 | 0.125 | 1 | 23 |
| ext1a | 0.659 | 0.117568085 | 0.161 | 0.125 | 1 | 23 |
| wtap | 0.65981 | 0.179335607 | 0.226 | 0.125 | 1 | 23 |
| zgc:56304 | 0.65998 | 0.201611637 | 0.419 | 0.25 | 1 | 23 |
| adh5 | 0.66023 | 0.241546574 | 0.645 | 0.5 | 1 | 23 |
| sptssa | 0.6615 | 0.196021274 | 0.226 | 0.125 | 1 | 23 |
| calm3b | 0.66157 | -0.21523101 | 0.581 | 0.75 | 1 | 23 |
| sae1 | 0.66225 | 0.179096902 | 0.258 | 0.125 | 1 | 23 |
| arf2a | 0.66305 | -0.19392864 | 0.161 | 0.25 | 1 | 23 |
| fosl1a | 0.66363 | -0.25599041 | 0.258 | 0.375 | 1 | 23 |
| hvcn1 | 0.66372 | 0.212856025 | 0.258 | 0.125 | 1 | 23 |
| psmd7 | 0.66424 | -0.15456252 | 0.452 | 0.5 | 1 | 23 |
| calcoco1a | 0.66436 | -0.22679114 | 0.226 | 0.375 | 1 | 23 |
| tmem134 | 0.66438 | -0.1565553 | 0.194 | 0.25 | 1 | 23 |
| znf503 | 0.66441 | -0.25933555 | 0.226 | 0.375 | 1 | 23 |
| homer3b | 0.66452 | -0.18470757 | 0.161 | 0.25 | 1 | 23 |
| pafah1b1a | 0.66593 | -0.23955322 | 0.129 | 0.25 | 1 | 23 |
| BX511129.1 | 0.66612 | 0.129151803 | 0.161 | 0.125 | 1 | 23 |
| atp1b3b | 0.66615 | 0.137901141 | 0.161 | 0.125 | 1 | 23 |
| atf4a | 0.66681 | -0.24741238 | 0.355 | 0.5 | 1 | 23 |
| zyx | 0.66699 | -0.16021461 | 0.129 | 0.25 | 1 | 23 |
| zbtb46 | 0.66707 | -0.24661749 | 0.129 | 0.25 | 1 | 23 |
| ppp2cb | 0.66731 | 0.129132753 | 0.516 | 0.375 | 1 | 23 |
| eif1axa | 0.66757 | 0.23059338 | 0.419 | 0.25 | 1 | 23 |
| si:ch73-138n13.1 | 0.66761 | -0.14030051 | 0.129 | 0.25 | 1 | 23 |
| prdx5 | 0.66807 | 0.197479837 | 0.581 | 0.5 | 1 | 23 |
| sec11a | 0.66842 | 0.124313666 | 0.387 | 0.375 | 1 | 23 |
| si:ch211-39i22.1 | 0.66904 | -0.24974712 | 0.452 | 0.625 | 1 | 23 |
| glod5 | 0.66917 | -0.21882396 | 0.387 | 0.5 | 1 | 23 |
| supt4h1 | 0.66957 | -0.19303039 | 0.129 | 0.25 | 1 | 23 |
| vaspb | 0.67044 | 0.118355357 | 0.161 | 0.125 | 1 | 23 |
| snip1 | 0.67051 | 0.19144376 | 0.226 | 0.125 | 1 | 23 |
| eif5a | 0.67084 | -0.11958518 | 0.871 | 0.875 | 1 | 23 |
| cbx1b | 0.6709 | 0.177585455 | 0.258 | 0.125 | 1 | 23 |
| smarce1 | 0.67094 | -0.19195801 | 0.581 | 0.75 | 1 | 23 |
| tefa | 0.67143 | 0.179222991 | 0.355 | 0.25 | 1 | 23 |
| wdr83os | 0.67162 | 0.18340593 | 0.258 | 0.125 | 1 | 23 |
| dbt | 0.6722 | -0.20818978 | 0.129 | 0.25 | 1 | 23 |
| fn1a | 0.67257 | -0.10362571 | 0.258 | 0.375 | 1 | 23 |
| cd99 | 0.67265 | -0.25429763 | 0.129 | 0.25 | 1 | 23 |
| sf3b3 | 0.67268 | -0.11653867 | 0.097 | 0.125 | 1 | 23 |

| | | | | | | |
|---|---|---|---|---|---|---|
| nudt12 | 0.67275 | -0.20407436 | 0.129 | 0.25 | 1 | 23 |
| ndfip1 | 0.6729 | 0.178427758 | 0.452 | 0.375 | 1 | 23 |
| itsn2a | 0.67313 | -0.19365496 | 0.161 | 0.25 | 1 | 23 |
| sipa1l2 | 0.67329 | -0.15527973 | 0.194 | 0.25 | 1 | 23 |
| tnk1 | 0.67331 | -0.11598371 | 0.097 | 0.125 | 1 | 23 |
| zgc:92744 | 0.67367 | 0.252568758 | 0.258 | 0.125 | 1 | 23 |
| dusp14 | 0.67464 | 0.154900994 | 0.161 | 0.125 | 1 | 23 |
| clptm1 | 0.675 | 0.176652835 | 0.226 | 0.125 | 1 | 23 |
| echs1 | 0.67526 | 0.124795626 | 0.226 | 0.125 | 1 | 23 |
| tm2d2 | 0.67534 | 0.216211191 | 0.387 | 0.25 | 1 | 23 |
| ap1m1 | 0.67557 | -0.24764047 | 0.226 | 0.375 | 1 | 23 |
| tab1 | 0.67574 | 0.157986596 | 0.258 | 0.125 | 1 | 23 |
| tma7 | 0.67576 | -0.19271634 | 0.677 | 0.75 | 1 | 23 |
| si:ch211-108c17.2 | 0.67606 | -0.11018483 | 0.097 | 0.125 | 1 | 23 |
| ammecr1 | 0.67686 | -0.21959727 | 0.161 | 0.25 | 1 | 23 |
| smc5 | 0.67742 | -0.19573724 | 0.129 | 0.25 | 1 | 23 |
| eci1 | 0.67792 | 0.16630708 | 0.258 | 0.125 | 1 | 23 |
| dpy30 | 0.67872 | -0.14549553 | 0.581 | 0.625 | 1 | 23 |
| frg1 | 0.67887 | -0.11324408 | 0.097 | 0.125 | 1 | 23 |
| emg1 | 0.67947 | 0.198019927 | 0.258 | 0.125 | 1 | 23 |
| ik | 0.67979 | 0.161869894 | 0.258 | 0.125 | 1 | 23 |
| sub1b | 0.68004 | -0.2269153 | 0.355 | 0.5 | 1 | 23 |
| mrpl30 | 0.68018 | -0.22440313 | 0.226 | 0.375 | 1 | 23 |
| ccnq | 0.68031 | 0.105953413 | 0.129 | 0.125 | 1 | 23 |
| fzd7b | 0.68101 | -0.19239083 | 0.194 | 0.25 | 1 | 23 |
| si:ch211-286o17.1 | 0.68133 | -0.20200306 | 0.226 | 0.375 | 1 | 23 |
| usf1l | 0.68179 | 0.194985854 | 0.258 | 0.125 | 1 | 23 |
| tcta | 0.68182 | 0.138012293 | 0.387 | 0.25 | 1 | 23 |
| sumo1 | 0.68253 | 0.209998875 | 0.548 | 0.375 | 1 | 23 |
| CU539058.1 | 0.68384 | -0.16701283 | 0.161 | 0.25 | 1 | 23 |
| atp5f1b | 0.6843 | -0.14909481 | 0.839 | 0.875 | 1 | 23 |
| tuba8l3 | 0.6843 | 0.130820474 | 0.161 | 0.125 | 1 | 23 |
| gmfb | 0.68444 | 0.161996252 | 0.387 | 0.25 | 1 | 23 |
| apmap | 0.68475 | 0.182675065 | 0.226 | 0.125 | 1 | 23 |
| aldh1a2 | 0.68559 | -0.21761222 | 0.613 | 0.75 | 1 | 23 |
| selenoh | 0.68566 | 0.104629207 | 0.355 | 0.375 | 1 | 23 |
| BX005340.1 | 0.68622 | -0.16528442 | 0.161 | 0.25 | 1 | 23 |
| mmp2 | 0.68632 | -0.1125461 | 0.484 | 0.5 | 1 | 23 |
| ccni | 0.68649 | 0.140371418 | 0.677 | 0.625 | 1 | 23 |
| fblim1 | 0.68746 | -0.24085985 | 0.226 | 0.375 | 1 | 23 |
| rgrb | 0.68981 | 0.101795965 | 0.387 | 0.25 | 1 | 23 |
| scp2b | 0.69005 | 0.200048455 | 0.258 | 0.125 | 1 | 23 |
| pnn | 0.69041 | -0.11062852 | 0.258 | 0.375 | 1 | 23 |
| tjp1a | 0.69117 | -0.15813775 | 0.323 | 0.375 | 1 | 23 |
| rrp36 | 0.69169 | 0.193335887 | 0.226 | 0.125 | 1 | 23 |
| enkur | 0.69181 | 0.180931968 | 0.194 | 0.125 | 1 | 23 |
| atp5mc3a | 0.6921 | 0.16819171 | 0.548 | 0.5 | 1 | 23 |
| atp6v0b | 0.69215 | -0.11063396 | 0.097 | 0.125 | 1 | 23 |

| | | | | | | |
|---|---|---|---|---|---|---|
| rfc2 | 0.69256 | -0.12425497 | 0.129 | 0.25 | 1 | 23 |
| zgc:136564 | 0.69265 | -0.11154308 | 0.097 | 0.125 | 1 | 23 |
| lsm7 | 0.69325 | 0.118695785 | 0.516 | 0.375 | 1 | 23 |
| pin4 | 0.69326 | 0.157334323 | 0.194 | 0.125 | 1 | 23 |
| ciarta | 0.69406 | 0.229717012 | 0.387 | 0.25 | 1 | 23 |
| mbnl1 | 0.69441 | -0.23787292 | 0.226 | 0.375 | 1 | 23 |
| CABZ01092746.1 | 0.6945 | 0.224252501 | 0.258 | 0.125 | 1 | 23 |
| smg5 | 0.69493 | -0.19807584 | 0.129 | 0.25 | 1 | 23 |
| zgc:55461 | 0.69516 | -0.22439303 | 0.129 | 0.25 | 1 | 23 |
| cxcl18b | 0.69569 | -0.12969782 | 0.355 | 0.375 | 1 | 23 |
| BX005183.3 | 0.69584 | -0.10186203 | 0.097 | 0.125 | 1 | 23 |
| gulp1a | 0.69584 | -0.1200794 | 0.226 | 0.25 | 1 | 23 |
| npepl1 | 0.69752 | -0.19468721 | 0.161 | 0.25 | 1 | 23 |
| tubb4b | 0.69758 | -0.15637764 | 0.839 | 0.875 | 1 | 23 |
| TST | 0.69771 | -0.18946193 | 0.161 | 0.25 | 1 | 23 |
| slc7a3a | 0.69803 | -0.21174919 | 0.129 | 0.25 | 1 | 23 |
| rbpms2a | 0.69836 | -0.14402559 | 0.129 | 0.25 | 1 | 23 |
| si:ch211-86h15.1 | 0.69877 | -0.11364554 | 0.097 | 0.125 | 1 | 23 |
| anp32b | 0.69894 | -0.19276141 | 0.419 | 0.5 | 1 | 23 |
| col27a1b | 0.6992 | 0.215528147 | 0.258 | 0.125 | 1 | 23 |
| march5l | 0.69953 | 0.153321328 | 0.194 | 0.125 | 1 | 23 |
| map3k4 | 0.69983 | -0.18708379 | 0.129 | 0.25 | 1 | 23 |
| chmp1a | 0.69987 | -0.15448582 | 0.129 | 0.25 | 1 | 23 |
| helz | 0.70067 | -0.18539758 | 0.129 | 0.25 | 1 | 23 |
| pip4k2ca | 0.70079 | 0.142924846 | 0.194 | 0.125 | 1 | 23 |
| ddt | 0.70092 | 0.205324968 | 0.258 | 0.125 | 1 | 23 |
| ewsr1a | 0.70094 | 0.200121057 | 0.226 | 0.125 | 1 | 23 |
| psmd5 | 0.70101 | -0.19959974 | 0.129 | 0.25 | 1 | 23 |
| wu:fb16f03 | 0.70127 | -0.20566057 | 0.129 | 0.25 | 1 | 23 |
| rfc4 | 0.70141 | -0.15433087 | 0.097 | 0.125 | 1 | 23 |
| bptf | 0.70235 | -0.11629183 | 0.226 | 0.25 | 1 | 23 |
| gpc4 | 0.70288 | -0.2266197 | 0.226 | 0.375 | 1 | 23 |
| mpc2 | 0.70291 | 0.212657772 | 0.258 | 0.125 | 1 | 23 |
| si:ch211-156j16.1 | 0.70299 | -0.23689559 | 0.484 | 0.625 | 1 | 23 |
| metrnla | 0.70304 | -0.15731096 | 0.129 | 0.25 | 1 | 23 |
| zgc:152791 | 0.70309 | -0.14248182 | 0.355 | 0.5 | 1 | 23 |
| zgc:174906 | 0.70322 | -0.23760936 | 0.226 | 0.375 | 1 | 23 |
| vapal | 0.70388 | 0.172599856 | 0.194 | 0.125 | 1 | 23 |
| spop | 0.70422 | -0.2104142 | 0.226 | 0.375 | 1 | 23 |
| psmd3 | 0.70456 | -0.1772406 | 0.387 | 0.5 | 1 | 23 |
| prkci | 0.70473 | -0.19055943 | 0.129 | 0.25 | 1 | 23 |
| fnbp4 | 0.70521 | -0.11035482 | 0.097 | 0.125 | 1 | 23 |
| atp5pb | 0.70551 | 0.139550487 | 0.774 | 0.625 | 1 | 23 |
| nckap1 | 0.70554 | -0.21352317 | 0.226 | 0.375 | 1 | 23 |
| si:ch211-147a11.3 | 0.70564 | 0.1547527 | 0.194 | 0.125 | 1 | 23 |
| tspan14 | 0.70582 | 0.195662999 | 0.226 | 0.125 | 1 | 23 |
| dynll2b | 0.70584 | -0.17954465 | 0.226 | 0.375 | 1 | 23 |
| idh3a | 0.70587 | -0.18655029 | 0.129 | 0.25 | 1 | 23 |

| | | | | | | |
|---|---|---|---|---|---|---|
| sept7b | 0.70596 | -0.18230669 | 0.226 | 0.375 | 1 | 23 |
| si:dkey-28b4.7 | 0.70684 | 0.172150176 | 0.774 | 0.625 | 1 | 23 |
| afdna | 0.70716 | -0.21058261 | 0.226 | 0.375 | 1 | 23 |
| npc2 | 0.70719 | -0.1087057 | 0.613 | 0.625 | 1 | 23 |
| cntrl | 0.70779 | -0.25385385 | 0.129 | 0.25 | 1 | 23 |
| pip4p2 | 0.70788 | 0.113009834 | 0.161 | 0.125 | 1 | 23 |
| ivd | 0.70805 | 0.158701839 | 0.258 | 0.125 | 1 | 23 |
| si:ch211-276i12.11 | 0.7085 | 0.196637027 | 0.387 | 0.25 | 1 | 23 |
| cnot8 | 0.70855 | 0.176510131 | 0.226 | 0.125 | 1 | 23 |
| phactr4b | 0.70883 | 0.176181593 | 0.194 | 0.125 | 1 | 23 |
| aamp | 0.70902 | 0.158697114 | 0.258 | 0.125 | 1 | 23 |
| degs1 | 0.70908 | 0.171821246 | 0.226 | 0.125 | 1 | 23 |
| vim | 0.70908 | 0.240919777 | 0.258 | 0.125 | 1 | 23 |
| dynlt3 | 0.70911 | -0.19507432 | 0.387 | 0.5 | 1 | 23 |
| gtf2a1 | 0.7093 | 0.204879046 | 0.258 | 0.125 | 1 | 23 |
| uqcrc2a | 0.70943 | 0.118866791 | 0.258 | 0.125 | 1 | 23 |
| arpp19b | 0.70943 | 0.1981218 | 0.258 | 0.125 | 1 | 23 |
| gstr | 0.70972 | 0.195973784 | 0.387 | 0.25 | 1 | 23 |
| myl9a | 0.71031 | -0.21499901 | 0.129 | 0.25 | 1 | 23 |
| emc7 | 0.71046 | 0.17473352 | 0.226 | 0.125 | 1 | 23 |
| sumo3a | 0.71064 | -0.11349694 | 0.839 | 0.875 | 1 | 23 |
| col1a2 | 0.71077 | 0.160801134 | 0.387 | 0.375 | 1 | 23 |
| psma3 | 0.71126 | 0.149800839 | 0.677 | 0.625 | 1 | 23 |
| prrg4 | 0.71148 | -0.13412949 | 0.129 | 0.25 | 1 | 23 |
| cast | 0.7118 | -0.19501855 | 0.129 | 0.25 | 1 | 23 |
| wasf2 | 0.71264 | -0.17367922 | 0.129 | 0.25 | 1 | 23 |
| ube2ka | 0.71322 | -0.16143302 | 0.161 | 0.25 | 1 | 23 |
| efnb2a | 0.71406 | -0.15371747 | 0.194 | 0.25 | 1 | 23 |
| trappc6bl | 0.71469 | -0.19849007 | 0.129 | 0.25 | 1 | 23 |
| nsfl1c | 0.71663 | 0.232143921 | 0.387 | 0.25 | 1 | 23 |
| tfr1b | 0.71683 | -0.12987287 | 0.097 | 0.125 | 1 | 23 |
| si:dkey-7j14.6 | 0.71692 | -0.13956091 | 0.581 | 0.625 | 1 | 23 |
| spred1 | 0.71746 | -0.14375435 | 0.129 | 0.25 | 1 | 23 |
| efcab2 | 0.71823 | -0.15817921 | 0.129 | 0.25 | 1 | 23 |
| c1d | 0.71829 | 0.11098626 | 0.161 | 0.125 | 1 | 23 |
| gtf3c6 | 0.71836 | -0.16553762 | 0.129 | 0.25 | 1 | 23 |
| atp6v1e1b | 0.71846 | -0.18765717 | 0.129 | 0.25 | 1 | 23 |
| twistnb | 0.71875 | -0.15360767 | 0.129 | 0.25 | 1 | 23 |
| abi2b | 0.71882 | -0.24010403 | 0.129 | 0.25 | 1 | 23 |
| pdcd6ip | 0.71933 | -0.13791607 | 0.129 | 0.25 | 1 | 23 |
| lima1a | 0.72008 | -0.12614066 | 0.097 | 0.125 | 1 | 23 |
| entpd1 | 0.72019 | -0.10766463 | 0.226 | 0.25 | 1 | 23 |
| gm2a | 0.7203 | 0.108702597 | 0.161 | 0.125 | 1 | 23 |
| csde1 | 0.72081 | -0.18367756 | 0.355 | 0.5 | 1 | 23 |
| lamb1a | 0.72124 | -0.17064706 | 0.129 | 0.25 | 1 | 23 |
| zgc:92907 | 0.72167 | -0.10816289 | 0.097 | 0.125 | 1 | 23 |
| tktb | 0.72238 | 0.162417395 | 0.226 | 0.125 | 1 | 23 |
| mcm6 | 0.72243 | -0.21528901 | 0.129 | 0.25 | 1 | 23 |

| | | | | | | |
|---|---|---|---|---|---|---|
| prkar1ab | 0.72268 | -0.16832974 | 0.129 | 0.25 | 1 | 23 |
| ndrg2 | 0.72269 | -0.1667554 | 0.161 | 0.25 | 1 | 23 |
| adam17a | 0.72292 | -0.11009165 | 0.097 | 0.125 | 1 | 23 |
| idh1 | 0.72324 | -0.11257811 | 0.097 | 0.125 | 1 | 23 |
| gemin2 | 0.72385 | -0.14933784 | 0.129 | 0.25 | 1 | 23 |
| hoxb7a | 0.72496 | 0.166744572 | 0.226 | 0.125 | 1 | 23 |
| tm9sf3 | 0.7252 | -0.19163245 | 0.194 | 0.25 | 1 | 23 |
| prkar2aa | 0.72546 | 0.166439567 | 0.194 | 0.125 | 1 | 23 |
| snx12 | 0.72619 | -0.17217329 | 0.129 | 0.25 | 1 | 23 |
| eif4ebp3l | 0.72638 | 0.201792203 | 0.387 | 0.25 | 1 | 23 |
| cited4a | 0.72647 | -0.16695249 | 0.129 | 0.25 | 1 | 23 |
| FP101882.1 | 0.72657 | -0.15286544 | 0.129 | 0.25 | 1 | 23 |
| efcab1 | 0.72682 | -0.15725001 | 0.129 | 0.25 | 1 | 23 |
| pxnb | 0.72694 | -0.14720635 | 0.129 | 0.25 | 1 | 23 |
| rab41 | 0.72743 | 0.114125796 | 0.226 | 0.125 | 1 | 23 |
| ctsa | 0.72763 | -0.18683333 | 0.129 | 0.25 | 1 | 23 |
| ctbp2a | 0.72764 | -0.20581612 | 0.129 | 0.25 | 1 | 23 |
| wipf2a | 0.72768 | -0.16110787 | 0.129 | 0.25 | 1 | 23 |
| tmem115 | 0.72772 | -0.18011866 | 0.129 | 0.25 | 1 | 23 |
| gse1 | 0.72773 | 0.113273881 | 0.161 | 0.125 | 1 | 23 |
| timm10b | 0.72796 | -0.10821556 | 0.097 | 0.125 | 1 | 23 |
| hspa12b | 0.72811 | 0.10529464 | 0.129 | 0.125 | 1 | 23 |
| rgs19 | 0.72827 | -0.10954005 | 0.097 | 0.125 | 1 | 23 |
| arrdc2 | 0.72853 | -0.10775651 | 0.097 | 0.125 | 1 | 23 |
| eif4a1b | 0.72891 | 0.182622341 | 0.71 | 0.625 | 1 | 23 |
| btr31 | 0.72899 | -0.10835966 | 0.097 | 0.125 | 1 | 23 |
| si:ch211-282b22.1 | 0.72949 | -0.13229873 | 0.194 | 0.25 | 1 | 23 |
| pank2 | 0.73284 | 0.160806254 | 0.194 | 0.125 | 1 | 23 |
| tgm1l1 | 0.73325 | -0.1063734 | 0.097 | 0.125 | 1 | 23 |
| psma6a | 0.73328 | 0.202052518 | 0.516 | 0.375 | 1 | 23 |
| reep5 | 0.73456 | 0.187634863 | 0.355 | 0.25 | 1 | 23 |
| psmc3 | 0.73506 | 0.159089638 | 0.387 | 0.25 | 1 | 23 |
| btg2 | 0.73521 | -0.11481561 | 0.742 | 0.75 | 1 | 23 |
| aldh6a1 | 0.73599 | -0.12378128 | 0.355 | 0.375 | 1 | 23 |
| zfyve19 | 0.73629 | -0.10626987 | 0.097 | 0.125 | 1 | 23 |
| sox6 | 0.73671 | -0.20848549 | 0.355 | 0.5 | 1 | 23 |
| ttl | 0.73868 | -0.10873567 | 0.097 | 0.125 | 1 | 23 |
| cdc42l | 0.73891 | -0.18878181 | 0.484 | 0.625 | 1 | 23 |
| hcfc1a | 0.73905 | 0.131590426 | 0.226 | 0.125 | 1 | 23 |
| pspc1 | 0.73943 | 0.106770912 | 0.161 | 0.125 | 1 | 23 |
| gspt1l | 0.73972 | 0.106435957 | 0.226 | 0.125 | 1 | 23 |
| sgpl1 | 0.73986 | -0.10628827 | 0.097 | 0.125 | 1 | 23 |
| rreb1a | 0.74014 | -0.10992849 | 0.097 | 0.125 | 1 | 23 |
| pdcd5 | 0.74034 | 0.158085494 | 0.387 | 0.25 | 1 | 23 |
| sumo2b | 0.74119 | 0.11765791 | 0.774 | 0.75 | 1 | 23 |
| grinab | 0.7413 | -0.15542265 | 0.355 | 0.5 | 1 | 23 |
| tusc2b | 0.74186 | -0.10711802 | 0.097 | 0.125 | 1 | 23 |
| atp5fa1 | 0.74209 | -0.11751865 | 0.613 | 0.75 | 1 | 23 |

| | | | | | | |
|---|---|---|---|---|---|---|
| march2 | 0.74306 | -0.10789711 | 0.097 | 0.125 | 1 | 23 |
| sacm1la | 0.74396 | -0.10541645 | 0.097 | 0.125 | 1 | 23 |
| perp | 0.74407 | 0.115930029 | 0.226 | 0.125 | 1 | 23 |
| acadm | 0.74537 | -0.10612315 | 0.355 | 0.375 | 1 | 23 |
| cdk2ap2 | 0.7456 | -0.15789617 | 0.194 | 0.25 | 1 | 23 |
| snrpc | 0.74598 | 0.155596888 | 0.387 | 0.25 | 1 | 23 |
| aebp2 | 0.74685 | 0.179310402 | 0.226 | 0.125 | 1 | 23 |
| mpzl2b | 0.74719 | 0.186805991 | 0.226 | 0.125 | 1 | 23 |
| si:ch211-269e2.1 | 0.74749 | 0.126523289 | 0.161 | 0.125 | 1 | 23 |
| ebna1bp2 | 0.74753 | -0.11648549 | 0.258 | 0.375 | 1 | 23 |
| ccnd2a | 0.74789 | 0.112609518 | 0.161 | 0.125 | 1 | 23 |
| si:ch211-213a13.1 | 0.748 | 0.170757789 | 0.226 | 0.125 | 1 | 23 |
| pcyox1 | 0.74807 | -0.17290295 | 0.29 | 0.375 | 1 | 23 |
| hars | 0.74855 | 0.134856992 | 0.194 | 0.125 | 1 | 23 |
| klf11b | 0.74858 | -0.13477838 | 0.452 | 0.5 | 1 | 23 |
| strap | 0.74916 | 0.146802196 | 0.194 | 0.125 | 1 | 23 |
| znf385a | 0.74922 | 0.179588486 | 0.226 | 0.125 | 1 | 23 |
| cldn11a | 0.75086 | -0.19966591 | 0.613 | 0.75 | 1 | 23 |
| mcm5 | 0.75092 | -0.13443642 | 0.097 | 0.125 | 1 | 23 |
| phf14 | 0.75169 | 0.11520676 | 0.161 | 0.125 | 1 | 23 |
| ppp3r1b | 0.75217 | 0.172072326 | 0.226 | 0.125 | 1 | 23 |
| rcc2 | 0.75235 | -0.15169487 | 0.323 | 0.375 | 1 | 23 |
| esd | 0.75296 | 0.212361511 | 0.516 | 0.375 | 1 | 23 |
| mrpl23 | 0.7533 | 0.148357202 | 0.194 | 0.125 | 1 | 23 |
| calr3a | 0.75449 | 0.185576007 | 0.387 | 0.25 | 1 | 23 |
| supt6h | 0.75453 | 0.173042127 | 0.194 | 0.125 | 1 | 23 |
| cald1b | 0.75454 | 0.140512745 | 0.161 | 0.125 | 1 | 23 |
| neo1a | 0.75563 | -0.17966072 | 0.194 | 0.25 | 1 | 23 |
| mrpl36 | 0.75575 | -0.17683694 | 0.355 | 0.5 | 1 | 23 |
| robo2 | 0.75616 | 0.163871601 | 0.226 | 0.125 | 1 | 23 |
| gabarapl2 | 0.75655 | 0.14312336 | 0.194 | 0.125 | 1 | 23 |
| srsf9 | 0.75721 | 0.189665277 | 0.387 | 0.25 | 1 | 23 |
| slc44a2 | 0.75735 | -0.16446156 | 0.161 | 0.25 | 1 | 23 |
| ube2ia | 0.75766 | 0.201728115 | 0.516 | 0.375 | 1 | 23 |
| hif1ab | 0.75782 | 0.134840131 | 0.419 | 0.375 | 1 | 23 |
| ppfibp1a | 0.75855 | -0.21666825 | 0.258 | 0.375 | 1 | 23 |
| CABZ01034698.2 | 0.75857 | -0.16088736 | 0.161 | 0.25 | 1 | 23 |
| rbm4.2 | 0.75949 | 0.137668966 | 0.323 | 0.25 | 1 | 23 |
| med19a | 0.75961 | 0.173345643 | 0.194 | 0.125 | 1 | 23 |
| krt94 | 0.75996 | -0.14796303 | 0.71 | 0.75 | 1 | 23 |
| si:ch211-14a17.11 | 0.76068 | 0.169215823 | 0.226 | 0.125 | 1 | 23 |
| STMP1 | 0.76151 | 0.165982656 | 0.355 | 0.25 | 1 | 23 |
| naa30 | 0.76263 | 0.121969734 | 0.226 | 0.125 | 1 | 23 |
| rdh10a | 0.76293 | -0.20722339 | 0.516 | 0.625 | 1 | 23 |
| tent4b | 0.76294 | 0.127147782 | 0.226 | 0.125 | 1 | 23 |
| ctgfa | 0.76303 | 0.119692033 | 0.355 | 0.25 | 1 | 23 |
| wdr93 | 0.76448 | 0.12879088 | 0.226 | 0.125 | 1 | 23 |
| ctsla | 0.76458 | -0.14950995 | 0.613 | 0.75 | 1 | 23 |

| | | | | | | |
|---|---|---|---|---|---|---|
| capza1a | 0.76494 | 0.185281977 | 0.387 | 0.25 | 1 | 23 |
| c6 | 0.76542 | 0.163358045 | 0.516 | 0.375 | 1 | 23 |
| wu:fi37e09 | 0.76641 | 0.103848169 | 0.355 | 0.25 | 1 | 23 |
| ncor2 | 0.76728 | -0.13620657 | 0.323 | 0.375 | 1 | 23 |
| sdf2 | 0.7675 | -0.12960319 | 0.194 | 0.25 | 1 | 23 |
| tmed7 | 0.76787 | 0.165634805 | 0.516 | 0.375 | 1 | 23 |
| mrpl33 | 0.76825 | 0.167488563 | 0.226 | 0.125 | 1 | 23 |
| myg1 | 0.76839 | 0.166721298 | 0.226 | 0.125 | 1 | 23 |
| scnm1 | 0.7687 | 0.140165743 | 0.161 | 0.125 | 1 | 23 |
| zgc:162339 | 0.76875 | 0.101912469 | 0.161 | 0.125 | 1 | 23 |
| pdap1a | 0.77068 | 0.151633752 | 0.226 | 0.125 | 1 | 23 |
| si:dkey-42i9.7 | 0.77146 | 0.18334323 | 0.323 | 0.25 | 1 | 23 |
| ncln | 0.77159 | 0.113590841 | 0.161 | 0.125 | 1 | 23 |
| tbc1d5 | 0.77334 | -0.10225868 | 0.097 | 0.125 | 1 | 23 |
| ndufa9a | 0.77345 | -0.15146715 | 0.161 | 0.25 | 1 | 23 |
| brk1 | 0.77578 | 0.172956919 | 0.516 | 0.375 | 1 | 23 |
| lamtor4 | 0.77609 | 0.122886552 | 0.226 | 0.125 | 1 | 23 |
| dspa | 0.77715 | -0.1028579 | 0.097 | 0.125 | 1 | 23 |
| eif4e2 | 0.77718 | -0.10659867 | 0.097 | 0.125 | 1 | 23 |
| srrm2 | 0.77726 | -0.16842471 | 0.194 | 0.25 | 1 | 23 |
| ndufb8 | 0.77761 | 0.165841804 | 0.581 | 0.5 | 1 | 23 |
| acad9 | 0.77785 | -0.12238877 | 0.097 | 0.125 | 1 | 23 |
| caprin1a | 0.77798 | -0.13291327 | 0.258 | 0.375 | 1 | 23 |
| spcs2 | 0.7784 | -0.11684748 | 0.355 | 0.375 | 1 | 23 |
| cdc40 | 0.77989 | 0.111307467 | 0.161 | 0.125 | 1 | 23 |
| mpzl1l | 0.7802 | 0.162921454 | 0.226 | 0.125 | 1 | 23 |
| bhlhe40 | 0.78053 | -0.10911309 | 0.226 | 0.25 | 1 | 23 |
| nkd2a | 0.781 | -0.21631808 | 0.161 | 0.25 | 1 | 23 |
| zgc:56493 | 0.78103 | 0.122920971 | 0.935 | 0.875 | 1 | 23 |
| ndufb7 | 0.78126 | 0.190408347 | 0.484 | 0.375 | 1 | 23 |
| golga7 | 0.78282 | 0.131600031 | 0.548 | 0.5 | 1 | 23 |
| ss18l2 | 0.78389 | 0.1372149 | 0.194 | 0.125 | 1 | 23 |
| npm3 | 0.78425 | 0.15222745 | 0.323 | 0.25 | 1 | 23 |
| lgals3b | 0.78442 | 0.110748348 | 0.355 | 0.25 | 1 | 23 |
| LEPROTL1 | 0.78501 | -0.18300142 | 0.29 | 0.375 | 1 | 23 |
| pgd | 0.78523 | -0.15504759 | 0.161 | 0.25 | 1 | 23 |
| dcun1d5 | 0.7857 | 0.136566722 | 0.226 | 0.125 | 1 | 23 |
| lef1 | 0.78872 | -0.10128795 | 0.097 | 0.125 | 1 | 23 |
| nfs1 | 0.78955 | -0.10305357 | 0.097 | 0.125 | 1 | 23 |
| brd9 | 0.78996 | -0.17561584 | 0.258 | 0.375 | 1 | 23 |
| uqcrc1 | 0.79274 | -0.1188833 | 0.258 | 0.375 | 1 | 23 |
| dnpep | 0.79302 | 0.146010231 | 0.226 | 0.125 | 1 | 23 |
| rbm4.1 | 0.79323 | 0.108588009 | 0.29 | 0.25 | 1 | 23 |
| arpc1a | 0.79436 | -0.13386542 | 0.258 | 0.375 | 1 | 23 |
| zranb2 | 0.79644 | -0.17321443 | 0.161 | 0.25 | 1 | 23 |
| srsf6b | 0.79657 | 0.110263642 | 0.29 | 0.25 | 1 | 23 |
| mknk2b | 0.79755 | -0.10828115 | 0.355 | 0.375 | 1 | 23 |
| AK6 | 0.79771 | -0.18303644 | 0.258 | 0.375 | 1 | 23 |

| | | | | | | |
|---|---|---|---|---|---|---|
| serinc2 | 0.79837 | 0.145863838 | 0.581 | 0.5 | 1 | 23 |
| pdlim3b | 0.79937 | -0.19966008 | 0.161 | 0.25 | 1 | 23 |
| ddah2 | 0.80025 | -0.12424717 | 0.323 | 0.375 | 1 | 23 |
| pum2 | 0.80073 | 0.123649398 | 0.226 | 0.125 | 1 | 23 |
| adrm1 | 0.80079 | 0.101671092 | 0.29 | 0.25 | 1 | 23 |
| tshz3b | 0.80179 | -0.13915709 | 0.258 | 0.375 | 1 | 23 |
| dnajc15 | 0.80205 | -0.1031232 | 0.097 | 0.125 | 1 | 23 |
| zgc:112982 | 0.80215 | 0.114889675 | 0.161 | 0.125 | 1 | 23 |
| tmem131 | 0.8024 | -0.10054667 | 0.097 | 0.125 | 1 | 23 |
| ccdc47 | 0.80331 | 0.183875908 | 0.226 | 0.125 | 1 | 23 |
| rrp8 | 0.80383 | 0.157461344 | 0.194 | 0.125 | 1 | 23 |
| mmgt1 | 0.80384 | 0.148566303 | 0.226 | 0.125 | 1 | 23 |
| sh3kbp1 | 0.80393 | 0.116779371 | 0.226 | 0.125 | 1 | 23 |
| elovl1b | 0.8051 | -0.12782655 | 0.452 | 0.5 | 1 | 23 |
| aes | 0.80543 | -0.17868567 | 0.161 | 0.25 | 1 | 23 |
| cwc25 | 0.80543 | 0.137449217 | 0.194 | 0.125 | 1 | 23 |
| si:dkeyp-27c8.2 | 0.80567 | 0.162376354 | 0.226 | 0.125 | 1 | 23 |
| hoxc9a | 0.80569 | -0.15358615 | 0.29 | 0.375 | 1 | 23 |
| CABZ01068251.1 | 0.80734 | 0.155936877 | 0.226 | 0.125 | 1 | 23 |
| appb | 0.80747 | 0.146640753 | 0.226 | 0.125 | 1 | 23 |
| immt | 0.80752 | 0.142752804 | 0.226 | 0.125 | 1 | 23 |
| iqgap1 | 0.80851 | 0.165523397 | 0.226 | 0.125 | 1 | 23 |
| ptk7a | 0.809 | -0.1614666 | 0.29 | 0.375 | 1 | 23 |
| pwp1 | 0.80963 | 0.131227845 | 0.226 | 0.125 | 1 | 23 |
| polr2i | 0.80983 | 0.13436143 | 0.226 | 0.125 | 1 | 23 |
| ube2ib | 0.81124 | -0.15512606 | 0.516 | 0.625 | 1 | 23 |
| ssuh2rs1 | 0.81195 | -0.15116613 | 0.258 | 0.375 | 1 | 23 |
| timp2a | 0.81249 | -0.14821142 | 0.258 | 0.375 | 1 | 23 |
| tmem88b | 0.81281 | 0.140181811 | 0.323 | 0.25 | 1 | 23 |
| chac1 | 0.81322 | 0.187665961 | 0.581 | 0.5 | 1 | 23 |
| fbxo28 | 0.81325 | -0.15917224 | 0.161 | 0.25 | 1 | 23 |
| ptprz1b | 0.81419 | 0.150616252 | 0.194 | 0.125 | 1 | 23 |
| gstm.3 | 0.81499 | 0.155162043 | 0.742 | 0.625 | 1 | 23 |
| tsc22d3 | 0.81507 | 0.107929337 | 0.29 | 0.25 | 1 | 23 |
| selenot2 | 0.81646 | -0.12910554 | 0.194 | 0.25 | 1 | 23 |
| rhoca | 0.81671 | -0.16575236 | 0.161 | 0.25 | 1 | 23 |
| hspa4b | 0.81702 | 0.156401007 | 0.355 | 0.25 | 1 | 23 |
| ndufv2 | 0.82037 | 0.107841004 | 0.161 | 0.125 | 1 | 23 |
| nudt21 | 0.82131 | -0.13434127 | 0.194 | 0.25 | 1 | 23 |
| manf | 0.82291 | -0.14878557 | 0.161 | 0.25 | 1 | 23 |
| skp1 | 0.82323 | 0.153845485 | 0.581 | 0.5 | 1 | 23 |
| mrpl14 | 0.82381 | 0.134033455 | 0.194 | 0.125 | 1 | 23 |
| lims1 | 0.82404 | 0.119645522 | 0.194 | 0.125 | 1 | 23 |
| trmt112 | 0.82658 | -0.11409592 | 0.452 | 0.5 | 1 | 23 |
| tgfb3 | 0.82698 | 0.118772227 | 0.29 | 0.25 | 1 | 23 |
| zgc:66433 | 0.82721 | -0.12647559 | 0.194 | 0.25 | 1 | 23 |
| vasnb | 0.82728 | 0.172016418 | 0.355 | 0.25 | 1 | 23 |
| kmt2bb | 0.82782 | -0.15313177 | 0.29 | 0.375 | 1 | 23 |

| | | | | | | |
|---|---|---|---|---|---|---|
| ifrd1 | 0.82882 | 0.142565666 | 0.194 | 0.125 | 1 | 23 |
| ergic2 | 0.82909 | -0.14549605 | 0.161 | 0.25 | 1 | 23 |
| ddx27 | 0.82996 | 0.186451367 | 0.355 | 0.25 | 1 | 23 |
| ralbp1 | 0.83028 | -0.13676742 | 0.161 | 0.25 | 1 | 23 |
| CABZ01087560.1 | 0.83058 | 0.105551922 | 0.484 | 0.375 | 1 | 23 |
| ewsr1b | 0.83283 | -0.15021888 | 0.387 | 0.5 | 1 | 23 |
| serpine2 | 0.83413 | -0.10101979 | 0.097 | 0.125 | 1 | 23 |
| plecb | 0.83423 | 0.104566585 | 0.161 | 0.125 | 1 | 23 |
| abcf1 | 0.83436 | -0.11876694 | 0.387 | 0.5 | 1 | 23 |
| si:ch211-114c12.2 | 0.83461 | -0.10194858 | 0.161 | 0.25 | 1 | 23 |
| ubtfl | 0.83501 | -0.13109854 | 0.194 | 0.25 | 1 | 23 |
| uqcc2 | 0.83535 | 0.136203434 | 0.194 | 0.125 | 1 | 23 |
| pphln1 | 0.83538 | -0.14816486 | 0.161 | 0.25 | 1 | 23 |
| zgc:174863 | 0.83584 | 0.129173318 | 0.355 | 0.25 | 1 | 23 |
| CABZ01086029.1 | 0.83774 | -0.11479276 | 0.323 | 0.375 | 1 | 23 |
| ruvbl1 | 0.83864 | -0.13757573 | 0.161 | 0.25 | 1 | 23 |
| pag1 | 0.83898 | -0.14374148 | 0.161 | 0.25 | 1 | 23 |
| myl6 | 0.83961 | -0.15797728 | 0.387 | 0.5 | 1 | 23 |
| ddx39aa | 0.83961 | -0.11912291 | 0.387 | 0.5 | 1 | 23 |
| nono | 0.84044 | -0.1284394 | 0.194 | 0.25 | 1 | 23 |
| rpa1 | 0.84067 | -0.10832194 | 0.161 | 0.25 | 1 | 23 |
| dkk3b | 0.84116 | -0.13625248 | 0.387 | 0.5 | 1 | 23 |
| setb | 0.84191 | 0.153647914 | 0.613 | 0.5 | 1 | 23 |
| dazap2 | 0.84191 | -0.12207134 | 0.194 | 0.25 | 1 | 23 |
| gtf2b | 0.8421 | 0.128608145 | 0.194 | 0.125 | 1 | 23 |
| psat1 | 0.84316 | -0.15115189 | 0.161 | 0.25 | 1 | 23 |
| rpf1 | 0.84319 | 0.135673878 | 0.323 | 0.25 | 1 | 23 |
| arf2b | 0.84481 | 0.153678882 | 0.323 | 0.25 | 1 | 23 |
| ube2e1 | 0.84504 | -0.10759324 | 0.452 | 0.5 | 1 | 23 |
| nup62l | 0.84776 | -0.10764722 | 0.161 | 0.25 | 1 | 23 |
| CU984600.2 | 0.84808 | -0.14322847 | 0.194 | 0.25 | 1 | 23 |
| hyi | 0.84813 | 0.127213541 | 0.194 | 0.125 | 1 | 23 |
| tmem18 | 0.84814 | 0.125277979 | 0.323 | 0.25 | 1 | 23 |
| rps6kb1b | 0.84869 | -0.13173052 | 0.161 | 0.25 | 1 | 23 |
| bmp4 | 0.84884 | 0.102907377 | 0.161 | 0.125 | 1 | 23 |
| vps28 | 0.8491 | 0.146187659 | 0.355 | 0.25 | 1 | 23 |
| psmc1a | 0.84994 | 0.14935802 | 0.484 | 0.375 | 1 | 23 |
| jagn1b | 0.85001 | 0.123201079 | 0.194 | 0.125 | 1 | 23 |
| srp72 | 0.85056 | -0.11863451 | 0.161 | 0.25 | 1 | 23 |
| pcdh1b | 0.85062 | 0.151562617 | 0.355 | 0.25 | 1 | 23 |
| bcap31 | 0.85175 | -0.14871993 | 0.161 | 0.25 | 1 | 23 |
| snx5 | 0.85198 | -0.13836383 | 0.29 | 0.375 | 1 | 23 |
| upf1 | 0.85212 | 0.104816206 | 0.161 | 0.125 | 1 | 23 |
| ap1s1 | 0.85298 | -0.10414547 | 0.161 | 0.25 | 1 | 23 |
| arl6ip1 | 0.85343 | 0.121351785 | 0.484 | 0.375 | 1 | 23 |
| tdp1 | 0.8543 | -0.12919457 | 0.161 | 0.25 | 1 | 23 |
| gpatch11 | 0.85492 | -0.11645538 | 0.161 | 0.25 | 1 | 23 |
| ppp2r5eb | 0.85497 | -0.13301487 | 0.161 | 0.25 | 1 | 23 |

| | | | | | | |
|---|---|---|---|---|---|---|
| fbxo25 | 0.855 | -0.12807985 | 0.161 | 0.25 | 1 | 23 |
| gas1a | 0.85507 | -0.13302141 | 0.194 | 0.25 | 1 | 23 |
| rars | 0.85599 | 0.131513614 | 0.194 | 0.125 | 1 | 23 |
| pop5 | 0.85795 | 0.118216126 | 0.194 | 0.125 | 1 | 23 |
| PRRC2B | 0.85831 | 0.102048462 | 0.161 | 0.125 | 1 | 23 |
| lhfpl2a | 0.85842 | 0.135275107 | 0.484 | 0.375 | 1 | 23 |
| igf2b | 0.85848 | -0.15414054 | 0.29 | 0.375 | 1 | 23 |
| ythdf1 | 0.86026 | 0.116535308 | 0.161 | 0.125 | 1 | 23 |
| atf5b | 0.86085 | 0.113365875 | 0.161 | 0.125 | 1 | 23 |
| tmem165 | 0.86142 | 0.118372172 | 0.194 | 0.125 | 1 | 23 |
| brms1la | 0.86273 | 0.118069312 | 0.194 | 0.125 | 1 | 23 |
| rnf216 | 0.86438 | 0.119583672 | 0.194 | 0.125 | 1 | 23 |
| slc9a3r1a | 0.86584 | 0.106974121 | 0.194 | 0.125 | 1 | 23 |
| ube2q2 | 0.86647 | -0.11792508 | 0.194 | 0.25 | 1 | 23 |
| ube2d2l | 0.86787 | -0.11188982 | 0.194 | 0.25 | 1 | 23 |
| gtf2e2 | 0.86819 | 0.116855436 | 0.194 | 0.125 | 1 | 23 |
| ap3d1 | 0.87096 | -0.10998071 | 0.194 | 0.25 | 1 | 23 |
| psmd14 | 0.87129 | 0.127751998 | 0.581 | 0.5 | 1 | 23 |
| creg2 | 0.87173 | 0.110973457 | 0.194 | 0.125 | 1 | 23 |
| gskip | 0.87185 | 0.11346219 | 0.194 | 0.125 | 1 | 23 |
| tfam | 0.87417 | 0.120457839 | 0.194 | 0.125 | 1 | 23 |
| igfbp7 | 0.87421 | 0.117209844 | 0.194 | 0.125 | 1 | 23 |
| znf318 | 0.87435 | 0.100021594 | 0.194 | 0.125 | 1 | 23 |
| si:dkey-67c22.2 | 0.87674 | -0.1373399 | 0.29 | 0.375 | 1 | 23 |
| h1f0 | 0.87918 | -0.1386325 | 0.548 | 0.625 | 1 | 23 |
| tmem50a | 0.87928 | 0.109235009 | 0.419 | 0.375 | 1 | 23 |
| ralba | 0.87942 | 0.113459662 | 0.194 | 0.125 | 1 | 23 |
| rab14 | 0.88035 | -0.11061914 | 0.194 | 0.25 | 1 | 23 |
| gtf2h5 | 0.88119 | 0.103656625 | 0.29 | 0.25 | 1 | 23 |
| si:dkey-16l2.16 | 0.8823 | 0.113756638 | 0.194 | 0.125 | 1 | 23 |
| nudt1 | 0.88242 | 0.108091353 | 0.194 | 0.125 | 1 | 23 |
| stat2 | 0.88297 | -0.15345363 | 0.29 | 0.375 | 1 | 23 |
| nop56 | 0.88487 | -0.11623088 | 0.548 | 0.625 | 1 | 23 |
| znf706 | 0.88802 | 0.117344188 | 0.452 | 0.375 | 1 | 23 |
| homezb | 0.88888 | 0.12171587 | 0.194 | 0.125 | 1 | 23 |
| mcm2 | 0.88925 | -0.1287277 | 0.226 | 0.25 | 1 | 23 |
| zgc:66388 | 0.8907 | -0.13574211 | 0.194 | 0.25 | 1 | 23 |
| gid8b | 0.89142 | 0.107163123 | 0.194 | 0.125 | 1 | 23 |
| eif2b1 | 0.89304 | 0.100024045 | 0.194 | 0.125 | 1 | 23 |
| LOC110439449 | 0.89354 | -0.11030699 | 0.097 | 0.125 | 1 | 23 |
| gnb1b | 0.89535 | 0.110172603 | 0.71 | 0.625 | 1 | 23 |
| BX548028.1 | 0.89631 | 0.114167778 | 0.194 | 0.125 | 1 | 23 |
| pet100 | 0.89755 | -0.10216116 | 0.29 | 0.375 | 1 | 23 |
| dachc | 0.89903 | 0.124515698 | 0.194 | 0.125 | 1 | 23 |
| rp9 | 0.89937 | 0.10362404 | 0.194 | 0.125 | 1 | 23 |
| si:dkey-261h17.1 | 0.89946 | 0.126699236 | 0.194 | 0.125 | 1 | 23 |
| dnajc8 | 0.89983 | 0.110958548 | 0.194 | 0.125 | 1 | 23 |
| vcp | 0.90033 | -0.12000668 | 0.29 | 0.375 | 1 | 23 |

| | | | | | | |
|---|---|---|---|---|---|---|
| otub1b | 0.90034 | -0.10466036 | 0.29 | 0.375 | 1 | 23 |
| nog1 | 0.90183 | 0.10182769 | 0.71 | 0.625 | 1 | 23 |
| fstl3 | 0.90185 | 0.108826186 | 0.29 | 0.25 | 1 | 23 |
| ptenb | 0.90186 | -0.12567116 | 0.29 | 0.375 | 1 | 23 |
| cldn5a | 0.90189 | -0.13848922 | 0.29 | 0.375 | 1 | 23 |
| cdh4 | 0.90514 | 0.131590022 | 0.323 | 0.25 | 1 | 23 |
| mibp2 | 0.90655 | 0.105941361 | 0.806 | 0.75 | 1 | 23 |
| morf4l1 | 0.90833 | 0.113817671 | 0.452 | 0.375 | 1 | 23 |
| sart1 | 0.90849 | -0.11090173 | 0.419 | 0.5 | 1 | 23 |
| map1lc3b | 0.91156 | 0.122692669 | 0.452 | 0.375 | 1 | 23 |
| copz1 | 0.91305 | 0.102912376 | 0.452 | 0.375 | 1 | 23 |
| si:dkey-222f8.3 | 0.91333 | -0.10467662 | 0.194 | 0.25 | 1 | 23 |
| si:ch211-262e15.1 | 0.9171 | -0.111501 | 0.194 | 0.25 | 1 | 23 |
| kank2 | 0.91849 | -0.10057851 | 0.097 | 0.125 | 1 | 23 |
| tubb6 | 0.91863 | 0.100048417 | 0.161 | 0.125 | 1 | 23 |
| smim15 | 0.91981 | 0.101335487 | 0.323 | 0.25 | 1 | 23 |
| cstf2 | 0.9228 | -0.11277371 | 0.194 | 0.25 | 1 | 23 |
| dido1 | 0.92342 | 0.109066511 | 0.452 | 0.375 | 1 | 23 |
| si:ch211-288g17.3 | 0.92688 | 0.110417892 | 0.677 | 0.625 | 1 | 23 |
| ccna2 | 0.00116 | -0.91621079 | 0 | 0.375 | 1 | 24 |
| timm8b | 0.00116 | -0.76566673 | 0 | 0.375 | 1 | 24 |
| dcn | 0.00371 | -0.71420971 | 0 | 0.312 | 1 | 24 |
| ndufb7 | 0.00417 | 0.842540849 | 0.333 | 0 | 1 | 24 |
| CR318588.1 | 0.00474 | -1.37101008 | 0.048 | 0.5 | 1 | 24 |
| calm2b | 0.00698 | -0.62796758 | 0.476 | 0.938 | 1 | 24 |
| psmb7 | 0.00727 | -0.96240052 | 0.095 | 0.562 | 1 | 24 |
| akirin1 | 0.00925 | 0.5884542 | 0.286 | 0 | 1 | 24 |
| eif4bb | 0.00925 | 0.692379377 | 0.286 | 0 | 1 | 24 |
| ndufa11 | 0.00925 | 0.605277158 | 0.286 | 0 | 1 | 24 |
| puf60a | 0.00925 | 0.654313158 | 0.286 | 0 | 1 | 24 |
| atp5md | 0.00957 | -0.82035675 | 0.333 | 0.812 | 1 | 24 |
| fundc2 | 0.0113 | -0.54680056 | 0 | 0.25 | 1 | 24 |
| utp15 | 0.0113 | -0.51478281 | 0 | 0.25 | 1 | 24 |
| arpc4 | 0.0113 | -0.6479708 | 0 | 0.25 | 1 | 24 |
| ctnnb2 | 0.0113 | -0.60691872 | 0 | 0.25 | 1 | 24 |
| kctd12.2 | 0.0113 | -0.69308983 | 0 | 0.25 | 1 | 24 |
| rab6a | 0.0113 | -0.5049319 | 0 | 0.25 | 1 | 24 |
| pim2 | 0.0113 | -0.62231732 | 0 | 0.25 | 1 | 24 |
| exosc2 | 0.0113 | -0.60954973 | 0 | 0.25 | 1 | 24 |
| cct6a | 0.01421 | -0.87634619 | 0.048 | 0.438 | 1 | 24 |
| ndufa8 | 0.01421 | -0.69904008 | 0.048 | 0.438 | 1 | 24 |
| ndufa12 | 0.01817 | -0.68484973 | 0.095 | 0.5 | 1 | 24 |
| mgst1.2 | 0.01997 | 0.594590262 | 0.238 | 0 | 1 | 24 |
| ncbp2 | 0.01997 | 0.488279702 | 0.238 | 0 | 1 | 24 |
| col4a1 | 0.01997 | 0.58015807 | 0.238 | 0 | 1 | 24 |
| ngdn | 0.01997 | 0.582994419 | 0.238 | 0 | 1 | 24 |
| sox4a | 0.01997 | 0.570288077 | 0.238 | 0 | 1 | 24 |
| arhgef1b | 0.01997 | 0.612523089 | 0.238 | 0 | 1 | 24 |

| | | | | | | |
|---|---|---|---|---|---|---|
| capza1b | 0.02297 | -0.81994897 | 0.143 | 0.562 | 1 | 24 |
| zgc:162730 | 0.02312 | -0.89020136 | 0.143 | 0.562 | 1 | 24 |
| marcksl1a | 0.02956 | 0.89305407 | 0.429 | 0.062 | 1 | 24 |
| polr2f | 0.03136 | 0.703897267 | 0.429 | 0.062 | 1 | 24 |
| srsf3b | 0.03208 | 0.847815199 | 0.524 | 0.125 | 1 | 24 |
| bmper | 0.03323 | -0.51907128 | 0 | 0.188 | 1 | 24 |
| ccdc93 | 0.03323 | -0.54207518 | 0 | 0.188 | 1 | 24 |
| cox20 | 0.03323 | -0.48025653 | 0 | 0.188 | 1 | 24 |
| crkl | 0.03323 | -0.536819 | 0 | 0.188 | 1 | 24 |
| zgc:152791 | 0.03323 | -0.64293117 | 0 | 0.188 | 1 | 24 |
| akt1s1 | 0.03323 | -0.33745193 | 0 | 0.188 | 1 | 24 |
| cfdp1 | 0.03323 | -0.3323137 | 0 | 0.188 | 1 | 24 |
| prkrip1 | 0.03323 | -0.3130048 | 0 | 0.188 | 1 | 24 |
| psmg3 | 0.03323 | -0.37674652 | 0 | 0.188 | 1 | 24 |
| tprkb | 0.03323 | -0.45905239 | 0 | 0.188 | 1 | 24 |
| wdr21 | 0.03323 | -0.47515425 | 0 | 0.188 | 1 | 24 |
| zgc:153372 | 0.03323 | -0.37749192 | 0 | 0.188 | 1 | 24 |
| PYURF | 0.03323 | -0.27898822 | 0 | 0.188 | 1 | 24 |
| afmid | 0.03323 | -0.49853514 | 0 | 0.188 | 1 | 24 |
| cops5 | 0.03323 | -0.28391019 | 0 | 0.188 | 1 | 24 |
| cox16 | 0.03323 | -0.40400016 | 0 | 0.188 | 1 | 24 |
| dhx38 | 0.03323 | -0.40402755 | 0 | 0.188 | 1 | 24 |
| ezh2 | 0.03323 | -0.28623479 | 0 | 0.188 | 1 | 24 |
| gstt1b | 0.03323 | -0.54257346 | 0 | 0.188 | 1 | 24 |
| med31 | 0.03323 | -0.44506631 | 0 | 0.188 | 1 | 24 |
| ppid | 0.03323 | -0.34290221 | 0 | 0.188 | 1 | 24 |
| sf3a3 | 0.03323 | -0.25603842 | 0 | 0.188 | 1 | 24 |
| timm10b | 0.03323 | -0.42076451 | 0 | 0.188 | 1 | 24 |
| tk1 | 0.03323 | -0.48048649 | 0 | 0.188 | 1 | 24 |
| tmem53 | 0.03323 | -0.4331447 | 0 | 0.188 | 1 | 24 |
| vaspb | 0.03323 | -0.41943614 | 0 | 0.188 | 1 | 24 |
| add1 | 0.03323 | -0.38384216 | 0 | 0.188 | 1 | 24 |
| akirin2 | 0.03323 | -0.40805484 | 0 | 0.188 | 1 | 24 |
| arl6ip4 | 0.03323 | -0.49222231 | 0 | 0.188 | 1 | 24 |
| cox8b | 0.03323 | -0.51884612 | 0 | 0.188 | 1 | 24 |
| dnmt3ab | 0.03323 | -0.33970605 | 0 | 0.188 | 1 | 24 |
| kifc1 | 0.03323 | -0.55386437 | 0 | 0.188 | 1 | 24 |
| mbd3a | 0.03323 | -0.44007346 | 0 | 0.188 | 1 | 24 |
| mrpl10 | 0.03323 | -0.49866861 | 0 | 0.188 | 1 | 24 |
| msi2a | 0.03323 | -0.39598281 | 0 | 0.188 | 1 | 24 |
| naa15a | 0.03323 | -0.38264205 | 0 | 0.188 | 1 | 24 |
| nr2f1a | 0.03323 | -0.49738253 | 0 | 0.188 | 1 | 24 |
| parp16 | 0.03323 | -0.29718448 | 0 | 0.188 | 1 | 24 |
| pdcd10a | 0.03323 | -0.41904282 | 0 | 0.188 | 1 | 24 |
| pole2 | 0.03323 | -0.45820693 | 0 | 0.188 | 1 | 24 |
| ppp2r5eb | 0.03323 | -0.39750418 | 0 | 0.188 | 1 | 24 |
| prdx1 | 0.03323 | -0.3816817 | 0 | 0.188 | 1 | 24 |
| rangap1a | 0.03323 | -0.40152717 | 0 | 0.188 | 1 | 24 |

| | | | | | | |
|---|---|---|---|---|---|---|
| rbp4 | 0.03323 | -0.5991905 | 0 | 0.188 | 1 | 24 |
| rcor1 | 0.03323 | -0.44331848 | 0 | 0.188 | 1 | 24 |
| rnaseh2b | 0.03323 | -0.40983154 | 0 | 0.188 | 1 | 24 |
| si:dkey-46a10.3 | 0.03323 | -0.39600551 | 0 | 0.188 | 1 | 24 |
| ssr2 | 0.03323 | -0.3967681 | 0 | 0.188 | 1 | 24 |
| stat2 | 0.03323 | -0.44403744 | 0 | 0.188 | 1 | 24 |
| uxt | 0.03323 | -0.47481189 | 0 | 0.188 | 1 | 24 |
| vapal | 0.03323 | -0.45967774 | 0 | 0.188 | 1 | 24 |
| zgc:77650 | 0.03323 | -0.35574999 | 0 | 0.188 | 1 | 24 |
| znf367 | 0.03323 | -0.52537775 | 0 | 0.188 | 1 | 24 |
| anapc16 | 0.03583 | -0.56518772 | 0.048 | 0.375 | 1 | 24 |
| elocb | 0.03704 | -0.61418956 | 0.048 | 0.375 | 1 | 24 |
| krt15 | 0.03728 | -0.71801018 | 0.048 | 0.375 | 1 | 24 |
| ppig | 0.0377 | -0.73602078 | 0.048 | 0.375 | 1 | 24 |
| etfb | 0.04226 | 0.378609252 | 0.19 | 0 | 1 | 24 |
| lamtor2 | 0.04226 | 0.461517734 | 0.19 | 0 | 1 | 24 |
| si:ch73-376l24.2 | 0.04226 | 0.464192056 | 0.19 | 0 | 1 | 24 |
| bhmt | 0.04226 | 0.429493396 | 0.19 | 0 | 1 | 24 |
| cebpb | 0.04226 | 0.380133105 | 0.19 | 0 | 1 | 24 |
| gcsha | 0.04226 | 0.509036771 | 0.19 | 0 | 1 | 24 |
| mrpl52 | 0.04226 | 0.517889572 | 0.19 | 0 | 1 | 24 |
| nanos1 | 0.04226 | 0.514130181 | 0.19 | 0 | 1 | 24 |
| tmem11 | 0.04226 | 0.483559146 | 0.19 | 0 | 1 | 24 |
| CDK2AP1 | 0.04226 | 0.456132114 | 0.19 | 0 | 1 | 24 |
| UBL4A | 0.04226 | 0.500376866 | 0.19 | 0 | 1 | 24 |
| elovl1b | 0.04226 | 0.43366034 | 0.19 | 0 | 1 | 24 |
| mrpl42 | 0.04226 | 0.513076076 | 0.19 | 0 | 1 | 24 |
| FQ323156.1 | 0.04226 | 0.362717747 | 0.19 | 0 | 1 | 24 |
| cald1a | 0.04226 | 0.488794971 | 0.19 | 0 | 1 | 24 |
| wdr83os | 0.04226 | 0.476499039 | 0.19 | 0 | 1 | 24 |
| nudt15 | 0.04226 | 0.399302211 | 0.19 | 0 | 1 | 24 |
| tpi1b | 0.04226 | 0.477326583 | 0.19 | 0 | 1 | 24 |
| ISCU (1 of many) | 0.04226 | 0.507112575 | 0.19 | 0 | 1 | 24 |
| aimp2 | 0.04226 | 0.508955311 | 0.19 | 0 | 1 | 24 |
| bhlhe40 | 0.04226 | 0.478271347 | 0.19 | 0 | 1 | 24 |
| si:ch73-37h15.2 | 0.04226 | 0.494121484 | 0.19 | 0 | 1 | 24 |
| spina | 0.04226 | 0.510897296 | 0.19 | 0 | 1 | 24 |
| eno3 | 0.04242 | -0.84852679 | 0.095 | 0.438 | 1 | 24 |
| krt94 | 0.05351 | -0.84317032 | 0.095 | 0.438 | 1 | 24 |
| gstm.3 | 0.05606 | -0.93364702 | 0.19 | 0.562 | 1 | 24 |
| si:ch211-156b7.4 | 0.05989 | -0.61512953 | 0.19 | 0.562 | 1 | 24 |
| tmem258 | 0.06127 | -0.60098672 | 0.143 | 0.5 | 1 | 24 |
| qkia | 0.06136 | 0.830807366 | 0.381 | 0.062 | 1 | 24 |
| h2afx1 | 0.07018 | -0.12393997 | 0.238 | 0.5 | 1 | 24 |
| vat1 | 0.07843 | -0.71893907 | 0.048 | 0.312 | 1 | 24 |
| gstm.1 | 0.0791 | -0.70261265 | 0.333 | 0.688 | 1 | 24 |
| syf2 | 0.07911 | -0.61787954 | 0.048 | 0.312 | 1 | 24 |
| uba2 | 0.0803 | -0.61835049 | 0.048 | 0.312 | 1 | 24 |

| | | | | | | |
|---|---|---|---|---|---|---|
| rbp5 | 0.08564 | -0.57134269 | 0.048 | 0.312 | 1 | 24 |
| s100a10b | 0.08583 | -0.52320214 | 0.333 | 0.688 | 1 | 24 |
| aldh6a1 | 0.08834 | 0.299446565 | 0.143 | 0 | 1 | 24 |
| gsto2 | 0.08834 | 0.306182203 | 0.143 | 0 | 1 | 24 |
| anapc11 | 0.08834 | 0.483316773 | 0.143 | 0 | 1 | 24 |
| ccnd1 | 0.08834 | 0.371382125 | 0.143 | 0 | 1 | 24 |
| ddx51 | 0.08834 | 0.328920137 | 0.143 | 0 | 1 | 24 |
| ddx54 | 0.08834 | 0.430762226 | 0.143 | 0 | 1 | 24 |
| gpd1c | 0.08834 | 0.306645462 | 0.143 | 0 | 1 | 24 |
| gstm.2 | 0.08834 | 0.369451182 | 0.143 | 0 | 1 | 24 |
| josd2 | 0.08834 | 0.261972117 | 0.143 | 0 | 1 | 24 |
| shmt1 | 0.08834 | 0.267745455 | 0.143 | 0 | 1 | 24 |
| si:ch211-246m6.4 | 0.08834 | 0.424005489 | 0.143 | 0 | 1 | 24 |
| si:ch211-276i12.11 | 0.08834 | 0.362329256 | 0.143 | 0 | 1 | 24 |
| si:dkey-33c12.4 | 0.08834 | 0.268050812 | 0.143 | 0 | 1 | 24 |
| CR847984.1 | 0.08834 | 0.329330392 | 0.143 | 0 | 1 | 24 |
| XLOC-008119 | 0.08834 | 0.365082852 | 0.143 | 0 | 1 | 24 |
| XLOC-043857 | 0.08834 | 0.247507862 | 0.143 | 0 | 1 | 24 |
| api5 | 0.08834 | 0.366000679 | 0.143 | 0 | 1 | 24 |
| arfip1 | 0.08834 | 0.274134273 | 0.143 | 0 | 1 | 24 |
| chd4a | 0.08834 | 0.377545689 | 0.143 | 0 | 1 | 24 |
| chek1 | 0.08834 | 0.406662086 | 0.143 | 0 | 1 | 24 |
| fnta | 0.08834 | 0.448211334 | 0.143 | 0 | 1 | 24 |
| gpx1b | 0.08834 | 0.422493478 | 0.143 | 0 | 1 | 24 |
| lsm1 | 0.08834 | 0.384818426 | 0.143 | 0 | 1 | 24 |
| mcrip1 | 0.08834 | 0.463415649 | 0.143 | 0 | 1 | 24 |
| mis12 | 0.08834 | 0.409147026 | 0.143 | 0 | 1 | 24 |
| mre11a | 0.08834 | 0.468720648 | 0.143 | 0 | 1 | 24 |
| ndufb10 | 0.08834 | 0.267403597 | 0.143 | 0 | 1 | 24 |
| nup37 | 0.08834 | 0.358727474 | 0.143 | 0 | 1 | 24 |
| phc2a | 0.08834 | 0.319131425 | 0.143 | 0 | 1 | 24 |
| polr3k | 0.08834 | 0.390344606 | 0.143 | 0 | 1 | 24 |
| psmc3 | 0.08834 | 0.316222649 | 0.143 | 0 | 1 | 24 |
| psmd13 | 0.08834 | 0.439453079 | 0.143 | 0 | 1 | 24 |
| psmf1 | 0.08834 | 0.354456803 | 0.143 | 0 | 1 | 24 |
| qdprb1 | 0.08834 | 0.286671764 | 0.143 | 0 | 1 | 24 |
| sema3bl | 0.08834 | 0.447353307 | 0.143 | 0 | 1 | 24 |
| si:ch211-140m22.7 | 0.08834 | 0.334986597 | 0.143 | 0 | 1 | 24 |
| si:dkey-14d8.7 | 0.08834 | 0.311372533 | 0.143 | 0 | 1 | 24 |
| utp11l | 0.08834 | 0.43412076 | 0.143 | 0 | 1 | 24 |
| wdr5 | 0.08834 | 0.296954409 | 0.143 | 0 | 1 | 24 |
| bsdc1 | 0.08834 | 0.381328236 | 0.143 | 0 | 1 | 24 |
| cluap1 | 0.08834 | 0.371132605 | 0.143 | 0 | 1 | 24 |
| ctdsplb | 0.08834 | 0.367669357 | 0.143 | 0 | 1 | 24 |
| dennd2da | 0.08834 | 0.33362594 | 0.143 | 0 | 1 | 24 |
| eif4a2 | 0.08834 | 0.362550714 | 0.143 | 0 | 1 | 24 |
| gpia | 0.08834 | 0.345792003 | 0.143 | 0 | 1 | 24 |
| id2a | 0.08834 | 0.412542212 | 0.143 | 0 | 1 | 24 |

| | | | | | | |
|---|---|---|---|---|---|---|
| rbbp6 | 0.08834 | 0.347695053 | 0.143 | 0 | 1 | 24 |
| si:ch211-39i22.1 | 0.08834 | 0.431090858 | 0.143 | 0 | 1 | 24 |
| MCUR1 | 0.08834 | 0.302286283 | 0.143 | 0 | 1 | 24 |
| adka | 0.08834 | 0.372301791 | 0.143 | 0 | 1 | 24 |
| ankrd11 | 0.08834 | 0.290219203 | 0.143 | 0 | 1 | 24 |
| ap1m1 | 0.08834 | 0.244901519 | 0.143 | 0 | 1 | 24 |
| arglu1b | 0.08834 | 0.378743084 | 0.143 | 0 | 1 | 24 |
| cdca7b | 0.08834 | 0.324635165 | 0.143 | 0 | 1 | 24 |
| ddx18 | 0.08834 | 0.516134051 | 0.143 | 0 | 1 | 24 |
| e2f4 | 0.08834 | 0.387559686 | 0.143 | 0 | 1 | 24 |
| ergic3 | 0.08834 | 0.275188249 | 0.143 | 0 | 1 | 24 |
| ip6k2a | 0.08834 | 0.27230736 | 0.143 | 0 | 1 | 24 |
| irf2a | 0.08834 | 0.313421127 | 0.143 | 0 | 1 | 24 |
| kdm1a | 0.08834 | 0.326798607 | 0.143 | 0 | 1 | 24 |
| lmnb2 | 0.08834 | 0.315924062 | 0.143 | 0 | 1 | 24 |
| map1lc3b | 0.08834 | 0.32770775 | 0.143 | 0 | 1 | 24 |
| mrpl35 | 0.08834 | 0.286220512 | 0.143 | 0 | 1 | 24 |
| ncapg | 0.08834 | 0.278799987 | 0.143 | 0 | 1 | 24 |
| npm1b | 0.08834 | 0.433510123 | 0.143 | 0 | 1 | 24 |
| pgam5 | 0.08834 | 0.423257779 | 0.143 | 0 | 1 | 24 |
| ppp1cb | 0.08834 | 0.282670657 | 0.143 | 0 | 1 | 24 |
| rap2c | 0.08834 | 0.361110169 | 0.143 | 0 | 1 | 24 |
| rpz5 | 0.08834 | 0.352741368 | 0.143 | 0 | 1 | 24 |
| serinc1 | 0.08834 | 0.466374741 | 0.143 | 0 | 1 | 24 |
| si:ch211-262e15.1 | 0.08834 | 0.336047446 | 0.143 | 0 | 1 | 24 |
| stag2b | 0.08834 | 0.378863904 | 0.143 | 0 | 1 | 24 |
| tcp11l2 | 0.08834 | 0.339142855 | 0.143 | 0 | 1 | 24 |
| tuba1a | 0.08834 | 0.352740111 | 0.143 | 0 | 1 | 24 |
| ube2s | 0.08834 | 0.419022769 | 0.143 | 0 | 1 | 24 |
| zc3h10 | 0.08834 | 0.270369878 | 0.143 | 0 | 1 | 24 |
| zgc:153317 | 0.08834 | 0.28282837 | 0.143 | 0 | 1 | 24 |
| zgc:56106 | 0.08834 | 0.292660473 | 0.143 | 0 | 1 | 24 |
| ube2d1b | 0.08871 | -0.51940384 | 0.048 | 0.312 | 1 | 24 |
| ppp2r2d | 0.08955 | -0.49202701 | 0.048 | 0.312 | 1 | 24 |
| si:ch211-22i13.2 | 0.09147 | 0.713723375 | 0.333 | 0.062 | 1 | 24 |
| sec11a | 0.09158 | -0.53257934 | 0.048 | 0.312 | 1 | 24 |
| dap1b | 0.0917 | -0.59431531 | 0.048 | 0.312 | 1 | 24 |
| mt-co2 | 0.09374 | 0.757034949 | 0.333 | 0.062 | 1 | 24 |
| ccdc50 | 0.09611 | -0.37637327 | 0 | 0.125 | 1 | 24 |
| chchd6a | 0.09611 | -0.27090232 | 0 | 0.125 | 1 | 24 |
| itfg2 | 0.09611 | -0.27212318 | 0 | 0.125 | 1 | 24 |
| mhc1laa | 0.09611 | -0.2712857 | 0 | 0.125 | 1 | 24 |
| ndufaf3 | 0.09611 | -0.38697848 | 0 | 0.125 | 1 | 24 |
| pbk | 0.09611 | -0.39115313 | 0 | 0.125 | 1 | 24 |
| CABZ01076667.1 | 0.09611 | -0.23464242 | 0 | 0.125 | 1 | 24 |
| CABZ01110881.1 | 0.09611 | -0.25317406 | 0 | 0.125 | 1 | 24 |
| COA4 | 0.09611 | -0.26791206 | 0 | 0.125 | 1 | 24 |
| FO082877.1 | 0.09611 | -0.29967819 | 0 | 0.125 | 1 | 24 |

| | | | | | | |
|---|---|---|---|---|---|---|
| LOC101882596 | 0.09611 | -0.33224794 | 0 | 0.125 | 1 | 24 |
| XLOC-029463 | 0.09611 | -0.23991567 | 0 | 0.125 | 1 | 24 |
| abi3a | 0.09611 | -0.45284776 | 0 | 0.125 | 1 | 24 |
| aldh2.1 | 0.09611 | -0.21025053 | 0 | 0.125 | 1 | 24 |
| apex2 | 0.09611 | -0.34030575 | 0 | 0.125 | 1 | 24 |
| arf4a | 0.09611 | -0.21751817 | 0 | 0.125 | 1 | 24 |
| arl8a | 0.09611 | -0.30235522 | 0 | 0.125 | 1 | 24 |
| ash1l | 0.09611 | -0.36377569 | 0 | 0.125 | 1 | 24 |
| atp6v1ba | 0.09611 | -0.47408832 | 0 | 0.125 | 1 | 24 |
| bcl6aa | 0.09611 | -0.19614219 | 0 | 0.125 | 1 | 24 |
| blcap | 0.09611 | -0.18288088 | 0 | 0.125 | 1 | 24 |
| calcoco2 | 0.09611 | -0.21244887 | 0 | 0.125 | 1 | 24 |
| cdca5 | 0.09611 | -0.42348998 | 0 | 0.125 | 1 | 24 |
| cenpp | 0.09611 | -0.3056944 | 0 | 0.125 | 1 | 24 |
| cfap36 | 0.09611 | -0.34527205 | 0 | 0.125 | 1 | 24 |
| cib1 | 0.09611 | -0.24851578 | 0 | 0.125 | 1 | 24 |
| cip2a | 0.09611 | -0.2822301 | 0 | 0.125 | 1 | 24 |
| cmc1 | 0.09611 | -0.35120887 | 0 | 0.125 | 1 | 24 |
| cmtm3 | 0.09611 | -0.3689408 | 0 | 0.125 | 1 | 24 |
| col6a3 | 0.09611 | -0.24167847 | 0 | 0.125 | 1 | 24 |
| cox5b2 | 0.09611 | -0.41190611 | 0 | 0.125 | 1 | 24 |
| cox7a1 | 0.09611 | -0.28779916 | 0 | 0.125 | 1 | 24 |
| crema | 0.09611 | -0.35194912 | 0 | 0.125 | 1 | 24 |
| ctnnbip1 | 0.09611 | -0.29830558 | 0 | 0.125 | 1 | 24 |
| cxcl18b | 0.09611 | -0.19078447 | 0 | 0.125 | 1 | 24 |
| dap3 | 0.09611 | -0.33769244 | 0 | 0.125 | 1 | 24 |
| dhdhl | 0.09611 | -0.33097061 | 0 | 0.125 | 1 | 24 |
| dnajc5ga | 0.09611 | -0.23164336 | 0 | 0.125 | 1 | 24 |
| dtd1 | 0.09611 | -0.32165342 | 0 | 0.125 | 1 | 24 |
| fam3c | 0.09611 | -0.23747019 | 0 | 0.125 | 1 | 24 |
| fbxo44 | 0.09611 | -0.33223277 | 0 | 0.125 | 1 | 24 |
| fbxw2 | 0.09611 | -0.19615373 | 0 | 0.125 | 1 | 24 |
| fkbp2 | 0.09611 | -0.13153424 | 0 | 0.125 | 1 | 24 |
| foxn3 | 0.09611 | -0.30887269 | 0 | 0.125 | 1 | 24 |
| gpr183a | 0.09611 | -0.22854017 | 0 | 0.125 | 1 | 24 |
| hdac8 | 0.09611 | -0.2324564 | 0 | 0.125 | 1 | 24 |
| hmbox1a | 0.09611 | -0.25807818 | 0 | 0.125 | 1 | 24 |
| hnrnpl | 0.09611 | -0.24525958 | 0 | 0.125 | 1 | 24 |
| hoxc8a | 0.09611 | -0.35705457 | 0 | 0.125 | 1 | 24 |
| idh3a | 0.09611 | -0.23428596 | 0 | 0.125 | 1 | 24 |
| ifngr2 | 0.09611 | -0.22915604 | 0 | 0.125 | 1 | 24 |
| igf2bp2b | 0.09611 | -0.25402174 | 0 | 0.125 | 1 | 24 |
| ints8 | 0.09611 | -0.29696139 | 0 | 0.125 | 1 | 24 |
| irf2 | 0.09611 | -0.19680995 | 0 | 0.125 | 1 | 24 |
| kat6a | 0.09611 | -0.2578321 | 0 | 0.125 | 1 | 24 |
| krt4 | 0.09611 | -0.37463978 | 0 | 0.125 | 1 | 24 |
| lap3 | 0.09611 | -0.30222577 | 0 | 0.125 | 1 | 24 |
| ldha | 0.09611 | -0.24694449 | 0 | 0.125 | 1 | 24 |

| | | | | | | |
|---|---|---|---|---|---|---|
| lgals3a | 0.09611 | -0.28944953 | 0 | 0.125 | 1 | 24 |
| lsg1 | 0.09611 | -0.20576981 | 0 | 0.125 | 1 | 24 |
| mcmbp | 0.09611 | -0.18185144 | 0 | 0.125 | 1 | 24 |
| mier1a | 0.09611 | -0.37025579 | 0 | 0.125 | 1 | 24 |
| mllt3 | 0.09611 | -0.32411888 | 0 | 0.125 | 1 | 24 |
| mrps18a | 0.09611 | -0.28854277 | 0 | 0.125 | 1 | 24 |
| nedd8l | 0.09611 | -0.17532886 | 0 | 0.125 | 1 | 24 |
| nudt3a | 0.09611 | -0.33838736 | 0 | 0.125 | 1 | 24 |
| pacsin3 | 0.09611 | -0.26275055 | 0 | 0.125 | 1 | 24 |
| pak1ip1 | 0.09611 | -0.3369783 | 0 | 0.125 | 1 | 24 |
| pak4 | 0.09611 | -0.33879048 | 0 | 0.125 | 1 | 24 |
| pdcd6ip | 0.09611 | -0.22864119 | 0 | 0.125 | 1 | 24 |
| pgr | 0.09611 | -0.34184252 | 0 | 0.125 | 1 | 24 |
| plk3 | 0.09611 | -0.2879451 | 0 | 0.125 | 1 | 24 |
| pmpcb | 0.09611 | -0.35003978 | 0 | 0.125 | 1 | 24 |
| ppp2r5ea | 0.09611 | -0.25370269 | 0 | 0.125 | 1 | 24 |
| ppp4cb | 0.09611 | -0.21333222 | 0 | 0.125 | 1 | 24 |
| prkacab | 0.09611 | -0.23236766 | 0 | 0.125 | 1 | 24 |
| prpf4 | 0.09611 | -0.34028578 | 0 | 0.125 | 1 | 24 |
| rbm25a | 0.09611 | -0.23498379 | 0 | 0.125 | 1 | 24 |
| samm50 | 0.09611 | -0.41478184 | 0 | 0.125 | 1 | 24 |
| selenom | 0.09611 | -0.18013642 | 0 | 0.125 | 1 | 24 |
| sept15 | 0.09611 | -0.37545741 | 0 | 0.125 | 1 | 24 |
| sh3glb1a | 0.09611 | -0.28218684 | 0 | 0.125 | 1 | 24 |
| shoc2 | 0.09611 | -0.30337539 | 0 | 0.125 | 1 | 24 |
| si:ch211-106e7.2 | 0.09611 | -0.33872518 | 0 | 0.125 | 1 | 24 |
| si:ch211-125a15.1 | 0.09611 | -0.17736245 | 0 | 0.125 | 1 | 24 |
| si:ch211-160f23.5 | 0.09611 | -0.20241816 | 0 | 0.125 | 1 | 24 |
| si:ch211-207i20.3 | 0.09611 | -0.29387485 | 0 | 0.125 | 1 | 24 |
| si:dkey-159f12.2 | 0.09611 | -0.19364745 | 0 | 0.125 | 1 | 24 |
| si:dkey-79d12.5 | 0.09611 | -0.2955537 | 0 | 0.125 | 1 | 24 |
| si:dkeyp-55f12.3 | 0.09611 | -0.45654875 | 0 | 0.125 | 1 | 24 |
| siah1 | 0.09611 | -0.25541884 | 0 | 0.125 | 1 | 24 |
| smfn | 0.09611 | -0.22128335 | 0 | 0.125 | 1 | 24 |
| smim4 | 0.09611 | -0.2357739 | 0 | 0.125 | 1 | 24 |
| smyd4 | 0.09611 | -0.35623285 | 0 | 0.125 | 1 | 24 |
| snap23.2 | 0.09611 | -0.35010061 | 0 | 0.125 | 1 | 24 |
| tbk1 | 0.09611 | -0.33444945 | 0 | 0.125 | 1 | 24 |
| tmem263 | 0.09611 | -0.33868172 | 0 | 0.125 | 1 | 24 |
| tox2 | 0.09611 | -0.20346545 | 0 | 0.125 | 1 | 24 |
| trub2 | 0.09611 | -0.30414105 | 0 | 0.125 | 1 | 24 |
| tsr3 | 0.09611 | -0.28517163 | 0 | 0.125 | 1 | 24 |
| ttc1 | 0.09611 | -0.25929124 | 0 | 0.125 | 1 | 24 |
| ube2e2 | 0.09611 | -0.30216045 | 0 | 0.125 | 1 | 24 |
| ubtfl | 0.09611 | -0.39664384 | 0 | 0.125 | 1 | 24 |
| ucmaa | 0.09611 | -0.38827755 | 0 | 0.125 | 1 | 24 |
| ufd1l | 0.09611 | -0.1750273 | 0 | 0.125 | 1 | 24 |
| urm1 | 0.09611 | -0.33037608 | 0 | 0.125 | 1 | 24 |

| | | | | | | |
|---|---|---|---|---|---|---|
| vps26b | 0.09611 | -0.19345563 | 0 | 0.125 | 1 | 24 |
| vps28 | 0.09611 | -0.41452662 | 0 | 0.125 | 1 | 24 |
| wasa | 0.09611 | -0.24071287 | 0 | 0.125 | 1 | 24 |
| wu:fj29h11 | 0.09611 | -0.22633271 | 0 | 0.125 | 1 | 24 |
| xpo7 | 0.09611 | -0.20152943 | 0 | 0.125 | 1 | 24 |
| zgc:171971 | 0.09611 | -0.34363078 | 0 | 0.125 | 1 | 24 |
| zgc:193812 | 0.09611 | -0.27899651 | 0 | 0.125 | 1 | 24 |
| zgc:194469 | 0.09611 | -0.37061778 | 0 | 0.125 | 1 | 24 |
| zgc:77929 | 0.09611 | -0.19116403 | 0 | 0.125 | 1 | 24 |
| znhit1 | 0.09611 | -0.22756312 | 0 | 0.125 | 1 | 24 |
| CABZ01078055.1 | 0.09611 | -0.45561729 | 0 | 0.125 | 1 | 24 |
| aebp2 | 0.09611 | -0.3093572 | 0 | 0.125 | 1 | 24 |
| ankrd12 | 0.09611 | -0.3209771 | 0 | 0.125 | 1 | 24 |
| ap3s1.1 | 0.09611 | -0.37073581 | 0 | 0.125 | 1 | 24 |
| atp2b1a | 0.09611 | -0.36925774 | 0 | 0.125 | 1 | 24 |
| ctdspl3 | 0.09611 | -0.30779846 | 0 | 0.125 | 1 | 24 |
| ctps1b | 0.09611 | -0.42127641 | 0 | 0.125 | 1 | 24 |
| ddx55 | 0.09611 | -0.25676858 | 0 | 0.125 | 1 | 24 |
| fam49ba | 0.09611 | -0.39780848 | 0 | 0.125 | 1 | 24 |
| grap2a | 0.09611 | -0.45330445 | 0 | 0.125 | 1 | 24 |
| hcst | 0.09611 | -0.31601119 | 0 | 0.125 | 1 | 24 |
| ice2 | 0.09611 | -0.35572173 | 0 | 0.125 | 1 | 24 |
| mis18bp1 | 0.09611 | -0.38450701 | 0 | 0.125 | 1 | 24 |
| msrb1b | 0.09611 | -0.25296297 | 0 | 0.125 | 1 | 24 |
| mvb12a | 0.09611 | -0.2907644 | 0 | 0.125 | 1 | 24 |
| nansa | 0.09611 | -0.36337093 | 0 | 0.125 | 1 | 24 |
| nedd1 | 0.09611 | -0.34930983 | 0 | 0.125 | 1 | 24 |
| nln | 0.09611 | -0.30442596 | 0 | 0.125 | 1 | 24 |
| nup107 | 0.09611 | -0.28690165 | 0 | 0.125 | 1 | 24 |
| p2ry10 | 0.09611 | -0.37179481 | 0 | 0.125 | 1 | 24 |
| prkaa1 | 0.09611 | -0.31917921 | 0 | 0.125 | 1 | 24 |
| psmd4a | 0.09611 | -0.25630242 | 0 | 0.125 | 1 | 24 |
| slc16a10 | 0.09611 | -0.41925741 | 0 | 0.125 | 1 | 24 |
| stoml2 | 0.09611 | -0.29121391 | 0 | 0.125 | 1 | 24 |
| timm50 | 0.09611 | -0.22245209 | 0 | 0.125 | 1 | 24 |
| vaspa | 0.09611 | -0.33434288 | 0 | 0.125 | 1 | 24 |
| zbed4 | 0.09611 | -0.29793121 | 0 | 0.125 | 1 | 24 |
| zgc:113411 | 0.09611 | -0.27573579 | 0 | 0.125 | 1 | 24 |
| zgc:174877 | 0.09611 | -0.29367378 | 0 | 0.125 | 1 | 24 |
| GNAZ | 0.09611 | -0.25055945 | 0 | 0.125 | 1 | 24 |
| LOC101884549 | 0.09611 | -0.27137397 | 0 | 0.125 | 1 | 24 |
| XLOC-039817 | 0.09611 | -0.19272307 | 0 | 0.125 | 1 | 24 |
| abcc5 | 0.09611 | -0.25794232 | 0 | 0.125 | 1 | 24 |
| abhd17b | 0.09611 | -0.30993468 | 0 | 0.125 | 1 | 24 |
| badb | 0.09611 | -0.3180473 | 0 | 0.125 | 1 | 24 |
| bahcc1b | 0.09611 | -0.30606161 | 0 | 0.125 | 1 | 24 |
| blnk | 0.09611 | -0.22718806 | 0 | 0.125 | 1 | 24 |
| ccnd2a | 0.09611 | -0.3740333 | 0 | 0.125 | 1 | 24 |

| | | | | | | |
|---|---|---|---|---|---|---|
| ccne1 | 0.09611 | -0.20444949 | 0 | 0.125 | 1 | 24 |
| cd7al | 0.09611 | -0.25652879 | 0 | 0.125 | 1 | 24 |
| cep104 | 0.09611 | -0.32884133 | 0 | 0.125 | 1 | 24 |
| cep350 | 0.09611 | -0.25602296 | 0 | 0.125 | 1 | 24 |
| commd5 | 0.09611 | -0.20128089 | 0 | 0.125 | 1 | 24 |
| csrnp1a | 0.09611 | -0.18171349 | 0 | 0.125 | 1 | 24 |
| ddx19 | 0.09611 | -0.20268148 | 0 | 0.125 | 1 | 24 |
| dhrs4 | 0.09611 | -0.25597916 | 0 | 0.125 | 1 | 24 |
| dhx15 | 0.09611 | -0.19544559 | 0 | 0.125 | 1 | 24 |
| dph1 | 0.09611 | -0.19913718 | 0 | 0.125 | 1 | 24 |
| dph5 | 0.09611 | -0.20357274 | 0 | 0.125 | 1 | 24 |
| dpp9 | 0.09611 | -0.32798996 | 0 | 0.125 | 1 | 24 |
| ech1 | 0.09611 | -0.18197197 | 0 | 0.125 | 1 | 24 |
| eif4g2a | 0.09611 | -0.20056238 | 0 | 0.125 | 1 | 24 |
| fbxl6 | 0.09611 | -0.32774083 | 0 | 0.125 | 1 | 24 |
| fkbp1b | 0.09611 | -0.25737235 | 0 | 0.125 | 1 | 24 |
| ftr82 | 0.09611 | -0.37942747 | 0 | 0.125 | 1 | 24 |
| gclm | 0.09611 | -0.37903555 | 0 | 0.125 | 1 | 24 |
| gemin7 | 0.09611 | -0.31886113 | 0 | 0.125 | 1 | 24 |
| gfi1ab | 0.09611 | -0.24605824 | 0 | 0.125 | 1 | 24 |
| gtf2h3 | 0.09611 | -0.30219092 | 0 | 0.125 | 1 | 24 |
| hibadhb | 0.09611 | -0.19501372 | 0 | 0.125 | 1 | 24 |
| kpna1 | 0.09611 | -0.20552227 | 0 | 0.125 | 1 | 24 |
| kras | 0.09611 | -0.33805435 | 0 | 0.125 | 1 | 24 |
| lcp2a | 0.09611 | -0.18298535 | 0 | 0.125 | 1 | 24 |
| lin37 | 0.09611 | -0.41054703 | 0 | 0.125 | 1 | 24 |
| lrrc8aa | 0.09611 | -0.27549972 | 0 | 0.125 | 1 | 24 |
| mad1l1 | 0.09611 | -0.20269996 | 0 | 0.125 | 1 | 24 |
| mbd3b | 0.09611 | -0.29964699 | 0 | 0.125 | 1 | 24 |
| mettl9 | 0.09611 | -0.19781834 | 0 | 0.125 | 1 | 24 |
| morc2 | 0.09611 | -0.20252755 | 0 | 0.125 | 1 | 24 |
| nap1l4b | 0.09611 | -0.19474901 | 0 | 0.125 | 1 | 24 |
| necap2 | 0.09611 | -0.32604415 | 0 | 0.125 | 1 | 24 |
| nfixb | 0.09611 | -0.25613678 | 0 | 0.125 | 1 | 24 |
| nob1 | 0.09611 | -0.30892801 | 0 | 0.125 | 1 | 24 |
| nolc1 | 0.09611 | -0.36293094 | 0 | 0.125 | 1 | 24 |
| pafah1b3 | 0.09611 | -0.23811517 | 0 | 0.125 | 1 | 24 |
| pdxka | 0.09611 | -0.27665341 | 0 | 0.125 | 1 | 24 |
| pepd | 0.09611 | -0.31149516 | 0 | 0.125 | 1 | 24 |
| phb2b | 0.09611 | -0.23017833 | 0 | 0.125 | 1 | 24 |
| pik3r1 | 0.09611 | -0.20230582 | 0 | 0.125 | 1 | 24 |
| ptena | 0.09611 | -0.2008876 | 0 | 0.125 | 1 | 24 |
| ptger4b | 0.09611 | -0.20471056 | 0 | 0.125 | 1 | 24 |
| pwp1 | 0.09611 | -0.23751831 | 0 | 0.125 | 1 | 24 |
| qars | 0.09611 | -0.30577234 | 0 | 0.125 | 1 | 24 |
| rab5ab | 0.09611 | -0.30519695 | 0 | 0.125 | 1 | 24 |
| rab8a | 0.09611 | -0.18566767 | 0 | 0.125 | 1 | 24 |
| rag1 | 0.09611 | -0.4642614 | 0 | 0.125 | 1 | 24 |

| | | | | | | |
|---|---|---|---|---|---|---|
| rfx5 | 0.09611 | -0.26825248 | 0 | 0.125 | 1 | 24 |
| rhogb | 0.09611 | -0.17353984 | 0 | 0.125 | 1 | 24 |
| ric8b | 0.09611 | -0.19954094 | 0 | 0.125 | 1 | 24 |
| rmnd5b | 0.09611 | -0.30456206 | 0 | 0.125 | 1 | 24 |
| scamp2 | 0.09611 | -0.30836775 | 0 | 0.125 | 1 | 24 |
| selenow2a | 0.09611 | -0.19821843 | 0 | 0.125 | 1 | 24 |
| sept9b | 0.09611 | -0.19121179 | 0 | 0.125 | 1 | 24 |
| sf1 | 0.09611 | -0.37896301 | 0 | 0.125 | 1 | 24 |
| sf3b2 | 0.09611 | -0.19831097 | 0 | 0.125 | 1 | 24 |
| si:ch211-11k18.4 | 0.09611 | -0.20232482 | 0 | 0.125 | 1 | 24 |
| si:dkey-217l24.1 | 0.09611 | -0.32574499 | 0 | 0.125 | 1 | 24 |
| si:dkey-34m19.3 | 0.09611 | -0.30403723 | 0 | 0.125 | 1 | 24 |
| si:rp71-68n21.9 | 0.09611 | -0.20099668 | 0 | 0.125 | 1 | 24 |
| sid1 | 0.09611 | -0.38012001 | 0 | 0.125 | 1 | 24 |
| slc25a43 | 0.09611 | -0.26403612 | 0 | 0.125 | 1 | 24 |
| slc29a1a | 0.09611 | -0.20929784 | 0 | 0.125 | 1 | 24 |
| slc43a1a | 0.09611 | -0.24899017 | 0 | 0.125 | 1 | 24 |
| slc6a6b | 0.09611 | -0.30513366 | 0 | 0.125 | 1 | 24 |
| smarca2 | 0.09611 | -0.26738967 | 0 | 0.125 | 1 | 24 |
| smco4 | 0.09611 | -0.3485989 | 0 | 0.125 | 1 | 24 |
| snai2 | 0.09611 | -0.32417934 | 0 | 0.125 | 1 | 24 |
| sntb2 | 0.09611 | -0.36521885 | 0 | 0.125 | 1 | 24 |
| spout1 | 0.09611 | -0.14075483 | 0 | 0.125 | 1 | 24 |
| sptssa | 0.09611 | -0.2793187 | 0 | 0.125 | 1 | 24 |
| ssuh2rs1 | 0.09611 | -0.36004164 | 0 | 0.125 | 1 | 24 |
| stat1a | 0.09611 | -0.20745104 | 0 | 0.125 | 1 | 24 |
| sypl2b | 0.09611 | -0.24503419 | 0 | 0.125 | 1 | 24 |
| taf2 | 0.09611 | -0.20979222 | 0 | 0.125 | 1 | 24 |
| tfr1b | 0.09611 | -0.18648919 | 0 | 0.125 | 1 | 24 |
| tmem39b | 0.09611 | -0.20100748 | 0 | 0.125 | 1 | 24 |
| tob1a | 0.09611 | -0.23796171 | 0 | 0.125 | 1 | 24 |
| tp53rk | 0.09611 | -0.30836533 | 0 | 0.125 | 1 | 24 |
| tsen15 | 0.09611 | -0.38210647 | 0 | 0.125 | 1 | 24 |
| tspan2a | 0.09611 | -0.28491982 | 0 | 0.125 | 1 | 24 |
| xpc | 0.09611 | -0.18632755 | 0 | 0.125 | 1 | 24 |
| zgc:194312 | 0.09611 | -0.24970121 | 0 | 0.125 | 1 | 24 |
| XLOC-042229 | 0.09611 | -0.42516005 | 0 | 0.125 | 1 | 24 |
| aaas | 0.09611 | -0.36378265 | 0 | 0.125 | 1 | 24 |
| aass | 0.09611 | -0.43205935 | 0 | 0.125 | 1 | 24 |
| aldh16a1 | 0.09611 | -0.26794544 | 0 | 0.125 | 1 | 24 |
| csk | 0.09611 | -0.26288696 | 0 | 0.125 | 1 | 24 |
| ctsf | 0.09611 | -0.43245953 | 0 | 0.125 | 1 | 24 |
| dnajc2 | 0.09611 | -0.29922612 | 0 | 0.125 | 1 | 24 |
| exoc8 | 0.09611 | -0.36820589 | 0 | 0.125 | 1 | 24 |
| med4 | 0.09611 | -0.35511419 | 0 | 0.125 | 1 | 24 |
| ncapd2 | 0.09611 | -0.29757253 | 0 | 0.125 | 1 | 24 |
| ndufv1 | 0.09611 | -0.25862015 | 0 | 0.125 | 1 | 24 |
| otud5b | 0.09611 | -0.32520683 | 0 | 0.125 | 1 | 24 |

| | | | | | | |
|---|---|---|---|---|---|---|
| pls3 | 0.09611 | -0.42433813 | 0 | 0.125 | 1 | 24 |
| rbpjb | 0.09611 | -0.33126215 | 0 | 0.125 | 1 | 24 |
| tra2b | 0.09611 | -0.25169735 | 0 | 0.125 | 1 | 24 |
| znf1015 | 0.09611 | -0.36932817 | 0 | 0.125 | 1 | 24 |
| sem1 | 0.10115 | 0.627743562 | 0.524 | 0.188 | 1 | 24 |
| eef1da | 0.10283 | -0.62889512 | 0.429 | 0.75 | 1 | 24 |
| rwdd1 | 0.11248 | -0.64875622 | 0.095 | 0.375 | 1 | 24 |
| ifi45 | 0.11945 | -0.70302294 | 0.095 | 0.375 | 1 | 24 |
| zgc:153713 | 0.1207 | -0.54074923 | 0.095 | 0.375 | 1 | 24 |
| eif4a1a | 0.12849 | -0.66531846 | 0.143 | 0.438 | 1 | 24 |
| vrk1 | 0.1331 | -0.58058414 | 0.143 | 0.438 | 1 | 24 |
| rad23b | 0.13364 | -0.4972906 | 0.143 | 0.438 | 1 | 24 |
| sh3bgrl3 | 0.13573 | -0.57177304 | 0.143 | 0.438 | 1 | 24 |
| atp5pb | 0.13574 | -0.50489895 | 0.429 | 0.75 | 1 | 24 |
| polr2d | 0.13724 | -0.52980666 | 0.19 | 0.5 | 1 | 24 |
| marcksb | 0.14527 | 0.374379387 | 0.286 | 0.062 | 1 | 24 |
| actb2 | 0.14572 | -0.40410885 | 0.762 | 0.938 | 1 | 24 |
| cirbpb | 0.14884 | -0.2262298 | 0.619 | 0.875 | 1 | 24 |
| atp5l | 0.15414 | 0.399982176 | 0.619 | 0.312 | 1 | 24 |
| rpl27a | 0.15562 | -0.60268038 | 0.429 | 0.688 | 1 | 24 |
| cox4i1 | 0.15702 | -0.56509191 | 0.381 | 0.688 | 1 | 24 |
| eif3f | 0.15712 | -0.3840234 | 0.381 | 0.688 | 1 | 24 |
| hmgb1a | 0.16089 | -0.55715083 | 0.381 | 0.688 | 1 | 24 |
| rps29 | 0.16303 | -0.2450722 | 0.905 | 1 | 1 | 24 |
| arhgef39 | 0.16767 | -0.61104644 | 0.048 | 0.25 | 1 | 24 |
| cfl1 | 0.16866 | -0.20210346 | 0.714 | 0.938 | 1 | 24 |
| bcas2 | 0.17102 | -0.60712368 | 0.048 | 0.25 | 1 | 24 |
| pfkfb4b | 0.17146 | 0.653461043 | 0.286 | 0.062 | 1 | 24 |
| eif1b | 0.17174 | 0.587645502 | 0.476 | 0.188 | 1 | 24 |
| nudt21 | 0.17486 | -0.55662285 | 0.048 | 0.25 | 1 | 24 |
| snrpb | 0.17647 | -0.45905787 | 0.381 | 0.688 | 1 | 24 |
| sap30l | 0.17766 | 0.586682301 | 0.286 | 0.062 | 1 | 24 |
| rpn1 | 0.18116 | 0.572009937 | 0.286 | 0.062 | 1 | 24 |
| anp32b | 0.18318 | -0.55505917 | 0.476 | 0.688 | 1 | 24 |
| coro1a | 0.18465 | -0.59304734 | 0.048 | 0.25 | 1 | 24 |
| PQLC1 | 0.1848 | 0.18506648 | 0.095 | 0 | 1 | 24 |
| caap1 | 0.1848 | 0.260343363 | 0.095 | 0 | 1 | 24 |
| crip2 | 0.1848 | 0.194937304 | 0.095 | 0 | 1 | 24 |
| pknox1.1 | 0.1848 | 0.194614655 | 0.095 | 0 | 1 | 24 |
| rabl3 | 0.1848 | 0.336620673 | 0.095 | 0 | 1 | 24 |
| ranbp9 | 0.1848 | 0.193209605 | 0.095 | 0 | 1 | 24 |
| actr6 | 0.1848 | 0.261119465 | 0.095 | 0 | 1 | 24 |
| atg13 | 0.1848 | 0.242744949 | 0.095 | 0 | 1 | 24 |
| chmp2a | 0.1848 | 0.245102274 | 0.095 | 0 | 1 | 24 |
| emp2 | 0.1848 | 0.335814601 | 0.095 | 0 | 1 | 24 |
| fhl2a | 0.1848 | 0.236843488 | 0.095 | 0 | 1 | 24 |
| ift46 | 0.1848 | 0.283590465 | 0.095 | 0 | 1 | 24 |
| il34 | 0.1848 | 0.306798347 | 0.095 | 0 | 1 | 24 |

| | | | | | | |
|---|---|---|---|---|---|---|
| map2k1 | 0.1848 | 0.310698326 | 0.095 | 0 | 1 | 24 |
| med22 | 0.1848 | 0.25837125 | 0.095 | 0 | 1 | 24 |
| ncaph | 0.1848 | 0.26119552 | 0.095 | 0 | 1 | 24 |
| pcgf1 | 0.1848 | 0.259654183 | 0.095 | 0 | 1 | 24 |
| pop4 | 0.1848 | 0.370406144 | 0.095 | 0 | 1 | 24 |
| rab23 | 0.1848 | 0.314628905 | 0.095 | 0 | 1 | 24 |
| rbpms2b | 0.1848 | 0.300550655 | 0.095 | 0 | 1 | 24 |
| sdhaf4 | 0.1848 | 0.32221244 | 0.095 | 0 | 1 | 24 |
| stxbp6 | 0.1848 | 0.22253003 | 0.095 | 0 | 1 | 24 |
| tdh | 0.1848 | 0.306055331 | 0.095 | 0 | 1 | 24 |
| tfdp1b | 0.1848 | 0.260328439 | 0.095 | 0 | 1 | 24 |
| ube2ka | 0.1848 | 0.260160171 | 0.095 | 0 | 1 | 24 |
| c1d | 0.1848 | 0.295829809 | 0.095 | 0 | 1 | 24 |
| ddx3xa | 0.1848 | 0.293830562 | 0.095 | 0 | 1 | 24 |
| gosr1 | 0.1848 | 0.29323597 | 0.095 | 0 | 1 | 24 |
| jupa | 0.1848 | 0.304556245 | 0.095 | 0 | 1 | 24 |
| mpped2 | 0.1848 | 0.252239943 | 0.095 | 0 | 1 | 24 |
| nuf2 | 0.1848 | 0.252434874 | 0.095 | 0 | 1 | 24 |
| prkcsh | 0.1848 | 0.252013952 | 0.095 | 0 | 1 | 24 |
| scamp4 | 0.1848 | 0.251179666 | 0.095 | 0 | 1 | 24 |
| zgc:153115 | 0.1848 | 0.253700709 | 0.095 | 0 | 1 | 24 |
| zgc:66160 | 0.1848 | 0.250149773 | 0.095 | 0 | 1 | 24 |
| znf576.2 | 0.1848 | 0.251073772 | 0.095 | 0 | 1 | 24 |
| BX957362.1 | 0.1848 | 0.241679389 | 0.095 | 0 | 1 | 24 |
| GNB4 | 0.1848 | 0.237755053 | 0.095 | 0 | 1 | 24 |
| ankrd10a | 0.1848 | 0.192774917 | 0.095 | 0 | 1 | 24 |
| aprt | 0.1848 | 0.306062503 | 0.095 | 0 | 1 | 24 |
| aqp12 | 0.1848 | 0.344647142 | 0.095 | 0 | 1 | 24 |
| baxb | 0.1848 | 0.263772059 | 0.095 | 0 | 1 | 24 |
| bcap31 | 0.1848 | 0.20827641 | 0.095 | 0 | 1 | 24 |
| bop1 | 0.1848 | 0.280427134 | 0.095 | 0 | 1 | 24 |
| cab39l | 0.1848 | 0.277093441 | 0.095 | 0 | 1 | 24 |
| cdc14b | 0.1848 | 0.207565892 | 0.095 | 0 | 1 | 24 |
| cenpf | 0.1848 | 0.22680462 | 0.095 | 0 | 1 | 24 |
| chd2 | 0.1848 | 0.163851096 | 0.095 | 0 | 1 | 24 |
| ciao1 | 0.1848 | 0.281192105 | 0.095 | 0 | 1 | 24 |
| clip2 | 0.1848 | 0.213639287 | 0.095 | 0 | 1 | 24 |
| cutc | 0.1848 | 0.247509026 | 0.095 | 0 | 1 | 24 |
| cybrd1 | 0.1848 | 0.218139603 | 0.095 | 0 | 1 | 24 |
| dlgap4a | 0.1848 | 0.287771006 | 0.095 | 0 | 1 | 24 |
| dnajb6a | 0.1848 | 0.20460228 | 0.095 | 0 | 1 | 24 |
| e2f3 | 0.1848 | 0.157368242 | 0.095 | 0 | 1 | 24 |
| ecd | 0.1848 | 0.168905364 | 0.095 | 0 | 1 | 24 |
| efemp2a | 0.1848 | 0.311817848 | 0.095 | 0 | 1 | 24 |
| emc10 | 0.1848 | 0.244244904 | 0.095 | 0 | 1 | 24 |
| hif1al | 0.1848 | 0.242796396 | 0.095 | 0 | 1 | 24 |
| hnrpkl | 0.1848 | 0.209590903 | 0.095 | 0 | 1 | 24 |
| hsbp1a | 0.1848 | 0.245572843 | 0.095 | 0 | 1 | 24 |

| | | | | | | |
|---|---|---|---|---|---|---|
| hspa5 | 0.1848 | 0.167188052 | 0.095 | 0 | 1 | 24 |
| irak1bp1 | 0.1848 | 0.275733371 | 0.095 | 0 | 1 | 24 |
| lama4 | 0.1848 | 0.289844962 | 0.095 | 0 | 1 | 24 |
| lamb1a | 0.1848 | 0.202580725 | 0.095 | 0 | 1 | 24 |
| lims1 | 0.1848 | 0.252004077 | 0.095 | 0 | 1 | 24 |
| mcfd2 | 0.1848 | 0.156394064 | 0.095 | 0 | 1 | 24 |
| med18 | 0.1848 | 0.162369605 | 0.095 | 0 | 1 | 24 |
| mrpl17 | 0.1848 | 0.166097659 | 0.095 | 0 | 1 | 24 |
| mrps28 | 0.1848 | 0.304621439 | 0.095 | 0 | 1 | 24 |
| mrps9 | 0.1848 | 0.308262854 | 0.095 | 0 | 1 | 24 |
| mybbp1a | 0.1848 | 0.291281691 | 0.095 | 0 | 1 | 24 |
| ncapg2 | 0.1848 | 0.363491523 | 0.095 | 0 | 1 | 24 |
| net1 | 0.1848 | 0.323001026 | 0.095 | 0 | 1 | 24 |
| nsd3 | 0.1848 | 0.296575148 | 0.095 | 0 | 1 | 24 |
| phlda2 | 0.1848 | 0.296408275 | 0.095 | 0 | 1 | 24 |
| ppp2r1bb | 0.1848 | 0.303646432 | 0.095 | 0 | 1 | 24 |
| prkacaa | 0.1848 | 0.155344839 | 0.095 | 0 | 1 | 24 |
| prrc1 | 0.1848 | 0.210049573 | 0.095 | 0 | 1 | 24 |
| psma6l | 0.1848 | 0.190945033 | 0.095 | 0 | 1 | 24 |
| ptp4a2a | 0.1848 | 0.266822655 | 0.095 | 0 | 1 | 24 |
| rhoca | 0.1848 | 0.301693152 | 0.095 | 0 | 1 | 24 |
| rnasekb | 0.1848 | 0.254106634 | 0.095 | 0 | 1 | 24 |
| rpf2 | 0.1848 | 0.280358169 | 0.095 | 0 | 1 | 24 |
| s100w | 0.1848 | 0.318611096 | 0.095 | 0 | 1 | 24 |
| si:ch211-121a2.2 | 0.1848 | 0.164067494 | 0.095 | 0 | 1 | 24 |
| si:ch211-25g7.5 | 0.1848 | 0.316405956 | 0.095 | 0 | 1 | 24 |
| si:ch73-308l14.2 | 0.1848 | 0.167196978 | 0.095 | 0 | 1 | 24 |
| si:ch73-376l24.3 | 0.1848 | 0.273965016 | 0.095 | 0 | 1 | 24 |
| spg21 | 0.1848 | 0.152619927 | 0.095 | 0 | 1 | 24 |
| ssbp3b | 0.1848 | 0.343231987 | 0.095 | 0 | 1 | 24 |
| tex10 | 0.1848 | 0.166136103 | 0.095 | 0 | 1 | 24 |
| tmem231 | 0.1848 | 0.212095981 | 0.095 | 0 | 1 | 24 |
| top2b | 0.1848 | 0.246786896 | 0.095 | 0 | 1 | 24 |
| tprb | 0.1848 | 0.1644434 | 0.095 | 0 | 1 | 24 |
| tubb5 | 0.1848 | 0.324560326 | 0.095 | 0 | 1 | 24 |
| u2af2a | 0.1848 | 0.251267088 | 0.095 | 0 | 1 | 24 |
| urod | 0.1848 | 0.158390559 | 0.095 | 0 | 1 | 24 |
| usp3 | 0.1848 | 0.245257575 | 0.095 | 0 | 1 | 24 |
| vps72a | 0.1848 | 0.168905364 | 0.095 | 0 | 1 | 24 |
| wars | 0.1848 | 0.272665696 | 0.095 | 0 | 1 | 24 |
| zgc:113336 | 0.1848 | 0.213639287 | 0.095 | 0 | 1 | 24 |
| BX470259.2 | 0.1848 | 0.274694637 | 0.095 | 0 | 1 | 24 |
| BX640522.2 | 0.1848 | 0.461650032 | 0.095 | 0 | 1 | 24 |
| CABZ01077217.1 | 0.1848 | 0.346531602 | 0.095 | 0 | 1 | 24 |
| CABZ01088109.1 | 0.1848 | 0.290806449 | 0.095 | 0 | 1 | 24 |
| CR293507.1 | 0.1848 | 0.247639344 | 0.095 | 0 | 1 | 24 |
| CT025742.1 | 0.1848 | 0.220162239 | 0.095 | 0 | 1 | 24 |
| CT573476.1 | 0.1848 | 0.333015822 | 0.095 | 0 | 1 | 24 |

| | | | | | | |
|---|---|---|---|---|---|---|
| LOC101882623 | 0.1848 | 0.263793722 | 0.095 | 0 | 1 | 24 |
| LOC108190702 | 0.1848 | 0.249827214 | 0.095 | 0 | 1 | 24 |
| MCMDC2 | 0.1848 | 0.272470741 | 0.095 | 0 | 1 | 24 |
| NAV1 (1 of many) | 0.1848 | 0.213650622 | 0.095 | 0 | 1 | 24 |
| XLOC-034235 | 0.1848 | 0.299676857 | 0.095 | 0 | 1 | 24 |
| ak1 | 0.1848 | 0.231431851 | 0.095 | 0 | 1 | 24 |
| alas1 | 0.1848 | 0.290617189 | 0.095 | 0 | 1 | 24 |
| aldh7a1 | 0.1848 | 0.252780406 | 0.095 | 0 | 1 | 24 |
| ap3s2 | 0.1848 | 0.211721239 | 0.095 | 0 | 1 | 24 |
| aqr | 0.1848 | 0.274037029 | 0.095 | 0 | 1 | 24 |
| arvcfb | 0.1848 | 0.235695861 | 0.095 | 0 | 1 | 24 |
| as3mt | 0.1848 | 0.189066441 | 0.095 | 0 | 1 | 24 |
| atp1b3a | 0.1848 | 0.27109448 | 0.095 | 0 | 1 | 24 |
| atp6v1d | 0.1848 | 0.175218236 | 0.095 | 0 | 1 | 24 |
| bnc2 | 0.1848 | 0.24607653 | 0.095 | 0 | 1 | 24 |
| bnip3la | 0.1848 | 0.296655063 | 0.095 | 0 | 1 | 24 |
| brdt | 0.1848 | 0.219979193 | 0.095 | 0 | 1 | 24 |
| brms1la | 0.1848 | 0.231039904 | 0.095 | 0 | 1 | 24 |
| calua | 0.1848 | 0.163842125 | 0.095 | 0 | 1 | 24 |
| capgb | 0.1848 | 0.204126674 | 0.095 | 0 | 1 | 24 |
| ccdc32 | 0.1848 | 0.287428749 | 0.095 | 0 | 1 | 24 |
| ccdc82 | 0.1848 | 0.2319918 | 0.095 | 0 | 1 | 24 |
| chchd1 | 0.1848 | 0.229413456 | 0.095 | 0 | 1 | 24 |
| cnn3b | 0.1848 | 0.313704695 | 0.095 | 0 | 1 | 24 |
| coil | 0.1848 | 0.193863954 | 0.095 | 0 | 1 | 24 |
| cpb1 | 0.1848 | 0.214985234 | 0.095 | 0 | 1 | 24 |
| cst14b.1 | 0.1848 | 0.122010246 | 0.095 | 0 | 1 | 24 |
| cybb | 0.1848 | 0.180837021 | 0.095 | 0 | 1 | 24 |
| dclre1c | 0.1848 | 0.184223688 | 0.095 | 0 | 1 | 24 |
| ddx20 | 0.1848 | 0.308218402 | 0.095 | 0 | 1 | 24 |
| dedd1 | 0.1848 | 0.310718045 | 0.095 | 0 | 1 | 24 |
| dhrs12 | 0.1848 | 0.166082205 | 0.095 | 0 | 1 | 24 |
| dhrs9 | 0.1848 | 0.12286373 | 0.095 | 0 | 1 | 24 |
| dlg5a | 0.1848 | 0.321994476 | 0.095 | 0 | 1 | 24 |
| dnajb1b | 0.1848 | 0.34480633 | 0.095 | 0 | 1 | 24 |
| dpf2 | 0.1848 | 0.229782469 | 0.095 | 0 | 1 | 24 |
| dub | 0.1848 | 0.196142265 | 0.095 | 0 | 1 | 24 |
| dync1li1 | 0.1848 | 0.308860164 | 0.095 | 0 | 1 | 24 |
| eif2b1 | 0.1848 | 0.165292804 | 0.095 | 0 | 1 | 24 |
| eif2b4 | 0.1848 | 0.197171301 | 0.095 | 0 | 1 | 24 |
| eif4ebp3 | 0.1848 | 0.315804366 | 0.095 | 0 | 1 | 24 |
| ero1b | 0.1848 | 0.199138573 | 0.095 | 0 | 1 | 24 |
| erp27 | 0.1848 | 0.171067912 | 0.095 | 0 | 1 | 24 |
| fam120c | 0.1848 | 0.177847359 | 0.095 | 0 | 1 | 24 |
| fbrs | 0.1848 | 0.245518271 | 0.095 | 0 | 1 | 24 |
| fbxl5 | 0.1848 | 0.209737655 | 0.095 | 0 | 1 | 24 |
| fcf1 | 0.1848 | 0.379972197 | 0.095 | 0 | 1 | 24 |
| flj11011l | 0.1848 | 0.200479247 | 0.095 | 0 | 1 | 24 |

| | | | | | | |
|---|---|---|---|---|---|---|
| ftr51 | 0.1848 | 0.325922624 | 0.095 | 0 | 1 | 24 |
| fxyd6l | 0.1848 | 0.203368971 | 0.095 | 0 | 1 | 24 |
| gemin8 | 0.1848 | 0.228408199 | 0.095 | 0 | 1 | 24 |
| gins4 | 0.1848 | 0.229501356 | 0.095 | 0 | 1 | 24 |
| glb1l | 0.1848 | 0.16519992 | 0.095 | 0 | 1 | 24 |
| gpr108 | 0.1848 | 0.166060509 | 0.095 | 0 | 1 | 24 |
| grpel1 | 0.1848 | 0.216514617 | 0.095 | 0 | 1 | 24 |
| hagh | 0.1848 | 0.350350614 | 0.095 | 0 | 1 | 24 |
| hectd1 | 0.1848 | 0.311904698 | 0.095 | 0 | 1 | 24 |
| ier5l | 0.1848 | 0.275193332 | 0.095 | 0 | 1 | 24 |
| igf2bp1 | 0.1848 | 0.157461971 | 0.095 | 0 | 1 | 24 |
| kif14 | 0.1848 | 0.324288954 | 0.095 | 0 | 1 | 24 |
| kif4 | 0.1848 | 0.278345127 | 0.095 | 0 | 1 | 24 |
| klf5l | 0.1848 | 0.259878325 | 0.095 | 0 | 1 | 24 |
| krt91 | 0.1848 | 0.393533721 | 0.095 | 0 | 1 | 24 |
| lrrc59 | 0.1848 | 0.245993835 | 0.095 | 0 | 1 | 24 |
| macf1a | 0.1848 | 0.324776408 | 0.095 | 0 | 1 | 24 |
| mob4 | 0.1848 | 0.364932587 | 0.095 | 0 | 1 | 24 |
| mrpl14 | 0.1848 | 0.281063815 | 0.095 | 0 | 1 | 24 |
| mrpl15 | 0.1848 | 0.364188864 | 0.095 | 0 | 1 | 24 |
| nat10 | 0.1848 | 0.249287549 | 0.095 | 0 | 1 | 24 |
| ndufaf4 | 0.1848 | 0.193772273 | 0.095 | 0 | 1 | 24 |
| nol4lb | 0.1848 | 0.196712478 | 0.095 | 0 | 1 | 24 |
| nt5c3a | 0.1848 | 0.145196757 | 0.095 | 0 | 1 | 24 |
| pcm1 | 0.1848 | 0.308393154 | 0.095 | 0 | 1 | 24 |
| pias4a | 0.1848 | 0.305913827 | 0.095 | 0 | 1 | 24 |
| pitrm1 | 0.1848 | 0.153192354 | 0.095 | 0 | 1 | 24 |
| pkig | 0.1848 | 0.27021207 | 0.095 | 0 | 1 | 24 |
| plgrkt | 0.1848 | 0.27096786 | 0.095 | 0 | 1 | 24 |
| pop7 | 0.1848 | 0.187933804 | 0.095 | 0 | 1 | 24 |
| ppp1r12a | 0.1848 | 0.295832552 | 0.095 | 0 | 1 | 24 |
| ppp6r3 | 0.1848 | 0.352832957 | 0.095 | 0 | 1 | 24 |
| prkab1a | 0.1848 | 0.238211393 | 0.095 | 0 | 1 | 24 |
| prpsap2 | 0.1848 | 0.165553237 | 0.095 | 0 | 1 | 24 |
| ptbp1b | 0.1848 | 0.143454955 | 0.095 | 0 | 1 | 24 |
| qtrt2 | 0.1848 | 0.30806525 | 0.095 | 0 | 1 | 24 |
| rab5if | 0.1848 | 0.254149734 | 0.095 | 0 | 1 | 24 |
| rab7a | 0.1848 | 0.235165925 | 0.095 | 0 | 1 | 24 |
| rars | 0.1848 | 0.266037723 | 0.095 | 0 | 1 | 24 |
| rgmd | 0.1848 | 0.347978518 | 0.095 | 0 | 1 | 24 |
| rnaseka | 0.1848 | 0.174770459 | 0.095 | 0 | 1 | 24 |
| rnmt | 0.1848 | 0.292082239 | 0.095 | 0 | 1 | 24 |
| rpp14 | 0.1848 | 0.34659948 | 0.095 | 0 | 1 | 24 |
| rprd1b | 0.1848 | 0.280107592 | 0.095 | 0 | 1 | 24 |
| rpusd4 | 0.1848 | 0.245003899 | 0.095 | 0 | 1 | 24 |
| sdr16c5b | 0.1848 | 0.255290177 | 0.095 | 0 | 1 | 24 |
| si:ch211-1o7.3 | 0.1848 | 0.123375751 | 0.095 | 0 | 1 | 24 |
| si:ch211-218c6.8 | 0.1848 | 0.14258793 | 0.095 | 0 | 1 | 24 |

| | | | | | | |
|---|---|---|---|---|---|---|
| si:dkey-14d8.6 | 0.1848 | 0.229027742 | 0.095 | 0 | 1 | 24 |
| si:dkey-222f8.3 | 0.1848 | 0.16074585 | 0.095 | 0 | 1 | 24 |
| si:dkey-226l10.6 | 0.1848 | 0.349299452 | 0.095 | 0 | 1 | 24 |
| si:dkeyp-113d7.1 | 0.1848 | 0.344807347 | 0.095 | 0 | 1 | 24 |
| slc48a1b | 0.1848 | 0.198713153 | 0.095 | 0 | 1 | 24 |
| smarcb1a | 0.1848 | 0.25873683 | 0.095 | 0 | 1 | 24 |
| snx17 | 0.1848 | 0.281564476 | 0.095 | 0 | 1 | 24 |
| snx3 | 0.1848 | 0.188785927 | 0.095 | 0 | 1 | 24 |
| srgap2 | 0.1848 | 0.143770711 | 0.095 | 0 | 1 | 24 |
| srrm1 | 0.1848 | 0.194142164 | 0.095 | 0 | 1 | 24 |
| srsf10b | 0.1848 | 0.273355144 | 0.095 | 0 | 1 | 24 |
| tcf7l2 | 0.1848 | 0.325922624 | 0.095 | 0 | 1 | 24 |
| ten1 | 0.1848 | 0.257592898 | 0.095 | 0 | 1 | 24 |
| tmeff1b | 0.1848 | 0.288733438 | 0.095 | 0 | 1 | 24 |
| tmem43 | 0.1848 | 0.141173598 | 0.095 | 0 | 1 | 24 |
| tnrc18 | 0.1848 | 0.332504393 | 0.095 | 0 | 1 | 24 |
| tp53inp2 | 0.1848 | 0.355503683 | 0.095 | 0 | 1 | 24 |
| trip12 | 0.1848 | 0.351783224 | 0.095 | 0 | 1 | 24 |
| tspan3a | 0.1848 | 0.202209961 | 0.095 | 0 | 1 | 24 |
| txlna | 0.1848 | 0.281068119 | 0.095 | 0 | 1 | 24 |
| uevld | 0.1848 | 0.273142242 | 0.095 | 0 | 1 | 24 |
| upf3b | 0.1848 | 0.311491975 | 0.095 | 0 | 1 | 24 |
| usp39 | 0.1848 | 0.163587692 | 0.095 | 0 | 1 | 24 |
| utp3 | 0.1848 | 0.151975422 | 0.095 | 0 | 1 | 24 |
| xaf1 | 0.1848 | 0.122317258 | 0.095 | 0 | 1 | 24 |
| yif1a | 0.1848 | 0.23587231 | 0.095 | 0 | 1 | 24 |
| yju2 | 0.1848 | 0.243891714 | 0.095 | 0 | 1 | 24 |
| ylpm1 | 0.1848 | 0.233233691 | 0.095 | 0 | 1 | 24 |
| zc3h7ba | 0.1848 | 0.308385954 | 0.095 | 0 | 1 | 24 |
| zgc:63831 | 0.1848 | 0.177003981 | 0.095 | 0 | 1 | 24 |
| zmp:0000000624 | 0.1848 | 0.291688491 | 0.095 | 0 | 1 | 24 |
| znf423 | 0.1848 | 0.323892251 | 0.095 | 0 | 1 | 24 |
| znf704 | 0.1848 | 0.318263663 | 0.095 | 0 | 1 | 24 |
| zw10 | 0.1848 | 0.19702802 | 0.095 | 0 | 1 | 24 |
| CABZ01113812.1 | 0.1848 | 0.257990442 | 0.095 | 0 | 1 | 24 |
| LOC110437782 | 0.1848 | 0.265163067 | 0.095 | 0 | 1 | 24 |
| acaa2 | 0.1848 | 0.194901049 | 0.095 | 0 | 1 | 24 |
| adcy9 | 0.1848 | 0.260066188 | 0.095 | 0 | 1 | 24 |
| cd2bp2 | 0.1848 | 0.246222016 | 0.095 | 0 | 1 | 24 |
| cdc20 | 0.1848 | 0.275008207 | 0.095 | 0 | 1 | 24 |
| fam129bb | 0.1848 | 0.263510104 | 0.095 | 0 | 1 | 24 |
| fermt2 | 0.1848 | 0.261029952 | 0.095 | 0 | 1 | 24 |
| gatad1 | 0.1848 | 0.195315142 | 0.095 | 0 | 1 | 24 |
| kat5b | 0.1848 | 0.198168449 | 0.095 | 0 | 1 | 24 |
| lrrtm1 | 0.1848 | 0.22811355 | 0.095 | 0 | 1 | 24 |
| map7d1b | 0.1848 | 0.311487592 | 0.095 | 0 | 1 | 24 |
| melk | 0.1848 | 0.266297649 | 0.095 | 0 | 1 | 24 |
| nufip2 | 0.1848 | 0.262774916 | 0.095 | 0 | 1 | 24 |

| | | | | | | |
|---|---|---|---|---|---|---|
| pmp22a | 0.1848 | 0.387385837 | 0.095 | 0 | 1 | 24 |
| rabl2 | 0.1848 | 0.2155837 | 0.095 | 0 | 1 | 24 |
| rbm14a | 0.1848 | 0.262432149 | 0.095 | 0 | 1 | 24 |
| rnf44 | 0.1848 | 0.238887494 | 0.095 | 0 | 1 | 24 |
| rybpa | 0.1848 | 0.194773624 | 0.095 | 0 | 1 | 24 |
| senp6a | 0.1848 | 0.23368184 | 0.095 | 0 | 1 | 24 |
| si:ch211-133n4.4 | 0.1848 | 0.278775218 | 0.095 | 0 | 1 | 24 |
| si:dkey-6e2.2 | 0.1848 | 0.269468669 | 0.095 | 0 | 1 | 24 |
| slc20a2 | 0.1848 | 0.300194962 | 0.095 | 0 | 1 | 24 |
| smarcd1 | 0.1848 | 0.196926064 | 0.095 | 0 | 1 | 24 |
| tacc3 | 0.1848 | 0.227989543 | 0.095 | 0 | 1 | 24 |
| tardbp | 0.1848 | 0.196650518 | 0.095 | 0 | 1 | 24 |
| uqcc2 | 0.1848 | 0.207599529 | 0.095 | 0 | 1 | 24 |
| zgc:161969 | 0.1848 | 0.214680018 | 0.095 | 0 | 1 | 24 |
| zgc:175264 | 0.1848 | 0.227196568 | 0.095 | 0 | 1 | 24 |
| znf503 | 0.1848 | 0.175149907 | 0.095 | 0 | 1 | 24 |
| znf800a | 0.1848 | 0.240605296 | 0.095 | 0 | 1 | 24 |
| BOLA2 | 0.1848 | 0.191840432 | 0.095 | 0 | 1 | 24 |
| CU074419.2 | 0.1848 | 0.108788516 | 0.095 | 0 | 1 | 24 |
| CU855930.1 | 0.1848 | 0.231736827 | 0.095 | 0 | 1 | 24 |
| FP236812.2 | 0.1848 | 0.182152206 | 0.095 | 0 | 1 | 24 |
| FQ976913.1 | 0.1848 | 0.244149869 | 0.095 | 0 | 1 | 24 |
| LOC110438237 | 0.1848 | 0.173553373 | 0.095 | 0 | 1 | 24 |
| XLOC-000946 | 0.1848 | 0.176272087 | 0.095 | 0 | 1 | 24 |
| aldob | 0.1848 | 0.262956475 | 0.095 | 0 | 1 | 24 |
| arfgef2 | 0.1848 | 0.125393759 | 0.095 | 0 | 1 | 24 |
| arl13b | 0.1848 | 0.257381195 | 0.095 | 0 | 1 | 24 |
| arl6ip5b | 0.1848 | 0.289878641 | 0.095 | 0 | 1 | 24 |
| arl8ba | 0.1848 | 0.128103771 | 0.095 | 0 | 1 | 24 |
| bahcc1a | 0.1848 | 0.227521884 | 0.095 | 0 | 1 | 24 |
| baxa | 0.1848 | 0.229586436 | 0.095 | 0 | 1 | 24 |
| ccng2 | 0.1848 | 0.309435607 | 0.095 | 0 | 1 | 24 |
| commd1 | 0.1848 | 0.138238795 | 0.095 | 0 | 1 | 24 |
| cxxc5a | 0.1848 | 0.236408786 | 0.095 | 0 | 1 | 24 |
| cyp20a1 | 0.1848 | 0.193600685 | 0.095 | 0 | 1 | 24 |
| edc4 | 0.1848 | 0.323077573 | 0.095 | 0 | 1 | 24 |
| eny2 | 0.1848 | 0.200778303 | 0.095 | 0 | 1 | 24 |
| ercc1 | 0.1848 | 0.239070293 | 0.095 | 0 | 1 | 24 |
| fbn2b | 0.1848 | 0.253101783 | 0.095 | 0 | 1 | 24 |
| fosb | 0.1848 | 0.284940922 | 0.095 | 0 | 1 | 24 |
| fstl1b | 0.1848 | 0.252953953 | 0.095 | 0 | 1 | 24 |
| gamt | 0.1848 | 0.19865795 | 0.095 | 0 | 1 | 24 |
| gigyf1b | 0.1848 | 0.137578765 | 0.095 | 0 | 1 | 24 |
| ifrd2 | 0.1848 | 0.248463704 | 0.095 | 0 | 1 | 24 |
| kpna4 | 0.1848 | 0.207515729 | 0.095 | 0 | 1 | 24 |
| kyat3 | 0.1848 | 0.167506996 | 0.095 | 0 | 1 | 24 |
| lmnb1 | 0.1848 | 0.167155748 | 0.095 | 0 | 1 | 24 |
| luc7l | 0.1848 | 0.274075325 | 0.095 | 0 | 1 | 24 |

| | | | | | | |
|---|---|---|---|---|---|---|
| matr3l1.1 | 0.1848 | 0.178731593 | 0.095 | 0 | 1 | 24 |
| mgaa | 0.1848 | 0.237009403 | 0.095 | 0 | 1 | 24 |
| mir21-1 | 0.1848 | 0.257865781 | 0.095 | 0 | 1 | 24 |
| mmp15b | 0.1848 | 0.144118734 | 0.095 | 0 | 1 | 24 |
| nccrp1 | 0.1848 | 0.199421164 | 0.095 | 0 | 1 | 24 |
| nktr | 0.1848 | 0.139033955 | 0.095 | 0 | 1 | 24 |
| pccb | 0.1848 | 0.246617415 | 0.095 | 0 | 1 | 24 |
| pcnp | 0.1848 | 0.144629787 | 0.095 | 0 | 1 | 24 |
| pdcd7 | 0.1848 | 0.308565004 | 0.095 | 0 | 1 | 24 |
| pdhx | 0.1848 | 0.165595812 | 0.095 | 0 | 1 | 24 |
| pdia4 | 0.1848 | 0.138208449 | 0.095 | 0 | 1 | 24 |
| pigu | 0.1848 | 0.149764322 | 0.095 | 0 | 1 | 24 |
| plxna2 | 0.1848 | 0.252788311 | 0.095 | 0 | 1 | 24 |
| polr2c | 0.1848 | 0.128064289 | 0.095 | 0 | 1 | 24 |
| ppp4r1 | 0.1848 | 0.251335889 | 0.095 | 0 | 1 | 24 |
| praf2 | 0.1848 | 0.168189265 | 0.095 | 0 | 1 | 24 |
| prpf4bb | 0.1848 | 0.245477639 | 0.095 | 0 | 1 | 24 |
| psmc1b | 0.1848 | 0.204825207 | 0.095 | 0 | 1 | 24 |
| quo | 0.1848 | 0.231736827 | 0.095 | 0 | 1 | 24 |
| rangap1b | 0.1848 | 0.144483221 | 0.095 | 0 | 1 | 24 |
| rasgef1ba | 0.1848 | 0.314478162 | 0.095 | 0 | 1 | 24 |
| rnf213a | 0.1848 | 0.234686004 | 0.095 | 0 | 1 | 24 |
| rptor | 0.1848 | 0.149764322 | 0.095 | 0 | 1 | 24 |
| scn3b | 0.1848 | 0.311044025 | 0.095 | 0 | 1 | 24 |
| sec61a1l | 0.1848 | 0.121587603 | 0.095 | 0 | 1 | 24 |
| setdb2 | 0.1848 | 0.218777678 | 0.095 | 0 | 1 | 24 |
| si:ch211-15d5.12 | 0.1848 | 0.160045202 | 0.095 | 0 | 1 | 24 |
| si:ch211-213a13.1 | 0.1848 | 0.214696588 | 0.095 | 0 | 1 | 24 |
| si:ch211-222e20.4 | 0.1848 | 0.275087111 | 0.095 | 0 | 1 | 24 |
| si:ch211-59h6.1 | 0.1848 | 0.247025964 | 0.095 | 0 | 1 | 24 |
| si:dkey-109l4.3 | 0.1848 | 0.140713275 | 0.095 | 0 | 1 | 24 |
| si:dkey-154b15.1 | 0.1848 | 0.225405952 | 0.095 | 0 | 1 | 24 |
| si:dkey-77f5.3 | 0.1848 | 0.230727373 | 0.095 | 0 | 1 | 24 |
| si:dkeyp-44a8.2 | 0.1848 | 0.22989389 | 0.095 | 0 | 1 | 24 |
| slc39a10 | 0.1848 | 0.171646688 | 0.095 | 0 | 1 | 24 |
| slka | 0.1848 | 0.311204991 | 0.095 | 0 | 1 | 24 |
| srd5a2a | 0.1848 | 0.216022772 | 0.095 | 0 | 1 | 24 |
| sumf1 | 0.1848 | 0.130758405 | 0.095 | 0 | 1 | 24 |
| tiprl | 0.1848 | 0.257982649 | 0.095 | 0 | 1 | 24 |
| tmem126a | 0.1848 | 0.24198202 | 0.095 | 0 | 1 | 24 |
| tnrc6b | 0.1848 | 0.228982697 | 0.095 | 0 | 1 | 24 |
| tp63 | 0.1848 | 0.14030898 | 0.095 | 0 | 1 | 24 |
| trim8a | 0.1848 | 0.154105946 | 0.095 | 0 | 1 | 24 |
| ubxn1 | 0.1848 | 0.165552829 | 0.095 | 0 | 1 | 24 |
| ythdf2 | 0.1848 | 0.310860683 | 0.095 | 0 | 1 | 24 |
| zbtb7a | 0.1848 | 0.309839024 | 0.095 | 0 | 1 | 24 |
| zfand5a | 0.1848 | 0.136379339 | 0.095 | 0 | 1 | 24 |
| znf131 | 0.1848 | 0.220948833 | 0.095 | 0 | 1 | 24 |

| | | | | | | |
|---|---|---|---|---|---|---|
| cnn3a | 0.1848 | 0.230931105 | 0.095 | 0 | 1 | 24 |
| fbxl14a | 0.1848 | 0.241027426 | 0.095 | 0 | 1 | 24 |
| gatc | 0.1848 | 0.239126221 | 0.095 | 0 | 1 | 24 |
| isy1 | 0.1848 | 0.239002654 | 0.095 | 0 | 1 | 24 |
| nr1d2a | 0.1848 | 0.229770246 | 0.095 | 0 | 1 | 24 |
| RASA2 | 0.1848 | 0.182414823 | 0.095 | 0 | 1 | 24 |
| fgf12b | 0.1848 | 0.187569177 | 0.095 | 0 | 1 | 24 |
| gatad2b | 0.1848 | 0.184620855 | 0.095 | 0 | 1 | 24 |
| gtse1 | 0.1848 | 0.213389065 | 0.095 | 0 | 1 | 24 |
| pdk2b | 0.1848 | 0.198306408 | 0.095 | 0 | 1 | 24 |
| si:ch211-155k24.9 | 0.1848 | 0.187569177 | 0.095 | 0 | 1 | 24 |
| sltm | 0.1848 | 0.1775183 | 0.095 | 0 | 1 | 24 |
| tnrc6a | 0.1848 | 0.177942394 | 0.095 | 0 | 1 | 24 |
| rpl5a | 0.18759 | 0.194539911 | 0.952 | 0.75 | 1 | 24 |
| cpsf6 | 0.18812 | -0.37197987 | 0.048 | 0.25 | 1 | 24 |
| cnbpa | 0.18872 | -0.28489796 | 0.238 | 0.5 | 1 | 24 |
| rps3 | 0.19069 | -0.25429217 | 0.714 | 0.938 | 1 | 24 |
| polr2b | 0.19097 | -0.43996354 | 0.048 | 0.25 | 1 | 24 |
| atp6v1g1 | 0.1919 | -0.41154074 | 0.048 | 0.25 | 1 | 24 |
| smim15 | 0.19492 | 0.440470441 | 0.286 | 0.062 | 1 | 24 |
| tuba8l4 | 0.19569 | -0.30579515 | 0.619 | 0.875 | 1 | 24 |
| ube2c | 0.19657 | 0.556815434 | 0.286 | 0.062 | 1 | 24 |
| CR383676.1 | 0.19663 | 0.866173942 | 0.429 | 0.188 | 1 | 24 |
| psma6a | 0.19755 | -0.49413089 | 0.048 | 0.25 | 1 | 24 |
| ptmab | 0.19941 | -0.27500372 | 0.714 | 0.938 | 1 | 24 |
| etf1b | 0.1995 | -0.41157229 | 0.048 | 0.25 | 1 | 24 |
| aldh1a2 | 0.2005 | -0.50790096 | 0.048 | 0.25 | 1 | 24 |
| zgc:77849 | 0.20088 | -0.41315156 | 0.048 | 0.25 | 1 | 24 |
| myl12.1 | 0.20112 | -0.42132425 | 0.333 | 0.625 | 1 | 24 |
| med10 | 0.20179 | -0.35856632 | 0.048 | 0.25 | 1 | 24 |
| chmp4bb | 0.20198 | -0.4803051 | 0.048 | 0.25 | 1 | 24 |
| CR753876.1 | 0.20239 | -0.40415773 | 0.048 | 0.25 | 1 | 24 |
| wu:fb55g09 | 0.20303 | -0.49151081 | 0.048 | 0.25 | 1 | 24 |
| calm1b | 0.20304 | -0.40833484 | 0.048 | 0.25 | 1 | 24 |
| sept7b | 0.20304 | -0.41943701 | 0.048 | 0.25 | 1 | 24 |
| sec61b | 0.21239 | 0.578308329 | 0.524 | 0.25 | 1 | 24 |
| polr2eb | 0.22297 | -0.58436392 | 0.095 | 0.312 | 1 | 24 |
| ppp1caa | 0.22349 | -0.39883223 | 0.095 | 0.312 | 1 | 24 |
| vamp3 | 0.22846 | -0.37479251 | 0.095 | 0.312 | 1 | 24 |
| rpl26 | 0.22986 | 0.18983164 | 1 | 0.875 | 1 | 24 |
| si:zfos-169g10.3 | 0.23075 | -0.59319064 | 0.143 | 0.375 | 1 | 24 |
| adssl | 0.23156 | -0.4338053 | 0.095 | 0.312 | 1 | 24 |
| nme2b.1 | 0.23958 | 0.178176897 | 1 | 1 | 1 | 24 |
| sdcbp2 | 0.24099 | -0.53129419 | 0.095 | 0.312 | 1 | 24 |
| cox6c | 0.24239 | -0.56254716 | 0.143 | 0.375 | 1 | 24 |
| eif5a | 0.24427 | 0.421677479 | 0.762 | 0.5 | 1 | 24 |
| rpl27 | 0.24463 | 0.128138717 | 1 | 0.875 | 1 | 24 |
| ddx6 | 0.24496 | -0.41806062 | 0.095 | 0.312 | 1 | 24 |

| | | | | | | |
|---|---|---|---|---|---|---|
| eif3ea | 0.2452 | -0.57022312 | 0.238 | 0.5 | 1 | 24 |
| arl6ip1 | 0.24611 | -0.54252713 | 0.095 | 0.312 | 1 | 24 |
| ahcy | 0.24662 | -0.37659987 | 0.143 | 0.375 | 1 | 24 |
| rhocb | 0.24715 | -0.48256444 | 0.095 | 0.312 | 1 | 24 |
| prkab1b | 0.24731 | -0.48554773 | 0.095 | 0.312 | 1 | 24 |
| orc4 | 0.24889 | -0.40592541 | 0.095 | 0.312 | 1 | 24 |
| kctd5a | 0.2489 | -0.47851159 | 0.095 | 0.312 | 1 | 24 |
| mcm4 | 0.2492 | -0.41024293 | 0.095 | 0.312 | 1 | 24 |
| atp5f1d | 0.24923 | -0.43109877 | 0.238 | 0.5 | 1 | 24 |
| rfc3 | 0.2497 | -0.47392379 | 0.095 | 0.312 | 1 | 24 |
| tmsb4x | 0.25067 | -0.22985054 | 0.905 | 1 | 1 | 24 |
| pomp | 0.2533 | 0.537982939 | 0.571 | 0.312 | 1 | 24 |
| eif1axb | 0.25395 | -0.53565117 | 0.19 | 0.438 | 1 | 24 |
| bzw1a | 0.25463 | 0.602636768 | 0.238 | 0.062 | 1 | 24 |
| naa10 | 0.25831 | -0.5222523 | 0.19 | 0.438 | 1 | 24 |
| nars | 0.25957 | -0.35736015 | 0.143 | 0.375 | 1 | 24 |
| arpc2 | 0.26078 | -0.52158033 | 0.19 | 0.438 | 1 | 24 |
| anxa1a | 0.26435 | -0.51889929 | 0.143 | 0.375 | 1 | 24 |
| hdac1 | 0.26484 | -0.42558001 | 0.19 | 0.438 | 1 | 24 |
| dck | 0.26588 | -0.48596878 | 0.143 | 0.375 | 1 | 24 |
| arhgdig | 0.26594 | -0.50760269 | 0.143 | 0.375 | 1 | 24 |
| polr2l | 0.26684 | -0.41994808 | 0.143 | 0.375 | 1 | 24 |
| gars | 0.26726 | -0.41897828 | 0.143 | 0.375 | 1 | 24 |
| metap2b | 0.27655 | 0.579793856 | 0.333 | 0.125 | 1 | 24 |
| AL935174.2 | 0.28371 | -0.21873615 | 0 | 0.062 | 1 | 24 |
| BX005085.4 | 0.28371 | -0.14247566 | 0 | 0.062 | 1 | 24 |
| BX005129.1 | 0.28371 | -0.22915862 | 0 | 0.062 | 1 | 24 |
| BX005461.3 | 0.28371 | -0.27785612 | 0 | 0.062 | 1 | 24 |
| BX284638.1 | 0.28371 | -0.21044265 | 0 | 0.062 | 1 | 24 |
| BX322661.2 | 0.28371 | -0.11235916 | 0 | 0.062 | 1 | 24 |
| BX511034.1 | 0.28371 | -0.21171263 | 0 | 0.062 | 1 | 24 |
| BX664614.1 | 0.28371 | -0.21388497 | 0 | 0.062 | 1 | 24 |
| BX855614.1 | 0.28371 | -0.11209045 | 0 | 0.062 | 1 | 24 |
| BX901883.3 | 0.28371 | -0.11395362 | 0 | 0.062 | 1 | 24 |
| CABZ01028771.1 | 0.28371 | -0.1293689 | 0 | 0.062 | 1 | 24 |
| CABZ01052576.1 | 0.28371 | -0.11366358 | 0 | 0.062 | 1 | 24 |
| CABZ01054965.1 | 0.28371 | -0.10715757 | 0 | 0.062 | 1 | 24 |
| CABZ01063556.1 | 0.28371 | -0.27981302 | 0 | 0.062 | 1 | 24 |
| CABZ01066926.1 | 0.28371 | -0.14451628 | 0 | 0.062 | 1 | 24 |
| CABZ01067348.1 | 0.28371 | -0.22294884 | 0 | 0.062 | 1 | 24 |
| CABZ01068366.1 | 0.28371 | -0.2091206 | 0 | 0.062 | 1 | 24 |
| CABZ01076275.1 | 0.28371 | -0.24279723 | 0 | 0.062 | 1 | 24 |
| CABZ01079663.1 | 0.28371 | -0.28131918 | 0 | 0.062 | 1 | 24 |
| CABZ01085658.1 | 0.28371 | -0.1128186 | 0 | 0.062 | 1 | 24 |
| CABZ01090890.1 | 0.28371 | -0.13857457 | 0 | 0.062 | 1 | 24 |
| CR352226.5 | 0.28371 | -0.1252822 | 0 | 0.062 | 1 | 24 |
| CR388132.2 | 0.28371 | -0.11283427 | 0 | 0.062 | 1 | 24 |
| CR450780.2 | 0.28371 | -0.22280482 | 0 | 0.062 | 1 | 24 |

| | | | | | | |
|---|---|---|---|---|---|---|
| CR925761.4 | 0.28371 | -0.13852083 | 0 | 0.062 | 1 | 24 |
| CT573793.2 | 0.28371 | -0.10763998 | 0 | 0.062 | 1 | 24 |
| CU462878.2 | 0.28371 | -0.10542595 | 0 | 0.062 | 1 | 24 |
| CU856358.1 | 0.28371 | -0.21075665 | 0 | 0.062 | 1 | 24 |
| CU929391.2 | 0.28371 | -0.16230395 | 0 | 0.062 | 1 | 24 |
| FRMD1 | 0.28371 | -0.1031652 | 0 | 0.062 | 1 | 24 |
| GPATCH2 | 0.28371 | -0.24312261 | 0 | 0.062 | 1 | 24 |
| LO018183.1 | 0.28371 | -0.21814485 | 0 | 0.062 | 1 | 24 |
| LOC100537138 | 0.28371 | -0.10754065 | 0 | 0.062 | 1 | 24 |
| LOC101882159 | 0.28371 | -0.28139612 | 0 | 0.062 | 1 | 24 |
| LOC101882264 | 0.28371 | -0.10406833 | 0 | 0.062 | 1 | 24 |
| LOC101883391 | 0.28371 | -0.14623801 | 0 | 0.062 | 1 | 24 |
| LOC101885677 | 0.28371 | -0.11395362 | 0 | 0.062 | 1 | 24 |
| LOC101886015 | 0.28371 | -0.11170614 | 0 | 0.062 | 1 | 24 |
| LOC101886030 | 0.28371 | -0.14154043 | 0 | 0.062 | 1 | 24 |
| LOC103909099 | 0.28371 | -0.26318467 | 0 | 0.062 | 1 | 24 |
| LOC103909928 | 0.28371 | -0.10407511 | 0 | 0.062 | 1 | 24 |
| LOC108190050 | 0.28371 | -0.28171799 | 0 | 0.062 | 1 | 24 |
| LOC108190056 | 0.28371 | -0.14920215 | 0 | 0.062 | 1 | 24 |
| LOC108191970 | 0.28371 | -0.21331761 | 0 | 0.062 | 1 | 24 |
| LOC110437842 | 0.28371 | -0.21534305 | 0 | 0.062 | 1 | 24 |
| LOC110439849 | 0.28371 | -0.22639534 | 0 | 0.062 | 1 | 24 |
| LOC570723 | 0.28371 | -0.23655828 | 0 | 0.062 | 1 | 24 |
| LOC794642 | 0.28371 | -0.10542595 | 0 | 0.062 | 1 | 24 |
| METTL18 | 0.28371 | -0.27507897 | 0 | 0.062 | 1 | 24 |
| MMP23B | 0.28371 | -0.14067267 | 0 | 0.062 | 1 | 24 |
| NDUFAF8 | 0.28371 | -0.1048814 | 0 | 0.062 | 1 | 24 |
| NHLRC3 | 0.28371 | -0.10797901 | 0 | 0.062 | 1 | 24 |
| PAXBP1 | 0.28371 | -0.11150302 | 0 | 0.062 | 1 | 24 |
| PDPR | 0.28371 | -0.11191679 | 0 | 0.062 | 1 | 24 |
| TMEM19 | 0.28371 | -0.15624873 | 0 | 0.062 | 1 | 24 |
| XLOC-002869 | 0.28371 | -0.14623801 | 0 | 0.062 | 1 | 24 |
| XLOC-005605 | 0.28371 | -0.11426752 | 0 | 0.062 | 1 | 24 |
| XLOC-014336 | 0.28371 | -0.11175686 | 0 | 0.062 | 1 | 24 |
| XLOC-019096 | 0.28371 | -0.10878534 | 0 | 0.062 | 1 | 24 |
| XLOC-034538 | 0.28371 | -0.20511094 | 0 | 0.062 | 1 | 24 |
| XLOC-043124 | 0.28371 | -0.1648955 | 0 | 0.062 | 1 | 24 |
| XRCC2 | 0.28371 | -0.1648955 | 0 | 0.062 | 1 | 24 |
| aagab | 0.28371 | -0.1423237 | 0 | 0.062 | 1 | 24 |
| aar2 | 0.28371 | -0.22719951 | 0 | 0.062 | 1 | 24 |
| abhd17ab | 0.28371 | -0.14089712 | 0 | 0.062 | 1 | 24 |
| acads | 0.28371 | -0.14475496 | 0 | 0.062 | 1 | 24 |
| acap1 | 0.28371 | -0.10568493 | 0 | 0.062 | 1 | 24 |
| acbd7 | 0.28371 | -0.22983317 | 0 | 0.062 | 1 | 24 |
| accs | 0.28371 | -0.11233506 | 0 | 0.062 | 1 | 24 |
| acin1b | 0.28371 | -0.10357267 | 0 | 0.062 | 1 | 24 |
| acy1 | 0.28371 | -0.14101855 | 0 | 0.062 | 1 | 24 |
| ada | 0.28371 | -0.22137909 | 0 | 0.062 | 1 | 24 |

| | | | | | | |
|---|---|---|---|---|---|---|
| adam19a | 0.28371 | -0.24401277 | 0 | 0.062 | 1 | 24 |
| adap2 | 0.28371 | -0.28146432 | 0 | 0.062 | 1 | 24 |
| adh8b | 0.28371 | -0.17534096 | 0 | 0.062 | 1 | 24 |
| adma | 0.28371 | -0.28171799 | 0 | 0.062 | 1 | 24 |
| adnpb | 0.28371 | -0.14375163 | 0 | 0.062 | 1 | 24 |
| adora2ab | 0.28371 | -0.11279285 | 0 | 0.062 | 1 | 24 |
| adora2b | 0.28371 | -0.10709576 | 0 | 0.062 | 1 | 24 |
| adsl | 0.28371 | -0.10212055 | 0 | 0.062 | 1 | 24 |
| aes | 0.28371 | -0.11244983 | 0 | 0.062 | 1 | 24 |
| agpat3 | 0.28371 | -0.24432137 | 0 | 0.062 | 1 | 24 |
| ahsa1b | 0.28371 | -0.10659241 | 0 | 0.062 | 1 | 24 |
| akt2 | 0.28371 | -0.10451173 | 0 | 0.062 | 1 | 24 |
| akt2l | 0.28371 | -0.13549106 | 0 | 0.062 | 1 | 24 |
| alcamb | 0.28371 | -0.23276486 | 0 | 0.062 | 1 | 24 |
| aldh3b1 | 0.28371 | -0.15686129 | 0 | 0.062 | 1 | 24 |
| amn1 | 0.28371 | -0.22805465 | 0 | 0.062 | 1 | 24 |
| ankrd10b | 0.28371 | -0.28068614 | 0 | 0.062 | 1 | 24 |
| ankrd52a | 0.28371 | -0.10389612 | 0 | 0.062 | 1 | 24 |
| anln | 0.28371 | -0.24268566 | 0 | 0.062 | 1 | 24 |
| aoc2 | 0.28371 | -0.1407675 | 0 | 0.062 | 1 | 24 |
| aplp1 | 0.28371 | -0.24284459 | 0 | 0.062 | 1 | 24 |
| apoeb | 0.28371 | -0.21003068 | 0 | 0.062 | 1 | 24 |
| arcn1b | 0.28371 | -0.2293082 | 0 | 0.062 | 1 | 24 |
| arfgap2 | 0.28371 | -0.10633263 | 0 | 0.062 | 1 | 24 |
| arhgef2 | 0.28371 | -0.24361209 | 0 | 0.062 | 1 | 24 |
| arid1aa | 0.28371 | -0.11251962 | 0 | 0.062 | 1 | 24 |
| arl8bb | 0.28371 | -0.11078328 | 0 | 0.062 | 1 | 24 |
| arl9 | 0.28371 | -0.14501909 | 0 | 0.062 | 1 | 24 |
| arnt2 | 0.28371 | -0.22096021 | 0 | 0.062 | 1 | 24 |
| arpc5la | 0.28371 | -0.22733119 | 0 | 0.062 | 1 | 24 |
| asap1b | 0.28371 | -0.11297395 | 0 | 0.062 | 1 | 24 |
| asb11 | 0.28371 | -0.22239799 | 0 | 0.062 | 1 | 24 |
| atad2b | 0.28371 | -0.17940432 | 0 | 0.062 | 1 | 24 |
| atf7ip2 | 0.28371 | -0.24307755 | 0 | 0.062 | 1 | 24 |
| atrxl | 0.28371 | -0.20156319 | 0 | 0.062 | 1 | 24 |
| babam1 | 0.28371 | -0.22104448 | 0 | 0.062 | 1 | 24 |
| bbx | 0.28371 | -0.10684706 | 0 | 0.062 | 1 | 24 |
| bcap29 | 0.28371 | -0.10136137 | 0 | 0.062 | 1 | 24 |
| bcl11aa | 0.28371 | -0.10544243 | 0 | 0.062 | 1 | 24 |
| bcl2l1 | 0.28371 | -0.10149809 | 0 | 0.062 | 1 | 24 |
| bgna | 0.28371 | -0.22192273 | 0 | 0.062 | 1 | 24 |
| bin2b | 0.28371 | -0.13421602 | 0 | 0.062 | 1 | 24 |
| blzf1 | 0.28371 | -0.24259859 | 0 | 0.062 | 1 | 24 |
| bmpr1aa | 0.28371 | -0.10191777 | 0 | 0.062 | 1 | 24 |
| bola3 | 0.28371 | -0.13285026 | 0 | 0.062 | 1 | 24 |
| brd1a | 0.28371 | -0.22252302 | 0 | 0.062 | 1 | 24 |
| bri3 | 0.28371 | -0.10853742 | 0 | 0.062 | 1 | 24 |
| bri3bp | 0.28371 | -0.10455679 | 0 | 0.062 | 1 | 24 |

| | | | | | | |
|---|---|---|---|---|---|---|
| brms1lb | 0.28371 | -0.26883557 | 0 | 0.062 | 1 | 24 |
| bscl2 | 0.28371 | -0.23261047 | 0 | 0.062 | 1 | 24 |
| bub3 | 0.28371 | -0.10932622 | 0 | 0.062 | 1 | 24 |
| c1ql3a | 0.28371 | -0.10544243 | 0 | 0.062 | 1 | 24 |
| c1qtnf6a | 0.28371 | -0.1532939 | 0 | 0.062 | 1 | 24 |
| ca8 | 0.28371 | -0.23195406 | 0 | 0.062 | 1 | 24 |
| cables2b | 0.28371 | -0.15083988 | 0 | 0.062 | 1 | 24 |
| cahz | 0.28371 | -0.223735 | 0 | 0.062 | 1 | 24 |
| camk2g2 | 0.28371 | -0.11280718 | 0 | 0.062 | 1 | 24 |
| cand1 | 0.28371 | -0.11143011 | 0 | 0.062 | 1 | 24 |
| casp7 | 0.28371 | -0.14272933 | 0 | 0.062 | 1 | 24 |
| casp8ap2 | 0.28371 | -0.11167552 | 0 | 0.062 | 1 | 24 |
| casp9 | 0.28371 | -0.11309583 | 0 | 0.062 | 1 | 24 |
| castor2 | 0.28371 | -0.10546014 | 0 | 0.062 | 1 | 24 |
| cav1 | 0.28371 | -0.2105379 | 0 | 0.062 | 1 | 24 |
| cav2 | 0.28371 | -0.10215797 | 0 | 0.062 | 1 | 24 |
| cbfa2t3 | 0.28371 | -0.1123466 | 0 | 0.062 | 1 | 24 |
| cbfb | 0.28371 | -0.11069158 | 0 | 0.062 | 1 | 24 |
| cbx8a | 0.28371 | -0.11217899 | 0 | 0.062 | 1 | 24 |
| cbx8b | 0.28371 | -0.10203124 | 0 | 0.062 | 1 | 24 |
| ccbe1 | 0.28371 | -0.1405453 | 0 | 0.062 | 1 | 24 |
| ccdc47 | 0.28371 | -0.11041879 | 0 | 0.062 | 1 | 24 |
| ccdc59 | 0.28371 | -0.13399864 | 0 | 0.062 | 1 | 24 |
| ccl33.3 | 0.28371 | -0.48125377 | 0 | 0.062 | 1 | 24 |
| ccl39.6 | 0.28371 | -0.20982686 | 0 | 0.062 | 1 | 24 |
| ccm2 | 0.28371 | -0.2813079 | 0 | 0.062 | 1 | 24 |
| cd28l | 0.28371 | -0.10130788 | 0 | 0.062 | 1 | 24 |
| cdc37l1 | 0.28371 | -0.23093888 | 0 | 0.062 | 1 | 24 |
| cdc6 | 0.28371 | -0.10845628 | 0 | 0.062 | 1 | 24 |
| cdc7 | 0.28371 | -0.22830482 | 0 | 0.062 | 1 | 24 |
| cdk11b | 0.28371 | -0.13804123 | 0 | 0.062 | 1 | 24 |
| cdon | 0.28371 | -0.12834632 | 0 | 0.062 | 1 | 24 |
| cep290 | 0.28371 | -0.11181485 | 0 | 0.062 | 1 | 24 |
| cep55l | 0.28371 | -0.27388895 | 0 | 0.062 | 1 | 24 |
| cep85l | 0.28371 | -0.10968857 | 0 | 0.062 | 1 | 24 |
| cercam | 0.28371 | -0.2308633 | 0 | 0.062 | 1 | 24 |
| cfap410 | 0.28371 | -0.21914565 | 0 | 0.062 | 1 | 24 |
| chd1l | 0.28371 | -0.23302361 | 0 | 0.062 | 1 | 24 |
| chn1 | 0.28371 | -0.22457524 | 0 | 0.062 | 1 | 24 |
| chordc1a | 0.28371 | -0.28117373 | 0 | 0.062 | 1 | 24 |
| chrdl2 | 0.28371 | -0.2145679 | 0 | 0.062 | 1 | 24 |
| chst1 | 0.28371 | -0.22712508 | 0 | 0.062 | 1 | 24 |
| cicb | 0.28371 | -0.10455302 | 0 | 0.062 | 1 | 24 |
| cldng | 0.28371 | -0.10716512 | 0 | 0.062 | 1 | 24 |
| clgn | 0.28371 | -0.10864044 | 0 | 0.062 | 1 | 24 |
| clptm1l | 0.28371 | -0.109119 | 0 | 0.062 | 1 | 24 |
| clpxa | 0.28371 | -0.23094062 | 0 | 0.062 | 1 | 24 |
| cltca | 0.28371 | -0.16217496 | 0 | 0.062 | 1 | 24 |

| | | | | | | |
|---|---|---|---|---|---|---|
| cmip | 0.28371 | -0.1126457 | 0 | 0.062 | 1 | 24 |
| coa7 | 0.28371 | -0.22950076 | 0 | 0.062 | 1 | 24 |
| col12a1b | 0.28371 | -0.10613971 | 0 | 0.062 | 1 | 24 |
| col18a1a | 0.28371 | -0.24198958 | 0 | 0.062 | 1 | 24 |
| col4a2 | 0.28371 | -0.22332824 | 0 | 0.062 | 1 | 24 |
| commd6 | 0.28371 | -0.16258758 | 0 | 0.062 | 1 | 24 |
| coro1cb | 0.28371 | -0.14957539 | 0 | 0.062 | 1 | 24 |
| cplane2 | 0.28371 | -0.21967762 | 0 | 0.062 | 1 | 24 |
| cpne3 | 0.28371 | -0.16339569 | 0 | 0.062 | 1 | 24 |
| cpsf2 | 0.28371 | -0.10760112 | 0 | 0.062 | 1 | 24 |
| crb3a | 0.28371 | -0.14247566 | 0 | 0.062 | 1 | 24 |
| crbn | 0.28371 | -0.1124709 | 0 | 0.062 | 1 | 24 |
| crebl2 | 0.28371 | -0.22842945 | 0 | 0.062 | 1 | 24 |
| crfb4 | 0.28371 | -0.10484479 | 0 | 0.062 | 1 | 24 |
| cry2 | 0.28371 | -0.14147645 | 0 | 0.062 | 1 | 24 |
| csgalnact1a | 0.28371 | -0.21953584 | 0 | 0.062 | 1 | 24 |
| csnk2a2b | 0.28371 | -0.22808233 | 0 | 0.062 | 1 | 24 |
| ctnnd1 | 0.28371 | -0.23084119 | 0 | 0.062 | 1 | 24 |
| cttn | 0.28371 | -0.21942366 | 0 | 0.062 | 1 | 24 |
| cul1a | 0.28371 | -0.2299957 | 0 | 0.062 | 1 | 24 |
| cul3b | 0.28371 | -0.10934051 | 0 | 0.062 | 1 | 24 |
| cul4a | 0.28371 | -0.11235041 | 0 | 0.062 | 1 | 24 |
| cx43.4 | 0.28371 | -0.15759952 | 0 | 0.062 | 1 | 24 |
| cxcl8b.1 | 0.28371 | -0.10348554 | 0 | 0.062 | 1 | 24 |
| cyfip2 | 0.28371 | -0.18609369 | 0 | 0.062 | 1 | 24 |
| cyldl | 0.28371 | -0.108591 | 0 | 0.062 | 1 | 24 |
| cyth1b | 0.28371 | -0.11090535 | 0 | 0.062 | 1 | 24 |
| cyth4a | 0.28371 | -0.18966666 | 0 | 0.062 | 1 | 24 |
| dachc | 0.28371 | -0.2279583 | 0 | 0.062 | 1 | 24 |
| daxx | 0.28371 | -0.21277055 | 0 | 0.062 | 1 | 24 |
| ddx1 | 0.28371 | -0.22655157 | 0 | 0.062 | 1 | 24 |
| ddx42 | 0.28371 | -0.14375404 | 0 | 0.062 | 1 | 24 |
| ddx52 | 0.28371 | -0.2625338 | 0 | 0.062 | 1 | 24 |
| def6c | 0.28371 | -0.10880696 | 0 | 0.062 | 1 | 24 |
| defbl1 | 0.28371 | -0.26002319 | 0 | 0.062 | 1 | 24 |
| depdc1a | 0.28371 | -0.22280482 | 0 | 0.062 | 1 | 24 |
| dffb | 0.28371 | -0.24433627 | 0 | 0.062 | 1 | 24 |
| dhdds | 0.28371 | -0.10640576 | 0 | 0.062 | 1 | 24 |
| dhps | 0.28371 | -0.15997585 | 0 | 0.062 | 1 | 24 |
| dhx29 | 0.28371 | -0.15854342 | 0 | 0.062 | 1 | 24 |
| dhx33 | 0.28371 | -0.22084642 | 0 | 0.062 | 1 | 24 |
| dhx34 | 0.28371 | -0.12031838 | 0 | 0.062 | 1 | 24 |
| dido1 | 0.28371 | -0.10417127 | 0 | 0.062 | 1 | 24 |
| dimt1l | 0.28371 | -0.10841708 | 0 | 0.062 | 1 | 24 |
| dio2 | 0.28371 | -0.1446903 | 0 | 0.062 | 1 | 24 |
| dixdc1b | 0.28371 | -0.1998878 | 0 | 0.062 | 1 | 24 |
| dkk3b | 0.28371 | -0.15801847 | 0 | 0.062 | 1 | 24 |
| dlc1 | 0.28371 | -0.21715347 | 0 | 0.062 | 1 | 24 |

| | | | | | | |
|---|---|---|---|---|---|---|
| dnaja1 | 0.28371 | -0.11180234 | 0 | 0.062 | 1 | 24 |
| dnajb1a | 0.28371 | -0.10322358 | 0 | 0.062 | 1 | 24 |
| dnajc11a | 0.28371 | -0.1407391 | 0 | 0.062 | 1 | 24 |
| dnajc24 | 0.28371 | -0.22092923 | 0 | 0.062 | 1 | 24 |
| dnajc5aa | 0.28371 | -0.10139348 | 0 | 0.062 | 1 | 24 |
| dnajc5ab | 0.28371 | -0.21119482 | 0 | 0.062 | 1 | 24 |
| dnase1l4.2 | 0.28371 | -0.10573611 | 0 | 0.062 | 1 | 24 |
| dnmt1 | 0.28371 | -0.10413148 | 0 | 0.062 | 1 | 24 |
| dnmt3aa | 0.28371 | -0.11395362 | 0 | 0.062 | 1 | 24 |
| dot1l | 0.28371 | -0.24456886 | 0 | 0.062 | 1 | 24 |
| dpf1 | 0.28371 | -0.11247833 | 0 | 0.062 | 1 | 24 |
| dph3 | 0.28371 | -0.1287818 | 0 | 0.062 | 1 | 24 |
| dynll2b | 0.28371 | -0.11707058 | 0 | 0.062 | 1 | 24 |
| e2f1 | 0.28371 | -0.1130649 | 0 | 0.062 | 1 | 24 |
| eaf1 | 0.28371 | -0.15147531 | 0 | 0.062 | 1 | 24 |
| ebp | 0.28371 | -0.14799715 | 0 | 0.062 | 1 | 24 |
| ecrg4a | 0.28371 | -0.22651725 | 0 | 0.062 | 1 | 24 |
| ect2 | 0.28371 | -0.24380812 | 0 | 0.062 | 1 | 24 |
| eef1akmt1 | 0.28371 | -0.2230048 | 0 | 0.062 | 1 | 24 |
| eefsec | 0.28371 | -0.21315672 | 0 | 0.062 | 1 | 24 |
| egfl6 | 0.28371 | -0.1252822 | 0 | 0.062 | 1 | 24 |
| ehbp1l1b | 0.28371 | -0.11018417 | 0 | 0.062 | 1 | 24 |
| eif3eb | 0.28371 | -0.22131473 | 0 | 0.062 | 1 | 24 |
| eipr1 | 0.28371 | -0.28099866 | 0 | 0.062 | 1 | 24 |
| elmsan1b | 0.28371 | -0.15859385 | 0 | 0.062 | 1 | 24 |
| emc8 | 0.28371 | -0.10844021 | 0 | 0.062 | 1 | 24 |
| emid1 | 0.28371 | -0.22944898 | 0 | 0.062 | 1 | 24 |
| emx2 | 0.28371 | -0.23886396 | 0 | 0.062 | 1 | 24 |
| enosf1 | 0.28371 | -0.10416901 | 0 | 0.062 | 1 | 24 |
| enpp4 | 0.28371 | -0.10878534 | 0 | 0.062 | 1 | 24 |
| epb41l5 | 0.28371 | -0.16064421 | 0 | 0.062 | 1 | 24 |
| epdl1 | 0.28371 | -0.243714 | 0 | 0.062 | 1 | 24 |
| erap1b | 0.28371 | -0.11139551 | 0 | 0.062 | 1 | 24 |
| eri1 | 0.28371 | -0.22067262 | 0 | 0.062 | 1 | 24 |
| esr2a | 0.28371 | -0.2380352 | 0 | 0.062 | 1 | 24 |
| eva1ba | 0.28371 | -0.13521058 | 0 | 0.062 | 1 | 24 |
| evla | 0.28371 | -0.23120253 | 0 | 0.062 | 1 | 24 |
| exd3 | 0.28371 | -0.13716447 | 0 | 0.062 | 1 | 24 |
| faap24 | 0.28371 | -0.14247566 | 0 | 0.062 | 1 | 24 |
| fam102ab | 0.28371 | -0.20592514 | 0 | 0.062 | 1 | 24 |
| fam114a2 | 0.28371 | -0.20887704 | 0 | 0.062 | 1 | 24 |
| fam122b | 0.28371 | -0.10152141 | 0 | 0.062 | 1 | 24 |
| fam135a | 0.28371 | -0.11219255 | 0 | 0.062 | 1 | 24 |
| fam173b | 0.28371 | -0.19121391 | 0 | 0.062 | 1 | 24 |
| fam177a1 | 0.28371 | -0.15715344 | 0 | 0.062 | 1 | 24 |
| fam208b | 0.28371 | -0.11179654 | 0 | 0.062 | 1 | 24 |
| fam20b | 0.28371 | -0.11182625 | 0 | 0.062 | 1 | 24 |
| fam210ab | 0.28371 | -0.22417616 | 0 | 0.062 | 1 | 24 |

| | | | | | | |
|---|---|---|---|---|---|---|
| farp2 | 0.28371 | -0.10264118 | 0 | 0.062 | 1 | 24 |
| farsa | 0.28371 | -0.1391545 | 0 | 0.062 | 1 | 24 |
| fbxl7 | 0.28371 | -0.13877382 | 0 | 0.062 | 1 | 24 |
| fbxo11a | 0.28371 | -0.11107632 | 0 | 0.062 | 1 | 24 |
| fbxw4 | 0.28371 | -0.11112445 | 0 | 0.062 | 1 | 24 |
| fcer1gl | 0.28371 | -0.27926151 | 0 | 0.062 | 1 | 24 |
| fgf24 | 0.28371 | -0.16037105 | 0 | 0.062 | 1 | 24 |
| fkbp8 | 0.28371 | -0.10858596 | 0 | 0.062 | 1 | 24 |
| fktn | 0.28371 | -0.10414403 | 0 | 0.062 | 1 | 24 |
| fli1a | 0.28371 | -0.14539118 | 0 | 0.062 | 1 | 24 |
| fndc3a | 0.28371 | -0.2221631 | 0 | 0.062 | 1 | 24 |
| foxg1c | 0.28371 | -0.2158708 | 0 | 0.062 | 1 | 24 |
| foxn2b | 0.28371 | -0.1024098 | 0 | 0.062 | 1 | 24 |
| foxo3a | 0.28371 | -0.22288071 | 0 | 0.062 | 1 | 24 |
| frmd4bb | 0.28371 | -0.22483636 | 0 | 0.062 | 1 | 24 |
| fzd2 | 0.28371 | -0.15889326 | 0 | 0.062 | 1 | 24 |
| gcat | 0.28371 | -0.22927232 | 0 | 0.062 | 1 | 24 |
| gga3a | 0.28371 | -0.21097194 | 0 | 0.062 | 1 | 24 |
| gig2g | 0.28371 | -0.158987 | 0 | 0.062 | 1 | 24 |
| gins1 | 0.28371 | -0.10784316 | 0 | 0.062 | 1 | 24 |
| glmp | 0.28371 | -0.2129826 | 0 | 0.062 | 1 | 24 |
| glrx5 | 0.28371 | -0.22163182 | 0 | 0.062 | 1 | 24 |
| glsb | 0.28371 | -0.10705112 | 0 | 0.062 | 1 | 24 |
| gnal2 | 0.28371 | -0.27996118 | 0 | 0.062 | 1 | 24 |
| gpr141 | 0.28371 | -0.22280482 | 0 | 0.062 | 1 | 24 |
| gpr184 | 0.28371 | -0.17876721 | 0 | 0.062 | 1 | 24 |
| gpr22a | 0.28371 | -0.22483636 | 0 | 0.062 | 1 | 24 |
| grhpra | 0.28371 | -0.21919201 | 0 | 0.062 | 1 | 24 |
| grk7b | 0.28371 | -0.23323932 | 0 | 0.062 | 1 | 24 |
| grm8a | 0.28371 | -0.23120429 | 0 | 0.062 | 1 | 24 |
| gstt1a | 0.28371 | -0.185371 | 0 | 0.062 | 1 | 24 |
| gtf2b | 0.28371 | -0.24325621 | 0 | 0.062 | 1 | 24 |
| gtf2h2 | 0.28371 | -0.11117624 | 0 | 0.062 | 1 | 24 |
| gtf2h4 | 0.28371 | -0.10979609 | 0 | 0.062 | 1 | 24 |
| gtf2h5 | 0.28371 | -0.21044617 | 0 | 0.062 | 1 | 24 |
| gtpbp1 | 0.28371 | -0.11356682 | 0 | 0.062 | 1 | 24 |
| hacl1 | 0.28371 | -0.11202735 | 0 | 0.062 | 1 | 24 |
| hic1 | 0.28371 | -0.13613796 | 0 | 0.062 | 1 | 24 |
| hint2 | 0.28371 | -0.19159289 | 0 | 0.062 | 1 | 24 |
| hist1h2a3 | 0.28371 | -0.10987728 | 0 | 0.062 | 1 | 24 |
| hk2 | 0.28371 | -0.12076194 | 0 | 0.062 | 1 | 24 |
| hlfa | 0.28371 | -0.14403248 | 0 | 0.062 | 1 | 24 |
| hmga2 | 0.28371 | -0.19237229 | 0 | 0.062 | 1 | 24 |
| hmgxb4a | 0.28371 | -0.11357285 | 0 | 0.062 | 1 | 24 |
| homezb | 0.28371 | -0.24005267 | 0 | 0.062 | 1 | 24 |
| hoxa9b | 0.28371 | -0.13852083 | 0 | 0.062 | 1 | 24 |
| hoxb6b | 0.28371 | -0.10878534 | 0 | 0.062 | 1 | 24 |
| hoxc10a | 0.28371 | -0.14623801 | 0 | 0.062 | 1 | 24 |

| | | | | | | |
|---|---|---|---|---|---|---|
| hps4 | 0.28371 | -0.1446901 | 0 | 0.062 | 1 | 24 |
| hsdl1 | 0.28371 | -0.14623801 | 0 | 0.062 | 1 | 24 |
| ide | 0.28371 | -0.163967 | 0 | 0.062 | 1 | 24 |
| igflr1 | 0.28371 | -0.21722815 | 0 | 0.062 | 1 | 24 |
| ikzf5 | 0.28371 | -0.11211296 | 0 | 0.062 | 1 | 24 |
| ilkap | 0.28371 | -0.2204149 | 0 | 0.062 | 1 | 24 |
| illr1 | 0.28371 | -0.24088812 | 0 | 0.062 | 1 | 24 |
| im:7145024 | 0.28371 | -0.11027682 | 0 | 0.062 | 1 | 24 |
| impact | 0.28371 | -0.10245624 | 0 | 0.062 | 1 | 24 |
| ing1 | 0.28371 | -0.22095594 | 0 | 0.062 | 1 | 24 |
| inpp5d | 0.28371 | -0.10237127 | 0 | 0.062 | 1 | 24 |
| ints12 | 0.28371 | -0.13722139 | 0 | 0.062 | 1 | 24 |
| iqgap3 | 0.28371 | -0.1648955 | 0 | 0.062 | 1 | 24 |
| ireb2 | 0.28371 | -0.10690475 | 0 | 0.062 | 1 | 24 |
| irf1b | 0.28371 | -0.28062318 | 0 | 0.062 | 1 | 24 |
| isg15 | 0.28371 | -0.13836474 | 0 | 0.062 | 1 | 24 |
| itga1 | 0.28371 | -0.28095292 | 0 | 0.062 | 1 | 24 |
| itga4 | 0.28371 | -0.10452935 | 0 | 0.062 | 1 | 24 |
| itgb1bp1 | 0.28371 | -0.23287752 | 0 | 0.062 | 1 | 24 |
| katnb1 | 0.28371 | -0.15950825 | 0 | 0.062 | 1 | 24 |
| kcnk5a | 0.28371 | -0.11395362 | 0 | 0.062 | 1 | 24 |
| kctd13 | 0.28371 | -0.10825153 | 0 | 0.062 | 1 | 24 |
| kctd3 | 0.28371 | -0.11395362 | 0 | 0.062 | 1 | 24 |
| kdelr3 | 0.28371 | -0.21223743 | 0 | 0.062 | 1 | 24 |
| kdm3b | 0.28371 | -0.10623441 | 0 | 0.062 | 1 | 24 |
| keap1b | 0.28371 | -0.11170547 | 0 | 0.062 | 1 | 24 |
| kif11 | 0.28371 | -0.22912428 | 0 | 0.062 | 1 | 24 |
| kif21b | 0.28371 | -0.10693478 | 0 | 0.062 | 1 | 24 |
| kif5ba | 0.28371 | -0.11221195 | 0 | 0.062 | 1 | 24 |
| klhdc10 | 0.28371 | -0.11119301 | 0 | 0.062 | 1 | 24 |
| kmt2ba | 0.28371 | -0.23040175 | 0 | 0.062 | 1 | 24 |
| knstrn | 0.28371 | -0.22895319 | 0 | 0.062 | 1 | 24 |
| kri1 | 0.28371 | -0.10935419 | 0 | 0.062 | 1 | 24 |
| l3mbtl2 | 0.28371 | -0.11292871 | 0 | 0.062 | 1 | 24 |
| lamtor5 | 0.28371 | -0.15312272 | 0 | 0.062 | 1 | 24 |
| lats1 | 0.28371 | -0.10032963 | 0 | 0.062 | 1 | 24 |
| leo1 | 0.28371 | -0.16322142 | 0 | 0.062 | 1 | 24 |
| lgals17 | 0.28371 | -0.1254404 | 0 | 0.062 | 1 | 24 |
| lig3 | 0.28371 | -0.14424904 | 0 | 0.062 | 1 | 24 |
| lipg | 0.28371 | -0.20639628 | 0 | 0.062 | 1 | 24 |
| lman1 | 0.28371 | -0.20761664 | 0 | 0.062 | 1 | 24 |
| lrrc47 | 0.28371 | -0.10213885 | 0 | 0.062 | 1 | 24 |
| lrwd1 | 0.28371 | -0.11037723 | 0 | 0.062 | 1 | 24 |
| lsm12a | 0.28371 | -0.14283342 | 0 | 0.062 | 1 | 24 |
| lztfl1 | 0.28371 | -0.11144225 | 0 | 0.062 | 1 | 24 |
| lztr1 | 0.28371 | -0.10131572 | 0 | 0.062 | 1 | 24 |
| maff | 0.28371 | -0.22723283 | 0 | 0.062 | 1 | 24 |
| man1a1 | 0.28371 | -0.2217704 | 0 | 0.062 | 1 | 24 |

| | | | | | | |
|---|---|---|---|---|---|---|
| map3k12 | 0.28371 | -0.11395362 | 0 | 0.062 | 1 | 24 |
| map7d3 | 0.28371 | -0.10489567 | 0 | 0.062 | 1 | 24 |
| mapk1 | 0.28371 | -0.11232541 | 0 | 0.062 | 1 | 24 |
| mapk6 | 0.28371 | -0.22112243 | 0 | 0.062 | 1 | 24 |
| maptb | 0.28371 | -0.10502131 | 0 | 0.062 | 1 | 24 |
| mdm2 | 0.28371 | -0.22122341 | 0 | 0.062 | 1 | 24 |
| med13b | 0.28371 | -0.11325843 | 0 | 0.062 | 1 | 24 |
| med19b | 0.28371 | -0.14341409 | 0 | 0.062 | 1 | 24 |
| med30 | 0.28371 | -0.208574 | 0 | 0.062 | 1 | 24 |
| mettl14 | 0.28371 | -0.11222054 | 0 | 0.062 | 1 | 24 |
| mettl17 | 0.28371 | -0.14070525 | 0 | 0.062 | 1 | 24 |
| mfng | 0.28371 | -0.28149006 | 0 | 0.062 | 1 | 24 |
| mfsd5 | 0.28371 | -0.10791299 | 0 | 0.062 | 1 | 24 |
| mgat1a | 0.28371 | -0.10998166 | 0 | 0.062 | 1 | 24 |
| mgat2 | 0.28371 | -0.21331761 | 0 | 0.062 | 1 | 24 |
| micu2 | 0.28371 | -0.10570931 | 0 | 0.062 | 1 | 24 |
| mier2 | 0.28371 | -0.11031357 | 0 | 0.062 | 1 | 24 |
| morc3a | 0.28371 | -0.10368577 | 0 | 0.062 | 1 | 24 |
| mphosph10 | 0.28371 | -0.10786806 | 0 | 0.062 | 1 | 24 |
| mrap2a | 0.28371 | -0.11395362 | 0 | 0.062 | 1 | 24 |
| mrgbp | 0.28371 | -0.15433462 | 0 | 0.062 | 1 | 24 |
| mri1 | 0.28371 | -0.15874224 | 0 | 0.062 | 1 | 24 |
| mrpl16 | 0.28371 | -0.20976596 | 0 | 0.062 | 1 | 24 |
| mrpl21 | 0.28371 | -0.22506572 | 0 | 0.062 | 1 | 24 |
| mrpl37 | 0.28371 | -0.24363094 | 0 | 0.062 | 1 | 24 |
| mrpl38 | 0.28371 | -0.15691215 | 0 | 0.062 | 1 | 24 |
| mrpl4 | 0.28371 | -0.28072045 | 0 | 0.062 | 1 | 24 |
| mrpl46 | 0.28371 | -0.10957884 | 0 | 0.062 | 1 | 24 |
| mrps31 | 0.28371 | -0.13275351 | 0 | 0.062 | 1 | 24 |
| mthfd2 | 0.28371 | -0.10490986 | 0 | 0.062 | 1 | 24 |
| mtnr1aa | 0.28371 | -0.23120429 | 0 | 0.062 | 1 | 24 |
| mto1 | 0.28371 | -0.11283254 | 0 | 0.062 | 1 | 24 |
| mtor | 0.28371 | -0.11259973 | 0 | 0.062 | 1 | 24 |
| mycbp2 | 0.28371 | -0.10622071 | 0 | 0.062 | 1 | 24 |
| myofl | 0.28371 | -0.19690601 | 0 | 0.062 | 1 | 24 |
| naa15b | 0.28371 | -0.10494368 | 0 | 0.062 | 1 | 24 |
| nab1b | 0.28371 | -0.11330732 | 0 | 0.062 | 1 | 24 |
| napga | 0.28371 | -0.2113488 | 0 | 0.062 | 1 | 24 |
| nat8l | 0.28371 | -0.2217704 | 0 | 0.062 | 1 | 24 |
| nav2b | 0.28371 | -0.10165106 | 0 | 0.062 | 1 | 24 |
| ncalda | 0.28371 | -0.14045852 | 0 | 0.062 | 1 | 24 |
| ncdn | 0.28371 | -0.10863601 | 0 | 0.062 | 1 | 24 |
| ndnl2 | 0.28371 | -0.1417097 | 0 | 0.062 | 1 | 24 |
| ndrg3b | 0.28371 | -0.14172255 | 0 | 0.062 | 1 | 24 |
| nelfb | 0.28371 | -0.22998044 | 0 | 0.062 | 1 | 24 |
| nfatc2ip | 0.28371 | -0.10896133 | 0 | 0.062 | 1 | 24 |
| nfatc3a | 0.28371 | -0.14584877 | 0 | 0.062 | 1 | 24 |
| nfe2l1b | 0.28371 | -0.11338451 | 0 | 0.062 | 1 | 24 |

| | | | | | | |
|---|---|---|---|---|---|---|
| nipa2 | 0.28371 | -0.11231669 | 0 | 0.062 | 1 | 24 |
| nlk2 | 0.28371 | -0.14127379 | 0 | 0.062 | 1 | 24 |
| nol12 | 0.28371 | -0.20897356 | 0 | 0.062 | 1 | 24 |
| nol6 | 0.28371 | -0.26491497 | 0 | 0.062 | 1 | 24 |
| nol9 | 0.28371 | -0.23167083 | 0 | 0.062 | 1 | 24 |
| nprl3 | 0.28371 | -0.14384441 | 0 | 0.062 | 1 | 24 |
| nr0b1 | 0.28371 | -0.15059478 | 0 | 0.062 | 1 | 24 |
| nr1d2b | 0.28371 | -0.11085252 | 0 | 0.062 | 1 | 24 |
| nr4a1 | 0.28371 | -0.21564181 | 0 | 0.062 | 1 | 24 |
| nras | 0.28371 | -0.10313254 | 0 | 0.062 | 1 | 24 |
| nrbf2b | 0.28371 | -0.22018514 | 0 | 0.062 | 1 | 24 |
| nsun4 | 0.28371 | -0.22639747 | 0 | 0.062 | 1 | 24 |
| nsun5 | 0.28371 | -0.15701411 | 0 | 0.062 | 1 | 24 |
| nub1 | 0.28371 | -0.24344318 | 0 | 0.062 | 1 | 24 |
| nudt9 | 0.28371 | -0.11053489 | 0 | 0.062 | 1 | 24 |
| nup35 | 0.28371 | -0.28071166 | 0 | 0.062 | 1 | 24 |
| ogfod1 | 0.28371 | -0.10115976 | 0 | 0.062 | 1 | 24 |
| oip5-as1 | 0.28371 | -0.15055726 | 0 | 0.062 | 1 | 24 |
| orc5 | 0.28371 | -0.11056714 | 0 | 0.062 | 1 | 24 |
| os9 | 0.28371 | -0.10964443 | 0 | 0.062 | 1 | 24 |
| otulina | 0.28371 | -0.13573693 | 0 | 0.062 | 1 | 24 |
| palld | 0.28371 | -0.13835198 | 0 | 0.062 | 1 | 24 |
| paplna | 0.28371 | -0.17940227 | 0 | 0.062 | 1 | 24 |
| parp4 | 0.28371 | -0.10878534 | 0 | 0.062 | 1 | 24 |
| pcyt2 | 0.28371 | -0.24430841 | 0 | 0.062 | 1 | 24 |
| pdcd2l | 0.28371 | -0.14097676 | 0 | 0.062 | 1 | 24 |
| pde6ha | 0.28371 | -0.16100772 | 0 | 0.062 | 1 | 24 |
| pdk4 | 0.28371 | -0.10512839 | 0 | 0.062 | 1 | 24 |
| pdlim1 | 0.28371 | -0.22503699 | 0 | 0.062 | 1 | 24 |
| per3 | 0.28371 | -0.11054246 | 0 | 0.062 | 1 | 24 |
| phrf1 | 0.28371 | -0.24443474 | 0 | 0.062 | 1 | 24 |
| phyhipla | 0.28371 | -0.11395362 | 0 | 0.062 | 1 | 24 |
| pi15a | 0.28371 | -0.10951567 | 0 | 0.062 | 1 | 24 |
| pigf | 0.28371 | -0.11176212 | 0 | 0.062 | 1 | 24 |
| pigp | 0.28371 | -0.21148111 | 0 | 0.062 | 1 | 24 |
| pik3c2b | 0.28371 | -0.11266916 | 0 | 0.062 | 1 | 24 |
| pik3r2 | 0.28371 | -0.13908573 | 0 | 0.062 | 1 | 24 |
| pitpnm3 | 0.28371 | -0.1569114 | 0 | 0.062 | 1 | 24 |
| pja2 | 0.28371 | -0.23067411 | 0 | 0.062 | 1 | 24 |
| pla2g6 | 0.28371 | -0.10673566 | 0 | 0.062 | 1 | 24 |
| plcd3a | 0.28371 | -0.10191777 | 0 | 0.062 | 1 | 24 |
| pnp5b | 0.28371 | -0.22174856 | 0 | 0.062 | 1 | 24 |
| pof1b | 0.28371 | -0.21796122 | 0 | 0.062 | 1 | 24 |
| polr1a | 0.28371 | -0.11152064 | 0 | 0.062 | 1 | 24 |
| polr1b | 0.28371 | -0.11251225 | 0 | 0.062 | 1 | 24 |
| pop5 | 0.28371 | -0.10866925 | 0 | 0.062 | 1 | 24 |
| pou2f1b | 0.28371 | -0.1125533 | 0 | 0.062 | 1 | 24 |
| ppfibp1a | 0.28371 | -0.15576447 | 0 | 0.062 | 1 | 24 |

| | | | | | | |
|---|---|---|---|---|---|---|
| ppfibp1b | 0.28371 | -0.12873964 | 0 | 0.062 | 1 | 24 |
| ppm1bb | 0.28371 | -0.11201676 | 0 | 0.062 | 1 | 24 |
| ppp1r32 | 0.28371 | -0.22712508 | 0 | 0.062 | 1 | 24 |
| ppp2cab | 0.28371 | -0.10544328 | 0 | 0.062 | 1 | 24 |
| prc1a | 0.28371 | -0.219418 | 0 | 0.062 | 1 | 24 |
| prcp | 0.28371 | -0.11100912 | 0 | 0.062 | 1 | 24 |
| prkag2a | 0.28371 | -0.11264192 | 0 | 0.062 | 1 | 24 |
| prkar1aa | 0.28371 | -0.12548421 | 0 | 0.062 | 1 | 24 |
| prkar1b | 0.28371 | -0.10764255 | 0 | 0.062 | 1 | 24 |
| prkn | 0.28371 | -0.13585955 | 0 | 0.062 | 1 | 24 |
| prpf6 | 0.28371 | -0.22782888 | 0 | 0.062 | 1 | 24 |
| prr11 | 0.28371 | -0.19695877 | 0 | 0.062 | 1 | 24 |
| prxl2b | 0.28371 | -0.21247843 | 0 | 0.062 | 1 | 24 |
| ptpn4a | 0.28371 | -0.21943429 | 0 | 0.062 | 1 | 24 |
| ptprja | 0.28371 | -0.10847005 | 0 | 0.062 | 1 | 24 |
| pum2 | 0.28371 | -0.24352115 | 0 | 0.062 | 1 | 24 |
| purab | 0.28371 | -0.11273645 | 0 | 0.062 | 1 | 24 |
| pus1 | 0.28371 | -0.10866913 | 0 | 0.062 | 1 | 24 |
| pwp2h | 0.28371 | -0.1421379 | 0 | 0.062 | 1 | 24 |
| pxdc1b | 0.28371 | -0.12938901 | 0 | 0.062 | 1 | 24 |
| pygb | 0.28371 | -0.11175399 | 0 | 0.062 | 1 | 24 |
| r3hcc1l | 0.28371 | -0.14451279 | 0 | 0.062 | 1 | 24 |
| rab1aa | 0.28371 | -0.243724 | 0 | 0.062 | 1 | 24 |
| rab3il1 | 0.28371 | -0.21360878 | 0 | 0.062 | 1 | 24 |
| rab5aa | 0.28371 | -0.10985025 | 0 | 0.062 | 1 | 24 |
| rab6bb | 0.28371 | -0.15621953 | 0 | 0.062 | 1 | 24 |
| rabgef1 | 0.28371 | -0.1013391 | 0 | 0.062 | 1 | 24 |
| rabggta | 0.28371 | -0.11039992 | 0 | 0.062 | 1 | 24 |
| rabggtb | 0.28371 | -0.1427117 | 0 | 0.062 | 1 | 24 |
| rac3b | 0.28371 | -0.11151535 | 0 | 0.062 | 1 | 24 |
| rag2 | 0.28371 | -0.11395362 | 0 | 0.062 | 1 | 24 |
| rb1 | 0.28371 | -0.15753891 | 0 | 0.062 | 1 | 24 |
| rb1cc1 | 0.28371 | -0.23613797 | 0 | 0.062 | 1 | 24 |
| rbbp8l | 0.28371 | -0.10368463 | 0 | 0.062 | 1 | 24 |
| rbm42 | 0.28371 | -0.13906758 | 0 | 0.062 | 1 | 24 |
| rbm45 | 0.28371 | -0.16094049 | 0 | 0.062 | 1 | 24 |
| rcor3 | 0.28371 | -0.22280482 | 0 | 0.062 | 1 | 24 |
| rex1bd | 0.28371 | -0.10570002 | 0 | 0.062 | 1 | 24 |
| rfxap | 0.28371 | -0.11003653 | 0 | 0.062 | 1 | 24 |
| rgcc | 0.28371 | -0.10268977 | 0 | 0.062 | 1 | 24 |
| rgs7a | 0.28371 | -0.24390313 | 0 | 0.062 | 1 | 24 |
| rgs7bpb | 0.28371 | -0.20363243 | 0 | 0.062 | 1 | 24 |
| rhebl1 | 0.28371 | -0.13782319 | 0 | 0.062 | 1 | 24 |
| rmi1 | 0.28371 | -0.21368693 | 0 | 0.062 | 1 | 24 |
| rnf170 | 0.28371 | -0.1098279 | 0 | 0.062 | 1 | 24 |
| rnf220a | 0.28371 | -0.24406248 | 0 | 0.062 | 1 | 24 |
| rnf40 | 0.28371 | -0.10453903 | 0 | 0.062 | 1 | 24 |
| rpp25l | 0.28371 | -0.18622 | 0 | 0.062 | 1 | 24 |

| | | | | | | |
|---|---|---|---|---|---|---|
| rpp38 | 0.28371 | -0.10001748 | 0 | 0.062 | 1 | 24 |
| rrp9 | 0.28371 | -0.22561883 | 0 | 0.062 | 1 | 24 |
| rsu1 | 0.28371 | -0.15212776 | 0 | 0.062 | 1 | 24 |
| rtn4ip1 | 0.28371 | -0.10897483 | 0 | 0.062 | 1 | 24 |
| rufy2 | 0.28371 | -0.16294972 | 0 | 0.062 | 1 | 24 |
| ruvbl1 | 0.28371 | -0.22793599 | 0 | 0.062 | 1 | 24 |
| s100a1 | 0.28371 | -0.2014057 | 0 | 0.062 | 1 | 24 |
| s100a10a | 0.28371 | -0.10172186 | 0 | 0.062 | 1 | 24 |
| s100b | 0.28371 | -0.21155729 | 0 | 0.062 | 1 | 24 |
| sapcd2 | 0.28371 | -0.23030443 | 0 | 0.062 | 1 | 24 |
| sart3 | 0.28371 | -0.21143344 | 0 | 0.062 | 1 | 24 |
| sbf1 | 0.28371 | -0.15776666 | 0 | 0.062 | 1 | 24 |
| sec22c | 0.28371 | -0.1124699 | 0 | 0.062 | 1 | 24 |
| selenoj | 0.28371 | -0.10007309 | 0 | 0.062 | 1 | 24 |
| sema3ab | 0.28371 | -0.22526544 | 0 | 0.062 | 1 | 24 |
| senp5 | 0.28371 | -0.22865529 | 0 | 0.062 | 1 | 24 |
| serinc2 | 0.28371 | -0.11112688 | 0 | 0.062 | 1 | 24 |
| serpinb1l1 | 0.28371 | -0.1420855 | 0 | 0.062 | 1 | 24 |
| setd4 | 0.28371 | -0.13546829 | 0 | 0.062 | 1 | 24 |
| sh2b1 | 0.28371 | -0.10364568 | 0 | 0.062 | 1 | 24 |
| sh2b3 | 0.28371 | -0.110569 | 0 | 0.062 | 1 | 24 |
| sh3glb2b | 0.28371 | -0.1113125 | 0 | 0.062 | 1 | 24 |
| shisa4 | 0.28371 | -0.11253582 | 0 | 0.062 | 1 | 24 |
| si:cabz01069012.2 | 0.28371 | -0.23125898 | 0 | 0.062 | 1 | 24 |
| si:cabz01102082.1 | 0.28371 | -0.11125698 | 0 | 0.062 | 1 | 24 |
| si:ch1073-272j17.3 | 0.28371 | -0.22058121 | 0 | 0.062 | 1 | 24 |
| si:ch211-106h4.12 | 0.28371 | -0.13584498 | 0 | 0.062 | 1 | 24 |
| si:ch211-108d22.2 | 0.28371 | -0.16288983 | 0 | 0.062 | 1 | 24 |
| si:ch211-10a23.2 | 0.28371 | -0.18876532 | 0 | 0.062 | 1 | 24 |
| si:ch211-114l13.9 | 0.28371 | -0.14247566 | 0 | 0.062 | 1 | 24 |
| si:ch211-120g10.1 | 0.28371 | -0.15258353 | 0 | 0.062 | 1 | 24 |
| si:ch211-154o6.3 | 0.28371 | -0.10888065 | 0 | 0.062 | 1 | 24 |
| si:ch211-161c3.6 | 0.28371 | -0.15214783 | 0 | 0.062 | 1 | 24 |
| si:ch211-183d21.1 | 0.28371 | -0.1648955 | 0 | 0.062 | 1 | 24 |
| si:ch211-214c7.4 | 0.28371 | -0.14247566 | 0 | 0.062 | 1 | 24 |
| si:ch211-223a10.1 | 0.28371 | -0.1963201 | 0 | 0.062 | 1 | 24 |
| si:ch211-238n5.4 | 0.28371 | -0.21220634 | 0 | 0.062 | 1 | 24 |
| si:ch211-243j20.2 | 0.28371 | -0.11142513 | 0 | 0.062 | 1 | 24 |
| si:ch211-244b2.3 | 0.28371 | -0.11031515 | 0 | 0.062 | 1 | 24 |
| si:ch211-256e16.3 | 0.28371 | -0.10542595 | 0 | 0.062 | 1 | 24 |
| si:ch211-261d7.3 | 0.28371 | -0.21310575 | 0 | 0.062 | 1 | 24 |
| si:ch211-271b14.1 | 0.28371 | -0.23013958 | 0 | 0.062 | 1 | 24 |
| si:ch211-66e2.3 | 0.28371 | -0.15790661 | 0 | 0.062 | 1 | 24 |
| si:ch73-100l22.3 | 0.28371 | -0.11114047 | 0 | 0.062 | 1 | 24 |
| si:ch73-234b20.5 | 0.28371 | -0.10416012 | 0 | 0.062 | 1 | 24 |
| si:ch73-264p11.1 | 0.28371 | -0.28147847 | 0 | 0.062 | 1 | 24 |
| si:ch73-27e22.1 | 0.28371 | -0.11253322 | 0 | 0.062 | 1 | 24 |
| si:dkey-103e21.5 | 0.28371 | -0.10919742 | 0 | 0.062 | 1 | 24 |

| | | | | | | |
|---|---|---|---|---|---|---|
| si:dkey-108k21.10 | 0.28371 | -0.16397554 | 0 | 0.062 | 1 | 24 |
| si:dkey-111k8.5 | 0.28371 | -0.11320582 | 0 | 0.062 | 1 | 24 |
| si:dkey-156m2.3 | 0.28371 | -0.14737693 | 0 | 0.062 | 1 | 24 |
| si:dkey-165n16.1 | 0.28371 | -0.11139402 | 0 | 0.062 | 1 | 24 |
| si:dkey-184a18.5 | 0.28371 | -0.15985321 | 0 | 0.062 | 1 | 24 |
| si:dkey-184p18.2 | 0.28371 | -0.14258612 | 0 | 0.062 | 1 | 24 |
| si:dkey-185e18.7 | 0.28371 | -0.14216771 | 0 | 0.062 | 1 | 24 |
| si:dkey-188i13.9 | 0.28371 | -0.13595561 | 0 | 0.062 | 1 | 24 |
| si:dkey-199f5.8 | 0.28371 | -0.15634004 | 0 | 0.062 | 1 | 24 |
| si:dkey-21e2.12 | 0.28371 | -0.23120429 | 0 | 0.062 | 1 | 24 |
| si:dkey-230p4.1 | 0.28371 | -0.10714683 | 0 | 0.062 | 1 | 24 |
| si:dkey-23a23.3 | 0.28371 | -0.14422464 | 0 | 0.062 | 1 | 24 |
| si:dkey-276j7.1 | 0.28371 | -0.2257606 | 0 | 0.062 | 1 | 24 |
| si:dkey-40c23.2 | 0.28371 | -0.24318513 | 0 | 0.062 | 1 | 24 |
| si:dkey-4e7.3 | 0.28371 | -0.16245189 | 0 | 0.062 | 1 | 24 |
| si:dkey-68o6.6 | 0.28371 | -0.14247566 | 0 | 0.062 | 1 | 24 |
| si:dkey-6n6.1 | 0.28371 | -0.21788567 | 0 | 0.062 | 1 | 24 |
| si:dkey-70b23.2 | 0.28371 | -0.16238109 | 0 | 0.062 | 1 | 24 |
| si:dkey-77f5.4 | 0.28371 | -0.21821405 | 0 | 0.062 | 1 | 24 |
| si:dkey-78k11.9 | 0.28371 | -0.22280482 | 0 | 0.062 | 1 | 24 |
| si:dkey-7n6.2 | 0.28371 | -0.22181267 | 0 | 0.062 | 1 | 24 |
| si:dkey-89b17.4 | 0.28371 | -0.11316154 | 0 | 0.062 | 1 | 24 |
| si:dkey-92i17.2 | 0.28371 | -0.27441076 | 0 | 0.062 | 1 | 24 |
| si:dkey-9i23.15 | 0.28371 | -0.12851743 | 0 | 0.062 | 1 | 24 |
| si:dkey-9i23.16 | 0.28371 | -0.10207734 | 0 | 0.062 | 1 | 24 |
| si:dkeyp-20g2.1 | 0.28371 | -0.10490251 | 0 | 0.062 | 1 | 24 |
| si:dkeyp-66d1.7 | 0.28371 | -0.21199012 | 0 | 0.062 | 1 | 24 |
| si:dkeyp-97b10.3 | 0.28371 | -0.11089569 | 0 | 0.062 | 1 | 24 |
| si:rp71-36a1.5 | 0.28371 | -0.11191789 | 0 | 0.062 | 1 | 24 |
| si:zfos-464b6.2 | 0.28371 | -0.11159095 | 0 | 0.062 | 1 | 24 |
| sinhcaf | 0.28371 | -0.11501912 | 0 | 0.062 | 1 | 24 |
| sirt3 | 0.28371 | -0.10639802 | 0 | 0.062 | 1 | 24 |
| six1a | 0.28371 | -0.23120429 | 0 | 0.062 | 1 | 24 |
| skap2 | 0.28371 | -0.22589892 | 0 | 0.062 | 1 | 24 |
| slc1a4 | 0.28371 | -0.14874294 | 0 | 0.062 | 1 | 24 |
| slc25a32a | 0.28371 | -0.15915563 | 0 | 0.062 | 1 | 24 |
| slc35f2 | 0.28371 | -0.10003114 | 0 | 0.062 | 1 | 24 |
| slc39a7 | 0.28371 | -0.11147858 | 0 | 0.062 | 1 | 24 |
| slc4a1a | 0.28371 | -0.22985356 | 0 | 0.062 | 1 | 24 |
| slc7a3b | 0.28371 | -0.11343786 | 0 | 0.062 | 1 | 24 |
| slc7a6os | 0.28371 | -0.11175604 | 0 | 0.062 | 1 | 24 |
| slmapa | 0.28371 | -0.22031435 | 0 | 0.062 | 1 | 24 |
| smad6a | 0.28371 | -0.14361486 | 0 | 0.062 | 1 | 24 |
| smarcc1a | 0.28371 | -0.10090894 | 0 | 0.062 | 1 | 24 |
| smarcc2 | 0.28371 | -0.16317909 | 0 | 0.062 | 1 | 24 |
| smc6 | 0.28371 | -0.19945647 | 0 | 0.062 | 1 | 24 |
| smcr8a | 0.28371 | -0.10775388 | 0 | 0.062 | 1 | 24 |
| smu1a | 0.28371 | -0.14512247 | 0 | 0.062 | 1 | 24 |

| | | | | | | |
|---|---|---|---|---|---|---|
| snapc1b | 0.28371 | -0.10972235 | 0 | 0.062 | 1 | 24 |
| snx5 | 0.28371 | -0.28039564 | 0 | 0.062 | 1 | 24 |
| socs3a | 0.28371 | -0.19676098 | 0 | 0.062 | 1 | 24 |
| soga1 | 0.28371 | -0.21248664 | 0 | 0.062 | 1 | 24 |
| sord | 0.28371 | -0.11139621 | 0 | 0.062 | 1 | 24 |
| soul5 | 0.28371 | -0.15742163 | 0 | 0.062 | 1 | 24 |
| sox12 | 0.28371 | -0.10878534 | 0 | 0.062 | 1 | 24 |
| sox8a | 0.28371 | -0.21388497 | 0 | 0.062 | 1 | 24 |
| sp4 | 0.28371 | -0.26289136 | 0 | 0.062 | 1 | 24 |
| spsb3a | 0.28371 | -0.24475203 | 0 | 0.062 | 1 | 24 |
| sptan1 | 0.28371 | -0.24438042 | 0 | 0.062 | 1 | 24 |
| sra1 | 0.28371 | -0.11050414 | 0 | 0.062 | 1 | 24 |
| srgap3 | 0.28371 | -0.20554047 | 0 | 0.062 | 1 | 24 |
| srpk1a | 0.28371 | -0.15824863 | 0 | 0.062 | 1 | 24 |
| srpra | 0.28371 | -0.10345203 | 0 | 0.062 | 1 | 24 |
| srrm4 | 0.28371 | -0.10696802 | 0 | 0.062 | 1 | 24 |
| srsf2b | 0.28371 | -0.10578655 | 0 | 0.062 | 1 | 24 |
| ssr1 | 0.28371 | -0.16585268 | 0 | 0.062 | 1 | 24 |
| stag2a | 0.28371 | -0.11295858 | 0 | 0.062 | 1 | 24 |
| stat1b | 0.28371 | -0.28069719 | 0 | 0.062 | 1 | 24 |
| stk17a | 0.28371 | -0.21534305 | 0 | 0.062 | 1 | 24 |
| stk38b | 0.28371 | -0.24088812 | 0 | 0.062 | 1 | 24 |
| stk38l | 0.28371 | -0.13646504 | 0 | 0.062 | 1 | 24 |
| stn1 | 0.28371 | -0.14143277 | 0 | 0.062 | 1 | 24 |
| strada | 0.28371 | -0.10337581 | 0 | 0.062 | 1 | 24 |
| strn3 | 0.28371 | -0.15855742 | 0 | 0.062 | 1 | 24 |
| strn4 | 0.28371 | -0.10226134 | 0 | 0.062 | 1 | 24 |
| stx2a | 0.28371 | -0.10390891 | 0 | 0.062 | 1 | 24 |
| stxbp3 | 0.28371 | -0.14429132 | 0 | 0.062 | 1 | 24 |
| stxbp4 | 0.28371 | -0.10581206 | 0 | 0.062 | 1 | 24 |
| suox | 0.28371 | -0.14550024 | 0 | 0.062 | 1 | 24 |
| surf6 | 0.28371 | -0.11133368 | 0 | 0.062 | 1 | 24 |
| sypl1 | 0.28371 | -0.14312033 | 0 | 0.062 | 1 | 24 |
| tab1 | 0.28371 | -0.16148638 | 0 | 0.062 | 1 | 24 |
| taf11 | 0.28371 | -0.10881153 | 0 | 0.062 | 1 | 24 |
| taf6l | 0.28371 | -0.11363461 | 0 | 0.062 | 1 | 24 |
| tagapa | 0.28371 | -0.22636071 | 0 | 0.062 | 1 | 24 |
| tango2 | 0.28371 | -0.11007993 | 0 | 0.062 | 1 | 24 |
| tango6 | 0.28371 | -0.11035925 | 0 | 0.062 | 1 | 24 |
| taok2a | 0.28371 | -0.24477004 | 0 | 0.062 | 1 | 24 |
| tarsl2 | 0.28371 | -0.24497379 | 0 | 0.062 | 1 | 24 |
| tbcc | 0.28371 | -0.20652876 | 0 | 0.062 | 1 | 24 |
| tbl2 | 0.28371 | -0.10400328 | 0 | 0.062 | 1 | 24 |
| tbp | 0.28371 | -0.10618435 | 0 | 0.062 | 1 | 24 |
| tbpl1 | 0.28371 | -0.11270633 | 0 | 0.062 | 1 | 24 |
| tcf23 | 0.28371 | -0.1648955 | 0 | 0.062 | 1 | 24 |
| tent4b | 0.28371 | -0.11093392 | 0 | 0.062 | 1 | 24 |
| tent5c | 0.28371 | -0.22963119 | 0 | 0.062 | 1 | 24 |

| | | | | | | |
|---|---|---|---|---|---|---|
| tjp2b | 0.28371 | -0.21904049 | 0 | 0.062 | 1 | 24 |
| tm2d3 | 0.28371 | -0.11112758 | 0 | 0.062 | 1 | 24 |
| tmco1 | 0.28371 | -0.15673115 | 0 | 0.062 | 1 | 24 |
| tmem129 | 0.28371 | -0.11160779 | 0 | 0.062 | 1 | 24 |
| tmem203 | 0.28371 | -0.21693729 | 0 | 0.062 | 1 | 24 |
| tmem246 | 0.28371 | -0.15258353 | 0 | 0.062 | 1 | 24 |
| tmem5 | 0.28371 | -0.22294884 | 0 | 0.062 | 1 | 24 |
| tmem59 | 0.28371 | -0.22893585 | 0 | 0.062 | 1 | 24 |
| tmem88b | 0.28371 | -0.13624128 | 0 | 0.062 | 1 | 24 |
| tmub1 | 0.28371 | -0.22971762 | 0 | 0.062 | 1 | 24 |
| tmx2b | 0.28371 | -0.10972665 | 0 | 0.062 | 1 | 24 |
| tnfrsf19 | 0.28371 | -0.13559489 | 0 | 0.062 | 1 | 24 |
| tnip1 | 0.28371 | -0.15492676 | 0 | 0.062 | 1 | 24 |
| tnk2b | 0.28371 | -0.23120429 | 0 | 0.062 | 1 | 24 |
| tox4a | 0.28371 | -0.1998878 | 0 | 0.062 | 1 | 24 |
| tpd52l1 | 0.28371 | -0.26342641 | 0 | 0.062 | 1 | 24 |
| tpd52l2a | 0.28371 | -0.22071964 | 0 | 0.062 | 1 | 24 |
| triqk | 0.28371 | -0.15910206 | 0 | 0.062 | 1 | 24 |
| trmt12 | 0.28371 | -0.11021329 | 0 | 0.062 | 1 | 24 |
| trmt1l | 0.28371 | -0.23209587 | 0 | 0.062 | 1 | 24 |
| trmt2a | 0.28371 | -0.21239104 | 0 | 0.062 | 1 | 24 |
| trpt1 | 0.28371 | -0.1438909 | 0 | 0.062 | 1 | 24 |
| tsg101a | 0.28371 | -0.1016692 | 0 | 0.062 | 1 | 24 |
| tsg101b | 0.28371 | -0.1072342 | 0 | 0.062 | 1 | 24 |
| tspan2b | 0.28371 | -0.21821405 | 0 | 0.062 | 1 | 24 |
| ttf1 | 0.28371 | -0.10980775 | 0 | 0.062 | 1 | 24 |
| ttyh2 | 0.28371 | -0.10559262 | 0 | 0.062 | 1 | 24 |
| ubald1a | 0.28371 | -0.10232636 | 0 | 0.062 | 1 | 24 |
| ubald2 | 0.28371 | -0.10661987 | 0 | 0.062 | 1 | 24 |
| ube2g1a | 0.28371 | -0.10773688 | 0 | 0.062 | 1 | 24 |
| ube2h | 0.28371 | -0.14288353 | 0 | 0.062 | 1 | 24 |
| ublcp1 | 0.28371 | -0.20994076 | 0 | 0.062 | 1 | 24 |
| ubqln4 | 0.28371 | -0.14067524 | 0 | 0.062 | 1 | 24 |
| ubxn8 | 0.28371 | -0.11023943 | 0 | 0.062 | 1 | 24 |
| ufsp2 | 0.28371 | -0.1002482 | 0 | 0.062 | 1 | 24 |
| ulk1a | 0.28371 | -0.10191777 | 0 | 0.062 | 1 | 24 |
| unc119a | 0.28371 | -0.10758111 | 0 | 0.062 | 1 | 24 |
| uraha | 0.28371 | -0.21284195 | 0 | 0.062 | 1 | 24 |
| usf2 | 0.28371 | -0.11088138 | 0 | 0.062 | 1 | 24 |
| usp22 | 0.28371 | -0.10548033 | 0 | 0.062 | 1 | 24 |
| usp4 | 0.28371 | -0.2312524 | 0 | 0.062 | 1 | 24 |
| usp48 | 0.28371 | -0.11205862 | 0 | 0.062 | 1 | 24 |
| uvrag | 0.28371 | -0.11218024 | 0 | 0.062 | 1 | 24 |
| vipr2 | 0.28371 | -0.11006021 | 0 | 0.062 | 1 | 24 |
| vkorc1l1 | 0.28371 | -0.15837531 | 0 | 0.062 | 1 | 24 |
| vps13c | 0.28371 | -0.10703637 | 0 | 0.062 | 1 | 24 |
| vps16 | 0.28371 | -0.10871408 | 0 | 0.062 | 1 | 24 |
| vps26c | 0.28371 | -0.26002146 | 0 | 0.062 | 1 | 24 |

| | | | | | | |
|---|---|---|---|---|---|---|
| vps33b | 0.28371 | -0.21258674 | 0 | 0.062 | 1 | 24 |
| vps37a | 0.28371 | -0.16105404 | 0 | 0.062 | 1 | 24 |
| vps45 | 0.28371 | -0.11093188 | 0 | 0.062 | 1 | 24 |
| vrk3 | 0.28371 | -0.23077686 | 0 | 0.062 | 1 | 24 |
| wars2 | 0.28371 | -0.23002656 | 0 | 0.062 | 1 | 24 |
| washc2c | 0.28371 | -0.10811966 | 0 | 0.062 | 1 | 24 |
| wdr3 | 0.28371 | -0.10631326 | 0 | 0.062 | 1 | 24 |
| wdr36 | 0.28371 | -0.11142634 | 0 | 0.062 | 1 | 24 |
| wdr45 | 0.28371 | -0.11158349 | 0 | 0.062 | 1 | 24 |
| wsb2 | 0.28371 | -0.10701481 | 0 | 0.062 | 1 | 24 |
| wu:fc75a09 | 0.28371 | -0.24488467 | 0 | 0.062 | 1 | 24 |
| wu:fi04e12 | 0.28371 | -0.243447 | 0 | 0.062 | 1 | 24 |
| wwp2 | 0.28371 | -0.10504009 | 0 | 0.062 | 1 | 24 |
| xkr8.3 | 0.28371 | -0.11030137 | 0 | 0.062 | 1 | 24 |
| xrcc4 | 0.28371 | -0.21300383 | 0 | 0.062 | 1 | 24 |
| xrcc5 | 0.28371 | -0.2215885 | 0 | 0.062 | 1 | 24 |
| yars | 0.28371 | -0.13898842 | 0 | 0.062 | 1 | 24 |
| ywhae2 | 0.28371 | -0.11165258 | 0 | 0.062 | 1 | 24 |
| yy1a | 0.28371 | -0.11076979 | 0 | 0.062 | 1 | 24 |
| zc3h4 | 0.28371 | -0.1115722 | 0 | 0.062 | 1 | 24 |
| zdhhc20a | 0.28371 | -0.10624618 | 0 | 0.062 | 1 | 24 |
| zdhhc6 | 0.28371 | -0.1107518 | 0 | 0.062 | 1 | 24 |
| zgc:101851 | 0.28371 | -0.23862679 | 0 | 0.062 | 1 | 24 |
| zgc:110425 | 0.28371 | -0.1614174 | 0 | 0.062 | 1 | 24 |
| zgc:110626 | 0.28371 | -0.16228481 | 0 | 0.062 | 1 | 24 |
| zgc:112052 | 0.28371 | -0.11147111 | 0 | 0.062 | 1 | 24 |
| zgc:112083 | 0.28371 | -0.16035636 | 0 | 0.062 | 1 | 24 |
| zgc:112982 | 0.28371 | -0.14100943 | 0 | 0.062 | 1 | 24 |
| zgc:114200 | 0.28371 | -0.2196181 | 0 | 0.062 | 1 | 24 |
| zgc:123105 | 0.28371 | -0.13913712 | 0 | 0.062 | 1 | 24 |
| zgc:136864 | 0.28371 | -0.22240567 | 0 | 0.062 | 1 | 24 |
| zgc:153738 | 0.28371 | -0.22763065 | 0 | 0.062 | 1 | 24 |
| zgc:154077 | 0.28371 | -0.10938216 | 0 | 0.062 | 1 | 24 |
| zgc:158517 | 0.28371 | -0.11284294 | 0 | 0.062 | 1 | 24 |
| zgc:158564 | 0.28371 | -0.10701091 | 0 | 0.062 | 1 | 24 |
| zgc:162816 | 0.28371 | -0.14881304 | 0 | 0.062 | 1 | 24 |
| zgc:162898 | 0.28371 | -0.18435046 | 0 | 0.062 | 1 | 24 |
| zgc:163077 | 0.28371 | -0.10191777 | 0 | 0.062 | 1 | 24 |
| zgc:171506 | 0.28371 | -0.15683543 | 0 | 0.062 | 1 | 24 |
| zgc:173619 | 0.28371 | -0.21887771 | 0 | 0.062 | 1 | 24 |
| zgc:174895 | 0.28371 | -0.15721735 | 0 | 0.062 | 1 | 24 |
| zgc:194989 | 0.28371 | -0.1569114 | 0 | 0.062 | 1 | 24 |
| zgc:195081 | 0.28371 | -0.26315105 | 0 | 0.062 | 1 | 24 |
| zgc:195175 | 0.28371 | -0.13365807 | 0 | 0.062 | 1 | 24 |
| zgc:64002 | 0.28371 | -0.13128415 | 0 | 0.062 | 1 | 24 |
| zgc:64022 | 0.28371 | -0.21887771 | 0 | 0.062 | 1 | 24 |
| zgc:76872 | 0.28371 | -0.11045806 | 0 | 0.062 | 1 | 24 |
| zgc:77056 | 0.28371 | -0.11016819 | 0 | 0.062 | 1 | 24 |

| | | | | | | |
|---|---|---|---|---|---|---|
| zgc:77118 | 0.28371 | -0.14307425 | 0 | 0.062 | 1 | 24 |
| zgc:77784 | 0.28371 | -0.10542595 | 0 | 0.062 | 1 | 24 |
| zgc:86839 | 0.28371 | -0.24420145 | 0 | 0.062 | 1 | 24 |
| zmat5 | 0.28371 | -0.1271172 | 0 | 0.062 | 1 | 24 |
| zmp:0000000521 | 0.28371 | -0.22386392 | 0 | 0.062 | 1 | 24 |
| znf296 | 0.28371 | -0.11053969 | 0 | 0.062 | 1 | 24 |
| znf598 | 0.28371 | -0.23017141 | 0 | 0.062 | 1 | 24 |
| znf609b | 0.28371 | -0.22172704 | 0 | 0.062 | 1 | 24 |
| znf770 | 0.28371 | -0.22926051 | 0 | 0.062 | 1 | 24 |
| znrf1 | 0.28371 | -0.28002155 | 0 | 0.062 | 1 | 24 |
| AL954696.1 | 0.28371 | -0.16509442 | 0 | 0.062 | 1 | 24 |
| BX571880.1 | 0.28371 | -0.10332134 | 0 | 0.062 | 1 | 24 |
| BX890608.1 | 0.28371 | -0.10332134 | 0 | 0.062 | 1 | 24 |
| CABZ01021532.1 | 0.28371 | -0.20899044 | 0 | 0.062 | 1 | 24 |
| CABZ01046432.1 | 0.28371 | -0.10004196 | 0 | 0.062 | 1 | 24 |
| CABZ01072487.1 | 0.28371 | -0.17431292 | 0 | 0.062 | 1 | 24 |
| CR385050.1 | 0.28371 | -0.10004196 | 0 | 0.062 | 1 | 24 |
| CT027702.1 | 0.28371 | -0.10332134 | 0 | 0.062 | 1 | 24 |
| CT583672.1 | 0.28371 | -0.10004196 | 0 | 0.062 | 1 | 24 |
| CU550719.1 | 0.28371 | -0.16841074 | 0 | 0.062 | 1 | 24 |
| CU633999.2 | 0.28371 | -0.16815332 | 0 | 0.062 | 1 | 24 |
| FAR2 | 0.28371 | -0.2088893 | 0 | 0.062 | 1 | 24 |
| HOOK3 | 0.28371 | -0.19827612 | 0 | 0.062 | 1 | 24 |
| LOC100151196 | 0.28371 | -0.1026794 | 0 | 0.062 | 1 | 24 |
| LOC101882070 | 0.28371 | -0.14695366 | 0 | 0.062 | 1 | 24 |
| LOC101884253 | 0.28371 | -0.10332134 | 0 | 0.062 | 1 | 24 |
| LOC101885675 | 0.28371 | -0.10332134 | 0 | 0.062 | 1 | 24 |
| LOC101886485 | 0.28371 | -0.16594458 | 0 | 0.062 | 1 | 24 |
| LOC108191465 | 0.28371 | -0.20604245 | 0 | 0.062 | 1 | 24 |
| LOC110439449 | 0.28371 | -0.16181782 | 0 | 0.062 | 1 | 24 |
| LOC110439500 | 0.28371 | -0.13953036 | 0 | 0.062 | 1 | 24 |
| NFX1 | 0.28371 | -0.17709285 | 0 | 0.062 | 1 | 24 |
| PDCL3 | 0.28371 | -0.1701965 | 0 | 0.062 | 1 | 24 |
| SBNO2 | 0.28371 | -0.20764706 | 0 | 0.062 | 1 | 24 |
| SMIM10 | 0.28371 | -0.24059141 | 0 | 0.062 | 1 | 24 |
| XLOC-013920 | 0.28371 | -0.16861128 | 0 | 0.062 | 1 | 24 |
| XLOC-018328 | 0.28371 | -0.16576655 | 0 | 0.062 | 1 | 24 |
| XLOC-043760 | 0.28371 | -0.1328754 | 0 | 0.062 | 1 | 24 |
| aak1b | 0.28371 | -0.2410401 | 0 | 0.062 | 1 | 24 |
| abcb8 | 0.28371 | -0.16807254 | 0 | 0.062 | 1 | 24 |
| acod1 | 0.28371 | -0.16572585 | 0 | 0.062 | 1 | 24 |
| adgrl3.1 | 0.28371 | -0.14937257 | 0 | 0.062 | 1 | 24 |
| adkb | 0.28371 | -0.14207162 | 0 | 0.062 | 1 | 24 |
| alms1 | 0.28371 | -0.21199423 | 0 | 0.062 | 1 | 24 |
| ankrd40 | 0.28371 | -0.20437834 | 0 | 0.062 | 1 | 24 |
| aox6 | 0.28371 | -0.16824934 | 0 | 0.062 | 1 | 24 |
| apbb1 | 0.28371 | -0.16816162 | 0 | 0.062 | 1 | 24 |
| aplp2 | 0.28371 | -0.2093157 | 0 | 0.062 | 1 | 24 |

| | | | | | | |
|---|---|---|---|---|---|---|
| arpin | 0.28371 | -0.16205864 | 0 | 0.062 | 1 | 24 |
| asip1 | 0.28371 | -0.15702794 | 0 | 0.062 | 1 | 24 |
| aspm | 0.28371 | -0.16598679 | 0 | 0.062 | 1 | 24 |
| atp2b2 | 0.28371 | -0.20945461 | 0 | 0.062 | 1 | 24 |
| atp8a1 | 0.28371 | -0.21291849 | 0 | 0.062 | 1 | 24 |
| atpaf2 | 0.28371 | -0.20190651 | 0 | 0.062 | 1 | 24 |
| b3galnt2 | 0.28371 | -0.21164514 | 0 | 0.062 | 1 | 24 |
| babam2 | 0.28371 | -0.15431933 | 0 | 0.062 | 1 | 24 |
| bmp8a | 0.28371 | -0.20241496 | 0 | 0.062 | 1 | 24 |
| bmt2 | 0.28371 | -0.10018709 | 0 | 0.062 | 1 | 24 |
| borcs5 | 0.28371 | -0.10094039 | 0 | 0.062 | 1 | 24 |
| brat1 | 0.28371 | -0.16206399 | 0 | 0.062 | 1 | 24 |
| c19h1orf109 | 0.28371 | -0.20049841 | 0 | 0.062 | 1 | 24 |
| cad | 0.28371 | -0.17451687 | 0 | 0.062 | 1 | 24 |
| castor1 | 0.28371 | -0.21071433 | 0 | 0.062 | 1 | 24 |
| cbx6b | 0.28371 | -0.20986289 | 0 | 0.062 | 1 | 24 |
| ccdc106a | 0.28371 | -0.21533329 | 0 | 0.062 | 1 | 24 |
| ccdc43 | 0.28371 | -0.21093374 | 0 | 0.062 | 1 | 24 |
| ccl32b.3 | 0.28371 | -0.20761432 | 0 | 0.062 | 1 | 24 |
| ccnb3 | 0.28371 | -0.16794533 | 0 | 0.062 | 1 | 24 |
| ccz1 | 0.28371 | -0.23556683 | 0 | 0.062 | 1 | 24 |
| cdc23 | 0.28371 | -0.21145934 | 0 | 0.062 | 1 | 24 |
| cdc73 | 0.28371 | -0.212974 | 0 | 0.062 | 1 | 24 |
| cdk5rap1 | 0.28371 | -0.15937296 | 0 | 0.062 | 1 | 24 |
| cep120 | 0.28371 | -0.10114141 | 0 | 0.062 | 1 | 24 |
| cep135 | 0.28371 | -0.10052313 | 0 | 0.062 | 1 | 24 |
| cfd | 0.28371 | -0.23219768 | 0 | 0.062 | 1 | 24 |
| chchd3b | 0.28371 | -0.16319928 | 0 | 0.062 | 1 | 24 |
| chpt1 | 0.28371 | -0.16011689 | 0 | 0.062 | 1 | 24 |
| chst14 | 0.28371 | -0.20352148 | 0 | 0.062 | 1 | 24 |
| clic2 | 0.28371 | -0.1675465 | 0 | 0.062 | 1 | 24 |
| cltcb | 0.28371 | -0.21110312 | 0 | 0.062 | 1 | 24 |
| cnot10 | 0.28371 | -0.21379224 | 0 | 0.062 | 1 | 24 |
| creb1b | 0.28371 | -0.10052313 | 0 | 0.062 | 1 | 24 |
| cript | 0.28371 | -0.23748973 | 0 | 0.062 | 1 | 24 |
| crot | 0.28371 | -0.24041453 | 0 | 0.062 | 1 | 24 |
| cspg5a | 0.28371 | -0.21388766 | 0 | 0.062 | 1 | 24 |
| ctdsp2 | 0.28371 | -0.21096164 | 0 | 0.062 | 1 | 24 |
| ctsa | 0.28371 | -0.1740051 | 0 | 0.062 | 1 | 24 |
| cxcl12a | 0.28371 | -0.15661795 | 0 | 0.062 | 1 | 24 |
| cxcr3.1 | 0.28371 | -0.17073272 | 0 | 0.062 | 1 | 24 |
| cxxc5b | 0.28371 | -0.17431292 | 0 | 0.062 | 1 | 24 |
| cygb1 | 0.28371 | -0.21582301 | 0 | 0.062 | 1 | 24 |
| cyp11c1 | 0.28371 | -0.27742078 | 0 | 0.062 | 1 | 24 |
| dclre1b | 0.28371 | -0.16825697 | 0 | 0.062 | 1 | 24 |
| dglucy | 0.28371 | -0.16698886 | 0 | 0.062 | 1 | 24 |
| dhrs1 | 0.28371 | -0.1668856 | 0 | 0.062 | 1 | 24 |
| dlg3 | 0.28371 | -0.24036982 | 0 | 0.062 | 1 | 24 |

| | | | | | | |
|---|---|---|---|---|---|---|
| dock8 | 0.28371 | -0.1670487 | 0 | 0.062 | 1 | 24 |
| dyrk1ab | 0.28371 | -0.22498846 | 0 | 0.062 | 1 | 24 |
| eapp | 0.28371 | -0.24029891 | 0 | 0.062 | 1 | 24 |
| efhd2 | 0.28371 | -0.20840565 | 0 | 0.062 | 1 | 24 |
| efna2b | 0.28371 | -0.21490115 | 0 | 0.062 | 1 | 24 |
| eif3hb | 0.28371 | -0.17762878 | 0 | 0.062 | 1 | 24 |
| emsy | 0.28371 | -0.21165961 | 0 | 0.062 | 1 | 24 |
| esr2b | 0.28371 | -0.21802177 | 0 | 0.062 | 1 | 24 |
| exoc3l4 | 0.28371 | -0.19909695 | 0 | 0.062 | 1 | 24 |
| fabp11a | 0.28371 | -0.37974957 | 0 | 0.062 | 1 | 24 |
| fam126a | 0.28371 | -0.10118982 | 0 | 0.062 | 1 | 24 |
| fam69c | 0.28371 | -0.16562432 | 0 | 0.062 | 1 | 24 |
| fbxl3a | 0.28371 | -0.10062252 | 0 | 0.062 | 1 | 24 |
| fbxo42 | 0.28371 | -0.17762487 | 0 | 0.062 | 1 | 24 |
| fhl3b | 0.28371 | -0.16916391 | 0 | 0.062 | 1 | 24 |
| fkbp7 | 0.28371 | -0.16657302 | 0 | 0.062 | 1 | 24 |
| fopnl | 0.28371 | -0.17481876 | 0 | 0.062 | 1 | 24 |
| foxc1b | 0.28371 | -0.130917 | 0 | 0.062 | 1 | 24 |
| fxyd1 | 0.28371 | -0.2048019 | 0 | 0.062 | 1 | 24 |
| g0s2 | 0.28371 | -0.12521343 | 0 | 0.062 | 1 | 24 |
| g6pc3 | 0.28371 | -0.23478926 | 0 | 0.062 | 1 | 24 |
| gatd1 | 0.28371 | -0.10101549 | 0 | 0.062 | 1 | 24 |
| get4 | 0.28371 | -0.21080132 | 0 | 0.062 | 1 | 24 |
| gid8b | 0.28371 | -0.21358668 | 0 | 0.062 | 1 | 24 |
| gig2i | 0.28371 | -0.10117026 | 0 | 0.062 | 1 | 24 |
| got1 | 0.28371 | -0.1662352 | 0 | 0.062 | 1 | 24 |
| gpank1 | 0.28371 | -0.10040758 | 0 | 0.062 | 1 | 24 |
| gpt2l | 0.28371 | -0.23148479 | 0 | 0.062 | 1 | 24 |
| grb10a | 0.28371 | -0.18122665 | 0 | 0.062 | 1 | 24 |
| grcc10 | 0.28371 | -0.1670218 | 0 | 0.062 | 1 | 24 |
| gstz1 | 0.28371 | -0.19160685 | 0 | 0.062 | 1 | 24 |
| gtf3c3 | 0.28371 | -0.14936661 | 0 | 0.062 | 1 | 24 |
| gzm3 | 0.28371 | -0.14695366 | 0 | 0.062 | 1 | 24 |
| hddc2 | 0.28371 | -0.16041782 | 0 | 0.062 | 1 | 24 |
| hexb | 0.28371 | -0.10042086 | 0 | 0.062 | 1 | 24 |
| hmg20a | 0.28371 | -0.17012909 | 0 | 0.062 | 1 | 24 |
| hnrnpua | 0.28371 | -0.1712149 | 0 | 0.062 | 1 | 24 |
| hoxb6a | 0.28371 | -0.20937302 | 0 | 0.062 | 1 | 24 |
| hsbp1l1 | 0.28371 | -0.17752912 | 0 | 0.062 | 1 | 24 |
| hspa12a | 0.28371 | -0.24029896 | 0 | 0.062 | 1 | 24 |
| igf2b | 0.28371 | -0.15691977 | 0 | 0.062 | 1 | 24 |
| il12rb2 | 0.28371 | -0.10332134 | 0 | 0.062 | 1 | 24 |
| ilk | 0.28371 | -0.21180865 | 0 | 0.062 | 1 | 24 |
| ints11 | 0.28371 | -0.24072534 | 0 | 0.062 | 1 | 24 |
| itpk1b | 0.28371 | -0.16435405 | 0 | 0.062 | 1 | 24 |
| itsn1 | 0.28371 | -0.17583738 | 0 | 0.062 | 1 | 24 |
| jagn1a | 0.28371 | -0.10332134 | 0 | 0.062 | 1 | 24 |
| kat7a | 0.28371 | -0.10126973 | 0 | 0.062 | 1 | 24 |

| | | | | | | |
|---|---|---|---|---|---|---|
| kif20a | 0.28371 | -0.16864827 | 0 | 0.062 | 1 | 24 |
| klhl29 | 0.28371 | -0.17826717 | 0 | 0.062 | 1 | 24 |
| klhl35 | 0.28371 | -0.21127279 | 0 | 0.062 | 1 | 24 |
| lamp2 | 0.28371 | -0.17213114 | 0 | 0.062 | 1 | 24 |
| loxl2a | 0.28371 | -0.15065479 | 0 | 0.062 | 1 | 24 |
| lrata | 0.28371 | -0.1698591 | 0 | 0.062 | 1 | 24 |
| lrrfip2 | 0.28371 | -0.16811118 | 0 | 0.062 | 1 | 24 |
| lyn | 0.28371 | -0.20654064 | 0 | 0.062 | 1 | 24 |
| mafa | 0.28371 | -0.10163725 | 0 | 0.062 | 1 | 24 |
| mapk15 | 0.28371 | -0.2066081 | 0 | 0.062 | 1 | 24 |
| mblac1 | 0.28371 | -0.1984154 | 0 | 0.062 | 1 | 24 |
| mccc2 | 0.28371 | -0.14157166 | 0 | 0.062 | 1 | 24 |
| mcee | 0.28371 | -0.17454906 | 0 | 0.062 | 1 | 24 |
| med1 | 0.28371 | -0.15242612 | 0 | 0.062 | 1 | 24 |
| med24 | 0.28371 | -0.16233567 | 0 | 0.062 | 1 | 24 |
| meis3 | 0.28371 | -0.15604599 | 0 | 0.062 | 1 | 24 |
| mettl23 | 0.28371 | -0.14100167 | 0 | 0.062 | 1 | 24 |
| mettl24 | 0.28371 | -0.17653591 | 0 | 0.062 | 1 | 24 |
| mettl3 | 0.28371 | -0.16855281 | 0 | 0.062 | 1 | 24 |
| mmadhc | 0.28371 | -0.21038868 | 0 | 0.062 | 1 | 24 |
| mms19 | 0.28371 | -0.2109313 | 0 | 0.062 | 1 | 24 |
| mospd2 | 0.28371 | -0.12807907 | 0 | 0.062 | 1 | 24 |
| mrpl24 | 0.28371 | -0.16555818 | 0 | 0.062 | 1 | 24 |
| mrpl3 | 0.28371 | -0.10003672 | 0 | 0.062 | 1 | 24 |
| mrps34 | 0.28371 | -0.16466897 | 0 | 0.062 | 1 | 24 |
| msrb3 | 0.28371 | -0.16156499 | 0 | 0.062 | 1 | 24 |
| mt-atp8 | 0.28371 | -0.17731782 | 0 | 0.062 | 1 | 24 |
| mtif3 | 0.28371 | -0.16761944 | 0 | 0.062 | 1 | 24 |
| mybl2a | 0.28371 | -0.18349056 | 0 | 0.062 | 1 | 24 |
| myd88 | 0.28371 | -0.12755083 | 0 | 0.062 | 1 | 24 |
| nadkb | 0.28371 | -0.16083881 | 0 | 0.062 | 1 | 24 |
| naxe | 0.28371 | -0.21303313 | 0 | 0.062 | 1 | 24 |
| nell2a | 0.28371 | -0.1675683 | 0 | 0.062 | 1 | 24 |
| nf1b | 0.28371 | -0.19467869 | 0 | 0.062 | 1 | 24 |
| nfkbie | 0.28371 | -0.14343158 | 0 | 0.062 | 1 | 24 |
| nme6 | 0.28371 | -0.21075815 | 0 | 0.062 | 1 | 24 |
| nme7 | 0.28371 | -0.21438392 | 0 | 0.062 | 1 | 24 |
| nog1 | 0.28371 | -0.18073462 | 0 | 0.062 | 1 | 24 |
| nrf1 | 0.28371 | -0.21094559 | 0 | 0.062 | 1 | 24 |
| nsdhl | 0.28371 | -0.16823842 | 0 | 0.062 | 1 | 24 |
| nsfb | 0.28371 | -0.21640508 | 0 | 0.062 | 1 | 24 |
| nudt17 | 0.28371 | -0.16688636 | 0 | 0.062 | 1 | 24 |
| nup98 | 0.28371 | -0.17753733 | 0 | 0.062 | 1 | 24 |
| ociad1 | 0.28371 | -0.17167256 | 0 | 0.062 | 1 | 24 |
| oip5 | 0.28371 | -0.14082393 | 0 | 0.062 | 1 | 24 |
| osbpl9 | 0.28371 | -0.2115098 | 0 | 0.062 | 1 | 24 |
| osgep | 0.28371 | -0.17203126 | 0 | 0.062 | 1 | 24 |
| patj | 0.28371 | -0.1771559 | 0 | 0.062 | 1 | 24 |

| | | | | | | |
|---|---|---|---|---|---|---|
| pbxip1b | 0.28371 | -0.1737234 | 0 | 0.062 | 1 | 24 |
| pde6h | 0.28371 | -0.20261552 | 0 | 0.062 | 1 | 24 |
| pde8a | 0.28371 | -0.10048699 | 0 | 0.062 | 1 | 24 |
| pdhb | 0.28371 | -0.17562896 | 0 | 0.062 | 1 | 24 |
| pemt | 0.28371 | -0.16886017 | 0 | 0.062 | 1 | 24 |
| phf14 | 0.28371 | -0.19677434 | 0 | 0.062 | 1 | 24 |
| phf23a | 0.28371 | -0.12573656 | 0 | 0.062 | 1 | 24 |
| phyhd1 | 0.28371 | -0.16553666 | 0 | 0.062 | 1 | 24 |
| pisd | 0.28371 | -0.16476558 | 0 | 0.062 | 1 | 24 |
| pitx2 | 0.28371 | -0.16572585 | 0 | 0.062 | 1 | 24 |
| pogza | 0.28371 | -0.17826717 | 0 | 0.062 | 1 | 24 |
| polr2j | 0.28371 | -0.17259496 | 0 | 0.062 | 1 | 24 |
| ppme1 | 0.28371 | -0.14240385 | 0 | 0.062 | 1 | 24 |
| ppp2r2bb | 0.28371 | -0.21533329 | 0 | 0.062 | 1 | 24 |
| ppp4r3b | 0.28371 | -0.24009709 | 0 | 0.062 | 1 | 24 |
| prickle2b | 0.28371 | -0.16878283 | 0 | 0.062 | 1 | 24 |
| psat1 | 0.28371 | -0.16697964 | 0 | 0.062 | 1 | 24 |
| psmc3ip | 0.28371 | -0.17039375 | 0 | 0.062 | 1 | 24 |
| ptcd3 | 0.28371 | -0.21079091 | 0 | 0.062 | 1 | 24 |
| ptp4a2b | 0.28371 | -0.1669184 | 0 | 0.062 | 1 | 24 |
| pts | 0.28371 | -0.17269739 | 0 | 0.062 | 1 | 24 |
| qki2 | 0.28371 | -0.16775435 | 0 | 0.062 | 1 | 24 |
| rabif | 0.28371 | -0.14061914 | 0 | 0.062 | 1 | 24 |
| rad1 | 0.28371 | -0.17628379 | 0 | 0.062 | 1 | 24 |
| rad9a | 0.28371 | -0.16870867 | 0 | 0.062 | 1 | 24 |
| rarab | 0.28371 | -0.17132455 | 0 | 0.062 | 1 | 24 |
| rdh10a | 0.28371 | -0.16877682 | 0 | 0.062 | 1 | 24 |
| rgs17 | 0.28371 | -0.16875457 | 0 | 0.062 | 1 | 24 |
| rgs5b | 0.28371 | -0.1676175 | 0 | 0.062 | 1 | 24 |
| rhoj | 0.28371 | -0.16653274 | 0 | 0.062 | 1 | 24 |
| rnd3b | 0.28371 | -0.23752517 | 0 | 0.062 | 1 | 24 |
| rnf8 | 0.28371 | -0.16762111 | 0 | 0.062 | 1 | 24 |
| rps6ka3a | 0.28371 | -0.12573656 | 0 | 0.062 | 1 | 24 |
| rps6ka4 | 0.28371 | -0.10162949 | 0 | 0.062 | 1 | 24 |
| rpz | 0.28371 | -0.21012942 | 0 | 0.062 | 1 | 24 |
| rragca | 0.28371 | -0.15823961 | 0 | 0.062 | 1 | 24 |
| rtraf | 0.28371 | -0.13400399 | 0 | 0.062 | 1 | 24 |
| rufy3 | 0.28371 | -0.17707645 | 0 | 0.062 | 1 | 24 |
| sass6 | 0.28371 | -0.12374833 | 0 | 0.062 | 1 | 24 |
| scaf4a | 0.28371 | -0.10092753 | 0 | 0.062 | 1 | 24 |
| scarb2a | 0.28371 | -0.17572955 | 0 | 0.062 | 1 | 24 |
| sec23a | 0.28371 | -0.20972052 | 0 | 0.062 | 1 | 24 |
| selenou1b | 0.28371 | -0.15514195 | 0 | 0.062 | 1 | 24 |
| serbp1b | 0.28371 | -0.20991082 | 0 | 0.062 | 1 | 24 |
| serpinb1 | 0.28371 | -0.13389276 | 0 | 0.062 | 1 | 24 |
| si:ch1073-153i20.4 | 0.28371 | -0.21179762 | 0 | 0.062 | 1 | 24 |
| si:ch1073-209e23.1 | 0.28371 | -0.16951636 | 0 | 0.062 | 1 | 24 |
| si:ch1073-357b18.4 | 0.28371 | -0.10221403 | 0 | 0.062 | 1 | 24 |

| | | | | | | |
|---|---|---|---|---|---|---|
| si:ch1073-385f13.3 | 0.28371 | -0.10332134 | 0 | 0.062 | 1 | 24 |
| si:ch211-113a14.18 | 0.28371 | -0.17814288 | 0 | 0.062 | 1 | 24 |
| si:ch211-134m17.9 | 0.28371 | -0.2036497 | 0 | 0.062 | 1 | 24 |
| si:ch211-147a11.3 | 0.28371 | -0.20664272 | 0 | 0.062 | 1 | 24 |
| si:ch211-162e15.3 | 0.28371 | -0.14117198 | 0 | 0.062 | 1 | 24 |
| si:ch211-207k7.4 | 0.28371 | -0.16755548 | 0 | 0.062 | 1 | 24 |
| si:ch211-25d12.7 | 0.28371 | -0.10004196 | 0 | 0.062 | 1 | 24 |
| si:ch211-262h13.3 | 0.28371 | -0.14504616 | 0 | 0.062 | 1 | 24 |
| si:ch211-284e13.9 | 0.28371 | -0.10118105 | 0 | 0.062 | 1 | 24 |
| si:ch73-127m5.2 | 0.28371 | -0.17783406 | 0 | 0.062 | 1 | 24 |
| si:ch73-238c9.1 | 0.28371 | -0.16568436 | 0 | 0.062 | 1 | 24 |
| si:ch73-27e22.8 | 0.28371 | -0.2410401 | 0 | 0.062 | 1 | 24 |
| si:dkey-147f3.8 | 0.28371 | -0.15775904 | 0 | 0.062 | 1 | 24 |
| si:dkey-183k8.2 | 0.28371 | -0.23262741 | 0 | 0.062 | 1 | 24 |
| si:dkey-21c19.3 | 0.28371 | -0.1767917 | 0 | 0.062 | 1 | 24 |
| si:dkey-222h21.2 | 0.28371 | -0.15554609 | 0 | 0.062 | 1 | 24 |
| si:dkey-22a18.2 | 0.28371 | -0.10004196 | 0 | 0.062 | 1 | 24 |
| si:dkey-23a13.1.1 | 0.28371 | -0.1698591 | 0 | 0.062 | 1 | 24 |
| si:dkey-23f9.4 | 0.28371 | -0.21314058 | 0 | 0.062 | 1 | 24 |
| si:dkey-25o16.4 | 0.28371 | -0.17577955 | 0 | 0.062 | 1 | 24 |
| si:dkey-264d12.4 | 0.28371 | -0.10074611 | 0 | 0.062 | 1 | 24 |
| si:dkey-33c9.6 | 0.28371 | -0.10004196 | 0 | 0.062 | 1 | 24 |
| si:dkey-44g23.5 | 0.28371 | -0.14537784 | 0 | 0.062 | 1 | 24 |
| si:dkey-69o16.5 | 0.28371 | -0.1646116 | 0 | 0.062 | 1 | 24 |
| si:dkeyp-68b7.5 | 0.28371 | -0.17826717 | 0 | 0.062 | 1 | 24 |
| si:dkeyp-84f3.5 | 0.28371 | -0.10039796 | 0 | 0.062 | 1 | 24 |
| slc10a7 | 0.28371 | -0.10154798 | 0 | 0.062 | 1 | 24 |
| slc25a1a | 0.28371 | -0.16763576 | 0 | 0.062 | 1 | 24 |
| slc38a5a | 0.28371 | -0.16940522 | 0 | 0.062 | 1 | 24 |
| slc6a7 | 0.28371 | -0.10218744 | 0 | 0.062 | 1 | 24 |
| slc7a8a | 0.28371 | -0.10140796 | 0 | 0.062 | 1 | 24 |
| slx4 | 0.28371 | -0.1698591 | 0 | 0.062 | 1 | 24 |
| smg5 | 0.28371 | -0.10154595 | 0 | 0.062 | 1 | 24 |
| snx2 | 0.28371 | -0.16913319 | 0 | 0.062 | 1 | 24 |
| snx6 | 0.28371 | -0.21276404 | 0 | 0.062 | 1 | 24 |
| spag7 | 0.28371 | -0.21345843 | 0 | 0.062 | 1 | 24 |
| spo11 | 0.28371 | -0.20587967 | 0 | 0.062 | 1 | 24 |
| stk17b | 0.28371 | -0.1643497 | 0 | 0.062 | 1 | 24 |
| stk25b | 0.28371 | -0.21385325 | 0 | 0.062 | 1 | 24 |
| strap | 0.28371 | -0.16080053 | 0 | 0.062 | 1 | 24 |
| stx10 | 0.28371 | -0.16594458 | 0 | 0.062 | 1 | 24 |
| sulf2b | 0.28371 | -0.21453445 | 0 | 0.062 | 1 | 24 |
| sybl1 | 0.28371 | -0.16650231 | 0 | 0.062 | 1 | 24 |
| syngr1a | 0.28371 | -0.20697614 | 0 | 0.062 | 1 | 24 |
| taf1 | 0.28371 | -0.2108599 | 0 | 0.062 | 1 | 24 |
| taf1a | 0.28371 | -0.1644702 | 0 | 0.062 | 1 | 24 |
| tbc1d16 | 0.28371 | -0.10103612 | 0 | 0.062 | 1 | 24 |
| tbl3 | 0.28371 | -0.17585193 | 0 | 0.062 | 1 | 24 |

| | | | | | | |
|---|---|---|---|---|---|---|
| tespa1 | 0.28371 | -0.10197799 | 0 | 0.062 | 1 | 24 |
| themis | 0.28371 | -0.10332134 | 0 | 0.062 | 1 | 24 |
| timm9 | 0.28371 | -0.23846628 | 0 | 0.062 | 1 | 24 |
| tmem42a | 0.28371 | -0.1546895 | 0 | 0.062 | 1 | 24 |
| tmpoa | 0.28371 | -0.16582509 | 0 | 0.062 | 1 | 24 |
| tnpo1 | 0.28371 | -0.10004196 | 0 | 0.062 | 1 | 24 |
| toe1 | 0.28371 | -0.16225476 | 0 | 0.062 | 1 | 24 |
| tonsl | 0.28371 | -0.10195119 | 0 | 0.062 | 1 | 24 |
| tpst1 | 0.28371 | -0.14195736 | 0 | 0.062 | 1 | 24 |
| tradd | 0.28371 | -0.10052376 | 0 | 0.062 | 1 | 24 |
| traf2a | 0.28371 | -0.16077266 | 0 | 0.062 | 1 | 24 |
| traip | 0.28371 | -0.20795658 | 0 | 0.062 | 1 | 24 |
| trappc6b | 0.28371 | -0.1733101 | 0 | 0.062 | 1 | 24 |
| trip4 | 0.28371 | -0.10173511 | 0 | 0.062 | 1 | 24 |
| trove2 | 0.28371 | -0.16583673 | 0 | 0.062 | 1 | 24 |
| tshz1 | 0.28371 | -0.16608084 | 0 | 0.062 | 1 | 24 |
| ttyh3a | 0.28371 | -0.21357392 | 0 | 0.062 | 1 | 24 |
| twnk | 0.28371 | -0.10092373 | 0 | 0.062 | 1 | 24 |
| txn | 0.28371 | -0.25924588 | 0 | 0.062 | 1 | 24 |
| txnl4b | 0.28371 | -0.10126111 | 0 | 0.062 | 1 | 24 |
| ubiad1 | 0.28371 | -0.21100296 | 0 | 0.062 | 1 | 24 |
| ugp2a | 0.28371 | -0.17381683 | 0 | 0.062 | 1 | 24 |
| unk | 0.28371 | -0.1678361 | 0 | 0.062 | 1 | 24 |
| uqcrb | 0.28371 | -0.21187923 | 0 | 0.062 | 1 | 24 |
| utp20 | 0.28371 | -0.16917207 | 0 | 0.062 | 1 | 24 |
| utp25 | 0.28371 | -0.19845257 | 0 | 0.062 | 1 | 24 |
| uxs1 | 0.28371 | -0.17477934 | 0 | 0.062 | 1 | 24 |
| vamp5 | 0.28371 | -0.16871701 | 0 | 0.062 | 1 | 24 |
| vegfc | 0.28371 | -0.1698591 | 0 | 0.062 | 1 | 24 |
| vps11 | 0.28371 | -0.10078626 | 0 | 0.062 | 1 | 24 |
| vps13a | 0.28371 | -0.10215865 | 0 | 0.062 | 1 | 24 |
| vps35l | 0.28371 | -0.13690613 | 0 | 0.062 | 1 | 24 |
| wacb | 0.28371 | -0.21090087 | 0 | 0.062 | 1 | 24 |
| waplb | 0.28371 | -0.15829591 | 0 | 0.062 | 1 | 24 |
| wbp1la | 0.28371 | -0.10005547 | 0 | 0.062 | 1 | 24 |
| xiap | 0.28371 | -0.17475345 | 0 | 0.062 | 1 | 24 |
| yipf5 | 0.28371 | -0.20519841 | 0 | 0.062 | 1 | 24 |
| zbtb3 | 0.28371 | -0.16736123 | 0 | 0.062 | 1 | 24 |
| zdhhc21 | 0.28371 | -0.20941285 | 0 | 0.062 | 1 | 24 |
| zfp36l1b | 0.28371 | -0.21266181 | 0 | 0.062 | 1 | 24 |
| zgc:109744 | 0.28371 | -0.21038916 | 0 | 0.062 | 1 | 24 |
| zgc:110269 | 0.28371 | -0.2116534 | 0 | 0.062 | 1 | 24 |
| zgc:110591 | 0.28371 | -0.16795708 | 0 | 0.062 | 1 | 24 |
| zgc:113149 | 0.28371 | -0.10206373 | 0 | 0.062 | 1 | 24 |
| zgc:113518 | 0.28371 | -0.21414524 | 0 | 0.062 | 1 | 24 |
| zgc:114045 | 0.28371 | -0.20362168 | 0 | 0.062 | 1 | 24 |
| zgc:114130 | 0.28371 | -0.16034385 | 0 | 0.062 | 1 | 24 |
| zgc:153012 | 0.28371 | -0.16909447 | 0 | 0.062 | 1 | 24 |

| | | | | | | |
|---|---|---|---|---|---|---|
| zgc:165555.3 | 0.28371 | -0.21127099 | 0 | 0.062 | 1 | 24 |
| zgc:172121 | 0.28371 | -0.14151154 | 0 | 0.062 | 1 | 24 |
| zgc:66474 | 0.28371 | -0.16182086 | 0 | 0.062 | 1 | 24 |
| zgc:86764 | 0.28371 | -0.17735959 | 0 | 0.062 | 1 | 24 |
| zgc:91890 | 0.28371 | -0.15070465 | 0 | 0.062 | 1 | 24 |
| zmp:0000000652 | 0.28371 | -0.21059314 | 0 | 0.062 | 1 | 24 |
| znf330 | 0.28371 | -0.14198531 | 0 | 0.062 | 1 | 24 |
| znf346 | 0.28371 | -0.16236892 | 0 | 0.062 | 1 | 24 |
| znf513a | 0.28371 | -0.16107524 | 0 | 0.062 | 1 | 24 |
| cox8a | 0.28567 | -0.25300792 | 0.381 | 0.625 | 1 | 24 |
| nfyba | 0.28715 | 0.327766429 | 0.333 | 0.125 | 1 | 24 |
| apex1 | 0.28898 | 0.522512594 | 0.238 | 0.062 | 1 | 24 |
| si:dkey-180p18.9 | 0.29028 | 0.537988729 | 0.238 | 0.062 | 1 | 24 |
| cwc25 | 0.29056 | 0.547180956 | 0.238 | 0.062 | 1 | 24 |
| rps27.1 | 0.29479 | 0.255147738 | 0.952 | 0.875 | 1 | 24 |
| si:dkey-188i13.10 | 0.2979 | -0.55671441 | 0.048 | 0.188 | 1 | 24 |
| rps15 | 0.30103 | -0.2056502 | 0.905 | 1 | 1 | 24 |
| ubc | 0.30633 | -0.48775338 | 0.381 | 0.625 | 1 | 24 |
| emd | 0.31152 | 0.521459325 | 0.238 | 0.062 | 1 | 24 |
| psmd10 | 0.31215 | 0.458118062 | 0.238 | 0.062 | 1 | 24 |
| sub1a | 0.31391 | -0.46896217 | 0.381 | 0.625 | 1 | 24 |
| h3f3a | 0.31522 | -0.23371774 | 0.667 | 0.875 | 1 | 24 |
| mrps11 | 0.3158 | 0.403052072 | 0.238 | 0.062 | 1 | 24 |
| tra2a | 0.32394 | 0.457026891 | 0.238 | 0.062 | 1 | 24 |
| tubb2b | 0.32633 | -0.27992757 | 0.667 | 0.875 | 1 | 24 |
| ap1s1 | 0.32645 | -0.52683109 | 0.048 | 0.188 | 1 | 24 |
| si:dkey-239h2.3 | 0.32805 | 0.43214987 | 0.238 | 0.062 | 1 | 24 |
| swi5 | 0.33075 | 0.476928457 | 0.238 | 0.062 | 1 | 24 |
| cacybp | 0.3308 | 0.331651331 | 0.238 | 0.062 | 1 | 24 |
| hmgb1b | 0.33599 | 0.373157646 | 0.619 | 0.375 | 1 | 24 |
| slc25a5 | 0.339 | -0.18724193 | 0.905 | 1 | 1 | 24 |
| ppp2r1ba | 0.34269 | -0.14729791 | 0.048 | 0.188 | 1 | 24 |
| sec61a1 | 0.34476 | -0.43532605 | 0.048 | 0.188 | 1 | 24 |
| ostc | 0.35441 | -0.43585268 | 0.048 | 0.188 | 1 | 24 |
| rpl39 | 0.35647 | -0.1696018 | 0.952 | 1 | 1 | 24 |
| rpl6 | 0.36137 | -0.26100227 | 0.81 | 0.938 | 1 | 24 |
| paf1 | 0.36582 | -0.41592175 | 0.048 | 0.188 | 1 | 24 |
| clpp | 0.36674 | -0.40949114 | 0.048 | 0.188 | 1 | 24 |
| gins3 | 0.36682 | -0.40460794 | 0.048 | 0.188 | 1 | 24 |
| BX005183.3 | 0.3687 | -0.25954549 | 0.048 | 0.188 | 1 | 24 |
| setb | 0.36968 | -0.51551853 | 0.238 | 0.438 | 1 | 24 |
| rplp2 | 0.3698 | -0.16601371 | 0.905 | 1 | 1 | 24 |
| ccnl1a | 0.37233 | -0.39091675 | 0.048 | 0.188 | 1 | 24 |
| rraga | 0.37281 | -0.4509846 | 0.048 | 0.188 | 1 | 24 |
| fam76b | 0.37426 | -0.49188403 | 0.048 | 0.188 | 1 | 24 |
| rps17 | 0.37433 | 0.107444018 | 0.762 | 0.812 | 1 | 24 |
| mrps21 | 0.37463 | -0.38495361 | 0.048 | 0.188 | 1 | 24 |
| tspo | 0.37468 | -0.46178611 | 0.048 | 0.188 | 1 | 24 |

| | | | | | | |
|---|---|---|---|---|---|---|
| eif3k | 0.37549 | 0.506636888 | 0.381 | 0.188 | 1 | 24 |
| tcf12 | 0.37611 | -0.23114228 | 0.048 | 0.188 | 1 | 24 |
| ywhabl | 0.37827 | -0.36862247 | 0.333 | 0.562 | 1 | 24 |
| zgc:113184 | 0.37954 | -0.24834564 | 0.048 | 0.188 | 1 | 24 |
| psmc1a | 0.38522 | -0.11726131 | 0.095 | 0.25 | 1 | 24 |
| lxn | 0.38677 | -0.36126268 | 0.048 | 0.188 | 1 | 24 |
| reep3b | 0.38788 | -0.19004067 | 0.048 | 0.188 | 1 | 24 |
| fabp3 | 0.38827 | 0.48967664 | 0.524 | 0.312 | 1 | 24 |
| ccdc12 | 0.38989 | -0.2424779 | 0.048 | 0.188 | 1 | 24 |
| mt-co1 | 0.39018 | 0.54281175 | 0.381 | 0.188 | 1 | 24 |
| trnau1apa | 0.39052 | -0.24188201 | 0.048 | 0.188 | 1 | 24 |
| imp4 | 0.39062 | -0.18288116 | 0.048 | 0.188 | 1 | 24 |
| arhgdia | 0.39399 | 0.461551181 | 0.524 | 0.312 | 1 | 24 |
| rcc2 | 0.39422 | 0.48554909 | 0.286 | 0.125 | 1 | 24 |
| CABZ01072614.1 | 0.39534 | 0.537902064 | 0.19 | 0.062 | 1 | 24 |
| trir | 0.39556 | -0.27210165 | 0.048 | 0.188 | 1 | 24 |
| ppp3r1b | 0.39606 | -0.2546835 | 0.048 | 0.188 | 1 | 24 |
| hspb1 | 0.39671 | -0.29457418 | 0.048 | 0.188 | 1 | 24 |
| serp1 | 0.3972 | -0.37691473 | 0.048 | 0.188 | 1 | 24 |
| ube2ia | 0.3974 | -0.46415653 | 0.143 | 0.312 | 1 | 24 |
| psmd8 | 0.39821 | -0.4088329 | 0.048 | 0.188 | 1 | 24 |
| tia1l | 0.39855 | -0.26610542 | 0.048 | 0.188 | 1 | 24 |
| tpm1 | 0.39905 | -0.27656714 | 0.095 | 0.25 | 1 | 24 |
| tgif1 | 0.39922 | -0.1966947 | 0.048 | 0.188 | 1 | 24 |
| eif4a1b | 0.39957 | -0.30204727 | 0.286 | 0.5 | 1 | 24 |
| armc1l | 0.39985 | -0.30829596 | 0.048 | 0.188 | 1 | 24 |
| supt16h | 0.4001 | -0.21861154 | 0.048 | 0.188 | 1 | 24 |
| wdr18 | 0.40045 | -0.23265167 | 0.048 | 0.188 | 1 | 24 |
| zgc:174268 | 0.40048 | -0.31230955 | 0.048 | 0.188 | 1 | 24 |
| gskip | 0.40074 | -0.29479504 | 0.048 | 0.188 | 1 | 24 |
| cox6b1 | 0.40084 | -0.38630264 | 0.048 | 0.188 | 1 | 24 |
| spdl1 | 0.40135 | -0.35023948 | 0.048 | 0.188 | 1 | 24 |
| uchl3 | 0.40214 | -0.23014388 | 0.048 | 0.188 | 1 | 24 |
| rps26l | 0.4022 | -0.14055763 | 0.905 | 1 | 1 | 24 |
| wdr12 | 0.4023 | -0.23696578 | 0.048 | 0.188 | 1 | 24 |
| asf1bb | 0.40263 | -0.30900132 | 0.048 | 0.188 | 1 | 24 |
| cpne1 | 0.40295 | -0.30423015 | 0.048 | 0.188 | 1 | 24 |
| snrpb2 | 0.40347 | -0.34105089 | 0.048 | 0.188 | 1 | 24 |
| psmb10 | 0.40363 | -0.25841233 | 0.048 | 0.188 | 1 | 24 |
| uba1 | 0.40371 | -0.25419035 | 0.048 | 0.188 | 1 | 24 |
| atp6v0b | 0.40384 | -0.26649586 | 0.048 | 0.125 | 1 | 24 |
| ckbb | 0.40384 | -0.31293056 | 0.048 | 0.125 | 1 | 24 |
| col1a1a | 0.40384 | -0.14563887 | 0.048 | 0.125 | 1 | 24 |
| commd8 | 0.40384 | -0.20360147 | 0.048 | 0.125 | 1 | 24 |
| faub | 0.40384 | -0.21554096 | 0.048 | 0.125 | 1 | 24 |
| kif15 | 0.40384 | -0.2033513 | 0.048 | 0.125 | 1 | 24 |
| lta4h | 0.40384 | -0.21294826 | 0.048 | 0.125 | 1 | 24 |
| mrpl43 | 0.40384 | -0.10525682 | 0.048 | 0.125 | 1 | 24 |

| | | | | | | |
|---|---|---|---|---|---|---|
| pef1 | 0.40384 | -0.26107956 | 0.048 | 0.125 | 1 | 24 |
| si:dkey-12j5.1 | 0.40384 | -0.17217583 | 0.048 | 0.125 | 1 | 24 |
| CABZ01087560.1 | 0.40384 | -0.20601353 | 0.048 | 0.125 | 1 | 24 |
| LOC103909972 | 0.40384 | -0.12751366 | 0.048 | 0.125 | 1 | 24 |
| aip | 0.40384 | -0.15752895 | 0.048 | 0.125 | 1 | 24 |
| ap2a1 | 0.40384 | -0.2030082 | 0.048 | 0.125 | 1 | 24 |
| bckdhb | 0.40384 | -0.215339 | 0.048 | 0.125 | 1 | 24 |
| cenpo | 0.40384 | -0.17462403 | 0.048 | 0.125 | 1 | 24 |
| dcp2 | 0.40384 | -0.10217773 | 0.048 | 0.125 | 1 | 24 |
| emg1 | 0.40384 | -0.12896371 | 0.048 | 0.125 | 1 | 24 |
| flot2a | 0.40384 | -0.31456736 | 0.048 | 0.125 | 1 | 24 |
| golgb1 | 0.40384 | -0.10315058 | 0.048 | 0.125 | 1 | 24 |
| incenp | 0.40384 | -0.28676978 | 0.048 | 0.125 | 1 | 24 |
| khdrbs1b | 0.40384 | -0.16104107 | 0.048 | 0.125 | 1 | 24 |
| klf6a | 0.40384 | -0.16276948 | 0.048 | 0.125 | 1 | 24 |
| llph | 0.40384 | -0.17387646 | 0.048 | 0.125 | 1 | 24 |
| lsm14aa | 0.40384 | -0.2034205 | 0.048 | 0.125 | 1 | 24 |
| pcmt | 0.40384 | -0.16002394 | 0.048 | 0.125 | 1 | 24 |
| pelo | 0.40384 | -0.2632462 | 0.048 | 0.125 | 1 | 24 |
| prpf38b | 0.40384 | -0.15411005 | 0.048 | 0.125 | 1 | 24 |
| sccpdhb | 0.40384 | -0.10145884 | 0.048 | 0.125 | 1 | 24 |
| sgta | 0.40384 | -0.20999706 | 0.048 | 0.125 | 1 | 24 |
| si:ch211-150o23.3 | 0.40384 | -0.28057795 | 0.048 | 0.125 | 1 | 24 |
| srrm2 | 0.40384 | -0.13702384 | 0.048 | 0.125 | 1 | 24 |
| stip1 | 0.40384 | -0.17913536 | 0.048 | 0.125 | 1 | 24 |
| stk35 | 0.40384 | -0.22017017 | 0.048 | 0.125 | 1 | 24 |
| supt4h1 | 0.40384 | -0.14433677 | 0.048 | 0.125 | 1 | 24 |
| zgc:92664 | 0.40384 | -0.14329314 | 0.048 | 0.125 | 1 | 24 |
| XLOC-041870 | 0.40384 | -0.17191832 | 0.048 | 0.125 | 1 | 24 |
| acaa1 | 0.40384 | -0.13401362 | 0.048 | 0.125 | 1 | 24 |
| actr1 | 0.40384 | -0.10303783 | 0.048 | 0.125 | 1 | 24 |
| atf1 | 0.40384 | -0.26102894 | 0.048 | 0.125 | 1 | 24 |
| aurkb | 0.40384 | -0.14694444 | 0.048 | 0.125 | 1 | 24 |
| blmh | 0.40384 | -0.2143959 | 0.048 | 0.125 | 1 | 24 |
| bod1 | 0.40384 | -0.16729786 | 0.048 | 0.125 | 1 | 24 |
| bola1 | 0.40384 | -0.18746898 | 0.048 | 0.125 | 1 | 24 |
| calm3a | 0.40384 | -0.28077792 | 0.048 | 0.125 | 1 | 24 |
| caprin1a | 0.40384 | -0.29027124 | 0.048 | 0.125 | 1 | 24 |
| ccdc58 | 0.40384 | -0.14045249 | 0.048 | 0.125 | 1 | 24 |
| cetn3 | 0.40384 | -0.16208711 | 0.048 | 0.125 | 1 | 24 |
| cnot3b | 0.40384 | -0.11171182 | 0.048 | 0.125 | 1 | 24 |
| coq7 | 0.40384 | -0.19834485 | 0.048 | 0.125 | 1 | 24 |
| csnk1g2b | 0.40384 | -0.14366937 | 0.048 | 0.125 | 1 | 24 |
| dpm1 | 0.40384 | -0.22268038 | 0.048 | 0.125 | 1 | 24 |
| elmo1 | 0.40384 | -0.10763699 | 0.048 | 0.125 | 1 | 24 |
| fam210aa | 0.40384 | -0.20572763 | 0.048 | 0.125 | 1 | 24 |
| fkbp5 | 0.40384 | -0.15463409 | 0.048 | 0.125 | 1 | 24 |
| gsdf | 0.40384 | -0.34854063 | 0.048 | 0.125 | 1 | 24 |

| | | | | | | |
|---|---|---|---|---|---|---|
| ing4 | 0.40384 | -0.23582972 | 0.048 | 0.125 | 1 | 24 |
| kat7b | 0.40384 | -0.13596591 | 0.048 | 0.125 | 1 | 24 |
| lgals9l3 | 0.40384 | -0.18422487 | 0.048 | 0.125 | 1 | 24 |
| mbnl1 | 0.40384 | -0.3994781 | 0.048 | 0.125 | 1 | 24 |
| mgst3b | 0.40384 | -0.27740715 | 0.048 | 0.125 | 1 | 24 |
| mlf2 | 0.40384 | -0.30164812 | 0.048 | 0.125 | 1 | 24 |
| myb | 0.40384 | -0.15313865 | 0.048 | 0.125 | 1 | 24 |
| myh10 | 0.40384 | -0.16900186 | 0.048 | 0.125 | 1 | 24 |
| ndc80 | 0.40384 | -0.22715282 | 0.048 | 0.125 | 1 | 24 |
| ndufaf2 | 0.40384 | -0.20359163 | 0.048 | 0.125 | 1 | 24 |
| ndufv3 | 0.40384 | -0.18778836 | 0.048 | 0.125 | 1 | 24 |
| npepps | 0.40384 | -0.11906815 | 0.048 | 0.125 | 1 | 24 |
| nudcd3 | 0.40384 | -0.22592527 | 0.048 | 0.125 | 1 | 24 |
| osr2 | 0.40384 | -0.17176916 | 0.048 | 0.125 | 1 | 24 |
| pex3 | 0.40384 | -0.20710668 | 0.048 | 0.125 | 1 | 24 |
| pmm2 | 0.40384 | -0.17723303 | 0.048 | 0.125 | 1 | 24 |
| ppp1r14bb | 0.40384 | -0.1262023 | 0.048 | 0.125 | 1 | 24 |
| psmd11a | 0.40384 | -0.15061243 | 0.048 | 0.125 | 1 | 24 |
| psph | 0.40384 | -0.13161181 | 0.048 | 0.125 | 1 | 24 |
| ptenb | 0.40384 | -0.16436703 | 0.048 | 0.125 | 1 | 24 |
| pum1 | 0.40384 | -0.12683769 | 0.048 | 0.125 | 1 | 24 |
| rras | 0.40384 | -0.27624358 | 0.048 | 0.125 | 1 | 24 |
| sec24a | 0.40384 | -0.17163085 | 0.048 | 0.125 | 1 | 24 |
| sephs1 | 0.40384 | -0.25757946 | 0.048 | 0.125 | 1 | 24 |
| sf3b3 | 0.40384 | -0.11411903 | 0.048 | 0.125 | 1 | 24 |
| si:ch1073-443f11.2 | 0.40384 | -0.16031124 | 0.048 | 0.125 | 1 | 24 |
| si:ch211-105c13.3 | 0.40384 | -0.17606619 | 0.048 | 0.125 | 1 | 24 |
| si:ch211-266i6.3 | 0.40384 | -0.12257563 | 0.048 | 0.125 | 1 | 24 |
| si:dkey-222b8.1 | 0.40384 | -0.11303569 | 0.048 | 0.125 | 1 | 24 |
| si:dkey-6n21.12 | 0.40384 | -0.19760116 | 0.048 | 0.125 | 1 | 24 |
| sirt7 | 0.40384 | -0.29350425 | 0.048 | 0.125 | 1 | 24 |
| slc30a7 | 0.40384 | -0.30580159 | 0.048 | 0.125 | 1 | 24 |
| spi1b | 0.40384 | -0.23434428 | 0.048 | 0.125 | 1 | 24 |
| spryd7a | 0.40384 | -0.15931483 | 0.048 | 0.125 | 1 | 24 |
| stx12 | 0.40384 | -0.24017793 | 0.048 | 0.125 | 1 | 24 |
| tmem176 | 0.40384 | -0.17732825 | 0.048 | 0.125 | 1 | 24 |
| trappc2l | 0.40384 | -0.19353191 | 0.048 | 0.125 | 1 | 24 |
| txndc17 | 0.40384 | -0.27094907 | 0.048 | 0.125 | 1 | 24 |
| uchl5 | 0.40384 | -0.14962669 | 0.048 | 0.125 | 1 | 24 |
| wipf1b | 0.40384 | -0.16359499 | 0.048 | 0.125 | 1 | 24 |
| wu:fb18f06 | 0.40384 | -0.24791923 | 0.048 | 0.125 | 1 | 24 |
| zbtb18 | 0.40384 | -0.20874927 | 0.048 | 0.125 | 1 | 24 |
| zgc:123258 | 0.40384 | -0.29690564 | 0.048 | 0.125 | 1 | 24 |
| zgc:153409 | 0.40384 | -0.15068654 | 0.048 | 0.125 | 1 | 24 |
| XLOC-013964 | 0.40384 | -0.18938115 | 0.048 | 0.125 | 1 | 24 |
| XLOC-025423 | 0.40384 | -0.31563044 | 0.048 | 0.125 | 1 | 24 |
| XLOC-042595 | 0.40384 | -0.1394488 | 0.048 | 0.125 | 1 | 24 |
| XLOC-043861 | 0.40384 | -0.17903457 | 0.048 | 0.125 | 1 | 24 |

| | | | | | | |
|---|---|---|---|---|---|---|
| aamp | 0.40384 | -0.19839003 | 0.048 | 0.125 | 1 | 24 |
| actl6a | 0.40384 | -0.25823269 | 0.048 | 0.125 | 1 | 24 |
| akap8l | 0.40384 | -0.18981623 | 0.048 | 0.125 | 1 | 24 |
| bms1 | 0.40384 | -0.15961619 | 0.048 | 0.125 | 1 | 24 |
| chmp1a | 0.40384 | -0.26197122 | 0.048 | 0.125 | 1 | 24 |
| ctdp1 | 0.40384 | -0.12714294 | 0.048 | 0.125 | 1 | 24 |
| dcps | 0.40384 | -0.32622365 | 0.048 | 0.125 | 1 | 24 |
| dctd | 0.40384 | -0.18737749 | 0.048 | 0.125 | 1 | 24 |
| dera | 0.40384 | -0.12250268 | 0.048 | 0.125 | 1 | 24 |
| dohh | 0.40384 | -0.12332672 | 0.048 | 0.125 | 1 | 24 |
| dusp14 | 0.40384 | -0.17950252 | 0.048 | 0.125 | 1 | 24 |
| eed | 0.40384 | -0.1108735 | 0.048 | 0.125 | 1 | 24 |
| fam13b | 0.40384 | -0.13080481 | 0.048 | 0.125 | 1 | 24 |
| fhit | 0.40384 | -0.18512428 | 0.048 | 0.125 | 1 | 24 |
| fignl1 | 0.40384 | -0.18331667 | 0.048 | 0.125 | 1 | 24 |
| fip1l1b | 0.40384 | -0.16066737 | 0.048 | 0.125 | 1 | 24 |
| foxj2 | 0.40384 | -0.22249967 | 0.048 | 0.125 | 1 | 24 |
| gnai2a | 0.40384 | -0.10386612 | 0.048 | 0.125 | 1 | 24 |
| gng12a | 0.40384 | -0.1715616 | 0.048 | 0.125 | 1 | 24 |
| gtf2a1 | 0.40384 | -0.16457887 | 0.048 | 0.125 | 1 | 24 |
| hdgfl2 | 0.40384 | -0.2854228 | 0.048 | 0.125 | 1 | 24 |
| jak1 | 0.40384 | -0.24141494 | 0.048 | 0.125 | 1 | 24 |
| jam2a | 0.40384 | -0.31184308 | 0.048 | 0.125 | 1 | 24 |
| lin7c | 0.40384 | -0.19339977 | 0.048 | 0.125 | 1 | 24 |
| map2k6 | 0.40384 | -0.32469132 | 0.048 | 0.125 | 1 | 24 |
| mapre1a | 0.40384 | -0.17094654 | 0.048 | 0.125 | 1 | 24 |
| mmp2 | 0.40384 | -0.24892842 | 0.048 | 0.125 | 1 | 24 |
| mrpl45 | 0.40384 | -0.17866999 | 0.048 | 0.125 | 1 | 24 |
| mtx2 | 0.40384 | -0.13228897 | 0.048 | 0.125 | 1 | 24 |
| mus81 | 0.40384 | -0.20942117 | 0.048 | 0.125 | 1 | 24 |
| paics | 0.40384 | -0.12621496 | 0.048 | 0.125 | 1 | 24 |
| pnp5a | 0.40384 | -0.14212072 | 0.048 | 0.125 | 1 | 24 |
| ppih | 0.40384 | -0.13157221 | 0.048 | 0.125 | 1 | 24 |
| ppt1 | 0.40384 | -0.18681554 | 0.048 | 0.125 | 1 | 24 |
| psmd1 | 0.40384 | -0.19924351 | 0.048 | 0.125 | 1 | 24 |
| pwwp2b | 0.40384 | -0.12624154 | 0.048 | 0.125 | 1 | 24 |
| rimkla | 0.40384 | -0.17614095 | 0.048 | 0.125 | 1 | 24 |
| rmdn1 | 0.40384 | -0.17185496 | 0.048 | 0.125 | 1 | 24 |
| rnf7 | 0.40384 | -0.14981951 | 0.048 | 0.125 | 1 | 24 |
| rrp15 | 0.40384 | -0.12608323 | 0.048 | 0.125 | 1 | 24 |
| sdccag8 | 0.40384 | -0.11838882 | 0.048 | 0.125 | 1 | 24 |
| selenok | 0.40384 | -0.25904081 | 0.048 | 0.125 | 1 | 24 |
| si:ch73-311h14.2 | 0.40384 | -0.25117572 | 0.048 | 0.125 | 1 | 24 |
| si:dkey-78l4.14 | 0.40384 | -0.13802283 | 0.048 | 0.125 | 1 | 24 |
| slc25a12 | 0.40384 | -0.12199847 | 0.048 | 0.125 | 1 | 24 |
| slc25a51b | 0.40384 | -0.23587147 | 0.048 | 0.125 | 1 | 24 |
| spryd4 | 0.40384 | -0.17994746 | 0.048 | 0.125 | 1 | 24 |
| srprb | 0.40384 | -0.23268723 | 0.048 | 0.125 | 1 | 24 |

| | | | | | | |
|---|---|---|---|---|---|---|
| tmem18 | 0.40384 | -0.14059049 | 0.048 | 0.125 | 1 | 24 |
| tp53i11a | 0.40384 | -0.12814922 | 0.048 | 0.125 | 1 | 24 |
| trim35-28 | 0.40384 | -0.23934593 | 0.048 | 0.125 | 1 | 24 |
| ube2a | 0.40384 | -0.13642997 | 0.048 | 0.125 | 1 | 24 |
| wasb | 0.40384 | -0.13433048 | 0.048 | 0.125 | 1 | 24 |
| zgc:152830 | 0.40384 | -0.21879278 | 0.048 | 0.125 | 1 | 24 |
| brd3b | 0.40384 | -0.18031197 | 0.048 | 0.125 | 1 | 24 |
| cenpk | 0.40384 | -0.10528719 | 0.048 | 0.125 | 1 | 24 |
| gosr2 | 0.40384 | -0.19157929 | 0.048 | 0.125 | 1 | 24 |
| rbm4.1 | 0.40384 | -0.10840893 | 0.048 | 0.125 | 1 | 24 |
| scp2b | 0.40384 | -0.15801811 | 0.048 | 0.125 | 1 | 24 |
| ssb | 0.40384 | -0.23747505 | 0.048 | 0.125 | 1 | 24 |
| tfdp1a | 0.40384 | -0.22637832 | 0.048 | 0.125 | 1 | 24 |
| cops2 | 0.40387 | -0.26249055 | 0.048 | 0.188 | 1 | 24 |
| prelid1a | 0.4041 | -0.22999162 | 0.048 | 0.188 | 1 | 24 |
| cdk2ap2 | 0.40423 | -0.24033671 | 0.048 | 0.188 | 1 | 24 |
| suv39h1b | 0.40441 | -0.36886898 | 0.048 | 0.188 | 1 | 24 |
| grpel2 | 0.40448 | -0.27458361 | 0.048 | 0.188 | 1 | 24 |
| zgc:86598 | 0.40454 | -0.27920639 | 0.048 | 0.188 | 1 | 24 |
| inka1a | 0.4046 | -0.36252083 | 0.048 | 0.188 | 1 | 24 |
| ddx27 | 0.40466 | -0.22903871 | 0.048 | 0.188 | 1 | 24 |
| XLOC-000431 | 0.40477 | -0.36818704 | 0.048 | 0.188 | 1 | 24 |
| sdhdb | 0.40484 | -0.2607618 | 0.048 | 0.188 | 1 | 24 |
| pum3 | 0.40487 | -0.25891758 | 0.048 | 0.188 | 1 | 24 |
| cd3eap | 0.40488 | -0.32418289 | 0.048 | 0.188 | 1 | 24 |
| ube2d2l | 0.40489 | -0.24837716 | 0.048 | 0.188 | 1 | 24 |
| wdr61 | 0.40634 | -0.23509125 | 0.095 | 0.25 | 1 | 24 |
| krt8 | 0.41104 | -0.42986933 | 0.286 | 0.5 | 1 | 24 |
| dek | 0.41325 | -0.44950446 | 0.095 | 0.25 | 1 | 24 |
| ucp2 | 0.41398 | -0.43166862 | 0.095 | 0.25 | 1 | 24 |
| h2afy2 | 0.41667 | 0.516282031 | 0.286 | 0.125 | 1 | 24 |
| wdr1 | 0.41711 | 0.500248993 | 0.19 | 0.062 | 1 | 24 |
| coa6 | 0.42192 | -0.38171714 | 0.095 | 0.25 | 1 | 24 |
| tbcb | 0.42599 | -0.20102128 | 0.095 | 0.25 | 1 | 24 |
| arpc3 | 0.42602 | -0.42091786 | 0.238 | 0.438 | 1 | 24 |
| tma7 | 0.42681 | -0.27320758 | 0.476 | 0.688 | 1 | 24 |
| nutf2l | 0.42767 | -0.39305788 | 0.238 | 0.438 | 1 | 24 |
| NC-002333.4 | 0.42845 | 0.419796887 | 0.381 | 0.188 | 1 | 24 |
| si:ch1073-325m22.2 | 0.42881 | 0.304925379 | 0.381 | 0.188 | 1 | 24 |
| eif1ad | 0.43044 | -0.43791743 | 0.095 | 0.25 | 1 | 24 |
| MZT1 | 0.43152 | -0.39445033 | 0.095 | 0.25 | 1 | 24 |
| myl9b | 0.43163 | -0.42547975 | 0.095 | 0.25 | 1 | 24 |
| odc1 | 0.43261 | 0.425412793 | 0.19 | 0.062 | 1 | 24 |
| srsf11 | 0.43356 | -0.39687323 | 0.19 | 0.375 | 1 | 24 |
| rpl8 | 0.4339 | 0.137023955 | 1 | 0.938 | 1 | 24 |
| tank | 0.43436 | -0.37040854 | 0.095 | 0.25 | 1 | 24 |
| tomm7 | 0.43534 | -0.44030876 | 0.143 | 0.312 | 1 | 24 |
| khsrp | 0.43573 | 0.207239876 | 0.286 | 0.125 | 1 | 24 |

| | | | | | | |
|---|---|---|---|---|---|---|
| htatsf1 | 0.43651 | -0.22965804 | 0.095 | 0.25 | 1 | 24 |
| prps1a | 0.43662 | 0.465907086 | 0.286 | 0.125 | 1 | 24 |
| tomm40 | 0.43667 | -0.37078613 | 0.095 | 0.25 | 1 | 24 |
| myg1 | 0.43765 | -0.3733385 | 0.095 | 0.25 | 1 | 24 |
| mcm7 | 0.43926 | -0.24511232 | 0.19 | 0.375 | 1 | 24 |
| lsm5 | 0.43971 | -0.24890546 | 0.19 | 0.375 | 1 | 24 |
| cd63 | 0.44041 | -0.25785815 | 0.095 | 0.25 | 1 | 24 |
| rpa3 | 0.44162 | -0.3670353 | 0.238 | 0.438 | 1 | 24 |
| prpf40a | 0.44436 | -0.37505181 | 0.095 | 0.25 | 1 | 24 |
| med21 | 0.44558 | -0.24252417 | 0.095 | 0.25 | 1 | 24 |
| actb1 | 0.44589 | -0.14748901 | 0.714 | 0.875 | 1 | 24 |
| higd2a | 0.44648 | -0.40751462 | 0.095 | 0.25 | 1 | 24 |
| denr | 0.44648 | -0.32174247 | 0.095 | 0.25 | 1 | 24 |
| eif4a3 | 0.44748 | -0.4074435 | 0.143 | 0.312 | 1 | 24 |
| ndufb9 | 0.44796 | 0.460759992 | 0.286 | 0.125 | 1 | 24 |
| phb | 0.44803 | -0.34470281 | 0.095 | 0.25 | 1 | 24 |
| psmd14 | 0.44897 | -0.39960749 | 0.143 | 0.312 | 1 | 24 |
| rab11ba | 0.44905 | -0.24516352 | 0.095 | 0.25 | 1 | 24 |
| ywhah | 0.44974 | -0.38613939 | 0.143 | 0.312 | 1 | 24 |
| aimp1 | 0.44998 | -0.4475943 | 0.095 | 0.25 | 1 | 24 |
| rpl22l1 | 0.45158 | 0.281477932 | 0.81 | 0.625 | 1 | 24 |
| spcs1 | 0.45173 | -0.38104336 | 0.19 | 0.375 | 1 | 24 |
| syncrip | 0.45205 | -0.30571058 | 0.19 | 0.375 | 1 | 24 |
| pfn2l | 0.45207 | -0.46302551 | 0.143 | 0.312 | 1 | 24 |
| eno1a | 0.45294 | -0.26832025 | 0.095 | 0.25 | 1 | 24 |
| st13 | 0.45306 | -0.39572885 | 0.095 | 0.25 | 1 | 24 |
| pim3 | 0.45316 | 0.412923215 | 0.19 | 0.062 | 1 | 24 |
| lgals3b | 0.45367 | -0.33551213 | 0.095 | 0.25 | 1 | 24 |
| si:dkey-16p21.8 | 0.45371 | -0.33048732 | 0.095 | 0.25 | 1 | 24 |
| drap1 | 0.45433 | -0.34495204 | 0.095 | 0.25 | 1 | 24 |
| mrps18c | 0.45453 | 0.373463698 | 0.19 | 0.062 | 1 | 24 |
| cdkn2aipnl | 0.45476 | -0.33200917 | 0.095 | 0.25 | 1 | 24 |
| rbm39a | 0.45505 | -0.28606521 | 0.095 | 0.25 | 1 | 24 |
| tmsb | 0.45546 | 0.31521118 | 0.286 | 0.125 | 1 | 24 |
| mcts1 | 0.45583 | -0.40722607 | 0.19 | 0.375 | 1 | 24 |
| tmed2 | 0.45731 | -0.33602451 | 0.19 | 0.375 | 1 | 24 |
| gins2 | 0.45967 | -0.41650244 | 0.143 | 0.312 | 1 | 24 |
| txnipa | 0.4607 | -0.32678737 | 0.19 | 0.375 | 1 | 24 |
| cdc42 | 0.46075 | -0.31785644 | 0.19 | 0.375 | 1 | 24 |
| rab1ab | 0.46076 | -0.36641918 | 0.19 | 0.375 | 1 | 24 |
| hnrnpa0l | 0.46078 | 0.242283325 | 0.857 | 0.688 | 1 | 24 |
| rpl14 | 0.4614 | -0.21527933 | 0.81 | 0.938 | 1 | 24 |
| arf5 | 0.4636 | -0.36472324 | 0.143 | 0.312 | 1 | 24 |
| sesn1 | 0.46628 | 0.433579535 | 0.19 | 0.062 | 1 | 24 |
| psme1 | 0.46679 | -0.41108817 | 0.143 | 0.312 | 1 | 24 |
| ppa1b | 0.46735 | 0.275020891 | 0.286 | 0.125 | 1 | 24 |
| gar1 | 0.46745 | -0.34081377 | 0.143 | 0.312 | 1 | 24 |
| mrpl11 | 0.46764 | -0.309246 | 0.143 | 0.312 | 1 | 24 |

| | | | | | | |
|---|---|---|---|---|---|---|
| si:rp71-45k5.4 | 0.46774 | -0.33600423 | 0.143 | 0.312 | 1 | 24 |
| fen1 | 0.46774 | -0.36860379 | 0.143 | 0.312 | 1 | 24 |
| eef1g | 0.46802 | -0.2104552 | 0.81 | 0.938 | 1 | 24 |
| dynll1 | 0.46975 | 0.303231243 | 0.619 | 0.562 | 1 | 24 |
| b2m | 0.47296 | 0.276054353 | 0.19 | 0.062 | 1 | 24 |
| timm23a | 0.47497 | 0.387297892 | 0.19 | 0.062 | 1 | 24 |
| tagln2 | 0.47642 | 0.130168442 | 0.333 | 0.188 | 1 | 24 |
| cct2 | 0.47687 | 0.368717632 | 0.571 | 0.375 | 1 | 24 |
| med11 | 0.47717 | 0.191785535 | 0.19 | 0.062 | 1 | 24 |
| ctbp2a | 0.47925 | 0.285617912 | 0.19 | 0.062 | 1 | 24 |
| nop53 | 0.47948 | 0.329786881 | 0.286 | 0.125 | 1 | 24 |
| taf12 | 0.48028 | 0.388313321 | 0.19 | 0.062 | 1 | 24 |
| uba52 | 0.48036 | -0.23050382 | 0.714 | 0.875 | 1 | 24 |
| slc25a11 | 0.48044 | 0.368239979 | 0.19 | 0.062 | 1 | 24 |
| eif4ebp2 | 0.48242 | 0.390084461 | 0.286 | 0.125 | 1 | 24 |
| sept7a | 0.48458 | 0.377951417 | 0.19 | 0.062 | 1 | 24 |
| gmfb | 0.48725 | 0.337179821 | 0.286 | 0.125 | 1 | 24 |
| pcbp2 | 0.48782 | 0.36513454 | 0.19 | 0.062 | 1 | 24 |
| seta | 0.48809 | -0.33900767 | 0.429 | 0.625 | 1 | 24 |
| mt-nd1 | 0.48841 | 0.356926894 | 0.286 | 0.125 | 1 | 24 |
| tomm6 | 0.48855 | 0.339407656 | 0.286 | 0.125 | 1 | 24 |
| lsm4 | 0.48874 | -0.42398625 | 0.286 | 0.375 | 1 | 24 |
| max | 0.48972 | 0.254810395 | 0.19 | 0.062 | 1 | 24 |
| pdcd5 | 0.4913 | 0.222287962 | 0.19 | 0.062 | 1 | 24 |
| smc4 | 0.49259 | 0.21486484 | 0.19 | 0.062 | 1 | 24 |
| ss18 | 0.49288 | 0.338116003 | 0.19 | 0.062 | 1 | 24 |
| rpa2 | 0.4935 | -0.29300981 | 0.429 | 0.625 | 1 | 24 |
| echs1 | 0.49361 | 0.216859673 | 0.19 | 0.062 | 1 | 24 |
| rps3a | 0.49386 | 0.128648072 | 1 | 0.938 | 1 | 24 |
| timm8a | 0.49603 | 0.251529035 | 0.19 | 0.062 | 1 | 24 |
| ptp4a1 | 0.4963 | 0.452337753 | 0.333 | 0.188 | 1 | 24 |
| ola1 | 0.49681 | 0.451475351 | 0.381 | 0.25 | 1 | 24 |
| nkap | 0.49748 | 0.233278309 | 0.19 | 0.062 | 1 | 24 |
| sf3b4 | 0.49759 | 0.388307175 | 0.19 | 0.062 | 1 | 24 |
| clns1a | 0.49977 | 0.300999555 | 0.19 | 0.062 | 1 | 24 |
| hcfc1a | 0.5002 | 0.325072908 | 0.19 | 0.062 | 1 | 24 |
| slc38a5b | 0.50115 | 0.274397491 | 0.19 | 0.062 | 1 | 24 |
| rps13 | 0.50245 | 0.137471264 | 0.952 | 0.938 | 1 | 24 |
| fam32a | 0.50448 | 0.3956259 | 0.429 | 0.25 | 1 | 24 |
| cks2 | 0.50458 | 0.310552565 | 0.19 | 0.062 | 1 | 24 |
| ncl | 0.50695 | 0.423594838 | 0.476 | 0.312 | 1 | 24 |
| tubb4b | 0.50766 | -0.17049243 | 0.81 | 0.938 | 1 | 24 |
| ctnnb1 | 0.50782 | 0.289355506 | 0.19 | 0.062 | 1 | 24 |
| slc25a55a | 0.50828 | 0.197239531 | 0.19 | 0.062 | 1 | 24 |
| bud31 | 0.50986 | 0.441662964 | 0.333 | 0.188 | 1 | 24 |
| pdrg1 | 0.51083 | 0.272469909 | 0.19 | 0.062 | 1 | 24 |
| cox14 | 0.51125 | 0.259886316 | 0.19 | 0.062 | 1 | 24 |
| adi1 | 0.51141 | 0.249934168 | 0.19 | 0.062 | 1 | 24 |

| | | | | | | |
|---|---|---|---|---|---|---|
| selenbp1 | 0.51161 | 0.27869653 | 0.19 | 0.062 | 1 | 24 |
| add3a | 0.51164 | 0.259164917 | 0.19 | 0.062 | 1 | 24 |
| hadhb | 0.51184 | 0.309114284 | 0.19 | 0.062 | 1 | 24 |
| sdad1 | 0.51201 | 0.287633819 | 0.19 | 0.062 | 1 | 24 |
| phf20b | 0.51208 | 0.276405165 | 0.19 | 0.062 | 1 | 24 |
| aars | 0.51226 | 0.278112471 | 0.19 | 0.062 | 1 | 24 |
| nop10 | 0.51333 | 0.428419688 | 0.333 | 0.25 | 1 | 24 |
| ccng1 | 0.5166 | 0.370662839 | 0.429 | 0.25 | 1 | 24 |
| pabpc1a | 0.51955 | 0.115137373 | 0.714 | 0.562 | 1 | 24 |
| vdac2 | 0.51999 | 0.354536849 | 0.429 | 0.25 | 1 | 24 |
| fkbp1aa | 0.52491 | -0.24372274 | 0.571 | 0.75 | 1 | 24 |
| tomm5 | 0.52512 | 0.340712588 | 0.429 | 0.25 | 1 | 24 |
| sumo3b | 0.52787 | 0.443102127 | 0.381 | 0.25 | 1 | 24 |
| rpl36a | 0.52787 | 0.112732548 | 1 | 1 | 1 | 24 |
| npm3 | 0.53493 | 0.417783422 | 0.238 | 0.188 | 1 | 24 |
| ranbp1 | 0.53518 | -0.31623204 | 0.524 | 0.625 | 1 | 24 |
| erh | 0.55698 | 0.364694368 | 0.476 | 0.375 | 1 | 24 |
| sod1 | 0.57107 | 0.359155054 | 0.476 | 0.375 | 1 | 24 |
| magoh | 0.57847 | 0.385865003 | 0.238 | 0.125 | 1 | 24 |
| si:dkey-102m7.3 | 0.57957 | 0.387912324 | 0.238 | 0.125 | 1 | 24 |
| zgc:153867 | 0.57988 | -0.18710372 | 0.524 | 0.688 | 1 | 24 |
| ywhabb | 0.58359 | -0.23119393 | 0.333 | 0.5 | 1 | 24 |
| ssr4 | 0.58563 | -0.1570292 | 0.286 | 0.438 | 1 | 24 |
| cnot8 | 0.5872 | 0.415890141 | 0.143 | 0.062 | 1 | 24 |
| stmn1a | 0.58733 | -0.22218082 | 0.762 | 0.875 | 1 | 24 |
| ndufs5 | 0.58746 | 0.357808267 | 0.333 | 0.188 | 1 | 24 |
| psma1 | 0.58883 | 0.361053542 | 0.333 | 0.188 | 1 | 24 |
| atp5mc1 | 0.58942 | -0.24927443 | 0.476 | 0.5 | 1 | 24 |
| rps10 | 0.58955 | -0.12356298 | 0.952 | 1 | 1 | 24 |
| pfn1 | 0.58973 | -0.29026924 | 0.333 | 0.5 | 1 | 24 |
| mrpl54 | 0.5947 | 0.387510567 | 0.238 | 0.125 | 1 | 24 |
| mycbp | 0.59612 | 0.369548764 | 0.238 | 0.125 | 1 | 24 |
| prdx5 | 0.59675 | 0.295360069 | 0.476 | 0.312 | 1 | 24 |
| uqcrh | 0.59954 | 0.35013947 | 0.333 | 0.188 | 1 | 24 |
| psmb1 | 0.60105 | -0.39195769 | 0.19 | 0.312 | 1 | 24 |
| mif | 0.60139 | 0.376674313 | 0.381 | 0.25 | 1 | 24 |
| fthl27 | 0.60165 | -0.23412307 | 0.524 | 0.688 | 1 | 24 |
| uqcrq | 0.60246 | 0.274999635 | 0.333 | 0.188 | 1 | 24 |
| psmb3 | 0.60405 | 0.337541148 | 0.333 | 0.188 | 1 | 24 |
| rpl36 | 0.60487 | -0.10581217 | 0.952 | 1 | 1 | 24 |
| sub1b | 0.60544 | -0.3612794 | 0.333 | 0.438 | 1 | 24 |
| ost4 | 0.60737 | -0.14545964 | 0.238 | 0.375 | 1 | 24 |
| si:dkey-51e6.1 | 0.60785 | -0.39844602 | 0.095 | 0.188 | 1 | 24 |
| prdx2 | 0.61334 | -0.10060535 | 0.619 | 0.75 | 1 | 24 |
| sdc4 | 0.61889 | 0.377967019 | 0.143 | 0.062 | 1 | 24 |
| tpm3 | 0.6238 | -0.31400603 | 0.286 | 0.438 | 1 | 24 |
| mcm3 | 0.62687 | 0.191164816 | 0.143 | 0.188 | 1 | 24 |
| tp53 | 0.63046 | -0.34770877 | 0.19 | 0.312 | 1 | 24 |

| | | | | | | |
|---|---|---|---|---|---|---|
| gapdhs | 0.63279 | -0.26882668 | 0.286 | 0.438 | 1 | 24 |
| lasp1 | 0.63411 | -0.23200005 | 0.286 | 0.438 | 1 | 24 |
| rap1aa | 0.63456 | 0.353659397 | 0.238 | 0.188 | 1 | 24 |
| abce1 | 0.63639 | -0.35550907 | 0.095 | 0.188 | 1 | 24 |
| ndufb5 | 0.63715 | -0.36515472 | 0.143 | 0.25 | 1 | 24 |
| mrps33 | 0.63715 | -0.36248725 | 0.143 | 0.25 | 1 | 24 |
| myl6 | 0.63798 | -0.17014527 | 0.19 | 0.312 | 1 | 24 |
| elavl1a | 0.63855 | -0.34927411 | 0.19 | 0.312 | 1 | 24 |
| rhoaa | 0.63956 | -0.16266255 | 0.19 | 0.312 | 1 | 24 |
| tia1 | 0.63994 | -0.34607509 | 0.095 | 0.188 | 1 | 24 |
| ccni | 0.64105 | -0.31628517 | 0.19 | 0.312 | 1 | 24 |
| cap1 | 0.64186 | 0.289617236 | 0.238 | 0.125 | 1 | 24 |
| gbp | 0.64416 | -0.10459664 | 0.143 | 0.25 | 1 | 24 |
| srp19 | 0.64472 | -0.38183997 | 0.095 | 0.188 | 1 | 24 |
| si:dkey-28b4.7 | 0.64482 | 0.162173446 | 0.238 | 0.125 | 1 | 24 |
| snrpe | 0.6466 | 0.295973958 | 0.524 | 0.375 | 1 | 24 |
| slirp | 0.64669 | 0.322709833 | 0.238 | 0.125 | 1 | 24 |
| rfc4 | 0.64912 | -0.18016245 | 0.238 | 0.375 | 1 | 24 |
| atf4b | 0.6493 | -0.36876035 | 0.19 | 0.312 | 1 | 24 |
| cct7 | 0.65037 | 0.329086323 | 0.381 | 0.312 | 1 | 24 |
| drg2 | 0.65057 | 0.337584605 | 0.238 | 0.125 | 1 | 24 |
| cdv3 | 0.65259 | -0.11730899 | 0.19 | 0.312 | 1 | 24 |
| gpatch4 | 0.65457 | 0.339359137 | 0.143 | 0.062 | 1 | 24 |
| brk1 | 0.65486 | -0.33358968 | 0.095 | 0.188 | 1 | 24 |
| si:dkey-42i9.4 | 0.65634 | 0.288846135 | 0.238 | 0.125 | 1 | 24 |
| csnk2b | 0.65675 | -0.3013964 | 0.238 | 0.375 | 1 | 24 |
| pin1 | 0.65862 | 0.185485066 | 0.238 | 0.125 | 1 | 24 |
| nudc | 0.66026 | -0.1128085 | 0.143 | 0.25 | 1 | 24 |
| dnaja2a | 0.66131 | 0.199329539 | 0.238 | 0.125 | 1 | 24 |
| mad2l1 | 0.66255 | -0.29286574 | 0.238 | 0.375 | 1 | 24 |
| ctsba | 0.66453 | 0.215864621 | 0.238 | 0.125 | 1 | 24 |
| ifi46 | 0.665 | 0.299460533 | 0.238 | 0.125 | 1 | 24 |
| ywhae1 | 0.66565 | -0.24163163 | 0.238 | 0.375 | 1 | 24 |
| krt18a.1 | 0.6659 | -0.23545664 | 0.19 | 0.312 | 1 | 24 |
| dut | 0.66621 | -0.22554477 | 0.476 | 0.625 | 1 | 24 |
| ppdpfb | 0.66658 | -0.21454333 | 0.476 | 0.625 | 1 | 24 |
| psmb4 | 0.66728 | -0.24066115 | 0.238 | 0.375 | 1 | 24 |
| nipsnap2 | 0.66786 | 0.334058269 | 0.143 | 0.062 | 1 | 24 |
| LOC101885938 | 0.66833 | -0.32058003 | 0.095 | 0.188 | 1 | 24 |
| cebpd | 0.66846 | -0.25538139 | 0.238 | 0.375 | 1 | 24 |
| ube2na | 0.66884 | -0.17298134 | 0.19 | 0.312 | 1 | 24 |
| si:dkey-7j14.6 | 0.66908 | -0.34551176 | 0.095 | 0.188 | 1 | 24 |
| selenof | 0.67003 | 0.335098664 | 0.333 | 0.25 | 1 | 24 |
| taf13 | 0.67064 | 0.263746762 | 0.238 | 0.125 | 1 | 24 |
| dpy30 | 0.67067 | 0.292803868 | 0.238 | 0.125 | 1 | 24 |
| si:ch73-281n10.2 | 0.67174 | 0.189996987 | 0.857 | 0.75 | 1 | 24 |
| rplp0 | 0.67197 | 0.119201098 | 0.952 | 0.938 | 1 | 24 |
| ndufa7 | 0.67208 | 0.276831095 | 0.238 | 0.125 | 1 | 24 |

| | | | | | | |
|---|---|---|---|---|---|---|
| gid8a | 0.67463 | -0.10692571 | 0.143 | 0.25 | 1 | 24 |
| vma21 | 0.67489 | 0.232592853 | 0.238 | 0.125 | 1 | 24 |
| h2afx | 0.67521 | -0.35387029 | 0.095 | 0.188 | 1 | 24 |
| hmgb3a | 0.67539 | 0.230116336 | 0.238 | 0.125 | 1 | 24 |
| fam50a | 0.6757 | -0.11029475 | 0.095 | 0.188 | 1 | 24 |
| eif2s1a | 0.6758 | 0.308315996 | 0.143 | 0.062 | 1 | 24 |
| snrpa | 0.67587 | -0.306697 | 0.095 | 0.188 | 1 | 24 |
| hells | 0.67621 | 0.234456719 | 0.238 | 0.125 | 1 | 24 |
| ndfip1 | 0.6765 | -0.28975136 | 0.095 | 0.188 | 1 | 24 |
| nfkbiab | 0.67669 | 0.184558734 | 0.238 | 0.125 | 1 | 24 |
| mcl1b | 0.67712 | 0.218322798 | 0.238 | 0.125 | 1 | 24 |
| uqcrfs1 | 0.6775 | 0.34703086 | 0.286 | 0.188 | 1 | 24 |
| hmgn3 | 0.67776 | 0.285648312 | 0.238 | 0.125 | 1 | 24 |
| tmpob | 0.67777 | -0.29228477 | 0.095 | 0.188 | 1 | 24 |
| cct8 | 0.67814 | -0.29371992 | 0.19 | 0.312 | 1 | 24 |
| arf6b | 0.67815 | 0.232031597 | 0.238 | 0.125 | 1 | 24 |
| entr1 | 0.67827 | 0.304991689 | 0.143 | 0.062 | 1 | 24 |
| hars | 0.67877 | 0.241217924 | 0.238 | 0.125 | 1 | 24 |
| id1 | 0.67879 | 0.309917429 | 0.238 | 0.125 | 1 | 24 |
| rnf185 | 0.67879 | 0.215920905 | 0.238 | 0.125 | 1 | 24 |
| dynlrb1 | 0.6788 | 0.293862707 | 0.381 | 0.25 | 1 | 24 |
| rpf1 | 0.67913 | 0.294871917 | 0.143 | 0.062 | 1 | 24 |
| psma6b | 0.67928 | -0.29145434 | 0.143 | 0.25 | 1 | 24 |
| hp1bp3 | 0.6798 | -0.2975798 | 0.19 | 0.312 | 1 | 24 |
| snrpf | 0.68003 | 0.272794406 | 0.619 | 0.5 | 1 | 24 |
| gpx4b | 0.68022 | 0.187060584 | 0.381 | 0.25 | 1 | 24 |
| mapre1b | 0.6804 | -0.12567394 | 0.143 | 0.25 | 1 | 24 |
| skp1 | 0.682 | 0.20495284 | 0.381 | 0.25 | 1 | 24 |
| rrm1 | 0.68275 | -0.29936523 | 0.143 | 0.25 | 1 | 24 |
| pycard | 0.68333 | -0.32321663 | 0.095 | 0.188 | 1 | 24 |
| lsm12b | 0.68411 | -0.1880386 | 0.19 | 0.312 | 1 | 24 |
| tmed7 | 0.68485 | -0.25283292 | 0.19 | 0.312 | 1 | 24 |
| twf1a | 0.68521 | -0.31565147 | 0.143 | 0.25 | 1 | 24 |
| ppp2cb | 0.68572 | -0.16748622 | 0.19 | 0.312 | 1 | 24 |
| si:ch1073-429i10.3.1 | 0.68629 | 0.210858655 | 0.381 | 0.25 | 1 | 24 |
| ftsj1 | 0.68635 | 0.289201587 | 0.143 | 0.062 | 1 | 24 |
| pole4 | 0.68661 | -0.13311628 | 0.143 | 0.25 | 1 | 24 |
| pfdn6 | 0.68776 | -0.20860348 | 0.19 | 0.312 | 1 | 24 |
| ddx39aa | 0.68867 | -0.20378066 | 0.19 | 0.312 | 1 | 24 |
| mrpl47 | 0.68899 | 0.288189138 | 0.143 | 0.062 | 1 | 24 |
| mcm5 | 0.69029 | -0.12284787 | 0.095 | 0.188 | 1 | 24 |
| slc25a3b | 0.69066 | -0.27020065 | 0.429 | 0.562 | 1 | 24 |
| chtf8 | 0.6912 | -0.27742299 | 0.143 | 0.25 | 1 | 24 |
| arl14ep | 0.69155 | -0.19122433 | 0.19 | 0.312 | 1 | 24 |
| CABZ01045617.1 | 0.69184 | 0.322065759 | 0.143 | 0.062 | 1 | 24 |
| calm1a | 0.69252 | -0.11033963 | 0.238 | 0.188 | 1 | 24 |
| med28 | 0.69311 | 0.280957954 | 0.143 | 0.062 | 1 | 24 |
| h2afva | 0.69319 | -0.2656134 | 0.19 | 0.312 | 1 | 24 |

| | | | | | | |
|---|---|---|---|---|---|---|
| acot13 | 0.69333 | 0.300322417 | 0.143 | 0.062 | 1 | 24 |
| nup50 | 0.69337 | -0.21436867 | 0.19 | 0.312 | 1 | 24 |
| nucks1b | 0.69355 | -0.19172974 | 0.143 | 0.25 | 1 | 24 |
| psmd9 | 0.69363 | 0.288473014 | 0.143 | 0.062 | 1 | 24 |
| mrpl53 | 0.69385 | 0.287459352 | 0.143 | 0.062 | 1 | 24 |
| ywhaqb | 0.69532 | -0.22207686 | 0.19 | 0.312 | 1 | 24 |
| csnk1a1 | 0.69538 | -0.27706916 | 0.143 | 0.25 | 1 | 24 |
| hmgn6 | 0.69542 | 0.146846555 | 0.81 | 0.688 | 1 | 24 |
| jun | 0.69673 | 0.276069768 | 0.381 | 0.25 | 1 | 24 |
| esd | 0.69772 | 0.25503985 | 0.143 | 0.062 | 1 | 24 |
| atp5if1a | 0.69777 | 0.243911081 | 0.381 | 0.25 | 1 | 24 |
| polr3h | 0.69823 | -0.10876684 | 0.095 | 0.188 | 1 | 24 |
| psenen | 0.69839 | 0.108654602 | 0.143 | 0.062 | 1 | 24 |
| pnisr | 0.6995 | 0.279686592 | 0.143 | 0.062 | 1 | 24 |
| ndufs6 | 0.70021 | -0.10550196 | 0.095 | 0.188 | 1 | 24 |
| wtap | 0.70025 | -0.12644751 | 0.143 | 0.25 | 1 | 24 |
| pkma | 0.70031 | -0.25005307 | 0.095 | 0.188 | 1 | 24 |
| adam10a | 0.70041 | 0.274123308 | 0.143 | 0.062 | 1 | 24 |
| ppm1aa | 0.70058 | 0.265759517 | 0.143 | 0.062 | 1 | 24 |
| commd2 | 0.70146 | 0.297989609 | 0.143 | 0.062 | 1 | 24 |
| cope | 0.70203 | -0.13776462 | 0.143 | 0.25 | 1 | 24 |
| atf4a | 0.70245 | -0.12155972 | 0.095 | 0.188 | 1 | 24 |
| zgc:114188 | 0.70275 | 0.10446827 | 0.952 | 0.875 | 1 | 24 |
| prpf39 | 0.70303 | 0.247445572 | 0.143 | 0.062 | 1 | 24 |
| kpna2 | 0.70325 | 0.172479882 | 0.143 | 0.062 | 1 | 24 |
| cox5ab | 0.70353 | -0.29792649 | 0.143 | 0.25 | 1 | 24 |
| naa40 | 0.70392 | 0.105581008 | 0.143 | 0.062 | 1 | 24 |
| park7 | 0.70409 | 0.267059158 | 0.143 | 0.062 | 1 | 24 |
| mt-nd2 | 0.70446 | -0.21468578 | 0.143 | 0.25 | 1 | 24 |
| tm9sf2 | 0.70459 | 0.261697369 | 0.143 | 0.062 | 1 | 24 |
| kif22 | 0.70514 | -0.2818368 | 0.095 | 0.188 | 1 | 24 |
| nek7 | 0.70515 | 0.293216095 | 0.143 | 0.062 | 1 | 24 |
| pttg1 | 0.70542 | -0.30364166 | 0.095 | 0.188 | 1 | 24 |
| mdka | 0.7066 | -0.32291163 | 0.095 | 0.188 | 1 | 24 |
| cmpk | 0.70735 | -0.10606951 | 0.095 | 0.188 | 1 | 24 |
| bnip3lb | 0.70737 | 0.258469521 | 0.143 | 0.062 | 1 | 24 |
| brd2a | 0.70763 | 0.267328188 | 0.143 | 0.062 | 1 | 24 |
| nup210 | 0.70769 | 0.263602604 | 0.143 | 0.062 | 1 | 24 |
| cenpn | 0.70781 | 0.249457851 | 0.143 | 0.062 | 1 | 24 |
| siva1 | 0.70794 | 0.303256185 | 0.286 | 0.188 | 1 | 24 |
| ubb | 0.70803 | -0.17172596 | 0.143 | 0.25 | 1 | 24 |
| pdia6 | 0.70806 | 0.11727819 | 0.143 | 0.062 | 1 | 24 |
| LOC103909284 | 0.70833 | 0.251985138 | 0.143 | 0.062 | 1 | 24 |
| rp9 | 0.70859 | 0.245494782 | 0.143 | 0.062 | 1 | 24 |
| fn1b | 0.7093 | 0.340588674 | 0.143 | 0.062 | 1 | 24 |
| acp1 | 0.70958 | -0.2667785 | 0.143 | 0.25 | 1 | 24 |
| u2surp | 0.70977 | 0.115299123 | 0.143 | 0.062 | 1 | 24 |
| hnrnpm | 0.70998 | 0.11969406 | 0.143 | 0.062 | 1 | 24 |

| | | | | | | |
|---|---|---|---|---|---|---|
| ogt.1 | 0.71002 | 0.243619972 | 0.143 | 0.062 | 1 | 24 |
| psmc2 | 0.71038 | 0.250127553 | 0.143 | 0.062 | 1 | 24 |
| psma2 | 0.71066 | -0.28355263 | 0.143 | 0.25 | 1 | 24 |
| COX3 | 0.71074 | 0.381317666 | 0.333 | 0.25 | 1 | 24 |
| ddx21 | 0.71105 | -0.11903997 | 0.095 | 0.188 | 1 | 24 |
| rer1 | 0.71118 | -0.13089644 | 0.095 | 0.188 | 1 | 24 |
| celf1 | 0.71124 | 0.280309304 | 0.143 | 0.062 | 1 | 24 |
| hspa9 | 0.71237 | 0.214150024 | 0.143 | 0.062 | 1 | 24 |
| nasp | 0.71279 | -0.24191392 | 0.143 | 0.25 | 1 | 24 |
| bud23 | 0.71281 | 0.238181116 | 0.143 | 0.062 | 1 | 24 |
| si:zfos-1192g2.3 | 0.71377 | 0.239451416 | 0.143 | 0.062 | 1 | 24 |
| ier3ip1 | 0.71398 | 0.161040561 | 0.143 | 0.062 | 1 | 24 |
| arpc1a | 0.71417 | -0.18896009 | 0.095 | 0.188 | 1 | 24 |
| sparc | 0.71472 | -0.30117539 | 0.143 | 0.25 | 1 | 24 |
| vps29 | 0.71486 | -0.21022434 | 0.095 | 0.188 | 1 | 24 |
| zgc:194981 | 0.71556 | 0.127891927 | 0.143 | 0.062 | 1 | 24 |
| eif4e1c | 0.71564 | -0.25706746 | 0.143 | 0.25 | 1 | 24 |
| mt-nd5 | 0.71588 | 0.226116635 | 0.143 | 0.062 | 1 | 24 |
| pabpc1b | 0.71601 | -0.26703432 | 0.095 | 0.188 | 1 | 24 |
| ncaph2 | 0.71639 | -0.21550507 | 0.095 | 0.188 | 1 | 24 |
| sdhb | 0.71641 | -0.20297691 | 0.143 | 0.25 | 1 | 24 |
| csde1 | 0.71654 | -0.19673125 | 0.143 | 0.25 | 1 | 24 |
| stau1 | 0.7166 | -0.24355686 | 0.143 | 0.25 | 1 | 24 |
| litaf | 0.71685 | -0.23755773 | 0.095 | 0.188 | 1 | 24 |
| smc3 | 0.71693 | -0.20488111 | 0.095 | 0.188 | 1 | 24 |
| calm3b | 0.71697 | -0.24213676 | 0.429 | 0.562 | 1 | 24 |
| mrpl27 | 0.71756 | -0.2071655 | 0.095 | 0.188 | 1 | 24 |
| dynlt3 | 0.71758 | 0.26644274 | 0.143 | 0.062 | 1 | 24 |
| nedd8 | 0.71784 | -0.17368918 | 0.095 | 0.188 | 1 | 24 |
| tardbpl | 0.71837 | -0.18364086 | 0.095 | 0.188 | 1 | 24 |
| btf3l4 | 0.71845 | -0.13255925 | 0.095 | 0.188 | 1 | 24 |
| cops9 | 0.7188 | -0.14203072 | 0.095 | 0.188 | 1 | 24 |
| chd3 | 0.71914 | -0.16217314 | 0.095 | 0.188 | 1 | 24 |
| rpl38 | 0.71963 | -0.12001679 | 0.857 | 0.938 | 1 | 24 |
| caprin1b | 0.71971 | -0.31850315 | 0.19 | 0.25 | 1 | 24 |
| abhd16a | 0.71997 | -0.18104139 | 0.095 | 0.188 | 1 | 24 |
| smarca4a | 0.72001 | 0.221912999 | 0.143 | 0.062 | 1 | 24 |
| cstf2 | 0.72044 | 0.12103111 | 0.095 | 0.062 | 1 | 24 |
| erp44 | 0.72044 | 0.115987673 | 0.095 | 0.062 | 1 | 24 |
| lrpap1 | 0.72044 | 0.194911505 | 0.095 | 0.062 | 1 | 24 |
| ndfip1l | 0.72044 | 0.115631861 | 0.095 | 0.062 | 1 | 24 |
| pde7a | 0.72044 | 0.11717699 | 0.095 | 0.062 | 1 | 24 |
| si:dkey-27i16.2 | 0.72044 | 0.190003663 | 0.095 | 0.062 | 1 | 24 |
| smn1 | 0.72044 | 0.152212347 | 0.095 | 0.062 | 1 | 24 |
| srsf6a | 0.72044 | 0.111248848 | 0.095 | 0.062 | 1 | 24 |
| atp6v0ca | 0.72044 | 0.160547596 | 0.095 | 0.062 | 1 | 24 |
| marcksa | 0.72044 | 0.113193636 | 0.095 | 0.062 | 1 | 24 |
| polr1c | 0.72044 | 0.127876493 | 0.095 | 0.062 | 1 | 24 |

| | | | | | | |
|---|---|---|---|---|---|---|
| racgap1 | 0.72044 | 0.143990051 | 0.095 | 0.062 | 1 | 24 |
| rgs4 | 0.72044 | 0.164923349 | 0.095 | 0.062 | 1 | 24 |
| si:dkey-177p2.6 | 0.72044 | 0.114441375 | 0.095 | 0.062 | 1 | 24 |
| ss18l2 | 0.72044 | 0.156748654 | 0.095 | 0.062 | 1 | 24 |
| timm17a | 0.72044 | 0.147159061 | 0.095 | 0.062 | 1 | 24 |
| CABZ01005379.1 | 0.72044 | 0.112981931 | 0.095 | 0.062 | 1 | 24 |
| FO082781.1 | 0.72044 | 0.132469899 | 0.095 | 0.062 | 1 | 24 |
| acadvl | 0.72044 | 0.131916532 | 0.095 | 0.062 | 1 | 24 |
| atp6v1e1a | 0.72044 | 0.232901793 | 0.095 | 0.062 | 1 | 24 |
| btbd10b | 0.72044 | 0.145561147 | 0.095 | 0.062 | 1 | 24 |
| capns1a | 0.72044 | 0.168307154 | 0.095 | 0.062 | 1 | 24 |
| ciarta | 0.72044 | 0.110086913 | 0.095 | 0.062 | 1 | 24 |
| csnk1db | 0.72044 | 0.114378915 | 0.095 | 0.062 | 1 | 24 |
| dctpp1 | 0.72044 | 0.208755517 | 0.095 | 0.062 | 1 | 24 |
| dpf2l | 0.72044 | 0.154094952 | 0.095 | 0.062 | 1 | 24 |
| enoph1 | 0.72044 | 0.210143485 | 0.095 | 0.062 | 1 | 24 |
| fbp1a | 0.72044 | 0.120536653 | 0.095 | 0.062 | 1 | 24 |
| foxk2 | 0.72044 | 0.197769902 | 0.095 | 0.062 | 1 | 24 |
| gnal | 0.72044 | -0.14008171 | 0.095 | 0.062 | 1 | 24 |
| guk1a | 0.72044 | 0.16940804 | 0.095 | 0.062 | 1 | 24 |
| hsd11b2 | 0.72044 | 0.127450613 | 0.095 | 0.062 | 1 | 24 |
| ino80c | 0.72044 | 0.165178049 | 0.095 | 0.062 | 1 | 24 |
| ivd | 0.72044 | 0.220320131 | 0.095 | 0.062 | 1 | 24 |
| larp4ab | 0.72044 | 0.253318882 | 0.095 | 0.062 | 1 | 24 |
| mapkapk2a | 0.72044 | 0.103660171 | 0.095 | 0.062 | 1 | 24 |
| midn | 0.72044 | 0.171217279 | 0.095 | 0.062 | 1 | 24 |
| mmab | 0.72044 | 0.127130894 | 0.095 | 0.062 | 1 | 24 |
| napab | 0.72044 | 0.234565499 | 0.095 | 0.062 | 1 | 24 |
| nfia | 0.72044 | 0.117287229 | 0.095 | 0.062 | 1 | 24 |
| parp1 | 0.72044 | 0.290859593 | 0.095 | 0.062 | 1 | 24 |
| pdcd10b | 0.72044 | 0.175998741 | 0.095 | 0.062 | 1 | 24 |
| pdcd6 | 0.72044 | 0.141879677 | 0.095 | 0.062 | 1 | 24 |
| pex5 | 0.72044 | -0.1263528 | 0.095 | 0.062 | 1 | 24 |
| prpf31 | 0.72044 | 0.173629752 | 0.095 | 0.062 | 1 | 24 |
| psap | 0.72044 | 0.107544865 | 0.095 | 0.062 | 1 | 24 |
| psmd12 | 0.72044 | 0.162090382 | 0.095 | 0.062 | 1 | 24 |
| rae1 | 0.72044 | 0.109625358 | 0.095 | 0.062 | 1 | 24 |
| rgrb | 0.72044 | 0.201320599 | 0.095 | 0.062 | 1 | 24 |
| rn7sk | 0.72044 | 0.102510571 | 0.095 | 0.062 | 1 | 24 |
| s100u | 0.72044 | 0.117862877 | 0.095 | 0.062 | 1 | 24 |
| serpinh1b | 0.72044 | 0.281111518 | 0.095 | 0.062 | 1 | 24 |
| sgk1 | 0.72044 | 0.255286627 | 0.095 | 0.062 | 1 | 24 |
| si:ch211-212k18.7 | 0.72044 | 0.152052252 | 0.095 | 0.062 | 1 | 24 |
| si:ch211-85n16.3 | 0.72044 | 0.156588477 | 0.095 | 0.062 | 1 | 24 |
| smim7 | 0.72044 | 0.225463809 | 0.095 | 0.062 | 1 | 24 |
| srp9 | 0.72044 | 0.130371343 | 0.095 | 0.062 | 1 | 24 |
| srsf1b | 0.72044 | 0.10880829 | 0.095 | 0.062 | 1 | 24 |
| tomm40l | 0.72044 | 0.173698177 | 0.095 | 0.062 | 1 | 24 |

| | | | | | | |
|---|---|---|---|---|---|---|
| trap1 | 0.72044 | 0.129380699 | 0.095 | 0.062 | 1 | 24 |
| trappc3 | 0.72044 | 0.146586644 | 0.095 | 0.062 | 1 | 24 |
| tuba1b | 0.72044 | 0.107450383 | 0.095 | 0.062 | 1 | 24 |
| tuba8l2 | 0.72044 | 0.170792765 | 0.095 | 0.062 | 1 | 24 |
| yrk | 0.72044 | -0.1463551 | 0.095 | 0.062 | 1 | 24 |
| zfp36l2 | 0.72044 | 0.102189315 | 0.095 | 0.062 | 1 | 24 |
| zgc:112496 | 0.72044 | 0.137564901 | 0.095 | 0.062 | 1 | 24 |
| zgc:174863 | 0.72044 | 0.108565175 | 0.095 | 0.062 | 1 | 24 |
| zgc:85936 | 0.72044 | 0.111972585 | 0.095 | 0.062 | 1 | 24 |
| actr2a | 0.72044 | 0.210590839 | 0.095 | 0.062 | 1 | 24 |
| angptl1a | 0.72044 | 0.191562717 | 0.095 | 0.062 | 1 | 24 |
| atad5a | 0.72044 | 0.107302278 | 0.095 | 0.062 | 1 | 24 |
| ccl25b | 0.72044 | 0.223138485 | 0.095 | 0.062 | 1 | 24 |
| chd6 | 0.72044 | 0.109482407 | 0.095 | 0.062 | 1 | 24 |
| coq9 | 0.72044 | 0.22445622 | 0.095 | 0.062 | 1 | 24 |
| ctsd | 0.72044 | 0.136481179 | 0.095 | 0.062 | 1 | 24 |
| dctn5 | 0.72044 | 0.18604395 | 0.095 | 0.062 | 1 | 24 |
| grinab | 0.72044 | 0.108335132 | 0.095 | 0.062 | 1 | 24 |
| gtf2f1 | 0.72044 | 0.211376349 | 0.095 | 0.062 | 1 | 24 |
| histh1l | 0.72044 | 0.118989353 | 0.095 | 0.062 | 1 | 24 |
| immp2l | 0.72044 | 0.271072579 | 0.095 | 0.062 | 1 | 24 |
| irf2bpl | 0.72044 | 0.126918411 | 0.095 | 0.062 | 1 | 24 |
| lancl1 | 0.72044 | 0.217179464 | 0.095 | 0.062 | 1 | 24 |
| lcmt1 | 0.72044 | 0.129322617 | 0.095 | 0.062 | 1 | 24 |
| lgals2a | 0.72044 | 0.276279901 | 0.095 | 0.062 | 1 | 24 |
| mrps25 | 0.72044 | 0.12164745 | 0.095 | 0.062 | 1 | 24 |
| mylipb | 0.72044 | 0.104783442 | 0.095 | 0.062 | 1 | 24 |
| ndc1 | 0.72044 | 0.144979483 | 0.095 | 0.062 | 1 | 24 |
| parp12b | 0.72044 | 0.180235448 | 0.095 | 0.062 | 1 | 24 |
| prkar2aa | 0.72044 | 0.14475593 | 0.095 | 0.062 | 1 | 24 |
| ptges3a | 0.72044 | 0.101665533 | 0.095 | 0.062 | 1 | 24 |
| rrp36 | 0.72044 | 0.151166084 | 0.095 | 0.062 | 1 | 24 |
| si:dkey-11f4.14 | 0.72044 | 0.135879114 | 0.095 | 0.062 | 1 | 24 |
| ska1 | 0.72044 | 0.150398988 | 0.095 | 0.062 | 1 | 24 |
| smim12 | 0.72044 | 0.137466369 | 0.095 | 0.062 | 1 | 24 |
| snapc3 | 0.72044 | 0.150396077 | 0.095 | 0.062 | 1 | 24 |
| spred2b | 0.72044 | 0.17926546 | 0.095 | 0.062 | 1 | 24 |
| stau2 | 0.72044 | 0.195396257 | 0.095 | 0.062 | 1 | 24 |
| timmdc1 | 0.72044 | 0.253221546 | 0.095 | 0.062 | 1 | 24 |
| xrn2 | 0.72044 | 0.15011481 | 0.095 | 0.062 | 1 | 24 |
| zcrb1 | 0.72044 | 0.232168134 | 0.095 | 0.062 | 1 | 24 |
| zgc:136564 | 0.72044 | 0.256945284 | 0.095 | 0.062 | 1 | 24 |
| kars | 0.72063 | 0.285831706 | 0.286 | 0.188 | 1 | 24 |
| ndufv2 | 0.72106 | -0.15266656 | 0.095 | 0.188 | 1 | 24 |
| cbx5 | 0.72112 | -0.22077471 | 0.429 | 0.562 | 1 | 24 |
| abcf1 | 0.7212 | -0.14706707 | 0.095 | 0.188 | 1 | 24 |
| pfn2 | 0.72165 | -0.21066134 | 0.429 | 0.562 | 1 | 24 |
| snrpd1 | 0.72212 | -0.24291709 | 0.524 | 0.625 | 1 | 24 |

| | | | | | | |
|---|---|---|---|---|---|---|
| serf2 | 0.72224 | -0.20729477 | 0.429 | 0.562 | 1 | 24 |
| si:ch211-197h24.9 | 0.72229 | 0.164504893 | 0.143 | 0.062 | 1 | 24 |
| ddt | 0.72235 | -0.23205605 | 0.095 | 0.188 | 1 | 24 |
| ash2l | 0.72249 | -0.23258128 | 0.095 | 0.188 | 1 | 24 |
| arf1 | 0.72318 | 0.172770707 | 0.143 | 0.062 | 1 | 24 |
| ctbp1 | 0.72398 | -0.17842566 | 0.095 | 0.188 | 1 | 24 |
| tktb | 0.72403 | 0.184468236 | 0.143 | 0.062 | 1 | 24 |
| ska2 | 0.72421 | 0.119420449 | 0.143 | 0.062 | 1 | 24 |
| clic1 | 0.72443 | -0.18829637 | 0.095 | 0.188 | 1 | 24 |
| mcm6 | 0.72463 | -0.22076196 | 0.095 | 0.188 | 1 | 24 |
| ctps1a | 0.7248 | -0.17703503 | 0.095 | 0.188 | 1 | 24 |
| mrpl57 | 0.72484 | -0.1782635 | 0.095 | 0.188 | 1 | 24 |
| nifk | 0.72486 | -0.21340425 | 0.095 | 0.188 | 1 | 24 |
| usp14 | 0.72523 | 0.247582142 | 0.143 | 0.062 | 1 | 24 |
| mid1ip1l | 0.72537 | 0.175629681 | 0.143 | 0.062 | 1 | 24 |
| tuba8l | 0.72569 | -0.10321806 | 0.571 | 0.688 | 1 | 24 |
| ube2e1 | 0.7258 | 0.304781448 | 0.286 | 0.188 | 1 | 24 |
| polr2gl | 0.72621 | 0.182752108 | 0.143 | 0.062 | 1 | 24 |
| mydgf | 0.72643 | 0.119855192 | 0.143 | 0.062 | 1 | 24 |
| hat1 | 0.72657 | 0.191816598 | 0.143 | 0.062 | 1 | 24 |
| usp5 | 0.72687 | 0.182511062 | 0.143 | 0.062 | 1 | 24 |
| kdm6bb | 0.72768 | 0.151117148 | 0.143 | 0.062 | 1 | 24 |
| smim13 | 0.72792 | 0.224947346 | 0.143 | 0.062 | 1 | 24 |
| zgc:175088 | 0.72798 | 0.185002406 | 0.143 | 0.062 | 1 | 24 |
| arpc5a | 0.72808 | 0.183986072 | 0.143 | 0.062 | 1 | 24 |
| dsn1 | 0.72859 | 0.267243649 | 0.143 | 0.062 | 1 | 24 |
| mzt2b | 0.72871 | 0.233973162 | 0.143 | 0.062 | 1 | 24 |
| lsm14ab | 0.72915 | 0.164548437 | 0.143 | 0.062 | 1 | 24 |
| minos1 | 0.72921 | 0.131180039 | 0.143 | 0.062 | 1 | 24 |
| cops8 | 0.72944 | 0.128405807 | 0.143 | 0.062 | 1 | 24 |
| arih2 | 0.72967 | 0.208420016 | 0.143 | 0.062 | 1 | 24 |
| zgc:194627 | 0.73027 | 0.225336654 | 0.143 | 0.062 | 1 | 24 |
| ndufa9a | 0.73092 | 0.124993069 | 0.143 | 0.062 | 1 | 24 |
| srsf1a | 0.73112 | 0.166530534 | 0.143 | 0.062 | 1 | 24 |
| pold1 | 0.73135 | 0.16419452 | 0.143 | 0.062 | 1 | 24 |
| sec22bb | 0.73145 | 0.161167493 | 0.143 | 0.062 | 1 | 24 |
| fdx2 | 0.73148 | 0.195593394 | 0.143 | 0.062 | 1 | 24 |
| cenps | 0.7318 | 0.205665874 | 0.143 | 0.062 | 1 | 24 |
| huwe1 | 0.73229 | 0.155170583 | 0.143 | 0.062 | 1 | 24 |
| chrac1 | 0.73239 | 0.182364491 | 0.143 | 0.062 | 1 | 24 |
| smim8 | 0.7324 | 0.204691988 | 0.143 | 0.062 | 1 | 24 |
| arf2b | 0.73246 | 0.248931349 | 0.143 | 0.062 | 1 | 24 |
| eif1axa | 0.73262 | 0.145480248 | 0.143 | 0.062 | 1 | 24 |
| pafah1b1a | 0.73267 | 0.184461078 | 0.143 | 0.062 | 1 | 24 |
| bub1bb | 0.73278 | 0.184784346 | 0.143 | 0.062 | 1 | 24 |
| psmd11b | 0.73286 | 0.197865479 | 0.143 | 0.062 | 1 | 24 |
| serinc5 | 0.73289 | 0.190094181 | 0.143 | 0.062 | 1 | 24 |
| gps2 | 0.73298 | 0.157237004 | 0.143 | 0.062 | 1 | 24 |

| | | | | | | |
|---|---|---|---|---|---|---|
| irf2bp2a | 0.73303 | 0.200989198 | 0.143 | 0.062 | 1 | 24 |
| exosc4 | 0.73303 | 0.131866354 | 0.143 | 0.062 | 1 | 24 |
| mt-cyb | 0.73861 | 0.320400686 | 0.286 | 0.188 | 1 | 24 |
| si:ch211-214p16.1 | 0.73888 | -0.20849137 | 0.19 | 0.188 | 1 | 24 |
| tomm20a | 0.74153 | 0.281054267 | 0.286 | 0.25 | 1 | 24 |
| nhp2 | 0.74298 | 0.306665298 | 0.19 | 0.125 | 1 | 24 |
| snrpa1 | 0.74486 | 0.284622652 | 0.286 | 0.188 | 1 | 24 |
| hnrnpa0b | 0.74549 | 0.223297367 | 0.619 | 0.5 | 1 | 24 |
| arpc1b | 0.74617 | -0.1884012 | 0.143 | 0.125 | 1 | 24 |
| prmt1 | 0.75938 | 0.240851669 | 0.429 | 0.312 | 1 | 24 |
| rtf1 | 0.76051 | 0.287387985 | 0.19 | 0.125 | 1 | 24 |
| tars | 0.76064 | 0.315628182 | 0.19 | 0.125 | 1 | 24 |
| eif2s1b | 0.76127 | 0.281045608 | 0.333 | 0.25 | 1 | 24 |
| sri | 0.76138 | 0.266549889 | 0.286 | 0.188 | 1 | 24 |
| nsa2 | 0.76371 | -0.16541593 | 0.381 | 0.5 | 1 | 24 |
| mknk2b | 0.76475 | 0.296621604 | 0.19 | 0.125 | 1 | 24 |
| psmb5 | 0.76745 | 0.296487458 | 0.19 | 0.125 | 1 | 24 |
| rbx1 | 0.76791 | -0.17197271 | 0.381 | 0.5 | 1 | 24 |
| cbx3a | 0.76841 | -0.21609708 | 0.381 | 0.5 | 1 | 24 |
| jpt1b | 0.77055 | -0.27662817 | 0.286 | 0.375 | 1 | 24 |
| si:dkey-88p24.11 | 0.77068 | 0.15085815 | 0.286 | 0.188 | 1 | 24 |
| bysl | 0.77256 | 0.235147605 | 0.286 | 0.188 | 1 | 24 |
| atp5f1e | 0.77457 | -0.19531894 | 0.524 | 0.562 | 1 | 24 |
| mcl1a | 0.7788 | 0.131237362 | 0.286 | 0.188 | 1 | 24 |
| ube2ib | 0.78034 | -0.11270776 | 0.333 | 0.438 | 1 | 24 |
| smc1al | 0.78148 | 0.226644942 | 0.286 | 0.188 | 1 | 24 |
| higd1a | 0.78178 | -0.27719162 | 0.143 | 0.188 | 1 | 24 |
| ube2l3b | 0.78376 | 0.215362483 | 0.286 | 0.188 | 1 | 24 |
| ptmaa | 0.78385 | -0.14148414 | 0.714 | 0.812 | 1 | 24 |
| arpc5b | 0.78666 | -0.2825607 | 0.095 | 0.125 | 1 | 24 |
| nfyc | 0.78872 | -0.17369587 | 0.143 | 0.125 | 1 | 24 |
| idh2 | 0.78985 | -0.22388077 | 0.381 | 0.438 | 1 | 24 |
| sept2 | 0.79016 | -0.113482 | 0.333 | 0.438 | 1 | 24 |
| si:ch211-282b22.1 | 0.79171 | 0.255499711 | 0.19 | 0.125 | 1 | 24 |
| carm1 | 0.79248 | 0.143931133 | 0.095 | 0.125 | 1 | 24 |
| dnajc9 | 0.79269 | 0.24426942 | 0.286 | 0.25 | 1 | 24 |
| chd4b | 0.79531 | -0.27619432 | 0.143 | 0.188 | 1 | 24 |
| rpl5b | 0.7955 | -0.1113486 | 0.619 | 0.625 | 1 | 24 |
| si:ch211-235e9.6 | 0.79608 | 0.211229595 | 0.333 | 0.312 | 1 | 24 |
| arf6a | 0.79691 | 0.20553662 | 0.19 | 0.188 | 1 | 24 |
| atp5fa1 | 0.79733 | -0.12876733 | 0.333 | 0.438 | 1 | 24 |
| nudcd2 | 0.79882 | 0.122273757 | 0.095 | 0.125 | 1 | 24 |
| eif3c | 0.79939 | -0.27003247 | 0.143 | 0.188 | 1 | 24 |
| rbm28 | 0.80259 | -0.27066107 | 0.095 | 0.125 | 1 | 24 |
| usf1l | 0.80309 | -0.27170862 | 0.095 | 0.125 | 1 | 24 |
| anp32a | 0.80332 | -0.15544777 | 0.333 | 0.438 | 1 | 24 |
| lsm7 | 0.80415 | 0.106783732 | 0.476 | 0.375 | 1 | 24 |
| eif3i | 0.80529 | -0.24225635 | 0.286 | 0.375 | 1 | 24 |

| | | | | | | |
|---|---|---|---|---|---|---|
| hnrnpa1a | 0.80578 | 0.242757047 | 0.143 | 0.125 | 1 | 24 |
| alyref | 0.80608 | 0.252705973 | 0.19 | 0.125 | 1 | 24 |
| suclg1 | 0.80848 | 0.24313234 | 0.143 | 0.125 | 1 | 24 |
| btf3 | 0.8136 | 0.129354036 | 0.714 | 0.688 | 1 | 24 |
| rps27l | 0.81374 | 0.245166997 | 0.19 | 0.125 | 1 | 24 |
| top2a | 0.81474 | -0.26261337 | 0.095 | 0.125 | 1 | 24 |
| sarnp | 0.81674 | 0.255512753 | 0.238 | 0.188 | 1 | 24 |
| sh3bgrl | 0.81681 | 0.23503901 | 0.19 | 0.125 | 1 | 24 |
| rab11a | 0.8169 | 0.241516505 | 0.19 | 0.125 | 1 | 24 |
| psma5 | 0.81833 | -0.24416779 | 0.238 | 0.312 | 1 | 24 |
| mrps10 | 0.81845 | -0.25354695 | 0.143 | 0.188 | 1 | 24 |
| eif2s2 | 0.81853 | -0.12457007 | 0.524 | 0.625 | 1 | 24 |
| cycsb | 0.82028 | -0.24555515 | 0.238 | 0.312 | 1 | 24 |
| asf1ba | 0.82085 | -0.24339128 | 0.238 | 0.312 | 1 | 24 |
| hsp90ab1 | 0.82166 | -0.10425471 | 0.857 | 0.875 | 1 | 24 |
| ndufb4 | 0.82166 | 0.23935211 | 0.238 | 0.188 | 1 | 24 |
| ndufab1b | 0.82168 | 0.211707461 | 0.19 | 0.125 | 1 | 24 |
| fkbp3 | 0.82307 | 0.158367046 | 0.238 | 0.25 | 1 | 24 |
| bcl2l10 | 0.82362 | 0.265607638 | 0.19 | 0.125 | 1 | 24 |
| srsf7a | 0.82418 | -0.22436133 | 0.286 | 0.375 | 1 | 24 |
| hnrnpa0a | 0.82443 | 0.224698244 | 0.19 | 0.125 | 1 | 24 |
| mrpl12 | 0.82454 | 0.25178862 | 0.19 | 0.125 | 1 | 24 |
| rbbp4 | 0.8254 | -0.22558635 | 0.286 | 0.375 | 1 | 24 |
| nsfl1c | 0.82571 | 0.245317745 | 0.238 | 0.188 | 1 | 24 |
| LEPROTL1 | 0.8262 | -0.14637203 | 0.143 | 0.125 | 1 | 24 |
| pcna | 0.82627 | 0.163243799 | 0.714 | 0.625 | 1 | 24 |
| ponzr1 | 0.82702 | 0.185281589 | 0.476 | 0.438 | 1 | 24 |
| dldh | 0.82937 | -0.24706454 | 0.095 | 0.125 | 1 | 24 |
| pabpn1 | 0.8324 | -0.21226479 | 0.286 | 0.375 | 1 | 24 |
| sf3b6 | 0.83563 | 0.212764885 | 0.19 | 0.125 | 1 | 24 |
| snw1 | 0.83602 | 0.197609839 | 0.19 | 0.125 | 1 | 24 |
| gdi2 | 0.8377 | -0.24399247 | 0.143 | 0.188 | 1 | 24 |
| si:ch211-66k16.27 | 0.83781 | 0.143305883 | 0.238 | 0.25 | 1 | 24 |
| adh5 | 0.83909 | 0.204116628 | 0.333 | 0.25 | 1 | 24 |
| ppil3 | 0.8393 | 0.211125585 | 0.19 | 0.125 | 1 | 24 |
| ppp6c | 0.83998 | 0.233816141 | 0.19 | 0.125 | 1 | 24 |
| psme2 | 0.84044 | -0.23451753 | 0.19 | 0.25 | 1 | 24 |
| sod2 | 0.84079 | -0.23993102 | 0.143 | 0.188 | 1 | 24 |
| mrpl20 | 0.84108 | 0.184934101 | 0.19 | 0.188 | 1 | 24 |
| aif1l | 0.84151 | -0.24386197 | 0.095 | 0.125 | 1 | 24 |
| psmd6 | 0.84244 | -0.23507133 | 0.143 | 0.188 | 1 | 24 |
| eif3ja | 0.84265 | 0.200542468 | 0.19 | 0.125 | 1 | 24 |
| nip7 | 0.84346 | 0.10212562 | 0.095 | 0.125 | 1 | 24 |
| msra | 0.84441 | -0.2398007 | 0.095 | 0.125 | 1 | 24 |
| si:ch73-42p12.2 | 0.84525 | 0.105735833 | 0.19 | 0.125 | 1 | 24 |
| selenow1 | 0.84657 | -0.20032316 | 0.429 | 0.5 | 1 | 24 |
| rcl1 | 0.84659 | -0.23743945 | 0.095 | 0.125 | 1 | 24 |
| ldhba | 0.84683 | -0.17677054 | 0.286 | 0.375 | 1 | 24 |

| | | | | | | |
|---|---|---|---|---|---|---|
| zgc:158803 | 0.84722 | -0.23361584 | 0.143 | 0.188 | 1 | 24 |
| ilf2 | 0.84817 | -0.18748535 | 0.286 | 0.375 | 1 | 24 |
| COX7A2 (1 of many) | 0.84843 | -0.14783451 | 0.286 | 0.375 | 1 | 24 |
| esco2 | 0.84858 | -0.23099351 | 0.143 | 0.188 | 1 | 24 |
| sars | 0.84864 | 0.209836955 | 0.19 | 0.125 | 1 | 24 |
| phf5a | 0.84865 | 0.194605416 | 0.333 | 0.25 | 1 | 24 |
| cox7b | 0.84877 | 0.186018178 | 0.524 | 0.438 | 1 | 24 |
| chchd7 | 0.84883 | -0.11889671 | 0.048 | 0.062 | 1 | 24 |
| ptk7a | 0.84883 | -0.10931408 | 0.048 | 0.062 | 1 | 24 |
| si:ch211-67e16.4 | 0.84883 | -0.1100701 | 0.048 | 0.062 | 1 | 24 |
| tufm | 0.84883 | -0.11679687 | 0.048 | 0.062 | 1 | 24 |
| wipi1 | 0.84883 | -0.11481377 | 0.048 | 0.062 | 1 | 24 |
| si:dkey-170l10.1 | 0.84883 | 0.146476187 | 0.048 | 0.062 | 1 | 24 |
| BX322530.1 | 0.84883 | -0.16532653 | 0.048 | 0.062 | 1 | 24 |
| CABZ01085201.1 | 0.84883 | -0.14115675 | 0.048 | 0.062 | 1 | 24 |
| DHDH | 0.84883 | -0.19095091 | 0.048 | 0.062 | 1 | 24 |
| SCYL2 | 0.84883 | -0.15950878 | 0.048 | 0.062 | 1 | 24 |
| adipor1b | 0.84883 | -0.18475842 | 0.048 | 0.062 | 1 | 24 |
| aktip | 0.84883 | 0.100014547 | 0.048 | 0.062 | 1 | 24 |
| anxa5b | 0.84883 | -0.10413519 | 0.048 | 0.062 | 1 | 24 |
| apool | 0.84883 | -0.11428878 | 0.048 | 0.062 | 1 | 24 |
| asb12a | 0.84883 | -0.18949778 | 0.048 | 0.062 | 1 | 24 |
| atg101 | 0.84883 | -0.11034677 | 0.048 | 0.062 | 1 | 24 |
| ccnb1 | 0.84883 | -0.23553591 | 0.048 | 0.062 | 1 | 24 |
| cdc37 | 0.84883 | -0.11512673 | 0.048 | 0.062 | 1 | 24 |
| cdca8 | 0.84883 | -0.11480242 | 0.048 | 0.062 | 1 | 24 |
| csnk1da | 0.84883 | -0.18303212 | 0.048 | 0.062 | 1 | 24 |
| cyba | 0.84883 | -0.21324673 | 0.048 | 0.062 | 1 | 24 |
| dicer1 | 0.84883 | -0.17949927 | 0.048 | 0.062 | 1 | 24 |
| dnm2a | 0.84883 | -0.17247197 | 0.048 | 0.062 | 1 | 24 |
| dynll2a | 0.84883 | -0.10295697 | 0.048 | 0.062 | 1 | 24 |
| eci1 | 0.84883 | -0.18965891 | 0.048 | 0.062 | 1 | 24 |
| elac1 | 0.84883 | -0.16074837 | 0.048 | 0.062 | 1 | 24 |
| eme1 | 0.84883 | -0.18107719 | 0.048 | 0.062 | 1 | 24 |
| erbin | 0.84883 | -0.14581499 | 0.048 | 0.062 | 1 | 24 |
| fcho2 | 0.84883 | -0.12793841 | 0.048 | 0.062 | 1 | 24 |
| glipr1a | 0.84883 | -0.16498666 | 0.048 | 0.062 | 1 | 24 |
| glmna | 0.84883 | -0.14779863 | 0.048 | 0.062 | 1 | 24 |
| hs1bp3 | 0.84883 | -0.16089092 | 0.048 | 0.062 | 1 | 24 |
| il12rb2l | 0.84883 | -0.11135264 | 0.048 | 0.062 | 1 | 24 |
| l2hgdh | 0.84883 | -0.16329853 | 0.048 | 0.062 | 1 | 24 |
| lamc1 | 0.84883 | -0.13899569 | 0.048 | 0.062 | 1 | 24 |
| map4k4 | 0.84883 | -0.13194606 | 0.048 | 0.062 | 1 | 24 |
| mbip | 0.84883 | -0.11785499 | 0.048 | 0.062 | 1 | 24 |
| mef2cb | 0.84883 | -0.16459877 | 0.048 | 0.062 | 1 | 24 |
| mettl7a | 0.84883 | -0.18194158 | 0.048 | 0.062 | 1 | 24 |
| myef2 | 0.84883 | -0.1255734 | 0.048 | 0.062 | 1 | 24 |
| naa38 | 0.84883 | -0.10055409 | 0.048 | 0.062 | 1 | 24 |

| | | | | | | |
|---|---|---|---|---|---|---|
| naxd | 0.84883 | -0.16004978 | 0.048 | 0.062 | 1 | 24 |
| nnt | 0.84883 | -0.15228136 | 0.048 | 0.062 | 1 | 24 |
| nucb2a | 0.84883 | -0.10887422 | 0.048 | 0.062 | 1 | 24 |
| nudt22 | 0.84883 | -0.11085301 | 0.048 | 0.062 | 1 | 24 |
| oaz2b | 0.84883 | -0.11147545 | 0.048 | 0.062 | 1 | 24 |
| pam16 | 0.84883 | -0.14596754 | 0.048 | 0.062 | 1 | 24 |
| pds5a | 0.84883 | -0.13253791 | 0.048 | 0.062 | 1 | 24 |
| phf11 | 0.84883 | -0.21883623 | 0.048 | 0.062 | 1 | 24 |
| plek | 0.84883 | -0.23596674 | 0.048 | 0.062 | 1 | 24 |
| rab5c | 0.84883 | -0.15046727 | 0.048 | 0.062 | 1 | 24 |
| rfxank | 0.84883 | -0.12802422 | 0.048 | 0.062 | 1 | 24 |
| rock2a | 0.84883 | -0.1882016 | 0.048 | 0.062 | 1 | 24 |
| rpp21 | 0.84883 | 0.126025821 | 0.048 | 0.062 | 1 | 24 |
| sec13 | 0.84883 | -0.1596516 | 0.048 | 0.062 | 1 | 24 |
| sept9a | 0.84883 | 0.100487111 | 0.048 | 0.062 | 1 | 24 |
| si:ch1073-314i13.4 | 0.84883 | -0.10009023 | 0.048 | 0.062 | 1 | 24 |
| si:ch211-206k20.5 | 0.84883 | -0.1814525 | 0.048 | 0.062 | 1 | 24 |
| si:dkey-112a7.4 | 0.84883 | 0.116673058 | 0.048 | 0.062 | 1 | 24 |
| si:dkey-287g12.6 | 0.84883 | -0.13009873 | 0.048 | 0.062 | 1 | 24 |
| si:dkeyp-27c8.2 | 0.84883 | -0.11622979 | 0.048 | 0.062 | 1 | 24 |
| slc20a1b | 0.84883 | -0.12638268 | 0.048 | 0.062 | 1 | 24 |
| snd1 | 0.84883 | 0.145453586 | 0.048 | 0.062 | 1 | 24 |
| snx7 | 0.84883 | -0.18435719 | 0.048 | 0.062 | 1 | 24 |
| stx4 | 0.84883 | -0.13346988 | 0.048 | 0.062 | 1 | 24 |
| sympk | 0.84883 | -0.15742621 | 0.048 | 0.062 | 1 | 24 |
| tefa | 0.84883 | -0.11676078 | 0.048 | 0.062 | 1 | 24 |
| tirap | 0.84883 | -0.24182854 | 0.048 | 0.062 | 1 | 24 |
| tnpo3 | 0.84883 | -0.10687222 | 0.048 | 0.062 | 1 | 24 |
| trnt1 | 0.84883 | -0.17898533 | 0.048 | 0.062 | 1 | 24 |
| unm-hu7910 | 0.84883 | -0.14081772 | 0.048 | 0.062 | 1 | 24 |
| uqcc3 | 0.84883 | -0.11059793 | 0.048 | 0.062 | 1 | 24 |
| utp23 | 0.84883 | -0.11490394 | 0.048 | 0.062 | 1 | 24 |
| wipf1a | 0.84883 | -0.2073719 | 0.048 | 0.062 | 1 | 24 |
| zgc:162972 | 0.84883 | -0.10512342 | 0.048 | 0.062 | 1 | 24 |
| zranb2 | 0.84883 | -0.12514682 | 0.048 | 0.062 | 1 | 24 |
| abt1 | 0.84883 | 0.109437349 | 0.048 | 0.062 | 1 | 24 |
| anxa11a | 0.84883 | -0.12705999 | 0.048 | 0.062 | 1 | 24 |
| appa | 0.84883 | -0.19842467 | 0.048 | 0.062 | 1 | 24 |
| atf7ip | 0.84883 | -0.13134681 | 0.048 | 0.062 | 1 | 24 |
| bcam | 0.84883 | -0.10156964 | 0.048 | 0.062 | 1 | 24 |
| bida | 0.84883 | -0.13145104 | 0.048 | 0.062 | 1 | 24 |
| cenpl | 0.84883 | -0.11143722 | 0.048 | 0.062 | 1 | 24 |
| cndp2 | 0.84883 | -0.16552861 | 0.048 | 0.062 | 1 | 24 |
| ddah2 | 0.84883 | -0.14407064 | 0.048 | 0.062 | 1 | 24 |
| dnajc1 | 0.84883 | -0.11542551 | 0.048 | 0.062 | 1 | 24 |
| elf2b | 0.84883 | -0.11948436 | 0.048 | 0.062 | 1 | 24 |
| entpd1 | 0.84883 | 0.107378608 | 0.048 | 0.062 | 1 | 24 |
| etv5a | 0.84883 | -0.16864357 | 0.048 | 0.062 | 1 | 24 |

| | | | | | | |
|---|---|---|---|---|---|---|
| fbxw11a | 0.84883 | -0.16568738 | 0.048 | 0.062 | 1 | 24 |
| fuca1.1 | 0.84883 | 0.117378369 | 0.048 | 0.062 | 1 | 24 |
| hacd2 | 0.84883 | -0.17920006 | 0.048 | 0.062 | 1 | 24 |
| helz | 0.84883 | -0.14444102 | 0.048 | 0.062 | 1 | 24 |
| hspa4a | 0.84883 | 0.117345241 | 0.048 | 0.062 | 1 | 24 |
| lamp1a | 0.84883 | -0.17518518 | 0.048 | 0.062 | 1 | 24 |
| lyrm2 | 0.84883 | -0.10298216 | 0.048 | 0.062 | 1 | 24 |
| lyrm7 | 0.84883 | -0.17877217 | 0.048 | 0.062 | 1 | 24 |
| mrps36 | 0.84883 | 0.125095372 | 0.048 | 0.062 | 1 | 24 |
| mtap | 0.84883 | -0.1211753 | 0.048 | 0.062 | 1 | 24 |
| mybl2b | 0.84883 | -0.14282095 | 0.048 | 0.062 | 1 | 24 |
| nckap1 | 0.84883 | -0.12891413 | 0.048 | 0.062 | 1 | 24 |
| ndufa10 | 0.84883 | -0.13311644 | 0.048 | 0.062 | 1 | 24 |
| otub1b | 0.84883 | -0.17536413 | 0.048 | 0.062 | 1 | 24 |
| parp3 | 0.84883 | 0.168888412 | 0.048 | 0.062 | 1 | 24 |
| pcid2 | 0.84883 | -0.16630889 | 0.048 | 0.062 | 1 | 24 |
| ptgr1 | 0.84883 | -0.12403569 | 0.048 | 0.062 | 1 | 24 |
| rab28 | 0.84883 | -0.14214967 | 0.048 | 0.062 | 1 | 24 |
| rassf2a | 0.84883 | -0.11019373 | 0.048 | 0.062 | 1 | 24 |
| sftpbb | 0.84883 | -0.10930078 | 0.048 | 0.062 | 1 | 24 |
| sostdc1a | 0.84883 | 0.157015677 | 0.048 | 0.062 | 1 | 24 |
| swap70b | 0.84883 | -0.21976939 | 0.048 | 0.062 | 1 | 24 |
| timeless | 0.84883 | -0.11790425 | 0.048 | 0.062 | 1 | 24 |
| tnfsf10 | 0.84883 | 0.108373219 | 0.048 | 0.062 | 1 | 24 |
| trit1 | 0.84883 | 0.104549572 | 0.048 | 0.062 | 1 | 24 |
| ube3a | 0.84883 | -0.12127414 | 0.048 | 0.062 | 1 | 24 |
| wasf2 | 0.84883 | -0.13089076 | 0.048 | 0.062 | 1 | 24 |
| zgc:193742 | 0.84883 | -0.17367021 | 0.048 | 0.062 | 1 | 24 |
| zgc:64051 | 0.84883 | -0.10403346 | 0.048 | 0.062 | 1 | 24 |
| UTP14C | 0.84883 | -0.11665117 | 0.048 | 0.062 | 1 | 24 |
| arrdc3b | 0.84883 | -0.10053553 | 0.048 | 0.062 | 1 | 24 |
| eloa | 0.84883 | -0.11507974 | 0.048 | 0.062 | 1 | 24 |
| pdk1 | 0.84883 | -0.10695846 | 0.048 | 0.062 | 1 | 24 |
| rpe | 0.84883 | -0.11676504 | 0.048 | 0.062 | 1 | 24 |
| si:ch211-153a8.4 | 0.84883 | -0.1096293 | 0.048 | 0.062 | 1 | 24 |
| si:ch211-199g17.2 | 0.84883 | -0.10115407 | 0.048 | 0.062 | 1 | 24 |
| surf4l | 0.84883 | -0.11348328 | 0.048 | 0.062 | 1 | 24 |
| wbp11 | 0.84883 | 0.114346426 | 0.048 | 0.062 | 1 | 24 |
| zgc:114181 | 0.84883 | -0.18943259 | 0.048 | 0.062 | 1 | 24 |
| plp2 | 0.84883 | -0.13788394 | 0.048 | 0.062 | 1 | 24 |
| rhov | 0.84883 | -0.14162539 | 0.048 | 0.062 | 1 | 24 |
| zgc:162944 | 0.84896 | -0.16649455 | 0.286 | 0.375 | 1 | 24 |
| fbl | 0.84907 | -0.16682046 | 0.286 | 0.375 | 1 | 24 |
| etfa | 0.85146 | 0.161839508 | 0.19 | 0.125 | 1 | 24 |
| gadd45ba | 0.85187 | 0.197467606 | 0.19 | 0.125 | 1 | 24 |
| atp5mf | 0.85279 | -0.23208938 | 0.19 | 0.25 | 1 | 24 |
| grb2b | 0.85349 | -0.2294834 | 0.095 | 0.125 | 1 | 24 |
| s100v2 | 0.85369 | -0.22633284 | 0.095 | 0.125 | 1 | 24 |

| | | | | | | |
|---|---|---|---|---|---|---|
| safb | 0.85447 | -0.2256035 | 0.095 | 0.125 | 1 | 24 |
| rnf10 | 0.85463 | 0.196919085 | 0.19 | 0.125 | 1 | 24 |
| jdp2b | 0.85613 | 0.172043812 | 0.19 | 0.188 | 1 | 24 |
| si:ch73-46j18.5 | 0.85629 | 0.189004231 | 0.333 | 0.25 | 1 | 24 |
| meaf6 | 0.85652 | 0.177292794 | 0.19 | 0.125 | 1 | 24 |
| snu13b | 0.85711 | 0.144607457 | 0.381 | 0.375 | 1 | 24 |
| COX5B | 0.8575 | -0.19048877 | 0.381 | 0.438 | 1 | 24 |
| ddx39ab | 0.85861 | 0.178379811 | 0.333 | 0.25 | 1 | 24 |
| psma3 | 0.85862 | 0.148660905 | 0.333 | 0.25 | 1 | 24 |
| snrpg | 0.85871 | 0.143563985 | 0.524 | 0.5 | 1 | 24 |
| rrp1 | 0.85894 | 0.210495387 | 0.143 | 0.125 | 1 | 24 |
| lsm6 | 0.85897 | 0.171550978 | 0.333 | 0.25 | 1 | 24 |
| nop56 | 0.85912 | 0.141641721 | 0.333 | 0.25 | 1 | 24 |
| taf15 | 0.85996 | -0.22343158 | 0.143 | 0.188 | 1 | 24 |
| chmp6b | 0.86111 | 0.178821093 | 0.19 | 0.125 | 1 | 24 |
| ier2a | 0.86167 | 0.169037268 | 0.19 | 0.188 | 1 | 24 |
| rfc2 | 0.86178 | 0.128628907 | 0.19 | 0.125 | 1 | 24 |
| cenpw | 0.86199 | 0.122390449 | 0.19 | 0.125 | 1 | 24 |
| polr2h | 0.86203 | -0.22187373 | 0.19 | 0.25 | 1 | 24 |
| ufm1 | 0.86204 | -0.21400543 | 0.143 | 0.188 | 1 | 24 |
| btg3 | 0.86271 | 0.205238981 | 0.143 | 0.125 | 1 | 24 |
| mt-nd4 | 0.86283 | 0.24956938 | 0.238 | 0.188 | 1 | 24 |
| rfc5 | 0.86323 | 0.167769318 | 0.19 | 0.125 | 1 | 24 |
| spcs3 | 0.86329 | 0.125134448 | 0.19 | 0.125 | 1 | 24 |
| prdx3 | 0.86342 | 0.121350168 | 0.19 | 0.125 | 1 | 24 |
| dtymk | 0.86366 | 0.157953302 | 0.19 | 0.125 | 1 | 24 |
| rab10 | 0.86382 | 0.188452705 | 0.19 | 0.125 | 1 | 24 |
| prdx4 | 0.86449 | 0.14407646 | 0.19 | 0.125 | 1 | 24 |
| snrnp27 | 0.86471 | 0.154257756 | 0.19 | 0.125 | 1 | 24 |
| CU694197.1 | 0.86484 | 0.157326074 | 0.19 | 0.125 | 1 | 24 |
| sdhc | 0.86496 | 0.142732294 | 0.19 | 0.125 | 1 | 24 |
| fam162a | 0.86505 | 0.122505138 | 0.19 | 0.125 | 1 | 24 |
| chtopa | 0.86522 | 0.154120239 | 0.19 | 0.125 | 1 | 24 |
| rbb4l | 0.86524 | 0.118718508 | 0.19 | 0.125 | 1 | 24 |
| ciao2b | 0.86549 | 0.110969824 | 0.19 | 0.125 | 1 | 24 |
| gtpbp4 | 0.86572 | 0.144419441 | 0.19 | 0.125 | 1 | 24 |
| ehd1b | 0.86574 | 0.146626155 | 0.19 | 0.125 | 1 | 24 |
| bzw1b | 0.86578 | 0.141731596 | 0.19 | 0.125 | 1 | 24 |
| ap2m1a | 0.86605 | -0.21543451 | 0.19 | 0.25 | 1 | 24 |
| p4hb | 0.86721 | -0.22024868 | 0.143 | 0.188 | 1 | 24 |
| gpx1a | 0.86927 | -0.22393113 | 0.19 | 0.25 | 1 | 24 |
| btg1 | 0.8703 | 0.120484795 | 0.238 | 0.25 | 1 | 24 |
| si:ch211-286b5.5 | 0.87086 | 0.211622387 | 0.238 | 0.188 | 1 | 24 |
| ndufc2 | 0.87131 | -0.21494831 | 0.143 | 0.188 | 1 | 24 |
| atp5meb | 0.87163 | -0.14899757 | 0.476 | 0.562 | 1 | 24 |
| cox6a1 | 0.87554 | 0.190914153 | 0.381 | 0.312 | 1 | 24 |
| si:ch211-196l7.4 | 0.87809 | -0.16591452 | 0.238 | 0.312 | 1 | 24 |
| lig1 | 0.87901 | 0.189200382 | 0.286 | 0.25 | 1 | 24 |

| | | | | | | |
|---|---|---|---|---|---|---|
| gabarapb | 0.87907 | -0.19612284 | 0.19 | 0.25 | 1 | 24 |
| taf9 | 0.87909 | -0.21013196 | 0.143 | 0.188 | 1 | 24 |
| mdh1aa | 0.87927 | -0.15954171 | 0.238 | 0.312 | 1 | 24 |
| rrm2 | 0.88065 | -0.14135241 | 0.524 | 0.562 | 1 | 24 |
| jpt2 | 0.88112 | -0.13609414 | 0.238 | 0.312 | 1 | 24 |
| u2af1 | 0.88154 | -0.1282553 | 0.238 | 0.312 | 1 | 24 |
| ndufb8 | 0.88166 | -0.12813704 | 0.238 | 0.312 | 1 | 24 |
| cdc26 | 0.88178 | 0.199489031 | 0.143 | 0.125 | 1 | 24 |
| mat2aa | 0.88271 | -0.20471169 | 0.143 | 0.188 | 1 | 24 |
| cox7c | 0.88273 | 0.102538994 | 0.571 | 0.562 | 1 | 24 |
| h3f3c | 0.88369 | -0.128151 | 0.429 | 0.438 | 1 | 24 |
| marcksl1b | 0.88465 | 0.206452754 | 0.238 | 0.188 | 1 | 24 |
| dad1 | 0.88553 | 0.181382457 | 0.286 | 0.25 | 1 | 24 |
| twistnb | 0.88591 | 0.187237688 | 0.143 | 0.125 | 1 | 24 |
| cyc1 | 0.886 | 0.178535272 | 0.286 | 0.25 | 1 | 24 |
| atp5mea | 0.8861 | -0.17547935 | 0.19 | 0.25 | 1 | 24 |
| tcf25 | 0.8869 | -0.20026449 | 0.095 | 0.125 | 1 | 24 |
| rbfox2 | 0.88698 | -0.1986451 | 0.095 | 0.125 | 1 | 24 |
| nop58 | 0.88799 | 0.179827319 | 0.238 | 0.188 | 1 | 24 |
| arid4a | 0.88981 | -0.19372775 | 0.095 | 0.125 | 1 | 24 |
| arl1 | 0.88996 | 0.183552432 | 0.143 | 0.125 | 1 | 24 |
| atf6 | 0.89365 | -0.19746845 | 0.095 | 0.125 | 1 | 24 |
| cd81a | 0.89462 | -0.18550164 | 0.143 | 0.188 | 1 | 24 |
| rtn4a | 0.89482 | -0.18857198 | 0.143 | 0.188 | 1 | 24 |
| si:ch211-51e12.7 | 0.89504 | -0.1704579 | 0.19 | 0.25 | 1 | 24 |
| psmb2 | 0.8964 | -0.16215993 | 0.286 | 0.312 | 1 | 24 |
| abracl | 0.89644 | 0.115879887 | 0.238 | 0.25 | 1 | 24 |
| gnb1a | 0.8968 | -0.18134923 | 0.143 | 0.188 | 1 | 24 |
| hnrnpub | 0.89756 | 0.181205625 | 0.238 | 0.188 | 1 | 24 |
| luc7l3 | 0.89774 | -0.15471476 | 0.19 | 0.25 | 1 | 24 |
| nsmce2 | 0.89775 | -0.18615944 | 0.095 | 0.125 | 1 | 24 |
| usp1 | 0.89811 | 0.176655325 | 0.143 | 0.125 | 1 | 24 |
| trmt112 | 0.89856 | -0.18719059 | 0.143 | 0.188 | 1 | 24 |
| srsf6b | 0.89917 | -0.17847102 | 0.143 | 0.188 | 1 | 24 |
| si:dkey-112e7.2 | 0.8997 | 0.176450317 | 0.143 | 0.125 | 1 | 24 |
| NDUFB1 | 0.90267 | 0.1072989 | 0.238 | 0.25 | 1 | 24 |
| LOC100330864 | 0.90274 | -0.18900082 | 0.095 | 0.125 | 1 | 24 |
| ppdpfa | 0.90325 | -0.10537279 | 0.143 | 0.125 | 1 | 24 |
| fkbp4 | 0.90327 | -0.1684431 | 0.143 | 0.188 | 1 | 24 |
| hnrnpabb | 0.90376 | 0.13439116 | 0.619 | 0.562 | 1 | 24 |
| celf2 | 0.90431 | -0.15521366 | 0.19 | 0.25 | 1 | 24 |
| eif4g2b | 0.90506 | 0.169956284 | 0.143 | 0.125 | 1 | 24 |
| npepl1 | 0.90591 | -0.18611815 | 0.095 | 0.125 | 1 | 24 |
| mob3a | 0.9068 | -0.18455088 | 0.095 | 0.125 | 1 | 24 |
| rbm8a | 0.90772 | -0.12484864 | 0.476 | 0.5 | 1 | 24 |
| CZQB01141835.1 | 0.90801 | 0.139191488 | 0.19 | 0.188 | 1 | 24 |
| mt-nd3 | 0.90819 | -0.18479763 | 0.095 | 0.125 | 1 | 24 |
| gadd45gip1 | 0.90898 | -0.16393318 | 0.143 | 0.188 | 1 | 24 |

| | | | | | | |
|---|---|---|---|---|---|---|
| dkc1 | 0.90899 | -0.12600816 | 0.19 | 0.25 | 1 | 24 |
| hnrnpk | 0.9094 | 0.172455255 | 0.286 | 0.25 | 1 | 24 |
| kmt5aa | 0.91017 | -0.1234031 | 0.19 | 0.25 | 1 | 24 |
| calm2a | 0.91031 | -0.14801176 | 0.19 | 0.25 | 1 | 24 |
| zgc:56676 | 0.91036 | -0.12605209 | 0.19 | 0.25 | 1 | 24 |
| tmem256 | 0.91078 | -0.1727561 | 0.095 | 0.125 | 1 | 24 |
| pdia3 | 0.91121 | -0.11868792 | 0.19 | 0.188 | 1 | 24 |
| eif3m | 0.91126 | 0.103553497 | 0.571 | 0.5 | 1 | 24 |
| elof1 | 0.91128 | 0.135378067 | 0.19 | 0.188 | 1 | 24 |
| atp5f1b | 0.91132 | -0.11048842 | 0.429 | 0.5 | 1 | 24 |
| CU929070.1 | 0.91144 | -0.17377881 | 0.095 | 0.125 | 1 | 24 |
| psmc4 | 0.91154 | 0.16918804 | 0.238 | 0.188 | 1 | 24 |
| si:ch211-251p5.5 | 0.91157 | -0.1788133 | 0.095 | 0.125 | 1 | 24 |
| zgc:110540 | 0.91252 | -0.13315654 | 0.238 | 0.25 | 1 | 24 |
| mych | 0.9128 | -0.18034738 | 0.143 | 0.188 | 1 | 24 |
| npm1a | 0.91343 | -0.1085198 | 0.429 | 0.438 | 1 | 24 |
| ebna1bp2 | 0.91479 | -0.15653205 | 0.143 | 0.188 | 1 | 24 |
| pde4ba | 0.91493 | -0.17361857 | 0.095 | 0.125 | 1 | 24 |
| rsf1b.1 | 0.91572 | -0.1735803 | 0.095 | 0.125 | 1 | 24 |
| eif5b | 0.91596 | 0.136238344 | 0.19 | 0.188 | 1 | 24 |
| sephs2 | 0.91741 | 0.160348724 | 0.143 | 0.125 | 1 | 24 |
| slc25a20 | 0.91743 | -0.1558461 | 0.095 | 0.125 | 1 | 24 |
| prps1b | 0.91753 | -0.164328 | 0.095 | 0.125 | 1 | 24 |
| vbp1 | 0.91808 | 0.13579671 | 0.19 | 0.188 | 1 | 24 |
| clic4 | 0.91847 | -0.16092468 | 0.143 | 0.188 | 1 | 24 |
| arf2a | 0.9193 | -0.13963472 | 0.143 | 0.188 | 1 | 24 |
| tpx2 | 0.92071 | -0.15286553 | 0.143 | 0.188 | 1 | 24 |
| si:dkey-67c22.2 | 0.92131 | 0.157072401 | 0.143 | 0.125 | 1 | 24 |
| cks1b | 0.92202 | -0.10599647 | 0.19 | 0.188 | 1 | 24 |
| hspd1 | 0.92239 | 0.126620502 | 0.19 | 0.188 | 1 | 24 |
| ptges3b | 0.92266 | 0.158302392 | 0.238 | 0.188 | 1 | 24 |
| slbp | 0.9227 | 0.155048579 | 0.286 | 0.25 | 1 | 24 |
| eif3ha | 0.92282 | 0.144166448 | 0.429 | 0.375 | 1 | 24 |
| srsf5b | 0.92486 | -0.15467909 | 0.095 | 0.125 | 1 | 24 |
| romo1 | 0.92549 | -0.11620756 | 0.476 | 0.5 | 1 | 24 |
| tmem107 | 0.92557 | 0.158190366 | 0.143 | 0.125 | 1 | 24 |
| atox1 | 0.92645 | -0.16182095 | 0.095 | 0.125 | 1 | 24 |
| g3bp1 | 0.92727 | -0.1404608 | 0.143 | 0.188 | 1 | 24 |
| hnrnpa1b | 0.92763 | -0.12365273 | 0.143 | 0.188 | 1 | 24 |
| pfdn1 | 0.92821 | 0.130797925 | 0.333 | 0.312 | 1 | 24 |
| ndufab1a | 0.9287 | 0.124333936 | 0.238 | 0.188 | 1 | 24 |
| pno1 | 0.92919 | 0.148177409 | 0.143 | 0.125 | 1 | 24 |
| hirip3 | 0.92936 | 0.148578915 | 0.143 | 0.125 | 1 | 24 |
| prdx6 | 0.92988 | -0.10616375 | 0.19 | 0.188 | 1 | 24 |
| si:ch211-217k17.7 | 0.93069 | -0.11281268 | 0.143 | 0.188 | 1 | 24 |
| syncripl | 0.93085 | -0.1448641 | 0.095 | 0.125 | 1 | 24 |
| capza1a | 0.93232 | 0.15237754 | 0.143 | 0.125 | 1 | 24 |
| golga7 | 0.93267 | 0.14939411 | 0.143 | 0.125 | 1 | 24 |

| | | | | | | |
|---|---|---|---|---|---|---|
| nono | 0.93318 | 0.149280787 | 0.143 | 0.125 | 1 | 24 |
| kpnb3 | 0.93319 | -0.11285374 | 0.143 | 0.188 | 1 | 24 |
| ddx5 | 0.93331 | 0.145088679 | 0.286 | 0.25 | 1 | 24 |
| rad21a | 0.93341 | 0.145294106 | 0.286 | 0.25 | 1 | 24 |
| si:ch211-39k3.2 | 0.93483 | -0.15419025 | 0.095 | 0.125 | 1 | 24 |
| ndufs7 | 0.93484 | -0.10337322 | 0.143 | 0.188 | 1 | 24 |
| STMP1 | 0.93494 | 0.138497875 | 0.286 | 0.25 | 1 | 24 |
| tyms | 0.93681 | -0.1032225 | 0.143 | 0.188 | 1 | 24 |
| myl12.2 | 0.93682 | -0.10879219 | 0.143 | 0.188 | 1 | 24 |
| XLOC-010899 | 0.94027 | -0.12597922 | 0.095 | 0.125 | 1 | 24 |
| rsl24d1 | 0.94043 | -0.1153251 | 0.381 | 0.438 | 1 | 24 |
| klf2a | 0.94047 | -0.12696077 | 0.095 | 0.125 | 1 | 24 |
| nck1b | 0.94173 | -0.129092 | 0.095 | 0.125 | 1 | 24 |
| CABZ01115113.1 | 0.94186 | 0.113590639 | 0.19 | 0.188 | 1 | 24 |
| pdcd4a | 0.94355 | -0.12374506 | 0.095 | 0.125 | 1 | 24 |
| mrpl30 | 0.94471 | 0.134328859 | 0.143 | 0.125 | 1 | 24 |
| ddost | 0.94489 | -0.12005734 | 0.095 | 0.125 | 1 | 24 |
| top1l | 0.94536 | 0.103041168 | 0.19 | 0.188 | 1 | 24 |
| ube2d3 | 0.94673 | 0.133363032 | 0.143 | 0.125 | 1 | 24 |
| mtch2 | 0.94724 | -0.12499071 | 0.095 | 0.125 | 1 | 24 |
| zgc:65873 | 0.94809 | -0.12406419 | 0.095 | 0.125 | 1 | 24 |
| rnaseh2c | 0.94847 | -0.11672864 | 0.095 | 0.125 | 1 | 24 |
| hsdl2 | 0.94864 | -0.12725809 | 0.095 | 0.125 | 1 | 24 |
| ndufs2 | 0.94881 | 0.125241652 | 0.143 | 0.125 | 1 | 24 |
| mycb | 0.95002 | -0.11771998 | 0.095 | 0.125 | 1 | 24 |
| med20 | 0.95012 | -0.11112936 | 0.095 | 0.125 | 1 | 24 |
| LOC101884451 | 0.95047 | -0.11384842 | 0.095 | 0.125 | 1 | 24 |
| krtcap2 | 0.95124 | 0.125574681 | 0.143 | 0.125 | 1 | 24 |
| ubr7 | 0.95139 | 0.129385108 | 0.143 | 0.125 | 1 | 24 |
| prim2 | 0.95175 | -0.12640616 | 0.095 | 0.125 | 1 | 24 |
| cxcr4b | 0.95377 | -0.11806772 | 0.095 | 0.125 | 1 | 24 |
| brd7 | 0.95406 | -0.10205327 | 0.095 | 0.125 | 1 | 24 |
| rhoac | 0.95482 | 0.12410773 | 0.143 | 0.125 | 1 | 24 |
| dlgap5 | 0.95567 | 0.121634623 | 0.143 | 0.125 | 1 | 24 |
| morf4l1 | 0.9562 | -0.10171465 | 0.238 | 0.25 | 1 | 24 |
| aldoaa | 0.95673 | -0.10472359 | 0.333 | 0.375 | 1 | 24 |
| capzb | 0.95727 | -0.10506171 | 0.238 | 0.25 | 1 | 24 |
| xbp1 | 0.9579 | 0.116919727 | 0.143 | 0.125 | 1 | 24 |
| sat1a.2 | 0.95875 | -0.10576602 | 0.095 | 0.125 | 1 | 24 |
| rnaseh2a | 0.95885 | 0.119811598 | 0.143 | 0.125 | 1 | 24 |
| eif4ea | 0.95935 | 0.116605209 | 0.143 | 0.125 | 1 | 24 |
| mrpl51 | 0.9636 | 0.111901903 | 0.143 | 0.125 | 1 | 24 |
| churc1 | 0.96502 | 0.104274723 | 0.143 | 0.125 | 1 | 24 |
| srsf3a | 0.96571 | 0.104704762 | 0.143 | 0.125 | 1 | 24 |
| myca | 0.96603 | 0.100625991 | 0.143 | 0.125 | 1 | 24 |
| atp1a1a.1 | 0.96834 | 0.105441359 | 0.143 | 0.125 | 1 | 24 |
| reep5 | 0.97157 | 0.100772988 | 0.143 | 0.125 | 1 | 24 |
| rps3 | 0.00253 | 1.573056498 | 0.767 | 0 | 1 | 25 |

| | | | | | | |
|---|---|---|---|---|---|---|
| ciao2a | 0.00417 | -1.16579666 | 0 | 0.5 | 1 | 25 |
| copg2 | 0.00417 | -1.16413485 | 0 | 0.5 | 1 | 25 |
| phf6 | 0.00417 | -1.16172226 | 0 | 0.5 | 1 | 25 |
| cndp2 | 0.00417 | -0.92790772 | 0 | 0.5 | 1 | 25 |
| csde1 | 0.00417 | -0.91849476 | 0 | 0.5 | 1 | 25 |
| csrnp1b | 0.00417 | -0.92842324 | 0 | 0.5 | 1 | 25 |
| fbln2 | 0.00417 | -0.8508485 | 0 | 0.5 | 1 | 25 |
| srsf11 | 0.00417 | -0.89767125 | 0 | 0.5 | 1 | 25 |
| tpst1l | 0.00417 | -0.97287344 | 0 | 0.5 | 1 | 25 |
| si:ch1073-325m22.2 | 0.00444 | -1.21514354 | 0.033 | 0.75 | 1 | 25 |
| zgc:101715 | 0.00452 | -1.39240152 | 0.033 | 0.75 | 1 | 25 |
| cfl1l | 0.00588 | 1.495093256 | 0.7 | 0 | 1 | 25 |
| rpl10a | 0.00588 | 1.444307865 | 0.7 | 0 | 1 | 25 |
| rpl14 | 0.00849 | 1.392994791 | 0.667 | 0 | 1 | 25 |
| rps5 | 0.01194 | 1.473201752 | 0.633 | 0 | 1 | 25 |
| CR354540.1 | 0.01204 | -1.24016711 | 0.067 | 0.75 | 1 | 25 |
| lasp1 | 0.01641 | 1.301897371 | 0.6 | 0 | 1 | 25 |
| rpl38 | 0.01641 | 1.316228799 | 0.6 | 0 | 1 | 25 |
| rps14 | 0.01641 | 1.348914116 | 0.6 | 0 | 1 | 25 |
| LOC101883999 | 0.01741 | -1.11340858 | 0.033 | 0.5 | 1 | 25 |
| desi1b | 0.01741 | -1.08483896 | 0.033 | 0.5 | 1 | 25 |
| myorg | 0.01741 | -1.1097167 | 0.033 | 0.5 | 1 | 25 |
| eif4g1a | 0.01741 | -0.82588706 | 0.033 | 0.5 | 1 | 25 |
| bckdk | 0.01741 | -0.78740919 | 0.033 | 0.5 | 1 | 25 |
| cmklr1 | 0.01741 | -0.84470439 | 0.033 | 0.5 | 1 | 25 |
| puf60b | 0.01741 | -0.92062099 | 0.033 | 0.5 | 1 | 25 |
| tmem176l.3a | 0.01741 | -0.91801633 | 0.033 | 0.5 | 1 | 25 |
| tspan37 | 0.01741 | -1.03825032 | 0.033 | 0.5 | 1 | 25 |
| CU571315.1 | 0.01741 | -0.86901015 | 0.033 | 0.5 | 1 | 25 |
| faf2 | 0.01741 | -1.06797897 | 0.033 | 0.5 | 1 | 25 |
| rab22a | 0.01741 | -0.96387825 | 0.033 | 0.5 | 1 | 25 |
| ncoa5 | 0.01741 | -1.08023196 | 0.033 | 0.5 | 1 | 25 |
| rpl17 | 0.0221 | 1.151479357 | 0.567 | 0 | 1 | 25 |
| rpl26 | 0.0221 | 1.546985928 | 0.567 | 0 | 1 | 25 |
| rpl7a | 0.0221 | 1.4518927 | 0.567 | 0 | 1 | 25 |
| rplp0 | 0.0221 | 1.324845123 | 0.567 | 0 | 1 | 25 |
| rps20 | 0.0221 | 1.492137927 | 0.567 | 0 | 1 | 25 |
| mt-nd2 | 0.02224 | -0.92597239 | 0.3 | 1 | 1 | 25 |
| abracl | 0.02926 | 1.122953287 | 0.533 | 0 | 1 | 25 |
| rps13 | 0.03817 | 1.195083623 | 0.5 | 0 | 1 | 25 |
| hnrnpa0l | 0.03817 | 1.049629956 | 0.5 | 0 | 1 | 25 |
| icn | 0.03817 | 1.456689592 | 0.5 | 0 | 1 | 25 |
| rpl12 | 0.03817 | 1.184438507 | 0.5 | 0 | 1 | 25 |
| rps10 | 0.03817 | 1.108170172 | 0.5 | 0 | 1 | 25 |
| cox6a2 | 0.04159 | -1.08040148 | 0.133 | 0.75 | 1 | 25 |
| rps16 | 0.04636 | 0.576239556 | 0.667 | 1 | 1 | 25 |
| rpl5a | 0.04915 | 0.942347721 | 0.467 | 0 | 1 | 25 |
| rpl10 | 0.05362 | 1.47214828 | 0.8 | 0.25 | 1 | 25 |

| | | | | | | |
|---|---|---|---|---|---|---|
| gpia | 0.06257 | 1.006684949 | 0.433 | 0 | 1 | 25 |
| gstp1 | 0.06257 | 0.9208919 | 0.433 | 0 | 1 | 25 |
| si:dkey-151g10.6 | 0.06257 | 1.386548383 | 0.433 | 0 | 1 | 25 |
| LOC101883382 | 0.06566 | -0.72916768 | 0 | 0.25 | 1 | 25 |
| alg2 | 0.06566 | -0.71850252 | 0 | 0.25 | 1 | 25 |
| arcn1b | 0.06566 | -0.71993819 | 0 | 0.25 | 1 | 25 |
| atg3 | 0.06566 | -0.72479302 | 0 | 0.25 | 1 | 25 |
| ctsc | 0.06566 | -0.72805239 | 0 | 0.25 | 1 | 25 |
| dtx4b | 0.06566 | -0.71668049 | 0 | 0.25 | 1 | 25 |
| fam3c | 0.06566 | -0.72324199 | 0 | 0.25 | 1 | 25 |
| gnpat | 0.06566 | -0.72556869 | 0 | 0.25 | 1 | 25 |
| hsd17b4 | 0.06566 | -0.72321067 | 0 | 0.25 | 1 | 25 |
| itprid2 | 0.06566 | -0.71991992 | 0 | 0.25 | 1 | 25 |
| klc1a | 0.06566 | -0.72957726 | 0 | 0.25 | 1 | 25 |
| lrmda | 0.06566 | -0.72957726 | 0 | 0.25 | 1 | 25 |
| map4k4 | 0.06566 | -0.72635036 | 0 | 0.25 | 1 | 25 |
| map7d3 | 0.06566 | -0.72670508 | 0 | 0.25 | 1 | 25 |
| mapk9 | 0.06566 | -0.72439309 | 0 | 0.25 | 1 | 25 |
| osbpl2a | 0.06566 | -0.72710799 | 0 | 0.25 | 1 | 25 |
| si:ch211-194k22.8 | 0.06566 | -0.72289244 | 0 | 0.25 | 1 | 25 |
| si:dkey-14d8.1 | 0.06566 | -0.41963395 | 0 | 0.25 | 1 | 25 |
| slc25a51b | 0.06566 | -0.70151104 | 0 | 0.25 | 1 | 25 |
| smg9 | 0.06566 | -0.56187027 | 0 | 0.25 | 1 | 25 |
| snapc3 | 0.06566 | -0.71923578 | 0 | 0.25 | 1 | 25 |
| spata13 | 0.06566 | -0.70139125 | 0 | 0.25 | 1 | 25 |
| ssh2b | 0.06566 | -0.72877915 | 0 | 0.25 | 1 | 25 |
| tnfrsf1b | 0.06566 | -0.7293303 | 0 | 0.25 | 1 | 25 |
| top1l | 0.06566 | -0.72329845 | 0 | 0.25 | 1 | 25 |
| top3b | 0.06566 | -0.70179838 | 0 | 0.25 | 1 | 25 |
| tspo | 0.06566 | -0.12868034 | 0 | 0.25 | 1 | 25 |
| zgc:56095 | 0.06566 | -0.7128034 | 0 | 0.25 | 1 | 25 |
| zmat5 | 0.06566 | -0.69764267 | 0 | 0.25 | 1 | 25 |
| znf1010 | 0.06566 | -0.72095693 | 0 | 0.25 | 1 | 25 |
| LOC108190761 | 0.06566 | -0.65742138 | 0 | 0.25 | 1 | 25 |
| XLOC-018356 | 0.06566 | -0.93418817 | 0 | 0.25 | 1 | 25 |
| acox1 | 0.06566 | -0.65898175 | 0 | 0.25 | 1 | 25 |
| alad | 0.06566 | -0.64847655 | 0 | 0.25 | 1 | 25 |
| btg2 | 0.06566 | -0.65434492 | 0 | 0.25 | 1 | 25 |
| casp7 | 0.06566 | -0.65012589 | 0 | 0.25 | 1 | 25 |
| ccnl1b | 0.06566 | -0.6516034 | 0 | 0.25 | 1 | 25 |
| cdc34a | 0.06566 | -0.65380732 | 0 | 0.25 | 1 | 25 |
| cfhl2 | 0.06566 | -0.65853868 | 0 | 0.25 | 1 | 25 |
| ctcf | 0.06566 | -0.64978773 | 0 | 0.25 | 1 | 25 |
| dennd5a | 0.06566 | -0.63934641 | 0 | 0.25 | 1 | 25 |
| dnajc8 | 0.06566 | -0.64445733 | 0 | 0.25 | 1 | 25 |
| eml3 | 0.06566 | -0.65974977 | 0 | 0.25 | 1 | 25 |
| exoc3 | 0.06566 | -0.65354482 | 0 | 0.25 | 1 | 25 |
| foxg1c | 0.06566 | -0.64085282 | 0 | 0.25 | 1 | 25 |

| | | | | | | |
|---|---|---|---|---|---|---|
| ftr50 | 0.06566 | -0.65848158 | 0 | 0.25 | 1 | 25 |
| habp2 | 0.06566 | -0.66140293 | 0 | 0.25 | 1 | 25 |
| insig1 | 0.06566 | -0.80001606 | 0 | 0.25 | 1 | 25 |
| itpka | 0.06566 | -0.65564778 | 0 | 0.25 | 1 | 25 |
| klf11a | 0.06566 | -0.66015536 | 0 | 0.25 | 1 | 25 |
| lyst | 0.06566 | -0.65922484 | 0 | 0.25 | 1 | 25 |
| mir22a | 0.06566 | -0.65788515 | 0 | 0.25 | 1 | 25 |
| moxd1 | 0.06566 | -0.66211141 | 0 | 0.25 | 1 | 25 |
| mrpl9 | 0.06566 | -0.65086301 | 0 | 0.25 | 1 | 25 |
| nav2b | 0.06566 | -0.44683854 | 0 | 0.25 | 1 | 25 |
| nfkb2 | 0.06566 | -0.65343737 | 0 | 0.25 | 1 | 25 |
| pnpla6 | 0.06566 | -0.65352359 | 0 | 0.25 | 1 | 25 |
| poldip2 | 0.06566 | -0.65220138 | 0 | 0.25 | 1 | 25 |
| polr2c | 0.06566 | -0.65315553 | 0 | 0.25 | 1 | 25 |
| prdm1a | 0.06566 | -0.6621957 | 0 | 0.25 | 1 | 25 |
| prkcba | 0.06566 | -0.66181903 | 0 | 0.25 | 1 | 25 |
| rad1 | 0.06566 | -0.64912763 | 0 | 0.25 | 1 | 25 |
| shc1 | 0.06566 | -0.65901412 | 0 | 0.25 | 1 | 25 |
| si:ch1073-127d16.1 | 0.06566 | -0.64982035 | 0 | 0.25 | 1 | 25 |
| si:ch211-108d22.2 | 0.06566 | -0.65586051 | 0 | 0.25 | 1 | 25 |
| si:ch211-165d12.4 | 0.06566 | -0.66309396 | 0 | 0.25 | 1 | 25 |
| si:ch211-215p11.1 | 0.06566 | -0.66309396 | 0 | 0.25 | 1 | 25 |
| si:ch73-223f5.1 | 0.06566 | -0.65969141 | 0 | 0.25 | 1 | 25 |
| si:dkey-23k10.5 | 0.06566 | -0.65765611 | 0 | 0.25 | 1 | 25 |
| sik1 | 0.06566 | -0.65972496 | 0 | 0.25 | 1 | 25 |
| slc2a3b | 0.06566 | -0.66277163 | 0 | 0.25 | 1 | 25 |
| smap2 | 0.06566 | -0.66209114 | 0 | 0.25 | 1 | 25 |
| sox4b | 0.06566 | -0.66166472 | 0 | 0.25 | 1 | 25 |
| ssbp3a | 0.06566 | -0.65778277 | 0 | 0.25 | 1 | 25 |
| strip1 | 0.06566 | -0.65197346 | 0 | 0.25 | 1 | 25 |
| szrd1 | 0.06566 | -0.6545786 | 0 | 0.25 | 1 | 25 |
| tob1a | 0.06566 | -0.7957536 | 0 | 0.25 | 1 | 25 |
| unc93b1 | 0.06566 | -0.66105414 | 0 | 0.25 | 1 | 25 |
| uxs1 | 0.06566 | -0.63378354 | 0 | 0.25 | 1 | 25 |
| ANKRD50 | 0.06566 | -0.64694062 | 0 | 0.25 | 1 | 25 |
| CABZ01088428.1 | 0.06566 | -0.67127947 | 0 | 0.25 | 1 | 25 |
| LOC100150882 | 0.06566 | -0.6693241 | 0 | 0.25 | 1 | 25 |
| atg5 | 0.06566 | -0.664685 | 0 | 0.25 | 1 | 25 |
| auh | 0.06566 | -0.51072348 | 0 | 0.25 | 1 | 25 |
| ccdc130 | 0.06566 | -0.65620675 | 0 | 0.25 | 1 | 25 |
| crtac1a | 0.06566 | -0.61032868 | 0 | 0.25 | 1 | 25 |
| dcun1d3 | 0.06566 | -0.66771979 | 0 | 0.25 | 1 | 25 |
| ero1a | 0.06566 | -0.67011234 | 0 | 0.25 | 1 | 25 |
| fbxo11a | 0.06566 | -0.66499142 | 0 | 0.25 | 1 | 25 |
| ggcta | 0.06566 | -0.66427249 | 0 | 0.25 | 1 | 25 |
| gpatch3 | 0.06566 | -0.66700088 | 0 | 0.25 | 1 | 25 |
| hck | 0.06566 | -0.67061445 | 0 | 0.25 | 1 | 25 |
| hhla2a.2 | 0.06566 | -0.67237999 | 0 | 0.25 | 1 | 25 |

| | | | | | | |
|---|---|---|---|---|---|---|
| kdm2aa | 0.06566 | -0.66593541 | 0 | 0.25 | 1 | 25 |
| klhl20 | 0.06566 | -0.66607398 | 0 | 0.25 | 1 | 25 |
| me1 | 0.06566 | -0.67208917 | 0 | 0.25 | 1 | 25 |
| metrnla | 0.06566 | -0.66521837 | 0 | 0.25 | 1 | 25 |
| mlec | 0.06566 | -0.66787387 | 0 | 0.25 | 1 | 25 |
| mob1ba | 0.06566 | -0.62934391 | 0 | 0.25 | 1 | 25 |
| mov10b.1 | 0.06566 | -0.6616052 | 0 | 0.25 | 1 | 25 |
| mpzl2b | 0.06566 | -0.65748865 | 0 | 0.25 | 1 | 25 |
| nedd9 | 0.06566 | -0.66667891 | 0 | 0.25 | 1 | 25 |
| net1 | 0.06566 | -0.6613853 | 0 | 0.25 | 1 | 25 |
| nucb1 | 0.06566 | -0.66854782 | 0 | 0.25 | 1 | 25 |
| olfml2ba | 0.06566 | -0.66467098 | 0 | 0.25 | 1 | 25 |
| prkdc | 0.06566 | -0.66334591 | 0 | 0.25 | 1 | 25 |
| pusl1 | 0.06566 | -0.36462388 | 0 | 0.25 | 1 | 25 |
| r3hdm1 | 0.06566 | -0.66660132 | 0 | 0.25 | 1 | 25 |
| rab1ba | 0.06566 | -0.66555427 | 0 | 0.25 | 1 | 25 |
| ric8b | 0.06566 | -0.66119862 | 0 | 0.25 | 1 | 25 |
| rnf220a | 0.06566 | -0.66661919 | 0 | 0.25 | 1 | 25 |
| sb:cb81 | 0.06566 | -0.6689998 | 0 | 0.25 | 1 | 25 |
| scp2b | 0.06566 | -0.66414277 | 0 | 0.25 | 1 | 25 |
| scrn3 | 0.06566 | -0.63966934 | 0 | 0.25 | 1 | 25 |
| si:cabz01102082.1 | 0.06566 | -0.67067583 | 0 | 0.25 | 1 | 25 |
| si:dkey-68o6.6 | 0.06566 | -0.67107359 | 0 | 0.25 | 1 | 25 |
| slc39a11 | 0.06566 | -0.66960411 | 0 | 0.25 | 1 | 25 |
| vps13a | 0.06566 | -0.67057326 | 0 | 0.25 | 1 | 25 |
| ywhag2 | 0.06566 | -0.66541944 | 0 | 0.25 | 1 | 25 |
| zgc:113411 | 0.06566 | -0.66845744 | 0 | 0.25 | 1 | 25 |
| BX537358.1 | 0.06566 | -0.34936654 | 0 | 0.25 | 1 | 25 |
| CABZ01005379.1 | 0.06566 | -0.39206429 | 0 | 0.25 | 1 | 25 |
| CABZ01055347.1 | 0.06566 | -0.40676798 | 0 | 0.25 | 1 | 25 |
| CABZ01087514.1 | 0.06566 | -0.41260013 | 0 | 0.25 | 1 | 25 |
| GNAZ | 0.06566 | -0.40896812 | 0 | 0.25 | 1 | 25 |
| NHLRC3 | 0.06566 | -0.35269028 | 0 | 0.25 | 1 | 25 |
| XLOC-022621 | 0.06566 | -0.40042883 | 0 | 0.25 | 1 | 25 |
| abcg2c | 0.06566 | -0.37471598 | 0 | 0.25 | 1 | 25 |
| abtb1 | 0.06566 | -0.38872704 | 0 | 0.25 | 1 | 25 |
| adsl | 0.06566 | -0.31073913 | 0 | 0.25 | 1 | 25 |
| amacr | 0.06566 | -0.32731457 | 0 | 0.25 | 1 | 25 |
| arl6ip5a | 0.06566 | -0.41285072 | 0 | 0.25 | 1 | 25 |
| ash1l | 0.06566 | -0.39654451 | 0 | 0.25 | 1 | 25 |
| atxn7l3 | 0.06566 | -0.40499279 | 0 | 0.25 | 1 | 25 |
| bloc1s4 | 0.06566 | -0.35498658 | 0 | 0.25 | 1 | 25 |
| cdc6 | 0.06566 | -0.37703738 | 0 | 0.25 | 1 | 25 |
| cratb | 0.06566 | -0.51255469 | 0 | 0.25 | 1 | 25 |
| det1 | 0.06566 | -0.36871618 | 0 | 0.25 | 1 | 25 |
| dhps | 0.06566 | -0.24122311 | 0 | 0.25 | 1 | 25 |
| dlat | 0.06566 | -0.39586856 | 0 | 0.25 | 1 | 25 |
| dyrk1b | 0.06566 | -0.37147584 | 0 | 0.25 | 1 | 25 |

| | | | | | | |
|---|---|---|---|---|---|---|
| fahd2a | 0.06566 | -0.3852781 | 0 | 0.25 | 1 | 25 |
| fgd4a | 0.06566 | -0.41095119 | 0 | 0.25 | 1 | 25 |
| galk2 | 0.06566 | -0.41169977 | 0 | 0.25 | 1 | 25 |
| galnt1 | 0.06566 | -0.38019289 | 0 | 0.25 | 1 | 25 |
| gpr137ba | 0.06566 | -0.41542801 | 0 | 0.25 | 1 | 25 |
| gpr174 | 0.06566 | -0.41497815 | 0 | 0.25 | 1 | 25 |
| hacd2 | 0.06566 | -0.40692125 | 0 | 0.25 | 1 | 25 |
| hgh1 | 0.06566 | -0.38167627 | 0 | 0.25 | 1 | 25 |
| hsp90b1 | 0.06566 | -0.37616813 | 0 | 0.25 | 1 | 25 |
| larp1b | 0.06566 | -0.39042463 | 0 | 0.25 | 1 | 25 |
| map3k1 | 0.06566 | -0.38712201 | 0 | 0.25 | 1 | 25 |
| mcoln1a | 0.06566 | -0.40160476 | 0 | 0.25 | 1 | 25 |
| mta3 | 0.06566 | -0.37779041 | 0 | 0.25 | 1 | 25 |
| naxd | 0.06566 | -0.40028451 | 0 | 0.25 | 1 | 25 |
| naxe | 0.06566 | -0.34374841 | 0 | 0.25 | 1 | 25 |
| ndnl2 | 0.06566 | -0.3104069 | 0 | 0.25 | 1 | 25 |
| ndufa11 | 0.06566 | -0.37081623 | 0 | 0.25 | 1 | 25 |
| nr2f6a | 0.06566 | -0.40864181 | 0 | 0.25 | 1 | 25 |
| phip | 0.06566 | -0.3907649 | 0 | 0.25 | 1 | 25 |
| pms1 | 0.06566 | -0.39056999 | 0 | 0.25 | 1 | 25 |
| pofut2 | 0.06566 | -0.32718674 | 0 | 0.25 | 1 | 25 |
| polr2i | 0.06566 | -0.39629921 | 0 | 0.25 | 1 | 25 |
| pop7 | 0.06566 | -0.35915246 | 0 | 0.25 | 1 | 25 |
| ppp3ccb | 0.06566 | -0.54732072 | 0 | 0.25 | 1 | 25 |
| pptc7a | 0.06566 | -0.40293187 | 0 | 0.25 | 1 | 25 |
| ptpn9a | 0.06566 | -0.41062321 | 0 | 0.25 | 1 | 25 |
| ptrh2 | 0.06566 | -0.37886765 | 0 | 0.25 | 1 | 25 |
| ralgps1 | 0.06566 | -0.41805488 | 0 | 0.25 | 1 | 25 |
| rhbdl2 | 0.06566 | -0.41133875 | 0 | 0.25 | 1 | 25 |
| ripor2 | 0.06566 | -0.38949291 | 0 | 0.25 | 1 | 25 |
| sdf2l1 | 0.06566 | -0.40167752 | 0 | 0.25 | 1 | 25 |
| si:ch211-286o17.1 | 0.06566 | -0.36141378 | 0 | 0.25 | 1 | 25 |
| si:ch73-170d6.3 | 0.06566 | -0.41480919 | 0 | 0.25 | 1 | 25 |
| si:dkey-19e4.5 | 0.06566 | -0.37903 | 0 | 0.25 | 1 | 25 |
| si:dkey-1j5.4 | 0.06566 | -0.41805488 | 0 | 0.25 | 1 | 25 |
| si:dkey-84k17.2 | 0.06566 | -0.32556659 | 0 | 0.25 | 1 | 25 |
| sinhcaf | 0.06566 | -0.38195987 | 0 | 0.25 | 1 | 25 |
| sorl1 | 0.06566 | -0.40651551 | 0 | 0.25 | 1 | 25 |
| srm | 0.06566 | -0.39925164 | 0 | 0.25 | 1 | 25 |
| sumo3b | 0.06566 | -0.35456627 | 0 | 0.25 | 1 | 25 |
| tapt1b | 0.06566 | -0.41182398 | 0 | 0.25 | 1 | 25 |
| tbcd | 0.06566 | -0.40532805 | 0 | 0.25 | 1 | 25 |
| tgfbr2a | 0.06566 | -0.55815721 | 0 | 0.25 | 1 | 25 |
| tle2c | 0.06566 | -0.40338989 | 0 | 0.25 | 1 | 25 |
| tlr22 | 0.06566 | -0.41386938 | 0 | 0.25 | 1 | 25 |
| tmem147 | 0.06566 | -0.37550642 | 0 | 0.25 | 1 | 25 |
| tmppe | 0.06566 | -0.36655683 | 0 | 0.25 | 1 | 25 |
| ube2q1 | 0.06566 | -0.38256122 | 0 | 0.25 | 1 | 25 |

| | | | | | | |
|---|---|---|---|---|---|---|
| ubxn7 | 0.06566 | -0.35566062 | 0 | 0.25 | 1 | 25 |
| uggt1 | 0.06566 | -0.40870445 | 0 | 0.25 | 1 | 25 |
| vps4b | 0.06566 | -0.39318442 | 0 | 0.25 | 1 | 25 |
| itgae.2 | 0.06578 | -1.01809467 | 0.167 | 0.75 | 1 | 25 |
| lcp1 | 0.06743 | -0.93540001 | 0.167 | 0.75 | 1 | 25 |
| aldh9a1a.1 | 0.07888 | 0.788399562 | 0.4 | 0 | 1 | 25 |
| f11r.1 | 0.07888 | 0.93999038 | 0.4 | 0 | 1 | 25 |
| hnrnpaba | 0.09858 | 0.704402392 | 0.367 | 0 | 1 | 25 |
| seta | 0.09858 | 0.747247267 | 0.367 | 0 | 1 | 25 |
| si:ch73-234b20.5 | 0.09858 | 0.783010305 | 0.367 | 0 | 1 | 25 |
| ncl | 0.09858 | 0.832103207 | 0.367 | 0 | 1 | 25 |
| nrros | 0.09858 | 0.887859937 | 0.367 | 0 | 1 | 25 |
| prdx5 | 0.09858 | 0.912153814 | 0.367 | 0 | 1 | 25 |
| rplp1 | 0.09858 | 1.22202952 | 0.367 | 0 | 1 | 25 |
| selenof | 0.09858 | 0.753278429 | 0.367 | 0 | 1 | 25 |
| tcnbb | 0.09858 | 1.204821069 | 0.367 | 0 | 1 | 25 |
| actn4 | 0.09858 | 0.821329739 | 0.367 | 0 | 1 | 25 |
| ak2 | 0.09858 | 0.807210978 | 0.367 | 0 | 1 | 25 |
| arl6ip1 | 0.09858 | 0.83147604 | 0.367 | 0 | 1 | 25 |
| lxn | 0.11462 | -0.55102653 | 0.5 | 1 | 1 | 25 |
| cnbpb | 0.11491 | -0.9061826 | 0.067 | 0.5 | 1 | 25 |
| rbm4.3 | 0.11614 | -0.85000778 | 0.067 | 0.5 | 1 | 25 |
| zbtb8os | 0.1179 | -0.85629061 | 0.067 | 0.5 | 1 | 25 |
| ctsa | 0.11814 | -0.68208388 | 0.067 | 0.5 | 1 | 25 |
| pik3r1 | 0.1198 | -0.79761698 | 0.067 | 0.5 | 1 | 25 |
| lpcat3 | 0.12169 | -0.80133198 | 0.067 | 0.5 | 1 | 25 |
| dnaja2a | 0.1223 | 0.715036563 | 0.333 | 0 | 1 | 25 |
| gsr | 0.1223 | 0.685570218 | 0.333 | 0 | 1 | 25 |
| pxna | 0.1223 | 0.731184137 | 0.333 | 0 | 1 | 25 |
| rab44 | 0.1223 | 0.874606416 | 0.333 | 0 | 1 | 25 |
| si:ch211-260e23.8 | 0.1223 | 0.766562862 | 0.333 | 0 | 1 | 25 |
| ywhaz | 0.1223 | 0.636917205 | 0.333 | 0 | 1 | 25 |
| cited4b | 0.1223 | 0.75137264 | 0.333 | 0 | 1 | 25 |
| ran | 0.1223 | 0.782425124 | 0.333 | 0 | 1 | 25 |
| amd1 | 0.15076 | 0.748459398 | 0.3 | 0 | 1 | 25 |
| atp5f1e | 0.15076 | 0.555161487 | 0.3 | 0 | 1 | 25 |
| rap1b | 0.15076 | 0.685419958 | 0.3 | 0 | 1 | 25 |
| RASSF5 | 0.15076 | 0.699939634 | 0.3 | 0 | 1 | 25 |
| anp32a | 0.15076 | 0.658060023 | 0.3 | 0 | 1 | 25 |
| atf3 | 0.15076 | 0.772691784 | 0.3 | 0 | 1 | 25 |
| cct5 | 0.15076 | 0.697121533 | 0.3 | 0 | 1 | 25 |
| cybc1 | 0.15076 | 0.631722586 | 0.3 | 0 | 1 | 25 |
| eif2s2 | 0.15076 | 0.621649929 | 0.3 | 0 | 1 | 25 |
| eif2s3 | 0.15076 | 0.616695773 | 0.3 | 0 | 1 | 25 |
| eif3f | 0.15076 | 0.709327984 | 0.3 | 0 | 1 | 25 |
| eif3i | 0.15076 | 0.569269292 | 0.3 | 0 | 1 | 25 |
| fkbp1aa | 0.15076 | 0.745565097 | 0.3 | 0 | 1 | 25 |
| fmo5 | 0.15076 | 0.696283961 | 0.3 | 0 | 1 | 25 |

| | | | | | | |
|---|---|---|---|---|---|---|
| hmga1a | 0.15076 | 0.544050683 | 0.3 | 0 | 1 | 25 |
| mob1a | 0.15076 | 0.636999707 | 0.3 | 0 | 1 | 25 |
| ndfip1 | 0.15076 | 0.68906391 | 0.3 | 0 | 1 | 25 |
| pak2a | 0.15076 | 0.629236579 | 0.3 | 0 | 1 | 25 |
| ponzr1 | 0.15076 | 0.596061628 | 0.3 | 0 | 1 | 25 |
| rhoab | 0.15076 | 0.638530735 | 0.3 | 0 | 1 | 25 |
| si:ch211-196l7.4 | 0.15076 | 0.631252913 | 0.3 | 0 | 1 | 25 |
| skap2 | 0.15076 | 0.740332376 | 0.3 | 0 | 1 | 25 |
| spcs1 | 0.15076 | 0.623914981 | 0.3 | 0 | 1 | 25 |
| vaspa | 0.15076 | 0.612165983 | 0.3 | 0 | 1 | 25 |
| ywhabb | 0.15076 | 0.615035181 | 0.3 | 0 | 1 | 25 |
| rps29 | 0.15934 | 1.300880149 | 0.667 | 0.25 | 1 | 25 |
| NC-002333.4 | 0.16334 | -0.71662168 | 0.567 | 1 | 1 | 25 |
| mt-cyb | 0.17649 | -0.60632364 | 0.567 | 1 | 1 | 25 |
| ybx1 | 0.18185 | 0.815589727 | 0.733 | 0.25 | 1 | 25 |
| CABZ01110881.1 | 0.18301 | -0.52832509 | 0.033 | 0.25 | 1 | 25 |
| dusp6 | 0.18301 | -0.52002448 | 0.033 | 0.25 | 1 | 25 |
| itm2cb | 0.18301 | -0.50719925 | 0.033 | 0.25 | 1 | 25 |
| mfng | 0.18301 | -0.6780314 | 0.033 | 0.25 | 1 | 25 |
| samsn1a | 0.18301 | -0.53782497 | 0.033 | 0.25 | 1 | 25 |
| FP074902.2 | 0.18301 | -0.59910833 | 0.033 | 0.25 | 1 | 25 |
| XLOC-014477 | 0.18301 | -0.54213176 | 0.033 | 0.25 | 1 | 25 |
| XLOC-042595 | 0.18301 | -0.33400836 | 0.033 | 0.25 | 1 | 25 |
| aktip | 0.18301 | -0.5976005 | 0.033 | 0.25 | 1 | 25 |
| ano10a | 0.18301 | -0.54437514 | 0.033 | 0.25 | 1 | 25 |
| ap5m1 | 0.18301 | -0.69500985 | 0.033 | 0.25 | 1 | 25 |
| babam1 | 0.18301 | -0.51297997 | 0.033 | 0.25 | 1 | 25 |
| c6ast1 | 0.18301 | -0.33564005 | 0.033 | 0.25 | 1 | 25 |
| camkk1a | 0.18301 | -0.62265363 | 0.033 | 0.25 | 1 | 25 |
| cbx1a | 0.18301 | -0.63362659 | 0.033 | 0.25 | 1 | 25 |
| ccdc12 | 0.18301 | -0.62687613 | 0.033 | 0.25 | 1 | 25 |
| ccndbp1 | 0.18301 | -0.50969952 | 0.033 | 0.25 | 1 | 25 |
| chd4a | 0.18301 | -0.58038166 | 0.033 | 0.25 | 1 | 25 |
| commd9 | 0.18301 | -0.62094429 | 0.033 | 0.25 | 1 | 25 |
| csnk1da | 0.18301 | -0.62357812 | 0.033 | 0.25 | 1 | 25 |
| cstf1 | 0.18301 | -0.68511243 | 0.033 | 0.25 | 1 | 25 |
| cutc | 0.18301 | -0.32447033 | 0.033 | 0.25 | 1 | 25 |
| dap1b | 0.18301 | -0.5929282 | 0.033 | 0.25 | 1 | 25 |
| dctn1a | 0.18301 | -0.31644158 | 0.033 | 0.25 | 1 | 25 |
| ddx21 | 0.18301 | -0.75628127 | 0.033 | 0.25 | 1 | 25 |
| dffb | 0.18301 | -0.62277988 | 0.033 | 0.25 | 1 | 25 |
| dram2b | 0.18301 | -0.60885976 | 0.033 | 0.25 | 1 | 25 |
| dtnbp1a | 0.18301 | -0.64132467 | 0.033 | 0.25 | 1 | 25 |
| gpr132b | 0.18301 | -0.54792262 | 0.033 | 0.25 | 1 | 25 |
| htatsf1 | 0.18301 | -0.69025816 | 0.033 | 0.25 | 1 | 25 |
| il12rb2l | 0.18301 | -0.57974244 | 0.033 | 0.25 | 1 | 25 |
| kptn | 0.18301 | -0.5314255 | 0.033 | 0.25 | 1 | 25 |
| lsm7 | 0.18301 | -0.5715681 | 0.033 | 0.25 | 1 | 25 |

| | | | | | | |
|---|---|---|---|---|---|---|
| lyn | 0.18301 | -0.55928215 | 0.033 | 0.25 | 1 | 25 |
| mapk3 | 0.18301 | -0.63022117 | 0.033 | 0.25 | 1 | 25 |
| mesd | 0.18301 | -0.36026968 | 0.033 | 0.25 | 1 | 25 |
| ncdn | 0.18301 | -0.60827042 | 0.033 | 0.25 | 1 | 25 |
| nlrc3l1 | 0.18301 | -0.57590576 | 0.033 | 0.25 | 1 | 25 |
| nsd2 | 0.18301 | -0.28090186 | 0.033 | 0.25 | 1 | 25 |
| orai1a | 0.18301 | -0.52372583 | 0.033 | 0.25 | 1 | 25 |
| pag1 | 0.18301 | -0.51246863 | 0.033 | 0.25 | 1 | 25 |
| ppp2r1ba | 0.18301 | -0.35005196 | 0.033 | 0.25 | 1 | 25 |
| prkcsh | 0.18301 | -0.30903467 | 0.033 | 0.25 | 1 | 25 |
| prpf18 | 0.18301 | -0.60721357 | 0.033 | 0.25 | 1 | 25 |
| psmg2 | 0.18301 | -0.32536133 | 0.033 | 0.25 | 1 | 25 |
| ptk6b | 0.18301 | -0.62258799 | 0.033 | 0.25 | 1 | 25 |
| rfc3 | 0.18301 | -0.31922986 | 0.033 | 0.25 | 1 | 25 |
| rsu1 | 0.18301 | -0.69947887 | 0.033 | 0.25 | 1 | 25 |
| samd11 | 0.18301 | -0.588846 | 0.033 | 0.25 | 1 | 25 |
| scaf11 | 0.18301 | -0.35468368 | 0.033 | 0.25 | 1 | 25 |
| scinla | 0.18301 | -0.62094981 | 0.033 | 0.25 | 1 | 25 |
| sel1l | 0.18301 | -0.55456851 | 0.033 | 0.25 | 1 | 25 |
| sephs2 | 0.18301 | -0.60527706 | 0.033 | 0.25 | 1 | 25 |
| serp1 | 0.18301 | -0.60902368 | 0.033 | 0.25 | 1 | 25 |
| sh3d21 | 0.18301 | -0.33232621 | 0.033 | 0.25 | 1 | 25 |
| si:ch1073-188e1.1 | 0.18301 | -0.52997565 | 0.033 | 0.25 | 1 | 25 |
| si:ch211-162e15.3 | 0.18301 | -0.62793053 | 0.033 | 0.25 | 1 | 25 |
| si:ch211-191c10.1 | 0.18301 | -0.62686427 | 0.033 | 0.25 | 1 | 25 |
| si:ch211-286b5.9 | 0.18301 | -0.53707232 | 0.033 | 0.25 | 1 | 25 |
| si:ch73-100l22.3 | 0.18301 | -0.49809009 | 0.033 | 0.25 | 1 | 25 |
| si:dkey-199f5.8 | 0.18301 | -0.2756937 | 0.033 | 0.25 | 1 | 25 |
| si:dkeyp-44a8.2 | 0.18301 | -0.61466978 | 0.033 | 0.25 | 1 | 25 |
| slc35b3 | 0.18301 | -0.30912979 | 0.033 | 0.25 | 1 | 25 |
| slc6a9 | 0.18301 | -0.31336336 | 0.033 | 0.25 | 1 | 25 |
| smim7 | 0.18301 | -0.34234134 | 0.033 | 0.25 | 1 | 25 |
| spata20 | 0.18301 | -0.37141294 | 0.033 | 0.25 | 1 | 25 |
| spice1 | 0.18301 | -0.50865645 | 0.033 | 0.25 | 1 | 25 |
| supt6h | 0.18301 | -0.62802895 | 0.033 | 0.25 | 1 | 25 |
| tet1 | 0.18301 | -0.24496393 | 0.033 | 0.25 | 1 | 25 |
| tmem120a | 0.18301 | -0.59283599 | 0.033 | 0.25 | 1 | 25 |
| tpm4a | 0.18301 | -0.62146554 | 0.033 | 0.25 | 1 | 25 |
| usp9 | 0.18301 | -0.36246003 | 0.033 | 0.25 | 1 | 25 |
| vmp1 | 0.18301 | -0.25821722 | 0.033 | 0.25 | 1 | 25 |
| wdr45 | 0.18301 | -0.64545832 | 0.033 | 0.25 | 1 | 25 |
| wdr82 | 0.18301 | -0.62141237 | 0.033 | 0.25 | 1 | 25 |
| zc3h7bb | 0.18301 | -0.5468226 | 0.033 | 0.25 | 1 | 25 |
| zfr2 | 0.18301 | -0.64150668 | 0.033 | 0.25 | 1 | 25 |
| zgc:153993 | 0.18301 | -0.62023912 | 0.033 | 0.25 | 1 | 25 |
| zgc:158564 | 0.18301 | -0.50010659 | 0.033 | 0.25 | 1 | 25 |
| zgc:163080 | 0.18301 | -0.40820699 | 0.033 | 0.25 | 1 | 25 |
| CABZ01109481.1 | 0.18301 | -0.61257835 | 0.033 | 0.25 | 1 | 25 |

| | | | | | | |
|---|---|---|---|---|---|---|
| adgre5a | 0.18301 | -0.26966924 | 0.033 | 0.25 | 1 | 25 |
| alyref | 0.18301 | -0.35506579 | 0.033 | 0.25 | 1 | 25 |
| arf2a | 0.18301 | -0.702062 | 0.033 | 0.25 | 1 | 25 |
| b3gnt2b | 0.18301 | -0.26106687 | 0.033 | 0.25 | 1 | 25 |
| cacybp | 0.18301 | -0.24778222 | 0.033 | 0.25 | 1 | 25 |
| decr1 | 0.18301 | -0.28724149 | 0.033 | 0.25 | 1 | 25 |
| erp44 | 0.18301 | -0.52341799 | 0.033 | 0.25 | 1 | 25 |
| fam120c | 0.18301 | -0.28863228 | 0.033 | 0.25 | 1 | 25 |
| fam172a | 0.18301 | -0.35212154 | 0.033 | 0.25 | 1 | 25 |
| fdx2 | 0.18301 | -0.20997924 | 0.033 | 0.25 | 1 | 25 |
| grasp | 0.18301 | -0.81398779 | 0.033 | 0.25 | 1 | 25 |
| hmgcl | 0.18301 | -0.26191517 | 0.033 | 0.25 | 1 | 25 |
| junba | 0.18301 | -0.52583228 | 0.033 | 0.25 | 1 | 25 |
| larp4ab | 0.18301 | -0.28049421 | 0.033 | 0.25 | 1 | 25 |
| map3k15 | 0.18301 | -0.35675788 | 0.033 | 0.25 | 1 | 25 |
| mapk12a | 0.18301 | -0.55287227 | 0.033 | 0.25 | 1 | 25 |
| pam | 0.18301 | -0.34737659 | 0.033 | 0.25 | 1 | 25 |
| pcsk7 | 0.18301 | -0.5482924 | 0.033 | 0.25 | 1 | 25 |
| phkg2 | 0.18301 | -0.61808846 | 0.033 | 0.25 | 1 | 25 |
| sfpq | 0.18301 | -0.55458562 | 0.033 | 0.25 | 1 | 25 |
| stag2b | 0.18301 | -0.29732341 | 0.033 | 0.25 | 1 | 25 |
| tm9sf3 | 0.18301 | -0.28264332 | 0.033 | 0.25 | 1 | 25 |
| usp4 | 0.18301 | -0.24219169 | 0.033 | 0.25 | 1 | 25 |
| vbp1 | 0.18301 | -0.6126783 | 0.033 | 0.25 | 1 | 25 |
| zgc:76872 | 0.18301 | -0.49154068 | 0.033 | 0.25 | 1 | 25 |
| CU856539.2 | 0.18301 | -0.57084905 | 0.033 | 0.25 | 1 | 25 |
| PDPR | 0.18301 | -0.63985743 | 0.033 | 0.25 | 1 | 25 |
| abhd18 | 0.18301 | -0.3852119 | 0.033 | 0.25 | 1 | 25 |
| adcy9 | 0.18301 | -0.60016457 | 0.033 | 0.25 | 1 | 25 |
| adipor1b | 0.18301 | -0.31108806 | 0.033 | 0.25 | 1 | 25 |
| aifm1 | 0.18301 | -0.10822917 | 0.033 | 0.25 | 1 | 25 |
| ap1ar | 0.18301 | -0.35329244 | 0.033 | 0.25 | 1 | 25 |
| ap2m1b | 0.18301 | -0.63941934 | 0.033 | 0.25 | 1 | 25 |
| btf3l4 | 0.18301 | -0.64153556 | 0.033 | 0.25 | 1 | 25 |
| cd82b | 0.18301 | -0.26847246 | 0.033 | 0.25 | 1 | 25 |
| cfap20 | 0.18301 | -0.26046601 | 0.033 | 0.25 | 1 | 25 |
| chd4b | 0.18301 | -0.70911045 | 0.033 | 0.25 | 1 | 25 |
| chic2 | 0.18301 | -0.55884956 | 0.033 | 0.25 | 1 | 25 |
| chid1 | 0.18301 | -0.61336902 | 0.033 | 0.25 | 1 | 25 |
| crkl | 0.18301 | -0.5208564 | 0.033 | 0.25 | 1 | 25 |
| ctsh | 0.18301 | -0.5407837 | 0.033 | 0.25 | 1 | 25 |
| ctss2.2 | 0.18301 | -0.5958552 | 0.033 | 0.25 | 1 | 25 |
| cybrd1 | 0.18301 | -0.37458385 | 0.033 | 0.25 | 1 | 25 |
| ddt | 0.18301 | -0.70329484 | 0.033 | 0.25 | 1 | 25 |
| dhdhl | 0.18301 | -0.52935704 | 0.033 | 0.25 | 1 | 25 |
| dusp7 | 0.18301 | -0.69788447 | 0.033 | 0.25 | 1 | 25 |
| fen1 | 0.18301 | -0.26293703 | 0.033 | 0.25 | 1 | 25 |
| glmp | 0.18301 | -0.52295393 | 0.033 | 0.25 | 1 | 25 |

| | | | | | | |
|---|---|---|---|---|---|---|
| hectd3 | 0.18301 | -0.61081843 | 0.033 | 0.25 | 1 | 25 |
| hexa | 0.18301 | -0.64587074 | 0.033 | 0.25 | 1 | 25 |
| hpse | 0.18301 | -0.63518459 | 0.033 | 0.25 | 1 | 25 |
| icmt | 0.18301 | -0.54098747 | 0.033 | 0.25 | 1 | 25 |
| maff | 0.18301 | -0.59245973 | 0.033 | 0.25 | 1 | 25 |
| map1lc3b | 0.18301 | -0.72504771 | 0.033 | 0.25 | 1 | 25 |
| march2 | 0.18301 | -0.5990829 | 0.033 | 0.25 | 1 | 25 |
| med20 | 0.18301 | -0.74111511 | 0.033 | 0.25 | 1 | 25 |
| mettl7a | 0.18301 | -0.33467584 | 0.033 | 0.25 | 1 | 25 |
| mrps5 | 0.18301 | -0.62348892 | 0.033 | 0.25 | 1 | 25 |
| noctb | 0.18301 | -0.6486094 | 0.033 | 0.25 | 1 | 25 |
| nucb2a | 0.18301 | -0.58842419 | 0.033 | 0.25 | 1 | 25 |
| parp8 | 0.18301 | -0.59088014 | 0.033 | 0.25 | 1 | 25 |
| pm20d1.2 | 0.18301 | -0.29446536 | 0.033 | 0.25 | 1 | 25 |
| prpf4ba | 0.18301 | -0.25657635 | 0.033 | 0.25 | 1 | 25 |
| prpf8 | 0.18301 | -0.59990918 | 0.033 | 0.25 | 1 | 25 |
| prss16 | 0.18301 | -0.34651866 | 0.033 | 0.25 | 1 | 25 |
| psmd8 | 0.18301 | -0.64604728 | 0.033 | 0.25 | 1 | 25 |
| ptprea | 0.18301 | -0.64255722 | 0.033 | 0.25 | 1 | 25 |
| rad23b | 0.18301 | -0.70712526 | 0.033 | 0.25 | 1 | 25 |
| retreg2 | 0.18301 | -0.58165523 | 0.033 | 0.25 | 1 | 25 |
| serinc5 | 0.18301 | -0.41960363 | 0.033 | 0.25 | 1 | 25 |
| sesn1 | 0.18301 | -0.55007766 | 0.033 | 0.25 | 1 | 25 |
| si:cabz01007802.1 | 0.18301 | -0.59693504 | 0.033 | 0.25 | 1 | 25 |
| si:ch211-114l13.12 | 0.18301 | -0.62975333 | 0.033 | 0.25 | 1 | 25 |
| si:ch211-14c7.2 | 0.18301 | -0.38471426 | 0.033 | 0.25 | 1 | 25 |
| si:ch211-277c7.6 | 0.18301 | -0.54921863 | 0.033 | 0.25 | 1 | 25 |
| si:dkey-206d17.12 | 0.18301 | -0.60721626 | 0.033 | 0.25 | 1 | 25 |
| si:dkey-24p1.7 | 0.18301 | -0.60516614 | 0.033 | 0.25 | 1 | 25 |
| slbp | 0.18301 | -0.20367497 | 0.033 | 0.25 | 1 | 25 |
| slc10a7 | 0.18301 | -0.54946907 | 0.033 | 0.25 | 1 | 25 |
| slc11a2 | 0.18301 | -0.60616974 | 0.033 | 0.25 | 1 | 25 |
| slc43a2a | 0.18301 | -0.54944096 | 0.033 | 0.25 | 1 | 25 |
| tomm70a | 0.18301 | -0.26453712 | 0.033 | 0.25 | 1 | 25 |
| trim71 | 0.18301 | -0.60385233 | 0.033 | 0.25 | 1 | 25 |
| ube2na | 0.18301 | -0.62990176 | 0.033 | 0.25 | 1 | 25 |
| unkl | 0.18301 | -0.41093956 | 0.033 | 0.25 | 1 | 25 |
| ylpm1 | 0.18301 | -0.55773308 | 0.033 | 0.25 | 1 | 25 |
| zgc:153935 | 0.18301 | -0.62868069 | 0.033 | 0.25 | 1 | 25 |
| zgc:66474 | 0.18301 | -0.59349573 | 0.033 | 0.25 | 1 | 25 |
| zmp:0000001132 | 0.18301 | -0.34727233 | 0.033 | 0.25 | 1 | 25 |
| zrsr2 | 0.18301 | -0.63730113 | 0.033 | 0.25 | 1 | 25 |
| zyg11 | 0.18301 | -0.69706357 | 0.033 | 0.25 | 1 | 25 |
| cyp2ae1 | 0.18301 | -0.47301765 | 0.033 | 0.25 | 1 | 25 |
| lmbrd1 | 0.18301 | -0.64893903 | 0.033 | 0.25 | 1 | 25 |
| mrps9 | 0.18301 | -0.66445119 | 0.033 | 0.25 | 1 | 25 |
| tmem150aa | 0.18301 | -0.68107552 | 0.033 | 0.25 | 1 | 25 |
| ubl3b | 0.18301 | -0.66679325 | 0.033 | 0.25 | 1 | 25 |

| | | | | | | |
|---|---|---|---|---|---|---|
| vps26a | 0.18301 | -0.62026339 | 0.033 | 0.25 | 1 | 25 |
| ndufa2 | 0.18482 | -0.98427346 | 0.1 | 0.5 | 1 | 25 |
| ndufb7 | 0.18486 | 0.558891657 | 0.267 | 0 | 1 | 25 |
| ccni | 0.18486 | 0.712224258 | 0.267 | 0 | 1 | 25 |
| eif4ebp2 | 0.18486 | 0.620441889 | 0.267 | 0 | 1 | 25 |
| klf6a | 0.18486 | 0.520966553 | 0.267 | 0 | 1 | 25 |
| parvg | 0.18486 | 0.6655399 | 0.267 | 0 | 1 | 25 |
| wasa | 0.18486 | 0.670974579 | 0.267 | 0 | 1 | 25 |
| acin1a | 0.18486 | 0.608440409 | 0.267 | 0 | 1 | 25 |
| adam10a | 0.18486 | 0.698339696 | 0.267 | 0 | 1 | 25 |
| atp5if1a | 0.18486 | 0.428711939 | 0.267 | 0 | 1 | 25 |
| ddx61 | 0.18486 | 0.640940249 | 0.267 | 0 | 1 | 25 |
| eif3c | 0.18486 | 0.48903812 | 0.267 | 0 | 1 | 25 |
| fam49ba | 0.18486 | 0.587377287 | 0.267 | 0 | 1 | 25 |
| gpr183a | 0.18486 | 0.708863445 | 0.267 | 0 | 1 | 25 |
| ldhba | 0.18486 | 0.663988947 | 0.267 | 0 | 1 | 25 |
| mdh2 | 0.18486 | 0.573622892 | 0.267 | 0 | 1 | 25 |
| mrtfab | 0.18486 | 0.610152973 | 0.267 | 0 | 1 | 25 |
| paip2b | 0.18486 | 0.530653924 | 0.267 | 0 | 1 | 25 |
| pfdn5 | 0.18486 | 0.627868632 | 0.267 | 0 | 1 | 25 |
| pgp | 0.18486 | 0.562051113 | 0.267 | 0 | 1 | 25 |
| psma1 | 0.18486 | 0.60495913 | 0.267 | 0 | 1 | 25 |
| rab11ba | 0.18486 | 0.616775242 | 0.267 | 0 | 1 | 25 |
| serinc2 | 0.18486 | 0.606705273 | 0.267 | 0 | 1 | 25 |
| tagapb | 0.18486 | 0.61606553 | 0.267 | 0 | 1 | 25 |
| triobpa | 0.18486 | 0.647526104 | 0.267 | 0 | 1 | 25 |
| txnipa | 0.18486 | 0.633512847 | 0.267 | 0 | 1 | 25 |
| ywhaba | 0.18486 | 0.625880292 | 0.267 | 0 | 1 | 25 |
| atp6v1g1 | 0.18486 | 0.55162977 | 0.267 | 0 | 1 | 25 |
| f8 | 0.18486 | 0.731869967 | 0.267 | 0 | 1 | 25 |
| lamtor5 | 0.18486 | 0.506250305 | 0.267 | 0 | 1 | 25 |
| lrp10 | 0.18486 | 0.563892771 | 0.267 | 0 | 1 | 25 |
| mtdha | 0.18486 | 0.491138475 | 0.267 | 0 | 1 | 25 |
| naga | 0.18486 | 0.578788949 | 0.267 | 0 | 1 | 25 |
| psma8 | 0.18486 | 0.503619715 | 0.267 | 0 | 1 | 25 |
| snrpf | 0.18486 | 0.50037029 | 0.267 | 0 | 1 | 25 |
| LOC100151196 | 0.18649 | -0.99120431 | 0.1 | 0.5 | 1 | 25 |
| si:dkey-27b3.2 | 0.18793 | -0.97871458 | 0.1 | 0.5 | 1 | 25 |
| b2m | 0.19039 | -0.94809048 | 0.1 | 0.5 | 1 | 25 |
| NAV1 (1 of many) | 0.19346 | -0.59579502 | 0.1 | 0.5 | 1 | 25 |
| TMEM19 | 0.19357 | -0.89725501 | 0.1 | 0.5 | 1 | 25 |
| si:ch73-269m14.2 | 0.19722 | -0.74368871 | 0.1 | 0.5 | 1 | 25 |
| rps6ka5 | 0.19845 | -0.75913416 | 0.1 | 0.5 | 1 | 25 |
| mia3 | 0.19845 | -0.86528859 | 0.1 | 0.5 | 1 | 25 |
| ryr3 | 0.19882 | -0.83614787 | 0.1 | 0.5 | 1 | 25 |
| mt-atp6 | 0.21353 | -0.52646701 | 0.6 | 1 | 1 | 25 |
| rps28 | 0.22031 | 1.227032201 | 0.633 | 0.25 | 1 | 25 |
| rpl34 | 0.22483 | 0.918229666 | 0.7 | 0.25 | 1 | 25 |

| | | | | | | |
|---|---|---|---|---|---|---|
| rpl4 | 0.22498 | 0.87115276 | 0.7 | 0.25 | 1 | 25 |
| glrx | 0.22574 | 0.452661059 | 0.233 | 0 | 1 | 25 |
| lman2 | 0.22574 | 0.583013133 | 0.233 | 0 | 1 | 25 |
| ndufb8 | 0.22574 | 0.42424205 | 0.233 | 0 | 1 | 25 |
| puraa | 0.22574 | 0.531621596 | 0.233 | 0 | 1 | 25 |
| ugp2b | 0.22574 | 0.503462739 | 0.233 | 0 | 1 | 25 |
| CABZ01111959.1 | 0.22574 | 0.652459938 | 0.233 | 0 | 1 | 25 |
| alox5a | 0.22574 | 0.580400394 | 0.233 | 0 | 1 | 25 |
| csrp1a | 0.22574 | 0.402401762 | 0.233 | 0 | 1 | 25 |
| eif3ha | 0.22574 | 0.463338374 | 0.233 | 0 | 1 | 25 |
| mtpn | 0.22574 | 0.548143542 | 0.233 | 0 | 1 | 25 |
| nt5dc2 | 0.22574 | 0.612859981 | 0.233 | 0 | 1 | 25 |
| rps6ka1 | 0.22574 | 0.525877145 | 0.233 | 0 | 1 | 25 |
| syf2 | 0.22574 | 0.515430084 | 0.233 | 0 | 1 | 25 |
| trnau1apb | 0.22574 | 0.535207411 | 0.233 | 0 | 1 | 25 |
| yrk | 0.22574 | 0.7033224 | 0.233 | 0 | 1 | 25 |
| ap2m1a | 0.22574 | 0.444553411 | 0.233 | 0 | 1 | 25 |
| atp5pd | 0.22574 | 0.41864334 | 0.233 | 0 | 1 | 25 |
| cib1 | 0.22574 | 0.509266738 | 0.233 | 0 | 1 | 25 |
| cybb | 0.22574 | 0.498943451 | 0.233 | 0 | 1 | 25 |
| gpx4b | 0.22574 | 0.55004039 | 0.233 | 0 | 1 | 25 |
| hmgb1a | 0.22574 | 0.342913194 | 0.233 | 0 | 1 | 25 |
| idh2 | 0.22574 | 0.539304411 | 0.233 | 0 | 1 | 25 |
| larp6a | 0.22574 | 0.648627683 | 0.233 | 0 | 1 | 25 |
| lgals3b | 0.22574 | 0.403936394 | 0.233 | 0 | 1 | 25 |
| pim2 | 0.22574 | 0.658515489 | 0.233 | 0 | 1 | 25 |
| ppifb | 0.22574 | 0.450031909 | 0.233 | 0 | 1 | 25 |
| ppp1caa | 0.22574 | 0.455360038 | 0.233 | 0 | 1 | 25 |
| psmb1 | 0.22574 | 0.406346568 | 0.233 | 0 | 1 | 25 |
| sdcbp2 | 0.22574 | 0.500007075 | 0.233 | 0 | 1 | 25 |
| si:dkey-42l23.4 | 0.22574 | 0.595435657 | 0.233 | 0 | 1 | 25 |
| smarce1 | 0.22574 | 0.382114729 | 0.233 | 0 | 1 | 25 |
| sumo2b | 0.22574 | 0.550808733 | 0.233 | 0 | 1 | 25 |
| tcp1 | 0.22574 | 0.434501782 | 0.233 | 0 | 1 | 25 |
| ubb | 0.22574 | 0.60308686 | 0.233 | 0 | 1 | 25 |
| STMP1 | 0.22574 | 0.482329722 | 0.233 | 0 | 1 | 25 |
| eif4ebp1 | 0.22574 | 0.462938046 | 0.233 | 0 | 1 | 25 |
| gnb1b | 0.22574 | 0.556627875 | 0.233 | 0 | 1 | 25 |
| jpt2 | 0.22574 | 0.528401187 | 0.233 | 0 | 1 | 25 |
| ndufa5 | 0.22574 | 0.370865594 | 0.233 | 0 | 1 | 25 |
| papss2b | 0.22574 | 0.506888277 | 0.233 | 0 | 1 | 25 |
| pde7a | 0.22574 | 0.518466282 | 0.233 | 0 | 1 | 25 |
| pgk1 | 0.22574 | 0.409654557 | 0.233 | 0 | 1 | 25 |
| si:dkey-185m8.2 | 0.22574 | 0.462097502 | 0.233 | 0 | 1 | 25 |
| sult6b1 | 0.22574 | 0.500094117 | 0.233 | 0 | 1 | 25 |
| myl12.1 | 0.22923 | -0.25853315 | 0.633 | 1 | 1 | 25 |
| ubc | 0.23044 | 0.811113846 | 0.7 | 0.25 | 1 | 25 |
| rps27a | 0.23126 | 0.656073212 | 0.7 | 0.25 | 1 | 25 |

| | | | | | | |
|---|---|---|---|---|---|---|
| cox7c | 0.23233 | -0.58691222 | 0.3 | 0.75 | 1 | 25 |
| rps24 | 0.2324 | 0.67983322 | 0.6 | 0.75 | 1 | 25 |
| CT030188.1 | 0.24348 | -0.33338699 | 0.633 | 1 | 1 | 25 |
| anxa11a | 0.25375 | -1.01692778 | 0.133 | 0.5 | 1 | 25 |
| xbp1 | 0.25417 | -0.3890425 | 0.633 | 1 | 1 | 25 |
| dnase1l1 | 0.26331 | -0.90687884 | 0.133 | 0.5 | 1 | 25 |
| LOC101884451 | 0.26507 | -0.91962875 | 0.133 | 0.5 | 1 | 25 |
| litaf | 0.26612 | -0.91588522 | 0.133 | 0.5 | 1 | 25 |
| rps9 | 0.27032 | 0.5605006 | 0.533 | 0.75 | 1 | 25 |
| zgc:100846 | 0.27077 | -0.87124548 | 0.133 | 0.5 | 1 | 25 |
| sptssa | 0.27208 | -0.84641877 | 0.133 | 0.5 | 1 | 25 |
| uqcrq | 0.27211 | -0.7914282 | 0.333 | 0.75 | 1 | 25 |
| rnf10 | 0.27315 | -0.8804995 | 0.133 | 0.5 | 1 | 25 |
| gde1 | 0.27443 | -0.75531299 | 0.133 | 0.5 | 1 | 25 |
| bud31 | 0.27485 | 0.383676036 | 0.2 | 0 | 1 | 25 |
| irf2 | 0.27485 | 0.4926117 | 0.2 | 0 | 1 | 25 |
| med10 | 0.27485 | 0.356516426 | 0.2 | 0 | 1 | 25 |
| slc40a1 | 0.27485 | 0.489406671 | 0.2 | 0 | 1 | 25 |
| calm2a | 0.27485 | 0.431712781 | 0.2 | 0 | 1 | 25 |
| capns1a | 0.27485 | 0.346448944 | 0.2 | 0 | 1 | 25 |
| cbr1l | 0.27485 | 0.458622031 | 0.2 | 0 | 1 | 25 |
| cyp3c1 | 0.27485 | 0.452761061 | 0.2 | 0 | 1 | 25 |
| ddx39ab | 0.27485 | 0.367064305 | 0.2 | 0 | 1 | 25 |
| eif3g | 0.27485 | 0.510742594 | 0.2 | 0 | 1 | 25 |
| hcls1 | 0.27485 | 0.488087717 | 0.2 | 0 | 1 | 25 |
| hnrnpk | 0.27485 | 0.369469401 | 0.2 | 0 | 1 | 25 |
| iqgap1 | 0.27485 | 0.484456725 | 0.2 | 0 | 1 | 25 |
| lsm12b | 0.27485 | 0.45343221 | 0.2 | 0 | 1 | 25 |
| psen1 | 0.27485 | 0.466703603 | 0.2 | 0 | 1 | 25 |
| rps6kb1b | 0.27485 | 0.477770042 | 0.2 | 0 | 1 | 25 |
| sept12 | 0.27485 | 0.39925879 | 0.2 | 0 | 1 | 25 |
| sgk1 | 0.27485 | 0.432713368 | 0.2 | 0 | 1 | 25 |
| si:dkey-27i16.2 | 0.27485 | 0.414489721 | 0.2 | 0 | 1 | 25 |
| tram1 | 0.27485 | 0.539538774 | 0.2 | 0 | 1 | 25 |
| u2af1 | 0.27485 | 0.33676631 | 0.2 | 0 | 1 | 25 |
| znf706 | 0.27485 | 0.376856476 | 0.2 | 0 | 1 | 25 |
| XLOC-011085 | 0.27485 | 0.419121605 | 0.2 | 0 | 1 | 25 |
| agtrap | 0.27485 | 0.402021266 | 0.2 | 0 | 1 | 25 |
| anxa4 | 0.27485 | 0.455448124 | 0.2 | 0 | 1 | 25 |
| capns1b | 0.27485 | 0.481113441 | 0.2 | 0 | 1 | 25 |
| ccdc88b | 0.27485 | 0.510426569 | 0.2 | 0 | 1 | 25 |
| ccnd2a | 0.27485 | 0.29209962 | 0.2 | 0 | 1 | 25 |
| cct4 | 0.27485 | 0.384620482 | 0.2 | 0 | 1 | 25 |
| cox17 | 0.27485 | 0.332821731 | 0.2 | 0 | 1 | 25 |
| eif5a2 | 0.27485 | 0.359840066 | 0.2 | 0 | 1 | 25 |
| erlin1 | 0.27485 | 0.502730573 | 0.2 | 0 | 1 | 25 |
| gnai1 | 0.27485 | 0.476459663 | 0.2 | 0 | 1 | 25 |
| gtf2f2a | 0.27485 | 0.47750878 | 0.2 | 0 | 1 | 25 |

| | | | | | | |
|---|---|---|---|---|---|---|
| h2afx1 | 0.27485 | 0.502334213 | 0.2 | 0 | 1 | 25 |
| hsd3b7 | 0.27485 | 0.332857939 | 0.2 | 0 | 1 | 25 |
| ifi30 | 0.27485 | 0.342444694 | 0.2 | 0 | 1 | 25 |
| ilk | 0.27485 | 0.475265511 | 0.2 | 0 | 1 | 25 |
| lpcat2 | 0.27485 | 0.446894342 | 0.2 | 0 | 1 | 25 |
| marcksl1a | 0.27485 | 0.523549129 | 0.2 | 0 | 1 | 25 |
| msra | 0.27485 | 0.3693238 | 0.2 | 0 | 1 | 25 |
| plek | 0.27485 | 0.436095091 | 0.2 | 0 | 1 | 25 |
| ppp1r14aa | 0.27485 | 0.339313162 | 0.2 | 0 | 1 | 25 |
| rbx1 | 0.27485 | 0.268319436 | 0.2 | 0 | 1 | 25 |
| rnaseka | 0.27485 | 0.422089452 | 0.2 | 0 | 1 | 25 |
| si:ch211-199g17.2 | 0.27485 | 0.502244902 | 0.2 | 0 | 1 | 25 |
| si:ch73-343l4.8 | 0.27485 | 0.53133535 | 0.2 | 0 | 1 | 25 |
| si:dkey-172h23.2 | 0.27485 | 0.438787735 | 0.2 | 0 | 1 | 25 |
| vamp3 | 0.27485 | 0.419649025 | 0.2 | 0 | 1 | 25 |
| vdrb | 0.27485 | 0.488575276 | 0.2 | 0 | 1 | 25 |
| vma21 | 0.27485 | 0.458044058 | 0.2 | 0 | 1 | 25 |
| zgc:153713 | 0.27485 | 0.422139626 | 0.2 | 0 | 1 | 25 |
| zgc:162944 | 0.27485 | 0.287756407 | 0.2 | 0 | 1 | 25 |
| abcc13 | 0.27485 | 0.435215923 | 0.2 | 0 | 1 | 25 |
| cox7a2a | 0.27485 | 0.303391153 | 0.2 | 0 | 1 | 25 |
| kdelr2b | 0.27485 | 0.382140879 | 0.2 | 0 | 1 | 25 |
| ndufb5 | 0.27485 | 0.380905295 | 0.2 | 0 | 1 | 25 |
| pfdn4 | 0.27485 | 0.422183219 | 0.2 | 0 | 1 | 25 |
| ptbp1a | 0.27485 | 0.39853207 | 0.2 | 0 | 1 | 25 |
| pycard | 0.27485 | 0.320189448 | 0.2 | 0 | 1 | 25 |
| qkia | 0.27485 | 0.3021126 | 0.2 | 0 | 1 | 25 |
| sarnp | 0.27485 | 0.304944897 | 0.2 | 0 | 1 | 25 |
| sf3b6 | 0.27485 | 0.371646245 | 0.2 | 0 | 1 | 25 |
| si:ch1073-429i10.3.1 | 0.27485 | 0.566580518 | 0.2 | 0 | 1 | 25 |
| snrpb | 0.27485 | 0.351381236 | 0.2 | 0 | 1 | 25 |
| prdx2 | 0.27485 | 0.466441814 | 0.2 | 0 | 1 | 25 |
| rpl28 | 0.27547 | 1.041887363 | 0.633 | 0.25 | 1 | 25 |
| rpsa | 0.27781 | 1.03733021 | 0.633 | 0.25 | 1 | 25 |
| banf1 | 0.27801 | -0.85282109 | 0.133 | 0.5 | 1 | 25 |
| ccdc28a | 0.28016 | -0.81183336 | 0.133 | 0.5 | 1 | 25 |
| mt-nd4 | 0.28079 | -0.25661675 | 0.667 | 1 | 1 | 25 |
| rpl9 | 0.28228 | 1.005668624 | 0.633 | 0.25 | 1 | 25 |
| mkrn1 | 0.28293 | -0.73682645 | 0.133 | 0.5 | 1 | 25 |
| b2ml | 0.28399 | -0.6244066 | 0.133 | 0.5 | 1 | 25 |
| CT027744.2 | 0.28608 | -0.70278219 | 0.133 | 0.5 | 1 | 25 |
| uqcr10 | 0.28616 | -0.61332179 | 0.133 | 0.5 | 1 | 25 |
| tmx1 | 0.28619 | -0.68143039 | 0.133 | 0.5 | 1 | 25 |
| rpl36a | 0.2906 | 0.726682927 | 0.667 | 0.25 | 1 | 25 |
| slc25a3b | 0.29083 | -0.53070926 | 0.333 | 0.75 | 1 | 25 |
| rpl15 | 0.30103 | 0.961425825 | 0.633 | 0.25 | 1 | 25 |
| rpl19 | 0.30481 | 0.720820302 | 0.733 | 0.75 | 1 | 25 |
| b3gnt5a | 0.3107 | -0.21194796 | 0.067 | 0.25 | 1 | 25 |

| | | | | | | |
|---|---|---|---|---|---|---|
| cdk13 | 0.3107 | -0.21535617 | 0.067 | 0.25 | 1 | 25 |
| cyth1b | 0.3107 | -0.27122336 | 0.067 | 0.25 | 1 | 25 |
| dnajc1 | 0.3107 | -0.51730052 | 0.067 | 0.25 | 1 | 25 |
| fkbp3 | 0.3107 | -0.470715 | 0.067 | 0.25 | 1 | 25 |
| foxo1a | 0.3107 | -0.4808237 | 0.067 | 0.25 | 1 | 25 |
| golga7 | 0.3107 | -0.42333342 | 0.067 | 0.25 | 1 | 25 |
| got2b | 0.3107 | -0.53166596 | 0.067 | 0.25 | 1 | 25 |
| mpc2 | 0.3107 | -0.30759904 | 0.067 | 0.25 | 1 | 25 |
| ndufb4 | 0.3107 | -0.46118139 | 0.067 | 0.25 | 1 | 25 |
| ppp1r2 | 0.3107 | -0.24047038 | 0.067 | 0.25 | 1 | 25 |
| tmem209 | 0.3107 | -0.2092869 | 0.067 | 0.25 | 1 | 25 |
| zgc:77056 | 0.3107 | -0.57009027 | 0.067 | 0.25 | 1 | 25 |
| CABZ01034698.2 | 0.3107 | -0.21481848 | 0.067 | 0.25 | 1 | 25 |
| LOC101886015 | 0.3107 | -0.45812696 | 0.067 | 0.25 | 1 | 25 |
| acap1 | 0.3107 | -0.27499835 | 0.067 | 0.25 | 1 | 25 |
| anp32e | 0.3107 | -0.53707412 | 0.067 | 0.25 | 1 | 25 |
| cd28 | 0.3107 | -0.43406313 | 0.067 | 0.25 | 1 | 25 |
| dachd | 0.3107 | -0.60955907 | 0.067 | 0.25 | 1 | 25 |
| derl1 | 0.3107 | -0.20609522 | 0.067 | 0.25 | 1 | 25 |
| fkbp5 | 0.3107 | -0.5077029 | 0.067 | 0.25 | 1 | 25 |
| fundc1 | 0.3107 | -0.49787373 | 0.067 | 0.25 | 1 | 25 |
| hnrpkl | 0.3107 | -0.14954316 | 0.067 | 0.25 | 1 | 25 |
| hyi | 0.3107 | -0.55953647 | 0.067 | 0.25 | 1 | 25 |
| ifngr2 | 0.3107 | -0.51469271 | 0.067 | 0.25 | 1 | 25 |
| kmt5aa | 0.3107 | -0.32908931 | 0.067 | 0.25 | 1 | 25 |
| runx3 | 0.3107 | -0.20824282 | 0.067 | 0.25 | 1 | 25 |
| serinc1 | 0.3107 | -0.40801783 | 0.067 | 0.25 | 1 | 25 |
| si:rp71-36a1.3 | 0.3107 | -0.50392907 | 0.067 | 0.25 | 1 | 25 |
| smim15 | 0.3107 | -0.22343032 | 0.067 | 0.25 | 1 | 25 |
| tmed9 | 0.3107 | -0.49691452 | 0.067 | 0.25 | 1 | 25 |
| tomm6 | 0.3107 | -0.42174681 | 0.067 | 0.25 | 1 | 25 |
| ubtd2 | 0.3107 | -0.29992231 | 0.067 | 0.25 | 1 | 25 |
| uqcrc2b | 0.3107 | -0.20317209 | 0.067 | 0.25 | 1 | 25 |
| ypel5 | 0.3107 | -0.6583392 | 0.067 | 0.25 | 1 | 25 |
| zbtb10 | 0.3107 | -0.25230238 | 0.067 | 0.25 | 1 | 25 |
| ccng2 | 0.3107 | -0.63847236 | 0.067 | 0.25 | 1 | 25 |
| glb1l | 0.3107 | -0.34136495 | 0.067 | 0.25 | 1 | 25 |
| hdac1 | 0.3107 | -0.25927332 | 0.067 | 0.25 | 1 | 25 |
| ino80c | 0.3107 | -0.46130408 | 0.067 | 0.25 | 1 | 25 |
| mctp2b | 0.3107 | -0.57121148 | 0.067 | 0.25 | 1 | 25 |
| ppp1r12c | 0.3107 | -0.14391401 | 0.067 | 0.25 | 1 | 25 |
| si:dkeyp-19e1.3 | 0.3107 | -0.44566176 | 0.067 | 0.25 | 1 | 25 |
| tec | 0.3107 | -0.68589679 | 0.067 | 0.25 | 1 | 25 |
| zgc:153441 | 0.3107 | -0.33765937 | 0.067 | 0.25 | 1 | 25 |
| BX649485.2 | 0.3107 | -0.41547937 | 0.067 | 0.25 | 1 | 25 |
| CR855311.1 | 0.3107 | -0.44613968 | 0.067 | 0.25 | 1 | 25 |
| akap17a | 0.3107 | -0.57618362 | 0.067 | 0.25 | 1 | 25 |
| bzw1b | 0.3107 | -0.52259191 | 0.067 | 0.25 | 1 | 25 |

| | | | | | | |
|---|---|---|---|---|---|---|
| cab39l1 | 0.3107 | -0.54537608 | 0.067 | 0.25 | 1 | 25 |
| crema | 0.3107 | -0.45981178 | 0.067 | 0.25 | 1 | 25 |
| dctn3 | 0.3107 | -0.24969828 | 0.067 | 0.25 | 1 | 25 |
| ddx5 | 0.3107 | -0.6171625 | 0.067 | 0.25 | 1 | 25 |
| dnase1l1l | 0.3107 | -0.26012939 | 0.067 | 0.25 | 1 | 25 |
| dok2 | 0.3107 | -0.55148784 | 0.067 | 0.25 | 1 | 25 |
| glud1a | 0.3107 | -0.60854538 | 0.067 | 0.25 | 1 | 25 |
| gnai2a | 0.3107 | -0.32299252 | 0.067 | 0.25 | 1 | 25 |
| gtf2b | 0.3107 | -0.6014431 | 0.067 | 0.25 | 1 | 25 |
| gusb | 0.3107 | -0.38933103 | 0.067 | 0.25 | 1 | 25 |
| hmgb3b | 0.3107 | -0.32598808 | 0.067 | 0.25 | 1 | 25 |
| ikzf1 | 0.3107 | -0.50738276 | 0.067 | 0.25 | 1 | 25 |
| lgals9l3 | 0.3107 | -0.48374752 | 0.067 | 0.25 | 1 | 25 |
| lsm8 | 0.3107 | -0.52333683 | 0.067 | 0.25 | 1 | 25 |
| nde1 | 0.3107 | -0.13686505 | 0.067 | 0.25 | 1 | 25 |
| ndfip1l | 0.3107 | -0.55587289 | 0.067 | 0.25 | 1 | 25 |
| ndufs6 | 0.3107 | -0.60077232 | 0.067 | 0.25 | 1 | 25 |
| ntan1 | 0.3107 | -0.34095594 | 0.067 | 0.25 | 1 | 25 |
| os9 | 0.3107 | -0.20834494 | 0.067 | 0.25 | 1 | 25 |
| plp2 | 0.3107 | -0.34209586 | 0.067 | 0.25 | 1 | 25 |
| plpp5 | 0.3107 | -0.50260719 | 0.067 | 0.25 | 1 | 25 |
| polr2f | 0.3107 | -0.53855457 | 0.067 | 0.25 | 1 | 25 |
| ppp1cab | 0.3107 | -0.62939332 | 0.067 | 0.25 | 1 | 25 |
| pstpip1a | 0.3107 | -0.29519511 | 0.067 | 0.25 | 1 | 25 |
| purba | 0.3107 | -0.48245455 | 0.067 | 0.25 | 1 | 25 |
| ranbp9 | 0.3107 | -0.42853244 | 0.067 | 0.25 | 1 | 25 |
| rcc2 | 0.3107 | -0.53976844 | 0.067 | 0.25 | 1 | 25 |
| rgs18 | 0.3107 | -0.4376219 | 0.067 | 0.25 | 1 | 25 |
| rnf146 | 0.3107 | -0.57270069 | 0.067 | 0.25 | 1 | 25 |
| sc:d217 | 0.3107 | -0.259168 | 0.067 | 0.25 | 1 | 25 |
| sh3glb1b | 0.3107 | -0.25725193 | 0.067 | 0.25 | 1 | 25 |
| si:ch211-137j23.6 | 0.3107 | -0.47885566 | 0.067 | 0.25 | 1 | 25 |
| si:ch211-195h23.3 | 0.3107 | -0.53729177 | 0.067 | 0.25 | 1 | 25 |
| si:dkey-177p2.6 | 0.3107 | -0.5671728 | 0.067 | 0.25 | 1 | 25 |
| si:dkey-239h2.3 | 0.3107 | -0.53302516 | 0.067 | 0.25 | 1 | 25 |
| socs6a | 0.3107 | -0.24319396 | 0.067 | 0.25 | 1 | 25 |
| syne2b | 0.3107 | -0.5920655 | 0.067 | 0.25 | 1 | 25 |
| tardbpl | 0.3107 | -0.55403943 | 0.067 | 0.25 | 1 | 25 |
| tax1bp3 | 0.3107 | -0.59032566 | 0.067 | 0.25 | 1 | 25 |
| tbc1d10b | 0.3107 | -0.30675349 | 0.067 | 0.25 | 1 | 25 |
| tm2d2 | 0.3107 | -0.4867846 | 0.067 | 0.25 | 1 | 25 |
| tm9sf1 | 0.3107 | -0.1872236 | 0.067 | 0.25 | 1 | 25 |
| tmc6b | 0.3107 | -0.50078476 | 0.067 | 0.25 | 1 | 25 |
| tubb2b | 0.3107 | -0.5201474 | 0.067 | 0.25 | 1 | 25 |
| zgc:101679 | 0.3107 | -0.5881287 | 0.067 | 0.25 | 1 | 25 |
| zgc:136439 | 0.3107 | -0.44150219 | 0.067 | 0.25 | 1 | 25 |
| zgc:92360 | 0.3107 | -0.64297882 | 0.067 | 0.25 | 1 | 25 |
| brd4 | 0.3107 | -0.52453352 | 0.067 | 0.25 | 1 | 25 |

| | | | | | | |
|---|---|---|---|---|---|---|
| flna | 0.3107 | -0.31808928 | 0.067 | 0.25 | 1 | 25 |
| ginm1 | 0.3107 | -0.57682924 | 0.067 | 0.25 | 1 | 25 |
| mcm4 | 0.3107 | -0.31398268 | 0.067 | 0.25 | 1 | 25 |
| med31 | 0.3107 | -0.30777597 | 0.067 | 0.25 | 1 | 25 |
| mydgf | 0.3107 | -0.3063158 | 0.067 | 0.25 | 1 | 25 |
| nadkb | 0.3107 | -0.57906413 | 0.067 | 0.25 | 1 | 25 |
| ndufc2 | 0.3107 | -0.55774269 | 0.067 | 0.25 | 1 | 25 |
| pcf11 | 0.3107 | -0.60635069 | 0.067 | 0.25 | 1 | 25 |
| pgam1a | 0.3107 | -0.6058431 | 0.067 | 0.25 | 1 | 25 |
| plscr3b | 0.3107 | -0.62852599 | 0.067 | 0.25 | 1 | 25 |
| ptp4a2b | 0.3107 | -0.54457344 | 0.067 | 0.25 | 1 | 25 |
| rab14 | 0.3107 | -0.61024602 | 0.067 | 0.25 | 1 | 25 |
| sema4e | 0.3107 | -0.30490259 | 0.067 | 0.25 | 1 | 25 |
| si:ch211-14a17.11 | 0.3107 | -0.57007391 | 0.067 | 0.25 | 1 | 25 |
| si:rp71-1g18.13 | 0.3107 | -0.32422094 | 0.067 | 0.25 | 1 | 25 |
| slmapa | 0.3107 | -0.60100009 | 0.067 | 0.25 | 1 | 25 |
| tmbim1a | 0.3107 | -0.49164619 | 0.067 | 0.25 | 1 | 25 |
| tmem14ca | 0.3107 | -0.55459683 | 0.067 | 0.25 | 1 | 25 |
| trappc2l | 0.3107 | -0.61311102 | 0.067 | 0.25 | 1 | 25 |
| zfand5a | 0.3107 | -0.58218842 | 0.067 | 0.25 | 1 | 25 |
| rpl18a | 0.31113 | 0.227245219 | 0.5 | 0.75 | 1 | 25 |
| rpl39 | 0.32922 | 0.944396011 | 0.533 | 0.5 | 1 | 25 |
| FAR2 | 0.33414 | 0.350095567 | 0.167 | 0 | 1 | 25 |
| LOC101885706 | 0.33414 | 0.383517895 | 0.167 | 0 | 1 | 25 |
| arhgef3l | 0.33414 | 0.393722418 | 0.167 | 0 | 1 | 25 |
| ctsba | 0.33414 | 0.441310022 | 0.167 | 0 | 1 | 25 |
| hopx | 0.33414 | 0.362860074 | 0.167 | 0 | 1 | 25 |
| ilf2 | 0.33414 | 0.420005708 | 0.167 | 0 | 1 | 25 |
| kdm4b | 0.33414 | 0.254841778 | 0.167 | 0 | 1 | 25 |
| mycbp | 0.33414 | 0.258973427 | 0.167 | 0 | 1 | 25 |
| ppp6c | 0.33414 | 0.266649341 | 0.167 | 0 | 1 | 25 |
| selenot2 | 0.33414 | 0.413681003 | 0.167 | 0 | 1 | 25 |
| si:ch211-214j24.10 | 0.33414 | 0.256417745 | 0.167 | 0 | 1 | 25 |
| si:rp71-36a1.1 | 0.33414 | 0.388752501 | 0.167 | 0 | 1 | 25 |
| stat1a | 0.33414 | 0.357350283 | 0.167 | 0 | 1 | 25 |
| tomm7 | 0.33414 | 0.234357035 | 0.167 | 0 | 1 | 25 |
| ube2v2 | 0.33414 | 0.226526075 | 0.167 | 0 | 1 | 25 |
| BX649411.1 | 0.33414 | 0.363206325 | 0.167 | 0 | 1 | 25 |
| CABZ01026681.1 | 0.33414 | 0.356335581 | 0.167 | 0 | 1 | 25 |
| ISCU (1 of many) | 0.33414 | 0.486118112 | 0.167 | 0 | 1 | 25 |
| abhd16a | 0.33414 | 0.246761488 | 0.167 | 0 | 1 | 25 |
| acsl4b | 0.33414 | 0.473196742 | 0.167 | 0 | 1 | 25 |
| aldh8a1 | 0.33414 | 0.352067799 | 0.167 | 0 | 1 | 25 |
| arf1 | 0.33414 | 0.313881556 | 0.167 | 0 | 1 | 25 |
| arf6a | 0.33414 | 0.284676142 | 0.167 | 0 | 1 | 25 |
| arf6b | 0.33414 | 0.319305582 | 0.167 | 0 | 1 | 25 |
| bbip1 | 0.33414 | 0.465209037 | 0.167 | 0 | 1 | 25 |
| btk | 0.33414 | 0.344906286 | 0.167 | 0 | 1 | 25 |

| | | | | | | |
|---|---|---|---|---|---|---|
| calm2b | 0.33414 | 0.361864292 | 0.167 | 0 | 1 | 25 |
| cbx3a | 0.33414 | 0.388461917 | 0.167 | 0 | 1 | 25 |
| ccng1 | 0.33414 | 0.409485013 | 0.167 | 0 | 1 | 25 |
| cct3 | 0.33414 | 0.226411262 | 0.167 | 0 | 1 | 25 |
| cct6a | 0.33414 | 0.285673852 | 0.167 | 0 | 1 | 25 |
| cdipt | 0.33414 | 0.360930709 | 0.167 | 0 | 1 | 25 |
| cmtm7 | 0.33414 | 0.359192982 | 0.167 | 0 | 1 | 25 |
| cops7a | 0.33414 | 0.330087532 | 0.167 | 0 | 1 | 25 |
| cox5ab | 0.33414 | 0.237812057 | 0.167 | 0 | 1 | 25 |
| cst14a.2 | 0.33414 | 0.345719089 | 0.167 | 0 | 1 | 25 |
| cycsb | 0.33414 | 0.331024094 | 0.167 | 0 | 1 | 25 |
| ddx39aa | 0.33414 | 0.358474065 | 0.167 | 0 | 1 | 25 |
| dlst | 0.33414 | 0.303143811 | 0.167 | 0 | 1 | 25 |
| dynlt3 | 0.33414 | 0.370462298 | 0.167 | 0 | 1 | 25 |
| elovl5 | 0.33414 | 0.310154528 | 0.167 | 0 | 1 | 25 |
| erh | 0.33414 | 0.247880344 | 0.167 | 0 | 1 | 25 |
| fam162a | 0.33414 | 0.280584904 | 0.167 | 0 | 1 | 25 |
| fam32a | 0.33414 | 0.30836635 | 0.167 | 0 | 1 | 25 |
| foxq1a | 0.33414 | 0.374852113 | 0.167 | 0 | 1 | 25 |
| gdi2 | 0.33414 | 0.29270836 | 0.167 | 0 | 1 | 25 |
| gtpbp4 | 0.33414 | 0.26128277 | 0.167 | 0 | 1 | 25 |
| gyg1a | 0.33414 | 0.440017663 | 0.167 | 0 | 1 | 25 |
| hif1al | 0.33414 | 0.450370987 | 0.167 | 0 | 1 | 25 |
| hmha1b | 0.33414 | 0.467531764 | 0.167 | 0 | 1 | 25 |
| id2a | 0.33414 | 0.348945625 | 0.167 | 0 | 1 | 25 |
| ier3ip1 | 0.33414 | 0.339939109 | 0.167 | 0 | 1 | 25 |
| ip6k2b | 0.33414 | 0.316415296 | 0.167 | 0 | 1 | 25 |
| klf2b | 0.33414 | 0.382358927 | 0.167 | 0 | 1 | 25 |
| lamtor4 | 0.33414 | 0.245498496 | 0.167 | 0 | 1 | 25 |
| lsm3 | 0.33414 | 0.308197204 | 0.167 | 0 | 1 | 25 |
| madd | 0.33414 | 0.35704639 | 0.167 | 0 | 1 | 25 |
| mettl5 | 0.33414 | 0.319438625 | 0.167 | 0 | 1 | 25 |
| mif | 0.33414 | 0.293246929 | 0.167 | 0 | 1 | 25 |
| mob3a | 0.33414 | 0.426569707 | 0.167 | 0 | 1 | 25 |
| mphosph8 | 0.33414 | 0.356541828 | 0.167 | 0 | 1 | 25 |
| mrpl57 | 0.33414 | 0.206061864 | 0.167 | 0 | 1 | 25 |
| msrb1b | 0.33414 | 0.325628554 | 0.167 | 0 | 1 | 25 |
| ndufa8 | 0.33414 | 0.371012595 | 0.167 | 0 | 1 | 25 |
| nedd8l | 0.33414 | 0.308634262 | 0.167 | 0 | 1 | 25 |
| nfyba | 0.33414 | 0.369087909 | 0.167 | 0 | 1 | 25 |
| nop53 | 0.33414 | 0.204329429 | 0.167 | 0 | 1 | 25 |
| nudc | 0.33414 | 0.239481501 | 0.167 | 0 | 1 | 25 |
| nupr1a | 0.33414 | 0.237890741 | 0.167 | 0 | 1 | 25 |
| ostf1 | 0.33414 | 0.3077958 | 0.167 | 0 | 1 | 25 |
| pcbp2 | 0.33414 | 0.24142414 | 0.167 | 0 | 1 | 25 |
| pef1 | 0.33414 | 0.389703854 | 0.167 | 0 | 1 | 25 |
| pfdn1 | 0.33414 | 0.280573247 | 0.167 | 0 | 1 | 25 |
| pla2g15 | 0.33414 | 0.367152952 | 0.167 | 0 | 1 | 25 |

| | | | | | | |
|---|---|---|---|---|---|---|
| prdx3 | 0.33414 | 0.198212138 | 0.167 | 0 | 1 | 25 |
| psenen | 0.33414 | 0.300648024 | 0.167 | 0 | 1 | 25 |
| psma2 | 0.33414 | 0.426169315 | 0.167 | 0 | 1 | 25 |
| ptpn6 | 0.33414 | 0.338904266 | 0.167 | 0 | 1 | 25 |
| pygb | 0.33414 | 0.350972897 | 0.167 | 0 | 1 | 25 |
| rab11fip1b | 0.33414 | 0.416750709 | 0.167 | 0 | 1 | 25 |
| rps27l | 0.33414 | 0.24442936 | 0.167 | 0 | 1 | 25 |
| samsn1b | 0.33414 | 0.43204125 | 0.167 | 0 | 1 | 25 |
| sc:d156 | 0.33414 | 0.552682929 | 0.167 | 0 | 1 | 25 |
| sdhc | 0.33414 | 0.279877562 | 0.167 | 0 | 1 | 25 |
| selenot1b | 0.33414 | 0.25096157 | 0.167 | 0 | 1 | 25 |
| sem1 | 0.33414 | 0.33810556 | 0.167 | 0 | 1 | 25 |
| si:dkey-11f4.14 | 0.33414 | 0.519524633 | 0.167 | 0 | 1 | 25 |
| si:dkey-46a10.3 | 0.33414 | 0.362483887 | 0.167 | 0 | 1 | 25 |
| si:dkeyp-68b7.12 | 0.33414 | 0.405169137 | 0.167 | 0 | 1 | 25 |
| si:zfos-169g10.3 | 0.33414 | 0.134289195 | 0.167 | 0 | 1 | 25 |
| sptlc2a | 0.33414 | 0.293895281 | 0.167 | 0 | 1 | 25 |
| sub1a | 0.33414 | 0.370224136 | 0.167 | 0 | 1 | 25 |
| tmed2 | 0.33414 | 0.325277971 | 0.167 | 0 | 1 | 25 |
| tmem131 | 0.33414 | 0.421888149 | 0.167 | 0 | 1 | 25 |
| tmem165 | 0.33414 | 0.411966901 | 0.167 | 0 | 1 | 25 |
| tomm20b | 0.33414 | 0.345472135 | 0.167 | 0 | 1 | 25 |
| tprg1 | 0.33414 | 0.412099611 | 0.167 | 0 | 1 | 25 |
| tradd | 0.33414 | 0.306668739 | 0.167 | 0 | 1 | 25 |
| tspan31 | 0.33414 | 0.411562086 | 0.167 | 0 | 1 | 25 |
| u2af2b | 0.33414 | 0.410541836 | 0.167 | 0 | 1 | 25 |
| ufm1 | 0.33414 | 0.312589851 | 0.167 | 0 | 1 | 25 |
| usf1l | 0.33414 | 0.178620419 | 0.167 | 0 | 1 | 25 |
| vps25 | 0.33414 | 0.361466661 | 0.167 | 0 | 1 | 25 |
| ywhaqa | 0.33414 | 0.322476327 | 0.167 | 0 | 1 | 25 |
| ywhaqb | 0.33414 | 0.357882866 | 0.167 | 0 | 1 | 25 |
| zgc:152863 | 0.33414 | 0.566001997 | 0.167 | 0 | 1 | 25 |
| zgc:158343 | 0.33414 | 0.425534805 | 0.167 | 0 | 1 | 25 |
| zgc:86598 | 0.33414 | 0.238187914 | 0.167 | 0 | 1 | 25 |
| ccdc124 | 0.33414 | 0.326296698 | 0.167 | 0 | 1 | 25 |
| FBXO48 | 0.33414 | 0.363173531 | 0.167 | 0 | 1 | 25 |
| arhgef5 | 0.33414 | 0.508896066 | 0.167 | 0 | 1 | 25 |
| atf4a | 0.33414 | 0.468994996 | 0.167 | 0 | 1 | 25 |
| atp1b1a | 0.33414 | 0.29214432 | 0.167 | 0 | 1 | 25 |
| bin2b | 0.33414 | 0.348672737 | 0.167 | 0 | 1 | 25 |
| chchd2 | 0.33414 | 0.330279299 | 0.167 | 0 | 1 | 25 |
| fam49bb | 0.33414 | 0.442529571 | 0.167 | 0 | 1 | 25 |
| fth1a | 0.33414 | 0.313349532 | 0.167 | 0 | 1 | 25 |
| gra | 0.33414 | 0.420726513 | 0.167 | 0 | 1 | 25 |
| jak1 | 0.33414 | 0.41937321 | 0.167 | 0 | 1 | 25 |
| lrrc8da | 0.33414 | 0.261287897 | 0.167 | 0 | 1 | 25 |
| ptbp3 | 0.33414 | 0.366011253 | 0.167 | 0 | 1 | 25 |
| rab1ab | 0.33414 | 0.366454417 | 0.167 | 0 | 1 | 25 |

| | | | | | | |
|---|---|---|---|---|---|---|
| rbm39a | 0.33414 | 0.403172225 | 0.167 | 0 | 1 | 25 |
| rnf44 | 0.33414 | 0.3954283 | 0.167 | 0 | 1 | 25 |
| sept7b | 0.33414 | 0.367047505 | 0.167 | 0 | 1 | 25 |
| si:ch1073-340j19.1 | 0.33414 | 0.447009627 | 0.167 | 0 | 1 | 25 |
| si:dkey-53k12.2 | 0.33414 | 0.38373998 | 0.167 | 0 | 1 | 25 |
| syncrip | 0.33414 | 0.448918882 | 0.167 | 0 | 1 | 25 |
| unc13d | 0.33414 | 0.309352706 | 0.167 | 0 | 1 | 25 |
| zgc:56304 | 0.33414 | 0.473856013 | 0.167 | 0 | 1 | 25 |
| ly6m5 | 0.33449 | 0.835873276 | 0.633 | 0.25 | 1 | 25 |
| npc2 | 0.3413 | -0.81329712 | 0.4 | 0.75 | 1 | 25 |
| zgc:114045 | 0.34333 | -0.86154235 | 0.167 | 0.5 | 1 | 25 |
| reep5 | 0.35141 | -0.58466778 | 0.367 | 0.75 | 1 | 25 |
| capzb | 0.35329 | -0.58234018 | 0.367 | 0.75 | 1 | 25 |
| ak1 | 0.3577 | -0.81447825 | 0.167 | 0.5 | 1 | 25 |
| csk | 0.3599 | -0.8153712 | 0.167 | 0.5 | 1 | 25 |
| rplp2 | 0.36197 | 0.284250399 | 0.567 | 0.75 | 1 | 25 |
| arpc1b | 0.36492 | -0.16201927 | 0.733 | 1 | 1 | 25 |
| si:rp71-68n21.9 | 0.37001 | -0.45079472 | 0.167 | 0.5 | 1 | 25 |
| hsbp1b | 0.37444 | -0.73209752 | 0.167 | 0.5 | 1 | 25 |
| hspa8 | 0.37603 | 0.405055404 | 0.833 | 0.5 | 1 | 25 |
| syngr2a | 0.37757 | -0.6707795 | 0.167 | 0.5 | 1 | 25 |
| calr3a | 0.37868 | -0.51453911 | 0.167 | 0.5 | 1 | 25 |
| si:dkey-69o16.5 | 0.37909 | -0.52492466 | 0.167 | 0.5 | 1 | 25 |
| rpl27 | 0.38767 | 0.819228719 | 0.6 | 0.25 | 1 | 25 |
| atp5meb | 0.3941 | -0.88299937 | 0.2 | 0.5 | 1 | 25 |
| ptmaa | 0.39544 | 0.832412538 | 0.6 | 0.25 | 1 | 25 |
| ppib | 0.40413 | -0.31047666 | 0.4 | 0.75 | 1 | 25 |
| eif3ja | 0.40637 | 0.284672133 | 0.133 | 0 | 1 | 25 |
| jakmip1 | 0.40637 | 0.399366053 | 0.133 | 0 | 1 | 25 |
| pbdc1 | 0.40637 | 0.324045741 | 0.133 | 0 | 1 | 25 |
| ube2v1 | 0.40637 | 0.321589625 | 0.133 | 0 | 1 | 25 |
| BX284696.1 | 0.40637 | 0.272112387 | 0.133 | 0 | 1 | 25 |
| BX323557.1 | 0.40637 | 0.323954611 | 0.133 | 0 | 1 | 25 |
| BX901883.3 | 0.40637 | 0.314887818 | 0.133 | 0 | 1 | 25 |
| CABZ01079663.1 | 0.40637 | 0.208677736 | 0.133 | 0 | 1 | 25 |
| CDPF1 | 0.40637 | 0.272363974 | 0.133 | 0 | 1 | 25 |
| CR855337.1 | 0.40637 | 0.404568357 | 0.133 | 0 | 1 | 25 |
| FERMT3 (1 of many) | 0.40637 | 0.39642732 | 0.133 | 0 | 1 | 25 |
| MRPS12 | 0.40637 | 0.251017486 | 0.133 | 0 | 1 | 25 |
| acin1b | 0.40637 | 0.302567743 | 0.133 | 0 | 1 | 25 |
| actn1 | 0.40637 | 0.306565369 | 0.133 | 0 | 1 | 25 |
| adam19a | 0.40637 | 0.33576948 | 0.133 | 0 | 1 | 25 |
| adam28 | 0.40637 | 0.442194283 | 0.133 | 0 | 1 | 25 |
| adipor2 | 0.40637 | 0.325772576 | 0.133 | 0 | 1 | 25 |
| ap1s1 | 0.40637 | 0.257429449 | 0.133 | 0 | 1 | 25 |
| arhgap15 | 0.40637 | 0.324344872 | 0.133 | 0 | 1 | 25 |
| atp2b1a | 0.40637 | 0.410507068 | 0.133 | 0 | 1 | 25 |
| atp5f1d | 0.40637 | 0.29064961 | 0.133 | 0 | 1 | 25 |

| | | | | | | |
|---|---|---|---|---|---|---|
| atp5md | 0.40637 | 0.287161652 | 0.133 | 0 | 1 | 25 |
| bbx | 0.40637 | 0.307795998 | 0.133 | 0 | 1 | 25 |
| bcas2 | 0.40637 | 0.28924221 | 0.133 | 0 | 1 | 25 |
| chd7 | 0.40637 | 0.416436025 | 0.133 | 0 | 1 | 25 |
| cited4a | 0.40637 | 0.403060941 | 0.133 | 0 | 1 | 25 |
| ckap5 | 0.40637 | 0.291071622 | 0.133 | 0 | 1 | 25 |
| cnksr1 | 0.40637 | 0.218791836 | 0.133 | 0 | 1 | 25 |
| cyb5a | 0.40637 | 0.274116576 | 0.133 | 0 | 1 | 25 |
| dazap2 | 0.40637 | 0.236636591 | 0.133 | 0 | 1 | 25 |
| dctn2 | 0.40637 | 0.271305614 | 0.133 | 0 | 1 | 25 |
| ddit3 | 0.40637 | 0.400803182 | 0.133 | 0 | 1 | 25 |
| dynll1 | 0.40637 | 0.314004575 | 0.133 | 0 | 1 | 25 |
| erg | 0.40637 | 0.340538807 | 0.133 | 0 | 1 | 25 |
| fubp1 | 0.40637 | 0.265094681 | 0.133 | 0 | 1 | 25 |
| g6pd | 0.40637 | 0.358112082 | 0.133 | 0 | 1 | 25 |
| ggctb | 0.40637 | 0.259933261 | 0.133 | 0 | 1 | 25 |
| gigyf1a | 0.40637 | 0.373863429 | 0.133 | 0 | 1 | 25 |
| glipr2 | 0.40637 | 0.248206797 | 0.133 | 0 | 1 | 25 |
| gmip | 0.40637 | 0.357721738 | 0.133 | 0 | 1 | 25 |
| hdac9b | 0.40637 | 0.303799854 | 0.133 | 0 | 1 | 25 |
| heca | 0.40637 | 0.240805343 | 0.133 | 0 | 1 | 25 |
| hist2h2l | 0.40637 | 0.226122246 | 0.133 | 0 | 1 | 25 |
| histh1l | 0.40637 | 0.404250099 | 0.133 | 0 | 1 | 25 |
| hk2 | 0.40637 | 0.26054865 | 0.133 | 0 | 1 | 25 |
| irf2bp2a | 0.40637 | 0.305980886 | 0.133 | 0 | 1 | 25 |
| itga1 | 0.40637 | 0.357550648 | 0.133 | 0 | 1 | 25 |
| itsn2a | 0.40637 | 0.264505108 | 0.133 | 0 | 1 | 25 |
| kars | 0.40637 | 0.273418079 | 0.133 | 0 | 1 | 25 |
| khsrp | 0.40637 | 0.186817753 | 0.133 | 0 | 1 | 25 |
| kif5ba | 0.40637 | 0.352977836 | 0.133 | 0 | 1 | 25 |
| klf7b | 0.40637 | 0.281158996 | 0.133 | 0 | 1 | 25 |
| krtcap2 | 0.40637 | 0.209746924 | 0.133 | 0 | 1 | 25 |
| lsm5 | 0.40637 | 0.243782317 | 0.133 | 0 | 1 | 25 |
| max | 0.40637 | 0.199221295 | 0.133 | 0 | 1 | 25 |
| minos1 | 0.40637 | 0.22153566 | 0.133 | 0 | 1 | 25 |
| mrpl18 | 0.40637 | 0.264354801 | 0.133 | 0 | 1 | 25 |
| mrps14 | 0.40637 | 0.271169304 | 0.133 | 0 | 1 | 25 |
| mvb12a | 0.40637 | 0.339475739 | 0.133 | 0 | 1 | 25 |
| nap1l4b | 0.40637 | 0.310680039 | 0.133 | 0 | 1 | 25 |
| ncalda | 0.40637 | 0.23232629 | 0.133 | 0 | 1 | 25 |
| ngdn | 0.40637 | 0.351982995 | 0.133 | 0 | 1 | 25 |
| otud5a | 0.40637 | 0.26132153 | 0.133 | 0 | 1 | 25 |
| pdk2b | 0.40637 | 0.356118406 | 0.133 | 0 | 1 | 25 |
| pik3r5 | 0.40637 | 0.335924175 | 0.133 | 0 | 1 | 25 |
| ppp1r9ba | 0.40637 | 0.256455991 | 0.133 | 0 | 1 | 25 |
| prkacab | 0.40637 | 0.29818483 | 0.133 | 0 | 1 | 25 |
| prkar1aa | 0.40637 | 0.207184909 | 0.133 | 0 | 1 | 25 |
| psma3 | 0.40637 | 0.192694708 | 0.133 | 0 | 1 | 25 |

| | | | | | | |
|---|---|---|---|---|---|---|
| psme1 | 0.40637 | 0.179786814 | 0.133 | 0 | 1 | 25 |
| rab27a | 0.40637 | 0.367266892 | 0.133 | 0 | 1 | 25 |
| rnf144b | 0.40637 | 0.356763698 | 0.133 | 0 | 1 | 25 |
| s1pr4 | 0.40637 | 0.356264634 | 0.133 | 0 | 1 | 25 |
| sccpdhb | 0.40637 | 0.312127928 | 0.133 | 0 | 1 | 25 |
| sec61a1 | 0.40637 | 0.268092212 | 0.133 | 0 | 1 | 25 |
| sec61a1l | 0.40637 | 0.321844642 | 0.133 | 0 | 1 | 25 |
| selenot1a | 0.40637 | 0.23346784 | 0.133 | 0 | 1 | 25 |
| si:ch211-191j22.3 | 0.40637 | 0.159938338 | 0.133 | 0 | 1 | 25 |
| si:ch211-8c17.2 | 0.40637 | 0.363453918 | 0.133 | 0 | 1 | 25 |
| si:dkey-163m14.6 | 0.40637 | 0.363455149 | 0.133 | 0 | 1 | 25 |
| si:dkey-180p18.9 | 0.40637 | 0.304162848 | 0.133 | 0 | 1 | 25 |
| si:dkey-187e18.1 | 0.40637 | 0.304290547 | 0.133 | 0 | 1 | 25 |
| si:dkey-286j15.3 | 0.40637 | 0.338205492 | 0.133 | 0 | 1 | 25 |
| skp1 | 0.40637 | 0.182224889 | 0.133 | 0 | 1 | 25 |
| slc16a3 | 0.40637 | 0.277693125 | 0.133 | 0 | 1 | 25 |
| slc38a5b | 0.40637 | 0.360924257 | 0.133 | 0 | 1 | 25 |
| smc1al | 0.40637 | 0.278470903 | 0.133 | 0 | 1 | 25 |
| smim29 | 0.40637 | 0.231104979 | 0.133 | 0 | 1 | 25 |
| snx12 | 0.40637 | 0.387643304 | 0.133 | 0 | 1 | 25 |
| sptlc3 | 0.40637 | 0.299705751 | 0.133 | 0 | 1 | 25 |
| srsf7a | 0.40637 | 0.278300895 | 0.133 | 0 | 1 | 25 |
| ssuh2.3 | 0.40637 | 0.303862472 | 0.133 | 0 | 1 | 25 |
| sypl2b | 0.40637 | 0.260483536 | 0.133 | 0 | 1 | 25 |
| taf13 | 0.40637 | 0.283890021 | 0.133 | 0 | 1 | 25 |
| tm9sf2 | 0.40637 | 0.344699485 | 0.133 | 0 | 1 | 25 |
| tspan3a | 0.40637 | 0.234699615 | 0.133 | 0 | 1 | 25 |
| txndc15 | 0.40637 | 0.202800713 | 0.133 | 0 | 1 | 25 |
| ube2al | 0.40637 | 0.274693768 | 0.133 | 0 | 1 | 25 |
| ube2e1 | 0.40637 | 0.299619651 | 0.133 | 0 | 1 | 25 |
| ube4b | 0.40637 | 0.376694769 | 0.133 | 0 | 1 | 25 |
| unm-hu7910 | 0.40637 | 0.365428326 | 0.133 | 0 | 1 | 25 |
| usp8 | 0.40637 | 0.280385018 | 0.133 | 0 | 1 | 25 |
| wu:fi37e09 | 0.40637 | 0.349882046 | 0.133 | 0 | 1 | 25 |
| zfp36l1a | 0.40637 | 0.285508712 | 0.133 | 0 | 1 | 25 |
| zgc:101016 | 0.40637 | 0.300922651 | 0.133 | 0 | 1 | 25 |
| zgc:113333 | 0.40637 | 0.235958576 | 0.133 | 0 | 1 | 25 |
| zgc:152891 | 0.40637 | 0.196769094 | 0.133 | 0 | 1 | 25 |
| zgc:194312 | 0.40637 | 0.348532392 | 0.133 | 0 | 1 | 25 |
| GNB4 | 0.40637 | 0.257090115 | 0.133 | 0 | 1 | 25 |
| LOC110437825 | 0.40637 | 0.276125273 | 0.133 | 0 | 1 | 25 |
| arl6ip5b | 0.40637 | 0.180873858 | 0.133 | 0 | 1 | 25 |
| arrb2a | 0.40637 | 0.242543347 | 0.133 | 0 | 1 | 25 |
| atf4b | 0.40637 | 0.218063585 | 0.133 | 0 | 1 | 25 |
| atp6v1d | 0.40637 | 0.171657823 | 0.133 | 0 | 1 | 25 |
| b3gnt5b | 0.40637 | 0.277694855 | 0.133 | 0 | 1 | 25 |
| blcap | 0.40637 | 0.325118596 | 0.133 | 0 | 1 | 25 |
| brd2a | 0.40637 | 0.255749931 | 0.133 | 0 | 1 | 25 |

| | | | | | | |
|---|---|---|---|---|---|---|
| cactin | 0.40637 | 0.284946291 | 0.133 | 0 | 1 | 25 |
| cd44b | 0.40637 | 0.263619486 | 0.133 | 0 | 1 | 25 |
| commd7 | 0.40637 | 0.222911351 | 0.133 | 0 | 1 | 25 |
| cs | 0.40637 | 0.272383138 | 0.133 | 0 | 1 | 25 |
| ctdsp1 | 0.40637 | 0.27618501 | 0.133 | 0 | 1 | 25 |
| def6a | 0.40637 | 0.170458096 | 0.133 | 0 | 1 | 25 |
| dhx15 | 0.40637 | 0.304880528 | 0.133 | 0 | 1 | 25 |
| dynlrb1 | 0.40637 | 0.206358795 | 0.133 | 0 | 1 | 25 |
| ebag9 | 0.40637 | 0.235873489 | 0.133 | 0 | 1 | 25 |
| eif2s1a | 0.40637 | 0.210086018 | 0.133 | 0 | 1 | 25 |
| eif3s10 | 0.40637 | 0.227010423 | 0.133 | 0 | 1 | 25 |
| eif3s6ip | 0.40637 | 0.194926382 | 0.133 | 0 | 1 | 25 |
| eif4bb | 0.40637 | 0.308741762 | 0.133 | 0 | 1 | 25 |
| emc6 | 0.40637 | 0.235265467 | 0.133 | 0 | 1 | 25 |
| fam126a | 0.40637 | 0.375765532 | 0.133 | 0 | 1 | 25 |
| fam49a | 0.40637 | 0.294507719 | 0.133 | 0 | 1 | 25 |
| fli1b | 0.40637 | 0.268064124 | 0.133 | 0 | 1 | 25 |
| fryl | 0.40637 | 0.311060756 | 0.133 | 0 | 1 | 25 |
| glod5 | 0.40637 | 0.330450675 | 0.133 | 0 | 1 | 25 |
| gm2a | 0.40637 | 0.321845435 | 0.133 | 0 | 1 | 25 |
| gpr183b | 0.40637 | 0.315412508 | 0.133 | 0 | 1 | 25 |
| h2afva | 0.40637 | 0.1926527 | 0.133 | 0 | 1 | 25 |
| hikeshi | 0.40637 | 0.280495345 | 0.133 | 0 | 1 | 25 |
| il10ra | 0.40637 | 0.25174561 | 0.133 | 0 | 1 | 25 |
| insra | 0.40637 | 0.288338452 | 0.133 | 0 | 1 | 25 |
| lsm4 | 0.40637 | 0.232149039 | 0.133 | 0 | 1 | 25 |
| mapk14a | 0.40637 | 0.267111852 | 0.133 | 0 | 1 | 25 |
| mbpb | 0.40637 | 0.358149571 | 0.133 | 0 | 1 | 25 |
| metap2b | 0.40637 | 0.169775872 | 0.133 | 0 | 1 | 25 |
| midn | 0.40637 | 0.293634137 | 0.133 | 0 | 1 | 25 |
| mknk2b | 0.40637 | 0.378377511 | 0.133 | 0 | 1 | 25 |
| mrpl36 | 0.40637 | 0.173192947 | 0.133 | 0 | 1 | 25 |
| naa10 | 0.40637 | 0.255465593 | 0.133 | 0 | 1 | 25 |
| nadka | 0.40637 | 0.193338863 | 0.133 | 0 | 1 | 25 |
| nfe2l2a | 0.40637 | 0.294419502 | 0.133 | 0 | 1 | 25 |
| nr1d2a | 0.40637 | 0.238735858 | 0.133 | 0 | 1 | 25 |
| ostc | 0.40637 | 0.334091994 | 0.133 | 0 | 1 | 25 |
| parm1 | 0.40637 | 0.283159052 | 0.133 | 0 | 1 | 25 |
| pcyox1 | 0.40637 | 0.232549588 | 0.133 | 0 | 1 | 25 |
| pde4ba | 0.40637 | 0.245242428 | 0.133 | 0 | 1 | 25 |
| pgm1 | 0.40637 | 0.226104295 | 0.133 | 0 | 1 | 25 |
| phf5a | 0.40637 | 0.216409112 | 0.133 | 0 | 1 | 25 |
| plekho2 | 0.40637 | 0.35462056 | 0.133 | 0 | 1 | 25 |
| rab3db | 0.40637 | 0.151163777 | 0.133 | 0 | 1 | 25 |
| rhocb | 0.40637 | 0.264326885 | 0.133 | 0 | 1 | 25 |
| rraga | 0.40637 | 0.214608884 | 0.133 | 0 | 1 | 25 |
| rsrp1 | 0.40637 | 0.229095764 | 0.133 | 0 | 1 | 25 |
| scamp2 | 0.40637 | 0.341870742 | 0.133 | 0 | 1 | 25 |

| | | | | | | |
|---|---|---|---|---|---|---|
| scarb2a | 0.40637 | 0.361496538 | 0.133 | 0 | 1 | 25 |
| sdhaf4 | 0.40637 | 0.221223047 | 0.133 | 0 | 1 | 25 |
| sema4aa | 0.40637 | 0.404033399 | 0.133 | 0 | 1 | 25 |
| sept9b | 0.40637 | 0.260041635 | 0.133 | 0 | 1 | 25 |
| si:ch211-197h24.9 | 0.40637 | 0.296939739 | 0.133 | 0 | 1 | 25 |
| si:ch211-235e9.6 | 0.40637 | 0.187172442 | 0.133 | 0 | 1 | 25 |
| si:dkey-112a7.4 | 0.40637 | 0.243349623 | 0.133 | 0 | 1 | 25 |
| slc25a11 | 0.40637 | 0.173161013 | 0.133 | 0 | 1 | 25 |
| slc25a28 | 0.40637 | 0.332157681 | 0.133 | 0 | 1 | 25 |
| slc25a55a | 0.40637 | 0.214454753 | 0.133 | 0 | 1 | 25 |
| srsf3b | 0.40637 | 0.248877592 | 0.133 | 0 | 1 | 25 |
| ssuh2rs1 | 0.40637 | 0.269155642 | 0.133 | 0 | 1 | 25 |
| stim1a | 0.40637 | 0.306200416 | 0.133 | 0 | 1 | 25 |
| stx11a | 0.40637 | 0.342979973 | 0.133 | 0 | 1 | 25 |
| suco | 0.40637 | 0.238171703 | 0.133 | 0 | 1 | 25 |
| swap70b | 0.40637 | 0.232206713 | 0.133 | 0 | 1 | 25 |
| taf12 | 0.40637 | 0.142488738 | 0.133 | 0 | 1 | 25 |
| taf9 | 0.40637 | 0.215501017 | 0.133 | 0 | 1 | 25 |
| tfr1b | 0.40637 | 0.284666358 | 0.133 | 0 | 1 | 25 |
| tma7 | 0.40637 | 0.243507756 | 0.133 | 0 | 1 | 25 |
| tmem18 | 0.40637 | 0.280210683 | 0.133 | 0 | 1 | 25 |
| tob1b | 0.40637 | 0.326282927 | 0.133 | 0 | 1 | 25 |
| tpmt.1 | 0.40637 | 0.213021004 | 0.133 | 0 | 1 | 25 |
| triqk | 0.40637 | 0.299312913 | 0.133 | 0 | 1 | 25 |
| uqcrfs1 | 0.40637 | 0.169853616 | 0.133 | 0 | 1 | 25 |
| usp39 | 0.40637 | 0.247555132 | 0.133 | 0 | 1 | 25 |
| vcp | 0.40637 | 0.249225224 | 0.133 | 0 | 1 | 25 |
| wsb1 | 0.40637 | 0.213388833 | 0.133 | 0 | 1 | 25 |
| zc4h2 | 0.40637 | 0.255245188 | 0.133 | 0 | 1 | 25 |
| zgc:165573 | 0.40637 | 0.17565801 | 0.133 | 0 | 1 | 25 |
| zgc:65873 | 0.40637 | 0.369658039 | 0.133 | 0 | 1 | 25 |
| znf451 | 0.40637 | 0.27241273 | 0.133 | 0 | 1 | 25 |
| CZQB01141835.1 | 0.40637 | 0.163720428 | 0.133 | 0 | 1 | 25 |
| LOC108181220 | 0.40637 | 0.291035406 | 0.133 | 0 | 1 | 25 |
| hdlbpa | 0.40637 | 0.314266806 | 0.133 | 0 | 1 | 25 |
| ncf4 | 0.40637 | 0.354269891 | 0.133 | 0 | 1 | 25 |
| ola1 | 0.40637 | 0.343238176 | 0.133 | 0 | 1 | 25 |
| thy1 | 0.41084 | -0.37145508 | 0.733 | 1 | 1 | 25 |
| si:ch211-9d9.8 | 0.41323 | -0.60096473 | 0.4 | 0.75 | 1 | 25 |
| nr4a1 | 0.41405 | -0.24141873 | 0.2 | 0.5 | 1 | 25 |
| rpl13a | 0.41552 | 0.522860005 | 0.6 | 0.25 | 1 | 25 |
| tpt1 | 0.43243 | 0.242205762 | 0.9 | 1 | 1 | 25 |
| srprb | 0.43825 | -0.77463044 | 0.2 | 0.5 | 1 | 25 |
| gabarapb | 0.43895 | -0.77373922 | 0.2 | 0.5 | 1 | 25 |
| si:ch211-132p1.3 | 0.44461 | -0.76212187 | 0.2 | 0.5 | 1 | 25 |
| rpl23 | 0.44861 | 1.002959238 | 0.467 | 0.25 | 1 | 25 |
| slc25a5 | 0.44873 | 0.601635915 | 0.9 | 0.75 | 1 | 25 |
| hmgb2a | 0.45545 | 0.802445524 | 0.7 | 0.5 | 1 | 25 |

| | | | | | | |
|---|---|---|---|---|---|---|
| mapk11 | 0.45956 | -0.36563742 | 0.2 | 0.5 | 1 | 25 |
| ndufa6 | 0.46199 | -0.69248144 | 0.2 | 0.5 | 1 | 25 |
| bdh1 | 0.46531 | -0.61075667 | 0.2 | 0.5 | 1 | 25 |
| gsto2 | 0.46682 | 0.914848828 | 0.5 | 0.25 | 1 | 25 |
| rpl35a | 0.46936 | 1.00857737 | 0.467 | 0.25 | 1 | 25 |
| ndufb2 | 0.47032 | -0.58279195 | 0.2 | 0.5 | 1 | 25 |
| BX005340.1 | 0.47441 | -0.45778318 | 0.2 | 0.5 | 1 | 25 |
| mcl1b | 0.47562 | -0.35190097 | 0.433 | 0.75 | 1 | 25 |
| rps25 | 0.47648 | 0.69430266 | 0.567 | 0.25 | 1 | 25 |
| BX908782.3 | 0.47761 | 0.483426566 | 0.933 | 0.75 | 1 | 25 |
| rpl30 | 0.47809 | 0.824573684 | 0.533 | 0.25 | 1 | 25 |
| rps11 | 0.48293 | 0.80100193 | 0.533 | 0.25 | 1 | 25 |
| rpl13 | 0.48353 | -0.37290398 | 0.433 | 0.75 | 1 | 25 |
| atp5fa1 | 0.49034 | -0.37296157 | 0.433 | 0.75 | 1 | 25 |
| eef1b2 | 0.49037 | 0.491282552 | 0.567 | 0.25 | 1 | 25 |
| rpl24 | 0.49078 | 0.606611079 | 0.567 | 0.25 | 1 | 25 |
| rpl11 | 0.49103 | 0.55882304 | 0.567 | 0.25 | 1 | 25 |
| si:ch211-193e13.5 | 0.49121 | 0.557182172 | 0.567 | 0.25 | 1 | 25 |
| itm2ba | 0.49127 | 0.78136961 | 0.533 | 0.25 | 1 | 25 |
| adam8a | 0.49167 | 0.595611094 | 0.567 | 0.25 | 1 | 25 |
| rps19 | 0.49171 | 0.597326725 | 0.567 | 0.25 | 1 | 25 |
| rack1 | 0.49468 | 0.623874201 | 0.767 | 0.5 | 1 | 25 |
| rps27.1 | 0.49554 | 0.735386747 | 0.7 | 0.5 | 1 | 25 |
| acp5a | 0.49575 | 0.265882232 | 0.1 | 0 | 1 | 25 |
| arhgef2 | 0.49575 | 0.276436489 | 0.1 | 0 | 1 | 25 |
| atp5mc1 | 0.49575 | 0.257348765 | 0.1 | 0 | 1 | 25 |
| atrx | 0.49575 | 0.247010032 | 0.1 | 0 | 1 | 25 |
| brms1 | 0.49575 | 0.294166157 | 0.1 | 0 | 1 | 25 |
| cdc123 | 0.49575 | 0.244409184 | 0.1 | 0 | 1 | 25 |
| cdkn1d | 0.49575 | 0.338057325 | 0.1 | 0 | 1 | 25 |
| cds2 | 0.49575 | 0.242534633 | 0.1 | 0 | 1 | 25 |
| col14a1a | 0.49575 | 0.29626875 | 0.1 | 0 | 1 | 25 |
| cops3 | 0.49575 | 0.286310414 | 0.1 | 0 | 1 | 25 |
| cpsf6 | 0.49575 | 0.297499967 | 0.1 | 0 | 1 | 25 |
| dnm1l | 0.49575 | 0.346078783 | 0.1 | 0 | 1 | 25 |
| eif1ad | 0.49575 | 0.269854224 | 0.1 | 0 | 1 | 25 |
| eif5 | 0.49575 | 0.236970921 | 0.1 | 0 | 1 | 25 |
| elof1 | 0.49575 | 0.182108755 | 0.1 | 0 | 1 | 25 |
| emc2 | 0.49575 | 0.141223574 | 0.1 | 0 | 1 | 25 |
| ergic2 | 0.49575 | 0.228236359 | 0.1 | 0 | 1 | 25 |
| esf1 | 0.49575 | 0.203677246 | 0.1 | 0 | 1 | 25 |
| fhit | 0.49575 | 0.246900403 | 0.1 | 0 | 1 | 25 |
| fmnl1a | 0.49575 | 0.29715166 | 0.1 | 0 | 1 | 25 |
| gbp | 0.49575 | 0.231224895 | 0.1 | 0 | 1 | 25 |
| gnptg | 0.49575 | 0.275250332 | 0.1 | 0 | 1 | 25 |
| gpatch8 | 0.49575 | 0.185180488 | 0.1 | 0 | 1 | 25 |
| gskip | 0.49575 | 0.298856929 | 0.1 | 0 | 1 | 25 |
| hgsnat | 0.49575 | 0.2719107 | 0.1 | 0 | 1 | 25 |

| | | | | | | |
|---|---|---|---|---|---|---|
| itgb1bp1 | 0.49575 | 0.221995991 | 0.1 | 0 | 1 | 25 |
| lactb | 0.49575 | 0.193483752 | 0.1 | 0 | 1 | 25 |
| larp4b | 0.49575 | 0.144971982 | 0.1 | 0 | 1 | 25 |
| lcmt1 | 0.49575 | 0.142049291 | 0.1 | 0 | 1 | 25 |
| lgals3a | 0.49575 | 0.192696023 | 0.1 | 0 | 1 | 25 |
| m6pr | 0.49575 | 0.25920761 | 0.1 | 0 | 1 | 25 |
| mcrs1 | 0.49575 | 0.115480198 | 0.1 | 0 | 1 | 25 |
| mkrn4 | 0.49575 | 0.26860481 | 0.1 | 0 | 1 | 25 |
| nagk | 0.49575 | 0.192959596 | 0.1 | 0 | 1 | 25 |
| ndufab1a | 0.49575 | 0.131746196 | 0.1 | 0 | 1 | 25 |
| ndufb9 | 0.49575 | 0.188213963 | 0.1 | 0 | 1 | 25 |
| nop58 | 0.49575 | 0.100437571 | 0.1 | 0 | 1 | 25 |
| ophn1 | 0.49575 | 0.22503904 | 0.1 | 0 | 1 | 25 |
| park7 | 0.49575 | 0.204311251 | 0.1 | 0 | 1 | 25 |
| pdcd6ip | 0.49575 | 0.245156065 | 0.1 | 0 | 1 | 25 |
| pggt1b | 0.49575 | 0.277998382 | 0.1 | 0 | 1 | 25 |
| prkar2aa | 0.49575 | 0.210672706 | 0.1 | 0 | 1 | 25 |
| ptpn22 | 0.49575 | 0.314111008 | 0.1 | 0 | 1 | 25 |
| pum1 | 0.49575 | 0.203613044 | 0.1 | 0 | 1 | 25 |
| rhbdf1a | 0.49575 | 0.313709062 | 0.1 | 0 | 1 | 25 |
| rsl1d1 | 0.49575 | 0.206715096 | 0.1 | 0 | 1 | 25 |
| si:ch211-15b10.6 | 0.49575 | 0.290576917 | 0.1 | 0 | 1 | 25 |
| si:ch73-359m17.2 | 0.49575 | 0.169552242 | 0.1 | 0 | 1 | 25 |
| si:zfos-464b6.2 | 0.49575 | 0.357043468 | 0.1 | 0 | 1 | 25 |
| slc12a9 | 0.49575 | 0.319314693 | 0.1 | 0 | 1 | 25 |
| smim20 | 0.49575 | 0.289385889 | 0.1 | 0 | 1 | 25 |
| spink2.1 | 0.49575 | 0.341066206 | 0.1 | 0 | 1 | 25 |
| srp19 | 0.49575 | 0.25297046 | 0.1 | 0 | 1 | 25 |
| tab1 | 0.49575 | 0.232755617 | 0.1 | 0 | 1 | 25 |
| tnfaip8l2b | 0.49575 | 0.225912131 | 0.1 | 0 | 1 | 25 |
| tspy | 0.49575 | 0.188594254 | 0.1 | 0 | 1 | 25 |
| yipf6 | 0.49575 | 0.224343781 | 0.1 | 0 | 1 | 25 |
| zc3h11a | 0.49575 | 0.240274164 | 0.1 | 0 | 1 | 25 |
| zgc:153759 | 0.49575 | 0.217514925 | 0.1 | 0 | 1 | 25 |
| BX005309.1 | 0.49575 | 0.170789718 | 0.1 | 0 | 1 | 25 |
| BX537342.1 | 0.49575 | 0.118497412 | 0.1 | 0 | 1 | 25 |
| CABZ01058650.1 | 0.49575 | 0.166052578 | 0.1 | 0 | 1 | 25 |
| CABZ01087560.1 | 0.49575 | 0.358778427 | 0.1 | 0 | 1 | 25 |
| LOC101884098 | 0.49575 | 0.248437299 | 0.1 | 0 | 1 | 25 |
| XLOC-004189 | 0.49575 | 0.265269594 | 0.1 | 0 | 1 | 25 |
| XLOC-039619 | 0.49575 | 0.147380812 | 0.1 | 0 | 1 | 25 |
| ablim1a | 0.49575 | 0.182633232 | 0.1 | 0 | 1 | 25 |
| adcy7 | 0.49575 | 0.305823821 | 0.1 | 0 | 1 | 25 |
| adgre5b.3 | 0.49575 | 0.257979216 | 0.1 | 0 | 1 | 25 |
| adora2b | 0.49575 | 0.211554543 | 0.1 | 0 | 1 | 25 |
| adssl | 0.49575 | 0.226772818 | 0.1 | 0 | 1 | 25 |
| afap1l1b | 0.49575 | 0.217582373 | 0.1 | 0 | 1 | 25 |
| aldocb | 0.49575 | 0.146354281 | 0.1 | 0 | 1 | 25 |

| | | | | | | |
|---|---|---|---|---|---|---|
| ankhd1 | 0.49575 | 0.208875142 | 0.1 | 0 | 1 | 25 |
| ap1g2 | 0.49575 | 0.197913576 | 0.1 | 0 | 1 | 25 |
| ap1m1 | 0.49575 | 0.20204391 | 0.1 | 0 | 1 | 25 |
| ap2b1 | 0.49575 | 0.252247518 | 0.1 | 0 | 1 | 25 |
| ap3d1 | 0.49575 | 0.176768836 | 0.1 | 0 | 1 | 25 |
| appbp2 | 0.49575 | 0.256859751 | 0.1 | 0 | 1 | 25 |
| arf5 | 0.49575 | 0.161773585 | 0.1 | 0 | 1 | 25 |
| arrdc3b | 0.49575 | 0.26595847 | 0.1 | 0 | 1 | 25 |
| asap1b | 0.49575 | 0.246071746 | 0.1 | 0 | 1 | 25 |
| atg101 | 0.49575 | 0.244723739 | 0.1 | 0 | 1 | 25 |
| atp6v1ba | 0.49575 | 0.132546622 | 0.1 | 0 | 1 | 25 |
| atp6v1c1a | 0.49575 | 0.13344857 | 0.1 | 0 | 1 | 25 |
| bada | 0.49575 | 0.18106106 | 0.1 | 0 | 1 | 25 |
| baz2ba | 0.49575 | 0.213054099 | 0.1 | 0 | 1 | 25 |
| bcl2l1 | 0.49575 | 0.169591591 | 0.1 | 0 | 1 | 25 |
| bola1 | 0.49575 | 0.210798442 | 0.1 | 0 | 1 | 25 |
| btbd10b | 0.49575 | 0.293799174 | 0.1 | 0 | 1 | 25 |
| casd1 | 0.49575 | 0.24949717 | 0.1 | 0 | 1 | 25 |
| cbx5 | 0.49575 | 0.234695072 | 0.1 | 0 | 1 | 25 |
| cdc26 | 0.49575 | 0.146882769 | 0.1 | 0 | 1 | 25 |
| cdc42se1 | 0.49575 | 0.164001582 | 0.1 | 0 | 1 | 25 |
| cdv3 | 0.49575 | 0.315329738 | 0.1 | 0 | 1 | 25 |
| chchd6b | 0.49575 | 0.181513845 | 0.1 | 0 | 1 | 25 |
| chmp1b | 0.49575 | 0.156937667 | 0.1 | 0 | 1 | 25 |
| ciao1 | 0.49575 | 0.375253494 | 0.1 | 0 | 1 | 25 |
| cltca | 0.49575 | 0.232525382 | 0.1 | 0 | 1 | 25 |
| coa5 | 0.49575 | 0.192167998 | 0.1 | 0 | 1 | 25 |
| cops9 | 0.49575 | 0.211798371 | 0.1 | 0 | 1 | 25 |
| cox6b1 | 0.49575 | 0.189665238 | 0.1 | 0 | 1 | 25 |
| cst14b.2 | 0.49575 | 0.197837554 | 0.1 | 0 | 1 | 25 |
| ctdspl2a | 0.49575 | 0.127149907 | 0.1 | 0 | 1 | 25 |
| cyb5r2 | 0.49575 | 0.139667067 | 0.1 | 0 | 1 | 25 |
| cyfip1 | 0.49575 | 0.273800528 | 0.1 | 0 | 1 | 25 |
| ddx27 | 0.49575 | 0.169172865 | 0.1 | 0 | 1 | 25 |
| dennd2da | 0.49575 | 0.111838292 | 0.1 | 0 | 1 | 25 |
| dnaja2b | 0.49575 | 0.296951988 | 0.1 | 0 | 1 | 25 |
| dock8 | 0.49575 | 0.166957693 | 0.1 | 0 | 1 | 25 |
| dop1b | 0.49575 | 0.205207949 | 0.1 | 0 | 1 | 25 |
| eef1db | 0.49575 | 0.166565749 | 0.1 | 0 | 1 | 25 |
| eef2k | 0.49575 | 0.268119804 | 0.1 | 0 | 1 | 25 |
| eif4a1a | 0.49575 | 0.258250555 | 0.1 | 0 | 1 | 25 |
| eif4a2 | 0.49575 | 0.212691262 | 0.1 | 0 | 1 | 25 |
| emc7 | 0.49575 | 0.312134147 | 0.1 | 0 | 1 | 25 |
| ero1b | 0.49575 | 0.201939027 | 0.1 | 0 | 1 | 25 |
| fam117ab | 0.49575 | 0.285105646 | 0.1 | 0 | 1 | 25 |
| fbxl14b | 0.49575 | 0.182479917 | 0.1 | 0 | 1 | 25 |
| fli1a | 0.49575 | 0.153969161 | 0.1 | 0 | 1 | 25 |
| flot1b | 0.49575 | 0.182437276 | 0.1 | 0 | 1 | 25 |

| | | | | | | |
|---|---|---|---|---|---|---|
| fnta | 0.49575 | 0.144354453 | 0.1 | 0 | 1 | 25 |
| gabpa | 0.49575 | 0.260613651 | 0.1 | 0 | 1 | 25 |
| gigyf1b | 0.49575 | 0.290218823 | 0.1 | 0 | 1 | 25 |
| gins2 | 0.49575 | 0.178848544 | 0.1 | 0 | 1 | 25 |
| gnao1a | 0.49575 | 0.232709596 | 0.1 | 0 | 1 | 25 |
| gramd1c | 0.49575 | 0.335682235 | 0.1 | 0 | 1 | 25 |
| grk6 | 0.49575 | 0.180492251 | 0.1 | 0 | 1 | 25 |
| grpel2 | 0.49575 | 0.174392577 | 0.1 | 0 | 1 | 25 |
| hectd1 | 0.49575 | 0.230026997 | 0.1 | 0 | 1 | 25 |
| hephl1a | 0.49575 | 0.191000932 | 0.1 | 0 | 1 | 25 |
| hif1ab | 0.49575 | 0.146048535 | 0.1 | 0 | 1 | 25 |
| higd1a | 0.49575 | 0.178424095 | 0.1 | 0 | 1 | 25 |
| higd2a | 0.49575 | 0.11329834 | 0.1 | 0 | 1 | 25 |
| hmgb2b | 0.49575 | 0.302559361 | 0.1 | 0 | 1 | 25 |
| hnrnpa0a | 0.49575 | 0.144619046 | 0.1 | 0 | 1 | 25 |
| ier5 | 0.49575 | 0.266919602 | 0.1 | 0 | 1 | 25 |
| ints13 | 0.49575 | 0.254503332 | 0.1 | 0 | 1 | 25 |
| kcnip3b | 0.49575 | 0.18278536 | 0.1 | 0 | 1 | 25 |
| khdrbs1b | 0.49575 | 0.132037439 | 0.1 | 0 | 1 | 25 |
| lmnb1 | 0.49575 | 0.258681016 | 0.1 | 0 | 1 | 25 |
| lmod1a | 0.49575 | 0.230490444 | 0.1 | 0 | 1 | 25 |
| lrmp | 0.49575 | 0.297414398 | 0.1 | 0 | 1 | 25 |
| man1a2 | 0.49575 | 0.332238236 | 0.1 | 0 | 1 | 25 |
| map2k6 | 0.49575 | 0.298637368 | 0.1 | 0 | 1 | 25 |
| mapkapk2a | 0.49575 | 0.29061538 | 0.1 | 0 | 1 | 25 |
| matr3l1.1 | 0.49575 | 0.208277644 | 0.1 | 0 | 1 | 25 |
| mbd3a | 0.49575 | 0.22365458 | 0.1 | 0 | 1 | 25 |
| mbd4 | 0.49575 | 0.179192622 | 0.1 | 0 | 1 | 25 |
| mboat7 | 0.49575 | 0.253792659 | 0.1 | 0 | 1 | 25 |
| mcts1 | 0.49575 | 0.183137044 | 0.1 | 0 | 1 | 25 |
| mgat5 | 0.49575 | 0.301636035 | 0.1 | 0 | 1 | 25 |
| mrps18c | 0.49575 | 0.169759271 | 0.1 | 0 | 1 | 25 |
| msl2b | 0.49575 | 0.205988719 | 0.1 | 0 | 1 | 25 |
| myb | 0.49575 | 0.115828572 | 0.1 | 0 | 1 | 25 |
| nabp1a | 0.49575 | 0.263557679 | 0.1 | 0 | 1 | 25 |
| ndrg3b | 0.49575 | 0.109933331 | 0.1 | 0 | 1 | 25 |
| ndufb10 | 0.49575 | 0.12130063 | 0.1 | 0 | 1 | 25 |
| nek7 | 0.49575 | 0.203360339 | 0.1 | 0 | 1 | 25 |
| nfatc2ip | 0.49575 | 0.214094855 | 0.1 | 0 | 1 | 25 |
| nup50 | 0.49575 | 0.11232377 | 0.1 | 0 | 1 | 25 |
| ociad1 | 0.49575 | 0.144114039 | 0.1 | 0 | 1 | 25 |
| opa1 | 0.49575 | 0.161931014 | 0.1 | 0 | 1 | 25 |
| p2ry13 | 0.49575 | 0.259985479 | 0.1 | 0 | 1 | 25 |
| pdcd10a | 0.49575 | 0.193543966 | 0.1 | 0 | 1 | 25 |
| pdcd10b | 0.49575 | 0.236514739 | 0.1 | 0 | 1 | 25 |
| pfn2 | 0.49575 | 0.229284821 | 0.1 | 0 | 1 | 25 |
| pitpnaa | 0.49575 | 0.212935795 | 0.1 | 0 | 1 | 25 |
| pitpnb | 0.49575 | 0.179758802 | 0.1 | 0 | 1 | 25 |

| | | | | | | |
|---|---|---|---|---|---|---|
| plcg2 | 0.49575 | 0.236104335 | 0.1 | 0 | 1 | 25 |
| pnn | 0.49575 | 0.231642853 | 0.1 | 0 | 1 | 25 |
| polr1d | 0.49575 | 0.225733941 | 0.1 | 0 | 1 | 25 |
| ppp2cb | 0.49575 | 0.219777041 | 0.1 | 0 | 1 | 25 |
| ppp4r1 | 0.49575 | 0.249931981 | 0.1 | 0 | 1 | 25 |
| ppp6r3 | 0.49575 | 0.275220016 | 0.1 | 0 | 1 | 25 |
| prkaa1 | 0.49575 | 0.212833651 | 0.1 | 0 | 1 | 25 |
| prrc2a | 0.49575 | 0.21986778 | 0.1 | 0 | 1 | 25 |
| psma5 | 0.49575 | 0.173149867 | 0.1 | 0 | 1 | 25 |
| psma6b | 0.49575 | 0.140928583 | 0.1 | 0 | 1 | 25 |
| psmd7 | 0.49575 | 0.208659738 | 0.1 | 0 | 1 | 25 |
| ptcd3 | 0.49575 | 0.124330768 | 0.1 | 0 | 1 | 25 |
| rabgap1l | 0.49575 | 0.304760833 | 0.1 | 0 | 1 | 25 |
| rbbp6 | 0.49575 | 0.163464638 | 0.1 | 0 | 1 | 25 |
| rbm4.2 | 0.49575 | 0.235185899 | 0.1 | 0 | 1 | 25 |
| rdh14b | 0.49575 | 0.19094181 | 0.1 | 0 | 1 | 25 |
| rhof | 0.49575 | 0.275832654 | 0.1 | 0 | 1 | 25 |
| rnf11a | 0.49575 | 0.268088623 | 0.1 | 0 | 1 | 25 |
| rtn4ip1 | 0.49575 | 0.163423126 | 0.1 | 0 | 1 | 25 |
| sars | 0.49575 | 0.293025872 | 0.1 | 0 | 1 | 25 |
| sdhdb | 0.49575 | 0.26584012 | 0.1 | 0 | 1 | 25 |
| selp | 0.49575 | 0.346618124 | 0.1 | 0 | 1 | 25 |
| sema4ab | 0.49575 | 0.241660763 | 0.1 | 0 | 1 | 25 |
| sept7a | 0.49575 | 0.100809714 | 0.1 | 0 | 1 | 25 |
| serpina10a | 0.49575 | 0.234599003 | 0.1 | 0 | 1 | 25 |
| serpinb14 | 0.49575 | 0.349245546 | 0.1 | 0 | 1 | 25 |
| sft2d1 | 0.49575 | 0.202521419 | 0.1 | 0 | 1 | 25 |
| sh3gl3a | 0.49575 | 0.169383535 | 0.1 | 0 | 1 | 25 |
| si:ch211-122f10.4 | 0.49575 | 0.20353103 | 0.1 | 0 | 1 | 25 |
| si:ch211-214k5.3 | 0.49575 | 0.160176815 | 0.1 | 0 | 1 | 25 |
| si:ch211-233a24.2 | 0.49575 | 0.305837425 | 0.1 | 0 | 1 | 25 |
| si:ch211-288g17.3 | 0.49575 | 0.225525012 | 0.1 | 0 | 1 | 25 |
| si:ch211-51e12.7 | 0.49575 | 0.276999847 | 0.1 | 0 | 1 | 25 |
| si:ch211-66k16.27 | 0.49575 | 0.146527828 | 0.1 | 0 | 1 | 25 |
| si:ch73-208g10.1 | 0.49575 | 0.271631527 | 0.1 | 0 | 1 | 25 |
| si:ch73-233k15.2 | 0.49575 | 0.305058547 | 0.1 | 0 | 1 | 25 |
| si:ch73-359m17.5 | 0.49575 | 0.228535573 | 0.1 | 0 | 1 | 25 |
| si:ch73-7i4.3 | 0.49575 | 0.294023877 | 0.1 | 0 | 1 | 25 |
| si:dkey-238m4.4 | 0.49575 | 0.29000605 | 0.1 | 0 | 1 | 25 |
| si:dkey-56i24.1 | 0.49575 | 0.118497412 | 0.1 | 0 | 1 | 25 |
| si:dkey-67c22.2 | 0.49575 | 0.213409242 | 0.1 | 0 | 1 | 25 |
| si:dkey-7j14.6 | 0.49575 | 0.236633784 | 0.1 | 0 | 1 | 25 |
| si:dkeyp-110e4.6 | 0.49575 | 0.340926401 | 0.1 | 0 | 1 | 25 |
| slc3a2b | 0.49575 | 0.216334692 | 0.1 | 0 | 1 | 25 |
| slc4a7 | 0.49575 | 0.278314041 | 0.1 | 0 | 1 | 25 |
| snrpe | 0.49575 | 0.132128089 | 0.1 | 0 | 1 | 25 |
| sntb1 | 0.49575 | 0.217140183 | 0.1 | 0 | 1 | 25 |
| spast | 0.49575 | 0.332024966 | 0.1 | 0 | 1 | 25 |

| | | | | | | |
|---|---|---|---|---|---|---|
| srebf2 | 0.49575 | 0.252229994 | 0.1 | 0 | 1 | 25 |
| srp72 | 0.49575 | 0.3356613 | 0.1 | 0 | 1 | 25 |
| st6galnac5b | 0.49575 | 0.289483777 | 0.1 | 0 | 1 | 25 |
| supt4h1 | 0.49575 | 0.104595494 | 0.1 | 0 | 1 | 25 |
| swi5 | 0.49575 | 0.176739167 | 0.1 | 0 | 1 | 25 |
| sybl1 | 0.49575 | 0.188470995 | 0.1 | 0 | 1 | 25 |
| tacc1 | 0.49575 | 0.276143098 | 0.1 | 0 | 1 | 25 |
| tax1bp1b | 0.49575 | 0.165335801 | 0.1 | 0 | 1 | 25 |
| tmc8 | 0.49575 | 0.293869528 | 0.1 | 0 | 1 | 25 |
| tmem214 | 0.49575 | 0.294713735 | 0.1 | 0 | 1 | 25 |
| tmpob | 0.49575 | 0.204174958 | 0.1 | 0 | 1 | 25 |
| tmub1 | 0.49575 | 0.287830624 | 0.1 | 0 | 1 | 25 |
| tnikb | 0.49575 | 0.155507415 | 0.1 | 0 | 1 | 25 |
| trim35-28 | 0.49575 | 0.21755161 | 0.1 | 0 | 1 | 25 |
| uba1 | 0.49575 | 0.317274058 | 0.1 | 0 | 1 | 25 |
| ubald2 | 0.49575 | 0.191163697 | 0.1 | 0 | 1 | 25 |
| ube2d4 | 0.49575 | 0.267867672 | 0.1 | 0 | 1 | 25 |
| ube2ib | 0.49575 | 0.185333124 | 0.1 | 0 | 1 | 25 |
| ube2l3a | 0.49575 | 0.160797899 | 0.1 | 0 | 1 | 25 |
| utp18 | 0.49575 | 0.335803285 | 0.1 | 0 | 1 | 25 |
| uvrag | 0.49575 | 0.19155958 | 0.1 | 0 | 1 | 25 |
| xpo6 | 0.49575 | 0.319969754 | 0.1 | 0 | 1 | 25 |
| yy1a | 0.49575 | 0.238775628 | 0.1 | 0 | 1 | 25 |
| zeb1a | 0.49575 | 0.219241876 | 0.1 | 0 | 1 | 25 |
| zgc:103510 | 0.49575 | 0.278246653 | 0.1 | 0 | 1 | 25 |
| zgc:111986 | 0.49575 | 0.252220708 | 0.1 | 0 | 1 | 25 |
| zgc:153733 | 0.49575 | 0.289544477 | 0.1 | 0 | 1 | 25 |
| zgc:173548 | 0.49575 | 0.184150412 | 0.1 | 0 | 1 | 25 |
| zgc:175088 | 0.49575 | 0.171872051 | 0.1 | 0 | 1 | 25 |
| zgc:55943 | 0.49575 | 0.267499478 | 0.1 | 0 | 1 | 25 |
| zgc:91910 | 0.49575 | 0.183962427 | 0.1 | 0 | 1 | 25 |
| zmp:0000000936 | 0.49575 | 0.243829038 | 0.1 | 0 | 1 | 25 |
| zmynd11 | 0.49575 | 0.228973263 | 0.1 | 0 | 1 | 25 |
| znhit3 | 0.49575 | 0.267503385 | 0.1 | 0 | 1 | 25 |
| zyx | 0.49575 | 0.223698871 | 0.1 | 0 | 1 | 25 |
| FQ323156.1 | 0.49575 | 0.18693237 | 0.1 | 0 | 1 | 25 |
| acot8 | 0.49575 | 0.214529112 | 0.1 | 0 | 1 | 25 |
| acss2 | 0.49575 | 0.223856393 | 0.1 | 0 | 1 | 25 |
| atp8a1 | 0.49575 | 0.173015951 | 0.1 | 0 | 1 | 25 |
| bloc1s6 | 0.49575 | 0.219159496 | 0.1 | 0 | 1 | 25 |
| bnip4 | 0.49575 | 0.133092165 | 0.1 | 0 | 1 | 25 |
| brk1 | 0.49575 | 0.14203012 | 0.1 | 0 | 1 | 25 |
| camk1b | 0.49575 | 0.220935499 | 0.1 | 0 | 1 | 25 |
| canx | 0.49575 | 0.182823193 | 0.1 | 0 | 1 | 25 |
| chmp2bb | 0.49575 | 0.183319717 | 0.1 | 0 | 1 | 25 |
| chtf8 | 0.49575 | 0.252412375 | 0.1 | 0 | 1 | 25 |
| csgalnact1a | 0.49575 | 0.235778771 | 0.1 | 0 | 1 | 25 |
| dpp7 | 0.49575 | 0.136392623 | 0.1 | 0 | 1 | 25 |

| | | | | | | |
|---|---|---|---|---|---|---|
| eif3k | 0.49575 | 0.151543969 | 0.1 | 0 | 1 | 25 |
| ep300b | 0.49575 | 0.208334637 | 0.1 | 0 | 1 | 25 |
| gadd45ba | 0.49575 | 0.234179146 | 0.1 | 0 | 1 | 25 |
| galnt7 | 0.49575 | 0.227724631 | 0.1 | 0 | 1 | 25 |
| gmfg | 0.49575 | 0.193311458 | 0.1 | 0 | 1 | 25 |
| ipmkb | 0.49575 | 0.17113211 | 0.1 | 0 | 1 | 25 |
| itgav | 0.49575 | 0.148959615 | 0.1 | 0 | 1 | 25 |
| kdm6bb | 0.49575 | 0.147273534 | 0.1 | 0 | 1 | 25 |
| lyar | 0.49575 | 0.112781718 | 0.1 | 0 | 1 | 25 |
| med19a | 0.49575 | 0.237868599 | 0.1 | 0 | 1 | 25 |
| mrpl11 | 0.49575 | 0.206273617 | 0.1 | 0 | 1 | 25 |
| mrpl30 | 0.49575 | 0.217101221 | 0.1 | 0 | 1 | 25 |
| osgep | 0.49575 | 0.211550582 | 0.1 | 0 | 1 | 25 |
| otub1b | 0.49575 | 0.223965024 | 0.1 | 0 | 1 | 25 |
| pak2b | 0.49575 | 0.207804818 | 0.1 | 0 | 1 | 25 |
| pcca | 0.49575 | 0.212522664 | 0.1 | 0 | 1 | 25 |
| pdhb | 0.49575 | 0.127578705 | 0.1 | 0 | 1 | 25 |
| plekho1b | 0.49575 | 0.224923121 | 0.1 | 0 | 1 | 25 |
| ppm1g | 0.49575 | 0.139554982 | 0.1 | 0 | 1 | 25 |
| prkab1b | 0.49575 | 0.19180014 | 0.1 | 0 | 1 | 25 |
| safb | 0.49575 | 0.18775963 | 0.1 | 0 | 1 | 25 |
| si:ch211-203d1.3 | 0.49575 | 0.27925493 | 0.1 | 0 | 1 | 25 |
| si:ch73-91k6.2 | 0.49575 | 0.276935343 | 0.1 | 0 | 1 | 25 |
| si:dkey-42i9.6 | 0.49575 | 0.175124871 | 0.1 | 0 | 1 | 25 |
| si:rp71-45k5.4 | 0.49575 | 0.119171005 | 0.1 | 0 | 1 | 25 |
| slc37a1 | 0.49575 | 0.142680902 | 0.1 | 0 | 1 | 25 |
| snrpc | 0.49575 | 0.233003148 | 0.1 | 0 | 1 | 25 |
| snu13b | 0.49575 | 0.169681217 | 0.1 | 0 | 1 | 25 |
| snx1a | 0.49575 | 0.105357186 | 0.1 | 0 | 1 | 25 |
| taf11 | 0.49575 | 0.175940124 | 0.1 | 0 | 1 | 25 |
| tbk1 | 0.49575 | 0.142787887 | 0.1 | 0 | 1 | 25 |
| tmem131l | 0.49575 | 0.206328168 | 0.1 | 0 | 1 | 25 |
| tmem256 | 0.49575 | 0.159966513 | 0.1 | 0 | 1 | 25 |
| tpp1 | 0.49575 | 0.188928693 | 0.1 | 0 | 1 | 25 |
| txnrd3 | 0.49575 | 0.140468558 | 0.1 | 0 | 1 | 25 |
| uap1l1 | 0.49575 | 0.222106868 | 0.1 | 0 | 1 | 25 |
| unc50 | 0.49575 | 0.192319231 | 0.1 | 0 | 1 | 25 |
| wtap | 0.49575 | 0.153669941 | 0.1 | 0 | 1 | 25 |
| zdhhc4 | 0.49575 | 0.213079209 | 0.1 | 0 | 1 | 25 |
| zfand5b | 0.49575 | 0.228247554 | 0.1 | 0 | 1 | 25 |
| zgc:153012 | 0.49575 | 0.1180872 | 0.1 | 0 | 1 | 25 |
| LO017700.1 | 0.49575 | 0.182254396 | 0.1 | 0 | 1 | 25 |
| XLOC-043857 | 0.49575 | 0.116559828 | 0.1 | 0 | 1 | 25 |
| afmid | 0.49575 | 0.177714994 | 0.1 | 0 | 1 | 25 |
| cdc73 | 0.49575 | 0.182440395 | 0.1 | 0 | 1 | 25 |
| cers2b | 0.49575 | 0.219804389 | 0.1 | 0 | 1 | 25 |
| clptm1l | 0.49575 | 0.200492172 | 0.1 | 0 | 1 | 25 |
| creb3l3l | 0.49575 | 0.23187685 | 0.1 | 0 | 1 | 25 |

| | | | | | | |
|---|---|---|---|---|---|---|
| ctnnb1 | 0.49575 | 0.136930155 | 0.1 | 0 | 1 | 25 |
| ctsz | 0.49575 | 0.236831427 | 0.1 | 0 | 1 | 25 |
| dctn5 | 0.49575 | 0.209530347 | 0.1 | 0 | 1 | 25 |
| emc10 | 0.49575 | 0.223214947 | 0.1 | 0 | 1 | 25 |
| galm | 0.49575 | 0.11489634 | 0.1 | 0 | 1 | 25 |
| hexb | 0.49575 | 0.241893463 | 0.1 | 0 | 1 | 25 |
| hic1l | 0.49575 | 0.145719549 | 0.1 | 0 | 1 | 25 |
| ino80e | 0.49575 | 0.153458648 | 0.1 | 0 | 1 | 25 |
| isg20 | 0.49575 | 0.107169193 | 0.1 | 0 | 1 | 25 |
| man2b1 | 0.49575 | 0.135619218 | 0.1 | 0 | 1 | 25 |
| mcm2 | 0.49575 | 0.151683135 | 0.1 | 0 | 1 | 25 |
| mir21-1 | 0.49575 | 0.194317419 | 0.1 | 0 | 1 | 25 |
| mob4 | 0.49575 | 0.188925553 | 0.1 | 0 | 1 | 25 |
| mroh1 | 0.49575 | 0.18722517 | 0.1 | 0 | 1 | 25 |
| myo18ab | 0.49575 | 0.207545208 | 0.1 | 0 | 1 | 25 |
| nat9 | 0.49575 | 0.191598109 | 0.1 | 0 | 1 | 25 |
| pgam5 | 0.49575 | 0.254105997 | 0.1 | 0 | 1 | 25 |
| pik3cg | 0.49575 | 0.13921787 | 0.1 | 0 | 1 | 25 |
| plcg1 | 0.49575 | 0.14692388 | 0.1 | 0 | 1 | 25 |
| ppid | 0.49575 | 0.119465899 | 0.1 | 0 | 1 | 25 |
| psmb3 | 0.49575 | 0.211866666 | 0.1 | 0 | 1 | 25 |
| ptrhd1 | 0.49575 | 0.104777014 | 0.1 | 0 | 1 | 25 |
| rasgef1ba | 0.49575 | 0.1880592 | 0.1 | 0 | 1 | 25 |
| rgs13 | 0.49575 | 0.264259082 | 0.1 | 0 | 1 | 25 |
| si:ch211-218c6.8 | 0.49575 | 0.110331147 | 0.1 | 0 | 1 | 25 |
| si:dkey-237j10.2 | 0.49575 | 0.206543793 | 0.1 | 0 | 1 | 25 |
| srsf9 | 0.49575 | 0.193669975 | 0.1 | 0 | 1 | 25 |
| susd6 | 0.49575 | 0.274666689 | 0.1 | 0 | 1 | 25 |
| taf15 | 0.49575 | 0.159784822 | 0.1 | 0 | 1 | 25 |
| tbl1xr1b | 0.49575 | 0.228968145 | 0.1 | 0 | 1 | 25 |
| tfam | 0.49575 | 0.256958088 | 0.1 | 0 | 1 | 25 |
| tia1l | 0.49575 | 0.183247617 | 0.1 | 0 | 1 | 25 |
| timm23a | 0.49575 | 0.160752922 | 0.1 | 0 | 1 | 25 |
| tm2d3 | 0.49575 | 0.135882335 | 0.1 | 0 | 1 | 25 |
| tmem39a | 0.49575 | 0.155201552 | 0.1 | 0 | 1 | 25 |
| trappc5 | 0.49575 | 0.158354992 | 0.1 | 0 | 1 | 25 |
| ube2d1b | 0.49575 | 0.232059512 | 0.1 | 0 | 1 | 25 |
| ube2l3b | 0.49575 | 0.202624679 | 0.1 | 0 | 1 | 25 |
| ufc1 | 0.49575 | 0.153137306 | 0.1 | 0 | 1 | 25 |
| usp33 | 0.49575 | 0.196899189 | 0.1 | 0 | 1 | 25 |
| wbp1 | 0.49575 | 0.13037513 | 0.1 | 0 | 1 | 25 |
| zgc:158403 | 0.49575 | 0.167852033 | 0.1 | 0 | 1 | 25 |
| ddx47 | 0.49575 | 0.251227173 | 0.1 | 0 | 1 | 25 |
| rps26 | 0.49819 | 0.899650672 | 0.467 | 0.25 | 1 | 25 |
| nccrp1 | 0.50414 | -0.11620349 | 0.8 | 1 | 1 | 25 |
| vamp8 | 0.52053 | 0.349349472 | 0.533 | 0.25 | 1 | 25 |
| rpl7 | 0.52411 | 0.761280051 | 0.6 | 0.5 | 1 | 25 |
| sec61g | 0.52875 | 0.701892693 | 0.533 | 0.25 | 1 | 25 |

| | | | | | | |
|---|---|---|---|---|---|---|
| si:ch211-67e16.3 | 0.53389 | 0.6970836 | 0.533 | 0.25 | 1 | 25 |
| tspan14 | 0.53932 | -0.67355703 | 0.233 | 0.5 | 1 | 25 |
| tmem30ab | 0.54622 | -0.63050577 | 0.233 | 0.5 | 1 | 25 |
| COX5B | 0.54763 | -0.78712791 | 0.3 | 0.5 | 1 | 25 |
| mt-co2 | 0.55053 | -0.26595533 | 0.8 | 1 | 1 | 25 |
| zgc:175264 | 0.55122 | -0.62354132 | 0.233 | 0.5 | 1 | 25 |
| ppiaa | 0.55141 | 0.438338252 | 0.733 | 0.75 | 1 | 25 |
| si:ch211-212k18.7 | 0.55143 | -0.59111682 | 0.233 | 0.5 | 1 | 25 |
| cirbpa | 0.55352 | -0.22988989 | 0.233 | 0.5 | 1 | 25 |
| p4hb | 0.55354 | -0.64281495 | 0.233 | 0.5 | 1 | 25 |
| rps2 | 0.55402 | 0.674451071 | 0.533 | 0.25 | 1 | 25 |
| zgc:64051 | 0.56083 | -0.34630001 | 0.467 | 0.75 | 1 | 25 |
| si:ch211-276a23.5 | 0.56113 | 0.735760565 | 0.5 | 0.25 | 1 | 25 |
| cnp | 0.56281 | 0.217526325 | 0.5 | 0.25 | 1 | 25 |
| cyp4f3 | 0.5664 | -0.37188889 | 0.233 | 0.5 | 1 | 25 |
| tpst1 | 0.56696 | -0.45524131 | 0.233 | 0.5 | 1 | 25 |
| si:ch211-222l21.1 | 0.56779 | 0.889904867 | 0.4 | 0.25 | 1 | 25 |
| rplp2l | 0.56877 | 0.751603152 | 0.5 | 0.25 | 1 | 25 |
| CU499336.2 | 0.56928 | -0.45152218 | 0.233 | 0.5 | 1 | 25 |
| dub | 0.56931 | -0.46507986 | 0.233 | 0.5 | 1 | 25 |
| BX908782.2 | 0.57988 | -0.31062226 | 0.933 | 1 | 1 | 25 |
| cebpb | 0.59399 | 0.270259896 | 0.933 | 0.75 | 1 | 25 |
| actb1 | 0.59776 | 0.24634575 | 0.933 | 0.75 | 1 | 25 |
| alox5ap | 0.60419 | -0.50104838 | 0.5 | 0.75 | 1 | 25 |
| cox4i1 | 0.60899 | 0.291705076 | 0.5 | 0.25 | 1 | 25 |
| znf644a | 0.60945 | 0.232826009 | 0.067 | 0 | 1 | 25 |
| FBXO46 | 0.60945 | 0.163305991 | 0.067 | 0 | 1 | 25 |
| UBL4A | 0.60945 | 0.139328454 | 0.067 | 0 | 1 | 25 |
| baz1a | 0.60945 | 0.149030732 | 0.067 | 0 | 1 | 25 |
| bptf | 0.60945 | 0.105496438 | 0.067 | 0 | 1 | 25 |
| cd164 | 0.60945 | 0.141312376 | 0.067 | 0 | 1 | 25 |
| cnppd1 | 0.60945 | 0.1122655 | 0.067 | 0 | 1 | 25 |
| ephx2 | 0.60945 | 0.1657753 | 0.067 | 0 | 1 | 25 |
| fdx1 | 0.60945 | 0.117094172 | 0.067 | 0 | 1 | 25 |
| gpat3 | 0.60945 | 0.164298841 | 0.067 | 0 | 1 | 25 |
| insig2 | 0.60945 | 0.167010413 | 0.067 | 0 | 1 | 25 |
| krit1 | 0.60945 | 0.129906809 | 0.067 | 0 | 1 | 25 |
| lonrf1l | 0.60945 | 0.111453506 | 0.067 | 0 | 1 | 25 |
| magoh | 0.60945 | 0.140432028 | 0.067 | 0 | 1 | 25 |
| map2k2a | 0.60945 | 0.150874186 | 0.067 | 0 | 1 | 25 |
| mat2aa | 0.60945 | 0.144216535 | 0.067 | 0 | 1 | 25 |
| mplkip | 0.60945 | 0.102843205 | 0.067 | 0 | 1 | 25 |
| nasp | 0.60945 | 0.186454533 | 0.067 | 0 | 1 | 25 |
| pacs1a | 0.60945 | 0.149499652 | 0.067 | 0 | 1 | 25 |
| pcmt | 0.60945 | 0.19417667 | 0.067 | 0 | 1 | 25 |
| pde4ca | 0.60945 | 0.156263814 | 0.067 | 0 | 1 | 25 |
| pqlc3 | 0.60945 | 0.110140456 | 0.067 | 0 | 1 | 25 |
| rdh12l | 0.60945 | 0.117261532 | 0.067 | 0 | 1 | 25 |

| | | | | | | |
|---|---|---|---|---|---|---|
| selenoh | 0.60945 | 0.217681602 | 0.067 | 0 | 1 | 25 |
| si:ch211-163c2.2 | 0.60945 | 0.154971382 | 0.067 | 0 | 1 | 25 |
| si:ch211-171h4.5 | 0.60945 | 0.156263814 | 0.067 | 0 | 1 | 25 |
| si:ch211-241f5.3 | 0.60945 | 0.150819218 | 0.067 | 0 | 1 | 25 |
| si:ch73-199g24.2 | 0.60945 | 0.154971382 | 0.067 | 0 | 1 | 25 |
| slc30a5 | 0.60945 | 0.101437872 | 0.067 | 0 | 1 | 25 |
| smim4 | 0.60945 | 0.104237333 | 0.067 | 0 | 1 | 25 |
| timmdc1 | 0.60945 | 0.141515963 | 0.067 | 0 | 1 | 25 |
| tom1l2a | 0.60945 | 0.103535101 | 0.067 | 0 | 1 | 25 |
| trap1 | 0.60945 | 0.105001494 | 0.067 | 0 | 1 | 25 |
| tsr1 | 0.60945 | 0.143777378 | 0.067 | 0 | 1 | 25 |
| ttpal | 0.60945 | 0.130683123 | 0.067 | 0 | 1 | 25 |
| vrk1 | 0.60945 | 0.145245127 | 0.067 | 0 | 1 | 25 |
| zer1 | 0.60945 | 0.154622456 | 0.067 | 0 | 1 | 25 |
| zgc:101663 | 0.60945 | 0.103535101 | 0.067 | 0 | 1 | 25 |
| zgc:113426 | 0.60945 | 0.150427362 | 0.067 | 0 | 1 | 25 |
| znf207b | 0.60945 | 0.104256787 | 0.067 | 0 | 1 | 25 |
| zw10 | 0.60945 | 0.103535101 | 0.067 | 0 | 1 | 25 |
| LEPROTL1 | 0.60945 | 0.124076424 | 0.067 | 0 | 1 | 25 |
| LOC101882213 | 0.60945 | 0.233275989 | 0.067 | 0 | 1 | 25 |
| RASA2 | 0.60945 | 0.189389803 | 0.067 | 0 | 1 | 25 |
| XLOC-002710 | 0.60945 | 0.223977996 | 0.067 | 0 | 1 | 25 |
| XLOC-010337 | 0.60945 | 0.198562948 | 0.067 | 0 | 1 | 25 |
| abca2 | 0.60945 | 0.148352448 | 0.067 | 0 | 1 | 25 |
| acadl | 0.60945 | 0.109004506 | 0.067 | 0 | 1 | 25 |
| adi1 | 0.60945 | 0.128016349 | 0.067 | 0 | 1 | 25 |
| agla | 0.60945 | 0.169618793 | 0.067 | 0 | 1 | 25 |
| ago2 | 0.60945 | 0.172526019 | 0.067 | 0 | 1 | 25 |
| akap10 | 0.60945 | 0.102214253 | 0.067 | 0 | 1 | 25 |
| alg5 | 0.60945 | 0.110980943 | 0.067 | 0 | 1 | 25 |
| ankrd44 | 0.60945 | 0.113768974 | 0.067 | 0 | 1 | 25 |
| ankrd52a | 0.60945 | 0.156586297 | 0.067 | 0 | 1 | 25 |
| ap3s1.1 | 0.60945 | 0.18387245 | 0.067 | 0 | 1 | 25 |
| api5 | 0.60945 | 0.157590344 | 0.067 | 0 | 1 | 25 |
| arglu1b | 0.60945 | 0.161705425 | 0.067 | 0 | 1 | 25 |
| arhgap27 | 0.60945 | 0.20159206 | 0.067 | 0 | 1 | 25 |
| arid2 | 0.60945 | 0.160086352 | 0.067 | 0 | 1 | 25 |
| atf1 | 0.60945 | 0.13962302 | 0.067 | 0 | 1 | 25 |
| atf6 | 0.60945 | 0.163692258 | 0.067 | 0 | 1 | 25 |
| atl3 | 0.60945 | 0.139913887 | 0.067 | 0 | 1 | 25 |
| blnk | 0.60945 | 0.24274236 | 0.067 | 0 | 1 | 25 |
| brd9 | 0.60945 | 0.169001473 | 0.067 | 0 | 1 | 25 |
| c1galt1b | 0.60945 | 0.229446894 | 0.067 | 0 | 1 | 25 |
| caspa | 0.60945 | 0.145823489 | 0.067 | 0 | 1 | 25 |
| cbfb | 0.60945 | 0.176830761 | 0.067 | 0 | 1 | 25 |
| cdkn1bb | 0.60945 | 0.133494394 | 0.067 | 0 | 1 | 25 |
| cfh | 0.60945 | 0.216804472 | 0.067 | 0 | 1 | 25 |
| clns1a | 0.60945 | 0.10303452 | 0.067 | 0 | 1 | 25 |

| | | | | | | |
|---|---|---|---|---|---|---|
| cnr2 | 0.60945 | 0.184085933 | 0.067 | 0 | 1 | 25 |
| coa6 | 0.60945 | 0.233668345 | 0.067 | 0 | 1 | 25 |
| commd3 | 0.60945 | 0.122941523 | 0.067 | 0 | 1 | 25 |
| coq8a | 0.60945 | 0.165982624 | 0.067 | 0 | 1 | 25 |
| cpsf3 | 0.60945 | 0.134931622 | 0.067 | 0 | 1 | 25 |
| crfb4 | 0.60945 | 0.203586656 | 0.067 | 0 | 1 | 25 |
| cul2 | 0.60945 | 0.110582042 | 0.067 | 0 | 1 | 25 |
| ddx46 | 0.60945 | 0.142632206 | 0.067 | 0 | 1 | 25 |
| dffa | 0.60945 | 0.214987366 | 0.067 | 0 | 1 | 25 |
| dpm2 | 0.60945 | 0.165280449 | 0.067 | 0 | 1 | 25 |
| eed | 0.60945 | 0.203653644 | 0.067 | 0 | 1 | 25 |
| ehmt1a | 0.60945 | 0.24005638 | 0.067 | 0 | 1 | 25 |
| eif4g3b | 0.60945 | 0.197852707 | 0.067 | 0 | 1 | 25 |
| eif5b | 0.60945 | 0.15908927 | 0.067 | 0 | 1 | 25 |
| eny2 | 0.60945 | 0.166398496 | 0.067 | 0 | 1 | 25 |
| fam207a | 0.60945 | 0.109329842 | 0.067 | 0 | 1 | 25 |
| far1 | 0.60945 | 0.201950991 | 0.067 | 0 | 1 | 25 |
| flvcr2b | 0.60945 | 0.221634053 | 0.067 | 0 | 1 | 25 |
| fundc2 | 0.60945 | 0.116658265 | 0.067 | 0 | 1 | 25 |
| g6pc3 | 0.60945 | 0.149469329 | 0.067 | 0 | 1 | 25 |
| gatc | 0.60945 | 0.168738385 | 0.067 | 0 | 1 | 25 |
| gclc | 0.60945 | 0.156675909 | 0.067 | 0 | 1 | 25 |
| gclm | 0.60945 | 0.147907606 | 0.067 | 0 | 1 | 25 |
| gfer | 0.60945 | 0.190415976 | 0.067 | 0 | 1 | 25 |
| gltpa | 0.60945 | 0.198833062 | 0.067 | 0 | 1 | 25 |
| hint2 | 0.60945 | 0.178404303 | 0.067 | 0 | 1 | 25 |
| hnrnpdl | 0.60945 | 0.160905369 | 0.067 | 0 | 1 | 25 |
| ifngr1l | 0.60945 | 0.156262565 | 0.067 | 0 | 1 | 25 |
| irf2bp1 | 0.60945 | 0.189184959 | 0.067 | 0 | 1 | 25 |
| jpt1b | 0.60945 | 0.138316184 | 0.067 | 0 | 1 | 25 |
| kdm5ba | 0.60945 | 0.217037199 | 0.067 | 0 | 1 | 25 |
| lnpep | 0.60945 | 0.162098602 | 0.067 | 0 | 1 | 25 |
| map2k4a | 0.60945 | 0.180529987 | 0.067 | 0 | 1 | 25 |
| med6 | 0.60945 | 0.127787786 | 0.067 | 0 | 1 | 25 |
| mocs3 | 0.60945 | 0.223013556 | 0.067 | 0 | 1 | 25 |
| mrpl28 | 0.60945 | 0.182893101 | 0.067 | 0 | 1 | 25 |
| mvp | 0.60945 | 0.177371621 | 0.067 | 0 | 1 | 25 |
| naa35 | 0.60945 | 0.11945466 | 0.067 | 0 | 1 | 25 |
| nipblb | 0.60945 | 0.158944587 | 0.067 | 0 | 1 | 25 |
| nkrf | 0.60945 | 0.166271089 | 0.067 | 0 | 1 | 25 |
| npc1 | 0.60945 | 0.190354804 | 0.067 | 0 | 1 | 25 |
| nup107 | 0.60945 | 0.129923826 | 0.067 | 0 | 1 | 25 |
| nup160 | 0.60945 | 0.11173615 | 0.067 | 0 | 1 | 25 |
| p2ry10 | 0.60945 | 0.16220921 | 0.067 | 0 | 1 | 25 |
| pcgf5b | 0.60945 | 0.201812559 | 0.067 | 0 | 1 | 25 |
| pdha1a | 0.60945 | 0.20536969 | 0.067 | 0 | 1 | 25 |
| pdlim5b | 0.60945 | 0.131797432 | 0.067 | 0 | 1 | 25 |
| phc2a | 0.60945 | 0.112432617 | 0.067 | 0 | 1 | 25 |

| | | | | | | |
|---|---|---|---|---|---|---|
| phf20b | 0.60945 | 0.168638663 | 0.067 | 0 | 1 | 25 |
| pla2g7 | 0.60945 | 0.12848984 | 0.067 | 0 | 1 | 25 |
| pleca | 0.60945 | 0.174889387 | 0.067 | 0 | 1 | 25 |
| ppdpfb | 0.60945 | 0.17889028 | 0.067 | 0 | 1 | 25 |
| preb | 0.60945 | 0.238860518 | 0.067 | 0 | 1 | 25 |
| ptger4b | 0.60945 | 0.172857029 | 0.067 | 0 | 1 | 25 |
| pudp | 0.60945 | 0.169270238 | 0.067 | 0 | 1 | 25 |
| rab8a | 0.60945 | 0.261831853 | 0.067 | 0 | 1 | 25 |
| rasal3 | 0.60945 | 0.177698103 | 0.067 | 0 | 1 | 25 |
| rbm5 | 0.60945 | 0.178775493 | 0.067 | 0 | 1 | 25 |
| rgs2 | 0.60945 | 0.258807472 | 0.067 | 0 | 1 | 25 |
| rinl | 0.60945 | 0.20159206 | 0.067 | 0 | 1 | 25 |
| rnf13 | 0.60945 | 0.157004586 | 0.067 | 0 | 1 | 25 |
| rnf145b | 0.60945 | 0.136481296 | 0.067 | 0 | 1 | 25 |
| rp2 | 0.60945 | 0.110216575 | 0.067 | 0 | 1 | 25 |
| sat1a.1 | 0.60945 | 0.214124542 | 0.067 | 0 | 1 | 25 |
| sco1 | 0.60945 | 0.208793589 | 0.067 | 0 | 1 | 25 |
| sec62 | 0.60945 | 0.168558236 | 0.067 | 0 | 1 | 25 |
| sec63 | 0.60945 | 0.151495954 | 0.067 | 0 | 1 | 25 |
| selenow2a | 0.60945 | 0.184657372 | 0.067 | 0 | 1 | 25 |
| sf3b5 | 0.60945 | 0.156154012 | 0.067 | 0 | 1 | 25 |
| shisa4 | 0.60945 | 0.189200103 | 0.067 | 0 | 1 | 25 |
| si:ch211-117m20.4 | 0.60945 | 0.223977996 | 0.067 | 0 | 1 | 25 |
| si:ch211-148l7.4 | 0.60945 | 0.105149861 | 0.067 | 0 | 1 | 25 |
| si:ch211-209f23.6 | 0.60945 | 0.175060018 | 0.067 | 0 | 1 | 25 |
| si:ch211-74f19.2 | 0.60945 | 0.211932984 | 0.067 | 0 | 1 | 25 |
| si:dkey-165a24.10 | 0.60945 | 0.157630962 | 0.067 | 0 | 1 | 25 |
| si:dkey-16l2.16 | 0.60945 | 0.15574674 | 0.067 | 0 | 1 | 25 |
| si:dkeyp-55f12.3 | 0.60945 | 0.132998926 | 0.067 | 0 | 1 | 25 |
| si:zfos-943e10.1 | 0.60945 | 0.14565408 | 0.067 | 0 | 1 | 25 |
| slc35a3a | 0.60945 | 0.209241729 | 0.067 | 0 | 1 | 25 |
| slc35d1a | 0.60945 | 0.12768246 | 0.067 | 0 | 1 | 25 |
| smdt1b | 0.60945 | 0.107748628 | 0.067 | 0 | 1 | 25 |
| snrnp70 | 0.60945 | 0.146722696 | 0.067 | 0 | 1 | 25 |
| snrpd1 | 0.60945 | 0.107898226 | 0.067 | 0 | 1 | 25 |
| snx13 | 0.60945 | 0.14569028 | 0.067 | 0 | 1 | 25 |
| spsb4b | 0.60945 | 0.174322881 | 0.067 | 0 | 1 | 25 |
| srsf3a | 0.60945 | 0.162980142 | 0.067 | 0 | 1 | 25 |
| stk35 | 0.60945 | 0.169324145 | 0.067 | 0 | 1 | 25 |
| sulf2b | 0.60945 | 0.25140863 | 0.067 | 0 | 1 | 25 |
| syne1b | 0.60945 | 0.104791437 | 0.067 | 0 | 1 | 25 |
| tdh | 0.60945 | 0.1510191 | 0.067 | 0 | 1 | 25 |
| tesk2 | 0.60945 | 0.122803396 | 0.067 | 0 | 1 | 25 |
| tmed1b | 0.60945 | 0.193626736 | 0.067 | 0 | 1 | 25 |
| tp53i11a | 0.60945 | 0.108272022 | 0.067 | 0 | 1 | 25 |
| trappc3 | 0.60945 | 0.180911085 | 0.067 | 0 | 1 | 25 |
| trpt1 | 0.60945 | 0.218263722 | 0.067 | 0 | 1 | 25 |
| ttc1 | 0.60945 | 0.156469694 | 0.067 | 0 | 1 | 25 |

| | | | | | | |
|---|---|---|---|---|---|---|
| ttc39c | 0.60945 | 0.149838428 | 0.067 | 0 | 1 | 25 |
| ube2ka | 0.60945 | 0.141766308 | 0.067 | 0 | 1 | 25 |
| ugt5b2 | 0.60945 | 0.190017491 | 0.067 | 0 | 1 | 25 |
| vamp5 | 0.60945 | 0.198503963 | 0.067 | 0 | 1 | 25 |
| vkorc1l1 | 0.60945 | 0.149841458 | 0.067 | 0 | 1 | 25 |
| waca | 0.60945 | 0.146951759 | 0.067 | 0 | 1 | 25 |
| washc2c | 0.60945 | 0.151222479 | 0.067 | 0 | 1 | 25 |
| wdfy4 | 0.60945 | 0.174312973 | 0.067 | 0 | 1 | 25 |
| wipf1a | 0.60945 | 0.17051788 | 0.067 | 0 | 1 | 25 |
| wwc1 | 0.60945 | 0.146630691 | 0.067 | 0 | 1 | 25 |
| xkr9 | 0.60945 | 0.250583494 | 0.067 | 0 | 1 | 25 |
| yars | 0.60945 | 0.139889764 | 0.067 | 0 | 1 | 25 |
| zbtb16a | 0.60945 | 0.240120026 | 0.067 | 0 | 1 | 25 |
| zdhhc5a | 0.60945 | 0.144329893 | 0.067 | 0 | 1 | 25 |
| zgc:112356 | 0.60945 | 0.1674032 | 0.067 | 0 | 1 | 25 |
| zgc:55558 | 0.60945 | 0.148321835 | 0.067 | 0 | 1 | 25 |
| znf131 | 0.60945 | 0.127306214 | 0.067 | 0 | 1 | 25 |
| LOC793824 | 0.60945 | 0.178461499 | 0.067 | 0 | 1 | 25 |
| arpp19a | 0.60945 | 0.147711655 | 0.067 | 0 | 1 | 25 |
| brd3b | 0.60945 | 0.175137971 | 0.067 | 0 | 1 | 25 |
| cpvl | 0.60945 | 0.195010146 | 0.067 | 0 | 1 | 25 |
| fbxo9 | 0.60945 | 0.172151251 | 0.067 | 0 | 1 | 25 |
| gps2 | 0.60945 | 0.230713653 | 0.067 | 0 | 1 | 25 |
| hnrnpabb | 0.60945 | 0.161837776 | 0.067 | 0 | 1 | 25 |
| lgals9l1 | 0.60945 | 0.177637341 | 0.067 | 0 | 1 | 25 |
| lpar5b | 0.60945 | 0.247881425 | 0.067 | 0 | 1 | 25 |
| mbnl3 | 0.60945 | 0.176190512 | 0.067 | 0 | 1 | 25 |
| mt-nd5 | 0.60945 | 0.203462188 | 0.067 | 0 | 1 | 25 |
| ppcdc | 0.60945 | 0.127983782 | 0.067 | 0 | 1 | 25 |
| si:dkey-32n7.4 | 0.60945 | 0.232014449 | 0.067 | 0 | 1 | 25 |
| si:dkeyp-28d2.4 | 0.60945 | 0.156715468 | 0.067 | 0 | 1 | 25 |
| tbc1d15 | 0.60945 | 0.151794394 | 0.067 | 0 | 1 | 25 |
| wdr43 | 0.60945 | 0.165579505 | 0.067 | 0 | 1 | 25 |
| zgc:153118 | 0.60945 | 0.213290927 | 0.067 | 0 | 1 | 25 |
| CABZ01076758.1 | 0.60945 | 0.218984788 | 0.067 | 0 | 1 | 25 |
| LOC103910438 | 0.60945 | 0.189856319 | 0.067 | 0 | 1 | 25 |
| adipor1a | 0.60945 | 0.148658368 | 0.067 | 0 | 1 | 25 |
| ahcyl1 | 0.60945 | 0.139353955 | 0.067 | 0 | 1 | 25 |
| aldh2.2 | 0.60945 | 0.12173909 | 0.067 | 0 | 1 | 25 |
| ankrd12 | 0.60945 | 0.15799319 | 0.067 | 0 | 1 | 25 |
| ap2a1 | 0.60945 | 0.127063866 | 0.067 | 0 | 1 | 25 |
| apaf1 | 0.60945 | 0.103026734 | 0.067 | 0 | 1 | 25 |
| arhgef18b | 0.60945 | 0.169361146 | 0.067 | 0 | 1 | 25 |
| arih2 | 0.60945 | 0.170437657 | 0.067 | 0 | 1 | 25 |
| arl5c | 0.60945 | 0.127414845 | 0.067 | 0 | 1 | 25 |
| atp6v1e1a | 0.60945 | 0.126312685 | 0.067 | 0 | 1 | 25 |
| atxn1a | 0.60945 | 0.15210267 | 0.067 | 0 | 1 | 25 |
| aup1 | 0.60945 | 0.132090731 | 0.067 | 0 | 1 | 25 |

| | | | | | | |
|---|---|---|---|---|---|---|
| aven | 0.60945 | 0.153944376 | 0.067 | 0 | 1 | 25 |
| bnipl | 0.60945 | 0.128001988 | 0.067 | 0 | 1 | 25 |
| cdc5l | 0.60945 | 0.219215037 | 0.067 | 0 | 1 | 25 |
| cflara | 0.60945 | 0.126044447 | 0.067 | 0 | 1 | 25 |
| copb2 | 0.60945 | 0.158503746 | 0.067 | 0 | 1 | 25 |
| cops8 | 0.60945 | 0.128711145 | 0.067 | 0 | 1 | 25 |
| cst3 | 0.60945 | 0.189527349 | 0.067 | 0 | 1 | 25 |
| dapp1 | 0.60945 | 0.243120636 | 0.067 | 0 | 1 | 25 |
| ech1 | 0.60945 | 0.134830241 | 0.067 | 0 | 1 | 25 |
| eif4a3 | 0.60945 | 0.116812833 | 0.067 | 0 | 1 | 25 |
| elmo1 | 0.60945 | 0.133811363 | 0.067 | 0 | 1 | 25 |
| etf1b | 0.60945 | 0.180071951 | 0.067 | 0 | 1 | 25 |
| ewsr1a | 0.60945 | 0.184802288 | 0.067 | 0 | 1 | 25 |
| fam84b | 0.60945 | 0.106959855 | 0.067 | 0 | 1 | 25 |
| fzr1a | 0.60945 | 0.176370639 | 0.067 | 0 | 1 | 25 |
| gmeb1 | 0.60945 | 0.182831047 | 0.067 | 0 | 1 | 25 |
| gspt1l | 0.60945 | 0.171999852 | 0.067 | 0 | 1 | 25 |
| hdgfl2 | 0.60945 | 0.215705095 | 0.067 | 0 | 1 | 25 |
| hnrnpa1b | 0.60945 | 0.112695982 | 0.067 | 0 | 1 | 25 |
| idh3b | 0.60945 | 0.125227928 | 0.067 | 0 | 1 | 25 |
| ifrd1 | 0.60945 | 0.249589029 | 0.067 | 0 | 1 | 25 |
| ift27 | 0.60945 | 0.143089687 | 0.067 | 0 | 1 | 25 |
| inpp5d | 0.60945 | 0.185578578 | 0.067 | 0 | 1 | 25 |
| itsn2b | 0.60945 | 0.123370836 | 0.067 | 0 | 1 | 25 |
| jmjd1cb | 0.60945 | 0.146436498 | 0.067 | 0 | 1 | 25 |
| lclat1 | 0.60945 | 0.1240111 | 0.067 | 0 | 1 | 25 |
| lin37 | 0.60945 | 0.144486497 | 0.067 | 0 | 1 | 25 |
| lpar2a | 0.60945 | 0.169209317 | 0.067 | 0 | 1 | 25 |
| manf | 0.60945 | 0.127851015 | 0.067 | 0 | 1 | 25 |
| mccc2 | 0.60945 | 0.178055027 | 0.067 | 0 | 1 | 25 |
| mcm6 | 0.60945 | 0.20854338 | 0.067 | 0 | 1 | 25 |
| med22 | 0.60945 | 0.165098872 | 0.067 | 0 | 1 | 25 |
| mff | 0.60945 | 0.227594454 | 0.067 | 0 | 1 | 25 |
| mycbp2 | 0.60945 | 0.230050392 | 0.067 | 0 | 1 | 25 |
| nol10 | 0.60945 | 0.147379629 | 0.067 | 0 | 1 | 25 |
| nosip | 0.60945 | 0.147849513 | 0.067 | 0 | 1 | 25 |
| npm3 | 0.60945 | 0.213204295 | 0.067 | 0 | 1 | 25 |
| pafah1b1a | 0.60945 | 0.123961769 | 0.067 | 0 | 1 | 25 |
| pbxip1b | 0.60945 | 0.177160628 | 0.067 | 0 | 1 | 25 |
| pex19 | 0.60945 | 0.168032265 | 0.067 | 0 | 1 | 25 |
| plekhg5b | 0.60945 | 0.155915821 | 0.067 | 0 | 1 | 25 |
| ppp2r2d | 0.60945 | 0.158826889 | 0.067 | 0 | 1 | 25 |
| psmc4 | 0.60945 | 0.137119242 | 0.067 | 0 | 1 | 25 |
| ptenb | 0.60945 | 0.132559153 | 0.067 | 0 | 1 | 25 |
| ptgr1 | 0.60945 | 0.124618903 | 0.067 | 0 | 1 | 25 |
| rhogd | 0.60945 | 0.171628171 | 0.067 | 0 | 1 | 25 |
| rpz5 | 0.60945 | 0.224382916 | 0.067 | 0 | 1 | 25 |
| scarf1 | 0.60945 | 0.17352069 | 0.067 | 0 | 1 | 25 |

| | | | | | | |
|---|---|---|---|---|---|---|
| sec22c | 0.60945 | 0.160612137 | 0.067 | 0 | 1 | 25 |
| sec24a | 0.60945 | 0.219721139 | 0.067 | 0 | 1 | 25 |
| si:ch211-121a2.2 | 0.60945 | 0.131574269 | 0.067 | 0 | 1 | 25 |
| si:ch73-281n10.2 | 0.60945 | 0.150194881 | 0.067 | 0 | 1 | 25 |
| si:dkey-53k12.1 | 0.60945 | 0.18224949 | 0.067 | 0 | 1 | 25 |
| slc30a7 | 0.60945 | 0.257708073 | 0.067 | 0 | 1 | 25 |
| smim19 | 0.60945 | 0.1313027 | 0.067 | 0 | 1 | 25 |
| snx27a | 0.60945 | 0.188670533 | 0.067 | 0 | 1 | 25 |
| sod2 | 0.60945 | 0.173389207 | 0.067 | 0 | 1 | 25 |
| stk24b | 0.60945 | 0.130149117 | 0.067 | 0 | 1 | 25 |
| taf1 | 0.60945 | 0.187625527 | 0.067 | 0 | 1 | 25 |
| tctex1d2 | 0.60945 | 0.18275798 | 0.067 | 0 | 1 | 25 |
| tfip11 | 0.60945 | 0.209577382 | 0.067 | 0 | 1 | 25 |
| tlr5a | 0.60945 | 0.233494714 | 0.067 | 0 | 1 | 25 |
| tspan4b | 0.60945 | 0.186770511 | 0.067 | 0 | 1 | 25 |
| ube2w | 0.60945 | 0.132067345 | 0.067 | 0 | 1 | 25 |
| wasf2 | 0.60945 | 0.109469338 | 0.067 | 0 | 1 | 25 |
| zbtb11 | 0.60945 | 0.17352069 | 0.067 | 0 | 1 | 25 |
| zdhhc6 | 0.60945 | 0.137559732 | 0.067 | 0 | 1 | 25 |
| zgc:103697 | 0.60945 | 0.135013177 | 0.067 | 0 | 1 | 25 |
| zgc:153044 | 0.60945 | 0.133626019 | 0.067 | 0 | 1 | 25 |
| znf622 | 0.60945 | 0.139177293 | 0.067 | 0 | 1 | 25 |
| BX649485.1 | 0.60945 | 0.26230231 | 0.067 | 0 | 1 | 25 |
| CABZ01045617.1 | 0.60945 | 0.240740622 | 0.067 | 0 | 1 | 25 |
| CABZ01078737.1 | 0.60945 | 0.227118341 | 0.067 | 0 | 1 | 25 |
| CABZ01115113.1 | 0.60945 | 0.132371281 | 0.067 | 0 | 1 | 25 |
| CHST13 | 0.60945 | 0.220674074 | 0.067 | 0 | 1 | 25 |
| COX7A2 (1 of many) | 0.60945 | 0.172466506 | 0.067 | 0 | 1 | 25 |
| LO018176.1 | 0.60945 | 0.245991131 | 0.067 | 0 | 1 | 25 |
| LOC100004944 | 0.60945 | 0.235106639 | 0.067 | 0 | 1 | 25 |
| NPC2 (1 of many) | 0.60945 | 0.103545979 | 0.067 | 0 | 1 | 25 |
| SBNO2 | 0.60945 | 0.158094494 | 0.067 | 0 | 1 | 25 |
| acsl3b | 0.60945 | 0.171782302 | 0.067 | 0 | 1 | 25 |
| acvr2aa | 0.60945 | 0.231640243 | 0.067 | 0 | 1 | 25 |
| apol1 | 0.60945 | 0.246892446 | 0.067 | 0 | 1 | 25 |
| arfip1 | 0.60945 | 0.15317867 | 0.067 | 0 | 1 | 25 |
| arl1 | 0.60945 | 0.201931907 | 0.067 | 0 | 1 | 25 |
| asns | 0.60945 | 0.101328622 | 0.067 | 0 | 1 | 25 |
| atp6v0b | 0.60945 | 0.135955556 | 0.067 | 0 | 1 | 25 |
| b4galt6 | 0.60945 | 0.171747808 | 0.067 | 0 | 1 | 25 |
| baz1b | 0.60945 | 0.232623575 | 0.067 | 0 | 1 | 25 |
| bloc1s5 | 0.60945 | 0.101257075 | 0.067 | 0 | 1 | 25 |
| bsdc1 | 0.60945 | 0.201148882 | 0.067 | 0 | 1 | 25 |
| c1qbp | 0.60945 | 0.122034021 | 0.067 | 0 | 1 | 25 |
| ccdc84 | 0.60945 | 0.23569655 | 0.067 | 0 | 1 | 25 |
| chico | 0.60945 | 0.120411651 | 0.067 | 0 | 1 | 25 |
| chmp5b | 0.60945 | 0.162637832 | 0.067 | 0 | 1 | 25 |
| ckbb | 0.60945 | 0.13963538 | 0.067 | 0 | 1 | 25 |

| | | | | | | |
|---|---|---|---|---|---|---|
| clcn7 | 0.60945 | 0.151676766 | 0.067 | 0 | 1 | 25 |
| clk4b | 0.60945 | 0.213621304 | 0.067 | 0 | 1 | 25 |
| copb1 | 0.60945 | 0.151776806 | 0.067 | 0 | 1 | 25 |
| crata | 0.60945 | 0.174382076 | 0.067 | 0 | 1 | 25 |
| ctps1a | 0.60945 | 0.102058867 | 0.067 | 0 | 1 | 25 |
| cxcr3.2 | 0.60945 | 0.150616626 | 0.067 | 0 | 1 | 25 |
| dars | 0.60945 | 0.27122549 | 0.067 | 0 | 1 | 25 |
| ddx6 | 0.60945 | 0.133195672 | 0.067 | 0 | 1 | 25 |
| dnmt1 | 0.60945 | 0.137465112 | 0.067 | 0 | 1 | 25 |
| drg1 | 0.60945 | 0.103845261 | 0.067 | 0 | 1 | 25 |
| e2f4 | 0.60945 | 0.216290184 | 0.067 | 0 | 1 | 25 |
| eif4g2a | 0.60945 | 0.155246434 | 0.067 | 0 | 1 | 25 |
| foxo4 | 0.60945 | 0.154987169 | 0.067 | 0 | 1 | 25 |
| foxp1b | 0.60945 | 0.235265795 | 0.067 | 0 | 1 | 25 |
| gadd45ab | 0.60945 | 0.232197009 | 0.067 | 0 | 1 | 25 |
| galns | 0.60945 | 0.178753666 | 0.067 | 0 | 1 | 25 |
| golgb1 | 0.60945 | 0.110421125 | 0.067 | 0 | 1 | 25 |
| gosr1 | 0.60945 | 0.175237474 | 0.067 | 0 | 1 | 25 |
| gosr2 | 0.60945 | 0.111830652 | 0.067 | 0 | 1 | 25 |
| gpd1c | 0.60945 | 0.137492021 | 0.067 | 0 | 1 | 25 |
| gys1 | 0.60945 | 0.152477003 | 0.067 | 0 | 1 | 25 |
| hadhaa | 0.60945 | 0.227208636 | 0.067 | 0 | 1 | 25 |
| hk1 | 0.60945 | 0.170572612 | 0.067 | 0 | 1 | 25 |
| hps1 | 0.60945 | 0.239809049 | 0.067 | 0 | 1 | 25 |
| igf2bp2a | 0.60945 | 0.230450789 | 0.067 | 0 | 1 | 25 |
| ipcef1 | 0.60945 | 0.101145647 | 0.067 | 0 | 1 | 25 |
| itgb3a | 0.60945 | 0.183286476 | 0.067 | 0 | 1 | 25 |
| kansl1b | 0.60945 | 0.210312156 | 0.067 | 0 | 1 | 25 |
| kdm1a | 0.60945 | 0.167424916 | 0.067 | 0 | 1 | 25 |
| kdm4aa | 0.60945 | 0.151028554 | 0.067 | 0 | 1 | 25 |
| lman1 | 0.60945 | 0.165640676 | 0.067 | 0 | 1 | 25 |
| lsm6 | 0.60945 | 0.179927122 | 0.067 | 0 | 1 | 25 |
| maco1a | 0.60945 | 0.212482468 | 0.067 | 0 | 1 | 25 |
| mkln1 | 0.60945 | 0.167232459 | 0.067 | 0 | 1 | 25 |
| myd88 | 0.60945 | 0.184342463 | 0.067 | 0 | 1 | 25 |
| ndufaf3 | 0.60945 | 0.178418131 | 0.067 | 0 | 1 | 25 |
| ndufv3 | 0.60945 | 0.139965201 | 0.067 | 0 | 1 | 25 |
| nfic | 0.60945 | 0.175660662 | 0.067 | 0 | 1 | 25 |
| nfkbiaa | 0.60945 | 0.216260248 | 0.067 | 0 | 1 | 25 |
| nfkbie | 0.60945 | 0.198757167 | 0.067 | 0 | 1 | 25 |
| nucks1b | 0.60945 | 0.117176539 | 0.067 | 0 | 1 | 25 |
| oga | 0.60945 | 0.126672614 | 0.067 | 0 | 1 | 25 |
| pdia4 | 0.60945 | 0.197452597 | 0.067 | 0 | 1 | 25 |
| per1b | 0.60945 | 0.233239344 | 0.067 | 0 | 1 | 25 |
| pik3r3a | 0.60945 | 0.221744676 | 0.067 | 0 | 1 | 25 |
| polr2d | 0.60945 | 0.120219582 | 0.067 | 0 | 1 | 25 |
| pom121 | 0.60945 | 0.190248835 | 0.067 | 0 | 1 | 25 |
| prelid3b | 0.60945 | 0.190575404 | 0.067 | 0 | 1 | 25 |

| | | | | | | |
|---|---|---|---|---|---|---|
| prkcaa | 0.60945 | 0.156858641 | 0.067 | 0 | 1 | 25 |
| prpsap2 | 0.60945 | 0.23992873 | 0.067 | 0 | 1 | 25 |
| ptgs1 | 0.60945 | 0.13657778 | 0.067 | 0 | 1 | 25 |
| rap2b | 0.60945 | 0.106731105 | 0.067 | 0 | 1 | 25 |
| rcan1a | 0.60945 | 0.255818095 | 0.067 | 0 | 1 | 25 |
| rnf111 | 0.60945 | 0.23758167 | 0.067 | 0 | 1 | 25 |
| rpa2 | 0.60945 | 0.154968218 | 0.067 | 0 | 1 | 25 |
| sftpbb | 0.60945 | 0.249090935 | 0.067 | 0 | 1 | 25 |
| shoc2 | 0.60945 | 0.165051568 | 0.067 | 0 | 1 | 25 |
| si:busm1-266f07.2 | 0.60945 | 0.266556974 | 0.067 | 0 | 1 | 25 |
| si:ch211-214p16.1 | 0.60945 | 0.212577999 | 0.067 | 0 | 1 | 25 |
| si:ch211-225b11.1 | 0.60945 | 0.198126336 | 0.067 | 0 | 1 | 25 |
| si:ch211-260e23.9 | 0.60945 | 0.148678349 | 0.067 | 0 | 1 | 25 |
| si:ch73-139j3.4 | 0.60945 | 0.190081293 | 0.067 | 0 | 1 | 25 |
| si:ch73-376l24.3 | 0.60945 | 0.237065503 | 0.067 | 0 | 1 | 25 |
| si:dkey-20d21.12 | 0.60945 | 0.224924712 | 0.067 | 0 | 1 | 25 |
| si:dkeyp-80c12.4 | 0.60945 | 0.113970091 | 0.067 | 0 | 1 | 25 |
| skiv2l | 0.60945 | 0.161071321 | 0.067 | 0 | 1 | 25 |
| ssrp1a | 0.60945 | 0.142672683 | 0.067 | 0 | 1 | 25 |
| stat5a | 0.60945 | 0.201161727 | 0.067 | 0 | 1 | 25 |
| stk11 | 0.60945 | 0.167801828 | 0.067 | 0 | 1 | 25 |
| sub1b | 0.60945 | 0.18882624 | 0.067 | 0 | 1 | 25 |
| sult2st3 | 0.60945 | 0.120411651 | 0.067 | 0 | 1 | 25 |
| tfg | 0.60945 | 0.148068545 | 0.067 | 0 | 1 | 25 |
| tmco1 | 0.60945 | 0.141023248 | 0.067 | 0 | 1 | 25 |
| tmem154 | 0.60945 | 0.192446487 | 0.067 | 0 | 1 | 25 |
| tomm22 | 0.60945 | 0.152625363 | 0.067 | 0 | 1 | 25 |
| trappc1 | 0.60945 | 0.154259725 | 0.067 | 0 | 1 | 25 |
| txndc9 | 0.60945 | 0.133413409 | 0.067 | 0 | 1 | 25 |
| ube2e3 | 0.60945 | 0.121154612 | 0.067 | 0 | 1 | 25 |
| ube2j2 | 0.60945 | 0.162084547 | 0.067 | 0 | 1 | 25 |
| ubr3 | 0.60945 | 0.18241927 | 0.067 | 0 | 1 | 25 |
| ubtfl | 0.60945 | 0.212096777 | 0.067 | 0 | 1 | 25 |
| unm-sa1614 | 0.60945 | 0.183289603 | 0.067 | 0 | 1 | 25 |
| vps39 | 0.60945 | 0.238888538 | 0.067 | 0 | 1 | 25 |
| yipf4 | 0.60945 | 0.221625259 | 0.067 | 0 | 1 | 25 |
| zgc:100829 | 0.60945 | 0.231847249 | 0.067 | 0 | 1 | 25 |
| zgc:100918 | 0.60945 | 0.109262192 | 0.067 | 0 | 1 | 25 |
| zgc:114119 | 0.60945 | 0.25162803 | 0.067 | 0 | 1 | 25 |
| zgc:114130 | 0.60945 | 0.139667984 | 0.067 | 0 | 1 | 25 |
| zgc:162396 | 0.60945 | 0.166614254 | 0.067 | 0 | 1 | 25 |
| zgc:92606 | 0.60945 | 0.189823325 | 0.067 | 0 | 1 | 25 |
| zmiz1a | 0.60945 | 0.158136626 | 0.067 | 0 | 1 | 25 |
| CABZ01079427.1 | 0.60945 | 0.150081677 | 0.067 | 0 | 1 | 25 |
| asah1b | 0.60945 | 0.224818293 | 0.067 | 0 | 1 | 25 |
| grb2b | 0.60945 | 0.151607999 | 0.067 | 0 | 1 | 25 |
| march7 | 0.60945 | 0.152159358 | 0.067 | 0 | 1 | 25 |
| plekhb2 | 0.60945 | 0.239897071 | 0.067 | 0 | 1 | 25 |

| | | | | | | |
|---|---|---|---|---|---|---|
| psmd13 | 0.60945 | 0.237964545 | 0.067 | 0 | 1 | 25 |
| rps6ka3a | 0.60945 | 0.223530777 | 0.067 | 0 | 1 | 25 |
| ubap2l | 0.60945 | 0.237342222 | 0.067 | 0 | 1 | 25 |
| zfr | 0.60945 | 0.145661551 | 0.067 | 0 | 1 | 25 |
| BX248113.1 | 0.60945 | 0.119222816 | 0.067 | 0 | 1 | 25 |
| BX296551.1 | 0.60945 | 0.18202637 | 0.067 | 0 | 1 | 25 |
| BX548028.1 | 0.60945 | 0.200040628 | 0.067 | 0 | 1 | 25 |
| CABZ01038697.1 | 0.60945 | 0.16232579 | 0.067 | 0 | 1 | 25 |
| CABZ01066694.1 | 0.60945 | 0.181433454 | 0.067 | 0 | 1 | 25 |
| CABZ01068251.1 | 0.60945 | 0.161859955 | 0.067 | 0 | 1 | 25 |
| CR293507.1 | 0.60945 | 0.138696734 | 0.067 | 0 | 1 | 25 |
| CU570889.1 | 0.60945 | 0.122942025 | 0.067 | 0 | 1 | 25 |
| FQ377903.3 | 0.60945 | 0.165515981 | 0.067 | 0 | 1 | 25 |
| FQ976913.1 | 0.60945 | 0.113909271 | 0.067 | 0 | 1 | 25 |
| LOC103909972 | 0.60945 | 0.185027097 | 0.067 | 0 | 1 | 25 |
| LOC103911430 | 0.60945 | 0.22631977 | 0.067 | 0 | 1 | 25 |
| MARF1 | 0.60945 | 0.136361579 | 0.067 | 0 | 1 | 25 |
| PDCL3 | 0.60945 | 0.154489981 | 0.067 | 0 | 1 | 25 |
| PYURF | 0.60945 | 0.161864299 | 0.067 | 0 | 1 | 25 |
| XLOC-000431 | 0.60945 | 0.136344009 | 0.067 | 0 | 1 | 25 |
| XLOC-012680 | 0.60945 | 0.164900513 | 0.067 | 0 | 1 | 25 |
| abca1a | 0.60945 | 0.167084458 | 0.067 | 0 | 1 | 25 |
| abce1 | 0.60945 | 0.194266494 | 0.067 | 0 | 1 | 25 |
| abi3a | 0.60945 | 0.204857386 | 0.067 | 0 | 1 | 25 |
| ada2a | 0.60945 | 0.225310417 | 0.067 | 0 | 1 | 25 |
| agps | 0.60945 | 0.146760912 | 0.067 | 0 | 1 | 25 |
| aldh2.1 | 0.60945 | 0.18319221 | 0.067 | 0 | 1 | 25 |
| ankrd10b | 0.60945 | 0.12382492 | 0.067 | 0 | 1 | 25 |
| ankrd22 | 0.60945 | 0.162344133 | 0.067 | 0 | 1 | 25 |
| apbb1ip | 0.60945 | 0.119222816 | 0.067 | 0 | 1 | 25 |
| aph1b | 0.60945 | 0.127363408 | 0.067 | 0 | 1 | 25 |
| arid3a | 0.60945 | 0.134811607 | 0.067 | 0 | 1 | 25 |
| arid4a | 0.60945 | 0.165437357 | 0.067 | 0 | 1 | 25 |
| arpin | 0.60945 | 0.160802249 | 0.067 | 0 | 1 | 25 |
| arpp19b | 0.60945 | 0.19827223 | 0.067 | 0 | 1 | 25 |
| atad1b | 0.60945 | 0.20981493 | 0.067 | 0 | 1 | 25 |
| atp10d | 0.60945 | 0.148679033 | 0.067 | 0 | 1 | 25 |
| carm1 | 0.60945 | 0.115639978 | 0.067 | 0 | 1 | 25 |
| casp8 | 0.60945 | 0.171044563 | 0.067 | 0 | 1 | 25 |
| cc2d1b | 0.60945 | 0.159422562 | 0.067 | 0 | 1 | 25 |
| cdca4 | 0.60945 | 0.182216388 | 0.067 | 0 | 1 | 25 |
| cdk2ap2 | 0.60945 | 0.173222639 | 0.067 | 0 | 1 | 25 |
| cep131 | 0.60945 | 0.216444635 | 0.067 | 0 | 1 | 25 |
| chtopa | 0.60945 | 0.168242925 | 0.067 | 0 | 1 | 25 |
| churc1 | 0.60945 | 0.125963238 | 0.067 | 0 | 1 | 25 |
| ciarta | 0.60945 | 0.160592376 | 0.067 | 0 | 1 | 25 |
| cldni | 0.60945 | 0.206470952 | 0.067 | 0 | 1 | 25 |
| cluap1 | 0.60945 | 0.152828944 | 0.067 | 0 | 1 | 25 |

| | | | | | | |
|---|---|---|---|---|---|---|
| copa | 0.60945 | 0.123495847 | 0.067 | 0 | 1 | 25 |
| cope | 0.60945 | 0.114780634 | 0.067 | 0 | 1 | 25 |
| copz1 | 0.60945 | 0.13583839 | 0.067 | 0 | 1 | 25 |
| ctbp1 | 0.60945 | 0.208530935 | 0.067 | 0 | 1 | 25 |
| ctdsplb | 0.60945 | 0.127431476 | 0.067 | 0 | 1 | 25 |
| cwc25 | 0.60945 | 0.138240068 | 0.067 | 0 | 1 | 25 |
| cwc27 | 0.60945 | 0.128511687 | 0.067 | 0 | 1 | 25 |
| cyp20a1 | 0.60945 | 0.17760928 | 0.067 | 0 | 1 | 25 |
| dcp2 | 0.60945 | 0.224665089 | 0.067 | 0 | 1 | 25 |
| dcun1d5 | 0.60945 | 0.123657367 | 0.067 | 0 | 1 | 25 |
| dhdds | 0.60945 | 0.112689422 | 0.067 | 0 | 1 | 25 |
| dhrs7 | 0.60945 | 0.199131908 | 0.067 | 0 | 1 | 25 |
| diaph2 | 0.60945 | 0.174521234 | 0.067 | 0 | 1 | 25 |
| dohh | 0.60945 | 0.15807148 | 0.067 | 0 | 1 | 25 |
| eaf1 | 0.60945 | 0.212087459 | 0.067 | 0 | 1 | 25 |
| eci2 | 0.60945 | 0.150930538 | 0.067 | 0 | 1 | 25 |
| efhd2 | 0.60945 | 0.121367918 | 0.067 | 0 | 1 | 25 |
| egr1 | 0.60945 | 0.225487725 | 0.067 | 0 | 1 | 25 |
| eif2ak3 | 0.60945 | 0.125404317 | 0.067 | 0 | 1 | 25 |
| elmo3 | 0.60945 | 0.144045698 | 0.067 | 0 | 1 | 25 |
| emd | 0.60945 | 0.11653624 | 0.067 | 0 | 1 | 25 |
| entpd1 | 0.60945 | 0.184911502 | 0.067 | 0 | 1 | 25 |
| fam50a | 0.60945 | 0.214720053 | 0.067 | 0 | 1 | 25 |
| fpgt | 0.60945 | 0.112410194 | 0.067 | 0 | 1 | 25 |
| gadd45bb | 0.60945 | 0.217004013 | 0.067 | 0 | 1 | 25 |
| gapdh | 0.60945 | 0.188534418 | 0.067 | 0 | 1 | 25 |
| gb:bc139872 | 0.60945 | 0.130450474 | 0.067 | 0 | 1 | 25 |
| gcnt4a | 0.60945 | 0.15226015 | 0.067 | 0 | 1 | 25 |
| gemin7 | 0.60945 | 0.103076334 | 0.067 | 0 | 1 | 25 |
| gpr141 | 0.60945 | 0.163721774 | 0.067 | 0 | 1 | 25 |
| gpr155a | 0.60945 | 0.160462083 | 0.067 | 0 | 1 | 25 |
| gstr | 0.60945 | 0.105149827 | 0.067 | 0 | 1 | 25 |
| hadh | 0.60945 | 0.159731118 | 0.067 | 0 | 1 | 25 |
| hint1 | 0.60945 | 0.237482257 | 0.067 | 0 | 1 | 25 |
| hnrnpul1l | 0.60945 | 0.106362503 | 0.067 | 0 | 1 | 25 |
| ift20 | 0.60945 | 0.145105564 | 0.067 | 0 | 1 | 25 |
| ino80db | 0.60945 | 0.124667397 | 0.067 | 0 | 1 | 25 |
| iscub | 0.60945 | 0.231564445 | 0.067 | 0 | 1 | 25 |
| jagn1b | 0.60945 | 0.187509635 | 0.067 | 0 | 1 | 25 |
| jak2b | 0.60945 | 0.174468534 | 0.067 | 0 | 1 | 25 |
| kat5b | 0.60945 | 0.150029902 | 0.067 | 0 | 1 | 25 |
| kdm7ab | 0.60945 | 0.143744581 | 0.067 | 0 | 1 | 25 |
| kif5bb | 0.60945 | 0.207940432 | 0.067 | 0 | 1 | 25 |
| klf11b | 0.60945 | 0.277964164 | 0.067 | 0 | 1 | 25 |
| klhl8 | 0.60945 | 0.137442433 | 0.067 | 0 | 1 | 25 |
| lamtor2 | 0.60945 | 0.217105781 | 0.067 | 0 | 1 | 25 |
| lig1 | 0.60945 | 0.163398869 | 0.067 | 0 | 1 | 25 |
| luc7l3 | 0.60945 | 0.104157554 | 0.067 | 0 | 1 | 25 |

| | | | | | | |
|---|---|---|---|---|---|---|
| manbal | 0.60945 | 0.141512625 | 0.067 | 0 | 1 | 25 |
| mapk12b | 0.60945 | 0.116014957 | 0.067 | 0 | 1 | 25 |
| mark2a | 0.60945 | 0.158837883 | 0.067 | 0 | 1 | 25 |
| mcm3 | 0.60945 | 0.133645095 | 0.067 | 0 | 1 | 25 |
| mcm5 | 0.60945 | 0.19824853 | 0.067 | 0 | 1 | 25 |
| mea1 | 0.60945 | 0.146488149 | 0.067 | 0 | 1 | 25 |
| med21 | 0.60945 | 0.192481563 | 0.067 | 0 | 1 | 25 |
| metap1 | 0.60945 | 0.119382944 | 0.067 | 0 | 1 | 25 |
| morf4l1 | 0.60945 | 0.223953264 | 0.067 | 0 | 1 | 25 |
| mrpl12 | 0.60945 | 0.193656198 | 0.067 | 0 | 1 | 25 |
| mrps16 | 0.60945 | 0.156698762 | 0.067 | 0 | 1 | 25 |
| mtap | 0.60945 | 0.16215069 | 0.067 | 0 | 1 | 25 |
| ndufa12 | 0.60945 | 0.111973096 | 0.067 | 0 | 1 | 25 |
| nfatc3a | 0.60945 | 0.116714254 | 0.067 | 0 | 1 | 25 |
| nfe2l3 | 0.60945 | 0.147998321 | 0.067 | 0 | 1 | 25 |
| nfil3-5 | 0.60945 | 0.237102997 | 0.067 | 0 | 1 | 25 |
| nfybb | 0.60945 | 0.163835359 | 0.067 | 0 | 1 | 25 |
| nipsnap2 | 0.60945 | 0.228857033 | 0.067 | 0 | 1 | 25 |
| npepl1 | 0.60945 | 0.114571265 | 0.067 | 0 | 1 | 25 |
| npepps | 0.60945 | 0.138691249 | 0.067 | 0 | 1 | 25 |
| otub1a | 0.60945 | 0.136149557 | 0.067 | 0 | 1 | 25 |
| papolg | 0.60945 | 0.150108923 | 0.067 | 0 | 1 | 25 |
| pcid2 | 0.60945 | 0.158240781 | 0.067 | 0 | 1 | 25 |
| pdpk1b | 0.60945 | 0.228132358 | 0.067 | 0 | 1 | 25 |
| phb | 0.60945 | 0.110769887 | 0.067 | 0 | 1 | 25 |
| pin4 | 0.60945 | 0.103039639 | 0.067 | 0 | 1 | 25 |
| pip5k1aa | 0.60945 | 0.153473141 | 0.067 | 0 | 1 | 25 |
| ppip5k1b | 0.60945 | 0.176944499 | 0.067 | 0 | 1 | 25 |
| ppm1aa | 0.60945 | 0.111088879 | 0.067 | 0 | 1 | 25 |
| ppp3r1b | 0.60945 | 0.160654505 | 0.067 | 0 | 1 | 25 |
| prdx6 | 0.60945 | 0.187548933 | 0.067 | 0 | 1 | 25 |
| prkx | 0.60945 | 0.116323329 | 0.067 | 0 | 1 | 25 |
| pstpip2 | 0.60945 | 0.174985358 | 0.067 | 0 | 1 | 25 |
| ranbp1 | 0.60945 | 0.258585492 | 0.067 | 0 | 1 | 25 |
| rapgef1a | 0.60945 | 0.131629242 | 0.067 | 0 | 1 | 25 |
| rasal2 | 0.60945 | 0.179283008 | 0.067 | 0 | 1 | 25 |
| rassf2b | 0.60945 | 0.111278936 | 0.067 | 0 | 1 | 25 |
| rbm28 | 0.60945 | 0.12353995 | 0.067 | 0 | 1 | 25 |
| rbm39b | 0.60945 | 0.112288898 | 0.067 | 0 | 1 | 25 |
| rcor1 | 0.60945 | 0.183797285 | 0.067 | 0 | 1 | 25 |
| rexo4 | 0.60945 | 0.154033419 | 0.067 | 0 | 1 | 25 |
| rffl | 0.60945 | 0.184471207 | 0.067 | 0 | 1 | 25 |
| rfxap | 0.60945 | 0.156161904 | 0.067 | 0 | 1 | 25 |
| rmdn1 | 0.60945 | 0.176492925 | 0.067 | 0 | 1 | 25 |
| rnaseh2c | 0.60945 | 0.19786893 | 0.067 | 0 | 1 | 25 |
| rnf11b | 0.60945 | 0.172789101 | 0.067 | 0 | 1 | 25 |
| rps6kb1a | 0.60945 | 0.230001735 | 0.067 | 0 | 1 | 25 |
| rtf2 | 0.60945 | 0.110633357 | 0.067 | 0 | 1 | 25 |

| | | | | | | |
|---|---|---|---|---|---|---|
| rwdd1 | 0.60945 | 0.177645061 | 0.067 | 0 | 1 | 25 |
| sbds | 0.60945 | 0.168516267 | 0.067 | 0 | 1 | 25 |
| sec23b | 0.60945 | 0.140192974 | 0.067 | 0 | 1 | 25 |
| serpinb1l1 | 0.60945 | 0.131261274 | 0.067 | 0 | 1 | 25 |
| si:ch211-193k19.1 | 0.60945 | 0.157863607 | 0.067 | 0 | 1 | 25 |
| si:ch211-197h24.6 | 0.60945 | 0.160029091 | 0.067 | 0 | 1 | 25 |
| si:ch211-51h9.7 | 0.60945 | 0.122537931 | 0.067 | 0 | 1 | 25 |
| si:ch73-42p12.2 | 0.60945 | 0.102260449 | 0.067 | 0 | 1 | 25 |
| si:dkey-172j4.3 | 0.60945 | 0.186594724 | 0.067 | 0 | 1 | 25 |
| si:dkey-188h10.3 | 0.60945 | 0.221713594 | 0.067 | 0 | 1 | 25 |
| si:dkey-21o19.2 | 0.60945 | 0.18524586 | 0.067 | 0 | 1 | 25 |
| si:dkey-237i9.1 | 0.60945 | 0.2041049 | 0.067 | 0 | 1 | 25 |
| si:dkey-48p11.3 | 0.60945 | 0.156656117 | 0.067 | 0 | 1 | 25 |
| si:zfos-2326c3.2 | 0.60945 | 0.119760716 | 0.067 | 0 | 1 | 25 |
| siah1 | 0.60945 | 0.174384213 | 0.067 | 0 | 1 | 25 |
| sirt2 | 0.60945 | 0.161168721 | 0.067 | 0 | 1 | 25 |
| slc25a14 | 0.60945 | 0.115729607 | 0.067 | 0 | 1 | 25 |
| slc25a15a | 0.60945 | 0.108325585 | 0.067 | 0 | 1 | 25 |
| slc25a20 | 0.60945 | 0.154786341 | 0.067 | 0 | 1 | 25 |
| slc30a9 | 0.60945 | 0.113080838 | 0.067 | 0 | 1 | 25 |
| slc37a4a | 0.60945 | 0.239360731 | 0.067 | 0 | 1 | 25 |
| slc38a10 | 0.60945 | 0.163438221 | 0.067 | 0 | 1 | 25 |
| slc38a5a | 0.60945 | 0.168471448 | 0.067 | 0 | 1 | 25 |
| slc45a4 | 0.60945 | 0.172671241 | 0.067 | 0 | 1 | 25 |
| sltm | 0.60945 | 0.182532884 | 0.067 | 0 | 1 | 25 |
| smarcb1a | 0.60945 | 0.1608257 | 0.067 | 0 | 1 | 25 |
| snd1 | 0.60945 | 0.140009997 | 0.067 | 0 | 1 | 25 |
| snx5 | 0.60945 | 0.138921134 | 0.067 | 0 | 1 | 25 |
| sqor | 0.60945 | 0.112016252 | 0.067 | 0 | 1 | 25 |
| srcap | 0.60945 | 0.187743888 | 0.067 | 0 | 1 | 25 |
| srsf4 | 0.60945 | 0.150072652 | 0.067 | 0 | 1 | 25 |
| ss18 | 0.60945 | 0.133361643 | 0.067 | 0 | 1 | 25 |
| stau2 | 0.60945 | 0.109505542 | 0.067 | 0 | 1 | 25 |
| stip1 | 0.60945 | 0.1418178 | 0.067 | 0 | 1 | 25 |
| strn4 | 0.60945 | 0.137421337 | 0.067 | 0 | 1 | 25 |
| stx12 | 0.60945 | 0.221854959 | 0.067 | 0 | 1 | 25 |
| stx5al | 0.60945 | 0.153380828 | 0.067 | 0 | 1 | 25 |
| syap1 | 0.60945 | 0.120791638 | 0.067 | 0 | 1 | 25 |
| tapbp.2 | 0.60945 | 0.104585133 | 0.067 | 0 | 1 | 25 |
| terfa | 0.60945 | 0.12132228 | 0.067 | 0 | 1 | 25 |
| tes | 0.60945 | 0.203124769 | 0.067 | 0 | 1 | 25 |
| tfe3b | 0.60945 | 0.186429492 | 0.067 | 0 | 1 | 25 |
| tiam1a | 0.60945 | 0.245645526 | 0.067 | 0 | 1 | 25 |
| tm9sf4 | 0.60945 | 0.214790502 | 0.067 | 0 | 1 | 25 |
| tmem39b | 0.60945 | 0.154418236 | 0.067 | 0 | 1 | 25 |
| tmem42a | 0.60945 | 0.165528743 | 0.067 | 0 | 1 | 25 |
| tmod2 | 0.60945 | 0.226509708 | 0.067 | 0 | 1 | 25 |
| tmub2 | 0.60945 | 0.145633228 | 0.067 | 0 | 1 | 25 |

| | | | | | | |
|---|---|---|---|---|---|---|
| tp53 | 0.60945 | 0.145209438 | 0.067 | 0 | 1 | 25 |
| ubald1a | 0.60945 | 0.175491133 | 0.067 | 0 | 1 | 25 |
| ulk1a | 0.60945 | 0.169738671 | 0.067 | 0 | 1 | 25 |
| unc119a | 0.60945 | 0.211808644 | 0.067 | 0 | 1 | 25 |
| vgll3 | 0.60945 | 0.170607734 | 0.067 | 0 | 1 | 25 |
| vps26b | 0.60945 | 0.149837872 | 0.067 | 0 | 1 | 25 |
| wbp1lb | 0.60945 | 0.152685834 | 0.067 | 0 | 1 | 25 |
| zbtb7b | 0.60945 | 0.111122417 | 0.067 | 0 | 1 | 25 |
| zgc:162952 | 0.60945 | 0.147285175 | 0.067 | 0 | 1 | 25 |
| zgc:171679 | 0.60945 | 0.20370278 | 0.067 | 0 | 1 | 25 |
| zgc:66388 | 0.60945 | 0.149466967 | 0.067 | 0 | 1 | 25 |
| zgc:85932 | 0.60945 | 0.155535146 | 0.067 | 0 | 1 | 25 |
| znf217 | 0.60945 | 0.174191068 | 0.067 | 0 | 1 | 25 |
| znf296 | 0.60945 | 0.164010601 | 0.067 | 0 | 1 | 25 |
| rpl32 | 0.61029 | 0.68789647 | 0.567 | 0.5 | 1 | 25 |
| HACD4 | 0.62149 | 0.565938708 | 0.5 | 0.25 | 1 | 25 |
| pdia3 | 0.62727 | -0.5865712 | 0.267 | 0.5 | 1 | 25 |
| zmp:0000001082 | 0.63131 | -0.21432861 | 0.267 | 0.5 | 1 | 25 |
| rpl36 | 0.63741 | 0.509541793 | 0.5 | 0.25 | 1 | 25 |
| uba52 | 0.63898 | 0.172417599 | 0.467 | 0.25 | 1 | 25 |
| zgc:55262 | 0.65208 | -0.49304026 | 0.267 | 0.5 | 1 | 25 |
| zgc:193541 | 0.65462 | -0.42493416 | 0.267 | 0.5 | 1 | 25 |
| ndufs3 | 0.65502 | -0.73546898 | 0.1 | 0.25 | 1 | 25 |
| si:dkey-262k9.4 | 0.65512 | -0.30188194 | 0.267 | 0.5 | 1 | 25 |
| cnksr2b | 0.65849 | -0.39405597 | 0.267 | 0.5 | 1 | 25 |
| arpc4l | 0.65868 | 0.191364162 | 0.467 | 0.25 | 1 | 25 |
| atp5l | 0.65914 | -0.34188834 | 0.267 | 0.5 | 1 | 25 |
| rac2 | 0.65916 | 0.356672564 | 0.733 | 0.5 | 1 | 25 |
| pim1 | 0.65923 | -0.4315818 | 0.267 | 0.5 | 1 | 25 |
| sec11a | 0.65938 | -0.36867696 | 0.267 | 0.5 | 1 | 25 |
| si:ch211-264f5.2 | 0.65939 | -0.4369093 | 0.267 | 0.5 | 1 | 25 |
| CT573383.1 | 0.66143 | -0.61678239 | 0.3 | 0.5 | 1 | 25 |
| rps3a | 0.66206 | 0.3531047 | 0.667 | 0.75 | 1 | 25 |
| id3 | 0.66376 | 0.673425423 | 0.433 | 0.25 | 1 | 25 |
| nme2b.1 | 0.66711 | 0.232553413 | 0.633 | 0.75 | 1 | 25 |
| mt-co1 | 0.66919 | -0.2718025 | 0.9 | 1 | 1 | 25 |
| si:dkey-222f2.7 | 0.67024 | 0.673570314 | 0.433 | 0.25 | 1 | 25 |
| sec61b | 0.67479 | 0.722912089 | 0.4 | 0.25 | 1 | 25 |
| tubb4b | 0.67768 | 0.493082876 | 0.7 | 0.5 | 1 | 25 |
| rps8a | 0.68368 | 0.542105102 | 0.533 | 0.5 | 1 | 25 |
| faua | 0.68379 | 0.704104969 | 0.4 | 0.25 | 1 | 25 |
| cfl1 | 0.68421 | 0.143058706 | 0.633 | 0.75 | 1 | 25 |
| cirbpb | 0.68494 | 0.545736464 | 0.533 | 0.5 | 1 | 25 |
| trib3 | 0.68992 | -0.6920669 | 0.1 | 0.25 | 1 | 25 |
| wdr1 | 0.69044 | 0.257675443 | 0.467 | 0.25 | 1 | 25 |
| rps21 | 0.6905 | 0.563657192 | 0.467 | 0.25 | 1 | 25 |
| twf1a | 0.6908 | 0.261645721 | 0.467 | 0.25 | 1 | 25 |
| lyrm4 | 0.6958 | -0.59240804 | 0.1 | 0.25 | 1 | 25 |

| | | | | | | |
|---|---|---|---|---|---|---|
| odc1 | 0.69926 | 0.260688568 | 0.467 | 0.25 | 1 | 25 |
| zgc:113184 | 0.6995 | -0.62228853 | 0.1 | 0.25 | 1 | 25 |
| rpl23a | 0.70036 | 0.549889976 | 0.567 | 0.5 | 1 | 25 |
| pabpc1a | 0.70125 | 0.485768365 | 0.467 | 0.25 | 1 | 25 |
| romo1 | 0.70153 | 0.655457104 | 0.4 | 0.25 | 1 | 25 |
| pomp | 0.70363 | -0.57314343 | 0.1 | 0.25 | 1 | 25 |
| slc35e1 | 0.70604 | -0.54751606 | 0.1 | 0.25 | 1 | 25 |
| atp1a1a.1 | 0.70637 | -0.5433878 | 0.1 | 0.25 | 1 | 25 |
| dhrs9 | 0.7073 | -0.33003519 | 0.533 | 0.75 | 1 | 25 |
| ncf1 | 0.70851 | -0.26194225 | 0.533 | 0.75 | 1 | 25 |
| BOLA2 | 0.70883 | -0.56347205 | 0.1 | 0.25 | 1 | 25 |
| rpl5b | 0.70917 | 0.414960283 | 0.467 | 0.25 | 1 | 25 |
| tmsb1 | 0.70926 | 0.425477132 | 0.467 | 0.25 | 1 | 25 |
| LOC101886073 | 0.71001 | -0.56395043 | 0.1 | 0.25 | 1 | 25 |
| ddost | 0.7101 | -0.53451867 | 0.1 | 0.25 | 1 | 25 |
| rpl22 | 0.71033 | 0.429999637 | 0.7 | 0.5 | 1 | 25 |
| zmat2 | 0.7104 | -0.55938074 | 0.1 | 0.25 | 1 | 25 |
| stk17b | 0.71112 | 0.594730389 | 0.433 | 0.25 | 1 | 25 |
| klf13 | 0.71127 | -0.58912034 | 0.1 | 0.25 | 1 | 25 |
| nop10 | 0.71167 | -0.52724784 | 0.1 | 0.25 | 1 | 25 |
| prdx1 | 0.7136 | -0.52562781 | 0.1 | 0.25 | 1 | 25 |
| psme2 | 0.71367 | -0.52087878 | 0.1 | 0.25 | 1 | 25 |
| XLOC-000946 | 0.71565 | -0.51727812 | 0.1 | 0.25 | 1 | 25 |
| crebzf | 0.71941 | -0.5663752 | 0.1 | 0.25 | 1 | 25 |
| clint1a | 0.72026 | -0.50195479 | 0.1 | 0.25 | 1 | 25 |
| myh11a | 0.72126 | -0.49667545 | 0.1 | 0.25 | 1 | 25 |
| ctss1 | 0.72152 | -0.47445872 | 0.3 | 0.5 | 1 | 25 |
| LOC103909115 | 0.72202 | -0.54477238 | 0.4 | 0.5 | 1 | 25 |
| edf1 | 0.72259 | -0.49049228 | 0.1 | 0.25 | 1 | 25 |
| sptan1 | 0.72296 | -0.49298848 | 0.1 | 0.25 | 1 | 25 |
| cnpy3 | 0.72298 | -0.55504276 | 0.1 | 0.25 | 1 | 25 |
| ca6 | 0.72326 | 0.172722421 | 0.433 | 0.25 | 1 | 25 |
| crip1 | 0.72476 | 0.573701672 | 0.433 | 0.25 | 1 | 25 |
| ip6k2a | 0.72565 | -0.48592505 | 0.1 | 0.25 | 1 | 25 |
| ndufs5 | 0.72652 | -0.47300907 | 0.1 | 0.25 | 1 | 25 |
| eif3m | 0.72774 | 0.537313712 | 0.433 | 0.25 | 1 | 25 |
| zgc:174863 | 0.72828 | -0.45525351 | 0.3 | 0.5 | 1 | 25 |
| napab | 0.72865 | -0.50105699 | 0.1 | 0.25 | 1 | 25 |
| thrap3b | 0.72923 | -0.49967879 | 0.1 | 0.25 | 1 | 25 |
| mych | 0.72984 | -0.41878396 | 0.1 | 0.25 | 1 | 25 |
| pisd | 0.73012 | -0.52759608 | 0.1 | 0.25 | 1 | 25 |
| srsf2a | 0.73086 | -0.49048739 | 0.1 | 0.25 | 1 | 25 |
| rps15 | 0.73092 | 0.23793275 | 0.667 | 0.75 | 1 | 25 |
| mpx | 0.73097 | -0.20577745 | 0.967 | 1 | 1 | 25 |
| sdf4 | 0.7314 | -0.41075142 | 0.1 | 0.25 | 1 | 25 |
| ndufa1 | 0.73144 | -0.45938353 | 0.1 | 0.25 | 1 | 25 |
| pdcl | 0.73148 | -0.52314801 | 0.1 | 0.25 | 1 | 25 |
| ccr9a | 0.73154 | -0.55423333 | 0.1 | 0.25 | 1 | 25 |

| | | | | | | |
|---|---|---|---|---|---|---|
| ptprr | 0.73156 | -0.4671096 | 0.1 | 0.25 | 1 | 25 |
| si:ch211-243a20.4 | 0.7316 | -0.13087538 | 0.1 | 0.25 | 1 | 25 |
| si:cabz01036022.1 | 0.73269 | -0.54990337 | 0.1 | 0.25 | 1 | 25 |
| fbxl6 | 0.73292 | -0.45805863 | 0.1 | 0.25 | 1 | 25 |
| hnrnpd | 0.73307 | -0.48222861 | 0.1 | 0.25 | 1 | 25 |
| si:dkey-21a6.5 | 0.73309 | -0.12598726 | 0.1 | 0.25 | 1 | 25 |
| tmem176l.1 | 0.7331 | -0.1201947 | 0.1 | 0.25 | 1 | 25 |
| ap1s2 | 0.73323 | -0.45191768 | 0.1 | 0.25 | 1 | 25 |
| zbtb2b | 0.73428 | -0.44849809 | 0.1 | 0.25 | 1 | 25 |
| CU570894.1 | 0.7359 | -0.46981924 | 0.1 | 0.25 | 1 | 25 |
| lman2lb | 0.73598 | -0.14565423 | 0.1 | 0.25 | 1 | 25 |
| rpl3 | 0.73638 | 0.462677427 | 0.667 | 0.5 | 1 | 25 |
| dpy30 | 0.73654 | -0.42319772 | 0.1 | 0.25 | 1 | 25 |
| BX323087.1 | 0.73654 | -0.41255842 | 0.1 | 0.25 | 1 | 25 |
| stt3b | 0.73693 | -0.31115414 | 0.1 | 0.25 | 1 | 25 |
| drap1 | 0.73708 | -0.43029322 | 0.1 | 0.25 | 1 | 25 |
| caprin1b | 0.73737 | -0.22572626 | 0.3 | 0.5 | 1 | 25 |
| dedd1 | 0.73763 | -0.13766035 | 0.1 | 0.25 | 1 | 25 |
| si:dkey-260g12.1 | 0.73774 | -0.29400181 | 0.1 | 0.25 | 1 | 25 |
| gmds | 0.73853 | -0.15985387 | 0.1 | 0.25 | 1 | 25 |
| kctd5a | 0.73872 | -0.42153903 | 0.1 | 0.25 | 1 | 25 |
| ctdsp2 | 0.73888 | -0.27653367 | 0.1 | 0.25 | 1 | 25 |
| hnrnpub | 0.7391 | -0.41640216 | 0.1 | 0.25 | 1 | 25 |
| cxcr4b | 0.73958 | 0.164106555 | 0.9 | 0.75 | 1 | 25 |
| CR381673.1 | 0.73991 | -0.25848531 | 0.3 | 0.5 | 1 | 25 |
| gadd45aa | 0.74002 | -0.3000226 | 0.1 | 0.25 | 1 | 25 |
| mrpl47 | 0.74005 | -0.15838655 | 0.1 | 0.25 | 1 | 25 |
| mt-nd1 | 0.74036 | -0.28466074 | 0.3 | 0.5 | 1 | 25 |
| ncor2 | 0.7404 | -0.44141745 | 0.1 | 0.25 | 1 | 25 |
| gnai2b | 0.74059 | -0.44031835 | 0.1 | 0.25 | 1 | 25 |
| chia.6 | 0.74079 | 0.510579485 | 0.433 | 0.25 | 1 | 25 |
| tuba8l | 0.74116 | -0.39790388 | 0.1 | 0.25 | 1 | 25 |
| si:ch211-217k17.7 | 0.74133 | -0.16393479 | 0.1 | 0.25 | 1 | 25 |
| atp5mc3a | 0.74144 | -0.14499294 | 0.1 | 0.25 | 1 | 25 |
| si:ch211-136m16.8 | 0.74145 | -0.38367781 | 0.3 | 0.5 | 1 | 25 |
| elovl1b | 0.74147 | -0.29872027 | 0.3 | 0.5 | 1 | 25 |
| b3gnt3.4 | 0.74167 | -0.32156553 | 0.1 | 0.25 | 1 | 25 |
| brdt | 0.74175 | -0.38927921 | 0.1 | 0.25 | 1 | 25 |
| thoc2 | 0.74191 | -0.32117432 | 0.1 | 0.25 | 1 | 25 |
| mlx | 0.74202 | -0.19428925 | 0.1 | 0.25 | 1 | 25 |
| vps29 | 0.74215 | -0.18269062 | 0.1 | 0.25 | 1 | 25 |
| hmgn7 | 0.74233 | -0.26812394 | 0.3 | 0.5 | 1 | 25 |
| hnrnpr | 0.74237 | -0.19098484 | 0.1 | 0.25 | 1 | 25 |
| st7 | 0.74237 | -0.32625116 | 0.1 | 0.25 | 1 | 25 |
| ehmt1b | 0.74254 | -0.19966263 | 0.1 | 0.25 | 1 | 25 |
| LOC110438237 | 0.74254 | -0.36621641 | 0.3 | 0.5 | 1 | 25 |
| f5 | 0.74258 | -0.33215693 | 0.1 | 0.25 | 1 | 25 |
| elf2b | 0.74276 | -0.38391746 | 0.1 | 0.25 | 1 | 25 |

| | | | | | | |
|---|---|---|---|---|---|---|
| socs3b | 0.74282 | -0.38234216 | 0.1 | 0.25 | 1 | 25 |
| nfkbiab | 0.74285 | -0.34858103 | 0.3 | 0.5 | 1 | 25 |
| usp1 | 0.7429 | -0.19415822 | 0.1 | 0.25 | 1 | 25 |
| tln1 | 0.74294 | -0.31621387 | 0.3 | 0.5 | 1 | 25 |
| ubl7b | 0.74296 | -0.37967122 | 0.1 | 0.25 | 1 | 25 |
| zgc:162698 | 0.7431 | -0.37832163 | 0.1 | 0.25 | 1 | 25 |
| mgst1.2 | 0.74319 | -0.32155528 | 0.1 | 0.25 | 1 | 25 |
| wdr83os | 0.74323 | -0.19661002 | 0.1 | 0.25 | 1 | 25 |
| slc7a11 | 0.74325 | -0.37679462 | 0.1 | 0.25 | 1 | 25 |
| oxa1l | 0.74328 | -0.19818918 | 0.1 | 0.25 | 1 | 25 |
| eif2s1b | 0.74332 | -0.19083914 | 0.1 | 0.25 | 1 | 25 |
| cyc1 | 0.74337 | -0.18763755 | 0.1 | 0.25 | 1 | 25 |
| lamp1a | 0.74338 | -0.21819871 | 0.1 | 0.25 | 1 | 25 |
| tmem199 | 0.74355 | -0.21969604 | 0.1 | 0.25 | 1 | 25 |
| LOC566587 | 0.74357 | -0.22527148 | 0.1 | 0.25 | 1 | 25 |
| rpl6 | 0.74617 | 0.325349081 | 0.7 | 0.75 | 1 | 25 |
| eif3ba | 0.74762 | 0.468596479 | 0.433 | 0.25 | 1 | 25 |
| si:ch211-66e2.3 | 0.74925 | 0.209895119 | 0.433 | 0.25 | 1 | 25 |
| tpi1b | 0.75578 | 0.571757703 | 0.4 | 0.25 | 1 | 25 |
| fam107b | 0.75722 | -0.50339216 | 0.333 | 0.5 | 1 | 25 |
| ssr2 | 0.75812 | 0.616253748 | 0.333 | 0.25 | 1 | 25 |
| il6r | 0.76123 | -0.16264023 | 0.567 | 0.75 | 1 | 25 |
| rpl21 | 0.76375 | 0.492309238 | 0.567 | 0.5 | 1 | 25 |
| rps18 | 0.76443 | 0.420192343 | 0.5 | 0.5 | 1 | 25 |
| rps12 | 0.76449 | 0.635662491 | 0.333 | 0.25 | 1 | 25 |
| dusp2 | 0.76771 | 0.432162471 | 0.433 | 0.25 | 1 | 25 |
| ctsd | 0.76848 | -0.48896786 | 0.333 | 0.5 | 1 | 25 |
| pfn1 | 0.77026 | 0.155912395 | 1 | 1 | 1 | 25 |
| rps23 | 0.77166 | 0.449108213 | 0.633 | 0.5 | 1 | 25 |
| cotl1 | 0.77248 | 0.308464467 | 0.833 | 0.75 | 1 | 25 |
| ndufa4l | 0.77333 | 0.376008137 | 0.433 | 0.25 | 1 | 25 |
| ier2a | 0.77464 | 0.541566081 | 0.4 | 0.25 | 1 | 25 |
| zgc:162730 | 0.77596 | -0.30175039 | 0.567 | 0.75 | 1 | 25 |
| itm2bb | 0.77604 | -0.28302734 | 0.567 | 0.75 | 1 | 25 |
| mir223 | 0.7765 | 0.287910058 | 0.433 | 0.25 | 1 | 25 |
| cox6a1 | 0.77696 | 0.281812748 | 0.433 | 0.25 | 1 | 25 |
| rhogb | 0.77841 | 0.331414191 | 0.433 | 0.25 | 1 | 25 |
| actb2 | 0.77919 | 0.121434329 | 1 | 1 | 1 | 25 |
| rps27.2 | 0.78321 | 0.588382845 | 0.3 | 0.25 | 1 | 25 |
| ipo7 | 0.78841 | -0.5607168 | 0.133 | 0.25 | 1 | 25 |
| cox7b | 0.78882 | -0.42728244 | 0.333 | 0.5 | 1 | 25 |
| capza1a | 0.79009 | 0.480655785 | 0.4 | 0.25 | 1 | 25 |
| ch25hl2 | 0.79253 | -0.124719 | 0.9 | 1 | 1 | 25 |
| eif3d | 0.79408 | 0.483492258 | 0.4 | 0.25 | 1 | 25 |
| sms | 0.79514 | 0.408606634 | 0.633 | 0.5 | 1 | 25 |
| arhgdig | 0.79815 | 0.157666679 | 0.667 | 0.5 | 1 | 25 |
| arhgdia | 0.80108 | -0.16688611 | 0.333 | 0.5 | 1 | 25 |
| FP085398.1 | 0.80165 | -0.51079768 | 0.133 | 0.25 | 1 | 25 |

| | | | | | | |
|---|---|---|---|---|---|---|
| mt-nd3 | 0.80323 | -0.38497628 | 0.333 | 0.5 | 1 | 25 |
| h2afvb | 0.80398 | 0.528377292 | 0.367 | 0.25 | 1 | 25 |
| rtn4a | 0.80493 | -0.50047032 | 0.133 | 0.25 | 1 | 25 |
| tegt | 0.80875 | -0.16155094 | 0.333 | 0.5 | 1 | 25 |
| itgb2 | 0.80924 | 0.184709933 | 0.667 | 0.5 | 1 | 25 |
| ndufs7 | 0.80967 | -0.46777985 | 0.133 | 0.25 | 1 | 25 |
| NDUFB1 | 0.81018 | -0.48628646 | 0.133 | 0.25 | 1 | 25 |
| ndufab1b | 0.81189 | -0.47082384 | 0.133 | 0.25 | 1 | 25 |
| rbmx | 0.81375 | -0.48006798 | 0.133 | 0.25 | 1 | 25 |
| ppt1 | 0.81376 | -0.21680247 | 0.333 | 0.5 | 1 | 25 |
| ssr4 | 0.81545 | -0.17923997 | 0.333 | 0.5 | 1 | 25 |
| ssr3 | 0.81755 | -0.25317405 | 0.333 | 0.5 | 1 | 25 |
| ppiab | 0.81889 | 0.31049572 | 0.5 | 0.5 | 1 | 25 |
| itpkcb | 0.82028 | -0.50033865 | 0.133 | 0.25 | 1 | 25 |
| csnk1a1 | 0.822 | -0.43469323 | 0.133 | 0.25 | 1 | 25 |
| COX3 | 0.82514 | -0.17400831 | 0.933 | 1 | 1 | 25 |
| rpl29 | 0.82816 | 0.204020226 | 0.4 | 0.25 | 1 | 25 |
| eif5a | 0.82891 | 0.483103928 | 0.367 | 0.25 | 1 | 25 |
| rnf130 | 0.83153 | -0.40932737 | 0.133 | 0.25 | 1 | 25 |
| eif1axb | 0.83228 | -0.4219803 | 0.133 | 0.25 | 1 | 25 |
| derl2 | 0.83241 | -0.40708409 | 0.133 | 0.25 | 1 | 25 |
| mbnl1 | 0.83265 | 0.170872505 | 0.4 | 0.25 | 1 | 25 |
| rps15a | 0.83328 | 0.400835498 | 0.6 | 0.5 | 1 | 25 |
| rer1 | 0.83449 | -0.40147005 | 0.133 | 0.25 | 1 | 25 |
| si:ch211-284o19.8 | 0.83526 | 0.46850292 | 0.367 | 0.25 | 1 | 25 |
| tuba8l4 | 0.83576 | -0.26610218 | 0.3 | 0.25 | 1 | 25 |
| CABZ01058046.1 | 0.83591 | -0.39600898 | 0.133 | 0.25 | 1 | 25 |
| ucp2 | 0.83612 | -0.39851181 | 0.367 | 0.5 | 1 | 25 |
| tmem50a | 0.83613 | -0.39610709 | 0.133 | 0.25 | 1 | 25 |
| setb | 0.83643 | -0.34371562 | 0.133 | 0.25 | 1 | 25 |
| eef1da | 0.83677 | 0.215836484 | 0.4 | 0.25 | 1 | 25 |
| dbi | 0.83689 | 0.192192041 | 0.4 | 0.25 | 1 | 25 |
| zgc:66427 | 0.83759 | -0.38903486 | 0.133 | 0.25 | 1 | 25 |
| echdc2 | 0.83782 | -0.38426816 | 0.133 | 0.25 | 1 | 25 |
| fermt3b | 0.83789 | -0.38928861 | 0.133 | 0.25 | 1 | 25 |
| nsa2 | 0.83932 | 0.215032406 | 0.4 | 0.25 | 1 | 25 |
| ppp1r18 | 0.83934 | -0.38156955 | 0.133 | 0.25 | 1 | 25 |
| zgc:153867 | 0.84005 | 0.333349792 | 0.4 | 0.25 | 1 | 25 |
| btf3 | 0.84153 | 0.310556413 | 0.4 | 0.25 | 1 | 25 |
| tmem59 | 0.84241 | 0.300662684 | 0.4 | 0.25 | 1 | 25 |
| tpm3 | 0.84244 | 0.283504051 | 0.4 | 0.25 | 1 | 25 |
| dhrs13a.1 | 0.84381 | -0.14676785 | 0.133 | 0.25 | 1 | 25 |
| mmp25b | 0.84571 | -0.10134411 | 0.133 | 0.25 | 1 | 25 |
| cyb5r3 | 0.84663 | -0.34147818 | 0.133 | 0.25 | 1 | 25 |
| rap1aa | 0.84749 | -0.48685998 | 0.167 | 0.25 | 1 | 25 |
| ywhae1 | 0.8496 | -0.3124872 | 0.133 | 0.25 | 1 | 25 |
| CABZ01090890.1 | 0.84992 | -0.18606805 | 0.133 | 0.25 | 1 | 25 |
| atg16l1 | 0.84994 | -0.22826887 | 0.133 | 0.25 | 1 | 25 |

| | | | | | | |
|---|---|---|---|---|---|---|
| alas1 | 0.85002 | -0.22614258 | 0.133 | 0.25 | 1 | 25 |
| hp1bp3 | 0.85009 | -0.12027519 | 0.133 | 0.25 | 1 | 25 |
| alox12 | 0.85017 | -0.13235445 | 0.133 | 0.25 | 1 | 25 |
| rab11fip4b | 0.85074 | -0.30740747 | 0.133 | 0.25 | 1 | 25 |
| h3f3c | 0.85078 | 0.420910104 | 0.367 | 0.25 | 1 | 25 |
| adam19b | 0.85082 | -0.29711146 | 0.133 | 0.25 | 1 | 25 |
| pabpn1 | 0.85093 | -0.3064987 | 0.133 | 0.25 | 1 | 25 |
| sptbn2 | 0.85128 | -0.14281251 | 0.133 | 0.25 | 1 | 25 |
| rab10 | 0.85135 | -0.13247591 | 0.133 | 0.25 | 1 | 25 |
| hacd3 | 0.85141 | -0.20472014 | 0.133 | 0.25 | 1 | 25 |
| tmem258 | 0.85159 | -0.19363664 | 0.133 | 0.25 | 1 | 25 |
| agr1 | 0.85174 | -0.13221544 | 0.133 | 0.25 | 1 | 25 |
| DHDH | 0.85204 | -0.25670721 | 0.133 | 0.25 | 1 | 25 |
| furina | 0.85217 | -0.15788063 | 0.133 | 0.25 | 1 | 25 |
| mgst3b | 0.85226 | -0.16049229 | 0.133 | 0.25 | 1 | 25 |
| CU694197.1 | 0.85231 | -0.16863509 | 0.133 | 0.25 | 1 | 25 |
| rab2a | 0.85233 | -0.26865788 | 0.133 | 0.25 | 1 | 25 |
| arpc3 | 0.85491 | 0.290941842 | 0.5 | 0.5 | 1 | 25 |
| si:ch1073-429i10.1 | 0.85574 | 0.323580949 | 0.567 | 0.5 | 1 | 25 |
| bsg | 0.85579 | -0.47388311 | 0.2 | 0.25 | 1 | 25 |
| ctss2.1 | 0.85713 | -0.35027855 | 0.367 | 0.5 | 1 | 25 |
| pnrc2 | 0.85983 | 0.257693306 | 0.833 | 0.75 | 1 | 25 |
| hbxb | 0.86001 | -0.45923218 | 0.167 | 0.25 | 1 | 25 |
| mmp9 | 0.86396 | -0.42621921 | 0.2 | 0.25 | 1 | 25 |
| hbegfa | 0.8651 | -0.2096158 | 0.3 | 0.25 | 1 | 25 |
| spi1b | 0.86537 | 0.287344122 | 0.633 | 0.5 | 1 | 25 |
| cox5aa | 0.86679 | 0.437626778 | 0.333 | 0.25 | 1 | 25 |
| ndufb6 | 0.86743 | -0.44376219 | 0.167 | 0.25 | 1 | 25 |
| rpl22l1 | 0.86771 | -0.40322182 | 0.233 | 0.25 | 1 | 25 |
| plxnc1 | 0.86896 | -0.31123912 | 0.367 | 0.5 | 1 | 25 |
| psmb6 | 0.87187 | -0.4303473 | 0.167 | 0.25 | 1 | 25 |
| rps4x | 0.87341 | 0.133997724 | 0.633 | 0.5 | 1 | 25 |
| rps7 | 0.87467 | 0.292370461 | 0.533 | 0.5 | 1 | 25 |
| tagln2 | 0.87483 | -0.35082451 | 0.4 | 0.5 | 1 | 25 |
| npsn | 0.87507 | -0.1039268 | 1 | 1 | 1 | 25 |
| rab32a | 0.87554 | -0.4190058 | 0.167 | 0.25 | 1 | 25 |
| il34 | 0.87557 | -0.15388902 | 0.367 | 0.5 | 1 | 25 |
| ncf2 | 0.87737 | -0.42436425 | 0.167 | 0.25 | 1 | 25 |
| timp2b | 0.87921 | -0.17759538 | 0.367 | 0.5 | 1 | 25 |
| atp5f1c | 0.87938 | -0.43803606 | 0.2 | 0.25 | 1 | 25 |
| add3a | 0.87971 | -0.17005323 | 0.367 | 0.5 | 1 | 25 |
| cct2 | 0.87979 | 0.323781976 | 0.367 | 0.25 | 1 | 25 |
| arpc2 | 0.88094 | -0.24069142 | 0.367 | 0.5 | 1 | 25 |
| antxr2a | 0.88098 | -0.19311099 | 0.367 | 0.5 | 1 | 25 |
| CABZ01073834.1 | 0.88112 | -0.21649103 | 0.367 | 0.5 | 1 | 25 |
| zgc:114188 | 0.88813 | 0.321046669 | 0.567 | 0.5 | 1 | 25 |
| birc2 | 0.88871 | -0.41538148 | 0.2 | 0.25 | 1 | 25 |
| mdh1aa | 0.88936 | -0.41724904 | 0.2 | 0.25 | 1 | 25 |

| | | | | | | |
|---|---|---|---|---|---|---|
| atp5mf | 0.89357 | -0.37294515 | 0.167 | 0.25 | 1 | 25 |
| si:ch73-376l24.2 | 0.89511 | -0.37808399 | 0.167 | 0.25 | 1 | 25 |
| socs3a | 0.89569 | 0.203177959 | 0.367 | 0.25 | 1 | 25 |
| spcs2 | 0.89627 | -0.39754866 | 0.2 | 0.25 | 1 | 25 |
| degs1 | 0.89707 | -0.36314983 | 0.167 | 0.25 | 1 | 25 |
| cdc42 | 0.89711 | 0.17858211 | 0.367 | 0.25 | 1 | 25 |
| scinlb | 0.89858 | -0.23628214 | 0.667 | 0.75 | 1 | 25 |
| slc9a3r1a | 0.89917 | 0.364524303 | 0.333 | 0.25 | 1 | 25 |
| rpn1 | 0.89949 | -0.35477973 | 0.167 | 0.25 | 1 | 25 |
| h3f3d | 0.9012 | 0.262711759 | 0.5 | 0.5 | 1 | 25 |
| zgc:56493 | 0.90429 | 0.23425885 | 0.5 | 0.5 | 1 | 25 |
| ywhabl | 0.90446 | -0.38765684 | 0.2 | 0.25 | 1 | 25 |
| rab5ab | 0.90716 | -0.32943119 | 0.167 | 0.25 | 1 | 25 |
| ptmab | 0.90773 | -0.26106768 | 0.467 | 0.5 | 1 | 25 |
| fthl27 | 0.91207 | -0.27349115 | 0.4 | 0.5 | 1 | 25 |
| zgc:172106 | 0.91221 | -0.32238374 | 0.167 | 0.25 | 1 | 25 |
| smox | 0.91244 | -0.31030855 | 0.167 | 0.25 | 1 | 25 |
| hpgd | 0.91293 | 0.258009317 | 0.2 | 0.25 | 1 | 25 |
| ost4 | 0.91391 | -0.36604829 | 0.2 | 0.25 | 1 | 25 |
| dusp1 | 0.91408 | -0.31596424 | 0.167 | 0.25 | 1 | 25 |
| hnrnpa0b | 0.91673 | 0.354128886 | 0.3 | 0.25 | 1 | 25 |
| gsto1 | 0.91694 | -0.29060809 | 0.167 | 0.25 | 1 | 25 |
| rnf213a | 0.91703 | -0.29197712 | 0.167 | 0.25 | 1 | 25 |
| si:dkey-102g19.3 | 0.91739 | 0.33674266 | 0.267 | 0.25 | 1 | 25 |
| LOC101885065 | 0.91759 | 0.312834616 | 0.333 | 0.25 | 1 | 25 |
| rad21a | 0.9177 | -0.28952342 | 0.167 | 0.25 | 1 | 25 |
| ampd3b | 0.9179 | -0.28559871 | 0.167 | 0.25 | 1 | 25 |
| lbr | 0.91936 | -0.29052641 | 0.167 | 0.25 | 1 | 25 |
| zgc:77849 | 0.91949 | -0.27582362 | 0.167 | 0.25 | 1 | 25 |
| tpd52l2b | 0.92035 | -0.27353671 | 0.167 | 0.25 | 1 | 25 |
| cct8 | 0.92286 | -0.34351649 | 0.2 | 0.25 | 1 | 25 |
| myo1f | 0.92378 | -0.34218826 | 0.2 | 0.25 | 1 | 25 |
| cyba | 0.9239 | -0.23494793 | 0.4 | 0.5 | 1 | 25 |
| sri | 0.9248 | 0.28226719 | 0.333 | 0.25 | 1 | 25 |
| slc7a3a | 0.92498 | -0.13377593 | 0.167 | 0.25 | 1 | 25 |
| ATP11B | 0.92623 | -0.23109638 | 0.167 | 0.25 | 1 | 25 |
| elf1 | 0.92632 | -0.23137949 | 0.167 | 0.25 | 1 | 25 |
| tnrc6c1 | 0.92665 | -0.15089488 | 0.167 | 0.25 | 1 | 25 |
| sh3kbp1 | 0.92678 | -0.13075169 | 0.3 | 0.25 | 1 | 25 |
| LO018102.1 | 0.9274 | -0.2111017 | 0.167 | 0.25 | 1 | 25 |
| vwa11 | 0.92742 | -0.2198637 | 0.167 | 0.25 | 1 | 25 |
| zgc:110239 | 0.92751 | -0.19142999 | 0.167 | 0.25 | 1 | 25 |
| spag1a | 0.92778 | -0.12694254 | 0.3 | 0.25 | 1 | 25 |
| pfkfb4b | 0.92797 | -0.16880098 | 0.167 | 0.25 | 1 | 25 |
| atp6v0ca | 0.92813 | -0.1017177 | 0.167 | 0.25 | 1 | 25 |
| txn | 0.92813 | -0.10434016 | 0.167 | 0.25 | 1 | 25 |
| elocb | 0.92833 | -0.11275661 | 0.167 | 0.25 | 1 | 25 |
| usp36 | 0.92843 | -0.33326186 | 0.2 | 0.25 | 1 | 25 |

| | | | | | | |
|---|---|---|---|---|---|---|
| h3f3a | 0.92845 | -0.30064456 | 0.233 | 0.25 | 1 | 25 |
| si:dkey-27h10.2 | 0.92947 | 0.246375579 | 0.233 | 0.25 | 1 | 25 |
| msna | 0.93168 | -0.13832166 | 0.4 | 0.5 | 1 | 25 |
| oaz1b | 0.93467 | 0.306694336 | 0.267 | 0.25 | 1 | 25 |
| actr2a | 0.93503 | 0.308033824 | 0.3 | 0.25 | 1 | 25 |
| sap30l | 0.93581 | -0.31231334 | 0.2 | 0.25 | 1 | 25 |
| capza1b | 0.93695 | 0.136049541 | 0.467 | 0.5 | 1 | 25 |
| eef2b | 0.93826 | 0.226684777 | 0.533 | 0.5 | 1 | 25 |
| si:ch211-213a13.1 | 0.9384 | -0.3054961 | 0.2 | 0.25 | 1 | 25 |
| sat1a.2 | 0.9387 | -0.28966715 | 0.233 | 0.25 | 1 | 25 |
| ube2d2 | 0.93918 | -0.30187361 | 0.2 | 0.25 | 1 | 25 |
| psmb4 | 0.93963 | -0.30345527 | 0.2 | 0.25 | 1 | 25 |
| prkcda | 0.94048 | -0.23813529 | 0.433 | 0.5 | 1 | 25 |
| si:ch211-102c2.4 | 0.94128 | 0.176640276 | 0.5 | 0.5 | 1 | 25 |
| rpl8 | 0.94163 | 0.245816072 | 0.533 | 0.5 | 1 | 25 |
| eno1a | 0.94221 | 0.106798058 | 0.333 | 0.25 | 1 | 25 |
| vat1 | 0.94305 | -0.27012852 | 0.233 | 0.25 | 1 | 25 |
| wbp2nl | 0.94322 | 0.288160073 | 0.3 | 0.25 | 1 | 25 |
| naca | 0.94342 | 0.213776351 | 0.567 | 0.5 | 1 | 25 |
| rsl24d1 | 0.94363 | 0.136091382 | 0.333 | 0.25 | 1 | 25 |
| cbx7a | 0.94406 | -0.26905776 | 0.233 | 0.25 | 1 | 25 |
| tbcb | 0.94461 | -0.29032814 | 0.2 | 0.25 | 1 | 25 |
| cox6b2 | 0.94503 | 0.189053122 | 0.333 | 0.25 | 1 | 25 |
| vsir | 0.94523 | -0.10598268 | 0.667 | 0.75 | 1 | 25 |
| si:ch73-248e21.5 | 0.94528 | 0.170050233 | 0.333 | 0.25 | 1 | 25 |
| gpx1a | 0.94532 | 0.282963236 | 0.3 | 0.25 | 1 | 25 |
| si:ch73-248e21.7 | 0.94558 | 0.152751526 | 0.767 | 0.75 | 1 | 25 |
| rpl27a | 0.9457 | -0.27944899 | 0.2 | 0.25 | 1 | 25 |
| plpp1a | 0.94868 | 0.266153304 | 0.3 | 0.25 | 1 | 25 |
| sumo3a | 0.94904 | 0.235311103 | 0.233 | 0.25 | 1 | 25 |
| mrps15 | 0.94905 | 0.267387792 | 0.3 | 0.25 | 1 | 25 |
| glo1 | 0.94921 | -0.27169147 | 0.2 | 0.25 | 1 | 25 |
| ndufb3 | 0.95008 | 0.113599236 | 0.2 | 0.25 | 1 | 25 |
| asah1a | 0.95118 | 0.262185693 | 0.267 | 0.25 | 1 | 25 |
| notch2 | 0.95172 | -0.16358125 | 0.267 | 0.25 | 1 | 25 |
| gnb1a | 0.9537 | -0.25846719 | 0.2 | 0.25 | 1 | 25 |
| magt1 | 0.95656 | 0.191361306 | 0.233 | 0.25 | 1 | 25 |
| hmgn2 | 0.959 | 0.124854201 | 0.767 | 0.75 | 1 | 25 |
| rgs19 | 0.95938 | -0.23394107 | 0.2 | 0.25 | 1 | 25 |
| CABZ01001434.1 | 0.96054 | 0.235917723 | 0.3 | 0.25 | 1 | 25 |
| nme2b.2 | 0.96291 | -0.22090935 | 0.233 | 0.25 | 1 | 25 |
| arpc5a | 0.96468 | -0.21209657 | 0.2 | 0.25 | 1 | 25 |
| chac1 | 0.96494 | -0.21078126 | 0.2 | 0.25 | 1 | 25 |
| cd7al | 0.96538 | -0.11958797 | 0.733 | 0.75 | 1 | 25 |
| cd81a | 0.96742 | 0.170463334 | 0.233 | 0.25 | 1 | 25 |
| atp5if1b | 0.96751 | -0.21234021 | 0.233 | 0.25 | 1 | 25 |
| arrb2b | 0.96823 | -0.21300172 | 0.233 | 0.25 | 1 | 25 |
| serbp1a | 0.96881 | -0.12864246 | 0.433 | 0.5 | 1 | 25 |

| | | | | | | |
|---|---|---|---|---|---|---|
| psap | 0.9689 | 0.215645398 | 0.267 | 0.25 | 1 | 25 |
| atp5po | 0.96921 | -0.14100901 | 0.267 | 0.25 | 1 | 25 |
| glipr1a | 0.96935 | 0.115775665 | 0.567 | 0.5 | 1 | 25 |
| lpar5a | 0.96959 | -0.10848635 | 0.433 | 0.5 | 1 | 25 |
| anxa11b | 0.96975 | -0.202744 | 0.233 | 0.25 | 1 | 25 |
| si:dkey-19b23.14 | 0.97028 | 0.188456206 | 0.3 | 0.25 | 1 | 25 |
| rgs14a | 0.97102 | -0.174073 | 0.2 | 0.25 | 1 | 25 |
| lrrfip1a | 0.97192 | -0.16763825 | 0.2 | 0.25 | 1 | 25 |
| rps17 | 0.97295 | 0.157586269 | 0.533 | 0.5 | 1 | 25 |
| XLOC-039817 | 0.97346 | -0.15123381 | 0.2 | 0.25 | 1 | 25 |
| nfil3-6 | 0.97544 | -0.10242895 | 0.2 | 0.25 | 1 | 25 |
| si:ch211-9d9.1 | 0.97547 | -0.10705826 | 0.2 | 0.25 | 1 | 25 |
| spns3 | 0.97548 | -0.12292953 | 0.2 | 0.25 | 1 | 25 |
| tpm1 | 0.97594 | -0.10667673 | 0.267 | 0.25 | 1 | 25 |
| sod1 | 0.97765 | -0.11482577 | 0.267 | 0.25 | 1 | 25 |
| farsa | 0.97792 | 0.134352909 | 0.233 | 0.25 | 1 | 25 |
| srd5a2a | 0.97853 | 0.179859213 | 0.267 | 0.25 | 1 | 25 |
| LOC101882843 | 0.97916 | 0.1024719 | 0.3 | 0.25 | 1 | 25 |
| calm3b | 0.97983 | -0.17482357 | 0.233 | 0.25 | 1 | 25 |
| aldoaa | 0.98049 | -0.16967408 | 0.233 | 0.25 | 1 | 25 |
| pkma | 0.98058 | 0.167929887 | 0.267 | 0.25 | 1 | 25 |
| ank3b | 0.98132 | 0.118220137 | 0.233 | 0.25 | 1 | 25 |
| atp5pf | 0.98354 | 0.126267732 | 0.233 | 0.25 | 1 | 25 |
| wu:fb55g09 | 0.98364 | 0.106790896 | 0.233 | 0.25 | 1 | 25 |
| h1fx | 0.98416 | -0.14650245 | 0.233 | 0.25 | 1 | 25 |
| eif3ea | 0.98546 | 0.104475607 | 0.233 | 0.25 | 1 | 25 |
| zgc:174904 | 0.98661 | -0.13076456 | 0.233 | 0.25 | 1 | 25 |
| eif1b | 0.98845 | -0.13049255 | 0.233 | 0.25 | 1 | 25 |
| hmgn6 | 0.98922 | -0.12346497 | 0.233 | 0.25 | 1 | 25 |
| pxk | 0.98928 | -0.12623415 | 0.233 | 0.25 | 1 | 25 |
| cox6c | 0.98995 | -0.12024951 | 0.233 | 0.25 | 1 | 25 |
| GCA | 0.99246 | -0.10366034 | 0.233 | 0.25 | 1 | 25 |
| mt-co2 | 0.00496 | 0.535293005 | 0.444 | 0.571 | 1 | 26 |
| eif5a2 | 0.0057 | -1.20765223 | 0 | 0.429 | 1 | 26 |
| cd81a | 0.00911 | 1.343372601 | 0.5 | 0 | 1 | 26 |
| si:dkey-203a12.9 | 0.01361 | -1.49657653 | 0.111 | 0.571 | 1 | 26 |
| eef1da | 0.01647 | 1.161331139 | 0.444 | 0 | 1 | 26 |
| cebpb | 0.0192 | -0.44941594 | 0.222 | 0.571 | 1 | 26 |
| rpl7 | 0.02427 | -0.67712253 | 0.667 | 1 | 1 | 26 |
| CR383676.1 | 0.02492 | 1.481473 | 0.556 | 0.143 | 1 | 26 |
| rpl13 | 0.02572 | -0.76525195 | 0.556 | 1 | 1 | 26 |
| rps9 | 0.02762 | -0.59228999 | 0.5 | 1 | 1 | 26 |
| nfkbiab | 0.02869 | 1.180858593 | 0.389 | 0 | 1 | 26 |
| cfp | 0.02869 | 1.103872366 | 0.389 | 0 | 1 | 26 |
| pdia2 | 0.02869 | 0.849098729 | 0.389 | 0 | 1 | 26 |
| atp5fa1 | 0.02869 | 0.998712676 | 0.389 | 0 | 1 | 26 |
| uqcrq | 0.03076 | -1.12377168 | 0.111 | 0.571 | 1 | 26 |
| si:ch1073-110a20.3 | 0.0318 | -0.68726641 | 0.556 | 0.857 | 1 | 26 |

| | | | | | | |
|---|---|---|---|---|---|---|
| ppp2cab | 0.03223 | -0.84118128 | 0 | 0.286 | 1 | 26 |
| h3f3c | 0.03223 | -0.85613666 | 0 | 0.286 | 1 | 26 |
| si:dkey-27h10.2 | 0.03223 | -0.89516711 | 0 | 0.286 | 1 | 26 |
| snrpf | 0.03223 | -0.88362986 | 0 | 0.286 | 1 | 26 |
| BX005340.1 | 0.03223 | -0.85857823 | 0 | 0.286 | 1 | 26 |
| fkbp4 | 0.03223 | -0.87071351 | 0 | 0.286 | 1 | 26 |
| imp3 | 0.03223 | -0.7865265 | 0 | 0.286 | 1 | 26 |
| nsfl1c | 0.03223 | -0.78353469 | 0 | 0.286 | 1 | 26 |
| ppp1cab | 0.03223 | -0.79373992 | 0 | 0.286 | 1 | 26 |
| psma2 | 0.03223 | -0.86611634 | 0 | 0.286 | 1 | 26 |
| psme1 | 0.03223 | -0.86721291 | 0 | 0.286 | 1 | 26 |
| rhof | 0.03223 | -0.8266937 | 0 | 0.286 | 1 | 26 |
| si:dkeyp-27c8.2 | 0.03223 | -0.79477446 | 0 | 0.286 | 1 | 26 |
| skp1 | 0.03223 | -0.78188307 | 0 | 0.286 | 1 | 26 |
| slc11a2 | 0.03223 | -0.79370178 | 0 | 0.286 | 1 | 26 |
| sub1b | 0.03223 | -0.7251331 | 0 | 0.286 | 1 | 26 |
| yy1a | 0.03223 | -0.85448194 | 0 | 0.286 | 1 | 26 |
| zgc:193541 | 0.03223 | -0.77856083 | 0 | 0.286 | 1 | 26 |
| LOC100536213 | 0.03223 | -0.83931442 | 0 | 0.286 | 1 | 26 |
| TSTD1 | 0.03223 | -0.827907 | 0 | 0.286 | 1 | 26 |
| cenpw | 0.03223 | -0.77888333 | 0 | 0.286 | 1 | 26 |
| cpne1 | 0.03223 | -0.89003747 | 0 | 0.286 | 1 | 26 |
| fybb | 0.03223 | -0.84891767 | 0 | 0.286 | 1 | 26 |
| h2afva | 0.03223 | -0.83262095 | 0 | 0.286 | 1 | 26 |
| mcts1 | 0.03223 | -0.80433728 | 0 | 0.286 | 1 | 26 |
| ndufa2 | 0.03223 | -0.78156802 | 0 | 0.286 | 1 | 26 |
| prpf40a | 0.03223 | -0.81523616 | 0 | 0.286 | 1 | 26 |
| rab5ab | 0.03223 | -0.79602841 | 0 | 0.286 | 1 | 26 |
| txndc17 | 0.03223 | -0.86174262 | 0 | 0.286 | 1 | 26 |
| sept7a | 0.03223 | -0.90620999 | 0 | 0.286 | 1 | 26 |
| yipf4 | 0.03223 | -0.90467344 | 0 | 0.286 | 1 | 26 |
| rpl30 | 0.03272 | 0.105087287 | 0.722 | 0.857 | 1 | 26 |
| crip1 | 0.04511 | -0.78572862 | 0.444 | 0.857 | 1 | 26 |
| pfn1 | 0.04656 | -0.37988096 | 0.833 | 1 | 1 | 26 |
| dub | 0.04833 | -1.05638263 | 0.111 | 0.571 | 1 | 26 |
| nfil3-6 | 0.04852 | 0.91613979 | 0.333 | 0 | 1 | 26 |
| socs3a | 0.04852 | 1.004286033 | 0.333 | 0 | 1 | 26 |
| rpl27a | 0.05274 | -0.98545624 | 0.222 | 0.571 | 1 | 26 |
| cox6b2 | 0.05476 | -1.06046841 | 0.111 | 0.571 | 1 | 26 |
| rps2 | 0.05556 | -0.32924018 | 0.611 | 1 | 1 | 26 |
| hmgn2 | 0.05647 | -0.90393478 | 0.444 | 0.857 | 1 | 26 |
| gng5 | 0.06871 | -1.02359582 | 0.111 | 0.571 | 1 | 26 |
| snx5 | 0.08011 | 0.871294929 | 0.278 | 0 | 1 | 26 |
| qkia | 0.08011 | 0.847875433 | 0.278 | 0 | 1 | 26 |
| atp5if1a | 0.08011 | 0.6983507 | 0.278 | 0 | 1 | 26 |
| c6ast3 | 0.08011 | 0.780927955 | 0.278 | 0 | 1 | 26 |
| cox7b | 0.08011 | 0.735636721 | 0.278 | 0 | 1 | 26 |
| ndufb3 | 0.08011 | 0.740831058 | 0.278 | 0 | 1 | 26 |

| | | | | | | |
|---|---|---|---|---|---|---|
| serpinb1l3 | 0.08011 | 0.739014858 | 0.278 | 0 | 1 | 26 |
| si:dkey-118j18.2 | 0.08011 | 0.743961273 | 0.278 | 0 | 1 | 26 |
| cox6c | 0.08011 | 0.739582558 | 0.278 | 0 | 1 | 26 |
| eif3ha | 0.08011 | 0.769680309 | 0.278 | 0 | 1 | 26 |
| wasa | 0.08011 | 0.812279469 | 0.278 | 0 | 1 | 26 |
| rps5 | 0.0813 | -0.4042907 | 0.611 | 1 | 1 | 26 |
| rpl18 | 0.08453 | -0.43605285 | 0.611 | 1 | 1 | 26 |
| zgc:114188 | 0.0876 | -0.48894217 | 0.778 | 1 | 1 | 26 |
| zgc:56493 | 0.09334 | -0.17085529 | 0.667 | 0.571 | 1 | 26 |
| eef1g | 0.09589 | 0.31850277 | 0.778 | 0.429 | 1 | 26 |
| ssr4 | 0.09591 | 0.401495514 | 0.444 | 0.143 | 1 | 26 |
| cfl1 | 0.09648 | -0.46481214 | 0.722 | 1 | 1 | 26 |
| ufm1 | 0.10281 | -0.9135785 | 0.056 | 0.429 | 1 | 26 |
| rpl23 | 0.10296 | -0.53775755 | 0.667 | 1 | 1 | 26 |
| alas1 | 0.10353 | -0.90216852 | 0.056 | 0.429 | 1 | 26 |
| ywhabb | 0.10354 | -0.8939403 | 0.056 | 0.429 | 1 | 26 |
| rps23 | 0.10684 | -0.4789037 | 0.444 | 0.857 | 1 | 26 |
| rpl39 | 0.10906 | -0.51531643 | 0.722 | 1 | 1 | 26 |
| rpl10 | 0.11205 | -0.42525948 | 0.778 | 1 | 1 | 26 |
| FO203432.1 | 0.11372 | -0.6103759 | 0.278 | 0.429 | 1 | 26 |
| rps26l | 0.11595 | -0.34384492 | 0.667 | 1 | 1 | 26 |
| rpl19 | 0.11763 | -0.52417595 | 0.722 | 1 | 1 | 26 |
| clca1 | 0.11946 | 0.849292279 | 0.556 | 0.143 | 1 | 26 |
| si:dkey-151g10.6 | 0.11957 | -0.16217468 | 0.778 | 1 | 1 | 26 |
| rps15 | 0.1233 | -0.51074748 | 0.667 | 1 | 1 | 26 |
| myl12.1 | 0.12887 | -0.82875522 | 0.278 | 0.714 | 1 | 26 |
| atp5po | 0.12995 | 0.711429339 | 0.222 | 0 | 1 | 26 |
| cdc42 | 0.12995 | 0.720599996 | 0.222 | 0 | 1 | 26 |
| cxcl8b.1 | 0.12995 | 0.78356469 | 0.222 | 0 | 1 | 26 |
| eif1axb | 0.12995 | 0.566738671 | 0.222 | 0 | 1 | 26 |
| eif2s2 | 0.12995 | 0.602833597 | 0.222 | 0 | 1 | 26 |
| fkbp5 | 0.12995 | 0.641631847 | 0.222 | 0 | 1 | 26 |
| itgb2 | 0.12995 | 0.663031638 | 0.222 | 0 | 1 | 26 |
| klf6a | 0.12995 | 0.648442528 | 0.222 | 0 | 1 | 26 |
| limd2 | 0.12995 | 0.758545119 | 0.222 | 0 | 1 | 26 |
| nr4a1 | 0.12995 | 0.75034174 | 0.222 | 0 | 1 | 26 |
| si:ch73-208g10.1 | 0.12995 | 0.777097948 | 0.222 | 0 | 1 | 26 |
| si:dkey-163m14.2 | 0.12995 | 0.542567112 | 0.222 | 0 | 1 | 26 |
| vaspa | 0.12995 | 0.731196464 | 0.222 | 0 | 1 | 26 |
| zmat2 | 0.12995 | 0.608911592 | 0.222 | 0 | 1 | 26 |
| adam8a | 0.12995 | 0.655618841 | 0.222 | 0 | 1 | 26 |
| atf6 | 0.12995 | 0.646528833 | 0.222 | 0 | 1 | 26 |
| atp5mc3b | 0.12995 | 0.618689192 | 0.222 | 0 | 1 | 26 |
| btg1 | 0.12995 | 0.641584539 | 0.222 | 0 | 1 | 26 |
| dnaja2a | 0.12995 | 0.649353985 | 0.222 | 0 | 1 | 26 |
| ehd1a | 0.12995 | 0.678163908 | 0.222 | 0 | 1 | 26 |
| eif3g | 0.12995 | 0.666426588 | 0.222 | 0 | 1 | 26 |
| furina | 0.12995 | 0.642578878 | 0.222 | 0 | 1 | 26 |

| | | | | | | |
|---|---|---|---|---|---|---|
| glipr1a | 0.12995 | 0.718134141 | 0.222 | 0 | 1 | 26 |
| ier2b | 0.12995 | 0.716302964 | 0.222 | 0 | 1 | 26 |
| prdx4 | 0.12995 | 0.568469908 | 0.222 | 0 | 1 | 26 |
| reep5 | 0.12995 | 0.648131285 | 0.222 | 0 | 1 | 26 |
| spcs1 | 0.12995 | 0.63695534 | 0.222 | 0 | 1 | 26 |
| spcs2 | 0.12995 | 0.567754863 | 0.222 | 0 | 1 | 26 |
| ssr1 | 0.12995 | 0.66335339 | 0.222 | 0 | 1 | 26 |
| tpm1 | 0.12995 | 0.678424217 | 0.222 | 0 | 1 | 26 |
| rps18 | 0.1353 | -0.46898044 | 0.667 | 1 | 1 | 26 |
| ppdpfb | 0.13546 | -0.78897947 | 0.278 | 0.571 | 1 | 26 |
| tubb4b | 0.1363 | -0.73123655 | 0.278 | 0.714 | 1 | 26 |
| cd63 | 0.14127 | -0.25590237 | 0.444 | 0.714 | 1 | 26 |
| rps7 | 0.14274 | -0.41788307 | 0.667 | 1 | 1 | 26 |
| LOC100329433 | 0.14295 | -0.70129274 | 0.056 | 0.286 | 1 | 26 |
| asb12a | 0.14295 | -0.62867142 | 0.056 | 0.286 | 1 | 26 |
| atp1b1a | 0.14295 | -0.61252515 | 0.056 | 0.286 | 1 | 26 |
| ccdc28a | 0.14295 | -0.60302838 | 0.056 | 0.286 | 1 | 26 |
| cip2a | 0.14295 | -0.64203158 | 0.056 | 0.286 | 1 | 26 |
| clta | 0.14295 | -0.6030782 | 0.056 | 0.286 | 1 | 26 |
| letm1 | 0.14295 | -0.71447973 | 0.056 | 0.286 | 1 | 26 |
| myd88 | 0.14295 | -0.59508765 | 0.056 | 0.286 | 1 | 26 |
| pfdn4 | 0.14295 | -0.58823634 | 0.056 | 0.286 | 1 | 26 |
| romo1 | 0.14295 | -0.59783134 | 0.056 | 0.286 | 1 | 26 |
| sf3a2 | 0.14295 | -0.63899734 | 0.056 | 0.286 | 1 | 26 |
| ube2d1b | 0.14295 | -0.6467073 | 0.056 | 0.286 | 1 | 26 |
| ywhabl | 0.14295 | -0.69048765 | 0.056 | 0.286 | 1 | 26 |
| zgc:65873 | 0.14295 | -0.58627642 | 0.056 | 0.286 | 1 | 26 |
| zgc:91860 | 0.14295 | -0.64640993 | 0.056 | 0.286 | 1 | 26 |
| eif1b | 0.14295 | -0.63607572 | 0.056 | 0.286 | 1 | 26 |
| glrx | 0.14295 | -0.59199226 | 0.056 | 0.286 | 1 | 26 |
| ncbp2 | 0.14295 | -0.7392001 | 0.056 | 0.286 | 1 | 26 |
| psme2 | 0.14295 | -0.65528743 | 0.056 | 0.286 | 1 | 26 |
| rpn1 | 0.14295 | -0.63780395 | 0.056 | 0.286 | 1 | 26 |
| si:dkey-51e6.1 | 0.14295 | -0.64169242 | 0.056 | 0.286 | 1 | 26 |
| zgc:64051 | 0.14295 | -0.68203539 | 0.056 | 0.286 | 1 | 26 |
| atp6v0ca | 0.14295 | -0.62072205 | 0.056 | 0.286 | 1 | 26 |
| pabpc1a | 0.15094 | 0.460224665 | 0.444 | 0.143 | 1 | 26 |
| ccng1 | 0.15211 | -0.75759591 | 0.333 | 0.714 | 1 | 26 |
| mt-cyb | 0.15566 | 0.456242395 | 0.389 | 0.429 | 1 | 26 |
| mt-atp6 | 0.15672 | 0.680741984 | 0.389 | 0.286 | 1 | 26 |
| si:ch211-214p16.1 | 0.15902 | -0.31396186 | 0.833 | 1 | 1 | 26 |
| BX322787.1 | 0.16545 | -0.25311245 | 0.944 | 1 | 1 | 26 |
| CR381673.1 | 0.16831 | -0.53224518 | 0 | 0.143 | 1 | 26 |
| DHDH | 0.16831 | -0.5294056 | 0 | 0.143 | 1 | 26 |
| XLOC-043857 | 0.16831 | -0.52926184 | 0 | 0.143 | 1 | 26 |
| add1 | 0.16831 | -0.53105072 | 0 | 0.143 | 1 | 26 |
| atp13a2 | 0.16831 | -0.53224618 | 0 | 0.143 | 1 | 26 |
| atp7a | 0.16831 | -0.53233385 | 0 | 0.143 | 1 | 26 |

| | | | | | | |
|---|---|---|---|---|---|---|
| bckdha | 0.16831 | -0.52967842 | 0 | 0.143 | 1 | 26 |
| cnot6a | 0.16831 | -0.53068153 | 0 | 0.143 | 1 | 26 |
| cs | 0.16831 | -0.53043737 | 0 | 0.143 | 1 | 26 |
| ctbp1 | 0.16831 | -0.52902495 | 0 | 0.143 | 1 | 26 |
| dok2 | 0.16831 | -0.53260785 | 0 | 0.143 | 1 | 26 |
| gnaia | 0.16831 | -0.52973394 | 0 | 0.143 | 1 | 26 |
| gpat4 | 0.16831 | -0.45587174 | 0 | 0.143 | 1 | 26 |
| meaf6 | 0.16831 | -0.52878817 | 0 | 0.143 | 1 | 26 |
| napab | 0.16831 | -0.52928059 | 0 | 0.143 | 1 | 26 |
| nat8l | 0.16831 | -0.38769177 | 0 | 0.143 | 1 | 26 |
| pisd | 0.16831 | -0.62443889 | 0 | 0.143 | 1 | 26 |
| polr2d | 0.16831 | -0.52932698 | 0 | 0.143 | 1 | 26 |
| rab1bb | 0.16831 | -0.30197698 | 0 | 0.143 | 1 | 26 |
| rae1 | 0.16831 | -0.53108856 | 0 | 0.143 | 1 | 26 |
| raf1b | 0.16831 | -0.53064277 | 0 | 0.143 | 1 | 26 |
| rhbdl2 | 0.16831 | -0.53207657 | 0 | 0.143 | 1 | 26 |
| sept2 | 0.16831 | -0.53097599 | 0 | 0.143 | 1 | 26 |
| sf1 | 0.16831 | -0.53087957 | 0 | 0.143 | 1 | 26 |
| sh3gl1a | 0.16831 | -0.53082217 | 0 | 0.143 | 1 | 26 |
| snx9b | 0.16831 | -0.53080296 | 0 | 0.143 | 1 | 26 |
| ssbp1 | 0.16831 | -0.5288404 | 0 | 0.143 | 1 | 26 |
| stard3nl | 0.16831 | -0.53153835 | 0 | 0.143 | 1 | 26 |
| surf2 | 0.16831 | -0.526839 | 0 | 0.143 | 1 | 26 |
| tmem106a | 0.16831 | -0.53081977 | 0 | 0.143 | 1 | 26 |
| ttc32 | 0.16831 | -0.48798257 | 0 | 0.143 | 1 | 26 |
| zgc:110626 | 0.16831 | -0.53031211 | 0 | 0.143 | 1 | 26 |
| LOC100151196 | 0.16831 | -0.51738045 | 0 | 0.143 | 1 | 26 |
| PRRC2B | 0.16831 | -0.5157551 | 0 | 0.143 | 1 | 26 |
| RNF14 (1 of many) | 0.16831 | -0.50923038 | 0 | 0.143 | 1 | 26 |
| XLOC-019062 | 0.16831 | -0.51455264 | 0 | 0.143 | 1 | 26 |
| anxa2a | 0.16831 | -0.49620721 | 0 | 0.143 | 1 | 26 |
| anxa4 | 0.16831 | -0.51252016 | 0 | 0.143 | 1 | 26 |
| arpc1a | 0.16831 | -0.51416211 | 0 | 0.143 | 1 | 26 |
| ccl35.1 | 0.16831 | -0.51520541 | 0 | 0.143 | 1 | 26 |
| cct6a | 0.16831 | -0.51413098 | 0 | 0.143 | 1 | 26 |
| ciao1 | 0.16831 | -0.51312953 | 0 | 0.143 | 1 | 26 |
| ets2 | 0.16831 | -0.51482986 | 0 | 0.143 | 1 | 26 |
| ewsr1b | 0.16831 | -0.51304209 | 0 | 0.143 | 1 | 26 |
| gtpbp4 | 0.16831 | -0.51150105 | 0 | 0.143 | 1 | 26 |
| ivns1abpb | 0.16831 | -0.51525789 | 0 | 0.143 | 1 | 26 |
| march8 | 0.16831 | -0.5149216 | 0 | 0.143 | 1 | 26 |
| mrps21 | 0.16831 | -0.51235785 | 0 | 0.143 | 1 | 26 |
| mvb12a | 0.16831 | -0.51490127 | 0 | 0.143 | 1 | 26 |
| ncalda | 0.16831 | -0.51662765 | 0 | 0.143 | 1 | 26 |
| nedd8l | 0.16831 | -0.51364139 | 0 | 0.143 | 1 | 26 |
| nup214 | 0.16831 | -0.51370218 | 0 | 0.143 | 1 | 26 |
| rnft1 | 0.16831 | -0.5169772 | 0 | 0.143 | 1 | 26 |
| shmt1 | 0.16831 | -0.51053931 | 0 | 0.143 | 1 | 26 |

| | | | | | | |
|---|---|---|---|---|---|---|
| si:ch211-212d10.2 | 0.16831 | -0.4881036 | 0 | 0.143 | 1 | 26 |
| si:ch211-246m6.4 | 0.16831 | -0.50888634 | 0 | 0.143 | 1 | 26 |
| si:dkey-21o19.2 | 0.16831 | -0.51686201 | 0 | 0.143 | 1 | 26 |
| snapc5 | 0.16831 | -0.51345224 | 0 | 0.143 | 1 | 26 |
| sult1st5 | 0.16831 | -0.51624324 | 0 | 0.143 | 1 | 26 |
| timm23a | 0.16831 | -0.50818627 | 0 | 0.143 | 1 | 26 |
| uchl3 | 0.16831 | -0.60668981 | 0 | 0.143 | 1 | 26 |
| vdac1 | 0.16831 | -0.47587428 | 0 | 0.143 | 1 | 26 |
| zcchc17 | 0.16831 | -0.51101896 | 0 | 0.143 | 1 | 26 |
| zgc:162952 | 0.16831 | -0.51608468 | 0 | 0.143 | 1 | 26 |
| CT027744.2 | 0.16831 | -0.55994419 | 0 | 0.143 | 1 | 26 |
| LOC100537613 | 0.16831 | -0.46336724 | 0 | 0.143 | 1 | 26 |
| agk | 0.16831 | -0.45716028 | 0 | 0.143 | 1 | 26 |
| ahr2 | 0.16831 | -0.46340452 | 0 | 0.143 | 1 | 26 |
| arf1 | 0.16831 | -0.46088256 | 0 | 0.143 | 1 | 26 |
| arf2a | 0.16831 | -0.4616997 | 0 | 0.143 | 1 | 26 |
| atrx | 0.16831 | -0.46431401 | 0 | 0.143 | 1 | 26 |
| brinp3a.1 | 0.16831 | -0.46517867 | 0 | 0.143 | 1 | 26 |
| btbd9 | 0.16831 | -0.46393928 | 0 | 0.143 | 1 | 26 |
| cbr1l | 0.16831 | -0.46066869 | 0 | 0.143 | 1 | 26 |
| cdc37 | 0.16831 | -0.45367237 | 0 | 0.143 | 1 | 26 |
| cenpx | 0.16831 | -0.45532486 | 0 | 0.143 | 1 | 26 |
| chchd1 | 0.16831 | -0.46196014 | 0 | 0.143 | 1 | 26 |
| cmc2 | 0.16831 | -0.46259275 | 0 | 0.143 | 1 | 26 |
| cops9 | 0.16831 | -0.45783902 | 0 | 0.143 | 1 | 26 |
| cryzl1 | 0.16831 | -0.46321679 | 0 | 0.143 | 1 | 26 |
| cybc1 | 0.16831 | -0.46410195 | 0 | 0.143 | 1 | 26 |
| dctn2 | 0.16831 | -0.46190195 | 0 | 0.143 | 1 | 26 |
| derl1 | 0.16831 | -0.46212312 | 0 | 0.143 | 1 | 26 |
| dync1i2a | 0.16831 | -0.46379318 | 0 | 0.143 | 1 | 26 |
| eif2b2 | 0.16831 | -0.46193115 | 0 | 0.143 | 1 | 26 |
| fbxw8 | 0.16831 | -0.44422437 | 0 | 0.143 | 1 | 26 |
| gars | 0.16831 | -0.46054155 | 0 | 0.143 | 1 | 26 |
| gtf3c5 | 0.16831 | -0.43885073 | 0 | 0.143 | 1 | 26 |
| gtpbp1 | 0.16831 | -0.46423617 | 0 | 0.143 | 1 | 26 |
| hnrnpd | 0.16831 | -0.46217358 | 0 | 0.143 | 1 | 26 |
| hnrnpl2 | 0.16831 | -0.4593116 | 0 | 0.143 | 1 | 26 |
| hp1bp3 | 0.16831 | -0.46244213 | 0 | 0.143 | 1 | 26 |
| hsf1 | 0.16831 | -0.46414999 | 0 | 0.143 | 1 | 26 |
| ino80b | 0.16831 | -0.45737746 | 0 | 0.143 | 1 | 26 |
| ino80e | 0.16831 | -0.4617787 | 0 | 0.143 | 1 | 26 |
| khk | 0.16831 | -0.46499178 | 0 | 0.143 | 1 | 26 |
| kmt5aa | 0.16831 | -0.46325585 | 0 | 0.143 | 1 | 26 |
| lig4 | 0.16831 | -0.4605522 | 0 | 0.143 | 1 | 26 |
| lsm3 | 0.16831 | -0.45149257 | 0 | 0.143 | 1 | 26 |
| lsm7 | 0.16831 | -0.45377636 | 0 | 0.143 | 1 | 26 |
| mccc2 | 0.16831 | -0.46219037 | 0 | 0.143 | 1 | 26 |
| mrps14 | 0.16831 | -0.45562037 | 0 | 0.143 | 1 | 26 |

| | | | | | | |
|---|---|---|---|---|---|---|
| namptb | 0.16831 | -0.46275797 | 0 | 0.143 | 1 | 26 |
| necap2 | 0.16831 | -0.46286902 | 0 | 0.143 | 1 | 26 |
| nfyal | 0.16831 | -0.46063935 | 0 | 0.143 | 1 | 26 |
| nhp2 | 0.16831 | -0.45819216 | 0 | 0.143 | 1 | 26 |
| nosip | 0.16831 | -0.46099776 | 0 | 0.143 | 1 | 26 |
| nprl2 | 0.16831 | -0.45221553 | 0 | 0.143 | 1 | 26 |
| nudcd3 | 0.16831 | -0.45486605 | 0 | 0.143 | 1 | 26 |
| osbpl2b | 0.16831 | -0.46565665 | 0 | 0.143 | 1 | 26 |
| oser1 | 0.16831 | -0.46522547 | 0 | 0.143 | 1 | 26 |
| pabpn1 | 0.16831 | -0.46088323 | 0 | 0.143 | 1 | 26 |
| pmpcb | 0.16831 | -0.46215571 | 0 | 0.143 | 1 | 26 |
| ppp1r9ba | 0.16831 | -0.46332721 | 0 | 0.143 | 1 | 26 |
| prkar1ab | 0.16831 | -0.55984107 | 0 | 0.143 | 1 | 26 |
| purba | 0.16831 | -0.46074698 | 0 | 0.143 | 1 | 26 |
| pygmb | 0.16831 | -0.39643747 | 0 | 0.143 | 1 | 26 |
| rab11fip4b | 0.16831 | -0.46541726 | 0 | 0.143 | 1 | 26 |
| rap2b | 0.16831 | -0.46537411 | 0 | 0.143 | 1 | 26 |
| rbm4.3 | 0.16831 | -0.45703673 | 0 | 0.143 | 1 | 26 |
| rpz5 | 0.16831 | -0.44730293 | 0 | 0.143 | 1 | 26 |
| rtn4ip1 | 0.16831 | -0.46221806 | 0 | 0.143 | 1 | 26 |
| samm50 | 0.16831 | -0.55729263 | 0 | 0.143 | 1 | 26 |
| sec23ip | 0.16831 | -0.4642733 | 0 | 0.143 | 1 | 26 |
| si:ch1073-325m22.2 | 0.16831 | -0.45440255 | 0 | 0.143 | 1 | 26 |
| si:ch211-136m16.8 | 0.16831 | -0.46404153 | 0 | 0.143 | 1 | 26 |
| si:ch211-191c10.1 | 0.16831 | -0.46411024 | 0 | 0.143 | 1 | 26 |
| si:dkey-78l4.14 | 0.16831 | -0.45868244 | 0 | 0.143 | 1 | 26 |
| smg6 | 0.16831 | -0.46240998 | 0 | 0.143 | 1 | 26 |
| spina | 0.16831 | -0.46034246 | 0 | 0.143 | 1 | 26 |
| srp72 | 0.16831 | -0.46028125 | 0 | 0.143 | 1 | 26 |
| st7 | 0.16831 | -0.4621693 | 0 | 0.143 | 1 | 26 |
| suclg1 | 0.16831 | -0.45468427 | 0 | 0.143 | 1 | 26 |
| sumo2b | 0.16831 | -0.45908065 | 0 | 0.143 | 1 | 26 |
| tbk1 | 0.16831 | -0.46469912 | 0 | 0.143 | 1 | 26 |
| triobpb | 0.16831 | -0.4605172 | 0 | 0.143 | 1 | 26 |
| ube2e3 | 0.16831 | -0.45904291 | 0 | 0.143 | 1 | 26 |
| yipf2 | 0.16831 | -0.55804466 | 0 | 0.143 | 1 | 26 |
| zcchc10 | 0.16831 | -0.55602095 | 0 | 0.143 | 1 | 26 |
| zgc:113921 | 0.16831 | -0.46329857 | 0 | 0.143 | 1 | 26 |
| zgc:153990 | 0.16831 | -0.46235881 | 0 | 0.143 | 1 | 26 |
| zgc:77849 | 0.16831 | -0.46213237 | 0 | 0.143 | 1 | 26 |
| BX005254.1 | 0.16831 | -0.3776556 | 0 | 0.143 | 1 | 26 |
| CABZ01064670.1 | 0.16831 | -0.43933557 | 0 | 0.143 | 1 | 26 |
| CABZ01068251.1 | 0.16831 | -0.43281399 | 0 | 0.143 | 1 | 26 |
| CABZ01087560.1 | 0.16831 | -0.42708694 | 0 | 0.143 | 1 | 26 |
| FBXO46 | 0.16831 | -0.47043695 | 0 | 0.143 | 1 | 26 |
| NC-002333.17 | 0.16831 | -0.41378527 | 0 | 0.143 | 1 | 26 |
| NDUFAF8 | 0.16831 | -0.43812244 | 0 | 0.143 | 1 | 26 |
| STMP1 | 0.16831 | -0.4616777 | 0 | 0.143 | 1 | 26 |

| | | | | | | |
|---|---|---|---|---|---|---|
| UBL4A | 0.16831 | -0.46396491 | 0 | 0.143 | 1 | 26 |
| XLOC-032112 | 0.16831 | -0.44132843 | 0 | 0.143 | 1 | 26 |
| aamdc | 0.16831 | -0.46921035 | 0 | 0.143 | 1 | 26 |
| acat1 | 0.16831 | -0.46111852 | 0 | 0.143 | 1 | 26 |
| adnp2a | 0.16831 | -0.46740281 | 0 | 0.143 | 1 | 26 |
| alox5ap | 0.16831 | -0.47075185 | 0 | 0.143 | 1 | 26 |
| ankrd13c | 0.16831 | -0.29138913 | 0 | 0.143 | 1 | 26 |
| ap2m1a | 0.16831 | -0.43363092 | 0 | 0.143 | 1 | 26 |
| api5 | 0.16831 | -0.43519253 | 0 | 0.143 | 1 | 26 |
| appl2 | 0.16831 | -0.42780054 | 0 | 0.143 | 1 | 26 |
| atg12 | 0.16831 | -0.46781731 | 0 | 0.143 | 1 | 26 |
| atg4c | 0.16831 | -0.46887916 | 0 | 0.143 | 1 | 26 |
| brd7 | 0.16831 | -0.53478686 | 0 | 0.143 | 1 | 26 |
| capns1b | 0.16831 | -0.45908258 | 0 | 0.143 | 1 | 26 |
| cdk4 | 0.16831 | -0.43845747 | 0 | 0.143 | 1 | 26 |
| cdkn2aipnl | 0.16831 | -0.43822001 | 0 | 0.143 | 1 | 26 |
| cfap20 | 0.16831 | -0.43595784 | 0 | 0.143 | 1 | 26 |
| clpp | 0.16831 | -0.43644609 | 0 | 0.143 | 1 | 26 |
| cltcb | 0.16831 | -0.44017934 | 0 | 0.143 | 1 | 26 |
| cnot7 | 0.16831 | -0.41341727 | 0 | 0.143 | 1 | 26 |
| cox17 | 0.16831 | -0.46436429 | 0 | 0.143 | 1 | 26 |
| csnk2a1 | 0.16831 | -0.46133041 | 0 | 0.143 | 1 | 26 |
| cul1a | 0.16831 | -0.43932869 | 0 | 0.143 | 1 | 26 |
| dctpp1 | 0.16831 | -0.42735184 | 0 | 0.143 | 1 | 26 |
| decr1 | 0.16831 | -0.46912154 | 0 | 0.143 | 1 | 26 |
| dnph1 | 0.16831 | -0.58621202 | 0 | 0.143 | 1 | 26 |
| dnttip2 | 0.16831 | -0.43952723 | 0 | 0.143 | 1 | 26 |
| ehhadh | 0.16831 | -0.34083693 | 0 | 0.143 | 1 | 26 |
| eif4e1c | 0.16831 | -0.43074997 | 0 | 0.143 | 1 | 26 |
| elovl1b | 0.16831 | -0.46935548 | 0 | 0.143 | 1 | 26 |
| fam162a | 0.16831 | -0.43513385 | 0 | 0.143 | 1 | 26 |
| fbxw4 | 0.16831 | -0.43955063 | 0 | 0.143 | 1 | 26 |
| flot2a | 0.16831 | -0.43623476 | 0 | 0.143 | 1 | 26 |
| gabpb2b | 0.16831 | -0.43883546 | 0 | 0.143 | 1 | 26 |
| galt | 0.16831 | -0.47004604 | 0 | 0.143 | 1 | 26 |
| gclm | 0.16831 | -0.43868062 | 0 | 0.143 | 1 | 26 |
| gcsha | 0.16831 | -0.41901504 | 0 | 0.143 | 1 | 26 |
| gins2 | 0.16831 | -0.46633776 | 0 | 0.143 | 1 | 26 |
| gins4 | 0.16831 | -0.43626542 | 0 | 0.143 | 1 | 26 |
| hddc3 | 0.16831 | -0.1772948 | 0 | 0.143 | 1 | 26 |
| hmgb2b | 0.16831 | -0.37600483 | 0 | 0.143 | 1 | 26 |
| hnrnpabb | 0.16831 | -0.41772234 | 0 | 0.143 | 1 | 26 |
| hypk | 0.16831 | -0.43125201 | 0 | 0.143 | 1 | 26 |
| idh3g | 0.16831 | -0.44937793 | 0 | 0.143 | 1 | 26 |
| ilf2 | 0.16831 | -0.43266145 | 0 | 0.143 | 1 | 26 |
| illr4 | 0.16831 | -0.46819683 | 0 | 0.143 | 1 | 26 |
| isca2 | 0.16831 | -0.43028578 | 0 | 0.143 | 1 | 26 |
| isy1 | 0.16831 | -0.46805505 | 0 | 0.143 | 1 | 26 |

| | | | | | | |
|---|---|---|---|---|---|---|
| keap1a | 0.16831 | -0.47053088 | 0 | 0.143 | 1 | 26 |
| kif5ba | 0.16831 | -0.44021482 | 0 | 0.143 | 1 | 26 |
| lcorl | 0.16831 | -0.46978276 | 0 | 0.143 | 1 | 26 |
| lcp1 | 0.16831 | -0.47084082 | 0 | 0.143 | 1 | 26 |
| lias | 0.16831 | -0.41799456 | 0 | 0.143 | 1 | 26 |
| lsm5 | 0.16831 | -0.42954571 | 0 | 0.143 | 1 | 26 |
| lyrm2 | 0.16831 | -0.43754133 | 0 | 0.143 | 1 | 26 |
| mcm6 | 0.16831 | -0.43990549 | 0 | 0.143 | 1 | 26 |
| med4 | 0.16831 | -0.43578952 | 0 | 0.143 | 1 | 26 |
| megf8l | 0.16831 | -0.47063307 | 0 | 0.143 | 1 | 26 |
| mfap1 | 0.16831 | -0.43019102 | 0 | 0.143 | 1 | 26 |
| mis12 | 0.16831 | -0.43330534 | 0 | 0.143 | 1 | 26 |
| mrpl38 | 0.16831 | -0.43585257 | 0 | 0.143 | 1 | 26 |
| mrps28 | 0.16831 | -0.43614471 | 0 | 0.143 | 1 | 26 |
| naa50 | 0.16831 | -0.43716698 | 0 | 0.143 | 1 | 26 |
| ndufa4 | 0.16831 | -0.47015694 | 0 | 0.143 | 1 | 26 |
| ndufa8 | 0.16831 | -0.42908919 | 0 | 0.143 | 1 | 26 |
| ndufab1a | 0.16831 | -0.43174027 | 0 | 0.143 | 1 | 26 |
| ndufb10 | 0.16831 | -0.46392373 | 0 | 0.143 | 1 | 26 |
| nectin3b | 0.16831 | -0.39383038 | 0 | 0.143 | 1 | 26 |
| nfyba | 0.16831 | -0.43435352 | 0 | 0.143 | 1 | 26 |
| nop10 | 0.16831 | -0.43334602 | 0 | 0.143 | 1 | 26 |
| nudcd1 | 0.16831 | -0.44041551 | 0 | 0.143 | 1 | 26 |
| nup62l | 0.16831 | -0.4360091 | 0 | 0.143 | 1 | 26 |
| pa2g4b | 0.16831 | -0.43577678 | 0 | 0.143 | 1 | 26 |
| papss1 | 0.16831 | -0.44180747 | 0 | 0.143 | 1 | 26 |
| parp12a | 0.16831 | -0.46808905 | 0 | 0.143 | 1 | 26 |
| pbdc1 | 0.16831 | -0.43436381 | 0 | 0.143 | 1 | 26 |
| pgam1a | 0.16831 | -0.43885546 | 0 | 0.143 | 1 | 26 |
| pgk1 | 0.16831 | -0.46669093 | 0 | 0.143 | 1 | 26 |
| phb2a | 0.16831 | -0.43234736 | 0 | 0.143 | 1 | 26 |
| pip4k2ca | 0.16831 | -0.46202367 | 0 | 0.143 | 1 | 26 |
| plaa | 0.16831 | -0.438765 | 0 | 0.143 | 1 | 26 |
| polr2c | 0.16831 | -0.43703232 | 0 | 0.143 | 1 | 26 |
| polr2f | 0.16831 | -0.43009213 | 0 | 0.143 | 1 | 26 |
| polr3h | 0.16831 | -0.43662844 | 0 | 0.143 | 1 | 26 |
| prdx3 | 0.16831 | -0.46030612 | 0 | 0.143 | 1 | 26 |
| prim1 | 0.16831 | -0.43753937 | 0 | 0.143 | 1 | 26 |
| prps1b | 0.16831 | -0.43932922 | 0 | 0.143 | 1 | 26 |
| psmb2 | 0.16831 | -0.46479217 | 0 | 0.143 | 1 | 26 |
| qsox1 | 0.16831 | -0.43924491 | 0 | 0.143 | 1 | 26 |
| rab5aa | 0.16831 | -0.46974616 | 0 | 0.143 | 1 | 26 |
| rad54l | 0.16831 | -0.44608691 | 0 | 0.143 | 1 | 26 |
| rbm24b | 0.16831 | -0.42970019 | 0 | 0.143 | 1 | 26 |
| rdh12l | 0.16831 | -0.47004083 | 0 | 0.143 | 1 | 26 |
| rnaseh2a | 0.16831 | -0.43440775 | 0 | 0.143 | 1 | 26 |
| rpa2 | 0.16831 | -0.42525714 | 0 | 0.143 | 1 | 26 |
| rpa3 | 0.16831 | -0.43023517 | 0 | 0.143 | 1 | 26 |

| | | | | | | |
|---|---|---|---|---|---|---|
| rpap3 | 0.16831 | -0.43673087 | 0 | 0.143 | 1 | 26 |
| rrm2 | 0.16831 | -0.3616214 | 0 | 0.143 | 1 | 26 |
| sart1 | 0.16831 | -0.43312886 | 0 | 0.143 | 1 | 26 |
| scube3 | 0.16831 | -0.4232555 | 0 | 0.143 | 1 | 26 |
| selenoh | 0.16831 | -0.42186243 | 0 | 0.143 | 1 | 26 |
| si:ch211-191j22.3 | 0.16831 | -0.46699407 | 0 | 0.143 | 1 | 26 |
| si:ch211-197h24.9 | 0.16831 | -0.46533088 | 0 | 0.143 | 1 | 26 |
| si:ch211-213a13.2 | 0.16831 | -0.44080288 | 0 | 0.143 | 1 | 26 |
| si:ch211-51e12.7 | 0.16831 | -0.43069425 | 0 | 0.143 | 1 | 26 |
| si:dkey-126g1.7 | 0.16831 | -0.47001419 | 0 | 0.143 | 1 | 26 |
| si:dkey-184a18.5 | 0.16831 | -0.46814605 | 0 | 0.143 | 1 | 26 |
| si:dkey-260j18.2 | 0.16831 | -0.4394335 | 0 | 0.143 | 1 | 26 |
| slc22a2 | 0.16831 | -0.46377458 | 0 | 0.143 | 1 | 26 |
| slc25a33 | 0.16831 | -0.46707042 | 0 | 0.143 | 1 | 26 |
| smad2 | 0.16831 | -0.43907719 | 0 | 0.143 | 1 | 26 |
| sptssa | 0.16831 | -0.46780837 | 0 | 0.143 | 1 | 26 |
| suclg2 | 0.16831 | -0.4670584 | 0 | 0.143 | 1 | 26 |
| taf13 | 0.16831 | -0.53343817 | 0 | 0.143 | 1 | 26 |
| timm50 | 0.16831 | -0.43489915 | 0 | 0.143 | 1 | 26 |
| tk1 | 0.16831 | -0.43900077 | 0 | 0.143 | 1 | 26 |
| tomm6 | 0.16831 | -0.43529365 | 0 | 0.143 | 1 | 26 |
| tyms | 0.16831 | -0.43527053 | 0 | 0.143 | 1 | 26 |
| uba3 | 0.16831 | -0.46779027 | 0 | 0.143 | 1 | 26 |
| ube2e2 | 0.16831 | -0.45765313 | 0 | 0.143 | 1 | 26 |
| vti1a | 0.16831 | -0.44173018 | 0 | 0.143 | 1 | 26 |
| wdr83 | 0.16831 | -0.43771036 | 0 | 0.143 | 1 | 26 |
| wtap | 0.16831 | -0.43683807 | 0 | 0.143 | 1 | 26 |
| zgc:153119 | 0.16831 | -0.4232555 | 0 | 0.143 | 1 | 26 |
| zgc:174263 | 0.16831 | -0.13997594 | 0 | 0.143 | 1 | 26 |
| zgc:194246 | 0.16831 | -0.36943072 | 0 | 0.143 | 1 | 26 |
| zmp:0000000650 | 0.16831 | -0.46563262 | 0 | 0.143 | 1 | 26 |
| CABZ01058650.1 | 0.16831 | -0.49707136 | 0 | 0.143 | 1 | 26 |
| CDPF1 | 0.16831 | -0.49137837 | 0 | 0.143 | 1 | 26 |
| CR855389.1 | 0.16831 | -0.49991624 | 0 | 0.143 | 1 | 26 |
| LOC101885769 | 0.16831 | -0.50057634 | 0 | 0.143 | 1 | 26 |
| LOC103910438 | 0.16831 | -0.49770622 | 0 | 0.143 | 1 | 26 |
| MZT1 | 0.16831 | -0.49669742 | 0 | 0.143 | 1 | 26 |
| abhd17b | 0.16831 | -0.49880791 | 0 | 0.143 | 1 | 26 |
| ankrd10b | 0.16831 | -0.49832627 | 0 | 0.143 | 1 | 26 |
| arl8a | 0.16831 | -0.49921164 | 0 | 0.143 | 1 | 26 |
| caprin1a | 0.16831 | -0.48962276 | 0 | 0.143 | 1 | 26 |
| chmp4bb | 0.16831 | -0.49819041 | 0 | 0.143 | 1 | 26 |
| commd2 | 0.16831 | -0.48327693 | 0 | 0.143 | 1 | 26 |
| csnk1a1 | 0.16831 | -0.49515045 | 0 | 0.143 | 1 | 26 |
| dynlt3 | 0.16831 | -0.49557642 | 0 | 0.143 | 1 | 26 |
| gmppb | 0.16831 | -0.49907603 | 0 | 0.143 | 1 | 26 |
| golga5 | 0.16831 | -0.4985648 | 0 | 0.143 | 1 | 26 |
| itgb1bp1 | 0.16831 | -0.4979783 | 0 | 0.143 | 1 | 26 |

| | | | | | | |
|---|---|---|---|---|---|---|
| mff | 0.16831 | -0.49814502 | 0 | 0.143 | 1 | 26 |
| mrpl33 | 0.16831 | -0.49750756 | 0 | 0.143 | 1 | 26 |
| mrpl47 | 0.16831 | -0.49636441 | 0 | 0.143 | 1 | 26 |
| pkp4 | 0.16831 | -0.48984618 | 0 | 0.143 | 1 | 26 |
| psmd11a | 0.16831 | -0.49680215 | 0 | 0.143 | 1 | 26 |
| rars | 0.16831 | -0.49837634 | 0 | 0.143 | 1 | 26 |
| rfc4 | 0.16831 | -0.49093912 | 0 | 0.143 | 1 | 26 |
| rtn4a | 0.16831 | -0.4976566 | 0 | 0.143 | 1 | 26 |
| si:ch211-255f4.9 | 0.16831 | -0.50030665 | 0 | 0.143 | 1 | 26 |
| si:ch211-85n16.4 | 0.16831 | -0.49805571 | 0 | 0.143 | 1 | 26 |
| si:dkey-165a24.10 | 0.16831 | -0.49961392 | 0 | 0.143 | 1 | 26 |
| slc35d1a | 0.16831 | -0.49564024 | 0 | 0.143 | 1 | 26 |
| smarcb1a | 0.16831 | -0.49676387 | 0 | 0.143 | 1 | 26 |
| snx17 | 0.16831 | -0.49805664 | 0 | 0.143 | 1 | 26 |
| tfcp2 | 0.16831 | -0.48586509 | 0 | 0.143 | 1 | 26 |
| tmem120a | 0.16831 | -0.50014946 | 0 | 0.143 | 1 | 26 |
| tmem18 | 0.16831 | -0.49669456 | 0 | 0.143 | 1 | 26 |
| ubl3b | 0.16831 | -0.49946972 | 0 | 0.143 | 1 | 26 |
| zgc:101715 | 0.16831 | -0.49953377 | 0 | 0.143 | 1 | 26 |
| zgc:153012 | 0.16831 | -0.50033377 | 0 | 0.143 | 1 | 26 |
| zgc:162339 | 0.16831 | -0.59273199 | 0 | 0.143 | 1 | 26 |
| zgc:195282 | 0.16831 | -0.49987822 | 0 | 0.143 | 1 | 26 |
| znrf1 | 0.16831 | -0.49650211 | 0 | 0.143 | 1 | 26 |
| rps25 | 0.17619 | -0.11639021 | 0.778 | 1 | 1 | 26 |
| lxn | 0.17752 | -0.81966891 | 0.111 | 0.429 | 1 | 26 |
| rps29 | 0.17953 | -0.39829139 | 0.833 | 1 | 1 | 26 |
| morf4l1 | 0.18242 | -0.83281297 | 0.111 | 0.429 | 1 | 26 |
| rpl29 | 0.18265 | -0.43077896 | 0.722 | 1 | 1 | 26 |
| rpl22 | 0.18721 | -0.61684211 | 0.5 | 0.857 | 1 | 26 |
| rpl14 | 0.18885 | 0.537640661 | 0.889 | 0.571 | 1 | 26 |
| spink2.3 | 0.20153 | -0.31711085 | 0.722 | 1 | 1 | 26 |
| LOC108179162 | 0.20312 | -0.79724318 | 0.222 | 0.571 | 1 | 26 |
| uba52 | 0.20582 | 0.182013833 | 0.667 | 0.714 | 1 | 26 |
| rpl21 | 0.20845 | -0.36863422 | 0.778 | 1 | 1 | 26 |
| CR392341.1 | 0.2086 | 0.473086727 | 0.167 | 0 | 1 | 26 |
| acadvl | 0.2086 | 0.579734812 | 0.167 | 0 | 1 | 26 |
| cdaa | 0.2086 | 0.56056695 | 0.167 | 0 | 1 | 26 |
| celf2 | 0.2086 | 0.567964086 | 0.167 | 0 | 1 | 26 |
| fundc2 | 0.2086 | 0.527199656 | 0.167 | 0 | 1 | 26 |
| ier3ip1 | 0.2086 | 0.431911131 | 0.167 | 0 | 1 | 26 |
| krt8 | 0.2086 | 0.444745148 | 0.167 | 0 | 1 | 26 |
| mbnl1 | 0.2086 | 0.488657478 | 0.167 | 0 | 1 | 26 |
| mctp2b | 0.2086 | 0.485218412 | 0.167 | 0 | 1 | 26 |
| myofl | 0.2086 | 0.536297845 | 0.167 | 0 | 1 | 26 |
| nbas | 0.2086 | 0.512331035 | 0.167 | 0 | 1 | 26 |
| nucb2b | 0.2086 | 0.501047432 | 0.167 | 0 | 1 | 26 |
| timp2b | 0.2086 | 0.505333856 | 0.167 | 0 | 1 | 26 |
| tmub1 | 0.2086 | 0.519422372 | 0.167 | 0 | 1 | 26 |

| | | | | | | |
|---|---|---|---|---|---|---|
| txnipa | 0.2086 | 0.531609231 | 0.167 | 0 | 1 | 26 |
| CR855337.1 | 0.2086 | 0.521276094 | 0.167 | 0 | 1 | 26 |
| NDUFB1 | 0.2086 | 0.418727132 | 0.167 | 0 | 1 | 26 |
| aimp1 | 0.2086 | 0.439767953 | 0.167 | 0 | 1 | 26 |
| calm3a | 0.2086 | 0.458331485 | 0.167 | 0 | 1 | 26 |
| camk2g1 | 0.2086 | 0.434743004 | 0.167 | 0 | 1 | 26 |
| cpb1 | 0.2086 | 0.499205434 | 0.167 | 0 | 1 | 26 |
| eif3c | 0.2086 | 0.481324189 | 0.167 | 0 | 1 | 26 |
| fabp3 | 0.2086 | 0.457135284 | 0.167 | 0 | 1 | 26 |
| fam32a | 0.2086 | 0.515775954 | 0.167 | 0 | 1 | 26 |
| far1 | 0.2086 | 0.541937453 | 0.167 | 0 | 1 | 26 |
| gpx4b | 0.2086 | 0.497055768 | 0.167 | 0 | 1 | 26 |
| icn | 0.2086 | 0.628268915 | 0.167 | 0 | 1 | 26 |
| itm2ba | 0.2086 | 0.429689684 | 0.167 | 0 | 1 | 26 |
| max | 0.2086 | 0.482860947 | 0.167 | 0 | 1 | 26 |
| ncl | 0.2086 | 0.460338587 | 0.167 | 0 | 1 | 26 |
| ndufa6 | 0.2086 | 0.590733562 | 0.167 | 0 | 1 | 26 |
| pfdn1 | 0.2086 | 0.496729683 | 0.167 | 0 | 1 | 26 |
| pla2g1b | 0.2086 | 0.226551506 | 0.167 | 0 | 1 | 26 |
| pomp | 0.2086 | 0.466681418 | 0.167 | 0 | 1 | 26 |
| ppifb | 0.2086 | 0.466931645 | 0.167 | 0 | 1 | 26 |
| prkcbb | 0.2086 | 0.58132347 | 0.167 | 0 | 1 | 26 |
| pstpip2 | 0.2086 | 0.594421024 | 0.167 | 0 | 1 | 26 |
| sept6 | 0.2086 | 0.537057145 | 0.167 | 0 | 1 | 26 |
| si:ch73-1a9.3 | 0.2086 | 0.549188392 | 0.167 | 0 | 1 | 26 |
| sort1b | 0.2086 | 0.600380894 | 0.167 | 0 | 1 | 26 |
| srpra | 0.2086 | 0.553704072 | 0.167 | 0 | 1 | 26 |
| syncrip | 0.2086 | 0.478248756 | 0.167 | 0 | 1 | 26 |
| syngr2a | 0.2086 | 0.405659126 | 0.167 | 0 | 1 | 26 |
| taf3 | 0.2086 | 0.479237216 | 0.167 | 0 | 1 | 26 |
| tmem30aa | 0.2086 | 0.371903197 | 0.167 | 0 | 1 | 26 |
| tpmt.1 | 0.2086 | 0.496750043 | 0.167 | 0 | 1 | 26 |
| uba1 | 0.2086 | 0.479733738 | 0.167 | 0 | 1 | 26 |
| vcp | 0.2086 | 0.56957824 | 0.167 | 0 | 1 | 26 |
| yaf2 | 0.2086 | 0.484640141 | 0.167 | 0 | 1 | 26 |
| zbtb20 | 0.2086 | 0.561470304 | 0.167 | 0 | 1 | 26 |
| LOC101883382 | 0.2086 | 0.523296532 | 0.167 | 0 | 1 | 26 |
| MYADM | 0.2086 | 0.548481431 | 0.167 | 0 | 1 | 26 |
| ap1m1 | 0.2086 | 0.560588873 | 0.167 | 0 | 1 | 26 |
| caprin1b | 0.2086 | 0.575868823 | 0.167 | 0 | 1 | 26 |
| cct7 | 0.2086 | 0.561937531 | 0.167 | 0 | 1 | 26 |
| cd7al | 0.2086 | 0.513272564 | 0.167 | 0 | 1 | 26 |
| gdi2 | 0.2086 | 0.570550841 | 0.167 | 0 | 1 | 26 |
| itm2bb | 0.2086 | 0.445536545 | 0.167 | 0 | 1 | 26 |
| junba | 0.2086 | 0.552994298 | 0.167 | 0 | 1 | 26 |
| krtcap2 | 0.2086 | 0.419889989 | 0.167 | 0 | 1 | 26 |
| lrrfip1a | 0.2086 | 0.452590928 | 0.167 | 0 | 1 | 26 |
| mir223 | 0.2086 | 0.572355479 | 0.167 | 0 | 1 | 26 |

| | | | | | | |
|---|---|---|---|---|---|---|
| prss59.1 | 0.2086 | 0.705938994 | 0.167 | 0 | 1 | 26 |
| rflnb | 0.2086 | 0.53448589 | 0.167 | 0 | 1 | 26 |
| rxrba | 0.2086 | 0.574456682 | 0.167 | 0 | 1 | 26 |
| slc2a3b | 0.2086 | 0.579479466 | 0.167 | 0 | 1 | 26 |
| tegt | 0.2086 | 0.581917529 | 0.167 | 0 | 1 | 26 |
| ctrl | 0.2086 | 0.763670917 | 0.167 | 0 | 1 | 26 |
| eif2s3 | 0.2086 | 0.554969497 | 0.167 | 0 | 1 | 26 |
| gpx1a | 0.2086 | 0.568966722 | 0.167 | 0 | 1 | 26 |
| hars | 0.2086 | 0.44637587 | 0.167 | 0 | 1 | 26 |
| rogdi | 0.2086 | 0.576242886 | 0.167 | 0 | 1 | 26 |
| tia1l | 0.2086 | 0.528706092 | 0.167 | 0 | 1 | 26 |
| pnrc2 | 0.21085 | 0.502682866 | 0.611 | 0.286 | 1 | 26 |
| si:dkey-102g19.3 | 0.21154 | 0.84555043 | 0.5 | 0.143 | 1 | 26 |
| tpt1 | 0.21322 | -0.36355511 | 0.722 | 1 | 1 | 26 |
| h3f3a | 0.21403 | -0.66528891 | 0.111 | 0.429 | 1 | 26 |
| rps12 | 0.21567 | -0.34232739 | 0.722 | 1 | 1 | 26 |
| tpm3 | 0.21599 | -0.78001981 | 0.111 | 0.429 | 1 | 26 |
| rpl24 | 0.22183 | -0.36472785 | 0.833 | 1 | 1 | 26 |
| ppp1r18 | 0.22653 | -0.70339393 | 0.111 | 0.429 | 1 | 26 |
| LOC103909115 | 0.23725 | -0.77205406 | 0.222 | 0.571 | 1 | 26 |
| cnp | 0.23884 | -0.7371815 | 0.111 | 0.429 | 1 | 26 |
| ywhaba | 0.24051 | -0.72708039 | 0.111 | 0.429 | 1 | 26 |
| atp5f1d | 0.24168 | -0.71293979 | 0.111 | 0.429 | 1 | 26 |
| cfi | 0.25248 | -0.7159756 | 0.222 | 0.571 | 1 | 26 |
| rps20 | 0.25747 | -0.29176208 | 0.889 | 1 | 1 | 26 |
| rpl23a | 0.26496 | 0.442820844 | 0.722 | 0.571 | 1 | 26 |
| mt-co1 | 0.26576 | 0.584527051 | 0.556 | 0.429 | 1 | 26 |
| xbp1 | 0.28502 | 0.370914154 | 0.556 | 0.286 | 1 | 26 |
| rpl32 | 0.29899 | -0.32870334 | 0.833 | 1 | 1 | 26 |
| zgc:153867 | 0.30461 | -0.11793089 | 0.333 | 0.286 | 1 | 26 |
| ppiaa | 0.30708 | -0.30376846 | 0.778 | 1 | 1 | 26 |
| eif4ebp1 | 0.30768 | 0.408755605 | 0.389 | 0.143 | 1 | 26 |
| rpl15 | 0.30808 | -0.29743875 | 0.778 | 1 | 1 | 26 |
| rpl36 | 0.30816 | -0.31098143 | 0.778 | 1 | 1 | 26 |
| actb1 | 0.30899 | -0.27928647 | 0.833 | 1 | 1 | 26 |
| rpl37 | 0.31033 | -0.25102994 | 0.778 | 1 | 1 | 26 |
| mt-nd1 | 0.3171 | 0.462095334 | 0.222 | 0.143 | 1 | 26 |
| rpl18a | 0.32088 | -0.39447147 | 0.722 | 0.857 | 1 | 26 |
| gapdhs | 0.32687 | -0.49622988 | 0.389 | 0.714 | 1 | 26 |
| rps26 | 0.33019 | -0.45119648 | 0.556 | 0.857 | 1 | 26 |
| eef1b2 | 0.33426 | 0.55230454 | 0.722 | 0.429 | 1 | 26 |
| BX908782.1 | 0.33482 | 0.21993791 | 0.111 | 0 | 1 | 26 |
| CABZ01090890.1 | 0.33482 | 0.301285495 | 0.111 | 0 | 1 | 26 |
| CELA1 (1 of many) | 0.33482 | 0.6914238 | 0.111 | 0 | 1 | 26 |
| LEPROTL1 | 0.33482 | 0.382033684 | 0.111 | 0 | 1 | 26 |
| LOC100148254 | 0.33482 | 0.381264207 | 0.111 | 0 | 1 | 26 |
| LOC100332329 | 0.33482 | 0.410689507 | 0.111 | 0 | 1 | 26 |
| LOC103909452 | 0.33482 | 0.345198386 | 0.111 | 0 | 1 | 26 |

| | | | | | | |
|---|---|---|---|---|---|---|
| RASSF5 | 0.33482 | 0.40677326 | 0.111 | 0 | 1 | 26 |
| XLOC-000946 | 0.33482 | 0.308230345 | 0.111 | 0 | 1 | 26 |
| XLOC-013964 | 0.33482 | 0.408734531 | 0.111 | 0 | 1 | 26 |
| acin1a | 0.33482 | 0.540212287 | 0.111 | 0 | 1 | 26 |
| acp5b | 0.33482 | 0.446523487 | 0.111 | 0 | 1 | 26 |
| ahcy | 0.33482 | 0.346520551 | 0.111 | 0 | 1 | 26 |
| anp32e | 0.33482 | 0.399585237 | 0.111 | 0 | 1 | 26 |
| ap3s1.1 | 0.33482 | 0.440437725 | 0.111 | 0 | 1 | 26 |
| apoda.2 | 0.33482 | 0.437166949 | 0.111 | 0 | 1 | 26 |
| arcn1b | 0.33482 | 0.391459897 | 0.111 | 0 | 1 | 26 |
| arf6a | 0.33482 | 0.381242575 | 0.111 | 0 | 1 | 26 |
| arhgef2 | 0.33482 | 0.291700268 | 0.111 | 0 | 1 | 26 |
| arrb2b | 0.33482 | 0.354902138 | 0.111 | 0 | 1 | 26 |
| atp2b2 | 0.33482 | 0.242875326 | 0.111 | 0 | 1 | 26 |
| atp5f1c | 0.33482 | 0.43712241 | 0.111 | 0 | 1 | 26 |
| atp5md | 0.33482 | 0.436383602 | 0.111 | 0 | 1 | 26 |
| atp6v1g1 | 0.33482 | 0.386530251 | 0.111 | 0 | 1 | 26 |
| bach1b | 0.33482 | 0.441309543 | 0.111 | 0 | 1 | 26 |
| banf1 | 0.33482 | 0.348082256 | 0.111 | 0 | 1 | 26 |
| bcas2 | 0.33482 | 0.376422212 | 0.111 | 0 | 1 | 26 |
| birc2 | 0.33482 | 0.387099621 | 0.111 | 0 | 1 | 26 |
| brd2a | 0.33482 | 0.423404309 | 0.111 | 0 | 1 | 26 |
| cbfa2t3 | 0.33482 | 0.382149904 | 0.111 | 0 | 1 | 26 |
| ccr9a | 0.33482 | 0.411107249 | 0.111 | 0 | 1 | 26 |
| cct4 | 0.33482 | 0.434354265 | 0.111 | 0 | 1 | 26 |
| cdkn3 | 0.33482 | 0.399676098 | 0.111 | 0 | 1 | 26 |
| cds1 | 0.33482 | 0.381664081 | 0.111 | 0 | 1 | 26 |
| cel.2 | 0.33482 | 0.336041432 | 0.111 | 0 | 1 | 26 |
| chd9 | 0.33482 | 0.384343677 | 0.111 | 0 | 1 | 26 |
| chmp2a | 0.33482 | 0.450817956 | 0.111 | 0 | 1 | 26 |
| clstn1 | 0.33482 | 0.335786887 | 0.111 | 0 | 1 | 26 |
| cnn2 | 0.33482 | 0.486312777 | 0.111 | 0 | 1 | 26 |
| copa | 0.33482 | 0.419700096 | 0.111 | 0 | 1 | 26 |
| cpa4 | 0.33482 | 0.185973189 | 0.111 | 0 | 1 | 26 |
| cry2 | 0.33482 | 0.40663618 | 0.111 | 0 | 1 | 26 |
| csde1 | 0.33482 | 0.388995191 | 0.111 | 0 | 1 | 26 |
| ctrb1 | 0.33482 | 0.64819629 | 0.111 | 0 | 1 | 26 |
| ctsa | 0.33482 | 0.437089064 | 0.111 | 0 | 1 | 26 |
| ctsba | 0.33482 | 0.279283763 | 0.111 | 0 | 1 | 26 |
| cyp4f3 | 0.33482 | 0.370121113 | 0.111 | 0 | 1 | 26 |
| ddx47 | 0.33482 | 0.405632152 | 0.111 | 0 | 1 | 26 |
| def6a | 0.33482 | 0.39352966 | 0.111 | 0 | 1 | 26 |
| dhrs1 | 0.33482 | 0.384545588 | 0.111 | 0 | 1 | 26 |
| dnajc19 | 0.33482 | 0.289352782 | 0.111 | 0 | 1 | 26 |
| dynlrb1 | 0.33482 | 0.366808999 | 0.111 | 0 | 1 | 26 |
| eif3ja | 0.33482 | 0.377797447 | 0.111 | 0 | 1 | 26 |
| ela2 | 0.33482 | 0.513359575 | 0.111 | 0 | 1 | 26 |
| elmod2 | 0.33482 | 0.282368167 | 0.111 | 0 | 1 | 26 |

| | | | | | | |
|---|---|---|---|---|---|---|
| endou | 0.33482 | 0.178520995 | 0.111 | 0 | 1 | 26 |
| ero1b | 0.33482 | 0.416988897 | 0.111 | 0 | 1 | 26 |
| etfb | 0.33482 | 0.375804643 | 0.111 | 0 | 1 | 26 |
| fkbp2 | 0.33482 | 0.296112708 | 0.111 | 0 | 1 | 26 |
| g6pd | 0.33482 | 0.399484169 | 0.111 | 0 | 1 | 26 |
| galnt12 | 0.33482 | 0.369285977 | 0.111 | 0 | 1 | 26 |
| gatc | 0.33482 | 0.387955327 | 0.111 | 0 | 1 | 26 |
| ggh | 0.33482 | 0.295591624 | 0.111 | 0 | 1 | 26 |
| golgb1 | 0.33482 | 0.437141154 | 0.111 | 0 | 1 | 26 |
| gpia | 0.33482 | 0.371105299 | 0.111 | 0 | 1 | 26 |
| gpkow | 0.33482 | 0.372597722 | 0.111 | 0 | 1 | 26 |
| grasp | 0.33482 | 0.372493129 | 0.111 | 0 | 1 | 26 |
| hdlbpa | 0.33482 | 0.412793848 | 0.111 | 0 | 1 | 26 |
| higd1a | 0.33482 | 0.361551401 | 0.111 | 0 | 1 | 26 |
| hm13 | 0.33482 | 0.386451594 | 0.111 | 0 | 1 | 26 |
| hnrnpa0b | 0.33482 | 0.387195787 | 0.111 | 0 | 1 | 26 |
| htatip2 | 0.33482 | 0.372145407 | 0.111 | 0 | 1 | 26 |
| ifrd1 | 0.33482 | 0.290757221 | 0.111 | 0 | 1 | 26 |
| ilk | 0.33482 | 0.418601687 | 0.111 | 0 | 1 | 26 |
| ipmkb | 0.33482 | 0.433999637 | 0.111 | 0 | 1 | 26 |
| irf2bp2b | 0.33482 | 0.367573018 | 0.111 | 0 | 1 | 26 |
| itga4 | 0.33482 | 0.4238596 | 0.111 | 0 | 1 | 26 |
| jpt2 | 0.33482 | 0.40999161 | 0.111 | 0 | 1 | 26 |
| jun | 0.33482 | 0.462659695 | 0.111 | 0 | 1 | 26 |
| kars | 0.33482 | 0.354839968 | 0.111 | 0 | 1 | 26 |
| kdm1a | 0.33482 | 0.36537329 | 0.111 | 0 | 1 | 26 |
| klf11a | 0.33482 | 0.42853155 | 0.111 | 0 | 1 | 26 |
| larp6a | 0.33482 | 0.387713853 | 0.111 | 0 | 1 | 26 |
| ldhba | 0.33482 | 0.408482391 | 0.111 | 0 | 1 | 26 |
| luc7l | 0.33482 | 0.268888892 | 0.111 | 0 | 1 | 26 |
| m17 | 0.33482 | 0.421464894 | 0.111 | 0 | 1 | 26 |
| maff | 0.33482 | 0.378097876 | 0.111 | 0 | 1 | 26 |
| marcksl1a | 0.33482 | 0.368977915 | 0.111 | 0 | 1 | 26 |
| mast2 | 0.33482 | 0.391318485 | 0.111 | 0 | 1 | 26 |
| mcm2 | 0.33482 | 0.373998402 | 0.111 | 0 | 1 | 26 |
| med10 | 0.33482 | 0.387823915 | 0.111 | 0 | 1 | 26 |
| mettl15 | 0.33482 | 0.396926979 | 0.111 | 0 | 1 | 26 |
| mfap3l | 0.33482 | 0.442269973 | 0.111 | 0 | 1 | 26 |
| mfsd6b | 0.33482 | 0.298928752 | 0.111 | 0 | 1 | 26 |
| mgst1.2 | 0.33482 | 0.407702641 | 0.111 | 0 | 1 | 26 |
| mgst3b | 0.33482 | 0.324604962 | 0.111 | 0 | 1 | 26 |
| mknk2b | 0.33482 | 0.305950762 | 0.111 | 0 | 1 | 26 |
| mob3a | 0.33482 | 0.392008944 | 0.111 | 0 | 1 | 26 |
| mrpl36 | 0.33482 | 0.364718844 | 0.111 | 0 | 1 | 26 |
| mrps2 | 0.33482 | 0.404545771 | 0.111 | 0 | 1 | 26 |
| ms4a17a.9 | 0.33482 | 0.402877702 | 0.111 | 0 | 1 | 26 |
| mtdha | 0.33482 | 0.371545457 | 0.111 | 0 | 1 | 26 |
| ndufa11 | 0.33482 | 0.350417114 | 0.111 | 0 | 1 | 26 |

| | | | | | | |
|---|---|---|---|---|---|---|
| ndufa12 | 0.33482 | 0.274123665 | 0.111 | 0 | 1 | 26 |
| ndufa3 | 0.33482 | 0.27481512 | 0.111 | 0 | 1 | 26 |
| ndufb8 | 0.33482 | 0.380514463 | 0.111 | 0 | 1 | 26 |
| nop56 | 0.33482 | 0.353164279 | 0.111 | 0 | 1 | 26 |
| npm1a | 0.33482 | 0.381503414 | 0.111 | 0 | 1 | 26 |
| nudc | 0.33482 | 0.392755622 | 0.111 | 0 | 1 | 26 |
| ostc | 0.33482 | 0.265175947 | 0.111 | 0 | 1 | 26 |
| pacsin2 | 0.33482 | 0.395622711 | 0.111 | 0 | 1 | 26 |
| peli3 | 0.33482 | 0.275439218 | 0.111 | 0 | 1 | 26 |
| piezo1 | 0.33482 | 0.392232429 | 0.111 | 0 | 1 | 26 |
| pkma | 0.33482 | 0.392031131 | 0.111 | 0 | 1 | 26 |
| plek | 0.33482 | 0.394651717 | 0.111 | 0 | 1 | 26 |
| ppm1k | 0.33482 | 0.391110877 | 0.111 | 0 | 1 | 26 |
| prdx6 | 0.33482 | 0.379224718 | 0.111 | 0 | 1 | 26 |
| prmt1 | 0.33482 | 0.385157464 | 0.111 | 0 | 1 | 26 |
| prss1 | 0.33482 | 0.597625185 | 0.111 | 0 | 1 | 26 |
| psma5 | 0.33482 | 0.431256289 | 0.111 | 0 | 1 | 26 |
| psma6a | 0.33482 | 0.400068893 | 0.111 | 0 | 1 | 26 |
| psma6b | 0.33482 | 0.403807506 | 0.111 | 0 | 1 | 26 |
| psmb3 | 0.33482 | 0.365239658 | 0.111 | 0 | 1 | 26 |
| psmb4 | 0.33482 | 0.287262902 | 0.111 | 0 | 1 | 26 |
| psmb7 | 0.33482 | 0.345310106 | 0.111 | 0 | 1 | 26 |
| ptges3b | 0.33482 | 0.381767015 | 0.111 | 0 | 1 | 26 |
| ptprc | 0.33482 | 0.378241455 | 0.111 | 0 | 1 | 26 |
| rab1ab | 0.33482 | 0.383136499 | 0.111 | 0 | 1 | 26 |
| rab27a | 0.33482 | 0.418832534 | 0.111 | 0 | 1 | 26 |
| rab5c | 0.33482 | 0.378769129 | 0.111 | 0 | 1 | 26 |
| rab6ba | 0.33482 | 0.439301998 | 0.111 | 0 | 1 | 26 |
| rbx1 | 0.33482 | 0.40849896 | 0.111 | 0 | 1 | 26 |
| rer1 | 0.33482 | 0.346071526 | 0.111 | 0 | 1 | 26 |
| rgs13 | 0.33482 | 0.387805885 | 0.111 | 0 | 1 | 26 |
| rhbdf1a | 0.33482 | 0.251815669 | 0.111 | 0 | 1 | 26 |
| rnf10 | 0.33482 | 0.383573806 | 0.111 | 0 | 1 | 26 |
| rpp14 | 0.33482 | 0.440052425 | 0.111 | 0 | 1 | 26 |
| rps6ka1 | 0.33482 | 0.405913858 | 0.111 | 0 | 1 | 26 |
| rsrp1 | 0.33482 | 0.436982062 | 0.111 | 0 | 1 | 26 |
| samsn1a | 0.33482 | 0.440683579 | 0.111 | 0 | 1 | 26 |
| scn4bb | 0.33482 | 0.404148668 | 0.111 | 0 | 1 | 26 |
| selenof | 0.33482 | 0.277698967 | 0.111 | 0 | 1 | 26 |
| sesn2 | 0.33482 | 0.432960891 | 0.111 | 0 | 1 | 26 |
| sf3b6 | 0.33482 | 0.39652454 | 0.111 | 0 | 1 | 26 |
| sgk1 | 0.33482 | 0.303106618 | 0.111 | 0 | 1 | 26 |
| sh3kbp1 | 0.33482 | 0.423525877 | 0.111 | 0 | 1 | 26 |
| si:ch1073-67j19.1 | 0.33482 | 0.425089977 | 0.111 | 0 | 1 | 26 |
| si:ch211-105j21.9 | 0.33482 | 0.397135255 | 0.111 | 0 | 1 | 26 |
| si:ch211-195b11.3 | 0.33482 | 0.1254247 | 0.111 | 0 | 1 | 26 |
| si:ch211-210b2.4 | 0.33482 | 0.386547766 | 0.111 | 0 | 1 | 26 |
| si:ch211-215e19.3 | 0.33482 | 0.263717245 | 0.111 | 0 | 1 | 26 |

| | | | | | | |
|---|---|---|---|---|---|---|
| si:ch211-240l19.8 | 0.33482 | 0.40677415 | 0.111 | 0 | 1 | 26 |
| si:ch211-270n8.1 | 0.33482 | 0.478606292 | 0.111 | 0 | 1 | 26 |
| si:ch73-233k15.2 | 0.33482 | 0.372899221 | 0.111 | 0 | 1 | 26 |
| si:ch73-237c6.1 | 0.33482 | 0.38879287 | 0.111 | 0 | 1 | 26 |
| si:ch73-281n10.2 | 0.33482 | 0.444620882 | 0.111 | 0 | 1 | 26 |
| si:dkey-11f4.14 | 0.33482 | 0.390043701 | 0.111 | 0 | 1 | 26 |
| si:dkey-172j4.3 | 0.33482 | 0.355661887 | 0.111 | 0 | 1 | 26 |
| si:dkey-73n10.1 | 0.33482 | 0.382334009 | 0.111 | 0 | 1 | 26 |
| si:dkey-7i4.24 | 0.33482 | 0.386904017 | 0.111 | 0 | 1 | 26 |
| si:dkeyp-75h12.5 | 0.33482 | 0.346417149 | 0.111 | 0 | 1 | 26 |
| si:rp71-36a1.5 | 0.33482 | 0.385507654 | 0.111 | 0 | 1 | 26 |
| sirt7 | 0.33482 | 0.268301634 | 0.111 | 0 | 1 | 26 |
| slc25a10 | 0.33482 | 0.360410469 | 0.111 | 0 | 1 | 26 |
| slc25a36a | 0.33482 | 0.423567245 | 0.111 | 0 | 1 | 26 |
| slc6a16b | 0.33482 | 0.301518168 | 0.111 | 0 | 1 | 26 |
| slirp | 0.33482 | 0.379660402 | 0.111 | 0 | 1 | 26 |
| smad3a | 0.33482 | 0.377191596 | 0.111 | 0 | 1 | 26 |
| smim8 | 0.33482 | 0.368147541 | 0.111 | 0 | 1 | 26 |
| snd1 | 0.33482 | 0.357706265 | 0.111 | 0 | 1 | 26 |
| snrpd3l | 0.33482 | 0.365701427 | 0.111 | 0 | 1 | 26 |
| sod1 | 0.33482 | 0.403808453 | 0.111 | 0 | 1 | 26 |
| spink4 | 0.33482 | 0.234614939 | 0.111 | 0 | 1 | 26 |
| spns3 | 0.33482 | 0.390031161 | 0.111 | 0 | 1 | 26 |
| srp19 | 0.33482 | 0.457503116 | 0.111 | 0 | 1 | 26 |
| srsf9 | 0.33482 | 0.398711169 | 0.111 | 0 | 1 | 26 |
| sult1st1 | 0.33482 | 0.344280039 | 0.111 | 0 | 1 | 26 |
| syap1 | 0.33482 | 0.343994714 | 0.111 | 0 | 1 | 26 |
| sycn.2 | 0.33482 | 0.454576234 | 0.111 | 0 | 1 | 26 |
| tbl2 | 0.33482 | 0.272074183 | 0.111 | 0 | 1 | 26 |
| tcf25 | 0.33482 | 0.380917812 | 0.111 | 0 | 1 | 26 |
| thoc2 | 0.33482 | 0.38695711 | 0.111 | 0 | 1 | 26 |
| tle3a | 0.33482 | 0.440672523 | 0.111 | 0 | 1 | 26 |
| tmed7 | 0.33482 | 0.383545115 | 0.111 | 0 | 1 | 26 |
| tmed8 | 0.33482 | 0.422017493 | 0.111 | 0 | 1 | 26 |
| tmem59 | 0.33482 | 0.298164105 | 0.111 | 0 | 1 | 26 |
| tmx1 | 0.33482 | 0.384821582 | 0.111 | 0 | 1 | 26 |
| tmx4 | 0.33482 | 0.37197481 | 0.111 | 0 | 1 | 26 |
| tomm5 | 0.33482 | 0.409981251 | 0.111 | 0 | 1 | 26 |
| tpi1b | 0.33482 | 0.387895955 | 0.111 | 0 | 1 | 26 |
| trappc11 | 0.33482 | 0.264921918 | 0.111 | 0 | 1 | 26 |
| tsc22d3 | 0.33482 | 0.414885261 | 0.111 | 0 | 1 | 26 |
| u2af1 | 0.33482 | 0.355652434 | 0.111 | 0 | 1 | 26 |
| ube2d2 | 0.33482 | 0.40843732 | 0.111 | 0 | 1 | 26 |
| ube2d4 | 0.33482 | 0.41891003 | 0.111 | 0 | 1 | 26 |
| ube2l3b | 0.33482 | 0.35986914 | 0.111 | 0 | 1 | 26 |
| uqcrfs1 | 0.33482 | 0.384906697 | 0.111 | 0 | 1 | 26 |
| vps29 | 0.33482 | 0.369110206 | 0.111 | 0 | 1 | 26 |
| yrk | 0.33482 | 0.392570919 | 0.111 | 0 | 1 | 26 |

| | | | | | | |
|---|---|---|---|---|---|---|
| ywhae2 | 0.33482 | 0.404941334 | 0.111 | 0 | 1 | 26 |
| zgc:112160 | 0.33482 | 0.454015005 | 0.111 | 0 | 1 | 26 |
| zgc:174904 | 0.33482 | 0.387270035 | 0.111 | 0 | 1 | 26 |
| zgc:66427 | 0.33482 | 0.405594263 | 0.111 | 0 | 1 | 26 |
| zgc:77112 | 0.33482 | 0.436274893 | 0.111 | 0 | 1 | 26 |
| zgc:85858 | 0.33482 | 0.393743674 | 0.111 | 0 | 1 | 26 |
| zgc:92744 | 0.33482 | 0.420021882 | 0.111 | 0 | 1 | 26 |
| zyx | 0.33482 | 0.434207042 | 0.111 | 0 | 1 | 26 |
| arl5c | 0.33482 | 0.466588845 | 0.111 | 0 | 1 | 26 |
| camk2g2 | 0.33482 | 0.465309811 | 0.111 | 0 | 1 | 26 |
| cited4b | 0.33482 | 0.363151031 | 0.111 | 0 | 1 | 26 |
| cyth1a | 0.33482 | 0.366748563 | 0.111 | 0 | 1 | 26 |
| viml | 0.33482 | 0.466817536 | 0.111 | 0 | 1 | 26 |
| CABZ01026681.1 | 0.33482 | 0.208175784 | 0.111 | 0 | 1 | 26 |
| CABZ01073834.1 | 0.33482 | 0.330701534 | 0.111 | 0 | 1 | 26 |
| CABZ01079427.1 | 0.33482 | 0.32638001 | 0.111 | 0 | 1 | 26 |
| FBXO48 | 0.33482 | 0.330581706 | 0.111 | 0 | 1 | 26 |
| XLOC-024867 | 0.33482 | 0.311734372 | 0.111 | 0 | 1 | 26 |
| acadm | 0.33482 | 0.289406398 | 0.111 | 0 | 1 | 26 |
| actn1 | 0.33482 | 0.33585085 | 0.111 | 0 | 1 | 26 |
| ada | 0.33482 | 0.304994956 | 0.111 | 0 | 1 | 26 |
| adam10a | 0.33482 | 0.328036021 | 0.111 | 0 | 1 | 26 |
| adgre13 | 0.33482 | 0.292517271 | 0.111 | 0 | 1 | 26 |
| ahcyl1 | 0.33482 | 0.31076265 | 0.111 | 0 | 1 | 26 |
| aldh9a1a.1 | 0.33482 | 0.261192636 | 0.111 | 0 | 1 | 26 |
| ampd3b | 0.33482 | 0.329659424 | 0.111 | 0 | 1 | 26 |
| atf4b | 0.33482 | 0.306427592 | 0.111 | 0 | 1 | 26 |
| b2m | 0.33482 | 0.29047296 | 0.111 | 0 | 1 | 26 |
| bhlhe40 | 0.33482 | 0.331992515 | 0.111 | 0 | 1 | 26 |
| carhsp1 | 0.33482 | 0.323774309 | 0.111 | 0 | 1 | 26 |
| ccni | 0.33482 | 0.287578522 | 0.111 | 0 | 1 | 26 |
| cct3 | 0.33482 | 0.315933386 | 0.111 | 0 | 1 | 26 |
| cd59 | 0.33482 | 0.357034656 | 0.111 | 0 | 1 | 26 |
| cfl1l | 0.33482 | 0.200231858 | 0.111 | 0 | 1 | 26 |
| cox6a2 | 0.33482 | 0.262720382 | 0.111 | 0 | 1 | 26 |
| cox8b | 0.33482 | 0.191372627 | 0.111 | 0 | 1 | 26 |
| dennd4c | 0.33482 | 0.305441519 | 0.111 | 0 | 1 | 26 |
| edf1 | 0.33482 | 0.31730916 | 0.111 | 0 | 1 | 26 |
| erlec1 | 0.33482 | 0.328466943 | 0.111 | 0 | 1 | 26 |
| fbp1a | 0.33482 | 0.306509439 | 0.111 | 0 | 1 | 26 |
| grk3 | 0.33482 | 0.303995243 | 0.111 | 0 | 1 | 26 |
| grk6 | 0.33482 | 0.368829205 | 0.111 | 0 | 1 | 26 |
| gsr | 0.33482 | 0.334721745 | 0.111 | 0 | 1 | 26 |
| hint1 | 0.33482 | 0.232620059 | 0.111 | 0 | 1 | 26 |
| hspa14 | 0.33482 | 0.3045468 | 0.111 | 0 | 1 | 26 |
| id3 | 0.33482 | 0.326656261 | 0.111 | 0 | 1 | 26 |
| il16 | 0.33482 | 0.367060494 | 0.111 | 0 | 1 | 26 |
| ipo7 | 0.33482 | 0.323719653 | 0.111 | 0 | 1 | 26 |

| | | | | | | |
|---|---|---|---|---|---|---|
| irf2bp2a | 0.33482 | 0.303464558 | 0.111 | 0 | 1 | 26 |
| itgb3a | 0.33482 | 0.304146383 | 0.111 | 0 | 1 | 26 |
| mbnl3 | 0.33482 | 0.34112837 | 0.111 | 0 | 1 | 26 |
| mmp25b | 0.33482 | 0.215948241 | 0.111 | 0 | 1 | 26 |
| mrps15 | 0.33482 | 0.330746025 | 0.111 | 0 | 1 | 26 |
| ndufs6 | 0.33482 | 0.319574564 | 0.111 | 0 | 1 | 26 |
| npepl1 | 0.33482 | 0.210683208 | 0.111 | 0 | 1 | 26 |
| peli1b | 0.33482 | 0.211617072 | 0.111 | 0 | 1 | 26 |
| plekhj1 | 0.33482 | 0.302537604 | 0.111 | 0 | 1 | 26 |
| purab | 0.33482 | 0.326775816 | 0.111 | 0 | 1 | 26 |
| rabgap1l | 0.33482 | 0.308879593 | 0.111 | 0 | 1 | 26 |
| rca2.1 | 0.33482 | 0.323477491 | 0.111 | 0 | 1 | 26 |
| rhogb | 0.33482 | 0.331252212 | 0.111 | 0 | 1 | 26 |
| rp2 | 0.33482 | 0.331064811 | 0.111 | 0 | 1 | 26 |
| selenow2a | 0.33482 | 0.390954437 | 0.111 | 0 | 1 | 26 |
| sesn1 | 0.33482 | 0.303308345 | 0.111 | 0 | 1 | 26 |
| si:ch211-102c2.4 | 0.33482 | 0.312606283 | 0.111 | 0 | 1 | 26 |
| si:dkey-172h23.2 | 0.33482 | 0.332192058 | 0.111 | 0 | 1 | 26 |
| si:dkey-238c7.16 | 0.33482 | 0.305862736 | 0.111 | 0 | 1 | 26 |
| si:dkey-24p1.7 | 0.33482 | 0.318016891 | 0.111 | 0 | 1 | 26 |
| slc25a28 | 0.33482 | 0.333310894 | 0.111 | 0 | 1 | 26 |
| slc6a6b | 0.33482 | 0.346606366 | 0.111 | 0 | 1 | 26 |
| stat1a | 0.33482 | 0.305442843 | 0.111 | 0 | 1 | 26 |
| stim1a | 0.33482 | 0.345501854 | 0.111 | 0 | 1 | 26 |
| supt4h1 | 0.33482 | 0.323813213 | 0.111 | 0 | 1 | 26 |
| syk | 0.33482 | 0.384010052 | 0.111 | 0 | 1 | 26 |
| taf11 | 0.33482 | 0.292977141 | 0.111 | 0 | 1 | 26 |
| tln1 | 0.33482 | 0.31246706 | 0.111 | 0 | 1 | 26 |
| tmem176l.3a | 0.33482 | 0.296429636 | 0.111 | 0 | 1 | 26 |
| trappc6bl | 0.33482 | 0.277236355 | 0.111 | 0 | 1 | 26 |
| tspan2b | 0.33482 | 0.300838197 | 0.111 | 0 | 1 | 26 |
| ttc9c | 0.33482 | 0.332157912 | 0.111 | 0 | 1 | 26 |
| tuba8l | 0.33482 | 0.29919868 | 0.111 | 0 | 1 | 26 |
| ube2z | 0.33482 | 0.305640306 | 0.111 | 0 | 1 | 26 |
| wasf2 | 0.33482 | 0.301253198 | 0.111 | 0 | 1 | 26 |
| ywhaqb | 0.33482 | 0.340859712 | 0.111 | 0 | 1 | 26 |
| zgc:63831 | 0.33482 | 0.329895746 | 0.111 | 0 | 1 | 26 |
| zmp:0000001082 | 0.33482 | 0.373377116 | 0.111 | 0 | 1 | 26 |
| frg1 | 0.33482 | 0.300770367 | 0.111 | 0 | 1 | 26 |
| msra | 0.33482 | 0.360307448 | 0.111 | 0 | 1 | 26 |
| npc2 | 0.33482 | 0.292345851 | 0.111 | 0 | 1 | 26 |
| si:dkey-14d8.7 | 0.33482 | 0.288952345 | 0.111 | 0 | 1 | 26 |
| rpl35a | 0.35122 | 0.265540415 | 0.833 | 0.571 | 1 | 26 |
| rps13 | 0.3544 | -0.14171126 | 0.889 | 0.857 | 1 | 26 |
| tmsb4x | 0.356 | -0.14347172 | 0.944 | 1 | 1 | 26 |
| actb2 | 0.36659 | -0.19543613 | 0.944 | 1 | 1 | 26 |
| nme2b.1 | 0.3676 | -0.2838431 | 0.833 | 1 | 1 | 26 |
| cox6b1 | 0.36811 | -0.51349592 | 0.167 | 0.286 | 1 | 26 |

| | | | | | | |
|---|---|---|---|---|---|---|
| MFAP4 (1 of many) | 0.36935 | -0.22639794 | 0.889 | 1 | 1 | 26 |
| COX3 | 0.37372 | 0.76203065 | 0.556 | 0.286 | 1 | 26 |
| zgc:162730 | 0.38379 | 0.710221245 | 0.389 | 0.143 | 1 | 26 |
| calm3b | 0.39632 | -0.59016963 | 0.167 | 0.429 | 1 | 26 |
| sumo3a | 0.3965 | -0.61146779 | 0.167 | 0.429 | 1 | 26 |
| rpl36a | 0.3975 | -0.14468833 | 0.833 | 1 | 1 | 26 |
| h1fx | 0.40368 | -0.5575554 | 0.278 | 0.571 | 1 | 26 |
| hmgn6 | 0.40452 | -0.64716405 | 0.167 | 0.429 | 1 | 26 |
| tuba8l4 | 0.41089 | -0.55465673 | 0.167 | 0.429 | 1 | 26 |
| csnk2b | 0.4164 | -0.55935574 | 0.167 | 0.429 | 1 | 26 |
| eif3ea | 0.41794 | -0.64310225 | 0.111 | 0.286 | 1 | 26 |
| setb | 0.41859 | -0.5670499 | 0.167 | 0.429 | 1 | 26 |
| faua | 0.42362 | -0.37442555 | 0.667 | 0.857 | 1 | 26 |
| ube2v1 | 0.42641 | -0.54412595 | 0.222 | 0.429 | 1 | 26 |
| cfbl | 0.43214 | 0.473107921 | 0.611 | 0.429 | 1 | 26 |
| cotl1 | 0.4541 | -0.32144496 | 0.611 | 0.857 | 1 | 26 |
| cox5aa | 0.4544 | 0.527969292 | 0.389 | 0.143 | 1 | 26 |
| rpl8 | 0.46483 | -0.25424739 | 0.778 | 0.857 | 1 | 26 |
| ptmaa | 0.46601 | -0.36032887 | 0.611 | 0.857 | 1 | 26 |
| mibp2 | 0.46682 | -0.3064059 | 0.389 | 0.429 | 1 | 26 |
| ssr2 | 0.46848 | -0.46248187 | 0.389 | 0.571 | 1 | 26 |
| serf2 | 0.46888 | -0.15505765 | 0.333 | 0.286 | 1 | 26 |
| slc25a5 | 0.47127 | -0.33824845 | 0.611 | 0.857 | 1 | 26 |
| pdia3 | 0.47127 | 0.559697619 | 0.389 | 0.143 | 1 | 26 |
| msna | 0.47144 | 0.612909793 | 0.389 | 0.143 | 1 | 26 |
| rpl9 | 0.47858 | 0.155126531 | 0.889 | 0.714 | 1 | 26 |
| CT030188.1 | 0.48014 | -0.54529743 | 0.222 | 0.429 | 1 | 26 |
| rpl27 | 0.48032 | -0.13858816 | 0.722 | 0.714 | 1 | 26 |
| atp5mf | 0.48219 | -0.51378184 | 0.111 | 0.286 | 1 | 26 |
| rplp0 | 0.49182 | 0.181995025 | 0.722 | 0.714 | 1 | 26 |
| rps27.2 | 0.49492 | -0.41090885 | 0.333 | 0.429 | 1 | 26 |
| cox7c | 0.50186 | 0.127620735 | 0.278 | 0.143 | 1 | 26 |
| GCA | 0.50832 | -0.21377267 | 0.056 | 0.143 | 1 | 26 |
| LOC558761 | 0.50832 | -0.2624323 | 0.056 | 0.143 | 1 | 26 |
| acp1 | 0.50832 | -0.26936972 | 0.056 | 0.143 | 1 | 26 |
| adh5 | 0.50832 | -0.28383839 | 0.056 | 0.143 | 1 | 26 |
| anp32a | 0.50832 | -0.2637003 | 0.056 | 0.143 | 1 | 26 |
| apooa | 0.50832 | -0.24404786 | 0.056 | 0.143 | 1 | 26 |
| atp5if1b | 0.50832 | -0.23618992 | 0.056 | 0.143 | 1 | 26 |
| cbx5 | 0.50832 | -0.225046 | 0.056 | 0.143 | 1 | 26 |
| copz1 | 0.50832 | -0.23729606 | 0.056 | 0.143 | 1 | 26 |
| ebna1bp2 | 0.50832 | -0.25463731 | 0.056 | 0.143 | 1 | 26 |
| ikzf2 | 0.50832 | -0.24802303 | 0.056 | 0.143 | 1 | 26 |
| lman1 | 0.50832 | -0.24174309 | 0.056 | 0.143 | 1 | 26 |
| mars | 0.50832 | -0.25994193 | 0.056 | 0.143 | 1 | 26 |
| nme2b.2 | 0.50832 | -0.2699336 | 0.056 | 0.143 | 1 | 26 |
| psmb1 | 0.50832 | -0.25950574 | 0.056 | 0.143 | 1 | 26 |
| psmc4 | 0.50832 | -0.21707603 | 0.056 | 0.143 | 1 | 26 |

| | | | | | | |
|---|---|---|---|---|---|---|
| rab35b | 0.50832 | -0.26370522 | 0.056 | 0.143 | 1 | 26 |
| selenok | 0.50832 | -0.21251589 | 0.056 | 0.143 | 1 | 26 |
| snrpb | 0.50832 | -0.24469442 | 0.056 | 0.143 | 1 | 26 |
| spi1b | 0.50832 | -0.24033181 | 0.056 | 0.143 | 1 | 26 |
| taf12 | 0.50832 | -0.26225758 | 0.056 | 0.143 | 1 | 26 |
| anxa11b | 0.50832 | -0.22215266 | 0.056 | 0.143 | 1 | 26 |
| arl1 | 0.50832 | -0.22074447 | 0.056 | 0.143 | 1 | 26 |
| cct2 | 0.50832 | -0.3554031 | 0.056 | 0.143 | 1 | 26 |
| chmp5b | 0.50832 | -0.26345929 | 0.056 | 0.143 | 1 | 26 |
| dhdhl | 0.50832 | -0.37725212 | 0.056 | 0.143 | 1 | 26 |
| dusp2 | 0.50832 | -0.2518264 | 0.056 | 0.143 | 1 | 26 |
| gabarapb | 0.50832 | -0.29642191 | 0.056 | 0.143 | 1 | 26 |
| ikbkg | 0.50832 | -0.22316864 | 0.056 | 0.143 | 1 | 26 |
| map4k6 | 0.50832 | -0.29630649 | 0.056 | 0.143 | 1 | 26 |
| nap1l1 | 0.50832 | -0.30235272 | 0.056 | 0.143 | 1 | 26 |
| ndufa5 | 0.50832 | -0.28692684 | 0.056 | 0.143 | 1 | 26 |
| ndufc2 | 0.50832 | -0.24142112 | 0.056 | 0.143 | 1 | 26 |
| nfe2 | 0.50832 | -0.32092323 | 0.056 | 0.143 | 1 | 26 |
| polr2l | 0.50832 | -0.23905432 | 0.056 | 0.143 | 1 | 26 |
| ptpn6 | 0.50832 | -0.29715891 | 0.056 | 0.143 | 1 | 26 |
| rab42b | 0.50832 | -0.21669302 | 0.056 | 0.143 | 1 | 26 |
| rbmx | 0.50832 | -0.21290827 | 0.056 | 0.143 | 1 | 26 |
| si:dkey-262k9.4 | 0.50832 | -0.25149876 | 0.056 | 0.143 | 1 | 26 |
| si:dkeyp-44a8.2 | 0.50832 | -0.22281089 | 0.056 | 0.143 | 1 | 26 |
| snrpd1 | 0.50832 | -0.2032524 | 0.056 | 0.143 | 1 | 26 |
| stt3b | 0.50832 | -0.29681174 | 0.056 | 0.143 | 1 | 26 |
| ube2l3a | 0.50832 | -0.29920261 | 0.056 | 0.143 | 1 | 26 |
| vps37a | 0.50832 | -0.37068168 | 0.056 | 0.143 | 1 | 26 |
| atp6ap1b | 0.50832 | -0.33540837 | 0.056 | 0.143 | 1 | 26 |
| brk1 | 0.50832 | -0.33178888 | 0.056 | 0.143 | 1 | 26 |
| echdc2 | 0.50832 | -0.41143777 | 0.056 | 0.143 | 1 | 26 |
| eif3k | 0.50832 | -0.41351155 | 0.056 | 0.143 | 1 | 26 |
| fam49ba | 0.50832 | -0.33682404 | 0.056 | 0.143 | 1 | 26 |
| gskip | 0.50832 | -0.30970814 | 0.056 | 0.143 | 1 | 26 |
| rgs14a | 0.50832 | -0.31341173 | 0.056 | 0.143 | 1 | 26 |
| si:ch211-74f19.2 | 0.50832 | -0.26277843 | 0.056 | 0.143 | 1 | 26 |
| tcp1 | 0.50832 | -0.33918811 | 0.056 | 0.143 | 1 | 26 |
| trir | 0.50832 | -0.31261242 | 0.056 | 0.143 | 1 | 26 |
| uqcr10 | 0.50832 | -0.31662921 | 0.056 | 0.143 | 1 | 26 |
| CABZ01072950.1 | 0.50832 | -0.26661064 | 0.056 | 0.143 | 1 | 26 |
| LOC101885065 | 0.50832 | -0.27537533 | 0.056 | 0.143 | 1 | 26 |
| TMEM208 | 0.50832 | -0.25049569 | 0.056 | 0.143 | 1 | 26 |
| ak1 | 0.50832 | -0.31147062 | 0.056 | 0.143 | 1 | 26 |
| arfip1 | 0.50832 | -0.2770167 | 0.056 | 0.143 | 1 | 26 |
| blcap | 0.50832 | -0.33603473 | 0.056 | 0.143 | 1 | 26 |
| cib1 | 0.50832 | -0.20805289 | 0.056 | 0.143 | 1 | 26 |
| clint1a | 0.50832 | -0.38177154 | 0.056 | 0.143 | 1 | 26 |
| cmtm7 | 0.50832 | -0.28519799 | 0.056 | 0.143 | 1 | 26 |

| | | | | | | |
|---|---|---|---|---|---|---|
| cox7a2a | 0.50832 | -0.3483604 | 0.056 | 0.143 | 1 | 26 |
| cyb5a | 0.50832 | -0.27131251 | 0.056 | 0.143 | 1 | 26 |
| dnase1l4.2 | 0.50832 | -0.42604839 | 0.056 | 0.143 | 1 | 26 |
| eif2s1b | 0.50832 | -0.26643605 | 0.056 | 0.143 | 1 | 26 |
| elf1 | 0.50832 | -0.33737787 | 0.056 | 0.143 | 1 | 26 |
| fam107b | 0.50832 | -0.29699294 | 0.056 | 0.143 | 1 | 26 |
| gnai1 | 0.50832 | -0.37223932 | 0.056 | 0.143 | 1 | 26 |
| gng2 | 0.50832 | -0.25135364 | 0.056 | 0.143 | 1 | 26 |
| hivep2a | 0.50832 | -0.27712126 | 0.056 | 0.143 | 1 | 26 |
| ier5 | 0.50832 | -0.25258823 | 0.056 | 0.143 | 1 | 26 |
| ndufaf3 | 0.50832 | -0.27562533 | 0.056 | 0.143 | 1 | 26 |
| pfdn5 | 0.50832 | -0.34028751 | 0.056 | 0.143 | 1 | 26 |
| plekho2 | 0.50832 | -0.25322962 | 0.056 | 0.143 | 1 | 26 |
| ppp6c | 0.50832 | -0.27581477 | 0.056 | 0.143 | 1 | 26 |
| psip1b | 0.50832 | -0.34717461 | 0.056 | 0.143 | 1 | 26 |
| sarnp | 0.50832 | -0.38514069 | 0.056 | 0.143 | 1 | 26 |
| scamp2 | 0.50832 | -0.27578411 | 0.056 | 0.143 | 1 | 26 |
| sec61a1 | 0.50832 | -0.29375558 | 0.056 | 0.143 | 1 | 26 |
| serpinb1l1 | 0.50832 | -0.25268369 | 0.056 | 0.143 | 1 | 26 |
| smim15 | 0.50832 | -0.20657399 | 0.056 | 0.143 | 1 | 26 |
| tagapb | 0.50832 | -0.22266471 | 0.056 | 0.143 | 1 | 26 |
| twf1a | 0.50832 | -0.33790042 | 0.056 | 0.143 | 1 | 26 |
| vamp3 | 0.50832 | -0.27943776 | 0.056 | 0.143 | 1 | 26 |
| CU694197.1 | 0.50832 | -0.24236691 | 0.056 | 0.143 | 1 | 26 |
| abi3a | 0.50832 | -0.36959959 | 0.056 | 0.143 | 1 | 26 |
| calr3a | 0.50832 | -0.1993763 | 0.056 | 0.143 | 1 | 26 |
| cst14a.2 | 0.50832 | -0.33577302 | 0.056 | 0.143 | 1 | 26 |
| dido1 | 0.50832 | -0.40697628 | 0.056 | 0.143 | 1 | 26 |
| erp44 | 0.50832 | -0.27942093 | 0.056 | 0.143 | 1 | 26 |
| gosr2 | 0.50832 | -0.23876368 | 0.056 | 0.143 | 1 | 26 |
| ilf3b | 0.50832 | -0.23864423 | 0.056 | 0.143 | 1 | 26 |
| pdia4 | 0.50832 | -0.30216734 | 0.056 | 0.143 | 1 | 26 |
| ppp1caa | 0.50832 | -0.40823446 | 0.056 | 0.143 | 1 | 26 |
| sap18 | 0.50832 | -0.27190287 | 0.056 | 0.143 | 1 | 26 |
| taf15 | 0.50832 | -0.22345223 | 0.056 | 0.143 | 1 | 26 |
| vamp8 | 0.50832 | -0.29148027 | 0.056 | 0.143 | 1 | 26 |
| vhl | 0.50832 | -0.23589123 | 0.056 | 0.143 | 1 | 26 |
| CABZ01058647.1 | 0.50832 | -0.41170449 | 0.056 | 0.143 | 1 | 26 |
| aspm | 0.50832 | -0.31317878 | 0.056 | 0.143 | 1 | 26 |
| crfb4 | 0.50832 | -0.27653755 | 0.056 | 0.143 | 1 | 26 |
| gbp | 0.50832 | -0.26234895 | 0.056 | 0.143 | 1 | 26 |
| lamtor4 | 0.50832 | -0.32021503 | 0.056 | 0.143 | 1 | 26 |
| litaf | 0.50832 | -0.28063751 | 0.056 | 0.143 | 1 | 26 |
| lsm4 | 0.50832 | -0.31479294 | 0.056 | 0.143 | 1 | 26 |
| med29 | 0.50832 | -0.20956253 | 0.056 | 0.143 | 1 | 26 |
| nat15 | 0.50832 | -0.33797645 | 0.056 | 0.143 | 1 | 26 |
| ndufs5 | 0.50832 | -0.28185642 | 0.056 | 0.143 | 1 | 26 |
| nfkbie | 0.50832 | -0.31481666 | 0.056 | 0.143 | 1 | 26 |

| | | | | | | |
|---|---|---|---|---|---|---|
| polr2j | 0.50832 | -0.27731539 | 0.056 | 0.143 | 1 | 26 |
| psmd8 | 0.50832 | -0.31554511 | 0.056 | 0.143 | 1 | 26 |
| rab14 | 0.50832 | -0.2988295 | 0.056 | 0.143 | 1 | 26 |
| sh3glb1a | 0.50832 | -0.30659329 | 0.056 | 0.143 | 1 | 26 |
| si:dkey-33c12.4 | 0.50832 | -0.40722999 | 0.056 | 0.143 | 1 | 26 |
| smco4 | 0.50832 | -0.43238476 | 0.056 | 0.143 | 1 | 26 |
| tmed9 | 0.50832 | -0.44696993 | 0.056 | 0.143 | 1 | 26 |
| tmem251 | 0.50832 | -0.34662994 | 0.056 | 0.143 | 1 | 26 |
| zgc:162396 | 0.50832 | -0.36697796 | 0.056 | 0.143 | 1 | 26 |
| rspry1 | 0.50853 | -0.55067828 | 0.111 | 0.286 | 1 | 26 |
| si:dkey-77g12.4 | 0.50856 | -0.22161259 | 0.944 | 1 | 1 | 26 |
| LOC110438970 | 0.51405 | 0.515785759 | 0.5 | 0.286 | 1 | 26 |
| capzb | 0.51576 | -0.38001284 | 0.333 | 0.571 | 1 | 26 |
| arpc5b | 0.517 | -0.50902983 | 0.222 | 0.429 | 1 | 26 |
| timp4.2 | 0.51824 | -0.51613102 | 0.111 | 0.286 | 1 | 26 |
| eef1db | 0.5217 | -0.51855765 | 0.111 | 0.286 | 1 | 26 |
| ak2 | 0.52681 | -0.46216423 | 0.278 | 0.429 | 1 | 26 |
| rplp2l | 0.53014 | 0.32621295 | 0.889 | 0.714 | 1 | 26 |
| mapre1a | 0.53356 | -0.51706717 | 0.111 | 0.286 | 1 | 26 |
| hsp90ab1 | 0.53415 | -0.10631862 | 0.778 | 0.714 | 1 | 26 |
| itln1 | 0.53476 | 0.178869775 | 0.889 | 0.714 | 1 | 26 |
| arpc3 | 0.54313 | -0.24461468 | 0.389 | 0.429 | 1 | 26 |
| BX000363.1 | 0.54902 | 0.131770998 | 0.056 | 0 | 1 | 26 |
| BX950187.1 | 0.54902 | 0.12728812 | 0.056 | 0 | 1 | 26 |
| CABZ01041494.1 | 0.54902 | 0.130448378 | 0.056 | 0 | 1 | 26 |
| CABZ01086293.1 | 0.54902 | 0.121710708 | 0.056 | 0 | 1 | 26 |
| CABZ01088428.1 | 0.54902 | 0.127770195 | 0.056 | 0 | 1 | 26 |
| CABZ01102170.1 | 0.54902 | 0.127260957 | 0.056 | 0 | 1 | 26 |
| CABZ01112656.1 | 0.54902 | 0.117069722 | 0.056 | 0 | 1 | 26 |
| DIPK2B | 0.54902 | 0.112116917 | 0.056 | 0 | 1 | 26 |
| LO018197.1 | 0.54902 | 0.129512238 | 0.056 | 0 | 1 | 26 |
| LOC101884451 | 0.54902 | 0.127454314 | 0.056 | 0 | 1 | 26 |
| LOC101886073 | 0.54902 | 0.129702433 | 0.056 | 0 | 1 | 26 |
| LOC103909069 | 0.54902 | 0.131770998 | 0.056 | 0 | 1 | 26 |
| LOC108190726 | 0.54902 | 0.131770998 | 0.056 | 0 | 1 | 26 |
| LOC110438237 | 0.54902 | 0.130892337 | 0.056 | 0 | 1 | 26 |
| LOC793824 | 0.54902 | 0.128231797 | 0.056 | 0 | 1 | 26 |
| RASA2 | 0.54902 | 0.130658212 | 0.056 | 0 | 1 | 26 |
| XLOC-034437 | 0.54902 | 0.126603165 | 0.056 | 0 | 1 | 26 |
| arhgef18b | 0.54902 | 0.131770998 | 0.056 | 0 | 1 | 26 |
| armc1l | 0.54902 | 0.168632404 | 0.056 | 0 | 1 | 26 |
| asxl2 | 0.54902 | 0.119464693 | 0.056 | 0 | 1 | 26 |
| atf1 | 0.54902 | 0.121029481 | 0.056 | 0 | 1 | 26 |
| birc6 | 0.54902 | 0.125365683 | 0.056 | 0 | 1 | 26 |
| brdt | 0.54902 | 0.129694304 | 0.056 | 0 | 1 | 26 |
| btbd10b | 0.54902 | 0.128334188 | 0.056 | 0 | 1 | 26 |
| btg2 | 0.54902 | 0.123219708 | 0.056 | 0 | 1 | 26 |
| calm2a | 0.54902 | 0.123262351 | 0.056 | 0 | 1 | 26 |

| | | | | | | |
|---|---|---|---|---|---|---|
| casp3a | 0.54902 | 0.129639118 | 0.056 | 0 | 1 | 26 |
| ccdc88b | 0.54902 | 0.130480686 | 0.056 | 0 | 1 | 26 |
| ccdc97 | 0.54902 | 0.118963416 | 0.056 | 0 | 1 | 26 |
| ccl39.7 | 0.54902 | 0.114568311 | 0.056 | 0 | 1 | 26 |
| ccr6a | 0.54902 | 0.131770998 | 0.056 | 0 | 1 | 26 |
| cdc26 | 0.54902 | 0.123972877 | 0.056 | 0 | 1 | 26 |
| cdc34b | 0.54902 | 0.122987079 | 0.056 | 0 | 1 | 26 |
| chordc1b | 0.54902 | 0.123285712 | 0.056 | 0 | 1 | 26 |
| cndp2 | 0.54902 | 0.127552258 | 0.056 | 0 | 1 | 26 |
| cnot3a | 0.54902 | 0.127467038 | 0.056 | 0 | 1 | 26 |
| cnsta | 0.54902 | 0.131770998 | 0.056 | 0 | 1 | 26 |
| coq4 | 0.54902 | 0.128252791 | 0.056 | 0 | 1 | 26 |
| cript | 0.54902 | 0.118728106 | 0.056 | 0 | 1 | 26 |
| crlf3 | 0.54902 | 0.129168613 | 0.056 | 0 | 1 | 26 |
| cul5a | 0.54902 | 0.131770998 | 0.056 | 0 | 1 | 26 |
| cxcr2 | 0.54902 | 0.127572288 | 0.056 | 0 | 1 | 26 |
| cxcr4b | 0.54902 | 0.123923309 | 0.056 | 0 | 1 | 26 |
| def6c | 0.54902 | 0.173735557 | 0.056 | 0 | 1 | 26 |
| dennd3b | 0.54902 | 0.131770998 | 0.056 | 0 | 1 | 26 |
| diablob | 0.54902 | 0.125287807 | 0.056 | 0 | 1 | 26 |
| eef2k | 0.54902 | 0.129840662 | 0.056 | 0 | 1 | 26 |
| egr1 | 0.54902 | 0.172210428 | 0.056 | 0 | 1 | 26 |
| elf2a | 0.54902 | 0.128653287 | 0.056 | 0 | 1 | 26 |
| elmo1 | 0.54902 | 0.128968104 | 0.056 | 0 | 1 | 26 |
| elovl6 | 0.54902 | 0.131770998 | 0.056 | 0 | 1 | 26 |
| epd | 0.54902 | 0.123756327 | 0.056 | 0 | 1 | 26 |
| epsti1 | 0.54902 | 0.1234221 | 0.056 | 0 | 1 | 26 |
| fam13b | 0.54902 | 0.129644973 | 0.056 | 0 | 1 | 26 |
| fgfr1op | 0.54902 | 0.126577444 | 0.056 | 0 | 1 | 26 |
| flna | 0.54902 | 0.116906312 | 0.056 | 0 | 1 | 26 |
| ftr14 | 0.54902 | 0.126740164 | 0.056 | 0 | 1 | 26 |
| gabarapa | 0.54902 | 0.124642414 | 0.056 | 0 | 1 | 26 |
| gfi1ab | 0.54902 | 0.129296503 | 0.056 | 0 | 1 | 26 |
| ghdc | 0.54902 | 0.131770998 | 0.056 | 0 | 1 | 26 |
| gmnn | 0.54902 | 0.12322514 | 0.056 | 0 | 1 | 26 |
| gucd1 | 0.54902 | 0.127169629 | 0.056 | 0 | 1 | 26 |
| hadh | 0.54902 | 0.108678694 | 0.056 | 0 | 1 | 26 |
| herc1 | 0.54902 | 0.105609926 | 0.056 | 0 | 1 | 26 |
| herc3 | 0.54902 | 0.131770998 | 0.056 | 0 | 1 | 26 |
| hmgn3 | 0.54902 | 0.122489198 | 0.056 | 0 | 1 | 26 |
| hnrnpub | 0.54902 | 0.124250134 | 0.056 | 0 | 1 | 26 |
| hnrnpul1 | 0.54902 | 0.127761316 | 0.056 | 0 | 1 | 26 |
| hnrnpul1l | 0.54902 | 0.124134959 | 0.056 | 0 | 1 | 26 |
| hs6st2 | 0.54902 | 0.130033176 | 0.056 | 0 | 1 | 26 |
| hsbp1b | 0.54902 | 0.120363571 | 0.056 | 0 | 1 | 26 |
| ikbkb | 0.54902 | 0.128796709 | 0.056 | 0 | 1 | 26 |
| ino80db | 0.54902 | 0.125009372 | 0.056 | 0 | 1 | 26 |
| ireb2 | 0.54902 | 0.123615425 | 0.056 | 0 | 1 | 26 |

| | | | | | | |
|---|---|---|---|---|---|---|
| itga1 | 0.54902 | 0.121544472 | 0.056 | 0 | 1 | 26 |
| itpk1b | 0.54902 | 0.130574238 | 0.056 | 0 | 1 | 26 |
| itpripl2 | 0.54902 | 0.126356137 | 0.056 | 0 | 1 | 26 |
| jak1 | 0.54902 | 0.174121758 | 0.056 | 0 | 1 | 26 |
| jak3 | 0.54902 | 0.131770998 | 0.056 | 0 | 1 | 26 |
| kat8 | 0.54902 | 0.10954412 | 0.056 | 0 | 1 | 26 |
| kcnq5b | 0.54902 | 0.125036554 | 0.056 | 0 | 1 | 26 |
| kdm4b | 0.54902 | 0.17252476 | 0.056 | 0 | 1 | 26 |
| keap1b | 0.54902 | 0.128684841 | 0.056 | 0 | 1 | 26 |
| lcp2a | 0.54902 | 0.128437783 | 0.056 | 0 | 1 | 26 |
| ldb1a | 0.54902 | 0.105582355 | 0.056 | 0 | 1 | 26 |
| lgals3a | 0.54902 | 0.122345314 | 0.056 | 0 | 1 | 26 |
| lpar5a | 0.54902 | 0.130828688 | 0.056 | 0 | 1 | 26 |
| lrrc59 | 0.54902 | 0.125024084 | 0.056 | 0 | 1 | 26 |
| ltb4r | 0.54902 | 0.130620691 | 0.056 | 0 | 1 | 26 |
| lypla2 | 0.54902 | 0.127285907 | 0.056 | 0 | 1 | 26 |
| man2b1 | 0.54902 | 0.131770998 | 0.056 | 0 | 1 | 26 |
| mbd1b | 0.54902 | 0.127243588 | 0.056 | 0 | 1 | 26 |
| mboat2a | 0.54902 | 0.130873778 | 0.056 | 0 | 1 | 26 |
| megf8 | 0.54902 | 0.1274924 | 0.056 | 0 | 1 | 26 |
| mief1 | 0.54902 | 0.124520425 | 0.056 | 0 | 1 | 26 |
| moxd1 | 0.54902 | 0.129546449 | 0.056 | 0 | 1 | 26 |
| mrpl39 | 0.54902 | 0.125395135 | 0.056 | 0 | 1 | 26 |
| mtmr6 | 0.54902 | 0.166625325 | 0.056 | 0 | 1 | 26 |
| ncaldb | 0.54902 | 0.130754355 | 0.056 | 0 | 1 | 26 |
| nr2c1 | 0.54902 | 0.121020586 | 0.056 | 0 | 1 | 26 |
| nrm | 0.54902 | 0.131770998 | 0.056 | 0 | 1 | 26 |
| numb | 0.54902 | 0.128918872 | 0.056 | 0 | 1 | 26 |
| otud5a | 0.54902 | 0.125804015 | 0.056 | 0 | 1 | 26 |
| p2ry13 | 0.54902 | 0.131770998 | 0.056 | 0 | 1 | 26 |
| pak2b | 0.54902 | 0.12751718 | 0.056 | 0 | 1 | 26 |
| pcif1 | 0.54902 | 0.174972798 | 0.056 | 0 | 1 | 26 |
| pcmt | 0.54902 | 0.127586825 | 0.056 | 0 | 1 | 26 |
| pdcd10b | 0.54902 | 0.171684522 | 0.056 | 0 | 1 | 26 |
| per3 | 0.54902 | 0.121226918 | 0.056 | 0 | 1 | 26 |
| pfdn2 | 0.54902 | 0.12648285 | 0.056 | 0 | 1 | 26 |
| pigx | 0.54902 | 0.126003534 | 0.056 | 0 | 1 | 26 |
| pik3cd | 0.54902 | 0.128770287 | 0.056 | 0 | 1 | 26 |
| pim1 | 0.54902 | 0.171209137 | 0.056 | 0 | 1 | 26 |
| pitpnc1b | 0.54902 | 0.130923097 | 0.056 | 0 | 1 | 26 |
| pkn1a | 0.54902 | 0.131770998 | 0.056 | 0 | 1 | 26 |
| plcb3 | 0.54902 | 0.123832263 | 0.056 | 0 | 1 | 26 |
| plpp1a | 0.54902 | 0.127208992 | 0.056 | 0 | 1 | 26 |
| polr2k | 0.54902 | 0.116241389 | 0.056 | 0 | 1 | 26 |
| ppp1r26 | 0.54902 | 0.131770998 | 0.056 | 0 | 1 | 26 |
| prkar2aa | 0.54902 | 0.125862313 | 0.056 | 0 | 1 | 26 |
| prkchb | 0.54902 | 0.121625952 | 0.056 | 0 | 1 | 26 |
| prkcsh | 0.54902 | 0.129126281 | 0.056 | 0 | 1 | 26 |

| | | | | | | |
|---|---|---|---|---|---|---|
| prpf3 | 0.54902 | 0.127023367 | 0.056 | 0 | 1 | 26 |
| psmc2 | 0.54902 | 0.123977674 | 0.056 | 0 | 1 | 26 |
| ptbp3 | 0.54902 | 0.127793051 | 0.056 | 0 | 1 | 26 |
| ptprr | 0.54902 | 0.131770998 | 0.056 | 0 | 1 | 26 |
| rab3ab | 0.54902 | 0.131770998 | 0.056 | 0 | 1 | 26 |
| rasgef1ba | 0.54902 | 0.124490828 | 0.056 | 0 | 1 | 26 |
| rbm14b | 0.54902 | 0.128959964 | 0.056 | 0 | 1 | 26 |
| rcc2 | 0.54902 | 0.121892561 | 0.056 | 0 | 1 | 26 |
| rffl | 0.54902 | 0.127241515 | 0.056 | 0 | 1 | 26 |
| rfx1b | 0.54902 | 0.128875176 | 0.056 | 0 | 1 | 26 |
| rgs2 | 0.54902 | 0.127744442 | 0.056 | 0 | 1 | 26 |
| rpz4 | 0.54902 | 0.12464141 | 0.056 | 0 | 1 | 26 |
| rtf2 | 0.54902 | 0.127111328 | 0.056 | 0 | 1 | 26 |
| s1pr2 | 0.54902 | 0.125622288 | 0.056 | 0 | 1 | 26 |
| scnm1 | 0.54902 | 0.127845716 | 0.056 | 0 | 1 | 26 |
| sec23a | 0.54902 | 0.131770998 | 0.056 | 0 | 1 | 26 |
| selenot1a | 0.54902 | 0.120850424 | 0.056 | 0 | 1 | 26 |
| selenot1b | 0.54902 | 0.118106289 | 0.056 | 0 | 1 | 26 |
| si:ch211-173p18.3 | 0.54902 | 0.123619503 | 0.056 | 0 | 1 | 26 |
| si:ch211-194m7.4 | 0.54902 | 0.129664748 | 0.056 | 0 | 1 | 26 |
| si:ch211-194m7.5 | 0.54902 | 0.1200493 | 0.056 | 0 | 1 | 26 |
| si:ch211-199g17.2 | 0.54902 | 0.12604343 | 0.056 | 0 | 1 | 26 |
| si:ch211-203d1.3 | 0.54902 | 0.131770998 | 0.056 | 0 | 1 | 26 |
| si:ch211-212c13.10 | 0.54902 | 0.128556958 | 0.056 | 0 | 1 | 26 |
| si:ch211-212k18.7 | 0.54902 | 0.127631704 | 0.056 | 0 | 1 | 26 |
| si:ch211-262e15.1 | 0.54902 | 0.125766347 | 0.056 | 0 | 1 | 26 |
| si:ch73-238c9.1 | 0.54902 | 0.120059954 | 0.056 | 0 | 1 | 26 |
| si:dkey-121a9.3 | 0.54902 | 0.131770998 | 0.056 | 0 | 1 | 26 |
| si:dkey-188i13.11 | 0.54902 | 0.103810543 | 0.056 | 0 | 1 | 26 |
| si:dkey-204f11.64 | 0.54902 | 0.11995426 | 0.056 | 0 | 1 | 26 |
| si:dkey-237i9.1 | 0.54902 | 0.127162626 | 0.056 | 0 | 1 | 26 |
| si:dkey-250d21.1 | 0.54902 | 0.171048792 | 0.056 | 0 | 1 | 26 |
| si:dkey-56i24.1 | 0.54902 | 0.131770998 | 0.056 | 0 | 1 | 26 |
| sirt2 | 0.54902 | 0.125999349 | 0.056 | 0 | 1 | 26 |
| slc30a7 | 0.54902 | 0.125570768 | 0.056 | 0 | 1 | 26 |
| slc35a4 | 0.54902 | 0.131047373 | 0.056 | 0 | 1 | 26 |
| slc48a1a | 0.54902 | 0.131770998 | 0.056 | 0 | 1 | 26 |
| smarca2 | 0.54902 | 0.125216628 | 0.056 | 0 | 1 | 26 |
| smchd1 | 0.54902 | 0.12163175 | 0.056 | 0 | 1 | 26 |
| smim14 | 0.54902 | 0.118876453 | 0.056 | 0 | 1 | 26 |
| sp3b | 0.54902 | 0.124968908 | 0.056 | 0 | 1 | 26 |
| sptbn2 | 0.54902 | 0.123268014 | 0.056 | 0 | 1 | 26 |
| sri | 0.54902 | 0.128166088 | 0.056 | 0 | 1 | 26 |
| srprb | 0.54902 | 0.123051331 | 0.056 | 0 | 1 | 26 |
| srrm2 | 0.54902 | 0.120090297 | 0.056 | 0 | 1 | 26 |
| stat5a | 0.54902 | 0.131093777 | 0.056 | 0 | 1 | 26 |
| stk35 | 0.54902 | 0.128271916 | 0.056 | 0 | 1 | 26 |
| stxbp2 | 0.54902 | 0.129201392 | 0.056 | 0 | 1 | 26 |

| | | | | | | |
|---|---|---|---|---|---|---|
| styx | 0.54902 | 0.121075984 | 0.056 | 0 | 1 | 26 |
| sult3st1 | 0.54902 | 0.131770998 | 0.056 | 0 | 1 | 26 |
| supt20 | 0.54902 | 0.124530741 | 0.056 | 0 | 1 | 26 |
| tab2 | 0.54902 | 0.123089036 | 0.056 | 0 | 1 | 26 |
| tcp11l2 | 0.54902 | 0.169979041 | 0.056 | 0 | 1 | 26 |
| terfa | 0.54902 | 0.122819283 | 0.056 | 0 | 1 | 26 |
| tet3 | 0.54902 | 0.12950224 | 0.056 | 0 | 1 | 26 |
| thap4 | 0.54902 | 0.12724848 | 0.056 | 0 | 1 | 26 |
| tiam2a | 0.54902 | 0.174972798 | 0.056 | 0 | 1 | 26 |
| tm7sf3 | 0.54902 | 0.127049669 | 0.056 | 0 | 1 | 26 |
| tmbim1a | 0.54902 | 0.131770998 | 0.056 | 0 | 1 | 26 |
| tmem131l | 0.54902 | 0.130716904 | 0.056 | 0 | 1 | 26 |
| tom1l2a | 0.54902 | 0.131770998 | 0.056 | 0 | 1 | 26 |
| tomm7 | 0.54902 | 0.12298436 | 0.056 | 0 | 1 | 26 |
| trim25 | 0.54902 | 0.127530664 | 0.056 | 0 | 1 | 26 |
| trim44 | 0.54902 | 0.126518815 | 0.056 | 0 | 1 | 26 |
| trip12 | 0.54902 | 0.124606967 | 0.056 | 0 | 1 | 26 |
| tspan14 | 0.54902 | 0.128531454 | 0.056 | 0 | 1 | 26 |
| ttc37 | 0.54902 | 0.131770998 | 0.056 | 0 | 1 | 26 |
| ubxn6 | 0.54902 | 0.131770998 | 0.056 | 0 | 1 | 26 |
| ugt1a7 | 0.54902 | 0.11108833 | 0.056 | 0 | 1 | 26 |
| usb1 | 0.54902 | 0.126809856 | 0.056 | 0 | 1 | 26 |
| usp37 | 0.54902 | 0.102997043 | 0.056 | 0 | 1 | 26 |
| usp39 | 0.54902 | 0.118846783 | 0.056 | 0 | 1 | 26 |
| usp7 | 0.54902 | 0.125185146 | 0.056 | 0 | 1 | 26 |
| vhll | 0.54902 | 0.173153118 | 0.056 | 0 | 1 | 26 |
| wdr82 | 0.54902 | 0.12298679 | 0.056 | 0 | 1 | 26 |
| wu:fi37e09 | 0.54902 | 0.172452366 | 0.056 | 0 | 1 | 26 |
| xrcc4 | 0.54902 | 0.127249017 | 0.056 | 0 | 1 | 26 |
| zbtb11 | 0.54902 | 0.131770998 | 0.056 | 0 | 1 | 26 |
| zc4h2 | 0.54902 | 0.120319174 | 0.056 | 0 | 1 | 26 |
| zgc:162964 | 0.54902 | 0.112448444 | 0.056 | 0 | 1 | 26 |
| zgc:92606 | 0.54902 | 0.106127275 | 0.056 | 0 | 1 | 26 |
| znf770 | 0.54902 | 0.122967619 | 0.056 | 0 | 1 | 26 |
| BX323555.3 | 0.54902 | 0.180312144 | 0.056 | 0 | 1 | 26 |
| BX511132.1 | 0.54902 | 0.223333834 | 0.056 | 0 | 1 | 26 |
| BX649485.1 | 0.54902 | 0.217753382 | 0.056 | 0 | 1 | 26 |
| CABZ01072614.1 | 0.54902 | 0.197009891 | 0.056 | 0 | 1 | 26 |
| CABZ01088367.1 | 0.54902 | 0.214424013 | 0.056 | 0 | 1 | 26 |
| CABZ01110881.1 | 0.54902 | 0.187547483 | 0.056 | 0 | 1 | 26 |
| CABZ01113192.1 | 0.54902 | 0.214514592 | 0.056 | 0 | 1 | 26 |
| CR293507.1 | 0.54902 | 0.216316142 | 0.056 | 0 | 1 | 26 |
| CR936442.1 | 0.54902 | 0.222475528 | 0.056 | 0 | 1 | 26 |
| CU856539.2 | 0.54902 | 0.233321707 | 0.056 | 0 | 1 | 26 |
| FQ377903.3 | 0.54902 | 0.222174553 | 0.056 | 0 | 1 | 26 |
| GNAZ | 0.54902 | 0.217968157 | 0.056 | 0 | 1 | 26 |
| ISCU (1 of many) | 0.54902 | 0.185284907 | 0.056 | 0 | 1 | 26 |
| LO018176.1 | 0.54902 | 0.232695742 | 0.056 | 0 | 1 | 26 |

| | | | | | | |
|---|---|---|---|---|---|---|
| LOC100329806 | 0.54902 | 0.19089453 | 0.056 | 0 | 1 | 26 |
| LOC101884921 | 0.54902 | 0.217627056 | 0.056 | 0 | 1 | 26 |
| LOC110437842 | 0.54902 | 0.223333834 | 0.056 | 0 | 1 | 26 |
| LOC796453 | 0.54902 | 0.216308118 | 0.056 | 0 | 1 | 26 |
| RSBN1 | 0.54902 | 0.234517178 | 0.056 | 0 | 1 | 26 |
| TENT5A | 0.54902 | 0.189012131 | 0.056 | 0 | 1 | 26 |
| USP34 | 0.54902 | 0.214295939 | 0.056 | 0 | 1 | 26 |
| XLOC-008185 | 0.54902 | 0.218206971 | 0.056 | 0 | 1 | 26 |
| XLOC-012620 | 0.54902 | 0.218206971 | 0.056 | 0 | 1 | 26 |
| XLOC-042229 | 0.54902 | 0.278763398 | 0.056 | 0 | 1 | 26 |
| abca2 | 0.54902 | 0.21770088 | 0.056 | 0 | 1 | 26 |
| acaa1 | 0.54902 | 0.229773799 | 0.056 | 0 | 1 | 26 |
| acaa2 | 0.54902 | 0.182940075 | 0.056 | 0 | 1 | 26 |
| aco2 | 0.54902 | 0.217585724 | 0.056 | 0 | 1 | 26 |
| actr1 | 0.54902 | 0.233535377 | 0.056 | 0 | 1 | 26 |
| agap2 | 0.54902 | 0.217942147 | 0.056 | 0 | 1 | 26 |
| agps | 0.54902 | 0.215157441 | 0.056 | 0 | 1 | 26 |
| aldh7a1 | 0.54902 | 0.166771998 | 0.056 | 0 | 1 | 26 |
| amdhd2 | 0.54902 | 0.189401253 | 0.056 | 0 | 1 | 26 |
| ankmy2a | 0.54902 | 0.215507254 | 0.056 | 0 | 1 | 26 |
| ano10a | 0.54902 | 0.233810962 | 0.056 | 0 | 1 | 26 |
| ap1s1 | 0.54902 | 0.186480025 | 0.056 | 0 | 1 | 26 |
| arcn1a | 0.54902 | 0.216800013 | 0.056 | 0 | 1 | 26 |
| arf4a | 0.54902 | 0.216989482 | 0.056 | 0 | 1 | 26 |
| arhgap15 | 0.54902 | 0.217657306 | 0.056 | 0 | 1 | 26 |
| arhgap25 | 0.54902 | 0.216885867 | 0.056 | 0 | 1 | 26 |
| arid4a | 0.54902 | 0.233595557 | 0.056 | 0 | 1 | 26 |
| arpp19b | 0.54902 | 0.178681907 | 0.056 | 0 | 1 | 26 |
| asap2b | 0.54902 | 0.217351977 | 0.056 | 0 | 1 | 26 |
| asna1 | 0.54902 | 0.233113871 | 0.056 | 0 | 1 | 26 |
| atox1 | 0.54902 | 0.231732259 | 0.056 | 0 | 1 | 26 |
| atp11c | 0.54902 | 0.217417699 | 0.056 | 0 | 1 | 26 |
| atp2a3 | 0.54902 | 0.217471817 | 0.056 | 0 | 1 | 26 |
| atp2b1a | 0.54902 | 0.222448683 | 0.056 | 0 | 1 | 26 |
| atp6v0d1 | 0.54902 | 0.216122093 | 0.056 | 0 | 1 | 26 |
| bahcc1b | 0.54902 | 0.223051079 | 0.056 | 0 | 1 | 26 |
| baz2a | 0.54902 | 0.216682676 | 0.056 | 0 | 1 | 26 |
| bbip1 | 0.54902 | 0.1631906 | 0.056 | 0 | 1 | 26 |
| bbox1 | 0.54902 | 0.234517178 | 0.056 | 0 | 1 | 26 |
| bbs1 | 0.54902 | 0.223333834 | 0.056 | 0 | 1 | 26 |
| bmi1b | 0.54902 | 0.231517566 | 0.056 | 0 | 1 | 26 |
| bmt2 | 0.54902 | 0.217440893 | 0.056 | 0 | 1 | 26 |
| brd4 | 0.54902 | 0.217936128 | 0.056 | 0 | 1 | 26 |
| brpf3a | 0.54902 | 0.215622642 | 0.056 | 0 | 1 | 26 |
| bysl | 0.54902 | 0.214090716 | 0.056 | 0 | 1 | 26 |
| cab39l1 | 0.54902 | 0.216171009 | 0.056 | 0 | 1 | 26 |
| calhm6 | 0.54902 | 0.217911341 | 0.056 | 0 | 1 | 26 |
| caskb | 0.54902 | 0.217215092 | 0.056 | 0 | 1 | 26 |

| | | | | | | |
|---|---|---|---|---|---|---|
| cbx8a | 0.54902 | 0.232853452 | 0.056 | 0 | 1 | 26 |
| ccnd1 | 0.54902 | 0.210372681 | 0.056 | 0 | 1 | 26 |
| ccnd2a | 0.54902 | 0.216617638 | 0.056 | 0 | 1 | 26 |
| ccnd2b | 0.54902 | 0.216726919 | 0.056 | 0 | 1 | 26 |
| cd8a | 0.54902 | 0.19089453 | 0.056 | 0 | 1 | 26 |
| cecr2 | 0.54902 | 0.221803338 | 0.056 | 0 | 1 | 26 |
| cers1 | 0.54902 | 0.217841713 | 0.056 | 0 | 1 | 26 |
| cers5 | 0.54902 | 0.222501654 | 0.056 | 0 | 1 | 26 |
| chmp4ba | 0.54902 | 0.180254097 | 0.056 | 0 | 1 | 26 |
| chrac1 | 0.54902 | 0.214247432 | 0.056 | 0 | 1 | 26 |
| chtopa | 0.54902 | 0.221539753 | 0.056 | 0 | 1 | 26 |
| cisd2 | 0.54902 | 0.216547645 | 0.056 | 0 | 1 | 26 |
| cltca | 0.54902 | 0.279368915 | 0.056 | 0 | 1 | 26 |
| cmc1 | 0.54902 | 0.186574878 | 0.056 | 0 | 1 | 26 |
| cnbpb | 0.54902 | 0.179733934 | 0.056 | 0 | 1 | 26 |
| cnksr1 | 0.54902 | 0.215755376 | 0.056 | 0 | 1 | 26 |
| cnot6b | 0.54902 | 0.217942147 | 0.056 | 0 | 1 | 26 |
| cnr2 | 0.54902 | 0.217942147 | 0.056 | 0 | 1 | 26 |
| coa5 | 0.54902 | 0.18044463 | 0.056 | 0 | 1 | 26 |
| copb1 | 0.54902 | 0.216932468 | 0.056 | 0 | 1 | 26 |
| cops8 | 0.54902 | 0.186091529 | 0.056 | 0 | 1 | 26 |
| coq6 | 0.54902 | 0.215762283 | 0.056 | 0 | 1 | 26 |
| cpt1ab | 0.54902 | 0.234517178 | 0.056 | 0 | 1 | 26 |
| crcp | 0.54902 | 0.179333285 | 0.056 | 0 | 1 | 26 |
| csnk1da | 0.54902 | 0.179878562 | 0.056 | 0 | 1 | 26 |
| csnk2a4 | 0.54902 | 0.218206971 | 0.056 | 0 | 1 | 26 |
| csrnp1b | 0.54902 | 0.222020423 | 0.056 | 0 | 1 | 26 |
| ctu2 | 0.54902 | 0.20663084 | 0.056 | 0 | 1 | 26 |
| cwc22 | 0.54902 | 0.188821978 | 0.056 | 0 | 1 | 26 |
| cyp3c1 | 0.54902 | 0.217407936 | 0.056 | 0 | 1 | 26 |
| ddc | 0.54902 | 0.19089453 | 0.056 | 0 | 1 | 26 |
| ddx23 | 0.54902 | 0.278480322 | 0.056 | 0 | 1 | 26 |
| dedd1 | 0.54902 | 0.233277391 | 0.056 | 0 | 1 | 26 |
| dennd2da | 0.54902 | 0.220290716 | 0.056 | 0 | 1 | 26 |
| dennd4a | 0.54902 | 0.217452775 | 0.056 | 0 | 1 | 26 |
| dldh | 0.54902 | 0.179551641 | 0.056 | 0 | 1 | 26 |
| dnajb11 | 0.54902 | 0.216902496 | 0.056 | 0 | 1 | 26 |
| dnajc10 | 0.54902 | 0.217120492 | 0.056 | 0 | 1 | 26 |
| dnajc18 | 0.54902 | 0.210952962 | 0.056 | 0 | 1 | 26 |
| dnajc3b | 0.54902 | 0.217207176 | 0.056 | 0 | 1 | 26 |
| dnajc7 | 0.54902 | 0.215486514 | 0.056 | 0 | 1 | 26 |
| dock8 | 0.54902 | 0.217255868 | 0.056 | 0 | 1 | 26 |
| dpy30 | 0.54902 | 0.224191705 | 0.056 | 0 | 1 | 26 |
| dr1 | 0.54902 | 0.221341062 | 0.056 | 0 | 1 | 26 |
| dusp1 | 0.54902 | 0.234221418 | 0.056 | 0 | 1 | 26 |
| edrf1 | 0.54902 | 0.216221698 | 0.056 | 0 | 1 | 26 |
| egr3 | 0.54902 | 0.233770144 | 0.056 | 0 | 1 | 26 |
| ehd4 | 0.54902 | 0.217744473 | 0.056 | 0 | 1 | 26 |

| | | | | | | |
|---|---|---|---|---|---|---|
| eif4g2b | 0.54902 | 0.216622454 | 0.056 | 0 | 1 | 26 |
| eif6 | 0.54902 | 0.214785645 | 0.056 | 0 | 1 | 26 |
| erfl3 | 0.54902 | 0.217111057 | 0.056 | 0 | 1 | 26 |
| ergic2 | 0.54902 | 0.145540744 | 0.056 | 0 | 1 | 26 |
| fam192a | 0.54902 | 0.185347884 | 0.056 | 0 | 1 | 26 |
| fam49bb | 0.54902 | 0.217668693 | 0.056 | 0 | 1 | 26 |
| fbxo11a | 0.54902 | 0.234033353 | 0.056 | 0 | 1 | 26 |
| fbxw7 | 0.54902 | 0.21648505 | 0.056 | 0 | 1 | 26 |
| fdx2 | 0.54902 | 0.217765045 | 0.056 | 0 | 1 | 26 |
| fli1b | 0.54902 | 0.222915115 | 0.056 | 0 | 1 | 26 |
| fmr1 | 0.54902 | 0.233095686 | 0.056 | 0 | 1 | 26 |
| fndc3a | 0.54902 | 0.217029912 | 0.056 | 0 | 1 | 26 |
| foxj3 | 0.54902 | 0.216994693 | 0.056 | 0 | 1 | 26 |
| ftr93 | 0.54902 | 0.223333834 | 0.056 | 0 | 1 | 26 |
| fus | 0.54902 | 0.223333834 | 0.056 | 0 | 1 | 26 |
| gb:bc139872 | 0.54902 | 0.189504783 | 0.056 | 0 | 1 | 26 |
| gbe1b | 0.54902 | 0.233242406 | 0.056 | 0 | 1 | 26 |
| gcat | 0.54902 | 0.214643853 | 0.056 | 0 | 1 | 26 |
| glod4 | 0.54902 | 0.232206556 | 0.056 | 0 | 1 | 26 |
| gna13b | 0.54902 | 0.217722432 | 0.056 | 0 | 1 | 26 |
| gnai2a | 0.54902 | 0.217479331 | 0.056 | 0 | 1 | 26 |
| gnb1b | 0.54902 | 0.232792817 | 0.056 | 0 | 1 | 26 |
| gnb4b | 0.54902 | 0.218206971 | 0.056 | 0 | 1 | 26 |
| gpcpd1 | 0.54902 | 0.218918011 | 0.056 | 0 | 1 | 26 |
| gpn2 | 0.54902 | 0.214898524 | 0.056 | 0 | 1 | 26 |
| gpr141 | 0.54902 | 0.222692497 | 0.056 | 0 | 1 | 26 |
| gpr155a | 0.54902 | 0.233115574 | 0.056 | 0 | 1 | 26 |
| gps2 | 0.54902 | 0.221839473 | 0.056 | 0 | 1 | 26 |
| gstt1a | 0.54902 | 0.221957825 | 0.056 | 0 | 1 | 26 |
| gtf2b | 0.54902 | 0.185931667 | 0.056 | 0 | 1 | 26 |
| h6pd | 0.54902 | 0.218206971 | 0.056 | 0 | 1 | 26 |
| haao | 0.54902 | 0.23303527 | 0.056 | 0 | 1 | 26 |
| hadhb | 0.54902 | 0.215104994 | 0.056 | 0 | 1 | 26 |
| hck | 0.54902 | 0.217368728 | 0.056 | 0 | 1 | 26 |
| hdhd2 | 0.54902 | 0.181707892 | 0.056 | 0 | 1 | 26 |
| heatr5a | 0.54902 | 0.232496886 | 0.056 | 0 | 1 | 26 |
| higd2a | 0.54902 | 0.186388714 | 0.056 | 0 | 1 | 26 |
| hipk3b | 0.54902 | 0.217440728 | 0.056 | 0 | 1 | 26 |
| hnrnph1l | 0.54902 | 0.221669807 | 0.056 | 0 | 1 | 26 |
| hnrpkl | 0.54902 | 0.221603589 | 0.056 | 0 | 1 | 26 |
| hspa4b | 0.54902 | 0.18358169 | 0.056 | 0 | 1 | 26 |
| hsph1 | 0.54902 | 0.218206971 | 0.056 | 0 | 1 | 26 |
| ifngr1l | 0.54902 | 0.231193057 | 0.056 | 0 | 1 | 26 |
| il12rb2l | 0.54902 | 0.217218406 | 0.056 | 0 | 1 | 26 |
| ing5a | 0.54902 | 0.233174234 | 0.056 | 0 | 1 | 26 |
| ip6k2b | 0.54902 | 0.232657776 | 0.056 | 0 | 1 | 26 |
| ipcef1 | 0.54902 | 0.217517579 | 0.056 | 0 | 1 | 26 |
| iqgap1 | 0.54902 | 0.217299707 | 0.056 | 0 | 1 | 26 |

| | | | | | | |
|---|---|---|---|---|---|---|
| irf2bpl | 0.54902 | 0.218725668 | 0.056 | 0 | 1 | 26 |
| ist1 | 0.54902 | 0.187723606 | 0.056 | 0 | 1 | 26 |
| itcha | 0.54902 | 0.223333834 | 0.056 | 0 | 1 | 26 |
| josd2 | 0.54902 | 0.215547761 | 0.056 | 0 | 1 | 26 |
| kansl1a | 0.54902 | 0.221972197 | 0.056 | 0 | 1 | 26 |
| kcnk1a | 0.54902 | 0.234517178 | 0.056 | 0 | 1 | 26 |
| kctd5a | 0.54902 | 0.184924515 | 0.056 | 0 | 1 | 26 |
| kdelr2b | 0.54902 | 0.220898313 | 0.056 | 0 | 1 | 26 |
| kdm2ba | 0.54902 | 0.234993744 | 0.056 | 0 | 1 | 26 |
| klhl6 | 0.54902 | 0.223333834 | 0.056 | 0 | 1 | 26 |
| kmt2ca | 0.54902 | 0.217624006 | 0.056 | 0 | 1 | 26 |
| kpnb3 | 0.54902 | 0.182346313 | 0.056 | 0 | 1 | 26 |
| krt18b | 0.54902 | 0.187725027 | 0.056 | 0 | 1 | 26 |
| ksr1b | 0.54902 | 0.216057511 | 0.056 | 0 | 1 | 26 |
| lclat1 | 0.54902 | 0.215411257 | 0.056 | 0 | 1 | 26 |
| lgmn | 0.54902 | 0.179357559 | 0.056 | 0 | 1 | 26 |
| lmod1a | 0.54902 | 0.217942147 | 0.056 | 0 | 1 | 26 |
| lnpep | 0.54902 | 0.217363674 | 0.056 | 0 | 1 | 26 |
| lrch3 | 0.54902 | 0.215067271 | 0.056 | 0 | 1 | 26 |
| lrrc20 | 0.54902 | 0.232708741 | 0.056 | 0 | 1 | 26 |
| lrrc47 | 0.54902 | 0.217400284 | 0.056 | 0 | 1 | 26 |
| manf | 0.54902 | 0.177287432 | 0.056 | 0 | 1 | 26 |
| map4k3b | 0.54902 | 0.218206971 | 0.056 | 0 | 1 | 26 |
| matr3l1.1 | 0.54902 | 0.216144039 | 0.056 | 0 | 1 | 26 |
| med12 | 0.54902 | 0.217375229 | 0.056 | 0 | 1 | 26 |
| med8 | 0.54902 | 0.186267592 | 0.056 | 0 | 1 | 26 |
| mia3 | 0.54902 | 0.214740434 | 0.056 | 0 | 1 | 26 |
| mid1ip1l | 0.54902 | 0.228906571 | 0.056 | 0 | 1 | 26 |
| mif | 0.54902 | 0.181835526 | 0.056 | 0 | 1 | 26 |
| mrps30 | 0.54902 | 0.184233158 | 0.056 | 0 | 1 | 26 |
| mrtfab | 0.54902 | 0.217263994 | 0.056 | 0 | 1 | 26 |
| mthfd2 | 0.54902 | 0.232891638 | 0.056 | 0 | 1 | 26 |
| mtmr4 | 0.54902 | 0.215951029 | 0.056 | 0 | 1 | 26 |
| mtmr8 | 0.54902 | 0.222695013 | 0.056 | 0 | 1 | 26 |
| mycb | 0.54902 | 0.229770841 | 0.056 | 0 | 1 | 26 |
| mylk5 | 0.54902 | 0.218206971 | 0.056 | 0 | 1 | 26 |
| nap1l4b | 0.54902 | 0.215938253 | 0.056 | 0 | 1 | 26 |
| nbeal2 | 0.54902 | 0.222446834 | 0.056 | 0 | 1 | 26 |
| ndfip1 | 0.54902 | 0.184893322 | 0.056 | 0 | 1 | 26 |
| ndfip1l | 0.54902 | 0.262690591 | 0.056 | 0 | 1 | 26 |
| ndfip2 | 0.54902 | 0.222610305 | 0.056 | 0 | 1 | 26 |
| ndrg1a | 0.54902 | 0.23296666 | 0.056 | 0 | 1 | 26 |
| ndufa10 | 0.54902 | 0.184016788 | 0.056 | 0 | 1 | 26 |
| ndufa7 | 0.54902 | 0.23143293 | 0.056 | 0 | 1 | 26 |
| ndufv1 | 0.54902 | 0.186502383 | 0.056 | 0 | 1 | 26 |
| nectin4b | 0.54902 | 0.217942147 | 0.056 | 0 | 1 | 26 |
| nedd8 | 0.54902 | 0.185026664 | 0.056 | 0 | 1 | 26 |
| nedd9 | 0.54902 | 0.223333834 | 0.056 | 0 | 1 | 26 |

| | | | | | | |
|---|---|---|---|---|---|---|
| nktr | 0.54902 | 0.231938784 | 0.056 | 0 | 1 | 26 |
| nlk2 | 0.54902 | 0.217701154 | 0.056 | 0 | 1 | 26 |
| nme4 | 0.54902 | 0.177671347 | 0.056 | 0 | 1 | 26 |
| nop53 | 0.54902 | 0.181885401 | 0.056 | 0 | 1 | 26 |
| nphp4 | 0.54902 | 0.218206971 | 0.056 | 0 | 1 | 26 |
| npl | 0.54902 | 0.221343831 | 0.056 | 0 | 1 | 26 |
| nr1d2a | 0.54902 | 0.221337452 | 0.056 | 0 | 1 | 26 |
| nr3c1 | 0.54902 | 0.215986133 | 0.056 | 0 | 1 | 26 |
| nsd1a | 0.54902 | 0.216028314 | 0.056 | 0 | 1 | 26 |
| nucb1 | 0.54902 | 0.189468563 | 0.056 | 0 | 1 | 26 |
| nupr1a | 0.54902 | 0.179711423 | 0.056 | 0 | 1 | 26 |
| otud4 | 0.54902 | 0.141187662 | 0.056 | 0 | 1 | 26 |
| oxr1a | 0.54902 | 0.213965965 | 0.056 | 0 | 1 | 26 |
| pcbp2 | 0.54902 | 0.218182638 | 0.056 | 0 | 1 | 26 |
| pcyox1 | 0.54902 | 0.234230736 | 0.056 | 0 | 1 | 26 |
| pdap1b | 0.54902 | 0.222144011 | 0.056 | 0 | 1 | 26 |
| pfkpb | 0.54902 | 0.262478962 | 0.056 | 0 | 1 | 26 |
| pgm1 | 0.54902 | 0.141099093 | 0.056 | 0 | 1 | 26 |
| phf8 | 0.54902 | 0.233448334 | 0.056 | 0 | 1 | 26 |
| pigu | 0.54902 | 0.217942147 | 0.056 | 0 | 1 | 26 |
| pik3r2 | 0.54902 | 0.216497954 | 0.056 | 0 | 1 | 26 |
| pim2 | 0.54902 | 0.233248699 | 0.056 | 0 | 1 | 26 |
| pip5k1aa | 0.54902 | 0.217269776 | 0.056 | 0 | 1 | 26 |
| pla2g4aa | 0.54902 | 0.263401744 | 0.056 | 0 | 1 | 26 |
| plcg1 | 0.54902 | 0.233489971 | 0.056 | 0 | 1 | 26 |
| plk3 | 0.54902 | 0.213930109 | 0.056 | 0 | 1 | 26 |
| plscr3b | 0.54902 | 0.18544018 | 0.056 | 0 | 1 | 26 |
| pnkd | 0.54902 | 0.217942147 | 0.056 | 0 | 1 | 26 |
| ppan | 0.54902 | 0.221278777 | 0.056 | 0 | 1 | 26 |
| ppid | 0.54902 | 0.215633122 | 0.056 | 0 | 1 | 26 |
| ppm1aa | 0.54902 | 0.216527093 | 0.056 | 0 | 1 | 26 |
| ppm1g | 0.54902 | 0.262496383 | 0.056 | 0 | 1 | 26 |
| ppp2r5d | 0.54902 | 0.263226788 | 0.056 | 0 | 1 | 26 |
| ppp3cca | 0.54902 | 0.215593064 | 0.056 | 0 | 1 | 26 |
| ppt1 | 0.54902 | 0.21568825 | 0.056 | 0 | 1 | 26 |
| pqlc3 | 0.54902 | 0.217256163 | 0.056 | 0 | 1 | 26 |
| prkacaa | 0.54902 | 0.216146034 | 0.056 | 0 | 1 | 26 |
| prkn | 0.54902 | 0.165120251 | 0.056 | 0 | 1 | 26 |
| prkx | 0.54902 | 0.217312772 | 0.056 | 0 | 1 | 26 |
| prpf8 | 0.54902 | 0.263202879 | 0.056 | 0 | 1 | 26 |
| psma1 | 0.54902 | 0.215286727 | 0.056 | 0 | 1 | 26 |
| psmd13 | 0.54902 | 0.187041036 | 0.056 | 0 | 1 | 26 |
| psme3 | 0.54902 | 0.215999537 | 0.056 | 0 | 1 | 26 |
| ptbp1a | 0.54902 | 0.217312772 | 0.056 | 0 | 1 | 26 |
| puf60a | 0.54902 | 0.232893232 | 0.056 | 0 | 1 | 26 |
| pxmp2 | 0.54902 | 0.218206971 | 0.056 | 0 | 1 | 26 |
| rab3da | 0.54902 | 0.234517178 | 0.056 | 0 | 1 | 26 |
| rab41 | 0.54902 | 0.216449637 | 0.056 | 0 | 1 | 26 |

| | | | | | | |
|---|---|---|---|---|---|---|
| rad23aa | 0.54902 | 0.217942147 | 0.056 | 0 | 1 | 26 |
| rad23b | 0.54902 | 0.185675907 | 0.056 | 0 | 1 | 26 |
| raf1a | 0.54902 | 0.222155767 | 0.056 | 0 | 1 | 26 |
| rapgef1b | 0.54902 | 0.212907722 | 0.056 | 0 | 1 | 26 |
| rassf2b | 0.54902 | 0.233521038 | 0.056 | 0 | 1 | 26 |
| rassf8a | 0.54902 | 0.19089453 | 0.056 | 0 | 1 | 26 |
| rbm27 | 0.54902 | 0.222323874 | 0.056 | 0 | 1 | 26 |
| rbm33a | 0.54902 | 0.214786836 | 0.056 | 0 | 1 | 26 |
| rdh1 | 0.54902 | 0.107706504 | 0.056 | 0 | 1 | 26 |
| relb | 0.54902 | 0.222390237 | 0.056 | 0 | 1 | 26 |
| rell1 | 0.54902 | 0.182744235 | 0.056 | 0 | 1 | 26 |
| rgs19 | 0.54902 | 0.217809568 | 0.056 | 0 | 1 | 26 |
| rmdn1 | 0.54902 | 0.212489821 | 0.056 | 0 | 1 | 26 |
| rnf111 | 0.54902 | 0.223333834 | 0.056 | 0 | 1 | 26 |
| rnf144b | 0.54902 | 0.234011846 | 0.056 | 0 | 1 | 26 |
| rnf145a | 0.54902 | 0.214181264 | 0.056 | 0 | 1 | 26 |
| rnf146 | 0.54902 | 0.187353878 | 0.056 | 0 | 1 | 26 |
| rnf170 | 0.54902 | 0.185827215 | 0.056 | 0 | 1 | 26 |
| rnf34a | 0.54902 | 0.175616926 | 0.056 | 0 | 1 | 26 |
| rprd1a | 0.54902 | 0.212554004 | 0.056 | 0 | 1 | 26 |
| rps6ka5 | 0.54902 | 0.217628405 | 0.056 | 0 | 1 | 26 |
| rps6kal | 0.54902 | 0.213854304 | 0.056 | 0 | 1 | 26 |
| rps6kb1b | 0.54902 | 0.179315796 | 0.056 | 0 | 1 | 26 |
| rrp9 | 0.54902 | 0.221037446 | 0.056 | 0 | 1 | 26 |
| rsu1 | 0.54902 | 0.221680057 | 0.056 | 0 | 1 | 26 |
| runx3 | 0.54902 | 0.217158271 | 0.056 | 0 | 1 | 26 |
| rxrbb | 0.54902 | 0.213541119 | 0.056 | 0 | 1 | 26 |
| sars | 0.54902 | 0.232409194 | 0.056 | 0 | 1 | 26 |
| sb:cb81 | 0.54902 | 0.179084738 | 0.056 | 0 | 1 | 26 |
| scly | 0.54902 | 0.218206971 | 0.056 | 0 | 1 | 26 |
| scpep1 | 0.54902 | 0.222487326 | 0.056 | 0 | 1 | 26 |
| sdhaf2 | 0.54902 | 0.18492469 | 0.056 | 0 | 1 | 26 |
| sdhaf4 | 0.54902 | 0.181316139 | 0.056 | 0 | 1 | 26 |
| sdhc | 0.54902 | 0.220218938 | 0.056 | 0 | 1 | 26 |
| sec24d | 0.54902 | 0.233670427 | 0.056 | 0 | 1 | 26 |
| sec63 | 0.54902 | 0.232836739 | 0.056 | 0 | 1 | 26 |
| selplg | 0.54902 | 0.21765361 | 0.056 | 0 | 1 | 26 |
| seta | 0.54902 | 0.184592267 | 0.056 | 0 | 1 | 26 |
| sfpq | 0.54902 | 0.220376909 | 0.056 | 0 | 1 | 26 |
| si:ch1073-392o20.2 | 0.54902 | 0.127555223 | 0.056 | 0 | 1 | 26 |
| si:ch1073-475a24.1 | 0.54902 | 0.131088603 | 0.056 | 0 | 1 | 26 |
| si:ch211-121a2.2 | 0.54902 | 0.230796673 | 0.056 | 0 | 1 | 26 |
| si:ch211-13c6.2 | 0.54902 | 0.215079968 | 0.056 | 0 | 1 | 26 |
| si:ch211-148l7.4 | 0.54902 | 0.217049157 | 0.056 | 0 | 1 | 26 |
| si:ch211-160o17.4 | 0.54902 | 0.216661155 | 0.056 | 0 | 1 | 26 |
| si:ch211-196l7.4 | 0.54902 | 0.173296001 | 0.056 | 0 | 1 | 26 |
| si:ch211-196p9.1 | 0.54902 | 0.234517178 | 0.056 | 0 | 1 | 26 |
| si:ch211-212k18.4 | 0.54902 | 0.215286836 | 0.056 | 0 | 1 | 26 |

| | | | | | | |
|---|---|---|---|---|---|---|
| si:ch211-217k17.7 | 0.54902 | 0.183437616 | 0.056 | 0 | 1 | 26 |
| si:ch211-241f5.3 | 0.54902 | 0.217297287 | 0.056 | 0 | 1 | 26 |
| si:ch211-281l24.3 | 0.54902 | 0.217718291 | 0.056 | 0 | 1 | 26 |
| si:ch211-286b5.5 | 0.54902 | 0.163836271 | 0.056 | 0 | 1 | 26 |
| si:ch211-288g17.3 | 0.54902 | 0.18092396 | 0.056 | 0 | 1 | 26 |
| si:ch211-39i2.2 | 0.54902 | 0.222786534 | 0.056 | 0 | 1 | 26 |
| si:ch211-86h15.1 | 0.54902 | 0.218206971 | 0.056 | 0 | 1 | 26 |
| si:ch211-87m7.2 | 0.54902 | 0.217687463 | 0.056 | 0 | 1 | 26 |
| si:ch73-138e16.6 | 0.54902 | 0.218206971 | 0.056 | 0 | 1 | 26 |
| si:ch73-147o17.1 | 0.54902 | 0.185629681 | 0.056 | 0 | 1 | 26 |
| si:ch73-236c18.8 | 0.54902 | 0.234517178 | 0.056 | 0 | 1 | 26 |
| si:ch73-359m17.5 | 0.54902 | 0.225736722 | 0.056 | 0 | 1 | 26 |
| si:ch73-42p12.2 | 0.54902 | 0.177918718 | 0.056 | 0 | 1 | 26 |
| si:ch73-60h1.1 | 0.54902 | 0.217942147 | 0.056 | 0 | 1 | 26 |
| si:dkey-180p18.9 | 0.54902 | 0.188091083 | 0.056 | 0 | 1 | 26 |
| si:dkey-33c9.6 | 0.54902 | 0.218206971 | 0.056 | 0 | 1 | 26 |
| si:dkey-42i9.4 | 0.54902 | 0.2151935 | 0.056 | 0 | 1 | 26 |
| si:dkeyp-82a1.6 | 0.54902 | 0.21760487 | 0.056 | 0 | 1 | 26 |
| si:dkeyp-87d8.8 | 0.54902 | 0.217942147 | 0.056 | 0 | 1 | 26 |
| slc16a3 | 0.54902 | 0.234110424 | 0.056 | 0 | 1 | 26 |
| slc25a51b | 0.54902 | 0.176794714 | 0.056 | 0 | 1 | 26 |
| slc25a55a | 0.54902 | 0.216353145 | 0.056 | 0 | 1 | 26 |
| slc30a9 | 0.54902 | 0.215993471 | 0.056 | 0 | 1 | 26 |
| slc38a5a | 0.54902 | 0.217606782 | 0.056 | 0 | 1 | 26 |
| smarce1 | 0.54902 | 0.216062955 | 0.056 | 0 | 1 | 26 |
| smim1 | 0.54902 | 0.205158882 | 0.056 | 0 | 1 | 26 |
| snrpa | 0.54902 | 0.183993206 | 0.056 | 0 | 1 | 26 |
| snx16 | 0.54902 | 0.232474035 | 0.056 | 0 | 1 | 26 |
| snx27a | 0.54902 | 0.1283788 | 0.056 | 0 | 1 | 26 |
| socs3b | 0.54902 | 0.220977112 | 0.056 | 0 | 1 | 26 |
| sptlc3 | 0.54902 | 0.19089453 | 0.056 | 0 | 1 | 26 |
| srp54 | 0.54902 | 0.232834374 | 0.056 | 0 | 1 | 26 |
| ssuh2.3 | 0.54902 | 0.233386199 | 0.056 | 0 | 1 | 26 |
| st3gal3a | 0.54902 | 0.176724296 | 0.056 | 0 | 1 | 26 |
| stk40 | 0.54902 | 0.232594753 | 0.056 | 0 | 1 | 26 |
| stx11a | 0.54902 | 0.222148808 | 0.056 | 0 | 1 | 26 |
| sv2 | 0.54902 | 0.215575069 | 0.056 | 0 | 1 | 26 |
| tars | 0.54902 | 0.215861108 | 0.056 | 0 | 1 | 26 |
| tbl1xr1a | 0.54902 | 0.216388883 | 0.056 | 0 | 1 | 26 |
| tec | 0.54902 | 0.234112016 | 0.056 | 0 | 1 | 26 |
| tefa | 0.54902 | 0.184922491 | 0.056 | 0 | 1 | 26 |
| tfe3b | 0.54902 | 0.223333834 | 0.056 | 0 | 1 | 26 |
| thoc1 | 0.54902 | 0.21592963 | 0.056 | 0 | 1 | 26 |
| thrab | 0.54902 | 0.217942147 | 0.056 | 0 | 1 | 26 |
| tm9sf4 | 0.54902 | 0.216144277 | 0.056 | 0 | 1 | 26 |
| tmed5 | 0.54902 | 0.222669432 | 0.056 | 0 | 1 | 26 |
| tmem267 | 0.54902 | 0.217942147 | 0.056 | 0 | 1 | 26 |
| tmpob | 0.54902 | 0.187791879 | 0.056 | 0 | 1 | 26 |

| | | | | | | |
|---|---|---|---|---|---|---|
| tnfaip3 | 0.54902 | 0.234517178 | 0.056 | 0 | 1 | 26 |
| tnfaip8l2a | 0.54902 | 0.217942147 | 0.056 | 0 | 1 | 26 |
| top2a | 0.54902 | 0.116262311 | 0.056 | 0 | 1 | 26 |
| tph1b | 0.54902 | 0.217942147 | 0.056 | 0 | 1 | 26 |
| tpra | 0.54902 | 0.184701606 | 0.056 | 0 | 1 | 26 |
| trim8b | 0.54902 | 0.217176069 | 0.056 | 0 | 1 | 26 |
| trpc4apb | 0.54902 | 0.201987538 | 0.056 | 0 | 1 | 26 |
| ttyh3a | 0.54902 | 0.221652639 | 0.056 | 0 | 1 | 26 |
| ubap2b | 0.54902 | 0.215693209 | 0.056 | 0 | 1 | 26 |
| ube2e1 | 0.54902 | 0.232285151 | 0.056 | 0 | 1 | 26 |
| ube2g1a | 0.54902 | 0.215981998 | 0.056 | 0 | 1 | 26 |
| ufsp2 | 0.54902 | 0.17228413 | 0.056 | 0 | 1 | 26 |
| ugt5c2 | 0.54902 | 0.234517178 | 0.056 | 0 | 1 | 26 |
| unc119b | 0.54902 | 0.175616926 | 0.056 | 0 | 1 | 26 |
| uso1 | 0.54902 | 0.216219902 | 0.056 | 0 | 1 | 26 |
| vsir | 0.54902 | 0.217295575 | 0.056 | 0 | 1 | 26 |
| waca | 0.54902 | 0.233989995 | 0.056 | 0 | 1 | 26 |
| wbp11 | 0.54902 | 0.216033163 | 0.056 | 0 | 1 | 26 |
| wdr26b | 0.54902 | 0.215162993 | 0.056 | 0 | 1 | 26 |
| wdr45 | 0.54902 | 0.187572528 | 0.056 | 0 | 1 | 26 |
| wdr61 | 0.54902 | 0.222226943 | 0.056 | 0 | 1 | 26 |
| xpo6 | 0.54902 | 0.218206971 | 0.056 | 0 | 1 | 26 |
| xpo7 | 0.54902 | 0.216660403 | 0.056 | 0 | 1 | 26 |
| yipf3 | 0.54902 | 0.18394466 | 0.056 | 0 | 1 | 26 |
| yipf6 | 0.54902 | 0.216945554 | 0.056 | 0 | 1 | 26 |
| yy1b | 0.54902 | 0.220603136 | 0.056 | 0 | 1 | 26 |
| zbtb10 | 0.54902 | 0.217835477 | 0.056 | 0 | 1 | 26 |
| zc3h11a | 0.54902 | 0.27837189 | 0.056 | 0 | 1 | 26 |
| zc3h7bb | 0.54902 | 0.217454597 | 0.056 | 0 | 1 | 26 |
| zcchc7 | 0.54902 | 0.221569898 | 0.056 | 0 | 1 | 26 |
| zdhhc5a | 0.54902 | 0.216835685 | 0.056 | 0 | 1 | 26 |
| zfp36l1a | 0.54902 | 0.21710443 | 0.056 | 0 | 1 | 26 |
| zfr2 | 0.54902 | 0.217329984 | 0.056 | 0 | 1 | 26 |
| zgc:100846 | 0.54902 | 0.217277332 | 0.056 | 0 | 1 | 26 |
| zgc:101559 | 0.54902 | 0.234517178 | 0.056 | 0 | 1 | 26 |
| zgc:103678 | 0.54902 | 0.118114684 | 0.056 | 0 | 1 | 26 |
| zgc:113184 | 0.54902 | 0.187813944 | 0.056 | 0 | 1 | 26 |
| zgc:113208 | 0.54902 | 0.217774551 | 0.056 | 0 | 1 | 26 |
| zgc:162193 | 0.54902 | 0.215157236 | 0.056 | 0 | 1 | 26 |
| zgc:171857 | 0.54902 | 0.234517178 | 0.056 | 0 | 1 | 26 |
| zgc:194312 | 0.54902 | 0.234203718 | 0.056 | 0 | 1 | 26 |
| zgc:56304 | 0.54902 | 0.215071676 | 0.056 | 0 | 1 | 26 |
| zgc:56676 | 0.54902 | 0.232028534 | 0.056 | 0 | 1 | 26 |
| zmiz1a | 0.54902 | 0.217720377 | 0.056 | 0 | 1 | 26 |
| znf410 | 0.54902 | 0.233171789 | 0.056 | 0 | 1 | 26 |
| zswim5 | 0.54902 | 0.222942667 | 0.056 | 0 | 1 | 26 |
| AL954142.1 | 0.54902 | 0.203511008 | 0.056 | 0 | 1 | 26 |
| BX511067.1 | 0.54902 | 0.226041426 | 0.056 | 0 | 1 | 26 |

| | | | | | | |
|---|---|---|---|---|---|---|
| BX548028.1 | 0.54902 | 0.200675425 | 0.056 | 0 | 1 | 26 |
| BX649485.2 | 0.54902 | 0.351348151 | 0.056 | 0 | 1 | 26 |
| CABZ01001434.1 | 0.54902 | 0.245909724 | 0.056 | 0 | 1 | 26 |
| CABZ01055347.1 | 0.54902 | 0.192115565 | 0.056 | 0 | 1 | 26 |
| CABZ01076758.1 | 0.54902 | 0.192202815 | 0.056 | 0 | 1 | 26 |
| CABZ01090373.1 | 0.54902 | 0.215462002 | 0.056 | 0 | 1 | 26 |
| COA4 | 0.54902 | 0.173656529 | 0.056 | 0 | 1 | 26 |
| CU467110.1 | 0.54902 | 0.217234822 | 0.056 | 0 | 1 | 26 |
| FO907087.1 | 0.54902 | 0.199800574 | 0.056 | 0 | 1 | 26 |
| HOOK3 | 0.54902 | 0.189536956 | 0.056 | 0 | 1 | 26 |
| LO017700.1 | 0.54902 | 0.249121047 | 0.056 | 0 | 1 | 26 |
| LOC100535743 | 0.54902 | 0.217866993 | 0.056 | 0 | 1 | 26 |
| LOC100536946 | 0.54902 | 0.248276741 | 0.056 | 0 | 1 | 26 |
| LOC101883751 | 0.54902 | 0.202860194 | 0.056 | 0 | 1 | 26 |
| LOC101883798 | 0.54902 | 0.202860194 | 0.056 | 0 | 1 | 26 |
| LOC103909972 | 0.54902 | 0.198528033 | 0.056 | 0 | 1 | 26 |
| LOC103910871 | 0.54902 | 0.200784515 | 0.056 | 0 | 1 | 26 |
| LOC107195253 | 0.54902 | 0.194055005 | 0.056 | 0 | 1 | 26 |
| LOC108179023 | 0.54902 | 0.195974677 | 0.056 | 0 | 1 | 26 |
| LOC110437920 | 0.54902 | 0.219588437 | 0.056 | 0 | 1 | 26 |
| LOC570723 | 0.54902 | 0.215155289 | 0.056 | 0 | 1 | 26 |
| PLBD2 | 0.54902 | 0.170992437 | 0.056 | 0 | 1 | 26 |
| PYURF | 0.54902 | 0.201766117 | 0.056 | 0 | 1 | 26 |
| UTP14C | 0.54902 | 0.246942221 | 0.056 | 0 | 1 | 26 |
| XLOC-010899 | 0.54902 | 0.191434427 | 0.056 | 0 | 1 | 26 |
| XLOC-011085 | 0.54902 | 0.248454944 | 0.056 | 0 | 1 | 26 |
| XLOC-014397 | 0.54902 | 0.196536482 | 0.056 | 0 | 1 | 26 |
| XLOC-016094 | 0.54902 | 0.182407285 | 0.056 | 0 | 1 | 26 |
| XLOC-018981 | 0.54902 | 0.194055005 | 0.056 | 0 | 1 | 26 |
| XLOC-025819 | 0.54902 | 0.223035611 | 0.056 | 0 | 1 | 26 |
| XLOC-026836 | 0.54902 | 0.200541837 | 0.056 | 0 | 1 | 26 |
| XLOC-029303 | 0.54902 | 0.202860194 | 0.056 | 0 | 1 | 26 |
| XLOC-035202 | 0.54902 | 0.202028039 | 0.056 | 0 | 1 | 26 |
| abcb6b | 0.54902 | 0.182360353 | 0.056 | 0 | 1 | 26 |
| abcb8 | 0.54902 | 0.194055005 | 0.056 | 0 | 1 | 26 |
| acad9 | 0.54902 | 0.216505331 | 0.056 | 0 | 1 | 26 |
| acadl | 0.54902 | 0.216762622 | 0.056 | 0 | 1 | 26 |
| adam23a | 0.54902 | 0.104823151 | 0.056 | 0 | 1 | 26 |
| alg12 | 0.54902 | 0.21786721 | 0.056 | 0 | 1 | 26 |
| alg5 | 0.54902 | 0.210746348 | 0.056 | 0 | 1 | 26 |
| alox5a | 0.54902 | 0.191757168 | 0.056 | 0 | 1 | 26 |
| amfra | 0.54902 | 0.193213551 | 0.056 | 0 | 1 | 26 |
| amy2a | 0.54902 | 0.242622418 | 0.056 | 0 | 1 | 26 |
| anapc16 | 0.54902 | 0.189417306 | 0.056 | 0 | 1 | 26 |
| angptl1a | 0.54902 | 0.215360151 | 0.056 | 0 | 1 | 26 |
| ano5b | 0.54902 | 0.199321482 | 0.056 | 0 | 1 | 26 |
| anxa1a | 0.54902 | 0.183452372 | 0.056 | 0 | 1 | 26 |
| aph1b | 0.54902 | 0.192705306 | 0.056 | 0 | 1 | 26 |

| | | | | | | |
|---|---|---|---|---|---|---|
| arfgap1 | 0.54902 | 0.188306275 | 0.056 | 0 | 1 | 26 |
| arfgap2 | 0.54902 | 0.189393082 | 0.056 | 0 | 1 | 26 |
| arfgef2 | 0.54902 | 0.191886903 | 0.056 | 0 | 1 | 26 |
| arglu1b | 0.54902 | 0.197590233 | 0.056 | 0 | 1 | 26 |
| arl14ep | 0.54902 | 0.201639392 | 0.056 | 0 | 1 | 26 |
| arl2 | 0.54902 | 0.216248108 | 0.056 | 0 | 1 | 26 |
| arl6ip5a | 0.54902 | 0.217064285 | 0.056 | 0 | 1 | 26 |
| asphd2 | 0.54902 | 0.218381918 | 0.056 | 0 | 1 | 26 |
| atl2 | 0.54902 | 0.186703747 | 0.056 | 0 | 1 | 26 |
| atp1a1a.1 | 0.54902 | 0.197713256 | 0.056 | 0 | 1 | 26 |
| atp1a3a | 0.54902 | 0.29377892 | 0.056 | 0 | 1 | 26 |
| atp5meb | 0.54902 | 0.219533964 | 0.056 | 0 | 1 | 26 |
| atp6v1c1a | 0.54902 | 0.218176209 | 0.056 | 0 | 1 | 26 |
| atxn7l2b | 0.54902 | 0.197939454 | 0.056 | 0 | 1 | 26 |
| b2ml | 0.54902 | 0.176002384 | 0.056 | 0 | 1 | 26 |
| b4galt1l | 0.54902 | 0.194055005 | 0.056 | 0 | 1 | 26 |
| b4galt4 | 0.54902 | 0.191204278 | 0.056 | 0 | 1 | 26 |
| bag1 | 0.54902 | 0.196162403 | 0.056 | 0 | 1 | 26 |
| batf3 | 0.54902 | 0.202860194 | 0.056 | 0 | 1 | 26 |
| bnip1a | 0.54902 | 0.215155289 | 0.056 | 0 | 1 | 26 |
| brms1la | 0.54902 | 0.189442538 | 0.056 | 0 | 1 | 26 |
| brpf3b | 0.54902 | 0.199897823 | 0.056 | 0 | 1 | 26 |
| bsdc1 | 0.54902 | 0.188247233 | 0.056 | 0 | 1 | 26 |
| btaf1 | 0.54902 | 0.188106073 | 0.056 | 0 | 1 | 26 |
| btf3l4 | 0.54902 | 0.194614598 | 0.056 | 0 | 1 | 26 |
| bzw1b | 0.54902 | 0.189936952 | 0.056 | 0 | 1 | 26 |
| c1ql3a | 0.54902 | 0.246142018 | 0.056 | 0 | 1 | 26 |
| c6ast4 | 0.54902 | 0.340731672 | 0.056 | 0 | 1 | 26 |
| calhm2 | 0.54902 | 0.219588437 | 0.056 | 0 | 1 | 26 |
| calm1a | 0.54902 | 0.246399123 | 0.056 | 0 | 1 | 26 |
| calm2b | 0.54902 | 0.211614094 | 0.056 | 0 | 1 | 26 |
| canx | 0.54902 | 0.238726866 | 0.056 | 0 | 1 | 26 |
| casc4 | 0.54902 | 0.192198252 | 0.056 | 0 | 1 | 26 |
| ccl33.3 | 0.54902 | 0.200650069 | 0.056 | 0 | 1 | 26 |
| ccr12b.2 | 0.54902 | 0.217077161 | 0.056 | 0 | 1 | 26 |
| cd44b | 0.54902 | 0.182360353 | 0.056 | 0 | 1 | 26 |
| cd74b | 0.54902 | 0.198495356 | 0.056 | 0 | 1 | 26 |
| cdc73 | 0.54902 | 0.247638996 | 0.056 | 0 | 1 | 26 |
| cdca4 | 0.54902 | 0.247525577 | 0.056 | 0 | 1 | 26 |
| cdipt | 0.54902 | 0.214546782 | 0.056 | 0 | 1 | 26 |
| cdon | 0.54902 | 0.157540322 | 0.056 | 0 | 1 | 26 |
| cel.1 | 0.54902 | 0.312833637 | 0.056 | 0 | 1 | 26 |
| cenph | 0.54902 | 0.217647508 | 0.056 | 0 | 1 | 26 |
| cep104 | 0.54902 | 0.162500711 | 0.056 | 0 | 1 | 26 |
| cfhl1 | 0.54902 | 0.189006922 | 0.056 | 0 | 1 | 26 |
| cflarb | 0.54902 | 0.202860194 | 0.056 | 0 | 1 | 26 |
| chaf1a | 0.54902 | 0.18747107 | 0.056 | 0 | 1 | 26 |
| chchd3a | 0.54902 | 0.223936754 | 0.056 | 0 | 1 | 26 |

| | | | | | | |
|---|---|---|---|---|---|---|
| chd7 | 0.54902 | 0.201121126 | 0.056 | 0 | 1 | 26 |
| chic1 | 0.54902 | 0.203949734 | 0.056 | 0 | 1 | 26 |
| chmp6b | 0.54902 | 0.198943551 | 0.056 | 0 | 1 | 26 |
| ciarta | 0.54902 | 0.247246209 | 0.056 | 0 | 1 | 26 |
| cks1b | 0.54902 | 0.199125357 | 0.056 | 0 | 1 | 26 |
| clec16a | 0.54902 | 0.201787992 | 0.056 | 0 | 1 | 26 |
| clgn | 0.54902 | 0.169159196 | 0.056 | 0 | 1 | 26 |
| clk2b | 0.54902 | 0.191752642 | 0.056 | 0 | 1 | 26 |
| clk4a | 0.54902 | 0.19927166 | 0.056 | 0 | 1 | 26 |
| clns1a | 0.54902 | 0.157540322 | 0.056 | 0 | 1 | 26 |
| clptm1l | 0.54902 | 0.181472464 | 0.056 | 0 | 1 | 26 |
| cluha | 0.54902 | 0.195904005 | 0.056 | 0 | 1 | 26 |
| cnbpa | 0.54902 | 0.204849256 | 0.056 | 0 | 1 | 26 |
| cnih1 | 0.54902 | 0.179915975 | 0.056 | 0 | 1 | 26 |
| cnot9 | 0.54902 | 0.186765571 | 0.056 | 0 | 1 | 26 |
| cnpy3 | 0.54902 | 0.193029763 | 0.056 | 0 | 1 | 26 |
| coa3a | 0.54902 | 0.192807986 | 0.056 | 0 | 1 | 26 |
| coa6 | 0.54902 | 0.192709561 | 0.056 | 0 | 1 | 26 |
| col4a3bpb | 0.54902 | 0.193890466 | 0.056 | 0 | 1 | 26 |
| copb2 | 0.54902 | 0.199064765 | 0.056 | 0 | 1 | 26 |
| cops3 | 0.54902 | 0.180325162 | 0.056 | 0 | 1 | 26 |
| cox20 | 0.54902 | 0.195417483 | 0.056 | 0 | 1 | 26 |
| cpa1 | 0.54902 | 0.188655814 | 0.056 | 0 | 1 | 26 |
| cpa5 | 0.54902 | 0.324504076 | 0.056 | 0 | 1 | 26 |
| creb3l1 | 0.54902 | 0.249121047 | 0.056 | 0 | 1 | 26 |
| ctdspl3 | 0.54902 | 0.171817744 | 0.056 | 0 | 1 | 26 |
| ctsla | 0.54902 | 0.224259856 | 0.056 | 0 | 1 | 26 |
| cyfip1 | 0.54902 | 0.199206886 | 0.056 | 0 | 1 | 26 |
| cyp20a1 | 0.54902 | 0.216133552 | 0.056 | 0 | 1 | 26 |
| dck | 0.54902 | 0.212048896 | 0.056 | 0 | 1 | 26 |
| ddx42 | 0.54902 | 0.214791942 | 0.056 | 0 | 1 | 26 |
| ddx46 | 0.54902 | 0.24726824 | 0.056 | 0 | 1 | 26 |
| deaf1 | 0.54902 | 0.202860194 | 0.056 | 0 | 1 | 26 |
| dennd6aa | 0.54902 | 0.194055005 | 0.056 | 0 | 1 | 26 |
| denr | 0.54902 | 0.163499411 | 0.056 | 0 | 1 | 26 |
| desi1b | 0.54902 | 0.192057609 | 0.056 | 0 | 1 | 26 |
| dgkzb | 0.54902 | 0.194055005 | 0.056 | 0 | 1 | 26 |
| dhrs13a.1 | 0.54902 | 0.178175472 | 0.056 | 0 | 1 | 26 |
| dhrs4 | 0.54902 | 0.162663189 | 0.056 | 0 | 1 | 26 |
| dlg1 | 0.54902 | 0.249121047 | 0.056 | 0 | 1 | 26 |
| dlst | 0.54902 | 0.215279526 | 0.056 | 0 | 1 | 26 |
| dnajb6a | 0.54902 | 0.21555165 | 0.056 | 0 | 1 | 26 |
| dnajc1 | 0.54902 | 0.197762562 | 0.056 | 0 | 1 | 26 |
| dnajc3a | 0.54902 | 0.207726501 | 0.056 | 0 | 1 | 26 |
| dnajc8 | 0.54902 | 0.198174968 | 0.056 | 0 | 1 | 26 |
| dnmt1 | 0.54902 | 0.200685041 | 0.056 | 0 | 1 | 26 |
| dock5 | 0.54902 | 0.239229532 | 0.056 | 0 | 1 | 26 |
| dohh | 0.54902 | 0.212822904 | 0.056 | 0 | 1 | 26 |

| | | | | | | |
|---|---|---|---|---|---|---|
| dpp9 | 0.54902 | 0.191274521 | 0.056 | 0 | 1 | 26 |
| dse | 0.54902 | 0.217738298 | 0.056 | 0 | 1 | 26 |
| dtnbp1a | 0.54902 | 0.202755898 | 0.056 | 0 | 1 | 26 |
| eef1akmt2 | 0.54902 | 0.193630685 | 0.056 | 0 | 1 | 26 |
| egln3 | 0.54902 | 0.194055005 | 0.056 | 0 | 1 | 26 |
| eif2b1 | 0.54902 | 0.18872242 | 0.056 | 0 | 1 | 26 |
| eif2s1a | 0.54902 | 0.220273929 | 0.056 | 0 | 1 | 26 |
| eif3i | 0.54902 | 0.199233856 | 0.056 | 0 | 1 | 26 |
| eif4h | 0.54902 | 0.189782009 | 0.056 | 0 | 1 | 26 |
| eif5 | 0.54902 | 0.245911633 | 0.056 | 0 | 1 | 26 |
| ela2l | 0.54902 | 0.198819571 | 0.056 | 0 | 1 | 26 |
| epc1b | 0.54902 | 0.247634171 | 0.056 | 0 | 1 | 26 |
| epg5 | 0.54902 | 0.247249805 | 0.056 | 0 | 1 | 26 |
| exoc6 | 0.54902 | 0.202259157 | 0.056 | 0 | 1 | 26 |
| eya3 | 0.54902 | 0.189015685 | 0.056 | 0 | 1 | 26 |
| ezh2 | 0.54902 | 0.19256548 | 0.056 | 0 | 1 | 26 |
| f3b | 0.54902 | 0.247994329 | 0.056 | 0 | 1 | 26 |
| fam172a | 0.54902 | 0.191636079 | 0.056 | 0 | 1 | 26 |
| fam173a | 0.54902 | 0.248074373 | 0.056 | 0 | 1 | 26 |
| fastkd2 | 0.54902 | 0.217972868 | 0.056 | 0 | 1 | 26 |
| fdx1b | 0.54902 | 0.20089549 | 0.056 | 0 | 1 | 26 |
| fetub | 0.54902 | 0.195012846 | 0.056 | 0 | 1 | 26 |
| fgfr1op2 | 0.54902 | 0.199711461 | 0.056 | 0 | 1 | 26 |
| fgl2a | 0.54902 | 0.247800842 | 0.056 | 0 | 1 | 26 |
| fkbp11 | 0.54902 | 0.155036542 | 0.056 | 0 | 1 | 26 |
| fkbp9 | 0.54902 | 0.201078779 | 0.056 | 0 | 1 | 26 |
| frmd8 | 0.54902 | 0.215729809 | 0.056 | 0 | 1 | 26 |
| ftr67 | 0.54902 | 0.200354775 | 0.056 | 0 | 1 | 26 |
| fubp1 | 0.54902 | 0.248270584 | 0.056 | 0 | 1 | 26 |
| fyco1b | 0.54902 | 0.202860194 | 0.056 | 0 | 1 | 26 |
| g3bp2 | 0.54902 | 0.189290434 | 0.056 | 0 | 1 | 26 |
| gabpb2a | 0.54902 | 0.203119954 | 0.056 | 0 | 1 | 26 |
| gadd45bb | 0.54902 | 0.200864848 | 0.056 | 0 | 1 | 26 |
| gatad2b | 0.54902 | 0.24830587 | 0.056 | 0 | 1 | 26 |
| gatd1 | 0.54902 | 0.217393766 | 0.056 | 0 | 1 | 26 |
| gatm | 0.54902 | 0.12534885 | 0.056 | 0 | 1 | 26 |
| gnpnat1 | 0.54902 | 0.189051188 | 0.056 | 0 | 1 | 26 |
| gnptg | 0.54902 | 0.220419854 | 0.056 | 0 | 1 | 26 |
| got2b | 0.54902 | 0.222698362 | 0.056 | 0 | 1 | 26 |
| gpd1c | 0.54902 | 0.180800393 | 0.056 | 0 | 1 | 26 |
| gpr137 | 0.54902 | 0.194055005 | 0.056 | 0 | 1 | 26 |
| grb2b | 0.54902 | 0.216792091 | 0.056 | 0 | 1 | 26 |
| grwd1 | 0.54902 | 0.224326055 | 0.056 | 0 | 1 | 26 |
| gstcd | 0.54902 | 0.145627899 | 0.056 | 0 | 1 | 26 |
| gstm.2 | 0.54902 | 0.224138439 | 0.056 | 0 | 1 | 26 |
| gtf2a2 | 0.54902 | 0.217251555 | 0.056 | 0 | 1 | 26 |
| gtf2e2 | 0.54902 | 0.173555156 | 0.056 | 0 | 1 | 26 |
| gtf2f1 | 0.54902 | 0.199529482 | 0.056 | 0 | 1 | 26 |

| | | | | | | |
|---|---|---|---|---|---|---|
| gzf1 | 0.54902 | 0.200652716 | 0.056 | 0 | 1 | 26 |
| h2afx1 | 0.54902 | 0.190520842 | 0.056 | 0 | 1 | 26 |
| hcfc1b | 0.54902 | 0.19175069 | 0.056 | 0 | 1 | 26 |
| helb | 0.54902 | 0.194055005 | 0.056 | 0 | 1 | 26 |
| hexim1 | 0.54902 | 0.211278194 | 0.056 | 0 | 1 | 26 |
| hif1al | 0.54902 | 0.200336933 | 0.056 | 0 | 1 | 26 |
| histh1l | 0.54902 | 0.169537249 | 0.056 | 0 | 1 | 26 |
| hmgb3a | 0.54902 | 0.175895834 | 0.056 | 0 | 1 | 26 |
| hmgcl | 0.54902 | 0.224089442 | 0.056 | 0 | 1 | 26 |
| hmha1b | 0.54902 | 0.21723737 | 0.056 | 0 | 1 | 26 |
| hnrnpc | 0.54902 | 0.19146322 | 0.056 | 0 | 1 | 26 |
| hnrnph3 | 0.54902 | 0.201380968 | 0.056 | 0 | 1 | 26 |
| hnrnpua | 0.54902 | 0.201442628 | 0.056 | 0 | 1 | 26 |
| hprt1 | 0.54902 | 0.215261644 | 0.056 | 0 | 1 | 26 |
| hsbp1a | 0.54902 | 0.187206775 | 0.056 | 0 | 1 | 26 |
| hsdl2 | 0.54902 | 0.195495435 | 0.056 | 0 | 1 | 26 |
| hspb8 | 0.54902 | 0.140866281 | 0.056 | 0 | 1 | 26 |
| hspe1 | 0.54902 | 0.181844064 | 0.056 | 0 | 1 | 26 |
| htatsf1 | 0.54902 | 0.218168926 | 0.056 | 0 | 1 | 26 |
| iars | 0.54902 | 0.189840218 | 0.056 | 0 | 1 | 26 |
| igf2r | 0.54902 | 0.198134414 | 0.056 | 0 | 1 | 26 |
| igfbp5b | 0.54902 | 0.258033728 | 0.056 | 0 | 1 | 26 |
| il1rl1 | 0.54902 | 0.202860194 | 0.056 | 0 | 1 | 26 |
| il4 | 0.54902 | 0.215284455 | 0.056 | 0 | 1 | 26 |
| ilf3a | 0.54902 | 0.179866927 | 0.056 | 0 | 1 | 26 |
| imp4 | 0.54902 | 0.201438199 | 0.056 | 0 | 1 | 26 |
| irf7 | 0.54902 | 0.179837245 | 0.056 | 0 | 1 | 26 |
| isg20 | 0.54902 | 0.198386721 | 0.056 | 0 | 1 | 26 |
| itm2cb | 0.54902 | 0.170377158 | 0.056 | 0 | 1 | 26 |
| itpa | 0.54902 | 0.201054471 | 0.056 | 0 | 1 | 26 |
| jac8 | 0.54902 | 0.191414371 | 0.056 | 0 | 1 | 26 |
| jac9 | 0.54902 | 0.24819012 | 0.056 | 0 | 1 | 26 |
| katna1 | 0.54902 | 0.182360353 | 0.056 | 0 | 1 | 26 |
| kctd12.2 | 0.54902 | 0.21880293 | 0.056 | 0 | 1 | 26 |
| khsrp | 0.54902 | 0.190186952 | 0.056 | 0 | 1 | 26 |
| klf2b | 0.54902 | 0.201614631 | 0.056 | 0 | 1 | 26 |
| kmt5c | 0.54902 | 0.201417646 | 0.056 | 0 | 1 | 26 |
| kpnb1 | 0.54902 | 0.199994455 | 0.056 | 0 | 1 | 26 |
| ktn1 | 0.54902 | 0.191972987 | 0.056 | 0 | 1 | 26 |
| lamp1a | 0.54902 | 0.248389373 | 0.056 | 0 | 1 | 26 |
| lamtor2 | 0.54902 | 0.19125811 | 0.056 | 0 | 1 | 26 |
| larp1 | 0.54902 | 0.247003723 | 0.056 | 0 | 1 | 26 |
| larp4ab | 0.54902 | 0.19690754 | 0.056 | 0 | 1 | 26 |
| leng1 | 0.54902 | 0.217068771 | 0.056 | 0 | 1 | 26 |
| lgals3b | 0.54902 | 0.190317168 | 0.056 | 0 | 1 | 26 |
| lifrb | 0.54902 | 0.249121047 | 0.056 | 0 | 1 | 26 |
| limk2 | 0.54902 | 0.206213826 | 0.056 | 0 | 1 | 26 |
| lims1 | 0.54902 | 0.21169241 | 0.056 | 0 | 1 | 26 |

| | | | | | | |
|---|---|---|---|---|---|---|
| lin52 | 0.54902 | 0.19946408 | 0.056 | 0 | 1 | 26 |
| llph | 0.54902 | 0.215111224 | 0.056 | 0 | 1 | 26 |
| lonrf1l | 0.54902 | 0.218161526 | 0.056 | 0 | 1 | 26 |
| lrmp | 0.54902 | 0.243526298 | 0.056 | 0 | 1 | 26 |
| lrrc40 | 0.54902 | 0.194683783 | 0.056 | 0 | 1 | 26 |
| lrrc8da | 0.54902 | 0.201236034 | 0.056 | 0 | 1 | 26 |
| lsm12b | 0.54902 | 0.176400874 | 0.056 | 0 | 1 | 26 |
| lsm6 | 0.54902 | 0.19805802 | 0.056 | 0 | 1 | 26 |
| map1lc3b | 0.54902 | 0.223127 | 0.056 | 0 | 1 | 26 |
| map2k2a | 0.54902 | 0.179451955 | 0.056 | 0 | 1 | 26 |
| map2k6 | 0.54902 | 0.200754617 | 0.056 | 0 | 1 | 26 |
| mapk1 | 0.54902 | 0.192441965 | 0.056 | 0 | 1 | 26 |
| mapk6 | 0.54902 | 0.202063075 | 0.056 | 0 | 1 | 26 |
| mapk8ip3 | 0.54902 | 0.193052133 | 0.056 | 0 | 1 | 26 |
| mcm3ap | 0.54902 | 0.247546324 | 0.056 | 0 | 1 | 26 |
| mcm4 | 0.54902 | 0.192415837 | 0.056 | 0 | 1 | 26 |
| mcm5 | 0.54902 | 0.196082048 | 0.056 | 0 | 1 | 26 |
| mcm7 | 0.54902 | 0.15598308 | 0.056 | 0 | 1 | 26 |
| mdh1aa | 0.54902 | 0.184846265 | 0.056 | 0 | 1 | 26 |
| mdkb | 0.54902 | 0.21593024 | 0.056 | 0 | 1 | 26 |
| mdm4 | 0.54902 | 0.185396591 | 0.056 | 0 | 1 | 26 |
| med21 | 0.54902 | 0.247977976 | 0.056 | 0 | 1 | 26 |
| mfap4 | 0.54902 | 0.248121201 | 0.056 | 0 | 1 | 26 |
| micall1a | 0.54902 | 0.191572592 | 0.056 | 0 | 1 | 26 |
| mios | 0.54902 | 0.214824328 | 0.056 | 0 | 1 | 26 |
| mlec | 0.54902 | 0.192534342 | 0.056 | 0 | 1 | 26 |
| mon2 | 0.54902 | 0.246711921 | 0.056 | 0 | 1 | 26 |
| mpc2 | 0.54902 | 0.204873496 | 0.056 | 0 | 1 | 26 |
| mroh1 | 0.54902 | 0.199594537 | 0.056 | 0 | 1 | 26 |
| mrpl20 | 0.54902 | 0.188455124 | 0.056 | 0 | 1 | 26 |
| mrpl23 | 0.54902 | 0.200486729 | 0.056 | 0 | 1 | 26 |
| mrpl51 | 0.54902 | 0.223653228 | 0.056 | 0 | 1 | 26 |
| mrps10 | 0.54902 | 0.189411872 | 0.056 | 0 | 1 | 26 |
| mrps34 | 0.54902 | 0.208771368 | 0.056 | 0 | 1 | 26 |
| mrps7 | 0.54902 | 0.214946509 | 0.056 | 0 | 1 | 26 |
| mrps9 | 0.54902 | 0.212326552 | 0.056 | 0 | 1 | 26 |
| mrrf | 0.54902 | 0.180036726 | 0.056 | 0 | 1 | 26 |
| msh2 | 0.54902 | 0.200493453 | 0.056 | 0 | 1 | 26 |
| msl1b | 0.54902 | 0.191468286 | 0.056 | 0 | 1 | 26 |
| mtfr1l | 0.54902 | 0.198719151 | 0.056 | 0 | 1 | 26 |
| mtmr10 | 0.54902 | 0.201652339 | 0.056 | 0 | 1 | 26 |
| mtmr1a | 0.54902 | 0.194055005 | 0.056 | 0 | 1 | 26 |
| mtpn | 0.54902 | 0.197729364 | 0.056 | 0 | 1 | 26 |
| mus81 | 0.54902 | 0.196749394 | 0.056 | 0 | 1 | 26 |
| mybbp1a | 0.54902 | 0.199332363 | 0.056 | 0 | 1 | 26 |
| myg1 | 0.54902 | 0.187957268 | 0.056 | 0 | 1 | 26 |
| myh9a | 0.54902 | 0.193005722 | 0.056 | 0 | 1 | 26 |
| myo18ab | 0.54902 | 0.198913623 | 0.056 | 0 | 1 | 26 |

| | | | | | | |
|---|---|---|---|---|---|---|
| mzt2b | 0.54902 | 0.19917029 | 0.056 | 0 | 1 | 26 |
| nars | 0.54902 | 0.213306263 | 0.056 | 0 | 1 | 26 |
| nat10 | 0.54902 | 0.217108711 | 0.056 | 0 | 1 | 26 |
| nbr1b | 0.54902 | 0.191282237 | 0.056 | 0 | 1 | 26 |
| ndufa9a | 0.54902 | 0.21051753 | 0.056 | 0 | 1 | 26 |
| ndufaf5 | 0.54902 | 0.210979516 | 0.056 | 0 | 1 | 26 |
| ndufb4 | 0.54902 | 0.221087398 | 0.056 | 0 | 1 | 26 |
| nek7 | 0.54902 | 0.202612571 | 0.056 | 0 | 1 | 26 |
| nfatc3a | 0.54902 | 0.248636932 | 0.056 | 0 | 1 | 26 |
| nif3l1 | 0.54902 | 0.200882732 | 0.056 | 0 | 1 | 26 |
| nifk | 0.54902 | 0.200390666 | 0.056 | 0 | 1 | 26 |
| nipa2 | 0.54902 | 0.248407535 | 0.056 | 0 | 1 | 26 |
| nkiras2 | 0.54902 | 0.203425934 | 0.056 | 0 | 1 | 26 |
| nkrf | 0.54902 | 0.178267478 | 0.056 | 0 | 1 | 26 |
| nme7 | 0.54902 | 0.181763953 | 0.056 | 0 | 1 | 26 |
| nnt | 0.54902 | 0.188664983 | 0.056 | 0 | 1 | 26 |
| nol11 | 0.54902 | 0.190636181 | 0.056 | 0 | 1 | 26 |
| notch2 | 0.54902 | 0.193090516 | 0.056 | 0 | 1 | 26 |
| npm3 | 0.54902 | 0.223897637 | 0.056 | 0 | 1 | 26 |
| nrf1 | 0.54902 | 0.190442474 | 0.056 | 0 | 1 | 26 |
| nudt22 | 0.54902 | 0.21742612 | 0.056 | 0 | 1 | 26 |
| nudt3b | 0.54902 | 0.193072926 | 0.056 | 0 | 1 | 26 |
| nup50 | 0.54902 | 0.191573642 | 0.056 | 0 | 1 | 26 |
| osbp | 0.54902 | 0.193413761 | 0.056 | 0 | 1 | 26 |
| osmr | 0.54902 | 0.194055005 | 0.056 | 0 | 1 | 26 |
| otulinb | 0.54902 | 0.197631291 | 0.056 | 0 | 1 | 26 |
| p2rx4b | 0.54902 | 0.192764978 | 0.056 | 0 | 1 | 26 |
| papola | 0.54902 | 0.190912264 | 0.056 | 0 | 1 | 26 |
| parp16 | 0.54902 | 0.216883286 | 0.056 | 0 | 1 | 26 |
| parp8 | 0.54902 | 0.19343414 | 0.056 | 0 | 1 | 26 |
| pcca | 0.54902 | 0.215177651 | 0.056 | 0 | 1 | 26 |
| pdap1a | 0.54902 | 0.194016883 | 0.056 | 0 | 1 | 26 |
| pdcd2 | 0.54902 | 0.222346198 | 0.056 | 0 | 1 | 26 |
| pdcd4a | 0.54902 | 0.148922139 | 0.056 | 0 | 1 | 26 |
| pdcd4b | 0.54902 | 0.247934971 | 0.056 | 0 | 1 | 26 |
| pdcd6 | 0.54902 | 0.178925449 | 0.056 | 0 | 1 | 26 |
| pef1 | 0.54902 | 0.225813574 | 0.056 | 0 | 1 | 26 |
| pex26 | 0.54902 | 0.19604068 | 0.056 | 0 | 1 | 26 |
| pgd | 0.54902 | 0.216523752 | 0.056 | 0 | 1 | 26 |
| pgs1 | 0.54902 | 0.245917545 | 0.056 | 0 | 1 | 26 |
| phf10 | 0.54902 | 0.180745965 | 0.056 | 0 | 1 | 26 |
| phip | 0.54902 | 0.201652647 | 0.056 | 0 | 1 | 26 |
| phlda2 | 0.54902 | 0.193399424 | 0.056 | 0 | 1 | 26 |
| pim3 | 0.54902 | 0.169219154 | 0.056 | 0 | 1 | 26 |
| pkn1b | 0.54902 | 0.182360353 | 0.056 | 0 | 1 | 26 |
| pmm2 | 0.54902 | 0.194561897 | 0.056 | 0 | 1 | 26 |
| pola1 | 0.54902 | 0.215424492 | 0.056 | 0 | 1 | 26 |
| polr1b | 0.54902 | 0.188044284 | 0.056 | 0 | 1 | 26 |

| | | | | | | |
|---|---|---|---|---|---|---|
| ponzr1 | 0.54902 | 0.211344054 | 0.056 | 0 | 1 | 26 |
| ppm1ba | 0.54902 | 0.24856992 | 0.056 | 0 | 1 | 26 |
| ppp1r12c | 0.54902 | 0.248399157 | 0.056 | 0 | 1 | 26 |
| ppp2r5cb | 0.54902 | 0.194055005 | 0.056 | 0 | 1 | 26 |
| ppp3r1a | 0.54902 | 0.248922752 | 0.056 | 0 | 1 | 26 |
| prelid1b | 0.54902 | 0.193218476 | 0.056 | 0 | 1 | 26 |
| prkaa1 | 0.54902 | 0.24743938 | 0.056 | 0 | 1 | 26 |
| prkcda | 0.54902 | 0.201141102 | 0.056 | 0 | 1 | 26 |
| prps1a | 0.54902 | 0.200577744 | 0.056 | 0 | 1 | 26 |
| prrc2c | 0.54902 | 0.189664147 | 0.056 | 0 | 1 | 26 |
| prss59.2 | 0.54902 | 0.335010102 | 0.056 | 0 | 1 | 26 |
| psma3 | 0.54902 | 0.214172992 | 0.056 | 0 | 1 | 26 |
| psma8 | 0.54902 | 0.198748244 | 0.056 | 0 | 1 | 26 |
| psmb6 | 0.54902 | 0.197587245 | 0.056 | 0 | 1 | 26 |
| psmc5 | 0.54902 | 0.199470012 | 0.056 | 0 | 1 | 26 |
| psmd7 | 0.54902 | 0.196869035 | 0.056 | 0 | 1 | 26 |
| ptgir | 0.54902 | 0.202228214 | 0.056 | 0 | 1 | 26 |
| ptprea | 0.54902 | 0.248233914 | 0.056 | 0 | 1 | 26 |
| puraa | 0.54902 | 0.192062828 | 0.056 | 0 | 1 | 26 |
| purg | 0.54902 | 0.246365947 | 0.056 | 0 | 1 | 26 |
| pwwp2a | 0.54902 | 0.196820532 | 0.056 | 0 | 1 | 26 |
| qars | 0.54902 | 0.20185865 | 0.056 | 0 | 1 | 26 |
| qpctla | 0.54902 | 0.182360353 | 0.056 | 0 | 1 | 26 |
| rab6a | 0.54902 | 0.189957896 | 0.056 | 0 | 1 | 26 |
| rab9b | 0.54902 | 0.194055005 | 0.056 | 0 | 1 | 26 |
| rabgef1 | 0.54902 | 0.178257431 | 0.056 | 0 | 1 | 26 |
| rabgef1l | 0.54902 | 0.194055005 | 0.056 | 0 | 1 | 26 |
| rasal3 | 0.54902 | 0.202860194 | 0.056 | 0 | 1 | 26 |
| rasgrp2 | 0.54902 | 0.202860194 | 0.056 | 0 | 1 | 26 |
| rasgrp3 | 0.54902 | 0.202860194 | 0.056 | 0 | 1 | 26 |
| rasgrp4 | 0.54902 | 0.248764269 | 0.056 | 0 | 1 | 26 |
| rbck1 | 0.54902 | 0.210710529 | 0.056 | 0 | 1 | 26 |
| rcor1 | 0.54902 | 0.201777449 | 0.056 | 0 | 1 | 26 |
| rfc5 | 0.54902 | 0.19919131 | 0.056 | 0 | 1 | 26 |
| rfxank | 0.54902 | 0.181405058 | 0.056 | 0 | 1 | 26 |
| rfxap | 0.54902 | 0.155336973 | 0.056 | 0 | 1 | 26 |
| rhoaa | 0.54902 | 0.180734686 | 0.056 | 0 | 1 | 26 |
| rida.1 | 0.54902 | 0.206213826 | 0.056 | 0 | 1 | 26 |
| rif1 | 0.54902 | 0.192063996 | 0.056 | 0 | 1 | 26 |
| rnaset2l | 0.54902 | 0.181437352 | 0.056 | 0 | 1 | 26 |
| rnf11b | 0.54902 | 0.187991903 | 0.056 | 0 | 1 | 26 |
| rnf216 | 0.54902 | 0.215155289 | 0.056 | 0 | 1 | 26 |
| rnf25 | 0.54902 | 0.191267428 | 0.056 | 0 | 1 | 26 |
| rnf44 | 0.54902 | 0.20205335 | 0.056 | 0 | 1 | 26 |
| rnf8 | 0.54902 | 0.181304899 | 0.056 | 0 | 1 | 26 |
| rpa1 | 0.54902 | 0.18691602 | 0.056 | 0 | 1 | 26 |
| rpe | 0.54902 | 0.173329928 | 0.056 | 0 | 1 | 26 |
| rpl7l1 | 0.54902 | 0.188075047 | 0.056 | 0 | 1 | 26 |

| | | | | | | |
|---|---|---|---|---|---|---|
| rtf1 | 0.54902 | 0.198998429 | 0.056 | 0 | 1 | 26 |
| rtn1b | 0.54902 | 0.155062634 | 0.056 | 0 | 1 | 26 |
| rtn3 | 0.54902 | 0.198222881 | 0.056 | 0 | 1 | 26 |
| ruvbl1 | 0.54902 | 0.195296695 | 0.056 | 0 | 1 | 26 |
| s100a10b | 0.54902 | 0.197591941 | 0.056 | 0 | 1 | 26 |
| sacs | 0.54902 | 0.202860194 | 0.056 | 0 | 1 | 26 |
| safb | 0.54902 | 0.201113508 | 0.056 | 0 | 1 | 26 |
| saraf | 0.54902 | 0.206763574 | 0.056 | 0 | 1 | 26 |
| sardh | 0.54902 | 0.215424492 | 0.056 | 0 | 1 | 26 |
| sbno1 | 0.54902 | 0.200811218 | 0.056 | 0 | 1 | 26 |
| sco1 | 0.54902 | 0.225551506 | 0.056 | 0 | 1 | 26 |
| scrib | 0.54902 | 0.199800574 | 0.056 | 0 | 1 | 26 |
| sdad1 | 0.54902 | 0.217995064 | 0.056 | 0 | 1 | 26 |
| sdcbp2 | 0.54902 | 0.190187058 | 0.056 | 0 | 1 | 26 |
| sec22ba | 0.54902 | 0.216063683 | 0.056 | 0 | 1 | 26 |
| selenos | 0.54902 | 0.20655926 | 0.056 | 0 | 1 | 26 |
| sf3b1 | 0.54902 | 0.198446516 | 0.056 | 0 | 1 | 26 |
| sh3bgrl | 0.54902 | 0.248493486 | 0.056 | 0 | 1 | 26 |
| shmt2 | 0.54902 | 0.11918688 | 0.056 | 0 | 1 | 26 |
| shoc2 | 0.54902 | 0.192107902 | 0.056 | 0 | 1 | 26 |
| si:ch1073-228j22.1 | 0.54902 | 0.221875098 | 0.056 | 0 | 1 | 26 |
| si:ch1073-314i13.4 | 0.54902 | 0.20395884 | 0.056 | 0 | 1 | 26 |
| si:ch211-113e8.11 | 0.54902 | 0.201091033 | 0.056 | 0 | 1 | 26 |
| si:ch211-133n4.4 | 0.54902 | 0.214533742 | 0.056 | 0 | 1 | 26 |
| si:ch211-161h7.5 | 0.54902 | 0.178257431 | 0.056 | 0 | 1 | 26 |
| si:ch211-166e11.5 | 0.54902 | 0.202860194 | 0.056 | 0 | 1 | 26 |
| si:ch211-198c19.1 | 0.54902 | 0.203949734 | 0.056 | 0 | 1 | 26 |
| si:ch211-213a13.1 | 0.54902 | 0.248463824 | 0.056 | 0 | 1 | 26 |
| si:ch211-214e3.5 | 0.54902 | 0.194055005 | 0.056 | 0 | 1 | 26 |
| si:ch211-235e9.6 | 0.54902 | 0.195614372 | 0.056 | 0 | 1 | 26 |
| si:ch211-262i1.6 | 0.54902 | 0.175742921 | 0.056 | 0 | 1 | 26 |
| si:ch211-276i12.11 | 0.54902 | 0.1987162 | 0.056 | 0 | 1 | 26 |
| si:ch211-59d17.3 | 0.54902 | 0.203949734 | 0.056 | 0 | 1 | 26 |
| si:ch73-103b9.2 | 0.54902 | 0.202860194 | 0.056 | 0 | 1 | 26 |
| si:ch73-299h12.8 | 0.54902 | 0.215424492 | 0.056 | 0 | 1 | 26 |
| si:dkey-112a7.4 | 0.54902 | 0.210939684 | 0.056 | 0 | 1 | 26 |
| si:dkey-177p2.6 | 0.54902 | 0.247985394 | 0.056 | 0 | 1 | 26 |
| si:dkey-17o15.2 | 0.54902 | 0.191565877 | 0.056 | 0 | 1 | 26 |
| si:dkey-185m8.2 | 0.54902 | 0.238402099 | 0.056 | 0 | 1 | 26 |
| si:dkey-19e4.5 | 0.54902 | 0.242026864 | 0.056 | 0 | 1 | 26 |
| si:dkey-239h2.3 | 0.54902 | 0.244697238 | 0.056 | 0 | 1 | 26 |
| si:dkey-253d23.4 | 0.54902 | 0.190636181 | 0.056 | 0 | 1 | 26 |
| si:dkey-261h17.1 | 0.54902 | 0.214417707 | 0.056 | 0 | 1 | 26 |
| si:dkey-28b4.7 | 0.54902 | 0.176036184 | 0.056 | 0 | 1 | 26 |
| si:dkey-42l23.2 | 0.54902 | 0.248568083 | 0.056 | 0 | 1 | 26 |
| si:dkey-79d12.5 | 0.54902 | 0.18860215 | 0.056 | 0 | 1 | 26 |
| si:dkey-93m18.3 | 0.54902 | 0.202860194 | 0.056 | 0 | 1 | 26 |
| si:dkeyp-113d7.1 | 0.54902 | 0.199120927 | 0.056 | 0 | 1 | 26 |

| | | | | | | |
|---|---|---|---|---|---|---|
| si:dkeyp-113d7.10 | 0.54902 | 0.218711554 | 0.056 | 0 | 1 | 26 |
| si:dkeyp-120h9.1 | 0.54902 | 0.186117021 | 0.056 | 0 | 1 | 26 |
| si:rp71-45k5.4 | 0.54902 | 0.209812055 | 0.056 | 0 | 1 | 26 |
| sirt1 | 0.54902 | 0.198953836 | 0.056 | 0 | 1 | 26 |
| siva1 | 0.54902 | 0.203973773 | 0.056 | 0 | 1 | 26 |
| slc17a5 | 0.54902 | 0.191295145 | 0.056 | 0 | 1 | 26 |
| slc25a14 | 0.54902 | 0.193169481 | 0.056 | 0 | 1 | 26 |
| slc25a40 | 0.54902 | 0.187016206 | 0.056 | 0 | 1 | 26 |
| slc35b4 | 0.54902 | 0.201078779 | 0.056 | 0 | 1 | 26 |
| slc3a2b | 0.54902 | 0.190363659 | 0.056 | 0 | 1 | 26 |
| smarca4a | 0.54902 | 0.191887013 | 0.056 | 0 | 1 | 26 |
| smdt1b | 0.54902 | 0.190205339 | 0.056 | 0 | 1 | 26 |
| smek1 | 0.54902 | 0.198105833 | 0.056 | 0 | 1 | 26 |
| smpd5 | 0.54902 | 0.202860194 | 0.056 | 0 | 1 | 26 |
| snrnp27 | 0.54902 | 0.177855808 | 0.056 | 0 | 1 | 26 |
| snrpb2 | 0.54902 | 0.201394326 | 0.056 | 0 | 1 | 26 |
| snrpe | 0.54902 | 0.210215394 | 0.056 | 0 | 1 | 26 |
| snrpg | 0.54902 | 0.212967947 | 0.056 | 0 | 1 | 26 |
| sod2 | 0.54902 | 0.187602919 | 0.056 | 0 | 1 | 26 |
| sostdc1a | 0.54902 | 0.190871688 | 0.056 | 0 | 1 | 26 |
| sqstm1 | 0.54902 | 0.181389404 | 0.056 | 0 | 1 | 26 |
| srd5a2a | 0.54902 | 0.198600618 | 0.056 | 0 | 1 | 26 |
| srsf11 | 0.54902 | 0.197444358 | 0.056 | 0 | 1 | 26 |
| srsf1b | 0.54902 | 0.188524512 | 0.056 | 0 | 1 | 26 |
| srsf4 | 0.54902 | 0.215451686 | 0.056 | 0 | 1 | 26 |
| srsf5a | 0.54902 | 0.196932279 | 0.056 | 0 | 1 | 26 |
| srsf7a | 0.54902 | 0.174552588 | 0.056 | 0 | 1 | 26 |
| sssca1 | 0.54902 | 0.198977711 | 0.056 | 0 | 1 | 26 |
| ssuh2.4 | 0.54902 | 0.21885039 | 0.056 | 0 | 1 | 26 |
| st13 | 0.54902 | 0.194530285 | 0.056 | 0 | 1 | 26 |
| st6galnac5b | 0.54902 | 0.210979516 | 0.056 | 0 | 1 | 26 |
| stab2 | 0.54902 | 0.202860194 | 0.056 | 0 | 1 | 26 |
| stam2 | 0.54902 | 0.195513867 | 0.056 | 0 | 1 | 26 |
| stard7 | 0.54902 | 0.21886398 | 0.056 | 0 | 1 | 26 |
| stk35l | 0.54902 | 0.246578316 | 0.056 | 0 | 1 | 26 |
| stx4 | 0.54902 | 0.192139131 | 0.056 | 0 | 1 | 26 |
| sub1a | 0.54902 | 0.191513917 | 0.056 | 0 | 1 | 26 |
| sucla2 | 0.54902 | 0.21473495 | 0.056 | 0 | 1 | 26 |
| sult2st1 | 0.54902 | 0.218584577 | 0.056 | 0 | 1 | 26 |
| supt3h | 0.54902 | 0.218685065 | 0.056 | 0 | 1 | 26 |
| svild | 0.54902 | 0.189970566 | 0.056 | 0 | 1 | 26 |
| syf2 | 0.54902 | 0.21428124 | 0.056 | 0 | 1 | 26 |
| taf10 | 0.54902 | 0.181225857 | 0.056 | 0 | 1 | 26 |
| tagln3b | 0.54902 | 0.197939454 | 0.056 | 0 | 1 | 26 |
| taok3a | 0.54902 | 0.191848756 | 0.056 | 0 | 1 | 26 |
| tapbpl | 0.54902 | 0.202860194 | 0.056 | 0 | 1 | 26 |
| tbc1d1 | 0.54902 | 0.246982168 | 0.056 | 0 | 1 | 26 |
| tbl1x | 0.54902 | 0.19169214 | 0.056 | 0 | 1 | 26 |

| | | | | | | |
|---|---|---|---|---|---|---|
| tgm1l1 | 0.54902 | 0.200868795 | 0.056 | 0 | 1 | 26 |
| thrap3b | 0.54902 | 0.197951263 | 0.056 | 0 | 1 | 26 |
| timm9 | 0.54902 | 0.196033721 | 0.056 | 0 | 1 | 26 |
| tiparp | 0.54902 | 0.2475318 | 0.056 | 0 | 1 | 26 |
| tmem106c | 0.54902 | 0.183131181 | 0.056 | 0 | 1 | 26 |
| tmem11 | 0.54902 | 0.198203215 | 0.056 | 0 | 1 | 26 |
| tmem131 | 0.54902 | 0.192351769 | 0.056 | 0 | 1 | 26 |
| tmem147 | 0.54902 | 0.18989506 | 0.056 | 0 | 1 | 26 |
| tmem14ca | 0.54902 | 0.21312301 | 0.056 | 0 | 1 | 26 |
| tmem176l.1 | 0.54902 | 0.201341898 | 0.056 | 0 | 1 | 26 |
| tmem181 | 0.54902 | 0.182360353 | 0.056 | 0 | 1 | 26 |
| tmem222b | 0.54902 | 0.202156722 | 0.056 | 0 | 1 | 26 |
| tmem263 | 0.54902 | 0.200021842 | 0.056 | 0 | 1 | 26 |
| tmem42a | 0.54902 | 0.202274059 | 0.056 | 0 | 1 | 26 |
| tmem50a | 0.54902 | 0.210440154 | 0.056 | 0 | 1 | 26 |
| tmod2 | 0.54902 | 0.201396647 | 0.056 | 0 | 1 | 26 |
| tnip1 | 0.54902 | 0.192023564 | 0.056 | 0 | 1 | 26 |
| tob1b | 0.54902 | 0.164289516 | 0.056 | 0 | 1 | 26 |
| top1l | 0.54902 | 0.191691969 | 0.056 | 0 | 1 | 26 |
| top2b | 0.54902 | 0.210979516 | 0.056 | 0 | 1 | 26 |
| tor4aa | 0.54902 | 0.202860194 | 0.056 | 0 | 1 | 26 |
| tp53 | 0.54902 | 0.186390507 | 0.056 | 0 | 1 | 26 |
| tp53rk | 0.54902 | 0.216366221 | 0.056 | 0 | 1 | 26 |
| tpd52 | 0.54902 | 0.192410987 | 0.056 | 0 | 1 | 26 |
| tpm2 | 0.54902 | 0.200810534 | 0.056 | 0 | 1 | 26 |
| tpp2 | 0.54902 | 0.190006235 | 0.056 | 0 | 1 | 26 |
| tra2a | 0.54902 | 0.215109372 | 0.056 | 0 | 1 | 26 |
| traf5 | 0.54902 | 0.194055005 | 0.056 | 0 | 1 | 26 |
| trappc5 | 0.54902 | 0.248385269 | 0.056 | 0 | 1 | 26 |
| trip11 | 0.54902 | 0.201241361 | 0.056 | 0 | 1 | 26 |
| triqk | 0.54902 | 0.192509831 | 0.056 | 0 | 1 | 26 |
| trnau1apa | 0.54902 | 0.198443784 | 0.056 | 0 | 1 | 26 |
| tuba8l2 | 0.54902 | 0.187857009 | 0.056 | 0 | 1 | 26 |
| txn2 | 0.54902 | 0.19620221 | 0.056 | 0 | 1 | 26 |
| txndc9 | 0.54902 | 0.211669411 | 0.056 | 0 | 1 | 26 |
| ubald2 | 0.54902 | 0.190086848 | 0.056 | 0 | 1 | 26 |
| ube4b | 0.54902 | 0.248165512 | 0.056 | 0 | 1 | 26 |
| ubtf | 0.54902 | 0.166310342 | 0.056 | 0 | 1 | 26 |
| uggt2 | 0.54902 | 0.181356037 | 0.056 | 0 | 1 | 26 |
| upf3b | 0.54902 | 0.19109386 | 0.056 | 0 | 1 | 26 |
| uqcrb | 0.54902 | 0.157280963 | 0.056 | 0 | 1 | 26 |
| ushbp1 | 0.54902 | 0.247120835 | 0.056 | 0 | 1 | 26 |
| usp4 | 0.54902 | 0.188650906 | 0.056 | 0 | 1 | 26 |
| vipas39 | 0.54902 | 0.182360353 | 0.056 | 0 | 1 | 26 |
| vma21 | 0.54902 | 0.200535287 | 0.056 | 0 | 1 | 26 |
| vps11 | 0.54902 | 0.178257431 | 0.056 | 0 | 1 | 26 |
| vps26bl | 0.54902 | 0.194055005 | 0.056 | 0 | 1 | 26 |
| vps26c | 0.54902 | 0.199900076 | 0.056 | 0 | 1 | 26 |

| | | | | | | |
|---|---|---|---|---|---|---|
| vps28 | 0.54902 | 0.178624878 | 0.056 | 0 | 1 | 26 |
| vrk1 | 0.54902 | 0.214785408 | 0.056 | 0 | 1 | 26 |
| wbp1 | 0.54902 | 0.189739975 | 0.056 | 0 | 1 | 26 |
| wdr75 | 0.54902 | 0.20186769 | 0.056 | 0 | 1 | 26 |
| wipf1b | 0.54902 | 0.192692477 | 0.056 | 0 | 1 | 26 |
| wnk1b | 0.54902 | 0.199369499 | 0.056 | 0 | 1 | 26 |
| wu:fc75a09 | 0.54902 | 0.246836291 | 0.056 | 0 | 1 | 26 |
| xiap | 0.54902 | 0.214728493 | 0.056 | 0 | 1 | 26 |
| ywhaqa | 0.54902 | 0.200513904 | 0.056 | 0 | 1 | 26 |
| zc3h13 | 0.54902 | 0.26172372 | 0.056 | 0 | 1 | 26 |
| zcrb1 | 0.54902 | 0.201456487 | 0.056 | 0 | 1 | 26 |
| zfpl1 | 0.54902 | 0.20143959 | 0.056 | 0 | 1 | 26 |
| zfx | 0.54902 | 0.200518923 | 0.056 | 0 | 1 | 26 |
| zgc:103700 | 0.54902 | 0.247688847 | 0.056 | 0 | 1 | 26 |
| zgc:136461 | 0.54902 | 0.236414212 | 0.056 | 0 | 1 | 26 |
| zgc:165582 | 0.54902 | 0.24859705 | 0.056 | 0 | 1 | 26 |
| zgc:171497 | 0.54902 | 0.199233698 | 0.056 | 0 | 1 | 26 |
| zgc:174268 | 0.54902 | 0.20119203 | 0.056 | 0 | 1 | 26 |
| zgc:174863 | 0.54902 | 0.2006536 | 0.056 | 0 | 1 | 26 |
| zgc:194627 | 0.54902 | 0.210229181 | 0.056 | 0 | 1 | 26 |
| zgc:56106 | 0.54902 | 0.202865123 | 0.056 | 0 | 1 | 26 |
| zgc:77929 | 0.54902 | 0.198727289 | 0.056 | 0 | 1 | 26 |
| zgc:85932 | 0.54902 | 0.221875098 | 0.056 | 0 | 1 | 26 |
| zmym2 | 0.54902 | 0.202860194 | 0.056 | 0 | 1 | 26 |
| zmynd11 | 0.54902 | 0.164204439 | 0.056 | 0 | 1 | 26 |
| znf576.2 | 0.54902 | 0.216249774 | 0.056 | 0 | 1 | 26 |
| znfl1l | 0.54902 | 0.248307811 | 0.056 | 0 | 1 | 26 |
| CABZ01109481.1 | 0.54902 | 0.257538057 | 0.056 | 0 | 1 | 26 |
| CR385067.1 | 0.54902 | 0.257736812 | 0.056 | 0 | 1 | 26 |
| FP074902.2 | 0.54902 | 0.255479784 | 0.056 | 0 | 1 | 26 |
| GMEB2 | 0.54902 | 0.256972049 | 0.056 | 0 | 1 | 26 |
| agla | 0.54902 | 0.257736812 | 0.056 | 0 | 1 | 26 |
| ccdc127b | 0.54902 | 0.256642181 | 0.056 | 0 | 1 | 26 |
| dnajb9b | 0.54902 | 0.301867012 | 0.056 | 0 | 1 | 26 |
| edem3 | 0.54902 | 0.257736812 | 0.056 | 0 | 1 | 26 |
| emc2 | 0.54902 | 0.255394994 | 0.056 | 0 | 1 | 26 |
| epn2 | 0.54902 | 0.257431221 | 0.056 | 0 | 1 | 26 |
| fam122b | 0.54902 | 0.256857646 | 0.056 | 0 | 1 | 26 |
| gtse1 | 0.54902 | 0.257736812 | 0.056 | 0 | 1 | 26 |
| jdp2b | 0.54902 | 0.255642396 | 0.056 | 0 | 1 | 26 |
| larp4aa | 0.54902 | 0.256165519 | 0.056 | 0 | 1 | 26 |
| mcrs1 | 0.54902 | 0.25561669 | 0.056 | 0 | 1 | 26 |
| mrtfbb | 0.54902 | 0.257736812 | 0.056 | 0 | 1 | 26 |
| ncf1 | 0.54902 | 0.256983438 | 0.056 | 0 | 1 | 26 |
| nfya | 0.54902 | 0.256824547 | 0.056 | 0 | 1 | 26 |
| npsn | 0.54902 | 0.240226619 | 0.056 | 0 | 1 | 26 |
| nsd3 | 0.54902 | 0.255623877 | 0.056 | 0 | 1 | 26 |
| nucks1a | 0.54902 | 0.2536325 | 0.056 | 0 | 1 | 26 |

| | | | | | | |
|---|---|---|---|---|---|---|
| prkar1aa | 0.54902 | 0.2573064 | 0.056 | 0 | 1 | 26 |
| rbbp6 | 0.54902 | 0.25628586 | 0.056 | 0 | 1 | 26 |
| rnf4 | 0.54902 | 0.252013572 | 0.056 | 0 | 1 | 26 |
| si:ch211-119e14.1 | 0.54902 | 0.256950744 | 0.056 | 0 | 1 | 26 |
| si:dkey-206d17.12 | 0.54902 | 0.25690295 | 0.056 | 0 | 1 | 26 |
| smg7 | 0.54902 | 0.256488493 | 0.056 | 0 | 1 | 26 |
| smim20 | 0.54902 | 0.256724122 | 0.056 | 0 | 1 | 26 |
| sptan1 | 0.54902 | 0.25498434 | 0.056 | 0 | 1 | 26 |
| tmem115 | 0.54902 | 0.256716082 | 0.056 | 0 | 1 | 26 |
| trnt1 | 0.54902 | 0.256602816 | 0.056 | 0 | 1 | 26 |
| tspan17 | 0.54902 | 0.235424188 | 0.056 | 0 | 1 | 26 |
| ubl7b | 0.54902 | 0.256686997 | 0.056 | 0 | 1 | 26 |
| wbp1la | 0.54902 | 0.256312759 | 0.056 | 0 | 1 | 26 |
| zdhhc13 | 0.54902 | 0.257219095 | 0.056 | 0 | 1 | 26 |
| zfhx3 | 0.54902 | 0.257736812 | 0.056 | 0 | 1 | 26 |
| slc25a3b | 0.55461 | -0.3889873 | 0.333 | 0.571 | 1 | 26 |
| ywhae1 | 0.55636 | -0.5051452 | 0.111 | 0.286 | 1 | 26 |
| CU075735.1 | 0.55699 | -0.49401398 | 0.111 | 0.286 | 1 | 26 |
| hcls1 | 0.55723 | -0.5570214 | 0.111 | 0.286 | 1 | 26 |
| rpl5a | 0.55971 | 0.37817637 | 0.667 | 0.429 | 1 | 26 |
| anxa11a | 0.56053 | -0.39313012 | 0.111 | 0.286 | 1 | 26 |
| zgc:162944 | 0.56474 | -0.4823256 | 0.111 | 0.286 | 1 | 26 |
| ccdc124 | 0.56643 | -0.4744541 | 0.111 | 0.286 | 1 | 26 |
| si:ch73-359m17.2 | 0.57022 | -0.19247613 | 0.889 | 1 | 1 | 26 |
| eif3m | 0.57259 | -0.48185219 | 0.111 | 0.286 | 1 | 26 |
| COX5B | 0.57409 | -0.23226009 | 0.278 | 0.286 | 1 | 26 |
| metap2b | 0.57449 | -0.50504782 | 0.111 | 0.286 | 1 | 26 |
| rpl22l1 | 0.57524 | 0.13278919 | 0.278 | 0.143 | 1 | 26 |
| atp5l | 0.57973 | -0.36778025 | 0.222 | 0.286 | 1 | 26 |
| arpc4l | 0.58063 | -0.42600627 | 0.111 | 0.286 | 1 | 26 |
| rps3a | 0.5873 | 0.345791307 | 0.778 | 0.571 | 1 | 26 |
| arf5 | 0.58868 | 0.41750949 | 0.333 | 0.143 | 1 | 26 |
| fkbp3 | 0.59116 | -0.47173644 | 0.111 | 0.286 | 1 | 26 |
| prkab1b | 0.59127 | -0.46163277 | 0.111 | 0.286 | 1 | 26 |
| rps19 | 0.5913 | 0.187477392 | 0.889 | 0.714 | 1 | 26 |
| si:ch211-212d10.1 | 0.59175 | -0.40163601 | 0.222 | 0.429 | 1 | 26 |
| grap2b | 0.59303 | 0.236438033 | 0.444 | 0.286 | 1 | 26 |
| hmgb2a | 0.59436 | 0.466094541 | 0.5 | 0.286 | 1 | 26 |
| eif3s6ip | 0.59481 | -0.47989658 | 0.111 | 0.286 | 1 | 26 |
| sept12 | 0.597 | -0.42084139 | 0.111 | 0.286 | 1 | 26 |
| cox5ab | 0.59979 | -0.4701382 | 0.111 | 0.286 | 1 | 26 |
| hacd3 | 0.60008 | -0.44649415 | 0.111 | 0.286 | 1 | 26 |
| agr2 | 0.6002 | 0.275285023 | 0.667 | 0.571 | 1 | 26 |
| eif4a1a | 0.60037 | -0.46147895 | 0.111 | 0.286 | 1 | 26 |
| ctsd | 0.60049 | -0.45958221 | 0.111 | 0.286 | 1 | 26 |
| vdac2 | 0.60071 | 0.387919153 | 0.333 | 0.143 | 1 | 26 |
| sh3bgrl3 | 0.60083 | -0.46918397 | 0.222 | 0.429 | 1 | 26 |
| elocb | 0.60087 | -0.45647272 | 0.111 | 0.286 | 1 | 26 |

| | | | | | | |
|---|---|---|---|---|---|---|
| capza1a | 0.60118 | -0.4424665 | 0.222 | 0.429 | 1 | 26 |
| arpc2 | 0.60264 | -0.23187375 | 0.389 | 0.571 | 1 | 26 |
| rpl35 | 0.60294 | -0.18405459 | 0.944 | 1 | 1 | 26 |
| rpl3 | 0.61381 | 0.131783177 | 0.722 | 0.714 | 1 | 26 |
| rps17 | 0.61402 | -0.15771003 | 0.889 | 1 | 1 | 26 |
| eno3 | 0.61457 | 0.426143322 | 0.333 | 0.143 | 1 | 26 |
| hsp90b1 | 0.61477 | 0.476178795 | 0.278 | 0.143 | 1 | 26 |
| wasb | 0.61487 | 0.474572143 | 0.333 | 0.143 | 1 | 26 |
| serbp1a | 0.61502 | 0.469268605 | 0.333 | 0.143 | 1 | 26 |
| gyg1a | 0.61689 | 0.480428895 | 0.333 | 0.143 | 1 | 26 |
| atp5pf | 0.61755 | -0.33770549 | 0.222 | 0.286 | 1 | 26 |
| hmga1a | 0.62891 | -0.40152447 | 0.167 | 0.286 | 1 | 26 |
| rpl34 | 0.63434 | -0.16070575 | 0.833 | 0.857 | 1 | 26 |
| myh11a | 0.63657 | 0.255443941 | 0.278 | 0.143 | 1 | 26 |
| si:ch73-248e21.7 | 0.6379 | -0.40832352 | 0.167 | 0.286 | 1 | 26 |
| si:ch211-14a17.10 | 0.64111 | 0.226651152 | 0.778 | 0.714 | 1 | 26 |
| ost4 | 0.64125 | -0.31358598 | 0.222 | 0.286 | 1 | 26 |
| hnrnpa1b | 0.64969 | -0.10185341 | 0.167 | 0.143 | 1 | 26 |
| arhgdia | 0.65371 | 0.348237369 | 0.222 | 0.143 | 1 | 26 |
| mcl1a | 0.658 | -0.21879707 | 0.389 | 0.429 | 1 | 26 |
| rps8a | 0.65947 | -0.19296529 | 0.778 | 0.857 | 1 | 26 |
| rpl38 | 0.65997 | -0.25352487 | 0.722 | 0.857 | 1 | 26 |
| cfhl2 | 0.66062 | 0.254348353 | 0.278 | 0.143 | 1 | 26 |
| rpl26 | 0.66718 | -0.14976268 | 0.944 | 1 | 1 | 26 |
| rpl11 | 0.66991 | -0.13406155 | 1 | 1 | 1 | 26 |
| hspa5 | 0.6705 | 0.447327298 | 0.278 | 0.143 | 1 | 26 |
| si:ch1073-429i10.3.1 | 0.67815 | -0.38589798 | 0.167 | 0.286 | 1 | 26 |
| ier2a | 0.68223 | 0.427141163 | 0.278 | 0.143 | 1 | 26 |
| lta4h | 0.68378 | -0.29246231 | 0.333 | 0.429 | 1 | 26 |
| cox6a1 | 0.68398 | 0.208464501 | 0.278 | 0.143 | 1 | 26 |
| fth1a | 0.68941 | -0.22948596 | 0.5 | 0.571 | 1 | 26 |
| rpl28 | 0.69997 | -0.13046578 | 0.722 | 0.857 | 1 | 26 |
| oaz1b | 0.70038 | -0.28934673 | 0.389 | 0.571 | 1 | 26 |
| cst14b.2 | 0.70098 | -0.17889385 | 0.444 | 0.429 | 1 | 26 |
| mt-nd4 | 0.70104 | 0.423391216 | 0.444 | 0.286 | 1 | 26 |
| erh | 0.70409 | -0.20836795 | 0.167 | 0.286 | 1 | 26 |
| LOC110437886 | 0.71474 | 0.305866911 | 0.611 | 0.429 | 1 | 26 |
| pdlim1 | 0.72145 | -0.36862089 | 0.167 | 0.286 | 1 | 26 |
| p4hb | 0.72176 | 0.424822792 | 0.278 | 0.143 | 1 | 26 |
| eif3ba | 0.73603 | 0.313084231 | 0.278 | 0.143 | 1 | 26 |
| calr3b | 0.73814 | -0.24223759 | 0.167 | 0.286 | 1 | 26 |
| c3a.4 | 0.7407 | -0.23049541 | 0.167 | 0.286 | 1 | 26 |
| rps6 | 0.74745 | -0.21565648 | 0.778 | 0.857 | 1 | 26 |
| atp5mc1 | 0.75125 | -0.26532846 | 0.222 | 0.286 | 1 | 26 |
| cirbpb | 0.75254 | 0.237229869 | 0.722 | 0.571 | 1 | 26 |
| rpl6 | 0.75306 | 0.173938129 | 0.722 | 0.571 | 1 | 26 |
| h3f3d | 0.75484 | 0.364773723 | 0.278 | 0.143 | 1 | 26 |
| prelid3b | 0.75609 | -0.36040334 | 0.167 | 0.286 | 1 | 26 |

| | | | | | | |
|---|---|---|---|---|---|---|
| arpc5a | 0.75694 | -0.33830067 | 0.278 | 0.429 | 1 | 26 |
| rps16 | 0.76014 | -0.19904304 | 0.722 | 0.857 | 1 | 26 |
| CABZ01112049.1 | 0.76186 | 0.307809133 | 0.444 | 0.286 | 1 | 26 |
| rpl4 | 0.76268 | -0.21738426 | 0.556 | 0.714 | 1 | 26 |
| mt-nd2 | 0.76268 | 0.331483205 | 0.444 | 0.286 | 1 | 26 |
| eno1a | 0.76401 | 0.3115955 | 0.444 | 0.286 | 1 | 26 |
| tmed2 | 0.76843 | 0.320715954 | 0.278 | 0.143 | 1 | 26 |
| cirbpa | 0.76868 | 0.346134948 | 0.278 | 0.143 | 1 | 26 |
| nfkbiaa | 0.76873 | 0.357971404 | 0.278 | 0.143 | 1 | 26 |
| rack1 | 0.76898 | -0.1754759 | 0.722 | 0.857 | 1 | 26 |
| dad1 | 0.77659 | -0.29521979 | 0.278 | 0.429 | 1 | 26 |
| aldoaa | 0.7766 | -0.33621137 | 0.167 | 0.286 | 1 | 26 |
| rhoab | 0.77912 | 0.108535924 | 0.222 | 0.143 | 1 | 26 |
| fkbp1aa | 0.78673 | -0.24024325 | 0.167 | 0.286 | 1 | 26 |
| ssr3 | 0.78724 | 0.191046415 | 0.833 | 0.714 | 1 | 26 |
| cox4i1 | 0.79017 | -0.24902057 | 0.167 | 0.286 | 1 | 26 |
| paip2b | 0.79107 | -0.24837143 | 0.167 | 0.286 | 1 | 26 |
| hnrnpa0l | 0.79158 | -0.25970638 | 0.444 | 0.571 | 1 | 26 |
| LOC110438722 | 0.80207 | 0.316694839 | 0.222 | 0.143 | 1 | 26 |
| rps27a | 0.80289 | -0.17895119 | 0.778 | 0.857 | 1 | 26 |
| cox8a | 0.80648 | -0.14978514 | 0.389 | 0.429 | 1 | 26 |
| srsf3b | 0.80687 | -0.26217571 | 0.167 | 0.286 | 1 | 26 |
| atp5mc3a | 0.80792 | -0.25833654 | 0.167 | 0.286 | 1 | 26 |
| actr2a | 0.81064 | -0.30258616 | 0.167 | 0.286 | 1 | 26 |
| dbi | 0.82341 | -0.2586319 | 0.444 | 0.571 | 1 | 26 |
| selenow1 | 0.82838 | 0.280078303 | 0.222 | 0.143 | 1 | 26 |
| pinx1 | 0.83324 | -0.17374639 | 0.111 | 0.143 | 1 | 26 |
| XLOC-000431 | 0.83324 | -0.12717383 | 0.111 | 0.143 | 1 | 26 |
| ca6 | 0.83324 | -0.34190843 | 0.111 | 0.143 | 1 | 26 |
| csrp1a | 0.83324 | -0.17511299 | 0.111 | 0.143 | 1 | 26 |
| dynll1 | 0.83324 | -0.13293748 | 0.111 | 0.143 | 1 | 26 |
| gata2a | 0.83324 | -0.11039222 | 0.111 | 0.143 | 1 | 26 |
| glo1 | 0.83324 | -0.22243963 | 0.111 | 0.143 | 1 | 26 |
| hdac1 | 0.83324 | -0.22739839 | 0.111 | 0.143 | 1 | 26 |
| pak2a | 0.83324 | -0.16205437 | 0.111 | 0.143 | 1 | 26 |
| ppp3r1b | 0.83324 | -0.17484374 | 0.111 | 0.143 | 1 | 26 |
| skap2 | 0.83324 | -0.1612748 | 0.111 | 0.143 | 1 | 26 |
| tbcb | 0.83324 | -0.10707315 | 0.111 | 0.143 | 1 | 26 |
| tes | 0.83324 | -0.19818084 | 0.111 | 0.143 | 1 | 26 |
| tnrc6c1 | 0.83324 | -0.14692708 | 0.111 | 0.143 | 1 | 26 |
| tomm20b | 0.83324 | -0.11179899 | 0.111 | 0.143 | 1 | 26 |
| tpd52l2b | 0.83324 | -0.13986272 | 0.111 | 0.143 | 1 | 26 |
| trib3 | 0.83324 | -0.10089346 | 0.111 | 0.143 | 1 | 26 |
| zgc:153713 | 0.83324 | -0.11860661 | 0.111 | 0.143 | 1 | 26 |
| caspa | 0.83324 | -0.16651687 | 0.111 | 0.143 | 1 | 26 |
| cmpk | 0.83324 | -0.19685129 | 0.111 | 0.143 | 1 | 26 |
| creg1 | 0.83324 | -0.17814667 | 0.111 | 0.143 | 1 | 26 |
| elof1 | 0.83324 | -0.12266328 | 0.111 | 0.143 | 1 | 26 |

| | | | | | | |
|---|---|---|---|---|---|---|
| gnb1a | 0.83324 | -0.14067089 | 0.111 | 0.143 | 1 | 26 |
| hmgb1a | 0.83324 | -0.10974037 | 0.111 | 0.143 | 1 | 26 |
| id2a | 0.83324 | -0.10394782 | 0.111 | 0.143 | 1 | 26 |
| pde4ba | 0.83324 | -0.18251523 | 0.111 | 0.143 | 1 | 26 |
| si:zfos-169g10.3 | 0.83324 | -0.19228526 | 0.111 | 0.143 | 1 | 26 |
| snu13b | 0.83324 | -0.10952095 | 0.111 | 0.143 | 1 | 26 |
| stk17b | 0.83324 | -0.17811198 | 0.111 | 0.143 | 1 | 26 |
| calr | 0.83324 | -0.11044261 | 0.111 | 0.143 | 1 | 26 |
| kdelr2a | 0.83324 | -0.18996714 | 0.111 | 0.143 | 1 | 26 |
| rps27l | 0.83324 | -0.15048287 | 0.111 | 0.143 | 1 | 26 |
| sdha | 0.83324 | -0.171352 | 0.111 | 0.143 | 1 | 26 |
| serpine2 | 0.83324 | -0.11332302 | 0.111 | 0.143 | 1 | 26 |
| si:dkey-222f8.3 | 0.83324 | -0.10767708 | 0.111 | 0.143 | 1 | 26 |
| wipf1a | 0.83324 | -0.15525045 | 0.111 | 0.143 | 1 | 26 |
| serp1 | 0.83624 | -0.23669543 | 0.333 | 0.429 | 1 | 26 |
| tmsb1 | 0.83848 | -0.13804121 | 0.556 | 0.571 | 1 | 26 |
| ndufa1 | 0.8398 | 0.110663314 | 0.222 | 0.143 | 1 | 26 |
| cebpa | 0.84652 | -0.21819709 | 0.222 | 0.286 | 1 | 26 |
| fosab | 0.84816 | 0.138319222 | 0.222 | 0.143 | 1 | 26 |
| capgb | 0.84931 | -0.2248575 | 0.444 | 0.571 | 1 | 26 |
| rps11 | 0.85136 | -0.14447744 | 0.667 | 0.714 | 1 | 26 |
| capza1b | 0.85429 | -0.22125404 | 0.222 | 0.286 | 1 | 26 |
| sem1 | 0.8562 | 0.253466565 | 0.222 | 0.143 | 1 | 26 |
| ywhaz | 0.86037 | 0.150916153 | 0.222 | 0.143 | 1 | 26 |
| si:dkey-203a12.4 | 0.86462 | -0.26218024 | 0.222 | 0.286 | 1 | 26 |
| ubc | 0.86521 | -0.1053914 | 0.278 | 0.286 | 1 | 26 |
| oaz1a | 0.87589 | 0.245913032 | 0.389 | 0.286 | 1 | 26 |
| antxr2a | 0.88475 | 0.187764628 | 0.222 | 0.143 | 1 | 26 |
| si:dkey-203a12.5 | 0.8884 | 0.170944983 | 0.667 | 0.571 | 1 | 26 |
| NC-002333.4 | 0.88865 | 0.242497642 | 0.389 | 0.286 | 1 | 26 |
| c3a.5 | 0.88887 | 0.19118305 | 0.222 | 0.143 | 1 | 26 |
| ran | 0.88888 | -0.2190002 | 0.333 | 0.429 | 1 | 26 |
| ddx5 | 0.89123 | 0.13703948 | 0.167 | 0.143 | 1 | 26 |
| h2afvb | 0.89223 | -0.16265247 | 0.333 | 0.429 | 1 | 26 |
| atf4a | 0.89298 | 0.230914702 | 0.222 | 0.143 | 1 | 26 |
| atp5f1e | 0.89421 | -0.22337588 | 0.333 | 0.429 | 1 | 26 |
| zgc:158343 | 0.90083 | 0.188287943 | 0.222 | 0.143 | 1 | 26 |
| tma7 | 0.90131 | 0.171382409 | 0.222 | 0.143 | 1 | 26 |
| abracl | 0.90354 | 0.197772962 | 0.222 | 0.143 | 1 | 26 |
| spink2.4 | 0.90372 | -0.10614943 | 0.778 | 0.857 | 1 | 26 |
| eif3f | 0.90486 | 0.161066607 | 0.333 | 0.286 | 1 | 26 |
| si:zfos-2330d3.1 | 0.90504 | 0.103762275 | 0.611 | 0.571 | 1 | 26 |
| rpl5b | 0.90544 | 0.172859613 | 0.5 | 0.429 | 1 | 26 |
| sec61b | 0.90755 | -0.19627534 | 0.333 | 0.429 | 1 | 26 |
| hspa8 | 0.91528 | -0.10494555 | 0.667 | 0.714 | 1 | 26 |
| si:ch1073-443f11.2 | 0.9179 | -0.18245515 | 0.222 | 0.286 | 1 | 26 |
| ucp2 | 0.923 | -0.11650804 | 0.222 | 0.286 | 1 | 26 |
| ddx61 | 0.93305 | 0.122322299 | 0.167 | 0.143 | 1 | 26 |

| | | | | | | |
|---|---|---|---|---|---|---|
| si:ch1073-376c22.1 | 0.93636 | -0.12542955 | 0.389 | 0.429 | 1 | 26 |
| rsl24d1 | 0.93801 | -0.12708105 | 0.222 | 0.286 | 1 | 26 |
| srsf2a | 0.9394 | 0.119266778 | 0.167 | 0.143 | 1 | 26 |
| rac2 | 0.94371 | -0.10122053 | 0.5 | 0.571 | 1 | 26 |
| tram1 | 0.9447 | 0.112886353 | 0.167 | 0.143 | 1 | 26 |
| idh2 | 0.94554 | -0.15832395 | 0.222 | 0.286 | 1 | 26 |
| khdrbs1a | 0.94619 | -0.15338019 | 0.222 | 0.286 | 1 | 26 |
| scinlb | 0.94805 | -0.14836882 | 0.222 | 0.286 | 1 | 26 |
| arhgdig | 0.94953 | 0.135683856 | 0.5 | 0.429 | 1 | 26 |
| sat1a.2 | 0.95933 | 0.108504107 | 0.167 | 0.143 | 1 | 26 |
| ndufb7 | 0.96208 | 0.100552048 | 0.167 | 0.143 | 1 | 26 |
| ola1 | 0.97276 | 0.105432474 | 0.167 | 0.143 | 1 | 26 |
| ckbb | 0.97336 | 0.107553391 | 0.333 | 0.286 | 1 | 26 |
| igic1s1 | 0.00158 | 1.527381879 | 0.905 | 0 | 1 | 27 |
| CU896602.1 | 0.00158 | 1.731599455 | 0.905 | 0 | 1 | 27 |
| romo1 | 0.00158 | 1.463067443 | 0.905 | 0 | 1 | 27 |
| rhof | 0.0023 | -1.38608106 | 0.048 | 1 | 1 | 27 |
| mt-co2 | 0.00232 | -1.68225467 | 0.048 | 1 | 1 | 27 |
| serbp1a | 0.00246 | 0.379850586 | 0.857 | 1 | 1 | 27 |
| mt-nd4 | 0.00247 | -1.03264275 | 0.048 | 1 | 1 | 27 |
| dync1i2a | 0.00386 | -1.0616317 | 0 | 0.667 | 1 | 27 |
| mt-nd3 | 0.00386 | -1.18605785 | 0 | 0.667 | 1 | 27 |
| nnt | 0.00386 | -1.04388033 | 0 | 0.667 | 1 | 27 |
| rbpjb | 0.00386 | -1.25880634 | 0 | 0.667 | 1 | 27 |
| si:ch73-335l21.4 | 0.00386 | -1.22124875 | 0 | 0.667 | 1 | 27 |
| si:dkey-118j18.2 | 0.00386 | -1.13189715 | 0 | 0.667 | 1 | 27 |
| si:rp71-36a1.3 | 0.00386 | -1.07642605 | 0 | 0.667 | 1 | 27 |
| slc12a7b | 0.00386 | -1.10720189 | 0 | 0.667 | 1 | 27 |
| slc9a3r1a | 0.00386 | -1.15647855 | 0 | 0.667 | 1 | 27 |
| spink2.1 | 0.00386 | -0.93178045 | 0 | 0.667 | 1 | 27 |
| sucla2 | 0.00386 | -1.14613261 | 0 | 0.667 | 1 | 27 |
| si:ch211-243a20.4 | 0.00525 | -1.30855238 | 0.095 | 1 | 1 | 27 |
| XLOC-014079 | 0.00578 | -1.62560626 | 0.095 | 1 | 1 | 27 |
| erp44 | 0.00603 | -1.2743335 | 0.095 | 1 | 1 | 27 |
| cebpb | 0.00641 | -1.35266796 | 0.095 | 1 | 1 | 27 |
| nudcd3 | 0.00656 | -1.18242345 | 0.095 | 1 | 1 | 27 |
| zfp36l1a | 0.01066 | -1.08888031 | 0.476 | 1 | 1 | 27 |
| pak2a | 0.01086 | -1.21046188 | 0.143 | 1 | 1 | 27 |
| XLOC-026705 | 0.01089 | 1.495003118 | 0.762 | 0 | 1 | 27 |
| hmga1a | 0.01089 | 1.647750098 | 0.762 | 0 | 1 | 27 |
| si:dkey-78k11.9 | 0.01116 | -1.21034673 | 0.143 | 1 | 1 | 27 |
| cox6b2 | 0.01479 | 1.188276466 | 0.952 | 0.333 | 1 | 27 |
| mt-cyb | 0.01597 | -1.07998868 | 0.048 | 0.667 | 1 | 27 |
| si:cabz01074946.1 | 0.01597 | -1.18036529 | 0.048 | 0.667 | 1 | 27 |
| pmaip1 | 0.01597 | -1.14577689 | 0.048 | 0.667 | 1 | 27 |
| mccc2 | 0.01597 | -1.03296107 | 0.048 | 0.667 | 1 | 27 |
| pigv | 0.01597 | -0.95384658 | 0.048 | 0.667 | 1 | 27 |
| sptbn2 | 0.01597 | -1.04013528 | 0.048 | 0.667 | 1 | 27 |

| | | | | | | |
|---|---|---|---|---|---|---|
| tlr19 | 0.01597 | -1.02287068 | 0.048 | 0.667 | 1 | 27 |
| XLOC-043857 | 0.01597 | -0.92672275 | 0.048 | 0.667 | 1 | 27 |
| hmgcl | 0.01597 | -1.01234792 | 0.048 | 0.667 | 1 | 27 |
| selenot1a | 0.01597 | -1.01837838 | 0.048 | 0.667 | 1 | 27 |
| tmem50a | 0.01597 | -1.14945839 | 0.048 | 0.667 | 1 | 27 |
| ddx61 | 0.01716 | 1.226245387 | 0.714 | 0 | 1 | 27 |
| tubb2b | 0.01716 | 1.562645877 | 0.714 | 0 | 1 | 27 |
| cox7a2a | 0.01716 | 1.073993981 | 0.714 | 0 | 1 | 27 |
| atp2a3 | 0.01927 | -1.12496686 | 0.19 | 1 | 1 | 27 |
| si:dkey-27i16.2 | 0.0205 | -1.20656315 | 0.238 | 1 | 1 | 27 |
| COX3 | 0.023 | -1.31807122 | 0.19 | 1 | 1 | 27 |
| rpl5b | 0.02332 | 0.429369269 | 0.952 | 1 | 1 | 27 |
| rplp2l | 0.02335 | 0.226766003 | 1 | 1 | 1 | 27 |
| pgam1a | 0.02419 | -1.05422882 | 0.19 | 1 | 1 | 27 |
| ccl35.1 | 0.02538 | -1.01297418 | 0.429 | 0.667 | 1 | 27 |
| mknk2b | 0.02563 | 1.113540791 | 0.667 | 0 | 1 | 27 |
| si:ch73-281n10.2 | 0.02563 | 1.393907552 | 0.667 | 0 | 1 | 27 |
| mpeg1.1 | 0.02702 | -1.16617029 | 0.238 | 1 | 1 | 27 |
| mt-nd1 | 0.02704 | -0.43289395 | 0.095 | 0.667 | 1 | 27 |
| ywhabl | 0.03676 | 1.005028318 | 0.619 | 0 | 1 | 27 |
| zgc:193541 | 0.03676 | 0.842112328 | 0.619 | 0 | 1 | 27 |
| ptprc | 0.03697 | -1.08189549 | 0.286 | 1 | 1 | 27 |
| id3 | 0.03903 | -1.01829512 | 0.524 | 1 | 1 | 27 |
| pax5 | 0.04111 | 0.391580127 | 0.857 | 0.333 | 1 | 27 |
| lcp1 | 0.04553 | -1.13980428 | 0.095 | 0.667 | 1 | 27 |
| rpl18 | 0.04863 | 0.24797735 | 1 | 1 | 1 | 27 |
| cct6a | 0.05105 | 0.857899873 | 0.571 | 0 | 1 | 27 |
| cyba | 0.05105 | 0.953713303 | 0.571 | 0 | 1 | 27 |
| ighv1-4 | 0.05105 | 1.267354426 | 0.571 | 0 | 1 | 27 |
| mdh2 | 0.05105 | 0.786996149 | 0.571 | 0 | 1 | 27 |
| selenow1 | 0.05105 | 0.902453534 | 0.571 | 0 | 1 | 27 |
| sh3bgrl3 | 0.05105 | 0.856366308 | 0.571 | 0 | 1 | 27 |
| uqcr10 | 0.05105 | 0.803375348 | 0.571 | 0 | 1 | 27 |
| cotl1 | 0.05296 | -1.00757557 | 0.524 | 1 | 1 | 27 |
| rps20 | 0.0535 | 0.158629083 | 1 | 1 | 1 | 27 |
| XLOC-043043 | 0.05841 | -1.14943228 | 0.286 | 1 | 1 | 27 |
| BX322618.1 | 0.06747 | -0.88434574 | 0 | 0.333 | 1 | 27 |
| BX465862.1 | 0.06747 | -0.57107448 | 0 | 0.333 | 1 | 27 |
| BX571825.4 | 0.06747 | -0.96032316 | 0 | 0.333 | 1 | 27 |
| BX908750.1 | 0.06747 | -0.66171474 | 0 | 0.333 | 1 | 27 |
| CABZ01021433.1 | 0.06747 | -0.61454244 | 0 | 0.333 | 1 | 27 |
| CABZ01072952.1 | 0.06747 | -0.63997695 | 0 | 0.333 | 1 | 27 |
| CABZ01076758.1 | 0.06747 | -0.61831611 | 0 | 0.333 | 1 | 27 |
| CABZ01080371.2 | 0.06747 | -0.46260909 | 0 | 0.333 | 1 | 27 |
| CT573344.1 | 0.06747 | -0.55077371 | 0 | 0.333 | 1 | 27 |
| CU570889.1 | 0.06747 | -0.66664069 | 0 | 0.333 | 1 | 27 |
| CU929070.1 | 0.06747 | -0.6177936 | 0 | 0.333 | 1 | 27 |
| LOC100002469 | 0.06747 | -0.61378834 | 0 | 0.333 | 1 | 27 |

| | | | | | | |
|---|---|---|---|---|---|---|
| LOC101884606 | 0.06747 | -0.65658618 | 0 | 0.333 | 1 | 27 |
| LOC101885938 | 0.06747 | -0.38972768 | 0 | 0.333 | 1 | 27 |
| LOC101887156 | 0.06747 | -0.66664069 | 0 | 0.333 | 1 | 27 |
| LOC110439835 | 0.06747 | -0.54361053 | 0 | 0.333 | 1 | 27 |
| LOC557729 | 0.06747 | -0.73009281 | 0 | 0.333 | 1 | 27 |
| MFAP4 (1 of many) | 0.06747 | -0.47217755 | 0 | 0.333 | 1 | 27 |
| RSBN1 | 0.06747 | -0.61126269 | 0 | 0.333 | 1 | 27 |
| SFMBT1 | 0.06747 | -0.5813844 | 0 | 0.333 | 1 | 27 |
| XLOC-002129 | 0.06747 | -0.66664069 | 0 | 0.333 | 1 | 27 |
| XLOC-004189 | 0.06747 | -0.63550177 | 0 | 0.333 | 1 | 27 |
| XLOC-007299 | 0.06747 | -0.60622603 | 0 | 0.333 | 1 | 27 |
| XLOC-032844 | 0.06747 | -0.5459439 | 0 | 0.333 | 1 | 27 |
| abca2 | 0.06747 | -0.61974369 | 0 | 0.333 | 1 | 27 |
| acadm | 0.06747 | -0.70923534 | 0 | 0.333 | 1 | 27 |
| akap11 | 0.06747 | -0.69103568 | 0 | 0.333 | 1 | 27 |
| ankmy2a | 0.06747 | -0.6044529 | 0 | 0.333 | 1 | 27 |
| ankra2 | 0.06747 | -0.57732998 | 0 | 0.333 | 1 | 27 |
| arhgef4 | 0.06747 | -0.60758918 | 0 | 0.333 | 1 | 27 |
| aspscr1 | 0.06747 | -0.59740848 | 0 | 0.333 | 1 | 27 |
| asxl1 | 0.06747 | -0.5969177 | 0 | 0.333 | 1 | 27 |
| atp6v1aa | 0.06747 | -0.72207245 | 0 | 0.333 | 1 | 27 |
| bada | 0.06747 | -0.62734333 | 0 | 0.333 | 1 | 27 |
| bckdk | 0.06747 | -0.59400711 | 0 | 0.333 | 1 | 27 |
| bpgm | 0.06747 | -0.66327425 | 0 | 0.333 | 1 | 27 |
| brox | 0.06747 | -0.58847911 | 0 | 0.333 | 1 | 27 |
| c3a.5 | 0.06747 | -0.7137781 | 0 | 0.333 | 1 | 27 |
| camk4 | 0.06747 | -0.66664069 | 0 | 0.333 | 1 | 27 |
| capns1a | 0.06747 | -0.61762109 | 0 | 0.333 | 1 | 27 |
| casp9 | 0.06747 | -0.73097713 | 0 | 0.333 | 1 | 27 |
| ccdc59 | 0.06747 | -0.59741953 | 0 | 0.333 | 1 | 27 |
| ccdc88c | 0.06747 | -0.7318652 | 0 | 0.333 | 1 | 27 |
| cdc42se1 | 0.06747 | -0.58828117 | 0 | 0.333 | 1 | 27 |
| cdkn1d | 0.06747 | -0.66163034 | 0 | 0.333 | 1 | 27 |
| chd2 | 0.06747 | -0.65222766 | 0 | 0.333 | 1 | 27 |
| ciao2b | 0.06747 | -0.61008338 | 0 | 0.333 | 1 | 27 |
| clasp2 | 0.06747 | -0.61066218 | 0 | 0.333 | 1 | 27 |
| cox18 | 0.06747 | -0.66806947 | 0 | 0.333 | 1 | 27 |
| cpeb4b | 0.06747 | -0.60849642 | 0 | 0.333 | 1 | 27 |
| cry5 | 0.06747 | -0.62584481 | 0 | 0.333 | 1 | 27 |
| csf2rb | 0.06747 | -0.79254753 | 0 | 0.333 | 1 | 27 |
| csrp1a | 0.06747 | -0.60459209 | 0 | 0.333 | 1 | 27 |
| cwc22 | 0.06747 | -0.56847865 | 0 | 0.333 | 1 | 27 |
| cxcl8b.3 | 0.06747 | -0.20311753 | 0 | 0.333 | 1 | 27 |
| cyp3c1 | 0.06747 | -0.6006249 | 0 | 0.333 | 1 | 27 |
| dcaf15 | 0.06747 | -0.59617162 | 0 | 0.333 | 1 | 27 |
| ddx18 | 0.06747 | -0.50814277 | 0 | 0.333 | 1 | 27 |
| dhdds | 0.06747 | -0.57381952 | 0 | 0.333 | 1 | 27 |
| dnaja1 | 0.06747 | -0.59798553 | 0 | 0.333 | 1 | 27 |

| | | | | | | |
|---|---|---|---|---|---|---|
| eif2ak3 | 0.06747 | -0.65185728 | 0 | 0.333 | 1 | 27 |
| eif4a2 | 0.06747 | -0.69665005 | 0 | 0.333 | 1 | 27 |
| eif4e2rs1 | 0.06747 | -0.58787775 | 0 | 0.333 | 1 | 27 |
| eif4g2a | 0.06747 | -0.72167177 | 0 | 0.333 | 1 | 27 |
| epsti1 | 0.06747 | -0.65969292 | 0 | 0.333 | 1 | 27 |
| exoc1 | 0.06747 | -0.58998738 | 0 | 0.333 | 1 | 27 |
| ezra | 0.06747 | -0.39939725 | 0 | 0.333 | 1 | 27 |
| faf1 | 0.06747 | -0.60641367 | 0 | 0.333 | 1 | 27 |
| fam114a1 | 0.06747 | -0.68063007 | 0 | 0.333 | 1 | 27 |
| fam219b | 0.06747 | -0.64757783 | 0 | 0.333 | 1 | 27 |
| fbxw11b | 0.06747 | -0.72099396 | 0 | 0.333 | 1 | 27 |
| fip1l1b | 0.06747 | -0.60209474 | 0 | 0.333 | 1 | 27 |
| gigyf1a | 0.06747 | -0.59252556 | 0 | 0.333 | 1 | 27 |
| git2a | 0.06747 | -0.6593323 | 0 | 0.333 | 1 | 27 |
| glmp | 0.06747 | -0.61667446 | 0 | 0.333 | 1 | 27 |
| glod5 | 0.06747 | -0.60506414 | 0 | 0.333 | 1 | 27 |
| glud1b | 0.06747 | -0.7031641 | 0 | 0.333 | 1 | 27 |
| gpd1c | 0.06747 | -0.65660283 | 0 | 0.333 | 1 | 27 |
| gpkow | 0.06747 | -0.69768421 | 0 | 0.333 | 1 | 27 |
| hacl1 | 0.06747 | -0.6432522 | 0 | 0.333 | 1 | 27 |
| hccsb | 0.06747 | -0.65025177 | 0 | 0.333 | 1 | 27 |
| helz2 | 0.06747 | -0.66664069 | 0 | 0.333 | 1 | 27 |
| hhex | 0.06747 | -0.60953975 | 0 | 0.333 | 1 | 27 |
| hivep2a | 0.06747 | -0.61013552 | 0 | 0.333 | 1 | 27 |
| hnrnpl | 0.06747 | -0.64708675 | 0 | 0.333 | 1 | 27 |
| hps4 | 0.06747 | -0.59276343 | 0 | 0.333 | 1 | 27 |
| igf2bp3 | 0.06747 | -0.60222985 | 0 | 0.333 | 1 | 27 |
| im:7147486 | 0.06747 | -0.5737071 | 0 | 0.333 | 1 | 27 |
| ints2 | 0.06747 | -0.60165005 | 0 | 0.333 | 1 | 27 |
| irak3 | 0.06747 | -0.66422598 | 0 | 0.333 | 1 | 27 |
| itgb1a | 0.06747 | -0.57862533 | 0 | 0.333 | 1 | 27 |
| itpr1a | 0.06747 | -0.71636042 | 0 | 0.333 | 1 | 27 |
| kat6b | 0.06747 | -0.79129437 | 0 | 0.333 | 1 | 27 |
| kazald3 | 0.06747 | -0.64219706 | 0 | 0.333 | 1 | 27 |
| kcnh5b | 0.06747 | -0.62069873 | 0 | 0.333 | 1 | 27 |
| kdelr2b | 0.06747 | -0.70956165 | 0 | 0.333 | 1 | 27 |
| kdrl | 0.06747 | -0.55058315 | 0 | 0.333 | 1 | 27 |
| kidins220b | 0.06747 | -0.72544832 | 0 | 0.333 | 1 | 27 |
| kif21b | 0.06747 | -0.66462884 | 0 | 0.333 | 1 | 27 |
| klf12a | 0.06747 | -0.5508004 | 0 | 0.333 | 1 | 27 |
| klf3 | 0.06747 | -0.51688145 | 0 | 0.333 | 1 | 27 |
| klhl36 | 0.06747 | -0.58597504 | 0 | 0.333 | 1 | 27 |
| kri1 | 0.06747 | -0.70830355 | 0 | 0.333 | 1 | 27 |
| krit1 | 0.06747 | -0.72121036 | 0 | 0.333 | 1 | 27 |
| lpar2a | 0.06747 | -0.58171605 | 0 | 0.333 | 1 | 27 |
| lpin1 | 0.06747 | -0.62658713 | 0 | 0.333 | 1 | 27 |
| man1a2 | 0.06747 | -0.59828537 | 0 | 0.333 | 1 | 27 |
| map3k12 | 0.06747 | -0.62069873 | 0 | 0.333 | 1 | 27 |

| | | | | | | |
|---|---|---|---|---|---|---|
| mbip | 0.06747 | -0.57146102 | 0 | 0.333 | 1 | 27 |
| mboat7 | 0.06747 | -0.61220742 | 0 | 0.333 | 1 | 27 |
| med17 | 0.06747 | -0.70604453 | 0 | 0.333 | 1 | 27 |
| mesd | 0.06747 | -0.60431634 | 0 | 0.333 | 1 | 27 |
| mindy2 | 0.06747 | -0.62069873 | 0 | 0.333 | 1 | 27 |
| mlh3 | 0.06747 | -0.72053771 | 0 | 0.333 | 1 | 27 |
| mob1ba | 0.06747 | -0.63575011 | 0 | 0.333 | 1 | 27 |
| mon1bb | 0.06747 | -0.66064782 | 0 | 0.333 | 1 | 27 |
| mpdu1b | 0.06747 | -0.60846787 | 0 | 0.333 | 1 | 27 |
| mpped2 | 0.06747 | -0.70785819 | 0 | 0.333 | 1 | 27 |
| mri1 | 0.06747 | -0.61753244 | 0 | 0.333 | 1 | 27 |
| mrps25 | 0.06747 | -0.55269292 | 0 | 0.333 | 1 | 27 |
| mt-nd2 | 0.06747 | -0.93357903 | 0 | 0.333 | 1 | 27 |
| mtmr10 | 0.06747 | -0.66664069 | 0 | 0.333 | 1 | 27 |
| mybbp1a | 0.06747 | -0.58946017 | 0 | 0.333 | 1 | 27 |
| napab | 0.06747 | -0.63810189 | 0 | 0.333 | 1 | 27 |
| ndrg3b | 0.06747 | -0.6991155 | 0 | 0.333 | 1 | 27 |
| nek6 | 0.06747 | -0.72615483 | 0 | 0.333 | 1 | 27 |
| nf1a | 0.06747 | -0.60342081 | 0 | 0.333 | 1 | 27 |
| nfs1 | 0.06747 | -0.56042481 | 0 | 0.333 | 1 | 27 |
| nocta | 0.06747 | -0.66360893 | 0 | 0.333 | 1 | 27 |
| nr2c2ap | 0.06747 | -0.54875237 | 0 | 0.333 | 1 | 27 |
| osbpl9 | 0.06747 | -0.6167946 | 0 | 0.333 | 1 | 27 |
| osgepl1 | 0.06747 | -0.55249092 | 0 | 0.333 | 1 | 27 |
| oxr1a | 0.06747 | -0.55379184 | 0 | 0.333 | 1 | 27 |
| pan2 | 0.06747 | -0.71623375 | 0 | 0.333 | 1 | 27 |
| pargl | 0.06747 | -0.60002536 | 0 | 0.333 | 1 | 27 |
| pcmtd2 | 0.06747 | -0.65463477 | 0 | 0.333 | 1 | 27 |
| pex5 | 0.06747 | -0.45098225 | 0 | 0.333 | 1 | 27 |
| pgk1 | 0.06747 | -0.71321132 | 0 | 0.333 | 1 | 27 |
| phactr4a | 0.06747 | -0.65254442 | 0 | 0.333 | 1 | 27 |
| piga | 0.06747 | -0.65658618 | 0 | 0.333 | 1 | 27 |
| pigb | 0.06747 | -0.61673361 | 0 | 0.333 | 1 | 27 |
| pik3ap1 | 0.06747 | -0.62069873 | 0 | 0.333 | 1 | 27 |
| pip4p1b | 0.06747 | -0.66664069 | 0 | 0.333 | 1 | 27 |
| pja2 | 0.06747 | -0.48546758 | 0 | 0.333 | 1 | 27 |
| plekhf2 | 0.06747 | -0.59034573 | 0 | 0.333 | 1 | 27 |
| ppp1r37 | 0.06747 | -0.57209349 | 0 | 0.333 | 1 | 27 |
| pstpip2 | 0.06747 | -0.61046877 | 0 | 0.333 | 1 | 27 |
| ptdss1a | 0.06747 | -0.65837274 | 0 | 0.333 | 1 | 27 |
| pus7 | 0.06747 | -0.60681758 | 0 | 0.333 | 1 | 27 |
| rab3il1 | 0.06747 | -0.61723009 | 0 | 0.333 | 1 | 27 |
| rab40c | 0.06747 | -0.39749888 | 0 | 0.333 | 1 | 27 |
| rab4a | 0.06747 | -0.71154954 | 0 | 0.333 | 1 | 27 |
| rabep1 | 0.06747 | -0.58323479 | 0 | 0.333 | 1 | 27 |
| rac1b | 0.06747 | -0.54797461 | 0 | 0.333 | 1 | 27 |
| rap2aa | 0.06747 | -0.65730901 | 0 | 0.333 | 1 | 27 |
| rasal2 | 0.06747 | -0.72739911 | 0 | 0.333 | 1 | 27 |

| | | | | | | |
|---|---|---|---|---|---|---|
| rfx1a | 0.06747 | -0.65658618 | 0 | 0.333 | 1 | 27 |
| rif1 | 0.06747 | -0.58537887 | 0 | 0.333 | 1 | 27 |
| rmdn2 | 0.06747 | -0.60595425 | 0 | 0.333 | 1 | 27 |
| rmdn3 | 0.06747 | -0.78436369 | 0 | 0.333 | 1 | 27 |
| rnpepl1 | 0.06747 | -0.72546825 | 0 | 0.333 | 1 | 27 |
| rorab | 0.06747 | -0.65977748 | 0 | 0.333 | 1 | 27 |
| rprd2b | 0.06747 | -0.78869565 | 0 | 0.333 | 1 | 27 |
| rtn3 | 0.06747 | -0.6069145 | 0 | 0.333 | 1 | 27 |
| scdb | 0.06747 | -0.7318652 | 0 | 0.333 | 1 | 27 |
| sdad1 | 0.06747 | -0.5746472 | 0 | 0.333 | 1 | 27 |
| selenoo1 | 0.06747 | -0.59589084 | 0 | 0.333 | 1 | 27 |
| sema3aa | 0.06747 | -0.7318652 | 0 | 0.333 | 1 | 27 |
| si:cabz01101003.1 | 0.06747 | -0.66276472 | 0 | 0.333 | 1 | 27 |
| si:ch211-132g1.1 | 0.06747 | -0.7318652 | 0 | 0.333 | 1 | 27 |
| si:ch211-212k18.9 | 0.06747 | -0.57753408 | 0 | 0.333 | 1 | 27 |
| si:ch211-214p16.2 | 0.06747 | -0.60884327 | 0 | 0.333 | 1 | 27 |
| si:ch211-275j6.5 | 0.06747 | -0.53257702 | 0 | 0.333 | 1 | 27 |
| si:ch73-158p21.3 | 0.06747 | -0.61380998 | 0 | 0.333 | 1 | 27 |
| si:ch73-285p12.4 | 0.06747 | -0.61671575 | 0 | 0.333 | 1 | 27 |
| si:ch73-382f3.1 | 0.06747 | -0.48319299 | 0 | 0.333 | 1 | 27 |
| si:dkey-171c9.3 | 0.06747 | -0.65986894 | 0 | 0.333 | 1 | 27 |
| si:dkey-29h14.10 | 0.06747 | -0.62069873 | 0 | 0.333 | 1 | 27 |
| si:dkey-53k12.2 | 0.06747 | -0.61899002 | 0 | 0.333 | 1 | 27 |
| si:dkey-89b17.4 | 0.06747 | -0.61329852 | 0 | 0.333 | 1 | 27 |
| si:dkeyp-2e4.6 | 0.06747 | -0.54361053 | 0 | 0.333 | 1 | 27 |
| slain2 | 0.06747 | -0.72775696 | 0 | 0.333 | 1 | 27 |
| slc27a4 | 0.06747 | -0.62069873 | 0 | 0.333 | 1 | 27 |
| slc2a13b | 0.06747 | -0.63140244 | 0 | 0.333 | 1 | 27 |
| slc50a1 | 0.06747 | -0.58381031 | 0 | 0.333 | 1 | 27 |
| slc9a6a | 0.06747 | -0.56812372 | 0 | 0.333 | 1 | 27 |
| smco4 | 0.06747 | -0.59079437 | 0 | 0.333 | 1 | 27 |
| smfn | 0.06747 | -0.57394677 | 0 | 0.333 | 1 | 27 |
| sntb1 | 0.06747 | -0.57978976 | 0 | 0.333 | 1 | 27 |
| snu13a | 0.06747 | -0.6382734 | 0 | 0.333 | 1 | 27 |
| spg11 | 0.06747 | -0.61478482 | 0 | 0.333 | 1 | 27 |
| st8sia1 | 0.06747 | -0.72459115 | 0 | 0.333 | 1 | 27 |
| stim1a | 0.06747 | -0.61649585 | 0 | 0.333 | 1 | 27 |
| stoml2 | 0.06747 | -0.70703041 | 0 | 0.333 | 1 | 27 |
| strap | 0.06747 | -0.57680982 | 0 | 0.333 | 1 | 27 |
| swt1 | 0.06747 | -0.5640406 | 0 | 0.333 | 1 | 27 |
| tbcd | 0.06747 | -0.60864997 | 0 | 0.333 | 1 | 27 |
| tbck | 0.06747 | -0.16618272 | 0 | 0.333 | 1 | 27 |
| tbl1x | 0.06747 | -0.71585509 | 0 | 0.333 | 1 | 27 |
| tbl1xr1b | 0.06747 | -0.64297642 | 0 | 0.333 | 1 | 27 |
| tbl2 | 0.06747 | -0.60560291 | 0 | 0.333 | 1 | 27 |
| tdg.2 | 0.06747 | -0.44208933 | 0 | 0.333 | 1 | 27 |
| thoc3 | 0.06747 | -0.5870908 | 0 | 0.333 | 1 | 27 |
| tmem168a | 0.06747 | -0.59569405 | 0 | 0.333 | 1 | 27 |

| | | | | | | |
|---|---|---|---|---|---|---|
| tmem173 | 0.06747 | -0.61945117 | 0 | 0.333 | 1 | 27 |
| tmem178b | 0.06747 | -0.60401132 | 0 | 0.333 | 1 | 27 |
| tnfaip8l1 | 0.06747 | -0.72553064 | 0 | 0.333 | 1 | 27 |
| tnfrsf1a | 0.06747 | -0.66352685 | 0 | 0.333 | 1 | 27 |
| tnpo3 | 0.06747 | -0.65232732 | 0 | 0.333 | 1 | 27 |
| tomm34 | 0.06747 | -0.6477935 | 0 | 0.333 | 1 | 27 |
| top2b | 0.06747 | -0.60885149 | 0 | 0.333 | 1 | 27 |
| trim25 | 0.06747 | -0.60541642 | 0 | 0.333 | 1 | 27 |
| tshz2 | 0.06747 | -0.58690975 | 0 | 0.333 | 1 | 27 |
| ttl | 0.06747 | -0.62826245 | 0 | 0.333 | 1 | 27 |
| uba7 | 0.06747 | -0.60830278 | 0 | 0.333 | 1 | 27 |
| ubald1b | 0.06747 | -0.60831685 | 0 | 0.333 | 1 | 27 |
| ube2j2 | 0.06747 | -0.71477211 | 0 | 0.333 | 1 | 27 |
| ube2kb | 0.06747 | -0.60133431 | 0 | 0.333 | 1 | 27 |
| ugt1b5 | 0.06747 | -0.62689776 | 0 | 0.333 | 1 | 27 |
| unc50 | 0.06747 | -0.64769328 | 0 | 0.333 | 1 | 27 |
| urm1 | 0.06747 | -0.59443799 | 0 | 0.333 | 1 | 27 |
| usp44 | 0.06747 | -0.64438655 | 0 | 0.333 | 1 | 27 |
| utp11l | 0.06747 | -0.54813207 | 0 | 0.333 | 1 | 27 |
| utrn | 0.06747 | -0.72164683 | 0 | 0.333 | 1 | 27 |
| wapla | 0.06747 | -0.59153077 | 0 | 0.333 | 1 | 27 |
| wdr3 | 0.06747 | -0.59098484 | 0 | 0.333 | 1 | 27 |
| wdsub1 | 0.06747 | -0.61485352 | 0 | 0.333 | 1 | 27 |
| wnk1b | 0.06747 | -0.72657972 | 0 | 0.333 | 1 | 27 |
| xpo6 | 0.06747 | -0.60729678 | 0 | 0.333 | 1 | 27 |
| zfr | 0.06747 | -0.72380687 | 0 | 0.333 | 1 | 27 |
| zgc:110269 | 0.06747 | -0.60086866 | 0 | 0.333 | 1 | 27 |
| zgc:123105 | 0.06747 | -0.62048367 | 0 | 0.333 | 1 | 27 |
| zgc:162255 | 0.06747 | -0.60793729 | 0 | 0.333 | 1 | 27 |
| zgc:64002 | 0.06747 | -0.63829781 | 0 | 0.333 | 1 | 27 |
| zgc:77650 | 0.06747 | -0.65873141 | 0 | 0.333 | 1 | 27 |
| zgc:92335 | 0.06747 | -0.58710744 | 0 | 0.333 | 1 | 27 |
| zgc:92818 | 0.06747 | -0.8113086 | 0 | 0.333 | 1 | 27 |
| zmat5 | 0.06747 | -0.63201867 | 0 | 0.333 | 1 | 27 |
| zmynd8 | 0.06747 | -0.53295723 | 0 | 0.333 | 1 | 27 |
| znf1158 | 0.06747 | -0.59435384 | 0 | 0.333 | 1 | 27 |
| znf1160 | 0.06747 | -0.72446934 | 0 | 0.333 | 1 | 27 |
| znf1164 | 0.06747 | -0.59803602 | 0 | 0.333 | 1 | 27 |
| znf131 | 0.06747 | -0.58925425 | 0 | 0.333 | 1 | 27 |
| znf318 | 0.06747 | -0.51673265 | 0 | 0.333 | 1 | 27 |
| znf592 | 0.06747 | -0.70678733 | 0 | 0.333 | 1 | 27 |
| gtpbp4 | 0.06908 | 0.842429309 | 0.524 | 0 | 1 | 27 |
| itm2ba | 0.06908 | 0.879061668 | 0.524 | 0 | 1 | 27 |
| nutf2l | 0.06908 | 1.029991613 | 0.524 | 0 | 1 | 27 |
| rbx1 | 0.06908 | 0.924963229 | 0.524 | 0 | 1 | 27 |
| tegt | 0.06908 | 0.835927028 | 0.524 | 0 | 1 | 27 |
| ywhaz | 0.06908 | 0.867030314 | 0.524 | 0 | 1 | 27 |
| cox6c | 0.06908 | 0.766675741 | 0.524 | 0 | 1 | 27 |

| | | | | | | |
|---|---|---|---|---|---|---|
| psma2 | 0.06908 | 0.806923044 | 0.524 | 0 | 1 | 27 |
| psma3 | 0.06908 | 0.931750756 | 0.524 | 0 | 1 | 27 |
| ptp4a3 | 0.06908 | 1.029688821 | 0.524 | 0 | 1 | 27 |
| skap2 | 0.06908 | 0.975820647 | 0.524 | 0 | 1 | 27 |
| tram1 | 0.06908 | 1.003581292 | 0.524 | 0 | 1 | 27 |
| ywhabb | 0.06908 | 0.823373726 | 0.524 | 0 | 1 | 27 |
| zgc:153713 | 0.06908 | 0.879761061 | 0.524 | 0 | 1 | 27 |
| abhd3 | 0.07273 | -0.96488456 | 0.095 | 0.667 | 1 | 27 |
| XLOC-000431 | 0.07338 | -0.49893815 | 0.952 | 1 | 1 | 27 |
| crip1 | 0.07479 | 0.293633818 | 0.952 | 1 | 1 | 27 |
| arpc1b | 0.07521 | -0.80109867 | 0.667 | 1 | 1 | 27 |
| usp7 | 0.07628 | -0.99611064 | 0.095 | 0.667 | 1 | 27 |
| ginm1 | 0.07704 | -0.94378095 | 0.095 | 0.667 | 1 | 27 |
| XLOC-023492 | 0.07889 | -1.10248573 | 0.095 | 0.667 | 1 | 27 |
| LOC101886979 | 0.08054 | -0.84667427 | 0.333 | 1 | 1 | 27 |
| ipo7 | 0.08114 | -1.06207893 | 0.143 | 0.667 | 1 | 27 |
| ilk | 0.08166 | -1.0122294 | 0.143 | 0.667 | 1 | 27 |
| BX120005.1 | 0.0847 | -1.08102806 | 0.095 | 0.667 | 1 | 27 |
| tomm7 | 0.08757 | -0.91542444 | 0.286 | 0.667 | 1 | 27 |
| arl6ip1 | 0.08846 | -0.8514028 | 0.429 | 1 | 1 | 27 |
| lbh | 0.08897 | -0.96730294 | 0.095 | 0.667 | 1 | 27 |
| ccng1 | 0.09153 | 0.745003766 | 0.476 | 0 | 1 | 27 |
| morf4l1 | 0.09153 | 0.824285685 | 0.476 | 0 | 1 | 27 |
| spi1b | 0.09153 | 0.831405883 | 0.476 | 0 | 1 | 27 |
| ubb | 0.09153 | 0.880008309 | 0.476 | 0 | 1 | 27 |
| BX571825.3 | 0.09153 | 1.130719508 | 0.476 | 0 | 1 | 27 |
| CU571382.4 | 0.09153 | 0.903746558 | 0.476 | 0 | 1 | 27 |
| fam32a | 0.09153 | 0.786148343 | 0.476 | 0 | 1 | 27 |
| hint1 | 0.09153 | 0.854356715 | 0.476 | 0 | 1 | 27 |
| hsbp1b | 0.09153 | 0.75553588 | 0.476 | 0 | 1 | 27 |
| lsm12b | 0.09153 | 0.732313789 | 0.476 | 0 | 1 | 27 |
| mdh1aa | 0.09153 | 0.824843426 | 0.476 | 0 | 1 | 27 |
| ndufb8 | 0.09153 | 0.774983482 | 0.476 | 0 | 1 | 27 |
| pacsin3 | 0.09153 | 0.930118105 | 0.476 | 0 | 1 | 27 |
| rgcc | 0.09153 | 0.810362418 | 0.476 | 0 | 1 | 27 |
| rhoab | 0.09153 | 0.795272603 | 0.476 | 0 | 1 | 27 |
| si:ch211-145b13.6 | 0.09153 | 0.933725884 | 0.476 | 0 | 1 | 27 |
| sub1a | 0.09153 | 0.745110152 | 0.476 | 0 | 1 | 27 |
| tp53 | 0.09153 | 0.765389355 | 0.476 | 0 | 1 | 27 |
| tent4b | 0.09185 | -0.90129863 | 0.095 | 0.667 | 1 | 27 |
| ebag9 | 0.09729 | -0.80267301 | 0.095 | 0.667 | 1 | 27 |
| pdpk1a | 0.10135 | -0.90184682 | 0.095 | 0.667 | 1 | 27 |
| mt-co1 | 0.10178 | -0.93994115 | 0.381 | 1 | 1 | 27 |
| mrap2a | 0.10317 | -0.97343052 | 0.095 | 0.667 | 1 | 27 |
| si:ch211-214p16.1 | 0.1047 | -0.77212953 | 0.619 | 1 | 1 | 27 |
| hcls1 | 0.10599 | -0.96407472 | 0.095 | 0.667 | 1 | 27 |
| zgc:56493 | 0.10616 | 0.731293299 | 0.952 | 0.667 | 1 | 27 |
| dusp6 | 0.10941 | -0.89648707 | 0.095 | 0.667 | 1 | 27 |

| | | | | | | |
|---|---|---|---|---|---|---|
| tmc6a | 0.10993 | -0.95536885 | 0.095 | 0.667 | 1 | 27 |
| ctsh | 0.11033 | -0.90087156 | 0.095 | 0.667 | 1 | 27 |
| irf4b | 0.1107 | -0.68775504 | 0.476 | 0.667 | 1 | 27 |
| mbnl1 | 0.11248 | 0.661901806 | 0.905 | 0.333 | 1 | 27 |
| s1pr4 | 0.11285 | -0.858531 | 0.095 | 0.667 | 1 | 27 |
| oaz1b | 0.115 | 0.471580084 | 0.857 | 0.333 | 1 | 27 |
| eef2b | 0.11647 | 0.105348906 | 0.857 | 1 | 1 | 27 |
| sumo2b | 0.11704 | -0.77949895 | 0.571 | 1 | 1 | 27 |
| gabarapb | 0.11783 | -0.72053522 | 0.714 | 1 | 1 | 27 |
| atp5mf | 0.11918 | 0.720859172 | 0.429 | 0 | 1 | 27 |
| atrx | 0.11918 | 0.789285768 | 0.429 | 0 | 1 | 27 |
| ilf2 | 0.11918 | 0.699925409 | 0.429 | 0 | 1 | 27 |
| ndufa6 | 0.11918 | 0.517560546 | 0.429 | 0 | 1 | 27 |
| pabpn1 | 0.11918 | 0.759373219 | 0.429 | 0 | 1 | 27 |
| pacsin2 | 0.11918 | 0.89688659 | 0.429 | 0 | 1 | 27 |
| rel | 0.11918 | 0.836170415 | 0.429 | 0 | 1 | 27 |
| sap18 | 0.11918 | 0.607645206 | 0.429 | 0 | 1 | 27 |
| sept2 | 0.11918 | 0.716614197 | 0.429 | 0 | 1 | 27 |
| spcs1 | 0.11918 | 0.710170126 | 0.429 | 0 | 1 | 27 |
| srsf9 | 0.11918 | 0.706039652 | 0.429 | 0 | 1 | 27 |
| swap70a | 0.11918 | 0.787136887 | 0.429 | 0 | 1 | 27 |
| tfdp2 | 0.11918 | 0.820956046 | 0.429 | 0 | 1 | 27 |
| tmem258 | 0.11918 | 0.777628761 | 0.429 | 0 | 1 | 27 |
| LOC103909452 | 0.11918 | 0.960626048 | 0.429 | 0 | 1 | 27 |
| adh5 | 0.11918 | 0.730551121 | 0.429 | 0 | 1 | 27 |
| psmb2 | 0.11918 | 0.631902286 | 0.429 | 0 | 1 | 27 |
| sp6 | 0.11918 | 0.760556246 | 0.429 | 0 | 1 | 27 |
| ube2na | 0.11918 | 0.598451741 | 0.429 | 0 | 1 | 27 |
| actb2 | 0.12626 | -0.23141878 | 1 | 1 | 1 | 27 |
| arpc4 | 0.13221 | -0.59159145 | 0.524 | 0.667 | 1 | 27 |
| lasp1 | 0.13291 | -0.70215152 | 0.524 | 1 | 1 | 27 |
| si:ch73-234b20.5 | 0.13508 | -0.7640729 | 0.429 | 1 | 1 | 27 |
| atf4a | 0.13796 | -0.70206436 | 0.524 | 1 | 1 | 27 |
| XLOC-018356 | 0.14109 | -0.83262594 | 0.429 | 1 | 1 | 27 |
| LOC100536213 | 0.14184 | -0.77214013 | 0.619 | 1 | 1 | 27 |
| gapdhs | 0.14377 | -0.78131455 | 0.381 | 0.667 | 1 | 27 |
| bcl2l10 | 0.14668 | -0.61262124 | 0.429 | 1 | 1 | 27 |
| capza1a | 0.14674 | -0.72671013 | 0.476 | 1 | 1 | 27 |
| gnb1b | 0.14739 | -0.62914356 | 0.429 | 1 | 1 | 27 |
| tagapa | 0.14856 | -0.71896022 | 0.429 | 1 | 1 | 27 |
| rps3a | 0.14861 | 0.224178119 | 1 | 1 | 1 | 27 |
| cpne1 | 0.14882 | -0.84264601 | 0.143 | 0.667 | 1 | 27 |
| zgc:111986 | 0.14969 | -0.88413473 | 0.143 | 0.667 | 1 | 27 |
| arf6b | 0.15183 | -0.7985112 | 0.286 | 0.667 | 1 | 27 |
| capga | 0.15301 | 0.888119704 | 0.381 | 0 | 1 | 27 |
| egr3 | 0.15301 | 0.73013005 | 0.381 | 0 | 1 | 27 |
| h2afy2 | 0.15301 | 0.606061945 | 0.381 | 0 | 1 | 27 |
| hopx | 0.15301 | 0.697083846 | 0.381 | 0 | 1 | 27 |

| | | | | | | |
|---|---|---|---|---|---|---|
| si:ch211-260e23.9 | 0.15301 | 0.707139779 | 0.381 | 0 | 1 | 27 |
| zfand5b | 0.15301 | 0.627015234 | 0.381 | 0 | 1 | 27 |
| cnbpa | 0.15301 | 0.70414378 | 0.381 | 0 | 1 | 27 |
| eif4a3 | 0.15301 | 0.532899284 | 0.381 | 0 | 1 | 27 |
| ndufa2 | 0.15301 | 0.542372515 | 0.381 | 0 | 1 | 27 |
| ndufs6 | 0.15301 | 0.561981982 | 0.381 | 0 | 1 | 27 |
| si:ch211-288g17.3 | 0.15301 | 0.838841122 | 0.381 | 0 | 1 | 27 |
| LOC108179343 | 0.15301 | 0.783104156 | 0.381 | 0 | 1 | 27 |
| STMP1 | 0.15301 | 0.606116728 | 0.381 | 0 | 1 | 27 |
| cnpy3 | 0.15301 | 0.714694203 | 0.381 | 0 | 1 | 27 |
| h2afx1 | 0.15301 | 0.781057171 | 0.381 | 0 | 1 | 27 |
| lmo2 | 0.15301 | 0.546840618 | 0.381 | 0 | 1 | 27 |
| psmg3 | 0.15301 | 0.60910176 | 0.381 | 0 | 1 | 27 |
| ptp4a1 | 0.15301 | 0.559463785 | 0.381 | 0 | 1 | 27 |
| stk35 | 0.15301 | 0.583270451 | 0.381 | 0 | 1 | 27 |
| syf2 | 0.15301 | 0.559107858 | 0.381 | 0 | 1 | 27 |
| hnrnpk | 0.15301 | 0.613641574 | 0.381 | 0 | 1 | 27 |
| snrpc | 0.15301 | 0.628400056 | 0.381 | 0 | 1 | 27 |
| taf12 | 0.15301 | 0.572649837 | 0.381 | 0 | 1 | 27 |
| twf1a | 0.15301 | 0.645783045 | 0.381 | 0 | 1 | 27 |
| zgc:91910 | 0.15301 | 0.644853978 | 0.381 | 0 | 1 | 27 |
| zmat2 | 0.15301 | 0.674216614 | 0.381 | 0 | 1 | 27 |
| lxn | 0.15365 | -0.82604121 | 0.143 | 0.667 | 1 | 27 |
| CR936442.1 | 0.15419 | -0.43868096 | 0.619 | 0.667 | 1 | 27 |
| LOC103910438 | 0.15641 | -0.25954183 | 0.476 | 0.333 | 1 | 27 |
| ywhae1 | 0.15722 | -0.20488672 | 0.429 | 0.333 | 1 | 27 |
| sec61b | 0.15855 | 0.869154846 | 0.81 | 0.333 | 1 | 27 |
| gtf2h5 | 0.15906 | -0.83599906 | 0.143 | 0.667 | 1 | 27 |
| rps6 | 0.15926 | 0.235588738 | 1 | 1 | 1 | 27 |
| galnt7 | 0.16301 | -0.88153226 | 0.143 | 0.667 | 1 | 27 |
| rnaset2l | 0.16539 | -0.90912344 | 0.143 | 0.667 | 1 | 27 |
| sod1 | 0.16605 | 0.181091616 | 0.667 | 0.333 | 1 | 27 |
| calm1b | 0.16701 | -0.65503677 | 0.476 | 1 | 1 | 27 |
| rplp0 | 0.1679 | 0.326384441 | 1 | 1 | 1 | 27 |
| rac2 | 0.16862 | -0.40825179 | 0.905 | 1 | 1 | 27 |
| nsa2 | 0.17976 | 0.632360177 | 0.857 | 0.333 | 1 | 27 |
| mob1a | 0.17994 | -0.64308573 | 0.476 | 1 | 1 | 27 |
| ccr2 | 0.18019 | -0.95657099 | 0.19 | 0.667 | 1 | 27 |
| slka | 0.18042 | -0.85312178 | 0.143 | 0.667 | 1 | 27 |
| ponzr1 | 0.18091 | -0.24649882 | 0.762 | 0.667 | 1 | 27 |
| cap1 | 0.18248 | -0.70974864 | 0.524 | 1 | 1 | 27 |
| ikzf1 | 0.18347 | -0.1593891 | 0.476 | 0.333 | 1 | 27 |
| map2k6 | 0.18444 | -0.84209368 | 0.143 | 0.667 | 1 | 27 |
| mhc2b | 0.18447 | -0.73236032 | 0.571 | 1 | 1 | 27 |
| zgc:86598 | 0.18608 | -0.74380652 | 0.143 | 0.667 | 1 | 27 |
| nccrp1 | 0.18873 | -0.59683375 | 0.048 | 0.333 | 1 | 27 |
| skap1 | 0.18873 | -0.50874988 | 0.048 | 0.333 | 1 | 27 |
| tmem176l.3a | 0.18873 | -0.52327766 | 0.048 | 0.333 | 1 | 27 |

| | | | | | | |
|---|---|---|---|---|---|---|
| tmx1 | 0.18873 | -0.51797647 | 0.048 | 0.333 | 1 | 27 |
| BX005340.1 | 0.18873 | -0.52630071 | 0.048 | 0.333 | 1 | 27 |
| CABZ01046432.1 | 0.18873 | -0.52959155 | 0.048 | 0.333 | 1 | 27 |
| CABZ01077217.1 | 0.18873 | -0.49774957 | 0.048 | 0.333 | 1 | 27 |
| CABZ01112656.1 | 0.18873 | -0.42759175 | 0.048 | 0.333 | 1 | 27 |
| CR847994.1 | 0.18873 | -0.63557586 | 0.048 | 0.333 | 1 | 27 |
| CR848667.1 | 0.18873 | -0.10265853 | 0.048 | 0.333 | 1 | 27 |
| CU550719.2 | 0.18873 | -0.57754943 | 0.048 | 0.333 | 1 | 27 |
| FP074902.2 | 0.18873 | -0.62520503 | 0.048 | 0.333 | 1 | 27 |
| LOC101884451 | 0.18873 | -0.50681232 | 0.048 | 0.333 | 1 | 27 |
| LOC108191614 | 0.18873 | -0.58722581 | 0.048 | 0.333 | 1 | 27 |
| XLOC-012680 | 0.18873 | -0.65307089 | 0.048 | 0.333 | 1 | 27 |
| abraxas1 | 0.18873 | -0.65154408 | 0.048 | 0.333 | 1 | 27 |
| actr2b | 0.18873 | -0.48428989 | 0.048 | 0.333 | 1 | 27 |
| adam19a | 0.18873 | -0.49484867 | 0.048 | 0.333 | 1 | 27 |
| adnpa | 0.18873 | -0.65315316 | 0.048 | 0.333 | 1 | 27 |
| alg12 | 0.18873 | -0.6273134 | 0.048 | 0.333 | 1 | 27 |
| apbb1 | 0.18873 | -0.52410236 | 0.048 | 0.333 | 1 | 27 |
| arf1 | 0.18873 | -0.43810136 | 0.048 | 0.333 | 1 | 27 |
| arrb2b | 0.18873 | -0.53874122 | 0.048 | 0.333 | 1 | 27 |
| babam2 | 0.18873 | -0.47152968 | 0.048 | 0.333 | 1 | 27 |
| bloc1s1 | 0.18873 | -0.51279027 | 0.048 | 0.333 | 1 | 27 |
| bloc1s5 | 0.18873 | -0.53238133 | 0.048 | 0.333 | 1 | 27 |
| bola1 | 0.18873 | -0.50092088 | 0.048 | 0.333 | 1 | 27 |
| bzw1b | 0.18873 | -0.58582847 | 0.048 | 0.333 | 1 | 27 |
| ccne1 | 0.18873 | -0.53820267 | 0.048 | 0.333 | 1 | 27 |
| cd9a | 0.18873 | -0.55833096 | 0.048 | 0.333 | 1 | 27 |
| cdk17 | 0.18873 | -0.56615055 | 0.048 | 0.333 | 1 | 27 |
| chst2a | 0.18873 | -0.29321634 | 0.048 | 0.333 | 1 | 27 |
| cldn12 | 0.18873 | -0.51788753 | 0.048 | 0.333 | 1 | 27 |
| commd1 | 0.18873 | -0.53825209 | 0.048 | 0.333 | 1 | 27 |
| copb1 | 0.18873 | -0.58119958 | 0.048 | 0.333 | 1 | 27 |
| coq3 | 0.18873 | -0.53963446 | 0.048 | 0.333 | 1 | 27 |
| cratb | 0.18873 | -0.44512848 | 0.048 | 0.333 | 1 | 27 |
| ddt | 0.18873 | -0.5038441 | 0.048 | 0.333 | 1 | 27 |
| ddx31 | 0.18873 | -0.51039888 | 0.048 | 0.333 | 1 | 27 |
| dffb | 0.18873 | -0.52982101 | 0.048 | 0.333 | 1 | 27 |
| dock4b | 0.18873 | -0.6590639 | 0.048 | 0.333 | 1 | 27 |
| dtnbp1a | 0.18873 | -0.52695815 | 0.048 | 0.333 | 1 | 27 |
| dyrk1ab | 0.18873 | -0.567075 | 0.048 | 0.333 | 1 | 27 |
| eif2b1 | 0.18873 | -0.48995988 | 0.048 | 0.333 | 1 | 27 |
| eif2b4 | 0.18873 | -0.56055124 | 0.048 | 0.333 | 1 | 27 |
| eif3jb | 0.18873 | -0.65261759 | 0.048 | 0.333 | 1 | 27 |
| enah | 0.18873 | -0.50381279 | 0.048 | 0.333 | 1 | 27 |
| evi5b | 0.18873 | -0.53489885 | 0.048 | 0.333 | 1 | 27 |
| faf2 | 0.18873 | -0.57134041 | 0.048 | 0.333 | 1 | 27 |
| fam162a | 0.18873 | -0.47417939 | 0.048 | 0.333 | 1 | 27 |
| fam210aa | 0.18873 | -0.51137749 | 0.048 | 0.333 | 1 | 27 |

| | | | | | | |
|---|---|---|---|---|---|---|
| fbxl14a | 0.18873 | -0.55731834 | 0.048 | 0.333 | 1 | 27 |
| fbxo25 | 0.18873 | -0.48060593 | 0.048 | 0.333 | 1 | 27 |
| gcnt4a | 0.18873 | -0.55442534 | 0.048 | 0.333 | 1 | 27 |
| ggh | 0.18873 | -0.50993234 | 0.048 | 0.333 | 1 | 27 |
| glrx5 | 0.18873 | -0.64650802 | 0.048 | 0.333 | 1 | 27 |
| gpr108 | 0.18873 | -0.54286203 | 0.048 | 0.333 | 1 | 27 |
| gpr137 | 0.18873 | -0.5867218 | 0.048 | 0.333 | 1 | 27 |
| gzf1 | 0.18873 | -0.39813414 | 0.048 | 0.333 | 1 | 27 |
| hck | 0.18873 | -0.54186413 | 0.048 | 0.333 | 1 | 27 |
| herpud2 | 0.18873 | -0.54289759 | 0.048 | 0.333 | 1 | 27 |
| hmgxb4a | 0.18873 | -0.58293253 | 0.048 | 0.333 | 1 | 27 |
| ift74 | 0.18873 | -0.53535255 | 0.048 | 0.333 | 1 | 27 |
| igflr1 | 0.18873 | -0.81286863 | 0.048 | 0.333 | 1 | 27 |
| il16 | 0.18873 | -0.60716036 | 0.048 | 0.333 | 1 | 27 |
| ing5a | 0.18873 | -0.53391477 | 0.048 | 0.333 | 1 | 27 |
| ints10 | 0.18873 | -0.41730923 | 0.048 | 0.333 | 1 | 27 |
| itga4 | 0.18873 | -0.84036993 | 0.048 | 0.333 | 1 | 27 |
| jkamp | 0.18873 | -0.58634728 | 0.048 | 0.333 | 1 | 27 |
| kif4 | 0.18873 | -0.5218565 | 0.048 | 0.333 | 1 | 27 |
| knop1 | 0.18873 | -0.51300118 | 0.048 | 0.333 | 1 | 27 |
| ldb1a | 0.18873 | -0.49976285 | 0.048 | 0.333 | 1 | 27 |
| ldha | 0.18873 | -0.4805458 | 0.048 | 0.333 | 1 | 27 |
| letm2 | 0.18873 | -0.65517679 | 0.048 | 0.333 | 1 | 27 |
| lpxn | 0.18873 | -0.45986504 | 0.048 | 0.333 | 1 | 27 |
| ltv1 | 0.18873 | -0.45526358 | 0.048 | 0.333 | 1 | 27 |
| malt3 | 0.18873 | -0.65376161 | 0.048 | 0.333 | 1 | 27 |
| man1a1 | 0.18873 | -0.6699447 | 0.048 | 0.333 | 1 | 27 |
| mfsd1 | 0.18873 | -0.56473239 | 0.048 | 0.333 | 1 | 27 |
| mipep | 0.18873 | -0.57708086 | 0.048 | 0.333 | 1 | 27 |
| mir223 | 0.18873 | -0.5705188 | 0.048 | 0.333 | 1 | 27 |
| mrpl19 | 0.18873 | -0.54664906 | 0.048 | 0.333 | 1 | 27 |
| mrpl4 | 0.18873 | -0.49821406 | 0.048 | 0.333 | 1 | 27 |
| mrtfab | 0.18873 | -0.50043552 | 0.048 | 0.333 | 1 | 27 |
| mxi1 | 0.18873 | -0.71489951 | 0.048 | 0.333 | 1 | 27 |
| n4bp1 | 0.18873 | -0.5518527 | 0.048 | 0.333 | 1 | 27 |
| nat9 | 0.18873 | -0.52102712 | 0.048 | 0.333 | 1 | 27 |
| ndrg1a | 0.18873 | -0.63728205 | 0.048 | 0.333 | 1 | 27 |
| nfatc3b | 0.18873 | -0.53778235 | 0.048 | 0.333 | 1 | 27 |
| nufip2 | 0.18873 | -0.48775598 | 0.048 | 0.333 | 1 | 27 |
| nuggc.1 | 0.18873 | -0.4985869 | 0.048 | 0.333 | 1 | 27 |
| nup160 | 0.18873 | -0.56459461 | 0.048 | 0.333 | 1 | 27 |
| ostf1 | 0.18873 | -0.67901403 | 0.048 | 0.333 | 1 | 27 |
| oxnad1 | 0.18873 | -0.52210532 | 0.048 | 0.333 | 1 | 27 |
| parp1 | 0.18873 | -0.45630208 | 0.048 | 0.333 | 1 | 27 |
| pbrm1l | 0.18873 | -0.67261341 | 0.048 | 0.333 | 1 | 27 |
| pcsk7 | 0.18873 | -0.5608161 | 0.048 | 0.333 | 1 | 27 |
| pggt1b | 0.18873 | -0.49426349 | 0.048 | 0.333 | 1 | 27 |
| pik3r2 | 0.18873 | -0.60251806 | 0.048 | 0.333 | 1 | 27 |

| | | | | | | |
|---|---|---|---|---|---|---|
| pla2g4aa | 0.18873 | -0.51469495 | 0.048 | 0.333 | 1 | 27 |
| poldip2 | 0.18873 | -0.63085784 | 0.048 | 0.333 | 1 | 27 |
| pom121 | 0.18873 | -0.55792949 | 0.048 | 0.333 | 1 | 27 |
| ppp1r12a | 0.18873 | -0.47353823 | 0.048 | 0.333 | 1 | 27 |
| ppp6c | 0.18873 | -0.57313039 | 0.048 | 0.333 | 1 | 27 |
| prelid1b | 0.18873 | -0.58673727 | 0.048 | 0.333 | 1 | 27 |
| prpf4bb | 0.18873 | -0.48840814 | 0.048 | 0.333 | 1 | 27 |
| prps1a | 0.18873 | -0.56496488 | 0.048 | 0.333 | 1 | 27 |
| ptenb | 0.18873 | -0.63819998 | 0.048 | 0.333 | 1 | 27 |
| ptprea | 0.18873 | -0.46606576 | 0.048 | 0.333 | 1 | 27 |
| ptrhd1 | 0.18873 | -0.59865995 | 0.048 | 0.333 | 1 | 27 |
| rab27a | 0.18873 | -0.49694724 | 0.048 | 0.333 | 1 | 27 |
| rnf145b | 0.18873 | -0.60425432 | 0.048 | 0.333 | 1 | 27 |
| rnpc3 | 0.18873 | -0.56052957 | 0.048 | 0.333 | 1 | 27 |
| rpf1 | 0.18873 | -0.44459806 | 0.048 | 0.333 | 1 | 27 |
| rps19bp1 | 0.18873 | -0.4438641 | 0.048 | 0.333 | 1 | 27 |
| rps6ka3a | 0.18873 | -0.65946162 | 0.048 | 0.333 | 1 | 27 |
| sccpdhb | 0.18873 | -0.54696708 | 0.048 | 0.333 | 1 | 27 |
| scinlb | 0.18873 | -0.54535367 | 0.048 | 0.333 | 1 | 27 |
| sdhc | 0.18873 | -0.48881152 | 0.048 | 0.333 | 1 | 27 |
| si:ch1073-340j19.1 | 0.18873 | -0.46272068 | 0.048 | 0.333 | 1 | 27 |
| si:ch211-108d22.2 | 0.18873 | -0.53672428 | 0.048 | 0.333 | 1 | 27 |
| si:ch211-162e15.3 | 0.18873 | -0.45637094 | 0.048 | 0.333 | 1 | 27 |
| si:ch211-191j22.3 | 0.18873 | -0.5719558 | 0.048 | 0.333 | 1 | 27 |
| si:ch211-197h24.6 | 0.18873 | -0.49758179 | 0.048 | 0.333 | 1 | 27 |
| si:ch211-253b8.2 | 0.18873 | -0.47284749 | 0.048 | 0.333 | 1 | 27 |
| si:ch73-208g10.1 | 0.18873 | -0.60414079 | 0.048 | 0.333 | 1 | 27 |
| si:ch73-340n8.4 | 0.18873 | -0.6281148 | 0.048 | 0.333 | 1 | 27 |
| si:dkey-204f11.64 | 0.18873 | -0.76225005 | 0.048 | 0.333 | 1 | 27 |
| si:dkey-222f8.3 | 0.18873 | -0.52730516 | 0.048 | 0.333 | 1 | 27 |
| si:dkey-23i12.5 | 0.18873 | -0.55749176 | 0.048 | 0.333 | 1 | 27 |
| si:dkey-51e6.1 | 0.18873 | -0.52921923 | 0.048 | 0.333 | 1 | 27 |
| si:dkey-57k2.7 | 0.18873 | -0.52311255 | 0.048 | 0.333 | 1 | 27 |
| si:dkey-72l14.4 | 0.18873 | -0.49007079 | 0.048 | 0.333 | 1 | 27 |
| si:dkeyp-68b7.7 | 0.18873 | -0.14652654 | 0.048 | 0.333 | 1 | 27 |
| si:dkeyp-69e1.8 | 0.18873 | -0.63718333 | 0.048 | 0.333 | 1 | 27 |
| si:rp71-36a1.1 | 0.18873 | -0.54385172 | 0.048 | 0.333 | 1 | 27 |
| slc16a10 | 0.18873 | -0.50163611 | 0.048 | 0.333 | 1 | 27 |
| slc20a1b | 0.18873 | -0.58821403 | 0.048 | 0.333 | 1 | 27 |
| slc30a4 | 0.18873 | -0.17229386 | 0.048 | 0.333 | 1 | 27 |
| slc35a1 | 0.18873 | -0.48262257 | 0.048 | 0.333 | 1 | 27 |
| smn1 | 0.18873 | -0.46895622 | 0.048 | 0.333 | 1 | 27 |
| snapc1b | 0.18873 | -0.53212676 | 0.048 | 0.333 | 1 | 27 |
| spns1 | 0.18873 | -0.55164909 | 0.048 | 0.333 | 1 | 27 |
| tada3l | 0.18873 | -0.5102486 | 0.048 | 0.333 | 1 | 27 |
| taf11 | 0.18873 | -0.56318682 | 0.048 | 0.333 | 1 | 27 |
| tarbp2 | 0.18873 | -0.4834926 | 0.048 | 0.333 | 1 | 27 |
| tbc1d15 | 0.18873 | -0.51844248 | 0.048 | 0.333 | 1 | 27 |

| | | | | | | |
|---|---|---|---|---|---|---|
| tdrd3 | 0.18873 | -0.64637921 | 0.048 | 0.333 | 1 | 27 |
| tecrb | 0.18873 | -0.58422153 | 0.048 | 0.333 | 1 | 27 |
| tespa1 | 0.18873 | -0.66316143 | 0.048 | 0.333 | 1 | 27 |
| tex10 | 0.18873 | -0.58216977 | 0.048 | 0.333 | 1 | 27 |
| thap12a | 0.18873 | -0.46214732 | 0.048 | 0.333 | 1 | 27 |
| themis2 | 0.18873 | -0.53599912 | 0.048 | 0.333 | 1 | 27 |
| tlk2 | 0.18873 | -0.48716017 | 0.048 | 0.333 | 1 | 27 |
| tlr22 | 0.18873 | -0.60532998 | 0.048 | 0.333 | 1 | 27 |
| tmem246 | 0.18873 | -0.5176093 | 0.048 | 0.333 | 1 | 27 |
| tmem42a | 0.18873 | -0.55684315 | 0.048 | 0.333 | 1 | 27 |
| trpc4apa | 0.18873 | -0.54184155 | 0.048 | 0.333 | 1 | 27 |
| tusc2b | 0.18873 | -0.49548235 | 0.048 | 0.333 | 1 | 27 |
| ubap1 | 0.18873 | -0.54276203 | 0.048 | 0.333 | 1 | 27 |
| uchl5 | 0.18873 | -0.62999217 | 0.048 | 0.333 | 1 | 27 |
| ugt1a7 | 0.18873 | -0.57441463 | 0.048 | 0.333 | 1 | 27 |
| upf1 | 0.18873 | -0.55116902 | 0.048 | 0.333 | 1 | 27 |
| vps11 | 0.18873 | -0.64294907 | 0.048 | 0.333 | 1 | 27 |
| vps37b | 0.18873 | -0.53392675 | 0.048 | 0.333 | 1 | 27 |
| waca | 0.18873 | -0.46654468 | 0.048 | 0.333 | 1 | 27 |
| wdr12 | 0.18873 | -0.47438715 | 0.048 | 0.333 | 1 | 27 |
| xiap | 0.18873 | -0.74132275 | 0.048 | 0.333 | 1 | 27 |
| zbtb1 | 0.18873 | -0.64712617 | 0.048 | 0.333 | 1 | 27 |
| zc3h15 | 0.18873 | -0.57649003 | 0.048 | 0.333 | 1 | 27 |
| zdhhc18b | 0.18873 | -0.53536311 | 0.048 | 0.333 | 1 | 27 |
| zdhhc23a | 0.18873 | -0.52713372 | 0.048 | 0.333 | 1 | 27 |
| zfand5a | 0.18873 | -0.5956043 | 0.048 | 0.333 | 1 | 27 |
| zgc:110843 | 0.18873 | -0.54611571 | 0.048 | 0.333 | 1 | 27 |
| zgc:112052 | 0.18873 | -0.69945545 | 0.048 | 0.333 | 1 | 27 |
| zgc:113149 | 0.18873 | -0.5446521 | 0.048 | 0.333 | 1 | 27 |
| zgc:114119 | 0.18873 | -0.52017808 | 0.048 | 0.333 | 1 | 27 |
| zgc:162200 | 0.18873 | -0.47133372 | 0.048 | 0.333 | 1 | 27 |
| zgc:56235 | 0.18873 | -0.46065061 | 0.048 | 0.333 | 1 | 27 |
| zgc:77056 | 0.18873 | -0.47170726 | 0.048 | 0.333 | 1 | 27 |
| zgc:92664 | 0.18873 | -0.52330757 | 0.048 | 0.333 | 1 | 27 |
| znf710b | 0.18873 | -0.46899803 | 0.048 | 0.333 | 1 | 27 |
| CABZ01034698.2 | 0.18873 | -0.47928207 | 0.048 | 0.333 | 1 | 27 |
| CR387933.1 | 0.18873 | -0.4837321 | 0.048 | 0.333 | 1 | 27 |
| FP236734.1 | 0.18873 | -0.59037984 | 0.048 | 0.333 | 1 | 27 |
| XLOC-032256 | 0.18873 | -0.52806275 | 0.048 | 0.333 | 1 | 27 |
| bin3 | 0.18873 | -0.59180154 | 0.048 | 0.333 | 1 | 27 |
| ephx2 | 0.18873 | -0.5243579 | 0.048 | 0.333 | 1 | 27 |
| farsb | 0.18873 | -0.51484165 | 0.048 | 0.333 | 1 | 27 |
| gbp | 0.18873 | -0.42544976 | 0.048 | 0.333 | 1 | 27 |
| gnal | 0.18873 | -0.65969156 | 0.048 | 0.333 | 1 | 27 |
| inpp5ka | 0.18873 | -0.47118586 | 0.048 | 0.333 | 1 | 27 |
| itpripl2 | 0.18873 | -0.55013009 | 0.048 | 0.333 | 1 | 27 |
| si:ch211-68a17.7 | 0.18873 | -0.48071089 | 0.048 | 0.333 | 1 | 27 |
| si:dkey-34m19.3 | 0.18873 | -0.52774251 | 0.048 | 0.333 | 1 | 27 |

| | | | | | | |
|---|---|---|---|---|---|---|
| st3gal5 | 0.18873 | -0.52981153 | 0.048 | 0.333 | 1 | 27 |
| tcp11l2 | 0.18873 | -0.50683927 | 0.048 | 0.333 | 1 | 27 |
| tmem251 | 0.18873 | -0.46826492 | 0.048 | 0.333 | 1 | 27 |
| triobpa | 0.18873 | -0.65068792 | 0.048 | 0.333 | 1 | 27 |
| trnau1apa | 0.18873 | -0.4688333 | 0.048 | 0.333 | 1 | 27 |
| BX901920.1 | 0.18873 | -0.65183392 | 0.048 | 0.333 | 1 | 27 |
| CR753876.1 | 0.18873 | -0.46339149 | 0.048 | 0.333 | 1 | 27 |
| CU929230.1 | 0.18873 | -0.53663601 | 0.048 | 0.333 | 1 | 27 |
| XLOC-032002 | 0.18873 | -0.72357752 | 0.048 | 0.333 | 1 | 27 |
| XLOC-041870 | 0.18873 | -0.86959811 | 0.048 | 0.333 | 1 | 27 |
| ano10b | 0.18873 | -0.52176025 | 0.048 | 0.333 | 1 | 27 |
| arl8bb | 0.18873 | -0.48749932 | 0.048 | 0.333 | 1 | 27 |
| ccnt1 | 0.18873 | -0.48473268 | 0.048 | 0.333 | 1 | 27 |
| ddx49 | 0.18873 | -0.58984963 | 0.048 | 0.333 | 1 | 27 |
| ech1 | 0.18873 | -0.48284638 | 0.048 | 0.333 | 1 | 27 |
| exoc6 | 0.18873 | -0.58753169 | 0.048 | 0.333 | 1 | 27 |
| fam3a | 0.18873 | -0.72648003 | 0.048 | 0.333 | 1 | 27 |
| golga5 | 0.18873 | -0.58786281 | 0.048 | 0.333 | 1 | 27 |
| ilf3b | 0.18873 | -0.52528684 | 0.048 | 0.333 | 1 | 27 |
| lonrf1 | 0.18873 | -0.45063429 | 0.048 | 0.333 | 1 | 27 |
| lyst | 0.18873 | -0.48457183 | 0.048 | 0.333 | 1 | 27 |
| os9 | 0.18873 | -0.49047676 | 0.048 | 0.333 | 1 | 27 |
| ptk2ba | 0.18873 | -0.47362726 | 0.048 | 0.333 | 1 | 27 |
| rab33ba | 0.18873 | -0.50180786 | 0.048 | 0.333 | 1 | 27 |
| sec61a1l | 0.18873 | -0.53267849 | 0.048 | 0.333 | 1 | 27 |
| serpinb1l1 | 0.18873 | -0.53945444 | 0.048 | 0.333 | 1 | 27 |
| si:dkey-211g8.4 | 0.18873 | -0.66868591 | 0.048 | 0.333 | 1 | 27 |
| syap1 | 0.18873 | -0.51827216 | 0.048 | 0.333 | 1 | 27 |
| xpot | 0.18873 | -0.52932335 | 0.048 | 0.333 | 1 | 27 |
| CU915256.1 | 0.18873 | -0.60512514 | 0.048 | 0.333 | 1 | 27 |
| LOC101885027 | 0.18873 | -0.59753497 | 0.048 | 0.333 | 1 | 27 |
| asb12a | 0.18873 | -0.58452774 | 0.048 | 0.333 | 1 | 27 |
| clk4b | 0.18873 | -0.62562542 | 0.048 | 0.333 | 1 | 27 |
| loxl2b | 0.18873 | -0.587376 | 0.048 | 0.333 | 1 | 27 |
| map2k4b | 0.18873 | -0.56189186 | 0.048 | 0.333 | 1 | 27 |
| med8 | 0.18873 | -0.55640005 | 0.048 | 0.333 | 1 | 27 |
| scamp2 | 0.18873 | -0.59951108 | 0.048 | 0.333 | 1 | 27 |
| zgc:112496 | 0.18873 | -0.75397626 | 0.048 | 0.333 | 1 | 27 |
| zgc:77262 | 0.18873 | -0.59071516 | 0.048 | 0.333 | 1 | 27 |
| bcam | 0.18873 | -0.66477542 | 0.048 | 0.333 | 1 | 27 |
| psmf1 | 0.18873 | -0.55217929 | 0.048 | 0.333 | 1 | 27 |
| si:ch211-166g5.4 | 0.18873 | -0.6301762 | 0.048 | 0.333 | 1 | 27 |
| slc11a2 | 0.18873 | -0.65061112 | 0.048 | 0.333 | 1 | 27 |
| stk24b | 0.18873 | -0.65979299 | 0.048 | 0.333 | 1 | 27 |
| zgc:101783 | 0.18873 | -0.56851436 | 0.048 | 0.333 | 1 | 27 |
| ctsla | 0.18873 | -0.3175434 | 0.048 | 0.333 | 1 | 27 |
| rps26 | 0.18935 | 0.318931952 | 1 | 1 | 1 | 27 |
| rps7 | 0.19261 | 0.189248612 | 1 | 1 | 1 | 27 |

| | | | | | | |
|---|---|---|---|---|---|---|
| slc46a2 | 0.19418 | 0.685878668 | 0.333 | 0 | 1 | 27 |
| baxa | 0.19418 | 0.670068865 | 0.333 | 0 | 1 | 27 |
| gadd45ba | 0.19418 | 0.633250242 | 0.333 | 0 | 1 | 27 |
| h2afx | 0.19418 | 0.781050208 | 0.333 | 0 | 1 | 27 |
| mapkapk2a | 0.19418 | 0.562344631 | 0.333 | 0 | 1 | 27 |
| ncf2 | 0.19418 | 0.615265484 | 0.333 | 0 | 1 | 27 |
| aimp1 | 0.19418 | 0.531834173 | 0.333 | 0 | 1 | 27 |
| akirin1 | 0.19418 | 0.602405709 | 0.333 | 0 | 1 | 27 |
| ankrd11 | 0.19418 | 0.603457978 | 0.333 | 0 | 1 | 27 |
| calr3a | 0.19418 | 0.514375362 | 0.333 | 0 | 1 | 27 |
| cks1b | 0.19418 | 0.607910178 | 0.333 | 0 | 1 | 27 |
| cnot9 | 0.19418 | 0.535973948 | 0.333 | 0 | 1 | 27 |
| cobll1a | 0.19418 | 0.591160153 | 0.333 | 0 | 1 | 27 |
| csnk2a1 | 0.19418 | 0.451502321 | 0.333 | 0 | 1 | 27 |
| cycsb | 0.19418 | 0.576655241 | 0.333 | 0 | 1 | 27 |
| dldh | 0.19418 | 0.464962501 | 0.333 | 0 | 1 | 27 |
| dtx4a | 0.19418 | 0.58141031 | 0.333 | 0 | 1 | 27 |
| dut | 0.19418 | 0.854710155 | 0.333 | 0 | 1 | 27 |
| dynll1 | 0.19418 | 0.756478132 | 0.333 | 0 | 1 | 27 |
| eif3ja | 0.19418 | 0.536175305 | 0.333 | 0 | 1 | 27 |
| eif5 | 0.19418 | 0.533517758 | 0.333 | 0 | 1 | 27 |
| iqgap1 | 0.19418 | 0.608631277 | 0.333 | 0 | 1 | 27 |
| itgb2 | 0.19418 | 0.60051068 | 0.333 | 0 | 1 | 27 |
| kif22 | 0.19418 | 0.50996269 | 0.333 | 0 | 1 | 27 |
| lamtor4 | 0.19418 | 0.65221631 | 0.333 | 0 | 1 | 27 |
| lbr | 0.19418 | 0.797468568 | 0.333 | 0 | 1 | 27 |
| lmnb2 | 0.19418 | 0.434627911 | 0.333 | 0 | 1 | 27 |
| lsm4 | 0.19418 | 0.523726028 | 0.333 | 0 | 1 | 27 |
| med19a | 0.19418 | 0.564532362 | 0.333 | 0 | 1 | 27 |
| mki67 | 0.19418 | 0.649916524 | 0.333 | 0 | 1 | 27 |
| pcna | 0.19418 | 0.80583903 | 0.333 | 0 | 1 | 27 |
| pfdn6 | 0.19418 | 0.472170749 | 0.333 | 0 | 1 | 27 |
| polr3glb | 0.19418 | 0.49829944 | 0.333 | 0 | 1 | 27 |
| prdx3 | 0.19418 | 0.449933321 | 0.333 | 0 | 1 | 27 |
| psap | 0.19418 | 0.565027713 | 0.333 | 0 | 1 | 27 |
| puf60b | 0.19418 | 0.560278452 | 0.333 | 0 | 1 | 27 |
| rab11ba | 0.19418 | 0.485907699 | 0.333 | 0 | 1 | 27 |
| rrm1 | 0.19418 | 0.593601807 | 0.333 | 0 | 1 | 27 |
| rrm2 | 0.19418 | 1.130006614 | 0.333 | 0 | 1 | 27 |
| rtn4a | 0.19418 | 0.522889262 | 0.333 | 0 | 1 | 27 |
| sf3b5 | 0.19418 | 0.554801557 | 0.333 | 0 | 1 | 27 |
| si:ch211-22i13.2 | 0.19418 | 0.600274742 | 0.333 | 0 | 1 | 27 |
| smc2 | 0.19418 | 0.500754735 | 0.333 | 0 | 1 | 27 |
| snrpa | 0.19418 | 0.560780311 | 0.333 | 0 | 1 | 27 |
| snx5 | 0.19418 | 0.60090866 | 0.333 | 0 | 1 | 27 |
| srsf1a | 0.19418 | 0.495927274 | 0.333 | 0 | 1 | 27 |
| stmn1a | 0.19418 | 1.030173194 | 0.333 | 0 | 1 | 27 |
| tomm20b | 0.19418 | 0.619064773 | 0.333 | 0 | 1 | 27 |

| | | | | | | |
|---|---|---|---|---|---|---|
| tp53inp1 | 0.19418 | 0.805249025 | 0.333 | 0 | 1 | 27 |
| uba2 | 0.19418 | 0.54153034 | 0.333 | 0 | 1 | 27 |
| cdc42 | 0.19418 | 0.466252889 | 0.333 | 0 | 1 | 27 |
| ddx39aa | 0.19418 | 0.54601136 | 0.333 | 0 | 1 | 27 |
| si:ch211-147a11.3 | 0.19418 | 0.54124872 | 0.333 | 0 | 1 | 27 |
| si:ch211-51e12.7 | 0.19418 | 0.462869577 | 0.333 | 0 | 1 | 27 |
| trappc6bl | 0.19418 | 0.460977268 | 0.333 | 0 | 1 | 27 |
| arpc4l | 0.19418 | 0.603537711 | 0.333 | 0 | 1 | 27 |
| atp6v1g1 | 0.19418 | 0.606439301 | 0.333 | 0 | 1 | 27 |
| csnk1a1 | 0.19418 | 0.485335039 | 0.333 | 0 | 1 | 27 |
| nfkbie | 0.19418 | 0.60527368 | 0.333 | 0 | 1 | 27 |
| pde7a | 0.19418 | 0.538391511 | 0.333 | 0 | 1 | 27 |
| psmc6 | 0.19418 | 0.527817408 | 0.333 | 0 | 1 | 27 |
| sem1 | 0.19418 | 0.556730574 | 0.333 | 0 | 1 | 27 |
| sept7b | 0.19418 | 0.485409819 | 0.333 | 0 | 1 | 27 |
| sh3kbp1 | 0.19418 | 0.48332538 | 0.333 | 0 | 1 | 27 |
| snx1a | 0.19418 | 0.609117641 | 0.333 | 0 | 1 | 27 |
| syngr2a | 0.19418 | 0.582553818 | 0.333 | 0 | 1 | 27 |
| xpc | 0.19418 | 0.645899586 | 0.333 | 0 | 1 | 27 |
| fthl27 | 0.19843 | -0.6618013 | 0.714 | 1 | 1 | 27 |
| zgc:114188 | 0.20182 | 0.523875071 | 0.905 | 0.667 | 1 | 27 |
| sftpba | 0.20193 | -0.73998581 | 0.571 | 1 | 1 | 27 |
| rplp1 | 0.20218 | 0.106051323 | 1 | 1 | 1 | 27 |
| xbp1 | 0.20637 | -0.86214975 | 0.333 | 0.667 | 1 | 27 |
| ncl | 0.21262 | -0.47147174 | 0.571 | 0.667 | 1 | 27 |
| tmed7 | 0.21536 | 0.463143441 | 0.81 | 0.333 | 1 | 27 |
| psma5 | 0.21716 | -0.16403848 | 0.476 | 0.333 | 1 | 27 |
| ppdpfb | 0.21818 | 0.307576166 | 0.905 | 1 | 1 | 27 |
| pfn1 | 0.21839 | -0.15448475 | 1 | 1 | 1 | 27 |
| kctd5a | 0.21925 | -0.51589715 | 0.762 | 1 | 1 | 27 |
| grb2b | 0.22074 | -0.78127534 | 0.238 | 0.667 | 1 | 27 |
| btf3 | 0.22475 | -0.24070599 | 1 | 1 | 1 | 27 |
| rps9 | 0.22478 | 0.188192393 | 1 | 1 | 1 | 27 |
| gpx1a | 0.22735 | -0.59590499 | 0.714 | 1 | 1 | 27 |
| btk | 0.23038 | -0.79656382 | 0.19 | 0.667 | 1 | 27 |
| rpl17 | 0.2322 | 0.205900325 | 0.952 | 1 | 1 | 27 |
| nmi | 0.23298 | -0.77700515 | 0.19 | 0.667 | 1 | 27 |
| prkcha | 0.23369 | -0.77416867 | 0.19 | 0.667 | 1 | 27 |
| nosip | 0.23438 | -0.70503489 | 0.238 | 0.667 | 1 | 27 |
| ube2ia | 0.23446 | -0.66216198 | 0.333 | 0.667 | 1 | 27 |
| ppiaa | 0.23493 | 0.155343882 | 1 | 1 | 1 | 27 |
| cox5ab | 0.24009 | -0.5877621 | 0.429 | 0.667 | 1 | 27 |
| zgc:112356 | 0.24207 | -0.81470076 | 0.238 | 0.667 | 1 | 27 |
| adssl | 0.24423 | 0.49583771 | 0.286 | 0 | 1 | 27 |
| ccna2 | 0.24423 | 0.545297301 | 0.286 | 0 | 1 | 27 |
| cox6b1 | 0.24423 | 0.342634701 | 0.286 | 0 | 1 | 27 |
| gemin8 | 0.24423 | 0.523861107 | 0.286 | 0 | 1 | 27 |
| gng2 | 0.24423 | 0.607503932 | 0.286 | 0 | 1 | 27 |

| | | | | | | |
|---|---|---|---|---|---|---|
| jdp2b | 0.24423 | 0.417398123 | 0.286 | 0 | 1 | 27 |
| jpt1b | 0.24423 | 0.412244554 | 0.286 | 0 | 1 | 27 |
| lsm3 | 0.24423 | 0.501062509 | 0.286 | 0 | 1 | 27 |
| lsm6 | 0.24423 | 0.53096829 | 0.286 | 0 | 1 | 27 |
| lsm7 | 0.24423 | 0.595042302 | 0.286 | 0 | 1 | 27 |
| meaf6 | 0.24423 | 0.470346877 | 0.286 | 0 | 1 | 27 |
| mier1a | 0.24423 | 0.49876615 | 0.286 | 0 | 1 | 27 |
| ndfip1 | 0.24423 | 0.464127592 | 0.286 | 0 | 1 | 27 |
| nop56 | 0.24423 | 0.483142075 | 0.286 | 0 | 1 | 27 |
| nop58 | 0.24423 | 0.511290365 | 0.286 | 0 | 1 | 27 |
| nup50 | 0.24423 | 0.39295613 | 0.286 | 0 | 1 | 27 |
| nup62l | 0.24423 | 0.445222825 | 0.286 | 0 | 1 | 27 |
| paip2b | 0.24423 | 0.514611808 | 0.286 | 0 | 1 | 27 |
| pdcd10b | 0.24423 | 0.406032517 | 0.286 | 0 | 1 | 27 |
| ppih | 0.24423 | 0.423013679 | 0.286 | 0 | 1 | 27 |
| psmb10 | 0.24423 | 0.457427791 | 0.286 | 0 | 1 | 27 |
| rhoac | 0.24423 | 0.528686157 | 0.286 | 0 | 1 | 27 |
| rps27l | 0.24423 | 0.533331869 | 0.286 | 0 | 1 | 27 |
| saraf | 0.24423 | 0.529714441 | 0.286 | 0 | 1 | 27 |
| selenow2a | 0.24423 | 0.471297628 | 0.286 | 0 | 1 | 27 |
| si:ch211-14a17.11 | 0.24423 | 0.526825287 | 0.286 | 0 | 1 | 27 |
| si:ch73-42p12.2 | 0.24423 | 0.439284746 | 0.286 | 0 | 1 | 27 |
| si:dkey-112a7.4 | 0.24423 | 0.347033613 | 0.286 | 0 | 1 | 27 |
| tspan3a | 0.24423 | 0.474365404 | 0.286 | 0 | 1 | 27 |
| ube2ib | 0.24423 | 0.513967769 | 0.286 | 0 | 1 | 27 |
| zgc:165555.3 | 0.24423 | 0.593561708 | 0.286 | 0 | 1 | 27 |
| RASA2 | 0.24423 | 0.55482568 | 0.286 | 0 | 1 | 27 |
| bcas2 | 0.24423 | 0.437517143 | 0.286 | 0 | 1 | 27 |
| cenpx | 0.24423 | 0.563777684 | 0.286 | 0 | 1 | 27 |
| dtymk | 0.24423 | 0.436902268 | 0.286 | 0 | 1 | 27 |
| mbpb | 0.24423 | 0.50211818 | 0.286 | 0 | 1 | 27 |
| mcts1 | 0.24423 | 0.44925752 | 0.286 | 0 | 1 | 27 |
| ndufs4 | 0.24423 | 0.450771866 | 0.286 | 0 | 1 | 27 |
| selenoh | 0.24423 | 0.468548807 | 0.286 | 0 | 1 | 27 |
| smchd1 | 0.24423 | 0.400338029 | 0.286 | 0 | 1 | 27 |
| wasf2 | 0.24423 | 0.584474599 | 0.286 | 0 | 1 | 27 |
| FP016018.1 | 0.24423 | 0.519533351 | 0.286 | 0 | 1 | 27 |
| LOC110438240 | 0.24423 | 0.5651885 | 0.286 | 0 | 1 | 27 |
| XLOC-028370 | 0.24423 | 0.663553956 | 0.286 | 0 | 1 | 27 |
| atp6v1ba | 0.24423 | 0.451240486 | 0.286 | 0 | 1 | 27 |
| brd2a | 0.24423 | 0.491824092 | 0.286 | 0 | 1 | 27 |
| brdt | 0.24423 | 0.547939137 | 0.286 | 0 | 1 | 27 |
| cbx1b | 0.24423 | 0.519895906 | 0.286 | 0 | 1 | 27 |
| cnn2 | 0.24423 | 0.476204149 | 0.286 | 0 | 1 | 27 |
| commd3 | 0.24423 | 0.558441594 | 0.286 | 0 | 1 | 27 |
| ddx21 | 0.24423 | 0.561291093 | 0.286 | 0 | 1 | 27 |
| eif2s2 | 0.24423 | 0.607140129 | 0.286 | 0 | 1 | 27 |
| elavl1a | 0.24423 | 0.490287843 | 0.286 | 0 | 1 | 27 |

| | | | | | | |
|---|---|---|---|---|---|---|
| fam49a | 0.24423 | 0.4423108 | 0.286 | 0 | 1 | 27 |
| gpatch8 | 0.24423 | 0.4266715 | 0.286 | 0 | 1 | 27 |
| hmgb1b | 0.24423 | 0.697268008 | 0.286 | 0 | 1 | 27 |
| hmha1b | 0.24423 | 0.500498919 | 0.286 | 0 | 1 | 27 |
| irf2bp2b | 0.24423 | 0.517566787 | 0.286 | 0 | 1 | 27 |
| khdrbs1b | 0.24423 | 0.574549171 | 0.286 | 0 | 1 | 27 |
| klf2a | 0.24423 | 0.562856796 | 0.286 | 0 | 1 | 27 |
| kpnb1 | 0.24423 | 0.473043127 | 0.286 | 0 | 1 | 27 |
| laptm4a | 0.24423 | 0.554406915 | 0.286 | 0 | 1 | 27 |
| lnpa | 0.24423 | 0.560124666 | 0.286 | 0 | 1 | 27 |
| mad2l1 | 0.24423 | 0.435938269 | 0.286 | 0 | 1 | 27 |
| mgst3a | 0.24423 | 0.430422797 | 0.286 | 0 | 1 | 27 |
| mibp | 0.24423 | 0.832708762 | 0.286 | 0 | 1 | 27 |
| mrps14 | 0.24423 | 0.423170552 | 0.286 | 0 | 1 | 27 |
| ndufb3 | 0.24423 | 0.546622543 | 0.286 | 0 | 1 | 27 |
| nkap | 0.24423 | 0.432918025 | 0.286 | 0 | 1 | 27 |
| pcbp2 | 0.24423 | 0.444935334 | 0.286 | 0 | 1 | 27 |
| phf5a | 0.24423 | 0.504500893 | 0.286 | 0 | 1 | 27 |
| psmd7 | 0.24423 | 0.386804084 | 0.286 | 0 | 1 | 27 |
| ptpn6 | 0.24423 | 0.464409724 | 0.286 | 0 | 1 | 27 |
| rer1 | 0.24423 | 0.511990682 | 0.286 | 0 | 1 | 27 |
| rpa1 | 0.24423 | 0.442806272 | 0.286 | 0 | 1 | 27 |
| si:dkey-24p1.1 | 0.24423 | 0.620077577 | 0.286 | 0 | 1 | 27 |
| snrpg | 0.24423 | 0.619003918 | 0.286 | 0 | 1 | 27 |
| stk17b | 0.24423 | 0.44070325 | 0.286 | 0 | 1 | 27 |
| top2a | 0.24423 | 0.484041654 | 0.286 | 0 | 1 | 27 |
| trim44 | 0.24423 | 0.553467163 | 0.286 | 0 | 1 | 27 |
| tspan14 | 0.24423 | 0.488424856 | 0.286 | 0 | 1 | 27 |
| uba1 | 0.24423 | 0.463955321 | 0.286 | 0 | 1 | 27 |
| ube2c | 0.24423 | 0.465648865 | 0.286 | 0 | 1 | 27 |
| uck2a | 0.24423 | 0.451127276 | 0.286 | 0 | 1 | 27 |
| vps29 | 0.24423 | 0.501969288 | 0.286 | 0 | 1 | 27 |
| zgc:153409 | 0.24423 | 0.476408137 | 0.286 | 0 | 1 | 27 |
| aip | 0.24423 | 0.525316337 | 0.286 | 0 | 1 | 27 |
| anp32e | 0.24423 | 0.497642256 | 0.286 | 0 | 1 | 27 |
| cad | 0.24423 | 0.56340246 | 0.286 | 0 | 1 | 27 |
| cdipt | 0.24423 | 0.489193363 | 0.286 | 0 | 1 | 27 |
| elf1 | 0.24423 | 0.50145142 | 0.286 | 0 | 1 | 27 |
| etfa | 0.24423 | 0.48833781 | 0.286 | 0 | 1 | 27 |
| kmt2a | 0.24423 | 0.474383788 | 0.286 | 0 | 1 | 27 |
| lin52 | 0.24423 | 0.400394564 | 0.286 | 0 | 1 | 27 |
| metap2b | 0.24423 | 0.406160812 | 0.286 | 0 | 1 | 27 |
| ndufb5 | 0.24423 | 0.422516789 | 0.286 | 0 | 1 | 27 |
| polr2l | 0.24423 | 0.371491912 | 0.286 | 0 | 1 | 27 |
| rangap1a | 0.24423 | 0.501083984 | 0.286 | 0 | 1 | 27 |
| rap1b | 0.24423 | 0.446475411 | 0.286 | 0 | 1 | 27 |
| sarnp | 0.24423 | 0.565734238 | 0.286 | 0 | 1 | 27 |
| skp1 | 0.24423 | 0.371212068 | 0.286 | 0 | 1 | 27 |

| | | | | | | |
|---|---|---|---|---|---|---|
| slc44a2 | 0.24423 | 0.496638015 | 0.286 | 0 | 1 | 27 |
| taf10 | 0.24423 | 0.523750843 | 0.286 | 0 | 1 | 27 |
| taf15 | 0.24423 | 0.491830948 | 0.286 | 0 | 1 | 27 |
| thoc5 | 0.24423 | 0.497136324 | 0.286 | 0 | 1 | 27 |
| tnrc6b | 0.24423 | 0.386574619 | 0.286 | 0 | 1 | 27 |
| vcp | 0.24423 | 0.45443709 | 0.286 | 0 | 1 | 27 |
| ywhaqa | 0.24423 | 0.589622537 | 0.286 | 0 | 1 | 27 |
| wipf1a | 0.24633 | -0.78121313 | 0.238 | 0.667 | 1 | 27 |
| coro1a | 0.24663 | -0.49844692 | 0.81 | 1 | 1 | 27 |
| si:ch211-132p1.3 | 0.25232 | -0.77983191 | 0.19 | 0.667 | 1 | 27 |
| psmb1 | 0.25574 | -0.20492978 | 0.429 | 0.333 | 1 | 27 |
| arpc2 | 0.25769 | 0.702393271 | 0.81 | 0.333 | 1 | 27 |
| sap30l | 0.26103 | -0.69629845 | 0.143 | 0.333 | 1 | 27 |
| hmgb2b | 0.2641 | 0.901625962 | 0.524 | 0.333 | 1 | 27 |
| sat1a.2 | 0.26503 | 0.266527491 | 0.714 | 0.333 | 1 | 27 |
| nt5dc2 | 0.26525 | -0.71327086 | 0.19 | 0.667 | 1 | 27 |
| si:dkey-183p4.10 | 0.26929 | -0.75756762 | 0.286 | 0.667 | 1 | 27 |
| sc:d0284 | 0.27152 | -0.66917531 | 0.19 | 0.667 | 1 | 27 |
| si:ch211-198c19.3 | 0.27182 | -1.08149411 | 0.19 | 0.667 | 1 | 27 |
| si:ch211-105j21.9 | 0.27211 | -0.67402464 | 0.19 | 0.667 | 1 | 27 |
| srprb | 0.27211 | -0.65535794 | 0.19 | 0.667 | 1 | 27 |
| slc25a3b | 0.27241 | 0.640749613 | 0.81 | 0.333 | 1 | 27 |
| rpl23 | 0.27366 | -0.19377781 | 0.952 | 1 | 1 | 27 |
| nop53 | 0.27433 | -0.34264708 | 0.667 | 0.667 | 1 | 27 |
| NDUFB1 | 0.27439 | -0.68260942 | 0.238 | 0.667 | 1 | 27 |
| cnbpb | 0.27723 | -0.71335626 | 0.333 | 0.667 | 1 | 27 |
| mtdha | 0.2776 | -0.69518996 | 0.238 | 0.667 | 1 | 27 |
| dcaf7 | 0.28232 | -0.68430304 | 0.238 | 0.667 | 1 | 27 |
| rps10 | 0.28327 | 0.1577845 | 1 | 1 | 1 | 27 |
| ncaldb | 0.2833 | -0.71161784 | 0.238 | 0.667 | 1 | 27 |
| COX5B | 0.28723 | 0.371417084 | 0.762 | 0.333 | 1 | 27 |
| vma21 | 0.30058 | -0.48165083 | 0.238 | 0.333 | 1 | 27 |
| LO018102.1 | 0.30516 | 0.402614751 | 0.238 | 0 | 1 | 27 |
| acadvl | 0.30516 | 0.34627513 | 0.238 | 0 | 1 | 27 |
| ago2 | 0.30516 | 0.421272992 | 0.238 | 0 | 1 | 27 |
| armc1l | 0.30516 | 0.44969106 | 0.238 | 0 | 1 | 27 |
| arrdc3b | 0.30516 | 0.351982342 | 0.238 | 0 | 1 | 27 |
| ash2l | 0.30516 | 0.32880542 | 0.238 | 0 | 1 | 27 |
| bcl11aa | 0.30516 | 0.421107436 | 0.238 | 0 | 1 | 27 |
| bri3bp | 0.30516 | 0.434857386 | 0.238 | 0 | 1 | 27 |
| cdc26 | 0.30516 | 0.474896605 | 0.238 | 0 | 1 | 27 |
| chaf1a | 0.30516 | 0.596723757 | 0.238 | 0 | 1 | 27 |
| chd3 | 0.30516 | 0.34704127 | 0.238 | 0 | 1 | 27 |
| chd4a | 0.30516 | 0.390117271 | 0.238 | 0 | 1 | 27 |
| crkl | 0.30516 | 0.356309657 | 0.238 | 0 | 1 | 27 |
| cul4b | 0.30516 | 0.44979109 | 0.238 | 0 | 1 | 27 |
| dars | 0.30516 | 0.355288072 | 0.238 | 0 | 1 | 27 |
| dazap2 | 0.30516 | 0.34562449 | 0.238 | 0 | 1 | 27 |

| | | | | | | |
|---|---|---|---|---|---|---|
| dnajc8 | 0.30516 | 0.445611419 | 0.238 | 0 | 1 | 27 |
| eed | 0.30516 | 0.449601581 | 0.238 | 0 | 1 | 27 |
| eif4g2b | 0.30516 | 0.357746879 | 0.238 | 0 | 1 | 27 |
| elf2b | 0.30516 | 0.41557277 | 0.238 | 0 | 1 | 27 |
| ewsr1a | 0.30516 | 0.360104573 | 0.238 | 0 | 1 | 27 |
| fen1 | 0.30516 | 0.42483912 | 0.238 | 0 | 1 | 27 |
| fgfr1op2 | 0.30516 | 0.368707475 | 0.238 | 0 | 1 | 27 |
| fosb | 0.30516 | 0.358740374 | 0.238 | 0 | 1 | 27 |
| gmps | 0.30516 | 0.376581388 | 0.238 | 0 | 1 | 27 |
| gsto2 | 0.30516 | 0.40253573 | 0.238 | 0 | 1 | 27 |
| hmbsa | 0.30516 | 0.434181757 | 0.238 | 0 | 1 | 27 |
| ivns1abpb | 0.30516 | 0.390434922 | 0.238 | 0 | 1 | 27 |
| jam2a | 0.30516 | 0.253840936 | 0.238 | 0 | 1 | 27 |
| lig1 | 0.30516 | 0.46965235 | 0.238 | 0 | 1 | 27 |
| luc7l3 | 0.30516 | 0.429687939 | 0.238 | 0 | 1 | 27 |
| m6pr | 0.30516 | 0.47848575 | 0.238 | 0 | 1 | 27 |
| manf | 0.30516 | 0.50526969 | 0.238 | 0 | 1 | 27 |
| marcksb | 0.30516 | 0.407540623 | 0.238 | 0 | 1 | 27 |
| mcph1 | 0.30516 | 0.488276381 | 0.238 | 0 | 1 | 27 |
| minos1 | 0.30516 | 0.373698725 | 0.238 | 0 | 1 | 27 |
| mrpl18 | 0.30516 | 0.453948356 | 0.238 | 0 | 1 | 27 |
| mrps36 | 0.30516 | 0.349371878 | 0.238 | 0 | 1 | 27 |
| mycbp | 0.30516 | 0.45380428 | 0.238 | 0 | 1 | 27 |
| nap1l4b | 0.30516 | 0.370457669 | 0.238 | 0 | 1 | 27 |
| nasp | 0.30516 | 0.4507543 | 0.238 | 0 | 1 | 27 |
| nrros | 0.30516 | 0.390793409 | 0.238 | 0 | 1 | 27 |
| nudc | 0.30516 | 0.406024255 | 0.238 | 0 | 1 | 27 |
| pa2g4a | 0.30516 | 0.372070034 | 0.238 | 0 | 1 | 27 |
| pagr1 | 0.30516 | 0.365022184 | 0.238 | 0 | 1 | 27 |
| pane1 | 0.30516 | 0.366933145 | 0.238 | 0 | 1 | 27 |
| pfkfb4b | 0.30516 | 0.4277151 | 0.238 | 0 | 1 | 27 |
| ppig | 0.30516 | 0.420329174 | 0.238 | 0 | 1 | 27 |
| ppp4r2a | 0.30516 | 0.422365057 | 0.238 | 0 | 1 | 27 |
| prdx6 | 0.30516 | 0.345016555 | 0.238 | 0 | 1 | 27 |
| prim2 | 0.30516 | 0.360109114 | 0.238 | 0 | 1 | 27 |
| psmc4 | 0.30516 | 0.415585667 | 0.238 | 0 | 1 | 27 |
| psmd8 | 0.30516 | 0.459361084 | 0.238 | 0 | 1 | 27 |
| psmd9 | 0.30516 | 0.413826797 | 0.238 | 0 | 1 | 27 |
| ptgr1 | 0.30516 | 0.351462205 | 0.238 | 0 | 1 | 27 |
| rab24 | 0.30516 | 0.295193986 | 0.238 | 0 | 1 | 27 |
| rabif | 0.30516 | 0.379797479 | 0.238 | 0 | 1 | 27 |
| rabl6b | 0.30516 | 0.430201523 | 0.238 | 0 | 1 | 27 |
| rbbp4 | 0.30516 | 0.546203044 | 0.238 | 0 | 1 | 27 |
| rpa3 | 0.30516 | 0.480012948 | 0.238 | 0 | 1 | 27 |
| sars | 0.30516 | 0.423632859 | 0.238 | 0 | 1 | 27 |
| sf3a3 | 0.30516 | 0.322233955 | 0.238 | 0 | 1 | 27 |
| sfpq | 0.30516 | 0.423520067 | 0.238 | 0 | 1 | 27 |
| si:ch211-276i12.11 | 0.30516 | 0.395350339 | 0.238 | 0 | 1 | 27 |

| | | | | | | |
|---|---|---|---|---|---|---|
| smc1al | 0.30516 | 0.40413855 | 0.238 | 0 | 1 | 27 |
| sptssa | 0.30516 | 0.373850357 | 0.238 | 0 | 1 | 27 |
| srrt | 0.30516 | 0.403110996 | 0.238 | 0 | 1 | 27 |
| stag2b | 0.30516 | 0.418827442 | 0.238 | 0 | 1 | 27 |
| sult1st1 | 0.30516 | 0.33675505 | 0.238 | 0 | 1 | 27 |
| sumo3b | 0.30516 | 0.523232547 | 0.238 | 0 | 1 | 27 |
| tfam | 0.30516 | 0.392609013 | 0.238 | 0 | 1 | 27 |
| tmc8 | 0.30516 | 0.53052456 | 0.238 | 0 | 1 | 27 |
| tmpob | 0.30516 | 0.451714868 | 0.238 | 0 | 1 | 27 |
| tyms | 0.30516 | 0.471359897 | 0.238 | 0 | 1 | 27 |
| wu:fi37e09 | 0.30516 | 0.424249555 | 0.238 | 0 | 1 | 27 |
| zgc:66388 | 0.30516 | 0.364028565 | 0.238 | 0 | 1 | 27 |
| CABZ01005379.1 | 0.30516 | 0.466078461 | 0.238 | 0 | 1 | 27 |
| CR855337.1 | 0.30516 | 0.470583478 | 0.238 | 0 | 1 | 27 |
| CR944667.2 | 0.30516 | 0.428302451 | 0.238 | 0 | 1 | 27 |
| actl6a | 0.30516 | 0.399647198 | 0.238 | 0 | 1 | 27 |
| adipor2 | 0.30516 | 0.437254163 | 0.238 | 0 | 1 | 27 |
| aldh6a1 | 0.30516 | 0.293297515 | 0.238 | 0 | 1 | 27 |
| ccnl1b | 0.30516 | 0.410482394 | 0.238 | 0 | 1 | 27 |
| cdca4 | 0.30516 | 0.462774805 | 0.238 | 0 | 1 | 27 |
| chtopa | 0.30516 | 0.468773304 | 0.238 | 0 | 1 | 27 |
| churc1 | 0.30516 | 0.379573268 | 0.238 | 0 | 1 | 27 |
| clint1a | 0.30516 | 0.450684879 | 0.238 | 0 | 1 | 27 |
| cmpk | 0.30516 | 0.368852825 | 0.238 | 0 | 1 | 27 |
| cry1ab | 0.30516 | 0.450355377 | 0.238 | 0 | 1 | 27 |
| ddx3xa | 0.30516 | 0.441447704 | 0.238 | 0 | 1 | 27 |
| dek | 0.30516 | 0.416149685 | 0.238 | 0 | 1 | 27 |
| egr1 | 0.30516 | 0.581443228 | 0.238 | 0 | 1 | 27 |
| eny2 | 0.30516 | 0.337363828 | 0.238 | 0 | 1 | 27 |
| fbxl3a | 0.30516 | 0.429516523 | 0.238 | 0 | 1 | 27 |
| fmnl1a | 0.30516 | 0.467595918 | 0.238 | 0 | 1 | 27 |
| gigyf1b | 0.30516 | 0.439124524 | 0.238 | 0 | 1 | 27 |
| im:7138535 | 0.30516 | 0.436985498 | 0.238 | 0 | 1 | 27 |
| ivd | 0.30516 | 0.36450754 | 0.238 | 0 | 1 | 27 |
| mbd4 | 0.30516 | 0.40216559 | 0.238 | 0 | 1 | 27 |
| mef2cb | 0.30516 | 0.416947037 | 0.238 | 0 | 1 | 27 |
| mvb12a | 0.30516 | 0.508532682 | 0.238 | 0 | 1 | 27 |
| ncoa5 | 0.30516 | 0.469217896 | 0.238 | 0 | 1 | 27 |
| ndufa5 | 0.30516 | 0.40664684 | 0.238 | 0 | 1 | 27 |
| ndufb2 | 0.30516 | 0.283388726 | 0.238 | 0 | 1 | 27 |
| nenf | 0.30516 | 0.400446064 | 0.238 | 0 | 1 | 27 |
| npm1a | 0.30516 | 0.351201599 | 0.238 | 0 | 1 | 27 |
| ost4 | 0.30516 | 0.34470802 | 0.238 | 0 | 1 | 27 |
| pi4k2b | 0.30516 | 0.367511248 | 0.238 | 0 | 1 | 27 |
| pinx1 | 0.30516 | 0.513538788 | 0.238 | 0 | 1 | 27 |
| psmb7 | 0.30516 | 0.490476293 | 0.238 | 0 | 1 | 27 |
| psmd6 | 0.30516 | 0.382898428 | 0.238 | 0 | 1 | 27 |
| rab2a | 0.30516 | 0.375161913 | 0.238 | 0 | 1 | 27 |

| | | | | | | |
|---|---|---|---|---|---|---|
| rad23b | 0.30516 | 0.422031454 | 0.238 | 0 | 1 | 27 |
| rassf2b | 0.30516 | 0.337132213 | 0.238 | 0 | 1 | 27 |
| rxrba | 0.30516 | 0.348912208 | 0.238 | 0 | 1 | 27 |
| sdhaf4 | 0.30516 | 0.408431229 | 0.238 | 0 | 1 | 27 |
| si:ch211-102c2.4 | 0.30516 | 0.518451419 | 0.238 | 0 | 1 | 27 |
| si:ch211-107o10.3 | 0.30516 | 0.358531755 | 0.238 | 0 | 1 | 27 |
| si:ch211-213a13.1 | 0.30516 | 0.37136716 | 0.238 | 0 | 1 | 27 |
| si:dkey-172h23.2 | 0.30516 | 0.44455588 | 0.238 | 0 | 1 | 27 |
| sqstm1 | 0.30516 | 0.433558846 | 0.238 | 0 | 1 | 27 |
| ss18 | 0.30516 | 0.455978338 | 0.238 | 0 | 1 | 27 |
| ss18l2 | 0.30516 | 0.381922183 | 0.238 | 0 | 1 | 27 |
| stk17al | 0.30516 | 0.532299786 | 0.238 | 0 | 1 | 27 |
| tab1 | 0.30516 | 0.480394961 | 0.238 | 0 | 1 | 27 |
| tm9sf2 | 0.30516 | 0.383663589 | 0.238 | 0 | 1 | 27 |
| tpst1l | 0.30516 | 0.453804308 | 0.238 | 0 | 1 | 27 |
| trappc3 | 0.30516 | 0.476192852 | 0.238 | 0 | 1 | 27 |
| ubl7b | 0.30516 | 0.454891765 | 0.238 | 0 | 1 | 27 |
| uqcrc2b | 0.30516 | 0.363242507 | 0.238 | 0 | 1 | 27 |
| vps4b | 0.30516 | 0.438595174 | 0.238 | 0 | 1 | 27 |
| zgc:174268 | 0.30516 | 0.377443746 | 0.238 | 0 | 1 | 27 |
| zgc:56304 | 0.30516 | 0.422964597 | 0.238 | 0 | 1 | 27 |
| zgc:92606 | 0.30516 | 0.401232061 | 0.238 | 0 | 1 | 27 |
| casc3 | 0.30516 | 0.443504536 | 0.238 | 0 | 1 | 27 |
| ccng2 | 0.30516 | 0.344810529 | 0.238 | 0 | 1 | 27 |
| creb1b | 0.30516 | 0.429527125 | 0.238 | 0 | 1 | 27 |
| cyc1 | 0.30516 | 0.302290275 | 0.238 | 0 | 1 | 27 |
| ehd4 | 0.30516 | 0.380743262 | 0.238 | 0 | 1 | 27 |
| eif4ebp2 | 0.30516 | 0.363204501 | 0.238 | 0 | 1 | 27 |
| evla | 0.30516 | 0.565973294 | 0.238 | 0 | 1 | 27 |
| fubp1 | 0.30516 | 0.369679555 | 0.238 | 0 | 1 | 27 |
| gars | 0.30516 | 0.304653887 | 0.238 | 0 | 1 | 27 |
| hspa5 | 0.30516 | 0.430204652 | 0.238 | 0 | 1 | 27 |
| ist1 | 0.30516 | 0.496043438 | 0.238 | 0 | 1 | 27 |
| naga | 0.30516 | 0.440629269 | 0.238 | 0 | 1 | 27 |
| npepl1 | 0.30516 | 0.517650342 | 0.238 | 0 | 1 | 27 |
| ppm1g | 0.30516 | 0.376757079 | 0.238 | 0 | 1 | 27 |
| prpf38b | 0.30516 | 0.394138735 | 0.238 | 0 | 1 | 27 |
| psma6b | 0.30516 | 0.50367714 | 0.238 | 0 | 1 | 27 |
| psmb3 | 0.30516 | 0.370844654 | 0.238 | 0 | 1 | 27 |
| reep5 | 0.30516 | 0.412621981 | 0.238 | 0 | 1 | 27 |
| stxbp2 | 0.30516 | 0.353753233 | 0.238 | 0 | 1 | 27 |
| terfa | 0.30516 | 0.392889234 | 0.238 | 0 | 1 | 27 |
| tmco1 | 0.30516 | 0.345959335 | 0.238 | 0 | 1 | 27 |
| tmem106a | 0.30516 | 0.446423176 | 0.238 | 0 | 1 | 27 |
| CABZ01115113.1 | 0.30516 | 0.429275785 | 0.238 | 0 | 1 | 27 |
| brd3b | 0.30516 | 0.365609918 | 0.238 | 0 | 1 | 27 |
| cenpn | 0.30516 | 0.324502837 | 0.238 | 0 | 1 | 27 |
| cks2 | 0.30516 | 0.468653158 | 0.238 | 0 | 1 | 27 |

| | | | | | | |
|---|---|---|---|---|---|---|
| hnrnpl2 | 0.30516 | 0.38973628 | 0.238 | 0 | 1 | 27 |
| klhl6 | 0.30516 | 0.375456349 | 0.238 | 0 | 1 | 27 |
| lsm12a | 0.30516 | 0.40720684 | 0.238 | 0 | 1 | 27 |
| ncapg | 0.30516 | 0.402352185 | 0.238 | 0 | 1 | 27 |
| ndufs5 | 0.30516 | 0.337120235 | 0.238 | 0 | 1 | 27 |
| rfc3 | 0.30516 | 0.422455746 | 0.238 | 0 | 1 | 27 |
| si:ch211-226m16.2 | 0.30516 | 0.434162735 | 0.238 | 0 | 1 | 27 |
| stx2a | 0.30516 | 0.325174525 | 0.238 | 0 | 1 | 27 |
| wdr82 | 0.30516 | 0.42610691 | 0.238 | 0 | 1 | 27 |
| zgc:110540 | 0.30516 | 0.553544444 | 0.238 | 0 | 1 | 27 |
| cxcr4a | 0.3073 | 0.850212031 | 0.714 | 0.333 | 1 | 27 |
| tnfaip8l2a | 0.30959 | -0.7058957 | 0.238 | 0.667 | 1 | 27 |
| si:ch1073-429i10.3.1 | 0.31227 | -0.52889537 | 0.667 | 1 | 1 | 27 |
| XLOC-042595 | 0.31905 | -0.46197803 | 0.857 | 1 | 1 | 27 |
| wdr1 | 0.32043 | -0.72562468 | 0.286 | 0.667 | 1 | 27 |
| si:dkeyp-68b7.12 | 0.32139 | -0.16166177 | 0.429 | 0.333 | 1 | 27 |
| CR847984.1 | 0.32179 | -0.53077755 | 0.095 | 0.333 | 1 | 27 |
| atp11a | 0.32179 | -0.41172042 | 0.095 | 0.333 | 1 | 27 |
| ciao1 | 0.32179 | -0.52360169 | 0.095 | 0.333 | 1 | 27 |
| dgkzb | 0.32179 | -0.44831815 | 0.095 | 0.333 | 1 | 27 |
| ero1b | 0.32179 | -0.53401922 | 0.095 | 0.333 | 1 | 27 |
| gpr55a | 0.32179 | -0.49984121 | 0.095 | 0.333 | 1 | 27 |
| hyal1 | 0.32179 | -0.40299447 | 0.095 | 0.333 | 1 | 27 |
| iah1 | 0.32179 | -0.33707709 | 0.095 | 0.333 | 1 | 27 |
| il12rb2l | 0.32179 | -0.4878036 | 0.095 | 0.333 | 1 | 27 |
| map1lc3b | 0.32179 | -0.47532124 | 0.095 | 0.333 | 1 | 27 |
| mrps2 | 0.32179 | -0.47845629 | 0.095 | 0.333 | 1 | 27 |
| napepld | 0.32179 | -0.20193874 | 0.095 | 0.333 | 1 | 27 |
| nedd8 | 0.32179 | -0.40806942 | 0.095 | 0.333 | 1 | 27 |
| nelfe | 0.32179 | -0.48814792 | 0.095 | 0.333 | 1 | 27 |
| si:ch211-218c6.8 | 0.32179 | -0.65855888 | 0.095 | 0.333 | 1 | 27 |
| stard3nl | 0.32179 | -0.52125304 | 0.095 | 0.333 | 1 | 27 |
| txnl4a | 0.32179 | -0.46029724 | 0.095 | 0.333 | 1 | 27 |
| ubr5 | 0.32179 | -0.45464019 | 0.095 | 0.333 | 1 | 27 |
| usp22 | 0.32179 | -0.53300126 | 0.095 | 0.333 | 1 | 27 |
| BX323082.1 | 0.32179 | -0.53753833 | 0.095 | 0.333 | 1 | 27 |
| BX548028.1 | 0.32179 | -0.5850552 | 0.095 | 0.333 | 1 | 27 |
| CR381673.1 | 0.32179 | -0.33336883 | 0.095 | 0.333 | 1 | 27 |
| LOC110437917 | 0.32179 | -0.51980564 | 0.095 | 0.333 | 1 | 27 |
| add1 | 0.32179 | -0.49289953 | 0.095 | 0.333 | 1 | 27 |
| agap2 | 0.32179 | -0.51824026 | 0.095 | 0.333 | 1 | 27 |
| akirin2 | 0.32179 | -0.41198541 | 0.095 | 0.333 | 1 | 27 |
| aldh7a1 | 0.32179 | -0.33496839 | 0.095 | 0.333 | 1 | 27 |
| angel2 | 0.32179 | -0.48995825 | 0.095 | 0.333 | 1 | 27 |
| arf4a | 0.32179 | -0.50280963 | 0.095 | 0.333 | 1 | 27 |
| arglu1a | 0.32179 | -0.52134369 | 0.095 | 0.333 | 1 | 27 |
| batf3 | 0.32179 | -0.48280363 | 0.095 | 0.333 | 1 | 27 |
| bnip4 | 0.32179 | -0.47082251 | 0.095 | 0.333 | 1 | 27 |

| | | | | | | |
|---|---|---|---|---|---|---|
| borcs7 | 0.32179 | -0.47285596 | 0.095 | 0.333 | 1 | 27 |
| btbd9 | 0.32179 | -0.45234836 | 0.095 | 0.333 | 1 | 27 |
| cbx1a | 0.32179 | -0.43314373 | 0.095 | 0.333 | 1 | 27 |
| cd81a | 0.32179 | -0.63805116 | 0.095 | 0.333 | 1 | 27 |
| cdk9 | 0.32179 | -0.43379825 | 0.095 | 0.333 | 1 | 27 |
| chd9 | 0.32179 | -0.44536205 | 0.095 | 0.333 | 1 | 27 |
| chordc1a | 0.32179 | -0.46877144 | 0.095 | 0.333 | 1 | 27 |
| clasrp | 0.32179 | -0.48968173 | 0.095 | 0.333 | 1 | 27 |
| cstf3 | 0.32179 | -0.43397273 | 0.095 | 0.333 | 1 | 27 |
| cyb561a3a | 0.32179 | -0.46006124 | 0.095 | 0.333 | 1 | 27 |
| dcun1d1 | 0.32179 | -0.53164693 | 0.095 | 0.333 | 1 | 27 |
| dgkg | 0.32179 | -0.3782794 | 0.095 | 0.333 | 1 | 27 |
| eaf1 | 0.32179 | -0.45783627 | 0.095 | 0.333 | 1 | 27 |
| ebf1a | 0.32179 | -0.50442701 | 0.095 | 0.333 | 1 | 27 |
| emc10 | 0.32179 | -0.43220514 | 0.095 | 0.333 | 1 | 27 |
| emc8 | 0.32179 | -0.46737752 | 0.095 | 0.333 | 1 | 27 |
| eno1a | 0.32179 | -0.57666313 | 0.095 | 0.333 | 1 | 27 |
| etfb | 0.32179 | -0.45046281 | 0.095 | 0.333 | 1 | 27 |
| fam120c | 0.32179 | -0.53174016 | 0.095 | 0.333 | 1 | 27 |
| fuca1.1 | 0.32179 | -0.33714162 | 0.095 | 0.333 | 1 | 27 |
| gabarapl2 | 0.32179 | -0.43258357 | 0.095 | 0.333 | 1 | 27 |
| gfi1ab | 0.32179 | -0.42973518 | 0.095 | 0.333 | 1 | 27 |
| glg1a | 0.32179 | -0.52993608 | 0.095 | 0.333 | 1 | 27 |
| gnpnat1 | 0.32179 | -0.54871423 | 0.095 | 0.333 | 1 | 27 |
| gpalpp1 | 0.32179 | -0.46163924 | 0.095 | 0.333 | 1 | 27 |
| hbegfa | 0.32179 | -0.46445157 | 0.095 | 0.333 | 1 | 27 |
| hdac9b | 0.32179 | -0.43787344 | 0.095 | 0.333 | 1 | 27 |
| hes6 | 0.32179 | -0.45497299 | 0.095 | 0.333 | 1 | 27 |
| inpp4aa | 0.32179 | -0.47440495 | 0.095 | 0.333 | 1 | 27 |
| jak2a | 0.32179 | -0.40419489 | 0.095 | 0.333 | 1 | 27 |
| kmt2ba | 0.32179 | -0.46299694 | 0.095 | 0.333 | 1 | 27 |
| lgals9l1 | 0.32179 | -0.58055744 | 0.095 | 0.333 | 1 | 27 |
| med13a | 0.32179 | -0.45390921 | 0.095 | 0.333 | 1 | 27 |
| mrpl21 | 0.32179 | -0.39257188 | 0.095 | 0.333 | 1 | 27 |
| mrpl41 | 0.32179 | -0.457008 | 0.095 | 0.333 | 1 | 27 |
| mrpl43 | 0.32179 | -0.52186949 | 0.095 | 0.333 | 1 | 27 |
| mrps15 | 0.32179 | -0.54078771 | 0.095 | 0.333 | 1 | 27 |
| mrps26 | 0.32179 | -0.49098282 | 0.095 | 0.333 | 1 | 27 |
| mt-nd5 | 0.32179 | -0.46474143 | 0.095 | 0.333 | 1 | 27 |
| naa50 | 0.32179 | -0.50466848 | 0.095 | 0.333 | 1 | 27 |
| nap1l4a | 0.32179 | -0.36312726 | 0.095 | 0.333 | 1 | 27 |
| ncf4 | 0.32179 | -0.44391259 | 0.095 | 0.333 | 1 | 27 |
| ndufa7 | 0.32179 | -0.41421741 | 0.095 | 0.333 | 1 | 27 |
| nrbp1 | 0.32179 | -0.54532875 | 0.095 | 0.333 | 1 | 27 |
| nsmce2 | 0.32179 | -0.35198473 | 0.095 | 0.333 | 1 | 27 |
| nsmce4a | 0.32179 | -0.5270948 | 0.095 | 0.333 | 1 | 27 |
| nuggc.3 | 0.32179 | -0.39618705 | 0.095 | 0.333 | 1 | 27 |
| nup58 | 0.32179 | -0.67531155 | 0.095 | 0.333 | 1 | 27 |

| | | | | | | |
|---|---|---|---|---|---|---|
| ociad1 | 0.32179 | -0.5929536 | 0.095 | 0.333 | 1 | 27 |
| ophn1 | 0.32179 | -0.50220825 | 0.095 | 0.333 | 1 | 27 |
| ormdl1 | 0.32179 | -0.36597693 | 0.095 | 0.333 | 1 | 27 |
| pank2 | 0.32179 | -0.47322871 | 0.095 | 0.333 | 1 | 27 |
| pcnp | 0.32179 | -0.52073808 | 0.095 | 0.333 | 1 | 27 |
| pdap1a | 0.32179 | -0.53664892 | 0.095 | 0.333 | 1 | 27 |
| pgam5 | 0.32179 | -0.54285041 | 0.095 | 0.333 | 1 | 27 |
| pgap2 | 0.32179 | -0.44354119 | 0.095 | 0.333 | 1 | 27 |
| phlda2 | 0.32179 | -0.50322531 | 0.095 | 0.333 | 1 | 27 |
| plrg1 | 0.32179 | -0.51509473 | 0.095 | 0.333 | 1 | 27 |
| pnn | 0.32179 | -0.58040596 | 0.095 | 0.333 | 1 | 27 |
| pou2f1b | 0.32179 | -0.38367962 | 0.095 | 0.333 | 1 | 27 |
| prpf3 | 0.32179 | -0.6938235 | 0.095 | 0.333 | 1 | 27 |
| rab1ba | 0.32179 | -0.58899747 | 0.095 | 0.333 | 1 | 27 |
| rexo4 | 0.32179 | -0.23886524 | 0.095 | 0.333 | 1 | 27 |
| ripor1.1 | 0.32179 | -0.34255041 | 0.095 | 0.333 | 1 | 27 |
| rpz2 | 0.32179 | -0.49714507 | 0.095 | 0.333 | 1 | 27 |
| sco1 | 0.32179 | -0.4898082 | 0.095 | 0.333 | 1 | 27 |
| sdc4 | 0.32179 | -0.39035878 | 0.095 | 0.333 | 1 | 27 |
| sh3bgrl | 0.32179 | -0.53840309 | 0.095 | 0.333 | 1 | 27 |
| sh3glb1b | 0.32179 | -0.47264135 | 0.095 | 0.333 | 1 | 27 |
| si:ch211-114l13.12 | 0.32179 | -0.39428681 | 0.095 | 0.333 | 1 | 27 |
| si:ch211-67e16.3 | 0.32179 | -0.36561685 | 0.095 | 0.333 | 1 | 27 |
| si:dkey-187e18.1 | 0.32179 | -0.5283095 | 0.095 | 0.333 | 1 | 27 |
| si:dkey-206d17.12 | 0.32179 | -0.39688279 | 0.095 | 0.333 | 1 | 27 |
| si:dkey-262k9.4 | 0.32179 | -0.41243088 | 0.095 | 0.333 | 1 | 27 |
| si:dkeyp-85e10.3 | 0.32179 | -0.38877792 | 0.095 | 0.333 | 1 | 27 |
| si:rp71-36a1.5 | 0.32179 | -0.40364117 | 0.095 | 0.333 | 1 | 27 |
| smad2 | 0.32179 | -0.45911903 | 0.095 | 0.333 | 1 | 27 |
| smim19 | 0.32179 | -0.37467364 | 0.095 | 0.333 | 1 | 27 |
| snu13b | 0.32179 | -0.60382552 | 0.095 | 0.333 | 1 | 27 |
| snx6 | 0.32179 | -0.39567918 | 0.095 | 0.333 | 1 | 27 |
| snx9b | 0.32179 | -0.40488354 | 0.095 | 0.333 | 1 | 27 |
| srsf4 | 0.32179 | -0.56964418 | 0.095 | 0.333 | 1 | 27 |
| stat2 | 0.32179 | -0.4360025 | 0.095 | 0.333 | 1 | 27 |
| sugt1 | 0.32179 | -0.46166837 | 0.095 | 0.333 | 1 | 27 |
| ten1 | 0.32179 | -0.43599621 | 0.095 | 0.333 | 1 | 27 |
| tfpt | 0.32179 | -0.33338685 | 0.095 | 0.333 | 1 | 27 |
| tm9sf4 | 0.32179 | -0.43389148 | 0.095 | 0.333 | 1 | 27 |
| tmem209 | 0.32179 | -0.3927087 | 0.095 | 0.333 | 1 | 27 |
| tmem263 | 0.32179 | -0.47756963 | 0.095 | 0.333 | 1 | 27 |
| ubald1a | 0.32179 | -0.48082753 | 0.095 | 0.333 | 1 | 27 |
| unm-hu7910 | 0.32179 | -0.31949699 | 0.095 | 0.333 | 1 | 27 |
| wdr45 | 0.32179 | -0.41010499 | 0.095 | 0.333 | 1 | 27 |
| yipf6 | 0.32179 | -0.43170423 | 0.095 | 0.333 | 1 | 27 |
| zc3h18 | 0.32179 | -0.45130919 | 0.095 | 0.333 | 1 | 27 |
| zgc:101583 | 0.32179 | -0.43945005 | 0.095 | 0.333 | 1 | 27 |
| zgc:114200 | 0.32179 | -0.41893283 | 0.095 | 0.333 | 1 | 27 |

| | | | | | | |
|---|---|---|---|---|---|---|
| zgc:152830 | 0.32179 | -0.42170609 | 0.095 | 0.333 | 1 | 27 |
| zyg11 | 0.32179 | -0.37845522 | 0.095 | 0.333 | 1 | 27 |
| BX649442.1 | 0.32179 | -0.38379762 | 0.095 | 0.333 | 1 | 27 |
| XLOC-005449 | 0.32179 | -0.44113292 | 0.095 | 0.333 | 1 | 27 |
| eef1db | 0.32179 | -0.40163526 | 0.095 | 0.333 | 1 | 27 |
| ikbkg | 0.32179 | -0.37150968 | 0.095 | 0.333 | 1 | 27 |
| ktn1 | 0.32179 | -0.37247106 | 0.095 | 0.333 | 1 | 27 |
| lrmp | 0.32179 | -0.3375845 | 0.095 | 0.333 | 1 | 27 |
| pes | 0.32179 | -0.44343538 | 0.095 | 0.333 | 1 | 27 |
| prkcbb | 0.32179 | -0.3469141 | 0.095 | 0.333 | 1 | 27 |
| si:ch211-212k18.4 | 0.32179 | -0.40266618 | 0.095 | 0.333 | 1 | 27 |
| si:ch211-214b16.4 | 0.32179 | -0.38696913 | 0.095 | 0.333 | 1 | 27 |
| sil1 | 0.32179 | -0.57522938 | 0.095 | 0.333 | 1 | 27 |
| socs3a | 0.32179 | -0.39140435 | 0.095 | 0.333 | 1 | 27 |
| zbtb8os | 0.32179 | -0.39573293 | 0.095 | 0.333 | 1 | 27 |
| zgc:152863 | 0.32179 | -0.38017494 | 0.095 | 0.333 | 1 | 27 |
| si:ch211-246m6.4 | 0.32179 | -0.48005133 | 0.095 | 0.333 | 1 | 27 |
| tomm5 | 0.32448 | -0.5827626 | 0.429 | 0.667 | 1 | 27 |
| ywhaba | 0.33283 | -0.32540306 | 0.619 | 1 | 1 | 27 |
| hmgb2a | 0.34326 | 0.12526723 | 0.762 | 1 | 1 | 27 |
| rps27.2 | 0.34497 | 0.174041252 | 0.952 | 1 | 1 | 27 |
| atp5l | 0.35425 | 0.309098322 | 0.714 | 0.333 | 1 | 27 |
| zgc:64051 | 0.35449 | 0.250890885 | 0.667 | 0.333 | 1 | 27 |
| bhlhe40 | 0.35947 | -0.67223941 | 0.238 | 0.667 | 1 | 27 |
| cox8a | 0.35996 | 0.539349205 | 0.762 | 0.333 | 1 | 27 |
| hnrnpa1a | 0.36143 | -0.67956357 | 0.333 | 0.667 | 1 | 27 |
| rps29 | 0.36183 | -0.11307065 | 1 | 1 | 1 | 27 |
| calm3a | 0.36303 | -0.56289029 | 0.238 | 0.667 | 1 | 27 |
| ahcy | 0.36305 | 0.5067015 | 0.333 | 0.333 | 1 | 27 |
| uqcrq | 0.36388 | -0.62462532 | 0.333 | 0.667 | 1 | 27 |
| cope | 0.36499 | -0.60430771 | 0.238 | 0.667 | 1 | 27 |
| tsc22d3 | 0.3657 | 0.595236475 | 0.762 | 0.333 | 1 | 27 |
| pou2f2a | 0.36654 | -0.55497105 | 0.429 | 0.667 | 1 | 27 |
| litaf | 0.37281 | -0.18956647 | 0.429 | 0.333 | 1 | 27 |
| nfkbiaa | 0.37875 | 0.15986856 | 0.619 | 0.333 | 1 | 27 |
| hspa8 | 0.37933 | -0.40041041 | 0.905 | 1 | 1 | 27 |
| coa5 | 0.37981 | 0.301665657 | 0.19 | 0 | 1 | 27 |
| lamtor5 | 0.37981 | 0.413866739 | 0.19 | 0 | 1 | 27 |
| mrps30 | 0.37981 | 0.34961936 | 0.19 | 0 | 1 | 27 |
| si:ch73-248e21.5 | 0.37981 | 0.40693536 | 0.19 | 0 | 1 | 27 |
| smim29 | 0.37981 | 0.334442205 | 0.19 | 0 | 1 | 27 |
| tbc1d22a | 0.37981 | 0.321095476 | 0.19 | 0 | 1 | 27 |
| CABZ01066694.1 | 0.37981 | 0.293521306 | 0.19 | 0 | 1 | 27 |
| atp5mea | 0.37981 | 0.407577253 | 0.19 | 0 | 1 | 27 |
| atp6v0ca | 0.37981 | 0.38812147 | 0.19 | 0 | 1 | 27 |
| bida | 0.37981 | 0.396595012 | 0.19 | 0 | 1 | 27 |
| casp8 | 0.37981 | 0.348745448 | 0.19 | 0 | 1 | 27 |
| cdc37 | 0.37981 | 0.319709242 | 0.19 | 0 | 1 | 27 |

| | | | | | | |
|---|---|---|---|---|---|---|
| ciao2a | 0.37981 | 0.415835965 | 0.19 | 0 | 1 | 27 |
| cstf1 | 0.37981 | 0.334226446 | 0.19 | 0 | 1 | 27 |
| dnajc19 | 0.37981 | 0.290578278 | 0.19 | 0 | 1 | 27 |
| fnbp4 | 0.37981 | 0.406062247 | 0.19 | 0 | 1 | 27 |
| gtf2f2a | 0.37981 | 0.353139328 | 0.19 | 0 | 1 | 27 |
| hcst | 0.37981 | 0.39352158 | 0.19 | 0 | 1 | 27 |
| inip | 0.37981 | 0.332770011 | 0.19 | 0 | 1 | 27 |
| isoc2 | 0.37981 | 0.360521019 | 0.19 | 0 | 1 | 27 |
| jmjd1cb | 0.37981 | 0.448657203 | 0.19 | 0 | 1 | 27 |
| lamtor3 | 0.37981 | 0.374958868 | 0.19 | 0 | 1 | 27 |
| ndufa1 | 0.37981 | 0.256644357 | 0.19 | 0 | 1 | 27 |
| parp16 | 0.37981 | 0.325951224 | 0.19 | 0 | 1 | 27 |
| pin4 | 0.37981 | 0.271164314 | 0.19 | 0 | 1 | 27 |
| ppp3r1a | 0.37981 | 0.424228416 | 0.19 | 0 | 1 | 27 |
| rab11fip2 | 0.37981 | 0.318222016 | 0.19 | 0 | 1 | 27 |
| rgs14a | 0.37981 | 0.338192163 | 0.19 | 0 | 1 | 27 |
| rtf2 | 0.37981 | 0.392975107 | 0.19 | 0 | 1 | 27 |
| si:ch211-1a19.3 | 0.37981 | 0.370783705 | 0.19 | 0 | 1 | 27 |
| si:ch211-286b5.5 | 0.37981 | 0.315730316 | 0.19 | 0 | 1 | 27 |
| si:ch73-269m14.2 | 0.37981 | 0.274434537 | 0.19 | 0 | 1 | 27 |
| si:dkey-46a10.3 | 0.37981 | 0.345600092 | 0.19 | 0 | 1 | 27 |
| srp68 | 0.37981 | 0.301719182 | 0.19 | 0 | 1 | 27 |
| tex30 | 0.37981 | 0.349013389 | 0.19 | 0 | 1 | 27 |
| tmbim4 | 0.37981 | 0.32025002 | 0.19 | 0 | 1 | 27 |
| tnrc6a | 0.37981 | 0.262370856 | 0.19 | 0 | 1 | 27 |
| ube2q2 | 0.37981 | 0.297690829 | 0.19 | 0 | 1 | 27 |
| vbp1 | 0.37981 | 0.308573112 | 0.19 | 0 | 1 | 27 |
| vmp1 | 0.37981 | 0.425888987 | 0.19 | 0 | 1 | 27 |
| zgc:110319 | 0.37981 | 0.292377032 | 0.19 | 0 | 1 | 27 |
| znf706 | 0.37981 | 0.28148618 | 0.19 | 0 | 1 | 27 |
| BX284696.1 | 0.37981 | 0.40137631 | 0.19 | 0 | 1 | 27 |
| CABZ01088428.1 | 0.37981 | 0.463463667 | 0.19 | 0 | 1 | 27 |
| CABZ01109481.1 | 0.37981 | 0.412426463 | 0.19 | 0 | 1 | 27 |
| FERMT3 (1 of many) | 0.37981 | 0.360509373 | 0.19 | 0 | 1 | 27 |
| KIF2A | 0.37981 | 0.298289504 | 0.19 | 0 | 1 | 27 |
| LOC100330864 | 0.37981 | 0.363014256 | 0.19 | 0 | 1 | 27 |
| SLC7A1 (1 of many) | 0.37981 | 0.32543251 | 0.19 | 0 | 1 | 27 |
| TMEM208 | 0.37981 | 0.378837868 | 0.19 | 0 | 1 | 27 |
| XLOC-042899 | 0.37981 | 0.313373714 | 0.19 | 0 | 1 | 27 |
| acin1b | 0.37981 | 0.400768674 | 0.19 | 0 | 1 | 27 |
| adka | 0.37981 | 0.257115158 | 0.19 | 0 | 1 | 27 |
| aldh9a1a.1 | 0.37981 | 0.297480709 | 0.19 | 0 | 1 | 27 |
| alyref | 0.37981 | 0.303995013 | 0.19 | 0 | 1 | 27 |
| anapc10 | 0.37981 | 0.324208668 | 0.19 | 0 | 1 | 27 |
| anxa11a | 0.37981 | 0.352981242 | 0.19 | 0 | 1 | 27 |
| ap1s2 | 0.37981 | 0.302698004 | 0.19 | 0 | 1 | 27 |
| ap2m1a | 0.37981 | 0.301884897 | 0.19 | 0 | 1 | 27 |
| araf | 0.37981 | 0.3612144 | 0.19 | 0 | 1 | 27 |

| | | | | | | |
|---|---|---|---|---|---|---|
| arcn1a | 0.37981 | 0.362894067 | 0.19 | 0 | 1 | 27 |
| arf2a | 0.37981 | 0.34074141 | 0.19 | 0 | 1 | 27 |
| arhgef1b | 0.37981 | 0.389187407 | 0.19 | 0 | 1 | 27 |
| asf1ba | 0.37981 | 0.251102253 | 0.19 | 0 | 1 | 27 |
| asf1bb | 0.37981 | 0.366555737 | 0.19 | 0 | 1 | 27 |
| atg3 | 0.37981 | 0.328158782 | 0.19 | 0 | 1 | 27 |
| atp6v1e1a | 0.37981 | 0.357734366 | 0.19 | 0 | 1 | 27 |
| baz1a | 0.37981 | 0.288087883 | 0.19 | 0 | 1 | 27 |
| baz1b | 0.37981 | 0.33926215 | 0.19 | 0 | 1 | 27 |
| birc2 | 0.37981 | 0.279389698 | 0.19 | 0 | 1 | 27 |
| birc5a | 0.37981 | 0.412059857 | 0.19 | 0 | 1 | 27 |
| brd3a | 0.37981 | 0.296860431 | 0.19 | 0 | 1 | 27 |
| brms1la | 0.37981 | 0.341040954 | 0.19 | 0 | 1 | 27 |
| cand1 | 0.37981 | 0.347071725 | 0.19 | 0 | 1 | 27 |
| caprin1b | 0.37981 | 0.322857402 | 0.19 | 0 | 1 | 27 |
| carm1 | 0.37981 | 0.305147887 | 0.19 | 0 | 1 | 27 |
| ccdc6b | 0.37981 | 0.396702168 | 0.19 | 0 | 1 | 27 |
| cdc27 | 0.37981 | 0.430024206 | 0.19 | 0 | 1 | 27 |
| cdv3 | 0.37981 | 0.320046964 | 0.19 | 0 | 1 | 27 |
| chrna1 | 0.37981 | 0.437241631 | 0.19 | 0 | 1 | 27 |
| clk4a | 0.37981 | 0.308987953 | 0.19 | 0 | 1 | 27 |
| clns1a | 0.37981 | 0.286057345 | 0.19 | 0 | 1 | 27 |
| cmc1 | 0.37981 | 0.316774536 | 0.19 | 0 | 1 | 27 |
| coa6 | 0.37981 | 0.263934471 | 0.19 | 0 | 1 | 27 |
| cops3 | 0.37981 | 0.335682226 | 0.19 | 0 | 1 | 27 |
| cops5 | 0.37981 | 0.32102597 | 0.19 | 0 | 1 | 27 |
| cox7a2l | 0.37981 | 0.323892986 | 0.19 | 0 | 1 | 27 |
| creb1a | 0.37981 | 0.387202634 | 0.19 | 0 | 1 | 27 |
| cst14b.1 | 0.37981 | 0.349436018 | 0.19 | 0 | 1 | 27 |
| ctdsp1 | 0.37981 | 0.358034405 | 0.19 | 0 | 1 | 27 |
| cwc25 | 0.37981 | 0.284366634 | 0.19 | 0 | 1 | 27 |
| cybb | 0.37981 | 0.448777926 | 0.19 | 0 | 1 | 27 |
| dapp1 | 0.37981 | 0.421721661 | 0.19 | 0 | 1 | 27 |
| dedd1 | 0.37981 | 0.369172549 | 0.19 | 0 | 1 | 27 |
| dlgap5 | 0.37981 | 0.351170209 | 0.19 | 0 | 1 | 27 |
| dlst | 0.37981 | 0.418500652 | 0.19 | 0 | 1 | 27 |
| dock8 | 0.37981 | 0.321895344 | 0.19 | 0 | 1 | 27 |
| elf2a | 0.37981 | 0.298240343 | 0.19 | 0 | 1 | 27 |
| elof1 | 0.37981 | 0.242337813 | 0.19 | 0 | 1 | 27 |
| epc2 | 0.37981 | 0.311051153 | 0.19 | 0 | 1 | 27 |
| ercc1 | 0.37981 | 0.2629573 | 0.19 | 0 | 1 | 27 |
| esf1 | 0.37981 | 0.397487446 | 0.19 | 0 | 1 | 27 |
| fahd2a | 0.37981 | 0.317065608 | 0.19 | 0 | 1 | 27 |
| fam133b | 0.37981 | 0.397728503 | 0.19 | 0 | 1 | 27 |
| fbxl6 | 0.37981 | 0.349641156 | 0.19 | 0 | 1 | 27 |
| fkbp5 | 0.37981 | 0.326672844 | 0.19 | 0 | 1 | 27 |
| fopnl | 0.37981 | 0.323200455 | 0.19 | 0 | 1 | 27 |
| foxm1 | 0.37981 | 0.310033523 | 0.19 | 0 | 1 | 27 |

| | | | | | | |
|---|---|---|---|---|---|---|
| g3bp2 | 0.37981 | 0.317997597 | 0.19 | 0 | 1 | 27 |
| glrx3 | 0.37981 | 0.365561572 | 0.19 | 0 | 1 | 27 |
| glud1a | 0.37981 | 0.291817844 | 0.19 | 0 | 1 | 27 |
| gnai1 | 0.37981 | 0.251183336 | 0.19 | 0 | 1 | 27 |
| gnaia | 0.37981 | 0.27333474 | 0.19 | 0 | 1 | 27 |
| gps2 | 0.37981 | 0.361684685 | 0.19 | 0 | 1 | 27 |
| grap2a | 0.37981 | 0.378192336 | 0.19 | 0 | 1 | 27 |
| gspt1l | 0.37981 | 0.307497829 | 0.19 | 0 | 1 | 27 |
| hadh | 0.37981 | 0.278152198 | 0.19 | 0 | 1 | 27 |
| hcfc1a | 0.37981 | 0.301253879 | 0.19 | 0 | 1 | 27 |
| hdlbpa | 0.37981 | 0.241142421 | 0.19 | 0 | 1 | 27 |
| hsdl2 | 0.37981 | 0.275458208 | 0.19 | 0 | 1 | 27 |
| hspe1 | 0.37981 | 0.223203898 | 0.19 | 0 | 1 | 27 |
| htatsf1 | 0.37981 | 0.371498016 | 0.19 | 0 | 1 | 27 |
| igf2bp1 | 0.37981 | 0.320698855 | 0.19 | 0 | 1 | 27 |
| illr1 | 0.37981 | 0.445754857 | 0.19 | 0 | 1 | 27 |
| irf2 | 0.37981 | 0.361930677 | 0.19 | 0 | 1 | 27 |
| irf5 | 0.37981 | 0.253650242 | 0.19 | 0 | 1 | 27 |
| isca2 | 0.37981 | 0.314163827 | 0.19 | 0 | 1 | 27 |
| itpka | 0.37981 | 0.303104398 | 0.19 | 0 | 1 | 27 |
| kdm1a | 0.37981 | 0.312848364 | 0.19 | 0 | 1 | 27 |
| klhdc3 | 0.37981 | 0.284593029 | 0.19 | 0 | 1 | 27 |
| lgals3a | 0.37981 | 0.347165717 | 0.19 | 0 | 1 | 27 |
| lrrc47 | 0.37981 | 0.326444948 | 0.19 | 0 | 1 | 27 |
| lrrfip1a | 0.37981 | 0.440857262 | 0.19 | 0 | 1 | 27 |
| maff | 0.37981 | 0.349640231 | 0.19 | 0 | 1 | 27 |
| magoh | 0.37981 | 0.384413668 | 0.19 | 0 | 1 | 27 |
| mbd1b | 0.37981 | 0.325440893 | 0.19 | 0 | 1 | 27 |
| mia3 | 0.37981 | 0.381785591 | 0.19 | 0 | 1 | 27 |
| mkrn1 | 0.37981 | 0.38349293 | 0.19 | 0 | 1 | 27 |
| mpc2 | 0.37981 | 0.287748991 | 0.19 | 0 | 1 | 27 |
| mrpl23 | 0.37981 | 0.32449575 | 0.19 | 0 | 1 | 27 |
| mrpl32 | 0.37981 | 0.35278351 | 0.19 | 0 | 1 | 27 |
| mrpl37 | 0.37981 | 0.27748335 | 0.19 | 0 | 1 | 27 |
| mrpl42 | 0.37981 | 0.353511718 | 0.19 | 0 | 1 | 27 |
| mrps6 | 0.37981 | 0.300796879 | 0.19 | 0 | 1 | 27 |
| mxd3 | 0.37981 | 0.29137943 | 0.19 | 0 | 1 | 27 |
| naxd | 0.37981 | 0.353555524 | 0.19 | 0 | 1 | 27 |
| ncoa4 | 0.37981 | 0.273503569 | 0.19 | 0 | 1 | 27 |
| ndufs8a | 0.37981 | 0.335110058 | 0.19 | 0 | 1 | 27 |
| ndufv3 | 0.37981 | 0.393185635 | 0.19 | 0 | 1 | 27 |
| nek9 | 0.37981 | 0.328351439 | 0.19 | 0 | 1 | 27 |
| nfyc | 0.37981 | 0.288458836 | 0.19 | 0 | 1 | 27 |
| nop10 | 0.37981 | 0.271972581 | 0.19 | 0 | 1 | 27 |
| npm3 | 0.37981 | 0.264250946 | 0.19 | 0 | 1 | 27 |
| nr3c1 | 0.37981 | 0.389732702 | 0.19 | 0 | 1 | 27 |
| nrbf2b | 0.37981 | 0.285466009 | 0.19 | 0 | 1 | 27 |
| nudt5 | 0.37981 | 0.343712062 | 0.19 | 0 | 1 | 27 |

| | | | | | | |
|---|---|---|---|---|---|---|
| nuf2 | 0.37981 | 0.335962605 | 0.19 | 0 | 1 | 27 |
| nusap1 | 0.37981 | 0.398059882 | 0.19 | 0 | 1 | 27 |
| nutf2 | 0.37981 | 0.295694121 | 0.19 | 0 | 1 | 27 |
| papola | 0.37981 | 0.381240846 | 0.19 | 0 | 1 | 27 |
| pbk | 0.37981 | 0.310423566 | 0.19 | 0 | 1 | 27 |
| pbxip1b | 0.37981 | 0.267248599 | 0.19 | 0 | 1 | 27 |
| pgd | 0.37981 | 0.306093507 | 0.19 | 0 | 1 | 27 |
| pmm2 | 0.37981 | 0.241044064 | 0.19 | 0 | 1 | 27 |
| pole4 | 0.37981 | 0.258261809 | 0.19 | 0 | 1 | 27 |
| polr2h | 0.37981 | 0.289966533 | 0.19 | 0 | 1 | 27 |
| ppp1cab | 0.37981 | 0.374590615 | 0.19 | 0 | 1 | 27 |
| ppp2r5ea | 0.37981 | 0.351617058 | 0.19 | 0 | 1 | 27 |
| prcp | 0.37981 | 0.413092135 | 0.19 | 0 | 1 | 27 |
| prkab1b | 0.37981 | 0.334447866 | 0.19 | 0 | 1 | 27 |
| psma6a | 0.37981 | 0.377099276 | 0.19 | 0 | 1 | 27 |
| psma6l | 0.37981 | 0.316641749 | 0.19 | 0 | 1 | 27 |
| psmc1a | 0.37981 | 0.292281078 | 0.19 | 0 | 1 | 27 |
| ptpn2b | 0.37981 | 0.384898129 | 0.19 | 0 | 1 | 27 |
| pttg1 | 0.37981 | 0.286784067 | 0.19 | 0 | 1 | 27 |
| puf60a | 0.37981 | 0.281860241 | 0.19 | 0 | 1 | 27 |
| rad51 | 0.37981 | 0.304359231 | 0.19 | 0 | 1 | 27 |
| rbbp5 | 0.37981 | 0.278958086 | 0.19 | 0 | 1 | 27 |
| rbm22 | 0.37981 | 0.319411294 | 0.19 | 0 | 1 | 27 |
| rbm38 | 0.37981 | 0.301288963 | 0.19 | 0 | 1 | 27 |
| rbp5 | 0.37981 | 0.38537187 | 0.19 | 0 | 1 | 27 |
| rfc2 | 0.37981 | 0.344896661 | 0.19 | 0 | 1 | 27 |
| rnaseka | 0.37981 | 0.345898825 | 0.19 | 0 | 1 | 27 |
| rps6kb1b | 0.37981 | 0.333975303 | 0.19 | 0 | 1 | 27 |
| ruvbl2 | 0.37981 | 0.329900094 | 0.19 | 0 | 1 | 27 |
| safb | 0.37981 | 0.362827877 | 0.19 | 0 | 1 | 27 |
| samd13 | 0.37981 | 0.320950598 | 0.19 | 0 | 1 | 27 |
| sar1b | 0.37981 | 0.267469908 | 0.19 | 0 | 1 | 27 |
| sde2 | 0.37981 | 0.380472942 | 0.19 | 0 | 1 | 27 |
| sdhdb | 0.37981 | 0.226140542 | 0.19 | 0 | 1 | 27 |
| sec62 | 0.37981 | 0.319810989 | 0.19 | 0 | 1 | 27 |
| selenot1b | 0.37981 | 0.388859876 | 0.19 | 0 | 1 | 27 |
| serbp1b | 0.37981 | 0.372739392 | 0.19 | 0 | 1 | 27 |
| sf3a1 | 0.37981 | 0.29787043 | 0.19 | 0 | 1 | 27 |
| sf3b1 | 0.37981 | 0.370263547 | 0.19 | 0 | 1 | 27 |
| sf3b2 | 0.37981 | 0.392253226 | 0.19 | 0 | 1 | 27 |
| shoc2 | 0.37981 | 0.313910119 | 0.19 | 0 | 1 | 27 |
| si:ch1073-322p19.1 | 0.37981 | 0.308114362 | 0.19 | 0 | 1 | 27 |
| si:ch211-156b7.4 | 0.37981 | 0.287738846 | 0.19 | 0 | 1 | 27 |
| si:ch211-161h7.4 | 0.37981 | 0.338551025 | 0.19 | 0 | 1 | 27 |
| si:ch211-271b14.1 | 0.37981 | 0.298514898 | 0.19 | 0 | 1 | 27 |
| si:ch211-93g21.1 | 0.37981 | 0.376325713 | 0.19 | 0 | 1 | 27 |
| si:ch73-71c20.5 | 0.37981 | 0.4309916 | 0.19 | 0 | 1 | 27 |
| si:dkey-159f12.2 | 0.37981 | 0.259692022 | 0.19 | 0 | 1 | 27 |

| | | | | | | |
|---|---|---|---|---|---|---|
| si:dkey-165a24.10 | 0.37981 | 0.302315776 | 0.19 | 0 | 1 | 27 |
| si:dkey-185m8.2 | 0.37981 | 0.422304103 | 0.19 | 0 | 1 | 27 |
| si:dkey-250d21.1 | 0.37981 | 0.320198098 | 0.19 | 0 | 1 | 27 |
| si:dkey-33c12.4 | 0.37981 | 0.264240716 | 0.19 | 0 | 1 | 27 |
| si:dkey-67c22.2 | 0.37981 | 0.331312041 | 0.19 | 0 | 1 | 27 |
| si:dkey-6i22.5 | 0.37981 | 0.28394203 | 0.19 | 0 | 1 | 27 |
| si:dkey-92i15.4 | 0.37981 | 0.377998237 | 0.19 | 0 | 1 | 27 |
| si:dkeyp-117h8.2 | 0.37981 | 0.435564422 | 0.19 | 0 | 1 | 27 |
| si:zfos-464b6.2 | 0.37981 | 0.268612298 | 0.19 | 0 | 1 | 27 |
| siva1 | 0.37981 | 0.275791682 | 0.19 | 0 | 1 | 27 |
| slbp | 0.37981 | 0.347292947 | 0.19 | 0 | 1 | 27 |
| slc15a4 | 0.37981 | 0.391029077 | 0.19 | 0 | 1 | 27 |
| slc25a51b | 0.37981 | 0.320487588 | 0.19 | 0 | 1 | 27 |
| slc30a7 | 0.37981 | 0.312046761 | 0.19 | 0 | 1 | 27 |
| slc3a2b | 0.37981 | 0.369788994 | 0.19 | 0 | 1 | 27 |
| smarca4a | 0.37981 | 0.299657532 | 0.19 | 0 | 1 | 27 |
| smarcd2 | 0.37981 | 0.277992849 | 0.19 | 0 | 1 | 27 |
| smndc1 | 0.37981 | 0.374225731 | 0.19 | 0 | 1 | 27 |
| snrpb2 | 0.37981 | 0.315245903 | 0.19 | 0 | 1 | 27 |
| snx18b | 0.37981 | 0.355080495 | 0.19 | 0 | 1 | 27 |
| sod2 | 0.37981 | 0.320200872 | 0.19 | 0 | 1 | 27 |
| spc24 | 0.37981 | 0.305809956 | 0.19 | 0 | 1 | 27 |
| spc25 | 0.37981 | 0.390055134 | 0.19 | 0 | 1 | 27 |
| spdl1 | 0.37981 | 0.358155983 | 0.19 | 0 | 1 | 27 |
| srrm1 | 0.37981 | 0.353985085 | 0.19 | 0 | 1 | 27 |
| st13 | 0.37981 | 0.332853099 | 0.19 | 0 | 1 | 27 |
| stat6 | 0.37981 | 0.32099069 | 0.19 | 0 | 1 | 27 |
| stip1 | 0.37981 | 0.280125665 | 0.19 | 0 | 1 | 27 |
| sub1b | 0.37981 | 0.26197739 | 0.19 | 0 | 1 | 27 |
| taf13 | 0.37981 | 0.313934592 | 0.19 | 0 | 1 | 27 |
| tmem30ab | 0.37981 | 0.399996545 | 0.19 | 0 | 1 | 27 |
| tmem33 | 0.37981 | 0.281422285 | 0.19 | 0 | 1 | 27 |
| tmsb | 0.37981 | 0.342808713 | 0.19 | 0 | 1 | 27 |
| tpx2 | 0.37981 | 0.427141593 | 0.19 | 0 | 1 | 27 |
| tradd | 0.37981 | 0.330610081 | 0.19 | 0 | 1 | 27 |
| trappc2l | 0.37981 | 0.282777231 | 0.19 | 0 | 1 | 27 |
| trappc5 | 0.37981 | 0.311857977 | 0.19 | 0 | 1 | 27 |
| trmt112 | 0.37981 | 0.319035745 | 0.19 | 0 | 1 | 27 |
| tspy | 0.37981 | 0.333454399 | 0.19 | 0 | 1 | 27 |
| u2af2a | 0.37981 | 0.287082373 | 0.19 | 0 | 1 | 27 |
| ubap2l | 0.37981 | 0.271948031 | 0.19 | 0 | 1 | 27 |
| ube2a | 0.37981 | 0.298067686 | 0.19 | 0 | 1 | 27 |
| ube2e3 | 0.37981 | 0.280645231 | 0.19 | 0 | 1 | 27 |
| ube2g1b | 0.37981 | 0.284803687 | 0.19 | 0 | 1 | 27 |
| ubtf | 0.37981 | 0.250795732 | 0.19 | 0 | 1 | 27 |
| uqcc3 | 0.37981 | 0.279054113 | 0.19 | 0 | 1 | 27 |
| uqcrc2a | 0.37981 | 0.286798509 | 0.19 | 0 | 1 | 27 |
| vav2 | 0.37981 | 0.361322502 | 0.19 | 0 | 1 | 27 |

| | | | | | | |
|---|---|---|---|---|---|---|
| vdac1 | 0.37981 | 0.335609432 | 0.19 | 0 | 1 | 27 |
| vdrb | 0.37981 | 0.320334158 | 0.19 | 0 | 1 | 27 |
| vrk1 | 0.37981 | 0.255940506 | 0.19 | 0 | 1 | 27 |
| wdr26b | 0.37981 | 0.377226249 | 0.19 | 0 | 1 | 27 |
| yeats4 | 0.37981 | 0.23834829 | 0.19 | 0 | 1 | 27 |
| zgc:100918 | 0.37981 | 0.350000972 | 0.19 | 0 | 1 | 27 |
| zgc:110591 | 0.37981 | 0.361878526 | 0.19 | 0 | 1 | 27 |
| zgc:136254 | 0.37981 | 0.409767065 | 0.19 | 0 | 1 | 27 |
| zgc:86839 | 0.37981 | 0.292103724 | 0.19 | 0 | 1 | 27 |
| zgpat | 0.37981 | 0.369008995 | 0.19 | 0 | 1 | 27 |
| aurkb | 0.37981 | 0.358310721 | 0.19 | 0 | 1 | 27 |
| ccdc25 | 0.37981 | 0.255184006 | 0.19 | 0 | 1 | 27 |
| cflara | 0.37981 | 0.364482073 | 0.19 | 0 | 1 | 27 |
| fam204a | 0.37981 | 0.306883118 | 0.19 | 0 | 1 | 27 |
| kat7b | 0.37981 | 0.288016804 | 0.19 | 0 | 1 | 27 |
| mapk3 | 0.37981 | 0.327232222 | 0.19 | 0 | 1 | 27 |
| mcm7 | 0.37981 | 0.331786782 | 0.19 | 0 | 1 | 27 |
| parp2 | 0.37981 | 0.41155279 | 0.19 | 0 | 1 | 27 |
| pds5b | 0.37981 | 0.2745973 | 0.19 | 0 | 1 | 27 |
| polr2a | 0.37981 | 0.344077825 | 0.19 | 0 | 1 | 27 |
| psmg2 | 0.37981 | 0.314684708 | 0.19 | 0 | 1 | 27 |
| setbp1 | 0.37981 | 0.353095974 | 0.19 | 0 | 1 | 27 |
| smc4 | 0.37981 | 0.298759415 | 0.19 | 0 | 1 | 27 |
| spg7 | 0.37981 | 0.298133334 | 0.19 | 0 | 1 | 27 |
| traf5 | 0.37981 | 0.299574133 | 0.19 | 0 | 1 | 27 |
| usf1l | 0.37981 | 0.339071974 | 0.19 | 0 | 1 | 27 |
| usp39 | 0.37981 | 0.352795803 | 0.19 | 0 | 1 | 27 |
| yy1b | 0.37981 | 0.312335221 | 0.19 | 0 | 1 | 27 |
| znf207b | 0.37981 | 0.278111149 | 0.19 | 0 | 1 | 27 |
| sec61g | 0.38085 | -0.52939261 | 0.667 | 1 | 1 | 27 |
| wasb | 0.38415 | -0.23380098 | 0.381 | 0.333 | 1 | 27 |
| nme2b.1 | 0.3842 | -0.17033132 | 0.952 | 1 | 1 | 27 |
| gnai2a | 0.3845 | -0.556173 | 0.429 | 0.667 | 1 | 27 |
| uqcrh | 0.38868 | -0.582814 | 0.286 | 0.667 | 1 | 27 |
| stat1a | 0.38951 | -0.55154692 | 0.19 | 0.333 | 1 | 27 |
| glo1 | 0.38995 | -0.25834304 | 0.667 | 0.667 | 1 | 27 |
| hmgn6 | 0.39286 | -0.43904642 | 0.714 | 1 | 1 | 27 |
| cd83 | 0.39409 | -0.41539684 | 0.333 | 0.333 | 1 | 27 |
| chac1 | 0.39436 | -0.65175605 | 0.333 | 0.667 | 1 | 27 |
| gdi2 | 0.39489 | -0.34771545 | 0.619 | 0.667 | 1 | 27 |
| vsir | 0.3954 | -0.34212097 | 0.333 | 0.333 | 1 | 27 |
| cacybp | 0.39679 | -0.46317484 | 0.238 | 0.333 | 1 | 27 |
| capzb | 0.39744 | -0.43452326 | 0.667 | 1 | 1 | 27 |
| gatc | 0.39757 | -0.5604613 | 0.19 | 0.333 | 1 | 27 |
| arl5c | 0.40155 | -0.7192493 | 0.143 | 0.333 | 1 | 27 |
| atp5mc3a | 0.40524 | -0.55131393 | 0.381 | 0.667 | 1 | 27 |
| lcp2a | 0.40858 | -0.55241159 | 0.19 | 0.333 | 1 | 27 |
| tubb4b | 0.41225 | 0.726530338 | 0.667 | 0.333 | 1 | 27 |

| | | | | | | |
|---|---|---|---|---|---|---|
| ctdsp2 | 0.41268 | -0.33173216 | 0.333 | 0.333 | 1 | 27 |
| tln1 | 0.41295 | -0.59275328 | 0.286 | 0.667 | 1 | 27 |
| ier2a | 0.41394 | -0.2175053 | 0.714 | 0.667 | 1 | 27 |
| eif3ea | 0.4141 | -0.40160943 | 0.714 | 1 | 1 | 27 |
| rps27a | 0.41453 | 0.18665233 | 1 | 1 | 1 | 27 |
| rbmx | 0.41587 | -0.37023397 | 0.286 | 0.333 | 1 | 27 |
| pnrc2 | 0.41633 | 0.311795264 | 0.952 | 0.667 | 1 | 27 |
| actb1 | 0.41785 | -0.44196316 | 0.762 | 1 | 1 | 27 |
| mt-atp6 | 0.418 | -0.58260123 | 0.19 | 0.333 | 1 | 27 |
| h3f3a | 0.41979 | 0.647676315 | 0.476 | 0.333 | 1 | 27 |
| khdrbs1a | 0.42129 | -0.41018603 | 0.714 | 1 | 1 | 27 |
| sumo3a | 0.42435 | 0.627775497 | 0.714 | 0.333 | 1 | 27 |
| tuba8l4 | 0.4247 | 0.1549324 | 0.476 | 0.667 | 1 | 27 |
| rpl28 | 0.42517 | 0.119522046 | 1 | 1 | 1 | 27 |
| cct8 | 0.42539 | -0.5801011 | 0.381 | 0.667 | 1 | 27 |
| cd74a | 0.43044 | -0.38539572 | 0.857 | 1 | 1 | 27 |
| ddit3 | 0.4309 | -0.59870535 | 0.429 | 0.667 | 1 | 27 |
| XLOC-010899 | 0.43165 | -0.44146152 | 0.857 | 1 | 1 | 27 |
| lpar5a | 0.43479 | -0.43765503 | 0.238 | 0.333 | 1 | 27 |
| ybx1 | 0.43485 | 0.116935683 | 0.857 | 1 | 1 | 27 |
| hnrnpa0b | 0.43785 | 0.511266982 | 0.524 | 0.333 | 1 | 27 |
| rpl31 | 0.43806 | 0.19747651 | 1 | 1 | 1 | 27 |
| tbca | 0.43906 | -0.2187437 | 0.381 | 0.333 | 1 | 27 |
| eif5a | 0.44245 | -0.30603252 | 0.857 | 1 | 1 | 27 |
| BX571825.5 | 0.45036 | -0.2137803 | 0.714 | 1 | 1 | 27 |
| gstm.2 | 0.45103 | -0.55966116 | 0.143 | 0.333 | 1 | 27 |
| CT573342.2 | 0.45374 | 0.651130612 | 0.667 | 0.333 | 1 | 27 |
| RASSF5 | 0.45774 | -0.54458234 | 0.333 | 0.667 | 1 | 27 |
| eif4bb | 0.45847 | -0.52370752 | 0.286 | 0.667 | 1 | 27 |
| spcs2 | 0.46042 | -0.50574201 | 0.429 | 0.667 | 1 | 27 |
| mapre1a | 0.46192 | -0.36255184 | 0.286 | 0.333 | 1 | 27 |
| cd7al | 0.46362 | -0.29222935 | 0.333 | 0.333 | 1 | 27 |
| tmsb4x | 0.465 | -0.13285252 | 1 | 1 | 1 | 27 |
| plk3 | 0.46722 | -0.23981159 | 0.381 | 0.333 | 1 | 27 |
| hnrnpabb | 0.46901 | -0.19536965 | 0.381 | 0.667 | 1 | 27 |
| tyk2 | 0.47063 | -0.5046381 | 0.143 | 0.333 | 1 | 27 |
| ier2b | 0.47119 | 0.623620745 | 0.667 | 0.333 | 1 | 27 |
| CABZ01044053.1 | 0.47255 | 0.244709606 | 0.143 | 0 | 1 | 27 |
| CT027980.1 | 0.47255 | 0.343630889 | 0.143 | 0 | 1 | 27 |
| CU570894.1 | 0.47255 | 0.27727169 | 0.143 | 0 | 1 | 27 |
| XLOC-016788 | 0.47255 | 0.343472827 | 0.143 | 0 | 1 | 27 |
| abcf1 | 0.47255 | 0.275447796 | 0.143 | 0 | 1 | 27 |
| abhd13 | 0.47255 | 0.248271896 | 0.143 | 0 | 1 | 27 |
| arhgef6 | 0.47255 | 0.318310724 | 0.143 | 0 | 1 | 27 |
| arid4a | 0.47255 | 0.212831044 | 0.143 | 0 | 1 | 27 |
| arpc1a | 0.47255 | 0.277035994 | 0.143 | 0 | 1 | 27 |
| atp6v0b | 0.47255 | 0.229644474 | 0.143 | 0 | 1 | 27 |
| bcap31 | 0.47255 | 0.183406628 | 0.143 | 0 | 1 | 27 |

| | | | | | | |
|---|---|---|---|---|---|---|
| bub3 | 0.47255 | 0.226256692 | 0.143 | 0 | 1 | 27 |
| ccz1 | 0.47255 | 0.257126857 | 0.143 | 0 | 1 | 27 |
| chek2 | 0.47255 | 0.277700473 | 0.143 | 0 | 1 | 27 |
| chordc1b | 0.47255 | 0.202063266 | 0.143 | 0 | 1 | 27 |
| ciita | 0.47255 | 0.326707751 | 0.143 | 0 | 1 | 27 |
| cops8 | 0.47255 | 0.257797524 | 0.143 | 0 | 1 | 27 |
| coq10b | 0.47255 | 0.327241703 | 0.143 | 0 | 1 | 27 |
| cox14 | 0.47255 | 0.225334361 | 0.143 | 0 | 1 | 27 |
| crfb4 | 0.47255 | 0.259746439 | 0.143 | 0 | 1 | 27 |
| ctcf | 0.47255 | 0.272193604 | 0.143 | 0 | 1 | 27 |
| dctd | 0.47255 | 0.257085677 | 0.143 | 0 | 1 | 27 |
| dnmt1 | 0.47255 | 0.210133037 | 0.143 | 0 | 1 | 27 |
| dnph1 | 0.47255 | 0.203308181 | 0.143 | 0 | 1 | 27 |
| dusp5 | 0.47255 | 0.282431267 | 0.143 | 0 | 1 | 27 |
| fam50a | 0.47255 | 0.220964874 | 0.143 | 0 | 1 | 27 |
| fbl | 0.47255 | 0.285906026 | 0.143 | 0 | 1 | 27 |
| gabarapa | 0.47255 | 0.262822149 | 0.143 | 0 | 1 | 27 |
| gadd45ab | 0.47255 | 0.346304855 | 0.143 | 0 | 1 | 27 |
| galnt1 | 0.47255 | 0.324712647 | 0.143 | 0 | 1 | 27 |
| hexim1 | 0.47255 | 0.21785969 | 0.143 | 0 | 1 | 27 |
| ing1 | 0.47255 | 0.303266125 | 0.143 | 0 | 1 | 27 |
| klf8 | 0.47255 | 0.28657368 | 0.143 | 0 | 1 | 27 |
| letm1 | 0.47255 | 0.342095601 | 0.143 | 0 | 1 | 27 |
| mbd2 | 0.47255 | 0.238988193 | 0.143 | 0 | 1 | 27 |
| mlec | 0.47255 | 0.288087387 | 0.143 | 0 | 1 | 27 |
| mrpl12 | 0.47255 | 0.234175565 | 0.143 | 0 | 1 | 27 |
| msantd1 | 0.47255 | 0.193061487 | 0.143 | 0 | 1 | 27 |
| msi2b | 0.47255 | 0.292301325 | 0.143 | 0 | 1 | 27 |
| mtmr11 | 0.47255 | 0.327142796 | 0.143 | 0 | 1 | 27 |
| mus81 | 0.47255 | 0.225991617 | 0.143 | 0 | 1 | 27 |
| pafah1b1a | 0.47255 | 0.257240779 | 0.143 | 0 | 1 | 27 |
| pcmtd1 | 0.47255 | 0.283695407 | 0.143 | 0 | 1 | 27 |
| phf20a | 0.47255 | 0.259691224 | 0.143 | 0 | 1 | 27 |
| pik3cd | 0.47255 | 0.309666513 | 0.143 | 0 | 1 | 27 |
| pitpnb | 0.47255 | 0.31820223 | 0.143 | 0 | 1 | 27 |
| pole3 | 0.47255 | 0.273817153 | 0.143 | 0 | 1 | 27 |
| ppil3 | 0.47255 | 0.256183385 | 0.143 | 0 | 1 | 27 |
| ppp4cb | 0.47255 | 0.285603368 | 0.143 | 0 | 1 | 27 |
| prpf19 | 0.47255 | 0.323111755 | 0.143 | 0 | 1 | 27 |
| psmd10 | 0.47255 | 0.251987701 | 0.143 | 0 | 1 | 27 |
| pudp | 0.47255 | 0.262126313 | 0.143 | 0 | 1 | 27 |
| rad18 | 0.47255 | 0.20649408 | 0.143 | 0 | 1 | 27 |
| rapgef1a | 0.47255 | 0.292309277 | 0.143 | 0 | 1 | 27 |
| rnf2 | 0.47255 | 0.219038508 | 0.143 | 0 | 1 | 27 |
| rnf213a | 0.47255 | 0.284823674 | 0.143 | 0 | 1 | 27 |
| scp2b | 0.47255 | 0.231472292 | 0.143 | 0 | 1 | 27 |
| scpep1 | 0.47255 | 0.232666525 | 0.143 | 0 | 1 | 27 |
| sf3b3 | 0.47255 | 0.217509226 | 0.143 | 0 | 1 | 27 |

| | | | | | | |
|---|---|---|---|---|---|---|
| si:ch211-195h23.3 | 0.47255 | 0.280767463 | 0.143 | 0 | 1 | 27 |
| si:ch73-376l24.2 | 0.47255 | 0.384226536 | 0.143 | 0 | 1 | 27 |
| si:dkey-6e2.2 | 0.47255 | 0.360619587 | 0.143 | 0 | 1 | 27 |
| si:dkeyp-55f12.3 | 0.47255 | 0.29172844 | 0.143 | 0 | 1 | 27 |
| smc3 | 0.47255 | 0.279535313 | 0.143 | 0 | 1 | 27 |
| smg7 | 0.47255 | 0.27017464 | 0.143 | 0 | 1 | 27 |
| snrnp27 | 0.47255 | 0.244042795 | 0.143 | 0 | 1 | 27 |
| snx3 | 0.47255 | 0.221266398 | 0.143 | 0 | 1 | 27 |
| thoc7 | 0.47255 | 0.243705914 | 0.143 | 0 | 1 | 27 |
| tia1l | 0.47255 | 0.242702756 | 0.143 | 0 | 1 | 27 |
| tmem11 | 0.47255 | 0.259802951 | 0.143 | 0 | 1 | 27 |
| tnfaip3 | 0.47255 | 0.225489876 | 0.143 | 0 | 1 | 27 |
| wdfy1 | 0.47255 | 0.283092042 | 0.143 | 0 | 1 | 27 |
| yaf2 | 0.47255 | 0.200637403 | 0.143 | 0 | 1 | 27 |
| ypel5 | 0.47255 | 0.253543602 | 0.143 | 0 | 1 | 27 |
| ywhah | 0.47255 | 0.247523553 | 0.143 | 0 | 1 | 27 |
| zbtb20 | 0.47255 | 0.348253907 | 0.143 | 0 | 1 | 27 |
| zfyve19 | 0.47255 | 0.264148156 | 0.143 | 0 | 1 | 27 |
| zgc:162025 | 0.47255 | 0.224958592 | 0.143 | 0 | 1 | 27 |
| zgc:77086 | 0.47255 | 0.180334244 | 0.143 | 0 | 1 | 27 |
| CABZ01068251.1 | 0.47255 | 0.242951192 | 0.143 | 0 | 1 | 27 |
| CABZ01073834.1 | 0.47255 | 0.230955945 | 0.143 | 0 | 1 | 27 |
| CABZ01075242.1 | 0.47255 | 0.205916374 | 0.143 | 0 | 1 | 27 |
| CABZ01087560.1 | 0.47255 | 0.252346768 | 0.143 | 0 | 1 | 27 |
| CR318588.1 | 0.47255 | 0.439424767 | 0.143 | 0 | 1 | 27 |
| CU019662.1 | 0.47255 | 0.221851421 | 0.143 | 0 | 1 | 27 |
| CU467110.1 | 0.47255 | 0.24330221 | 0.143 | 0 | 1 | 27 |
| FQ323156.1 | 0.47255 | 0.226531285 | 0.143 | 0 | 1 | 27 |
| FQ976913.1 | 0.47255 | 0.252408421 | 0.143 | 0 | 1 | 27 |
| GNAZ | 0.47255 | 0.370667826 | 0.143 | 0 | 1 | 27 |
| LEPROTL1 | 0.47255 | 0.256626885 | 0.143 | 0 | 1 | 27 |
| LO017938.1 | 0.47255 | 0.268064005 | 0.143 | 0 | 1 | 27 |
| LOC103911430 | 0.47255 | 0.298112626 | 0.143 | 0 | 1 | 27 |
| LOC108190616 | 0.47255 | 0.229940218 | 0.143 | 0 | 1 | 27 |
| LOC110438237 | 0.47255 | 0.286065754 | 0.143 | 0 | 1 | 27 |
| UBL4A | 0.47255 | 0.299083818 | 0.143 | 0 | 1 | 27 |
| XLOC-005448 | 0.47255 | 0.339029928 | 0.143 | 0 | 1 | 27 |
| XLOC-012410 | 0.47255 | 0.264173245 | 0.143 | 0 | 1 | 27 |
| abcf2a | 0.47255 | 0.217283382 | 0.143 | 0 | 1 | 27 |
| adcy7 | 0.47255 | 0.226467342 | 0.143 | 0 | 1 | 27 |
| adprhl2 | 0.47255 | 0.196003609 | 0.143 | 0 | 1 | 27 |
| ago4 | 0.47255 | 0.307115246 | 0.143 | 0 | 1 | 27 |
| aldh3b1 | 0.47255 | 0.25621612 | 0.143 | 0 | 1 | 27 |
| anapc11 | 0.47255 | 0.265367237 | 0.143 | 0 | 1 | 27 |
| anapc2 | 0.47255 | 0.208104059 | 0.143 | 0 | 1 | 27 |
| ankrd33ba | 0.47255 | 0.246433344 | 0.143 | 0 | 1 | 27 |
| anxa4 | 0.47255 | 0.272944018 | 0.143 | 0 | 1 | 27 |
| ap3s1.1 | 0.47255 | 0.258436049 | 0.143 | 0 | 1 | 27 |

| | | | | | | |
|---|---|---|---|---|---|---|
| arglu1b | 0.47255 | 0.173577157 | 0.143 | 0 | 1 | 27 |
| arih1 | 0.47255 | 0.310181649 | 0.143 | 0 | 1 | 27 |
| arpp19b | 0.47255 | 0.238921359 | 0.143 | 0 | 1 | 27 |
| atf6 | 0.47255 | 0.254011112 | 0.143 | 0 | 1 | 27 |
| atp10d | 0.47255 | 0.279370269 | 0.143 | 0 | 1 | 27 |
| atp6ap2 | 0.47255 | 0.277748497 | 0.143 | 0 | 1 | 27 |
| aup1 | 0.47255 | 0.168108569 | 0.143 | 0 | 1 | 27 |
| b3gnt3.4 | 0.47255 | 0.21749424 | 0.143 | 0 | 1 | 27 |
| bcl2a | 0.47255 | 0.255383435 | 0.143 | 0 | 1 | 27 |
| bcl6aa | 0.47255 | 0.262187855 | 0.143 | 0 | 1 | 27 |
| blcap | 0.47255 | 0.328736746 | 0.143 | 0 | 1 | 27 |
| bola3 | 0.47255 | 0.183222902 | 0.143 | 0 | 1 | 27 |
| brd8 | 0.47255 | 0.232677916 | 0.143 | 0 | 1 | 27 |
| bsdc1 | 0.47255 | 0.22648833 | 0.143 | 0 | 1 | 27 |
| btf3l4 | 0.47255 | 0.179375824 | 0.143 | 0 | 1 | 27 |
| btg3 | 0.47255 | 0.275062486 | 0.143 | 0 | 1 | 27 |
| cant1a | 0.47255 | 0.199869144 | 0.143 | 0 | 1 | 27 |
| canx | 0.47255 | 0.216891639 | 0.143 | 0 | 1 | 27 |
| capns1b | 0.47255 | 0.182303333 | 0.143 | 0 | 1 | 27 |
| caprin1a | 0.47255 | 0.23613319 | 0.143 | 0 | 1 | 27 |
| cars | 0.47255 | 0.216234881 | 0.143 | 0 | 1 | 27 |
| casp3a | 0.47255 | 0.250451365 | 0.143 | 0 | 1 | 27 |
| cbfb | 0.47255 | 0.247467025 | 0.143 | 0 | 1 | 27 |
| cdc14b | 0.47255 | 0.29818824 | 0.143 | 0 | 1 | 27 |
| cdc20 | 0.47255 | 0.22571989 | 0.143 | 0 | 1 | 27 |
| cdc25b | 0.47255 | 0.164735528 | 0.143 | 0 | 1 | 27 |
| cdc34a | 0.47255 | 0.356775576 | 0.143 | 0 | 1 | 27 |
| cdca8 | 0.47255 | 0.282548524 | 0.143 | 0 | 1 | 27 |
| cdk2ap2 | 0.47255 | 0.220851212 | 0.143 | 0 | 1 | 27 |
| cdkn2c | 0.47255 | 0.255523914 | 0.143 | 0 | 1 | 27 |
| cep55l | 0.47255 | 0.242082331 | 0.143 | 0 | 1 | 27 |
| cers1 | 0.47255 | 0.333091095 | 0.143 | 0 | 1 | 27 |
| ckap5 | 0.47255 | 0.226081628 | 0.143 | 0 | 1 | 27 |
| cltca | 0.47255 | 0.309396606 | 0.143 | 0 | 1 | 27 |
| cmtm8a | 0.47255 | 0.311541922 | 0.143 | 0 | 1 | 27 |
| cnih4 | 0.47255 | 0.250028565 | 0.143 | 0 | 1 | 27 |
| cnot8 | 0.47255 | 0.275780078 | 0.143 | 0 | 1 | 27 |
| commd2 | 0.47255 | 0.213577306 | 0.143 | 0 | 1 | 27 |
| commd7 | 0.47255 | 0.256149258 | 0.143 | 0 | 1 | 27 |
| copg2 | 0.47255 | 0.200113968 | 0.143 | 0 | 1 | 27 |
| coq7 | 0.47255 | 0.204812158 | 0.143 | 0 | 1 | 27 |
| crema | 0.47255 | 0.277411917 | 0.143 | 0 | 1 | 27 |
| cryl1 | 0.47255 | 0.289271095 | 0.143 | 0 | 1 | 27 |
| csde1 | 0.47255 | 0.238875643 | 0.143 | 0 | 1 | 27 |
| csnk1db | 0.47255 | 0.286015798 | 0.143 | 0 | 1 | 27 |
| csnk1g2b | 0.47255 | 0.283571546 | 0.143 | 0 | 1 | 27 |
| cxcr3.2 | 0.47255 | 0.209295468 | 0.143 | 0 | 1 | 27 |
| cyb5r3 | 0.47255 | 0.185167811 | 0.143 | 0 | 1 | 27 |

| | | | | | | |
|---|---|---|---|---|---|---|
| cyfip1 | 0.47255 | 0.267899914 | 0.143 | 0 | 1 | 27 |
| cyth1a | 0.47255 | 0.306714329 | 0.143 | 0 | 1 | 27 |
| cyth1b | 0.47255 | 0.246953994 | 0.143 | 0 | 1 | 27 |
| dclre1b | 0.47255 | 0.218315022 | 0.143 | 0 | 1 | 27 |
| ddost | 0.47255 | 0.327865302 | 0.143 | 0 | 1 | 27 |
| ddx23 | 0.47255 | 0.281007569 | 0.143 | 0 | 1 | 27 |
| dera | 0.47255 | 0.254805726 | 0.143 | 0 | 1 | 27 |
| dffa | 0.47255 | 0.203570182 | 0.143 | 0 | 1 | 27 |
| dhrs13a.1 | 0.47255 | 0.257267091 | 0.143 | 0 | 1 | 27 |
| dhx8 | 0.47255 | 0.207335914 | 0.143 | 0 | 1 | 27 |
| dmap1 | 0.47255 | 0.180529658 | 0.143 | 0 | 1 | 27 |
| dnajc1 | 0.47255 | 0.169360331 | 0.143 | 0 | 1 | 27 |
| dnajc4 | 0.47255 | 0.263255587 | 0.143 | 0 | 1 | 27 |
| dnajc7 | 0.47255 | 0.307972216 | 0.143 | 0 | 1 | 27 |
| dram2b | 0.47255 | 0.187018764 | 0.143 | 0 | 1 | 27 |
| drg2 | 0.47255 | 0.327185721 | 0.143 | 0 | 1 | 27 |
| dtnba | 0.47255 | 0.239296761 | 0.143 | 0 | 1 | 27 |
| dynlrb1 | 0.47255 | 0.187209308 | 0.143 | 0 | 1 | 27 |
| egr2a | 0.47255 | 0.332867757 | 0.143 | 0 | 1 | 27 |
| ehmt1b | 0.47255 | 0.222171916 | 0.143 | 0 | 1 | 27 |
| eif2b2 | 0.47255 | 0.205370131 | 0.143 | 0 | 1 | 27 |
| eif2b3 | 0.47255 | 0.203268019 | 0.143 | 0 | 1 | 27 |
| eif4g1a | 0.47255 | 0.210704507 | 0.143 | 0 | 1 | 27 |
| eif4h | 0.47255 | 0.229312254 | 0.143 | 0 | 1 | 27 |
| emc9 | 0.47255 | 0.219288095 | 0.143 | 0 | 1 | 27 |
| etf1b | 0.47255 | 0.272090195 | 0.143 | 0 | 1 | 27 |
| ethe1 | 0.47255 | 0.319500763 | 0.143 | 0 | 1 | 27 |
| fam76b | 0.47255 | 0.23929539 | 0.143 | 0 | 1 | 27 |
| farsa | 0.47255 | 0.233118232 | 0.143 | 0 | 1 | 27 |
| faub | 0.47255 | 0.205294652 | 0.143 | 0 | 1 | 27 |
| fbxl14b | 0.47255 | 0.217071078 | 0.143 | 0 | 1 | 27 |
| fkbp2 | 0.47255 | 0.291756546 | 0.143 | 0 | 1 | 27 |
| foxo3b | 0.47255 | 0.300532018 | 0.143 | 0 | 1 | 27 |
| gabpa | 0.47255 | 0.268375345 | 0.143 | 0 | 1 | 27 |
| gadd45aa | 0.47255 | 0.247489082 | 0.143 | 0 | 1 | 27 |
| gb:bc139872 | 0.47255 | 0.230007436 | 0.143 | 0 | 1 | 27 |
| ggnbp2 | 0.47255 | 0.221417546 | 0.143 | 0 | 1 | 27 |
| glipr1a | 0.47255 | 0.163116395 | 0.143 | 0 | 1 | 27 |
| gltpa | 0.47255 | 0.336950068 | 0.143 | 0 | 1 | 27 |
| glyr1 | 0.47255 | 0.166502633 | 0.143 | 0 | 1 | 27 |
| gmds | 0.47255 | 0.226297865 | 0.143 | 0 | 1 | 27 |
| gmeb1 | 0.47255 | 0.186169358 | 0.143 | 0 | 1 | 27 |
| gosr2 | 0.47255 | 0.231379519 | 0.143 | 0 | 1 | 27 |
| gstm.1 | 0.47255 | 0.210572445 | 0.143 | 0 | 1 | 27 |
| gtf2a1 | 0.47255 | 0.287247786 | 0.143 | 0 | 1 | 27 |
| gtf2b | 0.47255 | 0.231583695 | 0.143 | 0 | 1 | 27 |
| gtf2e2 | 0.47255 | 0.186816829 | 0.143 | 0 | 1 | 27 |
| hars | 0.47255 | 0.230695471 | 0.143 | 0 | 1 | 27 |

| | | | | | | |
|---|---|---|---|---|---|---|
| hiat1a | 0.47255 | 0.238110145 | 0.143 | 0 | 1 | 27 |
| hirip3 | 0.47255 | 0.221047825 | 0.143 | 0 | 1 | 27 |
| hmox2b | 0.47255 | 0.249209808 | 0.143 | 0 | 1 | 27 |
| hnrnpr | 0.47255 | 0.30122237 | 0.143 | 0 | 1 | 27 |
| hnrnpul1 | 0.47255 | 0.25180026 | 0.143 | 0 | 1 | 27 |
| hprt1 | 0.47255 | 0.184796001 | 0.143 | 0 | 1 | 27 |
| hps5 | 0.47255 | 0.193464253 | 0.143 | 0 | 1 | 27 |
| hspa4b | 0.47255 | 0.281773562 | 0.143 | 0 | 1 | 27 |
| huwe1 | 0.47255 | 0.204685329 | 0.143 | 0 | 1 | 27 |
| idh3b | 0.47255 | 0.252943836 | 0.143 | 0 | 1 | 27 |
| ighd | 0.47255 | 0.386814609 | 0.143 | 0 | 1 | 27 |
| ino80c | 0.47255 | 0.297146314 | 0.143 | 0 | 1 | 27 |
| insra | 0.47255 | 0.241068477 | 0.143 | 0 | 1 | 27 |
| ip6k2b | 0.47255 | 0.251245458 | 0.143 | 0 | 1 | 27 |
| jak1 | 0.47255 | 0.273308974 | 0.143 | 0 | 1 | 27 |
| jun | 0.47255 | 0.223782145 | 0.143 | 0 | 1 | 27 |
| kdm4b | 0.47255 | 0.246103436 | 0.143 | 0 | 1 | 27 |
| keap1b | 0.47255 | 0.234525862 | 0.143 | 0 | 1 | 27 |
| kmt2e | 0.47255 | 0.228248285 | 0.143 | 0 | 1 | 27 |
| krcp | 0.47255 | 0.29795557 | 0.143 | 0 | 1 | 27 |
| lclat1 | 0.47255 | 0.241949471 | 0.143 | 0 | 1 | 27 |
| lcorl | 0.47255 | 0.296608344 | 0.143 | 0 | 1 | 27 |
| ldlrad3 | 0.47255 | 0.245826845 | 0.143 | 0 | 1 | 27 |
| lgals3b | 0.47255 | 0.271108827 | 0.143 | 0 | 1 | 27 |
| lin37 | 0.47255 | 0.19741628 | 0.143 | 0 | 1 | 27 |
| lipea | 0.47255 | 0.355740066 | 0.143 | 0 | 1 | 27 |
| lman2 | 0.47255 | 0.254056628 | 0.143 | 0 | 1 | 27 |
| lnpep | 0.47255 | 0.331746903 | 0.143 | 0 | 1 | 27 |
| lonrf1l | 0.47255 | 0.295410335 | 0.143 | 0 | 1 | 27 |
| lpar5b | 0.47255 | 0.315913907 | 0.143 | 0 | 1 | 27 |
| lsm14aa | 0.47255 | 0.200716384 | 0.143 | 0 | 1 | 27 |
| ly6m2 | 0.47255 | 0.260821044 | 0.143 | 0 | 1 | 27 |
| lyrm2 | 0.47255 | 0.285295371 | 0.143 | 0 | 1 | 27 |
| lyrm4 | 0.47255 | 0.207011851 | 0.143 | 0 | 1 | 27 |
| macf1a | 0.47255 | 0.282836487 | 0.143 | 0 | 1 | 27 |
| map2k2a | 0.47255 | 0.294462438 | 0.143 | 0 | 1 | 27 |
| map3k14a | 0.47255 | 0.183639676 | 0.143 | 0 | 1 | 27 |
| march6 | 0.47255 | 0.286492777 | 0.143 | 0 | 1 | 27 |
| med11 | 0.47255 | 0.246878645 | 0.143 | 0 | 1 | 27 |
| med19b | 0.47255 | 0.233660804 | 0.143 | 0 | 1 | 27 |
| med20 | 0.47255 | 0.199709564 | 0.143 | 0 | 1 | 27 |
| med21 | 0.47255 | 0.284438957 | 0.143 | 0 | 1 | 27 |
| mettl14 | 0.47255 | 0.219368931 | 0.143 | 0 | 1 | 27 |
| mfng | 0.47255 | 0.279111093 | 0.143 | 0 | 1 | 27 |
| mlf2 | 0.47255 | 0.246021195 | 0.143 | 0 | 1 | 27 |
| mov10b.1 | 0.47255 | 0.206596819 | 0.143 | 0 | 1 | 27 |
| mrpl13 | 0.47255 | 0.268872133 | 0.143 | 0 | 1 | 27 |
| mrpl24 | 0.47255 | 0.213829986 | 0.143 | 0 | 1 | 27 |

| | | | | | | |
|---|---|---|---|---|---|---|
| mrpl28 | 0.47255 | 0.228463386 | 0.143 | 0 | 1 | 27 |
| mrpl34 | 0.47255 | 0.277272857 | 0.143 | 0 | 1 | 27 |
| mrpl39 | 0.47255 | 0.305554457 | 0.143 | 0 | 1 | 27 |
| mrpl47 | 0.47255 | 0.217472931 | 0.143 | 0 | 1 | 27 |
| mrps9 | 0.47255 | 0.299420972 | 0.143 | 0 | 1 | 27 |
| mtch2 | 0.47255 | 0.248036716 | 0.143 | 0 | 1 | 27 |
| mthfd2 | 0.47255 | 0.300344735 | 0.143 | 0 | 1 | 27 |
| mut | 0.47255 | 0.221597735 | 0.143 | 0 | 1 | 27 |
| mydgf | 0.47255 | 0.209364868 | 0.143 | 0 | 1 | 27 |
| myl12.2 | 0.47255 | 0.227759296 | 0.143 | 0 | 1 | 27 |
| naa38 | 0.47255 | 0.266715137 | 0.143 | 0 | 1 | 27 |
| nabp1a | 0.47255 | 0.355724369 | 0.143 | 0 | 1 | 27 |
| nars | 0.47255 | 0.306023848 | 0.143 | 0 | 1 | 27 |
| nbr1a | 0.47255 | 0.225237211 | 0.143 | 0 | 1 | 27 |
| ncaph2 | 0.47255 | 0.241459274 | 0.143 | 0 | 1 | 27 |
| ncbp2 | 0.47255 | 0.23428822 | 0.143 | 0 | 1 | 27 |
| ndfip2 | 0.47255 | 0.194606161 | 0.143 | 0 | 1 | 27 |
| ndufa12 | 0.47255 | 0.318414614 | 0.143 | 0 | 1 | 27 |
| ndufab1a | 0.47255 | 0.150593782 | 0.143 | 0 | 1 | 27 |
| ndufaf3 | 0.47255 | 0.286632492 | 0.143 | 0 | 1 | 27 |
| ndufv1 | 0.47255 | 0.286801972 | 0.143 | 0 | 1 | 27 |
| nfatc2ip | 0.47255 | 0.207484196 | 0.143 | 0 | 1 | 27 |
| nfybb | 0.47255 | 0.202174269 | 0.143 | 0 | 1 | 27 |
| nsfl1c | 0.47255 | 0.283389794 | 0.143 | 0 | 1 | 27 |
| nt5c2a | 0.47255 | 0.243388284 | 0.143 | 0 | 1 | 27 |
| nup107 | 0.47255 | 0.204682659 | 0.143 | 0 | 1 | 27 |
| nup93 | 0.47255 | 0.23346963 | 0.143 | 0 | 1 | 27 |
| nupr1a | 0.47255 | 0.381800675 | 0.143 | 0 | 1 | 27 |
| ogfrl2 | 0.47255 | 0.189901127 | 0.143 | 0 | 1 | 27 |
| ormdl2 | 0.47255 | 0.247028466 | 0.143 | 0 | 1 | 27 |
| oser1 | 0.47255 | 0.227369449 | 0.143 | 0 | 1 | 27 |
| osgn1 | 0.47255 | 0.249513979 | 0.143 | 0 | 1 | 27 |
| otub1a | 0.47255 | 0.272976246 | 0.143 | 0 | 1 | 27 |
| otud5a | 0.47255 | 0.313054736 | 0.143 | 0 | 1 | 27 |
| pag1 | 0.47255 | 0.202240526 | 0.143 | 0 | 1 | 27 |
| parn | 0.47255 | 0.248151838 | 0.143 | 0 | 1 | 27 |
| paxx | 0.47255 | 0.160352999 | 0.143 | 0 | 1 | 27 |
| pdhb | 0.47255 | 0.23847428 | 0.143 | 0 | 1 | 27 |
| pdpk1b | 0.47255 | 0.257337636 | 0.143 | 0 | 1 | 27 |
| pdxka | 0.47255 | 0.257121652 | 0.143 | 0 | 1 | 27 |
| pef1 | 0.47255 | 0.255051603 | 0.143 | 0 | 1 | 27 |
| pfdn1 | 0.47255 | 0.302211337 | 0.143 | 0 | 1 | 27 |
| pgrmc2 | 0.47255 | 0.194056832 | 0.143 | 0 | 1 | 27 |
| phax | 0.47255 | 0.237889549 | 0.143 | 0 | 1 | 27 |
| pigp | 0.47255 | 0.218435057 | 0.143 | 0 | 1 | 27 |
| pkma | 0.47255 | 0.250455943 | 0.143 | 0 | 1 | 27 |
| plcg1 | 0.47255 | 0.212334627 | 0.143 | 0 | 1 | 27 |
| plgrkt | 0.47255 | 0.286236991 | 0.143 | 0 | 1 | 27 |

| | | | | | | |
|---|---|---|---|---|---|---|
| polr2eb | 0.47255 | 0.188720649 | 0.143 | 0 | 1 | 27 |
| polr3k | 0.47255 | 0.251896044 | 0.143 | 0 | 1 | 27 |
| pou6f1 | 0.47255 | 0.224256803 | 0.143 | 0 | 1 | 27 |
| ppid | 0.47255 | 0.217486778 | 0.143 | 0 | 1 | 27 |
| ppp1r2 | 0.47255 | 0.227931128 | 0.143 | 0 | 1 | 27 |
| ppp2cb | 0.47255 | 0.290175402 | 0.143 | 0 | 1 | 27 |
| ppp4r3b | 0.47255 | 0.18282765 | 0.143 | 0 | 1 | 27 |
| primpol | 0.47255 | 0.209007445 | 0.143 | 0 | 1 | 27 |
| prpf40a | 0.47255 | 0.243981257 | 0.143 | 0 | 1 | 27 |
| psip1b | 0.47255 | 0.337413783 | 0.143 | 0 | 1 | 27 |
| psmd13 | 0.47255 | 0.190044236 | 0.143 | 0 | 1 | 27 |
| ptges3a | 0.47255 | 0.23807921 | 0.143 | 0 | 1 | 27 |
| ptp4a2b | 0.47255 | 0.302377542 | 0.143 | 0 | 1 | 27 |
| ptpn1 | 0.47255 | 0.275107899 | 0.143 | 0 | 1 | 27 |
| qdprb1 | 0.47255 | 0.186337053 | 0.143 | 0 | 1 | 27 |
| rab11fip4b | 0.47255 | 0.218458801 | 0.143 | 0 | 1 | 27 |
| rab43 | 0.47255 | 0.311333484 | 0.143 | 0 | 1 | 27 |
| rab5ab | 0.47255 | 0.202927608 | 0.143 | 0 | 1 | 27 |
| ramp2 | 0.47255 | 0.357727268 | 0.143 | 0 | 1 | 27 |
| ranbp3b | 0.47255 | 0.182724881 | 0.143 | 0 | 1 | 27 |
| rangrf | 0.47255 | 0.207287582 | 0.143 | 0 | 1 | 27 |
| rb1 | 0.47255 | 0.181971581 | 0.143 | 0 | 1 | 27 |
| rgs18 | 0.47255 | 0.244162231 | 0.143 | 0 | 1 | 27 |
| rgs3a | 0.47255 | 0.228298236 | 0.143 | 0 | 1 | 27 |
| rnf44 | 0.47255 | 0.257112745 | 0.143 | 0 | 1 | 27 |
| rpe | 0.47255 | 0.206124743 | 0.143 | 0 | 1 | 27 |
| rprd1b | 0.47255 | 0.205432801 | 0.143 | 0 | 1 | 27 |
| rsf1b.1 | 0.47255 | 0.200729732 | 0.143 | 0 | 1 | 27 |
| rsrc2 | 0.47255 | 0.261833135 | 0.143 | 0 | 1 | 27 |
| rtf1 | 0.47255 | 0.211639972 | 0.143 | 0 | 1 | 27 |
| rtn4rl2b | 0.47255 | 0.362122683 | 0.143 | 0 | 1 | 27 |
| rtraf | 0.47255 | 0.181575662 | 0.143 | 0 | 1 | 27 |
| sdhb | 0.47255 | 0.216043585 | 0.143 | 0 | 1 | 27 |
| sephs1 | 0.47255 | 0.282937396 | 0.143 | 0 | 1 | 27 |
| sf1 | 0.47255 | 0.312938092 | 0.143 | 0 | 1 | 27 |
| sgta | 0.47255 | 0.271393224 | 0.143 | 0 | 1 | 27 |
| sh2b3 | 0.47255 | 0.244352438 | 0.143 | 0 | 1 | 27 |
| si:ch211-150o23.2 | 0.47255 | 0.294732799 | 0.143 | 0 | 1 | 27 |
| si:ch211-197h24.8 | 0.47255 | 0.195668671 | 0.143 | 0 | 1 | 27 |
| si:ch211-199b20.3 | 0.47255 | 0.273297068 | 0.143 | 0 | 1 | 27 |
| si:ch211-25d12.7 | 0.47255 | 0.228599307 | 0.143 | 0 | 1 | 27 |
| si:ch73-248e21.1 | 0.47255 | 0.276161075 | 0.143 | 0 | 1 | 27 |
| si:ch73-91k6.2 | 0.47255 | 0.291574642 | 0.143 | 0 | 1 | 27 |
| si:dkey-21c19.3 | 0.47255 | 0.222319328 | 0.143 | 0 | 1 | 27 |
| si:dkey-222b8.1 | 0.47255 | 0.207234375 | 0.143 | 0 | 1 | 27 |
| si:dkey-237j10.2 | 0.47255 | 0.263264834 | 0.143 | 0 | 1 | 27 |
| si:dkey-88p24.11 | 0.47255 | 0.217931042 | 0.143 | 0 | 1 | 27 |
| si:dkeyp-113d7.10 | 0.47255 | 0.212258338 | 0.143 | 0 | 1 | 27 |

| | | | | | | |
|---|---|---|---|---|---|---|
| si:rp71-79p20.2 | 0.47255 | 0.23511247 | 0.143 | 0 | 1 | 27 |
| si:zfos-169g10.3 | 0.47255 | 0.279862766 | 0.143 | 0 | 1 | 27 |
| sidt2 | 0.47255 | 0.329477227 | 0.143 | 0 | 1 | 27 |
| sirt7 | 0.47255 | 0.209947213 | 0.143 | 0 | 1 | 27 |
| slc12a2 | 0.47255 | 0.258665745 | 0.143 | 0 | 1 | 27 |
| slc35f6 | 0.47255 | 0.330102603 | 0.143 | 0 | 1 | 27 |
| slc39a7 | 0.47255 | 0.269832911 | 0.143 | 0 | 1 | 27 |
| slc43a1a | 0.47255 | 0.260363961 | 0.143 | 0 | 1 | 27 |
| smarcb1b | 0.47255 | 0.196810659 | 0.143 | 0 | 1 | 27 |
| snd1 | 0.47255 | 0.186590199 | 0.143 | 0 | 1 | 27 |
| snrnp70 | 0.47255 | 0.232769375 | 0.143 | 0 | 1 | 27 |
| snx16 | 0.47255 | 0.17560639 | 0.143 | 0 | 1 | 27 |
| sowahd | 0.47255 | 0.272891909 | 0.143 | 0 | 1 | 27 |
| sp4 | 0.47255 | 0.195820217 | 0.143 | 0 | 1 | 27 |
| spast | 0.47255 | 0.213362931 | 0.143 | 0 | 1 | 27 |
| srp19 | 0.47255 | 0.213270133 | 0.143 | 0 | 1 | 27 |
| srsf5b | 0.47255 | 0.1902081 | 0.143 | 0 | 1 | 27 |
| ssbp1 | 0.47255 | 0.200055361 | 0.143 | 0 | 1 | 27 |
| ssbp3b | 0.47255 | 0.23168051 | 0.143 | 0 | 1 | 27 |
| stk11 | 0.47255 | 0.265853989 | 0.143 | 0 | 1 | 27 |
| stk19 | 0.47255 | 0.20549423 | 0.143 | 0 | 1 | 27 |
| stk35l | 0.47255 | 0.271017243 | 0.143 | 0 | 1 | 27 |
| stt3b | 0.47255 | 0.199690855 | 0.143 | 0 | 1 | 27 |
| stub1 | 0.47255 | 0.227072462 | 0.143 | 0 | 1 | 27 |
| suclg1 | 0.47255 | 0.22205102 | 0.143 | 0 | 1 | 27 |
| suclg2 | 0.47255 | 0.257992319 | 0.143 | 0 | 1 | 27 |
| swi5 | 0.47255 | 0.215185067 | 0.143 | 0 | 1 | 27 |
| tbk1 | 0.47255 | 0.232756453 | 0.143 | 0 | 1 | 27 |
| tcf4 | 0.47255 | 0.211477079 | 0.143 | 0 | 1 | 27 |
| tefb | 0.47255 | 0.340445907 | 0.143 | 0 | 1 | 27 |
| tet3 | 0.47255 | 0.322895641 | 0.143 | 0 | 1 | 27 |
| tfdp1a | 0.47255 | 0.197967247 | 0.143 | 0 | 1 | 27 |
| tfr1b | 0.47255 | 0.23156593 | 0.143 | 0 | 1 | 27 |
| thoc1 | 0.47255 | 0.278464918 | 0.143 | 0 | 1 | 27 |
| thrap3a | 0.47255 | 0.247610118 | 0.143 | 0 | 1 | 27 |
| timm23a | 0.47255 | 0.234039808 | 0.143 | 0 | 1 | 27 |
| tlr20.2 | 0.47255 | 0.297420591 | 0.143 | 0 | 1 | 27 |
| tmc6b | 0.47255 | 0.230462924 | 0.143 | 0 | 1 | 27 |
| tmed9 | 0.47255 | 0.304624109 | 0.143 | 0 | 1 | 27 |
| tmem110l | 0.47255 | 0.241516348 | 0.143 | 0 | 1 | 27 |
| tmem147 | 0.47255 | 0.21006109 | 0.143 | 0 | 1 | 27 |
| tmem176l.1 | 0.47255 | 0.213707448 | 0.143 | 0 | 1 | 27 |
| tmub1 | 0.47255 | 0.367853042 | 0.143 | 0 | 1 | 27 |
| tomm6 | 0.47255 | 0.187657728 | 0.143 | 0 | 1 | 27 |
| tomm70a | 0.47255 | 0.236190376 | 0.143 | 0 | 1 | 27 |
| trnt1 | 0.47255 | 0.264884031 | 0.143 | 0 | 1 | 27 |
| trrap | 0.47255 | 0.182764196 | 0.143 | 0 | 1 | 27 |
| tubg1 | 0.47255 | 0.184778503 | 0.143 | 0 | 1 | 27 |

| | | | | | | |
|---|---|---|---|---|---|---|
| txndc12 | 0.47255 | 0.238180936 | 0.143 | 0 | 1 | 27 |
| txndc15 | 0.47255 | 0.210732489 | 0.143 | 0 | 1 | 27 |
| ubap2b | 0.47255 | 0.27937274 | 0.143 | 0 | 1 | 27 |
| ube2g2 | 0.47255 | 0.165467862 | 0.143 | 0 | 1 | 27 |
| ube2t | 0.47255 | 0.194345706 | 0.143 | 0 | 1 | 27 |
| upf3b | 0.47255 | 0.20750033 | 0.143 | 0 | 1 | 27 |
| vps26b | 0.47255 | 0.304172772 | 0.143 | 0 | 1 | 27 |
| vps35 | 0.47255 | 0.238579969 | 0.143 | 0 | 1 | 27 |
| washc2c | 0.47255 | 0.280408922 | 0.143 | 0 | 1 | 27 |
| wdfy4 | 0.47255 | 0.312633106 | 0.143 | 0 | 1 | 27 |
| wdr61 | 0.47255 | 0.277919619 | 0.143 | 0 | 1 | 27 |
| xrn2 | 0.47255 | 0.267302014 | 0.143 | 0 | 1 | 27 |
| yipf4 | 0.47255 | 0.221137599 | 0.143 | 0 | 1 | 27 |
| yju2 | 0.47255 | 0.265344415 | 0.143 | 0 | 1 | 27 |
| zfand3 | 0.47255 | 0.39327034 | 0.143 | 0 | 1 | 27 |
| zfand6 | 0.47255 | 0.206802212 | 0.143 | 0 | 1 | 27 |
| zfhx3 | 0.47255 | 0.242488524 | 0.143 | 0 | 1 | 27 |
| zgc:110239 | 0.47255 | 0.220575216 | 0.143 | 0 | 1 | 27 |
| zgc:110339 | 0.47255 | 0.297647463 | 0.143 | 0 | 1 | 27 |
| zgc:113208 | 0.47255 | 0.260494656 | 0.143 | 0 | 1 | 27 |
| zgc:152774 | 0.47255 | 0.266886316 | 0.143 | 0 | 1 | 27 |
| zgc:162339 | 0.47255 | 0.219672995 | 0.143 | 0 | 1 | 27 |
| zgc:163080 | 0.47255 | 0.141294603 | 0.143 | 0 | 1 | 27 |
| zgc:56106 | 0.47255 | 0.290359192 | 0.143 | 0 | 1 | 27 |
| zgc:63470 | 0.47255 | 0.24537741 | 0.143 | 0 | 1 | 27 |
| zgc:66427 | 0.47255 | 0.258303246 | 0.143 | 0 | 1 | 27 |
| zgc:85789 | 0.47255 | 0.190745686 | 0.143 | 0 | 1 | 27 |
| znf865 | 0.47255 | 0.202762716 | 0.143 | 0 | 1 | 27 |
| zswim5 | 0.47255 | 0.263700564 | 0.143 | 0 | 1 | 27 |
| FAM72B | 0.47255 | 0.20978759 | 0.143 | 0 | 1 | 27 |
| FO082781.1 | 0.47255 | 0.207897705 | 0.143 | 0 | 1 | 27 |
| MZT1 | 0.47255 | 0.276026321 | 0.143 | 0 | 1 | 27 |
| akip1 | 0.47255 | 0.211575515 | 0.143 | 0 | 1 | 27 |
| ap2m1b | 0.47255 | 0.216502096 | 0.143 | 0 | 1 | 27 |
| bod1 | 0.47255 | 0.195130729 | 0.143 | 0 | 1 | 27 |
| cenpk | 0.47255 | 0.262801292 | 0.143 | 0 | 1 | 27 |
| cenpl | 0.47255 | 0.198961015 | 0.143 | 0 | 1 | 27 |
| cenpw | 0.47255 | 0.208518255 | 0.143 | 0 | 1 | 27 |
| cited4a | 0.47255 | 0.246183652 | 0.143 | 0 | 1 | 27 |
| clk2b | 0.47255 | 0.210476565 | 0.143 | 0 | 1 | 27 |
| ctdspl2b | 0.47255 | 0.175745229 | 0.143 | 0 | 1 | 27 |
| dctn2 | 0.47255 | 0.23303377 | 0.143 | 0 | 1 | 27 |
| diaph2 | 0.47255 | 0.204997136 | 0.143 | 0 | 1 | 27 |
| e2f8 | 0.47255 | 0.189622397 | 0.143 | 0 | 1 | 27 |
| flot1b | 0.47255 | 0.160217977 | 0.143 | 0 | 1 | 27 |
| gemin2 | 0.47255 | 0.20569446 | 0.143 | 0 | 1 | 27 |
| higd1a | 0.47255 | 0.219509298 | 0.143 | 0 | 1 | 27 |
| hint2 | 0.47255 | 0.19962126 | 0.143 | 0 | 1 | 27 |

| | | | | | | |
|---|---|---|---|---|---|---|
| hnrnpc | 0.47255 | 0.198767082 | 0.143 | 0 | 1 | 27 |
| hnrnpd | 0.47255 | 0.242944226 | 0.143 | 0 | 1 | 27 |
| kdelr2a | 0.47255 | 0.186516006 | 0.143 | 0 | 1 | 27 |
| kifc1 | 0.47255 | 0.334945654 | 0.143 | 0 | 1 | 27 |
| limk1a | 0.47255 | 0.23583235 | 0.143 | 0 | 1 | 27 |
| lmnb1 | 0.47255 | 0.290407679 | 0.143 | 0 | 1 | 27 |
| myb | 0.47255 | 0.220944379 | 0.143 | 0 | 1 | 27 |
| myef2 | 0.47255 | 0.236397162 | 0.143 | 0 | 1 | 27 |
| ncapd3 | 0.47255 | 0.19343794 | 0.143 | 0 | 1 | 27 |
| ndc1 | 0.47255 | 0.204577334 | 0.143 | 0 | 1 | 27 |
| ndc80 | 0.47255 | 0.1937824 | 0.143 | 0 | 1 | 27 |
| ndnl2 | 0.47255 | 0.187489293 | 0.143 | 0 | 1 | 27 |
| npm1b | 0.47255 | 0.232687019 | 0.143 | 0 | 1 | 27 |
| orc3 | 0.47255 | 0.209869068 | 0.143 | 0 | 1 | 27 |
| orc4 | 0.47255 | 0.178998791 | 0.143 | 0 | 1 | 27 |
| pdcd6 | 0.47255 | 0.254391 | 0.143 | 0 | 1 | 27 |
| phip | 0.47255 | 0.197436311 | 0.143 | 0 | 1 | 27 |
| polr2b | 0.47255 | 0.203695093 | 0.143 | 0 | 1 | 27 |
| polr2gl | 0.47255 | 0.231910177 | 0.143 | 0 | 1 | 27 |
| pop7 | 0.47255 | 0.172118382 | 0.143 | 0 | 1 | 27 |
| ppp2r1ba | 0.47255 | 0.225858718 | 0.143 | 0 | 1 | 27 |
| psmb5 | 0.47255 | 0.216193147 | 0.143 | 0 | 1 | 27 |
| rad1 | 0.47255 | 0.203399886 | 0.143 | 0 | 1 | 27 |
| rbm26 | 0.47255 | 0.272279088 | 0.143 | 0 | 1 | 27 |
| rdh10b | 0.47255 | 0.244592402 | 0.143 | 0 | 1 | 27 |
| rnf181 | 0.47255 | 0.239897566 | 0.143 | 0 | 1 | 27 |
| sdf2l1 | 0.47255 | 0.215440731 | 0.143 | 0 | 1 | 27 |
| si:ch211-114c12.2 | 0.47255 | 0.22621146 | 0.143 | 0 | 1 | 27 |
| si:dkey-185e18.7 | 0.47255 | 0.20978759 | 0.143 | 0 | 1 | 27 |
| si:dkey-19e4.5 | 0.47255 | 0.197151922 | 0.143 | 0 | 1 | 27 |
| slc25a43 | 0.47255 | 0.195844944 | 0.143 | 0 | 1 | 27 |
| srsf6b | 0.47255 | 0.250565723 | 0.143 | 0 | 1 | 27 |
| tab2 | 0.47255 | 0.235721154 | 0.143 | 0 | 1 | 27 |
| timmdc1 | 0.47255 | 0.204648587 | 0.143 | 0 | 1 | 27 |
| tk1 | 0.47255 | 0.268576686 | 0.143 | 0 | 1 | 27 |
| txnrd3 | 0.47255 | 0.250378917 | 0.143 | 0 | 1 | 27 |
| xrcc5 | 0.47255 | 0.208457591 | 0.143 | 0 | 1 | 27 |
| zgc:173587 | 0.47255 | 0.205309483 | 0.143 | 0 | 1 | 27 |
| zgc:77929 | 0.47255 | 0.185392445 | 0.143 | 0 | 1 | 27 |
| cisd2 | 0.47255 | 0.287706798 | 0.143 | 0 | 1 | 27 |
| rfx5 | 0.47255 | 0.255282771 | 0.143 | 0 | 1 | 27 |
| cxcr3.1 | 0.47255 | 0.304960095 | 0.143 | 0 | 1 | 27 |
| fcer1gl | 0.47255 | 0.25886648 | 0.143 | 0 | 1 | 27 |
| prpf6 | 0.47255 | 0.278395418 | 0.143 | 0 | 1 | 27 |
| si:ch211-239j9.1 | 0.47255 | 0.326869239 | 0.143 | 0 | 1 | 27 |
| slc25a16 | 0.47255 | 0.266578763 | 0.143 | 0 | 1 | 27 |
| snx12 | 0.47255 | 0.304051627 | 0.143 | 0 | 1 | 27 |
| srgn | 0.47255 | 0.284214901 | 0.143 | 0 | 1 | 27 |

| | | | | | | |
|---|---|---|---|---|---|---|
| mau2 | 0.47255 | 0.261277241 | 0.143 | 0 | 1 | 27 |
| si:ch1073-443f11.2 | 0.47255 | 0.189082917 | 0.143 | 0 | 1 | 27 |
| zgc:153631 | 0.47255 | 0.250025969 | 0.143 | 0 | 1 | 27 |
| si:ch73-1a9.3 | 0.47416 | -0.38917023 | 0.714 | 1 | 1 | 27 |
| mcl1b | 0.47509 | -0.51631409 | 0.19 | 0.333 | 1 | 27 |
| ndufs7 | 0.47558 | -0.45619558 | 0.19 | 0.333 | 1 | 27 |
| XLOC-013964 | 0.47928 | -0.27062149 | 0.952 | 1 | 1 | 27 |
| rps26l | 0.48064 | 0.146214922 | 1 | 1 | 1 | 27 |
| glrx | 0.48131 | 0.375372036 | 0.667 | 0.333 | 1 | 27 |
| mych | 0.48186 | 0.101787296 | 0.524 | 0.333 | 1 | 27 |
| dynlt3 | 0.48208 | -0.49712787 | 0.19 | 0.333 | 1 | 27 |
| zgc:63831 | 0.48547 | -0.5258229 | 0.333 | 0.667 | 1 | 27 |
| ak2 | 0.48823 | 0.307907636 | 0.667 | 0.333 | 1 | 27 |
| igl1c3 | 0.48939 | -0.37356978 | 0.714 | 1 | 1 | 27 |
| h3f3c | 0.48969 | 0.608216543 | 0.571 | 0.333 | 1 | 27 |
| LOC101886766 | 0.49143 | 0.183694211 | 0.952 | 1 | 1 | 27 |
| fam49ba | 0.49216 | -0.35793656 | 0.286 | 0.333 | 1 | 27 |
| atp5f1b | 0.49596 | -0.34942855 | 0.619 | 0.667 | 1 | 27 |
| rpl6 | 0.49749 | 0.184086535 | 1 | 1 | 1 | 27 |
| cirbpa | 0.49789 | 0.443064275 | 0.81 | 0.667 | 1 | 27 |
| baxb | 0.50214 | -0.51613989 | 0.143 | 0.333 | 1 | 27 |
| mibp2 | 0.50407 | -0.2928347 | 0.905 | 1 | 1 | 27 |
| psmc5 | 0.50485 | -0.45304297 | 0.19 | 0.333 | 1 | 27 |
| scfd1 | 0.50656 | -0.46665592 | 0.143 | 0.333 | 1 | 27 |
| mcl1a | 0.50817 | -0.27639821 | 0.952 | 1 | 1 | 27 |
| cox4i1 | 0.50978 | -0.33383422 | 0.762 | 1 | 1 | 27 |
| htatip2 | 0.51025 | -0.51204958 | 0.143 | 0.333 | 1 | 27 |
| ifi30 | 0.51379 | -0.47217634 | 0.524 | 0.667 | 1 | 27 |
| slc25a55a | 0.51815 | -0.5397544 | 0.333 | 0.667 | 1 | 27 |
| si:rp71-45k5.4 | 0.5185 | -0.18490632 | 0.381 | 0.333 | 1 | 27 |
| zgc:77849 | 0.51869 | -0.12565595 | 0.429 | 0.333 | 1 | 27 |
| rap1aa | 0.51944 | -0.44080094 | 0.429 | 0.667 | 1 | 27 |
| ran | 0.51948 | 0.199997757 | 1 | 1 | 1 | 27 |
| nap1l1 | 0.52205 | -0.46144148 | 0.333 | 0.667 | 1 | 27 |
| h2afvb | 0.52311 | 0.235908329 | 0.619 | 0.667 | 1 | 27 |
| irf2bp2a | 0.52554 | -0.30040327 | 0.286 | 0.333 | 1 | 27 |
| atp5meb | 0.52773 | -0.16355125 | 0.429 | 0.333 | 1 | 27 |
| NC-002333.4 | 0.52787 | -0.43016183 | 0.571 | 0.667 | 1 | 27 |
| fth1a | 0.5294 | 0.390600862 | 0.333 | 0.333 | 1 | 27 |
| mrpl33 | 0.52959 | -0.4509497 | 0.19 | 0.333 | 1 | 27 |
| qkia | 0.52986 | -0.51249489 | 0.333 | 0.667 | 1 | 27 |
| prelid3b | 0.53215 | -0.48882262 | 0.333 | 0.667 | 1 | 27 |
| pbdc1 | 0.53507 | -0.46480307 | 0.143 | 0.333 | 1 | 27 |
| tnrc6c1 | 0.53981 | -0.51169641 | 0.333 | 0.667 | 1 | 27 |
| h1fx | 0.54698 | -0.20310943 | 0.762 | 1 | 1 | 27 |
| bcl7bb | 0.54862 | -0.47602992 | 0.333 | 0.667 | 1 | 27 |
| atf4b | 0.54894 | 0.501725356 | 0.667 | 0.333 | 1 | 27 |
| inpp5d | 0.54944 | -0.32058051 | 0.286 | 0.333 | 1 | 27 |

| | | | | | | |
|---|---|---|---|---|---|---|
| ppp1r18 | 0.54952 | -0.52851234 | 0.143 | 0.333 | 1 | 27 |
| rps25 | 0.55056 | 0.199094359 | 1 | 1 | 1 | 27 |
| rbm8a | 0.55231 | 0.205214663 | 0.619 | 0.667 | 1 | 27 |
| cox5aa | 0.55373 | 0.14809144 | 0.619 | 0.333 | 1 | 27 |
| rps2 | 0.55678 | -0.10471176 | 1 | 1 | 1 | 27 |
| ncor2 | 0.55787 | -0.44564587 | 0.19 | 0.333 | 1 | 27 |
| ndufa3 | 0.55913 | -0.35090239 | 0.333 | 0.667 | 1 | 27 |
| atp5f1c | 0.55917 | 0.1108696 | 0.571 | 0.333 | 1 | 27 |
| ssr3 | 0.55999 | -0.42860781 | 0.333 | 0.667 | 1 | 27 |
| si:ch211-153b23.7 | 0.56012 | -0.45310373 | 0.19 | 0.333 | 1 | 27 |
| eif2s3 | 0.56069 | 0.418227223 | 0.667 | 0.333 | 1 | 27 |
| eif3k | 0.56158 | 0.438796451 | 0.667 | 0.333 | 1 | 27 |
| dusp1 | 0.56188 | -0.53211258 | 0.333 | 0.667 | 1 | 27 |
| cct5 | 0.56298 | 0.507854155 | 0.619 | 0.333 | 1 | 27 |
| pabpc1a | 0.56363 | -0.32328043 | 0.762 | 1 | 1 | 27 |
| wasa | 0.56439 | -0.44485134 | 0.476 | 0.667 | 1 | 27 |
| eif5b | 0.56717 | -0.38300407 | 0.238 | 0.333 | 1 | 27 |
| fkbp3 | 0.56766 | -0.37714128 | 0.238 | 0.333 | 1 | 27 |
| histh1l | 0.56767 | 0.557820843 | 0.571 | 0.333 | 1 | 27 |
| med29 | 0.57126 | -0.48205583 | 0.19 | 0.333 | 1 | 27 |
| ctss2.1 | 0.57263 | -0.43551165 | 0.238 | 0.333 | 1 | 27 |
| hsp90ab1 | 0.57277 | 0.126050431 | 1 | 1 | 1 | 27 |
| rhoaa | 0.57467 | -0.30323343 | 0.286 | 0.333 | 1 | 27 |
| max | 0.57636 | -0.39180704 | 0.238 | 0.333 | 1 | 27 |
| rhocb | 0.57794 | -0.43040898 | 0.19 | 0.333 | 1 | 27 |
| hnrnpaba | 0.58091 | 0.529831517 | 0.619 | 0.333 | 1 | 27 |
| parvg | 0.58499 | 0.52719056 | 0.619 | 0.333 | 1 | 27 |
| cox7c | 0.58774 | -0.2027453 | 0.667 | 0.667 | 1 | 27 |
| drap1 | 0.5885 | -0.44793529 | 0.143 | 0.333 | 1 | 27 |
| dbi | 0.59009 | -0.23517611 | 0.333 | 0.333 | 1 | 27 |
| tmem30aa | 0.59014 | -0.5206072 | 0.143 | 0.333 | 1 | 27 |
| FP101882.1 | 0.5908 | 0.138781721 | 0.095 | 0 | 1 | 27 |
| LO018493.1 | 0.5908 | 0.220809635 | 0.095 | 0 | 1 | 27 |
| LOC108190785 | 0.5908 | 0.104006954 | 0.095 | 0 | 1 | 27 |
| aagab | 0.5908 | 0.187247623 | 0.095 | 0 | 1 | 27 |
| ada2a | 0.5908 | 0.196401031 | 0.095 | 0 | 1 | 27 |
| aldh2.2 | 0.5908 | 0.181649068 | 0.095 | 0 | 1 | 27 |
| baz2ba | 0.5908 | 0.197836942 | 0.095 | 0 | 1 | 27 |
| cgas | 0.5908 | 0.195681876 | 0.095 | 0 | 1 | 27 |
| ctns | 0.5908 | 0.152873708 | 0.095 | 0 | 1 | 27 |
| cul4a | 0.5908 | 0.179748874 | 0.095 | 0 | 1 | 27 |
| cux1b | 0.5908 | 0.210901367 | 0.095 | 0 | 1 | 27 |
| cwf19l2 | 0.5908 | 0.145141814 | 0.095 | 0 | 1 | 27 |
| cyp20a1 | 0.5908 | 0.200036412 | 0.095 | 0 | 1 | 27 |
| egr2b | 0.5908 | 0.147785998 | 0.095 | 0 | 1 | 27 |
| fam129bb | 0.5908 | 0.232852084 | 0.095 | 0 | 1 | 27 |
| fbrs | 0.5908 | 0.199946914 | 0.095 | 0 | 1 | 27 |
| fhit | 0.5908 | 0.181739658 | 0.095 | 0 | 1 | 27 |

| | | | | | | |
|---|---|---|---|---|---|---|
| ftr50 | 0.5908 | 0.189461829 | 0.095 | 0 | 1 | 27 |
| fzr1a | 0.5908 | 0.189914652 | 0.095 | 0 | 1 | 27 |
| gsr | 0.5908 | 0.200969694 | 0.095 | 0 | 1 | 27 |
| hivep1 | 0.5908 | 0.142826414 | 0.095 | 0 | 1 | 27 |
| hyi | 0.5908 | 0.129501156 | 0.095 | 0 | 1 | 27 |
| icmt | 0.5908 | 0.174777893 | 0.095 | 0 | 1 | 27 |
| kansl1a | 0.5908 | 0.19712424 | 0.095 | 0 | 1 | 27 |
| mat2aa | 0.5908 | 0.206980482 | 0.095 | 0 | 1 | 27 |
| mrpl27 | 0.5908 | 0.140093277 | 0.095 | 0 | 1 | 27 |
| nck1b | 0.5908 | 0.22617096 | 0.095 | 0 | 1 | 27 |
| pdcl | 0.5908 | 0.20958958 | 0.095 | 0 | 1 | 27 |
| pelp1 | 0.5908 | 0.149513733 | 0.095 | 0 | 1 | 27 |
| phf8 | 0.5908 | 0.224787143 | 0.095 | 0 | 1 | 27 |
| plekha3 | 0.5908 | 0.188630559 | 0.095 | 0 | 1 | 27 |
| plscr3b | 0.5908 | 0.151606958 | 0.095 | 0 | 1 | 27 |
| rbbp6 | 0.5908 | 0.167823824 | 0.095 | 0 | 1 | 27 |
| rgs2 | 0.5908 | 0.253389523 | 0.095 | 0 | 1 | 27 |
| selenot2 | 0.5908 | 0.194783283 | 0.095 | 0 | 1 | 27 |
| si:ch211-212k18.7 | 0.5908 | 0.234230512 | 0.095 | 0 | 1 | 27 |
| si:ch211-217k17.7 | 0.5908 | 0.1748686 | 0.095 | 0 | 1 | 27 |
| si:ch211-281l24.3 | 0.5908 | 0.16006521 | 0.095 | 0 | 1 | 27 |
| si:ch73-37h15.2 | 0.5908 | 0.179294486 | 0.095 | 0 | 1 | 27 |
| si:dkey-286j15.3 | 0.5908 | 0.137919275 | 0.095 | 0 | 1 | 27 |
| si:dkeyp-19e1.3 | 0.5908 | 0.158312465 | 0.095 | 0 | 1 | 27 |
| slc25a20 | 0.5908 | 0.171381213 | 0.095 | 0 | 1 | 27 |
| snap23.1 | 0.5908 | 0.204368006 | 0.095 | 0 | 1 | 27 |
| sra1 | 0.5908 | 0.182983615 | 0.095 | 0 | 1 | 27 |
| supt4h1 | 0.5908 | 0.204500279 | 0.095 | 0 | 1 | 27 |
| tmem18 | 0.5908 | 0.193170616 | 0.095 | 0 | 1 | 27 |
| tmem243b | 0.5908 | 0.132593483 | 0.095 | 0 | 1 | 27 |
| tor1 | 0.5908 | 0.20728841 | 0.095 | 0 | 1 | 27 |
| tpd52l2b | 0.5908 | 0.180071549 | 0.095 | 0 | 1 | 27 |
| txn | 0.5908 | 0.171165728 | 0.095 | 0 | 1 | 27 |
| ube2nb | 0.5908 | 0.142231744 | 0.095 | 0 | 1 | 27 |
| utp23 | 0.5908 | 0.179074396 | 0.095 | 0 | 1 | 27 |
| vaspa | 0.5908 | 0.23143854 | 0.095 | 0 | 1 | 27 |
| zgc:103508 | 0.5908 | 0.147785998 | 0.095 | 0 | 1 | 27 |
| zgc:112982 | 0.5908 | 0.180404638 | 0.095 | 0 | 1 | 27 |
| zgc:112994 | 0.5908 | 0.234366413 | 0.095 | 0 | 1 | 27 |
| znf1001 | 0.5908 | 0.121320698 | 0.095 | 0 | 1 | 27 |
| znf16l | 0.5908 | 0.183754148 | 0.095 | 0 | 1 | 27 |
| ATP11B | 0.5908 | 0.154883569 | 0.095 | 0 | 1 | 27 |
| CABZ01063556.1 | 0.5908 | 0.147990289 | 0.095 | 0 | 1 | 27 |
| CABZ01074397.1 | 0.5908 | 0.132214776 | 0.095 | 0 | 1 | 27 |
| CABZ01074708.1 | 0.5908 | 0.149729789 | 0.095 | 0 | 1 | 27 |
| CU571074.1 | 0.5908 | 0.153748725 | 0.095 | 0 | 1 | 27 |
| CU694197.1 | 0.5908 | 0.134812403 | 0.095 | 0 | 1 | 27 |
| CU856344.1 | 0.5908 | 0.181404978 | 0.095 | 0 | 1 | 27 |

| | | | | | | |
|---|---|---|---|---|---|---|
| CU856539.2 | 0.5908 | 0.163576811 | 0.095 | 0 | 1 | 27 |
| LOC101885675 | 0.5908 | 0.264267834 | 0.095 | 0 | 1 | 27 |
| LOC110439840 | 0.5908 | 0.134424492 | 0.095 | 0 | 1 | 27 |
| MAP3K11 | 0.5908 | 0.182809291 | 0.095 | 0 | 1 | 27 |
| RPL37A | 0.5908 | 0.122372046 | 0.095 | 0 | 1 | 27 |
| aar2 | 0.5908 | 0.137346084 | 0.095 | 0 | 1 | 27 |
| afmid | 0.5908 | 0.178827852 | 0.095 | 0 | 1 | 27 |
| arfgef2 | 0.5908 | 0.206173773 | 0.095 | 0 | 1 | 27 |
| arhgef3l | 0.5908 | 0.186238531 | 0.095 | 0 | 1 | 27 |
| atp1a1a.1 | 0.5908 | 0.102939527 | 0.095 | 0 | 1 | 27 |
| atp2b1a | 0.5908 | 0.187102858 | 0.095 | 0 | 1 | 27 |
| aurkaip1 | 0.5908 | 0.159348584 | 0.095 | 0 | 1 | 27 |
| bcat2 | 0.5908 | 0.153698989 | 0.095 | 0 | 1 | 27 |
| borcs8 | 0.5908 | 0.168557946 | 0.095 | 0 | 1 | 27 |
| bysl | 0.5908 | 0.113420205 | 0.095 | 0 | 1 | 27 |
| bzw1a | 0.5908 | 0.155780978 | 0.095 | 0 | 1 | 27 |
| c1d | 0.5908 | 0.131839096 | 0.095 | 0 | 1 | 27 |
| capn2b | 0.5908 | 0.126239309 | 0.095 | 0 | 1 | 27 |
| ccdc12 | 0.5908 | 0.123422812 | 0.095 | 0 | 1 | 27 |
| ccdc9 | 0.5908 | 0.1271637 | 0.095 | 0 | 1 | 27 |
| ccnt2b | 0.5908 | 0.128411063 | 0.095 | 0 | 1 | 27 |
| cdca5 | 0.5908 | 0.146011641 | 0.095 | 0 | 1 | 27 |
| cdca7b | 0.5908 | 0.127089844 | 0.095 | 0 | 1 | 27 |
| cdk4 | 0.5908 | 0.147212609 | 0.095 | 0 | 1 | 27 |
| cenpi | 0.5908 | 0.132214776 | 0.095 | 0 | 1 | 27 |
| cenpp | 0.5908 | 0.154269196 | 0.095 | 0 | 1 | 27 |
| cep350 | 0.5908 | 0.185951155 | 0.095 | 0 | 1 | 27 |
| chchd3a | 0.5908 | 0.124092719 | 0.095 | 0 | 1 | 27 |
| chid1 | 0.5908 | 0.131516104 | 0.095 | 0 | 1 | 27 |
| chmp1a | 0.5908 | 0.188796878 | 0.095 | 0 | 1 | 27 |
| chmp1b | 0.5908 | 0.123250907 | 0.095 | 0 | 1 | 27 |
| cnpy1 | 0.5908 | 0.249098355 | 0.095 | 0 | 1 | 27 |
| copa | 0.5908 | 0.120527735 | 0.095 | 0 | 1 | 27 |
| csnk2a2a | 0.5908 | 0.181108017 | 0.095 | 0 | 1 | 27 |
| ddx47 | 0.5908 | 0.144531545 | 0.095 | 0 | 1 | 27 |
| desi2 | 0.5908 | 0.152581321 | 0.095 | 0 | 1 | 27 |
| dhodh | 0.5908 | 0.123334836 | 0.095 | 0 | 1 | 27 |
| dhx9 | 0.5908 | 0.141667447 | 0.095 | 0 | 1 | 27 |
| dock10 | 0.5908 | 0.160633248 | 0.095 | 0 | 1 | 27 |
| dot1l | 0.5908 | 0.173717175 | 0.095 | 0 | 1 | 27 |
| dpm3 | 0.5908 | 0.138178174 | 0.095 | 0 | 1 | 27 |
| dpp7 | 0.5908 | 0.113958036 | 0.095 | 0 | 1 | 27 |
| dtd1 | 0.5908 | 0.205574801 | 0.095 | 0 | 1 | 27 |
| dus3l | 0.5908 | 0.197187152 | 0.095 | 0 | 1 | 27 |
| ect2 | 0.5908 | 0.173328527 | 0.095 | 0 | 1 | 27 |
| eif1ad | 0.5908 | 0.173983358 | 0.095 | 0 | 1 | 27 |
| eif4ba | 0.5908 | 0.180448056 | 0.095 | 0 | 1 | 27 |
| elk1 | 0.5908 | 0.136739328 | 0.095 | 0 | 1 | 27 |

| | | | | | | |
|---|---|---|---|---|---|---|
| eme1 | 0.5908 | 0.129656066 | 0.095 | 0 | 1 | 27 |
| erlec1 | 0.5908 | 0.118740626 | 0.095 | 0 | 1 | 27 |
| exosc2 | 0.5908 | 0.115000118 | 0.095 | 0 | 1 | 27 |
| exosc8 | 0.5908 | 0.128353157 | 0.095 | 0 | 1 | 27 |
| fam126a | 0.5908 | 0.195413556 | 0.095 | 0 | 1 | 27 |
| far1 | 0.5908 | 0.160363342 | 0.095 | 0 | 1 | 27 |
| fibpa | 0.5908 | 0.184641522 | 0.095 | 0 | 1 | 27 |
| fnta | 0.5908 | 0.138970929 | 0.095 | 0 | 1 | 27 |
| ftr24 | 0.5908 | 0.116883546 | 0.095 | 0 | 1 | 27 |
| ggctb | 0.5908 | 0.129754253 | 0.095 | 0 | 1 | 27 |
| gkap1 | 0.5908 | 0.124522552 | 0.095 | 0 | 1 | 27 |
| gopc | 0.5908 | 0.106843132 | 0.095 | 0 | 1 | 27 |
| gosr1 | 0.5908 | 0.185552789 | 0.095 | 0 | 1 | 27 |
| hadhaa | 0.5908 | 0.139298544 | 0.095 | 0 | 1 | 27 |
| hax1 | 0.5908 | 0.122413115 | 0.095 | 0 | 1 | 27 |
| hddc3 | 0.5908 | 0.152591783 | 0.095 | 0 | 1 | 27 |
| hist2h2l | 0.5908 | 0.153276087 | 0.095 | 0 | 1 | 27 |
| hmg20b | 0.5908 | 0.14164849 | 0.095 | 0 | 1 | 27 |
| hsbp1a | 0.5908 | 0.157251574 | 0.095 | 0 | 1 | 27 |
| hsp90b1 | 0.5908 | 0.241903596 | 0.095 | 0 | 1 | 27 |
| ik | 0.5908 | 0.123376575 | 0.095 | 0 | 1 | 27 |
| ilf3a | 0.5908 | 0.127581457 | 0.095 | 0 | 1 | 27 |
| impa1 | 0.5908 | 0.129299118 | 0.095 | 0 | 1 | 27 |
| irf2bp1 | 0.5908 | 0.1279216 | 0.095 | 0 | 1 | 27 |
| irf2bpl | 0.5908 | 0.179828722 | 0.095 | 0 | 1 | 27 |
| itgb1bp1 | 0.5908 | 0.131191537 | 0.095 | 0 | 1 | 27 |
| kansl1b | 0.5908 | 0.143327086 | 0.095 | 0 | 1 | 27 |
| kif18a | 0.5908 | 0.157813332 | 0.095 | 0 | 1 | 27 |
| lman2la | 0.5908 | 0.125656768 | 0.095 | 0 | 1 | 27 |
| lrrc8aa | 0.5908 | 0.143234096 | 0.095 | 0 | 1 | 27 |
| med10 | 0.5908 | 0.118939049 | 0.095 | 0 | 1 | 27 |
| mfap1 | 0.5908 | 0.190822054 | 0.095 | 0 | 1 | 27 |
| mical1 | 0.5908 | 0.180391192 | 0.095 | 0 | 1 | 27 |
| mlx | 0.5908 | 0.218837383 | 0.095 | 0 | 1 | 27 |
| mrpl15 | 0.5908 | 0.121185636 | 0.095 | 0 | 1 | 27 |
| mrpl17 | 0.5908 | 0.127398792 | 0.095 | 0 | 1 | 27 |
| mrpl2 | 0.5908 | 0.124217836 | 0.095 | 0 | 1 | 27 |
| mrpl40 | 0.5908 | 0.136509166 | 0.095 | 0 | 1 | 27 |
| mrps18a | 0.5908 | 0.176879666 | 0.095 | 0 | 1 | 27 |
| mrps18b | 0.5908 | 0.183216405 | 0.095 | 0 | 1 | 27 |
| mrps7 | 0.5908 | 0.10177543 | 0.095 | 0 | 1 | 27 |
| mtif2 | 0.5908 | 0.167337105 | 0.095 | 0 | 1 | 27 |
| naa15b | 0.5908 | 0.128528135 | 0.095 | 0 | 1 | 27 |
| nab1a | 0.5908 | 0.192885183 | 0.095 | 0 | 1 | 27 |
| napaa | 0.5908 | 0.211156784 | 0.095 | 0 | 1 | 27 |
| naxe | 0.5908 | 0.161476311 | 0.095 | 0 | 1 | 27 |
| nedd8l | 0.5908 | 0.186501272 | 0.095 | 0 | 1 | 27 |
| ngdn | 0.5908 | 0.10397904 | 0.095 | 0 | 1 | 27 |

| | | | | | | |
|---|---|---|---|---|---|---|
| nop14 | 0.5908 | 0.131858668 | 0.095 | 0 | 1 | 27 |
| notch2 | 0.5908 | 0.139010595 | 0.095 | 0 | 1 | 27 |
| npepps | 0.5908 | 0.166228725 | 0.095 | 0 | 1 | 27 |
| nsfb | 0.5908 | 0.122346555 | 0.095 | 0 | 1 | 27 |
| nt5c2b | 0.5908 | 0.121334451 | 0.095 | 0 | 1 | 27 |
| nt5c3a | 0.5908 | 0.12098684 | 0.095 | 0 | 1 | 27 |
| ntan1 | 0.5908 | 0.146024639 | 0.095 | 0 | 1 | 27 |
| ntmt1 | 0.5908 | 0.171974095 | 0.095 | 0 | 1 | 27 |
| nubp2 | 0.5908 | 0.121210269 | 0.095 | 0 | 1 | 27 |
| nucb2a | 0.5908 | 0.150853516 | 0.095 | 0 | 1 | 27 |
| nup37 | 0.5908 | 0.12600494 | 0.095 | 0 | 1 | 27 |
| nup98 | 0.5908 | 0.158913093 | 0.095 | 0 | 1 | 27 |
| ogt.1 | 0.5908 | 0.208566561 | 0.095 | 0 | 1 | 27 |
| p2ry10 | 0.5908 | 0.155396296 | 0.095 | 0 | 1 | 27 |
| pdia6 | 0.5908 | 0.150596451 | 0.095 | 0 | 1 | 27 |
| pgp | 0.5908 | 0.177004815 | 0.095 | 0 | 1 | 27 |
| pigf | 0.5908 | 0.214386952 | 0.095 | 0 | 1 | 27 |
| pigl | 0.5908 | 0.146450171 | 0.095 | 0 | 1 | 27 |
| pknox1.1 | 0.5908 | 0.162011226 | 0.095 | 0 | 1 | 27 |
| polr2m | 0.5908 | 0.123282259 | 0.095 | 0 | 1 | 27 |
| polr3h | 0.5908 | 0.182899772 | 0.095 | 0 | 1 | 27 |
| ptger4b | 0.5908 | 0.152721794 | 0.095 | 0 | 1 | 27 |
| ptprja | 0.5908 | 0.192091702 | 0.095 | 0 | 1 | 27 |
| rchy1 | 0.5908 | 0.1968173 | 0.095 | 0 | 1 | 27 |
| reep3b | 0.5908 | 0.133545945 | 0.095 | 0 | 1 | 27 |
| rhbdf1a | 0.5908 | 0.188016808 | 0.095 | 0 | 1 | 27 |
| rnf144b | 0.5908 | 0.137975935 | 0.095 | 0 | 1 | 27 |
| rnf24 | 0.5908 | 0.13542751 | 0.095 | 0 | 1 | 27 |
| rnpep | 0.5908 | 0.13542751 | 0.095 | 0 | 1 | 27 |
| ruvbl1 | 0.5908 | 0.13823899 | 0.095 | 0 | 1 | 27 |
| sacm1lb | 0.5908 | 0.153659544 | 0.095 | 0 | 1 | 27 |
| si:ch1073-475a24.1 | 0.5908 | 0.132214776 | 0.095 | 0 | 1 | 27 |
| si:ch211-137a8.4 | 0.5908 | 0.118476134 | 0.095 | 0 | 1 | 27 |
| si:ch211-15d5.11 | 0.5908 | 0.194326868 | 0.095 | 0 | 1 | 27 |
| si:dkey-108k21.10 | 0.5908 | 0.11954106 | 0.095 | 0 | 1 | 27 |
| si:dkey-192g7.3 | 0.5908 | 0.239650123 | 0.095 | 0 | 1 | 27 |
| si:dkey-222h21.6 | 0.5908 | 0.136184995 | 0.095 | 0 | 1 | 27 |
| si:dkey-261h17.1 | 0.5908 | 0.114254134 | 0.095 | 0 | 1 | 27 |
| si:dkey-261l7.2 | 0.5908 | 0.156359044 | 0.095 | 0 | 1 | 27 |
| si:dkey-9k7.3 | 0.5908 | 0.198559765 | 0.095 | 0 | 1 | 27 |
| si:dkeyp-82a1.6 | 0.5908 | 0.128199907 | 0.095 | 0 | 1 | 27 |
| si:zfos-1192g2.3 | 0.5908 | 0.172990719 | 0.095 | 0 | 1 | 27 |
| slc35a4 | 0.5908 | 0.134650623 | 0.095 | 0 | 1 | 27 |
| slc5a11 | 0.5908 | 0.202601013 | 0.095 | 0 | 1 | 27 |
| slc5a8l | 0.5908 | 0.113486875 | 0.095 | 0 | 1 | 27 |
| smarcc1a | 0.5908 | 0.124278541 | 0.095 | 0 | 1 | 27 |
| smim20 | 0.5908 | 0.136888536 | 0.095 | 0 | 1 | 27 |
| smim4 | 0.5908 | 0.107512973 | 0.095 | 0 | 1 | 27 |

| | | | | | | |
|---|---|---|---|---|---|---|
| sp100.1 | 0.5908 | 0.147190912 | 0.095 | 0 | 1 | 27 |
| spryd7a | 0.5908 | 0.195081483 | 0.095 | 0 | 1 | 27 |
| spsb4b | 0.5908 | 0.127395842 | 0.095 | 0 | 1 | 27 |
| ssb | 0.5908 | 0.170464526 | 0.095 | 0 | 1 | 27 |
| ssr1 | 0.5908 | 0.195113643 | 0.095 | 0 | 1 | 27 |
| ssrp1a | 0.5908 | 0.130710964 | 0.095 | 0 | 1 | 27 |
| sssca1 | 0.5908 | 0.209616062 | 0.095 | 0 | 1 | 27 |
| st3gal4 | 0.5908 | 0.136041469 | 0.095 | 0 | 1 | 27 |
| stx5a | 0.5908 | 0.196260612 | 0.095 | 0 | 1 | 27 |
| sumf1 | 0.5908 | 0.162371782 | 0.095 | 0 | 1 | 27 |
| surf4l | 0.5908 | 0.134401299 | 0.095 | 0 | 1 | 27 |
| susd6 | 0.5908 | 0.185337542 | 0.095 | 0 | 1 | 27 |
| suz12a | 0.5908 | 0.134751833 | 0.095 | 0 | 1 | 27 |
| sv2 | 0.5908 | 0.1382056 | 0.095 | 0 | 1 | 27 |
| tasp1 | 0.5908 | 0.179016092 | 0.095 | 0 | 1 | 27 |
| tbl1xr1a | 0.5908 | 0.120447198 | 0.095 | 0 | 1 | 27 |
| thap11 | 0.5908 | 0.116749368 | 0.095 | 0 | 1 | 27 |
| thap12b | 0.5908 | 0.149729789 | 0.095 | 0 | 1 | 27 |
| timm10 | 0.5908 | 0.171153307 | 0.095 | 0 | 1 | 27 |
| timm44 | 0.5908 | 0.139615938 | 0.095 | 0 | 1 | 27 |
| tle3a | 0.5908 | 0.26223967 | 0.095 | 0 | 1 | 27 |
| tm2d2 | 0.5908 | 0.21175419 | 0.095 | 0 | 1 | 27 |
| tmed4 | 0.5908 | 0.142139598 | 0.095 | 0 | 1 | 27 |
| tpd52l2a | 0.5908 | 0.129453238 | 0.095 | 0 | 1 | 27 |
| trnau1apb | 0.5908 | 0.117460918 | 0.095 | 0 | 1 | 27 |
| tsen2 | 0.5908 | 0.147310297 | 0.095 | 0 | 1 | 27 |
| ube2r2 | 0.5908 | 0.20029221 | 0.095 | 0 | 1 | 27 |
| unk | 0.5908 | 0.180505982 | 0.095 | 0 | 1 | 27 |
| usp10 | 0.5908 | 0.140176934 | 0.095 | 0 | 1 | 27 |
| wdr76 | 0.5908 | 0.160313 | 0.095 | 0 | 1 | 27 |
| xab2 | 0.5908 | 0.121618107 | 0.095 | 0 | 1 | 27 |
| ypel1 | 0.5908 | 0.121844285 | 0.095 | 0 | 1 | 27 |
| zc3h13 | 0.5908 | 0.169062838 | 0.095 | 0 | 1 | 27 |
| zdhhc21 | 0.5908 | 0.13542751 | 0.095 | 0 | 1 | 27 |
| zfp36l2 | 0.5908 | 0.109456427 | 0.095 | 0 | 1 | 27 |
| zgc:101679 | 0.5908 | 0.122994924 | 0.095 | 0 | 1 | 27 |
| zgc:103697 | 0.5908 | 0.121010638 | 0.095 | 0 | 1 | 27 |
| zgc:153990 | 0.5908 | 0.125964043 | 0.095 | 0 | 1 | 27 |
| zgc:193742 | 0.5908 | 0.153314143 | 0.095 | 0 | 1 | 27 |
| zmynd11 | 0.5908 | 0.123779133 | 0.095 | 0 | 1 | 27 |
| znf410 | 0.5908 | 0.126214897 | 0.095 | 0 | 1 | 27 |
| znf516 | 0.5908 | 0.175925981 | 0.095 | 0 | 1 | 27 |
| AL954142.1 | 0.5908 | 0.128772287 | 0.095 | 0 | 1 | 27 |
| BX284666.1 | 0.5908 | 0.180602361 | 0.095 | 0 | 1 | 27 |
| BX323022.1 | 0.5908 | 0.136212374 | 0.095 | 0 | 1 | 27 |
| BX323557.1 | 0.5908 | 0.178576342 | 0.095 | 0 | 1 | 27 |
| CABZ01029938.1 | 0.5908 | 0.204934047 | 0.095 | 0 | 1 | 27 |
| CABZ01045617.1 | 0.5908 | 0.17992107 | 0.095 | 0 | 1 | 27 |

| | | | | | | |
|---|---|---|---|---|---|---|
| CABZ01055347.1 | 0.5908 | 0.138908533 | 0.095 | 0 | 1 | 27 |
| CABZ01058650.1 | 0.5908 | 0.133844183 | 0.095 | 0 | 1 | 27 |
| CABZ01076275.1 | 0.5908 | 0.201846106 | 0.095 | 0 | 1 | 27 |
| CABZ01079427.1 | 0.5908 | 0.225966675 | 0.095 | 0 | 1 | 27 |
| CABZ01084963.1 | 0.5908 | 0.199532258 | 0.095 | 0 | 1 | 27 |
| CABZ01088109.1 | 0.5908 | 0.128995454 | 0.095 | 0 | 1 | 27 |
| CABZ01088229.1 | 0.5908 | 0.123657382 | 0.095 | 0 | 1 | 27 |
| CR847936.1 | 0.5908 | 0.137403771 | 0.095 | 0 | 1 | 27 |
| CT027744.2 | 0.5908 | 0.202555406 | 0.095 | 0 | 1 | 27 |
| CU104709.1 | 0.5908 | 0.120917998 | 0.095 | 0 | 1 | 27 |
| LO018176.1 | 0.5908 | 0.260744669 | 0.095 | 0 | 1 | 27 |
| LOC100151196 | 0.5908 | 0.18846151 | 0.095 | 0 | 1 | 27 |
| LOC101885083 | 0.5908 | 0.137651158 | 0.095 | 0 | 1 | 27 |
| LOC101886015 | 0.5908 | 0.170125347 | 0.095 | 0 | 1 | 27 |
| LOC110438361 | 0.5908 | 0.184695431 | 0.095 | 0 | 1 | 27 |
| LOC110439849 | 0.5908 | 0.169145928 | 0.095 | 0 | 1 | 27 |
| LOC110439931 | 0.5908 | 0.181512501 | 0.095 | 0 | 1 | 27 |
| LOC558761 | 0.5908 | 0.131476611 | 0.095 | 0 | 1 | 27 |
| LOC566587 | 0.5908 | 0.239482456 | 0.095 | 0 | 1 | 27 |
| NFX1 | 0.5908 | 0.193212321 | 0.095 | 0 | 1 | 27 |
| PYURF | 0.5908 | 0.125407342 | 0.095 | 0 | 1 | 27 |
| XLOC-011085 | 0.5908 | 0.196856319 | 0.095 | 0 | 1 | 27 |
| XLOC-026761 | 0.5908 | 0.233789511 | 0.095 | 0 | 1 | 27 |
| XLOC-029426 | 0.5908 | 0.184668957 | 0.095 | 0 | 1 | 27 |
| XLOC-034437 | 0.5908 | 0.185774733 | 0.095 | 0 | 1 | 27 |
| abcc5 | 0.5908 | 0.236014477 | 0.095 | 0 | 1 | 27 |
| abtb1 | 0.5908 | 0.138464903 | 0.095 | 0 | 1 | 27 |
| acadl | 0.5908 | 0.130713309 | 0.095 | 0 | 1 | 27 |
| acap1 | 0.5908 | 0.221538591 | 0.095 | 0 | 1 | 27 |
| adat3 | 0.5908 | 0.186951619 | 0.095 | 0 | 1 | 27 |
| adgre5b.3 | 0.5908 | 0.262260728 | 0.095 | 0 | 1 | 27 |
| adi1 | 0.5908 | 0.130096579 | 0.095 | 0 | 1 | 27 |
| aes | 0.5908 | 0.132715892 | 0.095 | 0 | 1 | 27 |
| agpat2 | 0.5908 | 0.214427446 | 0.095 | 0 | 1 | 27 |
| ahsa1b | 0.5908 | 0.139062025 | 0.095 | 0 | 1 | 27 |
| ambra1a | 0.5908 | 0.158306858 | 0.095 | 0 | 1 | 27 |
| ammecr1 | 0.5908 | 0.142318991 | 0.095 | 0 | 1 | 27 |
| ankrd13a | 0.5908 | 0.154921653 | 0.095 | 0 | 1 | 27 |
| ankrd49 | 0.5908 | 0.137403771 | 0.095 | 0 | 1 | 27 |
| ankrd54 | 0.5908 | 0.136985827 | 0.095 | 0 | 1 | 27 |
| ap1m1 | 0.5908 | 0.13410356 | 0.095 | 0 | 1 | 27 |
| apbb1ip | 0.5908 | 0.237458081 | 0.095 | 0 | 1 | 27 |
| apopt1 | 0.5908 | 0.149662254 | 0.095 | 0 | 1 | 27 |
| aqr | 0.5908 | 0.166133464 | 0.095 | 0 | 1 | 27 |
| arfgap3 | 0.5908 | 0.247970919 | 0.095 | 0 | 1 | 27 |
| arhgap27 | 0.5908 | 0.237740763 | 0.095 | 0 | 1 | 27 |
| arhgap4a | 0.5908 | 0.186769588 | 0.095 | 0 | 1 | 27 |
| arhgef1a | 0.5908 | 0.243981862 | 0.095 | 0 | 1 | 27 |

| | | | | | | |
|---|---|---|---|---|---|---|
| arid2 | 0.5908 | 0.137191874 | 0.095 | 0 | 1 | 27 |
| asap1b | 0.5908 | 0.220528686 | 0.095 | 0 | 1 | 27 |
| asnsd1 | 0.5908 | 0.170821889 | 0.095 | 0 | 1 | 27 |
| aspm | 0.5908 | 0.155747803 | 0.095 | 0 | 1 | 27 |
| atp6v0a1a | 0.5908 | 0.19269927 | 0.095 | 0 | 1 | 27 |
| atrip | 0.5908 | 0.156152711 | 0.095 | 0 | 1 | 27 |
| aurka | 0.5908 | 0.157869332 | 0.095 | 0 | 1 | 27 |
| axin1 | 0.5908 | 0.201605597 | 0.095 | 0 | 1 | 27 |
| b4galt1l | 0.5908 | 0.204487425 | 0.095 | 0 | 1 | 27 |
| bcl10 | 0.5908 | 0.182764686 | 0.095 | 0 | 1 | 27 |
| blzf1 | 0.5908 | 0.1262736 | 0.095 | 0 | 1 | 27 |
| bmf2 | 0.5908 | 0.162575304 | 0.095 | 0 | 1 | 27 |
| brd1b | 0.5908 | 0.215121553 | 0.095 | 0 | 1 | 27 |
| brd4 | 0.5908 | 0.193395818 | 0.095 | 0 | 1 | 27 |
| brd9 | 0.5908 | 0.163137699 | 0.095 | 0 | 1 | 27 |
| brwd3 | 0.5908 | 0.172550894 | 0.095 | 0 | 1 | 27 |
| btbd7 | 0.5908 | 0.263894818 | 0.095 | 0 | 1 | 27 |
| bub1bb | 0.5908 | 0.15181464 | 0.095 | 0 | 1 | 27 |
| bxdc2 | 0.5908 | 0.157061902 | 0.095 | 0 | 1 | 27 |
| camlg | 0.5908 | 0.164050743 | 0.095 | 0 | 1 | 27 |
| casp2 | 0.5908 | 0.198586392 | 0.095 | 0 | 1 | 27 |
| casp8ap2 | 0.5908 | 0.210718014 | 0.095 | 0 | 1 | 27 |
| cbfa2t3 | 0.5908 | 0.238788196 | 0.095 | 0 | 1 | 27 |
| cbx8a | 0.5908 | 0.191572014 | 0.095 | 0 | 1 | 27 |
| ccdc115 | 0.5908 | 0.124405639 | 0.095 | 0 | 1 | 27 |
| ccdc167 | 0.5908 | 0.152178407 | 0.095 | 0 | 1 | 27 |
| ccl39.1 | 0.5908 | 0.28117186 | 0.095 | 0 | 1 | 27 |
| ccl39.3 | 0.5908 | 0.214374159 | 0.095 | 0 | 1 | 27 |
| ccl39.7 | 0.5908 | 0.189576349 | 0.095 | 0 | 1 | 27 |
| ccnb1 | 0.5908 | 0.150859843 | 0.095 | 0 | 1 | 27 |
| ccnc | 0.5908 | 0.163905219 | 0.095 | 0 | 1 | 27 |
| ccnd1 | 0.5908 | 0.158600714 | 0.095 | 0 | 1 | 27 |
| cd3eap | 0.5908 | 0.176938067 | 0.095 | 0 | 1 | 27 |
| cdc23 | 0.5908 | 0.134223459 | 0.095 | 0 | 1 | 27 |
| cdc34b | 0.5908 | 0.130960716 | 0.095 | 0 | 1 | 27 |
| cdc7 | 0.5908 | 0.151361123 | 0.095 | 0 | 1 | 27 |
| cdc73 | 0.5908 | 0.129195229 | 0.095 | 0 | 1 | 27 |
| cemip2 | 0.5908 | 0.130202965 | 0.095 | 0 | 1 | 27 |
| cenpe | 0.5908 | 0.186980815 | 0.095 | 0 | 1 | 27 |
| cenpf | 0.5908 | 0.181182562 | 0.095 | 0 | 1 | 27 |
| cenph | 0.5908 | 0.152776402 | 0.095 | 0 | 1 | 27 |
| cep135 | 0.5908 | 0.131151367 | 0.095 | 0 | 1 | 27 |
| cetn3 | 0.5908 | 0.150404071 | 0.095 | 0 | 1 | 27 |
| chd4b | 0.5908 | 0.128432285 | 0.095 | 0 | 1 | 27 |
| chek1 | 0.5908 | 0.158073496 | 0.095 | 0 | 1 | 27 |
| chic1 | 0.5908 | 0.248296761 | 0.095 | 0 | 1 | 27 |
| chmp4bb | 0.5908 | 0.218785669 | 0.095 | 0 | 1 | 27 |
| chmp5b | 0.5908 | 0.140719505 | 0.095 | 0 | 1 | 27 |

| | | | | | | |
|---|---|---|---|---|---|---|
| chmp6b | 0.5908 | 0.126361462 | 0.095 | 0 | 1 | 27 |
| cldn11b | 0.5908 | 0.131956192 | 0.095 | 0 | 1 | 27 |
| clp1 | 0.5908 | 0.223483139 | 0.095 | 0 | 1 | 27 |
| cmtm6 | 0.5908 | 0.213190617 | 0.095 | 0 | 1 | 27 |
| cnot6a | 0.5908 | 0.154397134 | 0.095 | 0 | 1 | 27 |
| coa3a | 0.5908 | 0.178683701 | 0.095 | 0 | 1 | 27 |
| coq4 | 0.5908 | 0.185038267 | 0.095 | 0 | 1 | 27 |
| coq8a | 0.5908 | 0.237403013 | 0.095 | 0 | 1 | 27 |
| cox10 | 0.5908 | 0.153087333 | 0.095 | 0 | 1 | 27 |
| cox19 | 0.5908 | 0.120161876 | 0.095 | 0 | 1 | 27 |
| cox8b | 0.5908 | 0.14752361 | 0.095 | 0 | 1 | 27 |
| creg1 | 0.5908 | 0.181945429 | 0.095 | 0 | 1 | 27 |
| crtc1a | 0.5908 | 0.162575304 | 0.095 | 0 | 1 | 27 |
| ctdspl3 | 0.5908 | 0.199991128 | 0.095 | 0 | 1 | 27 |
| ctr9 | 0.5908 | 0.134468339 | 0.095 | 0 | 1 | 27 |
| dbf4 | 0.5908 | 0.14464628 | 0.095 | 0 | 1 | 27 |
| dcps | 0.5908 | 0.131702645 | 0.095 | 0 | 1 | 27 |
| dcun1d5 | 0.5908 | 0.189170354 | 0.095 | 0 | 1 | 27 |
| dda1 | 0.5908 | 0.234538828 | 0.095 | 0 | 1 | 27 |
| ddit4 | 0.5908 | 0.187186581 | 0.095 | 0 | 1 | 27 |
| ddx27 | 0.5908 | 0.13660265 | 0.095 | 0 | 1 | 27 |
| ddx39b | 0.5908 | 0.201914304 | 0.095 | 0 | 1 | 27 |
| ddx42 | 0.5908 | 0.18568529 | 0.095 | 0 | 1 | 27 |
| ddx46 | 0.5908 | 0.143598665 | 0.095 | 0 | 1 | 27 |
| ddx6 | 0.5908 | 0.261448388 | 0.095 | 0 | 1 | 27 |
| dgkza | 0.5908 | 0.26658539 | 0.095 | 0 | 1 | 27 |
| dhrs13a.3 | 0.5908 | 0.145612194 | 0.095 | 0 | 1 | 27 |
| dicer1 | 0.5908 | 0.275239811 | 0.095 | 0 | 1 | 27 |
| dkc1 | 0.5908 | 0.225283858 | 0.095 | 0 | 1 | 27 |
| dlgap4a | 0.5908 | 0.140849889 | 0.095 | 0 | 1 | 27 |
| dmac2l | 0.5908 | 0.185576284 | 0.095 | 0 | 1 | 27 |
| dnajc18 | 0.5908 | 0.174126649 | 0.095 | 0 | 1 | 27 |
| dnajc2 | 0.5908 | 0.170474928 | 0.095 | 0 | 1 | 27 |
| dnajc5gb | 0.5908 | 0.179603377 | 0.095 | 0 | 1 | 27 |
| dnajc9 | 0.5908 | 0.159062564 | 0.095 | 0 | 1 | 27 |
| dnase1l4.2 | 0.5908 | 0.20208179 | 0.095 | 0 | 1 | 27 |
| dnm1l | 0.5908 | 0.137012762 | 0.095 | 0 | 1 | 27 |
| dusp22b | 0.5908 | 0.198638633 | 0.095 | 0 | 1 | 27 |
| dynll2b | 0.5908 | 0.122875091 | 0.095 | 0 | 1 | 27 |
| ecd | 0.5908 | 0.14464628 | 0.095 | 0 | 1 | 27 |
| egln1b | 0.5908 | 0.262716371 | 0.095 | 0 | 1 | 27 |
| eif2a | 0.5908 | 0.20296026 | 0.095 | 0 | 1 | 27 |
| elavl1b | 0.5908 | 0.177439485 | 0.095 | 0 | 1 | 27 |
| elp2 | 0.5908 | 0.154046505 | 0.095 | 0 | 1 | 27 |
| emc3 | 0.5908 | 0.166924101 | 0.095 | 0 | 1 | 27 |
| emd | 0.5908 | 0.20903544 | 0.095 | 0 | 1 | 27 |
| ep400 | 0.5908 | 0.17751428 | 0.095 | 0 | 1 | 27 |
| epc1b | 0.5908 | 0.17115885 | 0.095 | 0 | 1 | 27 |

| | | | | | | |
|---|---|---|---|---|---|---|
| esco2 | 0.5908 | 0.180491999 | 0.095 | 0 | 1 | 27 |
| esd | 0.5908 | 0.213336254 | 0.095 | 0 | 1 | 27 |
| etfdh | 0.5908 | 0.139447072 | 0.095 | 0 | 1 | 27 |
| etnk1 | 0.5908 | 0.26478075 | 0.095 | 0 | 1 | 27 |
| exosc5 | 0.5908 | 0.156250803 | 0.095 | 0 | 1 | 27 |
| ezh2 | 0.5908 | 0.170117729 | 0.095 | 0 | 1 | 27 |
| fam177a1 | 0.5908 | 0.213500575 | 0.095 | 0 | 1 | 27 |
| fam208ab | 0.5908 | 0.212803562 | 0.095 | 0 | 1 | 27 |
| fam210b | 0.5908 | 0.101355879 | 0.095 | 0 | 1 | 27 |
| fbxo3 | 0.5908 | 0.160010658 | 0.095 | 0 | 1 | 27 |
| fbxo5 | 0.5908 | 0.126998402 | 0.095 | 0 | 1 | 27 |
| fkbp4 | 0.5908 | 0.15740336 | 0.095 | 0 | 1 | 27 |
| flcn | 0.5908 | 0.172904203 | 0.095 | 0 | 1 | 27 |
| flot2a | 0.5908 | 0.191313945 | 0.095 | 0 | 1 | 27 |
| fosl2 | 0.5908 | 0.191297558 | 0.095 | 0 | 1 | 27 |
| foxn2b | 0.5908 | 0.172022381 | 0.095 | 0 | 1 | 27 |
| foxn3 | 0.5908 | 0.100049724 | 0.095 | 0 | 1 | 27 |
| frmd8 | 0.5908 | 0.160881291 | 0.095 | 0 | 1 | 27 |
| fundc2 | 0.5908 | 0.161926273 | 0.095 | 0 | 1 | 27 |
| g2e3 | 0.5908 | 0.151360955 | 0.095 | 0 | 1 | 27 |
| gadd45gip1 | 0.5908 | 0.123280506 | 0.095 | 0 | 1 | 27 |
| gatad2b | 0.5908 | 0.19154864 | 0.095 | 0 | 1 | 27 |
| golga7 | 0.5908 | 0.222927217 | 0.095 | 0 | 1 | 27 |
| golgb1 | 0.5908 | 0.199622349 | 0.095 | 0 | 1 | 27 |
| gpam | 0.5908 | 0.204706631 | 0.095 | 0 | 1 | 27 |
| grk6 | 0.5908 | 0.205734332 | 0.095 | 0 | 1 | 27 |
| grwd1 | 0.5908 | 0.191107886 | 0.095 | 0 | 1 | 27 |
| gse1 | 0.5908 | 0.19614877 | 0.095 | 0 | 1 | 27 |
| gtpbp1 | 0.5908 | 0.163433568 | 0.095 | 0 | 1 | 27 |
| guk1a | 0.5908 | 0.176007683 | 0.095 | 0 | 1 | 27 |
| hadhab | 0.5908 | 0.163111291 | 0.095 | 0 | 1 | 27 |
| haus7 | 0.5908 | 0.160547452 | 0.095 | 0 | 1 | 27 |
| hcfc1b | 0.5908 | 0.20165977 | 0.095 | 0 | 1 | 27 |
| hecw2b | 0.5908 | 0.264842041 | 0.095 | 0 | 1 | 27 |
| helz | 0.5908 | 0.159381027 | 0.095 | 0 | 1 | 27 |
| hexb | 0.5908 | 0.155416423 | 0.095 | 0 | 1 | 27 |
| hgs | 0.5908 | 0.130972799 | 0.095 | 0 | 1 | 27 |
| hikeshi | 0.5908 | 0.182399905 | 0.095 | 0 | 1 | 27 |
| hlx1 | 0.5908 | 0.168591153 | 0.095 | 0 | 1 | 27 |
| hmces | 0.5908 | 0.12770585 | 0.095 | 0 | 1 | 27 |
| hmga2 | 0.5908 | 0.103401212 | 0.095 | 0 | 1 | 27 |
| hmgb3b | 0.5908 | 0.194761421 | 0.095 | 0 | 1 | 27 |
| hnrnpm | 0.5908 | 0.153050459 | 0.095 | 0 | 1 | 27 |
| hnrnpua | 0.5908 | 0.199574561 | 0.095 | 0 | 1 | 27 |
| hpf1 | 0.5908 | 0.128313452 | 0.095 | 0 | 1 | 27 |
| hs1bp3 | 0.5908 | 0.199155234 | 0.095 | 0 | 1 | 27 |
| ifih1 | 0.5908 | 0.174702467 | 0.095 | 0 | 1 | 27 |
| ift20 | 0.5908 | 0.183553144 | 0.095 | 0 | 1 | 27 |

| | | | | | | |
|---|---|---|---|---|---|---|
| ikzf5 | 0.5908 | 0.146795304 | 0.095 | 0 | 1 | 27 |
| il12rb2 | 0.5908 | 0.232598408 | 0.095 | 0 | 1 | 27 |
| il6r | 0.5908 | 0.205067911 | 0.095 | 0 | 1 | 27 |
| im:7145024 | 0.5908 | 0.160316859 | 0.095 | 0 | 1 | 27 |
| ipcef1 | 0.5908 | 0.133947754 | 0.095 | 0 | 1 | 27 |
| ipo8 | 0.5908 | 0.123112902 | 0.095 | 0 | 1 | 27 |
| irgf1 | 0.5908 | 0.185614345 | 0.095 | 0 | 1 | 27 |
| isoc1 | 0.5908 | 0.233339096 | 0.095 | 0 | 1 | 27 |
| itgav | 0.5908 | 0.196427804 | 0.095 | 0 | 1 | 27 |
| itpkca | 0.5908 | 0.158652878 | 0.095 | 0 | 1 | 27 |
| jade1 | 0.5908 | 0.140379856 | 0.095 | 0 | 1 | 27 |
| jagn1b | 0.5908 | 0.179638099 | 0.095 | 0 | 1 | 27 |
| kat5b | 0.5908 | 0.193027201 | 0.095 | 0 | 1 | 27 |
| kat7a | 0.5908 | 0.174777338 | 0.095 | 0 | 1 | 27 |
| katnb1 | 0.5908 | 0.126490369 | 0.095 | 0 | 1 | 27 |
| kbtbd4 | 0.5908 | 0.160057686 | 0.095 | 0 | 1 | 27 |
| kctd2 | 0.5908 | 0.158677147 | 0.095 | 0 | 1 | 27 |
| kdm2ba | 0.5908 | 0.190074247 | 0.095 | 0 | 1 | 27 |
| kdm3b | 0.5908 | 0.153224059 | 0.095 | 0 | 1 | 27 |
| kdm6a | 0.5908 | 0.118201934 | 0.095 | 0 | 1 | 27 |
| keap1a | 0.5908 | 0.149713108 | 0.095 | 0 | 1 | 27 |
| klhl11 | 0.5908 | 0.18397847 | 0.095 | 0 | 1 | 27 |
| knstrn | 0.5908 | 0.1305979 | 0.095 | 0 | 1 | 27 |
| kpna4 | 0.5908 | 0.138130114 | 0.095 | 0 | 1 | 27 |
| kras | 0.5908 | 0.213623173 | 0.095 | 0 | 1 | 27 |
| larp4aa | 0.5908 | 0.19531545 | 0.095 | 0 | 1 | 27 |
| lin54 | 0.5908 | 0.1353731 | 0.095 | 0 | 1 | 27 |
| lrp10 | 0.5908 | 0.246126462 | 0.095 | 0 | 1 | 27 |
| lrrfip2 | 0.5908 | 0.13122536 | 0.095 | 0 | 1 | 27 |
| lsm1 | 0.5908 | 0.127496106 | 0.095 | 0 | 1 | 27 |
| lsm11 | 0.5908 | 0.160293749 | 0.095 | 0 | 1 | 27 |
| ly6m5 | 0.5908 | 0.203662196 | 0.095 | 0 | 1 | 27 |
| lyrm5a | 0.5908 | 0.178810661 | 0.095 | 0 | 1 | 27 |
| mad1l1 | 0.5908 | 0.157970949 | 0.095 | 0 | 1 | 27 |
| maip1 | 0.5908 | 0.229063489 | 0.095 | 0 | 1 | 27 |
| malt2 | 0.5908 | 0.207395328 | 0.095 | 0 | 1 | 27 |
| map2k4a | 0.5908 | 0.280680917 | 0.095 | 0 | 1 | 27 |
| map3k4 | 0.5908 | 0.197344352 | 0.095 | 0 | 1 | 27 |
| march5l | 0.5908 | 0.138960793 | 0.095 | 0 | 1 | 27 |
| mark3a | 0.5908 | 0.125014332 | 0.095 | 0 | 1 | 27 |
| mat2b | 0.5908 | 0.13419339 | 0.095 | 0 | 1 | 27 |
| mbnl3 | 0.5908 | 0.246468534 | 0.095 | 0 | 1 | 27 |
| mcee | 0.5908 | 0.170020482 | 0.095 | 0 | 1 | 27 |
| mcm4 | 0.5908 | 0.216381734 | 0.095 | 0 | 1 | 27 |
| mcm6 | 0.5908 | 0.19677732 | 0.095 | 0 | 1 | 27 |
| mcoln2 | 0.5908 | 0.149524317 | 0.095 | 0 | 1 | 27 |
| med27 | 0.5908 | 0.163853969 | 0.095 | 0 | 1 | 27 |
| med31 | 0.5908 | 0.153442319 | 0.095 | 0 | 1 | 27 |

| | | | | | | |
|---|---|---|---|---|---|---|
| melk | 0.5908 | 0.13952144 | 0.095 | 0 | 1 | 27 |
| metap1 | 0.5908 | 0.125197522 | 0.095 | 0 | 1 | 27 |
| mgat3b | 0.5908 | 0.103609034 | 0.095 | 0 | 1 | 27 |
| micall1a | 0.5908 | 0.221671786 | 0.095 | 0 | 1 | 27 |
| midn | 0.5908 | 0.193416442 | 0.095 | 0 | 1 | 27 |
| mir21-1 | 0.5908 | 0.265157227 | 0.095 | 0 | 1 | 27 |
| mis12 | 0.5908 | 0.178750673 | 0.095 | 0 | 1 | 27 |
| mis18a | 0.5908 | 0.15213496 | 0.095 | 0 | 1 | 27 |
| mob3c | 0.5908 | 0.207415405 | 0.095 | 0 | 1 | 27 |
| morc3a | 0.5908 | 0.137403771 | 0.095 | 0 | 1 | 27 |
| mospd1 | 0.5908 | 0.183480244 | 0.095 | 0 | 1 | 27 |
| mplkip | 0.5908 | 0.162359948 | 0.095 | 0 | 1 | 27 |
| mrpl22 | 0.5908 | 0.122813046 | 0.095 | 0 | 1 | 27 |
| mrpl53 | 0.5908 | 0.13789039 | 0.095 | 0 | 1 | 27 |
| mrps16 | 0.5908 | 0.169108504 | 0.095 | 0 | 1 | 27 |
| mrps35 | 0.5908 | 0.156936615 | 0.095 | 0 | 1 | 27 |
| mta2 | 0.5908 | 0.151118309 | 0.095 | 0 | 1 | 27 |
| mtm1 | 0.5908 | 0.14898154 | 0.095 | 0 | 1 | 27 |
| myca | 0.5908 | 0.190919631 | 0.095 | 0 | 1 | 27 |
| nckipsd | 0.5908 | 0.184917967 | 0.095 | 0 | 1 | 27 |
| ncoa3 | 0.5908 | 0.21230508 | 0.095 | 0 | 1 | 27 |
| ndel1a | 0.5908 | 0.123474211 | 0.095 | 0 | 1 | 27 |
| ndfip1l | 0.5908 | 0.113222383 | 0.095 | 0 | 1 | 27 |
| neil1 | 0.5908 | 0.238121492 | 0.095 | 0 | 1 | 27 |
| neil3 | 0.5908 | 0.162771358 | 0.095 | 0 | 1 | 27 |
| nemp1 | 0.5908 | 0.133436909 | 0.095 | 0 | 1 | 27 |
| nfya | 0.5908 | 0.152558837 | 0.095 | 0 | 1 | 27 |
| nhp2 | 0.5908 | 0.210415536 | 0.095 | 0 | 1 | 27 |
| nifk | 0.5908 | 0.220345141 | 0.095 | 0 | 1 | 27 |
| nit1 | 0.5908 | 0.160253031 | 0.095 | 0 | 1 | 27 |
| nol10 | 0.5908 | 0.167915904 | 0.095 | 0 | 1 | 27 |
| nol7 | 0.5908 | 0.132397848 | 0.095 | 0 | 1 | 27 |
| nrp2b | 0.5908 | 0.182377605 | 0.095 | 0 | 1 | 27 |
| nsd2 | 0.5908 | 0.147411124 | 0.095 | 0 | 1 | 27 |
| nsl1 | 0.5908 | 0.159172102 | 0.095 | 0 | 1 | 27 |
| nup205 | 0.5908 | 0.151274714 | 0.095 | 0 | 1 | 27 |
| nupl2 | 0.5908 | 0.15833382 | 0.095 | 0 | 1 | 27 |
| nxt2 | 0.5908 | 0.171893774 | 0.095 | 0 | 1 | 27 |
| orai1a | 0.5908 | 0.17193645 | 0.095 | 0 | 1 | 27 |
| orc6 | 0.5908 | 0.138685566 | 0.095 | 0 | 1 | 27 |
| pank4 | 0.5908 | 0.175380553 | 0.095 | 0 | 1 | 27 |
| parp12b | 0.5908 | 0.116770529 | 0.095 | 0 | 1 | 27 |
| paxip1 | 0.5908 | 0.156732059 | 0.095 | 0 | 1 | 27 |
| pdcd7 | 0.5908 | 0.124544237 | 0.095 | 0 | 1 | 27 |
| pds5a | 0.5908 | 0.201571186 | 0.095 | 0 | 1 | 27 |
| pdxdc1 | 0.5908 | 0.177513475 | 0.095 | 0 | 1 | 27 |
| pex12 | 0.5908 | 0.141595371 | 0.095 | 0 | 1 | 27 |
| pfn2 | 0.5908 | 0.153591789 | 0.095 | 0 | 1 | 27 |

| | | | | | | |
|---|---|---|---|---|---|---|
| phb | 0.5908 | 0.142028335 | 0.095 | 0 | 1 | 27 |
| phc2a | 0.5908 | 0.207329024 | 0.095 | 0 | 1 | 27 |
| phf10 | 0.5908 | 0.251392248 | 0.095 | 0 | 1 | 27 |
| phf14 | 0.5908 | 0.169089623 | 0.095 | 0 | 1 | 27 |
| pias4a | 0.5908 | 0.141113813 | 0.095 | 0 | 1 | 27 |
| pih1d1 | 0.5908 | 0.160647962 | 0.095 | 0 | 1 | 27 |
| pik3cg | 0.5908 | 0.187593755 | 0.095 | 0 | 1 | 27 |
| pip5k1aa | 0.5908 | 0.204532758 | 0.095 | 0 | 1 | 27 |
| pitpnaa | 0.5908 | 0.193584034 | 0.095 | 0 | 1 | 27 |
| pitpnc1b | 0.5908 | 0.145827141 | 0.095 | 0 | 1 | 27 |
| plekho1b | 0.5908 | 0.251038026 | 0.095 | 0 | 1 | 27 |
| plk4 | 0.5908 | 0.133436909 | 0.095 | 0 | 1 | 27 |
| pno1 | 0.5908 | 0.13736263 | 0.095 | 0 | 1 | 27 |
| ppil2 | 0.5908 | 0.131679841 | 0.095 | 0 | 1 | 27 |
| ppp1r11 | 0.5908 | 0.148038646 | 0.095 | 0 | 1 | 27 |
| ppp1r12c | 0.5908 | 0.177262548 | 0.095 | 0 | 1 | 27 |
| ppp1r8b | 0.5908 | 0.168311534 | 0.095 | 0 | 1 | 27 |
| ppp2r2d | 0.5908 | 0.17287287 | 0.095 | 0 | 1 | 27 |
| ppp2r3c | 0.5908 | 0.135598046 | 0.095 | 0 | 1 | 27 |
| ppp6r3 | 0.5908 | 0.124601389 | 0.095 | 0 | 1 | 27 |
| prex1 | 0.5908 | 0.13210169 | 0.095 | 0 | 1 | 27 |
| prkar1ab | 0.5908 | 0.181169284 | 0.095 | 0 | 1 | 27 |
| prkcba | 0.5908 | 0.162622224 | 0.095 | 0 | 1 | 27 |
| prkcda | 0.5908 | 0.151587724 | 0.095 | 0 | 1 | 27 |
| prkcsh | 0.5908 | 0.124522371 | 0.095 | 0 | 1 | 27 |
| prkrip1 | 0.5908 | 0.226237681 | 0.095 | 0 | 1 | 27 |
| prmt1 | 0.5908 | 0.181191169 | 0.095 | 0 | 1 | 27 |
| prmt2 | 0.5908 | 0.111361367 | 0.095 | 0 | 1 | 27 |
| prodha | 0.5908 | 0.23718893 | 0.095 | 0 | 1 | 27 |
| prpf38a | 0.5908 | 0.182108646 | 0.095 | 0 | 1 | 27 |
| prpsap2 | 0.5908 | 0.191820289 | 0.095 | 0 | 1 | 27 |
| prr11 | 0.5908 | 0.164105875 | 0.095 | 0 | 1 | 27 |
| prune2 | 0.5908 | 0.230234569 | 0.095 | 0 | 1 | 27 |
| psip1a | 0.5908 | 0.194211195 | 0.095 | 0 | 1 | 27 |
| psmd11a | 0.5908 | 0.173636129 | 0.095 | 0 | 1 | 27 |
| ptbp2b | 0.5908 | 0.13479394 | 0.095 | 0 | 1 | 27 |
| ptcd1 | 0.5908 | 0.183087295 | 0.095 | 0 | 1 | 27 |
| ptgir | 0.5908 | 0.134651354 | 0.095 | 0 | 1 | 27 |
| purba | 0.5908 | 0.185021593 | 0.095 | 0 | 1 | 27 |
| pwp1 | 0.5908 | 0.151886843 | 0.095 | 0 | 1 | 27 |
| qars | 0.5908 | 0.167208461 | 0.095 | 0 | 1 | 27 |
| qsox1 | 0.5908 | 0.275341142 | 0.095 | 0 | 1 | 27 |
| rab44 | 0.5908 | 0.226599717 | 0.095 | 0 | 1 | 27 |
| rab4b | 0.5908 | 0.215614344 | 0.095 | 0 | 1 | 27 |
| rab5aa | 0.5908 | 0.208944494 | 0.095 | 0 | 1 | 27 |
| rab8a | 0.5908 | 0.156110854 | 0.095 | 0 | 1 | 27 |
| rabep2 | 0.5908 | 0.137403771 | 0.095 | 0 | 1 | 27 |
| rae1 | 0.5908 | 0.142656277 | 0.095 | 0 | 1 | 27 |

| | | | | | | |
|---|---|---|---|---|---|---|
| ralbp1 | 0.5908 | 0.127295936 | 0.095 | 0 | 1 | 27 |
| rap2c | 0.5908 | 0.171562697 | 0.095 | 0 | 1 | 27 |
| rbm14a | 0.5908 | 0.127772399 | 0.095 | 0 | 1 | 27 |
| rbm14b | 0.5908 | 0.222480948 | 0.095 | 0 | 1 | 27 |
| rbm19 | 0.5908 | 0.225202554 | 0.095 | 0 | 1 | 27 |
| rbm25b | 0.5908 | 0.184015186 | 0.095 | 0 | 1 | 27 |
| rbm39b | 0.5908 | 0.189253845 | 0.095 | 0 | 1 | 27 |
| recql | 0.5908 | 0.136581527 | 0.095 | 0 | 1 | 27 |
| relch | 0.5908 | 0.163041264 | 0.095 | 0 | 1 | 27 |
| rmi1 | 0.5908 | 0.129614999 | 0.095 | 0 | 1 | 27 |
| rnf114 | 0.5908 | 0.217837976 | 0.095 | 0 | 1 | 27 |
| rnf121 | 0.5908 | 0.171708941 | 0.095 | 0 | 1 | 27 |
| rnf146 | 0.5908 | 0.1515751 | 0.095 | 0 | 1 | 27 |
| rnf185 | 0.5908 | 0.164742095 | 0.095 | 0 | 1 | 27 |
| rnf40 | 0.5908 | 0.19110617 | 0.095 | 0 | 1 | 27 |
| rnf7 | 0.5908 | 0.206908098 | 0.095 | 0 | 1 | 27 |
| rnmt | 0.5908 | 0.201169594 | 0.095 | 0 | 1 | 27 |
| rpp21 | 0.5908 | 0.182927197 | 0.095 | 0 | 1 | 27 |
| rps6ka1 | 0.5908 | 0.195639111 | 0.095 | 0 | 1 | 27 |
| rraga | 0.5908 | 0.184608421 | 0.095 | 0 | 1 | 27 |
| rwdd1 | 0.5908 | 0.17458001 | 0.095 | 0 | 1 | 27 |
| sae1 | 0.5908 | 0.218752677 | 0.095 | 0 | 1 | 27 |
| samd11 | 0.5908 | 0.197943781 | 0.095 | 0 | 1 | 27 |
| sart1 | 0.5908 | 0.146805202 | 0.095 | 0 | 1 | 27 |
| scamp3 | 0.5908 | 0.178097558 | 0.095 | 0 | 1 | 27 |
| seh1l | 0.5908 | 0.187656685 | 0.095 | 0 | 1 | 27 |
| selplg | 0.5908 | 0.267091983 | 0.095 | 0 | 1 | 27 |
| sema4aa | 0.5908 | 0.261395907 | 0.095 | 0 | 1 | 27 |
| sept9b | 0.5908 | 0.156704262 | 0.095 | 0 | 1 | 27 |
| serpine2 | 0.5908 | 0.165043521 | 0.095 | 0 | 1 | 27 |
| setdb1b | 0.5908 | 0.218768127 | 0.095 | 0 | 1 | 27 |
| sfswap | 0.5908 | 0.110506784 | 0.095 | 0 | 1 | 27 |
| sgo1 | 0.5908 | 0.160088123 | 0.095 | 0 | 1 | 27 |
| shc1 | 0.5908 | 0.184397634 | 0.095 | 0 | 1 | 27 |
| si:ch1073-157b13.1 | 0.5908 | 0.281619588 | 0.095 | 0 | 1 | 27 |
| si:ch211-122f10.4 | 0.5908 | 0.237840676 | 0.095 | 0 | 1 | 27 |
| si:ch211-132b12.8 | 0.5908 | 0.271382312 | 0.095 | 0 | 1 | 27 |
| si:ch211-167j9.5 | 0.5908 | 0.164072019 | 0.095 | 0 | 1 | 27 |
| si:ch211-195d17.2 | 0.5908 | 0.145077901 | 0.095 | 0 | 1 | 27 |
| si:ch211-262e15.1 | 0.5908 | 0.182552746 | 0.095 | 0 | 1 | 27 |
| si:ch211-284b7.3 | 0.5908 | 0.212110227 | 0.095 | 0 | 1 | 27 |
| si:ch211-67e16.4 | 0.5908 | 0.206054149 | 0.095 | 0 | 1 | 27 |
| si:ch211-69g19.2 | 0.5908 | 0.155481233 | 0.095 | 0 | 1 | 27 |
| si:ch73-308l14.2 | 0.5908 | 0.200054741 | 0.095 | 0 | 1 | 27 |
| si:ch73-376l24.6 | 0.5908 | 0.145003494 | 0.095 | 0 | 1 | 27 |
| si:ch73-62l21.1 | 0.5908 | 0.170469803 | 0.095 | 0 | 1 | 27 |
| si:dkey-100n23.3 | 0.5908 | 0.175096544 | 0.095 | 0 | 1 | 27 |
| si:dkey-102m7.3 | 0.5908 | 0.109201368 | 0.095 | 0 | 1 | 27 |

| | | | | | | |
|---|---|---|---|---|---|---|
| si:dkey-109l4.3 | 0.5908 | 0.128363845 | 0.095 | 0 | 1 | 27 |
| si:dkey-11f4.14 | 0.5908 | 0.25314595 | 0.095 | 0 | 1 | 27 |
| si:dkey-177p2.6 | 0.5908 | 0.147637998 | 0.095 | 0 | 1 | 27 |
| si:dkey-234h16.7 | 0.5908 | 0.197148423 | 0.095 | 0 | 1 | 27 |
| si:dkey-24p1.7 | 0.5908 | 0.170883659 | 0.095 | 0 | 1 | 27 |
| si:dkey-253d23.4 | 0.5908 | 0.108047379 | 0.095 | 0 | 1 | 27 |
| si:dkey-25e12.3 | 0.5908 | 0.232437014 | 0.095 | 0 | 1 | 27 |
| si:dkey-56i24.1 | 0.5908 | 0.170469803 | 0.095 | 0 | 1 | 27 |
| si:dkey-7j14.6 | 0.5908 | 0.185236855 | 0.095 | 0 | 1 | 27 |
| si:dkeyp-110e4.6 | 0.5908 | 0.157993537 | 0.095 | 0 | 1 | 27 |
| si:dkeyp-97b10.3 | 0.5908 | 0.144527701 | 0.095 | 0 | 1 | 27 |
| si:zfos-1011f11.2 | 0.5908 | 0.158677147 | 0.095 | 0 | 1 | 27 |
| ska1 | 0.5908 | 0.143556315 | 0.095 | 0 | 1 | 27 |
| skp2 | 0.5908 | 0.135862472 | 0.095 | 0 | 1 | 27 |
| slc10a7 | 0.5908 | 0.156861871 | 0.095 | 0 | 1 | 27 |
| slc25a26 | 0.5908 | 0.180058671 | 0.095 | 0 | 1 | 27 |
| slc25a46 | 0.5908 | 0.195101493 | 0.095 | 0 | 1 | 27 |
| slc35a3a | 0.5908 | 0.188351377 | 0.095 | 0 | 1 | 27 |
| slc38a2 | 0.5908 | 0.21567688 | 0.095 | 0 | 1 | 27 |
| slc39a9 | 0.5908 | 0.180652329 | 0.095 | 0 | 1 | 27 |
| slc6a22.1 | 0.5908 | 0.247143091 | 0.095 | 0 | 1 | 27 |
| slc7a3b | 0.5908 | 0.159617737 | 0.095 | 0 | 1 | 27 |
| slx1b | 0.5908 | 0.194846782 | 0.095 | 0 | 1 | 27 |
| smarca5 | 0.5908 | 0.170710551 | 0.095 | 0 | 1 | 27 |
| smarcd1 | 0.5908 | 0.162657784 | 0.095 | 0 | 1 | 27 |
| smek1 | 0.5908 | 0.128302503 | 0.095 | 0 | 1 | 27 |
| smg5 | 0.5908 | 0.20815961 | 0.095 | 0 | 1 | 27 |
| smim7 | 0.5908 | 0.108128969 | 0.095 | 0 | 1 | 27 |
| smu1a | 0.5908 | 0.111178119 | 0.095 | 0 | 1 | 27 |
| snapc3 | 0.5908 | 0.147071702 | 0.095 | 0 | 1 | 27 |
| snn | 0.5908 | 0.219872339 | 0.095 | 0 | 1 | 27 |
| snrnp25 | 0.5908 | 0.153627171 | 0.095 | 0 | 1 | 27 |
| snrnp40 | 0.5908 | 0.169535668 | 0.095 | 0 | 1 | 27 |
| snw1 | 0.5908 | 0.179463575 | 0.095 | 0 | 1 | 27 |
| spag1a | 0.5908 | 0.166410089 | 0.095 | 0 | 1 | 27 |
| spop | 0.5908 | 0.186106027 | 0.095 | 0 | 1 | 27 |
| spout1 | 0.5908 | 0.161940592 | 0.095 | 0 | 1 | 27 |
| sptan1 | 0.5908 | 0.151754758 | 0.095 | 0 | 1 | 27 |
| srgap2 | 0.5908 | 0.135407342 | 0.095 | 0 | 1 | 27 |
| srm | 0.5908 | 0.172716217 | 0.095 | 0 | 1 | 27 |
| srpk1a | 0.5908 | 0.168197146 | 0.095 | 0 | 1 | 27 |
| ssh2b | 0.5908 | 0.195732991 | 0.095 | 0 | 1 | 27 |
| st3gal3a | 0.5908 | 0.237280501 | 0.095 | 0 | 1 | 27 |
| stimate | 0.5908 | 0.260968014 | 0.095 | 0 | 1 | 27 |
| stn1 | 0.5908 | 0.187608092 | 0.095 | 0 | 1 | 27 |
| stx12 | 0.5908 | 0.118473134 | 0.095 | 0 | 1 | 27 |
| supt6h | 0.5908 | 0.152874965 | 0.095 | 0 | 1 | 27 |
| susd1 | 0.5908 | 0.193912692 | 0.095 | 0 | 1 | 27 |

| | | | | | | |
|---|---|---|---|---|---|---|
| suv39h1b | 0.5908 | 0.181810069 | 0.095 | 0 | 1 | 27 |
| swap70b | 0.5908 | 0.165879114 | 0.095 | 0 | 1 | 27 |
| taf3 | 0.5908 | 0.168765184 | 0.095 | 0 | 1 | 27 |
| taf7 | 0.5908 | 0.145729173 | 0.095 | 0 | 1 | 27 |
| tardbp | 0.5908 | 0.20702372 | 0.095 | 0 | 1 | 27 |
| tax1bp3 | 0.5908 | 0.172351647 | 0.095 | 0 | 1 | 27 |
| tbc1d17 | 0.5908 | 0.148698282 | 0.095 | 0 | 1 | 27 |
| tbcb | 0.5908 | 0.182584273 | 0.095 | 0 | 1 | 27 |
| tcf25 | 0.5908 | 0.233341586 | 0.095 | 0 | 1 | 27 |
| tcirg1b | 0.5908 | 0.178395285 | 0.095 | 0 | 1 | 27 |
| tec | 0.5908 | 0.138223093 | 0.095 | 0 | 1 | 27 |
| tescb | 0.5908 | 0.18269885 | 0.095 | 0 | 1 | 27 |
| tfcp2 | 0.5908 | 0.20950899 | 0.095 | 0 | 1 | 27 |
| tfdp1b | 0.5908 | 0.126003371 | 0.095 | 0 | 1 | 27 |
| thap4 | 0.5908 | 0.148363065 | 0.095 | 0 | 1 | 27 |
| tia1 | 0.5908 | 0.196194729 | 0.095 | 0 | 1 | 27 |
| ticam1 | 0.5908 | 0.24744802 | 0.095 | 0 | 1 | 27 |
| timeless | 0.5908 | 0.141821927 | 0.095 | 0 | 1 | 27 |
| tiparp | 0.5908 | 0.290065053 | 0.095 | 0 | 1 | 27 |
| tmem107 | 0.5908 | 0.114136282 | 0.095 | 0 | 1 | 27 |
| tmem165 | 0.5908 | 0.153108599 | 0.095 | 0 | 1 | 27 |
| tmem192 | 0.5908 | 0.126139241 | 0.095 | 0 | 1 | 27 |
| tmem256 | 0.5908 | 0.139542229 | 0.095 | 0 | 1 | 27 |
| tmod2 | 0.5908 | 0.213086263 | 0.095 | 0 | 1 | 27 |
| tollip | 0.5908 | 0.240727656 | 0.095 | 0 | 1 | 27 |
| tomm40 | 0.5908 | 0.161137913 | 0.095 | 0 | 1 | 27 |
| tprkb | 0.5908 | 0.173938415 | 0.095 | 0 | 1 | 27 |
| traf2b | 0.5908 | 0.137403771 | 0.095 | 0 | 1 | 27 |
| traip | 0.5908 | 0.14464628 | 0.095 | 0 | 1 | 27 |
| trappc4 | 0.5908 | 0.252299624 | 0.095 | 0 | 1 | 27 |
| trdmt1 | 0.5908 | 0.185411401 | 0.095 | 0 | 1 | 27 |
| triap1 | 0.5908 | 0.14985798 | 0.095 | 0 | 1 | 27 |
| trim35-28 | 0.5908 | 0.149699351 | 0.095 | 0 | 1 | 27 |
| tsr2 | 0.5908 | 0.137449351 | 0.095 | 0 | 1 | 27 |
| tsr3 | 0.5908 | 0.201896161 | 0.095 | 0 | 1 | 27 |
| tvp23b | 0.5908 | 0.212241866 | 0.095 | 0 | 1 | 27 |
| txndc11 | 0.5908 | 0.19681655 | 0.095 | 0 | 1 | 27 |
| txnl1 | 0.5908 | 0.133503444 | 0.095 | 0 | 1 | 27 |
| ube2d1a | 0.5908 | 0.145726306 | 0.095 | 0 | 1 | 27 |
| ube2d2l | 0.5908 | 0.152854253 | 0.095 | 0 | 1 | 27 |
| ube2f | 0.5908 | 0.15208388 | 0.095 | 0 | 1 | 27 |
| ube2z | 0.5908 | 0.147129073 | 0.095 | 0 | 1 | 27 |
| ublcp1 | 0.5908 | 0.147758838 | 0.095 | 0 | 1 | 27 |
| ubn2b | 0.5908 | 0.154046089 | 0.095 | 0 | 1 | 27 |
| ubqln4 | 0.5908 | 0.156044264 | 0.095 | 0 | 1 | 27 |
| ubxn7 | 0.5908 | 0.197842272 | 0.095 | 0 | 1 | 27 |
| ugt5b2 | 0.5908 | 0.246109763 | 0.095 | 0 | 1 | 27 |
| unc119.1 | 0.5908 | 0.15975454 | 0.095 | 0 | 1 | 27 |

| | | | | | | |
|---|---|---|---|---|---|---|
| unc13d | 0.5908 | 0.202834862 | 0.095 | 0 | 1 | 27 |
| uqcc1 | 0.5908 | 0.165754106 | 0.095 | 0 | 1 | 27 |
| usf1 | 0.5908 | 0.151721323 | 0.095 | 0 | 1 | 27 |
| usf2 | 0.5908 | 0.223328641 | 0.095 | 0 | 1 | 27 |
| usp3 | 0.5908 | 0.158788695 | 0.095 | 0 | 1 | 27 |
| uxs1 | 0.5908 | 0.168412544 | 0.095 | 0 | 1 | 27 |
| vgll4a | 0.5908 | 0.171547357 | 0.095 | 0 | 1 | 27 |
| vkorc1l1 | 0.5908 | 0.160191364 | 0.095 | 0 | 1 | 27 |
| vps28 | 0.5908 | 0.155882422 | 0.095 | 0 | 1 | 27 |
| vps39 | 0.5908 | 0.292254837 | 0.095 | 0 | 1 | 27 |
| vps4a | 0.5908 | 0.158306858 | 0.095 | 0 | 1 | 27 |
| vrk2 | 0.5908 | 0.233713124 | 0.095 | 0 | 1 | 27 |
| waplb | 0.5908 | 0.14023586 | 0.095 | 0 | 1 | 27 |
| wdr41 | 0.5908 | 0.165939246 | 0.095 | 0 | 1 | 27 |
| wtap | 0.5908 | 0.201541659 | 0.095 | 0 | 1 | 27 |
| xrcc3 | 0.5908 | 0.151105825 | 0.095 | 0 | 1 | 27 |
| ythdc1 | 0.5908 | 0.230464734 | 0.095 | 0 | 1 | 27 |
| zbtb11 | 0.5908 | 0.210525832 | 0.095 | 0 | 1 | 27 |
| zbtb41 | 0.5908 | 0.227809844 | 0.095 | 0 | 1 | 27 |
| zc3h11a | 0.5908 | 0.151773259 | 0.095 | 0 | 1 | 27 |
| zc3h14 | 0.5908 | 0.163827643 | 0.095 | 0 | 1 | 27 |
| zfp36l1b | 0.5908 | 0.243193852 | 0.095 | 0 | 1 | 27 |
| zfx | 0.5908 | 0.174285342 | 0.095 | 0 | 1 | 27 |
| zgc:101577 | 0.5908 | 0.160010658 | 0.095 | 0 | 1 | 27 |
| zgc:101715 | 0.5908 | 0.225586983 | 0.095 | 0 | 1 | 27 |
| zgc:101851 | 0.5908 | 0.181848842 | 0.095 | 0 | 1 | 27 |
| zgc:103692 | 0.5908 | 0.190374089 | 0.095 | 0 | 1 | 27 |
| zgc:110224 | 0.5908 | 0.160557877 | 0.095 | 0 | 1 | 27 |
| zgc:113184 | 0.5908 | 0.176645437 | 0.095 | 0 | 1 | 27 |
| zgc:136439 | 0.5908 | 0.118731108 | 0.095 | 0 | 1 | 27 |
| zgc:153293 | 0.5908 | 0.143787272 | 0.095 | 0 | 1 | 27 |
| zgc:153659 | 0.5908 | 0.200701032 | 0.095 | 0 | 1 | 27 |
| zgc:158803 | 0.5908 | 0.140305054 | 0.095 | 0 | 1 | 27 |
| zgc:162698 | 0.5908 | 0.196752191 | 0.095 | 0 | 1 | 27 |
| zgc:174863 | 0.5908 | 0.172739622 | 0.095 | 0 | 1 | 27 |
| zgc:194981 | 0.5908 | 0.201522986 | 0.095 | 0 | 1 | 27 |
| zgc:76872 | 0.5908 | 0.165461613 | 0.095 | 0 | 1 | 27 |
| zgc:86764 | 0.5908 | 0.162175333 | 0.095 | 0 | 1 | 27 |
| zmp:0000001082 | 0.5908 | 0.181328833 | 0.095 | 0 | 1 | 27 |
| zmp:0000001228 | 0.5908 | 0.253389204 | 0.095 | 0 | 1 | 27 |
| znf740b | 0.5908 | 0.163905219 | 0.095 | 0 | 1 | 27 |
| zranb2 | 0.5908 | 0.190607685 | 0.095 | 0 | 1 | 27 |
| zswim8 | 0.5908 | 0.18756577 | 0.095 | 0 | 1 | 27 |
| zwilch | 0.5908 | 0.151530486 | 0.095 | 0 | 1 | 27 |
| zyx | 0.5908 | 0.142635155 | 0.095 | 0 | 1 | 27 |
| BX530097.1 | 0.5908 | 0.210551835 | 0.095 | 0 | 1 | 27 |
| NC-002333.17 | 0.5908 | 0.126191526 | 0.095 | 0 | 1 | 27 |
| XLOC-026836 | 0.5908 | 0.252627947 | 0.095 | 0 | 1 | 27 |

| | | | | | | |
|---|---|---|---|---|---|---|
| acbd4 | 0.5908 | 0.18379772 | 0.095 | 0 | 1 | 27 |
| aco2 | 0.5908 | 0.142475859 | 0.095 | 0 | 1 | 27 |
| adora2b | 0.5908 | 0.232784379 | 0.095 | 0 | 1 | 27 |
| akr1a1b | 0.5908 | 0.149715572 | 0.095 | 0 | 1 | 27 |
| ank3a | 0.5908 | 0.219274012 | 0.095 | 0 | 1 | 27 |
| ap3s2 | 0.5908 | 0.105275524 | 0.095 | 0 | 1 | 27 |
| arl5a | 0.5908 | 0.202008525 | 0.095 | 0 | 1 | 27 |
| asph | 0.5908 | 0.112762441 | 0.095 | 0 | 1 | 27 |
| atox1 | 0.5908 | 0.129790195 | 0.095 | 0 | 1 | 27 |
| atp6v1e1b | 0.5908 | 0.151178222 | 0.095 | 0 | 1 | 27 |
| bach2b | 0.5908 | 0.218231695 | 0.095 | 0 | 1 | 27 |
| btbd10b | 0.5908 | 0.140353343 | 0.095 | 0 | 1 | 27 |
| caap1 | 0.5908 | 0.144709623 | 0.095 | 0 | 1 | 27 |
| cblb | 0.5908 | 0.233829576 | 0.095 | 0 | 1 | 27 |
| ccdc90b | 0.5908 | 0.146190448 | 0.095 | 0 | 1 | 27 |
| cdc6 | 0.5908 | 0.144153083 | 0.095 | 0 | 1 | 27 |
| cdkn2aipnl | 0.5908 | 0.199807432 | 0.095 | 0 | 1 | 27 |
| cmip | 0.5908 | 0.143017293 | 0.095 | 0 | 1 | 27 |
| cnot4a | 0.5908 | 0.180049457 | 0.095 | 0 | 1 | 27 |
| cops7a | 0.5908 | 0.211057577 | 0.095 | 0 | 1 | 27 |
| cpt2 | 0.5908 | 0.125983338 | 0.095 | 0 | 1 | 27 |
| cylda | 0.5908 | 0.13063434 | 0.095 | 0 | 1 | 27 |
| dpf2 | 0.5908 | 0.210479486 | 0.095 | 0 | 1 | 27 |
| dpp3 | 0.5908 | 0.146310234 | 0.095 | 0 | 1 | 27 |
| dscc1 | 0.5908 | 0.173787496 | 0.095 | 0 | 1 | 27 |
| ebna1bp2 | 0.5908 | 0.17817778 | 0.095 | 0 | 1 | 27 |
| echdc1 | 0.5908 | 0.134207063 | 0.095 | 0 | 1 | 27 |
| ell2 | 0.5908 | 0.131996197 | 0.095 | 0 | 1 | 27 |
| exoc4 | 0.5908 | 0.149756912 | 0.095 | 0 | 1 | 27 |
| fam122b | 0.5908 | 0.124970561 | 0.095 | 0 | 1 | 27 |
| fam49bb | 0.5908 | 0.187344592 | 0.095 | 0 | 1 | 27 |
| ftr67 | 0.5908 | 0.198307113 | 0.095 | 0 | 1 | 27 |
| gar1 | 0.5908 | 0.194562748 | 0.095 | 0 | 1 | 27 |
| gatad1 | 0.5908 | 0.183775615 | 0.095 | 0 | 1 | 27 |
| gmppb | 0.5908 | 0.113423459 | 0.095 | 0 | 1 | 27 |
| gnptg | 0.5908 | 0.183139053 | 0.095 | 0 | 1 | 27 |
| grinab | 0.5908 | 0.141108222 | 0.095 | 0 | 1 | 27 |
| gtf2f1 | 0.5908 | 0.162312677 | 0.095 | 0 | 1 | 27 |
| hdgfl2 | 0.5908 | 0.141314679 | 0.095 | 0 | 1 | 27 |
| josd2 | 0.5908 | 0.160056468 | 0.095 | 0 | 1 | 27 |
| kmt5aa | 0.5908 | 0.181972671 | 0.095 | 0 | 1 | 27 |
| las1l | 0.5908 | 0.149756912 | 0.095 | 0 | 1 | 27 |
| lin9 | 0.5908 | 0.169519602 | 0.095 | 0 | 1 | 27 |
| man2a2 | 0.5908 | 0.12865484 | 0.095 | 0 | 1 | 27 |
| man2c1 | 0.5908 | 0.117982223 | 0.095 | 0 | 1 | 27 |
| map4k4 | 0.5908 | 0.121142365 | 0.095 | 0 | 1 | 27 |
| map4k5 | 0.5908 | 0.137842961 | 0.095 | 0 | 1 | 27 |
| med12 | 0.5908 | 0.165487041 | 0.095 | 0 | 1 | 27 |

| | | | | | | |
|---|---|---|---|---|---|---|
| mgat1a | 0.5908 | 0.180893245 | 0.095 | 0 | 1 | 27 |
| mgmt | 0.5908 | 0.111174996 | 0.095 | 0 | 1 | 27 |
| mmadhc | 0.5908 | 0.164343322 | 0.095 | 0 | 1 | 27 |
| mpnd | 0.5908 | 0.127437249 | 0.095 | 0 | 1 | 27 |
| mrps17 | 0.5908 | 0.127027641 | 0.095 | 0 | 1 | 27 |
| msi2a | 0.5908 | 0.212821555 | 0.095 | 0 | 1 | 27 |
| msrb1b | 0.5908 | 0.146285327 | 0.095 | 0 | 1 | 27 |
| myo18ab | 0.5908 | 0.180444254 | 0.095 | 0 | 1 | 27 |
| ncapd2 | 0.5908 | 0.162416415 | 0.095 | 0 | 1 | 27 |
| nudt12 | 0.5908 | 0.164434305 | 0.095 | 0 | 1 | 27 |
| paf1 | 0.5908 | 0.159314693 | 0.095 | 0 | 1 | 27 |
| pbx4 | 0.5908 | 0.134142577 | 0.095 | 0 | 1 | 27 |
| pde12 | 0.5908 | 0.134207063 | 0.095 | 0 | 1 | 27 |
| pdia4 | 0.5908 | 0.148309774 | 0.095 | 0 | 1 | 27 |
| pdk2b | 0.5908 | 0.13165739 | 0.095 | 0 | 1 | 27 |
| pdk4 | 0.5908 | 0.10880288 | 0.095 | 0 | 1 | 27 |
| pigx | 0.5908 | 0.115912728 | 0.095 | 0 | 1 | 27 |
| ppp1r9ba | 0.5908 | 0.202807812 | 0.095 | 0 | 1 | 27 |
| prdm8b | 0.5908 | 0.196250385 | 0.095 | 0 | 1 | 27 |
| prkacaa | 0.5908 | 0.126089062 | 0.095 | 0 | 1 | 27 |
| prkcbp1l | 0.5908 | 0.215373469 | 0.095 | 0 | 1 | 27 |
| prps1b | 0.5908 | 0.136828393 | 0.095 | 0 | 1 | 27 |
| prrc2a | 0.5908 | 0.123731556 | 0.095 | 0 | 1 | 27 |
| psmd5 | 0.5908 | 0.140003844 | 0.095 | 0 | 1 | 27 |
| rassf3 | 0.5908 | 0.194760047 | 0.095 | 0 | 1 | 27 |
| rfc4 | 0.5908 | 0.206064239 | 0.095 | 0 | 1 | 27 |
| rtca | 0.5908 | 0.158728052 | 0.095 | 0 | 1 | 27 |
| rufy2 | 0.5908 | 0.118403561 | 0.095 | 0 | 1 | 27 |
| sbds | 0.5908 | 0.215869233 | 0.095 | 0 | 1 | 27 |
| serac1 | 0.5908 | 0.134289272 | 0.095 | 0 | 1 | 27 |
| shmt1 | 0.5908 | 0.193188649 | 0.095 | 0 | 1 | 27 |
| si:ch211-39i2.2 | 0.5908 | 0.224261021 | 0.095 | 0 | 1 | 27 |
| si:dkey-103i16.6 | 0.5908 | 0.134207063 | 0.095 | 0 | 1 | 27 |
| si:dkey-1h24.6 | 0.5908 | 0.149756912 | 0.095 | 0 | 1 | 27 |
| si:dkey-56d12.4 | 0.5908 | 0.192851404 | 0.095 | 0 | 1 | 27 |
| si:dkey-78l4.14 | 0.5908 | 0.150877976 | 0.095 | 0 | 1 | 27 |
| sirt1 | 0.5908 | 0.193438988 | 0.095 | 0 | 1 | 27 |
| sirt2 | 0.5908 | 0.19664245 | 0.095 | 0 | 1 | 27 |
| slc25a36a | 0.5908 | 0.122959823 | 0.095 | 0 | 1 | 27 |
| smarca2 | 0.5908 | 0.16336141 | 0.095 | 0 | 1 | 27 |
| smg6 | 0.5908 | 0.141603056 | 0.095 | 0 | 1 | 27 |
| sntb2 | 0.5908 | 0.158112236 | 0.095 | 0 | 1 | 27 |
| srbd1 | 0.5908 | 0.145857931 | 0.095 | 0 | 1 | 27 |
| ssu72 | 0.5908 | 0.178709856 | 0.095 | 0 | 1 | 27 |
| stag2a | 0.5908 | 0.134207063 | 0.095 | 0 | 1 | 27 |
| strn3 | 0.5908 | 0.158572401 | 0.095 | 0 | 1 | 27 |
| supt16h | 0.5908 | 0.165359874 | 0.095 | 0 | 1 | 27 |
| sympk | 0.5908 | 0.217563149 | 0.095 | 0 | 1 | 27 |

| | | | | | | |
|---|---|---|---|---|---|---|
| taf6 | 0.5908 | 0.124547917 | 0.095 | 0 | 1 | 27 |
| tbc1d5 | 0.5908 | 0.135974273 | 0.095 | 0 | 1 | 27 |
| tfg | 0.5908 | 0.121244242 | 0.095 | 0 | 1 | 27 |
| tm2d3 | 0.5908 | 0.145304617 | 0.095 | 0 | 1 | 27 |
| tmem203 | 0.5908 | 0.145384403 | 0.095 | 0 | 1 | 27 |
| tnfaip8l2b | 0.5908 | 0.237711234 | 0.095 | 0 | 1 | 27 |
| triobpb | 0.5908 | 0.183754159 | 0.095 | 0 | 1 | 27 |
| trmt2a | 0.5908 | 0.150318911 | 0.095 | 0 | 1 | 27 |
| ube3a | 0.5908 | 0.138299897 | 0.095 | 0 | 1 | 27 |
| ubr4 | 0.5908 | 0.113822965 | 0.095 | 0 | 1 | 27 |
| xpo7 | 0.5908 | 0.1406614 | 0.095 | 0 | 1 | 27 |
| yipf5 | 0.5908 | 0.188730341 | 0.095 | 0 | 1 | 27 |
| zbtb18 | 0.5908 | 0.180193014 | 0.095 | 0 | 1 | 27 |
| zc3h4 | 0.5908 | 0.23541733 | 0.095 | 0 | 1 | 27 |
| zcchc17 | 0.5908 | 0.145586615 | 0.095 | 0 | 1 | 27 |
| zgc:153654 | 0.5908 | 0.164957705 | 0.095 | 0 | 1 | 27 |
| zgc:55733 | 0.5908 | 0.144505239 | 0.095 | 0 | 1 | 27 |
| zgc:77739 | 0.5908 | 0.129054357 | 0.095 | 0 | 1 | 27 |
| znf277 | 0.5908 | 0.126990562 | 0.095 | 0 | 1 | 27 |
| znf362b | 0.5908 | 0.134157423 | 0.095 | 0 | 1 | 27 |
| AK6 | 0.5908 | 0.140807715 | 0.095 | 0 | 1 | 27 |
| CZQB01141835.1 | 0.5908 | 0.173089867 | 0.095 | 0 | 1 | 27 |
| LOC103909811 | 0.5908 | 0.112905387 | 0.095 | 0 | 1 | 27 |
| aamp | 0.5908 | 0.209266715 | 0.095 | 0 | 1 | 27 |
| abcb7 | 0.5908 | 0.170667833 | 0.095 | 0 | 1 | 27 |
| acsl3b | 0.5908 | 0.144089775 | 0.095 | 0 | 1 | 27 |
| ankrd13c | 0.5908 | 0.138436582 | 0.095 | 0 | 1 | 27 |
| arhgap11a | 0.5908 | 0.138976429 | 0.095 | 0 | 1 | 27 |
| arhgap4b | 0.5908 | 0.203228038 | 0.095 | 0 | 1 | 27 |
| arl14ep | 0.5908 | 0.120480875 | 0.095 | 0 | 1 | 27 |
| arl8a | 0.5908 | 0.106660449 | 0.095 | 0 | 1 | 27 |
| atad2 | 0.5908 | 0.136129792 | 0.095 | 0 | 1 | 27 |
| atad2b | 0.5908 | 0.170275846 | 0.095 | 0 | 1 | 27 |
| atg5 | 0.5908 | 0.177200531 | 0.095 | 0 | 1 | 27 |
| atp6v1h | 0.5908 | 0.112781451 | 0.095 | 0 | 1 | 27 |
| bap1 | 0.5908 | 0.176217147 | 0.095 | 0 | 1 | 27 |
| birc6 | 0.5908 | 0.245615785 | 0.095 | 0 | 1 | 27 |
| camk2b1 | 0.5908 | 0.141554186 | 0.095 | 0 | 1 | 27 |
| cbr1l | 0.5908 | 0.106298314 | 0.095 | 0 | 1 | 27 |
| ccdc14 | 0.5908 | 0.123250297 | 0.095 | 0 | 1 | 27 |
| ccne2 | 0.5908 | 0.199329199 | 0.095 | 0 | 1 | 27 |
| ccnl1a | 0.5908 | 0.135143121 | 0.095 | 0 | 1 | 27 |
| cdc45 | 0.5908 | 0.121747129 | 0.095 | 0 | 1 | 27 |
| cdk2 | 0.5908 | 0.115927213 | 0.095 | 0 | 1 | 27 |
| cep170aa | 0.5908 | 0.210870326 | 0.095 | 0 | 1 | 27 |
| chchd7 | 0.5908 | 0.130083906 | 0.095 | 0 | 1 | 27 |
| ciartb | 0.5908 | 0.172899441 | 0.095 | 0 | 1 | 27 |
| cltcb | 0.5908 | 0.133198565 | 0.095 | 0 | 1 | 27 |

| | | | | | | |
|---|---|---|---|---|---|---|
| commd5 | 0.5908 | 0.23404256 | 0.095 | 0 | 1 | 27 |
| copb2 | 0.5908 | 0.180967519 | 0.095 | 0 | 1 | 27 |
| ctbp1 | 0.5908 | 0.134355968 | 0.095 | 0 | 1 | 27 |
| ctdspl2a | 0.5908 | 0.146205172 | 0.095 | 0 | 1 | 27 |
| dclre1a | 0.5908 | 0.108010901 | 0.095 | 0 | 1 | 27 |
| ddx3xb | 0.5908 | 0.14300198 | 0.095 | 0 | 1 | 27 |
| decr1 | 0.5908 | 0.108005837 | 0.095 | 0 | 1 | 27 |
| dnaja2b | 0.5908 | 0.214526629 | 0.095 | 0 | 1 | 27 |
| dok2 | 0.5908 | 0.233664097 | 0.095 | 0 | 1 | 27 |
| dusp23b | 0.5908 | 0.194443313 | 0.095 | 0 | 1 | 27 |
| eftud2 | 0.5908 | 0.117671565 | 0.095 | 0 | 1 | 27 |
| eif1axa | 0.5908 | 0.176448562 | 0.095 | 0 | 1 | 27 |
| eif4g3b | 0.5908 | 0.172179518 | 0.095 | 0 | 1 | 27 |
| emc7 | 0.5908 | 0.129216226 | 0.095 | 0 | 1 | 27 |
| ets1 | 0.5908 | 0.219190429 | 0.095 | 0 | 1 | 27 |
| faah2a | 0.5908 | 0.180622447 | 0.095 | 0 | 1 | 27 |
| fdx1 | 0.5908 | 0.177037951 | 0.095 | 0 | 1 | 27 |
| gemin7 | 0.5908 | 0.131431986 | 0.095 | 0 | 1 | 27 |
| gins1 | 0.5908 | 0.140191229 | 0.095 | 0 | 1 | 27 |
| gnpda2 | 0.5908 | 0.207900235 | 0.095 | 0 | 1 | 27 |
| got2a | 0.5908 | 0.180375948 | 0.095 | 0 | 1 | 27 |
| grcc10 | 0.5908 | 0.169056433 | 0.095 | 0 | 1 | 27 |
| grpel2 | 0.5908 | 0.106882292 | 0.095 | 0 | 1 | 27 |
| hat1 | 0.5908 | 0.110829765 | 0.095 | 0 | 1 | 27 |
| haus1 | 0.5908 | 0.13780084 | 0.095 | 0 | 1 | 27 |
| hnrnpa0a | 0.5908 | 0.185621663 | 0.095 | 0 | 1 | 27 |
| ifi35 | 0.5908 | 0.185065537 | 0.095 | 0 | 1 | 27 |
| ighv5-1 | 0.5908 | 0.35562301 | 0.095 | 0 | 1 | 27 |
| ipmkb | 0.5908 | 0.220023797 | 0.095 | 0 | 1 | 27 |
| klf11a | 0.5908 | 0.175728064 | 0.095 | 0 | 1 | 27 |
| l2hgdh | 0.5908 | 0.117534941 | 0.095 | 0 | 1 | 27 |
| limd2 | 0.5908 | 0.164085297 | 0.095 | 0 | 1 | 27 |
| map2k7 | 0.5908 | 0.18105223 | 0.095 | 0 | 1 | 27 |
| mapk6 | 0.5908 | 0.282356209 | 0.095 | 0 | 1 | 27 |
| matr3l1.1 | 0.5908 | 0.209345629 | 0.095 | 0 | 1 | 27 |
| mettl5 | 0.5908 | 0.186598778 | 0.095 | 0 | 1 | 27 |
| mmab | 0.5908 | 0.178827845 | 0.095 | 0 | 1 | 27 |
| mtg1 | 0.5908 | 0.121050284 | 0.095 | 0 | 1 | 27 |
| myg1 | 0.5908 | 0.134832831 | 0.095 | 0 | 1 | 27 |
| naa20 | 0.5908 | 0.134627146 | 0.095 | 0 | 1 | 27 |
| ndufab1b | 0.5908 | 0.1270108 | 0.095 | 0 | 1 | 27 |
| nktr | 0.5908 | 0.237494998 | 0.095 | 0 | 1 | 27 |
| notch1a | 0.5908 | 0.205558012 | 0.095 | 0 | 1 | 27 |
| nub1 | 0.5908 | 0.198031588 | 0.095 | 0 | 1 | 27 |
| nubpl | 0.5908 | 0.157634463 | 0.095 | 0 | 1 | 27 |
| nup155 | 0.5908 | 0.166176275 | 0.095 | 0 | 1 | 27 |
| pcid2 | 0.5908 | 0.111983147 | 0.095 | 0 | 1 | 27 |
| pex16 | 0.5908 | 0.130255744 | 0.095 | 0 | 1 | 27 |

| | | | | | | |
|---|---|---|---|---|---|---|
| pgam1b | 0.5908 | 0.197692085 | 0.095 | 0 | 1 | 27 |
| pmepa1 | 0.5908 | 0.210870326 | 0.095 | 0 | 1 | 27 |
| pms2 | 0.5908 | 0.117515639 | 0.095 | 0 | 1 | 27 |
| pold3 | 0.5908 | 0.105142866 | 0.095 | 0 | 1 | 27 |
| polr2i | 0.5908 | 0.111988903 | 0.095 | 0 | 1 | 27 |
| ppm1aa | 0.5908 | 0.198319269 | 0.095 | 0 | 1 | 27 |
| ppp1r10 | 0.5908 | 0.102331726 | 0.095 | 0 | 1 | 27 |
| pqlc3 | 0.5908 | 0.162302553 | 0.095 | 0 | 1 | 27 |
| prkag1 | 0.5908 | 0.205648768 | 0.095 | 0 | 1 | 27 |
| psmc3ip | 0.5908 | 0.115033076 | 0.095 | 0 | 1 | 27 |
| psmd12 | 0.5908 | 0.102639592 | 0.095 | 0 | 1 | 27 |
| ptena | 0.5908 | 0.191965101 | 0.095 | 0 | 1 | 27 |
| rabggtb | 0.5908 | 0.140697116 | 0.095 | 0 | 1 | 27 |
| rad50 | 0.5908 | 0.144089775 | 0.095 | 0 | 1 | 27 |
| rhot1a | 0.5908 | 0.129372827 | 0.095 | 0 | 1 | 27 |
| sephs2 | 0.5908 | 0.134457066 | 0.095 | 0 | 1 | 27 |
| setd2 | 0.5908 | 0.105593956 | 0.095 | 0 | 1 | 27 |
| setdb2 | 0.5908 | 0.194227781 | 0.095 | 0 | 1 | 27 |
| si:ch211-166e11.5 | 0.5908 | 0.210870326 | 0.095 | 0 | 1 | 27 |
| si:ch211-189a15.5 | 0.5908 | 0.200912946 | 0.095 | 0 | 1 | 27 |
| si:ch211-251b21.1 | 0.5908 | 0.19895712 | 0.095 | 0 | 1 | 27 |
| si:ch211-284e13.9 | 0.5908 | 0.121747129 | 0.095 | 0 | 1 | 27 |
| si:dkey-230p4.1 | 0.5908 | 0.137956559 | 0.095 | 0 | 1 | 27 |
| si:dkey-82o10.4 | 0.5908 | 0.248992941 | 0.095 | 0 | 1 | 27 |
| smap1 | 0.5908 | 0.139040017 | 0.095 | 0 | 1 | 27 |
| sp100.2 | 0.5908 | 0.144089775 | 0.095 | 0 | 1 | 27 |
| srp54 | 0.5908 | 0.183500928 | 0.095 | 0 | 1 | 27 |
| srp9 | 0.5908 | 0.102551453 | 0.095 | 0 | 1 | 27 |
| syne3 | 0.5908 | 0.121747129 | 0.095 | 0 | 1 | 27 |
| synj1 | 0.5908 | 0.180622447 | 0.095 | 0 | 1 | 27 |
| tmem243a | 0.5908 | 0.222967016 | 0.095 | 0 | 1 | 27 |
| tmx4 | 0.5908 | 0.142595611 | 0.095 | 0 | 1 | 27 |
| tprb | 0.5908 | 0.164502401 | 0.095 | 0 | 1 | 27 |
| trioa | 0.5908 | 0.116408883 | 0.095 | 0 | 1 | 27 |
| tubgcp3 | 0.5908 | 0.151397487 | 0.095 | 0 | 1 | 27 |
| tufm | 0.5908 | 0.224505739 | 0.095 | 0 | 1 | 27 |
| ubr7 | 0.5908 | 0.109376002 | 0.095 | 0 | 1 | 27 |
| uqcrc1 | 0.5908 | 0.128339403 | 0.095 | 0 | 1 | 27 |
| wdr43 | 0.5908 | 0.164447144 | 0.095 | 0 | 1 | 27 |
| wrb | 0.5908 | 0.183603476 | 0.095 | 0 | 1 | 27 |
| zbed4 | 0.5908 | 0.190035657 | 0.095 | 0 | 1 | 27 |
| zgc:136564 | 0.5908 | 0.168217604 | 0.095 | 0 | 1 | 27 |
| zgc:158564 | 0.5908 | 0.190882204 | 0.095 | 0 | 1 | 27 |
| zgc:171506 | 0.5908 | 0.121747129 | 0.095 | 0 | 1 | 27 |
| zgc:171971 | 0.5908 | 0.134046598 | 0.095 | 0 | 1 | 27 |
| zgc:77287 | 0.5908 | 0.140220589 | 0.095 | 0 | 1 | 27 |
| znf1035 | 0.5908 | 0.275331908 | 0.095 | 0 | 1 | 27 |
| cab39l | 0.59391 | -0.44278824 | 0.143 | 0.333 | 1 | 27 |

| | | | | | | |
|---|---|---|---|---|---|---|
| psmc2 | 0.59498 | -0.47805106 | 0.143 | 0.333 | 1 | 27 |
| bpifcl | 0.59641 | -0.26863797 | 0.333 | 0.333 | 1 | 27 |
| psmd1 | 0.59641 | -0.48143599 | 0.143 | 0.333 | 1 | 27 |
| mrpl57 | 0.59955 | -0.47457409 | 0.143 | 0.333 | 1 | 27 |
| ranbp1 | 0.60023 | -0.40826575 | 0.429 | 0.667 | 1 | 27 |
| b2ml | 0.60078 | -0.45261117 | 0.381 | 0.667 | 1 | 27 |
| ucp2 | 0.6043 | 0.229938776 | 0.905 | 0.667 | 1 | 27 |
| rab14 | 0.60522 | -0.42385974 | 0.143 | 0.333 | 1 | 27 |
| dctn5 | 0.60682 | -0.41523415 | 0.143 | 0.333 | 1 | 27 |
| gstr | 0.60954 | -0.4243548 | 0.143 | 0.333 | 1 | 27 |
| COX7A2 (1 of many) | 0.61002 | -0.40425012 | 0.381 | 0.667 | 1 | 27 |
| copz1 | 0.61226 | -0.28026181 | 0.286 | 0.333 | 1 | 27 |
| exosc1 | 0.61294 | -0.44336815 | 0.143 | 0.333 | 1 | 27 |
| dusp2 | 0.61387 | -0.53312873 | 0.381 | 0.667 | 1 | 27 |
| snrpd2 | 0.61423 | 0.483528201 | 0.571 | 0.333 | 1 | 27 |
| crebzf | 0.61484 | -0.43933323 | 0.143 | 0.333 | 1 | 27 |
| creb3l3l | 0.61789 | -0.50733147 | 0.143 | 0.333 | 1 | 27 |
| ap2s1 | 0.61885 | -0.41438363 | 0.143 | 0.333 | 1 | 27 |
| dub | 0.61941 | -0.46527062 | 0.381 | 0.667 | 1 | 27 |
| ube2e1 | 0.61949 | 0.277824568 | 0.619 | 0.333 | 1 | 27 |
| eif4e1c | 0.62024 | -0.48885733 | 0.143 | 0.333 | 1 | 27 |
| gpia | 0.62235 | -0.41621854 | 0.381 | 0.667 | 1 | 27 |
| dpy30 | 0.62322 | -0.37619838 | 0.429 | 0.667 | 1 | 27 |
| ufd1l | 0.62343 | -0.42216107 | 0.143 | 0.333 | 1 | 27 |
| psmc3 | 0.62367 | -0.35824961 | 0.238 | 0.333 | 1 | 27 |
| myd88 | 0.62417 | -0.4394523 | 0.143 | 0.333 | 1 | 27 |
| LOC103909115 | 0.62487 | 0.31398063 | 0.619 | 0.333 | 1 | 27 |
| gng5 | 0.62495 | 0.478833282 | 0.619 | 0.333 | 1 | 27 |
| COA4 | 0.62811 | -0.41998183 | 0.143 | 0.333 | 1 | 27 |
| wdr5 | 0.63121 | -0.41646157 | 0.143 | 0.333 | 1 | 27 |
| wdr44 | 0.63393 | -0.43395516 | 0.143 | 0.333 | 1 | 27 |
| e2f4 | 0.63681 | -0.46657527 | 0.143 | 0.333 | 1 | 27 |
| cfap20 | 0.63782 | -0.40083747 | 0.143 | 0.333 | 1 | 27 |
| atp5fa1 | 0.63858 | 0.411300579 | 0.619 | 0.333 | 1 | 27 |
| atp5md | 0.64119 | -0.36145098 | 0.381 | 0.667 | 1 | 27 |
| mbd3a | 0.64134 | -0.40222822 | 0.381 | 0.667 | 1 | 27 |
| aldoaa | 0.6422 | 0.321267772 | 0.619 | 0.333 | 1 | 27 |
| fosab | 0.64263 | 0.479085315 | 0.619 | 0.333 | 1 | 27 |
| hnrnpub | 0.64446 | -0.34472895 | 0.238 | 0.333 | 1 | 27 |
| igl4v8 | 0.64459 | -0.17792689 | 1 | 1 | 1 | 27 |
| phf11 | 0.64687 | -0.42149522 | 0.143 | 0.333 | 1 | 27 |
| nudt21 | 0.6484 | -0.31889224 | 0.238 | 0.333 | 1 | 27 |
| mphosph8 | 0.64851 | -0.39554363 | 0.381 | 0.667 | 1 | 27 |
| tmem167a | 0.64854 | -0.38915092 | 0.143 | 0.333 | 1 | 27 |
| sumo1 | 0.65008 | -0.20439566 | 0.333 | 0.333 | 1 | 27 |
| tmed10 | 0.6541 | -0.42056172 | 0.19 | 0.333 | 1 | 27 |
| rbm4.3 | 0.65453 | -0.10652462 | 0.476 | 0.667 | 1 | 27 |
| cst14a.2 | 0.65526 | -0.3845071 | 0.381 | 0.667 | 1 | 27 |

| | | | | | | |
|---|---|---|---|---|---|---|
| pfdn5 | 0.65604 | -0.34463343 | 0.238 | 0.333 | 1 | 27 |
| cd74b | 0.65626 | -0.18227137 | 1 | 1 | 1 | 27 |
| cops2 | 0.6563 | -0.35352348 | 0.19 | 0.333 | 1 | 27 |
| dnaja2a | 0.65702 | 0.385388249 | 0.619 | 0.333 | 1 | 27 |
| atp5pf | 0.65709 | -0.40921094 | 0.429 | 0.667 | 1 | 27 |
| ptges3b | 0.65735 | -0.27509054 | 0.286 | 0.333 | 1 | 27 |
| cops9 | 0.65757 | -0.45384633 | 0.143 | 0.333 | 1 | 27 |
| sinhcafl | 0.65763 | -0.19088022 | 0.333 | 0.333 | 1 | 27 |
| eif3d | 0.65802 | 0.399816233 | 0.619 | 0.333 | 1 | 27 |
| atp5f1d | 0.65838 | 0.355703935 | 0.619 | 0.333 | 1 | 27 |
| derl2 | 0.66089 | 0.105510544 | 0.524 | 0.333 | 1 | 27 |
| ube2l3b | 0.66367 | -0.42324277 | 0.19 | 0.333 | 1 | 27 |
| pde4ba | 0.66446 | -0.21515445 | 0.429 | 0.667 | 1 | 27 |
| mff | 0.66458 | -0.39506137 | 0.143 | 0.333 | 1 | 27 |
| wu:fb55g09 | 0.66551 | -0.26911977 | 0.905 | 1 | 1 | 27 |
| eif3c | 0.66599 | -0.20888157 | 0.667 | 0.667 | 1 | 27 |
| kdm6bb | 0.66715 | -0.23354013 | 0.143 | 0.333 | 1 | 27 |
| CU207241.1 | 0.66735 | -0.27473703 | 0.286 | 0.333 | 1 | 27 |
| ndufv2 | 0.66746 | -0.38056279 | 0.143 | 0.333 | 1 | 27 |
| dbnlb | 0.6695 | -0.45637631 | 0.143 | 0.333 | 1 | 27 |
| fli1b | 0.66961 | -0.19131277 | 0.333 | 0.333 | 1 | 27 |
| ncf1 | 0.66984 | 0.296507116 | 0.762 | 0.667 | 1 | 27 |
| mob4 | 0.67079 | -0.41946332 | 0.143 | 0.333 | 1 | 27 |
| plp2 | 0.67123 | -0.36030293 | 0.19 | 0.333 | 1 | 27 |
| myl12.1 | 0.67131 | 0.126666794 | 0.857 | 0.667 | 1 | 27 |
| srsf5a | 0.67137 | 0.403061202 | 0.524 | 0.333 | 1 | 27 |
| srrm4 | 0.67298 | -0.26270326 | 0.143 | 0.333 | 1 | 27 |
| brms1 | 0.67557 | -0.31214966 | 0.238 | 0.333 | 1 | 27 |
| pdha1a | 0.67816 | -0.39097492 | 0.143 | 0.333 | 1 | 27 |
| cul1a | 0.6811 | -0.38792504 | 0.143 | 0.333 | 1 | 27 |
| pdcd10a | 0.68118 | -0.33215903 | 0.238 | 0.333 | 1 | 27 |
| dap1b | 0.68264 | 0.136489596 | 0.857 | 0.667 | 1 | 27 |
| ndufa8 | 0.68534 | -0.37732992 | 0.429 | 0.667 | 1 | 27 |
| bptf | 0.68541 | -0.45538687 | 0.143 | 0.333 | 1 | 27 |
| npc2 | 0.68607 | -0.38023295 | 0.19 | 0.333 | 1 | 27 |
| cnp | 0.68758 | -0.34881127 | 0.19 | 0.333 | 1 | 27 |
| eif2s1a | 0.68771 | -0.39953638 | 0.143 | 0.333 | 1 | 27 |
| rbm4.2 | 0.68884 | -0.3405031 | 0.238 | 0.333 | 1 | 27 |
| arpc3 | 0.68915 | -0.15263961 | 0.667 | 0.667 | 1 | 27 |
| tomm22 | 0.68973 | -0.43538932 | 0.143 | 0.333 | 1 | 27 |
| dnmt3ab | 0.69128 | -0.3267364 | 0.238 | 0.333 | 1 | 27 |
| rab5c | 0.69421 | -0.24900748 | 0.143 | 0.333 | 1 | 27 |
| znf296 | 0.69451 | -0.28252627 | 0.286 | 0.333 | 1 | 27 |
| srrm2 | 0.69644 | -0.34379629 | 0.143 | 0.333 | 1 | 27 |
| thoc2 | 0.69853 | -0.38733162 | 0.19 | 0.333 | 1 | 27 |
| ndufa9a | 0.70053 | -0.37428322 | 0.143 | 0.333 | 1 | 27 |
| cd63 | 0.70101 | -0.38383706 | 0.429 | 0.667 | 1 | 27 |
| spg21 | 0.70106 | -0.40781495 | 0.143 | 0.333 | 1 | 27 |

| | | | | | | |
|---|---|---|---|---|---|---|
| tspan2a | 0.70119 | 0.408890687 | 0.571 | 0.333 | 1 | 27 |
| banf1 | 0.70314 | 0.252699459 | 0.381 | 0.333 | 1 | 27 |
| rpl13a | 0.70331 | -0.13884522 | 1 | 1 | 1 | 27 |
| rad21a | 0.70483 | 0.426558551 | 0.571 | 0.333 | 1 | 27 |
| MCMDC2 | 0.70581 | -0.38158506 | 0.143 | 0.333 | 1 | 27 |
| prkar2aa | 0.70605 | -0.37367373 | 0.143 | 0.333 | 1 | 27 |
| atp1b1a | 0.70629 | -0.37454741 | 0.429 | 0.667 | 1 | 27 |
| vamp8 | 0.70635 | -0.3053964 | 0.238 | 0.333 | 1 | 27 |
| ppib | 0.70692 | 0.463679558 | 0.524 | 0.333 | 1 | 27 |
| snrpf | 0.70715 | 0.35206788 | 0.476 | 0.333 | 1 | 27 |
| bccip | 0.7082 | -0.34954711 | 0.143 | 0.333 | 1 | 27 |
| rnaseh2b | 0.70823 | -0.38663877 | 0.143 | 0.333 | 1 | 27 |
| naa15a | 0.7083 | -0.36022969 | 0.143 | 0.333 | 1 | 27 |
| arhgdia | 0.70941 | -0.35889445 | 0.476 | 0.667 | 1 | 27 |
| srsf7a | 0.70959 | -0.26495911 | 0.571 | 0.667 | 1 | 27 |
| ola1 | 0.70964 | 0.43063634 | 0.571 | 0.333 | 1 | 27 |
| cox6a1 | 0.71201 | 0.129936775 | 0.524 | 0.333 | 1 | 27 |
| anln | 0.7124 | -0.37573418 | 0.143 | 0.333 | 1 | 27 |
| uqcrfs1 | 0.71251 | -0.24910926 | 0.286 | 0.333 | 1 | 27 |
| seta | 0.71283 | 0.23223476 | 0.714 | 0.667 | 1 | 27 |
| arih2 | 0.71294 | -0.36642614 | 0.143 | 0.333 | 1 | 27 |
| znhit1 | 0.71297 | -0.15005322 | 0.143 | 0.333 | 1 | 27 |
| LOC110437938 | 0.71415 | -0.39949663 | 0.143 | 0.333 | 1 | 27 |
| ndufa4l | 0.71473 | 0.38113219 | 0.571 | 0.333 | 1 | 27 |
| tmem59 | 0.71535 | -0.39096218 | 0.143 | 0.333 | 1 | 27 |
| usp1 | 0.71549 | -0.36591014 | 0.143 | 0.333 | 1 | 27 |
| psmb6 | 0.71702 | 0.38196028 | 0.571 | 0.333 | 1 | 27 |
| hnrpkl | 0.71881 | -0.35498132 | 0.19 | 0.333 | 1 | 27 |
| rhoh | 0.71973 | -0.42664545 | 0.143 | 0.333 | 1 | 27 |
| psme4b | 0.71988 | -0.37337821 | 0.143 | 0.333 | 1 | 27 |
| tcf12 | 0.72022 | -0.40721743 | 0.143 | 0.333 | 1 | 27 |
| mtap | 0.721 | -0.36971671 | 0.19 | 0.333 | 1 | 27 |
| rack1 | 0.72146 | 0.100958559 | 1 | 1 | 1 | 27 |
| jpt2 | 0.72239 | -0.25565204 | 0.571 | 0.667 | 1 | 27 |
| myo1f | 0.72382 | -0.42166824 | 0.143 | 0.333 | 1 | 27 |
| zcrb1 | 0.72524 | -0.32290057 | 0.143 | 0.333 | 1 | 27 |
| snrpb | 0.72648 | 0.368158292 | 0.571 | 0.333 | 1 | 27 |
| dnajb11 | 0.72761 | -0.28806742 | 0.143 | 0.333 | 1 | 27 |
| XLOC-039757 | 0.72789 | -0.49146304 | 0.143 | 0.333 | 1 | 27 |
| ankrd52a | 0.72811 | -0.38365049 | 0.143 | 0.333 | 1 | 27 |
| uprt | 0.72879 | -0.36335441 | 0.143 | 0.333 | 1 | 27 |
| prelid1a | 0.72957 | -0.32752278 | 0.19 | 0.333 | 1 | 27 |
| calm3b | 0.72996 | -0.1392844 | 0.476 | 0.667 | 1 | 27 |
| pomp | 0.73263 | 0.38161362 | 0.571 | 0.333 | 1 | 27 |
| slu7 | 0.73371 | -0.35418219 | 0.143 | 0.333 | 1 | 27 |
| gnai2b | 0.73448 | -0.36086317 | 0.143 | 0.333 | 1 | 27 |
| zgc:162944 | 0.73489 | 0.308068228 | 0.571 | 0.333 | 1 | 27 |
| dhx15 | 0.73565 | -0.34972065 | 0.143 | 0.333 | 1 | 27 |

| | | | | | | |
|---|---|---|---|---|---|---|
| tacc3 | 0.73594 | -0.23176279 | 0.143 | 0.333 | 1 | 27 |
| cptp | 0.73678 | -0.35291958 | 0.143 | 0.333 | 1 | 27 |
| tma7 | 0.73704 | 0.396818704 | 0.476 | 0.333 | 1 | 27 |
| gpr183a | 0.73724 | -0.31304943 | 0.238 | 0.333 | 1 | 27 |
| gm2a | 0.73756 | -0.36064357 | 0.429 | 0.667 | 1 | 27 |
| cited4b | 0.73836 | 0.316699357 | 0.571 | 0.333 | 1 | 27 |
| igl3v5 | 0.73848 | -0.18175624 | 0.952 | 1 | 1 | 27 |
| cstf2 | 0.73885 | -0.37141806 | 0.143 | 0.333 | 1 | 27 |
| si:ch211-195b13.1 | 0.73961 | -0.2790404 | 0.143 | 0.333 | 1 | 27 |
| xirp2a | 0.74034 | -0.24163878 | 0.143 | 0.333 | 1 | 27 |
| tpp1 | 0.7404 | -0.35819665 | 0.143 | 0.333 | 1 | 27 |
| rsrp1 | 0.74078 | -0.27817181 | 0.143 | 0.333 | 1 | 27 |
| u2af2b | 0.74092 | -0.25720479 | 0.286 | 0.333 | 1 | 27 |
| ankrd12 | 0.74126 | -0.40206243 | 0.19 | 0.333 | 1 | 27 |
| zgc:103700 | 0.74221 | 0.369059194 | 0.571 | 0.333 | 1 | 27 |
| prdx5 | 0.74241 | 0.274650482 | 0.571 | 0.333 | 1 | 27 |
| akap8l | 0.74245 | -0.34500752 | 0.19 | 0.333 | 1 | 27 |
| si:ch211-214j24.10 | 0.74375 | -0.25859135 | 0.286 | 0.333 | 1 | 27 |
| ccr9b | 0.74429 | -0.40814756 | 0.143 | 0.333 | 1 | 27 |
| unc93b1 | 0.74484 | -0.35409067 | 0.143 | 0.333 | 1 | 27 |
| ppp3ccb | 0.74498 | -0.39955298 | 0.19 | 0.333 | 1 | 27 |
| si:ch211-196l7.4 | 0.74525 | -0.30028173 | 0.524 | 0.667 | 1 | 27 |
| atf7ip | 0.74559 | -0.11495362 | 0.19 | 0.333 | 1 | 27 |
| hif1al | 0.74586 | -0.2806779 | 0.143 | 0.333 | 1 | 27 |
| gskip | 0.74631 | -0.36081777 | 0.143 | 0.333 | 1 | 27 |
| atp5if1a | 0.74679 | -0.26570059 | 0.429 | 0.667 | 1 | 27 |
| ube2d2 | 0.74746 | -0.28854977 | 0.429 | 0.667 | 1 | 27 |
| dpm2 | 0.74759 | -0.30966592 | 0.143 | 0.333 | 1 | 27 |
| ppp1caa | 0.74843 | -0.31570714 | 0.429 | 0.667 | 1 | 27 |
| sec11a | 0.74877 | 0.317464734 | 0.571 | 0.333 | 1 | 27 |
| prdx4 | 0.74884 | -0.20625511 | 0.143 | 0.333 | 1 | 27 |
| cab39l1 | 0.74905 | -0.34490853 | 0.19 | 0.333 | 1 | 27 |
| edf1 | 0.74906 | -0.28819798 | 0.429 | 0.667 | 1 | 27 |
| acin1a | 0.74908 | 0.351067246 | 0.571 | 0.333 | 1 | 27 |
| pik3r5 | 0.74914 | -0.39888628 | 0.143 | 0.333 | 1 | 27 |
| ndufb9 | 0.74915 | -0.19301792 | 0.333 | 0.333 | 1 | 27 |
| otud5b | 0.74949 | -0.31741038 | 0.19 | 0.333 | 1 | 27 |
| ciapin1 | 0.75026 | -0.27570604 | 0.143 | 0.333 | 1 | 27 |
| oaz1a | 0.75133 | -0.1067716 | 0.714 | 0.667 | 1 | 27 |
| fuca2 | 0.75245 | -0.28992372 | 0.143 | 0.333 | 1 | 27 |
| psenen | 0.75282 | -0.26011184 | 0.286 | 0.333 | 1 | 27 |
| chmp2bb | 0.75285 | -0.35565204 | 0.143 | 0.333 | 1 | 27 |
| nudcd2 | 0.75318 | -0.31715127 | 0.143 | 0.333 | 1 | 27 |
| pkn1b | 0.75365 | -0.36236438 | 0.143 | 0.333 | 1 | 27 |
| ier5 | 0.75511 | 0.181053923 | 0.524 | 0.333 | 1 | 27 |
| srp72 | 0.75521 | -0.35282467 | 0.143 | 0.333 | 1 | 27 |
| arf2b | 0.75564 | -0.34543842 | 0.143 | 0.333 | 1 | 27 |
| serinc1 | 0.75583 | -0.34985042 | 0.143 | 0.333 | 1 | 27 |

| | | | | | | |
|---|---|---|---|---|---|---|
| kpna2 | 0.75708 | -0.26440146 | 0.143 | 0.333 | 1 | 27 |
| rasal3 | 0.75725 | -0.34359271 | 0.19 | 0.333 | 1 | 27 |
| si:ch73-248e21.7 | 0.75751 | -0.35519132 | 0.143 | 0.333 | 1 | 27 |
| cd40 | 0.75766 | -0.35484894 | 0.143 | 0.333 | 1 | 27 |
| arhgdig | 0.75781 | -0.12906589 | 0.857 | 1 | 1 | 27 |
| ufc1 | 0.75792 | -0.31617955 | 0.143 | 0.333 | 1 | 27 |
| atp5mc3b | 0.75828 | 0.196635383 | 0.857 | 0.667 | 1 | 27 |
| nfkb2 | 0.75837 | -0.18094977 | 0.333 | 0.333 | 1 | 27 |
| idh1 | 0.75915 | -0.32133724 | 0.143 | 0.333 | 1 | 27 |
| iscub | 0.7593 | -0.31710388 | 0.143 | 0.333 | 1 | 27 |
| snrpd1 | 0.75983 | 0.284467753 | 0.429 | 0.333 | 1 | 27 |
| CU207301.2 | 0.75992 | -0.33567662 | 0.143 | 0.333 | 1 | 27 |
| pim1 | 0.76022 | -0.31773708 | 0.143 | 0.333 | 1 | 27 |
| clgn | 0.76029 | -0.27217154 | 0.143 | 0.333 | 1 | 27 |
| cdk11b | 0.7603 | -0.30015531 | 0.143 | 0.333 | 1 | 27 |
| smad3a | 0.76035 | -0.34155688 | 0.143 | 0.333 | 1 | 27 |
| sbno1 | 0.76037 | -0.31813518 | 0.143 | 0.333 | 1 | 27 |
| nfkbiab | 0.7604 | 0.233047426 | 0.857 | 0.667 | 1 | 27 |
| SIPA1 | 0.76041 | -0.32359828 | 0.143 | 0.333 | 1 | 27 |
| mrpl20 | 0.76057 | -0.30660425 | 0.143 | 0.333 | 1 | 27 |
| btg1 | 0.76059 | 0.231368061 | 0.857 | 0.667 | 1 | 27 |
| sftpbb | 0.76091 | 0.187858496 | 0.524 | 0.333 | 1 | 27 |
| blmh | 0.76149 | -0.32630031 | 0.19 | 0.333 | 1 | 27 |
| si:dkey-112e7.2 | 0.76217 | 0.164790351 | 0.524 | 0.333 | 1 | 27 |
| rpl27a | 0.76271 | -0.17343159 | 0.857 | 1 | 1 | 27 |
| wdr83os | 0.76392 | -0.37966389 | 0.19 | 0.333 | 1 | 27 |
| XLOC-025423 | 0.76693 | 0.391784225 | 0.381 | 0.333 | 1 | 27 |
| tardbpl | 0.76869 | -0.28488533 | 0.238 | 0.333 | 1 | 27 |
| celf2 | 0.76937 | -0.129864 | 0.857 | 1 | 1 | 27 |
| grasp | 0.77139 | -0.29567144 | 0.238 | 0.333 | 1 | 27 |
| ezrb | 0.7724 | -0.26761635 | 0.238 | 0.333 | 1 | 27 |
| fam107b | 0.77377 | -0.20578549 | 0.333 | 0.333 | 1 | 27 |
| rab5if | 0.77444 | -0.33078789 | 0.19 | 0.333 | 1 | 27 |
| zgc:194312 | 0.77573 | -0.34149461 | 0.19 | 0.333 | 1 | 27 |
| eif3s6ip | 0.77814 | 0.16777332 | 0.524 | 0.333 | 1 | 27 |
| naa10 | 0.77994 | -0.32025868 | 0.19 | 0.333 | 1 | 27 |
| fam43a | 0.7803 | -0.35976003 | 0.19 | 0.333 | 1 | 27 |
| ube2d1b | 0.78416 | -0.346404 | 0.19 | 0.333 | 1 | 27 |
| ldhba | 0.78509 | -0.19556721 | 0.476 | 0.667 | 1 | 27 |
| ssr4 | 0.78572 | 0.382768731 | 0.524 | 0.333 | 1 | 27 |
| gpx4b | 0.78626 | -0.28190471 | 0.238 | 0.333 | 1 | 27 |
| eif3ba | 0.78686 | 0.119024585 | 0.476 | 0.333 | 1 | 27 |
| spcs3 | 0.78957 | 0.31051468 | 0.429 | 0.333 | 1 | 27 |
| psma4 | 0.79209 | -0.17096639 | 0.333 | 0.333 | 1 | 27 |
| GCA | 0.79301 | -0.3021358 | 0.238 | 0.333 | 1 | 27 |
| si:dkey-180p18.9 | 0.7933 | -0.1536491 | 0.19 | 0.333 | 1 | 27 |
| il10ra | 0.7936 | -0.32983198 | 0.19 | 0.333 | 1 | 27 |
| plcg2 | 0.79428 | -0.22536822 | 0.286 | 0.333 | 1 | 27 |

| | | | | | | |
|---|---|---|---|---|---|---|
| snrpd3l | 0.79431 | -0.21206162 | 0.286 | 0.333 | 1 | 27 |
| si:ch211-235e9.6 | 0.79452 | -0.25691136 | 0.238 | 0.333 | 1 | 27 |
| srsf11 | 0.79488 | 0.187957205 | 0.524 | 0.333 | 1 | 27 |
| pcf11 | 0.79667 | -0.33904685 | 0.19 | 0.333 | 1 | 27 |
| tnip2 | 0.79796 | -0.32267463 | 0.19 | 0.333 | 1 | 27 |
| hnrnpa0l | 0.79858 | 0.265317001 | 0.81 | 0.667 | 1 | 27 |
| si:busm1-266f07.2 | 0.79878 | -0.18563384 | 0.905 | 1 | 1 | 27 |
| irf2a | 0.80046 | -0.3081102 | 0.19 | 0.333 | 1 | 27 |
| txnipa | 0.80112 | -0.29302678 | 0.476 | 0.667 | 1 | 27 |
| pik3r3a | 0.80355 | -0.28629736 | 0.19 | 0.333 | 1 | 27 |
| mrpl30 | 0.80424 | -0.21943248 | 0.286 | 0.333 | 1 | 27 |
| calr | 0.80597 | -0.23158204 | 0.286 | 0.333 | 1 | 27 |
| psma8 | 0.80615 | 0.217043909 | 0.524 | 0.333 | 1 | 27 |
| ISCU (1 of many) | 0.80705 | -0.27126365 | 0.238 | 0.333 | 1 | 27 |
| sept12 | 0.80759 | -0.2784578 | 0.238 | 0.333 | 1 | 27 |
| srsf2a | 0.81018 | -0.27737176 | 0.476 | 0.667 | 1 | 27 |
| anapc13 | 0.81337 | -0.30274727 | 0.19 | 0.333 | 1 | 27 |
| lsm5 | 0.81427 | -0.27223131 | 0.476 | 0.667 | 1 | 27 |
| rab7a | 0.8149 | -0.32238117 | 0.19 | 0.333 | 1 | 27 |
| bin2b | 0.81558 | -0.29744806 | 0.19 | 0.333 | 1 | 27 |
| calm2b | 0.81564 | -0.24818473 | 0.476 | 0.667 | 1 | 27 |
| dad1 | 0.8185 | -0.23917268 | 0.286 | 0.333 | 1 | 27 |
| rhogb | 0.82041 | 0.316896532 | 0.524 | 0.333 | 1 | 27 |
| ndufb4 | 0.82155 | 0.194888786 | 0.524 | 0.333 | 1 | 27 |
| ubc | 0.82161 | -0.15844637 | 0.952 | 1 | 1 | 27 |
| si:ch73-7i4.3 | 0.82207 | -0.35082089 | 0.19 | 0.333 | 1 | 27 |
| tcp1 | 0.82315 | 0.199989094 | 0.524 | 0.333 | 1 | 27 |
| anp32a | 0.82403 | -0.28551239 | 0.476 | 0.667 | 1 | 27 |
| arf6a | 0.82541 | -0.2548611 | 0.476 | 0.667 | 1 | 27 |
| ube2v2 | 0.82706 | -0.26385823 | 0.476 | 0.667 | 1 | 27 |
| si:ch211-222l21.1 | 0.82806 | 0.34813234 | 0.524 | 0.333 | 1 | 27 |
| mzt2b | 0.82898 | -0.29761675 | 0.19 | 0.333 | 1 | 27 |
| fam53b | 0.82965 | -0.2477207 | 0.476 | 0.667 | 1 | 27 |
| u2af1 | 0.82981 | 0.239305726 | 0.524 | 0.333 | 1 | 27 |
| si:dkey-42i9.4 | 0.83005 | 0.259038748 | 0.524 | 0.333 | 1 | 27 |
| syncrip | 0.83018 | 0.27944208 | 0.524 | 0.333 | 1 | 27 |
| idh2 | 0.8304 | 0.228436558 | 0.524 | 0.333 | 1 | 27 |
| smdt1b | 0.83049 | -0.12866356 | 0.333 | 0.333 | 1 | 27 |
| erh | 0.83064 | 0.280062342 | 0.524 | 0.333 | 1 | 27 |
| eif3g | 0.83066 | 0.263109047 | 0.524 | 0.333 | 1 | 27 |
| si:ch211-198c19.1 | 0.83104 | -0.21962052 | 0.286 | 0.333 | 1 | 27 |
| ugp2b | 0.83148 | -0.29021545 | 0.19 | 0.333 | 1 | 27 |
| calr3b | 0.83167 | -0.2859074 | 0.19 | 0.333 | 1 | 27 |
| tmem14ca | 0.83378 | -0.19695789 | 0.286 | 0.333 | 1 | 27 |
| dck | 0.83453 | 0.116340263 | 0.333 | 0.333 | 1 | 27 |
| csk | 0.83481 | -0.26403077 | 0.238 | 0.333 | 1 | 27 |
| psmd3 | 0.83548 | -0.24396015 | 0.238 | 0.333 | 1 | 27 |
| pin1 | 0.83633 | -0.20458393 | 0.286 | 0.333 | 1 | 27 |

| | | | | | | |
|---|---|---|---|---|---|---|
| zc4h2 | 0.84163 | -0.27883996 | 0.19 | 0.333 | 1 | 27 |
| u2surp | 0.84176 | -0.24151649 | 0.238 | 0.333 | 1 | 27 |
| imp4 | 0.84191 | -0.21035154 | 0.286 | 0.333 | 1 | 27 |
| CT573231.1 | 0.8421 | -0.1608999 | 0.667 | 0.667 | 1 | 27 |
| hadhb | 0.84439 | -0.26633204 | 0.19 | 0.333 | 1 | 27 |
| ip6k2a | 0.84491 | -0.27097172 | 0.19 | 0.333 | 1 | 27 |
| si:dkeyp-113d7.1 | 0.84623 | -0.26735265 | 0.19 | 0.333 | 1 | 27 |
| DDX17 | 0.84851 | -0.30712576 | 0.19 | 0.333 | 1 | 27 |
| lypla2 | 0.84979 | -0.29021121 | 0.19 | 0.333 | 1 | 27 |
| calm2a | 0.85105 | 0.144736979 | 0.476 | 0.333 | 1 | 27 |
| ndufa11 | 0.85374 | -0.23241172 | 0.238 | 0.333 | 1 | 27 |
| zgc:85858 | 0.85399 | -0.26591495 | 0.19 | 0.333 | 1 | 27 |
| rnaseh2a | 0.85438 | -0.25596198 | 0.19 | 0.333 | 1 | 27 |
| ndufb7 | 0.85484 | 0.276408927 | 0.476 | 0.333 | 1 | 27 |
| clta | 0.85516 | -0.25453794 | 0.19 | 0.333 | 1 | 27 |
| smarcb1a | 0.85606 | -0.25757705 | 0.19 | 0.333 | 1 | 27 |
| snrpa1 | 0.85611 | -0.26533215 | 0.19 | 0.333 | 1 | 27 |
| rnf216 | 0.86014 | -0.23824634 | 0.19 | 0.333 | 1 | 27 |
| hmgb1a | 0.8602 | 0.259556246 | 0.476 | 0.333 | 1 | 27 |
| mrpl36 | 0.86031 | -0.19133679 | 0.19 | 0.333 | 1 | 27 |
| gadd45ga | 0.86172 | -0.25602636 | 0.19 | 0.333 | 1 | 27 |
| ctsba | 0.86247 | -0.23464282 | 0.238 | 0.333 | 1 | 27 |
| sf3a2 | 0.86258 | -0.23504222 | 0.19 | 0.333 | 1 | 27 |
| ndufb10 | 0.86279 | -0.17922846 | 0.286 | 0.333 | 1 | 27 |
| sh3d21 | 0.86496 | -0.25230605 | 0.19 | 0.333 | 1 | 27 |
| ccr9a | 0.86499 | -0.11150209 | 0.905 | 1 | 1 | 27 |
| dhfr | 0.86509 | -0.22742708 | 0.19 | 0.333 | 1 | 27 |
| ptbp1a | 0.86527 | -0.23121473 | 0.238 | 0.333 | 1 | 27 |
| itm2bb | 0.86546 | -0.12788223 | 0.667 | 0.667 | 1 | 27 |
| rpl22l1 | 0.86603 | 0.143526555 | 0.81 | 0.667 | 1 | 27 |
| ada | 0.86631 | -0.24781354 | 0.19 | 0.333 | 1 | 27 |
| MRPS12 | 0.86641 | -0.24185057 | 0.19 | 0.333 | 1 | 27 |
| calm1a | 0.8675 | -0.20040393 | 0.19 | 0.333 | 1 | 27 |
| nfyba | 0.86795 | 0.271295046 | 0.476 | 0.333 | 1 | 27 |
| zgc:194275 | 0.86805 | -0.1400184 | 0.952 | 1 | 1 | 27 |
| yy1a | 0.86807 | -0.24977336 | 0.238 | 0.333 | 1 | 27 |
| eif3i | 0.86843 | -0.17023027 | 0.619 | 0.667 | 1 | 27 |
| LOC103909972 | 0.86907 | -0.2273672 | 0.19 | 0.333 | 1 | 27 |
| cib1 | 0.86924 | -0.23860184 | 0.19 | 0.333 | 1 | 27 |
| foxp1b | 0.86958 | -0.23199313 | 0.19 | 0.333 | 1 | 27 |
| ciarta | 0.86961 | -0.21556977 | 0.19 | 0.333 | 1 | 27 |
| emc2 | 0.86976 | -0.2243889 | 0.19 | 0.333 | 1 | 27 |
| top1l | 0.87014 | -0.23871987 | 0.19 | 0.333 | 1 | 27 |
| hsf2 | 0.87026 | -0.22278983 | 0.19 | 0.333 | 1 | 27 |
| rnaset2 | 0.87027 | -0.23014951 | 0.19 | 0.333 | 1 | 27 |
| serinc2 | 0.87028 | -0.22320614 | 0.19 | 0.333 | 1 | 27 |
| antxr2a | 0.87245 | -0.14214964 | 0.333 | 0.333 | 1 | 27 |
| zgc:64065 | 0.87653 | -0.25688669 | 0.238 | 0.333 | 1 | 27 |

| | | | | | | |
|---|---|---|---|---|---|---|
| si:ch1073-325m22.2 | 0.87761 | 0.113365613 | 0.476 | 0.333 | 1 | 27 |
| tob1a | 0.87943 | 0.216619101 | 0.381 | 0.333 | 1 | 27 |
| selenof | 0.88005 | 0.242292658 | 0.476 | 0.333 | 1 | 27 |
| sept7a | 0.88052 | -0.2477559 | 0.238 | 0.333 | 1 | 27 |
| ppiab | 0.88162 | 0.102748849 | 1 | 1 | 1 | 27 |
| tuba8l | 0.88301 | 0.256930973 | 0.381 | 0.333 | 1 | 27 |
| hnrnpa1b | 0.88449 | 0.175654803 | 0.476 | 0.333 | 1 | 27 |
| clic1 | 0.88659 | -0.2100834 | 0.524 | 0.667 | 1 | 27 |
| cct4 | 0.88758 | 0.229348921 | 0.476 | 0.333 | 1 | 27 |
| rcc2 | 0.89064 | 0.222221816 | 0.476 | 0.333 | 1 | 27 |
| rpn1 | 0.89186 | -0.16483193 | 0.286 | 0.333 | 1 | 27 |
| ifrd1 | 0.89302 | -0.18882654 | 0.286 | 0.333 | 1 | 27 |
| h2afva | 0.89363 | 0.166397172 | 0.381 | 0.333 | 1 | 27 |
| arhgap15 | 0.8955 | -0.19348635 | 0.571 | 0.667 | 1 | 27 |
| XLOC-042596 | 0.89572 | 0.185112636 | 0.476 | 0.333 | 1 | 27 |
| eno3 | 0.89615 | -0.18457461 | 0.524 | 0.667 | 1 | 27 |
| chmp2a | 0.89695 | 0.219980461 | 0.476 | 0.333 | 1 | 27 |
| anp32b | 0.89787 | 0.119133731 | 0.333 | 0.333 | 1 | 27 |
| magt1 | 0.89846 | 0.214566845 | 0.476 | 0.333 | 1 | 27 |
| sdcbp2 | 0.89879 | 0.192956626 | 0.476 | 0.333 | 1 | 27 |
| pycard | 0.89962 | -0.1768617 | 0.524 | 0.667 | 1 | 27 |
| atp5pb | 0.89965 | -0.15932969 | 0.524 | 0.667 | 1 | 27 |
| denr | 0.90023 | -0.2027888 | 0.238 | 0.333 | 1 | 27 |
| capza1b | 0.9005 | -0.18154071 | 0.571 | 0.667 | 1 | 27 |
| cytip | 0.90155 | -0.19459376 | 0.286 | 0.333 | 1 | 27 |
| ndufs3 | 0.90177 | -0.20311417 | 0.238 | 0.333 | 1 | 27 |
| cox17 | 0.90399 | -0.21563394 | 0.238 | 0.333 | 1 | 27 |
| rnf10 | 0.90697 | -0.17887129 | 0.286 | 0.333 | 1 | 27 |
| sri | 0.90754 | -0.17225093 | 0.571 | 0.667 | 1 | 27 |
| selenok | 0.90806 | -0.22948066 | 0.238 | 0.333 | 1 | 27 |
| mmgt1 | 0.90826 | -0.2135724 | 0.238 | 0.333 | 1 | 27 |
| atp5po | 0.91004 | -0.18531279 | 0.571 | 0.667 | 1 | 27 |
| zgc:195282 | 0.91151 | -0.21597876 | 0.238 | 0.333 | 1 | 27 |
| polr2d | 0.91379 | -0.19574909 | 0.238 | 0.333 | 1 | 27 |
| crlf3 | 0.9169 | -0.20116245 | 0.238 | 0.333 | 1 | 27 |
| snrpe | 0.91723 | 0.188705893 | 0.429 | 0.333 | 1 | 27 |
| ccdc124 | 0.91753 | -0.19570581 | 0.238 | 0.333 | 1 | 27 |
| psma1 | 0.91769 | -0.16410393 | 0.571 | 0.667 | 1 | 27 |
| mrps33 | 0.91851 | -0.19273179 | 0.238 | 0.333 | 1 | 27 |
| ddx5 | 0.91908 | -0.10355419 | 0.238 | 0.333 | 1 | 27 |
| lrrc59 | 0.91989 | -0.22356771 | 0.238 | 0.333 | 1 | 27 |
| ddx39ab | 0.92257 | 0.199700668 | 0.429 | 0.333 | 1 | 27 |
| LOC103908723 | 0.92525 | -0.12585949 | 0.952 | 1 | 1 | 27 |
| tpm1 | 0.92559 | -0.15158139 | 0.286 | 0.333 | 1 | 27 |
| rsl1d1 | 0.92593 | -0.20323174 | 0.238 | 0.333 | 1 | 27 |
| eif4ebp1 | 0.92815 | -0.19221469 | 0.238 | 0.333 | 1 | 27 |
| anxa11b | 0.92818 | -0.15760384 | 0.571 | 0.667 | 1 | 27 |
| serinc5 | 0.93097 | -0.14684422 | 0.286 | 0.333 | 1 | 27 |

| | | | | | | |
|---|---|---|---|---|---|---|
| echs1 | 0.9313 | -0.14920177 | 0.286 | 0.333 | 1 | 27 |
| slirp | 0.93202 | -0.17006196 | 0.238 | 0.333 | 1 | 27 |
| tra2a | 0.9322 | -0.13954417 | 0.238 | 0.333 | 1 | 27 |
| prpf39 | 0.93394 | -0.17154342 | 0.238 | 0.333 | 1 | 27 |
| rab1ab | 0.93525 | -0.14182194 | 0.286 | 0.333 | 1 | 27 |
| tnip1 | 0.93696 | -0.17999181 | 0.238 | 0.333 | 1 | 27 |
| yrk | 0.93856 | -0.15820849 | 0.286 | 0.333 | 1 | 27 |
| anapc16 | 0.93923 | -0.12063381 | 0.238 | 0.333 | 1 | 27 |
| znf704 | 0.93966 | -0.12817228 | 0.238 | 0.333 | 1 | 27 |
| elocb | 0.93971 | 0.1683539 | 0.429 | 0.333 | 1 | 27 |
| tspan36 | 0.93976 | -0.16615589 | 0.238 | 0.333 | 1 | 27 |
| ptbp3 | 0.94 | -0.14327329 | 0.286 | 0.333 | 1 | 27 |
| cct2 | 0.94033 | 0.166724158 | 0.429 | 0.333 | 1 | 27 |
| chrac1 | 0.94205 | -0.13975973 | 0.238 | 0.333 | 1 | 27 |
| vamp3 | 0.94285 | -0.13909418 | 0.286 | 0.333 | 1 | 27 |
| pfdn2 | 0.94344 | -0.1327551 | 0.238 | 0.333 | 1 | 27 |
| si:ch73-376l24.3 | 0.94351 | -0.15572966 | 0.238 | 0.333 | 1 | 27 |
| cct7 | 0.94352 | -0.14823839 | 0.571 | 0.667 | 1 | 27 |
| ube2w | 0.94385 | -0.146355 | 0.238 | 0.333 | 1 | 27 |
| ube2l3a | 0.94427 | -0.14177595 | 0.238 | 0.333 | 1 | 27 |
| actr2a | 0.94556 | -0.15289828 | 0.286 | 0.333 | 1 | 27 |
| abracl | 0.94655 | 0.101784256 | 0.429 | 0.333 | 1 | 27 |
| eif2s1b | 0.94841 | -0.12995425 | 0.571 | 0.667 | 1 | 27 |
| atp5pd | 0.94861 | -0.14092673 | 0.571 | 0.667 | 1 | 27 |
| ywhaqb | 0.94931 | 0.120478195 | 0.429 | 0.333 | 1 | 27 |
| srsf3b | 0.95136 | 0.155072035 | 0.429 | 0.333 | 1 | 27 |
| si:ch211-119e14.1 | 0.95185 | 0.120796464 | 0.714 | 0.667 | 1 | 27 |
| prdx2 | 0.95199 | 0.138339044 | 0.429 | 0.333 | 1 | 27 |
| ctss2.2 | 0.95308 | -0.13355265 | 0.286 | 0.333 | 1 | 27 |
| atp5mc1 | 0.95329 | 0.110690659 | 0.429 | 0.333 | 1 | 27 |
| mhc2dab | 0.95345 | -0.14031814 | 0.286 | 0.333 | 1 | 27 |
| pa2g4b | 0.95412 | -0.12791191 | 0.286 | 0.333 | 1 | 27 |
| serf2 | 0.95861 | -0.11268432 | 0.619 | 0.667 | 1 | 27 |
| sesn1 | 0.9594 | -0.12945174 | 0.619 | 0.667 | 1 | 27 |
| mtpn | 0.95947 | -0.12091404 | 0.286 | 0.333 | 1 | 27 |
| vdac2 | 0.96657 | -0.11081036 | 0.619 | 0.667 | 1 | 27 |
| atp5if1b | 0.96891 | -0.11280988 | 0.286 | 0.333 | 1 | 27 |
| tpm3 | 0.97219 | 0.111678715 | 0.381 | 0.333 | 1 | 27 |
| tagapb | 0.97679 | 0.106380991 | 0.381 | 0.333 | 1 | 27 |
| pim2 | 0.98047 | 0.100725977 | 0.381 | 0.333 | 1 | 27 |
