## Supplementary material for "Direct Male Development in Chromosomally ZZ Zebrafish": SuppTable S4 30dpf Cell Counts per Cluster

| Cluster | ZW cell number | ZZ cell number |
|---|---|---|
| 0 | 845 | 0 |
| 1 | 770 | 10 |
| 2 | 644 | 0 |
| 3 | 550 | 0 |
| 4 | 454 | 0 |
| 5 | 360 | 52 |
| 6 | 284 | 99 |
| 7 | 151 | 230 |
| 8 | 265 | 60 |
| 9 | 33 | 255 |
| 10 | 174 | 113 |
| 11 | 17 | 269 |
| 12 | 110 | 165 |
| 13 | 263 | 0 |
| 14 | 168 | 86 |
| 15 | 244 | 1 |
| 16 | 155 | 76 |
| 17 | 162 | 68 |
| 18 | 149 | 79 |
| 19 | 114 | 87 |
| 20 | 197 | 1 |
| 21 | 103 | 90 |
| 22 | 0 | 181 |
| 23 | 113 | 66 |
| 24 | 169 | 8 |
| 25 | 78 | 98 |
| 26 | 49 | 126 |
| 27 | 167 | 0 |
| 28 | 161 | 0 |
| 29 | 149 | 0 |
| 30 | 55 | 69 |
| 31 | 91 | 27 |
| 32 | 49 | 59 |
| 33 | 76 | 17 |
| 34 | 88 | 0 |
| 35 | 2 | 86 |
| 36 | 46 | 34 |
| 37 | 48 | 29 |
| 38 | 69 | 0 |
| 39 | 0 | 65 |
| 40 | 0 | 62 |
| 41 | 27 | 34 |
| 42 | 56 | 0 |
| 43 | 56 | 0 |
| 44 | 0 | 47 |
| 45 | 0 | 43 |

| | | | |
|---|---|---|---|
| 46 | 0 | 41 | |
| 47 | 26 | 10 | |
| | 7787 | 2843 | 10630 |
