## Supplementary material for "Direct Male Development in Chromosomally ZZ Zebrafish": SuppTable S5 30dpf_AllMarkers_minpct10

| | p_val | avg_log2FC | pct.1 | pct.2 | p_val_adj | cluster |
|---|---|---|---|---|---|---|
| zgc:153499 | 4.51502832360698e-195 | 0.818360267 | 0.96 | 0.478 | 1.13638747876864e-190 | 0 |
| cldnd | 7.78404847566281e-185 | 0.748834732 | 0.992 | 0.575 | 1.95916716083957e-180 | 0 |
| h1m | 1.44815420929968e-181 | 0.734724714 | 0.985 | 0.572 | 3.64485932938637e-177 | 0 |
| h2af1al | 2.80208953830733e-181 | 0.745382156 | 0.987 | 0.562 | 7.05257915896573e-177 | 0 |
| si:dkey-241l7.2 | 1.60645421936916e-178 | 0.716893974 | 0.996 | 0.614 | 4.04328462473024e-174 | 0 |
| zp3 | 1.67497654996596e-171 | 0.734709839 | 0.972 | 0.555 | 4.21574847860931e-167 | 0 |
| mt2 | 2.14548764790514e-168 | 0.755906411 | 0.963 | 0.501 | 5.39997786101244e-164 | 0 |
| si:dkey-240e12.6 | 9.18272121688273e-168 | 0.744144458 | 0.969 | 0.538 | 2.31119910307722e-163 | 0 |
| zp2.5 | 2.56927025092836e-166 | 0.74506964 | 0.962 | 0.533 | 6.4665962945616e-162 | 0 |
| zp3.2 | 4.72762327939029e-163 | 0.722574616 | 0.928 | 0.53 | 1.18989550318974e-158 | 0 |
| acp5a | 3.61340763328987e-162 | 0.723099176 | 0.916 | 0.511 | 9.09458567222727e-158 | 0 |
| si:ch73-189n23.1 | 4.73343923831302e-161 | 0.750392259 | 0.93 | 0.494 | 1.191359321891e-156 | 0 |
| zp2.3 | 4.79705344516535e-160 | 0.745007416 | 0.921 | 0.488 | 1.20737038161367e-155 | 0 |
| si:ch211-226h8.14 | 7.10465956544729e-160 | 0.716648491 | 0.938 | 0.538 | 1.78817176602743e-155 | 0 |
| zp2.6 | 7.0156888335846e-158 | 0.74951939 | 0.898 | 0.471 | 1.76577872252491e-153 | 0 |
| tmsb5 | 2.46263956185304e-155 | 0.755032818 | 0.903 | 0.457 | 6.19821751322791e-151 | 0 |
| si:dkeyp-98a7.9 | 6.14408131834753e-154 | 0.739380016 | 0.907 | 0.482 | 1.54640382701489e-149 | 0 |
| ccna1 | 6.70593223066934e-154 | 0.742119999 | 0.857 | 0.447 | 1.68781608313717e-149 | 0 |
| zp2.2 | 2.65387237826702e-153 | 0.733655022 | 0.933 | 0.499 | 6.67953138886026e-149 | 0 |
| si:ch211-125e6.12 | 5.35443800630678e-152 | 0.735399201 | 0.882 | 0.472 | 1.34765850180735e-147 | 0 |
| cldng | 1.31944915292783e-151 | 0.726420962 | 0.893 | 0.485 | 3.32092157300406e-147 | 0 |
| si:dkey-90l23.1 | 1.54107918412487e-150 | 0.750967898 | 0.83 | 0.418 | 3.87874219852388e-146 | 0 |
| si:dkey-241l7.5 | 7.2598250383891e-149 | 0.684397612 | 0.938 | 0.557 | 1.82722536391215e-144 | 0 |
| BX649337.1 | 4.86937848698389e-146 | 0.74191598 | 0.821 | 0.409 | 1.22557387138898e-141 | 0 |
| zgc:171776 | 5.66768658132893e-146 | 0.722149073 | 0.871 | 0.467 | 1.42650003565468e-141 | 0 |
| si:zfos-1505d6.3 | 1.43569151425798e-145 | 0.719056229 | 0.893 | 0.485 | 3.6134919722359e-141 | 0 |
| crp2 | 2.24294410196698e-145 | 0.730706081 | 0.885 | 0.459 | 5.6452660102407e-141 | 0 |
| zp2.1 | 1.27780038723482e-144 | 0.710342766 | 0.91 | 0.506 | 3.21609579463132e-140 | 0 |
| zgc:77118 | 1.52846424384703e-142 | 0.728605321 | 0.831 | 0.429 | 3.8469916553386e-138 | 0 |
| zgc:175135 | 7.82348966446329e-142 | 0.731257855 | 0.875 | 0.45 | 1.96909411364877e-137 | 0 |
| zgc:171779 | 2.70494312118449e-141 | 0.707296467 | 0.877 | 0.485 | 6.80807134170924e-137 | 0 |
| si:dkey-241l7.4 | 1.67512972067235e-135 | 0.646311213 | 0.949 | 0.583 | 4.21613399396024e-131 | 0 |
| si:ch211-226h8.4 | 3.94516753051219e-135 | 0.721376966 | 0.809 | 0.414 | 9.92959215754613e-131 | 0 |
| CR790368.1 | 5.20120899834925e-135 | 0.686471727 | 0.879 | 0.493 | 1.30909229279452e-130 | 0 |

| | | | | | | |
|---|---|---|---|---|---|---|
| si:dkey-241l7.6 | 1.32388439210131e-128 | 0.665763101 | 0.857 | 0.502 | 3.33208462647978e-124 | 0 |
| si:dkey-229d11.3 | 1.17170417027519e-125 | 0.709791867 | 0.801 | 0.411 | 2.94906222616562e-121 | 0 |
| rrm2 | 7.44384722527322e-124 | 0.663704928 | 0.844 | 0.472 | 1.87354190812902e-119 | 0 |
| lamp2 | 6.87477812148881e-123 | 0.635236096 | 0.843 | 0.533 | 1.73031290539752e-118 | 0 |
| epcam | 1.47249033518649e-122 | 0.666668134 | 0.854 | 0.479 | 3.70611092463087e-118 | 0 |
| si:ch211-12h2.8 | 1.92885855648933e-120 | 0.635842458 | 0.86 | 0.524 | 4.854744100828e-116 | 0 |
| CU571163.1 | 4.508728575394e-120 | 0.70923584 | 0.757 | 0.375 | 1.13480189514091e-115 | 0 |
| ccnb2 | 6.4468059525407e-120 | 0.703253862 | 0.728 | 0.358 | 1.62259659019497e-115 | 0 |
| si:dkeyp-98a7.8 | 1.82747919014482e-118 | 0.677566417 | 0.8 | 0.432 | 4.59958237367549e-114 | 0 |
| LOC110437821 | 4.92342310146245e-117 | 0.699986883 | 0.753 | 0.39 | 1.23917636040708e-112 | 0 |
| zar1l | 9.41298783104742e-117 | 0.688139478 | 0.73 | 0.373 | 2.36915490719633e-112 | 0 |
| qdprb2 | 1.84740422605273e-111 | 0.666599705 | 0.753 | 0.409 | 4.64973169655211e-107 | 0 |
| thy1 | 7.94485723810921e-111 | 0.650156795 | 0.754 | 0.428 | 1.99964111825971e-106 | 0 |
| mid1ip1l | 4.17733401853214e-110 | 0.603244659 | 0.826 | 0.522 | 1.05139319912435e-105 | 0 |
| si:dkeyp-20e4.8 | 5.00702539880651e-106 | 0.675113927 | 0.727 | 0.384 | 1.26021822262561e-101 | 0 |
| si:ch211-12h2.6 | 5.12595307360012e-106 | 0.633639576 | 0.778 | 0.452 | 1.29015112909441e-101 | 0 |
| si:dkeyp-46h3.6 | 2.30027434278499e-103 | 0.653946491 | 0.72 | 0.388 | 5.78956049335555e-99 | 0 |
| zp3a.2 | 2.83486940934097e-101 | 0.618872702 | 0.801 | 0.468 | 7.13508281637028e-97 | 0 |
| hbba1 | 6.20850324437445e-101 | 0.676256351 | 0.703 | 0.33 | 1.5626181815766e-96 | 0 |
| si:dkeyp-98a7.4 | 1.15190133672589e-100 | 0.638616851 | 0.74 | 0.402 | 2.8992204744054e-96 | 0 |
| si:ch211-14a17.7 | 1.33576970664055e-100 | 0.647501936 | 0.717 | 0.385 | 3.3619987746436e-96 | 0 |
| zp3b | 1.52162559418798e-99 | 0.65196914 | 0.707 | 0.364 | 3.82977945801174e-95 | 0 |
| zar1 | 4.99949126378266e-98 | 0.626659197 | 0.738 | 0.418 | 1.25832195618146e-93 | 0 |
| zgc:195245 | 1.41818767450572e-94 | 0.630981831 | 0.686 | 0.366 | 3.56943655796344e-90 | 0 |
| si:dkey-229d11.5 | 1.76062878122169e-94 | 0.633840372 | 0.672 | 0.369 | 4.43132657945687e-90 | 0 |
| tuba4l | 5.37550579313195e-94 | 0.626254026 | 0.696 | 0.383 | 1.35296105307338e-89 | 0 |
| ctss2.1 | 8.60775761863308e-94 | 0.556972205 | 0.796 | 0.515 | 2.16648651503376e-89 | 0 |
| LOC100537499 | 4.73385034726699e-93 | 0.644987473 | 0.614 | 0.301 | 1.19146279390363e-88 | 0 |
| nanog | 1.58864408834621e-91 | 0.627798215 | 0.646 | 0.339 | 3.99845830595857e-87 | 0 |
| si:dkeyp-50f7.2 | 8.68603373076548e-90 | 0.615802296 | 0.698 | 0.381 | 2.18618782969636e-85 | 0 |
| slbp2 | 2.71270084762934e-89 | 0.614571617 | 0.688 | 0.376 | 6.8275967633983e-85 | 0 |
| LOC100535162 | 2.16516573439143e-87 | 0.636906667 | 0.595 | 0.288 | 5.44950563688979e-83 | 0 |
| zgc:173443 | 3.76883460998141e-87 | 0.624669189 | 0.606 | 0.31 | 9.4857798298622e-83 | 0 |
| ccnb1 | 1.30901098921829e-86 | 0.606237352 | 0.664 | 0.374 | 3.29464975876353e-82 | 0 |
| gtf3ab | 1.90214519053089e-86 | 0.622428643 | 0.594 | 0.3 | 4.78750923004721e-82 | 0 |

| | | | | | | |
|---|---|---|---|---|---|---|
| siva1 | 2.40156973060168e-86 | 0.594295799 | 0.707 | 0.408 | 6.04451085495137e-82 | 0 |
| btg4 | 6.77776075199132e-86 | 0.613081718 | 0.643 | 0.349 | 1.70589460366869e-81 | 0 |
| cth1 | 1.19301339667993e-85 | 0.605022394 | 0.656 | 0.359 | 3.00269541810372e-81 | 0 |
| zpcx | 5.92279104649362e-83 | 0.614078013 | 0.611 | 0.319 | 1.49070727849198e-78 | 0 |
| si:ch211-5k11.8 | 8.83472422742269e-83 | 0.551213157 | 0.683 | 0.334 | 2.22361174080002e-78 | 0 |
| si:dkeyp-46h3.8 | 1.32809256488599e-82 | 0.610998135 | 0.632 | 0.331 | 3.34267617656154e-78 | 0 |
| AL929032.1 | 1.60009111800973e-79 | 0.611097569 | 0.595 | 0.297 | 4.02726933491869e-75 | 0 |
| si:ch211-132b12.8 | 3.70759264004633e-79 | 0.598451717 | 0.605 | 0.323 | 9.33163991573259e-75 | 0 |
| hnrnpabb | 1.26744240752838e-78 | 0.472900753 | 0.774 | 0.584 | 3.19002579550817e-74 | 0 |
| kpna7 | 1.44663989308277e-76 | 0.594969304 | 0.586 | 0.31 | 3.64104794690003e-72 | 0 |
| s100a1 | 2.79798164312559e-76 | 0.581303759 | 0.625 | 0.345 | 7.04223999758281e-72 | 0 |
| zgc:171517 | 1.24469947328668e-75 | 0.581371266 | 0.644 | 0.37 | 3.13278410431524e-71 | 0 |
| npm2b | 1.60749652793732e-74 | 0.570123381 | 0.649 | 0.378 | 4.04590801116543e-70 | 0 |
| hbaa1 | 7.73966238231529e-73 | 0.585043022 | 0.557 | 0.261 | 1.94799562500494e-68 | 0 |
| zgc:136254 | 2.9249386515097e-68 | 0.566075013 | 0.575 | 0.313 | 7.36177809198475e-64 | 0 |
| si:dkey-208k4.2 | 7.10607812613387e-66 | 0.561081375 | 0.525 | 0.28 | 1.78852880356663e-61 | 0 |
| si:dkeyp-46h3.2 | 7.68188490063019e-66 | 0.537401143 | 0.622 | 0.389 | 1.93345361063961e-61 | 0 |
| cpeb1b | 1.96928479412746e-65 | 0.563862409 | 0.495 | 0.254 | 4.95649289833941e-61 | 0 |
| si:dkey-90l23.2 | 2.60617525258277e-65 | 0.557759291 | 0.582 | 0.328 | 6.55948249322557e-61 | 0 |
| gapdh | 2.45825508409672e-64 | 0.527049085 | 0.653 | 0.403 | 6.18718222116304e-60 | 0 |
| wu:fi42e03 | 3.38597551586908e-64 | 0.547516699 | 0.583 | 0.333 | 8.52216177589088e-60 | 0 |
| uchl1 | 1.74266983144346e-63 | 0.544362218 | 0.567 | 0.329 | 4.38612569876003e-59 | 0 |
| si:dkeyp-46h3.5 | 1.25253578621041e-62 | 0.551911776 | 0.511 | 0.274 | 3.15250732031297e-58 | 0 |
| lsm14b | 3.43257781491205e-62 | 0.55001617 | 0.52 | 0.284 | 8.63945510235215e-58 | 0 |
| nasp | 7.35387935044633e-62 | 0.54460614 | 0.56 | 0.324 | 1.85089789371384e-57 | 0 |
| ca15b | 9.46588132003139e-61 | 0.544897994 | 0.465 | 0.234 | 2.3824676694387e-56 | 0 |
| si:ch211-146l10.8 | 2.15002920415081e-60 | 0.545497504 | 0.524 | 0.291 | 5.41140850392718e-56 | 0 |
| si:ch211-226l4.6 | 3.3153672924769e-59 | 0.537796824 | 0.511 | 0.277 | 8.34444793843512e-55 | 0 |
| avd | 1.44220594916199e-58 | 0.536095087 | 0.479 | 0.25 | 3.62988815344582e-54 | 0 |
| si:ch73-90k17.1 | 1.63744765866694e-58 | 0.539909245 | 0.457 | 0.234 | 4.12129201209881e-54 | 0 |
| si:dkeyp-98a7.7 | 4.5572027568817e-58 | 0.526892462 | 0.396 | 0.181 | 1.14700236187955e-53 | 0 |
| arhgef10la | 1.23001202116043e-55 | 0.527033073 | 0.478 | 0.261 | 3.09581725605868e-51 | 0 |
| zgc:56699 | 2.66764533305877e-55 | 0.53203981 | 0.453 | 0.237 | 6.71419653877563e-51 | 0 |
| si:dkey-37o8.1 | 3.10998750773622e-50 | 0.501820061 | 0.503 | 0.294 | 7.82752755822129e-46 | 0 |
| si:ch211-251f6.6 | 8.15430404473384e-50 | 0.49624098 | 0.525 | 0.318 | 2.05235678501906e-45 | 0 |

| | | | | | | |
|---|---|---|---|---|---|---|
| si:dkey-152b24.6 | 1.38579606613139e-49 | 0.499430125 | 0.502 | 0.284 | 3.48791011884609e-45 | 0 |
| si:ch73-14h10.2 | 3.32711038417666e-48 | 0.495973075 | 0.467 | 0.258 | 8.37400412593423e-44 | 0 |
| zp3a.1 | 5.23983127128714e-48 | 0.474725451 | 0.569 | 0.369 | 1.31881313267026e-43 | 0 |
| zgc:171474 | 2.49024347728125e-47 | 0.486005745 | 0.498 | 0.291 | 6.26769380796918e-43 | 0 |
| pigrl2.3 | 2.67335524937536e-47 | 0.493357088 | 0.381 | 0.191 | 6.72856782715284e-43 | 0 |
| si:ch211-250e5.16 | 3.42404261049923e-47 | 0.481592631 | 0.534 | 0.334 | 8.6179728463655e-43 | 0 |
| LOC101885477 | 4.85324448237517e-47 | 0.492088177 | 0.466 | 0.267 | 1.22151310376901e-42 | 0 |
| si:ch211-145c1.1 | 8.24562079293039e-45 | 0.483420557 | 0.417 | 0.221 | 2.07534029737265e-40 | 0 |
| zgc:55461 | 1.25755845334801e-44 | 0.47579013 | 0.472 | 0.276 | 3.1651488712316e-40 | 0 |
| pabpn1l | 2.37318412463213e-44 | 0.477789984 | 0.462 | 0.272 | 5.97306712328662e-40 | 0 |
| birc5b | 2.07303314261919e-43 | 0.470589022 | 0.44 | 0.247 | 5.21761711665825e-39 | 0 |
| zgc:56231 | 3.3312396666022e-43 | 0.474122264 | 0.398 | 0.216 | 8.38439711687107e-39 | 0 |
| sinup | 7.09263431640356e-43 | 0.470688387 | 0.37 | 0.192 | 1.78514513109561e-38 | 0 |
| elovl7b | 8.54271019187079e-43 | 0.468904579 | 0.424 | 0.236 | 2.15011472819196e-38 | 0 |
| gmnn | 1.33016219954787e-42 | 0.468839397 | 0.476 | 0.291 | 3.34788524004203e-38 | 0 |
| si:ch1073-263o8.2 | 4.21278786189479e-39 | 0.457292814 | 0.43 | 0.253 | 1.0603165769603e-34 | 0 |
| haus6 | 1.44502091185066e-38 | 0.453408239 | 0.368 | 0.199 | 3.63697313303693e-34 | 0 |
| ccna2 | 2.60059858000697e-37 | 0.44158884 | 0.445 | 0.274 | 6.54544656601955e-33 | 0 |
| mkrn4 | 5.00904185672441e-36 | 0.440205422 | 0.382 | 0.218 | 1.26072574491897e-31 | 0 |
| zgc:113424 | 9.1580476151415e-35 | 0.428321343 | 0.301 | 0.152 | 2.30498900425496e-30 | 0 |
| rcc1 | 1.98734391674113e-34 | 0.435651491 | 0.348 | 0.193 | 5.00194590404576e-30 | 0 |
| zp3c | 9.47660971572975e-34 | 0.429281194 | 0.33 | 0.18 | 2.38516789935202e-29 | 0 |
| buc | 1.5083621620588e-32 | 0.415459091 | 0.318 | 0.17 | 3.7963967256858e-28 | 0 |
| slc16a3 | 1.59003614009008e-32 | 0.422546856 | 0.394 | 0.24 | 4.00196196099273e-28 | 0 |
| gdf3 | 1.8109328776179e-32 | 0.418647776 | 0.374 | 0.214 | 4.55793695967649e-28 | 0 |
| map1lc3c | 2.02536800764768e-32 | 0.424277349 | 0.343 | 0.193 | 5.09764873844844e-28 | 0 |
| zgc:92591 | 3.13326988893225e-32 | 0.416645345 | 0.349 | 0.196 | 7.88612698345357e-28 | 0 |
| rbpms2b | 6.78403656868274e-32 | 0.419466705 | 0.362 | 0.212 | 1.70747416397176e-27 | 0 |
| si:ch1073-190k2.1 | 9.28909236174986e-32 | 0.416716619 | 0.327 | 0.179 | 2.33797165652882e-27 | 0 |
| si:dkey-19b23.11 | 1.36908275761359e-31 | 0.411149474 | 0.434 | 0.269 | 3.44584439263764e-27 | 0 |
| ctsba | 2.27076646116112e-31 | 0.393983651 | 0.502 | 0.369 | 5.71529210609643e-27 | 0 |
| spink2.1 | 3.32992367929925e-31 | 0.370830816 | 0.587 | 0.427 | 8.38108490842828e-27 | 0 |
| sybu | 9.75159472009673e-31 | 0.39909847 | 0.301 | 0.158 | 2.45437887510115e-26 | 0 |
| zgc:173856 | 1.27928911730941e-30 | 0.407309643 | 0.362 | 0.211 | 3.21984277935605e-26 | 0 |
| zgc:113886 | 1.94918037576688e-30 | 0.392740419 | 0.267 | 0.133 | 4.90589208776765e-26 | 0 |

| | | | | | | |
|---|---|---|---|---|---|---|
| si:dkey-188i13.11 | 1.38817282222852e-29 | 0.410994604 | 0.444 | 0.308 | 3.49389217626695e-25 | 0 |
| rbm4.3 | 1.40220437552815e-29 | 0.357876857 | 0.535 | 0.439 | 3.52920819276681e-25 | 0 |
| AL929192.1 | 5.04819020596399e-29 | 0.380064093 | 0.296 | 0.156 | 1.27057899293908e-24 | 0 |
| snu13b | 5.97691150304452e-29 | 0.382100155 | 0.507 | 0.386 | 1.50432885620127e-24 | 0 |
| slbp | 6.02952669673926e-29 | 0.401907058 | 0.374 | 0.225 | 1.5175715743023e-24 | 0 |
| gyg1a | 6.21755074248315e-29 | 0.389963444 | 0.489 | 0.363 | 1.56489534637558e-24 | 0 |
| h3f3a | 8.73907581508509e-29 | 0.335507606 | 0.589 | 0.501 | 2.19953799189877e-24 | 0 |
| cdca7a | 9.31564560300058e-29 | 0.401630457 | 0.356 | 0.214 | 2.34465484181921e-24 | 0 |
| larp6a | 1.28500234945395e-28 | 0.406365194 | 0.38 | 0.242 | 3.23422241334064e-24 | 0 |
| LOC100536860 | 2.80408628583118e-28 | 0.37850198 | 0.479 | 0.343 | 7.0576047728085e-24 | 0 |
| crp3 | 7.82881330348257e-27 | 0.385692389 | 0.349 | 0.21 | 1.97043402035353e-22 | 0 |
| zgc:103482 | 1.8310641457226e-26 | 0.387142968 | 0.311 | 0.181 | 4.6086053483692e-22 | 0 |
| elavl2 | 2.64741874762095e-26 | 0.387515062 | 0.353 | 0.216 | 6.66328824588716e-22 | 0 |
| zgc:162945 | 2.90822462348496e-26 | 0.37656174 | 0.282 | 0.155 | 7.31971055484929e-22 | 0 |
| larp6b | 2.95741425646233e-26 | 0.382655368 | 0.322 | 0.188 | 7.44351594209004e-22 | 0 |
| nfr | 2.99135763465566e-26 | 0.350130798 | 0.217 | 0.104 | 7.52894803066484e-22 | 0 |
| inka1b | 3.73768384975044e-26 | 0.383286733 | 0.305 | 0.177 | 9.40737648143687e-22 | 0 |
| LOC101883897 | 7.24152437718996e-26 | 0.368602073 | 0.283 | 0.157 | 1.82261927049494e-21 | 0 |
| CR383662.2 | 1.0974061157376e-25 | 0.323445538 | 0.182 | 0.081 | 2.76206145269996e-21 | 0 |
| zglp1 | 4.60568317986594e-25 | 0.368906675 | 0.323 | 0.191 | 1.15920439954046e-20 | 0 |
| tram2 | 1.23345493771347e-24 | 0.374701235 | 0.238 | 0.126 | 3.10448273273103e-20 | 0 |
| gbgt1l4 | 3.09695936898071e-24 | 0.3542791 | 0.263 | 0.143 | 7.79473703578755e-20 | 0 |
| tph1a | 6.36990841264357e-24 | 0.355087924 | 0.25 | 0.135 | 1.60324224837826e-19 | 0 |
| si:dkey-42i9.8 | 6.65189693471999e-24 | 0.34868977 | 0.233 | 0.121 | 1.67421593949967e-19 | 0 |
| mibp | 6.92793661835253e-24 | 0.373796639 | 0.342 | 0.216 | 1.74369236747315e-19 | 0 |
| zgc:162879 | 7.74719851652665e-24 | 0.362676294 | 0.299 | 0.174 | 1.94989239462459e-19 | 0 |
| org | 8.8048055641422e-24 | 0.362724274 | 0.364 | 0.233 | 2.21608151243895e-19 | 0 |
| tac3b | 9.44531788169996e-24 | 0.33705986 | 0.225 | 0.114 | 2.37729205764506e-19 | 0 |
| CU467646.2 | 1.94024783643837e-23 | 0.333362453 | 0.231 | 0.119 | 4.88340977953174e-19 | 0 |
| patl2 | 1.193040286622e-22 | 0.352050083 | 0.286 | 0.166 | 3.00276309739891e-18 | 0 |
| dynll1 | 1.71932224729572e-22 | 0.314716044 | 0.528 | 0.451 | 4.32736216421861e-18 | 0 |
| pttg1 | 1.7734959451487e-22 | 0.356605555 | 0.266 | 0.152 | 4.46371194434478e-18 | 0 |
| hsd17b12a | 1.81141161565492e-22 | 0.351182763 | 0.275 | 0.159 | 4.55914189544187e-18 | 0 |
| cdc20 | 2.37169935413873e-22 | 0.346339591 | 0.26 | 0.146 | 5.96933010443177e-18 | 0 |
| retsatl | 2.70284646357419e-22 | 0.357779246 | 0.318 | 0.197 | 6.80279426416987e-18 | 0 |

| | | | | | | |
|---|---|---|---|---|---|---|
| plk1 | 3.2517554315847e-22 | 0.35545595 | 0.285 | 0.171 | 8.18434324575552e-18 | 0 |
| elovl7a | 1.51890755195173e-21 | 0.341549012 | 0.296 | 0.176 | 3.82293841750732e-17 | 0 |
| gstp1 | 1.56281329992707e-21 | 0.331706198 | 0.459 | 0.36 | 3.93344479458644e-17 | 0 |
| dctpp1 | 3.80356237200403e-21 | 0.345847456 | 0.282 | 0.169 | 9.57318613409694e-17 | 0 |
| si:ch73-78o10.1 | 1.37228118998934e-20 | 0.328192526 | 0.232 | 0.127 | 3.45389452708418e-16 | 0 |
| hspa5 | 4.15235654897545e-20 | 0.336268966 | 0.408 | 0.309 | 1.04510661981163e-15 | 0 |
| ankrd22 | 9.33731162358506e-20 | 0.328063456 | 0.24 | 0.137 | 2.35010796254012e-15 | 0 |
| ddx19 | 2.51536748537988e-19 | 0.336656997 | 0.246 | 0.146 | 6.33092842395261e-15 | 0 |
| si:ch211-146l10.7 | 4.9941518615064e-19 | 0.322202242 | 0.295 | 0.183 | 1.25697808202255e-14 | 0 |
| si:dkey-46g23.2 | 6.65561761001558e-19 | 0.320946936 | 0.24 | 0.139 | 1.67515239626482e-14 | 0 |
| tmem54a | 7.90181412562458e-19 | 0.318072406 | 0.238 | 0.135 | 1.98880759727845e-14 | 0 |
| CR847984.1 | 1.62058207268505e-18 | 0.324957535 | 0.262 | 0.158 | 4.078843018741e-14 | 0 |
| zp2l1 | 1.86896438316678e-18 | 0.322865126 | 0.262 | 0.158 | 4.70399645599246e-14 | 0 |
| tmprss4b | 3.3055507961184e-18 | 0.261717858 | 0.142 | 0.065 | 8.3197407987504e-14 | 0 |
| si:dkey-241l7.3 | 4.44085881542755e-18 | 0.321514975 | 0.458 | 0.36 | 1.11771975525496e-13 | 0 |
| lpcat4 | 6.24881834155657e-18 | 0.294748507 | 0.192 | 0.102 | 1.57276508838637e-13 | 0 |
| LOC103909972 | 8.00778158224836e-18 | 0.318217694 | 0.436 | 0.364 | 2.01547854643609e-13 | 0 |
| cldn7b | 8.83973558661014e-18 | 0.317337177 | 0.256 | 0.154 | 2.22487304979391e-13 | 0 |
| snu13a | 1.44924848013732e-17 | 0.292179939 | 0.182 | 0.096 | 3.64761349965761e-13 | 0 |
| BX571780.1 | 1.4715034752131e-17 | 0.266793589 | 0.157 | 0.077 | 3.70362709676384e-13 | 0 |
| ppp1r3b | 2.08366827179683e-17 | 0.295590396 | 0.193 | 0.105 | 5.24438467328544e-13 | 0 |
| h2afva | 2.60261872715333e-17 | 0.324318408 | 0.404 | 0.319 | 6.55053107437223e-13 | 0 |
| cox17 | 2.73045577212422e-17 | 0.301283597 | 0.445 | 0.374 | 6.87228413285945e-13 | 0 |
| CABZ01059627.2 | 3.44658203087939e-17 | 0.306983736 | 0.221 | 0.129 | 8.67470231352033e-13 | 0 |
| sox19b | 3.70948561185906e-17 | 0.274326385 | 0.179 | 0.093 | 9.33640433648807e-13 | 0 |
| cx44.2 | 4.47318704362925e-17 | 0.297617705 | 0.202 | 0.112 | 1.12585644701105e-12 | 0 |
| zgc:173742 | 4.50476768199147e-17 | 0.3117634 | 0.249 | 0.15 | 1.13380497788043e-12 | 0 |
| degs2 | 5.24177865676911e-17 | 0.290624563 | 0.188 | 0.102 | 1.31930327012222e-12 | 0 |
| eif4a1a | 6.05239170326291e-17 | 0.280905515 | 0.477 | 0.423 | 1.52332646779424e-12 | 0 |
| zgc:165539 | 7.76760802604803e-17 | 0.288750781 | 0.206 | 0.115 | 1.95502926407603e-12 | 0 |
| BX001030.1 | 8.45190624539224e-17 | 0.298978993 | 0.207 | 0.118 | 2.12726028290277e-12 | 0 |
| zgc:152986 | 1.09351826734282e-16 | 0.276829337 | 0.166 | 0.085 | 2.75227612707514e-12 | 0 |
| zgc:56676 | 1.41813714757644e-16 | 0.325083777 | 0.379 | 0.291 | 3.56930938673514e-12 | 0 |
| pou5f3 | 1.47404183850088e-16 | 0.305496057 | 0.236 | 0.142 | 3.71001590332287e-12 | 0 |
| orc6 | 1.54071066908789e-16 | 0.293915969 | 0.187 | 0.103 | 3.8778146830273e-12 | 0 |

| | | | | | | |
|---|---|---|---|---|---|---|
| CDPF1 | 2.84379207518967e-16 | 0.323787727 | 0.357 | 0.264 | 7.15754027404488e-12 | 0 |
| BX537276.1 | 3.58018794600845e-16 | 0.286449299 | 0.206 | 0.117 | 9.01097504130867e-12 | 0 |
| zc3h11a | 3.96964476968032e-16 | 0.328842385 | 0.307 | 0.212 | 9.99119892080839e-12 | 0 |
| nanos3 | 4.65625111131976e-16 | 0.28594888 | 0.181 | 0.098 | 1.17193184220807e-11 | 0 |
| gpr160 | 7.11845475010368e-16 | 0.257479664 | 0.153 | 0.077 | 1.7916438760536e-11 | 0 |
| si:dkeyp-114g9.1 | 8.11630555102147e-16 | 0.28493166 | 0.182 | 0.1 | 2.0427929441366e-11 | 0 |
| si:ch211-14a17.6 | 1.12044509233605e-15 | 0.26109747 | 0.159 | 0.082 | 2.82004825290061e-11 | 0 |
| si:ch73-160p18.3 | 1.16646713924442e-15 | 0.312288693 | 0.331 | 0.236 | 2.93588114276427e-11 | 0 |
| rnf4 | 1.51543313382552e-15 | 0.301237283 | 0.215 | 0.13 | 3.81419365452545e-11 | 0 |
| zgc:113984 | 1.57620814452953e-15 | 0.271484556 | 0.179 | 0.097 | 3.96715827896637e-11 | 0 |
| ccne1 | 1.71329784921972e-15 | 0.289096249 | 0.189 | 0.107 | 4.31219935670111e-11 | 0 |
| cdca8 | 2.79936612641435e-15 | 0.299463728 | 0.251 | 0.16 | 7.04572460357228e-11 | 0 |
| zgc:171750 | 4.62797258107848e-15 | 0.295814043 | 0.264 | 0.17 | 1.16481441893164e-10 | 0 |
| CU655961.4 | 5.11413426836861e-15 | 0.255741696 | 0.161 | 0.084 | 1.28717645400569e-10 | 0 |
| zgc:114123 | 6.18870880406972e-15 | 0.273528609 | 0.179 | 0.099 | 1.55763611889631e-10 | 0 |
| ctsla | 6.19148410707846e-15 | 0.280155769 | 0.451 | 0.384 | 1.55833463491058e-10 | 0 |
| si:dkey-68o6.5 | 8.33771551982264e-15 | 0.28407077 | 0.202 | 0.118 | 2.09851961918416e-10 | 0 |
| tmem144b | 1.1187314755454e-14 | 0.2579607 | 0.167 | 0.09 | 2.81573525080021e-10 | 0 |
| srsf2a | 1.34063751148482e-14 | 0.295485451 | 0.42 | 0.365 | 3.37425055265613e-10 | 0 |
| zgc:66024 | 1.45719885433528e-14 | 0.277744014 | 0.225 | 0.137 | 3.66762379647646e-10 | 0 |
| chia.4 | 1.53014819680254e-14 | 0.283245042 | 0.231 | 0.142 | 3.85122999653232e-10 | 0 |
| ccnh | 1.87725776799401e-14 | 0.28410681 | 0.202 | 0.121 | 4.72487007626412e-10 | 0 |
| CABZ01059627.1 | 4.2712450386412e-14 | 0.277226704 | 0.217 | 0.132 | 1.0750296637756e-09 | 0 |
| prrg2 | 4.60711491890049e-14 | 0.254796735 | 0.159 | 0.085 | 1.15956475393806e-09 | 0 |
| aurka | 4.93794576954042e-14 | 0.272376422 | 0.194 | 0.115 | 1.24283157073563e-09 | 0 |
| il13ra2 | 7.84717218237674e-14 | 0.270957484 | 0.182 | 0.106 | 1.9750547665824e-09 | 0 |
| rpz2 | 1.40698535762509e-13 | 0.306190935 | 0.259 | 0.177 | 3.54124144660658e-09 | 0 |
| itm2cb | 1.5436498331327e-13 | 0.287597248 | 0.213 | 0.134 | 3.88521226501169e-09 | 0 |
| CT009596.1 | 2.00918303641194e-13 | 0.252475888 | 0.157 | 0.087 | 5.0569127843452e-09 | 0 |
| parn | 2.19960500432003e-13 | 0.292984302 | 0.222 | 0.144 | 5.53618583537309e-09 | 0 |
| si:ch211-69g19.2 | 2.29627925638897e-13 | 0.268252838 | 0.196 | 0.118 | 5.77950526040539e-09 | 0 |
| vstm2l | 2.38894066307654e-13 | 0.250315894 | 0.164 | 0.092 | 6.01272475489734e-09 | 0 |
| mad2l1 | 2.8475405626229e-13 | 0.281302507 | 0.267 | 0.182 | 7.16697484206557e-09 | 0 |
| g3bp1 | 3.22437787730788e-13 | 0.30300436 | 0.285 | 0.207 | 8.1154366793962e-09 | 0 |
| BX284697.1 | 5.55396554976609e-13 | 0.258880848 | 0.187 | 0.11 | 1.39787758922063e-08 | 0 |

| | | | | | | |
|---|---|---|---|---|---|---|
| brf2 | 6.20609295909531e-13 | 0.250161134 | 0.162 | 0.091 | 1.5620115368747e-08 | 0 |
| pabpc1l | 6.46205431048097e-13 | 0.260870874 | 0.221 | 0.138 | 1.62643444940495e-08 | 0 |
| prr11 | 6.91802908225515e-13 | 0.25552685 | 0.168 | 0.097 | 1.7411987397128e-08 | 0 |
| prdx1 | 8.67716980078156e-13 | 0.289777848 | 0.244 | 0.163 | 2.18395686715871e-08 | 0 |
| figla | 9.09787672110387e-13 | 0.261862478 | 0.161 | 0.092 | 2.28984459193463e-08 | 0 |
| rasl11b | 1.92992395577969e-12 | 0.264070125 | 0.207 | 0.13 | 4.8574256043019e-08 | 0 |
| cpm | 2.80617333597021e-12 | 0.257108043 | 0.176 | 0.105 | 7.06285766930342e-08 | 0 |
| ddx39ab | 4.83577621514433e-12 | 0.290312136 | 0.353 | 0.287 | 1.21711651558968e-07 | 0 |
| chtopa | 7.94826845787179e-12 | 0.29121253 | 0.289 | 0.218 | 2.00049968816175e-07 | 0 |
| fdx1 | 1.07269852274216e-11 | 0.28748053 | 0.291 | 0.218 | 2.69987491188975e-07 | 0 |
| hells | 2.71788818603626e-11 | 0.264997966 | 0.2 | 0.13 | 6.84065277543466e-07 | 0 |
| tpx2 | 5.14041545671631e-11 | 0.25002259 | 0.192 | 0.123 | 1.29379116630093e-06 | 0 |
| kpna2 | 6.71633263367615e-11 | 0.25046936 | 0.23 | 0.155 | 1.69043376056995e-06 | 0 |
| imp3 | 7.61138019894736e-11 | 0.261734883 | 0.2 | 0.132 | 1.91570828227306e-06 | 0 |
| faub | 1.59986507237359e-10 | 0.271221641 | 0.267 | 0.197 | 4.02670040065709e-06 | 0 |
| tent5c | 1.60705219544204e-10 | 0.254532734 | 0.205 | 0.137 | 4.04478967070807e-06 | 0 |
| rdh10b | 3.30141717056177e-10 | 0.25726782 | 0.246 | 0.175 | 8.30933687658692e-06 | 0 |
| nifk | 1.59036703849629e-09 | 0.250650883 | 0.18 | 0.119 | 4.00279479919131e-05 | 0 |
| nutf2l | 1.06574030031367e-08 | 0.261025719 | 0.284 | 0.232 | 0.000268236 | 0 |
| anp32b | 1.35256116853114e-08 | 0.25626858 | 0.304 | 0.254 | 0.000340426 | 0 |
| rab5if | 3.88691154790422e-08 | 0.251631587 | 0.181 | 0.128 | 0.000978297 | 0 |
| si:dkey-241l7.21 | 1.76302670960865e-180 | 0.71723755 | 0.986 | 0.618 | 4.43736192541401e-176 | 1 |
| crp21 | 8.98783481645588e-169 | 0.790301958 | 0.926 | 0.458 | 2.26214814495378e-164 | 1 |
| si:dkey-229d11.31 | 1.63154753187118e-167 | 0.814851048 | 0.887 | 0.407 | 4.10644198296657e-163 | 1 |
| zp31 | 8.8692530881346e-167 | 0.740290588 | 0.972 | 0.557 | 2.2323023097526e-162 | 1 |
| zp2.21 | 9.28472542958634e-167 | 0.76981472 | 0.953 | 0.5 | 2.33687254337259e-162 | 1 |
| si:ch73-189n23.11 | 2.87926843081656e-166 | 0.772642812 | 0.942 | 0.496 | 7.2468307135222e-162 | 1 |
| si:dkey-240e12.61 | 5.31275140493405e-165 | 0.754078001 | 0.968 | 0.541 | 1.33716640110785e-160 | 1 |
| zp3.21 | 5.46682487940541e-163 | 0.740104872 | 0.946 | 0.531 | 1.37594515389755e-158 | 1 |
| si:ch211-125e6.121 | 1.17687361570036e-162 | 0.781525674 | 0.933 | 0.471 | 2.96207320335623e-158 | 1 |
| tmsb51 | 2.18097734726772e-161 | 0.791431854 | 0.942 | 0.457 | 5.48930188533813e-157 | 1 |
| zp2.51 | 8.29999465146986e-161 | 0.741846429 | 0.959 | 0.536 | 2.08902565382845e-156 | 1 |
| zp2.11 | 9.8023632484186e-161 | 0.759700824 | 0.942 | 0.507 | 2.46715680599448e-156 | 1 |
| zp2.61 | 2.95267430953892e-158 | 0.769912308 | 0.921 | 0.472 | 7.4315859696785e-154 | 1 |
| si:dkeyp-98a7.91 | 8.46453009391341e-158 | 0.762741146 | 0.936 | 0.483 | 2.13043757933707e-153 | 1 |

| | | | | | | |
|---|---|---|---|---|---|---|
| si:ch211-226h8.141 | 2.18643329994129e-157 | 0.732819925 | 0.959 | 0.539 | 5.50303397262222e-153 | 1 |
| cldnd1 | 2.75512507960194e-157 | 0.723501417 | 0.977 | 0.579 | 6.93437431285013e-153 | 1 |
| si:zfos-1505d6.31 | 4.00507992515949e-157 | 0.75718299 | 0.922 | 0.485 | 1.00803856636339e-152 | 1 |
| zgc:1534991 | 3.40444497304554e-156 | 0.776986087 | 0.955 | 0.482 | 8.56864755265832e-152 | 1 |
| si:dkey-241l7.51 | 3.47968985015446e-155 | 0.710244078 | 0.955 | 0.558 | 8.75803138385377e-151 | 1 |
| BX649337.11 | 1.47922964573942e-154 | 0.783503021 | 0.888 | 0.406 | 3.72307309536155e-150 | 1 |
| mt21 | 2.06261368875455e-153 | 0.746294322 | 0.956 | 0.505 | 5.19139239322632e-149 | 1 |
| h2af1al1 | 9.47680342774155e-150 | 0.715258905 | 0.972 | 0.566 | 2.38521665472827e-145 | 1 |
| h1m1 | 1.95304045329806e-149 | 0.702442856 | 0.969 | 0.576 | 4.91560751690588e-145 | 1 |
| zp2.31 | 6.96201798164806e-148 | 0.742577182 | 0.931 | 0.49 | 1.752270305801e-143 | 1 |
| zp3a.21 | 1.44690160512705e-146 | 0.734825214 | 0.876 | 0.465 | 3.64170664994428e-142 | 1 |
| si:ch211-12h2.81 | 3.00549463783305e-144 | 0.703877691 | 0.919 | 0.522 | 7.56452945396201e-140 | 1 |
| LOC1104378211 | 5.25807646521445e-142 | 0.774309743 | 0.828 | 0.387 | 1.32340526552982e-137 | 1 |
| cldng1 | 6.27026809791604e-142 | 0.733785263 | 0.921 | 0.486 | 1.57816377756449e-137 | 1 |
| zgc:1751351 | 3.69102351361797e-141 | 0.749980957 | 0.897 | 0.451 | 9.28993708142506e-137 | 1 |
| si:ch211-226h8.41 | 8.55279088910301e-140 | 0.753987775 | 0.856 | 0.413 | 2.15265193887834e-135 | 1 |
| zgc:1717791 | 1.93822392181649e-138 | 0.72098808 | 0.903 | 0.485 | 4.87831578881992e-134 | 1 |
| acp5a1 | 4.16274437844577e-137 | 0.714690925 | 0.954 | 0.511 | 1.04772113261102e-132 | 1 |
| si:dkeyp-20e4.81 | 4.98517748359124e-137 | 0.764538311 | 0.808 | 0.38 | 1.25471932084508e-132 | 1 |
| rrm21 | 5.80638237426606e-136 | 0.710528802 | 0.897 | 0.471 | 1.46140837977903e-131 | 1 |
| zgc:1717761 | 6.26573605948657e-136 | 0.726074238 | 0.906 | 0.467 | 1.57702310881217e-131 | 1 |
| si:dkey-241l7.41 | 5.56576650023318e-134 | 0.655279902 | 0.954 | 0.585 | 1.40084777044369e-129 | 1 |
| si:dkey-90l23.11 | 3.12686593669021e-132 | 0.73083868 | 0.838 | 0.42 | 7.8700088760556e-128 | 1 |
| CR790368.11 | 1.35726946450162e-129 | 0.697172725 | 0.903 | 0.494 | 3.41611151520412e-125 | 1 |
| zgc:771181 | 3.23275662958778e-129 | 0.723965141 | 0.864 | 0.429 | 8.13652516100949e-125 | 1 |
| si:dkeyp-98a7.81 | 4.65763494886952e-129 | 0.723463745 | 0.858 | 0.43 | 1.17228014028097e-124 | 1 |
| epcam1 | 1.19820052067877e-128 | 0.696161475 | 0.883 | 0.479 | 3.01575089049639e-124 | 1 |
| si:dkeyp-98a7.41 | 4.03689329979388e-128 | 0.721213271 | 0.818 | 0.398 | 1.01604567462512e-123 | 1 |
| si:ch211-12h2.61 | 4.36978417255214e-128 | 0.704118616 | 0.854 | 0.448 | 1.09983097838965e-123 | 1 |
| LOC1005351621 | 3.28480426020437e-127 | 0.748861691 | 0.708 | 0.281 | 8.26752384250839e-123 | 1 |
| si:dkey-229d11.51 | 5.00320047551112e-127 | 0.730257276 | 0.774 | 0.363 | 1.25925552768139e-122 | 1 |
| CU571163.11 | 5.92054874712588e-123 | 0.730638188 | 0.791 | 0.375 | 1.49014291416411e-118 | 1 |
| si:dkeyp-46h3.81 | 2.76565532491172e-122 | 0.732475149 | 0.746 | 0.324 | 6.9608778872703e-118 | 1 |
| si:dkey-241l7.61 | 3.74810572645411e-121 | 0.662896722 | 0.859 | 0.504 | 9.43360730291235e-117 | 1 |
| si:dkeyp-50f7.21 | 4.16279854694205e-120 | 0.713305666 | 0.782 | 0.376 | 1.04773476627984e-115 | 1 |

| | | | | | | |
|---|---|---|---|---|---|---|
| ccnb21 | 3.06228217946179e-119 | 0.71709604 | 0.772 | 0.357 | 7.70745801748738e-115 | 1 |
| si:ch211-14a17.71 | 1.20447070297964e-118 | 0.710585796 | 0.786 | 0.382 | 3.03153231232946e-114 | 1 |
| kpna71 | 1.80503334905064e-118 | 0.725984784 | 0.706 | 0.302 | 4.54308843622555e-114 | 1 |
| zp3b1 | 2.38334463132556e-118 | 0.716092978 | 0.769 | 0.362 | 5.99864010258329e-114 | 1 |
| qdprb21 | 4.24553869465912e-118 | 0.70541434 | 0.818 | 0.406 | 1.06855963405875e-113 | 1 |
| zar11 | 8.39064832254755e-118 | 0.697319146 | 0.814 | 0.414 | 2.11184227630199e-113 | 1 |
| zgc:1362541 | 1.25788639637984e-117 | 0.72115483 | 0.709 | 0.304 | 3.16597427104843e-113 | 1 |
| zgc:1715171 | 5.53829436538862e-117 | 0.710347397 | 0.754 | 0.363 | 1.39393330882466e-112 | 1 |
| zgc:1734431 | 2.93470209218711e-116 | 0.716117501 | 0.714 | 0.303 | 7.38635169582573e-112 | 1 |
| zgc:1952451 | 1.86300583150894e-115 | 0.703363826 | 0.774 | 0.362 | 4.68899937732485e-111 | 1 |
| slbp21 | 2.76672006304275e-115 | 0.70229821 | 0.787 | 0.37 | 6.9635577266723e-111 | 1 |
| siva11 | 1.56722926513941e-114 | 0.688840097 | 0.806 | 0.402 | 3.94455933742939e-110 | 1 |
| si:dkey-152b24.61 | 1.05663719832784e-113 | 0.71140878 | 0.66 | 0.273 | 2.65945016447134e-109 | 1 |
| zar1l1 | 5.8228686764545e-113 | 0.698309782 | 0.776 | 0.372 | 1.46555781717683e-108 | 1 |
| AL929032.11 | 1.063394457446e-112 | 0.717283465 | 0.686 | 0.292 | 2.67645750994583e-108 | 1 |
| ddx4 | 2.18380794537879e-112 | 0.623977607 | 0.373 | 0.098 | 5.49642621772389e-108 | 1 |
| mid1ip1l1 | 2.33125142779925e-112 | 0.630871728 | 0.877 | 0.52 | 5.86752671862794e-108 | 1 |
| ccna11 | 3.30775168799216e-112 | 0.681057961 | 0.864 | 0.449 | 8.32528022350747e-108 | 1 |
| thy11 | 9.86520518092331e-112 | 0.676508664 | 0.822 | 0.425 | 2.48297349198659e-107 | 1 |
| s100a11 | 4.91476974179255e-111 | 0.696841556 | 0.735 | 0.338 | 1.23699839631177e-106 | 1 |
| ccnb11 | 8.09506595807763e-111 | 0.687909315 | 0.765 | 0.368 | 2.03744715098856e-106 | 1 |
| si:dkeyp-46h3.61 | 3.15074681434304e-110 | 0.687037733 | 0.768 | 0.386 | 7.93011465702001e-106 | 1 |
| nasp1 | 1.18770946369151e-108 | 0.698223333 | 0.686 | 0.316 | 2.98934594916517e-104 | 1 |
| hnrnpabb1 | 4.12471963990047e-108 | 0.565864541 | 0.856 | 0.579 | 1.03815068616655e-103 | 1 |
| tuba4l1 | 2.07167127339452e-107 | 0.684316903 | 0.776 | 0.379 | 5.21418942800666e-103 | 1 |
| npm2b1 | 5.05194202160117e-106 | 0.677859487 | 0.767 | 0.37 | 1.2715232874168e-101 | 1 |
| nanog1 | 1.7538603631738e-102 | 0.678519799 | 0.722 | 0.335 | 4.41429114807215e-98 | 1 |
| lamp21 | 3.08437305923382e-102 | 0.609819853 | 0.873 | 0.533 | 7.7630585527856e-98 | 1 |
| si:dkeyp-46h3.21 | 3.94349693860457e-102 | 0.664677258 | 0.751 | 0.381 | 9.92538744477384e-98 | 1 |
| zpcx1 | 7.09456452292564e-102 | 0.681652002 | 0.695 | 0.315 | 1.78563094477515e-97 | 1 |
| avd1 | 3.76609783065111e-100 | 0.674432204 | 0.606 | 0.242 | 9.47889162996577e-96 | 1 |
| uchl11 | 6.24683335961936e-100 | 0.665335642 | 0.697 | 0.32 | 1.5722654882826e-95 | 1 |
| si:ch211-132b12.81 | 5.65451105250153e-99 | 0.66799727 | 0.7 | 0.317 | 1.42318388680411e-94 | 1 |
| si:ch211-226l4.61 | 1.2442989024613e-98 | 0.663590734 | 0.633 | 0.269 | 3.13177590760485e-94 | 1 |
| cth11 | 1.50155768042472e-97 | 0.657151951 | 0.736 | 0.355 | 3.77927052586098e-93 | 1 |

| | | | | | | |
|---|---|---|---|---|---|---|
| si:ch211-145c1.11 | 6.63711571736461e-97 | 0.663973539 | 0.554 | 0.212 | 1.6704956549035e-92 | 1 |
| stmn1a | 1.01416227170063e-96 | 0.637977311 | 0.499 | 0.185 | 2.55254502164331e-92 | 1 |
| wu:fi42e031 | 1.47057899476614e-95 | 0.659823083 | 0.688 | 0.327 | 3.70130027192689e-91 | 1 |
| ccna21 | 3.57864392201109e-93 | 0.654070898 | 0.61 | 0.262 | 9.00708888730971e-89 | 1 |
| LOC1005374991 | 1.57419431858489e-92 | 0.658136958 | 0.663 | 0.299 | 3.96208968044631e-88 | 1 |
| si:dkey-90l23.21 | 4.07291055096119e-92 | 0.651673296 | 0.676 | 0.322 | 1.02511085657142e-87 | 1 |
| zgc:1714741 | 1.15117660830589e-91 | 0.650447114 | 0.636 | 0.281 | 2.89739640544509e-87 | 1 |
| si:dkeyp-46h3.51 | 4.01608033773146e-91 | 0.646173123 | 0.613 | 0.267 | 1.01080726020363e-86 | 1 |
| gmnn1 | 1.38117788421172e-90 | 0.644142544 | 0.633 | 0.28 | 3.47628661677248e-86 | 1 |
| gtf3ab1 | 3.60894645136197e-90 | 0.634961152 | 0.668 | 0.297 | 9.08335732343294e-86 | 1 |
| zgc:566991 | 7.37274518924546e-90 | 0.630356189 | 0.568 | 0.23 | 1.85564623668119e-85 | 1 |
| si:dkey-19b23.111 | 7.69120145893315e-90 | 0.646913475 | 0.591 | 0.257 | 1.93579849519888e-85 | 1 |
| pigrl2.31 | 1.22610156602405e-88 | 0.616716077 | 0.495 | 0.183 | 3.08597503152594e-84 | 1 |
| si:ch211-146l10.81 | 1.27957395599064e-87 | 0.645920267 | 0.626 | 0.285 | 3.22055968983285e-83 | 1 |
| ctss2.11 | 1.23492339829775e-86 | 0.556602972 | 0.824 | 0.515 | 3.1081787011756e-82 | 1 |
| si:ch73-90k17.11 | 1.38025962632474e-86 | 0.632482827 | 0.553 | 0.228 | 3.47397545349674e-82 | 1 |
| zp3a.11 | 4.53313578149568e-84 | 0.619770689 | 0.69 | 0.36 | 1.14094494484465e-79 | 1 |
| si:ch73-14h10.21 | 5.16671772240042e-84 | 0.62816938 | 0.582 | 0.25 | 1.30041118355096e-79 | 1 |
| btg41 | 8.88080649570187e-84 | 0.619607159 | 0.696 | 0.347 | 2.2352101869032e-79 | 1 |
| si:ch211-251f6.61 | 4.69455723616641e-83 | 0.617994055 | 0.64 | 0.31 | 1.18157311077072e-78 | 1 |
| gapdh1 | 5.29728180093991e-81 | 0.598664404 | 0.746 | 0.398 | 1.33327285647857e-76 | 1 |
| arhgef10la1 | 6.05337590561986e-81 | 0.61409851 | 0.578 | 0.254 | 1.52357418168546e-76 | 1 |
| ca15b1 | 7.72598014471387e-81 | 0.605882963 | 0.549 | 0.229 | 1.94455194262303e-76 | 1 |
| si:dkey-208k4.21 | 3.39524333813053e-80 | 0.608321176 | 0.615 | 0.274 | 8.54548795774074e-76 | 1 |
| AL929192.11 | 6.82315056110008e-79 | 0.56479868 | 0.418 | 0.147 | 1.71731876472328e-74 | 1 |
| si:dkey-37o8.11 | 8.72843775244593e-79 | 0.593737651 | 0.619 | 0.286 | 2.19686049791312e-74 | 1 |
| LOC1018854771 | 2.01252867452655e-78 | 0.610771136 | 0.577 | 0.26 | 5.06533342091588e-74 | 1 |
| zgc:562311 | 7.48906663634784e-78 | 0.591020173 | 0.518 | 0.208 | 1.88492318170239e-73 | 1 |
| zgc:554611 | 5.04430322806725e-77 | 0.602085328 | 0.588 | 0.268 | 1.26960067947225e-72 | 1 |
| mkrn41 | 9.42629732314915e-77 | 0.581535634 | 0.512 | 0.209 | 2.37250477326341e-72 | 1 |
| CU467646.21 | 1.13896553616821e-76 | 0.523504216 | 0.35 | 0.111 | 2.86666235798177e-72 | 1 |
| elovl7b1 | 4.45315046718463e-76 | 0.590707035 | 0.54 | 0.228 | 1.1208134410857e-71 | 1 |
| si:dkeyp-98a7.71 | 8.67179329461102e-76 | 0.575880733 | 0.464 | 0.177 | 2.18260365432065e-71 | 1 |
| gdf31 | 4.05889476084337e-75 | 0.578160481 | 0.501 | 0.205 | 1.02158322235667e-70 | 1 |
| CABZ01059627.11 | 4.06792779110683e-74 | 0.530558628 | 0.36 | 0.121 | 1.02385674574368e-69 | 1 |

| | | | | | | |
|---|---|---|---|---|---|---|
| tmem54a1 | 5.62442185672528e-74 | 0.529514217 | 0.371 | 0.126 | 1.41561073711919e-69 | 1 |
| cpeb1b1 | 5.28869334103484e-73 | 0.589080133 | 0.567 | 0.25 | 1.33111122700506e-68 | 1 |
| lsm14b1 | 1.52090903630201e-71 | 0.574981508 | 0.596 | 0.28 | 3.82797595346852e-67 | 1 |
| org1 | 1.29381848114307e-69 | 0.574060607 | 0.504 | 0.223 | 3.25641173518899e-65 | 1 |
| zgc:1737421 | 3.30280840519012e-69 | 0.532363099 | 0.385 | 0.14 | 8.312838475023e-65 | 1 |
| si:ch211-250e5.161 | 3.93767498301841e-68 | 0.563940296 | 0.626 | 0.328 | 9.91073416475904e-64 | 1 |
| larp6b1 | 8.09722241325715e-68 | 0.540529014 | 0.449 | 0.179 | 2.03798990919269e-63 | 1 |
| zgc:660241 | 1.12686966533136e-67 | 0.508887391 | 0.36 | 0.127 | 2.8362182606725e-63 | 1 |
| crp31 | 1.44148442304228e-67 | 0.564129002 | 0.474 | 0.201 | 3.62807214435512e-63 | 1 |
| LOC1018838971 | 1.54365637930065e-67 | 0.518669634 | 0.403 | 0.148 | 3.8852287410618e-63 | 1 |
| chia.41 | 2.82290647156615e-67 | 0.508352333 | 0.371 | 0.132 | 7.10497329828485e-63 | 1 |
| patl21 | 2.84357764054102e-67 | 0.524146329 | 0.413 | 0.157 | 7.1570005634777e-63 | 1 |
| si:ch1073-263o8.21 | 1.21199955358173e-66 | 0.567765878 | 0.531 | 0.246 | 3.05048167640986e-62 | 1 |
| pabpn1l1 | 6.26529257056049e-66 | 0.547114026 | 0.571 | 0.264 | 1.57691148708437e-61 | 1 |
| birc5b1 | 9.93509912294357e-65 | 0.551777544 | 0.535 | 0.241 | 2.50056509825367e-60 | 1 |
| sinup1 | 1.36404780935134e-64 | 0.538099195 | 0.451 | 0.187 | 3.4331719313564e-60 | 1 |
| rbpms2b1 | 1.93580112318675e-64 | 0.54132639 | 0.476 | 0.204 | 4.87221784694872e-60 | 1 |
| zgc:1738561 | 2.26304189796578e-64 | 0.554843237 | 0.464 | 0.204 | 5.69585015299007e-60 | 1 |
| zgc:1034821 | 1.64206232982168e-63 | 0.525158432 | 0.428 | 0.173 | 4.13290667792818e-59 | 1 |
| elovl7a1 | 7.51143249672898e-63 | 0.515818073 | 0.423 | 0.167 | 1.89055244510172e-58 | 1 |
| zgc:1628791 | 2.68494059027492e-62 | 0.522143893 | 0.414 | 0.166 | 6.75772697166295e-58 | 1 |
| zp3c1 | 3.97600764748205e-62 | 0.514743922 | 0.428 | 0.173 | 1.00072136479476e-57 | 1 |
| map1lc3c1 | 7.25520524584892e-62 | 0.523564023 | 0.445 | 0.186 | 1.82606260832772e-57 | 1 |
| snu13b1 | 9.55839227329046e-62 | 0.532223581 | 0.641 | 0.376 | 2.40575175126448e-57 | 1 |
| slc16a31 | 2.86095074179107e-61 | 0.54187936 | 0.506 | 0.232 | 7.20072692201396e-57 | 1 |
| zgc:111868 | 3.80137635989059e-61 | 0.499576683 | 0.356 | 0.133 | 9.56768416020863e-57 | 1 |
| zgc:1629451 | 6.81893271902165e-61 | 0.500927411 | 0.385 | 0.148 | 1.71625717605056e-56 | 1 |
| zgc:925911 | 8.36059092942905e-61 | 0.51974313 | 0.45 | 0.189 | 2.104277131028e-56 | 1 |
| retsatl1 | 1.4977783373013e-60 | 0.529093668 | 0.436 | 0.188 | 3.76975829715365e-56 | 1 |
| hsp90aa1.2 | 1.5009804344907e-60 | 0.380066391 | 0.162 | 0.036 | 3.77781765556964e-56 | 1 |
| larp6a1 | 1.99889149289901e-60 | 0.542328322 | 0.494 | 0.234 | 5.03100999847753e-56 | 1 |
| si:dkey-188i13.111 | 4.38076493431648e-60 | 0.54489675 | 0.56 | 0.3 | 1.10259472631811e-55 | 1 |
| dctpp11 | 4.58487293028699e-60 | 0.498017902 | 0.401 | 0.16 | 1.15396666782393e-55 | 1 |
| haus61 | 6.07093387794489e-60 | 0.515388667 | 0.453 | 0.193 | 1.52799334773995e-55 | 1 |
| hsd17b12a1 | 8.28482887274919e-60 | 0.496004395 | 0.385 | 0.151 | 2.08520857898224e-55 | 1 |

| | | | | | | |
|---|---|---|---|---|---|---|
| top2a | 1.7760117043682e-59 | 0.352583284 | 0.153 | 0.033 | 4.47004385872433e-55 | 1 |
| pttg11 | 2.94658310634719e-59 | 0.492793189 | 0.372 | 0.144 | 7.41625502036524e-55 | 1 |
| rcc11 | 4.16171478397645e-58 | 0.520990224 | 0.433 | 0.187 | 1.04746199397903e-53 | 1 |
| zgc:114130 | 5.73875597627412e-58 | 0.364074409 | 0.229 | 0.065 | 1.44438749166843e-53 | 1 |
| gbgt1l41 | 1.10739836881675e-57 | 0.479924332 | 0.36 | 0.136 | 2.78721095447489e-53 | 1 |
| zgc:1134241 | 6.13336135202544e-57 | 0.469150317 | 0.381 | 0.147 | 1.54370571869128e-52 | 1 |
| sybu1 | 1.79834348972174e-56 | 0.485841009 | 0.382 | 0.152 | 4.52625072928064e-52 | 1 |
| zgc:171977 | 3.7726377723977e-56 | 0.516065007 | 0.468 | 0.214 | 9.49535200934776e-52 | 1 |
| buc1 | 7.23303817545101e-56 | 0.492991434 | 0.397 | 0.164 | 1.82048337837926e-51 | 1 |
| si:ch211-146l10.71 | 1.40240476807562e-55 | 0.494058219 | 0.412 | 0.174 | 3.52971256076953e-51 | 1 |
| mad2l11 | 5.64548718296316e-55 | 0.486156587 | 0.405 | 0.171 | 1.42091266908e-50 | 1 |
| inka1b1 | 1.07047275333486e-54 | 0.484667703 | 0.404 | 0.17 | 2.69427287286851e-50 | 1 |
| tac3b1 | 4.53114701687009e-54 | 0.445803101 | 0.303 | 0.108 | 1.14044439267603e-49 | 1 |
| cdca7a1 | 2.2626793062994e-53 | 0.497036168 | 0.453 | 0.207 | 5.69493754602496e-49 | 1 |
| zglp11 | 2.86744647893729e-53 | 0.476424004 | 0.427 | 0.183 | 7.21707604283728e-49 | 1 |
| CR847984.11 | 7.88884642729801e-53 | 0.472618518 | 0.371 | 0.15 | 1.98554375728664e-48 | 1 |
| zgc:101744 | 8.99930100933998e-53 | 0.437241814 | 0.323 | 0.121 | 2.26503407104078e-48 | 1 |
| zgc:1138861 | 2.43402568490838e-51 | 0.443672371 | 0.335 | 0.129 | 6.12619924634591e-47 | 1 |
| cdc6 | 2.69424303854427e-51 | 0.400013531 | 0.268 | 0.091 | 6.78114030371207e-47 | 1 |
| zgc:1717501 | 2.86471040411443e-51 | 0.47514636 | 0.379 | 0.161 | 7.21018961611562e-47 | 1 |
| sox19b1 | 3.66356921658129e-51 | 0.386458176 | 0.262 | 0.087 | 9.22083736121344e-47 | 1 |
| slbp1 | 5.58909425898107e-51 | 0.48926756 | 0.465 | 0.219 | 1.40671913404294e-46 | 1 |
| gyg1a1 | 1.54876335975575e-50 | 0.49597722 | 0.6 | 0.355 | 3.89808250016924e-46 | 1 |
| spink2.11 | 6.45887757041075e-50 | 0.474558482 | 0.699 | 0.419 | 1.62563489569668e-45 | 1 |
| si:ch211-14a17.61 | 8.1379029874766e-50 | 0.379590423 | 0.237 | 0.076 | 2.04822880291799e-45 | 1 |
| LOC1005368601 | 1.24494150947211e-49 | 0.487908882 | 0.583 | 0.336 | 3.13339328519036e-45 | 1 |
| si:ch73-78o10.11 | 2.86391158116207e-49 | 0.420132967 | 0.318 | 0.121 | 7.20817905862681e-45 | 1 |
| si:ch1073-190k2.11 | 3.6248788891636e-49 | 0.474655791 | 0.396 | 0.175 | 9.12345767613587e-45 | 1 |
| CDPF11 | 4.75351320081319e-49 | 0.49280118 | 0.491 | 0.254 | 1.19641173751267e-44 | 1 |
| pabpc1l1 | 8.49996115717482e-49 | 0.43042303 | 0.331 | 0.13 | 2.13935522364933e-44 | 1 |
| cdc201 | 2.66208089413063e-48 | 0.433006426 | 0.347 | 0.14 | 6.70019140243739e-44 | 1 |
| zgc:171426 | 3.43967047834467e-48 | 0.397518903 | 0.272 | 0.097 | 8.65730662694571e-44 | 1 |
| CU655961.3 | 7.1382199193757e-48 | 0.382248119 | 0.258 | 0.089 | 1.79661857150767e-43 | 1 |
| zgc:92287 | 1.89396614905597e-47 | 0.416280382 | 0.313 | 0.121 | 4.76692340055897e-43 | 1 |
| mcm3l | 2.87862585176319e-47 | 0.374846826 | 0.236 | 0.078 | 7.24521340630276e-43 | 1 |

| | | | | | | |
|---|---|---|---|---|---|---|
| elavl21 | 4.8321082503418e-47 | 0.46907035 | 0.44 | 0.21 | 1.21619332552853e-42 | 1 |
| h3f3a1 | 9.25066072311096e-47 | 0.430402789 | 0.703 | 0.493 | 2.3282987973998e-42 | 1 |
| aurka1 | 1.24258754794331e-46 | 0.409964915 | 0.288 | 0.109 | 3.12746859941853e-42 | 1 |
| zgc:1139841 | 1.32550938614801e-46 | 0.375189979 | 0.26 | 0.091 | 3.33617457399592e-42 | 1 |
| si:dkey-216e24.3 | 1.72329225841196e-46 | 0.407922694 | 0.294 | 0.112 | 4.33735428519707e-42 | 1 |
| rpain | 4.35691287175754e-46 | 0.409458238 | 0.286 | 0.109 | 1.09659140069266e-41 | 1 |
| pou5f31 | 1.05064318256706e-45 | 0.42448403 | 0.332 | 0.135 | 2.64436382620304e-41 | 1 |
| BX548000.1 | 1.26339295076611e-45 | 0.286958755 | 0.144 | 0.035 | 3.17983371778322e-41 | 1 |
| BX537276.11 | 2.55629421474129e-45 | 0.403731745 | 0.291 | 0.111 | 6.43393690908234e-41 | 1 |
| zp2l11 | 5.4625697476071e-45 | 0.43562827 | 0.355 | 0.152 | 1.37487417977523e-40 | 1 |
| si:dkey-46g23.21 | 1.08321692923476e-44 | 0.425827763 | 0.326 | 0.133 | 2.72634868919097e-40 | 1 |
| BX284697.11 | 1.11563725745333e-44 | 0.389539354 | 0.277 | 0.104 | 2.8079474132843e-40 | 1 |
| buc2l | 1.39161378334702e-44 | 0.348519298 | 0.219 | 0.071 | 3.50255273130611e-40 | 1 |
| zgc:566761 | 2.3876143190383e-44 | 0.471199558 | 0.504 | 0.282 | 6.00938647958751e-40 | 1 |
| gstp11 | 3.25210186072111e-44 | 0.45400393 | 0.574 | 0.352 | 8.18521517324895e-40 | 1 |
| hmgb2a | 5.54284360359228e-44 | 0.348792401 | 0.794 | 0.61 | 1.39507830658814e-39 | 1 |
| rasl11b1 | 6.70701988678678e-44 | 0.405267189 | 0.308 | 0.123 | 1.68808983530537e-39 | 1 |
| rdh10b1 | 7.05190366780768e-44 | 0.446343734 | 0.365 | 0.166 | 1.77489363415051e-39 | 1 |
| ubc | 8.16553137628172e-44 | 0.309727123 | 0.771 | 0.651 | 2.05518259209635e-39 | 1 |
| plk11 | 9.82795402706368e-44 | 0.433438228 | 0.372 | 0.165 | 2.47359774907166e-39 | 1 |
| parn1 | 1.11250406593007e-43 | 0.424559812 | 0.323 | 0.136 | 2.80006148353939e-39 | 1 |
| si:ch73-160p18.31 | 1.44247668268361e-43 | 0.46743094 | 0.45 | 0.227 | 3.63056956264638e-39 | 1 |
| ypel3 | 1.88289890622163e-43 | 0.397978851 | 0.295 | 0.116 | 4.73906825706923e-39 | 1 |
| prr111 | 3.61390373347968e-43 | 0.383987828 | 0.249 | 0.091 | 9.09583430679502e-39 | 1 |
| LOC110437779 | 3.76580078701164e-43 | 0.350989767 | 0.222 | 0.075 | 9.4781440008296e-39 | 1 |
| snx10a | 4.23334646233998e-43 | 0.405546547 | 0.291 | 0.116 | 1.06549097110635e-38 | 1 |
| kpna21 | 6.12591071990428e-43 | 0.417876464 | 0.341 | 0.146 | 1.54183046909271e-38 | 1 |
| BX001030.11 | 8.19623489353118e-43 | 0.387368836 | 0.287 | 0.112 | 2.06291036035286e-38 | 1 |
| zgc:152652 | 8.81916093253932e-43 | 0.394285246 | 0.272 | 0.104 | 2.21969461511082e-38 | 1 |
| evla | 1.327910143059e-42 | 0.388157632 | 0.249 | 0.093 | 3.34221703906519e-38 | 1 |
| si:ch73-15b2.5 | 1.42329796944405e-42 | 0.360710632 | 0.241 | 0.085 | 3.58229865929373e-38 | 1 |
| eif4e1b | 1.66369257962038e-42 | 0.352205266 | 0.224 | 0.077 | 4.18734785364654e-38 | 1 |
| rbm4.31 | 1.66978982925664e-42 | 0.424787577 | 0.636 | 0.432 | 4.20269402125604e-38 | 1 |
| zgc:1141231 | 3.75840396655899e-42 | 0.379593916 | 0.254 | 0.094 | 9.45952694343231e-38 | 1 |
| CABZ01059627.21 | 3.98351792320792e-42 | 0.39959488 | 0.303 | 0.123 | 1.0026116260922e-37 | 1 |

| | | | | | | |
|---|---|---|---|---|---|---|
| brf21 | 6.65288297474224e-42 | 0.368604162 | 0.238 | 0.086 | 1.67446411591288e-37 | 1 |
| CR383662.21 | 8.66023351239921e-42 | 0.337172675 | 0.227 | 0.078 | 2.17969417273576e-37 | 1 |
| si:dkey-241l7.31 | 9.25126033969163e-42 | 0.455216972 | 0.581 | 0.351 | 2.32844971489699e-37 | 1 |
| zgc:86839 | 1.99441957984322e-41 | 0.383976888 | 0.262 | 0.1 | 5.0197546405074e-37 | 1 |
| zgc:1655391 | 5.70808683126959e-41 | 0.387421733 | 0.278 | 0.11 | 1.43666837456224e-36 | 1 |
| tph1a1 | 8.27516758032718e-41 | 0.395464836 | 0.313 | 0.131 | 2.08277692829255e-36 | 1 |
| pif1 | 1.28929691589185e-40 | 0.365454773 | 0.258 | 0.097 | 3.24503140760819e-36 | 1 |
| cx44.21 | 4.72023376216455e-40 | 0.371651265 | 0.273 | 0.107 | 1.18803563559919e-35 | 1 |
| dazl | 1.00093975407695e-39 | 0.371104213 | 0.254 | 0.097 | 2.51926526703629e-35 | 1 |
| degs21 | 1.70584380039413e-39 | 0.368469147 | 0.254 | 0.097 | 4.29343826121197e-35 | 1 |
| cpm1 | 6.81486715299911e-39 | 0.358676982 | 0.256 | 0.099 | 1.71523391373835e-34 | 1 |
| si:dkeyp-114g9.11 | 9.45760371841633e-39 | 0.351971194 | 0.25 | 0.095 | 2.38038427988821e-34 | 1 |
| tpx21 | 1.00143219156608e-38 | 0.38119703 | 0.281 | 0.116 | 2.52050468295266e-34 | 1 |
| thop1 | 2.10752973321027e-38 | 0.366864577 | 0.267 | 0.108 | 5.30444158551694e-34 | 1 |
| vstm2l1 | 3.40956468783617e-38 | 0.34131561 | 0.235 | 0.087 | 8.58153336281487e-34 | 1 |
| il13ra21 | 3.88484649786675e-38 | 0.360553795 | 0.256 | 0.101 | 9.77777015048081e-34 | 1 |
| exd2 | 7.89406929702565e-38 | 0.338300211 | 0.228 | 0.085 | 1.98685830136839e-33 | 1 |
| zgc:1529861 | 8.26448908124919e-38 | 0.333917298 | 0.223 | 0.081 | 2.08008925685961e-33 | 1 |
| yrdc | 8.96327303925497e-38 | 0.364793963 | 0.278 | 0.113 | 2.25596619125008e-33 | 1 |
| tmem144b1 | 9.95780845445936e-38 | 0.335859448 | 0.231 | 0.086 | 2.50628080990288e-33 | 1 |
| orc61 | 1.84187762025374e-37 | 0.357413505 | 0.251 | 0.098 | 4.63582178241663e-33 | 1 |
| BX571780.11 | 2.05841061190653e-37 | 0.326308633 | 0.209 | 0.073 | 5.18081366910755e-33 | 1 |
| srsf2a1 | 2.18763271323355e-37 | 0.424450935 | 0.549 | 0.355 | 5.50605277593752e-33 | 1 |
| st8sia7.1 | 2.37554737498635e-37 | 0.306341939 | 0.191 | 0.064 | 5.97901518810313e-33 | 1 |
| cdca81 | 5.10550532332335e-37 | 0.395492887 | 0.337 | 0.154 | 1.28500463482725e-32 | 1 |
| dnd1 | 5.27338455914899e-37 | 0.354774411 | 0.254 | 0.101 | 1.32725815969221e-32 | 1 |
| ankrd221 | 8.2094742907231e-37 | 0.386719805 | 0.304 | 0.133 | 2.0662425842321e-32 | 1 |
| nfr1 | 1.15872365070767e-36 | 0.34849267 | 0.256 | 0.102 | 2.91639155646613e-32 | 1 |
| krcp | 1.73571525492918e-36 | 0.317094815 | 0.195 | 0.068 | 4.36862172513126e-32 | 1 |
| piwil1 | 1.8388631743801e-36 | 0.285351563 | 0.136 | 0.038 | 4.62823472359729e-32 | 1 |
| cldn7b1 | 2.00125044585121e-36 | 0.375823708 | 0.333 | 0.149 | 5.0369472471629e-32 | 1 |
| si:ch73-303b9.1 | 6.42495551738727e-36 | 0.300698374 | 0.203 | 0.071 | 1.6170970541712e-31 | 1 |
| mos | 7.20212536336816e-36 | 0.381192143 | 0.292 | 0.127 | 1.81270293270613e-31 | 1 |
| aurkb | 9.01211242827524e-36 | 0.368488331 | 0.278 | 0.119 | 2.26825857707259e-31 | 1 |
| si:ch211-286b5.4 | 1.19118222942696e-35 | 0.297321099 | 0.186 | 0.063 | 2.99808655324472e-31 | 1 |

| | | | | | | |
|---|---|---|---|---|---|---|
| ddx191 | 1.28809657224527e-35 | 0.376828148 | 0.313 | 0.141 | 3.24201026268413e-31 | 1 |
| rogdi | 1.58338823830303e-35 | 0.365257612 | 0.259 | 0.107 | 3.98522985698489e-31 | 1 |
| si:ch211-69g19.21 | 3.21231656844251e-35 | 0.361462453 | 0.269 | 0.113 | 8.08507957111295e-31 | 1 |
| LOC1039099721 | 3.37339987588215e-35 | 0.416095183 | 0.538 | 0.356 | 8.49051014760778e-31 | 1 |
| coil | 3.38082971236471e-35 | 0.350140469 | 0.245 | 0.099 | 8.50921030305074e-31 | 1 |
| CU326366.2 | 4.57028112059266e-35 | 0.277230469 | 0.162 | 0.051 | 1.15029405524197e-30 | 1 |
| tram21 | 7.44613325639021e-35 | 0.359576268 | 0.285 | 0.123 | 1.87411727930085e-30 | 1 |
| hells1 | 2.19699120369827e-34 | 0.371535261 | 0.281 | 0.124 | 5.52960716058818e-30 | 1 |
| zfand6 | 4.09946619545642e-34 | 0.368824624 | 0.269 | 0.117 | 1.03179464673443e-29 | 1 |
| depdc7b | 5.98781403172018e-34 | 0.283082623 | 0.177 | 0.06 | 1.50707291364365e-29 | 1 |
| BX005461.1 | 6.25149410219782e-34 | 0.295597481 | 0.2 | 0.072 | 1.57343855058217e-29 | 1 |
| si:dkey-43p13.5 | 8.23312727845073e-34 | 0.307286165 | 0.206 | 0.076 | 2.07219580471326e-29 | 1 |
| si:ch211-113a14.27 | 1.08508544565124e-33 | 0.310855266 | 0.195 | 0.071 | 2.7310515581596e-29 | 1 |
| ndc1 | 2.09999497314065e-33 | 0.330287776 | 0.224 | 0.089 | 5.28547734789771e-29 | 1 |
| CT009596.11 | 2.23517108469713e-33 | 0.313410408 | 0.217 | 0.082 | 5.6257021030742e-29 | 1 |
| zgc:113425 | 3.57423798120498e-33 | 0.296942037 | 0.178 | 0.062 | 8.99599957489482e-29 | 1 |
| znf395a | 3.86508004731556e-33 | 0.341052851 | 0.223 | 0.089 | 9.72801997108854e-29 | 1 |
| zgc:163030 | 4.20458242556963e-33 | 0.359853056 | 0.267 | 0.115 | 1.05825135069162e-28 | 1 |
| rpa3 | 4.70699892529875e-33 | 0.38597791 | 0.317 | 0.154 | 1.18470455950844e-28 | 1 |
| tatdn2 | 5.05719685053525e-33 | 0.29528234 | 0.186 | 0.066 | 1.27284587531122e-28 | 1 |
| znf1173 | 1.55427806065242e-32 | 0.256456356 | 0.128 | 0.037 | 3.91196245085608e-28 | 1 |
| prdx11 | 2.56572831190663e-32 | 0.378134311 | 0.324 | 0.157 | 6.4576815882378e-28 | 1 |
| tmem44 | 4.62763665050294e-32 | 0.265888406 | 0.163 | 0.055 | 1.16472986856509e-27 | 1 |
| gpx8 | 6.73651005066626e-32 | 0.312802855 | 0.2 | 0.077 | 1.69551221465219e-27 | 1 |
| lpcat41 | 7.29329560997509e-32 | 0.320332275 | 0.241 | 0.099 | 1.83564957207463e-27 | 1 |
| umps | 7.92981281560461e-32 | 0.299216085 | 0.19 | 0.07 | 1.99585458755953e-27 | 1 |
| zgc:55888 | 8.20269625654961e-32 | 0.304613159 | 0.205 | 0.078 | 2.06453662081097e-27 | 1 |
| zgc:173544 | 1.00278517629713e-31 | 0.293487021 | 0.19 | 0.069 | 2.52391001022224e-27 | 1 |
| gpr1601 | 1.05289374558199e-31 | 0.30064957 | 0.197 | 0.074 | 2.65002826825531e-27 | 1 |
| prc1a | 1.33255893772959e-31 | 0.273989413 | 0.173 | 0.061 | 3.3539175903716e-27 | 1 |
| cdca9 | 1.75568590221002e-31 | 0.299479095 | 0.205 | 0.078 | 4.4188858472724e-27 | 1 |
| mibp1 | 3.38392062257732e-31 | 0.385057184 | 0.399 | 0.212 | 8.51698981496486e-27 | 1 |
| zgc:101016 | 3.41057479017018e-31 | 0.320031971 | 0.219 | 0.089 | 8.58407568937933e-27 | 1 |
| nek2 | 3.83841793505173e-31 | 0.282621962 | 0.174 | 0.062 | 9.66091410073171e-27 | 1 |
| si:dkey-42i9.81 | 4.8384969592487e-31 | 0.329387154 | 0.271 | 0.119 | 1.2178012996733e-26 | 1 |

| | | | | | | |
|---|---|---|---|---|---|---|
| ddx39ab1 | 6.37149725326208e-31 | 0.400070556 | 0.467 | 0.278 | 1.60364214367353e-26 | 1 |
| h2afva1 | 6.93369453680466e-31 | 0.393323159 | 0.499 | 0.312 | 1.74514157796836e-26 | 1 |
| ctsba1 | 1.22462657767914e-30 | 0.38161664 | 0.559 | 0.365 | 3.08226263336063e-26 | 1 |
| tspan13a | 1.22621252561285e-30 | 0.272887662 | 0.163 | 0.056 | 3.08625430571499e-26 | 1 |
| lmnl3 | 1.97192365887558e-30 | 0.289271772 | 0.186 | 0.069 | 4.96313465702395e-26 | 1 |
| rbbp4 | 2.15660878300467e-30 | 0.374182286 | 0.322 | 0.161 | 5.42796864594444e-26 | 1 |
| metap1d | 2.30546858150978e-30 | 0.311536172 | 0.208 | 0.083 | 5.80263387280198e-26 | 1 |
| ttc32 | 2.74290030393726e-30 | 0.314167208 | 0.229 | 0.096 | 6.90360577497968e-26 | 1 |
| dynll2a | 3.15566135573034e-30 | 0.343563523 | 0.279 | 0.128 | 7.94248406623769e-26 | 1 |
| selenoh | 6.38240121632784e-30 | 0.385640779 | 0.408 | 0.234 | 1.60638656213755e-25 | 1 |
| si:dkey-189g17.2 | 7.27551399363717e-30 | 0.261315435 | 0.151 | 0.051 | 1.83117411705854e-25 | 1 |
| cx43.4 | 9.98404674117162e-30 | 0.346965235 | 0.255 | 0.116 | 2.51288472428548e-25 | 1 |
| ccnh1 | 1.06383049799859e-29 | 0.343231331 | 0.259 | 0.117 | 2.67755498041266e-25 | 1 |
| mrps18b | 1.17515881937545e-29 | 0.345772187 | 0.258 | 0.118 | 2.95775723248606e-25 | 1 |
| zgc:92313 | 1.65772568182851e-29 | 0.334176279 | 0.262 | 0.118 | 4.17232976859418e-25 | 1 |
| fbxo32 | 2.17025795258344e-29 | 0.306998621 | 0.229 | 0.097 | 5.46232224085725e-25 | 1 |
| hspa51 | 3.94356137598301e-29 | 0.388995825 | 0.486 | 0.304 | 9.92554962721164e-25 | 1 |
| bmb | 1.64731141993606e-28 | 0.284850284 | 0.204 | 0.081 | 4.14611811283707e-24 | 1 |
| zc3h11a1 | 1.66561330915136e-28 | 0.3727653 | 0.377 | 0.207 | 4.19218213780306e-24 | 1 |
| ccne11 | 1.69130045788467e-28 | 0.302411476 | 0.24 | 0.103 | 4.25683412244992e-24 | 1 |
| fdx11 | 2.08113058560667e-28 | 0.378239774 | 0.378 | 0.211 | 5.23799757091343e-24 | 1 |
| npm2a | 2.56454440186391e-28 | 0.277489079 | 0.187 | 0.072 | 6.45470180505127e-24 | 1 |
| snu13a1 | 2.69790752115886e-28 | 0.308930589 | 0.222 | 0.094 | 6.79036344000473e-24 | 1 |
| BX957362.1 | 2.74424988027978e-28 | 0.335285376 | 0.267 | 0.125 | 6.90700252367617e-24 | 1 |
| cited1 | 3.07416505929567e-28 | 0.253737177 | 0.162 | 0.058 | 7.73736603774126e-24 | 1 |
| lrp13 | 3.7565560490485e-28 | 0.277310974 | 0.171 | 0.064 | 9.45487591985018e-24 | 1 |
| pcna | 4.33826051058479e-28 | 0.338888822 | 0.277 | 0.135 | 1.09189678790908e-23 | 1 |
| dynll11 | 4.61375753762797e-28 | 0.351757475 | 0.603 | 0.445 | 1.16123663464558e-23 | 1 |
| si:dkey-87o1.2 | 4.62791288345314e-28 | 0.272810715 | 0.191 | 0.075 | 1.16479939363632e-23 | 1 |
| BX005461.3 | 5.01575128730461e-28 | 0.261704025 | 0.167 | 0.061 | 1.2624144415017e-23 | 1 |
| LOC100538023 | 6.7585651150372e-28 | 0.265645824 | 0.183 | 0.07 | 1.70106325380371e-23 | 1 |
| itm2cb1 | 7.66067342332628e-28 | 0.33805386 | 0.273 | 0.13 | 1.92811489391699e-23 | 1 |
| lrrc59 | 1.13020533202962e-27 | 0.345233985 | 0.277 | 0.135 | 2.84461380018534e-23 | 1 |
| exd1 | 1.31155404101333e-27 | 0.26582522 | 0.153 | 0.055 | 3.30105036582645e-23 | 1 |
| wu:fa19b12 | 2.19804117094388e-27 | 0.254938591 | 0.169 | 0.063 | 5.53224982314865e-23 | 1 |

| | | | | | | |
|---|---|---|---|---|---|---|
| zgc:175136 | 2.86355505704791e-27 | 0.316680757 | 0.227 | 0.1 | 7.20728172308388e-23 | 1 |
| cox171 | 2.92770078577695e-27 | 0.361715008 | 0.531 | 0.368 | 7.36873010772202e-23 | 1 |
| dcun1d1 | 3.29003300341061e-27 | 0.309160041 | 0.213 | 0.091 | 8.28068406628416e-23 | 1 |
| ddx41 | 3.86591244614256e-27 | 0.258772685 | 0.168 | 0.063 | 9.73011503569621e-23 | 1 |
| bzw1b | 4.04020855730618e-27 | 0.358939213 | 0.551 | 0.393 | 1.01688009178839e-22 | 1 |
| CU655961.41 | 4.14920702965288e-27 | 0.291766138 | 0.2 | 0.082 | 1.04431391729333e-22 | 1 |
| si:ch211-157c3.4 | 4.62070673370338e-27 | 0.283080522 | 0.203 | 0.083 | 1.1629856778058e-22 | 1 |
| pias4b | 5.40530184318992e-27 | 0.264792401 | 0.16 | 0.059 | 1.36046042091247e-22 | 1 |
| snrpb2 | 7.64223803147604e-27 | 0.347679338 | 0.283 | 0.141 | 1.9234748901422e-22 | 1 |
| g3bp11 | 1.17281838295864e-26 | 0.366123154 | 0.359 | 0.202 | 2.95186658806861e-22 | 1 |
| si:ch73-340n8.4 | 1.28135799128228e-26 | 0.32419726 | 0.255 | 0.12 | 3.22504992825837e-22 | 1 |
| mylipa | 1.39498199025073e-26 | 0.322130773 | 0.237 | 0.108 | 3.51103017126206e-22 | 1 |
| cdt1 | 1.83962587294557e-26 | 0.287965599 | 0.196 | 0.081 | 4.63015435961671e-22 | 1 |
| sec61a1l | 2.23520864741105e-26 | 0.321787627 | 0.251 | 0.119 | 5.62579664466888e-22 | 1 |
| chrac1 | 2.28037826429504e-26 | 0.320943804 | 0.255 | 0.121 | 5.73948405340418e-22 | 1 |
| gemin7 | 2.76473793134665e-26 | 0.339065634 | 0.288 | 0.147 | 6.95856889940639e-22 | 1 |
| ccdc58 | 2.90508130419379e-26 | 0.3080097 | 0.228 | 0.103 | 7.31179913452535e-22 | 1 |
| rpp30 | 3.45227586960842e-26 | 0.256237674 | 0.155 | 0.058 | 8.68903313621743e-22 | 1 |
| aif1l | 3.9288057265629e-26 | 0.33933149 | 0.337 | 0.178 | 9.88841113318615e-22 | 1 |
| ctsla1 | 4.28713998313865e-26 | 0.354216303 | 0.549 | 0.377 | 1.07903026235617e-21 | 1 |
| si:ch73-281n10.2 | 4.683730214913e-26 | 0.349216625 | 0.542 | 0.383 | 1.17884805779145e-21 | 1 |
| bokb | 4.93665333834881e-26 | 0.250049535 | 0.151 | 0.055 | 1.24250627872901e-21 | 1 |
| lman1 | 5.38844001494488e-26 | 0.299347572 | 0.203 | 0.088 | 1.35621646736148e-21 | 1 |
| eif4a1a1 | 6.21267615428986e-26 | 0.340058902 | 0.568 | 0.416 | 1.56366846127321e-21 | 1 |
| timm13 | 6.33670396073042e-26 | 0.296532747 | 0.212 | 0.093 | 1.59488501987624e-21 | 1 |
| cfl1l | 7.06239375686527e-26 | 0.320183727 | 0.291 | 0.144 | 1.77753388466542e-21 | 1 |
| zgc:153284 | 9.47167458828726e-26 | 0.296071655 | 0.214 | 0.094 | 2.38392577712602e-21 | 1 |
| marcksb | 1.04154690393985e-25 | 0.338052642 | 0.319 | 0.167 | 2.6214694025262e-21 | 1 |
| mrpl22 | 1.07819543095734e-25 | 0.265803966 | 0.165 | 0.065 | 2.71371008017654e-21 | 1 |
| si:dkey-68o6.51 | 1.08060404517782e-25 | 0.308024077 | 0.249 | 0.115 | 2.71977232130805e-21 | 1 |
| ube2c | 1.32265441605574e-25 | 0.314169731 | 0.273 | 0.132 | 3.32898889977068e-21 | 1 |
| hmga1a | 1.83084227870384e-25 | 0.331973072 | 0.594 | 0.442 | 4.60804693126971e-21 | 1 |
| hmgb3b | 1.96084016905932e-25 | 0.350524222 | 0.314 | 0.169 | 4.93523862150541e-21 | 1 |
| CABZ01069022.2 | 1.96865300429233e-25 | 0.252980677 | 0.151 | 0.056 | 4.95490274650338e-21 | 1 |
| slc29a2 | 4.39888178942244e-25 | 0.265384916 | 0.172 | 0.069 | 1.10715455757973e-20 | 1 |

| | | | | | | |
|---|---|---|---|---|---|---|
| nfs1 | 4.43430348179141e-25 | 0.277618014 | 0.186 | 0.078 | 1.11606984333208e-20 | 1 |
| si:ch211-210c8.7 | 6.0611517134923e-25 | 0.316271515 | 0.25 | 0.12 | 1.52553127476888e-20 | 1 |
| spsb1 | 6.90759351951936e-25 | 0.277821526 | 0.201 | 0.086 | 1.73857221292783e-20 | 1 |
| chchd10 | 7.53314260607041e-25 | 0.325750447 | 0.285 | 0.144 | 1.89601666252186e-20 | 1 |
| tmprss4b1 | 8.38959322157119e-25 | 0.251490902 | 0.164 | 0.064 | 2.11157671793725e-20 | 1 |
| gar1 | 1.16545118757148e-24 | 0.310161227 | 0.235 | 0.112 | 2.93332409399866e-20 | 1 |
| vps9d1 | 1.8121306084036e-24 | 0.275776857 | 0.206 | 0.09 | 4.56095152829102e-20 | 1 |
| kmt5ab | 2.40304964088456e-24 | 0.273668494 | 0.195 | 0.083 | 6.04823564114235e-20 | 1 |
| golt1bb | 2.71126968053926e-24 | 0.274903708 | 0.187 | 0.08 | 6.82399465894928e-20 | 1 |
| lipg | 3.5120425151018e-24 | 0.253194115 | 0.187 | 0.078 | 8.83945980625972e-20 | 1 |
| ncbp2 | 4.01765041205246e-24 | 0.33149646 | 0.303 | 0.162 | 1.01120243220948e-19 | 1 |
| ppp1r3b1 | 4.21353597091814e-24 | 0.292119687 | 0.224 | 0.103 | 1.06050486852039e-19 | 1 |
| spryd4 | 5.19721867220321e-24 | 0.259459743 | 0.176 | 0.072 | 1.30808796760683e-19 | 1 |
| mcfd2 | 5.29254283262127e-24 | 0.297188802 | 0.227 | 0.107 | 1.33208010554245e-19 | 1 |
| nanos31 | 5.72127056830813e-24 | 0.278567143 | 0.217 | 0.096 | 1.43998658933747e-19 | 1 |
| mcm7 | 6.26328088882502e-24 | 0.328994242 | 0.286 | 0.15 | 1.57640516690837e-19 | 1 |
| trub2 | 6.46286780635693e-24 | 0.251821675 | 0.144 | 0.055 | 1.62663919818197e-19 | 1 |
| rpz21 | 6.60703954266563e-24 | 0.333781739 | 0.319 | 0.173 | 1.66292578249351e-19 | 1 |
| dnajc9 | 7.15216837891544e-24 | 0.270525826 | 0.195 | 0.085 | 1.80012925928923e-19 | 1 |
| COQ10A | 7.21747385326178e-24 | 0.26686716 | 0.183 | 0.077 | 1.81656599412746e-19 | 1 |
| mrpl48 | 8.10872021146309e-24 | 0.271704486 | 0.177 | 0.074 | 2.04088379002314e-19 | 1 |
| faub1 | 1.3254828185252e-23 | 0.347062402 | 0.338 | 0.192 | 3.33610770594608e-19 | 1 |
| figla1 | 1.40688062454104e-23 | 0.275659298 | 0.203 | 0.089 | 3.54097784390734e-19 | 1 |
| fen1 | 1.54075060638289e-23 | 0.263222811 | 0.187 | 0.081 | 3.87791520120508e-19 | 1 |
| rbm7 | 1.70258148155866e-23 | 0.298267766 | 0.236 | 0.113 | 4.28522733093498e-19 | 1 |
| casd1 | 2.17604738064424e-23 | 0.25178549 | 0.173 | 0.071 | 5.4768936523435e-19 | 1 |
| tmem70 | 2.66508012010451e-23 | 0.279761792 | 0.217 | 0.098 | 6.70774015429105e-19 | 1 |
| h2afx | 3.02164060070609e-23 | 0.29767997 | 0.264 | 0.131 | 7.60516722791715e-19 | 1 |
| polr2k | 3.08187508151437e-23 | 0.338526657 | 0.326 | 0.183 | 7.75677139266352e-19 | 1 |
| pop4 | 3.29818234949778e-23 | 0.302959722 | 0.232 | 0.111 | 8.30119515545097e-19 | 1 |
| cnih1 | 3.47687323292754e-23 | 0.298209627 | 0.229 | 0.11 | 8.75094223995532e-19 | 1 |
| gtf2a1 | 7.68662946874332e-23 | 0.315403899 | 0.281 | 0.148 | 1.93464777098801e-18 | 1 |
| srsf7a | 1.20713990403429e-22 | 0.345803435 | 0.404 | 0.262 | 3.03825042446391e-18 | 1 |
| zgc:110239 | 1.32365326271121e-22 | 0.336446887 | 0.35 | 0.206 | 3.33150289691785e-18 | 1 |
| zgc:112980 | 1.45215139702752e-22 | 0.260506706 | 0.196 | 0.086 | 3.65491985117857e-18 | 1 |

| | | | | | | |
|---|---|---|---|---|---|---|
| asf1ba | 1.86311148796249e-22 | 0.272863506 | 0.201 | 0.091 | 4.68926530405278e-18 | 1 |
| snrpg | 2.5844607868163e-22 | 0.343864873 | 0.405 | 0.26 | 6.50482935433794e-18 | 1 |
| arpp19b | 3.02934432661111e-22 | 0.307749037 | 0.253 | 0.13 | 7.6245567356475e-18 | 1 |
| tent5c1 | 3.19014157622547e-22 | 0.29834687 | 0.263 | 0.133 | 8.0292673332019e-18 | 1 |
| eif5a2 | 5.34467051010436e-22 | 0.319151881 | 0.527 | 0.396 | 1.34520012068817e-17 | 1 |
| fbxw11b | 6.40902563711209e-22 | 0.30756201 | 0.265 | 0.138 | 1.61308766260474e-17 | 1 |
| ckbb | 6.64830991217456e-22 | 0.334709762 | 0.415 | 0.257 | 1.67331312179522e-17 | 1 |
| rbm4.2 | 8.16634378783406e-22 | 0.324508531 | 0.291 | 0.161 | 2.05538706795995e-17 | 1 |
| cnep1r1 | 8.57509024838061e-22 | 0.301892199 | 0.235 | 0.117 | 2.15826446461492e-17 | 1 |
| si:ch211-226h8.8 | 1.215056681028e-21 | 0.266837245 | 0.2 | 0.091 | 3.05817616047938e-17 | 1 |
| akirin1 | 1.29414173594894e-21 | 0.308375126 | 0.254 | 0.133 | 3.25722533520987e-17 | 1 |
| atp5mc1 | 1.494188391165e-21 | 0.332709385 | 0.463 | 0.324 | 3.7607227617232e-17 | 1 |
| hsp90b1 | 2.12220982796239e-21 | 0.330940813 | 0.353 | 0.212 | 5.34138991599854e-17 | 1 |
| rac1a | 2.20404335156051e-21 | 0.305904728 | 0.267 | 0.141 | 5.54735671154264e-17 | 1 |
| tomm20a | 7.08937870296609e-21 | 0.329449342 | 0.362 | 0.224 | 1.78432572574953e-16 | 1 |
| nutf2l1 | 1.04159857483079e-20 | 0.32640217 | 0.364 | 0.226 | 2.62159945299162e-16 | 1 |
| ppm1g | 1.24158532585524e-20 | 0.308224505 | 0.249 | 0.132 | 3.12494610664506e-16 | 1 |
| eno3 | 1.26524092312223e-20 | 0.310418326 | 0.505 | 0.375 | 3.18448487940633e-16 | 1 |
| snrpa1 | 1.39464241209159e-20 | 0.283871843 | 0.229 | 0.116 | 3.51017548699332e-16 | 1 |
| gfer | 1.45417929966185e-20 | 0.260457454 | 0.199 | 0.093 | 3.66002387931891e-16 | 1 |
| anp32b1 | 1.90522054627255e-20 | 0.330618234 | 0.386 | 0.248 | 4.79524959291337e-16 | 1 |
| imp31 | 1.98001773448418e-20 | 0.286671433 | 0.247 | 0.128 | 4.98350663592322e-16 | 1 |
| tacc3 | 3.51831113568127e-20 | 0.258948985 | 0.192 | 0.09 | 8.85523729739618e-16 | 1 |
| dkc1 | 4.10492872472242e-20 | 0.316341029 | 0.292 | 0.168 | 1.03316951072539e-15 | 1 |
| top1l | 4.15314871922129e-20 | 0.328461447 | 0.35 | 0.221 | 1.04530600114081e-15 | 1 |
| cbx3a | 5.30163739733165e-20 | 0.328805363 | 0.426 | 0.289 | 1.3343691165344e-15 | 1 |
| cstf1 | 6.95039156387169e-20 | 0.257145022 | 0.194 | 0.091 | 1.74934405271086e-15 | 1 |
| mpzl2b | 1.45627829796957e-19 | 0.262339921 | 0.188 | 0.089 | 3.6653068481596e-15 | 1 |
| pon3.2 | 1.88234285685626e-19 | 0.259898881 | 0.191 | 0.091 | 4.73766873642152e-15 | 1 |
| nap1l1 | 1.9259194866909e-19 | 0.311375922 | 0.478 | 0.355 | 4.84734675605233e-15 | 1 |
| glulb | 2.1421906691293e-19 | 0.288178715 | 0.265 | 0.145 | 5.39167969513153e-15 | 1 |
| txnl4a | 2.29409735392853e-19 | 0.266851092 | 0.214 | 0.107 | 5.77401363010272e-15 | 1 |
| mrps18c | 2.91132086002655e-19 | 0.29030696 | 0.224 | 0.118 | 7.32750347260083e-15 | 1 |
| ptgr1 | 4.30081880704571e-19 | 0.25927606 | 0.205 | 0.102 | 1.08247308554534e-14 | 1 |
| fbl | 4.32873762261303e-19 | 0.31732358 | 0.322 | 0.198 | 1.08949997223547e-14 | 1 |

| | | | | | | |
|---|---|---|---|---|---|---|
| mat2aa | 4.73670851919592e-19 | 0.305813342 | 0.3 | 0.177 | 1.19218216719642e-14 | 1 |
| ebna1bp2 | 4.94329226067708e-19 | 0.287420324 | 0.244 | 0.131 | 1.24417722908982e-14 | 1 |
| ivns1abpb | 5.71430864344511e-19 | 0.264831134 | 0.218 | 0.111 | 1.4382343424687e-14 | 1 |
| exosc6 | 6.6162855277104e-19 | 0.250449365 | 0.176 | 0.081 | 1.66525290446943e-14 | 1 |
| nifk1 | 7.40996371940756e-19 | 0.275884284 | 0.224 | 0.116 | 1.86501376853769e-14 | 1 |
| selenom | 1.48756390016797e-18 | 0.262403359 | 0.215 | 0.11 | 3.74404958033277e-14 | 1 |
| hprt1 | 2.31518313689784e-18 | 0.26966595 | 0.222 | 0.115 | 5.82708443725817e-14 | 1 |
| ncl | 2.51065944001847e-18 | 0.284899794 | 0.524 | 0.414 | 6.31907874458249e-14 | 1 |
| paip2b | 3.07460149250896e-18 | 0.317261101 | 0.415 | 0.298 | 7.7384644964958e-14 | 1 |
| btg3 | 3.12202066265828e-18 | 0.303568153 | 0.292 | 0.172 | 7.85781380584462e-14 | 1 |
| pim3 | 3.46242887908525e-18 | 0.260530063 | 0.233 | 0.123 | 8.71458724576968e-14 | 1 |
| zgc:77112 | 4.00387850392506e-18 | 0.257301554 | 0.223 | 0.114 | 1.0077361806529e-13 | 1 |
| emc6 | 6.14039562373326e-18 | 0.279639405 | 0.232 | 0.126 | 1.54547617453742e-13 | 1 |
| f11r.1 | 1.11684567475327e-17 | 0.290401473 | 0.292 | 0.171 | 2.81098887878651e-13 | 1 |
| dut | 1.50063579127221e-17 | 0.262042733 | 0.223 | 0.118 | 3.77695022305304e-13 | 1 |
| khsrp | 2.40941362262111e-17 | 0.299752481 | 0.31 | 0.193 | 6.06425314677507e-13 | 1 |
| spcs3 | 3.10774952858257e-17 | 0.305920404 | 0.379 | 0.261 | 7.82189478848948e-13 | 1 |
| sumo3b | 5.31729717452058e-17 | 0.277231489 | 0.222 | 0.12 | 1.33831052585509e-12 | 1 |
| ube2al | 7.03047942572831e-17 | 0.2849263 | 0.264 | 0.155 | 1.76950136666156e-12 | 1 |
| UBL4A | 7.72371879793037e-17 | 0.262481787 | 0.221 | 0.119 | 1.9439827842511e-12 | 1 |
| si:dkey-33c12.4 | 1.00133132609739e-16 | 0.295220899 | 0.287 | 0.176 | 2.52025081465451e-12 | 1 |
| uqcr10 | 1.17868802427707e-16 | 0.29150279 | 0.465 | 0.351 | 2.96663988830295e-12 | 1 |
| hnrnpaba | 1.86124181282318e-16 | 0.257137951 | 0.546 | 0.458 | 4.68455951869467e-12 | 1 |
| ube2g1b | 5.65843470310306e-16 | 0.278694197 | 0.235 | 0.135 | 1.42417143042401e-11 | 1 |
| lsm1 | 8.05409382896425e-16 | 0.261996688 | 0.233 | 0.131 | 2.02713487581201e-11 | 1 |
| thoc7 | 8.25485291425626e-16 | 0.255106676 | 0.206 | 0.111 | 2.07766392998916e-11 | 1 |
| cbx5 | 1.35971211104194e-15 | 0.292674851 | 0.308 | 0.197 | 3.42225941228145e-11 | 1 |
| lsm12a | 1.59220358566506e-15 | 0.2541879 | 0.219 | 0.121 | 4.0074172047604e-11 | 1 |
| snrpd2 | 1.9320975765444e-15 | 0.293518969 | 0.367 | 0.256 | 4.8628963904046e-11 | 1 |
| STMP1 | 2.33229186005244e-15 | 0.296491376 | 0.406 | 0.3 | 5.87014538256599e-11 | 1 |
| sub1b | 2.64680121053477e-15 | 0.285110194 | 0.331 | 0.217 | 6.66173396679495e-11 | 1 |
| npm1a | 3.95601265385928e-15 | 0.285996508 | 0.409 | 0.297 | 9.95688824849843e-11 | 1 |
| dap1b | 4.72900718235956e-15 | 0.279485182 | 0.36 | 0.245 | 1.19024381772808e-10 | 1 |
| rtn4a | 4.73245477257861e-15 | 0.279500019 | 0.346 | 0.234 | 1.19111154171031e-10 | 1 |
| aktip | 8.81891611579639e-15 | 0.256210661 | 0.233 | 0.134 | 2.21963299718479e-10 | 1 |

| | | | | | | |
|---|---|---|---|---|---|---|
| tuba8l4 | 9.52113238814487e-15 | 0.266113165 | 0.5 | 0.399 | 2.39637381077218e-10 | 1 |
| snrpe | 1.23414183254621e-14 | 0.288617875 | 0.391 | 0.284 | 3.10621157833555e-10 | 1 |
| tubb2b | 1.26337962978993e-14 | 0.272372079 | 0.363 | 0.243 | 3.17980019021829e-10 | 1 |
| lsm6 | 5.09189513558018e-14 | 0.282792145 | 0.335 | 0.227 | 1.28157908667417e-09 | 1 |
| pcf11 | 6.75077999891433e-14 | 0.276854235 | 0.341 | 0.234 | 1.69910381792675e-09 | 1 |
| slc25a25a | 7.02038523812714e-14 | 0.266684279 | 0.283 | 0.181 | 1.76696076058422e-09 | 1 |
| snrpc | 9.51329052709518e-14 | 0.265276266 | 0.278 | 0.176 | 2.39440009276458e-09 | 1 |
| cycsb | 1.29069990641249e-13 | 0.264673666 | 0.429 | 0.321 | 3.24856259444959e-09 | 1 |
| rpf1 | 1.5317556785599e-13 | 0.256009143 | 0.222 | 0.131 | 3.85527586736742e-09 | 1 |
| dad1 | 8.31798862967312e-13 | 0.253597475 | 0.44 | 0.347 | 2.09355455820243e-08 | 1 |
| snrpb | 1.33306963601963e-12 | 0.259737661 | 0.433 | 0.344 | 3.3552029668978e-08 | 1 |
| chtopa1 | 1.43734395649883e-12 | 0.251473771 | 0.322 | 0.216 | 3.61765100411192e-08 | 1 |
| hspd1 | 3.20479555075173e-12 | 0.251567826 | 0.251 | 0.161 | 8.06614992168703e-08 | 1 |
| setb | 5.2306829449881e-12 | 0.255456646 | 0.441 | 0.357 | 1.31651059042405e-07 | 1 |
| prmt1 | 5.4439552148292e-12 | 0.25322785 | 0.315 | 0.217 | 1.37018908802036e-07 | 1 |
| zfand5b | 1.25761944407191e-11 | 0.25426902 | 0.312 | 0.217 | 3.16530237878459e-07 | 1 |
| si:dkey-240e12.62 | 9.67393619729668e-144 | 0.758504766 | 0.983 | 0.546 | 2.4348330014976e-139 | 2 |
| zgc:1534992 | 5.16447299085507e-143 | 0.797790833 | 0.967 | 0.487 | 1.29984620706831e-138 | 2 |
| h2af1al2 | 6.83365957858977e-140 | 0.736974182 | 0.991 | 0.57 | 1.71996377933526e-135 | 2 |
| zp2.22 | 1.06880113607439e-137 | 0.757763147 | 0.949 | 0.506 | 2.69006557938563e-133 | 2 |
| cldnd2 | 3.03026425300478e-137 | 0.735228482 | 0.995 | 0.583 | 7.62687209838774e-133 | 2 |
| si:dkey-241l7.22 | 4.60461444243054e-137 | 0.699294429 | 0.997 | 0.622 | 1.15893540901534e-132 | 2 |
| si:zfos-1505d6.32 | 2.10585145093996e-134 | 0.763546724 | 0.943 | 0.49 | 5.30021751687079e-130 | 2 |
| zp32 | 5.23416580701753e-132 | 0.719516664 | 0.972 | 0.563 | 1.31738719196824e-127 | 2 |
| zp2.62 | 1.89817160492937e-130 | 0.757984663 | 0.924 | 0.478 | 4.77750811244674e-126 | 2 |
| h1m2 | 9.43713866315362e-130 | 0.709665419 | 0.988 | 0.58 | 2.37523343012913e-125 | 2 |
| si:dkey-241l7.52 | 1.88857686443831e-129 | 0.702502107 | 0.955 | 0.564 | 4.75335911010477e-125 | 2 |
| zp2.12 | 7.19213934260632e-129 | 0.738325666 | 0.936 | 0.513 | 1.81018955114058e-124 | 2 |
| zp2.52 | 1.20667852951273e-128 | 0.724969745 | 0.958 | 0.542 | 3.0370891909306e-124 | 2 |
| si:ch211-226h8.142 | 5.81499966405216e-128 | 0.719323306 | 0.961 | 0.545 | 1.46357726544529e-123 | 2 |
| cldng2 | 3.90727644817432e-127 | 0.740927681 | 0.921 | 0.492 | 9.83422409240996e-123 | 2 |
| si:dkey-229d11.32 | 2.34358066914498e-126 | 0.779949367 | 0.874 | 0.414 | 5.898558186171e-122 | 2 |
| si:ch73-189n23.12 | 7.84914782255654e-125 | 0.741886281 | 0.944 | 0.502 | 1.97555201545926e-120 | 2 |
| crp22 | 8.48104384732334e-125 | 0.753459594 | 0.915 | 0.466 | 2.13459392593281e-120 | 2 |
| tmsb52 | 2.0530635553991e-124 | 0.758683341 | 0.93 | 0.464 | 5.16735566258398e-120 | 2 |

| | | | | | | |
|---|---|---|---|---|---|---|
| zp3.22 | 2.53147038241561e-124 | 0.709978628 | 0.938 | 0.537 | 6.37145780550184e-120 | 2 |
| si:dkeyp-98a7.92 | 1.94049050590686e-123 | 0.736582284 | 0.915 | 0.49 | 4.88402055431697e-119 | 2 |
| si:ch211-125e6.122 | 7.2376770403314e-123 | 0.746069289 | 0.918 | 0.478 | 1.82165093428101e-118 | 2 |
| zp2.32 | 1.07471370336871e-121 | 0.727854068 | 0.921 | 0.497 | 2.70494692000869e-117 | 2 |
| acp5a2 | 3.23274678721255e-121 | 0.717222616 | 0.949 | 0.518 | 8.13650038873526e-117 | 2 |
| mt22 | 3.38289882616531e-121 | 0.727554627 | 0.958 | 0.511 | 8.51441805557547e-117 | 2 |
| zgc:1751352 | 2.29676674124066e-119 | 0.741948954 | 0.888 | 0.457 | 5.78073221102861e-115 | 2 |
| zgc:1717762 | 6.36344413008944e-119 | 0.731239934 | 0.902 | 0.473 | 1.60161525310221e-114 | 2 |
| si:dkey-241l7.42 | 1.5588260151691e-117 | 0.664045338 | 0.967 | 0.589 | 3.9234091975791e-113 | 2 |
| si:dkey-90l23.12 | 1.00764462997587e-115 | 0.741050314 | 0.845 | 0.426 | 2.53614076918626e-111 | 2 |
| BX649337.12 | 3.34216918901555e-114 | 0.740264233 | 0.854 | 0.415 | 8.41190563183324e-110 | 2 |
| ccna12 | 4.69677407384255e-114 | 0.721844082 | 0.876 | 0.454 | 1.18213106664543e-109 | 2 |
| CR790368.12 | 2.9427961619764e-113 | 0.703170945 | 0.911 | 0.499 | 7.40672366007839e-109 | 2 |
| si:ch211-12h2.82 | 1.05659923007448e-111 | 0.680711024 | 0.915 | 0.528 | 2.65935460217445e-107 | 2 |
| si:ch211-226h8.42 | 1.6759071201114e-111 | 0.735960944 | 0.851 | 0.419 | 4.21809063060838e-107 | 2 |
| zgc:1717792 | 2.69462011412796e-110 | 0.70223329 | 0.893 | 0.492 | 6.78208936524865e-106 | 2 |
| epcam2 | 2.95038152112418e-108 | 0.698255556 | 0.898 | 0.484 | 7.42581525051744e-104 | 2 |
| si:dkey-241l7.62 | 4.77500819913573e-108 | 0.674730216 | 0.87 | 0.508 | 1.20182181364047e-103 | 2 |
| si:dkeyp-98a7.82 | 3.28339370722085e-107 | 0.711260208 | 0.834 | 0.437 | 8.26397362170416e-103 | 2 |
| rrm22 | 1.50878193709811e-105 | 0.680941913 | 0.877 | 0.478 | 3.79745325748224e-101 | 2 |
| mid1ip1l2 | 2.0349956531536e-103 | 0.647388363 | 0.88 | 0.525 | 5.1218805594223e-99 | 2 |
| si:dkeyp-98a7.42 | 1.23468059203888e-102 | 0.705601879 | 0.809 | 0.404 | 3.10756758210266e-98 | 2 |
| LOC1104378212 | 8.65482835389873e-102 | 0.724935926 | 0.795 | 0.395 | 2.17833374839277e-97 | 2 |
| qdprb22 | 2.31050559528046e-101 | 0.704426395 | 0.801 | 0.413 | 5.81531153276138e-97 | 2 |
| ccnb22 | 1.92143386448831e-100 | 0.716620439 | 0.759 | 0.364 | 4.83605689353063e-96 | 2 |
| CU571163.12 | 5.94946710671184e-99 | 0.71736564 | 0.787 | 0.38 | 1.4974213760883e-94 | 2 |
| zgc:771182 | 6.89147746495638e-98 | 0.684125575 | 0.828 | 0.437 | 1.73451596315487e-93 | 2 |
| si:dkeyp-20e4.82 | 3.48231035875606e-95 | 0.709657474 | 0.778 | 0.388 | 8.76462694195313e-91 | 2 |
| zp3b2 | 5.83293182832274e-95 | 0.699900157 | 0.756 | 0.368 | 1.46809061187055e-90 | 2 |
| zar12 | 7.1702828487875e-95 | 0.679729784 | 0.797 | 0.42 | 1.80468849021133e-90 | 2 |
| si:dkeyp-46h3.62 | 1.02615064978283e-94 | 0.693334206 | 0.776 | 0.391 | 2.58271857043841e-90 | 2 |
| zp3a.22 | 1.03968265209866e-94 | 0.656659043 | 0.837 | 0.473 | 2.61677726706711e-90 | 2 |
| zgc:1952452 | 1.1385562906127e-92 | 0.689665364 | 0.752 | 0.369 | 2.86563232784311e-88 | 2 |
| npm2b2 | 1.90444551801595e-92 | 0.687270242 | 0.758 | 0.376 | 4.79329892429434e-88 | 2 |
| si:dkey-90l23.22 | 3.67883395244549e-92 | 0.700104779 | 0.689 | 0.326 | 9.25925717491006e-88 | 2 |

| | | | | | | |
|---|---|---|---|---|---|---|
| si:dkeyp-50f7.22 | 1.21945989024019e-91 | 0.679468968 | 0.755 | 0.383 | 3.06925859774554e-87 | 2 |
| si:ch211-12h2.62 | 4.17360247863911e-91 | 0.652001255 | 0.807 | 0.456 | 1.05045400784868e-86 | 2 |
| tuba4l2 | 1.13843550737403e-87 | 0.668809003 | 0.75 | 0.386 | 2.86532832850969e-83 | 2 |
| LOC1005374992 | 3.67615369884304e-87 | 0.68837256 | 0.672 | 0.303 | 9.25251124461805e-83 | 2 |
| ccnb12 | 5.5319802554992e-86 | 0.666182317 | 0.736 | 0.375 | 1.39234411050659e-81 | 2 |
| zar1l2 | 6.12128061556952e-86 | 0.670170974 | 0.748 | 0.379 | 1.54066511813269e-81 | 2 |
| lamp22 | 7.71755857112927e-86 | 0.603055004 | 0.856 | 0.538 | 1.94243231676753e-81 | 2 |
| LOC1005351622 | 1.54713423976727e-85 | 0.688270726 | 0.66 | 0.29 | 3.89398216807024e-81 | 2 |
| thy12 | 1.29003441789625e-84 | 0.645078931 | 0.793 | 0.432 | 3.24688762640307e-80 | 2 |
| slbp22 | 1.48434306832806e-84 | 0.662708252 | 0.747 | 0.378 | 3.73594306867489e-80 | 2 |
| s100a12 | 8.30615665382334e-83 | 0.660235018 | 0.702 | 0.346 | 2.0905765682008e-78 | 2 |
| siva12 | 2.7234527006883e-82 | 0.638314719 | 0.752 | 0.411 | 6.85465810236239e-78 | 2 |
| gapdh2 | 2.81160488801116e-81 | 0.637660138 | 0.75 | 0.402 | 7.07652834263528e-77 | 2 |
| si:ch211-14a17.72 | 3.30135064142429e-81 | 0.650642925 | 0.745 | 0.39 | 8.3091694294008e-77 | 2 |
| si:dkey-229d11.52 | 1.53406286785273e-80 | 0.653628144 | 0.733 | 0.371 | 3.86108283209853e-76 | 2 |
| ctss2.12 | 5.95142895085217e-78 | 0.568839133 | 0.823 | 0.519 | 1.49791515263998e-73 | 2 |
| AL929032.12 | 5.17765873617397e-77 | 0.660143687 | 0.64 | 0.3 | 1.30316492730763e-72 | 2 |
| nanog2 | 7.15306675491057e-77 | 0.645482697 | 0.691 | 0.342 | 1.80035537154344e-72 | 2 |
| si:dkeyp-46h3.82 | 1.04813393266157e-75 | 0.647813817 | 0.666 | 0.335 | 2.63804829511591e-71 | 2 |
| zpcx2 | 1.31405560998166e-74 | 0.643904861 | 0.658 | 0.322 | 3.30734656476285e-70 | 2 |
| cth12 | 1.44333372429e-74 | 0.628638053 | 0.693 | 0.363 | 3.6327266506655e-70 | 2 |
| zgc:1715172 | 4.76649155737859e-73 | 0.627330219 | 0.696 | 0.372 | 1.19967826007662e-68 | 2 |
| zgc:1362542 | 6.61027862614893e-73 | 0.636891287 | 0.638 | 0.314 | 1.66374102741542e-68 | 2 |
| btg42 | 2.91740813793764e-72 | 0.628511031 | 0.691 | 0.352 | 7.34282454237524e-68 | 2 |
| zgc:1734432 | 2.54199421718527e-70 | 0.627034745 | 0.641 | 0.314 | 6.39794524523361e-66 | 2 |
| si:dkeyp-46h3.22 | 5.29160904302035e-70 | 0.607375097 | 0.702 | 0.389 | 1.33184508003779e-65 | 2 |
| si:ch211-132b12.82 | 7.04946793254966e-70 | 0.623539029 | 0.649 | 0.326 | 1.77428058394342e-65 | 2 |
| hnrnpabb2 | 4.46002383111029e-68 | 0.493963672 | 0.806 | 0.586 | 1.12254339805215e-63 | 2 |
| gtf3ab2 | 8.01501605349825e-68 | 0.61119842 | 0.629 | 0.304 | 2.01729939050498e-63 | 2 |
| uchl12 | 2.10682875761776e-67 | 0.607549054 | 0.644 | 0.329 | 5.30267730004813e-63 | 2 |
| si:ch211-146l10.82 | 1.26474845451988e-66 | 0.622063385 | 0.589 | 0.292 | 3.18324538518108e-62 | 2 |
| si:ch73-90k17.12 | 4.61339010921609e-66 | 0.615624711 | 0.53 | 0.234 | 1.1611441565886e-61 | 2 |
| zgc:1714742 | 2.55351581628514e-62 | 0.602977704 | 0.578 | 0.29 | 6.42694395800807e-58 | 2 |
| nasp2 | 2.66275279636995e-61 | 0.591508854 | 0.618 | 0.325 | 6.70188251318353e-57 | 2 |
| wu:fi42e032 | 3.47907901434865e-60 | 0.582841153 | 0.632 | 0.335 | 8.75649397121411e-56 | 2 |

| | | | | | | |
|---|---|---|---|---|---|---|
| avd2 | 6.85721151845575e-60 | 0.5906515 | 0.533 | 0.251 | 1.72589156708013e-55 | 2 |
| zgc:554612 | 7.82549746571377e-59 | 0.588495266 | 0.554 | 0.275 | 1.9695994571455e-54 | 2 |
| si:ch73-14h10.22 | 1.20719956449572e-57 | 0.579452676 | 0.534 | 0.258 | 3.03840058387928e-53 | 2 |
| kpna72 | 1.74400507399637e-57 | 0.576512821 | 0.612 | 0.314 | 4.38948637074146e-53 | 2 |
| arhgef10la2 | 1.8495014803839e-57 | 0.578447186 | 0.543 | 0.261 | 4.65501027597823e-53 | 2 |
| si:dkey-37o8.12 | 3.1079405251313e-57 | 0.575127355 | 0.575 | 0.294 | 7.82237550770297e-53 | 2 |
| si:ch211-145c1.12 | 1.65949895110745e-56 | 0.577163554 | 0.492 | 0.22 | 4.17679291004235e-52 | 2 |
| birc5b2 | 3.08358912314011e-56 | 0.574509719 | 0.519 | 0.246 | 7.76108546403135e-52 | 2 |
| si:dkey-152b24.62 | 8.95737331254619e-56 | 0.57136694 | 0.562 | 0.284 | 2.25448128903475e-51 | 2 |
| LOC1018854772 | 6.5760923712379e-54 | 0.568285725 | 0.536 | 0.267 | 1.65513668891687e-49 | 2 |
| si:ch211-250e5.162 | 1.2705396561112e-52 | 0.549588523 | 0.609 | 0.333 | 3.19782126046628e-48 | 2 |
| spink2.12 | 3.23212508088696e-52 | 0.5153348 | 0.702 | 0.423 | 8.13493561608439e-48 | 2 |
| si:dkeyp-46h3.52 | 1.81858497144005e-51 | 0.554567772 | 0.536 | 0.277 | 4.57719651461747e-47 | 2 |
| si:dkey-208k4.22 | 2.79084701118955e-51 | 0.54674011 | 0.559 | 0.282 | 7.02428284246299e-47 | 2 |
| cd9b | 8.62316308682613e-50 | 0.424172967 | 0.207 | 0.058 | 2.17036391732327e-45 | 2 |
| si:dkeyp-98a7.72 | 1.30938466510917e-48 | 0.527241806 | 0.421 | 0.184 | 3.29559026361328e-44 | 2 |
| si:ch211-226l4.62 | 1.82090712653239e-48 | 0.534579758 | 0.543 | 0.28 | 4.58304114676938e-44 | 2 |
| hbba11 | 2.37760573987692e-47 | 0.534862715 | 0.632 | 0.342 | 5.98419588669621e-43 | 2 |
| gmnn2 | 9.22986188806596e-47 | 0.529165865 | 0.542 | 0.291 | 2.32306393860732e-42 | 2 |
| slc16a32 | 1.52788309756335e-46 | 0.533565918 | 0.478 | 0.237 | 3.84552896825719e-42 | 2 |
| si:dkey-19b23.112 | 2.64621588692967e-46 | 0.528260135 | 0.516 | 0.267 | 6.66026076581328e-42 | 2 |
| cpeb1b2 | 2.72476607109826e-46 | 0.528444372 | 0.512 | 0.258 | 6.85796372434721e-42 | 2 |
| mkrn42 | 3.59874709340175e-46 | 0.5281325 | 0.449 | 0.217 | 9.05768655938287e-42 | 2 |
| pabpn1l2 | 4.13095985396419e-46 | 0.524600117 | 0.517 | 0.272 | 1.03972128564425e-41 | 2 |
| zp3a.12 | 1.14573086712772e-45 | 0.507231111 | 0.615 | 0.37 | 2.88369001947375e-41 | 2 |
| si:ch1073-263o8.22 | 2.4554706074657e-45 | 0.52639277 | 0.494 | 0.252 | 6.18017397193043e-41 | 2 |
| zgc:1738562 | 1.01331529546775e-44 | 0.515419126 | 0.436 | 0.209 | 2.55041326716278e-40 | 2 |
| elovl7b2 | 1.22697427529439e-43 | 0.507633842 | 0.475 | 0.237 | 3.08817155348844e-39 | 2 |
| si:ch211-5k11.81 | 1.37914226422606e-43 | 0.444354113 | 0.643 | 0.344 | 3.47116316483058e-39 | 2 |
| haus62 | 1.6241794398409e-43 | 0.510242854 | 0.424 | 0.198 | 4.08789723213556e-39 | 2 |
| h3f3a2 | 2.63178348814796e-43 | 0.449910383 | 0.685 | 0.497 | 6.6239358613196e-39 | 2 |
| elovl7a2 | 1.88448719522366e-42 | 0.499817044 | 0.38 | 0.173 | 4.74306582165843e-38 | 2 |
| si:ch211-251f6.62 | 2.33523696726594e-42 | 0.503775411 | 0.554 | 0.32 | 5.87755792291164e-38 | 2 |
| lsm14b2 | 3.22468575503346e-42 | 0.505980587 | 0.528 | 0.288 | 8.11621157684371e-38 | 2 |
| ccna22 | 1.69511608096613e-41 | 0.501201354 | 0.508 | 0.273 | 4.26643766418365e-37 | 2 |

| | | | | | | |
|---|---|---|---|---|---|---|
| ca15b2 | 4.35020227490406e-41 | 0.494192217 | 0.472 | 0.238 | 1.0949024105706e-36 | 2 |
| larp6b2 | 3.47234698151689e-40 | 0.489515564 | 0.396 | 0.186 | 8.73955011777986e-36 | 2 |
| si:ch211-157c3.41 | 3.74624613762187e-40 | 0.411013469 | 0.238 | 0.082 | 9.42892690378047e-36 | 2 |
| zp3c2 | 1.16231901794094e-39 | 0.476331066 | 0.385 | 0.179 | 2.92544073625555e-35 | 2 |
| crp32 | 1.33013988584543e-39 | 0.489549749 | 0.424 | 0.208 | 3.34782907868437e-35 | 2 |
| zgc:1719771 | 5.67010362728593e-39 | 0.492278646 | 0.429 | 0.22 | 1.4271083819516e-34 | 2 |
| dhrs13a.2 | 8.20387730159164e-39 | 0.348219758 | 0.163 | 0.045 | 2.0648338780376e-34 | 2 |
| map1lc3c2 | 1.21406414492227e-38 | 0.479009697 | 0.399 | 0.192 | 3.05567804635486e-34 | 2 |
| AL929192.12 | 2.53181479671367e-38 | 0.461172759 | 0.349 | 0.155 | 6.37232466184863e-34 | 2 |
| zgc:1134242 | 3.20793229062515e-38 | 0.46412512 | 0.346 | 0.152 | 8.07404478227443e-34 | 2 |
| hbaa11 | 1.34008380330965e-37 | 0.474486977 | 0.511 | 0.27 | 3.37285692455006e-33 | 2 |
| org2 | 2.11447110779942e-37 | 0.484135343 | 0.441 | 0.231 | 5.32191233122036e-33 | 2 |
| zglp12 | 7.62668623989393e-37 | 0.469767093 | 0.391 | 0.189 | 1.9195606597189e-32 | 2 |
| zgc:562312 | 3.19353366583836e-36 | 0.469872095 | 0.425 | 0.218 | 8.03780488354856e-32 | 2 |
| zp2l12 | 3.49067344678551e-36 | 0.463897243 | 0.337 | 0.156 | 8.78567599821444e-32 | 2 |
| gdf32 | 6.56081787603808e-36 | 0.466309842 | 0.421 | 0.214 | 1.65129225122002e-31 | 2 |
| pou5f32 | 1.66053556854631e-35 | 0.445167123 | 0.315 | 0.139 | 4.17940197247421e-31 | 2 |
| zgc:925912 | 2.40513331973587e-35 | 0.461695341 | 0.398 | 0.196 | 6.05348005244322e-31 | 2 |
| si:ch1073-190k2.12 | 3.96503568598315e-35 | 0.458868869 | 0.37 | 0.18 | 9.979598318051e-31 | 2 |
| sinup2 | 4.50511130956069e-35 | 0.463339112 | 0.393 | 0.194 | 1.13389146550333e-30 | 2 |
| pigrl2.32 | 5.3497518917703e-35 | 0.458198985 | 0.394 | 0.194 | 1.34647905363967e-30 | 2 |
| patl22 | 1.1498609524946e-34 | 0.45024644 | 0.351 | 0.165 | 2.89408503133366e-30 | 2 |
| rcc12 | 1.20308690155176e-34 | 0.459192936 | 0.39 | 0.193 | 3.02804942251563e-30 | 2 |
| gbgt1l42 | 2.14200033958657e-34 | 0.437349683 | 0.318 | 0.142 | 5.39120065470543e-30 | 2 |
| si:ch73-160p18.32 | 2.14989653230479e-34 | 0.471945679 | 0.429 | 0.232 | 5.41107458215794e-30 | 2 |
| sybu2 | 1.32357955541974e-33 | 0.44655976 | 0.339 | 0.158 | 3.33131738303593e-29 | 2 |
| zgc:566992 | 5.15031277956746e-33 | 0.452002557 | 0.446 | 0.242 | 1.29628222348933e-28 | 2 |
| rbm4.32 | 6.07180573287374e-33 | 0.412041971 | 0.592 | 0.438 | 1.52821278490699e-28 | 2 |
| slbp3 | 7.1191184652245e-33 | 0.459192154 | 0.419 | 0.225 | 1.79181092651235e-28 | 2 |
| rbpms2b2 | 1.49028261324398e-32 | 0.452739716 | 0.399 | 0.213 | 3.75089230927378e-28 | 2 |
| CU467646.22 | 2.08399428514066e-32 | 0.398260771 | 0.281 | 0.118 | 5.24520521627053e-28 | 2 |
| LOC1018838972 | 3.66397503110224e-32 | 0.427165619 | 0.334 | 0.156 | 9.22185875578122e-28 | 2 |
| zgc:1628792 | 4.64762920711609e-32 | 0.434079736 | 0.354 | 0.173 | 1.16976179513905e-27 | 2 |
| LOC1005368602 | 4.97538612597397e-32 | 0.437883515 | 0.54 | 0.342 | 1.25225493404639e-27 | 2 |
| snu13b2 | 1.65839683757842e-31 | 0.427396086 | 0.564 | 0.384 | 4.17401900050112e-27 | 2 |

| | | | | | | |
|---|---|---|---|---|---|---|
| chia.42 | 4.95281555889435e-31 | 0.420962605 | 0.303 | 0.139 | 1.24657414801812e-26 | 2 |
| si:dkey-188i13.112 | 4.90526173521068e-30 | 0.444415182 | 0.489 | 0.308 | 1.23460532613518e-25 | 2 |
| rasl11b2 | 5.98849270790086e-30 | 0.399203059 | 0.284 | 0.127 | 1.50724372965157e-25 | 2 |
| CR847984.12 | 6.16088841787534e-30 | 0.416337511 | 0.325 | 0.156 | 1.55063400589504e-25 | 2 |
| gyg1a2 | 1.04832028309524e-29 | 0.427095399 | 0.534 | 0.363 | 2.63851732052241e-25 | 2 |
| h2afva2 | 1.16523886939027e-29 | 0.436105617 | 0.484 | 0.316 | 2.93278971036837e-25 | 2 |
| inka1b2 | 1.35235350505008e-29 | 0.424139234 | 0.351 | 0.177 | 3.40373853686055e-25 | 2 |
| plk12 | 1.92110772138495e-29 | 0.426814763 | 0.339 | 0.17 | 4.83523602395379e-25 | 2 |
| zgc:1138862 | 3.7380592571196e-29 | 0.394702019 | 0.293 | 0.134 | 9.40832134424433e-25 | 2 |
| cdc202 | 4.15992542420466e-29 | 0.407803765 | 0.307 | 0.145 | 1.04701163001807e-24 | 2 |
| cldn7b2 | 4.92163964092186e-29 | 0.406997413 | 0.318 | 0.152 | 1.23872748122362e-24 | 2 |
| si:dkey-241l7.32 | 5.38304438569951e-29 | 0.424664064 | 0.547 | 0.356 | 1.35485844143671e-24 | 2 |
| ubc1 | 6.25876264961423e-29 | 0.266351439 | 0.727 | 0.655 | 1.57526797128141e-24 | 2 |
| mibp2 | 5.07315966332048e-28 | 0.418186141 | 0.391 | 0.215 | 1.27686355566113e-23 | 2 |
| si:dkey-46g23.22 | 7.20457840110216e-28 | 0.398197305 | 0.292 | 0.138 | 1.8133203377734e-23 | 2 |
| tph1a2 | 8.77712849748463e-28 | 0.388923018 | 0.289 | 0.135 | 2.20911547153191e-23 | 2 |
| ddx39ab2 | 1.41345287619777e-27 | 0.423911138 | 0.452 | 0.282 | 3.55751954410216e-23 | 2 |
| pabpc1l2 | 1.44099730811007e-27 | 0.395997564 | 0.286 | 0.135 | 3.62684612478223e-23 | 2 |
| CABZ01059627.12 | 3.74039115822285e-27 | 0.386164859 | 0.278 | 0.13 | 9.4141905061311e-23 | 2 |
| buc2 | 4.90731301309622e-27 | 0.403627528 | 0.337 | 0.171 | 1.23512161226619e-22 | 2 |
| cdca7a2 | 1.27767865529816e-26 | 0.417245637 | 0.384 | 0.215 | 3.21578940751993e-22 | 2 |
| larp6a2 | 1.29831718045179e-26 | 0.422018495 | 0.416 | 0.242 | 3.26773451147912e-22 | 2 |
| CABZ01059627.22 | 2.08702776257999e-26 | 0.38533638 | 0.272 | 0.127 | 5.25284017563758e-22 | 2 |
| dctpp12 | 2.79777623578326e-26 | 0.406128496 | 0.326 | 0.168 | 7.0417230078429e-22 | 2 |
| vstm2l2 | 3.20676654751644e-26 | 0.343376827 | 0.217 | 0.09 | 8.07111072344412e-22 | 2 |
| zgc:1034822 | 3.20761786449597e-26 | 0.406181899 | 0.345 | 0.182 | 8.07325340314991e-22 | 2 |
| hspa52 | 3.76011821989096e-26 | 0.408345595 | 0.469 | 0.307 | 9.46384154764356e-22 | 2 |
| hsd17b12a2 | 3.99321212579468e-26 | 0.392258256 | 0.317 | 0.158 | 1.00505155994126e-21 | 2 |
| zgc:1526521 | 5.06351072342996e-26 | 0.364075116 | 0.244 | 0.108 | 1.27443501398009e-21 | 2 |
| retsatl2 | 8.79128756077717e-26 | 0.408547318 | 0.362 | 0.196 | 2.212679166172e-21 | 2 |
| rdh10b2 | 9.02358174279602e-26 | 0.408740748 | 0.325 | 0.171 | 2.27114528884433e-21 | 2 |
| zgc:1630301 | 1.13697758553579e-25 | 0.369885274 | 0.258 | 0.118 | 2.86165888503502e-21 | 2 |
| si:dkey-43p13.51 | 1.84069362245961e-25 | 0.337829501 | 0.196 | 0.078 | 4.63284177836859e-21 | 2 |
| mad2l12 | 1.85874575448508e-25 | 0.394634788 | 0.337 | 0.179 | 4.6782771894635e-21 | 2 |
| si:ch73-15b2.51 | 3.47157535228806e-25 | 0.343172832 | 0.213 | 0.089 | 8.73760800417382e-21 | 2 |

| | | | | | | |
|---|---|---|---|---|---|---|
| BX537276.12 | 4.40179320412478e-25 | 0.367021556 | 0.252 | 0.116 | 1.10788733154617e-20 | 2 |
| tmem54a2 | 1.003273786919e-24 | 0.376452986 | 0.278 | 0.135 | 2.52513979429642e-20 | 2 |
| zgc:1655392 | 1.49588750677679e-24 | 0.357124519 | 0.248 | 0.114 | 3.76499926580651e-20 | 2 |
| trim35-29 | 1.55913485227237e-24 | 0.267496515 | 0.135 | 0.044 | 3.92418650968432e-20 | 2 |
| zgc:923131 | 2.27254896792383e-24 | 0.366592253 | 0.255 | 0.12 | 5.71977849736748e-20 | 2 |
| faub2 | 3.69031586498444e-24 | 0.403489513 | 0.345 | 0.194 | 9.28815600057935e-20 | 2 |
| zgc:1629452 | 3.88940030217673e-24 | 0.376512243 | 0.306 | 0.156 | 9.78923162054862e-20 | 2 |
| si:ch211-146l10.72 | 1.06843288184216e-23 | 0.38790254 | 0.335 | 0.182 | 2.68913872030853e-19 | 2 |
| zgc:1017441 | 2.36210861530509e-23 | 0.370231055 | 0.261 | 0.128 | 5.94519117386138e-19 | 2 |
| tmem701 | 2.94157133025865e-23 | 0.344657499 | 0.222 | 0.1 | 7.40364088112799e-19 | 2 |
| zc3h11a2 | 4.20890671144495e-23 | 0.39324144 | 0.362 | 0.211 | 1.05933973020358e-18 | 2 |
| gstp12 | 4.47223811786939e-23 | 0.375779155 | 0.505 | 0.359 | 1.12561761188655e-18 | 2 |
| degs22 | 4.56276810310748e-23 | 0.340815727 | 0.225 | 0.101 | 1.14840310387112e-18 | 2 |
| prrg21 | 5.50914321453435e-23 | 0.324439624 | 0.199 | 0.084 | 1.38659625566615e-18 | 2 |
| zgc:660242 | 1.55941662512961e-22 | 0.355103202 | 0.273 | 0.136 | 3.92489570378871e-18 | 2 |
| tac3b2 | 1.72316748949686e-22 | 0.347154983 | 0.244 | 0.115 | 4.33704025431464e-18 | 2 |
| CDPF12 | 2.82396677776638e-22 | 0.397538767 | 0.415 | 0.262 | 7.10764198296021e-18 | 2 |
| lpcat42 | 4.3329493430448e-22 | 0.33211118 | 0.224 | 0.102 | 1.09056002015095e-17 | 2 |
| ctsla2 | 6.22079769156519e-22 | 0.361947017 | 0.52 | 0.381 | 1.56571257099004e-17 | 2 |
| CR383662.22 | 6.32032648845259e-22 | 0.302420704 | 0.194 | 0.082 | 1.59076297387863e-17 | 2 |
| si:ch73-78o10.12 | 9.83505247456448e-22 | 0.344033718 | 0.259 | 0.128 | 2.47538435732313e-17 | 2 |
| dynll12 | 1.17418691690922e-21 | 0.340465472 | 0.567 | 0.45 | 2.95531105116882e-17 | 2 |
| chchd101 | 1.39697880262116e-21 | 0.365902117 | 0.278 | 0.146 | 3.51605594831719e-17 | 2 |
| BX001030.12 | 1.6283870730845e-21 | 0.334479831 | 0.244 | 0.117 | 4.09848742424637e-17 | 2 |
| pif11 | 1.81466586535642e-21 | 0.333035594 | 0.22 | 0.102 | 4.56733251651558e-17 | 2 |
| pttg12 | 1.82639888938812e-21 | 0.351290204 | 0.292 | 0.153 | 4.59686336470095e-17 | 2 |
| CU655961.31 | 2.32999314857834e-21 | 0.306268958 | 0.211 | 0.094 | 5.86435975565682e-17 | 2 |
| mos1 | 6.79972656736118e-21 | 0.343703176 | 0.259 | 0.131 | 1.71142317973913e-16 | 2 |
| zgc:1737422 | 6.81474034958457e-21 | 0.34546027 | 0.289 | 0.15 | 1.71520199858694e-16 | 2 |
| ctsba2 | 7.95254099389944e-21 | 0.359357508 | 0.514 | 0.371 | 2.00157504275455e-16 | 2 |
| zgc:1118681 | 1.07009983747863e-20 | 0.344158253 | 0.273 | 0.142 | 2.69333428094995e-16 | 2 |
| srsf2a2 | 2.17321911471187e-20 | 0.36068787 | 0.488 | 0.362 | 5.46977518981829e-16 | 2 |
| cx43.41 | 5.92217044174382e-20 | 0.327639665 | 0.239 | 0.118 | 1.4905510784825e-15 | 2 |
| BX649350.1 | 7.83732473165756e-20 | 0.258973922 | 0.137 | 0.052 | 1.97257626171089e-15 | 2 |
| zgc:1717502 | 8.13289360891984e-20 | 0.353652961 | 0.304 | 0.169 | 2.04696799242904e-15 | 2 |

| | | | | | | |
|---|---|---|---|---|---|---|
| kpna22 | 1.28414889952605e-19 | 0.345543037 | 0.283 | 0.153 | 3.23207436521711e-15 | 2 |
| BX571780.12 | 1.28672654911515e-19 | 0.287615341 | 0.179 | 0.077 | 3.23856205146793e-15 | 2 |
| nanos32 | 1.97362188087955e-19 | 0.316017805 | 0.208 | 0.098 | 4.96740891198574e-15 | 2 |
| aurka2 | 2.7102133849138e-19 | 0.311339827 | 0.233 | 0.115 | 6.82133606848954e-15 | 2 |
| buc2l1 | 2.7971149604661e-19 | 0.287710221 | 0.175 | 0.076 | 7.04005864399712e-15 | 2 |
| zgc:113293 | 5.67665046911949e-19 | 0.25079107 | 0.132 | 0.05 | 1.42875615657269e-14 | 2 |
| cx44.22 | 6.16148866852177e-19 | 0.317375426 | 0.228 | 0.112 | 1.55078508298025e-14 | 2 |
| dph3 | 7.51245515354244e-19 | 0.280624831 | 0.166 | 0.071 | 1.8908098375951e-14 | 2 |
| zgc:922871 | 8.84439516088993e-19 | 0.321296706 | 0.248 | 0.128 | 2.22604581804439e-14 | 2 |
| sox19b2 | 2.42057559027208e-18 | 0.285677429 | 0.2 | 0.093 | 6.09234670315579e-14 | 2 |
| cdca82 | 2.83123722560741e-18 | 0.336043255 | 0.287 | 0.16 | 7.12594097313129e-14 | 2 |
| ccnh2 | 3.82512862560684e-18 | 0.319234265 | 0.234 | 0.121 | 9.62746623778986e-14 | 2 |
| il13ra22 | 6.0373713065214e-18 | 0.306190987 | 0.214 | 0.106 | 1.51954598413837e-13 | 2 |
| yrdc1 | 6.47215820547483e-18 | 0.314883756 | 0.231 | 0.118 | 1.62897749873596e-13 | 2 |
| ttc321 | 7.07509545511411e-18 | 0.298102666 | 0.205 | 0.099 | 1.78073077509767e-13 | 2 |
| ypel31 | 8.44070833805953e-18 | 0.318408437 | 0.236 | 0.123 | 2.1244418816062e-13 | 2 |
| zgc:1139842 | 1.01715464335839e-17 | 0.295895804 | 0.202 | 0.097 | 2.56007652186874e-13 | 2 |
| BX284697.12 | 1.12920468682495e-17 | 0.306706734 | 0.22 | 0.11 | 2.84209527626971e-13 | 2 |
| hsd3b2 | 1.22258120922395e-17 | 0.250091732 | 0.14 | 0.057 | 3.07711464549577e-13 | 2 |
| mcm3l1 | 1.4417103666323e-17 | 0.271451287 | 0.183 | 0.084 | 3.62864082177683e-13 | 2 |
| dynll2a1 | 1.75726356835489e-17 | 0.316907649 | 0.248 | 0.132 | 4.42285667519242e-13 | 2 |
| zgc:1714261 | 1.76705739423869e-17 | 0.297218231 | 0.21 | 0.103 | 4.44750675555936e-13 | 2 |
| tmprss4b2 | 1.77655051914332e-17 | 0.266928196 | 0.154 | 0.066 | 4.47140000163181e-13 | 2 |
| abcf2a | 1.90977571725967e-17 | 0.341783464 | 0.258 | 0.146 | 4.80671450277086e-13 | 2 |
| rpain1 | 2.09625605624747e-17 | 0.303619716 | 0.225 | 0.115 | 5.27606686796926e-13 | 2 |
| e2f5 | 3.34620281215482e-17 | 0.272158655 | 0.157 | 0.068 | 8.42205785791247e-13 | 2 |
| si:dkey-216e24.31 | 3.74923409689089e-17 | 0.306938858 | 0.23 | 0.119 | 9.43644729846468e-13 | 2 |
| tatdn21 | 3.88748814260132e-17 | 0.261739573 | 0.158 | 0.069 | 9.78441890611327e-13 | 2 |
| phex | 6.75247263261671e-17 | 0.281163397 | 0.177 | 0.083 | 1.6995298369033e-12 | 2 |
| si:dkey-68o6.52 | 9.98735183640829e-17 | 0.30695362 | 0.228 | 0.118 | 2.5137165837056e-12 | 2 |
| rnf41 | 1.03918524527021e-16 | 0.312719057 | 0.242 | 0.13 | 2.61552534382059e-12 | 2 |
| eif4a1a2 | 1.04943375531641e-16 | 0.310599108 | 0.514 | 0.421 | 2.64131981875588e-12 | 2 |
| ube2c1 | 1.28703375873902e-16 | 0.31050519 | 0.248 | 0.136 | 3.23933526737025e-12 | 2 |
| si:dkeyp-114g9.12 | 1.61933761148415e-16 | 0.288979688 | 0.202 | 0.1 | 4.07571083434446e-12 | 2 |
| snx10a1 | 1.65954810751214e-16 | 0.30030596 | 0.233 | 0.122 | 4.1769166317973e-12 | 2 |

| | | | | | | |
|---|---|---|---|---|---|---|
| gpr1602 | 1.71483152486462e-16 | 0.266355151 | 0.169 | 0.077 | 4.31605946493176e-12 | 2 |
| spint2 | 2.19854133409965e-16 | 0.269971958 | 0.163 | 0.074 | 5.53350868379542e-12 | 2 |
| zgc:558881 | 2.30705774394893e-16 | 0.267448735 | 0.175 | 0.082 | 5.80663363574506e-12 | 2 |
| elavl22 | 2.50356357274685e-16 | 0.333364358 | 0.349 | 0.219 | 6.30121915624654e-12 | 2 |
| nfr2 | 3.01417968102567e-16 | 0.27975123 | 0.213 | 0.107 | 7.58638883917352e-12 | 2 |
| dck | 3.33964529926916e-16 | 0.282072361 | 0.177 | 0.085 | 8.40555325373054e-12 | 2 |
| si:dkey-42i9.82 | 4.14310237304522e-16 | 0.291167633 | 0.234 | 0.123 | 1.04277743627175e-11 | 2 |
| itm2cb2 | 4.43999208144487e-16 | 0.309727376 | 0.242 | 0.134 | 1.11750160697886e-11 | 2 |
| prkrip1 | 5.28068613154718e-16 | 0.276771982 | 0.166 | 0.078 | 1.32909589244911e-11 | 2 |
| prdx12 | 6.41006639933073e-16 | 0.316024735 | 0.281 | 0.162 | 1.61334961204755e-11 | 2 |
| h2afx1 | 7.09865528187058e-16 | 0.310516868 | 0.244 | 0.134 | 1.78666054789401e-11 | 2 |
| evla1 | 8.17816467499349e-16 | 0.287941527 | 0.196 | 0.099 | 2.05836226704911e-11 | 2 |
| zgc:1129801 | 8.31563119948793e-16 | 0.272765981 | 0.183 | 0.089 | 2.09296121659912e-11 | 2 |
| znf395a1 | 9.59741026933884e-16 | 0.273922913 | 0.189 | 0.093 | 2.41557219068989e-11 | 2 |
| si:ch211-69g19.22 | 9.96004121456481e-16 | 0.29357164 | 0.222 | 0.118 | 2.50684277329382e-11 | 2 |
| umps1 | 1.19246916173591e-15 | 0.258010552 | 0.16 | 0.074 | 3.00132563317311e-11 | 2 |
| bzw1b1 | 1.29446401076525e-15 | 0.305384188 | 0.491 | 0.399 | 3.25803646869505e-11 | 2 |
| lmnl31 | 1.97339540447179e-15 | 0.253164328 | 0.158 | 0.073 | 4.96683889351505e-11 | 2 |
| cdca91 | 2.30770603106728e-15 | 0.260570381 | 0.172 | 0.082 | 5.80826530959323e-11 | 2 |
| selenoh1 | 2.34074406925449e-15 | 0.337816494 | 0.354 | 0.24 | 5.89141874790663e-11 | 2 |
| si:ch211-226h8.6 | 2.94038047886946e-15 | 0.252829917 | 0.161 | 0.075 | 7.40064362726655e-11 | 2 |
| rogdi1 | 5.48937042067172e-15 | 0.29191506 | 0.211 | 0.112 | 1.38161964117887e-10 | 2 |
| ppp1r3b2 | 7.96149726667566e-15 | 0.285657876 | 0.202 | 0.106 | 2.0038292470496e-10 | 2 |
| si:ch73-303b9.11 | 1.08555994342938e-14 | 0.251484911 | 0.16 | 0.076 | 2.73224582161741e-10 | 2 |
| lipg1 | 1.14303357127484e-14 | 0.252031325 | 0.168 | 0.081 | 2.87690119554164e-10 | 2 |
| spryd41 | 1.20793792311117e-14 | 0.26156991 | 0.157 | 0.075 | 3.04025895867851e-10 | 2 |
| thop11 | 1.40573487606564e-14 | 0.290754672 | 0.21 | 0.113 | 3.53809410956961e-10 | 2 |
| lsm12a1 | 1.54460874878971e-14 | 0.298218125 | 0.22 | 0.122 | 3.88762575982882e-10 | 2 |
| nifk2 | 1.56872451963362e-14 | 0.294439136 | 0.216 | 0.118 | 3.94832274346586e-10 | 2 |
| h2afvb | 1.7398740136921e-14 | 0.266323042 | 0.557 | 0.467 | 4.37908890506164e-10 | 2 |
| cox172 | 1.93797678971004e-14 | 0.300350119 | 0.47 | 0.374 | 4.87769378202121e-10 | 2 |
| ccne12 | 2.24758173636225e-14 | 0.280184387 | 0.203 | 0.107 | 5.65693847225013e-10 | 2 |
| zgc:1735441 | 2.79064228064071e-14 | 0.252010867 | 0.155 | 0.073 | 7.0237675561446e-10 | 2 |
| mrps18b1 | 3.95622168678325e-14 | 0.291037686 | 0.219 | 0.122 | 9.95741436346477e-10 | 2 |
| bmb1 | 4.56078601828892e-14 | 0.263377132 | 0.171 | 0.085 | 1.14790423294314e-09 | 2 |

| | | | | | | |
|---|---|---|---|---|---|---|
| ddx192 | 4.71616097327125e-14 | 0.301505455 | 0.25 | 0.148 | 1.18701055536264e-09 | 2 |
| tram22 | 4.95301045292758e-14 | 0.294468412 | 0.228 | 0.129 | 1.24662320089734e-09 | 2 |
| rpz22 | 5.20034006586492e-14 | 0.321950425 | 0.283 | 0.178 | 1.30887359117754e-09 | 2 |
| tmem144b2 | 5.24808353260728e-14 | 0.254924925 | 0.18 | 0.091 | 1.32089014432193e-09 | 2 |
| BX957362.11 | 9.24636157531334e-14 | 0.292912638 | 0.227 | 0.129 | 2.32721674489062e-09 | 2 |
| snu13a2 | 1.76873691760708e-13 | 0.261927999 | 0.186 | 0.098 | 4.45173394792526e-09 | 2 |
| psme3 | 1.88538799264064e-13 | 0.298963724 | 0.222 | 0.128 | 4.74533303867723e-09 | 2 |
| ncbp21 | 2.3106117721243e-13 | 0.307777395 | 0.267 | 0.166 | 5.81557876925964e-09 | 2 |
| ankrd222 | 2.9556384618974e-13 | 0.282479341 | 0.241 | 0.139 | 7.43904644474956e-09 | 2 |
| brf22 | 3.22599226771485e-13 | 0.26410447 | 0.177 | 0.092 | 8.1194999386115e-09 | 2 |
| tpx22 | 3.98146321643465e-13 | 0.27100186 | 0.217 | 0.122 | 1.00209447694444e-08 | 2 |
| fdx12 | 3.98554893872215e-13 | 0.310857914 | 0.323 | 0.217 | 1.00312281238698e-08 | 2 |
| mylipa1 | 3.99817040944907e-13 | 0.27404302 | 0.203 | 0.112 | 1.00629951035424e-08 | 2 |
| cfl1l1 | 6.16273808963322e-13 | 0.278819936 | 0.25 | 0.149 | 1.55109954977979e-08 | 2 |
| marcksb1 | 6.80736807935114e-13 | 0.295983981 | 0.273 | 0.172 | 1.71334647189189e-08 | 2 |
| dnd11 | 7.18126674371783e-13 | 0.257025976 | 0.197 | 0.106 | 1.80745302672634e-08 | 2 |
| gfer1 | 7.34547963096159e-13 | 0.250742456 | 0.18 | 0.095 | 1.84878376831672e-08 | 2 |
| parn2 | 7.66671520590141e-13 | 0.280172356 | 0.244 | 0.144 | 1.92963555017332e-08 | 2 |
| vps9d11 | 8.6824543984916e-13 | 0.257180624 | 0.177 | 0.093 | 2.18528694755635e-08 | 2 |
| cdt11 | 8.73597981923198e-13 | 0.255590043 | 0.165 | 0.085 | 2.1987587607025e-08 | 2 |
| cnot8 | 9.47783843074535e-13 | 0.26242791 | 0.175 | 0.093 | 2.3854771546343e-08 | 2 |
| si:ch73-281n10.21 | 1.20116572444023e-12 | 0.277832374 | 0.481 | 0.389 | 3.0232140118436e-08 | 2 |
| rbbp41 | 1.57892783577227e-12 | 0.292351227 | 0.266 | 0.167 | 3.97400346985523e-08 | 2 |
| zgc:1141232 | 1.61374045950732e-12 | 0.251307386 | 0.188 | 0.1 | 4.06162336253397e-08 | 2 |
| ckbb1 | 1.62659507935679e-12 | 0.293677045 | 0.37 | 0.262 | 4.09397715523312e-08 | 2 |
| hmga1a1 | 1.63457956795461e-12 | 0.258193361 | 0.525 | 0.448 | 4.11407331458496e-08 | 2 |
| zgc:566762 | 2.7005874361144e-12 | 0.307915601 | 0.39 | 0.292 | 6.79710851795633e-08 | 2 |
| g3bp12 | 2.96719905934556e-12 | 0.29988858 | 0.306 | 0.207 | 7.46814331246684e-08 | 2 |
| hells2 | 2.97486769086656e-12 | 0.276798065 | 0.22 | 0.13 | 7.48744449114204e-08 | 2 |
| idh1 | 4.44444412843616e-12 | 0.279115834 | 0.21 | 0.123 | 1.1186221426861e-07 | 2 |
| btg31 | 5.12781905381174e-12 | 0.298681728 | 0.27 | 0.175 | 1.29062077765388e-07 | 2 |
| tuba8l41 | 7.63801657288376e-12 | 0.263994656 | 0.475 | 0.402 | 1.92241239122911e-07 | 2 |
| zgc:771121 | 8.24913499796456e-12 | 0.263516629 | 0.203 | 0.117 | 2.0762247876377e-07 | 2 |
| gemin8 | 1.01526994784774e-11 | 0.265556279 | 0.179 | 0.1 | 2.55533293173798e-07 | 2 |
| zgc:868391 | 1.81984015236691e-11 | 0.254163043 | 0.189 | 0.107 | 4.58035567949227e-07 | 2 |

| | | | | | | |
|---|---|---|---|---|---|---|
| zfand61 | 1.8823630544664e-11 | 0.260276636 | 0.21 | 0.123 | 4.73771957178647e-07 | 2 |
| snrpb21 | 1.88331622664931e-11 | 0.282670706 | 0.236 | 0.146 | 4.74011861085365e-07 | 2 |
| ube2g1a | 2.12082207184947e-11 | 0.263029574 | 0.174 | 0.097 | 5.33789707263792e-07 | 2 |
| ebna1bp21 | 3.12534442900939e-11 | 0.277139462 | 0.219 | 0.134 | 7.86617939337374e-07 | 2 |
| chtopa2 | 4.6449837952708e-11 | 0.290393348 | 0.311 | 0.218 | 1.16909597143171e-06 | 2 |
| alyref | 4.94467292732576e-11 | 0.299794057 | 0.36 | 0.274 | 1.24452472907862e-06 | 2 |
| mphosph6 | 6.15627963793829e-11 | 0.255631894 | 0.174 | 0.098 | 1.54947402207269e-06 | 2 |
| dap1b1 | 8.44737569956311e-11 | 0.291303661 | 0.331 | 0.248 | 2.12611998982304e-06 | 2 |
| imp32 | 1.34184927768907e-10 | 0.260416328 | 0.216 | 0.132 | 3.37730044701561e-06 | 2 |
| gemin71 | 1.43006199432655e-10 | 0.271862704 | 0.236 | 0.152 | 3.5993230335205e-06 | 2 |
| zgc:1102391 | 1.62466903462416e-10 | 0.287333218 | 0.298 | 0.211 | 4.08912949324556e-06 | 2 |
| f11r.11 | 2.34577636291485e-10 | 0.269922317 | 0.264 | 0.175 | 5.90408452782038e-06 | 2 |
| selenom1 | 2.82688619022065e-10 | 0.25637192 | 0.189 | 0.113 | 7.11498985216634e-06 | 2 |
| atp5mc11 | 3.40283585463495e-10 | 0.267346899 | 0.405 | 0.329 | 8.56459756253071e-06 | 2 |
| tmco1 | 6.87467353548188e-10 | 0.259047227 | 0.193 | 0.118 | 1.73028658214544e-05 | 2 |
| fbxw11b1 | 7.1165416203646e-10 | 0.251292061 | 0.225 | 0.143 | 1.79116236042957e-05 | 2 |
| cbx51 | 9.29859413714104e-10 | 0.274025852 | 0.283 | 0.2 | 2.34036315837703e-05 | 2 |
| sap18 | 1.16289792939469e-09 | 0.277946885 | 0.295 | 0.216 | 2.9268977984935e-05 | 2 |
| top1l1 | 1.38686872197212e-09 | 0.283048426 | 0.303 | 0.226 | 3.49060988633164e-05 | 2 |
| ilf2 | 1.42262112517214e-09 | 0.279449023 | 0.32 | 0.242 | 3.58059510994576e-05 | 2 |
| ctsd | 5.64888867063579e-09 | 0.255226806 | 0.205 | 0.134 | 0.000142177 | 2 |
| rtn4a1 | 7.22693674782388e-09 | 0.273740044 | 0.309 | 0.238 | 0.000181895 | 2 |
| cbx3a1 | 2.31385410943925e-08 | 0.253797188 | 0.366 | 0.295 | 0.000582374 | 2 |
| etf1b | 2.76874438519469e-08 | 0.262233859 | 0.275 | 0.205 | 0.000696865 | 2 |
| snrpg1 | 3.40300062541217e-07 | 0.254289748 | 0.326 | 0.267 | 0.008565012 | 2 |
| h2af1al3 | 4.24754850729568e-112 | 0.717609715 | 0.991 | 0.574 | 1.06906548380125e-107 | 3 |
| h1m3 | 4.19535925932338e-110 | 0.699232898 | 0.98 | 0.584 | 1.0559299719791e-105 | 3 |
| cldnd3 | 5.91071230320667e-108 | 0.706668471 | 0.991 | 0.587 | 1.48766717959409e-103 | 3 |
| tmsb53 | 4.77311091644473e-107 | 0.75236811 | 0.924 | 0.469 | 1.20134428655997e-102 | 3 |
| zp2.53 | 1.63382735526097e-106 | 0.711285692 | 0.953 | 0.546 | 4.11218007045634e-102 | 3 |
| zgc:1534993 | 3.59243677429271e-106 | 0.754435567 | 0.947 | 0.493 | 9.04180411721731e-102 | 3 |
| si:dkey-241l7.23 | 4.8982621159675e-106 | 0.67679798 | 0.995 | 0.626 | 1.23284359196786e-101 | 3 |
| si:ch211-226h8.143 | 1.27874821195765e-100 | 0.691472005 | 0.945 | 0.55 | 3.21848137467621e-96 | 3 |
| si:zfos-1505d6.33 | 5.8169695437417e-100 | 0.717244284 | 0.924 | 0.495 | 1.46407306446435e-95 | 3 |
| si:dkey-241l7.43 | 7.50773189915137e-100 | 0.644910453 | 0.942 | 0.594 | 1.88962104169741e-95 | 3 |

| | | | | | | |
|---|---|---|---|---|---|---|
| si:dkey-240e12.63 | 2.30389888718227e-99 | 0.706285085 | 0.965 | 0.551 | 5.79868310914906e-95 | 3 |
| acp5a3 | 1.23829759055516e-96 | 0.690190935 | 0.935 | 0.522 | 3.11667120566827e-92 | 3 |
| zp2.63 | 3.00604814325303e-96 | 0.710265029 | 0.889 | 0.484 | 7.56592257175354e-92 | 3 |
| si:ch73-189n23.13 | 1.12687730809779e-95 | 0.698983511 | 0.913 | 0.507 | 2.83623749675133e-91 | 3 |
| zp2.13 | 2.59722028403669e-95 | 0.693820356 | 0.913 | 0.518 | 6.53694373289195e-91 | 3 |
| zp33 | 3.64950186389152e-95 | 0.678291056 | 0.965 | 0.567 | 9.18543124122858e-91 | 3 |
| zp2.33 | 3.03645877004917e-94 | 0.701163156 | 0.915 | 0.501 | 7.64246307833675e-90 | 3 |
| mt23 | 1.33932359681986e-93 | 0.68735354 | 0.931 | 0.517 | 3.37094356083591e-89 | 3 |
| si:dkeyp-98a7.93 | 2.73611217535622e-93 | 0.700365554 | 0.905 | 0.495 | 6.88652073415407e-89 | 3 |
| zp3.23 | 3.42232431893249e-91 | 0.667541553 | 0.918 | 0.542 | 8.61364807832117e-87 | 3 |
| zgc:1751353 | 5.84763872249497e-91 | 0.706031188 | 0.88 | 0.462 | 1.47179219006476e-86 | 3 |
| si:ch211-125e6.123 | 6.64875550983952e-91 | 0.698974502 | 0.9 | 0.483 | 1.67342527427151e-86 | 3 |
| zgc:1717763 | 2.75754941187114e-90 | 0.688563673 | 0.875 | 0.479 | 6.94047611473847e-86 | 3 |
| hbaa12 | 5.57175870171142e-90 | 0.760479586 | 0.658 | 0.264 | 1.40235594763375e-85 | 3 |
| crp23 | 5.80730227535227e-90 | 0.701628141 | 0.895 | 0.471 | 1.46163990968341e-85 | 3 |
| CR790368.13 | 2.43637783782546e-88 | 0.669304245 | 0.887 | 0.504 | 6.1321193800229e-84 | 3 |
| cldng3 | 6.11185353989346e-88 | 0.675792246 | 0.88 | 0.498 | 1.53829241745578e-83 | 3 |
| si:dkey-241l7.63 | 2.76030842184367e-86 | 0.652258113 | 0.867 | 0.512 | 6.94742026693834e-82 | 3 |
| ccna13 | 1.60163603328502e-85 | 0.678645382 | 0.845 | 0.46 | 4.03115773217506e-81 | 3 |
| hbba12 | 2.53905106143174e-85 | 0.735971914 | 0.747 | 0.338 | 6.39053761651754e-81 | 3 |
| zgc:1717793 | 1.25951104977023e-84 | 0.660995276 | 0.867 | 0.497 | 3.17006336116669e-80 | 3 |
| zgc:771183 | 4.29910101232388e-84 | 0.682337639 | 0.833 | 0.441 | 1.0820407337918e-79 | 3 |
| si:dkey-241l7.53 | 5.40679240535756e-84 | 0.635374566 | 0.931 | 0.569 | 1.36083558050444e-79 | 3 |
| zp3a.23 | 1.29627519072122e-83 | 0.661556851 | 0.844 | 0.476 | 3.26259502752625e-79 | 3 |
| si:ch211-12h2.83 | 5.28948511091553e-83 | 0.635639589 | 0.891 | 0.533 | 1.33131050756633e-78 | 3 |
| si:ch211-226h8.43 | 1.62442149712031e-82 | 0.690888968 | 0.833 | 0.424 | 4.0885064661021e-78 | 3 |
| si:dkeyp-75b4.10 | 2.43520685456662e-80 | 0.640948403 | 0.753 | 0.423 | 6.12917213225874e-76 | 3 |
| lamp23 | 7.26530873790927e-80 | 0.613287642 | 0.845 | 0.542 | 1.82860555624438e-75 | 3 |
| zp2.23 | 3.9761712632624e-79 | 0.645689428 | 0.893 | 0.514 | 1.00076254525051e-74 | 3 |
| si:dkey-229d11.33 | 2.75831675640628e-78 | 0.676024379 | 0.805 | 0.422 | 6.94240744419897e-74 | 3 |
| BX649337.13 | 1.064077415197e-77 | 0.672493452 | 0.824 | 0.421 | 2.67817644630933e-73 | 3 |
| si:ch211-5k11.82 | 2.06291328392029e-76 | 0.667740233 | 0.715 | 0.342 | 5.19214644429897e-72 | 3 |
| mid1ip1l3 | 2.33303788323045e-76 | 0.600594703 | 0.844 | 0.53 | 5.87202304830272e-72 | 3 |
| epcam3 | 7.22606518963741e-74 | 0.630576004 | 0.86 | 0.49 | 1.81872834757984e-69 | 3 |
| si:dkeyp-98a7.83 | 2.55790246442898e-72 | 0.63880174 | 0.796 | 0.443 | 6.43798471272129e-68 | 3 |

| | | | | | | |
|---|---|---|---|---|---|---|
| qdprb23 | 4.17023806596444e-72 | 0.647078085 | 0.775 | 0.418 | 1.04960721882259e-67 | 3 |
| si:dkeyp-98a7.43 | 2.57594194408945e-68 | 0.630709854 | 0.747 | 0.411 | 6.48338827907874e-64 | 3 |
| rrm23 | 5.43422663087494e-68 | 0.598775534 | 0.824 | 0.484 | 1.36774050072491e-63 | 3 |
| si:dkeyp-20e4.83 | 3.63214449672244e-67 | 0.648835559 | 0.74 | 0.394 | 9.14174448380071e-63 | 3 |
| CU571163.13 | 1.96638117448524e-66 | 0.641811577 | 0.735 | 0.387 | 4.94918477806191e-62 | 3 |
| LOC1104378213 | 5.60443334863182e-65 | 0.634041716 | 0.729 | 0.402 | 1.41057982951714e-60 | 3 |
| si:dkeyp-46h3.83 | 6.87715892758746e-62 | 0.626706849 | 0.656 | 0.339 | 1.73091213048449e-57 | 3 |
| si:ch211-14a17.73 | 1.28088285337348e-61 | 0.612898372 | 0.722 | 0.395 | 3.22385405365572e-57 | 3 |
| zar13 | 2.36096882660586e-60 | 0.595120402 | 0.736 | 0.427 | 5.9423224396843e-56 | 3 |
| si:dkeyp-50f7.23 | 4.28173417818529e-60 | 0.605205146 | 0.716 | 0.389 | 1.07766967530745e-55 | 3 |
| si:ch211-12h2.63 | 6.35042251070335e-60 | 0.577357743 | 0.765 | 0.462 | 1.59833784171893e-55 | 3 |
| zgc:1952453 | 9.47223643369764e-60 | 0.602719715 | 0.695 | 0.375 | 2.38406718799736e-55 | 3 |
| zar1l3 | 2.6168321260828e-59 | 0.601193105 | 0.704 | 0.385 | 6.58630477813779e-55 | 3 |
| si:dkey-229d11.53 | 4.7083154123445e-59 | 0.605474129 | 0.689 | 0.377 | 1.18503590613299e-54 | 3 |
| slbp23 | 1.29396441217956e-58 | 0.597571978 | 0.696 | 0.385 | 3.25677902901474e-54 | 3 |
| siva13 | 6.15379240676134e-57 | 0.581582756 | 0.713 | 0.416 | 1.54884801085776e-52 | 3 |
| si:dkey-90l23.13 | 4.72969528454021e-56 | 0.579772754 | 0.747 | 0.435 | 1.19041700616593e-51 | 3 |
| npm2b3 | 6.15533308826958e-56 | 0.592061798 | 0.684 | 0.384 | 1.54923578498657e-51 | 3 |
| nanog3 | 1.51656124802916e-55 | 0.594130369 | 0.645 | 0.348 | 3.81703300516459e-51 | 3 |
| btg43 | 1.51024918045133e-54 | 0.587294214 | 0.651 | 0.357 | 3.80114616227796e-50 | 3 |
| s100a13 | 2.00531660186041e-54 | 0.584964668 | 0.64 | 0.352 | 5.04718135522246e-50 | 3 |
| zpcx3 | 2.23953577002094e-54 | 0.597677613 | 0.622 | 0.327 | 5.6366875795657e-50 | 3 |
| AL929032.13 | 3.14101776075215e-54 | 0.604219123 | 0.607 | 0.305 | 7.90562760203708e-50 | 3 |
| ctss2.13 | 9.15710420397037e-54 | 0.509717011 | 0.782 | 0.524 | 2.3047515570973e-49 | 3 |
| si:dkeyp-46h3.23 | 9.81623261892442e-54 | 0.572371919 | 0.678 | 0.393 | 2.47064758785709e-49 | 3 |
| spink2.13 | 1.66172879492485e-53 | 0.555234053 | 0.718 | 0.425 | 4.18240520394636e-49 | 3 |
| LOC1005351623 | 2.68221156443851e-53 | 0.599663713 | 0.593 | 0.297 | 6.75085828653529e-49 | 3 |
| zgc:1362543 | 5.72045305643554e-53 | 0.58961079 | 0.615 | 0.318 | 1.43978082977426e-48 | 3 |
| ccnb13 | 1.11013844265735e-52 | 0.571944228 | 0.671 | 0.382 | 2.79410744632429e-48 | 3 |
| gapdh3 | 6.93666371562531e-52 | 0.557544923 | 0.682 | 0.409 | 1.74588889058573e-47 | 3 |
| si:dkey-203a12.2 | 2.37966758917238e-51 | 0.558902708 | 0.496 | 0.236 | 5.98938535518797e-47 | 3 |
| zp3b3 | 6.40406422666139e-51 | 0.572947442 | 0.675 | 0.376 | 1.61183892520841e-46 | 3 |
| tuba4l3 | 1.24202879651454e-50 | 0.561591155 | 0.678 | 0.393 | 3.12606227794744e-46 | 3 |
| zgc:1734433 | 6.40198750741644e-50 | 0.575222804 | 0.604 | 0.319 | 1.61131623574164e-45 | 3 |
| kpna73 | 3.73040815688609e-49 | 0.57470181 | 0.598 | 0.317 | 9.38906429006661e-45 | 3 |

| | | | | | | |
|---|---|---|---|---|---|---|
| LOC1005374993 | 1.29499705829662e-48 | 0.571145123 | 0.593 | 0.311 | 3.25937809602676e-44 | 3 |
| thy13 | 3.17269348032838e-48 | 0.534406227 | 0.711 | 0.44 | 7.9853522206385e-44 | 3 |
| cth13 | 3.67426604270845e-48 | 0.552289295 | 0.642 | 0.369 | 9.24776020289289e-44 | 3 |
| gtf3ab3 | 9.13128119698186e-48 | 0.565237266 | 0.589 | 0.309 | 2.29825216446837e-43 | 3 |
| si:ch211-132b12.83 | 6.693772450288e-47 | 0.561220886 | 0.609 | 0.331 | 1.68475558801299e-42 | 3 |
| ccnb23 | 2.64350676912737e-46 | 0.549432394 | 0.653 | 0.373 | 6.65344218721668e-42 | 3 |
| si:dkeyp-46h3.63 | 4.58003681692233e-45 | 0.540642467 | 0.678 | 0.4 | 1.15274946645118e-40 | 3 |
| si:dkey-90l23.23 | 5.78592053808795e-45 | 0.548942306 | 0.593 | 0.335 | 1.45625834023136e-40 | 3 |
| uchl13 | 1.94131478599325e-44 | 0.543944945 | 0.595 | 0.334 | 4.88609518486641e-40 | 3 |
| zgc:1715173 | 1.30610300677376e-42 | 0.531727987 | 0.638 | 0.378 | 3.28733065774888e-38 | 3 |
| zp3a.13 | 2.17146745303638e-42 | 0.530292299 | 0.615 | 0.372 | 5.46536643254727e-38 | 3 |
| avd3 | 2.54802041686323e-41 | 0.5384521 | 0.5 | 0.256 | 6.41311258720307e-37 | 3 |
| si:ch73-14h10.23 | 3.61082656498446e-41 | 0.536511044 | 0.509 | 0.262 | 9.08808938140939e-37 | 3 |
| wu:fi42e033 | 5.44019723312396e-41 | 0.529092194 | 0.587 | 0.341 | 1.36924324160497e-36 | 3 |
| spink2.2 | 6.98066866466242e-39 | 0.466567278 | 0.315 | 0.124 | 1.75696449620888e-34 | 3 |
| hnrnpabb3 | 2.03638738840824e-38 | 0.407027043 | 0.749 | 0.591 | 5.12538341788469e-34 | 3 |
| si:ch211-226l4.63 | 1.82379020630511e-37 | 0.50937228 | 0.515 | 0.284 | 4.59029757024932e-33 | 3 |
| si:ch211-146l10.83 | 3.96609607792733e-37 | 0.513080998 | 0.538 | 0.297 | 9.98226721853529e-33 | 3 |
| ccna23 | 8.81755479261657e-37 | 0.512824561 | 0.502 | 0.275 | 2.21929036575366e-32 | 3 |
| lsm14b3 | 1.76487638018282e-36 | 0.506868615 | 0.518 | 0.291 | 4.44201736128215e-32 | 3 |
| si:dkey-152b24.63 | 1.82287723122444e-35 | 0.501189366 | 0.522 | 0.289 | 4.5879997032688e-31 | 3 |
| si:dkeyp-98a7.73 | 2.13112301948532e-35 | 0.495879357 | 0.396 | 0.188 | 5.3638235277426e-31 | 3 |
| zgc:1714743 | 1.33698159288611e-34 | 0.494803007 | 0.524 | 0.296 | 3.36504897113506e-30 | 3 |
| si:dkey-37o8.13 | 1.9611164428555e-34 | 0.492481034 | 0.515 | 0.299 | 4.93593397502302e-30 | 3 |
| birc5b3 | 7.01423669374539e-34 | 0.496385355 | 0.458 | 0.252 | 1.76541323344878e-29 | 3 |
| slbp4 | 1.4384434355656e-33 | 0.495951116 | 0.429 | 0.226 | 3.62041828297505e-29 | 3 |
| si:ch211-250e5.163 | 2.2557725973733e-33 | 0.478760819 | 0.549 | 0.339 | 5.67755405032886e-29 | 3 |
| cpeb1b3 | 2.55523758420869e-33 | 0.489628095 | 0.475 | 0.263 | 6.43127747569486e-29 | 3 |
| zgc:554613 | 1.59197797539847e-32 | 0.485857308 | 0.495 | 0.28 | 4.0068493662804e-28 | 3 |
| pabpn1l3 | 1.87924050122239e-32 | 0.491457913 | 0.487 | 0.276 | 4.72986041752663e-28 | 3 |
| gmnn3 | 1.96514441503256e-32 | 0.484297357 | 0.507 | 0.295 | 4.94607197819544e-28 | 3 |
| LOC1018854773 | 5.96548101977666e-32 | 0.482837281 | 0.482 | 0.272 | 1.50145191786759e-27 | 3 |
| LOC1005368603 | 3.84754973782892e-31 | 0.462632775 | 0.535 | 0.344 | 9.68389793514162e-27 | 3 |
| si:dkey-241l7.33 | 5.79129744872697e-31 | 0.47739606 | 0.545 | 0.358 | 1.45761165487009e-26 | 3 |
| nasp3 | 5.25270835239014e-30 | 0.462209855 | 0.538 | 0.332 | 1.32205416521307e-25 | 3 |

| | | | | | | |
|---|---|---|---|---|---|---|
| zmp:0000001323 | 1.36731023217539e-29 | 0.412103386 | 0.309 | 0.138 | 3.44138312336224e-25 | 3 |
| BX322787.1 | 2.17164745893458e-29 | 0.42927446 | 0.296 | 0.13 | 5.46581948939243e-25 | 3 |
| h3f3a3 | 5.94379860039052e-29 | 0.400315587 | 0.644 | 0.501 | 1.49599466973229e-24 | 3 |
| si:ch1073-263o8.23 | 6.03675706912665e-29 | 0.461057175 | 0.449 | 0.257 | 1.51939138672849e-24 | 3 |
| si:ch211-14a17.10 | 7.63008415245857e-28 | 0.345085999 | 0.222 | 0.084 | 1.9204158803323e-23 | 3 |
| crp33 | 8.70355876166065e-28 | 0.452770167 | 0.396 | 0.212 | 2.19059870472237e-23 | 3 |
| ca15b3 | 9.37919218000584e-28 | 0.452536935 | 0.435 | 0.242 | 2.36064887978567e-23 | 3 |
| elovl7b3 | 1.10938940601546e-27 | 0.457148588 | 0.427 | 0.242 | 2.79222219600032e-23 | 3 |
| arhgef10la3 | 2.49019157014165e-27 | 0.449155009 | 0.462 | 0.268 | 6.26756316288952e-23 | 3 |
| pigrl2.33 | 4.37056617337486e-27 | 0.447682806 | 0.378 | 0.197 | 1.10002780017672e-22 | 3 |
| hsd17b12a3 | 5.98634098330338e-27 | 0.433294867 | 0.325 | 0.159 | 1.50670216208763e-22 | 3 |
| map1lc3c3 | 1.88381392271488e-26 | 0.447968038 | 0.364 | 0.196 | 4.74137126208108e-22 | 3 |
| si:ch211-251f6.63 | 3.42122275335805e-26 | 0.437210636 | 0.511 | 0.325 | 8.61087554792689e-22 | 3 |
| slc16a33 | 6.14397024577306e-26 | 0.4368717 | 0.424 | 0.243 | 1.54637587115862e-21 | 3 |
| buc3 | 8.75241088883943e-26 | 0.433354391 | 0.34 | 0.173 | 2.202894296612e-21 | 3 |
| si:dkey-208k4.23 | 1.28585852451608e-25 | 0.433724493 | 0.478 | 0.289 | 3.23637732035452e-21 | 3 |
| si:ch73-359m17.2 | 1.68476411226422e-25 | 0.412135014 | 0.353 | 0.183 | 4.24038279415782e-21 | 3 |
| zgc:925913 | 7.61362889739085e-25 | 0.422501017 | 0.373 | 0.2 | 1.9162742571843e-20 | 3 |
| si:dkey-203a12.9 | 8.43837800070176e-25 | 0.401425138 | 0.285 | 0.134 | 2.12385535899663e-20 | 3 |
| spink2.4 | 1.97266471527215e-24 | 0.311751907 | 0.184 | 0.067 | 4.96499982186847e-20 | 3 |
| si:dkey-19b23.113 | 2.23911213262157e-24 | 0.430190038 | 0.451 | 0.273 | 5.63562132659523e-20 | 3 |
| haus63 | 2.87562230177525e-24 | 0.420324055 | 0.376 | 0.203 | 7.23765377133813e-20 | 3 |
| si:ch211-145c1.13 | 3.24832708707335e-24 | 0.430126907 | 0.404 | 0.228 | 8.1757144454549e-20 | 3 |
| si:dkeyp-46h3.53 | 3.62001330194965e-24 | 0.427122108 | 0.458 | 0.284 | 9.11121147967709e-20 | 3 |
| mibp3 | 7.95196623864413e-24 | 0.432706712 | 0.376 | 0.217 | 2.00143038260434e-19 | 3 |
| rbpms2b3 | 2.2246401241543e-23 | 0.420965505 | 0.38 | 0.215 | 5.59919672848395e-19 | 3 |
| elavl23 | 3.04207566851958e-23 | 0.423416117 | 0.382 | 0.218 | 7.65660025009694e-19 | 3 |
| si:ch73-90k17.13 | 4.90182490159963e-23 | 0.419404649 | 0.415 | 0.243 | 1.23374030948361e-18 | 3 |
| zgc:562313 | 1.01040913757834e-22 | 0.416074176 | 0.387 | 0.222 | 2.54309875837092e-18 | 3 |
| retsatl3 | 1.18813405075292e-22 | 0.423611334 | 0.353 | 0.198 | 2.99041459234003e-18 | 3 |
| gyg1a3 | 1.95458164751433e-22 | 0.407225183 | 0.513 | 0.365 | 4.91948654862882e-18 | 3 |
| gdf33 | 2.55371930668719e-22 | 0.413937295 | 0.38 | 0.218 | 6.427456123001e-18 | 3 |
| org3 | 8.10848614965844e-22 | 0.417820475 | 0.393 | 0.235 | 2.04082487900753e-17 | 3 |
| zgc:1719772 | 2.84043074421223e-20 | 0.407510892 | 0.376 | 0.225 | 7.14908014010777e-16 | 3 |
| zglp13 | 8.62132590887306e-20 | 0.390818004 | 0.338 | 0.194 | 2.16990151800426e-15 | 3 |

| | | | | | | |
|---|---|---|---|---|---|---|
| zgc:1134243 | 2.72086285731419e-19 | 0.374713201 | 0.295 | 0.157 | 6.84813972557408e-15 | 3 |
| ypel32 | 7.26882193124684e-19 | 0.369749724 | 0.244 | 0.123 | 1.82948979187552e-14 | 3 |
| BX001030.13 | 1.12212261375372e-18 | 0.355342909 | 0.24 | 0.118 | 2.82427040655673e-14 | 3 |
| sybu3 | 1.32973335743126e-18 | 0.372871246 | 0.298 | 0.162 | 3.34680588731874e-14 | 3 |
| zgc:566993 | 1.54370160403293e-18 | 0.376965238 | 0.398 | 0.247 | 3.88534256719049e-14 | 3 |
| bmb2 | 1.71650379266289e-18 | 0.329955348 | 0.191 | 0.085 | 4.32026839575324e-14 | 3 |
| gbgt1l43 | 1.93783579491723e-18 | 0.367138956 | 0.276 | 0.146 | 4.87733891222717e-14 | 3 |
| zgc:1138863 | 6.01003536826578e-18 | 0.355998865 | 0.264 | 0.138 | 1.51266580183881e-13 | 3 |
| CDPF13 | 1.02122481399667e-17 | 0.386717956 | 0.395 | 0.265 | 2.57032073434822e-13 | 3 |
| CR847984.13 | 1.20513810864249e-17 | 0.364331289 | 0.289 | 0.16 | 3.03321210564227e-13 | 3 |
| cldn7b3 | 1.29873013800308e-17 | 0.364033391 | 0.282 | 0.156 | 3.26877388433995e-13 | 3 |
| si:ch73-160p18.33 | 1.33533533416201e-17 | 0.38032078 | 0.375 | 0.236 | 3.36090550255237e-13 | 3 |
| snu13b3 | 1.73341272048999e-17 | 0.350698475 | 0.511 | 0.389 | 4.36282647620125e-13 | 3 |
| si:ch211-146l10.73 | 2.20644116801386e-17 | 0.367775491 | 0.318 | 0.185 | 5.55339177577409e-13 | 3 |
| zgc:1717503 | 3.30680386237767e-17 | 0.368739147 | 0.298 | 0.171 | 8.32289464121836e-13 | 3 |
| zgc:922872 | 3.38005891151757e-17 | 0.351088147 | 0.245 | 0.129 | 8.50727027439858e-13 | 3 |
| zgc:1738563 | 4.7039570446419e-17 | 0.372185458 | 0.349 | 0.216 | 1.18393894856592e-12 | 3 |
| zgc:1629453 | 6.96290769324317e-17 | 0.352998613 | 0.285 | 0.158 | 1.75249423731237e-12 | 3 |
| elovl7a3 | 1.65785999753842e-16 | 0.35293366 | 0.309 | 0.179 | 4.17266782780445e-12 | 3 |
| larp6a3 | 1.98668458541803e-16 | 0.370939319 | 0.375 | 0.246 | 5.00028643303865e-12 | 3 |
| tph1a3 | 2.56673644599893e-16 | 0.341145988 | 0.256 | 0.138 | 6.4602189609347e-12 | 3 |
| zgc:1118682 | 2.63939237160727e-16 | 0.342946851 | 0.264 | 0.144 | 6.64308666009833e-12 | 3 |
| kpna23 | 3.63520309254837e-16 | 0.34892401 | 0.275 | 0.154 | 9.149442663635e-12 | 3 |
| AL929192.13 | 4.12191883671891e-16 | 0.335050674 | 0.289 | 0.16 | 1.03744575201378e-11 | 3 |
| si:ch211-226h8.61 | 6.1433694714154e-16 | 0.307386832 | 0.167 | 0.075 | 1.54622466226054e-11 | 3 |
| nfr3 | 7.84033860388018e-16 | 0.324525248 | 0.213 | 0.108 | 1.9733348232106e-11 | 3 |
| CR383662.23 | 1.17280498278184e-15 | 0.292574648 | 0.182 | 0.084 | 2.95183286116362e-11 | 3 |
| gstp13 | 1.17304448908331e-15 | 0.344084276 | 0.467 | 0.363 | 2.95243567457379e-11 | 3 |
| mkrn43 | 1.20833843840516e-15 | 0.348026038 | 0.358 | 0.224 | 3.04126701562194e-11 | 3 |
| si:ch73-78o10.13 | 1.38651210344734e-15 | 0.328513406 | 0.244 | 0.13 | 3.4897123131666e-11 | 3 |
| rasl11b3 | 2.78593019174932e-15 | 0.327503984 | 0.244 | 0.131 | 7.01190769961385e-11 | 3 |
| cdca7a3 | 2.86121973435138e-15 | 0.356568272 | 0.344 | 0.219 | 7.20140394938898e-11 | 3 |
| sox19b3 | 1.13483314947475e-14 | 0.300394817 | 0.193 | 0.095 | 2.85626155391301e-10 | 3 |
| rdh10b3 | 1.34611371751891e-14 | 0.341732235 | 0.291 | 0.175 | 3.38803361562334e-10 | 3 |
| pou5f33 | 1.67624919344982e-14 | 0.326493014 | 0.256 | 0.143 | 4.21895159499386e-10 | 3 |

| | | | | | | |
|---|---|---|---|---|---|---|
| si:dkey-46g23.23 | 1.80068630168456e-14 | 0.329952571 | 0.251 | 0.142 | 4.53214735270988e-10 | 3 |
| zp3c3 | 2.0931980253768e-14 | 0.336190556 | 0.305 | 0.186 | 5.26837011007086e-10 | 3 |
| mad2l13 | 2.70338695620052e-14 | 0.332560668 | 0.3 | 0.183 | 6.80415463006108e-10 | 3 |
| CT009596.12 | 2.74414686651175e-14 | 0.30017269 | 0.178 | 0.088 | 6.90674324832342e-10 | 3 |
| ctsla3 | 3.10906195538271e-14 | 0.317455098 | 0.48 | 0.384 | 7.82519803550275e-10 | 3 |
| zp2l13 | 3.34946474804464e-14 | 0.334392005 | 0.273 | 0.161 | 8.43026782435356e-10 | 3 |
| cdca83 | 4.27215023806854e-14 | 0.335644774 | 0.273 | 0.162 | 1.07525749341947e-09 | 3 |
| ctsba3 | 4.58899626687844e-14 | 0.320674893 | 0.485 | 0.374 | 1.15500447041063e-09 | 3 |
| BX284697.13 | 6.17018045463723e-14 | 0.307171422 | 0.211 | 0.111 | 1.55297271862764e-09 | 3 |
| si:ch1073-190k2.13 | 1.26056449719696e-13 | 0.330948404 | 0.3 | 0.185 | 3.17271478299503e-09 | 3 |
| zgc:1628793 | 1.42746792970704e-13 | 0.324202702 | 0.295 | 0.178 | 3.59279403227966e-09 | 3 |
| h2afva3 | 1.97948736773267e-13 | 0.334097431 | 0.42 | 0.321 | 4.98217175584636e-09 | 3 |
| si:dkey-68o6.53 | 2.15499361832741e-13 | 0.320667386 | 0.218 | 0.12 | 5.42390343796825e-09 | 3 |
| cx43.42 | 2.49796016427252e-13 | 0.315224377 | 0.218 | 0.121 | 6.28711593745751e-09 | 3 |
| inka1b3 | 2.67935321379406e-13 | 0.319661381 | 0.298 | 0.181 | 6.74366410379827e-09 | 3 |
| dynll13 | 2.68326303941387e-13 | 0.290159896 | 0.529 | 0.453 | 6.75350474390078e-09 | 3 |
| zgc:1034823 | 2.91560091478722e-13 | 0.322075693 | 0.3 | 0.186 | 7.33827594242795e-09 | 3 |
| pabpc1l3 | 2.93584332488317e-13 | 0.316595244 | 0.242 | 0.139 | 7.38922406439844e-09 | 3 |
| rcc13 | 2.97184142627266e-13 | 0.326699095 | 0.315 | 0.199 | 7.47982768578565e-09 | 3 |
| tent5c2 | 3.39562830657909e-13 | 0.321906437 | 0.238 | 0.137 | 8.5464568848289e-09 | 3 |
| yrdc2 | 3.9606507633172e-13 | 0.310906187 | 0.218 | 0.12 | 9.96856190619306e-09 | 3 |
| cdc203 | 4.5579405903161e-13 | 0.312976514 | 0.255 | 0.15 | 1.14718806717666e-08 | 3 |
| CABZ01059627.23 | 5.10883430906817e-13 | 0.309086706 | 0.231 | 0.131 | 1.28584250724937e-08 | 3 |
| sinup3 | 5.72504500971192e-13 | 0.319852719 | 0.315 | 0.2 | 1.44093657849439e-08 | 3 |
| fdx13 | 6.62266430528854e-13 | 0.338467405 | 0.324 | 0.218 | 1.66685837899807e-08 | 3 |
| zgc:1737423 | 9.11810962523026e-13 | 0.31358115 | 0.258 | 0.153 | 2.2949370115742e-08 | 3 |
| degs23 | 1.11387577762441e-12 | 0.285886259 | 0.196 | 0.104 | 2.80351394470287e-08 | 3 |
| pttg1ipb | 1.12838688477276e-12 | 0.304282634 | 0.204 | 0.111 | 2.84003695028456e-08 | 3 |
| ddx39ab3 | 1.29146876020977e-12 | 0.335033722 | 0.385 | 0.287 | 3.25049772257198e-08 | 3 |
| si:dkey-216e24.32 | 1.56620735160896e-12 | 0.297958709 | 0.216 | 0.121 | 3.9419872832646e-08 | 3 |
| mcm3l2 | 3.13888698702914e-12 | 0.265288389 | 0.169 | 0.085 | 7.90026465765363e-08 | 3 |
| CU467646.23 | 3.64118745711472e-12 | 0.287680703 | 0.22 | 0.123 | 9.16450471081205e-08 | 3 |
| znf395a2 | 3.83372512934989e-12 | 0.282896632 | 0.18 | 0.095 | 9.64910277806073e-08 | 3 |
| dynll2a2 | 4.61159775170541e-12 | 0.296257829 | 0.231 | 0.134 | 1.16069303812673e-07 | 3 |
| larp6b3 | 4.75587352453384e-12 | 0.305143654 | 0.304 | 0.193 | 1.19700580738992e-07 | 3 |

| | | | | | | |
|---|---|---|---|---|---|---|
| BX537276.13 | 5.06181738994386e-12 | 0.282145688 | 0.215 | 0.119 | 1.27400881887497e-07 | 3 |
| zgc:1532841 | 6.32159898372412e-12 | 0.282412134 | 0.184 | 0.098 | 1.59108324821352e-07 | 3 |
| zgc:868392 | 6.89425701946808e-12 | 0.287574066 | 0.196 | 0.107 | 1.73521554922992e-07 | 3 |
| BX005461.11 | 8.59705197963146e-12 | 0.257442819 | 0.156 | 0.077 | 2.16379201275344e-07 | 3 |
| patl23 | 1.49212241480589e-11 | 0.299806042 | 0.273 | 0.171 | 3.75552290582495e-07 | 3 |
| prdx13 | 2.43524900884497e-11 | 0.304328337 | 0.262 | 0.164 | 6.1292782303619e-07 | 3 |
| LOC1018838973 | 2.58751639300368e-11 | 0.289048237 | 0.262 | 0.162 | 6.51252000955096e-07 | 3 |
| dazl1 | 3.57294878132656e-11 | 0.266661367 | 0.191 | 0.104 | 8.99275478772083e-07 | 3 |
| zgc:1655393 | 4.97069502888818e-11 | 0.282191936 | 0.205 | 0.118 | 1.25107423182087e-06 | 3 |
| tacc31 | 5.51300060448308e-11 | 0.271228798 | 0.173 | 0.093 | 1.38756712214235e-06 | 3 |
| tac3b3 | 5.91212887332597e-11 | 0.274844666 | 0.207 | 0.118 | 1.48802371612741e-06 | 3 |
| dctpp13 | 7.12610283762011e-11 | 0.299776888 | 0.267 | 0.173 | 1.7935688232006e-06 | 3 |
| si:dkeyp-114g9.13 | 7.29021407159583e-11 | 0.268657598 | 0.185 | 0.102 | 1.83487397967995e-06 | 3 |
| pttg13 | 8.69662804104427e-11 | 0.294339575 | 0.249 | 0.156 | 2.18885431165043e-06 | 3 |
| plk13 | 1.03864842448309e-10 | 0.285643903 | 0.273 | 0.175 | 2.6141742195815e-06 | 3 |
| pif12 | 1.26731231905414e-10 | 0.26712293 | 0.187 | 0.105 | 3.18969837582737e-06 | 3 |
| zgc:1141233 | 1.28801416078228e-10 | 0.25897173 | 0.184 | 0.101 | 3.24180284127292e-06 | 3 |
| prrg22 | 1.49869063012364e-10 | 0.257859636 | 0.164 | 0.087 | 3.77205444695818e-06 | 3 |
| hspa53 | 1.5479795298863e-10 | 0.295640366 | 0.396 | 0.313 | 3.89610967877083e-06 | 3 |
| faub3 | 1.66200070153589e-10 | 0.308452947 | 0.291 | 0.198 | 4.18308956569568e-06 | 3 |
| si:ch211-226h8.81 | 2.15085455949421e-10 | 0.272550503 | 0.173 | 0.095 | 5.41348584079097e-06 | 3 |
| mos2 | 2.2034293021882e-10 | 0.277011224 | 0.224 | 0.134 | 5.54581121067748e-06 | 3 |
| nanos33 | 2.31549099438563e-10 | 0.259896103 | 0.182 | 0.101 | 5.82785928376918e-06 | 3 |
| rbm4.33 | 2.66743055619787e-10 | 0.25639702 | 0.502 | 0.444 | 6.71365596689442e-06 | 3 |
| zgc:1526522 | 4.72384430748007e-10 | 0.261865062 | 0.195 | 0.112 | 1.18894437374966e-05 | 3 |
| fbxo321 | 7.82023567387231e-10 | 0.263829175 | 0.18 | 0.102 | 1.96827511675692e-05 | 3 |
| zgc:660243 | 8.94447851097218e-10 | 0.267718733 | 0.227 | 0.14 | 2.25123579642659e-05 | 3 |
| si:ch211-69g19.23 | 1.183496079094e-09 | 0.264144961 | 0.202 | 0.12 | 2.97874128147169e-05 | 3 |
| chia.43 | 1.19282796571074e-09 | 0.26948047 | 0.233 | 0.145 | 3.00222870689736e-05 | 3 |
| si:dkey-188i13.113 | 1.32723582075226e-09 | 0.287178155 | 0.398 | 0.315 | 3.34051983725137e-05 | 3 |
| eif4a1a3 | 1.44740242430154e-09 | 0.254424224 | 0.469 | 0.425 | 3.64296716172455e-05 | 3 |
| parn3 | 1.6210320949178e-09 | 0.282044283 | 0.229 | 0.146 | 4.07997567969862e-05 | 3 |
| cox173 | 2.05121834923308e-09 | 0.266233123 | 0.44 | 0.376 | 5.16271146318475e-05 | 3 |
| si:ch73-340n8.41 | 2.2242335027238e-09 | 0.275038911 | 0.205 | 0.126 | 5.59817330300553e-05 | 3 |
| si:dkey-42i9.83 | 3.1832529325638e-09 | 0.254479318 | 0.207 | 0.125 | 8.01192930596983e-05 | 3 |

| | | | | | | |
|---|---|---|---|---|---|---|
| zgc:1630302 | 3.86979847916411e-09 | 0.259080333 | 0.202 | 0.122 | 9.73989579220816e-05 | 3 |
| g3bp13 | 4.08753151669331e-09 | 0.297436164 | 0.289 | 0.209 | 0.000102879 | 3 |
| cx44.23 | 4.14418894014155e-09 | 0.253230658 | 0.193 | 0.115 | 0.000104305 | 3 |
| si:ch211-210c8.71 | 4.42844446094016e-09 | 0.279996688 | 0.202 | 0.126 | 0.00011146 | 3 |
| rnf42 | 4.75544169669082e-09 | 0.270523341 | 0.211 | 0.132 | 0.00011969 | 3 |
| tram23 | 5.89262311155757e-09 | 0.27370916 | 0.209 | 0.131 | 0.000148311 | 3 |
| cstf11 | 9.84942463466169e-09 | 0.253346865 | 0.164 | 0.095 | 0.0002479 | 3 |
| aurkb1 | 1.14469583586104e-08 | 0.259078391 | 0.202 | 0.126 | 0.000288108 | 3 |
| f11r.12 | 1.20425944447968e-08 | 0.286114225 | 0.255 | 0.176 | 0.0003031 | 3 |
| ankrd223 | 1.48244802671067e-08 | 0.258884599 | 0.222 | 0.141 | 0.000373117 | 3 |
| zc3h11a3 | 1.48778601119826e-08 | 0.282968726 | 0.295 | 0.216 | 0.000374461 | 3 |
| marcksb2 | 2.89560364752412e-08 | 0.274131233 | 0.251 | 0.175 | 0.000728794 | 3 |
| chchd102 | 1.08719445293786e-07 | 0.259197152 | 0.224 | 0.15 | 0.00273636 | 3 |
| BX957362.12 | 1.53180805546752e-07 | 0.254314475 | 0.2 | 0.132 | 0.003855408 | 3 |
| snrpb22 | 1.57783452704881e-07 | 0.256784638 | 0.218 | 0.148 | 0.003971252 | 3 |
| hprt11 | 2.53017361686013e-07 | 0.252248987 | 0.184 | 0.119 | 0.006368194 | 3 |
| ddx193 | 2.6247584944059e-07 | 0.253945459 | 0.22 | 0.15 | 0.006606255 | 3 |
| cbx3a2 | 3.99356856733738e-07 | 0.256534307 | 0.351 | 0.296 | 0.010051413 | 3 |
| zgc:1102392 | 4.86378300610101e-07 | 0.266889578 | 0.278 | 0.213 | 0.012241655 | 3 |
| selenoh2 | 6.52697010233787e-07 | 0.25703765 | 0.307 | 0.244 | 0.016427731 | 3 |
| zgc:566763 | 8.81926629510699e-07 | 0.254872453 | 0.355 | 0.295 | 0.022197211 | 3 |
| drap1 | 1.73282952132478e-05 | 0.256011297 | 0.264 | 0.217 | 0.436135862 | 3 |
| hbba13 | 2.51335052928978e-132 | 0.951828551 | 0.879 | 0.336 | 6.32585194716946e-128 | 4 |
| si:ch211-5k11.83 | 2.65966887420281e-123 | 0.88951102 | 0.837 | 0.341 | 6.69412058948105e-119 | 4 |
| hbaa13 | 1.43883074887077e-107 | 0.883984487 | 0.729 | 0.265 | 3.62139311183283e-103 | 4 |
| si:dkey-240e12.64 | 1.63089911607273e-98 | 0.733317288 | 0.976 | 0.554 | 4.10480998524345e-94 | 4 |
| zgc:1534994 | 1.16664311301778e-97 | 0.774379757 | 0.963 | 0.496 | 2.93632405115446e-93 | 4 |
| h1m4 | 7.550168895886e-92 | 0.689757214 | 0.976 | 0.588 | 1.90030200940555e-87 | 4 |
| h2af1al4 | 1.40031936394207e-90 | 0.702300977 | 0.991 | 0.578 | 3.52446380710579e-86 | 4 |
| cldnd4 | 1.75451986161481e-89 | 0.696583669 | 0.987 | 0.591 | 4.41595103969832e-85 | 4 |
| si:dkey-241l7.24 | 6.11756250693229e-89 | 0.670341364 | 0.993 | 0.629 | 1.53972930736979e-84 | 4 |
| mt24 | 3.14209589251693e-87 | 0.708694152 | 0.943 | 0.52 | 7.90834115187585e-83 | 4 |
| si:dkey-241l7.54 | 2.7053645926453e-86 | 0.678605504 | 0.958 | 0.571 | 6.80913214322897e-82 | 4 |
| tmsb54 | 3.70503700160812e-86 | 0.731524944 | 0.903 | 0.474 | 9.32520762934747e-82 | 4 |
| zp34 | 8.24252946176897e-85 | 0.687194416 | 0.965 | 0.571 | 2.07456224023263e-80 | 4 |

| | | | | | | |
|---|---|---|---|---|---|---|
| zgc:1717764 | 1.1295463152362e-84 | 0.714810862 | 0.883 | 0.482 | 2.842955120818e-80 | 4 |
| zp2.24 | 2.05784027530987e-81 | 0.692672371 | 0.919 | 0.516 | 5.17937818892741e-77 | 4 |
| si:ch211-226h8.144 | 2.06305782840781e-81 | 0.680117551 | 0.943 | 0.554 | 5.19251024831961e-77 | 4 |
| si:ch73-189n23.14 | 2.93220315003893e-81 | 0.708508421 | 0.936 | 0.51 | 7.38006210833298e-77 | 4 |
| crp24 | 8.19096203721882e-81 | 0.713225182 | 0.899 | 0.475 | 2.0615832351476e-76 | 4 |
| cldng4 | 1.64595085596967e-78 | 0.688301969 | 0.89 | 0.501 | 4.14269370939007e-74 | 4 |
| si:dkeyp-98a7.94 | 7.6361645305358e-77 | 0.691851038 | 0.907 | 0.498 | 1.92194625069056e-72 | 4 |
| zp2.54 | 6.10702899030948e-76 | 0.672716665 | 0.945 | 0.55 | 1.53707812657099e-71 | 4 |
| zp3.24 | 7.33300731359028e-76 | 0.664631979 | 0.923 | 0.545 | 1.84564461075754e-71 | 4 |
| zp2.34 | 8.48691544124161e-76 | 0.684435626 | 0.901 | 0.506 | 2.1360717474061e-71 | 4 |
| zp2.14 | 2.00060716490047e-75 | 0.680107843 | 0.921 | 0.522 | 5.03532817333799e-71 | 4 |
| acp5a4 | 9.74108633505105e-75 | 0.661569776 | 0.907 | 0.527 | 2.451734019669e-70 | 4 |
| zgc:1751354 | 1.80999749801476e-74 | 0.696779748 | 0.872 | 0.466 | 4.55558270275335e-70 | 4 |
| si:ch211-12h2.84 | 2.42982100651177e-72 | 0.641585243 | 0.888 | 0.536 | 6.11561649128948e-68 | 4 |
| zgc:771184 | 1.27449489982539e-70 | 0.671927386 | 0.808 | 0.445 | 3.20777621337053e-66 | 4 |
| zgc:1717794 | 1.4311552398487e-70 | 0.658390328 | 0.868 | 0.5 | 3.60207462317518e-66 | 4 |
| CR790368.14 | 2.10500872141168e-68 | 0.647533869 | 0.883 | 0.508 | 5.29809645092106e-64 | 4 |
| si:dkey-241l7.44 | 5.1299395755598e-68 | 0.600705366 | 0.934 | 0.598 | 1.29115449177265e-63 | 4 |
| si:dkey-229d11.34 | 1.20980821091252e-66 | 0.684694765 | 0.824 | 0.425 | 3.04496628604572e-62 | 4 |
| si:zfos-1505d6.34 | 1.2352649422525e-66 | 0.656069025 | 0.896 | 0.5 | 3.10903833315531e-62 | 4 |
| BX649337.14 | 6.8040000061987e-66 | 0.668210068 | 0.802 | 0.425 | 1.71249876156015e-61 | 4 |
| zp2.64 | 4.53184596799264e-65 | 0.648455576 | 0.859 | 0.489 | 1.14062031168407e-60 | 4 |
| epcam4 | 1.09780128125725e-64 | 0.629459387 | 0.839 | 0.494 | 2.76305604479637e-60 | 4 |
| rrm24 | 4.19645652073656e-63 | 0.618745567 | 0.824 | 0.488 | 1.05620614170418e-58 | 4 |
| zar1l4 | 1.38191508249239e-62 | 0.661703521 | 0.727 | 0.387 | 3.47814207112509e-58 | 4 |
| si:ch211-125e6.124 | 2.00248363636471e-62 | 0.646992666 | 0.87 | 0.488 | 5.04005106436633e-58 | 4 |
| ccna14 | 1.47428676054809e-61 | 0.633579607 | 0.815 | 0.465 | 3.71063234762348e-57 | 4 |
| CU571163.14 | 6.0409035439317e-59 | 0.652305448 | 0.727 | 0.391 | 1.52043501297217e-54 | 4 |
| lamp24 | 7.95336189604026e-59 | 0.580636439 | 0.83 | 0.545 | 2.00178165561437e-54 | 4 |
| si:dkeyp-98a7.44 | 6.90164995769649e-58 | 0.631573421 | 0.76 | 0.414 | 1.73707627785263e-53 | 4 |
| zp3b4 | 1.57855359665671e-57 | 0.644371744 | 0.714 | 0.377 | 3.97306154742528e-53 | 4 |
| si:dkeyp-98a7.84 | 1.79618006523152e-56 | 0.624972892 | 0.793 | 0.447 | 4.5208056061812e-52 | 4 |
| zar14 | 5.18296306798939e-56 | 0.622350263 | 0.762 | 0.429 | 1.30449997458225e-51 | 4 |
| si:ch211-226h8.44 | 1.49299169939164e-55 | 0.625973781 | 0.778 | 0.431 | 3.75771080819881e-51 | 4 |
| si:dkey-241l7.64 | 1.7096400236828e-54 | 0.584698281 | 0.833 | 0.517 | 4.30299297560723e-50 | 4 |

| | | | | | | |
|---|---|---|---|---|---|---|
| mid1ip1l4 | 2.45267765036489e-54 | 0.558535882 | 0.808 | 0.535 | 6.1731443782034e-50 | 4 |
| tuba4l4 | 5.75469493177448e-54 | 0.616572653 | 0.703 | 0.395 | 1.44839916737832e-49 | 4 |
| LOC1104378214 | 2.09249611773413e-53 | 0.632753729 | 0.736 | 0.405 | 5.26660347872504e-49 | 4 |
| npm2b4 | 5.63297460527769e-53 | 0.619025054 | 0.694 | 0.386 | 1.41776337840234e-48 | 4 |
| zp3a.24 | 5.40337340303771e-52 | 0.583993868 | 0.795 | 0.481 | 1.35997505181056e-47 | 4 |
| zgc:1952454 | 1.0012795030951e-50 | 0.609037422 | 0.685 | 0.379 | 2.52012038134005e-46 | 4 |
| si:ch211-12h2.64 | 1.50132585626795e-50 | 0.579217476 | 0.769 | 0.465 | 3.77868704764081e-46 | 4 |
| si:dkeyp-20e4.84 | 1.80135407533481e-50 | 0.619353722 | 0.731 | 0.397 | 4.53382807221018e-46 | 4 |
| spink2.14 | 8.71296765826253e-50 | 0.574642755 | 0.72 | 0.427 | 2.1929668299081e-45 | 4 |
| slbp24 | 1.32037200535196e-49 | 0.602071367 | 0.696 | 0.387 | 3.32324430027035e-45 | 4 |
| si:dkey-90l23.14 | 4.60971945576385e-49 | 0.588927069 | 0.753 | 0.438 | 1.1602202898212e-44 | 4 |
| thy14 | 1.865818457287e-48 | 0.571812143 | 0.731 | 0.441 | 4.69607847514564e-44 | 4 |
| si:ch211-14a17.74 | 4.22321718304134e-48 | 0.591084899 | 0.687 | 0.399 | 1.06294153279968e-43 | 4 |
| qdprb24 | 1.00597672670703e-47 | 0.585148541 | 0.714 | 0.424 | 2.53194282344891e-43 | 4 |
| siva14 | 2.9404836990242e-47 | 0.573179604 | 0.711 | 0.419 | 7.400903422074e-43 | 4 |
| cth14 | 5.72390806862726e-45 | 0.576833472 | 0.654 | 0.371 | 1.44065042179279e-40 | 4 |
| AL929032.14 | 9.02849742508201e-45 | 0.600309214 | 0.581 | 0.309 | 2.27238251691889e-40 | 4 |
| btg44 | 1.87572589809743e-44 | 0.578718496 | 0.641 | 0.36 | 4.72101451292142e-40 | 4 |
| gapdh4 | 7.93772919999214e-43 | 0.555305288 | 0.676 | 0.412 | 1.99784706234602e-38 | 4 |
| si:dkeyp-46h3.84 | 1.07106991204868e-42 | 0.581482934 | 0.623 | 0.343 | 2.69577586163533e-38 | 4 |
| si:dkey-229d11.54 | 1.22051453819108e-42 | 0.566866477 | 0.659 | 0.381 | 3.07191304117313e-38 | 4 |
| nanog4 | 1.61328277332618e-42 | 0.564253931 | 0.628 | 0.351 | 4.06047141218466e-38 | 4 |
| zgc:1362544 | 1.08659297573942e-41 | 0.569125246 | 0.601 | 0.322 | 2.73484586063855e-37 | 4 |
| zgc:1715174 | 1.32774904127351e-41 | 0.564194719 | 0.648 | 0.38 | 3.3418115619813e-37 | 4 |
| zgc:1734434 | 2.44914089673373e-41 | 0.564365572 | 0.601 | 0.322 | 6.16424272298912e-37 | 4 |
| ccnb24 | 3.0222403649294e-41 | 0.558760522 | 0.645 | 0.376 | 7.60667677449081e-37 | 4 |
| si:dkeyp-50f7.24 | 3.52065145444023e-41 | 0.546129922 | 0.665 | 0.394 | 8.86112764568062e-37 | 4 |
| LOC1005374994 | 4.64346978416826e-41 | 0.575394107 | 0.588 | 0.314 | 1.16871490997731e-36 | 4 |
| LOC1005351624 | 6.37417534151365e-40 | 0.568761796 | 0.577 | 0.3 | 1.60431619170557e-35 | 4 |
| gtf3ab4 | 1.108455780809e-39 | 0.562797851 | 0.577 | 0.312 | 2.78987235471817e-35 | 4 |
| si:dkeyp-46h3.24 | 2.59212164221767e-39 | 0.541736511 | 0.65 | 0.397 | 6.52411096129766e-35 | 4 |
| si:dkeyp-46h3.64 | 4.65854494405084e-39 | 0.542658522 | 0.674 | 0.402 | 1.17250917696816e-34 | 4 |
| ctss2.14 | 7.74591968017151e-39 | 0.47477192 | 0.749 | 0.528 | 1.94957052430237e-34 | 4 |
| si:ch211-132b12.84 | 1.56776455517403e-38 | 0.549325139 | 0.588 | 0.335 | 3.94590660891753e-34 | 4 |
| kpna74 | 5.97852961695193e-38 | 0.551140287 | 0.579 | 0.321 | 1.50473611929063e-33 | 4 |

| | | | | | | |
|---|---|---|---|---|---|---|
| si:dkeyp-75b4.101 | 1.0157155481481e-37 | 0.489141844 | 0.667 | 0.43 | 2.55645446313395e-33 | 4 |
| si:dkey-152b24.64 | 5.33320245443426e-37 | 0.547222053 | 0.535 | 0.291 | 1.34231372575656e-32 | 4 |
| haus64 | 2.89926870860394e-36 | 0.54937357 | 0.421 | 0.203 | 7.29716941268525e-32 | 4 |
| s100a14 | 1.14706404042582e-35 | 0.526465066 | 0.612 | 0.356 | 2.88704548334776e-31 | 4 |
| si:dkey-90l23.24 | 3.02501635255491e-35 | 0.53042445 | 0.584 | 0.338 | 7.61366365774545e-31 | 4 |
| wu:fi42e034 | 7.96645072562057e-35 | 0.525423539 | 0.579 | 0.343 | 2.00507598313144e-30 | 4 |
| zpcx4 | 9.78716583511893e-35 | 0.522351539 | 0.577 | 0.332 | 2.46333176904108e-30 | 4 |
| si:dkey-208k4.24 | 9.99458628348805e-34 | 0.527687768 | 0.515 | 0.289 | 2.51553742169111e-29 | 4 |
| ccnb14 | 3.71244515296808e-32 | 0.495644733 | 0.621 | 0.387 | 9.34385320550535e-28 | 4 |
| hnrnpabb4 | 3.71081967819063e-31 | 0.394415397 | 0.733 | 0.593 | 9.339762048038e-27 | 4 |
| ca15b4 | 4.8889764046804e-30 | 0.505676731 | 0.452 | 0.244 | 1.23050647129401e-25 | 4 |
| zgc:1719773 | 5.24454010814021e-30 | 0.520560706 | 0.423 | 0.224 | 1.31999829981781e-25 | 4 |
| si:ch211-145c1.14 | 9.78530979844446e-30 | 0.514880265 | 0.434 | 0.228 | 2.46286462317048e-25 | 4 |
| gmnn4 | 2.383541902124e-29 | 0.498668598 | 0.502 | 0.297 | 5.99913661345589e-25 | 4 |
| avd4 | 1.33439346405489e-28 | 0.491765801 | 0.471 | 0.259 | 3.35853490967976e-24 | 4 |
| ccna24 | 2.33408974816106e-28 | 0.493048106 | 0.476 | 0.279 | 5.87467048714658e-24 | 4 |
| zgc:1714744 | 3.54016955923935e-28 | 0.489021809 | 0.507 | 0.298 | 8.91025276364951e-24 | 4 |
| pigrl2.34 | 5.66514531931278e-28 | 0.497065716 | 0.388 | 0.198 | 1.42586042541783e-23 | 4 |
| larp6b4 | 5.67688385791936e-28 | 0.496248796 | 0.374 | 0.191 | 1.42881489819972e-23 | 4 |
| cpeb1b4 | 6.03390217991496e-28 | 0.486656427 | 0.469 | 0.265 | 1.5186728396628e-23 | 4 |
| gdf34 | 1.00918919839652e-27 | 0.487659811 | 0.412 | 0.218 | 2.54002829344421e-23 | 4 |
| lsm14b4 | 3.49429138052885e-27 | 0.482583112 | 0.491 | 0.295 | 8.79478197565307e-23 | 4 |
| zgc:925914 | 8.92518584588147e-26 | 0.47481425 | 0.383 | 0.201 | 2.24638002554991e-21 | 4 |
| arhgef10la4 | 4.45759414744415e-25 | 0.477744864 | 0.454 | 0.27 | 1.12193187097022e-20 | 4 |
| zglp14 | 6.07470325315106e-25 | 0.473341771 | 0.366 | 0.194 | 1.52894206178559e-20 | 4 |
| si:ch211-250e5.164 | 6.8581312038607e-25 | 0.46145528 | 0.524 | 0.342 | 1.7261230426997e-20 | 4 |
| si:ch211-251f6.64 | 9.33315631492725e-25 | 0.457952373 | 0.5 | 0.327 | 2.34906211290404e-20 | 4 |
| nasp4 | 1.46418801019385e-24 | 0.454391106 | 0.529 | 0.335 | 3.68521480285691e-20 | 4 |
| si:dkey-37o8.14 | 2.09777816642757e-24 | 0.461867136 | 0.491 | 0.302 | 5.27989786708155e-20 | 4 |
| si:ch73-14h10.24 | 2.87439395891244e-24 | 0.460771002 | 0.458 | 0.266 | 7.23456215518671e-20 | 4 |
| uchl14 | 5.71839952143946e-24 | 0.442010331 | 0.533 | 0.34 | 1.4392639755511e-19 | 4 |
| birc5b4 | 2.85283912223358e-23 | 0.452048218 | 0.434 | 0.255 | 7.18031078674968e-19 | 4 |
| slbp5 | 6.17589215894733e-23 | 0.462630158 | 0.396 | 0.23 | 1.55441029748545e-18 | 4 |
| pabpn1l4 | 7.93268393765933e-23 | 0.44748263 | 0.46 | 0.279 | 1.99657722026948e-18 | 4 |
| h3f3a4 | 1.53927329137787e-22 | 0.380554699 | 0.619 | 0.503 | 3.87419694706897e-18 | 4 |

| | | | | | | |
|---|---|---|---|---|---|---|
| elovl7b4 | 1.85196473937584e-21 | 0.438882714 | 0.416 | 0.244 | 4.66121005253506e-17 | 4 |
| si:ch211-146l10.84 | 1.82757057059386e-20 | 0.426590339 | 0.476 | 0.302 | 4.59981236912768e-16 | 4 |
| zp3a.14 | 2.1923643757184e-20 | 0.412214794 | 0.537 | 0.378 | 5.51796189724565e-16 | 4 |
| si:dkeyp-98a7.74 | 5.45318460716383e-20 | 0.42556548 | 0.35 | 0.192 | 1.37251203377706e-15 | 4 |
| zgc:554614 | 6.24395292166426e-20 | 0.418956786 | 0.452 | 0.284 | 1.57154051085368e-15 | 4 |
| LOC1018854774 | 1.60831785775042e-19 | 0.420843286 | 0.441 | 0.276 | 4.04797521617203e-15 | 4 |
| si:dkey-241l7.34 | 4.99127272783713e-19 | 0.406184649 | 0.509 | 0.361 | 1.25625343286933e-14 | 4 |
| mkrn44 | 5.22068936570703e-19 | 0.427057364 | 0.374 | 0.225 | 1.3139953064548e-14 | 4 |
| si:ch73-90k17.14 | 5.47567419507939e-19 | 0.413934301 | 0.407 | 0.245 | 1.37817243815953e-14 | 4 |
| si:dkeyp-46h3.54 | 8.0157241147025e-19 | 0.407793582 | 0.447 | 0.286 | 2.01747760242947e-14 | 4 |
| si:dkey-19b23.114 | 2.14710022900478e-18 | 0.411286305 | 0.434 | 0.275 | 5.40403656638213e-14 | 4 |
| si:ch211-146l10.74 | 4.49327925844036e-18 | 0.408769399 | 0.326 | 0.186 | 1.13091345655685e-13 | 4 |
| si:ch1073-263o8.24 | 5.35183261560242e-18 | 0.410023432 | 0.412 | 0.26 | 1.34700275102097e-13 | 4 |
| si:ch211-226l4.64 | 5.73138497754253e-18 | 0.405884273 | 0.452 | 0.289 | 1.44253228499768e-13 | 4 |
| AL929192.14 | 6.11451222097552e-18 | 0.395943534 | 0.302 | 0.161 | 1.53896158089733e-13 | 4 |
| org4 | 8.52137085747312e-18 | 0.413196435 | 0.385 | 0.237 | 2.14474383111741e-13 | 4 |
| zgc:562314 | 3.24638535304948e-17 | 0.393897183 | 0.372 | 0.225 | 8.17082729509023e-13 | 4 |
| inka1b4 | 5.57612280826532e-17 | 0.398429079 | 0.317 | 0.182 | 1.4034543496123e-12 | 4 |
| slc16a34 | 2.68746952750073e-16 | 0.389940089 | 0.388 | 0.246 | 6.76409205376658e-12 | 4 |
| LOC1018838974 | 3.26718178997578e-16 | 0.383494007 | 0.291 | 0.161 | 8.22316984719003e-12 | 4 |
| elavl24 | 1.04735435342502e-15 | 0.385149501 | 0.361 | 0.221 | 2.63608617213544e-11 | 4 |
| zgc:1655394 | 1.04759059789577e-15 | 0.360821249 | 0.233 | 0.117 | 2.63668077584387e-11 | 4 |
| rbpms2b4 | 1.15719974363283e-15 | 0.389412155 | 0.35 | 0.218 | 2.91255603474946e-11 | 4 |
| sinup4 | 1.33841918995057e-15 | 0.385449401 | 0.333 | 0.2 | 3.36866725918659e-11 | 4 |
| mibp4 | 1.59455368072216e-15 | 0.384435842 | 0.35 | 0.22 | 4.01333215900961e-11 | 4 |
| zgc:566994 | 1.94444937585184e-15 | 0.380161133 | 0.385 | 0.249 | 4.8939846340815e-11 | 4 |
| cldn7b4 | 2.02493645275616e-15 | 0.372876186 | 0.282 | 0.157 | 5.09656255794198e-11 | 4 |
| tph1a4 | 2.28907780556512e-15 | 0.364284202 | 0.26 | 0.139 | 5.76137992882686e-11 | 4 |
| zgc:1526523 | 2.66867633830307e-15 | 0.353360263 | 0.225 | 0.111 | 6.71679147587499e-11 | 4 |
| zgc:1738564 | 2.75254407326955e-15 | 0.378627648 | 0.352 | 0.217 | 6.92787817801213e-11 | 4 |
| rasl11b4 | 2.85780866753163e-15 | 0.359751158 | 0.251 | 0.131 | 7.19281863531036e-11 | 4 |
| si:dkey-203a12.21 | 6.06684365392192e-15 | 0.325876459 | 0.383 | 0.243 | 1.52696387925561e-10 | 4 |
| zgc:1717504 | 9.25448763933248e-15 | 0.377116943 | 0.295 | 0.172 | 2.32926199394359e-10 | 4 |
| zgc:1628794 | 3.45761502925393e-14 | 0.365024122 | 0.302 | 0.179 | 8.70247126712922e-10 | 4 |
| BX537276.14 | 5.87975609661293e-14 | 0.345571145 | 0.227 | 0.119 | 1.47987581195651e-09 | 4 |

| | | | | | | |
|---|---|---|---|---|---|---|
| si:dkey-68o6.54 | 8.53493118819754e-14 | 0.347702028 | 0.229 | 0.12 | 2.14815683075744e-09 | 4 |
| pabpc1l4 | 1.22584469304937e-13 | 0.344296175 | 0.253 | 0.14 | 3.08532850793596e-09 | 4 |
| zp3c4 | 1.73199323331969e-13 | 0.352199725 | 0.308 | 0.187 | 4.35925376894232e-09 | 4 |
| rbm4.34 | 3.64439456582763e-13 | 0.311899558 | 0.515 | 0.444 | 9.17257668273157e-09 | 4 |
| rcc14 | 8.73321132700906e-13 | 0.352135153 | 0.317 | 0.2 | 2.19806195889491e-08 | 4 |
| crp34 | 9.75877666566248e-13 | 0.350628769 | 0.335 | 0.216 | 2.45618649898059e-08 | 4 |
| si:ch73-15b2.52 | 1.01430839778721e-12 | 0.312635345 | 0.187 | 0.093 | 2.55291280639063e-08 | 4 |
| ddx39ab4 | 1.27755548649357e-12 | 0.356371337 | 0.392 | 0.287 | 3.21547940395568e-08 | 4 |
| snu13b4 | 1.2881303819392e-12 | 0.325183427 | 0.485 | 0.391 | 3.24209535830278e-08 | 4 |
| zgc:1034824 | 1.39582405831949e-12 | 0.344184396 | 0.304 | 0.187 | 3.51314957238432e-08 | 4 |
| zgc:1737424 | 1.93924482806231e-12 | 0.340682119 | 0.262 | 0.153 | 4.88088530775003e-08 | 4 |
| buc4 | 2.34876558971115e-12 | 0.34281097 | 0.291 | 0.176 | 5.91160811274398e-08 | 4 |
| chia.44 | 2.57086769426445e-12 | 0.336239335 | 0.251 | 0.145 | 6.4706168996942e-08 | 4 |
| pou5f34 | 3.08518570020648e-12 | 0.334767326 | 0.251 | 0.145 | 7.76510388884969e-08 | 4 |
| zgc:660244 | 3.150705284974e-12 | 0.328454238 | 0.244 | 0.14 | 7.93001013175107e-08 | 4 |
| CR847984.14 | 3.24883318446363e-12 | 0.328118684 | 0.273 | 0.162 | 8.1769882419765e-08 | 4 |
| si:ch73-78o10.14 | 3.82094152468589e-12 | 0.319700201 | 0.236 | 0.131 | 9.61692772348193e-08 | 4 |
| mos3 | 5.44749358967519e-12 | 0.339894603 | 0.236 | 0.135 | 1.37107966158535e-07 | 4 |
| si:ch73-359m17.21 | 5.8994366253231e-12 | 0.311331064 | 0.304 | 0.187 | 1.48482920422757e-07 | 4 |
| CABZ01075618.1 | 6.48837027748265e-12 | 0.258397017 | 0.128 | 0.054 | 1.63305791513961e-07 | 4 |
| zgc:1141301 | 1.10086410534394e-11 | 0.278360966 | 0.156 | 0.074 | 2.77076486674016e-07 | 4 |
| dynll2a3 | 1.38744406675468e-11 | 0.324866838 | 0.233 | 0.135 | 3.49205797161485e-07 | 4 |
| ankrd224 | 1.50564780291998e-11 | 0.329162841 | 0.242 | 0.141 | 3.78956495516931e-07 | 4 |
| LOC1005368604 | 1.63557981677478e-11 | 0.318533109 | 0.449 | 0.35 | 4.11659084084045e-07 | 4 |
| kmt5ab1 | 1.84651194823274e-11 | 0.300510141 | 0.174 | 0.088 | 4.64748592250699e-07 | 4 |
| retsatl4 | 1.85159761510371e-11 | 0.338022736 | 0.311 | 0.202 | 4.66028603745454e-07 | 4 |
| zgc:566764 | 2.0990653184079e-11 | 0.34893462 | 0.388 | 0.294 | 5.28313749990084e-07 | 4 |
| patl24 | 2.28337930137936e-11 | 0.330809324 | 0.278 | 0.171 | 5.74703736364172e-07 | 4 |
| CDPF14 | 2.36582931865277e-11 | 0.347828022 | 0.368 | 0.267 | 5.95455581211717e-07 | 4 |
| CU655961.42 | 2.57150082284966e-11 | 0.301299209 | 0.172 | 0.087 | 6.4722104210303e-07 | 4 |
| si:ch73-160p18.34 | 3.32878516697425e-11 | 0.335979908 | 0.348 | 0.239 | 8.3782193867575e-07 | 4 |
| cdt12 | 3.38630877319354e-11 | 0.296658033 | 0.17 | 0.086 | 8.52300055125082e-07 | 4 |
| lpcat43 | 5.4794943320903e-11 | 0.301276487 | 0.196 | 0.105 | 1.37913392844381e-06 | 4 |
| cdca7a4 | 5.85409835763113e-11 | 0.335439119 | 0.322 | 0.221 | 1.47341801563218e-06 | 4 |
| si:ch73-281n10.22 | 8.56269979466951e-11 | 0.308284169 | 0.456 | 0.392 | 2.15514591132037e-06 | 4 |

| | | | | | | |
|---|---|---|---|---|---|---|
| tac3b4 | 1.03573046147859e-10 | 0.300941542 | 0.211 | 0.119 | 2.60682999849547e-06 | 4 |
| nfr4 | 1.21393246484475e-10 | 0.298838747 | 0.198 | 0.109 | 3.05534662076775e-06 | 4 |
| tmem54a3 | 1.22849802351004e-10 | 0.308340655 | 0.236 | 0.139 | 3.09200667537242e-06 | 4 |
| si:ch211-202f5.3 | 1.31748412203136e-10 | 0.291338859 | 0.172 | 0.089 | 3.31597578674073e-06 | 4 |
| hsd17b12a4 | 1.34285817545805e-10 | 0.315050028 | 0.264 | 0.164 | 3.37983974181036e-06 | 4 |
| dynll14 | 1.36434839669564e-10 | 0.277936083 | 0.513 | 0.455 | 3.43392847964324e-06 | 4 |
| map1lc3c4 | 1.64110348648852e-10 | 0.323925255 | 0.304 | 0.2 | 4.13049336514294e-06 | 4 |
| snx10a2 | 2.07067864601729e-10 | 0.305396325 | 0.216 | 0.125 | 5.21169108416091e-06 | 4 |
| elovl7a4 | 3.5001033132654e-10 | 0.308020571 | 0.282 | 0.181 | 8.80941002915768e-06 | 4 |
| dctpp14 | 5.16194639511018e-10 | 0.3218351 | 0.267 | 0.174 | 1.29921028818528e-05 | 4 |
| gyg1a4 | 5.94110992343951e-10 | 0.306155093 | 0.447 | 0.37 | 1.49531795663049e-05 | 4 |
| ypel33 | 6.73233683431639e-10 | 0.303508829 | 0.214 | 0.126 | 1.69446185782909e-05 | 4 |
| zgc:1134244 | 1.11988841688571e-09 | 0.297324969 | 0.256 | 0.16 | 2.81864715645965e-05 | 4 |
| oclna | 1.46740706563101e-09 | 0.26014408 | 0.152 | 0.078 | 3.69331684348669e-05 | 4 |
| vstm2l3 | 1.65819588007446e-09 | 0.265593738 | 0.174 | 0.094 | 4.17351321055941e-05 | 4 |
| si:ch1073-190k2.14 | 1.79236368725275e-09 | 0.298582871 | 0.286 | 0.187 | 4.51120016444645e-05 | 4 |
| si:ch211-157c3.42 | 1.97876107464786e-09 | 0.269255481 | 0.165 | 0.088 | 4.9803437487812e-05 | 4 |
| zgc:1138864 | 2.67457772714601e-09 | 0.292087746 | 0.229 | 0.14 | 6.7316446814538e-05 | 4 |
| cx44.24 | 4.17285999097276e-09 | 0.2766729 | 0.2 | 0.116 | 0.000105027 | 4 |
| sybu4 | 6.33678494549581e-09 | 0.287601535 | 0.258 | 0.165 | 0.000159491 | 4 |
| ctsla4 | 6.44056577277115e-09 | 0.275979955 | 0.458 | 0.386 | 0.000162103 | 4 |
| sox19b4 | 6.82086309475025e-09 | 0.260535861 | 0.174 | 0.097 | 0.000171674 | 4 |
| phex1 | 6.84011396000526e-09 | 0.260916579 | 0.159 | 0.085 | 0.000172159 | 4 |
| mad2l14 | 7.27717536956064e-09 | 0.300627255 | 0.273 | 0.185 | 0.000183159 | 4 |
| h2afx2 | 8.36514359101313e-09 | 0.294134416 | 0.22 | 0.137 | 0.000210542 | 4 |
| rnf43 | 8.82322933073609e-09 | 0.295655193 | 0.214 | 0.133 | 0.000222072 | 4 |
| dnd12 | 9.29177865559213e-09 | 0.27415203 | 0.187 | 0.108 | 0.000233865 | 4 |
| zgc:1141234 | 1.05831089038582e-08 | 0.26401224 | 0.181 | 0.102 | 0.000266366 | 4 |
| zgc:1017442 | 1.23954752040233e-08 | 0.287009793 | 0.214 | 0.132 | 0.000311982 | 4 |
| cdc204 | 1.71365429830447e-08 | 0.280234926 | 0.238 | 0.152 | 0.00043131 | 4 |
| cox174 | 1.91298897007754e-08 | 0.270147773 | 0.427 | 0.378 | 0.00048148 | 4 |
| zgc:922873 | 1.92332760306114e-08 | 0.273639767 | 0.214 | 0.132 | 0.000484082 | 4 |
| snrpb23 | 2.05741483652021e-08 | 0.300893257 | 0.229 | 0.148 | 0.000517831 | 4 |
| aif1l1 | 2.1187683539211e-08 | 0.299161678 | 0.269 | 0.187 | 0.000533273 | 4 |
| bzw1b2 | 2.2485363412611e-08 | 0.266478942 | 0.452 | 0.403 | 0.000565934 | 4 |

| | | | | | | |
|---|---|---|---|---|---|---|
| si:dkey-42i9.84 | 2.69390750918057e-08 | 0.282127265 | 0.205 | 0.126 | 0.00067803 | 4 |
| zgc:1129802 | 2.95944867470515e-08 | 0.257515664 | 0.163 | 0.091 | 0.000744864 | 4 |
| zgc:1630303 | 3.04991879620494e-08 | 0.266314578 | 0.203 | 0.123 | 0.000767634 | 4 |
| si:dkey-46g23.24 | 3.63985691322206e-08 | 0.275978861 | 0.225 | 0.144 | 0.000916116 | 4 |
| cpm2 | 3.95788060321547e-08 | 0.260159882 | 0.183 | 0.108 | 0.000996159 | 4 |
| si:dkey-92i17.2 | 4.23634777330895e-08 | 0.270364826 | 0.161 | 0.091 | 0.001066246 | 4 |
| fbxo322 | 4.32128844357035e-08 | 0.266577565 | 0.176 | 0.103 | 0.001087625 | 4 |
| pttg14 | 7.1624579041022e-08 | 0.276181113 | 0.238 | 0.158 | 0.001802719 | 4 |
| CABZ01059627.24 | 1.17774912488594e-07 | 0.266645802 | 0.209 | 0.133 | 0.002964277 | 4 |
| smn1 | 1.56670351731542e-07 | 0.255976877 | 0.159 | 0.092 | 0.003943236 | 4 |
| larp6a4 | 2.15716568253077e-07 | 0.277868939 | 0.328 | 0.249 | 0.00542937 | 4 |
| pif13 | 2.2638346595601e-07 | 0.252097983 | 0.176 | 0.106 | 0.005697845 | 4 |
| si:dkey-203a12.91 | 3.39661812988172e-07 | 0.25043766 | 0.216 | 0.139 | 0.008548948 | 4 |
| zp2l14 | 4.22126106622162e-07 | 0.261003859 | 0.24 | 0.163 | 0.010624492 | 4 |
| si:dkey-188i13.114 | 5.00030579495717e-07 | 0.280973958 | 0.379 | 0.316 | 0.01258527 | 4 |
| CABZ01059627.13 | 5.08157737199574e-07 | 0.254606088 | 0.209 | 0.136 | 0.012789822 | 4 |
| tent5c3 | 5.4335875812219e-07 | 0.264610828 | 0.209 | 0.139 | 0.013675797 | 4 |
| zgc:1629454 | 6.25586542907288e-07 | 0.259290953 | 0.236 | 0.162 | 0.015745388 | 4 |
| zgc:868393 | 7.63894330489616e-07 | 0.257156201 | 0.174 | 0.109 | 0.019226456 | 4 |
| BX284697.14 | 1.00236138173853e-06 | 0.250102203 | 0.181 | 0.114 | 0.025228434 | 4 |
| cnih11 | 1.43909603288697e-06 | 0.258645179 | 0.181 | 0.116 | 0.036220608 | 4 |
| aurkb2 | 2.3349822057617e-06 | 0.253082055 | 0.192 | 0.128 | 0.058769167 | 4 |
| pcf111 | 4.55776797529843e-06 | 0.265453237 | 0.295 | 0.239 | 0.114714462 | 4 |
| aktip1 | 4.60663498781632e-06 | 0.264865182 | 0.2 | 0.139 | 0.115944396 | 4 |
| fdx14 | 4.00062703407913e-05 | 0.251016342 | 0.273 | 0.222 | 1 | 4 |
| anp32b2 | 4.02905887904096e-05 | 0.252087639 | 0.3 | 0.257 | 1 | 4 |
| ccl36.1 | 0 | 1.929860379 | 0.905 | 0.072 | 0 | 5 |
| XLOC-042595 | 0 | 1.81899206 | 0.966 | 0.135 | 0 | 5 |
| XLOC-000431 | 0 | 1.811956113 | 0.99 | 0.252 | 0 | 5 |
| ccl38.6 | 0 | 1.771962088 | 0.893 | 0.081 | 0 | 5 |
| XLOC-013964 | 0 | 1.727910541 | 0.968 | 0.145 | 0 | 5 |
| XLOC-010899 | 0 | 1.695543749 | 0.976 | 0.232 | 0 | 5 |
| LOC103910871 | 0 | 1.653956476 | 0.951 | 0.142 | 0 | 5 |
| si:ch211-214p16.1 | 0 | 1.596676135 | 0.99 | 0.231 | 0 | 5 |
| CR936442.1 | 0 | 1.545501831 | 0.939 | 0.141 | 0 | 5 |

| | | | | | | |
|---|---|---|---|---|---|---|
| dusp2 | 0 | 1.395741238 | 0.881 | 0.153 | 0 | 5 |
| XLOC-028370 | 0 | 1.377998162 | 0.801 | 0.117 | 0 | 5 |
| XLOC-025423 | 0 | 1.331948302 | 0.837 | 0.145 | 0 | 5 |
| XLOC-020301 | 0 | 1.310011085 | 0.692 | 0.063 | 0 | 5 |
| XLOC-026836 | 0 | 1.303813984 | 0.636 | 0.058 | 0 | 5 |
| ccl34b.4 | 0 | 1.290444168 | 0.573 | 0.018 | 0 | 5 |
| LOC100536213 | 0 | 1.226308433 | 0.738 | 0.083 | 0 | 5 |
| CR753876.1 | 0 | 1.226256506 | 0.716 | 0.093 | 0 | 5 |
| ccl32b.3 | 0 | 1.200876101 | 0.575 | 0.011 | 0 | 5 |
| LOC108190717 | 0 | 1.200590589 | 0.682 | 0.075 | 0 | 5 |
| BX005105.1 | 0 | 1.095943303 | 0.585 | 0.024 | 0 | 5 |
| XLOC-032002 | 0 | 1.045654459 | 0.5 | 0.03 | 0 | 5 |
| si:dkey-183i3.6 | 0 | 1.041530637 | 0.505 | 0.029 | 0 | 5 |
| si:ch73-223p23.2 | 0 | 0.891904714 | 0.464 | 0.016 | 0 | 5 |
| LOC101882070 | 0 | 0.887431335 | 0.432 | 0.007 | 0 | 5 |
| si:dkey-109a10.2 | 0 | 0.880459279 | 0.454 | 0.017 | 0 | 5 |
| CR848749.2 | 0 | 0.820709603 | 0.386 | 0.014 | 0 | 5 |
| LOC100004978 | 0 | 0.782268556 | 0.364 | 0.003 | 0 | 5 |
| si:dkey-183p4.10 | 0 | 0.781962579 | 0.4 | 0.022 | 0 | 5 |
| LOC108185170 | 0 | 0.717730755 | 0.337 | 0.003 | 0 | 5 |
| si:dkey-260g12.1 | 0 | 0.706698762 | 0.376 | 0.021 | 0 | 5 |
| FP016106.1 | 0 | 0.682452947 | 0.337 | 0.014 | 0 | 5 |
| cxcr3.1 | 0 | 0.603815653 | 0.301 | 0.011 | 0 | 5 |
| si:ch211-93g21.1 | 2.00235937872583e-302 | 0.732296389 | 0.371 | 0.023 | 5.03973832031504e-298 | 5 |
| XLOC-043760 | 1.09682108195273e-290 | 0.881003387 | 0.444 | 0.037 | 2.76058898116682e-286 | 5 |
| si:ch73-285p12.4 | 1.18882684216079e-279 | 0.515218323 | 0.252 | 0.009 | 2.9921582790345e-275 | 5 |
| zgc:136605 | 8.07878246769812e-268 | 0.604106423 | 0.277 | 0.013 | 2.03334875929494e-263 | 5 |
| si:rp71-81e14.2 | 2.24882361601566e-258 | 0.696916821 | 0.347 | 0.024 | 5.66006415914983e-254 | 5 |
| tnfsf14 | 1.90339218277759e-257 | 0.503159492 | 0.231 | 0.008 | 4.79064778483292e-253 | 5 |
| faslg | 5.24808216709753e-254 | 0.496348063 | 0.233 | 0.009 | 1.32088980063678e-249 | 5 |
| XLOC-023492 | 5.29754474968477e-246 | 0.984353164 | 0.57 | 0.078 | 1.33333903804816e-241 | 5 |
| c3a.5 | 5.72593177764875e-244 | 0.872961031 | 0.485 | 0.053 | 1.44115976911641e-239 | 5 |
| gzm3 | 1.31376762616433e-241 | 0.364706836 | 0.165 | 0.003 | 3.306621738293e-237 | 5 |
| wasb | 2.85815276197016e-241 | 1.076244231 | 0.784 | 0.156 | 7.19368468660269e-237 | 5 |
| si:ch211-119e14.1 | 1.31477548248227e-237 | 0.849215175 | 0.495 | 0.059 | 3.30915841185963e-233 | 5 |

| | | | | | | |
|---|---|---|---|---|---|---|
| si:ch211-167j9.5 | 1.27370110952424e-235 | 0.577634696 | 0.291 | 0.018 | 3.20577832256156e-231 | 5 |
| gpr183a | 7.72056109217364e-232 | 0.821701128 | 0.454 | 0.051 | 1.94318802128918e-227 | 5 |
| FP236356.1 | 9.31547519835032e-226 | 0.752569202 | 0.391 | 0.037 | 2.34461195267279e-221 | 5 |
| si:ch73-226l13.2 | 8.94625891927912e-225 | 0.473315948 | 0.223 | 0.01 | 2.25168390739336e-220 | 5 |
| ptpn6 | 8.10016373669739e-224 | 0.89762099 | 0.534 | 0.075 | 2.03873021088937e-219 | 5 |
| wasa | 4.66526841196867e-222 | 0.861956858 | 0.541 | 0.077 | 1.1742014066084e-217 | 5 |
| BX005183.3 | 1.28994588547042e-218 | 0.859062482 | 0.5 | 0.066 | 3.2466647991405e-214 | 5 |
| LOC103910136 | 4.12296274432651e-213 | 0.60705853 | 0.299 | 0.022 | 1.03770849311954e-208 | 5 |
| si:ch211-132g1.1 | 1.68550259818455e-208 | 0.494299841 | 0.26 | 0.016 | 4.2422414893707e-204 | 5 |
| gzmk | 5.28134238309775e-208 | 0.450135019 | 0.206 | 0.009 | 1.32926106440187e-203 | 5 |
| sla2 | 2.09271769955591e-206 | 0.826027711 | 0.451 | 0.056 | 5.26716117801227e-202 | 5 |
| LOC103909107 | 5.42401246276465e-205 | 0.586897469 | 0.265 | 0.017 | 1.36516969675323e-200 | 5 |
| id3 | 3.96617962475056e-204 | 1.092048772 | 0.784 | 0.19 | 9.98247749753469e-200 | 5 |
| LOC100535806 | 4.94127366615562e-197 | 0.622773589 | 0.303 | 0.025 | 1.24366916903471e-192 | 5 |
| XLOC-008993 | 3.11557169884402e-196 | 0.585616502 | 0.296 | 0.024 | 7.84158240882052e-192 | 5 |
| si:ch73-266m14.1 | 9.89900229104827e-196 | 0.288783123 | 0.112 | 0.001 | 2.49147988663394e-191 | 5 |
| lpar6b | 1.01651217511534e-190 | 0.434986222 | 0.214 | 0.011 | 2.5584594935478e-186 | 5 |
| ccl38a.5 | 4.86101310562433e-189 | 0.469981482 | 0.17 | 0.006 | 1.22346838855459e-184 | 5 |
| si:dkey-27i16.2 | 2.83053580778896e-184 | 0.82038599 | 0.5 | 0.077 | 7.12417557462402e-180 | 5 |
| mrps15 | 8.4988384458506e-183 | 0.889534323 | 0.585 | 0.108 | 2.13907264843614e-178 | 5 |
| kctd5a | 8.59316393364092e-181 | 0.963846373 | 0.675 | 0.158 | 2.16281343045808e-176 | 5 |
| arpc1b | 5.65473593201371e-180 | 1.073306073 | 0.925 | 0.287 | 1.42324048672853e-175 | 5 |
| cabz01093075.1 | 1.95775994683973e-178 | 0.451574298 | 0.218 | 0.013 | 4.92748601020092e-174 | 5 |
| btg1 | 3.49420087666868e-177 | 1.016970634 | 0.9 | 0.312 | 8.79455418648739e-173 | 5 |
| cd247l | 1.27112990244977e-173 | 0.436859699 | 0.221 | 0.014 | 3.19930685147582e-169 | 5 |
| XLOC-043761 | 2.55140148851393e-173 | 0.514191781 | 0.223 | 0.014 | 6.42162240644072e-169 | 5 |
| il2rb | 5.24048770848084e-170 | 0.456327021 | 0.218 | 0.014 | 1.31897835134754e-165 | 5 |
| si:ch73-46n24.1 | 1.65918084810817e-166 | 0.260184187 | 0.1 | 0.001 | 4.17599227660345e-162 | 5 |
| ncaldb | 2.35243619918404e-166 | 0.701637173 | 0.43 | 0.062 | 5.92084666972632e-162 | 5 |
| XLOC-033137 | 7.85070331922249e-165 | 0.571466002 | 0.284 | 0.027 | 1.97594351841511e-160 | 5 |
| si:dkeyp-85e10.3 | 8.00123320155895e-162 | 0.443699656 | 0.226 | 0.016 | 2.01383038450037e-157 | 5 |
| ccl38a.4 | 4.75065543252374e-160 | 0.511607897 | 0.163 | 0.007 | 1.1956924658119e-155 | 5 |
| LOC110439931 | 9.87553260642157e-160 | 0.450777442 | 0.226 | 0.017 | 2.48557280171024e-155 | 5 |
| coro1a | 2.98088063396817e-157 | 0.946983803 | 0.786 | 0.211 | 7.5025784676345e-153 | 5 |
| cyth4a | 8.65650383115506e-157 | 0.410010593 | 0.204 | 0.013 | 2.17875544926342e-152 | 5 |

| | | | | | | |
|---|---|---|---|---|---|---|
| rgs13 | 6.9861542830255e-156 | 0.893424924 | 0.566 | 0.117 | 1.75834517149469e-151 | 5 |
| pfn1 | 1.12433500781927e-154 | 0.979816059 | 0.995 | 0.508 | 2.82983878118031e-150 | 5 |
| rplp2 | 7.33620052780193e-150 | 0.55879395 | 0.993 | 0.75 | 1.84644831084247e-145 | 5 |
| LOC101885675 | 1.67698730062465e-149 | 0.345159574 | 0.155 | 0.007 | 4.22080933694219e-145 | 5 |
| entpd3 | 5.50374687991259e-149 | 0.419505545 | 0.197 | 0.013 | 1.3852380522052e-144 | 5 |
| si:ch211-196f2.6 | 1.80544241538368e-145 | 0.36217536 | 0.184 | 0.011 | 4.54411801527917e-141 | 5 |
| CU499336.2 | 3.36232471642822e-144 | 0.507165944 | 0.272 | 0.028 | 8.46263507877819e-140 | 5 |
| XLOC-042596 | 1.50093685537872e-143 | 0.582111511 | 0.294 | 0.033 | 3.77770797130269e-139 | 5 |
| hcst | 2.80740866302905e-140 | 0.576899073 | 0.325 | 0.042 | 7.06596686397781e-136 | 5 |
| ptprc | 2.92992515832654e-137 | 0.766430469 | 0.534 | 0.113 | 7.37432863099208e-133 | 5 |
| nitr2b | 7.96497509306101e-134 | 0.343711895 | 0.102 | 0.003 | 2.00470458117253e-129 | 5 |
| nitr10a | 1.72611127305931e-133 | 0.273025054 | 0.1 | 0.002 | 4.34444946316299e-129 | 5 |
| si:ch211-226m16.2 | 4.65771343565953e-133 | 0.63566012 | 0.371 | 0.058 | 1.17229989462115e-128 | 5 |
| rplp1 | 7.40659322543889e-131 | 0.414247655 | 1 | 0.873 | 1.86416544891072e-126 | 5 |
| FP101882.1 | 2.94261948205305e-130 | 0.480262724 | 0.255 | 0.027 | 7.40627897437933e-126 | 5 |
| rac2 | 6.11833540100598e-130 | 0.850804606 | 0.709 | 0.204 | 1.53992383707919e-125 | 5 |
| rpl18a | 1.49554061418838e-129 | 0.499106931 | 0.993 | 0.778 | 3.76412617185073e-125 | 5 |
| rplp2l | 2.86940839363947e-129 | 0.519495506 | 0.995 | 0.767 | 7.22201398595118e-125 | 5 |
| rpl37 | 1.0278489435983e-128 | 0.437387383 | 1 | 0.845 | 2.58699300614256e-124 | 5 |
| si:dkey-171c9.3 | 1.26936918382868e-128 | 0.473627006 | 0.252 | 0.027 | 3.1948752987784e-124 | 5 |
| CR855337.1 | 5.0882189524077e-128 | 0.486144521 | 0.25 | 0.027 | 1.28065382813149e-123 | 5 |
| themis | 2.04626104329182e-127 | 0.282931956 | 0.141 | 0.007 | 5.15023441986119e-123 | 5 |
| rps20 | 2.83113712488074e-127 | 0.345799215 | 0.993 | 0.914 | 7.12568902961232e-123 | 5 |
| si:ch211-25d12.7 | 4.26914134849865e-127 | 0.351138152 | 0.177 | 0.013 | 1.07450018600363e-122 | 5 |
| h1fx | 1.87198525579668e-126 | 0.922001837 | 0.614 | 0.168 | 4.71159969031466e-122 | 5 |
| rps27.1 | 3.41865156985537e-126 | 0.602160569 | 0.983 | 0.669 | 8.60440413616899e-122 | 5 |
| rps29 | 3.63719213166063e-126 | 0.431995535 | 0.998 | 0.827 | 9.15444887617665e-122 | 5 |
| laptm5 | 1.1198108530041e-125 | 0.769433104 | 0.624 | 0.154 | 2.81845193592602e-121 | 5 |
| rps2 | 2.284788184153e-125 | 0.471192331 | 0.99 | 0.824 | 5.75058338069468e-121 | 5 |
| tagapb | 1.12827814769759e-124 | 0.596106312 | 0.359 | 0.058 | 2.83976326994005e-120 | 5 |
| rps6 | 8.35428545961306e-124 | 0.480762563 | 0.995 | 0.782 | 2.10269010733001e-119 | 5 |
| sh2d1ab | 2.17972039755251e-122 | 0.374901811 | 0.158 | 0.01 | 5.48613826859992e-118 | 5 |
| itk | 1.00607072837891e-121 | 0.312509401 | 0.146 | 0.008 | 2.53217941625688e-117 | 5 |
| wdr19 | 2.80782538941818e-120 | 0.486279811 | 0.238 | 0.026 | 7.06701572262663e-116 | 5 |
| sept12 | 2.04332498940335e-119 | 0.574911867 | 0.333 | 0.052 | 5.1428446658293e-115 | 5 |

| | | | | | | |
|---|---|---|---|---|---|---|
| rps17 | 2.86474402595269e-119 | 0.478813015 | 0.976 | 0.76 | 7.21027423892034e-115 | 5 |
| capgb | 1.40998211997287e-118 | 0.699192718 | 0.478 | 0.102 | 3.54878399775972e-114 | 5 |
| arhgdig | 1.01391054525866e-117 | 0.823069342 | 0.714 | 0.227 | 2.55191145136152e-113 | 5 |
| slc6a15 | 7.03804357795795e-117 | 0.314235303 | 0.138 | 0.008 | 1.77140518813624e-112 | 5 |
| rpl11 | 1.2244414655577e-116 | 0.454784426 | 0.998 | 0.808 | 3.08179672466216e-112 | 5 |
| rps15a | 1.23710981652643e-114 | 0.503126001 | 0.995 | 0.773 | 3.11368169721537e-110 | 5 |
| rps26l | 8.41994257839547e-114 | 0.590831294 | 0.978 | 0.654 | 2.11921534755636e-109 | 5 |
| rpl36 | 1.39328359050255e-113 | 0.474254163 | 0.99 | 0.783 | 3.50675546893587e-109 | 5 |
| wu:fb55g09 | 1.63381175110807e-113 | 0.828927397 | 0.714 | 0.234 | 4.11214079636391e-109 | 5 |
| rps5 | 2.1834572760988e-112 | 0.549896482 | 0.978 | 0.692 | 5.49554361821308e-108 | 5 |
| zgc:64051 | 3.35076144367494e-112 | 0.714749092 | 0.583 | 0.148 | 8.43353147758545e-108 | 5 |
| calm1b | 1.83990904289432e-111 | 0.620744409 | 0.42 | 0.084 | 4.6308670700607e-107 | 5 |
| rpl19 | 3.90674571195703e-110 | 0.388868254 | 0.995 | 0.847 | 9.83288828242464e-106 | 5 |
| XLOC-011085 | 5.16640280145296e-110 | 0.56661269 | 0.325 | 0.054 | 1.3003319210977e-105 | 5 |
| si:dkey-78o7.3 | 6.19089166331769e-110 | 0.426636981 | 0.197 | 0.019 | 1.55818552274043e-105 | 5 |
| rpl34 | 2.52592618784456e-109 | 0.470379108 | 0.993 | 0.777 | 6.35750362218597e-105 | 5 |
| rps21 | 5.57621979236525e-109 | 0.5085059 | 0.99 | 0.742 | 1.40347875954041e-104 | 5 |
| adgre5b.3 | 1.03259944090047e-108 | 0.465299887 | 0.252 | 0.032 | 2.59894953280239e-104 | 5 |
| samsn1b | 1.18569600654093e-108 | 0.378638915 | 0.189 | 0.018 | 2.98427827886287e-104 | 5 |
| rpl15 | 1.85243761141126e-108 | 0.502261259 | 0.983 | 0.736 | 4.66240022416101e-104 | 5 |
| rpl32 | 4.30090864964059e-108 | 0.441368119 | 0.99 | 0.807 | 1.08249569802804e-103 | 5 |
| cxcr4b | 1.8677459041274e-107 | 0.810503225 | 0.609 | 0.169 | 4.70092966609826e-103 | 5 |
| rps16 | 1.11909592162821e-106 | 0.508369613 | 0.99 | 0.729 | 2.81665252514605e-102 | 5 |
| BX511079.2 | 1.2672972325729e-106 | 0.301310381 | 0.141 | 0.009 | 3.18966040466273e-102 | 5 |
| cd40lg | 6.27711224900726e-106 | 0.276658136 | 0.129 | 0.008 | 1.57988638195264e-101 | 5 |
| skap1 | 1.79367406923613e-105 | 0.376077735 | 0.189 | 0.018 | 4.51449826486042e-101 | 5 |
| ptmaa | 2.51017713767319e-105 | 0.8019073 | 0.891 | 0.378 | 6.31786483780965e-101 | 5 |
| grap2a | 6.30450597893248e-105 | 0.486978303 | 0.272 | 0.039 | 1.58678110983752e-100 | 5 |
| rsl24d1 | 6.56833970832621e-105 | 0.75317803 | 0.883 | 0.429 | 1.65318542118862e-100 | 5 |
| rpl7a | 4.80413808523116e-104 | 0.427004531 | 0.993 | 0.804 | 1.20915351467183e-99 | 5 |
| zgc:174945 | 1.7078835161983e-103 | 0.378078124 | 0.197 | 0.02 | 4.29857202191951e-99 | 5 |
| csk | 3.55259765514562e-103 | 0.600311481 | 0.425 | 0.09 | 8.94153303823602e-99 | 5 |
| hmgn2 | 1.33358993088837e-102 | 0.86658183 | 0.886 | 0.427 | 3.35651249705294e-98 | 5 |
| rpl35 | 1.02834548623873e-101 | 0.422697775 | 0.99 | 0.798 | 2.58824275431426e-97 | 5 |
| rpl8 | 3.17337182005204e-101 | 0.45551316 | 0.99 | 0.773 | 7.98705953388899e-97 | 5 |

| | | | | | | |
|---|---|---|---|---|---|---|
| rps7 | 1.06906866095521e-100 | 0.414024211 | 0.993 | 0.815 | 2.69073891275817e-96 | 5 |
| tox | 1.47715537096235e-100 | 0.349627518 | 0.17 | 0.015 | 3.71785235317513e-96 | 5 |
| rpl26 | 5.50607133861806e-100 | 0.38597935 | 0.993 | 0.836 | 1.38582309521678e-95 | 5 |
| si:dkey-185m8.2 | 7.33721435241584e-100 | 0.491924939 | 0.269 | 0.041 | 1.84670348035954e-95 | 5 |
| srgn | 1.6487357060967e-99 | 0.738691415 | 0.699 | 0.221 | 4.14970289867478e-95 | 5 |
| XLOC-005449 | 1.78601800246709e-99 | 0.537883753 | 0.289 | 0.046 | 4.49522871040941e-95 | 5 |
| tspan33a | 2.20611730305123e-99 | 0.294001314 | 0.15 | 0.012 | 5.55257664004963e-95 | 5 |
| si:ch211-132g1.4 | 2.22574174197464e-99 | 0.341859796 | 0.17 | 0.015 | 5.60196939037598e-95 | 5 |
| rpl27 | 2.32152177673027e-99 | 0.460560595 | 0.985 | 0.755 | 5.84303815985241e-95 | 5 |
| syk | 4.46091816080108e-99 | 0.572871015 | 0.359 | 0.069 | 1.12276849189202e-94 | 5 |
| cebpb | 1.40402314850712e-97 | 0.829763963 | 0.852 | 0.336 | 3.53378586247756e-93 | 5 |
| rpl18 | 5.17371995697729e-97 | 0.458554786 | 0.993 | 0.767 | 1.30217357597161e-92 | 5 |
| pik3r1 | 7.40240609211952e-97 | 0.602323481 | 0.376 | 0.08 | 1.86311158932556e-92 | 5 |
| BX901920.1 | 2.43144419222088e-95 | 0.58571187 | 0.34 | 0.065 | 6.11970188740072e-91 | 5 |
| sh2b3 | 4.98300390451406e-95 | 0.512866856 | 0.303 | 0.053 | 1.25417225272714e-90 | 5 |
| rps28 | 8.93603786970908e-95 | 0.40240274 | 0.993 | 0.821 | 2.24911137142708e-90 | 5 |
| tox2 | 1.25009430710589e-94 | 0.274175283 | 0.133 | 0.01 | 3.14636236155483e-90 | 5 |
| si:ch73-264p11.1 | 8.43604228900164e-94 | 0.393032804 | 0.192 | 0.021 | 2.12326748371882e-89 | 5 |
| rps14 | 1.09968910529411e-92 | 0.456940238 | 0.99 | 0.751 | 2.76780750911475e-88 | 5 |
| rpl23 | 2.24487058150066e-91 | 0.322427273 | 0.988 | 0.869 | 5.65011476657902e-87 | 5 |
| tagapa | 4.33111069263015e-91 | 0.659553473 | 0.41 | 0.095 | 1.09009725022808e-86 | 5 |
| rps9 | 3.66164005276777e-90 | 0.435414771 | 0.988 | 0.77 | 9.2159818488112e-86 | 5 |
| cytip | 1.36486665784609e-89 | 0.266833747 | 0.141 | 0.012 | 3.43523289113283e-85 | 5 |
| rpl10 | 5.05154268448357e-89 | 0.429281207 | 0.981 | 0.766 | 1.27142277825767e-84 | 5 |
| il12rb2l | 1.74286521070548e-88 | 0.448830582 | 0.243 | 0.037 | 4.38661744882462e-84 | 5 |
| rpl10a | 2.57826923581123e-88 | 0.43574506 | 0.978 | 0.758 | 6.4892458396133e-84 | 5 |
| rpsa | 2.98034201375652e-88 | 0.465955656 | 0.995 | 0.744 | 7.50122281442378e-84 | 5 |
| rps12 | 1.32114592917403e-87 | 0.377428803 | 0.993 | 0.835 | 3.32519218913812e-83 | 5 |
| rps19 | 2.46131726461972e-87 | 0.345908639 | 0.993 | 0.859 | 6.19488942332137e-83 | 5 |
| nitr1m | 2.58907748327945e-86 | 0.333447319 | 0.119 | 0.008 | 6.51644911766604e-82 | 5 |
| slc25a5 | 2.63908406351341e-86 | 0.389547785 | 0.985 | 0.801 | 6.6423106794569e-82 | 5 |
| si:dkey-211g8.4 | 6.0505116870794e-86 | 0.257966827 | 0.126 | 0.01 | 1.52285328652101e-81 | 5 |
| cmah | 2.58991795694857e-85 | 0.287108825 | 0.143 | 0.013 | 6.51856450584386e-81 | 5 |
| LOC103909115 | 4.33413807636359e-85 | 0.617726663 | 0.524 | 0.153 | 1.09085921243995e-80 | 5 |
| tmsb4x | 6.38613809085596e-85 | 0.638740226 | 1 | 0.676 | 1.60732709608754e-80 | 5 |

| | | | | | | |
|---|---|---|---|---|---|---|
| rpl22 | 4.40678528124578e-84 | 0.506695966 | 0.981 | 0.684 | 1.10914378743675e-79 | 5 |
| rpl38 | 1.28222694424352e-82 | 0.47814872 | 0.988 | 0.719 | 3.22723699596652e-78 | 5 |
| rplp0 | 1.835238330843e-82 | 0.347238544 | 0.983 | 0.847 | 4.61911135489875e-78 | 5 |
| si:ch211-105j21.9 | 1.89188525201186e-82 | 0.38637471 | 0.209 | 0.029 | 4.76168599078865e-78 | 5 |
| rpl28 | 1.49661507083417e-81 | 0.43803805 | 0.981 | 0.742 | 3.76683047178252e-77 | 5 |
| rpl21 | 2.08299835241828e-81 | 0.433226602 | 0.988 | 0.749 | 5.24269855320156e-77 | 5 |
| rps3 | 2.56027666061228e-81 | 0.480376466 | 0.973 | 0.691 | 6.44396032709506e-77 | 5 |
| rpl7 | 3.02269612774456e-81 | 0.386673217 | 0.978 | 0.799 | 7.60782388392028e-77 | 5 |
| rps15 | 4.33096943549045e-81 | 0.512237256 | 0.976 | 0.675 | 1.09006169721859e-76 | 5 |
| ptger4b | 5.69514838752159e-81 | 0.305507476 | 0.148 | 0.015 | 1.43341189765531e-76 | 5 |
| CR786582.2 | 9.51588670910665e-81 | 0.256178247 | 0.117 | 0.009 | 2.39505352581505e-76 | 5 |
| sesn1 | 4.04450178073346e-80 | 0.689924237 | 0.502 | 0.157 | 1.0179606531928e-75 | 5 |
| CABZ01079663.1 | 5.12023737988893e-80 | 0.410368619 | 0.221 | 0.033 | 1.28871254614425e-75 | 5 |
| zfp36l1a | 5.17316834149503e-80 | 0.695831642 | 0.646 | 0.235 | 1.30203473987088e-75 | 5 |
| faua | 5.49421261477282e-80 | 0.501832347 | 0.983 | 0.68 | 1.38283837301217e-75 | 5 |
| rpl39 | 5.98041245375407e-80 | 0.335772501 | 0.99 | 0.87 | 1.50521001048536e-75 | 5 |
| rps13 | 7.85777847711749e-80 | 0.468877863 | 0.988 | 0.716 | 1.9777242649057e-75 | 5 |
| si:ch211-284f22.3 | 1.4646859319412e-79 | 0.382339422 | 0.201 | 0.028 | 3.68646802210279e-75 | 5 |
| rps24 | 5.78883427296125e-79 | 0.428289118 | 0.981 | 0.751 | 1.45699169816162e-74 | 5 |
| naca | 1.64465769202505e-78 | 0.551542025 | 0.956 | 0.607 | 4.13943894505785e-74 | 5 |
| gzm3.3 | 4.86878999658033e-78 | 0.261021859 | 0.119 | 0.009 | 1.2254257542393e-73 | 5 |
| cotl1 | 1.32507727523182e-77 | 0.709796558 | 0.743 | 0.292 | 3.33508699403098e-73 | 5 |
| rpl9 | 3.42677247252726e-77 | 0.557012957 | 0.973 | 0.618 | 8.62484363610387e-73 | 5 |
| rpl29 | 2.74219733724411e-76 | 0.389993933 | 0.993 | 0.803 | 6.90183647810969e-72 | 5 |
| rpl35a | 4.01732268241912e-76 | 0.426131026 | 0.983 | 0.752 | 1.01111994593807e-71 | 5 |
| b2m | 8.49813283298561e-76 | 0.693239241 | 0.684 | 0.279 | 2.13889505273415e-71 | 5 |
| rps4x | 2.17609654995117e-75 | 0.46883361 | 0.971 | 0.702 | 5.47701740657209e-71 | 5 |
| rps26 | 4.92780376222729e-75 | 0.439558062 | 0.983 | 0.725 | 1.24027892891499e-70 | 5 |
| LOC100151196 | 6.20209601526443e-75 | 0.393944577 | 0.218 | 0.035 | 1.56100554608191e-70 | 5 |
| mbnl1 | 7.32914150290117e-75 | 0.622944612 | 0.529 | 0.172 | 1.8446716248652e-70 | 5 |
| rpl30 | 2.57693889964352e-74 | 0.465443732 | 0.976 | 0.699 | 6.48589751651277e-70 | 5 |
| rps18 | 1.14516300841178e-73 | 0.457155607 | 0.971 | 0.717 | 2.88226077587161e-69 | 5 |
| CR385050.1 | 1.30823904068302e-73 | 0.254358023 | 0.121 | 0.011 | 3.29270684149508e-69 | 5 |
| scinlb | 1.69560413339498e-73 | 0.478958559 | 0.333 | 0.075 | 4.26766604334183e-69 | 5 |
| si:ch211-191c10.1 | 4.00995053831476e-73 | 0.442586619 | 0.255 | 0.048 | 1.00926445098844e-68 | 5 |

| | | | | | | |
|---|---|---|---|---|---|---|
| rpl36a | 4.60407384315573e-73 | 0.372873005 | 0.993 | 0.803 | 1.15879934558387e-68 | 5 |
| hsp90ab1 | 1.27985754347093e-72 | 0.383124496 | 0.988 | 0.784 | 3.22127345116198e-68 | 5 |
| CR381673.1 | 6.62795314213042e-72 | 0.438864948 | 0.262 | 0.05 | 1.6681895263428e-67 | 5 |
| pip5k1ba | 1.57536270652462e-71 | 0.256839604 | 0.124 | 0.012 | 3.96503039605183e-67 | 5 |
| rps10 | 8.01960369387932e-71 | 0.467800648 | 0.973 | 0.685 | 2.01845405371249e-66 | 5 |
| ms4a17a.6 | 8.46558098243842e-71 | 0.405859682 | 0.238 | 0.043 | 2.13070207746993e-66 | 5 |
| mfng | 9.90420266818568e-71 | 0.387646601 | 0.218 | 0.037 | 2.49278876955565e-66 | 5 |
| gnai2b | 1.51515643497524e-70 | 0.446480647 | 0.286 | 0.06 | 3.81349723118919e-66 | 5 |
| actb2 | 8.00899524570636e-70 | 0.38237417 | 0.998 | 0.864 | 2.01578401339183e-65 | 5 |
| lcp1 | 4.64919587700419e-69 | 0.47877001 | 0.335 | 0.08 | 1.17015611028318e-64 | 5 |
| rps25 | 9.86416099075114e-69 | 0.435226971 | 0.985 | 0.716 | 2.48271067976216e-64 | 5 |
| rpl24 | 3.39835688739378e-68 | 0.401446479 | 0.985 | 0.769 | 8.5533244498814e-64 | 5 |
| tpt1 | 9.73292669793482e-68 | 0.361200754 | 0.983 | 0.795 | 2.44968032060321e-63 | 5 |
| map2k6 | 1.4791259890883e-67 | 0.491946895 | 0.328 | 0.081 | 3.72281220193634e-63 | 5 |
| si:dkeyp-68b7.12 | 2.17388110275936e-66 | 0.441490723 | 0.267 | 0.056 | 5.47144134753504e-62 | 5 |
| rasal3 | 2.87861434058732e-66 | 0.309891404 | 0.167 | 0.023 | 7.24518443382422e-62 | 5 |
| arhgef3l | 6.57893549449446e-66 | 0.318719268 | 0.18 | 0.027 | 1.65585227460931e-61 | 5 |
| ccni | 7.13653827569959e-66 | 0.636477724 | 0.653 | 0.282 | 1.79619531861083e-61 | 5 |
| il34 | 9.20547153171769e-66 | 0.413979892 | 0.233 | 0.043 | 2.31692512981802e-61 | 5 |
| si:ch1073-220m6.1 | 1.51591314517998e-65 | 0.266236202 | 0.143 | 0.017 | 3.81540179510348e-61 | 5 |
| lbh | 2.63547678750645e-65 | 0.566935597 | 0.512 | 0.161 | 6.63323152647499e-61 | 5 |
| crip1 | 1.20430873660989e-64 | 0.624055791 | 0.672 | 0.257 | 3.03112465917343e-60 | 5 |
| nsa2 | 2.8608978923519e-64 | 0.619135187 | 0.767 | 0.349 | 7.20059390526049e-60 | 5 |
| si:ch211-121a2.2 | 7.31761130023382e-64 | 0.333189596 | 0.187 | 0.03 | 1.84176958815585e-59 | 5 |
| XLOC-014079 | 8.89417679744758e-64 | 0.337194359 | 0.184 | 0.028 | 2.23857535814958e-59 | 5 |
| rps3a | 1.27332268433649e-63 | 0.373137275 | 0.976 | 0.772 | 3.20482586420652e-59 | 5 |
| si:ch211-102c2.4 | 1.34917544073599e-63 | 0.561505195 | 0.442 | 0.133 | 3.39573966678842e-59 | 5 |
| si:dkey-262k9.2 | 2.2513972321529e-63 | 0.336703034 | 0.187 | 0.03 | 5.66654169360564e-59 | 5 |
| ikzf1 | 3.53760409431956e-63 | 0.308756247 | 0.182 | 0.028 | 8.90379574499291e-59 | 5 |
| unc119b | 4.84610717384285e-63 | 0.37345616 | 0.209 | 0.036 | 1.21971671458451e-58 | 5 |
| rps23 | 1.66956085099006e-62 | 0.421395835 | 0.968 | 0.715 | 4.20211770585689e-58 | 5 |
| rpl13 | 3.32020475319927e-62 | 0.345809671 | 0.983 | 0.799 | 8.35662334332724e-58 | 5 |
| eif3f | 3.61627011029133e-62 | 0.604815366 | 0.835 | 0.418 | 9.10179024059224e-58 | 5 |
| rpl23a | 6.94110946966342e-61 | 0.49408323 | 0.947 | 0.624 | 1.74700784241959e-56 | 5 |
| arrdc3b | 1.8451804450892e-60 | 0.354694613 | 0.189 | 0.032 | 4.644134662245e-56 | 5 |

| | | | | | | |
|---|---|---|---|---|---|---|
| cd74a | 3.54361590693202e-60 | 0.47506932 | 0.607 | 0.206 | 8.9189268761572e-56 | 5 |
| gpr132b | 6.69994086349093e-60 | 0.350254034 | 0.184 | 0.031 | 1.68630811593203e-55 | 5 |
| si:ch211-230g15.5 | 1.55663345771461e-59 | 0.288869 | 0.141 | 0.018 | 3.9178907497219e-55 | 5 |
| prkacba | 5.57628419562289e-59 | 0.26380636 | 0.133 | 0.017 | 1.40349496919632e-54 | 5 |
| zbtb32 | 1.04291924438146e-58 | 0.366007597 | 0.218 | 0.041 | 2.62492344618371e-54 | 5 |
| sept9b | 2.63458482643373e-58 | 0.375722178 | 0.216 | 0.042 | 6.63098654965106e-54 | 5 |
| rps8a | 7.95691734077184e-58 | 0.36462817 | 0.985 | 0.781 | 2.00267652549886e-53 | 5 |
| dok2 | 2.32697234124469e-57 | 0.341459605 | 0.189 | 0.033 | 5.85675668567877e-53 | 5 |
| phf11 | 2.84320408394711e-57 | 0.265579431 | 0.141 | 0.019 | 7.15606035888649e-53 | 5 |
| RASSF5 | 3.23748332572844e-57 | 0.382868797 | 0.243 | 0.052 | 8.1484217825259e-53 | 5 |
| prf1.9 | 6.38707056363155e-57 | 0.264227468 | 0.126 | 0.015 | 1.60756179016042e-52 | 5 |
| lgals17 | 1.55639170113638e-56 | 0.409746412 | 0.226 | 0.047 | 3.91728227259016e-52 | 5 |
| fli1b | 1.70489816407765e-56 | 0.316786866 | 0.177 | 0.03 | 4.29105818916705e-52 | 5 |
| ptpn4b | 1.99666624367578e-56 | 0.281673299 | 0.155 | 0.023 | 5.02540926870758e-52 | 5 |
| nme2b.1 | 2.14251152875014e-56 | 0.28694095 | 0.995 | 0.853 | 5.39248726671123e-52 | 5 |
| msna | 2.30820975416491e-56 | 0.58572528 | 0.563 | 0.231 | 5.80953313025767e-52 | 5 |
| XLOC-041870 | 1.12244517573865e-55 | 0.34396147 | 0.158 | 0.024 | 2.82508226281662e-51 | 5 |
| wipf1a | 1.37327318589883e-55 | 0.405677907 | 0.257 | 0.059 | 3.45639128158876e-51 | 5 |
| rpl27a | 1.42788395945432e-55 | 0.570503072 | 0.891 | 0.482 | 3.59384113755059e-51 | 5 |
| itga4 | 1.57653125675635e-55 | 0.413931958 | 0.255 | 0.059 | 3.96797152013005e-51 | 5 |
| gfi1ab | 1.60058438620769e-55 | 0.269401528 | 0.141 | 0.019 | 4.02851084164614e-51 | 5 |
| rps11 | 2.1821696111639e-55 | 0.357961103 | 0.964 | 0.759 | 5.49230269433843e-51 | 5 |
| myl12.1 | 3.61255443417088e-55 | 0.580922755 | 0.767 | 0.379 | 9.09243825536469e-51 | 5 |
| nkl.2 | 3.63541412276459e-55 | 0.525802456 | 0.451 | 0.15 | 9.14997380558621e-51 | 5 |
| slc44a2 | 6.98980506496324e-55 | 0.363788088 | 0.218 | 0.045 | 1.7592640368006e-50 | 5 |
| CR318588.1 | 1.3450128244986e-54 | 0.847760193 | 0.49 | 0.199 | 3.38526277798052e-50 | 5 |
| fam49a | 2.26301573392384e-54 | 0.362897941 | 0.221 | 0.046 | 5.69578430071291e-50 | 5 |
| ccl33.3 | 2.64123189810981e-54 | 0.363473792 | 0.233 | 0.049 | 6.64771656435257e-50 | 5 |
| glipr1a | 2.94685803812406e-54 | 0.398827346 | 0.25 | 0.058 | 7.41694699615445e-50 | 5 |
| XLOC-043043 | 4.54381992111191e-54 | 0.330425118 | 0.201 | 0.038 | 1.14363403594466e-49 | 5 |
| p2ry10 | 4.6590779398357e-54 | 0.255646506 | 0.131 | 0.018 | 1.17264332667725e-49 | 5 |
| FERMT3 (1 of many) | 1.39897757066016e-53 | 0.338679761 | 0.192 | 0.037 | 3.52108664759455e-49 | 5 |
| tmc8 | 9.1084344125143e-53 | 0.302808983 | 0.175 | 0.031 | 2.29250185728572e-48 | 5 |
| zgc:152791 | 1.79329653241602e-52 | 0.430488832 | 0.26 | 0.062 | 4.51354804243787e-48 | 5 |
| arpc3 | 1.90924314598679e-52 | 0.565201953 | 0.658 | 0.299 | 4.80537407413415e-48 | 5 |

| | | | | | | |
|---|---|---|---|---|---|---|
| si:ch211-132p1.3 | 6.12068596208527e-52 | 0.447754913 | 0.296 | 0.082 | 1.54051544979724e-47 | 5 |
| tefa | 2.1498078564857e-51 | 0.493870074 | 0.388 | 0.13 | 5.41085139398887e-47 | 5 |
| rpl3 | 2.24341355102476e-51 | 0.264940471 | 0.99 | 0.872 | 5.64644756657422e-47 | 5 |
| arhgap27 | 2.25117492604006e-51 | 0.259221586 | 0.136 | 0.02 | 5.66598217135023e-47 | 5 |
| il21r.1 | 3.55276752491173e-50 | 0.258059286 | 0.136 | 0.02 | 8.94196058345033e-46 | 5 |
| ifi45 | 4.31224008226002e-50 | 0.507116587 | 0.379 | 0.121 | 1.08534770630403e-45 | 5 |
| rpl31 | 4.71561752296977e-50 | 0.440354997 | 0.939 | 0.645 | 1.18687377435626e-45 | 5 |
| ccr9a | 4.84996578536161e-50 | 0.42946971 | 0.299 | 0.081 | 1.22068788851766e-45 | 5 |
| rpl5a | 4.43879439445308e-49 | 0.537263988 | 0.867 | 0.479 | 1.11720016113989e-44 | 5 |
| uba52 | 1.23761127651899e-48 | 0.451845971 | 0.927 | 0.62 | 3.11494382187066e-44 | 5 |
| rpl6 | 1.34356034478854e-48 | 0.437553033 | 0.954 | 0.658 | 3.38160703179828e-44 | 5 |
| zgc:114188 | 1.40778482007264e-48 | 0.494540628 | 0.966 | 0.616 | 3.54325361364082e-44 | 5 |
| CR855389.1 | 1.51606571466925e-48 | 0.27848012 | 0.15 | 0.025 | 3.81578579725104e-44 | 5 |
| si:ch211-196l7.4 | 1.74830356954827e-48 | 0.543210106 | 0.646 | 0.303 | 4.40030525419603e-44 | 5 |
| fcer1g | 2.08219758280307e-48 | 0.332845192 | 0.189 | 0.038 | 5.24068309615704e-44 | 5 |
| lpar5a | 4.8830460812788e-48 | 0.336428516 | 0.184 | 0.037 | 1.22901386819706e-43 | 5 |
| ppiaa | 5.05156959829348e-48 | 0.320129753 | 0.976 | 0.798 | 1.27142955219449e-43 | 5 |
| histh1l | 8.00879434359676e-48 | 0.473857956 | 0.333 | 0.103 | 2.01573344833987e-43 | 5 |
| arpc4 | 5.91303927978549e-47 | 0.505973522 | 0.532 | 0.216 | 1.48825285632921e-42 | 5 |
| sik1 | 6.55698220068877e-47 | 0.478969383 | 0.34 | 0.109 | 1.65032685009136e-42 | 5 |
| rack1 | 1.21497482026982e-46 | 0.351016641 | 0.951 | 0.738 | 3.0579701251371e-42 | 5 |
| rpl13a | 2.49080016157686e-46 | 0.334441048 | 0.961 | 0.758 | 6.2690949266728e-42 | 5 |
| bcl2l10 | 3.45755133043361e-46 | 0.515347332 | 0.522 | 0.205 | 8.70231094356837e-42 | 5 |
| sat1a.2 | 4.7776299964515e-46 | 0.529520316 | 0.624 | 0.288 | 1.20248169380688e-41 | 5 |
| pycard | 8.31003558956613e-46 | 0.388982659 | 0.252 | 0.067 | 2.0915528575379e-41 | 5 |
| mcl1a | 1.17465315926785e-45 | 0.456064149 | 0.908 | 0.615 | 2.95648453656126e-41 | 5 |
| hmgn3 | 1.62339900771926e-45 | 0.385090811 | 0.226 | 0.056 | 4.08593296252861e-41 | 5 |
| ikzf2 | 1.73371554284844e-45 | 0.261280178 | 0.15 | 0.026 | 4.36358864979524e-41 | 5 |
| FQ323156.1 | 2.48017411236713e-45 | 0.454872352 | 0.456 | 0.161 | 6.24235022341683e-41 | 5 |
| notch2 | 2.54993869090223e-45 | 0.453405809 | 0.337 | 0.113 | 6.41794069113182e-41 | 5 |
| eef1g | 1.82515499483012e-44 | 0.449843244 | 0.915 | 0.611 | 4.59373260648793e-40 | 5 |
| vaspa | 2.39216363201388e-44 | 0.379743979 | 0.26 | 0.07 | 6.02083664541572e-40 | 5 |
| btf3 | 2.58893413300398e-43 | 0.461936631 | 0.908 | 0.577 | 6.51608831935772e-39 | 5 |
| lasp1 | 5.3947655242247e-43 | 0.484066261 | 0.527 | 0.227 | 1.35780853479211e-38 | 5 |
| si:ch73-234b20.5 | 5.73254901558661e-43 | 0.393738016 | 0.303 | 0.094 | 1.44282526173299e-38 | 5 |

| | | | | | | |
|---|---|---|---|---|---|---|
| vsir | 7.90378108649155e-43 | 0.385524119 | 0.274 | 0.08 | 1.98930266165906e-38 | 5 |
| tnip1 | 1.76013302890817e-42 | 0.396199867 | 0.265 | 0.076 | 4.43007882045898e-38 | 5 |
| hmgn6 | 1.97194575226407e-42 | 0.500796053 | 0.561 | 0.256 | 4.96319026387344e-38 | 5 |
| si:ch211-241f5.3 | 6.25949521961097e-42 | 0.385445055 | 0.235 | 0.063 | 1.57545235182388e-37 | 5 |
| tnrc6c1 | 1.2628284326671e-41 | 0.452234557 | 0.35 | 0.126 | 3.17841288217982e-37 | 5 |
| pnrc2 | 1.49106209849788e-41 | 0.491582306 | 0.932 | 0.555 | 3.75285419570933e-37 | 5 |
| zgc:162944 | 1.73595145279616e-41 | 0.504252298 | 0.532 | 0.244 | 4.36921621154266e-37 | 5 |
| cd74b | 2.42814960937244e-41 | 0.408297729 | 0.558 | 0.213 | 6.1114097518295e-37 | 5 |
| si:rp71-68n21.9 | 3.69346353934917e-41 | 0.297989071 | 0.18 | 0.04 | 9.29607838218794e-37 | 5 |
| nfkbie | 5.17322806217012e-41 | 0.394258323 | 0.286 | 0.088 | 1.3020497709676e-36 | 5 |
| eef1da | 5.59256065268393e-41 | 0.483933441 | 0.823 | 0.481 | 1.40759159067402e-36 | 5 |
| rpl12 | 1.89486024579403e-40 | 0.303753855 | 0.973 | 0.793 | 4.76917375263899e-36 | 5 |
| celf2 | 3.50490783290493e-40 | 0.473255475 | 0.529 | 0.233 | 8.82150252463841e-36 | 5 |
| elf1 | 4.21324032573412e-40 | 0.389658286 | 0.303 | 0.098 | 1.06043045758402e-35 | 5 |
| rps27a | 1.4167796647569e-39 | 0.346201166 | 0.966 | 0.745 | 3.56589273822664e-35 | 5 |
| sept2 | 4.41656542905316e-39 | 0.439341249 | 0.403 | 0.158 | 1.11160535283839e-34 | 5 |
| zgc:152863 | 6.62542423911323e-39 | 0.301692137 | 0.177 | 0.04 | 1.66755302674241e-34 | 5 |
| spry2 | 7.31793304086037e-39 | 0.331641508 | 0.218 | 0.059 | 1.84185056705415e-34 | 5 |
| serbp1a | 1.06477695626299e-38 | 0.47938259 | 0.811 | 0.471 | 2.67993712121831e-34 | 5 |
| nptna | 2.31573797643416e-38 | 0.346237533 | 0.24 | 0.066 | 5.82848091288714e-34 | 5 |
| abi3a | 2.39756365340032e-38 | 0.274007341 | 0.175 | 0.04 | 6.03442795924327e-34 | 5 |
| ube2e3 | 2.65079568819696e-38 | 0.43249601 | 0.381 | 0.149 | 6.67178766762294e-34 | 5 |
| rps27.2 | 3.20139416938369e-38 | 0.489056537 | 0.888 | 0.505 | 8.05758898492181e-34 | 5 |
| si:dkey-165a24.10 | 1.5776410418031e-37 | 0.257481085 | 0.146 | 0.03 | 3.97076473811423e-33 | 5 |
| arl11 | 2.75111360715511e-37 | 0.272926911 | 0.136 | 0.026 | 6.9242778378487e-33 | 5 |
| skia | 5.20808894429818e-37 | 0.306503105 | 0.17 | 0.04 | 1.31082390639041e-32 | 5 |
| ptmab | 7.59145547865453e-37 | 0.471938047 | 0.852 | 0.53 | 1.91069342942256e-32 | 5 |
| ccr2 | 9.25476273407471e-37 | 0.296643904 | 0.18 | 0.043 | 2.32933123253926e-32 | 5 |
| cfl1 | 9.82836382620666e-37 | 0.451319195 | 0.883 | 0.567 | 2.47370089141796e-32 | 5 |
| mknk2b | 2.36091031520133e-36 | 0.450810143 | 0.444 | 0.19 | 5.94217517233023e-32 | 5 |
| cox6b1 | 2.41436161701512e-36 | 0.380244155 | 0.323 | 0.114 | 6.07670675386536e-32 | 5 |
| si:ch73-248e21.7 | 8.25746702346399e-36 | 0.26570574 | 0.172 | 0.041 | 2.07832187513565e-31 | 5 |
| cish | 8.40386555438058e-36 | 0.311230187 | 0.194 | 0.051 | 2.11516892138205e-31 | 5 |
| clic1 | 2.20027832251449e-35 | 0.372791255 | 0.306 | 0.108 | 5.53788050993672e-31 | 5 |
| arpc2 | 2.39648059032773e-35 | 0.452487009 | 0.544 | 0.259 | 6.03170199779587e-31 | 5 |

| | | | | | | |
|---|---|---|---|---|---|---|
| mob1a | 1.78746030637009e-34 | 0.405662101 | 0.391 | 0.159 | 4.49885884510288e-30 | 5 |
| il16 | 3.10223186350289e-34 | 0.288520295 | 0.18 | 0.046 | 7.80800737725042e-30 | 5 |
| rpl17 | 3.31580665361193e-34 | 0.331419664 | 0.961 | 0.739 | 8.34555376647586e-30 | 5 |
| si:dkey-112e7.2 | 4.72392954529654e-34 | 0.433811794 | 0.444 | 0.199 | 1.18896582725569e-29 | 5 |
| litaf | 9.1656935330484e-34 | 0.408378294 | 0.333 | 0.129 | 2.30691340533295e-29 | 5 |
| pgam1a | 2.01317744909745e-33 | 0.409550691 | 0.374 | 0.151 | 5.06696632163338e-29 | 5 |
| BX248082.1 | 2.46745287116304e-33 | 0.325355648 | 0.163 | 0.039 | 6.21033213143026e-29 | 5 |
| acap1 | 4.22417628348253e-33 | 0.251773184 | 0.158 | 0.037 | 1.06318292878972e-28 | 5 |
| ercc1 | 2.25871940810053e-32 | 0.253911861 | 0.197 | 0.053 | 5.68497087824823e-28 | 5 |
| fscn1a | 5.44254368999125e-32 | 0.296407083 | 0.194 | 0.054 | 1.3698338213339e-27 | 5 |
| rbpjb | 6.81802131850757e-32 | 0.314203348 | 0.192 | 0.054 | 1.71602778565517e-27 | 5 |
| zgc:162730 | 2.96714819345775e-31 | 0.466631286 | 0.743 | 0.38 | 7.46801528811381e-27 | 5 |
| socs1a | 3.26010757414938e-31 | 0.277383847 | 0.172 | 0.045 | 8.20536475337658e-27 | 5 |
| tmem131l | 1.73757951020271e-30 | 0.283243849 | 0.189 | 0.054 | 4.3733138692292e-26 | 5 |
| rps6ka1 | 2.6476116826017e-30 | 0.294776987 | 0.223 | 0.069 | 6.66377384394023e-26 | 5 |
| wipf1b | 3.88636745764106e-30 | 0.290660052 | 0.199 | 0.059 | 9.78159825413679e-26 | 5 |
| cap1 | 9.34025345757637e-30 | 0.430506505 | 0.473 | 0.226 | 2.3508483927374e-25 | 5 |
| cd63 | 1.2491374713418e-29 | 0.420370639 | 0.643 | 0.331 | 3.14395410162017e-25 | 5 |
| gapdhs | 1.29835778481312e-29 | 0.416168007 | 0.556 | 0.28 | 3.26783670859613e-25 | 5 |
| sumo2b | 5.87891666245369e-29 | 0.405327499 | 0.515 | 0.262 | 1.47966453477297e-24 | 5 |
| si:dkey-187e18.1 | 1.35194163910051e-28 | 0.30231934 | 0.206 | 0.065 | 3.40270191145206e-24 | 5 |
| ponzr1 | 1.8335154558797e-28 | 0.424098086 | 0.532 | 0.265 | 4.61477505090361e-24 | 5 |
| isg15 | 2.0700362981925e-28 | 0.298903347 | 0.209 | 0.064 | 5.2100743589207e-24 | 5 |
| ppp1r18 | 2.54870358537708e-28 | 0.264766735 | 0.177 | 0.05 | 6.41483205403558e-24 | 5 |
| rnf213a | 2.85868599183313e-28 | 0.334006057 | 0.24 | 0.085 | 7.19502677284482e-24 | 5 |
| fam49ba | 4.31616598590568e-28 | 0.313226633 | 0.274 | 0.099 | 1.0863358169926e-23 | 5 |
| CR383676.1 | 6.30865665690296e-28 | 0.569754607 | 0.983 | 0.718 | 1.58782579397591e-23 | 5 |
| rnf44 | 7.45796937549285e-28 | 0.330842979 | 0.24 | 0.085 | 1.8770963121178e-23 | 5 |
| rpl4 | 7.47627106072523e-28 | 0.367665224 | 0.893 | 0.619 | 1.88170266327393e-23 | 5 |
| arpc5a | 5.82041897876261e-27 | 0.328973559 | 0.284 | 0.11 | 1.46494125276476e-22 | 5 |
| GCA | 8.87067710063673e-27 | 0.353775458 | 0.386 | 0.175 | 2.23266071945926e-22 | 5 |
| hmha1b | 1.28084721258873e-26 | 0.267024469 | 0.216 | 0.071 | 3.22376434936456e-22 | 5 |
| stat2 | 1.68617168383729e-26 | 0.273272037 | 0.187 | 0.058 | 4.24392551105006e-22 | 5 |
| psme2 | 2.16735040373855e-26 | 0.324288333 | 0.248 | 0.093 | 5.45500423116955e-22 | 5 |
| pabpc1a | 5.71753710209078e-26 | 0.400335461 | 0.85 | 0.53 | 1.43904691322523e-21 | 5 |

| | | | | | | |
|---|---|---|---|---|---|---|
| cdkn1bb | 5.78155263221346e-26 | 0.347572291 | 0.333 | 0.143 | 1.45515898200181e-21 | 5 |
| romo1 | 7.69274320198784e-26 | 0.400129691 | 0.658 | 0.378 | 1.93618653650832e-21 | 5 |
| pam16 | 9.26961716859227e-26 | 0.298717274 | 0.265 | 0.099 | 2.33306994516299e-21 | 5 |
| eif3m | 1.83342817446901e-25 | 0.396604722 | 0.617 | 0.35 | 4.61455537232105e-21 | 5 |
| psme1 | 5.18529769181083e-25 | 0.292441067 | 0.233 | 0.086 | 1.30508757605187e-20 | 5 |
| ier2b | 8.2801970098069e-25 | 0.392716542 | 0.434 | 0.218 | 2.0840427853983e-20 | 5 |
| nr4a1 | 1.14690450185502e-24 | 0.373960562 | 0.362 | 0.162 | 2.88664394071889e-20 | 5 |
| dnmt3ab | 1.94375881653458e-24 | 0.305458314 | 0.211 | 0.076 | 4.89224656533589e-20 | 5 |
| tnfrsf9a | 2.55790380031723e-24 | 0.299057379 | 0.165 | 0.049 | 6.43798807501845e-20 | 5 |
| nrros | 2.92880778019958e-24 | 0.250439456 | 0.155 | 0.045 | 7.37151630198433e-20 | 5 |
| irf2bp2a | 4.30862277007845e-24 | 0.346399085 | 0.369 | 0.171 | 1.08443726500104e-19 | 5 |
| eif3ba | 4.69551057639799e-24 | 0.351001585 | 0.415 | 0.204 | 1.18181305697361e-19 | 5 |
| hmgb1a | 7.5136407697903e-24 | 0.362375874 | 0.379 | 0.184 | 1.89110824534852e-19 | 5 |
| txnipa | 1.11522457837455e-23 | 0.376484907 | 0.415 | 0.205 | 2.80690874131091e-19 | 5 |
| actr2a | 9.38363784086012e-23 | 0.302827387 | 0.296 | 0.125 | 2.36176780816608e-18 | 5 |
| tpi1b | 1.03794779646702e-22 | 0.32351135 | 0.35 | 0.162 | 2.61241080892785e-18 | 5 |
| si:dkey-262k9.4 | 1.1067882009494e-22 | 0.298335381 | 0.228 | 0.087 | 2.78567522296954e-18 | 5 |
| rpl14 | 1.33251180221327e-22 | 0.342818241 | 0.896 | 0.627 | 3.35379895499058e-18 | 5 |
| elmo1 | 1.85598208465271e-22 | 0.268016494 | 0.194 | 0.068 | 4.67132130886241e-18 | 5 |
| fabp3 | 2.59211631169189e-22 | 0.37041126 | 0.451 | 0.236 | 6.52409754489731e-18 | 5 |
| dtnbp1a | 2.71804830631687e-22 | 0.27539405 | 0.223 | 0.083 | 6.84105578216893e-18 | 5 |
| plk3 | 3.71589843229565e-22 | 0.292627099 | 0.286 | 0.116 | 9.35254476424493e-18 | 5 |
| cox7b | 5.728968047168e-22 | 0.369954243 | 0.689 | 0.408 | 1.44192396779171e-17 | 5 |
| ran | 6.87750027736237e-22 | 0.326244422 | 0.891 | 0.636 | 1.73099804480933e-17 | 5 |
| arhgdia | 9.2146013185723e-22 | 0.357309752 | 0.427 | 0.224 | 2.31922300587146e-17 | 5 |
| ucp2 | 3.85586919553898e-21 | 0.367613871 | 0.709 | 0.407 | 9.70483717825205e-17 | 5 |
| mt-co1 | 3.87708860043052e-21 | 0.373583703 | 0.973 | 0.647 | 9.75824429842357e-17 | 5 |
| serf2 | 7.52827793580876e-21 | 0.354734448 | 0.575 | 0.336 | 1.89479227366371e-16 | 5 |
| rpl22l1 | 1.14068082629821e-20 | 0.370270182 | 0.653 | 0.384 | 2.87097957170996e-16 | 5 |
| si:ch211-222l21.1 | 1.4405536142055e-20 | 0.367844412 | 0.498 | 0.269 | 3.62572939159382e-16 | 5 |
| eif3ea | 3.39818010286516e-20 | 0.326913582 | 0.493 | 0.271 | 8.55287950090133e-16 | 5 |
| mt-co2 | 4.28321130444169e-20 | 0.364428305 | 0.976 | 0.712 | 1.07804145321493e-15 | 5 |
| gtpbp4 | 5.10809148572697e-20 | 0.347225084 | 0.425 | 0.232 | 1.28565554604262e-15 | 5 |
| eno1a | 5.3208648613219e-20 | 0.285044953 | 0.291 | 0.13 | 1.33920847694611e-15 | 5 |
| mibp21 | 5.77923932459503e-20 | 0.366802792 | 0.774 | 0.483 | 1.45457674560732e-15 | 5 |

| | | | | | | |
|---|---|---|---|---|---|---|
| slc25a3b | 7.61991375647563e-20 | 0.351048182 | 0.704 | 0.429 | 1.91785609336735e-15 | 5 |
| rpl5b | 2.67564282003414e-19 | 0.322544359 | 0.874 | 0.622 | 6.73432541374393e-15 | 5 |
| gng5 | 2.90384855676134e-19 | 0.338103344 | 0.471 | 0.266 | 7.30869643251262e-15 | 5 |
| nop53 | 8.39714790570328e-19 | 0.340155506 | 0.437 | 0.246 | 2.11347815638646e-14 | 5 |
| cdc42l | 1.60434788433346e-18 | 0.305136135 | 0.4 | 0.211 | 4.03798319007888e-14 | 5 |
| sh3bgrl3 | 2.90865891647918e-18 | 0.276725079 | 0.303 | 0.143 | 7.32080362688645e-14 | 5 |
| arpc4l | 4.82086895074442e-18 | 0.295311505 | 0.371 | 0.192 | 1.21336450621286e-13 | 5 |
| zgc:56493 | 4.83003756978791e-18 | 0.325644105 | 0.82 | 0.565 | 1.21567215593992e-13 | 5 |
| ddx21 | 1.51714552971592e-17 | 0.318277556 | 0.376 | 0.203 | 3.81850358374201e-13 | 5 |
| gnai2a | 2.59487671927338e-17 | 0.273978669 | 0.291 | 0.14 | 6.53104521473917e-13 | 5 |
| capzb | 6.34865389816837e-17 | 0.292753801 | 0.444 | 0.248 | 1.59789269963e-12 | 5 |
| b2ml | 8.32362496837769e-17 | 0.312918044 | 0.459 | 0.262 | 2.09497316829098e-12 | 5 |
| COX3 | 1.15932879935352e-16 | 0.340233861 | 0.983 | 0.691 | 2.91791465509288e-12 | 5 |
| rhoab | 1.83523239949659e-16 | 0.294755057 | 0.347 | 0.185 | 4.61909642629298e-12 | 5 |
| arl5c | 7.79106212181066e-16 | 0.257973727 | 0.24 | 0.109 | 1.96093242543853e-11 | 5 |
| tomm20b | 1.33107991242063e-15 | 0.265507573 | 0.383 | 0.21 | 3.35019503157149e-11 | 5 |
| oaz1b | 1.78600651127016e-15 | 0.288063638 | 0.536 | 0.317 | 4.49519978821587e-11 | 5 |
| zgc:153713 | 1.92677473973038e-15 | 0.256396462 | 0.294 | 0.149 | 4.8494993424274e-11 | 5 |
| cox4i1 | 2.01150941295271e-15 | 0.305568507 | 0.624 | 0.385 | 5.06276804146067e-11 | 5 |
| si:dkey-188i13.10 | 4.00070322278102e-15 | 0.253191132 | 0.221 | 0.101 | 1.00693699414176e-10 | 5 |
| stat1a | 4.55316275909157e-15 | 0.253813279 | 0.245 | 0.117 | 1.14598553483576e-10 | 5 |
| cct2 | 4.77715134685388e-15 | 0.275544618 | 0.393 | 0.223 | 1.20236122248965e-10 | 5 |
| eif3g | 5.3726434079555e-15 | 0.291107583 | 0.464 | 0.277 | 1.35224061934832e-10 | 5 |
| eif4ebp2 | 1.13802837587964e-14 | 0.255880927 | 0.323 | 0.171 | 2.86430361925146e-10 | 5 |
| ywhabl | 1.21453903945614e-14 | 0.274277417 | 0.466 | 0.273 | 3.05687330840716e-10 | 5 |
| capza1a | 1.58885150754115e-14 | 0.265075017 | 0.354 | 0.196 | 3.99898035933033e-10 | 5 |
| glrx | 1.63595260807333e-14 | 0.28592385 | 0.51 | 0.306 | 4.11752911925975e-10 | 5 |
| crema | 1.74233671291775e-14 | 0.285831832 | 0.325 | 0.175 | 4.3852872727427e-10 | 5 |
| eif3s10 | 4.94963257310221e-14 | 0.25696479 | 0.301 | 0.162 | 1.24577302232409e-09 | 5 |
| si:ch73-1a9.3 | 7.89729470206933e-14 | 0.301762367 | 0.434 | 0.261 | 1.98767010356383e-09 | 5 |
| cox8a | 1.62970097282125e-13 | 0.263326019 | 0.383 | 0.224 | 4.1017943784938e-09 | 5 |
| eif3c | 1.63686579900265e-13 | 0.251634459 | 0.303 | 0.166 | 4.11982752950977e-09 | 5 |
| pde4ba | 1.94332176786867e-13 | 0.269100148 | 0.265 | 0.137 | 4.89114655754866e-09 | 5 |
| mt-atp6 | 2.79752116855549e-13 | 0.300905949 | 0.891 | 0.581 | 7.04108102913732e-09 | 5 |
| eif5a | 6.1892246384414e-13 | 0.292142907 | 0.714 | 0.472 | 1.55776594924932e-08 | 5 |

| | | | | | | |
|---|---|---|---|---|---|---|
| eif3i | 6.93484643121691e-13 | 0.257409239 | 0.444 | 0.269 | 1.74543149827298e-08 | 5 |
| atp5l | 3.4520606280056e-11 | 0.258476328 | 0.658 | 0.443 | 8.68849139462729e-07 | 5 |
| h2afx11 | 8.70184601318842e-11 | 0.259885197 | 0.325 | 0.196 | 2.19016762305939e-06 | 5 |
| NC-002333.4 | 7.84327636797595e-10 | 0.271309208 | 0.903 | 0.614 | 1.97407422905587e-05 | 5 |
| grn1 | 0 | 1.750155959 | 0.812 | 0.062 | 0 | 6 |
| cd74a1 | 0 | 1.740126307 | 0.93 | 0.195 | 0 | 6 |
| lgals2a | 0 | 1.580141238 | 0.817 | 0.102 | 0 | 6 |
| c1qb | 0 | 1.554587967 | 0.721 | 0.048 | 0 | 6 |
| c1qc | 0 | 1.498864582 | 0.668 | 0.046 | 0 | 6 |
| si:dkey-5n18.1 | 0 | 1.386898394 | 0.595 | 0.04 | 0 | 6 |
| lgals3bpb | 0 | 1.208849598 | 0.428 | 0.028 | 0 | 6 |
| ccl34b.1 | 0 | 1.054912724 | 0.363 | 0.019 | 0 | 6 |
| si:busm1-266f07.2 | 2.28725858037108e-305 | 1.482221585 | 0.775 | 0.125 | 5.75680112093597e-301 | 6 |
| lygl1 | 2.82133211090658e-301 | 1.313223057 | 0.632 | 0.079 | 7.10101078994078e-297 | 6 |
| cd74b1 | 9.44072431802048e-274 | 1.482743189 | 0.88 | 0.202 | 2.37613590360257e-269 | 6 |
| zgc:153118 | 2.49480142004422e-260 | 1.053492826 | 0.368 | 0.026 | 6.27916569410931e-256 | 6 |
| fcer1gl | 9.47419745859615e-242 | 1.284131069 | 0.77 | 0.149 | 2.38456075835406e-237 | 6 |
| NPC2 (1 of many) | 8.21450981438158e-227 | 0.892592927 | 0.332 | 0.025 | 2.0675099751817e-222 | 6 |
| c1qa | 1.01640643441568e-226 | 0.970690098 | 0.407 | 0.04 | 2.55819335478082e-222 | 6 |
| CU499330.1 | 4.20219955550695e-179 | 0.99812395 | 0.439 | 0.059 | 1.05765160612554e-174 | 6 |
| marco | 1.69273027403692e-170 | 0.756732161 | 0.295 | 0.027 | 4.26043282672351e-166 | 6 |
| si:ch211-147m6.1 | 1.24997000303968e-162 | 0.728924677 | 0.277 | 0.025 | 3.14604950065057e-158 | 6 |
| tmsb4x1 | 6.82370059176061e-162 | 0.810883877 | 0.997 | 0.677 | 1.71745720194023e-157 | 6 |
| LOC100537504 | 8.54145006751607e-156 | 0.692897806 | 0.243 | 0.019 | 2.14979756749312e-151 | 6 |
| mhc2dab | 1.03750530546852e-145 | 1.016655213 | 0.467 | 0.085 | 2.61129710333372e-141 | 6 |
| actb21 | 2.27448154140008e-143 | 0.585360293 | 0.997 | 0.864 | 5.72464259154987e-139 | 6 |
| ms4a17a.10 | 2.10413328924684e-127 | 0.771228577 | 0.319 | 0.044 | 5.29589307570538e-123 | 6 |
| pfn11 | 3.99859282664721e-127 | 0.93234886 | 0.979 | 0.51 | 1.00640582853884e-122 | 6 |
| si:dkey-7i4.24 | 3.20532200935735e-124 | 0.678491116 | 0.201 | 0.016 | 8.06747496535152e-120 | 6 |
| CU929150.1 | 1.47835415573849e-118 | 0.610553399 | 0.209 | 0.019 | 3.72086957457822e-114 | 6 |
| si:ch211-283g2.2 | 2.53328664766934e-117 | 0.544807462 | 0.201 | 0.018 | 6.37602916351897e-113 | 6 |
| slc3a2a | 4.24587201420434e-116 | 0.64700125 | 0.243 | 0.028 | 1.06864352725509e-111 | 6 |
| grna | 1.47268199472649e-114 | 0.730079918 | 0.313 | 0.048 | 3.70659331252711e-110 | 6 |
| slc40a1 | 6.62144025851819e-99 | 0.615945723 | 0.235 | 0.03 | 1.66655029866644e-94 | 6 |
| zgc:103700 | 8.70523985429011e-99 | 0.652545646 | 0.248 | 0.034 | 2.19102181892628e-94 | 6 |

| | | | | | | |
|---|---|---|---|---|---|---|
| tmsb1 | 1.18224024482765e-91 | 0.831000714 | 0.517 | 0.145 | 2.97558047220672e-87 | 6 |
| havcr1 | 1.42273999385576e-90 | 0.572963208 | 0.217 | 0.028 | 3.58089429053557e-86 | 6 |
| psap | 2.07204117799284e-87 | 0.879920477 | 0.653 | 0.279 | 5.21512044089019e-83 | 6 |
| krt18b | 3.20903288602122e-87 | 0.864396723 | 0.475 | 0.141 | 8.0768148708268e-83 | 6 |
| sftpbb | 6.63539821565818e-82 | 0.695533663 | 0.303 | 0.059 | 1.67006337689901e-77 | 6 |
| ccl35.1 | 2.62919994863409e-79 | 0.825117756 | 0.342 | 0.075 | 6.61743335071714e-75 | 6 |
| grn2 | 1.8683640249385e-73 | 0.497638409 | 0.154 | 0.017 | 4.70248541436771e-69 | 6 |
| si:ch211-194m7.3 | 2.00717016308828e-73 | 0.466542303 | 0.157 | 0.018 | 5.05184658347688e-69 | 6 |
| zgc:174904 | 5.89549138076646e-70 | 0.617531712 | 0.238 | 0.043 | 1.48383622562511e-65 | 6 |
| cotl11 | 2.10752694756066e-67 | 0.756669918 | 0.642 | 0.297 | 5.30443457431543e-63 | 6 |
| si:ch211-165i18.2 | 1.14614542927205e-66 | 0.522271332 | 0.198 | 0.031 | 2.88473343093482e-62 | 6 |
| BX649485.2 | 4.15567591255986e-66 | 0.540086311 | 0.185 | 0.027 | 1.04594207043219e-61 | 6 |
| rnaset2l | 4.7653858526676e-66 | 0.533996027 | 0.217 | 0.038 | 1.19939996525791e-61 | 6 |
| ccl34b.8 | 3.78294233155957e-63 | 0.391256993 | 0.136 | 0.015 | 9.52128755430228e-59 | 6 |
| LOC100536946 | 8.20724540359258e-63 | 0.40516081 | 0.131 | 0.014 | 2.06568159563022e-58 | 6 |
| ifi30 | 5.7061384528097e-62 | 0.780091277 | 0.454 | 0.167 | 1.43617798718767e-57 | 6 |
| CR318588.11 | 2.72406508929847e-61 | 0.950210818 | 0.512 | 0.199 | 6.85619942325532e-57 | 6 |
| rbp2a | 3.51348526355385e-59 | 0.3451032 | 0.11 | 0.011 | 8.84309105983868e-55 | 6 |
| rplp21 | 2.00634270826384e-56 | 0.411592063 | 0.958 | 0.752 | 5.04976396242927e-52 | 6 |
| txn | 1.29232991797372e-54 | 0.543504802 | 0.206 | 0.04 | 3.25266517054806e-50 | 6 |
| rps201 | 2.21216214205357e-54 | 0.254761784 | 0.982 | 0.915 | 5.56779089533463e-50 | 6 |
| cmklr1 | 2.42533024683962e-54 | 0.453617433 | 0.167 | 0.027 | 6.10431369827064e-50 | 6 |
| rps15a1 | 1.30626121083333e-52 | 0.388898094 | 0.958 | 0.775 | 3.2877288415464e-48 | 6 |
| rpl391 | 5.33201009188634e-52 | 0.28463403 | 0.966 | 0.871 | 1.34201362002687e-47 | 6 |
| cfp | 6.07877330332493e-51 | 0.556489918 | 0.277 | 0.07 | 1.52996645271385e-46 | 6 |
| spi1a | 6.64965277451432e-51 | 0.445383282 | 0.178 | 0.032 | 1.67365110681751e-46 | 6 |
| gpx1a | 1.57536314881412e-50 | 0.685960782 | 0.501 | 0.216 | 3.96503150925025e-46 | 6 |
| mhc2b | 8.84829982028925e-49 | 0.414116606 | 0.157 | 0.026 | 2.2270285817686e-44 | 6 |
| itgae.1 | 1.44061730283714e-48 | 0.443483744 | 0.167 | 0.03 | 3.6258896895108e-44 | 6 |
| lta4h | 1.85577101544813e-48 | 0.539618692 | 0.258 | 0.067 | 4.67079006878139e-44 | 6 |
| gng10 | 8.67402934814232e-48 | 0.517381619 | 0.217 | 0.05 | 2.18316644663394e-43 | 6 |
| ppiab | 2.37812366874304e-45 | 0.397809165 | 0.82 | 0.652 | 5.98549946185935e-41 | 6 |
| rps22 | 3.98445380209002e-42 | 0.328967881 | 0.969 | 0.825 | 1.00284717744804e-37 | 6 |
| rps291 | 1.16229832573728e-40 | 0.29906249 | 0.971 | 0.828 | 2.92538865604815e-36 | 6 |
| ctss2.2 | 3.16530468430117e-40 | 0.438868236 | 0.175 | 0.039 | 7.96675535991761e-36 | 6 |

| | | | | | | |
|---|---|---|---|---|---|---|
| fabp11a | 4.25800761190003e-40 | 0.708398538 | 0.264 | 0.079 | 1.07169793583912e-35 | 6 |
| scpep1 | 3.14135970568994e-39 | 0.564879292 | 0.321 | 0.114 | 7.90648824325101e-35 | 6 |
| rps27.11 | 1.18212806010314e-38 | 0.40286297 | 0.906 | 0.673 | 2.97529811447359e-34 | 6 |
| arpc1b1 | 4.49981348915509e-38 | 0.594121179 | 0.601 | 0.301 | 1.13255805708544e-33 | 6 |
| mfap4 | 4.55915236507702e-38 | 0.345879433 | 0.125 | 0.022 | 1.14749305876623e-33 | 6 |
| rpl35a1 | 4.24534912259057e-37 | 0.31840425 | 0.914 | 0.755 | 1.06851192066482e-32 | 6 |
| rpl18a1 | 8.5939007136934e-37 | 0.311860693 | 0.93 | 0.781 | 2.16299887062949e-32 | 6 |
| cndp2 | 3.01405199720475e-36 | 0.500763767 | 0.214 | 0.06 | 7.58606747176464e-32 | 6 |
| rpl221 | 1.41655108899403e-35 | 0.349699292 | 0.864 | 0.69 | 3.56531743588906e-31 | 6 |
| zgc:64065 | 1.52919193520148e-35 | 0.308295529 | 0.104 | 0.016 | 3.8488231817086e-31 | 6 |
| rpl361 | 1.81279808224818e-34 | 0.315422268 | 0.943 | 0.785 | 4.56263149321044e-30 | 6 |
| rpl291 | 4.39974894775388e-33 | 0.287535482 | 0.95 | 0.805 | 1.10737281266017e-28 | 6 |
| rps211 | 4.82942917220209e-33 | 0.330057417 | 0.916 | 0.745 | 1.21551902835154e-28 | 6 |
| slc25a51 | 6.24304978592362e-33 | 0.252601263 | 0.909 | 0.804 | 1.57131320061912e-28 | 6 |
| rpl351 | 3.32669133093383e-32 | 0.265118485 | 0.916 | 0.802 | 8.37294941082735e-28 | 6 |
| rpl111 | 5.87807746338089e-32 | 0.284476015 | 0.943 | 0.81 | 1.47945331675834e-27 | 6 |
| rps281 | 1.64358118497617e-31 | 0.268960652 | 0.953 | 0.823 | 4.13672948446651e-27 | 6 |
| rpl181 | 3.54836837699538e-31 | 0.306607019 | 0.935 | 0.769 | 8.93088836805966e-27 | 6 |
| dpp7 | 1.72563191561305e-30 | 0.392027712 | 0.17 | 0.045 | 4.34324296840649e-26 | 6 |
| rpl241 | 5.20731021152876e-30 | 0.276571063 | 0.906 | 0.773 | 1.31062790713967e-25 | 6 |
| rps241 | 5.65664636192193e-30 | 0.302297466 | 0.924 | 0.754 | 1.42372132283213e-25 | 6 |
| rpl321 | 1.50912263318293e-29 | 0.273988303 | 0.937 | 0.81 | 3.79831075545811e-25 | 6 |
| si:ch211-1o7.3 | 1.98853576133179e-29 | 0.274194367 | 0.102 | 0.018 | 5.00494565769599e-25 | 6 |
| rpl371 | 4.10168065876142e-29 | 0.27415159 | 0.984 | 0.846 | 1.03235200500366e-24 | 6 |
| rps91 | 1.99073335500166e-28 | 0.280016731 | 0.916 | 0.773 | 5.01047678120367e-24 | 6 |
| rpl171 | 4.7136638527894e-28 | 0.268240436 | 0.869 | 0.743 | 1.18638205510856e-23 | 6 |
| rps51 | 1.36351560426123e-27 | 0.318589039 | 0.864 | 0.697 | 3.4318324243651e-23 | 6 |
| rpl151 | 2.27981153498546e-27 | 0.29406534 | 0.893 | 0.74 | 5.7380576524049e-23 | 6 |
| rpl81 | 4.66930942808993e-27 | 0.279865638 | 0.924 | 0.776 | 1.17521848995595e-22 | 6 |
| f13a1b | 1.15408647328226e-26 | 0.309411948 | 0.125 | 0.029 | 2.90472024460411e-22 | 6 |
| rpl36a1 | 1.18846833322066e-26 | 0.252596585 | 0.935 | 0.806 | 2.99125594788309e-22 | 6 |
| arpc5b | 2.17449282119539e-26 | 0.491827287 | 0.269 | 0.11 | 5.47298098166668e-22 | 6 |
| atp6v1g1 | 2.43571308624375e-26 | 0.495317821 | 0.298 | 0.129 | 6.1304462667669e-22 | 6 |
| faua1 | 2.92539007586766e-26 | 0.317119003 | 0.862 | 0.685 | 7.36291428195132e-22 | 6 |
| actb1 | 4.76813638812012e-26 | 0.256461237 | 0.869 | 0.783 | 1.20009224752595e-21 | 6 |

| | | | | | | |
|---|---|---|---|---|---|---|
| rpl381 | 2.65828322984157e-25 | 0.287454847 | 0.875 | 0.724 | 6.69063306118824e-21 | 6 |
| rps161 | 3.07226723487444e-25 | 0.309804938 | 0.916 | 0.733 | 7.73258940345547e-21 | 6 |
| mrc1b | 3.15936657683513e-25 | 0.344580933 | 0.149 | 0.04 | 7.95180973723634e-21 | 6 |
| anxa3b | 7.92161917261018e-25 | 0.320803054 | 0.125 | 0.031 | 1.99379232955426e-20 | 6 |
| si:ch211-15b10.6 | 8.10618549030556e-25 | 0.42689101 | 0.198 | 0.069 | 2.04024582605501e-20 | 6 |
| rps151 | 9.40795062308063e-25 | 0.319423337 | 0.864 | 0.68 | 2.36788709232316e-20 | 6 |
| zgc:152830 | 2.70781412961483e-24 | 0.409506633 | 0.188 | 0.063 | 6.81529738282757e-20 | 6 |
| tuba8l2 | 3.55605980564256e-24 | 0.382597675 | 0.18 | 0.058 | 8.95024692482175e-20 | 6 |
| sec61g | 6.1200601896419e-24 | 0.271033909 | 0.775 | 0.677 | 1.54035794913097e-19 | 6 |
| ckbb2 | 7.90516569494888e-24 | 0.487835296 | 0.465 | 0.261 | 1.98965115376168e-19 | 6 |
| rpsa1 | 1.140750613068e-23 | 0.274889346 | 0.896 | 0.748 | 2.87115521803084e-19 | 6 |
| rpl211 | 4.24529989783221e-23 | 0.259230142 | 0.888 | 0.753 | 1.06849953128539e-18 | 6 |
| atp6v0ca | 1.89013585891139e-22 | 0.478919791 | 0.366 | 0.195 | 4.75728294329407e-18 | 6 |
| LOC110437731 | 2.511350099553e-22 | 0.473779313 | 0.23 | 0.087 | 6.32081706556494e-18 | 6 |
| naga | 2.96285997924788e-22 | 0.37197286 | 0.154 | 0.048 | 7.45722228176899e-18 | 6 |
| rps141 | 3.69851786652467e-22 | 0.253546682 | 0.883 | 0.756 | 9.30879961825594e-18 | 6 |
| rpl341 | 7.02132588944025e-22 | 0.265881287 | 0.932 | 0.78 | 1.76719751311322e-17 | 6 |
| tpp1 | 7.20995329800756e-22 | 0.398752815 | 0.193 | 0.069 | 1.81467314557552e-17 | 6 |
| npc2 | 1.35965925717411e-21 | 0.470904027 | 0.525 | 0.355 | 3.42212638438153e-17 | 6 |
| laptm51 | 2.64314112642058e-21 | 0.449106062 | 0.332 | 0.167 | 6.65252190108796e-17 | 6 |
| rpl91 | 3.18952748826748e-21 | 0.322234107 | 0.807 | 0.625 | 8.02772173522043e-17 | 6 |
| rpl281 | 4.43822458924478e-21 | 0.260973414 | 0.893 | 0.746 | 1.11705674686702e-16 | 6 |
| ybx1 | 5.81129207831988e-21 | 0.265950253 | 0.786 | 0.667 | 1.46264410319233e-16 | 6 |
| kctd12.2 | 2.85432078660835e-20 | 0.39412364 | 0.198 | 0.077 | 7.18403998781455e-16 | 6 |
| itm2bb | 3.05982683457194e-20 | 0.451507226 | 0.384 | 0.219 | 7.70127815993411e-16 | 6 |
| zgc:1141881 | 5.67167147213656e-20 | 0.311902477 | 0.794 | 0.624 | 1.42750299282205e-15 | 6 |
| ctsba4 | 8.09218026219176e-20 | 0.4542023 | 0.54 | 0.373 | 2.03672085019104e-15 | 6 |
| arl6ip1 | 1.03055369246496e-19 | 0.42742244 | 0.522 | 0.367 | 2.59380058856506e-15 | 6 |
| rplp2l1 | 1.11370524420377e-19 | 0.265162759 | 0.924 | 0.77 | 2.80308472913646e-15 | 6 |
| spi1b | 2.16356252702995e-19 | 0.353693371 | 0.167 | 0.059 | 5.44547052428168e-15 | 6 |
| mdh1aa | 3.4332456003347e-19 | 0.457751691 | 0.352 | 0.203 | 8.6411358514824e-15 | 6 |
| naca1 | 6.41953085635984e-19 | 0.290653019 | 0.762 | 0.615 | 1.61573172123721e-14 | 6 |
| rpl301 | 6.67150438832556e-19 | 0.260477665 | 0.843 | 0.705 | 1.67915093949766e-14 | 6 |
| cxcr3.2 | 1.22270502380261e-18 | 0.278183192 | 0.115 | 0.033 | 3.07742627440878e-14 | 6 |
| gm2a | 1.23801291026881e-18 | 0.415204985 | 0.256 | 0.118 | 3.11595469385557e-14 | 6 |

| | | | | | | |
|---|---|---|---|---|---|---|
| wasb1 | 1.36549465725606e-18 | 0.435780987 | 0.326 | 0.174 | 3.43681350284777e-14 | 6 |
| si:ch211-212k18.7 | 1.90321147959916e-18 | 0.422840112 | 0.277 | 0.136 | 4.79019297300313e-14 | 6 |
| gstt1a | 2.09667143373825e-18 | 0.276040662 | 0.123 | 0.036 | 5.27711233157581e-14 | 6 |
| atp6v1e1b | 9.482114329237e-18 | 0.372246009 | 0.196 | 0.08 | 2.38655335552566e-13 | 6 |
| tcima | 3.07226308112255e-17 | 0.353789596 | 0.175 | 0.066 | 7.73257894887734e-13 | 6 |
| rpl61 | 3.1726261865027e-17 | 0.256898031 | 0.796 | 0.665 | 7.98518284880864e-13 | 6 |
| rpl23a1 | 1.55776683004573e-16 | 0.26230821 | 0.76 | 0.632 | 3.92074333454209e-12 | 6 |
| rpl27a1 | 5.28770497217272e-16 | 0.307753077 | 0.642 | 0.493 | 1.33086246444615e-11 | 6 |
| crip11 | 8.38568651490216e-16 | 0.393534708 | 0.42 | 0.268 | 2.11059343893572e-11 | 6 |
| coro1a1 | 4.55576784957695e-15 | 0.37036462 | 0.389 | 0.228 | 1.14664121006002e-10 | 6 |
| CR855311.1 | 9.1581039584873e-15 | 0.333622429 | 0.196 | 0.084 | 2.30500318531167e-10 | 6 |
| mapre1a | 1.46732962578134e-14 | 0.38102434 | 0.334 | 0.195 | 3.69312193512907e-10 | 6 |
| pepd | 2.98020841614249e-14 | 0.327717938 | 0.164 | 0.069 | 7.50088656258902e-10 | 6 |
| XLOC-0234921 | 3.49311897469032e-14 | 0.362439016 | 0.201 | 0.093 | 8.79183114739807e-10 | 6 |
| rps26l1 | 4.1397828038427e-14 | 0.25230036 | 0.802 | 0.661 | 1.04194193389917e-09 | 6 |
| ctsd1 | 5.54356449542636e-14 | 0.395420704 | 0.256 | 0.134 | 1.39525974785386e-09 | 6 |
| pycard1 | 7.85377615784066e-14 | 0.323605337 | 0.164 | 0.071 | 1.97671692116692e-09 | 6 |
| pdia3 | 1.65784168838378e-13 | 0.375081285 | 0.399 | 0.282 | 4.17262174549314e-09 | 6 |
| rac21 | 3.40893376867939e-13 | 0.361555361 | 0.35 | 0.219 | 8.57994540238916e-09 | 6 |
| fgl2a | 4.22595646039492e-13 | 0.312114494 | 0.154 | 0.064 | 1.0636309815168e-08 | 6 |
| zgc:153317 | 1.18133407009352e-12 | 0.278764683 | 0.112 | 0.041 | 2.97329972101837e-08 | 6 |
| ppdpfb | 1.4177118245207e-12 | 0.307059894 | 0.574 | 0.519 | 3.56823889113614e-08 | 6 |
| gabarapb | 1.43089802971344e-12 | 0.385909245 | 0.373 | 0.267 | 3.60142725098576e-08 | 6 |
| arpc21 | 1.53602768662891e-12 | 0.367704867 | 0.379 | 0.266 | 3.86602808447631e-08 | 6 |
| gapdhs1 | 2.30531012496536e-12 | 0.366743987 | 0.394 | 0.287 | 5.8022350535253e-08 | 6 |
| BX248082.11 | 4.88096347045913e-12 | 0.268345056 | 0.112 | 0.041 | 1.22848969587986e-07 | 6 |
| cst14b.1 | 7.70603307957528e-12 | 0.295517744 | 0.125 | 0.05 | 1.9395314657983e-07 | 6 |
| cd631 | 1.09597661716157e-11 | 0.335176088 | 0.452 | 0.339 | 2.75846354773396e-07 | 6 |
| eef1db | 1.16165523174041e-11 | 0.357322061 | 0.379 | 0.273 | 2.92377005276744e-07 | 6 |
| ctsa | 1.17849058238494e-11 | 0.324341803 | 0.18 | 0.088 | 2.96614294680467e-07 | 6 |
| zgc:1627301 | 1.63377744408481e-11 | 0.321476704 | 0.577 | 0.388 | 4.11205444901705e-07 | 6 |
| clic2 | 1.70355132261874e-11 | 0.269343491 | 0.138 | 0.059 | 4.28766832389912e-07 | 6 |
| cebpa | 4.34382397851965e-11 | 0.276104465 | 0.133 | 0.057 | 1.09329705715361e-06 | 6 |
| cap11 | 1.06917599947482e-10 | 0.356960872 | 0.337 | 0.232 | 2.69100907307817e-06 | 6 |
| arpc4l1 | 1.57207750003139e-10 | 0.351132618 | 0.298 | 0.195 | 3.95676185982901e-06 | 6 |

| | | | | | | |
|---|---|---|---|---|---|---|
| btg11 | 2.06156231383145e-10 | 0.326532046 | 0.433 | 0.332 | 5.18874618768238e-06 | 6 |
| xaf1 | 3.82280960485872e-10 | 0.270144134 | 0.133 | 0.06 | 9.62162949446892e-06 | 6 |
| rap1b | 4.34546740129102e-10 | 0.356123931 | 0.298 | 0.201 | 1.09371069023094e-05 | 6 |
| si:ch211-66e2.3 | 5.25336819205792e-10 | 0.264177438 | 0.131 | 0.058 | 1.32222024025906e-05 | 6 |
| ptmaa1 | 6.82108350380605e-10 | 0.28346449 | 0.525 | 0.393 | 1.71679850707295e-05 | 6 |
| ms4a17a.61 | 2.49712116643915e-09 | 0.25277671 | 0.112 | 0.048 | 6.2850042638107e-05 | 6 |
| eif4ebp1 | 3.01173154512384e-09 | 0.323130672 | 0.342 | 0.245 | 7.5802271259222e-05 | 6 |
| ctsh | 4.52577577019859e-09 | 0.260328112 | 0.117 | 0.053 | 0.000113909 | 6 |
| cox7b1 | 8.50575733164989e-09 | 0.261801187 | 0.483 | 0.417 | 0.000214081 | 6 |
| ctsk | 1.56881856877504e-08 | 0.292101762 | 0.128 | 0.062 | 0.000394856 | 6 |
| prdx5 | 3.25399031527174e-08 | 0.316071593 | 0.24 | 0.158 | 0.000818997 | 6 |
| smox | 1.16991020516379e-07 | 0.267841462 | 0.154 | 0.084 | 0.002944547 | 6 |
| COX7A2 (1 of many) | 1.68289519659224e-07 | 0.283164992 | 0.342 | 0.26 | 0.004235679 | 6 |
| arpc31 | 2.09020331973448e-07 | 0.271740783 | 0.386 | 0.31 | 0.005260833 | 6 |
| suclg1 | 2.80729665719777e-07 | 0.273212317 | 0.183 | 0.109 | 0.007065685 | 6 |
| si:dkey-180p18.9 | 7.76982231249315e-07 | 0.257394608 | 0.167 | 0.099 | 0.019555866 | 6 |
| nccrp1 | 8.90093880621824e-07 | 0.26044131 | 0.282 | 0.193 | 0.022402773 | 6 |
| zgc:640511 | 1.77794700632322e-06 | 0.252690143 | 0.24 | 0.162 | 0.044749148 | 6 |
| glrx1 | 1.82715430148444e-06 | 0.25687887 | 0.379 | 0.312 | 0.045987647 | 6 |
| ctsz | 2.34504771737231e-06 | 0.270179099 | 0.149 | 0.087 | 0.059022506 | 6 |
| vdac3 | 3.88429421459961e-06 | 0.25936864 | 0.225 | 0.154 | 0.097763801 | 6 |
| slc6a6b | 1.00511140916029e-05 | 0.251140885 | 0.164 | 0.103 | 0.252976491 | 6 |
| si:dkey-112e7.21 | 5.27329234390925e-05 | 0.254975587 | 0.261 | 0.206 | 1 | 6 |
| igfbp5b | 0 | 1.906908155 | 0.814 | 0.047 | 0 | 7 |
| si:dkey-261h17.1 | 0 | 1.782637649 | 0.819 | 0.066 | 0 | 7 |
| pmp22a | 0 | 1.653823111 | 0.906 | 0.113 | 0 | 7 |
| rbp4 | 0 | 1.640238612 | 0.832 | 0.104 | 0 | 7 |
| krt94 | 0 | 1.591381511 | 0.84 | 0.105 | 0 | 7 |
| endouc | 0 | 1.495967277 | 0.69 | 0.041 | 0 | 7 |
| col1a1b | 0 | 1.447449246 | 0.837 | 0.124 | 0 | 7 |
| col1a2 | 0 | 1.427783685 | 0.843 | 0.133 | 0 | 7 |
| dcn | 0 | 1.411797697 | 0.803 | 0.096 | 0 | 7 |
| pcolcea | 0 | 1.365953697 | 0.64 | 0.031 | 0 | 7 |
| si:ch211-105c13.3 | 0 | 1.32496397 | 0.651 | 0.04 | 0 | 7 |
| plpp3 | 0 | 1.308555565 | 0.646 | 0.035 | 0 | 7 |

| | | | | | | | |
|---|---|---|---|---|---|---|---|
| fstl1b | 0 | 1.276994841 | 0.727 | 0.105 | | 0 | 7 |
| fn1a | 0 | 1.23668567 | 0.612 | 0.046 | | 0 | 7 |
| CABZ01092746.1 | 0 | 1.206275631 | 0.58 | 0.05 | | 0 | 7 |
| neu3.5 | 0 | 1.181229113 | 0.514 | 0.002 | | 0 | 7 |
| spon2b | 0 | 1.12227883 | 0.425 | 0.01 | | 0 | 7 |
| htra1b | 0 | 1.103195054 | 0.467 | 0.025 | | 0 | 7 |
| AL732488.1 | 0 | 1.084400608 | 0.491 | 0.025 | | 0 | 7 |
| thbs3a | 0 | 1.07575335 | 0.462 | 0.004 | | 0 | 7 |
| mfap5 | 0 | 1.041538719 | 0.488 | 0.03 | | 0 | 7 |
| mgp | 0 | 1.013505998 | 0.367 | 0.008 | | 0 | 7 |
| emilin2a | 0 | 1.004343951 | 0.459 | 0.009 | | 0 | 7 |
| si:ch73-173h19.3 | 0 | 0.959734093 | 0.423 | 0.008 | | 0 | 7 |
| CABZ01039096.1 | 0 | 0.945054464 | 0.417 | 0.008 | | 0 | 7 |
| im:7151449 | 0 | 0.937653902 | 0.404 | 0.015 | | 0 | 7 |
| slit3 | 0 | 0.931306945 | 0.433 | 0.013 | | 0 | 7 |
| BX663503.3 | 0 | 0.927412497 | 0.409 | 0.017 | | 0 | 7 |
| osr1 | 0 | 0.880301753 | 0.391 | 0.015 | | 0 | 7 |
| efnb1 | 0 | 0.872145851 | 0.388 | 0.022 | | 0 | 7 |
| crhbp | 0 | 0.831782541 | 0.31 | 0.007 | | 0 | 7 |
| adamts15b | 0 | 0.830428738 | 0.36 | 0.012 | | 0 | 7 |
| flnb | 0 | 0.786117511 | 0.331 | 0.011 | | 0 | 7 |
| mustn1a | 0 | 0.778787815 | 0.341 | 0.012 | | 0 | 7 |
| plvapb | 0 | 0.759907081 | 0.294 | 0.005 | | 0 | 7 |
| crabp2b | 0 | 0.725920302 | 0.304 | 0.009 | | 0 | 7 |
| dock9b | 0 | 0.722283129 | 0.328 | 0.013 | | 0 | 7 |
| hapln3 | 0 | 0.695221437 | 0.289 | 0.008 | | 0 | 7 |
| itih5 | 0 | 0.651748163 | 0.268 | 0.001 | | 0 | 7 |
| tspan35 | 0 | 0.647816479 | 0.278 | 0.009 | | 0 | 7 |
| tgfb2 | 0 | 0.637876751 | 0.27 | 0.006 | | 0 | 7 |
| crabp2a | 0 | 0.63244019 | 0.241 | 0.003 | | 0 | 7 |
| slc6a4a | 0 | 0.561415162 | 0.231 | 0.005 | | 0 | 7 |
| adgrg6 | 0 | 0.557071545 | 0.22 | 0.005 | | 0 | 7 |
| tll1 | 0 | 0.49018455 | 0.189 | 0.002 | | 0 | 7 |
| col1a1a | 1.10041504418903e-306 | 1.323599562 | 0.79 | 0.118 | 2.76963462471937e-302 | | 7 |
| emid1 | 4.73588656720351e-305 | 0.704901243 | 0.299 | 0.012 | 1.19197529009945e-300 | | 7 |

| | | | | | | |
|---|---|---|---|---|---|---|
| si:ch211-227m13.1 | 3.41079393353351e-304 | 0.531455747 | 0.236 | 0.006 | 8.5846272513105e-300 | 7 |
| plpp1a | 1.71813971168267e-303 | 1.076505671 | 0.535 | 0.053 | 4.3243858403341e-299 | 7 |
| fbln2 | 8.2487629725559e-302 | 0.705161816 | 0.32 | 0.015 | 2.0761311525626e-297 | 7 |
| abca1a | 4.37117715021066e-298 | 1.09195639 | 0.528 | 0.053 | 1.10018157693652e-293 | 7 |
| mfge8b | 4.19484387389444e-296 | 0.907255801 | 0.373 | 0.023 | 1.05580025462049e-291 | 7 |
| LOC108180269 | 1.21465774423587e-292 | 1.00957754 | 0.478 | 0.042 | 3.05717207646727e-288 | 7 |
| fbln5 | 1.29419606968914e-288 | 0.84990813 | 0.391 | 0.027 | 3.25736208780059e-284 | 7 |
| gfpt2 | 1.50693224102597e-282 | 0.509656645 | 0.186 | 0.003 | 3.79279775743827e-278 | 7 |
| htra1a | 2.2454821369059e-272 | 0.64676568 | 0.273 | 0.011 | 5.65165399037845e-268 | 7 |
| lrp1aa | 3.5571532992235e-270 | 0.905303501 | 0.457 | 0.042 | 8.95299913881563e-266 | 7 |
| XLOC-016094 | 1.0713569696761e-267 | 1.044296012 | 0.551 | 0.064 | 2.69649835697777e-263 | 7 |
| ablim1b | 3.38306637039844e-264 | 0.784138819 | 0.367 | 0.026 | 8.51483974765585e-260 | 7 |
| nid1a | 2.69053708685398e-263 | 0.785938898 | 0.375 | 0.027 | 6.77181279390278e-259 | 7 |
| tgfbr3 | 5.27623382510922e-263 | 0.873842507 | 0.396 | 0.031 | 1.32797529144174e-258 | 7 |
| CR855311.11 | 9.41257143082498e-262 | 1.217197956 | 0.564 | 0.07 | 2.36905010342434e-257 | 7 |
| si:ch211-67e16.11 | 1.43468245505794e-258 | 0.471052497 | 0.171 | 0.003 | 3.61095227113532e-254 | 7 |
| vasna | 1.0478170247248e-257 | 0.782447576 | 0.354 | 0.024 | 2.63725066952985e-253 | 7 |
| kirrel1a | 4.24286434231922e-256 | 0.704648463 | 0.312 | 0.018 | 1.06788652631832e-251 | 7 |
| crabp1b | 4.8558479856981e-254 | 0.422947911 | 0.165 | 0.003 | 1.22216837952036e-249 | 7 |
| si:ch211-198c19.3 | 9.99650650094139e-246 | 1.011377109 | 0.472 | 0.05 | 2.51602072122194e-241 | 7 |
| kcne4 | 1.26499153447943e-242 | 0.578393977 | 0.213 | 0.007 | 3.18385719313127e-238 | 7 |
| dpysl3 | 1.20923062789113e-241 | 0.548753481 | 0.252 | 0.011 | 3.04351256733918e-237 | 7 |
| fstb | 1.01320787107382e-240 | 0.677007713 | 0.268 | 0.013 | 2.55014289070571e-236 | 7 |
| tgfbr2b | 5.00827556876023e-239 | 0.601075795 | 0.257 | 0.012 | 1.26053287790126e-234 | 7 |
| si:ch211-39i22.1 | 6.89517131522913e-238 | 0.987891354 | 0.493 | 0.058 | 1.73544566833002e-233 | 7 |
| gpc6a | 5.17253735842722e-237 | 0.441535001 | 0.189 | 0.005 | 1.30187592774255e-232 | 7 |
| id2b | 3.13556038251957e-235 | 0.846551634 | 0.336 | 0.024 | 7.8918919267635e-231 | 7 |
| slc6a2 | 1.36345305904976e-233 | 0.363354369 | 0.129 | 0.001 | 3.43167500432234e-229 | 7 |
| neu3.4 | 1.1863143291046e-231 | 0.366093649 | 0.113 | 0 | 2.98583453492337e-227 | 7 |
| ca10b | 1.42509815820944e-229 | 0.423378389 | 0.171 | 0.004 | 3.58682955439735e-225 | 7 |
| nptx1l | 1.45752240180296e-229 | 0.437324205 | 0.171 | 0.004 | 3.66843813309787e-225 | 7 |
| ccl25b | 1.09196101551695e-228 | 1.199027657 | 0.507 | 0.063 | 2.74835667995462e-224 | 7 |
| nova2 | 1.14470268345248e-225 | 0.698594744 | 0.312 | 0.021 | 2.88110218398155e-221 | 7 |
| tmem100a | 7.86421583484525e-223 | 0.60262944 | 0.255 | 0.013 | 1.9793444834722e-218 | 7 |
| add3a | 5.76509839500301e-219 | 1.082844193 | 0.622 | 0.108 | 1.45101761503831e-214 | 7 |

| | | | | | | |
|---|---|---|---|---|---|---|
| col5a2a | 3.54036919451208e-218 | 0.985081979 | 0.525 | 0.072 | 8.91075522566747e-214 | 7 |
| col5a1 | 4.92881812572071e-218 | 1.036523987 | 0.612 | 0.096 | 1.24053423406265e-213 | 7 |
| marcksl1a | 1.6881671534988e-217 | 1.301551006 | 0.785 | 0.196 | 4.24894790864113e-213 | 7 |
| icn | 7.73949406993401e-216 | 1.349471358 | 0.895 | 0.254 | 1.94795326246169e-211 | 7 |
| cbln2b | 4.46622544334842e-215 | 0.33082263 | 0.118 | 0.001 | 1.12410428183636e-210 | 7 |
| hapln1b | 1.58327065405007e-213 | 0.416643195 | 0.168 | 0.004 | 3.98493390917863e-209 | 7 |
| tgm1l1 | 7.9262939486646e-211 | 0.777801698 | 0.339 | 0.028 | 1.99496892393939e-206 | 7 |
| tsku | 3.58955929266135e-210 | 0.633330876 | 0.297 | 0.021 | 9.03456178369935e-206 | 7 |
| eva1ba | 1.68956015118508e-204 | 0.875869941 | 0.441 | 0.053 | 4.25245394451772e-200 | 7 |
| june | 2.67419274891666e-204 | 0.677794474 | 0.299 | 0.022 | 6.73067572974834e-200 | 7 |
| bag2 | 1.57495265602066e-202 | 0.462977014 | 0.192 | 0.007 | 3.96399833993839e-198 | 7 |
| pamr1 | 1.71944355830926e-199 | 0.433374963 | 0.178 | 0.006 | 4.32766749190858e-195 | 7 |
| akap12b | 1.28043846891137e-197 | 0.823578504 | 0.42 | 0.048 | 3.22273558240303e-193 | 7 |
| LOC100537342 | 2.94393287594163e-195 | 0.384128535 | 0.157 | 0.004 | 7.4095846554575e-191 | 7 |
| aplp1 | 6.00237233054866e-193 | 0.479512088 | 0.207 | 0.01 | 1.51073709187579e-188 | 7 |
| ebf3a | 6.56101070317087e-192 | 0.373258601 | 0.15 | 0.004 | 1.65134078388108e-187 | 7 |
| zgc:195173 | 1.38434216810609e-191 | 0.963404873 | 0.504 | 0.078 | 3.48425080290622e-187 | 7 |
| s100a10b | 2.84679401614368e-191 | 1.245767293 | 0.835 | 0.237 | 7.16509585923203e-187 | 7 |
| gpr146 | 6.57438661028829e-191 | 0.537463028 | 0.223 | 0.012 | 1.65470736594346e-186 | 7 |
| nfixb | 1.29320468824106e-190 | 0.822344966 | 0.428 | 0.054 | 3.25486687983391e-186 | 7 |
| emp2 | 2.5964717895142e-190 | 0.775154441 | 0.367 | 0.039 | 6.53505984702829e-186 | 7 |
| has2 | 1.83650176764744e-186 | 0.341493931 | 0.129 | 0.002 | 4.62229129899183e-182 | 7 |
| cyp2n13 | 6.48761399531143e-184 | 0.68286393 | 0.315 | 0.028 | 1.63286756647993e-179 | 7 |
| zcchc24 | 1.74841016929313e-183 | 0.633262196 | 0.291 | 0.024 | 4.40057355509388e-179 | 7 |
| cx43 | 5.11280355898644e-183 | 0.863866347 | 0.381 | 0.044 | 1.2868415277613e-178 | 7 |
| CR383676.11 | 1.05103779599605e-182 | 0.939656454 | 0.982 | 0.718 | 2.64535702874245e-178 | 7 |
| colec12 | 3.37926635302175e-182 | 0.757175006 | 0.381 | 0.043 | 8.50527548392045e-178 | 7 |
| angptl2a | 1.71344152381716e-179 | 0.26679985 | 0.1 | 0.001 | 4.31256097129541e-175 | 7 |
| tnfsf10l | 2.11247076735003e-179 | 0.571113699 | 0.26 | 0.019 | 5.31687767434329e-175 | 7 |
| anxa1a | 9.00377323578983e-179 | 1.105054438 | 0.625 | 0.127 | 2.26615968571594e-174 | 7 |
| xkr9 | 1.23309034486535e-178 | 0.491504107 | 0.207 | 0.011 | 3.10356508899161e-174 | 7 |
| fgl2b | 3.56809907323782e-178 | 0.455936518 | 0.194 | 0.009 | 8.98054855743226e-174 | 7 |
| peli1a | 4.58243089874943e-178 | 0.438567228 | 0.168 | 0.006 | 1.15335203290624e-173 | 7 |
| mdka | 1.2357001127102e-177 | 1.000358363 | 0.501 | 0.08 | 3.11013361368031e-173 | 7 |
| ddr2l | 1.23740249948688e-177 | 0.578374608 | 0.257 | 0.019 | 3.11441835095852e-173 | 7 |

| | | | | | | |
|---|---|---|---|---|---|---|
| krt8 | 2.34777518933709e-176 | 1.150193571 | 0.861 | 0.257 | 5.90911537404251e-172 | 7 |
| ppap2d | 1.08372512250909e-173 | 0.630815484 | 0.304 | 0.028 | 2.72762776084312e-169 | 7 |
| itga5 | 7.24825317626069e-173 | 0.632757202 | 0.27 | 0.022 | 1.82431284193305e-168 | 7 |
| pmp22b | 3.37698016374562e-171 | 0.715862518 | 0.339 | 0.036 | 8.49952137413135e-167 | 7 |
| c7a | 3.61008264213328e-166 | 0.716867166 | 0.333 | 0.036 | 9.08621700198526e-162 | 7 |
| galr1b | 6.36978120936841e-162 | 0.285427572 | 0.105 | 0.002 | 1.60321023258593e-157 | 7 |
| c6 | 8.60177143513266e-159 | 0.962161239 | 0.486 | 0.083 | 2.16497985250854e-154 | 7 |
| ackr3b | 6.35114671774083e-158 | 0.591526447 | 0.234 | 0.017 | 1.59852011738819e-153 | 7 |
| angptl1a | 1.75858155494312e-156 | 0.602399924 | 0.249 | 0.02 | 4.42617391563634e-152 | 7 |
| crabp1a | 2.72180074975375e-153 | 0.32383999 | 0.126 | 0.004 | 6.85050030705522e-149 | 7 |
| arrdc3a | 4.15284786995768e-153 | 0.83306758 | 0.412 | 0.061 | 1.04523028038965e-148 | 7 |
| angptl4 | 1.29713095376587e-150 | 0.717164339 | 0.336 | 0.04 | 3.26474889753331e-146 | 7 |
| BX640522.2 | 2.02180532028897e-149 | 0.654797207 | 0.244 | 0.02 | 5.0886818106353e-145 | 7 |
| mast2 | 9.52664487357793e-147 | 0.654845297 | 0.304 | 0.034 | 2.39776124823083e-142 | 7 |
| sema3d | 1.53334869064573e-143 | 0.441974646 | 0.178 | 0.01 | 3.85928531948624e-139 | 7 |
| lpar1 | 1.25370080455816e-142 | 0.770931011 | 0.375 | 0.055 | 3.15543955499243e-138 | 7 |
| znf385a | 1.46015267410056e-142 | 0.661685256 | 0.315 | 0.038 | 3.6750582654437e-138 | 7 |
| bicc1b | 1.66625870471933e-141 | 0.478305087 | 0.21 | 0.016 | 4.19380653390808e-137 | 7 |
| kank2 | 1.74361921107211e-140 | 0.69018361 | 0.339 | 0.045 | 4.3885151923474e-136 | 7 |
| adamts15a | 1.5826021060074e-136 | 0.682182628 | 0.318 | 0.04 | 3.98325124061003e-132 | 7 |
| klf2a | 5.47143667889791e-134 | 0.839104342 | 0.425 | 0.075 | 1.37710589771181e-129 | 7 |
| mfap2 | 5.6699696848599e-134 | 0.809250469 | 0.459 | 0.085 | 1.42707466998239e-129 | 7 |
| c1qtnf2 | 8.6147636626695e-133 | 0.461636819 | 0.207 | 0.016 | 2.16824986625729e-128 | 7 |
| ramp2 | 4.42322672142678e-132 | 0.667341321 | 0.289 | 0.034 | 1.11328193351591e-127 | 7 |
| clec19a | 1.08553525922265e-131 | 0.349326441 | 0.126 | 0.005 | 2.73218369393749e-127 | 7 |
| col6a3 | 1.72087559068591e-131 | 0.820480485 | 0.47 | 0.089 | 4.33127177419738e-127 | 7 |
| rnd3b | 5.58757037094712e-131 | 0.578969141 | 0.286 | 0.033 | 1.40633558666368e-126 | 7 |
| rgs5a | 3.92837762196541e-130 | 0.538687357 | 0.231 | 0.021 | 9.88733363672475e-126 | 7 |
| cyp2ad3 | 3.53845228530874e-129 | 0.641978111 | 0.323 | 0.043 | 8.90593055689357e-125 | 7 |
| ifitm1 | 1.17598734122167e-128 | 0.332066219 | 0.102 | 0.003 | 2.95984253912081e-124 | 7 |
| tnfaip2a | 1.8262138210033e-128 | 0.502629862 | 0.215 | 0.018 | 4.59639756608322e-124 | 7 |
| tns2b | 1.14010849325133e-126 | 0.347844492 | 0.15 | 0.008 | 2.86953906666426e-122 | 7 |
| fkbp1ab | 3.3687815748765e-126 | 0.718565691 | 0.365 | 0.055 | 8.47888634580667e-122 | 7 |
| adrb3a | 5.34684029573093e-125 | 0.319529797 | 0.129 | 0.006 | 1.34574623403252e-120 | 7 |
| antxr2a | 1.74737279310592e-123 | 0.869816606 | 0.556 | 0.141 | 4.39796258296828e-119 | 7 |

| | | | | | | |
|---|---|---|---|---|---|---|
| cyr61 | 3.09877924011721e-123 | 0.814883922 | 0.367 | 0.059 | 7.799317469451e-119 | 7 |
| ahnak | 3.05374735391419e-122 | 0.859254437 | 0.504 | 0.115 | 7.68597671506662e-118 | 7 |
| cyp2p6 | 4.67927516537191e-121 | 0.461149272 | 0.21 | 0.019 | 1.17772676637246e-116 | 7 |
| slc7a1 | 3.88163778870349e-119 | 0.553243171 | 0.262 | 0.031 | 9.76969415038782e-115 | 7 |
| eda | 8.60559625389881e-118 | 0.371052689 | 0.136 | 0.007 | 2.16594252114379e-113 | 7 |
| sparc | 9.36114315492231e-118 | 0.843931832 | 0.591 | 0.145 | 2.3561061206624e-113 | 7 |
| wscd2 | 1.90176696624091e-117 | 0.336872933 | 0.136 | 0.007 | 4.78655727733174e-113 | 7 |
| dio2 | 1.99762608334248e-116 | 0.26636844 | 0.105 | 0.004 | 5.02782508916469e-112 | 7 |
| si:dkey-240h12.3 | 3.94095054662192e-116 | 0.441691725 | 0.192 | 0.016 | 9.9189784307927e-112 | 7 |
| jund | 9.59853946481288e-116 | 0.758003721 | 0.417 | 0.083 | 2.41585639789875e-111 | 7 |
| ca5a | 2.55492221187053e-115 | 0.510061136 | 0.231 | 0.024 | 6.43048371505695e-111 | 7 |
| pdgfra | 2.93966978699955e-115 | 0.590886032 | 0.297 | 0.04 | 7.39885488689917e-111 | 7 |
| bgnb | 1.01243371458444e-114 | 0.490330514 | 0.218 | 0.021 | 2.54819441623758e-110 | 7 |
| abhd15a | 1.23952634395566e-114 | 0.536393185 | 0.255 | 0.029 | 3.119763855102e-110 | 7 |
| ank3b | 3.37342313423493e-114 | 0.565182106 | 0.26 | 0.031 | 8.49056868655589e-110 | 7 |
| arhgap21a | 3.90063031351903e-114 | 0.335305563 | 0.139 | 0.008 | 9.81749643609603e-110 | 7 |
| sema4ba | 9.17538835088929e-113 | 0.598726902 | 0.286 | 0.039 | 2.30935349403532e-108 | 7 |
| dhrs3b | 2.06182850383021e-110 | 0.407663976 | 0.16 | 0.011 | 5.18941616129025e-106 | 7 |
| ip6k2b | 9.68586301366923e-110 | 0.813185512 | 0.47 | 0.112 | 2.43783486191041e-105 | 7 |
| itpkcb | 1.74794506021257e-109 | 0.685869016 | 0.373 | 0.068 | 4.39940292204902e-105 | 7 |
| rgcc | 6.46771623037691e-109 | 0.689772394 | 0.357 | 0.062 | 1.62785949802356e-104 | 7 |
| sost | 7.37408141879393e-109 | 0.30876149 | 0.126 | 0.007 | 1.85598255229624e-104 | 7 |
| socs3a | 1.10214589279354e-108 | 0.983501353 | 0.764 | 0.284 | 2.77399099757207e-104 | 7 |
| ptx3a | 2.59693768422076e-108 | 0.560826981 | 0.252 | 0.03 | 6.53623245741523e-104 | 7 |
| hspb1 | 4.98394731629292e-106 | 0.654158848 | 0.333 | 0.055 | 1.25440970003776e-101 | 7 |
| abi3bpa | 5.051541768875e-106 | 0.351975596 | 0.144 | 0.009 | 1.27142254780815e-101 | 7 |
| si:ch73-46j18.5 | 6.4090154060852e-106 | 0.844263625 | 0.58 | 0.163 | 1.61308508755758e-101 | 7 |
| synm | 4.34433919566343e-105 | 0.294337719 | 0.115 | 0.006 | 1.09342673215653e-100 | 7 |
| itgb1a | 5.47599899794746e-105 | 0.577759977 | 0.278 | 0.04 | 1.3782541877934e-100 | 7 |
| nectin1b | 8.77346624674652e-105 | 0.561592811 | 0.265 | 0.036 | 2.20819371964363e-100 | 7 |
| fam107b | 9.34766439535115e-104 | 0.731166428 | 0.43 | 0.094 | 2.35271365166593e-99 | 7 |
| fgfr2 | 1.72944834879123e-103 | 0.463442844 | 0.207 | 0.022 | 4.35284854907265e-99 | 7 |
| rem1 | 3.39837419749385e-103 | 0.526285403 | 0.234 | 0.027 | 8.55336801767228e-99 | 7 |
| cebpd | 8.20020913437278e-103 | 0.794153872 | 0.488 | 0.118 | 2.06391063703029e-98 | 7 |
| tmsb11 | 2.36335560184389e-102 | 0.873699577 | 0.541 | 0.144 | 5.94832971428089e-98 | 7 |

| | | | | | | |
|---|---|---|---|---|---|---|
| ackr4b | 3.12582012624502e-100 | 0.302437401 | 0.113 | 0.006 | 7.8673766757461e-96 | 7 |
| cxcl12a | 4.18192340159731e-100 | 0.827646188 | 0.378 | 0.074 | 1.05254830094803e-95 | 7 |
| junba | 1.74993117533316e-99 | 0.914731545 | 0.756 | 0.295 | 4.40440177519603e-95 | 7 |
| ugt5g1 | 2.66933539335318e-99 | 0.302544931 | 0.129 | 0.008 | 6.71845025153063e-95 | 7 |
| loxl1 | 5.54261008898853e-98 | 0.427184569 | 0.184 | 0.018 | 1.39501953329752e-93 | 7 |
| anxa5b | 1.64666906012778e-97 | 0.718030407 | 0.391 | 0.085 | 4.1445013574356e-93 | 7 |
| tns1b | 1.83235266168214e-97 | 0.495927312 | 0.228 | 0.028 | 4.61184841418777e-93 | 7 |
| cyp2ad6 | 1.94883278528981e-97 | 0.362335 | 0.157 | 0.013 | 4.90501723729593e-93 | 7 |
| agtr2 | 2.61832755479395e-96 | 0.410368104 | 0.163 | 0.014 | 6.59006862266088e-92 | 7 |
| si:ch1073-459j12.1 | 5.27840926016949e-96 | 0.632754719 | 0.325 | 0.058 | 1.32852282669206e-91 | 7 |
| fbn2b | 1.83219552023963e-94 | 0.432844554 | 0.194 | 0.021 | 4.61145290489112e-90 | 7 |
| scara3 | 4.72856669698299e-94 | 0.462981489 | 0.215 | 0.026 | 1.19013295196365e-89 | 7 |
| NC-002333.41 | 4.86374364406831e-93 | 0.68565547 | 0.969 | 0.612 | 1.22415563777555e-88 | 7 |
| si:ch211-212k18.8 | 8.98546504779858e-92 | 0.477354001 | 0.252 | 0.037 | 2.26155169788043e-87 | 7 |
| fabp11a1 | 1.08264162047905e-91 | 0.716305109 | 0.373 | 0.075 | 2.72490069458372e-87 | 7 |
| prelp | 7.10168840298131e-91 | 0.297380107 | 0.121 | 0.008 | 1.78742395414637e-86 | 7 |
| s100v2 | 1.70725799332542e-89 | 0.736214938 | 0.441 | 0.112 | 4.29699764340075e-85 | 7 |
| vim | 7.48704897661418e-89 | 0.438938525 | 0.223 | 0.029 | 1.88441535692402e-84 | 7 |
| nfia | 3.97928916929302e-88 | 0.634685189 | 0.346 | 0.072 | 1.00154729101936e-83 | 7 |
| klf3 | 6.31026126525563e-88 | 0.453949128 | 0.223 | 0.03 | 1.58822965785219e-83 | 7 |
| emilin1b | 9.48400313633107e-87 | 0.295959796 | 0.134 | 0.01 | 2.38702874938317e-82 | 7 |
| lhfpl6 | 3.01597222627924e-86 | 0.355935708 | 0.165 | 0.016 | 7.59090049632223e-82 | 7 |
| lrp1ab | 1.55429206086339e-85 | 0.61208491 | 0.32 | 0.062 | 3.91199768798706e-81 | 7 |
| hspb8 | 4.86084255595842e-85 | 0.39338329 | 0.178 | 0.019 | 1.22342546290917e-80 | 7 |
| klf6a | 8.43813695628866e-85 | 0.747233389 | 0.52 | 0.157 | 2.12379469052829e-80 | 7 |
| cd81a | 2.07762700273477e-84 | 0.784372071 | 0.591 | 0.195 | 5.22917940318315e-80 | 7 |
| tshz2 | 2.08110882931039e-84 | 0.44815059 | 0.192 | 0.023 | 5.23794281249132e-80 | 7 |
| LOC103909284 | 4.65935380416447e-84 | 0.563831488 | 0.302 | 0.057 | 1.17271275897016e-79 | 7 |
| vat1 | 5.69975299038043e-84 | 0.72995672 | 0.459 | 0.125 | 1.43457083014885e-79 | 7 |
| grb10a | 9.70543357720931e-84 | 0.485304296 | 0.231 | 0.033 | 2.44276057704781e-79 | 7 |
| fosab | 1.01773093822475e-83 | 0.917753517 | 0.743 | 0.337 | 2.56152699841787e-79 | 7 |
| cyp2p7 | 2.37884658088572e-83 | 0.398727384 | 0.181 | 0.021 | 5.98731895943126e-79 | 7 |
| tmem176 | 1.32799091070738e-82 | 0.57432921 | 0.307 | 0.059 | 3.3424203231594e-78 | 7 |
| iqsec1b | 4.65898233472513e-80 | 0.295336814 | 0.126 | 0.01 | 1.17261926382697e-75 | 7 |
| prss59.2 | 7.41460971540523e-80 | 0.355976653 | 0.152 | 0.015 | 1.86618311927034e-75 | 7 |

| | | | | | | |
|---|---|---|---|---|---|---|
| zfp36l1b | 9.85465243971721e-80 | 0.722554599 | 0.446 | 0.12 | 2.48031747255242e-75 | 7 |
| rpz5 | 3.53690381972332e-79 | 0.636063952 | 0.386 | 0.094 | 8.90203322386161e-75 | 7 |
| plod2 | 3.85985772538451e-78 | 0.38554227 | 0.171 | 0.02 | 9.71487590902027e-74 | 7 |
| clu | 3.95494508760951e-78 | 0.409384197 | 0.181 | 0.022 | 9.95420129100437e-74 | 7 |
| lix1l | 1.71713267093696e-77 | 0.295625034 | 0.136 | 0.012 | 4.32185121948122e-73 | 7 |
| tril | 1.53593956939088e-76 | 0.376576525 | 0.16 | 0.017 | 3.86580630219989e-72 | 7 |
| s100w | 3.7508944151282e-76 | 0.334667986 | 0.147 | 0.015 | 9.44062615343616e-72 | 7 |
| pdgfrb | 2.62308218783156e-75 | 0.484046712 | 0.255 | 0.044 | 6.60203555855324e-71 | 7 |
| pabpc1b | 7.69879364458746e-75 | 0.558569772 | 0.299 | 0.062 | 1.93770937240622e-70 | 7 |
| lama4 | 4.9394964797643e-74 | 0.412316791 | 0.213 | 0.032 | 1.24322186899188e-69 | 7 |
| si:ch211-106h4.12 | 8.71146005686259e-74 | 0.608066585 | 0.278 | 0.052 | 2.19258738171174e-69 | 7 |
| tlr5b | 1.18533634159636e-73 | 0.29146057 | 0.131 | 0.012 | 2.98337303816389e-69 | 7 |
| COX31 | 1.47458053216584e-73 | 0.570372359 | 0.979 | 0.692 | 3.71137174140821e-69 | 7 |
| tuft1a | 2.46450779839232e-72 | 0.417755925 | 0.199 | 0.029 | 6.20291967777364e-68 | 7 |
| prss59.1 | 6.20272220731665e-72 | 0.259438489 | 0.113 | 0.009 | 1.56116315235953e-67 | 7 |
| slc38a5b | 7.96960612555977e-72 | 0.642713083 | 0.391 | 0.107 | 2.00587016574214e-67 | 7 |
| col6a2 | 1.59177533996336e-70 | 0.522401646 | 0.286 | 0.059 | 4.00633935315377e-66 | 7 |
| lxn | 1.0570241176876e-68 | 0.687380704 | 0.446 | 0.139 | 2.66042400180791e-64 | 7 |
| klf13 | 1.48074042251127e-68 | 0.639010017 | 0.365 | 0.097 | 3.72687556941861e-64 | 7 |
| col4a1 | 2.25690322122629e-68 | 0.543047117 | 0.268 | 0.053 | 5.68039971750445e-64 | 7 |
| mmp14a | 2.56809894430973e-68 | 0.457285269 | 0.249 | 0.047 | 6.46364823293315e-64 | 7 |
| mt-atp61 | 3.28879160287271e-68 | 0.619866092 | 0.948 | 0.58 | 8.27755958527032e-64 | 7 |
| magi3a | 6.99624569159964e-68 | 0.298198489 | 0.131 | 0.013 | 1.76088507811871e-63 | 7 |
| XLOC-040020 | 1.57664581718156e-67 | 0.267783348 | 0.105 | 0.008 | 3.96825985726427e-63 | 7 |
| htra3b | 1.95880763751654e-67 | 0.287715319 | 0.123 | 0.012 | 4.93012294286537e-63 | 7 |
| sesn3 | 5.87139213268078e-67 | 0.373492045 | 0.163 | 0.021 | 1.47777068587442e-62 | 7 |
| XLOC-042917 | 7.42346346575828e-67 | 0.270920952 | 0.113 | 0.01 | 1.8684115196967e-62 | 7 |
| zgc:114181 | 7.8344293161829e-67 | 0.252345416 | 0.115 | 0.01 | 1.97184751459007e-62 | 7 |
| sult1st6 | 1.1073558513701e-66 | 0.297572971 | 0.129 | 0.013 | 2.78710394231339e-62 | 7 |
| ddit3 | 3.7336478649091e-66 | 0.715230337 | 0.488 | 0.167 | 9.39721831118972e-62 | 7 |
| LOC566587 | 1.05272849843958e-65 | 0.491476208 | 0.234 | 0.043 | 2.64961235772258e-61 | 7 |
| dub | 1.07898904305857e-64 | 0.563301288 | 0.286 | 0.065 | 2.71570752247413e-60 | 7 |
| dedd1 | 2.8348634849187e-64 | 0.670498998 | 0.428 | 0.138 | 7.13506790519187e-60 | 7 |
| hic1 | 7.27748269528222e-64 | 0.476229795 | 0.257 | 0.052 | 1.83166961957558e-59 | 7 |
| LO018340.1 | 7.70481937105829e-64 | 0.387356564 | 0.163 | 0.022 | 1.93922598750166e-59 | 7 |

| | | | | | | |
|---|---|---|---|---|---|---|
| zgc:1627302 | 1.03324290454691e-63 | 0.737787779 | 0.782 | 0.38 | 2.60056906645412e-59 | 7 |
| neu3.2 | 1.32134249351581e-63 | 0.374688806 | 0.189 | 0.028 | 3.32568692192994e-59 | 7 |
| gadd45ba | 3.0869637253226e-63 | 0.74723887 | 0.501 | 0.181 | 7.76957900026444e-59 | 7 |
| rras2 | 4.08107701717239e-63 | 0.321700446 | 0.147 | 0.018 | 1.02716627445212e-58 | 7 |
| cald1a | 5.32212848413599e-63 | 0.569448134 | 0.349 | 0.092 | 1.33952651817219e-58 | 7 |
| dhrs3a | 1.48436992353709e-62 | 0.375700181 | 0.176 | 0.026 | 3.73601066055049e-58 | 7 |
| si:ch1073-291c23.2 | 9.30602142100185e-61 | 0.474352125 | 0.26 | 0.056 | 2.34223253145195e-56 | 7 |
| myl6 | 4.78110570633413e-60 | 0.564733904 | 0.346 | 0.094 | 1.20335649522724e-55 | 7 |
| CABZ01039304.1 | 1.58530143300243e-59 | 0.274923538 | 0.118 | 0.012 | 3.99004517672382e-55 | 7 |
| adh8a | 2.30961971073552e-59 | 0.486041474 | 0.218 | 0.04 | 5.81308184995023e-55 | 7 |
| rbp5 | 3.98528939358319e-59 | 0.597174195 | 0.331 | 0.088 | 1.00305748747095e-54 | 7 |
| sostdc1a | 7.91917888400972e-59 | 0.272551736 | 0.1 | 0.009 | 1.99317813331641e-54 | 7 |
| il15 | 8.94307313778669e-58 | 0.466854295 | 0.244 | 0.051 | 2.25088207804953e-53 | 7 |
| cxcl8b.1 | 1.39732799291456e-57 | 0.471040342 | 0.186 | 0.031 | 3.51693482536665e-53 | 7 |
| ccl19a.1 | 4.30256518419025e-57 | 0.287887505 | 0.1 | 0.009 | 1.08291263120884e-52 | 7 |
| ptges | 4.67616925118694e-57 | 0.347041569 | 0.147 | 0.02 | 1.17694503883124e-52 | 7 |
| mt-cyb | 1.03237153057817e-56 | 0.552120407 | 0.945 | 0.614 | 2.5983759053122e-52 | 7 |
| lpin1 | 1.81494215365344e-56 | 0.288071858 | 0.121 | 0.013 | 4.56802790653034e-52 | 7 |
| capn2a | 3.71061178400824e-56 | 0.434590146 | 0.231 | 0.048 | 9.33923879917035e-52 | 7 |
| mpzl1l | 2.05097953106669e-55 | 0.356954321 | 0.171 | 0.027 | 5.16211038174174e-51 | 7 |
| col6a1 | 3.02870593638314e-55 | 0.546482712 | 0.315 | 0.083 | 7.62294997128274e-51 | 7 |
| si:ch211-1a19.3 | 3.84014186733731e-55 | 0.452836255 | 0.231 | 0.048 | 9.66525306590129e-51 | 7 |
| mmp14b | 1.24394959415174e-54 | 0.454679702 | 0.239 | 0.052 | 3.13089673352052e-50 | 7 |
| kitlga | 1.86983937444946e-54 | 0.26370728 | 0.115 | 0.013 | 4.70619872155184e-50 | 7 |
| myo1ca | 2.12368448119139e-54 | 0.322095194 | 0.147 | 0.021 | 5.34510147071061e-50 | 7 |
| igf2b | 2.5291229476884e-54 | 0.474160784 | 0.244 | 0.054 | 6.36554954703694e-50 | 7 |
| mt-co21 | 2.99788223092728e-54 | 0.504333999 | 0.984 | 0.712 | 7.54536978702086e-50 | 7 |
| boc | 6.84870163446407e-54 | 0.414953131 | 0.218 | 0.044 | 1.72374971437826e-49 | 7 |
| tagln2 | 1.23287881221032e-53 | 0.611418627 | 0.478 | 0.18 | 3.10303268245215e-49 | 7 |
| cldn5a | 4.51889870434118e-53 | 0.464770877 | 0.218 | 0.045 | 1.13736161489563e-48 | 7 |
| btg2 | 5.87896599007786e-53 | 0.682279839 | 0.661 | 0.365 | 1.4796769500427e-48 | 7 |
| stom | 1.42046994299543e-51 | 0.303584313 | 0.131 | 0.018 | 3.5751807995252e-47 | 7 |
| celf21 | 4.12463500362295e-51 | 0.631214323 | 0.53 | 0.234 | 1.03812938406186e-46 | 7 |
| egfra | 4.40233769736381e-51 | 0.339473852 | 0.157 | 0.025 | 1.1080243750495e-46 | 7 |
| uap1 | 8.18634100819174e-51 | 0.363965462 | 0.192 | 0.036 | 2.06042016835178e-46 | 7 |

| | | | | | | |
|---|---|---|---|---|---|---|
| fgfr1a | 1.20819821608771e-50 | 0.288849579 | 0.131 | 0.018 | 3.04091409007115e-46 | 7 |
| loxl2a | 3.62805575899458e-50 | 0.39869061 | 0.194 | 0.038 | 9.13145353981346e-46 | 7 |
| CABZ01072614.1 | 1.32822201914263e-49 | 0.53819923 | 0.278 | 0.075 | 3.34300199998008e-45 | 7 |
| il11ra | 3.17089070574702e-49 | 0.259073036 | 0.108 | 0.012 | 7.98081481729468e-45 | 7 |
| sema3b | 1.21666665489492e-48 | 0.295252696 | 0.134 | 0.019 | 3.06222830370501e-44 | 7 |
| ctdspla | 1.11100358912937e-47 | 0.348228566 | 0.173 | 0.032 | 2.79628493347971e-43 | 7 |
| sc:d0202 | 1.87973063747249e-47 | 0.382021695 | 0.207 | 0.043 | 4.73109404145451e-43 | 7 |
| tcf12 | 1.97082993916365e-47 | 0.408053776 | 0.228 | 0.053 | 4.96038187388099e-43 | 7 |
| sptan1 | 2.86250494005805e-47 | 0.403727292 | 0.21 | 0.046 | 7.20463868363211e-43 | 7 |
| lpl | 8.01804982825469e-47 | 0.315400759 | 0.123 | 0.017 | 2.01806296127342e-42 | 7 |
| cxcl18b | 2.00251947223949e-46 | 0.557720676 | 0.336 | 0.101 | 5.04014125967957e-42 | 7 |
| cdon | 7.91724200982782e-46 | 0.420557802 | 0.223 | 0.053 | 1.99269064145356e-41 | 7 |
| dlc1 | 1.20669912218636e-45 | 0.324944611 | 0.157 | 0.028 | 3.03714102063085e-41 | 7 |
| hdac4 | 6.37765938718266e-45 | 0.328228016 | 0.155 | 0.027 | 1.60519309116e-40 | 7 |
| cdh6 | 1.86594858938596e-44 | 0.255195831 | 0.121 | 0.017 | 4.69640600462551e-40 | 7 |
| ldlrad4a | 2.67638992396457e-44 | 0.327019231 | 0.157 | 0.029 | 6.73620579962642e-40 | 7 |
| rbms2a | 5.39390015056272e-44 | 0.3036955 | 0.15 | 0.026 | 1.35759072889513e-39 | 7 |
| krt4 | 1.07181422444424e-43 | 0.349095723 | 0.184 | 0.037 | 2.69764922150371e-39 | 7 |
| paplna | 1.45178486529021e-43 | 0.297832545 | 0.131 | 0.021 | 3.65399732744892e-39 | 7 |
| si:ch211-212k18.9 | 1.17590913195004e-42 | 0.322640711 | 0.168 | 0.033 | 2.95964569420506e-38 | 7 |
| diaph2 | 1.56928990863266e-42 | 0.334619934 | 0.16 | 0.031 | 3.94974577103755e-38 | 7 |
| fxyd1 | 2.24728898674726e-42 | 0.450235333 | 0.265 | 0.076 | 5.65620165074419e-38 | 7 |
| jun | 2.35885708182455e-42 | 0.592536771 | 0.525 | 0.22 | 5.93700738924422e-38 | 7 |
| mgst1.2 | 4.29372165322662e-42 | 0.477139395 | 0.31 | 0.097 | 1.08068680290061e-37 | 7 |
| zgc:158463 | 6.65097868480428e-42 | 0.480918408 | 0.249 | 0.069 | 1.67398482517839e-37 | 7 |
| lamb1b | 6.85279010324923e-42 | 0.290789034 | 0.139 | 0.024 | 1.7247787410868e-37 | 7 |
| rdh10a | 8.15146310121611e-42 | 0.34660462 | 0.178 | 0.037 | 2.05164174794508e-37 | 7 |
| ghra | 9.64807650066773e-42 | 0.323200816 | 0.15 | 0.027 | 2.42832437445306e-37 | 7 |
| tns2a | 9.67310994679623e-42 | 0.329053505 | 0.168 | 0.034 | 2.43462504250914e-37 | 7 |
| ndfip1l | 2.15377113121887e-41 | 0.439316789 | 0.234 | 0.062 | 5.42082656016477e-37 | 7 |
| nfil3-6 | 5.18418162231117e-41 | 0.555206456 | 0.349 | 0.125 | 1.3048066725195e-36 | 7 |
| crispld2 | 5.86149919806337e-41 | 0.269668916 | 0.11 | 0.016 | 1.47528073316057e-36 | 7 |
| jam3b | 9.54702282442177e-41 | 0.280906955 | 0.139 | 0.024 | 2.40289017467872e-36 | 7 |
| tuba8l3 | 2.02283152854963e-40 | 0.361334094 | 0.189 | 0.043 | 5.09126467420657e-36 | 7 |
| crip12 | 3.11381350945547e-40 | 0.637380758 | 0.517 | 0.264 | 7.83715722194846e-36 | 7 |

| | | | | | | |
|---|---|---|---|---|---|---|
| mych | 4.92250534114963e-40 | 0.449363842 | 0.249 | 0.07 | 1.23894536931395e-35 | 7 |
| llgl1 | 5.43521138135909e-40 | 0.300283649 | 0.147 | 0.027 | 1.36798835257427e-35 | 7 |
| XLOC-000946 | 6.05975687604164e-40 | 0.356728843 | 0.181 | 0.04 | 1.52518020813092e-35 | 7 |
| ehd1b | 7.15532780797059e-40 | 0.348424085 | 0.194 | 0.044 | 1.80092445598812e-35 | 7 |
| anxa13 | 1.07912565527227e-39 | 0.366593802 | 0.194 | 0.046 | 2.71605136175478e-35 | 7 |
| sik2b | 2.54804627483753e-39 | 0.348362997 | 0.168 | 0.036 | 6.41317766913859e-35 | 7 |
| anxa2a | 2.65658596261659e-39 | 0.527565729 | 0.307 | 0.101 | 6.68636120930971e-35 | 7 |
| plekho2 | 2.75994833651732e-39 | 0.415439854 | 0.247 | 0.07 | 6.94651396818045e-35 | 7 |
| qki2 | 4.2199503402386e-39 | 0.300686801 | 0.136 | 0.024 | 1.06211930113465e-34 | 7 |
| krt15 | 7.57527670022823e-39 | 0.441228894 | 0.276 | 0.08 | 1.90662139268044e-34 | 7 |
| fbxo34 | 7.61796790554855e-39 | 0.35549544 | 0.194 | 0.045 | 1.91736634214751e-34 | 7 |
| dusp8a | 1.24099229447816e-38 | 0.30147247 | 0.136 | 0.025 | 3.12345350597208e-34 | 7 |
| ulk2 | 1.30004420471629e-38 | 0.344740485 | 0.189 | 0.044 | 3.27208125885044e-34 | 7 |
| gulp1a | 3.09184565142081e-38 | 0.302513132 | 0.147 | 0.028 | 7.78186632006103e-34 | 7 |
| mir199-3 | 1.07856209418644e-37 | 0.25457142 | 0.108 | 0.016 | 2.71463293485784e-33 | 7 |
| hsd11b2 | 1.41842281511801e-37 | 0.515441395 | 0.349 | 0.117 | 3.57002838337052e-33 | 7 |
| ccdc85b | 1.60977641251719e-37 | 0.329516582 | 0.189 | 0.045 | 4.05164625266451e-33 | 7 |
| pim1 | 4.98527430291412e-37 | 0.550981875 | 0.399 | 0.165 | 1.25474368930045e-32 | 7 |
| glis2a | 6.64006865776965e-37 | 0.299460119 | 0.131 | 0.024 | 1.67123888047404e-32 | 7 |
| b2ml1 | 7.04976275592972e-37 | 0.536208384 | 0.512 | 0.26 | 1.77435478803995e-32 | 7 |
| CR847566.1 | 7.47663465940838e-37 | 0.333119277 | 0.155 | 0.032 | 1.88179417742649e-32 | 7 |
| dcbld2 | 1.05778927303039e-36 | 0.254418825 | 0.118 | 0.02 | 2.66234982129018e-32 | 7 |
| mef2aa | 1.09787312507532e-36 | 0.381482515 | 0.231 | 0.065 | 2.76323686850208e-32 | 7 |
| timp2a | 1.44854703208593e-36 | 0.37044317 | 0.205 | 0.053 | 3.64584802505707e-32 | 7 |
| mt-nd4 | 2.71168243198188e-36 | 0.498920427 | 0.751 | 0.448 | 6.8250335130552e-32 | 7 |
| appa | 6.52377220397436e-36 | 0.407437483 | 0.244 | 0.073 | 1.64196822601831e-31 | 7 |
| flna | 9.2553372078532e-36 | 0.439991374 | 0.312 | 0.109 | 2.32947582184457e-31 | 7 |
| mt-co11 | 1.10733574358766e-35 | 0.442820307 | 0.948 | 0.649 | 2.78705333303578e-31 | 7 |
| pdlim1 | 2.86388876352375e-35 | 0.353859941 | 0.197 | 0.051 | 7.20812162891293e-31 | 7 |
| svila | 3.77919621825019e-35 | 0.295269645 | 0.134 | 0.026 | 9.5118589617139e-31 | 7 |
| TCIM (1 of many) | 5.26750382113282e-35 | 0.406380594 | 0.173 | 0.04 | 1.32577803674092e-30 | 7 |
| glula | 7.89392908297402e-35 | 0.499465814 | 0.346 | 0.13 | 1.98682301089373e-30 | 7 |
| spon1b | 8.23885453373407e-35 | 0.357872652 | 0.173 | 0.042 | 2.07363729759553e-30 | 7 |
| BX322530.1 | 2.59147598775388e-34 | 0.348548816 | 0.178 | 0.044 | 6.52248591357774e-30 | 7 |
| rbms3 | 2.88697225408979e-34 | 0.282773456 | 0.147 | 0.031 | 7.2662204663186e-30 | 7 |

| | | | | | | |
|---|---|---|---|---|---|---|
| wu:fj16a03 | 3.35828139192341e-34 | 0.378605993 | 0.192 | 0.048 | 8.45245843533203e-30 | 7 |
| spry4 | 3.40511283937793e-34 | 0.343203066 | 0.178 | 0.044 | 8.57032850543031e-30 | 7 |
| CABZ01074363.1 | 3.73728043147092e-34 | 0.268105717 | 0.134 | 0.026 | 9.40636111796917e-30 | 7 |
| appb | 4.38782351312407e-34 | 0.31365111 | 0.16 | 0.037 | 1.1043713000182e-29 | 7 |
| reep3a | 3.01457805684212e-33 | 0.265628455 | 0.126 | 0.024 | 7.58739151126592e-29 | 7 |
| arl6ip5b | 3.13826729066549e-33 | 0.386200268 | 0.273 | 0.092 | 7.89870494387596e-29 | 7 |
| cast | 1.33540546522835e-32 | 0.375098331 | 0.223 | 0.066 | 3.36108201543324e-28 | 7 |
| svild | 3.60153311176071e-32 | 0.387181932 | 0.244 | 0.077 | 9.06469868899053e-28 | 7 |
| tspan14 | 4.51535316850973e-32 | 0.433816918 | 0.318 | 0.119 | 1.13646923898221e-27 | 7 |
| fzd2 | 4.73935414350842e-32 | 0.265949746 | 0.129 | 0.026 | 1.19284804437964e-27 | 7 |
| si:ch73-237c6.1 | 4.98594040893646e-32 | 0.31009548 | 0.163 | 0.039 | 1.25491134152522e-27 | 7 |
| hopx | 5.87564482438515e-32 | 0.370389993 | 0.213 | 0.061 | 1.4788410458495e-27 | 7 |
| krt18b1 | 1.05164016247929e-31 | 0.448327113 | 0.367 | 0.145 | 2.64687312494412e-27 | 7 |
| foxp4 | 1.34050415005615e-31 | 0.348683997 | 0.181 | 0.048 | 3.37391489527632e-27 | 7 |
| cst3 | 1.58755489367126e-31 | 0.299396033 | 0.165 | 0.041 | 3.99571691188119e-27 | 7 |
| lrrc17 | 1.6229560961404e-31 | 0.293864462 | 0.144 | 0.032 | 4.08481819837577e-27 | 7 |
| cd28l | 1.66695033184604e-31 | 0.274679611 | 0.152 | 0.035 | 4.19554729022331e-27 | 7 |
| pnpla6 | 5.05145369221971e-31 | 0.252492981 | 0.11 | 0.02 | 1.27140037979478e-26 | 7 |
| mt-nd2 | 5.73753559753871e-31 | 0.474168605 | 0.669 | 0.393 | 1.44408033454452e-26 | 7 |
| chd3 | 7.18463523104233e-31 | 0.421510141 | 0.276 | 0.1 | 1.80830084130105e-26 | 7 |
| abcc5 | 1.24418868569208e-30 | 0.404491493 | 0.26 | 0.09 | 3.13149850301839e-26 | 7 |
| mknk2b1 | 1.30426210320301e-30 | 0.490997531 | 0.412 | 0.192 | 3.28269728755165e-26 | 7 |
| nrp1a | 7.36603100790996e-30 | 0.36394053 | 0.186 | 0.052 | 1.85395634438086e-25 | 7 |
| lamc1 | 1.0698515160579e-29 | 0.32546245 | 0.186 | 0.052 | 2.69270928076614e-25 | 7 |
| lamb1a | 1.17082346579298e-29 | 0.281941697 | 0.15 | 0.036 | 2.94684558105436e-25 | 7 |
| ckap4 | 1.35577877767182e-29 | 0.302125259 | 0.16 | 0.041 | 3.41235960552221e-25 | 7 |
| zmiz2 | 2.0033922626982e-29 | 0.251533647 | 0.121 | 0.025 | 5.04233798598509e-25 | 7 |
| cish1 | 2.46586871757049e-29 | 0.328271366 | 0.184 | 0.052 | 6.20634497525316e-25 | 7 |
| si:ch73-335l21.4 | 2.65620653647417e-29 | 0.565487789 | 0.344 | 0.144 | 6.68540623165183e-25 | 7 |
| tshz3b | 3.9949092999479e-29 | 0.254645011 | 0.126 | 0.027 | 1.00547872170389e-24 | 7 |
| foxo3b | 5.00938273206269e-29 | 0.364983776 | 0.205 | 0.063 | 1.26081153983286e-24 | 7 |
| si:ch211-133n4.4 | 6.53667711192261e-29 | 0.267782755 | 0.163 | 0.042 | 1.6452162622998e-24 | 7 |
| sdc4 | 1.66110866662136e-28 | 0.458474346 | 0.517 | 0.25 | 4.1808444030193e-24 | 7 |
| adam10a | 1.72300769772783e-28 | 0.368582426 | 0.255 | 0.09 | 4.33663807441118e-24 | 7 |
| ubl3a | 1.95442076573257e-28 | 0.303412718 | 0.16 | 0.042 | 4.9190816252723e-24 | 7 |

| | | | | | | |
|---|---|---|---|---|---|---|
| sntb2 | 4.25911911479412e-28 | 0.287102828 | 0.163 | 0.043 | 1.07197769000253e-23 | 7 |
| c4 | 4.55595353850507e-28 | 0.322362747 | 0.152 | 0.039 | 1.14668794610634e-23 | 7 |
| grk4 | 1.23128965620959e-27 | 0.27044376 | 0.126 | 0.028 | 3.09903293571392e-23 | 7 |
| tmod2 | 1.86209500721346e-27 | 0.298732785 | 0.163 | 0.044 | 4.68670692365555e-23 | 7 |
| gli3 | 2.21422964483236e-27 | 0.260927059 | 0.129 | 0.029 | 5.57299459307856e-23 | 7 |
| krt91 | 4.44320698024288e-27 | 0.305370965 | 0.213 | 0.065 | 1.11831076485733e-22 | 7 |
| etnk1 | 8.00874153226105e-27 | 0.286982938 | 0.134 | 0.032 | 2.01572015625478e-22 | 7 |
| XLOC-024750 | 1.16017990596083e-26 | 0.368378421 | 0.215 | 0.072 | 2.9200568053128e-22 | 7 |
| sesn11 | 1.36858715869427e-26 | 0.438608817 | 0.365 | 0.163 | 3.44459701971761e-22 | 7 |
| eif4ebp3l | 1.38719890110828e-26 | 0.457087375 | 0.423 | 0.201 | 3.49144091419943e-22 | 7 |
| prss1 | 1.46134753167837e-26 | 0.252022393 | 0.108 | 0.021 | 3.6780656024813e-22 | 7 |
| mt-nd1 | 2.85314784004799e-26 | 0.442325237 | 0.656 | 0.405 | 7.18108779861678e-22 | 7 |
| fkbp5 | 3.78014596162266e-26 | 0.412027575 | 0.289 | 0.118 | 9.51424937080808e-22 | 7 |
| ptp4a3 | 4.35285909192816e-26 | 0.303122355 | 0.171 | 0.049 | 1.0955711048474e-21 | 7 |
| mcl1b | 5.24294457893313e-26 | 0.456648838 | 0.459 | 0.237 | 1.31959672107168e-21 | 7 |
| perp | 7.70527706965994e-26 | 0.298520077 | 0.176 | 0.052 | 1.93934118566271e-21 | 7 |
| eppk1 | 8.96680356994162e-26 | 0.28829012 | 0.147 | 0.038 | 2.25685479051861e-21 | 7 |
| rtn3 | 2.40518506940612e-25 | 0.337900979 | 0.22 | 0.075 | 6.05361030118826e-21 | 7 |
| si:dkey-238c7.16 | 3.36261608577778e-25 | 0.299779451 | 0.192 | 0.059 | 8.46336842629408e-21 | 7 |
| FO082781.1 | 1.64662718634363e-24 | 0.289974732 | 0.157 | 0.045 | 4.14439596530827e-20 | 7 |
| si:dkey-9i23.16 | 2.48167908941445e-24 | 0.265129977 | 0.142 | 0.038 | 6.24613810014723e-20 | 7 |
| cebpa1 | 2.66533779893276e-24 | 0.310536377 | 0.178 | 0.055 | 6.70838870613385e-20 | 7 |
| tjp1a | 2.85661270494322e-24 | 0.277670871 | 0.157 | 0.045 | 7.18980851707159e-20 | 7 |
| pbx1a | 3.01300920626712e-24 | 0.286144765 | 0.147 | 0.04 | 7.58344287125372e-20 | 7 |
| junbb | 5.71884082894192e-24 | 0.325217609 | 0.202 | 0.07 | 1.43937504823639e-19 | 7 |
| nfic | 1.04560832621865e-23 | 0.31647016 | 0.21 | 0.073 | 2.63169159625971e-19 | 7 |
| hdlbpa | 2.02689871406024e-23 | 0.400163568 | 0.336 | 0.155 | 5.10150137341823e-19 | 7 |
| slc43a2a | 5.95383615689798e-23 | 0.377064555 | 0.249 | 0.099 | 1.49852102232965e-18 | 7 |
| f3b | 8.04125311301909e-23 | 0.393593025 | 0.252 | 0.098 | 2.02390299601577e-18 | 7 |
| ash1l | 2.03145456763746e-22 | 0.26584064 | 0.142 | 0.04 | 5.11296800128671e-18 | 7 |
| rbpjb1 | 3.87109585720844e-22 | 0.321416324 | 0.171 | 0.055 | 9.74316116300791e-18 | 7 |
| rab5aa | 4.10083932024372e-22 | 0.333374289 | 0.205 | 0.074 | 1.03214024851214e-17 | 7 |
| si:dkeyp-19e1.3 | 4.13661052987797e-22 | 0.333033958 | 0.189 | 0.065 | 1.04114350426499e-17 | 7 |
| mt-nd3 | 4.40183348387908e-22 | 0.404828565 | 0.622 | 0.39 | 1.10789746955753e-17 | 7 |
| egr1 | 6.29238627318913e-22 | 0.471170385 | 0.362 | 0.186 | 1.58373070109897e-17 | 7 |

| | | | | | | |
|---|---|---|---|---|---|---|
| fosl1a | 7.74261108220018e-22 | 0.390966192 | 0.228 | 0.089 | 1.94873778327896e-17 | 7 |
| itm2ba | 1.02113303314444e-21 | 0.411992603 | 0.467 | 0.269 | 2.57008973112124e-17 | 7 |
| dbn1 | 1.39347054727082e-21 | 0.28151481 | 0.165 | 0.053 | 3.50722602042593e-17 | 7 |
| zgc:113208 | 1.40551278637287e-21 | 0.277166712 | 0.144 | 0.043 | 3.53753513202188e-17 | 7 |
| stau2 | 8.00024130912802e-21 | 0.314671412 | 0.215 | 0.083 | 2.01358073509443e-16 | 7 |
| mrtfab | 8.17644822512006e-21 | 0.267226041 | 0.16 | 0.051 | 2.05793025378047e-16 | 7 |
| zfp36l2 | 8.50736700743606e-21 | 0.337643724 | 0.265 | 0.107 | 2.14121920210158e-16 | 7 |
| rerea | 1.09908957794014e-20 | 0.255672211 | 0.157 | 0.049 | 2.76629855871753e-16 | 7 |
| gpx4a | 4.53550127486015e-20 | 0.270117976 | 0.197 | 0.072 | 1.14154031586955e-15 | 7 |
| pdk2b | 5.41588155764482e-20 | 0.359103148 | 0.234 | 0.097 | 1.36312322924363e-15 | 7 |
| cyp3c1 | 6.69247829356727e-20 | 0.28996881 | 0.171 | 0.058 | 1.68442986170795e-15 | 7 |
| myl9b | 8.79160488137539e-20 | 0.266256753 | 0.207 | 0.077 | 2.21275903259337e-15 | 7 |
| ptprga | 1.19640523000316e-19 | 0.304052368 | 0.178 | 0.064 | 3.01123232339496e-15 | 7 |
| sulf2b | 1.2051331586467e-19 | 0.251731817 | 0.129 | 0.037 | 3.03319964699789e-15 | 7 |
| sdcbp2 | 1.36307063062227e-19 | 0.391758849 | 0.378 | 0.205 | 3.43071247021319e-15 | 7 |
| pdcd6 | 2.23550959676869e-19 | 0.277332141 | 0.194 | 0.072 | 5.62655410410711e-15 | 7 |
| calm2b | 4.25578155440219e-19 | 0.400634949 | 0.381 | 0.207 | 1.07113765942749e-14 | 7 |
| fxyd6l | 5.02281560114505e-19 | 0.321293757 | 0.26 | 0.11 | 1.2641924586522e-14 | 7 |
| plecb | 1.84607948418882e-18 | 0.278502539 | 0.147 | 0.048 | 4.64639745375484e-14 | 7 |
| csrnp1b | 5.9975004018569e-18 | 0.363497312 | 0.339 | 0.176 | 1.50951087614336e-13 | 7 |
| cd632 | 6.07243857898731e-18 | 0.365878739 | 0.538 | 0.336 | 1.52837206594532e-13 | 7 |
| rab10 | 8.25207764937996e-18 | 0.354203122 | 0.36 | 0.191 | 2.07696542357244e-13 | 7 |
| ier2b1 | 2.15980039516932e-17 | 0.372608319 | 0.394 | 0.22 | 5.43600161460166e-13 | 7 |
| sdc2 | 2.99647397746793e-17 | 0.284868115 | 0.189 | 0.074 | 7.54182535388904e-13 | 7 |
| clic21 | 6.49299693943443e-17 | 0.285794607 | 0.16 | 0.058 | 1.63422239968625e-12 | 7 |
| aoc2 | 6.5342520363771e-17 | 0.315981149 | 0.171 | 0.064 | 1.64460589503575e-12 | 7 |
| ppib | 1.14785967478473e-16 | 0.359442758 | 0.514 | 0.329 | 2.88904801546569e-12 | 7 |
| itgav | 2.08588594390182e-16 | 0.257970408 | 0.155 | 0.056 | 5.24996633220649e-12 | 7 |
| si:ch211-196l7.41 | 5.93182107946311e-16 | 0.345254529 | 0.488 | 0.31 | 1.49298004749007e-11 | 7 |
| tmem176l.1 | 8.4330131880309e-16 | 0.324796989 | 0.336 | 0.177 | 2.1225050892955e-11 | 7 |
| atp2b2 | 8.64240687543786e-16 | 0.262930486 | 0.176 | 0.07 | 2.17520738647895e-11 | 7 |
| ywhaqb | 3.75084075659628e-15 | 0.311895105 | 0.26 | 0.129 | 9.44049110027717e-11 | 7 |
| h2afx12 | 6.97096424005309e-15 | 0.326676677 | 0.352 | 0.195 | 1.75452198957896e-10 | 7 |
| fam168a | 7.28486292183902e-15 | 0.250510567 | 0.15 | 0.057 | 1.83352714879766e-10 | 7 |
| rbm39a | 7.31882114952127e-15 | 0.320149756 | 0.273 | 0.14 | 1.84207409512301e-10 | 7 |

| | | | | | | |
|---|---|---|---|---|---|---|
| tmsb4x2 | 1.0859240089055e-14 | 0.378147541 | 0.971 | 0.678 | 2.73316213801425e-10 | 7 |
| per3 | 1.8059376549064e-14 | 0.282678909 | 0.181 | 0.077 | 4.54536448363391e-10 | 7 |
| txnipa1 | 1.38743389281425e-13 | 0.326821223 | 0.354 | 0.208 | 3.49203236482417e-09 | 7 |
| ciarta | 1.99153834725179e-13 | 0.275595515 | 0.215 | 0.102 | 5.01250286619803e-09 | 7 |
| marcksl1b | 2.54224716785027e-13 | 0.31283814 | 0.234 | 0.114 | 6.39858189676236e-09 | 7 |
| pnrc21 | 7.70567959012576e-13 | 0.287055082 | 0.759 | 0.563 | 1.93944249603875e-08 | 7 |
| klf11b | 9.02269111516524e-13 | 0.254522708 | 0.176 | 0.078 | 2.27092112677594e-08 | 7 |
| rhoaa | 1.51750604983941e-12 | 0.296541822 | 0.273 | 0.152 | 3.81941097684081e-08 | 7 |
| socs3b | 1.71612455403774e-12 | 0.301849364 | 0.283 | 0.153 | 4.31931389005758e-08 | 7 |
| aldh9a1a.1 | 2.02026362718196e-12 | 0.302355455 | 0.289 | 0.162 | 5.08480152325428e-08 | 7 |
| wls | 3.23904256869281e-12 | 0.266111989 | 0.239 | 0.121 | 8.15234624114294e-08 | 7 |
| slc3a2b | 2.55187117623822e-11 | 0.270437596 | 0.223 | 0.116 | 6.42280456347398e-07 | 7 |
| tob1b | 3.02288347922238e-11 | 0.284915517 | 0.299 | 0.171 | 7.60829542885482e-07 | 7 |
| notch21 | 3.81010278752672e-11 | 0.261210463 | 0.223 | 0.118 | 9.589647705926e-07 | 7 |
| acin1a | 3.86067814395166e-11 | 0.303819429 | 0.344 | 0.214 | 9.71694082051193e-07 | 7 |
| arf2a | 6.12731278445291e-11 | 0.252467966 | 0.236 | 0.125 | 1.54218335471895e-06 | 7 |
| higd1a | 1.14545095664859e-10 | 0.27998141 | 0.457 | 0.307 | 2.88298551278883e-06 | 7 |
| zfp36l1a1 | 1.39322656641989e-10 | 0.29682428 | 0.383 | 0.246 | 3.50661194502222e-06 | 7 |
| si:dkey-7j14.6 | 4.40058235776574e-10 | 0.252928181 | 0.294 | 0.173 | 1.10758257362606e-05 | 7 |
| tcima1 | 3.05666359987358e-09 | 0.257787402 | 0.144 | 0.067 | 7.69331661452182e-05 | 7 |
| h3f3c | 3.24314125692419e-09 | 0.255058218 | 0.383 | 0.254 | 8.16266222955249e-05 | 7 |
| ier2a | 1.50211374373889e-06 | 0.255296908 | 0.354 | 0.261 | 0.037806701 | 7 |
| krt941 | 7.95861674982502e-11 | 0.390091529 | 0.243 | 0.128 | 2.00310424976346e-06 | 8 |
| icn1 | 1.19166433111641e-10 | 0.476222494 | 0.412 | 0.273 | 2.9992999549869e-06 | 8 |
| epcam5 | 6.59689954453042e-10 | 0.307376039 | 0.668 | 0.504 | 1.66037364636286e-05 | 8 |
| krt81 | 1.80588666190096e-09 | 0.340287064 | 0.415 | 0.275 | 4.54523613933851e-05 | 8 |
| si:ch211-39i22.11 | 2.55759784325099e-08 | 0.280096969 | 0.148 | 0.071 | 0.000643722 | 8 |
| BX649337.15 | 3.98087942764146e-08 | 0.292891937 | 0.609 | 0.436 | 0.001001948 | 8 |
| gtf3ab5 | 7.70129639869203e-08 | 0.289945437 | 0.465 | 0.319 | 0.001938339 | 8 |
| cldnd5 | 8.36867078521613e-08 | 0.275380485 | 0.751 | 0.603 | 0.002106311 | 8 |
| acp5a5 | 2.15107616943157e-07 | 0.264538181 | 0.68 | 0.539 | 0.005414044 | 8 |
| si:ch211-5k11.84 | 4.83718318466757e-07 | 0.267844378 | 0.489 | 0.358 | 0.012174706 | 8 |
| hbba14 | 5.57677808399889e-07 | 0.275641827 | 0.492 | 0.355 | 0.014036193 | 8 |
| CR790368.15 | 5.89640958147661e-07 | 0.255903891 | 0.643 | 0.52 | 0.014840673 | 8 |
| h1m5 | 8.29126030581296e-07 | 0.258840374 | 0.732 | 0.601 | 0.020868273 | 8 |

| | | | | | | |
|---|---|---|---|---|---|---|
| si:dkey-241l7.55 | 9.50539541536134e-07 | 0.257901366 | 0.726 | 0.583 | 0.02392413 | 8 |
| zp3.25 | 1.49279404577455e-06 | 0.251886223 | 0.686 | 0.557 | 0.037572133 | 8 |
| zp2.55 | 2.27944014189857e-06 | 0.261324193 | 0.708 | 0.563 | 0.057371229 | 8 |
| zp35 | 2.72918441949788e-06 | 0.263649829 | 0.748 | 0.583 | 0.068690843 | 8 |
| h2af1al5 | 2.77029939930574e-06 | 0.254726538 | 0.729 | 0.592 | 0.069725666 | 8 |
| hbaa14 | 3.06927698926485e-06 | 0.266475923 | 0.397 | 0.281 | 0.077250633 | 8 |
| zgc:1751355 | 3.34036159881412e-06 | 0.25406712 | 0.612 | 0.479 | 0.084073561 | 8 |
| zp2.15 | 5.18408646545096e-06 | 0.250066828 | 0.671 | 0.534 | 0.130478272 | 8 |
| si:ch73-189n23.15 | 6.72811044848986e-06 | 0.251921404 | 0.658 | 0.524 | 0.169339812 | 8 |
| cxcl18b1 | 0 | 1.724643582 | 0.83 | 0.089 | 0 | 9 |
| tagln | 0 | 1.659935085 | 0.83 | 0.043 | 0 | 9 |
| rbp41 | 0 | 1.575477923 | 0.92 | 0.108 | 0 | 9 |
| hsd11b21 | 0 | 1.53302498 | 0.92 | 0.104 | 0 | 9 |
| mmp16a | 0 | 1.500483492 | 0.84 | 0.05 | 0 | 9 |
| ccl25b1 | 0 | 1.471232722 | 0.694 | 0.061 | 0 | 9 |
| fn1b | 0 | 1.442972054 | 0.865 | 0.068 | 0 | 9 |
| wu:fj16a031 | 0 | 1.39368333 | 0.667 | 0.036 | 0 | 9 |
| nid2a | 0 | 1.321155558 | 0.795 | 0.035 | 0 | 9 |
| col6a11 | 0 | 1.296898407 | 0.84 | 0.07 | 0 | 9 |
| aoc21 | 0 | 1.220109224 | 0.656 | 0.052 | 0 | 9 |
| col4a11 | 0 | 1.205756673 | 0.753 | 0.041 | 0 | 9 |
| cald1a1 | 0 | 1.159274387 | 0.767 | 0.082 | 0 | 9 |
| mfap21 | 0 | 1.124812178 | 0.76 | 0.08 | 0 | 9 |
| si:ch1073-291c23.21 | 0 | 1.074446915 | 0.708 | 0.045 | 0 | 9 |
| cpn1 | 0 | 1.069444494 | 0.674 | 0.04 | 0 | 9 |
| cd82a | 0 | 0.9934 | 0.639 | 0.042 | 0 | 9 |
| tmem1761 | 0 | 0.9830279 | 0.656 | 0.052 | 0 | 9 |
| inhbaa | 0 | 0.922269628 | 0.503 | 0.029 | 0 | 9 |
| si:dkey-23i12.7 | 0 | 0.901102854 | 0.458 | 0.021 | 0 | 9 |
| adamts15a1 | 0 | 0.860729502 | 0.538 | 0.036 | 0 | 9 |
| dkk3b | 0 | 0.834510018 | 0.497 | 0.03 | 0 | 9 |
| rnd1b | 0 | 0.830849902 | 0.438 | 0.013 | 0 | 9 |
| cd248a | 0 | 0.813566583 | 0.503 | 0.022 | 0 | 9 |
| mdkb | 0 | 0.709444133 | 0.344 | 0.01 | 0 | 9 |
| cyp1c1 | 0 | 0.695824429 | 0.323 | 0.008 | 0 | 9 |

| | | | | | | |
|---|---|---|---|---|---|---|
| fhl2a | 0 | 0.680831921 | 0.368 | 0.008 | 0 | 9 |
| si:ch211-270g19.5 | 0 | 0.659562598 | 0.351 | 0.014 | 0 | 9 |
| fank1 | 0 | 0.647043552 | 0.375 | 0.011 | 0 | 9 |
| ccdc3b | 0 | 0.61942223 | 0.375 | 0.016 | 0 | 9 |
| si:ch211-106j24.1 | 0 | 0.563638503 | 0.326 | 0.01 | 0 | 9 |
| mmrn2a | 0 | 0.511435992 | 0.292 | 0.008 | 0 | 9 |
| myocd | 0 | 0.440183724 | 0.212 | 0.003 | 0 | 9 |
| rxfp3.3b | 0 | 0.429411739 | 0.208 | 0.002 | 0 | 9 |
| rbp51 | 2.3580823160261e-307 | 1.337191591 | 0.719 | 0.079 | 5.93505738120609e-303 | 9 |
| myl9b1 | 9.20745756860079e-300 | 1.054207997 | 0.663 | 0.065 | 2.31742499544113e-295 | 9 |
| angptl2b | 5.44197567822243e-298 | 0.37834915 | 0.222 | 0.004 | 1.3696908584518e-293 | 9 |
| loxl2a1 | 7.75599619995877e-298 | 0.792553429 | 0.483 | 0.031 | 1.95210668356762e-293 | 9 |
| selenbp1 | 8.37289536026593e-295 | 0.942089981 | 0.618 | 0.056 | 2.10737403322533e-290 | 9 |
| cyp2ad31 | 1.72383359377593e-293 | 0.871681284 | 0.535 | 0.04 | 4.33871677217464e-289 | 9 |
| dcn1 | 1.59609270895868e-291 | 1.183480465 | 0.84 | 0.102 | 4.01720573917809e-287 | 9 |
| slco2b1 | 9.07354632686059e-286 | 0.964763522 | 0.594 | 0.052 | 2.28372087500754e-281 | 9 |
| col6a4a | 9.9054340014256e-285 | 0.839988703 | 0.5 | 0.035 | 2.49309868381881e-280 | 9 |
| nrp1a1 | 1.52842970194353e-282 | 0.84102638 | 0.545 | 0.044 | 3.84690471682167e-278 | 9 |
| krt18a.1 | 7.31539841135718e-282 | 1.350111061 | 0.819 | 0.11 | 1.84121262615449e-277 | 9 |
| csrp1b | 1.19042619912705e-281 | 0.551151554 | 0.188 | 0.003 | 2.99618370058288e-277 | 9 |
| acta2 | 2.55984606494008e-281 | 0.930320338 | 0.271 | 0.008 | 6.4428765608477e-277 | 9 |
| asic1b | 1.31269081869854e-277 | 0.463377262 | 0.257 | 0.007 | 3.30391152158236e-273 | 9 |
| c61 | 7.92670475955069e-277 | 1.003135552 | 0.733 | 0.079 | 1.99507232093131e-272 | 9 |
| lrrc171 | 2.79481937044986e-276 | 0.679202315 | 0.424 | 0.025 | 7.03428087348524e-272 | 9 |
| col1a21 | 1.65843094426787e-275 | 1.291426563 | 0.938 | 0.136 | 4.17410484362781e-271 | 9 |
| zfp36l1b1 | 6.24293462891136e-273 | 1.303337428 | 0.802 | 0.114 | 1.5712842167507e-268 | 9 |
| prss23 | 6.75550985466034e-272 | 0.66630647 | 0.396 | 0.022 | 1.70029427531946e-267 | 9 |
| atp1a3b | 7.51555888020684e-272 | 0.647410156 | 0.385 | 0.021 | 1.89159101455926e-267 | 9 |
| aebp1 | 1.21369127453022e-270 | 0.700594904 | 0.444 | 0.029 | 3.05473956886511e-266 | 9 |
| pdgfra1 | 4.7583023679624e-266 | 0.805809637 | 0.493 | 0.037 | 1.19761712299246e-261 | 9 |
| si:ch1073-459j12.11 | 1.36304190428681e-264 | 0.881269813 | 0.583 | 0.054 | 3.43064016889948e-260 | 9 |
| col18a1a | 9.70265093079487e-264 | 0.873209179 | 0.451 | 0.032 | 2.44206021277176e-259 | 9 |
| col6a31 | 1.43791019589501e-261 | 1.006400273 | 0.733 | 0.085 | 3.61907617204815e-257 | 9 |
| ntn1a | 6.72145236381985e-259 | 0.435547018 | 0.24 | 0.007 | 1.69172234544982e-254 | 9 |
| cyp1c2 | 9.3496512370577e-256 | 0.564605313 | 0.26 | 0.009 | 2.35321371985505e-251 | 9 |

| | | | | | | |
|---|---|---|---|---|---|---|
| cygb1 | 3.12482137799919e-253 | 0.953819913 | 0.507 | 0.043 | 7.86486292628617e-249 | 9 |
| col1a1a1 | 1.04456770812588e-252 | 1.119041895 | 0.875 | 0.122 | 2.62907246458202e-248 | 9 |
| igf2b1 | 1.27688175662781e-251 | 0.901406845 | 0.531 | 0.048 | 3.21378369325653e-247 | 9 |
| nid1b | 3.35067663201103e-250 | 0.847561491 | 0.517 | 0.044 | 8.43331801510857e-246 | 9 |
| mylkb | 6.81477616452758e-248 | 0.374790498 | 0.16 | 0.002 | 1.71521101284995e-243 | 9 |
| tenm3 | 3.35870974738769e-245 | 0.726657681 | 0.462 | 0.035 | 8.45353656320008e-241 | 9 |
| zgc:158343 | 5.90222838074433e-245 | 1.504666444 | 0.83 | 0.15 | 1.48553186114954e-240 | 9 |
| col6a21 | 6.06878560745922e-245 | 0.816097548 | 0.559 | 0.053 | 1.52745264954141e-240 | 9 |
| col5a11 | 2.03952222554427e-244 | 1.008556126 | 0.757 | 0.097 | 5.13327348947238e-240 | 9 |
| lama41 | 6.76666144980989e-238 | 0.695608539 | 0.406 | 0.028 | 1.70310102030265e-233 | 9 |
| si:ch73-46j18.51 | 9.77755925050989e-236 | 1.201362655 | 0.896 | 0.158 | 2.46091388776083e-231 | 9 |
| lamc11 | 1.97748676663849e-234 | 0.754867768 | 0.5 | 0.044 | 4.97713644295241e-230 | 9 |
| pmp22a1 | 2.24706222497011e-233 | 1.101173016 | 0.83 | 0.123 | 5.65563091402727e-229 | 9 |
| cdon1 | 1.97849268408504e-229 | 0.765508541 | 0.503 | 0.047 | 4.97966823657365e-225 | 9 |
| cd81a1 | 4.42156068040797e-229 | 1.280979739 | 0.92 | 0.19 | 1.11286260765188e-224 | 9 |
| socs3b1 | 6.59693867464898e-228 | 1.185504267 | 0.816 | 0.139 | 1.6603834950224e-223 | 9 |
| col1a1b1 | 4.46371728375463e-227 | 1.067933297 | 0.858 | 0.13 | 1.1234730031482e-222 | 9 |
| spon1b1 | 3.17722982068078e-226 | 0.687625837 | 0.441 | 0.035 | 7.99676973567145e-222 | 9 |
| myl9a | 9.89433930394553e-226 | 0.699030686 | 0.365 | 0.023 | 2.49030625941005e-221 | 9 |
| cyp3c11 | 1.58037924063636e-224 | 0.784179257 | 0.514 | 0.05 | 3.97765651075766e-220 | 9 |
| col12a1b | 1.28263268091726e-223 | 0.648733555 | 0.368 | 0.024 | 3.22825819460065e-219 | 9 |
| sfrp1b | 4.87968582444227e-223 | 0.459171089 | 0.247 | 0.009 | 1.22816812515388e-218 | 9 |
| irf8 | 4.17346311255793e-217 | 0.902303908 | 0.434 | 0.037 | 1.0504189307997e-212 | 9 |
| zgc:1951731 | 9.14945088524178e-216 | 0.893810209 | 0.628 | 0.078 | 2.3028252933065e-211 | 9 |
| vcam1b | 2.52951948779216e-215 | 0.582999189 | 0.302 | 0.016 | 6.36654759882408e-211 | 9 |
| mdka1 | 4.28444046210435e-215 | 0.956667646 | 0.639 | 0.08 | 1.07835081990704e-210 | 9 |
| rgs5b | 3.10964366716266e-213 | 1.175810231 | 0.552 | 0.065 | 7.8266621458817e-209 | 9 |
| cxcl12a1 | 4.28329035816883e-208 | 1.011063918 | 0.594 | 0.07 | 1.07806135024751e-203 | 9 |
| gstm.3 | 3.81676116432824e-204 | 1.159567745 | 0.632 | 0.088 | 9.60640617449775e-200 | 9 |
| snx17 | 2.07425848234849e-201 | 0.734530289 | 0.479 | 0.048 | 5.22070117422292e-197 | 9 |
| socs3a1 | 4.52013464016889e-201 | 1.366305192 | 0.997 | 0.282 | 1.13767268758411e-196 | 9 |
| fosl1a1 | 1.24997522747413e-200 | 0.850467939 | 0.615 | 0.079 | 3.14606265002965e-196 | 9 |
| vegfaa | 3.28108134624311e-198 | 0.443211831 | 0.264 | 0.013 | 8.25815364035929e-194 | 9 |
| cfd | 9.6680888180677e-196 | 0.996778828 | 0.51 | 0.057 | 2.43336127461946e-191 | 9 |
| si:ch211-160j14.3 | 5.25366030980057e-194 | 0.331804625 | 0.174 | 0.005 | 1.32229376337371e-189 | 9 |

| | | | | | | |
|---|---|---|---|---|---|---|
| tpm4a | 1.49826278008692e-193 | 0.909774734 | 0.583 | 0.077 | 3.77097759120077e-189 | 9 |
| robo2 | 7.08506579356654e-193 | 0.671510313 | 0.42 | 0.038 | 1.78324020958276e-188 | 9 |
| fstl1b1 | 8.48140109512559e-193 | 0.926145505 | 0.722 | 0.11 | 2.13468384163216e-188 | 9 |
| TCIM (1 of many)1 | 2.42279763222654e-192 | 0.947490112 | 0.399 | 0.035 | 6.09793936055098e-188 | 9 |
| CABZ01068367.1 | 2.8744980660855e-192 | 0.440689691 | 0.264 | 0.014 | 7.2348241825306e-188 | 9 |
| sparc1 | 2.94859294068824e-192 | 1.015891263 | 0.833 | 0.142 | 7.42131357241822e-188 | 9 |
| CU459094.2 | 4.41993878984119e-192 | 0.35849173 | 0.198 | 0.007 | 1.11245439401513e-187 | 9 |
| cdh2 | 6.64248991650796e-192 | 0.709757847 | 0.458 | 0.046 | 1.67184828708589e-187 | 9 |
| si:dkey-9i23.161 | 2.43069987810075e-191 | 0.588124154 | 0.385 | 0.032 | 6.11782852319177e-187 | 9 |
| jun1 | 7.41270699589918e-191 | 1.244257049 | 0.931 | 0.211 | 1.86570422379787e-186 | 9 |
| col4a2 | 8.78363799176537e-189 | 0.354294494 | 0.212 | 0.008 | 2.21075384614743e-184 | 9 |
| olfml3b | 6.16311343440479e-187 | 0.65274815 | 0.406 | 0.037 | 1.55119402030534e-182 | 9 |
| si:ch211-212k18.91 | 5.12924645355825e-186 | 0.549272533 | 0.365 | 0.029 | 1.29098003989607e-181 | 9 |
| cnn1b | 1.43484492730418e-185 | 0.310331441 | 0.139 | 0.003 | 3.61136119753189e-181 | 9 |
| bag3 | 2.42736337021996e-184 | 0.616005544 | 0.358 | 0.029 | 6.10943086650662e-180 | 9 |
| mxra8a | 1.34903934393444e-183 | 0.620028202 | 0.406 | 0.037 | 3.39539712474859e-179 | 9 |
| eva1ba1 | 9.42851182374365e-183 | 0.72246138 | 0.493 | 0.055 | 2.37306214091804e-178 | 9 |
| im:7152348 | 6.21630307672627e-182 | 0.44599763 | 0.233 | 0.011 | 1.56458132138123e-177 | 9 |
| spry41 | 6.27649044422374e-182 | 0.726291261 | 0.406 | 0.038 | 1.57972987990667e-177 | 9 |
| gadd45ba1 | 4.61870074357315e-181 | 1.140684585 | 0.83 | 0.175 | 1.16248079014993e-176 | 9 |
| si:dkey-226m8.10 | 1.00139310810431e-180 | 0.340520096 | 0.191 | 0.007 | 2.52040631378774e-176 | 9 |
| col5a2a1 | 1.70309546157219e-180 | 0.768462016 | 0.576 | 0.074 | 4.28652096723103e-176 | 9 |
| mgst1.1 | 4.74935699838116e-179 | 0.58772144 | 0.378 | 0.033 | 1.19536566292255e-174 | 9 |
| BX088707.3 | 6.57670878386645e-179 | 0.367269899 | 0.153 | 0.004 | 1.65529183381135e-174 | 9 |
| cyp27c1 | 1.65903907711969e-178 | 0.357128635 | 0.208 | 0.009 | 4.17563545320255e-174 | 9 |
| rdh10a1 | 3.58842441400002e-176 | 0.636132291 | 0.375 | 0.033 | 9.03170540759665e-172 | 9 |
| krt82 | 6.40174694527786e-176 | 1.22304223 | 0.972 | 0.26 | 1.61125568865698e-171 | 9 |
| si:dkey-261h17.11 | 2.09874541048766e-174 | 0.834816175 | 0.587 | 0.079 | 5.28233232365638e-170 | 9 |
| cebpd1 | 7.26819944617721e-173 | 0.954684468 | 0.694 | 0.116 | 1.82933311860834e-168 | 9 |
| nrp2a | 1.84210901338659e-172 | 0.403681232 | 0.24 | 0.013 | 4.63640417579271e-168 | 9 |
| zgc:1627303 | 2.37627346044068e-171 | 1.236144189 | 0.99 | 0.378 | 5.98084267258316e-167 | 9 |
| c1qtnf5 | 7.51538263656016e-170 | 0.479441725 | 0.288 | 0.019 | 1.89154665579583e-165 | 9 |
| sdc41 | 1.1138737745261e-168 | 1.160137376 | 0.899 | 0.242 | 2.80350890310475e-164 | 9 |
| pdgfrb1 | 4.19547301636463e-167 | 0.664859223 | 0.41 | 0.042 | 1.05595860348881e-162 | 9 |
| zgc:1527911 | 5.70126950569886e-166 | 0.841035097 | 0.479 | 0.059 | 1.43495252188935e-161 | 9 |

| | | | | | | |
|---|---|---|---|---|---|---|
| kif26ba | 1.60553169255892e-165 | 0.317422514 | 0.17 | 0.006 | 4.04096271700155e-161 | 9 |
| si:ch211-212k18.81 | 2.25009310360394e-164 | 0.551035392 | 0.375 | 0.035 | 5.66325933246075e-160 | 9 |
| f3b1 | 7.93795003808826e-162 | 0.919462844 | 0.587 | 0.09 | 1.99790264508643e-157 | 9 |
| lxn1 | 1.22793911573267e-161 | 0.959935998 | 0.708 | 0.135 | 3.09059996038755e-157 | 9 |
| junba1 | 2.68526910581872e-161 | 1.177841766 | 0.993 | 0.293 | 6.75855381243514e-157 | 9 |
| pcdh18b | 1.68060006718861e-155 | 0.374281958 | 0.219 | 0.012 | 4.22990230910702e-151 | 9 |
| cbx7a | 8.97237411776646e-154 | 0.685765158 | 0.403 | 0.045 | 2.25825684170064e-149 | 9 |
| svild1 | 1.40957968269934e-153 | 0.720086567 | 0.514 | 0.071 | 3.54777110338598e-149 | 9 |
| LOC103909099 | 2.0039510000423e-153 | 0.261945323 | 0.153 | 0.005 | 5.04374427200646e-149 | 9 |
| fosab1 | 1.37710545210429e-152 | 1.232522327 | 0.983 | 0.334 | 3.46603671240129e-148 | 9 |
| CABZ01068358.1 | 2.58936278036118e-152 | 0.755968566 | 0.444 | 0.056 | 6.51716718189106e-148 | 9 |
| pbx1a1 | 1.55375337606933e-151 | 0.585343924 | 0.358 | 0.035 | 3.91064187222889e-147 | 9 |
| lamb1a1 | 3.98514884723048e-151 | 0.525882796 | 0.34 | 0.031 | 1.00302211335944e-146 | 9 |
| tmem47 | 1.20407905480305e-149 | 0.50223828 | 0.33 | 0.03 | 3.03054657303379e-145 | 9 |
| pim11 | 1.8701411354461e-149 | 0.994227075 | 0.736 | 0.158 | 4.70695822380429e-145 | 9 |
| fosb | 5.65233106715784e-149 | 0.994917814 | 0.632 | 0.114 | 1.42263520629296e-144 | 9 |
| gadd45ab | 1.71750535039434e-148 | 0.726101384 | 0.472 | 0.063 | 4.3227892164075e-144 | 9 |
| nme3 | 8.53667004144238e-148 | 0.658821289 | 0.483 | 0.066 | 2.14859448273063e-143 | 9 |
| cyp2p61 | 4.75856695937409e-147 | 0.453131355 | 0.264 | 0.019 | 1.19768371800486e-142 | 9 |
| si:ch73-86n18.1 | 2.36958841065819e-146 | 0.364151609 | 0.205 | 0.011 | 5.96401707078561e-142 | 9 |
| f10 | 2.76916774283185e-146 | 0.559665132 | 0.382 | 0.041 | 6.96971829193349e-142 | 9 |
| si:ch211-178n15.1 | 2.84955848903414e-145 | 0.502008026 | 0.295 | 0.025 | 7.17205376105004e-141 | 9 |
| cldn11a | 5.49982044008207e-145 | 0.710352032 | 0.434 | 0.054 | 1.38424980656426e-140 | 9 |
| igfbp7 | 9.58026649329977e-145 | 0.459731441 | 0.274 | 0.021 | 2.41125727369862e-140 | 9 |
| cplx2l | 3.22514025429862e-144 | 0.343887147 | 0.219 | 0.013 | 8.11735550604419e-140 | 9 |
| rasl11a | 1.03259596226554e-143 | 0.306825327 | 0.181 | 0.008 | 2.59894077742614e-139 | 9 |
| si:ch73-335l21.41 | 8.32488889510136e-143 | 1.133538959 | 0.663 | 0.136 | 2.09529128600806e-138 | 9 |
| MMP23B | 4.03339239861689e-142 | 0.355386687 | 0.222 | 0.014 | 1.01516453280789e-137 | 9 |
| anos1a | 9.17018968508076e-142 | 0.354976473 | 0.205 | 0.011 | 2.30804504183798e-137 | 9 |
| gchfr | 2.4591886008906e-141 | 0.588007456 | 0.323 | 0.031 | 6.18953178958156e-137 | 9 |
| gstt1b | 2.28019318327771e-139 | 0.632223285 | 0.465 | 0.064 | 5.73901822299167e-135 | 9 |
| si:ch211-191i18.2 | 3.50004772548836e-139 | 0.403806166 | 0.236 | 0.016 | 8.80927012028165e-135 | 9 |
| cpox | 3.63279119417266e-138 | 0.637676573 | 0.375 | 0.043 | 9.14337215661317e-134 | 9 |
| bhmt | 1.13075770285562e-136 | 0.56890653 | 0.354 | 0.038 | 2.84600406231732e-132 | 9 |
| sema4ba1 | 4.0028830272528e-136 | 0.531784012 | 0.361 | 0.039 | 1.00748562912926e-131 | 9 |

| | | | | | | |
|---|---|---|---|---|---|---|
| il13ra1 | 6.49524367704338e-136 | 0.468392699 | 0.288 | 0.025 | 1.63478788107505e-131 | 9 |
| chd31 | 8.10238475933599e-136 | 0.705430035 | 0.559 | 0.094 | 2.03928922007728e-131 | 9 |
| csrnp1b1 | 1.56477838919692e-135 | 0.894189065 | 0.736 | 0.166 | 3.93839072776973e-131 | 9 |
| lpar11 | 3.35954608632199e-135 | 0.5838523 | 0.434 | 0.056 | 8.45564154466382e-131 | 9 |
| cpxm1a | 3.07489074084371e-134 | 0.347188882 | 0.184 | 0.009 | 7.73919250562954e-130 | 9 |
| mafba | 4.36363187005703e-134 | 0.651923807 | 0.451 | 0.064 | 1.09828250537465e-129 | 9 |
| nog1 | 1.18562362280498e-133 | 0.273125954 | 0.139 | 0.005 | 2.98409609623786e-129 | 9 |
| esr2a | 1.54485411610442e-133 | 0.46500059 | 0.292 | 0.026 | 3.88824332482321e-129 | 9 |
| lrp1ab1 | 2.46318256129567e-133 | 0.634938106 | 0.448 | 0.061 | 6.19958418852508e-129 | 9 |
| s100v21 | 7.01253618434727e-133 | 0.723082966 | 0.618 | 0.11 | 1.76498523223836e-128 | 9 |
| aldh9a1a.11 | 7.71818536659341e-133 | 0.842704468 | 0.698 | 0.152 | 1.94259007491789e-128 | 9 |
| ms4a17a.11 | 2.29915416452873e-132 | 0.490134646 | 0.319 | 0.032 | 5.78674111670237e-128 | 9 |
| boc1 | 7.39431937717858e-132 | 0.544692655 | 0.365 | 0.041 | 1.86107624404208e-127 | 9 |
| caln2 | 1.20884807469344e-131 | 0.283723811 | 0.17 | 0.008 | 3.04254971919593e-127 | 9 |
| pbx1b | 5.85719626374861e-131 | 0.422226455 | 0.267 | 0.022 | 1.47419772762289e-126 | 9 |
| aldh1a2 | 9.22531751568926e-131 | 0.68229736 | 0.479 | 0.069 | 2.32192016552383e-126 | 9 |
| col16a1 | 3.74597133044431e-130 | 0.451158611 | 0.295 | 0.027 | 9.42823524159528e-126 | 9 |
| tpm4b | 4.85737247830696e-130 | 0.324564031 | 0.184 | 0.01 | 1.22255207906508e-125 | 9 |
| ephb3 | 3.07329418016543e-129 | 0.498332833 | 0.33 | 0.035 | 7.73517412205837e-125 | 9 |
| mmp2 | 7.23195924009888e-129 | 0.657894048 | 0.517 | 0.082 | 1.82021182114049e-124 | 9 |
| cnn3a | 7.54317808312189e-129 | 0.562458804 | 0.389 | 0.049 | 1.89854249174095e-124 | 9 |
| si:dkey-188i13.101 | 2.22914792066609e-128 | 0.706447226 | 0.545 | 0.094 | 5.61054240152447e-124 | 9 |
| ier2a1 | 2.35173196586636e-128 | 0.999718874 | 0.861 | 0.248 | 5.91907418488905e-124 | 9 |
| gsta.1 | 4.73720950828587e-128 | 0.502776675 | 0.33 | 0.035 | 1.19230826114047e-123 | 9 |
| adh8a1 | 1.14461583276612e-127 | 0.70881727 | 0.34 | 0.038 | 2.88088358948905e-123 | 9 |
| si:dkey-7j14.61 | 1.74629616778351e-127 | 0.826827334 | 0.722 | 0.162 | 4.39525282469431e-123 | 9 |
| zgc:112437 | 3.34574524664275e-126 | 0.358860259 | 0.205 | 0.013 | 8.42090621127514e-122 | 9 |
| jdp2b | 1.08621997457298e-125 | 1.026832819 | 0.684 | 0.158 | 2.73390705400275e-121 | 9 |
| CABZ01092746.11 | 2.90949213867163e-125 | 0.626995223 | 0.431 | 0.059 | 7.32290076382261e-121 | 9 |
| mmp14a1 | 2.72791920996864e-124 | 0.546554528 | 0.368 | 0.045 | 6.86589985957007e-120 | 9 |
| h2afy2 | 4.58197332349172e-124 | 0.654156565 | 0.514 | 0.088 | 1.15323686578963e-119 | 9 |
| nr2f1a | 6.13996912476182e-124 | 0.522649476 | 0.34 | 0.039 | 1.5453688290113e-119 | 9 |
| flna1 | 8.25693255688028e-124 | 0.761970069 | 0.559 | 0.104 | 2.0781873552412e-119 | 9 |
| cox4i2 | 4.77214783582837e-122 | 0.449853124 | 0.285 | 0.027 | 1.20110188879964e-117 | 9 |
| c1qtnf6a | 8.70556662886829e-122 | 0.362213427 | 0.215 | 0.015 | 2.19110406481986e-117 | 9 |

| | | | | | | |
|---|---|---|---|---|---|---|
| pik3r2 | 3.52243788005711e-121 | 0.570655462 | 0.392 | 0.053 | 8.86562390031573e-117 | 9 |
| c4b | 6.49388424444833e-121 | 0.427097802 | 0.215 | 0.015 | 1.6344457254852e-116 | 9 |
| tshz3b1 | 1.46219033930452e-120 | 0.454933895 | 0.264 | 0.024 | 3.68018686499556e-116 | 9 |
| nfia1 | 1.60301810669463e-119 | 0.621587342 | 0.458 | 0.071 | 4.03463627273972e-115 | 9 |
| nfil3-61 | 1.6933242374162e-119 | 0.812151715 | 0.594 | 0.12 | 4.26192777315284e-115 | 9 |
| s100a10b1 | 3.62017424627698e-119 | 0.955485206 | 0.903 | 0.24 | 9.11161656045454e-115 | 9 |
| zeb2a | 4.69705361529114e-119 | 0.546636482 | 0.33 | 0.038 | 1.18220142443263e-114 | 9 |
| col15a1a | 5.72671247827992e-119 | 0.345872022 | 0.226 | 0.017 | 1.44135626365827e-114 | 9 |
| gpc1b | 1.58708364299892e-118 | 0.4466463 | 0.288 | 0.029 | 3.99453082106397e-114 | 9 |
| CABZ01068366.1 | 6.2020289129394e-117 | 0.412492476 | 0.264 | 0.024 | 1.56098865709772e-112 | 9 |
| cldn5a1 | 1.60505623187781e-116 | 0.589594086 | 0.347 | 0.043 | 4.03976603001325e-112 | 9 |
| marcksl1a1 | 6.45178284599891e-116 | 0.872416716 | 0.792 | 0.202 | 1.62384922450947e-111 | 9 |
| xirp2a | 1.06677038302263e-115 | 0.388739014 | 0.243 | 0.021 | 2.68495437702966e-111 | 9 |
| dclk1a | 4.42269455780994e-115 | 0.463108902 | 0.302 | 0.033 | 1.11314799325518e-110 | 9 |
| atf3 | 2.29194769874105e-114 | 0.794843069 | 0.476 | 0.082 | 5.76860316296134e-110 | 9 |
| myl61 | 3.46193999327468e-114 | 0.810485159 | 0.5 | 0.092 | 8.71335676907303e-110 | 9 |
| emilin1a | 5.19073756808521e-114 | 0.35823908 | 0.208 | 0.015 | 1.30645673851137e-109 | 9 |
| colec121 | 2.16396458807842e-113 | 0.537863155 | 0.361 | 0.047 | 5.44648247173458e-109 | 9 |
| tnfsf12 | 2.18180221920139e-113 | 0.374287536 | 0.226 | 0.018 | 5.49137800550798e-109 | 9 |
| mtss1la | 2.7033544163599e-113 | 0.380993403 | 0.233 | 0.02 | 6.80407273053622e-109 | 9 |
| crim1 | 5.57966321905797e-113 | 0.458695587 | 0.288 | 0.031 | 1.4043454356047e-108 | 9 |
| gas1a | 6.14140807411305e-113 | 0.48497597 | 0.316 | 0.037 | 1.54573099817351e-108 | 9 |
| il151 | 1.26735987471632e-112 | 0.530613251 | 0.372 | 0.049 | 3.18981806867352e-108 | 9 |
| ifi451 | 2.07669065971907e-112 | 0.970321437 | 0.562 | 0.119 | 5.22682272144693e-108 | 9 |
| mrc2 | 1.68015190172303e-111 | 0.410766906 | 0.236 | 0.021 | 4.22877432144669e-107 | 9 |
| rgl1 | 1.03494309424629e-110 | 0.446729102 | 0.285 | 0.03 | 2.60484827390848e-106 | 9 |
| lgals9l3 | 1.0821657400045e-110 | 0.519547942 | 0.392 | 0.056 | 2.72370295101734e-106 | 9 |
| tns2a1 | 3.5878822620061e-110 | 0.416029006 | 0.292 | 0.032 | 9.03034086524315e-106 | 9 |
| fstl3 | 1.84253828426241e-109 | 0.40291261 | 0.229 | 0.02 | 4.63748460766005e-105 | 9 |
| anxa1a1 | 3.40329968010861e-108 | 0.796867441 | 0.618 | 0.132 | 8.56576496486536e-104 | 9 |
| spry21 | 4.27881353491226e-108 | 0.53266757 | 0.385 | 0.056 | 1.07693457860207e-103 | 9 |
| tmem176l.11 | 6.23388818049697e-108 | 0.779151556 | 0.698 | 0.169 | 1.56900731614928e-103 | 9 |
| nfic1 | 1.73516935179972e-107 | 0.575191756 | 0.427 | 0.068 | 4.3672477415447e-103 | 9 |
| sh3gl1b | 6.03644418553202e-106 | 0.406700334 | 0.281 | 0.031 | 1.51931263705655e-101 | 9 |
| slc7a11 | 1.03323508324652e-105 | 0.434968651 | 0.288 | 0.032 | 2.60054938102317e-101 | 9 |

| | | | | | | |
|---|---|---|---|---|---|---|
| ar | 1.92713848828431e-104 | 0.497388529 | 0.368 | 0.052 | 4.85041486116278e-100 | 9 |
| ms4a17a.9 | 2.55511830002921e-104 | 0.493553212 | 0.354 | 0.049 | 6.43097724934353e-100 | 9 |
| LOC1039092841 | 4.75160711716609e-104 | 0.541715614 | 0.382 | 0.057 | 1.19593199531953e-99 | 9 |
| postnb | 5.03454386213976e-104 | 0.621976839 | 0.438 | 0.07 | 1.26714434466196e-99 | 9 |
| h3f3c1 | 7.54380171929816e-104 | 0.833153494 | 0.809 | 0.244 | 1.89869945473015e-99 | 9 |
| CABZ01080702.1 | 9.91829886986749e-104 | 0.616501397 | 0.444 | 0.076 | 2.49633664255695e-99 | 9 |
| grb10a1 | 9.95243693382508e-104 | 0.449033429 | 0.292 | 0.033 | 2.50492885187444e-99 | 9 |
| znf385a1 | 1.43308336867826e-103 | 0.463102904 | 0.319 | 0.04 | 3.60692753062631e-99 | 9 |
| zgc:154058 | 5.78983045496403e-103 | 0.275150347 | 0.17 | 0.011 | 1.4572424272099e-98 | 9 |
| si:ch211-106h4.121 | 2.17574664669992e-102 | 0.719315118 | 0.361 | 0.052 | 5.47613673507902e-98 | 9 |
| meis2b | 4.42387702614488e-102 | 0.29453007 | 0.177 | 0.012 | 1.11344560871041e-97 | 9 |
| zgc:92744 | 4.83764329423633e-102 | 0.908950474 | 0.878 | 0.341 | 1.21758644072634e-97 | 9 |
| slc38a5b1 | 2.02971784418556e-100 | 0.626462394 | 0.528 | 0.106 | 5.10859684203063e-96 | 9 |
| palm1a | 4.15255113488373e-99 | 0.292152123 | 0.181 | 0.013 | 1.04515559513889e-94 | 9 |
| col27a1b | 9.24710637878485e-99 | 0.317786158 | 0.194 | 0.016 | 2.32740420447636e-94 | 9 |
| ece1 | 1.2803110360046e-98 | 0.420936459 | 0.257 | 0.028 | 3.22241484651999e-94 | 9 |
| meis1b | 1.90718319357388e-98 | 0.516616426 | 0.347 | 0.05 | 4.80018937990611e-94 | 9 |
| inka1a | 2.06578346846966e-98 | 0.345461726 | 0.194 | 0.016 | 5.19937041179129e-94 | 9 |
| cyp2n131 | 3.01471031598451e-98 | 0.419281035 | 0.278 | 0.032 | 7.58772439430141e-94 | 9 |
| nr2f2 | 2.9629403601472e-96 | 0.441707698 | 0.302 | 0.038 | 7.4574245924545e-92 | 9 |
| fkbp7 | 6.19751875918614e-96 | 0.371311465 | 0.267 | 0.031 | 1.55985349649956e-91 | 9 |
| rbp7b | 8.34104418345774e-96 | 0.297128988 | 0.111 | 0.005 | 2.09935741053448e-91 | 9 |
| p3h2 | 1.4261251125267e-95 | 0.287421129 | 0.174 | 0.013 | 3.58941429571844e-91 | 9 |
| gamt | 2.16637962505951e-95 | 0.435685101 | 0.299 | 0.039 | 5.45256087831227e-91 | 9 |
| junbb1 | 1.28401928935723e-94 | 0.522454264 | 0.392 | 0.066 | 3.23174814938322e-90 | 9 |
| klf11b1 | 1.43774896002849e-94 | 0.553790594 | 0.413 | 0.073 | 3.61867035749571e-90 | 9 |
| cebpb1 | 2.38248210096731e-94 | 0.943244151 | 0.899 | 0.341 | 5.99646919992463e-90 | 9 |
| s1pr2 | 2.96174567660839e-94 | 0.386587458 | 0.25 | 0.027 | 7.45441769345566e-90 | 9 |
| cyp2p71 | 7.4031200058436e-94 | 0.384950173 | 0.219 | 0.021 | 1.86329127427077e-89 | 9 |
| lrp1aa1 | 1.68613558317534e-93 | 0.500445604 | 0.337 | 0.049 | 4.24383464929402e-89 | 9 |
| BX322530.11 | 3.54909397706545e-93 | 0.438679251 | 0.309 | 0.042 | 8.93271463087603e-89 | 9 |
| tmem88b | 8.6426011918803e-93 | 0.389744976 | 0.271 | 0.032 | 2.17525629398435e-88 | 9 |
| itga9 | 2.71188544330255e-92 | 0.380516826 | 0.243 | 0.026 | 6.82554447224818e-88 | 9 |
| b2ml2 | 7.22179006180994e-92 | 0.785072489 | 0.802 | 0.255 | 1.81765234065694e-87 | 9 |
| aldob | 5.39285194886971e-91 | 0.376521665 | 0.271 | 0.033 | 1.35732690701102e-86 | 9 |

| | | | | | | |
|---|---|---|---|---|---|---|
| socs2 | 6.44997227391032e-91 | 0.353370076 | 0.222 | 0.022 | 1.62339352162049e-86 | 9 |
| plekha6 | 2.09461361325303e-90 | 0.310692947 | 0.181 | 0.015 | 5.27193300319655e-86 | 9 |
| FP102052.2 | 4.01850645377665e-90 | 0.294045883 | 0.184 | 0.015 | 1.01141788935105e-85 | 9 |
| si:ch211-286o17.1 | 2.12117189585595e-89 | 0.375970751 | 0.267 | 0.032 | 5.33877754467984e-85 | 9 |
| clec11a | 2.17817592696562e-89 | 0.297172994 | 0.17 | 0.013 | 5.48225099057976e-85 | 9 |
| spred2b | 3.88294579068254e-89 | 0.484894027 | 0.368 | 0.06 | 9.77298626056889e-85 | 9 |
| deptor | 6.44136359039851e-89 | 0.383699199 | 0.267 | 0.033 | 1.6212268020674e-84 | 9 |
| tmeff1b | 8.53549516272653e-89 | 0.343850709 | 0.233 | 0.025 | 2.14829877750664e-84 | 9 |
| nfil3 | 7.91987561731807e-88 | 0.393497685 | 0.247 | 0.029 | 1.99335349412278e-83 | 9 |
| CR855311.12 | 1.67883147180724e-87 | 0.74101432 | 0.417 | 0.078 | 4.22545093139164e-83 | 9 |
| ier5 | 4.88128420317504e-87 | 0.639902737 | 0.545 | 0.127 | 1.22857042109713e-82 | 9 |
| osr2 | 6.15117634513432e-87 | 0.445102916 | 0.295 | 0.04 | 1.54818957430686e-82 | 9 |
| hmgn61 | 1.72989430469641e-86 | 0.770350149 | 0.781 | 0.253 | 4.35397097549039e-82 | 9 |
| nav3 | 2.45379000922468e-86 | 0.306758724 | 0.177 | 0.015 | 6.17594407421759e-82 | 9 |
| kank21 | 3.28071529763826e-86 | 0.477772081 | 0.319 | 0.048 | 8.25723233262573e-82 | 9 |
| gng7 | 5.18854704250225e-86 | 0.428822292 | 0.319 | 0.048 | 1.30590540512739e-81 | 9 |
| rhoua | 7.19147215518318e-86 | 0.325313579 | 0.205 | 0.02 | 1.81002162673805e-81 | 9 |
| sfrp1a | 1.65795261173455e-85 | 0.387087655 | 0.226 | 0.024 | 4.17290092847469e-81 | 9 |
| ptk2aa | 1.7719885520034e-85 | 0.279391667 | 0.17 | 0.014 | 4.45991798653736e-81 | 9 |
| lgals171 | 3.65671347007044e-85 | 0.522484346 | 0.312 | 0.047 | 9.2035821328203e-81 | 9 |
| LOC553316 | 9.85178018074066e-85 | 0.263562522 | 0.153 | 0.011 | 2.47959455369062e-80 | 9 |
| ghrb | 1.21902794051111e-84 | 0.403788656 | 0.278 | 0.037 | 3.06817142347242e-80 | 9 |
| prss35 | 1.52476077672081e-84 | 0.326845962 | 0.188 | 0.017 | 3.83767039892861e-80 | 9 |
| hmgb1b | 1.61490195163402e-84 | 0.608337115 | 0.521 | 0.118 | 4.06454672206767e-80 | 9 |
| c41 | 1.88224288828329e-84 | 0.428357057 | 0.274 | 0.036 | 4.73741712552022e-80 | 9 |
| CR383676.12 | 2.2477655407181e-84 | 0.795529893 | 0.997 | 0.72 | 5.65740108943339e-80 | 9 |
| ppib1 | 9.80492978192047e-84 | 0.790007746 | 0.872 | 0.32 | 2.46780277681156e-79 | 9 |
| adh5 | 1.55149795244091e-83 | 0.696107421 | 0.719 | 0.21 | 3.90496519649852e-79 | 9 |
| actn1 | 4.01987995177153e-83 | 0.432166209 | 0.281 | 0.039 | 1.01176358506138e-78 | 9 |
| maff | 5.01165577890014e-83 | 0.520840218 | 0.403 | 0.076 | 1.26138364299138e-78 | 9 |
| fblim1 | 5.32605785744417e-83 | 0.316785834 | 0.212 | 0.022 | 1.34051550214012e-78 | 9 |
| cox8b | 6.63027231649872e-83 | 0.530353761 | 0.441 | 0.088 | 1.66877323933956e-78 | 9 |
| fhl3b | 2.30721232494682e-82 | 0.260171037 | 0.146 | 0.01 | 5.80702270065865e-78 | 9 |
| atf4b | 3.31968417388283e-82 | 0.72920138 | 0.788 | 0.252 | 8.35531309724569e-78 | 9 |
| irf2bp2b | 1.96577242873128e-81 | 0.517619172 | 0.396 | 0.076 | 4.94765262587377e-77 | 9 |

| | | | | | | |
|---|---|---|---|---|---|---|
| egr11 | 1.41713562321424e-80 | 0.725028168 | 0.649 | 0.18 | 3.56678865006793e-76 | 9 |
| hic11 | 1.68641553868969e-79 | 0.479209693 | 0.323 | 0.053 | 4.24453926932809e-75 | 9 |
| cdh11 | 4.28169393220042e-79 | 0.352551548 | 0.208 | 0.023 | 1.07765954579552e-74 | 9 |
| ptges1 | 5.98722271789385e-79 | 0.329164555 | 0.194 | 0.02 | 1.5069240858667e-74 | 9 |
| agtr21 | 1.38442210172371e-78 | 0.330346516 | 0.17 | 0.015 | 3.4844519878284e-74 | 9 |
| thbs1a | 1.68491216383056e-78 | 0.281594662 | 0.146 | 0.011 | 4.24075542514514e-74 | 9 |
| krt911 | 3.3959031039535e-78 | 0.601217678 | 0.354 | 0.062 | 8.54714852234057e-74 | 9 |
| ppp1cbl | 3.43518709457033e-78 | 0.478153187 | 0.347 | 0.062 | 8.64602239832407e-74 | 9 |
| zgc:1132081 | 6.22831643551072e-78 | 0.387361829 | 0.278 | 0.04 | 1.56760496365369e-73 | 9 |
| yap1 | 6.34275074081452e-78 | 0.398426904 | 0.295 | 0.045 | 1.59640693395561e-73 | 9 |
| gadd45bb | 9.79783914005097e-78 | 0.557860315 | 0.403 | 0.081 | 2.46601813315943e-73 | 9 |
| b2m1 | 2.18193993510822e-77 | 0.743651023 | 0.819 | 0.28 | 5.49172462267387e-73 | 9 |
| rpz51 | 2.98613710640257e-77 | 0.571298694 | 0.444 | 0.095 | 7.51580848310462e-73 | 9 |
| ppap2d1 | 3.20122029927122e-77 | 0.376491991 | 0.247 | 0.032 | 8.05715137123574e-73 | 9 |
| gstm.1 | 5.43803018117965e-77 | 0.744991532 | 0.76 | 0.249 | 1.36869781630111e-72 | 9 |
| srpx | 1.01786982547576e-76 | 0.305911738 | 0.222 | 0.026 | 2.56187656373994e-72 | 9 |
| si:dkeyp-75h12.5 | 2.87485161354494e-76 | 0.351750415 | 0.253 | 0.034 | 7.23571402613126e-72 | 9 |
| mgst1.21 | 9.73855924703036e-76 | 0.552770614 | 0.444 | 0.095 | 2.45109797688507e-71 | 9 |
| slc43a2a1 | 1.44862477650657e-75 | 0.531638536 | 0.444 | 0.095 | 3.64604369998938e-71 | 9 |
| rnd3b1 | 3.46723192429689e-75 | 0.381976928 | 0.26 | 0.036 | 8.72667603026284e-71 | 9 |
| fam168a1 | 6.92561928111378e-75 | 0.410810108 | 0.316 | 0.053 | 1.74310911686353e-70 | 9 |
| si:dkey-163m14.2 | 9.4421709932158e-75 | 0.302643816 | 0.167 | 0.015 | 2.37650001728249e-70 | 9 |
| fat1a | 1.08867435998106e-74 | 0.330836119 | 0.208 | 0.024 | 2.74008449663633e-70 | 9 |
| loxl2b | 1.33174848913369e-74 | 0.402602719 | 0.26 | 0.037 | 3.35187777230058e-70 | 9 |
| irf1b | 1.44592965235293e-74 | 0.411474766 | 0.25 | 0.034 | 3.63926034200708e-70 | 9 |
| asip2b | 2.04752080677812e-74 | 0.400378487 | 0.167 | 0.015 | 5.15340511857986e-70 | 9 |
| si:dkey-51e6.1 | 3.53118567238768e-74 | 0.524260057 | 0.472 | 0.108 | 8.88764121883255e-70 | 9 |
| appbp2 | 1.24516745193348e-73 | 0.389725913 | 0.26 | 0.037 | 3.13396195977137e-69 | 9 |
| zgc:1141882 | 1.56936240606852e-73 | 0.602932987 | 0.983 | 0.62 | 3.94992823983385e-69 | 9 |
| crispld21 | 2.65043983871262e-73 | 0.285431668 | 0.163 | 0.015 | 6.67089203005579e-69 | 9 |
| ckap41 | 2.03043224695988e-72 | 0.370327879 | 0.264 | 0.039 | 5.11039492237331e-68 | 9 |
| pdgfrl | 6.41124162489507e-72 | 0.311801623 | 0.205 | 0.024 | 1.61364540456984e-67 | 9 |
| selenow1 | 6.88255739429162e-72 | 0.736602609 | 0.906 | 0.373 | 1.73227087056926e-67 | 9 |
| notch3 | 1.09734738346753e-71 | 0.292491843 | 0.184 | 0.02 | 2.76191362944944e-67 | 9 |
| ybx11 | 1.33253017093797e-71 | 0.537820101 | 0.969 | 0.663 | 3.35384518723378e-67 | 9 |

| | | | | | | |
|---|---|---|---|---|---|---|
| hmgb3a | 1.67787262036715e-70 | 0.401119489 | 0.278 | 0.044 | 4.22303759820207e-66 | 9 |
| si:ch73-237c6.11 | 1.78143378470866e-70 | 0.363630276 | 0.257 | 0.038 | 4.48369069273323e-66 | 9 |
| tshz21 | 2.13897604057977e-70 | 0.290814648 | 0.205 | 0.024 | 5.38358879653522e-66 | 9 |
| CU459056.1 | 2.17988182844739e-70 | 0.285226841 | 0.184 | 0.02 | 5.48654457401925e-66 | 9 |
| tmem119b | 2.81755946373695e-70 | 0.254921667 | 0.156 | 0.014 | 7.09151541427954e-66 | 9 |
| tmem100a1 | 4.12512901203635e-70 | 0.279391908 | 0.174 | 0.018 | 1.03825372103943e-65 | 9 |
| si:dkey-58f10.11 | 4.30310675789472e-70 | 0.307776159 | 0.194 | 0.022 | 1.08304893989452e-65 | 9 |
| qkia | 7.49320837928012e-70 | 0.585506591 | 0.58 | 0.159 | 1.88596561698101e-65 | 9 |
| rpl92 | 7.84006024533402e-70 | 0.601615623 | 0.997 | 0.621 | 1.97326476314812e-65 | 9 |
| gas6 | 8.40774425110864e-70 | 0.302989682 | 0.188 | 0.021 | 2.11614515056153e-65 | 9 |
| krt942 | 1.23070147218321e-69 | 0.591612138 | 0.507 | 0.121 | 3.09755253533791e-65 | 9 |
| mvp | 1.40876385112697e-69 | 0.388416445 | 0.292 | 0.048 | 3.54571773690147e-65 | 9 |
| zeb1b | 1.5073986402828e-69 | 0.316773473 | 0.198 | 0.023 | 3.79397163772779e-65 | 9 |
| snai2 | 6.81431846550016e-69 | 0.300638028 | 0.194 | 0.023 | 1.71509581458174e-64 | 9 |
| cebpa2 | 1.51815524715445e-68 | 0.467544001 | 0.302 | 0.053 | 3.82104494156305e-64 | 9 |
| mgst3b | 2.23326162748726e-68 | 0.584080215 | 0.58 | 0.163 | 5.62089619022269e-64 | 9 |
| pgrmc1 | 4.40538868955974e-68 | 0.51493448 | 0.528 | 0.132 | 1.10879227927529e-63 | 9 |
| clic22 | 5.57199402443457e-68 | 0.443910807 | 0.309 | 0.055 | 1.40241517600994e-63 | 9 |
| grinab | 6.70104031933408e-68 | 0.502815163 | 0.451 | 0.107 | 1.6865848379732e-63 | 9 |
| higd1a1 | 1.03126894269011e-67 | 0.69727122 | 0.812 | 0.298 | 2.59560080185673e-63 | 9 |
| sh3pxd2aa | 1.05891031304915e-67 | 0.341407678 | 0.215 | 0.028 | 2.6651713669134e-63 | 9 |
| sult1st1 | 1.56350597799168e-67 | 0.32581541 | 0.184 | 0.021 | 3.93518819600725e-63 | 9 |
| ctdsp2 | 2.83653440106962e-67 | 0.424529796 | 0.326 | 0.061 | 7.13927343405212e-63 | 9 |
| fzd7b | 4.22437827977329e-67 | 0.336861529 | 0.24 | 0.034 | 1.06323376923614e-62 | 9 |
| mxra8b | 9.99191013983023e-67 | 0.273072992 | 0.181 | 0.02 | 2.51486386309387e-62 | 9 |
| tagln21 | 2.60429930269468e-66 | 0.621677841 | 0.608 | 0.179 | 6.55476091495224e-62 | 9 |
| celf22 | 2.90671153997645e-66 | 0.67799672 | 0.691 | 0.232 | 7.31590227496672e-62 | 9 |
| pabpc1b1 | 3.00073963984243e-66 | 0.417533458 | 0.333 | 0.063 | 7.55256159951942e-62 | 9 |
| ifi46 | 3.95518796129844e-66 | 0.552430137 | 0.465 | 0.114 | 9.95481257979204e-62 | 9 |
| dhrs13l1 | 1.35483999513898e-65 | 0.311521149 | 0.215 | 0.029 | 3.40999678376531e-61 | 9 |
| si:ch211-74m13.3 | 2.38760887763691e-65 | 0.415515273 | 0.33 | 0.062 | 6.00937278412435e-61 | 9 |
| ptbp1b | 2.47825175303828e-65 | 0.317379437 | 0.226 | 0.031 | 6.23751183722206e-61 | 9 |
| rps27.21 | 3.6124786007628e-65 | 0.650350028 | 0.951 | 0.508 | 9.0922473902599e-61 | 9 |
| hoxc3a | 3.68532016651532e-65 | 0.262698883 | 0.17 | 0.018 | 9.27558232710241e-61 | 9 |
| si:dkey-239h2.3 | 6.0626103922729e-65 | 0.467035109 | 0.382 | 0.083 | 1.52589840963117e-60 | 9 |

| | | | | | | |
|---|---|---|---|---|---|---|
| vdac31 | 8.19495341603126e-65 | 0.545128558 | 0.538 | 0.146 | 2.06258782528091e-60 | 9 |
| grb10b | 3.69792669620461e-64 | 0.325968287 | 0.226 | 0.032 | 9.30731170167739e-60 | 9 |
| fxyd11 | 7.14204664164118e-64 | 0.504688649 | 0.354 | 0.075 | 1.79758171923467e-59 | 9 |
| si:ch211-133n4.41 | 2.15175815739824e-63 | 0.362764022 | 0.253 | 0.041 | 5.41576010635564e-59 | 9 |
| timp2a1 | 3.35767593409444e-63 | 0.39704164 | 0.292 | 0.052 | 8.4509345585223e-59 | 9 |
| ca10a | 6.99199154551007e-63 | 0.279482072 | 0.174 | 0.02 | 1.75981435208943e-58 | 9 |
| CABZ01074363.11 | 1.05737307249963e-62 | 0.290654764 | 0.198 | 0.025 | 2.66130228617432e-58 | 9 |
| ppdpfa | 1.13773203101857e-62 | 0.490291553 | 0.344 | 0.071 | 2.86355774887065e-58 | 9 |
| fgfr21 | 1.16738569400401e-62 | 0.302796061 | 0.191 | 0.024 | 2.93819305323869e-58 | 9 |
| CR318588.12 | 1.68164075271636e-62 | 0.914147481 | 0.59 | 0.2 | 4.23252161051182e-58 | 9 |
| hsp70.3 | 3.70961876179473e-62 | 0.290213608 | 0.17 | 0.019 | 9.33673946156116e-58 | 9 |
| ptenb | 4.33254027933187e-62 | 0.437625802 | 0.354 | 0.074 | 1.09045706290504e-57 | 9 |
| vim1 | 6.75873336604206e-62 | 0.336668855 | 0.219 | 0.031 | 1.70110560089913e-57 | 9 |
| CR847953.1 | 9.70478880363098e-62 | 0.321708264 | 0.17 | 0.02 | 2.44259829398588e-57 | 9 |
| fermt2 | 1.31627224693899e-61 | 0.294467264 | 0.201 | 0.027 | 3.31292561832073e-57 | 9 |
| arrdc3a1 | 1.3870780809768e-61 | 0.490506112 | 0.33 | 0.067 | 3.49113682201051e-57 | 9 |
| cyr611 | 1.79532034371192e-61 | 0.537554487 | 0.319 | 0.063 | 4.51864177308852e-57 | 9 |
| angptl41 | 2.7907095926997e-61 | 0.442437119 | 0.264 | 0.045 | 7.02393697386587e-57 | 9 |
| ghra1 | 3.47384409764494e-61 | 0.30554053 | 0.201 | 0.027 | 8.74331820936255e-57 | 9 |
| arhgap31 | 5.50437719723919e-61 | 0.275167707 | 0.198 | 0.026 | 1.38539669677313e-56 | 9 |
| tspan4a | 1.09659734635197e-60 | 0.301977438 | 0.208 | 0.029 | 2.76002586103327e-56 | 9 |
| ptgs2a | 1.25424604463815e-60 | 0.265027549 | 0.118 | 0.009 | 3.15681186974977e-56 | 9 |
| creg2 | 1.28404633532543e-60 | 0.290290301 | 0.198 | 0.026 | 3.23181622138056e-56 | 9 |
| btg21 | 1.37855919064837e-60 | 0.707193199 | 0.858 | 0.362 | 3.46969562694289e-56 | 9 |
| ctdspla1 | 1.96298536221622e-60 | 0.31879089 | 0.219 | 0.032 | 4.94063785816202e-56 | 9 |
| antxr2a1 | 2.24550285912394e-60 | 0.539637441 | 0.524 | 0.146 | 5.65170614612905e-56 | 9 |
| si:dkey-79d12.5 | 6.77334259692197e-60 | 0.473487699 | 0.333 | 0.071 | 1.70478259821929e-55 | 9 |
| mycb | 7.80120671965206e-60 | 0.63071707 | 0.635 | 0.202 | 1.96348571926923e-55 | 9 |
| eif4a2 | 1.24516804678259e-59 | 0.434373765 | 0.361 | 0.079 | 3.1339634569471e-55 | 9 |
| si:ch211-204c21.1 | 1.61825250465645e-59 | 0.405558084 | 0.333 | 0.068 | 4.07297972896982e-55 | 9 |
| ier5l | 1.83976551433693e-59 | 0.260522027 | 0.156 | 0.017 | 4.63050582303462e-55 | 9 |
| rps27 | 2.2890007558318e-59 | 0.393424488 | 0.969 | 0.826 | 5.76118600235305e-55 | 9 |
| klf6a1 | 2.66826511807852e-59 | 0.577975307 | 0.545 | 0.159 | 6.71575647569182e-55 | 9 |
| map1lc3b | 2.77664110608214e-59 | 0.469627317 | 0.41 | 0.1 | 6.98852799989814e-55 | 9 |
| si:ch73-27e22.2 | 2.96370658689185e-59 | 0.290867275 | 0.156 | 0.017 | 7.4593531085481e-55 | 9 |

| | | | | | | |
|---|---|---|---|---|---|---|
| BX569787.1 | 3.93228358524672e-59 | 0.271648862 | 0.174 | 0.021 | 9.89716455570747e-55 | 9 |
| serinc2 | 8.10220848424399e-59 | 0.452594103 | 0.427 | 0.105 | 2.03924485339937e-54 | 9 |
| edil3a | 1.46451604216475e-58 | 0.280903671 | 0.17 | 0.02 | 3.68604042652447e-54 | 9 |
| tsku1 | 2.39800523494031e-58 | 0.280925336 | 0.194 | 0.026 | 6.03553937582125e-54 | 9 |
| cux1a | 3.13984100351727e-58 | 0.356174266 | 0.247 | 0.041 | 7.90266582175262e-54 | 9 |
| rpl27a2 | 4.21819669717575e-58 | 0.64424905 | 0.917 | 0.487 | 1.06167792671217e-53 | 9 |
| tuba8l31 | 5.97820314020486e-58 | 0.361844511 | 0.25 | 0.043 | 1.50465394835816e-53 | 9 |
| tpm1 | 8.70385758234082e-58 | 0.66747648 | 0.597 | 0.199 | 2.19067391489936e-53 | 9 |
| rps152 | 1.62805451306407e-57 | 0.494475474 | 0.965 | 0.679 | 4.09765040393095e-53 | 9 |
| foxo3b1 | 2.09188691060351e-57 | 0.398449379 | 0.306 | 0.061 | 5.26507016529797e-53 | 9 |
| ctgfa | 3.65507242588665e-57 | 0.44835376 | 0.184 | 0.025 | 9.19945178871412e-53 | 9 |
| rps92 | 4.20741506711348e-57 | 0.410245137 | 0.979 | 0.772 | 1.05896429824179e-52 | 9 |
| tns1b1 | 6.64213572282621e-57 | 0.327412966 | 0.208 | 0.031 | 1.67175914007813e-52 | 9 |
| tnfrsf1a | 1.76357795315847e-56 | 0.382001169 | 0.267 | 0.05 | 4.43874935030454e-52 | 9 |
| vasnb | 2.15437682822192e-56 | 0.303073694 | 0.198 | 0.028 | 5.42235103895175e-52 | 9 |
| arhgap5 | 2.76522081785338e-56 | 0.28558622 | 0.184 | 0.025 | 6.95978427645516e-52 | 9 |
| c7a1 | 3.34036262466556e-56 | 0.363886899 | 0.243 | 0.041 | 8.40735869002076e-52 | 9 |
| cnbpb | 3.44742088339883e-56 | 0.51977818 | 0.569 | 0.176 | 8.67681362142653e-52 | 9 |
| itm2ba1 | 4.68295265806397e-56 | 0.607037397 | 0.719 | 0.263 | 1.17865235450812e-51 | 9 |
| tspan9b | 5.84189754796034e-56 | 0.322646952 | 0.24 | 0.04 | 1.47034719384614e-51 | 9 |
| si:ch211-197h24.9 | 7.82633910113955e-56 | 0.385788489 | 0.309 | 0.063 | 1.96981128836581e-51 | 9 |
| abl1 | 1.17335056734478e-55 | 0.298510289 | 0.201 | 0.029 | 2.95320604295008e-51 | 9 |
| rpl292 | 1.52764473642241e-55 | 0.387911902 | 0.993 | 0.805 | 3.84492903710155e-51 | 9 |
| sgce | 1.60273519869353e-55 | 0.270625744 | 0.181 | 0.024 | 4.03392422159174e-51 | 9 |
| rps282 | 1.83409849416113e-55 | 0.377920706 | 0.997 | 0.823 | 4.61624249995416e-51 | 9 |
| homeza | 1.9011907150291e-55 | 0.360387089 | 0.295 | 0.057 | 4.78510691065675e-51 | 9 |
| rps181 | 1.96398305789106e-55 | 0.46557789 | 0.979 | 0.719 | 4.943148958406e-51 | 9 |
| pi4kb | 3.62635138822121e-55 | 0.298778981 | 0.215 | 0.034 | 9.12716380901396e-51 | 9 |
| cxcl8a | 3.71977277341973e-55 | 0.287107172 | 0.132 | 0.013 | 9.36229609342011e-51 | 9 |
| ccdc85b1 | 5.13818533238326e-55 | 0.367979958 | 0.25 | 0.045 | 1.29322986630754e-50 | 9 |
| ldlrad4a1 | 6.33756995934008e-55 | 0.280220766 | 0.198 | 0.029 | 1.59510298306631e-50 | 9 |
| mprip | 9.6473241379711e-55 | 0.263174067 | 0.17 | 0.022 | 2.42813501228595e-50 | 9 |
| jpt1b | 1.35714788563275e-54 | 0.592991911 | 0.736 | 0.26 | 3.41580551334908e-50 | 9 |
| raraa | 2.03631993243699e-54 | 0.278058314 | 0.194 | 0.028 | 5.12521363795066e-50 | 9 |
| ywhaz | 2.33174741847529e-54 | 0.620936826 | 0.712 | 0.272 | 5.86877507756046e-50 | 9 |

| | | | | | | |
|---|---|---|---|---|---|---|
| zcchc241 | 2.67476114296364e-54 | 0.277129459 | 0.198 | 0.029 | 6.73210632072519e-50 | 9 |
| rpl362 | 3.39811030786789e-54 | 0.422868531 | 0.993 | 0.785 | 8.55270383387269e-50 | 9 |
| SBNO2 | 3.91968730484639e-54 | 0.302220669 | 0.201 | 0.03 | 9.86546097756788e-50 | 9 |
| mob2a | 4.58129704701038e-54 | 0.291302216 | 0.222 | 0.036 | 1.15306665376204e-49 | 9 |
| si:ch73-1a9.31 | 5.52574080625475e-54 | 0.602101007 | 0.708 | 0.256 | 1.39077370352626e-49 | 9 |
| atp10a | 8.11565394747709e-54 | 0.376104852 | 0.271 | 0.052 | 2.04262894204051e-49 | 9 |
| XLOC-041269 | 9.7237834289135e-54 | 0.280066556 | 0.167 | 0.021 | 2.44737905122324e-49 | 9 |
| adh8b | 3.45044131055588e-53 | 0.30250747 | 0.215 | 0.035 | 8.68441573453809e-49 | 9 |
| eif1b | 3.49517930878082e-53 | 0.630511235 | 0.865 | 0.372 | 8.79701680227045e-49 | 9 |
| fbn2b1 | 4.08042863566517e-53 | 0.260773334 | 0.174 | 0.023 | 1.02700308331057e-48 | 9 |
| itgb1a1 | 4.26842675201229e-53 | 0.329484468 | 0.243 | 0.043 | 1.07432032921397e-48 | 9 |
| si:ch211-1a19.31 | 7.11726682990761e-53 | 0.402062678 | 0.26 | 0.049 | 1.79134488841945e-48 | 9 |
| tgfb1a | 7.18909091953443e-53 | 0.251119632 | 0.16 | 0.02 | 1.80942229353762e-48 | 9 |
| rps162 | 9.77357412194927e-53 | 0.445936794 | 0.979 | 0.733 | 2.45991087075341e-48 | 9 |
| rpl62 | 1.02079274399631e-52 | 0.504167903 | 0.986 | 0.661 | 2.56923325736432e-48 | 9 |
| wu:fb16f03 | 1.21550825945558e-52 | 0.433796978 | 0.347 | 0.082 | 3.05931273822375e-48 | 9 |
| dhrs3a1 | 1.56381738397739e-52 | 0.27460637 | 0.188 | 0.027 | 3.93597197373268e-48 | 9 |
| gstz1 | 1.6298388435818e-52 | 0.278080761 | 0.194 | 0.029 | 4.10214138541102e-48 | 9 |
| rplp11 | 2.21156722223171e-52 | 0.334552531 | 0.997 | 0.875 | 5.56629354163499e-48 | 9 |
| hmgn7 | 2.46335382396027e-52 | 0.592555343 | 0.722 | 0.27 | 6.20001523952559e-48 | 9 |
| plpp31 | 3.00303849925937e-52 | 0.376519481 | 0.267 | 0.051 | 7.5583475987859e-48 | 9 |
| nsa21 | 3.29200999673266e-52 | 0.638761851 | 0.83 | 0.352 | 8.28565996077643e-48 | 9 |
| ckba | 4.75514291684114e-52 | 0.466210986 | 0.288 | 0.061 | 1.19682192073975e-47 | 9 |
| rps131 | 5.70479340339457e-52 | 0.439859724 | 0.969 | 0.719 | 1.43583945170038e-47 | 9 |
| pcolcea1 | 7.9177912826862e-52 | 0.404739208 | 0.253 | 0.047 | 1.99282888793929e-47 | 9 |
| rpl33 | 1.28436129535737e-51 | 0.291970218 | 0.993 | 0.873 | 3.23260894428497e-47 | 9 |
| tnfsf10 | 2.2580156904678e-51 | 0.27012233 | 0.174 | 0.024 | 5.6831996913384e-47 | 9 |
| crtap | 2.77777475326915e-51 | 0.264083874 | 0.188 | 0.028 | 6.99138127650314e-47 | 9 |
| rbms31 | 4.04595981935013e-51 | 0.282691208 | 0.198 | 0.03 | 1.01832762693223e-46 | 9 |
| si:ch211-222l21.11 | 6.19380799575542e-51 | 0.618049424 | 0.712 | 0.266 | 1.55891953445168e-46 | 9 |
| pnrc22 | 1.10482461362337e-50 | 0.596087199 | 0.965 | 0.559 | 2.78073307002865e-46 | 9 |
| rps191 | 1.12982877497626e-50 | 0.325506586 | 1 | 0.86 | 2.84366604373776e-46 | 9 |
| clic4 | 1.27352470859222e-50 | 0.296381742 | 0.247 | 0.045 | 3.20533433905575e-46 | 9 |
| sumo2b1 | 1.64942019317328e-50 | 0.563388479 | 0.694 | 0.26 | 4.15142568419782e-46 | 9 |
| rpsa2 | 1.7384274789362e-50 | 0.43041548 | 0.986 | 0.747 | 4.37544812173451e-46 | 9 |

| | | | | | | |
|---|---|---|---|---|---|---|
| sptlc2a | 2.57353309811882e-50 | 0.306909011 | 0.219 | 0.037 | 6.47732545465526e-46 | 9 |
| rpl392 | 3.67222493926215e-50 | 0.309753936 | 0.983 | 0.872 | 9.2426229496289e-46 | 9 |
| cavin2b | 4.91961358768439e-50 | 0.36134967 | 0.292 | 0.061 | 1.23821754388428e-45 | 9 |
| rpl342 | 1.26830188414762e-49 | 0.40388522 | 0.99 | 0.779 | 3.19218901221114e-45 | 9 |
| ptprga1 | 1.41593757848379e-49 | 0.377420341 | 0.288 | 0.062 | 3.56377329128585e-45 | 9 |
| mych1 | 1.43461218056639e-49 | 0.432680594 | 0.309 | 0.07 | 3.61077539726754e-45 | 9 |
| isg151 | 2.01078838588546e-49 | 0.491046986 | 0.292 | 0.064 | 5.06095328843512e-45 | 9 |
| rps8a1 | 2.09582230941499e-49 | 0.376038917 | 0.983 | 0.784 | 5.2749751705666e-45 | 9 |
| rhoab1 | 2.54149152596815e-49 | 0.532308364 | 0.545 | 0.182 | 6.39668002170924e-45 | 9 |
| rpl322 | 5.55506910740098e-49 | 0.366819738 | 0.976 | 0.81 | 1.39815534364175e-44 | 9 |
| FO082781.11 | 6.6768009695602e-49 | 0.31919625 | 0.236 | 0.044 | 1.68048403602861e-44 | 9 |
| tax1bp3 | 6.78346402691388e-49 | 0.359997973 | 0.347 | 0.084 | 1.70733006093395e-44 | 9 |
| stxbp6 | 7.93583476037022e-49 | 0.279687964 | 0.226 | 0.039 | 1.99737025083758e-44 | 9 |
| rpl7a1 | 1.12355409240133e-48 | 0.365028383 | 0.986 | 0.806 | 2.82787329516492e-44 | 9 |
| sept21 | 1.23773118543649e-48 | 0.487166244 | 0.507 | 0.158 | 3.1152456206251e-44 | 9 |
| mcl1b1 | 1.57564489608007e-48 | 0.552222769 | 0.66 | 0.233 | 3.96574063894393e-44 | 9 |
| her6 | 1.68867792268528e-48 | 0.272431329 | 0.153 | 0.02 | 4.25023346360659e-44 | 9 |
| myo1ca1 | 3.5159111757798e-48 | 0.250926728 | 0.16 | 0.022 | 8.84919683832019e-44 | 9 |
| aste1a | 3.89676535594017e-48 | 0.257138954 | 0.153 | 0.02 | 9.80776872436581e-44 | 9 |
| pdk2a | 3.98057417748522e-48 | 0.257638311 | 0.174 | 0.025 | 1.00187071473125e-43 | 9 |
| mmp14b1 | 5.08093372438542e-48 | 0.363427336 | 0.26 | 0.053 | 1.27882020909057e-43 | 9 |
| stat1a1 | 6.66639709124519e-48 | 0.444490491 | 0.41 | 0.114 | 1.6778654838955e-43 | 9 |
| angptl1a1 | 8.85351007853533e-48 | 0.282691573 | 0.17 | 0.024 | 2.22833995166656e-43 | 9 |
| ablim1b1 | 1.01565374705895e-47 | 0.299838337 | 0.201 | 0.033 | 2.55629891597266e-43 | 9 |
| slc3a2b1 | 1.10139099024241e-47 | 0.457540401 | 0.403 | 0.112 | 2.77209098334113e-43 | 9 |
| CU984600.2 | 1.12910274561693e-47 | 0.25329088 | 0.163 | 0.023 | 2.84183870044326e-43 | 9 |
| eef1db1 | 1.45625198473996e-47 | 0.574519235 | 0.684 | 0.265 | 3.66524062039201e-43 | 9 |
| il6st | 1.49686963226218e-47 | 0.333338385 | 0.243 | 0.048 | 3.76747117744068e-43 | 9 |
| igfbp5b1 | 1.88342846736178e-47 | 0.37397385 | 0.306 | 0.068 | 4.74040110950286e-43 | 9 |
| si:ch211-286b5.5 | 2.14360987963069e-47 | 0.410373896 | 0.451 | 0.13 | 5.39525170604248e-43 | 9 |
| rbfox2 | 3.09503523025282e-47 | 0.270151876 | 0.198 | 0.033 | 7.78989417102332e-43 | 9 |
| proza | 6.14598580825759e-47 | 0.251809235 | 0.188 | 0.03 | 1.54688316808035e-42 | 9 |
| ets1 | 1.00700692619906e-46 | 0.2965688 | 0.201 | 0.034 | 2.53453573255042e-42 | 9 |
| rps71 | 1.0915238920129e-46 | 0.348288914 | 0.979 | 0.817 | 2.74725648380726e-42 | 9 |
| ndufa4l | 1.56841605994499e-46 | 0.591624361 | 0.854 | 0.385 | 3.94754638127555e-42 | 9 |

| | | | | | | |
|---|---|---|---|---|---|---|
| rpl261 | 1.98195857716766e-46 | 0.331558572 | 0.99 | 0.838 | 4.98839154287329e-42 | 9 |
| rpl131 | 2.9883005149869e-46 | 0.351525542 | 0.993 | 0.801 | 7.52125356617053e-42 | 9 |
| rpl372 | 6.6637986134706e-46 | 0.348008462 | 0.993 | 0.847 | 1.67721147302441e-41 | 9 |
| myh11a | 6.86859482458471e-46 | 0.481741356 | 0.274 | 0.062 | 1.72875663139973e-41 | 9 |
| foxp41 | 1.61324121089359e-45 | 0.326367731 | 0.24 | 0.048 | 4.06036680369808e-41 | 9 |
| akap12b1 | 1.65039798922836e-45 | 0.321909415 | 0.267 | 0.056 | 4.15388669908887e-41 | 9 |
| stau21 | 1.86194234360637e-45 | 0.369167554 | 0.326 | 0.081 | 4.68632268462286e-41 | 9 |
| gli31 | 2.65444276590528e-45 | 0.256694541 | 0.181 | 0.029 | 6.680966997507e-41 | 9 |
| si:ch211-198c19.31 | 5.00711478801676e-45 | 0.389021272 | 0.274 | 0.059 | 1.26024072099594e-40 | 9 |
| rpl242 | 5.10123445698892e-45 | 0.385279038 | 0.983 | 0.772 | 1.28392970047954e-40 | 9 |
| rgs5a1 | 5.27559787995064e-45 | 0.276304493 | 0.167 | 0.025 | 1.32781523040478e-40 | 9 |
| ndrg2 | 5.33597376588397e-45 | 0.299712624 | 0.205 | 0.037 | 1.34301123713534e-40 | 9 |
| ptmaa2 | 5.41086315872463e-45 | 0.613995971 | 0.875 | 0.385 | 1.3618601484194e-40 | 9 |
| enc1 | 8.62067548904381e-45 | 0.260133545 | 0.177 | 0.028 | 2.16973781383744e-40 | 9 |
| rpl36a2 | 1.05101559995564e-44 | 0.347159403 | 0.986 | 0.806 | 2.64530116352836e-40 | 9 |
| rpl23a2 | 2.2166010681429e-44 | 0.493285254 | 0.948 | 0.628 | 5.57896322840887e-40 | 9 |
| rpl10a1 | 2.49858470167336e-44 | 0.383910986 | 0.976 | 0.76 | 6.28868783564167e-40 | 9 |
| rpl112 | 2.57936565248398e-44 | 0.375492112 | 0.997 | 0.81 | 6.49200541073694e-40 | 9 |
| rps242 | 3.72902318534484e-44 | 0.390280963 | 0.976 | 0.754 | 9.38557845519442e-40 | 9 |
| lims1 | 5.3223219898361e-44 | 0.287702022 | 0.219 | 0.041 | 1.33957522162185e-39 | 9 |
| ccnl1a | 6.12828225167238e-44 | 0.335764541 | 0.264 | 0.058 | 1.54242735992342e-39 | 9 |
| ponzr11 | 7.8661667240178e-44 | 0.589491541 | 0.66 | 0.265 | 1.97983550276804e-39 | 9 |
| rps27a1 | 8.22684232978378e-44 | 0.392554887 | 0.983 | 0.747 | 2.07061394598328e-39 | 9 |
| anxa131 | 8.69061095000365e-44 | 0.287228891 | 0.233 | 0.046 | 2.18733987000642e-39 | 9 |
| sulf2b1 | 1.10940205801765e-43 | 0.271034582 | 0.201 | 0.036 | 2.79225403982462e-39 | 9 |
| rpl282 | 1.12805914813193e-43 | 0.396794178 | 0.976 | 0.745 | 2.83921206993326e-39 | 9 |
| h2afx13 | 1.30852897758284e-43 | 0.525437414 | 0.542 | 0.192 | 3.29343658367826e-39 | 9 |
| s100u | 1.74806866458408e-43 | 0.265427805 | 0.191 | 0.033 | 4.39971402189168e-39 | 9 |
| si:dkey-42i9.6 | 2.75897886346801e-43 | 0.352847566 | 0.295 | 0.07 | 6.94407390146263e-39 | 9 |
| tpm2 | 3.30258752431128e-43 | 0.275869666 | 0.139 | 0.018 | 8.31228253993906e-39 | 9 |
| neo1a | 4.17786107616371e-43 | 0.255434894 | 0.163 | 0.025 | 1.05152585425964e-38 | 9 |
| hspb11 | 5.75819035026406e-43 | 0.33122014 | 0.271 | 0.06 | 1.44927892925796e-38 | 9 |
| zgc:174863 | 7.06254251087207e-43 | 0.372645113 | 0.358 | 0.097 | 1.77757132456139e-38 | 9 |
| aldh2.2 | 9.60279814002513e-43 | 0.342742692 | 0.312 | 0.078 | 2.41692826386292e-38 | 9 |
| rplp2l2 | 1.56416894491196e-42 | 0.406061777 | 0.99 | 0.77 | 3.93685681744892e-38 | 9 |

| | | | | | | |
|---|---|---|---|---|---|---|
| pdia31 | 2.19361711867958e-42 | 0.526927269 | 0.701 | 0.275 | 5.52111492600464e-38 | 9 |
| sh3pxd2b | 2.19371067149791e-42 | 0.294099061 | 0.215 | 0.042 | 5.5213503890931e-38 | 9 |
| col12a1a | 2.4364348721062e-42 | 0.254652224 | 0.16 | 0.024 | 6.1322629296041e-38 | 9 |
| rps142 | 3.45693696536098e-42 | 0.399232209 | 0.99 | 0.754 | 8.70076464811705e-38 | 9 |
| ctdsp1 | 3.63513218775754e-42 | 0.278643859 | 0.233 | 0.047 | 9.14926420336695e-38 | 9 |
| rps15a2 | 4.94432377028308e-42 | 0.403003661 | 0.979 | 0.776 | 1.24443684974255e-37 | 9 |
| ptena | 5.43525780362929e-42 | 0.290778387 | 0.233 | 0.048 | 1.36800003659546e-37 | 9 |
| rpl182 | 1.14237091535998e-41 | 0.385241366 | 0.983 | 0.77 | 2.87523335686954e-37 | 9 |
| flot2a | 2.08565843835441e-41 | 0.320362072 | 0.25 | 0.056 | 5.2493937234942e-37 | 9 |
| bgnb1 | 2.3227844420492e-41 | 0.285338123 | 0.16 | 0.025 | 5.84621616219364e-37 | 9 |
| rps101 | 4.03907989161465e-41 | 0.444446506 | 0.976 | 0.688 | 1.01659601792049e-36 | 9 |
| eppk11 | 5.89723777152162e-41 | 0.288049368 | 0.201 | 0.038 | 1.48427577471428e-36 | 9 |
| ddit31 | 6.22012243010011e-41 | 0.492164087 | 0.497 | 0.17 | 1.5655426144319e-36 | 9 |
| rpl82 | 1.27220751476039e-40 | 0.360433115 | 0.965 | 0.777 | 3.20201909390041e-36 | 9 |
| rps4x1 | 1.82482561137993e-40 | 0.428539239 | 0.976 | 0.705 | 4.59290358128215e-36 | 9 |
| si:dkey-28b4.7 | 3.91993831882584e-40 | 0.348939973 | 0.326 | 0.087 | 9.86609275465276e-36 | 9 |
| rpl35a2 | 4.67922403906276e-40 | 0.377996286 | 0.965 | 0.755 | 1.17771389839171e-35 | 9 |
| appb1 | 6.40822560398582e-40 | 0.278610684 | 0.194 | 0.037 | 1.61288630226719e-35 | 9 |
| capns1a | 7.21853715820355e-40 | 0.397539992 | 0.438 | 0.137 | 1.81683361734825e-35 | 9 |
| rps61 | 8.82685412582608e-40 | 0.357766975 | 0.979 | 0.785 | 2.22163091492917e-35 | 9 |
| ptx3a1 | 1.90080306122839e-39 | 0.316953113 | 0.188 | 0.034 | 4.78413122480574e-35 | 9 |
| nfixb1 | 1.92145826476786e-39 | 0.323188346 | 0.264 | 0.061 | 4.83611830659423e-35 | 9 |
| rpl22l11 | 2.26665446913413e-39 | 0.571832814 | 0.802 | 0.383 | 5.70494263336369e-35 | 9 |
| faua2 | 2.88587312506415e-39 | 0.435582616 | 0.965 | 0.684 | 7.26345406847397e-35 | 9 |
| sept10 | 2.99739683770812e-39 | 0.276152678 | 0.215 | 0.044 | 7.54414810082758e-35 | 9 |
| gnl3 | 3.04252575389676e-39 | 0.293932026 | 0.25 | 0.057 | 7.65773306998275e-35 | 9 |
| dedd11 | 5.29818561745509e-39 | 0.427745802 | 0.434 | 0.14 | 1.33350033805727e-34 | 9 |
| rps111 | 6.90241598333491e-39 | 0.366317617 | 0.986 | 0.761 | 1.73726907884556e-34 | 9 |
| rpl382 | 7.80937392408967e-39 | 0.418682269 | 0.979 | 0.723 | 1.96554132295413e-34 | 9 |
| rpl172 | 7.93232007292678e-39 | 0.372988344 | 0.972 | 0.742 | 1.99648563915494e-34 | 9 |
| b4galt6 | 8.70800540640276e-39 | 0.324672657 | 0.264 | 0.063 | 2.19171788073751e-34 | 9 |
| abcc51 | 2.01825015723146e-38 | 0.360784315 | 0.326 | 0.09 | 5.07973382073586e-34 | 9 |
| gpx4a1 | 2.11776982365994e-38 | 0.320945281 | 0.281 | 0.07 | 5.33021486916971e-34 | 9 |
| eef1g1 | 2.28908141908143e-38 | 0.472888952 | 0.934 | 0.614 | 5.76138902368604e-34 | 9 |
| rps212 | 3.27037621150846e-38 | 0.411492768 | 0.972 | 0.745 | 8.23120988674566e-34 | 9 |

| | | | | | | |
|---|---|---|---|---|---|---|
| emp21 | 4.23246574303623e-38 | 0.26953432 | 0.219 | 0.046 | 1.06526930286479e-33 | 9 |
| tnrc6c11 | 5.91777784961004e-38 | 0.421648854 | 0.399 | 0.128 | 1.48944550696835e-33 | 9 |
| rpl5a1 | 6.9404315576337e-38 | 0.555612545 | 0.91 | 0.483 | 1.74683721874083e-33 | 9 |
| tpm3 | 7.19905874052209e-38 | 0.479329975 | 0.625 | 0.247 | 1.811931094402e-33 | 9 |
| si:ch211-15b10.61 | 7.87934000904945e-38 | 0.340843903 | 0.271 | 0.068 | 1.98315108687766e-33 | 9 |
| ahnak1 | 7.95367500783308e-38 | 0.378939357 | 0.399 | 0.121 | 2.00186046272151e-33 | 9 |
| rps26l2 | 9.79969682675262e-38 | 0.46183554 | 0.962 | 0.658 | 2.46648569432537e-33 | 9 |
| rpl302 | 1.14030087272851e-37 | 0.416147221 | 0.969 | 0.703 | 2.87002326657039e-33 | 9 |
| rpl71 | 1.57713957687589e-37 | 0.316911538 | 0.962 | 0.802 | 3.96950260103894e-33 | 9 |
| rps3a1 | 1.65030023768402e-37 | 0.353977368 | 0.986 | 0.774 | 4.1536406682269e-33 | 9 |
| capns1b | 5.03170594135178e-37 | 0.356641519 | 0.382 | 0.117 | 1.26643006837883e-32 | 9 |
| eif4a1b | 5.84903538856468e-37 | 0.492409748 | 0.66 | 0.272 | 1.47214371694784e-32 | 9 |
| rps261 | 6.14919217516852e-37 | 0.380153155 | 0.965 | 0.729 | 1.54769017856816e-32 | 9 |
| calm2a | 1.10314448532761e-36 | 0.461718207 | 0.535 | 0.196 | 2.77650435512106e-32 | 9 |
| add3a1 | 1.13240675588255e-36 | 0.393673715 | 0.385 | 0.119 | 2.85015456388079e-32 | 9 |
| anxa4 | 1.85086178387576e-36 | 0.387405469 | 0.406 | 0.131 | 4.65843402383689e-32 | 9 |
| rps292 | 2.04681088144391e-36 | 0.326964678 | 0.997 | 0.829 | 5.15161830750617e-32 | 9 |
| pno1 | 2.73108333846756e-36 | 0.395888997 | 0.476 | 0.165 | 6.873863654589e-32 | 9 |
| mt-nd21 | 2.77089028023924e-36 | 0.520097626 | 0.847 | 0.391 | 6.97405374633415e-32 | 9 |
| rpl18a2 | 4.56105632676327e-36 | 0.354937281 | 0.969 | 0.781 | 1.14797226688305e-31 | 9 |
| fosl2 | 5.19676556549287e-36 | 0.392309835 | 0.375 | 0.117 | 1.3079739251789e-31 | 9 |
| gapdhs2 | 5.23963621097473e-36 | 0.485563085 | 0.688 | 0.28 | 1.31876403794023e-31 | 9 |
| snai1a | 5.32654880733665e-36 | 0.275105538 | 0.236 | 0.054 | 1.34063906931856e-31 | 9 |
| dbi | 1.21664574465131e-35 | 0.452030862 | 0.542 | 0.206 | 3.06217567471288e-31 | 9 |
| cldnc | 1.47632021382401e-35 | 0.27883273 | 0.153 | 0.026 | 3.71575034617364e-31 | 9 |
| tgif1 | 1.64955223986066e-35 | 0.322005121 | 0.312 | 0.087 | 4.1517580325053e-31 | 9 |
| tmsb4x3 | 1.8322195981471e-35 | 0.545941591 | 1 | 0.68 | 4.61151350657644e-31 | 9 |
| hdlbpa1 | 3.15449190822149e-35 | 0.397173532 | 0.448 | 0.154 | 7.93954068380266e-31 | 9 |
| CABZ01072614.11 | 5.38309050598512e-35 | 0.335252645 | 0.285 | 0.077 | 1.35487004945139e-30 | 9 |
| pleca | 5.70691597138049e-35 | 0.284617187 | 0.236 | 0.056 | 1.43637368083676e-30 | 9 |
| pdk2b1 | 5.86532896073616e-35 | 0.421329121 | 0.319 | 0.096 | 1.47624464612768e-30 | 9 |
| uba521 | 6.88135789119012e-35 | 0.467806466 | 0.965 | 0.623 | 1.73196896763364e-30 | 9 |
| eif3g1 | 7.06156108379561e-35 | 0.454714245 | 0.667 | 0.274 | 1.77732430918052e-30 | 9 |
| serf21 | 1.06328402395416e-34 | 0.495936857 | 0.75 | 0.334 | 2.67617955989022e-30 | 9 |
| serpinf1 | 1.15032759778843e-34 | 0.275544944 | 0.194 | 0.041 | 2.8952595308737e-30 | 9 |

| | | | | | | |
|---|---|---|---|---|---|---|
| cflara | 1.60738613015899e-34 | 0.299260216 | 0.243 | 0.06 | 4.04563015099716e-30 | 9 |
| rpl222 | 1.67260028659509e-34 | 0.403770227 | 0.941 | 0.689 | 4.20976766133118e-30 | 9 |
| romo11 | 2.18690888596327e-34 | 0.50986965 | 0.812 | 0.377 | 5.50423097508094e-30 | 9 |
| rpl152 | 2.2169441511593e-34 | 0.385435916 | 0.976 | 0.74 | 5.57982673405283e-30 | 9 |
| serinc5 | 2.22371073970237e-34 | 0.308845795 | 0.292 | 0.079 | 5.5968575607569e-30 | 9 |
| rpl271 | 2.663954167844e-34 | 0.357519276 | 0.969 | 0.758 | 6.70490624504657e-30 | 9 |
| dnmt3ab1 | 5.63970797357145e-34 | 0.322255816 | 0.278 | 0.075 | 1.4194580998682e-29 | 9 |
| jund1 | 1.11607452249387e-33 | 0.368297302 | 0.306 | 0.089 | 2.80904796566483e-29 | 9 |
| rps52 | 1.21164123404497e-33 | 0.409120828 | 0.962 | 0.696 | 3.0495798219678e-29 | 9 |
| tob1b1 | 1.38488154035399e-33 | 0.386493171 | 0.476 | 0.168 | 3.48560834891696e-29 | 9 |
| arl6ip5b1 | 1.4078309274762e-33 | 0.323605153 | 0.316 | 0.092 | 3.54336966136485e-29 | 9 |
| krt18b2 | 1.58483526632425e-33 | 0.371596993 | 0.438 | 0.145 | 3.9888718818115e-29 | 9 |
| rplp01 | 1.76071962870058e-33 | 0.284387216 | 0.986 | 0.848 | 4.43155523347649e-29 | 9 |
| ywhaqb1 | 1.94777159100954e-33 | 0.384371326 | 0.382 | 0.127 | 4.90234631741192e-29 | 9 |
| flot1b | 1.99819803125973e-33 | 0.298838689 | 0.285 | 0.078 | 5.02926462487761e-29 | 9 |
| h1f0 | 2.13597733506249e-33 | 0.322300247 | 0.24 | 0.059 | 5.37604135461877e-29 | 9 |
| baz2ba | 3.23170025399039e-33 | 0.26493943 | 0.226 | 0.054 | 8.13386636926842e-29 | 9 |
| hmgb1a1 | 3.23457429111219e-33 | 0.414956781 | 0.49 | 0.183 | 8.14110003330028e-29 | 9 |
| ube2e31 | 5.11563047792248e-33 | 0.390324845 | 0.427 | 0.15 | 1.28755303498831e-28 | 9 |
| lgals3b | 6.28953764235273e-33 | 0.286626398 | 0.274 | 0.075 | 1.58301372920376e-28 | 9 |
| tomm5 | 6.80788196348065e-33 | 0.404090074 | 0.535 | 0.208 | 1.71347581138844e-28 | 9 |
| rpl352 | 1.97557467065997e-32 | 0.317188491 | 0.972 | 0.801 | 4.97232388858407e-28 | 9 |
| rpl121 | 3.0658087804568e-32 | 0.315463214 | 0.983 | 0.795 | 7.71633411953172e-28 | 9 |
| tspan7 | 3.1559560459314e-32 | 0.27165591 | 0.278 | 0.075 | 7.94322577200474e-28 | 9 |
| txnipa2 | 4.53077172278932e-32 | 0.446825529 | 0.521 | 0.204 | 1.14034993490884e-27 | 9 |
| map1lc3a | 5.66581217536979e-32 | 0.287147959 | 0.253 | 0.067 | 1.42602826641882e-27 | 9 |
| prkab1b | 5.8634812104557e-32 | 0.326309511 | 0.361 | 0.117 | 1.47577958585959e-27 | 9 |
| dbn11 | 6.43656942296871e-32 | 0.263041876 | 0.219 | 0.053 | 1.62002015806699e-27 | 9 |
| pim2 | 1.2102398814792e-31 | 0.304114135 | 0.295 | 0.086 | 3.046052757695e-27 | 9 |
| rps121 | 1.51018804024724e-31 | 0.292269387 | 0.986 | 0.837 | 3.80099227849827e-27 | 9 |
| rpl212 | 1.67008775285257e-31 | 0.353594644 | 0.976 | 0.752 | 4.20344386515464e-27 | 9 |
| naca2 | 2.87495469125533e-31 | 0.444936581 | 0.927 | 0.612 | 7.23597346242054e-27 | 9 |
| ctnnb1 | 3.67278561244084e-31 | 0.338441047 | 0.42 | 0.144 | 9.24403410795236e-27 | 9 |
| pfn2l | 3.68911965441161e-31 | 0.382771352 | 0.538 | 0.203 | 9.28514525818859e-27 | 9 |
| rps27l | 4.06583789663007e-31 | 0.392655366 | 0.51 | 0.197 | 1.02333074020282e-26 | 9 |

| | | | | | | |
|---|---|---|---|---|---|---|
| rps31 | 4.54529214063031e-31 | 0.385094775 | 0.948 | 0.695 | 1.14400457887524e-26 | 9 |
| emd | 4.63773014708867e-31 | 0.255589741 | 0.215 | 0.053 | 1.16727030072075e-26 | 9 |
| bhlhe40 | 1.38859367604689e-30 | 0.37096822 | 0.438 | 0.155 | 3.49495142324243e-26 | 9 |
| si:ch211-140m22.7 | 2.37280316947758e-30 | 0.277428035 | 0.24 | 0.063 | 5.97210829725812e-26 | 9 |
| rps251 | 3.0070466471543e-30 | 0.376949279 | 0.979 | 0.72 | 7.56843570622265e-26 | 9 |
| XLOC-0160941 | 3.51693195272427e-30 | 0.272801397 | 0.274 | 0.076 | 8.85176603181172e-26 | 9 |
| cry2 | 4.46121354652162e-30 | 0.300091371 | 0.271 | 0.077 | 1.12284283752403e-25 | 9 |
| ddt | 8.28541993039644e-30 | 0.299492238 | 0.309 | 0.096 | 2.08535734228148e-25 | 9 |
| serbp1a1 | 8.44098195433405e-30 | 0.502835969 | 0.882 | 0.473 | 2.12451074808634e-25 | 9 |
| atrx | 8.9306844028093e-30 | 0.312569047 | 0.319 | 0.101 | 2.24776395734307e-25 | 9 |
| nop531 | 9.54894814952907e-30 | 0.418928106 | 0.583 | 0.244 | 2.40337475975497e-25 | 9 |
| entpd1 | 1.16054940321345e-29 | 0.266951244 | 0.25 | 0.067 | 2.92098679294792e-25 | 9 |
| foxo1a | 1.39747882357871e-29 | 0.253038904 | 0.174 | 0.038 | 3.51731445106526e-25 | 9 |
| icn2 | 1.96154356907937e-29 | 0.399785515 | 0.656 | 0.267 | 4.93700900901587e-25 | 9 |
| zbtb20 | 3.22758544534613e-29 | 0.325159783 | 0.302 | 0.095 | 8.12350980739167e-25 | 9 |
| cct21 | 4.1790005927775e-29 | 0.377855241 | 0.549 | 0.221 | 1.05181265919617e-24 | 9 |
| nop56 | 4.28487275078871e-29 | 0.379469977 | 0.438 | 0.166 | 1.07845962264601e-24 | 9 |
| btg12 | 4.62739875973837e-29 | 0.440425202 | 0.736 | 0.324 | 1.16466999383855e-24 | 9 |
| fabp31 | 7.68431784677798e-29 | 0.391126577 | 0.576 | 0.235 | 1.93406595885555e-24 | 9 |
| zgc:153867 | 1.51075120372137e-28 | 0.483566477 | 0.788 | 0.429 | 3.80240970464631e-24 | 9 |
| stat21 | 1.83002952110589e-28 | 0.265177429 | 0.222 | 0.058 | 4.60600130167141e-24 | 9 |
| appa1 | 2.93434587901156e-28 | 0.29353989 | 0.257 | 0.074 | 7.3854551428842e-24 | 9 |
| rpl101 | 3.93082540826076e-28 | 0.323903124 | 0.965 | 0.769 | 9.89349447005151e-24 | 9 |
| rpl311 | 5.02037836903053e-28 | 0.429257639 | 0.972 | 0.648 | 1.2635790317013e-23 | 9 |
| eif4ebp3l1 | 5.63729612452036e-28 | 0.403999386 | 0.507 | 0.2 | 1.41885106158053e-23 | 9 |
| crebzf | 8.12772068571591e-28 | 0.312209903 | 0.306 | 0.098 | 2.04566601938784e-23 | 9 |
| pin1 | 8.75087553070783e-28 | 0.30173738 | 0.392 | 0.139 | 2.20250786232385e-23 | 9 |
| metap1 | 9.00952326591396e-28 | 0.250623592 | 0.264 | 0.077 | 2.26760691079788e-23 | 9 |
| eif3m1 | 2.96276346138861e-27 | 0.44299989 | 0.743 | 0.35 | 7.45697935596898e-23 | 9 |
| msna1 | 3.02455320187337e-27 | 0.401280976 | 0.556 | 0.236 | 7.61249795379508e-23 | 9 |
| ndfip1l1 | 5.33077917556683e-27 | 0.283773346 | 0.229 | 0.064 | 1.34170381069842e-22 | 9 |
| slc20a1b | 6.71701431422755e-27 | 0.325052929 | 0.302 | 0.098 | 1.69060533274793e-22 | 9 |
| cdc42 | 8.8959200984626e-27 | 0.368574898 | 0.503 | 0.207 | 2.23901412958205e-22 | 9 |
| mt-nd11 | 9.75116570902812e-27 | 0.438724072 | 0.799 | 0.403 | 2.45427089730529e-22 | 9 |
| wls1 | 1.07539176722049e-26 | 0.305074489 | 0.347 | 0.119 | 2.70665353891726e-22 | 9 |

| | | | | | | |
|---|---|---|---|---|---|---|
| LOC101884451 | 1.37700900297776e-26 | 0.305366866 | 0.267 | 0.082 | 3.46579395959472e-22 | 9 |
| zgc:1749041 | 1.49094291907881e-26 | 0.26110682 | 0.188 | 0.046 | 3.75255423302945e-22 | 9 |
| usp36 | 1.85112457037556e-26 | 0.276416585 | 0.243 | 0.072 | 4.65909543117825e-22 | 9 |
| cxcl8b.11 | 2.50875153179661e-26 | 0.300390048 | 0.153 | 0.033 | 6.31427673037888e-22 | 9 |
| gstr | 4.18119086076628e-26 | 0.290569628 | 0.333 | 0.115 | 1.05236392774627e-21 | 9 |
| ppiaa1 | 6.10861274961598e-26 | 0.302477302 | 0.99 | 0.8 | 1.53747674295085e-21 | 9 |
| zgc:110339 | 1.01846491229027e-25 | 0.250295448 | 0.292 | 0.093 | 2.56337433774338e-21 | 9 |
| zfp36l1a2 | 1.28391980700379e-25 | 0.397409347 | 0.562 | 0.242 | 3.23149776224785e-21 | 9 |
| marcksl1b1 | 1.5282717524419e-25 | 0.34124356 | 0.326 | 0.112 | 3.84650717372103e-21 | 9 |
| tpt11 | 2.1726841491376e-25 | 0.287888253 | 0.976 | 0.797 | 5.46842873496443e-21 | 9 |
| hnrnpa0b | 2.45850222160371e-25 | 0.405163936 | 0.663 | 0.304 | 6.18780424155437e-21 | 9 |
| si:ch211-132p1.31 | 2.77206126999877e-25 | 0.279372448 | 0.271 | 0.085 | 6.9770010104599e-21 | 9 |
| skp1 | 2.94778012379434e-25 | 0.354156867 | 0.569 | 0.245 | 7.41926779357798e-21 | 9 |
| larp4ab | 4.14091932788404e-25 | 0.259341899 | 0.233 | 0.068 | 1.04222798563513e-20 | 9 |
| rhoaa1 | 4.16350493508296e-25 | 0.317708427 | 0.392 | 0.149 | 1.04791255711103e-20 | 9 |
| rpl13a1 | 7.16920907674962e-25 | 0.30928259 | 0.969 | 0.76 | 1.80441823252711e-20 | 9 |
| rplp22 | 9.19924537338617e-25 | 0.374586556 | 0.983 | 0.753 | 2.31535806802757e-20 | 9 |
| zfand5a | 1.02308443722139e-24 | 0.260772381 | 0.243 | 0.073 | 2.57500122004253e-20 | 9 |
| vdac1 | 1.09637503762693e-24 | 0.270628437 | 0.438 | 0.165 | 2.75946633220321e-20 | 9 |
| fn1a1 | 1.12325586738831e-24 | 0.279283408 | 0.219 | 0.062 | 2.82712269262963e-20 | 9 |
| anxa11b | 1.44642004769871e-24 | 0.30035167 | 0.385 | 0.146 | 3.64049461805287e-20 | 9 |
| sdcbp21 | 1.49696081459994e-24 | 0.338149179 | 0.49 | 0.203 | 3.76770067426658e-20 | 9 |
| rps27.12 | 2.70223196951613e-24 | 0.408862358 | 0.962 | 0.673 | 6.80124764407514e-20 | 9 |
| cd82b | 3.25902998516194e-24 | 0.250193071 | 0.24 | 0.072 | 8.2026525696541e-20 | 9 |
| eif3ha | 3.91885660780989e-24 | 0.322579336 | 0.49 | 0.203 | 9.86337019619671e-20 | 9 |
| midn | 3.9742222417307e-24 | 0.334345075 | 0.361 | 0.136 | 1.0002719960212e-19 | 9 |
| fxyd6l1 | 4.7008270764145e-24 | 0.256286071 | 0.323 | 0.11 | 1.18315116686277e-19 | 9 |
| rbm28 | 6.4373853947177e-24 | 0.258486419 | 0.264 | 0.085 | 1.6202255299965e-19 | 9 |
| ier2b2 | 9.95250168745099e-24 | 0.368802474 | 0.507 | 0.218 | 2.50494514971454e-19 | 9 |
| cd633 | 1.14694796923106e-23 | 0.399168363 | 0.712 | 0.333 | 2.88675334375765e-19 | 9 |
| ciarta1 | 2.13804660914973e-23 | 0.27154122 | 0.295 | 0.101 | 5.38124951056896e-19 | 9 |
| lgals2a1 | 2.32720426281297e-23 | 0.267361451 | 0.347 | 0.122 | 5.85734040907397e-19 | 9 |
| eif3i1 | 6.7952244945292e-23 | 0.351583967 | 0.59 | 0.267 | 1.71029005302805e-18 | 9 |
| mt-nd41 | 6.84473589751926e-23 | 0.414159865 | 0.872 | 0.447 | 1.72275157804662e-18 | 9 |
| sesn12 | 9.48131353391097e-23 | 0.33940143 | 0.406 | 0.164 | 2.38635180335005e-18 | 9 |

| | | | | | | |
|---|---|---|---|---|---|---|
| NDUFB1 | 1.21889118847011e-22 | 0.313573148 | 0.458 | 0.193 | 3.06782723226042e-18 | 9 |
| pfn2 | 2.0046260859492e-22 | 0.330600253 | 0.51 | 0.221 | 5.04544339572554e-18 | 9 |
| rack11 | 2.01402349155617e-22 | 0.312268981 | 0.951 | 0.74 | 5.06909572589772e-18 | 9 |
| cnn2 | 2.32815227423426e-22 | 0.313864412 | 0.344 | 0.131 | 5.85972645902021e-18 | 9 |
| rps231 | 4.44969160278162e-22 | 0.336673743 | 0.958 | 0.718 | 1.11994287950411e-17 | 9 |
| tcp1 | 6.33353085203506e-22 | 0.285656201 | 0.521 | 0.22 | 1.5940863801487e-17 | 9 |
| prdx51 | 8.07307932966166e-22 | 0.277893416 | 0.392 | 0.155 | 2.03191333648254e-17 | 9 |
| acin1a1 | 2.02158521694419e-21 | 0.333488291 | 0.479 | 0.212 | 5.08812783252684e-17 | 9 |
| tegt | 2.16588323363787e-21 | 0.30646356 | 0.476 | 0.207 | 5.45131151074316e-17 | 9 |
| eif3f1 | 2.9672768009199e-21 | 0.440180725 | 0.819 | 0.423 | 7.46833898023529e-17 | 9 |
| rpl41 | 1.06220525278994e-20 | 0.392030437 | 0.941 | 0.621 | 2.67346440074701e-16 | 9 |
| mtpn | 1.32614562722546e-20 | 0.272351919 | 0.358 | 0.143 | 3.33777592916375e-16 | 9 |
| COX32 | 3.31945588690533e-20 | 0.413481212 | 0.997 | 0.694 | 8.35473852175201e-16 | 9 |
| tomm20b1 | 4.46968512841707e-20 | 0.306218459 | 0.469 | 0.21 | 1.12497504997129e-15 | 9 |
| cct8 | 5.22666016543244e-20 | 0.296829653 | 0.451 | 0.197 | 1.31549809703769e-15 | 9 |
| ppdpfb1 | 5.22906105717928e-20 | 0.407837127 | 0.844 | 0.512 | 1.31610237748145e-15 | 9 |
| rsl1d1 | 7.21031587139257e-20 | 0.266472828 | 0.424 | 0.176 | 1.8147644016708e-15 | 9 |
| irf2bp2a1 | 8.3975899121687e-20 | 0.295023141 | 0.406 | 0.172 | 2.11358940499374e-15 | 9 |
| laptm4a | 8.48741644912863e-20 | 0.260514665 | 0.285 | 0.106 | 2.13619784608119e-15 | 9 |
| eif3s101 | 1.03523838623829e-19 | 0.274393783 | 0.385 | 0.161 | 2.60559149432315e-15 | 9 |
| srsf5a | 1.09553229810573e-19 | 0.319809647 | 0.601 | 0.282 | 2.75734524110231e-15 | 9 |
| pabpc1a1 | 1.25611406103773e-19 | 0.411365972 | 0.875 | 0.533 | 3.16151348022586e-15 | 9 |
| mt-atp62 | 2.46367517282705e-19 | 0.412813247 | 0.955 | 0.583 | 6.20082404248841e-15 | 9 |
| si:dkey-42i9.4 | 3.175652912397e-19 | 0.255350619 | 0.354 | 0.143 | 7.992800815212e-15 | 9 |
| eif2s3 | 3.60532289179015e-19 | 0.3251351 | 0.583 | 0.279 | 9.07423718634663e-15 | 9 |
| eif2s2 | 3.70412405728165e-19 | 0.328344755 | 0.597 | 0.288 | 9.32290983977218e-15 | 9 |
| COX7A2 (1 of many)1 | 5.05926858290764e-19 | 0.299803967 | 0.549 | 0.255 | 1.27336730963202e-14 | 9 |
| gabarapb1 | 7.38778860576852e-19 | 0.300061037 | 0.566 | 0.263 | 1.85943251418588e-14 | 9 |
| crema1 | 8.94129587418605e-19 | 0.283250711 | 0.406 | 0.174 | 2.25043475857389e-14 | 9 |
| mibp22 | 9.31034722192525e-19 | 0.4090452 | 0.844 | 0.485 | 2.34332129228637e-14 | 9 |
| sh3bgrl | 1.86931728260863e-18 | 0.253982096 | 0.271 | 0.102 | 4.70488466859766e-14 | 9 |
| rpl5b1 | 3.88014616910788e-18 | 0.383786824 | 0.948 | 0.623 | 9.76593989302762e-14 | 9 |
| caprin1b | 5.58144899132515e-18 | 0.29235011 | 0.514 | 0.242 | 1.40479489662663e-13 | 9 |
| btf31 | 7.94554618549378e-18 | 0.405852976 | 0.938 | 0.58 | 1.99981451942693e-13 | 9 |
| mphosph8 | 1.6188122075098e-17 | 0.260069299 | 0.292 | 0.116 | 4.07438844508141e-13 | 9 |

| | | | | | | |
|---|---|---|---|---|---|---|
| ip6k2b1 | 1.77011159445247e-17 | 0.286049202 | 0.295 | 0.12 | 4.45519387207743e-13 | 9 |
| ak2 | 3.84385956271512e-17 | 0.285897465 | 0.479 | 0.228 | 9.67461013339769e-13 | 9 |
| hsp90ab11 | 4.10302769834136e-17 | 0.268555125 | 0.976 | 0.787 | 1.03269104139554e-12 | 9 |
| rsl24d11 | 5.74240960572437e-17 | 0.391034323 | 0.788 | 0.437 | 1.44530707366477e-12 | 9 |
| eif3c1 | 2.91907551537598e-16 | 0.260144415 | 0.368 | 0.166 | 7.34702116464981e-12 | 9 |
| uqcrh | 2.95386766439768e-16 | 0.258915969 | 0.514 | 0.245 | 7.43458952452253e-12 | 9 |
| rpl141 | 3.36939005759791e-16 | 0.371463248 | 0.948 | 0.628 | 8.48041783596817e-12 | 9 |
| prdx2 | 3.60798612307661e-16 | 0.346907806 | 0.743 | 0.394 | 9.08094027317153e-12 | 9 |
| ddx211 | 3.94917761086649e-16 | 0.263849324 | 0.438 | 0.203 | 9.93968512878988e-12 | 9 |
| rab101 | 4.03175096434353e-16 | 0.258308636 | 0.413 | 0.191 | 1.01475140021562e-11 | 9 |
| eif3ea1 | 6.28432256974381e-16 | 0.288903911 | 0.549 | 0.272 | 1.58170114757882e-11 | 9 |
| mt-nd31 | 1.12758444868825e-15 | 0.299940461 | 0.743 | 0.389 | 2.83801729890345e-11 | 9 |
| eif3d | 1.14119595609371e-15 | 0.26586342 | 0.51 | 0.248 | 2.87227610189227e-11 | 9 |
| oaz1b1 | 1.27238601458778e-15 | 0.295941886 | 0.625 | 0.317 | 3.202468360116e-11 | 9 |
| ccni1 | 1.430117507639e-15 | 0.284780909 | 0.576 | 0.289 | 3.59946275497659e-11 | 9 |
| tmem258 | 2.06115419720549e-15 | 0.270514443 | 0.556 | 0.277 | 5.1877189989465e-11 | 9 |
| ddx5 | 2.64038476592415e-15 | 0.326323983 | 0.653 | 0.351 | 6.64558441735449e-11 | 9 |
| mknk2b2 | 4.65487620377892e-15 | 0.262452902 | 0.41 | 0.194 | 1.17158579172912e-10 | 9 |
| syncrip | 7.26304660580745e-14 | 0.254928575 | 0.583 | 0.295 | 1.82803620021568e-09 | 9 |
| si:ch211-196l7.42 | 8.35055441662639e-14 | 0.286476838 | 0.576 | 0.31 | 2.1017510411207e-09 | 9 |
| NC-002333.42 | 1.06896098029467e-13 | 0.342186598 | 0.944 | 0.616 | 2.69046789130366e-09 | 9 |
| eif5a1 | 2.90642254494333e-13 | 0.352649254 | 0.809 | 0.472 | 7.31517490336787e-09 | 9 |
| ubb | 5.27233849725800e-13 | 0.255415821 | 0.594 | 0.311 | 1.32699487637487e-08 | 9 |
| mcl1a1 | 1.07382280325774e-12 | 0.321188869 | 0.92 | 0.618 | 2.7027046135194e-08 | 9 |
| eef1da1 | 1.20026810598783e-12 | 0.353444149 | 0.837 | 0.485 | 3.02095479596077e-08 | 9 |
| eef2b | 1.61847279947841e-11 | 0.317346966 | 0.899 | 0.608 | 4.07353418900722e-07 | 9 |
| fkbp1aa | 2.1310674661127e-11 | 0.277876957 | 0.688 | 0.382 | 5.36368370545905e-07 | 9 |
| mt-cyb1 | 4.16686175460035e-10 | 0.293474899 | 0.948 | 0.617 | 1.04875743501536e-05 | 9 |
| sec61b | 4.57398257370209e-09 | 0.266571002 | 0.774 | 0.448 | 0.000115123 | 9 |
| mt-co12 | 1.52298572315086e-08 | 0.316886075 | 0.983 | 0.651 | 0.00038332 | 9 |
| zgc:564931 | 2.22991672436593e-08 | 0.290650577 | 0.854 | 0.567 | 0.000561248 | 9 |
| ptmab1 | 3.92500973281871e-08 | 0.282936004 | 0.823 | 0.535 | 0.000987886 | 9 |
| mt-co22 | 6.90176244214925e-08 | 0.308697653 | 0.993 | 0.715 | 0.001737105 | 9 |
| si:ch211-14a17.101 | 0 | 2.442187302 | 0.993 | 0.066 | 0 | 10 |
| spink2.21 | 0 | 2.175369385 | 0.934 | 0.112 | 0 | 10 |

| | | | | | | |
|---|---|---|---|---|---|---|
| spink2.3 | 0 | 2.169960595 | 0.909 | 0.05 | 0 | 10 |
| spink2.41 | 0 | 2.160547754 | 0.885 | 0.05 | 0 | 10 |
| si:dkey-203a12.22 | 0 | 2.146976088 | 0.997 | 0.228 | 0 | 10 |
| zmp:00000013231 | 0 | 2.144795159 | 0.958 | 0.124 | 0 | 10 |
| si:ch211-194m7.8 | 0 | 2.121137437 | 0.92 | 0.037 | 0 | 10 |
| ccl34b.9 | 0 | 2.056041534 | 0.882 | 0.03 | 0 | 10 |
| LOC110437886 | 0 | 2.044892271 | 0.892 | 0.015 | 0 | 10 |
| itln1 | 0 | 2.035823323 | 0.902 | 0.039 | 0 | 10 |
| si:dkey-77g12.4 | 0 | 2.018465115 | 0.826 | 0.029 | 0 | 10 |
| BX322787.11 | 0 | 2.016849274 | 0.934 | 0.117 | 0 | 10 |
| MFAP4 (1 of many) | 0 | 2.010790104 | 0.885 | 0.044 | 0 | 10 |
| si:ch73-359m17.22 | 0 | 1.972037109 | 0.976 | 0.17 | 0 | 10 |
| si:ch1073-110a20.3 | 0 | 1.964954935 | 0.868 | 0.03 | 0 | 10 |
| clca1 | 0 | 1.95376674 | 0.861 | 0.012 | 0 | 10 |
| si:dkey-203a12.5 | 0 | 1.90047866 | 0.794 | 0.032 | 0 | 10 |
| cfbl | 0 | 1.885305644 | 0.861 | 0.046 | 0 | 10 |
| LOC110438970 | 0 | 1.833059632 | 0.801 | 0.012 | 0 | 10 |
| LOC1104377311 | 0 | 1.799581244 | 0.854 | 0.071 | 0 | 10 |
| CABZ01112049.1 | 0 | 1.74192134 | 0.756 | 0.008 | 0 | 10 |
| FO203432.1 | 0 | 1.699001848 | 0.76 | 0.024 | 0 | 10 |
| cfp1 | 0 | 1.548150104 | 0.753 | 0.059 | 0 | 10 |
| cfi | 0 | 1.50912578 | 0.693 | 0.005 | 0 | 10 |
| si:ch1073-376c22.1 | 0 | 1.482463815 | 0.648 | 0.013 | 0 | 10 |
| c3a.51 | 0 | 1.473578878 | 0.655 | 0.054 | 0 | 10 |
| cfhl2 | 0 | 1.399427227 | 0.624 | 0.007 | 0 | 10 |
| BX908782.3 | 0 | 1.322928276 | 0.554 | 0.023 | 0 | 10 |
| agr2 | 0 | 1.295352516 | 0.564 | 0.015 | 0 | 10 |
| spink2.5 | 0 | 1.290562087 | 0.383 | 0.012 | 0 | 10 |
| si:zfos-2330d3.1 | 0 | 1.277877848 | 0.491 | 0.015 | 0 | 10 |
| c3a.4 | 0 | 1.256359456 | 0.568 | 0.004 | 0 | 10 |
| si:dkey-203a12.4 | 0 | 1.211042556 | 0.345 | 0.01 | 0 | 10 |
| si:dkey-203a12.7 | 0 | 1.187546544 | 0.422 | 0.007 | 0 | 10 |
| ca6 | 0 | 1.100082166 | 0.488 | 0.008 | 0 | 10 |
| LOC108179162 | 0 | 0.994344881 | 0.422 | 0.006 | 0 | 10 |
| LOC101883382 | 0 | 0.867182676 | 0.369 | 0.003 | 0 | 10 |

| | | | | | | |
|---|---|---|---|---|---|---|
| cfh | 0 | 0.856528805 | 0.362 | 0.007 | 0 | 10 |
| timp4.2 | 0 | 0.850903796 | 0.355 | 0.006 | 0 | 10 |
| LOC103909452 | 0 | 0.829871073 | 0.345 | 0.008 | 0 | 10 |
| cd59 | 0 | 0.827465155 | 0.338 | 0.006 | 0 | 10 |
| pdia2 | 0 | 0.746947375 | 0.324 | 0.009 | 0 | 10 |
| nfe2 | 0 | 0.715657544 | 0.31 | 0.006 | 0 | 10 |
| LOC110438722 | 0 | 0.608594571 | 0.226 | 0.001 | 0 | 10 |
| osmr | 0 | 0.600736921 | 0.23 | 0.003 | 0 | 10 |
| LOC110437920 | 0 | 0.55404501 | 0.213 | 0.001 | 0 | 10 |
| BX323555.3 | 0 | 0.472136705 | 0.188 | 0.002 | 0 | 10 |
| cfhl1 | 0 | 0.453150399 | 0.181 | 0.001 | 0 | 10 |
| si:dkey-102g19.3 | 3.09700025661234e-300 | 1.057107574 | 0.509 | 0.036 | 7.79483994586759e-296 | 10 |
| viml | 5.33728387440873e-299 | 0.524101263 | 0.209 | 0.003 | 1.34334097834993e-294 | 10 |
| itgb3a | 8.96624267993687e-285 | 0.653640521 | 0.275 | 0.008 | 2.25671362011331e-280 | 10 |
| si:ch211-212d10.1 | 2.86099385873393e-284 | 0.843048148 | 0.348 | 0.015 | 7.20083544304743e-280 | 10 |
| si:dkey-27h10.2 | 8.97753098897632e-278 | 0.837643389 | 0.373 | 0.019 | 2.25955477461545e-273 | 10 |
| cdaa | 1.82381015480703e-252 | 0.90438186 | 0.401 | 0.026 | 4.5903477786338e-248 | 10 |
| si:dkey-203a12.92 | 1.00198595842129e-243 | 1.591122454 | 0.753 | 0.125 | 2.52189845875054e-239 | 10 |
| h6pd | 8.89234060712347e-243 | 0.626372317 | 0.251 | 0.009 | 2.23811320740691e-238 | 10 |
| si:ch211-214p16.2 | 2.24692658572812e-236 | 1.130868862 | 0.561 | 0.061 | 5.6552895236191e-232 | 10 |
| si:dkeyp-66d1.7 | 1.9700869773487e-235 | 0.510321096 | 0.195 | 0.004 | 4.95851191328895e-231 | 10 |
| cxcr2 | 1.0012158404133e-226 | 0.398503921 | 0.157 | 0.002 | 2.51996014873623e-222 | 10 |
| si:dkeyp-75b4.102 | 1.60799956161762e-217 | 1.714156742 | 0.997 | 0.424 | 4.04717409663538e-213 | 10 |
| CR848841.1 | 1.56910602180376e-209 | 0.650000963 | 0.261 | 0.012 | 3.94928294627787e-205 | 10 |
| nfkbiaa | 8.89707809737172e-196 | 1.331282543 | 0.721 | 0.139 | 2.23930558632749e-191 | 10 |
| si:ch211-194m7.5 | 8.99967983409728e-190 | 0.365805781 | 0.143 | 0.003 | 2.26512941744395e-185 | 10 |
| adgre13 | 1.30195696470258e-188 | 0.453988212 | 0.167 | 0.004 | 3.27689548445993e-184 | 10 |
| clstn1 | 1.83129937447514e-175 | 0.628950797 | 0.268 | 0.016 | 4.60919739561647e-171 | 10 |
| si:ch73-160i9.2 | 2.77292570836669e-171 | 0.27544666 | 0.105 | 0.001 | 6.97917671538813e-167 | 10 |
| cnr2 | 2.01612015763618e-165 | 0.303623456 | 0.118 | 0.002 | 5.07437282475451e-161 | 10 |
| si:ch211-214p16.11 | 1.36436054358763e-163 | 1.305230414 | 0.93 | 0.242 | 3.4339590521557e-159 | 10 |
| spink2.15 | 1.128779299994e-154 | 1.234709193 | 0.934 | 0.426 | 2.84102462015489e-150 | 10 |
| si:ch211-270n8.1 | 2.54529002762451e-152 | 0.335655303 | 0.132 | 0.003 | 6.40624047052814e-148 | 10 |
| si:ch211-276a23.5 | 4.68102693789401e-147 | 0.422595629 | 0.171 | 0.007 | 1.17816766999854e-142 | 10 |
| myofl | 4.13534678806854e-146 | 0.553842828 | 0.233 | 0.015 | 1.04082543308897e-141 | 10 |

| | | | | | | |
|---|---|---|---|---|---|---|
| cxcl8b.12 | 4.17965792707097e-140 | 0.723171989 | 0.31 | 0.029 | 1.05197810366449e-135 | 10 |
| gata2a | 6.60842063433033e-138 | 0.51020287 | 0.213 | 0.013 | 1.6632733894546e-133 | 10 |
| adam8a | 2.78923685130623e-136 | 0.752614518 | 0.376 | 0.044 | 7.02023023105265e-132 | 10 |
| BX936393.1 | 2.20419358893427e-131 | 0.355387363 | 0.15 | 0.006 | 5.54773484398865e-127 | 10 |
| ank3b1 | 6.83883090573389e-126 | 0.678205691 | 0.307 | 0.032 | 1.72126535066416e-121 | 10 |
| ltb4r | 7.13084807751361e-108 | 0.550070387 | 0.244 | 0.023 | 1.7947631526294e-103 | 10 |
| qsox1 | 7.61206653268958e-108 | 0.564506233 | 0.258 | 0.026 | 1.91588102561264e-103 | 10 |
| cyp4f3 | 1.11758270406692e-99 | 0.404948657 | 0.174 | 0.012 | 2.81284390786602e-95 | 10 |
| sult5a1 | 1.74146668138942e-99 | 0.348068725 | 0.125 | 0.006 | 4.38309749038902e-95 | 10 |
| si:ch211-74f19.2 | 1.41290677533282e-98 | 0.57627413 | 0.251 | 0.027 | 3.55614506283517e-94 | 10 |
| srgn1 | 7.19189986933591e-97 | 0.980289145 | 0.7 | 0.227 | 1.81012927811316e-92 | 10 |
| myh11a1 | 9.2196494823424e-97 | 0.709645441 | 0.366 | 0.059 | 2.32049357821076e-92 | 10 |
| itgb2 | 5.85276458541762e-95 | 0.821022836 | 0.446 | 0.091 | 1.47308231850376e-90 | 10 |
| cd81a2 | 5.06320038113407e-91 | 0.952828597 | 0.641 | 0.197 | 1.27435690392763e-86 | 10 |
| mmp25b | 3.94759110487015e-85 | 0.306750138 | 0.129 | 0.008 | 9.93569205184767e-81 | 10 |
| fcer1gl1 | 1.75635420611214e-83 | 0.841506038 | 0.624 | 0.159 | 4.42056790136365e-79 | 10 |
| lrrc8da | 2.50735082147763e-83 | 0.442615135 | 0.192 | 0.019 | 6.31075128257705e-79 | 10 |
| alox5a | 1.46524979081678e-82 | 0.540997679 | 0.237 | 0.029 | 3.68788719850675e-78 | 10 |
| cd634 | 4.15777152276936e-81 | 0.884211871 | 0.742 | 0.332 | 1.04646951456582e-76 | 10 |
| cd7al | 7.27291664150322e-81 | 0.466880518 | 0.206 | 0.022 | 1.83052038949995e-76 | 10 |
| furina | 5.56255727554273e-80 | 0.638145712 | 0.324 | 0.055 | 1.40004004068135e-75 | 10 |
| selp | 1.33354512696631e-79 | 0.337611094 | 0.132 | 0.009 | 3.3563997300615e-75 | 10 |
| si:cabz01036022.1 | 2.4800223259046e-79 | 0.332287161 | 0.132 | 0.009 | 6.2419681920693e-75 | 10 |
| slc7a3a | 4.52273315191129e-78 | 0.581027303 | 0.265 | 0.038 | 1.13832670700455e-73 | 10 |
| atp1a3a | 3.996278222549e-73 | 0.423869848 | 0.174 | 0.018 | 1.00582326583336e-68 | 10 |
| mctp2b | 4.21737393702683e-73 | 0.415405497 | 0.171 | 0.017 | 1.06147084621028e-68 | 10 |
| dhrs13a.1 | 2.11388217122935e-72 | 0.438362672 | 0.181 | 0.019 | 5.32043003676714e-68 | 10 |
| alox5ap | 7.19150300167063e-72 | 0.383683027 | 0.178 | 0.018 | 1.81002939049048e-67 | 10 |
| illr4 | 1.58796623135807e-69 | 0.303507691 | 0.118 | 0.008 | 3.99675220770512e-65 | 10 |
| pfn12 | 4.4231516476479e-69 | 0.809012388 | 0.951 | 0.516 | 1.1132630381965e-64 | 10 |
| slc6a6b1 | 3.87877979477505e-67 | 0.721613203 | 0.401 | 0.097 | 9.76250086546933e-63 | 10 |
| lpar5b | 4.48975278795627e-67 | 0.435090833 | 0.185 | 0.022 | 1.13002587920071e-62 | 10 |
| LOC100148254 | 1.95842348371366e-65 | 0.411457124 | 0.178 | 0.02 | 4.9291560661589e-61 | 10 |
| spns3 | 4.08597315467604e-65 | 0.310166963 | 0.132 | 0.011 | 1.02839858330041e-60 | 10 |
| si:ch73-248e21.71 | 7.55857180211802e-64 | 0.501589415 | 0.247 | 0.04 | 1.90241693687508e-59 | 10 |

| | | | | | | |
|---|---|---|---|---|---|---|
| rgcc1 | 2.59467974315322e-63 | 0.627405606 | 0.321 | 0.065 | 6.53054944554233e-59 | 10 |
| si:ch73-237c6.12 | 1.3491358360311e-61 | 0.477708324 | 0.237 | 0.038 | 3.39563998570667e-57 | 10 |
| mt-co13 | 1.98896748218105e-61 | 0.588549024 | 0.934 | 0.652 | 5.00603225590148e-57 | 10 |
| mt-atp63 | 4.57255785729332e-61 | 0.657207871 | 0.899 | 0.584 | 1.15086708710216e-56 | 10 |
| def6c | 1.18755471714074e-60 | 0.281978402 | 0.118 | 0.01 | 2.98895646757153e-56 | 10 |
| coro1a2 | 6.74147002343718e-60 | 0.785037906 | 0.603 | 0.224 | 1.6967605901989e-55 | 10 |
| mt-co23 | 1.54114396742815e-59 | 0.550384271 | 0.948 | 0.716 | 3.87890525161991e-55 | 10 |
| ehd1a | 2.6361670671001e-57 | 0.415391574 | 0.199 | 0.029 | 6.63496889118425e-53 | 10 |
| limd2 | 1.46991836093321e-56 | 0.580359489 | 0.303 | 0.065 | 3.6996375226328e-52 | 10 |
| si:ch211-136m16.8 | 6.4040220165837e-56 | 0.409627705 | 0.188 | 0.026 | 1.61182830135395e-51 | 10 |
| si:dkey-206d17.12 | 1.67939383322542e-55 | 0.54797532 | 0.254 | 0.049 | 4.22686633884506e-51 | 10 |
| samsn1a | 1.38978904584046e-54 | 0.468063705 | 0.206 | 0.033 | 3.49796004947585e-50 | 10 |
| capgb1 | 1.84564220821936e-54 | 0.641532771 | 0.397 | 0.109 | 4.6452968738673e-50 | 10 |
| sema4aa | 1.35969518778762e-53 | 0.286512215 | 0.115 | 0.01 | 3.42221681814265e-49 | 10 |
| CT030188.1 | 2.7384389664118e-53 | 0.502610243 | 0.251 | 0.048 | 6.89237703456186e-49 | 10 |
| glipr1a1 | 9.96595376520905e-53 | 0.540148564 | 0.279 | 0.06 | 2.50833090316547e-48 | 10 |
| LOC108190702 | 1.00185329419734e-52 | 0.269370081 | 0.101 | 0.008 | 2.52156455616528e-48 | 10 |
| anks1b | 5.53395824248862e-52 | 0.348503437 | 0.136 | 0.015 | 1.39284195005196e-47 | 10 |
| pygb | 5.68128384811135e-52 | 0.448531994 | 0.23 | 0.041 | 1.42992233173115e-47 | 10 |
| mt-nd42 | 6.45541760601121e-52 | 0.671272662 | 0.791 | 0.449 | 1.62476405725696e-47 | 10 |
| tagapa1 | 3.65567407141085e-51 | 0.66528804 | 0.369 | 0.1 | 9.20096607033397e-47 | 10 |
| zmp:0000001082 | 5.30617805992337e-51 | 0.485011521 | 0.261 | 0.053 | 1.33551195590211e-46 | 10 |
| actb22 | 6.96663814706113e-51 | 0.397436501 | 0.983 | 0.866 | 1.75343315523382e-46 | 10 |
| mt-cyb2 | 2.4912803093286e-50 | 0.573008807 | 0.885 | 0.619 | 6.27030341054916e-46 | 10 |
| eno1a1 | 8.02014350762172e-50 | 0.692661172 | 0.408 | 0.129 | 2.01858991943331e-45 | 10 |
| cyba | 1.22471154036649e-49 | 0.533151718 | 0.296 | 0.069 | 3.08247647594842e-45 | 10 |
| pla2g4aa | 4.5997264426072e-49 | 0.297757212 | 0.129 | 0.014 | 1.15770514833981e-44 | 10 |
| zgc:1583431 | 1.33491534497096e-48 | 0.765495388 | 0.463 | 0.16 | 3.35984843175741e-44 | 10 |
| MYADM | 5.57108635310086e-48 | 0.421689559 | 0.195 | 0.033 | 1.40218672421195e-43 | 10 |
| pim21 | 5.61893494368113e-48 | 0.633528722 | 0.321 | 0.085 | 1.4142297359751e-43 | 10 |
| COX33 | 5.61756282374238e-47 | 0.512609248 | 0.923 | 0.696 | 1.41388438710772e-42 | 10 |
| calm1b1 | 2.5134976970405e-46 | 0.57219733 | 0.334 | 0.09 | 6.32622235368124e-42 | 10 |
| grap2b | 3.28352447632549e-46 | 0.620818773 | 0.366 | 0.108 | 8.26430275446362e-42 | 10 |
| nr4a11 | 5.69372329900525e-45 | 0.652549473 | 0.463 | 0.161 | 1.43305321712663e-40 | 10 |
| dicp3.1 | 2.2042488326674e-44 | 0.288001799 | 0.125 | 0.015 | 5.54787388694058e-40 | 10 |

| | | | | | | |
|---|---|---|---|---|---|---|
| ncf1 | 2.82673261658539e-44 | 0.439204505 | 0.181 | 0.031 | 7.11460332268377e-40 | 10 |
| cahz | 1.90450261352738e-43 | 0.39267455 | 0.202 | 0.037 | 4.79344262798707e-39 | 10 |
| xbp1 | 1.16840993743409e-42 | 0.665635697 | 0.638 | 0.338 | 2.94077097152787e-38 | 10 |
| m17 | 3.75496158614478e-42 | 0.312639706 | 0.111 | 0.013 | 9.45086281616781e-38 | 10 |
| si:ch1073-443f11.2 | 3.86261773100134e-42 | 0.37032679 | 0.185 | 0.033 | 9.72182256715728e-38 | 10 |
| grasp | 1.68169508397527e-41 | 0.54067146 | 0.282 | 0.073 | 4.23265835685737e-37 | 10 |
| moxd1 | 6.619863811234e-41 | 0.276587902 | 0.108 | 0.012 | 1.66615352264949e-36 | 10 |
| myh9a | 2.87199594856487e-40 | 0.436122358 | 0.233 | 0.051 | 7.22852660294292e-36 | 10 |
| hacd3 | 1.28984223208672e-39 | 0.353220803 | 0.181 | 0.033 | 3.24640391393906e-35 | 10 |
| CR383676.13 | 3.72071786282137e-39 | 0.609975404 | 0.948 | 0.722 | 9.3646747889351e-35 | 10 |
| mir223 | 5.11342363292268e-39 | 0.387470586 | 0.164 | 0.028 | 1.28699759417031e-34 | 10 |
| vsir1 | 1.02268358991916e-38 | 0.548192129 | 0.289 | 0.082 | 2.57399232746753e-34 | 10 |
| hcar1-4 | 8.02003962358875e-37 | 0.372512659 | 0.174 | 0.033 | 2.01856377286105e-32 | 10 |
| mbnl11 | 1.12624849759904e-36 | 0.611345644 | 0.446 | 0.179 | 2.83465484360702e-32 | 10 |
| rgs19 | 4.3931359277606e-36 | 0.375293473 | 0.167 | 0.032 | 1.10570838165806e-31 | 10 |
| unc13d | 1.05091004506489e-35 | 0.344020718 | 0.153 | 0.027 | 2.64503549242382e-31 | 10 |
| camk1b | 1.76275185285831e-35 | 0.311659104 | 0.129 | 0.02 | 4.43667013845907e-31 | 10 |
| si:dkey-24p1.7 | 1.87816978676699e-35 | 0.255086711 | 0.101 | 0.012 | 4.72716553631383e-31 | 10 |
| vaspa1 | 1.6791339156233e-34 | 0.449798531 | 0.265 | 0.072 | 4.22621215223229e-30 | 10 |
| ssr3 | 1.0651859845555e-33 | 0.62586372 | 0.526 | 0.268 | 2.68096660452774e-29 | 10 |
| syngr2a | 7.60773426903965e-33 | 0.387590571 | 0.216 | 0.053 | 1.91479063817459e-28 | 10 |
| lta4h1 | 1.07875517977435e-32 | 0.42849152 | 0.254 | 0.069 | 2.71511891197406e-28 | 10 |
| cebpb2 | 1.2415767980779e-32 | 0.601302817 | 0.697 | 0.347 | 3.12492464308227e-28 | 10 |
| ikzf21 | 2.27166322490568e-32 | 0.316713819 | 0.15 | 0.028 | 5.71754917076511e-28 | 10 |
| si:zfos-741a10.3 | 3.72847268297134e-32 | 0.304969686 | 0.139 | 0.025 | 9.38419289577056e-28 | 10 |
| si:dkey-73n10.1 | 1.83464095364403e-31 | 0.283274065 | 0.122 | 0.019 | 4.61760781622667e-27 | 10 |
| slc2a3b | 1.8512808840303e-31 | 0.329231703 | 0.146 | 0.027 | 4.65948885701587e-27 | 10 |
| pnp5a | 4.73490490743971e-31 | 0.514844968 | 0.254 | 0.075 | 1.1917282161535e-26 | 10 |
| si:ch211-215e19.3 | 8.39815616412976e-31 | 0.285788513 | 0.122 | 0.02 | 2.11373192494982e-26 | 10 |
| relb | 1.14253885854178e-30 | 0.393268092 | 0.178 | 0.04 | 2.87565605306382e-26 | 10 |
| camk2g2 | 1.45330750102038e-29 | 0.34128903 | 0.16 | 0.034 | 3.65782964931818e-25 | 10 |
| cxcr4b1 | 2.68384724789338e-29 | 0.534742457 | 0.446 | 0.178 | 6.75497513822284e-25 | 10 |
| p4hb | 5.42196505384299e-28 | 0.578517378 | 0.449 | 0.211 | 1.36465438440174e-23 | 10 |
| gsr | 9.14235206299114e-28 | 0.33542814 | 0.153 | 0.033 | 2.30103859073424e-23 | 10 |
| ptpn61 | 1.71994305588055e-27 | 0.4423662 | 0.272 | 0.088 | 4.32892467734575e-23 | 10 |

| | | | | | | |
|---|---|---|---|---|---|---|
| msna2 | 2.75577795571929e-27 | 0.55005468 | 0.474 | 0.238 | 6.93601753674989e-23 | 10 |
| arl5c1 | 8.68999875681953e-27 | 0.506856431 | 0.3 | 0.109 | 2.18718578710391e-22 | 10 |
| hck | 1.03504669091308e-26 | 0.301196745 | 0.146 | 0.031 | 2.60510901635914e-22 | 10 |
| XLOC-0160942 | 1.25621064073981e-26 | 0.40423613 | 0.251 | 0.076 | 3.16175656167802e-22 | 10 |
| CABZ01073834.1 | 3.73769789414176e-26 | 0.317503382 | 0.167 | 0.04 | 9.4074118297654e-22 | 10 |
| rab11fip4b | 1.58033862166392e-25 | 0.336389313 | 0.146 | 0.033 | 3.97755427686593e-21 | 10 |
| mt-nd12 | 1.95165320036528e-25 | 0.505617222 | 0.645 | 0.407 | 4.91211593999937e-21 | 10 |
| LOC101884267 | 3.39457121862097e-25 | 0.37086336 | 0.202 | 0.056 | 8.54379630014711e-21 | 10 |
| smox1 | 3.69582647294424e-25 | 0.429950771 | 0.251 | 0.082 | 9.30202564975335e-21 | 10 |
| tnfaip3 | 3.98774345160614e-25 | 0.46393303 | 0.209 | 0.063 | 1.00367514933475e-20 | 10 |
| mt-nd22 | 4.04173940697501e-25 | 0.496049643 | 0.641 | 0.396 | 1.01726539134154e-20 | 10 |
| plxnc1 | 4.18928007190898e-25 | 0.339129024 | 0.167 | 0.042 | 1.05439990129877e-20 | 10 |
| atp2a3 | 1.48917913222385e-24 | 0.298044398 | 0.139 | 0.031 | 3.7481149578942e-20 | 10 |
| cap12 | 2.25164594759585e-24 | 0.522786202 | 0.456 | 0.23 | 5.66716768550399e-20 | 10 |
| mrtfab1 | 4.2985493509583e-24 | 0.383704652 | 0.185 | 0.051 | 1.0819018861427e-19 | 10 |
| cotl12 | 1.23617337652778e-23 | 0.512389968 | 0.557 | 0.302 | 3.11132477138276e-19 | 10 |
| rasal31 | 1.43392384587288e-23 | 0.254137982 | 0.125 | 0.026 | 3.60904292767744e-19 | 10 |
| antxr2a2 | 2.39973563540316e-23 | 0.475775534 | 0.352 | 0.151 | 6.0398946207462e-19 | 10 |
| itga41 | 1.7242827021003e-22 | 0.407804218 | 0.202 | 0.063 | 4.33984713291625e-18 | 10 |
| marcksl1b2 | 1.9561382571973e-22 | 0.513780839 | 0.289 | 0.114 | 4.92340437953988e-18 | 10 |
| far1 | 2.22376494706366e-22 | 0.293388528 | 0.143 | 0.034 | 5.59699399526452e-18 | 10 |
| LOC103911430 | 4.76769754080644e-22 | 0.304534723 | 0.146 | 0.036 | 1.19998179404557e-17 | 10 |
| birc2 | 1.17764678998635e-21 | 0.427368877 | 0.272 | 0.105 | 2.96401920571666e-17 | 10 |
| ccr9a1 | 1.71939162022054e-21 | 0.410786424 | 0.244 | 0.085 | 4.32753676893309e-17 | 10 |
| yrk | 4.45631955863182e-21 | 0.332944319 | 0.199 | 0.061 | 1.12161106971204e-16 | 10 |
| sort1b | 6.64001049110449e-21 | 0.261868992 | 0.129 | 0.03 | 1.67122424050609e-16 | 10 |
| actn11 | 1.65456447735464e-18 | 0.289042019 | 0.15 | 0.043 | 4.16437333305389e-14 | 10 |
| si:ch73-208g10.1 | 2.40955065131409e-18 | 0.286380228 | 0.146 | 0.041 | 6.06459803429244e-14 | 10 |
| gapdhs3 | 3.35186432118297e-18 | 0.443552844 | 0.481 | 0.286 | 8.43630730998542e-14 | 10 |
| si:dkey-118j18.2 | 5.56061915897791e-18 | 0.297023642 | 0.167 | 0.052 | 1.39955223612315e-13 | 10 |
| rab44 | 8.15335246174657e-18 | 0.32067333 | 0.181 | 0.059 | 2.05211728109699e-13 | 10 |
| cd74b2 | 8.28660918532331e-18 | 0.385242178 | 0.46 | 0.22 | 2.08565666585402e-13 | 10 |
| wdr1 | 1.03981929824141e-17 | 0.409719134 | 0.307 | 0.139 | 2.61712119174381e-13 | 10 |
| fybb | 1.38967366456507e-17 | 0.256628669 | 0.143 | 0.04 | 3.49766964634383e-13 | 10 |
| itgae.2 | 1.4533263163615e-17 | 0.270010089 | 0.129 | 0.034 | 3.65787700565025e-13 | 10 |

| | | | | | | |
|---|---|---|---|---|---|---|
| rac22 | 2.39446012921228e-17 | 0.412991441 | 0.425 | 0.218 | 6.02661669921439e-13 | 10 |
| cox6a2 | 3.36519832233077e-17 | 0.257347714 | 0.129 | 0.035 | 8.46986765747432e-13 | 10 |
| rhogb | 5.55626478189721e-17 | 0.372626874 | 0.237 | 0.095 | 1.39845628295571e-12 | 10 |
| mt-nd32 | 1.05139682337161e-16 | 0.423561136 | 0.578 | 0.393 | 2.646260664744e-12 | 10 |
| arpc1b2 | 1.68612532189451e-16 | 0.428537272 | 0.557 | 0.305 | 4.2438088226763e-12 | 10 |
| pnrc23 | 2.78960789606114e-16 | 0.358504741 | 0.78 | 0.564 | 7.02116411359629e-12 | 10 |
| ccng2 | 4.25867523141964e-16 | 0.376462741 | 0.213 | 0.084 | 1.07186596899601e-11 | 10 |
| arpc5a1 | 7.72336930455228e-16 | 0.380675749 | 0.258 | 0.113 | 1.94389482026276e-11 | 10 |
| ptprc1 | 1.03033125068016e-15 | 0.357634933 | 0.282 | 0.125 | 2.59324072483689e-11 | 10 |
| zbtb201 | 3.058557480353e-15 | 0.39096872 | 0.226 | 0.097 | 7.69808332230046e-11 | 10 |
| itgav1 | 6.71013292203535e-15 | 0.298367474 | 0.164 | 0.057 | 1.68887335514708e-10 | 10 |
| pgd | 1.08689304616564e-14 | 0.268125331 | 0.174 | 0.062 | 2.73560110789429e-10 | 10 |
| cmtm7 | 1.50976395094938e-14 | 0.259022102 | 0.143 | 0.046 | 3.79992488814449e-10 | 10 |
| ahr2 | 2.75255529865156e-14 | 0.255487003 | 0.143 | 0.047 | 6.92790643117611e-10 | 10 |
| arrb2b | 8.35345041662605e-14 | 0.341664676 | 0.213 | 0.09 | 2.10247993536061e-09 | 10 |
| nfkbiab | 9.57685958611368e-14 | 0.381189302 | 0.338 | 0.179 | 2.41039978922895e-09 | 10 |
| laptm52 | 9.71310602235809e-14 | 0.358393992 | 0.331 | 0.168 | 2.44469165476731e-09 | 10 |
| tram1 | 1.23743930527783e-13 | 0.321872326 | 0.233 | 0.101 | 3.11451098745377e-09 | 10 |
| n4bp1 | 1.60628950984346e-13 | 0.254567147 | 0.136 | 0.044 | 4.042870067325e-09 | 10 |
| dbnlb | 1.65147740451636e-13 | 0.254510414 | 0.129 | 0.041 | 4.15660347942724e-09 | 10 |
| zgc:66427 | 4.70270604814767e-13 | 0.282364616 | 0.199 | 0.081 | 1.18362408525829e-08 | 10 |
| parvg | 1.78534936608686e-12 | 0.271397368 | 0.171 | 0.067 | 4.49354581950403e-08 | 10 |
| cyth1a | 4.45654873583381e-12 | 0.287892414 | 0.167 | 0.067 | 1.12166875132201e-07 | 10 |
| fam107b1 | 6.54755049147678e-12 | 0.295576027 | 0.226 | 0.102 | 1.64795298319979e-07 | 10 |
| sec11a | 1.45633792054859e-11 | 0.338867426 | 0.321 | 0.176 | 3.66545691222876e-07 | 10 |
| arhgdig1 | 1.85814461433226e-11 | 0.354090727 | 0.397 | 0.241 | 4.67676417981286e-07 | 10 |
| stk17b | 4.46463183155683e-10 | 0.293090118 | 0.223 | 0.11 | 1.12370318568454e-05 | 10 |
| tmsb12 | 7.84540492737232e-10 | 0.264607979 | 0.3 | 0.154 | 1.97460996617034e-05 | 10 |
| sh3kbp1 | 1.27904802108603e-09 | 0.263939626 | 0.16 | 0.07 | 3.21923596427144e-05 | 10 |
| actr2a1 | 1.35699702860434e-09 | 0.292280291 | 0.244 | 0.129 | 3.41542582129427e-05 | 10 |
| tagln22 | 1.60440022391957e-09 | 0.307483956 | 0.317 | 0.187 | 4.03811492358316e-05 | 10 |
| maff1 | 3.74960975510447e-09 | 0.279762649 | 0.174 | 0.083 | 9.43739279262245e-05 | 10 |
| tmsb4x4 | 1.6778568864334e-08 | 0.317454333 | 0.934 | 0.681 | 0.0004223 | 10 |
| cox6b11 | 1.7563598677555e-08 | 0.287261576 | 0.22 | 0.12 | 0.000442058 | 10 |
| gyg1a5 | 2.00221147614395e-08 | 0.307231698 | 0.523 | 0.369 | 0.000503937 | 10 |

| | | | | | | |
|---|---|---|---|---|---|---|
| socs3a2 | 5.34910275727701e-08 | 0.264681832 | 0.477 | 0.296 | 0.001346316 | 10 |
| capzb1 | 4.88949182430606e-07 | 0.282550526 | 0.359 | 0.253 | 0.012306362 | 10 |
| fosl21 | 6.96829184620251e-06 | 0.277194237 | 0.199 | 0.121 | 0.175384937 | 10 |
| ddit32 | 7.29282433033039e-06 | 0.253854813 | 0.268 | 0.176 | 0.183553096 | 10 |
| rps27l1 | 0.000378406 | 0.254400824 | 0.265 | 0.204 | 1 | 10 |
| cyp17a1 | 0 | 2.274840838 | 0.983 | 0.051 | 0 | 11 |
| cyp11c1 | 0 | 1.96580855 | 0.885 | 0.002 | 0 | 11 |
| rgs16 | 0 | 1.844346576 | 0.794 | 0.005 | 0 | 11 |
| star | 0 | 1.843854628 | 0.741 | 0.005 | 0 | 11 |
| hsd3b1 | 0 | 1.693626195 | 0.853 | 0.001 | 0 | 11 |
| pi15a | 0 | 1.560333179 | 0.797 | 0.027 | 0 | 11 |
| XLOC-018328 | 0 | 1.549397764 | 0.818 | 0.001 | 0 | 11 |
| ptx3a2 | 0 | 1.44932407 | 0.71 | 0.02 | 0 | 11 |
| cyp11a2 | 0 | 1.425512294 | 0.766 | 0.001 | 0 | 11 |
| prkag2a | 0 | 1.342922596 | 0.769 | 0.024 | 0 | 11 |
| ece2b | 0 | 1.255329984 | 0.703 | 0.031 | 0 | 11 |
| pik3r3b | 0 | 1.206011581 | 0.664 | 0.034 | 0 | 11 |
| fdx1b | 0 | 1.191818667 | 0.626 | 0.004 | 0 | 11 |
| osr21 | 0 | 1.086870222 | 0.626 | 0.031 | 0 | 11 |
| fam184a | 0 | 1.076601061 | 0.587 | 0.003 | 0 | 11 |
| esr2b | 0 | 1.018426956 | 0.598 | 0.021 | 0 | 11 |
| nr2f1a1 | 0 | 1.01142542 | 0.654 | 0.03 | 0 | 11 |
| robo21 | 0 | 0.98541483 | 0.619 | 0.033 | 0 | 11 |
| fdxr | 0 | 0.98001377 | 0.549 | 0.013 | 0 | 11 |
| nell2a | 0 | 0.945187399 | 0.549 | 0.008 | 0 | 11 |
| islr2 | 0 | 0.881814345 | 0.458 | 0.001 | 0 | 11 |
| mctp1b | 0 | 0.876016707 | 0.5 | 0.013 | 0 | 11 |
| CABZ01074994.1 | 0 | 0.841911715 | 0.427 | 0.001 | 0 | 11 |
| nr5a1b | 0 | 0.821753797 | 0.479 | 0.002 | 0 | 11 |
| meis3 | 0 | 0.741951978 | 0.388 | 0.005 | 0 | 11 |
| tspan9a | 0 | 0.671785189 | 0.378 | 0.013 | 0 | 11 |
| nol4lb | 0 | 0.668871687 | 0.409 | 0.016 | 0 | 11 |
| CABZ01079281.1 | 0 | 0.606255171 | 0.36 | 0.005 | 0 | 11 |
| arhgap20b | 0 | 0.590655047 | 0.367 | 0.013 | 0 | 11 |
| CABZ01038709.1 | 0 | 0.584746663 | 0.304 | 0.004 | 0 | 11 |

| | | | | | | |
|---|---|---|---|---|---|---|
| prox1a | 0 | 0.560279399 | 0.336 | 0.011 | 0 | 11 |
| nlgn4xa | 0 | 0.450808105 | 0.266 | 0.005 | 0 | 11 |
| camkvl | 0 | 0.449976324 | 0.269 | 0.004 | 0 | 11 |
| inhbab | 0 | 0.431112909 | 0.234 | 0.004 | 0 | 11 |
| si:dkey-183j2.10 | 0 | 0.375982082 | 0.196 | 0.002 | 0 | 11 |
| stmn2b | 0 | 0.371812529 | 0.203 | 0.001 | 0 | 11 |
| pcdh8 | 0 | 0.369826248 | 0.199 | 0 | 0 | 11 |
| zgc:153395 | 0 | 0.367983816 | 0.206 | 0.001 | 0 | 11 |
| XLOC-001115 | 0 | 0.359953427 | 0.189 | 0.001 | 0 | 11 |
| unc5db | 0 | 0.352165703 | 0.196 | 0 | 0 | 11 |
| cyp4t8 | 0 | 0.344411678 | 0.192 | 0.001 | 0 | 11 |
| pklr | 0 | 0.330141347 | 0.171 | 0.001 | 0 | 11 |
| rprm3 | 0 | 0.318993377 | 0.164 | 0.001 | 0 | 11 |
| chrnb3a | 0 | 0.281811263 | 0.157 | 0 | 0 | 11 |
| celf4 | 1.68217881288988e-303 | 0.333659789 | 0.171 | 0.001 | 4.23387585416253e-299 | 11 |
| sypl1 | 9.13049072646785e-292 | 0.578037106 | 0.346 | 0.015 | 2.29805321094469e-287 | 11 |
| BX901962.2 | 2.09862140801909e-287 | 0.737737583 | 0.483 | 0.031 | 5.28202022184325e-283 | 11 |
| nrxn1a | 1.6100427251524e-286 | 0.39675792 | 0.238 | 0.005 | 4.05231653493607e-282 | 11 |
| cyp17a2 | 2.79025982716079e-282 | 0.321346396 | 0.189 | 0.003 | 7.022804958981e-278 | 11 |
| lrrtm1 | 1.12045643693749e-281 | 0.320859284 | 0.168 | 0.002 | 2.82007680612796e-277 | 11 |
| hmgb3a1 | 8.15816626355255e-280 | 0.75572949 | 0.507 | 0.038 | 2.05332886687354e-275 | 11 |
| insl3 | 6.0345171342687e-278 | 0.349244678 | 0.126 | 0 | 1.51882761752409e-273 | 11 |
| cox7a1 | 5.44813337308912e-272 | 0.659300837 | 0.395 | 0.022 | 1.3712406886728e-267 | 11 |
| hsd11b22 | 6.42616858066148e-269 | 1.515605689 | 0.773 | 0.108 | 1.61740237006669e-264 | 11 |
| oprd1a | 1.99360471194124e-262 | 0.281236005 | 0.122 | 0 | 5.01770369948491e-258 | 11 |
| pex5lb | 3.79203811369638e-261 | 0.300635598 | 0.154 | 0.001 | 9.54418072836241e-257 | 11 |
| ar1 | 1.32551266034164e-258 | 0.848493411 | 0.538 | 0.048 | 3.33618281481388e-254 | 11 |
| hs6st1b | 2.02568062108599e-247 | 0.472522655 | 0.266 | 0.01 | 5.09843555521132e-243 | 11 |
| ift172 | 8.04322915164893e-246 | 0.346930955 | 0.206 | 0.005 | 2.02440034517852e-241 | 11 |
| col18a1a1 | 1.47885137217679e-243 | 0.691981289 | 0.444 | 0.032 | 3.72212101863176e-239 | 11 |
| ppdpfa1 | 2.95654486145948e-241 | 0.910894704 | 0.601 | 0.064 | 7.44132776180735e-237 | 11 |
| brinp2 | 1.54050023704704e-238 | 0.294295495 | 0.157 | 0.002 | 3.8772850466237e-234 | 11 |
| tmsb2 | 1.95990865319095e-237 | 0.307414602 | 0.175 | 0.003 | 4.93289408921629e-233 | 11 |
| dachd | 4.43333935783273e-234 | 0.386311369 | 0.227 | 0.007 | 1.11582718297292e-229 | 11 |
| jag1b | 8.52609347083649e-232 | 0.531133642 | 0.332 | 0.018 | 2.14593246567484e-227 | 11 |

| | | | | | | |
|---|---|---|---|---|---|---|
| dab1a | 2.52334104617492e-228 | 0.286552121 | 0.15 | 0.002 | 6.35099707911766e-224 | 11 |
| LOC101882765 | 9.66637074141596e-228 | 0.281289737 | 0.168 | 0.003 | 2.43292885190698e-223 | 11 |
| nme31 | 1.11440147790857e-222 | 0.813832124 | 0.573 | 0.064 | 2.80483707974807e-218 | 11 |
| sipa1l2.1 | 4.67383849223555e-220 | 0.321613797 | 0.189 | 0.005 | 1.17635841011077e-215 | 11 |
| atf31 | 3.21710371498273e-219 | 1.115844385 | 0.615 | 0.078 | 8.09712834024003e-215 | 11 |
| skib | 4.21883513625306e-214 | 0.431273058 | 0.266 | 0.012 | 1.06183861544353e-209 | 11 |
| CR318588.3 | 1.11355986390874e-206 | 0.913469838 | 0.423 | 0.036 | 2.80271882147191e-202 | 11 |
| dclk1a1 | 1.53773075511502e-203 | 0.591373852 | 0.392 | 0.03 | 3.870314537549e-199 | 11 |
| atp6ap1lb | 2.10481760992487e-193 | 0.282588894 | 0.171 | 0.004 | 5.29761544241991e-189 | 11 |
| mtbl | 6.24256912083187e-188 | 0.414826859 | 0.238 | 0.011 | 1.57119222202217e-183 | 11 |
| fam222a | 1.05450385954808e-187 | 0.366266484 | 0.234 | 0.01 | 2.65408076409657e-183 | 11 |
| cox6a21 | 1.11497223993505e-184 | 0.574551497 | 0.36 | 0.028 | 2.80627363069252e-180 | 11 |
| vamp2 | 5.30594696153374e-181 | 0.258372394 | 0.15 | 0.003 | 1.33545379074843e-176 | 11 |
| zgc:927441 | 1.33028585626494e-180 | 1.25635216 | 0.93 | 0.34 | 3.34819647163322e-176 | 11 |
| ctnnd2a | 5.48065476890917e-180 | 0.338536744 | 0.192 | 0.007 | 1.37942599878675e-175 | 11 |
| atp1a3b1 | 1.14250772379612e-177 | 0.539733804 | 0.318 | 0.023 | 2.87557769002246e-173 | 11 |
| cited4a | 3.55711830728238e-177 | 0.648098155 | 0.381 | 0.034 | 8.95291106759902e-173 | 11 |
| nr2f21 | 8.73413327732927e-176 | 0.640367623 | 0.392 | 0.036 | 2.198294004571e-171 | 11 |
| fundc2 | 5.77177651114593e-172 | 0.752988433 | 0.563 | 0.078 | 1.45269843009032e-167 | 11 |
| hmgb1b1 | 5.21636396306949e-170 | 0.864481283 | 0.678 | 0.113 | 1.31290664586496e-165 | 11 |
| inhbaa1 | 7.79004764239404e-167 | 0.609025766 | 0.367 | 0.033 | 1.96067709111416e-162 | 11 |
| b4galt61 | 3.37527179696674e-165 | 0.773321522 | 0.469 | 0.057 | 8.4952215857856e-161 | 11 |
| socs21 | 2.64899578481017e-164 | 0.500842259 | 0.29 | 0.021 | 6.66725749078872e-160 | 11 |
| atp10a1 | 7.16717483192627e-162 | 0.648296675 | 0.43 | 0.048 | 1.80390623344752e-157 | 11 |
| ckba1 | 1.36866913282535e-161 | 0.661775217 | 0.469 | 0.056 | 3.44480334040811e-157 | 11 |
| irs2b | 4.3027634602568e-160 | 0.415223316 | 0.238 | 0.014 | 1.08296253531203e-155 | 11 |
| bag31 | 1.82382976381592e-155 | 0.584097807 | 0.332 | 0.029 | 4.59039713254829e-151 | 11 |
| acsbg2 | 2.93690713122221e-155 | 0.525825632 | 0.346 | 0.031 | 7.39190155857318e-151 | 11 |
| cxcl18b2 | 6.72365973546309e-154 | 0.893840589 | 0.608 | 0.095 | 1.6922779188187e-149 | 11 |
| tbx2b | 2.70124377134028e-153 | 0.381829715 | 0.206 | 0.01 | 6.79876044808634e-149 | 11 |
| cldn11a1 | 3.15636923669309e-153 | 0.754919072 | 0.444 | 0.053 | 7.94426573183284e-149 | 11 |
| cdon2 | 4.80538233461558e-152 | 0.625340962 | 0.423 | 0.049 | 1.20946667979939e-147 | 11 |
| pgr | 6.53198295351266e-151 | 0.366784567 | 0.231 | 0.014 | 1.6440347895696e-146 | 11 |
| meis1b1 | 9.525986743269e-149 | 0.612499955 | 0.416 | 0.049 | 2.39759560341337e-144 | 11 |
| pcdh11 | 4.61702908478554e-148 | 0.370518436 | 0.21 | 0.011 | 1.16206005034967e-143 | 11 |

| | | | | | | |
|---|---|---|---|---|---|---|
| pgrmc11 | 1.37677973803126e-145 | 0.910848331 | 0.661 | 0.129 | 3.46521692265087e-141 | 11 |
| acss1 | 2.68145429204601e-145 | 0.501260834 | 0.318 | 0.029 | 6.74895230765061e-141 | 11 |
| dhcr7 | 7.35608611514361e-144 | 0.334473041 | 0.217 | 0.012 | 1.8514533143205e-139 | 11 |
| pde4ba1 | 1.21852086940298e-139 | 0.871467354 | 0.654 | 0.128 | 3.06689517620036e-135 | 11 |
| robo1 | 1.46695919637412e-139 | 0.384087755 | 0.248 | 0.018 | 3.69218960135401e-135 | 11 |
| gpr27 | 1.54511160306289e-134 | 0.357954002 | 0.21 | 0.013 | 3.888891393749e-130 | 11 |
| socs3a3 | 1.02619143242865e-133 | 1.159294536 | 0.899 | 0.284 | 2.58282121627966e-129 | 11 |
| jun2 | 5.97566815000384e-133 | 1.068249461 | 0.829 | 0.214 | 1.50401591667447e-128 | 11 |
| tmsb | 1.5354200017856e-130 | 0.501512868 | 0.269 | 0.023 | 3.86449860249418e-126 | 11 |
| hspe1 | 2.77100946568535e-130 | 1.051476031 | 0.878 | 0.281 | 6.97435372418346e-126 | 11 |
| gramd1a | 6.20626483236756e-128 | 0.304910969 | 0.185 | 0.01 | 1.56205479565859e-123 | 11 |
| zgc:92606 | 2.87441649536336e-127 | 0.647889505 | 0.49 | 0.078 | 7.23461887718004e-123 | 11 |
| fam89b | 4.73472167066541e-127 | 0.348959716 | 0.206 | 0.013 | 1.19168209728978e-122 | 11 |
| tp53inp1 | 9.02626266904281e-127 | 0.590717302 | 0.381 | 0.048 | 2.27182005117139e-122 | 11 |
| edil3a1 | 1.844874286159e-125 | 0.372904826 | 0.241 | 0.019 | 4.6433640908336e-121 | 11 |
| si:dkey-151g10.6 | 1.03801847731148e-124 | 0.268277232 | 1 | 0.992 | 2.61258870554526e-120 | 11 |
| cux1a1 | 1.43906809767685e-124 | 0.497634329 | 0.343 | 0.039 | 3.62199049504286e-120 | 11 |
| nt5c2b | 1.51524280883065e-124 | 0.529250148 | 0.339 | 0.038 | 3.81371462554587e-120 | 11 |
| sox12 | 1.56570006566692e-119 | 0.304543759 | 0.164 | 0.008 | 3.94071049527708e-115 | 11 |
| hist2h2l | 7.15073722116335e-119 | 0.665255577 | 0.448 | 0.07 | 1.7997690511946e-114 | 11 |
| asph | 9.94699990043225e-119 | 0.451298307 | 0.294 | 0.03 | 2.50356040493979e-114 | 11 |
| FO082781.12 | 1.09344836274206e-117 | 0.561101913 | 0.339 | 0.041 | 2.7521001841855e-113 | 11 |
| ghrb1 | 2.15960667061242e-117 | 0.469509905 | 0.322 | 0.036 | 5.43551402926439e-113 | 11 |
| dnmt3ab2 | 1.65622861006051e-116 | 0.590339115 | 0.451 | 0.071 | 4.16856178866131e-112 | 11 |
| socs3b2 | 2.89630936873243e-115 | 0.823358365 | 0.654 | 0.144 | 7.28972105016265e-111 | 11 |
| pde8a | 6.26747971467623e-114 | 0.260796734 | 0.161 | 0.009 | 1.57746196938686e-109 | 11 |
| ptprda | 6.30850752376168e-114 | 0.277753531 | 0.164 | 0.009 | 1.58778825865558e-109 | 11 |
| si:ch73-46j18.52 | 6.62349368902823e-114 | 0.812272074 | 0.72 | 0.163 | 1.66706712659152e-109 | 11 |
| syngr1a | 7.23607099697501e-114 | 0.263030928 | 0.164 | 0.009 | 1.82124670922864e-109 | 11 |
| slc4a4a | 3.20420044694034e-113 | 0.291568944 | 0.154 | 0.008 | 8.06465210490413e-109 | 11 |
| msi2a | 3.64018256949504e-112 | 0.459006723 | 0.332 | 0.039 | 9.16197550916207e-108 | 11 |
| hipk2 | 2.88551973779165e-110 | 0.344374883 | 0.217 | 0.017 | 7.26256462804781e-106 | 11 |
| irf2bpl | 1.83266458080482e-109 | 0.474641858 | 0.318 | 0.038 | 4.61263348342764e-105 | 11 |
| slc51a | 4.41664176223031e-108 | 0.264438758 | 0.168 | 0.01 | 1.11162456513575e-103 | 11 |
| junba2 | 1.04582644371739e-107 | 1.005616954 | 0.885 | 0.296 | 2.63224057619229e-103 | 11 |

| | | | | | | |
|---|---|---|---|---|---|---|
| mtus1b | 1.1175366617371e-107 | 0.340577237 | 0.224 | 0.019 | 2.81272802392611e-103 | 11 |
| tpst1 | 4.33958931090586e-106 | 0.393241839 | 0.276 | 0.029 | 1.0922312336619e-101 | 11 |
| gfra1a | 2.63559172384204e-102 | 0.274297851 | 0.168 | 0.011 | 6.63352080973803e-98 | 11 |
| marcksl1b3 | 3.48219620668984e-101 | 0.690971388 | 0.535 | 0.107 | 8.76433963261766e-97 | 11 |
| nfic2 | 2.56065931043687e-100 | 0.560571671 | 0.416 | 0.069 | 6.44492341843856e-96 | 11 |
| ifi452 | 8.5256927797161e-98 | 0.878050387 | 0.542 | 0.119 | 2.14583161572675e-93 | 11 |
| klf11b2 | 1.74877127195416e-97 | 0.569382474 | 0.42 | 0.072 | 4.40148241438143e-93 | 11 |
| aldh2.1 | 1.35839699229796e-96 | 0.36346929 | 0.255 | 0.028 | 3.41894938991474e-92 | 11 |
| cplx2l1 | 1.14695646114108e-95 | 0.287108102 | 0.182 | 0.014 | 2.88677471704598e-91 | 11 |
| dusp5 | 3.64124558025736e-94 | 0.667541779 | 0.388 | 0.066 | 9.16465100094975e-90 | 11 |
| atf4b1 | 3.79528214661598e-94 | 0.818916901 | 0.783 | 0.252 | 9.55234563481777e-90 | 11 |
| ifi461 | 5.51318356930152e-94 | 0.701304309 | 0.524 | 0.112 | 1.3876131725575e-89 | 11 |
| fosl1a2 | 1.5281176950229e-93 | 0.613373399 | 0.451 | 0.084 | 3.84611942660314e-89 | 11 |
| CABZ01086041.1 | 1.74770477172707e-93 | 0.253182027 | 0.147 | 0.009 | 4.39879813995987e-89 | 11 |
| rps283 | 4.77468273513005e-93 | 0.4444619 | 1 | 0.823 | 1.20173989760488e-88 | 11 |
| cbx7a1 | 6.13799066364133e-93 | 0.556300318 | 0.325 | 0.047 | 1.54487087013189e-88 | 11 |
| lifrb | 2.10420164608131e-92 | 0.25913853 | 0.168 | 0.012 | 5.29606512302206e-88 | 11 |
| rpl343 | 2.79525253599529e-92 | 0.495535724 | 1 | 0.779 | 7.03537110784655e-88 | 11 |
| rpl93 | 6.48147487971858e-92 | 0.651301613 | 0.997 | 0.621 | 1.63132241247637e-87 | 11 |
| gbp | 7.52426711557433e-92 | 0.590198513 | 0.472 | 0.093 | 1.8937827903189e-87 | 11 |
| rpl323 | 7.72533237465773e-92 | 0.464392798 | 0.99 | 0.81 | 1.94438890537761e-87 | 11 |
| mdka2 | 9.23692771064069e-91 | 0.615076915 | 0.455 | 0.086 | 2.32484233549116e-86 | 11 |
| cadm2a | 2.61643090947293e-90 | 0.276454025 | 0.171 | 0.013 | 6.58529495605242e-86 | 11 |
| tgfbr31 | 5.3306707435045e-90 | 0.416643306 | 0.29 | 0.038 | 1.34167651943265e-85 | 11 |
| gadd45ga | 2.56442399168588e-89 | 0.530935269 | 0.322 | 0.047 | 6.45439874467419e-85 | 11 |
| enc11 | 1.89756927783824e-88 | 0.390904061 | 0.238 | 0.026 | 4.77599211539107e-84 | 11 |
| slc38a2 | 1.17730508909073e-87 | 0.457019342 | 0.297 | 0.041 | 2.96315917873246e-83 | 11 |
| rpl373 | 1.35340262801421e-87 | 0.425128452 | 1 | 0.846 | 3.40637907444896e-83 | 11 |
| h1f01 | 1.40205775060286e-87 | 0.57199691 | 0.35 | 0.056 | 3.52883915249234e-83 | 11 |
| rps72 | 4.72931139226488e-87 | 0.433072149 | 0.979 | 0.817 | 1.19032038431915e-82 | 11 |
| prkar2aa | 1.05198613904939e-85 | 0.551835229 | 0.455 | 0.091 | 2.64774391337342e-81 | 11 |
| crim11 | 1.68352754962117e-84 | 0.385447796 | 0.255 | 0.032 | 4.23727048964153e-80 | 11 |
| zgc:1141883 | 1.23658082741939e-83 | 0.627633326 | 0.979 | 0.62 | 3.11235028453187e-79 | 11 |
| si:dkey-188i13.102 | 1.59685338319711e-83 | 0.627500243 | 0.455 | 0.096 | 4.01912028016881e-79 | 11 |
| nt5dc2 | 2.69044660145097e-83 | 0.492440508 | 0.392 | 0.071 | 6.77158505119194e-79 | 11 |

| | | | | | | |
|---|---|---|---|---|---|---|
| rps262 | 3.82354475813101e-83 | 0.493632387 | 0.976 | 0.729 | 9.62347980173994e-79 | 11 |
| mafba1 | 3.89362664634039e-83 | 0.552515081 | 0.371 | 0.066 | 9.79986890617412e-79 | 11 |
| rpl183 | 4.48159830898972e-83 | 0.484941979 | 0.997 | 0.769 | 1.12797347838962e-78 | 11 |
| fmnl3 | 4.517677381243e-83 | 0.263446393 | 0.182 | 0.016 | 1.13705422008505e-78 | 11 |
| si:dkey-44g23.5 | 6.01599822101067e-83 | 0.520968406 | 0.385 | 0.067 | 1.51416659224617e-78 | 11 |
| NC-002333.43 | 9.65357881125497e-83 | 0.709578377 | 0.909 | 0.617 | 2.42970925100476e-78 | 11 |
| fosab2 | 3.03143127452982e-81 | 0.909271915 | 0.878 | 0.337 | 7.62980937486412e-77 | 11 |
| znf503 | 3.65001263356578e-81 | 0.287242212 | 0.189 | 0.018 | 9.18671679742171e-77 | 11 |
| ptp4a31 | 4.76661688992023e-81 | 0.412914677 | 0.308 | 0.046 | 1.19970980502402e-76 | 11 |
| msi2b | 4.9248649831958e-81 | 0.525463669 | 0.437 | 0.085 | 1.23953926762055e-76 | 11 |
| hmgn62 | 5.67827210290009e-81 | 0.763392641 | 0.759 | 0.254 | 1.42916430557892e-76 | 11 |
| rpl363 | 1.00020134414888e-80 | 0.473433944 | 1 | 0.785 | 2.51740676308831e-76 | 11 |
| zbtb4 | 4.78224416767768e-80 | 0.299043477 | 0.189 | 0.018 | 1.2036430345628e-75 | 11 |
| rpl22l12 | 1.02461642238788e-79 | 0.778310595 | 0.892 | 0.38 | 2.57885707350805e-75 | 11 |
| rps163 | 1.29371806506209e-79 | 0.50782577 | 0.993 | 0.732 | 3.25615899795478e-75 | 11 |
| rps93 | 3.53482066426101e-79 | 0.462579338 | 0.99 | 0.772 | 8.89679012987854e-75 | 11 |
| bnip3lb | 7.6400921555564e-79 | 0.432665454 | 0.343 | 0.058 | 1.92293479463199e-74 | 11 |
| rps27.13 | 2.48080019796045e-77 | 0.569524937 | 0.983 | 0.673 | 6.24392601824666e-73 | 11 |
| rpl27a3 | 1.64545496302275e-76 | 0.702180217 | 0.913 | 0.487 | 4.14144559643195e-72 | 11 |
| rpl293 | 2.07604212940507e-76 | 0.431671894 | 0.997 | 0.805 | 5.22519043549961e-72 | 11 |
| b2m2 | 3.06431671220978e-76 | 0.774005231 | 0.783 | 0.281 | 7.71257873296079e-72 | 11 |
| tubb5 | 8.12983006864518e-76 | 0.433112729 | 0.311 | 0.05 | 2.04619692997731e-71 | 11 |
| nlk2 | 1.3666985018493e-75 | 0.251537366 | 0.164 | 0.015 | 3.4398434593045e-71 | 11 |
| rplp02 | 3.85659828368671e-75 | 0.385545457 | 0.993 | 0.848 | 9.70667222021107e-71 | 11 |
| adh51 | 4.45879824000766e-75 | 0.693948085 | 0.682 | 0.211 | 1.12223492902753e-70 | 11 |
| rplp2l3 | 2.9027403387051e-74 | 0.477017703 | 0.99 | 0.77 | 7.30590715848686e-70 | 11 |
| rps192 | 6.1093580188252e-74 | 0.362448889 | 0.993 | 0.861 | 1.53766431975812e-69 | 11 |
| tbc1d4 | 7.62106395643226e-74 | 0.275547207 | 0.171 | 0.016 | 1.91814558719444e-69 | 11 |
| cox8a1 | 1.43040107660514e-73 | 0.67260754 | 0.692 | 0.217 | 3.60017646970748e-69 | 11 |
| nupr1b | 1.93910486355079e-73 | 0.465805008 | 0.28 | 0.043 | 4.88053303107099e-69 | 11 |
| plxna4 | 2.10705510230499e-73 | 0.283944203 | 0.185 | 0.019 | 5.30324698699143e-69 | 11 |
| rpl113 | 2.2579582154504e-73 | 0.436476829 | 1 | 0.81 | 5.6830550324671e-69 | 11 |
| rps182 | 3.41954647370105e-73 | 0.506033556 | 0.983 | 0.719 | 8.60665651965817e-69 | 11 |
| CR318588.13 | 3.46934453543779e-73 | 0.909321621 | 0.643 | 0.198 | 8.73199326124338e-69 | 11 |
| faua3 | 8.87453969008492e-73 | 0.525124884 | 0.965 | 0.684 | 2.23363289459747e-68 | 11 |

| | | | | | | |
|---|---|---|---|---|---|---|
| rpl283 | 1.16124691560507e-72 | 0.467144842 | 0.986 | 0.744 | 2.92274236188641e-68 | 11 |
| rpl243 | 2.77158994847587e-72 | 0.451633295 | 0.993 | 0.772 | 6.97581474131891e-68 | 11 |
| aoc22 | 3.92156810408169e-72 | 0.456503329 | 0.339 | 0.061 | 9.87019476116322e-68 | 11 |
| rps8a2 | 1.7012551487338e-71 | 0.422116228 | 0.979 | 0.784 | 4.2818890838481e-67 | 11 |
| rps252 | 2.02300209047322e-71 | 0.477931513 | 0.979 | 0.72 | 5.09169396151206e-67 | 11 |
| rps122 | 3.52775962724738e-70 | 0.382261455 | 0.983 | 0.837 | 8.87901820581892e-66 | 11 |
| fndc3a | 7.05350164422794e-70 | 0.255481082 | 0.157 | 0.015 | 1.77529582883573e-65 | 11 |
| znfx1 | 2.09714635410159e-69 | 0.356437184 | 0.21 | 0.026 | 5.27830765863829e-65 | 11 |
| fat1a1 | 1.13671954618864e-68 | 0.279891847 | 0.203 | 0.024 | 2.8610094258022e-64 | 11 |
| rpl63 | 2.9264898835489e-68 | 0.537394188 | 0.979 | 0.661 | 7.36568238790422e-64 | 11 |
| rps153 | 5.37321377851963e-68 | 0.52835561 | 0.979 | 0.678 | 1.35238417591561e-63 | 11 |
| maff2 | 1.24328107518599e-67 | 0.460034887 | 0.374 | 0.077 | 3.12921413813561e-63 | 11 |
| lgals172 | 2.83586885554741e-67 | 0.481344804 | 0.283 | 0.047 | 7.13759832252727e-63 | 11 |
| rpl36a3 | 3.24106441234748e-67 | 0.395432941 | 0.99 | 0.806 | 8.15743501943737e-63 | 11 |
| cx431 | 4.30029916866397e-67 | 0.392538131 | 0.294 | 0.049 | 1.08234229776103e-62 | 11 |
| rpl7a2 | 5.62281953518922e-67 | 0.404655091 | 0.983 | 0.806 | 1.41520744881177e-62 | 11 |
| rpl383 | 1.00634876285782e-66 | 0.486818765 | 0.979 | 0.723 | 2.53287920123684e-62 | 11 |
| rps102 | 1.60325871821928e-66 | 0.509873406 | 0.986 | 0.688 | 4.03524186788611e-62 | 11 |
| rpl262 | 2.32524773853244e-66 | 0.372441129 | 0.99 | 0.838 | 5.85241603311229e-62 | 11 |
| rps27.22 | 3.20698997274893e-66 | 0.645968021 | 0.92 | 0.509 | 8.07167306241179e-62 | 11 |
| rps132 | 8.92621010554915e-66 | 0.483714257 | 0.986 | 0.719 | 2.24663782146567e-61 | 11 |
| si:dkey-28b4.71 | 1.57349218048531e-65 | 0.44401177 | 0.395 | 0.085 | 3.96032246906349e-61 | 11 |
| rps26l3 | 1.85334298230833e-65 | 0.539793238 | 0.962 | 0.658 | 4.66467895217184e-61 | 11 |
| si:ch211-286b5.51 | 2.07032775964523e-65 | 0.526046056 | 0.5 | 0.129 | 5.21080793825108e-61 | 11 |
| rps213 | 2.43144634411817e-65 | 0.482573008 | 0.993 | 0.745 | 6.11970730351101e-61 | 11 |
| rpl353 | 3.23724755182564e-65 | 0.399589713 | 0.983 | 0.801 | 8.14782836318996e-61 | 11 |
| rpl35a3 | 9.23725933991595e-65 | 0.449015649 | 0.986 | 0.755 | 2.32492580326345e-60 | 11 |
| cd99l2 | 1.28823599523484e-64 | 0.278811275 | 0.178 | 0.02 | 3.24236117640657e-60 | 11 |
| rplp12 | 2.56413394287978e-64 | 0.359019943 | 0.997 | 0.875 | 6.45366872083412e-60 | 11 |
| cyb5a | 3.03260220865694e-64 | 0.376780249 | 0.297 | 0.053 | 7.63275649896865e-60 | 11 |
| rps3a2 | 3.86568786031505e-64 | 0.413324706 | 0.979 | 0.774 | 9.72954977562694e-60 | 11 |
| CR383676.14 | 8.51021644225563e-64 | 0.691016081 | 0.927 | 0.722 | 2.14193637635132e-59 | 11 |
| rpl272 | 1.20874935999249e-63 | 0.429555595 | 0.969 | 0.759 | 3.0423012641651e-59 | 11 |
| si:ch211-178n15.11 | 1.64402105593741e-63 | 0.315620968 | 0.206 | 0.027 | 4.13783659568887e-59 | 11 |
| ptprga2 | 4.45851414295096e-63 | 0.380856807 | 0.322 | 0.061 | 1.12216342463933e-58 | 11 |

| | | | | | | |
|---|---|---|---|---|---|---|
| rps4x2 | 4.59771855937022e-63 | 0.48732805 | 0.976 | 0.705 | 1.15719978420789e-58 | 11 |
| wu:fc75a09 | 7.55368208008698e-63 | 0.293396673 | 0.175 | 0.02 | 1.90118624273709e-58 | 11 |
| rps243 | 2.43643063643081e-62 | 0.437604309 | 0.979 | 0.754 | 6.13225226883271e-58 | 11 |
| slc25a52 | 3.304178882924e-62 | 0.38593513 | 0.983 | 0.803 | 8.31628783043143e-58 | 11 |
| XLOC-0412691 | 5.93927061123975e-62 | 0.289308257 | 0.178 | 0.021 | 1.49485502014293e-57 | 11 |
| stat1a2 | 7.99549699910303e-62 | 0.541794262 | 0.444 | 0.113 | 2.01238663970424e-57 | 11 |
| eif1b1 | 1.24686214333395e-61 | 0.694032309 | 0.839 | 0.373 | 3.13822732855722e-57 | 11 |
| rpl312 | 1.5570053106867e-61 | 0.533918502 | 0.979 | 0.648 | 3.91882666646735e-57 | 11 |
| elmsan1b | 4.31378545010203e-61 | 0.360776071 | 0.252 | 0.041 | 1.08573665993618e-56 | 11 |
| mt-nd13 | 8.4591265599289e-61 | 0.707440891 | 0.836 | 0.402 | 2.1290775638685e-56 | 11 |
| eef1g2 | 8.97607512139694e-61 | 0.55342749 | 0.944 | 0.614 | 2.25918834730439e-56 | 11 |
| rpl303 | 1.72347470342291e-60 | 0.485354781 | 0.979 | 0.703 | 4.33781348104513e-56 | 11 |
| rps112 | 2.82997273073597e-60 | 0.406020294 | 0.972 | 0.762 | 7.12275836598937e-56 | 11 |
| rps143 | 3.9115166718211e-60 | 0.440896611 | 0.99 | 0.754 | 9.84489631130652e-56 | 11 |
| h3f3c2 | 4.6193659489827e-60 | 0.637051907 | 0.706 | 0.247 | 1.16264821569946e-55 | 11 |
| rpl18a3 | 6.70888266982672e-60 | 0.416966239 | 0.969 | 0.782 | 1.68855867916869e-55 | 11 |
| kif5bb | 7.44892422255766e-60 | 0.258158339 | 0.185 | 0.023 | 1.87481973757554e-55 | 11 |
| rgs4 | 8.93864912623952e-60 | 0.406959751 | 0.241 | 0.038 | 2.24976859858323e-55 | 11 |
| c42 | 2.25961986117328e-59 | 0.339760483 | 0.238 | 0.037 | 5.68723722858703e-55 | 11 |
| si:dkey-23i12.71 | 5.31650625142423e-59 | 0.362055047 | 0.203 | 0.028 | 1.33811145842096e-54 | 11 |
| atp1b1a | 6.31656977205304e-59 | 0.632540515 | 0.612 | 0.205 | 1.58981744592803e-54 | 11 |
| fam168a2 | 1.18821352135015e-58 | 0.369406145 | 0.287 | 0.054 | 2.9906146118862e-54 | 11 |
| anxa132 | 1.39223800035628e-58 | 0.345177962 | 0.262 | 0.045 | 3.50412382309671e-54 | 11 |
| nipsnap2 | 1.85074797359236e-58 | 0.386342141 | 0.364 | 0.079 | 4.65814757473462e-54 | 11 |
| mvp1 | 7.35027385630366e-58 | 0.387502177 | 0.269 | 0.049 | 1.84999042689307e-53 | 11 |
| hspd11 | 7.85665511481582e-58 | 0.569033432 | 0.542 | 0.157 | 1.97744152584799e-53 | 11 |
| mt-nd23 | 8.89081660664297e-58 | 0.683580635 | 0.843 | 0.391 | 2.23772963172597e-53 | 11 |
| rps293 | 9.02086426339517e-58 | 0.366141409 | 0.986 | 0.829 | 2.27046132645393e-53 | 11 |
| antxr1c | 1.03505839721972e-57 | 0.319466613 | 0.22 | 0.034 | 2.60513847996231e-53 | 11 |
| diaph21 | 1.17870975289346e-57 | 0.29381895 | 0.21 | 0.031 | 2.96669457705755e-53 | 11 |
| rps62 | 1.46815503129067e-57 | 0.405389895 | 0.983 | 0.785 | 3.69519939825548e-53 | 11 |
| rpl23a3 | 3.16128163135159e-57 | 0.526481562 | 0.937 | 0.628 | 7.95662973794881e-53 | 11 |
| eef1a1l1 | 3.47118599408997e-57 | 0.276914445 | 1 | 0.914 | 8.73662802852504e-53 | 11 |
| si:ch73-1a9.32 | 5.8321598825884e-57 | 0.648571757 | 0.699 | 0.256 | 1.46789632084867e-52 | 11 |
| zgc:1527912 | 6.82014037617531e-57 | 0.544831019 | 0.308 | 0.064 | 1.71656113127956e-52 | 11 |

| | | | | | | |
|---|---|---|---|---|---|---|
| gas1a1 | 8.94719180108967e-57 | 0.325071455 | 0.238 | 0.039 | 2.25191870441626e-52 | 11 |
| rps15a3 | 3.67452727635627e-56 | 0.437084065 | 0.979 | 0.776 | 9.2484177018611e-52 | 11 |
| rpl10a2 | 3.98425866663605e-56 | 0.413420387 | 0.972 | 0.76 | 1.00279806380563e-51 | 11 |
| rpl231 | 2.20303200468408e-55 | 0.300918621 | 0.993 | 0.87 | 5.54481125258935e-51 | 11 |
| rpl83 | 6.70916133670496e-55 | 0.398386216 | 0.962 | 0.777 | 1.68862881683527e-50 | 11 |
| ier2a2 | 1.49317864716626e-54 | 0.674209245 | 0.678 | 0.253 | 3.75818133705277e-50 | 11 |
| zgc:1132082 | 5.02266990498664e-54 | 0.349910809 | 0.238 | 0.041 | 1.26415578838609e-49 | 11 |
| mt-co24 | 7.0758147509463e-54 | 0.519453897 | 0.934 | 0.716 | 1.78091181466567e-49 | 11 |
| nfia2 | 2.37330164612759e-53 | 0.397146323 | 0.336 | 0.075 | 5.97336291313853e-49 | 11 |
| flot2a1 | 2.37814625157174e-53 | 0.33436947 | 0.28 | 0.055 | 5.98555630058091e-49 | 11 |
| sept101 | 2.69523843202878e-53 | 0.325499933 | 0.245 | 0.043 | 6.78364560957325e-49 | 11 |
| mgst1.11 | 4.36014152189905e-53 | 0.283181472 | 0.227 | 0.038 | 1.09740401964677e-48 | 11 |
| rpl102 | 5.57429200931885e-53 | 0.398071066 | 0.972 | 0.768 | 1.40299355582546e-48 | 11 |
| isg152 | 8.3632648159723e-53 | 0.559450845 | 0.297 | 0.064 | 2.10495012153207e-48 | 11 |
| rpl132 | 8.50433567493716e-53 | 0.359818261 | 0.983 | 0.801 | 2.14045624602493e-48 | 11 |
| rpl122 | 1.35378999820351e-52 | 0.369241786 | 0.986 | 0.795 | 3.40735404647842e-48 | 11 |
| rpl153 | 2.81221259029785e-52 | 0.436908434 | 0.976 | 0.74 | 7.07805786852067e-48 | 11 |
| fam102ab | 5.86136231838139e-52 | 0.28941709 | 0.199 | 0.03 | 1.47524628191341e-47 | 11 |
| zbtb202 | 8.90930932904737e-52 | 0.424547565 | 0.374 | 0.093 | 2.24238406502793e-47 | 11 |
| junbb2 | 1.20995170981496e-51 | 0.372628448 | 0.311 | 0.068 | 3.04532745843327e-47 | 11 |
| rps27a2 | 1.5746729049906e-51 | 0.401629696 | 0.965 | 0.747 | 3.96329423457083e-47 | 11 |
| higd1a2 | 2.44213701679474e-51 | 0.607876133 | 0.762 | 0.3 | 6.14661465757068e-47 | 11 |
| ewsr1a | 4.93278157097225e-51 | 0.279968535 | 0.224 | 0.038 | 1.241531793598e-46 | 11 |
| echs1 | 1.08524765110755e-50 | 0.345503892 | 0.325 | 0.073 | 2.73145981307258e-46 | 11 |
| eif4ebp11 | 1.28146627538192e-50 | 0.580469446 | 0.661 | 0.237 | 3.22532246850876e-46 | 11 |
| rps210 | 3.55959642886308e-50 | 0.382778247 | 0.979 | 0.826 | 8.95914825180548e-46 | 11 |
| pdia32 | 1.02916938871912e-49 | 0.599470096 | 0.706 | 0.275 | 2.59031643446715e-45 | 11 |
| rpl5b2 | 1.12667133278329e-49 | 0.492546424 | 0.927 | 0.624 | 2.83571907748225e-45 | 11 |
| COX34 | 2.07991825211767e-49 | 0.514487447 | 0.934 | 0.696 | 5.23494624875495e-45 | 11 |
| rpl13a2 | 2.69043818960192e-49 | 0.374832652 | 0.965 | 0.76 | 6.77156387940908e-45 | 11 |
| il4r.1 | 3.48253810904781e-49 | 0.312530682 | 0.224 | 0.04 | 8.76520016666243e-45 | 11 |
| egr12 | 5.24004493744843e-49 | 0.614007292 | 0.538 | 0.183 | 1.3188669103064e-44 | 11 |
| rack12 | 9.81525468579578e-49 | 0.406991565 | 0.986 | 0.739 | 2.47040145186794e-44 | 11 |
| rpl393 | 1.12530314987244e-48 | 0.308595728 | 0.983 | 0.872 | 2.83227549791394e-44 | 11 |
| rpl310 | 1.21271995988808e-48 | 0.290218012 | 0.993 | 0.873 | 3.0522948670423e-44 | 11 |

| | | | | | | |
|---|---|---|---|---|---|---|
| rpl173 | 1.27722827810653e-48 | 0.402535519 | 0.962 | 0.742 | 3.21465585316632e-44 | 11 |
| hnrnpa0a | 1.34702181486173e-48 | 0.350266513 | 0.297 | 0.065 | 3.3903192058255e-44 | 11 |
| hmgb1a2 | 2.64659486965463e-48 | 0.525238946 | 0.545 | 0.181 | 6.66121462743374e-44 | 11 |
| ywhaqb2 | 3.76970818901881e-48 | 0.467222048 | 0.434 | 0.126 | 9.48797854094145e-44 | 11 |
| uba522 | 4.53916737843319e-48 | 0.497677876 | 0.944 | 0.624 | 1.14246303747785e-43 | 11 |
| sfpq | 5.96994271015435e-48 | 0.402210446 | 0.388 | 0.102 | 1.50257488071875e-43 | 11 |
| rsl24d12 | 1.03008038560679e-47 | 0.621897252 | 0.878 | 0.435 | 2.59260932253374e-43 | 11 |
| serinc1 | 2.2771234575986e-47 | 0.331847668 | 0.329 | 0.077 | 5.73129203042992e-43 | 11 |
| mt-co14 | 2.48520626762138e-47 | 0.519062966 | 0.909 | 0.653 | 6.25501565497625e-43 | 11 |
| nop532 | 2.81916580369105e-47 | 0.558576691 | 0.647 | 0.243 | 7.09555841131001e-43 | 11 |
| sema4ba2 | 3.1231349490289e-47 | 0.325663002 | 0.231 | 0.043 | 7.86061835321084e-43 | 11 |
| porb | 3.50458436374418e-47 | 0.291707929 | 0.22 | 0.039 | 8.82068838510772e-43 | 11 |
| pkma | 1.2429844497693e-46 | 0.463610297 | 0.535 | 0.17 | 3.12846756162436e-42 | 11 |
| h2afx14 | 2.06174898299585e-46 | 0.590650798 | 0.538 | 0.192 | 5.18921601530225e-42 | 11 |
| zgc:1951732 | 1.03742112266657e-45 | 0.383223035 | 0.346 | 0.086 | 2.6110852236395e-41 | 11 |
| ybx12 | 1.28036625630178e-45 | 0.479839292 | 0.972 | 0.663 | 3.22255383048596e-41 | 11 |
| naca3 | 4.21165729293032e-45 | 0.497103501 | 0.92 | 0.613 | 1.06003202405763e-40 | 11 |
| irf1b1 | 4.55931060901991e-45 | 0.320172641 | 0.203 | 0.036 | 1.14753288718422e-40 | 11 |
| rps27l2 | 9.53947571216806e-45 | 0.486808091 | 0.566 | 0.196 | 2.40099064199558e-40 | 11 |
| arhgef4 | 1.15272807620285e-44 | 0.267291973 | 0.189 | 0.031 | 2.90130129499495e-40 | 11 |
| phactr4a | 2.3018568168299e-44 | 0.297286959 | 0.196 | 0.034 | 5.79354342227917e-40 | 11 |
| si:dkey-42i9.41 | 3.47497736735457e-44 | 0.451532913 | 0.455 | 0.141 | 8.74617053589471e-40 | 11 |
| tmem243b | 3.91498394309404e-44 | 0.253769021 | 0.238 | 0.046 | 9.8536230863734e-40 | 11 |
| gabarapl2 | 1.44836284748212e-43 | 0.323147964 | 0.297 | 0.07 | 3.64538445082775e-39 | 11 |
| il6st1 | 1.54335933534046e-43 | 0.280740081 | 0.238 | 0.048 | 3.88448111111839e-39 | 11 |
| hp1bp3 | 3.56246458046527e-43 | 0.372069411 | 0.35 | 0.093 | 8.96636710257304e-39 | 11 |
| mt-nd33 | 5.61179662080296e-43 | 0.578770176 | 0.811 | 0.387 | 1.4124330914899e-38 | 11 |
| jam2a | 6.1112439629865e-43 | 0.257523559 | 0.196 | 0.034 | 1.53813899304407e-38 | 11 |
| rpl191 | 7.25659769304007e-43 | 0.300569135 | 0.976 | 0.849 | 1.82641307336125e-38 | 11 |
| rps202 | 9.74145217629452e-43 | 0.266093261 | 1 | 0.915 | 2.45182609825157e-38 | 11 |
| rpl142 | 1.12958876817621e-42 | 0.492501606 | 0.972 | 0.628 | 2.8430619706227e-38 | 11 |
| mt-nd43 | 1.16808664824838e-42 | 0.586591678 | 0.839 | 0.448 | 2.93995728497634e-38 | 11 |
| b2ml3 | 1.75741200863617e-42 | 0.543082048 | 0.65 | 0.259 | 4.42323028453638e-38 | 11 |
| parp12b | 1.96043461705466e-42 | 0.257135481 | 0.171 | 0.028 | 4.93421788766488e-38 | 11 |
| rps232 | 6.45575577792451e-42 | 0.391538007 | 0.937 | 0.719 | 1.62484917174582e-37 | 11 |

| | | | | | | |
|---|---|---|---|---|---|---|
| stat22 | 9.50176451043977e-42 | 0.333123161 | 0.259 | 0.058 | 2.39149910963259e-37 | 11 |
| rps32 | 1.14561944885673e-41 | 0.418491115 | 0.948 | 0.695 | 2.88340959082751e-37 | 11 |
| DHDH | 1.49209463458575e-41 | 0.293224588 | 0.259 | 0.057 | 3.75545298578887e-37 | 11 |
| eif3f2 | 1.62071135861506e-41 | 0.586554173 | 0.836 | 0.423 | 4.07916841849825e-37 | 11 |
| hnrnpa0b1 | 1.64421645950562e-41 | 0.547982603 | 0.72 | 0.303 | 4.13832840692969e-37 | 11 |
| oaz1b2 | 1.80473539762028e-41 | 0.541074452 | 0.745 | 0.314 | 4.54233852227047e-37 | 11 |
| hipk3b | 1.8825860306459e-41 | 0.271864951 | 0.203 | 0.038 | 4.73828078053267e-37 | 11 |
| irs1 | 3.20830223263585e-41 | 0.251675504 | 0.154 | 0.023 | 8.07497588932118e-37 | 11 |
| mibp23 | 3.78471025131575e-41 | 0.554656964 | 0.853 | 0.485 | 9.52573723153661e-37 | 11 |
| map1lc3a1 | 4.32705426397631e-41 | 0.324993982 | 0.28 | 0.067 | 1.0890762877002e-36 | 11 |
| aldh7a1 | 9.61057428348133e-41 | 0.298419346 | 0.262 | 0.059 | 2.41888544140942e-36 | 11 |
| tjp2b | 2.08141525617465e-40 | 0.286299272 | 0.259 | 0.057 | 5.23871405826596e-36 | 11 |
| bcl2l1 | 3.24127417634034e-40 | 0.320892547 | 0.301 | 0.075 | 8.15796297443101e-36 | 11 |
| rpl213 | 8.40916618132657e-40 | 0.373797158 | 0.969 | 0.752 | 2.11650303617808e-35 | 11 |
| mknk2a | 2.57729814568042e-39 | 0.270491507 | 0.227 | 0.047 | 6.48680170286304e-35 | 11 |
| rpl5a2 | 2.72293780657293e-39 | 0.562252712 | 0.899 | 0.483 | 6.8533621653634e-35 | 11 |
| CABZ01080702.11 | 2.78807447735442e-39 | 0.333803378 | 0.311 | 0.08 | 7.01730465205334e-35 | 11 |
| fabp32 | 8.73560333254134e-39 | 0.482649506 | 0.615 | 0.234 | 2.19866400276733e-34 | 11 |
| rps53 | 2.93941222987922e-38 | 0.429563425 | 0.965 | 0.696 | 7.398206641383e-34 | 11 |
| rpsa3 | 5.66021209350797e-38 | 0.404170489 | 0.99 | 0.747 | 1.42461878181502e-33 | 11 |
| rpl223 | 6.72826540556296e-38 | 0.428206027 | 0.955 | 0.689 | 1.69343711992614e-33 | 11 |
| eef1db2 | 9.8910885213897e-38 | 0.503543053 | 0.647 | 0.266 | 2.48948806994857e-33 | 11 |
| si:dkey-51e6.11 | 1.31224316748307e-37 | 0.374190044 | 0.371 | 0.111 | 3.30278482823814e-33 | 11 |
| aldh6a1 | 2.03121996900719e-37 | 0.29989235 | 0.308 | 0.081 | 5.11237753999419e-33 | 11 |
| chd32 | 2.11165874405441e-37 | 0.362473802 | 0.346 | 0.099 | 5.31483389291055e-33 | 11 |
| eif2s1b | 6.07564753160262e-37 | 0.401447092 | 0.483 | 0.167 | 1.52917972722906e-32 | 11 |
| ddx51 | 9.9010546414048e-37 | 0.522803136 | 0.78 | 0.348 | 2.49199644269517e-32 | 11 |
| cish2 | 1.26429882046568e-36 | 0.275973426 | 0.231 | 0.051 | 3.18211370123006e-32 | 11 |
| nsa22 | 2.11142580951374e-36 | 0.549537376 | 0.741 | 0.355 | 5.31424761996514e-32 | 11 |
| rbbp6 | 3.18998837078091e-36 | 0.34295525 | 0.346 | 0.1 | 8.02888173041846e-32 | 11 |
| brd3b | 7.12907463381495e-36 | 0.281450589 | 0.241 | 0.057 | 1.79431679458489e-31 | 11 |
| hsp90ab12 | 8.58766646977685e-36 | 0.338095348 | 0.99 | 0.786 | 2.16142977377813e-31 | 11 |
| ccnl1a1 | 1.15492626371884e-35 | 0.294072551 | 0.245 | 0.059 | 2.90683391315394e-31 | 11 |
| si:ch211-222l21.12 | 1.90739445364253e-35 | 0.508844813 | 0.657 | 0.267 | 4.80072110037289e-31 | 11 |
| CABZ01072614.12 | 3.56205789015038e-35 | 0.334121487 | 0.287 | 0.077 | 8.9653435037195e-31 | 11 |

| | | | | | | |
|---|---|---|---|---|---|---|
| mphosph81 | 4.5793556520905e-35 | 0.352255682 | 0.371 | 0.114 | 1.15257802407466e-30 | 11 |
| NDUFB11 | 4.67756519896229e-35 | 0.431514338 | 0.51 | 0.192 | 1.17729638492682e-30 | 11 |
| mid1ip1b | 5.49712954429389e-35 | 0.26914936 | 0.22 | 0.049 | 1.38357253500333e-30 | 11 |
| rpl72 | 6.71019403010383e-35 | 0.314025059 | 0.962 | 0.802 | 1.68888873543683e-30 | 11 |
| txndc9 | 7.36291748652297e-35 | 0.323363456 | 0.367 | 0.112 | 1.85317270218297e-30 | 11 |
| ptp4a1 | 8.00614403107924e-35 | 0.341545162 | 0.367 | 0.113 | 2.01506639118234e-30 | 11 |
| ube2e32 | 2.68437422972189e-34 | 0.387139333 | 0.437 | 0.15 | 6.75630149878702e-30 | 11 |
| uqcrc1 | 6.67874500927806e-34 | 0.313896877 | 0.311 | 0.089 | 1.6809733313852e-29 | 11 |
| ccnd1 | 7.04160842995342e-34 | 0.258496799 | 0.161 | 0.03 | 1.77230242573498e-29 | 11 |
| rab11fip1b | 1.26085707482701e-33 | 0.262879054 | 0.231 | 0.054 | 3.17345117163209e-29 | 11 |
| atrx1 | 1.39640103728654e-33 | 0.326057463 | 0.336 | 0.1 | 3.5146017707465e-29 | 11 |
| rnf213a1 | 2.17791131889934e-33 | 0.349811292 | 0.297 | 0.085 | 5.48158499853775e-29 | 11 |
| tomm51 | 4.84279937608163e-33 | 0.412316515 | 0.535 | 0.208 | 1.21888417496599e-28 | 11 |
| laptm4a1 | 6.99596436881815e-33 | 0.314962379 | 0.343 | 0.104 | 1.76081427198784e-28 | 11 |
| agrn | 7.55821160919875e-33 | 0.25348623 | 0.21 | 0.047 | 1.90232627991923e-28 | 11 |
| mt-nd5 | 8.37520613206369e-33 | 0.33629061 | 0.364 | 0.115 | 2.10795563137911e-28 | 11 |
| si:ch211-196l7.43 | 1.64248448601265e-32 | 0.470331499 | 0.696 | 0.306 | 4.13396920284524e-28 | 11 |
| ndufc2 | 2.58561050605101e-32 | 0.344793708 | 0.392 | 0.13 | 6.50772308267978e-28 | 11 |
| si:dkeyp-44a8.2 | 2.83939344099239e-32 | 0.254028093 | 0.224 | 0.054 | 7.14646935163374e-28 | 11 |
| eef2b1 | 4.40997636529602e-32 | 0.454576359 | 0.937 | 0.607 | 1.10994695138136e-27 | 11 |
| gtpbp41 | 5.2129413555827e-32 | 0.435378508 | 0.566 | 0.231 | 1.31204520978661e-27 | 11 |
| serpine1 | 6.3756534897547e-32 | 0.290451084 | 0.192 | 0.042 | 1.60468822683636e-27 | 11 |
| cct22 | 7.73783705925373e-32 | 0.438432711 | 0.542 | 0.221 | 1.94753620944357e-27 | 11 |
| gadd45ab1 | 8.50657566875671e-32 | 0.316256604 | 0.259 | 0.069 | 2.14102003006938e-27 | 11 |
| btf32 | 8.56356012488252e-32 | 0.480386563 | 0.934 | 0.58 | 2.15536244783168e-27 | 11 |
| gng71 | 1.59094377460186e-31 | 0.260558443 | 0.213 | 0.051 | 4.00424638629542e-27 | 11 |
| dldh | 1.85797934991217e-31 | 0.304416672 | 0.29 | 0.084 | 4.67634822579393e-27 | 11 |
| anxa41 | 2.77229173354744e-31 | 0.358652525 | 0.388 | 0.132 | 6.97758106416554e-27 | 11 |
| atp5l1 | 3.07418153710926e-31 | 0.502196009 | 0.815 | 0.442 | 7.73740751075029e-27 | 11 |
| zgc:1583432 | 8.48301343239485e-31 | 0.477867331 | 0.43 | 0.161 | 2.13508965079946e-26 | 11 |
| stau22 | 1.73373183410674e-30 | 0.300654438 | 0.283 | 0.082 | 4.36362965326325e-26 | 11 |
| cbx1b | 1.76403454687808e-30 | 0.257874419 | 0.192 | 0.044 | 4.43989855103743e-26 | 11 |
| appa2 | 3.30151496384775e-30 | 0.282970115 | 0.266 | 0.074 | 8.3095830125084e-26 | 11 |
| cct7 | 4.76961093281903e-30 | 0.398475297 | 0.514 | 0.203 | 1.20046337568122e-25 | 11 |
| rnf7 | 6.46947079869037e-30 | 0.299064467 | 0.311 | 0.095 | 1.62830110532238e-25 | 11 |

| | | | | | | |
|---|---|---|---|---|---|---|
| pno11 | 7.98679207785206e-30 | 0.361768465 | 0.448 | 0.166 | 2.01019569807458e-25 | 11 |
| elocb | 1.22223793525358e-29 | 0.368446938 | 0.444 | 0.166 | 3.07625065923972e-25 | 11 |
| eif3m2 | 1.49647940320507e-29 | 0.479084643 | 0.727 | 0.351 | 3.76648900992683e-25 | 11 |
| grinab1 | 1.93861960582464e-29 | 0.303449005 | 0.339 | 0.11 | 4.87931168590003e-25 | 11 |
| cnbpb1 | 2.63293713199565e-29 | 0.352554406 | 0.472 | 0.179 | 6.62683946751986e-25 | 11 |
| sesn13 | 3.55738924892765e-29 | 0.425124095 | 0.43 | 0.163 | 8.953593000626e-25 | 11 |
| ppib2 | 5.32091543176376e-29 | 0.466808837 | 0.72 | 0.325 | 1.33922120502062e-24 | 11 |
| irf2bp2b1 | 6.72805848791517e-29 | 0.268643419 | 0.273 | 0.079 | 1.69338504082337e-24 | 11 |
| h2afy21 | 7.39629650939706e-29 | 0.303635596 | 0.301 | 0.094 | 1.86157386845015e-24 | 11 |
| mt-atp64 | 9.05551553129972e-29 | 0.467823245 | 0.881 | 0.585 | 2.27918270407283e-24 | 11 |
| srsf5a1 | 9.53240934608901e-29 | 0.425230739 | 0.636 | 0.281 | 2.39921210831714e-24 | 11 |
| add3a2 | 2.49717438883788e-28 | 0.303113474 | 0.36 | 0.12 | 6.28513821926606e-24 | 11 |
| rnf10 | 2.70283962624003e-28 | 0.287510998 | 0.318 | 0.102 | 6.80277705528354e-24 | 11 |
| eif3i2 | 3.08135703508938e-28 | 0.421293984 | 0.601 | 0.266 | 7.75546752161647e-24 | 11 |
| eif4a1b1 | 3.79641490970359e-28 | 0.432609768 | 0.608 | 0.274 | 9.55519668623297e-24 | 11 |
| cct81 | 4.74653117594263e-28 | 0.361211643 | 0.497 | 0.196 | 1.194654431673e-23 | 11 |
| eif3c2 | 8.77273044596966e-28 | 0.355473084 | 0.43 | 0.164 | 2.2080085259461e-23 | 11 |
| ppiaa2 | 2.15184687004421e-27 | 0.306422199 | 0.986 | 0.8 | 5.41598338721427e-23 | 11 |
| MRPS12 | 5.28896984570281e-27 | 0.258996714 | 0.269 | 0.081 | 1.33118082046494e-22 | 11 |
| rpl42 | 6.69941094618079e-27 | 0.43171007 | 0.948 | 0.621 | 1.68617474104424e-22 | 11 |
| pim22 | 1.49757184606992e-26 | 0.258657113 | 0.28 | 0.086 | 3.76923857937337e-22 | 11 |
| etfa | 1.86822117947001e-26 | 0.253933879 | 0.276 | 0.085 | 4.70212588660807e-22 | 11 |
| ndufa5 | 8.60579794320393e-26 | 0.308005128 | 0.378 | 0.138 | 2.165993284325e-21 | 11 |
| eif3ea2 | 1.09076735643787e-25 | 0.391615638 | 0.605 | 0.27 | 2.74535235941848e-21 | 11 |
| pfn21 | 1.56408328293178e-25 | 0.358046824 | 0.528 | 0.22 | 3.936641214811e-21 | 11 |
| grpel2 | 1.95122939564329e-25 | 0.272405199 | 0.297 | 0.097 | 4.91104926589459e-21 | 11 |
| romo12 | 1.95214513784323e-25 | 0.440608424 | 0.752 | 0.378 | 4.91335409743762e-21 | 11 |
| ola1 | 2.3035071865024e-25 | 0.347676759 | 0.531 | 0.224 | 5.79769723770788e-21 | 11 |
| atp5f1e | 2.9622102170538e-25 | 0.4540665 | 0.822 | 0.476 | 7.45558689530272e-21 | 11 |
| zgc:111986 | 3.26063487014901e-25 | 0.31395691 | 0.441 | 0.172 | 8.20669190467804e-21 | 11 |
| cox6b2 | 6.3390232143772e-25 | 0.436993143 | 0.853 | 0.527 | 1.5954687528266e-20 | 11 |
| tcp11 | 1.26105916586534e-24 | 0.352678367 | 0.517 | 0.22 | 3.17395981456647e-20 | 11 |
| idh2 | 1.93173174114577e-24 | 0.338202453 | 0.472 | 0.194 | 4.86197561928979e-20 | 11 |
| smc1al | 2.26114035109231e-24 | 0.273476746 | 0.28 | 0.091 | 5.69106414966422e-20 | 11 |
| eif3ba1 | 2.32596273115986e-24 | 0.361672731 | 0.483 | 0.205 | 5.85421559805625e-20 | 11 |

| | | | | | | |
|---|---|---|---|---|---|---|
| mt-cyb3 | 3.52614466314438e-24 | 0.415369986 | 0.878 | 0.619 | 8.87495350266808e-20 | 11 |
| u2surp | 3.79867913532357e-24 | 0.270077301 | 0.28 | 0.092 | 9.56089551569589e-20 | 11 |
| tktb | 3.80584210620722e-24 | 0.304963404 | 0.434 | 0.168 | 9.57892399711296e-20 | 11 |
| atp5mf | 9.71375292328796e-24 | 0.392460202 | 0.598 | 0.282 | 2.44485447326235e-19 | 11 |
| esd | 2.15152196417569e-23 | 0.260485772 | 0.287 | 0.096 | 5.41516563163378e-19 | 11 |
| ndufb8 | 2.17608238519491e-23 | 0.349370251 | 0.545 | 0.242 | 5.47698175529706e-19 | 11 |
| cyc1 | 2.31477636926974e-23 | 0.284023011 | 0.357 | 0.133 | 5.826060643815e-19 | 11 |
| si:dkey-79d12.51 | 2.51878343989849e-23 | 0.267441662 | 0.238 | 0.074 | 6.3395260398805e-19 | 11 |
| prdx3 | 2.71532850362218e-23 | 0.288787149 | 0.353 | 0.131 | 6.83421031076665e-19 | 11 |
| rhoaa2 | 3.86329745624423e-23 | 0.304104607 | 0.385 | 0.15 | 9.72353336762111e-19 | 11 |
| gadd45bb1 | 4.01312903911763e-23 | 0.299419023 | 0.259 | 0.085 | 1.01006444785552e-18 | 11 |
| eif3g2 | 5.46399652633082e-23 | 0.380664253 | 0.587 | 0.276 | 1.3752332857122e-18 | 11 |
| atp5mc3a | 8.4161406451182e-23 | 0.335857172 | 0.503 | 0.218 | 2.1182584389698e-18 | 11 |
| cebpd2 | 1.903465273299e-22 | 0.350033262 | 0.336 | 0.126 | 4.79083174636626e-18 | 11 |
| map1lc3b1 | 2.07950413467444e-22 | 0.277561948 | 0.294 | 0.104 | 5.23390395656209e-18 | 11 |
| sumo2b2 | 2.75794314674699e-22 | 0.354883872 | 0.573 | 0.263 | 6.94146710604751e-18 | 11 |
| caprin1b1 | 3.21521479475009e-22 | 0.333823755 | 0.542 | 0.241 | 8.09237411690649e-18 | 11 |
| ostc | 4.14157354970623e-22 | 0.279295448 | 0.434 | 0.175 | 1.04239264672556e-17 | 11 |
| eef1b2 | 4.88557566128842e-22 | 0.292163263 | 0.909 | 0.731 | 1.22965053818968e-17 | 11 |
| hmgn71 | 5.71897237152051e-22 | 0.357431393 | 0.594 | 0.273 | 1.439408156188e-17 | 11 |
| rpz52 | 9.66070601930653e-22 | 0.254766362 | 0.287 | 0.099 | 2.43150309799926e-17 | 11 |
| selenok | 1.00196513158732e-21 | 0.256908099 | 0.332 | 0.124 | 2.52184603969212e-17 | 11 |
| tegt1 | 6.3306494905822e-21 | 0.325876152 | 0.465 | 0.207 | 1.59336117028463e-16 | 11 |
| rnf441 | 7.41265752995156e-21 | 0.253365942 | 0.255 | 0.087 | 1.86569177371351e-16 | 11 |
| cox6b12 | 1.40331735791575e-20 | 0.258564849 | 0.315 | 0.117 | 3.53200945813815e-16 | 11 |
| cirbpb | 2.27141685333988e-20 | 0.302434859 | 0.962 | 0.754 | 5.71692907817114e-16 | 11 |
| mycb1 | 2.88259741308068e-20 | 0.322169912 | 0.476 | 0.207 | 7.25520942898275e-16 | 11 |
| eif2s31 | 3.38548376923928e-20 | 0.350369478 | 0.577 | 0.279 | 8.52092409879834e-16 | 11 |
| prdx52 | 3.50603777743431e-20 | 0.262664289 | 0.385 | 0.155 | 8.82434648202441e-16 | 11 |
| si:dkey-7j14.62 | 4.84947280683475e-20 | 0.287647793 | 0.409 | 0.171 | 1.22056381075224e-15 | 11 |
| qkia1 | 9.74775025256274e-20 | 0.293312895 | 0.392 | 0.164 | 2.45341126106752e-15 | 11 |
| prdx21 | 1.09260196329562e-19 | 0.408227163 | 0.696 | 0.396 | 2.74996988141874e-15 | 11 |
| pfn2l1 | 1.70201527340282e-19 | 0.271231071 | 0.476 | 0.204 | 4.28380224162757e-15 | 11 |
| hnrnpa1a | 4.45702205144507e-19 | 0.287538991 | 0.371 | 0.156 | 1.12178788012821e-14 | 11 |
| cct4 | 6.650896951041e-19 | 0.289849078 | 0.49 | 0.222 | 1.67396425360751e-14 | 11 |

| | | | | | | |
|---|---|---|---|---|---|---|
| atf4a | 9.57343657043483e-19 | 0.25855725 | 0.416 | 0.178 | 2.40953825041274e-14 | 11 |
| ndufa4l1 | 9.97340980291e-19 | 0.391845123 | 0.699 | 0.389 | 2.51020751329442e-14 | 11 |
| eif3k | 2.91549666780719e-18 | 0.276412416 | 0.493 | 0.225 | 7.3380135632039e-14 | 11 |
| dynlrb1 | 3.08388343233256e-18 | 0.251070918 | 0.455 | 0.2 | 7.76182621083782e-14 | 11 |
| tomm20b2 | 4.92617450134615e-18 | 0.292782223 | 0.458 | 0.21 | 1.23986886024381e-13 | 11 |
| ndufb9 | 5.97259660829854e-18 | 0.250068905 | 0.399 | 0.172 | 1.50324284034266e-13 | 11 |
| ptmab2 | 6.15180236301952e-18 | 0.391862908 | 0.857 | 0.534 | 1.54834713674838e-13 | 11 |
| atp5pf | 7.61764585122901e-18 | 0.334757528 | 0.58 | 0.293 | 1.91728528429583e-13 | 11 |
| jpt1b1 | 1.06425705301009e-17 | 0.29767887 | 0.563 | 0.265 | 2.67862857672109e-13 | 11 |
| itm2ba2 | 1.40013964582287e-17 | 0.296543867 | 0.559 | 0.268 | 3.52401147457157e-13 | 11 |
| eif3ha1 | 1.48311068315213e-17 | 0.262217142 | 0.451 | 0.204 | 3.7328412784256e-13 | 11 |
| ldhba | 1.48745234083618e-17 | 0.299202887 | 0.507 | 0.243 | 3.74376879665059e-13 | 11 |
| atp5pd | 2.79542838597148e-17 | 0.310250064 | 0.524 | 0.257 | 7.03581370465161e-13 | 11 |
| rplp23 | 3.08586492768986e-17 | 0.338258945 | 0.979 | 0.753 | 7.76681343650261e-13 | 11 |
| hmgb2b | 3.79207126202438e-17 | 0.298231565 | 0.647 | 0.313 | 9.54426415938916e-13 | 11 |
| gabarapb2 | 4.17350029018122e-17 | 0.323786179 | 0.524 | 0.264 | 1.05042828803571e-12 | 11 |
| tnrc6c12 | 5.94403178312576e-17 | 0.25194743 | 0.315 | 0.13 | 1.49605335949492e-12 | 11 |
| rbm39a1 | 1.46400191874803e-16 | 0.25384047 | 0.329 | 0.14 | 3.6847464292969e-12 | 11 |
| ywhaz1 | 1.62952020903584e-16 | 0.300334891 | 0.559 | 0.276 | 4.1013394141223e-12 | 11 |
| rgcc2 | 1.99883474870743e-16 | 0.251471323 | 0.199 | 0.069 | 5.03086717902174e-12 | 11 |
| eif3d1 | 2.40551593451382e-16 | 0.274730068 | 0.514 | 0.248 | 6.05444305557784e-12 | 11 |
| tma7 | 2.42592878726354e-16 | 0.305688964 | 0.587 | 0.299 | 6.1058201646636e-12 | 11 |
| tpt12 | 3.06206612419919e-16 | 0.255044239 | 0.969 | 0.798 | 7.70691422799695e-12 | 11 |
| cct5 | 3.98421756713818e-16 | 0.273816853 | 0.507 | 0.246 | 1.00278771947301e-11 | 11 |
| sdc42 | 4.72448237918229e-16 | 0.296013126 | 0.528 | 0.253 | 1.18910497001639e-11 | 11 |
| midn1 | 1.2034180164631e-15 | 0.295713045 | 0.311 | 0.138 | 3.02888280563597e-11 | 11 |
| atp5f1d | 1.26363963189617e-15 | 0.282257324 | 0.51 | 0.254 | 3.18045458951946e-11 | 11 |
| nfil3-62 | 4.05986881241125e-15 | 0.26982343 | 0.297 | 0.128 | 1.02182838139579e-10 | 11 |
| gnb1b | 9.9082656034227e-15 | 0.262845164 | 0.5 | 0.25 | 2.49381136972546e-10 | 11 |
| eif2s21 | 1.58677416220837e-14 | 0.276499403 | 0.559 | 0.289 | 3.99375188886224e-10 | 11 |
| pnrc24 | 3.9219500920536e-14 | 0.364262239 | 0.881 | 0.561 | 9.8711561866897e-10 | 11 |
| vdac2 | 1.31754477192696e-13 | 0.264614634 | 0.559 | 0.29 | 3.31612843646298e-09 | 11 |
| fkbp1aa1 | 2.84324014014883e-13 | 0.311145036 | 0.689 | 0.382 | 7.15615110874059e-09 | 11 |
| serbp1a2 | 4.63705464682858e-13 | 0.356439313 | 0.79 | 0.476 | 1.16710028406029e-08 | 11 |
| serf22 | 1.72209087006624e-12 | 0.281734315 | 0.615 | 0.338 | 4.33433051086972e-08 | 11 |

| | | | | | | |
|---|---|---|---|---|---|---|
| eif5a3 | 3.72493497491825e-11 | 0.32395609 | 0.78 | 0.473 | 9.37528883837175e-07 | 11 |
| slc25a3b1 | 5.37076604427869e-11 | 0.302360283 | 0.731 | 0.432 | 1.3517681056845e-06 | 11 |
| eef1da2 | 8.25842591716592e-11 | 0.328654194 | 0.808 | 0.486 | 2.07856321909149e-06 | 11 |
| cox4i11 | 3.55817269894419e-10 | 0.267190992 | 0.664 | 0.387 | 8.95556486597264e-06 | 11 |
| cox7c | 1.56242900870463e-08 | 0.260144675 | 0.745 | 0.476 | 0.000393248 | 11 |
| si:ch211-132g1.41 | 0 | 1.015319382 | 0.487 | 0.009 | 0 | 12 |
| si:dkey-109a10.21 | 0 | 0.943231275 | 0.476 | 0.022 | 0 | 12 |
| zgc:1366051 | 0 | 0.866432451 | 0.415 | 0.013 | 0 | 12 |
| si:ch211-196f2.61 | 0 | 0.807631958 | 0.364 | 0.009 | 0 | 12 |
| foxp3a | 0 | 0.571328338 | 0.171 | 0.001 | 0 | 12 |
| si:ch73-361p23.3 | 0 | 0.566653712 | 0.244 | 0.002 | 0 | 12 |
| tcf7 | 3.8480431523573e-307 | 0.7239215 | 0.375 | 0.016 | 9.68513981016809e-303 | 12 |
| XLOC-0425951 | 3.01966861813652e-293 | 1.649678692 | 0.924 | 0.147 | 7.6002039449878e-289 | 12 |
| zap70 | 3.38147319632768e-291 | 0.507565904 | 0.251 | 0.006 | 8.51082988783714e-287 | 12 |
| si:ch211-67e16.3 | 3.272608997721e-290 | 0.664583962 | 0.309 | 0.011 | 8.23682958636399e-286 | 12 |
| XLOC-0139641 | 1.55157952930295e-285 | 1.621057406 | 0.938 | 0.156 | 3.90517051730259e-281 | 12 |
| sla21 | 6.80593953347241e-269 | 1.087464721 | 0.604 | 0.057 | 1.71298692117967e-264 | 12 |
| LOC1005358061 | 7.65557812831969e-264 | 0.811277599 | 0.418 | 0.026 | 1.92683245911678e-259 | 12 |
| LOC1039108711 | 1.00233795828352e-247 | 1.437003342 | 0.895 | 0.154 | 2.52278440720378e-243 | 12 |
| CR936442.11 | 1.7974730826033e-245 | 1.394332659 | 0.891 | 0.153 | 4.52406000160424e-241 | 12 |
| dusp21 | 1.35511744524218e-241 | 1.401551672 | 0.887 | 0.163 | 3.41069509793004e-237 | 12 |
| si:dkey-27i16.21 | 3.98552776137557e-239 | 1.115576705 | 0.655 | 0.078 | 1.00311748226062e-234 | 12 |
| tnfrsf9b | 1.16038950194308e-235 | 0.597877525 | 0.276 | 0.011 | 2.92058433744055e-231 | 12 |
| XLOC-0004311 | 3.54829332894077e-225 | 1.703340925 | 0.985 | 0.262 | 8.93069947961102e-221 | 12 |
| XLOC-0108991 | 1.94493665042873e-222 | 1.588974298 | 0.964 | 0.242 | 4.89521105546407e-218 | 12 |
| cd28 | 6.1386433424171e-222 | 0.366914465 | 0.138 | 0.001 | 1.54503514285296e-217 | 12 |
| gpr184 | 4.53772221590618e-220 | 0.352422956 | 0.164 | 0.003 | 1.14209930452143e-215 | 12 |
| si:dkey-260g12.11 | 9.76813218680181e-220 | 0.717193525 | 0.382 | 0.026 | 2.45854119009615e-215 | 12 |
| si:dkey-266f7.4 | 1.09624036635129e-209 | 0.312696064 | 0.135 | 0.002 | 2.75912737806957e-205 | 12 |
| FP236356.11 | 4.42758461919616e-204 | 0.863827301 | 0.447 | 0.04 | 1.11437877280548e-199 | 12 |
| gzmk1 | 4.99179796153623e-203 | 0.539783927 | 0.247 | 0.01 | 1.25638562893905e-198 | 12 |
| tagapa2 | 1.04230922397919e-196 | 1.099827384 | 0.651 | 0.093 | 2.62338808583323e-192 | 12 |
| rgs131 | 1.23592678033781e-191 | 1.128918562 | 0.72 | 0.119 | 3.11070411343224e-187 | 12 |
| si:ch211-119e14.11 | 7.0388755931695e-184 | 0.859921464 | 0.535 | 0.063 | 1.77161459804483e-179 | 12 |
| XLOC-0425961 | 4.23961803780383e-179 | 0.73273063 | 0.389 | 0.034 | 1.06706946393484e-174 | 12 |

| | | | | | | |
|---|---|---|---|---|---|---|
| cxcr4a | 2.60554969022922e-178 | 0.569262479 | 0.244 | 0.012 | 6.55790801533794e-174 | 12 |
| il7r | 4.88608722645032e-173 | 0.505438638 | 0.244 | 0.013 | 1.22977929402528e-168 | 12 |
| si:ch1073-220m6.11 | 1.23309884758678e-170 | 0.487743451 | 0.265 | 0.015 | 3.10358648949116e-166 | 12 |
| tnfrsf18 | 2.38748374705888e-163 | 0.580839934 | 0.215 | 0.01 | 6.00905784297249e-159 | 12 |
| LOC1018856751 | 2.09623572840865e-162 | 0.397527179 | 0.196 | 0.008 | 5.27601570483173e-158 | 12 |
| b3gnt5a | 2.50687795352793e-162 | 0.637762671 | 0.316 | 0.025 | 6.30956112123444e-158 | 12 |
| si:ch211-214p16.12 | 1.38558580537947e-161 | 1.308088755 | 0.945 | 0.242 | 3.48738091355959e-157 | 12 |
| si:ch211-93g21.11 | 4.20863785432853e-161 | 0.637459492 | 0.338 | 0.028 | 1.05927206155595e-156 | 12 |
| FP016106.11 | 1.95294900202276e-157 | 0.557520033 | 0.284 | 0.02 | 4.9153773431911e-153 | 12 |
| CR848749.21 | 3.43881392821577e-149 | 0.58531165 | 0.287 | 0.022 | 8.65515077592628e-145 | 12 |
| zfp36l1a3 | 5.75525008675998e-148 | 1.116077393 | 0.862 | 0.234 | 1.44853889433662e-143 | 12 |
| nab1a | 1.36444569572235e-147 | 0.571014356 | 0.251 | 0.017 | 3.43417337156359e-143 | 12 |
| tnfrsf9a1 | 9.54678409588818e-146 | 0.866906983 | 0.396 | 0.045 | 2.4028300890941e-141 | 12 |
| si:dkey-183i3.61 | 3.08731388418726e-145 | 0.823902006 | 0.371 | 0.039 | 7.77046031511092e-141 | 12 |
| sh2d1ab1 | 4.60876788768279e-145 | 0.424465725 | 0.207 | 0.011 | 1.15998078965088e-140 | 12 |
| skap11 | 8.31693791954139e-143 | 0.486282579 | 0.262 | 0.019 | 2.09329010496937e-138 | 12 |
| cxcr4b2 | 9.53191103378253e-143 | 1.117612684 | 0.756 | 0.171 | 2.39908668809272e-138 | 12 |
| c3a.52 | 1.21124902222535e-142 | 0.801258333 | 0.462 | 0.059 | 3.04859266403899e-138 | 12 |
| si:ch73-40i7.2 | 2.03955883828085e-139 | 0.332974996 | 0.167 | 0.007 | 5.13336564006908e-135 | 12 |
| si:ch73-223p23.21 | 4.93959518795696e-131 | 0.564030447 | 0.295 | 0.027 | 1.24324671285689e-126 | 12 |
| si:dkey-262k9.21 | 1.78643982991239e-127 | 0.554980213 | 0.302 | 0.029 | 4.49629040790648e-123 | 12 |
| cebpb3 | 1.21847574505251e-125 | 1.122763049 | 0.931 | 0.341 | 3.06678160272265e-121 | 12 |
| s1pr5b | 1.89106841264072e-125 | 0.331073033 | 0.138 | 0.005 | 4.75963008777542e-121 | 12 |
| irf4b | 8.12674642501549e-123 | 0.507215562 | 0.247 | 0.02 | 2.04542080771215e-118 | 12 |
| ncaldb1 | 8.53621239151942e-122 | 0.696458525 | 0.451 | 0.066 | 2.14847929682152e-117 | 12 |
| samsn1b1 | 1.22105960944918e-121 | 0.456779722 | 0.24 | 0.019 | 3.07328493102265e-117 | 12 |
| LOC1005362131 | 1.88611206933806e-121 | 0.835658672 | 0.542 | 0.096 | 4.74715546731695e-117 | 12 |
| si:dkeyp-85e10.31 | 4.19738604633451e-119 | 0.467992088 | 0.236 | 0.019 | 1.05644009400193e-114 | 12 |
| ccr9a2 | 1.21981495753193e-118 | 0.881471993 | 0.476 | 0.079 | 3.07015226661211e-114 | 12 |
| wasb2 | 7.49529302147477e-118 | 0.918797295 | 0.709 | 0.166 | 1.88649030057499e-113 | 12 |
| adgre5b.31 | 1.197244714166e-117 | 0.565768677 | 0.313 | 0.034 | 3.01334522108439e-113 | 12 |
| ptpn4b1 | 3.13014628605467e-116 | 0.48817748 | 0.255 | 0.022 | 7.87826518737101e-112 | 12 |
| si:ch211-132g1.11 | 3.47568276848018e-116 | 0.455631353 | 0.24 | 0.02 | 8.74794595998777e-112 | 12 |
| zgc:640512 | 2.88901460439748e-113 | 0.845028491 | 0.676 | 0.151 | 7.27136085780801e-109 | 12 |
| cd4-1 | 1.33542822617257e-110 | 0.365336327 | 0.131 | 0.005 | 3.36113930245374e-106 | 12 |

| | | | | | | |
|---|---|---|---|---|---|---|
| ptpn11b | 3.74354859789962e-110 | 0.417255541 | 0.225 | 0.018 | 9.42213746605357e-106 | 12 |
| si:dkey-183p4.101 | 6.59738832839201e-109 | 0.5201459 | 0.284 | 0.03 | 1.66049666837298e-104 | 12 |
| itk1 | 1.20453028732142e-108 | 0.345123987 | 0.167 | 0.01 | 3.03168228015928e-104 | 12 |
| arhgdig2 | 1.49301965714768e-105 | 0.906873 | 0.793 | 0.231 | 3.757781175075e-101 | 12 |
| gpr55a | 2.73083365972262e-105 | 0.273748591 | 0.127 | 0.005 | 6.87323523815586e-101 | 12 |
| pfn13 | 6.81597915572811e-105 | 0.971196576 | 0.996 | 0.515 | 1.71551379370521e-100 | 12 |
| hcst1 | 6.37935595097831e-104 | 0.585435079 | 0.342 | 0.045 | 1.60562009930173e-99 | 12 |
| btg13 | 7.26184057569854e-104 | 0.946108868 | 0.88 | 0.321 | 1.82773265449757e-99 | 12 |
| myca | 1.51007199407412e-103 | 0.54217565 | 0.28 | 0.031 | 3.80070020188515e-99 | 12 |
| wu:fb55g091 | 2.0015374434921e-102 | 0.962244867 | 0.76 | 0.239 | 5.03766959152528e-98 | 12 |
| abo | 4.0651064816313e-102 | 0.416139658 | 0.204 | 0.016 | 1.02314665036178e-97 | 12 |
| coro1a3 | 6.92520428320873e-101 | 0.918596924 | 0.778 | 0.219 | 1.7430046660408e-96 | 12 |
| srgn2 | 8.00826288532223e-101 | 0.896418777 | 0.789 | 0.225 | 2.01559968560675e-96 | 12 |
| rplp13 | 3.59653659910632e-100 | 0.427865884 | 1 | 0.875 | 9.0521229662907e-96 | 12 |
| rps27.14 | 6.49311842930089e-97 | 0.621292411 | 0.989 | 0.673 | 1.63425297747074e-92 | 12 |
| sik11 | 1.62227741388045e-96 | 0.75644733 | 0.516 | 0.108 | 4.08311002299569e-92 | 12 |
| rps54 | 5.28714477346988e-96 | 0.584365707 | 0.978 | 0.696 | 1.33072146803463e-91 | 12 |
| rpl18a4 | 6.7936516638255e-96 | 0.511731939 | 0.996 | 0.781 | 1.70989418726824e-91 | 12 |
| cxcr3.11 | 4.20570894085711e-94 | 0.407889192 | 0.204 | 0.018 | 1.05853488332433e-89 | 12 |
| rplp2l4 | 5.92161372155667e-93 | 0.527650195 | 0.993 | 0.77 | 1.4904109575786e-88 | 12 |
| si:ch211-25d12.71 | 1.13690625336369e-92 | 0.344247454 | 0.185 | 0.015 | 2.86147934909107e-88 | 12 |
| si:ch211-105j21.91 | 4.96447571381309e-92 | 0.462312949 | 0.262 | 0.03 | 1.24950889240962e-87 | 12 |
| cytip1 | 2.05166085114335e-91 | 0.336980168 | 0.171 | 0.013 | 5.16382519624269e-87 | 12 |
| rps171 | 2.0767821382243e-91 | 0.502702153 | 0.982 | 0.763 | 5.22705296369673e-87 | 12 |
| rplp24 | 2.18214812560837e-91 | 0.539789836 | 1 | 0.753 | 5.49224861734371e-87 | 12 |
| si:dkeyp-68b7.121 | 2.37779092997359e-91 | 0.575709853 | 0.36 | 0.057 | 5.98466199165053e-87 | 12 |
| rac23 | 2.93897712431407e-91 | 0.841229261 | 0.731 | 0.21 | 7.39711152418608e-87 | 12 |
| rps63 | 5.21107494330813e-91 | 0.480764696 | 0.978 | 0.785 | 1.31157545248122e-86 | 12 |
| tagapb1 | 1.02555003431336e-90 | 0.650273407 | 0.371 | 0.061 | 2.58120688136329e-86 | 12 |
| pde4ba2 | 1.97370513730581e-90 | 0.836899158 | 0.542 | 0.131 | 4.96761846008499e-86 | 12 |
| rpl324 | 7.40644899331898e-90 | 0.470967811 | 0.996 | 0.81 | 1.86412914712845e-85 | 12 |
| rps123 | 2.52725841036918e-89 | 0.430646126 | 0.989 | 0.837 | 6.3608566930582e-85 | 12 |
| rpl374 | 1.7926349592124e-88 | 0.434333513 | 0.996 | 0.847 | 4.51188292884168e-84 | 12 |
| cremb | 3.80680795934871e-88 | 0.42700667 | 0.193 | 0.017 | 9.58135495288477e-84 | 12 |
| rpl7a3 | 5.68065006987187e-88 | 0.458153303 | 0.996 | 0.806 | 1.42976281608605e-83 | 12 |

| | | | | | | |
|---|---|---|---|---|---|---|
| mxd | 1.98950735965749e-87 | 0.309909531 | 0.175 | 0.014 | 5.00739107352193e-83 | 12 |
| rps203 | 2.4350662736686e-87 | 0.346531063 | 0.993 | 0.915 | 6.1288183041965e-83 | 12 |
| rpl114 | 5.78541865188119e-87 | 0.462813018 | 0.996 | 0.81 | 1.45613202049198e-82 | 12 |
| rps214 | 7.59254795638508e-87 | 0.471648276 | 0.996 | 0.826 | 1.91096839514256e-82 | 12 |
| si:ch73-285p12.41 | 1.08385481076939e-86 | 0.342167497 | 0.178 | 0.014 | 2.72795417322549e-82 | 12 |
| wasa1 | 2.1561485263367e-86 | 0.630817464 | 0.444 | 0.086 | 5.42681022593683e-82 | 12 |
| rps215 | 3.27602099621385e-86 | 0.526294709 | 0.993 | 0.745 | 8.24541724537063e-82 | 12 |
| rpl154 | 3.63282883591083e-86 | 0.522637659 | 0.982 | 0.74 | 9.14346689710397e-82 | 12 |
| rpl84 | 6.02278665622498e-86 | 0.482864759 | 0.989 | 0.776 | 1.51587517350527e-81 | 12 |
| arpc1b3 | 1.32779299517018e-85 | 0.888945997 | 0.884 | 0.296 | 3.34192218954383e-81 | 12 |
| rpl364 | 1.40739292473736e-85 | 0.489275555 | 0.996 | 0.786 | 3.54226725227145e-81 | 12 |
| tpt13 | 2.41247693611049e-85 | 0.451699671 | 0.993 | 0.797 | 6.0719632004965e-81 | 12 |
| si:ch211-226m16.21 | 2.93759684945793e-84 | 0.623523595 | 0.364 | 0.062 | 7.39363751040066e-80 | 12 |
| rpl73 | 3.87298711466685e-84 | 0.445556469 | 0.985 | 0.801 | 9.747921268905e-80 | 12 |
| ccl38.61 | 1.15359659023582e-83 | 0.958931787 | 0.473 | 0.102 | 2.90348725796454e-79 | 12 |
| ptprc2 | 1.70953960114821e-83 | 0.693224554 | 0.524 | 0.119 | 4.30274022212994e-79 | 12 |
| rsl24d13 | 6.03124360353056e-83 | 0.79255827 | 0.909 | 0.434 | 1.51800370257261e-78 | 12 |
| sept9b1 | 6.59516408018184e-83 | 0.546639881 | 0.295 | 0.043 | 1.65993684734097e-78 | 12 |
| LOC101882375 | 6.63255669934924e-83 | 0.326417676 | 0.16 | 0.012 | 1.66934819565921e-78 | 12 |
| rps26l4 | 2.62010060848122e-81 | 0.599684985 | 0.993 | 0.658 | 6.59453122148639e-77 | 12 |
| rpl273 | 3.13496026416069e-81 | 0.486842786 | 0.985 | 0.758 | 7.89038148886604e-77 | 12 |
| rpl344 | 3.7759852756316e-81 | 0.474441281 | 0.989 | 0.78 | 9.50377734023718e-77 | 12 |
| cyth4a1 | 1.48228645524466e-80 | 0.345896912 | 0.182 | 0.016 | 3.73076677920528e-76 | 12 |
| rps73 | 1.22495483881314e-79 | 0.432700616 | 0.989 | 0.817 | 3.0830888338088e-75 | 12 |
| rpl184 | 1.33838671955522e-79 | 0.479794765 | 0.989 | 0.77 | 3.36858553444854e-75 | 12 |
| rps164 | 2.67967489526224e-79 | 0.516450997 | 0.989 | 0.733 | 6.74447374388553e-75 | 12 |
| ikzf22 | 3.01302171414515e-79 | 0.440774412 | 0.225 | 0.026 | 7.58347435233194e-75 | 12 |
| il12rb2l1 | 1.89501575449356e-78 | 0.457163185 | 0.276 | 0.039 | 4.76956515248485e-74 | 12 |
| rps15a4 | 2.17757935795292e-78 | 0.493322578 | 0.985 | 0.776 | 5.48074948603171e-74 | 12 |
| h1fx1 | 5.63153201622841e-78 | 0.867436526 | 0.604 | 0.175 | 1.41740029316453e-73 | 12 |
| LOC1039091151 | 1.0268317131272e-77 | 0.721173924 | 0.578 | 0.157 | 2.58443273876986e-73 | 12 |
| XLOC-0110851 | 1.09451529516303e-77 | 0.553255414 | 0.335 | 0.057 | 2.75478554639584e-73 | 12 |
| LOC1104399311 | 2.48918956611171e-77 | 0.374268464 | 0.196 | 0.02 | 6.26504121894656e-73 | 12 |
| rps294 | 1.32685583293832e-76 | 0.407213091 | 0.993 | 0.829 | 3.33956344592246e-72 | 12 |
| rpl192 | 1.85463668972884e-76 | 0.389593644 | 0.993 | 0.849 | 4.66793508437853e-72 | 12 |

| | | | | | | |
|---|---|---|---|---|---|---|
| themis1 | 2.26096334165703e-76 | 0.252175152 | 0.135 | 0.009 | 5.69061863461658e-72 | 12 |
| rpl10a3 | 6.14336706781803e-76 | 0.470944316 | 0.989 | 0.76 | 1.54622405729912e-71 | 12 |
| rpl103 | 1.08092199691132e-74 | 0.453337723 | 0.985 | 0.768 | 2.72057257402611e-70 | 12 |
| laptm53 | 1.53399713408252e-74 | 0.708087551 | 0.604 | 0.161 | 3.86091738677229e-70 | 12 |
| rps193 | 2.00400703056581e-74 | 0.368968188 | 0.993 | 0.861 | 5.04388529523109e-70 | 12 |
| rps4x3 | 2.96582478646378e-74 | 0.519920399 | 0.967 | 0.705 | 7.46468440505068e-70 | 12 |
| si:dkey-42i9.7 | 7.09724010015914e-74 | 0.412120267 | 0.185 | 0.019 | 1.78630436080905e-69 | 12 |
| BX005105.11 | 9.28258057280965e-74 | 0.498968291 | 0.273 | 0.039 | 2.33633270437046e-69 | 12 |
| rps94 | 4.0464232800023e-73 | 0.453474249 | 0.985 | 0.773 | 1.01844427534378e-68 | 12 |
| sept121 | 1.39368838288191e-72 | 0.509843022 | 0.324 | 0.056 | 3.50777429087549e-68 | 12 |
| si:ch73-208g10.11 | 4.66084871483769e-72 | 0.425007069 | 0.265 | 0.038 | 1.1730890130375e-67 | 12 |
| rplp03 | 2.10463713315116e-71 | 0.377882746 | 0.985 | 0.849 | 5.29716120042816e-67 | 12 |
| il21r.11 | 2.49113985951048e-71 | 0.366839218 | 0.189 | 0.02 | 6.26994991240194e-67 | 12 |
| nr4a12 | 3.82287500870145e-71 | 0.769546354 | 0.564 | 0.159 | 9.62179410940067e-67 | 12 |
| rpl354 | 2.55432777326489e-70 | 0.428029713 | 0.993 | 0.801 | 6.4289875725304e-66 | 12 |
| slc25a53 | 3.25149043017504e-70 | 0.404868963 | 0.985 | 0.803 | 8.18367626370757e-66 | 12 |
| rpl384 | 3.36526790321442e-70 | 0.504333634 | 0.985 | 0.723 | 8.47004278560037e-66 | 12 |
| LOC1039091071 | 5.35223952756964e-70 | 0.433021537 | 0.196 | 0.022 | 1.347105166694e-65 | 12 |
| rpl263 | 9.88649038936369e-70 | 0.385991969 | 0.993 | 0.838 | 2.48833076609895e-65 | 12 |
| calm1b2 | 1.11306006677095e-69 | 0.565874932 | 0.415 | 0.088 | 2.8014608820558e-65 | 12 |
| rpsa4 | 1.42214751558184e-69 | 0.474604214 | 0.985 | 0.747 | 3.57940308196794e-65 | 12 |
| rel | 1.70473718755164e-69 | 0.45136417 | 0.236 | 0.032 | 4.29065302734872e-65 | 12 |
| dgkaa | 4.17221583711872e-69 | 0.253227363 | 0.131 | 0.01 | 1.05010500404441e-64 | 12 |
| rps144 | 1.75493542903601e-68 | 0.470075007 | 1 | 0.754 | 4.41699698134072e-64 | 12 |
| faua4 | 1.84961334316355e-68 | 0.528238939 | 0.982 | 0.684 | 4.65529182340835e-64 | 12 |
| BX511079.21 | 1.10937072808076e-67 | 0.273611701 | 0.138 | 0.011 | 2.79217518550646e-63 | 12 |
| rps183 | 2.47680590262258e-67 | 0.499456115 | 0.982 | 0.72 | 6.23387277631077e-63 | 12 |
| csrnp1a | 4.19320606144324e-67 | 0.517019775 | 0.284 | 0.047 | 1.05538803360465e-62 | 12 |
| rpl232 | 1.62061571083245e-66 | 0.33135264 | 0.989 | 0.87 | 4.0789276825942e-62 | 12 |
| rpl94 | 2.41944454060811e-66 | 0.591808004 | 0.982 | 0.622 | 6.08949996425655e-62 | 12 |
| hsp90ab13 | 3.14370140815912e-66 | 0.418991724 | 0.989 | 0.787 | 7.91238207419568e-62 | 12 |
| rps133 | 2.56588200156019e-65 | 0.488067791 | 0.985 | 0.719 | 6.45806840972685e-61 | 12 |
| ptmaa3 | 5.73584063947773e-65 | 0.762324478 | 0.895 | 0.385 | 1.44365373055015e-60 | 12 |
| agap2 | 1.52307754728043e-64 | 0.295756659 | 0.156 | 0.015 | 3.83343387875011e-60 | 12 |
| rpl35a4 | 2.01348095226546e-64 | 0.447508626 | 0.978 | 0.755 | 5.06773020875693e-60 | 12 |

| | | | | | | |
|---|---|---|---|---|---|---|
| rps33 | 6.09022135130464e-64 | 0.495996666 | 0.978 | 0.694 | 1.53284781190986e-59 | 12 |
| CU207241.1 | 6.43325726819231e-64 | 0.321228283 | 0.145 | 0.013 | 1.61918652183132e-59 | 12 |
| ms4a17a.62 | 2.21102298018067e-63 | 0.455151761 | 0.269 | 0.045 | 5.56492373881673e-59 | 12 |
| rps233 | 4.4315719344406e-63 | 0.476079222 | 0.982 | 0.718 | 1.11538234017935e-58 | 12 |
| vsir2 | 5.05045609724645e-62 | 0.536331997 | 0.367 | 0.08 | 1.27114929511596e-57 | 12 |
| si:dkey-262k9.41 | 6.20742731062026e-62 | 0.548742928 | 0.378 | 0.085 | 1.56234737981001e-57 | 12 |
| wipf1a1 | 1.68373716359725e-61 | 0.477887535 | 0.316 | 0.06 | 4.23779806705793e-57 | 12 |
| rpl284 | 1.95822542558336e-61 | 0.450839521 | 0.985 | 0.745 | 4.92865757365076e-57 | 12 |
| CR855337.11 | 2.13289197531053e-61 | 0.386132456 | 0.218 | 0.031 | 5.36827581265906e-57 | 12 |
| tnip11 | 2.36547143082846e-61 | 0.569226709 | 0.353 | 0.076 | 5.95365504425215e-57 | 12 |
| rps154 | 3.09114989636635e-61 | 0.522492268 | 0.982 | 0.679 | 7.78011517416446e-57 | 12 |
| rpl133 | 4.72371352268556e-61 | 0.385902022 | 0.989 | 0.801 | 1.18891145652473e-56 | 12 |
| lpar5b1 | 6.19878488235167e-61 | 0.367362828 | 0.182 | 0.022 | 1.56017216703909e-56 | 12 |
| RASA2 | 7.01181169707151e-61 | 0.427439225 | 0.247 | 0.04 | 1.76480288603593e-56 | 12 |
| capgb2 | 1.49187466011214e-60 | 0.613406974 | 0.433 | 0.108 | 3.75489933203626e-56 | 12 |
| rpl244 | 2.33183514230758e-60 | 0.437014882 | 1 | 0.772 | 5.86899586967394e-56 | 12 |
| rpl313 | 2.46935184673695e-60 | 0.316534571 | 0.996 | 0.873 | 6.21511166305223e-56 | 12 |
| cabz01093075.11 | 2.45113636706032e-59 | 0.311148964 | 0.16 | 0.017 | 6.16926512225411e-55 | 12 |
| rpl304 | 2.76038914888971e-59 | 0.487936517 | 0.978 | 0.703 | 6.94762344884052e-55 | 12 |
| prkacba1 | 3.49857855958541e-59 | 0.30907843 | 0.16 | 0.017 | 8.80557237662053e-55 | 12 |
| litaf1 | 5.79644201639342e-59 | 0.58908132 | 0.465 | 0.128 | 1.45890649110606e-54 | 12 |
| rpl394 | 8.34326459929703e-59 | 0.338761728 | 0.989 | 0.872 | 2.09991626699707e-54 | 12 |
| rps284 | 1.09125134878448e-58 | 0.389844546 | 0.996 | 0.823 | 2.74657051975566e-54 | 12 |
| nfkbie1 | 1.41625516872974e-58 | 0.577490201 | 0.375 | 0.088 | 3.56457263417587e-54 | 12 |
| rps244 | 1.79927769243565e-58 | 0.437264236 | 0.985 | 0.754 | 4.52860202409129e-54 | 12 |
| LOC1001511961 | 1.22260829171313e-57 | 0.403696508 | 0.233 | 0.037 | 3.07718280941277e-53 | 12 |
| irf2bp2a2 | 1.63330298774524e-57 | 0.655676961 | 0.535 | 0.169 | 4.110860289856e-53 | 12 |
| rpl214 | 1.8690643439559e-57 | 0.428880935 | 0.978 | 0.752 | 4.7042480473026e-53 | 12 |
| tnfaip31 | 4.77311096615507e-57 | 0.487562947 | 0.302 | 0.061 | 1.20134429907157e-52 | 12 |
| rpl36a4 | 5.24303899416144e-57 | 0.38650293 | 0.993 | 0.806 | 1.31962048444049e-52 | 12 |
| mcl1a2 | 7.46825506564847e-57 | 0.572385785 | 0.956 | 0.617 | 1.87968511747306e-52 | 12 |
| rps103 | 4.63526209756813e-56 | 0.491870245 | 0.985 | 0.688 | 1.16664911733692e-51 | 12 |
| si:dkey-53k12.2 | 9.89127398478256e-56 | 0.388999049 | 0.244 | 0.041 | 2.48953474922992e-51 | 12 |
| si:dkey-171c9.31 | 1.37226607257546e-55 | 0.355597111 | 0.211 | 0.031 | 3.45385647806518e-51 | 12 |
| nr4a3 | 1.69564624255756e-55 | 0.498673829 | 0.269 | 0.05 | 4.26777202789312e-51 | 12 |

| | | | | | | |
|---|---|---|---|---|---|---|
| fam49a1 | 1.81212533844785e-55 | 0.439466417 | 0.262 | 0.047 | 4.56093826433938e-51 | 12 |
| eef1a1l11 | 3.71745464827369e-55 | 0.275712348 | 0.996 | 0.914 | 9.35646160424005e-51 | 12 |
| wipf1b1 | 8.53616558586978e-55 | 0.458413317 | 0.291 | 0.059 | 2.14846751630757e-50 | 12 |
| naca4 | 1.27533935684596e-54 | 0.543908569 | 0.942 | 0.612 | 3.2099016272456e-50 | 12 |
| ets2 | 3.58791796643815e-54 | 0.5786732 | 0.353 | 0.085 | 9.03043072972817e-50 | 12 |
| hmha1b1 | 3.69535359193975e-54 | 0.47275749 | 0.324 | 0.07 | 9.30083545555316e-50 | 12 |
| sat1a.21 | 4.62720612857748e-54 | 0.681032219 | 0.713 | 0.29 | 1.16462151050167e-49 | 12 |
| rps3a3 | 5.38091326799741e-54 | 0.393187759 | 0.971 | 0.775 | 1.35432206042227e-49 | 12 |
| acap11 | 1.25503088792638e-53 | 0.373883133 | 0.225 | 0.037 | 3.15878724182191e-49 | 12 |
| zgc:195282 | 2.67238623099076e-53 | 0.261061901 | 0.131 | 0.013 | 6.72612890478065e-49 | 12 |
| rps8a3 | 3.16970858753002e-53 | 0.390220286 | 0.978 | 0.784 | 7.9778395439543e-49 | 12 |
| cd28l1 | 3.81078902445473e-53 | 0.423612172 | 0.215 | 0.034 | 9.59137489565011e-49 | 12 |
| b2m3 | 4.32541780145482e-53 | 0.710813416 | 0.698 | 0.284 | 1.08866440644816e-48 | 12 |
| rpl294 | 8.48774624056843e-53 | 0.385658494 | 0.989 | 0.805 | 2.13628085128867e-48 | 12 |
| si:ch211-256a21.4 | 1.82310972040917e-52 | 0.275988396 | 0.109 | 0.009 | 4.58858485529785e-48 | 12 |
| si:rp71-81e14.21 | 2.54097217205345e-52 | 0.380128393 | 0.207 | 0.032 | 6.39537285984132e-48 | 12 |
| ccr21 | 2.86972755851179e-52 | 0.399159301 | 0.244 | 0.043 | 7.22281729201834e-48 | 12 |
| pnrc25 | 5.23375809989664e-52 | 0.608868245 | 0.956 | 0.56 | 1.31728457616299e-47 | 12 |
| ptpn62 | 7.75125471485562e-52 | 0.528942649 | 0.356 | 0.086 | 1.95091329918201e-47 | 12 |
| ldha | 1.92411530183444e-51 | 0.459537838 | 0.313 | 0.068 | 4.84280580318709e-47 | 12 |
| p2ry101 | 5.3328288560527e-51 | 0.289072579 | 0.153 | 0.018 | 1.3422196947799e-46 | 12 |
| gpr183a1 | 1.56665443693918e-50 | 0.50097357 | 0.287 | 0.061 | 3.94311255233222e-46 | 12 |
| rack13 | 2.82060965408285e-50 | 0.41751683 | 0.982 | 0.74 | 7.09919243836112e-46 | 12 |
| sh2b31 | 4.1916829124829e-50 | 0.456102431 | 0.276 | 0.057 | 1.05500467224282e-45 | 12 |
| traf1 | 5.78847307360197e-50 | 0.302202309 | 0.142 | 0.016 | 1.45690078789488e-45 | 12 |
| runx3 | 7.20541118061179e-50 | 0.333404626 | 0.149 | 0.018 | 1.81352994004818e-45 | 12 |
| arhgef3l1 | 1.61830791608517e-49 | 0.326669247 | 0.189 | 0.028 | 4.07311919399476e-45 | 12 |
| si:ch73-226l13.21 | 1.61870598186747e-49 | 0.313485525 | 0.135 | 0.015 | 4.07412108576225e-45 | 12 |
| rps253 | 1.93459091604045e-49 | 0.434901139 | 0.978 | 0.72 | 4.8691718765822e-45 | 12 |
| si:ch211-167j9.51 | 4.55174277831955e-49 | 0.324795903 | 0.175 | 0.025 | 1.14562813987525e-44 | 12 |
| rpl123 | 4.92236312750244e-49 | 0.361295977 | 0.978 | 0.795 | 1.23890957556109e-44 | 12 |
| ccni2 | 2.64121615138576e-48 | 0.639127752 | 0.684 | 0.286 | 6.64767693142283e-44 | 12 |
| rps263 | 4.77605920578343e-48 | 0.415072343 | 0.967 | 0.729 | 1.20208634150363e-43 | 12 |
| abi3a1 | 8.16167583446112e-48 | 0.346834028 | 0.225 | 0.04 | 2.05421219077552e-43 | 12 |
| zgc:152774 | 9.56080962779189e-47 | 0.255354069 | 0.135 | 0.016 | 2.40636017521894e-42 | 12 |

| | | | | | | |
|---|---|---|---|---|---|---|
| kctd5a1 | 1.16776832379297e-46 | 0.595506575 | 0.498 | 0.169 | 2.93915609415453e-42 | 12 |
| rpl224 | 2.02156164947263e-46 | 0.48134274 | 0.993 | 0.688 | 5.08806851555765e-42 | 12 |
| ucp21 | 2.59235507106253e-46 | 0.625445514 | 0.829 | 0.408 | 6.52469847835729e-42 | 12 |
| btf33 | 5.09512061083492e-46 | 0.529784923 | 0.92 | 0.581 | 1.28239090654104e-41 | 12 |
| rasal32 | 5.1651360869608e-46 | 0.296978625 | 0.171 | 0.025 | 1.30001310172716e-41 | 12 |
| tnip2 | 5.20714488019388e-46 | 0.273995088 | 0.142 | 0.018 | 1.310586294896e-41 | 12 |
| rpl23a4 | 5.60344423049937e-46 | 0.51634398 | 0.96 | 0.628 | 1.41033087837439e-41 | 12 |
| ikzf11 | 8.5647659040158e-46 | 0.300841552 | 0.189 | 0.03 | 2.15566593037816e-41 | 12 |
| XLOC-0089931 | 9.4238178079588e-46 | 0.319516309 | 0.189 | 0.03 | 2.37188070408515e-41 | 12 |
| mbnl12 | 1.72898709744617e-45 | 0.580118492 | 0.516 | 0.177 | 4.35168762556226e-41 | 12 |
| eef1g3 | 2.00942731828318e-45 | 0.498695162 | 0.924 | 0.615 | 5.05752761738693e-41 | 12 |
| rnf442 | 2.36249689732094e-45 | 0.477972914 | 0.331 | 0.085 | 5.94616844086708e-41 | 12 |
| fmnl1a | 4.21683111257242e-45 | 0.34083037 | 0.204 | 0.035 | 1.06133422272335e-40 | 12 |
| ppp1r181 | 5.30687887869476e-45 | 0.385977531 | 0.247 | 0.05 | 1.33568834497869e-40 | 12 |
| ets11 | 5.95646854752184e-45 | 0.343724321 | 0.2 | 0.035 | 1.49918356872577e-40 | 12 |
| id31 | 6.04808776034482e-45 | 0.640880481 | 0.564 | 0.203 | 1.52224320840119e-40 | 12 |
| lbh1 | 7.00056366861773e-45 | 0.553610635 | 0.52 | 0.165 | 1.7619718697544e-40 | 12 |
| nsa23 | 8.45986938260919e-45 | 0.620100712 | 0.767 | 0.354 | 2.12926452490891e-40 | 12 |
| cd83 | 1.45757064468662e-44 | 0.641735895 | 0.36 | 0.099 | 3.66855955561174e-40 | 12 |
| CU499336.21 | 1.65142758894327e-44 | 0.334103683 | 0.196 | 0.033 | 4.15647809861132e-40 | 12 |
| chmp2a | 2.28797338858991e-44 | 0.561193327 | 0.411 | 0.129 | 5.75860022174194e-40 | 12 |
| CR392341.1 | 2.86698026367321e-44 | 0.398590821 | 0.262 | 0.056 | 7.2159026256391e-40 | 12 |
| si:dkey-185m8.21 | 3.57140935828507e-44 | 0.375552684 | 0.229 | 0.045 | 8.98888021386769e-40 | 12 |
| AL954142.1 | 4.59173904297566e-44 | 0.27015818 | 0.153 | 0.021 | 1.15569479972654e-39 | 12 |
| tnfsf141 | 6.2789761926274e-44 | 0.275763912 | 0.124 | 0.014 | 1.58035551792239e-39 | 12 |
| rps113 | 1.25612119476981e-43 | 0.37226536 | 0.967 | 0.762 | 3.16153143511614e-39 | 12 |
| rpl314 | 1.90812106501775e-43 | 0.479506581 | 0.964 | 0.648 | 4.80254990854318e-39 | 12 |
| lcp2a | 1.91526189745677e-43 | 0.332627121 | 0.193 | 0.033 | 4.82052266970895e-39 | 12 |
| gtpbp42 | 9.94570741848564e-43 | 0.573390121 | 0.589 | 0.23 | 2.50323510015865e-38 | 12 |
| mob1a1 | 1.00844644960745e-42 | 0.524172204 | 0.476 | 0.16 | 2.53815886901699e-38 | 12 |
| si:ch211-196l7.44 | 1.01348405717415e-42 | 0.599167658 | 0.705 | 0.306 | 2.55083802350162e-38 | 12 |
| hmgn21 | 4.63922919539826e-42 | 0.673671122 | 0.829 | 0.435 | 1.16764759618979e-37 | 12 |
| tefa1 | 5.90095672575601e-42 | 0.52907796 | 0.418 | 0.133 | 1.48521179830553e-37 | 12 |
| selplg | 2.1442595353688e-41 | 0.384758953 | 0.218 | 0.044 | 5.39688682456973e-37 | 12 |
| uba523 | 2.37942964971726e-41 | 0.489444164 | 0.953 | 0.624 | 5.98878648537337e-37 | 12 |

| | | | | | | |
|---|---|---|---|---|---|---|
| scinlb1 | 2.81652193326566e-41 | 0.426310757 | 0.313 | 0.079 | 7.08890405383634e-37 | 12 |
| rpl13a3 | 3.11915833143972e-41 | 0.367737479 | 0.975 | 0.76 | 7.85060960440064e-37 | 12 |
| CR381673.11 | 3.61123278326657e-41 | 0.39394863 | 0.247 | 0.054 | 9.08911179220363e-37 | 12 |
| si:ch73-248e21.5 | 4.96077261280284e-41 | 0.354375107 | 0.215 | 0.042 | 1.24857685891635e-36 | 12 |
| CU929418.1 | 6.34380367958058e-41 | 0.279026589 | 0.153 | 0.023 | 1.59667194811364e-36 | 12 |
| rpl64 | 6.68435643718959e-41 | 0.467062456 | 0.964 | 0.662 | 1.68238567167625e-36 | 12 |
| stk17al | 8.70815725684917e-41 | 0.501886296 | 0.396 | 0.115 | 2.19175609997637e-36 | 12 |
| crema2 | 1.81072052104285e-40 | 0.600575913 | 0.473 | 0.173 | 4.55740247941276e-36 | 12 |
| vaspa2 | 2.02218286427993e-40 | 0.399040837 | 0.295 | 0.071 | 5.08963205110615e-36 | 12 |
| il2rb1 | 2.10744702744431e-40 | 0.261990245 | 0.138 | 0.019 | 5.30423342337459e-36 | 12 |
| rpl27a4 | 8.27051619225532e-40 | 0.583400881 | 0.905 | 0.487 | 2.08160622042874e-35 | 12 |
| si:ch211-191c10.11 | 1.3127828214898e-39 | 0.375695854 | 0.236 | 0.051 | 3.30414308340767e-35 | 12 |
| rpl5a3 | 1.70306018034565e-39 | 0.567215808 | 0.895 | 0.484 | 4.28643216791196e-35 | 12 |
| CABZ01079663.11 | 2.35199128331282e-39 | 0.308905744 | 0.196 | 0.037 | 5.91972686097004e-35 | 12 |
| msna3 | 4.82745671218876e-39 | 0.570086752 | 0.578 | 0.235 | 1.21502257989079e-34 | 12 |
| si:ch211-132p1.32 | 7.72562188383844e-39 | 0.451171813 | 0.309 | 0.085 | 1.9444617719433e-34 | 12 |
| gnai2b1 | 1.45504654188539e-38 | 0.394535244 | 0.265 | 0.064 | 3.66220664127134e-34 | 12 |
| celf23 | 2.65186485768272e-38 | 0.551444814 | 0.585 | 0.236 | 6.67447866030163e-34 | 12 |
| igfbp1a | 3.04038949964887e-38 | 0.440116984 | 0.124 | 0.016 | 7.65235633166625e-34 | 12 |
| sept22 | 3.98673026348584e-38 | 0.498317165 | 0.458 | 0.16 | 1.00342014001675e-33 | 12 |
| actb23 | 1.10384954053502e-37 | 0.343633175 | 0.996 | 0.865 | 2.77827890857259e-33 | 12 |
| arhgap15 | 2.46160766786431e-37 | 0.275591913 | 0.16 | 0.027 | 6.19562033924769e-33 | 12 |
| tox1 | 4.36315966683406e-37 | 0.263116603 | 0.131 | 0.019 | 1.09816365654546e-32 | 12 |
| lpar5a1 | 6.19580066942301e-37 | 0.328414094 | 0.196 | 0.039 | 1.55942107048708e-32 | 12 |
| mrps151 | 8.22700327797562e-37 | 0.471266933 | 0.385 | 0.12 | 2.07065445503368e-32 | 12 |
| mknk2b3 | 1.16038378220647e-36 | 0.529032896 | 0.505 | 0.191 | 2.92056994143547e-32 | 12 |
| lcp11 | 1.48601410481134e-36 | 0.406732234 | 0.309 | 0.084 | 3.74014890039965e-32 | 12 |
| spint1a | 4.41170819962172e-36 | 0.333136067 | 0.236 | 0.053 | 1.11038283676279e-31 | 12 |
| eif3f3 | 6.08462526809161e-36 | 0.555171555 | 0.825 | 0.424 | 1.53143933372598e-31 | 12 |
| cd247l1 | 6.64427437347053e-36 | 0.255316772 | 0.131 | 0.019 | 1.6722974170588e-31 | 12 |
| myl12.11 | 7.93203131774669e-36 | 0.556735095 | 0.796 | 0.384 | 1.99641296236366e-31 | 12 |
| si:ch211-284f22.31 | 8.77029671347573e-36 | 0.30518188 | 0.171 | 0.031 | 2.20739597981471e-31 | 12 |
| grap2a1 | 3.17356060582056e-35 | 0.317297425 | 0.207 | 0.044 | 7.98753468878977e-31 | 12 |
| rps27a3 | 3.71696138172654e-35 | 0.372338785 | 0.978 | 0.747 | 9.35522010166752e-31 | 12 |
| ddb2 | 7.48039856746478e-35 | 0.398953227 | 0.291 | 0.079 | 1.88274151544521e-30 | 12 |

| | | | | | | |
|---|---|---|---|---|---|---|
| ppiaa3 | 7.80880274339598e-35 | 0.324762645 | 0.978 | 0.801 | 1.96539756248534e-30 | 12 |
| LOC1018820701 | 8.8263694724296e-35 | 0.263067096 | 0.135 | 0.02 | 2.2215089325158e-30 | 12 |
| dnajb9b | 1.65164171650335e-34 | 0.26117878 | 0.138 | 0.022 | 4.15701703626727e-30 | 12 |
| dtnbp1a1 | 1.97392190263037e-34 | 0.402451913 | 0.298 | 0.083 | 4.96816403673038e-30 | 12 |
| arhgap4b | 4.18573258190835e-34 | 0.278786112 | 0.156 | 0.028 | 1.05350703354051e-29 | 12 |
| tes | 4.73025768208332e-34 | 0.344457033 | 0.244 | 0.06 | 1.19055855600355e-29 | 12 |
| eef1da3 | 5.07010440781162e-34 | 0.518390176 | 0.829 | 0.486 | 1.27609457840211e-29 | 12 |
| cd74a2 | 5.10403789688379e-34 | 0.414195561 | 0.578 | 0.212 | 1.28463529826668e-29 | 12 |
| slc44a21 | 1.90003241357357e-33 | 0.337008196 | 0.211 | 0.048 | 4.78219158172331e-29 | 12 |
| tnfb | 2.41951054565756e-33 | 0.404225295 | 0.218 | 0.05 | 6.08966609236551e-29 | 12 |
| prex1 | 2.66763934565395e-33 | 0.262609155 | 0.145 | 0.025 | 6.71418146907642e-29 | 12 |
| rpl22l13 | 3.03946316651004e-33 | 0.549880137 | 0.731 | 0.385 | 7.65002484378912e-29 | 12 |
| si:ch211-150o23.2 | 3.33128664949118e-33 | 0.386245401 | 0.262 | 0.068 | 8.38451536810436e-29 | 12 |
| cotl13 | 6.27732583499105e-33 | 0.552250481 | 0.68 | 0.299 | 1.5799401394089e-28 | 12 |
| csk1 | 7.91510701495772e-33 | 0.416463143 | 0.324 | 0.098 | 1.99215328459471e-28 | 12 |
| serbp1a3 | 8.26736268401133e-33 | 0.51731496 | 0.833 | 0.475 | 2.08081251393881e-28 | 12 |
| n4bp11 | 2.38077146004593e-32 | 0.303280969 | 0.196 | 0.043 | 5.9921636877896e-28 | 12 |
| tmsb4x5 | 2.88158981049078e-32 | 0.53680068 | 0.996 | 0.68 | 7.25267339402424e-28 | 12 |
| XLOC-0254231 | 3.132977194687e-32 | 0.573043034 | 0.444 | 0.164 | 7.8853903013077e-28 | 12 |
| si:dkey-25e12.3 | 4.74226975143505e-32 | 0.370581611 | 0.207 | 0.049 | 1.19358187373869e-27 | 12 |
| si:ch73-234b20.51 | 4.84921081992744e-32 | 0.397452156 | 0.316 | 0.097 | 1.22049787126754e-27 | 12 |
| si:dkey-187e18.11 | 1.64959162376192e-31 | 0.37815057 | 0.247 | 0.066 | 4.15185715784637e-27 | 12 |
| arl5c2 | 3.05914605959787e-31 | 0.448607929 | 0.335 | 0.109 | 7.69956471740187e-27 | 12 |
| si:cabz01074946.1 | 3.80064820044824e-31 | 0.315323186 | 0.229 | 0.056 | 9.56585145570817e-27 | 12 |
| batf3 | 3.8228800639998e-31 | 0.297488557 | 0.189 | 0.041 | 9.6218068330811e-27 | 12 |
| tiparp | 5.65599431709088e-31 | 0.320805437 | 0.189 | 0.042 | 1.4235572096686e-26 | 12 |
| si:ch211-121a2.21 | 7.71387346670716e-31 | 0.270526454 | 0.164 | 0.033 | 1.94150481283552e-26 | 12 |
| FERMT3 (1 of many)1 | 8.34962596848424e-31 | 0.277430662 | 0.182 | 0.039 | 2.1015173600078e-26 | 12 |
| crip13 | 1.0336616906781e-30 | 0.508757519 | 0.622 | 0.264 | 2.60162310926772e-26 | 12 |
| pgam1a1 | 1.14234459585114e-30 | 0.457297827 | 0.415 | 0.152 | 2.87516711329774e-26 | 12 |
| CR383676.15 | 2.11824174612377e-30 | 0.630810744 | 0.993 | 0.721 | 5.33140265081893e-26 | 12 |
| ddx212 | 4.21168105818609e-30 | 0.479826674 | 0.487 | 0.202 | 1.06003800553486e-25 | 12 |
| sept6 | 1.09792610853206e-29 | 0.44082648 | 0.502 | 0.2 | 2.76337022256434e-25 | 12 |
| mfng1 | 1.16617006668301e-29 | 0.281425365 | 0.182 | 0.04 | 2.93513344083447e-25 | 12 |
| elf11 | 2.36190781545242e-29 | 0.400777618 | 0.313 | 0.1 | 5.94468578071219e-25 | 12 |

| | | | | | | |
|---|---|---|---|---|---|---|
| kmt2bb | 2.90587206635627e-29 | 0.381232118 | 0.273 | 0.083 | 7.31378940381211e-25 | 12 |
| XLOC-0247501 | 3.4333462616172e-29 | 0.366247791 | 0.255 | 0.073 | 8.64138920586432e-25 | 12 |
| aldocb | 7.90087418352468e-29 | 0.356220549 | 0.236 | 0.064 | 1.98857102325133e-24 | 12 |
| zgc:1749451 | 8.96396602226313e-29 | 0.252651799 | 0.135 | 0.024 | 2.25614060814341e-24 | 12 |
| FP101882.11 | 9.91616551925383e-29 | 0.277016076 | 0.16 | 0.033 | 2.495799699541e-24 | 12 |
| tmc81 | 2.24320576969542e-28 | 0.258528611 | 0.16 | 0.033 | 5.64592460174639e-24 | 12 |
| fli1b1 | 4.2497819746171e-28 | 0.279422781 | 0.156 | 0.032 | 1.06962762519138e-23 | 12 |
| myh11a2 | 4.88116441603941e-28 | 0.304767291 | 0.236 | 0.063 | 1.22854027187296e-23 | 12 |
| cdkn1bb1 | 4.95094297136493e-28 | 0.44497699 | 0.382 | 0.144 | 1.24610283646284e-23 | 12 |
| hmgb1a3 | 5.52493370363776e-28 | 0.446102763 | 0.451 | 0.184 | 1.39057056386859e-23 | 12 |
| rpl174 | 6.17991858419075e-28 | 0.339466255 | 0.956 | 0.742 | 1.55542370845497e-23 | 12 |
| cd74b3 | 7.27770567791013e-28 | 0.424923023 | 0.549 | 0.218 | 1.8317257420732e-23 | 12 |
| si:dkey-118j18.21 | 9.15282772751412e-28 | 0.298752862 | 0.204 | 0.051 | 2.30367521073803e-23 | 12 |
| psme11 | 2.53355680852631e-27 | 0.349759971 | 0.276 | 0.087 | 6.37670913137987e-23 | 12 |
| ccl36.11 | 2.65140968673139e-27 | 0.611942014 | 0.305 | 0.099 | 6.67333304053423e-23 | 12 |
| rbp52 | 2.90692261478786e-27 | 0.375151791 | 0.291 | 0.091 | 7.31643352915956e-23 | 12 |
| cd635 | 3.42144098093071e-27 | 0.482667023 | 0.68 | 0.334 | 8.6114248049045e-23 | 12 |
| CABZ01084575.1 | 3.56885620103785e-27 | 0.291679566 | 0.178 | 0.041 | 8.98245417239216e-23 | 12 |
| ier2b3 | 3.91706991065844e-27 | 0.463796697 | 0.509 | 0.219 | 9.85887325813623e-23 | 12 |
| arg2 | 9.43074917563919e-27 | 0.251630988 | 0.116 | 0.02 | 2.37362526001663e-22 | 12 |
| rpl5b3 | 1.00124760231728e-26 | 0.407514358 | 0.902 | 0.625 | 2.52004009027237e-22 | 12 |
| il15l | 1.31800358510916e-26 | 0.297658258 | 0.142 | 0.028 | 3.31728322336126e-22 | 12 |
| pabpc1a2 | 3.55362463211289e-26 | 0.447876727 | 0.865 | 0.534 | 8.94411783656492e-22 | 12 |
| hmgn31 | 1.71074535259763e-25 | 0.355080777 | 0.211 | 0.059 | 4.30577497795299e-21 | 12 |
| nop533 | 1.9610996635092e-25 | 0.436719872 | 0.538 | 0.246 | 4.93589174308631e-21 | 12 |
| serpinb1l1 | 2.08606459414578e-25 | 0.265431123 | 0.182 | 0.045 | 5.25041597700552e-21 | 12 |
| tpi1b1 | 2.52051472970423e-25 | 0.39675347 | 0.407 | 0.163 | 6.34388352319257e-21 | 12 |
| eif3m3 | 3.79152594058929e-25 | 0.458976258 | 0.687 | 0.352 | 9.54289163986919e-21 | 12 |
| gapdhs4 | 1.98441624213438e-24 | 0.448922976 | 0.589 | 0.283 | 4.99457723982801e-20 | 12 |
| arpc22 | 3.59139053883304e-24 | 0.433517515 | 0.56 | 0.262 | 9.03917084718889e-20 | 12 |
| unc119b1 | 4.96285048578835e-24 | 0.253544192 | 0.167 | 0.04 | 1.24909983876807e-19 | 12 |
| zgc:1629441 | 5.88309659593655e-24 | 0.436618906 | 0.527 | 0.248 | 1.48071658223127e-19 | 12 |
| dub1 | 7.57782104445495e-24 | 0.323052118 | 0.229 | 0.069 | 1.90726177867887e-19 | 12 |
| pik3r11 | 1.38564127352639e-23 | 0.356894746 | 0.262 | 0.087 | 3.48752052133857e-19 | 12 |
| rpl143 | 2.15888179805199e-23 | 0.393142249 | 0.92 | 0.629 | 5.43368959751705e-19 | 12 |

| | | | | | | |
|---|---|---|---|---|---|---|
| ccnd2a | 2.37308187004764e-23 | 0.362345443 | 0.255 | 0.084 | 5.97280975872291e-19 | 12 |
| rpl43 | 2.87289253955915e-23 | 0.387017105 | 0.898 | 0.622 | 7.23078323281643e-19 | 12 |
| fam49ba1 | 3.52145799132026e-23 | 0.32783246 | 0.295 | 0.1 | 8.86315761835397e-19 | 12 |
| ptmab3 | 3.66173857933665e-23 | 0.446872042 | 0.822 | 0.535 | 9.2162298303324e-19 | 12 |
| eno1a2 | 4.40375078319576e-23 | 0.351081898 | 0.345 | 0.131 | 1.10838003462254e-18 | 12 |
| eif3ea3 | 4.96100355327813e-23 | 0.412425241 | 0.567 | 0.272 | 1.24863498432457e-18 | 12 |
| si:dkey-112e7.22 | 1.42777310386869e-22 | 0.392877338 | 0.455 | 0.202 | 3.5935621251271e-18 | 12 |
| grap2b1 | 1.49858062375781e-22 | 0.358147149 | 0.305 | 0.11 | 3.77177757193603e-18 | 12 |
| tnrc6c13 | 1.8635813630839e-22 | 0.378532406 | 0.331 | 0.13 | 4.69044793274586e-18 | 12 |
| zgc:1627304 | 2.01046058848499e-22 | 0.467119605 | 0.76 | 0.385 | 5.06012825515787e-18 | 12 |
| sumo2b3 | 2.39523785289798e-22 | 0.42618052 | 0.535 | 0.265 | 6.02857415195892e-18 | 12 |
| arpc5a2 | 2.5065029354359e-22 | 0.359199955 | 0.302 | 0.112 | 6.30861723819861e-18 | 12 |
| BX005183.31 | 2.62173784617688e-22 | 0.342210986 | 0.244 | 0.078 | 6.59865198504259e-18 | 12 |
| slc25a33 | 4.33761798047395e-22 | 0.323025568 | 0.28 | 0.096 | 1.09173506950549e-17 | 12 |
| dusp1 | 5.19420692539405e-22 | 0.391977136 | 0.382 | 0.156 | 1.30732994105243e-17 | 12 |
| elmo11 | 7.7050386300686e-22 | 0.293231931 | 0.222 | 0.068 | 1.93928117280196e-17 | 12 |
| ponzr12 | 8.28050582815437e-22 | 0.426246864 | 0.56 | 0.268 | 2.08412051188817e-17 | 12 |
| si:rp71-68n21.91 | 9.43014081120378e-22 | 0.259857415 | 0.164 | 0.042 | 2.37347214077188e-17 | 12 |
| si:dkey-172h23.2 | 1.25585281977835e-21 | 0.27149611 | 0.156 | 0.039 | 3.16085596210014e-17 | 12 |
| si:dkey-112a7.4 | 1.47332620036851e-21 | 0.329800099 | 0.316 | 0.118 | 3.70821471370749e-17 | 12 |
| b2ml4 | 1.54864432592637e-21 | 0.410225813 | 0.538 | 0.262 | 3.89778290392409e-17 | 12 |
| rgs5b1 | 1.57805019336058e-21 | 0.345692557 | 0.233 | 0.074 | 3.97179453166923e-17 | 12 |
| bcl2l101 | 1.58082487582992e-21 | 0.442065943 | 0.465 | 0.21 | 3.97877812997633e-17 | 12 |
| nfkbiab1 | 1.82854050576896e-21 | 0.401492183 | 0.407 | 0.178 | 4.6022535989699e-17 | 12 |
| nfkbiaa1 | 2.90372332705255e-21 | 0.445195915 | 0.356 | 0.149 | 7.30838124185857e-17 | 12 |
| clic11 | 3.55820960830791e-21 | 0.319598179 | 0.298 | 0.111 | 8.95565776315017e-17 | 12 |
| mbpb | 4.27683484674153e-21 | 0.259421975 | 0.196 | 0.058 | 1.07643656257638e-16 | 12 |
| psme21 | 6.82021293909291e-21 | 0.312741609 | 0.265 | 0.094 | 1.7165793946403e-16 | 12 |
| eif5a4 | 1.0453564544473e-20 | 0.42432027 | 0.815 | 0.472 | 2.6310576601984e-16 | 12 |
| eif3g3 | 1.33866997657131e-20 | 0.396766811 | 0.553 | 0.277 | 3.36929846403233e-16 | 12 |
| arpc32 | 1.69583542958178e-20 | 0.407314169 | 0.607 | 0.305 | 4.26824819271439e-16 | 12 |
| cap13 | 1.98496990536278e-20 | 0.414609489 | 0.48 | 0.229 | 4.99597075480759e-16 | 12 |
| prkar1ab | 2.12606712434993e-20 | 0.288732324 | 0.215 | 0.069 | 5.35109834527635e-16 | 12 |
| birc21 | 3.87319095523196e-20 | 0.333222866 | 0.28 | 0.104 | 9.74843431522332e-16 | 12 |
| romo13 | 5.04994374555535e-20 | 0.414438144 | 0.705 | 0.38 | 1.27102034131883e-15 | 12 |

| | | | | | | |
|---|---|---|---|---|---|---|
| ccl32b.31 | 5.36753460427298e-20 | 0.268525018 | 0.131 | 0.031 | 1.35095478454947e-15 | 12 |
| oaz1b3 | 1.08664003276902e-19 | 0.406299774 | 0.596 | 0.318 | 2.73496429847636e-15 | 12 |
| cdc42l1 | 1.18868496483016e-19 | 0.362804019 | 0.455 | 0.212 | 2.99180118798102e-15 | 12 |
| higd1a3 | 2.11274507953031e-19 | 0.415757963 | 0.571 | 0.305 | 5.31756809066983e-15 | 12 |
| eif3ba2 | 2.83660042614228e-19 | 0.366091289 | 0.436 | 0.206 | 7.1394396125575e-15 | 12 |
| cfl11 | 3.29775685503934e-19 | 0.397050092 | 0.855 | 0.572 | 8.30012422844851e-15 | 12 |
| mibp24 | 3.29793076513317e-19 | 0.418353738 | 0.825 | 0.486 | 8.30056194276367e-15 | 12 |
| XLOC-0234922 | 3.37635628887816e-19 | 0.336559563 | 0.258 | 0.093 | 8.49795114347743e-15 | 12 |
| zgc:1141884 | 4.40210720179173e-19 | 0.411962483 | 0.945 | 0.621 | 1.10796636161896e-14 | 12 |
| hcls1 | 6.49764995141173e-19 | 0.271164222 | 0.211 | 0.069 | 1.63539351627082e-14 | 12 |
| XLOC-0054491 | 7.26948569565427e-19 | 0.255273103 | 0.178 | 0.052 | 1.82965685473922e-14 | 12 |
| prelid3b | 7.5423728854705e-19 | 0.372941155 | 0.48 | 0.234 | 1.89833983154407e-14 | 12 |
| tsc22d3 | 1.63420155534902e-18 | 0.376094567 | 0.327 | 0.141 | 4.11312189465796e-14 | 12 |
| si:busm1-266f07.21 | 3.09122886256011e-18 | 0.30720864 | 0.356 | 0.143 | 7.78031392417755e-14 | 12 |
| kdm6bb | 7.2754795366055e-18 | 0.263508293 | 0.167 | 0.05 | 1.83116544456824e-13 | 12 |
| dok21 | 8.71798170757934e-18 | 0.250800045 | 0.138 | 0.037 | 2.19422881598064e-13 | 12 |
| rhoab2 | 9.8944206825241e-18 | 0.356052696 | 0.396 | 0.186 | 2.49032674158449e-13 | 12 |
| capza1a1 | 1.34746107286481e-17 | 0.342474406 | 0.415 | 0.196 | 3.39142477429345e-13 | 12 |
| ccl34b.41 | 1.68859072060164e-17 | 0.312546137 | 0.138 | 0.037 | 4.25001398468228e-13 | 12 |
| midn2 | 5.62571746664065e-17 | 0.32611044 | 0.32 | 0.138 | 1.41593682917879e-12 | 12 |
| serf23 | 2.30436655103426e-16 | 0.362279896 | 0.615 | 0.338 | 5.79986017229812e-12 | 12 |
| lasp11 | 3.87445105582481e-16 | 0.342612297 | 0.458 | 0.233 | 9.75160586240546e-12 | 12 |
| gng51 | 7.16999439205471e-16 | 0.342192579 | 0.509 | 0.268 | 1.80461588853625e-11 | 12 |
| sesn14 | 7.41200060813412e-16 | 0.358752681 | 0.353 | 0.165 | 1.86552643306128e-11 | 12 |
| rnf213a2 | 8.06637436592431e-16 | 0.296780608 | 0.225 | 0.087 | 2.03022576415949e-11 | 12 |
| pycard2 | 9.67940790827765e-16 | 0.253250628 | 0.2 | 0.071 | 2.4362101764344e-11 | 12 |
| si:dkey-11f4.14 | 1.47321622649259e-15 | 0.315877271 | 0.258 | 0.107 | 3.70793792045921e-11 | 12 |
| histh1l1 | 1.49800293794292e-15 | 0.327248714 | 0.262 | 0.108 | 3.77032359450853e-11 | 12 |
| rps27.23 | 3.77226893519144e-15 | 0.379190939 | 0.822 | 0.512 | 9.49442368298334e-11 | 12 |
| rhogb1 | 9.81737566201617e-15 | 0.256435586 | 0.24 | 0.095 | 2.47093528037285e-10 | 12 |
| arf6b | 1.24438356442371e-14 | 0.262796281 | 0.244 | 0.099 | 3.13198899329803e-10 | 12 |
| NC-002333.44 | 1.42351259962265e-14 | 0.358198393 | 0.927 | 0.617 | 3.58283886199024e-10 | 12 |
| sh3bgrl1 | 2.80542865871915e-14 | 0.262484398 | 0.247 | 0.103 | 7.06098339113024e-10 | 12 |
| egr3 | 3.24291521554889e-14 | 0.255060288 | 0.167 | 0.057 | 8.16209330601501e-10 | 12 |
| anxa11a | 5.4410708502245e-14 | 0.260126745 | 0.247 | 0.104 | 1.369463122293e-09 | 12 |

| | | | | | | |
|---|---|---|---|---|---|---|
| fosl22 | 5.75224862422593e-14 | 0.309293056 | 0.269 | 0.12 | 1.44778345623142e-09 | 12 |
| ankrd11 | 9.65747781687124e-14 | 0.2656908 | 0.218 | 0.089 | 2.43069059172832e-09 | 12 |
| slc25a3b2 | 1.18824846473352e-13 | 0.346547004 | 0.716 | 0.432 | 2.99070256088779e-09 | 12 |
| itm2bb1 | 1.54647419669939e-13 | 0.294818961 | 0.429 | 0.219 | 3.89232090567269e-09 | 12 |
| cct82 | 1.97365550877201e-13 | 0.281030976 | 0.393 | 0.199 | 4.96749355002826e-09 | 12 |
| chac1 | 2.8104109441695e-13 | 0.344724516 | 0.651 | 0.394 | 7.07352330538023e-09 | 12 |
| atf4b2 | 4.34406315863215e-13 | 0.319738281 | 0.473 | 0.261 | 1.09335725639613e-08 | 12 |
| csrnp1b2 | 4.54388057777701e-13 | 0.307316773 | 0.353 | 0.177 | 1.1436493026207e-08 | 12 |
| ddit33 | 9.52683487806203e-13 | 0.315046179 | 0.345 | 0.174 | 2.39780907045943e-08 | 12 |
| eif3i3 | 1.15275191474939e-12 | 0.297438848 | 0.487 | 0.27 | 2.90136129423274e-08 | 12 |
| eif3s102 | 1.4223756679139e-12 | 0.270063198 | 0.327 | 0.163 | 3.5799773185725e-08 | 12 |
| GCA1 | 1.52075326110276e-12 | 0.290006566 | 0.349 | 0.179 | 3.82758388286955e-08 | 12 |
| idh21 | 1.58671055456855e-12 | 0.278291311 | 0.382 | 0.197 | 3.99359179479359e-08 | 12 |
| XLOC-0283701 | 2.88124238515321e-12 | 0.313251329 | 0.295 | 0.139 | 7.25179895919212e-08 | 12 |
| gnb1a | 3.51414778812145e-12 | 0.293780376 | 0.36 | 0.19 | 8.84475856792288e-08 | 12 |
| cnbpb2 | 7.28770162240942e-12 | 0.270967339 | 0.349 | 0.182 | 1.83424162134423e-07 | 12 |
| notch22 | 7.91536316251258e-12 | 0.25868549 | 0.255 | 0.118 | 1.99221775437279e-07 | 12 |
| nop561 | 9.31880398833918e-12 | 0.255906479 | 0.335 | 0.169 | 2.34544977582509e-07 | 12 |
| tob1a | 1.23165305513799e-11 | 0.262211542 | 0.269 | 0.128 | 3.0999475744768e-07 | 12 |
| si:ch211-222l21.13 | 1.55756039097301e-11 | 0.340578184 | 0.465 | 0.273 | 3.92022374803996e-07 | 12 |
| eif4ebp21 | 2.3675000261142e-11 | 0.259655886 | 0.335 | 0.172 | 5.95876081572683e-07 | 12 |
| ppt1 | 4.97430827996472e-11 | 0.281344192 | 0.313 | 0.162 | 1.25198365098432e-06 | 12 |
| arpc4l2 | 5.73081253002057e-11 | 0.253620492 | 0.367 | 0.195 | 1.44238820568088e-06 | 12 |
| eif3c3 | 7.45224131910577e-11 | 0.266079979 | 0.316 | 0.167 | 1.87565461760573e-06 | 12 |
| eef2b2 | 1.51856724099568e-10 | 0.308015734 | 0.873 | 0.609 | 3.82208188886203e-06 | 12 |
| hnrnpa0b2 | 1.90509195642801e-10 | 0.290296449 | 0.513 | 0.309 | 4.79492594513366e-06 | 12 |
| arhgdia1 | 1.95406670526873e-10 | 0.268154984 | 0.404 | 0.227 | 4.91819049049087e-06 | 12 |
| srsf5a2 | 2.20667799547185e-10 | 0.276815941 | 0.48 | 0.286 | 5.5539878468031e-06 | 12 |
| eif3d2 | 2.6403875098007e-10 | 0.254583268 | 0.44 | 0.251 | 6.64559132341738e-06 | 12 |
| ywhaba | 3.2894725095301e-10 | 0.266077935 | 0.538 | 0.318 | 8.27927335923632e-06 | 12 |
| arpc41 | 4.44876043533487e-10 | 0.256134009 | 0.404 | 0.224 | 1.11970851396943e-05 | 12 |
| ier2a3 | 4.65423720454709e-10 | 0.256737225 | 0.462 | 0.26 | 1.17142496201246e-05 | 12 |
| hmgn63 | 1.349762583141e-09 | 0.278228912 | 0.44 | 0.263 | 3.39721744550757e-05 | 12 |
| zgc:564932 | 1.26821449875957e-08 | 0.280593842 | 0.815 | 0.568 | 0.000319197 | 12 |
| txnipa3 | 3.09269773523682e-08 | 0.255442451 | 0.353 | 0.209 | 0.000778401 | 12 |

| | | | | | | |
|---|---|---|---|---|---|---|
| ran1 | 1.1519265191104e-07 | 0.252238568 | 0.862 | 0.64 | 0.002899284 | 12 |
| CR318588.14 | 1.2273668481105e-06 | 0.381707134 | 0.316 | 0.207 | 0.030891596 | 12 |
| mt-co25 | 3.82731786610884e-05 | 0.268761459 | 0.971 | 0.715 | 0.963297634 | 12 |
| mt-co15 | 5.85802125396645e-05 | 0.258447214 | 0.949 | 0.652 | 1 | 12 |
| COX35 | 6.82065684919815e-05 | 0.252999259 | 0.964 | 0.695 | 1 | 12 |
| si:ch211-226h8.145 | 9.9857660504243e-58 | 0.71097331 | 0.966 | 0.56 | 2.51331745723129e-53 | 13 |
| si:dkey-240e12.65 | 1.26047878139172e-56 | 0.71694376 | 0.977 | 0.562 | 3.17249904488482e-52 | 13 |
| CU571163.15 | 1.79026349337391e-56 | 0.780795951 | 0.837 | 0.394 | 4.5059141864728e-52 | 13 |
| h2af1al6 | 2.59251458192896e-56 | 0.695126861 | 0.981 | 0.586 | 6.525099951257e-52 | 13 |
| si:ch211-125e6.125 | 1.34277182790846e-55 | 0.732391049 | 0.92 | 0.494 | 3.37962241366282e-51 | 13 |
| cldnd6 | 1.64389395831417e-54 | 0.691605155 | 0.996 | 0.598 | 4.13751670368093e-50 | 13 |
| zp36 | 3.03407719412219e-54 | 0.691686747 | 0.981 | 0.578 | 7.63646888988614e-50 | 13 |
| si:dkey-241l7.25 | 6.94207011905367e-54 | 0.660550778 | 0.992 | 0.636 | 1.74724962826462e-49 | 13 |
| mt25 | 5.145412553384e-53 | 0.713636962 | 0.966 | 0.527 | 1.29504888556122e-48 | 13 |
| acp5a6 | 8.04507915128914e-53 | 0.691610848 | 0.928 | 0.534 | 2.02486597158796e-48 | 13 |
| zgc:1534995 | 5.04351529074522e-52 | 0.737441746 | 0.966 | 0.505 | 1.26940236352767e-47 | 13 |
| zgc:1751356 | 6.45666464999812e-52 | 0.721290839 | 0.89 | 0.473 | 1.62507792575803e-47 | 13 |
| si:dkey-241l7.56 | 9.96821137186109e-52 | 0.674006048 | 0.973 | 0.578 | 2.50889912018372e-47 | 13 |
| tmsb55 | 1.04259886535176e-51 | 0.721455124 | 0.913 | 0.482 | 2.62411708420384e-47 | 13 |
| zgc:1717795 | 1.28799164894225e-51 | 0.693772972 | 0.882 | 0.507 | 3.24174618122275e-47 | 13 |
| si:zfos-1505d6.35 | 3.2297292138609e-51 | 0.705457475 | 0.916 | 0.507 | 8.12890545836649e-47 | 13 |
| si:dkey-229d11.35 | 7.02504428771477e-51 | 0.749632453 | 0.886 | 0.431 | 1.76813339677493e-46 | 13 |
| crp25 | 3.46840113754453e-49 | 0.705615499 | 0.901 | 0.482 | 8.72961882308584e-45 | 13 |
| si:ch73-189n23.16 | 4.25298441454086e-49 | 0.696632054 | 0.932 | 0.518 | 1.07043364729579e-44 | 13 |
| zp2.16 | 1.41439782316099e-48 | 0.687682074 | 0.928 | 0.529 | 3.55989788111389e-44 | 13 |
| h1m6 | 2.46331048177028e-48 | 0.654821367 | 0.97 | 0.596 | 6.19990615156762e-44 | 13 |
| zp2.56 | 2.73921854517919e-48 | 0.671147021 | 0.943 | 0.558 | 6.89433915636149e-44 | 13 |
| zp2.35 | 6.52347219099537e-48 | 0.704243719 | 0.947 | 0.512 | 1.64189271575162e-43 | 13 |
| zp2.25 | 1.74102451429971e-47 | 0.688295284 | 0.947 | 0.523 | 4.38198460004093e-43 | 13 |
| cldng5 | 2.07264549691358e-46 | 0.692545132 | 0.935 | 0.507 | 5.2166414511818e-42 | 13 |
| si:dkeyp-98a7.95 | 1.78748698992506e-45 | 0.684204391 | 0.92 | 0.506 | 4.49892600494239e-41 | 13 |
| si:dkey-241l7.45 | 1.97730814959045e-45 | 0.617859501 | 0.947 | 0.603 | 4.97668688170421e-41 | 13 |
| BX649337.16 | 9.6658528578689e-45 | 0.698159489 | 0.852 | 0.431 | 2.43279850579702e-40 | 13 |
| zgc:1717765 | 2.28536634397316e-44 | 0.674572013 | 0.882 | 0.489 | 5.75203855114604e-40 | 13 |
| si:ch211-226h8.45 | 2.81379859161345e-44 | 0.694580689 | 0.84 | 0.435 | 7.08204967523188e-40 | 13 |

| | | | | | | |
|---|---|---|---|---|---|---|
| si:dkeyp-46h3.65 | 5.44579375720213e-44 | 0.699272962 | 0.787 | 0.405 | 1.37065183075021e-39 | 13 |
| zp3.26 | 5.93668015177823e-44 | 0.655704407 | 0.947 | 0.551 | 1.49420302740106e-39 | 13 |
| slbp25 | 2.39310501209989e-43 | 0.686773171 | 0.764 | 0.391 | 6.02320600495422e-39 | 13 |
| lamp25 | 4.23706196879622e-43 | 0.631744192 | 0.897 | 0.549 | 1.06642612692632e-38 | 13 |
| rrm25 | 8.37866960986371e-43 | 0.652850677 | 0.886 | 0.492 | 2.1088273541066e-38 | 13 |
| si:dkey-90l23.15 | 1.26989499870387e-42 | 0.67612582 | 0.821 | 0.442 | 3.19619872223776e-38 | 13 |
| CR790368.16 | 1.724454179466e-42 | 0.639308667 | 0.878 | 0.515 | 4.34027872429798e-38 | 13 |
| si:ch211-12h2.85 | 7.43146888164369e-42 | 0.626740408 | 0.894 | 0.542 | 1.8704264028209e-37 | 13 |
| zp2.65 | 3.80640007791532e-41 | 0.660413838 | 0.886 | 0.496 | 9.58032835610507e-37 | 13 |
| qdprb25 | 7.08207131215801e-41 | 0.67126269 | 0.81 | 0.427 | 1.78248652855705e-36 | 13 |
| si:dkeyp-20e4.85 | 7.53054484776127e-41 | 0.688359302 | 0.776 | 0.402 | 1.89536283273303e-36 | 13 |
| si:dkeyp-98a7.85 | 2.53403524180243e-40 | 0.664295336 | 0.84 | 0.452 | 6.37791330009253e-36 | 13 |
| ccna15 | 3.1062566627887e-40 | 0.653288627 | 0.859 | 0.47 | 7.81813739457287e-36 | 13 |
| si:ch211-12h2.65 | 4.83321270995516e-39 | 0.636497948 | 0.817 | 0.469 | 1.21647130696861e-34 | 13 |
| zp3a.25 | 5.69700396642752e-39 | 0.630941898 | 0.837 | 0.486 | 1.43387892831014e-34 | 13 |
| LOC1104378215 | 7.8115336876879e-39 | 0.674433474 | 0.791 | 0.41 | 1.96608491385417e-34 | 13 |
| siva15 | 8.63102222740015e-39 | 0.650946421 | 0.787 | 0.423 | 2.17234198441434e-34 | 13 |
| zgc:1715175 | 9.39232485754221e-39 | 0.665834784 | 0.715 | 0.383 | 2.3639542433948e-34 | 13 |
| zar1l5 | 2.35523899251843e-37 | 0.650721392 | 0.745 | 0.393 | 5.92790102026964e-33 | 13 |
| mid1ip1l5 | 1.04815399798305e-36 | 0.582329449 | 0.844 | 0.539 | 2.63809879752354e-32 | 13 |
| epcam6 | 6.15107238739796e-35 | 0.604592398 | 0.844 | 0.5 | 1.54816340918419e-30 | 13 |
| zp3b5 | 2.35993133310643e-34 | 0.636837498 | 0.722 | 0.383 | 5.93971117229557e-30 | 13 |
| si:ch211-14a17.75 | 2.97903667062428e-34 | 0.628533898 | 0.741 | 0.403 | 7.49793739629424e-30 | 13 |
| uchl15 | 3.68488675020685e-34 | 0.641529028 | 0.669 | 0.34 | 9.27449146159562e-30 | 13 |
| gtf3ab6 | 6.78859517281702e-34 | 0.63598513 | 0.669 | 0.315 | 1.70862151904632e-29 | 13 |
| zgc:1952455 | 1.72940775173916e-33 | 0.623160561 | 0.715 | 0.384 | 4.35274637035229e-29 | 13 |
| wu:fi42e035 | 4.72395480446766e-33 | 0.634734946 | 0.677 | 0.345 | 1.18897218473647e-28 | 13 |
| si:dkeyp-46h3.85 | 8.15077522712064e-33 | 0.63587841 | 0.684 | 0.347 | 2.05146861691399e-28 | 13 |
| zgc:771185 | 1.0479926594422e-32 | 0.601878544 | 0.791 | 0.452 | 2.63769272455007e-28 | 13 |
| si:dkeyp-50f7.25 | 1.09559232371121e-31 | 0.608403171 | 0.726 | 0.398 | 2.75749631954873e-27 | 13 |
| thy15 | 4.01382731978975e-31 | 0.588406096 | 0.768 | 0.446 | 1.01024019811788e-26 | 13 |
| si:dkey-229d11.55 | 6.28764892463414e-30 | 0.599064631 | 0.7 | 0.385 | 1.58253835784117e-25 | 13 |
| AL929032.15 | 6.35620161787537e-30 | 0.621755932 | 0.635 | 0.313 | 1.59979238520305e-25 | 13 |
| si:dkey-241l7.65 | 2.31781721681337e-29 | 0.551379275 | 0.821 | 0.523 | 5.83371415299758e-25 | 13 |
| btg45 | 5.57563077839531e-29 | 0.596131159 | 0.684 | 0.364 | 1.40333051061432e-24 | 13 |

| | | | | | | |
|---|---|---|---|---|---|---|
| s100a15 | 1.0160514828761e-28 | 0.589157344 | 0.681 | 0.359 | 2.55729997725086e-24 | 13 |
| ccnb25 | 1.83782489807556e-28 | 0.585854429 | 0.696 | 0.38 | 4.62562148596638e-24 | 13 |
| si:dkeyp-98a7.45 | 2.46538543388363e-28 | 0.57433584 | 0.734 | 0.421 | 6.20512859854172e-24 | 13 |
| tuba4l5 | 3.36692662105754e-28 | 0.573693479 | 0.7 | 0.401 | 8.47421761253972e-24 | 13 |
| hbba15 | 4.99415736667582e-28 | 0.595400115 | 0.696 | 0.351 | 1.25697946761864e-23 | 13 |
| zgc:554615 | 7.35886128699392e-28 | 0.589971662 | 0.582 | 0.284 | 1.8521517973235e-23 | 13 |
| npm2b5 | 7.50727271795355e-28 | 0.577379971 | 0.7 | 0.392 | 1.88950547038173e-23 | 13 |
| ccnb15 | 7.75491899278221e-28 | 0.576607857 | 0.677 | 0.39 | 1.95183556129335e-23 | 13 |
| si:dkey-152b24.65 | 1.41707613396718e-27 | 0.591271683 | 0.593 | 0.294 | 3.566638921582e-23 | 13 |
| ctss2.15 | 5.61429835526583e-27 | 0.507625759 | 0.798 | 0.531 | 1.41306275303686e-22 | 13 |
| si:ch211-132b12.85 | 9.87500238770717e-27 | 0.577279333 | 0.639 | 0.338 | 2.48543935096202e-22 | 13 |
| zar15 | 1.19158797718552e-26 | 0.556403288 | 0.734 | 0.436 | 2.99910777977824e-22 | 13 |
| bambia | 1.74944237879624e-26 | 0.3165726 | 0.114 | 0.019 | 4.40317152319226e-22 | 13 |
| nanog5 | 2.77095676889653e-26 | 0.570773843 | 0.662 | 0.355 | 6.97422109163568e-22 | 13 |
| zp3a.15 | 6.27451356776689e-26 | 0.563392482 | 0.654 | 0.378 | 1.57923231987125e-21 | 13 |
| zgc:562315 | 1.18255364021781e-25 | 0.570655129 | 0.487 | 0.224 | 2.9763692570642e-21 | 13 |
| zgc:1734435 | 1.28336384311343e-25 | 0.56804685 | 0.616 | 0.326 | 3.2300984567322e-21 | 13 |
| avd5 | 1.82459318659975e-25 | 0.572373525 | 0.536 | 0.262 | 4.59231859135292e-21 | 13 |
| zpcx5 | 4.08011886820396e-25 | 0.563614968 | 0.624 | 0.335 | 1.02692511793825e-20 | 13 |
| cth15 | 1.09363692630815e-24 | 0.549459739 | 0.662 | 0.376 | 2.75257477982498e-20 | 13 |
| LOC1005374995 | 1.51609175261554e-24 | 0.562058564 | 0.605 | 0.318 | 3.81585133215804e-20 | 13 |
| LOC1005351625 | 1.94265989197977e-24 | 0.566627169 | 0.586 | 0.305 | 4.88948068212389e-20 | 13 |
| zglp15 | 9.16975633533387e-24 | 0.548501141 | 0.433 | 0.195 | 2.30793597204018e-19 | 13 |
| si:ch1073-263o8.25 | 1.33088010164807e-23 | 0.563226603 | 0.51 | 0.261 | 3.34969212783802e-19 | 13 |
| si:ch73-14h10.25 | 1.61698430073705e-23 | 0.555495015 | 0.525 | 0.268 | 4.06978778652507e-19 | 13 |
| si:ch211-5k11.85 | 2.69807713516421e-23 | 0.471154915 | 0.681 | 0.354 | 6.7907903414948e-19 | 13 |
| si:ch211-226l4.65 | 3.51680703554352e-23 | 0.548337567 | 0.555 | 0.289 | 8.85145162775949e-19 | 13 |
| cpeb1b5 | 5.2207485496837e-23 | 0.545575039 | 0.532 | 0.267 | 1.31401020246989e-18 | 13 |
| si:dkeyp-46h3.25 | 6.84610064096484e-23 | 0.525361584 | 0.65 | 0.402 | 1.72309507032444e-18 | 13 |
| zgc:1362545 | 2.6952567553037e-22 | 0.540463993 | 0.601 | 0.327 | 6.78369172742388e-18 | 13 |
| zgc:1714745 | 2.91327961495772e-22 | 0.538679787 | 0.555 | 0.301 | 7.33243346288707e-18 | 13 |
| LOC1018854775 | 3.12931369581431e-22 | 0.537697553 | 0.548 | 0.276 | 7.87616964099503e-18 | 13 |
| birc5b5 | 7.67582837713579e-22 | 0.531146416 | 0.51 | 0.256 | 1.93192924424131e-17 | 13 |
| si:dkeyp-46h3.55 | 2.52573415043809e-21 | 0.529377886 | 0.536 | 0.287 | 6.35702028323762e-17 | 13 |
| arhgef10la5 | 3.53050031763582e-21 | 0.530875764 | 0.521 | 0.272 | 8.8859162494576e-17 | 13 |

| | | | | | | |
|---|---|---|---|---|---|---|
| si:ch211-146l10.85 | 9.85108102032652e-21 | 0.529900526 | 0.555 | 0.304 | 2.47941858200598e-16 | 13 |
| ttll12 | 1.01309623970008e-20 | 0.292418421 | 0.122 | 0.026 | 2.54986192570113e-16 | 13 |
| CU467646.24 | 1.94792135611082e-20 | 0.48300199 | 0.312 | 0.124 | 4.90272326119533e-16 | 13 |
| si:ch211-145c1.15 | 5.74421205687358e-20 | 0.518292409 | 0.468 | 0.231 | 1.44576073259451e-15 | 13 |
| elovl7b5 | 8.7312176265984e-20 | 0.511683227 | 0.479 | 0.245 | 2.19756016443855e-15 | 13 |
| gapdh5 | 1.12718886502531e-19 | 0.488964138 | 0.669 | 0.417 | 2.8370216543822e-15 | 13 |
| hnrnpabb5 | 1.28812881681476e-19 | 0.404607012 | 0.768 | 0.595 | 3.24209141904106e-15 | 13 |
| kpna75 | 1.89333804030489e-19 | 0.510687473 | 0.57 | 0.326 | 4.76534251364337e-15 | 13 |
| pigrl2.35 | 2.97170144110013e-19 | 0.498117559 | 0.422 | 0.201 | 7.47947535710492e-15 | 13 |
| pabpn1l5 | 3.17780642643884e-19 | 0.50643443 | 0.513 | 0.281 | 7.99822099470391e-15 | 13 |
| LOC1005368605 | 4.16850599390348e-19 | 0.490490216 | 0.578 | 0.348 | 1.04917127360557e-14 | 13 |
| si:dkey-90l23.25 | 1.01567477258007e-18 | 0.499717795 | 0.582 | 0.342 | 2.55635183510678e-14 | 13 |
| si:ch211-250e5.165 | 4.67168939937874e-18 | 0.48754979 | 0.567 | 0.344 | 1.17581750492964e-13 | 13 |
| AL929192.15 | 4.82809377023919e-18 | 0.474419684 | 0.361 | 0.162 | 1.2151829210315e-13 | 13 |
| si:dkey-19b23.115 | 8.72755117683781e-18 | 0.490262501 | 0.506 | 0.276 | 2.19663735569831e-13 | 13 |
| gdf35 | 9.64580765659834e-18 | 0.489749342 | 0.433 | 0.221 | 2.42775332908924e-13 | 13 |
| zgc:925915 | 2.30648659246012e-17 | 0.478692726 | 0.411 | 0.203 | 5.80519610456288e-13 | 13 |
| sybu5 | 4.88593662828907e-17 | 0.471004046 | 0.354 | 0.164 | 1.22974138997407e-12 | 13 |
| si:dkey-208k4.25 | 1.1472159626949e-16 | 0.470086192 | 0.513 | 0.294 | 2.8874278565068e-12 | 13 |
| hbaa15 | 1.28577377273982e-16 | 0.455233127 | 0.521 | 0.279 | 3.23616400860885e-12 | 13 |
| zgc:566995 | 1.97780263480769e-16 | 0.470515729 | 0.456 | 0.249 | 4.97793145154749e-12 | 13 |
| zgc:173545 | 2.71112227493966e-16 | 0.338060879 | 0.156 | 0.047 | 6.82362365379562e-12 | 13 |
| si:dkey-46g23.25 | 3.14338093579e-16 | 0.42869505 | 0.323 | 0.143 | 7.91157547728985e-12 | 13 |
| nasp5 | 4.03102275286403e-16 | 0.468424629 | 0.548 | 0.338 | 1.01456811666835e-11 | 13 |
| cdca7a5 | 4.42273149633006e-16 | 0.476480721 | 0.414 | 0.22 | 1.11315729031131e-11 | 13 |
| lsm14b5 | 5.14311032347886e-16 | 0.462902764 | 0.51 | 0.298 | 1.29446943731639e-11 | 13 |
| gmnn5 | 3.33436337269645e-15 | 0.457182845 | 0.513 | 0.301 | 8.39225917273969e-11 | 13 |
| chchd103 | 4.0340355707965e-15 | 0.45189796 | 0.316 | 0.15 | 1.01532641281377e-10 | 13 |
| snu13b5 | 8.55200311228962e-15 | 0.43106515 | 0.563 | 0.391 | 2.15245366333217e-10 | 13 |
| org5 | 1.08511418241063e-14 | 0.462952553 | 0.426 | 0.239 | 2.73112388570931e-10 | 13 |
| ccna25 | 1.23061523788603e-14 | 0.448173171 | 0.487 | 0.282 | 3.09733549223535e-10 | 13 |
| si:dkeyp-98a7.75 | 2.17189506479581e-14 | 0.43432058 | 0.38 | 0.194 | 5.46644268858458e-10 | 13 |
| si:ch211-146l10.75 | 2.28386196399443e-14 | 0.436363877 | 0.365 | 0.187 | 5.74825217717759e-10 | 13 |
| si:dkey-37o8.15 | 4.16608727635295e-14 | 0.439644074 | 0.51 | 0.305 | 1.04856250658528e-09 | 13 |
| zgc:1138865 | 4.34281831548107e-14 | 0.425512687 | 0.3 | 0.14 | 1.09304394182343e-09 | 13 |

| | | | | | | |
|---|---|---|---|---|---|---|
| imp33 | 4.78936445787928e-14 | 0.428814653 | 0.285 | 0.133 | 1.20543514040364e-09 | 13 |
| crp35 | 4.8507928572273e-14 | 0.439551517 | 0.399 | 0.217 | 1.22089605423554e-09 | 13 |
| si:ch73-90k17.15 | 5.18329402873276e-14 | 0.44149848 | 0.441 | 0.247 | 1.30458327409175e-09 | 13 |
| slbp6 | 5.80862131450586e-14 | 0.441720773 | 0.418 | 0.232 | 1.46197189864798e-09 | 13 |
| elavl25 | 7.00302351965785e-14 | 0.442042506 | 0.403 | 0.222 | 1.76259098966268e-09 | 13 |
| yrdc3 | 1.12044419782879e-13 | 0.406614986 | 0.27 | 0.122 | 2.82004600151528e-09 | 13 |
| ccne13 | 2.71033219662866e-13 | 0.387896043 | 0.251 | 0.11 | 6.82163510569468e-09 | 13 |
| cyp11a1 | 2.81858340105359e-13 | 0.336701386 | 0.183 | 0.067 | 7.09409256211178e-09 | 13 |
| ca15b5 | 2.83758611660301e-13 | 0.421958783 | 0.437 | 0.248 | 7.14192049687812e-09 | 13 |
| spink2.16 | 3.9891974734903e-13 | 0.393952975 | 0.639 | 0.435 | 1.00404111210277e-08 | 13 |
| slc16a35 | 5.4149583275851e-13 | 0.430221575 | 0.422 | 0.248 | 1.36289086146989e-08 | 13 |
| sinup5 | 6.00536019384064e-13 | 0.414265543 | 0.38 | 0.202 | 1.51148910718775e-08 | 13 |
| adat3 | 6.53970176345284e-13 | 0.293264753 | 0.133 | 0.042 | 1.64597753684345e-08 | 13 |
| kif23 | 7.91863228543656e-13 | 0.299397314 | 0.152 | 0.052 | 1.99304055992153e-08 | 13 |
| h3f3a5 | 1.36788693182551e-12 | 0.368112139 | 0.643 | 0.505 | 3.44283461871162e-08 | 13 |
| patl25 | 1.43294629612578e-12 | 0.408707697 | 0.331 | 0.172 | 3.60658253271898e-08 | 13 |
| larp6a5 | 1.45778200330766e-12 | 0.418007569 | 0.422 | 0.248 | 3.66909152412505e-08 | 13 |
| rcc15 | 2.06919852161696e-12 | 0.409330331 | 0.373 | 0.201 | 5.20796575905774e-08 | 13 |
| LOC1018838975 | 2.11351235392798e-12 | 0.391625761 | 0.323 | 0.163 | 5.31949924360132e-08 | 13 |
| pou5f35 | 2.57312522955723e-12 | 0.388810751 | 0.297 | 0.146 | 6.47629889027258e-08 | 13 |
| pabpc1l5 | 4.43056401293216e-12 | 0.380573136 | 0.289 | 0.141 | 1.1151286564149e-07 | 13 |
| trmt9b | 7.86020796391594e-12 | 0.260823715 | 0.125 | 0.04 | 1.978335742438e-07 | 13 |
| depdc7b1 | 1.4772170885699e-11 | 0.312754892 | 0.171 | 0.066 | 3.71800769022157e-07 | 13 |
| cax1 | 1.68821060690697e-11 | 0.300704887 | 0.156 | 0.058 | 4.24905727652415e-07 | 13 |
| si:ch1073-190k2.15 | 3.13623122280557e-11 | 0.381590571 | 0.346 | 0.187 | 7.89358036467935e-07 | 13 |
| si:ch211-69g19.24 | 3.40272370363633e-11 | 0.356450548 | 0.255 | 0.121 | 8.56431528968228e-07 | 13 |
| si:dkey-216e24.33 | 6.45778372902158e-11 | 0.372791522 | 0.251 | 0.122 | 1.62535958675744e-06 | 13 |
| cx44.25 | 9.23821382888587e-11 | 0.352493093 | 0.243 | 0.116 | 2.32516603859228e-06 | 13 |
| COQ10A1 | 1.02983653130214e-10 | 0.33470784 | 0.19 | 0.082 | 2.59199556563435e-06 | 13 |
| buc5 | 1.05385186135547e-10 | 0.377397417 | 0.327 | 0.178 | 2.65243974984557e-06 | 13 |
| zgc:1129803 | 1.21194011258836e-10 | 0.339930139 | 0.205 | 0.092 | 3.05033206937364e-06 | 13 |
| haus65 | 1.42431198858694e-10 | 0.38040664 | 0.365 | 0.208 | 3.58485084407446e-06 | 13 |
| si:ch73-78o10.15 | 1.45718679196686e-10 | 0.358226711 | 0.266 | 0.132 | 3.66759343670138e-06 | 13 |
| dctpp15 | 1.56397046064919e-10 | 0.388376576 | 0.312 | 0.174 | 3.93635725240795e-06 | 13 |
| tmem54a4 | 2.96194543329575e-10 | 0.353398632 | 0.274 | 0.14 | 7.45492046106206e-06 | 13 |

| | | | | | | |
|---|---|---|---|---|---|---|
| zgc:1738565 | 3.09831854572643e-10 | 0.385798054 | 0.365 | 0.219 | 7.79815794773884e-06 | 13 |
| zgc:1132931 | 4.00264028895985e-10 | 0.269080939 | 0.141 | 0.053 | 1.0074245343283e-05 | 13 |
| BX571780.13 | 4.45846122111162e-10 | 0.307601771 | 0.186 | 0.08 | 1.12215010474158e-05 | 13 |
| nek21 | 5.96568405118232e-10 | 0.299274717 | 0.163 | 0.068 | 1.50150301884208e-05 | 13 |
| ypel34 | 6.72718149850946e-10 | 0.358372113 | 0.247 | 0.126 | 1.69316431135985e-05 | 13 |
| zgc:1630304 | 7.36375083149337e-10 | 0.343414228 | 0.247 | 0.123 | 1.85338244677857e-05 | 13 |
| vstm2l4 | 7.39241623324578e-10 | 0.305464866 | 0.209 | 0.095 | 1.86059724174563e-05 | 13 |
| ddx39ab5 | 7.6222093308892e-10 | 0.38634475 | 0.43 | 0.288 | 1.9184338664915e-05 | 13 |
| rbm4.35 | 9.02238849843322e-10 | 0.332545707 | 0.555 | 0.444 | 2.27084496117066e-05 | 13 |
| ccnh3 | 9.94605705537786e-10 | 0.342150469 | 0.247 | 0.124 | 2.50332310026805e-05 | 13 |
| tph1b | 1.02166688372795e-09 | 0.260950576 | 0.114 | 0.04 | 2.57143337965489e-05 | 13 |
| zgc:1034825 | 1.45733527041939e-09 | 0.361492427 | 0.327 | 0.188 | 3.66796714211855e-05 | 13 |
| BX537276.15 | 1.53474222970712e-09 | 0.324437644 | 0.243 | 0.121 | 3.86279271794986e-05 | 13 |
| zgc:1134245 | 1.69135724659333e-09 | 0.350598942 | 0.297 | 0.161 | 4.25697705395076e-05 | 13 |
| zgc:1118683 | 1.69712707844022e-09 | 0.344647706 | 0.278 | 0.147 | 4.27149914372619e-05 | 13 |
| zgc:1628795 | 1.84078836502573e-09 | 0.35366594 | 0.323 | 0.181 | 4.63308023593325e-05 | 13 |
| tph1a5 | 1.93443264849963e-09 | 0.344302825 | 0.266 | 0.141 | 4.86877353300873e-05 | 13 |
| itm2cb3 | 2.15701910719128e-09 | 0.349672024 | 0.259 | 0.137 | 5.42900139088972e-05 | 13 |
| si:ch211-251f6.65 | 2.1711111711486e-09 | 0.362996793 | 0.479 | 0.331 | 5.46446970666392e-05 | 13 |
| larp6b5 | 2.28529612590355e-09 | 0.348755103 | 0.342 | 0.195 | 5.75186181928663e-05 | 13 |
| zgc:1017443 | 2.28634675413078e-09 | 0.3278756 | 0.259 | 0.133 | 5.75450614547176e-05 | 13 |
| snx10a3 | 2.35177868857088e-09 | 0.336856803 | 0.247 | 0.125 | 5.91919178126404e-05 | 13 |
| gyg1a6 | 2.6152232160528e-09 | 0.357396973 | 0.498 | 0.37 | 6.5822553124833e-05 | 13 |
| CABZ01059627.14 | 2.71681710174653e-09 | 0.33501128 | 0.259 | 0.136 | 6.83795696338584e-05 | 13 |
| kpna24 | 2.97050940419565e-09 | 0.347103824 | 0.285 | 0.158 | 7.47647511942002e-05 | 13 |
| si:ch211-286b5.41 | 3.01406773272871e-09 | 0.284398612 | 0.163 | 0.07 | 7.58610707650488e-05 | 13 |
| si:dkey-42i9.85 | 3.32752213573962e-09 | 0.330999035 | 0.247 | 0.127 | 8.37504046344305e-05 | 13 |
| tatdn22 | 3.67791255345386e-09 | 0.283328276 | 0.167 | 0.072 | 9.25693810578802e-05 | 13 |
| zgc:1629455 | 3.89370755916712e-09 | 0.334140092 | 0.297 | 0.162 | 9.80007255566772e-05 | 13 |
| elovl7a5 | 3.9127945139672e-09 | 0.348327521 | 0.316 | 0.182 | 9.84811251220406e-05 | 13 |
| dynll15 | 3.93724207760153e-09 | 0.317419044 | 0.57 | 0.454 | 9.9096445851153e-05 | 13 |
| retsatl5 | 4.57338176854109e-09 | 0.358712583 | 0.338 | 0.203 | 0.000115107 | 13 |
| si:ch211-157c3.43 | 6.02291253207358e-09 | 0.282708428 | 0.194 | 0.089 | 0.000151591 | 13 |
| si:ch211-14a17.62 | 6.48443354505383e-09 | 0.302362739 | 0.186 | 0.086 | 0.000163207 | 13 |
| mkrn45 | 7.38353386120623e-09 | 0.351623212 | 0.369 | 0.228 | 0.000185836 | 13 |

| | | | | | | |
|---|---|---|---|---|---|---|
| CT009596.13 | 7.56303998805172e-09 | 0.308029862 | 0.19 | 0.09 | 0.000190354 | 13 |
| CR847984.15 | 7.80389975486492e-09 | 0.333618201 | 0.297 | 0.163 | 0.000196416 | 13 |
| ankrd225 | 8.09495422745306e-09 | 0.330345528 | 0.266 | 0.142 | 0.000203742 | 13 |
| pop41 | 8.32389361404757e-09 | 0.33031013 | 0.228 | 0.117 | 0.000209504 | 13 |
| gbgt1l44 | 8.62926253498989e-09 | 0.337500195 | 0.274 | 0.15 | 0.00021719 | 13 |
| lipg2 | 1.00584677918655e-08 | 0.279224428 | 0.183 | 0.083 | 0.000253162 | 13 |
| rasl11b5 | 1.04492627753913e-08 | 0.328077896 | 0.251 | 0.134 | 0.000262997 | 13 |
| ctsba5 | 1.1316776502218e-08 | 0.339327414 | 0.502 | 0.376 | 0.000284832 | 13 |
| hsd17b12a5 | 1.16285161807135e-08 | 0.337998137 | 0.289 | 0.165 | 0.000292678 | 13 |
| map1lc3c5 | 1.22537863702236e-08 | 0.345959825 | 0.335 | 0.201 | 0.000308416 | 13 |
| rpain2 | 1.44135459498938e-08 | 0.330192245 | 0.228 | 0.119 | 0.000362775 | 13 |
| pif14 | 2.10274823697447e-08 | 0.304536912 | 0.213 | 0.106 | 0.000529241 | 13 |
| zgc:1737425 | 2.51238266083315e-08 | 0.335285593 | 0.274 | 0.155 | 0.000632342 | 13 |
| oclna1 | 2.56680888276779e-08 | 0.282960948 | 0.171 | 0.078 | 0.00064604 | 13 |
| si:dkey-68o6.55 | 3.08558246246479e-08 | 0.314285403 | 0.232 | 0.122 | 0.00077661 | 13 |
| si:ch73-281n10.23 | 3.21082961218526e-08 | 0.318134104 | 0.506 | 0.392 | 0.000808134 | 13 |
| zgc:558882 | 3.46680615413014e-08 | 0.288224504 | 0.179 | 0.085 | 0.00087256 | 13 |
| zp3c5 | 5.14640084928238e-08 | 0.330965954 | 0.312 | 0.189 | 0.001295298 | 13 |
| nfr5 | 5.62630893798792e-08 | 0.306758179 | 0.213 | 0.11 | 0.001416086 | 13 |
| gstp14 | 6.08451681610255e-08 | 0.327535822 | 0.479 | 0.365 | 0.001531412 | 13 |
| rbpms2b5 | 6.12720872519758e-08 | 0.342357964 | 0.346 | 0.221 | 0.001542157 | 13 |
| anp32b3 | 7.36858914616096e-08 | 0.351432096 | 0.369 | 0.256 | 0.0018546 | 13 |
| cdc205 | 8.99581612160998e-08 | 0.32028751 | 0.266 | 0.152 | 0.002264157 | 13 |
| tbcc | 9.02328801223358e-08 | 0.259369813 | 0.125 | 0.052 | 0.002271071 | 13 |
| rdh10b4 | 9.05674995229848e-08 | 0.323432351 | 0.297 | 0.178 | 0.002279493 | 13 |
| cdca92 | 9.29105857763977e-08 | 0.27205922 | 0.179 | 0.085 | 0.002338467 | 13 |
| kif22 | 1.02315134655898e-07 | 0.286298889 | 0.179 | 0.087 | 0.00257517 | 13 |
| si:ch211-202f5.31 | 1.12536364044764e-07 | 0.290043558 | 0.183 | 0.09 | 0.002832428 | 13 |
| pttg15 | 1.15120791392751e-07 | 0.300850264 | 0.278 | 0.158 | 0.002897475 | 13 |
| orc62 | 1.20908482468308e-07 | 0.299238608 | 0.205 | 0.107 | 0.003043146 | 13 |
| snrpc1 | 1.33504007613428e-07 | 0.350805612 | 0.289 | 0.181 | 0.003360162 | 13 |
| si:ch211-241j12.3 | 1.38730090275228e-07 | 0.250491214 | 0.129 | 0.055 | 0.003491698 | 13 |
| cpm3 | 1.54664599735664e-07 | 0.291179731 | 0.205 | 0.108 | 0.003892753 | 13 |
| zgc:922874 | 1.60959859737776e-07 | 0.294966553 | 0.24 | 0.132 | 0.004051199 | 13 |
| zgc:1139843 | 1.9318406184645e-07 | 0.279987111 | 0.198 | 0.101 | 0.00486225 | 13 |

| | | | | | | |
|---|---|---|---|---|---|---|
| akip1 | 2.17634662195245e-07 | 0.255631411 | 0.137 | 0.061 | 0.005477647 | 13 |
| si:dkey-87o1.21 | 2.98545949491922e-07 | 0.2613773 | 0.167 | 0.081 | 0.007514103 | 13 |
| prrg23 | 3.49950719811189e-07 | 0.268857448 | 0.179 | 0.089 | 0.00880791 | 13 |
| zgc:1717505 | 3.64683560740903e-07 | 0.308844966 | 0.289 | 0.174 | 0.009178721 | 13 |
| BX005461.12 | 3.78224202545413e-07 | 0.26202755 | 0.163 | 0.079 | 0.009519525 | 13 |
| LOC1005380231 | 4.34988978812751e-07 | 0.25770417 | 0.16 | 0.076 | 0.010948238 | 13 |
| zgc:566765 | 4.52415155005386e-07 | 0.329033286 | 0.403 | 0.296 | 0.011386837 | 13 |
| si:ch73-160p18.35 | 4.74269101574822e-07 | 0.323412207 | 0.357 | 0.241 | 0.011936879 | 13 |
| chia.45 | 4.74765810698937e-07 | 0.291759226 | 0.255 | 0.147 | 0.011949381 | 13 |
| tac3b5 | 4.92088700998568e-07 | 0.272947715 | 0.221 | 0.12 | 0.012385381 | 13 |
| BX001030.14 | 5.06002586751106e-07 | 0.294427873 | 0.221 | 0.122 | 0.012735579 | 13 |
| LOC1104377791 | 6.62283616416208e-07 | 0.268273667 | 0.167 | 0.083 | 0.016669016 | 13 |
| cdca84 | 6.73992765971919e-07 | 0.311507661 | 0.27 | 0.165 | 0.016963724 | 13 |
| kmt5ab2 | 7.0850984446001e-07 | 0.279163165 | 0.175 | 0.089 | 0.017832484 | 13 |
| LOC1039099722 | 7.24777061080403e-07 | 0.309358018 | 0.46 | 0.367 | 0.018241914 | 13 |
| si:ch211-113a14.271 | 7.27702441644844e-07 | 0.2531373 | 0.16 | 0.078 | 0.018315543 | 13 |
| tent5c4 | 7.88229509576388e-07 | 0.298142124 | 0.24 | 0.14 | 0.019838949 | 13 |
| si:ch73-15b2.53 | 7.98537658589403e-07 | 0.269309469 | 0.183 | 0.095 | 0.020098394 | 13 |
| cx43.43 | 8.0151365794171e-07 | 0.287922263 | 0.221 | 0.123 | 0.020173297 | 13 |
| zgc:660245 | 8.10346210064741e-07 | 0.280224703 | 0.247 | 0.142 | 0.020395604 | 13 |
| dazl2 | 8.88058924975084e-07 | 0.256035968 | 0.202 | 0.106 | 0.022351555 | 13 |
| id2a | 1.16186124426964e-06 | 0.305204565 | 0.27 | 0.17 | 0.029242886 | 13 |
| ttc322 | 1.22222168935715e-06 | 0.286728299 | 0.19 | 0.103 | 0.030762098 | 13 |
| zgc:1719774 | 1.47488150081632e-06 | 0.307345472 | 0.346 | 0.23 | 0.037121292 | 13 |
| slc48a1b | 1.6593379818758e-06 | 0.26030749 | 0.167 | 0.086 | 0.041763878 | 13 |
| mibp5 | 1.8899406348207e-06 | 0.301154345 | 0.335 | 0.223 | 0.047567916 | 13 |
| fdx15 | 1.90243798348196e-06 | 0.30304646 | 0.327 | 0.221 | 0.047882462 | 13 |
| nanos34 | 1.91176230615173e-06 | 0.270200161 | 0.19 | 0.103 | 0.048117145 | 13 |
| plk14 | 1.99918003710773e-06 | 0.279959005 | 0.289 | 0.177 | 0.050317362 | 13 |
| vps9d12 | 2.07755193188204e-06 | 0.26107958 | 0.183 | 0.096 | 0.052289905 | 13 |
| ube2c2 | 2.15221300227783e-06 | 0.290635747 | 0.236 | 0.14 | 0.054169049 | 13 |
| zgc:923132 | 2.37513831814126e-06 | 0.27074229 | 0.221 | 0.126 | 0.059779856 | 13 |
| rbbp42 | 2.56861299559807e-06 | 0.295380317 | 0.27 | 0.171 | 0.06464942 | 13 |
| zgc:1010161 | 2.65755687396657e-06 | 0.278600649 | 0.179 | 0.097 | 0.066888049 | 13 |
| mrps18b2 | 2.69415306868582e-06 | 0.284166533 | 0.217 | 0.126 | 0.067809139 | 13 |

| | | | | | | |
|---|---|---|---|---|---|---|
| figla2 | 2.74180316974686e-06 | 0.266021464 | 0.179 | 0.095 | 0.069008444 | 13 |
| sumo3a | 2.94729508132933e-06 | 0.295370738 | 0.437 | 0.37 | 0.07418047 | 13 |
| eno31 | 3.10624151992152e-06 | 0.277054343 | 0.456 | 0.383 | 0.078180993 | 13 |
| aurkb3 | 3.30460908281071e-06 | 0.274022298 | 0.221 | 0.128 | 0.083173706 | 13 |
| degs24 | 3.43693561850051e-06 | 0.263321265 | 0.194 | 0.106 | 0.086504233 | 13 |
| prdx14 | 3.54393974843506e-06 | 0.294518412 | 0.266 | 0.167 | 0.08919742 | 13 |
| chtopa3 | 3.96900650857517e-06 | 0.306872967 | 0.319 | 0.222 | 0.099895925 | 13 |
| parn4 | 4.06430911833962e-06 | 0.283479766 | 0.243 | 0.148 | 0.102294596 | 13 |
| rab5if1 | 4.40642935052996e-06 | 0.294754398 | 0.217 | 0.13 | 0.11090542 | 13 |
| fbxo323 | 4.42685392765668e-06 | 0.254629945 | 0.19 | 0.104 | 0.111419487 | 13 |
| dck1 | 4.44555805671423e-06 | 0.255138672 | 0.167 | 0.089 | 0.111890251 | 13 |
| rnf45 | 4.90770367932609e-06 | 0.284115862 | 0.224 | 0.134 | 0.123521994 | 13 |
| faub4 | 6.08465225246837e-06 | 0.298193396 | 0.3 | 0.201 | 0.153144613 | 13 |
| CABZ01059627.25 | 6.28289050891365e-06 | 0.257535285 | 0.228 | 0.134 | 0.158134071 | 13 |
| si:dkey-241l7.35 | 6.40961545834941e-06 | 0.284569451 | 0.468 | 0.365 | 0.161323611 | 13 |
| zgc:1526524 | 6.80021074548916e-06 | 0.254401644 | 0.202 | 0.114 | 0.171154504 | 13 |
| zc3h11a4 | 8.15145041250668e-06 | 0.299449919 | 0.312 | 0.217 | 0.205163855 | 13 |
| ctsla5 | 8.41415933065058e-06 | 0.270242766 | 0.456 | 0.388 | 0.211775976 | 13 |
| dynll2a4 | 8.88480157378283e-06 | 0.265496806 | 0.228 | 0.136 | 0.223621571 | 13 |
| cldn7b5 | 1.08867643974013e-05 | 0.280008879 | 0.251 | 0.16 | 0.274008973 | 13 |
| snrpb24 | 1.27544426833292e-05 | 0.277931632 | 0.24 | 0.149 | 0.321016568 | 13 |
| mos4 | 1.33121214360981e-05 | 0.25379504 | 0.228 | 0.137 | 0.335052784 | 13 |
| cnep1r11 | 1.53456012060749e-05 | 0.271660282 | 0.205 | 0.124 | 0.386233437 | 13 |
| evla2 | 1.79998893022563e-05 | 0.260415196 | 0.179 | 0.103 | 0.453039214 | 13 |
| hspa54 | 2.04475836307499e-05 | 0.273571679 | 0.411 | 0.315 | 0.514645232 | 13 |
| si:dkey-188i13.115 | 2.12740138373757e-05 | 0.276724182 | 0.418 | 0.317 | 0.535445654 | 13 |
| tpx23 | 2.13084190893465e-05 | 0.254803759 | 0.209 | 0.126 | 0.5363116 | 13 |
| ckbb3 | 2.14242188938574e-05 | 0.266579061 | 0.365 | 0.266 | 0.539226165 | 13 |
| mrgbp | 2.32740840066276e-05 | 0.250954214 | 0.167 | 0.094 | 0.58578542 | 13 |
| CDPF15 | 2.59351629518727e-05 | 0.28307756 | 0.361 | 0.269 | 0.652762116 | 13 |
| bzw1b3 | 2.70380510760975e-05 | 0.25513133 | 0.464 | 0.403 | 0.680520708 | 13 |
| si:ch73-340n8.42 | 3.38394563799756e-05 | 0.253276606 | 0.209 | 0.128 | 0.851705278 | 13 |
| inka1b5 | 3.50954207303509e-05 | 0.260203685 | 0.281 | 0.185 | 0.883316644 | 13 |
| rpa31 | 3.60089647115831e-05 | 0.277981083 | 0.247 | 0.163 | 0.906309633 | 13 |
| adipor1a | 3.77849640049096e-05 | 0.251185111 | 0.171 | 0.1 | 0.951009759 | 13 |

| | | | | | | | |
|---|---|---|---|---|---|---|---|
| h2afva4 | 3.9890229307843e-05 | 0.273591498 | 0.403 | 0.324 | | 1 | 13 |
| zp2l15 | 4.469080492054e-05 | 0.251910933 | 0.255 | 0.164 | | 1 | 13 |
| ivns1abpb1 | 4.78371723118034e-05 | 0.261950839 | 0.19 | 0.117 | | 1 | 13 |
| BX957362.13 | 5.3325502620206e-05 | 0.25006989 | 0.213 | 0.133 | | 1 | 13 |
| gemin72 | 8.6198364353584e-05 | 0.267405901 | 0.232 | 0.155 | | 1 | 13 |
| g3bp14 | 0.000117987 | 0.260871416 | 0.293 | 0.211 | | 1 | 13 |
| aktip2 | 0.000125439 | 0.25630575 | 0.213 | 0.14 | | 1 | 13 |
| sub1b1 | 0.000168879 | 0.270829905 | 0.297 | 0.223 | | 1 | 13 |
| tia1l | 0.000212908 | 0.250778899 | 0.221 | 0.149 | | 1 | 13 |
| tspan36 | 0.000231192 | 0.258135973 | 0.228 | 0.158 | | 1 | 13 |
| btg32 | 0.000353875 | 0.251482527 | 0.251 | 0.179 | | 1 | 13 |
| ccl36.12 | 0 | 2.067974882 | 0.992 | 0.082 | | 0 | 14 |
| si:dkey-21e2.12 | 0 | 1.699931509 | 0.752 | 0.004 | | 0 | 14 |
| si:dkey-21e2.11 | 0 | 1.420149098 | 0.563 | 0.002 | | 0 | 14 |
| ncaldb2 | 0 | 1.370171671 | 0.772 | 0.059 | | 0 | 14 |
| zbtb321 | 0 | 1.367522249 | 0.732 | 0.032 | | 0 | 14 |
| ca2 | 0 | 1.314396813 | 0.661 | 0.043 | | 0 | 14 |
| si:ch211-277c7.6 | 0 | 1.246206897 | 0.591 | 0.026 | | 0 | 14 |
| il341 | 0 | 1.135993708 | 0.594 | 0.037 | | 0 | 14 |
| si:ch211-207n23.2 | 0 | 1.135851517 | 0.587 | 0.016 | | 0 | 14 |
| si:dkey-78l4.4 | 0 | 1.135348318 | 0.567 | 0.022 | | 0 | 14 |
| ccr22 | 0 | 1.06825278 | 0.594 | 0.035 | | 0 | 14 |
| si:rp71-81e14.22 | 0 | 1.000541066 | 0.492 | 0.025 | | 0 | 14 |
| blnk | 0 | 0.834852356 | 0.398 | 0.017 | | 0 | 14 |
| dicp1.5-6 | 0 | 0.703934797 | 0.358 | 0.011 | | 0 | 14 |
| BX000438.2 | 0 | 0.66938427 | 0.276 | 0.001 | | 0 | 14 |
| fam167b | 0 | 0.599770078 | 0.291 | 0.006 | | 0 | 14 |
| XLOC-026336 | 0 | 0.442885644 | 0.201 | 0.002 | | 0 | 14 |
| scinlb2 | 5.46835206595685e-303 | 1.175380539 | 0.713 | 0.07 | 1.37632953148068e-298 | | 14 |
| CABZ01084575.11 | 2.31164012640274e-291 | 0.967939876 | 0.508 | 0.033 | 5.81816703414305e-287 | | 14 |
| si:ch211-102c2.41 | 2.58645268505815e-286 | 1.423552403 | 0.89 | 0.127 | 6.50984276302285e-282 | | 14 |
| ccl38.62 | 9.62938568202072e-271 | 1.526712964 | 0.783 | 0.096 | 2.42362008230779e-266 | | 14 |
| ponzr6 | 1.5820209301618e-266 | 0.727779404 | 0.362 | 0.017 | 3.98178847912423e-262 | | 14 |
| LOC1081907171 | 6.2515362912844e-266 | 1.291616671 | 0.732 | 0.083 | 1.57344916915337e-261 | | 14 |
| znrf1 | 1.28018482628342e-263 | 0.575134653 | 0.291 | 0.01 | 3.22209718927274e-259 | | 14 |

| | | | | | | |
|---|---|---|---|---|---|---|
| obsl1a | 8.05970146510063e-262 | 0.422679298 | 0.189 | 0.003 | 2.02854626175118e-257 | 14 |
| si:ch211-241f5.31 | 3.78972791973383e-249 | 1.041093728 | 0.594 | 0.056 | 9.53836620117808e-245 | 14 |
| XLOC-0139642 | 1.66976358898116e-246 | 1.568909795 | 0.917 | 0.158 | 4.20262797710667e-242 | 14 |
| LOC1039108712 | 1.52044517332174e-244 | 1.482529917 | 0.913 | 0.155 | 3.82680845673348e-240 | 14 |
| mctp1a | 6.16992485931324e-233 | 0.492052483 | 0.236 | 0.007 | 1.55290838784055e-228 | 14 |
| si:ch73-40a2.1 | 2.838262613475e-230 | 0.448046397 | 0.224 | 0.006 | 7.14362317185522e-226 | 14 |
| si:ch211-214p16.13 | 6.00736643929263e-230 | 1.636603213 | 0.988 | 0.243 | 1.51199405910556e-225 | 14 |
| CR936442.12 | 2.17407442212005e-224 | 1.45255305 | 0.866 | 0.155 | 5.47192791303395e-220 | 14 |
| LOC103910169 | 6.13770620703535e-224 | 0.351495354 | 0.169 | 0.003 | 1.54479927524873e-219 | 14 |
| XLOC-0425952 | 2.58259690106077e-223 | 1.447377344 | 0.882 | 0.149 | 6.50013814027984e-219 | 14 |
| tnxba | 3.30200899731996e-219 | 0.407797008 | 0.197 | 0.005 | 8.31082644535462e-215 | 14 |
| itpr1b | 1.71334330480165e-215 | 0.563436584 | 0.264 | 0.011 | 4.31231376385527e-211 | 14 |
| sla22 | 4.28446270581386e-211 | 1.010143987 | 0.563 | 0.059 | 1.07835641842629e-206 | 14 |
| nitr1m1 | 7.80928048367725e-204 | 0.533115727 | 0.224 | 0.008 | 1.96551780493673e-199 | 14 |
| XLOC-0203011 | 5.55348091813195e-202 | 1.110617252 | 0.614 | 0.074 | 1.39775561228463e-197 | 14 |
| si:ch211-191c10.12 | 1.13578845515189e-201 | 0.878478538 | 0.48 | 0.045 | 2.85866596277179e-197 | 14 |
| CT573231.1 | 7.18441587996571e-200 | 0.655428152 | 0.327 | 0.019 | 1.80824563282857e-195 | 14 |
| creb5b | 1.09251711036034e-194 | 0.581423557 | 0.283 | 0.014 | 2.74975631506593e-190 | 14 |
| tnfb1 | 1.65828008796072e-194 | 1.003217962 | 0.469 | 0.044 | 4.17372515338835e-190 | 14 |
| XLOC-0425962 | 5.09739633029136e-194 | 0.805761789 | 0.417 | 0.034 | 1.28296368237103e-189 | 14 |
| XLOC-0268361 | 7.25503694039931e-186 | 1.122788434 | 0.563 | 0.069 | 1.8260202475291e-181 | 14 |
| XLOC-0108992 | 7.06535424051594e-185 | 1.451816731 | 0.957 | 0.244 | 1.77827900879546e-180 | 14 |
| XLOC-0283702 | 3.02510313829468e-183 | 1.263325989 | 0.744 | 0.128 | 7.61388208877388e-179 | 14 |
| dusp22 | 6.5059914387153e-181 | 1.221350413 | 0.843 | 0.165 | 1.63749298521025e-176 | 14 |
| XLOC-0320021 | 1.50222705014494e-178 | 0.843042516 | 0.425 | 0.039 | 3.78095526250979e-174 | 14 |
| XLOC-0004312 | 5.13481193046421e-176 | 1.540427599 | 0.961 | 0.264 | 1.29238081477854e-171 | 14 |
| CR392341.11 | 1.16045347171402e-173 | 0.846828981 | 0.472 | 0.051 | 2.92074534295702e-169 | 14 |
| XLOC-0254232 | 7.04693933351628e-168 | 1.322395107 | 0.78 | 0.157 | 1.77364416085271e-163 | 14 |
| syk1 | 1.76919139636232e-166 | 0.905641972 | 0.539 | 0.069 | 4.45287782550432e-162 | 14 |
| arpc1b4 | 1.26537108947639e-160 | 1.248503121 | 0.972 | 0.295 | 3.18481249510313e-156 | 14 |
| stim1a | 7.2750797858065e-160 | 0.842260457 | 0.496 | 0.06 | 1.83106483128964e-155 | 14 |
| si:dkey-183i3.62 | 1.83286578270787e-157 | 0.742746261 | 0.406 | 0.039 | 4.61313988849743e-153 | 14 |
| LOC1005362132 | 2.25040095370492e-153 | 0.933468403 | 0.618 | 0.095 | 5.66403416037992e-149 | 14 |
| zgc:640513 | 2.92832845556746e-153 | 1.080464429 | 0.744 | 0.151 | 7.37030988981774e-149 | 14 |
| CU914617.1 | 6.85885439922666e-153 | 0.260071374 | 0.102 | 0.001 | 1.72630506374136e-148 | 14 |

| | | | | | | |
|---|---|---|---|---|---|---|
| prkcba | 1.8500251721666e-152 | 0.505392281 | 0.248 | 0.015 | 4.65632835582611e-148 | 14 |
| gramd2aa | 3.98821869982537e-149 | 0.481876194 | 0.244 | 0.014 | 1.00379476455905e-144 | 14 |
| BX901920.11 | 8.54810522228027e-149 | 0.890772037 | 0.496 | 0.065 | 2.15147260339572e-144 | 14 |
| orai1a | 3.6067857388298e-144 | 0.5283677 | 0.276 | 0.019 | 9.07791902606073e-140 | 14 |
| mbnl13 | 8.96245097041978e-144 | 1.04635715 | 0.772 | 0.172 | 2.25575928474495e-139 | 14 |
| XLOC-0430431 | 2.5032581288826e-143 | 0.750397683 | 0.37 | 0.037 | 6.30045038458461e-139 | 14 |
| rgs132 | 1.21017325360547e-140 | 0.991825008 | 0.665 | 0.121 | 3.04588506199962e-136 | 14 |
| laptm54 | 4.65332165293374e-138 | 1.048540594 | 0.736 | 0.159 | 1.17119452682689e-133 | 14 |
| il2rb2 | 5.95889626264519e-137 | 0.508625362 | 0.248 | 0.016 | 1.49979460034517e-132 | 14 |
| XLOC-0437601 | 2.50876922543323e-136 | 0.802292221 | 0.394 | 0.044 | 6.3143212634929e-132 | 14 |
| ttyh2l | 3.10578435871616e-136 | 0.376478691 | 0.181 | 0.008 | 7.81694865245271e-132 | 14 |
| si:ch211-214p16.21 | 3.63868931641981e-136 | 0.833673079 | 0.48 | 0.064 | 9.15821714049701e-132 | 14 |
| gfi1ab1 | 3.08956832068002e-135 | 0.503299459 | 0.26 | 0.018 | 7.77613450631955e-131 | 14 |
| wdr191 | 1.27072811299292e-134 | 0.622006585 | 0.311 | 0.027 | 3.19829558759189e-130 | 14 |
| gpx1a1 | 1.18897375616187e-133 | 1.298454625 | 0.764 | 0.213 | 2.99252804688381e-129 | 14 |
| calm1b3 | 2.46355333234174e-133 | 0.847163545 | 0.543 | 0.086 | 6.20051738217092e-129 | 14 |
| CABZ01117603.1 | 3.72225034120043e-132 | 0.45365503 | 0.228 | 0.014 | 9.36853188376736e-128 | 14 |
| CABZ01044746.1 | 2.45797715603946e-130 | 0.905866488 | 0.575 | 0.095 | 6.18648270403571e-126 | 14 |
| XLOC-0234923 | 4.86187154234708e-125 | 0.931525772 | 0.524 | 0.086 | 1.22368444849334e-120 | 14 |
| CR753876.11 | 1.21086981428891e-122 | 0.974049546 | 0.583 | 0.106 | 3.04763823558375e-118 | 14 |
| srgn3 | 1.50114777959584e-122 | 1.024933533 | 0.846 | 0.225 | 3.77823884646478e-118 | 14 |
| slc41a1 | 5.9570075449749e-121 | 0.384819505 | 0.189 | 0.01 | 1.49931922899473e-116 | 14 |
| CU972454.4 | 8.53141182864536e-121 | 0.473283873 | 0.248 | 0.019 | 2.14727104315175e-116 | 14 |
| cd44b | 1.79431793738282e-117 | 0.583534921 | 0.303 | 0.03 | 4.51611881659881e-113 | 14 |
| mef2d | 6.40224478091669e-117 | 0.748699787 | 0.441 | 0.066 | 1.61138098890892e-112 | 14 |
| id32 | 7.12879691796395e-117 | 1.053624089 | 0.76 | 0.199 | 1.79424689628235e-112 | 14 |
| camk1db | 4.78443352766077e-116 | 0.424088804 | 0.22 | 0.015 | 1.20419407457694e-111 | 14 |
| bhlhe401 | 1.00388162542707e-114 | 0.963071165 | 0.665 | 0.151 | 2.52666966303739e-110 | 14 |
| gzm3.4 | 1.61696086587126e-114 | 0.331568024 | 0.138 | 0.005 | 4.06972880331138e-110 | 14 |
| atp2b1a | 2.83503092690203e-114 | 0.761246895 | 0.449 | 0.066 | 7.13548933991971e-110 | 14 |
| sik12 | 2.27297768455827e-113 | 0.844823923 | 0.567 | 0.107 | 5.72085753426471e-109 | 14 |
| arhgdig3 | 8.91350714771074e-113 | 0.983690528 | 0.811 | 0.232 | 2.24344061400732e-108 | 14 |
| fcer1g1 | 6.61392672842369e-112 | 0.588700607 | 0.331 | 0.037 | 1.66465921827696e-107 | 14 |
| adam19b | 4.38212349534693e-111 | 0.388788644 | 0.185 | 0.011 | 1.10293666254387e-106 | 14 |
| FP236356.12 | 2.48991000814602e-109 | 0.654536089 | 0.354 | 0.043 | 6.26685449950271e-105 | 14 |

| | | | | | | |
|---|---|---|---|---|---|---|
| dock10 | 3.5183577719782e-109 | 0.523242363 | 0.276 | 0.027 | 8.85535467629193e-105 | 14 |
| LOC1039091072 | 2.86569822461282e-108 | 0.526796236 | 0.248 | 0.021 | 7.212675861528e-104 | 14 |
| XLOC-0418701 | 1.03321496211085e-107 | 0.589162608 | 0.26 | 0.024 | 2.6004987381368e-103 | 14 |
| irf4b1 | 9.05811958414759e-107 | 0.511513038 | 0.24 | 0.02 | 2.27983811813411e-102 | 14 |
| zgc:153018 | 1.1399639935112e-106 | 0.491866927 | 0.232 | 0.019 | 2.86917537526835e-102 | 14 |
| kdm2ba | 2.47850620074325e-106 | 0.537766284 | 0.299 | 0.032 | 6.23815225665069e-102 | 14 |
| BX322618.1 | 1.68622473095079e-105 | 0.590662503 | 0.26 | 0.024 | 4.24405902533003e-101 | 14 |
| ptprc3 | 2.2171548958765e-104 | 0.813196342 | 0.583 | 0.118 | 5.58035715743157e-100 | 14 |
| tagapa3 | 6.34007629311014e-104 | 0.857995847 | 0.52 | 0.098 | 1.59573380221289e-99 | 14 |
| wasa2 | 1.15025105538806e-103 | 0.713083398 | 0.492 | 0.085 | 2.89506688130621e-99 | 14 |
| si:ch211-167j9.52 | 1.27213789017569e-103 | 0.466216775 | 0.252 | 0.023 | 3.2018438557832e-99 | 14 |
| XLOC-0140791 | 1.66694193569031e-103 | 0.561572685 | 0.28 | 0.029 | 4.19552615793893e-99 | 14 |
| tmsb4x6 | 1.19178990097125e-102 | 0.762726807 | 1 | 0.681 | 2.99961600175453e-98 | 14 |
| plcg2 | 6.81156142364633e-102 | 0.630890615 | 0.346 | 0.045 | 1.71440189471754e-97 | 14 |
| capgb3 | 1.52608596448438e-101 | 0.799346665 | 0.543 | 0.106 | 3.84100576401075e-97 | 14 |
| hcls11 | 3.02892066342009e-101 | 0.662619285 | 0.413 | 0.064 | 7.62349041776203e-97 | 14 |
| plp2 | 5.34415669053573e-100 | 0.583158357 | 0.343 | 0.045 | 1.34507079744094e-95 | 14 |
| XLOC-039757 | 5.47938642587924e-100 | 0.516302369 | 0.244 | 0.022 | 1.37910676952955e-95 | 14 |
| CABZ01111959.1 | 4.66938765503174e-98 | 0.440688679 | 0.24 | 0.022 | 1.17523817889494e-93 | 14 |
| pik3r12 | 2.53352844195363e-97 | 0.709406372 | 0.461 | 0.083 | 6.37663773555308e-93 | 14 |
| ptmaa4 | 4.2517098889411e-97 | 0.939779574 | 0.933 | 0.385 | 1.07011286194758e-92 | 14 |
| BX005183.32 | 2.73172776799675e-95 | 0.703518763 | 0.437 | 0.074 | 6.87548561927101e-91 | 14 |
| caspa | 3.42963696237711e-95 | 0.426355803 | 0.224 | 0.02 | 8.63205327060695e-91 | 14 |
| XLOC-0437611 | 7.0691509531367e-95 | 0.462820612 | 0.213 | 0.018 | 1.77923460339498e-90 | 14 |
| atp11c | 4.42508108743379e-94 | 0.512484596 | 0.268 | 0.029 | 1.11374865889621e-89 | 14 |
| gsdmea | 1.80447874591163e-93 | 0.437367793 | 0.22 | 0.02 | 4.54169255558498e-89 | 14 |
| cd247l2 | 3.61435450344072e-93 | 0.387057708 | 0.209 | 0.017 | 9.09696884970995e-89 | 14 |
| grapa | 1.97563552697321e-92 | 0.478704487 | 0.24 | 0.024 | 4.97247705783887e-88 | 14 |
| coro1a4 | 1.69714430432911e-91 | 0.907413229 | 0.776 | 0.221 | 4.27154249956592e-87 | 14 |
| tmc6a | 5.82538231871445e-91 | 0.332816075 | 0.157 | 0.01 | 1.46619047579724e-86 | 14 |
| pxna | 1.24132056479544e-89 | 0.705850629 | 0.461 | 0.084 | 3.12427972953365e-85 | 14 |
| cd40lg1 | 1.35041549011873e-89 | 0.303817295 | 0.15 | 0.009 | 3.39886074707984e-85 | 14 |
| BX908750.1 | 4.7161924212188e-89 | 0.488279451 | 0.256 | 0.028 | 1.18701847049656e-84 | 14 |
| si:dkey-183p4.102 | 1.0435548451078e-87 | 0.474395778 | 0.268 | 0.031 | 2.62652318965182e-83 | 14 |
| sla1 | 2.14865031673941e-87 | 0.365424826 | 0.205 | 0.018 | 5.40793798220143e-83 | 14 |

| | | | | | | |
|---|---|---|---|---|---|---|
| lck | 2.62865714046181e-87 | 0.298077141 | 0.142 | 0.008 | 6.61606715682834e-83 | 14 |
| atp5if1b | 3.19355127841631e-87 | 0.827069901 | 0.657 | 0.182 | 8.03784921264601e-83 | 14 |
| tmc82 | 1.08036538133124e-85 | 0.476314335 | 0.264 | 0.031 | 2.71917162827261e-81 | 14 |
| aldocb1 | 1.54848299104982e-84 | 0.594033824 | 0.374 | 0.061 | 3.89737684017328e-80 | 14 |
| cotl14 | 1.73628196620537e-84 | 0.903204746 | 0.839 | 0.296 | 4.3700480807423e-80 | 14 |
| si:ch73-208g10.12 | 1.75061779511719e-84 | 0.525477046 | 0.291 | 0.038 | 4.40612992853045e-80 | 14 |
| ptmab4 | 8.27418890355335e-84 | 0.776538154 | 0.941 | 0.533 | 2.08253060513534e-79 | 14 |
| rplp25 | 3.16961504452936e-83 | 0.536294485 | 0.996 | 0.753 | 7.97760410557594e-79 | 14 |
| XLOC-0054492 | 6.13443671185026e-83 | 0.62709037 | 0.327 | 0.049 | 1.54397637600559e-78 | 14 |
| si:dkey-1c7.1 | 6.63892042514209e-83 | 0.373163408 | 0.213 | 0.021 | 1.67094988180401e-78 | 14 |
| tagapb2 | 2.41955891155825e-82 | 0.61843901 | 0.37 | 0.062 | 6.08978782450097e-78 | 14 |
| LOC1104399312 | 4.00056114734248e-82 | 0.407863666 | 0.209 | 0.02 | 1.00690123517463e-77 | 14 |
| fscn1a1 | 4.54689204524282e-82 | 0.602127532 | 0.339 | 0.052 | 1.14440725886717e-77 | 14 |
| si:ch211-93g21.12 | 5.24722160042106e-81 | 0.441185724 | 0.26 | 0.031 | 1.32067320460998e-76 | 14 |
| si:dkey-53k12.21 | 2.19477636592344e-80 | 0.52933 | 0.291 | 0.04 | 5.5240326353927e-76 | 14 |
| arpc5a3 | 2.47366766501478e-80 | 0.702100303 | 0.488 | 0.108 | 6.22597414607571e-76 | 14 |
| lasp12 | 9.57589755018379e-80 | 0.811356598 | 0.713 | 0.227 | 2.41015765440576e-75 | 14 |
| tnip12 | 3.39441017020235e-79 | 0.703930277 | 0.398 | 0.076 | 8.54339095738228e-75 | 14 |
| zgc:1749452 | 9.06508705024404e-79 | 0.403308108 | 0.217 | 0.022 | 2.28159175967592e-74 | 14 |
| cox6b13 | 1.313064276639e-78 | 0.6993133 | 0.496 | 0.113 | 3.30485147787271e-74 | 14 |
| mef2ca | 2.24398199862578e-78 | 0.282310364 | 0.134 | 0.008 | 5.64787829234124e-74 | 14 |
| vaspb | 5.20022374321108e-78 | 0.553526164 | 0.303 | 0.045 | 1.3088443139288e-73 | 14 |
| CR381673.12 | 2.02969781647357e-77 | 0.539398621 | 0.331 | 0.052 | 5.10854643428234e-73 | 14 |
| myl12.12 | 2.31919589661318e-77 | 0.810513434 | 0.87 | 0.383 | 5.83718415218572e-73 | 14 |
| si:rp71-68n21.92 | 6.05585113688963e-77 | 0.493737336 | 0.283 | 0.04 | 1.52419717264375e-72 | 14 |
| ponzr13 | 8.59073074813962e-77 | 0.863873872 | 0.74 | 0.264 | 2.16220102199926e-72 | 14 |
| elf12 | 1.03942081289964e-76 | 0.687650812 | 0.453 | 0.097 | 2.61611824398711e-72 | 14 |
| rac24 | 4.49427000626148e-76 | 0.794814642 | 0.717 | 0.211 | 1.13116281787595e-71 | 14 |
| si:ch211-119e14.12 | 1.631893037534e-75 | 0.624948357 | 0.378 | 0.068 | 4.10731158616931e-71 | 14 |
| hmgn22 | 1.22993211267135e-74 | 0.95160086 | 0.878 | 0.435 | 3.09561613438251e-70 | 14 |
| ikzf12 | 3.08016905135568e-74 | 0.411367973 | 0.24 | 0.029 | 7.75247748535711e-70 | 14 |
| unc13d1 | 4.08707203986162e-74 | 0.391581603 | 0.224 | 0.026 | 1.02867516171277e-69 | 14 |
| CABZ01054965.1 | 5.57232446105261e-74 | 0.355655013 | 0.161 | 0.013 | 1.40249834360233e-69 | 14 |
| fpr1 | 1.17635944862409e-72 | 0.36940255 | 0.181 | 0.017 | 2.96077909624196e-68 | 14 |
| CR392341.4 | 1.94581121130951e-72 | 0.268660475 | 0.122 | 0.007 | 4.8974122377449e-68 | 14 |

| | | | | | | |
|---|---|---|---|---|---|---|
| si:dkey-151g10.3 | 3.13839117763471e-72 | 0.325169366 | 0.165 | 0.014 | 7.89901675498881e-68 | 14 |
| casz1 | 4.13389827032106e-72 | 0.36630178 | 0.189 | 0.019 | 1.04046085565711e-67 | 14 |
| dok22 | 1.43325575330007e-71 | 0.448638164 | 0.256 | 0.034 | 3.60736140548095e-67 | 14 |
| si:ch211-222l21.14 | 1.75067410992242e-71 | 0.831333006 | 0.756 | 0.266 | 4.40627166726373e-67 | 14 |
| crema3 | 1.86859625205873e-71 | 0.799885969 | 0.587 | 0.171 | 4.70306990680661e-67 | 14 |
| LOC110438754 | 2.48987989043283e-71 | 0.274873982 | 0.15 | 0.012 | 6.26677869623038e-67 | 14 |
| XLOC-0110852 | 5.07151691909788e-71 | 0.538284959 | 0.335 | 0.058 | 1.27645009336775e-66 | 14 |
| RASSF51 | 6.01483563148982e-71 | 0.55429627 | 0.319 | 0.053 | 1.51387398008967e-66 | 14 |
| yrk1 | 3.50785687595895e-70 | 0.530291989 | 0.339 | 0.058 | 8.82892497110109e-66 | 14 |
| arpc23 | 5.100935017513e-69 | 0.776896542 | 0.724 | 0.259 | 1.28385433455785e-64 | 14 |
| arpc33 | 5.57417097553668e-69 | 0.791894867 | 0.78 | 0.302 | 1.40296309283283e-64 | 14 |
| XLOC-0331371 | 7.17927088227506e-69 | 0.47211634 | 0.24 | 0.032 | 1.80695068835981e-64 | 14 |
| nptna1 | 1.88182793823322e-67 | 0.57013074 | 0.358 | 0.066 | 4.73637273773919e-63 | 14 |
| vaspa3 | 3.48326952307286e-67 | 0.574735008 | 0.366 | 0.07 | 8.76704106262208e-63 | 14 |
| cap14 | 7.55423412026904e-67 | 0.748686819 | 0.693 | 0.225 | 1.90132518573052e-62 | 14 |
| wasb3 | 8.91254763453422e-67 | 0.74633019 | 0.594 | 0.17 | 2.24319911413592e-62 | 14 |
| fybb1 | 1.09870140932318e-66 | 0.459905385 | 0.26 | 0.038 | 2.7653215771255e-62 | 14 |
| si:ch211-132g1.12 | 1.63514627406593e-66 | 0.386535769 | 0.193 | 0.021 | 4.11549965719655e-62 | 14 |
| pycard3 | 4.70708248705066e-66 | 0.557497854 | 0.35 | 0.067 | 1.18472559116578e-61 | 14 |
| si:cabz01074946.11 | 5.59411985562998e-66 | 0.521955236 | 0.315 | 0.055 | 1.40798402646351e-61 | 14 |
| entpd31 | 1.01649135725408e-65 | 0.35631436 | 0.169 | 0.016 | 2.55840709707278e-61 | 14 |
| rps295 | 1.76849904350895e-65 | 0.40034202 | 0.988 | 0.829 | 4.45113524260767e-61 | 14 |
| fmnl1a1 | 6.45086974913021e-65 | 0.464230269 | 0.244 | 0.035 | 1.62361940715858e-60 | 14 |
| si:dkeyp-19e1.31 | 3.62042478781573e-64 | 0.545149623 | 0.335 | 0.063 | 9.1122471484534e-60 | 14 |
| cyth4b | 4.43975215492722e-64 | 0.329215035 | 0.165 | 0.016 | 1.11744121987363e-59 | 14 |
| si:dkey-260g12.12 | 4.67486851385041e-64 | 0.426844791 | 0.228 | 0.03 | 1.17661765625101e-59 | 14 |
| gcnt4a | 7.22340657520133e-64 | 0.3080186 | 0.146 | 0.013 | 1.81805920091242e-59 | 14 |
| itga42 | 1.67133402672359e-63 | 0.526209095 | 0.327 | 0.06 | 4.2065806118606e-59 | 14 |
| h1fx2 | 1.72545429656324e-63 | 0.824594164 | 0.579 | 0.176 | 4.34279591902001e-59 | 14 |
| rplp14 | 5.88657669286411e-63 | 0.371806526 | 0.996 | 0.875 | 1.48159248782697e-58 | 14 |
| wipf1a2 | 9.0522310203665e-63 | 0.53310505 | 0.327 | 0.061 | 2.27835602551604e-58 | 14 |
| si:ch211-215e19.31 | 1.46407821447001e-62 | 0.342830012 | 0.177 | 0.019 | 3.68493845799956e-58 | 14 |
| strbp | 2.3606678786685e-62 | 0.306425265 | 0.161 | 0.016 | 5.94156498382076e-58 | 14 |
| samd11 | 1.32327192747611e-61 | 0.332544767 | 0.161 | 0.016 | 3.33054311426462e-57 | 14 |
| CABZ01088229.1 | 1.99583337575612e-61 | 0.265724074 | 0.134 | 0.011 | 5.02331302344059e-57 | 14 |

| | | | | | | |
|---|---|---|---|---|---|---|
| si:dkeyp-68b7.122 | 2.48921240824624e-61 | 0.499571271 | 0.315 | 0.058 | 6.26509871031497e-57 | 14 |
| arpc42 | 3.19506147323075e-61 | 0.721558113 | 0.654 | 0.218 | 8.04165022197449e-57 | 14 |
| cers1 | 6.9312548937698e-61 | 0.346743891 | 0.169 | 0.018 | 1.74452754421292e-56 | 14 |
| LOC1039101361 | 3.50004612532756e-60 | 0.403585618 | 0.213 | 0.028 | 8.80926609283693e-56 | 14 |
| bcl2l102 | 5.50824279980549e-60 | 0.749675714 | 0.63 | 0.207 | 1.38636963028304e-55 | 14 |
| csf2rb | 1.08168701290938e-59 | 0.461560059 | 0.24 | 0.036 | 2.72249804279162e-55 | 14 |
| rps6ka11 | 1.22700621370574e-59 | 0.530264501 | 0.343 | 0.069 | 3.08825193927597e-55 | 14 |
| pip5k1ba1 | 2.68995556441044e-59 | 0.29064539 | 0.142 | 0.013 | 6.77034916006464e-55 | 14 |
| dtnbp1a2 | 3.385784112238e-59 | 0.528309027 | 0.378 | 0.081 | 8.52168003209183e-55 | 14 |
| sox5 | 4.12616847615512e-59 | 0.26890075 | 0.126 | 0.01 | 1.03851534376348e-54 | 14 |
| XLOC-0089932 | 1.19336417232539e-58 | 0.406043129 | 0.217 | 0.03 | 3.00357828532577e-54 | 14 |
| zgc:195175 | 1.29143219667787e-58 | 0.407892734 | 0.224 | 0.032 | 3.25040569581852e-54 | 14 |
| pfn14 | 4.23970570846799e-58 | 0.798213511 | 0.984 | 0.516 | 1.06709152976431e-53 | 14 |
| rin2 | 3.90730374528162e-57 | 0.357786797 | 0.173 | 0.02 | 9.83429279649931e-53 | 14 |
| rps216 | 6.72600178967196e-57 | 0.478258209 | 0.984 | 0.746 | 1.69286739044254e-52 | 14 |
| LOC1018823751 | 1.41293255802062e-55 | 0.302741416 | 0.138 | 0.013 | 3.5562099552821e-51 | 14 |
| rpl115 | 1.49800570003578e-55 | 0.407611442 | 0.984 | 0.811 | 3.77033054642004e-51 | 14 |
| rps27.15 | 2.67237361006021e-55 | 0.529358924 | 0.972 | 0.674 | 6.72609713916054e-51 | 14 |
| rpl375 | 1.1886889530612e-54 | 0.376070397 | 0.988 | 0.847 | 2.99181122595974e-50 | 14 |
| msna4 | 7.68238323669241e-54 | 0.697199001 | 0.638 | 0.235 | 1.93357903684311e-49 | 14 |
| arhgdia2 | 1.71119942703324e-53 | 0.67724972 | 0.622 | 0.222 | 4.30691783789997e-49 | 14 |
| actb24 | 1.21247258325341e-52 | 0.417481073 | 1 | 0.866 | 3.05167224479052e-48 | 14 |
| arl111 | 2.63579417274592e-52 | 0.362889291 | 0.193 | 0.026 | 6.63403035338419e-48 | 14 |
| clic12 | 1.1826588009896e-51 | 0.570203191 | 0.409 | 0.108 | 2.97663393621072e-47 | 14 |
| pim23 | 1.66669718920115e-51 | 0.573984967 | 0.354 | 0.085 | 4.19491015550036e-47 | 14 |
| cmah1 | 3.19540872662799e-51 | 0.274174927 | 0.142 | 0.015 | 8.04252422404998e-47 | 14 |
| ldha1 | 1.54274402186756e-50 | 0.485426754 | 0.319 | 0.069 | 3.88293242863846e-46 | 14 |
| megf8l | 5.16614287725622e-50 | 0.263974589 | 0.13 | 0.013 | 1.30026650077662e-45 | 14 |
| arhgef3l2 | 5.28349737947553e-50 | 0.327484855 | 0.197 | 0.029 | 1.3298034554402e-45 | 14 |
| rpl193 | 7.05633416987457e-50 | 0.343689234 | 0.988 | 0.849 | 1.77600874721573e-45 | 14 |
| gapdhs5 | 1.1207059642055e-49 | 0.662540536 | 0.705 | 0.281 | 2.82070484130883e-45 | 14 |
| cebpb4 | 2.65773434572202e-49 | 0.743283636 | 0.811 | 0.345 | 6.68925157474775e-45 | 14 |
| cyfip2 | 9.35243027232629e-49 | 0.3064493 | 0.173 | 0.023 | 2.3539131752418e-44 | 14 |
| rpl274 | 3.30628353728337e-48 | 0.422696669 | 0.98 | 0.759 | 8.32158503498851e-44 | 14 |
| inpp5d | 9.26089553248639e-48 | 0.367014378 | 0.201 | 0.031 | 2.3308747965715e-43 | 14 |

| | | | | | | |
|---|---|---|---|---|---|---|
| si:dkey-262k9.42 | 1.44232677321637e-47 | 0.518195672 | 0.35 | 0.086 | 3.63019225550828e-43 | 14 |
| cd636 | 2.44030773756662e-47 | 0.673545907 | 0.752 | 0.333 | 6.14201054468143e-43 | 14 |
| dub2 | 4.96021320077524e-47 | 0.442851175 | 0.307 | 0.067 | 1.24843606050312e-42 | 14 |
| rps172 | 7.33060128699946e-47 | 0.407761318 | 0.976 | 0.763 | 1.84503903792489e-42 | 14 |
| LOC1001511962 | 8.93167768091139e-47 | 0.376978702 | 0.22 | 0.038 | 2.24801395550859e-42 | 14 |
| ptpn22 | 9.80866118138775e-47 | 0.309435091 | 0.161 | 0.021 | 2.46874193274348e-42 | 14 |
| rps285 | 4.12556989047318e-46 | 0.363130019 | 0.98 | 0.824 | 1.03836468573319e-41 | 14 |
| celf24 | 4.21668647459672e-46 | 0.632263893 | 0.622 | 0.236 | 1.06129781879125e-41 | 14 |
| rpl345 | 5.76395562815993e-46 | 0.411300718 | 0.992 | 0.78 | 1.45072999205157e-41 | 14 |
| CU571074.1 | 2.42206103187095e-45 | 0.330678367 | 0.189 | 0.029 | 6.096085411116e-41 | 14 |
| rps26l5 | 3.45205390825816e-45 | 0.504450348 | 0.961 | 0.659 | 8.68847448169496e-41 | 14 |
| prkcaa | 5.44639349037401e-45 | 0.275972294 | 0.142 | 0.017 | 1.37080277759223e-40 | 14 |
| plxnc11 | 1.3247030613905e-44 | 0.386230816 | 0.224 | 0.041 | 3.33414513521374e-40 | 14 |
| mctp1b1 | 1.74340430902702e-44 | 0.312385855 | 0.165 | 0.023 | 4.38797430539012e-40 | 14 |
| rps155 | 2.07358792503772e-44 | 0.479710421 | 0.961 | 0.68 | 5.21901344852743e-40 | 14 |
| si:ch211-226m16.22 | 2.12182013638835e-44 | 0.466488173 | 0.291 | 0.065 | 5.34040910127583e-40 | 14 |
| tefa2 | 2.3185778229205e-44 | 0.559433816 | 0.437 | 0.133 | 5.8356285225086e-40 | 14 |
| egr31 | 6.64584794012613e-44 | 0.456890612 | 0.264 | 0.055 | 1.67269346805035e-39 | 14 |
| mt-co16 | 7.87922137649073e-44 | 0.555071144 | 0.996 | 0.651 | 1.98312122824895e-39 | 14 |
| rps217 | 1.37409942355883e-43 | 0.390171961 | 0.992 | 0.826 | 3.45847083915523e-39 | 14 |
| myo1f | 1.4852289724216e-43 | 0.398328457 | 0.228 | 0.043 | 3.73817280068793e-39 | 14 |
| arf6b1 | 1.74774906292047e-43 | 0.484427443 | 0.362 | 0.097 | 4.39890961646454e-39 | 14 |
| nfkbiaa2 | 4.66803066388054e-43 | 0.673563255 | 0.453 | 0.147 | 1.17489663779209e-38 | 14 |
| rps15a5 | 6.58574515302282e-43 | 0.420577029 | 0.976 | 0.777 | 1.65756619756431e-38 | 14 |
| skap2 | 8.42536984313744e-43 | 0.433868922 | 0.303 | 0.071 | 2.12058133581926e-38 | 14 |
| pgam1a2 | 9.93038427874376e-43 | 0.558053623 | 0.472 | 0.152 | 2.49937841911702e-38 | 14 |
| BX120005.1 | 3.33322329980612e-42 | 0.442808012 | 0.236 | 0.046 | 8.38938972328202e-38 | 14 |
| lrrfip1a | 3.51117755484338e-42 | 0.399252091 | 0.26 | 0.055 | 8.8372827877853e-38 | 14 |
| rplp2l5 | 4.00330799504394e-42 | 0.415751683 | 0.988 | 0.771 | 1.00759258927261e-37 | 14 |
| kif21b | 1.04125560717609e-41 | 0.304980478 | 0.157 | 0.022 | 2.6207362377015e-37 | 14 |
| syne1b | 1.12017823738307e-41 | 0.300781467 | 0.157 | 0.022 | 2.81937660566945e-37 | 14 |
| mrps152 | 1.36937442240828e-41 | 0.537922328 | 0.409 | 0.12 | 3.4465784837594e-37 | 14 |
| rhoab3 | 1.44737541165428e-41 | 0.580977762 | 0.512 | 0.184 | 3.64289917359266e-37 | 14 |
| cox8a2 | 2.86933885769329e-41 | 0.599343887 | 0.567 | 0.221 | 7.22183897092824e-37 | 14 |
| dusp11 | 7.46273398591039e-41 | 0.599653707 | 0.465 | 0.155 | 1.87829551691379e-36 | 14 |

| | | | | | | |
|---|---|---|---|---|---|---|
| tagln23 | 8.94660236176331e-41 | 0.594388819 | 0.512 | 0.183 | 2.25177034843221e-36 | 14 |
| rps204 | 1.41659186317439e-40 | 0.261789589 | 0.984 | 0.915 | 3.56542006042363e-36 | 14 |
| tln1 | 2.87078919963941e-40 | 0.505544832 | 0.402 | 0.123 | 7.22548933657243e-36 | 14 |
| ankrd44 | 3.6756829152464e-40 | 0.313242431 | 0.169 | 0.027 | 9.25132632938365e-36 | 14 |
| selplg1 | 3.71437475766431e-40 | 0.357260987 | 0.224 | 0.044 | 9.34870982756529e-36 | 14 |
| grap2a2 | 4.16808320700132e-40 | 0.372265746 | 0.224 | 0.044 | 1.04906486237016e-35 | 14 |
| irf2bp2a3 | 4.1830079177312e-40 | 0.570442278 | 0.488 | 0.171 | 1.05282126281377e-35 | 14 |
| si:dkey-171c9.32 | 4.57799963670343e-40 | 0.332375762 | 0.189 | 0.032 | 1.15223672856189e-35 | 14 |
| capzb2 | 4.71503645922075e-40 | 0.594052283 | 0.602 | 0.247 | 1.18672752642127e-35 | 14 |
| nkl.21 | 7.29987858435124e-40 | 0.59353678 | 0.469 | 0.154 | 1.83730644089536e-35 | 14 |
| rpl18a5 | 1.38978375557342e-39 | 0.379026262 | 0.965 | 0.782 | 3.49794673440273e-35 | 14 |
| pnrc26 | 1.72799434041703e-39 | 0.564286074 | 0.949 | 0.561 | 4.34918895539563e-35 | 14 |
| crip14 | 3.04574666639044e-39 | 0.615668515 | 0.654 | 0.264 | 7.6658397846381e-35 | 14 |
| sort1b1 | 3.19549505579408e-39 | 0.327573504 | 0.177 | 0.029 | 8.04274150592811e-35 | 14 |
| rpl355 | 9.85314081492159e-39 | 0.355188217 | 0.98 | 0.802 | 2.47993701170761e-34 | 14 |
| rpl365 | 1.35364113421726e-38 | 0.396697657 | 0.988 | 0.786 | 3.40697937071143e-34 | 14 |
| fli1b2 | 1.88903330345211e-38 | 0.315919686 | 0.185 | 0.032 | 4.75450792145861e-34 | 14 |
| btg14 | 4.32019977012051e-38 | 0.615224823 | 0.713 | 0.326 | 1.08735108014163e-33 | 14 |
| CR855389.11 | 1.54781910016321e-37 | 0.302191202 | 0.165 | 0.027 | 3.89570589320078e-33 | 14 |
| sept23 | 2.78884393672286e-37 | 0.514489768 | 0.465 | 0.16 | 7.01924130433777e-33 | 14 |
| rpl385 | 3.37409474762877e-37 | 0.43139263 | 0.976 | 0.724 | 8.49225907030686e-33 | 14 |
| si:ch73-223p23.22 | 3.89021296210927e-37 | 0.286190858 | 0.177 | 0.03 | 9.79127700433282e-33 | 14 |
| edaradd | 3.95236014585879e-37 | 0.270364849 | 0.15 | 0.022 | 9.94769525111199e-33 | 14 |
| rpl7a4 | 7.28088743885612e-37 | 0.34665304 | 0.984 | 0.807 | 1.8325265594857e-32 | 14 |
| cfl12 | 1.07802658475861e-36 | 0.535208866 | 0.913 | 0.571 | 2.71328511117895e-32 | 14 |
| rpl233 | 2.21034489681867e-36 | 0.270752623 | 0.988 | 0.871 | 5.5632170708029e-32 | 14 |
| gng52 | 2.22632145756509e-36 | 0.573224814 | 0.614 | 0.266 | 5.60342847654558e-32 | 14 |
| si:ch73-264p11.11 | 2.33132295432253e-36 | 0.287998062 | 0.157 | 0.025 | 5.86770674373439e-32 | 14 |
| pik3cg | 6.68529384367518e-36 | 0.270650909 | 0.154 | 0.024 | 1.68262160751461e-31 | 14 |
| si:ch211-196l7.45 | 9.55392421960378e-36 | 0.580516667 | 0.673 | 0.308 | 2.40462718683208e-31 | 14 |
| nr4a13 | 1.06888286539513e-35 | 0.551152056 | 0.469 | 0.162 | 2.69027128391301e-31 | 14 |
| rps64 | 1.15700810085662e-35 | 0.344866765 | 0.953 | 0.786 | 2.91207368904604e-31 | 14 |
| sh2b32 | 1.39652651607284e-35 | 0.42846742 | 0.248 | 0.059 | 3.51491758830373e-31 | 14 |
| rps104 | 1.42639743307026e-35 | 0.439998879 | 0.972 | 0.689 | 3.59009969929455e-31 | 14 |
| si:dkey-185m8.22 | 1.66749175984173e-35 | 0.345049304 | 0.217 | 0.045 | 4.19691001034565e-31 | 14 |

| | | | | | | |
|---|---|---|---|---|---|---|
| rps165 | 1.86158774912086e-35 | 0.41845735 | 0.984 | 0.733 | 4.68543020576229e-31 | 14 |
| prelid1b | 2.09594879384962e-35 | 0.262674542 | 0.142 | 0.021 | 5.2752935192401e-31 | 14 |
| rpl264 | 2.32447558486829e-35 | 0.314777887 | 0.984 | 0.839 | 5.850472599555e-31 | 14 |
| rpl395 | 1.47101066898853e-34 | 0.293812225 | 0.988 | 0.872 | 3.70238675277722e-30 | 14 |
| kctd5a2 | 2.92965875599641e-34 | 0.540668618 | 0.461 | 0.171 | 7.37365812296736e-30 | 14 |
| rpl85 | 3.40723171815714e-34 | 0.362367176 | 0.972 | 0.777 | 8.57566151142971e-30 | 14 |
| serf24 | 7.76118104411308e-34 | 0.544156503 | 0.72 | 0.336 | 1.95341165699282e-29 | 14 |
| rpsa5 | 8.15434483608477e-34 | 0.393214809 | 0.976 | 0.748 | 2.05236705179417e-29 | 14 |
| carhsp1 | 1.08756162316964e-33 | 0.380673664 | 0.268 | 0.067 | 2.73728384935566e-29 | 14 |
| rpl185 | 1.37291180355299e-33 | 0.37372312 | 0.976 | 0.77 | 3.45548171836251e-29 | 14 |
| dock8 | 2.39419142713081e-33 | 0.294604219 | 0.185 | 0.036 | 6.02594040294554e-29 | 14 |
| krt18b3 | 2.8953539163986e-33 | 0.538306112 | 0.425 | 0.146 | 7.28731627218363e-29 | 14 |
| rpl10a4 | 3.27221134028435e-33 | 0.365103683 | 0.969 | 0.761 | 8.23582872236168e-29 | 14 |
| rpl155 | 3.46494274632933e-33 | 0.392244844 | 0.965 | 0.741 | 8.72091439823628e-29 | 14 |
| sdcbp22 | 3.47243855132802e-33 | 0.535991836 | 0.508 | 0.204 | 8.73978058983749e-29 | 14 |
| rps194 | 3.73032323295826e-33 | 0.293279012 | 0.988 | 0.861 | 9.38885054503265e-29 | 14 |
| il161 | 6.05737469105879e-33 | 0.355040897 | 0.213 | 0.047 | 1.52458063599259e-28 | 14 |
| rpl225 | 1.087335265861e-32 | 0.428804688 | 0.957 | 0.69 | 2.73671413064556e-28 | 14 |
| dusp51 | 1.21618634672621e-32 | 0.431654818 | 0.268 | 0.07 | 3.06101941607521e-28 | 14 |
| ankrd12 | 1.41483435458595e-32 | 0.366057563 | 0.264 | 0.068 | 3.56099658705739e-28 | 14 |
| cd831 | 1.53960018945464e-32 | 0.533061303 | 0.335 | 0.1 | 3.87501971683837e-28 | 14 |
| bcar1 | 1.82821531868905e-32 | 0.252427054 | 0.126 | 0.019 | 4.60143513560848e-28 | 14 |
| histh1l2 | 2.52433745063788e-32 | 0.516393715 | 0.339 | 0.106 | 6.35350492951049e-28 | 14 |
| rps184 | 2.95957278337005e-32 | 0.402202287 | 0.961 | 0.721 | 7.44894873846407e-28 | 14 |
| BX248082.12 | 3.05152425292793e-32 | 0.417429816 | 0.193 | 0.04 | 7.6803813921943e-28 | 14 |
| ccdc88b | 3.57671299893366e-32 | 0.260965551 | 0.165 | 0.03 | 9.00222894701614e-28 | 14 |
| actr2a2 | 3.85760523417428e-32 | 0.459653921 | 0.382 | 0.126 | 9.70920661389324e-28 | 14 |
| elovl1b | 4.15099784650472e-32 | 0.451542356 | 0.354 | 0.113 | 1.04476464798677e-27 | 14 |
| grap2b2 | 4.98801230423729e-32 | 0.435445816 | 0.354 | 0.109 | 1.25543281685348e-27 | 14 |
| capza1a2 | 6.08575963377928e-32 | 0.497896061 | 0.496 | 0.195 | 1.53172484222591e-27 | 14 |
| rps74 | 6.24346665821487e-32 | 0.31147112 | 0.961 | 0.818 | 1.5714181232061e-27 | 14 |
| cmtm71 | 6.70136858910659e-32 | 0.330013249 | 0.205 | 0.045 | 1.68666746019224e-27 | 14 |
| unm-sa1614 | 8.88723776507078e-32 | 0.267606625 | 0.126 | 0.019 | 2.23682887309066e-27 | 14 |
| zgc:564933 | 9.8674040616614e-32 | 0.473219991 | 0.862 | 0.568 | 2.48352692827956e-27 | 14 |
| reep5 | 3.18787030633366e-31 | 0.478266506 | 0.421 | 0.154 | 8.02355077401118e-27 | 14 |

| | | | | | | |
|---|---|---|---|---|---|---|
| rps245 | 4.76105461152441e-31 | 0.366893031 | 0.972 | 0.755 | 1.19830983517458e-26 | 14 |
| rps134 | 6.02954767935546e-31 | 0.40625359 | 0.984 | 0.72 | 1.51757685541698e-26 | 14 |
| rps234 | 6.16741981653093e-31 | 0.384453181 | 0.961 | 0.719 | 1.55227789362267e-26 | 14 |
| rps145 | 7.1520769535106e-31 | 0.36739536 | 0.972 | 0.755 | 1.80010624842908e-26 | 14 |
| zgc:1141885 | 1.07367248416357e-30 | 0.482578311 | 0.953 | 0.622 | 2.70232627539129e-26 | 14 |
| CABZ01079663.12 | 3.86917803197336e-30 | 0.293860244 | 0.181 | 0.037 | 9.73833418867374e-26 | 14 |
| vsir3 | 4.19417934247083e-30 | 0.398759835 | 0.287 | 0.082 | 1.05563299870648e-25 | 14 |
| rpl215 | 4.30331886847891e-30 | 0.350679344 | 0.957 | 0.753 | 1.08310232600746e-25 | 14 |
| rps34 | 4.41489206332265e-30 | 0.406789331 | 0.965 | 0.695 | 1.11118418341768e-25 | 14 |
| rpl245 | 4.81839697209512e-30 | 0.351911874 | 0.972 | 0.773 | 1.21274233390662e-25 | 14 |
| rpl5a4 | 5.72185391333605e-30 | 0.522542932 | 0.862 | 0.485 | 1.44013341144755e-25 | 14 |
| si:dkey-78o7.31 | 8.12331448301741e-30 | 0.269129045 | 0.138 | 0.023 | 2.04455702223065e-25 | 14 |
| rpl325 | 8.44304583768673e-30 | 0.336781804 | 0.98 | 0.811 | 2.12503020688737e-25 | 14 |
| ywhabl1 | 8.76777050596159e-30 | 0.504507516 | 0.606 | 0.273 | 2.20676015864547e-25 | 14 |
| rpl104 | 1.16732944758943e-29 | 0.347630511 | 0.972 | 0.769 | 2.93805148663785e-25 | 14 |
| rps95 | 1.97081986867749e-29 | 0.349179342 | 0.976 | 0.773 | 4.96035652747438e-25 | 14 |
| rps55 | 3.19981416140228e-29 | 0.401644905 | 0.949 | 0.697 | 8.0536122628334e-25 | 14 |
| si:dkey-177p2.6 | 6.06566975107593e-29 | 0.363911107 | 0.26 | 0.072 | 1.5266684196483e-24 | 14 |
| pabpc1a3 | 1.10417834420295e-28 | 0.475242193 | 0.862 | 0.534 | 2.7791064745244e-24 | 14 |
| erbin | 1.13776067476505e-28 | 0.337303953 | 0.217 | 0.053 | 2.86362984231614e-24 | 14 |
| lcp12 | 1.18873567354153e-28 | 0.370018324 | 0.291 | 0.085 | 2.99192881673667e-24 | 14 |
| mt-atp65 | 1.46685038535926e-28 | 0.499533672 | 0.949 | 0.584 | 3.69191573491072e-24 | 14 |
| capza1b | 1.61246552443792e-28 | 0.479630509 | 0.5 | 0.208 | 4.0584144784578e-24 | 14 |
| hcst2 | 1.66994059200857e-28 | 0.304371329 | 0.209 | 0.049 | 4.20307347602637e-24 | 14 |
| eno1a3 | 1.94770556959741e-28 | 0.433822878 | 0.374 | 0.131 | 4.90218014811972e-24 | 14 |
| nfkbie2 | 2.41861055860705e-28 | 0.387968141 | 0.299 | 0.091 | 6.08740091495808e-24 | 14 |
| rpl35a5 | 2.68877427199435e-28 | 0.363053366 | 0.976 | 0.756 | 6.76737596518257e-24 | 14 |
| LOC1039091152 | 3.19775795150724e-28 | 0.4487355 | 0.429 | 0.161 | 8.04843698814858e-24 | 14 |
| zgc:1537131 | 3.47771385135233e-28 | 0.449059404 | 0.398 | 0.149 | 8.75305799246868e-24 | 14 |
| psme22 | 3.71360765307631e-28 | 0.393487155 | 0.299 | 0.094 | 9.34677910202775e-24 | 14 |
| bida | 6.65447127407132e-28 | 0.328911204 | 0.252 | 0.068 | 1.67486387497101e-23 | 14 |
| rasal33 | 1.06739325481465e-27 | 0.252636066 | 0.142 | 0.026 | 2.686522083043e-23 | 14 |
| fam49ba2 | 1.99999513245648e-27 | 0.383321109 | 0.319 | 0.1 | 5.03378774887971e-23 | 14 |
| cdc42l2 | 2.07046691091848e-27 | 0.473591006 | 0.5 | 0.211 | 5.21115816809072e-23 | 14 |
| arl6ip5b2 | 2.4367592144194e-27 | 0.394888085 | 0.295 | 0.093 | 6.1330792667722e-23 | 14 |

| | | | | | | |
|---|---|---|---|---|---|---|
| nab1a1 | 2.64387944632097e-27 | 0.275186597 | 0.122 | 0.02 | 6.65438017844524e-23 | 14 |
| si:ch211-105j21.92 | 2.89587278423974e-27 | 0.261801546 | 0.161 | 0.033 | 7.288622210653e-23 | 14 |
| mob1a2 | 3.55041569373625e-27 | 0.428343428 | 0.425 | 0.162 | 8.93604125956477e-23 | 14 |
| rpl144 | 3.72000755601395e-27 | 0.442816338 | 0.969 | 0.629 | 9.36288701773151e-23 | 14 |
| fam13b | 6.04470076456239e-27 | 0.252115692 | 0.154 | 0.031 | 1.52139073543271e-22 | 14 |
| rps124 | 6.43334213684675e-27 | 0.289712233 | 0.984 | 0.838 | 1.61920788242296e-22 | 14 |
| hmgn64 | 1.04823929986391e-26 | 0.490655268 | 0.563 | 0.261 | 2.63831349382747e-22 | 14 |
| ccni3 | 1.22009967170349e-26 | 0.495563924 | 0.602 | 0.289 | 3.07086886371052e-22 | 14 |
| XLOC-005448 | 1.41981656837499e-26 | 0.28480151 | 0.142 | 0.027 | 3.57353632094302e-22 | 14 |
| lpar5a2 | 1.62798945406611e-26 | 0.287643841 | 0.177 | 0.04 | 4.09748665693899e-22 | 14 |
| actn4 | 1.66145024854536e-26 | 0.374148405 | 0.291 | 0.091 | 4.18170413056381e-22 | 14 |
| rpl36a5 | 1.94754804293219e-26 | 0.306719564 | 0.972 | 0.807 | 4.90178366925604e-22 | 14 |
| gnai2a1 | 2.14302520442914e-26 | 0.415257812 | 0.382 | 0.14 | 5.39378013702771e-22 | 14 |
| rpl95 | 2.20843946005308e-26 | 0.454528584 | 0.941 | 0.624 | 5.5584212770076e-22 | 14 |
| actb11 | 2.42865471985912e-26 | 0.324894065 | 0.933 | 0.783 | 6.11268106441342e-22 | 14 |
| glrx2 | 2.83908548507621e-26 | 0.484407923 | 0.638 | 0.306 | 7.14569425738832e-22 | 14 |
| per31 | 3.13833791011313e-26 | 0.355947218 | 0.26 | 0.077 | 7.89888268596373e-22 | 14 |
| rel1 | 5.81309956765825e-26 | 0.291870786 | 0.161 | 0.035 | 1.46309903018391e-21 | 14 |
| zgc:1629442 | 6.01517115405493e-26 | 0.469640787 | 0.547 | 0.248 | 1.51395842776408e-21 | 14 |
| tpi1b2 | 6.44135573047233e-26 | 0.424897406 | 0.417 | 0.163 | 1.62122482380258e-21 | 14 |
| rpl315 | 8.22027387827573e-26 | 0.413959197 | 0.949 | 0.649 | 2.06896073242322e-21 | 14 |
| grasp1 | 9.18177537773776e-26 | 0.347849211 | 0.256 | 0.074 | 2.31096104482282e-21 | 14 |
| sh3kbp11 | 2.62862501113554e-25 | 0.33826173 | 0.24 | 0.069 | 6.61598629052703e-21 | 14 |
| rpl305 | 5.14528821896504e-25 | 0.377406064 | 0.953 | 0.704 | 1.29501759183131e-20 | 14 |
| rpl295 | 7.61452219043593e-25 | 0.319673741 | 0.984 | 0.806 | 1.91649909011082e-20 | 14 |
| slc25a54 | 8.80019527950267e-25 | 0.286844165 | 0.961 | 0.804 | 2.21492114989803e-20 | 14 |
| hsp90ab14 | 1.08643651587339e-24 | 0.308504186 | 0.972 | 0.787 | 2.73445206680174e-20 | 14 |
| eif3f4 | 1.11235952916097e-24 | 0.485844245 | 0.768 | 0.426 | 2.79969769894524e-20 | 14 |
| plekho21 | 1.62355915480085e-24 | 0.346446749 | 0.244 | 0.072 | 4.08633603671825e-20 | 14 |
| iqgap1 | 3.0905303875698e-24 | 0.318770572 | 0.232 | 0.067 | 7.77855593247442e-20 | 14 |
| chmp2a1 | 4.44265682186175e-24 | 0.400847389 | 0.35 | 0.131 | 1.11817229549438e-19 | 14 |
| tmem30aa | 5.06024197256031e-24 | 0.328179815 | 0.213 | 0.059 | 1.2736123020737e-19 | 14 |
| litaf2 | 5.74398045964446e-24 | 0.405637728 | 0.35 | 0.132 | 1.44570244188791e-19 | 14 |
| rpl285 | 7.73558392576103e-24 | 0.345923013 | 0.969 | 0.746 | 1.94696911827479e-19 | 14 |
| arrb2b1 | 9.08931320616441e-24 | 0.344712967 | 0.276 | 0.089 | 2.28768924085952e-19 | 14 |

| | | | | | | |
|---|---|---|---|---|---|---|
| gnai2b2 | 1.03102438229065e-23 | 0.306358783 | 0.228 | 0.065 | 2.59498526778735e-19 | 14 |
| rab8b | 2.45469844670104e-23 | 0.347914991 | 0.248 | 0.077 | 6.17823052050185e-19 | 14 |
| mt-co26 | 3.25853728595689e-23 | 0.436677 | 0.988 | 0.716 | 8.20141249502488e-19 | 14 |
| si:dkey-25e12.31 | 3.93008541734258e-23 | 0.311075385 | 0.189 | 0.049 | 9.89163198690954e-19 | 14 |
| LOC1018844511 | 5.77321347080301e-23 | 0.35057596 | 0.26 | 0.083 | 1.45306009846641e-18 | 14 |
| faua5 | 6.51364096106975e-23 | 0.381380827 | 0.945 | 0.685 | 1.63941829349164e-18 | 14 |
| mcl1a3 | 7.4171146298491e-23 | 0.406320704 | 0.894 | 0.619 | 1.86681358118672e-18 | 14 |
| cmip | 7.9928544978616e-23 | 0.326648795 | 0.217 | 0.063 | 2.01172154856679e-18 | 14 |
| rsl24d14 | 8.75517569761437e-23 | 0.4760799 | 0.783 | 0.438 | 2.20359017133256e-18 | 14 |
| glipr1a2 | 8.76764886153223e-23 | 0.317459495 | 0.217 | 0.062 | 2.20672954195905e-18 | 14 |
| rps114 | 2.44033082792265e-22 | 0.312167148 | 0.961 | 0.763 | 6.14206866079851e-18 | 14 |
| zdhhc18b | 2.87608772871649e-22 | 0.314686218 | 0.244 | 0.073 | 7.23882520440654e-18 | 14 |
| uba524 | 6.74176180864111e-22 | 0.405994346 | 0.929 | 0.625 | 1.69683402961688e-17 | 14 |
| si:ch73-234b20.52 | 7.68479125225354e-22 | 0.346514683 | 0.283 | 0.098 | 1.93418511027969e-17 | 14 |
| mob3a | 1.0666740968523e-21 | 0.360165337 | 0.386 | 0.151 | 2.68471203436755e-17 | 14 |
| rps254 | 1.64032210221971e-21 | 0.340904399 | 0.949 | 0.721 | 4.1285266990768e-17 | 14 |
| c3a.53 | 1.64092307049915e-21 | 0.334883272 | 0.224 | 0.066 | 4.1300392761393e-17 | 14 |
| nfkbiab2 | 1.96801187313557e-21 | 0.441160676 | 0.409 | 0.178 | 4.95328908349492e-17 | 14 |
| tgif11 | 4.30150975751919e-21 | 0.326819347 | 0.264 | 0.089 | 1.08264699087001e-16 | 14 |
| cpne1 | 9.46734368547118e-21 | 0.348907405 | 0.319 | 0.121 | 2.38283573219624e-16 | 14 |
| ccr9a3 | 9.67541889560452e-21 | 0.36839303 | 0.256 | 0.085 | 2.4352061818347e-16 | 14 |
| slc9a3r1a | 1.27970284573283e-20 | 0.252878674 | 0.177 | 0.047 | 3.22088409242495e-16 | 14 |
| rin3 | 1.47370889782427e-20 | 0.25383353 | 0.142 | 0.033 | 3.70917792493392e-16 | 14 |
| fosl23 | 2.80394131436721e-20 | 0.399034618 | 0.307 | 0.119 | 7.05723989413083e-16 | 14 |
| CR383676.16 | 3.04971668362716e-20 | 0.577182505 | 0.988 | 0.721 | 7.6758319210212e-16 | 14 |
| zgc:1528631 | 3.35038180019176e-20 | 0.254408926 | 0.165 | 0.042 | 8.43257595290263e-16 | 14 |
| rps4x4 | 3.45419825291021e-20 | 0.342419713 | 0.933 | 0.707 | 8.69387158274971e-16 | 14 |
| rps3a4 | 9.85319648061197e-20 | 0.290449891 | 0.957 | 0.775 | 2.47995102220523e-15 | 14 |
| cox7b2 | 1.39340732490681e-19 | 0.43027626 | 0.72 | 0.412 | 3.50706689605796e-15 | 14 |
| cox7a2a | 1.67177618958695e-19 | 0.382272196 | 0.508 | 0.238 | 4.20769349157139e-15 | 14 |
| anxa11a1 | 1.78921920301177e-19 | 0.336725851 | 0.28 | 0.104 | 4.50328581206032e-15 | 14 |
| tnfaip32 | 2.18606908224629e-19 | 0.316857769 | 0.205 | 0.063 | 5.50211727310569e-15 | 14 |
| rps8a4 | 3.03702717538832e-19 | 0.279938556 | 0.949 | 0.785 | 7.64389369773487e-15 | 14 |
| rab441 | 3.42737126338934e-19 | 0.301521563 | 0.197 | 0.059 | 8.62635073282462e-15 | 14 |
| cib1 | 4.47292684988486e-19 | 0.271369976 | 0.232 | 0.076 | 1.12579095884752e-14 | 14 |

| | | | | | | |
|---|---|---|---|---|---|---|
| tnrc6c14 | 7.77285788463273e-19 | 0.37230864 | 0.319 | 0.131 | 1.95635060098321e-14 | 14 |
| lbh2 | 8.57749529333266e-19 | 0.341369594 | 0.413 | 0.168 | 2.1588697903789e-14 | 14 |
| oaz1b4 | 9.72098108202688e-19 | 0.426633214 | 0.575 | 0.319 | 2.44667372853535e-14 | 14 |
| zgc:63831 | 1.51492309475582e-18 | 0.312096474 | 0.232 | 0.079 | 3.81290993719092e-14 | 14 |
| rpl74 | 1.69993600715576e-18 | 0.258005028 | 0.945 | 0.803 | 4.27856893641034e-14 | 14 |
| mef2aa1 | 2.00930894150259e-18 | 0.312397916 | 0.209 | 0.067 | 5.05722967486788e-14 | 14 |
| scamp2 | 2.51371434807893e-18 | 0.262584156 | 0.213 | 0.068 | 6.32676764267986e-14 | 14 |
| kdm6bb1 | 2.72438890258276e-18 | 0.265847952 | 0.173 | 0.05 | 6.85701442891054e-14 | 14 |
| arpc5b1 | 3.08694595556871e-18 | 0.32823533 | 0.291 | 0.111 | 7.76953427557089e-14 | 14 |
| COX36 | 3.53379602384644e-18 | 0.400212335 | 0.976 | 0.695 | 8.8942112124191e-14 | 14 |
| zmynd11 | 4.983550507123e-18 | 0.292514221 | 0.189 | 0.057 | 1.25430982713779e-13 | 14 |
| sh3bgrl31 | 5.442047515857e-18 | 0.337747116 | 0.346 | 0.145 | 1.36970893926605e-13 | 14 |
| arpc4l3 | 6.61110428905977e-18 | 0.36001446 | 0.421 | 0.194 | 1.66394883851345e-13 | 14 |
| sh3bgrl2 | 7.91231967570785e-18 | 0.336469648 | 0.268 | 0.102 | 1.99145173917891e-13 | 14 |
| rps264 | 9.21239749731065e-18 | 0.302089651 | 0.929 | 0.731 | 2.31866832609812e-13 | 14 |
| zgc:1627305 | 9.71134300181223e-18 | 0.419520013 | 0.768 | 0.385 | 2.44424792012612e-13 | 14 |
| si:dkey-112e7.23 | 1.0454285065524e-17 | 0.378693062 | 0.425 | 0.203 | 2.63123900814174e-13 | 14 |
| ndufa4l2 | 1.07445094645063e-17 | 0.40683497 | 0.705 | 0.39 | 2.70428558712158e-13 | 14 |
| nr4a31 | 1.08074900098744e-17 | 0.297535755 | 0.177 | 0.053 | 2.72013716058529e-13 | 14 |
| pde4ba3 | 1.35960736452208e-17 | 0.377835569 | 0.327 | 0.137 | 3.42199577576561e-13 | 14 |
| lmnb2 | 1.37040979137649e-17 | 0.256790893 | 0.189 | 0.059 | 3.44918440391548e-13 | 14 |
| ywhaba1 | 1.40675098449381e-17 | 0.402963562 | 0.583 | 0.317 | 3.54065155287246e-13 | 14 |
| gnb1a1 | 2.24522343232378e-17 | 0.369063092 | 0.402 | 0.19 | 5.65100285681573e-13 | 14 |
| mtpn1 | 3.56467559569221e-17 | 0.336106298 | 0.335 | 0.144 | 8.97193200679773e-13 | 14 |
| ier2b4 | 4.15770357087773e-17 | 0.405971883 | 0.445 | 0.221 | 1.04645241175422e-12 | 14 |
| mt-cyb4 | 5.10577634677717e-17 | 0.386970943 | 0.913 | 0.619 | 1.28507284872034e-12 | 14 |
| sub1a | 7.27325798739553e-17 | 0.359004089 | 0.472 | 0.229 | 1.83060630284758e-12 | 14 |
| cry1aa | 9.47372804398212e-17 | 0.260993107 | 0.181 | 0.057 | 2.38444261138986e-12 | 14 |
| slc25a55a | 1.51924809019362e-16 | 0.264829961 | 0.197 | 0.065 | 3.82379551820833e-12 | 14 |
| zgc:1538671 | 1.84829881257955e-16 | 0.394167509 | 0.717 | 0.432 | 4.65198328138147e-12 | 14 |
| csk2 | 2.21423304325563e-16 | 0.289720681 | 0.264 | 0.099 | 5.57300314657009e-12 | 14 |
| rnf443 | 2.24020428922566e-16 | 0.303927666 | 0.236 | 0.088 | 5.63837017555207e-12 | 14 |
| abracl | 2.8114031154657e-16 | 0.356939999 | 0.461 | 0.218 | 7.07602050131563e-12 | 14 |
| ccl39.7 | 3.59770445763554e-16 | 0.344084723 | 0.181 | 0.058 | 9.0550623494229e-12 | 14 |
| gnb1b1 | 4.15174283445551e-16 | 0.361940986 | 0.488 | 0.251 | 1.04495215400411e-11 | 14 |

| | | | | | | |
|---|---|---|---|---|---|---|
| uqcrh1 | 5.97109425883677e-16 | 0.350055487 | 0.484 | 0.247 | 1.50286471400663e-11 | 14 |
| pkma1 | 5.97260295025368e-16 | 0.348873157 | 0.378 | 0.175 | 1.50324443654935e-11 | 14 |
| naca5 | 7.47863452871839e-16 | 0.367500889 | 0.894 | 0.614 | 1.88229752453313e-11 | 14 |
| hmgb1a4 | 8.70959170401895e-16 | 0.342965398 | 0.39 | 0.186 | 2.19211713598453e-11 | 14 |
| ppiab1 | 1.05758102413413e-15 | 0.326685863 | 0.89 | 0.653 | 2.6618256796432e-11 | 14 |
| tpt14 | 1.14160640879746e-15 | 0.256962099 | 0.969 | 0.798 | 2.87330917030232e-11 | 14 |
| si:ch211-132p1.33 | 1.22435108230851e-15 | 0.287754258 | 0.232 | 0.087 | 3.0815692390623e-11 | 14 |
| GCA2 | 1.48616147757233e-15 | 0.322109596 | 0.382 | 0.178 | 3.74051982290179e-11 | 14 |
| sept7b | 1.77183981008872e-15 | 0.260994919 | 0.26 | 0.101 | 4.45954361801229e-11 | 14 |
| atp5f1e1 | 3.25448889308579e-15 | 0.376621718 | 0.787 | 0.478 | 8.19122309500762e-11 | 14 |
| atp5pf1 | 3.26005608327943e-15 | 0.351508215 | 0.551 | 0.295 | 8.20523515600601e-11 | 14 |
| arf6a | 3.29137665586183e-15 | 0.309432367 | 0.362 | 0.161 | 8.28406590513864e-11 | 14 |
| jak1 | 3.35071419537281e-15 | 0.304122487 | 0.24 | 0.094 | 8.43341255833382e-11 | 14 |
| h2afx15 | 3.48404555236889e-15 | 0.356600248 | 0.402 | 0.196 | 8.76899425075727e-11 | 14 |
| ppiaa4 | 4.12373411881626e-15 | 0.252954579 | 0.961 | 0.801 | 1.03790264036486e-10 | 14 |
| spcs2 | 5.97745457241781e-15 | 0.304819986 | 0.35 | 0.158 | 1.50446554133184e-10 | 14 |
| mrpl33 | 6.53761805482356e-15 | 0.262578101 | 0.248 | 0.096 | 1.64545308821854e-10 | 14 |
| cox4i12 | 7.17541162862579e-15 | 0.367513986 | 0.665 | 0.388 | 1.80597935280883e-10 | 14 |
| rps27a4 | 8.08689985310018e-15 | 0.290591037 | 0.953 | 0.748 | 2.03539182402678e-10 | 14 |
| gadd45bb2 | 1.03786029588905e-14 | 0.307035336 | 0.224 | 0.086 | 2.61219057872315e-10 | 14 |
| ap1s1 | 1.51390110625489e-14 | 0.252105979 | 0.248 | 0.096 | 3.81033769433294e-10 | 14 |
| rpl27a5 | 1.57423599949478e-14 | 0.395706092 | 0.791 | 0.491 | 3.96219458712842e-10 | 14 |
| rpl23a5 | 2.24339010391025e-14 | 0.356151618 | 0.906 | 0.63 | 5.64638855253171e-10 | 14 |
| cox6a1 | 2.76723480777897e-14 | 0.355885245 | 0.654 | 0.369 | 6.9648532876989e-10 | 14 |
| ywhabb | 3.67430040979243e-14 | 0.315730466 | 0.441 | 0.224 | 9.24784670140658e-10 | 14 |
| atp5pd1 | 4.56088540683419e-14 | 0.324881245 | 0.492 | 0.259 | 1.1479292480461e-09 | 14 |
| ip6k2b2 | 6.19523490670448e-14 | 0.296401871 | 0.28 | 0.121 | 1.55927867366845e-09 | 14 |
| rpl13a4 | 6.78813548669735e-14 | 0.267891706 | 0.957 | 0.761 | 1.70850582064686e-09 | 14 |
| mvb12a | 6.90879344443054e-14 | 0.25471078 | 0.205 | 0.076 | 1.73887422202872e-09 | 14 |
| elmo12 | 7.67025618744281e-14 | 0.254925206 | 0.193 | 0.069 | 1.93052677981748e-09 | 14 |
| brk1 | 7.93377341525083e-14 | 0.295531938 | 0.374 | 0.179 | 1.99685143088448e-09 | 14 |
| twf1a | 8.1772863496927e-14 | 0.281111481 | 0.331 | 0.149 | 2.05814120135416e-09 | 14 |
| ptpn63 | 1.32520023824296e-13 | 0.276736993 | 0.228 | 0.09 | 3.33539647963371e-09 | 14 |
| edf1 | 1.56894444676194e-13 | 0.344473038 | 0.48 | 0.266 | 3.94887627805513e-09 | 14 |
| rps27.24 | 1.6899002517482e-13 | 0.377125584 | 0.823 | 0.513 | 4.25330994362505e-09 | 14 |

| | | | | | | |
|---|---|---|---|---|---|---|
| NC-002333.45 | 1.89280561881096e-13 | 0.351627841 | 0.929 | 0.617 | 4.7640024619853e-09 | 14 |
| psme12 | 2.05581569535978e-13 | 0.265316173 | 0.22 | 0.089 | 5.17428252365104e-09 | 14 |
| ucp22 | 2.14568080663653e-13 | 0.364143082 | 0.661 | 0.413 | 5.40046402222348e-09 | 14 |
| COX5B | 2.6772274613845e-13 | 0.340272267 | 0.61 | 0.354 | 6.73831379755864e-09 | 14 |
| tpm11 | 2.94363195546467e-13 | 0.269735713 | 0.429 | 0.204 | 7.40882726870902e-09 | 14 |
| rhoac | 3.22097173207606e-13 | 0.28611478 | 0.327 | 0.154 | 8.10686375246223e-09 | 14 |
| hmgn72 | 3.29915624953004e-13 | 0.329017019 | 0.508 | 0.276 | 8.30364636444215e-09 | 14 |
| hnrnpa0b3 | 4.07607958997267e-13 | 0.338582328 | 0.547 | 0.309 | 1.02590847200022e-08 | 14 |
| cdkn1bb2 | 7.10519018057214e-13 | 0.303929719 | 0.311 | 0.147 | 1.7883053165482e-08 | 14 |
| itgb21 | 7.89323445665985e-13 | 0.277253864 | 0.236 | 0.098 | 1.98664818039672e-08 | 14 |
| notch23 | 8.23882313664448e-13 | 0.268708541 | 0.268 | 0.118 | 2.07362939526205e-08 | 14 |
| rack14 | 8.47092667258553e-13 | 0.278297177 | 0.949 | 0.741 | 2.13204753422305e-08 | 14 |
| sumo2b4 | 9.64272926549823e-13 | 0.332920023 | 0.472 | 0.267 | 2.42697852883325e-08 | 14 |
| mibp25 | 1.28446215001945e-12 | 0.354130816 | 0.78 | 0.488 | 3.23286278538394e-08 | 14 |
| NDUFB12 | 1.55656089570998e-12 | 0.301133479 | 0.378 | 0.196 | 3.91770811841246e-08 | 14 |
| b2m4 | 2.170800504404e-12 | 0.332519988 | 0.516 | 0.289 | 5.46368778953443e-08 | 14 |
| atp5f1d1 | 2.18383060579383e-12 | 0.314445103 | 0.465 | 0.256 | 5.4964832517225e-08 | 14 |
| ppt11 | 2.78932543256171e-12 | 0.313978192 | 0.327 | 0.162 | 7.02045318121458e-08 | 14 |
| cox7c1 | 3.24913966050265e-12 | 0.331564532 | 0.717 | 0.478 | 8.17775961151911e-08 | 14 |
| eif4ebp22 | 3.41855569446708e-12 | 0.28659909 | 0.346 | 0.173 | 8.6041628274042e-08 | 14 |
| selenof | 4.758999439245e-12 | 0.299514798 | 0.437 | 0.234 | 1.19779256886357e-07 | 14 |
| cxcr4b3 | 5.17883879111686e-12 | 0.299844553 | 0.374 | 0.181 | 1.3034619353362e-07 | 14 |
| mt-nd44 | 6.51775936008846e-12 | 0.334165688 | 0.728 | 0.452 | 1.64045485334066e-07 | 14 |
| btg22 | 6.76687267465821e-12 | 0.334522407 | 0.634 | 0.369 | 1.70315418348472e-07 | 14 |
| rhoaa3 | 8.81771534889373e-12 | 0.286320226 | 0.307 | 0.152 | 2.21933077616306e-07 | 14 |
| hmgb2a1 | 9.31759264771098e-12 | 0.338844662 | 0.815 | 0.619 | 2.34514489350238e-07 | 14 |
| atp5meb | 9.51895587610153e-12 | 0.306794257 | 0.547 | 0.319 | 2.39582600445599e-07 | 14 |
| mt-nd14 | 1.16795635987561e-11 | 0.310986551 | 0.685 | 0.407 | 2.93962936217093e-07 | 14 |
| serbp1a4 | 1.23773505501175e-11 | 0.334859189 | 0.732 | 0.478 | 3.11525535995907e-07 | 14 |
| si:ch1073-325m22.2 | 1.58883355313155e-11 | 0.296980609 | 0.413 | 0.228 | 3.9989351698768e-07 | 14 |
| ube2v2 | 2.21366891296143e-11 | 0.300291611 | 0.472 | 0.267 | 5.57158328703261e-07 | 14 |
| txnipa4 | 2.22292480121946e-11 | 0.320469225 | 0.386 | 0.209 | 5.59487943218927e-07 | 14 |
| romo14 | 2.74992717813222e-11 | 0.321865733 | 0.626 | 0.383 | 6.92129171464098e-07 | 14 |
| tomm20b3 | 3.1460098971921e-11 | 0.293714496 | 0.39 | 0.213 | 7.9181923102428e-07 | 14 |
| pomp | 3.7221959300519e-11 | 0.262005381 | 0.429 | 0.228 | 9.36839493634762e-07 | 14 |

| | | | | | | |
|---|---|---|---|---|---|---|
| atrx2 | 5.24176112726063e-11 | 0.273328518 | 0.228 | 0.104 | 1.31929885812023e-06 | 14 |
| rpl65 | 6.16423730266073e-11 | 0.316692865 | 0.925 | 0.663 | 1.55147688670668e-06 | 14 |
| uqcrq | 8.03432616481458e-11 | 0.28593377 | 0.476 | 0.269 | 2.02215955242218e-06 | 14 |
| tmed7 | 8.74450346121959e-11 | 0.271322305 | 0.382 | 0.205 | 2.20090407615436e-06 | 14 |
| sri | 1.46202299916349e-10 | 0.257600334 | 0.299 | 0.149 | 3.67976568659458e-06 | 14 |
| aldoaa | 1.69038799038012e-10 | 0.280859018 | 0.445 | 0.248 | 4.25453753298773e-06 | 14 |
| eif3i4 | 1.8514284524749e-10 | 0.279842646 | 0.472 | 0.271 | 4.65986027203408e-06 | 14 |
| ywhae1 | 2.15350264645239e-10 | 0.277332369 | 0.492 | 0.28 | 5.42015081085601e-06 | 14 |
| hnrnpa1a1 | 3.30639103047597e-10 | 0.253622234 | 0.311 | 0.158 | 8.32185558460498e-06 | 14 |
| egr13 | 4.50174828522839e-10 | 0.318422852 | 0.346 | 0.189 | 1.13304502590913e-05 | 14 |
| b2ml5 | 6.84562980411784e-10 | 0.277000759 | 0.461 | 0.265 | 1.72297656539842e-05 | 14 |
| atp5l2 | 8.15223264121167e-10 | 0.305099273 | 0.673 | 0.446 | 2.05183543346656e-05 | 14 |
| fkbp1aa2 | 8.19651304740512e-10 | 0.296402794 | 0.614 | 0.385 | 2.0629803689014e-05 | 14 |
| rap1b1 | 1.26640216697564e-09 | 0.254563851 | 0.366 | 0.201 | 3.18740761406098e-05 | 14 |
| rpl44 | 1.38618101997943e-09 | 0.298018181 | 0.874 | 0.624 | 3.48887900918624e-05 | 14 |
| btf34 | 1.64856778528712e-09 | 0.31581472 | 0.858 | 0.583 | 4.14928025878914e-05 | 14 |
| tegt2 | 1.77393257905354e-09 | 0.252812524 | 0.374 | 0.21 | 4.46481090821986e-05 | 14 |
| rpl5b4 | 2.00051866424544e-09 | 0.301576029 | 0.882 | 0.626 | 5.03510542603936e-05 | 14 |
| xbp11 | 2.1798677127476e-09 | 0.294921208 | 0.555 | 0.341 | 5.48650904621444e-05 | 14 |
| eif3ea4 | 3.04104591796856e-09 | 0.266136948 | 0.461 | 0.275 | 7.65400847093506e-05 | 14 |
| nsa24 | 3.13908277839387e-09 | 0.291936055 | 0.579 | 0.36 | 7.90075744493954e-05 | 14 |
| mt-nd34 | 4.21095867278555e-09 | 0.275561177 | 0.618 | 0.393 | 0.000105986 | 14 |
| eef1g4 | 5.16814698228635e-09 | 0.305793141 | 0.882 | 0.616 | 0.000130077 | 14 |
| anp32a | 5.76300037337941e-09 | 0.255016722 | 0.311 | 0.168 | 0.000145049 | 14 |
| calm3b | 6.40171791640423e-09 | 0.263509399 | 0.433 | 0.255 | 0.000161125 | 14 |
| ybx13 | 7.93068447434643e-09 | 0.281204876 | 0.913 | 0.666 | 0.000199607 | 14 |
| chac11 | 1.07684333706346e-08 | 0.276399939 | 0.622 | 0.396 | 0.000271031 | 14 |
| vdac21 | 1.28964451827838e-08 | 0.26240824 | 0.476 | 0.293 | 0.000324591 | 14 |
| cox5aa | 3.7009521160637e-08 | 0.260445644 | 0.551 | 0.35 | 0.000931493 | 14 |
| slc25a3b3 | 2.66113284279367e-07 | 0.256807443 | 0.657 | 0.434 | 0.006697805 | 14 |
| ctrb1 | 5.20598821582452e-92 | 0.510595318 | 0.2 | 0.016 | 1.31029517404087e-87 | 15 |
| cldnd7 | 4.49101247954632e-55 | 0.704515092 | 0.996 | 0.599 | 1.13034293097701e-50 | 15 |
| hbba16 | 6.19874509622108e-51 | 0.803669399 | 0.8 | 0.349 | 1.56016215326788e-46 | 15 |
| zgc:1751357 | 6.60781380733926e-51 | 0.738121717 | 0.91 | 0.473 | 1.66312065716922e-46 | 15 |
| mt26 | 7.655971308039e-51 | 0.711768318 | 0.951 | 0.528 | 1.92693141852034e-46 | 15 |

| | | | | | | |
|---|---|---|---|---|---|---|
| h2af1al7 | 8.46051837114197e-50 | 0.681805298 | 0.971 | 0.587 | 2.12942786883272e-45 | 15 |
| h1m7 | 5.2629505780532e-49 | 0.664706061 | 0.976 | 0.596 | 1.32463203099021e-44 | 15 |
| rrm26 | 8.99146446080814e-48 | 0.693066045 | 0.894 | 0.493 | 2.2630616901408e-43 | 15 |
| si:dkey-241l7.46 | 2.13096757834219e-47 | 0.639401287 | 0.955 | 0.604 | 5.36343229792947e-43 | 15 |
| si:dkey-240e12.66 | 1.41480770551903e-46 | 0.700798599 | 0.976 | 0.563 | 3.56092951402084e-42 | 15 |
| zp37 | 2.27161336329783e-46 | 0.67105197 | 0.971 | 0.579 | 5.71742367408431e-42 | 15 |
| CR790368.17 | 2.3474785610387e-46 | 0.682476836 | 0.91 | 0.515 | 5.90836879027831e-42 | 15 |
| zp2.57 | 1.36181049040318e-45 | 0.680077233 | 0.951 | 0.558 | 3.42754082329576e-41 | 15 |
| prss59.21 | 5.79098521990612e-45 | 0.378000561 | 0.143 | 0.017 | 1.45753306999817e-40 | 15 |
| si:zfos-1505d6.36 | 1.16620681514364e-44 | 0.696175386 | 0.927 | 0.507 | 2.93522593303504e-40 | 15 |
| zp2.36 | 1.74140312800837e-44 | 0.695224532 | 0.927 | 0.513 | 4.38293753288427e-40 | 15 |
| acp5a7 | 2.48158523706632e-43 | 0.664419171 | 0.922 | 0.535 | 6.24590188317223e-39 | 15 |
| si:dkeyp-98a7.96 | 4.63451161759393e-43 | 0.684952273 | 0.914 | 0.506 | 1.16646022903222e-38 | 15 |
| zp2.26 | 2.14902767669925e-42 | 0.663663338 | 0.91 | 0.524 | 5.40888775948435e-38 | 15 |
| si:dkey-241l7.57 | 3.38815122144366e-42 | 0.638775337 | 0.943 | 0.579 | 8.52763780925154e-38 | 15 |
| si:dkey-241l7.26 | 2.9428192703369e-41 | 0.631800463 | 0.992 | 0.636 | 7.40678182151095e-37 | 15 |
| zp2.17 | 8.00679185080288e-41 | 0.65688656 | 0.91 | 0.53 | 2.01522944092858e-36 | 15 |
| hbaa16 | 1.08329072486384e-40 | 0.743089509 | 0.645 | 0.276 | 2.72653442540979e-36 | 15 |
| zp2.66 | 1.58565215876374e-40 | 0.677177923 | 0.898 | 0.496 | 3.99092791839246e-36 | 15 |
| si:ch211-226h8.146 | 1.36265090941274e-39 | 0.643900519 | 0.943 | 0.561 | 3.42965607390092e-35 | 15 |
| ccna16 | 1.5007002135309e-39 | 0.662243132 | 0.853 | 0.471 | 3.77711236743592e-35 | 15 |
| crp26 | 2.60987121359589e-39 | 0.670699849 | 0.886 | 0.483 | 6.5687848574995e-35 | 15 |
| si:ch211-5k11.86 | 2.88011796160404e-39 | 0.684482387 | 0.739 | 0.353 | 7.2489688975612e-35 | 15 |
| si:dkey-229d11.36 | 3.26692861282856e-39 | 0.693498228 | 0.845 | 0.433 | 8.2225326256282e-35 | 15 |
| si:ch73-189n23.17 | 3.39915298472955e-39 | 0.656702782 | 0.906 | 0.519 | 8.55532814726581e-35 | 15 |
| prss11 | 4.77963081963523e-39 | 0.424587064 | 0.151 | 0.022 | 1.20298528099399e-34 | 15 |
| si:ch211-125e6.126 | 2.56069549383748e-38 | 0.656855901 | 0.878 | 0.496 | 6.44501448843954e-34 | 15 |
| zgc:1717766 | 2.74716257921717e-38 | 0.648304072 | 0.857 | 0.491 | 6.91433349563168e-34 | 15 |
| zgc:771186 | 6.76738292505448e-38 | 0.65888303 | 0.829 | 0.452 | 1.70328260840696e-33 | 15 |
| zgc:1534996 | 1.37305814987532e-37 | 0.671833571 | 0.927 | 0.507 | 3.4558500574212e-33 | 15 |
| tmsb56 | 3.20381849868548e-37 | 0.664993182 | 0.898 | 0.483 | 8.06369077934148e-33 | 15 |
| tuba4l6 | 6.51063866116145e-37 | 0.659353238 | 0.735 | 0.4 | 1.63866264462773e-32 | 15 |
| zar16 | 7.57000564279639e-36 | 0.648220123 | 0.788 | 0.435 | 1.90529472023542e-31 | 15 |
| BX649337.17 | 1.14289584211592e-35 | 0.653315918 | 0.812 | 0.433 | 2.87655454502156e-31 | 15 |
| cldng6 | 1.88619445424926e-35 | 0.637608018 | 0.894 | 0.509 | 4.74736282189997e-31 | 15 |

| | | | | | | |
|---|---|---|---|---|---|---|
| ela2l | 3.57609401073382e-35 | 0.274222551 | 0.102 | 0.011 | 9.00067101561595e-31 | 15 |
| zgc:1717796 | 9.8938794288559e-35 | 0.628207155 | 0.878 | 0.507 | 2.49019051344874e-30 | 15 |
| si:ch211-12h2.66 | 1.51140167264126e-34 | 0.621930923 | 0.808 | 0.47 | 3.80404686987078e-30 | 15 |
| si:ch211-12h2.86 | 1.99372392004618e-34 | 0.594218862 | 0.865 | 0.544 | 5.01800373436423e-30 | 15 |
| si:dkeyp-98a7.86 | 4.42610786266314e-34 | 0.632496169 | 0.808 | 0.453 | 1.11400708795369e-29 | 15 |
| cth16 | 1.2051365207809e-32 | 0.629135355 | 0.706 | 0.375 | 3.03320810915344e-28 | 15 |
| epcam7 | 1.68881291739389e-32 | 0.602820147 | 0.849 | 0.501 | 4.25057323178868e-28 | 15 |
| si:ch211-226h8.46 | 6.60419833721359e-32 | 0.622547966 | 0.788 | 0.437 | 1.66221067949329e-27 | 15 |
| si:dkeyp-50f7.26 | 5.40384702449475e-31 | 0.613535026 | 0.727 | 0.398 | 1.36009425759508e-26 | 15 |
| zp3a.26 | 8.82423368367728e-31 | 0.58775998 | 0.796 | 0.488 | 2.22097137584474e-26 | 15 |
| zp3b6 | 1.35034306589662e-30 | 0.624296106 | 0.698 | 0.384 | 3.39867846255521e-26 | 15 |
| mid1ip1l6 | 1.55094631061394e-30 | 0.551862852 | 0.82 | 0.54 | 3.90357676918423e-26 | 15 |
| hnrnpabb6 | 2.44348700030316e-30 | 0.506012788 | 0.816 | 0.594 | 6.15001243106301e-26 | 15 |
| lamp26 | 2.22422541778238e-29 | 0.543391766 | 0.816 | 0.551 | 5.59815295401648e-25 | 15 |
| gapdh6 | 2.50589911146694e-29 | 0.595355727 | 0.727 | 0.416 | 6.30709747365115e-25 | 15 |
| ccnb16 | 2.97130061152465e-28 | 0.594055848 | 0.682 | 0.39 | 7.47846650914639e-24 | 15 |
| siva16 | 3.68653806182352e-28 | 0.58206421 | 0.731 | 0.425 | 9.27864764780361e-24 | 15 |
| si:dkeyp-98a7.46 | 7.54201418422356e-28 | 0.588102853 | 0.751 | 0.421 | 1.89824955002723e-23 | 15 |
| zp3.27 | 2.25472685808275e-27 | 0.557729295 | 0.878 | 0.554 | 5.67492202910848e-23 | 15 |
| si:dkey-241l7.36 | 3.52023322391022e-27 | 0.612875739 | 0.629 | 0.362 | 8.86007500125963e-23 | 15 |
| si:dkey-241l7.66 | 5.80289132481744e-27 | 0.543904061 | 0.816 | 0.524 | 1.4605297175433e-22 | 15 |
| slbp26 | 7.98839782042635e-27 | 0.579523607 | 0.686 | 0.394 | 2.01059984742311e-22 | 15 |
| btg46 | 2.75656478550104e-26 | 0.584178072 | 0.657 | 0.366 | 6.93799790862757e-22 | 15 |
| zgc:1952456 | 2.93150839497746e-26 | 0.575190273 | 0.694 | 0.385 | 7.37831347931877e-22 | 15 |
| si:dkey-90l23.16 | 3.81737344480123e-26 | 0.572827116 | 0.767 | 0.444 | 9.60794722322021e-22 | 15 |
| s100a16 | 1.08676642011576e-25 | 0.579442305 | 0.649 | 0.361 | 2.73528240278935e-21 | 15 |
| CU571163.16 | 1.76703003262457e-24 | 0.56759481 | 0.698 | 0.398 | 4.44743788911278e-20 | 15 |
| si:dkeyp-46h3.66 | 9.04576618604476e-24 | 0.554737057 | 0.69 | 0.408 | 2.27672889136561e-19 | 15 |
| wu:fi42e036 | 1.81601480687441e-23 | 0.564857294 | 0.604 | 0.347 | 4.57072766742221e-19 | 15 |
| si:dkeyp-46h3.86 | 2.70160496614124e-23 | 0.56247565 | 0.629 | 0.349 | 6.7996695392809e-19 | 15 |
| zpcx6 | 3.17854732872675e-23 | 0.561177483 | 0.608 | 0.336 | 8.00008577167235e-19 | 15 |
| si:ch211-14a17.76 | 1.01915504271947e-22 | 0.542252743 | 0.686 | 0.405 | 2.56511132702064e-18 | 15 |
| zp3a.16 | 1.60835468656199e-22 | 0.550288994 | 0.62 | 0.379 | 4.04806791060788e-18 | 15 |
| zar1l6 | 5.57203545646514e-22 | 0.537125274 | 0.678 | 0.395 | 1.40242560403771e-17 | 15 |
| npm2b6 | 7.02021880529459e-22 | 0.538175539 | 0.673 | 0.393 | 1.7669188711046e-17 | 15 |

| | | | | | | |
|---|---|---|---|---|---|---|
| ccnb26 | 7.41328482604934e-22 | 0.540173177 | 0.661 | 0.381 | 1.86584965786836e-17 | 15 |
| si:dkey-152b24.66 | 1.41020232908397e-21 | 0.5463577 | 0.551 | 0.295 | 3.54933824207143e-17 | 15 |
| qdprb26 | 1.52423572121741e-21 | 0.530522077 | 0.71 | 0.43 | 3.83634888673209e-17 | 15 |
| si:dkey-19b23.116 | 8.57420046257645e-21 | 0.555317581 | 0.518 | 0.276 | 2.15804051442587e-16 | 15 |
| birc5b6 | 1.74247740994007e-20 | 0.541686141 | 0.494 | 0.257 | 4.38564139307816e-16 | 15 |
| si:ch211-132b12.86 | 1.77525876817059e-20 | 0.523641993 | 0.584 | 0.34 | 4.46814879360855e-16 | 15 |
| nanog6 | 1.87689998009912e-20 | 0.522638651 | 0.616 | 0.357 | 4.72396955991148e-16 | 15 |
| si:ch211-240l19.8 | 1.94078267678859e-20 | 0.276237401 | 0.106 | 0.02 | 4.8847559192092e-16 | 15 |
| zgc:1715176 | 5.81656603435118e-20 | 0.520117122 | 0.649 | 0.386 | 1.46397150518585e-15 | 15 |
| ca15b6 | 9.44908685305581e-20 | 0.540274319 | 0.473 | 0.247 | 2.37824067004562e-15 | 15 |
| gmnn6 | 1.08403109916878e-19 | 0.522136745 | 0.543 | 0.301 | 2.72839787349791e-15 | 15 |
| spink2.17 | 2.0273911922676e-19 | 0.481619019 | 0.669 | 0.434 | 5.10274089181832e-15 | 15 |
| zgc:1362546 | 2.35089353392e-19 | 0.5147047 | 0.584 | 0.328 | 5.91696393552326e-15 | 15 |
| lsm14b6 | 5.21162121152744e-19 | 0.523867243 | 0.527 | 0.298 | 1.31171294272934e-14 | 15 |
| kpna76 | 9.84626287900102e-19 | 0.512882966 | 0.571 | 0.326 | 2.47820590401577e-14 | 15 |
| zgc:1734436 | 1.35276114183123e-18 | 0.51050525 | 0.58 | 0.328 | 3.40476451787502e-14 | 15 |
| gtf3ab7 | 1.385617505066e-18 | 0.507100366 | 0.551 | 0.318 | 3.48746069850061e-14 | 15 |
| si:ch73-90k17.16 | 2.04681997703203e-18 | 0.518274908 | 0.469 | 0.247 | 5.15164120019192e-14 | 15 |
| LOC1104378216 | 2.59528081159655e-18 | 0.501739774 | 0.669 | 0.413 | 6.53206227470735e-14 | 15 |
| hnrnpa0l | 3.43374907586539e-18 | 0.265344782 | 0.837 | 0.757 | 8.64240304904559e-14 | 15 |
| zgc:554616 | 1.6188530792517e-17 | 0.501222071 | 0.514 | 0.286 | 4.07449131516859e-13 | 15 |
| si:dkeyp-20e4.86 | 9.14403122170845e-17 | 0.48473181 | 0.653 | 0.406 | 2.3014612181918e-12 | 15 |
| thy16 | 1.87870511842967e-16 | 0.450227892 | 0.661 | 0.449 | 4.72851291257564e-12 | 15 |
| avd6 | 2.86582799513911e-16 | 0.481440529 | 0.486 | 0.263 | 7.21300248096563e-12 | 15 |
| si:dkey-90l23.26 | 3.17655441346564e-16 | 0.478485399 | 0.567 | 0.343 | 7.99506980325168e-12 | 15 |
| si:ch73-14h10.26 | 4.58586197426837e-16 | 0.489575981 | 0.473 | 0.27 | 1.1542156003036e-11 | 15 |
| zgc:1134246 | 7.01532974381156e-16 | 0.474819133 | 0.339 | 0.16 | 1.76568834321993e-11 | 15 |
| pabpn1l6 | 8.96887827353647e-16 | 0.478084301 | 0.498 | 0.282 | 2.25737697266639e-11 | 15 |
| zgc:566996 | 1.12784459349992e-15 | 0.482419021 | 0.445 | 0.25 | 2.83867205737996e-11 | 15 |
| h3f3a6 | 2.06923465857871e-15 | 0.408207005 | 0.649 | 0.505 | 5.20805671217676e-11 | 15 |
| AL929192.16 | 2.08901989185065e-15 | 0.451523978 | 0.347 | 0.163 | 5.25785416579891e-11 | 15 |
| gbgt1l45 | 2.30877921300552e-15 | 0.450224672 | 0.327 | 0.149 | 5.81096640121359e-11 | 15 |
| si:dkey-229d11.56 | 2.35801865249095e-15 | 0.460461998 | 0.616 | 0.387 | 5.93489714645446e-11 | 15 |
| si:ch211-146l10.76 | 2.43476997511549e-15 | 0.473687122 | 0.371 | 0.187 | 6.12807255036818e-11 | 15 |
| si:ch211-145c1.16 | 2.51072197222679e-15 | 0.481706156 | 0.429 | 0.232 | 6.31923613189761e-11 | 15 |

| | | | | | | |
|---|---|---|---|---|---|---|
| haus66 | 9.08414597591562e-15 | 0.465228495 | 0.396 | 0.208 | 2.2863887006782e-10 | 15 |
| zgc:1629456 | 1.58929075921003e-14 | 0.45311849 | 0.335 | 0.161 | 4.00008591185572e-10 | 15 |
| elovl7b6 | 2.02896365421793e-14 | 0.460445306 | 0.445 | 0.247 | 5.10669862130112e-10 | 15 |
| snu13b6 | 2.55643601301699e-14 | 0.437223489 | 0.555 | 0.391 | 6.43429380116245e-10 | 15 |
| gdf36 | 2.63585474403172e-14 | 0.470956445 | 0.408 | 0.222 | 6.63418280525343e-10 | 15 |
| si:ch211-251f6.66 | 2.96588764907346e-14 | 0.456273026 | 0.518 | 0.33 | 7.46484262395298e-10 | 15 |
| si:ch211-146l10.86 | 3.38119569120611e-14 | 0.453972847 | 0.502 | 0.305 | 8.51013143519665e-10 | 15 |
| AL929032.16 | 3.5232499355087e-14 | 0.449153979 | 0.535 | 0.316 | 8.86766776268185e-10 | 15 |
| uchl16 | 3.64413254223924e-14 | 0.444580459 | 0.547 | 0.343 | 9.17191719556196e-10 | 15 |
| LOC1005351626 | 4.29504337101538e-14 | 0.45295813 | 0.522 | 0.307 | 1.08101946605086e-09 | 15 |
| ddx39ab6 | 5.53503739896781e-14 | 0.466244233 | 0.457 | 0.288 | 1.39311356294621e-09 | 15 |
| si:dkey-208k4.26 | 6.20299650673898e-14 | 0.450260569 | 0.486 | 0.295 | 1.56123219078113e-09 | 15 |
| si:dkey-37o8.16 | 1.72395415610713e-13 | 0.447298292 | 0.494 | 0.306 | 4.33902021550603e-09 | 15 |
| LOC1005374996 | 1.82269155786431e-13 | 0.4428443 | 0.535 | 0.321 | 4.58753238198869e-09 | 15 |
| si:dkeyp-46h3.26 | 3.08425235779723e-13 | 0.426050929 | 0.592 | 0.404 | 7.76275475933984e-09 | 15 |
| ctss2.16 | 3.64389071548294e-13 | 0.373739807 | 0.702 | 0.534 | 9.17130854179902e-09 | 15 |
| slc16a36 | 8.04335631045701e-13 | 0.442538051 | 0.424 | 0.248 | 2.02443234977892e-08 | 15 |
| si:ch73-160p18.36 | 1.14237797930703e-12 | 0.446763025 | 0.412 | 0.24 | 2.87525113611786e-08 | 15 |
| ccna26 | 1.3547015430791e-12 | 0.43327001 | 0.465 | 0.283 | 3.40964831377578e-08 | 15 |
| rcc16 | 1.59798896038756e-12 | 0.444660965 | 0.363 | 0.201 | 4.02197841439945e-08 | 15 |
| slbp7 | 1.68118149605628e-12 | 0.443002842 | 0.404 | 0.233 | 4.23136570742405e-08 | 15 |
| si:dkeyp-75b4.103 | 1.79647488214054e-12 | 0.371527597 | 0.612 | 0.436 | 4.52154763085952e-08 | 15 |
| crp36 | 2.87729625630735e-12 | 0.431778138 | 0.384 | 0.218 | 7.24186694749997e-08 | 15 |
| cpeb1b6 | 5.17488008086306e-12 | 0.412616348 | 0.457 | 0.269 | 1.30246556755242e-07 | 15 |
| cdc206 | 6.73985367645358e-12 | 0.412603671 | 0.298 | 0.152 | 1.6963537718266e-07 | 15 |
| LOC1018854776 | 9.95457392474797e-12 | 0.42208734 | 0.457 | 0.279 | 2.50546671111982e-07 | 15 |
| larp6b6 | 1.71434103105948e-11 | 0.415604875 | 0.355 | 0.195 | 4.3148249410736e-07 | 15 |
| arhgef10la6 | 2.0274827152117e-11 | 0.412707145 | 0.445 | 0.274 | 5.10297124591632e-07 | 15 |
| si:ch73-78o10.16 | 4.10976961159512e-11 | 0.388013393 | 0.269 | 0.132 | 1.03438791354238e-06 | 15 |
| si:ch211-250e5.166 | 6.69098742276567e-11 | 0.39781074 | 0.514 | 0.346 | 1.68405462443589e-06 | 15 |
| cx44.26 | 6.77333301014182e-11 | 0.380509745 | 0.245 | 0.116 | 1.70478018532259e-06 | 15 |
| inka1b6 | 8.97984903135721e-11 | 0.402458359 | 0.331 | 0.184 | 2.2601382027023e-06 | 15 |
| zgc:562316 | 1.07135295161644e-10 | 0.39960422 | 0.388 | 0.227 | 2.69648824392341e-06 | 15 |
| org6 | 1.10406713145991e-10 | 0.407720021 | 0.396 | 0.24 | 2.77882656317144e-06 | 15 |
| zgc:1714746 | 2.40865425059522e-10 | 0.391426423 | 0.478 | 0.303 | 6.0623418833231e-06 | 15 |

| | | | | | | |
|---|---|---|---|---|---|---|
| si:dkeyp-46h3.56 | 3.11048498762582e-10 | 0.394317486 | 0.445 | 0.289 | 7.82877966535543e-06 | 15 |
| zgc:1738566 | 3.84806665970438e-10 | 0.391486081 | 0.376 | 0.219 | 9.68519897580996e-06 | 15 |
| zgc:925916 | 6.0331730735978e-10 | 0.382717535 | 0.355 | 0.205 | 1.51848933089383e-05 | 15 |
| nasp6 | 6.49503162658785e-10 | 0.380966042 | 0.482 | 0.34 | 1.6347345100959e-05 | 15 |
| CDPF16 | 1.12859756786976e-09 | 0.390136484 | 0.416 | 0.268 | 2.84056721857139e-05 | 15 |
| LOC1005368606 | 1.20214937834307e-09 | 0.369274521 | 0.486 | 0.351 | 3.02568977035167e-05 | 15 |
| rdh10b5 | 1.53830282630429e-09 | 0.393025845 | 0.31 | 0.177 | 3.87175438352527e-05 | 15 |
| zgc:1628796 | 1.54171606096972e-09 | 0.375243875 | 0.318 | 0.181 | 3.88034515385469e-05 | 15 |
| larp6a6 | 1.97655470816402e-09 | 0.395477312 | 0.388 | 0.249 | 4.97479054497802e-05 | 15 |
| LOC1018838976 | 2.05864635281053e-09 | 0.372504415 | 0.298 | 0.164 | 5.18140700538883e-05 | 15 |
| srsf2a3 | 2.26355303693367e-09 | 0.373089146 | 0.478 | 0.367 | 5.69713663865835e-05 | 15 |
| nanos35 | 3.27064664064036e-09 | 0.35123929 | 0.212 | 0.102 | 8.23189052982773e-05 | 15 |
| st6galnac1.2 | 3.81310055951445e-09 | 0.274844019 | 0.139 | 0.054 | 9.59719279824191e-05 | 15 |
| si:dkeyp-98a7.76 | 4.49396863476042e-09 | 0.359870537 | 0.339 | 0.195 | 0.000113109 | 15 |
| si:ch211-226l4.66 | 4.81242862170855e-09 | 0.364187249 | 0.437 | 0.293 | 0.000121124 | 15 |
| dynll16 | 4.87381800270557e-09 | 0.331984735 | 0.547 | 0.455 | 0.000122669 | 15 |
| lin28a | 6.58392521679961e-09 | 0.305936962 | 0.163 | 0.07 | 0.000165711 | 15 |
| mos5 | 6.79994601732054e-09 | 0.357171987 | 0.257 | 0.136 | 0.000171148 | 15 |
| si:ch211-226h8.82 | 1.10693750934925e-08 | 0.353643382 | 0.2 | 0.097 | 0.000278605 | 15 |
| si:dkey-42i9.86 | 1.39593406187717e-08 | 0.348493703 | 0.241 | 0.127 | 0.000351343 | 15 |
| LOC1104377792 | 1.430340849398e-08 | 0.29847997 | 0.184 | 0.083 | 0.000360002 | 15 |
| cbx3a3 | 1.86477596543714e-08 | 0.366202909 | 0.412 | 0.297 | 0.000469345 | 15 |
| eif4a1a4 | 2.30167581584288e-08 | 0.326015162 | 0.506 | 0.425 | 0.000579309 | 15 |
| zgc:1034826 | 2.32569160448741e-08 | 0.351475673 | 0.314 | 0.189 | 0.000585353 | 15 |
| pabpc1l6 | 2.33010362281387e-08 | 0.347648324 | 0.257 | 0.142 | 0.000586464 | 15 |
| si:dkey-46g23.26 | 2.34380086909765e-08 | 0.336031073 | 0.265 | 0.145 | 0.000589911 | 15 |
| zgc:660246 | 3.20841764260564e-08 | 0.338286468 | 0.257 | 0.141 | 0.000807527 | 15 |
| lhx8a | 3.68989267787111e-08 | 0.287967404 | 0.139 | 0.058 | 0.000928709 | 15 |
| orc63 | 3.82807576511068e-08 | 0.322558862 | 0.212 | 0.107 | 0.000963488 | 15 |
| patl26 | 4.53980894167258e-08 | 0.352490719 | 0.294 | 0.173 | 0.001142625 | 15 |
| zp2l16 | 4.5830740614159e-08 | 0.344493507 | 0.282 | 0.164 | 0.001153514 | 15 |
| plk15 | 5.34071598617565e-08 | 0.343993569 | 0.298 | 0.177 | 0.001344205 | 15 |
| elovl7a6 | 7.146527069089e-08 | 0.336138414 | 0.31 | 0.182 | 0.001798709 | 15 |
| si:dkey-216e24.34 | 1.01685727883544e-07 | 0.326991449 | 0.229 | 0.123 | 0.002559328 | 15 |
| cdca85 | 1.0237963382239e-07 | 0.337511785 | 0.282 | 0.165 | 0.002576793 | 15 |

| | | | | | | |
|---|---|---|---|---|---|---|
| zgc:1630305 | 1.16811009932202e-07 | 0.333026905 | 0.229 | 0.124 | 0.002940016 | 15 |
| rbm4.36 | 1.17495544988721e-07 | 0.302170502 | 0.522 | 0.445 | 0.002957245 | 15 |
| sybu6 | 1.31216760655233e-07 | 0.330370564 | 0.286 | 0.166 | 0.003302595 | 15 |
| yrdc4 | 1.35278492665224e-07 | 0.341790809 | 0.224 | 0.123 | 0.003404824 | 15 |
| map1lc3c6 | 1.54854724449022e-07 | 0.335845671 | 0.322 | 0.202 | 0.003897539 | 15 |
| dazl3 | 1.57598245556346e-07 | 0.316257358 | 0.204 | 0.106 | 0.00396659 | 15 |
| BX005461.13 | 1.89205960316685e-07 | 0.285559902 | 0.167 | 0.079 | 0.004762125 | 15 |
| rbpms2b6 | 2.00939049230893e-07 | 0.333430453 | 0.351 | 0.221 | 0.005057435 | 15 |
| buc6 | 2.64470930745726e-07 | 0.335527494 | 0.294 | 0.179 | 0.006656469 | 15 |
| retsatl6 | 3.09833440102914e-07 | 0.337492854 | 0.322 | 0.203 | 0.007798198 | 15 |
| ankrd226 | 3.10836687487956e-07 | 0.332902443 | 0.249 | 0.143 | 0.007823449 | 15 |
| ddx411 | 3.60665054716784e-07 | 0.253191844 | 0.151 | 0.069 | 0.009077579 | 15 |
| buc2l2 | 4.06401528276047e-07 | 0.270004609 | 0.167 | 0.08 | 0.01022872 | 15 |
| snx10a4 | 4.08896718303817e-07 | 0.308007038 | 0.229 | 0.126 | 0.010291522 | 15 |
| si:ch211-202f5.32 | 6.23483550562455e-07 | 0.287823844 | 0.18 | 0.091 | 0.015692457 | 15 |
| hspa55 | 6.38780737390874e-07 | 0.32705349 | 0.416 | 0.315 | 0.016077472 | 15 |
| kpna25 | 8.01721662607101e-07 | 0.314744841 | 0.265 | 0.158 | 0.020178533 | 15 |
| aurka3 | 8.15581617770458e-07 | 0.300638942 | 0.216 | 0.119 | 0.020527374 | 15 |
| hsd17b12a6 | 9.14451992643262e-07 | 0.310301555 | 0.273 | 0.165 | 0.023015842 | 15 |
| si:ch73-132k15.2 | 9.38548107098318e-07 | 0.266657831 | 0.139 | 0.063 | 0.023622317 | 15 |
| BX537276.16 | 9.42228636986906e-07 | 0.293705763 | 0.22 | 0.122 | 0.023714953 | 15 |
| snu13a3 | 1.2425369215874e-06 | 0.284497536 | 0.192 | 0.101 | 0.031273412 | 15 |
| rbbp43 | 1.2553256342173e-06 | 0.329791552 | 0.273 | 0.171 | 0.031595291 | 15 |
| dynll2a5 | 1.48403325243187e-06 | 0.317055151 | 0.233 | 0.137 | 0.037351633 | 15 |
| tent5c5 | 1.77998118336247e-06 | 0.316544672 | 0.237 | 0.14 | 0.044800346 | 15 |
| si:dkey-68o6.56 | 2.11502586675831e-06 | 0.304597098 | 0.216 | 0.122 | 0.053233086 | 15 |
| faub5 | 2.22582752341602e-06 | 0.325728486 | 0.302 | 0.201 | 0.056021853 | 15 |
| hmgb3b1 | 2.25158393931333e-06 | 0.3362274 | 0.273 | 0.178 | 0.056670116 | 15 |
| ctsla6 | 2.27610161521508e-06 | 0.294685259 | 0.465 | 0.387 | 0.057287202 | 15 |
| bmb3 | 2.35788253123106e-06 | 0.269485448 | 0.171 | 0.088 | 0.059345545 | 15 |
| zgc:1655395 | 2.69115308727256e-06 | 0.283412822 | 0.216 | 0.12 | 0.067733632 | 15 |
| si:ch73-303b9.12 | 2.71933789593652e-06 | 0.258728206 | 0.159 | 0.079 | 0.068443016 | 15 |
| pou5f36 | 2.78260861437657e-06 | 0.302283953 | 0.245 | 0.147 | 0.070035476 | 15 |
| rpain3 | 3.04635293930494e-06 | 0.301973842 | 0.208 | 0.12 | 0.076673657 | 15 |
| LOC1005380232 | 3.20439870922936e-06 | 0.25169279 | 0.155 | 0.077 | 0.080651511 | 15 |

| | | | | | | |
|---|---|---|---|---|---|---|
| zfand62 | 3.41996523382207e-06 | 0.308258259 | 0.216 | 0.126 | 0.086077105 | 15 |
| rpf11 | 3.51386799633809e-06 | 0.322564861 | 0.224 | 0.136 | 0.088440544 | 15 |
| CR847984.16 | 3.75233774499431e-06 | 0.29537374 | 0.265 | 0.164 | 0.094442589 | 15 |
| ccne14 | 4.08374584069778e-06 | 0.283193286 | 0.2 | 0.111 | 0.102783799 | 15 |
| CABZ01059627.26 | 4.14812068215213e-06 | 0.285282351 | 0.229 | 0.134 | 0.104404049 | 15 |
| gyg1a7 | 4.89529700846856e-06 | 0.300879927 | 0.465 | 0.371 | 0.12320973 | 15 |
| zgc:1751361 | 5.86070941475128e-06 | 0.288600801 | 0.192 | 0.108 | 0.147508195 | 15 |
| prdx15 | 6.67334011753815e-06 | 0.306622572 | 0.261 | 0.167 | 0.167961297 | 15 |
| elavl26 | 7.65708868578339e-06 | 0.300188169 | 0.335 | 0.224 | 0.192721265 | 15 |
| alyref1 | 8.72439939920057e-06 | 0.318688822 | 0.363 | 0.277 | 0.219584408 | 15 |
| mibp6 | 1.0782450101781e-05 | 0.306600622 | 0.322 | 0.223 | 0.271383487 | 15 |
| marcksb3 | 1.09774831999803e-05 | 0.297467903 | 0.269 | 0.176 | 0.276292275 | 15 |
| pttg16 | 1.26645508034382e-05 | 0.295935416 | 0.249 | 0.159 | 0.318754079 | 15 |
| ube2c3 | 1.34651251544975e-05 | 0.269376395 | 0.233 | 0.14 | 0.338903735 | 15 |
| nabp1a | 1.46165431851141e-05 | 0.28953896 | 0.196 | 0.115 | 0.367883775 | 15 |
| gfer2 | 1.81444376145547e-05 | 0.264452782 | 0.176 | 0.099 | 0.45667735 | 15 |
| zgc:1719775 | 1.98499545966941e-05 | 0.29239442 | 0.331 | 0.231 | 0.499603507 | 15 |
| si:ch1073-190k2.16 | 2.05256026568264e-05 | 0.284416041 | 0.286 | 0.189 | 0.516608893 | 15 |
| h2afx3 | 2.66777356262914e-05 | 0.27289825 | 0.224 | 0.139 | 0.671451928 | 15 |
| CU467646.25 | 2.81858648905407e-05 | 0.254484503 | 0.212 | 0.126 | 0.709410033 | 15 |
| si:ch1073-263o8.26 | 2.96737701281551e-05 | 0.285357981 | 0.367 | 0.264 | 0.74685912 | 15 |
| zgc:1010162 | 2.98829635802128e-05 | 0.267943263 | 0.171 | 0.097 | 0.75212431 | 15 |
| imp34 | 3.3653327901075e-05 | 0.280175621 | 0.216 | 0.135 | 0.84702061 | 15 |
| sinup6 | 3.37195534130665e-05 | 0.28287731 | 0.298 | 0.204 | 0.84868744 | 15 |
| lsm11 | 4.06287054661773e-05 | 0.288215544 | 0.216 | 0.137 | 1 | 15 |
| mylipa2 | 4.14997181123308e-05 | 0.275156099 | 0.192 | 0.116 | 1 | 15 |
| cdca7a6 | 4.23658233954994e-05 | 0.289693801 | 0.31 | 0.223 | 1 | 15 |
| thop12 | 5.18811690870695e-05 | 0.256694025 | 0.196 | 0.117 | 1 | 15 |
| cdc61 | 5.30892130664267e-05 | 0.26016409 | 0.176 | 0.103 | 1 | 15 |
| tph1a6 | 5.5784348420658e-05 | 0.250951128 | 0.229 | 0.142 | 1 | 15 |
| snrpc2 | 6.80081090791033e-05 | 0.303186265 | 0.257 | 0.182 | 1 | 15 |
| chchd104 | 7.17183399039201e-05 | 0.275765015 | 0.233 | 0.152 | 1 | 15 |
| zp3c6 | 7.58929948935825e-05 | 0.260404328 | 0.282 | 0.19 | 1 | 15 |
| parn5 | 7.60882771133002e-05 | 0.271315934 | 0.229 | 0.148 | 1 | 15 |
| zgc:1717506 | 7.69004043689308e-05 | 0.269138493 | 0.261 | 0.175 | 1 | 15 |

| | | | | | | |
|---|---|---|---|---|---|---|
| dctpp16 | 8.80405365488443e-05 | 0.273141273 | 0.257 | 0.176 | 1 | 15 |
| chia.46 | 9.03095948924328e-05 | 0.26030556 | 0.229 | 0.148 | 1 | 15 |
| pigrl2.36 | 0.000108322 | 0.253841613 | 0.298 | 0.204 | 1 | 15 |
| tomm20a1 | 0.000134056 | 0.299430613 | 0.302 | 0.233 | 1 | 15 |
| zgc:566766 | 0.000192221 | 0.279820511 | 0.363 | 0.297 | 1 | 15 |
| zglp16 | 0.000198687 | 0.25148997 | 0.286 | 0.199 | 1 | 15 |
| rpa32 | 0.000236768 | 0.26200394 | 0.237 | 0.164 | 1 | 15 |
| mcm71 | 0.000359776 | 0.254788919 | 0.229 | 0.159 | 1 | 15 |
| chtopa4 | 0.000389193 | 0.273709757 | 0.286 | 0.223 | 1 | 15 |
| si:dkey-188i13.116 | 0.000416415 | 0.254915972 | 0.384 | 0.318 | 1 | 15 |
| zc3h11a5 | 0.000684737 | 0.26352764 | 0.286 | 0.218 | 1 | 15 |
| ebna1bp22 | 0.00090368 | 0.250898516 | 0.2 | 0.138 | 1 | 15 |
| gemin73 | 0.001040296 | 0.254684913 | 0.216 | 0.156 | 1 | 15 |
| ncbp22 | 0.001164359 | 0.253962747 | 0.233 | 0.171 | 1 | 15 |
| ddx3xb | 0.00211564 | 0.256022514 | 0.2 | 0.146 | 1 | 15 |
| CU499330.11 | 0 | 1.693498455 | 0.965 | 0.053 | 0 | 16 |
| LOC1104377312 | 0 | 1.667566186 | 0.848 | 0.076 | 0 | 16 |
| grn11 | 0 | 1.618110018 | 0.9 | 0.071 | 0 | 16 |
| lygl11 | 0 | 1.61210071 | 0.939 | 0.08 | 0 | 16 |
| BX649485.21 | 0 | 1.605428152 | 0.766 | 0.017 | 0 | 16 |
| c1qb1 | 0 | 1.602878013 | 0.926 | 0.053 | 0 | 16 |
| lgals2a2 | 0 | 1.590102647 | 0.965 | 0.109 | 0 | 16 |
| mrc1b1 | 0 | 1.490941299 | 0.87 | 0.026 | 0 | 16 |
| c1qc1 | 0 | 1.409788701 | 0.896 | 0.05 | 0 | 16 |
| cfp2 | 0 | 1.334073299 | 0.827 | 0.061 | 0 | 16 |
| CR855311.13 | 0 | 1.31166662 | 0.779 | 0.072 | 0 | 16 |
| si:ch211-165i18.21 | 0 | 1.303242252 | 0.723 | 0.022 | 0 | 16 |
| c1qa1 | 0 | 1.243147153 | 0.823 | 0.036 | 0 | 16 |
| si:dkey-5n18.11 | 0 | 1.220613139 | 0.684 | 0.046 | 0 | 16 |
| ccl34b.81 | 0 | 1.109142035 | 0.446 | 0.01 | 0 | 16 |
| cx32.2 | 0 | 1.086358648 | 0.714 | 0.016 | 0 | 16 |
| ms4a17a.101 | 0 | 1.07831232 | 0.701 | 0.039 | 0 | 16 |
| ccl34a.4 | 0 | 1.077901273 | 0.55 | 0.01 | 0 | 16 |
| si:dkey-102g19.31 | 0 | 1.059965218 | 0.675 | 0.035 | 0 | 16 |
| LOC1005369461 | 0 | 0.954881914 | 0.42 | 0.01 | 0 | 16 |

| | | | | | | |
|---|---|---|---|---|---|---|
| spi1a1 | 0 | 0.903837212 | 0.602 | 0.025 | 0 | 16 |
| itgae.11 | 0 | 0.885957621 | 0.593 | 0.023 | 0 | 16 |
| grna1 | 0 | 0.869276255 | 0.645 | 0.044 | 0 | 16 |
| timp2b | 0 | 0.859270948 | 0.524 | 0.028 | 0 | 16 |
| si:ch73-343l4.8 | 0 | 0.790584355 | 0.494 | 0.014 | 0 | 16 |
| cxcr3.21 | 0 | 0.788499791 | 0.58 | 0.024 | 0 | 16 |
| CABZ01001434.1 | 0 | 0.775135994 | 0.325 | 0.007 | 0 | 16 |
| cdaa1 | 0 | 0.74534321 | 0.511 | 0.026 | 0 | 16 |
| samsn1a1 | 0 | 0.731209343 | 0.524 | 0.026 | 0 | 16 |
| dhrs9 | 0 | 0.716413187 | 0.524 | 0.025 | 0 | 16 |
| moxd11 | 0 | 0.632503509 | 0.398 | 0.006 | 0 | 16 |
| ccr12b.2 | 0 | 0.594947598 | 0.355 | 0.004 | 0 | 16 |
| LOC564660 | 0 | 0.580184628 | 0.225 | 0.002 | 0 | 16 |
| jac3 | 0 | 0.565685861 | 0.216 | 0.002 | 0 | 16 |
| wu:fb64b08 | 0 | 0.443443029 | 0.273 | 0.003 | 0 | 16 |
| si:ch73-343l4.2 | 0 | 0.390429532 | 0.273 | 0.004 | 0 | 16 |
| si:ch211-264f5.2 | 4.22840149534015e-305 | 0.457021173 | 0.303 | 0.008 | 1.06424637236216e-300 | 16 |
| cybb | 1.16108880452927e-302 | 0.508661369 | 0.368 | 0.013 | 2.92234441211971e-298 | 16 |
| sftpbb1 | 2.27021999185874e-298 | 0.951440058 | 0.693 | 0.054 | 5.71391669750926e-294 | 16 |
| cd22 | 1.27328122036743e-295 | 0.384789105 | 0.255 | 0.005 | 3.20472150354278e-291 | 16 |
| zgc:1531181 | 1.04523192249921e-289 | 1.064013641 | 0.498 | 0.028 | 2.63074422573827e-285 | 16 |
| selp1 | 1.08560110130476e-287 | 0.393363188 | 0.273 | 0.006 | 2.73234941187394e-283 | 16 |
| clic23 | 1.75701345065994e-283 | 0.950635883 | 0.628 | 0.049 | 4.422227153966e-279 | 16 |
| cst14b.11 | 3.0181343174164e-278 | 0.757842172 | 0.58 | 0.041 | 7.59634226350533e-274 | 16 |
| si:ch211-147m6.11 | 3.79931240218541e-272 | 0.750884611 | 0.455 | 0.025 | 9.56248938506047e-268 | 16 |
| NPC2 (1 of many)1 | 9.87989518890157e-268 | 0.863586041 | 0.463 | 0.026 | 2.48667082009464e-263 | 16 |
| si:ch211-194m7.31 | 9.35694290421073e-267 | 0.666347597 | 0.364 | 0.015 | 2.3550489595608e-262 | 16 |
| si:busm1-266f07.22 | 3.77957939815948e-260 | 1.424429577 | 0.952 | 0.131 | 9.51282338722759e-256 | 16 |
| grn21 | 4.02371624873544e-260 | 0.859999787 | 0.351 | 0.015 | 1.01272914264422e-255 | 16 |
| fcer1gl2 | 1.26132712646724e-259 | 1.507616984 | 0.983 | 0.153 | 3.17463424460539e-255 | 16 |
| spi1b1 | 2.76154343112279e-259 | 0.790599927 | 0.623 | 0.051 | 6.95052866179295e-255 | 16 |
| krt18b4 | 9.96745881584816e-255 | 1.359314566 | 0.918 | 0.136 | 2.50870970936082e-250 | 16 |
| mfap41 | 1.67770243498427e-254 | 0.611186379 | 0.377 | 0.018 | 4.2226092586119e-250 | 16 |
| slc3a2a1 | 4.8441853766496e-251 | 0.625346885 | 0.45 | 0.026 | 1.21923301744894e-246 | 16 |
| ms4a17a.63 | 4.75314243511772e-249 | 0.711370489 | 0.541 | 0.04 | 1.19631841949478e-244 | 16 |

| | | | | | | |
|---|---|---|---|---|---|---|
| si:ch211-283g2.21 | 6.89182233786885e-249 | 0.570309738 | 0.368 | 0.017 | 1.73460276421821e-244 | 16 |
| jac9 | 3.08094585268983e-243 | 0.34427307 | 0.152 | 0.001 | 7.75443261663503e-239 | 16 |
| zgc:1528301 | 7.61251733541169e-241 | 0.824430005 | 0.619 | 0.055 | 1.91599448814977e-236 | 16 |
| lgals3bpb1 | 3.5136634598053e-240 | 0.986931646 | 0.485 | 0.033 | 8.84353956198396e-236 | 16 |
| BX649485.1 | 9.05929876146827e-237 | 0.645808954 | 0.368 | 0.018 | 2.28013490527395e-232 | 16 |
| marco1 | 1.1256876689939e-229 | 0.750838291 | 0.437 | 0.028 | 2.83324329409076e-225 | 16 |
| mir21-1 | 2.26087381136453e-227 | 1.024989649 | 0.766 | 0.095 | 5.69039329582339e-223 | 16 |
| calhm6 | 5.30869561158089e-226 | 0.386128487 | 0.281 | 0.01 | 1.33614559847879e-221 | 16 |
| cxcl8b.13 | 1.57126486605454e-224 | 0.994271746 | 0.424 | 0.028 | 3.95471654137268e-220 | 16 |
| smox2 | 4.60104010352392e-223 | 0.884773698 | 0.684 | 0.073 | 1.15803578365594e-218 | 16 |
| gpr84 | 9.12476831665679e-223 | 0.393424887 | 0.247 | 0.008 | 2.29661293761935e-218 | 16 |
| mhc2dab1 | 1.03530050083779e-221 | 1.104785536 | 0.732 | 0.085 | 2.60574783055864e-217 | 16 |
| fgl2a1 | 2.7072508827463e-220 | 0.903529175 | 0.593 | 0.056 | 6.81387974678415e-216 | 16 |
| LO018493.1 | 3.63558608461885e-220 | 0.489860343 | 0.355 | 0.018 | 9.15040661637718e-216 | 16 |
| ccl39.1 | 6.85275736980424e-220 | 0.709906401 | 0.221 | 0.006 | 1.72477050240603e-215 | 16 |
| zgc:136870 | 2.47503125191373e-217 | 0.422641061 | 0.299 | 0.013 | 6.22940615794166e-213 | 16 |
| ctss2.21 | 2.05002420790345e-216 | 0.66090401 | 0.468 | 0.034 | 5.1597059288722e-212 | 16 |
| si:dkey-56f14.7 | 8.15284880821166e-212 | 0.348176986 | 0.251 | 0.009 | 2.05199051653879e-207 | 16 |
| sb:cb81 | 1.78826390729307e-211 | 0.510799023 | 0.364 | 0.02 | 4.50088142826593e-207 | 16 |
| si:ch211-9d9.8 | 5.73756720714361e-211 | 0.379560614 | 0.234 | 0.007 | 1.44408829036597e-206 | 16 |
| cyba1 | 2.76950400253809e-208 | 0.807208866 | 0.615 | 0.063 | 6.97056462398813e-204 | 16 |
| ltb4r1 | 9.66685498338644e-208 | 0.484611231 | 0.372 | 0.022 | 2.43305073076853e-203 | 16 |
| ncf2 | 9.5368372937503e-205 | 0.471663939 | 0.325 | 0.017 | 2.40032657846401e-200 | 16 |
| havcr11 | 6.3347357955008e-204 | 0.63855229 | 0.403 | 0.026 | 1.5943896523696e-199 | 16 |
| tcnbb | 1.32430012055153e-202 | 0.618493891 | 0.277 | 0.012 | 3.33313097341614e-198 | 16 |
| tmsb13 | 1.67892684412732e-202 | 1.228840621 | 0.887 | 0.142 | 4.22569097398406e-198 | 16 |
| enpp1 | 2.69404435280536e-201 | 0.626225944 | 0.437 | 0.032 | 6.78064023157582e-197 | 16 |
| mapk11 | 4.64683322470384e-201 | 0.383814229 | 0.29 | 0.013 | 1.16956145432571e-196 | 16 |
| abca1b | 1.57699426479882e-200 | 0.453403594 | 0.29 | 0.013 | 3.96913686507214e-196 | 16 |
| LOC100536983 | 4.64219179522311e-199 | 0.360349488 | 0.221 | 0.007 | 1.16839325293971e-194 | 16 |
| zgc:640651 | 4.19116806161159e-197 | 0.481935945 | 0.29 | 0.013 | 1.05487508942702e-192 | 16 |
| ppp1r14ab | 1.70068610531643e-194 | 0.273303963 | 0.165 | 0.003 | 4.28045685847093e-190 | 16 |
| hlx1 | 5.37399241706367e-193 | 0.3515923 | 0.238 | 0.009 | 1.35258015145076e-188 | 16 |
| FP015862.2 | 1.80626343670508e-190 | 0.416911486 | 0.268 | 0.012 | 4.54618444384301e-186 | 16 |
| pnp5a1 | 1.82140411763309e-189 | 0.891596364 | 0.602 | 0.068 | 4.58429202367073e-185 | 16 |

| | | | | | | |
|---|---|---|---|---|---|---|
| f13a1b1 | 1.32857385735954e-186 | 0.557587697 | 0.372 | 0.025 | 3.34388754158822e-182 | 16 |
| lta4h2 | 1.38542011275823e-186 | 0.748552585 | 0.589 | 0.062 | 3.48696388180119e-182 | 16 |
| nrros1 | 3.4690902392422e-185 | 0.633372294 | 0.468 | 0.04 | 8.73135322314869e-181 | 16 |
| cd74b4 | 3.58105255197132e-185 | 1.368470569 | 0.987 | 0.209 | 9.0131511680566e-181 | 16 |
| m171 | 5.3409899353688e-184 | 0.445800267 | 0.247 | 0.01 | 1.34427375683297e-179 | 16 |
| si:ch211-152p11.4 | 7.77742862601057e-182 | 0.485607374 | 0.368 | 0.025 | 1.9575010108806e-177 | 16 |
| itgb22 | 1.17665008631398e-181 | 0.850373071 | 0.684 | 0.088 | 2.96151060224366e-177 | 16 |
| plekho1b | 1.5470039586277e-180 | 0.585566523 | 0.424 | 0.034 | 3.89365426347006e-176 | 16 |
| gpx1a2 | 2.27015844007104e-179 | 1.29081148 | 0.926 | 0.211 | 5.71376177781481e-175 | 16 |
| cmklr11 | 4.9201192634574e-177 | 0.554579832 | 0.364 | 0.025 | 1.23834481741959e-172 | 16 |
| cd832 | 4.11694808951378e-175 | 1.081963199 | 0.675 | 0.093 | 1.03619466464972e-170 | 16 |
| slc16a9a | 2.99118420962676e-174 | 0.347787898 | 0.203 | 0.007 | 7.5285115372096e-170 | 16 |
| hdac9b | 1.00887417429803e-173 | 0.466584737 | 0.316 | 0.019 | 2.53923540929071e-169 | 16 |
| asah1b | 2.02840432633109e-170 | 0.624928839 | 0.476 | 0.046 | 5.10529084894272e-166 | 16 |
| cd74a3 | 1.96736004150977e-168 | 1.341560122 | 0.957 | 0.205 | 4.95164848847593e-164 | 16 |
| fosl24 | 2.22769209269277e-168 | 0.931461289 | 0.723 | 0.11 | 5.60687822809843e-164 | 16 |
| ccl35.11 | 2.40484245346209e-168 | 1.129547387 | 0.589 | 0.073 | 6.05274797111873e-164 | 16 |
| ms4a17a.91 | 1.26371728767583e-167 | 0.667332419 | 0.481 | 0.048 | 3.1806500413513e-163 | 16 |
| si:dkey-262k9.43 | 7.00333490814302e-166 | 0.727132998 | 0.632 | 0.08 | 1.76266936303052e-161 | 16 |
| atp6v1e1b1 | 1.73125910731245e-164 | 0.718736879 | 0.597 | 0.073 | 4.35740604719471e-160 | 16 |
| AL928650.1 | 1.75210363088169e-162 | 0.458781103 | 0.203 | 0.008 | 4.40986962856612e-158 | 16 |
| ccl39.3 | 4.390311307325e-162 | 0.479978659 | 0.152 | 0.004 | 1.10499745294063e-157 | 16 |
| il1b | 2.57188167669793e-160 | 0.470524882 | 0.307 | 0.02 | 6.47316899208103e-156 | 16 |
| xpnpep2 | 8.10362927095461e-159 | 0.27740623 | 0.195 | 0.007 | 2.03960245120657e-154 | 16 |
| slc6a6b2 | 8.89275378552212e-159 | 0.821411743 | 0.662 | 0.093 | 2.23821720027806e-154 | 16 |
| slc2a6 | 8.29880668160434e-158 | 0.25659557 | 0.165 | 0.005 | 2.088726653693e-153 | 16 |
| si:ch211-193e13.5 | 3.39288313467684e-157 | 0.299053511 | 0.203 | 0.008 | 8.53954756166814e-153 | 16 |
| BX324132.2 | 2.24234139268271e-156 | 0.261994788 | 0.169 | 0.005 | 5.6437490512431e-152 | 16 |
| lect2l | 8.35722263145588e-156 | 0.602659967 | 0.355 | 0.028 | 2.10342936411113e-151 | 16 |
| ly6m5 | 9.51233009619588e-156 | 0.299339756 | 0.203 | 0.008 | 2.39415836191154e-151 | 16 |
| anxa5b1 | 6.41963809062801e-155 | 0.761973945 | 0.619 | 0.084 | 1.61575871103016e-150 | 16 |
| pmaip1 | 4.2167196247375e-153 | 0.956383428 | 0.636 | 0.095 | 1.06130616235018e-148 | 16 |
| naga1 | 4.63995607805901e-153 | 0.54496435 | 0.442 | 0.043 | 1.16783054528667e-148 | 16 |
| prdx53 | 4.8278579506368e-151 | 0.982517426 | 0.771 | 0.148 | 1.21512356759578e-146 | 16 |
| ncf4 | 6.90649806217654e-151 | 0.342342329 | 0.255 | 0.014 | 1.73829649726921e-146 | 16 |

| | | | | | | |
|---|---|---|---|---|---|---|
| txn1 | 1.31493170267952e-150 | 0.848591259 | 0.403 | 0.039 | 3.30955160247408e-146 | 16 |
| glb1l | 1.01637365222016e-148 | 0.437260852 | 0.286 | 0.018 | 2.55811084527292e-144 | 16 |
| zmp:0000000936 | 3.87513447676391e-148 | 0.368630285 | 0.255 | 0.014 | 9.75332596456708e-144 | 16 |
| FP102158.1 | 1.2990904253979e-147 | 0.334941272 | 0.242 | 0.013 | 3.26968069168398e-143 | 16 |
| atp1a3a1 | 4.17254801504587e-145 | 0.355741525 | 0.268 | 0.016 | 1.0501886099069e-140 | 16 |
| snx10b | 1.19518661715047e-144 | 0.29604809 | 0.19 | 0.008 | 3.00816519670601e-140 | 16 |
| si:dkey-7i4.241 | 3.2937528692874e-144 | 0.830460887 | 0.273 | 0.018 | 8.29004659670945e-140 | 16 |
| si:ch211-74m13.31 | 4.32278819473269e-143 | 0.762125031 | 0.494 | 0.06 | 1.08800256073227e-138 | 16 |
| XLOC-0054481 | 1.35187690694745e-142 | 0.564920637 | 0.312 | 0.023 | 3.40253898709605e-138 | 16 |
| LOC1005375041 | 2.12357006464735e-141 | 0.565933323 | 0.299 | 0.021 | 5.34481349571093e-137 | 16 |
| tcima2 | 1.18501068089723e-138 | 0.747705405 | 0.498 | 0.061 | 2.98255338275024e-134 | 16 |
| plbd1 | 1.23679415178058e-138 | 0.411380111 | 0.294 | 0.021 | 3.11288720061654e-134 | 16 |
| stat3 | 1.36634958538528e-138 | 0.547176167 | 0.429 | 0.045 | 3.43896527145622e-134 | 16 |
| myo1f1 | 1.99089692841313e-138 | 0.489381544 | 0.403 | 0.04 | 5.01088847912301e-134 | 16 |
| tgfbi | 2.25140495890138e-138 | 0.396921595 | 0.294 | 0.021 | 5.66656114105888e-134 | 16 |
| il4r.11 | 2.7785357347486e-137 | 0.597102814 | 0.381 | 0.037 | 6.99329659078875e-133 | 16 |
| si:dkey-11f4.141 | 3.89232394290367e-136 | 0.861433102 | 0.619 | 0.099 | 9.79659013189424e-132 | 16 |
| adgrg3 | 4.30474002403748e-136 | 0.394577974 | 0.281 | 0.02 | 1.08346001664999e-131 | 16 |
| il10ra | 4.33617003818444e-136 | 0.497513255 | 0.372 | 0.035 | 1.09137063691064e-131 | 16 |
| rnaset2l1 | 3.49881346169704e-135 | 0.484638378 | 0.385 | 0.037 | 8.80616360174528e-131 | 16 |
| laptm55 | 1.55637109032037e-134 | 0.903455284 | 0.827 | 0.158 | 3.91723039722735e-130 | 16 |
| vsir4 | 2.62146138644273e-134 | 0.661735712 | 0.558 | 0.077 | 6.59795616353771e-130 | 16 |
| parvg1 | 4.47606468232146e-134 | 0.619047886 | 0.489 | 0.06 | 1.12658071989349e-129 | 16 |
| LOC100150882 | 5.60499828771919e-128 | 0.350689616 | 0.177 | 0.008 | 1.41072201903604e-123 | 16 |
| trib3 | 1.71098136220737e-127 | 0.750372769 | 0.606 | 0.097 | 4.30636899053972e-123 | 16 |
| LOC1018842671 | 1.2434873852687e-124 | 0.522779893 | 0.442 | 0.051 | 3.12973339998278e-120 | 16 |
| si:ch211-15b10.62 | 8.16483846479183e-124 | 0.594139511 | 0.485 | 0.064 | 2.05500819320346e-119 | 16 |
| cebpa3 | 1.58793902662521e-122 | 0.598273695 | 0.429 | 0.051 | 3.99668373611299e-118 | 16 |
| cxcr4b4 | 1.70853930341032e-122 | 0.980558891 | 0.818 | 0.172 | 4.30022257275344e-118 | 16 |
| atp6v1ba | 6.05800536383572e-122 | 0.511457254 | 0.403 | 0.045 | 1.52473937002381e-117 | 16 |
| nr4a14 | 1.35195311612536e-121 | 0.937479244 | 0.753 | 0.156 | 3.40273079797591e-117 | 16 |
| rasgrp4 | 1.4240763854968e-121 | 0.405564426 | 0.299 | 0.025 | 3.5842578546569e-117 | 16 |
| s100a10b2 | 2.40583142657676e-121 | 1.184324242 | 0.883 | 0.244 | 6.05523711755104e-117 | 16 |
| ccl39.2 | 1.00862422090668e-120 | 0.402992559 | 0.126 | 0.004 | 2.53860630160002e-116 | 16 |
| entpd11 | 3.34839607581906e-120 | 0.634366664 | 0.476 | 0.063 | 8.42757808322898e-116 | 16 |

| | | | | | | |
|---|---|---|---|---|---|---|
| scpep11 | 1.97937471929664e-119 | 0.714429628 | 0.649 | 0.11 | 4.98188823099772e-115 | 16 |
| ctsh1 | 4.89298299189101e-118 | 0.550315468 | 0.403 | 0.047 | 1.23151488922905e-113 | 16 |
| pgd1 | 5.62779404034892e-118 | 0.541806475 | 0.446 | 0.057 | 1.41645948201542e-113 | 16 |
| atp6v1g11 | 2.12506881869861e-116 | 0.712523482 | 0.68 | 0.123 | 5.34858570978253e-112 | 16 |
| si:cabz01074946.12 | 4.16365721871769e-116 | 0.605225711 | 0.424 | 0.053 | 1.04795088537906e-111 | 16 |
| dpp71 | 1.20080278202995e-115 | 0.455938357 | 0.381 | 0.043 | 3.02230052209117e-111 | 16 |
| prcp | 1.01299492651073e-114 | 0.321868624 | 0.221 | 0.014 | 2.54960693053486e-110 | 16 |
| hexb | 2.57221399847199e-113 | 0.377147014 | 0.312 | 0.029 | 6.47400541275414e-109 | 16 |
| slc7a111 | 2.07754719749981e-112 | 0.286106785 | 0.165 | 0.008 | 5.22897854138727e-108 | 16 |
| zgc:638311 | 3.00907241000212e-112 | 0.548798517 | 0.506 | 0.073 | 7.57353434873433e-108 | 16 |
| icn3 | 4.42965386035673e-112 | 1.180796362 | 0.892 | 0.263 | 1.11489958011319e-107 | 16 |
| dnase2 | 5.46113510213031e-112 | 0.464471278 | 0.359 | 0.039 | 1.37451309385518e-107 | 16 |
| klf2b | 9.28765533916389e-112 | 0.349879816 | 0.238 | 0.017 | 2.33760997231416e-107 | 16 |
| zgc:1749042 | 2.23849325173335e-111 | 0.606136362 | 0.372 | 0.043 | 5.63406366528768e-107 | 16 |
| unc93b1 | 6.77813362112683e-111 | 0.341832087 | 0.247 | 0.019 | 1.70598845110141e-106 | 16 |
| crema4 | 4.9662624702604e-110 | 0.860762315 | 0.753 | 0.168 | 1.24995860113984e-105 | 16 |
| il4r.2 | 1.08386055371499e-109 | 0.392497671 | 0.195 | 0.012 | 2.72796862764526e-105 | 16 |
| plin2 | 2.74151391753383e-109 | 0.447789759 | 0.286 | 0.026 | 6.9001163790409e-105 | 16 |
| tcirg1b | 3.03198485173885e-109 | 0.463437626 | 0.394 | 0.047 | 7.63120267334151e-105 | 16 |
| BX248082.13 | 3.80227474552578e-109 | 0.74356928 | 0.338 | 0.037 | 9.56994530701383e-105 | 16 |
| ets21 | 6.79862092174982e-109 | 0.597618402 | 0.528 | 0.083 | 1.71114489979521e-104 | 16 |
| zgc:640514 | 1.09745192856973e-108 | 0.784221355 | 0.745 | 0.152 | 2.76217675901716e-104 | 16 |
| slc7a7 | 1.13910040231965e-108 | 0.329339917 | 0.199 | 0.012 | 2.86700180259832e-104 | 16 |
| ctsa1 | 1.42381721701108e-108 | 0.550059734 | 0.528 | 0.081 | 3.58360555349519e-104 | 16 |
| ccng21 | 1.72236895852633e-108 | 0.602023005 | 0.506 | 0.078 | 4.33503043171493e-104 | 16 |
| acsl4b | 8.52455791147409e-108 | 0.474342933 | 0.403 | 0.05 | 2.14554598073891e-103 | 16 |
| atp6v0ca1 | 1.00187697740136e-107 | 0.861319847 | 0.805 | 0.188 | 2.52162416442149e-103 | 16 |
| si:ch73-248e21.51 | 1.67307560604196e-107 | 0.461834615 | 0.355 | 0.04 | 4.210963992847e-103 | 16 |
| bach1b | 5.08296823311319e-107 | 0.372172508 | 0.255 | 0.021 | 1.27933227459226e-102 | 16 |
| LOC1001482541 | 7.89518594345003e-105 | 0.349678731 | 0.247 | 0.02 | 1.98713935010694e-100 | 16 |
| cndp21 | 1.44198157154461e-103 | 0.488566774 | 0.429 | 0.058 | 3.62932341742063e-99 | 16 |
| ugt5b2 | 1.91119135014925e-103 | 0.338722958 | 0.281 | 0.026 | 4.81027750919065e-99 | 16 |
| XLOC-024516 | 2.59558504536506e-103 | 0.258274709 | 0.173 | 0.01 | 6.53282800067933e-99 | 16 |
| gnpda1 | 7.0124613303222e-103 | 0.398773416 | 0.346 | 0.039 | 1.7649663922288e-98 | 16 |
| rab442 | 7.60919210080659e-103 | 0.496796857 | 0.416 | 0.055 | 1.91515755985201e-98 | 16 |

| | | | | | | |
|---|---|---|---|---|---|---|
| ptgis | 1.29785295780365e-102 | 0.305826085 | 0.19 | 0.012 | 3.26656610949601e-98 | 16 |
| cebpb5 | 2.66195704672606e-102 | 1.080264629 | 0.926 | 0.343 | 6.69987969090482e-98 | 16 |
| fuca2 | 2.83701293233686e-102 | 0.487413499 | 0.424 | 0.058 | 7.14047784939865e-98 | 16 |
| aplp2 | 4.50177251433118e-102 | 0.561011429 | 0.45 | 0.066 | 1.13305112413202e-97 | 16 |
| sec61a1 | 7.69574385852675e-102 | 0.822970233 | 0.749 | 0.179 | 1.9369417717526e-97 | 16 |
| si:dkeyp-23e4.3 | 9.71325354373972e-102 | 0.257037337 | 0.177 | 0.01 | 2.44472878442385e-97 | 16 |
| tpp11 | 2.35934338474987e-100 | 0.501293136 | 0.455 | 0.065 | 5.93823136507695e-96 | 16 |
| cd637 | 1.31045288047639e-99 | 0.981307792 | 0.935 | 0.33 | 3.29827885487103e-95 | 16 |
| lxn2 | 5.44545023831659e-99 | 0.74956412 | 0.667 | 0.139 | 1.3705653704819e-94 | 16 |
| glipr1a3 | 4.95648364524512e-98 | 0.447366979 | 0.42 | 0.058 | 1.24749736867174e-93 | 16 |
| rhogb2 | 5.16552448826136e-98 | 0.583943989 | 0.528 | 0.089 | 1.3001108584505e-93 | 16 |
| si:ch211-66e2.31 | 2.17178353858922e-97 | 0.482675818 | 0.398 | 0.053 | 5.46616198827521e-93 | 16 |
| dicp3.11 | 5.88553155480354e-96 | 0.276116277 | 0.199 | 0.014 | 1.4813294370285e-91 | 16 |
| ssr31 | 6.79294752743012e-96 | 0.883545221 | 0.887 | 0.261 | 1.70971696317889e-91 | 16 |
| arl5c3 | 1.02483472456748e-95 | 0.63102023 | 0.567 | 0.104 | 2.57940651826388e-91 | 16 |
| XLOC-0234924 | 2.20573844592186e-95 | 0.680651511 | 0.511 | 0.088 | 5.55162309454073e-91 | 16 |
| csf2rb1 | 4.27949831847844e-95 | 0.459440616 | 0.312 | 0.035 | 1.07710693177784e-90 | 16 |
| XLOC-0054493 | 9.6679284219125e-94 | 0.553369278 | 0.368 | 0.048 | 2.43332090451116e-89 | 16 |
| gpr183a2 | 1.59680510701534e-93 | 0.46630176 | 0.416 | 0.059 | 4.01899877384692e-89 | 16 |
| lpar5a3 | 3.15533612253714e-93 | 0.410945834 | 0.32 | 0.037 | 7.94166548681374e-89 | 16 |
| si:dkey-206d17.121 | 4.1490251138081e-93 | 0.440035251 | 0.364 | 0.047 | 1.04426813089436e-88 | 16 |
| zmp:00000010821 | 2.20009585470769e-92 | 0.463851918 | 0.381 | 0.052 | 5.53742125671378e-88 | 16 |
| nccrp11 | 3.91867969902721e-92 | 0.824759844 | 0.749 | 0.184 | 9.86292493448158e-88 | 16 |
| cotl15 | 5.11409277866663e-91 | 0.93828877 | 0.931 | 0.295 | 1.2871660114626e-86 | 16 |
| pdia33 | 1.15530874678382e-90 | 0.882810858 | 0.866 | 0.273 | 2.90779658478019e-86 | 16 |
| fermt3b | 2.30062465325443e-90 | 0.280042854 | 0.203 | 0.016 | 5.79044218977608e-86 | 16 |
| tp53i11b | 5.09536503680594e-90 | 0.251020957 | 0.225 | 0.019 | 1.28245242611369e-85 | 16 |
| zgc:101715 | 1.27192693783389e-89 | 0.263798932 | 0.229 | 0.02 | 3.20131290983412e-85 | 16 |
| si:ch211-1o7.31 | 1.81071592931039e-89 | 0.302345799 | 0.212 | 0.017 | 4.55739092248133e-85 | 16 |
| ubl3b | 1.38702569352743e-88 | 0.453610558 | 0.433 | 0.066 | 3.49100496803919e-84 | 16 |
| myofl1 | 2.87110352745761e-88 | 0.312320305 | 0.208 | 0.017 | 7.22628046825806e-84 | 16 |
| gnsb | 4.31440001506774e-88 | 0.400952611 | 0.299 | 0.034 | 1.0858913397924e-83 | 16 |
| anxa2a1 | 4.57009256541619e-88 | 0.632491389 | 0.532 | 0.099 | 1.1502465977896e-83 | 16 |
| actb25 | 1.71542147069153e-87 | 0.558892651 | 1 | 0.866 | 4.31754429958351e-83 | 16 |
| si:ch1073-429i10.3.1 | 1.80036133550356e-87 | 0.722912115 | 0.766 | 0.192 | 4.53132944532892e-83 | 16 |

| | | | | | | |
|---|---|---|---|---|---|---|
| si:zfos-1069f5.1 | 1.18941300623566e-86 | 0.257467885 | 0.165 | 0.01 | 2.99363359539454e-82 | 16 |
| BX901920.12 | 5.43272671030325e-86 | 0.622734184 | 0.42 | 0.068 | 1.36736298571622e-81 | 16 |
| snd1 | 1.41286278040248e-85 | 0.561952564 | 0.519 | 0.098 | 3.556034331995e-81 | 16 |
| atp7a | 3.47698666551556e-85 | 0.261660895 | 0.19 | 0.015 | 8.75122773843612e-81 | 16 |
| mef2cb | 3.98910978659936e-85 | 0.36861632 | 0.234 | 0.022 | 1.00401904218919e-80 | 16 |
| tmsb4x7 | 1.75207526577692e-84 | 0.739143778 | 1 | 0.681 | 4.40979823643394e-80 | 16 |
| rgs2 | 2.1923144828391e-84 | 0.396981239 | 0.242 | 0.024 | 5.51783632185772e-80 | 16 |
| si:ch211-204c21.11 | 8.13190173259187e-84 | 0.604654003 | 0.411 | 0.068 | 2.04671834707605e-79 | 16 |
| XLOC-0004313 | 1.12148982888935e-83 | 0.863741291 | 0.931 | 0.267 | 2.8226777503316e-79 | 16 |
| gm2a1 | 1.2000674931184e-83 | 0.568266723 | 0.576 | 0.113 | 3.0204498734297e-79 | 16 |
| atp6v1aa | 1.74158642830275e-83 | 0.376764766 | 0.286 | 0.033 | 4.38339888139519e-79 | 16 |
| arpc5b2 | 5.52993632589274e-83 | 0.522505743 | 0.55 | 0.106 | 1.39182967386394e-78 | 16 |
| cap15 | 9.02631675591193e-83 | 0.779902974 | 0.81 | 0.223 | 2.27183366429547e-78 | 16 |
| antxr2a3 | 9.68606579442351e-83 | 0.751031242 | 0.623 | 0.146 | 2.43788589979845e-78 | 16 |
| plxnc12 | 1.04244404306298e-82 | 0.374396715 | 0.312 | 0.039 | 2.62372741198522e-78 | 16 |
| echdc2 | 6.30419417454856e-82 | 0.367713065 | 0.273 | 0.031 | 1.58670263179213e-77 | 16 |
| nfkbiaa3 | 6.94662554919721e-82 | 0.785666507 | 0.615 | 0.144 | 1.74839618447745e-77 | 16 |
| gapdhs6 | 1.65293063954037e-81 | 0.828787959 | 0.861 | 0.278 | 4.16026112665917e-77 | 16 |
| junba3 | 3.8529126135665e-81 | 0.904849048 | 0.939 | 0.298 | 9.69739575708552e-77 | 16 |
| derl2 | 9.30931859377555e-81 | 0.540318144 | 0.515 | 0.098 | 2.34306239686737e-76 | 16 |
| pepd1 | 2.28345372741675e-80 | 0.411510165 | 0.407 | 0.065 | 5.74722468653521e-76 | 16 |
| ppiaa5 | 4.88764489946972e-80 | 0.461222879 | 1 | 0.801 | 1.23017134474753e-75 | 16 |
| psap1 | 3.19814109608227e-79 | 0.818701064 | 0.896 | 0.279 | 8.04940132472946e-75 | 16 |
| nfil3-63 | 3.42145247649364e-79 | 0.653832654 | 0.567 | 0.123 | 8.61145373808683e-75 | 16 |
| tram11 | 4.07236503936953e-79 | 0.492611773 | 0.506 | 0.095 | 1.02497355675892e-74 | 16 |
| tpi1b3 | 1.03075982670732e-78 | 0.62436993 | 0.667 | 0.158 | 2.59431940783964e-74 | 16 |
| ifngr1l | 1.46782395839729e-78 | 0.316746422 | 0.251 | 0.027 | 3.69436612089013e-74 | 16 |
| rgs191 | 1.96519875955277e-78 | 0.330625462 | 0.264 | 0.03 | 4.94620875791837e-74 | 16 |
| dram2b | 5.766999474154e-78 | 0.279394299 | 0.234 | 0.024 | 1.45149609764982e-73 | 16 |
| coro1a5 | 8.4245078845207e-78 | 0.767876776 | 0.844 | 0.22 | 2.12036438945501e-73 | 16 |
| itm2bb2 | 2.42866878938858e-77 | 0.727527971 | 0.775 | 0.212 | 6.11271647601212e-73 | 16 |
| slc2a3b1 | 3.1437789306262e-77 | 0.337081177 | 0.242 | 0.026 | 7.91257719049308e-73 | 16 |
| si:ch211-102c2.42 | 3.19418876170029e-77 | 0.613607242 | 0.615 | 0.134 | 8.03945369432345e-73 | 16 |
| rps15a6 | 7.09882936047877e-77 | 0.517819017 | 1 | 0.776 | 1.7867043617389e-72 | 16 |
| gsdmea1 | 1.01667765106671e-76 | 0.339964432 | 0.212 | 0.02 | 2.55887597996979e-72 | 16 |

| | | | | | | |
|---|---|---|---|---|---|---|
| ctsd2 | 1.15626076065264e-76 | 0.606687291 | 0.593 | 0.129 | 2.91019270848662e-72 | 16 |
| tspan37 | 1.23338313742255e-76 | 0.277159535 | 0.234 | 0.024 | 3.10430201857881e-72 | 16 |
| arhgef5 | 2.33138136429234e-76 | 0.315310389 | 0.238 | 0.025 | 5.86785375578738e-72 | 16 |
| stx11a | 2.59670341694435e-76 | 0.290622069 | 0.238 | 0.025 | 6.53564283010724e-72 | 16 |
| mboat2a | 4.61489193650134e-76 | 0.292918233 | 0.221 | 0.022 | 1.16152215149802e-71 | 16 |
| nfe2l2a | 4.85188836231095e-76 | 0.472388286 | 0.385 | 0.063 | 1.22117178191004e-71 | 16 |
| zgc:1627306 | 1.31735055107288e-75 | 0.929755291 | 0.944 | 0.382 | 3.31563960199532e-71 | 16 |
| tuba1b | 2.11030630757036e-75 | 0.328484784 | 0.247 | 0.028 | 5.31142994552385e-71 | 16 |
| srgn4 | 4.63473962734838e-75 | 0.761374951 | 0.805 | 0.227 | 1.16651761680731e-70 | 16 |
| arpc24 | 5.27948688898224e-75 | 0.745341912 | 0.848 | 0.257 | 1.32879405508794e-70 | 16 |
| dbi1 | 6.16940232964929e-75 | 0.689439438 | 0.745 | 0.204 | 1.55277687234943e-70 | 16 |
| CT030188.11 | 1.88174013567564e-74 | 0.38540751 | 0.329 | 0.047 | 4.73615174748202e-70 | 16 |
| lyn | 2.96787774157197e-74 | 0.356687959 | 0.303 | 0.041 | 7.4698514877625e-70 | 16 |
| si:zfos-741a10.31 | 4.26381394882787e-74 | 0.300079567 | 0.225 | 0.023 | 1.07315933278049e-69 | 16 |
| tuba8l21 | 8.64085820588539e-74 | 0.407698463 | 0.359 | 0.056 | 2.17481760183929e-69 | 16 |
| itgae.21 | 1.01342422814124e-73 | 0.34974339 | 0.264 | 0.032 | 2.55068743980868e-69 | 16 |
| slc43a2a2 | 2.00377709841666e-73 | 0.55310847 | 0.481 | 0.096 | 5.04330657900489e-69 | 16 |
| rab11fip4b1 | 2.32235939633076e-73 | 0.294741394 | 0.26 | 0.031 | 5.84514636462489e-69 | 16 |
| canx | 3.29247640779283e-73 | 0.477049595 | 0.442 | 0.082 | 8.28683387077376e-69 | 16 |
| si:ch1073-443f11.21 | 3.77545377445955e-73 | 0.286500258 | 0.264 | 0.032 | 9.50243960493724e-69 | 16 |
| crip15 | 5.00489892487882e-73 | 0.800005388 | 0.853 | 0.26 | 1.25968301040275e-68 | 16 |
| anxa42 | 5.44240736177723e-73 | 0.577494542 | 0.567 | 0.129 | 1.36979950888571e-68 | 16 |
| abcc3 | 1.72616372233959e-72 | 0.254944927 | 0.203 | 0.02 | 4.3445814727565e-68 | 16 |
| ehd4 | 2.38105056148982e-72 | 0.302474402 | 0.234 | 0.026 | 5.99286615821374e-68 | 16 |
| stat5a | 1.91665998979891e-71 | 0.387824863 | 0.294 | 0.041 | 4.82404152832487e-67 | 16 |
| tfe3a | 1.98623725520195e-71 | 0.31692054 | 0.273 | 0.035 | 4.9991605476178e-67 | 16 |
| gnao1a | 4.15650184302281e-71 | 0.275678201 | 0.203 | 0.02 | 1.04614994887041e-66 | 16 |
| si:ch73-248e21.72 | 6.64178782311976e-71 | 0.366860348 | 0.294 | 0.04 | 1.67167157720101e-66 | 16 |
| bhlhe402 | 7.27881188923418e-71 | 0.658408504 | 0.619 | 0.153 | 1.83200416440135e-66 | 16 |
| dbn12 | 7.92432107616621e-71 | 0.388972313 | 0.333 | 0.051 | 1.99447237166027e-66 | 16 |
| klf131 | 8.08568506437075e-71 | 0.506600864 | 0.485 | 0.099 | 2.03508607385147e-66 | 16 |
| wdr11 | 1.06716561119238e-70 | 0.564027595 | 0.58 | 0.134 | 2.68594912681009e-66 | 16 |
| si:dkey-112e7.24 | 2.52718646716277e-70 | 0.675028119 | 0.706 | 0.197 | 6.36067561920198e-66 | 16 |
| nme2b.11 | 2.66258174920218e-70 | 0.35764594 | 0.991 | 0.855 | 6.70145200456697e-66 | 16 |
| si:dkey-25e12.32 | 4.61422028825375e-70 | 0.407661217 | 0.316 | 0.047 | 1.16135310435059e-65 | 16 |

| | | | | | | |
|---|---|---|---|---|---|---|
| csrnp1b3 | 6.77940394206812e-70 | 0.660719054 | 0.654 | 0.171 | 1.70630817817912e-65 | 16 |
| birc22 | 8.89241828730213e-70 | 0.525153555 | 0.485 | 0.101 | 2.23813275873107e-65 | 16 |
| si:ch211-39i2.2 | 2.41500638132408e-69 | 0.431264768 | 0.368 | 0.061 | 6.07832956115457e-65 | 16 |
| nocta | 2.60756678879654e-69 | 0.357838448 | 0.294 | 0.041 | 6.562984850722e-65 | 16 |
| irf81 | 5.44759016799989e-69 | 0.377374442 | 0.299 | 0.042 | 1.37110396938389e-64 | 16 |
| triqk | 9.63702344872538e-69 | 0.343503148 | 0.312 | 0.046 | 2.42554243180969e-64 | 16 |
| rpl36a6 | 2.38963603170281e-68 | 0.427826457 | 0.996 | 0.807 | 6.0144749281928e-64 | 16 |
| si:ch211-212k18.71 | 3.04215169199669e-68 | 0.553988418 | 0.571 | 0.132 | 7.65679159358647e-64 | 16 |
| BX908782.2 | 3.1154695405431e-68 | 0.419550933 | 0.216 | 0.023 | 7.84132528659292e-64 | 16 |
| zgc:1037001 | 3.60569411508657e-68 | 0.419601932 | 0.273 | 0.036 | 9.0751715182614e-64 | 16 |
| xbp12 | 4.35399049465196e-68 | 0.806626275 | 0.879 | 0.334 | 1.09585586759895e-63 | 16 |
| atp6ap2 | 1.03593132653545e-67 | 0.472710424 | 0.468 | 0.095 | 2.60733555575707e-63 | 16 |
| rassf2a | 1.32525081127714e-67 | 0.271594737 | 0.19 | 0.019 | 3.33552376690344e-63 | 16 |
| cflarb | 1.46783126456437e-67 | 0.258158841 | 0.177 | 0.016 | 3.69438450978206e-63 | 16 |
| atp6v1d | 2.04139641898057e-67 | 0.260139218 | 0.208 | 0.022 | 5.1379906469322e-63 | 16 |
| ssr4 | 2.14819955714453e-67 | 0.752604959 | 0.857 | 0.297 | 5.40680346537706e-63 | 16 |
| rpl246 | 6.04154387497867e-67 | 0.464161986 | 0.996 | 0.773 | 1.52059617789338e-62 | 16 |
| lcp13 | 8.56596096394929e-67 | 0.484544035 | 0.429 | 0.082 | 2.1559667150164e-62 | 16 |
| rap1b2 | 1.79953562167677e-66 | 0.607447378 | 0.71 | 0.193 | 4.52925120619825e-62 | 16 |
| rps146 | 1.8544683361562e-66 | 0.480008118 | 0.991 | 0.755 | 4.66751135527153e-62 | 16 |
| arpc1b5 | 2.91042850089354e-66 | 0.830590051 | 0.913 | 0.298 | 7.32525749389895e-62 | 16 |
| wasb4 | 3.85857142255002e-66 | 0.589381558 | 0.675 | 0.169 | 9.71163841341615e-62 | 16 |
| gadd45bb3 | 7.84079958169599e-66 | 0.494055373 | 0.42 | 0.082 | 1.97345084671706e-61 | 16 |
| wipf1b2 | 8.85349044121342e-66 | 0.363622898 | 0.351 | 0.058 | 2.22833500914901e-61 | 16 |
| rabgap1l | 9.2360747855055e-66 | 0.28808054 | 0.255 | 0.033 | 2.32462766276388e-61 | 16 |
| si:ch73-237c6.13 | 1.31346996023127e-65 | 0.321645626 | 0.277 | 0.038 | 3.30587254290608e-61 | 16 |
| grap2b3 | 2.79266064953045e-65 | 0.467455503 | 0.502 | 0.106 | 7.02884758880319e-61 | 16 |
| suclg11 | 1.2482855134531e-64 | 0.47486905 | 0.481 | 0.103 | 3.1418098088101e-60 | 16 |
| BX323596.1 | 4.26933281307577e-64 | 0.288384183 | 0.126 | 0.008 | 1.07454837572304e-59 | 16 |
| rpl156 | 4.4894462415081e-64 | 0.505781709 | 0.996 | 0.74 | 1.12994872452517e-59 | 16 |
| elovl1b1 | 5.77749666563046e-64 | 0.46780084 | 0.502 | 0.11 | 1.45413813577253e-59 | 16 |
| rps96 | 1.30319011298202e-63 | 0.453510848 | 0.983 | 0.774 | 3.27999919536444e-59 | 16 |
| sec61g1 | 2.21494118937585e-63 | 0.552310475 | 0.991 | 0.674 | 5.57478547954008e-59 | 16 |
| rps218 | 2.59226220144396e-63 | 0.501660428 | 0.991 | 0.746 | 6.52446473481429e-59 | 16 |
| gng101 | 3.35905726300352e-63 | 0.36806933 | 0.316 | 0.05 | 8.45441122525355e-59 | 16 |

| | | | | | | |
|---|---|---|---|---|---|---|
| rplp2l6 | 4.2398241109868e-63 | 0.481339824 | 0.996 | 0.771 | 1.06712133049427e-58 | 16 |
| rpl396 | 5.11510294233204e-63 | 0.368363619 | 1 | 0.872 | 1.28742025955555e-58 | 16 |
| rnf144b | 6.74564855287051e-63 | 0.271596813 | 0.225 | 0.027 | 1.69781228427198e-58 | 16 |
| pfn15 | 9.13288355552378e-63 | 0.856968781 | 1 | 0.517 | 2.29865546208978e-58 | 16 |
| ifi301 | 1.07798144620085e-62 | 0.679154487 | 0.615 | 0.168 | 2.71317150194292e-58 | 16 |
| ahr21 | 4.44455736814386e-62 | 0.334942266 | 0.29 | 0.044 | 1.11865064398813e-57 | 16 |
| flna2 | 4.65447234680096e-62 | 0.470703722 | 0.489 | 0.108 | 1.17148414496633e-57 | 16 |
| atf32 | 4.68665635649931e-62 | 0.476666678 | 0.424 | 0.086 | 1.17958453836731e-57 | 16 |
| FP074902.2 | 5.28163792977162e-62 | 0.409518179 | 0.39 | 0.075 | 1.32933545054422e-57 | 16 |
| rps205 | 1.28668761370725e-61 | 0.323516314 | 1 | 0.915 | 3.23846405493979e-57 | 16 |
| cpox1 | 1.69464932110963e-61 | 0.332721855 | 0.299 | 0.046 | 4.26526287630083e-57 | 16 |
| vma21 | 2.83810998357374e-61 | 0.463338168 | 0.463 | 0.101 | 7.14323901765675e-57 | 16 |
| si:ch1073-392o20.2 | 2.85346151830183e-61 | 0.351455155 | 0.307 | 0.049 | 7.18187729541388e-57 | 16 |
| btg15 | 5.83207871217306e-61 | 0.726874065 | 0.896 | 0.323 | 1.46787589106684e-56 | 16 |
| rps195 | 7.13126423755579e-61 | 0.364062366 | 1 | 0.861 | 1.79486789595042e-56 | 16 |
| rpl35a6 | 8.80020392492813e-61 | 0.462503608 | 0.987 | 0.756 | 2.21492332586516e-56 | 16 |
| ppiab2 | 2.21656020346224e-60 | 0.554635239 | 0.978 | 0.651 | 5.57886037609412e-56 | 16 |
| rpl366 | 2.34698006761392e-60 | 0.455849795 | 0.991 | 0.787 | 5.90711413217749e-56 | 16 |
| hcls12 | 2.7500114439274e-60 | 0.362118048 | 0.364 | 0.066 | 6.92150380322088e-56 | 16 |
| gdi2 | 3.16391382435915e-60 | 0.57067315 | 0.641 | 0.177 | 7.96325470452955e-56 | 16 |
| pkma2 | 4.18167207519105e-60 | 0.56187339 | 0.632 | 0.17 | 1.05248504460483e-55 | 16 |
| rpl296 | 1.09484790713245e-59 | 0.42002679 | 0.996 | 0.806 | 2.75562269746167e-55 | 16 |
| pim12 | 1.15668689457427e-59 | 0.635881089 | 0.602 | 0.164 | 2.91126524495397e-55 | 16 |
| rac25 | 1.46422184777634e-59 | 0.647918485 | 0.745 | 0.212 | 3.68529996866827e-55 | 16 |
| celf25 | 1.66649531172543e-59 | 0.640925746 | 0.758 | 0.233 | 4.19440205008172e-55 | 16 |
| osbp | 2.03134581204268e-59 | 0.275829136 | 0.251 | 0.035 | 5.11269427433023e-55 | 16 |
| zgc:1141886 | 2.97081078112404e-59 | 0.59868846 | 0.978 | 0.622 | 7.47723365501109e-55 | 16 |
| ccr9a4 | 4.30228760549529e-59 | 0.469947822 | 0.407 | 0.082 | 1.08284276742711e-54 | 16 |
| smdt1b | 5.24135898530442e-59 | 0.467862501 | 0.524 | 0.127 | 1.31919764301127e-54 | 16 |
| dnase1l1 | 8.39807216879514e-59 | 0.340072737 | 0.273 | 0.042 | 2.11371078416405e-54 | 16 |
| wbp2nl | 8.65869264697389e-59 | 0.405147905 | 0.403 | 0.082 | 2.17930635231686e-54 | 16 |
| ostf1 | 1.50196224341754e-58 | 0.379688063 | 0.299 | 0.049 | 3.78028877045761e-54 | 16 |
| rpl376 | 1.84170647666149e-58 | 0.400710699 | 1 | 0.847 | 4.63539103110929e-54 | 16 |
| ehbp1l1b | 1.9455885200912e-58 | 0.251988257 | 0.182 | 0.019 | 4.89685174621755e-54 | 16 |
| cfbl1 | 3.24747082189982e-58 | 0.261662865 | 0.351 | 0.062 | 8.17355931163966e-54 | 16 |

| | | | | | | |
|---|---|---|---|---|---|---|
| rpl186 | 3.26492121393894e-58 | 0.459638367 | 0.996 | 0.77 | 8.21748020336291e-54 | 16 |
| eef1a1l12 | 4.79055868492763e-58 | 0.297733351 | 1 | 0.914 | 1.20573571540943e-53 | 16 |
| rpl234 | 5.23250089935871e-58 | 0.331731246 | 1 | 0.871 | 1.31696815135959e-53 | 16 |
| tagln24 | 5.42976498616213e-58 | 0.590787528 | 0.645 | 0.18 | 1.36661754936715e-53 | 16 |
| XLOC-0254233 | 5.68528898433252e-58 | 0.765356261 | 0.58 | 0.163 | 1.43093038446665e-53 | 16 |
| rpl386 | 9.71070856214275e-58 | 0.501182561 | 1 | 0.724 | 2.44408823800571e-53 | 16 |
| prdx6 | 1.06874501813872e-57 | 0.571112603 | 0.632 | 0.174 | 2.68992433615334e-53 | 16 |
| metrnla | 1.35396268876215e-57 | 0.422603808 | 0.42 | 0.088 | 3.40778869134545e-53 | 16 |
| pycard4 | 1.96586288812328e-57 | 0.385163977 | 0.359 | 0.068 | 4.94788030311748e-53 | 16 |
| arpc4l4 | 2.06523935090439e-57 | 0.588508719 | 0.649 | 0.189 | 5.19800092229126e-53 | 16 |
| si:dkey-180p18.91 | 2.1517675281573e-57 | 0.390855081 | 0.437 | 0.094 | 5.4157836916191e-53 | 16 |
| rps27.16 | 2.50965150198865e-57 | 0.55261111 | 0.987 | 0.674 | 6.31654186535523e-53 | 16 |
| cox6b14 | 3.27237934664235e-57 | 0.45475023 | 0.489 | 0.114 | 8.23625157756414e-53 | 16 |
| CABZ01073834.11 | 4.81326791949056e-57 | 0.296487201 | 0.26 | 0.038 | 1.21145140265658e-52 | 16 |
| rgs18 | 1.16823464458924e-56 | 0.280057587 | 0.229 | 0.031 | 2.94032977696665e-52 | 16 |
| rpl27a6 | 1.31387447530967e-56 | 0.681723499 | 0.935 | 0.489 | 3.30689066690691e-52 | 16 |
| rps26l6 | 1.32077741635077e-56 | 0.551807454 | 0.974 | 0.659 | 3.32426467921324e-52 | 16 |
| LOC1039091153 | 1.64297838173682e-56 | 0.503894383 | 0.602 | 0.158 | 4.1352122889934e-52 | 16 |
| eef1db3 | 1.99085805961814e-56 | 0.661905687 | 0.779 | 0.265 | 5.01079065025289e-52 | 16 |
| znf827 | 2.00787501992255e-56 | 0.279274263 | 0.212 | 0.027 | 5.05362063764307e-52 | 16 |
| rplp15 | 4.26185712225573e-56 | 0.367508106 | 1 | 0.875 | 1.07266681910054e-51 | 16 |
| rps235 | 5.89892066459223e-56 | 0.477990621 | 0.983 | 0.719 | 1.48469934207122e-51 | 16 |
| clic13 | 6.42410066150549e-56 | 0.463167812 | 0.463 | 0.108 | 1.61688189549432e-51 | 16 |
| hmha1b2 | 7.8593475757361e-56 | 0.361040584 | 0.364 | 0.07 | 1.97811919133702e-51 | 16 |
| rps166 | 8.21316179545775e-56 | 0.484701406 | 0.991 | 0.734 | 2.06717069229876e-51 | 16 |
| ckbb4 | 2.39966637562577e-55 | 0.692430558 | 0.801 | 0.257 | 6.03972030081249e-51 | 16 |
| rps125 | 2.83256202661931e-55 | 0.383353842 | 1 | 0.837 | 7.12927536479814e-51 | 16 |
| DHDH1 | 3.21505533730358e-55 | 0.312459772 | 0.32 | 0.057 | 8.09197277845938e-51 | 16 |
| ppp1r182 | 3.59841882433613e-55 | 0.297612126 | 0.299 | 0.05 | 9.0568603389716e-51 | 16 |
| rps156 | 5.28251502464982e-55 | 0.529711509 | 0.987 | 0.68 | 1.32955620655411e-50 | 16 |
| rab8b1 | 7.00692254521119e-55 | 0.385584697 | 0.372 | 0.075 | 1.7635723354042e-50 | 16 |
| bri3 | 9.80877798619909e-55 | 0.339124972 | 0.316 | 0.057 | 2.46877133134645e-50 | 16 |
| sdcbp23 | 1.02838501514744e-54 | 0.575327745 | 0.667 | 0.201 | 2.5883422446246e-50 | 16 |
| rps105 | 2.21232777341379e-54 | 0.51076335 | 0.991 | 0.689 | 5.56820777290516e-50 | 16 |
| ppt12 | 2.7429127299983e-54 | 0.503692529 | 0.58 | 0.156 | 6.90363705013271e-50 | 16 |

| | | | | | | |
|---|---|---|---|---|---|---|
| rpl18a6 | 3.69947007593386e-54 | 0.444641826 | 1 | 0.782 | 9.31119623411792e-50 | 16 |
| grinab2 | 3.7078335773378e-54 | 0.442506061 | 0.459 | 0.108 | 9.33224633080152e-50 | 16 |
| skap21 | 5.1829170297659e-54 | 0.346494912 | 0.359 | 0.07 | 1.30448838722178e-49 | 16 |
| mafbb | 1.04734913779413e-53 | 0.259962231 | 0.173 | 0.019 | 2.63607304491405e-49 | 16 |
| rpl306 | 1.55247035139832e-53 | 0.49440787 | 0.991 | 0.704 | 3.90741262743443e-49 | 16 |
| pdk2b2 | 1.59773867770141e-53 | 0.487031033 | 0.416 | 0.095 | 4.02134847790669e-49 | 16 |
| rps296 | 3.0408699839012e-53 | 0.385011909 | 1 | 0.83 | 7.65356566248094e-49 | 16 |
| arhgap151 | 4.4211627211944e-53 | 0.252098411 | 0.203 | 0.026 | 1.11276244529742e-48 | 16 |
| mbd2 | 5.14347964787265e-53 | 0.265741003 | 0.225 | 0.032 | 1.29456239257307e-48 | 16 |
| rpl286 | 5.55266403083213e-53 | 0.448343886 | 0.987 | 0.746 | 1.39755000992014e-48 | 16 |
| rps56 | 5.73693256884336e-53 | 0.504667874 | 0.978 | 0.697 | 1.44392855825219e-48 | 16 |
| jak11 | 8.06837653258067e-53 | 0.403574924 | 0.411 | 0.091 | 2.03072968948523e-48 | 16 |
| CR936442.13 | 9.28231751368591e-53 | 0.478278713 | 0.61 | 0.162 | 2.33626649501961e-48 | 16 |
| rps286 | 1.2104302272154e-52 | 0.397402363 | 1 | 0.824 | 3.04653183887843e-48 | 16 |
| rpl265 | 1.51945885427261e-52 | 0.373007964 | 1 | 0.839 | 3.82432599031874e-48 | 16 |
| mrtfab2 | 2.7083047851196e-52 | 0.345733652 | 0.286 | 0.049 | 6.81653231366752e-48 | 16 |
| rpl116 | 5.51140794936057e-52 | 0.414989385 | 1 | 0.811 | 1.38716626677456e-47 | 16 |
| rpl23a6 | 6.5793268715425e-52 | 0.560826019 | 0.987 | 0.629 | 1.65595078029853e-47 | 16 |
| mdh1aa1 | 1.03893316520627e-51 | 0.53445587 | 0.658 | 0.198 | 2.61489088350766e-47 | 16 |
| zgc:110591 | 1.06711242191765e-51 | 0.310668118 | 0.277 | 0.047 | 2.68581525472454e-47 | 16 |
| rpl96 | 1.24318047978961e-51 | 0.569920698 | 0.97 | 0.624 | 3.12896094958247e-47 | 16 |
| rpl326 | 1.39263838189285e-51 | 0.411835568 | 1 | 0.81 | 3.50513154338612e-47 | 16 |
| atp6v1h | 1.83324336229185e-51 | 0.264837375 | 0.26 | 0.042 | 4.61409021855236e-47 | 16 |
| fybb2 | 3.24386824767706e-51 | 0.310012118 | 0.247 | 0.039 | 8.1644919925784e-47 | 16 |
| si:ch211-74f19.21 | 3.38333576429729e-51 | 0.271684376 | 0.212 | 0.029 | 8.51551778515986e-47 | 16 |
| ddit34 | 3.58962515949858e-51 | 0.551667061 | 0.589 | 0.17 | 9.03472756394198e-47 | 16 |
| kdm7aa | 3.68613748407476e-51 | 0.254199152 | 0.208 | 0.028 | 9.27763943366777e-47 | 16 |
| rpl175 | 3.91050982525478e-51 | 0.428252632 | 0.974 | 0.743 | 9.84236217918376e-47 | 16 |
| dusp12 | 4.07982932153365e-51 | 0.572844973 | 0.545 | 0.154 | 1.0268522419368e-46 | 16 |
| npl | 4.91734438516565e-51 | 0.305637645 | 0.229 | 0.034 | 1.23764640830234e-46 | 16 |
| rpl105 | 1.05605201072859e-50 | 0.416261535 | 0.978 | 0.769 | 2.65797730580278e-46 | 16 |
| dennd4a | 2.29353187379113e-50 | 0.356331983 | 0.225 | 0.034 | 5.77259037314489e-46 | 16 |
| oaz1b5 | 6.19659157934451e-50 | 0.636349979 | 0.827 | 0.314 | 1.55962013460522e-45 | 16 |
| si:dkey-27i16.22 | 8.30020255929818e-50 | 0.486127733 | 0.385 | 0.086 | 2.08907798214976e-45 | 16 |
| pak2a | 1.12729594196798e-49 | 0.321556335 | 0.307 | 0.058 | 2.8372911563392e-45 | 16 |

| | | | | | | |
|---|---|---|---|---|---|---|
| rplp26 | 1.90367332193452e-49 | 0.47005991 | 0.996 | 0.754 | 4.791355383977e-45 | 16 |
| ufm1 | 2.7579101301186e-49 | 0.317692868 | 0.385 | 0.084 | 6.94138400649551e-45 | 16 |
| XLOC-0425953 | 3.88993066535444e-49 | 0.38227896 | 0.593 | 0.157 | 9.79056649163059e-45 | 16 |
| ssr1 | 4.22251938143781e-49 | 0.467116167 | 0.619 | 0.181 | 1.06276590311408e-44 | 16 |
| krtcap2 | 4.89676398404607e-49 | 0.476222446 | 0.589 | 0.17 | 1.23246652714455e-44 | 16 |
| rps255 | 5.29281047847599e-49 | 0.460305504 | 0.991 | 0.721 | 1.33214746932762e-44 | 16 |
| CR392341.12 | 6.83426403621151e-49 | 0.357266775 | 0.299 | 0.056 | 1.72011591527407e-44 | 16 |
| rps35 | 9.56614468960918e-49 | 0.474361752 | 0.97 | 0.696 | 2.40770295692774e-44 | 16 |
| rpl194 | 9.98148255493565e-49 | 0.352027563 | 1 | 0.85 | 2.51223934425175e-44 | 16 |
| aldoaa1 | 1.2706649212252e-48 | 0.582361295 | 0.723 | 0.243 | 3.1981365402317e-44 | 16 |
| rps8a5 | 1.92773206120469e-48 | 0.401043226 | 0.991 | 0.785 | 4.85190882484608e-44 | 16 |
| rpl10a5 | 2.34565416242987e-48 | 0.427510774 | 0.991 | 0.761 | 5.90377696141975e-44 | 16 |
| slc25a55 | 2.83636279977791e-48 | 0.374132497 | 0.991 | 0.804 | 7.13884153076102e-44 | 16 |
| myh9a1 | 3.10644715657813e-48 | 0.261527646 | 0.286 | 0.051 | 7.8186168483915e-44 | 16 |
| rpl275 | 3.39920314846673e-48 | 0.433001216 | 0.996 | 0.759 | 8.55545440437591e-44 | 16 |
| rps65 | 3.45186319643606e-48 | 0.413234139 | 0.996 | 0.786 | 8.68799447910991e-44 | 16 |
| mob1a3 | 3.81464703745404e-48 | 0.424372677 | 0.576 | 0.159 | 9.60108512856807e-44 | 16 |
| abi2b | 3.85661492177143e-48 | 0.344445741 | 0.273 | 0.049 | 9.70671409660651e-44 | 16 |
| psme23 | 4.29033194916762e-48 | 0.390336867 | 0.398 | 0.092 | 1.079833648286e-43 | 16 |
| arpc5a4 | 4.64833454440721e-48 | 0.412541207 | 0.446 | 0.11 | 1.16993932148185e-43 | 16 |
| ncf11 | 4.99104769233451e-48 | 0.269745344 | 0.212 | 0.031 | 1.25619679368367e-43 | 16 |
| arhgdia3 | 7.07614903703435e-48 | 0.539512544 | 0.688 | 0.222 | 1.78099595113117e-43 | 16 |
| FERMT3 (1 of many)2 | 1.00024757273764e-47 | 0.288626138 | 0.238 | 0.038 | 2.51752311582336e-43 | 16 |
| CABZ01068358.11 | 1.14940546618933e-47 | 0.381738851 | 0.307 | 0.061 | 2.89293861785194e-43 | 16 |
| gnai1 | 1.31645763454035e-47 | 0.319332248 | 0.294 | 0.056 | 3.3133922203746e-43 | 16 |
| rps75 | 1.38390056387393e-47 | 0.380838339 | 0.996 | 0.818 | 3.48313932921429e-43 | 16 |
| rpl66 | 1.45447783316043e-47 | 0.517044769 | 0.991 | 0.662 | 3.66077525828148e-43 | 16 |
| rplp04 | 1.5513890658011e-47 | 0.348846673 | 0.991 | 0.849 | 3.90469113971479e-43 | 16 |
| rpl86 | 1.94209098355782e-47 | 0.417824381 | 0.987 | 0.777 | 4.88804879651667e-43 | 16 |
| m6pr | 2.43520382438665e-47 | 0.354558318 | 0.398 | 0.091 | 6.12916450559875e-43 | 16 |
| krt83 | 2.45260216703603e-47 | 0.592701051 | 0.81 | 0.267 | 6.17295439421299e-43 | 16 |
| cox8a3 | 2.54274975932044e-47 | 0.535281662 | 0.68 | 0.22 | 6.39984686923362e-43 | 16 |
| rpn1 | 3.13276054245887e-47 | 0.391788062 | 0.528 | 0.142 | 7.88484500931473e-43 | 16 |
| sri1 | 3.30494671652985e-47 | 0.469486021 | 0.519 | 0.145 | 8.31822039083397e-43 | 16 |
| bloc1s6 | 4.72068668064431e-47 | 0.315692449 | 0.355 | 0.076 | 1.18814963065137e-42 | 16 |

| | | | | | | |
|---|---|---|---|---|---|---|
| rab1ab | 5.34767796659709e-47 | 0.39236098 | 0.459 | 0.117 | 1.34595706741282e-42 | 16 |
| slc7a3a1 | 7.06139009993948e-47 | 0.278343174 | 0.242 | 0.04 | 1.77728127425377e-42 | 16 |
| COX5B1 | 7.27941054333324e-47 | 0.632160496 | 0.866 | 0.349 | 1.83215483965154e-42 | 16 |
| rps27a5 | 8.15012343202712e-47 | 0.420847306 | 0.987 | 0.748 | 2.0513045666069e-42 | 16 |
| rps246 | 1.28609972157247e-46 | 0.430285994 | 0.996 | 0.755 | 3.23698438922574e-42 | 16 |
| rpl316 | 1.66047236258322e-46 | 0.516544825 | 0.987 | 0.649 | 4.1792428893857e-42 | 16 |
| vdac32 | 1.75320985264289e-46 | 0.423149565 | 0.537 | 0.148 | 4.41265387811689e-42 | 16 |
| cfl13 | 2.58849019672884e-46 | 0.612295022 | 0.965 | 0.571 | 6.51497097614681e-42 | 16 |
| rpl346 | 5.45077483686948e-46 | 0.419855523 | 0.996 | 0.78 | 1.37190551869168e-41 | 16 |
| rpsa6 | 7.5759520505373e-46 | 0.439658543 | 0.991 | 0.748 | 1.90679137159973e-41 | 16 |
| si:dkey-53k12.22 | 9.14384509679352e-46 | 0.270251869 | 0.247 | 0.042 | 2.30141437241196e-41 | 16 |
| slc20a1b1 | 1.17911325257574e-45 | 0.365671365 | 0.407 | 0.097 | 2.96771014540789e-41 | 16 |
| smim15 | 1.33602549933286e-45 | 0.402324837 | 0.506 | 0.138 | 3.36264257927087e-41 | 16 |
| rpl226 | 1.46304608196257e-45 | 0.482249822 | 0.974 | 0.69 | 3.68234068369159e-41 | 16 |
| syngr2a1 | 1.53029029797426e-45 | 0.258980318 | 0.281 | 0.052 | 3.85158765097141e-41 | 16 |
| rps4x5 | 1.70154148864738e-45 | 0.47090708 | 0.978 | 0.706 | 4.2826097727766e-41 | 16 |
| si:ch1073-325m22.2 | 1.80992072886799e-45 | 0.505074634 | 0.68 | 0.222 | 4.55538948248786e-41 | 16 |
| tln11 | 2.53203374384061e-45 | 0.415711881 | 0.463 | 0.122 | 6.37287572987243e-41 | 16 |
| rpl7a5 | 2.85111248528002e-45 | 0.381049633 | 0.991 | 0.807 | 7.17596501420128e-41 | 16 |
| capzb3 | 3.41528718871427e-45 | 0.534394351 | 0.727 | 0.245 | 8.59593632527494e-41 | 16 |
| egr14 | 4.09198827689594e-45 | 0.550107561 | 0.593 | 0.183 | 1.02991252941194e-40 | 16 |
| gstm.2 | 4.40301379271154e-45 | 0.279172714 | 0.26 | 0.047 | 1.10819454148757e-40 | 16 |
| rpl216 | 4.46857812004186e-45 | 0.417096271 | 0.983 | 0.753 | 1.12469642703334e-40 | 16 |
| mapre1a1 | 5.58481284379839e-45 | 0.449889158 | 0.641 | 0.19 | 1.40564154465562e-40 | 16 |
| gnai2a2 | 6.82468230342803e-45 | 0.438261141 | 0.498 | 0.138 | 1.7177042889498e-40 | 16 |
| rps3a5 | 7.19992133854389e-45 | 0.39508525 | 0.983 | 0.775 | 1.81214820169811e-40 | 16 |
| rpl124 | 8.2034973719962e-45 | 0.371355156 | 0.991 | 0.795 | 2.06473825355772e-40 | 16 |
| rps135 | 8.21191451586318e-45 | 0.458981309 | 0.991 | 0.72 | 2.0668567644976e-40 | 16 |
| hsp90ab15 | 8.21923021212417e-45 | 0.381969791 | 0.991 | 0.787 | 2.06869805208953e-40 | 16 |
| lgals3b1 | 9.54560676127839e-45 | 0.350402036 | 0.338 | 0.075 | 2.40253376574616e-40 | 16 |
| plp21 | 1.07933787963629e-44 | 0.252867719 | 0.264 | 0.048 | 2.71658550925658e-40 | 16 |
| faua6 | 1.27417266740495e-44 | 0.488071575 | 0.987 | 0.685 | 3.20696518659152e-40 | 16 |
| rps219 | 1.42648661159271e-44 | 0.403521363 | 0.996 | 0.827 | 3.5903241527177e-40 | 16 |
| csrp1a | 1.75334209279782e-44 | 0.297289797 | 0.312 | 0.064 | 4.41298671336285e-40 | 16 |
| cited4a1 | 2.16041525799511e-44 | 0.279495987 | 0.234 | 0.04 | 5.43754916284789e-40 | 16 |

| | | | | | | |
|---|---|---|---|---|---|---|
| sucla2 | 2.97748428075762e-44 | 0.277123315 | 0.277 | 0.053 | 7.49403018623885e-40 | 16 |
| ppib3 | 4.30102262117096e-44 | 0.615358699 | 0.853 | 0.324 | 1.08252438352252e-39 | 16 |
| gpia | 6.62601124024533e-44 | 0.349330568 | 0.455 | 0.117 | 1.66770076905735e-39 | 16 |
| atp6v0b | 8.10165850906448e-44 | 0.362862327 | 0.407 | 0.1 | 2.03910643014644e-39 | 16 |
| txnipa5 | 1.00401584765292e-43 | 0.566623892 | 0.61 | 0.204 | 2.52700748695765e-39 | 16 |
| dub3 | 1.26829786423191e-43 | 0.313472224 | 0.32 | 0.067 | 3.1921788944853e-39 | 16 |
| mt-nd51 | 1.41575502375147e-43 | 0.403095269 | 0.437 | 0.115 | 3.56331381928007e-39 | 16 |
| zgc:1629443 | 1.64229896833566e-43 | 0.526072934 | 0.714 | 0.245 | 4.13350227340401e-39 | 16 |
| vdac22 | 1.8584109198699e-43 | 0.577724784 | 0.762 | 0.287 | 4.67743444422055e-39 | 16 |
| capza1b1 | 3.31941815800552e-43 | 0.483565591 | 0.641 | 0.206 | 8.35464356188409e-39 | 16 |
| csde1 | 3.57627358416483e-43 | 0.447177306 | 0.636 | 0.201 | 9.00112298398447e-39 | 16 |
| zgc:664271 | 5.36153008358326e-43 | 0.340300849 | 0.346 | 0.078 | 1.34944350673707e-38 | 16 |
| si:dkey-238c7.161 | 6.4044475756242e-43 | 0.273943616 | 0.294 | 0.059 | 1.61193541030885e-38 | 16 |
| mt-atp66 | 8.66346081090865e-43 | 0.613151756 | 0.965 | 0.585 | 2.1805064514976e-38 | 16 |
| ssr2 | 9.3849439764797e-43 | 0.595934948 | 0.814 | 0.313 | 2.36209654944018e-38 | 16 |
| arpc34 | 1.01431695910165e-42 | 0.587981296 | 0.801 | 0.302 | 2.55293435436295e-38 | 16 |
| rack15 | 1.11075683637217e-42 | 0.417445798 | 0.996 | 0.74 | 2.79566388146511e-38 | 16 |
| rnaseka | 2.10977189978755e-42 | 0.410348872 | 0.459 | 0.124 | 5.31008489457529e-38 | 16 |
| tsc22d31 | 3.27128928245692e-42 | 0.441804235 | 0.485 | 0.139 | 8.23350799501581e-38 | 16 |
| XLOC-010897 | 3.32244175264901e-42 | 0.287488144 | 0.264 | 0.051 | 8.36225364724228e-38 | 16 |
| lamp1a | 3.50050350683203e-42 | 0.335779625 | 0.368 | 0.087 | 8.81041727634553e-38 | 16 |
| cited4b | 3.66150840365553e-42 | 0.393772787 | 0.472 | 0.126 | 9.21565050116061e-38 | 16 |
| mfng2 | 3.93453694526253e-42 | 0.272196229 | 0.229 | 0.04 | 9.90283603753128e-38 | 16 |
| arhgdig4 | 2.19826568617753e-41 | 0.530367946 | 0.697 | 0.236 | 5.53281490554023e-37 | 16 |
| calr3b | 2.19995316387737e-41 | 0.357989084 | 0.325 | 0.073 | 5.53706211816296e-37 | 16 |
| COX7A2 (1 of many)2 | 2.51671137895247e-41 | 0.510093137 | 0.719 | 0.253 | 6.33431086968547e-37 | 16 |
| pcyox1 | 2.54416304619313e-41 | 0.297030313 | 0.251 | 0.047 | 6.40340397096349e-37 | 16 |
| atp2b21 | 3.52284054384635e-41 | 0.313166459 | 0.312 | 0.068 | 8.86663736480688e-37 | 16 |
| eef1g5 | 5.22163879795595e-41 | 0.525562751 | 0.961 | 0.615 | 1.31423426905753e-36 | 16 |
| eno1a4 | 5.9347004303606e-41 | 0.409660824 | 0.463 | 0.129 | 1.49370475131746e-36 | 16 |
| ldhba1 | 1.13058884491883e-40 | 0.504462006 | 0.688 | 0.24 | 2.84557906377621e-36 | 16 |
| zgc:1537132 | 1.19302606645913e-40 | 0.404585553 | 0.502 | 0.147 | 3.00272730667097e-36 | 16 |
| ncor2 | 2.38016761973244e-40 | 0.323620149 | 0.312 | 0.069 | 5.99064388210459e-36 | 16 |
| ddost | 5.37603736155502e-40 | 0.300723883 | 0.381 | 0.094 | 1.35309484352978e-35 | 16 |
| rpl356 | 5.755880466435e-40 | 0.364528312 | 0.987 | 0.802 | 1.44869755459703e-35 | 16 |

| | | | | | | |
|---|---|---|---|---|---|---|
| ier3ip1 | 6.29890791040943e-40 | 0.363549407 | 0.506 | 0.147 | 1.58537213197095e-35 | 16 |
| msna5 | 6.9625495717669e-40 | 0.509901871 | 0.675 | 0.235 | 1.75240410171801e-35 | 16 |
| ap2m1a | 1.90618796836973e-39 | 0.362418809 | 0.481 | 0.137 | 4.79768449758977e-35 | 16 |
| uqcrq1 | 2.60784640714328e-39 | 0.495979941 | 0.736 | 0.264 | 6.56368862213891e-35 | 16 |
| mbnl3 | 1.02744596565973e-38 | 0.310813047 | 0.26 | 0.053 | 2.58597875096898e-34 | 16 |
| ahnak2 | 1.18960240458117e-38 | 0.400597134 | 0.437 | 0.122 | 2.99411029209033e-34 | 16 |
| socs3b3 | 1.21050116793819e-38 | 0.448257709 | 0.502 | 0.15 | 3.04671038958364e-34 | 16 |
| atp5pd2 | 1.65077111554948e-38 | 0.48301466 | 0.714 | 0.254 | 4.1548258207265e-34 | 16 |
| mt-nd35 | 1.79262371896995e-38 | 0.559137263 | 0.9 | 0.387 | 4.51185463827548e-34 | 16 |
| hdlbpa2 | 1.90953200109925e-38 | 0.397917903 | 0.511 | 0.154 | 4.80610109356671e-34 | 16 |
| ptprc4 | 2.13334966037404e-38 | 0.369777866 | 0.446 | 0.123 | 5.36942776019543e-34 | 16 |
| spcs1 | 2.31264893086897e-38 | 0.43427075 | 0.61 | 0.203 | 5.82070609410411e-34 | 16 |
| rpl134 | 3.43806363732039e-38 | 0.346755615 | 0.983 | 0.802 | 8.65326236877168e-34 | 16 |
| pde4ba4 | 4.79392971938516e-38 | 0.415752042 | 0.463 | 0.135 | 1.20658417107205e-33 | 16 |
| aoc23 | 8.12607740854138e-38 | 0.280957925 | 0.29 | 0.063 | 2.04525242295578e-33 | 16 |
| COX37 | 1.09580904851129e-37 | 0.524340023 | 0.991 | 0.696 | 2.75804179419807e-33 | 16 |
| spg21 | 1.18896636935546e-37 | 0.267768246 | 0.26 | 0.053 | 2.99250945503076e-33 | 16 |
| adipor2 | 1.23348803545455e-37 | 0.29607523 | 0.342 | 0.083 | 3.10456603643555e-33 | 16 |
| BX322618.11 | 1.32585508173878e-37 | 0.329899756 | 0.173 | 0.027 | 3.33704465522833e-33 | 16 |
| adh52 | 1.37375838803003e-37 | 0.450749166 | 0.632 | 0.215 | 3.45761248683277e-33 | 16 |
| etfb | 1.39208002900897e-37 | 0.30176832 | 0.351 | 0.087 | 3.50372622501269e-33 | 16 |
| rps185 | 1.3980124494033e-37 | 0.445735893 | 0.991 | 0.721 | 3.51865753390317e-33 | 16 |
| gnb1a2 | 1.76009190490291e-37 | 0.442790089 | 0.567 | 0.187 | 4.42997531545013e-33 | 16 |
| capza1a3 | 1.76063100105726e-37 | 0.438465944 | 0.584 | 0.194 | 4.43133216656101e-33 | 16 |
| ccl39.71 | 1.98914178880714e-37 | 0.57457657 | 0.26 | 0.057 | 5.00647096824868e-33 | 16 |
| rps115 | 2.31229061981592e-37 | 0.381181854 | 0.991 | 0.762 | 5.8198042610147e-33 | 16 |
| gabarapb3 | 2.72621346538394e-37 | 0.494308254 | 0.71 | 0.261 | 6.86160667102484e-33 | 16 |
| ifi462 | 3.05548265233749e-37 | 0.326313949 | 0.429 | 0.117 | 7.69034428766824e-33 | 16 |
| uba525 | 4.4515046878344e-37 | 0.510488674 | 0.991 | 0.624 | 1.12039921488104e-32 | 16 |
| si:ch73-42p12.2 | 7.81210470859827e-37 | 0.297921998 | 0.377 | 0.098 | 1.9662286341071e-32 | 16 |
| GCA3 | 8.37128141783334e-37 | 0.415770673 | 0.545 | 0.175 | 2.10696782005447e-32 | 16 |
| arrb2b2 | 8.53066981271251e-37 | 0.303566462 | 0.351 | 0.088 | 2.14708428516161e-32 | 16 |
| anxa1a2 | 9.58348993491619e-37 | 0.408956782 | 0.472 | 0.138 | 2.41206858171906e-32 | 16 |
| mt-co27 | 1.17476690174589e-36 | 0.506653709 | 0.978 | 0.716 | 2.95677081500424e-32 | 16 |
| rpl5a5 | 1.18475670882813e-36 | 0.593243894 | 0.939 | 0.485 | 2.98191416044952e-32 | 16 |

| | | | | | | |
|---|---|---|---|---|---|---|
| arf6b2 | 1.27342506332216e-36 | 0.354624476 | 0.368 | 0.097 | 3.20508354187553e-32 | 16 |
| si:dkey-79d12.52 | 1.42394162295343e-36 | 0.327952927 | 0.307 | 0.073 | 3.58391867081148e-32 | 16 |
| zgc:1584631 | 1.4248175850643e-36 | 0.299371966 | 0.303 | 0.071 | 3.58612337984834e-32 | 16 |
| mrps153 | 1.60644464669046e-36 | 0.371401391 | 0.429 | 0.12 | 4.04326053125522e-32 | 16 |
| rpl75 | 2.06892088810746e-36 | 0.350300012 | 0.987 | 0.802 | 5.20726698327767e-32 | 16 |
| jdp2b1 | 2.1209251732665e-36 | 0.479642424 | 0.511 | 0.165 | 5.33815656859446e-32 | 16 |
| eif4ebp12 | 2.32299943433458e-36 | 0.469479844 | 0.675 | 0.239 | 5.8467572762767e-32 | 16 |
| naca6 | 2.75302439273568e-36 | 0.489790522 | 0.939 | 0.614 | 6.92908709407643e-32 | 16 |
| actr2a3 | 3.19458509372255e-36 | 0.385775877 | 0.433 | 0.125 | 8.04045122239029e-32 | 16 |
| zbtb16a | 3.61572664723393e-36 | 0.265532486 | 0.234 | 0.046 | 9.10042239842308e-32 | 16 |
| atp1b1a1 | 4.56051397465142e-36 | 0.444670624 | 0.606 | 0.208 | 1.14783576228002e-31 | 16 |
| tmem2581 | 4.76537012130707e-36 | 0.471023876 | 0.74 | 0.275 | 1.19939600583178e-31 | 16 |
| mrpl331 | 5.91152193446676e-36 | 0.341661485 | 0.359 | 0.094 | 1.48787095568594e-31 | 16 |
| ctsc | 5.95824688043229e-36 | 0.31138872 | 0.299 | 0.069 | 1.499631157336e-31 | 16 |
| eif3k1 | 6.67271010007191e-36 | 0.431221749 | 0.641 | 0.223 | 1.6794544050871e-31 | 16 |
| fkbp51 | 7.19955354143875e-36 | 0.358678103 | 0.416 | 0.118 | 1.81205563084472e-31 | 16 |
| capgb4 | 7.57068278348528e-36 | 0.333929122 | 0.407 | 0.11 | 1.90546514977541e-31 | 16 |
| tpt15 | 7.87098463238631e-36 | 0.341915606 | 0.978 | 0.798 | 1.98104812212531e-31 | 16 |
| zeb2a1 | 8.15559214088618e-36 | 0.263992679 | 0.221 | 0.042 | 2.05268098593964e-31 | 16 |
| NDUFB13 | 1.28429016527408e-35 | 0.381398878 | 0.584 | 0.192 | 3.23242991697834e-31 | 16 |
| nt5dc21 | 1.29483822674193e-35 | 0.267602757 | 0.312 | 0.074 | 3.25897833288677e-31 | 16 |
| mt-nd15 | 1.3501708201768e-35 | 0.558932465 | 0.87 | 0.404 | 3.39824493730299e-31 | 16 |
| ier2b5 | 1.36309853705099e-35 | 0.465306108 | 0.623 | 0.217 | 3.43078270790364e-31 | 16 |
| fkbp2 | 2.15475213927817e-35 | 0.313302932 | 0.45 | 0.13 | 5.42329565934923e-31 | 16 |
| irf2 | 3.12363796609577e-35 | 0.314857157 | 0.281 | 0.065 | 7.86188439686645e-31 | 16 |
| eif3m4 | 3.49376131392068e-35 | 0.555958451 | 0.818 | 0.351 | 8.79344785100695e-31 | 16 |
| commd7 | 3.50622076208966e-35 | 0.261779222 | 0.307 | 0.073 | 8.82480703610348e-31 | 16 |
| eif3s6ip | 4.18399338835868e-35 | 0.358818101 | 0.472 | 0.144 | 1.053069295916e-30 | 16 |
| cox4i13 | 4.39820089307253e-35 | 0.56141196 | 0.874 | 0.384 | 1.10698318277743e-30 | 16 |
| eif3c4 | 4.83114192402627e-35 | 0.388547567 | 0.511 | 0.164 | 1.21595011085817e-30 | 16 |
| psme13 | 7.64675988645768e-35 | 0.300068832 | 0.338 | 0.086 | 1.92461299582253e-30 | 16 |
| ier2a4 | 8.46447075087862e-35 | 0.476150475 | 0.697 | 0.255 | 2.13042264328864e-30 | 16 |
| XLOC-0108993 | 9.49863449558275e-35 | 0.358711345 | 0.706 | 0.251 | 2.39071131619322e-30 | 16 |
| midn3 | 1.63494055687301e-34 | 0.39462607 | 0.446 | 0.136 | 4.11498188759367e-30 | 16 |
| eif2s1b1 | 1.95556028474884e-34 | 0.379788348 | 0.519 | 0.168 | 4.92194968068435e-30 | 16 |

| | | | | | | |
|---|---|---|---|---|---|---|
| selenof1 | 2.20516998325531e-34 | 0.422964783 | 0.649 | 0.23 | 5.5501923308553e-30 | 16 |
| eif3s103 | 3.33298269502199e-34 | 0.381071725 | 0.498 | 0.16 | 8.38878414510084e-30 | 16 |
| chmp2a2 | 3.949613210387e-34 | 0.334473573 | 0.437 | 0.129 | 9.94078148922305e-30 | 16 |
| ak21 | 4.74204218086824e-34 | 0.446603676 | 0.623 | 0.226 | 1.19352459650273e-29 | 16 |
| rasgef1ba | 5.10728610404038e-34 | 0.331211878 | 0.273 | 0.063 | 1.28545283952592e-29 | 16 |
| ccni4 | 7.52220188501441e-34 | 0.477253356 | 0.745 | 0.287 | 1.89326299243928e-29 | 16 |
| zgc:1583433 | 8.08442640604147e-34 | 0.415858384 | 0.498 | 0.161 | 2.03476928213658e-29 | 16 |
| lasp13 | 1.11769944518759e-33 | 0.432666558 | 0.641 | 0.23 | 2.81313773359265e-29 | 16 |
| rab5c | 1.21590728405743e-33 | 0.332368857 | 0.403 | 0.116 | 3.06031704324414e-29 | 16 |
| rpl22l14 | 1.79890410306794e-33 | 0.582120128 | 0.84 | 0.384 | 4.5276617370117e-29 | 16 |
| cbr1l | 2.12039621401878e-33 | 0.280526835 | 0.359 | 0.097 | 5.33682523106386e-29 | 16 |
| atp5pf2 | 2.4975098882783e-33 | 0.47632446 | 0.745 | 0.291 | 6.28598263780766e-29 | 16 |
| snx1a | 2.89141713824328e-33 | 0.276728588 | 0.394 | 0.11 | 7.2774077952445e-29 | 16 |
| rhoab4 | 3.67976039781519e-33 | 0.415542839 | 0.537 | 0.184 | 9.26158894526105e-29 | 16 |
| rsl24d15 | 3.78452070977372e-33 | 0.575405635 | 0.879 | 0.437 | 9.52526017442947e-29 | 16 |
| litaf3 | 4.29757095424102e-33 | 0.343870896 | 0.433 | 0.13 | 1.08165563347292e-28 | 16 |
| arpc43 | 5.20630192027563e-33 | 0.404103398 | 0.632 | 0.219 | 1.31037413031417e-28 | 16 |
| twf1a1 | 5.422710175388e-33 | 0.320786193 | 0.476 | 0.146 | 1.36484192404341e-28 | 16 |
| zfp36l1a4 | 5.98259930161242e-33 | 0.467200332 | 0.654 | 0.242 | 1.50576041822283e-28 | 16 |
| egr32 | 6.96971608844881e-33 | 0.29493583 | 0.251 | 0.056 | 1.75420784230168e-28 | 16 |
| ndufa2 | 8.51748658668912e-33 | 0.321300978 | 0.459 | 0.141 | 2.14376619900378e-28 | 16 |
| rps265 | 1.42649053031607e-32 | 0.400059102 | 0.991 | 0.73 | 3.59033401575253e-28 | 16 |
| lgals9l31 | 1.92375487607376e-32 | 0.333563085 | 0.26 | 0.061 | 4.84189864759005e-28 | 16 |
| ybx14 | 1.94250332440603e-32 | 0.434131751 | 0.952 | 0.665 | 4.88908661719754e-28 | 16 |
| b2ml6 | 2.1828372841241e-32 | 0.449333851 | 0.688 | 0.26 | 5.49398316041194e-28 | 16 |
| ptmaa5 | 2.36103325125044e-32 | 0.583183913 | 0.874 | 0.387 | 5.94248459007224e-28 | 16 |
| zgc:77486 | 3.20313719112181e-32 | 0.269893236 | 0.281 | 0.068 | 8.06197599633448e-28 | 16 |
| cox7b3 | 3.24556949084187e-32 | 0.548284738 | 0.874 | 0.409 | 8.1687738514999e-28 | 16 |
| CU929150.11 | 4.66099006280177e-32 | 0.389549174 | 0.147 | 0.023 | 1.17312458890658e-27 | 16 |
| mibp26 | 6.77363042085188e-32 | 0.543828676 | 0.887 | 0.486 | 1.70485504062421e-27 | 16 |
| XLOC-0283703 | 9.92023916324715e-32 | 0.469914237 | 0.429 | 0.137 | 2.49682499499767e-27 | 16 |
| ywhaba2 | 1.59252148169286e-31 | 0.465324874 | 0.788 | 0.313 | 4.00821731727276e-27 | 16 |
| ctsz1 | 1.76561235092851e-31 | 0.264596621 | 0.32 | 0.084 | 4.44386972605197e-27 | 16 |
| cox5ab | 1.82725203282697e-31 | 0.340587863 | 0.576 | 0.196 | 4.59901064142221e-27 | 16 |
| glo1 | 2.05085041828158e-31 | 0.368038964 | 0.554 | 0.191 | 5.1617854177729e-27 | 16 |

| | | | | | | |
|---|---|---|---|---|---|---|
| cyfip1 | 2.12908348447191e-31 | 0.257826623 | 0.242 | 0.054 | 5.35869022206734e-27 | 16 |
| ddx213 | 2.34765648093698e-31 | 0.372549143 | 0.58 | 0.201 | 5.90881659687029e-27 | 16 |
| mcl1b2 | 3.29863780028193e-31 | 0.431472339 | 0.641 | 0.236 | 8.30234147952958e-27 | 16 |
| tnfrsf1a1 | 4.35902249931851e-31 | 0.251620746 | 0.234 | 0.052 | 1.09712237285348e-26 | 16 |
| si:ch73-1a9.33 | 5.58513051096255e-31 | 0.449767874 | 0.684 | 0.259 | 1.40572149830416e-26 | 16 |
| socs3a4 | 6.04278262542866e-31 | 0.497741943 | 0.745 | 0.291 | 1.52090795899414e-26 | 16 |
| mt-cyb5 | 6.23330340946016e-31 | 0.511885277 | 0.961 | 0.619 | 1.56886013512703e-26 | 16 |
| atp5f1d2 | 7.33516883612022e-31 | 0.400474834 | 0.68 | 0.251 | 1.8461886443631e-26 | 16 |
| BX005183.33 | 9.91332164287704e-31 | 0.351847928 | 0.299 | 0.078 | 2.49508392429572e-26 | 16 |
| ccl34b.11 | 1.11126430609654e-30 | 0.284625724 | 0.165 | 0.028 | 2.79694113201438e-26 | 16 |
| cyth1a1 | 2.01847341707177e-30 | 0.258212922 | 0.268 | 0.065 | 5.08029574342795e-26 | 16 |
| lamtor5 | 3.07908918478602e-30 | 0.296935021 | 0.429 | 0.131 | 7.74975956918793e-26 | 16 |
| aimp1 | 3.23675211313371e-30 | 0.271510863 | 0.329 | 0.09 | 8.14658139354623e-26 | 16 |
| calr3a | 3.92637854531901e-30 | 0.300553103 | 0.45 | 0.141 | 9.88230216071343e-26 | 16 |
| eef2b3 | 6.06320471538714e-30 | 0.466527915 | 0.944 | 0.609 | 1.52604799481579e-25 | 16 |
| pgam1a3 | 6.18162780528806e-30 | 0.318741538 | 0.472 | 0.152 | 1.55585390231295e-25 | 16 |
| rpl145 | 8.97974373240214e-30 | 0.457923467 | 0.97 | 0.63 | 2.26011170000829e-25 | 16 |
| ola11 | 1.07379208852732e-29 | 0.393107371 | 0.602 | 0.224 | 2.7026273076144e-25 | 16 |
| sumo2b5 | 1.41286564618824e-29 | 0.405677911 | 0.688 | 0.262 | 3.55604154489119e-25 | 16 |
| cct23 | 1.85139903913112e-29 | 0.383296341 | 0.602 | 0.221 | 4.65978624158912e-25 | 16 |
| eif3ha2 | 1.96472136472771e-29 | 0.382569071 | 0.558 | 0.203 | 4.94500720288318e-25 | 16 |
| chmp1b | 3.15785523518678e-29 | 0.280610886 | 0.39 | 0.118 | 7.94800584144161e-25 | 16 |
| eif3f5 | 3.17044542626178e-29 | 0.554105664 | 0.87 | 0.424 | 7.97969409335827e-25 | 16 |
| cmip1 | 3.52825783290454e-29 | 0.252113395 | 0.255 | 0.062 | 8.88027213963743e-25 | 16 |
| reep51 | 4.62711988155904e-29 | 0.309708329 | 0.468 | 0.154 | 1.16459980298959e-24 | 16 |
| mt-co17 | 4.65477101692218e-29 | 0.487702256 | 0.974 | 0.653 | 1.17155931724914e-24 | 16 |
| cnn21 | 5.25402028705257e-29 | 0.360674887 | 0.407 | 0.131 | 1.32238436604826e-24 | 16 |
| rpl45 | 5.49192430198844e-29 | 0.461827334 | 0.961 | 0.622 | 1.38226242756747e-24 | 16 |
| btf35 | 7.66097741913055e-29 | 0.485559755 | 0.944 | 0.582 | 1.92819140662097e-24 | 16 |
| rpl317 | 9.85583641044981e-29 | 0.258834797 | 1 | 0.873 | 2.48061546614611e-24 | 16 |
| fosab3 | 1.02319371355113e-28 | 0.4764344 | 0.818 | 0.341 | 2.57527625763684e-24 | 16 |
| sec61b1 | 1.31516426793932e-28 | 0.5319814 | 0.874 | 0.448 | 3.31013694597646e-24 | 16 |
| gpx4b | 1.69620469102394e-28 | 0.325588883 | 0.558 | 0.198 | 4.26917758683815e-24 | 16 |
| selenok1 | 1.72855880183177e-28 | 0.279597472 | 0.398 | 0.123 | 4.35060964833039e-24 | 16 |
| atp5mf1 | 1.84911971149754e-28 | 0.389126514 | 0.719 | 0.281 | 4.65404940186816e-24 | 16 |

| | | | | | | |
|---|---|---|---|---|---|---|
| atp5pb | 1.90371821089668e-28 | 0.416853224 | 0.736 | 0.293 | 4.79146836500586e-24 | 16 |
| eif3i5 | 2.43195140694081e-28 | 0.426646045 | 0.667 | 0.267 | 6.12097849612933e-24 | 16 |
| atf4b3 | 2.75585123011211e-28 | 0.417020922 | 0.667 | 0.258 | 6.93620196106916e-24 | 16 |
| si:dkey-42i9.61 | 3.06312444920757e-28 | 0.259818485 | 0.277 | 0.072 | 7.70957792621052e-24 | 16 |
| mt-nd24 | 3.38999720210201e-28 | 0.488200142 | 0.853 | 0.393 | 8.53228395797054e-24 | 16 |
| mknk2b4 | 4.14528936891768e-28 | 0.402992256 | 0.524 | 0.192 | 1.04332788126289e-23 | 16 |
| cox6c | 4.48757279424563e-28 | 0.498742452 | 0.892 | 0.409 | 1.12947719658368e-23 | 16 |
| stk17b1 | 5.09992427898205e-28 | 0.261441529 | 0.364 | 0.107 | 1.28359994177699e-23 | 16 |
| aldh9a1a.12 | 5.62230232522951e-28 | 0.3272265 | 0.472 | 0.16 | 1.41507727223702e-23 | 16 |
| dad11 | 6.49146257816186e-28 | 0.460368404 | 0.805 | 0.344 | 1.63383621629756e-23 | 16 |
| mgst1.22 | 6.92191909993113e-28 | 0.2799408 | 0.342 | 0.1 | 1.74217781826167e-23 | 16 |
| cox6b21 | 7.34029574274051e-28 | 0.506311157 | 0.926 | 0.527 | 1.84747903549036e-23 | 16 |
| ube2v21 | 1.2134410434822e-27 | 0.406099263 | 0.667 | 0.263 | 3.05410976234035e-23 | 16 |
| sec11a1 | 1.37979059585069e-27 | 0.281725334 | 0.506 | 0.172 | 3.4727949506966e-23 | 16 |
| mtdha | 1.63027241184786e-27 | 0.295746686 | 0.416 | 0.135 | 4.10323263337988e-23 | 16 |
| eif3ba3 | 1.78818315502745e-27 | 0.376420365 | 0.55 | 0.205 | 4.5006781828886e-23 | 16 |
| magt1 | 2.10874254510532e-27 | 0.281349961 | 0.364 | 0.111 | 5.30749411177557e-23 | 16 |
| sh3bgrl32 | 2.1289730381339e-27 | 0.292277623 | 0.437 | 0.143 | 5.35841223967921e-23 | 16 |
| nsa25 | 2.59877100280883e-27 | 0.51326777 | 0.784 | 0.356 | 6.54084673696954e-23 | 16 |
| tomm52 | 2.60492251116496e-27 | 0.341788907 | 0.567 | 0.209 | 6.55632946835108e-23 | 16 |
| myl12.13 | 2.90382927630731e-27 | 0.507859284 | 0.844 | 0.384 | 7.30864790553787e-23 | 16 |
| nr4a32 | 3.29697685527212e-27 | 0.293235058 | 0.221 | 0.052 | 8.29816104703441e-23 | 16 |
| elf13 | 3.33338440617381e-27 | 0.270811729 | 0.342 | 0.101 | 8.38979521189885e-23 | 16 |
| ndufb3 | 3.37810518400901e-27 | 0.343000401 | 0.571 | 0.212 | 8.50235293763229e-23 | 16 |
| psmb2 | 3.91774287384532e-27 | 0.288812732 | 0.468 | 0.158 | 9.8605670391813e-23 | 16 |
| alas1 | 3.98714422135099e-27 | 0.284555488 | 0.42 | 0.135 | 1.00352432907183e-22 | 16 |
| ost4 | 4.40767579660081e-27 | 0.346385673 | 0.658 | 0.251 | 1.10936792124646e-22 | 16 |
| eif3ea5 | 5.46634420209343e-27 | 0.424611289 | 0.662 | 0.271 | 1.3758241722249e-22 | 16 |
| glrx3 | 5.60533751308333e-27 | 0.443910547 | 0.723 | 0.305 | 1.41080739866794e-22 | 16 |
| atp5l3 | 6.51642193062442e-27 | 0.509251462 | 0.892 | 0.442 | 1.64011823571886e-22 | 16 |
| vps29 | 6.94804243149412e-27 | 0.273652257 | 0.381 | 0.12 | 1.74875279958275e-22 | 16 |
| rps173 | 8.85654242129127e-27 | 0.348649099 | 0.991 | 0.764 | 2.2291031620148e-22 | 16 |
| rnf101 | 1.18003922816306e-26 | 0.258761989 | 0.342 | 0.103 | 2.9700407333636e-22 | 16 |
| aimp2 | 1.19138652061032e-26 | 0.251325297 | 0.381 | 0.119 | 2.99860073372411e-22 | 16 |
| cox5aa1 | 1.3122130385172e-26 | 0.457469166 | 0.788 | 0.345 | 3.30270899664393e-22 | 16 |

| | | | | | | |
|---|---|---|---|---|---|---|
| tmed2 | 1.74162645772407e-26 | 0.373666731 | 0.619 | 0.242 | 4.38349963144571e-22 | 16 |
| pfdn4 | 4.17940084496507e-26 | 0.301429002 | 0.476 | 0.167 | 1.05191339866926e-21 | 16 |
| cct71 | 4.63781790863104e-26 | 0.321658182 | 0.554 | 0.204 | 1.16729238942335e-21 | 16 |
| cox6a11 | 5.1851274563414e-26 | 0.465207316 | 0.818 | 0.366 | 1.30504472948657e-21 | 16 |
| pnrc27 | 5.21176267412336e-26 | 0.504428359 | 0.957 | 0.561 | 1.31174854745011e-21 | 16 |
| pomp1 | 6.19927804479894e-26 | 0.341902895 | 0.589 | 0.225 | 1.56029629109545e-21 | 16 |
| pgk1 | 7.00208903170756e-26 | 0.251871502 | 0.39 | 0.124 | 1.76235578839047e-21 | 16 |
| anxa11b1 | 8.70096968109313e-26 | 0.299960859 | 0.429 | 0.146 | 2.18994705903433e-21 | 16 |
| atf4a1 | 1.1959895432529e-25 | 0.297854617 | 0.498 | 0.177 | 3.01018608141323e-21 | 16 |
| itm2ba3 | 1.31867043803333e-25 | 0.394141706 | 0.662 | 0.267 | 3.3189616254861e-21 | 16 |
| cct83 | 2.3503736118598e-25 | 0.327544031 | 0.532 | 0.197 | 5.91565534368994e-21 | 16 |
| elmo13 | 2.64995997515918e-25 | 0.265004781 | 0.255 | 0.068 | 6.66968426147814e-21 | 16 |
| hnrnpa1a2 | 2.75225754707833e-25 | 0.301710943 | 0.446 | 0.155 | 6.92715702024146e-21 | 16 |
| metap2b | 3.30661398985991e-25 | 0.317552117 | 0.545 | 0.205 | 8.32241675107841e-21 | 16 |
| rnf213a3 | 4.64027301069647e-25 | 0.274332379 | 0.294 | 0.086 | 1.16791031406219e-20 | 16 |
| mt-nd45 | 4.74860519565052e-25 | 0.480188249 | 0.896 | 0.449 | 1.19517644169328e-20 | 16 |
| tnfb2 | 5.28427200472327e-25 | 0.264997139 | 0.212 | 0.051 | 1.3299984208688e-20 | 16 |
| uqcrh2 | 5.47595862282895e-25 | 0.357708306 | 0.615 | 0.244 | 1.37824402577982e-20 | 16 |
| mcl1a4 | 5.63953178468329e-25 | 0.461618536 | 0.974 | 0.618 | 1.41941375488694e-20 | 16 |
| cox7a2l | 5.83528489539484e-25 | 0.258731651 | 0.515 | 0.183 | 1.46868285532193e-20 | 16 |
| tomm20b4 | 5.97331338597423e-25 | 0.332249743 | 0.55 | 0.21 | 1.50342324611585e-20 | 16 |
| ndufa4l3 | 6.62204025332647e-25 | 0.476069378 | 0.835 | 0.388 | 1.66670131135974e-20 | 16 |
| anxa3b1 | 1.15532917460438e-24 | 0.251997576 | 0.156 | 0.031 | 2.90784799956175e-20 | 16 |
| odc1 | 1.33951700339574e-24 | 0.296150799 | 0.368 | 0.119 | 3.37143034584674e-20 | 16 |
| ponzr14 | 1.74227484897966e-24 | 0.40498756 | 0.658 | 0.267 | 4.3851315673969e-20 | 16 |
| dusp52 | 2.1029583119274e-24 | 0.290595504 | 0.255 | 0.071 | 5.29293577529006e-20 | 16 |
| ipo7 | 3.08862539580284e-24 | 0.264507986 | 0.338 | 0.107 | 7.77376125869616e-20 | 16 |
| gtpbp43 | 3.63635641384581e-24 | 0.362779908 | 0.58 | 0.232 | 9.15234545800851e-20 | 16 |
| fam120c | 9.44003114532898e-24 | 0.264720048 | 0.312 | 0.096 | 2.37596143896785e-19 | 16 |
| ndufa3 | 1.40535310721877e-23 | 0.252271839 | 0.494 | 0.179 | 3.53713323555891e-19 | 16 |
| rpl13a5 | 1.46615547704575e-23 | 0.324906484 | 0.983 | 0.761 | 3.69016672017645e-19 | 16 |
| pdia6 | 1.98265650246989e-23 | 0.291914761 | 0.351 | 0.115 | 4.99014815106646e-19 | 16 |
| b2m5 | 2.24729481828288e-23 | 0.386164736 | 0.684 | 0.286 | 5.65621632813618e-19 | 16 |
| si:dkey-7j14.63 | 3.04978553617536e-23 | 0.262140122 | 0.476 | 0.17 | 7.67600521599977e-19 | 16 |
| si:ch211-196l7.46 | 3.23226356938546e-23 | 0.406835877 | 0.697 | 0.308 | 8.13528417778627e-19 | 16 |

| | | | | | | |
|---|---|---|---|---|---|---|
| p4hb1 | 6.69209587902039e-23 | 0.32659983 | 0.528 | 0.211 | 1.68433361179064e-18 | 16 |
| psma2 | 8.28660855695757e-23 | 0.262523422 | 0.42 | 0.148 | 2.08565650770065e-18 | 16 |
| sat1a.22 | 1.01101643635888e-22 | 0.408856075 | 0.671 | 0.293 | 2.54462726867166e-18 | 16 |
| prdx22 | 1.80408781569492e-22 | 0.451031879 | 0.81 | 0.395 | 4.54070862332253e-18 | 16 |
| rsl1d11 | 2.40164557955758e-22 | 0.279736633 | 0.476 | 0.177 | 6.04470175918846e-18 | 16 |
| rps27.25 | 3.10903424265584e-22 | 0.508143655 | 0.944 | 0.511 | 7.82512828534047e-18 | 16 |
| zgc:564934 | 3.35189372416709e-22 | 0.46610775 | 0.944 | 0.566 | 8.43638131435615e-18 | 16 |
| zgc:193541 | 3.57921800351811e-22 | 0.427468481 | 0.762 | 0.362 | 9.00853379305473e-18 | 16 |
| eif3ja | 4.41089205478584e-22 | 0.259286189 | 0.494 | 0.186 | 1.11017742126905e-17 | 16 |
| nop562 | 4.90579668021374e-22 | 0.279733029 | 0.45 | 0.167 | 1.234739966443e-17 | 16 |
| rps27l3 | 5.5856526280974e-22 | 0.261704456 | 0.524 | 0.199 | 1.40585290996583e-17 | 16 |
| cct51 | 5.652679010833e-22 | 0.331363569 | 0.597 | 0.246 | 1.42272278023656e-17 | 16 |
| map3k14a | 6.09775745954054e-22 | 0.264432751 | 0.147 | 0.031 | 1.53474457499176e-17 | 16 |
| psma8 | 6.18290674054493e-22 | 0.284749585 | 0.519 | 0.202 | 1.55617579752775e-17 | 16 |
| csnk2b | 6.24526559945027e-22 | 0.278419211 | 0.515 | 0.198 | 1.57187089872564e-17 | 16 |
| tmed10 | 7.13491126051987e-22 | 0.271086622 | 0.45 | 0.168 | 1.79578581516025e-17 | 16 |
| arl6ip11 | 8.66031019388057e-22 | 0.424924714 | 0.801 | 0.363 | 2.1797134726978e-17 | 16 |
| serbp1a5 | 1.54829305827982e-21 | 0.483422223 | 0.9 | 0.475 | 3.89689879838449e-17 | 16 |
| eif3g4 | 1.56284781036455e-21 | 0.367730138 | 0.641 | 0.277 | 3.93353165390654e-17 | 16 |
| eif4a1b2 | 5.28938290353957e-21 | 0.350873223 | 0.645 | 0.275 | 1.33128478299187e-16 | 16 |
| atp5mc3a1 | 6.19754855952861e-21 | 0.26933006 | 0.55 | 0.218 | 1.55986099694776e-16 | 16 |
| arf2a1 | 7.87420005975939e-21 | 0.255265981 | 0.355 | 0.124 | 1.98185741304084e-16 | 16 |
| atp5po | 7.95661573554874e-21 | 0.306412712 | 0.632 | 0.264 | 2.00260061448026e-16 | 16 |
| abracl1 | 1.21554042092599e-20 | 0.26726208 | 0.554 | 0.217 | 3.05939368542861e-16 | 16 |
| cox7a2a1 | 1.33349825720358e-20 | 0.290121336 | 0.58 | 0.237 | 3.35628176355569e-16 | 16 |
| eif3d3 | 1.34578905359135e-20 | 0.320576783 | 0.593 | 0.248 | 3.38721646898408e-16 | 16 |
| idh22 | 1.41897285961745e-20 | 0.290746247 | 0.494 | 0.195 | 3.57141279037117e-16 | 16 |
| cd9b1 | 1.93847055346258e-20 | 0.330272476 | 0.221 | 0.063 | 4.87893653600996e-16 | 16 |
| eif2s32 | 2.27829916660712e-20 | 0.344963008 | 0.641 | 0.279 | 5.73425117243347e-16 | 16 |
| cct3 | 3.51201055592303e-20 | 0.277337458 | 0.506 | 0.202 | 8.83937936820266e-16 | 16 |
| ywhabb1 | 3.03672085105895e-19 | 0.27537312 | 0.537 | 0.222 | 7.64312271003028e-15 | 16 |
| serf25 | 3.9099229895014e-19 | 0.365438356 | 0.745 | 0.337 | 9.84088517227607e-15 | 16 |
| atp5md | 6.71209973987609e-19 | 0.357378906 | 0.762 | 0.353 | 1.68936838352941e-14 | 16 |
| rpl5b5 | 9.89088746039338e-19 | 0.40088474 | 0.944 | 0.625 | 2.48943746490641e-14 | 16 |
| nop534 | 1.70149371058554e-18 | 0.320763189 | 0.563 | 0.247 | 4.28248952017274e-14 | 16 |

| | | | | | | |
|---|---|---|---|---|---|---|
| gng53 | 2.00416760569876e-18 | 0.276029727 | 0.623 | 0.266 | 5.04428944678322e-14 | 16 |
| cox7c2 | 2.36822702918497e-18 | 0.420770108 | 0.883 | 0.475 | 5.96059060975565e-14 | 16 |
| NC-002333.46 | 2.42116904120636e-18 | 0.431993551 | 0.974 | 0.617 | 6.0938403598123e-14 | 16 |
| atp5f1c | 4.01016227963842e-18 | 0.268787769 | 0.584 | 0.25 | 1.00931774416219e-13 | 16 |
| tcp12 | 5.971180981639e-18 | 0.267433674 | 0.524 | 0.222 | 1.50288654126872e-13 | 16 |
| slc25a3b4 | 1.01844641617059e-17 | 0.416013886 | 0.823 | 0.431 | 2.56332778485977e-13 | 16 |
| tegt3 | 1.79144218883927e-17 | 0.269061759 | 0.489 | 0.208 | 4.50888084508956e-13 | 16 |
| romo15 | 3.35601115937323e-17 | 0.391874947 | 0.771 | 0.38 | 8.44674448702648e-13 | 16 |
| eef1b21 | 4.80338920233551e-17 | 0.305201135 | 0.948 | 0.732 | 1.20896502833582e-12 | 16 |
| CR383676.17 | 5.30297722858387e-17 | 0.555705884 | 0.97 | 0.722 | 1.33470633866227e-12 | 16 |
| hspa8 | 2.39235006481198e-16 | 0.260416394 | 1 | 0.826 | 6.02130587812527e-12 | 16 |
| atp5mc3b | 3.07854231972313e-16 | 0.37397723 | 0.887 | 0.599 | 7.74838316451114e-12 | 16 |
| pabpc1a4 | 3.09857025768457e-16 | 0.424895024 | 0.922 | 0.534 | 7.79879148156629e-12 | 16 |
| ywhaz2 | 3.39484981440492e-15 | 0.266717443 | 0.606 | 0.277 | 8.54449749787573e-11 | 16 |
| eif2s22 | 4.84010782637013e-15 | 0.263979011 | 0.623 | 0.289 | 1.2182067388191e-10 | 16 |
| atp5fa1 | 9.40341032827621e-15 | 0.28923395 | 0.732 | 0.351 | 2.36674434552384e-10 | 16 |
| atp5meb1 | 1.06177696980974e-14 | 0.270571434 | 0.675 | 0.317 | 2.67238645531413e-10 | 16 |
| npc21 | 1.5084512788444e-14 | 0.282288662 | 0.745 | 0.353 | 3.79662102372348e-10 | 16 |
| histh1l3 | 2.58580158139513e-14 | 0.261257536 | 0.277 | 0.108 | 6.5082040002134e-10 | 16 |
| fkbp1aa3 | 4.01739699400808e-14 | 0.329518393 | 0.775 | 0.381 | 1.01113864942189e-09 | 16 |
| ecm1b | 9.02402207354912e-14 | 0.401343846 | 0.165 | 0.053 | 2.27125611569158e-09 | 16 |
| eif1b2 | 1.97188819610424e-13 | 0.304126715 | 0.771 | 0.377 | 4.96304540077475e-09 | 16 |
| higd1a4 | 2.15114148754542e-13 | 0.259530151 | 0.636 | 0.305 | 5.41420801000306e-09 | 16 |
| ppdpfb2 | 9.58764499976496e-13 | 0.355168315 | 0.857 | 0.514 | 2.41311436999084e-08 | 16 |
| chac12 | 2.11000449024072e-12 | 0.310812796 | 0.74 | 0.393 | 5.31067030148687e-08 | 16 |
| eif5a5 | 1.92643980574712e-10 | 0.341161396 | 0.835 | 0.473 | 4.84865634708492e-06 | 16 |
| CR318588.15 | 1.28151845986403e-09 | 0.291417464 | 0.39 | 0.206 | 3.22545381163178e-05 | 16 |
| eef1da4 | 1.66457871642256e-09 | 0.327545184 | 0.827 | 0.487 | 4.18957817136394e-05 | 16 |
| ucp23 | 3.30871751919692e-09 | 0.265628799 | 0.771 | 0.411 | 8.32771112406672e-05 | 16 |
| ptmab5 | 1.81298120187825e-08 | 0.317842711 | 0.879 | 0.535 | 0.000456309 | 16 |
| dcn2 | 0 | 1.53264155 | 0.909 | 0.104 | 0 | 17 |
| col6a4a1 | 0 | 1.107253591 | 0.578 | 0.036 | 0 | 17 |
| asip2b1 | 0 | 1.042705732 | 0.378 | 0.012 | 0 | 17 |
| c62 | 4.05770617935954e-296 | 1.40463366 | 0.791 | 0.082 | 1.021284068283e-291 | 17 |
| hsd11b23 | 1.65662645947962e-280 | 1.654568166 | 0.861 | 0.109 | 4.16956313586426e-276 | 17 |

| | | | | | | |
|---|---|---|---|---|---|---|
| col1a1a2 | 6.56020446456135e-272 | 1.593246625 | 0.913 | 0.125 | 1.65113786168545e-267 | 17 |
| col1a22 | 6.752843782864e-264 | 1.617818894 | 0.948 | 0.141 | 1.69962325170904e-259 | 17 |
| rbp42 | 5.72759647275969e-258 | 1.569804332 | 0.857 | 0.114 | 1.44157875622889e-253 | 17 |
| si:ch211-106j24.11 | 1.75657740601246e-246 | 0.641079487 | 0.309 | 0.012 | 4.42112967319275e-242 | 17 |
| col6a32 | 2.64318524525699e-223 | 1.207331943 | 0.722 | 0.089 | 6.65263294378731e-219 | 17 |
| col5a12 | 5.2044703811443e-217 | 1.22781189 | 0.757 | 0.1 | 1.30991315023021e-212 | 17 |
| fn1b1 | 6.25834708506999e-207 | 1.208145005 | 0.657 | 0.077 | 1.57516337784127e-202 | 17 |
| postnb1 | 7.90932592575873e-206 | 1.163238076 | 0.622 | 0.068 | 1.99069824225422e-201 | 17 |
| c1qtnf51 | 1.03342491572575e-204 | 0.809219567 | 0.343 | 0.019 | 2.60102717039014e-200 | 17 |
| mmp16a1 | 5.41046939987879e-200 | 1.089464801 | 0.57 | 0.06 | 1.36176104325549e-195 | 17 |
| col1a1b2 | 3.63383065147538e-193 | 1.35738896 | 0.822 | 0.135 | 9.14598836669838e-189 | 17 |
| meis2b1 | 2.13146164983601e-189 | 0.555407244 | 0.261 | 0.011 | 5.36467582647225e-185 | 17 |
| zgc:113531 | 1.58067921619443e-184 | 0.56004724 | 0.261 | 0.012 | 3.97841151923976e-180 | 17 |
| crispld22 | 1.72323587977698e-177 | 0.689554555 | 0.27 | 0.013 | 4.33721238581067e-173 | 17 |
| si:ch1073-291c23.22 | 6.00178425120847e-176 | 0.876873285 | 0.504 | 0.053 | 1.51058907818666e-171 | 17 |
| mfap22 | 4.4551518784388e-175 | 1.067071501 | 0.635 | 0.086 | 1.12131717628426e-170 | 17 |
| si:ch1073-459j12.12 | 5.05499057399494e-175 | 0.970149379 | 0.522 | 0.058 | 1.27229057756879e-170 | 17 |
| pbx1a2 | 1.54905117285562e-169 | 0.733952168 | 0.413 | 0.036 | 3.89880689696032e-165 | 17 |
| col6a12 | 1.64948169488115e-167 | 0.995440471 | 0.609 | 0.08 | 4.15158047784638e-163 | 17 |
| pmp22a2 | 1.33866231558021e-166 | 1.16897178 | 0.761 | 0.128 | 3.36927918208384e-162 | 17 |
| pdgfra2 | 1.99528503288011e-158 | 0.809497794 | 0.426 | 0.041 | 5.02193289925596e-154 | 17 |
| si:ch73-46j18.53 | 3.64677810035637e-156 | 1.163163749 | 0.813 | 0.163 | 9.17857580078694e-152 | 17 |
| fstl1b2 | 4.11205184332322e-150 | 1.059945799 | 0.683 | 0.115 | 1.03496232844602e-145 | 17 |
| si:ch211-270g19.51 | 1.36853668391872e-147 | 0.592686616 | 0.274 | 0.017 | 3.44446997975502e-143 | 17 |
| tshz3b2 | 2.715626241999e-147 | 0.629740193 | 0.317 | 0.024 | 6.83495968848729e-143 | 17 |
| col6a22 | 5.0571139246347e-147 | 0.843114475 | 0.483 | 0.058 | 1.27282500369131e-142 | 17 |
| wfdc1 | 2.06686426242287e-146 | 0.305783767 | 0.157 | 0.005 | 5.20209066209213e-142 | 17 |
| XLOC-0412692 | 2.46146775026807e-146 | 0.633882514 | 0.287 | 0.019 | 6.1952681806497e-142 | 17 |
| loxl2a2 | 9.86402447906134e-146 | 0.724068366 | 0.383 | 0.036 | 2.48267632113495e-141 | 17 |
| BX640522.21 | 7.51654929502005e-145 | 0.754597571 | 0.304 | 0.022 | 1.8918402920636e-140 | 17 |
| col4a12 | 1.10479579114144e-139 | 0.862918385 | 0.448 | 0.052 | 2.78066052672388e-135 | 17 |
| si:ch211-106h4.122 | 1.71558340690159e-137 | 1.059158925 | 0.443 | 0.052 | 4.31795187683062e-133 | 17 |
| nrp1a2 | 4.18227879908901e-137 | 0.792369466 | 0.43 | 0.049 | 1.05263775094271e-132 | 17 |
| mxra8a1 | 2.97073790743695e-135 | 0.688292339 | 0.387 | 0.04 | 7.47705023922805e-131 | 17 |
| nid1b1 | 5.22299421765956e-135 | 0.791958149 | 0.43 | 0.049 | 1.31457541464273e-130 | 17 |

| | | | | | | |
|---|---|---|---|---|---|---|
| lrp1aa2 | 9.98527380179731e-134 | 0.748568438 | 0.426 | 0.048 | 2.51319356317437e-129 | 17 |
| igf2b2 | 3.79211467643546e-132 | 0.864405327 | 0.435 | 0.052 | 9.54437342912041e-128 | 17 |
| nid2a1 | 5.87947439509379e-128 | 0.788641215 | 0.413 | 0.048 | 1.47980491050116e-123 | 17 |
| aebp11 | 1.27898816259566e-127 | 0.614792303 | 0.348 | 0.034 | 3.21908530643702e-123 | 17 |
| sparc2 | 1.8124375853124e-125 | 1.116461755 | 0.722 | 0.148 | 4.56172415847279e-121 | 17 |
| ccl25b2 | 2.60449551494226e-123 | 0.982738597 | 0.496 | 0.069 | 6.55525476155817e-119 | 17 |
| col5a2a2 | 7.1274619638432e-123 | 0.828480253 | 0.526 | 0.078 | 1.7939109016797e-118 | 17 |
| prss231 | 5.22804011053875e-122 | 0.640413059 | 0.3 | 0.026 | 1.3158454154215e-117 | 17 |
| adamts10 | 8.86372368075483e-121 | 0.349163119 | 0.152 | 0.006 | 2.23091061320918e-116 | 17 |
| pdgfrb2 | 2.77665145836892e-120 | 0.693500115 | 0.387 | 0.045 | 6.98855405556874e-116 | 17 |
| gas1a2 | 2.11175914629241e-119 | 0.647373091 | 0.352 | 0.038 | 5.31508659530336e-115 | 17 |
| cebpd3 | 3.12474194537235e-119 | 0.961111124 | 0.639 | 0.12 | 7.86466300230767e-115 | 17 |
| slco2b11 | 4.91248768105814e-118 | 0.806152057 | 0.439 | 0.058 | 1.23642402444552e-113 | 17 |
| tmem119b1 | 2.48225435890099e-117 | 0.484084648 | 0.217 | 0.014 | 6.24758599591791e-113 | 17 |
| cyp2ad32 | 2.18529863706606e-116 | 0.724733988 | 0.387 | 0.046 | 5.50017813963157e-112 | 17 |
| cxcl18b3 | 2.40110670917258e-114 | 1.031933287 | 0.57 | 0.099 | 6.04334547631647e-110 | 17 |
| cd81a3 | 1.48034514009479e-107 | 1.024834214 | 0.77 | 0.197 | 3.72588068310458e-103 | 17 |
| si:dkey-261h17.12 | 2.55560124573041e-107 | 0.886765068 | 0.513 | 0.084 | 6.43219277537886e-103 | 17 |
| CABZ01092746.12 | 3.1502358671532e-107 | 0.818143403 | 0.43 | 0.061 | 7.92882865403789e-103 | 17 |
| ptges2 | 8.88292900373245e-106 | 0.528863956 | 0.243 | 0.02 | 2.23574440094942e-101 | 17 |
| abi3bpb | 3.74286216673099e-105 | 0.381255425 | 0.165 | 0.008 | 9.42040978744524e-101 | 17 |
| tmem1762 | 7.80090449697698e-105 | 0.685987364 | 0.426 | 0.06 | 1.96340965284414e-100 | 17 |
| col16a11 | 1.37505019674088e-103 | 0.557583869 | 0.291 | 0.029 | 3.46086384017712e-99 | 17 |
| mdka3 | 5.27590281495849e-102 | 0.881212517 | 0.5 | 0.087 | 1.3278919794969e-97 | 17 |
| cldn11a2 | 7.38475873626016e-102 | 0.775818531 | 0.404 | 0.056 | 1.85866992632932e-97 | 17 |
| cpn11 | 3.37232035574035e-101 | 0.696931333 | 0.374 | 0.05 | 8.48779310336288e-97 | 17 |
| fosab4 | 3.88432719843483e-99 | 1.151752778 | 0.9 | 0.339 | 9.77646312574062e-95 | 17 |
| vim2 | 2.44981940687066e-98 | 0.576924144 | 0.291 | 0.031 | 6.16595046515276e-94 | 17 |
| adh8a2 | 6.10171536809688e-98 | 0.826844421 | 0.33 | 0.04 | 1.5357407409963e-93 | 17 |
| zgc:1951733 | 8.76264141157395e-97 | 0.749348862 | 0.487 | 0.084 | 2.20546921687905e-92 | 17 |
| cald1a2 | 1.58690203527317e-96 | 0.769815758 | 0.509 | 0.092 | 3.99407373257904e-92 | 17 |
| rhoua1 | 4.66281533437618e-95 | 0.482234467 | 0.235 | 0.02 | 1.17358399150914e-90 | 17 |
| tagln1 | 1.13539564279249e-94 | 0.744380128 | 0.396 | 0.057 | 2.85767729334441e-90 | 17 |
| BX569787.11 | 3.195823117619e-94 | 0.462594114 | 0.235 | 0.02 | 8.04356720473527e-90 | 17 |
| socs3a5 | 1.17998109564847e-93 | 1.087488167 | 0.865 | 0.288 | 2.96989441963763e-89 | 17 |

| | | | | | | |
|---|---|---|---|---|---|---|
| col12a1a1 | 1.2506447196137e-93 | 0.512553968 | 0.248 | 0.023 | 3.14774769479572e-89 | 17 |
| zgc:1124371 | 8.58631206717935e-93 | 0.445426829 | 0.196 | 0.014 | 2.16108888418837e-88 | 17 |
| jun3 | 1.51270418170928e-92 | 1.059245373 | 0.761 | 0.219 | 3.80732515494408e-88 | 17 |
| ece11 | 1.73969090521835e-92 | 0.513051377 | 0.274 | 0.028 | 4.37862803934408e-88 | 17 |
| lrp1ab2 | 2.10440732298127e-92 | 0.693545849 | 0.413 | 0.064 | 5.29658279121155e-88 | 17 |
| mmp21 | 1.17051443504737e-89 | 0.793902824 | 0.47 | 0.086 | 2.94606778157072e-85 | 17 |
| bhmt1 | 1.17291543324036e-89 | 0.591262504 | 0.322 | 0.04 | 2.95211085392266e-85 | 17 |
| serpinf11 | 1.89955759834731e-89 | 0.584547589 | 0.313 | 0.039 | 4.78099651928035e-85 | 17 |
| nfixb2 | 1.94333145436225e-89 | 0.661536175 | 0.391 | 0.06 | 4.89117093748434e-85 | 17 |
| mrc21 | 4.2746070605779e-89 | 0.463216892 | 0.235 | 0.022 | 1.07587585107685e-84 | 17 |
| dhrs3a2 | 5.34694776811464e-89 | 0.515948988 | 0.257 | 0.026 | 1.34577328375677e-84 | 17 |
| angptl1a2 | 5.4107340644008e-88 | 0.526409706 | 0.243 | 0.024 | 1.36182765666904e-83 | 17 |
| CR383676.18 | 1.00679100212941e-87 | 0.84862817 | 0.991 | 0.722 | 2.5339922732595e-83 | 17 |
| mfap51 | 7.05782058428583e-87 | 0.590730705 | 0.317 | 0.04 | 1.7763828628589e-82 | 17 |
| lxn3 | 1.95841660677765e-86 | 0.854679341 | 0.596 | 0.14 | 4.92913875759867e-82 | 17 |
| nfia3 | 5.16046226906032e-86 | 0.711828821 | 0.426 | 0.074 | 1.29883674849979e-81 | 17 |
| pcdh18b1 | 4.40826597024278e-85 | 0.368594685 | 0.183 | 0.014 | 1.1095164620504e-80 | 17 |
| igfbp5b2 | 2.89299204448927e-84 | 0.701803657 | 0.413 | 0.067 | 7.28137167677505e-80 | 17 |
| nr2f22 | 1.44053385847164e-83 | 0.548596454 | 0.309 | 0.04 | 3.62567966838726e-79 | 17 |
| tenm31 | 4.353613050555e-83 | 0.552321266 | 0.313 | 0.041 | 1.09576086869419e-78 | 17 |
| s100a10b3 | 4.51769382306076e-83 | 0.981524319 | 0.809 | 0.246 | 1.13705835832616e-78 | 17 |
| inhbaa2 | 5.11091781920183e-83 | 0.628735187 | 0.291 | 0.036 | 1.28636690591491e-78 | 17 |
| lrrc172 | 6.50430541072755e-83 | 0.503996062 | 0.27 | 0.031 | 1.63706862882602e-78 | 17 |
| foxo3b2 | 3.7425906173206e-82 | 0.635184451 | 0.378 | 0.061 | 9.41972632473421e-78 | 17 |
| colec122 | 8.93862881033751e-81 | 0.583862764 | 0.339 | 0.049 | 2.24976348527385e-76 | 17 |
| adamts15a2 | 1.48226430847195e-79 | 0.586829244 | 0.317 | 0.044 | 3.73071103799305e-75 | 17 |
| col12a1b1 | 4.42493153793362e-79 | 0.55378576 | 0.252 | 0.028 | 1.11371101878251e-74 | 17 |
| slc38a5b2 | 5.61708139420708e-79 | 0.721308866 | 0.504 | 0.109 | 1.41376321610798e-74 | 17 |
| pcolcea2 | 1.9804663077106e-78 | 0.657812625 | 0.326 | 0.046 | 4.9846356498768e-74 | 17 |
| cdon3 | 3.67309043560485e-78 | 0.595515062 | 0.343 | 0.053 | 9.24480131737384e-74 | 17 |
| junba4 | 3.85663913035104e-78 | 0.985998882 | 0.826 | 0.3 | 9.70677502718055e-74 | 17 |
| p3h21 | 8.34911887581311e-78 | 0.369109504 | 0.174 | 0.014 | 2.1013897298534e-73 | 17 |
| marcksl1a2 | 4.21028485808673e-76 | 0.890373913 | 0.7 | 0.207 | 1.05968659593185e-71 | 17 |
| ms4a17a.8 | 7.94852880800922e-76 | 0.30230088 | 0.157 | 0.011 | 2.00056521568784e-71 | 17 |
| mmrn2a1 | 1.11243354937491e-74 | 0.343021282 | 0.165 | 0.013 | 2.79988400042171e-70 | 17 |

| | | | | | | |
|---|---|---|---|---|---|---|
| mir199-31 | 3.55986478230437e-74 | 0.386018882 | 0.183 | 0.016 | 8.95982367058187e-70 | 17 |
| si:dkey-163m14.21 | 6.10049068999744e-74 | 0.385269325 | 0.183 | 0.016 | 1.53543250176546e-69 | 17 |
| sult1st61 | 2.82347905056085e-73 | 0.341618308 | 0.17 | 0.014 | 7.1064144223566e-69 | 17 |
| c1qtnf6a1 | 2.94506559389893e-72 | 0.374761529 | 0.187 | 0.017 | 7.41243559328421e-68 | 17 |
| boc2 | 6.61031283256163e-72 | 0.540386105 | 0.304 | 0.045 | 1.66374963682744e-67 | 17 |
| si:dkey-9i23.162 | 2.97758619238056e-71 | 0.440389441 | 0.274 | 0.036 | 7.49428668760263e-67 | 17 |
| ier2a5 | 3.26409863169916e-71 | 0.948169478 | 0.709 | 0.255 | 8.21540984612361e-67 | 17 |
| rbp53 | 6.5893003996242e-71 | 0.733204933 | 0.435 | 0.089 | 1.65846101758142e-66 | 17 |
| mmp14a2 | 2.07346397498462e-70 | 0.53698347 | 0.313 | 0.048 | 5.2187014786388e-66 | 17 |
| cfd1 | 6.61939077145799e-70 | 0.73709946 | 0.361 | 0.063 | 1.66603446326826e-65 | 17 |
| BX322530.12 | 7.86797271886197e-70 | 0.514383755 | 0.296 | 0.043 | 1.98029005361037e-65 | 17 |
| fibinb | 8.72770073698283e-70 | 0.346761799 | 0.161 | 0.013 | 2.19667499849121e-65 | 17 |
| inka1a1 | 1.16768055365309e-69 | 0.397257557 | 0.183 | 0.017 | 2.93893518548947e-65 | 17 |
| slc7a12 | 1.86591449834054e-69 | 0.494565165 | 0.261 | 0.034 | 4.6963202008733e-65 | 17 |
| mmp14b2 | 5.45808321511889e-69 | 0.558133502 | 0.326 | 0.053 | 1.37374496441327e-64 | 17 |
| chd33 | 7.44532415944724e-69 | 0.667241646 | 0.452 | 0.098 | 1.87391363769128e-64 | 17 |
| lpar12 | 7.52147678996509e-69 | 0.560873043 | 0.352 | 0.06 | 1.89308049326631e-64 | 17 |
| cd82a1 | 1.22918928247855e-68 | 0.554679065 | 0.322 | 0.052 | 3.09374650507025e-64 | 17 |
| si:ch211-198c19.32 | 1.5575315515274e-68 | 0.63750551 | 0.348 | 0.059 | 3.92015116203931e-64 | 17 |
| FP102783.1 | 1.59375959608975e-68 | 0.309579391 | 0.117 | 0.007 | 4.0113335273983e-64 | 17 |
| anxa1a3 | 2.81220083217677e-68 | 0.763681551 | 0.557 | 0.136 | 7.07802827450571e-64 | 17 |
| eva1ba2 | 5.24554606585445e-68 | 0.57034888 | 0.352 | 0.061 | 1.32025148931491e-63 | 17 |
| ptprga3 | 6.17237782169833e-68 | 0.612039574 | 0.348 | 0.062 | 1.55352577394325e-63 | 17 |
| fblim11 | 7.31982491206948e-66 | 0.394468348 | 0.209 | 0.023 | 1.84232673211877e-61 | 17 |
| cyp3c12 | 1.12003657458246e-65 | 0.557087241 | 0.33 | 0.056 | 2.8190200545666e-61 | 17 |
| gpr271 | 2.00937451934404e-65 | 0.34795201 | 0.165 | 0.015 | 5.05739472773702e-61 | 17 |
| c43 | 4.56835925670461e-65 | 0.529668554 | 0.265 | 0.038 | 1.14981034131998e-60 | 17 |
| cyp2p72 | 6.0876300608293e-65 | 0.389581797 | 0.204 | 0.022 | 1.53219561001013e-60 | 17 |
| cyp2n132 | 7.97575836561297e-65 | 0.467542784 | 0.252 | 0.034 | 2.00741862304113e-60 | 17 |
| col15a1a1 | 1.29835083035402e-64 | 0.390625412 | 0.187 | 0.019 | 3.26781920491804e-60 | 17 |
| tsku2 | 2.74040352295486e-64 | 0.385652867 | 0.222 | 0.027 | 6.8973216269251e-60 | 17 |
| gadd45ba2 | 4.70917678062672e-64 | 0.84588796 | 0.609 | 0.184 | 1.18525270391594e-59 | 17 |
| tmem176l.12 | 6.50919384177756e-62 | 0.731669314 | 0.596 | 0.174 | 1.63829899803699e-57 | 17 |
| ephb31 | 7.68048222232062e-62 | 0.456530707 | 0.261 | 0.038 | 1.93310057053588e-57 | 17 |
| cldn5a2 | 1.37414815236248e-61 | 0.55586939 | 0.287 | 0.046 | 3.45859348468112e-57 | 17 |

| | | | | | | |
|---|---|---|---|---|---|---|
| cyp2p62 | 2.55405508927284e-61 | 0.374712125 | 0.196 | 0.022 | 6.4283012541908e-57 | 17 |
| svild2 | 5.9978418420951e-61 | 0.592311077 | 0.378 | 0.077 | 1.50959681323691e-56 | 17 |
| cyp27c11 | 8.1762789051139e-61 | 0.290138699 | 0.139 | 0.011 | 2.05788763762812e-56 | 17 |
| icn4 | 1.33077694215989e-60 | 0.896025205 | 0.757 | 0.267 | 3.34943248572222e-56 | 17 |
| hoxc9a | 1.62385892668151e-60 | 0.26361219 | 0.135 | 0.01 | 4.0870905325647e-56 | 17 |
| clec11a1 | 2.589898376664e-60 | 0.319363622 | 0.157 | 0.014 | 6.51851522422561e-56 | 17 |
| loxl2b1 | 2.88848645408417e-59 | 0.470168689 | 0.257 | 0.038 | 7.27003155628444e-55 | 17 |
| wu:fj16a032 | 3.01205397529937e-59 | 0.618549384 | 0.291 | 0.048 | 7.58103865043099e-55 | 17 |
| emp22 | 2.55558469474652e-58 | 0.50583721 | 0.278 | 0.046 | 6.43215111820751e-54 | 17 |
| tshz22 | 6.24327951119073e-58 | 0.40285224 | 0.204 | 0.025 | 1.57137102017159e-53 | 17 |
| dkk3b1 | 6.73989321423073e-58 | 0.447331351 | 0.252 | 0.038 | 1.69636372308973e-53 | 17 |
| tnfsf101 | 9.17401597306282e-58 | 0.383093676 | 0.2 | 0.024 | 2.30900808026018e-53 | 17 |
| nupr1b1 | 6.74670605128302e-57 | 0.454708759 | 0.274 | 0.045 | 1.69807844604742e-52 | 17 |
| lama42 | 1.4912229091166e-56 | 0.459655587 | 0.235 | 0.034 | 3.75325893995558e-52 | 17 |
| zfp36l1b2 | 2.04207953674604e-56 | 0.722742388 | 0.478 | 0.125 | 5.1397099860361e-52 | 17 |
| zgc:1627307 | 5.06382330980886e-56 | 0.8323667 | 0.857 | 0.384 | 1.27451368884579e-51 | 17 |
| tril1 | 6.88799503164279e-55 | 0.353944129 | 0.174 | 0.019 | 1.73363946951417e-50 | 17 |
| tnfrsf19 | 7.50835567125007e-55 | 0.271954817 | 0.143 | 0.013 | 1.88977803889693e-50 | 17 |
| dedd12 | 1.81781862793306e-54 | 0.69448354 | 0.496 | 0.14 | 4.57526770464473e-50 | 17 |
| fosb1 | 2.07163306206157e-54 | 0.734248404 | 0.465 | 0.12 | 5.21409325390275e-50 | 17 |
| col2a1b | 4.03831657342109e-54 | 0.303103194 | 0.139 | 0.013 | 1.01640389836435e-49 | 17 |
| LOC1039092842 | 8.22145791824538e-54 | 0.517784609 | 0.313 | 0.06 | 2.06925874344318e-49 | 17 |
| rgs7bpb | 1.11004568953165e-53 | 0.271241383 | 0.148 | 0.014 | 2.79387399598222e-49 | 17 |
| meis1b2 | 1.8503487727346e-53 | 0.488075063 | 0.291 | 0.053 | 4.65714282609571e-49 | 17 |
| b2ml7 | 2.36938222553291e-53 | 0.711551423 | 0.696 | 0.26 | 5.96349812344378e-49 | 17 |
| pabpc1b2 | 2.98928968423356e-53 | 0.50613174 | 0.326 | 0.065 | 7.52374320624745e-49 | 17 |
| timp2a2 | 3.38103066578263e-53 | 0.518937817 | 0.291 | 0.054 | 8.50971608270831e-49 | 17 |
| ccbe1 | 4.14753731341146e-53 | 0.3155985 | 0.174 | 0.02 | 1.04389366641253e-48 | 17 |
| krt84 | 4.38880809329947e-53 | 0.759103404 | 0.761 | 0.268 | 1.10461910900254e-48 | 17 |
| f3b2 | 6.64837304838408e-53 | 0.654103538 | 0.409 | 0.097 | 1.67332901254779e-48 | 17 |
| myl9b2 | 1.16088377921024e-52 | 0.524661864 | 0.357 | 0.075 | 2.92182838389426e-48 | 17 |
| fam198a | 1.17973153637803e-52 | 0.289141574 | 0.139 | 0.013 | 2.96926630390986e-48 | 17 |
| ip6k2b3 | 3.63458523146003e-52 | 0.654257828 | 0.443 | 0.117 | 9.14788756906176e-48 | 17 |
| efnb11 | 8.32566304337526e-52 | 0.410094286 | 0.217 | 0.031 | 2.09548613138712e-47 | 17 |
| dhrs3b1 | 1.10768734504805e-51 | 0.302555631 | 0.143 | 0.014 | 2.78793827875144e-47 | 17 |

| | | | | | | |
|---|---|---|---|---|---|---|
| hic12 | 2.66322464155819e-51 | 0.495961373 | 0.291 | 0.055 | 6.7030701003378e-47 | 17 |
| cnn3a1 | 2.72153847917539e-51 | 0.454080939 | 0.287 | 0.053 | 6.84984019823654e-47 | 17 |
| endouc1 | 2.71155436067833e-50 | 0.563726167 | 0.304 | 0.059 | 6.82471117039129e-46 | 17 |
| sox4a | 9.671233114815e-50 | 0.454253518 | 0.248 | 0.041 | 2.43415266266779e-45 | 17 |
| si:ch211-212k18.92 | 1.00769643971616e-48 | 0.355836573 | 0.222 | 0.034 | 2.53627116912159e-44 | 17 |
| fzd7b1 | 1.34124611706861e-48 | 0.405202974 | 0.226 | 0.036 | 3.37578235205e-44 | 17 |
| ldlrad4a2 | 1.40969925999649e-48 | 0.354956435 | 0.204 | 0.029 | 3.54807206748517e-44 | 17 |
| foxp42 | 2.2687911235158e-48 | 0.440514469 | 0.265 | 0.048 | 5.71032037877691e-44 | 17 |
| lgals2a3 | 5.00038502995497e-48 | 0.70335626 | 0.452 | 0.121 | 1.25854690818937e-43 | 17 |
| pbx1b1 | 8.483983046581e-48 | 0.340124367 | 0.187 | 0.025 | 2.13533369299397e-43 | 17 |
| ahnak3 | 5.10624611746763e-47 | 0.60505033 | 0.443 | 0.122 | 1.28519108530543e-42 | 17 |
| mgst1.12 | 6.49389188342212e-47 | 0.40199336 | 0.23 | 0.039 | 1.63444764813851e-42 | 17 |
| id1 | 6.6725696287185e-47 | 0.613044818 | 0.4 | 0.101 | 1.67941904985216e-42 | 17 |
| tns2a2 | 1.47851590594736e-46 | 0.401006694 | 0.217 | 0.035 | 3.72127668367891e-42 | 17 |
| olfml3b1 | 2.49815742472517e-46 | 0.404786984 | 0.243 | 0.043 | 6.28761242229077e-42 | 17 |
| sema3d1 | 3.91940365763275e-46 | 0.262406467 | 0.135 | 0.014 | 9.86474706589586e-42 | 17 |
| col18a1a2 | 5.28137617181107e-46 | 0.475244581 | 0.23 | 0.039 | 1.32926956868313e-41 | 17 |
| yap11 | 9.54571911327197e-46 | 0.389034287 | 0.257 | 0.047 | 2.40256204361942e-41 | 17 |
| ntn1a1 | 1.07344773295188e-45 | 0.259006588 | 0.117 | 0.011 | 2.70176059906658e-41 | 17 |
| krt912 | 1.65512457671248e-45 | 0.480646208 | 0.313 | 0.065 | 4.16578304712765e-41 | 17 |
| ppib4 | 3.11241887041199e-45 | 0.683974063 | 0.739 | 0.326 | 7.83364705493993e-41 | 17 |
| aoc24 | 1.11269927346324e-44 | 0.472571429 | 0.3 | 0.063 | 2.80055280137964e-40 | 17 |
| mmp15a | 3.63390870860587e-44 | 0.26514684 | 0.135 | 0.015 | 9.14618482869011e-40 | 17 |
| tnfsf121 | 1.19813054789668e-43 | 0.334021569 | 0.161 | 0.021 | 3.01557477600115e-39 | 17 |
| nfil3-64 | 1.89533603004445e-43 | 0.588434709 | 0.439 | 0.126 | 4.77037125401889e-39 | 17 |
| vcam1b1 | 4.29382079060281e-43 | 0.323496904 | 0.161 | 0.021 | 1.08071175478682e-38 | 17 |
| cygb11 | 6.94077125392999e-43 | 0.487104487 | 0.261 | 0.051 | 1.74692271690164e-38 | 17 |
| map1lc3a2 | 9.59126341450866e-43 | 0.459400885 | 0.3 | 0.067 | 2.41402508879768e-38 | 17 |
| s100v22 | 1.98978165054543e-42 | 0.546976764 | 0.422 | 0.117 | 5.00808143625779e-38 | 17 |
| vdrb | 2.62671901394606e-42 | 0.346243899 | 0.204 | 0.033 | 6.61118908620084e-38 | 17 |
| lamb1a2 | 2.75968231024734e-42 | 0.367038763 | 0.213 | 0.036 | 6.94584440666154e-38 | 17 |
| pdk2a1 | 1.03049049078554e-41 | 0.315409413 | 0.178 | 0.026 | 2.59364151625811e-37 | 17 |
| si:dkey-7j14.64 | 2.09242214242292e-41 | 0.614403783 | 0.504 | 0.17 | 5.26641729026425e-37 | 17 |
| bgnb2 | 2.4599962295996e-41 | 0.375992729 | 0.174 | 0.025 | 6.19156451027923e-37 | 17 |
| ar2 | 4.2642562297623e-41 | 0.425095368 | 0.27 | 0.056 | 1.07327065046887e-36 | 17 |

| | | | | | | |
|---|---|---|---|---|---|---|
| aqp3a | 8.38407999009507e-41 | 0.451141663 | 0.178 | 0.027 | 2.11018909270703e-36 | 17 |
| pdgfrl1 | 1.13280170535309e-40 | 0.330335063 | 0.174 | 0.026 | 2.85114861220319e-36 | 17 |
| rgs5a2 | 1.93778396375e-40 | 0.332806773 | 0.174 | 0.025 | 4.87720845836238e-36 | 17 |
| f101 | 3.06951529461821e-40 | 0.396218708 | 0.239 | 0.046 | 7.72566304502458e-36 | 17 |
| frzb | 6.46209856691493e-40 | 0.292382863 | 0.139 | 0.017 | 1.62644558830682e-35 | 17 |
| cd248a1 | 8.83971601699331e-40 | 0.363054442 | 0.191 | 0.031 | 2.22486812431705e-35 | 17 |
| XLOC-0160943 | 1.05390207895828e-39 | 0.512560569 | 0.317 | 0.076 | 2.65256614253009e-35 | 17 |
| rerea1 | 2.35307625578428e-39 | 0.435635997 | 0.243 | 0.049 | 5.92245762818346e-35 | 17 |
| sostdc1a1 | 2.384242887425e-39 | 0.266913246 | 0.104 | 0.01 | 6.00090092335998e-35 | 17 |
| atoh8 | 2.85792431193239e-39 | 0.252758213 | 0.139 | 0.017 | 7.19310970070262e-35 | 17 |
| mdkb1 | 3.44434599904968e-39 | 0.323209152 | 0.135 | 0.016 | 8.66907444500813e-35 | 17 |
| spry22 | 3.45309004688984e-39 | 0.424919757 | 0.274 | 0.06 | 8.69108233901705e-35 | 17 |
| itgb1a2 | 3.64277838478236e-39 | 0.401711713 | 0.23 | 0.045 | 9.16850891665873e-35 | 17 |
| spred2b1 | 7.85595975497647e-39 | 0.417049059 | 0.283 | 0.064 | 1.97726651073003e-34 | 17 |
| rbms32 | 8.56129758617643e-39 | 0.333574908 | 0.191 | 0.032 | 2.15479298946474e-34 | 17 |
| krt943 | 1.06173157917811e-38 | 0.595977227 | 0.43 | 0.125 | 2.67227221163339e-34 | 17 |
| aldh9a1a.13 | 1.09420955007766e-38 | 0.571827077 | 0.478 | 0.159 | 2.75401601659046e-34 | 17 |
| hoxc3a1 | 1.43372522538523e-38 | 0.261024396 | 0.148 | 0.02 | 3.60854301977208e-34 | 17 |
| antxr2a4 | 3.40486771629385e-38 | 0.550613954 | 0.465 | 0.149 | 8.56971155513999e-34 | 17 |
| cox4i21 | 6.54473490943476e-38 | 0.317289623 | 0.187 | 0.031 | 1.64724432935564e-33 | 17 |
| spon1b2 | 8.41697208433904e-38 | 0.382283616 | 0.222 | 0.043 | 2.11846770390729e-33 | 17 |
| si:ch211-212k18.82 | 8.64962747214834e-38 | 0.345932947 | 0.217 | 0.041 | 2.17702473846502e-33 | 17 |
| deptor1 | 1.01739928391577e-37 | 0.368064198 | 0.2 | 0.036 | 2.56069225768761e-33 | 17 |
| tpm4a1 | 2.95894036183161e-37 | 0.47672308 | 0.33 | 0.085 | 7.44735699669399e-33 | 17 |
| smo | 4.1459532149891e-37 | 0.280991175 | 0.148 | 0.02 | 1.04349496468061e-32 | 17 |
| fzd21 | 4.61739872437624e-37 | 0.306142219 | 0.17 | 0.027 | 1.16215308493825e-32 | 17 |
| gstt1b1 | 8.61395433139056e-37 | 0.4292955 | 0.291 | 0.07 | 2.16804616566769e-32 | 17 |
| CU929237.1 | 8.8322286431738e-37 | 0.29828111 | 0.122 | 0.014 | 2.22298362720041e-32 | 17 |
| pid1 | 1.0007029690104e-36 | 0.277673339 | 0.122 | 0.014 | 2.51866930270228e-32 | 17 |
| si:ch211-133n4.42 | 1.00576041825338e-36 | 0.37997842 | 0.217 | 0.043 | 2.53139839670194e-32 | 17 |
| slit31 | 1.89063025862795e-36 | 0.348410006 | 0.165 | 0.025 | 4.75852729794069e-32 | 17 |
| ciarta2 | 2.11036994527778e-36 | 0.490817586 | 0.357 | 0.101 | 5.31159011526964e-32 | 17 |
| ghra2 | 5.2492056515131e-36 | 0.347898478 | 0.174 | 0.029 | 1.32117257042933e-31 | 17 |
| BX663516.2 | 6.1102298923005e-36 | 0.258259103 | 0.117 | 0.014 | 1.53788376159311e-31 | 17 |
| grb10b1 | 1.10594655012571e-35 | 0.326407178 | 0.191 | 0.034 | 2.7835568720114e-31 | 17 |

| | | | | | | |
|---|---|---|---|---|---|---|
| ctdsp21 | 1.56171772620802e-35 | 0.431300526 | 0.27 | 0.064 | 3.93068734509297e-31 | 17 |
| pik3r21 | 1.96105514832418e-35 | 0.391740718 | 0.257 | 0.058 | 4.93577970281714e-31 | 17 |
| ch25h | 2.51015144676525e-35 | 0.295261937 | 0.148 | 0.021 | 6.31780017636346e-31 | 17 |
| egr15 | 2.51511977384264e-35 | 0.604588142 | 0.509 | 0.185 | 6.33030495878455e-31 | 17 |
| hmgn65 | 3.29537083245882e-35 | 0.588312596 | 0.617 | 0.26 | 8.2941188482156e-31 | 17 |
| LOC101882334 | 4.60038522606708e-35 | 0.255725636 | 0.122 | 0.015 | 1.15787095754882e-30 | 17 |
| pfkfb4b | 5.85880126191422e-35 | 0.452509807 | 0.339 | 0.093 | 1.47460168961119e-30 | 17 |
| grb10a2 | 7.48262100726977e-35 | 0.324740925 | 0.2 | 0.037 | 1.88330088131973e-30 | 17 |
| edil3a2 | 1.37600722863216e-34 | 0.290230037 | 0.148 | 0.022 | 3.46327259374429e-30 | 17 |
| tcf121 | 1.48656753187205e-34 | 0.401996831 | 0.248 | 0.055 | 3.74154182096877e-30 | 17 |
| nr2f1a2 | 6.3193253130547e-34 | 0.37645485 | 0.213 | 0.043 | 1.59051098804274e-29 | 17 |
| cxcl12a2 | 7.49841644830904e-34 | 0.533078965 | 0.309 | 0.08 | 1.8872764358749e-29 | 17 |
| grhl1 | 1.60353188422756e-33 | 0.309788077 | 0.157 | 0.025 | 4.03592939941234e-29 | 17 |
| scube1 | 3.43860439996023e-33 | 0.251807588 | 0.13 | 0.018 | 8.6546234142599e-29 | 17 |
| zcchc242 | 4.40621762463524e-33 | 0.296792104 | 0.174 | 0.03 | 1.10900091394444e-28 | 17 |
| ccdc3b1 | 9.85384976025094e-33 | 0.268083569 | 0.148 | 0.023 | 2.48011544615756e-28 | 17 |
| stau23 | 3.02061455643161e-32 | 0.406554969 | 0.304 | 0.083 | 7.60258477708271e-28 | 17 |
| cdh21 | 3.55560795908896e-32 | 0.39249962 | 0.235 | 0.053 | 8.94910967223101e-28 | 17 |
| celf26 | 3.563417385143e-32 | 0.553851879 | 0.578 | 0.238 | 8.96876521666642e-28 | 17 |
| kirrel1a1 | 3.94908723535548e-32 | 0.307780238 | 0.157 | 0.026 | 9.93945766266621e-28 | 17 |
| egfra1 | 3.97089069008955e-32 | 0.295219113 | 0.161 | 0.027 | 9.99433477788639e-28 | 17 |
| tns1b2 | 4.39844800854066e-32 | 0.304048306 | 0.178 | 0.032 | 1.1070453792696e-27 | 17 |
| h1f02 | 4.40843575809327e-32 | 0.422188487 | 0.252 | 0.06 | 1.10955919595449e-27 | 17 |
| arrdc3a2 | 4.44410314989254e-32 | 0.463122497 | 0.274 | 0.069 | 1.11853632179645e-27 | 17 |
| vasna1 | 7.4417976310002e-32 | 0.369476096 | 0.178 | 0.033 | 1.87302604574644e-27 | 17 |
| FO744854.1 | 8.23924363176048e-32 | 0.26464855 | 0.122 | 0.016 | 2.07373522967779e-27 | 17 |
| myl62 | 8.30620894967147e-32 | 0.445431627 | 0.339 | 0.098 | 2.09058973054281e-27 | 17 |
| txnipa6 | 1.29942753401471e-31 | 0.574208813 | 0.517 | 0.206 | 3.27052916036162e-27 | 17 |
| rbfox1 | 1.72830058808532e-31 | 0.285858231 | 0.148 | 0.024 | 4.34995975015195e-27 | 17 |
| ier51 | 2.48732622104421e-31 | 0.495876285 | 0.396 | 0.133 | 6.26035136574617e-27 | 17 |
| SCARF2 | 2.6713386820391e-31 | 0.260840845 | 0.143 | 0.022 | 6.72349232882421e-27 | 17 |
| qkia2 | 3.48609864555346e-31 | 0.526989573 | 0.452 | 0.164 | 8.77416168099351e-27 | 17 |
| ackr3b1 | 4.50004368781102e-31 | 0.278102591 | 0.143 | 0.022 | 1.13261599578516e-26 | 17 |
| nfic3 | 6.62271767020165e-31 | 0.41166268 | 0.278 | 0.074 | 1.66687181041305e-26 | 17 |
| ulk21 | 7.66530167709773e-31 | 0.33310928 | 0.213 | 0.046 | 1.92927977910873e-26 | 17 |

| | | | | | | |
|---|---|---|---|---|---|---|
| ptenb1 | 8.22162136327474e-31 | 0.426869574 | 0.287 | 0.077 | 2.06929988092262e-26 | 17 |
| zgc:1141887 | 1.79034970956864e-30 | 0.476192461 | 0.926 | 0.623 | 4.50613118401331e-26 | 17 |
| zgc:171679 | 2.27471509952677e-30 | 0.283041057 | 0.152 | 0.026 | 5.72523043399892e-26 | 17 |
| h3f3c3 | 2.38359062048415e-30 | 0.559861128 | 0.574 | 0.252 | 5.99925923269656e-26 | 17 |
| zgc:1583434 | 2.66730957341976e-30 | 0.613038829 | 0.443 | 0.162 | 6.71335146534019e-26 | 17 |
| plpp32 | 2.97454657751275e-30 | 0.399800299 | 0.23 | 0.053 | 7.48663628094184e-26 | 17 |
| krt18b5 | 2.99319335557308e-30 | 0.470297789 | 0.439 | 0.146 | 7.53356835664189e-26 | 17 |
| selenow11 | 3.05232007799452e-30 | 0.56742834 | 0.726 | 0.38 | 7.68238440430441e-26 | 17 |
| si:ch73-335l21.42 | 3.3337477329728e-30 | 0.697329969 | 0.404 | 0.145 | 8.39070966911925e-26 | 17 |
| btg23 | 4.67629034666185e-30 | 0.613588987 | 0.691 | 0.369 | 1.17697551735132e-25 | 17 |
| mcl1b3 | 7.76257379566907e-30 | 0.568949753 | 0.561 | 0.238 | 1.95376219863195e-25 | 17 |
| entpd12 | 8.6062267639103e-30 | 0.384638421 | 0.265 | 0.068 | 2.16610121420858e-25 | 17 |
| mgst1.23 | 8.7492506201957e-30 | 0.469164087 | 0.33 | 0.1 | 2.20209888859705e-25 | 17 |
| bnc2 | 1.63636913666839e-29 | 0.28836957 | 0.165 | 0.03 | 4.11857748008067e-25 | 17 |
| fxyd12 | 1.94525859069433e-29 | 0.429005206 | 0.283 | 0.079 | 4.89602134691857e-25 | 17 |
| lamc12 | 2.06685449124437e-29 | 0.36473242 | 0.226 | 0.053 | 5.20206606901295e-25 | 17 |
| robo22 | 3.18892560376984e-29 | 0.369508015 | 0.204 | 0.045 | 8.02620685212831e-25 | 17 |
| socs3b4 | 1.00372622061573e-28 | 0.529223399 | 0.422 | 0.152 | 2.52627852466773e-24 | 17 |
| ddit35 | 1.21335185722932e-28 | 0.560479322 | 0.448 | 0.173 | 3.05388528946047e-24 | 17 |
| si:ch211-39i22.12 | 1.63000149660959e-28 | 0.374652328 | 0.265 | 0.069 | 4.10255076681668e-24 | 17 |
| ip6k2a | 1.99162214170522e-28 | 0.339852984 | 0.235 | 0.058 | 5.01271376845786e-24 | 17 |
| junbb3 | 2.51638226789846e-28 | 0.374905123 | 0.261 | 0.07 | 6.33348253007364e-24 | 17 |
| znf385a2 | 3.45004271722229e-28 | 0.35253798 | 0.2 | 0.044 | 8.68341251497678e-24 | 17 |
| osr22 | 3.67627382489025e-28 | 0.333976738 | 0.2 | 0.044 | 9.25281358986627e-24 | 17 |
| myl9a1 | 3.98598832744426e-28 | 0.2821485 | 0.161 | 0.03 | 1.00323340213445e-23 | 17 |
| rnd3b2 | 4.31677105335826e-28 | 0.309717956 | 0.187 | 0.039 | 1.08648810641974e-23 | 17 |
| ppap2d2 | 4.36764853716236e-28 | 0.322686995 | 0.174 | 0.035 | 1.0992934603184e-23 | 17 |
| CABZ01074363.12 | 5.45341354118855e-28 | 0.274738235 | 0.152 | 0.027 | 1.37256965418175e-23 | 17 |
| itm2ba4 | 6.89996780556475e-28 | 0.524093826 | 0.591 | 0.269 | 1.73665289698259e-23 | 17 |
| cyr612 | 7.62367660475095e-28 | 0.540312984 | 0.248 | 0.066 | 1.91880316464977e-23 | 17 |
| sesn15 | 8.73362614770735e-28 | 0.506451076 | 0.439 | 0.164 | 2.19816636511646e-23 | 17 |
| CR847566.11 | 1.53687819772105e-27 | 0.311079859 | 0.17 | 0.034 | 3.86816873584412e-23 | 17 |
| fkbp52 | 1.6325158190386e-27 | 0.465199665 | 0.352 | 0.119 | 4.10887906493824e-23 | 17 |
| rps27.26 | 1.78533185443404e-27 | 0.490996251 | 0.822 | 0.513 | 4.49350174442504e-23 | 17 |
| FO082781.13 | 3.60331219913249e-27 | 0.324658063 | 0.2 | 0.046 | 9.06917647399656e-23 | 17 |

| | | | | | | |
|---|---|---|---|---|---|---|
| zeb1b1 | 4.42574733904934e-27 | 0.264936152 | 0.143 | 0.025 | 1.11391634776533e-22 | 17 |
| mt-nd46 | 6.03099689125083e-27 | 0.520521919 | 0.839 | 0.45 | 1.51794160755892e-22 | 17 |
| appb2 | 9.91881159923292e-27 | 0.303880542 | 0.178 | 0.038 | 2.49646569141093e-22 | 17 |
| si:ch73-1a9.34 | 2.27057354999256e-26 | 0.515381457 | 0.583 | 0.261 | 5.71480656797628e-22 | 17 |
| sema4ba3 | 2.58410742862714e-26 | 0.330179518 | 0.196 | 0.045 | 6.50393998711164e-22 | 17 |
| klf2a1 | 3.6434837447292e-26 | 0.428916388 | 0.283 | 0.083 | 9.17028423710892e-22 | 17 |
| si:rp71-79p20.2 | 5.1461636043294e-26 | 0.318157008 | 0.174 | 0.037 | 1.29523791757367e-21 | 17 |
| jdp2b2 | 8.08616249212462e-26 | 0.568383914 | 0.426 | 0.167 | 2.03520623764284e-21 | 17 |
| gstm.31 | 2.12047573146477e-25 | 0.460030435 | 0.313 | 0.098 | 5.33702536852369e-21 | 17 |
| creg21 | 2.23866470275456e-25 | 0.270434208 | 0.148 | 0.029 | 5.63449519036294e-21 | 17 |
| TCIM (1 of many)2 | 3.21773042182554e-25 | 0.507379826 | 0.183 | 0.042 | 8.09870569869269e-21 | 17 |
| gpx4a2 | 4.82588481363193e-25 | 0.324771374 | 0.257 | 0.072 | 1.21462694874302e-20 | 17 |
| rpl97 | 7.38024766695353e-25 | 0.439498339 | 0.917 | 0.625 | 1.85753453529553e-20 | 17 |
| crim12 | 9.47493560764211e-25 | 0.2508455 | 0.165 | 0.035 | 2.38474654308744e-20 | 17 |
| c7a2 | 1.22721649004782e-24 | 0.336347432 | 0.187 | 0.043 | 3.08878118380135e-20 | 17 |
| CABZ01080702.12 | 2.01053016457679e-24 | 0.416283836 | 0.27 | 0.082 | 5.06030337122333e-20 | 17 |
| adh53 | 4.45412591514948e-24 | 0.463615511 | 0.504 | 0.218 | 1.12105895158397e-19 | 17 |
| nme32 | 4.52664742212813e-24 | 0.35336701 | 0.252 | 0.074 | 1.13931188967543e-19 | 17 |
| cavin2b1 | 4.93677544292365e-24 | 0.338302331 | 0.235 | 0.064 | 1.24253701122945e-19 | 17 |
| serinc51 | 6.20979201707851e-24 | 0.357918487 | 0.27 | 0.081 | 1.56294255277849e-19 | 17 |
| COX38 | 6.42784951812295e-24 | 0.457188234 | 0.991 | 0.696 | 1.61782544521637e-19 | 17 |
| add3a3 | 1.66124292787997e-23 | 0.432400874 | 0.343 | 0.121 | 4.1811823251811e-19 | 17 |
| ptprk | 1.6961687104825e-23 | 0.2734032 | 0.157 | 0.033 | 4.26908702741341e-19 | 17 |
| dbpb | 2.1898831206905e-23 | 0.274995855 | 0.174 | 0.04 | 5.51171682646592e-19 | 17 |
| pnrc28 | 2.38078791161418e-23 | 0.46626113 | 0.865 | 0.564 | 5.99220509474173e-19 | 17 |
| vasnb1 | 2.81609799817851e-23 | 0.263015346 | 0.148 | 0.03 | 7.08783705161549e-19 | 17 |
| ccdc85b2 | 3.00970975651475e-23 | 0.297437965 | 0.191 | 0.047 | 7.57513848617198e-19 | 17 |
| rpl367 | 3.95045127564588e-23 | 0.319719747 | 0.948 | 0.788 | 9.94289081567311e-19 | 17 |
| cux1a2 | 4.26627296245351e-23 | 0.301333108 | 0.183 | 0.044 | 1.07377824191992e-18 | 17 |
| si:ch211-222l21.15 | 5.14705145887619e-23 | 0.50979358 | 0.578 | 0.271 | 1.29546138168455e-18 | 17 |
| flot2a2 | 6.43777104477167e-23 | 0.305204103 | 0.213 | 0.057 | 1.62032259425858e-18 | 17 |
| rpl297 | 7.48375996452564e-23 | 0.292838193 | 0.957 | 0.807 | 1.88358754547146e-18 | 17 |
| s1pr21 | 8.86382202652887e-23 | 0.256948103 | 0.148 | 0.031 | 2.23093536585705e-18 | 17 |
| si:ch211-132b12.7 | 1.12574991923584e-22 | 0.302225886 | 0.196 | 0.05 | 2.83339997172467e-18 | 17 |
| hspb12 | 1.80262426735298e-22 | 0.316198966 | 0.226 | 0.062 | 4.53702501850073e-18 | 17 |

| | | | | | | |
|---|---|---|---|---|---|---|
| gng72 | 2.3367602923276e-22 | 0.27891213 | 0.2 | 0.052 | 5.88139197975933e-18 | 17 |
| mknk2b5 | 2.50082816026104e-22 | 0.439228127 | 0.461 | 0.194 | 6.294334396561e-18 | 17 |
| mt-nd25 | 3.34240095260825e-22 | 0.4727941 | 0.735 | 0.396 | 8.41248895761971e-18 | 17 |
| reck | 4.85001871787146e-22 | 0.256123851 | 0.13 | 0.026 | 1.22070121110107e-17 | 17 |
| cd28l2 | 9.92324994738572e-22 | 0.251524091 | 0.161 | 0.036 | 2.49758277925751e-17 | 17 |
| map1lc3b2 | 1.99257034206369e-21 | 0.367439501 | 0.3 | 0.105 | 5.0151002939401e-17 | 17 |
| selenbp11 | 2.22115872737289e-21 | 0.332809322 | 0.23 | 0.068 | 5.59043440092483e-17 | 17 |
| klf11b3 | 2.36088764273118e-21 | 0.369118382 | 0.248 | 0.078 | 5.9421181079901e-17 | 17 |
| ablim1b2 | 2.82962206931998e-21 | 0.256362256 | 0.157 | 0.035 | 7.12187578627146e-17 | 17 |
| rpsa7 | 3.1640129016117e-21 | 0.342206612 | 0.961 | 0.749 | 7.9635040720665e-17 | 17 |
| mt-atp67 | 3.22326584999848e-21 | 0.454722757 | 0.913 | 0.586 | 8.11263781786117e-17 | 17 |
| BX663503.31 | 4.74213781663705e-21 | 0.25327544 | 0.139 | 0.029 | 1.19354866706938e-16 | 17 |
| rps167 | 7.92058188554195e-21 | 0.346209961 | 0.948 | 0.735 | 1.99353125477205e-16 | 17 |
| rplp16 | 8.270502761608e-21 | 0.265903102 | 1 | 0.875 | 2.08160284006912e-16 | 17 |
| mgst3b1 | 1.2206575380149e-20 | 0.380875567 | 0.413 | 0.169 | 3.0722729574297e-16 | 17 |
| h2afx16 | 4.15605266435355e-20 | 0.421006161 | 0.448 | 0.196 | 1.04603689509114e-15 | 17 |
| htra1b1 | 5.20592374546973e-20 | 0.263237937 | 0.161 | 0.038 | 1.31027894749728e-15 | 17 |
| il152 | 6.50472655996456e-20 | 0.273074873 | 0.2 | 0.055 | 1.63717462787748e-15 | 17 |
| zgc:927442 | 7.41853646322407e-20 | 0.483958868 | 0.643 | 0.35 | 1.86717144242887e-15 | 17 |
| tnrc6c15 | 7.69216033394911e-20 | 0.390970026 | 0.335 | 0.131 | 1.93603983445165e-15 | 17 |
| rbm39a2 | 7.96266221460863e-20 | 0.376317935 | 0.357 | 0.14 | 2.00412245279484e-15 | 17 |
| h2afy22 | 8.01262043702818e-20 | 0.354391482 | 0.274 | 0.096 | 2.01669643779562e-15 | 17 |
| rps287 | 9.85159508397953e-20 | 0.262130613 | 0.948 | 0.825 | 2.47954796668681e-15 | 17 |
| klf6a2 | 9.98430717485418e-20 | 0.421085359 | 0.396 | 0.165 | 2.51295027283905e-15 | 17 |
| zgc:1132083 | 1.02265368908312e-19 | 0.284549599 | 0.17 | 0.044 | 2.57391707005331e-15 | 17 |
| tgfbi1 | 1.08446396901303e-19 | 0.272282639 | 0.122 | 0.025 | 2.7294873636089e-15 | 17 |
| mafba2 | 1.76077171589942e-19 | 0.343328136 | 0.226 | 0.071 | 4.43168633174725e-15 | 17 |
| rpl347 | 1.79063325018938e-19 | 0.308430459 | 0.961 | 0.781 | 4.50684482740165e-15 | 17 |
| ybx15 | 1.86072633289603e-19 | 0.347333598 | 0.883 | 0.667 | 4.68326210726602e-15 | 17 |
| NC-002333.47 | 1.89307120854112e-19 | 0.43018601 | 0.926 | 0.618 | 4.76467092477714e-15 | 17 |
| mt-cyb6 | 2.48004441980996e-19 | 0.415188421 | 0.943 | 0.619 | 6.24202380021968e-15 | 17 |
| rpl27a7 | 2.62468571153225e-19 | 0.442362036 | 0.778 | 0.492 | 6.60607146735552e-15 | 17 |
| hmgb1b2 | 3.5279068545956e-19 | 0.368308542 | 0.326 | 0.124 | 8.87938876233166e-15 | 17 |
| mt-nd16 | 6.59722645613862e-19 | 0.426992814 | 0.739 | 0.406 | 1.66045592674553e-14 | 17 |
| flna3 | 6.75760167107922e-19 | 0.380216656 | 0.3 | 0.112 | 1.70082076459393e-14 | 17 |

| | | | | | | |
|---|---|---|---|---|---|---|
| abcc52 | 7.25920544688394e-19 | 0.345882397 | 0.265 | 0.093 | 1.82706941892622e-14 | 17 |
| mt-co28 | 9.29614036614993e-19 | 0.412271853 | 0.983 | 0.716 | 2.33974556875628e-14 | 17 |
| lum | 9.31256168740789e-19 | 0.278856937 | 0.1 | 0.018 | 2.34387865110369e-14 | 17 |
| CABZ01068358.12 | 1.36223759116078e-18 | 0.309209202 | 0.209 | 0.063 | 3.42861579319257e-14 | 17 |
| nsa26 | 1.48642608962488e-18 | 0.445797279 | 0.643 | 0.359 | 3.74118582497686e-14 | 17 |
| slc43a2a3 | 1.58124930753051e-18 | 0.363416724 | 0.278 | 0.1 | 3.97984638212355e-14 | 17 |
| si:dkey-51e6.12 | 1.88759771477825e-18 | 0.333569753 | 0.304 | 0.114 | 4.75089468832538e-14 | 17 |
| sh3pxd2b1 | 2.64795797764899e-18 | 0.274334016 | 0.165 | 0.044 | 6.66464543394475e-14 | 17 |
| tsc22d32 | 2.87367339406635e-18 | 0.376950926 | 0.348 | 0.142 | 7.23274856552561e-14 | 17 |
| rpl247 | 3.11438716322134e-18 | 0.289160644 | 0.939 | 0.774 | 7.83860105111179e-14 | 17 |
| wls2 | 3.84896969898649e-18 | 0.345970402 | 0.317 | 0.121 | 9.6874718353791e-14 | 17 |
| si:ch211-204c21.12 | 5.29848401169091e-18 | 0.346792997 | 0.222 | 0.072 | 1.33357544090249e-13 | 17 |
| hmgb1a5 | 5.36936666161434e-18 | 0.385301485 | 0.417 | 0.186 | 1.35141589506171e-13 | 17 |
| rpl67 | 9.1755929068866e-18 | 0.351624817 | 0.896 | 0.665 | 2.30940497873429e-13 | 17 |
| rps157 | 9.41752425877281e-18 | 0.338951196 | 0.891 | 0.682 | 2.37029668069053e-13 | 17 |
| rps136 | 1.1403600086204e-17 | 0.309802812 | 0.909 | 0.722 | 2.8701721056967e-13 | 17 |
| anxa43 | 1.44715352850459e-17 | 0.332493703 | 0.335 | 0.134 | 3.64234071589321e-13 | 17 |
| ghrb2 | 1.88748322852344e-17 | 0.260779909 | 0.157 | 0.041 | 4.75060653787065e-13 | 17 |
| zgc:1748631 | 2.21357407058272e-17 | 0.312404701 | 0.274 | 0.1 | 5.57134457824965e-13 | 17 |
| cnbpb3 | 2.85803668664411e-17 | 0.367685972 | 0.404 | 0.182 | 7.19339253661455e-13 | 17 |
| itpkcb1 | 3.67476419107603e-17 | 0.27650491 | 0.23 | 0.076 | 9.24901399251926e-13 | 17 |
| spry42 | 4.3344702642552e-17 | 0.290892909 | 0.165 | 0.046 | 1.09094282081039e-12 | 17 |
| si:dkey-28b4.72 | 4.4250938683966e-17 | 0.312815165 | 0.252 | 0.09 | 1.11375187573674e-12 | 17 |
| b2m6 | 5.29245533820816e-17 | 0.42643611 | 0.552 | 0.289 | 1.33205808407361e-12 | 17 |
| eif4ebp3 | 5.3880895638345e-17 | 0.43488084 | 0.348 | 0.148 | 1.35612826232151e-12 | 17 |
| si:ch211-196l7.47 | 5.61862723531291e-17 | 0.403371192 | 0.596 | 0.311 | 1.41415228885591e-12 | 17 |
| rps220 | 5.83499606804071e-17 | 0.253985939 | 0.935 | 0.828 | 1.46861016036517e-12 | 17 |
| pik3r3a | 7.48831485321158e-17 | 0.276732942 | 0.239 | 0.081 | 1.88473396540482e-12 | 17 |
| pgrmc12 | 7.7045615768318e-17 | 0.327555174 | 0.343 | 0.139 | 1.93916110327279e-12 | 17 |
| rps2110 | 7.71199386320792e-17 | 0.322178685 | 0.939 | 0.747 | 1.9410317354308e-12 | 17 |
| rps186 | 1.26746596575191e-16 | 0.308021592 | 0.909 | 0.722 | 3.19008508920099e-12 | 17 |
| ms4a17a.92 | 1.2765103518851e-16 | 0.253720003 | 0.183 | 0.055 | 3.21284890465961e-12 | 17 |
| rpl7a6 | 1.42439526173123e-16 | 0.251290998 | 0.939 | 0.808 | 3.58506043425133e-12 | 17 |
| jpt1b2 | 1.45653068068511e-16 | 0.395371863 | 0.539 | 0.268 | 3.66594207021634e-12 | 17 |
| rpl287 | 1.72568761020253e-16 | 0.294420469 | 0.93 | 0.747 | 4.34338314611874e-12 | 17 |

| | | | | | | |
|---|---|---|---|---|---|---|
| eif4ebp3l2 | 1.84535972524744e-16 | 0.387692674 | 0.448 | 0.203 | 4.64458589247529e-12 | 17 |
| arl6ip5b3 | 2.16192245238911e-16 | 0.281691736 | 0.261 | 0.095 | 5.44134262041814e-12 | 17 |
| si:ch211-105c13.31 | 2.26792315492606e-16 | 0.266150681 | 0.196 | 0.059 | 5.70813578863341e-12 | 17 |
| rsrp1 | 2.66411996365279e-16 | 0.304611626 | 0.304 | 0.117 | 6.70532353651772e-12 | 17 |
| sdc43 | 3.09421295425573e-16 | 0.415638305 | 0.517 | 0.254 | 7.78782458456624e-12 | 17 |
| krt151 | 3.60581777884781e-16 | 0.296629235 | 0.243 | 0.083 | 9.07548276758206e-12 | 17 |
| krt18a.11 | 4.53191192191391e-16 | 0.362133966 | 0.317 | 0.125 | 1.14063691162651e-11 | 17 |
| fam168a3 | 7.27344807609932e-16 | 0.282817659 | 0.183 | 0.058 | 1.83065414627344e-11 | 17 |
| tomm53 | 8.50531253720333e-16 | 0.357774327 | 0.439 | 0.212 | 2.14070211248870e-11 | 17 |
| rplp27 | 9.05889894584263e-16 | 0.334245463 | 0.97 | 0.755 | 2.28003427567913e-11 | 17 |
| hmgn73 | 1.40211614645966e-15 | 0.386647718 | 0.53 | 0.276 | 3.52898612902432e-11 | 17 |
| gstm.11 | 1.46885650180161e-15 | 0.390025782 | 0.513 | 0.257 | 3.69696492938446e-11 | 17 |
| angptl42 | 1.60879333304508e-15 | 0.27387837 | 0.165 | 0.049 | 4.04917193994116e-11 | 17 |
| per1b | 1.6618312375072e-15 | 0.274456106 | 0.2 | 0.066 | 4.18266304168188e-11 | 17 |
| hist2h2l1 | 2.06692521427096e-15 | 0.272530738 | 0.222 | 0.077 | 5.20224407179857e-11 | 17 |
| hdlbpa3 | 2.13270457576134e-15 | 0.338999569 | 0.357 | 0.157 | 5.36780414673373e-11 | 17 |
| eif4a1b3 | 2.24632012540674e-15 | 0.381415208 | 0.526 | 0.277 | 5.65376312363624e-11 | 17 |
| rps27l4 | 2.2859652736913e-15 | 0.355324748 | 0.422 | 0.201 | 5.75354599735363e-11 | 17 |
| rpz53 | 2.59931068510455e-15 | 0.303596908 | 0.265 | 0.101 | 6.54220506333963e-11 | 17 |
| eef1g6 | 2.74477462440101e-15 | 0.359854814 | 0.87 | 0.617 | 6.90832325215492e-11 | 17 |
| wu:fb16f031 | 3.56446624790807e-15 | 0.302433253 | 0.235 | 0.086 | 8.97140509935983e-11 | 17 |
| rgs5b2 | 3.68759787295862e-15 | 0.406003945 | 0.213 | 0.075 | 9.28131508644955e-11 | 17 |
| rpl117 | 6.38909698085664e-15 | 0.271099636 | 0.97 | 0.812 | 1.60807181911181e-10 | 17 |
| XLOC-0004314 | 7.40658801683223e-15 | 0.302753574 | 0.574 | 0.275 | 1.8641641379565e-10 | 17 |
| atf4b4 | 8.76155298521787e-15 | 0.386437945 | 0.496 | 0.262 | 2.20519527084948e-10 | 17 |
| si:ch211-132p1.34 | 9.61442048076237e-15 | 0.297161734 | 0.235 | 0.087 | 2.41985349080308e-10 | 17 |
| rps106 | 1.07035461664923e-14 | 0.317325851 | 0.904 | 0.691 | 2.69397553464444e-10 | 17 |
| cd82b1 | 1.10121545530675e-14 | 0.251346665 | 0.213 | 0.073 | 2.77164917946156e-10 | 17 |
| rps247 | 1.13806308547379e-14 | 0.277242437 | 0.93 | 0.756 | 2.86439097982899e-10 | 17 |
| rps266 | 1.18926958927657e-14 | 0.284348165 | 0.917 | 0.731 | 2.9932726292502e-10 | 17 |
| sdc21 | 1.27330944843282e-14 | 0.276792221 | 0.213 | 0.075 | 3.20479255076055e-10 | 17 |
| rps4x6 | 1.4790588287293e-14 | 0.315642608 | 0.926 | 0.707 | 3.72264316602877e-10 | 17 |
| gbp1 | 2.76155087849097e-14 | 0.280599597 | 0.257 | 0.099 | 6.95054740607393e-10 | 17 |
| akap12b2 | 3.05266974817103e-14 | 0.250509785 | 0.183 | 0.059 | 7.68326448917166e-10 | 17 |
| zbtb203 | 3.38819245826337e-14 | 0.30202633 | 0.248 | 0.097 | 8.52774159820308e-10 | 17 |

| | | | | | | |
|---|---|---|---|---|---|---|
| sfrp1a1 | 3.85281733283996e-14 | 0.253194828 | 0.113 | 0.028 | 9.69715594502489e-10 | 17 |
| rps27a6 | 3.96938410533551e-14 | 0.267239314 | 0.922 | 0.749 | 9.99054285471895e-10 | 17 |
| si:ch211-140m22.71 | 4.57254533748921e-14 | 0.260366851 | 0.191 | 0.065 | 1.15086393599266e-09 | 17 |
| rps147 | 5.36987474040882e-14 | 0.271582362 | 0.922 | 0.757 | 1.3515437734135e-09 | 17 |
| cry21 | 6.6160214641541e-14 | 0.275604934 | 0.217 | 0.08 | 1.66518644231295e-09 | 17 |
| slc3a2b2 | 6.705213006788e-14 | 0.347924727 | 0.274 | 0.117 | 1.68763506167847e-09 | 17 |
| si:dkey-11f4.142 | 8.2952587654282e-14 | 0.289850809 | 0.265 | 0.107 | 2.08783367867062e-09 | 17 |
| rpl23a7 | 1.09695855903658e-13 | 0.328785139 | 0.852 | 0.632 | 2.76093499723917e-09 | 17 |
| CABZ01072614.13 | 1.16763495166982e-13 | 0.307313798 | 0.213 | 0.08 | 2.93882040985777e-09 | 17 |
| mt-co18 | 1.80290538393618e-13 | 0.377633312 | 0.97 | 0.653 | 4.53773256082897e-09 | 17 |
| adam10a1 | 4.04086804005748e-13 | 0.281527851 | 0.235 | 0.093 | 1.01704607700207e-08 | 17 |
| rplp2l7 | 4.56099196757019e-13 | 0.292883095 | 0.97 | 0.771 | 1.14795606831774e-08 | 17 |
| rpl187 | 8.20311939682441e-13 | 0.267795418 | 0.943 | 0.772 | 2.06464312098674e-08 | 17 |
| ier2b6 | 9.8098598075698e-13 | 0.369712038 | 0.417 | 0.222 | 2.46904361496724e-08 | 17 |
| rpl176 | 1.68474858390548e-12 | 0.255376843 | 0.913 | 0.744 | 4.2403437108317e-08 | 17 |
| CR855311.14 | 2.12348996142694e-12 | 0.28444106 | 0.222 | 0.085 | 5.34461188391547e-08 | 17 |
| rpl22l15 | 3.59533012713058e-12 | 0.358163729 | 0.639 | 0.389 | 9.04908639697497e-08 | 17 |
| rpl307 | 3.7017064282417e-12 | 0.290942495 | 0.909 | 0.706 | 9.31682490924154e-08 | 17 |
| si:dkey-188i13.103 | 4.42998128094784e-12 | 0.259231847 | 0.248 | 0.103 | 1.11498198860176e-07 | 17 |
| fxyd6l2 | 4.73506321228639e-12 | 0.278834347 | 0.265 | 0.112 | 1.19176805990036e-07 | 17 |
| serinc11 | 5.12673723080535e-12 | 0.254593447 | 0.209 | 0.081 | 1.2903484936214e-07 | 17 |
| capns1b1 | 5.9274689885443e-12 | 0.25846982 | 0.274 | 0.121 | 1.49188466972672e-07 | 17 |
| rpl146 | 6.60797807148479e-12 | 0.320338669 | 0.878 | 0.632 | 1.66316200081201e-07 | 17 |
| si:dkeyp-19e1.32 | 6.84035687417288e-12 | 0.250413382 | 0.183 | 0.067 | 1.72164942166057e-07 | 17 |
| uba526 | 7.40692715409999e-12 | 0.330025799 | 0.878 | 0.627 | 1.86424949541543e-07 | 17 |
| rpl318 | 7.50367691552274e-12 | 0.319711103 | 0.9 | 0.651 | 1.88860044286792e-07 | 17 |
| rpl387 | 8.12985063086727e-12 | 0.268596659 | 0.891 | 0.726 | 2.04620210528298e-07 | 17 |
| vdac33 | 9.51610953919234e-12 | 0.255244181 | 0.33 | 0.152 | 2.39510960991932e-07 | 17 |
| btg16 | 1.03753384031622e-11 | 0.327159571 | 0.591 | 0.33 | 2.6113689226919e-07 | 17 |
| irf2bp2a4 | 1.06073164380237e-11 | 0.305280818 | 0.352 | 0.175 | 2.66975547428619e-07 | 17 |
| marcksl1b4 | 1.13037754297928e-11 | 0.289576939 | 0.265 | 0.115 | 2.84504723792455e-07 | 17 |
| rps36 | 1.43944498695766e-11 | 0.275817259 | 0.891 | 0.698 | 3.62293908767373e-07 | 17 |
| zgc:1119861 | 2.19376629460461e-11 | 0.307473535 | 0.348 | 0.175 | 5.52149038689035e-07 | 17 |
| romo16 | 2.51605753625873e-11 | 0.337557296 | 0.617 | 0.383 | 6.3326652130096e-07 | 17 |
| bhlhe403 | 2.57358192233796e-11 | 0.300410991 | 0.33 | 0.159 | 6.47744834033242e-07 | 17 |

| | | | | | | |
|---|---|---|---|---|---|---|
| pim13 | 2.82408989420978e-11 | 0.333334192 | 0.339 | 0.17 | 7.10795185473659e-07 | 17 |
| serf26 | 3.27269551804794e-11 | 0.335460299 | 0.57 | 0.34 | 8.23704734937486e-07 | 17 |
| cebpb6 | 3.46441321441359e-11 | 0.355866418 | 0.617 | 0.35 | 8.71958161935757e-07 | 17 |
| ppdpfb3 | 4.06855382428723e-11 | 0.324463687 | 0.743 | 0.516 | 1.02401431203485e-06 | 17 |
| eif1b3 | 5.73956460038378e-11 | 0.333474971 | 0.622 | 0.38 | 1.44459101427059e-06 | 17 |
| jund2 | 7.04765781566387e-11 | 0.278381714 | 0.217 | 0.092 | 1.77382499562444e-06 | 17 |
| tmsb4x8 | 7.06935660553431e-11 | 0.408508291 | 0.996 | 0.681 | 1.77928636404693e-06 | 17 |
| vat11 | 9.53490302587496e-11 | 0.31303593 | 0.283 | 0.134 | 2.39983974258247e-06 | 17 |
| mycb2 | 1.3107906166803e-10 | 0.30149252 | 0.404 | 0.21 | 3.29912890312264e-06 | 17 |
| pdk2b3 | 2.36639238593164e-10 | 0.262197975 | 0.226 | 0.099 | 5.95597299615134e-06 | 17 |
| tob1b2 | 3.7490588341051e-10 | 0.267307847 | 0.343 | 0.172 | 9.43600617955913e-06 | 17 |
| pabpc1a5 | 3.82611285581403e-10 | 0.327354472 | 0.8 | 0.537 | 9.62994344679833e-06 | 17 |
| rps26l7 | 3.82870311159956e-10 | 0.308716425 | 0.891 | 0.661 | 9.63646286158494e-06 | 17 |
| mir21-11 | 5.41499810410311e-10 | 0.268985898 | 0.235 | 0.107 | 1.36290087282171e-05 | 17 |
| si:ch73-376l24.3 | 6.08147560459393e-10 | 0.254256777 | 0.248 | 0.113 | 1.53064659492025e-05 | 17 |
| rps256 | 6.92417487261743e-10 | 0.253246904 | 0.904 | 0.723 | 1.74274557368908e-05 | 17 |
| tpm12 | 9.13008338283877e-10 | 0.302155336 | 0.383 | 0.206 | 2.29795068662669e-05 | 17 |
| rpl5a6 | 9.47423300331413e-10 | 0.328000168 | 0.743 | 0.489 | 2.38456970460413e-05 | 17 |
| pno12 | 2.03495665351554e-09 | 0.25715839 | 0.326 | 0.17 | 5.12178240123327e-05 | 17 |
| mt-nd36 | 2.58853222321071e-09 | 0.296519725 | 0.617 | 0.393 | 6.51507675259904e-05 | 17 |
| pdia34 | 6.65492219098422e-09 | 0.288913548 | 0.474 | 0.282 | 0.000167498 | 17 |
| klf132 | 8.3184530880173e-09 | 0.256359984 | 0.222 | 0.105 | 0.000209367 | 17 |
| nop535 | 9.66283138259171e-09 | 0.267981173 | 0.43 | 0.25 | 0.000243204 | 17 |
| eif3d4 | 1.18934509567314e-08 | 0.257268107 | 0.435 | 0.252 | 0.000299346 | 17 |
| nfkbiab3 | 1.20040168190713e-08 | 0.257008484 | 0.335 | 0.18 | 0.000302129 | 17 |
| eef1db4 | 1.22342068943066e-08 | 0.266308565 | 0.465 | 0.272 | 0.000307923 | 17 |
| naca7 | 2.18554233039986e-08 | 0.277878757 | 0.826 | 0.616 | 0.000550079 | 17 |
| serbp1a6 | 2.61782855888306e-08 | 0.301622718 | 0.713 | 0.479 | 0.000658881 | 17 |
| ccni5 | 3.28792246893206e-08 | 0.257547632 | 0.487 | 0.292 | 0.000827537 | 17 |
| rpl5b6 | 3.87798087406591e-08 | 0.278290926 | 0.843 | 0.627 | 0.000976049 | 17 |
| glula1 | 4.74221989658213e-08 | 0.280641314 | 0.261 | 0.135 | 0.001193569 | 17 |
| ywhaz3 | 8.96131047070925e-08 | 0.269042773 | 0.452 | 0.28 | 0.002255472 | 17 |
| eef2b4 | 1.05610101612743e-07 | 0.263807439 | 0.813 | 0.612 | 0.002658101 | 17 |
| acin1a2 | 1.96177779902546e-07 | 0.250715697 | 0.365 | 0.216 | 0.004937599 | 17 |
| csrnp1b4 | 2.08060298057006e-07 | 0.270402909 | 0.309 | 0.179 | 0.00523667 | 17 |

| | | | | | | |
|---|---|---|---|---|---|---|
| eef1da5 | 5.22032053594908e-07 | 0.27157481 | 0.713 | 0.49 | 0.013139025 | 17 |
| tcima3 | 1.59461625712674e-06 | 0.266395652 | 0.148 | 0.068 | 0.040134897 | 17 |
| btf36 | 2.12490832822865e-06 | 0.266084341 | 0.813 | 0.585 | 0.053481818 | 17 |
| rsl24d16 | 2.38937925695565e-06 | 0.258005795 | 0.648 | 0.442 | 0.060138287 | 17 |
| ccl33.31 | 0 | 2.599760945 | 0.89 | 0.038 | 0 | 18 |
| sftpba | 0 | 1.336316727 | 0.588 | 0.016 | 0 | 18 |
| npsn | 0 | 1.076071373 | 0.509 | 0.027 | 0 | 18 |
| si:ch211-220f21.3 | 0 | 0.560243158 | 0.197 | 0.001 | 0 | 18 |
| FO393424.2 | 1.03448518935581e-301 | 0.562274983 | 0.211 | 0.003 | 2.60369577308965e-297 | 18 |
| si:dkey-222f2.7 | 3.85887311242609e-290 | 1.025182579 | 0.443 | 0.023 | 9.71239773666524e-286 | 18 |
| XLOC-018356 | 3.5383161338277e-252 | 0.953875137 | 0.399 | 0.021 | 8.90558787723094e-248 | 18 |
| fcer1g2 | 9.00757790483267e-236 | 0.977585392 | 0.478 | 0.035 | 2.26711728286733e-231 | 18 |
| CABZ01046427.2 | 1.12270990069828e-216 | 0.449996293 | 0.189 | 0.004 | 2.82574854906749e-212 | 18 |
| vaspa4 | 3.20858127229188e-212 | 1.157085777 | 0.601 | 0.065 | 8.07567820423143e-208 | 18 |
| dicp1.1 | 2.78967290305373e-204 | 0.564407058 | 0.241 | 0.008 | 7.02132772969594e-200 | 18 |
| CT573231.11 | 1.57442425949907e-201 | 0.777815594 | 0.342 | 0.019 | 3.96266841873321e-197 | 18 |
| si:ch73-380l3.4 | 3.07816467576148e-186 | 0.520093768 | 0.202 | 0.006 | 7.74743267242407e-182 | 18 |
| ccl35.12 | 2.27923675930525e-177 | 1.264158736 | 0.596 | 0.073 | 5.73661099949538e-173 | 18 |
| si:ch211-102c2.43 | 6.63208861453877e-177 | 1.304621222 | 0.75 | 0.132 | 1.66923038339326e-172 | 18 |
| fcer1gl3 | 1.05126068825331e-162 | 1.335531893 | 0.798 | 0.158 | 2.64591802626476e-158 | 18 |
| rnaset2l2 | 7.19116585439105e-141 | 0.783087709 | 0.382 | 0.037 | 1.80994453389168e-136 | 18 |
| si:ch211-214k5.3 | 1.82331263465912e-137 | 0.421674772 | 0.184 | 0.008 | 4.58909557017355e-133 | 18 |
| arpc1b6 | 1.14426163870456e-135 | 1.259766366 | 0.934 | 0.298 | 2.87999211845552e-131 | 18 |
| CABZ01046432.1 | 5.1743920932108e-134 | 0.514333486 | 0.228 | 0.013 | 1.30234274594023e-129 | 18 |
| coro1a6 | 7.50076545603876e-132 | 1.188269433 | 0.838 | 0.221 | 1.88786765763039e-127 | 18 |
| pfn16 | 1.74981915459157e-128 | 1.148158656 | 1 | 0.517 | 4.40411983019153e-124 | 18 |
| si:dkey-172o19.2 | 1.50912620044265e-125 | 0.358617212 | 0.14 | 0.004 | 3.7983197338941e-121 | 18 |
| pou2f2a | 1.52881685017995e-124 | 0.355939821 | 0.149 | 0.005 | 3.84787913021792e-120 | 18 |
| LOC101882805 | 1.70878639428401e-123 | 0.272815632 | 0.114 | 0.003 | 4.30084447577342e-119 | 18 |
| lcp2b | 6.73929433587369e-122 | 0.443630303 | 0.189 | 0.009 | 1.69621299139605e-117 | 18 |
| gpr132b1 | 4.96073596048557e-117 | 0.724414362 | 0.316 | 0.031 | 1.24856763389461e-112 | 18 |
| ephx2 | 5.97594308180977e-117 | 0.696005677 | 0.32 | 0.031 | 1.5040851142607e-112 | 18 |
| spi1b2 | 2.23260532666375e-113 | 0.794959968 | 0.417 | 0.055 | 5.61924434667998e-109 | 18 |
| kctd12.21 | 3.47243481220662e-112 | 0.887619804 | 0.469 | 0.073 | 8.73977117884284e-108 | 18 |
| si:dkey-33i11.9 | 2.85030665546962e-105 | 0.569501433 | 0.237 | 0.018 | 7.17393682115149e-101 | 18 |

| | | | | | | |
|---|---|---|---|---|---|---|
| CU972454.41 | 5.97890685500303e-104 | 0.54868303 | 0.241 | 0.02 | 1.50483106633571e-99 | 18 |
| actb26 | 6.5590060732592e-100 | 0.638166589 | 0.996 | 0.866 | 1.65083623857861e-95 | 18 |
| mef2b | 8.87422729715031e-100 | 0.371786245 | 0.158 | 0.008 | 2.23355426841976e-95 | 18 |
| cotl16 | 4.86962452861808e-98 | 1.06932757 | 0.838 | 0.298 | 1.22563579760789e-93 | 18 |
| bpifcl | 8.48682377612264e-98 | 0.389685906 | 0.175 | 0.01 | 2.13604867621231e-93 | 18 |
| LOC1005362133 | 5.39337631308216e-95 | 0.877663226 | 0.518 | 0.099 | 1.35745888423965e-90 | 18 |
| wipf1a3 | 5.54349345867907e-95 | 0.743617851 | 0.399 | 0.06 | 1.39524186861493e-90 | 18 |
| si:ch211-214p16.14 | 1.41154077607483e-93 | 1.156722776 | 0.816 | 0.248 | 3.55270697930273e-89 | 18 |
| cd74a4 | 2.33865361985148e-92 | 1.042204546 | 0.768 | 0.21 | 5.88615729580418e-88 | 18 |
| cd74b5 | 5.18336273441131e-91 | 1.05692075 | 0.759 | 0.214 | 1.30460056662398e-86 | 18 |
| XLOC-0283704 | 9.32587797167254e-90 | 0.971504788 | 0.592 | 0.133 | 2.34723022669026e-85 | 18 |
| tmsb4x9 | 1.78549338288481e-89 | 0.77556488 | 0.996 | 0.682 | 4.49390829538278e-85 | 18 |
| LOC100535743 | 1.75249243200199e-88 | 0.279544522 | 0.11 | 0.004 | 4.4108482021058e-84 | 18 |
| arpc5b3 | 2.09289043775645e-85 | 0.910411537 | 0.5 | 0.107 | 5.2675959427892e-81 | 18 |
| abi3a2 | 2.25388540230941e-83 | 0.624501473 | 0.303 | 0.039 | 5.67280416907255e-79 | 18 |
| itga1 | 6.95542610714183e-83 | 0.679434418 | 0.285 | 0.035 | 1.75061119690653e-78 | 18 |
| zdhhc23a | 9.41255077709892e-80 | 0.298307886 | 0.132 | 0.007 | 2.36904490508803e-75 | 18 |
| XLOC-0203012 | 2.06460289022893e-78 | 0.819993464 | 0.43 | 0.08 | 5.1963990144172e-74 | 18 |
| ptmaa6 | 5.73189299873491e-78 | 0.918769687 | 0.86 | 0.388 | 1.44266014885159e-73 | 18 |
| si:busm1-266f07.23 | 1.05266049332734e-77 | 0.922926772 | 0.588 | 0.139 | 2.64944119565557e-73 | 18 |
| CU855821.1 | 1.37802036428171e-74 | 0.422362755 | 0.158 | 0.012 | 3.46833945486063e-70 | 18 |
| yrk2 | 1.87421781094503e-72 | 0.666426025 | 0.351 | 0.059 | 4.71721880836756e-68 | 18 |
| grap2b4 | 1.3338100924054e-69 | 0.782959036 | 0.469 | 0.107 | 3.35706662157516e-65 | 18 |
| si:ch73-264p11.12 | 1.80011348872554e-67 | 0.457512514 | 0.215 | 0.024 | 4.53070563977332e-63 | 18 |
| fam49ba3 | 2.87806907583667e-62 | 0.722430207 | 0.43 | 0.098 | 7.24381205697331e-58 | 18 |
| LOC1081907172 | 7.52120171062184e-62 | 0.743485866 | 0.421 | 0.091 | 1.89301125854641e-57 | 18 |
| rac26 | 9.61892472797991e-61 | 0.804087394 | 0.649 | 0.214 | 2.42098716478526e-56 | 18 |
| si:ch73-380l3.2 | 1.89865094288264e-60 | 0.452937249 | 0.193 | 0.022 | 4.77871455814133e-56 | 18 |
| arpc35 | 5.70394655773531e-59 | 0.828933887 | 0.724 | 0.304 | 1.4356263091164e-54 | 18 |
| lcp14 | 9.12321181648543e-59 | 0.681811336 | 0.382 | 0.084 | 2.29622118209122e-54 | 18 |
| zgc:640515 | 1.7789613449929e-58 | 0.820524199 | 0.526 | 0.157 | 4.47746780921264e-54 | 18 |
| wasa3 | 2.42800814854196e-57 | 0.664618052 | 0.395 | 0.089 | 6.11105370906525e-53 | 18 |
| arpc25 | 9.28988611219249e-57 | 0.810558744 | 0.667 | 0.261 | 2.33817143557773e-52 | 18 |
| cnpy3 | 3.89423563028217e-56 | 0.538037953 | 0.276 | 0.046 | 9.80140165785719e-52 | 18 |
| BX120005.11 | 5.56447059314253e-55 | 0.604657353 | 0.272 | 0.046 | 1.40052160358804e-50 | 18 |

| | | | | | | |
|---|---|---|---|---|---|---|
| hcls13 | 9.86693527011315e-55 | 0.608862667 | 0.329 | 0.067 | 2.48340893813478e-50 | 18 |
| ptpn64 | 2.78351404008236e-54 | 0.689550041 | 0.377 | 0.087 | 7.0058264874833e-50 | 18 |
| sftpbb2 | 2.25918397039844e-53 | 0.62273201 | 0.32 | 0.062 | 5.68614013509583e-49 | 18 |
| ppp1r183 | 9.2637101933427e-53 | 0.514641362 | 0.281 | 0.05 | 2.33158321856242e-48 | 18 |
| arpc4l5 | 2.31816599054972e-52 | 0.782442835 | 0.553 | 0.191 | 5.8345919816146e-48 | 18 |
| ptprc5 | 7.47401700124347e-52 | 0.715074247 | 0.447 | 0.123 | 1.88113533904297e-47 | 18 |
| arpc44 | 1.38869865054618e-51 | 0.77524517 | 0.596 | 0.22 | 3.49521563355968e-47 | 18 |
| actb12 | 4.13609976578088e-50 | 0.471610071 | 0.947 | 0.783 | 1.04101495004939e-45 | 18 |
| il7r1 | 1.19674738888989e-49 | 0.317720295 | 0.149 | 0.016 | 3.01209350309697e-45 | 18 |
| mhc2dab2 | 1.3727048286136e-48 | 0.618624561 | 0.39 | 0.093 | 3.45496078313758e-44 | 18 |
| si:dkey-9i23.4 | 1.46070894473784e-48 | 0.497819773 | 0.154 | 0.017 | 3.67645834301067e-44 | 18 |
| arl112 | 3.76114107140957e-48 | 0.437396349 | 0.193 | 0.027 | 9.46641596263074e-44 | 18 |
| dtnbp1a3 | 6.70631104379238e-47 | 0.627093274 | 0.346 | 0.083 | 1.6879114266121e-42 | 18 |
| CR381673.13 | 9.92122280979805e-46 | 0.542061591 | 0.272 | 0.054 | 2.49707256899807e-41 | 18 |
| fgd4a | 3.39224744819722e-45 | 0.298059117 | 0.14 | 0.015 | 8.53794760236757e-41 | 18 |
| XLOC-0268362 | 4.50664472526926e-45 | 0.647829295 | 0.329 | 0.075 | 1.13427741090302e-40 | 18 |
| plekho22 | 3.43540339816476e-44 | 0.549984362 | 0.311 | 0.071 | 8.64656681284088e-40 | 18 |
| arhgdig5 | 1.16862799494053e-43 | 0.70762756 | 0.61 | 0.238 | 2.94131980046581e-39 | 18 |
| rplp28 | 8.06404037501489e-43 | 0.442763906 | 0.974 | 0.755 | 2.0296383219875e-38 | 18 |
| rhogb3 | 5.72784721988171e-42 | 0.582994639 | 0.355 | 0.093 | 1.44164186677203e-37 | 18 |
| fam49al | 8.9653620118802e-42 | 0.26994983 | 0.123 | 0.013 | 2.25649196477013e-37 | 18 |
| myl12.14 | 2.16542185524052e-41 | 0.672742681 | 0.732 | 0.387 | 5.45015026745487e-37 | 18 |
| CR753876.12 | 6.77044305446498e-40 | 0.65674391 | 0.39 | 0.111 | 1.70405281237829e-35 | 18 |
| capzb4 | 5.59212828472207e-39 | 0.667346518 | 0.592 | 0.248 | 1.4074827679817e-34 | 18 |
| rgs133 | 1.75911212218897e-38 | 0.645882618 | 0.417 | 0.128 | 4.42750930033741e-34 | 18 |
| capza1b2 | 2.21801368227632e-38 | 0.668490153 | 0.526 | 0.208 | 5.58251863692127e-34 | 18 |
| XLOC-0254234 | 2.42787330664371e-38 | 0.690650333 | 0.496 | 0.164 | 6.11071432549155e-34 | 18 |
| cfl14 | 5.45272354508609e-38 | 0.566953937 | 0.851 | 0.573 | 1.37239598906272e-33 | 18 |
| si:ch211-243a20.4 | 2.15904096034039e-37 | 0.273924757 | 0.132 | 0.016 | 5.43409019308074e-33 | 18 |
| washc2c | 3.3145582138554e-37 | 0.395494053 | 0.193 | 0.033 | 8.34241156845266e-33 | 18 |
| si:ch211-214p16.22 | 3.52023880221333e-37 | 0.56519184 | 0.289 | 0.07 | 8.86008904129073e-33 | 18 |
| b3gnt2l | 3.54863741335256e-37 | 0.264099002 | 0.14 | 0.018 | 8.93156550566706e-33 | 18 |
| LOC101885445 | 1.11792593159648e-36 | 0.267622365 | 0.11 | 0.012 | 2.81370777723517e-32 | 18 |
| ipcef1 | 1.68371767280428e-36 | 0.341980989 | 0.154 | 0.022 | 4.23774901068108e-32 | 18 |
| AL954142.11 | 9.45393766520458e-36 | 0.349263755 | 0.149 | 0.022 | 2.37946157095534e-31 | 18 |

| | | | | | | |
|---|---|---|---|---|---|---|
| XLOC-0320022 | 6.08764311697022e-35 | 0.463766369 | 0.219 | 0.044 | 1.53219889611023e-30 | 18 |
| tln12 | 2.72827803104433e-34 | 0.564962257 | 0.382 | 0.124 | 6.86680297633548e-30 | 18 |
| wasb5 | 1.52129538371841e-33 | 0.595384858 | 0.482 | 0.173 | 3.82894835128086e-29 | 18 |
| apbb1ip | 8.0810777615663e-33 | 0.311080123 | 0.158 | 0.026 | 2.03392646180862e-28 | 18 |
| cnn22 | 1.60349532351733e-32 | 0.584432671 | 0.386 | 0.131 | 4.03583737976078e-28 | 18 |
| zgc:1627308 | 2.37405541331721e-32 | 0.67060934 | 0.746 | 0.387 | 5.97526006977808e-28 | 18 |
| eomesa | 3.19773608061371e-32 | 0.381849388 | 0.197 | 0.038 | 8.04838194129666e-28 | 18 |
| syk2 | 4.80445444013785e-32 | 0.462065047 | 0.289 | 0.076 | 1.2092331380383e-27 | 18 |
| ppiab3 | 1.04090116945383e-31 | 0.435567568 | 0.89 | 0.653 | 2.61984415339835e-27 | 18 |
| mir2231 | 1.07143693477636e-31 | 0.325999888 | 0.167 | 0.029 | 2.69669962113861e-27 | 18 |
| rab443 | 3.17859448590281e-31 | 0.417301417 | 0.246 | 0.059 | 8.00020446156877e-27 | 18 |
| scgn | 1.8326136644151e-30 | 0.296412655 | 0.14 | 0.022 | 4.61250533196635e-26 | 18 |
| mpeg1.1 | 3.43857488092489e-30 | 0.260458912 | 0.114 | 0.015 | 8.65454911779985e-26 | 18 |
| laptm56 | 4.17424619097955e-30 | 0.57797019 | 0.447 | 0.167 | 1.05061602380764e-25 | 18 |
| si:ch211-66e2.32 | 1.81488879701127e-29 | 0.455656482 | 0.232 | 0.057 | 4.56789361319766e-25 | 18 |
| hck1 | 5.58693349742148e-29 | 0.321544164 | 0.167 | 0.031 | 1.40617529196601e-24 | 18 |
| cers11 | 8.15421414276522e-29 | 0.263459977 | 0.127 | 0.019 | 2.05233415759258e-24 | 18 |
| LOC1039091154 | 1.04913590060801e-28 | 0.5369981 | 0.43 | 0.162 | 2.64057014824029e-24 | 18 |
| itm2bb3 | 1.14678928180803e-27 | 0.579487192 | 0.5 | 0.218 | 2.88635394338263e-23 | 18 |
| si:ch211-153b23.7 | 1.29472033809341e-27 | 0.281052783 | 0.132 | 0.021 | 3.25868161894729e-23 | 18 |
| ccni6 | 2.51573073982239e-27 | 0.580187147 | 0.575 | 0.291 | 6.33184269905898e-23 | 18 |
| ccr23 | 3.74575667786039e-27 | 0.391017453 | 0.197 | 0.045 | 9.42769498250681e-23 | 18 |
| mrtfaa | 4.89729518669855e-27 | 0.279958834 | 0.136 | 0.023 | 1.23260022554016e-22 | 18 |
| ptmab6 | 1.23078663708821e-26 | 0.489649627 | 0.825 | 0.536 | 3.09776688688733e-22 | 18 |
| naca8 | 1.29262472539962e-26 | 0.441482386 | 0.882 | 0.615 | 3.2534071713583e-22 | 18 |
| ckbb5 | 1.35527864912694e-26 | 0.608386069 | 0.557 | 0.262 | 3.4111008319876e-22 | 18 |
| cyba2 | 1.40538638325162e-26 | 0.441497809 | 0.254 | 0.071 | 3.53721698800601e-22 | 18 |
| gm2a2 | 1.49628631552073e-26 | 0.494671263 | 0.351 | 0.118 | 3.76600302753413e-22 | 18 |
| si:ch211-277c7.61 | 2.61598660596659e-26 | 0.385304809 | 0.175 | 0.037 | 6.58417668855732e-22 | 18 |
| eif3f6 | 2.89242320127777e-26 | 0.513422623 | 0.715 | 0.428 | 7.27993995529602e-22 | 18 |
| grasp2 | 3.10791348514769e-26 | 0.424883542 | 0.263 | 0.074 | 7.82230745076823e-22 | 18 |
| rpl266 | 3.1306112886232e-26 | 0.29704459 | 0.978 | 0.839 | 7.87943555233574e-22 | 18 |
| stk17b2 | 6.99915877347754e-26 | 0.458424394 | 0.329 | 0.108 | 1.76161827169656e-21 | 18 |
| RASSF52 | 2.48296821588227e-25 | 0.376263216 | 0.219 | 0.056 | 6.24938270255409e-21 | 18 |
| il10ra1 | 1.08567862839621e-24 | 0.320580192 | 0.175 | 0.039 | 2.73254453981042e-20 | 18 |

| | | | | | | |
|---|---|---|---|---|---|---|
| mob1a4 | 1.60338526511868e-24 | 0.514928599 | 0.404 | 0.163 | 4.0355603737772e-20 | 18 |
| selplg2 | 5.86910412301776e-24 | 0.315196489 | 0.189 | 0.045 | 1.47719481672234e-19 | 18 |
| mcl1a5 | 9.12490834174259e-24 | 0.413114903 | 0.829 | 0.622 | 2.29664818053319e-19 | 18 |
| rps297 | 1.26722407403205e-23 | 0.299717679 | 0.982 | 0.83 | 3.18947627193127e-19 | 18 |
| dok23 | 1.30970834843029e-23 | 0.302535191 | 0.167 | 0.037 | 3.2964049421642e-19 | 18 |
| XLOC-0437602 | 1.3176518864042e-23 | 0.314672905 | 0.202 | 0.049 | 3.31639803289072e-19 | 18 |
| gabarapb4 | 1.65716713884239e-23 | 0.551464685 | 0.518 | 0.266 | 4.17092397175242e-19 | 18 |
| tagapa4 | 2.00874168107655e-23 | 0.481385154 | 0.307 | 0.103 | 5.05580193710157e-19 | 18 |
| rgs181 | 2.66094510141451e-23 | 0.308156196 | 0.154 | 0.033 | 6.69733272575019e-19 | 18 |
| zgc:1749453 | 2.91435260476033e-23 | 0.277895868 | 0.132 | 0.025 | 7.33513407092127e-19 | 18 |
| mapre1a2 | 3.92464413997907e-23 | 0.51050028 | 0.456 | 0.194 | 9.87793683591332e-19 | 18 |
| si:dkey-262k9.44 | 6.38619296346531e-23 | 0.403874645 | 0.276 | 0.088 | 1.60734090697459e-18 | 18 |
| myh11a3 | 1.40528566440847e-22 | 0.359560622 | 0.228 | 0.064 | 3.53696348874967e-18 | 18 |
| ucp24 | 1.44748788628982e-22 | 0.491527528 | 0.68 | 0.413 | 3.64318226100285e-18 | 18 |
| srgn5 | 2.79285943693684e-22 | 0.497369494 | 0.518 | 0.234 | 7.02934791682632e-18 | 18 |
| rps27.17 | 4.06488943558962e-22 | 0.380395753 | 0.895 | 0.677 | 1.02309202204355e-17 | 18 |
| tagapb3 | 6.59983343800317e-22 | 0.358549939 | 0.228 | 0.066 | 1.66111207801102e-17 | 18 |
| cd833 | 2.90940726462125e-21 | 0.506249355 | 0.294 | 0.101 | 7.32268714432524e-17 | 18 |
| ptprea | 3.3206846861665e-21 | 0.261720852 | 0.123 | 0.024 | 8.35783128661247e-17 | 18 |
| rps221 | 6.52655872816331e-21 | 0.315776004 | 0.987 | 0.827 | 1.64266956629142e-16 | 18 |
| lasp14 | 7.14969039606574e-21 | 0.514806153 | 0.469 | 0.234 | 1.79950557578579e-16 | 18 |
| skap22 | 8.17723257639568e-21 | 0.36743835 | 0.237 | 0.073 | 2.05812766715303e-16 | 18 |
| si:ch211-119e14.13 | 2.10186990807322e-20 | 0.362164434 | 0.237 | 0.072 | 5.29019637162948e-16 | 18 |
| ppiaa6 | 3.390526597739e-20 | 0.26749733 | 0.939 | 0.802 | 8.53361639384929e-16 | 18 |
| pnrc29 | 3.83112854665341e-20 | 0.426558557 | 0.811 | 0.565 | 9.64256743907198e-16 | 18 |
| h1fx3 | 4.32502242223743e-20 | 0.537592191 | 0.408 | 0.181 | 1.08856489345294e-15 | 18 |
| crip16 | 5.87442083011611e-20 | 0.49743816 | 0.535 | 0.267 | 1.47853297873192e-15 | 18 |
| itgb23 | 1.04461377168571e-19 | 0.450395935 | 0.272 | 0.097 | 2.62918840195576e-15 | 18 |
| abracl2 | 2.14901792105904e-19 | 0.475099951 | 0.465 | 0.219 | 5.40886320551349e-15 | 18 |
| rpl227 | 2.61352136193935e-19 | 0.34638815 | 0.899 | 0.692 | 6.57797191586515e-15 | 18 |
| si:ch211-105j21.93 | 2.78582027033175e-19 | 0.265530617 | 0.145 | 0.034 | 7.01163103839799e-15 | 18 |
| hsp90ab16 | 2.9805367911247e-19 | 0.268159349 | 0.921 | 0.789 | 7.50171304958175e-15 | 18 |
| alox5a1 | 9.80239127716008e-19 | 0.250959071 | 0.14 | 0.032 | 2.46716386054842e-14 | 18 |
| LOC1018842672 | 1.2818008670089e-18 | 0.322137482 | 0.197 | 0.057 | 3.22616460217471e-14 | 18 |
| nr4a15 | 1.7088651326334e-18 | 0.487069248 | 0.377 | 0.165 | 4.30104265232501e-14 | 18 |

| | | | | | | |
|---|---|---|---|---|---|---|
| fybb3 | 2.89544806035889e-18 | 0.282689115 | 0.158 | 0.041 | 7.2875532231173e-14 | 18 |
| myo1f2 | 3.17387638711671e-18 | 0.300036768 | 0.167 | 0.045 | 7.98832947873406e-14 | 18 |
| hmgn23 | 9.4903064922739e-18 | 0.501867844 | 0.697 | 0.44 | 2.38861524104042e-13 | 18 |
| gapdhs7 | 1.89555389563709e-17 | 0.456763509 | 0.522 | 0.286 | 4.770919599929e-13 | 18 |
| rpl327 | 2.05307229152333e-17 | 0.284306176 | 0.961 | 0.811 | 5.16737765053506e-13 | 18 |
| CABZ01109481.1 | 3.21837585405588e-17 | 0.254257016 | 0.123 | 0.028 | 8.10033018707325e-13 | 18 |
| rps187 | 3.57271618853058e-17 | 0.341236274 | 0.952 | 0.721 | 8.99216937491261e-13 | 18 |
| rpl388 | 4.46304261696029e-17 | 0.316354306 | 0.912 | 0.726 | 1.12330319626274e-12 | 18 |
| calm1b4 | 5.09771036186875e-17 | 0.358969432 | 0.259 | 0.093 | 1.28304272097875e-12 | 18 |
| rps107 | 1.06190115408236e-16 | 0.342588345 | 0.921 | 0.691 | 2.6726990147099e-12 | 18 |
| btg17 | 1.21569055252734e-16 | 0.473756542 | 0.548 | 0.331 | 3.05977155165605e-12 | 18 |
| tpi1b4 | 1.58357035680755e-16 | 0.444480942 | 0.355 | 0.165 | 3.98568823104893e-12 | 18 |
| rpl157 | 1.68167108915893e-16 | 0.312781471 | 0.93 | 0.742 | 4.2325979643041e-12 | 18 |
| rap1b3 | 2.57340017762725e-16 | 0.425759604 | 0.408 | 0.2 | 6.47699090707004e-12 | 18 |
| rpl368 | 3.30613667486502e-16 | 0.308373653 | 0.974 | 0.787 | 8.32121539696777e-12 | 18 |
| elf14 | 3.31943783548836e-16 | 0.349735814 | 0.268 | 0.102 | 8.35469308814066e-12 | 18 |
| rpl348 | 3.95442226837734e-16 | 0.296327119 | 0.961 | 0.781 | 9.95288540727893e-12 | 18 |
| csk3 | 4.01727457171796e-16 | 0.37171958 | 0.263 | 0.1 | 1.01110783695569e-11 | 18 |
| BX005183.34 | 4.96146649389545e-16 | 0.390782068 | 0.224 | 0.08 | 1.24875150184855e-11 | 18 |
| si:dkey-112e7.25 | 6.32368613559171e-16 | 0.429261377 | 0.404 | 0.204 | 1.59160856346708e-11 | 18 |
| mbnl14 | 7.41693194015855e-16 | 0.429523164 | 0.382 | 0.182 | 1.8667676000185e-11 | 18 |
| rpl35a7 | 1.07338448278804e-15 | 0.273299759 | 0.899 | 0.758 | 2.70160140472922e-11 | 18 |
| arhgap152 | 1.07558911761028e-15 | 0.253985221 | 0.118 | 0.028 | 2.70715025011331e-11 | 18 |
| rsl24d17 | 1.51658386812019e-15 | 0.415993875 | 0.68 | 0.441 | 3.81708993767171e-11 | 18 |
| mfng3 | 1.73617100119887e-15 | 0.25676663 | 0.149 | 0.041 | 4.36976879291744e-11 | 18 |
| rpl87 | 2.30784846111897e-15 | 0.2897945 | 0.961 | 0.778 | 5.80862379179034e-11 | 18 |
| cap16 | 3.13310855762899e-15 | 0.441435769 | 0.439 | 0.231 | 7.8857209286964e-11 | 18 |
| rps2111 | 3.75989219929076e-15 | 0.320736447 | 0.947 | 0.747 | 9.46327267639492e-11 | 18 |
| rps15a7 | 3.91502312294534e-15 | 0.302268951 | 0.947 | 0.778 | 9.85372169814114e-11 | 18 |
| cdkn1bb3 | 4.42807536027579e-15 | 0.402302716 | 0.325 | 0.147 | 1.11450228742781e-10 | 18 |
| zgc:1538672 | 4.75243850188516e-15 | 0.414913359 | 0.645 | 0.434 | 1.19614124653948e-10 | 18 |
| nptna2 | 9.60931769920503e-15 | 0.310753242 | 0.206 | 0.07 | 2.41856917171291e-10 | 18 |
| si:ch211-213a13.1 | 1.44023185498561e-14 | 0.356986453 | 0.228 | 0.087 | 3.62491955581329e-10 | 18 |
| rps174 | 1.51103085885911e-14 | 0.259966395 | 0.912 | 0.765 | 3.80311356866249e-10 | 18 |
| rps6ka12 | 5.47612570413431e-14 | 0.278447747 | 0.206 | 0.072 | 1.37828607847356e-09 | 18 |

| | | | | | | |
|---|---|---|---|---|---|---|
| rpsa8 | 5.71343607910408e-14 | 0.286379867 | 0.934 | 0.75 | 1.43801472674971e-09 | 18 |
| cox8a4 | 6.00588392176917e-14 | 0.413913676 | 0.412 | 0.226 | 1.51162092427008e-09 | 18 |
| idh23 | 8.61693195481176e-14 | 0.418164133 | 0.377 | 0.198 | 2.16879560370657e-09 | 18 |
| arl5c4 | 2.09362494498848e-13 | 0.35105883 | 0.263 | 0.111 | 5.26944462404152e-09 | 18 |
| cox8b1 | 3.05098210229778e-13 | 0.357251065 | 0.232 | 0.094 | 7.67901685327327e-09 | 18 |
| rps168 | 4.79509512027922e-13 | 0.308286093 | 0.943 | 0.735 | 1.20687749082308e-08 | 18 |
| rpl147 | 4.85038366709705e-13 | 0.317413563 | 0.842 | 0.632 | 1.22079306517166e-08 | 18 |
| rpl106 | 4.97322572553603e-13 | 0.252868815 | 0.925 | 0.771 | 1.25171118286016e-08 | 18 |
| XLOC-0234925 | 5.06392346427259e-13 | 0.331275049 | 0.237 | 0.094 | 1.27453889672277e-08 | 18 |
| rps57 | 5.98754533289668e-13 | 0.295725935 | 0.886 | 0.699 | 1.50700528483677e-08 | 18 |
| rpl118 | 9.83015991668675e-13 | 0.254899683 | 0.961 | 0.812 | 2.47415294943089e-08 | 18 |
| vsir5 | 1.02854875175199e-12 | 0.275353916 | 0.219 | 0.084 | 2.58875435328459e-08 | 18 |
| romo17 | 1.26735797112756e-12 | 0.36752058 | 0.596 | 0.384 | 3.18981327753095e-08 | 18 |
| limd21 | 1.35424181363834e-12 | 0.305134483 | 0.189 | 0.069 | 3.40849122074633e-08 | 18 |
| parvg2 | 1.38535722529905e-12 | 0.303127058 | 0.184 | 0.067 | 3.48680560035517e-08 | 18 |
| actr2a4 | 1.72112464665165e-12 | 0.360480503 | 0.281 | 0.129 | 4.33189862315755e-08 | 18 |
| cd638 | 1.83590152887557e-12 | 0.374253948 | 0.561 | 0.338 | 4.62078055802692e-08 | 18 |
| lrrfip1a1 | 2.7570851026841e-12 | 0.255724509 | 0.167 | 0.057 | 6.9393074949456e-08 | 18 |
| wdr12 | 3.62175372869842e-12 | 0.367001789 | 0.294 | 0.14 | 9.11559195976105e-08 | 18 |
| fgl2a2 | 5.16074890390267e-12 | 0.295042871 | 0.18 | 0.065 | 1.29890889162326e-07 | 18 |
| arhgdia4 | 5.62067728850897e-12 | 0.361580195 | 0.412 | 0.228 | 1.41466826674482e-07 | 18 |
| gng54 | 9.38469565797334e-12 | 0.378454804 | 0.452 | 0.27 | 2.36203405015531e-07 | 18 |
| plekhj1 | 1.49019295085137e-11 | 0.264242547 | 0.154 | 0.053 | 3.75066663799782e-07 | 18 |
| twf1a2 | 2.33709796764873e-11 | 0.335839699 | 0.307 | 0.15 | 5.8822418747751e-07 | 18 |
| zgc:1527913 | 5.56004994196353e-11 | 0.356250443 | 0.175 | 0.068 | 1.3994089698928e-06 | 18 |
| rps26l8 | 5.84717751692123e-11 | 0.310675816 | 0.877 | 0.662 | 1.4716761092339e-06 | 18 |
| rpl98 | 1.16710156266379e-10 | 0.30959444 | 0.838 | 0.627 | 2.93747792306848e-06 | 18 |
| zgc:927443 | 2.55334353424085e-10 | 0.35176124 | 0.539 | 0.352 | 6.42651034133079e-06 | 18 |
| prkar2aa1 | 2.69428905865427e-10 | 0.267333533 | 0.224 | 0.098 | 6.78125613172694e-06 | 18 |
| rps158 | 3.45370664284597e-10 | 0.284690211 | 0.877 | 0.682 | 8.69263424937902e-06 | 18 |
| rpl18a7 | 4.25659788838722e-10 | 0.254416226 | 0.952 | 0.783 | 1.07134312252818e-05 | 18 |
| clic14 | 6.15111100915595e-10 | 0.309346292 | 0.237 | 0.113 | 1.54817312989446e-05 | 18 |
| eif4ebp13 | 6.50282458430177e-10 | 0.338934369 | 0.417 | 0.245 | 1.63669591962291e-05 | 18 |
| rps148 | 6.65421171630574e-10 | 0.250345644 | 0.925 | 0.757 | 1.67479854687699e-05 | 18 |
| litaf4 | 8.32013874270368e-10 | 0.281136413 | 0.272 | 0.134 | 2.09409572015109e-05 | 18 |

| | | | | | | |
|---|---|---|---|---|---|---|
| arrb2b3 | 9.18216884486462e-10 | 0.279215112 | 0.206 | 0.091 | 2.31106007656398e-05 | 18 |
| arf6b3 | 1.12763891923323e-09 | 0.28726735 | 0.219 | 0.101 | 2.83815439581813e-05 | 18 |
| rpl68 | 1.44962816996097e-09 | 0.268994292 | 0.851 | 0.666 | 3.64856914097476e-05 | 18 |
| rpl308 | 1.8675124795316e-09 | 0.265852255 | 0.89 | 0.706 | 4.70034215973307e-05 | 18 |
| cxcr4b5 | 2.39246928297612e-09 | 0.360966744 | 0.338 | 0.182 | 6.02160593832261e-05 | 18 |
| si:ch1073-429i10.3.1 | 3.1170485640917e-09 | 0.329287797 | 0.351 | 0.201 | 7.8452995309624e-05 | 18 |
| sik13 | 3.53386011026095e-09 | 0.252774194 | 0.246 | 0.115 | 8.89437251151577e-05 | 18 |
| ak1 | 4.66398331496092e-09 | 0.259630086 | 0.175 | 0.075 | 0.000117388 | 18 |
| scinlb3 | 6.27900601749739e-09 | 0.266844547 | 0.189 | 0.083 | 0.000158036 | 18 |
| rplp2l8 | 6.31095854420402e-09 | 0.253291044 | 0.952 | 0.772 | 0.000158841 | 18 |
| pde4ba5 | 7.68239364460939e-09 | 0.311405771 | 0.268 | 0.139 | 0.000193358 | 18 |
| ifi453 | 8.81036171543739e-09 | 0.370929542 | 0.25 | 0.128 | 0.000221748 | 18 |
| rpl319 | 1.00302475051013e-08 | 0.252351146 | 0.811 | 0.653 | 0.000252451 | 18 |
| cox6b15 | 1.60219830704824e-08 | 0.25925224 | 0.241 | 0.12 | 0.000403257 | 18 |
| sumo2b6 | 1.84566919371486e-08 | 0.330880104 | 0.408 | 0.269 | 0.000464536 | 18 |
| sept61 | 1.96321815324453e-08 | 0.297186699 | 0.355 | 0.205 | 0.000494122 | 18 |
| pabpc1a6 | 2.682382016669e-08 | 0.276955538 | 0.702 | 0.539 | 0.000675129 | 18 |
| msna6 | 2.77499158240808e-08 | 0.307674711 | 0.39 | 0.241 | 0.000698438 | 18 |
| cdc42l3 | 2.82849850029319e-08 | 0.299016357 | 0.36 | 0.215 | 0.000711905 | 18 |
| zgc:1141888 | 3.30730796557255e-08 | 0.292335472 | 0.842 | 0.625 | 0.000832416 | 18 |
| capza1a4 | 5.1749776917654e-08 | 0.289244961 | 0.338 | 0.199 | 0.00130249 | 18 |
| btf37 | 8.00182561836468e-08 | 0.260556806 | 0.759 | 0.586 | 0.002013979 | 18 |
| rps27.27 | 9.99995056820364e-08 | 0.283164251 | 0.697 | 0.516 | 0.002516888 | 18 |
| serf27 | 1.76165298886631e-07 | 0.284032984 | 0.496 | 0.342 | 0.004433904 | 18 |
| hif1ab | 1.76730894609461e-07 | 0.254658863 | 0.219 | 0.113 | 0.00444814 | 18 |
| atp6v1g12 | 1.9615035109886e-07 | 0.258186259 | 0.25 | 0.132 | 0.004936908 | 18 |
| cox4i14 | 2.16542964765096e-07 | 0.282111078 | 0.535 | 0.391 | 0.00545017 | 18 |
| mtpn2 | 3.45964991490073e-07 | 0.268391502 | 0.259 | 0.146 | 0.008707593 | 18 |
| zfp36l1a5 | 1.04884741977367e-06 | 0.304186068 | 0.373 | 0.248 | 0.026398441 | 18 |
| oaz1b6 | 1.10913141347761e-06 | 0.272163535 | 0.465 | 0.322 | 0.027915729 | 18 |
| nsa27 | 1.53079088284008e-06 | 0.266497945 | 0.509 | 0.362 | 0.038528476 | 18 |
| cdc421 | 2.19848592260973e-06 | 0.262971298 | 0.333 | 0.212 | 0.055333692 | 18 |
| glo11 | 3.85640241031095e-06 | 0.264329918 | 0.311 | 0.197 | 0.097061792 | 18 |
| CR318588.16 | 0.00013983 | 0.355305578 | 0.298 | 0.208 | 1 | 18 |
| c1qb2 | 0 | 1.91417798 | 0.945 | 0.055 | 0 | 19 |

| | | | | | | |
|---|---|---|---|---|---|---|
| c1qc2 | 0 | 1.794197013 | 0.93 | 0.051 | 0 | 19 |
| si:dkey-5n18.12 | 0 | 1.776220193 | 0.915 | 0.044 | 0 | 19 |
| CU499330.12 | 0 | 1.765232567 | 0.905 | 0.056 | 0 | 19 |
| grn12 | 0 | 1.7110642 | 0.905 | 0.074 | 0 | 19 |
| lygl12 | 0 | 1.684706588 | 0.955 | 0.082 | 0 | 19 |
| c1qa2 | 0 | 1.669546382 | 0.896 | 0.037 | 0 | 19 |
| ccl35.13 | 0 | 1.642210067 | 0.831 | 0.07 | 0 | 19 |
| mhc2dab3 | 0 | 1.639095785 | 0.881 | 0.084 | 0 | 19 |
| ms4a17a.102 | 0 | 1.499358576 | 0.826 | 0.039 | 0 | 19 |
| cmklr12 | 0 | 1.461861596 | 0.791 | 0.017 | 0 | 19 |
| lgals3bpb2 | 0 | 1.457645268 | 0.652 | 0.031 | 0 | 19 |
| zgc:1749043 | 0 | 1.425712829 | 0.721 | 0.037 | 0 | 19 |
| havcr12 | 0 | 1.377282699 | 0.756 | 0.021 | 0 | 19 |
| mrc1b2 | 0 | 1.342553875 | 0.761 | 0.03 | 0 | 19 |
| sftpbb3 | 0 | 1.335032977 | 0.746 | 0.055 | 0 | 19 |
| itgae.12 | 0 | 1.326424551 | 0.761 | 0.021 | 0 | 19 |
| grna2 | 0 | 1.311885697 | 0.781 | 0.043 | 0 | 19 |
| marco2 | 0 | 1.284819883 | 0.687 | 0.024 | 0 | 19 |
| si:ch211-165i18.22 | 0 | 1.248275351 | 0.672 | 0.025 | 0 | 19 |
| LOC1005375042 | 0 | 1.246595876 | 0.587 | 0.016 | 0 | 19 |
| NPC2 (1 of many)2 | 0 | 1.222701673 | 0.647 | 0.024 | 0 | 19 |
| cx32.21 | 0 | 1.216386863 | 0.687 | 0.019 | 0 | 19 |
| si:ch211-147m6.12 | 0 | 1.163699944 | 0.627 | 0.022 | 0 | 19 |
| ccl34b.12 | 0 | 1.107524366 | 0.502 | 0.022 | 0 | 19 |
| slc3a2a2 | 0 | 1.09108458 | 0.632 | 0.024 | 0 | 19 |
| ctss2.22 | 0 | 1.049459967 | 0.602 | 0.033 | 0 | 19 |
| ccl34a.41 | 0 | 0.995632058 | 0.468 | 0.013 | 0 | 19 |
| CU929150.12 | 0 | 0.977252776 | 0.448 | 0.018 | 0 | 19 |
| zgc:640652 | 0 | 0.918314085 | 0.438 | 0.011 | 0 | 19 |
| si:ch211-194m7.32 | 0 | 0.897418191 | 0.458 | 0.014 | 0 | 19 |
| mafbb1 | 0 | 0.873994026 | 0.413 | 0.015 | 0 | 19 |
| si:ch211-283g2.22 | 0 | 0.805011337 | 0.468 | 0.016 | 0 | 19 |
| slc7a71 | 0 | 0.738966034 | 0.378 | 0.009 | 0 | 19 |
| abca1b1 | 0 | 0.674123971 | 0.383 | 0.012 | 0 | 19 |
| CABZ01076275.1 | 4.44786763121365e-300 | 0.802323098 | 0.393 | 0.014 | 1.11948380410016e-295 | 19 |

| | | | | | | |
|---|---|---|---|---|---|---|
| zgc:1531182 | 2.87141200632338e-293 | 1.182226505 | 0.532 | 0.029 | 7.22705687871532e-289 | 19 |
| mfap42 | 1.35006906702524e-289 | 0.874827925 | 0.423 | 0.018 | 3.39798883479582e-285 | 19 |
| lgals2a4 | 3.44168805060089e-289 | 1.614892977 | 0.945 | 0.112 | 8.66238465455738e-285 | 19 |
| si:ch73-158p21.3 | 8.00136751800499e-289 | 0.80291657 | 0.448 | 0.02 | 2.01386419060668e-284 | 19 |
| si:busm1-266f07.24 | 3.72456523691046e-287 | 1.812693916 | 0.97 | 0.133 | 9.37435824477993e-283 | 19 |
| adgrg31 | 2.45129797139514e-275 | 0.729495422 | 0.413 | 0.018 | 6.16967186420443e-271 | 19 |
| cfp3 | 9.86483988217334e-273 | 1.249056698 | 0.741 | 0.065 | 2.48288154994421e-268 | 19 |
| si:zfos-2330d3.3 | 1.18581097624693e-271 | 0.505759412 | 0.229 | 0.004 | 2.98456764611589e-267 | 19 |
| zgc:1037002 | 2.12107302281716e-258 | 1.048653972 | 0.517 | 0.032 | 5.3385286911285e-254 | 19 |
| si:zfos-1069f5.11 | 2.47210623441185e-256 | 0.492491594 | 0.294 | 0.008 | 6.22204418139118e-252 | 19 |
| rnaset2l3 | 9.2292787540287e-251 | 0.893623659 | 0.532 | 0.035 | 2.32291716960148e-246 | 19 |
| naga2 | 9.3774510243474e-250 | 0.91721082 | 0.572 | 0.042 | 2.360210648318e-245 | 19 |
| spi1a2 | 1.37971222059201e-247 | 0.795118644 | 0.488 | 0.029 | 3.47259768800803e-243 | 19 |
| cd834 | 1.54425430147562e-244 | 1.391738021 | 0.811 | 0.092 | 3.88673365138399e-240 | 19 |
| tcima4 | 4.44698646366017e-243 | 1.191741629 | 0.662 | 0.059 | 1.11926202303863e-238 | 19 |
| zgc:1368701 | 2.90513946718993e-233 | 0.537973049 | 0.328 | 0.013 | 7.31194552497033e-229 | 19 |
| LO018493.11 | 1.46165458681621e-230 | 0.682123059 | 0.383 | 0.019 | 3.67883842955773e-226 | 19 |
| si:ch211-74m13.32 | 1.24414471219644e-225 | 1.050746744 | 0.632 | 0.059 | 3.13138782612721e-221 | 19 |
| aatka | 1.57781519212169e-225 | 0.472683768 | 0.259 | 0.007 | 3.97120305705109e-221 | 19 |
| cxcr3.22 | 5.47936989313312e-224 | 0.769332007 | 0.448 | 0.028 | 1.37910260840267e-219 | 19 |
| il10ra2 | 1.52817812019585e-222 | 0.810123545 | 0.488 | 0.033 | 3.84627151072093e-218 | 19 |
| fgl2a3 | 1.48709441620858e-220 | 1.10836616 | 0.617 | 0.057 | 3.74286793615537e-216 | 19 |
| asah1b1 | 6.69393475180463e-219 | 0.952830831 | 0.552 | 0.046 | 1.68479643768171e-214 | 19 |
| fcer1gl4 | 2.96531207471192e-218 | 1.489616825 | 0.965 | 0.156 | 7.46339396084244e-214 | 19 |
| cd74a5 | 4.67766096741526e-210 | 1.858195511 | 0.98 | 0.207 | 1.17732048888875e-205 | 19 |
| zmp:00000009361 | 1.10115100785571e-208 | 0.559914531 | 0.318 | 0.014 | 2.77148697167203e-204 | 19 |
| zgc:1528302 | 7.67233979455229e-206 | 0.945706222 | 0.602 | 0.057 | 1.93105120289087e-201 | 19 |
| slc6a6b3 | 4.65201695338009e-205 | 1.147586504 | 0.751 | 0.093 | 1.17086614699623e-200 | 19 |
| cd74b6 | 9.41009562637222e-205 | 1.692760235 | 0.975 | 0.212 | 2.36842696820162e-200 | 19 |
| stoml3b | 1.1728733241936e-201 | 0.566508577 | 0.279 | 0.011 | 2.95200486966286e-197 | 19 |
| plin21 | 4.41537391680458e-201 | 0.800452935 | 0.398 | 0.024 | 1.11130546112055e-196 | 19 |
| BX649485.11 | 3.17814675538446e-197 | 0.712062776 | 0.358 | 0.02 | 7.99907756862715e-193 | 19 |
| plekho1b1 | 1.16432704511279e-194 | 0.722948229 | 0.463 | 0.035 | 2.93049473984438e-190 | 19 |
| slc40a11 | 1.73747059266412e-194 | 0.82667037 | 0.433 | 0.03 | 4.37303973467632e-190 | 19 |
| tmem163a | 2.45673223691519e-194 | 0.299924104 | 0.134 | 0.001 | 6.18334936709184e-190 | 19 |

| | | | | | | |
|---|---|---|---|---|---|---|
| pmaip11 | 1.31688281270891e-192 | 1.246433868 | 0.721 | 0.095 | 3.31446235130705e-188 | 19 |
| ctsh2 | 2.10097396208585e-189 | 0.866972003 | 0.517 | 0.046 | 5.28794136517388e-185 | 19 |
| mhc2b1 | 1.41746414608925e-188 | 0.861508948 | 0.383 | 0.024 | 3.56761550929204e-184 | 19 |
| tpp12 | 4.67452167772032e-182 | 0.94561474 | 0.602 | 0.064 | 1.17653036106543e-177 | 19 |
| slc16a9a1 | 3.75626284142431e-177 | 0.419469472 | 0.219 | 0.007 | 9.45413794558085e-173 | 19 |
| dnase21 | 3.78344684932318e-177 | 0.745827304 | 0.463 | 0.038 | 9.52255737506151e-173 | 19 |
| CR759843.2 | 1.68677078127072e-175 | 0.386895757 | 0.179 | 0.004 | 4.24543337938028e-171 | 19 |
| csf2rb2 | 6.84872226432345e-174 | 0.69693853 | 0.433 | 0.033 | 1.72375490670757e-169 | 19 |
| grn22 | 2.10752194583878e-169 | 0.907687791 | 0.303 | 0.016 | 5.30442198548163e-165 | 19 |
| si:busm1-194e12.11 | 2.37074227938132e-164 | 0.576159044 | 0.289 | 0.015 | 5.96692124297484e-160 | 19 |
| smox3 | 2.92260874448979e-162 | 0.908255758 | 0.622 | 0.076 | 7.35591394900636e-158 | 19 |
| FP015862.21 | 2.18746020846297e-161 | 0.490994568 | 0.264 | 0.013 | 5.50561859868046e-157 | 19 |
| plaua | 3.58332196574391e-161 | 0.300733941 | 0.139 | 0.002 | 9.01886305558084e-157 | 19 |
| si:cabz01074946.13 | 8.07943845461419e-157 | 0.883190201 | 0.502 | 0.052 | 2.03351386464184e-152 | 19 |
| rbp2a1 | 2.01303819091487e-155 | 0.507613729 | 0.234 | 0.01 | 5.06661582271363e-151 | 19 |
| mpeg1.11 | 3.45783240808142e-155 | 0.486307373 | 0.259 | 0.012 | 8.70301838790012e-151 | 19 |
| gng102 | 1.67918675576064e-153 | 0.773814115 | 0.483 | 0.048 | 4.22634514557396e-149 | 19 |
| si:dkey-56f14.71 | 4.12105192049035e-153 | 0.391701628 | 0.229 | 0.01 | 1.03722755786822e-148 | 19 |
| krt18b6 | 3.44769539388949e-149 | 1.178489059 | 0.786 | 0.141 | 8.67750453688047e-145 | 19 |
| nrros2 | 4.24702820625184e-149 | 0.739268366 | 0.443 | 0.042 | 1.06893452923153e-144 | 19 |
| psap2 | 1.94842413054829e-148 | 1.294845471 | 0.98 | 0.28 | 4.90398869417698e-144 | 19 |
| si:dkey-27i16.23 | 4.5380483858174e-148 | 0.954050294 | 0.622 | 0.083 | 1.14218139822638e-143 | 19 |
| atp7a1 | 5.06454003123967e-145 | 0.44719378 | 0.259 | 0.014 | 1.27469408046271e-140 | 19 |
| cndp22 | 6.25063097123356e-145 | 0.795490309 | 0.512 | 0.057 | 1.57322130914977e-140 | 19 |
| si:ch211-39i2.21 | 2.58986922891459e-144 | 0.82634384 | 0.522 | 0.059 | 6.51844186225512e-140 | 19 |
| fuca21 | 1.97994643079609e-143 | 0.769138125 | 0.507 | 0.057 | 4.98332717167068e-139 | 19 |
| cxcr4b6 | 9.2835352393676e-143 | 1.225627367 | 0.861 | 0.173 | 2.33657298439643e-138 | 19 |
| clic24 | 2.46402672745991e-141 | 0.78278656 | 0.488 | 0.053 | 6.20170887034384e-137 | 19 |
| LOC560447 | 6.64665384251522e-141 | 0.27273736 | 0.159 | 0.005 | 1.67289630562265e-136 | 19 |
| ifi302 | 1.54333358499152e-140 | 1.184596191 | 0.811 | 0.165 | 3.88441630006516e-136 | 19 |
| si:dkeyp-23e4.31 | 3.24384541130634e-140 | 0.427959004 | 0.219 | 0.01 | 8.16443451571692e-136 | 19 |
| gnsb1 | 3.10722217581657e-138 | 0.66206664 | 0.383 | 0.033 | 7.82056749431272e-134 | 19 |
| ms4a17a.64 | 1.28559996146635e-137 | 0.698024916 | 0.433 | 0.043 | 3.23572654301466e-133 | 19 |
| slc2a61 | 1.35612222757228e-137 | 0.331202008 | 0.164 | 0.005 | 3.41322403457666e-133 | 19 |
| nr4a16 | 2.19921333218473e-136 | 1.127425491 | 0.801 | 0.157 | 5.53520003577575e-132 | 19 |

| | | | | | | |
|---|---|---|---|---|---|---|
| p2rx7 | 2.63574495629253e-136 | 0.398994746 | 0.224 | 0.011 | 6.63390648049267e-132 | 19 |
| robo4 | 7.52212602593809e-136 | 0.277759901 | 0.134 | 0.003 | 1.89324389946836e-131 | 19 |
| calhm61 | 5.12941140876024e-135 | 0.423790487 | 0.234 | 0.012 | 1.29102155747086e-130 | 19 |
| scpep12 | 1.01741311450041e-134 | 0.963375066 | 0.692 | 0.111 | 2.56072706788608e-130 | 19 |
| AL928650.11 | 1.31592783451839e-134 | 0.430381754 | 0.199 | 0.008 | 3.31205876669933e-130 | 19 |
| si:ch211-66e2.33 | 3.83349312150269e-131 | 0.761412191 | 0.468 | 0.053 | 9.64851883751012e-127 | 19 |
| spi1b3 | 4.58532199956468e-131 | 0.709026982 | 0.483 | 0.055 | 1.15407969407043e-126 | 19 |
| slc43a2a4 | 1.69395977590676e-129 | 0.913913206 | 0.617 | 0.094 | 4.26352735997973e-125 | 19 |
| lipf | 1.85843183735121e-128 | 0.682086018 | 0.383 | 0.036 | 4.67748709142927e-124 | 19 |
| gm2a3 | 7.62313494080216e-128 | 0.921969156 | 0.687 | 0.112 | 1.9186668332505e-123 | 19 |
| npl1 | 5.25604333302398e-126 | 0.639772229 | 0.358 | 0.032 | 1.3228935464888e-121 | 19 |
| tmsb14 | 6.58473644605334e-125 | 1.026076683 | 0.791 | 0.146 | 1.65731231610717e-120 | 19 |
| gpr841 | 2.64767748658712e-124 | 0.376616812 | 0.199 | 0.009 | 6.66393946599112e-120 | 19 |
| mir21-12 | 2.95838865762587e-124 | 0.967500473 | 0.612 | 0.1 | 7.44596841237856e-120 | 19 |
| ctsa2 | 2.10377003390016e-123 | 0.822719298 | 0.567 | 0.082 | 5.29497879832331e-119 | 19 |
| enpp11 | 2.29381815231316e-122 | 0.617135128 | 0.368 | 0.035 | 5.773310907557e-118 | 19 |
| mpp1 | 3.43773810312592e-122 | 0.538519047 | 0.318 | 0.026 | 8.65244303175763e-118 | 19 |
| zgc:1533171 | 1.28855029733749e-121 | 0.602793692 | 0.378 | 0.037 | 3.24315224336873e-117 | 19 |
| ccng22 | 2.34852244112528e-120 | 0.81085275 | 0.542 | 0.079 | 5.91099613206822e-116 | 19 |
| si:dkey-11f4.143 | 3.69759548950306e-119 | 0.934012792 | 0.607 | 0.101 | 9.30647808753024e-115 | 19 |
| sb:cb811 | 2.84008117680424e-118 | 0.506600969 | 0.294 | 0.023 | 7.14820031389859e-114 | 19 |
| CABZ01056054.1 | 5.59448496113608e-118 | 0.343747086 | 0.184 | 0.008 | 1.40807591986834e-113 | 19 |
| ctsk1 | 9.1258730277771e-117 | 0.788697268 | 0.453 | 0.057 | 2.29689098236122e-112 | 19 |
| si:dkey-23a13.11 | 2.01737606982478e-116 | 0.263480398 | 0.134 | 0.004 | 5.077533830142e-112 | 19 |
| BX324132.21 | 1.16445876246409e-113 | 0.326070623 | 0.154 | 0.006 | 2.93082625924588e-109 | 19 |
| crema5 | 1.42200177790988e-113 | 0.978108325 | 0.781 | 0.169 | 3.57903627482137e-109 | 19 |
| nfil3-65 | 6.26179916330913e-113 | 0.965425986 | 0.657 | 0.123 | 1.57603223141328e-108 | 19 |
| sowahd | 2.45819266190176e-112 | 0.366680924 | 0.204 | 0.011 | 6.18702511074053e-108 | 19 |
| tnfb3 | 8.46519292282728e-112 | 0.75665626 | 0.413 | 0.048 | 2.1306044067464e-107 | 19 |
| abi2b1 | 1.21195908514105e-111 | 0.671871021 | 0.408 | 0.047 | 3.05037982139152e-107 | 19 |
| rassf2a1 | 3.66830606504611e-111 | 0.42151765 | 0.254 | 0.018 | 9.23275953511454e-107 | 19 |
| laptm57 | 4.92368066789583e-111 | 0.925936929 | 0.791 | 0.161 | 1.2392411873027e-106 | 19 |
| CR855311.15 | 1.34847766742422e-110 | 0.930664783 | 0.532 | 0.079 | 3.39398344114001e-106 | 19 |
| arl5c5 | 1.316243026863e-109 | 0.879081503 | 0.602 | 0.105 | 3.31285207431148e-105 | 19 |
| plbd11 | 3.33860310820196e-109 | 0.484346662 | 0.279 | 0.022 | 8.40293016303353e-105 | 19 |

| | | | | | | |
|---|---|---|---|---|---|---|
| fosl25 | 8.02157395150043e-108 | 0.889777157 | 0.622 | 0.114 | 2.01894994785314e-103 | 19 |
| si:ch73-236c18.7 | 1.15548873897756e-107 | 0.284356947 | 0.144 | 0.005 | 2.90824960713261e-103 | 19 |
| ctsd3 | 1.864883145716e-107 | 1.044017476 | 0.657 | 0.129 | 4.6937243894526e-103 | 19 |
| ccr9a5 | 2.74955190317993e-107 | 0.819062249 | 0.532 | 0.081 | 6.92034718511356e-103 | 19 |
| cybb1 | 3.73167925150143e-107 | 0.450109464 | 0.239 | 0.016 | 9.39226350810394e-103 | 19 |
| dpp72 | 4.32509083502847e-107 | 0.609241625 | 0.383 | 0.043 | 1.08858211226832e-102 | 19 |
| ncf21 | 1.59937633882008e-106 | 0.448436418 | 0.254 | 0.019 | 4.02547030717627e-102 | 19 |
| lta4h3 | 6.39414378034382e-106 | 0.694379477 | 0.483 | 0.066 | 1.60934204807473e-101 | 19 |
| si:ch211-212k18.72 | 7.99592564026617e-106 | 0.926092114 | 0.667 | 0.131 | 2.01249452439859e-101 | 19 |
| LOC108190616 | 2.29266485886137e-105 | 0.282657428 | 0.159 | 0.007 | 5.77040818326819e-101 | 19 |
| rab444 | 4.40146289833792e-104 | 0.66252161 | 0.433 | 0.056 | 1.10780419688267e-99 | 19 |
| anxa3b2 | 5.58584857318219e-104 | 0.562834376 | 0.308 | 0.029 | 1.40590222738423e-99 | 19 |
| cebpb7 | 6.06503475005561e-104 | 1.146657229 | 0.96 | 0.344 | 1.5265085962415e-99 | 19 |
| pepd2 | 9.82453760629563e-104 | 0.644269791 | 0.468 | 0.065 | 2.47273787012855e-99 | 19 |
| XLOC-0245161 | 4.4048944889861e-103 | 0.319187781 | 0.184 | 0.01 | 1.10866789393291e-98 | 19 |
| cd221 | 5.73496979240605e-103 | 0.295607418 | 0.164 | 0.008 | 1.44343454705068e-98 | 19 |
| atp6v1e1b2 | 4.37910789801396e-102 | 0.703646125 | 0.507 | 0.076 | 1.10217766685113e-97 | 19 |
| si:ch211-1o7.32 | 1.09597769310012e-101 | 0.399740487 | 0.239 | 0.017 | 2.75846625576369e-97 | 19 |
| ccl38.1 | 7.98906267215653e-101 | 0.272683105 | 0.144 | 0.006 | 2.01076718395508e-96 | 19 |
| samsn1a2 | 7.23851170654072e-100 | 0.546317276 | 0.318 | 0.032 | 1.82186101141923e-95 | 19 |
| dusp13 | 1.06344278560969e-99 | 0.994042906 | 0.687 | 0.152 | 2.67657914710103e-95 | 19 |
| rab8b2 | 5.67609410452964e-99 | 0.713180024 | 0.483 | 0.074 | 1.42861612516906e-94 | 19 |
| mertka | 9.39966504734068e-99 | 0.359420831 | 0.194 | 0.012 | 2.36580169576517e-94 | 19 |
| atp6v0ca2 | 3.3626101689139e-98 | 0.990502238 | 0.766 | 0.19 | 8.4633535341394e-94 | 19 |
| mbnl31 | 4.59880590140695e-98 | 0.646398231 | 0.398 | 0.051 | 1.15747345732512e-93 | 19 |
| slc43a2b | 6.81966213084416e-98 | 0.327019117 | 0.149 | 0.007 | 1.71644076171217e-93 | 19 |
| itgae.22 | 1.5888114056968e-97 | 0.537904184 | 0.313 | 0.032 | 3.99887942699827e-93 | 19 |
| timp4.3 | 1.82417343019262e-97 | 0.295008119 | 0.144 | 0.006 | 4.59126210645182e-93 | 19 |
| snx27a | 2.6189064864514e-97 | 0.531991368 | 0.333 | 0.036 | 6.59152573574954e-93 | 19 |
| wdfy4 | 8.1369667156661e-97 | 0.316147183 | 0.184 | 0.01 | 2.047993152666e-92 | 19 |
| tmsb4x10 | 5.81649655339677e-95 | 0.807288545 | 1 | 0.682 | 1.46395401752443e-90 | 19 |
| si:ch211-8c17.3 | 1.01233871843145e-93 | 0.468312142 | 0.289 | 0.028 | 2.54795532042013e-89 | 19 |
| glb1l1 | 2.25571782539983e-93 | 0.443337145 | 0.244 | 0.02 | 5.67741619474883e-89 | 19 |
| si:ch211-204c21.13 | 8.71780164307932e-93 | 0.690345069 | 0.453 | 0.068 | 2.19418349554663e-88 | 19 |
| CABZ01078737.1 | 2.346018881686e-92 | 0.297981202 | 0.164 | 0.009 | 5.90469492331549e-88 | 19 |

| | | | | | | |
|---|---|---|---|---|---|---|
| itm2bb4 | 6.56533162991722e-92 | 0.924031964 | 0.811 | 0.213 | 1.65242831793386e-87 | 19 |
| LOC101885706 | 5.10882764862487e-90 | 0.333739564 | 0.159 | 0.008 | 1.28584083088239e-85 | 19 |
| bach1b1 | 1.63681372618922e-89 | 0.437078741 | 0.249 | 0.022 | 4.11969646744564e-85 | 19 |
| ctsc1 | 1.33258561195487e-87 | 0.677271354 | 0.443 | 0.067 | 3.35398472672922e-83 | 19 |
| zgc:1627309 | 1.41122108572394e-87 | 1.050458774 | 0.98 | 0.383 | 3.55190235065859e-83 | 19 |
| f13a1b2 | 4.66264468797237e-87 | 0.473260555 | 0.279 | 0.028 | 1.17354104151577e-82 | 19 |
| slc43a3b | 2.0248375526869e-86 | 0.610269077 | 0.244 | 0.022 | 5.09631363635767e-82 | 19 |
| pdk2b4 | 5.59836424524522e-86 | 0.81896237 | 0.507 | 0.094 | 1.40905229688577e-81 | 19 |
| slc36a1 | 5.60503497640678e-86 | 0.281713187 | 0.169 | 0.01 | 1.41073125321182e-81 | 19 |
| cebpa4 | 9.22119730701067e-86 | 0.653380503 | 0.383 | 0.053 | 2.32088315020152e-81 | 19 |
| si:ch73-343l4.81 | 3.63367018640769e-85 | 0.447546818 | 0.234 | 0.02 | 9.1455844921695e-81 | 19 |
| kctd12.22 | 4.10495015521795e-85 | 0.663243742 | 0.458 | 0.074 | 1.03317490456681e-80 | 19 |
| LOC101885065 | 5.39318377028873e-85 | 0.324979442 | 0.184 | 0.012 | 1.35741042314397e-80 | 19 |
| trpv1 | 1.65689418772306e-84 | 0.323421227 | 0.179 | 0.012 | 4.17023698108017e-80 | 19 |
| XLOC-0234926 | 7.0481304975438e-84 | 0.77406264 | 0.502 | 0.089 | 1.7739439649268e-79 | 19 |
| mapk111 | 1.07283822571333e-83 | 0.345009721 | 0.204 | 0.015 | 2.70022653029789e-79 | 19 |
| tnfrsf21 | 1.30393610563815e-83 | 0.25741194 | 0.139 | 0.007 | 3.28187678428067e-79 | 19 |
| LOC1001482542 | 1.46191229888093e-82 | 0.41866556 | 0.234 | 0.021 | 3.67948706505343e-78 | 19 |
| tuba8l22 | 2.13900815592611e-82 | 0.543649569 | 0.393 | 0.056 | 5.38366962765042e-78 | 19 |
| rgs192 | 4.13518059107787e-82 | 0.439124869 | 0.284 | 0.03 | 1.04078360296839e-77 | 19 |
| gpx1a3 | 4.77670828409968e-82 | 0.895001342 | 0.796 | 0.216 | 1.20224970802505e-77 | 19 |
| dbn13 | 4.85330577381216e-82 | 0.54749918 | 0.368 | 0.051 | 1.22152853021078e-77 | 19 |
| atp6v1d1 | 2.30750565768063e-81 | 0.383067678 | 0.239 | 0.022 | 5.80776098981637e-77 | 19 |
| cadm4 | 1.28060340021935e-80 | 0.389685202 | 0.229 | 0.02 | 3.22315069801207e-76 | 19 |
| FERMT3 (1 of many)3 | 1.8846833322937e-80 | 0.463167841 | 0.313 | 0.037 | 4.74355947905002e-76 | 19 |
| tnfrsf181 | 4.95240966241844e-80 | 0.369685975 | 0.179 | 0.012 | 1.2464719879341e-75 | 19 |
| zgc:640516 | 5.32137065957952e-80 | 0.820353546 | 0.672 | 0.155 | 1.33933578130957e-75 | 19 |
| XLOC-0054494 | 7.51815639206056e-80 | 0.635312671 | 0.358 | 0.05 | 1.89224478231772e-75 | 19 |
| slc7a8a | 1.5810456409753e-79 | 0.401202184 | 0.254 | 0.025 | 3.97933377377073e-75 | 19 |
| cyba3 | 1.01919768708694e-78 | 0.62477868 | 0.423 | 0.068 | 2.56521865862913e-74 | 19 |
| tnfrsf1b | 1.96022773382763e-78 | 0.362621603 | 0.209 | 0.017 | 4.93369718327076e-74 | 19 |
| lgmn | 2.51090955435023e-78 | 0.703817477 | 0.373 | 0.056 | 6.31970825734409e-74 | 19 |
| rxrga | 2.61820359690158e-78 | 0.358343262 | 0.204 | 0.016 | 6.58975663304158e-74 | 19 |
| BX511021.2 | 6.25184325323634e-78 | 0.276156445 | 0.154 | 0.009 | 1.57352642840705e-73 | 19 |
| timp2b1 | 6.68612943641897e-78 | 0.5644235 | 0.289 | 0.034 | 1.68283191785229e-73 | 19 |

| | | | | | | |
|---|---|---|---|---|---|---|
| gstt1a1 | 5.89099009070223e-77 | 0.477318746 | 0.294 | 0.034 | 1.48270329592884e-72 | 19 |
| atp6v1g13 | 1.03305293369335e-76 | 0.731869222 | 0.587 | 0.126 | 2.60009092881279e-72 | 19 |
| actb27 | 1.19345700577296e-75 | 0.5602167 | 1 | 0.866 | 3.00381193782996e-71 | 19 |
| si:dkey-24p1.71 | 2.86611758565276e-75 | 0.283680249 | 0.169 | 0.012 | 7.21373135132943e-71 | 19 |
| si:zfos-741a10.32 | 7.85043457837078e-75 | 0.39267094 | 0.239 | 0.024 | 1.97587587903014e-70 | 19 |
| ets22 | 1.1123123911728e-74 | 0.621314092 | 0.468 | 0.085 | 2.79957905734281e-70 | 19 |
| grapa1 | 1.27098981810776e-74 | 0.412818672 | 0.244 | 0.025 | 3.19895427319542e-70 | 19 |
| wasb6 | 6.05107928828541e-74 | 0.753782693 | 0.711 | 0.17 | 1.52299614606856e-69 | 19 |
| ehbp1l1b1 | 6.77805399416428e-74 | 0.362517361 | 0.214 | 0.019 | 1.70596840979121e-69 | 19 |
| ncf41 | 9.56741728916435e-74 | 0.322162595 | 0.194 | 0.016 | 2.40802325750978e-69 | 19 |
| gnpda11 | 1.26849981876883e-73 | 0.469534906 | 0.313 | 0.04 | 3.19268719385926e-69 | 19 |
| tmem86a | 1.83850289260333e-73 | 0.297419084 | 0.164 | 0.011 | 4.62732793039332e-69 | 19 |
| slc30a1a | 2.94778601975845e-73 | 0.389404422 | 0.219 | 0.021 | 7.41928263313004e-69 | 19 |
| sdr16c5b | 3.63097057590751e-73 | 0.489483706 | 0.294 | 0.037 | 9.13878984250162e-69 | 19 |
| tspan371 | 1.13113665316646e-72 | 0.411687203 | 0.239 | 0.025 | 2.84695784235467e-68 | 19 |
| zgc:638312 | 1.8891718309466e-72 | 0.597834989 | 0.433 | 0.076 | 4.7548565813095e-68 | 19 |
| bri31 | 2.55426060815515e-72 | 0.521516616 | 0.368 | 0.056 | 6.42881852466569e-68 | 19 |
| si:dkey-262k9.45 | 3.30056893218678e-72 | 0.613253783 | 0.463 | 0.085 | 8.3072019454209e-68 | 19 |
| cflarb1 | 4.19002661976241e-72 | 0.328713443 | 0.194 | 0.016 | 1.054587799928e-67 | 19 |
| mir22a | 7.70394086577838e-72 | 0.276174115 | 0.159 | 0.011 | 1.93900487650776e-67 | 19 |
| zfp36l1a6 | 1.06571904740574e-71 | 0.901902155 | 0.771 | 0.24 | 2.68230827041551e-67 | 19 |
| entpd2a.1 | 9.43174033058831e-71 | 0.437759471 | 0.284 | 0.034 | 2.37387472380577e-66 | 19 |
| ctsz2 | 2.90725407257004e-70 | 0.736389558 | 0.438 | 0.082 | 7.31726777525154e-66 | 19 |
| ehd41 | 3.57520870508997e-70 | 0.381692055 | 0.244 | 0.026 | 8.99844278984095e-66 | 19 |
| arpc5b4 | 1.56732721545049e-69 | 0.630450807 | 0.522 | 0.107 | 3.94480586856733e-65 | 19 |
| dennd4a1 | 2.10926273734855e-69 | 0.475402641 | 0.274 | 0.034 | 5.30880338363256e-65 | 19 |
| si:ch211-15b10.63 | 3.12559610944853e-69 | 0.581267815 | 0.393 | 0.067 | 7.866812847871e-65 | 19 |
| zgc:123107 | 4.5051187594813e-69 | 0.256025171 | 0.154 | 0.011 | 1.13389334057385e-64 | 19 |
| BX649485.22 | 6.61351771509722e-69 | 0.527394789 | 0.254 | 0.029 | 1.66455627371282e-64 | 19 |
| ifngr1l1 | 8.1449878971766e-69 | 0.408777973 | 0.249 | 0.028 | 2.05001200384038e-64 | 19 |
| rnaseka1 | 8.61941702898428e-69 | 0.692285872 | 0.557 | 0.123 | 2.16942107202505e-64 | 19 |
| si:ch211-152p11.41 | 2.13792313100326e-68 | 0.413974892 | 0.249 | 0.028 | 5.3809387284221e-64 | 19 |
| csrnp1b5 | 3.04947461898369e-68 | 0.776172824 | 0.652 | 0.173 | 7.67522266852005e-64 | 19 |
| cd639 | 6.87473102133617e-68 | 0.873625048 | 0.866 | 0.333 | 1.7303010507601e-63 | 19 |
| FP102158.11 | 7.10678751693012e-68 | 0.289435541 | 0.179 | 0.015 | 1.78870735013614e-63 | 19 |

| | | | | | | |
|---|---|---|---|---|---|---|
| tspan361 | 7.87402403033071e-68 | 0.797879537 | 0.617 | 0.151 | 1.98181310819394e-63 | 19 |
| m172 | 1.07461177604213e-67 | 0.336640293 | 0.164 | 0.012 | 2.70469037912044e-63 | 19 |
| dicp3.12 | 3.8026752517409e-67 | 0.304961482 | 0.179 | 0.015 | 9.57095334110667e-63 | 19 |
| git2b | 5.02842766778366e-67 | 0.372650275 | 0.224 | 0.023 | 1.26560495970447e-62 | 19 |
| wipf1b3 | 1.18656125387263e-66 | 0.51393273 | 0.363 | 0.059 | 2.98645601987202e-62 | 19 |
| mgat4a | 8.43093488535982e-66 | 0.288784687 | 0.169 | 0.014 | 2.12198200129621e-61 | 19 |
| pcyox11 | 9.11222638172538e-66 | 0.477555945 | 0.318 | 0.047 | 2.29345625801646e-61 | 19 |
| CABZ01068358.13 | 2.91490226159992e-65 | 0.569056988 | 0.363 | 0.06 | 7.33651750222085e-61 | 19 |
| arhgef12b | 8.13072608081758e-65 | 0.332827964 | 0.189 | 0.017 | 2.04642244728098e-60 | 19 |
| mfsd1 | 1.91581499962743e-64 | 0.376691319 | 0.214 | 0.022 | 4.82191477256227e-60 | 19 |
| vsir6 | 8.19276155506146e-64 | 0.596078855 | 0.423 | 0.081 | 2.06203615579342e-59 | 19 |
| XLOC-0004315 | 8.8093866051418e-64 | 0.836597181 | 0.876 | 0.27 | 2.21723451464814e-59 | 19 |
| xaf11 | 1.11935581054596e-63 | 0.526655689 | 0.348 | 0.057 | 2.81730663956312e-59 | 19 |
| junba5 | 1.41495959893006e-63 | 0.876786577 | 0.9 | 0.3 | 3.56131181454707e-59 | 19 |
| rac27 | 2.27051119039373e-63 | 0.766773627 | 0.761 | 0.213 | 5.71464961510198e-59 | 19 |
| nck1a | 2.28147684798531e-63 | 0.408854227 | 0.239 | 0.028 | 5.74224907869423e-59 | 19 |
| cds1 | 4.08092110630793e-63 | 0.250774899 | 0.114 | 0.006 | 1.02712703324664e-58 | 19 |
| nocta1 | 4.44921200823187e-63 | 0.491066518 | 0.294 | 0.042 | 1.11982217035188e-58 | 19 |
| txnipa7 | 4.80160324877947e-63 | 0.823252066 | 0.677 | 0.204 | 1.2085155216853e-58 | 19 |
| si:dkey-206d17.122 | 7.57126530958212e-63 | 0.506702172 | 0.318 | 0.049 | 1.90561176576872e-58 | 19 |
| BX248082.14 | 9.16208998241089e-63 | 0.695640643 | 0.279 | 0.039 | 2.306006427673e-58 | 19 |
| rgs21 | 9.24967466100938e-63 | 0.458818643 | 0.224 | 0.025 | 2.32805061542945e-58 | 19 |
| lyn1 | 2.38872697849267e-62 | 0.443153544 | 0.294 | 0.042 | 6.01218693216819e-58 | 19 |
| mt-co19 | 2.88241656860592e-62 | 0.662753691 | 0.995 | 0.653 | 7.25475426152424e-58 | 19 |
| aplp21 | 4.71111070780175e-62 | 0.519657642 | 0.383 | 0.068 | 1.18573945404662e-57 | 19 |
| metrnla1 | 5.12801726287812e-62 | 0.588831835 | 0.443 | 0.088 | 1.29067066489379e-57 | 19 |
| egr33 | 1.42487255827387e-61 | 0.505357584 | 0.338 | 0.055 | 3.5862617419195e-57 | 19 |
| pfn17 | 1.43430149633195e-61 | 0.887965991 | 0.995 | 0.518 | 3.60999343611788e-57 | 19 |
| klf133 | 1.48860593489764e-61 | 0.642870475 | 0.463 | 0.1 | 3.74667227754387e-57 | 19 |
| irf82 | 1.76282424139095e-61 | 0.448351784 | 0.299 | 0.043 | 4.43685233315688e-57 | 19 |
| rbm47 | 1.87870150878332e-61 | 0.332851358 | 0.164 | 0.014 | 4.72850382745674e-57 | 19 |
| dhrs91 | 6.07306240283529e-61 | 0.413516511 | 0.249 | 0.031 | 1.52852907616961e-56 | 19 |
| flvcr2b | 1.04228401486966e-60 | 0.275361571 | 0.169 | 0.015 | 2.62332463702545e-56 | 19 |
| LOC101886114 | 1.37813270679377e-60 | 0.292893115 | 0.159 | 0.013 | 3.46862220972923e-56 | 19 |
| nfe2l2a1 | 2.00449495337327e-60 | 0.522619142 | 0.363 | 0.065 | 5.04511334814517e-56 | 19 |

| | | | | | | |
|---|---|---|---|---|---|---|
| si:ch73-248e21.52 | 4.34463574786936e-60 | 0.442986727 | 0.289 | 0.042 | 1.09350137138124e-55 | 19 |
| cotl17 | 6.84047971108992e-60 | 0.846451401 | 0.831 | 0.299 | 1.72168033848422e-55 | 19 |
| acsl4b1 | 9.90583827512435e-60 | 0.52105801 | 0.323 | 0.053 | 2.49320043546605e-55 | 19 |
| slc1a3a | 1.17372047069978e-59 | 0.307266562 | 0.174 | 0.016 | 2.95413705270427e-55 | 19 |
| atp6v1ba1 | 1.46742517661659e-59 | 0.476753329 | 0.308 | 0.048 | 3.69336242702631e-55 | 19 |
| COX39 | 1.61183849619237e-59 | 0.635218179 | 0.995 | 0.696 | 4.05683631106658e-55 | 19 |
| CABZ01001434.11 | 2.9821154306061e-59 | 0.354056919 | 0.144 | 0.011 | 7.50568632729249e-55 | 19 |
| klf6a3 | 4.44546078269466e-59 | 0.751318714 | 0.592 | 0.162 | 1.11887802439642e-54 | 19 |
| ddit36 | 5.4586872452595e-58 | 0.757773812 | 0.607 | 0.17 | 1.37389699275936e-53 | 19 |
| tfe3a1 | 1.4486005256709e-57 | 0.414756258 | 0.259 | 0.036 | 3.6459826630611e-53 | 19 |
| me2 | 3.67439348796679e-57 | 0.322392515 | 0.209 | 0.024 | 9.24808096986363e-53 | 19 |
| si:dkey-25e12.33 | 5.77208679118504e-57 | 0.448450113 | 0.303 | 0.048 | 1.45277652447336e-52 | 19 |
| ccl34b.82 | 8.36689770049909e-57 | 0.377736334 | 0.174 | 0.017 | 2.10586448223862e-52 | 19 |
| coro1a7 | 1.25749714577136e-56 | 0.75132163 | 0.761 | 0.224 | 3.16499456619193e-52 | 19 |
| nfkbiaa4 | 1.49585881781915e-56 | 0.719069034 | 0.562 | 0.147 | 3.76492705856901e-52 | 19 |
| abcc53 | 1.80842602991404e-56 | 0.544424285 | 0.428 | 0.09 | 4.55162747469066e-52 | 19 |
| rnf130 | 3.06634658911594e-56 | 0.380311673 | 0.289 | 0.044 | 7.7176877301459e-52 | 19 |
| mcl1b4 | 7.71414964308986e-56 | 0.797519039 | 0.706 | 0.236 | 1.94157432366929e-51 | 19 |
| gabarapb5 | 1.11696922112326e-55 | 0.737361108 | 0.751 | 0.262 | 2.81129983264514e-51 | 19 |
| btg18 | 2.31953720127241e-55 | 0.789352133 | 0.836 | 0.326 | 5.83804318188253e-51 | 19 |
| si:ch211-102c2.44 | 3.62467291939547e-55 | 0.656289102 | 0.552 | 0.137 | 9.12293927082645e-51 | 19 |
| abca1a1 | 7.08180159446652e-55 | 0.502499565 | 0.353 | 0.065 | 1.78241864331128e-50 | 19 |
| LOC1018842673 | 4.40530621251559e-54 | 0.455525295 | 0.323 | 0.055 | 1.10877152062805e-49 | 19 |
| LO018340.11 | 4.43980159294466e-54 | 0.30143227 | 0.204 | 0.024 | 1.11745366292824e-49 | 19 |
| si:ch1073-392o20.21 | 1.32088628438066e-53 | 0.424902419 | 0.303 | 0.05 | 3.32453868915769e-49 | 19 |
| fybb4 | 1.69079729343349e-53 | 0.404670105 | 0.264 | 0.039 | 4.25556770784275e-49 | 19 |
| LOC1005369462 | 1.79046514706234e-53 | 0.317903977 | 0.164 | 0.016 | 4.5064217286412e-49 | 19 |
| mfng4 | 6.14042826230233e-53 | 0.415688264 | 0.264 | 0.039 | 1.54548438933887e-48 | 19 |
| rnf144b1 | 8.1923080357099e-53 | 0.348631354 | 0.219 | 0.028 | 2.06192200950782e-48 | 19 |
| cap17 | 2.64370817255853e-52 | 0.693665595 | 0.716 | 0.226 | 6.65394909951257e-48 | 19 |
| grap2b5 | 3.62295861410382e-52 | 0.564126929 | 0.468 | 0.108 | 9.11862453583791e-48 | 19 |
| bpifcl1 | 3.63931471006467e-52 | 0.258717402 | 0.139 | 0.012 | 9.15979119376177e-48 | 19 |
| itgb24 | 7.9973906640922e-52 | 0.558400991 | 0.428 | 0.095 | 2.01286325624537e-47 | 19 |
| il4r.12 | 9.79855428622755e-52 | 0.394904074 | 0.264 | 0.04 | 2.46619812830061e-47 | 19 |
| sdcbp24 | 1.32054084406343e-51 | 0.702775075 | 0.637 | 0.203 | 3.32366925042326e-47 | 19 |

| | | | | | | |
|---|---|---|---|---|---|---|
| arrb2b4 | 1.90367640606349e-51 | 0.52940126 | 0.403 | 0.087 | 4.79136314642119e-47 | 19 |
| slc20a1b2 | 2.30546019565689e-51 | 0.585627799 | 0.428 | 0.098 | 5.80261276644884e-47 | 19 |
| cst14b.12 | 2.35668419327858e-51 | 0.456803905 | 0.289 | 0.049 | 5.93153844606285e-47 | 19 |
| b2ml8 | 3.66065779094281e-51 | 0.68817264 | 0.756 | 0.26 | 9.21350959402395e-47 | 19 |
| txn2 | 5.80483209538188e-51 | 0.475186162 | 0.269 | 0.042 | 1.46101819008667e-46 | 19 |
| mboat2a1 | 6.78715161043502e-51 | 0.334324818 | 0.194 | 0.023 | 1.70825818883039e-46 | 19 |
| stat31 | 8.60228837014381e-51 | 0.462130149 | 0.289 | 0.049 | 2.1651099598815e-46 | 19 |
| irf21 | 1.55971750024016e-50 | 0.458380134 | 0.338 | 0.065 | 3.92565297635446e-46 | 19 |
| atp2b22 | 2.99190055504093e-50 | 0.455276378 | 0.348 | 0.068 | 7.53031450698251e-46 | 19 |
| slc39a1 | 8.6066936783657e-50 | 0.287172017 | 0.169 | 0.018 | 2.16621873190786e-45 | 19 |
| pparg | 9.87845731217002e-50 | 0.312754132 | 0.154 | 0.015 | 2.48630892090007e-45 | 19 |
| si:ch211-243a20.41 | 1.69147986264374e-49 | 0.267005544 | 0.159 | 0.016 | 4.25728566628803e-45 | 19 |
| gapdhs8 | 2.14938545640945e-49 | 0.715276961 | 0.766 | 0.282 | 5.40978825523695e-45 | 19 |
| tcirg1b1 | 5.2171930395862e-49 | 0.398194184 | 0.294 | 0.05 | 1.31311531613345e-44 | 19 |
| crip17 | 8.37151264323545e-49 | 0.752803978 | 0.746 | 0.264 | 2.10702601717593e-44 | 19 |
| si:dkey-222f2.71 | 1.12232315596093e-48 | 0.31676942 | 0.214 | 0.028 | 2.82477515123806e-44 | 19 |
| CABZ01073834.12 | 3.34998336156364e-48 | 0.363728939 | 0.254 | 0.039 | 8.43157312271953e-44 | 19 |
| mt-co29 | 3.723622963351e-48 | 0.582599026 | 1 | 0.717 | 9.37198663645812e-44 | 19 |
| rgs182 | 1.17560105235782e-47 | 0.324121344 | 0.224 | 0.032 | 2.9588702886794e-43 | 19 |
| sat1a.23 | 1.20529096621063e-47 | 0.700815518 | 0.791 | 0.291 | 3.03359683285554e-43 | 19 |
| si:dkey-112e7.26 | 2.50998896918492e-47 | 0.62900454 | 0.632 | 0.2 | 6.31739123654152e-43 | 19 |
| cd2ap | 3.73149517517476e-47 | 0.266619575 | 0.154 | 0.016 | 9.39180020639735e-43 | 19 |
| me1 | 4.69241182378252e-47 | 0.250853934 | 0.149 | 0.015 | 1.18103313192782e-42 | 19 |
| zbtb16a1 | 4.69929199301955e-47 | 0.374313639 | 0.274 | 0.046 | 1.18276480172309e-42 | 19 |
| si:ch211-212g7.6 | 4.77973275963971e-47 | 0.250111809 | 0.144 | 0.014 | 1.20301093827372e-42 | 19 |
| tacc1 | 5.70540191750564e-47 | 0.371678873 | 0.229 | 0.034 | 1.43599260861699e-42 | 19 |
| jdp2b3 | 1.8833377869809e-46 | 0.755380476 | 0.547 | 0.165 | 4.74017287605223e-42 | 19 |
| rasgrp41 | 2.95101703319153e-46 | 0.341582811 | 0.204 | 0.028 | 7.42741477083976e-42 | 19 |
| mt-cyb7 | 3.03868121780915e-46 | 0.62978693 | 0.99 | 0.619 | 7.64805675710384e-42 | 19 |
| abcc31 | 3.78638745774303e-46 | 0.295419932 | 0.174 | 0.021 | 9.52995859239344e-42 | 19 |
| FBXO48 | 3.94857749133058e-46 | 0.422688547 | 0.303 | 0.057 | 9.93817468792993e-42 | 19 |
| arpc1b7 | 4.34159811570521e-46 | 0.742121299 | 0.856 | 0.301 | 1.09273682974184e-41 | 19 |
| il1b1 | 7.11780896834784e-46 | 0.346903026 | 0.184 | 0.023 | 1.79148133924347e-41 | 19 |
| lcp15 | 8.69517779024985e-46 | 0.485240466 | 0.383 | 0.084 | 2.18848929802798e-41 | 19 |
| celf27 | 1.17938831068499e-45 | 0.666302936 | 0.687 | 0.236 | 2.96840243916305e-41 | 19 |

| | | | | | | |
|---|---|---|---|---|---|---|
| CABZ01020840.1 | 1.98269517768416e-45 | 0.335387124 | 0.179 | 0.022 | 4.99024549271327e-41 | 19 |
| selplg3 | 9.62989603944306e-45 | 0.405456646 | 0.259 | 0.044 | 2.42374853416742e-40 | 19 |
| dnase1l11 | 1.24886859309495e-44 | 0.365553244 | 0.254 | 0.043 | 3.14327736196068e-40 | 19 |
| arpc26 | 1.93097406210349e-44 | 0.666040453 | 0.716 | 0.261 | 4.86006861690827e-40 | 19 |
| b3gnt3.4 | 2.00684438800931e-44 | 0.33790385 | 0.229 | 0.035 | 5.05102664018064e-40 | 19 |
| kdm7aa1 | 3.06104835121294e-44 | 0.362485904 | 0.204 | 0.029 | 7.70435259516785e-40 | 19 |
| plek | 6.09777326137245e-44 | 0.314306575 | 0.179 | 0.023 | 1.53474855215483e-39 | 19 |
| cul5a | 6.65881609108921e-44 | 0.289910301 | 0.184 | 0.023 | 1.67595742196624e-39 | 19 |
| ltb4r2 | 7.73083014631275e-44 | 0.313264129 | 0.194 | 0.026 | 1.94577263952545e-39 | 19 |
| adipor21 | 8.31209130233901e-44 | 0.50886391 | 0.368 | 0.083 | 2.0920702598857e-39 | 19 |
| hexb1 | 1.08161682059342e-43 | 0.337239049 | 0.214 | 0.032 | 2.72232137575158e-39 | 19 |
| anxa44 | 1.46779035163973e-43 | 0.543481057 | 0.478 | 0.132 | 3.69428153604205e-39 | 19 |
| rhogb4 | 2.22828509167987e-43 | 0.461868162 | 0.398 | 0.093 | 5.60837074724906e-39 | 19 |
| clic15 | 2.53904065118723e-43 | 0.490111936 | 0.433 | 0.11 | 6.39051141497314e-39 | 19 |
| egr16 | 4.44275154741103e-43 | 0.670768804 | 0.577 | 0.185 | 1.11819613696788e-38 | 19 |
| slc15a4 | 4.93049908119703e-43 | 0.276614104 | 0.169 | 0.021 | 1.24095731374648e-38 | 19 |
| lamp1a1 | 8.77073079918015e-43 | 0.487189375 | 0.378 | 0.087 | 2.20750523484565e-38 | 19 |
| klf2a2 | 1.04288216488519e-42 | 0.565617004 | 0.358 | 0.082 | 2.62483012079953e-38 | 19 |
| pfkfb4b1 | 1.46995627124297e-42 | 0.535086528 | 0.383 | 0.093 | 3.69973293909143e-38 | 19 |
| si:dkey-53k12.23 | 2.0582287874724e-42 | 0.369289625 | 0.249 | 0.042 | 5.18035603518928e-38 | 19 |
| xpr1b | 3.41118797881907e-42 | 0.339257157 | 0.224 | 0.036 | 8.58561902388971e-38 | 19 |
| sdc44 | 8.01827651465263e-42 | 0.700414114 | 0.692 | 0.252 | 2.01812001597292e-37 | 19 |
| LOC1104377313 | 1.08904753871911e-41 | 0.570972019 | 0.378 | 0.087 | 2.74102375020214e-37 | 19 |
| ndfip1l2 | 2.79095234528785e-41 | 0.433103359 | 0.308 | 0.064 | 7.024547957855e-37 | 19 |
| CR318588.17 | 3.60003594969097e-41 | 0.879693134 | 0.582 | 0.203 | 9.06093048177719e-37 | 19 |
| CR936442.14 | 5.49875532910116e-41 | 0.53883999 | 0.572 | 0.164 | 1.38398172878147e-36 | 19 |
| ctsba6 | 1.10367345755441e-40 | 0.740425625 | 0.811 | 0.371 | 2.7778357253187e-36 | 19 |
| plxnc13 | 1.32765303881297e-40 | 0.374968636 | 0.239 | 0.041 | 3.34156993338836e-36 | 19 |
| elovl1b2 | 1.84786780678593e-40 | 0.490430139 | 0.428 | 0.112 | 4.65089848289951e-36 | 19 |
| si:dkey-79d12.53 | 2.44164918930162e-40 | 0.456985411 | 0.328 | 0.073 | 6.14538684455324e-36 | 19 |
| b2m7 | 3.01260452700244e-40 | 0.640052316 | 0.756 | 0.286 | 7.58242433401244e-36 | 19 |
| unc93b11 | 3.78381162745394e-40 | 0.284918113 | 0.164 | 0.021 | 9.52347548513883e-36 | 19 |
| tnfaip33 | 9.8085061024948e-40 | 0.430721136 | 0.299 | 0.062 | 2.46870290093692e-35 | 19 |
| fabp11a2 | 1.38243815916378e-39 | 0.825776025 | 0.338 | 0.081 | 3.47945860279933e-35 | 19 |
| tsc22d33 | 1.78700933392675e-39 | 0.539043377 | 0.478 | 0.14 | 4.49772379256023e-35 | 19 |

| | | | | | | |
|---|---|---|---|---|---|---|
| marcksl1a3 | 2.86198591397456e-39 | 0.592345078 | 0.632 | 0.21 | 7.20333234688257e-35 | 19 |
| unm-sa16141 | 3.02470602336176e-39 | 0.259524535 | 0.154 | 0.019 | 7.61288259019922e-35 | 19 |
| chmp4ba | 3.42760257141713e-39 | 0.378378449 | 0.274 | 0.054 | 8.62693291199978e-35 | 19 |
| ch25h1 | 4.25501506427284e-39 | 0.30920213 | 0.164 | 0.021 | 1.07094474152683e-34 | 19 |
| rasgef1ba1 | 5.00573377593154e-39 | 0.463317064 | 0.299 | 0.063 | 1.25989313406421e-34 | 19 |
| myo1f3 | 6.06090779885687e-39 | 0.333367814 | 0.244 | 0.044 | 1.52546988389429e-34 | 19 |
| npc22 | 4.15005901959829e-38 | 0.681577045 | 0.801 | 0.353 | 1.04452835464269e-33 | 19 |
| skap23 | 5.26387150114057e-38 | 0.385261845 | 0.323 | 0.071 | 1.32486381812207e-33 | 19 |
| ugt5b21 | 6.00015821728609e-38 | 0.285955966 | 0.189 | 0.029 | 1.51017982170874e-33 | 19 |
| rplp29 | 6.93609988117387e-38 | 0.453996678 | 0.995 | 0.755 | 1.74574697909265e-33 | 19 |
| arpc4l6 | 7.79808494643577e-38 | 0.56542503 | 0.572 | 0.192 | 1.96270000016842e-33 | 19 |
| midn4 | 1.09289540806788e-37 | 0.53381028 | 0.463 | 0.136 | 2.75070845256606e-33 | 19 |
| zgc:774861 | 1.48750455236662e-37 | 0.401596815 | 0.308 | 0.068 | 3.74390020785154e-33 | 19 |
| si:dkey-69o16.5 | 1.94045696717121e-37 | 0.270174555 | 0.184 | 0.027 | 4.88393614067321e-33 | 19 |
| sh3pxd2b2 | 3.11113536828145e-37 | 0.365392126 | 0.234 | 0.043 | 7.83041660842757e-33 | 19 |
| jund3 | 3.15663596061913e-37 | 0.4726528 | 0.358 | 0.09 | 7.94493704928229e-33 | 19 |
| BX120005.12 | 5.686974448716e-37 | 0.392168994 | 0.249 | 0.047 | 1.43135459899733e-32 | 19 |
| nr4a33 | 8.02028891555326e-37 | 0.454649435 | 0.259 | 0.052 | 2.0186265171556e-32 | 19 |
| BX901920.13 | 8.68756220169891e-37 | 0.468158645 | 0.313 | 0.071 | 2.1865725305456e-32 | 19 |
| CR383676.19 | 9.37635597049585e-37 | 0.690423057 | 0.995 | 0.723 | 2.3599350342141e-32 | 19 |
| gnao1a1 | 1.20935376005715e-36 | 0.275286991 | 0.159 | 0.021 | 3.04382247868785e-32 | 19 |
| elmo14 | 2.65460686682643e-36 | 0.36241427 | 0.308 | 0.068 | 6.68138002311543e-32 | 19 |
| gadd45bb4 | 2.88638111699317e-36 | 0.459130002 | 0.343 | 0.085 | 7.26473263336012e-32 | 19 |
| arhgef51 | 3.09637006119876e-36 | 0.319954976 | 0.179 | 0.027 | 7.79325380703117e-32 | 19 |
| atp6ap21 | 3.65163213640982e-36 | 0.442277092 | 0.378 | 0.097 | 9.19079292412988e-32 | 19 |
| si:ch73-237c6.14 | 5.41279712492435e-36 | 0.324936916 | 0.224 | 0.04 | 1.36234690837221e-31 | 19 |
| TPGS1 | 5.77097474206699e-36 | 0.324083425 | 0.214 | 0.037 | 1.45249663283084e-31 | 19 |
| prss16 | 6.1143301198368e-36 | 0.262991403 | 0.169 | 0.024 | 1.53891574786172e-31 | 19 |
| nt5dc22 | 8.62479043386786e-36 | 0.414048536 | 0.318 | 0.075 | 2.1707735043002e-31 | 19 |
| ptprc6 | 1.60335579996532e-35 | 0.489353391 | 0.438 | 0.124 | 4.03548621293272e-31 | 19 |
| gpr183a3 | 1.82830108424941e-35 | 0.391021572 | 0.289 | 0.063 | 4.60165099894735e-31 | 19 |
| ypel1 | 2.48631097440806e-35 | 0.286010414 | 0.179 | 0.027 | 6.25779609148764e-31 | 19 |
| mgrn1b | 2.50731847831613e-35 | 0.275186941 | 0.179 | 0.027 | 6.31066987807387e-31 | 19 |
| pnrc210 | 2.85193491662393e-35 | 0.59108245 | 0.97 | 0.562 | 7.17803499165078e-31 | 19 |
| mt-atp68 | 4.9436572404005e-35 | 0.58992336 | 0.97 | 0.586 | 1.2442690908364e-30 | 19 |

| | | | | | | |
|---|---|---|---|---|---|---|
| slc12a2 | 5.2198322067211e-35 | 0.267390027 | 0.164 | 0.023 | 1.31377956810963e-30 | 19 |
| mcl1a6 | 1.15115816808266e-34 | 0.516308184 | 0.91 | 0.62 | 2.89734999324726e-30 | 19 |
| rap1b4 | 2.13876799537235e-34 | 0.597220725 | 0.542 | 0.198 | 5.38306516755267e-30 | 19 |
| LOC1039101362 | 2.22289190477317e-34 | 0.339713357 | 0.184 | 0.03 | 5.5947966351236e-30 | 19 |
| ucp25 | 5.56899165311171e-34 | 0.620785151 | 0.871 | 0.41 | 1.40165950917169e-29 | 19 |
| mknk2b6 | 7.15821825205458e-34 | 0.541975051 | 0.557 | 0.193 | 1.80165195185962e-29 | 19 |
| mef2cb1 | 7.52474982220588e-34 | 0.267649356 | 0.164 | 0.024 | 1.893904282751e-29 | 19 |
| hdac9b1 | 8.24968906475347e-34 | 0.260118642 | 0.159 | 0.023 | 2.0763642407078e-29 | 19 |
| eif3jb | 9.78312360570142e-34 | 0.462818044 | 0.378 | 0.104 | 2.46231438031899e-29 | 19 |
| bhlhe404 | 1.27463259480605e-33 | 0.559779495 | 0.483 | 0.157 | 3.20812277786734e-29 | 19 |
| ms4a17a.93 | 1.75383739754981e-33 | 0.372282111 | 0.254 | 0.054 | 4.41423334589311e-29 | 19 |
| lgals9l1 | 5.30965619769468e-33 | 0.352258405 | 0.154 | 0.022 | 1.33638736839777e-28 | 19 |
| osbp1 | 7.31704561372408e-33 | 0.291066544 | 0.204 | 0.036 | 1.84162721051821e-28 | 19 |
| litaf5 | 9.67786750842069e-33 | 0.480790137 | 0.428 | 0.131 | 2.4358224731944e-28 | 19 |
| lpar5b2 | 1.24789712899469e-32 | 0.265887631 | 0.159 | 0.023 | 3.14083228396674e-28 | 19 |
| snx9b | 1.31900252935754e-32 | 0.342640384 | 0.264 | 0.057 | 3.31979746613999e-28 | 19 |
| eif4ebp14 | 1.35842416642291e-32 | 0.548976665 | 0.642 | 0.241 | 3.41901778446982e-28 | 19 |
| si:dkey-7i4.242 | 1.44557788684266e-32 | 0.347243584 | 0.149 | 0.021 | 3.63837498339429e-28 | 19 |
| zmp:00000010822 | 1.61730381492971e-32 | 0.332268157 | 0.259 | 0.055 | 4.0705919717966e-28 | 19 |
| mdh1aa2 | 2.06428887284924e-32 | 0.509331676 | 0.567 | 0.202 | 5.19560866407425e-28 | 19 |
| dusp53 | 2.92566046710739e-32 | 0.370157936 | 0.299 | 0.07 | 7.3635948296626e-28 | 19 |
| fosab5 | 4.53671730326572e-32 | 0.643189223 | 0.796 | 0.343 | 1.14184637805895e-27 | 19 |
| si:ch211-136m16.81 | 6.74861996045971e-32 | 0.269135221 | 0.174 | 0.028 | 1.69856015784811e-27 | 19 |
| trib31 | 1.08859842505358e-31 | 0.475656986 | 0.363 | 0.103 | 2.73989337601735e-27 | 19 |
| arl6ip12 | 2.03380189004852e-31 | 0.599380848 | 0.796 | 0.364 | 5.11887597706313e-27 | 19 |
| CABZ01055347.1 | 2.06989786356395e-31 | 0.294130377 | 0.179 | 0.03 | 5.2097259328041e-27 | 19 |
| LOC1039114301 | 3.23328370912206e-31 | 0.273066303 | 0.199 | 0.036 | 8.1378517674893e-27 | 19 |
| mt-nd47 | 3.53487998975513e-31 | 0.58274535 | 0.881 | 0.45 | 8.8969394462147e-27 | 19 |
| mt-nd37 | 4.24360247810996e-31 | 0.585635829 | 0.801 | 0.391 | 1.06807230771549e-26 | 19 |
| parvg3 | 5.37675437095975e-31 | 0.375030173 | 0.279 | 0.065 | 1.35327530762686e-26 | 19 |
| efhd2 | 5.91439618674722e-31 | 0.35207412 | 0.259 | 0.058 | 1.48859437624241e-26 | 19 |
| ppt13 | 8.07010108520435e-31 | 0.490909642 | 0.478 | 0.16 | 2.03116374213508e-26 | 19 |
| npc1 | 8.08722078337059e-31 | 0.267352956 | 0.164 | 0.026 | 2.03547259896654e-26 | 19 |
| rab5aa1 | 9.88761320253118e-31 | 0.386577411 | 0.299 | 0.075 | 2.48861336694507e-26 | 19 |
| rps15a8 | 1.35102143804348e-30 | 0.414415418 | 0.995 | 0.777 | 3.40038585741163e-26 | 19 |

| | | | | | | |
|---|---|---|---|---|---|---|
| pycard5 | 1.78620202277355e-30 | 0.365022201 | 0.289 | 0.07 | 4.49569187111875e-26 | 19 |
| CR392341.13 | 2.34192200805812e-30 | 0.320457976 | 0.259 | 0.058 | 5.89438350208149e-26 | 19 |
| atp6v0b1 | 2.88439620061278e-30 | 0.401617139 | 0.363 | 0.102 | 7.25973679732232e-26 | 19 |
| ubl3b1 | 3.03257920416728e-30 | 0.316760865 | 0.294 | 0.07 | 7.63269859896862e-26 | 19 |
| si:ch1073-429i10.3.1 | 3.21217278289435e-30 | 0.49740238 | 0.547 | 0.198 | 8.0847176772668e-26 | 19 |
| si:dkeyp-68b7.123 | 4.99777725106314e-30 | 0.342534481 | 0.264 | 0.061 | 1.25789055632008e-25 | 19 |
| rnf213a4 | 6.04507352904619e-30 | 0.427034744 | 0.318 | 0.086 | 1.52148455652564e-25 | 19 |
| bloc1s61 | 6.38312274732726e-30 | 0.371147406 | 0.303 | 0.078 | 1.6065681642748e-25 | 19 |
| actr2a5 | 6.68787278057021e-30 | 0.440227323 | 0.413 | 0.126 | 1.68327070014172e-25 | 19 |
| sik14 | 7.06636707613547e-30 | 0.423128709 | 0.388 | 0.113 | 1.77853392939254e-25 | 19 |
| zgc:1583435 | 8.64885829062532e-30 | 0.530827636 | 0.478 | 0.162 | 2.17683114316749e-25 | 19 |
| tmem131 | 8.77663250197754e-30 | 0.3392451 | 0.264 | 0.061 | 2.20899063442273e-25 | 19 |
| si:ch211-106a19.1 | 1.21136577665707e-29 | 0.277892957 | 0.199 | 0.037 | 3.04888652326817e-25 | 19 |
| gnai2a3 | 2.03120781703993e-29 | 0.43341271 | 0.438 | 0.14 | 5.11234695470779e-25 | 19 |
| NC-002333.48 | 2.13608132084374e-29 | 0.530959569 | 0.97 | 0.618 | 5.37630307643161e-25 | 19 |
| chmp2a3 | 3.75111362110184e-29 | 0.430470029 | 0.413 | 0.131 | 9.44117787295123e-25 | 19 |
| cpox2 | 3.94963928606429e-29 | 0.303043554 | 0.229 | 0.049 | 9.94084711909522e-25 | 19 |
| dusp23 | 4.69821284781255e-29 | 0.49612267 | 0.512 | 0.175 | 1.18249319166594e-24 | 19 |
| pnp5a2 | 4.79015869384782e-29 | 0.416924026 | 0.294 | 0.075 | 1.20563504165456e-24 | 19 |
| tram12 | 4.95104604156379e-29 | 0.402566527 | 0.353 | 0.1 | 1.24612877820119e-24 | 19 |
| si:dkey-102g19.32 | 5.29031085758448e-29 | 0.280115287 | 0.224 | 0.046 | 1.33151833974544e-24 | 19 |
| XLOC-0108971 | 6.74219719743262e-29 | 0.327878387 | 0.234 | 0.052 | 1.69694361262182e-24 | 19 |
| arhgdig6 | 7.32915158203557e-29 | 0.486334126 | 0.647 | 0.238 | 1.84467416168253e-24 | 19 |
| si:ch211-215e19.32 | 7.76059128305671e-29 | 0.25099759 | 0.139 | 0.021 | 1.95326322003254e-24 | 19 |
| wdr13 | 1.34333282109045e-28 | 0.44090951 | 0.428 | 0.138 | 3.38103437740256e-24 | 19 |
| CABZ01080702.13 | 1.81991201090314e-28 | 0.382556393 | 0.308 | 0.082 | 4.58053654024212e-24 | 19 |
| pim14 | 2.22634930453811e-28 | 0.480741033 | 0.488 | 0.167 | 5.60349856459196e-24 | 19 |
| hcls14 | 2.6742841128292e-28 | 0.328351112 | 0.279 | 0.068 | 6.73090568357981e-24 | 19 |
| lxn4 | 3.63692876551425e-28 | 0.430858314 | 0.448 | 0.144 | 9.15378600992282e-24 | 19 |
| BX005183.35 | 4.47199205726272e-28 | 0.416687365 | 0.299 | 0.079 | 1.12555568089245e-23 | 19 |
| atp1b1a2 | 4.92072203243233e-28 | 0.481808089 | 0.557 | 0.21 | 1.23849652834289e-23 | 19 |
| prdx54 | 6.93884986032375e-28 | 0.454303636 | 0.453 | 0.156 | 1.74643912134489e-23 | 19 |
| chmp4bb | 8.63058382514454e-28 | 0.363109788 | 0.303 | 0.081 | 2.17223164295063e-23 | 19 |
| mgst1.24 | 8.92965553921021e-28 | 0.396215335 | 0.348 | 0.1 | 2.24750500266382e-23 | 19 |
| grasp3 | 1.12876119681785e-27 | 0.362608223 | 0.289 | 0.074 | 2.84097905627086e-23 | 19 |

| | | | | | | |
|---|---|---|---|---|---|---|
| histh1l4 | 1.20342174262959e-27 | 0.493238446 | 0.353 | 0.107 | 3.02889218402442e-23 | 19 |
| lpar5a4 | 1.67877549996791e-27 | 0.287633495 | 0.199 | 0.04 | 4.22531005586924e-23 | 19 |
| XLOC-0108994 | 2.04831739650522e-27 | 0.415082157 | 0.657 | 0.253 | 5.155410055264e-23 | 19 |
| abi3a3 | 2.07926365800358e-27 | 0.282538432 | 0.204 | 0.042 | 5.2332987008292e-23 | 19 |
| mapre1a3 | 2.46222831381003e-27 | 0.459193008 | 0.542 | 0.193 | 6.19718244302847e-23 | 19 |
| rhoab5 | 2.93936057545973e-27 | 0.489231972 | 0.498 | 0.186 | 7.39807663237459e-23 | 19 |
| smdt1b1 | 4.15165244601233e-27 | 0.418214689 | 0.398 | 0.131 | 1.04492940413684e-22 | 19 |
| rps27.18 | 4.90403922188107e-27 | 0.471269637 | 0.99 | 0.675 | 1.23429763175525e-22 | 19 |
| wbp2nl1 | 6.06994948601874e-27 | 0.33420601 | 0.308 | 0.085 | 1.52774558613606e-22 | 19 |
| tln13 | 1.0263820643161e-26 | 0.410536316 | 0.388 | 0.125 | 2.58330101767719e-22 | 19 |
| ppiaa7 | 1.6458609911646e-26 | 0.327470705 | 0.99 | 0.802 | 4.14246752866219e-22 | 19 |
| mbd21 | 2.20761809015131e-26 | 0.277385048 | 0.174 | 0.033 | 5.55635397110184e-22 | 19 |
| cfl15 | 2.64141187125386e-26 | 0.514847262 | 0.94 | 0.572 | 6.64816953875883e-22 | 19 |
| glmp | 2.68370050698534e-26 | 0.283045227 | 0.189 | 0.038 | 6.7546058060314e-22 | 19 |
| rpl397 | 4.11431355644798e-26 | 0.294502095 | 1 | 0.872 | 1.03553157902239e-21 | 19 |
| arhgdia5 | 4.38396622936715e-26 | 0.461990919 | 0.577 | 0.225 | 1.10340046026942e-21 | 19 |
| zgc:1748632 | 4.5392964422703e-26 | 0.38217908 | 0.333 | 0.099 | 1.14249552155501e-21 | 19 |
| rps206 | 5.54072721400138e-26 | 0.250657082 | 1 | 0.915 | 1.39454563249201e-21 | 19 |
| itm2ba5 | 6.73375283484399e-26 | 0.502156289 | 0.627 | 0.269 | 1.69481825100188e-21 | 19 |
| oaz1b7 | 7.12850459370688e-26 | 0.522326111 | 0.692 | 0.318 | 1.79417332119008e-21 | 19 |
| mob1a5 | 9.03798702529628e-26 | 0.414050114 | 0.463 | 0.163 | 2.27477095439682e-21 | 19 |
| itga43 | 1.11476973436404e-25 | 0.321062841 | 0.254 | 0.063 | 2.80576394442085e-21 | 19 |
| cited4b1 | 1.24746601325723e-25 | 0.406260781 | 0.403 | 0.128 | 3.13974720876711e-21 | 19 |
| nfkbie3 | 1.58962680231065e-25 | 0.358616973 | 0.318 | 0.091 | 4.00093169873568e-21 | 19 |
| mtss1la1 | 1.85664199479784e-25 | 0.253092331 | 0.139 | 0.023 | 4.67298223670669e-21 | 19 |
| pkma3 | 2.12101778144893e-25 | 0.42892364 | 0.483 | 0.174 | 5.33838965412882e-21 | 19 |
| BX322618.12 | 4.24566777100148e-25 | 0.277117051 | 0.154 | 0.028 | 1.06859212128336e-20 | 19 |
| rpl369 | 5.29723896942078e-25 | 0.381245492 | 1 | 0.787 | 1.33326207621352e-20 | 19 |
| capza1a5 | 5.54546662242501e-25 | 0.427382377 | 0.517 | 0.196 | 1.39573849419815e-20 | 19 |
| mt-nd17 | 6.07906817705537e-25 | 0.509215507 | 0.816 | 0.406 | 1.53004066948307e-20 | 19 |
| wasf2 | 8.57706573733632e-25 | 0.334820323 | 0.269 | 0.072 | 2.15876167543018e-20 | 19 |
| ier2b7 | 8.91341759625201e-25 | 0.497628962 | 0.542 | 0.22 | 2.24341807480067e-20 | 19 |
| XLOC-0254235 | 9.50355310802789e-25 | 0.652929358 | 0.433 | 0.167 | 2.39194928175954e-20 | 19 |
| m6pr1 | 1.44469562274489e-24 | 0.351514074 | 0.318 | 0.094 | 3.63615441288662e-20 | 19 |
| fosb2 | 1.64889032826531e-24 | 0.479109924 | 0.373 | 0.123 | 4.15009206721096e-20 | 19 |

| | | | | | | |
|---|---|---|---|---|---|---|
| znf8271 | 2.0090704143371e-24 | 0.258630797 | 0.154 | 0.029 | 5.05662932584506e-20 | 19 |
| eef1db5 | 2.18184185303093e-24 | 0.485432789 | 0.627 | 0.27 | 5.49147775989355e-20 | 19 |
| rpl377 | 3.07686579541512e-24 | 0.321919135 | 0.995 | 0.848 | 7.74416352048031e-20 | 19 |
| ifi463 | 3.82094689147594e-24 | 0.394379724 | 0.368 | 0.119 | 9.6169412311558e-20 | 19 |
| gdi21 | 6.10004040240214e-24 | 0.41648806 | 0.483 | 0.181 | 1.5353191688806e-19 | 19 |
| rpl389 | 6.74862406275664e-24 | 0.410840838 | 0.995 | 0.725 | 1.69856119035522e-19 | 19 |
| arl8bb | 8.71560619398469e-24 | 0.258731175 | 0.184 | 0.039 | 2.19363092296401e-19 | 19 |
| XLOC-0331372 | 9.30602408894798e-24 | 0.291011094 | 0.169 | 0.034 | 2.34223320294732e-19 | 19 |
| BX908750.11 | 9.83553826550769e-24 | 0.289083321 | 0.159 | 0.031 | 2.47550662604563e-19 | 19 |
| tpi1b5 | 1.09193419343727e-23 | 0.4109798 | 0.448 | 0.164 | 2.74828917146227e-19 | 19 |
| si:ch211-198c19.33 | 1.33563663550823e-23 | 0.33375071 | 0.244 | 0.062 | 3.36166384791067e-19 | 19 |
| gnai11 | 1.44754284952791e-23 | 0.311360816 | 0.229 | 0.058 | 3.64332059797679e-19 | 19 |
| ube2v22 | 1.87733774053593e-23 | 0.46827818 | 0.612 | 0.265 | 4.72507135915487e-19 | 19 |
| zgc:1105911 | 1.92099871514077e-23 | 0.265132028 | 0.209 | 0.049 | 4.83496166613781e-19 | 19 |
| pdcd6ip | 3.41771673008842e-23 | 0.27062503 | 0.204 | 0.047 | 8.60205123795956e-19 | 19 |
| psme24 | 4.49214343687248e-23 | 0.355029388 | 0.308 | 0.095 | 1.13062758162644e-18 | 19 |
| ostf11 | 4.74185395936401e-23 | 0.281184738 | 0.214 | 0.052 | 1.19347722303233e-18 | 19 |
| rpl228 | 4.78886073846724e-23 | 0.424251235 | 0.985 | 0.69 | 1.20530835926482e-18 | 19 |
| ahnak4 | 5.33771503278931e-23 | 0.413010307 | 0.368 | 0.124 | 1.34344949660274e-18 | 19 |
| slc35f6 | 6.2330637414857e-23 | 0.29087952 | 0.259 | 0.069 | 1.56879981309454e-18 | 19 |
| zfyve21 | 7.15619562402893e-23 | 0.298352053 | 0.224 | 0.056 | 1.80114287661184e-18 | 19 |
| itpkcb2 | 8.36176893289894e-23 | 0.331784043 | 0.269 | 0.076 | 2.10457362272134e-18 | 19 |
| rpl248 | 1.23186679559874e-22 | 0.358038253 | 0.995 | 0.774 | 3.10048553784246e-18 | 19 |
| syngr2a2 | 1.25274952601398e-22 | 0.271948641 | 0.219 | 0.054 | 3.15304528202458e-18 | 19 |
| COX7A2 (1 of many)3 | 1.66025874022199e-22 | 0.458759356 | 0.587 | 0.257 | 4.17870522326473e-18 | 19 |
| tagln25 | 1.76691240277416e-22 | 0.467127899 | 0.468 | 0.185 | 4.44714182654227e-18 | 19 |
| igf2b3 | 1.80755842576666e-22 | 0.274404623 | 0.229 | 0.057 | 4.54944380181212e-18 | 19 |
| dusp6 | 1.84128080177643e-22 | 0.308179596 | 0.194 | 0.046 | 4.63431964999109e-18 | 19 |
| krt85 | 2.032925656559e-22 | 0.483940235 | 0.647 | 0.272 | 5.11667058499335e-18 | 19 |
| si:ch211-196l7.48 | 2.28140155575739e-22 | 0.48258299 | 0.667 | 0.31 | 5.74205957568577e-18 | 19 |
| chmp1b1 | 2.87437763296215e-22 | 0.373808438 | 0.353 | 0.119 | 7.23452106440243e-18 | 19 |
| XLOC-0283705 | 2.95484399090036e-22 | 0.397523352 | 0.403 | 0.138 | 7.43704684069712e-18 | 19 |
| derl21 | 3.08328694532238e-22 | 0.340235199 | 0.328 | 0.103 | 7.76032491268191e-18 | 19 |
| gabarapa | 3.33563299240694e-22 | 0.374394263 | 0.433 | 0.158 | 8.39545467858904e-18 | 19 |
| ptenb2 | 3.60732208277512e-22 | 0.296423797 | 0.274 | 0.078 | 9.07926895013669e-18 | 19 |

| | | | | | | |
|---|---|---|---|---|---|---|
| elf15 | 3.87132513791669e-22 | 0.338890243 | 0.323 | 0.102 | 9.74373823962252e-18 | 19 |
| sec61a11 | 4.75924833610316e-22 | 0.40208859 | 0.478 | 0.186 | 1.1978552137138e-17 | 19 |
| LOC1039091155 | 5.77362789043272e-22 | 0.381348987 | 0.443 | 0.162 | 1.45316440374301e-17 | 19 |
| rap2b | 5.96017001243713e-22 | 0.27258802 | 0.204 | 0.049 | 1.5001151904303e-17 | 19 |
| zgc:1141889 | 6.34632198350024e-22 | 0.481434366 | 0.985 | 0.623 | 1.59730578002718e-17 | 19 |
| si:dkey-56m19.5 | 7.70681571088971e-22 | 0.269207288 | 0.194 | 0.046 | 1.93972844627383e-17 | 19 |
| rps2112 | 8.69782857805557e-22 | 0.397175749 | 0.995 | 0.747 | 2.18915647481081e-17 | 19 |
| rpl35a8 | 9.57421438286702e-22 | 0.368728635 | 0.995 | 0.757 | 2.4097340180238e-17 | 19 |
| irf7 | 1.27541643376492e-21 | 0.261628299 | 0.154 | 0.031 | 3.21009562214292e-17 | 19 |
| arpc36 | 2.33358844348658e-21 | 0.474476313 | 0.662 | 0.306 | 5.87340875341138e-17 | 19 |
| si:rp71-36a1.5 | 3.24440580273214e-21 | 0.250203313 | 0.194 | 0.046 | 8.16584496489653e-17 | 19 |
| nck1b | 3.55790260284928e-21 | 0.28104476 | 0.204 | 0.051 | 8.95488506111134e-17 | 19 |
| sqstm1 | 3.58122006933317e-21 | 0.351210173 | 0.259 | 0.075 | 9.01357279250465e-17 | 19 |
| rarga | 4.46962076492812e-21 | 0.278511986 | 0.214 | 0.055 | 1.12495885032476e-16 | 19 |
| xbp13 | 4.67761757241221e-21 | 0.516517553 | 0.672 | 0.34 | 1.17730956680043e-16 | 19 |
| stk35 | 6.06715560316111e-21 | 0.289625432 | 0.209 | 0.054 | 1.52704239375962e-16 | 19 |
| hif1ab1 | 6.80699523116624e-21 | 0.323368528 | 0.333 | 0.111 | 1.71325262973223e-16 | 19 |
| pdia35 | 8.76318691029591e-21 | 0.451048545 | 0.622 | 0.28 | 2.20560651345238e-16 | 19 |
| si:ch211-260e23.9 | 1.00256620787625e-20 | 0.260531231 | 0.199 | 0.049 | 2.52335888860372e-16 | 19 |
| cyfip11 | 1.26622177783927e-20 | 0.263075685 | 0.214 | 0.055 | 3.18695359264365e-16 | 19 |
| ppiab4 | 1.48980624556932e-20 | 0.405039983 | 0.955 | 0.653 | 3.74969333947342e-16 | 19 |
| cdc42l4 | 1.67780125075503e-20 | 0.40131333 | 0.512 | 0.212 | 4.22285796802532e-16 | 19 |
| rps298 | 1.83748605133322e-20 | 0.304067398 | 0.995 | 0.83 | 4.62476864260058e-16 | 19 |
| rps222 | 1.9397618617633e-20 | 0.329381037 | 0.99 | 0.827 | 4.88218662987204e-16 | 19 |
| sec61g2 | 2.29913747479636e-20 | 0.394144156 | 0.945 | 0.675 | 5.78669911031495e-16 | 19 |
| rpl18a8 | 2.90833174298836e-20 | 0.34843032 | 0.99 | 0.783 | 7.31998016392739e-16 | 19 |
| myl12.15 | 3.51091119430394e-20 | 0.481333019 | 0.761 | 0.387 | 8.8366123849436e-16 | 19 |
| glula2 | 4.98383321915256e-20 | 0.373887938 | 0.378 | 0.133 | 1.25438098292851e-15 | 19 |
| mt-nd52 | 5.31680097815578e-20 | 0.309464522 | 0.343 | 0.118 | 1.33818563819203e-15 | 19 |
| rab1ab1 | 5.45360435619675e-20 | 0.356562708 | 0.338 | 0.12 | 1.37261768041116e-15 | 19 |
| jak12 | 5.89074418383427e-20 | 0.354439622 | 0.289 | 0.094 | 1.48264140362925e-15 | 19 |
| fxyd13 | 6.57797384889844e-20 | 0.29907237 | 0.264 | 0.079 | 1.65561023802925e-15 | 19 |
| rpl119 | 7.44180281718016e-20 | 0.332302067 | 1 | 0.812 | 1.87302735105607e-15 | 19 |
| sptlc2b | 9.77042016538704e-20 | 0.265647639 | 0.184 | 0.045 | 2.45911705142626e-15 | 19 |
| cox8a5 | 1.05749888402777e-19 | 0.422340175 | 0.512 | 0.224 | 2.66161894120949e-15 | 19 |

| | | | | | | |
|---|---|---|---|---|---|---|
| lamtor51 | 1.14580772431971e-19 | 0.323258481 | 0.378 | 0.133 | 2.88388346134028e-15 | 19 |
| uqcrq2 | 1.70257896149403e-19 | 0.418628463 | 0.592 | 0.268 | 4.28522098818433e-15 | 19 |
| si:ch1073-325m22.2 | 1.82438340368133e-19 | 0.385961172 | 0.527 | 0.226 | 4.59179058872553e-15 | 19 |
| hmha1b3 | 2.4763221984927e-19 | 0.28333979 | 0.249 | 0.073 | 6.23265534138629e-15 | 19 |
| GCA4 | 3.07727825561435e-19 | 0.354993102 | 0.448 | 0.178 | 7.74520164155576e-15 | 19 |
| XLOC-0425954 | 4.1013488342301e-19 | 0.277198562 | 0.443 | 0.162 | 1.03226848808737e-14 | 19 |
| anxa5b2 | 4.63319069274944e-19 | 0.327354429 | 0.284 | 0.092 | 1.16612776545811e-14 | 19 |
| DHDH2 | 5.70580288399726e-19 | 0.278817836 | 0.214 | 0.06 | 1.43609352787327e-14 | 19 |
| rps97 | 6.47883013499796e-19 | 0.339237095 | 1 | 0.774 | 1.63065675667764e-14 | 19 |
| nme2b.12 | 8.18464389361978e-19 | 0.251527168 | 1 | 0.855 | 2.05999302158516e-14 | 19 |
| tmem59 | 8.66397111800079e-19 | 0.333854041 | 0.303 | 0.105 | 2.18063489068962e-14 | 19 |
| glud1b | 8.71569549166126e-19 | 0.276825688 | 0.239 | 0.07 | 2.19365339829622e-14 | 19 |
| rps26l9 | 9.23532381761632e-19 | 0.423761795 | 0.965 | 0.66 | 2.32443865165585e-14 | 19 |
| fkbp53 | 1.04086184626236e-18 | 0.349037294 | 0.333 | 0.12 | 2.61974518085773e-14 | 19 |
| srgn6 | 1.1786913208481e-18 | 0.386007411 | 0.557 | 0.233 | 2.96664818544257e-14 | 19 |
| grb2b | 1.22048448050371e-18 | 0.256248756 | 0.244 | 0.072 | 3.07183738897979e-14 | 19 |
| twf1a3 | 1.24202396022213e-18 | 0.342707903 | 0.393 | 0.149 | 3.12605010548307e-14 | 19 |
| slc37a2 | 1.43598039755718e-18 | 0.285207801 | 0.284 | 0.09 | 3.61421906261167e-14 | 19 |
| si:ch211-191c10.13 | 1.90094003435858e-18 | 0.256263229 | 0.199 | 0.053 | 4.78447597247711e-14 | 19 |
| msna7 | 2.03345325231463e-18 | 0.407777177 | 0.537 | 0.239 | 5.1179984907507e-14 | 19 |
| cd9b2 | 2.15560334647875e-18 | 0.479793689 | 0.219 | 0.064 | 5.42543806275238e-14 | 19 |
| arpc45 | 2.70162784962944e-18 | 0.355913795 | 0.532 | 0.222 | 6.79972713473233e-14 | 19 |
| vat12 | 2.85870918145318e-18 | 0.398846756 | 0.353 | 0.133 | 7.19508513879952e-14 | 19 |
| rpl298 | 4.42581787986527e-18 | 0.312420429 | 0.99 | 0.807 | 1.11393410218329e-13 | 19 |
| rps236 | 4.83754042541811e-18 | 0.362401916 | 0.98 | 0.72 | 1.21756054967348e-13 | 19 |
| faua7 | 9.43699178054411e-18 | 0.375504093 | 0.95 | 0.687 | 2.37519646124515e-13 | 19 |
| si:dkey-180p18.92 | 9.79963282909589e-18 | 0.284710305 | 0.289 | 0.098 | 2.46646958675515e-13 | 19 |
| dync1i2a | 9.8354462647743e-18 | 0.260125217 | 0.194 | 0.053 | 2.47548347038104e-13 | 19 |
| rps149 | 1.19137774635329e-17 | 0.338458384 | 0.985 | 0.756 | 2.99857864979659e-13 | 19 |
| ktn1 | 1.89975967066808e-17 | 0.254235413 | 0.204 | 0.058 | 4.7815051151045e-13 | 19 |
| rpl36a7 | 1.98061868229857e-17 | 0.301610828 | 0.995 | 0.807 | 4.98501916147728e-13 | 19 |
| stat23 | 2.34307628371812e-17 | 0.252381878 | 0.209 | 0.06 | 5.89728869849015e-13 | 19 |
| rpl357 | 2.39306210904558e-17 | 0.297988452 | 0.985 | 0.802 | 6.02309802225683e-13 | 19 |
| rps159 | 2.49781980643778e-17 | 0.397264293 | 0.97 | 0.681 | 6.28676267082324e-13 | 19 |
| tegt4 | 2.73102863950606e-17 | 0.36161452 | 0.478 | 0.209 | 6.8737259827728e-13 | 19 |

| | | | | | | |
|---|---|---|---|---|---|---|
| rpl188 | 2.79877981754808e-17 | 0.335638314 | 0.985 | 0.771 | 7.04424892278676e-13 | 19 |
| rps58 | 2.91770564924732e-17 | 0.381396095 | 0.97 | 0.698 | 7.34357334859058e-13 | 19 |
| tagapb4 | 3.11950806963699e-17 | 0.25238267 | 0.224 | 0.066 | 7.85148986046934e-13 | 19 |
| rpl27a8 | 3.28043761955304e-17 | 0.480618937 | 0.881 | 0.491 | 8.25653344465305e-13 | 19 |
| commd3 | 3.42422243541222e-17 | 0.262304858 | 0.308 | 0.107 | 8.61842544768901e-13 | 19 |
| coq10b | 3.65919937747631e-17 | 0.27702583 | 0.199 | 0.056 | 9.20983891317013e-13 | 19 |
| rab5c1 | 3.9139321487906e-17 | 0.303158407 | 0.323 | 0.119 | 9.85097582529106e-13 | 19 |
| rpl158 | 3.95362493198296e-17 | 0.354721557 | 0.99 | 0.741 | 9.95087859130792e-13 | 19 |
| kctd5a3 | 6.16425589937276e-17 | 0.35395384 | 0.418 | 0.173 | 1.55148156731313e-12 | 19 |
| vdac34 | 6.87847362411768e-17 | 0.308853789 | 0.388 | 0.152 | 1.73124302645418e-12 | 19 |
| ptmaa7 | 8.63567749137593e-17 | 0.450258395 | 0.786 | 0.39 | 2.17351366780441e-12 | 19 |
| ier2a6 | 1.15691410196664e-16 | 0.431588797 | 0.542 | 0.259 | 2.91183710323983e-12 | 19 |
| hif1al | 1.27909990607578e-16 | 0.302920134 | 0.333 | 0.124 | 3.21936655360213e-12 | 19 |
| rgs5b3 | 1.50553558148994e-16 | 0.283432787 | 0.239 | 0.075 | 3.78928250505202e-12 | 19 |
| pgd2 | 2.02496569762308e-16 | 0.251861829 | 0.209 | 0.062 | 5.09663616434752e-12 | 19 |
| rpl328 | 2.13742661010731e-16 | 0.306290174 | 0.99 | 0.811 | 5.37968903497909e-12 | 19 |
| si:ch73-1a9.35 | 2.99980729298356e-16 | 0.371020999 | 0.572 | 0.262 | 7.55021497571032e-12 | 19 |
| ppdpfb4 | 3.60347543210323e-16 | 0.428723194 | 0.811 | 0.516 | 9.06958731506063e-12 | 19 |
| maff3 | 3.84816073918708e-16 | 0.308734339 | 0.244 | 0.082 | 9.68543576445996e-12 | 19 |
| rps288 | 3.88341612593544e-16 | 0.297347502 | 0.995 | 0.824 | 9.7741700473669e-12 | 19 |
| rps108 | 4.41643252942737e-16 | 0.38482763 | 0.975 | 0.691 | 1.11157190333157e-11 | 19 |
| tob1a1 | 6.51798871693752e-16 | 0.334423078 | 0.328 | 0.128 | 1.64051258016601e-11 | 19 |
| ip6k2b4 | 7.02377224305235e-16 | 0.313784818 | 0.318 | 0.121 | 1.76781323585385e-11 | 19 |
| gnb1a3 | 7.36493764137491e-16 | 0.332992577 | 0.438 | 0.19 | 1.85368115495765e-11 | 19 |
| wee2 | 7.82070049180162e-16 | 0.251657175 | 0.274 | 0.092 | 1.96839210678155e-11 | 19 |
| capzb5 | 7.89170852312579e-16 | 0.365967811 | 0.532 | 0.25 | 1.98626411818553e-11 | 19 |
| slc25a56 | 9.64373735708111e-16 | 0.290401221 | 1 | 0.804 | 2.42723225540374e-11 | 19 |
| romo18 | 1.34483780015254e-15 | 0.410752427 | 0.716 | 0.382 | 3.38482225920392e-11 | 19 |
| zgc:1629444 | 1.56670403963261e-15 | 0.358637348 | 0.537 | 0.249 | 3.94323739735133e-11 | 19 |
| zgc:1537133 | 1.78374624301388e-15 | 0.302250057 | 0.368 | 0.151 | 4.48951091904163e-11 | 19 |
| zgc:153911 | 1.98238323570202e-15 | 0.25016743 | 0.1 | 0.019 | 4.98946036593841e-11 | 19 |
| higd1a5 | 2.69218193105896e-15 | 0.398861487 | 0.607 | 0.307 | 6.7759527022823e-11 | 19 |
| rpl309 | 3.51034084178322e-15 | 0.361862364 | 0.97 | 0.705 | 8.83517686468418e-11 | 19 |
| grinab3 | 3.825364107289e-15 | 0.282488476 | 0.299 | 0.112 | 9.62805892163569e-11 | 19 |
| snd11 | 3.84439332509036e-15 | 0.269151638 | 0.284 | 0.104 | 9.67595355991993e-11 | 19 |

| | | | | | | |
|---|---|---|---|---|---|---|
| cnn23 | 5.24644474446556e-15 | 0.306776132 | 0.333 | 0.133 | 1.32047767773454e-10 | 19 |
| ptp4a2b | 6.97223158832371e-15 | 0.307228126 | 0.363 | 0.148 | 1.75484096846519e-10 | 19 |
| rpl99 | 9.87388424303033e-15 | 0.425016116 | 0.97 | 0.625 | 2.4851579251283e-10 | 19 |
| notch24 | 1.26572889976532e-14 | 0.294010061 | 0.303 | 0.118 | 3.18571306781935e-10 | 19 |
| sh3bgrl33 | 1.5550443706966e-14 | 0.304660628 | 0.353 | 0.146 | 3.91389117660628e-10 | 19 |
| rps188 | 1.58166253714846e-14 | 0.361305532 | 0.995 | 0.721 | 3.98088643974895e-10 | 19 |
| capza1b3 | 1.62185929796577e-14 | 0.301217289 | 0.473 | 0.21 | 4.08205766705006e-10 | 19 |
| atf4b5 | 1.62797724079647e-14 | 0.368433377 | 0.537 | 0.261 | 4.09745591736064e-10 | 19 |
| csde11 | 1.64141211726043e-14 | 0.319621047 | 0.453 | 0.205 | 4.13127015793278e-10 | 19 |
| wu:fi37e09 | 1.91430306311922e-14 | 0.275741287 | 0.303 | 0.116 | 4.81810937956475e-10 | 19 |
| antxr2a5 | 2.05676712320795e-14 | 0.32619591 | 0.363 | 0.152 | 5.17667717240208e-10 | 19 |
| dbi2 | 2.13338125835453e-14 | 0.34583996 | 0.453 | 0.211 | 5.36950728915252e-10 | 19 |
| mt-nd26 | 2.50137777438498e-14 | 0.386052963 | 0.731 | 0.397 | 6.29571772034955e-10 | 19 |
| rplp2l9 | 2.9520733009326e-14 | 0.33176898 | 0.995 | 0.772 | 7.43007329111727e-10 | 19 |
| s100a10b4 | 2.99383990931181e-14 | 0.395607637 | 0.532 | 0.253 | 7.53519566774688e-10 | 19 |
| rpl23a8 | 3.35055031727606e-14 | 0.40053347 | 0.955 | 0.63 | 8.43300009355211e-10 | 19 |
| cox7b4 | 3.93182411279328e-14 | 0.392995961 | 0.746 | 0.413 | 9.8960081094894e-10 | 19 |
| rpl288 | 4.82844517962019e-14 | 0.331916483 | 0.99 | 0.746 | 1.2152713672586e-09 | 19 |
| atp5pf3 | 5.06315887381054e-14 | 0.356322624 | 0.587 | 0.296 | 1.27434645694937e-09 | 19 |
| icn5 | 6.00256054853407e-14 | 0.374249172 | 0.562 | 0.272 | 1.51078446446054e-09 | 19 |
| COX5B2 | 6.26683530043796e-14 | 0.363582572 | 0.687 | 0.354 | 1.57729977676723e-09 | 19 |
| rpl3110 | 7.78829120680557e-14 | 0.388056341 | 0.975 | 0.65 | 1.96023501384089e-09 | 19 |
| rpl7a7 | 8.54106190963166e-14 | 0.275397008 | 0.985 | 0.808 | 2.14969987203519e-09 | 19 |
| vma211 | 1.01115172177798e-13 | 0.253351053 | 0.279 | 0.105 | 2.544967768543e-09 | 19 |
| ccni7 | 1.13149736984619e-13 | 0.350310316 | 0.577 | 0.291 | 2.84786573016587e-09 | 19 |
| slc3a2b3 | 1.21888962530917e-13 | 0.291512503 | 0.294 | 0.117 | 3.06782329794064e-09 | 19 |
| rps196 | 1.24352097956606e-13 | 0.250523593 | 1 | 0.862 | 3.12981795346982e-09 | 19 |
| klf11b4 | 1.46672834036105e-13 | 0.261338928 | 0.224 | 0.079 | 3.69160855985473e-09 | 19 |
| glrx4 | 1.73856586928463e-13 | 0.356488205 | 0.602 | 0.308 | 4.37579643640248e-09 | 19 |
| ndufb31 | 1.85428796602187e-13 | 0.287574479 | 0.468 | 0.215 | 4.66705738168045e-09 | 19 |
| rps27a7 | 2.56782087577389e-13 | 0.305259158 | 0.975 | 0.749 | 6.4629483622353e-09 | 19 |
| arf6b4 | 2.9701322512257e-13 | 0.258133985 | 0.264 | 0.1 | 7.47552586310996e-09 | 19 |
| rps169 | 3.25894011709236e-13 | 0.333597779 | 0.97 | 0.735 | 8.20242638070976e-09 | 19 |
| cox4i15 | 3.85759046062568e-13 | 0.372342702 | 0.721 | 0.388 | 9.70916943034878e-09 | 19 |
| nccrp12 | 4.03260253129165e-13 | 0.290205355 | 0.433 | 0.192 | 1.0149657311008e-08 | 19 |

| | | | | | | |
|---|---|---|---|---|---|---|
| rhoaa4 | 4.70419595427143e-13 | 0.290535512 | 0.348 | 0.152 | 1.18399907973058e-08 | 19 |
| eif3f7 | 7.7353570415073e-13 | 0.393436175 | 0.766 | 0.428 | 1.94691201377697e-08 | 19 |
| eif3m5 | 1.49187679240922e-12 | 0.354562966 | 0.672 | 0.355 | 3.75490469881475e-08 | 19 |
| tnfrsf9a2 | 1.87125622143478e-12 | 0.323217534 | 0.164 | 0.051 | 4.70976478372919e-08 | 19 |
| btg24 | 2.11845301049911e-12 | 0.369123377 | 0.677 | 0.37 | 5.33193438212521e-08 | 19 |
| bsg | 2.27855520572417e-12 | 0.357354951 | 0.701 | 0.376 | 5.73489559728716e-08 | 19 |
| alas11 | 2.84588589959673e-12 | 0.255148628 | 0.328 | 0.138 | 7.16281022069502e-08 | 19 |
| vmp1 | 3.14528060949685e-12 | 0.269475617 | 0.199 | 0.069 | 7.91635676604261e-08 | 19 |
| rpl349 | 3.50577770851343e-12 | 0.301010035 | 0.99 | 0.781 | 8.82369191455746e-08 | 19 |
| rps27l5 | 5.14704208362235e-12 | 0.32349846 | 0.413 | 0.202 | 1.29545902202691e-07 | 19 |
| mafba3 | 5.75335989076575e-12 | 0.26855707 | 0.199 | 0.072 | 1.44806315090683e-07 | 19 |
| rps27.28 | 6.09543073542639e-12 | 0.408552372 | 0.896 | 0.513 | 1.53415896179947e-07 | 19 |
| socs3a6 | 8.22062695626861e-12 | 0.316301516 | 0.592 | 0.295 | 2.06904959862325e-07 | 19 |
| naca9 | 8.75218277950243e-12 | 0.372399952 | 0.925 | 0.615 | 2.20283688377297e-07 | 19 |
| rps257 | 1.03645531597047e-11 | 0.324036468 | 0.98 | 0.722 | 2.60865438476608e-07 | 19 |
| ckbb6 | 2.17811153614889e-11 | 0.316379599 | 0.537 | 0.263 | 5.48208892533314e-07 | 19 |
| nsa28 | 2.81746530289143e-11 | 0.347716643 | 0.677 | 0.359 | 7.09127842084743e-07 | 19 |
| ssr32 | 6.01830558399911e-11 | 0.292691069 | 0.527 | 0.27 | 1.51474733243674e-06 | 19 |
| rpl217 | 6.7688543696722e-11 | 0.277965327 | 0.96 | 0.754 | 1.70365295630279e-06 | 19 |
| chac13 | 7.07675665062106e-11 | 0.339402445 | 0.692 | 0.395 | 1.78114888139482e-06 | 19 |
| rps4x7 | 7.20885624255612e-11 | 0.33192129 | 0.98 | 0.707 | 1.81439702768895e-06 | 19 |
| ndufa4l4 | 8.46134807336503e-11 | 0.341848001 | 0.706 | 0.392 | 2.12963669658524e-06 | 19 |
| rpsa9 | 1.10204083057134e-10 | 0.314248244 | 0.98 | 0.749 | 2.773726566465e-06 | 19 |
| rpl88 | 1.10320919908989e-10 | 0.276756018 | 0.975 | 0.778 | 2.77666723318933e-06 | 19 |
| ywhaba3 | 1.10910952613204e-10 | 0.300665556 | 0.597 | 0.318 | 2.79151776632174e-06 | 19 |
| rpl276 | 1.15164875813642e-10 | 0.299115886 | 0.995 | 0.76 | 2.89858475935356e-06 | 19 |
| rps175 | 1.57035045287465e-10 | 0.278418158 | 0.98 | 0.764 | 3.9524150548402e-06 | 19 |
| rpl69 | 1.68315004088976e-10 | 0.356990155 | 0.97 | 0.664 | 4.23632033791544e-06 | 19 |
| fthl27 | 1.69846526543434e-10 | 0.307332767 | 0.915 | 0.693 | 4.2748672265717e-06 | 19 |
| atp5l4 | 3.33460882778916e-10 | 0.332907053 | 0.746 | 0.446 | 8.39287695866252e-06 | 19 |
| tpm31 | 4.28397724239572e-10 | 0.280299467 | 0.478 | 0.253 | 1.07823423213858e-05 | 19 |
| rps267 | 8.07163178743079e-10 | 0.304187492 | 0.98 | 0.731 | 2.03154900457846e-05 | 19 |
| rpl177 | 9.11949758249349e-10 | 0.286589493 | 0.975 | 0.743 | 2.29528634653779e-05 | 19 |
| uba527 | 1.45212291896772e-09 | 0.349820794 | 0.94 | 0.626 | 3.65484817474985e-05 | 19 |
| rpl107 | 1.9399007354435e-09 | 0.258961002 | 0.96 | 0.77 | 4.88253616103775e-05 | 19 |

| | | | | | | |
|---|---|---|---|---|---|---|
| NDUFB14 | 2.37892619411918e-09 | 0.251542129 | 0.383 | 0.197 | 5.98751933797857e-05 | 19 |
| zgc:1935411 | 2.54170902921796e-09 | 0.289598689 | 0.647 | 0.365 | 6.39722745563869e-05 | 19 |
| rps248 | 4.93590308092063e-09 | 0.282080887 | 0.98 | 0.756 | 0.000124232 | 19 |
| cox6c1 | 7.49100421085025e-09 | 0.298776543 | 0.716 | 0.413 | 0.000188541 | 19 |
| zgc:564935 | 8.88209500687685e-09 | 0.331280203 | 0.866 | 0.569 | 0.000223553 | 19 |
| zgc:927444 | 2.00928923807812e-08 | 0.257312699 | 0.632 | 0.351 | 0.000505718 | 19 |
| ppib5 | 2.33415186509389e-08 | 0.267307905 | 0.602 | 0.33 | 0.000587483 | 19 |
| eif1b4 | 2.8046530248578e-08 | 0.28573002 | 0.657 | 0.38 | 0.000705903 | 19 |
| rps116 | 2.92630382204677e-08 | 0.269033458 | 0.985 | 0.763 | 0.000736521 | 19 |
| rps137 | 3.12218535110779e-08 | 0.311740958 | 0.995 | 0.721 | 0.000785823 | 19 |
| rps37 | 5.43986478827131e-08 | 0.297191109 | 0.95 | 0.697 | 0.00136916 | 19 |
| btf38 | 1.05321062975234e-07 | 0.315340473 | 0.886 | 0.584 | 0.002650826 | 19 |
| rpl10a6 | 2.12104138692488e-07 | 0.25711786 | 0.97 | 0.762 | 0.005338449 | 19 |
| rpl5a7 | 3.58515519734831e-07 | 0.318286563 | 0.796 | 0.489 | 0.009023477 | 19 |
| h1fx4 | 5.05018121492576e-07 | 0.253017764 | 0.333 | 0.183 | 0.012710801 | 19 |
| pabpc1a7 | 6.16016326496659e-07 | 0.315457621 | 0.856 | 0.536 | 0.015504515 | 19 |
| rsl24d18 | 1.20558029878317e-06 | 0.289324472 | 0.756 | 0.441 | 0.030343251 | 19 |
| rpl148 | 1.36294140315159e-06 | 0.317193476 | 0.955 | 0.631 | 0.034303872 | 19 |
| cox6b22 | 3.25795752930965e-06 | 0.274553071 | 0.801 | 0.531 | 0.081999533 | 19 |
| mibp27 | 4.49738990473564e-06 | 0.276562845 | 0.776 | 0.489 | 0.113194807 | 19 |
| ctss2.17 | 1.31414150786961e-05 | 0.273290445 | 0.801 | 0.532 | 0.330756276 | 19 |
| serbp1a7 | 2.71027058331431e-05 | 0.262913325 | 0.746 | 0.479 | 0.682148003 | 19 |
| eef1g7 | 0.002102563 | 0.255313729 | 0.9 | 0.617 | 1 | 19 |
| cxcl12a3 | 0 | 1.990612824 | 0.97 | 0.068 | 0 | 20 |
| podxl | 0 | 1.674437181 | 0.879 | 0.033 | 0 | 20 |
| krt152 | 0 | 1.663518322 | 0.864 | 0.072 | 0 | 20 |
| nid1b2 | 0 | 1.651540667 | 0.879 | 0.041 | 0 | 20 |
| aldh1a21 | 0 | 1.597023719 | 0.884 | 0.065 | 0 | 20 |
| c63 | 0 | 1.562505619 | 0.904 | 0.082 | 0 | 20 |
| tmem88b1 | 0 | 1.525190739 | 0.798 | 0.024 | 0 | 20 |
| postnb2 | 0 | 1.512362118 | 0.828 | 0.066 | 0 | 20 |
| slco2b12 | 0 | 1.476565246 | 0.818 | 0.052 | 0 | 20 |
| XLOC-0160944 | 0 | 1.365915528 | 0.823 | 0.067 | 0 | 20 |
| il153 | 0 | 1.340662576 | 0.737 | 0.045 | 0 | 20 |
| akap12b3 | 0 | 1.300852398 | 0.712 | 0.049 | 0 | 20 |

| | | | | | | |
|---|---|---|---|---|---|---|
| hspb13 | 0 | 1.2835404 | 0.763 | 0.052 | 0 | 20 |
| ndnf | 0 | 1.281010304 | 0.687 | 0.019 | 0 | 20 |
| ace | 0 | 1.147561855 | 0.591 | 0.019 | 0 | 20 |
| jam2b | 0 | 1.144937703 | 0.606 | 0.01 | 0 | 20 |
| slc29a1a | 0 | 1.108762348 | 0.621 | 0.029 | 0 | 20 |
| col12a1b2 | 0 | 1.100609159 | 0.591 | 0.023 | 0 | 20 |
| c44 | 0 | 1.079961865 | 0.611 | 0.032 | 0 | 20 |
| gpr17 | 0 | 1.056879678 | 0.566 | 0.006 | 0 | 20 |
| wt1a | 0 | 0.983294333 | 0.551 | 0.025 | 0 | 20 |
| ptprk1 | 0 | 0.965190589 | 0.561 | 0.026 | 0 | 20 |
| grhl11 | 0 | 0.951681736 | 0.525 | 0.018 | 0 | 20 |
| tmsb3 | 0 | 0.949234262 | 0.485 | 0.021 | 0 | 20 |
| si:ch211-191i18.21 | 0 | 0.949234075 | 0.439 | 0.014 | 0 | 20 |
| cd151 | 0 | 0.742410842 | 0.414 | 0.01 | 0 | 20 |
| gria3b | 0 | 0.635502835 | 0.318 | 0.003 | 0 | 20 |
| vegfd | 0 | 0.538951737 | 0.288 | 0.004 | 0 | 20 |
| foxl2a | 4.21470020370711e-305 | 0.561261303 | 0.288 | 0.006 | 1.06079789427104e-300 | 20 |
| phc2a | 2.15850947235701e-302 | 1.483195386 | 0.833 | 0.081 | 5.43275249097536e-298 | 20 |
| ctgfa1 | 4.36063431324978e-299 | 0.994984948 | 0.465 | 0.021 | 1.09752805030184e-294 | 20 |
| arxa | 9.59669853081742e-296 | 0.36997857 | 0.182 | 0.001 | 2.41539305322144e-291 | 20 |
| CR786577.1 | 5.99933431081176e-294 | 0.372911522 | 0.172 | 0.001 | 1.50997245268821e-289 | 20 |
| steap4 | 8.85840769393279e-289 | 0.6590476 | 0.313 | 0.008 | 2.22957263248594e-284 | 20 |
| cyp19a1a | 1.00395027032838e-285 | 0.60808255 | 0.212 | 0.003 | 2.5268424353895e-281 | 20 |
| krt18a.12 | 5.02664232111223e-280 | 1.730677639 | 0.909 | 0.114 | 1.26515560580074e-275 | 20 |
| csf1a | 4.46250245464036e-274 | 0.478925384 | 0.247 | 0.005 | 1.12316724280843e-269 | 20 |
| entpd13 | 7.85158956528188e-274 | 1.259175494 | 0.697 | 0.061 | 1.9761665776858e-269 | 20 |
| CR751602.2 | 7.60172495506892e-272 | 0.555683559 | 0.273 | 0.006 | 1.9132781539413e-267 | 20 |
| s1pr3a | 1.19807792544174e-265 | 0.361537838 | 0.162 | 0.001 | 3.01544233054432e-261 | 20 |
| fn1b2 | 1.94397040424884e-265 | 1.371872609 | 0.783 | 0.076 | 4.89277911045391e-261 | 20 |
| tspan9b1 | 2.78292359221034e-257 | 0.886873678 | 0.545 | 0.036 | 7.00434038923419e-253 | 20 |
| lpar13 | 5.03634612469833e-255 | 1.103210714 | 0.657 | 0.055 | 1.26759795612532e-250 | 20 |
| FO744854.11 | 1.21360249021061e-254 | 0.641431269 | 0.343 | 0.012 | 3.05451610761108e-250 | 20 |
| bnc21 | 2.36973407745085e-252 | 0.820927831 | 0.46 | 0.025 | 5.96438369953605e-248 | 20 |
| cldn5a3 | 2.24915512827047e-243 | 0.994314235 | 0.566 | 0.041 | 5.66089854234394e-239 | 20 |
| spon1b3 | 3.89252291559573e-238 | 0.900428148 | 0.53 | 0.037 | 9.7970909262629e-234 | 20 |

| | | | | | | |
|---|---|---|---|---|---|---|
| c7a3 | 5.31241347422113e-231 | 0.959911086 | 0.525 | 0.037 | 1.33708134732672e-226 | 20 |
| wt1b | 8.47123646464642e-227 | 0.66766479 | 0.364 | 0.017 | 2.13212550578686e-222 | 20 |
| anxa2a2 | 6.11008031840263e-223 | 1.413763591 | 0.768 | 0.096 | 1.53784611533876e-218 | 20 |
| cebpd4 | 2.1104451995935e-221 | 1.347137485 | 0.859 | 0.118 | 5.31177952285688e-217 | 20 |
| sh3pxd2aa1 | 8.95235394265272e-221 | 0.820580632 | 0.429 | 0.026 | 2.25321796382626e-216 | 20 |
| si:ch211-204c21.14 | 7.07238104088257e-219 | 1.076945705 | 0.652 | 0.064 | 1.78004758417973e-214 | 20 |
| itga91 | 2.30345408242024e-215 | 0.788505158 | 0.419 | 0.025 | 5.79756358004351e-211 | 20 |
| rbms33 | 5.32640907161411e-215 | 0.768950088 | 0.439 | 0.027 | 1.34060389923455e-210 | 20 |
| mao | 5.9075421906569e-214 | 0.554456677 | 0.283 | 0.01 | 1.48686929396643e-209 | 20 |
| vasnb2 | 9.4007564139578e-210 | 0.735114909 | 0.419 | 0.025 | 2.36607638182904e-205 | 20 |
| lrp1ab3 | 4.31602160601866e-205 | 1.088467396 | 0.616 | 0.061 | 1.08629947801884e-200 | 20 |
| col6a33 | 2.2896410924689e-201 | 1.136869871 | 0.753 | 0.09 | 5.76279766563498e-197 | 20 |
| BX322530.13 | 6.95685687427503e-200 | 0.836010482 | 0.505 | 0.04 | 1.75097130668628e-195 | 20 |
| tmem98 | 2.06697783856354e-199 | 0.480903718 | 0.258 | 0.009 | 5.20237652188057e-195 | 20 |
| phactr4a1 | 2.62428487791667e-197 | 0.812099201 | 0.439 | 0.031 | 6.60506260922847e-193 | 20 |
| adma | 5.77609629774372e-193 | 0.689363376 | 0.303 | 0.014 | 1.45378567717912e-188 | 20 |
| cdh22 | 2.40721560439363e-190 | 0.877453966 | 0.535 | 0.048 | 6.05872095469833e-186 | 20 |
| eppk12 | 6.11813600874764e-189 | 0.856295659 | 0.455 | 0.034 | 1.53987365204169e-184 | 20 |
| krt944 | 3.61856712426244e-188 | 1.349407549 | 0.813 | 0.119 | 9.10757159505614e-184 | 20 |
| s1pr22 | 4.15693542957076e-185 | 0.695258567 | 0.399 | 0.026 | 1.04625907826866e-180 | 20 |
| si:ch211-197h24.91 | 1.91703707083271e-182 | 0.986929536 | 0.576 | 0.06 | 4.82499060357885e-178 | 20 |
| CABZ01080702.14 | 1.07613725257257e-181 | 0.998689036 | 0.646 | 0.076 | 2.7085298509999e-177 | 20 |
| gabra5 | 3.25987355669827e-181 | 0.30879432 | 0.152 | 0.002 | 8.20477575485387e-177 | 20 |
| si:ch211-106h4.123 | 6.35887465876091e-179 | 1.077497568 | 0.54 | 0.051 | 1.60046516286353e-174 | 20 |
| vcam1b2 | 9.61092023229049e-178 | 0.683979029 | 0.328 | 0.018 | 2.41897251326519e-173 | 20 |
| ugt1b5 | 8.99349092877143e-177 | 0.681547591 | 0.369 | 0.024 | 2.26357173186248e-172 | 20 |
| pdgfc | 4.64262333242961e-172 | 0.338493894 | 0.187 | 0.005 | 1.16850186653921e-167 | 20 |
| gstm.32 | 2.16660399059767e-169 | 1.25041335 | 0.682 | 0.092 | 5.45312558393527e-165 | 20 |
| met | 1.68768047456313e-168 | 0.389083183 | 0.187 | 0.005 | 4.24772298642793e-164 | 20 |
| pbx1b2 | 2.79168913442724e-168 | 0.641073 | 0.354 | 0.023 | 7.02640238243991e-164 | 20 |
| si:dkey-56m19.51 | 1.59504459903769e-166 | 0.804455196 | 0.46 | 0.041 | 4.01456775131796e-162 | 20 |
| hsd11b24 | 1.74804124021762e-165 | 1.239063034 | 0.783 | 0.113 | 4.39964499750373e-161 | 20 |
| tmem1763 | 2.82809582626749e-165 | 0.866870978 | 0.556 | 0.059 | 7.11803438513264e-161 | 20 |
| cd81a4 | 7.90755266955323e-164 | 1.340852142 | 0.904 | 0.196 | 1.99025193139985e-159 | 20 |
| lama5 | 6.32072438642598e-163 | 0.621933146 | 0.323 | 0.019 | 1.59086312081956e-158 | 20 |

| | | | | | | |
|---|---|---|---|---|---|---|
| BX569787.12 | 1.20359154445394e-156 | 0.603803899 | 0.318 | 0.02 | 3.02931955823613e-152 | 20 |
| kirrel1b | 6.24843029282849e-156 | 0.353546421 | 0.162 | 0.004 | 1.572667420402e-151 | 20 |
| ghrb3 | 1.95999701871048e-155 | 0.71632178 | 0.424 | 0.037 | 4.93311649639241e-151 | 20 |
| mmp16a2 | 1.55690576908979e-154 | 0.911025725 | 0.556 | 0.062 | 3.91857613022209e-150 | 20 |
| CR749775.2 | 8.724036652706e-154 | 0.417896346 | 0.187 | 0.006 | 2.19575278511957e-149 | 20 |
| krt86 | 1.35830985946709e-151 | 1.429072572 | 0.965 | 0.266 | 3.41873008529273e-147 | 20 |
| cygb12 | 2.37524877526899e-151 | 0.781437887 | 0.485 | 0.048 | 5.97826364247453e-147 | 20 |
| cxcl14 | 5.68295891723591e-149 | 0.45294255 | 0.237 | 0.011 | 1.43034392987911e-144 | 20 |
| paplna1 | 1.73556802967301e-148 | 0.543468307 | 0.308 | 0.019 | 4.36825117388399e-144 | 20 |
| vat13 | 1.10061841962399e-147 | 1.114382963 | 0.747 | 0.125 | 2.77014650035163e-143 | 20 |
| nid2a2 | 4.49661072251181e-146 | 0.787265191 | 0.475 | 0.048 | 1.131751952749e-141 | 20 |
| mmp22 | 1.8703824054217e-144 | 0.938187291 | 0.616 | 0.084 | 4.70756547620588e-140 | 20 |
| pleca1 | 1.90953624921718e-144 | 0.834500545 | 0.485 | 0.053 | 4.80611178565471e-140 | 20 |
| nr2f1a3 | 1.04775559881394e-143 | 0.731728004 | 0.424 | 0.04 | 2.63709606665482e-139 | 20 |
| wnt11r | 7.57400262132365e-143 | 0.340930289 | 0.172 | 0.005 | 1.90630071976095e-138 | 20 |
| cavin2b2 | 2.45957793233945e-142 | 0.893386196 | 0.51 | 0.059 | 6.19051169790516e-138 | 20 |
| itm2bb5 | 6.1852868987833e-138 | 1.183359947 | 0.884 | 0.212 | 1.55677485955477e-133 | 20 |
| sox4a1 | 1.69531686956324e-135 | 0.758568316 | 0.409 | 0.039 | 4.26694302900372e-131 | 20 |
| abl2 | 2.59099919655975e-135 | 0.49195926 | 0.253 | 0.014 | 6.52128587782124e-131 | 20 |
| si:ch211-198c19.34 | 1.22968746567277e-134 | 0.897797126 | 0.495 | 0.057 | 3.0950003823518e-130 | 20 |
| c4b1 | 3.64471950930014e-133 | 0.531422196 | 0.268 | 0.016 | 9.17339453295752e-129 | 20 |
| fstl1b3 | 4.36686570825725e-133 | 0.996924033 | 0.707 | 0.116 | 1.09909643011127e-128 | 20 |
| cd248a2 | 4.49189228181775e-132 | 0.631571151 | 0.348 | 0.029 | 1.13056436841071e-127 | 20 |
| AL954191.1 | 2.6409857439901e-130 | 0.459454616 | 0.227 | 0.012 | 6.64709701904869e-126 | 20 |
| fabp11a3 | 7.95224375884097e-128 | 1.080437515 | 0.556 | 0.077 | 2.00150023166268e-123 | 20 |
| zeb2b | 6.31045414868832e-127 | 0.371046469 | 0.182 | 0.007 | 1.58827820468336e-122 | 20 |
| ptprga4 | 1.31582948863657e-125 | 0.804149496 | 0.485 | 0.06 | 3.3118112399494e-121 | 20 |
| bmp3 | 8.17839650597499e-124 | 0.444784133 | 0.202 | 0.009 | 2.05842061658884e-119 | 20 |
| adamts2 | 1.70485552075417e-123 | 0.553986769 | 0.288 | 0.021 | 4.29095086018617e-119 | 20 |
| rgs7bpb1 | 1.73629278550698e-122 | 0.42993179 | 0.232 | 0.013 | 4.37007531184252e-118 | 20 |
| ftr82 | 2.51479759492072e-122 | 0.517021118 | 0.278 | 0.019 | 6.32949406665595e-118 | 20 |
| pabpc1b3 | 2.91947782910939e-122 | 0.800651815 | 0.49 | 0.062 | 7.34803374808543e-118 | 20 |
| gdpd4a | 9.62820976411607e-122 | 0.426550622 | 0.187 | 0.008 | 2.42332411553037e-117 | 20 |
| col6a13 | 1.2294544269995e-121 | 0.8618334 | 0.576 | 0.082 | 3.09441384731505e-117 | 20 |
| ahnak5 | 2.4836969882786e-121 | 0.979741288 | 0.672 | 0.118 | 6.2512169497984e-117 | 20 |

| | | | | | | |
|---|---|---|---|---|---|---|
| clu1 | 1.4906491808728e-120 | 0.560125553 | 0.298 | 0.022 | 3.75181492333875e-116 | 20 |
| fhl3b1 | 6.45583857496396e-120 | 0.387955514 | 0.207 | 0.01 | 1.62487001093268e-115 | 20 |
| col5a13 | 7.23116279132438e-120 | 0.90696162 | 0.652 | 0.104 | 1.82001136294843e-115 | 20 |
| tfpia | 7.27140469075485e-120 | 0.502176258 | 0.263 | 0.017 | 1.83013984661609e-115 | 20 |
| zfp36l1b3 | 7.74002206098177e-118 | 0.996248631 | 0.682 | 0.122 | 1.9480861525285e-113 | 20 |
| add3a4 | 9.36832618652565e-118 | 0.970682783 | 0.652 | 0.116 | 2.35791401788664e-113 | 20 |
| gna12a | 1.89044402672451e-115 | 0.504550187 | 0.278 | 0.021 | 4.75805857086292e-111 | 20 |
| tns2a3 | 4.22061176818667e-113 | 0.609846895 | 0.343 | 0.033 | 1.0622857759349e-108 | 20 |
| arhgap42b | 2.31640683788852e-112 | 0.321291246 | 0.172 | 0.007 | 5.83016437028161e-108 | 20 |
| myl9b3 | 1.62903048958261e-111 | 0.804016971 | 0.51 | 0.073 | 4.10010683923048e-107 | 20 |
| gas61 | 3.30541953646433e-111 | 0.506688518 | 0.273 | 0.021 | 8.31941043132706e-107 | 20 |
| col6a23 | 6.36895490936844e-110 | 0.719725539 | 0.46 | 0.059 | 1.60300226113894e-105 | 20 |
| dkk3b2 | 2.7181339945476e-109 | 0.57370623 | 0.359 | 0.036 | 6.84127145087685e-105 | 20 |
| si:ch211-136a13.1 | 8.323494688469e-108 | 0.326744882 | 0.162 | 0.007 | 2.09494037814076e-103 | 20 |
| tmem176l.13 | 1.18694904798846e-107 | 0.99733727 | 0.758 | 0.172 | 2.98743205888214e-103 | 20 |
| anxa1a4 | 3.99852518829561e-107 | 1.023816966 | 0.687 | 0.135 | 1.00638880464212e-102 | 20 |
| rasgef1bb | 5.48435964565075e-107 | 0.585300946 | 0.308 | 0.028 | 1.38035847921384e-102 | 20 |
| jcada | 1.29696303469251e-106 | 0.377223817 | 0.207 | 0.012 | 3.26432626201758e-102 | 20 |
| clic41 | 1.38986450419957e-106 | 0.637359719 | 0.389 | 0.044 | 3.4981499706199e-102 | 20 |
| slc43a3b1 | 1.52621172900562e-106 | 0.505498481 | 0.273 | 0.021 | 3.84132230073425e-102 | 20 |
| cfd2 | 1.36028179601936e-105 | 0.889841112 | 0.46 | 0.062 | 3.42369325240113e-101 | 20 |
| ext1a | 1.10238461169905e-104 | 0.408656378 | 0.222 | 0.014 | 2.77459182918534e-100 | 20 |
| bmp4 | 7.00656899839357e-104 | 0.362442675 | 0.202 | 0.012 | 1.76348335120568e-99 | 20 |
| f3b3 | 7.36210742459046e-103 | 0.868066182 | 0.571 | 0.095 | 1.85296881769517e-98 | 20 |
| zbtb16b | 9.68819859223518e-103 | 0.499292837 | 0.268 | 0.022 | 2.43842270367967e-98 | 20 |
| mtss1la2 | 1.03129789055158e-102 | 0.482283928 | 0.263 | 0.021 | 2.59567366072927e-98 | 20 |
| gpc4 | 1.6784391035434e-102 | 0.472396974 | 0.283 | 0.024 | 4.22446337970838e-98 | 20 |
| aadac | 7.87523795254347e-102 | 0.288754292 | 0.141 | 0.005 | 1.98211864027566e-97 | 20 |
| pard6a | 1.86705608965812e-101 | 0.263897573 | 0.141 | 0.005 | 4.69919347206051e-97 | 20 |
| ywhag1 | 1.87692252867811e-101 | 0.271140072 | 0.162 | 0.007 | 4.72402631242993e-97 | 20 |
| antxr1c1 | 3.69352269710625e-101 | 0.529672339 | 0.328 | 0.033 | 9.29622727634673e-97 | 20 |
| slc7a6 | 8.79868460275525e-101 | 0.42700569 | 0.222 | 0.015 | 2.21454092766747e-96 | 20 |
| f102 | 1.15675132323861e-100 | 0.610459898 | 0.379 | 0.044 | 2.91142740545925e-96 | 20 |
| sorbs1 | 3.88673619144061e-100 | 0.431142369 | 0.232 | 0.016 | 9.78252632023687e-96 | 20 |
| mxra8a2 | 4.65806198209189e-100 | 0.601838126 | 0.364 | 0.041 | 1.17238762027271e-95 | 20 |

| | | | | | | |
|---|---|---|---|---|---|---|
| acvrl1 | 1.54270225163127e-99 | 0.383918502 | 0.202 | 0.012 | 3.88282729713075e-95 | 20 |
| aebp12 | 3.17215725906377e-99 | 0.564132118 | 0.333 | 0.035 | 7.98400260533761e-95 | 20 |
| sparc3 | 4.16550919017191e-99 | 0.949809769 | 0.727 | 0.15 | 1.04841700807437e-94 | 20 |
| eva1ba3 | 7.22234642417245e-99 | 0.708353669 | 0.439 | 0.06 | 1.81779237149996e-94 | 20 |
| nr2f23 | 2.69820313102383e-98 | 0.627114556 | 0.354 | 0.04 | 6.79110746047389e-94 | 20 |
| pgr1 | 6.45728531877087e-97 | 0.389707503 | 0.222 | 0.016 | 1.62523414188144e-92 | 20 |
| wu:fc27b11 | 3.34716587749627e-96 | 0.372931026 | 0.187 | 0.011 | 8.42448179707037e-92 | 20 |
| adamts8a | 5.84321943420436e-96 | 0.258747741 | 0.111 | 0.003 | 1.47067989939489e-91 | 20 |
| tagln3 | 1.4618289202692e-95 | 0.696340941 | 0.429 | 0.058 | 3.67927720942556e-91 | 20 |
| CU539058.1 | 4.33224879917397e-95 | 0.394366226 | 0.212 | 0.014 | 1.0903837002641e-90 | 20 |
| hdac41 | 5.23280115785931e-95 | 0.51756298 | 0.288 | 0.027 | 1.31704372342161e-90 | 20 |
| slc43a2a5 | 2.85645184732561e-94 | 0.831334707 | 0.54 | 0.096 | 7.18940365453383e-90 | 20 |
| si:ch73-46j18.54 | 3.38849261198907e-94 | 0.998378932 | 0.712 | 0.167 | 8.52849705511529e-90 | 20 |
| pdgfrb3 | 5.06387382939461e-94 | 0.63418034 | 0.374 | 0.046 | 1.27452640412033e-89 | 20 |
| hspg2 | 7.49867020052765e-94 | 0.43834293 | 0.258 | 0.022 | 1.8873403027708e-89 | 20 |
| rbp43 | 8.62299490162991e-94 | 0.842751504 | 0.657 | 0.12 | 2.17032158679123e-89 | 20 |
| cald1a3 | 4.03393814973416e-93 | 0.793394827 | 0.535 | 0.093 | 1.01530189290659e-88 | 20 |
| col1a1a3 | 1.2391323587141e-92 | 0.840746162 | 0.682 | 0.132 | 3.11877223364751e-88 | 20 |
| tnfaip6 | 2.37336695266303e-92 | 0.370428402 | 0.192 | 0.012 | 5.97352728315757e-88 | 20 |
| atp1a3b2 | 2.68183948537353e-92 | 0.515322476 | 0.278 | 0.026 | 6.74992180073663e-88 | 20 |
| lxn5 | 5.16914536950101e-92 | 0.942602534 | 0.641 | 0.141 | 1.30102219804971e-87 | 20 |
| col27a1b1 | 7.69802015217171e-92 | 0.427633402 | 0.222 | 0.017 | 1.9375146921001e-87 | 20 |
| ppdpfa2 | 3.35371598028492e-91 | 0.698699309 | 0.465 | 0.071 | 8.44096775077913e-87 | 20 |
| jun4 | 3.80763161381286e-91 | 1.066293893 | 0.813 | 0.22 | 9.58342800880558e-87 | 20 |
| svild3 | 6.83612639758973e-91 | 0.743152116 | 0.475 | 0.076 | 1.72058465300936e-86 | 20 |
| dag1 | 5.35814029862533e-90 | 0.609918309 | 0.364 | 0.045 | 1.34859033176101e-85 | 20 |
| dchs1a | 6.83439012083101e-90 | 0.298541612 | 0.157 | 0.008 | 1.72014764951196e-85 | 20 |
| cxcl18b4 | 9.99686721230063e-90 | 0.846378285 | 0.566 | 0.101 | 2.51611150866395e-85 | 20 |
| grb10b2 | 2.1307269032964e-88 | 0.514976368 | 0.303 | 0.032 | 5.3628265429067e-84 | 20 |
| rhobtb4 | 4.63931299438276e-88 | 0.415305605 | 0.222 | 0.017 | 1.1676686875562e-83 | 20 |
| CABZ01068366.11 | 5.17479476139474e-88 | 0.487335902 | 0.273 | 0.026 | 1.30244409349544e-83 | 20 |
| ptprua | 8.18572321095618e-88 | 0.267272252 | 0.141 | 0.007 | 2.06026467496556e-83 | 20 |
| junba6 | 1.39740562212307e-87 | 1.076821768 | 0.899 | 0.301 | 3.51713021032155e-83 | 20 |
| greb1l | 2.1984841342982e-87 | 0.2786047 | 0.136 | 0.006 | 5.53336471761513e-83 | 20 |
| dcn3 | 3.19522745282805e-87 | 0.771805945 | 0.611 | 0.112 | 8.04206797602292e-83 | 20 |

| | | | | | | |
|---|---|---|---|---|---|---|
| nr0b1 | 7.99582511328584e-87 | 0.310026915 | 0.167 | 0.009 | 2.01246922276291e-82 | 20 |
| socs3a7 | 2.51212929492131e-86 | 1.124186997 | 0.869 | 0.29 | 6.32277822238745e-82 | 20 |
| gstt1b2 | 1.39598591966651e-85 | 0.636726238 | 0.439 | 0.067 | 3.51355696120865e-81 | 20 |
| pmp22a3 | 1.99934724365712e-85 | 0.8445376 | 0.646 | 0.132 | 5.03215707756061e-81 | 20 |
| col1a1b3 | 5.38882496569359e-85 | 0.806710098 | 0.677 | 0.139 | 1.35631335561542e-80 | 20 |
| clec11a2 | 1.06354281315638e-84 | 0.361415796 | 0.197 | 0.014 | 2.6768309064333e-80 | 20 |
| fosab6 | 5.4124057844477e-84 | 1.114620915 | 0.919 | 0.341 | 1.36224841188764e-79 | 20 |
| ms4a17a.81 | 7.54116755248812e-84 | 0.287606125 | 0.177 | 0.011 | 1.89803646128574e-79 | 20 |
| si:ch73-22o12.1 | 8.69799269789989e-84 | 0.47430712 | 0.247 | 0.023 | 2.18919778213442e-79 | 20 |
| scarb2a | 2.0967096940395e-83 | 0.626258342 | 0.404 | 0.059 | 5.27720862892803e-79 | 20 |
| xirp2a1 | 2.45187432341606e-83 | 0.42157117 | 0.247 | 0.023 | 6.17112248460588e-79 | 20 |
| col18a1a3 | 2.97243051375924e-83 | 0.64379539 | 0.318 | 0.038 | 7.48131036008063e-79 | 20 |
| id11 | 1.16684755588355e-82 | 0.90726448 | 0.52 | 0.099 | 2.9368386134033e-78 | 20 |
| stard13b | 1.56261660823068e-82 | 0.281120625 | 0.152 | 0.008 | 3.93294974125579e-78 | 20 |
| tspan4a1 | 2.22008177414529e-82 | 0.47292608 | 0.278 | 0.029 | 5.58772381734628e-78 | 20 |
| eif4ebp3l3 | 7.61300854310636e-82 | 0.914193344 | 0.758 | 0.198 | 1.91611812021444e-77 | 20 |
| col1a23 | 2.63572161460178e-81 | 0.836643637 | 0.692 | 0.148 | 6.63384773179122e-77 | 20 |
| s100a10b5 | 2.13355786660197e-79 | 1.024631186 | 0.833 | 0.247 | 5.3699517944505e-75 | 20 |
| foxo3b3 | 4.38782064701827e-79 | 0.636550931 | 0.399 | 0.062 | 1.10437057864803e-74 | 20 |
| si:dkey-7j14.65 | 4.50985886885283e-79 | 0.866850225 | 0.662 | 0.168 | 1.13508637870157e-74 | 20 |
| rab6ba | 6.58484924944543e-79 | 0.329882596 | 0.172 | 0.011 | 1.65734070759292e-74 | 20 |
| FP016056.1 | 7.98409385038922e-78 | 0.279410031 | 0.141 | 0.008 | 2.00951658120446e-73 | 20 |
| mcl1b5 | 3.43238854139556e-76 | 0.917830006 | 0.783 | 0.234 | 8.63897871983849e-72 | 20 |
| eif4ebp31 | 7.93038191127046e-76 | 0.934303405 | 0.596 | 0.144 | 1.99599782324766e-71 | 20 |
| cav1 | 2.33355053295994e-75 | 0.540465852 | 0.318 | 0.04 | 5.87331333640687e-71 | 20 |
| desi1a | 2.79453539224434e-75 | 0.425718599 | 0.237 | 0.023 | 7.03356612873979e-71 | 20 |
| mir199-32 | 4.07480653987362e-75 | 0.403965222 | 0.197 | 0.016 | 1.02558805802079e-70 | 20 |
| si:dkeyp-9d4.2 | 1.29763234523422e-74 | 0.31400821 | 0.172 | 0.012 | 3.26601084972002e-70 | 20 |
| s100v23 | 5.26919569670686e-74 | 0.768197285 | 0.545 | 0.116 | 1.32620386490415e-69 | 20 |
| mfap23 | 9.57854004525289e-74 | 0.682264648 | 0.49 | 0.091 | 2.4108227439897e-69 | 20 |
| arhgef52 | 1.13086740794023e-73 | 0.450556118 | 0.247 | 0.026 | 2.84628017904475e-69 | 20 |
| colec123 | 3.98278107118118e-73 | 0.555768575 | 0.348 | 0.05 | 1.00242616780559e-68 | 20 |
| si:dkey-28b4.73 | 4.7854345771549e-73 | 0.654155422 | 0.46 | 0.087 | 1.20444602872412e-68 | 20 |
| si:ch211-204a13.2 | 1.03278407710115e-72 | 0.253377327 | 0.131 | 0.007 | 2.59941424365589e-68 | 20 |
| sema4ba4 | 1.18639996708606e-72 | 0.56917028 | 0.318 | 0.043 | 2.98605007715891e-68 | 20 |

| | | | | | | |
|---|---|---|---|---|---|---|
| agrn1 | 4.12412866575139e-72 | 0.530454614 | 0.333 | 0.046 | 1.03800194388297e-67 | 20 |
| gata6 | 1.52771646704056e-71 | 0.367443536 | 0.197 | 0.017 | 3.84510957589438e-67 | 20 |
| selenbp12 | 2.73259502709633e-71 | 0.601505314 | 0.394 | 0.065 | 6.87766842369874e-67 | 20 |
| enc12 | 3.42259114889789e-71 | 0.468605973 | 0.253 | 0.028 | 8.61431966266109e-67 | 20 |
| grip1 | 1.14863058314788e-70 | 0.275599682 | 0.136 | 0.008 | 2.89098831472489e-66 | 20 |
| zgc:110182 | 1.61744380554708e-70 | 0.632429325 | 0.465 | 0.083 | 4.07094431418145e-66 | 20 |
| loxl2a3 | 2.2359269043616e-70 | 0.53544651 | 0.298 | 0.039 | 5.62760442558772e-66 | 20 |
| col21a1 | 4.98926857405099e-70 | 0.384125496 | 0.187 | 0.016 | 1.25574900740289e-65 | 20 |
| ugt5b22 | 8.85682171847495e-70 | 0.440399188 | 0.247 | 0.028 | 2.22917345832296e-65 | 20 |
| XLOC-024248 | 9.04556663385552e-70 | 0.365621212 | 0.212 | 0.02 | 2.2766786660751e-65 | 20 |
| lipf1 | 2.00936130671366e-69 | 0.512470886 | 0.293 | 0.038 | 5.05736147286761e-65 | 20 |
| tcf122 | 2.14343919636156e-69 | 0.548688986 | 0.354 | 0.054 | 5.39482211332241e-65 | 20 |
| dachd1 | 6.45323699584171e-69 | 0.281679937 | 0.152 | 0.01 | 1.6242152194834e-64 | 20 |
| tmem204 | 1.25764218357977e-68 | 0.26326699 | 0.126 | 0.007 | 3.16535961185193e-64 | 20 |
| mknk2b7 | 2.4576934265485e-68 | 0.821089487 | 0.682 | 0.19 | 6.18576858527993e-64 | 20 |
| si:ch211-250c4.4 | 2.60922707668682e-68 | 0.301601817 | 0.172 | 0.013 | 6.56716362931307e-64 | 20 |
| plod21 | 3.20067484766012e-68 | 0.392208397 | 0.217 | 0.021 | 8.05577852407577e-64 | 20 |
| CR383676.110 | 7.08976379723616e-68 | 0.821524598 | 1 | 0.723 | 1.78442265012637e-63 | 20 |
| slc38a5b3 | 2.60621722684624e-67 | 0.674927046 | 0.51 | 0.11 | 6.55958813824929e-63 | 20 |
| cnn24 | 3.50887094186181e-67 | 0.7263417 | 0.551 | 0.129 | 8.83147727357199e-63 | 20 |
| robo23 | 5.88641304564542e-67 | 0.472002711 | 0.313 | 0.044 | 1.48155129945849e-62 | 20 |
| dlc11 | 9.30879174859932e-67 | 0.44897126 | 0.247 | 0.028 | 2.34292979520496e-62 | 20 |
| ptprsa | 1.14004113913365e-66 | 0.40402903 | 0.242 | 0.027 | 2.86936954308548e-62 | 20 |
| CABZ01068367.11 | 1.17645803993949e-66 | 0.332204219 | 0.192 | 0.017 | 2.9610272407237e-62 | 20 |
| pdgfrl2 | 1.59731335319277e-66 | 0.39419198 | 0.232 | 0.025 | 4.02027797865087e-62 | 20 |
| ywhaz4 | 2.3090051790044e-66 | 0.881264402 | 0.758 | 0.275 | 5.81153513503616e-62 | 20 |
| si:ch211-222l21.16 | 2.3821175330445e-66 | 0.876387386 | 0.803 | 0.268 | 5.99555161891971e-62 | 20 |
| pfkfb4b2 | 1.75784669033219e-65 | 0.660595072 | 0.455 | 0.091 | 4.42432433489708e-61 | 20 |
| nudt4a | 4.33811270802918e-65 | 0.357638788 | 0.187 | 0.017 | 1.09185958748387e-60 | 20 |
| gsta.11 | 6.36406456396053e-65 | 0.446225332 | 0.288 | 0.039 | 1.60177141010323e-60 | 20 |
| tnfsf122 | 8.04301769837631e-65 | 0.357045902 | 0.207 | 0.021 | 2.02434712450433e-60 | 20 |
| znf385a3 | 1.1652044975895e-64 | 0.500078263 | 0.303 | 0.043 | 2.93270319998302e-60 | 20 |
| phldb2a | 2.24424775378716e-64 | 0.262739515 | 0.152 | 0.011 | 5.6485471715069e-60 | 20 |
| neo1a1 | 3.88811186268169e-64 | 0.404062594 | 0.227 | 0.025 | 9.78598874718355e-60 | 20 |
| col5a2a3 | 3.97677331510273e-64 | 0.604035718 | 0.434 | 0.081 | 1.00091407567821e-59 | 20 |

| | | | | | | |
|---|---|---|---|---|---|---|
| CR848833.1 | 2.71493584962345e-63 | 0.254813335 | 0.111 | 0.006 | 6.83322203991725e-59 | 20 |
| lrp1aa3 | 1.80498416243422e-62 | 0.551040964 | 0.328 | 0.051 | 4.54296463843069e-58 | 20 |
| hdlbpa4 | 4.20325061190336e-62 | 0.724309937 | 0.591 | 0.153 | 1.05791614650996e-57 | 20 |
| frzb1 | 5.44306422412878e-62 | 0.376618027 | 0.182 | 0.016 | 1.36996483457097e-57 | 20 |
| cavin1b | 5.88730533199924e-62 | 0.486005295 | 0.278 | 0.037 | 1.48177587901089e-57 | 20 |
| zgc:1951751 | 1.45126502946197e-61 | 0.420814605 | 0.258 | 0.033 | 3.65268895265282e-57 | 20 |
| mgst1.25 | 1.94206031077667e-61 | 0.652959615 | 0.465 | 0.098 | 4.88797159619379e-57 | 20 |
| rhpn2 | 2.01110213467791e-61 | 0.416727614 | 0.258 | 0.032 | 5.06174296277083e-57 | 20 |
| fam171a2a | 7.17573084345646e-61 | 0.311409048 | 0.167 | 0.014 | 1.80605969598956e-56 | 20 |
| epb41l5 | 8.63226579635288e-61 | 0.445459861 | 0.278 | 0.038 | 2.17265497828406e-56 | 20 |
| CR318588.31 | 9.1045293343354e-61 | 0.579314374 | 0.293 | 0.042 | 2.29151898815888e-56 | 20 |
| XLOC-034538 | 8.75695910732941e-59 | 0.309905719 | 0.162 | 0.014 | 2.20403903772374e-54 | 20 |
| anos1a1 | 9.75668868037781e-59 | 0.297064879 | 0.162 | 0.014 | 2.45566097396429e-54 | 20 |
| nrg2a | 9.80938915419748e-59 | 0.283318266 | 0.152 | 0.012 | 2.46892515621996e-54 | 20 |
| tle2b | 5.44707167005653e-58 | 0.29056768 | 0.152 | 0.012 | 1.37097346863653e-53 | 20 |
| rasgef1ba2 | 1.62829485900311e-57 | 0.534672077 | 0.354 | 0.062 | 4.09825533062492e-53 | 20 |
| nrxn1a1 | 3.49060243269468e-57 | 0.275251722 | 0.131 | 0.009 | 8.78549726284924e-53 | 20 |
| egfra2 | 3.74782738246701e-57 | 0.366743427 | 0.222 | 0.026 | 9.43290673893121e-53 | 20 |
| il6st2 | 5.08453390485028e-57 | 0.473828463 | 0.303 | 0.048 | 1.27972633851177e-52 | 20 |
| gulp1a1 | 7.87016047502777e-57 | 0.382229292 | 0.232 | 0.029 | 1.98084068995974e-52 | 20 |
| zgc:16273010 | 7.90396956328322e-57 | 0.882085281 | 0.904 | 0.385 | 1.98935009938275e-52 | 20 |
| chd34 | 8.03726038605515e-57 | 0.611019397 | 0.449 | 0.1 | 2.02289806656622e-52 | 20 |
| si:ch73-86n18.11 | 9.10737962731742e-57 | 0.301424134 | 0.157 | 0.013 | 2.29223637839952e-52 | 20 |
| cntfr | 9.34294070784478e-57 | 0.29751894 | 0.162 | 0.014 | 2.35152474675745e-52 | 20 |
| lamb1a3 | 1.23569827393654e-56 | 0.45804205 | 0.258 | 0.036 | 3.11012898567088e-52 | 20 |
| gadd45ba3 | 2.44184887564182e-56 | 0.765679477 | 0.641 | 0.184 | 6.14588943510289e-52 | 20 |
| olfml3b2 | 3.56881239153855e-56 | 0.447202393 | 0.283 | 0.042 | 8.98234390826339e-52 | 20 |
| stxbp61 | 5.79048527577699e-56 | 0.405982495 | 0.278 | 0.04 | 1.45740723906031e-51 | 20 |
| robo11 | 1.18583839753729e-55 | 0.336720171 | 0.192 | 0.021 | 2.98463666276161e-51 | 20 |
| deptor2 | 2.90167877547865e-55 | 0.428122725 | 0.253 | 0.035 | 7.30323531000221e-51 | 20 |
| si:ch211-1a19.32 | 5.60188222207389e-55 | 0.483469406 | 0.308 | 0.05 | 1.40993773647378e-50 | 20 |
| rhoj | 2.34086888558138e-54 | 0.28701024 | 0.152 | 0.013 | 5.89173289811978e-50 | 20 |
| si:ch211-74m13.33 | 4.77793799726856e-54 | 0.532255139 | 0.348 | 0.064 | 1.20255921453252e-49 | 20 |
| ksr1b | 7.12597590732184e-54 | 0.282519338 | 0.146 | 0.012 | 1.79353687611383e-49 | 20 |
| phactr4b | 7.96160640260486e-54 | 0.334770133 | 0.187 | 0.02 | 2.00385671547162e-49 | 20 |

| | | | | | | |
|---|---|---|---|---|---|---|
| marcksl1b5 | 9.31028583656603e-54 | 0.694434797 | 0.47 | 0.112 | 2.3433058422053e-49 | 20 |
| pi15a1 | 2.84864006347393e-53 | 0.474223035 | 0.283 | 0.043 | 7.16974217575753e-49 | 20 |
| wbp1lb | 3.4879289124901e-53 | 0.458877436 | 0.283 | 0.045 | 8.77876827984632e-49 | 20 |
| appa3 | 6.63780476746709e-53 | 0.508455587 | 0.374 | 0.073 | 1.67066908192379e-48 | 20 |
| hipk21 | 9.41594453722941e-53 | 0.351735289 | 0.182 | 0.02 | 2.36989908057527e-48 | 20 |
| tjp2b1 | 1.68080489359857e-52 | 0.468380113 | 0.328 | 0.058 | 4.23041783669824e-48 | 20 |
| ppp1r14bb | 1.70720091850073e-52 | 0.376319589 | 0.232 | 0.031 | 4.29685399177448e-48 | 20 |
| mir21-13 | 2.35726192559941e-52 | 0.615328761 | 0.444 | 0.103 | 5.93299254054116e-48 | 20 |
| itga11 | 5.83629936623774e-52 | 0.434334734 | 0.253 | 0.037 | 1.46893818748838e-47 | 20 |
| ephb32 | 6.31531390536375e-52 | 0.453774717 | 0.258 | 0.039 | 1.589501356841e-47 | 20 |
| kank3 | 1.14800499112025e-51 | 0.274037448 | 0.162 | 0.016 | 2.88941376215055e-47 | 20 |
| cdc42bpb | 1.85080791839225e-51 | 0.328933976 | 0.207 | 0.025 | 4.65829844980145e-47 | 20 |
| si:dkey-9i23.163 | 2.93323295727747e-51 | 0.42697157 | 0.253 | 0.037 | 7.38265403017166e-47 | 20 |
| usp43b | 3.6994059509414e-51 | 0.312522189 | 0.162 | 0.016 | 9.3110348379244e-47 | 20 |
| ar3 | 4.18663265838092e-51 | 0.505017242 | 0.313 | 0.056 | 1.05373357378789e-46 | 20 |
| mdka4 | 6.25437341544698e-51 | 0.645509509 | 0.409 | 0.09 | 1.57416324493385e-46 | 20 |
| hbp1 | 7.65649143003809e-51 | 0.465859531 | 0.298 | 0.051 | 1.92706232802629e-46 | 20 |
| pik3r22 | 2.29098439418385e-50 | 0.505482891 | 0.313 | 0.057 | 5.76617862172134e-46 | 20 |
| mxra8b1 | 3.13697074542096e-50 | 0.315603518 | 0.187 | 0.021 | 7.89544166915002e-46 | 20 |
| LOC1018844512 | 3.81700510132449e-50 | 0.591106102 | 0.379 | 0.082 | 9.60702013952362e-46 | 20 |
| si:dkey-58f10.111 | 4.04847784403145e-50 | 0.330681993 | 0.197 | 0.024 | 1.01896138856427e-45 | 20 |
| ccdc85b3 | 4.32488062290536e-50 | 0.46427356 | 0.278 | 0.046 | 1.08852920397905e-45 | 20 |
| tiam1a | 8.19198521094738e-50 | 0.265412853 | 0.126 | 0.01 | 2.06184075774335e-45 | 20 |
| b2ml9 | 3.92012653521225e-49 | 0.723151611 | 0.717 | 0.261 | 9.86656647647571e-45 | 20 |
| gamt1 | 4.64263582138344e-49 | 0.389292871 | 0.263 | 0.042 | 1.168505009884e-44 | 20 |
| si:dkeyp-75h12.51 | 1.41583976852909e-48 | 0.395642595 | 0.242 | 0.036 | 3.56352711341085e-44 | 20 |
| prss351 | 2.42382453215259e-48 | 0.323383999 | 0.172 | 0.019 | 6.10052396497485e-44 | 20 |
| nr1d2a | 3.53523977429746e-48 | 0.535502167 | 0.359 | 0.075 | 8.89784498792928e-44 | 20 |
| NAV1 (1 of many) | 3.55587725434536e-48 | 0.33658947 | 0.182 | 0.021 | 8.94978746146183e-44 | 20 |
| hspb81 | 5.4530680781527e-48 | 0.313892884 | 0.187 | 0.022 | 1.37248270459025e-43 | 20 |
| ifi464 | 8.04088473923287e-48 | 0.579231461 | 0.47 | 0.117 | 2.02381028001752e-43 | 20 |
| txnipa8 | 9.01524488431827e-48 | 0.723837051 | 0.621 | 0.205 | 2.26904698493406e-43 | 20 |
| mogat2 | 1.43336749906516e-47 | 0.315540725 | 0.152 | 0.015 | 3.6076426583971e-43 | 20 |
| mob2a1 | 1.88515742901642e-47 | 0.408937954 | 0.242 | 0.037 | 4.74475273309142e-43 | 20 |
| COX310 | 1.04254747796226e-46 | 0.594789265 | 0.995 | 0.696 | 2.62398774728322e-42 | 20 |

| | | | | | | |
|---|---|---|---|---|---|---|
| pgrmc13 | 1.81256884171468e-46 | 0.621436299 | 0.5 | 0.136 | 4.56205451771168e-42 | 20 |
| pkp3b | 2.95032663707138e-46 | 0.406792507 | 0.253 | 0.04 | 7.42567711284495e-42 | 20 |
| kank22 | 3.13770552808959e-46 | 0.411135836 | 0.288 | 0.051 | 7.89729104364868e-42 | 20 |
| calcoco1a | 3.42689865626938e-46 | 0.362153127 | 0.212 | 0.03 | 8.62516122796441e-42 | 20 |
| itgb5 | 4.68443520264076e-46 | 0.254176409 | 0.146 | 0.014 | 1.17902549615265e-41 | 20 |
| zgc:1951734 | 1.04261823866482e-45 | 0.526485022 | 0.389 | 0.087 | 2.62416584489548e-41 | 20 |
| mt-nd18 | 2.46786241559192e-45 | 0.700100695 | 0.889 | 0.405 | 6.2113629138033e-41 | 20 |
| sox6 | 3.10778246887824e-45 | 0.338401465 | 0.197 | 0.026 | 7.82197769591965e-41 | 20 |
| dpp3 | 3.30598350077517e-45 | 0.389183873 | 0.242 | 0.038 | 8.32082987310103e-41 | 20 |
| fkbp54 | 9.67645168987333e-45 | 0.577838343 | 0.449 | 0.118 | 2.43546612582422e-40 | 20 |
| ptpn21 | 1.25872437870058e-44 | 0.345215137 | 0.217 | 0.032 | 3.1680833887515e-40 | 20 |
| adam10a2 | 1.71968376701822e-44 | 0.499566449 | 0.389 | 0.09 | 4.32827207320815e-40 | 20 |
| mt-co210 | 1.74728931597165e-44 | 0.582523774 | 0.995 | 0.717 | 4.39775247936905e-40 | 20 |
| si:ch211-212k18.83 | 2.52121146710119e-44 | 0.388742377 | 0.247 | 0.041 | 6.34563714154698e-40 | 20 |
| mmp14a3 | 3.73610640734009e-44 | 0.402572007 | 0.278 | 0.05 | 9.40340621663428e-40 | 20 |
| rpz54 | 8.03456591009401e-44 | 0.568749369 | 0.404 | 0.099 | 2.02221989391156e-39 | 20 |
| si:dkey-11f4.144 | 1.1226421571289e-43 | 0.621711297 | 0.409 | 0.105 | 2.82557804527772e-39 | 20 |
| cybrd1 | 1.52856478700203e-43 | 0.402113505 | 0.278 | 0.05 | 3.8472447124054e-39 | 20 |
| egr17 | 1.54880091879558e-43 | 0.671525109 | 0.586 | 0.185 | 3.8981770325166e-39 | 20 |
| FO082781.14 | 2.01444440127442e-43 | 0.444366962 | 0.258 | 0.045 | 5.0701551135676e-39 | 20 |
| nfil3-66 | 2.19283318873618e-43 | 0.611444776 | 0.465 | 0.127 | 5.5191418527301e-39 | 20 |
| mpp5a | 3.68651709366959e-43 | 0.268495864 | 0.141 | 0.014 | 9.27859487305698e-39 | 20 |
| mt-nd48 | 3.73140967803653e-43 | 0.69336645 | 0.904 | 0.45 | 9.39158501865014e-39 | 20 |
| tcf4 | 2.02004418654585e-42 | 0.294676053 | 0.167 | 0.02 | 5.08424921311724e-38 | 20 |
| plxnb2a | 2.84571649099453e-42 | 0.324875635 | 0.172 | 0.021 | 7.16238383618413e-38 | 20 |
| fzd22 | 4.4561590676973e-42 | 0.314317434 | 0.192 | 0.027 | 1.12157067574873e-37 | 20 |
| actn12 | 6.71343125347419e-42 | 0.45345921 | 0.242 | 0.042 | 1.68970351218692e-37 | 20 |
| ctdsp22 | 7.9508425418891e-42 | 0.448676005 | 0.308 | 0.064 | 2.00114755936807e-37 | 20 |
| tnfrsfa | 1.12807315698472e-41 | 0.318334794 | 0.192 | 0.027 | 2.83924732881485e-37 | 20 |
| calm2b1 | 7.7123059307115e-41 | 0.638019009 | 0.606 | 0.205 | 1.94111027970078e-36 | 20 |
| mt-atp69 | 9.0221500288676e-41 | 0.634014305 | 0.965 | 0.586 | 2.27078494076569e-36 | 20 |
| mt-nd27 | 1.44110208600096e-40 | 0.67888423 | 0.854 | 0.394 | 3.62710984025581e-36 | 20 |
| nt5c2b1 | 1.63793152895932e-40 | 0.419684329 | 0.242 | 0.043 | 4.12250986523772e-36 | 20 |
| si:ch211-140m22.72 | 3.93777214241164e-40 | 0.450384324 | 0.303 | 0.063 | 9.91097870523585e-36 | 20 |
| lrrc1 | 4.38526947528123e-40 | 0.291338914 | 0.177 | 0.023 | 1.10372847423353e-35 | 20 |

| | | | | | | |
|---|---|---|---|---|---|---|
| pdk2a2 | 4.93775428664312e-40 | 0.327275406 | 0.187 | 0.026 | 1.24278337640521e-35 | 20 |
| creg22 | 1.3008709377735e-39 | 0.316590642 | 0.192 | 0.028 | 3.27416206328213e-35 | 20 |
| dpyda.1 | 1.31332228519441e-39 | 0.250399543 | 0.131 | 0.014 | 3.30550085960581e-35 | 20 |
| fkbp71 | 1.64727253260551e-39 | 0.317120896 | 0.212 | 0.034 | 4.1460202373148e-35 | 20 |
| BX663516.21 | 2.25577683163403e-39 | 0.251477612 | 0.131 | 0.014 | 5.6775647075397e-35 | 20 |
| tspan141 | 2.52601729816794e-39 | 0.536109062 | 0.439 | 0.121 | 6.3577329377589e-35 | 20 |
| mt-cyb8 | 3.64054359246377e-39 | 0.605935411 | 0.995 | 0.619 | 9.16288416787207e-35 | 20 |
| sptan11 | 3.74168536481841e-39 | 0.393800356 | 0.258 | 0.048 | 9.41744789471145e-35 | 20 |
| sik2b1 | 4.21664223016535e-39 | 0.362960292 | 0.222 | 0.037 | 1.06128668291032e-34 | 20 |
| itgav2 | 4.40863871344653e-39 | 0.42372579 | 0.278 | 0.055 | 1.10961027778736e-34 | 20 |
| serinc12 | 1.16422632720576e-38 | 0.465431605 | 0.338 | 0.079 | 2.93024124294417e-34 | 20 |
| fbn2b2 | 1.16679526532162e-38 | 0.2927577 | 0.177 | 0.024 | 2.93670700328799e-34 | 20 |
| zbtb46 | 1.19353774903383e-38 | 0.308959699 | 0.172 | 0.023 | 3.00401516054326e-34 | 20 |
| si:dkey-6n21.12 | 1.28972663599786e-38 | 0.329783656 | 0.207 | 0.033 | 3.24611297014301e-34 | 20 |
| rbpms2a | 1.29964949342961e-38 | 0.441760827 | 0.379 | 0.088 | 3.27108781001299e-34 | 20 |
| tpm4a2 | 1.67498458393117e-38 | 0.463571545 | 0.359 | 0.085 | 4.21576869929635e-34 | 20 |
| plpp1a1 | 2.06970009937628e-38 | 0.442861695 | 0.308 | 0.066 | 5.20922818012015e-34 | 20 |
| sh2d3cb | 2.08464440390652e-38 | 0.256452282 | 0.121 | 0.012 | 5.24684150019233e-34 | 20 |
| si:dkey-101k6.5 | 2.3527631273014e-38 | 0.327306516 | 0.152 | 0.019 | 5.92166951510491e-34 | 20 |
| myo1cb | 4.2857507700763e-38 | 0.34954217 | 0.207 | 0.033 | 1.0786806113205e-33 | 20 |
| nrp2a1 | 4.6477549763705e-38 | 0.268813823 | 0.141 | 0.016 | 1.16979345000269e-33 | 20 |
| atoh81 | 5.19638395900962e-38 | 0.259137637 | 0.146 | 0.018 | 1.30787787864313e-33 | 20 |
| kmt2cb | 7.70252963280732e-38 | 0.406845887 | 0.263 | 0.051 | 1.93864968328127e-33 | 20 |
| cyp2p73 | 9.33441250386902e-38 | 0.310670562 | 0.172 | 0.024 | 2.34937828309879e-33 | 20 |
| cgnl1 | 1.13123439247775e-37 | 0.275092225 | 0.167 | 0.022 | 2.84720384242726e-33 | 20 |
| rab11fip1b1 | 1.4485531895676e-37 | 0.432193755 | 0.273 | 0.055 | 3.64586352282269e-33 | 20 |
| reep3a1 | 1.61327636071257e-37 | 0.279320312 | 0.177 | 0.025 | 4.06045527227748e-33 | 20 |
| si:dkey-188i13.104 | 1.7856132230706e-37 | 0.477592847 | 0.389 | 0.101 | 4.49420992114638e-33 | 20 |
| erbb2 | 1.79427085930383e-37 | 0.307525062 | 0.187 | 0.028 | 4.51600032578181e-33 | 20 |
| cyp2ad33 | 2.25180061907062e-37 | 0.40801209 | 0.258 | 0.05 | 5.66755697813885e-33 | 20 |
| mt-nd38 | 2.34768624203015e-37 | 0.650479085 | 0.843 | 0.39 | 5.90889150256568e-33 | 20 |
| grinab4 | 2.72610361457764e-37 | 0.504811522 | 0.404 | 0.11 | 6.86133018753047e-33 | 20 |
| ralgps2 | 3.47500478831883e-37 | 0.262515988 | 0.157 | 0.02 | 8.74623955171966e-33 | 20 |
| b2m8 | 7.36536030054841e-37 | 0.656787256 | 0.697 | 0.287 | 1.85378753404503e-32 | 20 |
| cbx7a2 | 8.29935536494347e-37 | 0.413909711 | 0.258 | 0.051 | 2.08886475180262e-32 | 20 |

| | | | | | | |
|---|---|---|---|---|---|---|
| wls3 | 1.95342782004663e-36 | 0.523242509 | 0.424 | 0.12 | 4.91658248027537e-32 | 20 |
| cyp3c13 | 2.85521709116491e-36 | 0.404210309 | 0.278 | 0.058 | 7.18629589675295e-32 | 20 |
| palm2 | 3.19305143007207e-36 | 0.252754089 | 0.136 | 0.016 | 8.03659114434839e-32 | 20 |
| hmgn66 | 3.45894808554866e-36 | 0.619347365 | 0.662 | 0.26 | 8.70582643651742e-32 | 20 |
| junbb4 | 6.94842519673571e-36 | 0.448552823 | 0.303 | 0.07 | 1.74884913776641e-31 | 20 |
| mt-co110 | 7.67343590195455e-36 | 0.556156448 | 0.995 | 0.653 | 1.93132708216294e-31 | 20 |
| cyr613 | 7.89997252202674e-36 | 0.498696875 | 0.298 | 0.066 | 1.98834408406891e-31 | 20 |
| itm2ba6 | 1.07514449520548e-35 | 0.611748358 | 0.667 | 0.268 | 2.70603117998266e-31 | 20 |
| atp10a2 | 1.46902778296608e-35 | 0.406643369 | 0.263 | 0.054 | 3.69739602694732e-31 | 20 |
| serinc52 | 1.93969275999841e-35 | 0.426150349 | 0.333 | 0.08 | 4.88201270764e-31 | 20 |
| tjp1a1 | 2.94506975793964e-35 | 0.379102411 | 0.237 | 0.045 | 7.41244607375828e-31 | 20 |
| mphosph82 | 8.39938346681183e-35 | 0.511215067 | 0.404 | 0.115 | 2.11404082476187e-30 | 20 |
| ngdn | 1.71688126445235e-34 | 0.476171143 | 0.348 | 0.09 | 4.32121845450013e-30 | 20 |
| si:rp71-79p20.21 | 3.73203933155389e-34 | 0.380479823 | 0.207 | 0.037 | 9.39316979358799e-30 | 20 |
| ier2a7 | 7.55128257884964e-34 | 0.680174079 | 0.621 | 0.258 | 1.90058231227067e-29 | 20 |
| si:ch211-74f19.22 | 8.3737263118705e-34 | 0.330448753 | 0.187 | 0.03 | 2.10758317543469e-29 | 20 |
| si:ch211-212k18.93 | 1.05153046838303e-33 | 0.307432994 | 0.202 | 0.035 | 2.64659703587324e-29 | 20 |
| btg25 | 1.60286895666394e-33 | 0.654713029 | 0.768 | 0.368 | 4.03426087702747e-29 | 20 |
| pard3bb | 1.61765053461356e-33 | 0.28724073 | 0.167 | 0.025 | 4.07146463056888e-29 | 20 |
| nid1a1 | 1.90353138142854e-33 | 0.31597119 | 0.207 | 0.036 | 4.7909981339175e-29 | 20 |
| lamc13 | 3.75914354329809e-33 | 0.390866601 | 0.253 | 0.053 | 9.46138838412696e-29 | 20 |
| sypl2a | 4.48334081045518e-33 | 0.346671215 | 0.247 | 0.05 | 1.12841204858346e-28 | 20 |
| afdna | 4.76069171221525e-33 | 0.281613827 | 0.152 | 0.021 | 1.19821849704746e-28 | 20 |
| si:ch73-237c6.15 | 4.82401706670692e-33 | 0.320166327 | 0.217 | 0.04 | 1.21415685551946e-28 | 20 |
| pdlim11 | 5.33870497336479e-33 | 0.34440919 | 0.253 | 0.052 | 1.34369865474618e-28 | 20 |
| pbx4 | 7.16556758590217e-33 | 0.404535059 | 0.288 | 0.066 | 1.80350170569572e-28 | 20 |
| si:ch73-1a9.36 | 7.7134856208903e-33 | 0.620204899 | 0.636 | 0.261 | 1.94140719592188e-28 | 20 |
| si:ch211-196l7.49 | 7.89576887161259e-33 | 0.611743642 | 0.702 | 0.309 | 1.98728606729617e-28 | 20 |
| gng73 | 1.04349777227596e-32 | 0.347020559 | 0.247 | 0.051 | 2.62637954304136e-28 | 20 |
| ccl25b3 | 1.30722236503296e-32 | 0.545771237 | 0.308 | 0.074 | 3.29014797055146e-28 | 20 |
| znf609b | 1.74145026780401e-32 | 0.276935856 | 0.152 | 0.022 | 4.38305617903592e-28 | 20 |
| pabpn1 | 2.89812389029732e-32 | 0.588267876 | 0.672 | 0.269 | 7.29428801948934e-28 | 20 |
| itgb1a3 | 3.70071858575769e-32 | 0.357521929 | 0.227 | 0.045 | 9.31433860849354e-28 | 20 |
| celf28 | 4.33895341734443e-32 | 0.584942251 | 0.611 | 0.238 | 1.09207118561142e-27 | 20 |
| zbtb41 | 5.18259144050565e-32 | 0.272240804 | 0.146 | 0.021 | 1.30440643966087e-27 | 20 |

| | | | | | | |
|---|---|---|---|---|---|---|
| h1f03 | 9.93605469411635e-32 | 0.503433831 | 0.263 | 0.06 | 2.50080560596214e-27 | 20 |
| net1 | 1.07300575468974e-31 | 0.273518658 | 0.146 | 0.021 | 2.70064818397862e-27 | 20 |
| NC-002333.49 | 1.28608056082588e-31 | 0.554489628 | 0.98 | 0.618 | 3.23693616354265e-27 | 20 |
| jak13 | 2.24244246407126e-31 | 0.476415702 | 0.338 | 0.093 | 5.64400343782096e-27 | 20 |
| zgc:66433 | 2.34244501894292e-31 | 0.279039355 | 0.172 | 0.028 | 5.89569986817743e-27 | 20 |
| hcar1-41 | 3.22644627633997e-31 | 0.306841414 | 0.192 | 0.034 | 8.12064263292007e-27 | 20 |
| si:dkey-42i9.71 | 4.03977657577451e-31 | 0.28112129 | 0.146 | 0.021 | 1.01677136635669e-26 | 20 |
| si:ch211-199g17.2 | 5.84045544976711e-31 | 0.411287686 | 0.273 | 0.064 | 1.46998423215188e-26 | 20 |
| si:ch211-166a6.5 | 7.35859598171233e-31 | 0.293607783 | 0.182 | 0.031 | 1.85208502263718e-26 | 20 |
| ctbp2a | 9.37752804827438e-31 | 0.283452467 | 0.192 | 0.034 | 2.36023003447018e-26 | 20 |
| si:ch211-132p1.35 | 1.08291998551269e-30 | 0.439471687 | 0.323 | 0.086 | 2.72560131153689e-26 | 20 |
| zgc:1748633 | 1.45172506914315e-30 | 0.428622398 | 0.354 | 0.099 | 3.65384682652638e-26 | 20 |
| serinc21 | 1.56014497448381e-30 | 0.455210992 | 0.374 | 0.109 | 3.9267288862783e-26 | 20 |
| si:ch211-15b10.64 | 2.19284315203614e-30 | 0.411285538 | 0.283 | 0.069 | 5.51916692935977e-26 | 20 |
| rsrp11 | 3.0146060152367e-30 | 0.458723469 | 0.394 | 0.116 | 7.58746187974926e-26 | 20 |
| tfe3a2 | 3.46005278023033e-30 | 0.316033834 | 0.197 | 0.037 | 8.70860684256171e-26 | 20 |
| marcksl1a4 | 6.48662161245409e-30 | 0.538483996 | 0.576 | 0.211 | 1.63261779363857e-25 | 20 |
| ier2b8 | 6.99941873767513e-30 | 0.577087715 | 0.561 | 0.22 | 1.76168370208545e-25 | 20 |
| map1lc3a3 | 9.53046751664901e-30 | 0.394225291 | 0.278 | 0.068 | 2.39872336926539e-25 | 20 |
| crebzf1 | 1.43285405588607e-29 | 0.46944331 | 0.343 | 0.099 | 3.60635037325964e-25 | 20 |
| h3f3c4 | 2.23013361750242e-29 | 0.548413144 | 0.621 | 0.252 | 5.61302330189184e-25 | 20 |
| rsu1 | 2.60258222454837e-29 | 0.340917037 | 0.212 | 0.043 | 6.5504392009658e-25 | 20 |
| si:zfos-464b6.2 | 5.57785071444106e-29 | 0.316886877 | 0.217 | 0.044 | 1.40388924631767e-24 | 20 |
| mgst1.13 | 6.4752034061185e-29 | 0.30371518 | 0.202 | 0.04 | 1.62974394528597e-24 | 20 |
| arhgef1b | 1.07527500447723e-28 | 0.396562722 | 0.278 | 0.069 | 2.70635965876874e-24 | 20 |
| ddit37 | 1.75317103451472e-28 | 0.562263266 | 0.475 | 0.173 | 4.41255617677009e-24 | 20 |
| tspan18b | 1.95658281337926e-28 | 0.313225786 | 0.187 | 0.035 | 4.92452328299426e-24 | 20 |
| igfbp71 | 2.26656235294357e-28 | 0.250726844 | 0.157 | 0.025 | 5.70471078612366e-24 | 20 |
| klf6a4 | 2.2701616353097e-28 | 0.536909549 | 0.465 | 0.164 | 5.71376981991099e-24 | 20 |
| mfap52 | 2.52796848406508e-28 | 0.328743909 | 0.212 | 0.043 | 6.36264387754339e-24 | 20 |
| ifi454 | 2.95523546244429e-28 | 0.534850989 | 0.399 | 0.126 | 7.43803213542603e-24 | 20 |
| cast1 | 3.40216644617395e-28 | 0.397540761 | 0.273 | 0.068 | 8.56291272837522e-24 | 20 |
| fermt21 | 4.60210536546364e-28 | 0.28280247 | 0.167 | 0.029 | 1.15830389943354e-23 | 20 |
| lama43 | 5.01865925722048e-28 | 0.31384616 | 0.187 | 0.035 | 1.26314634844982e-23 | 20 |
| aldh9a1a.14 | 5.11461138218996e-28 | 0.504192827 | 0.455 | 0.161 | 1.28729653878339e-23 | 20 |

| | | | | | | |
|---|---|---|---|---|---|---|
| fam168a4 | 7.64949700603886e-28 | 0.344103535 | 0.242 | 0.057 | 1.92530190144992e-23 | 20 |
| stat1a3 | 1.05066066323124e-27 | 0.456390304 | 0.374 | 0.117 | 2.64440782328672e-23 | 20 |
| selenow12 | 1.12636327674969e-27 | 0.574971132 | 0.753 | 0.381 | 2.8349437312513e-23 | 20 |
| sh3pxd2b3 | 1.15570529166562e-27 | 0.341164081 | 0.207 | 0.043 | 2.90879464859319e-23 | 20 |
| anxa45 | 1.19584501355948e-27 | 0.440611236 | 0.409 | 0.133 | 3.00982231462784e-23 | 20 |
| oip5-as1 | 1.54169047338921e-27 | 0.256531207 | 0.157 | 0.026 | 3.88028075247331e-23 | 20 |
| si:ch1073-291c23.23 | 2.15227946003817e-27 | 0.331297713 | 0.253 | 0.059 | 5.41707217297008e-23 | 20 |
| ppfibp1a | 3.19865564574487e-27 | 0.280117917 | 0.207 | 0.042 | 8.05069639477527e-23 | 20 |
| arhgef41 | 3.8766967771018e-27 | 0.274900756 | 0.177 | 0.033 | 9.75725811828751e-23 | 20 |
| wwtr1 | 4.44638119396492e-27 | 0.257682282 | 0.146 | 0.024 | 1.11910968270903e-22 | 20 |
| tp53inp11 | 4.47017556551233e-27 | 0.35212019 | 0.232 | 0.053 | 1.1250984880838e-22 | 20 |
| tmsb4x11 | 6.78866557441442e-27 | 0.546431167 | 0.99 | 0.683 | 1.70863923842437e-22 | 20 |
| flna4 | 8.08765175452954e-27 | 0.425935527 | 0.364 | 0.112 | 2.03558107009754e-22 | 20 |
| si:ch73-335l21.43 | 1.17095372191513e-26 | 0.639905958 | 0.414 | 0.146 | 2.94717342268818e-22 | 20 |
| ulk22 | 1.5075195514831e-26 | 0.342129815 | 0.212 | 0.047 | 3.7942759591278e-22 | 20 |
| hmgb1b3 | 1.73625277968225e-26 | 0.445394926 | 0.384 | 0.124 | 4.36997462118225e-22 | 20 |
| si:dkey-23i12.72 | 2.3835850367654e-26 | 0.315207171 | 0.167 | 0.03 | 5.99924517903483e-22 | 20 |
| krt18b7 | 3.55764886363873e-26 | 0.444220254 | 0.444 | 0.147 | 8.95424642489233e-22 | 20 |
| nfia4 | 6.30470051444292e-26 | 0.420575014 | 0.283 | 0.078 | 1.58683007248014e-21 | 20 |
| stk35l | 1.18220061886679e-25 | 0.282327741 | 0.177 | 0.035 | 2.97548073762583e-21 | 20 |
| ecm1b1 | 1.98659554206663e-25 | 0.309903785 | 0.227 | 0.053 | 5.00006231982751e-21 | 20 |
| atf4b6 | 1.9904875497301e-25 | 0.517107913 | 0.606 | 0.26 | 5.00985811391568e-21 | 20 |
| dhrs13l11 | 2.10092590822628e-25 | 0.259400511 | 0.167 | 0.032 | 5.28782041841472e-21 | 20 |
| map1lc3b3 | 2.94495264393784e-25 | 0.392440215 | 0.338 | 0.104 | 7.41215130952714e-21 | 20 |
| socs3b5 | 5.2990052730871e-25 | 0.5172529 | 0.424 | 0.153 | 1.33370663718329e-20 | 20 |
| si:ch211-286b5.52 | 6.21354728115425e-25 | 0.409563876 | 0.399 | 0.134 | 1.56388771519371e-20 | 20 |
| sdcbp25 | 6.65906000533149e-25 | 0.447023445 | 0.53 | 0.205 | 1.67601881274188e-20 | 20 |
| SBNO21 | 9.10583732269355e-25 | 0.29700109 | 0.167 | 0.032 | 2.29184819574874e-20 | 20 |
| pi4kb1 | 1.68153537059267e-24 | 0.278372715 | 0.177 | 0.036 | 4.2322563742447e-20 | 20 |
| tefa3 | 2.3900107216406e-24 | 0.432766078 | 0.394 | 0.135 | 6.01541798529724e-20 | 20 |
| fbln51 | 2.87730263295225e-24 | 0.253364871 | 0.182 | 0.037 | 7.24188299687751e-20 | 20 |
| rnd3b3 | 3.48082682259517e-24 | 0.296589249 | 0.187 | 0.04 | 8.76089302978979e-20 | 20 |
| CR847566.12 | 3.63491440451277e-24 | 0.274801161 | 0.172 | 0.034 | 9.1487160647182e-20 | 20 |
| meis1b3 | 3.67808979433579e-24 | 0.312423875 | 0.227 | 0.055 | 9.25738420336375e-20 | 20 |
| irf2bp2b2 | 3.99302640723151e-24 | 0.365283362 | 0.283 | 0.08 | 1.0050048164361e-19 | 20 |

| | | | | | | |
|---|---|---|---|---|---|---|
| cflara1 | 4.30631909161756e-24 | 0.308265772 | 0.242 | 0.061 | 1.08385745216922e-19 | 20 |
| dedd13 | 4.74942017682479e-24 | 0.482975688 | 0.399 | 0.143 | 1.19538156430503e-19 | 20 |
| CR318588.18 | 5.83038142554195e-24 | 0.674179798 | 0.495 | 0.205 | 1.46744870099465e-19 | 20 |
| timp2a3 | 6.0155956715847e-24 | 0.314698076 | 0.227 | 0.056 | 1.51406527458115e-19 | 20 |
| gpx1a4 | 6.54499637751636e-24 | 0.456782974 | 0.561 | 0.22 | 1.64731013825709e-19 | 20 |
| fundc21 | 7.14316090630471e-24 | 0.346810826 | 0.298 | 0.087 | 1.79786216850783e-19 | 20 |
| qkia3 | 8.32284623122623e-24 | 0.453430925 | 0.444 | 0.165 | 2.09477716793733e-19 | 20 |
| ca5a1 | 8.67655002937511e-24 | 0.250568757 | 0.157 | 0.03 | 2.18380087689342e-19 | 20 |
| anxa5b3 | 1.00278236827187e-23 | 0.374631914 | 0.308 | 0.092 | 2.52390294270346e-19 | 20 |
| klf11b5 | 1.92159324916807e-23 | 0.379795852 | 0.273 | 0.078 | 4.83645804883111e-19 | 20 |
| cebpb8 | 3.71040389743794e-23 | 0.563911475 | 0.747 | 0.349 | 9.33871556946156e-19 | 20 |
| ppap2d3 | 5.06773102142198e-23 | 0.265419383 | 0.172 | 0.035 | 1.2754972207817e-18 | 20 |
| zmiz1a | 5.58285169062038e-23 | 0.264562821 | 0.177 | 0.037 | 1.40514794201224e-18 | 20 |
| map2k61 | 8.22237519195122e-23 | 0.351703259 | 0.293 | 0.087 | 2.0694896120622e-18 | 20 |
| si:ch211-133n4.43 | 9.07414996754402e-23 | 0.275845363 | 0.192 | 0.044 | 2.28387280533115e-18 | 20 |
| scara31 | 9.16632627145091e-23 | 0.25689918 | 0.157 | 0.03 | 2.30707265926148e-18 | 20 |
| tmem176l.3a | 1.06415643932496e-22 | 0.269346585 | 0.177 | 0.038 | 2.67837534213699e-18 | 20 |
| boc3 | 1.35873734446819e-22 | 0.295700189 | 0.202 | 0.047 | 3.41980602229199e-18 | 20 |
| esr2b1 | 1.73456424894304e-22 | 0.270025772 | 0.167 | 0.035 | 4.36572475816473e-18 | 20 |
| mvp2 | 1.74904396314368e-22 | 0.289880085 | 0.212 | 0.052 | 4.40216875083633e-18 | 20 |
| nupr1b2 | 1.79088614722916e-22 | 0.352612938 | 0.197 | 0.047 | 4.50748134396107e-18 | 20 |
| pnrc211 | 2.92145635711586e-22 | 0.489097393 | 0.899 | 0.564 | 7.3530135052249e-18 | 20 |
| si:dkey-17m8.1 | 3.48776700026442e-22 | 0.264001918 | 0.177 | 0.038 | 8.77836076296553e-18 | 20 |
| appbp21 | 4.81280857893078e-22 | 0.29123613 | 0.182 | 0.041 | 1.21133579123109e-17 | 20 |
| cnn3a2 | 5.22940091296157e-22 | 0.311451404 | 0.217 | 0.055 | 1.3161879157833e-17 | 20 |
| pdk2b5 | 5.84548281624534e-22 | 0.390084975 | 0.308 | 0.098 | 1.47124957002079e-17 | 20 |
| zbtb204 | 8.05569916413416e-22 | 0.393950372 | 0.303 | 0.096 | 2.02753892262093e-17 | 20 |
| rnf19a | 9.79156767839452e-22 | 0.253517155 | 0.192 | 0.044 | 2.46443966897512e-17 | 20 |
| emd1 | 1.15639340160725e-21 | 0.297253201 | 0.212 | 0.054 | 2.91052655250529e-17 | 20 |
| si:ch1073-459j12.13 | 1.2929326830758e-21 | 0.309696122 | 0.242 | 0.065 | 3.25418227003349e-17 | 20 |
| gstm.12 | 1.34103702652613e-21 | 0.466525306 | 0.596 | 0.257 | 3.37525609206361e-17 | 20 |
| nr3c1 | 1.62394013640427e-21 | 0.267944769 | 0.182 | 0.041 | 4.0872949293159e-17 | 20 |
| wu:fj16a033 | 1.77330987257546e-21 | 0.330053627 | 0.207 | 0.05 | 4.46324361828518e-17 | 20 |
| cebpg | 2.69524911711119e-21 | 0.305419579 | 0.232 | 0.062 | 6.78367250285715e-17 | 20 |
| arl4d | 3.85340258041413e-21 | 0.266873787 | 0.202 | 0.049 | 9.69862895464432e-17 | 20 |

| | | | | | | |
|---|---|---|---|---|---|---|
| hspe11 | 4.81975887192515e-21 | 0.485933826 | 0.646 | 0.29 | 1.21308511047484e-16 | 20 |
| serpinf12 | 7.79380943167654e-21 | 0.277274291 | 0.182 | 0.042 | 1.96162389585867e-16 | 20 |
| anxa133 | 8.75260762861286e-21 | 0.274385034 | 0.197 | 0.049 | 2.20294381404557e-16 | 20 |
| osr23 | 8.96158365060415e-21 | 0.316188356 | 0.187 | 0.045 | 2.25554098902056e-16 | 20 |
| jdp2b4 | 9.93981737725225e-21 | 0.507377334 | 0.424 | 0.168 | 2.50175263568062e-16 | 20 |
| cpn12 | 1.23680441915419e-20 | 0.268790687 | 0.212 | 0.054 | 3.11291304256919e-16 | 20 |
| wu:fi37e091 | 1.56064994024306e-20 | 0.342037922 | 0.343 | 0.116 | 3.92799983459775e-16 | 20 |
| crim13 | 1.59434963024493e-20 | 0.268608835 | 0.162 | 0.035 | 4.01281858436348e-16 | 20 |
| stk351 | 1.71288417749984e-20 | 0.314310529 | 0.207 | 0.054 | 4.31115818634935e-16 | 20 |
| XLOC-043861 | 1.89833726514873e-20 | 0.277161508 | 0.192 | 0.047 | 4.77792506265284e-16 | 20 |
| ccng23 | 1.96896547582805e-20 | 0.345565405 | 0.273 | 0.084 | 4.95568920611161e-16 | 20 |
| ier52 | 2.51558303335877e-20 | 0.413815548 | 0.364 | 0.134 | 6.33147093666068e-16 | 20 |
| ndfip1l3 | 2.62511670314633e-20 | 0.333642718 | 0.232 | 0.065 | 6.60715623014901e-16 | 20 |
| klf2a3 | 4.42870081803972e-20 | 0.363417372 | 0.273 | 0.084 | 1.11465970889242e-15 | 20 |
| tnfrsf1a2 | 6.91446677355378e-20 | 0.295388079 | 0.202 | 0.053 | 1.74030214223575e-15 | 20 |
| zgc:1132084 | 7.36128370204157e-20 | 0.268375831 | 0.182 | 0.044 | 1.85276149496684e-15 | 20 |
| ppp1cb | 9.64329189731992e-20 | 0.293576339 | 0.227 | 0.063 | 2.42712013763645e-15 | 20 |
| vdrb1 | 1.57818140137759e-19 | 0.262491401 | 0.157 | 0.035 | 3.97212476912726e-15 | 20 |
| ak11 | 1.59454245421158e-19 | 0.305389145 | 0.247 | 0.074 | 4.01330390300513e-15 | 20 |
| nckap1 | 1.61616493296131e-19 | 0.252652635 | 0.172 | 0.04 | 4.06772551977032e-15 | 20 |
| tax1bp31 | 1.88004278286658e-19 | 0.30815751 | 0.278 | 0.087 | 4.73187968019691e-15 | 20 |
| ube2e33 | 2.34219180301602e-19 | 0.392918218 | 0.394 | 0.153 | 5.89506254901102e-15 | 20 |
| CABZ01072614.14 | 2.53837525234573e-19 | 0.335091734 | 0.258 | 0.079 | 6.38883667262897e-15 | 20 |
| rhoab6 | 4.73619065706648e-19 | 0.420815996 | 0.444 | 0.187 | 1.19205182647706e-14 | 20 |
| ponzr15 | 4.82491290597349e-19 | 0.449811546 | 0.596 | 0.27 | 1.21438232930447e-14 | 20 |
| macf1a | 4.98712935107721e-19 | 0.286245806 | 0.197 | 0.052 | 1.25521058637262e-14 | 20 |
| hopx1 | 6.29891718178462e-19 | 0.283599866 | 0.227 | 0.064 | 1.58537446548337e-14 | 20 |
| si:dkey-51e6.13 | 6.44521337428478e-19 | 0.352037698 | 0.323 | 0.114 | 1.62219575417374e-14 | 20 |
| CABZ01068358.14 | 7.65403933385479e-19 | 0.315505706 | 0.222 | 0.063 | 1.92644515993791e-14 | 20 |
| ppdpfb5 | 9.03498935644961e-19 | 0.451661885 | 0.808 | 0.516 | 2.2740164711248e-14 | 20 |
| sdc45 | 9.19160257616983e-19 | 0.445451385 | 0.576 | 0.254 | 2.31343445239619e-14 | 20 |
| wu:fb16f032 | 1.48943561329102e-18 | 0.323360983 | 0.268 | 0.086 | 3.74876049509217e-14 | 20 |
| flot2a3 | 2.10502664632005e-18 | 0.289306532 | 0.207 | 0.058 | 5.29814156612294e-14 | 20 |
| ecrg4a | 2.44802228107253e-18 | 0.258294468 | 0.141 | 0.03 | 6.16142727923144e-14 | 20 |
| capns1b2 | 3.35448513044186e-18 | 0.349871061 | 0.328 | 0.12 | 8.44290362480912e-14 | 20 |

| | | | | | | |
|---|---|---|---|---|---|---|
| yap12 | 4.23606197447738e-18 | 0.263252378 | 0.187 | 0.049 | 1.06617443835621e-13 | 20 |
| csrp1a1 | 4.47951145952394e-18 | 0.263405565 | 0.227 | 0.066 | 1.12744823924758e-13 | 20 |
| rhoaa5 | 5.57095002831336e-18 | 0.381869474 | 0.379 | 0.152 | 1.40215241262619e-13 | 20 |
| ctnnb11 | 6.30425737410633e-18 | 0.351231173 | 0.384 | 0.148 | 1.58671853848882e-13 | 20 |
| pik3r3a1 | 7.11493557514881e-18 | 0.296906468 | 0.258 | 0.081 | 1.7907581349092e-13 | 20 |
| slc7a13 | 7.68680377257637e-18 | 0.259485212 | 0.157 | 0.037 | 1.93469164151975e-13 | 20 |
| gmfb | 9.17765669372179e-18 | 0.281784026 | 0.298 | 0.1 | 2.30992441324284e-13 | 20 |
| mafba4 | 1.12155190864113e-17 | 0.314447766 | 0.232 | 0.071 | 2.82283399885886e-13 | 20 |
| adamts15a3 | 1.25909439517522e-17 | 0.292978232 | 0.182 | 0.047 | 3.1690146832165e-13 | 20 |
| rbm39a3 | 1.33923866531265e-17 | 0.383351676 | 0.359 | 0.141 | 3.37072979672541e-13 | 20 |
| si:ch211-213a13.11 | 2.3994826877516e-17 | 0.303429231 | 0.263 | 0.087 | 6.03925797680201e-13 | 20 |
| nt5dc23 | 2.65407671017076e-17 | 0.300730944 | 0.242 | 0.076 | 6.68004567182879e-13 | 20 |
| rnf213a5 | 3.08765298212487e-17 | 0.323798194 | 0.263 | 0.087 | 7.7713137907101e-13 | 20 |
| tagln26 | 3.15104764478782e-17 | 0.37709387 | 0.444 | 0.186 | 7.93087181716647e-13 | 20 |
| si:ch211-214p16.23 | 5.32630471831752e-17 | 0.269085985 | 0.237 | 0.071 | 1.34057763455334e-12 | 20 |
| si:ch73-376l24.2 | 5.3280061706338e-17 | 0.353554342 | 0.313 | 0.114 | 1.34100587308682e-12 | 20 |
| zgc:11418810 | 7.69023853508434e-17 | 0.400465923 | 0.894 | 0.625 | 1.93555613689538e-12 | 20 |
| ppib6 | 1.17466329944675e-16 | 0.441306227 | 0.636 | 0.33 | 2.95651005837754e-12 | 20 |
| capn2a1 | 1.20829783448181e-16 | 0.26717008 | 0.187 | 0.052 | 3.04116481960727e-12 | 20 |
| irf22 | 1.27524360591687e-16 | 0.300328001 | 0.217 | 0.067 | 3.20966063173217e-12 | 20 |
| fxyd14 | 1.27905217535631e-16 | 0.277093685 | 0.247 | 0.08 | 3.2192464201543e-12 | 20 |
| mt-nd53 | 2.72119038623359e-16 | 0.337853694 | 0.313 | 0.118 | 6.84896408311132e-12 | 20 |
| psme25 | 5.31469712562488e-16 | 0.289953363 | 0.273 | 0.096 | 1.33765611954853e-11 | 20 |
| tsc22d34 | 5.40470275977991e-16 | 0.372680179 | 0.348 | 0.142 | 1.360309637609e-11 | 20 |
| tnrc6a | 6.33308655955922e-16 | 0.273236006 | 0.227 | 0.072 | 1.59397455617546e-11 | 20 |
| rerea2 | 7.61771558778637e-16 | 0.255284757 | 0.182 | 0.051 | 1.91730283628995e-11 | 20 |
| si:ch211-260e23.91 | 1.10463747204771e-15 | 0.273666577 | 0.177 | 0.05 | 2.78026205339689e-11 | 20 |
| acin1a3 | 1.74778292700294e-15 | 0.369375073 | 0.47 | 0.214 | 4.39899484897369e-11 | 20 |
| abca1a2 | 1.98348397734043e-15 | 0.253185855 | 0.217 | 0.067 | 4.99223082256813e-11 | 20 |
| stau24 | 1.99173845258576e-15 | 0.282530707 | 0.247 | 0.084 | 5.01300651131309e-11 | 20 |
| XLOC-0247502 | 2.3415897797077e-15 | 0.290451527 | 0.227 | 0.074 | 5.8935473165463e-11 | 20 |
| lrpap1 | 3.1261526005668e-15 | 0.257965589 | 0.187 | 0.055 | 7.86821348036658e-11 | 20 |
| slc3a2b4 | 3.26391129691915e-15 | 0.330087142 | 0.303 | 0.117 | 8.21493834321582e-11 | 20 |
| aplp22 | 3.72523954551865e-15 | 0.263757444 | 0.222 | 0.071 | 9.37605541211589e-11 | 20 |
| notch25 | 4.93969954301193e-15 | 0.326793029 | 0.303 | 0.118 | 1.24327297798067e-10 | 20 |

| | | | | | | |
|---|---|---|---|---|---|---|
| mycb3 | 5.69204633877578e-15 | 0.366268219 | 0.47 | 0.209 | 1.43263114300648e-10 | 20 |
| ptp4a11 | 6.82961496442546e-15 | 0.300164133 | 0.303 | 0.116 | 1.71894579039624e-10 | 20 |
| capns1a1 | 7.27883351401378e-15 | 0.317872081 | 0.348 | 0.142 | 1.83200960714213e-10 | 20 |
| jpt1b3 | 1.20279901034952e-14 | 0.357593796 | 0.566 | 0.268 | 3.02732482914871e-10 | 20 |
| gpatch8 | 3.82098684678029e-14 | 0.266386839 | 0.177 | 0.053 | 9.6170417946613e-10 | 20 |
| rplp2l10 | 4.13569326064117e-14 | 0.301981736 | 0.949 | 0.772 | 1.04091263677078e-09 | 20 |
| hif1al1 | 5.40339785471053e-14 | 0.302871677 | 0.313 | 0.125 | 1.35998120605209e-09 | 20 |
| stat24 | 6.02525872056803e-14 | 0.254032831 | 0.192 | 0.06 | 1.51649736737977e-09 | 20 |
| srsf5a3 | 9.34245265315688e-14 | 0.353785348 | 0.566 | 0.286 | 2.35140190827305e-09 | 20 |
| dynlt3 | 9.68565819250356e-14 | 0.295963595 | 0.343 | 0.145 | 2.43778331047122e-09 | 20 |
| rps289 | 1.00125701647213e-13 | 0.253978477 | 0.96 | 0.825 | 2.52006378475871e-09 | 20 |
| mgst3b2 | 1.31932780156966e-13 | 0.315643379 | 0.384 | 0.171 | 3.32061614377068e-09 | 20 |
| sumo2b7 | 1.3431042796502e-13 | 0.347935975 | 0.535 | 0.267 | 3.38045916145158e-09 | 20 |
| ubb1 | 1.80104836052929e-13 | 0.365423182 | 0.601 | 0.313 | 4.53305861861617e-09 | 20 |
| prkab1b1 | 2.16919101068495e-13 | 0.278941712 | 0.298 | 0.12 | 5.45963685479295e-09 | 20 |
| atrx3 | 2.37406614398197e-13 | 0.29238174 | 0.268 | 0.104 | 5.97528707778821e-09 | 20 |
| snx171 | 2.50102515997265e-13 | 0.262275787 | 0.182 | 0.058 | 6.29483022513516e-09 | 20 |
| jund4 | 3.15245829400984e-13 | 0.279139558 | 0.247 | 0.092 | 7.93442228019337e-09 | 20 |
| rps138 | 3.98020953462358e-13 | 0.278997132 | 0.889 | 0.723 | 1.00177893776941e-08 | 20 |
| rab5aa2 | 4.57772109323092e-13 | 0.265164061 | 0.217 | 0.076 | 1.15216662195529e-08 | 20 |
| fkbp1aa4 | 8.79250306650651e-13 | 0.377533036 | 0.672 | 0.385 | 2.21298509680902e-08 | 20 |
| smdt1b2 | 1.4208470625242e-12 | 0.266086802 | 0.313 | 0.132 | 3.57612997166715e-08 | 20 |
| tpm13 | 1.86375908612445e-12 | 0.35114888 | 0.429 | 0.205 | 4.69089524386662e-08 | 20 |
| tomm54 | 2.11651723940319e-12 | 0.306683679 | 0.439 | 0.212 | 5.32706223985389e-08 | 20 |
| gapdhs9 | 2.19356552557848e-12 | 0.343601088 | 0.561 | 0.286 | 5.52098507132849e-08 | 20 |
| si:dkey-42i9.42 | 2.68871040901436e-12 | 0.313096154 | 0.328 | 0.146 | 6.76721522844825e-08 | 20 |
| fam120c1 | 5.03030816143792e-12 | 0.266593855 | 0.247 | 0.098 | 1.26607826115231e-07 | 20 |
| plk31 | 5.23682608349942e-12 | 0.251222489 | 0.298 | 0.119 | 1.31805675695597e-07 | 20 |
| pim15 | 5.306389188481e-12 | 0.330808177 | 0.369 | 0.17 | 1.33556509484878e-07 | 20 |
| ywhaqb3 | 5.68925684005591e-12 | 0.343488957 | 0.293 | 0.131 | 1.43192905407367e-07 | 20 |
| eif1b5 | 7.06173136939138e-12 | 0.359821501 | 0.667 | 0.38 | 1.77736716836212e-07 | 20 |
| pfn2l2 | 8.54458581658755e-12 | 0.278498369 | 0.444 | 0.207 | 2.15058680417692e-07 | 20 |
| sh3bgrl4 | 1.33058314481843e-11 | 0.253442671 | 0.258 | 0.104 | 3.3489447171935e-07 | 20 |
| btg19 | 1.44740851347245e-11 | 0.336112839 | 0.616 | 0.33 | 3.64298248755882e-07 | 20 |
| si:ch211-191j22.3 | 1.6153130686562e-11 | 0.25195244 | 0.273 | 0.113 | 4.06558146250079e-07 | 20 |

| | | | | | | |
|---|---|---|---|---|---|---|
| atp1b1a3 | 2.2146488553624e-11 | 0.320375899 | 0.424 | 0.212 | 5.57404970406163e-07 | 20 |
| hmgb1a6 | 2.3762854764307e-11 | 0.323707839 | 0.379 | 0.188 | 5.98087291562844e-07 | 20 |
| rplp210 | 2.97098642621842e-11 | 0.306616279 | 0.955 | 0.756 | 7.47767573614915e-07 | 20 |
| zgc:1119862 | 3.18469334612204e-11 | 0.291991123 | 0.369 | 0.175 | 8.01555468285456e-07 | 20 |
| rps27l6 | 3.50319733526076e-11 | 0.299796233 | 0.409 | 0.202 | 8.81719737311781e-07 | 20 |
| si:ch211-214j24.10 | 3.7168912138867e-11 | 0.255836365 | 0.348 | 0.157 | 9.35504349623143e-07 | 20 |
| antxr2a6 | 4.99101615619911e-11 | 0.265811494 | 0.338 | 0.153 | 1.25618885635376e-06 | 20 |
| glula3 | 5.55120613739443e-11 | 0.258084985 | 0.313 | 0.134 | 1.3971830727208e-06 | 20 |
| rps249 | 5.9383937805385e-11 | 0.257928556 | 0.924 | 0.757 | 1.49463433062373e-06 | 20 |
| cldn11a3 | 7.87025135556469e-11 | 0.268188671 | 0.177 | 0.062 | 1.98086356368208e-06 | 20 |
| dbi3 | 8.68589950360065e-11 | 0.297863604 | 0.419 | 0.212 | 2.18615404606125e-06 | 20 |
| calm1a | 1.06107669182e-10 | 0.259531751 | 0.354 | 0.163 | 2.67062392564175e-06 | 20 |
| hnrnpa0a1 | 1.12417494861163e-10 | 0.254660597 | 0.187 | 0.069 | 2.82943592816062e-06 | 20 |
| ciarta3 | 1.31367923174097e-10 | 0.257951175 | 0.247 | 0.104 | 3.30639925836885e-06 | 20 |
| nfkbiaa5 | 1.46999726467734e-10 | 0.287341681 | 0.328 | 0.151 | 3.6998361154664e-06 | 20 |
| tpm32 | 1.60198875884819e-10 | 0.310488164 | 0.475 | 0.253 | 4.032045507145e-06 | 20 |
| rpl3610 | 1.8059258514904e-10 | 0.261295687 | 0.944 | 0.788 | 4.54533477561618e-06 | 20 |
| chac14 | 1.93134952112162e-10 | 0.335595754 | 0.677 | 0.396 | 4.86101360971101e-06 | 20 |
| higd1a6 | 2.13904849337976e-10 | 0.317658317 | 0.566 | 0.307 | 5.38377115298752e-06 | 20 |
| pomp2 | 2.86100115350008e-10 | 0.281291203 | 0.444 | 0.229 | 7.20085380324435e-06 | 20 |
| rab102 | 2.91690175889171e-10 | 0.291498902 | 0.384 | 0.194 | 7.34155003695454e-06 | 20 |
| sept24 | 4.34265773805359e-10 | 0.265281491 | 0.343 | 0.164 | 1.09300352609071e-05 | 20 |
| anxa11b2 | 4.61907499077231e-10 | 0.250572492 | 0.318 | 0.149 | 1.16257498442748e-05 | 20 |
| capza1a6 | 4.69665530392597e-10 | 0.274717884 | 0.394 | 0.198 | 1.18210117344513e-05 | 20 |
| rap1b5 | 5.37156560114321e-10 | 0.259338255 | 0.404 | 0.201 | 1.35196934615173e-05 | 20 |
| fabp33 | 6.59916564193918e-10 | 0.25863449 | 0.475 | 0.24 | 1.66094400041967e-05 | 20 |
| bhlhe405 | 6.99427377615734e-10 | 0.283388217 | 0.333 | 0.16 | 1.76038876672104e-05 | 20 |
| tegt5 | 7.39940524678041e-10 | 0.256900586 | 0.414 | 0.21 | 1.86235630656216e-05 | 20 |
| ndufa4l5 | 7.5158527979878e-10 | 0.328262528 | 0.652 | 0.393 | 1.89166499072555e-05 | 20 |
| fosb3 | 7.85677348642877e-10 | 0.29772862 | 0.278 | 0.125 | 1.97747131879926e-05 | 20 |
| tmem2582 | 7.95830121192118e-10 | 0.293806678 | 0.51 | 0.281 | 2.00302483202844e-05 | 20 |
| ywhabl2 | 8.29236629472143e-10 | 0.284252337 | 0.515 | 0.276 | 2.08710567271844e-05 | 20 |
| tnrc6c16 | 9.34901292575033e-10 | 0.254401623 | 0.288 | 0.132 | 2.3530530632821e-05 | 20 |
| zgc:1538673 | 1.18910812701459e-09 | 0.328402702 | 0.707 | 0.434 | 2.99286624488303e-05 | 20 |
| eef1db6 | 1.25915221184246e-09 | 0.295474964 | 0.5 | 0.272 | 3.16916020198628e-05 | 20 |

| | | | | | | |
|---|---|---|---|---|---|---|
| ptmab7 | 1.56932502857643e-09 | 0.33620627 | 0.813 | 0.537 | 3.94983416442401e-05 | 20 |
| cd6310 | 1.85063455424377e-09 | 0.309513736 | 0.591 | 0.338 | 4.65786210957616e-05 | 20 |
| rpl910 | 1.88444828051683e-09 | 0.346452693 | 0.899 | 0.626 | 4.7429678772328e-05 | 20 |
| hmgn74 | 6.50714376832002e-09 | 0.286129842 | 0.495 | 0.278 | 0.000163778 | 20 |
| zgc:1584632 | 6.59416296633332e-09 | 0.271236718 | 0.182 | 0.074 | 0.000165968 | 20 |
| hmgn24 | 9.96822017543687e-09 | 0.282247841 | 0.717 | 0.44 | 0.00025089 | 20 |
| rps2113 | 1.00398600781829e-08 | 0.268949517 | 0.934 | 0.748 | 0.000252693 | 20 |
| LOC103910438 | 1.11154928295928e-08 | 0.269163479 | 0.323 | 0.164 | 0.000279766 | 20 |
| zgc:927445 | 1.51554227283096e-08 | 0.28469073 | 0.606 | 0.351 | 0.000381447 | 20 |
| rps189 | 6.37364926640094e-08 | 0.2669565 | 0.929 | 0.723 | 0.001604184 | 20 |
| glrx5 | 6.59880420187022e-08 | 0.254959171 | 0.54 | 0.31 | 0.001660853 | 20 |
| gnb1b2 | 9.47300323415183e-08 | 0.258020179 | 0.439 | 0.253 | 0.00238426 | 20 |
| rps109 | 1.18310134972112e-07 | 0.284207088 | 0.909 | 0.692 | 0.002977748 | 20 |
| rpl149 | 1.68895846485459e-07 | 0.297387954 | 0.889 | 0.632 | 0.00425094 | 20 |
| oaz1b8 | 1.7593782894939e-07 | 0.263855134 | 0.535 | 0.321 | 0.004428179 | 20 |
| rps27.29 | 1.93536516812139e-07 | 0.303178703 | 0.742 | 0.516 | 0.004871121 | 20 |
| ptmaa8 | 2.21557614176709e-07 | 0.292058923 | 0.662 | 0.393 | 0.005576384 | 20 |
| rpsa10 | 2.50827639599448e-07 | 0.27046086 | 0.955 | 0.75 | 0.006313081 | 20 |
| serbp1a8 | 4.47123020643355e-07 | 0.303018071 | 0.742 | 0.48 | 0.011253639 | 20 |
| ucp26 | 6.45169551755033e-07 | 0.27650833 | 0.641 | 0.415 | 0.016238272 | 20 |
| rpl22l16 | 9.06784319925625e-07 | 0.275445687 | 0.646 | 0.389 | 0.022822855 | 20 |
| eef1da6 | 1.40792510400357e-06 | 0.276821116 | 0.712 | 0.491 | 0.035436067 | 20 |
| rpl23a9 | 1.61079626909319e-06 | 0.277823775 | 0.854 | 0.632 | 0.040542131 | 20 |
| rpl3111 | 2.36923016346495e-06 | 0.268609484 | 0.879 | 0.652 | 0.059631154 | 20 |
| rpl27a9 | 9.8055782512018e-06 | 0.271194186 | 0.707 | 0.494 | 0.246796599 | 20 |
| pabpc1a8 | 1.0922614523979e-05 | 0.27776203 | 0.803 | 0.537 | 0.274911285 | 20 |
| uba528 | 1.11011822148591e-05 | 0.277391248 | 0.874 | 0.628 | 0.279405655 | 20 |
| rpl5a8 | 1.88550105705097e-05 | 0.279485217 | 0.763 | 0.489 | 0.474561761 | 20 |
| si:ch73-167f10.1 | 0 | 1.145359654 | 0.513 | 0.007 | 0 | 21 |
| sycp1 | 0 | 0.971999665 | 0.383 | 0.007 | 0 | 21 |
| stmn1b | 0 | 0.852259939 | 0.389 | 0.007 | 0 | 21 |
| si:dkey-40g16.5 | 0 | 0.832727355 | 0.326 | 0.004 | 0 | 21 |
| tuba7l | 0 | 0.762951565 | 0.321 | 0.003 | 0 | 21 |
| smc1b | 0 | 0.518129739 | 0.212 | 0.002 | 0 | 21 |
| LOC100149647 | 0 | 0.489531919 | 0.176 | 0 | 0 | 21 |

| | | | | | | |
|---|---|---|---|---|---|---|
| CU929391.1 | 0 | 0.432161766 | 0.176 | 0.001 | 0 | 21 |
| hsp90aa1.21 | 4.50216504435428e-290 | 1.180698268 | 0.58 | 0.035 | 1.13314992001353e-285 | 21 |
| hormad1 | 1.78050462853909e-273 | 0.630200535 | 0.269 | 0.006 | 4.48135209957004e-269 | 21 |
| sycp3 | 8.54862010453324e-271 | 0.666073359 | 0.254 | 0.005 | 2.15160219410997e-266 | 21 |
| ddx42 | 4.69691299302171e-214 | 1.477933174 | 0.793 | 0.105 | 1.18216603121363e-209 | 21 |
| hnrnpa3 | 2.21917781553848e-208 | 0.506843071 | 0.207 | 0.005 | 5.5854486439288e-204 | 21 |
| si:ch211-176l24.4 | 7.92148976346129e-184 | 0.530719859 | 0.192 | 0.005 | 1.99375975856557e-179 | 21 |
| rnf212b | 1.83357883555851e-183 | 0.307386965 | 0.114 | 0.001 | 4.61493457121722e-179 | 21 |
| pcp4l1 | 2.66378158712644e-183 | 0.455455343 | 0.187 | 0.004 | 6.70447187663853e-179 | 21 |
| stag3 | 3.57064067445587e-179 | 0.548233212 | 0.238 | 0.008 | 8.98694551353797e-175 | 21 |
| LOC101886261 | 5.9705603443606e-167 | 0.334175751 | 0.119 | 0.001 | 1.50273033307212e-162 | 21 |
| parp1 | 2.91411948903263e-164 | 0.784498667 | 0.358 | 0.024 | 7.33454734194622e-160 | 21 |
| CCNB1IP1 | 4.47296808151715e-147 | 0.342535942 | 0.145 | 0.003 | 1.12580133643705e-142 | 21 |
| si:ch211-241j8.2 | 9.61049461789866e-147 | 0.366279174 | 0.135 | 0.003 | 2.41886539037891e-142 | 21 |
| ddx39b | 1.62833484679128e-143 | 0.691682762 | 0.332 | 0.023 | 4.09835597588898e-139 | 21 |
| cox5b2 | 1.47204076977744e-142 | 0.496001244 | 0.218 | 0.009 | 3.70497941345284e-138 | 21 |
| zgc:193690 | 4.75793654214771e-140 | 0.750407549 | 0.378 | 0.031 | 1.19752504829316e-135 | 21 |
| zte38 | 9.12759024553686e-139 | 0.314993869 | 0.109 | 0.002 | 2.29732318889917e-134 | 21 |
| spata22 | 1.87529454349308e-130 | 0.402890626 | 0.176 | 0.006 | 4.71992883651773e-126 | 21 |
| tdrd1 | 3.63822766859972e-123 | 0.715929043 | 0.358 | 0.031 | 9.15705521909864e-119 | 21 |
| chtopb | 9.69971727893633e-122 | 0.73433508 | 0.373 | 0.034 | 2.44132184193549e-117 | 21 |
| piwil11 | 1.06554308548144e-115 | 0.830151016 | 0.383 | 0.039 | 2.68186539184823e-111 | 21 |
| CU927934.3 | 5.16599100730818e-114 | 0.563404201 | 0.161 | 0.006 | 1.3002282766294e-109 | 21 |
| zgc:194246 | 3.36341365331257e-106 | 0.764801268 | 0.352 | 0.036 | 8.46537582402242e-102 | 21 |
| gcshb | 1.72465243628673e-100 | 0.450192952 | 0.202 | 0.012 | 4.34077771689006e-96 | 21 |
| hmgb2b1 | 1.94502297362756e-98 | 1.175263386 | 0.855 | 0.312 | 4.89542832232322e-94 | 21 |
| adad1 | 3.23332938484468e-94 | 0.53374593 | 0.259 | 0.021 | 8.13796672871558e-90 | 21 |
| mrto4 | 6.74001671318156e-94 | 0.692814451 | 0.342 | 0.039 | 1.69639480654067e-89 | 21 |
| mgmt | 2.25665321258604e-93 | 0.926519678 | 0.539 | 0.095 | 5.67977047075782e-89 | 21 |
| CABZ01080371.2 | 1.64863619259119e-92 | 0.328853586 | 0.135 | 0.005 | 4.14945243313276e-88 | 21 |
| tdrd9 | 7.98577687405913e-92 | 0.581053878 | 0.28 | 0.025 | 2.00994018143194e-87 | 21 |
| dpy30 | 8.71877575328831e-92 | 0.99855412 | 0.642 | 0.145 | 2.19442866934513e-87 | 21 |
| dmrt1 | 1.01389296240833e-91 | 0.498830037 | 0.228 | 0.017 | 2.55186719708553e-87 | 21 |
| si:ch211-277c7.7 | 2.69556115026195e-91 | 0.30400661 | 0.114 | 0.004 | 6.7844578590943e-87 | 21 |
| dut1 | 1.402211095067e-87 | 0.955308918 | 0.585 | 0.117 | 3.52922510517413e-83 | 21 |

| | | | | | | |
|---|---|---|---|---|---|---|
| stmn1a1 | 8.61223814629071e-87 | 1.048526419 | 0.741 | 0.198 | 2.16761421903991e-82 | 21 |
| si:dkeyp-93d12.1 | 4.6093451028687e-85 | 0.504562747 | 0.249 | 0.022 | 1.16012606894102e-80 | 21 |
| syce2 | 2.2525730627814e-84 | 0.542351686 | 0.223 | 0.018 | 5.66950114171451e-80 | 21 |
| dnph1 | 5.95877157509989e-84 | 0.530212384 | 0.254 | 0.023 | 1.49976321773689e-79 | 21 |
| si:ch211-173m16.2 | 6.41069607999448e-82 | 0.259564379 | 0.119 | 0.005 | 1.61350809637381e-77 | 21 |
| tuba1c | 8.87865121187786e-81 | 0.300568597 | 0.13 | 0.006 | 2.23466772351754e-76 | 21 |
| calm2b2 | 1.09063898798633e-77 | 1.032662553 | 0.684 | 0.204 | 2.74502926886278e-73 | 21 |
| iqck | 1.01380238564455e-75 | 0.408549155 | 0.181 | 0.013 | 2.55163922442876e-71 | 21 |
| fanci | 1.28179716531223e-75 | 0.32594762 | 0.13 | 0.006 | 3.22615528537435e-71 | 21 |
| si:dkey-276j7.1 | 6.53595157386256e-75 | 0.575028508 | 0.275 | 0.031 | 1.64503365162547e-70 | 21 |
| ddx43 | 6.99501079075888e-75 | 0.328856201 | 0.145 | 0.008 | 1.7605742659261e-70 | 21 |
| terb1 | 8.72557652498538e-75 | 0.371835664 | 0.166 | 0.011 | 2.19614035557357e-70 | 21 |
| top2a1 | 1.68335209947112e-73 | 0.617993236 | 0.301 | 0.037 | 4.23682889915887e-69 | 21 |
| pld6 | 9.94452792183214e-73 | 0.651633629 | 0.337 | 0.045 | 2.50293823264593e-68 | 21 |
| piwil2 | 1.26037234143862e-72 | 0.54557859 | 0.28 | 0.032 | 3.17223114616687e-68 | 21 |
| rnf17 | 3.54676932985095e-72 | 0.530561614 | 0.254 | 0.027 | 8.92686372630186e-68 | 21 |
| CABZ01083945.1 | 7.73827852927613e-72 | 0.422316925 | 0.176 | 0.013 | 1.94764732303351e-67 | 21 |
| si:ch211-222l21.17 | 4.7941127844967e-71 | 1.001431489 | 0.756 | 0.269 | 1.20663024672997e-66 | 21 |
| mycbp | 3.27909144453546e-68 | 0.962823625 | 0.637 | 0.189 | 8.25314525675131e-64 | 21 |
| chd4a | 5.06962217330502e-68 | 0.753023692 | 0.44 | 0.086 | 1.27597320479914e-63 | 21 |
| lsm61 | 1.04222162203865e-67 | 0.965830214 | 0.71 | 0.226 | 2.62316760050909e-63 | 21 |
| lsm7 | 5.19471969008652e-67 | 0.933673516 | 0.653 | 0.192 | 1.30745899879788e-62 | 21 |
| h2afx17 | 2.6239225182119e-65 | 0.932029316 | 0.632 | 0.193 | 6.60415058608753e-61 | 21 |
| mcm3 | 3.3130986070809e-65 | 0.497709646 | 0.244 | 0.028 | 8.33873788416191e-61 | 21 |
| gpd1b | 5.2656290793798e-63 | 0.336495907 | 0.161 | 0.012 | 1.3253061829891e-58 | 21 |
| iqcb1 | 2.11723744788282e-62 | 0.269075076 | 0.109 | 0.005 | 5.32887493257627e-58 | 21 |
| sub1b2 | 1.06504047261826e-61 | 0.911501726 | 0.679 | 0.217 | 2.68060036553289e-57 | 21 |
| nsmce1 | 1.89794889196209e-61 | 0.495645595 | 0.233 | 0.027 | 4.77694756617938e-57 | 21 |
| llph | 2.63250801701503e-61 | 0.616567149 | 0.311 | 0.048 | 6.62575942802512e-57 | 21 |
| zgc:161973 | 9.58989153003525e-61 | 0.336803824 | 0.15 | 0.011 | 2.41367979919457e-56 | 21 |
| odf3l2a | 8.0397273962126e-60 | 0.354323834 | 0.161 | 0.013 | 2.02351898835275e-55 | 21 |
| snrpd1 | 1.3274638364849e-59 | 0.918124419 | 0.674 | 0.229 | 3.34109373004885e-55 | 21 |
| adad2 | 4.41920377461576e-59 | 0.354577005 | 0.181 | 0.017 | 1.11226939803304e-54 | 21 |
| morc3a | 2.30650855950536e-58 | 0.423669977 | 0.181 | 0.017 | 5.80525139341903e-54 | 21 |
| trmt112 | 1.82539036444827e-57 | 0.718568152 | 0.435 | 0.094 | 4.59432500827984e-53 | 21 |

| | | | | | | |
|---|---|---|---|---|---|---|
| isg153 | 4.25736397034921e-57 | 0.765596999 | 0.352 | 0.065 | 1.07153593769719e-52 | 21 |
| srsf2b | 1.0815716717204e-56 | 0.497776705 | 0.259 | 0.035 | 2.72220774055308e-52 | 21 |
| si:dkey-4p15.3 | 1.78433780205468e-56 | 0.25784874 | 0.109 | 0.006 | 4.49099981399143e-52 | 21 |
| zgc:162025 | 2.65678312264304e-56 | 0.618124704 | 0.316 | 0.052 | 6.68685744138028e-52 | 21 |
| esco2 | 1.01091521086012e-55 | 0.54950053 | 0.269 | 0.038 | 2.54437249421383e-51 | 21 |
| acot13 | 1.98030936768264e-55 | 0.386801554 | 0.181 | 0.018 | 4.98424064752043e-51 | 21 |
| chaf1a | 3.49501358882962e-54 | 0.57150371 | 0.347 | 0.062 | 8.79659970172528e-50 | 21 |
| minos1 | 4.75618606317548e-54 | 0.799795671 | 0.554 | 0.159 | 1.19708447024064e-49 | 21 |
| hmgb1b4 | 5.13974120439959e-54 | 0.816020878 | 0.472 | 0.122 | 1.29362146373533e-49 | 21 |
| rbx1 | 3.95588684149187e-53 | 0.824661543 | 0.699 | 0.26 | 9.95657159135089e-49 | 21 |
| snrpf | 4.8062738771254e-53 | 0.855083048 | 0.668 | 0.244 | 1.20969107213369e-48 | 21 |
| stk31 | 4.31961065826249e-52 | 0.347768121 | 0.161 | 0.015 | 1.08720280657809e-47 | 21 |
| CR382383.2 | 3.1277875049232e-51 | 0.460013725 | 0.254 | 0.036 | 7.87232837114121e-47 | 21 |
| snrpg2 | 1.16281624662662e-50 | 0.844985231 | 0.699 | 0.262 | 2.92669221113455e-46 | 21 |
| sms | 2.88718191482687e-50 | 0.470816612 | 0.244 | 0.035 | 7.26674816142775e-46 | 21 |
| gnl31 | 3.05591940176493e-50 | 0.6057054 | 0.311 | 0.057 | 7.69144354230215e-46 | 21 |
| zgc:153681 | 1.69152401909478e-49 | 0.368556045 | 0.171 | 0.018 | 4.25739680365966e-45 | 21 |
| srsf7b | 1.09664521213228e-48 | 0.29925798 | 0.145 | 0.013 | 2.76014633441574e-44 | 21 |
| tdrkh | 1.08383155785097e-47 | 0.495676585 | 0.249 | 0.038 | 2.72789564795511e-43 | 21 |
| rpa2 | 1.40443159768499e-47 | 0.628103473 | 0.378 | 0.081 | 3.53481388821335e-43 | 21 |
| hirip3 | 1.58646663210716e-47 | 0.66491256 | 0.373 | 0.081 | 3.99297786635051e-43 | 21 |
| pole4 | 2.18624504256643e-47 | 0.586334213 | 0.358 | 0.074 | 5.50256014763545e-43 | 21 |
| nub1 | 4.42882418296707e-47 | 0.362394646 | 0.161 | 0.017 | 1.11469075861098e-42 | 21 |
| parp3 | 7.67492064511948e-47 | 0.383474302 | 0.166 | 0.018 | 1.93170077717012e-42 | 21 |
| smfn | 1.02700049325087e-46 | 0.613565599 | 0.363 | 0.076 | 2.58485754146312e-42 | 21 |
| snrpd3l | 3.13138801552993e-46 | 0.76603882 | 0.544 | 0.167 | 7.88139049628729e-42 | 21 |
| cmss1 | 3.17359879894768e-46 | 0.465501478 | 0.275 | 0.046 | 7.98763081707141e-42 | 21 |
| ssrp1a | 5.1121322476446e-46 | 0.582014809 | 0.321 | 0.062 | 1.28667256540967e-41 | 21 |
| ppih | 6.51204988694816e-46 | 0.572528216 | 0.352 | 0.073 | 1.63901783604598e-41 | 21 |
| si:dkey-119f1.1 | 2.30063661069411e-45 | 0.289087196 | 0.124 | 0.01 | 5.79047228545601e-41 | 21 |
| auh | 5.13336884103112e-45 | 0.418671147 | 0.207 | 0.029 | 1.29201760359912e-40 | 21 |
| ccnd3 | 7.21552958944565e-45 | 0.339074074 | 0.161 | 0.017 | 1.81607664236757e-40 | 21 |
| hells3 | 7.92329295519418e-45 | 0.697226087 | 0.487 | 0.129 | 1.99421360389282e-40 | 21 |
| sart1 | 1.75210495652632e-44 | 0.615835792 | 0.378 | 0.088 | 4.4098729650811e-40 | 21 |
| lsm4 | 2.3082493490293e-44 | 0.777146388 | 0.591 | 0.207 | 5.80963278657185e-40 | 21 |

| | | | | | | |
|---|---|---|---|---|---|---|
| cpsf2 | 2.97991398020462e-44 | 0.475135693 | 0.254 | 0.042 | 7.500145496777e-40 | 21 |
| tdrd6 | 3.34387804660007e-44 | 0.392263396 | 0.176 | 0.021 | 8.41620665548772e-40 | 21 |
| odc11 | 6.41422970190697e-44 | 0.692360318 | 0.446 | 0.119 | 1.61439747367297e-39 | 21 |
| dlgap5 | 1.73832186848658e-43 | 0.422619677 | 0.238 | 0.037 | 4.37518231079386e-39 | 21 |
| psmb7 | 3.08655499964815e-43 | 0.710334625 | 0.487 | 0.144 | 7.76855027861442e-39 | 21 |
| ndufb6 | 4.5930203342163e-43 | 0.65409746 | 0.409 | 0.107 | 1.1560172879189e-38 | 21 |
| dcun1d5 | 5.70121088682942e-43 | 0.603104319 | 0.368 | 0.083 | 1.4349377681061e-38 | 21 |
| lrrc20 | 1.07919456186025e-42 | 0.262672932 | 0.13 | 0.012 | 2.71622479274607e-38 | 21 |
| hpf1 | 1.71375728441671e-42 | 0.585076512 | 0.311 | 0.063 | 4.31335570914843e-38 | 21 |
| npm1a1 | 1.76593621892253e-42 | 0.796841614 | 0.679 | 0.298 | 4.44468486940612e-38 | 21 |
| gra | 3.42092320945116e-42 | 0.488925016 | 0.275 | 0.049 | 8.61012162586762e-38 | 21 |
| timm10 | 6.44315934542997e-42 | 0.516967487 | 0.29 | 0.056 | 1.62167877565127e-37 | 21 |
| hint1 | 7.64399126819467e-42 | 0.756956737 | 0.58 | 0.213 | 1.92391616229192e-37 | 21 |
| pcna1 | 7.7046376247146e-42 | 0.709917765 | 0.477 | 0.139 | 1.93918024376442e-37 | 21 |
| supt3h | 1.21334306663013e-41 | 0.41901315 | 0.207 | 0.031 | 3.05386316440138e-37 | 21 |
| ppa1b | 1.76541921669526e-41 | 0.621677389 | 0.352 | 0.081 | 4.44338362650031e-37 | 21 |
| dynll2b | 1.89141306124983e-41 | 0.700823519 | 0.456 | 0.129 | 4.7604975338597e-37 | 21 |
| fkbp3 | 1.94802487992447e-41 | 0.7214616 | 0.497 | 0.159 | 4.9029838202819e-37 | 21 |
| rabif | 5.23626710973274e-41 | 0.489323073 | 0.254 | 0.045 | 1.31791606884863e-36 | 21 |
| h2afvb1 | 7.2740973827216e-41 | 0.688834418 | 0.829 | 0.466 | 1.8308175702572e-36 | 21 |
| lin52 | 7.5254421414613e-41 | 0.495608111 | 0.259 | 0.047 | 1.8940785325844e-36 | 21 |
| tubb2b1 | 9.49097101711295e-41 | 0.778534617 | 0.632 | 0.244 | 2.38878249529716e-36 | 21 |
| mcm6 | 1.0122494063693e-40 | 0.452798887 | 0.218 | 0.034 | 2.5477305308909e-36 | 21 |
| wdr82 | 2.82319527379506e-40 | 0.582277639 | 0.358 | 0.083 | 7.10570018461479e-36 | 21 |
| cenpx | 3.1839189408318e-40 | 0.578713415 | 0.352 | 0.081 | 8.01360558217955e-36 | 21 |
| ube2t | 7.07695537363541e-40 | 0.46715559 | 0.264 | 0.048 | 1.7811988979903e-35 | 21 |
| hmgn25 | 1.06765360439072e-39 | 0.722356456 | 0.803 | 0.439 | 2.68717735689101e-35 | 21 |
| polr2l | 6.26837267058686e-39 | 0.7054906 | 0.56 | 0.196 | 1.57768671746001e-34 | 21 |
| henmt1 | 7.86971269630747e-39 | 0.330425743 | 0.161 | 0.02 | 1.98072798853363e-34 | 21 |
| setb1 | 1.66866988452919e-38 | 0.729371684 | 0.725 | 0.356 | 4.19987523237151e-34 | 21 |
| si:ch211-212d10.2 | 8.96055678269086e-38 | 0.294680656 | 0.14 | 0.016 | 2.25528253663546e-33 | 21 |
| dnajc91 | 1.38756745804288e-37 | 0.581705346 | 0.363 | 0.088 | 3.49236853514812e-33 | 21 |
| mki67 | 4.03894467220335e-37 | 0.46990451 | 0.228 | 0.04 | 1.01656198454686e-32 | 21 |
| ppid | 4.0583399295954e-37 | 0.561085048 | 0.352 | 0.088 | 1.02144357687987e-32 | 21 |
| efcab1 | 5.09408283574973e-37 | 0.287130683 | 0.13 | 0.014 | 1.28212970892985e-32 | 21 |

| | | | | | | |
|---|---|---|---|---|---|---|
| atp5mc3b1 | 6.87915481947649e-37 | 0.538196939 | 0.865 | 0.6 | 1.73141447651404e-32 | 21 |
| nars | 9.28522500322185e-37 | 0.562472312 | 0.301 | 0.069 | 2.33699828106091e-32 | 21 |
| eif4e1b1 | 9.61442187182659e-37 | 0.553955593 | 0.352 | 0.083 | 2.41985384092003e-32 | 21 |
| cirbpa | 1.52039818340888e-36 | 0.537085768 | 0.855 | 0.615 | 3.8266901878218e-32 | 21 |
| ascl1b | 3.02607508822183e-36 | 0.261206202 | 0.114 | 0.011 | 7.61632838954553e-32 | 21 |
| txndc17 | 3.84373013983675e-36 | 0.454548675 | 0.259 | 0.051 | 9.67428438895513e-32 | 21 |
| med28 | 4.31982256605423e-36 | 0.572895282 | 0.342 | 0.085 | 1.08725614165019e-31 | 21 |
| tomm20a2 | 4.38413421331864e-36 | 0.701387339 | 0.596 | 0.228 | 1.10344274015017e-31 | 21 |
| wdr76 | 8.04592299973545e-36 | 0.337809562 | 0.171 | 0.024 | 2.02507835980342e-31 | 21 |
| sub1a1 | 1.1214540455975e-35 | 0.72381491 | 0.57 | 0.229 | 2.82258768736434e-31 | 21 |
| cenps | 1.13478274244366e-35 | 0.393681719 | 0.212 | 0.035 | 2.85613468445645e-31 | 21 |
| ubtf | 1.41789631815769e-35 | 0.465658129 | 0.244 | 0.048 | 3.56870324317109e-31 | 21 |
| ss18l2 | 1.81630135576826e-35 | 0.444537162 | 0.264 | 0.054 | 4.57144888233313e-31 | 21 |
| psmb3 | 2.37528249175876e-35 | 0.639688808 | 0.43 | 0.133 | 5.97834850350763e-31 | 21 |
| hibadha | 2.45732898616931e-35 | 0.295209825 | 0.13 | 0.014 | 6.18485132528953e-31 | 21 |
| msh6 | 7.44338579526808e-35 | 0.339276439 | 0.181 | 0.027 | 1.87342577081102e-30 | 21 |
| lsm5 | 1.03089410312367e-34 | 0.704555668 | 0.585 | 0.233 | 2.59465736815197e-30 | 21 |
| ndufa21 | 1.66806765994827e-34 | 0.642952714 | 0.44 | 0.143 | 4.19835949332381e-30 | 21 |
| ccdc25 | 1.99008158169552e-34 | 0.559752518 | 0.332 | 0.086 | 5.00883633296945e-30 | 21 |
| nutf2l2 | 3.04062946664972e-34 | 0.678730324 | 0.596 | 0.229 | 7.65296030461069e-30 | 21 |
| nup107 | 5.32485823703796e-34 | 0.352173072 | 0.176 | 0.027 | 1.34021356968008e-29 | 21 |
| tubb4b | 5.83168679791259e-34 | 0.568508054 | 0.845 | 0.582 | 1.46777725016662e-29 | 21 |
| rbm8a | 8.91083190275575e-34 | 0.719149798 | 0.637 | 0.309 | 2.2427672816046e-29 | 21 |
| gstr1 | 2.53105793903842e-33 | 0.614803736 | 0.383 | 0.116 | 6.37041972676579e-29 | 21 |
| psmc5 | 3.72297857031248e-33 | 0.507651156 | 0.326 | 0.084 | 9.37036476361949e-29 | 21 |
| pfdn6 | 3.83305619285918e-33 | 0.58232502 | 0.373 | 0.106 | 9.64741913180726e-29 | 21 |
| chek1 | 4.02685206441167e-33 | 0.284948437 | 0.15 | 0.02 | 1.01351839609177e-28 | 21 |
| nudc | 4.62664242138503e-33 | 0.62701213 | 0.456 | 0.148 | 1.1644796310384e-28 | 21 |
| cks2 | 4.81367420136613e-33 | 0.457469724 | 0.223 | 0.042 | 1.21155365974184e-28 | 21 |
| snrnp27 | 7.14771821244264e-33 | 0.588263754 | 0.409 | 0.123 | 1.79900919688969e-28 | 21 |
| atp5mea | 7.21922195485157e-33 | 0.643552303 | 0.44 | 0.145 | 1.81700597381659e-28 | 21 |
| snrnp200 | 9.88371347863966e-33 | 0.396152585 | 0.207 | 0.038 | 2.48763184543882e-28 | 21 |
| edc4 | 1.10172226808578e-32 | 0.373286831 | 0.192 | 0.032 | 2.7729247765451e-28 | 21 |
| alkbh3 | 1.10214315527145e-32 | 0.2862865 | 0.13 | 0.016 | 2.7739841075027e-28 | 21 |
| snrpe1 | 1.25333670881495e-32 | 0.687087426 | 0.642 | 0.285 | 3.15452316241634e-28 | 21 |

| | | | | | | |
|---|---|---|---|---|---|---|
| zgc:56106 | 1.7981068971889e-32 | 0.391736885 | 0.223 | 0.043 | 4.52565524953475e-28 | 21 |
| erh | 1.98498223283328e-32 | 0.6704132 | 0.674 | 0.338 | 4.99600178181808e-28 | 21 |
| dnd13 | 2.42222313846377e-32 | 0.587117057 | 0.383 | 0.107 | 6.09649341719947e-28 | 21 |
| mybbp1a | 4.68933097555784e-32 | 0.345121253 | 0.166 | 0.025 | 1.18025771323815e-27 | 21 |
| atp5pd3 | 4.86849149844752e-32 | 0.668863071 | 0.585 | 0.258 | 1.22535062524426e-27 | 21 |
| zcchc2 | 7.70649909700756e-32 | 0.30095188 | 0.119 | 0.014 | 1.93964875772583e-27 | 21 |
| pomp3 | 1.06225716340545e-31 | 0.664086099 | 0.544 | 0.227 | 2.67359505457519e-27 | 21 |
| si:dkey-148h10.5 | 1.36952170236286e-31 | 0.251820938 | 0.119 | 0.014 | 3.44694917267708e-27 | 21 |
| mif | 1.49126588699926e-31 | 0.656936622 | 0.492 | 0.184 | 3.75336711098844e-27 | 21 |
| rad21l1 | 1.86268005004858e-31 | 0.281725536 | 0.124 | 0.015 | 4.68817941796726e-27 | 21 |
| eftud2 | 2.10109293475235e-31 | 0.318776655 | 0.171 | 0.027 | 5.2882408074782e-27 | 21 |
| vrk1 | 3.01740745252899e-31 | 0.434145641 | 0.233 | 0.048 | 7.59451281727021e-27 | 21 |
| nelfcd | 3.16680552122948e-31 | 0.284405953 | 0.161 | 0.024 | 7.97053281638247e-27 | 21 |
| dazl4 | 3.36803798938918e-31 | 0.551230429 | 0.378 | 0.104 | 8.47701481549363e-27 | 21 |
| seta | 4.13257327035375e-31 | 0.673006373 | 0.617 | 0.281 | 1.04012736641534e-26 | 21 |
| cct72 | 6.69862074167342e-31 | 0.649100696 | 0.513 | 0.206 | 1.68597585447178e-26 | 21 |
| hnrnpul1 | 8.97722126994079e-31 | 0.304543602 | 0.155 | 0.023 | 2.2594768214314e-26 | 21 |
| smc3 | 9.70871295292212e-31 | 0.443570945 | 0.233 | 0.049 | 2.44358596312097e-26 | 21 |
| si:ch211-156b7.4 | 1.90770214291627e-30 | 0.350963636 | 0.192 | 0.034 | 4.80149552350596e-26 | 21 |
| baz1a | 2.18360708321066e-30 | 0.481263267 | 0.295 | 0.074 | 5.4959206677329e-26 | 21 |
| ap3s2 | 2.27643927133689e-30 | 0.394702303 | 0.238 | 0.05 | 5.72957000202783e-26 | 21 |
| dhfr | 2.63422087543671e-30 | 0.400731118 | 0.238 | 0.049 | 6.63007052138666e-26 | 21 |
| nuf2 | 4.18965821080518e-30 | 0.352508587 | 0.176 | 0.029 | 1.05449507507756e-25 | 21 |
| exd11 | 4.92670464835959e-30 | 0.394730212 | 0.264 | 0.058 | 1.24000229294563e-25 | 21 |
| mbd3b | 5.05405223321843e-30 | 0.442004921 | 0.244 | 0.053 | 1.27205440657875e-25 | 21 |
| rad51 | 5.25608340380787e-30 | 0.365709495 | 0.212 | 0.041 | 1.3229036319044e-25 | 21 |
| nup93 | 7.4402216489114e-30 | 0.37288698 | 0.192 | 0.035 | 1.87262938681451e-25 | 21 |
| emd2 | 1.15827347466259e-29 | 0.454666082 | 0.238 | 0.054 | 2.91525850837828e-25 | 21 |
| zgc:66472 | 2.00206486975556e-29 | 0.348507161 | 0.197 | 0.036 | 5.03899707068776e-25 | 21 |
| ift46 | 2.61035244759368e-29 | 0.271965084 | 0.13 | 0.017 | 6.56999607534855e-25 | 21 |
| aaas | 2.65275960835483e-29 | 0.252483733 | 0.124 | 0.016 | 6.67673065826828e-25 | 21 |
| sf3b5 | 3.30032446110632e-29 | 0.558812301 | 0.435 | 0.146 | 8.30658663615851e-25 | 21 |
| sf3a2 | 3.34114854185855e-29 | 0.565003462 | 0.399 | 0.128 | 8.40933676500377e-25 | 21 |
| nudt21 | 3.72809671854967e-29 | 0.461293414 | 0.28 | 0.069 | 9.38324663091767e-25 | 21 |
| psmc1a | 3.79272453366241e-29 | 0.45244812 | 0.275 | 0.067 | 9.54590837877492e-25 | 21 |

| | | | | | | |
|---|---|---|---|---|---|---|
| srsf1b | 3.9361367651895e-29 | 0.439194795 | 0.259 | 0.061 | 9.90686262430545e-25 | 21 |
| txlna | 3.94457411471599e-29 | 0.41787049 | 0.233 | 0.05 | 9.92809858932869e-25 | 21 |
| lsm3 | 4.35344442337366e-29 | 0.610296153 | 0.461 | 0.164 | 1.09571842691892e-24 | 21 |
| birc5a | 4.50769475688853e-29 | 0.526012351 | 0.347 | 0.097 | 1.13454169336127e-24 | 21 |
| ndufc21 | 7.18649748477897e-29 | 0.590292108 | 0.394 | 0.133 | 1.80876955194402e-24 | 21 |
| eif4a3 | 8.19546395829413e-29 | 0.58490235 | 0.425 | 0.147 | 2.06271632366305e-24 | 21 |
| tomm6 | 8.77759665581511e-29 | 0.560181504 | 0.352 | 0.107 | 2.2092333023021e-24 | 21 |
| spata6l | 9.05480781439174e-29 | 0.403978256 | 0.202 | 0.039 | 2.27900457880426e-24 | 21 |
| psma5 | 9.52647786352215e-29 | 0.586927017 | 0.44 | 0.155 | 2.39771921346989e-24 | 21 |
| ndufs5 | 1.35718505789212e-28 | 0.60356915 | 0.42 | 0.149 | 3.41589907220867e-24 | 21 |
| mphosph83 | 1.4193887494672e-28 | 0.549188472 | 0.368 | 0.116 | 3.57245954353398e-24 | 21 |
| mrps25 | 1.939641997912e-28 | 0.477415875 | 0.301 | 0.08 | 4.88188494454472e-24 | 21 |
| ndufa8 | 2.06281648749779e-28 | 0.558486901 | 0.394 | 0.13 | 5.19190281738319e-24 | 21 |
| zgc:158564 | 4.89567181038311e-28 | 0.480875118 | 0.29 | 0.077 | 1.23219163795533e-23 | 21 |
| dusp22b | 6.2813579967676e-28 | 0.36892221 | 0.176 | 0.032 | 1.58095499420644e-23 | 21 |
| cenph | 7.36432725354174e-28 | 0.300310444 | 0.145 | 0.022 | 1.85352752644392e-23 | 21 |
| tfdp1a | 7.61056947972711e-28 | 0.362341493 | 0.192 | 0.037 | 1.91550423235252e-23 | 21 |
| nop16 | 8.76180419518816e-28 | 0.529191733 | 0.363 | 0.111 | 2.20525849788691e-23 | 21 |
| phb2b | 1.45807091825311e-27 | 0.389700217 | 0.233 | 0.051 | 3.66981869415126e-23 | 21 |
| hddc2 | 1.51638248153919e-27 | 0.379105591 | 0.233 | 0.052 | 3.81658306778598e-23 | 21 |
| hdhd2 | 1.62134427522126e-27 | 0.262329647 | 0.119 | 0.016 | 4.08076140630438e-23 | 21 |
| nsmce2 | 1.67217060557739e-27 | 0.353512322 | 0.187 | 0.036 | 4.20868619717774e-23 | 21 |
| gtf2f2a | 1.6911888066019e-27 | 0.457749708 | 0.244 | 0.058 | 4.25655310733632e-23 | 21 |
| zgc:110540 | 1.71076620591016e-27 | 0.25675109 | 0.119 | 0.016 | 4.30582746365529e-23 | 21 |
| ndufs7 | 2.22480974290948e-27 | 0.542487012 | 0.358 | 0.114 | 5.59962364192887e-23 | 21 |
| rbmx2 | 2.8601994731119e-27 | 0.330496724 | 0.166 | 0.029 | 7.19883605387534e-23 | 21 |
| mcm72 | 3.01761633618151e-27 | 0.572554834 | 0.446 | 0.155 | 7.59503855653524e-23 | 21 |
| calm3b1 | 3.53806337588257e-27 | 0.641277099 | 0.549 | 0.253 | 8.90495171075884e-23 | 21 |
| si:ch73-41e3.7 | 4.19531699874821e-27 | 0.290492569 | 0.155 | 0.025 | 1.05591933541494e-22 | 21 |
| dgcr6 | 6.07033374587338e-27 | 0.33967434 | 0.176 | 0.033 | 1.52784230049887e-22 | 21 |
| tcp13 | 9.90288384204631e-27 | 0.606543841 | 0.518 | 0.223 | 2.49245683420464e-22 | 21 |
| ppie | 9.91011039889492e-27 | 0.343433035 | 0.197 | 0.039 | 2.49427568629786e-22 | 21 |
| hddc3 | 1.41521349993944e-26 | 0.353699462 | 0.176 | 0.033 | 3.56195085799758e-22 | 21 |
| cox5aa2 | 1.73703360248083e-26 | 0.603168867 | 0.653 | 0.349 | 4.37193987408401e-22 | 21 |
| ilf21 | 1.81293899757503e-26 | 0.609271157 | 0.565 | 0.241 | 4.5629861629966e-22 | 21 |

| | | | | | | |
|---|---|---|---|---|---|---|
| exosc2 | 2.25364447242183e-26 | 0.386296077 | 0.212 | 0.046 | 5.6721977726385e-22 | 21 |
| gtf2h5 | 2.38215500949125e-26 | 0.520216731 | 0.358 | 0.111 | 5.99564594338852e-22 | 21 |
| eif2s23 | 4.30567815897789e-26 | 0.603724252 | 0.596 | 0.291 | 1.08369613583315e-21 | 21 |
| ccdc137 | 4.43879880274452e-26 | 0.333660339 | 0.161 | 0.028 | 1.11720127066277e-21 | 21 |
| tbcb | 5.95176233646339e-26 | 0.471896261 | 0.259 | 0.068 | 1.49799906246447e-21 | 21 |
| mtrf1 | 7.29729430982654e-26 | 0.283347341 | 0.135 | 0.021 | 1.83665600484024e-21 | 21 |
| myl63 | 1.03465695718081e-25 | 0.493978028 | 0.326 | 0.099 | 2.60412809552837e-21 | 21 |
| nedd8 | 1.52858636029662e-25 | 0.464754792 | 0.275 | 0.076 | 3.84729901023056e-21 | 21 |
| ubxn1 | 1.65872476944662e-25 | 0.375519642 | 0.202 | 0.044 | 4.1748443722202e-21 | 21 |
| yeats4 | 2.35212858352003e-25 | 0.403416482 | 0.218 | 0.05 | 5.92007243186157e-21 | 21 |
| eif4e1c | 2.40780420482454e-25 | 0.466153548 | 0.337 | 0.1 | 6.06020240312289e-21 | 21 |
| cops3 | 3.26059143617888e-25 | 0.399679815 | 0.238 | 0.058 | 8.20658258571861e-21 | 21 |
| xrcc5 | 4.14396347266103e-25 | 0.250890858 | 0.119 | 0.017 | 1.04299416643405e-20 | 21 |
| ube2c4 | 4.26981229319652e-25 | 0.525583784 | 0.415 | 0.137 | 1.07466905607463e-20 | 21 |
| rbb4l | 4.34056219558879e-25 | 0.446500317 | 0.285 | 0.079 | 1.09247609900774e-20 | 21 |
| ndufab1b | 4.50582478459319e-25 | 0.539371794 | 0.425 | 0.156 | 1.13407104003426e-20 | 21 |
| fip1l1b | 8.29458615013426e-25 | 0.355614048 | 0.202 | 0.044 | 2.08766438812729e-20 | 21 |
| pin11 | 8.39091632427952e-25 | 0.53778276 | 0.394 | 0.141 | 2.11190972965791e-20 | 21 |
| ebna1bp23 | 1.15408537742949e-24 | 0.530205184 | 0.394 | 0.135 | 2.90471748645228e-20 | 21 |
| crnkl1 | 1.36476267609341e-24 | 0.272798137 | 0.155 | 0.028 | 3.43497117945951e-20 | 21 |
| taf15 | 1.45441315058518e-24 | 0.557211238 | 0.466 | 0.182 | 3.66061245870783e-20 | 21 |
| isoc2 | 1.59386918669072e-24 | 0.340701649 | 0.176 | 0.036 | 4.01160935598187e-20 | 21 |
| pbdc1 | 1.95563605490444e-24 | 0.518197333 | 0.363 | 0.125 | 4.92214038658899e-20 | 21 |
| ppig | 2.20166601792264e-24 | 0.390216693 | 0.244 | 0.061 | 5.54137320050948e-20 | 21 |
| ndnl2 | 2.54146291600215e-24 | 0.313567239 | 0.181 | 0.037 | 6.39660801328582e-20 | 21 |
| psmd8 | 3.3715395746494e-24 | 0.492429145 | 0.347 | 0.112 | 8.48582795543508e-20 | 21 |
| tuba8l | 3.76898572374781e-24 | 0.597859156 | 0.451 | 0.186 | 9.48616016810086e-20 | 21 |
| ndufab1a | 3.87333331970254e-24 | 0.546532672 | 0.373 | 0.132 | 9.74879263235932e-20 | 21 |
| rfc4 | 5.14108163190723e-24 | 0.423963551 | 0.275 | 0.075 | 1.29395883593473e-19 | 21 |
| ptgr11 | 5.2582253520315e-24 | 0.489422973 | 0.337 | 0.105 | 1.32344273885281e-19 | 21 |
| ruvbl1 | 5.54773245510786e-24 | 0.42332785 | 0.301 | 0.085 | 1.3963087816261e-19 | 21 |
| denr | 6.45599281315044e-24 | 0.524758748 | 0.368 | 0.128 | 1.62490883114183e-19 | 21 |
| hmgb1a7 | 6.86937988248755e-24 | 0.637682556 | 0.44 | 0.187 | 1.72895422262329e-19 | 21 |
| luc7l3 | 8.93360751454725e-24 | 0.354282954 | 0.218 | 0.052 | 2.2484996753364e-19 | 21 |
| psmd4b | 9.46812383061309e-24 | 0.397611561 | 0.223 | 0.055 | 2.38303208692701e-19 | 21 |

| | | | | | | |
|---|---|---|---|---|---|---|
| gtf2e2 | 9.8448886641468e-24 | 0.421892855 | 0.259 | 0.07 | 2.47786002787911e-19 | 21 |
| cdk11b | 1.030257696785e-23 | 0.372532716 | 0.202 | 0.046 | 2.59305559703816e-19 | 21 |
| COX7A2 (1 of many)4 | 1.11544514122977e-23 | 0.591484061 | 0.539 | 0.258 | 2.8074638759612e-19 | 21 |
| ifi455 | 1.15962705593714e-23 | 0.54922452 | 0.368 | 0.126 | 2.91866533708818e-19 | 21 |
| metap2b1 | 1.23214408261433e-23 | 0.588142499 | 0.477 | 0.208 | 3.10118344153201e-19 | 21 |
| psma81 | 1.36556707519611e-23 | 0.539917331 | 0.487 | 0.204 | 3.43699577156109e-19 | 21 |
| psmc3ip | 1.40548274683774e-23 | 0.304534886 | 0.155 | 0.029 | 3.5374595255159e-19 | 21 |
| snrpa11 | 1.4446838145201e-23 | 0.505278879 | 0.363 | 0.12 | 3.63612469276564e-19 | 21 |
| hnrnpd | 1.48951594116758e-23 | 0.388117025 | 0.223 | 0.055 | 3.74896267232467e-19 | 21 |
| elavl27 | 1.71410847308736e-23 | 0.574650887 | 0.544 | 0.221 | 4.31423961591358e-19 | 21 |
| timm23a | 2.01748210755254e-23 | 0.49533443 | 0.337 | 0.111 | 5.07780071649898e-19 | 21 |
| CR762420.1 | 2.10069108671183e-23 | 0.270156854 | 0.114 | 0.017 | 5.28722939614501e-19 | 21 |
| ddx18 | 2.26314121661262e-23 | 0.500333525 | 0.326 | 0.104 | 5.6961001280923e-19 | 21 |
| fundc1 | 4.55093227698212e-23 | 0.438723671 | 0.269 | 0.076 | 1.14542414479363e-18 | 21 |
| dmap1 | 5.76121710409904e-23 | 0.263485017 | 0.135 | 0.023 | 1.45004073293069e-18 | 21 |
| eif4eb | 6.38773497696271e-23 | 0.377398896 | 0.212 | 0.051 | 1.60772901635174e-18 | 21 |
| utp11l | 6.52401699299252e-23 | 0.378148533 | 0.244 | 0.063 | 1.64202983696629e-18 | 21 |
| srrm2 | 7.85026661842469e-23 | 0.444518271 | 0.275 | 0.08 | 1.97583360519131e-18 | 21 |
| si:dkey-6i22.5 | 1.00425908748942e-22 | 0.336104339 | 0.176 | 0.037 | 2.52761969730213e-18 | 21 |
| nup50 | 1.25824538642493e-22 | 0.439931433 | 0.316 | 0.099 | 3.16687781309291e-18 | 21 |
| otub1b | 1.27585377978277e-22 | 0.286123817 | 0.15 | 0.028 | 3.21119637833524e-18 | 21 |
| rcc2 | 1.99921703435504e-22 | 0.513162063 | 0.383 | 0.141 | 5.03182935376819e-18 | 21 |
| aifm1 | 2.10560028771852e-22 | 0.277914268 | 0.135 | 0.024 | 5.29958536415875e-18 | 21 |
| srsf3b | 2.30885429927743e-22 | 0.552656191 | 0.534 | 0.243 | 5.81115538585136e-18 | 21 |
| si:ch211-235e9.6 | 2.62549214970457e-22 | 0.526568441 | 0.389 | 0.149 | 6.60810119159142e-18 | 21 |
| psmd7 | 2.95672719708109e-22 | 0.46750286 | 0.321 | 0.105 | 7.4417866823334e-18 | 21 |
| mtx2 | 3.31685102870347e-22 | 0.368747906 | 0.192 | 0.044 | 8.34818235414377e-18 | 21 |
| psmb1 | 3.41823243968656e-22 | 0.550797036 | 0.415 | 0.168 | 8.60334922744709e-18 | 21 |
| ndufa12 | 3.96326856793776e-22 | 0.473738719 | 0.337 | 0.115 | 9.97515065864254e-18 | 21 |
| snip1 | 4.09607821409562e-22 | 0.316686715 | 0.166 | 0.034 | 1.03094192570573e-17 | 21 |
| tyms | 4.70347015021668e-22 | 0.371982812 | 0.259 | 0.07 | 1.18381640210804e-17 | 21 |
| bcl7a | 5.64439859504205e-22 | 0.287148093 | 0.161 | 0.032 | 1.42063868238613e-17 | 21 |
| etfa1 | 7.58134713970921e-22 | 0.429747092 | 0.28 | 0.086 | 1.90814926159341e-17 | 21 |
| CABZ01112656.1 | 7.59744163250584e-22 | 0.414519755 | 0.218 | 0.057 | 1.91220008448539e-17 | 21 |
| unga | 7.65610620456142e-22 | 0.401177198 | 0.269 | 0.075 | 1.92696537062606e-17 | 21 |

| | | | | | | |
|---|---|---|---|---|---|---|
| si:dkey-170l10.1 | 9.24795989644883e-22 | 0.281325383 | 0.145 | 0.027 | 2.32761902633721e-17 | 21 |
| si:dkey-57a22.11 | 9.36603301554173e-22 | 0.255506894 | 0.119 | 0.019 | 2.3573368496817e-17 | 21 |
| psmd1 | 9.54544154647634e-22 | 0.378242365 | 0.259 | 0.074 | 2.40249218283263e-17 | 21 |
| polr2f | 1.17678852335471e-21 | 0.509366294 | 0.352 | 0.127 | 2.96185903443146e-17 | 21 |
| asf1bb | 1.44785296832112e-21 | 0.370757156 | 0.228 | 0.059 | 3.64410113596744e-17 | 21 |
| hnrnpa0b4 | 1.50975822134796e-21 | 0.578804265 | 0.565 | 0.31 | 3.79991046731067e-17 | 21 |
| gcsha | 1.64684996785276e-21 | 0.394129914 | 0.249 | 0.068 | 4.14495668408861e-17 | 21 |
| psmd11b | 1.64897327366391e-21 | 0.344974829 | 0.192 | 0.045 | 4.15030083248469e-17 | 21 |
| ptges3b | 1.68072413507727e-21 | 0.544064239 | 0.43 | 0.173 | 4.23021457557597e-17 | 21 |
| usp1 | 1.95658166792092e-21 | 0.328420905 | 0.187 | 0.042 | 4.92452039999015e-17 | 21 |
| zpr1 | 2.06860247611028e-21 | 0.279941872 | 0.15 | 0.029 | 5.20646557212197e-17 | 21 |
| ndufa6 | 2.20516829319955e-21 | 0.555883533 | 0.518 | 0.246 | 5.55018807715395e-17 | 21 |
| gemin2 | 2.57955239387854e-21 | 0.358240346 | 0.207 | 0.051 | 6.4924754201529e-17 | 21 |
| psmb5 | 2.66286408683177e-21 | 0.37093097 | 0.238 | 0.065 | 6.70216262014687e-17 | 21 |
| si:ch211-22i13.2 | 2.71870048003683e-21 | 0.398523256 | 0.218 | 0.057 | 6.84269723820469e-17 | 21 |
| prpf6 | 3.30332029624953e-21 | 0.289544343 | 0.155 | 0.032 | 8.31412685363043e-17 | 21 |
| atp6v0cb | 3.4717473422843e-21 | 0.265858646 | 0.119 | 0.02 | 8.73804088579536e-17 | 21 |
| mrpl38 | 3.49410884337853e-21 | 0.341342931 | 0.176 | 0.039 | 8.79432254789942e-17 | 21 |
| ciapin1 | 3.68958044856899e-21 | 0.306037164 | 0.176 | 0.039 | 9.28630503100328e-17 | 21 |
| spc24 | 5.33234747893924e-21 | 0.379184169 | 0.249 | 0.068 | 1.34209853697422e-16 | 21 |
| drg2 | 6.21467414452231e-21 | 0.393655124 | 0.254 | 0.073 | 1.56417133543482e-16 | 21 |
| snu13b7 | 6.59595304599112e-21 | 0.551071035 | 0.705 | 0.389 | 1.66013542214551e-16 | 21 |
| ufc1 | 6.73039321870734e-21 | 0.428642893 | 0.275 | 0.084 | 1.69397266921645e-16 | 21 |
| ttf1 | 7.54239647251414e-21 | 0.364424661 | 0.228 | 0.06 | 1.89834576816708e-16 | 21 |
| eif1ad | 7.9437330757061e-21 | 0.443749591 | 0.264 | 0.081 | 1.99935817782447e-16 | 21 |
| pqbp1 | 8.30845032792775e-21 | 0.253738199 | 0.14 | 0.026 | 2.09115386303613e-16 | 21 |
| cops5 | 8.78697330978e-21 | 0.349581233 | 0.197 | 0.048 | 2.21159331233853e-16 | 21 |
| ruvbl2 | 9.8768269036312e-21 | 0.308091249 | 0.202 | 0.049 | 2.48589856337494e-16 | 21 |
| sf3b4 | 1.16726198633817e-20 | 0.363060515 | 0.254 | 0.072 | 2.93788169341455e-16 | 21 |
| tsn | 1.20745465730525e-20 | 0.333007642 | 0.207 | 0.051 | 3.03904262697159e-16 | 21 |
| magoh | 1.29222333633826e-20 | 0.492488785 | 0.342 | 0.123 | 3.25239691522976e-16 | 21 |
| ankrd10a | 1.39157638973272e-20 | 0.306152771 | 0.212 | 0.053 | 3.50245861531828e-16 | 21 |
| hp1bp31 | 1.4495835993872e-20 | 0.438198832 | 0.29 | 0.096 | 3.64845696129764e-16 | 21 |
| rpa33 | 1.49426743516639e-20 | 0.475657228 | 0.425 | 0.161 | 3.76092170757029e-16 | 21 |
| pcmt | 1.54946359928618e-20 | 0.297829956 | 0.161 | 0.034 | 3.89984493304339e-16 | 21 |

| | | | | | | |
|---|---|---|---|---|---|---|
| ik | 1.70625301829988e-20 | 0.309858314 | 0.187 | 0.045 | 4.29446822175898e-16 | 21 |
| apex1 | 2.29577112926828e-20 | 0.391337926 | 0.249 | 0.071 | 5.77822635525533e-16 | 21 |
| pfdn41 | 2.60315180704933e-20 | 0.495466614 | 0.409 | 0.169 | 6.55187278316245e-16 | 21 |
| ppil1 | 2.60768538461944e-20 | 0.367439506 | 0.264 | 0.077 | 6.56328334454867e-16 | 21 |
| esf1 | 3.2929843672377e-20 | 0.362761089 | 0.181 | 0.043 | 8.28811235390058e-16 | 21 |
| cox7a2a2 | 3.63763080753478e-20 | 0.537611516 | 0.503 | 0.24 | 9.15555297948428e-16 | 21 |
| fabp7a | 4.18168306018026e-20 | 0.287805813 | 0.135 | 0.026 | 1.05248780941677e-15 | 21 |
| dhps | 4.36420839489335e-20 | 0.307533193 | 0.145 | 0.029 | 1.09842761091071e-15 | 21 |
| gkap1 | 4.62220359864142e-20 | 0.450539971 | 0.332 | 0.116 | 1.16336242374206e-15 | 21 |
| cnot1 | 4.69722922674762e-20 | 0.365122177 | 0.223 | 0.061 | 1.18224562408011e-15 | 21 |
| dhx16 | 5.16721838184367e-20 | 0.332801172 | 0.207 | 0.053 | 1.30053719452623e-15 | 21 |
| snrpd21 | 5.19839155759974e-20 | 0.5483575 | 0.534 | 0.259 | 1.30838317113228e-15 | 21 |
| pcm1 | 7.41378698786669e-20 | 0.27431951 | 0.124 | 0.023 | 1.86597604697617e-15 | 21 |
| top1mt | 8.51575164976581e-20 | 0.282945211 | 0.192 | 0.045 | 2.14332953272956e-15 | 21 |
| rpa1 | 1.0320842015164e-19 | 0.397312743 | 0.254 | 0.076 | 2.59765272679664e-15 | 21 |
| phf20b | 1.04869131489747e-19 | 0.336153607 | 0.187 | 0.046 | 2.63945117046544e-15 | 21 |
| zgc:112496 | 1.12051561868767e-19 | 0.264652034 | 0.15 | 0.031 | 2.820225760675e-15 | 21 |
| cox8b2 | 1.14285920576558e-19 | 0.473735298 | 0.28 | 0.094 | 2.8764623349914e-15 | 21 |
| mcts1 | 1.16576505326727e-19 | 0.455331651 | 0.306 | 0.105 | 2.93411406256839e-15 | 21 |
| rsl1d12 | 1.17302804989084e-19 | 0.504752459 | 0.42 | 0.179 | 2.95239429877027e-15 | 21 |
| larp7 | 1.20575581170647e-19 | 0.352031895 | 0.233 | 0.065 | 3.034766802484e-15 | 21 |
| prdx31 | 1.29832040586566e-19 | 0.482020891 | 0.352 | 0.133 | 3.26774262952327e-15 | 21 |
| dus3l | 1.30712508553798e-19 | 0.410414845 | 0.233 | 0.067 | 3.28990312779053e-15 | 21 |
| cct24 | 1.33224323408632e-19 | 0.526706776 | 0.472 | 0.225 | 3.35312299587187e-15 | 21 |
| psmb6 | 1.35221706759544e-19 | 0.481051012 | 0.404 | 0.165 | 3.40339513743095e-15 | 21 |
| tomm34 | 1.45298734770677e-19 | 0.282041995 | 0.166 | 0.037 | 3.65702385544317e-15 | 21 |
| g3bp2 | 1.51255802072527e-19 | 0.304677571 | 0.161 | 0.035 | 3.80695728236343e-15 | 21 |
| si:ch211-68a17.7 | 1.61787468588556e-19 | 0.326030716 | 0.187 | 0.046 | 4.07202879690537e-15 | 21 |
| prpf8 | 1.80784085126474e-19 | 0.362576268 | 0.207 | 0.056 | 4.55015463854822e-15 | 21 |
| phb | 1.99752848400268e-19 | 0.360986016 | 0.218 | 0.06 | 5.02757944138634e-15 | 21 |
| nolc1 | 2.06114241790669e-19 | 0.398395848 | 0.28 | 0.088 | 5.18768935162935e-15 | 21 |
| hspa4a | 2.14767152152113e-19 | 0.308509013 | 0.192 | 0.047 | 5.40547445251654e-15 | 21 |
| ralaa | 2.88718875015085e-19 | 0.250495536 | 0.124 | 0.023 | 7.26676536525466e-15 | 21 |
| eif5a21 | 3.19067818127645e-19 | 0.514445488 | 0.663 | 0.401 | 8.03061791445471e-15 | 21 |
| ube2f | 3.51402289834575e-19 | 0.300211862 | 0.145 | 0.031 | 8.84444423284642e-15 | 21 |

| | | | | | | |
|---|---|---|---|---|---|---|
| mrps33 | 3.66610667901364e-19 | 0.370748417 | 0.259 | 0.078 | 9.22722390040942e-15 | 21 |
| supt16h | 4.01389275354254e-19 | 0.39091247 | 0.249 | 0.075 | 1.01025666713912e-14 | 21 |
| prmt11 | 5.07539824712131e-19 | 0.505093253 | 0.487 | 0.22 | 1.27742698481796e-14 | 21 |
| nip7 | 5.90192081661076e-19 | 0.370521729 | 0.202 | 0.054 | 1.48545445033276e-14 | 21 |
| selenou1a | 6.33096487084954e-19 | 0.313998728 | 0.161 | 0.037 | 1.59344054834412e-14 | 21 |
| cox7b5 | 6.74803982544385e-19 | 0.489015427 | 0.653 | 0.415 | 1.69841414366596e-14 | 21 |
| atp5f1d3 | 6.84082025698121e-19 | 0.515588524 | 0.513 | 0.256 | 1.7217660504796e-14 | 21 |
| si:ch211-226l4.67 | 7.94099918020946e-19 | 0.533742177 | 0.622 | 0.29 | 1.99867008366692e-14 | 21 |
| krr1 | 8.64802197639636e-19 | 0.268230567 | 0.15 | 0.032 | 2.1766206512392e-14 | 21 |
| si:ch211-51e12.7 | 8.92558347880234e-19 | 0.476287357 | 0.425 | 0.175 | 2.24648010577976e-14 | 21 |
| cbx3a4 | 9.28481646816199e-19 | 0.518646955 | 0.58 | 0.294 | 2.33689545687169e-14 | 21 |
| mrps14 | 1.04875701918732e-18 | 0.396309511 | 0.28 | 0.092 | 2.63961654159257e-14 | 21 |
| hypk | 1.08068046437134e-18 | 0.50070919 | 0.456 | 0.208 | 2.71996466077623e-14 | 21 |
| psmc6 | 1.15907865733022e-18 | 0.361300453 | 0.264 | 0.083 | 2.91728507263442e-14 | 21 |
| zswim5 | 1.38731522104488e-18 | 0.337292986 | 0.171 | 0.042 | 3.49173367984785e-14 | 21 |
| cox7c3 | 1.45964899999489e-18 | 0.467884478 | 0.694 | 0.48 | 3.67379056808714e-14 | 21 |
| rplp05 | 1.50356528706999e-18 | 0.255576761 | 0.964 | 0.85 | 3.78432347102646e-14 | 21 |
| LOC110438491 | 1.74500561563309e-18 | 0.260842496 | 0.14 | 0.029 | 4.39200463398691e-14 | 21 |
| pole3 | 1.92306565786448e-18 | 0.299387249 | 0.181 | 0.045 | 4.84016395427911e-14 | 21 |
| nap1l11 | 1.92622385060033e-18 | 0.508414071 | 0.637 | 0.359 | 4.84811280957597e-14 | 21 |
| caprin1a | 2.13134131444839e-18 | 0.350752405 | 0.192 | 0.051 | 5.36437295433516e-14 | 21 |
| rpsa11 | 2.16786413618736e-18 | 0.336097456 | 0.948 | 0.75 | 5.45629724436996e-14 | 21 |
| CABZ01068251.1 | 3.04758007340833e-18 | 0.263553193 | 0.171 | 0.041 | 7.67045428676143e-14 | 21 |
| mrpl21 | 3.10679354600378e-18 | 0.300648268 | 0.207 | 0.055 | 7.81948867593691e-14 | 21 |
| ndufb5 | 3.697691230571e-18 | 0.465248686 | 0.358 | 0.142 | 9.30671905822415e-14 | 21 |
| gins1 | 3.87980450674682e-18 | 0.256159338 | 0.135 | 0.027 | 9.76507996303106e-14 | 21 |
| zgc:1737426 | 4.69339986598288e-18 | 0.431265392 | 0.409 | 0.153 | 1.18128181226923e-13 | 21 |
| csnk2b1 | 5.21462702515174e-18 | 0.492778996 | 0.435 | 0.201 | 1.31246947596044e-13 | 21 |
| ddx23 | 5.75073560328602e-18 | 0.311910367 | 0.192 | 0.051 | 1.44740264399106e-13 | 21 |
| sf3b6 | 6.23487104976191e-18 | 0.51106901 | 0.487 | 0.244 | 1.56925469451458e-13 | 21 |
| trip13 | 6.24751322880759e-18 | 0.278527121 | 0.155 | 0.035 | 1.57243660455858e-13 | 21 |
| ndufa9a | 7.96183836792802e-18 | 0.40271912 | 0.238 | 0.075 | 2.0039150988238e-13 | 21 |
| alkbh5 | 1.07798755524896e-17 | 0.302928457 | 0.176 | 0.044 | 2.71318687780612e-13 | 21 |
| mkln1 | 1.25061030078988e-17 | 0.296122397 | 0.161 | 0.038 | 3.14766106605805e-13 | 21 |
| ccdc106b | 1.54403449352859e-17 | 0.326017642 | 0.176 | 0.044 | 3.8861804167621e-13 | 21 |

| | | | | | | |
|---|---|---|---|---|---|---|
| msh2 | 1.81649503336343e-17 | 0.266559773 | 0.15 | 0.034 | 4.57193634947241e-13 | 21 |
| zgc:922875 | 1.85964589656701e-17 | 0.398740576 | 0.363 | 0.131 | 4.68054275706951e-13 | 21 |
| zgc:101851 | 2.14060076577668e-17 | 0.368702363 | 0.275 | 0.09 | 5.38767806738333e-13 | 21 |
| si:ch211-286b5.53 | 2.15122074607475e-17 | 0.434940938 | 0.342 | 0.135 | 5.41440749579555e-13 | 21 |
| cks1b | 2.3371283684784e-17 | 0.322473235 | 0.223 | 0.065 | 5.88231839062329e-13 | 21 |
| psma3 | 2.35623144817036e-17 | 0.495682398 | 0.394 | 0.175 | 5.93039893189998e-13 | 21 |
| cope | 2.36104669018141e-17 | 0.429727684 | 0.342 | 0.132 | 5.94251841451759e-13 | 21 |
| polr2d | 2.3833412264565e-17 | 0.445582525 | 0.332 | 0.13 | 5.99863153286835e-13 | 21 |
| zgc:63470 | 2.66433617498955e-17 | 0.311922995 | 0.171 | 0.043 | 6.7058677188312e-13 | 21 |
| cdk2 | 3.26016956816103e-17 | 0.271701372 | 0.166 | 0.04 | 8.20552078610451e-13 | 21 |
| sdhc | 3.65164472388892e-17 | 0.420549991 | 0.352 | 0.14 | 9.19082460555602e-13 | 21 |
| dynlrb11 | 3.78678158643577e-17 | 0.480281734 | 0.44 | 0.203 | 9.53095057490019e-13 | 21 |
| zgc:103697 | 4.19670284527482e-17 | 0.32797783 | 0.197 | 0.055 | 1.05626813912722e-12 | 21 |
| dnajc8 | 4.46477441510646e-17 | 0.363009695 | 0.223 | 0.068 | 1.12373907253814e-12 | 21 |
| cdk1 | 4.8940719521288e-17 | 0.375648452 | 0.275 | 0.09 | 1.2317889696313e-12 | 21 |
| mrpl30 | 6.42829243235255e-17 | 0.409800853 | 0.295 | 0.108 | 1.61793692229881e-12 | 21 |
| atp5mf2 | 6.93142832286093e-17 | 0.489674161 | 0.534 | 0.286 | 1.74457119458087e-12 | 21 |
| eif5b | 7.39361902501695e-17 | 0.434592277 | 0.347 | 0.141 | 1.86089997240652e-12 | 21 |
| ilf3b | 9.82768788508137e-17 | 0.39102419 | 0.306 | 0.112 | 2.47353076379613e-12 | 21 |
| phax | 1.11292548116457e-16 | 0.333597767 | 0.218 | 0.064 | 2.80112214354311e-12 | 21 |
| uchl3 | 1.36743308980715e-16 | 0.318455069 | 0.207 | 0.06 | 3.44169234373562e-12 | 21 |
| swt1 | 1.53048537183647e-16 | 0.313074741 | 0.161 | 0.041 | 3.8520786323752e-12 | 21 |
| tuba1b1 | 1.66107107710723e-16 | 0.288756 | 0.135 | 0.03 | 4.18074979397118e-12 | 21 |
| prpf39 | 1.78068147330163e-16 | 0.349550045 | 0.207 | 0.062 | 4.48179720015287e-12 | 21 |
| pfdn1 | 1.84042420438062e-16 | 0.480948936 | 0.342 | 0.145 | 4.63216368000557e-12 | 21 |
| uqcc2 | 2.15365517883545e-16 | 0.344723342 | 0.207 | 0.062 | 5.42053471961095e-12 | 21 |
| fam162a | 2.22167539068382e-16 | 0.351408344 | 0.249 | 0.082 | 5.5917347908121e-12 | 21 |
| chchd2 | 2.44877901807466e-16 | 0.357876093 | 0.223 | 0.07 | 6.16333191059212e-12 | 21 |
| sephs2 | 3.6362102969696e-16 | 0.296041746 | 0.176 | 0.048 | 9.15197769644278e-12 | 21 |
| skp11 | 3.79813346104906e-16 | 0.495620159 | 0.477 | 0.249 | 9.55952210811439e-12 | 21 |
| pcgf6 | 4.69544679905966e-16 | 0.267611711 | 0.166 | 0.042 | 1.18179700485533e-11 | 21 |
| cse1l | 5.1797815944948e-16 | 0.36619463 | 0.238 | 0.076 | 1.3036992295184e-11 | 21 |
| coil1 | 5.24477932093018e-16 | 0.367492978 | 0.301 | 0.106 | 1.32005850728492e-11 | 21 |
| mphosph10 | 6.15239049940679e-16 | 0.270451541 | 0.145 | 0.035 | 1.5484951647957e-11 | 21 |
| gtf2b | 6.44154611306123e-16 | 0.348745723 | 0.228 | 0.073 | 1.62127274119638e-11 | 21 |

| | | | | | | |
|---|---|---|---|---|---|---|
| chmp5b | 6.77266059894268e-16 | 0.410113167 | 0.295 | 0.113 | 1.70461094614788e-11 | 21 |
| ndufb7 | 7.00295355423928e-16 | 0.44889639 | 0.378 | 0.169 | 1.76257338006648e-11 | 21 |
| fen11 | 7.23844663677042e-16 | 0.38313856 | 0.254 | 0.086 | 1.82184463400875e-11 | 21 |
| pes | 7.55714305599405e-16 | 0.316768842 | 0.166 | 0.044 | 1.90205733576314e-11 | 21 |
| npm3 | 8.28950824209657e-16 | 0.411397755 | 0.29 | 0.111 | 2.08638632945329e-11 | 21 |
| cript | 1.0757120032349e-15 | 0.256489981 | 0.155 | 0.039 | 2.70745954094193e-11 | 21 |
| ISCU (1 of many) | 1.14433437114077e-15 | 0.397512406 | 0.29 | 0.11 | 2.88017517872421e-11 | 21 |
| mrpl46 | 1.16588341406896e-15 | 0.300732036 | 0.15 | 0.038 | 2.93441196487016e-11 | 21 |
| dynll17 | 1.19958898324011e-15 | 0.461205795 | 0.699 | 0.453 | 3.01924551191703e-11 | 21 |
| polr1c | 1.24048113384062e-15 | 0.271982266 | 0.155 | 0.039 | 3.12216696576346e-11 | 21 |
| idh3g | 1.33044346221359e-15 | 0.296390873 | 0.155 | 0.04 | 3.34859315004539e-11 | 21 |
| si:dkey-67c22.2 | 1.33384420501893e-15 | 0.356259185 | 0.254 | 0.089 | 3.35715247961215e-11 | 21 |
| flot1b1 | 1.3547452587769e-15 | 0.34958983 | 0.238 | 0.081 | 3.40975834181558e-11 | 21 |
| pop5 | 1.46632093707126e-15 | 0.254096315 | 0.171 | 0.046 | 3.69058316651466e-11 | 21 |
| rchy1 | 1.62764457481427e-15 | 0.333838831 | 0.233 | 0.075 | 4.09661863035004e-11 | 21 |
| sod1 | 1.6370316480857e-15 | 0.461385409 | 0.565 | 0.326 | 4.1202449550669e-11 | 21 |
| dpp31 | 1.95284754268795e-15 | 0.259249661 | 0.155 | 0.04 | 4.91512198019129e-11 | 21 |
| fam133b | 1.98732410860823e-15 | 0.429670388 | 0.342 | 0.143 | 5.00189604895604e-11 | 21 |
| eny2 | 2.02701968139368e-15 | 0.383588779 | 0.28 | 0.103 | 5.10180583609974e-11 | 21 |
| bud31 | 2.03762416025011e-15 | 0.464691575 | 0.461 | 0.226 | 5.12849624893349e-11 | 21 |
| mrps21 | 2.03767257619157e-15 | 0.345483566 | 0.254 | 0.087 | 5.12861810701656e-11 | 21 |
| maco1a | 2.0830158516636e-15 | 0.303426679 | 0.161 | 0.043 | 5.2427425970521e-11 | 21 |
| si:ch211-284b7.3 | 2.09731162985336e-15 | 0.327172548 | 0.176 | 0.05 | 5.27872364117792e-11 | 21 |
| selenoh3 | 2.15849789668632e-15 | 0.434925963 | 0.508 | 0.242 | 5.4327233561698e-11 | 21 |
| gins2 | 2.33324249302786e-15 | 0.344720612 | 0.218 | 0.067 | 5.87253803070183e-11 | 21 |
| nsfl1c | 2.35450883494517e-15 | 0.367229795 | 0.212 | 0.068 | 5.9260632866735e-11 | 21 |
| nup37 | 2.41194105280743e-15 | 0.273908036 | 0.155 | 0.04 | 6.07061443581102e-11 | 21 |
| pane1 | 2.43373496984606e-15 | 0.251456831 | 0.14 | 0.033 | 6.12546754560554e-11 | 21 |
| polr3h | 2.45817386608072e-15 | 0.265283577 | 0.14 | 0.034 | 6.18697780353857e-11 | 21 |
| trnau1apa | 2.58237473477175e-15 | 0.286403675 | 0.187 | 0.053 | 6.49957896994701e-11 | 21 |
| psma6a | 2.83057545017708e-15 | 0.346274614 | 0.28 | 0.101 | 7.12427535055069e-11 | 21 |
| baz1b | 2.84795441522788e-15 | 0.344154433 | 0.207 | 0.065 | 7.16801646768704e-11 | 21 |
| smim8 | 2.87700675033726e-15 | 0.273679444 | 0.15 | 0.038 | 7.24113828992386e-11 | 21 |
| rfc2 | 3.17473383044245e-15 | 0.282695407 | 0.166 | 0.044 | 7.99048757784059e-11 | 21 |
| tma71 | 3.35675409859946e-15 | 0.475984093 | 0.534 | 0.303 | 8.44861439076498e-11 | 21 |

| | | | | | | |
|---|---|---|---|---|---|---|
| trappc1 | 3.45072903901566e-15 | 0.401239757 | 0.264 | 0.099 | 8.68513991829852e-11 | 21 |
| si:dkey-92i17.21 | 3.47875623083545e-15 | 0.365048416 | 0.264 | 0.091 | 8.75568155738975e-11 | 21 |
| pdrg1 | 4.23328143430838e-15 | 0.259417132 | 0.15 | 0.038 | 1.06547460420108e-10 | 21 |
| atp5md1 | 4.47530290337898e-15 | 0.480177397 | 0.565 | 0.358 | 1.12638898775145e-10 | 21 |
| tuba8l23 | 5.31082426912901e-15 | 0.309042715 | 0.197 | 0.06 | 1.33668136029708e-10 | 21 |
| UTP14C | 5.6455541435388e-15 | 0.260109682 | 0.155 | 0.041 | 1.42092952238728e-10 | 21 |
| rrp36 | 5.70213234981253e-15 | 0.287752525 | 0.166 | 0.045 | 1.43516969112432e-10 | 21 |
| ing5b | 5.74598748656183e-15 | 0.258474862 | 0.181 | 0.05 | 1.44620759049275e-10 | 21 |
| si:dkeyp-55f12.3 | 5.75414145571759e-15 | 0.290583728 | 0.187 | 0.055 | 1.44825986298956e-10 | 21 |
| cenpw | 5.95106772963079e-15 | 0.256310944 | 0.145 | 0.036 | 1.49782423687077e-10 | 21 |
| hnrnpa0l1 | 6.60994325458789e-15 | 0.296435233 | 0.907 | 0.756 | 1.66365661774723e-10 | 21 |
| gnpda12 | 6.69842209809644e-15 | 0.289114764 | 0.161 | 0.043 | 1.68592585786989e-10 | 21 |
| mtap | 7.65263171064192e-15 | 0.37248518 | 0.238 | 0.083 | 1.92609087525147e-10 | 21 |
| pho | 8.09527744767415e-15 | 0.399791322 | 0.249 | 0.09 | 2.03750038080511e-10 | 21 |
| eme1 | 8.33755998438437e-15 | 0.254644036 | 0.135 | 0.032 | 2.0984804724697e-10 | 21 |
| trir | 8.36577923909176e-15 | 0.322782611 | 0.181 | 0.054 | 2.10558297668701e-10 | 21 |
| med20 | 8.53308482451905e-15 | 0.258902001 | 0.145 | 0.037 | 2.1476921194832e-10 | 21 |
| cct31 | 9.57641047268955e-15 | 0.422660186 | 0.425 | 0.205 | 2.41028675187123e-10 | 21 |
| c1qbp | 9.85972699404748e-15 | 0.443662311 | 0.378 | 0.17 | 2.48159468713181e-10 | 21 |
| atp5f1c1 | 1.02009880528083e-14 | 0.472984326 | 0.472 | 0.253 | 2.56748668301131e-10 | 21 |
| abcf1 | 1.21733083681027e-14 | 0.353936188 | 0.254 | 0.092 | 3.06389998316777e-10 | 21 |
| ndufs4 | 1.22965661756614e-14 | 0.367971454 | 0.264 | 0.098 | 3.09492274075221e-10 | 21 |
| ddx61 | 1.25007429507591e-14 | 0.479251371 | 0.42 | 0.214 | 3.14631199327655e-10 | 21 |
| suclg12 | 1.36354912135129e-14 | 0.383083569 | 0.28 | 0.108 | 3.43191678352907e-10 | 21 |
| si:dkey-189g17.21 | 1.48164750372687e-14 | 0.279430191 | 0.192 | 0.056 | 3.72915860213016e-10 | 21 |
| nkap | 1.52843412318637e-14 | 0.37855173 | 0.29 | 0.113 | 3.84691584464776e-10 | 21 |
| atp5l5 | 1.75404253833864e-14 | 0.40998683 | 0.663 | 0.448 | 4.41474966474451e-10 | 21 |
| bida1 | 1.84331176558713e-14 | 0.34988974 | 0.212 | 0.07 | 4.63943138280624e-10 | 21 |
| ube2d1b | 1.93993903535955e-14 | 0.408317279 | 0.311 | 0.13 | 4.88263255809645e-10 | 21 |
| psmd14 | 2.14936910397535e-14 | 0.355745222 | 0.264 | 0.097 | 5.40974709779555e-10 | 21 |
| bola3 | 2.17223113759395e-14 | 0.273642513 | 0.218 | 0.07 | 5.46728855021021e-10 | 21 |
| ndufa1 | 2.24323400584275e-14 | 0.457836259 | 0.44 | 0.229 | 5.64599566930561e-10 | 21 |
| parga | 2.72270511474752e-14 | 0.273440041 | 0.119 | 0.027 | 6.85277650330803e-10 | 21 |
| ndufa31 | 2.72923718031466e-14 | 0.421873866 | 0.389 | 0.182 | 6.86921705913397e-10 | 21 |
| snw1 | 3.41952570617786e-14 | 0.334513114 | 0.212 | 0.071 | 8.60660424987905e-10 | 21 |

| | | | | | | |
|---|---|---|---|---|---|---|
| mrpl13 | 3.64975842815996e-14 | 0.3222172 | 0.181 | 0.055 | 9.18607698783580e-10 | 21 |
| safb | 3.80483174545362e-14 | 0.307796804 | 0.176 | 0.053 | 9.57638102013222e-10 | 21 |
| cops4 | 4.34196744774522e-14 | 0.356946841 | 0.223 | 0.077 | 1.09282978692299e-09 | 21 |
| si:zfos-169g10.3 | 4.40410739892537e-14 | 0.38056733 | 0.249 | 0.094 | 1.10846979123553e-09 | 21 |
| hspe12 | 4.4288605159516e-14 | 0.461101164 | 0.539 | 0.293 | 1.11469990325986e-09 | 21 |
| mrpl14 | 4.76366359667083e-14 | 0.3429595 | 0.269 | 0.1 | 1.19896649064608e-09 | 21 |
| csnk2a1 | 4.93421289809251e-14 | 0.466114097 | 0.378 | 0.182 | 1.2418920443209e-09 | 21 |
| sap181 | 5.35305567258561e-14 | 0.448330831 | 0.435 | 0.217 | 1.34731058223307e-09 | 21 |
| polr2h | 5.5091191174818e-14 | 0.296974516 | 0.212 | 0.068 | 1.386590190679e-09 | 21 |
| orc4 | 5.69948867977603e-14 | 0.265239366 | 0.155 | 0.042 | 1.43450430581283e-09 | 21 |
| rpf12 | 6.00264576980601e-14 | 0.388790087 | 0.326 | 0.135 | 1.51080591380247e-09 | 21 |
| atp5mc3a2 | 6.99241216293682e-14 | 0.447714125 | 0.43 | 0.222 | 1.75992021728957e-09 | 21 |
| myg1 | 8.21621862627354e-14 | 0.317933971 | 0.197 | 0.064 | 2.06794006604679e-09 | 21 |
| atp5pf4 | 8.27272891505614e-14 | 0.451927736 | 0.503 | 0.297 | 2.08216314063048e-09 | 21 |
| rps19bp1 | 8.30449037237358e-14 | 0.347623046 | 0.254 | 0.093 | 2.09015718182271e-09 | 21 |
| fam168b | 8.64282417386226e-14 | 0.314936375 | 0.228 | 0.078 | 2.17531241631939e-09 | 21 |
| taf11 | 8.67245815996593e-14 | 0.258667908 | 0.155 | 0.043 | 2.18277099428182e-09 | 21 |
| rbm25b | 9.07207395427228e-14 | 0.290452281 | 0.171 | 0.051 | 2.28335029355079e-09 | 21 |
| mcm2 | 9.64943426568288e-14 | 0.327928034 | 0.233 | 0.081 | 2.42866611032972e-09 | 21 |
| rpl23a10 | 1.03538884816995e-13 | 0.339959704 | 0.834 | 0.633 | 2.60597019195893e-09 | 21 |
| si:ch211-276i12.9 | 1.08666679432242e-13 | 0.284137383 | 0.197 | 0.062 | 2.73503165463011e-09 | 21 |
| h3f3c5 | 1.10575576176393e-13 | 0.437499646 | 0.472 | 0.255 | 2.78307667678364e-09 | 21 |
| sem1 | 1.10901764124242e-13 | 0.446591131 | 0.466 | 0.251 | 2.79128650124305e-09 | 21 |
| cstf2 | 1.52887633859178e-13 | 0.296283669 | 0.207 | 0.069 | 3.84802885660166e-09 | 21 |
| prpf31 | 1.60000503002629e-13 | 0.300963471 | 0.238 | 0.082 | 4.02705266007316e-09 | 21 |
| mrpl11 | 1.7775043236668e-13 | 0.329542169 | 0.197 | 0.066 | 4.47380063223697e-09 | 21 |
| prdx16 | 1.78044667471098e-13 | 0.418315349 | 0.378 | 0.165 | 4.48120623558007e-09 | 21 |
| u2af2a | 1.85462618546528e-13 | 0.28103597 | 0.181 | 0.056 | 4.66790864619756e-09 | 21 |
| fbl1 | 1.86798637666981e-13 | 0.439098271 | 0.415 | 0.203 | 4.70153491144024e-09 | 21 |
| cct52 | 2.06279566160099e-13 | 0.455954154 | 0.456 | 0.25 | 5.19185040068354e-09 | 21 |
| dynll2a6 | 2.29118908387608e-13 | 0.369283441 | 0.332 | 0.135 | 5.76669380520772e-09 | 21 |
| srsf3a | 2.300787043161e-13 | 0.254784987 | 0.15 | 0.042 | 5.79085090893193e-09 | 21 |
| s100u1 | 2.31212407154419e-13 | 0.262138099 | 0.135 | 0.036 | 5.81938507566958e-09 | 21 |
| hdac1 | 2.49248636006844e-13 | 0.433279199 | 0.425 | 0.217 | 6.27333891965627e-09 | 21 |
| naa20 | 2.55483854621291e-13 | 0.293878119 | 0.181 | 0.058 | 6.43027313696328e-09 | 21 |

| | | | | | | |
|---|---|---|---|---|---|---|
| gstm.13 | 2.62569032245143e-13 | 0.463406204 | 0.466 | 0.259 | 6.60859997257799e-09 | 21 |
| ndufb10 | 2.76828500081187e-13 | 0.410767333 | 0.326 | 0.147 | 6.96749651854339e-09 | 21 |
| zgc:566997 | 3.17764271202698e-13 | 0.439770978 | 0.513 | 0.25 | 7.9978089419007e-09 | 21 |
| alg13 | 4.52598944393867e-13 | 0.292217668 | 0.202 | 0.065 | 1.13914628314492e-08 | 21 |
| cox5ab1 | 4.74345906698941e-13 | 0.415984993 | 0.399 | 0.201 | 1.19388121257057e-08 | 21 |
| hmgb2a2 | 4.9042509289151e-13 | 0.355326609 | 0.793 | 0.62 | 1.23435091629864e-08 | 21 |
| idh3b | 5.56557229487391e-13 | 0.250260647 | 0.187 | 0.059 | 1.40079889089681e-08 | 21 |
| uba1 | 5.94525065389612e-13 | 0.349590484 | 0.269 | 0.108 | 1.49636013707912e-08 | 21 |
| lyar | 6.15375139419033e-13 | 0.258318644 | 0.171 | 0.052 | 1.54883768840376e-08 | 21 |
| slirp | 6.56513482253658e-13 | 0.368311322 | 0.29 | 0.12 | 1.65237878348423e-08 | 21 |
| ncaph | 6.64904218049089e-13 | 0.250462892 | 0.14 | 0.037 | 1.67349742640775e-08 | 21 |
| zgc:56304 | 7.31168841873111e-13 | 0.349023674 | 0.275 | 0.111 | 1.84027885811043e-08 | 21 |
| mfap1 | 7.77043314652074e-13 | 0.276509756 | 0.171 | 0.053 | 1.95574031864781e-08 | 21 |
| rbbp44 | 7.88174968022985e-13 | 0.390076274 | 0.378 | 0.169 | 1.98375757701705e-08 | 21 |
| zgc:1034827 | 7.97502353972299e-13 | 0.377702314 | 0.425 | 0.187 | 2.00723367471288e-08 | 21 |
| ptp4a12 | 8.20186496098102e-13 | 0.351984699 | 0.28 | 0.117 | 2.06432739202931e-08 | 21 |
| prpf3 | 8.25583944754848e-13 | 0.272148082 | 0.14 | 0.038 | 2.07791223055348e-08 | 21 |
| gpn2 | 9.15885770386893e-13 | 0.28179133 | 0.176 | 0.055 | 2.30519289548677e-08 | 21 |
| srpk1a | 9.54366130687643e-13 | 0.292114709 | 0.202 | 0.068 | 2.40204411432773e-08 | 21 |
| gtf2f1 | 9.65046248144705e-13 | 0.284525284 | 0.192 | 0.064 | 2.42892490195541e-08 | 21 |
| mrps11 | 1.02861867229961e-12 | 0.317819799 | 0.197 | 0.067 | 2.58893033631088e-08 | 21 |
| COX5B3 | 1.16270703726559e-12 | 0.415540911 | 0.565 | 0.357 | 2.92641734209375e-08 | 21 |
| fhit | 1.25641102388132e-12 | 0.280189939 | 0.161 | 0.048 | 3.16226090600688e-08 | 21 |
| rexo4 | 1.51252127945844e-12 | 0.276994338 | 0.202 | 0.068 | 3.80686480826894e-08 | 21 |
| uqcrc11 | 1.57736982480305e-12 | 0.341372756 | 0.238 | 0.092 | 3.97008211204681e-08 | 21 |
| si:ch73-1a9.37 | 1.78343004197537e-12 | 0.441858416 | 0.456 | 0.265 | 4.4887150726478e-08 | 21 |
| ythdf2 | 1.86378023534922e-12 | 0.335907353 | 0.228 | 0.085 | 4.69094847435044e-08 | 21 |
| lsm14ab | 2.00549388687875e-12 | 0.29341181 | 0.207 | 0.073 | 5.04762756388514e-08 | 21 |
| hars | 2.28928923800891e-12 | 0.256240301 | 0.14 | 0.04 | 5.76191208314463e-08 | 21 |
| exosc8 | 2.29113905904733e-12 | 0.282365836 | 0.207 | 0.071 | 5.76656789771622e-08 | 21 |
| ndufb2 | 2.29999635585729e-12 | 0.419528404 | 0.373 | 0.185 | 5.78886082805721e-08 | 21 |
| atp5mc12 | 2.55690476788766e-12 | 0.422465341 | 0.554 | 0.33 | 6.43547361029645e-08 | 21 |
| atp5fa11 | 2.73095496205863e-12 | 0.420343759 | 0.565 | 0.355 | 6.87354054400537e-08 | 21 |
| tufm | 2.78631197608525e-12 | 0.292424979 | 0.228 | 0.084 | 7.01286861260897e-08 | 21 |
| hmgn32 | 3.05632463678056e-12 | 0.278522128 | 0.181 | 0.06 | 7.69246347831299e-08 | 21 |

| | | | | | | |
|---|---|---|---|---|---|---|
| btr04 | 3.18951273808946e-12 | 0.259570215 | 0.192 | 0.063 | 8.02768461049736e-08 | 21 |
| morc2 | 3.48425035117013e-12 | 0.272277428 | 0.15 | 0.044 | 8.76950970886011e-08 | 21 |
| eif3ja1 | 3.59608480588225e-12 | 0.423283646 | 0.373 | 0.189 | 9.05098584792503e-08 | 21 |
| clns1a | 3.84428152102305e-12 | 0.287161145 | 0.207 | 0.072 | 9.67567216026293e-08 | 21 |
| txndc91 | 4.49029179199156e-12 | 0.371712721 | 0.269 | 0.116 | 1.13016154112636e-07 | 21 |
| cnot9 | 4.77017874098972e-12 | 0.285005004 | 0.181 | 0.061 | 1.2006062873197e-07 | 21 |
| prmt5 | 4.80582093344759e-12 | 0.253160818 | 0.155 | 0.047 | 1.20957707073942e-07 | 21 |
| khdrbs1a | 5.32931427481726e-12 | 0.352707795 | 0.741 | 0.563 | 1.34133510982876e-07 | 21 |
| si:dkey-180p18.93 | 6.14871620858776e-12 | 0.354344996 | 0.244 | 0.099 | 1.54757038253945e-07 | 21 |
| atp5meb2 | 6.15249629425086e-12 | 0.42235396 | 0.513 | 0.321 | 1.5485217923e-07 | 21 |
| pfn2l3 | 7.70579243989203e-12 | 0.378666346 | 0.415 | 0.208 | 1.93947089919643e-07 | 21 |
| uqcrh3 | 8.24319501935396e-12 | 0.39652349 | 0.451 | 0.249 | 2.0747297544212e-07 | 21 |
| chrac11 | 8.26796087473299e-12 | 0.364000698 | 0.301 | 0.128 | 2.08096307256155e-07 | 21 |
| pcnp | 8.46799068402084e-12 | 0.261124789 | 0.161 | 0.05 | 2.13130857526121e-07 | 21 |
| brk11 | 9.04746895293656e-12 | 0.414489774 | 0.358 | 0.181 | 2.2771574607646e-07 | 21 |
| pgam5 | 9.46722273838429e-12 | 0.325354829 | 0.264 | 0.107 | 2.38280529102394e-07 | 21 |
| srsf5a4 | 1.00382758588447e-11 | 0.402192866 | 0.487 | 0.287 | 2.52653365091262e-07 | 21 |
| psmd3 | 1.01821964045672e-11 | 0.276731687 | 0.202 | 0.072 | 2.56275701306552e-07 | 21 |
| cox14 | 1.06917092023264e-11 | 0.316147088 | 0.254 | 0.103 | 2.69099628913354e-07 | 21 |
| rnaseh2a | 1.12072669456076e-11 | 0.253918348 | 0.155 | 0.048 | 2.82075701753999e-07 | 21 |
| rad23b | 1.12267436055972e-11 | 0.38644604 | 0.337 | 0.162 | 2.82565909809276e-07 | 21 |
| ndufv2 | 1.15616980010209e-11 | 0.276097492 | 0.207 | 0.075 | 2.90996376987694e-07 | 21 |
| psma21 | 1.31500024613817e-11 | 0.371577335 | 0.321 | 0.151 | 3.30972411950517e-07 | 21 |
| nop2 | 1.35850988852128e-11 | 0.274121068 | 0.15 | 0.047 | 3.41923353841921e-07 | 21 |
| esd1 | 1.41397470331288e-11 | 0.33617019 | 0.244 | 0.099 | 3.55883293076818e-07 | 21 |
| cct6a | 1.68515206263324e-11 | 0.399084423 | 0.389 | 0.207 | 4.24135922644161e-07 | 21 |
| smndc1 | 1.70663311574144e-11 | 0.342928949 | 0.259 | 0.109 | 4.29542488900962e-07 | 21 |
| kpna26 | 1.81045020697199e-11 | 0.346797948 | 0.352 | 0.157 | 4.55672212592781e-07 | 21 |
| cox6c2 | 2.00619848460886e-11 | 0.392512027 | 0.606 | 0.416 | 5.04940096591205e-07 | 21 |
| tomm22 | 2.03881645347002e-11 | 0.295605325 | 0.228 | 0.088 | 5.1314971317387e-07 | 21 |
| htatsf1 | 2.1407378974314e-11 | 0.251406366 | 0.171 | 0.056 | 5.3880232140451e-07 | 21 |
| mrpl53 | 2.49128858264831e-11 | 0.312531592 | 0.233 | 0.09 | 6.27032423366754e-07 | 21 |
| cfap20 | 2.60045313433405e-11 | 0.272692887 | 0.166 | 0.054 | 6.54508049380536e-07 | 21 |
| ndufb91 | 2.74928822373347e-11 | 0.369058615 | 0.352 | 0.175 | 6.91968353031478e-07 | 21 |
| thrap3b | 2.85533246247765e-11 | 0.311391789 | 0.223 | 0.087 | 7.18658627481e-07 | 21 |

| | | | | | | |
|---|---|---|---|---|---|---|
| lyrm9 | 2.92761708533889e-11 | 0.254673229 | 0.181 | 0.061 | 7.36851944208945e-07 | 21 |
| ndufa11 | 3.00417383300497e-11 | 0.369705691 | 0.321 | 0.153 | 7.56120512029022e-07 | 21 |
| taf7 | 4.07615266975038e-11 | 0.260883222 | 0.155 | 0.05 | 1.02592686544947e-06 | 21 |
| dctpp17 | 4.40960690163105e-11 | 0.3392668 | 0.378 | 0.174 | 1.10985396107152e-06 | 21 |
| zgc:554617 | 4.79579159084809e-11 | 0.400490152 | 0.544 | 0.287 | 1.20705278550056e-06 | 21 |
| lig1 | 4.86933154584700e-11 | 0.264540047 | 0.166 | 0.055 | 1.22556205677423e-06 | 21 |
| mrps18c1 | 5.03849813338636e-11 | 0.323779035 | 0.285 | 0.123 | 1.26813959519201e-06 | 21 |
| rrs1 | 5.17533596232598e-11 | 0.295086333 | 0.218 | 0.083 | 1.30258030835782e-06 | 21 |
| smu1a | 5.40418227692409e-11 | 0.258511161 | 0.197 | 0.071 | 1.36017863727902e-06 | 21 |
| pabpc1b4 | 5.44056628234995e-11 | 0.308704121 | 0.187 | 0.068 | 1.36933612760466e-06 | 21 |
| polr2eb | 5.64656149161842e-11 | 0.306391205 | 0.207 | 0.079 | 1.42118306182544e-06 | 21 |
| tnrc6b | 5.7117206900573e-11 | 0.308995528 | 0.218 | 0.085 | 1.43758298048052e-06 | 21 |
| si:dkey-188i13.117 | 6.05455879074393e-11 | 0.39806694 | 0.56 | 0.315 | 1.52387190204234e-06 | 21 |
| txn21 | 6.14078366756064e-11 | 0.313705568 | 0.223 | 0.089 | 1.54557384128834e-06 | 21 |
| phf5a | 6.21414734738382e-11 | 0.352198942 | 0.337 | 0.161 | 1.56403874586303e-06 | 21 |
| mpc1 | 6.52725044445798e-11 | 0.289111104 | 0.197 | 0.074 | 1.64284366436563e-06 | 21 |
| psip1b | 6.85777517853391e-11 | 0.283256212 | 0.176 | 0.062 | 1.7260334346852e-06 | 21 |
| rrp8 | 6.97045367072605e-11 | 0.289573976 | 0.187 | 0.067 | 1.75439348438504e-06 | 21 |
| psma1 | 7.23309357867221e-11 | 0.371411521 | 0.446 | 0.249 | 1.82049732281601e-06 | 21 |
| ndufs2 | 7.32635588704707e-11 | 0.294166704 | 0.218 | 0.086 | 1.84397051321088e-06 | 21 |
| timm17a | 7.46302495184029e-11 | 0.285541773 | 0.187 | 0.068 | 1.87836875012868e-06 | 21 |
| hexim1 | 8.09660028578944e-11 | 0.260266758 | 0.223 | 0.085 | 2.03783332593034e-06 | 21 |
| rbm19 | 8.77561520075024e-11 | 0.270155551 | 0.15 | 0.049 | 2.20873458987683e-06 | 21 |
| rrp9 | 8.8465436421224e-11 | 0.253297165 | 0.181 | 0.063 | 2.22658656928579e-06 | 21 |
| hspd12 | 9.65437881497615e-11 | 0.35605619 | 0.342 | 0.164 | 2.42991060394135e-06 | 21 |
| ubap2b | 9.86952896431536e-11 | 0.294566044 | 0.218 | 0.085 | 2.48406174502853e-06 | 21 |
| psmd13 | 1.02979213920221e-10 | 0.2905203 | 0.218 | 0.085 | 2.59188383515805e-06 | 21 |
| ndufa10 | 1.12435918927323e-10 | 0.261385704 | 0.207 | 0.079 | 2.82989964348179e-06 | 21 |
| rplp2l11 | 1.22658716875086e-10 | 0.279989642 | 0.953 | 0.772 | 3.08719724502905e-06 | 21 |
| strap | 1.28602966811317e-10 | 0.312494098 | 0.238 | 0.098 | 3.23680807167403e-06 | 21 |
| znf593 | 1.31752030630646e-10 | 0.30771062 | 0.275 | 0.119 | 3.31606685894273e-06 | 21 |
| vcp | 1.38992660547495e-10 | 0.355711333 | 0.285 | 0.13 | 3.49830627331991e-06 | 21 |
| seh1l | 1.52766025734818e-10 | 0.250047638 | 0.155 | 0.051 | 3.84496810171964e-06 | 21 |
| iscub | 1.59382262414743e-10 | 0.255601775 | 0.181 | 0.066 | 4.01149216271666e-06 | 21 |
| ywhabb2 | 1.76474975051747e-10 | 0.386377964 | 0.404 | 0.226 | 4.44169864707742e-06 | 21 |

| | | | | | | |
|---|---|---|---|---|---|---|
| pa2g4a | 1.84813429914185e-10 | 0.283759655 | 0.28 | 0.121 | 4.65156921751013e-06 | 21 |
| mad2l15 | 2.03115638645709e-10 | 0.319582786 | 0.394 | 0.185 | 5.11221750907384e-06 | 21 |
| imp35 | 2.1816303722351e-10 | 0.329918592 | 0.301 | 0.134 | 5.49094548387852e-06 | 21 |
| bccip | 2.22626696187871e-10 | 0.356789313 | 0.28 | 0.133 | 5.60329131635252e-06 | 21 |
| vps4b | 2.56048904417511e-10 | 0.317791822 | 0.233 | 0.099 | 6.44449487528434e-06 | 21 |
| u2af2b | 2.5787860182904e-10 | 0.334887535 | 0.264 | 0.118 | 6.49054652943512e-06 | 21 |
| mrpl52 | 2.80076551640893e-10 | 0.271310844 | 0.197 | 0.074 | 7.04924672824964e-06 | 21 |
| park7 | 2.90109143640867e-10 | 0.293597063 | 0.207 | 0.082 | 7.30175703629698e-06 | 21 |
| pfdn2 | 2.98072594664296e-10 | 0.341312961 | 0.28 | 0.128 | 7.50218913510566e-06 | 21 |
| sf3b2 | 3.0276268297183e-10 | 0.295340038 | 0.197 | 0.076 | 7.62023396771798e-06 | 21 |
| rwdd1 | 3.07163447755038e-10 | 0.333663476 | 0.275 | 0.125 | 7.73099681654655e-06 | 21 |
| ppp2r2d | 3.12255483008656e-10 | 0.255310577 | 0.155 | 0.053 | 7.85915825184487e-06 | 21 |
| rpia | 3.24886978162609e-10 | 0.25149302 | 0.176 | 0.063 | 8.1770803533747e-06 | 21 |
| pdcd4a | 3.50360250736142e-10 | 0.255376009 | 0.161 | 0.056 | 8.81821715077796e-06 | 21 |
| psmd12 | 3.68323096468149e-10 | 0.269461617 | 0.176 | 0.063 | 9.27032401500684e-06 | 21 |
| aimp11 | 3.80478371901616e-10 | 0.305521576 | 0.223 | 0.093 | 9.57626014239179e-06 | 21 |
| dkc11 | 4.17495656790919e-10 | 0.351428964 | 0.352 | 0.174 | 1.05079481857706e-05 | 21 |
| ddx27 | 4.24541797512298e-10 | 0.268805362 | 0.197 | 0.076 | 1.0685292501587e-05 | 21 |
| cct41 | 4.70738651705222e-10 | 0.381215994 | 0.399 | 0.226 | 1.18480211247687e-05 | 21 |
| rtca | 4.893670555893e-10 | 0.302567461 | 0.233 | 0.097 | 1.23168794221271e-05 | 21 |
| atp5pb1 | 4.92768440772059e-10 | 0.384612353 | 0.477 | 0.299 | 1.2402488885792e-05 | 21 |
| ndufb32 | 5.01274134115753e-10 | 0.388558321 | 0.389 | 0.217 | 1.26165686815594e-05 | 21 |
| ran2 | 5.45520526965694e-10 | 0.266788032 | 0.777 | 0.643 | 1.37302061431996e-05 | 21 |
| psma4 | 5.58932288279346e-10 | 0.357415659 | 0.337 | 0.174 | 1.40677667637029e-05 | 21 |
| taf10 | 5.68537824687536e-10 | 0.284246158 | 0.187 | 0.071 | 1.43095285095606e-05 | 21 |
| atox1 | 5.85562048695347e-10 | 0.290746629 | 0.249 | 0.109 | 1.47380112036132e-05 | 21 |
| sod2 | 6.03080172511494e-10 | 0.313934565 | 0.29 | 0.133 | 1.51789248619418e-05 | 21 |
| copa | 6.40832076187263e-10 | 0.258872679 | 0.176 | 0.065 | 1.61291025255572e-05 | 21 |
| e2f51 | 7.60400801055607e-10 | 0.270981699 | 0.192 | 0.072 | 1.91385277617686e-05 | 21 |
| ndufb81 | 8.10236359403765e-10 | 0.363842533 | 0.43 | 0.247 | 2.03928389298334e-05 | 21 |
| psmd6 | 8.53989763867949e-10 | 0.259954581 | 0.202 | 0.079 | 2.14940683667924e-05 | 21 |
| nhp2 | 9.56343363912539e-10 | 0.334632672 | 0.373 | 0.192 | 2.40702061263147e-05 | 21 |
| nop10 | 1.05514560839573e-09 | 0.372646754 | 0.42 | 0.24 | 2.6556959817712e-05 | 21 |
| ranbp1 | 1.07733986319622e-09 | 0.381228091 | 0.461 | 0.277 | 2.71155670167856e-05 | 21 |
| arl14ep | 1.12060997943065e-09 | 0.260120289 | 0.197 | 0.076 | 2.82046325722901e-05 | 21 |

| | | | | | | |
|---|---|---|---|---|---|---|
| cnbpa | 1.17410593675443e-09 | 0.358352571 | 0.389 | 0.211 | 2.95510723221724e-05 | 21 |
| smn11 | 1.18643960820794e-09 | 0.269787662 | 0.228 | 0.093 | 2.98614984989857e-05 | 21 |
| ube2a | 1.19605415382236e-09 | 0.264477314 | 0.192 | 0.074 | 3.01034869975549e-05 | 21 |
| txnl4a1 | 1.43785874302638e-09 | 0.27510759 | 0.259 | 0.113 | 3.61894667032309e-05 | 21 |
| atrx4 | 1.4559874119303e-09 | 0.320327339 | 0.233 | 0.104 | 3.66457471708738e-05 | 21 |
| matr3l1.1 | 1.48759068560034e-09 | 0.285240084 | 0.197 | 0.079 | 3.74411699658749e-05 | 21 |
| ndufa51 | 1.52010279581573e-09 | 0.360808415 | 0.285 | 0.142 | 3.82594672678861e-05 | 21 |
| sumo3b1 | 1.79518067098077e-09 | 0.317110805 | 0.275 | 0.125 | 4.5182902307915e-05 | 21 |
| srrm1 | 2.03248036929705e-09 | 0.301509695 | 0.259 | 0.119 | 5.11554984148375e-05 | 21 |
| si:ch211-282b22.1 | 2.32640962886915e-09 | 0.291621497 | 0.238 | 0.103 | 5.85534039490077e-05 | 21 |
| h2afy23 | 2.35424743622953e-09 | 0.327378815 | 0.218 | 0.097 | 5.9254053722461e-05 | 21 |
| ddx39aa | 2.55549530793329e-09 | 0.328325919 | 0.321 | 0.162 | 6.43192614053731e-05 | 21 |
| lsm8 | 2.81314477510526e-09 | 0.321589433 | 0.316 | 0.155 | 7.08040408446242e-05 | 21 |
| hnrnpl2 | 2.82132221547898e-09 | 0.307107002 | 0.259 | 0.118 | 7.10098588413904e-05 | 21 |
| mrpl32 | 2.9318886781599e-09 | 0.27944025 | 0.228 | 0.098 | 7.37927061406066e-05 | 21 |
| mcm5 | 3.06148027689215e-09 | 0.280727428 | 0.228 | 0.096 | 7.70543970890985e-05 | 21 |
| slc25a3b5 | 3.07029169764154e-09 | 0.328873248 | 0.611 | 0.437 | 7.72761717379399e-05 | 21 |
| wdr46 | 3.20568840541477e-09 | 0.270983082 | 0.187 | 0.073 | 8.06839714758843e-05 | 21 |
| ube2e1 | 3.64464806768114e-09 | 0.341651678 | 0.368 | 0.2 | 9.17321472154667e-05 | 21 |
| UBL4A1 | 3.80762325564777e-09 | 0.310222718 | 0.269 | 0.124 | 9.58340697213987e-05 | 21 |
| psmc3 | 3.87455770437495e-09 | 0.256733372 | 0.238 | 0.103 | 9.75187428614131e-05 | 21 |
| adh54 | 4.4950453289182e-09 | 0.376524234 | 0.378 | 0.221 | 0.000113136 | 21 |
| eif3k2 | 4.52264260555986e-09 | 0.360571761 | 0.389 | 0.229 | 0.00011383 | 21 |
| si:zfos-1192g2.3 | 4.52374804650896e-09 | 0.282172371 | 0.212 | 0.091 | 0.000113858 | 21 |
| nxt2 | 4.74896301050631e-09 | 0.270414798 | 0.212 | 0.088 | 0.000119527 | 21 |
| uqcrq3 | 4.79253388732254e-09 | 0.344633809 | 0.446 | 0.271 | 0.000120623 | 21 |
| snrpb1 | 4.8449667366987e-09 | 0.367146141 | 0.528 | 0.347 | 0.000121943 | 21 |
| ost41 | 5.07410933803419e-09 | 0.343573606 | 0.44 | 0.256 | 0.00012771 | 21 |
| hmgn75 | 5.24668352712581e-09 | 0.369558642 | 0.44 | 0.279 | 0.000132054 | 21 |
| nfr6 | 5.28923947790307e-09 | 0.250870511 | 0.259 | 0.11 | 0.000133125 | 21 |
| psmb4 | 5.536238392009e-09 | 0.349917392 | 0.326 | 0.175 | 0.000139342 | 21 |
| pprc1 | 6.2202138255715e-09 | 0.258545164 | 0.218 | 0.091 | 0.000156557 | 21 |
| mrpl57 | 6.95949609320614e-09 | 0.304127571 | 0.254 | 0.118 | 0.000175164 | 21 |
| prdx23 | 8.18380586188768e-09 | 0.340745679 | 0.585 | 0.4 | 0.000205978 | 21 |
| myl12.2 | 8.25604088091512e-09 | 0.291111739 | 0.264 | 0.123 | 0.000207796 | 21 |

| | | | | | | |
|---|---|---|---|---|---|---|
| COA4 | 8.39210267211753e-09 | 0.274760089 | 0.202 | 0.085 | 0.000211221 | 21 |
| naa10 | 8.75999700637364e-09 | 0.367100181 | 0.378 | 0.222 | 0.00022048 | 21 |
| lyrm4 | 9.58248977099352e-09 | 0.29587602 | 0.228 | 0.102 | 0.000241182 | 21 |
| ube2ia | 9.77622234363894e-09 | 0.308619959 | 0.311 | 0.156 | 0.000246058 | 21 |
| polr2k1 | 1.07486444945865e-08 | 0.34420074 | 0.358 | 0.19 | 0.000270533 | 21 |
| cox4i16 | 1.1508781119252e-08 | 0.325856867 | 0.565 | 0.391 | 0.000289665 | 21 |
| tktb1 | 1.21504631874104e-08 | 0.29957845 | 0.337 | 0.172 | 0.000305815 | 21 |
| nol12 | 1.35344743115426e-08 | 0.250195219 | 0.192 | 0.078 | 0.000340649 | 21 |
| grpel21 | 1.65232052596054e-08 | 0.266463642 | 0.223 | 0.1 | 0.000415873 | 21 |
| api5 | 1.67319600434527e-08 | 0.265072757 | 0.207 | 0.088 | 0.000421127 | 21 |
| si:dkey-46a10.3 | 1.87178556215949e-08 | 0.270352685 | 0.187 | 0.078 | 0.00047111 | 21 |
| adi1 | 1.87663674328434e-08 | 0.255253174 | 0.192 | 0.08 | 0.000472331 | 21 |
| mapre1a4 | 2.35540392402643e-08 | 0.328622002 | 0.363 | 0.197 | 0.000592832 | 21 |
| eif6 | 2.40603559826382e-08 | 0.327365705 | 0.264 | 0.131 | 0.000605575 | 21 |
| zmat2 | 2.58410542869032e-08 | 0.294497628 | 0.275 | 0.138 | 0.000650393 | 21 |
| ube2v23 | 2.58650084855022e-08 | 0.339726989 | 0.435 | 0.269 | 0.000650996 | 21 |
| ppp1cab | 3.06585190621816e-08 | 0.269546134 | 0.212 | 0.095 | 0.000771644 | 21 |
| h2afva5 | 3.31830170334594e-08 | 0.329461404 | 0.523 | 0.323 | 0.000835183 | 21 |
| ywhae11 | 3.37547741545524e-08 | 0.334577535 | 0.446 | 0.282 | 0.000849574 | 21 |
| si:rp71-45k5.4 | 3.41096734533931e-08 | 0.304159482 | 0.228 | 0.108 | 0.000858506 | 21 |
| lamtor52 | 4.23504098097553e-08 | 0.310136507 | 0.269 | 0.135 | 0.001065917 | 21 |
| gpx4b1 | 5.14091379204715e-08 | 0.32953343 | 0.352 | 0.203 | 0.001293917 | 21 |
| supt4h1 | 5.15888412106889e-08 | 0.282172897 | 0.202 | 0.091 | 0.00129844 | 21 |
| si:dkey-28b4.74 | 5.92862306748119e-08 | 0.276560947 | 0.202 | 0.092 | 0.001492175 | 21 |
| gapdhs10 | 6.47100667833523e-08 | 0.360648223 | 0.43 | 0.289 | 0.001628688 | 21 |
| rps1010 | 6.73880684991093e-08 | 0.254188919 | 0.86 | 0.693 | 0.00169609 | 21 |
| zgc:86598 | 8.42547548469823e-08 | 0.257263539 | 0.228 | 0.107 | 0.002120608 | 21 |
| tuba8l42 | 8.78937289446057e-08 | 0.330617707 | 0.585 | 0.404 | 0.002212197 | 21 |
| si:ch211-217k17.7 | 9.07424359232298e-08 | 0.262448594 | 0.228 | 0.107 | 0.002283896 | 21 |
| sumo3a1 | 9.10039340059235e-08 | 0.326768226 | 0.539 | 0.369 | 0.002290478 | 21 |
| ewsr1b | 1.07786946965642e-07 | 0.313793187 | 0.358 | 0.203 | 0.00271289 | 21 |
| atp5f1b | 1.10474175108365e-07 | 0.299892442 | 0.632 | 0.488 | 0.002780525 | 21 |
| ndufa4l6 | 1.1632828712255e-07 | 0.313842981 | 0.539 | 0.395 | 0.002927867 | 21 |
| eif4h | 1.17697823246895e-07 | 0.261601408 | 0.259 | 0.128 | 0.002962337 | 21 |
| chchd105 | 1.44303915127712e-07 | 0.274043399 | 0.301 | 0.151 | 0.003631985 | 21 |

| | | | | | | |
|---|---|---|---|---|---|---|
| mrpl12 | 1.54738644820444e-07 | 0.263893859 | 0.264 | 0.131 | 0.003894617 | 21 |
| med10 | 1.76006056932033e-07 | 0.268638094 | 0.29 | 0.152 | 0.004429896 | 21 |
| tbca | 1.81574354303573e-07 | 0.310626238 | 0.337 | 0.195 | 0.004570045 | 21 |
| sap30l | 1.94388877946914e-07 | 0.283413092 | 0.264 | 0.139 | 0.004892574 | 21 |
| fubp1 | 2.17120747018425e-07 | 0.28196866 | 0.238 | 0.12 | 0.005464712 | 21 |
| zgc:66388 | 2.76195133570294e-07 | 0.29855157 | 0.238 | 0.121 | 0.006951555 | 21 |
| ube2d2 | 3.12033975805738e-07 | 0.299165937 | 0.326 | 0.187 | 0.007853583 | 21 |
| cdv3 | 3.20664301573543e-07 | 0.292830916 | 0.259 | 0.137 | 0.0080708 | 21 |
| st13 | 3.37377441126035e-07 | 0.265129871 | 0.249 | 0.125 | 0.008491453 | 21 |
| rps2114 | 3.83270985791721e-07 | 0.256274393 | 0.922 | 0.748 | 0.009646547 | 21 |
| hnrnpabb7 | 4.18472704445138e-07 | 0.28343469 | 0.751 | 0.597 | 0.010532539 | 21 |
| pgk11 | 5.11272748654073e-07 | 0.259657639 | 0.249 | 0.128 | 0.012868224 | 21 |
| nop58 | 5.51105117699436e-07 | 0.300300878 | 0.311 | 0.175 | 0.013870765 | 21 |
| znf706 | 5.56101701049117e-07 | 0.271442693 | 0.326 | 0.182 | 0.013996524 | 21 |
| ifi465 | 5.70742946785776e-07 | 0.267851166 | 0.238 | 0.121 | 0.014365029 | 21 |
| zgc:1935412 | 5.77851838385876e-07 | 0.300071996 | 0.523 | 0.368 | 0.014543953 | 21 |
| elocb1 | 5.85135182187899e-07 | 0.284023504 | 0.301 | 0.171 | 0.014727267 | 21 |
| kpnb1 | 6.61395981212882e-07 | 0.264523209 | 0.254 | 0.131 | 0.016646675 | 21 |
| paip2b1 | 7.37596486479934e-07 | 0.307490806 | 0.466 | 0.304 | 0.018564566 | 21 |
| meaf6 | 9.23971984260133e-07 | 0.267079793 | 0.269 | 0.145 | 0.023255451 | 21 |
| eif3d5 | 9.39084266015925e-07 | 0.280806556 | 0.404 | 0.253 | 0.023635812 | 21 |
| syncrip1 | 9.78191605771073e-07 | 0.293259474 | 0.456 | 0.3 | 0.024620105 | 21 |
| rad21a | 9.78273440309186e-07 | 0.25473879 | 0.244 | 0.127 | 0.024622164 | 21 |
| adssl | 1.07433874737234e-06 | 0.251600584 | 0.218 | 0.107 | 0.027040032 | 21 |
| dohh | 1.08352381053507e-06 | 0.25399384 | 0.202 | 0.098 | 0.027271211 | 21 |
| eif4a21 | 1.17795269057023e-06 | 0.250544377 | 0.181 | 0.085 | 0.029647891 | 21 |
| psmb21 | 1.34737262728004e-06 | 0.290153061 | 0.285 | 0.163 | 0.033912022 | 21 |
| ahcy | 1.51761142999776e-06 | 0.281098958 | 0.295 | 0.167 | 0.038196762 | 21 |
| serbp1b | 1.63768685702352e-06 | 0.260402234 | 0.212 | 0.108 | 0.041218941 | 21 |
| oaz1a | 2.07884026997293e-06 | 0.264524342 | 0.617 | 0.48 | 0.052322331 | 21 |
| hnrnpa1a3 | 2.31406851081472e-06 | 0.271946843 | 0.28 | 0.16 | 0.05824279 | 21 |
| u2surp1 | 2.77618131244255e-06 | 0.253692489 | 0.192 | 0.095 | 0.069873707 | 21 |
| si:ch1073-325m22.2: | 2.88490655342001e-06 | 0.321102819 | 0.352 | 0.23 | 0.072610213 | 21 |
| mdh1aa3 | 4.63773322982652e-06 | 0.282913337 | 0.332 | 0.206 | 0.116727108 | 21 |
| eif3g5 | 4.97399185157314e-06 | 0.29064536 | 0.404 | 0.282 | 0.125190401 | 21 |

| | | | | | | |
|---|---|---|---|---|---|---|
| atp5if1a | 5.35229104762885e-06 | 0.27819961 | 0.497 | 0.339 | 0.134711813 | 21 |
| cct84 | 5.74023033566966e-06 | 0.289863865 | 0.321 | 0.202 | 0.144475857 | 21 |
| eef1da7 | 6.08433197374911e-06 | 0.252870432 | 0.637 | 0.492 | 0.153136551 | 21 |
| si:ch211-214j24.101 | 6.30563542498901e-06 | 0.262522959 | 0.275 | 0.159 | 0.158706538 | 21 |
| atp5po1 | 7.92721968516798e-06 | 0.268981837 | 0.409 | 0.269 | 0.199520192 | 21 |
| cox6a12 | 9.06632009800304e-06 | 0.264344996 | 0.518 | 0.373 | 0.228190211 | 21 |
| h3f3a7 | 1.01705659717179e-05 | 0.266760579 | 0.674 | 0.505 | 0.255982975 | 21 |
| zgc:1119863 | 1.1429519509295e-05 | 0.266390192 | 0.29 | 0.177 | 0.287669577 | 21 |
| pabpn11 | 1.42890461422494e-05 | 0.2691319 | 0.409 | 0.274 | 0.359641002 | 21 |
| ube2v1 | 1.44392018537989e-05 | 0.26320395 | 0.321 | 0.202 | 0.363420271 | 21 |
| tmem2583 | 1.66496551851945e-05 | 0.277656144 | 0.404 | 0.283 | 0.419055171 | 21 |
| idh24 | 1.72978285792909e-05 | 0.255568108 | 0.316 | 0.199 | 0.435369048 | 21 |
| fam32a | 1.80480599175431e-05 | 0.274230482 | 0.415 | 0.283 | 0.45425162 | 21 |
| nop563 | 2.61376776076245e-05 | 0.250143134 | 0.28 | 0.171 | 0.657859208 | 21 |
| ywhaqa | 2.78644319373876e-05 | 0.256882896 | 0.321 | 0.203 | 0.701319887 | 21 |
| rps27l7 | 3.00961825736808e-05 | 0.283739676 | 0.311 | 0.204 | 0.757490819 | 21 |
| ak22 | 3.15771560084645e-05 | 0.251222444 | 0.352 | 0.233 | 0.79476544 | 21 |
| eif2s1b2 | 3.40705108373504e-05 | 0.27761822 | 0.275 | 0.174 | 0.857520687 | 21 |
| uqcr101 | 5.38770177417536e-05 | 0.253263531 | 0.492 | 0.357 | 1 | 21 |
| vdac23 | 7.18300642614222e-05 | 0.250590098 | 0.409 | 0.296 | 1 | 21 |
| u2af1 | 0.000116354 | 0.2647706 | 0.285 | 0.187 | 1 | 21 |
| si:dkey-12l12.1 | 0 | 1.388589645 | 0.608 | 0.014 | 0 | 22 |
| lum1 | 0 | 1.240677494 | 0.552 | 0.011 | 0 | 22 |
| ecrg4a1 | 0 | 1.190222094 | 0.525 | 0.024 | 0 | 22 |
| hpdb | 0 | 1.136322976 | 0.519 | 0.007 | 0 | 22 |
| asip1 | 0 | 1.136021259 | 0.564 | 0.011 | 0 | 22 |
| tgfbi2 | 0 | 1.038309446 | 0.486 | 0.019 | 0 | 22 |
| scara32 | 0 | 1.001253623 | 0.547 | 0.024 | 0 | 22 |
| zmp:0000001268 | 0 | 0.97172719 | 0.221 | 0.001 | 0 | 22 |
| ctgfb | 0 | 0.791753984 | 0.331 | 0.007 | 0 | 22 |
| si:ch211-243a20.3 | 0 | 0.762874457 | 0.348 | 0.008 | 0 | 22 |
| rhag | 0 | 0.746633077 | 0.326 | 0.006 | 0 | 22 |
| si:dkey-282h22.5 | 0 | 0.678120786 | 0.265 | 0.002 | 0 | 22 |
| tat | 0 | 0.623324928 | 0.232 | 0.001 | 0 | 22 |
| hoxb8a | 0 | 0.519440933 | 0.265 | 0.004 | 0 | 22 |

| | | | | | | |
|---|---|---|---|---|---|---|
| ttc36 | 0 | 0.510511266 | 0.215 | 0.001 | 0 | 22 |
| prrx1b | 2.60621729834419e-302 | 0.534015299 | 0.249 | 0.004 | 6.5595883182025e-298 | 22 |
| tnfrsf191 | 2.9664737323445e-302 | 0.647350382 | 0.359 | 0.01 | 7.46631773693787e-298 | 22 |
| vcana | 2.04281595134146e-301 | 0.610521203 | 0.221 | 0.002 | 5.14156346793133e-297 | 22 |
| hgd | 1.9232652280602e-292 | 0.645316229 | 0.287 | 0.006 | 4.84066625250472e-288 | 22 |
| aqp3b | 2.6430292534727e-291 | 0.366455562 | 0.127 | 0 | 6.65224032806544e-287 | 22 |
| hyal4 | 5.21667554445718e-284 | 0.560417185 | 0.199 | 0.002 | 1.31298506778443e-279 | 22 |
| FQ323119.1 | 2.4404220607265e-271 | 0.444702639 | 0.221 | 0.003 | 6.14229828464254e-267 | 22 |
| tpbga | 8.59579767431374e-257 | 0.6054072 | 0.282 | 0.007 | 2.16347631664802e-252 | 22 |
| sp9 | 5.39997070872565e-255 | 0.311608207 | 0.122 | 0 | 1.35911862767916e-250 | 22 |
| col5a2a4 | 1.15747592073267e-250 | 1.338816576 | 0.773 | 0.076 | 2.91325114489207e-246 | 22 |
| clu2 | 3.11302181313131e-235 | 0.98265922 | 0.42 | 0.021 | 7.83516460147019e-231 | 22 |
| mfap24 | 1.65285537773244e-233 | 1.324765624 | 0.79 | 0.086 | 4.16007170021477e-229 | 22 |
| fgfr3 | 8.22472773347655e-231 | 0.70535171 | 0.331 | 0.012 | 2.07008172323871e-226 | 22 |
| bhmt2 | 1.06042985839022e-226 | 1.112716769 | 0.536 | 0.038 | 2.66899591058234e-222 | 22 |
| aspn | 7.21973388185441e-222 | 0.417804608 | 0.127 | 0.001 | 1.81713482072394e-217 | 22 |
| smoc2 | 1.82855116055726e-220 | 0.531936302 | 0.276 | 0.008 | 4.60228041600658e-216 | 22 |
| CU929237.11 | 1.8306346798765e-220 | 0.658213403 | 0.315 | 0.011 | 4.60752442578117e-216 | 22 |
| dpt | 2.81011554992197e-217 | 0.807904231 | 0.304 | 0.011 | 7.07277982759861e-213 | 22 |
| twist1a | 2.83902559896377e-217 | 0.566014381 | 0.304 | 0.011 | 7.1455435300319e-213 | 22 |
| cxcl8a1 | 8.04676023475723e-217 | 0.74192897 | 0.309 | 0.011 | 2.02528908348605e-212 | 22 |
| ccdc80 | 4.10285282081748e-216 | 0.744073268 | 0.398 | 0.02 | 1.03264702647155e-211 | 22 |
| twist2 | 1.74797019505807e-215 | 0.583526095 | 0.271 | 0.008 | 4.39946618394165e-211 | 22 |
| igfbp3 | 4.37216453394452e-205 | 0.656936106 | 0.282 | 0.01 | 1.1004300915485e-200 | 22 |
| sparc4 | 5.04733840386394e-200 | 1.553157265 | 0.917 | 0.148 | 1.27036460286852e-195 | 22 |
| ehd2b | 7.4452863354587e-200 | 0.795999703 | 0.42 | 0.025 | 1.8739041177716e-195 | 22 |
| col5a14 | 1.72050851410472e-199 | 1.361217874 | 0.796 | 0.103 | 4.33034787915017e-195 | 22 |
| cyr614 | 1.87397903999186e-198 | 1.214049262 | 0.63 | 0.06 | 4.71661784575551e-194 | 22 |
| col1a24 | 4.23831683524187e-198 | 1.662804956 | 0.912 | 0.145 | 1.06674196426203e-193 | 22 |
| mmp14b3 | 1.01364719394006e-197 | 1.044630311 | 0.569 | 0.05 | 2.55124862242774e-193 | 22 |
| col1a1b4 | 2.65191330372999e-197 | 1.564162777 | 0.895 | 0.136 | 6.67460059415802e-193 | 22 |
| mmp11b | 4.99023653043217e-197 | 0.50031544 | 0.16 | 0.002 | 1.25599263234447e-192 | 22 |
| si:dkey-85k7.7 | 4.7799129299977e-196 | 0.439408081 | 0.199 | 0.004 | 1.20305628535112e-191 | 22 |
| nradd | 2.01340608488715e-195 | 0.479596327 | 0.238 | 0.007 | 5.06754177505246e-191 | 22 |
| CABZ01092746.13 | 3.66490702093169e-194 | 1.190618489 | 0.613 | 0.059 | 9.22420448098297e-190 | 22 |

| | | | | | | |
|---|---|---|---|---|---|---|
| col6a34 | 1.15444858670942e-190 | 1.238996781 | 0.746 | 0.091 | 2.90563164788895e-186 | 22 |
| filip1l | 8.34528356840393e-190 | 0.598572665 | 0.271 | 0.01 | 2.10042442133159e-185 | 22 |
| col6a14 | 1.46321136725428e-188 | 1.228977215 | 0.696 | 0.081 | 3.68275669024231e-184 | 22 |
| si:ch211-39i22.13 | 2.89806672654332e-188 | 1.135599642 | 0.624 | 0.064 | 7.29414414403687e-184 | 22 |
| fmoda | 3.0171148103577e-185 | 0.339384646 | 0.138 | 0.002 | 7.59377626618929e-181 | 22 |
| lhfpl61 | 1.12779254878496e-183 | 0.642314403 | 0.331 | 0.016 | 2.83854106603686e-179 | 22 |
| col1a1a4 | 2.82674457144044e-183 | 1.504118725 | 0.856 | 0.13 | 7.11463341185845e-179 | 22 |
| hoxb9a | 1.12037149612729e-182 | 0.399889544 | 0.193 | 0.004 | 2.81986301860277e-178 | 22 |
| pmp22a4 | 3.23333844179904e-182 | 1.337222575 | 0.867 | 0.129 | 8.137989524164e-178 | 22 |
| twist3 | 5.35618794313802e-181 | 0.727567156 | 0.403 | 0.026 | 1.34809894340841e-176 | 22 |
| cldn19 | 1.7478107550468e-180 | 0.449583764 | 0.193 | 0.005 | 4.39906488937729e-176 | 22 |
| htra1a1 | 1.57992649994077e-178 | 0.677843005 | 0.32 | 0.016 | 3.97651700770093e-174 | 22 |
| ccl44 | 1.60832302024217e-178 | 0.418412465 | 0.144 | 0.002 | 4.04798820964753e-174 | 22 |
| lpl1 | 1.98522613306796e-178 | 0.670640241 | 0.32 | 0.016 | 4.99661565431876e-174 | 22 |
| sod3b | 2.36785040964334e-178 | 0.451091101 | 0.249 | 0.009 | 5.95964269603133e-174 | 22 |
| si:dkey-178e17.3 | 6.54629562094483e-177 | 0.560455017 | 0.282 | 0.012 | 1.6476371448356e-172 | 22 |
| CABZ01074363.13 | 1.25811834919936e-174 | 0.789447188 | 0.381 | 0.024 | 3.16655807309987e-170 | 22 |
| ms4a17a.7 | 6.76838553255442e-172 | 0.428177269 | 0.199 | 0.005 | 1.70353495468862e-167 | 22 |
| hhipl2 | 5.63525995777232e-169 | 0.277074014 | 0.127 | 0.002 | 1.41833857877171e-164 | 22 |
| col2a1b1 | 5.15955968175078e-168 | 0.53328961 | 0.265 | 0.011 | 1.29860957629985e-163 | 22 |
| cdh111 | 7.27897658385931e-168 | 0.761585083 | 0.359 | 0.022 | 1.83204561639155e-163 | 22 |
| mmp14a4 | 1.59658819952897e-167 | 0.853338554 | 0.508 | 0.046 | 4.01845283939446e-163 | 22 |
| zgc:162612 | 3.20831785104615e-167 | 0.390782068 | 0.177 | 0.004 | 8.07501519929805e-163 | 22 |
| postna | 1.70087784679625e-166 | 0.997796634 | 0.348 | 0.021 | 4.28093945260148e-162 | 22 |
| fah | 1.38842420200704e-165 | 0.378568859 | 0.193 | 0.005 | 3.49452487403152e-161 | 22 |
| epha4b | 5.25692636119228e-165 | 0.456917244 | 0.215 | 0.007 | 1.32311579584849e-160 | 22 |
| sertad4 | 5.97049220722435e-164 | 0.479910278 | 0.265 | 0.011 | 1.5027131836363e-159 | 22 |
| krt913 | 3.91943859021573e-162 | 1.245179125 | 0.575 | 0.061 | 9.86483498771398e-158 | 22 |
| crabp2b1 | 5.11699956468911e-161 | 0.60571239 | 0.298 | 0.015 | 1.2878976204366e-156 | 22 |
| ier5l1 | 9.03563518263532e-161 | 0.609387597 | 0.304 | 0.016 | 2.27417901911748e-156 | 22 |
| fstl1b4 | 3.09565729053932e-160 | 1.180706478 | 0.773 | 0.116 | 7.79145983455841e-156 | 22 |
| LO018188.1 | 1.91435803669535e-159 | 0.54487929 | 0.276 | 0.013 | 4.81824774255852e-155 | 22 |
| mmp23 | 5.42135538324193e-158 | 1.080775748 | 0.657 | 0.084 | 1.36450093640816e-153 | 22 |
| s100v24 | 5.3000320882634e-154 | 1.146770815 | 0.746 | 0.113 | 1.33396507629502e-149 | 22 |
| fxyd15 | 7.60910262820126e-153 | 0.962207731 | 0.608 | 0.074 | 1.91513504049198e-148 | 22 |

| | | | | | | |
|---|---|---|---|---|---|---|
| si:ch211-286o17.11 | 1.26098090649406e-151 | 0.78678885 | 0.409 | 0.032 | 3.17376284355489e-147 | 22 |
| c1qtnf21 | 1.66294648201401e-147 | 0.590664388 | 0.309 | 0.018 | 4.18547000058105e-143 | 22 |
| jun5 | 1.20438433872517e-145 | 1.457370585 | 0.928 | 0.219 | 3.03131494213737e-141 | 22 |
| rarga1 | 1.13223437283445e-143 | 0.844958782 | 0.492 | 0.051 | 2.84972069298702e-139 | 22 |
| ms4a17a.94 | 2.38665100096927e-143 | 0.772227339 | 0.492 | 0.05 | 6.00696190433956e-139 | 22 |
| atoh82 | 9.41108781708509e-138 | 0.534689308 | 0.276 | 0.016 | 2.36867669268215e-133 | 22 |
| prrx1a | 4.40928078562147e-137 | 0.385092669 | 0.204 | 0.008 | 1.10977188093307e-132 | 22 |
| f3b4 | 5.3906265281849e-137 | 1.259757065 | 0.635 | 0.094 | 1.35676679087886e-132 | 22 |
| fosl1a3 | 6.6171055344985e-136 | 1.103335865 | 0.608 | 0.085 | 1.66545929197793e-131 | 22 |
| si:ch73-335l21.44 | 1.44998927754526e-134 | 1.447047362 | 0.751 | 0.14 | 3.64947801265367e-130 | 22 |
| alx4a | 3.97098365905947e-134 | 0.301035187 | 0.138 | 0.003 | 9.99456877148678e-130 | 22 |
| zgc:56585 | 1.36174518354157e-132 | 0.413939855 | 0.204 | 0.008 | 3.42737645245577e-128 | 22 |
| si:ch211-170d8.8 | 5.51385126960903e-132 | 0.475865546 | 0.227 | 0.011 | 1.3877812260479e-127 | 22 |
| heyl | 1.25185243080297e-130 | 0.268091956 | 0.11 | 0.002 | 3.150787383088e-126 | 22 |
| fgl2b1 | 8.51901706666553e-129 | 0.534873335 | 0.238 | 0.012 | 2.14415140550905e-124 | 22 |
| sema3aa | 1.81028686710124e-128 | 0.270525687 | 0.133 | 0.003 | 4.55631101580711e-124 | 22 |
| thbs1b | 4.65326342960517e-128 | 0.48617207 | 0.204 | 0.009 | 1.17117987259732e-123 | 22 |
| rnd3b4 | 7.84800752830803e-125 | 0.740849283 | 0.392 | 0.036 | 1.97526501479985e-120 | 22 |
| serpine11 | 9.19546410527334e-125 | 0.920384881 | 0.409 | 0.04 | 2.31440636065625e-120 | 22 |
| enpp2 | 9.66244285915672e-125 | 0.27778572 | 0.149 | 0.004 | 2.43194024322115e-120 | 22 |
| bcl11ba | 4.8504157905031e-124 | 0.518630261 | 0.238 | 0.013 | 1.22080115031172e-119 | 22 |
| gpm6bb | 1.92530584467791e-123 | 0.516071692 | 0.293 | 0.02 | 4.84580228046984e-119 | 22 |
| olfml1 | 2.35039796511626e-123 | 0.337625291 | 0.155 | 0.005 | 5.91571663840113e-119 | 22 |
| XLOC-0400201 | 2.73626100087456e-123 | 0.376289881 | 0.199 | 0.008 | 6.88689531310118e-119 | 22 |
| adh8a3 | 3.96061630628823e-123 | 1.06639208 | 0.403 | 0.04 | 9.96847518129683e-119 | 22 |
| fosab7 | 2.10902421828604e-122 | 1.422790261 | 0.967 | 0.341 | 5.30820305500413e-118 | 22 |
| ms4a17a.111 | 5.94021718896779e-122 | 0.643430728 | 0.376 | 0.034 | 1.4950932642913e-117 | 22 |
| loxl11 | 1.63207022072212e-121 | 0.542662286 | 0.287 | 0.019 | 4.1077575385355e-117 | 22 |
| dcn4 | 1.92280397835089e-121 | 1.095809012 | 0.685 | 0.112 | 4.83950533311134e-117 | 22 |
| sema3e | 2.84508462793791e-121 | 0.431904516 | 0.193 | 0.008 | 7.16079350005692e-117 | 22 |
| s100a10b6 | 1.27999503526977e-120 | 1.354003024 | 0.928 | 0.246 | 3.22161950427048e-116 | 22 |
| fosb4 | 1.3974648946946e-120 | 1.179489105 | 0.685 | 0.118 | 3.51727939345685e-116 | 22 |
| cntfr1 | 3.68339909394441e-120 | 0.572839615 | 0.238 | 0.013 | 9.27074717954869e-116 | 22 |
| pigr | 1.68829979433926e-118 | 0.468679113 | 0.265 | 0.017 | 4.24928175237248e-114 | 22 |
| anxa1a5 | 2.59928529915642e-117 | 1.164745144 | 0.724 | 0.135 | 6.54214116944679e-113 | 22 |

| | | | | | | |
|---|---|---|---|---|---|---|
| gstz11 | 2.81398128610481e-117 | 0.619267854 | 0.337 | 0.028 | 7.08250949899719e-113 | 22 |
| pcdh18b2 | 3.73204793260605e-117 | 0.441433328 | 0.238 | 0.013 | 9.39319144157617e-113 | 22 |
| sox11a | 5.19251427084313e-117 | 0.611535325 | 0.265 | 0.017 | 1.30690391682851e-112 | 22 |
| capn2b | 6.86494501450745e-115 | 0.52025938 | 0.287 | 0.021 | 1.72783801070138e-110 | 22 |
| si:ch73-46j18.55 | 1.03330386939426e-114 | 1.107784928 | 0.801 | 0.167 | 2.60072250887842e-110 | 22 |
| apln | 1.07001731734739e-114 | 0.301249151 | 0.138 | 0.004 | 2.69312658603164e-110 | 22 |
| sdc46 | 3.3666969090984e-114 | 1.222181111 | 0.906 | 0.249 | 8.47363945050977e-110 | 22 |
| zgc:154093 | 6.07802586054364e-113 | 0.407790556 | 0.221 | 0.012 | 1.52977832884023e-108 | 22 |
| postnb3 | 2.44859159093031e-112 | 1.136427511 | 0.519 | 0.073 | 6.1628601752125e-108 | 22 |
| ier2a8 | 7.03973580512622e-112 | 1.258499136 | 0.89 | 0.254 | 1.77183110479222e-107 | 22 |
| ctsk2 | 1.07071682432284e-111 | 0.704009953 | 0.47 | 0.057 | 2.69488717513816e-107 | 22 |
| vmo1a | 5.91402976501008e-111 | 0.34264612 | 0.149 | 0.005 | 1.48850215155539e-106 | 22 |
| tcf123 | 1.639212985592e-110 | 0.716455794 | 0.448 | 0.053 | 4.12573516343651e-106 | 22 |
| junba7 | 1.86566645992033e-110 | 1.26995089 | 0.95 | 0.301 | 4.69569591297347e-106 | 22 |
| fstb1 | 2.87707770566692e-110 | 0.631087269 | 0.265 | 0.018 | 7.24131687739308e-106 | 22 |
| egr18 | 1.96755822547013e-109 | 1.138376331 | 0.796 | 0.182 | 4.95214729768576e-105 | 22 |
| lix1l1 | 3.02113935658965e-109 | 0.436621989 | 0.227 | 0.013 | 7.60390564660049e-105 | 22 |
| bgnb3 | 1.7530249148464e-107 | 0.593833287 | 0.298 | 0.024 | 4.4121884081769e-103 | 22 |
| ecm2 | 3.70834079818557e-107 | 0.333259057 | 0.16 | 0.006 | 9.33352295495326e-103 | 22 |
| spon2b1 | 2.65804405002251e-106 | 0.649677906 | 0.276 | 0.02 | 6.69003106950166e-102 | 22 |
| rgma | 3.36260663768287e-106 | 0.331908129 | 0.166 | 0.007 | 8.46334464638401e-102 | 22 |
| ier2b9 | 3.45890104538287e-106 | 1.150503211 | 0.834 | 0.216 | 8.70570804112414e-102 | 22 |
| serpinf13 | 4.45604459800475e-106 | 0.683199283 | 0.376 | 0.039 | 1.12154186487182e-101 | 22 |
| vwa1 | 3.04463204981461e-104 | 0.513762623 | 0.26 | 0.019 | 7.6630344061784e-100 | 22 |
| junbb5 | 3.13480285688964e-103 | 0.731746262 | 0.486 | 0.067 | 7.88998531050553e-99 | 22 |
| lrrc173 | 7.89734431725574e-103 | 0.61079004 | 0.331 | 0.031 | 1.9876825912101e-98 | 22 |
| klf6a5 | 2.93153364130463e-102 | 1.141739977 | 0.713 | 0.16 | 7.37837702179963e-98 | 22 |
| socs3a8 | 4.82106300231156e-102 | 1.251530051 | 0.934 | 0.29 | 1.2134133470518e-97 | 22 |
| marcksl1a5 | 1.52609346366578e-100 | 1.168980953 | 0.818 | 0.207 | 3.84102463870041e-96 | 22 |
| dlx5a | 2.26754986375709e-100 | 0.286318174 | 0.127 | 0.004 | 5.70719625209021e-96 | 22 |
| col6a24 | 5.15862226480713e-100 | 0.791844268 | 0.453 | 0.06 | 1.29837363782931e-95 | 22 |
| qdpra | 6.68702479407015e-100 | 0.350012694 | 0.171 | 0.008 | 1.68305727041952e-95 | 22 |
| mxra8b2 | 1.79150196543563e-99 | 0.484266419 | 0.265 | 0.02 | 4.50903129680495e-95 | 22 |
| mamdc2b | 7.90951499008789e-98 | 0.272417693 | 0.11 | 0.003 | 1.99074582785522e-93 | 22 |
| col5a3a | 1.27421383428563e-96 | 0.366946204 | 0.177 | 0.009 | 3.2070687995135e-92 | 22 |

| | | | | | | |
|---|---|---|---|---|---|---|
| pcolcea3 | 1.34320631150689e-96 | 0.756941628 | 0.398 | 0.046 | 3.3807159654317e-92 | 22 |
| jdp2b5 | 4.75386492361687e-96 | 1.254207246 | 0.702 | 0.163 | 1.19650026262513e-91 | 22 |
| colec124 | 1.81183577676991e-95 | 0.69802381 | 0.403 | 0.049 | 4.56020946655218e-91 | 22 |
| myo1ca2 | 4.21939943689355e-95 | 0.477252123 | 0.265 | 0.021 | 1.06198064427174e-90 | 22 |
| cd99 | 1.57947924021591e-94 | 0.694699938 | 0.398 | 0.049 | 3.97539129969943e-90 | 22 |
| cspg4 | 8.56979530847876e-94 | 0.287011432 | 0.133 | 0.005 | 2.15693178119102e-89 | 22 |
| serpinh1b | 8.79396795632799e-94 | 0.528993196 | 0.254 | 0.02 | 2.21335379492819e-89 | 22 |
| nfixb3 | 1.28357157608181e-93 | 0.738839487 | 0.442 | 0.06 | 3.23062129984031e-89 | 22 |
| cyp26b1 | 3.25657998689835e-92 | 0.331977636 | 0.127 | 0.004 | 8.19648616902446e-88 | 22 |
| zeb2a2 | 7.16539178288348e-92 | 0.734382781 | 0.354 | 0.041 | 1.80345745783394e-87 | 22 |
| si:ch211-133n4.44 | 9.0346062124939e-92 | 0.584151648 | 0.359 | 0.041 | 2.27392003762259e-87 | 22 |
| spock2 | 9.94452611483254e-92 | 0.327847417 | 0.122 | 0.004 | 2.5029377778422e-87 | 22 |
| klf15 | 4.9538231537531e-91 | 0.289153572 | 0.166 | 0.008 | 1.24682774956812e-86 | 22 |
| osr24 | 1.55794264802681e-89 | 0.675612929 | 0.359 | 0.042 | 3.92118585081868e-85 | 22 |
| zgc:16273011 | 2.05348205533782e-89 | 1.173834601 | 0.939 | 0.385 | 5.16840898507977e-85 | 22 |
| icn6 | 2.16546455632641e-89 | 1.28799303 | 0.845 | 0.267 | 5.45025774181794e-85 | 22 |
| apcdd1l | 2.82721005346066e-88 | 0.30611983 | 0.144 | 0.006 | 7.11580498355515e-84 | 22 |
| hspg21 | 5.87380605063123e-88 | 0.477200585 | 0.26 | 0.022 | 1.47837824488338e-83 | 22 |
| tcf71 | 6.60383294352115e-88 | 0.537851632 | 0.254 | 0.021 | 1.66211871355484e-83 | 22 |
| ccdc3b2 | 3.03064635197467e-86 | 0.454993022 | 0.254 | 0.022 | 7.62783380328505e-82 | 22 |
| MDFI | 1.40394863751515e-85 | 0.399816809 | 0.221 | 0.016 | 3.53359832576187e-81 | 22 |
| endouc2 | 2.23031203709121e-85 | 0.780507207 | 0.42 | 0.058 | 5.61347236615486e-81 | 22 |
| krt87 | 2.38736462250719e-85 | 1.104787762 | 0.89 | 0.268 | 6.00875801838835e-81 | 22 |
| ckap42 | 2.65671682614288e-85 | 0.553423569 | 0.343 | 0.04 | 6.68669057971902e-81 | 22 |
| fap | 3.77453509034685e-85 | 0.415976857 | 0.177 | 0.01 | 9.50012736889398e-81 | 22 |
| srpx2 | 4.55615269900282e-85 | 0.304596189 | 0.144 | 0.007 | 1.14673807281202e-80 | 22 |
| dlx3b | 6.20369662395569e-85 | 0.369555158 | 0.155 | 0.008 | 1.56140840328341e-80 | 22 |
| vim3 | 1.41657694705375e-84 | 0.569870403 | 0.304 | 0.032 | 3.56538251803958e-80 | 22 |
| anxa5b4 | 4.87807917300397e-84 | 0.809333722 | 0.508 | 0.089 | 1.22776374705337e-79 | 22 |
| plpp33 | 6.22020226307992e-84 | 0.650606523 | 0.392 | 0.051 | 1.56556270759459e-79 | 22 |
| jag1b1 | 1.16028339411142e-83 | 0.501389743 | 0.254 | 0.023 | 2.92031727463902e-79 | 22 |
| pleca2 | 4.453682065183e-83 | 0.67950891 | 0.398 | 0.055 | 1.12094723898591e-78 | 22 |
| antxr2a7 | 1.51351061126541e-82 | 0.920791935 | 0.652 | 0.147 | 3.8093548574939e-78 | 22 |
| atf33 | 1.61409753970997e-81 | 0.91086842 | 0.492 | 0.086 | 4.06252209769602e-77 | 22 |
| cdo1 | 2.03120077794292e-81 | 0.304638711 | 0.149 | 0.007 | 5.11232923800453e-77 | 22 |

| | | | | | | |
|---|---|---|---|---|---|---|
| FAT4 | 4.46575155560101e-81 | 0.344455004 | 0.204 | 0.015 | 1.12398500902922e-76 | 22 |
| rgs5b4 | 4.63752782529006e-81 | 1.03648083 | 0.442 | 0.071 | 1.16721937834726e-76 | 22 |
| tcf41 | 1.8851954536948e-80 | 0.429556705 | 0.232 | 0.019 | 4.74484843740444e-76 | 22 |
| h3f3c6 | 2.3116842686321e-80 | 0.943347973 | 0.823 | 0.249 | 5.81827813572014e-76 | 22 |
| rbp44 | 2.38217526392412e-80 | 1.135362182 | 0.586 | 0.122 | 5.99569692177062e-76 | 22 |
| emilin2a1 | 2.46115222972778e-80 | 0.50571587 | 0.243 | 0.021 | 6.19447404700185e-76 | 22 |
| cxcl18b5 | 4.01433471286052e-80 | 1.049886449 | 0.536 | 0.102 | 1.01036790387986e-75 | 22 |
| cd81a5 | 5.25861184098665e-80 | 0.935258126 | 0.785 | 0.199 | 1.32354001425793e-75 | 22 |
| gpc1a | 1.67789319622674e-79 | 0.314217468 | 0.138 | 0.007 | 4.22308938558308e-75 | 22 |
| slc16a10 | 2.38261104548939e-79 | 0.375474341 | 0.204 | 0.015 | 5.99679374039226e-75 | 22 |
| rbfox11 | 3.44569447220716e-79 | 0.45525067 | 0.249 | 0.022 | 8.6724684170982e-75 | 22 |
| fstl31 | 3.73035809753916e-79 | 0.482044401 | 0.243 | 0.022 | 9.38893829569632e-75 | 22 |
| sfrp1b1 | 4.29712406558799e-79 | 0.419865646 | 0.188 | 0.013 | 1.08154315606784e-74 | 22 |
| ism1 | 5.84363370720356e-79 | 0.369505986 | 0.149 | 0.008 | 1.47078416776606e-74 | 22 |
| si:dkey-261h17.13 | 5.98973460174131e-79 | 0.889685696 | 0.497 | 0.086 | 1.50755630191227e-74 | 22 |
| grb10b3 | 7.07001569493679e-79 | 0.543151715 | 0.298 | 0.033 | 1.77945225025864e-74 | 22 |
| pmp22b1 | 9.93324401144887e-79 | 0.659041792 | 0.337 | 0.042 | 2.50009818524157e-74 | 22 |
| akap12b4 | 1.06816732040567e-78 | 0.729330106 | 0.392 | 0.056 | 2.68847032872903e-74 | 22 |
| im:7142702 | 1.07866221539753e-78 | 0.265925269 | 0.144 | 0.007 | 2.71488492993405e-74 | 22 |
| efna5b | 3.08230183338734e-78 | 0.281167746 | 0.144 | 0.007 | 7.75784548445258e-74 | 22 |
| timp2a4 | 6.2670775138003e-78 | 0.736651481 | 0.376 | 0.053 | 1.5773607394484e-73 | 22 |
| irx1a | 1.16460145324675e-77 | 0.252784758 | 0.11 | 0.004 | 2.93118539767674e-73 | 22 |
| slc25a48 | 1.51092790108291e-77 | 0.282953947 | 0.144 | 0.007 | 3.80285443423558e-73 | 22 |
| cspg5a | 1.32397190875868e-76 | 0.275173133 | 0.138 | 0.007 | 3.33230489715473e-72 | 22 |
| scara5 | 1.50881898025691e-76 | 0.263275307 | 0.127 | 0.006 | 3.79754649140861e-72 | 22 |
| c1qtnf52 | 1.51708473792735e-76 | 0.637862639 | 0.243 | 0.023 | 3.81835057688935e-72 | 22 |
| ddit38 | 2.39842613930858e-76 | 0.97208311 | 0.68 | 0.17 | 6.03659875002576e-72 | 22 |
| si:busm1-57f23.1 | 1.24765102063213e-75 | 0.567337814 | 0.271 | 0.028 | 3.14021285382901e-71 | 22 |
| zcchc243 | 4.59839774309395e-75 | 0.489137596 | 0.276 | 0.029 | 1.15737072795932e-70 | 22 |
| bmp16 | 3.29478043650317e-74 | 0.292480092 | 0.116 | 0.005 | 8.29263288063484e-70 | 22 |
| ablim1b3 | 3.29858642544106e-74 | 0.58740325 | 0.293 | 0.033 | 8.3022121741926e-70 | 22 |
| plxdc2 | 8.49640725131335e-74 | 0.311442653 | 0.149 | 0.008 | 2.13846074108306e-69 | 22 |
| fkbp72 | 8.94675066632069e-74 | 0.517312374 | 0.287 | 0.033 | 2.25180767520626e-69 | 22 |
| angptl1a3 | 3.71254782813371e-73 | 0.567078898 | 0.249 | 0.024 | 9.34411162862974e-69 | 22 |
| cox4i22 | 5.52245970738803e-73 | 0.476016726 | 0.276 | 0.03 | 1.38994788375249e-68 | 22 |

| | | | | | | |
|---|---|---|---|---|---|---|
| smad1 | 8.74733224189643e-73 | 0.37180376 | 0.204 | 0.016 | 2.20161605196291e-68 | 22 |
| klf2a4 | 9.84121826525507e-73 | 0.862889388 | 0.453 | 0.081 | 2.47693622518205e-68 | 22 |
| btg26 | 1.06015204391173e-72 | 0.984986133 | 0.906 | 0.366 | 2.66829667932144e-68 | 22 |
| zfp36l1b4 | 6.48753057636516e-72 | 0.960712016 | 0.558 | 0.125 | 1.63284657076535e-67 | 22 |
| meis1a | 8.32590239380519e-72 | 0.346297208 | 0.182 | 0.013 | 2.09554637349683e-67 | 22 |
| ecm1b2 | 2.39038311627348e-71 | 0.592588129 | 0.359 | 0.051 | 6.01635526534871e-67 | 22 |
| tnfaip2a1 | 1.15090515657189e-70 | 0.448325015 | 0.232 | 0.022 | 2.89671318857578e-66 | 22 |
| tgfbr32 | 2.75580105482886e-70 | 0.515545429 | 0.315 | 0.04 | 6.93607567489876e-66 | 22 |
| si:ch211-178n15.12 | 1.15390138180233e-69 | 0.489180324 | 0.26 | 0.028 | 2.90425438785828e-65 | 22 |
| nupr1b3 | 3.7813307638014e-69 | 0.693909055 | 0.326 | 0.045 | 9.51723139941174e-65 | 22 |
| id12 | 4.38054548926637e-69 | 0.933698404 | 0.497 | 0.1 | 1.10253949419345e-64 | 22 |
| spred2b2 | 1.14516119930418e-68 | 0.613224556 | 0.392 | 0.063 | 2.8822562225287e-64 | 22 |
| zgc:1533172 | 1.43409556350047e-68 | 0.555676708 | 0.304 | 0.039 | 3.60947512377433e-64 | 22 |
| ppap2d4 | 3.08972715520438e-68 | 0.538480239 | 0.282 | 0.034 | 7.7765342769339e-64 | 22 |
| cnn3a3 | 5.7713852949979e-68 | 0.534118231 | 0.359 | 0.053 | 1.45259996489802e-63 | 22 |
| entpd5a | 6.63973686768541e-68 | 0.505918247 | 0.105 | 0.004 | 1.67115537222774e-63 | 22 |
| ramp21 | 7.75826599391126e-68 | 0.553404426 | 0.304 | 0.038 | 1.95267796800753e-63 | 22 |
| lfng | 9.88267102272459e-68 | 0.348407053 | 0.127 | 0.007 | 2.48736946970955e-63 | 22 |
| f13a1b3 | 4.1985352096678e-67 | 0.442710861 | 0.26 | 0.029 | 1.05672932692129e-62 | 22 |
| mex3b | 1.03555895837811e-66 | 0.273966464 | 0.138 | 0.008 | 2.60639834234187e-62 | 22 |
| gas1a3 | 1.70712325931699e-66 | 0.520216952 | 0.304 | 0.04 | 4.29665853137492e-62 | 22 |
| fbn2b3 | 5.79119757414546e-66 | 0.457251634 | 0.232 | 0.024 | 1.45758651743667e-61 | 22 |
| itgb4 | 1.89936964616863e-65 | 0.404493662 | 0.182 | 0.015 | 4.78052346244182e-61 | 22 |
| hoxb7a | 1.39651532804127e-64 | 0.338189578 | 0.171 | 0.013 | 3.51488942914707e-60 | 22 |
| tuba8l32 | 2.25119455142023e-64 | 0.525297192 | 0.315 | 0.044 | 5.66603156646958e-60 | 22 |
| lmna | 2.674514295354e-64 | 0.335539228 | 0.171 | 0.013 | 6.73148502997647e-60 | 22 |
| atf4b7 | 3.03622242327811e-64 | 0.885762676 | 0.762 | 0.258 | 7.64186821714868e-60 | 22 |
| BX640522.22 | 5.31889084495933e-64 | 0.408513024 | 0.238 | 0.025 | 1.33871163676781e-59 | 22 |
| vstm4a | 5.88389080075977e-64 | 0.288232949 | 0.149 | 0.01 | 1.48091647564323e-59 | 22 |
| pcsk6 | 8.41647610634263e-64 | 0.2746219 | 0.138 | 0.008 | 2.11834287120538e-59 | 22 |
| cd82a2 | 1.06647990541339e-63 | 0.538024862 | 0.348 | 0.053 | 2.68422327393496e-59 | 22 |
| tenm32 | 1.24265910334253e-63 | 0.546169603 | 0.309 | 0.042 | 3.12764869720281e-59 | 22 |
| tpm4a3 | 1.43745851879089e-63 | 0.670263811 | 0.448 | 0.084 | 3.61793934594479e-59 | 22 |
| aldob1 | 1.46060571547337e-63 | 0.43949588 | 0.282 | 0.035 | 3.67619852527493e-59 | 22 |
| ghra3 | 2.81493332192515e-63 | 0.440156339 | 0.249 | 0.028 | 7.08490567795341e-59 | 22 |

| | | | | | | |
|---|---|---|---|---|---|---|
| ier53 | 2.95519116930921e-63 | 0.809158558 | 0.547 | 0.131 | 7.43792065403436e-59 | 22 |
| her61 | 4.57552388241666e-63 | 0.456173471 | 0.21 | 0.02 | 1.15161360596545e-58 | 22 |
| XLOC-0429171 | 1.70260804849465e-62 | 0.314240785 | 0.155 | 0.011 | 4.28529419725619e-58 | 22 |
| arhgap21a1 | 1.93110132108963e-62 | 0.306416635 | 0.149 | 0.01 | 4.86038891505048e-58 | 22 |
| rgcc3 | 2.20253770347376e-62 | 0.74241431 | 0.387 | 0.067 | 5.54356714587311e-58 | 22 |
| CR383676.111 | 2.25252424041141e-62 | 0.816308204 | 0.989 | 0.723 | 5.66937826069147e-58 | 22 |
| socs3b6 | 2.45267660626401e-62 | 0.853171611 | 0.591 | 0.15 | 6.1731417503059e-58 | 22 |
| abi3bpa1 | 3.39670601807964e-62 | 0.305659239 | 0.16 | 0.012 | 8.54916937690464e-58 | 22 |
| col6a4a2 | 8.03402216027625e-62 | 0.584201243 | 0.309 | 0.043 | 2.02208303751993e-57 | 22 |
| cebpd5 | 9.25675284810285e-62 | 0.838940531 | 0.536 | 0.124 | 2.32983212433901e-57 | 22 |
| wnt5b | 5.63870688903643e-61 | 0.302637376 | 0.149 | 0.01 | 1.41920613690158e-56 | 22 |
| lsp1 | 6.9737378717575e-61 | 0.327685173 | 0.166 | 0.013 | 1.75522008494264e-56 | 22 |
| fgl2a4 | 8.083931314037e-61 | 0.671348779 | 0.37 | 0.062 | 2.03464467242997e-56 | 22 |
| gng74 | 1.01361007465756e-60 | 0.497053274 | 0.331 | 0.05 | 2.55115519690562e-56 | 22 |
| fkbp14 | 1.37497556505191e-60 | 0.458840705 | 0.287 | 0.038 | 3.46067599967915e-56 | 22 |
| mtmr11 | 2.42033083980389e-60 | 0.279326256 | 0.149 | 0.011 | 6.09173069070241e-56 | 22 |
| ahnak6 | 4.44216751466683e-60 | 0.749766568 | 0.53 | 0.122 | 1.11804914176649e-55 | 22 |
| sertad2a | 6.54487015725479e-60 | 0.374523384 | 0.21 | 0.021 | 1.64727836987946e-55 | 22 |
| BX663503.32 | 7.65386909673385e-60 | 0.428912484 | 0.243 | 0.028 | 1.92640231295694e-55 | 22 |
| nova21 | 8.73668328711737e-60 | 0.457970424 | 0.243 | 0.028 | 2.19893581653457e-55 | 22 |
| im:71523481 | 1.17781340649296e-59 | 0.353951029 | 0.171 | 0.014 | 2.96443856280212e-55 | 22 |
| efnb2a | 2.13616273292495e-59 | 0.300545294 | 0.138 | 0.009 | 5.3765079824988e-55 | 22 |
| rgs5a3 | 5.80946162181273e-59 | 0.54007675 | 0.227 | 0.025 | 1.46218339559405e-54 | 22 |
| tuft1a1 | 1.27665123949129e-58 | 0.4369866 | 0.254 | 0.031 | 3.21320350467564e-54 | 22 |
| LOC101883507 | 2.19563156960153e-58 | 0.32989872 | 0.182 | 0.016 | 5.5261850975301e-54 | 22 |
| s100w1 | 1.07579002905386e-57 | 0.35605509 | 0.182 | 0.017 | 2.70765592412565e-53 | 22 |
| csrnp1b6 | 1.70332314166146e-57 | 0.790556589 | 0.624 | 0.174 | 4.28709401524774e-53 | 22 |
| cd248a3 | 1.05432167065423e-56 | 0.434611509 | 0.249 | 0.031 | 2.65362221286963e-52 | 22 |
| htra3b1 | 1.15169459853249e-56 | 0.342561963 | 0.16 | 0.013 | 2.89870013504642e-52 | 22 |
| col11a1b | 3.79543854469252e-56 | 0.346736799 | 0.155 | 0.012 | 9.5527392731366e-52 | 22 |
| hmbox1a | 4.80488871345086e-56 | 0.43215101 | 0.238 | 0.029 | 1.20934244028845e-51 | 22 |
| fpr11 | 5.1157807963558e-56 | 0.35933523 | 0.188 | 0.018 | 1.28759086863479e-51 | 22 |
| chd35 | 5.53459927949778e-56 | 0.618762339 | 0.464 | 0.1 | 1.3930032926568e-51 | 22 |
| SCARF21 | 3.88803455178054e-55 | 0.39969519 | 0.204 | 0.022 | 9.78579416337644e-51 | 22 |
| si:ch73-1a9.38 | 8.30710623800753e-54 | 0.804014215 | 0.757 | 0.26 | 2.09081556904412e-49 | 22 |

| | | | | | | |
|---|---|---|---|---|---|---|
| edil3a3 | 1.61949642248784e-53 | 0.42557899 | 0.199 | 0.022 | 4.07611054575964e-49 | 22 |
| vat14 | 5.69670975477659e-53 | 0.69915092 | 0.53 | 0.13 | 1.43380487817972e-48 | 22 |
| zgc:1716791 | 7.79394028064025e-53 | 0.381558524 | 0.215 | 0.025 | 1.96165682923434e-48 | 22 |
| si:ch211-251b21.1 | 9.5796006477104e-53 | 0.269376328 | 0.11 | 0.007 | 2.41108968702223e-48 | 22 |
| dpysl31 | 1.2673666789632e-52 | 0.350661261 | 0.177 | 0.017 | 3.18983519428248e-48 | 22 |
| tnfsf10l1 | 2.27646652738728e-52 | 0.410569465 | 0.21 | 0.024 | 5.72963860278104e-48 | 22 |
| marcksl1b6 | 3.35714708143511e-52 | 0.820078094 | 0.47 | 0.112 | 8.44960348926404e-48 | 22 |
| kank23 | 4.42680899988119e-52 | 0.541118448 | 0.309 | 0.051 | 1.1141835571801e-47 | 22 |
| pdgfrl3 | 6.63731828647282e-52 | 0.382922516 | 0.215 | 0.026 | 1.67054663952234e-47 | 22 |
| anxa2a3 | 7.81899946123059e-52 | 0.686724786 | 0.459 | 0.102 | 1.96796397439713e-47 | 22 |
| krt18b8 | 7.89513107892104e-52 | 0.746709216 | 0.558 | 0.146 | 1.98712554125364e-47 | 22 |
| gadd45ba4 | 1.6599692404944e-51 | 0.865535368 | 0.613 | 0.186 | 4.17797658140034e-47 | 22 |
| fkbp9 | 4.74661489263444e-51 | 0.377383871 | 0.204 | 0.024 | 1.19467550232716e-46 | 22 |
| angptl43 | 5.88977561022874e-51 | 0.57084607 | 0.293 | 0.047 | 1.48239762333847e-46 | 22 |
| si:ch73-138n13.1 | 6.36460637778597e-51 | 0.455983638 | 0.254 | 0.036 | 1.60190777922495e-46 | 22 |
| pdgfrb4 | 6.98380793984398e-51 | 0.483546661 | 0.298 | 0.048 | 1.75775462037933e-46 | 22 |
| nectin1b1 | 8.41913675629951e-51 | 0.47149908 | 0.271 | 0.04 | 2.11901253019302e-46 | 22 |
| inhbaa3 | 1.16596796568801e-50 | 0.587445504 | 0.26 | 0.038 | 2.93462477284016e-46 | 22 |
| ppib7 | 1.2428237082429e-50 | 0.785711582 | 0.818 | 0.327 | 3.12806299127656e-46 | 22 |
| hbegfa | 1.52634425425984e-50 | 0.750368351 | 0.459 | 0.108 | 3.84165585354659e-46 | 22 |
| tmem1764 | 1.92655068373457e-50 | 0.522806689 | 0.348 | 0.063 | 4.84893541589154e-46 | 22 |
| ednraa | 3.04211468120204e-50 | 0.304943126 | 0.16 | 0.015 | 7.65669844111743e-46 | 22 |
| si:ch211-212k18.84 | 3.59656422948311e-50 | 0.43395059 | 0.271 | 0.041 | 9.05219250918603e-46 | 22 |
| fxyd6l3 | 5.83604457312974e-50 | 0.625668377 | 0.475 | 0.11 | 1.46887405861102e-45 | 22 |
| loxl2b2 | 6.10706414161522e-50 | 0.439436685 | 0.265 | 0.039 | 1.53708697380313e-45 | 22 |
| sfrp1a2 | 9.35405797951133e-50 | 0.480894335 | 0.215 | 0.027 | 2.35432285286321e-45 | 22 |
| kirrel1a2 | 6.60248594925431e-49 | 0.37699707 | 0.21 | 0.025 | 1.66177968856782e-44 | 22 |
| pdgfab | 6.66223514254027e-49 | 0.308895095 | 0.149 | 0.013 | 1.67681796302596e-44 | 22 |
| si:dkey-101k6.51 | 1.16042335360245e-48 | 0.318189819 | 0.177 | 0.018 | 2.92066953868202e-44 | 22 |
| FP102018.1 | 1.17611046329448e-48 | 0.30528505 | 0.166 | 0.016 | 2.96015242506589e-44 | 22 |
| s1pr5a | 1.21021927910431e-48 | 0.270612538 | 0.122 | 0.009 | 3.04600090357764e-44 | 22 |
| gpc41 | 1.97855650883964e-48 | 0.341495354 | 0.21 | 0.026 | 4.97982887709849e-44 | 22 |
| sept102 | 1.97937641085324e-48 | 0.422248608 | 0.282 | 0.045 | 4.98189248847653e-44 | 22 |
| sh3pxd2b4 | 2.93438350199253e-48 | 0.478802652 | 0.271 | 0.043 | 7.38554983616501e-44 | 22 |
| fhl1b | 3.260180174097e-48 | 0.315633377 | 0.16 | 0.015 | 8.20554748018474e-44 | 22 |

| | | | | | | |
|---|---|---|---|---|---|---|
| si:dkey-9i23.164 | 3.75723946630233e-48 | 0.440897006 | 0.254 | 0.038 | 9.45659601273633e-44 | 22 |
| tmeff1b1 | 4.65000259341062e-48 | 0.364735842 | 0.215 | 0.028 | 1.17035915273552e-43 | 22 |
| kalrnb | 5.5899994246341e-48 | 0.283253386 | 0.122 | 0.009 | 1.40694695518616e-43 | 22 |
| LOC1018861141 | 7.37139653739541e-48 | 0.31693671 | 0.149 | 0.014 | 1.85530679449705e-43 | 22 |
| pdgfra3 | 1.16569583146499e-47 | 0.511297065 | 0.282 | 0.046 | 2.93393983821423e-43 | 22 |
| rnd3a | 1.4142410347066e-47 | 0.409350566 | 0.221 | 0.029 | 3.55950326025305e-43 | 22 |
| si:ch1073-459j12.14 | 2.10183790494548e-47 | 0.554322942 | 0.337 | 0.063 | 5.29011582295729e-43 | 22 |
| jund5 | 2.51392357578533e-47 | 0.612657149 | 0.403 | 0.089 | 6.3272942478941e-43 | 22 |
| capns1b3 | 3.17450665566494e-47 | 0.614794005 | 0.47 | 0.118 | 7.98991580164308e-43 | 22 |
| capn2a2 | 1.71005751985046e-46 | 0.479129826 | 0.293 | 0.05 | 4.30404377171162e-42 | 22 |
| ip6k2b5 | 1.91799134906791e-46 | 0.651355219 | 0.47 | 0.118 | 4.82739242646902e-42 | 22 |
| gli32 | 2.17961148730441e-46 | 0.359594017 | 0.221 | 0.03 | 5.48586415239647e-42 | 22 |
| meis1b4 | 4.84100010956234e-46 | 0.482116577 | 0.304 | 0.054 | 1.21843131757574e-41 | 22 |
| CR847953.11 | 1.18686221593578e-45 | 0.355643362 | 0.182 | 0.021 | 2.98721351128876e-41 | 22 |
| mycb4 | 1.32283263555441e-45 | 0.763356974 | 0.635 | 0.207 | 3.32943746042689e-41 | 22 |
| plecb1 | 1.48140034116802e-45 | 0.476835567 | 0.282 | 0.048 | 3.72853651868578e-41 | 22 |
| sdc22 | 1.75129349896648e-45 | 0.516444806 | 0.359 | 0.073 | 4.40783060754873e-41 | 22 |
| serpine2 | 1.93267134965181e-45 | 0.534546649 | 0.32 | 0.06 | 4.86434051993864e-41 | 22 |
| lamb1b1 | 3.88762464178069e-45 | 0.366451929 | 0.199 | 0.025 | 9.78476246089783e-41 | 22 |
| nkd1 | 4.18576035679731e-45 | 0.283310944 | 0.144 | 0.013 | 1.05351402420232e-40 | 22 |
| tagln27 | 1.08618829018543e-44 | 0.784056929 | 0.575 | 0.184 | 2.73382730756772e-40 | 22 |
| prelp1 | 3.36768605696315e-44 | 0.338026497 | 0.122 | 0.01 | 8.47612903677056e-40 | 22 |
| si:dkey-226m8.101 | 5.29428721820705e-44 | 0.256397433 | 0.122 | 0.01 | 1.33251914995053e-39 | 22 |
| tmem100a2 | 1.09474375144563e-43 | 0.343324508 | 0.171 | 0.019 | 2.75536054801351e-39 | 22 |
| mcl1b6 | 1.22054263273448e-43 | 0.742991081 | 0.685 | 0.237 | 3.07198375232942e-39 | 22 |
| VIT | 1.26909371101716e-43 | 0.276976895 | 0.11 | 0.008 | 3.1941819612591e-39 | 22 |
| cygb13 | 2.38418592288314e-43 | 0.472783884 | 0.293 | 0.052 | 6.00075754930459e-39 | 22 |
| sox61 | 2.68591818003425e-43 | 0.43182225 | 0.199 | 0.026 | 6.7601874673282e-39 | 22 |
| CR847566.13 | 2.95267596863472e-43 | 0.454047942 | 0.227 | 0.033 | 7.43159014545673e-39 | 22 |
| cxcl141 | 4.69032065816411e-43 | 0.273580278 | 0.138 | 0.013 | 1.18050680645333e-38 | 22 |
| pcbd1 | 5.01612908502979e-43 | 0.49867771 | 0.315 | 0.059 | 1.26250952941115e-38 | 22 |
| frmd6 | 6.71509712355506e-43 | 0.276582366 | 0.16 | 0.017 | 1.69012279502757e-38 | 22 |
| ptprga5 | 7.31532292452747e-43 | 0.508380462 | 0.32 | 0.063 | 1.84119362687432e-38 | 22 |
| rhoua2 | 2.22360336824543e-42 | 0.323802807 | 0.182 | 0.022 | 5.59658731753693e-38 | 22 |
| cavin2b3 | 2.76953645232824e-42 | 0.504025377 | 0.32 | 0.063 | 6.97064629686494e-38 | 22 |

| | | | | | | |
|---|---|---|---|---|---|---|
| eva1ba4 | 2.90053202042431e-42 | 0.495892034 | 0.32 | 0.063 | 7.30034904220595e-38 | 22 |
| selenop | 3.55369207853133e-42 | 0.295424533 | 0.16 | 0.018 | 8.9442875924555e-38 | 22 |
| ogna | 4.32753678768505e-42 | 0.329081788 | 0.11 | 0.008 | 1.08919773409245e-37 | 22 |
| bckdhbl | 4.4461330602987e-42 | 0.341549128 | 0.204 | 0.028 | 1.11904722994658e-37 | 22 |
| s100u2 | 5.88095641794879e-42 | 0.370156053 | 0.227 | 0.034 | 1.48017792083353e-37 | 22 |
| mych2 | 6.29017207647166e-42 | 0.537606207 | 0.343 | 0.072 | 1.58317340992715e-37 | 22 |
| anxa46 | 6.9276013199024e-42 | 0.623408761 | 0.475 | 0.133 | 1.74360797620623e-37 | 22 |
| gulp1a2 | 9.43342902627681e-42 | 0.364864002 | 0.21 | 0.03 | 2.37429975162361e-37 | 22 |
| midn5 | 1.11526802119598e-41 | 0.651810123 | 0.486 | 0.136 | 2.80701808254817e-37 | 22 |
| ccl25b4 | 1.41019267728626e-41 | 0.938275197 | 0.337 | 0.074 | 3.5493139494618e-37 | 22 |
| CABZ01039096.11 | 1.45792808813254e-41 | 0.358293923 | 0.171 | 0.02 | 3.66945920502079e-37 | 22 |
| cd28l3 | 1.98958189332033e-41 | 0.396054849 | 0.232 | 0.036 | 5.00757866729794e-37 | 22 |
| wls4 | 2.5235090875096e-41 | 0.579209786 | 0.459 | 0.119 | 6.35142002235291e-37 | 22 |
| ccdc85b4 | 3.48646422906352e-41 | 0.445674185 | 0.265 | 0.047 | 8.77508181812998e-37 | 22 |
| krt41 | 1.0321741386729e-40 | 0.426074361 | 0.243 | 0.039 | 2.59787908962583e-36 | 22 |
| si:ch211-284f22.32 | 2.26485817605263e-40 | 0.363242817 | 0.215 | 0.032 | 5.70042154330686e-36 | 22 |
| reck1 | 4.95941853429712e-40 | 0.332089361 | 0.188 | 0.025 | 1.24823605089724e-35 | 22 |
| dhrs13l12 | 6.31372654798844e-40 | 0.346778818 | 0.21 | 0.031 | 1.58910183486321e-35 | 22 |
| sh3gl1b1 | 1.54967295649593e-39 | 0.377881218 | 0.221 | 0.034 | 3.90037186420462e-35 | 22 |
| itgb1a4 | 2.41505680348245e-39 | 0.445471021 | 0.254 | 0.045 | 6.07845646868497e-35 | 22 |
| sulf1 | 3.15628402422908e-39 | 0.325041298 | 0.155 | 0.018 | 7.94405126058216e-35 | 22 |
| bicc1b1 | 3.22277728487374e-39 | 0.305064147 | 0.166 | 0.02 | 8.11140814829872e-35 | 22 |
| hoxc3a2 | 4.44787613116573e-39 | 0.264667831 | 0.166 | 0.02 | 1.1194859434531e-34 | 22 |
| lrp1aa4 | 6.89882263989583e-39 | 0.418741278 | 0.282 | 0.053 | 1.73636467023538e-34 | 22 |
| myl9b4 | 8.40806559907216e-39 | 0.481873032 | 0.348 | 0.077 | 2.11622603063047e-34 | 22 |
| si:ch211-105c13.32 | 1.7619023544253e-38 | 0.474252846 | 0.298 | 0.058 | 4.43453203585304e-34 | 22 |
| calua | 1.84170375814296e-38 | 0.302875585 | 0.177 | 0.023 | 4.63538418887003e-34 | 22 |
| myl64 | 4.87287420839039e-38 | 0.56437109 | 0.392 | 0.098 | 1.22645370950978e-33 | 22 |
| olfml3b3 | 6.03908103241009e-38 | 0.37691332 | 0.249 | 0.043 | 1.5199763050473e-33 | 22 |
| crtap1 | 7.3102179598841e-38 | 0.31554538 | 0.199 | 0.029 | 1.83990875832323e-33 | 22 |
| nfia5 | 8.34830840583946e-38 | 0.484511611 | 0.343 | 0.077 | 2.10118574266573e-33 | 22 |
| phactr4b1 | 8.74670619989165e-38 | 0.297282581 | 0.166 | 0.021 | 2.20145848345073e-33 | 22 |
| wu:fi04e12 | 9.37904774331465e-38 | 0.268348953 | 0.16 | 0.019 | 2.36061252651486e-33 | 22 |
| itga51 | 1.69276059194083e-37 | 0.3404346 | 0.193 | 0.028 | 4.26050913385586e-33 | 22 |
| aoc25 | 5.61442594361605e-37 | 0.51145567 | 0.304 | 0.064 | 1.41309486574872e-32 | 22 |

| | | | | | | |
|---|---|---|---|---|---|---|
| spry43 | 7.56149906861499e-37 | 0.43224253 | 0.249 | 0.045 | 1.90315370057971e-32 | 22 |
| mrc22 | 1.27392568726033e-36 | 0.275122667 | 0.177 | 0.024 | 3.20634356226553e-32 | 22 |
| CABZ01039304.11 | 1.92485205060512e-36 | 0.252495922 | 0.133 | 0.014 | 4.84466012616802e-32 | 22 |
| mafba5 | 2.27383540257459e-36 | 0.48226485 | 0.315 | 0.07 | 5.72301632473998e-32 | 22 |
| XLOC-0004316 | 2.51854362807437e-36 | 0.640322165 | 0.773 | 0.273 | 6.33892245750037e-32 | 22 |
| foxo1a1 | 4.80086558913091e-36 | 0.359918049 | 0.227 | 0.039 | 1.20832986012836e-31 | 22 |
| hcar1-42 | 9.97877484089983e-36 | 0.36770698 | 0.21 | 0.034 | 2.51155783970608e-31 | 22 |
| slc16a1b | 1.20987025497188e-35 | 0.287132097 | 0.155 | 0.019 | 3.04512244473872e-31 | 22 |
| rbfox21 | 2.05034516500265e-35 | 0.357272285 | 0.21 | 0.034 | 5.16051374579517e-31 | 22 |
| zgc:110158 | 2.36658154199565e-35 | 0.34178537 | 0.193 | 0.03 | 5.95644908304884e-31 | 22 |
| eif1b6 | 2.70809760605544e-35 | 0.65632013 | 0.801 | 0.378 | 6.81601086468093e-31 | 22 |
| pard3bb1 | 5.76106622401339e-35 | 0.295182093 | 0.177 | 0.025 | 1.45000275792193e-30 | 22 |
| col12a1a2 | 8.0482959355315e-35 | 0.377474267 | 0.177 | 0.025 | 2.02567560401392e-30 | 22 |
| nfil31 | 1.13404706675173e-34 | 0.388662844 | 0.199 | 0.032 | 2.85428306230744e-30 | 22 |
| ciarta4 | 1.7410702232974e-34 | 0.527063341 | 0.381 | 0.102 | 4.38209964501724e-30 | 22 |
| osr11 | 3.10048929800129e-34 | 0.36184808 | 0.177 | 0.026 | 7.80362151413945e-30 | 22 |
| lzts2a | 4.13213913036972e-34 | 0.271927305 | 0.155 | 0.02 | 1.04001809772276e-29 | 22 |
| htra1b2 | 5.24272274556731e-34 | 0.372460963 | 0.221 | 0.038 | 1.31954088783184e-29 | 22 |
| tcf7l2 | 6.2484095107434e-34 | 0.264568873 | 0.133 | 0.015 | 1.57266218975901e-29 | 22 |
| ctnnb12 | 1.10384712787994e-33 | 0.513697427 | 0.486 | 0.146 | 2.77827283616102e-29 | 22 |
| dedd14 | 1.26223567213894e-33 | 0.569911281 | 0.464 | 0.143 | 3.17692096320651e-29 | 22 |
| hic13 | 1.38461639289118e-33 | 0.428940036 | 0.271 | 0.056 | 3.48494099926782e-29 | 22 |
| vasna2 | 1.67513796993664e-33 | 0.312850646 | 0.204 | 0.033 | 4.21615475653353e-29 | 22 |
| mgp1 | 1.85978984769232e-33 | 0.345811911 | 0.149 | 0.019 | 4.6809050676568e-29 | 22 |
| znf5031 | 1.89920500522636e-33 | 0.258365814 | 0.155 | 0.02 | 4.78010907765423e-29 | 22 |
| hmgn67 | 2.30506998255033e-33 | 0.612066632 | 0.674 | 0.261 | 5.80163063908092e-29 | 22 |
| wwtr11 | 3.67570534590265e-33 | 0.312709383 | 0.166 | 0.024 | 9.25138278510239e-29 | 22 |
| spry23 | 4.84520829923076e-33 | 0.421364475 | 0.282 | 0.061 | 1.21949047683339e-28 | 22 |
| higd1a7 | 4.92932761799186e-33 | 0.664599047 | 0.702 | 0.305 | 1.24066246817237e-28 | 22 |
| tspan4b | 5.24651620309748e-33 | 0.427000774 | 0.293 | 0.064 | 1.3204956631576e-28 | 22 |
| mfap53 | 1.44395451089912e-32 | 0.389044605 | 0.232 | 0.043 | 3.63428910848198e-28 | 22 |
| gsna | 1.55240310992904e-32 | 0.32082755 | 0.182 | 0.028 | 3.9072433873804e-28 | 22 |
| enam | 2.30632309863136e-32 | 0.475842286 | 0.116 | 0.012 | 5.80478460694527e-28 | 22 |
| zgc:153372 | 2.3550173269049e-32 | 0.275189567 | 0.182 | 0.028 | 5.92734311008694e-28 | 22 |
| si:dkey-6n21.121 | 2.62449397737584e-32 | 0.312862564 | 0.199 | 0.033 | 6.60558889165724e-28 | 22 |

| | | | | | | |
|---|---|---|---|---|---|---|
| antxr1c2 | 4.20539926993908e-32 | 0.37070097 | 0.204 | 0.036 | 1.05845694225097e-27 | 22 |
| pdk2b6 | 4.54191775347724e-32 | 0.495671197 | 0.365 | 0.097 | 1.14315527937269e-27 | 22 |
| nanos1 | 7.26768992728267e-32 | 0.293459824 | 0.144 | 0.019 | 1.82920487779778e-27 | 22 |
| ntn1a2 | 8.46722799232012e-32 | 0.255806003 | 0.11 | 0.011 | 2.13111661338705e-27 | 22 |
| pim16 | 1.86449927693603e-31 | 0.614356182 | 0.497 | 0.168 | 4.6927582301203e-27 | 22 |
| flna5 | 2.07486312783405e-31 | 0.515277412 | 0.392 | 0.112 | 5.22222300644553e-27 | 22 |
| aqp3a1 | 2.15359120797165e-31 | 0.377102406 | 0.177 | 0.027 | 5.42037371134385e-27 | 22 |
| nsa29 | 3.25826649521402e-31 | 0.628286583 | 0.773 | 0.358 | 8.20073094180417e-27 | 22 |
| si:dkey-26m3.3 | 7.05539683288147e-31 | 0.253768072 | 0.127 | 0.015 | 1.77577282886794e-26 | 22 |
| ndrg21 | 7.38263067508241e-31 | 0.33665774 | 0.21 | 0.038 | 1.85813431461149e-26 | 22 |
| LOC1039092843 | 1.37762112594318e-30 | 0.40437177 | 0.276 | 0.062 | 3.46733461188639e-26 | 22 |
| proza1 | 1.59558085830143e-30 | 0.293184945 | 0.188 | 0.031 | 4.01591746225887e-26 | 22 |
| nme33 | 1.81063485014197e-30 | 0.408936677 | 0.304 | 0.074 | 4.55718685432232e-26 | 22 |
| adgrl3.1 | 1.90690370825337e-30 | 0.326076011 | 0.16 | 0.024 | 4.7994859433029e-26 | 22 |
| fbln21 | 2.00116010900633e-30 | 0.286544818 | 0.16 | 0.023 | 5.03671987835802e-26 | 22 |
| yap13 | 2.38910135880553e-30 | 0.364896132 | 0.238 | 0.048 | 6.01312920997763e-26 | 22 |
| cav11 | 4.03283141222091e-30 | 0.408925762 | 0.221 | 0.043 | 1.01502333814188e-25 | 22 |
| si:ch211-212k18.94 | 4.14039819865047e-30 | 0.315968089 | 0.199 | 0.035 | 1.04209682261834e-25 | 22 |
| col16a12 | 4.40462434373283e-30 | 0.342732547 | 0.188 | 0.032 | 1.10859990107412e-25 | 22 |
| adamts15a4 | 5.48155958061693e-30 | 0.423056015 | 0.232 | 0.047 | 1.37965373084548e-25 | 22 |
| nid1b3 | 7.03348087607267e-30 | 0.393049187 | 0.254 | 0.053 | 1.77025680169873e-25 | 22 |
| sgce1 | 9.56901498080645e-30 | 0.310947726 | 0.166 | 0.026 | 2.40842538051918e-25 | 22 |
| cyp2n133 | 1.17821000801102e-29 | 0.361431091 | 0.199 | 0.036 | 2.96543676916294e-25 | 22 |
| phlda2 | 2.23842577678589e-29 | 0.506624705 | 0.238 | 0.05 | 5.6338938375924e-25 | 22 |
| id2b1 | 2.647687591953e-29 | 0.420075516 | 0.188 | 0.033 | 6.66396490018652e-25 | 22 |
| dpysl2b | 3.15266610292358e-29 | 0.269157688 | 0.138 | 0.019 | 7.93494531444836e-25 | 22 |
| atp2b23 | 4.08741437381527e-29 | 0.424573962 | 0.287 | 0.07 | 1.02876132374557e-24 | 22 |
| tspan142 | 4.28056240356312e-29 | 0.503470501 | 0.403 | 0.122 | 1.0773747513528e-24 | 22 |
| appb3 | 4.29262001023352e-29 | 0.3215043 | 0.204 | 0.038 | 1.08040953037568e-24 | 22 |
| ubtd2 | 4.30485156788183e-29 | 0.276916268 | 0.16 | 0.025 | 1.08348809112018e-24 | 22 |
| si:dkey-7j14.66 | 5.21534271459153e-29 | 0.504830867 | 0.508 | 0.171 | 1.31264960783554e-24 | 22 |
| agtr22 | 5.65098026280712e-29 | 0.282335491 | 0.133 | 0.017 | 1.42229522234593e-24 | 22 |
| h1f04 | 6.11687832021618e-29 | 0.451654013 | 0.265 | 0.061 | 1.53955710441521e-24 | 22 |
| glud1a | 7.04330263792292e-29 | 0.42060927 | 0.337 | 0.089 | 1.77272884093882e-24 | 22 |
| cnn25 | 9.29455160740202e-29 | 0.510464154 | 0.42 | 0.132 | 2.33934569406701e-24 | 22 |

| | | | | | | |
|---|---|---|---|---|---|---|
| ak4 | 1.30817158092548e-28 | 0.277099759 | 0.177 | 0.03 | 3.29253705203134e-24 | 22 |
| AL732488.11 | 1.62170372730741e-28 | 0.386729955 | 0.204 | 0.039 | 4.08166611126002e-24 | 22 |
| si:ch211-227m13.11 | 1.63168495029997e-28 | 0.272097547 | 0.11 | 0.013 | 4.10678785140998e-24 | 22 |
| calm2b3 | 2.64189905800179e-28 | 0.589001272 | 0.541 | 0.207 | 6.64939573908471e-24 | 22 |
| klhl24b | 2.85446854342596e-28 | 0.264954325 | 0.122 | 0.015 | 7.18441187694881e-24 | 22 |
| scpp7 | 4.65001083800952e-28 | 0.29460057 | 0.166 | 0.027 | 1.17036122781862e-23 | 22 |
| glis2a1 | 7.17077252092489e-28 | 0.287004716 | 0.16 | 0.025 | 1.80481173579159e-23 | 22 |
| fbp1a | 8.307419699669e-28 | 0.319681496 | 0.199 | 0.038 | 2.09089446420969e-23 | 22 |
| zgc:1132085 | 1.00122243575634e-27 | 0.328306856 | 0.215 | 0.043 | 2.51997674855512e-23 | 22 |
| tead1b | 1.02901466043308e-27 | 0.285802567 | 0.166 | 0.027 | 2.58992699884403e-23 | 22 |
| col21a11 | 1.23465794529051e-27 | 0.261551292 | 0.127 | 0.017 | 3.10751058250167e-23 | 22 |
| eif4a1b4 | 1.4173394753941e-27 | 0.573426156 | 0.641 | 0.276 | 3.56730172561941e-23 | 22 |
| parvaa | 1.74309934920046e-27 | 0.303192922 | 0.166 | 0.028 | 4.38720675200265e-23 | 22 |
| histh1l5 | 2.76609165253762e-27 | 0.54714235 | 0.359 | 0.107 | 6.96197608027194e-23 | 22 |
| ehd1b1 | 3.08181174287668e-27 | 0.322069697 | 0.227 | 0.047 | 7.75661197564632e-23 | 22 |
| im:71514491 | 4.1024173958716e-27 | 0.36186942 | 0.16 | 0.026 | 1.03253743436692e-22 | 22 |
| neo1a2 | 1.58813846729355e-26 | 0.266737899 | 0.16 | 0.027 | 3.99718570833113e-22 | 22 |
| ybx16 | 2.57324089946333e-26 | 0.437659647 | 0.934 | 0.667 | 6.47659001985926e-22 | 22 |
| irf2bp2b3 | 4.19942885938066e-26 | 0.421264363 | 0.298 | 0.08 | 1.05695424961752e-21 | 22 |
| serf28 | 7.3955523562978e-26 | 0.558214571 | 0.724 | 0.339 | 1.86138657255659e-21 | 22 |
| igf2b4 | 9.70353480267296e-26 | 0.35533893 | 0.249 | 0.057 | 2.44228267448476e-21 | 22 |
| boc4 | 9.9431973327762e-26 | 0.320996768 | 0.221 | 0.047 | 2.50260333668644e-21 | 22 |
| acsl1a | 1.11656604989739e-25 | 0.258949168 | 0.133 | 0.019 | 2.81028509098673e-21 | 22 |
| ptbp1b1 | 1.2937826629739e-25 | 0.277290013 | 0.182 | 0.034 | 3.25632158443902e-21 | 22 |
| notch26 | 2.98461710128438e-25 | 0.457735991 | 0.37 | 0.117 | 7.51198278222264e-21 | 22 |
| rem11 | 4.10981751101895e-25 | 0.300842998 | 0.177 | 0.032 | 1.03439996934836e-20 | 22 |
| vdac35 | 4.11393496100798e-25 | 0.468607919 | 0.442 | 0.151 | 1.0354362903361e-20 | 22 |
| cdkn1a | 5.55380602746934e-25 | 0.295187506 | 0.171 | 0.031 | 1.39783743905376e-20 | 22 |
| cbfb | 5.84906038088769e-25 | 0.347279548 | 0.193 | 0.04 | 1.47215000726562e-20 | 22 |
| tuba1b2 | 6.45568581968996e-25 | 0.313730177 | 0.166 | 0.03 | 1.62483156395777e-20 | 22 |
| hmgn76 | 8.35216036962487e-25 | 0.566958419 | 0.613 | 0.276 | 2.10215524343088e-20 | 22 |
| adma1 | 1.18708586352384e-24 | 0.327515187 | 0.122 | 0.017 | 2.98777640990316e-20 | 22 |
| svild4 | 1.20449499163896e-24 | 0.416259611 | 0.293 | 0.08 | 3.0315934444561e-20 | 22 |
| add3a5 | 1.21245402415504e-24 | 0.467901435 | 0.381 | 0.122 | 3.05162553339581e-20 | 22 |
| ctdsp23 | 1.38665832895944e-24 | 0.351755061 | 0.26 | 0.065 | 3.49008034815801e-20 | 22 |

| | | | | | | |
|---|---|---|---|---|---|---|
| rgs41 | 1.44944159153832e-24 | 0.376526206 | 0.199 | 0.041 | 3.6480995417428e-20 | 22 |
| nfil3-67 | 1.45838191472684e-24 | 0.488127556 | 0.392 | 0.128 | 3.67060144117599e-20 | 22 |
| myh10 | 1.51917721478952e-24 | 0.267897974 | 0.182 | 0.035 | 3.82361713190375e-20 | 22 |
| cavin1b1 | 1.79521486800901e-24 | 0.342061245 | 0.193 | 0.039 | 4.51837630129187e-20 | 22 |
| pbx1a3 | 2.3395136660725e-24 | 0.29908315 | 0.199 | 0.041 | 5.88832194613788e-20 | 22 |
| mef2aa2 | 9.64936070745867e-24 | 0.398380569 | 0.26 | 0.067 | 2.42864759646027e-19 | 22 |
| perp1 | 1.07358515592528e-23 | 0.341062188 | 0.227 | 0.053 | 2.70210647894833e-19 | 22 |
| col15a1a2 | 1.11721353236439e-23 | 0.26446721 | 0.133 | 0.021 | 2.81191473960793e-19 | 22 |
| zgc:1749044 | 2.18776201681124e-23 | 0.396964683 | 0.21 | 0.047 | 5.5063782201122e-19 | 22 |
| gadd45ga1 | 3.92559757767654e-23 | 0.341162867 | 0.221 | 0.051 | 9.88033654325408e-19 | 22 |
| calr | 4.64347396204716e-23 | 0.339443479 | 0.215 | 0.05 | 1.16871596150765e-18 | 22 |
| cish3 | 4.97282766568055e-23 | 0.406086488 | 0.221 | 0.053 | 1.25161099517514e-18 | 22 |
| fam168a5 | 5.67977049769261e-23 | 0.3251871 | 0.232 | 0.057 | 1.42954143656425e-18 | 22 |
| ackr3b2 | 7.11592630240417e-23 | 0.299558537 | 0.138 | 0.023 | 1.79100749105211e-18 | 22 |
| dnase1l12 | 8.36301760322204e-23 | 0.303593867 | 0.199 | 0.044 | 2.10488790055496e-18 | 22 |
| DDX17 | 9.2895654886382e-23 | 0.308597553 | 0.238 | 0.059 | 2.33809073783535e-18 | 22 |
| tmem176l.14 | 1.05839165343223e-22 | 0.456998448 | 0.481 | 0.178 | 2.66386595252357e-18 | 22 |
| dusp8a1 | 1.11161218469659e-22 | 0.259362755 | 0.149 | 0.027 | 2.79781670766286e-18 | 22 |
| ephb33 | 1.5956430255688e-22 | 0.320118386 | 0.188 | 0.04 | 4.01607393105412e-18 | 22 |
| flot2a4 | 1.64387323965109e-22 | 0.330599407 | 0.232 | 0.058 | 4.13746455687783e-18 | 22 |
| klf31 | 2.72272496258602e-22 | 0.302753765 | 0.171 | 0.035 | 6.85282645833275e-18 | 22 |
| mgst1.14 | 2.78585281209533e-22 | 0.271961508 | 0.188 | 0.04 | 7.01171294276275e-18 | 22 |
| prdx55 | 3.95359977043677e-22 | 0.480735457 | 0.42 | 0.157 | 9.95081526221229e-18 | 22 |
| slc3a2b5 | 5.05612470554032e-22 | 0.416034667 | 0.354 | 0.116 | 1.27257602713744e-17 | 22 |
| mgst3b3 | 5.19453259207062e-22 | 0.457486731 | 0.448 | 0.17 | 1.30741190809825e-17 | 22 |
| slc38a5b4 | 5.70614382024509e-22 | 0.412748326 | 0.348 | 0.114 | 1.43617933811749e-17 | 22 |
| plod3 | 7.14757193073649e-22 | 0.268443395 | 0.166 | 0.033 | 1.79897237924707e-17 | 22 |
| fzd23 | 8.2682917810351e-22 | 0.255710114 | 0.149 | 0.028 | 2.08104635836872e-17 | 22 |
| palm1b | 8.57715405099845e-22 | 0.29231071 | 0.232 | 0.057 | 2.1587839030958e-17 | 22 |
| sema4ba5 | 8.8688343254066e-22 | 0.324412939 | 0.199 | 0.045 | 2.23219691136159e-17 | 22 |
| slc7a14 | 1.43323474969627e-21 | 0.257054661 | 0.177 | 0.037 | 3.60730854151055e-17 | 22 |
| emp23 | 1.54416982823489e-21 | 0.318644987 | 0.204 | 0.048 | 3.8865210406844e-17 | 22 |
| zgc:926061 | 1.57553021351915e-21 | 0.367562133 | 0.287 | 0.085 | 3.96545199440636e-17 | 22 |
| anxa134 | 1.62664866229132e-21 | 0.329463423 | 0.204 | 0.049 | 4.09411201812101e-17 | 22 |
| nr1d2a1 | 1.77411287862126e-21 | 0.368819765 | 0.271 | 0.077 | 4.46526470420184e-17 | 22 |

| | | | | | | |
|---|---|---|---|---|---|---|
| celf29 | 2.25596600856971e-21 | 0.540794998 | 0.536 | 0.24 | 5.67804084696909e-17 | 22 |
| dag11 | 2.37902354351577e-21 | 0.346278688 | 0.204 | 0.049 | 5.98776435667483e-17 | 22 |
| daam1a | 2.4676608818678e-21 | 0.25782347 | 0.155 | 0.03 | 6.21085567357307e-17 | 22 |
| FO082781.15 | 3.24933893451392e-21 | 0.311517397 | 0.199 | 0.046 | 8.17826116427808e-17 | 22 |
| CT030188.12 | 6.43164246130443e-21 | 0.314237419 | 0.21 | 0.05 | 1.61878009108571e-16 | 22 |
| nfe2l2a2 | 6.75622875414335e-21 | 0.310963614 | 0.249 | 0.067 | 1.70047521513034e-16 | 22 |
| rhoab7 | 7.79347628407291e-21 | 0.486865423 | 0.459 | 0.187 | 1.96154004593831e-16 | 22 |
| rps223 | 7.91102670525568e-21 | 0.320358074 | 0.967 | 0.828 | 1.9911263114458e-16 | 22 |
| dbn14 | 8.31529132842889e-21 | 0.334894344 | 0.215 | 0.054 | 2.09287567445227e-16 | 22 |
| hsp70.31 | 8.96380281086425e-21 | 0.254089254 | 0.127 | 0.022 | 2.25609952946642e-16 | 22 |
| eef1da8 | 1.05639697617339e-20 | 0.491713128 | 0.801 | 0.489 | 2.6588455493308e-16 | 22 |
| si:ch1073-291c23.24 | 1.13214235024261e-20 | 0.327232964 | 0.232 | 0.06 | 2.84948908132562e-16 | 22 |
| anxa11b3 | 1.14191430988672e-20 | 0.411000116 | 0.403 | 0.148 | 2.87408412655387e-16 | 22 |
| selenow13 | 2.07115241996521e-20 | 0.513549685 | 0.735 | 0.382 | 5.21288352581045e-16 | 22 |
| tpt16 | 2.53685947862312e-20 | 0.291989013 | 0.945 | 0.8 | 6.38502162174654e-16 | 22 |
| arl6ip5b4 | 2.70277264557057e-20 | 0.392436128 | 0.298 | 0.095 | 6.80260847163657e-16 | 22 |
| rpz55 | 2.90972625802806e-20 | 0.50472196 | 0.304 | 0.101 | 7.32349001883083e-16 | 22 |
| rpl329 | 4.92984991431026e-20 | 0.317156335 | 0.972 | 0.812 | 1.24079392493275e-15 | 22 |
| reep52 | 6.04240045879244e-20 | 0.404056109 | 0.414 | 0.156 | 1.52081177147347e-15 | 22 |
| pnrc212 | 7.3423350753584e-20 | 0.478762947 | 0.884 | 0.565 | 1.84799231511695e-15 | 22 |
| tns1b3 | 1.02921021447536e-19 | 0.254750667 | 0.16 | 0.033 | 2.59041918881303e-15 | 22 |
| lgals3b2 | 1.17417221035844e-19 | 0.354571506 | 0.26 | 0.077 | 2.95527403625115e-15 | 22 |
| XLOC-0108995 | 1.3724647881448e-19 | 0.341920356 | 0.619 | 0.255 | 3.45435662528166e-15 | 22 |
| si:dkey-28b4.75 | 1.87161244080908e-19 | 0.359314806 | 0.287 | 0.09 | 4.71066135227237e-15 | 22 |
| zgc:11418811 | 1.93232868235701e-19 | 0.452378699 | 0.945 | 0.624 | 4.86347806062436e-15 | 22 |
| HACD4 | 2.42393098992624e-19 | 0.259491937 | 0.199 | 0.048 | 6.10079190854535e-15 | 22 |
| CABZ01068358.15 | 2.45899786339655e-19 | 0.327565212 | 0.232 | 0.063 | 6.18905172238277e-15 | 22 |
| col4a13 | 2.53970497794499e-19 | 0.334170483 | 0.221 | 0.058 | 6.39218345898974e-15 | 22 |
| si:ch211-196l7.410 | 3.17066791974955e-19 | 0.490194021 | 0.635 | 0.311 | 7.98025408721765e-15 | 22 |
| LOC108179036 | 3.32324291002658e-19 | 0.257005118 | 0.16 | 0.034 | 8.36427008024589e-15 | 22 |
| tnrc6c17 | 3.36986320035981e-19 | 0.385194152 | 0.365 | 0.131 | 8.4816086889856e-15 | 22 |
| mdka5 | 4.37625291667123e-19 | 0.472172677 | 0.287 | 0.092 | 1.10145909659698e-14 | 22 |
| nfic4 | 4.51459412071242e-19 | 0.347199112 | 0.254 | 0.075 | 1.13627819424211e-14 | 22 |
| slit32 | 6.2807415003734e-19 | 0.254285567 | 0.138 | 0.027 | 1.58079982822898e-14 | 22 |
| klf11b6 | 7.85058399855105e-19 | 0.34800861 | 0.26 | 0.079 | 1.97591348659531e-14 | 22 |

| | | | | | | |
|---|---|---|---|---|---|---|
| pdcd61 | 8.01813357780853e-19 | 0.346964367 | 0.249 | 0.073 | 2.01808404019863e-14 | 22 |
| maff4 | 9.53246763651991e-19 | 0.353621053 | 0.265 | 0.082 | 2.3992267794357e-14 | 22 |
| cnbpb4 | 1.01621368219612e-18 | 0.414307935 | 0.448 | 0.182 | 2.55770821671941e-14 | 22 |
| krt18a.13 | 1.1599312794122e-18 | 0.478311586 | 0.354 | 0.125 | 2.91943103715256e-14 | 22 |
| cald1a4 | 2.51006342378418e-18 | 0.373319819 | 0.298 | 0.097 | 6.31757863132241e-14 | 22 |
| si:ch211-106h4.124 | 4.67085855085289e-18 | 0.525822595 | 0.21 | 0.058 | 1.17560838866416e-13 | 22 |
| tspan71 | 4.99300860710962e-18 | 0.302267916 | 0.26 | 0.077 | 1.25669033632342e-13 | 22 |
| creb3l3l | 5.59203282669939e-18 | 0.278730772 | 0.199 | 0.052 | 1.40745874215197e-13 | 22 |
| trib32 | 6.17057573887218e-18 | 0.403812424 | 0.304 | 0.104 | 1.55307220771674e-13 | 22 |
| june1 | 6.79779101959907e-18 | 0.262381569 | 0.144 | 0.03 | 1.71093602172289e-13 | 22 |
| tnrc18 | 7.99606729864886e-18 | 0.26006918 | 0.171 | 0.041 | 2.01253017839693e-13 | 22 |
| rpl911 | 8.51102571144628e-18 | 0.428849185 | 0.917 | 0.627 | 2.14214006131391e-13 | 22 |
| tmod21 | 1.27355421046472e-17 | 0.286465966 | 0.182 | 0.045 | 3.20540859231866e-13 | 22 |
| itm2ba7 | 1.32241696367959e-17 | 0.445667496 | 0.575 | 0.271 | 3.32839125588516e-13 | 22 |
| aldh9a1a.15 | 1.47678144465823e-17 | 0.414606674 | 0.403 | 0.162 | 3.7169112180603e-13 | 22 |
| irf2bpl1 | 1.69364167952138e-17 | 0.25851042 | 0.177 | 0.043 | 4.26272674318735e-13 | 22 |
| adam10a3 | 1.7664631394499e-17 | 0.33494898 | 0.282 | 0.093 | 4.44601107568146e-13 | 22 |
| hdlbpa5 | 2.6747367131337e-17 | 0.372517201 | 0.403 | 0.157 | 6.73204483328621e-13 | 22 |
| dnmt3ab3 | 3.01499966736629e-17 | 0.341236215 | 0.249 | 0.078 | 7.58845266279421e-13 | 22 |
| map1lc3a4 | 8.05525824774656e-17 | 0.291829879 | 0.232 | 0.069 | 2.02742794837533e-12 | 22 |
| hopx2 | 9.39327695663815e-17 | 0.335040814 | 0.221 | 0.064 | 2.36419387721626e-12 | 22 |
| hmgb1b5 | 1.0115997513441e-16 | 0.344753933 | 0.343 | 0.125 | 2.54609541415797e-12 | 22 |
| h2afy24 | 1.03896186182781e-16 | 0.336729841 | 0.282 | 0.096 | 2.61496311003443e-12 | 22 |
| ube2e34 | 1.2726339561123e-16 | 0.347567115 | 0.392 | 0.154 | 3.20309240413906e-12 | 22 |
| smdt1b3 | 1.52056593119495e-16 | 0.377536681 | 0.343 | 0.132 | 3.82711239222458e-12 | 22 |
| atox11 | 2.28980309960509e-16 | 0.300177581 | 0.309 | 0.108 | 5.76320542139606e-12 | 22 |
| snai1a1 | 2.72291108434845e-16 | 0.257772578 | 0.204 | 0.056 | 6.85329490819662e-12 | 22 |
| rps1610 | 3.11433040991092e-16 | 0.350942139 | 0.956 | 0.736 | 7.83845820870479e-12 | 22 |
| si:dkey-56m19.52 | 3.24614466069786e-16 | 0.288178669 | 0.177 | 0.046 | 8.17022149651045e-12 | 22 |
| cebpb9 | 3.47238803462985e-16 | 0.51731144 | 0.646 | 0.351 | 8.73965344435986e-12 | 22 |
| cd82b2 | 3.9353828984485e-16 | 0.315044934 | 0.238 | 0.073 | 9.90496521710503e-12 | 22 |
| cdc422 | 4.91589712893731e-16 | 0.384346424 | 0.475 | 0.21 | 1.23728214838223e-11 | 22 |
| pdia36 | 5.03191345341831e-16 | 0.435338969 | 0.575 | 0.281 | 1.26648229709086e-11 | 22 |
| stau25 | 6.01571245637589e-16 | 0.332187856 | 0.254 | 0.085 | 1.51409466814525e-11 | 22 |
| sec61a12 | 6.72317678843785e-16 | 0.406933006 | 0.431 | 0.187 | 1.69215636588192e-11 | 22 |

| | | | | | | |
|---|---|---|---|---|---|---|
| rps1810 | 7.09338018770492e-16 | 0.340969755 | 0.939 | 0.723 | 1.78533285944345e-11 | 22 |
| mmp9 | 7.85366380804314e-16 | 0.315977692 | 0.105 | 0.019 | 1.97668864384638e-11 | 22 |
| COX311 | 1.1360388284304e-15 | 0.425107248 | 0.994 | 0.697 | 2.85929612727647e-11 | 22 |
| mmp16a3 | 1.35154778608077e-15 | 0.31432587 | 0.227 | 0.069 | 3.40171062278669e-11 | 22 |
| cd6311 | 1.58112308875235e-15 | 0.422875959 | 0.685 | 0.337 | 3.97952870208079e-11 | 22 |
| lxn6 | 1.64511100013989e-15 | 0.345440107 | 0.376 | 0.146 | 4.14057987625209e-11 | 22 |
| si:dkey-79d12.54 | 2.20134948361937e-15 | 0.345415788 | 0.232 | 0.075 | 5.54057651532161e-11 | 22 |
| faua8 | 2.20823238294653e-15 | 0.368918603 | 0.945 | 0.687 | 5.55790008463813e-11 | 22 |
| rps1510 | 2.38236359503599e-15 | 0.378756499 | 0.939 | 0.682 | 5.99617093234609e-11 | 22 |
| pgrmc14 | 2.40862660577241e-15 | 0.342253821 | 0.359 | 0.139 | 6.06227230406859e-11 | 22 |
| tob1b3 | 2.8606289271182e-15 | 0.38783273 | 0.409 | 0.172 | 7.1999169466638e-11 | 22 |
| sar1b | 3.14935216280923e-15 | 0.293981966 | 0.243 | 0.079 | 7.92660445857454e-11 | 22 |
| rpl229 | 3.30931518121255e-15 | 0.367786912 | 0.945 | 0.692 | 8.32921537959387e-11 | 22 |
| rpl289 | 4.47129782922229e-15 | 0.306011758 | 0.939 | 0.748 | 1.12538095063696e-10 | 22 |
| fosl26 | 4.81539631868915e-15 | 0.369181764 | 0.315 | 0.12 | 1.21198709945087e-10 | 22 |
| ptp4a13 | 5.65805923305622e-15 | 0.327938976 | 0.309 | 0.116 | 1.42407692836792e-10 | 22 |
| rpl3611 | 6.88048765169169e-15 | 0.322847547 | 0.972 | 0.788 | 1.73174993705428e-10 | 22 |
| ppiaa8 | 9.2648978714851e-15 | 0.268844146 | 0.961 | 0.803 | 2.33188214527408e-10 | 22 |
| glo12 | 1.56409585749857e-14 | 0.3335743 | 0.448 | 0.195 | 3.93667286373815e-10 | 22 |
| col12a1b3 | 1.7886374663759e-14 | 0.339599865 | 0.133 | 0.031 | 4.50182163912151e-10 | 22 |
| fkbp1aa5 | 1.90795168008878e-14 | 0.417398807 | 0.724 | 0.384 | 4.80212358361544e-10 | 22 |
| hp1bp32 | 1.96999764530281e-14 | 0.29909683 | 0.271 | 0.097 | 4.95828707346263e-10 | 22 |
| mxra8a3 | 2.12651484619382e-14 | 0.274091972 | 0.166 | 0.045 | 5.35222521638524e-10 | 22 |
| ptgdsb.2 | 2.7561344906142e-14 | 0.307912204 | 0.16 | 0.042 | 6.93691489942689e-10 | 22 |
| abcc54 | 3.78237601101717e-14 | 0.283484914 | 0.265 | 0.094 | 9.5198621821291e-10 | 22 |
| irf1b2 | 4.2178541667725e-14 | 0.250702692 | 0.149 | 0.038 | 1.06159171523497e-09 | 22 |
| pim24 | 1.00921413071738e-13 | 0.257511181 | 0.254 | 0.089 | 2.54009104560258e-09 | 22 |
| rpl189 | 1.48532359977084e-13 | 0.294030371 | 0.95 | 0.772 | 3.73841096826323e-09 | 22 |
| rps27.210 | 1.67648029294251e-13 | 0.42196129 | 0.851 | 0.514 | 4.21953324930702e-09 | 22 |
| rpl610 | 1.79183523378473e-13 | 0.354340395 | 0.923 | 0.665 | 4.50987009991278e-09 | 22 |
| gabarapb6 | 2.15795113453736e-13 | 0.364943276 | 0.541 | 0.266 | 5.43134721051707e-09 | 22 |
| rpl18a9 | 2.47321709888015e-13 | 0.285087081 | 0.939 | 0.784 | 6.22484011617144e-09 | 22 |
| tsc22d35 | 2.62681531954273e-13 | 0.350213037 | 0.337 | 0.143 | 6.6114314777571e-09 | 22 |
| gatad2b | 3.79064694987575e-13 | 0.254030561 | 0.188 | 0.058 | 9.54067930814227e-09 | 22 |
| rpl5a9 | 4.52447745791704e-13 | 0.428926883 | 0.84 | 0.488 | 1.13876573138314e-08 | 22 |

| | | | | | | |
|---|---|---|---|---|---|---|
| qkia4 | 5.28302986840571e-13 | 0.330869616 | 0.381 | 0.167 | 1.32968578757903e-08 | 22 |
| grpel22 | 6.78624587493534e-13 | 0.29283063 | 0.265 | 0.1 | 1.70803022426247e-08 | 22 |
| btg110 | 7.06878718775301e-13 | 0.382545638 | 0.63 | 0.33 | 1.77914304728555e-08 | 22 |
| slco2b13 | 1.35102492005622e-12 | 0.280634166 | 0.199 | 0.064 | 3.4003946212895e-08 | 22 |
| eef2b5 | 1.43455284691078e-12 | 0.360948685 | 0.878 | 0.611 | 3.61062606038973e-08 | 22 |
| sdcbp26 | 1.45089003317685e-12 | 0.330716278 | 0.442 | 0.207 | 3.65174512450282e-08 | 22 |
| h2afx18 | 1.587964372745e-12 | 0.37959365 | 0.414 | 0.197 | 3.9967475297619e-08 | 22 |
| ssr41 | 1.85280265703057e-12 | 0.377117889 | 0.58 | 0.304 | 4.66331900748023e-08 | 22 |
| rps4x8 | 2.49056828703658e-12 | 0.320061596 | 0.928 | 0.708 | 6.26851132164236e-08 | 22 |
| cebpg1 | 2.89741481927137e-12 | 0.255791216 | 0.193 | 0.063 | 7.29250335862411e-08 | 22 |
| rpl108 | 3.57105749148681e-12 | 0.269928569 | 0.934 | 0.771 | 8.98799460032314e-08 | 22 |
| map1lc3b4 | 3.60036408327273e-12 | 0.276457503 | 0.271 | 0.106 | 9.06175636118914e-08 | 22 |
| CABZ01072614.15 | 3.66511611063489e-12 | 0.339729035 | 0.221 | 0.08 | 9.22473073885696e-08 | 22 |
| si:ch211-286b5.54 | 4.70723818517037e-12 | 0.299261845 | 0.32 | 0.136 | 1.18476477882553e-07 | 22 |
| zgc:1583436 | 5.3362431028393e-12 | 0.419526783 | 0.359 | 0.165 | 1.34307902655362e-07 | 22 |
| metrnla2 | 6.4523342571265e-12 | 0.266670738 | 0.249 | 0.092 | 1.62398800917617e-07 | 22 |
| rab2a | 6.92239112953731e-12 | 0.296166393 | 0.381 | 0.171 | 1.74229662339325e-07 | 22 |
| rpl23a11 | 7.30859364666112e-12 | 0.360957689 | 0.912 | 0.632 | 1.83949993492814e-07 | 22 |
| aebp13 | 7.76144793721108e-12 | 0.25795972 | 0.138 | 0.039 | 1.95347883131666e-07 | 22 |
| sltm | 8.73777609957015e-12 | 0.264006623 | 0.182 | 0.06 | 2.19921086650081e-07 | 22 |
| rhoaa6 | 9.11979025035216e-12 | 0.305233579 | 0.343 | 0.153 | 2.29536000811114e-07 | 22 |
| mknk2b8 | 1.09113204548019e-11 | 0.351402363 | 0.409 | 0.196 | 2.7462702452691e-07 | 22 |
| rpl1110 | 1.17043792460981e-11 | 0.261934387 | 0.95 | 0.813 | 2.94587521245042e-07 | 22 |
| taf12 | 1.44538010570007e-11 | 0.263045254 | 0.282 | 0.115 | 3.63787718803649e-07 | 22 |
| tpm33 | 1.8389903641021e-11 | 0.346604348 | 0.492 | 0.253 | 4.62855484740858e-07 | 22 |
| rpl249 | 1.84158181621481e-11 | 0.268585123 | 0.939 | 0.775 | 4.63507727323105e-07 | 22 |
| ppdpfb6 | 1.95752581616277e-11 | 0.37661635 | 0.812 | 0.516 | 4.92689672670007e-07 | 22 |
| rps15a9 | 2.26340423052762e-11 | 0.308262077 | 0.972 | 0.778 | 5.69676210781497e-07 | 22 |
| zfp36l1a7 | 2.84043242562905e-11 | 0.347362687 | 0.492 | 0.246 | 7.14908437206575e-07 | 22 |
| serinc53 | 3.38483546834471e-11 | 0.283045682 | 0.221 | 0.083 | 8.51929239027681e-07 | 22 |
| capns1a2 | 4.70463130983389e-11 | 0.305652732 | 0.32 | 0.142 | 1.18410865437209e-06 | 22 |
| XLOC-0139643 | 5.18539736451436e-11 | 0.256808546 | 0.392 | 0.173 | 1.30511266267462e-06 | 22 |
| adh55 | 5.7673193324065e-11 | 0.307459307 | 0.448 | 0.22 | 1.45157660277339e-06 | 22 |
| cpne11 | 6.00431765713944e-11 | 0.289802568 | 0.287 | 0.123 | 1.51122671112543e-06 | 22 |
| rpl3410 | 6.23906353406952e-11 | 0.284245279 | 0.961 | 0.782 | 1.57030990088996e-06 | 22 |

| | | | | | | |
|---|---|---|---|---|---|---|
| mibp28 | 6.55344905161594e-11 | 0.419534034 | 0.768 | 0.49 | 1.64943759180122e-06 | 22 |
| sec61g3 | 7.75004416074833e-11 | 0.311008184 | 0.912 | 0.676 | 1.95060861481875e-06 | 22 |
| rab103 | 8.38881560955888e-11 | 0.3307025 | 0.392 | 0.194 | 2.11138100076987e-06 | 22 |
| glula4 | 9.23191129134954e-11 | 0.334139273 | 0.309 | 0.135 | 2.32357975291977e-06 | 22 |
| rps66 | 9.58743134302956e-11 | 0.253692723 | 0.945 | 0.788 | 2.41306059472711e-06 | 22 |
| anp32a1 | 1.75690780661297e-10 | 0.278648226 | 0.365 | 0.168 | 4.42196125846419e-06 | 22 |
| rps2115 | 1.847836053919e-10 | 0.308008986 | 0.945 | 0.748 | 4.65081856410874e-06 | 22 |
| arhgef1b1 | 1.85078120989727e-10 | 0.2856791 | 0.193 | 0.071 | 4.65823122719044e-06 | 22 |
| rpl89 | 2.14753611533446e-10 | 0.258002121 | 0.934 | 0.779 | 5.4051336486853e-06 | 22 |
| eef1g8 | 3.18703190355442e-10 | 0.332369449 | 0.862 | 0.618 | 8.02144059805613e-06 | 22 |
| rps1011 | 3.19992628648935e-10 | 0.334391978 | 0.95 | 0.692 | 8.05389447046503e-06 | 22 |
| rps2410 | 3.46505400202867e-10 | 0.276013333 | 0.95 | 0.757 | 8.72119441770596e-06 | 22 |
| romo19 | 3.80494538043256e-10 | 0.340942685 | 0.68 | 0.383 | 9.5766670280107e-06 | 22 |
| eif4ebp3l4 | 5.5378407640693e-10 | 0.290368536 | 0.425 | 0.205 | 1.3938191419086e-05 | 22 |
| tax1bp32 | 6.48899336348606e-10 | 0.27630425 | 0.221 | 0.088 | 1.63321473965581e-05 | 22 |
| mt-co111 | 6.60458398393886e-10 | 0.360600632 | 0.994 | 0.654 | 1.66230774291757e-05 | 22 |
| rpl27a10 | 6.65027915127124e-10 | 0.383109153 | 0.796 | 0.493 | 1.67380875958346e-05 | 22 |
| hmgb1a8 | 1.13700460220232e-09 | 0.319587099 | 0.37 | 0.188 | 2.86172688328302e-05 | 22 |
| prelid3b1 | 1.59206419498429e-09 | 0.294709132 | 0.453 | 0.237 | 4.00706637235596e-05 | 22 |
| b2ml10 | 1.69828566060575e-09 | 0.312395199 | 0.486 | 0.266 | 4.2744151791786e-05 | 22 |
| si:dkey-11f4.145 | 3.23890520345522e-09 | 0.275761702 | 0.249 | 0.108 | 8.15200050657643e-05 | 22 |
| tegt6 | 3.79812764090379e-09 | 0.258863219 | 0.414 | 0.21 | 9.55950745939075e-05 | 22 |
| rpl277 | 5.01555434813009e-09 | 0.269523637 | 0.95 | 0.761 | 0.000126236 | 22 |
| mt-atp610 | 5.01729953738195e-09 | 0.328171853 | 0.901 | 0.588 | 0.00012628 | 22 |
| dusp14 | 7.49705389386357e-09 | 0.293459334 | 0.326 | 0.159 | 0.000188693 | 22 |
| rps139 | 8.8531497192922e-09 | 0.269605243 | 0.917 | 0.723 | 0.000222825 | 22 |
| eif2s33 | 9.16064465685887e-09 | 0.283027214 | 0.514 | 0.283 | 0.000230564 | 22 |
| ppt14 | 1.12771142095147e-08 | 0.263264505 | 0.331 | 0.163 | 0.000283834 | 22 |
| naca10 | 1.48195950014045e-08 | 0.309648107 | 0.856 | 0.617 | 0.000372994 | 22 |
| eif3i6 | 2.87333776038132e-08 | 0.280044126 | 0.486 | 0.272 | 0.00072319 | 22 |
| gbp2 | 2.93878870786165e-08 | 0.255847775 | 0.227 | 0.101 | 0.000739664 | 22 |
| rpl3010 | 3.1615924521322e-08 | 0.284648361 | 0.928 | 0.706 | 0.000795741 | 22 |
| NC-002333.410 | 4.27043580837716e-08 | 0.315716589 | 0.906 | 0.62 | 0.001074826 | 22 |
| eif3ea6 | 4.89745785494647e-08 | 0.274557856 | 0.486 | 0.276 | 0.001232641 | 22 |
| eif3m6 | 5.86534503853055e-08 | 0.294609187 | 0.608 | 0.356 | 0.001476249 | 22 |

| | | | | | | |
|---|---|---|---|---|---|---|
| acin1a4 | 8.60512506252486e-08 | 0.264642465 | 0.398 | 0.216 | 0.002165824 | 22 |
| rps27a8 | 8.86276716633213e-08 | 0.255509743 | 0.934 | 0.75 | 0.00223067 | 22 |
| ubb2 | 1.37660093575877e-07 | 0.257465626 | 0.552 | 0.314 | 0.003464767 | 22 |
| rps27.19 | 1.67658514735512e-07 | 0.30889354 | 0.923 | 0.677 | 0.004219797 | 22 |
| mt-nd28 | 2.19204824033487e-07 | 0.281033225 | 0.657 | 0.399 | 0.005517166 | 22 |
| serbp1a9 | 2.33803539517451e-07 | 0.310485684 | 0.74 | 0.48 | 0.005884601 | 22 |
| rpl35a9 | 2.94099039931264e-07 | 0.256905846 | 0.95 | 0.758 | 0.007402179 | 22 |
| rsl24d19 | 3.28286243639344e-07 | 0.317055741 | 0.724 | 0.442 | 0.008262636 | 22 |
| bsg1 | 3.31089936999155e-07 | 0.26931671 | 0.646 | 0.378 | 0.008333203 | 22 |
| mt-nd19 | 4.91374302184317e-07 | 0.274601238 | 0.652 | 0.41 | 0.0123674 | 22 |
| eif3f8 | 7.18613979555301e-07 | 0.306180057 | 0.729 | 0.429 | 0.018086795 | 22 |
| rpl22l17 | 9.94784505357749e-07 | 0.29088632 | 0.646 | 0.39 | 0.025037731 | 22 |
| rpsa12 | 1.16407953263302e-06 | 0.259084189 | 0.95 | 0.75 | 0.029298718 | 22 |
| eif5a6 | 1.56025924784933e-06 | 0.291846083 | 0.74 | 0.477 | 0.039270165 | 22 |
| fabp34 | 2.27583876159085e-06 | 0.252491193 | 0.409 | 0.241 | 0.057280586 | 22 |
| sec61b2 | 3.14239480545088e-06 | 0.271596975 | 0.691 | 0.453 | 0.079090935 | 22 |
| myl12.16 | 3.60277872095478e-06 | 0.262622708 | 0.63 | 0.39 | 0.090678338 | 22 |
| tubb4b1 | 8.62186437450418e-06 | 0.268096069 | 0.823 | 0.583 | 0.217003704 | 22 |
| rplp211 | 1.07654106299468e-05 | 0.265341234 | 0.95 | 0.756 | 0.27095462 | 22 |
| mt-cyb9 | 1.20049087897705e-05 | 0.261900754 | 0.912 | 0.621 | 0.302151549 | 22 |
| bhlhe406 | 1.89257555686952e-05 | 0.263931931 | 0.276 | 0.161 | 0.476342342 | 22 |
| rpl3810 | 3.38376088737228e-05 | 0.254907474 | 0.939 | 0.726 | 0.851658778 | 22 |
| tmsb4x12 | 0.000152409 | 0.322293734 | 0.978 | 0.683 | 1 | 22 |
| rps26l10 | 0.000194519 | 0.272442344 | 0.912 | 0.662 | 1 | 22 |
| CR855311.16 | 0.00036058 | 0.261113727 | 0.16 | 0.086 | 1 | 22 |
| mt-co211 | 0.001524015 | 0.254526162 | 0.972 | 0.718 | 1 | 22 |
| prf1.91 | 3.52790844523476e-287 | 1.24359562 | 0.385 | 0.013 | 8.87939276581138e-283 | 23 |
| ccl36.13 | 3.53165452702624e-177 | 1.609120603 | 0.732 | 0.093 | 8.88882127907234e-173 | 23 |
| ccl38.63 | 9.33566991348145e-109 | 1.265640459 | 0.615 | 0.103 | 2.34969476052415e-104 | 23 |
| si:ch211-214p16.15 | 6.06714035299985e-87 | 1.222948109 | 0.866 | 0.25 | 1.52703855544653e-82 | 23 |
| FP236356.13 | 1.29597475564411e-83 | 0.734935174 | 0.363 | 0.046 | 3.26183886248067e-79 | 23 |
| prf1.7 | 6.94993059813088e-80 | 0.527434648 | 0.212 | 0.016 | 1.74922803224356e-75 | 23 |
| LOC1039108713 | 1.84979339401217e-72 | 1.081828305 | 0.654 | 0.165 | 4.65574499338922e-68 | 23 |
| si:dkey-151g10.61 | 1.58200245458701e-64 | 0.258566446 | 0.994 | 0.992 | 3.98174197795005e-60 | 23 |
| CR936442.15 | 5.70632940560458e-64 | 0.994439532 | 0.626 | 0.164 | 1.43622604809662e-59 | 23 |

| | | | | | | |
|---|---|---|---|---|---|---|
| rps299 | 4.237501342925e-63 | 0.461228039 | 0.972 | 0.831 | 1.06653671300079e-58 | 23 |
| zbtb322 | 5.20624678072623e-58 | 0.62362275 | 0.302 | 0.044 | 1.31036025224098e-53 | 23 |
| rplp212 | 5.47735131132046e-58 | 0.521846389 | 0.966 | 0.756 | 1.37859455154625e-53 | 23 |
| rplp2l12 | 6.84825364964166e-57 | 0.50863118 | 0.944 | 0.773 | 1.72363696107831e-52 | 23 |
| il342 | 6.92230703511396e-56 | 0.650342907 | 0.302 | 0.046 | 1.74227545766783e-51 | 23 |
| rps2810 | 1.40905798592812e-55 | 0.421006419 | 0.961 | 0.825 | 3.5464580447825e-51 | 23 |
| arpc1b8 | 7.30908450268936e-55 | 0.942078028 | 0.788 | 0.304 | 1.83962347848188e-50 | 23 |
| nkl.22 | 1.09841221591537e-54 | 1.039333055 | 0.553 | 0.155 | 2.76459370623739e-50 | 23 |
| XLOC-0004317 | 1.775576439658e-54 | 1.013422371 | 0.782 | 0.273 | 4.46894834097522e-50 | 23 |
| XLOC-0108996 | 6.21597270244575e-53 | 0.97297795 | 0.721 | 0.253 | 1.56449816947857e-48 | 23 |
| rpl358 | 8.70337027674975e-53 | 0.442627502 | 0.966 | 0.803 | 2.19055126495514e-48 | 23 |
| si:dkey-183i3.63 | 3.05632796615651e-51 | 0.611975417 | 0.285 | 0.044 | 7.69247185801931e-47 | 23 |
| rps207 | 5.38790559566019e-51 | 0.3307012 | 0.978 | 0.916 | 1.35608195937171e-46 | 23 |
| rps2116 | 1.80502092689002e-50 | 0.496586403 | 0.95 | 0.748 | 4.5430571708895e-46 | 23 |
| rps15a10 | 4.09432119821017e-50 | 0.478601741 | 0.961 | 0.778 | 1.03049970237752e-45 | 23 |
| CR753876.13 | 4.92382488356443e-50 | 0.795858964 | 0.464 | 0.111 | 1.23927748494433e-45 | 23 |
| rplp17 | 3.89169971802298e-48 | 0.379697568 | 0.983 | 0.876 | 9.79501902029203e-44 | 23 |
| scgn1 | 4.58794868939909e-48 | 0.461621431 | 0.19 | 0.022 | 1.15474080563486e-43 | 23 |
| LOC1081907173 | 1.66347915211738e-47 | 0.7541899 | 0.413 | 0.093 | 4.18681067796424e-43 | 23 |
| coro1a8 | 2.92622745794462e-47 | 0.845255485 | 0.659 | 0.226 | 7.36502188890082e-43 | 23 |
| syk3 | 3.27479523237688e-47 | 0.670531565 | 0.363 | 0.076 | 8.24233212036937e-43 | 23 |
| rpl1111 | 7.84376845950775e-47 | 0.417187439 | 0.961 | 0.813 | 1.97419808357351e-42 | 23 |
| ncaldb3 | 1.11740118576189e-46 | 0.640170111 | 0.352 | 0.071 | 2.81238704444411e-42 | 23 |
| rpl3811 | 1.76349046106547e-46 | 0.493688888 | 0.927 | 0.726 | 4.43852914145568e-42 | 23 |
| rpl378 | 2.19841394738446e-46 | 0.381405476 | 0.961 | 0.849 | 5.53318806417196e-42 | 23 |
| BX005105.12 | 9.94785348330638e-46 | 0.557368753 | 0.263 | 0.042 | 2.50377524321338e-41 | 23 |
| rpl398 | 1.68956911742089e-44 | 0.353603553 | 0.966 | 0.873 | 4.25247651163664e-40 | 23 |
| XLOC-0139644 | 1.75282383993993e-43 | 0.826340987 | 0.57 | 0.17 | 4.41168232274482e-39 | 23 |
| rps27.110 | 2.30259117107569e-43 | 0.53830454 | 0.95 | 0.677 | 5.7953917184804e-39 | 23 |
| tmsb4x13 | 3.58305713108343e-43 | 0.640072827 | 0.983 | 0.683 | 9.0181964932239e-39 | 23 |
| rpl3411 | 7.49028175437048e-43 | 0.434429485 | 0.939 | 0.782 | 1.88522901475751e-38 | 23 |
| rps26l11 | 1.38051089277055e-42 | 0.541374414 | 0.939 | 0.662 | 3.47460786601419e-38 | 23 |
| pfn18 | 8.92361092905861e-42 | 0.789961448 | 0.944 | 0.52 | 2.24598363473476e-37 | 23 |
| rpl278 | 1.54881298540446e-41 | 0.428872912 | 0.939 | 0.761 | 3.89820740296449e-37 | 23 |
| rpl1810 | 1.23091313871511e-40 | 0.419755521 | 0.944 | 0.772 | 3.09808527883205e-36 | 23 |

| | | | | | | |
|---|---|---|---|---|---|---|
| rpl299 | 6.74834980357669e-40 | 0.377149644 | 0.927 | 0.808 | 1.69849216206222e-35 | 23 |
| rps1811 | 3.38265997101821e-39 | 0.469410133 | 0.955 | 0.723 | 8.51381688105574e-35 | 23 |
| CABZ01084575.12 | 1.25328419101826e-38 | 0.514543345 | 0.24 | 0.041 | 3.15439098037385e-34 | 23 |
| CU855821.11 | 1.98489066434998e-37 | 0.305515701 | 0.128 | 0.013 | 4.99577131310247e-33 | 23 |
| rps176 | 3.13024653827408e-37 | 0.396808125 | 0.899 | 0.766 | 7.87851751218204e-33 | 23 |
| si:ch211-241f5.32 | 3.57512275111857e-37 | 0.604473525 | 0.302 | 0.065 | 8.99822645229033e-33 | 23 |
| ca21 | 1.19412729157234e-36 | 0.604551695 | 0.274 | 0.055 | 3.00549898015842e-32 | 23 |
| si:dkey-78l4.41 | 2.32357039583584e-36 | 0.426108134 | 0.207 | 0.032 | 5.84819432927922e-32 | 23 |
| sla23 | 3.91402669348553e-36 | 0.5810883 | 0.307 | 0.067 | 9.85121378483373e-32 | 23 |
| rpl235 | 1.67185575442498e-35 | 0.29738162 | 0.966 | 0.872 | 4.20789374831222e-31 | 23 |
| rpl2210 | 1.74369941027401e-35 | 0.453588948 | 0.894 | 0.693 | 4.38871704571867e-31 | 23 |
| uba529 | 1.92861391267219e-35 | 0.513217827 | 0.883 | 0.628 | 4.85412835680464e-31 | 23 |
| rps197 | 7.03197885304e-35 | 0.303481006 | 0.95 | 0.863 | 1.76987875752164e-30 | 23 |
| rpl3612 | 1.07040054164118e-34 | 0.401294026 | 0.961 | 0.788 | 2.69409112325669e-30 | 23 |
| rpl195 | 6.43423226106803e-34 | 0.296972028 | 0.916 | 0.852 | 1.61943191778821e-29 | 23 |
| alox5a2 | 2.15035956927036e-33 | 0.447864945 | 0.196 | 0.032 | 5.41223999989657e-29 | 23 |
| rpl10a7 | 2.64378931445642e-33 | 0.392951318 | 0.944 | 0.763 | 6.65415332555536e-29 | 23 |
| dicp1.5-61 | 4.23172295253518e-33 | 0.311279247 | 0.14 | 0.017 | 1.06508234992358e-28 | 23 |
| rpl36a8 | 8.80828722229061e-33 | 0.347806427 | 0.955 | 0.808 | 2.21695781097832e-28 | 23 |
| rps126 | 1.0463052230757e-32 | 0.327320594 | 0.944 | 0.839 | 2.63344561595923e-28 | 23 |
| rps1410 | 2.57080210727015e-31 | 0.363518991 | 0.899 | 0.758 | 6.47045182378825e-27 | 23 |
| ela3l | 3.12696634130672e-31 | 0.317500407 | 0.128 | 0.015 | 7.87026158443487e-27 | 23 |
| cotl18 | 3.21333591666683e-31 | 0.714457714 | 0.67 | 0.303 | 8.08764516865876e-27 | 23 |
| actb28 | 4.02700237749007e-31 | 0.385419513 | 0.983 | 0.867 | 1.01355622839047e-26 | 23 |
| fscn1a2 | 4.86415549065046e-31 | 0.526367249 | 0.257 | 0.056 | 1.22425929544181e-26 | 23 |
| rps2411 | 8.40653104804191e-31 | 0.380447345 | 0.933 | 0.757 | 2.11583979948167e-26 | 23 |
| ptpn65 | 1.20931441447441e-30 | 0.604989497 | 0.33 | 0.089 | 3.04372344979065e-26 | 23 |
| mbnl15 | 2.25209810421272e-30 | 0.710076469 | 0.48 | 0.181 | 5.668305718493e-26 | 23 |
| rps224 | 2.64160065031565e-30 | 0.364767866 | 0.966 | 0.828 | 6.64864467677946e-26 | 23 |
| cmah2 | 8.12374856522066e-30 | 0.297256134 | 0.128 | 0.016 | 2.04466627638039e-25 | 23 |
| si:ch211-277c7.62 | 6.24971535536863e-29 | 0.450285173 | 0.201 | 0.037 | 1.57299085779273e-24 | 23 |
| rpl267 | 1.48683583528212e-28 | 0.307687669 | 0.95 | 0.84 | 3.74221711382156e-24 | 23 |
| id33 | 2.13213425108852e-28 | 0.689711553 | 0.52 | 0.208 | 5.36636869656468e-24 | 23 |
| XLOC-0425955 | 3.92322317710168e-28 | 0.684237312 | 0.475 | 0.162 | 9.87436041444722e-24 | 23 |
| rps117 | 7.45444636292324e-28 | 0.355721942 | 0.933 | 0.764 | 1.87620960508415e-23 | 23 |

| | | | | | | |
|---|---|---|---|---|---|---|
| dock101 | 1.63143283745832e-27 | 0.369127333 | 0.173 | 0.03 | 4.10615330859884e-23 | 23 |
| rps1310 | 2.10753244717684e-27 | 0.373264037 | 0.888 | 0.723 | 5.3044484162994e-23 | 23 |
| ptmaa9 | 2.75227808713697e-27 | 0.648243908 | 0.698 | 0.393 | 6.92720871751503e-23 | 23 |
| rps1511 | 6.70811310163808e-27 | 0.438290314 | 0.916 | 0.683 | 1.68836498655129e-22 | 23 |
| rpl159 | 1.31410312123933e-26 | 0.391015371 | 0.939 | 0.743 | 3.30746614584727e-22 | 23 |
| rps258 | 1.90414374901401e-26 | 0.382479056 | 0.922 | 0.723 | 4.79253940189336e-22 | 23 |
| rpl3011 | 2.46817144902558e-26 | 0.411076025 | 0.916 | 0.707 | 6.21214072005247e-22 | 23 |
| rgs134 | 8.88437331251754e-26 | 0.597600219 | 0.391 | 0.13 | 2.23610791902754e-21 | 23 |
| rpl18a10 | 1.60140723002653e-25 | 0.34423012 | 0.922 | 0.784 | 4.03058185725376e-21 | 23 |
| dusp24 | 1.83830561983764e-25 | 0.663747221 | 0.464 | 0.177 | 4.62683141456936e-21 | 23 |
| hsp90ab17 | 2.40243729185396e-25 | 0.320015516 | 0.927 | 0.789 | 6.04669441986723e-21 | 23 |
| rpl218 | 2.78023521504249e-25 | 0.353435872 | 0.922 | 0.755 | 6.99757401274044e-21 | 23 |
| rpl3112 | 4.03603562862831e-25 | 0.421910714 | 0.872 | 0.653 | 1.01582980736946e-20 | 23 |
| rps268 | 4.18603811551077e-25 | 0.359120275 | 0.899 | 0.733 | 1.05358393329291e-20 | 23 |
| rps237 | 4.58370700820508e-25 | 0.380919018 | 0.922 | 0.721 | 1.15367321689514e-20 | 23 |
| rpl2410 | 1.26874649592161e-24 | 0.331113471 | 0.916 | 0.775 | 3.19330805558509e-20 | 23 |
| rpl7a8 | 3.46715017528007e-24 | 0.319240091 | 0.944 | 0.809 | 8.72647027616242e-20 | 23 |
| entpd32 | 3.53577207821421e-23 | 0.277678939 | 0.123 | 0.018 | 8.89918474365734e-19 | 23 |
| rpl135 | 5.31350994222831e-23 | 0.309937499 | 0.961 | 0.803 | 1.33735731735944e-18 | 23 |
| grap2a3 | 9.71608176305423e-23 | 0.382420378 | 0.201 | 0.046 | 2.44544061894312e-18 | 23 |
| rps98 | 2.32950676997647e-22 | 0.312865499 | 0.905 | 0.776 | 5.86313558935378e-18 | 23 |
| rpl109 | 2.98977016142537e-22 | 0.325728027 | 0.922 | 0.771 | 7.5249525192915e-18 | 23 |
| rps59 | 5.57514698298414e-22 | 0.384254892 | 0.894 | 0.7 | 1.40320874414728e-17 | 23 |
| si:rp71-81e14.23 | 1.33885605218826e-21 | 0.366921038 | 0.168 | 0.034 | 3.36976679775264e-17 | 23 |
| rpsa13 | 1.4068709289499e-21 | 0.339871384 | 0.911 | 0.751 | 3.540953441074e-17 | 23 |
| rps1012 | 1.82578534916859e-21 | 0.385296879 | 0.899 | 0.692 | 4.59531914532242e-17 | 23 |
| arhgdig7 | 1.99702594775423e-21 | 0.611959245 | 0.508 | 0.241 | 5.02631460790261e-17 | 23 |
| si:ch211-119e14.14 | 2.26930333613716e-21 | 0.463986691 | 0.257 | 0.073 | 5.71160956672362e-17 | 23 |
| rac28 | 3.97500278876576e-21 | 0.554737107 | 0.503 | 0.218 | 1.00046845190445e-16 | 23 |
| zgc:11418812 | 3.99620343940301e-21 | 0.436070809 | 0.877 | 0.626 | 1.00580444366334e-16 | 23 |
| rpl35a10 | 4.33579892016688e-21 | 0.313261381 | 0.888 | 0.759 | 1.0912772302168e-16 | 23 |
| rpl23a12 | 5.14683478627883e-21 | 0.404466045 | 0.844 | 0.633 | 1.29540684735852e-16 | 23 |
| srgn7 | 5.47706350254147e-21 | 0.570801237 | 0.525 | 0.235 | 1.37852211295466e-16 | 23 |
| rpl1410 | 9.44458744619854e-21 | 0.391297629 | 0.821 | 0.634 | 2.37710821433371e-16 | 23 |
| ppiaa9 | 2.79882915598684e-20 | 0.285441591 | 0.933 | 0.803 | 7.04437310270328e-16 | 23 |

| | | | | | | |
|---|---|---|---|---|---|---|
| XLOC-0254236 | 4.205907523752e-20 | 0.608810462 | 0.425 | 0.167 | 1.05858486465314e-15 | 23 |
| il162 | 7.93700358165006e-20 | 0.379076417 | 0.196 | 0.048 | 1.9976644314655e-15 | 23 |
| capgb5 | 9.68382366995057e-20 | 0.502706529 | 0.324 | 0.113 | 2.43732157948986e-15 | 23 |
| ptprc7 | 1.19645339048159e-19 | 0.500565002 | 0.346 | 0.126 | 3.01135353850312e-15 | 23 |
| faua9 | 1.32436718593179e-19 | 0.353073253 | 0.866 | 0.689 | 3.33329977027173e-15 | 23 |
| rps3a6 | 2.32033548875408e-19 | 0.301203868 | 0.927 | 0.777 | 5.84005239164515e-15 | 23 |
| rps27a9 | 3.01085958728506e-19 | 0.307687698 | 0.905 | 0.751 | 7.57803249523776e-15 | 23 |
| rpl5b7 | 4.61509437719585e-19 | 0.359065939 | 0.788 | 0.63 | 1.16157310379642e-14 | 23 |
| rpl3210 | 6.82876107918479e-19 | 0.302830395 | 0.95 | 0.812 | 1.71873087602002e-14 | 23 |
| rps38 | 2.20988878881853e-18 | 0.335157059 | 0.866 | 0.699 | 5.56206909257737e-14 | 23 |
| si:ch211-102c2.45 | 3.21107873865843e-18 | 0.498903009 | 0.369 | 0.141 | 8.0819640773294e-14 | 23 |
| creb5b1 | 4.05753731777444e-18 | 0.258989419 | 0.112 | 0.019 | 1.02124156751065e-13 | 23 |
| si:ch211-191c10.14 | 5.1854517558196e-18 | 0.339426615 | 0.201 | 0.053 | 1.30512635242224e-13 | 23 |
| rpl611 | 1.05679655930106e-17 | 0.370047337 | 0.877 | 0.666 | 2.65985126010484e-13 | 23 |
| ptmab8 | 1.45201782632227e-17 | 0.462204486 | 0.765 | 0.539 | 3.65458366707051e-13 | 23 |
| ldha2 | 2.02386421989193e-17 | 0.419132215 | 0.235 | 0.072 | 5.09386385504601e-13 | 23 |
| ccl34b.42 | 2.10946772265969e-17 | 0.373098829 | 0.162 | 0.037 | 5.30931931116218e-13 | 23 |
| rps76 | 3.06317854913229e-17 | 0.25280956 | 0.905 | 0.82 | 7.70971409031105e-13 | 23 |
| rsl24d110 | 3.49962232570229e-17 | 0.483736999 | 0.67 | 0.443 | 8.8081994315601e-13 | 23 |
| rpl13a6 | 3.76194443805985e-17 | 0.282203466 | 0.905 | 0.763 | 9.46843795615285e-13 | 23 |
| rpl810 | 4.177573820873e-17 | 0.289430434 | 0.905 | 0.78 | 1.05145355497552e-12 | 23 |
| eif3f9 | 5.71861615630739e-17 | 0.46796952 | 0.665 | 0.43 | 1.43931850038101e-12 | 23 |
| rps1611 | 9.71729148333452e-17 | 0.31704871 | 0.883 | 0.737 | 2.44574509344047e-12 | 23 |
| yrk3 | 2.63046863151056e-16 | 0.375197965 | 0.212 | 0.063 | 6.62062649864893e-12 | 23 |
| rpl5a10 | 2.9912958813421e-16 | 0.427048419 | 0.693 | 0.491 | 7.52879260374993e-12 | 23 |
| itga44 | 4.37910190897831e-16 | 0.389728952 | 0.212 | 0.064 | 1.10217615947075e-11 | 23 |
| wasb7 | 8.70322482338016e-16 | 0.515469969 | 0.397 | 0.176 | 2.19051465579655e-11 | 23 |
| edaradd1 | 1.32763042572529e-15 | 0.262197474 | 0.117 | 0.024 | 3.34151301850797e-11 | 23 |
| myl12.17 | 3.69725893570364e-15 | 0.482216304 | 0.626 | 0.39 | 9.30563101527249e-11 | 23 |
| tagapb5 | 3.83204922776384e-15 | 0.381767563 | 0.212 | 0.067 | 9.64488470135881e-11 | 23 |
| scinlb4 | 4.01181919825373e-15 | 0.428686888 | 0.24 | 0.082 | 1.00973477400848e-10 | 23 |
| si:ch73-264p11.13 | 5.55074670839212e-15 | 0.267533529 | 0.123 | 0.026 | 1.39706743903521e-10 | 23 |
| arpc37 | 6.70935784199233e-15 | 0.489583928 | 0.542 | 0.309 | 1.68867827525105e-10 | 23 |
| si:dkeyp-68b7.124 | 8.48386187926089e-15 | 0.35684024 | 0.201 | 0.062 | 2.13530319639117e-10 | 23 |
| elf16 | 9.38515425989737e-15 | 0.421779818 | 0.274 | 0.103 | 2.36214947567357e-10 | 23 |

| | | | | | | |
|---|---|---|---|---|---|---|
| wipf1a4 | 1.05919509202693e-14 | 0.359599662 | 0.207 | 0.065 | 2.66588812712257e-10 | 23 |
| arpc5a5 | 1.19722889922902e-14 | 0.472017772 | 0.285 | 0.114 | 3.01330541646951e-10 | 23 |
| rps67 | 1.22631804299743e-14 | 0.259196343 | 0.888 | 0.789 | 3.08651988242023e-10 | 23 |
| cfl16 | 1.68044026282417e-14 | 0.40085721 | 0.782 | 0.576 | 4.22950009750214e-10 | 23 |
| wdr192 | 3.10588545677123e-14 | 0.315920884 | 0.134 | 0.032 | 7.81720310614752e-10 | 23 |
| tagapa5 | 3.88651334438067e-14 | 0.476190473 | 0.274 | 0.105 | 9.78196543647171e-10 | 23 |
| wasa4 | 4.27719554522582e-14 | 0.423186965 | 0.251 | 0.092 | 1.07652734677789e-09 | 23 |
| XLOC-0425963 | 4.65708911276775e-14 | 0.290124586 | 0.156 | 0.041 | 1.17214275879251e-09 | 23 |
| rpl912 | 5.65443797662234e-14 | 0.370811173 | 0.849 | 0.628 | 1.42316549433608e-09 | 23 |
| hmgn26 | 5.90868014084296e-14 | 0.553689097 | 0.631 | 0.442 | 1.48715570464877e-09 | 23 |
| pabpc1a9 | 1.41639899932113e-13 | 0.391174532 | 0.765 | 0.538 | 3.56493464139135e-09 | 23 |
| rpl2810 | 1.67858545385512e-13 | 0.291046594 | 0.911 | 0.748 | 4.22483172880796e-09 | 23 |
| tefa4 | 3.27040907538156e-13 | 0.439651907 | 0.313 | 0.137 | 8.23129260182785e-09 | 23 |
| arhgdia6 | 3.93135519810124e-13 | 0.495699031 | 0.419 | 0.229 | 9.89482789810102e-09 | 23 |
| ponzr16 | 4.00761309746961e-13 | 0.468199663 | 0.492 | 0.272 | 1.00867614050213e-08 | 23 |
| kctd5a4 | 4.77099064954271e-13 | 0.444954472 | 0.363 | 0.174 | 1.2008106365834e-08 | 23 |
| si:dkey-183p4.103 | 9.11531890064019e-13 | 0.260454271 | 0.134 | 0.035 | 2.29423461410213e-08 | 23 |
| XLOC-0430432 | 2.47897727411632e-12 | 0.299987051 | 0.151 | 0.043 | 6.23933790122336e-08 | 23 |
| sik15 | 3.76850952239126e-12 | 0.427973504 | 0.274 | 0.116 | 9.48496161690655e-08 | 23 |
| rpl46 | 5.51768363891328e-12 | 0.305368712 | 0.771 | 0.627 | 1.38874579507808e-07 | 23 |
| CR381673.14 | 8.40782340387128e-12 | 0.335560071 | 0.173 | 0.057 | 2.11616507252036e-07 | 23 |
| calm1b5 | 1.33140274315299e-11 | 0.355129985 | 0.24 | 0.094 | 3.35100756424177e-07 | 23 |
| XLOC-0437603 | 1.63321923859258e-11 | 0.318194716 | 0.162 | 0.051 | 4.11064950161366e-07 | 23 |
| dusp15 | 1.79062587456977e-11 | 0.451396914 | 0.33 | 0.159 | 4.50682626370465e-07 | 23 |
| zgc:564936 | 2.17647562531506e-11 | 0.332795306 | 0.721 | 0.572 | 5.47797150135548e-07 | 23 |
| lasp15 | 2.37194894891455e-11 | 0.423560427 | 0.425 | 0.236 | 5.96995830952304e-07 | 23 |
| CU499336.22 | 3.26479351179713e-11 | 0.281250949 | 0.128 | 0.036 | 8.21715878984219e-07 | 23 |
| RASSF53 | 3.42025858401084e-11 | 0.302897617 | 0.173 | 0.058 | 8.60844883009689e-07 | 23 |
| mrps154 | 5.89466150372439e-11 | 0.370156569 | 0.285 | 0.124 | 1.48362735387239e-06 | 23 |
| msna8 | 6.33654920677552e-11 | 0.437298205 | 0.419 | 0.241 | 1.59484606985333e-06 | 23 |
| laptm58 | 7.2636886225673e-11 | 0.384744446 | 0.352 | 0.169 | 1.82819778941396e-06 | 23 |
| h1fx5 | 7.75924991283625e-11 | 0.452358256 | 0.363 | 0.183 | 1.95292561056175e-06 | 23 |
| LOC1005362134 | 9.06939556956597e-11 | 0.381685101 | 0.251 | 0.105 | 2.28267617090406e-06 | 23 |
| XLOC-0283706 | 1.01169020679983e-10 | 0.421427513 | 0.307 | 0.14 | 2.54632308149449e-06 | 23 |
| aldocb2 | 1.50903349334584e-10 | 0.323587012 | 0.184 | 0.067 | 3.79808639940214e-06 | 23 |

| | | | | | | |
|---|---|---|---|---|---|---|
| mt-co112 | 1.61725634999038e-10 | 0.3001352 | 0.782 | 0.657 | 4.07047250729078e-06 | 23 |
| si:ch211-93g21.13 | 1.81682801766888e-10 | 0.270192298 | 0.123 | 0.035 | 4.57277443767081e-06 | 23 |
| dtnbp1a4 | 2.66637190456285e-10 | 0.329970582 | 0.218 | 0.086 | 6.71099144659425e-06 | 23 |
| cap18 | 3.73587535999227e-10 | 0.414196283 | 0.408 | 0.233 | 9.40282469356455e-06 | 23 |
| ccr24 | 4.17141072872243e-10 | 0.271849439 | 0.145 | 0.046 | 1.04990236631215e-05 | 23 |
| BX901920.14 | 6.23665210524086e-10 | 0.292008226 | 0.196 | 0.073 | 1.56970296836807e-05 | 23 |
| atp5if1b1 | 6.40021877008866e-10 | 0.418222458 | 0.352 | 0.19 | 1.61087106224362e-05 | 23 |
| zgc:640517 | 8.83573384241049e-10 | 0.365105163 | 0.33 | 0.162 | 2.2238658507963e-05 | 23 |
| fcer1g3 | 1.8015907150794e-09 | 0.252350061 | 0.134 | 0.042 | 4.53442367078334e-05 | 23 |
| rpl22l18 | 3.06621245564477e-09 | 0.359547755 | 0.553 | 0.391 | 7.71735012961231e-05 | 23 |
| si:ch211-222l21.18 | 3.15400749345574e-09 | 0.411990979 | 0.447 | 0.275 | 7.93832146027876e-05 | 23 |
| gapdhs11 | 5.05538789215025e-09 | 0.380687186 | 0.458 | 0.288 | 0.000127239 | 23 |
| XLOC-0203013 | 6.98358725057454e-09 | 0.342822236 | 0.207 | 0.085 | 0.00017577 | 23 |
| CABZ01079663.13 | 7.7914256757792e-09 | 0.257554609 | 0.123 | 0.039 | 0.000196102 | 23 |
| gpr183a4 | 2.47723452600154e-08 | 0.267800545 | 0.168 | 0.065 | 0.000623495 | 23 |
| dok24 | 2.63877782888168e-08 | 0.250302156 | 0.117 | 0.038 | 0.000664154 | 23 |
| csk4 | 2.75919534368658e-08 | 0.320665653 | 0.223 | 0.101 | 0.000694462 | 23 |
| fcer1gl5 | 3.19397255703834e-08 | 0.319633385 | 0.346 | 0.168 | 0.000803891 | 23 |
| stim1a1 | 3.53740526276997e-08 | 0.268472493 | 0.173 | 0.069 | 0.00089033 | 23 |
| pde4ba6 | 3.94861803292199e-08 | 0.369843585 | 0.268 | 0.14 | 0.000993828 | 23 |
| ppp1r184 | 4.84144124333425e-08 | 0.259067588 | 0.145 | 0.054 | 0.001218542 | 23 |
| lamp1a2 | 1.10606883430599e-07 | 0.324699763 | 0.201 | 0.091 | 0.002783865 | 23 |
| cebpb10 | 1.2487964797351e-07 | 0.353727582 | 0.57 | 0.352 | 0.003143096 | 23 |
| vaspa5 | 1.91185241564799e-07 | 0.262775044 | 0.179 | 0.075 | 0.004811941 | 23 |
| rps6ka13 | 2.31422871998839e-07 | 0.266502021 | 0.173 | 0.074 | 0.005824682 | 23 |
| cox6b16 | 2.88886583141915e-07 | 0.3356984 | 0.235 | 0.12 | 0.007270986 | 23 |
| XLOC-0268363 | 3.26273909409032e-07 | 0.265180712 | 0.184 | 0.079 | 0.008211988 | 23 |
| COX312 | 3.39723217078483e-07 | 0.25484375 | 0.838 | 0.7 | 0.008550494 | 23 |
| mef2d1 | 4.30402364405541e-07 | 0.278913055 | 0.168 | 0.073 | 0.010832797 | 23 |
| si:ch73-234b20.53 | 6.19756595846122e-07 | 0.298942196 | 0.207 | 0.101 | 0.015598654 | 23 |
| pik3r13 | 7.22653048397021e-07 | 0.29526286 | 0.19 | 0.09 | 0.018188455 | 23 |
| mt-atp611 | 1.28582356459133e-06 | 0.269020223 | 0.709 | 0.591 | 0.032362893 | 23 |
| arpc46 | 1.33673787937445e-06 | 0.337533235 | 0.358 | 0.226 | 0.033644356 | 23 |
| actr2a6 | 2.46759996969076e-06 | 0.305769895 | 0.24 | 0.13 | 0.062107024 | 23 |
| tnip13 | 3.299704974218e-06 | 0.276137626 | 0.173 | 0.082 | 0.083050274 | 23 |

| | | | | | | |
|---|---|---|---|---|---|---|
| rpl27a11 | 4.50664790317461e-06 | 0.270174653 | 0.648 | 0.496 | 0.113427821 | 23 |
| sept25 | 4.69894225540923e-06 | 0.314412805 | 0.279 | 0.166 | 0.118267678 | 23 |
| pxna1 | 5.99447755274593e-06 | 0.253107417 | 0.19 | 0.092 | 0.150875006 | 23 |
| crip18 | 6.29141518455288e-06 | 0.28757347 | 0.419 | 0.271 | 0.158348629 | 23 |
| arpc27 | 9.44963281225999e-06 | 0.315131463 | 0.385 | 0.268 | 0.237837808 | 23 |
| ywhaba4 | 2.2387688174833e-05 | 0.287545379 | 0.436 | 0.322 | 0.563475724 | 23 |
| ccni8 | 6.47217632165958e-05 | 0.2875924 | 0.397 | 0.295 | 1 | 23 |
| capzb6 | 6.80448110612641e-05 | 0.293174933 | 0.352 | 0.254 | 1 | 23 |
| celf210 | 7.69433179709324e-05 | 0.287722453 | 0.346 | 0.243 | 1 | 23 |
| hmgn68 | 9.42128378065722e-05 | 0.274274003 | 0.369 | 0.266 | 1 | 23 |
| bcl2l103 | 0.000104562 | 0.259879822 | 0.33 | 0.215 | 1 | 23 |
| bhlhe407 | 0.000146923 | 0.305118303 | 0.251 | 0.162 | 1 | 23 |
| gsdf | 0 | 1.880529625 | 0.763 | 0.04 | 0 | 24 |
| bcam | 0 | 1.459986643 | 0.689 | 0.02 | 0 | 24 |
| alcama | 0 | 1.13121002 | 0.531 | 0.02 | 0 | 24 |
| igf3 | 0 | 1.087250454 | 0.446 | 0.009 | 0 | 24 |
| LOC103910903 | 0 | 0.908573775 | 0.35 | 0.007 | 0 | 24 |
| lamb2l | 0 | 0.476380763 | 0.215 | 0.002 | 0 | 24 |
| smoc21 | 6.62485475978222e-301 | 0.79658214 | 0.322 | 0.008 | 1.66740969448959e-296 | 24 |
| map1aa | 7.59561910039118e-278 | 0.51186551 | 0.226 | 0.003 | 1.91174137137746e-273 | 24 |
| foxl2b | 1.54988531850344e-268 | 0.662090861 | 0.282 | 0.006 | 3.90090635814132e-264 | 24 |
| LOC110438024 | 3.91038523309664e-262 | 0.663530493 | 0.266 | 0.006 | 9.84204859318093e-258 | 24 |
| bmp6 | 2.65695544475498e-238 | 0.690501223 | 0.294 | 0.009 | 6.68729115890382e-234 | 24 |
| si:dkey-194e6.1 | 4.52023912003668e-236 | 0.352420592 | 0.141 | 0.001 | 1.13769898412203e-231 | 24 |
| hsd17b1 | 2.55695706778507e-231 | 0.359974405 | 0.136 | 0.001 | 6.43560524390825e-227 | 24 |
| cdh4 | 8.03304443239638e-226 | 0.521021397 | 0.226 | 0.005 | 2.02183695318984e-221 | 24 |
| emx2 | 2.08183515184607e-215 | 0.69641622 | 0.328 | 0.013 | 5.23977089368137e-211 | 24 |
| nectin1b2 | 6.70138084201088e-211 | 1.035012981 | 0.508 | 0.036 | 1.68667054412572e-206 | 24 |
| metrn | 3.4211531453603e-208 | 0.635740577 | 0.277 | 0.009 | 8.61070035155734e-204 | 24 |
| slc4a4a1 | 5.19186884135078e-203 | 0.621696051 | 0.254 | 0.008 | 1.30674146867958e-198 | 24 |
| col4a5 | 6.56091457821808e-202 | 0.480954325 | 0.209 | 0.005 | 1.65131659019171e-197 | 24 |
| jam2a1 | 4.32473101973857e-198 | 1.01747804 | 0.463 | 0.031 | 1.088491550358e-193 | 24 |
| cldn11a4 | 2.12116632193653e-197 | 1.279975001 | 0.593 | 0.055 | 5.33876351568205e-193 | 24 |
| cx34.5 | 5.28185694365351e-189 | 0.431625212 | 0.169 | 0.003 | 1.32939057414815e-184 | 24 |
| lhx9 | 7.61751094817291e-187 | 0.416805779 | 0.147 | 0.002 | 1.91725133054564e-182 | 24 |

| | | | | | | |
|---|---|---|---|---|---|---|
| col4a6 | 6.82003764200869e-184 | 0.510401954 | 0.226 | 0.007 | 1.71653527411717e-179 | 24 |
| sema3ab | 3.45785638920029e-183 | 0.622005973 | 0.249 | 0.008 | 8.7030787459782e-179 | 24 |
| ccdc3a | 6.0315353245383e-177 | 0.271162807 | 0.113 | 0.001 | 1.51807712583305e-172 | 24 |
| ecrg4a2 | 6.82255376470548e-177 | 0.993057483 | 0.401 | 0.026 | 1.71716855703872e-172 | 24 |
| znf536 | 7.43792321141328e-176 | 0.450973844 | 0.203 | 0.005 | 1.87205089308061e-171 | 24 |
| foxl2a1 | 4.59063308473886e-174 | 0.480853685 | 0.232 | 0.007 | 1.15541644109792e-169 | 24 |
| inha | 5.32489150081147e-171 | 0.48015007 | 0.198 | 0.005 | 1.34022194183924e-166 | 24 |
| aldh1a22 | 6.01603917452041e-169 | 1.217470446 | 0.627 | 0.071 | 1.51417689983504e-164 | 24 |
| agrn2 | 3.5799372021598e-167 | 0.96096268 | 0.497 | 0.044 | 9.010343944116e-163 | 24 |
| cemip2 | 5.03391317414304e-165 | 0.729632823 | 0.322 | 0.017 | 1.26698560680006e-160 | 24 |
| itga3b | 2.57833521900437e-162 | 0.292644302 | 0.136 | 0.002 | 6.4894119127121e-158 | 24 |
| si:ch211-207d6.2 | 4.37828071416588e-154 | 0.575955061 | 0.249 | 0.01 | 1.10196947294841e-149 | 24 |
| rpz56 | 2.77113043790523e-152 | 1.246318018 | 0.667 | 0.095 | 6.97465819916367e-148 | 24 |
| lama51 | 4.70838139400694e-151 | 0.68148256 | 0.328 | 0.02 | 1.18505251305761e-146 | 24 |
| si:dkey-6n6.1 | 3.77306214991465e-149 | 0.361761716 | 0.147 | 0.003 | 9.49642012512019e-145 | 24 |
| bmpr2a | 7.31473370841637e-138 | 0.592942181 | 0.277 | 0.015 | 1.84104532707132e-133 | 24 |
| si:dkey-42i9.72 | 1.34309690567705e-134 | 0.625826371 | 0.299 | 0.019 | 3.38044060189857e-130 | 24 |
| itm2bb6 | 6.90259632168192e-121 | 1.381953651 | 0.831 | 0.214 | 1.73731446820412e-116 | 24 |
| thbs3b | 1.08144878673999e-120 | 0.33113392 | 0.13 | 0.003 | 2.72189845134587e-116 | 24 |
| pi15a2 | 1.52873311905907e-120 | 0.857454644 | 0.418 | 0.041 | 3.84766838735977e-116 | 24 |
| chka | 1.66696008030759e-118 | 0.604269901 | 0.305 | 0.022 | 4.19557182612618e-114 | 24 |
| trim35-33 | 1.23692264693345e-115 | 0.261127691 | 0.136 | 0.004 | 3.11321061006681e-111 | 24 |
| f103 | 2.65328084981693e-110 | 0.796207641 | 0.407 | 0.044 | 6.67804257090424e-106 | 24 |
| nr0b11 | 1.05620640974436e-105 | 0.441622205 | 0.192 | 0.009 | 2.65836591268557e-101 | 24 |
| podxl1 | 1.98560567943201e-105 | 0.782130313 | 0.401 | 0.042 | 4.99757093456243e-101 | 24 |
| XLOC-005605 | 1.23224716017871e-100 | 0.475775183 | 0.22 | 0.013 | 3.10144287745378e-96 | 24 |
| plxna41 | 7.35055325806567e-98 | 0.529674694 | 0.26 | 0.02 | 1.85006074952255e-93 | 24 |
| wu:fc27b111 | 3.0151189605793e-97 | 0.414698077 | 0.198 | 0.011 | 7.58875291188204e-93 | 24 |
| igsf3 | 2.72792613886104e-96 | 0.304694686 | 0.13 | 0.004 | 6.86591729889936e-92 | 24 |
| scn4bb | 8.48103740868715e-96 | 0.286270103 | 0.136 | 0.005 | 2.13459230539247e-91 | 24 |
| slc44a5b | 3.46574830606153e-95 | 0.392246398 | 0.186 | 0.01 | 8.72294191152627e-91 | 24 |
| ivns1abpa | 1.28566187838227e-94 | 0.648554958 | 0.299 | 0.028 | 3.23588238170033e-90 | 24 |
| nog11 | 9.88645970526674e-86 | 0.365602965 | 0.141 | 0.006 | 2.48832304321859e-81 | 24 |
| emilin1a1 | 1.62605422395631e-85 | 0.480314082 | 0.226 | 0.017 | 4.09261587627565e-81 | 24 |
| negr1 | 3.08985203526431e-85 | 0.302267042 | 0.13 | 0.005 | 7.77684858755675e-81 | 24 |

| | | | | | | |
|---|---|---|---|---|---|---|
| CABZ01021532.1 | 1.04461088519202e-83 | 0.470943748 | 0.203 | 0.014 | 2.62918113693979e-79 | 24 |
| cldnc1 | 1.20635782140378e-81 | 0.577494614 | 0.271 | 0.025 | 3.03628200069117e-77 | 24 |
| sparc5 | 2.37073029451142e-81 | 1.026844002 | 0.667 | 0.152 | 5.96689107825578e-77 | 24 |
| rab11fip1b2 | 3.14745806772423e-81 | 0.728176031 | 0.39 | 0.054 | 7.9218372106551e-77 | 24 |
| XLOC-009815 | 1.16942999366242e-80 | 0.383727264 | 0.153 | 0.008 | 2.94333835104895e-76 | 24 |
| cahz1 | 1.37265447342572e-80 | 0.69475702 | 0.322 | 0.037 | 3.45483404416518e-76 | 24 |
| ccdc187 | 4.00326669100579e-80 | 0.544062596 | 0.277 | 0.027 | 1.00758219345925e-75 | 24 |
| gstt1b3 | 1.15325969111636e-78 | 0.833029779 | 0.429 | 0.068 | 2.90263931657078e-74 | 24 |
| cadm2a1 | 6.83571254093086e-78 | 0.433601823 | 0.198 | 0.014 | 1.72048048942689e-73 | 24 |
| ptn | 2.86280525629871e-77 | 0.299671706 | 0.141 | 0.007 | 7.20539454957823e-73 | 24 |
| appa4 | 4.51630207500973e-77 | 0.806514547 | 0.441 | 0.073 | 1.1367080692592e-72 | 24 |
| ace1 | 4.83752944249822e-76 | 0.55928993 | 0.26 | 0.025 | 1.21755778538238e-71 | 24 |
| si:ch211-117c9.5 | 1.92516924511972e-74 | 0.426759842 | 0.203 | 0.016 | 4.84545847304181e-70 | 24 |
| st5 | 2.27085793943803e-74 | 0.338349308 | 0.164 | 0.01 | 5.71552234777158e-70 | 24 |
| large2 | 1.24234617161431e-73 | 0.291117062 | 0.124 | 0.006 | 3.12686107933605e-69 | 24 |
| hpn | 1.34515547755929e-73 | 0.310818249 | 0.13 | 0.006 | 3.38562182146897e-69 | 24 |
| ar4 | 1.84507004145628e-70 | 0.628573761 | 0.373 | 0.056 | 4.6438567873413e-66 | 24 |
| nos1apa | 1.43420892632588e-66 | 0.391802229 | 0.209 | 0.019 | 3.60976044666962e-62 | 24 |
| CABZ01068358.16 | 2.61047876462699e-66 | 0.698424196 | 0.379 | 0.061 | 6.57031400268968e-62 | 24 |
| slc7a3a2 | 7.73959185192029e-66 | 0.600502942 | 0.305 | 0.04 | 1.94797787320982e-61 | 24 |
| F7 (1 of many) | 8.47268999533526e-66 | 0.257941947 | 0.113 | 0.005 | 2.13249134492593e-61 | 24 |
| pgr2 | 5.13573912742878e-64 | 0.385054933 | 0.192 | 0.016 | 1.29261418098255e-59 | 24 |
| sdk2b | 8.3481906692111e-64 | 0.252019266 | 0.107 | 0.005 | 2.10115610953374e-59 | 24 |
| si:dkey-188i13.9 | 1.69514427907179e-63 | 0.475975968 | 0.22 | 0.022 | 4.26650863599578e-59 | 24 |
| colec11 | 1.74317848643221e-62 | 0.32455054 | 0.13 | 0.007 | 4.38740593250123e-58 | 24 |
| ifi456 | 1.33989932251051e-61 | 0.982697214 | 0.525 | 0.124 | 3.37239260482671e-57 | 24 |
| bambia1 | 3.04266742739087e-60 | 0.443598249 | 0.198 | 0.018 | 7.65808964800008e-56 | 24 |
| ncam1a | 4.77738654910207e-59 | 0.40929473 | 0.186 | 0.017 | 1.2024204205435e-54 | 24 |
| hsbp1a | 6.81441271989525e-59 | 0.419891263 | 0.215 | 0.022 | 1.71511953747044e-54 | 24 |
| pcdh1b | 8.27228023549231e-59 | 0.271362572 | 0.124 | 0.007 | 2.08205021247106e-54 | 24 |
| cx432 | 1.05451967070525e-58 | 0.67698494 | 0.328 | 0.051 | 2.65412055919803e-54 | 24 |
| fgfr31 | 1.23858716242537e-58 | 0.434376116 | 0.175 | 0.015 | 3.11740002910842e-54 | 24 |
| htra1b3 | 4.85653466708374e-58 | 0.603976639 | 0.277 | 0.037 | 1.22234121035831e-53 | 24 |
| phlda21 | 3.13776157582084e-57 | 0.692298959 | 0.316 | 0.049 | 7.89743211018347e-53 | 24 |
| hdac42 | 7.32364342147929e-56 | 0.461866733 | 0.237 | 0.029 | 1.84328781275212e-51 | 24 |

| | | | | | | |
|---|---|---|---|---|---|---|
| cdh1 | 1.57476064567218e-55 | 0.625958441 | 0.339 | 0.055 | 3.9635150690923e-51 | 24 |
| nanos11 | 1.67026223158304e-55 | 0.422699071 | 0.186 | 0.018 | 4.20388301067136e-51 | 24 |
| atp1b3b | 2.39989151220102e-55 | 0.637330491 | 0.35 | 0.059 | 6.04028694705876e-51 | 24 |
| ttll121 | 6.53835523676134e-55 | 0.51343403 | 0.22 | 0.025 | 1.64563862954046e-50 | 24 |
| cav12 | 7.3407572176125e-55 | 0.546380135 | 0.288 | 0.042 | 1.84759518410089e-50 | 24 |
| zgc:123010 | 2.70604236293794e-54 | 0.473049603 | 0.26 | 0.034 | 6.81083802327851e-50 | 24 |
| gata61 | 3.55702209644073e-53 | 0.333385679 | 0.181 | 0.017 | 8.95266891453167e-49 | 24 |
| cxcl142 | 7.27026908694504e-53 | 0.396934583 | 0.153 | 0.013 | 1.8298540264932e-48 | 24 |
| b2m9 | 7.74065979553715e-53 | 0.89822747 | 0.718 | 0.287 | 1.94824666393875e-48 | 24 |
| wt1a1 | 3.01091813956385e-52 | 0.479682213 | 0.243 | 0.032 | 7.57817986546826e-48 | 24 |
| ccdc125 | 4.38973262534011e-52 | 0.324955535 | 0.153 | 0.013 | 1.10485180447185e-47 | 24 |
| jam3b1 | 9.43634169995707e-52 | 0.42669594 | 0.215 | 0.025 | 2.3750328424622e-47 | 24 |
| vasnb3 | 1.3103119499796e-51 | 0.455580516 | 0.232 | 0.029 | 3.29792414690364e-47 | 24 |
| mpzl2b1 | 1.41111786060448e-51 | 0.716844184 | 0.429 | 0.091 | 3.55164254335542e-47 | 24 |
| CR318588.32 | 1.32064360718895e-48 | 0.598736461 | 0.277 | 0.043 | 3.32392789493388e-44 | 24 |
| krt18a.14 | 2.38684083994494e-48 | 0.807425566 | 0.492 | 0.123 | 6.00743971005743e-44 | 24 |
| zgc:1141811 | 2.40002961991863e-48 | 0.298315441 | 0.141 | 0.012 | 6.04063455037321e-44 | 24 |
| isg154 | 4.58967719093734e-48 | 0.651590326 | 0.345 | 0.065 | 1.15517585218702e-43 | 24 |
| ifi466 | 1.00453338227809e-47 | 0.746721116 | 0.463 | 0.118 | 2.52831006985572e-43 | 24 |
| ponzr17 | 1.1474174621328e-47 | 0.867612457 | 0.672 | 0.269 | 2.88793501044205e-43 | 24 |
| gpx1b | 5.38150167027846e-47 | 0.504760445 | 0.203 | 0.025 | 1.35447015539238e-42 | 24 |
| schip1 | 7.43002468268204e-47 | 0.358576122 | 0.175 | 0.019 | 1.87006291238424e-42 | 24 |
| alcamb | 1.17057535700145e-46 | 0.30218111 | 0.147 | 0.013 | 2.94622111603696e-42 | 24 |
| parp12a | 5.74852251661254e-46 | 0.361014594 | 0.192 | 0.023 | 1.44684563220621e-41 | 24 |
| si:ch211-156j16.1 | 2.94326991910148e-45 | 0.372909383 | 0.175 | 0.019 | 7.40791605938651e-41 | 24 |
| bmpr1bb | 4.00421979887675e-45 | 0.259462837 | 0.124 | 0.01 | 1.00782208117929e-40 | 24 |
| mmp24 | 8.13550572747615e-45 | 0.623908081 | 0.395 | 0.089 | 2.04762543654847e-40 | 24 |
| cyp26a1 | 1.11324160803755e-44 | 0.345435094 | 0.119 | 0.009 | 2.80191780326971e-40 | 24 |
| gstm.33 | 1.06760757168614e-43 | 0.744712744 | 0.412 | 0.098 | 2.68706149717684e-39 | 24 |
| hey2 | 4.09133184963213e-43 | 0.264412568 | 0.124 | 0.01 | 1.02974731323391e-38 | 24 |
| s100v25 | 1.32224784242282e-42 | 0.678197411 | 0.452 | 0.118 | 3.327965594594e-38 | 24 |
| fndc3ba | 6.39861786043965e-42 | 0.302239359 | 0.147 | 0.015 | 1.61046812929406e-37 | 24 |
| mbnl32 | 2.90981229708689e-41 | 0.503924004 | 0.288 | 0.053 | 7.32370657053799e-37 | 24 |
| atp11a | 3.14988412918389e-40 | 0.261903293 | 0.124 | 0.011 | 7.92794336474294e-36 | 24 |
| phactr4a2 | 7.38356862043421e-40 | 0.424135143 | 0.226 | 0.035 | 1.85837038607709e-35 | 24 |

| | | | | | | |
|---|---|---|---|---|---|---|
| cox5b21 | 7.62718844203806e-40 | 0.270872871 | 0.124 | 0.011 | 1.91968705897656e-35 | 24 |
| dag12 | 7.91168899096888e-40 | 0.490473353 | 0.266 | 0.048 | 1.99129300213696e-35 | 24 |
| aplp23 | 1.06734051116109e-39 | 0.547057164 | 0.328 | 0.07 | 2.68638933254134e-35 | 24 |
| zgc:927446 | 1.28927730362504e-39 | 0.764832264 | 0.751 | 0.349 | 3.24498204549387e-35 | 24 |
| ece2b1 | 1.41810588077533e-39 | 0.467898489 | 0.26 | 0.045 | 3.56923069132342e-35 | 24 |
| cd99l21 | 1.4553922680293e-39 | 0.331769457 | 0.175 | 0.022 | 3.66307679940295e-35 | 24 |
| tfr1a | 6.06333206249975e-39 | 0.395054864 | 0.164 | 0.02 | 1.52608004681056e-34 | 24 |
| CABZ01068366.12 | 7.21700508117824e-39 | 0.383855395 | 0.198 | 0.028 | 1.81644800888175e-34 | 24 |
| myl9a2 | 9.2161585772081e-38 | 0.375683896 | 0.203 | 0.03 | 2.31961495229751e-33 | 24 |
| tp53inp12 | 1.01156103453332e-37 | 0.583415504 | 0.271 | 0.053 | 2.5459979678169e-33 | 24 |
| tspan9b2 | 1.01770447543387e-37 | 0.436764405 | 0.243 | 0.042 | 2.5614603942195e-33 | 24 |
| fkbp1b | 3.26697780826682e-37 | 0.255007492 | 0.13 | 0.013 | 8.22265644562676e-33 | 24 |
| ptprfb | 6.45667644369147e-37 | 0.29668568 | 0.13 | 0.013 | 1.6250808941127e-32 | 24 |
| wu:fj16a034 | 1.15422508751554e-36 | 0.688012469 | 0.26 | 0.05 | 2.90506912276786e-32 | 24 |
| myh101 | 2.08282047018848e-36 | 0.422932862 | 0.215 | 0.035 | 5.24225084141738e-32 | 24 |
| h1f05 | 1.49072806192145e-35 | 0.539545099 | 0.288 | 0.06 | 3.75201345905011e-31 | 24 |
| lamb1a4 | 1.64696554334413e-35 | 0.39294187 | 0.22 | 0.037 | 4.14524757604284e-31 | 24 |
| atp1b1a4 | 2.48212713941393e-35 | 0.731143108 | 0.548 | 0.211 | 6.24726579719092e-31 | 24 |
| slc38a21 | 2.69351746703157e-35 | 0.452173429 | 0.243 | 0.045 | 6.77931411277175e-31 | 24 |
| lamc14 | 7.57780342616295e-35 | 0.48609213 | 0.266 | 0.053 | 1.90725734433095e-30 | 24 |
| si:ch73-1a9.39 | 8.85454128603029e-35 | 0.735305959 | 0.621 | 0.262 | 2.22859949628096e-30 | 24 |
| PDE3B | 4.06844129760866e-34 | 0.29025803 | 0.147 | 0.018 | 1.02398599019512e-29 | 24 |
| si:ch211-264f5.6 | 1.49841210103367e-33 | 0.264985852 | 0.119 | 0.012 | 3.77135341709165e-29 | 24 |
| arhgef18a | 1.54606982654904e-33 | 0.271297687 | 0.147 | 0.018 | 3.89130314644127e-29 | 24 |
| id13 | 2.1788042318227e-33 | 0.66375524 | 0.379 | 0.103 | 5.48383237107455e-29 | 24 |
| foxp43 | 5.20192646008935e-33 | 0.468125898 | 0.249 | 0.049 | 1.30927287073989e-28 | 24 |
| parvaa1 | 7.77988840945549e-33 | 0.321719455 | 0.181 | 0.027 | 1.95812011377585e-28 | 24 |
| arhgef53 | 8.06068962925919e-33 | 0.337420243 | 0.181 | 0.027 | 2.02879497278825e-28 | 24 |
| enc13 | 2.17336101576368e-32 | 0.364769789 | 0.186 | 0.029 | 5.4701323405756e-28 | 24 |
| rbp54 | 2.26974548897213e-32 | 0.632513299 | 0.35 | 0.092 | 5.71272242119394e-28 | 24 |
| man1b1b | 1.24941501992104e-31 | 0.341875421 | 0.175 | 0.027 | 3.14465266363926e-27 | 24 |
| chpfa | 2.52561301111056e-31 | 0.254163108 | 0.113 | 0.012 | 6.35671538766418e-27 | 24 |
| pwwp2b | 2.93054459031799e-31 | 0.33931246 | 0.203 | 0.035 | 7.37588767937135e-27 | 24 |
| fkbp1aa6 | 1.26014677939035e-30 | 0.636858806 | 0.734 | 0.384 | 3.17166342904757e-26 | 24 |
| COX313 | 2.76661649620116e-30 | 0.50557131 | 0.904 | 0.699 | 6.96329705928871e-26 | 24 |

| | | | | | | |
|---|---|---|---|---|---|---|
| spred2b3 | 3.1948263381447e-30 | 0.512950594 | 0.277 | 0.065 | 8.0410584104764e-26 | 24 |
| si:dkey-178e17.31 | 3.67628467888494e-30 | 0.255720085 | 0.124 | 0.015 | 9.2528409082855e-26 | 24 |
| tjp1a2 | 4.1868237530638e-30 | 0.387139166 | 0.232 | 0.046 | 1.05378167040863e-25 | 24 |
| CR318588.19 | 5.17919953255064e-30 | 0.811121996 | 0.542 | 0.205 | 1.30355273034767e-25 | 24 |
| si:dkey-73n10.11 | 1.82006856207748e-29 | 0.253084938 | 0.147 | 0.02 | 4.5809305638928e-25 | 24 |
| si:ch211-225b11.1 | 1.82764550666377e-29 | 0.396515588 | 0.186 | 0.032 | 4.60000097572205e-25 | 24 |
| svild5 | 2.50058534322704e-29 | 0.505919375 | 0.311 | 0.079 | 6.29372325036815e-25 | 24 |
| cavin1b2 | 3.20057436269356e-29 | 0.410917343 | 0.209 | 0.039 | 8.05552561346341e-25 | 24 |
| mt-co212 | 5.02134132625386e-29 | 0.49020257 | 0.915 | 0.719 | 1.26382139840483e-24 | 24 |
| ecm1b3 | 1.76198897968487e-28 | 0.489022319 | 0.243 | 0.053 | 4.43475006296886e-24 | 24 |
| si:ch211-286b5.55 | 2.53008874496971e-28 | 0.543222773 | 0.418 | 0.134 | 6.36798036221428e-24 | 24 |
| rdh10a2 | 1.21795149262336e-27 | 0.429163373 | 0.203 | 0.039 | 3.06546211178373e-23 | 24 |
| mt-co113 | 2.01055065489631e-27 | 0.498613645 | 0.876 | 0.656 | 5.06035494330851e-23 | 24 |
| hsd11b25 | 5.84940079201494e-27 | 0.66088882 | 0.395 | 0.121 | 1.47223568534224e-22 | 24 |
| c7a4 | 1.04575219575933e-26 | 0.379636419 | 0.215 | 0.043 | 2.63205370150665e-22 | 24 |
| uxs1 | 1.3391876861719e-26 | 0.273381442 | 0.169 | 0.029 | 3.37060148732606e-22 | 24 |
| cox6a22 | 1.40587755575231e-26 | 0.368701064 | 0.186 | 0.035 | 3.53845322007298e-22 | 24 |
| prkar2aa2 | 1.87960657730807e-26 | 0.506325487 | 0.333 | 0.096 | 4.73078179442669e-22 | 24 |
| sptbn2 | 2.58492012455769e-26 | 0.350058724 | 0.198 | 0.039 | 6.50598546149925e-22 | 24 |
| s100a10b7 | 4.75566988145644e-26 | 0.635537061 | 0.621 | 0.252 | 1.19695455246377e-21 | 24 |
| edem1 | 4.92909516517165e-26 | 0.304190621 | 0.175 | 0.031 | 1.24060396212205e-21 | 24 |
| cgnl11 | 1.03381217979246e-25 | 0.273060665 | 0.147 | 0.023 | 2.60200187531964e-21 | 24 |
| oaz1b9 | 1.21251812992308e-25 | 0.609869713 | 0.638 | 0.32 | 3.0517868812034e-21 | 24 |
| cebpd6 | 1.67654838284288e-25 | 0.518511083 | 0.401 | 0.127 | 4.21970462477724e-21 | 24 |
| krt88 | 1.82924071419099e-25 | 0.651283399 | 0.633 | 0.273 | 4.60401595354731e-21 | 24 |
| oip5-as11 | 2.02514756970515e-25 | 0.264950916 | 0.158 | 0.026 | 5.09709391819089e-21 | 24 |
| abhd15a1 | 2.1510052609101e-25 | 0.323483658 | 0.186 | 0.035 | 5.41386514118463e-21 | 24 |
| mt-cyb10 | 5.31263719258706e-25 | 0.516175718 | 0.87 | 0.622 | 1.33713765500224e-20 | 24 |
| stat1a4 | 5.65048289784445e-25 | 0.495010148 | 0.367 | 0.118 | 1.42217004055847e-20 | 24 |
| adam10a4 | 5.84406987529428e-25 | 0.479550303 | 0.316 | 0.092 | 1.47089394691282e-20 | 24 |
| glulb1 | 6.51447061872624e-25 | 0.666194579 | 0.412 | 0.149 | 1.63962711002721e-20 | 24 |
| zgc:100829 | 1.4591755212479e-24 | 0.312020301 | 0.181 | 0.035 | 3.67259886942885e-20 | 24 |
| pnp5a3 | 5.92306334622689e-24 | 0.526888327 | 0.277 | 0.076 | 1.49077581361185e-19 | 24 |
| mphosph84 | 1.46741376797309e-23 | 0.511333752 | 0.356 | 0.117 | 3.69333371261147e-19 | 24 |
| slc2a1b | 1.95961304666758e-23 | 0.279440409 | 0.124 | 0.018 | 4.93215007715764e-19 | 24 |

| | | | | | | |
|---|---|---|---|---|---|---|
| CABZ01068367.12 | 2.22833935732512e-23 | 0.278557296 | 0.124 | 0.019 | 5.60850732845159e-19 | 24 |
| ap2m1a1 | 2.43334033230769e-23 | 0.521279157 | 0.395 | 0.14 | 6.12447428238521e-19 | 24 |
| afdna1 | 3.39665501631373e-23 | 0.254650227 | 0.136 | 0.022 | 8.54904101056002e-19 | 24 |
| tmem176l.15 | 6.59108048070016e-23 | 0.543605207 | 0.458 | 0.178 | 1.65890904618742e-18 | 24 |
| mt-atp612 | 7.85886115173263e-23 | 0.51093882 | 0.814 | 0.589 | 1.97799676327958e-18 | 24 |
| lgals9l32 | 2.90487933154106e-22 | 0.369929327 | 0.243 | 0.062 | 7.31129078955569e-18 | 24 |
| prss12 | 4.2952075487083e-22 | 0.339831305 | 0.136 | 0.023 | 1.08106078793439e-17 | 24 |
| yap14 | 4.93870149596508e-22 | 0.350226259 | 0.209 | 0.049 | 1.24302177951945e-17 | 24 |
| pde4ba7 | 5.28212474128922e-22 | 0.528849978 | 0.384 | 0.138 | 1.32945797613508e-17 | 24 |
| txnipa9 | 5.56791559213042e-22 | 0.567011026 | 0.492 | 0.208 | 1.4013886753833e-17 | 24 |
| serpinf14 | 6.78059392903352e-22 | 0.339491986 | 0.192 | 0.042 | 1.70660768599845e-17 | 24 |
| cdh23 | 1.09055882218685e-21 | 0.384279671 | 0.22 | 0.054 | 2.74482749956208e-17 | 24 |
| glod5 | 1.29700156559444e-21 | 0.307551113 | 0.164 | 0.033 | 3.26442324044465e-17 | 24 |
| anxa2a4 | 1.42597947620616e-21 | 0.546524861 | 0.322 | 0.104 | 3.58904774366329e-17 | 24 |
| msi2a1 | 2.4381450794736e-21 | 0.3421764 | 0.198 | 0.045 | 6.1365673505271e-17 | 24 |
| NC-002333.411 | 2.44434755618952e-21 | 0.457841543 | 0.825 | 0.621 | 6.15217836417341e-17 | 24 |
| gulp1a3 | 2.7387255640129e-21 | 0.274749254 | 0.158 | 0.031 | 6.89309837206407e-17 | 24 |
| ahnak7 | 3.5081236974445e-21 | 0.491598097 | 0.362 | 0.125 | 8.82959653409807e-17 | 24 |
| si:ch211-222l21.19 | 3.68616328795142e-21 | 0.57437817 | 0.576 | 0.273 | 9.27770437944492e-17 | 24 |
| si:dkey-7j14.67 | 3.76050604008226e-21 | 0.529846654 | 0.435 | 0.173 | 9.46481765228305e-17 | 24 |
| atp1a1a.1 | 4.11187970671931e-21 | 0.521630499 | 0.395 | 0.146 | 1.03491900338418e-16 | 24 |
| myl9b5 | 5.50342257379755e-21 | 0.412437503 | 0.271 | 0.078 | 1.38515642759911e-16 | 24 |
| si:dkey-188i13.105 | 1.00064719494664e-20 | 0.447926748 | 0.316 | 0.102 | 2.5185289249612e-16 | 24 |
| nfic5 | 1.00200129773818e-20 | 0.42253362 | 0.26 | 0.075 | 2.52193706627722e-16 | 24 |
| ptprga6 | 1.39025810075321e-20 | 0.341209725 | 0.243 | 0.065 | 3.49914061378576e-16 | 24 |
| zgc:1951735 | 1.55191260680678e-20 | 0.419275121 | 0.294 | 0.09 | 3.90600884007197e-16 | 24 |
| tuba8l33 | 2.55009743457912e-20 | 0.381299386 | 0.192 | 0.046 | 6.41834023309219e-16 | 24 |
| cd74b7 | 2.64096391729405e-20 | 0.598031632 | 0.514 | 0.221 | 6.64704208343739e-16 | 24 |
| hmgb3a2 | 3.96507532310589e-20 | 0.34926612 | 0.198 | 0.048 | 9.9796980807252e-16 | 24 |
| cald1a5 | 4.22385615004056e-20 | 0.448502669 | 0.305 | 0.097 | 1.06310235440371e-15 | 24 |
| atp10a3 | 4.50828613927213e-20 | 0.379644177 | 0.215 | 0.055 | 1.1346905383934e-15 | 24 |
| arf2b | 6.07695891020651e-20 | 0.48452469 | 0.418 | 0.16 | 1.52950978810988e-15 | 24 |
| zbtb205 | 7.95897617197065e-20 | 0.4218691 | 0.299 | 0.097 | 2.00319471272329e-15 | 24 |
| gli33 | 1.10850135097693e-19 | 0.279319942 | 0.153 | 0.031 | 2.78998705027384e-15 | 24 |
| selenof2 | 1.1589151869879e-19 | 0.51296678 | 0.514 | 0.234 | 2.91687363412984e-15 | 24 |

| | | | | | | |
|---|---|---|---|---|---|---|
| zgc:1748634 | 4.11860231942421e-19 | 0.446541313 | 0.299 | 0.101 | 1.03661101777588e-14 | 24 |
| mt-nd29 | 4.9692357419422e-19 | 0.526874265 | 0.667 | 0.398 | 1.25070694388943e-14 | 24 |
| h2afx19 | 7.57343358979071e-19 | 0.515701428 | 0.458 | 0.197 | 1.90615750021442e-14 | 24 |
| rnf213a6 | 1.03474224658362e-18 | 0.386481835 | 0.277 | 0.087 | 2.60434276042632e-14 | 24 |
| cox7a11 | 1.20341341060234e-18 | 0.290883131 | 0.147 | 0.03 | 3.02887121314504e-14 | 24 |
| ndrg22 | 1.34415428778097e-18 | 0.312147938 | 0.169 | 0.039 | 3.38310192691594e-14 | 24 |
| rn7sk | 1.44101290557308e-18 | 0.399083488 | 0.169 | 0.039 | 3.62688538203689e-14 | 24 |
| si:ch211-199g17.21 | 1.52988937781259e-18 | 0.3550346 | 0.232 | 0.065 | 3.85057857501651e-14 | 24 |
| CABZ01080702.15 | 3.66375757491103e-18 | 0.387400956 | 0.266 | 0.083 | 9.22131144029358e-14 | 24 |
| mknk2b9 | 4.26210922907506e-18 | 0.526527022 | 0.441 | 0.195 | 1.0727302718659e-13 | 24 |
| map1lc3a5 | 6.58522335608165e-18 | 0.338471401 | 0.237 | 0.069 | 1.65743486649219e-13 | 24 |
| cd164 | 7.84437430769672e-18 | 0.367256466 | 0.237 | 0.071 | 1.97435056950419e-13 | 24 |
| cux1a3 | 7.91931060802565e-18 | 0.303633669 | 0.181 | 0.045 | 1.99321128693398e-13 | 24 |
| mt-nd39 | 7.94474359230917e-18 | 0.51376641 | 0.667 | 0.394 | 1.99961251474829e-13 | 24 |
| fam168a6 | 9.50153380334564e-18 | 0.333653608 | 0.209 | 0.058 | 2.39144104296406e-13 | 24 |
| itm2ba8 | 1.08263927049567e-17 | 0.51189698 | 0.525 | 0.272 | 2.72489477991054e-13 | 24 |
| gng75 | 2.01474743948628e-17 | 0.317027764 | 0.198 | 0.053 | 5.07091783044303e-13 | 24 |
| kmt2cb1 | 2.0654329766559e-17 | 0.336733458 | 0.198 | 0.053 | 5.19848825894523e-13 | 24 |
| ccdc85b5 | 2.58538080654632e-17 | 0.315997935 | 0.186 | 0.048 | 6.50714495199643e-13 | 24 |
| rpia1 | 4.31263682051723e-17 | 0.348770409 | 0.22 | 0.063 | 1.08544756135598e-12 | 24 |
| abl11 | 4.68570399538784e-17 | 0.262524544 | 0.147 | 0.032 | 1.17934483859917e-12 | 24 |
| mt-nd49 | 1.49456046836426e-16 | 0.483795102 | 0.684 | 0.455 | 3.76165924282601e-12 | 24 |
| atp5mea1 | 1.53740527047905e-16 | 0.446330164 | 0.367 | 0.147 | 3.86949532526871e-12 | 24 |
| si:dkey-28b4.76 | 2.53649932001431e-16 | 0.399678691 | 0.266 | 0.091 | 6.38411513854402e-12 | 24 |
| ctnnb13 | 6.13505455126772e-16 | 0.423297916 | 0.367 | 0.148 | 1.54413188000857e-11 | 24 |
| phc2a1 | 1.11851694076683e-15 | 0.351983276 | 0.277 | 0.092 | 2.81519528821605e-11 | 24 |
| tmem88b2 | 1.7838008550757e-15 | 0.264173062 | 0.153 | 0.036 | 4.48964837214002e-11 | 24 |
| gpx4a3 | 1.84592049461512e-15 | 0.342985897 | 0.232 | 0.073 | 4.64599729289681e-11 | 24 |
| hmgb1a9 | 1.88526277900129e-15 | 0.465394541 | 0.407 | 0.188 | 4.74501788846834e-11 | 24 |
| calm2b4 | 2.01720618656859e-15 | 0.456322301 | 0.452 | 0.209 | 5.07710625097448e-11 | 24 |
| mt-nd110 | 4.16047200762244e-15 | 0.476174385 | 0.638 | 0.41 | 1.04714919959849e-10 | 24 |
| atp8a1 | 4.38238537250274e-15 | 0.281837635 | 0.153 | 0.037 | 1.10300257440521e-10 | 24 |
| LOC1039092844 | 4.47119345562528e-15 | 0.320390467 | 0.209 | 0.063 | 1.12535468084633e-10 | 24 |
| trappc11 | 5.33912052234608e-15 | 0.352969009 | 0.277 | 0.099 | 1.34380324426928e-10 | 24 |
| hbp11 | 7.55762335393452e-15 | 0.321551049 | 0.186 | 0.054 | 1.90217822195178e-10 | 24 |

| | | | | | | |
|---|---|---|---|---|---|---|
| zfp36l1b5 | 8.05873416207111e-15 | 0.394336846 | 0.333 | 0.129 | 2.02830280125168e-10 | 24 |
| ywhaqb4 | 8.39500797683604e-15 | 0.381141568 | 0.328 | 0.131 | 2.11293955768986e-10 | 24 |
| cad | 8.77200140982452e-15 | 0.269828388 | 0.124 | 0.027 | 2.20782503483873e-10 | 24 |
| rgs22 | 8.88399320676159e-15 | 0.286383059 | 0.124 | 0.027 | 2.23601225020982e-10 | 24 |
| sptan12 | 9.08926338200861e-15 | 0.25820635 | 0.181 | 0.05 | 2.28767670061775e-10 | 24 |
| cavin2b4 | 9.79330987476287e-15 | 0.283669695 | 0.215 | 0.065 | 2.46487816237907e-10 | 24 |
| LOC1018844513 | 1.07871347344829e-14 | 0.338781152 | 0.249 | 0.085 | 2.715013941322e-10 | 24 |
| hspe13 | 1.29639736746244e-14 | 0.496639535 | 0.531 | 0.293 | 3.26290253416622e-10 | 24 |
| b2ml11 | 1.4139039753973e-14 | 0.466446726 | 0.497 | 0.266 | 3.55865491567746e-10 | 24 |
| ywhaz5 | 1.59262300958354e-14 | 0.470326657 | 0.514 | 0.28 | 4.00847285282082e-10 | 24 |
| purab | 2.86677126264356e-14 | 0.256442251 | 0.175 | 0.049 | 7.21537659094758e-10 | 24 |
| zgc:1101821 | 3.0957490753455e-14 | 0.32327756 | 0.26 | 0.087 | 7.7916908477371e-10 | 24 |
| sox4a2 | 4.01195362604821e-14 | 0.273784003 | 0.164 | 0.044 | 1.00976860814007e-09 | 24 |
| wu:fb16f033 | 4.07960502461048e-14 | 0.3285827 | 0.249 | 0.086 | 1.02679578864421e-09 | 24 |
| hist2h2l2 | 4.53625469553333e-14 | 0.387258772 | 0.226 | 0.078 | 1.14172994431878e-09 | 24 |
| hmgn69 | 5.41543478722014e-14 | 0.464444608 | 0.486 | 0.264 | 1.36301078159544e-09 | 24 |
| robo24 | 6.2265928228049e-14 | 0.250214028 | 0.169 | 0.047 | 1.56717114757177e-09 | 24 |
| lgals173 | 6.22799800913189e-14 | 0.275042085 | 0.181 | 0.052 | 1.5675248189184e-09 | 24 |
| pgrmc15 | 6.25374749314867e-14 | 0.386245738 | 0.339 | 0.14 | 1.57400570655059e-09 | 24 |
| pfn22 | 7.14775017799533e-14 | 0.440831747 | 0.458 | 0.225 | 1.79901724229964e-09 | 24 |
| hmgb1b6 | 1.06102445166612e-13 | 0.380345823 | 0.311 | 0.126 | 2.67049244239847e-09 | 24 |
| ncor1 | 1.3129938625999e-13 | 0.28348522 | 0.186 | 0.056 | 3.30467425277768e-09 | 24 |
| pfn2l4 | 1.35903649922532e-13 | 0.407122181 | 0.446 | 0.208 | 3.42055896490021e-09 | 24 |
| hsd3b7 | 1.44159379132253e-13 | 0.360976455 | 0.249 | 0.086 | 3.62834741337967e-09 | 24 |
| smarca2 | 2.16549592548954e-13 | 0.284982424 | 0.158 | 0.044 | 5.45033669486462e-09 | 24 |
| srpra | 2.24662849371246e-13 | 0.314671707 | 0.192 | 0.059 | 5.65453925582489e-09 | 24 |
| socs3a9 | 3.11236591773972e-13 | 0.495307858 | 0.559 | 0.297 | 7.8335137783591e-09 | 24 |
| XLOC-0160945 | 3.57452258085187e-13 | 0.317304875 | 0.232 | 0.079 | 8.99671588374608e-09 | 24 |
| ppib8 | 3.93148337188368e-13 | 0.439592894 | 0.571 | 0.331 | 9.89515049869404e-09 | 24 |
| stat25 | 4.30902420283049e-13 | 0.302422531 | 0.192 | 0.061 | 1.08453830161041e-08 | 24 |
| nucb2a | 4.36618424305173e-13 | 0.282706439 | 0.164 | 0.048 | 1.09892491213369e-08 | 24 |
| angptl44 | 4.81208233755305e-13 | 0.283157219 | 0.169 | 0.049 | 1.21115300353873e-08 | 24 |
| serinc13 | 4.86473834614715e-13 | 0.319144788 | 0.232 | 0.081 | 1.22440599434178e-08 | 24 |
| tpm4a4 | 7.28504176492694e-13 | 0.336662127 | 0.243 | 0.088 | 1.83357216181446e-08 | 24 |
| ctnnd1 | 7.5209357836347e-13 | 0.272405514 | 0.192 | 0.06 | 1.89294432738302e-08 | 24 |

| | | | | | | |
|---|---|---|---|---|---|---|
| psme14 | 8.05590883193836e-13 | 0.326687671 | 0.243 | 0.089 | 2.02759169391057e-08 | 24 |
| chchd6b | 9.5448845471476e-13 | 0.345849834 | 0.237 | 0.086 | 2.40235199167158e-08 | 24 |
| si:ch211-197h24.92 | 1.68461263734095e-12 | 0.28763137 | 0.203 | 0.068 | 4.24000154692343e-08 | 24 |
| gabarapb7 | 3.02901468158239e-12 | 0.411904994 | 0.486 | 0.267 | 7.62372705207472e-08 | 24 |
| slc38a5b5 | 3.53054364374115e-12 | 0.336349856 | 0.282 | 0.115 | 8.8860252969321e-08 | 24 |
| fn1b3 | 4.085536984047e-12 | 0.389171693 | 0.237 | 0.087 | 1.02828880351479e-07 | 24 |
| tnfrsf9a3 | 4.59423171198108e-12 | 0.325493968 | 0.169 | 0.052 | 1.15632217958852e-07 | 24 |
| flot1b2 | 4.8559213496105e-12 | 0.299557038 | 0.226 | 0.082 | 1.22218684448347e-07 | 24 |
| eif4ebp15 | 6.54378901114383e-12 | 0.414239566 | 0.463 | 0.245 | 1.64700625621479e-07 | 24 |
| fam120c2 | 6.84346526404297e-12 | 0.295818905 | 0.254 | 0.098 | 1.72243177230698e-07 | 24 |
| si:ch211-1a19.33 | 8.84211281088949e-12 | 0.289369917 | 0.169 | 0.053 | 2.22547137337278e-07 | 24 |
| krt945 | 9.52510123246247e-12 | 0.336622609 | 0.316 | 0.129 | 2.39737272919848e-07 | 24 |
| hp1bp33 | 1.44161062385955e-11 | 0.318240721 | 0.249 | 0.097 | 3.62838977919211e-07 | 24 |
| oaz1a1 | 4.05275680949704e-11 | 0.378247661 | 0.65 | 0.48 | 1.02003836138231e-06 | 24 |
| atrx5 | 5.39545366073418e-11 | 0.31893542 | 0.254 | 0.104 | 1.35798173187018e-06 | 24 |
| fthl271 | 6.2426828833895e-11 | 0.289744276 | 0.836 | 0.695 | 1.5712208549203e-06 | 24 |
| atp6v1e1a | 8.12749251285195e-11 | 0.253429516 | 0.153 | 0.048 | 2.04560859055971e-06 | 24 |
| atp5pb2 | 8.92737957939409e-11 | 0.376614337 | 0.52 | 0.299 | 2.2469321663377e-06 | 24 |
| add3a6 | 9.35335753939667e-11 | 0.320854386 | 0.288 | 0.123 | 2.35414655909075e-06 | 24 |
| si:ch73-376l24.31 | 1.15299790159254e-10 | 0.310900516 | 0.271 | 0.113 | 2.90198041851828e-06 | 24 |
| dynlt31 | 1.20858462094981e-10 | 0.338055809 | 0.316 | 0.146 | 3.04188663246858e-06 | 24 |
| marcksl1b7 | 1.40074045546384e-10 | 0.36268276 | 0.271 | 0.116 | 3.52552365235695e-06 | 24 |
| ndufb92 | 1.40084655692893e-10 | 0.339328027 | 0.362 | 0.175 | 3.52579069913442e-06 | 24 |
| cd74a6 | 1.50522882863889e-10 | 0.381349511 | 0.429 | 0.218 | 3.78851043880123e-06 | 24 |
| hmgb2b2 | 1.8998794538836e-10 | 0.396492041 | 0.542 | 0.318 | 4.78180659747962e-06 | 24 |
| dynll2b1 | 3.01767792594298e-10 | 0.295803368 | 0.305 | 0.132 | 7.5951935718059e-06 | 24 |
| nucks1a | 3.865477710059e-10 | 0.361622851 | 0.362 | 0.181 | 9.72902084844749e-06 | 24 |
| mvp3 | 5.48492066412543e-10 | 0.25596107 | 0.158 | 0.053 | 1.38049968195373e-05 | 24 |
| mcl1b7 | 5.76911425584065e-10 | 0.370756161 | 0.446 | 0.241 | 1.45202836705253e-05 | 24 |
| ndufa4l7 | 6.18095179501882e-10 | 0.37099662 | 0.582 | 0.395 | 1.55568375728829e-05 | 24 |
| cnn3a4 | 6.47991937170431e-10 | 0.251278664 | 0.164 | 0.056 | 1.63093090666426e-05 | 24 |
| glud1a1 | 6.89979124182705e-10 | 0.287294125 | 0.226 | 0.091 | 1.73660845765545e-05 | 24 |
| arrdc3a3 | 8.93291891867696e-10 | 0.296117982 | 0.192 | 0.072 | 2.24832636264181e-05 | 24 |
| iscub1 | 9.62521115710145e-10 | 0.268079542 | 0.181 | 0.066 | 2.42256939613086e-05 | 24 |
| calm1a1 | 1.47575982064562e-09 | 0.337417436 | 0.333 | 0.164 | 3.71433989258295e-05 | 24 |

| | | | | | | |
|---|---|---|---|---|---|---|
| ucp27 | 1.64582814888435e-09 | 0.359713337 | 0.61 | 0.416 | 4.14238486792701e-05 | 24 |
| h2afy25 | 2.78482886647214e-09 | 0.302586856 | 0.226 | 0.097 | 7.00913577402373e-05 | 24 |
| metrnla3 | 3.05303748548086e-09 | 0.264763349 | 0.226 | 0.093 | 7.68419004720678e-05 | 24 |
| ndufa52 | 3.62080121379822e-09 | 0.333672764 | 0.294 | 0.142 | 9.11319457500875e-05 | 24 |
| cdv31 | 4.54726449811401e-09 | 0.279215634 | 0.294 | 0.137 | 0.00011445 | 24 |
| zgc:1527914 | 5.07122259561497e-09 | 0.282205564 | 0.181 | 0.068 | 0.000127638 | 24 |
| cdc423 | 5.45242340071708e-09 | 0.331056535 | 0.39 | 0.212 | 0.000137232 | 24 |
| hnrnpa1a4 | 5.51567113788084e-09 | 0.334579136 | 0.316 | 0.159 | 0.000138824 | 24 |
| tp53 | 5.78645097984142e-09 | 0.291522117 | 0.237 | 0.103 | 0.000145639 | 24 |
| ndufab1a1 | 6.0113357010325e-09 | 0.280535535 | 0.288 | 0.134 | 0.000151299 | 24 |
| sdcbp27 | 6.17813084133128e-09 | 0.327501471 | 0.384 | 0.208 | 0.000155497 | 24 |
| rhoaa7 | 7.68375328523392e-09 | 0.317301457 | 0.305 | 0.154 | 0.000193392 | 24 |
| si:ch73-376l24.21 | 8.51016040138995e-09 | 0.267005304 | 0.26 | 0.115 | 0.000214192 | 24 |
| tnrc6c18 | 8.73731928528007e-09 | 0.304147996 | 0.277 | 0.133 | 0.00021991 | 24 |
| chd4a1 | 1.02553664977826e-08 | 0.265828504 | 0.215 | 0.091 | 0.000258117 | 24 |
| myl65 | 1.12240126234795e-08 | 0.291605851 | 0.232 | 0.101 | 0.000282497 | 24 |
| chd4b | 1.46111322460533e-08 | 0.280254152 | 0.232 | 0.102 | 0.000367748 | 24 |
| ier3ip11 | 2.18458219054298e-08 | 0.290998856 | 0.305 | 0.153 | 0.000549837 | 24 |
| ndufc22 | 2.62419573938753e-08 | 0.258100234 | 0.282 | 0.135 | 0.000660484 | 24 |
| hif1ab2 | 3.0405695419399e-08 | 0.28925317 | 0.243 | 0.113 | 0.000765281 | 24 |
| si:zfos-169g10.31 | 3.13919425035172e-08 | 0.283273478 | 0.215 | 0.095 | 0.000790104 | 24 |
| ndufa13 | 3.3035034428658e-08 | 0.317181314 | 0.407 | 0.23 | 0.000831459 | 24 |
| psme26 | 3.64493007381377e-08 | 0.250026746 | 0.22 | 0.097 | 0.000917392 | 24 |
| ptmaa10 | 3.77234036674566e-08 | 0.352011226 | 0.593 | 0.395 | 0.00094946 | 24 |
| notch27 | 4.3310923134589e-08 | 0.268457645 | 0.254 | 0.119 | 0.001090093 | 24 |
| rab104 | 4.60340247400325e-08 | 0.293529243 | 0.362 | 0.194 | 0.00115863 | 24 |
| gnb1b3 | 5.4087219843449e-08 | 0.323310471 | 0.429 | 0.254 | 0.001361321 | 24 |
| eef1db7 | 5.87240511564679e-08 | 0.334044764 | 0.446 | 0.274 | 0.001478026 | 24 |
| rab5c2 | 6.81748856650999e-08 | 0.261304323 | 0.254 | 0.12 | 0.001715894 | 24 |
| h3f3c7 | 9.36819656758097e-08 | 0.32007135 | 0.429 | 0.256 | 0.002357881 | 24 |
| jun6 | 9.54232181422946e-08 | 0.314563081 | 0.418 | 0.228 | 0.002401707 | 24 |
| ifi303 | 9.58697352152057e-08 | 0.314291428 | 0.328 | 0.175 | 0.002412945 | 24 |
| ndufa32 | 1.12016085467518e-07 | 0.273996526 | 0.345 | 0.183 | 0.002819333 | 24 |
| gpx4b2 | 1.12555955942717e-07 | 0.294180631 | 0.367 | 0.203 | 0.002832921 | 24 |
| si:ch73-46j18.56 | 1.64011465988492e-07 | 0.285136707 | 0.339 | 0.175 | 0.004128005 | 24 |

| | | | | | | |
|---|---|---|---|---|---|---|
| abcc55 | 1.66324904618668e-07 | 0.255646449 | 0.209 | 0.095 | 0.004186232 | 24 |
| tmem591 | 2.04453568441456e-07 | 0.262188812 | 0.226 | 0.107 | 0.005145892 | 24 |
| dbi4 | 2.53083159875791e-07 | 0.316261652 | 0.367 | 0.213 | 0.00636985 | 24 |
| mt-nd54 | 2.70737860730666e-07 | 0.287267323 | 0.243 | 0.12 | 0.006814201 | 24 |
| cnn26 | 2.82246630525345e-07 | 0.276388217 | 0.266 | 0.135 | 0.007103865 | 24 |
| rpl27a12 | 2.97578021031528e-07 | 0.302839532 | 0.695 | 0.495 | 0.007489741 | 24 |
| atf4b8 | 3.35150771094635e-07 | 0.287739862 | 0.441 | 0.264 | 0.00843541 | 24 |
| selenok2 | 3.55030657042446e-07 | 0.260592201 | 0.254 | 0.127 | 0.008935767 | 24 |
| atp5pf5 | 3.73320141048483e-07 | 0.315780367 | 0.463 | 0.298 | 0.009396095 | 24 |
| dpy301 | 4.04768509491232e-07 | 0.25566455 | 0.294 | 0.152 | 0.010187619 | 24 |
| cox6b23 | 4.17245495625947e-07 | 0.276530916 | 0.706 | 0.533 | 0.010501652 | 24 |
| midn6 | 6.02556293412577e-07 | 0.265633603 | 0.271 | 0.14 | 0.015165739 | 24 |
| edf11 | 6.04121348826335e-07 | 0.296246658 | 0.435 | 0.268 | 0.01520513 | 24 |
| si:ch73-42p12.21 | 7.16127640490623e-07 | 0.250757992 | 0.215 | 0.102 | 0.018024217 | 24 |
| si:ch211-191j22.31 | 7.163094978675e-07 | 0.250578597 | 0.232 | 0.114 | 0.018028794 | 24 |
| tegt7 | 7.69713811763478e-07 | 0.295609239 | 0.356 | 0.211 | 0.019372927 | 24 |
| CR383676.112 | 8.10656542982575e-07 | 0.280611257 | 0.797 | 0.727 | 0.020403415 | 24 |
| cox7c4 | 8.30252603071131e-07 | 0.286923895 | 0.644 | 0.481 | 0.020896628 | 24 |
| sub1a2 | 8.97560335141853e-07 | 0.290014623 | 0.39 | 0.233 | 0.022590696 | 24 |
| atp5mc3a3 | 1.0440485428262e-06 | 0.295998558 | 0.373 | 0.223 | 0.026277658 | 24 |
| fosab8 | 1.50881086186046e-06 | 0.367193069 | 0.508 | 0.349 | 0.037975261 | 24 |
| odc12 | 1.70691228305632e-06 | 0.306936712 | 0.237 | 0.123 | 0.042961275 | 24 |
| eif1b7 | 1.82923680183053e-06 | 0.282143551 | 0.565 | 0.382 | 0.046040061 | 24 |
| cfl1l2 | 1.88460136485514e-06 | 0.27474016 | 0.294 | 0.152 | 0.047433532 | 24 |
| fam133b1 | 2.34224410718871e-06 | 0.261651891 | 0.271 | 0.145 | 0.058951942 | 24 |
| cox5ab2 | 2.63020591900725e-06 | 0.252656936 | 0.35 | 0.202 | 0.066199653 | 24 |
| rps27l8 | 2.96876076883145e-06 | 0.288404132 | 0.339 | 0.203 | 0.07472074 | 24 |
| zgc:11418813 | 3.05889502441677e-06 | 0.268196581 | 0.808 | 0.627 | 0.076989329 | 24 |
| dynlrb12 | 3.59066344661881e-06 | 0.260870511 | 0.35 | 0.204 | 0.090373408 | 24 |
| hmgn27 | 3.99397782765677e-06 | 0.265480271 | 0.61 | 0.442 | 0.100524428 | 24 |
| mif1 | 4.02259204617687e-06 | 0.272726687 | 0.322 | 0.188 | 0.101244619 | 24 |
| tma72 | 4.63002823300601e-06 | 0.278514681 | 0.463 | 0.305 | 0.116533181 | 24 |
| vat15 | 5.0211711150603e-06 | 0.252784837 | 0.254 | 0.135 | 0.126377856 | 24 |
| atp5pd4 | 5.27911694325102e-06 | 0.28554339 | 0.407 | 0.262 | 0.132870094 | 24 |
| psap3 | 5.37838230093573e-06 | 0.261362645 | 0.469 | 0.29 | 0.135368504 | 24 |

| | | | | | | |
|---|---|---|---|---|---|---|
| atp5f1c2 | 6.84052202517216e-06 | 0.2639586 | 0.407 | 0.254 | 0.172169099 | 24 |
| ier2a9 | 7.02248105735809e-06 | 0.298431814 | 0.407 | 0.262 | 0.176748826 | 24 |
| ost42 | 7.63220596626529e-06 | 0.271876004 | 0.407 | 0.257 | 0.192094992 | 24 |
| si:dkey-42i9.43 | 1.12132772821458e-05 | 0.259032068 | 0.26 | 0.147 | 0.282226976 | 24 |
| ier2b10 | 1.58404619336878e-05 | 0.260278832 | 0.362 | 0.224 | 0.398688586 | 24 |
| atp5f1e2 | 2.57682717124697e-05 | 0.253305559 | 0.644 | 0.482 | 0.648561631 | 24 |
| rbx11 | 3.28895724529761e-05 | 0.253248447 | 0.407 | 0.265 | 0.827797649 | 24 |
| pomp4 | 9.36516437475473e-05 | 0.255813342 | 0.345 | 0.231 | 1 | 24 |
| tmsb4x14 | 0.000116417 | 0.299637505 | 0.938 | 0.684 | 1 | 24 |
| rpl913 | 0.000303361 | 0.250567507 | 0.831 | 0.628 | 1 | 24 |
| nkl.4 | 0 | 1.796117934 | 0.795 | 0.012 | 0 | 25 |
| prf1.2 | 0 | 1.743031285 | 0.79 | 0.007 | 0 | 25 |
| LOC1018842674 | 0 | 1.703944706 | 0.778 | 0.048 | 0 | 25 |
| cpa5 | 0 | 1.518024486 | 0.648 | 0.029 | 0 | 25 |
| prf1.71 | 0 | 1.507352744 | 0.665 | 0.008 | 0 | 25 |
| XLOC-0430433 | 0 | 1.414649168 | 0.636 | 0.035 | 0 | 25 |
| XLOC-0397571 | 0 | 1.145494593 | 0.511 | 0.02 | 0 | 25 |
| tspan2b | 0 | 0.859617276 | 0.381 | 0.007 | 0 | 25 |
| nkl.3 | 0 | 0.827357252 | 0.318 | 0.004 | 0 | 25 |
| si:dkey-21e2.3 | 7.89457339575129e-306 | 0.533672232 | 0.244 | 0.003 | 1.98698517797664e-301 | 25 |
| CABZ01084575.13 | 9.00958543136866e-278 | 1.149011735 | 0.585 | 0.035 | 2.26762255722118e-273 | 25 |
| nkl.23 | 1.11662668024064e-271 | 2.336479289 | 0.994 | 0.148 | 2.81043769149768e-267 | 25 |
| XLOC-0140792 | 5.74214504279917e-250 | 1.111890594 | 0.489 | 0.027 | 1.44524048582212e-245 | 25 |
| BX248410.3 | 2.34507480854999e-237 | 0.333031494 | 0.142 | 0.001 | 5.90231878563947e-233 | 25 |
| si:ch211-173d10.4 | 3.59279288209241e-219 | 0.466761113 | 0.205 | 0.004 | 9.04270040493839e-215 | 25 |
| csf2rb3 | 5.17083225905503e-219 | 1.044748657 | 0.5 | 0.033 | 1.30144677128156e-214 | 25 |
| BX120005.13 | 1.25095082791162e-214 | 1.223520455 | 0.551 | 0.042 | 3.14851813877075e-210 | 25 |
| LOC1081907174 | 2.43662592824689e-206 | 1.379961575 | 0.761 | 0.087 | 6.13274379880459e-202 | 25 |
| si:ch211-277c7.63 | 1.1472799972695e-205 | 1.086228165 | 0.483 | 0.032 | 2.8875890251276e-201 | 25 |
| mef2b1 | 4.11897748322214e-201 | 0.567314727 | 0.25 | 0.007 | 1.03670544275218e-196 | 25 |
| CABZ01088229.11 | 3.77633681330056e-178 | 0.620511105 | 0.261 | 0.01 | 9.50466212539619e-174 | 25 |
| ca22 | 4.98809855812993e-172 | 1.126755808 | 0.54 | 0.05 | 1.25545452609572e-167 | 25 |
| si:dkey-78l4.42 | 1.5779015372104e-163 | 0.839856473 | 0.409 | 0.028 | 3.97142037900485e-159 | 25 |
| hck2 | 7.93221822209199e-161 | 0.813647506 | 0.398 | 0.028 | 1.99646000431833e-156 | 25 |
| alox5a3 | 1.07932653061887e-158 | 0.842123921 | 0.398 | 0.028 | 2.71655694491464e-154 | 25 |

| | | | | | | |
|---|---|---|---|---|---|---|
| ptk2ba | 1.57507884742041e-158 | 0.699232702 | 0.324 | 0.018 | 3.96431595107242e-154 | 25 |
| stim1a2 | 7.4564272379107e-157 | 1.063404599 | 0.568 | 0.062 | 1.87670817150974e-152 | 25 |
| CR753876.14 | 8.82075121020504e-154 | 1.255063961 | 0.733 | 0.107 | 2.22009487209651e-149 | 25 |
| CABZ01073834.13 | 4.3044218708832e-148 | 0.867997934 | 0.432 | 0.037 | 1.08337994068259e-143 | 25 |
| zbtb323 | 3.16474945931399e-143 | 0.898804126 | 0.455 | 0.042 | 7.96535791414737e-139 | 25 |
| creb5b2 | 1.55974709807142e-136 | 0.632063291 | 0.284 | 0.016 | 3.92572747113595e-132 | 25 |
| sort1b2 | 1.0325829944496e-134 | 0.7474351 | 0.358 | 0.027 | 2.59890813873021e-130 | 25 |
| gcnt7 | 1.87562320556146e-131 | 0.315450251 | 0.131 | 0.003 | 4.72075604607764e-127 | 25 |
| edaradd2 | 6.7693781352548e-130 | 0.712236274 | 0.307 | 0.02 | 1.70378478286228e-125 | 25 |
| prr5a | 2.57566890826277e-127 | 0.6712068 | 0.301 | 0.02 | 6.48270107520656e-123 | 25 |
| gzm3.31 | 7.21089092811438e-127 | 0.516332641 | 0.222 | 0.01 | 1.81490913769711e-122 | 25 |
| srgn8 | 9.98211532726947e-123 | 1.334314043 | 0.881 | 0.229 | 2.51239860672045e-118 | 25 |
| efhd21 | 2.74464489550543e-122 | 0.954952909 | 0.472 | 0.054 | 6.90799673749761e-118 | 25 |
| ponzr61 | 1.39088169470221e-115 | 0.6477018 | 0.29 | 0.02 | 3.500710137396e-111 | 25 |
| arl113 | 3.62106096924433e-114 | 0.74528622 | 0.318 | 0.025 | 9.11384835349105e-110 | 25 |
| illr1 | 8.16902602982607e-113 | 0.521853786 | 0.233 | 0.013 | 2.05606216144692e-108 | 25 |
| BX005105.13 | 3.04256387094803e-110 | 0.768249942 | 0.392 | 0.04 | 7.65782900678909e-106 | 25 |
| sftpba1 | 1.10677328929881e-108 | 0.657118207 | 0.307 | 0.024 | 2.78563769183618e-104 | 25 |
| slc41a11 | 1.83647379690073e-105 | 0.455111643 | 0.21 | 0.011 | 4.62222089941945e-101 | 25 |
| XLOC-0437604 | 6.52634558045822e-102 | 0.883778047 | 0.403 | 0.047 | 1.64261591914553e-97 | 25 |
| arpc1b9 | 1.69017866969024e-101 | 1.23582998 | 0.903 | 0.302 | 4.25401069374337e-97 | 25 |
| LOC110440216 | 2.92533035754645e-100 | 0.370121164 | 0.159 | 0.007 | 7.36276397690866e-96 | 25 |
| si:ch211-102c2.46 | 3.82740019624088e-100 | 1.09353991 | 0.676 | 0.136 | 9.63318355391867e-96 | 25 |
| plcg21 | 2.14313284874589e-95 | 0.767475081 | 0.392 | 0.047 | 5.39405106700852e-91 | 25 |
| rps6ka14 | 4.46245089766459e-95 | 0.874224914 | 0.472 | 0.069 | 1.1231542664332e-90 | 25 |
| cyfip21 | 6.38636553539427e-95 | 0.574925994 | 0.273 | 0.022 | 1.60738434160338e-90 | 25 |
| mrtfaa1 | 7.71170428768765e-95 | 0.562957872 | 0.267 | 0.021 | 1.9409588521681e-90 | 25 |
| si:dkey-21o19.2 | 1.62757881533544e-93 | 0.445129219 | 0.21 | 0.013 | 4.09645312031777e-89 | 25 |
| rbbp8l | 1.83632566017441e-88 | 0.36381671 | 0.165 | 0.008 | 4.62184805409297e-84 | 25 |
| XLOC-0054482 | 5.35425771973581e-88 | 0.696517365 | 0.278 | 0.025 | 1.34761312548031e-83 | 25 |
| ephx21 | 7.76447920900975e-88 | 0.627868146 | 0.318 | 0.033 | 1.95424177211566e-83 | 25 |
| LOC110437825 | 2.61690361562597e-87 | 0.343742987 | 0.142 | 0.006 | 6.58648471016901e-83 | 25 |
| znrf11 | 3.90812228300588e-87 | 0.485126619 | 0.205 | 0.013 | 9.83635297409749e-83 | 25 |
| coro1a9 | 1.31937888639787e-84 | 1.091444521 | 0.784 | 0.225 | 3.32074471917481e-80 | 25 |
| adam19b1 | 2.68062579524576e-84 | 0.394831757 | 0.193 | 0.012 | 6.74686706405405e-80 | 25 |

| | | | | | | |
|---|---|---|---|---|---|---|
| tbx21 | 3.06744195043315e-82 | 0.354623977 | 0.142 | 0.007 | 7.72044464504519e-78 | 25 |
| pfn19 | 6.46066025982807e-81 | 1.027569574 | 0.989 | 0.52 | 1.62608358079613e-76 | 25 |
| ccl39.9 | 2.15708995620107e-80 | 0.395673977 | 0.114 | 0.004 | 5.42917971076246e-76 | 25 |
| il12rb2 | 4.19925497795892e-80 | 0.322593993 | 0.142 | 0.007 | 1.05691048540248e-75 | 25 |
| myh11a4 | 1.07774683953067e-79 | 0.782078871 | 0.415 | 0.062 | 2.71258102041475e-75 | 25 |
| syk4 | 1.29955903046272e-79 | 0.827462873 | 0.455 | 0.074 | 3.27086012377163e-75 | 25 |
| wipf1a5 | 1.68784294350593e-79 | 0.737774582 | 0.415 | 0.061 | 4.24813190451007e-75 | 25 |
| fa2h | 1.56438322493472e-78 | 0.351725642 | 0.165 | 0.009 | 3.93739613883819e-74 | 25 |
| fcer1gl6 | 2.32400193742318e-78 | 1.006880159 | 0.722 | 0.162 | 5.84928047630041e-74 | 25 |
| strbp1 | 5.49376521011079e-78 | 0.470688019 | 0.21 | 0.016 | 1.38272576573278e-73 | 25 |
| zgc:640518 | 6.87199357939916e-78 | 0.960675137 | 0.659 | 0.156 | 1.72961206399897e-73 | 25 |
| si:ch211-241f5.33 | 2.6094615393545e-77 | 0.789505022 | 0.415 | 0.064 | 6.56775374840134e-73 | 25 |
| gramd2aa1 | 4.06229465398204e-75 | 0.420475659 | 0.21 | 0.017 | 1.02243894146074e-70 | 25 |
| zgc:152891 | 6.30921602335568e-75 | 0.413165521 | 0.182 | 0.012 | 1.58796658091839e-70 | 25 |
| mbnl16 | 2.63298958239153e-74 | 1.033098866 | 0.659 | 0.178 | 6.62697147992123e-70 | 25 |
| nr4a34 | 7.73423827076573e-73 | 0.775584347 | 0.358 | 0.051 | 1.94663043036903e-68 | 25 |
| atp11c1 | 1.380258125978e-72 | 0.583731226 | 0.278 | 0.031 | 3.47397167727403e-68 | 25 |
| BX901920.15 | 8.78127862041092e-72 | 0.820180338 | 0.42 | 0.069 | 2.21016001597123e-67 | 25 |
| vaspa6 | 5.57978378542167e-71 | 0.707500049 | 0.432 | 0.071 | 1.40437578095278e-66 | 25 |
| ly75 | 8.33034942291989e-71 | 0.523199764 | 0.25 | 0.025 | 2.09666564625471e-66 | 25 |
| gfi1ab2 | 1.37618270820764e-70 | 0.479998322 | 0.227 | 0.021 | 3.4637142582878e-66 | 25 |
| hyal3 | 2.84678533274525e-69 | 0.25775794 | 0.102 | 0.004 | 7.16507400398651e-65 | 25 |
| bhlhe408 | 1.4811903834954e-65 | 0.958637011 | 0.597 | 0.156 | 3.72800807621957e-61 | 25 |
| stat4 | 4.14470777189041e-65 | 0.320493858 | 0.136 | 0.008 | 1.0431814991071e-60 | 25 |
| pstpip2 | 5.28248377336146e-65 | 0.470137591 | 0.21 | 0.019 | 1.32954834091735e-60 | 25 |
| si:ch73-208g10.13 | 1.05927717352356e-64 | 0.591789198 | 0.301 | 0.04 | 2.66609471804146e-60 | 25 |
| actb29 | 2.21075799878424e-64 | 0.563146738 | 1 | 0.867 | 5.56425680714005e-60 | 25 |
| ptprja | 9.98260230948699e-64 | 0.347699223 | 0.153 | 0.01 | 2.51252117527478e-59 | 25 |
| plat | 2.48298239096261e-63 | 0.317320585 | 0.148 | 0.01 | 6.24941837981381e-59 | 25 |
| egr34 | 3.19667307155754e-63 | 0.697848539 | 0.352 | 0.055 | 8.04570645380317e-59 | 25 |
| arhgdig8 | 6.31341533015281e-63 | 0.996271325 | 0.705 | 0.238 | 1.58902350444616e-58 | 25 |
| elf17 | 1.46260484615098e-62 | 0.79241288 | 0.472 | 0.1 | 3.6812301372774e-58 | 25 |
| sh3kbp12 | 1.63297042623622e-61 | 0.679091654 | 0.386 | 0.068 | 4.11002326579393e-57 | 25 |
| BX322618.13 | 1.74939421138249e-61 | 0.597527564 | 0.239 | 0.026 | 4.4030502906286e-57 | 25 |
| laptm59 | 6.03439051602666e-59 | 0.865634461 | 0.614 | 0.165 | 1.51879574897875e-54 | 25 |

| | | | | | | |
|---|---|---|---|---|---|---|
| sh3bp2 | 7.29871330442506e-59 | 0.378461816 | 0.182 | 0.016 | 1.83701315159074e-54 | 25 |
| dapp1 | 1.41523466069944e-58 | 0.296284093 | 0.125 | 0.007 | 3.56200411751441e-54 | 25 |
| actr2a7 | 2.69325050839576e-56 | 0.797693775 | 0.511 | 0.125 | 6.7786422045813e-52 | 25 |
| dicp1.11 | 4.17631487420947e-56 | 0.368160847 | 0.148 | 0.011 | 1.05113669068978e-51 | 25 |
| uba5210 | 3.62045861295818e-55 | 0.661343045 | 0.909 | 0.628 | 9.11233228295445e-51 | 25 |
| gab2 | 5.81250583525412e-55 | 0.366815034 | 0.165 | 0.014 | 1.46294959367511e-50 | 25 |
| pla2g15 | 1.20001026211118e-54 | 0.584156379 | 0.307 | 0.047 | 3.02030582870764e-50 | 25 |
| tent5ab | 1.29650493981563e-54 | 0.311415375 | 0.125 | 0.008 | 3.26317328302196e-50 | 25 |
| wasb8 | 1.88462860222753e-54 | 0.845781809 | 0.608 | 0.173 | 4.74342172894648e-50 | 25 |
| sla11 | 3.54481158155776e-54 | 0.395338118 | 0.193 | 0.02 | 8.92193626962273e-50 | 25 |
| tagapa6 | 4.42705657763987e-54 | 0.804511747 | 0.455 | 0.102 | 1.11424587002618e-49 | 25 |
| eef1a1l13 | 4.53471708992018e-54 | 0.330318332 | 0.989 | 0.915 | 1.14134294436201e-49 | 25 |
| itga45 | 6.70099610965373e-54 | 0.677784745 | 0.347 | 0.062 | 1.68657371083875e-49 | 25 |
| si:ch211-214p16.16 | 4.53254337313203e-53 | 1.000153356 | 0.756 | 0.252 | 1.1407958415836e-48 | 25 |
| dok25 | 7.53455241193641e-53 | 0.485478335 | 0.261 | 0.036 | 1.89637149656028e-48 | 25 |
| ptprc8 | 3.4711574310892e-52 | 0.785721654 | 0.494 | 0.123 | 8.73655613830841e-48 | 25 |
| cables2b | 4.35640137397097e-52 | 0.362004825 | 0.153 | 0.013 | 1.09646266181475e-47 | 25 |
| sik16 | 7.7150519239649e-52 | 0.858119667 | 0.46 | 0.112 | 1.94180141874273e-47 | 25 |
| ndfip2 | 2.31745101952589e-51 | 0.570276237 | 0.312 | 0.053 | 5.83279247104471e-47 | 25 |
| BX908750.12 | 6.08055218296631e-51 | 0.48331409 | 0.233 | 0.03 | 1.53041417893079e-46 | 25 |
| XLOC-0283707 | 9.97316198490879e-51 | 0.809564593 | 0.534 | 0.137 | 2.51014513998169e-46 | 25 |
| pabpc1a10 | 1.02309904212763e-50 | 0.703269431 | 0.869 | 0.537 | 2.57503797913103e-46 | 25 |
| dicp1.5-62 | 1.7445193288027e-50 | 0.391968248 | 0.17 | 0.016 | 4.39078069866351e-46 | 25 |
| camk1b1 | 2.83161476082159e-50 | 0.372116133 | 0.188 | 0.02 | 7.12689119151185e-46 | 25 |
| dock102 | 3.67966094447983e-50 | 0.480141997 | 0.227 | 0.029 | 9.26133863116127e-46 | 25 |
| stk17b3 | 5.88229810375272e-50 | 0.723057493 | 0.455 | 0.107 | 1.48051560973352e-45 | 25 |
| tec | 4.57273468380758e-49 | 0.47652566 | 0.227 | 0.03 | 1.15091159256753e-44 | 25 |
| rpl47 | 5.39309697558681e-49 | 0.596975004 | 0.881 | 0.625 | 1.35738857778544e-44 | 25 |
| dock81 | 1.25870798034267e-48 | 0.484073727 | 0.25 | 0.036 | 3.16804211572448e-44 | 25 |
| LOC1039108714 | 5.43470969180445e-48 | 0.827920731 | 0.591 | 0.167 | 1.36786208233026e-43 | 25 |
| wdr193 | 2.78373170753216e-47 | 0.515049401 | 0.227 | 0.031 | 7.00637433468769e-43 | 25 |
| yrk4 | 3.37611993680812e-47 | 0.569934657 | 0.33 | 0.061 | 8.49735626895237e-43 | 25 |
| aldocb3 | 3.82924877900843e-47 | 0.617554018 | 0.335 | 0.064 | 9.63783625188632e-43 | 25 |
| XLOC-0254237 | 6.88625600225467e-47 | 0.809122589 | 0.585 | 0.165 | 1.73320177320748e-42 | 25 |
| sla24 | 8.29419374077944e-46 | 0.652811166 | 0.341 | 0.066 | 2.08756562261678e-41 | 25 |

| | | | | | | |
|---|---|---|---|---|---|---|
| rps1013 | 4.23036793121836e-44 | 0.507026406 | 0.955 | 0.692 | 1.06474130460835e-39 | 25 |
| hsp90ab18 | 5.15287838303321e-44 | 0.418388815 | 0.966 | 0.789 | 1.29692796022563e-39 | 25 |
| irak1 | 9.49398620900533e-44 | 0.419046544 | 0.193 | 0.024 | 2.38954138894455e-39 | 25 |
| prkcha | 8.58086021422402e-43 | 0.36217733 | 0.165 | 0.018 | 2.15971670731804e-38 | 25 |
| eef2b6 | 8.79323907872796e-43 | 0.589530956 | 0.886 | 0.611 | 2.21317034372504e-38 | 25 |
| cotl19 | 8.95079180889862e-43 | 0.814988873 | 0.727 | 0.302 | 2.25282479038169e-38 | 25 |
| rps1812 | 3.47984794826978e-42 | 0.46991033 | 0.926 | 0.723 | 8.7584293010002e-38 | 25 |
| rpl3113 | 4.19415118418166e-42 | 0.519205858 | 0.886 | 0.653 | 1.05562591154668e-37 | 25 |
| si:zfos-741a10.33 | 4.4463919624755e-42 | 0.411882655 | 0.193 | 0.025 | 1.11911239303546e-37 | 25 |
| tmsb4x15 | 4.68828063752216e-41 | 0.632896914 | 0.989 | 0.683 | 1.17999335365795e-36 | 25 |
| tyk2 | 8.31325314531822e-41 | 0.401379807 | 0.193 | 0.026 | 2.09236268414514e-36 | 25 |
| rpl612 | 2.46614057026938e-40 | 0.513036175 | 0.92 | 0.665 | 6.207029201311e-36 | 25 |
| si:dkey-222f2.72 | 1.18653078913851e-39 | 0.387334609 | 0.205 | 0.029 | 2.98637934318271e-35 | 25 |
| eomesa1 | 1.37337671380847e-39 | 0.487156818 | 0.239 | 0.038 | 3.45665185098455e-35 | 25 |
| CABZ01079663.14 | 3.97864986536287e-39 | 0.437987302 | 0.233 | 0.037 | 1.00138638461318e-34 | 25 |
| CR392341.14 | 6.55451886178808e-39 | 0.545499632 | 0.29 | 0.057 | 1.64970685232344e-34 | 25 |
| si:ch211-226m16.23 | 7.06226424208789e-39 | 0.586904039 | 0.312 | 0.066 | 1.7775012870911e-34 | 25 |
| rps26l12 | 8.96308973060162e-39 | 0.506315658 | 0.903 | 0.662 | 2.25592005429512e-34 | 25 |
| actn41 | 1.03811491111603e-38 | 0.588086005 | 0.375 | 0.091 | 2.61283141978794e-34 | 25 |
| hcls15 | 1.26237530710775e-38 | 0.564696579 | 0.318 | 0.068 | 3.17727241045949e-34 | 25 |
| si:rp71-36a1.51 | 1.53733977978188e-38 | 0.481090254 | 0.256 | 0.046 | 3.86933049173301e-34 | 25 |
| gnsb2 | 1.60948350314921e-38 | 0.478087441 | 0.227 | 0.037 | 4.05090902907625e-34 | 25 |
| slc25a57 | 5.57471477263727e-38 | 0.362317677 | 0.943 | 0.806 | 1.40309996112507e-33 | 25 |
| cap19 | 4.39683596425638e-37 | 0.748980516 | 0.597 | 0.23 | 1.10663964384369e-32 | 25 |
| sh2d3ca | 8.91878017322938e-37 | 0.364911677 | 0.165 | 0.021 | 2.2447677818001e-32 | 25 |
| XLOC-0108997 | 9.68140302432313e-37 | 0.796814645 | 0.659 | 0.254 | 2.43671232719189e-32 | 25 |
| scgn2 | 1.53043407231286e-36 | 0.340199399 | 0.17 | 0.022 | 3.85194951660423e-32 | 25 |
| rpl1411 | 1.89164824502015e-36 | 0.521054534 | 0.858 | 0.633 | 4.76108946789122e-32 | 25 |
| atp2b1a1 | 3.54491920395056e-36 | 0.546509066 | 0.324 | 0.071 | 8.92220714442318e-32 | 25 |
| rpl219 | 4.01960834746986e-36 | 0.395893843 | 0.92 | 0.755 | 1.01169522497469e-31 | 25 |
| cfl17 | 7.87337650458724e-36 | 0.599253914 | 0.835 | 0.575 | 1.98165013243956e-31 | 25 |
| rps4x9 | 8.45088395650297e-36 | 0.462793954 | 0.949 | 0.708 | 2.12700298301223e-31 | 25 |
| zgc:11418814 | 9.74761417249277e-36 | 0.536264526 | 0.92 | 0.625 | 2.45337701107471e-31 | 25 |
| btk | 1.84827475547352e-35 | 0.343691136 | 0.17 | 0.023 | 4.6519227320513e-31 | 25 |
| ponzr18 | 2.42468217667288e-35 | 0.728090985 | 0.653 | 0.269 | 6.10268257046798e-31 | 25 |

| | | | | | | |
|---|---|---|---|---|---|---|
| rpl5a11 | 3.03309878469887e-35 | 0.622366774 | 0.83 | 0.489 | 7.63400633120858e-31 | 25 |
| msna9 | 1.29857856663208e-34 | 0.724978026 | 0.585 | 0.239 | 3.26839239435629e-30 | 25 |
| st3gal4 | 1.56720461885508e-34 | 0.342238199 | 0.159 | 0.021 | 3.94449730519635e-30 | 25 |
| wdr14 | 1.7659338598952e-34 | 0.645998143 | 0.449 | 0.139 | 4.44467893197023e-30 | 25 |
| rnf19b | 1.36164202853733e-33 | 0.3330211 | 0.165 | 0.023 | 3.42711682162562e-29 | 25 |
| lamp1a3 | 1.70050138813365e-33 | 0.600503045 | 0.347 | 0.088 | 4.27999194379359e-29 | 25 |
| ubash3ba | 1.76351558282639e-33 | 0.355393758 | 0.165 | 0.023 | 4.43859237041574e-29 | 25 |
| rpl36a9 | 1.80393033069065e-33 | 0.363308824 | 0.972 | 0.808 | 4.54031224931529e-29 | 25 |
| il163 | 1.89042209948486e-33 | 0.482590084 | 0.244 | 0.048 | 4.75800338219344e-29 | 25 |
| CR936442.16 | 2.05047112449403e-33 | 0.711401351 | 0.511 | 0.166 | 5.16083077323901e-29 | 25 |
| slc7a3b | 3.02458794089228e-33 | 0.375720777 | 0.188 | 0.029 | 7.61258538843179e-29 | 25 |
| si:ch73-264p11.14 | 3.59270136826731e-33 | 0.348629874 | 0.176 | 0.026 | 9.04247007379199e-29 | 25 |
| rps238 | 5.88021238360162e-33 | 0.414605924 | 0.909 | 0.721 | 1.47999065482869e-28 | 25 |
| abo1 | 8.28723867320233e-33 | 0.345119534 | 0.148 | 0.019 | 2.0858151016583e-28 | 25 |
| per32 | 8.8839096486783e-33 | 0.522995554 | 0.318 | 0.077 | 2.23599121947584e-28 | 25 |
| il343 | 1.38666130538764e-32 | 0.476601172 | 0.244 | 0.047 | 3.49008783953015e-28 | 25 |
| samd111 | 2.51111742220576e-32 | 0.275293074 | 0.142 | 0.018 | 6.32023143994968e-28 | 25 |
| ptpn221 | 7.74231700043043e-32 | 0.332264699 | 0.159 | 0.022 | 1.94866376583834e-27 | 25 |
| plp22 | 8.78218334921694e-32 | 0.436852059 | 0.244 | 0.049 | 2.21038772716441e-27 | 25 |
| si:dkey-10h3.2 | 4.40097093607195e-31 | 0.293991749 | 0.136 | 0.017 | 1.10768037489995e-26 | 25 |
| capzb7 | 5.01055448805382e-31 | 0.671624437 | 0.585 | 0.25 | 1.26110645909827e-26 | 25 |
| far11 | 5.16185961649903e-31 | 0.410198872 | 0.199 | 0.034 | 1.29918844687664e-26 | 25 |
| tnfrsf9a4 | 9.25449549290498e-31 | 0.456822549 | 0.25 | 0.05 | 2.32926397060925e-26 | 25 |
| csk5 | 2.39978545109075e-30 | 0.569955989 | 0.358 | 0.099 | 6.0400200018503e-26 | 25 |
| pptc7b | 5.41982556750918e-30 | 0.366017037 | 0.188 | 0.032 | 1.36411589708639e-25 | 25 |
| tmc83 | 6.57359056959638e-30 | 0.398404579 | 0.193 | 0.034 | 1.65450701046171e-25 | 25 |
| wdfy2 | 6.62416065690219e-30 | 0.276623283 | 0.125 | 0.015 | 1.66723499573571e-25 | 25 |
| RASSF54 | 1.08279673860721e-29 | 0.490658518 | 0.256 | 0.057 | 2.72529111140048e-25 | 25 |
| zgc:1530181 | 1.23968193311312e-29 | 0.323403178 | 0.153 | 0.022 | 3.12015545745242e-25 | 25 |
| adcy7 | 2.04229229942129e-29 | 0.344730358 | 0.165 | 0.025 | 5.14024548841344e-25 | 25 |
| edf12 | 2.07689431358172e-29 | 0.671801445 | 0.597 | 0.265 | 5.22733529785384e-25 | 25 |
| megf8l1 | 3.71804874133961e-29 | 0.254304471 | 0.119 | 0.014 | 9.35795687707766e-25 | 25 |
| mir2232 | 8.20418890627372e-29 | 0.418753671 | 0.176 | 0.029 | 2.06491230582003e-24 | 25 |
| dock11 | 1.67311111959283e-28 | 0.325952105 | 0.159 | 0.024 | 4.21105337690319e-24 | 25 |
| XLOC-0203014 | 1.99553454397391e-28 | 0.52313969 | 0.324 | 0.083 | 5.02256089372794e-24 | 25 |

| | | | | | | |
|---|---|---|---|---|---|---|
| rpsa14 | 2.56953309693853e-28 | 0.385311903 | 0.932 | 0.751 | 6.46725785168458e-24 | 25 |
| grap2b6 | 3.55545233587299e-28 | 0.572047909 | 0.369 | 0.11 | 8.94871798415872e-24 | 25 |
| arhgdia7 | 6.88343575075065e-28 | 0.645799739 | 0.54 | 0.227 | 1.73249194410643e-23 | 25 |
| XLOC-019150 | 7.93308609822039e-28 | 0.315912508 | 0.153 | 0.023 | 1.99667844006109e-23 | 25 |
| aif1l2 | 8.04150075416371e-28 | 0.597749824 | 0.534 | 0.184 | 2.02396532481546e-23 | 25 |
| si:dkey-151g10.31 | 8.64881309398774e-28 | 0.307744175 | 0.125 | 0.016 | 2.17681976762577e-23 | 25 |
| ikzf13 | 9.88700816978278e-28 | 0.339854507 | 0.182 | 0.032 | 2.48846108625263e-23 | 25 |
| limd22 | 1.05815397752784e-27 | 0.482448428 | 0.278 | 0.068 | 2.66326774603982e-23 | 25 |
| orai1a1 | 1.38150180226499e-27 | 0.316135364 | 0.153 | 0.023 | 3.47710188612075e-23 | 25 |
| rpl914 | 1.90077741490859e-27 | 0.479906296 | 0.898 | 0.627 | 4.78406667558344e-23 | 25 |
| CABZ01111959.11 | 2.02894506538338e-27 | 0.307639068 | 0.159 | 0.025 | 5.10665183506343e-23 | 25 |
| cmip2 | 2.29912237944493e-27 | 0.500297379 | 0.261 | 0.063 | 5.78666111682495e-23 | 25 |
| tmem30aa1 | 2.42197290444448e-27 | 0.417504374 | 0.256 | 0.059 | 6.0958636031963e-23 | 25 |
| rgs135 | 3.49078514615464e-27 | 0.606994569 | 0.403 | 0.13 | 8.78595713435661e-23 | 25 |
| rpl2411 | 3.60346136228789e-27 | 0.367676399 | 0.96 | 0.775 | 9.06955190274239e-23 | 25 |
| si:cabz01074946.14 | 4.47733016398173e-27 | 0.47206713 | 0.25 | 0.058 | 1.12689922897256e-22 | 25 |
| lcp16 | 5.03325374784551e-27 | 0.523947234 | 0.312 | 0.086 | 1.26681963579524e-22 | 25 |
| zeb1b2 | 7.38241960123878e-27 | 0.316682693 | 0.159 | 0.026 | 1.85808118943579e-22 | 25 |
| ldha3 | 1.26899330732568e-26 | 0.498835457 | 0.278 | 0.071 | 3.19392925520801e-22 | 25 |
| slc1a5 | 2.74942953875635e-26 | 0.256715827 | 0.131 | 0.018 | 6.92003920609586e-22 | 25 |
| rps1411 | 8.45969191387149e-26 | 0.368532468 | 0.938 | 0.758 | 2.12921985780232e-21 | 25 |
| ap1b1 | 1.04112735233146e-25 | 0.400664179 | 0.227 | 0.05 | 2.62041343308306e-21 | 25 |
| nptna3 | 1.78776517995427e-25 | 0.441945996 | 0.278 | 0.07 | 4.4996261814269e-21 | 25 |
| dtnba | 1.90182095241193e-25 | 0.355399271 | 0.182 | 0.034 | 4.7866931551256e-21 | 25 |
| fryl | 1.91381115695754e-25 | 0.35759678 | 0.182 | 0.034 | 4.81687130094643e-21 | 25 |
| scinlb5 | 2.05324756323999e-25 | 0.455516424 | 0.301 | 0.081 | 5.16781879191874e-21 | 25 |
| ptmaa11 | 3.63908716621494e-25 | 0.620002793 | 0.727 | 0.392 | 9.15921848864637e-21 | 25 |
| susd6 | 3.90317365885423e-25 | 0.421316428 | 0.239 | 0.056 | 9.82389778197022e-21 | 25 |
| pnp5b | 6.85681313712124e-25 | 0.2987931 | 0.148 | 0.024 | 1.72579129848204e-20 | 25 |
| icn7 | 1.52540993290225e-24 | 0.70879212 | 0.608 | 0.272 | 3.83930426012168e-20 | 25 |
| ccr25 | 1.62538721840036e-24 | 0.391162364 | 0.21 | 0.045 | 4.09093708999187e-20 | 25 |
| si:dkey-262k9.46 | 1.75995253686904e-24 | 0.497445874 | 0.307 | 0.089 | 4.42962454004569e-20 | 25 |
| prf1.92 | 2.03785000114848e-24 | 0.304017058 | 0.125 | 0.018 | 5.12906466789061e-20 | 25 |
| rps1311 | 2.51577974912143e-24 | 0.384331818 | 0.92 | 0.723 | 6.33196605056373e-20 | 25 |
| slc6a6b4 | 2.62421891941368e-24 | 0.502104998 | 0.335 | 0.101 | 6.60489659827229e-20 | 25 |

| | | | | | | |
|---|---|---|---|---|---|---|
| atp5if1b2 | 3.17838661425488e-24 | 0.582752145 | 0.472 | 0.188 | 7.9996812694181e-20 | 25 |
| rac29 | 3.97119534856764e-24 | 0.566737908 | 0.545 | 0.218 | 9.9951015728099e-20 | 25 |
| btf39 | 4.16059370856713e-24 | 0.472942987 | 0.841 | 0.585 | 1.04717983050926e-19 | 25 |
| rab27a | 4.5857006736224e-24 | 0.351481152 | 0.21 | 0.046 | 1.15417500254402e-19 | 25 |
| ptp4a2b1 | 4.74565884843618e-24 | 0.553394022 | 0.415 | 0.148 | 1.1944348755629e-19 | 25 |
| rpl1510 | 5.39774397958182e-24 | 0.370987946 | 0.926 | 0.743 | 1.35855818222095e-19 | 25 |
| aars | 6.11643014730482e-24 | 0.426904035 | 0.267 | 0.07 | 1.53944430377515e-19 | 25 |
| myo1f4 | 1.01897428555996e-23 | 0.373617056 | 0.205 | 0.045 | 2.56465637932585e-19 | 25 |
| si:ch211-136m16.82 | 1.40786930529514e-23 | 0.338724238 | 0.159 | 0.029 | 3.54346625449733e-19 | 25 |
| ccdc171 | 1.4365861779964e-23 | 0.267318101 | 0.131 | 0.02 | 3.61574375139915e-19 | 25 |
| rpl136 | 1.70325991673542e-23 | 0.294469364 | 0.926 | 0.804 | 4.28693488443138e-19 | 25 |
| lcp2a1 | 3.68275422126726e-23 | 0.334761529 | 0.176 | 0.035 | 9.26912409950757e-19 | 25 |
| rplp18 | 6.57629104210201e-23 | 0.285181184 | 0.977 | 0.876 | 1.65518669238665e-18 | 25 |
| itgb25 | 6.87018677222138e-23 | 0.496922222 | 0.318 | 0.097 | 1.7291573087004e-18 | 25 |
| rpl7a9 | 7.76500820424909e-23 | 0.304680994 | 0.938 | 0.809 | 1.95437491492745e-18 | 25 |
| acap12 | 1.13196215497988e-22 | 0.326428772 | 0.188 | 0.039 | 2.84903554786886e-18 | 25 |
| ccni9 | 1.26606245478237e-22 | 0.589036686 | 0.585 | 0.292 | 3.18655259244175e-18 | 25 |
| adam17b | 1.40777379469613e-22 | 0.285073203 | 0.131 | 0.021 | 3.54322586387068e-18 | 25 |
| cox8a6 | 1.48502638396872e-22 | 0.58217819 | 0.5 | 0.225 | 3.73766290581087e-18 | 25 |
| CU972454.42 | 2.21483889805887e-22 | 0.266205563 | 0.136 | 0.022 | 5.57452802252436e-18 | 25 |
| mthfd2 | 2.31868117975397e-22 | 0.295877286 | 0.153 | 0.028 | 5.83588866132277e-18 | 25 |
| arpc5a6 | 2.43872725612626e-22 | 0.49097446 | 0.341 | 0.113 | 6.13803263094418e-18 | 25 |
| agpat3 | 2.58274845249536e-22 | 0.316402855 | 0.159 | 0.03 | 6.50051958008557e-18 | 25 |
| BX005183.36 | 3.4138615817337e-22 | 0.47278267 | 0.278 | 0.079 | 8.59234821506555e-18 | 25 |
| mt-atp613 | 3.95592006715335e-22 | 0.508255618 | 0.864 | 0.588 | 9.95665521701828e-18 | 25 |
| mibp29 | 4.65476861483044e-22 | 0.502514561 | 0.761 | 0.49 | 1.17155871266667e-17 | 25 |
| adam8a1 | 7.85631315075459e-22 | 0.364479115 | 0.216 | 0.051 | 1.97735545691342e-17 | 25 |
| rps2811 | 8.62596245341899e-22 | 0.306620265 | 0.966 | 0.825 | 2.17106848990102e-17 | 25 |
| rps2910 | 8.90636864562542e-22 | 0.286235159 | 0.943 | 0.831 | 2.24164392441746e-17 | 25 |
| XLOC-0004318 | 1.02728977226722e-21 | 0.536246704 | 0.653 | 0.275 | 2.58558562781936e-17 | 25 |
| arhgap1 | 1.1202192506093e-21 | 0.300402358 | 0.153 | 0.029 | 2.81947983185854e-17 | 25 |
| jak14 | 1.13030735193777e-21 | 0.479882909 | 0.301 | 0.094 | 2.84487057409216e-17 | 25 |
| hspa81 | 1.28885725676155e-21 | 0.270068453 | 0.926 | 0.828 | 3.24392482954314e-17 | 25 |
| glipr1a4 | 2.10372159889927e-21 | 0.399701017 | 0.239 | 0.063 | 5.29485689226957e-17 | 25 |
| kif21b1 | 2.20609022493162e-21 | 0.299694625 | 0.136 | 0.024 | 5.55250848713039e-17 | 25 |

| | | | | | | |
|---|---|---|---|---|---|---|
| arpc5b5 | 4.03022875964355e-21 | 0.483634601 | 0.335 | 0.112 | 1.01436827651469e-16 | 25 |
| arhgef6 | 7.51003456812858e-21 | 0.267935366 | 0.142 | 0.026 | 1.89020060045228e-16 | 25 |
| rps39 | 1.02367944211785e-20 | 0.367615022 | 0.903 | 0.698 | 2.57649878786643e-16 | 25 |
| zmp:0000000521 | 2.38678990060351e-20 | 0.29476356 | 0.125 | 0.021 | 6.00731150082899e-16 | 25 |
| cdaa2 | 2.41351866307194e-20 | 0.341953339 | 0.165 | 0.034 | 6.07458512308577e-16 | 25 |
| rpl2211 | 2.45837452169375e-20 | 0.372899796 | 0.886 | 0.693 | 6.18748283365101e-16 | 25 |
| actb13 | 2.77556583471896e-20 | 0.315455969 | 0.909 | 0.784 | 6.98582164940416e-16 | 25 |
| si:ch211-200p22.4 | 3.05041342254698e-20 | 0.28238471 | 0.136 | 0.025 | 7.67758554320849e-16 | 25 |
| XLOC-0139645 | 4.18549395048608e-20 | 0.51753968 | 0.455 | 0.172 | 1.05344697239784e-15 | 25 |
| mt-co114 | 4.62739722221265e-20 | 0.446178222 | 0.909 | 0.655 | 1.1646696068587e-15 | 25 |
| zyx | 6.9309401899603e-20 | 0.441850479 | 0.233 | 0.065 | 1.74444833641111e-15 | 25 |
| cd6312 | 7.2155419941627e-20 | 0.545285665 | 0.642 | 0.338 | 1.81607976451081e-15 | 25 |
| rel2 | 9.41531810939925e-20 | 0.354051819 | 0.165 | 0.035 | 2.3697414149547e-15 | 25 |
| zmp:00000010823 | 9.75225788677879e-20 | 0.388243791 | 0.216 | 0.056 | 2.45454578752335e-15 | 25 |
| elmo3 | 1.65157696801828e-19 | 0.305644802 | 0.148 | 0.029 | 4.15685407080521e-15 | 25 |
| arpc28 | 2.18090123180965e-19 | 0.539788959 | 0.54 | 0.265 | 5.48911031034171e-15 | 25 |
| rpl268 | 2.18456950179279e-19 | 0.269492553 | 0.955 | 0.84 | 5.49834297906227e-15 | 25 |
| XLOC-0418702 | 2.86085530741872e-19 | 0.335826388 | 0.142 | 0.028 | 7.20048672324217e-15 | 25 |
| rps1612 | 3.7649038203195e-19 | 0.34197747 | 0.915 | 0.736 | 9.47588642536216e-15 | 25 |
| ccl36.14 | 4.19785112008443e-19 | 0.467127535 | 0.324 | 0.1 | 1.05655714841405e-14 | 25 |
| rps2117 | 4.80678568409492e-19 | 0.335532739 | 0.909 | 0.749 | 1.20981988882985e-14 | 25 |
| rpl5b8 | 4.91910882651377e-19 | 0.403846832 | 0.858 | 0.628 | 1.23809050054525e-14 | 25 |
| rpl3812 | 5.02041811168443e-19 | 0.34675613 | 0.915 | 0.727 | 1.26358903452985e-14 | 25 |
| taok3a | 1.95255012417876e-18 | 0.299860849 | 0.148 | 0.031 | 4.91437340754552e-14 | 25 |
| si:ch211-8c17.31 | 2.45433494419827e-18 | 0.272167057 | 0.148 | 0.031 | 6.17731562105262e-14 | 25 |
| si:dkeyp-68b7.125 | 2.81248539686824e-18 | 0.37873613 | 0.222 | 0.062 | 7.07874449537769e-14 | 25 |
| rpl2910 | 3.51655886008843e-18 | 0.297762642 | 0.96 | 0.807 | 8.85082699495658e-14 | 25 |
| nbeal2 | 3.6433586684521e-18 | 0.26951343 | 0.125 | 0.023 | 9.1699694326271e-14 | 25 |
| pde4ba8 | 4.43826238935583e-18 | 0.489288477 | 0.358 | 0.138 | 1.11706626077697e-13 | 25 |
| fam120c3 | 4.63078870610356e-18 | 0.406674699 | 0.29 | 0.098 | 1.1655232094392e-13 | 25 |
| dusp16 | 4.65064038354833e-18 | 0.54773266 | 0.386 | 0.159 | 1.17051967813528e-13 | 25 |
| lrrfip1a2 | 5.03345127755242e-18 | 0.359298866 | 0.21 | 0.057 | 1.26686935204717e-13 | 25 |
| tpt17 | 6.07911746848425e-18 | 0.26559976 | 0.903 | 0.801 | 1.5300530756428e-13 | 25 |
| pik3cg1 | 1.11723103790711e-17 | 0.261490846 | 0.131 | 0.026 | 2.81195879930841e-13 | 25 |
| mef2d2 | 1.42513945116283e-17 | 0.374267659 | 0.239 | 0.072 | 3.58693348463172e-13 | 25 |

| | | | | | | |
|---|---|---|---|---|---|---|
| nsa210 | 1.5751390026737e-17 | 0.508895723 | 0.625 | 0.361 | 3.96446735582943e-13 | 25 |
| arpc38 | 1.6896475586832e-17 | 0.50777729 | 0.585 | 0.308 | 4.25267394044975e-13 | 25 |
| ppp1r185 | 2.01550892395157e-17 | 0.334992802 | 0.199 | 0.053 | 5.07283441069372e-13 | 25 |
| kdm2ba1 | 3.29561628116994e-17 | 0.299435136 | 0.159 | 0.037 | 8.29473661807663e-13 | 25 |
| rpl1811 | 3.49918966604417e-17 | 0.3001886 | 0.92 | 0.773 | 8.80711047046658e-13 | 25 |
| CR381673.15 | 6.65067047372358e-17 | 0.3386606 | 0.205 | 0.056 | 1.67390725153149e-12 | 25 |
| rack16 | 6.68437766079726e-17 | 0.278117559 | 0.875 | 0.744 | 1.68239101344606e-12 | 25 |
| si:dkeyp-110e4.6 | 6.91814499416113e-17 | 0.250899045 | 0.119 | 0.022 | 1.74122791358041e-12 | 25 |
| ipo71 | 8.35613464599865e-17 | 0.42997881 | 0.301 | 0.108 | 2.1031555290514e-12 | 25 |
| CABZ01044746.11 | 8.70370984599479e-17 | 0.419584455 | 0.301 | 0.103 | 2.19063673113843e-12 | 25 |
| id34 | 1.02588702471525e-16 | 0.52698091 | 0.46 | 0.209 | 2.5820550525058e-12 | 25 |
| serbp1a10 | 1.32868740929181e-16 | 0.448394767 | 0.756 | 0.48 | 3.34417334044655e-12 | 25 |
| inpp5d1 | 1.71882711465794e-16 | 0.281141681 | 0.148 | 0.033 | 4.32611596488256e-12 | 25 |
| CABZ01072614.16 | 4.44716003344196e-16 | 0.410540768 | 0.244 | 0.08 | 1.11930570881701e-11 | 25 |
| stxbp2 | 4.91118686093988e-16 | 0.329687115 | 0.199 | 0.056 | 1.23609662102996e-11 | 25 |
| cxcr4b7 | 4.91139878604181e-16 | 0.458012413 | 0.438 | 0.181 | 1.23614996045886e-11 | 25 |
| vdac24 | 5.70747177439809e-16 | 0.491247408 | 0.54 | 0.294 | 1.43651357089825e-11 | 25 |
| oaz1b10 | 6.48569445974131e-16 | 0.480154158 | 0.568 | 0.321 | 1.63238443857229e-11 | 25 |
| sat1a.24 | 8.78218528713443e-16 | 0.503021686 | 0.54 | 0.297 | 2.21038821491886e-11 | 25 |
| ankrd441 | 1.02872708876126e-15 | 0.257047901 | 0.131 | 0.028 | 2.58920320970321e-11 | 25 |
| il10ra3 | 1.36082859075112e-15 | 0.317145014 | 0.159 | 0.04 | 3.42506948006148e-11 | 25 |
| COX314 | 1.42590070687637e-15 | 0.401528905 | 0.943 | 0.698 | 3.58884948913713e-11 | 25 |
| myo18ab | 2.09521234306817e-15 | 0.25657539 | 0.148 | 0.035 | 5.27343994626829e-11 | 25 |
| abracl3 | 2.20031741205388e-15 | 0.465943756 | 0.472 | 0.22 | 5.53797889439841e-11 | 25 |
| plekhj11 | 3.01957946502641e-15 | 0.317507485 | 0.188 | 0.053 | 7.59997955552498e-11 | 25 |
| ostf12 | 3.53958627696796e-15 | 0.293357141 | 0.188 | 0.053 | 8.90878470050066e-11 | 25 |
| wasa5 | 3.70967005406175e-15 | 0.367744476 | 0.267 | 0.092 | 9.33686855906802e-11 | 25 |
| rps27.111 | 5.32108329558478e-15 | 0.364498985 | 0.898 | 0.678 | 1.33926345466573e-10 | 25 |
| fam49ba4 | 5.55089176631625e-15 | 0.385988963 | 0.284 | 0.102 | 1.39710394866414e-10 | 25 |
| reep53 | 7.33325591036253e-15 | 0.42054506 | 0.364 | 0.157 | 1.84570718007914e-10 | 25 |
| dnajc5gb | 8.43293780538904e-15 | 0.251904268 | 0.136 | 0.031 | 2.12248611623837e-10 | 25 |
| casc4 | 8.6004792770161e-15 | 0.276940647 | 0.153 | 0.038 | 2.16465462923218e-10 | 25 |
| iqgap11 | 1.01156477556104e-14 | 0.33566255 | 0.216 | 0.068 | 2.54600738360959e-10 | 25 |
| tomm20b5 | 1.20040828680827e-14 | 0.447553126 | 0.438 | 0.213 | 3.02130761706774e-10 | 25 |
| rplp213 | 1.38034871760615e-14 | 0.334951533 | 0.955 | 0.756 | 3.47419968734291e-10 | 25 |

| | | | | | | |
|---|---|---|---|---|---|---|
| selplg4 | 1.79932322910786e-14 | 0.264992402 | 0.17 | 0.046 | 4.52871663534157e-10 | 25 |
| NC-002333.412 | 2.42522737643368e-14 | 0.398069344 | 0.869 | 0.621 | 6.10405478374594e-10 | 25 |
| prkar1aa | 2.58020292381543e-14 | 0.332584953 | 0.193 | 0.058 | 6.49411273895105e-10 | 25 |
| snx27a1 | 3.20831693168409e-14 | 0.263199192 | 0.153 | 0.04 | 8.07501288535569e-10 | 25 |
| metrnla4 | 9.01793589537247e-14 | 0.315017025 | 0.261 | 0.092 | 2.2697242855063e-09 | 25 |
| XLOC-0268364 | 1.12689073486718e-13 | 0.37788236 | 0.233 | 0.078 | 2.83627129058721e-09 | 25 |
| eef1db8 | 1.54683467227783e-13 | 0.468325129 | 0.489 | 0.273 | 3.89322818665607e-09 | 25 |
| rps510 | 1.7791577584298e-13 | 0.320570146 | 0.886 | 0.7 | 4.47796216219197e-09 | 25 |
| arpc47 | 1.89867956176865e-13 | 0.434773499 | 0.455 | 0.225 | 4.77878658901551e-09 | 25 |
| ncaldb4 | 2.3196829592834e-13 | 0.316792509 | 0.222 | 0.074 | 5.83841004022039e-09 | 25 |
| calr3b1 | 2.97762316291639e-13 | 0.3798269 | 0.222 | 0.076 | 7.49437973874425e-09 | 25 |
| rps1512 | 2.98528894662532e-13 | 0.330626513 | 0.881 | 0.683 | 7.51367374976127e-09 | 25 |
| rsl24d111 | 3.7572510578141e-13 | 0.420833625 | 0.676 | 0.443 | 9.45662518741232e-09 | 25 |
| rps27a10 | 4.62417084641152e-13 | 0.277536034 | 0.903 | 0.751 | 1.16385756033332e-08 | 25 |
| cct85 | 5.3254187094438e-13 | 0.41684987 | 0.409 | 0.201 | 1.34035463497991e-08 | 25 |
| calm1b6 | 6.76095945745755e-13 | 0.343819983 | 0.256 | 0.094 | 1.70166588584749e-08 | 25 |
| skap24 | 7.82959140106436e-13 | 0.329649949 | 0.216 | 0.074 | 1.97062985973389e-08 | 25 |
| cbfb1 | 9.33627064253194e-13 | 0.270092426 | 0.148 | 0.041 | 2.34984595801887e-08 | 25 |
| zmynd111 | 1.02905928613456e-12 | 0.298284543 | 0.188 | 0.058 | 2.59003931727208e-08 | 25 |
| sh3bgrl5 | 1.30851787533801e-12 | 0.382682825 | 0.261 | 0.104 | 3.29340864043823e-08 | 25 |
| NC-002333.17 | 1.38721653901204e-12 | 0.340465829 | 0.176 | 0.055 | 3.49148530703941e-08 | 25 |
| hmha1b4 | 1.52085750432543e-12 | 0.313532102 | 0.216 | 0.074 | 3.82784625263666e-08 | 25 |
| grb2b1 | 1.5651042222883e-12 | 0.360631674 | 0.21 | 0.073 | 3.93921081707742e-08 | 25 |
| ero1b | 2.17798502958884e-12 | 0.268984028 | 0.142 | 0.038 | 5.48177052097215e-08 | 25 |
| rpl35a11 | 2.62187664267875e-12 | 0.260097965 | 0.886 | 0.759 | 6.59900132195815e-08 | 25 |
| nr4a17 | 2.9427461762606e-12 | 0.40682594 | 0.364 | 0.166 | 7.40659785103031e-08 | 25 |
| eif4ebp23 | 3.01150610271984e-12 | 0.409907845 | 0.364 | 0.174 | 7.57965970993557e-08 | 25 |
| celf211 | 4.3576971860219e-12 | 0.426904548 | 0.449 | 0.241 | 1.09678880474985e-07 | 25 |
| plekho23 | 4.58067224511978e-12 | 0.322212743 | 0.21 | 0.074 | 1.1529093973742e-07 | 25 |
| cox6b17 | 5.27923066808512e-12 | 0.374158498 | 0.284 | 0.12 | 1.32872956685034e-07 | 25 |
| myl12.18 | 5.52466491456238e-12 | 0.415284137 | 0.602 | 0.391 | 1.39050291234621e-07 | 25 |
| clic16 | 5.84959023372007e-12 | 0.371026284 | 0.273 | 0.113 | 1.472283365925e-07 | 25 |
| dub4 | 5.88288826818282e-12 | 0.307452753 | 0.205 | 0.071 | 1.48066414821893e-07 | 25 |
| XLOC-0425964 | 9.11165167895258e-12 | 0.252766112 | 0.148 | 0.042 | 2.29331161107557e-07 | 25 |
| crlf3 | 9.41154404446707e-12 | 0.254253498 | 0.17 | 0.053 | 2.36879152055192e-07 | 25 |

| | | | | | | |
|---|---|---|---|---|---|---|
| rpl3412 | 9.91592413089125e-12 | 0.254103047 | 0.909 | 0.783 | 2.49573894450402e-07 | 25 |
| tpi1b6 | 1.4442532354841e-11 | 0.390267103 | 0.347 | 0.166 | 3.63504096838992e-07 | 25 |
| LOC110438237 | 1.66916347917683e-11 | 0.26070114 | 0.17 | 0.053 | 4.20111756074016e-07 | 25 |
| eno1a5 | 1.68749146714357e-11 | 0.347571797 | 0.307 | 0.134 | 4.24724727365365e-07 | 25 |
| per1b1 | 1.72989574861005e-11 | 0.304599124 | 0.193 | 0.067 | 4.35397460967664e-07 | 25 |
| elmo15 | 2.74322804672802e-11 | 0.308185591 | 0.199 | 0.07 | 6.90443067080974e-07 | 25 |
| mt-co213 | 3.10474039634543e-11 | 0.35835946 | 0.949 | 0.718 | 7.81432110356181e-07 | 25 |
| larp4ab1 | 3.30033517156703e-11 | 0.284866781 | 0.199 | 0.07 | 8.30661359331706e-07 | 25 |
| rps177 | 3.367297188769e-11 | 0.25549135 | 0.909 | 0.766 | 8.4751502944127e-07 | 25 |
| tnfaip34 | 3.39468979423542e-11 | 0.301807272 | 0.188 | 0.065 | 8.54409474311113e-07 | 25 |
| vsir7 | 4.21646839644323e-11 | 0.342534335 | 0.222 | 0.085 | 1.0612429307008e-06 | 25 |
| abca1a3 | 4.30091178298648e-11 | 0.320463584 | 0.193 | 0.068 | 1.08249648665987e-06 | 25 |
| atp10a4 | 4.6921794305055e-11 | 0.279239458 | 0.17 | 0.056 | 1.18097464086393e-06 | 25 |
| tnip14 | 5.4898597557529e-11 | 0.318475887 | 0.216 | 0.081 | 1.38174280192545e-06 | 25 |
| pum1 | 6.06340626691395e-11 | 0.257776501 | 0.182 | 0.061 | 1.52609872331957e-06 | 25 |
| birc23 | 7.05419173787309e-11 | 0.342109666 | 0.256 | 0.107 | 1.77546951850528e-06 | 25 |
| arpc4l7 | 7.45013042512347e-11 | 0.409349089 | 0.375 | 0.196 | 1.87512332669933e-06 | 25 |
| eef1g9 | 1.080038113305e-10 | 0.327000495 | 0.824 | 0.619 | 2.71834792737735e-06 | 25 |
| elovl1b3 | 1.18172359169439e-10 | 0.346271723 | 0.267 | 0.116 | 2.9742801079356e-06 | 25 |
| rpl23a13 | 1.259470924719e-10 | 0.29811402 | 0.807 | 0.633 | 3.16996237042525e-06 | 25 |
| rpl3613 | 1.32607288165485e-10 | 0.267693089 | 0.943 | 0.789 | 3.3375928358371e-06 | 25 |
| rplp2l13 | 1.34016148989589e-10 | 0.270141784 | 0.932 | 0.773 | 3.37305245391897e-06 | 25 |
| tefb | 1.37813405430823e-10 | 0.282814146 | 0.188 | 0.067 | 3.46862560128838e-06 | 25 |
| hnrnpa1a5 | 1.40375837634783e-10 | 0.396231828 | 0.324 | 0.159 | 3.53311945742985e-06 | 25 |
| eif3d6 | 1.80587703276103e-10 | 0.394114173 | 0.443 | 0.252 | 4.54521190375623e-06 | 25 |
| si:ch211-222l21.110 | 1.85147351758009e-10 | 0.398300307 | 0.489 | 0.274 | 4.65997369639734e-06 | 25 |
| si:dkeyp-19e1.33 | 2.20277721486378e-10 | 0.297255646 | 0.188 | 0.067 | 5.54416997209066e-06 | 25 |
| arrb2b5 | 2.52400031601644e-10 | 0.311282921 | 0.227 | 0.091 | 6.35265639538179e-06 | 25 |
| XLOC-0425956 | 3.08524495035392e-10 | 0.297687534 | 0.364 | 0.164 | 7.76525301554578e-06 | 25 |
| chac15 | 3.31033056480769e-10 | 0.400379515 | 0.574 | 0.398 | 8.33177099856447e-06 | 25 |
| rpl2811 | 3.49653654113119e-10 | 0.256981106 | 0.903 | 0.748 | 8.8004328203731e-06 | 25 |
| capza1b4 | 6.46303817233951e-10 | 0.374550383 | 0.392 | 0.212 | 1.62668207759613e-05 | 25 |
| zgc:638313 | 7.35012901669771e-10 | 0.318189082 | 0.205 | 0.081 | 1.84995397221265e-05 | 25 |
| pdia37 | 1.12929287377573e-09 | 0.391151028 | 0.472 | 0.283 | 2.84231723400613e-05 | 25 |
| fosl27 | 1.22363757732832e-09 | 0.345638125 | 0.267 | 0.121 | 3.07977341837766e-05 | 25 |

| | | | | | | |
|---|---|---|---|---|---|---|
| dusp25 | 1.35386768030041e-09 | 0.405439525 | 0.352 | 0.179 | 3.40754956454811e-05 | 25 |
| xbp14 | 1.58857765349088e-09 | 0.380401323 | 0.545 | 0.343 | 3.99829109607119e-05 | 25 |
| eif3c5 | 1.63961910172724e-09 | 0.344443137 | 0.33 | 0.168 | 4.12675731713728e-05 | 25 |
| hif1ab3 | 1.81089099243776e-09 | 0.344202457 | 0.25 | 0.113 | 4.5578315388666e-05 | 25 |
| grap2a4 | 2.18354739830496e-09 | 0.255793409 | 0.142 | 0.047 | 5.49577044679375e-05 | 25 |
| pik3r14 | 3.05457832804905e-09 | 0.295174895 | 0.216 | 0.09 | 7.68806819386665e-05 | 25 |
| tefa5 | 5.00819507584588e-09 | 0.369250906 | 0.278 | 0.138 | 0.000126051 | 25 |
| si:ch211-191c10.15 | 5.45860531793591e-09 | 0.267166422 | 0.153 | 0.054 | 0.000137388 | 25 |
| cct6a1 | 5.95570480610117e-09 | 0.336745827 | 0.381 | 0.207 | 0.000149899 | 25 |
| zgc:1538674 | 6.4142147231944e-09 | 0.344507834 | 0.614 | 0.436 | 0.000161439 | 25 |
| tmem1311 | 6.84319117581035e-09 | 0.25321383 | 0.17 | 0.063 | 0.000172236 | 25 |
| CABZ01090890.1 | 7.21784984290552e-09 | 0.256181109 | 0.153 | 0.055 | 0.000181666 | 25 |
| pgam1a4 | 9.27759811906606e-09 | 0.329117174 | 0.312 | 0.157 | 0.000233508 | 25 |
| rab445 | 9.49605258577367e-09 | 0.267609279 | 0.165 | 0.061 | 0.000239006 | 25 |
| sh2b33 | 1.11732908158552e-08 | 0.251663042 | 0.165 | 0.061 | 0.000281221 | 25 |
| rhogb5 | 1.63663258416876e-08 | 0.286157467 | 0.222 | 0.097 | 0.000411924 | 25 |
| lasp16 | 1.86152746155359e-08 | 0.334775579 | 0.409 | 0.236 | 0.000468528 | 25 |
| rhoac1 | 1.93658264206383e-08 | 0.336075792 | 0.301 | 0.155 | 0.000487418 | 25 |
| pnrc213 | 2.04183298461035e-08 | 0.334103843 | 0.824 | 0.566 | 0.000513909 | 25 |
| mapre1a5 | 3.17728088112985e-08 | 0.309367057 | 0.375 | 0.197 | 0.00079969 | 25 |
| tln14 | 5.59997548358305e-08 | 0.281588324 | 0.261 | 0.128 | 0.001409458 | 25 |
| mt-nd410 | 7.00691642474339e-08 | 0.321635565 | 0.67 | 0.455 | 0.001763571 | 25 |
| eif3s104 | 7.86351581643598e-08 | 0.32053707 | 0.307 | 0.165 | 0.001979168 | 25 |
| sh3bgrl34 | 9.47143268365682e-08 | 0.290290852 | 0.29 | 0.147 | 0.002383865 | 25 |
| crema6 | 1.43327643883306e-07 | 0.334868463 | 0.324 | 0.178 | 0.003607413 | 25 |
| eif2s1b3 | 1.49050509840635e-07 | 0.292771656 | 0.318 | 0.173 | 0.003751452 | 25 |
| mrps155 | 1.80893428428473e-07 | 0.28911134 | 0.256 | 0.125 | 0.004552907 | 25 |
| psap4 | 1.98751393187879e-07 | 0.315069058 | 0.483 | 0.29 | 0.005002374 | 25 |
| pkma4 | 4.19633687901393e-07 | 0.324224298 | 0.312 | 0.178 | 0.01056176 | 25 |
| ptmab9 | 4.32013194071174e-07 | 0.277325806 | 0.699 | 0.54 | 0.01087334 | 25 |
| cox4i17 | 5.29785220948282e-07 | 0.300779312 | 0.568 | 0.391 | 0.013334164 | 25 |
| eif3i7 | 5.60649719553344e-07 | 0.325554188 | 0.42 | 0.273 | 0.014110993 | 25 |
| eif4a1b5 | 9.51401723363929e-07 | 0.307119571 | 0.432 | 0.28 | 0.02394583 | 25 |
| ist1 | 9.59634234234952e-07 | 0.265676964 | 0.205 | 0.096 | 0.024153034 | 25 |
| gapdhs12 | 2.43041426925192e-06 | 0.290309195 | 0.443 | 0.289 | 0.061171097 | 25 |

| | | | | | | |
|---|---|---|---|---|---|---|
| BX248082.15 | 2.51235725096791e-06 | 0.263396807 | 0.114 | 0.042 | 0.06323352 | 25 |
| slc3a2b6 | 3.82151400423241e-06 | 0.269338978 | 0.227 | 0.118 | 0.096183686 | 25 |
| ip6k2b6 | 4.18075656149565e-06 | 0.264111829 | 0.233 | 0.123 | 0.105225462 | 25 |
| ola12 | 4.4301806730078e-06 | 0.28693603 | 0.364 | 0.23 | 0.111503217 | 25 |
| eif3ba4 | 4.73601456494012e-06 | 0.280091001 | 0.341 | 0.21 | 0.119200751 | 25 |
| atf4a2 | 5.43932033215706e-06 | 0.270400144 | 0.312 | 0.182 | 0.136902253 | 25 |
| rnf444 | 5.9612015280774e-06 | 0.26183709 | 0.182 | 0.09 | 0.150037481 | 25 |
| cct25 | 8.13496677330073e-06 | 0.268007836 | 0.364 | 0.227 | 0.204748979 | 25 |
| cdc42l5 | 1.08009957338562e-05 | 0.252429444 | 0.352 | 0.216 | 0.271850262 | 25 |
| idh25 | 1.62582897495819e-05 | 0.259148836 | 0.324 | 0.199 | 0.409204895 | 25 |
| eif3ea7 | 1.80087448969408e-05 | 0.290147778 | 0.403 | 0.277 | 0.4532621 | 25 |
| mt-nd111 | 2.30202576361648e-05 | 0.270447975 | 0.568 | 0.411 | 0.579396864 | 25 |
| ywhabl3 | 2.4920589606762e-05 | 0.272085382 | 0.403 | 0.279 | 0.62722632 | 25 |
| csde12 | 3.30056976586838e-05 | 0.274466103 | 0.324 | 0.208 | 0.830720404 | 25 |
| serf29 | 4.43366901026052e-05 | 0.263547667 | 0.472 | 0.343 | 1 | 25 |
| mt-nd310 | 6.36374004827784e-05 | 0.259458716 | 0.534 | 0.396 | 1 | 25 |
| h1fx6 | 0.000125544 | 0.256642174 | 0.295 | 0.184 | 1 | 25 |
| ccl39.72 | 0.000456399 | 0.347215412 | 0.119 | 0.06 | 1 | 25 |
| CR383676.113 | 0.000760821 | 0.310118869 | 0.835 | 0.726 | 1 | 25 |
| cavin1b3 | 0 | 1.406794986 | 0.703 | 0.031 | 0 | 26 |
| podxl2 | 0 | 1.401874327 | 0.669 | 0.038 | 0 | 26 |
| cldnc2 | 0 | 1.391427372 | 0.606 | 0.02 | 0 | 26 |
| s100a10a | 0 | 1.247805186 | 0.463 | 0.005 | 0 | 26 |
| gcga | 0 | 1.169485096 | 0.44 | 0.004 | 0 | 26 |
| krt153 | 9.76369561214168e-302 | 1.820237152 | 0.829 | 0.074 | 2.45742454861994e-297 | 26 |
| krt42 | 1.89601756564401e-301 | 1.386004149 | 0.594 | 0.033 | 4.7720866109694e-297 | 26 |
| gstm.34 | 5.85217217004663e-285 | 1.78360787 | 0.874 | 0.09 | 1.47293321347904e-280 | 26 |
| ucmaa | 7.01401078854285e-271 | 0.769561748 | 0.303 | 0.007 | 1.76535637536835e-266 | 26 |
| scn4ba | 4.70646424721372e-248 | 0.391419921 | 0.177 | 0.002 | 1.18456998638122e-243 | 26 |
| si:ch211-156j16.11 | 5.15827062783738e-247 | 0.769359249 | 0.389 | 0.016 | 1.29828513432039e-242 | 26 |
| cyp2ad2 | 2.66610514368758e-244 | 0.504317833 | 0.206 | 0.003 | 6.71032003614727e-240 | 26 |
| fgf24 | 3.27584926766364e-224 | 0.304814714 | 0.149 | 0.001 | 8.24498502178262e-220 | 26 |
| slc29a1b | 8.55041173230172e-207 | 0.550274099 | 0.229 | 0.006 | 2.15205312890302e-202 | 26 |
| fn1b4 | 3.75947134201389e-195 | 1.369232708 | 0.709 | 0.079 | 9.46221342071476e-191 | 26 |
| itgb41 | 5.69395582342758e-186 | 0.639429766 | 0.303 | 0.013 | 1.43311174119849e-181 | 26 |

| | | | | | | |
|---|---|---|---|---|---|---|
| fsta | 8.63800025040583e-181 | 0.334494029 | 0.149 | 0.002 | 2.17409828302464e-176 | 26 |
| rap1gap | 2.5575437579237e-179 | 0.488827998 | 0.217 | 0.006 | 6.43708188431816e-175 | 26 |
| pdlim3b | 4.99595622924002e-176 | 0.351749381 | 0.171 | 0.003 | 1.25743222333742e-171 | 26 |
| eppk13 | 4.43410405425905e-175 | 0.907819369 | 0.463 | 0.035 | 1.11601964941646e-170 | 26 |
| frya | 1.80649558181396e-172 | 0.476830995 | 0.217 | 0.006 | 4.54676872986757e-168 | 26 |
| sccpdha | 3.27834201392506e-169 | 0.366004834 | 0.166 | 0.003 | 8.25125901484799e-165 | 26 |
| cav13 | 4.49950108746546e-169 | 0.963880766 | 0.474 | 0.038 | 1.13247942870418e-164 | 26 |
| aldh1a23 | 9.22966364488624e-161 | 1.103869678 | 0.629 | 0.071 | 2.32301404278142e-156 | 26 |
| LOC110438169 | 7.99590682597683e-160 | 0.283557824 | 0.126 | 0.002 | 2.01248978903011e-155 | 26 |
| sox62 | 1.21557557221953e-158 | 0.747709863 | 0.366 | 0.024 | 3.05948215771932e-154 | 26 |
| postnb4 | 9.86047050487369e-157 | 1.251002774 | 0.611 | 0.071 | 2.48178182137166e-152 | 26 |
| bnc22 | 3.39315700164812e-156 | 0.732771219 | 0.389 | 0.027 | 8.54023685744815e-152 | 26 |
| krt18a.15 | 4.73762127633382e-152 | 1.338180916 | 0.766 | 0.118 | 1.19241189904046e-147 | 26 |
| krt946 | 1.00153035155881e-149 | 1.338638478 | 0.771 | 0.121 | 2.52075174183836e-145 | 26 |
| wt1a2 | 3.90597266802764e-149 | 0.718059122 | 0.394 | 0.029 | 9.83094260815878e-145 | 26 |
| cygb14 | 1.34154590435194e-148 | 0.972925497 | 0.497 | 0.048 | 3.37653688666339e-144 | 26 |
| si:ch211-105c13.33 | 6.82571578619274e-147 | 0.943216054 | 0.531 | 0.055 | 1.71796440622685e-142 | 26 |
| si:ch211-137i24.10 | 6.10606607725609e-146 | 0.460874026 | 0.206 | 0.007 | 1.53683577098459e-141 | 26 |
| col6a4a3 | 4.44597818943221e-145 | 0.85952033 | 0.457 | 0.041 | 1.11900825049819e-140 | 26 |
| cavin2b5 | 8.05164745856822e-145 | 1.020925268 | 0.537 | 0.06 | 2.02651914884704e-140 | 26 |
| krt89 | 4.29767090881807e-139 | 1.467236487 | 0.966 | 0.267 | 1.08168079104042e-134 | 26 |
| sfrp1a3 | 3.6353783177938e-138 | 0.696841699 | 0.349 | 0.025 | 9.14988368805522e-134 | 26 |
| krt914 | 5.49206634315996e-136 | 1.222429182 | 0.537 | 0.062 | 1.38229817790993e-131 | 26 |
| FP016056.11 | 1.95551185455695e-133 | 0.365153934 | 0.194 | 0.007 | 4.92182778673439e-129 | 26 |
| kcnq5a | 2.49781638817893e-133 | 0.351624946 | 0.154 | 0.004 | 6.28675406740755e-129 | 26 |
| anxa1a6 | 4.87012529028424e-131 | 1.352728846 | 0.743 | 0.135 | 1.22576183431164e-126 | 26 |
| plod22 | 2.7131391260045e-130 | 0.605313197 | 0.309 | 0.02 | 6.82869986624072e-126 | 26 |
| cfd3 | 4.89919540074106e-130 | 1.142243051 | 0.526 | 0.062 | 1.23307849041252e-125 | 26 |
| si:ch211-250c4.41 | 1.2736811822041e-129 | 0.442307255 | 0.246 | 0.012 | 3.20572816748949e-125 | 26 |
| usp43b1 | 7.77120371742541e-129 | 0.509783127 | 0.263 | 0.014 | 1.9559342636388e-124 | 26 |
| frzb2 | 3.84418117594047e-128 | 0.635438996 | 0.269 | 0.015 | 9.67541960172458e-124 | 26 |
| tmsb15 | 6.6901605083094e-123 | 1.316545808 | 0.76 | 0.148 | 1.68384649833639e-118 | 26 |
| hspb14 | 1.11924965201107e-121 | 0.882844011 | 0.497 | 0.058 | 2.81703944914666e-117 | 26 |
| si:ch211-106h4.125 | 9.18926752192685e-121 | 1.153569374 | 0.469 | 0.053 | 2.31284674259377e-116 | 26 |
| CU984579.1 | 3.84014906219876e-118 | 0.566990487 | 0.246 | 0.014 | 9.66527117464805e-114 | 26 |

| | | | | | | |
|---|---|---|---|---|---|---|
| dkk3b3 | 7.99413980850577e-118 | 0.694122222 | 0.389 | 0.036 | 2.01204504840282e-113 | 26 |
| paplna2 | 1.06790992945247e-117 | 0.572538084 | 0.291 | 0.02 | 2.68782250143892e-113 | 26 |
| si:ch211-63p21.1 | 2.34123182305027e-115 | 0.351323887 | 0.154 | 0.005 | 5.89264637543521e-111 | 26 |
| anxa2a5 | 2.81729122708518e-113 | 1.097261548 | 0.617 | 0.1 | 7.09084028945069e-109 | 26 |
| c1qtnf12 | 6.21119379333123e-112 | 0.27945026 | 0.143 | 0.004 | 1.56329536584354e-107 | 26 |
| tmem981 | 6.04265267655924e-110 | 0.388869041 | 0.206 | 0.01 | 1.5208752521632e-105 | 26 |
| slc43a3b2 | 2.31713911841442e-109 | 0.583551034 | 0.291 | 0.022 | 5.83200744713726e-105 | 26 |
| s100a10b8 | 5.79714742951404e-109 | 1.374326279 | 0.874 | 0.248 | 1.45908403653439e-104 | 26 |
| mmp25 | 1.74534608601277e-105 | 0.91942047 | 0.56 | 0.086 | 4.39286156388555e-101 | 26 |
| lurap1 | 6.49381984855987e-104 | 0.278631132 | 0.143 | 0.005 | 1.63442951768403e-99 | 26 |
| cldn5a4 | 2.80135028566189e-103 | 0.753171023 | 0.406 | 0.045 | 7.05071853398241e-99 | 26 |
| col1a1b5 | 4.77246758605047e-103 | 1.16228753 | 0.709 | 0.14 | 1.20118236673304e-98 | 26 |
| slc43a1b | 2.8032011445174e-102 | 0.413848322 | 0.206 | 0.011 | 7.05537696063586e-98 | 26 |
| col1a25 | 3.04867058364993e-102 | 1.157106223 | 0.743 | 0.148 | 7.67319899198852e-98 | 26 |
| si:ch73-46j18.57 | 3.16345518279144e-102 | 1.128240617 | 0.749 | 0.168 | 7.96210034956777e-98 | 26 |
| jam2b1 | 1.15648737559805e-101 | 0.477123563 | 0.251 | 0.017 | 2.91076307564274e-97 | 26 |
| col1a1a5 | 4.00577700469717e-101 | 1.122977162 | 0.697 | 0.133 | 1.00821401431223e-96 | 26 |
| cebpd7 | 4.61552959191868e-101 | 1.060152055 | 0.651 | 0.123 | 1.16168264299001e-96 | 26 |
| ehd2b1 | 1.24775766854633e-97 | 0.542181203 | 0.309 | 0.027 | 3.14048127596427e-93 | 26 |
| c64 | 4.85148712516327e-97 | 0.901737289 | 0.566 | 0.089 | 1.22107079453234e-92 | 26 |
| im:71514492 | 1.63275080264752e-96 | 0.509899213 | 0.291 | 0.024 | 4.10947049518354e-92 | 26 |
| zfp36l1b6 | 1.0465665700938e-95 | 1.025391757 | 0.646 | 0.124 | 2.63410340026908e-91 | 26 |
| ptprq | 2.57445044478635e-95 | 0.282408062 | 0.114 | 0.003 | 6.47963432448276e-91 | 26 |
| ppdpfa3 | 2.43063915056681e-93 | 0.851499005 | 0.486 | 0.071 | 6.1176756780616e-89 | 26 |
| adh8a4 | 2.88320246077678e-92 | 0.809505051 | 0.366 | 0.041 | 7.25673227352907e-88 | 26 |
| col6a35 | 1.62043911404031e-91 | 0.926321454 | 0.56 | 0.095 | 4.07848320612805e-87 | 26 |
| col4a14 | 2.27728816636927e-91 | 0.766720416 | 0.423 | 0.054 | 5.73170658593481e-87 | 26 |
| endouc3 | 1.42599593998352e-88 | 0.777417927 | 0.434 | 0.058 | 3.58908918134451e-84 | 26 |
| icn8 | 3.14972004435036e-87 | 1.2198711 | 0.88 | 0.267 | 7.92753037962543e-83 | 26 |
| gata62 | 3.5943078239466e-87 | 0.422317502 | 0.229 | 0.017 | 9.04651336209121e-83 | 26 |
| hsd11b26 | 2.35472365105964e-86 | 1.025880922 | 0.629 | 0.117 | 5.92660395735202e-82 | 26 |
| efna1b | 1.55814327084025e-84 | 0.407211366 | 0.217 | 0.015 | 3.92169079837782e-80 | 26 |
| bmp2b | 1.97892984737765e-83 | 0.263187262 | 0.137 | 0.006 | 4.98076853286481e-79 | 26 |
| nfia6 | 7.14301440768301e-83 | 0.744972566 | 0.474 | 0.075 | 1.79782529626974e-78 | 26 |
| cdh24 | 1.71335387015959e-82 | 0.682596384 | 0.389 | 0.052 | 4.31234035580467e-78 | 26 |

| | | | | | | |
|---|---|---|---|---|---|---|
| zgc:1124372 | 7.0243568306535e-82 | 0.354791708 | 0.211 | 0.015 | 1.76796037070718e-77 | 26 |
| smoc1 | 2.81472004157943e-81 | 0.278290044 | 0.143 | 0.007 | 7.08436887265127e-77 | 26 |
| sparc6 | 2.89131305388048e-81 | 0.977516851 | 0.686 | 0.152 | 7.27714582531178e-77 | 26 |
| emilin1a2 | 1.91213893380375e-80 | 0.400006469 | 0.223 | 0.017 | 4.81266248249067e-76 | 26 |
| pleca3 | 2.67598263734407e-80 | 0.714047072 | 0.394 | 0.055 | 6.73518069993128e-76 | 26 |
| sema3d2 | 1.19449774892316e-77 | 0.385763938 | 0.194 | 0.013 | 3.00643138426469e-73 | 26 |
| f3b5 | 1.76239460080156e-77 | 0.924514701 | 0.52 | 0.097 | 4.43577097075745e-73 | 26 |
| daam1b | 5.3695390198203e-77 | 0.356272152 | 0.177 | 0.011 | 1.35145927589857e-72 | 26 |
| myo1cb1 | 1.61376445465324e-76 | 0.510565797 | 0.297 | 0.032 | 4.06168375591675e-72 | 26 |
| col6a15 | 6.32060015508068e-76 | 0.797477675 | 0.491 | 0.085 | 1.59083185303226e-71 | 26 |
| ncam1a1 | 7.30273427812148e-76 | 0.415751472 | 0.211 | 0.016 | 1.8380251904604e-71 | 26 |
| bag32 | 7.43796670342383e-76 | 0.622716973 | 0.297 | 0.033 | 1.87206183958474e-71 | 26 |
| XLOC-0412693 | 7.67614479560224e-75 | 0.492400052 | 0.24 | 0.022 | 1.93200888360513e-70 | 26 |
| junba8 | 8.13294449540215e-75 | 1.100724746 | 0.851 | 0.303 | 2.04698080004777e-70 | 26 |
| cyp1c21 | 1.12288533971022e-73 | 0.437840341 | 0.183 | 0.013 | 2.82619011151665e-69 | 26 |
| AL954191.11 | 2.35102495025492e-73 | 0.443008285 | 0.183 | 0.013 | 5.91729469729661e-69 | 26 |
| socs3a10 | 1.41938810718267e-72 | 1.116848133 | 0.84 | 0.292 | 3.57245792696807e-68 | 26 |
| cxcl18b6 | 2.86340175166617e-72 | 1.025385261 | 0.52 | 0.102 | 7.20689586876859e-68 | 26 |
| fosb5 | 4.92934456302924e-72 | 1.058289601 | 0.554 | 0.121 | 1.24066673306883e-67 | 26 |
| cdh61 | 5.07925374542768e-72 | 0.480322392 | 0.211 | 0.017 | 1.27839737518669e-67 | 26 |
| CABZ01080702.16 | 9.29075565877165e-72 | 0.735446148 | 0.457 | 0.08 | 2.33839029175624e-67 | 26 |
| fosab9 | 9.51343504873595e-72 | 1.144233635 | 0.869 | 0.343 | 2.39443646741635e-67 | 26 |
| itgb3b | 1.57531647419321e-71 | 0.342250526 | 0.154 | 0.009 | 3.9649140338969e-67 | 26 |
| sost1 | 1.37260516976351e-70 | 0.33695852 | 0.149 | 0.009 | 3.45470995177778e-66 | 26 |
| f104 | 1.52100531170559e-69 | 0.597943887 | 0.337 | 0.046 | 3.8282182690318e-65 | 26 |
| mical2a | 7.32349840224946e-69 | 0.270557718 | 0.126 | 0.006 | 1.84325131286217e-64 | 26 |
| nrp1a3 | 1.65399382032282e-68 | 0.624909226 | 0.36 | 0.052 | 4.16293704637051e-64 | 26 |
| nanos12 | 1.85149643232587e-68 | 0.521626784 | 0.206 | 0.018 | 4.66003137052097e-64 | 26 |
| snrkb | 1.85991934778755e-68 | 0.366731221 | 0.2 | 0.016 | 4.68123100644649e-64 | 26 |
| thbs1a1 | 4.35726268419571e-68 | 0.379032055 | 0.171 | 0.012 | 1.09667944498522e-63 | 26 |
| jun7 | 8.08303604084313e-68 | 0.994574425 | 0.771 | 0.222 | 2.03441934111981e-63 | 26 |
| dcn5 | 1.15186065907238e-66 | 0.894764997 | 0.543 | 0.115 | 2.89911809281928e-62 | 26 |
| tjp1a3 | 2.66559779467379e-66 | 0.566525985 | 0.326 | 0.044 | 6.70904308941446e-62 | 26 |
| col6a25 | 3.56239694967027e-66 | 0.637311034 | 0.389 | 0.061 | 8.96619688262509e-62 | 26 |
| flnb1 | 2.13302512586261e-65 | 0.406668502 | 0.211 | 0.019 | 5.3686109392836e-61 | 26 |

| | | | | | | |
|---|---|---|---|---|---|---|
| LOC1018823341 | 3.97353910076499e-65 | 0.401256238 | 0.183 | 0.014 | 1.00010005627154e-60 | 26 |
| vim4 | 4.2039873391847e-65 | 0.523129146 | 0.274 | 0.032 | 1.0581015733994e-60 | 26 |
| id14 | 3.28745401981939e-64 | 0.917905913 | 0.486 | 0.101 | 8.27419302248342e-60 | 26 |
| LO018188.11 | 1.55749392699538e-63 | 0.323307023 | 0.183 | 0.015 | 3.92005646485468e-59 | 26 |
| cyp27c12 | 4.62541416252098e-62 | 0.296263135 | 0.16 | 0.011 | 1.1641704905649e-57 | 26 |
| edil3a4 | 6.08546358009988e-62 | 0.389832346 | 0.217 | 0.021 | 1.53165032847534e-57 | 26 |
| rnd1b1 | 3.87286510166708e-61 | 0.442756348 | 0.217 | 0.022 | 9.74761417438588e-57 | 26 |
| plekha61 | 8.75420460122866e-61 | 0.361196597 | 0.189 | 0.016 | 2.20334575608324e-56 | 26 |
| sh3bgrl21 | 1.21684646543867e-60 | 0.264708364 | 0.149 | 0.01 | 3.0626808688626e-56 | 26 |
| gadd45ba5 | 2.71161957570035e-60 | 0.950485671 | 0.64 | 0.185 | 6.82487531008022e-56 | 26 |
| rhpn21 | 3.53909895799568e-60 | 0.459641217 | 0.269 | 0.033 | 8.90755816737932e-56 | 26 |
| zgc:1540581 | 1.03279746477428e-59 | 0.312311633 | 0.166 | 0.013 | 2.59944793909038e-55 | 26 |
| znf385a4 | 4.89761116715197e-58 | 0.561814348 | 0.303 | 0.044 | 1.23267975466048e-53 | 26 |
| colec111 | 1.73729265107208e-57 | 0.288248064 | 0.126 | 0.008 | 4.37259187348331e-53 | 26 |
| eif4ebp3l5 | 3.16799973979177e-57 | 0.934509822 | 0.651 | 0.201 | 7.97353854508191e-53 | 26 |
| cyp2n134 | 4.60737084800475e-57 | 0.495670361 | 0.269 | 0.035 | 1.15962916873432e-52 | 26 |
| s100v26 | 5.34310599645021e-57 | 0.73717746 | 0.514 | 0.117 | 1.34480634824655e-52 | 26 |
| tpm4a5 | 2.07806145407332e-56 | 0.671215759 | 0.429 | 0.085 | 5.23027287375713e-52 | 26 |
| si:ch211-198c19.35 | 2.28720541686505e-56 | 0.641085712 | 0.36 | 0.06 | 5.75666731370763e-52 | 26 |
| loxl2a4 | 4.12751566943154e-56 | 0.482266999 | 0.286 | 0.039 | 1.03885441883922e-51 | 26 |
| hdlbpa6 | 5.76985328482012e-56 | 0.820314436 | 0.571 | 0.155 | 1.45221437325638e-51 | 26 |
| LOC1039092845 | 3.12192287110318e-55 | 0.582545608 | 0.354 | 0.061 | 7.85756767427959e-51 | 26 |
| mob2a2 | 2.16039761960224e-54 | 0.53458532 | 0.269 | 0.037 | 5.43750476877689e-50 | 26 |
| ctgfa2 | 8.39518170213334e-54 | 0.511659764 | 0.223 | 0.026 | 2.11298328260994e-49 | 26 |
| tgm1l11 | 8.16403578032794e-53 | 0.497165935 | 0.263 | 0.036 | 2.05480616555074e-48 | 26 |
| si:ch211-1a19.34 | 1.80455873522265e-52 | 0.560027755 | 0.314 | 0.051 | 4.5418938806819e-48 | 26 |
| wt1b1 | 4.69760614585713e-52 | 0.383243506 | 0.194 | 0.02 | 1.18234049085078e-47 | 26 |
| fap1 | 8.41926263683524e-52 | 0.261528445 | 0.143 | 0.011 | 2.11904421306506e-47 | 26 |
| cyp2ad34 | 1.69863581553097e-51 | 0.526888231 | 0.309 | 0.049 | 4.27529648410991e-47 | 26 |
| ghra4 | 5.14460184974476e-51 | 0.430776186 | 0.229 | 0.028 | 1.29484483956226e-46 | 26 |
| reep3a2 | 2.51640784931804e-50 | 0.390837226 | 0.211 | 0.025 | 6.33354691594856e-46 | 26 |
| tagln4 | 2.71296045310651e-50 | 0.568528799 | 0.343 | 0.06 | 6.82825016442376e-46 | 26 |
| dsc2l | 2.80300541274431e-50 | 0.350872388 | 0.206 | 0.023 | 7.05488432333615e-46 | 26 |
| igf2b5 | 4.59136525614239e-50 | 0.582755886 | 0.326 | 0.056 | 1.15560072131848e-45 | 26 |
| col5a15 | 4.89260167045075e-50 | 0.672044772 | 0.48 | 0.108 | 1.23141891443575e-45 | 26 |

| | | | | | | |
|---|---|---|---|---|---|---|
| lamb1b2 | 5.34806127764547e-50 | 0.372687677 | 0.211 | 0.025 | 1.34605354297059e-45 | 26 |
| cd81a6 | 1.40247032192933e-48 | 0.823366935 | 0.646 | 0.202 | 3.52987755326392e-44 | 26 |
| ahnak8 | 1.9836972106366e-48 | 0.709124501 | 0.491 | 0.123 | 4.99276750945125e-44 | 26 |
| lrp1ab4 | 4.20474937212305e-48 | 0.559432773 | 0.354 | 0.067 | 1.05829336946965e-43 | 26 |
| gpm6ab | 4.49914453654648e-48 | 0.303058228 | 0.166 | 0.016 | 1.13238968840338e-43 | 26 |
| tnfsf10l2 | 5.98893982769272e-48 | 0.37380404 | 0.206 | 0.024 | 1.50735626523198e-43 | 26 |
| esr2a1 | 8.78390075125722e-48 | 0.423235508 | 0.229 | 0.03 | 2.21081998008393e-43 | 26 |
| si:ch1073-459j12.15 | 9.99376415259457e-48 | 0.561874733 | 0.343 | 0.063 | 2.51533049956653e-43 | 26 |
| si:dkey-16l2.20 | 1.87417335480014e-47 | 0.338015537 | 0.171 | 0.018 | 4.71710691669647e-43 | 26 |
| prss232 | 3.36075765979623e-47 | 0.427576193 | 0.223 | 0.029 | 8.45869095394112e-43 | 26 |
| pik3r23 | 4.82814749116482e-47 | 0.531965022 | 0.32 | 0.058 | 1.21519644205127e-42 | 26 |
| ftr821 | 3.56805853091444e-46 | 0.334266436 | 0.189 | 0.021 | 8.98044651645856e-42 | 26 |
| slco2b14 | 4.22855644724438e-46 | 0.523783753 | 0.337 | 0.062 | 1.06428537220694e-41 | 26 |
| fstl1b5 | 1.29560483627023e-45 | 0.672383993 | 0.486 | 0.121 | 3.26090781240854e-41 | 26 |
| igfbp5b3 | 1.6187818268178e-45 | 0.559982988 | 0.36 | 0.07 | 4.07431197991772e-41 | 26 |
| LOC101885758 | 2.64450956025312e-45 | 0.310937431 | 0.149 | 0.014 | 6.65596611220109e-41 | 26 |
| pbx3b | 5.65557720328382e-45 | 0.26364851 | 0.131 | 0.011 | 1.42345222629451e-40 | 26 |
| phactr4a3 | 1.00428004559922e-44 | 0.42277993 | 0.24 | 0.035 | 2.52767244676867e-40 | 26 |
| fosl1a4 | 1.0654774730048e-44 | 0.671439031 | 0.394 | 0.089 | 2.68170025180578e-40 | 26 |
| zgc:16273012 | 1.22648757730667e-44 | 0.85413614 | 0.846 | 0.387 | 3.08694658332315e-40 | 26 |
| BX322530.14 | 1.61558379458382e-44 | 0.465954495 | 0.274 | 0.045 | 4.06626285258801e-40 | 26 |
| nfil3-68 | 3.96209423436943e-44 | 0.698407333 | 0.48 | 0.127 | 9.97219497848443e-40 | 26 |
| nfic6 | 1.04436794561648e-43 | 0.543832088 | 0.354 | 0.074 | 2.62856968232211e-39 | 26 |
| ptprsa1 | 1.59758927039908e-43 | 0.366064006 | 0.211 | 0.028 | 4.02097243466744e-39 | 26 |
| cdon4 | 2.05315323468731e-43 | 0.474883293 | 0.303 | 0.055 | 5.1675813763845e-39 | 26 |
| deptor3 | 2.08230063082358e-43 | 0.400466397 | 0.24 | 0.036 | 5.24094245771987e-39 | 26 |
| zgc:1584633 | 2.16743216203946e-43 | 0.625206709 | 0.343 | 0.071 | 5.45521000863712e-39 | 26 |
| ptges3 | 4.9239443753852e-43 | 0.400648898 | 0.183 | 0.022 | 1.2393075598407e-38 | 26 |
| c7a5 | 5.70526525404914e-43 | 0.497621596 | 0.263 | 0.043 | 1.43595821179163e-38 | 26 |
| cyp2p63 | 6.25924790878642e-43 | 0.346859477 | 0.189 | 0.023 | 1.57539010616245e-38 | 26 |
| jdp2b6 | 8.04221909615469e-43 | 0.899107975 | 0.531 | 0.166 | 2.02414612431117e-38 | 26 |
| ugt1b51 | 9.01158791679093e-43 | 0.348357343 | 0.206 | 0.027 | 2.26812656277711e-38 | 26 |
| slc12a7b | 2.85777624332556e-42 | 0.31252374 | 0.177 | 0.021 | 7.1927370268261e-38 | 26 |
| cav2 | 2.97285634240003e-42 | 0.257590969 | 0.131 | 0.012 | 7.48238212818663e-38 | 26 |
| h1f06 | 7.33815540919627e-42 | 0.595976707 | 0.309 | 0.06 | 1.84694033494061e-37 | 26 |

| | | | | | | |
|---|---|---|---|---|---|---|
| sult1st62 | 1.51741925030368e-41 | 0.250804293 | 0.149 | 0.015 | 3.81919251108933e-37 | 26 |
| pim17 | 1.62593937025633e-41 | 0.800527102 | 0.526 | 0.167 | 4.09232680099815e-37 | 26 |
| aoc26 | 4.69261311337205e-41 | 0.573205962 | 0.32 | 0.064 | 1.18108379450461e-36 | 26 |
| boc5 | 5.23437336335786e-41 | 0.481744057 | 0.269 | 0.047 | 1.31743943182354e-36 | 26 |
| fstb2 | 1.44499888776578e-40 | 0.319868959 | 0.171 | 0.02 | 3.63691770061768e-36 | 26 |
| xirp2a2 | 3.04924145852146e-40 | 0.308602963 | 0.189 | 0.024 | 7.67463582695267e-36 | 26 |
| pmp22a5 | 4.31345078599448e-40 | 0.626265038 | 0.509 | 0.136 | 1.08565242832695e-35 | 26 |
| cldn11a5 | 3.83105333510264e-39 | 0.477847667 | 0.309 | 0.06 | 9.64237813911984e-35 | 26 |
| appa5 | 4.99813116543407e-39 | 0.514589431 | 0.343 | 0.075 | 1.2579796330281e-34 | 26 |
| pdk2a3 | 7.29600223508335e-39 | 0.347419176 | 0.194 | 0.027 | 1.83633080254813e-34 | 26 |
| meis1b5 | 1.20855701045599e-38 | 0.460346696 | 0.286 | 0.055 | 3.04181713961668e-34 | 26 |
| btg27 | 1.65758083254475e-38 | 0.780166468 | 0.737 | 0.37 | 4.17196519743188e-34 | 26 |
| epb41l51 | 2.3822041955304e-38 | 0.368278973 | 0.24 | 0.039 | 5.99576973973047e-34 | 26 |
| cyr615 | 3.80151014749754e-38 | 0.632403967 | 0.314 | 0.066 | 9.56802089023655e-34 | 26 |
| ptpn211 | 1.48012714188917e-37 | 0.371375821 | 0.211 | 0.032 | 3.72533200342084e-33 | 26 |
| epha3 | 2.23316257516707e-37 | 0.256078369 | 0.137 | 0.014 | 5.620646885438e-33 | 26 |
| chka1 | 2.37900734927019e-37 | 0.313385595 | 0.183 | 0.024 | 5.98772359737813e-33 | 26 |
| rpz57 | 4.23828823054269e-37 | 0.535079877 | 0.4 | 0.099 | 1.06673476474529e-32 | 26 |
| tspan4a2 | 1.0524112041398e-36 | 0.321278899 | 0.206 | 0.031 | 2.64881375969946e-32 | 26 |
| slc7a15 | 1.14158688588452e-36 | 0.395058339 | 0.223 | 0.036 | 2.87326003308274e-32 | 26 |
| zgc:1101822 | 5.51638957059335e-36 | 0.493876079 | 0.371 | 0.085 | 1.38842009102264e-31 | 26 |
| stau26 | 1.37725312508861e-35 | 0.495951362 | 0.349 | 0.083 | 3.46640839053553e-31 | 26 |
| aldh9a1a.16 | 1.47826553176297e-35 | 0.656867466 | 0.497 | 0.161 | 3.72064651689421e-31 | 26 |
| BX663516.22 | 3.88503119293553e-35 | 0.273780069 | 0.131 | 0.014 | 9.77823500949944e-31 | 26 |
| CR383676.114 | 5.81889260152885e-35 | 0.638108175 | 0.903 | 0.725 | 1.4645570788788e-30 | 26 |
| gchfr1 | 6.57064817357694e-35 | 0.390168776 | 0.217 | 0.036 | 1.65376643880758e-30 | 26 |
| si:ch211-166a6.51 | 1.54116381834773e-34 | 0.374480034 | 0.2 | 0.031 | 3.87895521439939e-30 | 26 |
| LOC110439416 | 9.30949595606826e-34 | 0.253142335 | 0.126 | 0.013 | 2.34310703718282e-29 | 26 |
| ctsf | 1.09875193010005e-33 | 0.290649932 | 0.177 | 0.025 | 2.76544873286883e-29 | 26 |
| tspan9b3 | 1.14987570045287e-33 | 0.388413633 | 0.234 | 0.043 | 2.89412215046982e-29 | 26 |
| gamt2 | 1.87883367860936e-33 | 0.367295493 | 0.234 | 0.042 | 4.72883648569189e-29 | 26 |
| lmbr1l | 2.07201580909708e-33 | 0.258070444 | 0.131 | 0.015 | 5.21505658991645e-29 | 26 |
| lama52 | 3.2751777439315e-33 | 0.306816889 | 0.166 | 0.023 | 8.2432948637012e-29 | 26 |
| rhoua3 | 4.33964475560169e-33 | 0.28933425 | 0.166 | 0.023 | 1.09224518853739e-28 | 26 |
| klf2a5 | 4.51603670784013e-33 | 0.565629104 | 0.337 | 0.083 | 1.13664127899628e-28 | 26 |

| | | | | | | |
|---|---|---|---|---|---|---|
| vasna3 | 5.13534657826839e-33 | 0.328893788 | 0.206 | 0.033 | 1.29251538028437e-28 | 26 |
| scarb2a1 | 6.5396215578283e-33 | 0.476781041 | 0.286 | 0.062 | 1.64595734988981e-28 | 26 |
| pnrc214 | 6.9821520466247e-33 | 0.617655767 | 0.891 | 0.565 | 1.75733784861497e-28 | 26 |
| sostdc1a2 | 1.00804140198103e-32 | 0.250414312 | 0.109 | 0.01 | 2.53713940464606e-28 | 26 |
| si:ch211-66e2.34 | 1.07938354240212e-32 | 0.448174007 | 0.274 | 0.057 | 2.71670043787189e-28 | 26 |
| lxn7 | 1.39836329593816e-32 | 0.623761976 | 0.463 | 0.145 | 3.51954057954676e-28 | 26 |
| ier2a10 | 2.87639336741645e-32 | 0.726511715 | 0.623 | 0.259 | 7.23959446645046e-28 | 26 |
| pkp3b1 | 2.9544867234372e-32 | 0.34438094 | 0.229 | 0.041 | 7.43614763421908e-28 | 26 |
| dedd15 | 3.18973046621601e-32 | 0.628014172 | 0.451 | 0.143 | 8.02823261041909e-28 | 26 |
| foxp44 | 3.72374367260823e-32 | 0.392061421 | 0.251 | 0.049 | 9.37229044958766e-28 | 26 |
| rtn31 | 1.0578529093973e-31 | 0.467930767 | 0.32 | 0.077 | 2.66250998766208e-27 | 26 |
| cebpb11 | 1.61291801596721e-31 | 0.744623287 | 0.737 | 0.35 | 4.05955335438786e-27 | 26 |
| bcam1 | 1.94430726299561e-31 | 0.384481966 | 0.183 | 0.029 | 4.89362695023364e-27 | 26 |
| mknk2b10 | 2.90912599745997e-31 | 0.630612706 | 0.537 | 0.194 | 7.321979223007e-27 | 26 |
| pawr | 3.60422562330292e-31 | 0.255616938 | 0.143 | 0.018 | 9.07147547129112e-27 | 26 |
| clic42 | 6.11148860907255e-31 | 0.424993383 | 0.24 | 0.048 | 1.53820056801747e-26 | 26 |
| egr19 | 9.58108687718802e-31 | 0.692954127 | 0.509 | 0.187 | 2.41146375611945e-26 | 26 |
| gas1a4 | 9.63285234360991e-31 | 0.343002426 | 0.223 | 0.041 | 2.42449260636318e-26 | 26 |
| cald1a6 | 1.12535170063052e-30 | 0.484874307 | 0.366 | 0.096 | 2.83239769531695e-26 | 26 |
| col5a2a5 | 1.6482613605233e-30 | 0.466272154 | 0.337 | 0.084 | 4.14850901830109e-26 | 26 |
| cgnl12 | 2.22719858478652e-30 | 0.256395547 | 0.16 | 0.023 | 5.6056361180492e-26 | 26 |
| dag13 | 2.84336297960643e-30 | 0.394303859 | 0.24 | 0.048 | 7.15646028337141e-26 | 26 |
| cdkn1a1 | 3.25752745853263e-30 | 0.331427319 | 0.189 | 0.031 | 8.19887086038078e-26 | 26 |
| csrnp1b7 | 5.12846174727584e-30 | 0.631839821 | 0.497 | 0.176 | 1.29078253717186e-25 | 26 |
| gpr1461 | 7.62173683342552e-30 | 0.276161739 | 0.137 | 0.018 | 1.91831494360487e-25 | 26 |
| tmem1765 | 8.12913556580808e-30 | 0.412392781 | 0.286 | 0.065 | 2.04602213055824e-25 | 26 |
| hecw2b | 9.10294508232432e-30 | 0.26099534 | 0.137 | 0.018 | 2.29112024777021e-25 | 26 |
| plecb2 | 2.55607198378664e-29 | 0.357038284 | 0.24 | 0.049 | 6.4333775759926e-25 | 26 |
| cast2 | 3.46186503094389e-29 | 0.411326956 | 0.291 | 0.068 | 8.71316809638266e-25 | 26 |
| pabpc1b5 | 4.16020009805695e-29 | 0.425523225 | 0.286 | 0.067 | 1.04708076267995e-24 | 26 |
| tmem88b3 | 6.13480293774121e-29 | 0.341450447 | 0.2 | 0.036 | 1.54406855140009e-24 | 26 |
| myl9a3 | 9.69712368774456e-29 | 0.28240565 | 0.183 | 0.03 | 2.44066906096843e-24 | 26 |
| selenbp13 | 1.34216536627002e-28 | 0.424080295 | 0.286 | 0.067 | 3.378096010365e-24 | 26 |
| afdna2 | 3.21449527080375e-28 | 0.298041775 | 0.149 | 0.021 | 8.09056314708596e-24 | 26 |
| si:ch73-138n13.11 | 4.04113311032182e-28 | 0.335465607 | 0.2 | 0.037 | 1.0171127925369e-23 | 26 |

| | | | | | | |
|---|---|---|---|---|---|---|
| rsu11 | 1.06272373938663e-27 | 0.368087249 | 0.217 | 0.043 | 2.67476937966221e-23 | 26 |
| slc38a5b6 | 1.48774850613105e-27 | 0.506289098 | 0.377 | 0.113 | 3.74451421508123e-23 | 26 |
| itgav3 | 1.99323464775566e-27 | 0.395704337 | 0.251 | 0.056 | 5.01677228493623e-23 | 26 |
| rdh10a3 | 2.17583798919028e-27 | 0.365367403 | 0.206 | 0.039 | 5.47636663499302e-23 | 26 |
| olfml3b4 | 2.3736749236393e-27 | 0.38348273 | 0.217 | 0.044 | 5.97430241530776e-23 | 26 |
| gstt1b4 | 2.9733018023106e-27 | 0.44750203 | 0.286 | 0.071 | 7.48350330623555e-23 | 26 |
| pfkfb4b3 | 3.65636438569497e-27 | 0.432128689 | 0.343 | 0.094 | 9.20270352235567e-23 | 26 |
| si:ch211-232b12.5 | 1.54119344336908e-26 | 0.251654076 | 0.131 | 0.018 | 3.87902977761563e-22 | 26 |
| cyp2p74 | 3.46062683546491e-26 | 0.262627235 | 0.154 | 0.024 | 8.71005168218163e-22 | 26 |
| ulk23 | 4.09489577921941e-26 | 0.332058542 | 0.223 | 0.047 | 1.03064431867173e-21 | 26 |
| ptprga7 | 4.98648303585364e-26 | 0.445348892 | 0.263 | 0.064 | 1.255047915294e-21 | 26 |
| si:dkey-51e6.14 | 5.16864215651203e-26 | 0.486086713 | 0.371 | 0.114 | 1.30089554437251e-21 | 26 |
| CABZ01072614.17 | 5.22867337944802e-26 | 0.462461974 | 0.297 | 0.079 | 1.31600480287327e-21 | 26 |
| LOC1018844514 | 6.94352355129444e-26 | 0.445053257 | 0.309 | 0.084 | 1.7476154426253e-21 | 26 |
| irf2bp2b4 | 1.05605083793423e-25 | 0.472484546 | 0.297 | 0.081 | 2.65797435399666e-21 | 26 |
| map1lc3a6 | 1.93170383276861e-25 | 0.379705705 | 0.274 | 0.069 | 4.86190537669532e-21 | 26 |
| tns2a4 | 2.80426936037034e-25 | 0.306557235 | 0.189 | 0.036 | 7.05806555311612e-21 | 26 |
| gulp1a4 | 3.26481522709976e-25 | 0.281866453 | 0.171 | 0.03 | 8.21721344508739e-21 | 26 |
| chd36 | 5.08604964682205e-25 | 0.455307445 | 0.343 | 0.102 | 1.28010783560864e-20 | 26 |
| sdc47 | 6.07886443655071e-25 | 0.646864521 | 0.577 | 0.255 | 1.52998939003545e-20 | 26 |
| txnipa10 | 9.04271896207803e-25 | 0.575082301 | 0.52 | 0.208 | 2.27596193556542e-20 | 26 |
| actn13 | 1.09367781219971e-24 | 0.337326893 | 0.206 | 0.043 | 2.75267768552546e-20 | 26 |
| nfixb4 | 1.18703354241344e-24 | 0.424008982 | 0.257 | 0.064 | 2.98764472290039e-20 | 26 |
| aldob2 | 1.30889693840861e-24 | 0.314456676 | 0.189 | 0.037 | 3.29436270428062e-20 | 26 |
| perp2 | 1.32889081465834e-24 | 0.338473779 | 0.234 | 0.053 | 3.34468529141357e-20 | 26 |
| si:ch211-222l21.111 | 3.82230410061743e-24 | 0.610559845 | 0.611 | 0.272 | 9.62035719084402e-20 | 26 |
| tshz3b3 | 7.64858883474012e-24 | 0.277243311 | 0.16 | 0.028 | 1.92507332381574e-19 | 26 |
| irf83 | 9.94948914469024e-24 | 0.466153829 | 0.206 | 0.045 | 2.50418692282709e-19 | 26 |
| ephb34 | 1.1380246788739e-23 | 0.338982458 | 0.194 | 0.04 | 2.86429431425771e-19 | 26 |
| gli34 | 1.20195780207367e-23 | 0.312717266 | 0.166 | 0.031 | 3.02520759203921e-19 | 26 |
| rpsa15 | 1.23879306510492e-23 | 0.370601613 | 0.943 | 0.75 | 3.11791826556257e-19 | 26 |
| FO744854.12 | 1.24835504466216e-23 | 0.250608474 | 0.12 | 0.017 | 3.1419848119102e-19 | 26 |
| zbtb206 | 1.36742433439679e-23 | 0.421402002 | 0.326 | 0.097 | 3.44167030724328e-19 | 26 |
| midn7 | 1.78179967136302e-23 | 0.5490781 | 0.394 | 0.138 | 4.48461159285359e-19 | 26 |
| gpc42 | 2.29052170654341e-23 | 0.25698915 | 0.154 | 0.027 | 5.76501408319912e-19 | 26 |

| | | | | | | |
|---|---|---|---|---|---|---|
| pdk2b7 | 2.56697315012311e-23 | 0.445455678 | 0.326 | 0.098 | 6.46081472154487e-19 | 26 |
| mycb5 | 2.64715643342492e-23 | 0.5715551 | 0.514 | 0.209 | 6.66262802728717e-19 | 26 |
| fkbp55 | 5.34692656223927e-23 | 0.498170498 | 0.36 | 0.121 | 1.34576794645e-18 | 26 |
| selenow14 | 6.80834838872466e-23 | 0.57233806 | 0.697 | 0.383 | 1.71359320595811e-18 | 26 |
| fxyd16 | 1.00192298219802e-22 | 0.396431398 | 0.286 | 0.08 | 2.52173995389421e-18 | 26 |
| foxo3b4 | 1.24824052047281e-22 | 0.393052615 | 0.251 | 0.065 | 3.141696565978e-18 | 26 |
| myh102 | 1.580025909791e-22 | 0.2963718 | 0.177 | 0.035 | 3.97676721235296e-18 | 26 |
| per33 | 1.7204630027083e-22 | 0.406183181 | 0.28 | 0.078 | 4.33023333151652e-18 | 26 |
| col12a1a3 | 1.76905704295443e-22 | 0.277631024 | 0.149 | 0.026 | 4.452539671412e-18 | 26 |
| socs3b7 | 2.08631757209839e-22 | 0.556202195 | 0.417 | 0.153 | 5.25105269721444e-18 | 26 |
| zgc:1583437 | 2.35057838220056e-22 | 0.6353445 | 0.429 | 0.164 | 5.9161707301606e-18 | 26 |
| ier2b11 | 2.58558214863979e-22 | 0.567888563 | 0.514 | 0.221 | 6.50765170991148e-18 | 26 |
| NC-002333.413 | 3.55383122947034e-22 | 0.490037931 | 0.817 | 0.622 | 8.9446378214539e-18 | 26 |
| abcc56 | 5.52798001484015e-22 | 0.409609773 | 0.309 | 0.093 | 1.39133728993512e-17 | 26 |
| nfil32 | 8.44549953415229e-22 | 0.291023013 | 0.166 | 0.033 | 2.12564777775079e-17 | 26 |
| myl9b6 | 1.12605089831811e-21 | 0.379684681 | 0.28 | 0.078 | 2.83415750597685e-17 | 26 |
| CABZ01068366.13 | 1.26415761224613e-21 | 0.287049601 | 0.154 | 0.029 | 3.18175829426229e-17 | 26 |
| taok1b | 1.53078922673208e-21 | 0.261149083 | 0.137 | 0.023 | 3.85284340476197e-17 | 26 |
| cpn13 | 1.87471465764009e-21 | 0.334100895 | 0.223 | 0.054 | 4.71846932181435e-17 | 26 |
| ctdspla2 | 2.5525756771009e-21 | 0.275580627 | 0.171 | 0.035 | 6.42457772169526e-17 | 26 |
| XLOC-0247503 | 3.12075572469795e-21 | 0.352620808 | 0.269 | 0.074 | 7.85463008349228e-17 | 26 |
| klf11b7 | 3.48838842593841e-21 | 0.385759502 | 0.274 | 0.079 | 8.77992482924438e-17 | 26 |
| atp10a5 | 7.28688823201611e-21 | 0.328899901 | 0.223 | 0.055 | 1.83403689911613e-16 | 26 |
| gstm.14 | 1.07348534173486e-20 | 0.533540666 | 0.577 | 0.258 | 2.70185525661246e-16 | 26 |
| si:dkey-28b4.77 | 1.23157889908372e-20 | 0.424424406 | 0.291 | 0.09 | 3.09976093110382e-16 | 26 |
| tenm33 | 2.06130708833923e-20 | 0.315574618 | 0.194 | 0.045 | 5.18810381064101e-16 | 26 |
| fgl2a5 | 2.36866319520415e-20 | 0.337179341 | 0.246 | 0.065 | 5.96168839600932e-16 | 26 |
| lgals3b3 | 2.48682814343977e-20 | 0.391015763 | 0.263 | 0.077 | 6.25909775422356e-16 | 26 |
| dbpb1 | 3.39755606700153e-20 | 0.273289939 | 0.183 | 0.04 | 8.55130886503616e-16 | 26 |
| ghrb4 | 6.30429191845342e-20 | 0.314712319 | 0.183 | 0.041 | 1.58672723295554e-15 | 26 |
| b2ml12 | 8.87625165836581e-20 | 0.529754377 | 0.549 | 0.265 | 2.23406377989409e-15 | 26 |
| entpd14 | 9.48980190090145e-20 | 0.341176805 | 0.251 | 0.069 | 2.38848824043789e-15 | 26 |
| dclk1a2 | 1.21138151788641e-19 | 0.351588826 | 0.171 | 0.038 | 3.04892614236831e-15 | 26 |
| si:dkey-23i12.73 | 1.45542787141805e-19 | 0.286719394 | 0.154 | 0.031 | 3.66316640957208e-15 | 26 |
| itpkcb3 | 1.7553750654539e-19 | 0.362863465 | 0.263 | 0.076 | 4.41810350224093e-15 | 26 |

| | | | | | | |
|---|---|---|---|---|---|---|
| rgs5b5 | 1.76895916817679e-19 | 0.494504107 | 0.257 | 0.075 | 4.45229333038416e-15 | 26 |
| ccdc85b6 | 1.98423927465637e-19 | 0.265328559 | 0.2 | 0.048 | 4.99413183038262e-15 | 26 |
| sort1a | 4.78775063386563e-19 | 0.263478911 | 0.131 | 0.024 | 1.20502895703764e-14 | 26 |
| pmp22b2 | 4.83181336103442e-19 | 0.304608491 | 0.189 | 0.044 | 1.21611910483875e-14 | 26 |
| stat1a5 | 6.22355165051454e-19 | 0.417383468 | 0.337 | 0.119 | 1.566405714918e-14 | 26 |
| si:ch211-204c21.15 | 6.85834162266815e-19 | 0.358916029 | 0.251 | 0.072 | 1.72617600300935e-14 | 26 |
| hmgb1a10 | 8.99707072335869e-19 | 0.45924635 | 0.451 | 0.187 | 2.26447273036215e-14 | 26 |
| mfap25 | 1.53264409802458e-18 | 0.364104102 | 0.303 | 0.095 | 3.85751193031806e-14 | 26 |
| cd991 | 1.79623576687177e-18 | 0.292703393 | 0.206 | 0.052 | 4.52094580163957e-14 | 26 |
| si:dkey-56m19.53 | 1.83269998188853e-18 | 0.318510784 | 0.189 | 0.046 | 4.61272258441524e-14 | 26 |
| pdgfrb5 | 1.9580445629807e-18 | 0.280538259 | 0.2 | 0.05 | 4.92820236056613e-14 | 26 |
| fam120c4 | 2.50933673900379e-18 | 0.36920434 | 0.297 | 0.098 | 6.31574963839863e-14 | 26 |
| tuft1a2 | 3.20561023943502e-18 | 0.286152217 | 0.154 | 0.033 | 8.068200411634e-14 | 26 |
| lpar14 | 3.81561452727922e-18 | 0.349872395 | 0.229 | 0.063 | 9.60352020370907e-14 | 26 |
| si:ch73-1a9.310 | 3.84772032045581e-18 | 0.522076456 | 0.543 | 0.263 | 9.68432727455524e-14 | 26 |
| fat1a2 | 6.11654257840082e-18 | 0.268884187 | 0.137 | 0.027 | 1.5394726015577e-13 | 26 |
| timp2a5 | 7.02673309409515e-18 | 0.296678716 | 0.211 | 0.056 | 1.76855845245281e-13 | 26 |
| ldlrad4a3 | 7.33519847684285e-18 | 0.262814178 | 0.149 | 0.031 | 1.84619610463658e-13 | 26 |
| fzd7b2 | 1.08118662433989e-17 | 0.263258862 | 0.166 | 0.038 | 2.72123861480108e-13 | 26 |
| cbx7a3 | 1.37040326723417e-17 | 0.317513124 | 0.2 | 0.052 | 3.44916798330169e-13 | 26 |
| calm2b5 | 1.56444042931105e-17 | 0.455801015 | 0.486 | 0.208 | 3.93754011653297e-13 | 26 |
| ptmaa12 | 2.1425273911619e-17 | 0.512067942 | 0.726 | 0.392 | 5.39252719081538e-13 | 26 |
| spry44 | 2.25623708164041e-17 | 0.360416433 | 0.183 | 0.046 | 5.67872311078076e-13 | 26 |
| ndrg23 | 2.58666831465022e-17 | 0.295143392 | 0.166 | 0.039 | 6.51038548114314e-13 | 26 |
| adamts15a5 | 2.80123975123908e-17 | 0.306780675 | 0.189 | 0.048 | 7.05044032989365e-13 | 26 |
| fosl28 | 3.1499623981621e-17 | 0.420452975 | 0.331 | 0.12 | 7.9281403599342e-13 | 26 |
| mt-nd210 | 3.15682556171659e-17 | 0.5190309 | 0.646 | 0.399 | 7.94541425628449e-13 | 26 |
| cnn3a5 | 3.47867690600794e-17 | 0.297397062 | 0.206 | 0.056 | 8.75548190473138e-13 | 26 |
| cox7a12 | 5.02456145015084e-17 | 0.250498198 | 0.143 | 0.03 | 1.26463187138846e-12 | 26 |
| si:ch211-286o17.12 | 5.75018727928555e-17 | 0.281021804 | 0.16 | 0.037 | 1.44726463632338e-12 | 26 |
| CR847566.14 | 6.64011884602465e-17 | 0.276987144 | 0.154 | 0.035 | 1.67125151235594e-12 | 26 |
| qkia5 | 7.42571094409579e-17 | 0.409873072 | 0.411 | 0.166 | 1.86897718751947e-12 | 26 |
| ifi457 | 9.17854797274652e-17 | 0.476394108 | 0.343 | 0.127 | 2.31014873926057e-12 | 26 |
| atf34 | 1.09473457523097e-16 | 0.463096463 | 0.269 | 0.09 | 2.75533745239882e-12 | 26 |
| mir21-14 | 1.20649071405895e-16 | 0.391500196 | 0.303 | 0.107 | 3.03661647821497e-12 | 26 |

| | | | | | | |
|---|---|---|---|---|---|---|
| mt-nd112 | 1.25202892336833e-16 | 0.497672028 | 0.674 | 0.409 | 3.15123159722575e-12 | 26 |
| rbp55 | 1.7654314114875e-16 | 0.32872455 | 0.286 | 0.093 | 4.44341431957289e-12 | 26 |
| atf4b9 | 1.85224983818823e-16 | 0.46496705 | 0.537 | 0.262 | 4.66192761773595e-12 | 26 |
| rbp45 | 2.53839838994337e-16 | 0.500192628 | 0.343 | 0.126 | 6.38889490764848e-12 | 26 |
| flna6 | 4.84696235741533e-16 | 0.367418841 | 0.314 | 0.113 | 1.21993195573786e-11 | 26 |
| si:ch1073-291c23.25 | 1.01434473539457e-15 | 0.279965014 | 0.211 | 0.06 | 2.55300426451459e-11 | 26 |
| ncor11 | 1.2601244087379e-15 | 0.269179325 | 0.2 | 0.056 | 3.17160712435243e-11 | 26 |
| emp24 | 1.41610857734577e-15 | 0.257923657 | 0.183 | 0.048 | 3.56420367832157e-11 | 26 |
| h3f3c8 | 1.89803887898296e-15 | 0.443715672 | 0.52 | 0.255 | 4.77717405451221e-11 | 26 |
| tagln28 | 2.05122023839962e-15 | 0.403910452 | 0.434 | 0.186 | 5.16271621802801e-11 | 26 |
| myl66 | 2.07038930247372e-15 | 0.361402792 | 0.286 | 0.1 | 5.21096283539609e-11 | 26 |
| ywhaqb5 | 2.30115382217304e-15 | 0.366092263 | 0.337 | 0.131 | 5.79177405502732e-11 | 26 |
| rps2812 | 2.44642821236129e-15 | 0.250351246 | 0.92 | 0.826 | 6.15741516769213e-11 | 26 |
| tmsb4x16 | 3.08197487792433e-15 | 0.469503763 | 0.977 | 0.683 | 7.75702257024774e-11 | 26 |
| zgc:1951736 | 4.73796656075634e-15 | 0.315686534 | 0.269 | 0.09 | 1.19249880367676e-10 | 26 |
| myh11a5 | 7.10185283312582e-15 | 0.286671543 | 0.217 | 0.065 | 1.78746533956944e-10 | 26 |
| adh56 | 7.74161953195443e-15 | 0.423477352 | 0.469 | 0.22 | 1.94848821999761e-10 | 26 |
| kank24 | 9.92904631241006e-15 | 0.274138834 | 0.189 | 0.053 | 2.49904166637049e-10 | 26 |
| cry22 | 1.06993292438274e-14 | 0.283105464 | 0.246 | 0.08 | 2.69291417737892e-10 | 26 |
| aebp14 | 1.07621033328893e-14 | 0.251925448 | 0.154 | 0.038 | 2.70871378785491e-10 | 26 |
| mcl1b8 | 1.11017182304036e-14 | 0.449895247 | 0.497 | 0.24 | 2.79419146141029e-10 | 26 |
| ube2e35 | 1.74167699814541e-14 | 0.387572266 | 0.366 | 0.154 | 4.38362683663219e-10 | 26 |
| gadd45bb5 | 2.01798465019621e-14 | 0.367212357 | 0.251 | 0.087 | 5.07906556607884e-10 | 26 |
| cebpa5 | 2.8820962751151e-14 | 0.299581045 | 0.194 | 0.057 | 7.25394811483718e-10 | 26 |
| sesn16 | 3.17630497219572e-14 | 0.427361722 | 0.383 | 0.167 | 7.99444198451941e-10 | 26 |
| si:ch211-140m22.73 | 3.94331111090473e-14 | 0.278425013 | 0.211 | 0.065 | 9.92491973503612e-10 | 26 |
| rpl2911 | 6.36516483495575e-14 | 0.252958415 | 0.926 | 0.808 | 1.60204833731001e-09 | 26 |
| ctdsp24 | 6.43140718031167e-14 | 0.259656229 | 0.211 | 0.066 | 1.61872087321265e-09 | 26 |
| capns1b4 | 7.19061974389622e-14 | 0.33766151 | 0.309 | 0.121 | 1.80980708334124e-09 | 26 |
| glud1a2 | 7.41134880522004e-14 | 0.325739007 | 0.257 | 0.09 | 1.86536238078583e-09 | 26 |
| irf1b3 | 8.42467476507952e-14 | 0.309644107 | 0.149 | 0.038 | 2.12040639162286e-09 | 26 |
| rbm5 | 9.02335199957832e-14 | 0.261618098 | 0.217 | 0.068 | 2.27108746477387e-09 | 26 |
| rpl613 | 1.52691726870397e-13 | 0.343277865 | 0.874 | 0.666 | 3.84309807360102e-09 | 26 |
| rpl915 | 1.806438547867e-13 | 0.39037171 | 0.891 | 0.627 | 4.54662518112645e-09 | 26 |
| zgc:11418815 | 1.96495382172839e-13 | 0.388671151 | 0.891 | 0.625 | 4.94559227390818e-09 | 26 |

| | | | | | | |
|---|---|---|---|---|---|---|
| wu:fb16f034 | 2.28047153460834e-13 | 0.316355539 | 0.246 | 0.086 | 5.73971880545573e-09 | 26 |
| cish4 | 2.77961341382622e-13 | 0.262141669 | 0.183 | 0.054 | 6.99600900125922e-09 | 26 |
| cflara2 | 3.27213071433012e-13 | 0.260815116 | 0.2 | 0.062 | 8.23562579489748e-09 | 26 |
| rps1014 | 3.7179401653603e-13 | 0.321831195 | 0.891 | 0.693 | 9.35768360219535e-09 | 26 |
| mt-atp614 | 4.13558129682816e-13 | 0.407214085 | 0.777 | 0.59 | 1.04088445659868e-08 | 26 |
| rpl5a12 | 5.16393357066753e-13 | 0.414474082 | 0.789 | 0.49 | 1.29971044040131e-08 | 26 |
| tefa6 | 5.7042263797267e-13 | 0.367377505 | 0.326 | 0.137 | 1.43569673751341e-08 | 26 |
| kmt2bb1 | 6.73387226483554e-13 | 0.299023837 | 0.24 | 0.085 | 1.69484831033646e-08 | 26 |
| rpl3614 | 8.06243830821107e-13 | 0.266056506 | 0.914 | 0.789 | 2.02923509779364e-08 | 26 |
| ppib9 | 9.61863702039439e-13 | 0.414203839 | 0.6 | 0.331 | 2.42091475166306e-08 | 26 |
| cnn27 | 1.11230646452785e-12 | 0.317189874 | 0.326 | 0.134 | 2.79956414057015e-08 | 26 |
| hmgn610 | 1.51607038707711e-12 | 0.41965706 | 0.497 | 0.264 | 3.81579755723438e-08 | 26 |
| nr1d2a2 | 1.61025438563114e-12 | 0.308811249 | 0.223 | 0.078 | 4.05284926319501e-08 | 26 |
| rps27.211 | 1.69338863399312e-12 | 0.395149146 | 0.789 | 0.516 | 4.2620898528973e-08 | 26 |
| jund6 | 1.87670055832541e-12 | 0.29663612 | 0.251 | 0.092 | 4.72346763524921e-08 | 26 |
| eif4ebp32 | 2.12025597305399e-12 | 0.398961213 | 0.343 | 0.149 | 5.33647225857958e-08 | 26 |
| si:ch211-196l7.411 | 2.17716789118331e-12 | 0.420666407 | 0.549 | 0.313 | 5.47971386531928e-08 | 26 |
| rplp214 | 2.77834279680325e-12 | 0.317235926 | 0.949 | 0.756 | 6.99281098527409e-08 | 26 |
| maff5 | 2.79292235534166e-12 | 0.31606318 | 0.229 | 0.083 | 7.02950627615943e-08 | 26 |
| BX640522.23 | 3.81303190600426e-12 | 0.266830183 | 0.114 | 0.027 | 9.59702000422211e-08 | 26 |
| slc20a1b3 | 3.8946597000317e-12 | 0.389462262 | 0.257 | 0.101 | 9.80246899900978e-08 | 26 |
| COX315 | 4.0813539542456e-12 | 0.330050002 | 0.817 | 0.7 | 1.02723597674407e-07 | 26 |
| ddit39 | 4.19274081829151e-12 | 0.42253543 | 0.371 | 0.176 | 1.05527093655579e-07 | 26 |
| lamc15 | 4.59722053712285e-12 | 0.252604887 | 0.177 | 0.055 | 1.15707443698845e-07 | 26 |
| nme34 | 6.75638860258511e-12 | 0.251131272 | 0.217 | 0.075 | 1.70051544738465e-07 | 26 |
| si:ch73-335l21.45 | 8.52007822153216e-12 | 0.50143013 | 0.326 | 0.148 | 2.14441848757743e-07 | 26 |
| mt-cyb11 | 1.26126283760908e-11 | 0.365123731 | 0.789 | 0.623 | 3.17447243597829e-07 | 26 |
| capns1a3 | 1.43852284303269e-11 | 0.308177808 | 0.331 | 0.142 | 3.62061814362899e-07 | 26 |
| ndufa4l8 | 2.42367923589439e-11 | 0.381373977 | 0.651 | 0.393 | 6.10015826882259e-07 | 26 |
| elovl1b4 | 2.6096952290028e-11 | 0.285081744 | 0.286 | 0.116 | 6.56834192187714e-07 | 26 |
| hmgn77 | 3.60934927241389e-11 | 0.388438204 | 0.509 | 0.278 | 9.08437118373852e-07 | 26 |
| mt-nd411 | 3.75742033424704e-11 | 0.404281011 | 0.663 | 0.455 | 9.45705123926638e-07 | 26 |
| ctnnb14 | 4.24569860803703e-11 | 0.309828029 | 0.337 | 0.149 | 1.06859988265684e-06 | 26 |
| tmem176l.16 | 4.52426395640213e-11 | 0.321823309 | 0.389 | 0.18 | 1.13871199518685e-06 | 26 |
| rbm39a4 | 4.86993332782694e-11 | 0.325712071 | 0.32 | 0.142 | 1.22571351928076e-06 | 26 |

| | | | | | | |
|---|---|---|---|---|---|---|
| vat16 | 5.48857130072674e-11 | 0.309243778 | 0.314 | 0.134 | 1.38141851067991e-06 | 26 |
| pfn23 | 5.95435643562923e-11 | 0.340866216 | 0.451 | 0.225 | 1.49865197128352e-06 | 26 |
| rpl3012 | 9.52608394119632e-11 | 0.289889127 | 0.886 | 0.707 | 2.3976200671597e-06 | 26 |
| rpl1412 | 1.26887314283913e-10 | 0.316245128 | 0.834 | 0.634 | 3.19362681321181e-06 | 26 |
| anxa5b5 | 2.05565805679331e-10 | 0.30952531 | 0.234 | 0.093 | 5.17388576314309e-06 | 26 |
| ybx17 | 2.20136170715601e-10 | 0.288288216 | 0.84 | 0.669 | 5.54060728074096e-06 | 26 |
| tsc22d36 | 2.54657818018088e-10 | 0.291668058 | 0.32 | 0.143 | 6.40948262169726e-06 | 26 |
| usp361 | 3.34174794581453e-10 | 0.265847992 | 0.2 | 0.074 | 8.4108454048206e-06 | 26 |
| add3a7 | 4.58320957207675e-10 | 0.291925801 | 0.286 | 0.123 | 1.153548017196e-05 | 26 |
| h2afx110 | 4.90644497009421e-10 | 0.3217491 | 0.4 | 0.198 | 1.23490313452301e-05 | 26 |
| ip6k2b7 | 5.46440141401491e-10 | 0.300745805 | 0.28 | 0.122 | 1.37533519189341e-05 | 26 |
| h2afy26 | 6.03568680474975e-10 | 0.253014491 | 0.24 | 0.097 | 1.51912201188746e-05 | 26 |
| acin1a5 | 7.26721233144143e-10 | 0.332534802 | 0.417 | 0.215 | 1.82908467170049e-05 | 26 |
| hopx3 | 7.72330886259128e-10 | 0.25116438 | 0.183 | 0.065 | 1.9438796076256e-05 | 26 |
| klf6a6 | 1.02749823569794e-09 | 0.315009518 | 0.349 | 0.167 | 2.58611030942816e-05 | 26 |
| rplp2l14 | 1.11226379971738e-09 | 0.262416279 | 0.931 | 0.773 | 2.79945675750867e-05 | 26 |
| rps1613 | 1.56747621841921e-09 | 0.261080666 | 0.886 | 0.737 | 3.94518089413931e-05 | 26 |
| mmp16a4 | 2.24918479336394e-09 | 0.261497204 | 0.189 | 0.069 | 5.6609732064177e-05 | 26 |
| si:dkey-7j14.68 | 2.93812734330442e-09 | 0.297770965 | 0.354 | 0.174 | 7.3949727103629e-05 | 26 |
| eef1db9 | 3.55672258711126e-09 | 0.329365436 | 0.486 | 0.273 | 8.95191507950034e-05 | 26 |
| b2m10 | 4.09999308016939e-09 | 0.336183334 | 0.514 | 0.291 | 0.000103193 | 26 |
| zgc:1119864 | 4.5970819941406e-09 | 0.290407499 | 0.354 | 0.176 | 0.000115704 | 26 |
| tpm34 | 5.52500365840644e-09 | 0.322865874 | 0.457 | 0.254 | 0.000139059 | 26 |
| crebzf2 | 7.50776993105674e-09 | 0.277528667 | 0.234 | 0.102 | 0.000188963 | 26 |
| atrx6 | 8.25460323596056e-09 | 0.267726511 | 0.24 | 0.104 | 0.00020776 | 26 |
| rps1312 | 1.09874904547579e-08 | 0.250210006 | 0.874 | 0.724 | 0.000276544 | 26 |
| mt-nd55 | 2.00816726851789e-08 | 0.282737049 | 0.257 | 0.12 | 0.000505436 | 26 |
| mt-co214 | 2.09595392756752e-08 | 0.275202408 | 0.823 | 0.72 | 0.000527531 | 26 |
| fabp11a4 | 2.54931791644305e-08 | 0.27506203 | 0.206 | 0.084 | 0.000641638 | 26 |
| arrdc3a4 | 2.81339223114313e-08 | 0.256454273 | 0.183 | 0.072 | 0.000708103 | 26 |
| bhlhe409 | 3.00438116983879e-08 | 0.276374679 | 0.326 | 0.16 | 0.000756173 | 26 |
| uba5211 | 3.28939573581254e-08 | 0.275772615 | 0.806 | 0.629 | 0.000827908 | 26 |
| marcksl1a6 | 4.2744841442963e-08 | 0.329207932 | 0.394 | 0.215 | 0.001075845 | 26 |
| dbi5 | 6.24005502319717e-08 | 0.306486413 | 0.383 | 0.213 | 0.001570559 | 26 |
| rpl27a13 | 1.00994693318798e-07 | 0.30580941 | 0.703 | 0.495 | 0.002541935 | 26 |

| | | | | | | |
|---|---|---|---|---|---|---|
| itm2ba9 | 1.12479561609399e-07 | 0.308572112 | 0.457 | 0.273 | 0.002830998 | 26 |
| rhoaa8 | 1.17742841145069e-07 | 0.25130414 | 0.303 | 0.154 | 0.00296347 | 26 |
| zfp36l1a8 | 1.60055934650036e-07 | 0.302642403 | 0.434 | 0.247 | 0.004028448 | 26 |
| rps27l9 | 1.80770238467209e-07 | 0.278050322 | 0.371 | 0.203 | 0.004549806 | 26 |
| mdka6 | 3.45329528684509e-07 | 0.293360106 | 0.206 | 0.094 | 0.008691599 | 26 |
| btf310 | 3.68714721020467e-07 | 0.292455544 | 0.811 | 0.586 | 0.009280181 | 26 |
| si:dkey-11f4.146 | 3.80011246141633e-07 | 0.26252538 | 0.229 | 0.109 | 0.009564503 | 26 |
| crip19 | 4.96781088519915e-07 | 0.340737057 | 0.429 | 0.27 | 0.012503483 | 26 |
| celf212 | 9.85591578995283e-07 | 0.281045533 | 0.411 | 0.242 | 0.024806354 | 26 |
| serbp1a11 | 1.0383709891443e-06 | 0.2989823 | 0.703 | 0.481 | 0.026134759 | 26 |
| rpl23a14 | 1.56338022806486e-06 | 0.252582316 | 0.811 | 0.633 | 0.039348717 | 26 |
| srsf5a5 | 2.00461607941075e-06 | 0.271676017 | 0.463 | 0.288 | 0.050454182 | 26 |
| romo110 | 2.11394219340235e-06 | 0.287517993 | 0.577 | 0.385 | 0.053205811 | 26 |
| cnbpb5 | 2.55792940744005e-06 | 0.266821629 | 0.32 | 0.184 | 0.064380525 | 26 |
| mt-co115 | 2.65659410103056e-06 | 0.253335948 | 0.8 | 0.657 | 0.066863817 | 26 |
| XLOC-0254238 | 2.91806168866995e-06 | 0.271468557 | 0.314 | 0.169 | 0.073444695 | 26 |
| ptmab10 | 4.99108396308546e-06 | 0.268873713 | 0.709 | 0.54 | 0.125620592 | 26 |
| eif4a1b6 | 7.45055792503464e-06 | 0.264574474 | 0.44 | 0.28 | 0.187523092 | 26 |
| mt-nd311 | 1.71777620179038e-05 | 0.256617821 | 0.566 | 0.396 | 0.432347092 | 26 |
| igl4v8 | 2.65501663639222e-115 | 0.849825764 | 0.347 | 0.029 | 6.68241137213559e-111 | 27 |
| LOC101886766 | 5.32776101647377e-95 | 0.57801734 | 0.216 | 0.013 | 1.34094417023628e-90 | 27 |
| zp38 | 1.87705455394289e-40 | 0.71374663 | 0.982 | 0.581 | 4.72435860681885e-36 | 27 |
| cldnd8 | 1.09240347551193e-38 | 0.709104791 | 1 | 0.602 | 2.74947030751599e-34 | 27 |
| si:dkey-241l7.27 | 2.188068226615e-38 | 0.674812231 | 0.994 | 0.639 | 5.50714891956729e-34 | 27 |
| zgc:1534997 | 9.75449838048361e-38 | 0.754214262 | 0.958 | 0.509 | 2.45510969738392e-33 | 27 |
| si:ch211-125e6.127 | 2.81598323575621e-37 | 0.738610242 | 0.94 | 0.498 | 7.0875482060748e-33 | 27 |
| si:dkeyp-98a7.97 | 4.28540888226146e-36 | 0.726301073 | 0.94 | 0.509 | 1.07859456157639e-31 | 27 |
| mt27 | 8.77994895475638e-36 | 0.706634788 | 0.952 | 0.532 | 2.20982535242263e-31 | 27 |
| zp2.37 | 1.72294071782845e-35 | 0.71568276 | 0.922 | 0.516 | 4.33646949270243e-31 | 27 |
| zp2.58 | 2.36941769052688e-35 | 0.692886112 | 0.958 | 0.561 | 5.96358738528709e-31 | 27 |
| zp3.28 | 2.35860299161182e-34 | 0.683195974 | 0.94 | 0.555 | 5.93636786958779e-30 | 27 |
| h2af1al8 | 8.95223629613685e-34 | 0.687181239 | 0.994 | 0.59 | 2.25318835337468e-29 | 27 |
| si:ch73-189n23.18 | 1.13858652591974e-32 | 0.705702138 | 0.946 | 0.522 | 2.86570842708741e-28 | 27 |
| zp2.67 | 1.22250871927157e-32 | 0.711152225 | 0.922 | 0.499 | 3.07693219553462e-28 | 27 |
| tmsb57 | 1.22436931748159e-32 | 0.720329398 | 0.928 | 0.486 | 3.08161513516941e-28 | 27 |

| | | | | | | |
|---|---|---|---|---|---|---|
| si:zfos-1505d6.37 | 5.02906134826253e-32 | 0.710566263 | 0.958 | 0.51 | 1.2657644507442e-27 | 27 |
| h1m8 | 3.88058339481332e-31 | 0.652280102 | 0.97 | 0.599 | 9.76704034640565e-27 | 27 |
| si:dkey-241l7.47 | 4.15305947340741e-31 | 0.622115799 | 0.946 | 0.607 | 1.04528353886191e-26 | 27 |
| si:ch211-12h2.87 | 4.2277351422737e-31 | 0.651272446 | 0.904 | 0.546 | 1.06407865795887e-26 | 27 |
| si:ch211-226h8.147 | 1.10282203893231e-30 | 0.660416904 | 0.952 | 0.564 | 2.77569278978872e-26 | 27 |
| zp3a.27 | 1.87847098908926e-29 | 0.663705431 | 0.856 | 0.489 | 4.72792363243877e-25 | 27 |
| cldng7 | 1.8838052824209e-29 | 0.677022734 | 0.916 | 0.511 | 4.74134951532517e-25 | 27 |
| si:dkey-240e12.67 | 1.92416309755315e-29 | 0.658536984 | 0.94 | 0.567 | 4.84292610023152e-25 | 27 |
| si:dkey-229d11.37 | 2.5786116455064e-29 | 0.691669593 | 0.808 | 0.436 | 6.49010765057507e-25 | 27 |
| zgc:1751358 | 3.10135851259463e-29 | 0.690864829 | 0.898 | 0.477 | 7.80580924034942e-25 | 27 |
| si:dkey-241l7.58 | 4.85018195191645e-29 | 0.630494375 | 0.94 | 0.582 | 1.22074229547785e-24 | 27 |
| acp5a8 | 5.06076321627763e-29 | 0.660904251 | 0.952 | 0.537 | 1.27374349390492e-24 | 27 |
| zp2.18 | 8.09276176255397e-29 | 0.663276199 | 0.922 | 0.532 | 2.03686720801721e-24 | 27 |
| si:ch211-14a17.77 | 2.36061921302169e-28 | 0.689741082 | 0.79 | 0.406 | 5.94144249725429e-24 | 27 |
| zgc:1717797 | 3.45698647449555e-28 | 0.661741098 | 0.904 | 0.51 | 8.70088925765786e-24 | 27 |
| ccna17 | 7.77478892394912e-28 | 0.6645379 | 0.868 | 0.474 | 1.95683662426875e-23 | 27 |
| zp2.27 | 1.05949889116274e-27 | 0.644575448 | 0.904 | 0.527 | 2.66665275916751e-23 | 27 |
| crp27 | 2.65769150948862e-27 | 0.665224046 | 0.88 | 0.487 | 6.68914376023192e-23 | 27 |
| si:dkeyp-46h3.67 | 1.58443899235914e-26 | 0.670656281 | 0.772 | 0.408 | 3.98787449986871e-22 | 27 |
| CR790368.18 | 1.59279018588434e-26 | 0.634026243 | 0.886 | 0.518 | 4.00889361885228e-22 | 27 |
| si:dkeyp-98a7.87 | 3.36478677958847e-26 | 0.65392664 | 0.826 | 0.456 | 8.46883184554623e-22 | 27 |
| si:ch211-226h8.47 | 3.57297537183946e-26 | 0.652971943 | 0.79 | 0.44 | 8.99282171338273e-22 | 27 |
| zgc:771187 | 5.96511646699393e-26 | 0.650869533 | 0.832 | 0.455 | 1.5013601635777e-21 | 27 |
| zgc:1717767 | 3.10243725488699e-25 | 0.632123534 | 0.862 | 0.493 | 7.80852432682507e-21 | 27 |
| mid1ip1l7 | 3.21633539781639e-25 | 0.597806419 | 0.886 | 0.541 | 8.09519456276408e-21 | 27 |
| si:dkeyp-98a7.47 | 3.75993419425644e-25 | 0.642989307 | 0.772 | 0.423 | 9.46337837352404e-21 | 27 |
| epcam8 | 4.05612070222717e-25 | 0.63058084 | 0.874 | 0.503 | 1.02088501954356e-20 | 27 |
| nanog7 | 1.11438422803454e-24 | 0.662795078 | 0.719 | 0.357 | 2.80479366354015e-20 | 27 |
| cth17 | 4.04061502013004e-24 | 0.645799896 | 0.731 | 0.377 | 1.01698239441653e-19 | 27 |
| si:dkey-229d11.57 | 5.77422574196639e-24 | 0.648396875 | 0.731 | 0.387 | 1.45331487699552e-19 | 27 |
| si:dkey-19b23.117 | 7.57123039879008e-24 | 0.679914593 | 0.587 | 0.277 | 1.90560297907148e-19 | 27 |
| si:dkeyp-20e4.87 | 7.94938691130878e-24 | 0.657170021 | 0.76 | 0.406 | 2.00078119170731e-19 | 27 |
| LOC1104378217 | 1.55192425324101e-23 | 0.647289252 | 0.731 | 0.414 | 3.90603815298229e-19 | 27 |
| zgc:1734437 | 1.04476267471596e-22 | 0.649529803 | 0.671 | 0.328 | 2.62956317599259e-18 | 27 |
| si:dkeyp-50f7.27 | 3.75448049703214e-22 | 0.617027392 | 0.749 | 0.4 | 9.44965196298018e-18 | 27 |

| | | | | | | |
|---|---|---|---|---|---|---|
| BX649337.18 | 4.01536755337938e-22 | 0.621064049 | 0.802 | 0.436 | 1.01062785951006e-17 | 27 |
| LOC1005374997 | 4.5187032770422e-22 | 0.635322299 | 0.641 | 0.32 | 1.13731242779875e-17 | 27 |
| zgc:1715177 | 2.04125431777241e-21 | 0.618347898 | 0.707 | 0.387 | 5.13763299240137e-17 | 27 |
| btg47 | 3.91535536042715e-21 | 0.615862436 | 0.701 | 0.367 | 9.85455790665908e-17 | 27 |
| lamp27 | 4.28417482537443e-21 | 0.550973771 | 0.82 | 0.553 | 1.07828396179849e-16 | 27 |
| zar1l7 | 4.43244854421661e-21 | 0.612232083 | 0.749 | 0.396 | 1.11560297409388e-16 | 27 |
| npm2b7 | 7.57422693296913e-21 | 0.608253635 | 0.707 | 0.395 | 1.906357176759e-16 | 27 |
| AL929032.17 | 7.99913547424508e-21 | 0.63079336 | 0.653 | 0.315 | 2.01330240751275e-16 | 27 |
| slbp27 | 9.85255956791593e-21 | 0.602305401 | 0.731 | 0.395 | 2.47979071764876e-16 | 27 |
| zp3b7 | 1.20715764363656e-20 | 0.611363538 | 0.725 | 0.386 | 3.03829507326887e-16 | 27 |
| si:dkey-90l23.17 | 2.99528666117876e-20 | 0.591159438 | 0.772 | 0.446 | 7.53883699752083e-16 | 27 |
| rrm27 | 5.73304085389541e-20 | 0.561999823 | 0.814 | 0.497 | 1.44294905251694e-15 | 27 |
| zgc:1952457 | 7.26052838956536e-20 | 0.594736641 | 0.701 | 0.387 | 1.82740239036971e-15 | 27 |
| wu:fi42e037 | 1.20997048587276e-19 | 0.599193547 | 0.653 | 0.349 | 3.04537471589316e-15 | 27 |
| si:dkey-37o8.17 | 1.41333595721716e-19 | 0.606442731 | 0.599 | 0.306 | 3.55722527071987e-15 | 27 |
| si:dkey-241l7.67 | 1.56523911853158e-19 | 0.551998351 | 0.82 | 0.526 | 3.93955033743212e-15 | 27 |
| ccnb27 | 1.93386954704734e-19 | 0.595765223 | 0.701 | 0.383 | 4.86735626296344e-15 | 27 |
| si:ch1073-263o8.27 | 4.12413429169137e-19 | 0.606938066 | 0.545 | 0.262 | 1.0380033598758e-14 | 27 |
| si:ch211-132b12.87 | 1.14225251468751e-18 | 0.590249565 | 0.629 | 0.341 | 2.87493535421699e-14 | 27 |
| ccnb17 | 1.91720270905353e-18 | 0.575204493 | 0.689 | 0.392 | 4.82540749841683e-14 | 27 |
| qdprb27 | 2.22488924565576e-18 | 0.567323259 | 0.731 | 0.432 | 5.59982374239099e-14 | 27 |
| zgc:1630306 | 2.34265753787136e-18 | 0.550757002 | 0.341 | 0.123 | 5.89623475706844e-14 | 27 |
| LOC1018854777 | 2.58585971688741e-18 | 0.603274722 | 0.551 | 0.279 | 6.50835032143392e-14 | 27 |
| kpna77 | 2.62846268405507e-18 | 0.591931985 | 0.617 | 0.327 | 6.6155777294982e-14 | 27 |
| CU571163.17 | 5.17486954393625e-18 | 0.577527113 | 0.719 | 0.4 | 1.30246291551331e-13 | 27 |
| LOC1005351627 | 5.44332933942336e-18 | 0.58749756 | 0.599 | 0.308 | 1.37003156143947e-13 | 27 |
| s100a17 | 8.71997201952946e-18 | 0.572915074 | 0.677 | 0.362 | 2.19472975759537e-13 | 27 |
| tuba4l7 | 1.14976992888627e-17 | 0.567030236 | 0.719 | 0.403 | 2.89385593401384e-13 | 27 |
| hnrnpabb8 | 1.25489255057282e-17 | 0.466130346 | 0.802 | 0.596 | 3.15843906053673e-13 | 27 |
| siva17 | 2.20321003683985e-17 | 0.552023939 | 0.713 | 0.427 | 5.54525934172222e-13 | 27 |
| si:ch211-5k11.87 | 1.9295000221176e-16 | 0.504811491 | 0.677 | 0.357 | 4.85635860566778e-12 | 27 |
| zar17 | 2.79731804732176e-16 | 0.526383393 | 0.713 | 0.439 | 7.04056979330413e-12 | 27 |
| si:ch211-12h2.67 | 5.91035589773457e-16 | 0.524709381 | 0.766 | 0.473 | 1.48757747590081e-11 | 27 |
| zgc:1526525 | 7.85808576602502e-16 | 0.52957556 | 0.305 | 0.113 | 1.97780160645084e-11 | 27 |
| hbba17 | 1.48678035284846e-15 | 0.534838513 | 0.677 | 0.354 | 3.74207747008429e-11 | 27 |

| | | | | | | |
|---|---|---|---|---|---|---|
| arhgef10la7 | 1.76490619113457e-15 | 0.549824405 | 0.539 | 0.274 | 4.44209239246659e-11 | 27 |
| si:dkey-208k4.27 | 2.53158569936369e-15 | 0.540295394 | 0.557 | 0.295 | 6.37174804672847e-11 | 27 |
| si:ch211-226l4.68 | 2.70742110293419e-15 | 0.550139073 | 0.551 | 0.292 | 6.81430817397506e-11 | 27 |
| gdf37 | 4.69278640378296e-15 | 0.546320789 | 0.461 | 0.223 | 1.18112740996813e-10 | 27 |
| si:dkey-152b24.67 | 6.19401707655845e-15 | 0.542460385 | 0.563 | 0.297 | 1.558972157999e-10 | 27 |
| zgc:1362547 | 8.75549810708372e-15 | 0.532971691 | 0.599 | 0.33 | 2.2036713185719e-10 | 27 |
| ca15b7 | 1.10542996175573e-14 | 0.541220379 | 0.497 | 0.248 | 2.78225667074299e-10 | 27 |
| zgc:1034828 | 1.17897477147774e-14 | 0.529312937 | 0.413 | 0.188 | 2.96736160233231e-10 | 27 |
| gbgt1l46 | 1.23661931658167e-14 | 0.51493117 | 0.359 | 0.149 | 3.11244715790442e-10 | 27 |
| cpeb1b7 | 1.82372877281507e-14 | 0.528072193 | 0.533 | 0.269 | 4.59014294829825e-10 | 27 |
| si:dkey-90l23.27 | 2.09142506260342e-14 | 0.531923385 | 0.611 | 0.344 | 5.26390774006656e-10 | 27 |
| LOC1018838977 | 2.2953947659321e-14 | 0.512198478 | 0.377 | 0.164 | 5.77727908637450e-10 | 27 |
| si:dkeyp-46h3.87 | 3.89480892508883e-14 | 0.530274581 | 0.629 | 0.351 | 9.80284458355608e-10 | 27 |
| hbaa17 | 4.43366653468363e-14 | 0.547407943 | 0.533 | 0.281 | 1.11590953011452e-09 | 27 |
| si:dkeyp-46h3.27 | 1.78117839654028e-13 | 0.495090686 | 0.659 | 0.404 | 4.48304790625224e-09 | 27 |
| si:ch211-146l10.87 | 1.8580303679751e-13 | 0.516266115 | 0.545 | 0.306 | 4.67647663315653e-09 | 27 |
| zgc:1714747 | 2.04472681539668e-13 | 0.515697955 | 0.539 | 0.304 | 5.14637292167192e-09 | 27 |
| uchl17 | 2.16688497319589e-13 | 0.510073644 | 0.599 | 0.344 | 5.45383278903673e-09 | 27 |
| zpcx7 | 2.18527309543047e-13 | 0.51479754 | 0.587 | 0.339 | 5.50011385388895e-09 | 27 |
| zgc:554618 | 4.80902115314084e-13 | 0.505358925 | 0.527 | 0.288 | 1.21038253403402e-08 | 27 |
| ctss2.18 | 7.4822550379064e-13 | 0.435138764 | 0.76 | 0.534 | 1.88320877049066e-08 | 27 |
| spink2.18 | 1.66634172963977e-12 | 0.463597513 | 0.677 | 0.436 | 4.19401549933035e-08 | 27 |
| zp3a.17 | 2.34243374099981e-12 | 0.483173634 | 0.611 | 0.381 | 5.89567148272243e-08 | 27 |
| gapdh7 | 5.26269612439067e-12 | 0.467243223 | 0.647 | 0.42 | 1.32456798754789e-07 | 27 |
| avd7 | 6.62338224952973e-12 | 0.482598423 | 0.491 | 0.265 | 1.66703907838414e-07 | 27 |
| mos6 | 8.1439380537664e-12 | 0.471608291 | 0.311 | 0.136 | 2.04974776875247e-07 | 27 |
| gmnn7 | 9.41240076214747e-12 | 0.487092839 | 0.527 | 0.303 | 2.3690071478249e-07 | 27 |
| si:dkeyp-98a7.77 | 1.08934091916848e-11 | 0.471190047 | 0.407 | 0.195 | 2.74176215945514e-07 | 27 |
| gtf3ab8 | 1.22472149355778e-11 | 0.473935354 | 0.557 | 0.32 | 3.08250152713558e-07 | 27 |
| si:ch211-286b5.42 | 1.87429160363746e-11 | 0.388327329 | 0.204 | 0.07 | 4.71740453719514e-07 | 27 |
| plk16 | 2.09154877999375e-11 | 0.47244506 | 0.365 | 0.177 | 5.26421912436628e-07 | 27 |
| thy17 | 3.10681222126488e-11 | 0.447380385 | 0.695 | 0.45 | 7.81953567970159e-07 | 27 |
| si:ch211-251f6.67 | 3.39004541572654e-11 | 0.471205466 | 0.533 | 0.331 | 8.53240530684212e-07 | 27 |
| zgc:1737427 | 4.15815659256154e-11 | 0.475869114 | 0.329 | 0.155 | 1.04656643278182e-06 | 27 |
| bzw1b4 | 5.47207419021033e-11 | 0.44406826 | 0.563 | 0.402 | 1.37726635293404e-06 | 27 |

| | | | | | | |
|---|---|---|---|---|---|---|
| zgc:1139844 | 7.11609746495936e-11 | 0.41644812 | 0.251 | 0.101 | 1.79105057095562e-06 | 27 |
| AL929192.17 | 1.42284326528543e-10 | 0.441549629 | 0.347 | 0.164 | 3.58115421439689e-06 | 27 |
| si:ch211-146l10.77 | 1.46532593927837e-10 | 0.462217734 | 0.371 | 0.189 | 3.68807885656974e-06 | 27 |
| LOC1005368607 | 1.865447934809e-10 | 0.445588118 | 0.551 | 0.351 | 4.69514590712077e-06 | 27 |
| tmem702 | 2.30768356959736e-10 | 0.42148996 | 0.251 | 0.105 | 5.8082087763196e-06 | 27 |
| eif4e1b2 | 3.98365856018377e-10 | 0.383560309 | 0.222 | 0.086 | 1.00264702301265e-05 | 27 |
| pigrl2.37 | 5.31529525367016e-10 | 0.435298253 | 0.395 | 0.203 | 1.33780666239624e-05 | 27 |
| nasp7 | 5.42192500177387e-10 | 0.446585196 | 0.545 | 0.34 | 1.36464430369647e-05 | 27 |
| srsf2a4 | 6.72474558438414e-10 | 0.434295975 | 0.527 | 0.367 | 1.69255121613364e-05 | 27 |
| sybu7 | 6.87039341234687e-10 | 0.443449327 | 0.335 | 0.166 | 1.72920931795358e-05 | 27 |
| mkrn46 | 7.65324488501276e-10 | 0.445225597 | 0.413 | 0.228 | 1.92624520510886e-05 | 27 |
| si:ch73-14h10.27 | 9.1294455648187e-10 | 0.440256398 | 0.473 | 0.271 | 2.29779015420922e-05 | 27 |
| zglp17 | 9.16675394786074e-10 | 0.436763189 | 0.377 | 0.198 | 2.30718030113707e-05 | 27 |
| zgc:566998 | 9.40277633672597e-10 | 0.439639577 | 0.449 | 0.251 | 2.36658477619056e-05 | 27 |
| pabpn1l7 | 1.11651236215222e-09 | 0.436179091 | 0.485 | 0.284 | 2.81014996430092e-05 | 27 |
| si:dkeyp-46h3.57 | 1.21133129844537e-09 | 0.442681592 | 0.485 | 0.29 | 3.04879974505716e-05 | 27 |
| si:ch211-250e5.167 | 1.42183741675767e-09 | 0.430661803 | 0.539 | 0.346 | 3.57862259423737e-05 | 27 |
| sox19b5 | 1.79661309881482e-09 | 0.397771181 | 0.234 | 0.098 | 4.52189550840703e-05 | 27 |
| cdca86 | 1.9331166238377e-09 | 0.430002948 | 0.329 | 0.165 | 4.86546123053711e-05 | 27 |
| mibp7 | 2.25365289360275e-09 | 0.440682079 | 0.401 | 0.223 | 5.67221896790876e-05 | 27 |
| setd4 | 3.35907079328698e-09 | 0.349183338 | 0.186 | 0.07 | 8.454445279624e-05 | 27 |
| b4galnt2.2 | 4.32955257958593e-09 | 0.289627731 | 0.132 | 0.041 | 0.000108971 | 27 |
| crp37 | 6.09917874869116e-09 | 0.42209114 | 0.395 | 0.219 | 0.00015351 | 27 |
| fdx16 | 6.85300817791002e-09 | 0.428553022 | 0.383 | 0.221 | 0.000172483 | 27 |
| st8sia7.11 | 7.88902754959537e-09 | 0.345860604 | 0.186 | 0.071 | 0.000198559 | 27 |
| birc5b7 | 7.97204610558632e-09 | 0.424961864 | 0.443 | 0.26 | 0.000200648 | 27 |
| bach2a | 8.58004327076243e-09 | 0.320374214 | 0.15 | 0.051 | 0.000215951 | 27 |
| CDPF17 | 1.13705475800763e-08 | 0.423263995 | 0.443 | 0.269 | 0.000286185 | 27 |
| elovl7a7 | 1.34043610797829e-08 | 0.411256838 | 0.347 | 0.183 | 0.000337374 | 27 |
| cldn7b6 | 1.44359314189621e-08 | 0.412046624 | 0.311 | 0.16 | 0.000363338 | 27 |
| zgc:562317 | 1.58470317975826e-08 | 0.418364809 | 0.401 | 0.228 | 0.000398854 | 27 |
| zgc:1134247 | 1.67578893040416e-08 | 0.396720472 | 0.317 | 0.162 | 0.000421779 | 27 |
| yrdc5 | 2.15352916940474e-08 | 0.374657678 | 0.263 | 0.123 | 0.000542022 | 27 |
| ppfibp1a1 | 3.35018917879954e-08 | 0.271039493 | 0.132 | 0.043 | 0.000843209 | 27 |
| zgc:1738567 | 3.45643600521181e-08 | 0.405334902 | 0.383 | 0.22 | 0.00086995 | 27 |

| | | | | | | |
|---|---|---|---|---|---|---|
| si:dkey-188i13.118 | 4.26865551599844e-08 | 0.412450126 | 0.479 | 0.317 | 0.001074378 | 27 |
| si:ch73-160p18.37 | 4.7125666845442e-08 | 0.431138029 | 0.395 | 0.241 | 0.001186106 | 27 |
| suv39h1b | 8.60855000378503e-08 | 0.295022289 | 0.132 | 0.046 | 0.002166686 | 27 |
| si:dkey-42i9.87 | 1.01198059871515e-07 | 0.363970541 | 0.263 | 0.128 | 0.002547054 | 27 |
| zp2l17 | 1.02036227968496e-07 | 0.392993964 | 0.305 | 0.164 | 0.00256815 | 27 |
| gstp15 | 1.14107438479017e-07 | 0.38587734 | 0.509 | 0.366 | 0.00287197 | 27 |
| cdca93 | 1.15134737797857e-07 | 0.350202605 | 0.198 | 0.086 | 0.002897826 | 27 |
| zgc:1138866 | 1.1732030837663e-07 | 0.373314095 | 0.281 | 0.142 | 0.002952835 | 27 |
| LOC1104377793 | 1.19070412022315e-07 | 0.330119145 | 0.198 | 0.084 | 0.002996883 | 27 |
| larp6a7 | 1.24661956381803e-07 | 0.404237574 | 0.407 | 0.25 | 0.003137617 | 27 |
| lsm14b7 | 1.56118856095898e-07 | 0.388479973 | 0.473 | 0.3 | 0.003929355 | 27 |
| rcc17 | 1.85197688312544e-07 | 0.3946791 | 0.353 | 0.203 | 0.004661241 | 27 |
| snx10a5 | 2.02271788189415e-07 | 0.35619317 | 0.257 | 0.126 | 0.005090979 | 27 |
| elavl28 | 2.0453158922656e-07 | 0.394331285 | 0.377 | 0.224 | 0.005147856 | 27 |
| dynll18 | 2.07890672192051e-07 | 0.344943347 | 0.587 | 0.455 | 0.0052324 | 27 |
| rbm4.37 | 2.34647374758296e-07 | 0.34594099 | 0.551 | 0.445 | 0.00590584 | 27 |
| spsb11 | 2.47294471061857e-07 | 0.361026217 | 0.204 | 0.093 | 0.006224155 | 27 |
| pou5f37 | 2.54938739509173e-07 | 0.363383645 | 0.287 | 0.147 | 0.006416553 | 27 |
| rbpms2b7 | 2.62097908224221e-07 | 0.387527312 | 0.371 | 0.222 | 0.006596742 | 27 |
| zgc:1628797 | 2.75012951975472e-07 | 0.384002541 | 0.329 | 0.182 | 0.006921801 | 27 |
| slbp8 | 2.86704750061122e-07 | 0.385207274 | 0.389 | 0.234 | 0.007216072 | 27 |
| tac3b6 | 2.8856771451546e-07 | 0.341736426 | 0.251 | 0.121 | 0.007262961 | 27 |
| snu13b8 | 3.19036341316629e-07 | 0.373834271 | 0.533 | 0.393 | 0.008029826 | 27 |
| zgc:925917 | 3.88689213833128e-07 | 0.375063155 | 0.359 | 0.206 | 0.009782919 | 27 |
| si:dkeyp-75b4.104 | 3.99007126045621e-07 | 0.317944659 | 0.605 | 0.437 | 0.01004261 | 27 |
| h2afva6 | 4.68030995936035e-07 | 0.383367165 | 0.455 | 0.324 | 0.011779872 | 27 |
| ddx412 | 4.68127359050659e-07 | 0.302761209 | 0.168 | 0.069 | 0.011782297 | 27 |
| ccna27 | 4.69536249218462e-07 | 0.373168831 | 0.449 | 0.285 | 0.011817758 | 27 |
| slc16a37 | 5.47128181573041e-07 | 0.375035796 | 0.413 | 0.249 | 0.013770669 | 27 |
| LOC100334439 | 5.83396327039071e-07 | 0.269717473 | 0.144 | 0.055 | 0.014683502 | 27 |
| hspa56 | 6.06358302357516e-07 | 0.374440217 | 0.461 | 0.315 | 0.015261432 | 27 |
| zgc:1629457 | 6.57640906438797e-07 | 0.361138694 | 0.299 | 0.163 | 0.016552164 | 27 |
| zgc:868394 | 6.75365199290641e-07 | 0.342700877 | 0.228 | 0.11 | 0.016998267 | 27 |
| h3f3a8 | 7.58074285424373e-07 | 0.319797195 | 0.605 | 0.507 | 0.019079972 | 27 |
| dctpp18 | 7.76816728167471e-07 | 0.367026603 | 0.311 | 0.176 | 0.0195517 | 27 |

| | | | | | | |
|---|---|---|---|---|---|---|
| zgc:660247 | 9.17852993209754e-07 | 0.349828714 | 0.269 | 0.142 | 0.023101442 | 27 |
| BX005461.31 | 9.65933595742249e-07 | 0.298099863 | 0.162 | 0.067 | 0.024311583 | 27 |
| si:ch1073-190k2.17 | 1.09995408040484e-06 | 0.374728999 | 0.323 | 0.189 | 0.027684744 | 27 |
| CABZ01059627.15 | 1.14218361443783e-06 | 0.336239797 | 0.263 | 0.137 | 0.028747619 | 27 |
| g3bp15 | 1.14875880962917e-06 | 0.374680712 | 0.347 | 0.211 | 0.02891311 | 27 |
| degs25 | 1.36545827575247e-06 | 0.320407394 | 0.222 | 0.107 | 0.034367219 | 27 |
| rpz23 | 1.46461470198454e-06 | 0.373847782 | 0.311 | 0.182 | 0.036862887 | 27 |
| si:ch73-90k17.17 | 1.57783898763445e-06 | 0.357176775 | 0.407 | 0.249 | 0.039712629 | 27 |
| zp3c7 | 1.74731331054323e-06 | 0.362455572 | 0.323 | 0.19 | 0.043978129 | 27 |
| map1lc3c7 | 2.0622006037476e-06 | 0.344402224 | 0.347 | 0.202 | 0.051903527 | 27 |
| cat | 2.13460646973226e-06 | 0.301634532 | 0.174 | 0.078 | 0.05372591 | 27 |
| si:dkey-241l7.37 | 2.20458510917804e-06 | 0.357060158 | 0.515 | 0.365 | 0.055487203 | 27 |
| sinup7 | 2.22952405251137e-06 | 0.362245523 | 0.341 | 0.204 | 0.056114891 | 27 |
| zgc:1655396 | 2.25002925834362e-06 | 0.350863846 | 0.234 | 0.12 | 0.056630986 | 27 |
| tmem54a5 | 2.75703044847538e-06 | 0.335596064 | 0.263 | 0.142 | 0.069391699 | 27 |
| hsd17b12a7 | 2.97998484633271e-06 | 0.34787548 | 0.293 | 0.166 | 0.075003239 | 27 |
| org7 | 3.298100855542e-06 | 0.37614904 | 0.377 | 0.241 | 0.0830099 | 27 |
| zgc:1714262 | 3.36547020873392e-06 | 0.326097591 | 0.216 | 0.108 | 0.08470552 | 27 |
| si:ch211-210c8.72 | 3.9156343637856e-06 | 0.342942447 | 0.24 | 0.128 | 0.098552601 | 27 |
| si:ch211-202f5.33 | 3.99312942542053e-06 | 0.311668684 | 0.192 | 0.091 | 0.100503075 | 27 |
| si:ch211-113a14.272 | 4.14057153807703e-06 | 0.285340344 | 0.174 | 0.078 | 0.104214045 | 27 |
| larp6b7 | 4.3544504221386e-06 | 0.35181497 | 0.329 | 0.197 | 0.109597163 | 27 |
| mad2l16 | 4.59387677528866e-06 | 0.347812795 | 0.311 | 0.187 | 0.115623285 | 27 |
| aktip3 | 5.09521293347486e-06 | 0.353716792 | 0.251 | 0.14 | 0.128241414 | 27 |
| zgc:1717507 | 5.24596890830008e-06 | 0.34854216 | 0.299 | 0.175 | 0.132035791 | 27 |
| CR847984.17 | 5.27698596642209e-06 | 0.344081138 | 0.287 | 0.165 | 0.13281646 | 27 |
| mcm3l3 | 6.17185001577432e-06 | 0.297195152 | 0.186 | 0.088 | 0.155339293 | 27 |
| si:ch211-157c3.44 | 6.32281259166147e-06 | 0.287003146 | 0.192 | 0.09 | 0.15913887 | 27 |
| BX571780.14 | 6.46193614661248e-06 | 0.269662074 | 0.18 | 0.082 | 0.162640471 | 27 |
| si:dkey-81e3.2 | 6.60779072226198e-06 | 0.271309009 | 0.15 | 0.064 | 0.166311485 | 27 |
| cdc207 | 6.87452429488618e-06 | 0.327394624 | 0.275 | 0.153 | 0.173024902 | 27 |
| tmem441 | 7.01716992066746e-06 | 0.270066262 | 0.144 | 0.061 | 0.17661515 | 27 |
| retsatl7 | 7.38525941553015e-06 | 0.35338231 | 0.335 | 0.204 | 0.185879594 | 27 |
| si:dkey-189g17.22 | 7.50187095870266e-06 | 0.251953703 | 0.138 | 0.057 | 0.18881459 | 27 |
| unga1 | 7.67864259674699e-06 | 0.289633545 | 0.168 | 0.077 | 0.193263756 | 27 |

| | | | | | | |
|---|---|---|---|---|---|---|
| zgc:1719776 | 7.70140013573853e-06 | 0.361813872 | 0.365 | 0.231 | 0.19383654 | 27 |
| asz1 | 7.81288080158621e-06 | 0.283835427 | 0.162 | 0.073 | 0.196642397 | 27 |
| asf1ba1 | 8.57876996573981e-06 | 0.302620934 | 0.198 | 0.098 | 0.215919061 | 27 |
| CU467646.26 | 8.99886614610457e-06 | 0.31199193 | 0.24 | 0.127 | 0.226492462 | 27 |
| ppp4r2b | 9.99986637919034e-06 | 0.29226356 | 0.162 | 0.075 | 0.251686637 | 27 |
| BX284697.15 | 1.00762587537878e-05 | 0.312956998 | 0.222 | 0.115 | 0.253609357 | 27 |
| hmga1a2 | 1.10526111409534e-05 | 0.286751534 | 0.539 | 0.451 | 0.27818317 | 27 |
| tmco11 | 1.18113112975011e-05 | 0.336246679 | 0.222 | 0.121 | 0.297278894 | 27 |
| slc29a21 | 1.25482796960034e-05 | 0.285625972 | 0.162 | 0.075 | 0.315827652 | 27 |
| eif4a1a5 | 1.27249780339768e-05 | 0.305541511 | 0.527 | 0.425 | 0.320274972 | 27 |
| chia.47 | 1.31798483012864e-05 | 0.316564015 | 0.263 | 0.148 | 0.331723602 | 27 |
| gpr1603 | 1.39331798715987e-05 | 0.275769868 | 0.174 | 0.082 | 0.350684204 | 27 |
| CABZ01059627.27 | 1.52432922396821e-05 | 0.315971522 | 0.246 | 0.134 | 0.383658422 | 27 |
| si:dkey-46g23.27 | 1.54058853547058e-05 | 0.315327034 | 0.257 | 0.146 | 0.387750728 | 27 |
| si:dkey-203a12.23 | 1.58564601760057e-05 | 0.285100231 | 0.377 | 0.247 | 0.399091246 | 27 |
| pttg17 | 1.75116790880299e-05 | 0.319453982 | 0.275 | 0.159 | 0.440751451 | 27 |
| cx44.27 | 1.88850154660848e-05 | 0.29978512 | 0.222 | 0.118 | 0.475316954 | 27 |
| cpm4 | 1.94919434659141e-05 | 0.292113687 | 0.21 | 0.109 | 0.490592725 | 27 |
| patl27 | 2.0926742829379e-05 | 0.313638279 | 0.293 | 0.174 | 0.52670519 | 27 |
| phex2 | 2.14042862922194e-05 | 0.272734661 | 0.18 | 0.087 | 0.538724482 | 27 |
| pabpc1l7 | 2.32840452455103e-05 | 0.314148732 | 0.251 | 0.143 | 0.586036135 | 27 |
| tuba8l43 | 2.34137151215502e-05 | 0.302608443 | 0.491 | 0.406 | 0.589299796 | 27 |
| rnf46 | 2.5699951310282e-05 | 0.31443436 | 0.24 | 0.135 | 0.646842075 | 27 |
| si:dkey-216e24.35 | 2.82398053273649e-05 | 0.298974038 | 0.228 | 0.124 | 0.71076766 | 27 |
| tmem144b3 | 2.87704417121357e-05 | 0.293344125 | 0.186 | 0.095 | 0.724123247 | 27 |
| zgc:566767 | 2.88485494340592e-05 | 0.34044957 | 0.407 | 0.296 | 0.726089141 | 27 |
| zgc:922876 | 3.11270273092773e-05 | 0.302195614 | 0.24 | 0.133 | 0.78343615 | 27 |
| ccne15 | 3.13872855587993e-05 | 0.299631237 | 0.21 | 0.112 | 0.78998659 | 27 |
| zgc:1118684 | 3.52227498962757e-05 | 0.31132062 | 0.257 | 0.148 | 0.886521392 | 27 |
| rdh10b6 | 3.6928282149911e-05 | 0.316300851 | 0.293 | 0.179 | 0.929447933 | 27 |
| rpf13 | 3.92137565008638e-05 | 0.338812374 | 0.234 | 0.137 | 0.986971037 | 27 |
| h2afvb2 | 4.5819192858127e-05 | 0.272059559 | 0.551 | 0.471 | 1 | 27 |
| ube2c5 | 4.7241132201926e-05 | 0.29689608 | 0.246 | 0.141 | 1 | 27 |
| orc64 | 4.75257802029345e-05 | 0.282855973 | 0.204 | 0.108 | 1 | 27 |
| ruvbl11 | 5.08671836362206e-05 | 0.285352555 | 0.174 | 0.088 | 1 | 27 |

| | | | | | | | |
|---|---|---|---|---|---|---|---|
| spryd42 | 5.16457089856435e-05 | 0.259510845 | 0.162 | 0.078 | | 1 | 27 |
| ndc11 | 5.17025804371743e-05 | 0.289987848 | 0.186 | 0.098 | | 1 | 27 |
| ccnh4 | 5.39379030052168e-05 | 0.290881847 | 0.228 | 0.126 | | 1 | 27 |
| thoc71 | 6.59674629714533e-05 | 0.304427017 | 0.21 | 0.117 | | 1 | 27 |
| ankrd227 | 6.65315443936364e-05 | 0.308043008 | 0.246 | 0.144 | | 1 | 27 |
| CT573476.1 | 6.87594871618148e-05 | 0.257160808 | 0.132 | 0.06 | | 1 | 27 |
| cdca7a7 | 7.2085904518913e-05 | 0.319791548 | 0.335 | 0.223 | | 1 | 27 |
| slc35f2 | 7.26598714504943e-05 | 0.258010191 | 0.156 | 0.076 | | 1 | 27 |
| st3gal8 | 7.31317178662511e-05 | 0.276399171 | 0.156 | 0.078 | | 1 | 27 |
| si:ch211-145c1.17 | 7.33171500235392e-05 | 0.316070033 | 0.353 | 0.235 | | 1 | 27 |
| elovl7b7 | 8.04860795576597e-05 | 0.315113672 | 0.371 | 0.249 | | 1 | 27 |
| zgc:1017444 | 0.000100826 | 0.284164284 | 0.234 | 0.134 | | 1 | 27 |
| spc241 | 0.000116728 | 0.262539106 | 0.144 | 0.07 | | 1 | 27 |
| si:ch211-69g19.25 | 0.000123032 | 0.283479261 | 0.216 | 0.123 | | 1 | 27 |
| fbxo324 | 0.000138327 | 0.269595568 | 0.192 | 0.105 | | 1 | 27 |
| ivns1abpb2 | 0.000148083 | 0.29748314 | 0.204 | 0.117 | | 1 | 27 |
| aurka4 | 0.000149339 | 0.289554615 | 0.21 | 0.12 | | 1 | 27 |
| ypel35 | 0.00015016 | 0.277546632 | 0.222 | 0.128 | | 1 | 27 |
| si:dkey-68o6.57 | 0.0001503 | 0.296740054 | 0.216 | 0.123 | | 1 | 27 |
| rangrf | 0.000153799 | 0.26674419 | 0.15 | 0.076 | | 1 | 27 |
| faub6 | 0.000179463 | 0.301960531 | 0.305 | 0.201 | | 1 | 27 |
| gyg1a8 | 0.0001825 | 0.300824399 | 0.467 | 0.371 | | 1 | 27 |
| lipg3 | 0.000198793 | 0.262428932 | 0.162 | 0.085 | | 1 | 27 |
| tph1a7 | 0.000199931 | 0.274231348 | 0.24 | 0.143 | | 1 | 27 |
| BX537276.17 | 0.000200044 | 0.26466845 | 0.216 | 0.122 | | 1 | 27 |
| c1d | 0.000210094 | 0.314089723 | 0.216 | 0.129 | | 1 | 27 |
| tomm40 | 0.00023087 | 0.296529867 | 0.186 | 0.107 | | 1 | 27 |
| pop42 | 0.000237834 | 0.290588379 | 0.204 | 0.118 | | 1 | 27 |
| chtopa5 | 0.000253477 | 0.308712158 | 0.317 | 0.222 | | 1 | 27 |
| fbxw11b2 | 0.000315209 | 0.296083341 | 0.234 | 0.146 | | 1 | 27 |
| kpna27 | 0.000327363 | 0.284317873 | 0.251 | 0.159 | | 1 | 27 |
| snrpb25 | 0.000357592 | 0.293385371 | 0.24 | 0.15 | | 1 | 27 |
| hells4 | 0.000390046 | 0.274169156 | 0.222 | 0.134 | | 1 | 27 |
| mcm51 | 0.000398845 | 0.260100939 | 0.174 | 0.097 | | 1 | 27 |
| rpain4 | 0.000416711 | 0.274597375 | 0.204 | 0.12 | | 1 | 27 |

| | | | | | | |
|---|---|---|---|---|---|---|
| ctsla7 | 0.000479605 | 0.264512404 | 0.473 | 0.388 | 1 | 27 |
| rac1a1 | 0.000503067 | 0.285709225 | 0.234 | 0.149 | 1 | 27 |
| prdx17 | 0.000518276 | 0.287023339 | 0.257 | 0.168 | 1 | 27 |
| rps19bp11 | 0.000531382 | 0.26078196 | 0.168 | 0.094 | 1 | 27 |
| sec61a1l1 | 0.000566608 | 0.2766809 | 0.21 | 0.128 | 1 | 27 |
| ctsba7 | 0.00058899 | 0.264389799 | 0.467 | 0.378 | 1 | 27 |
| si:ch73-281n10.24 | 0.000597804 | 0.257783198 | 0.473 | 0.394 | 1 | 27 |
| gar11 | 0.000632781 | 0.278550911 | 0.198 | 0.12 | 1 | 27 |
| chchd106 | 0.000670425 | 0.276347081 | 0.24 | 0.153 | 1 | 27 |
| dynll2a7 | 0.000773444 | 0.266277507 | 0.222 | 0.137 | 1 | 27 |
| cfl1l3 | 0.000855687 | 0.263120678 | 0.24 | 0.153 | 1 | 27 |
| buc7 | 0.000937126 | 0.262357133 | 0.275 | 0.18 | 1 | 27 |
| BX001030.15 | 0.000970441 | 0.2528569 | 0.204 | 0.123 | 1 | 27 |
| mrpl18 | 0.000971481 | 0.27735364 | 0.18 | 0.109 | 1 | 27 |
| rbm71 | 0.000992459 | 0.25968005 | 0.198 | 0.121 | 1 | 27 |
| pcf112 | 0.001243072 | 0.288748807 | 0.317 | 0.24 | 1 | 27 |
| nop161 | 0.001275169 | 0.260591451 | 0.186 | 0.114 | 1 | 27 |
| mcfd21 | 0.001398921 | 0.259957369 | 0.186 | 0.114 | 1 | 27 |
| prps1a | 0.001458933 | 0.271339593 | 0.192 | 0.121 | 1 | 27 |
| zgc:1102393 | 0.001482078 | 0.27692779 | 0.293 | 0.216 | 1 | 27 |
| marcksb4 | 0.001540205 | 0.251063391 | 0.263 | 0.177 | 1 | 27 |
| mrps16 | 0.001636784 | 0.251389509 | 0.144 | 0.082 | 1 | 27 |
| inka1b7 | 0.001773142 | 0.271532452 | 0.269 | 0.186 | 1 | 27 |
| zc3h11a6 | 0.001831515 | 0.256516129 | 0.305 | 0.218 | 1 | 27 |
| snrpc3 | 0.001960977 | 0.267893544 | 0.257 | 0.183 | 1 | 27 |
| bud311 | 0.002199873 | 0.280807712 | 0.299 | 0.229 | 1 | 27 |
| hmgb3b2 | 0.002498223 | 0.279433921 | 0.251 | 0.179 | 1 | 27 |
| slc25a25a1 | 0.003103495 | 0.268012227 | 0.257 | 0.188 | 1 | 27 |
| eif4g2a | 0.004541796 | 0.250851259 | 0.198 | 0.134 | 1 | 27 |
| pcbp2 | 0.005508235 | 0.266307767 | 0.281 | 0.221 | 1 | 27 |
| fbl2 | 0.006195377 | 0.255786043 | 0.269 | 0.206 | 1 | 27 |
| phospho1 | 8.50072845043177e-146 | 0.720074589 | 0.267 | 0.012 | 2.13954834368917e-141 | 28 |
| slc8a4b | 2.87205870925564e-77 | 0.596518028 | 0.211 | 0.015 | 7.22868456532551e-73 | 28 |
| si:ch73-189n23.19 | 8.09112885544136e-39 | 0.752484123 | 0.963 | 0.522 | 2.03645622162604e-34 | 28 |
| sgms2a | 3.78773771275384e-38 | 0.345305312 | 0.124 | 0.011 | 9.53335704923013e-34 | 28 |

| | | | | | | |
|---|---|---|---|---|---|---|
| h1m9 | 5.26440255653112e-38 | 0.694450804 | 0.981 | 0.599 | 1.32499747945332e-33 | 28 |
| zgc:1534998 | 5.93046907480378e-38 | 0.761341066 | 0.963 | 0.51 | 1.49263976143736e-33 | 28 |
| si:dkey-240e12.68 | 4.98296332259074e-37 | 0.718730365 | 0.988 | 0.566 | 1.25416203866286e-32 | 28 |
| zp2.38 | 7.73916991982142e-37 | 0.739596543 | 0.957 | 0.516 | 1.94787167711985e-32 | 28 |
| zp2.59 | 8.67635000029196e-36 | 0.707315025 | 0.981 | 0.561 | 2.18375053157348e-31 | 28 |
| tmsb58 | 2.59555155837605e-35 | 0.74435716 | 0.938 | 0.486 | 6.53274371727667e-31 | 28 |
| zp2.28 | 3.49242605025671e-35 | 0.730780956 | 0.975 | 0.526 | 8.79008712589112e-31 | 28 |
| CR790368.19 | 3.3847729969284e-34 | 0.7111611 | 0.944 | 0.517 | 8.51913515596908e-30 | 28 |
| zgc:1717768 | 3.99748996090119e-34 | 0.722266158 | 0.919 | 0.493 | 1.00612824825922e-29 | 28 |
| h2af1al9 | 1.32753788289113e-33 | 0.682504984 | 0.988 | 0.59 | 3.34128009744868e-29 | 28 |
| si:ch211-226h8.48 | 2.14648516893983e-33 | 0.731498723 | 0.851 | 0.439 | 5.40248852170465e-29 | 28 |
| si:dkey-241l7.28 | 2.43437423372882e-33 | 0.659820815 | 0.994 | 0.639 | 6.12707650887207e-29 | 28 |
| acp5a9 | 6.09464118153908e-33 | 0.68874439 | 0.944 | 0.537 | 1.53396023898157e-28 | 28 |
| crp28 | 1.61759999958034e-32 | 0.718607226 | 0.919 | 0.486 | 4.07133743894376e-28 | 28 |
| cldnd9 | 2.4643054216293e-32 | 0.680888461 | 0.988 | 0.602 | 6.20241031569879e-28 | 28 |
| cldng8 | 2.76054222006625e-32 | 0.70610957 | 0.938 | 0.511 | 6.94800871368473e-28 | 28 |
| si:zfos-1505d6.38 | 8.17951753023048e-32 | 0.710139658 | 0.944 | 0.51 | 2.05870276718371e-27 | 28 |
| mt28 | 1.03563843299505e-31 | 0.693979404 | 0.963 | 0.532 | 2.60659837200524e-27 | 28 |
| si:ch211-125e6.128 | 1.53372144277663e-31 | 0.712261077 | 0.932 | 0.498 | 3.86022349932449e-27 | 28 |
| si:ch211-12h2.88 | 1.67984504784307e-31 | 0.66476213 | 0.919 | 0.546 | 4.22800200091623e-27 | 28 |
| si:dkeyp-20e4.88 | 2.3299599139744e-31 | 0.748813219 | 0.845 | 0.405 | 5.86427610748217e-27 | 28 |
| si:dkey-241l7.48 | 8.37792777339061e-31 | 0.641383574 | 0.975 | 0.606 | 2.10864064128468e-26 | 28 |
| BX649337.19 | 9.86935201359937e-31 | 0.717709752 | 0.876 | 0.435 | 2.48401720830282e-26 | 28 |
| zp3.29 | 1.47845588096193e-30 | 0.672564331 | 0.95 | 0.555 | 3.72112560679309e-26 | 28 |
| si:ch211-226h8.148 | 3.22787785928846e-30 | 0.666595907 | 0.969 | 0.564 | 8.12424578404313e-26 | 28 |
| si:ch211-12h2.68 | 4.05413537459197e-30 | 0.686116125 | 0.851 | 0.472 | 1.02038533243105e-25 | 28 |
| zp39 | 6.13470896366823e-30 | 0.671064839 | 0.975 | 0.582 | 1.54404489906566e-25 | 28 |
| si:dkeyp-98a7.98 | 6.8644422755052e-30 | 0.690792303 | 0.925 | 0.51 | 1.7277114763219e-25 | 28 |
| epcam9 | 9.32838088475692e-30 | 0.676970312 | 0.901 | 0.503 | 2.34786018488447e-25 | 28 |
| rrm28 | 1.09383081444729e-29 | 0.675867843 | 0.913 | 0.496 | 2.75306277688238e-25 | 28 |
| ccna18 | 1.25853200219726e-29 | 0.689446271 | 0.888 | 0.473 | 3.16759919633028e-25 | 28 |
| zgc:1714748 | 1.73737325507281e-29 | 0.737485952 | 0.677 | 0.302 | 4.37279474569274e-25 | 28 |
| si:dkey-241l7.59 | 1.89874181365005e-29 | 0.644613619 | 0.944 | 0.582 | 4.7789432707758e-25 | 28 |
| zp2.19 | 3.0403297208399e-29 | 0.672720422 | 0.925 | 0.533 | 7.65220587438195e-25 | 28 |
| zar1l8 | 5.4145967196748e-29 | 0.706990431 | 0.807 | 0.395 | 1.36279984837495e-24 | 28 |

| | | | | | | |
|---|---|---|---|---|---|---|
| si:dkeyp-98a7.48 | 8.8159469611903e-29 | 0.697649624 | 0.826 | 0.423 | 2.21888569066199e-24 | 28 |
| si:dkey-241l7.68 | 9.4156240800642e-29 | 0.658691912 | 0.901 | 0.525 | 2.36981842471136e-24 | 28 |
| zp3a.28 | 1.59921704065093e-28 | 0.672929623 | 0.882 | 0.489 | 4.02506936961434e-24 | 28 |
| si:ch211-14a17.78 | 4.17549299888003e-28 | 0.69768838 | 0.795 | 0.406 | 1.05092983288811e-23 | 28 |
| zp2.68 | 5.8955119924766e-28 | 0.675450216 | 0.913 | 0.499 | 1.48384141338644e-23 | 28 |
| si:dkey-229d11.38 | 6.58349473732867e-28 | 0.700747757 | 0.851 | 0.436 | 1.65699979043825e-23 | 28 |
| zgc:1717798 | 8.73712308729433e-28 | 0.664789232 | 0.901 | 0.51 | 2.19904650984111e-23 | 28 |
| si:dkey-90l23.18 | 2.88160246275907e-27 | 0.689143346 | 0.857 | 0.445 | 7.25270523851832e-23 | 28 |
| ccnb28 | 2.85857262964411e-26 | 0.689107614 | 0.789 | 0.381 | 7.19474145155126e-22 | 28 |
| si:dkeyp-46h3.68 | 3.38734435417581e-26 | 0.678113902 | 0.783 | 0.408 | 8.52560700502511e-22 | 28 |
| CU571163.18 | 7.3604313240667e-26 | 0.690279667 | 0.789 | 0.399 | 1.85254695995435e-21 | 28 |
| zgc:1751359 | 7.53801471300292e-26 | 0.669778448 | 0.888 | 0.477 | 1.8972429231157e-21 | 28 |
| zgc:1715178 | 1.50980005672841e-25 | 0.676172408 | 0.764 | 0.386 | 3.80001576277974e-21 | 28 |
| LOC1104378218 | 1.89709495056501e-25 | 0.676529974 | 0.795 | 0.413 | 4.77479828107707e-21 | 28 |
| si:dkeyp-98a7.88 | 5.00280317065431e-25 | 0.658900129 | 0.851 | 0.455 | 1.25915553002198e-20 | 28 |
| si:dkey-208k4.28 | 5.55086650746299e-25 | 0.683439115 | 0.658 | 0.294 | 1.39709759126336e-20 | 28 |
| zar18 | 5.53189877372784e-24 | 0.63936975 | 0.807 | 0.438 | 1.39232360235956e-19 | 28 |
| slbp28 | 9.40231408465305e-24 | 0.652239072 | 0.77 | 0.395 | 2.36646843196633e-19 | 28 |
| si:ch211-250e5.168 | 1.11263340240086e-23 | 0.659411367 | 0.689 | 0.344 | 2.80038701050272e-19 | 28 |
| thy18 | 1.22922746448322e-23 | 0.629369696 | 0.814 | 0.448 | 3.09384260535782e-19 | 28 |
| zpcx8 | 1.73730421710375e-23 | 0.666480821 | 0.696 | 0.337 | 4.37262098402844e-19 | 28 |
| qdprb28 | 2.98868310855705e-23 | 0.637292249 | 0.783 | 0.431 | 7.52221651592724e-19 | 28 |
| si:dkey-90l23.28 | 3.73165263137541e-23 | 0.655722517 | 0.696 | 0.343 | 9.39219650790878e-19 | 28 |
| LOC1005374998 | 4.33552197111713e-23 | 0.658873643 | 0.677 | 0.32 | 1.09120752491047e-18 | 28 |
| LOC1005351628 | 1.2165272752085e-22 | 0.666867542 | 0.658 | 0.307 | 3.06187749897228e-18 | 28 |
| hnrnpabb9 | 1.30473357859915e-22 | 0.533775704 | 0.863 | 0.595 | 3.28388394397621e-18 | 28 |
| nasp8 | 1.82889575516998e-22 | 0.653633883 | 0.689 | 0.338 | 4.60314772618732e-18 | 28 |
| btg48 | 2.35395591598966e-22 | 0.643515703 | 0.714 | 0.367 | 5.92467164495438e-18 | 28 |
| lamp28 | 3.57883451917848e-22 | 0.57124659 | 0.863 | 0.553 | 9.0075686013203e-18 | 28 |
| zp3b8 | 4.58459902083619e-22 | 0.64344038 | 0.764 | 0.386 | 1.15389772755426e-17 | 28 |
| s100a18 | 8.49739278015395e-22 | 0.636599827 | 0.72 | 0.362 | 2.13870878883695e-17 | 28 |
| avd8 | 9.11872246604923e-22 | 0.656925792 | 0.59 | 0.264 | 2.29509125747993e-17 | 28 |
| nanog8 | 1.06793769649698e-21 | 0.635157651 | 0.714 | 0.358 | 2.68789238831325e-17 | 28 |
| mid1ip1l8 | 1.59961208024008e-21 | 0.561693762 | 0.839 | 0.542 | 4.02606364475627e-17 | 28 |
| cpeb1b8 | 4.2835433309566e-21 | 0.635782134 | 0.59 | 0.269 | 1.07812502096847e-16 | 28 |

| | | | | | | |
|---|---|---|---|---|---|---|
| zgc:771188 | 5.24520660290917e-21 | 0.60847959 | 0.814 | 0.455 | 1.32016604988621e-16 | 28 |
| si:ch211-132b12.88 | 6.65960007973239e-21 | 0.630600218 | 0.671 | 0.34 | 1.67615474406784e-16 | 28 |
| npm2b8 | 1.22262516746533e-20 | 0.613212386 | 0.727 | 0.394 | 3.07722528399348e-16 | 28 |
| si:dkeyp-50f7.28 | 2.04455379044836e-20 | 0.605145062 | 0.727 | 0.401 | 5.14593743517948e-16 | 28 |
| gapdh8 | 7.57847926827668e-20 | 0.592985279 | 0.758 | 0.418 | 1.90742744703256e-15 | 28 |
| si:dkeyp-46h3.28 | 8.11973015360441e-20 | 0.6013121 | 0.727 | 0.403 | 2.04365488236069e-15 | 28 |
| elovl7b8 | 5.43100295267352e-19 | 0.613072359 | 0.547 | 0.247 | 1.3669291331584e-14 | 28 |
| cth18 | 6.91170717536241e-19 | 0.590101409 | 0.696 | 0.378 | 1.73960757896696e-14 | 28 |
| bgna | 6.97971756064284e-19 | 0.307306177 | 0.106 | 0.016 | 1.7567251128382e-14 | 28 |
| si:dkeyp-98a7.78 | 1.36350937596449e-18 | 0.593689701 | 0.478 | 0.194 | 3.43181674836504e-14 | 28 |
| AL929032.18 | 1.54884075676327e-18 | 0.604456938 | 0.627 | 0.316 | 3.89827730069746e-14 | 28 |
| zgc:1734438 | 3.04560384855666e-18 | 0.591384746 | 0.671 | 0.328 | 7.66548032643226e-14 | 28 |
| si:dkeyp-46h3.88 | 4.40944100648655e-18 | 0.593316395 | 0.665 | 0.35 | 1.1098122069226e-13 | 28 |
| gtf3ab9 | 1.4217854504118e-17 | 0.58050543 | 0.621 | 0.319 | 3.57849180014146e-13 | 28 |
| si:ch211-146l10.88 | 1.76785758068824e-17 | 0.591158852 | 0.602 | 0.305 | 4.44952074483424e-13 | 28 |
| siva18 | 1.97696825624706e-17 | 0.554351901 | 0.72 | 0.427 | 4.97583140414823e-13 | 28 |
| ccna28 | 3.49356754299922e-17 | 0.574746099 | 0.571 | 0.283 | 8.79296014897474e-13 | 28 |
| tuba4l8 | 4.51141060643094e-17 | 0.56745422 | 0.714 | 0.403 | 1.1354769355326e-12 | 28 |
| LOC1018854778 | 5.62401304194378e-17 | 0.584250967 | 0.559 | 0.279 | 1.41550784252683e-12 | 28 |
| si:ch211-145c1.18 | 6.50073843877927e-17 | 0.58915021 | 0.503 | 0.233 | 1.63617085765635e-12 | 28 |
| uchl18 | 3.55739634475463e-16 | 0.552319167 | 0.627 | 0.343 | 8.95361086011294e-12 | 28 |
| si:dkey-229d11.58 | 4.58961325597813e-16 | 0.555349151 | 0.671 | 0.388 | 1.15515976039714e-11 | 28 |
| zgc:1952458 | 2.11717396622692e-15 | 0.534668834 | 0.665 | 0.388 | 5.32871515559652e-11 | 28 |
| kpna78 | 3.89563526209664e-15 | 0.540599806 | 0.609 | 0.327 | 9.80492439117103e-11 | 28 |
| hbba18 | 4.49368099907646e-15 | 0.545490689 | 0.665 | 0.355 | 1.13101457065755e-10 | 28 |
| chia.48 | 4.76861587026332e-15 | 0.52635985 | 0.36 | 0.146 | 1.20021292838657e-10 | 28 |
| si:ch73-90k17.18 | 5.04496669508447e-15 | 0.544064489 | 0.522 | 0.248 | 1.26976766748581e-10 | 28 |
| ccne16 | 5.81745413371859e-15 | 0.496743819 | 0.304 | 0.11 | 1.46419503091563e-10 | 28 |
| CDPF18 | 6.51111150137965e-15 | 0.558874175 | 0.516 | 0.268 | 1.63878165378224e-10 | 28 |
| zgc:660248 | 6.61022210411104e-15 | 0.512516555 | 0.354 | 0.141 | 1.66372680138371e-10 | 28 |
| arhgef10la8 | 1.26875965663872e-14 | 0.550387257 | 0.528 | 0.274 | 3.193341179794e-10 | 28 |
| wu:fi42e038 | 1.276707723696e-14 | 0.539724787 | 0.627 | 0.349 | 3.21334566977046e-10 | 28 |
| zgc:1362548 | 4.47912940386259e-14 | 0.526012796 | 0.602 | 0.33 | 1.12735207965818e-09 | 28 |
| si:dkey-19b23.118 | 7.266723028355e-14 | 0.535505334 | 0.528 | 0.278 | 1.82896151900667e-09 | 28 |
| ctss2.19 | 1.43045532715884e-13 | 0.454945746 | 0.783 | 0.534 | 3.60031301292609e-09 | 28 |

| | | | | | | |
|---|---|---|---|---|---|---|
| ddx413 | 2.19422049535165e-13 | 0.402480157 | 0.217 | 0.068 | 5.52263356475056e-09 | 28 |
| cdca87 | 2.56931065361987e-13 | 0.490683325 | 0.379 | 0.164 | 6.46669798409585e-09 | 28 |
| rcc18 | 8.80445587081879e-13 | 0.505296045 | 0.422 | 0.202 | 2.21599349812638e-08 | 28 |
| BX284697.16 | 9.85920616145108e-13 | 0.470516067 | 0.292 | 0.114 | 2.48146359877562e-08 | 28 |
| si:dkey-241l7.38 | 1.36112057652418e-12 | 0.515947897 | 0.59 | 0.364 | 3.42580437905372e-08 | 28 |
| si:dkeyp-46h3.58 | 2.01119590347468e-12 | 0.507133431 | 0.528 | 0.289 | 5.06197896945543e-08 | 28 |
| h3f3a9 | 3.18934076280876e-12 | 0.434125221 | 0.708 | 0.505 | 8.02725176591336e-08 | 28 |
| plk17 | 4.09381375705548e-12 | 0.490683685 | 0.379 | 0.177 | 1.03037198451329e-07 | 28 |
| buc8 | 4.98383168427243e-12 | 0.49021575 | 0.379 | 0.178 | 1.25438059661453e-07 | 28 |
| LOC1018838978 | 5.34311137080706e-12 | 0.464685912 | 0.373 | 0.164 | 1.34480770091843e-07 | 28 |
| si:dkey-152b24.68 | 1.21365693421872e-11 | 0.488277957 | 0.534 | 0.297 | 3.05465313773509e-07 | 28 |
| si:ch211-59o9.10 | 1.45901183186703e-11 | 0.318943804 | 0.143 | 0.038 | 3.67218687962612e-07 | 28 |
| spink2.19 | 1.55432139963234e-11 | 0.450210079 | 0.683 | 0.436 | 3.91207153073464e-07 | 28 |
| ccnb18 | 1.56249149842482e-11 | 0.474128573 | 0.64 | 0.393 | 3.93263485238543e-07 | 28 |
| larp6b8 | 1.66897197618276e-11 | 0.47953376 | 0.41 | 0.196 | 4.2006355668544e-07 | 28 |
| larp6a8 | 2.00755913480457e-11 | 0.488808754 | 0.466 | 0.249 | 5.05282558638963e-07 | 28 |
| si:dkey-37o8.18 | 2.10210663905423e-11 | 0.474126513 | 0.547 | 0.307 | 5.29079219983559e-07 | 28 |
| LOC1039099723 | 3.25641665863256e-11 | 0.464290381 | 0.559 | 0.367 | 8.1960750881123e-07 | 28 |
| crp38 | 1.3939371846665e-10 | 0.467928099 | 0.422 | 0.218 | 3.50840050008712e-06 | 28 |
| elovl7a8 | 1.64649057654683e-10 | 0.469911715 | 0.366 | 0.183 | 4.14405213211072e-06 | 28 |
| zgc:554619 | 1.66106814142053e-10 | 0.466387101 | 0.516 | 0.288 | 4.18074240514132e-06 | 28 |
| zgc:925918 | 1.87065539957524e-10 | 0.458428867 | 0.41 | 0.205 | 4.70825257519092e-06 | 28 |
| mkrn47 | 1.95925529063726e-10 | 0.464957354 | 0.429 | 0.228 | 4.93124964100492e-06 | 28 |
| birc5b8 | 2.34435286170004e-10 | 0.463712273 | 0.46 | 0.26 | 5.90050171761284e-06 | 28 |
| si:ch1073-190k2.18 | 2.46516212080779e-10 | 0.45814608 | 0.379 | 0.188 | 6.20456654186114e-06 | 28 |
| pabpn1l8 | 2.6226723008477e-10 | 0.465587926 | 0.497 | 0.284 | 6.60100391400357e-06 | 28 |
| LOC1005368608 | 3.16424325158492e-10 | 0.450593792 | 0.547 | 0.351 | 7.96408383991408e-06 | 28 |
| si:ch73-14h10.28 | 5.82425240333608e-10 | 0.455422196 | 0.478 | 0.271 | 1.46590608739566e-05 | 28 |
| gmnn8 | 6.73940734528051e-10 | 0.451335905 | 0.522 | 0.303 | 1.69624143473365e-05 | 28 |
| rbm72 | 7.62085579517051e-10 | 0.424225141 | 0.273 | 0.12 | 1.91809319508647e-05 | 28 |
| zgc:1141302 | 8.59427762481658e-10 | 0.337069379 | 0.205 | 0.075 | 2.16309373539009e-05 | 28 |
| si:ch211-5k11.88 | 1.07527166914223e-09 | 0.421303256 | 0.59 | 0.358 | 2.70635126406409e-05 | 28 |
| gfer3 | 1.15432208634644e-09 | 0.373528176 | 0.242 | 0.098 | 2.90531325912536e-05 | 28 |
| zgc:1629458 | 1.65191732009183e-09 | 0.438617069 | 0.329 | 0.162 | 4.15771070293913e-05 | 28 |
| gdf38 | 2.19098853512957e-09 | 0.448519002 | 0.41 | 0.224 | 5.51449904406761e-05 | 28 |

| | | | | | | |
|---|---|---|---|---|---|---|
| si:ch211-226l4.69 | 2.22632864540211e-09 | 0.435094106 | 0.497 | 0.293 | 5.60344656761257e-05 | 28 |
| haus67 | 2.23922866653932e-09 | 0.438949172 | 0.391 | 0.209 | 5.63591463081281e-05 | 28 |
| BX571780.15 | 2.31876032178477e-09 | 0.356100028 | 0.211 | 0.081 | 5.83608785390008e-05 | 28 |
| zgc:566999 | 2.58111296746799e-09 | 0.444783369 | 0.453 | 0.251 | 6.49640322782019e-05 | 28 |
| CR383662.24 | 2.76221210471423e-09 | 0.35136846 | 0.224 | 0.087 | 6.95221164635526e-05 | 28 |
| COQ10A2 | 3.07420457687273e-09 | 0.375143478 | 0.211 | 0.083 | 7.73746549953097e-05 | 28 |
| elavl29 | 3.10782551890185e-09 | 0.431675599 | 0.416 | 0.224 | 7.82208604852406e-05 | 28 |
| zp3c8 | 3.48893845324969e-09 | 0.418940158 | 0.373 | 0.189 | 8.78130919298416e-05 | 28 |
| CR847984.18 | 3.6181304844847e-09 | 0.421631068 | 0.335 | 0.164 | 9.10647261639955e-05 | 28 |
| tph1a8 | 4.13310152300752e-09 | 0.401085582 | 0.304 | 0.142 | 0.000104026 | 28 |
| skp2 | 4.49655503243934e-09 | 0.285719444 | 0.155 | 0.051 | 0.000113174 | 28 |
| wu:fa19b121 | 4.70539839422385e-09 | 0.351363981 | 0.186 | 0.069 | 0.00011843 | 28 |
| tpx24 | 4.70960876044054e-09 | 0.388839925 | 0.28 | 0.126 | 0.000118536 | 28 |
| si:dkey-42i9.88 | 4.90482634812339e-09 | 0.404075359 | 0.28 | 0.127 | 0.00012345 | 28 |
| zgc:1141235 | 4.97314639842429e-09 | 0.386994723 | 0.242 | 0.104 | 0.000125169 | 28 |
| zp3a.18 | 5.13764263279173e-09 | 0.424962184 | 0.571 | 0.382 | 0.000129309 | 28 |
| CABZ01069022.21 | 5.38616608878604e-09 | 0.318876775 | 0.174 | 0.061 | 0.000135564 | 28 |
| zgc:562318 | 5.46543799303283e-09 | 0.4323297 | 0.416 | 0.228 | 0.00013756 | 28 |
| si:ch211-251f6.68 | 7.20260812899889e-09 | 0.421952607 | 0.528 | 0.332 | 0.000181282 | 28 |
| nanos36 | 7.77178473001389e-09 | 0.374521489 | 0.242 | 0.103 | 0.000195608 | 28 |
| cdca7a8 | 7.8224628824016e-09 | 0.425204946 | 0.41 | 0.222 | 0.000196884 | 28 |
| hbaa18 | 8.20103664770037e-09 | 0.417103659 | 0.497 | 0.281 | 0.000206412 | 28 |
| pif15 | 8.58852346377903e-09 | 0.374155117 | 0.248 | 0.107 | 0.000216165 | 28 |
| dynll19 | 1.00299096961299e-08 | 0.379290043 | 0.615 | 0.455 | 0.000252443 | 28 |
| ctsla8 | 1.17632883761262e-08 | 0.410526987 | 0.553 | 0.387 | 0.00029607 | 28 |
| zgc:1529862 | 1.40591044580583e-08 | 0.358749116 | 0.217 | 0.09 | 0.000353854 | 28 |
| pabpc1l8 | 1.50217768770064e-08 | 0.405901152 | 0.292 | 0.142 | 0.000378083 | 28 |
| pou5f38 | 1.71637098778773e-08 | 0.394993247 | 0.304 | 0.147 | 0.000431993 | 28 |
| sinup8 | 2.2892459948292e-08 | 0.413146584 | 0.379 | 0.203 | 0.00057618 | 28 |
| slc16a38 | 2.34696467813527e-08 | 0.413083904 | 0.435 | 0.249 | 0.000590708 | 28 |
| CU856358.1 | 2.45975077456621e-08 | 0.257864249 | 0.124 | 0.038 | 0.000619095 | 28 |
| lpcat44 | 2.5077364534083e-08 | 0.376493659 | 0.242 | 0.107 | 0.000631172 | 28 |
| BX571701.1 | 3.05331438058505e-08 | 0.289538881 | 0.149 | 0.051 | 0.000768489 | 28 |
| zgc:1138867 | 3.29376580814606e-08 | 0.390349188 | 0.292 | 0.142 | 0.000829008 | 28 |
| ypel36 | 3.56297585293241e-08 | 0.388991658 | 0.267 | 0.127 | 0.000896765 | 28 |

| | | | | | | |
|---|---|---|---|---|---|---|
| vps9d13 | 4.15482419700626e-08 | 0.35836084 | 0.224 | 0.097 | 0.001045728 | 28 |
| ddx194 | 6.11408211614598e-08 | 0.39588817 | 0.298 | 0.152 | 0.001538853 | 28 |
| sybu8 | 6.440271537214e-08 | 0.384901051 | 0.329 | 0.167 | 0.001620952 | 28 |
| zgc:1628798 | 6.71518686394012e-08 | 0.385335283 | 0.348 | 0.182 | 0.001690145 | 28 |
| rasl11b6 | 6.76807529394661e-08 | 0.370733237 | 0.28 | 0.134 | 0.001703457 | 28 |
| zglp18 | 6.83066967740556e-08 | 0.395137677 | 0.366 | 0.199 | 0.001719211 | 28 |
| si:ch211-69g19.26 | 7.68410388273343e-08 | 0.366614561 | 0.261 | 0.122 | 0.001934012 | 28 |
| gstp16 | 8.64844087354713e-08 | 0.38590025 | 0.522 | 0.366 | 0.002176726 | 28 |
| zgc:112146 | 8.9524161626795e-08 | 0.262399866 | 0.124 | 0.04 | 0.002253234 | 28 |
| h2afx4 | 9.08908924163964e-08 | 0.383106482 | 0.28 | 0.139 | 0.002287633 | 28 |
| imp36 | 9.22517164312004e-08 | 0.385595873 | 0.273 | 0.135 | 0.002321883 | 28 |
| si:dkey-81e3.21 | 9.28615650140979e-08 | 0.309339797 | 0.168 | 0.064 | 0.002337233 | 28 |
| dnd14 | 9.47081613600246e-08 | 0.353208954 | 0.242 | 0.11 | 0.00238371 | 28 |
| prdx18 | 9.98853276918401e-08 | 0.392037226 | 0.317 | 0.167 | 0.002514014 | 28 |
| prrg24 | 1.03229034139065e-07 | 0.324898703 | 0.211 | 0.089 | 0.002598172 | 28 |
| zgc:1719777 | 1.0576028557691e-07 | 0.39346958 | 0.398 | 0.23 | 0.002661881 | 28 |
| buc2l3 | 1.12465431507331e-07 | 0.344892223 | 0.193 | 0.08 | 0.002830642 | 28 |
| org8 | 1.17313299802936e-07 | 0.410574321 | 0.41 | 0.241 | 0.002952658 | 28 |
| si:dkey-43p13.52 | 1.20496551453667e-07 | 0.338305006 | 0.199 | 0.084 | 0.003032778 | 28 |
| inka1b8 | 1.36614944985178e-07 | 0.388224902 | 0.342 | 0.185 | 0.003438462 | 28 |
| rpp21 | 1.60021918141331e-07 | 0.343373523 | 0.193 | 0.082 | 0.004027592 | 28 |
| mylipa3 | 1.65594073706681e-07 | 0.35006743 | 0.248 | 0.116 | 0.004167837 | 28 |
| dynll2a8 | 1.65746190447081e-07 | 0.37457483 | 0.273 | 0.137 | 0.004171666 | 28 |
| use1 | 1.8478334171735e-07 | 0.295196929 | 0.137 | 0.049 | 0.004650812 | 28 |
| h2afva7 | 1.86171658125787e-07 | 0.400387947 | 0.478 | 0.324 | 0.004685754 | 28 |
| tent5c6 | 1.87370940481404e-07 | 0.378766371 | 0.28 | 0.14 | 0.004715939 | 28 |
| map1lc3c8 | 2.01073554257847e-07 | 0.379680375 | 0.373 | 0.202 | 0.00506082 | 28 |
| bmb4 | 2.25837088946507e-07 | 0.328841022 | 0.205 | 0.089 | 0.005684094 | 28 |
| zgc:1034829 | 2.40905102296884e-07 | 0.376517205 | 0.348 | 0.189 | 0.006063341 | 28 |
| tacc32 | 2.56478211163098e-07 | 0.348725684 | 0.211 | 0.096 | 0.0064553 | 28 |
| tmem33 | 2.66251646155173e-07 | 0.329578376 | 0.193 | 0.083 | 0.006701288 | 28 |
| selenoh4 | 2.92644169779002e-07 | 0.40404227 | 0.391 | 0.245 | 0.007365561 | 28 |
| ankrd228 | 2.98473304043164e-07 | 0.360466409 | 0.286 | 0.143 | 0.007512275 | 28 |
| BX537276.18 | 3.01931715695004e-07 | 0.343867102 | 0.255 | 0.122 | 0.007599319 | 28 |
| rdh10b7 | 3.13475368457815e-07 | 0.381202518 | 0.329 | 0.178 | 0.007889862 | 28 |

| | | | | | | |
|---|---|---|---|---|---|---|
| BX649350.11 | 3.46584534377327e-07 | 0.267534039 | 0.149 | 0.055 | 0.008723186 | 28 |
| slbp9 | 3.46813153656103e-07 | 0.394630894 | 0.391 | 0.235 | 0.00872894 | 28 |
| birc5a1 | 3.6216958395863e-07 | 0.344603593 | 0.217 | 0.099 | 0.009115446 | 28 |
| si:ch73-340n8.43 | 3.76718872713294e-07 | 0.356197212 | 0.261 | 0.128 | 0.009481637 | 28 |
| CU929143.1 | 3.94976716126049e-07 | 0.335527012 | 0.205 | 0.091 | 0.009941169 | 28 |
| si:dkey-68o6.58 | 3.99914362012193e-07 | 0.37526041 | 0.248 | 0.123 | 0.010065445 | 28 |
| zgc:1134248 | 4.11130969901144e-07 | 0.364308703 | 0.311 | 0.162 | 0.010347755 | 28 |
| si:ch73-132k15.21 | 4.22437242376462e-07 | 0.309783631 | 0.161 | 0.064 | 0.010632323 | 28 |
| si:dkey-216e24.36 | 4.5008578806787e-07 | 0.339400024 | 0.255 | 0.124 | 0.011328209 | 28 |
| CU019646.1 | 4.56672069110124e-07 | 0.2743657 | 0.143 | 0.053 | 0.011493979 | 28 |
| ca15b8 | 5.31099991876163e-07 | 0.389774026 | 0.41 | 0.25 | 0.013367256 | 28 |
| zgc:923133 | 5.42920253891792e-07 | 0.350790827 | 0.255 | 0.127 | 0.01366476 | 28 |
| lsm14b8 | 6.23648460300234e-07 | 0.382642351 | 0.472 | 0.3 | 0.015696608 | 28 |
| cldn7b7 | 6.48861617815548e-07 | 0.370904572 | 0.298 | 0.16 | 0.016331198 | 28 |
| hspa57 | 7.31096633259486e-07 | 0.37289953 | 0.46 | 0.315 | 0.018400971 | 28 |
| CT009596.14 | 7.54589120687241e-07 | 0.303834391 | 0.205 | 0.09 | 0.018992254 | 28 |
| pttg18 | 7.81092104682781e-07 | 0.361336235 | 0.298 | 0.159 | 0.019659307 | 28 |
| tac3b7 | 8.38980312098245e-07 | 0.330552801 | 0.248 | 0.121 | 0.021116295 | 28 |
| snu13b9 | 1.03998655760214e-06 | 0.355036863 | 0.553 | 0.393 | 0.026175422 | 28 |
| zgc:566768 | 1.05844076248251e-06 | 0.376162845 | 0.441 | 0.296 | 0.026639896 | 28 |
| mos7 | 1.13816724307424e-06 | 0.352368069 | 0.267 | 0.137 | 0.028646531 | 28 |
| gbgt1l47 | 1.15302732033583e-06 | 0.35066969 | 0.286 | 0.151 | 0.029020545 | 28 |
| BX001030.16 | 1.2048871906115e-06 | 0.329760383 | 0.248 | 0.123 | 0.030325806 | 28 |
| si:ch211-266i6.3 | 1.21361559518518e-06 | 0.283681084 | 0.161 | 0.066 | 0.030545491 | 28 |
| ccnh5 | 1.26952479370518e-06 | 0.324821056 | 0.255 | 0.126 | 0.03195267 | 28 |
| si:dkey-188i13.119 | 1.33902635242265e-06 | 0.386507613 | 0.453 | 0.317 | 0.033701954 | 28 |
| si:dkeyp-75b4.105 | 1.37877954923557e-06 | 0.29555319 | 0.602 | 0.437 | 0.034702502 | 28 |
| zc3h11a7 | 1.42855331447372e-06 | 0.377975227 | 0.36 | 0.218 | 0.035955258 | 28 |
| zgc:1655397 | 1.44536973189802e-06 | 0.340161532 | 0.242 | 0.12 | 0.036378511 | 28 |
| degs26 | 1.55545867615298e-06 | 0.326225816 | 0.224 | 0.107 | 0.039149339 | 28 |
| zgc:174268 | 1.60272467500185e-06 | 0.310946692 | 0.186 | 0.083 | 0.040338977 | 28 |
| zgc:1102394 | 1.73528311756962e-06 | 0.377876652 | 0.348 | 0.215 | 0.043675341 | 28 |
| patl28 | 1.76707172097451e-06 | 0.33393336 | 0.323 | 0.174 | 0.044475428 | 28 |
| ctsba8 | 1.83903526882492e-06 | 0.344355678 | 0.528 | 0.377 | 0.046286679 | 28 |
| si:ch211-286b5.43 | 2.06443076476819e-06 | 0.27102864 | 0.168 | 0.07 | 0.051959658 | 28 |

| | | | | | | |
|---|---|---|---|---|---|---|
| CABZ01059627.28 | 2.37025787320127e-06 | 0.328183548 | 0.261 | 0.134 | 0.05965702 | 28 |
| si:ch73-303b9.13 | 2.61664775623499e-06 | 0.277403148 | 0.18 | 0.079 | 0.065858407 | 28 |
| pim31 | 2.76533385236494e-06 | 0.33587171 | 0.248 | 0.129 | 0.069600688 | 28 |
| AL929192.18 | 2.90489975980694e-06 | 0.345004839 | 0.298 | 0.165 | 0.073113422 | 28 |
| zgc:1134251 | 3.23820330159265e-06 | 0.287143204 | 0.161 | 0.069 | 0.081502339 | 28 |
| cx43.44 | 3.50358039680482e-06 | 0.326878058 | 0.242 | 0.124 | 0.088181615 | 28 |
| si:ch211-14a17.63 | 3.57687121766498e-06 | 0.273542662 | 0.193 | 0.087 | 0.090026272 | 28 |
| zgc:1717508 | 3.75249890085926e-06 | 0.352355193 | 0.304 | 0.175 | 0.094446645 | 28 |
| asz11 | 4.23026479050651e-06 | 0.281805426 | 0.168 | 0.073 | 0.106471535 | 28 |
| st3gal81 | 4.57354534935308e-06 | 0.286327098 | 0.174 | 0.077 | 0.115111563 | 28 |
| zp2l18 | 5.36939985560129e-06 | 0.347826778 | 0.292 | 0.165 | 0.135142425 | 28 |
| zgc:1630307 | 5.37279456722067e-06 | 0.315862308 | 0.242 | 0.125 | 0.135227866 | 28 |
| BX005461.14 | 5.44814152422996e-06 | 0.263262209 | 0.18 | 0.08 | 0.137124274 | 28 |
| si:ch211-11k18.4 | 5.79908417386581e-06 | 0.255019333 | 0.137 | 0.055 | 0.14595715 | 28 |
| spryd43 | 6.33244538921366e-06 | 0.28309472 | 0.174 | 0.078 | 0.159381318 | 28 |
| cdc208 | 6.55703138261035e-06 | 0.335764957 | 0.28 | 0.153 | 0.165033923 | 28 |
| ctnnbip1 | 6.82190359598866e-06 | 0.30844845 | 0.155 | 0.069 | 0.171700492 | 28 |
| vstm2l5 | 7.37704145259612e-06 | 0.309373809 | 0.199 | 0.096 | 0.185672756 | 28 |
| zgc:922877 | 8.26494316740144e-06 | 0.306608347 | 0.255 | 0.133 | 0.208020355 | 28 |
| brf23 | 8.86465168684859e-06 | 0.290302811 | 0.199 | 0.095 | 0.223114418 | 28 |
| cdt13 | 9.51721381392193e-06 | 0.286364663 | 0.186 | 0.088 | 0.239538754 | 28 |
| fbxw11b3 | 1.07744387662592e-05 | 0.331803984 | 0.261 | 0.146 | 0.271181849 | 28 |
| CU655961.43 | 1.09943271500749e-05 | 0.297061501 | 0.186 | 0.089 | 0.27671622 | 28 |
| si:ch211-146l10.78 | 1.22925644810287e-05 | 0.339799438 | 0.317 | 0.19 | 0.309391555 | 28 |
| si:dkeyp-114g9.14 | 1.24017814896481e-05 | 0.300732161 | 0.211 | 0.105 | 0.312140438 | 28 |
| parn6 | 1.35946191575819e-05 | 0.319095846 | 0.267 | 0.148 | 0.34216297 | 28 |
| zgc:1139845 | 1.4749182325447e-05 | 0.292136381 | 0.205 | 0.102 | 0.37122217 | 28 |
| BX323557.1 | 1.49969658722695e-05 | 0.298838163 | 0.186 | 0.091 | 0.377458634 | 28 |
| rac1a2 | 1.66800221544485e-05 | 0.330252666 | 0.261 | 0.149 | 0.419819478 | 28 |
| ndc12 | 1.76549949348195e-05 | 0.310313807 | 0.193 | 0.098 | 0.444358568 | 28 |
| pigrl2.38 | 1.82043497898499e-05 | 0.336308766 | 0.335 | 0.204 | 0.45818528 | 28 |
| eif5a22 | 1.88458656024249e-05 | 0.30642841 | 0.497 | 0.404 | 0.474331591 | 28 |
| c1d1 | 1.94981963934746e-05 | 0.322708166 | 0.236 | 0.129 | 0.490750105 | 28 |
| eif4a1a6 | 2.01279954706132e-05 | 0.307310084 | 0.516 | 0.426 | 0.506601518 | 28 |
| zgc:1526526 | 2.1552864209228e-05 | 0.278928884 | 0.224 | 0.115 | 0.542464039 | 28 |

| | | | | | | |
|---|---|---|---|---|---|---|
| tatdn23 | 2.16975657303732e-05 | 0.262418302 | 0.161 | 0.073 | 0.546106032 | 28 |
| zgc:1118685 | 2.23544112877603e-05 | 0.321091927 | 0.261 | 0.148 | 0.562638178 | 28 |
| zgc:1017445 | 2.27724304458622e-05 | 0.301329133 | 0.248 | 0.134 | 0.573159302 | 28 |
| gyg1a9 | 2.35614946310944e-05 | 0.325501367 | 0.491 | 0.371 | 0.593019258 | 28 |
| hsd17b12a8 | 2.3594435931226e-05 | 0.309929065 | 0.286 | 0.166 | 0.593848358 | 28 |
| ube2l3a | 2.43875297035524e-05 | 0.339186924 | 0.267 | 0.16 | 0.613809735 | 28 |
| itm2cb4 | 2.49551140911767e-05 | 0.313894017 | 0.248 | 0.139 | 0.628095267 | 28 |
| bzw1b5 | 2.49661423019778e-05 | 0.310591645 | 0.503 | 0.403 | 0.628372836 | 28 |
| trmt61a | 2.60516464003292e-05 | 0.302561032 | 0.168 | 0.082 | 0.655693888 | 28 |
| phex3 | 2.67880242431963e-05 | 0.268287491 | 0.18 | 0.087 | 0.674227782 | 28 |
| si:ch73-160p18.38 | 2.79918947325219e-05 | 0.325590973 | 0.373 | 0.242 | 0.704527999 | 28 |
| ddx39ab7 | 2.88150627117273e-05 | 0.335312425 | 0.41 | 0.29 | 0.725246313 | 28 |
| tmco12 | 3.10836944354371e-05 | 0.307549019 | 0.224 | 0.121 | 0.782345505 | 28 |
| gemin81 | 3.15916427020637e-05 | 0.292376914 | 0.199 | 0.103 | 0.795130055 | 28 |
| lipg4 | 3.43565428834328e-05 | 0.27014747 | 0.174 | 0.085 | 0.864719828 | 28 |
| CABZ01059627.16 | 3.49248097355695e-05 | 0.291479159 | 0.248 | 0.137 | 0.879022536 | 28 |
| mibp8 | 4.03679142335565e-05 | 0.314834106 | 0.354 | 0.224 | 1 | 28 |
| zgc:1737428 | 4.85718806734709e-05 | 0.321251866 | 0.267 | 0.156 | 1 | 28 |
| e2f52 | 5.01524204573056e-05 | 0.25708152 | 0.155 | 0.073 | 1 | 28 |
| tmem54a6 | 5.19072752181319e-05 | 0.308802533 | 0.248 | 0.142 | 1 | 28 |
| mad2l17 | 5.59518690343788e-05 | 0.310686993 | 0.304 | 0.187 | 1 | 28 |
| zgc:1735442 | 6.13143801157602e-05 | 0.254034384 | 0.161 | 0.077 | 1 | 28 |
| dazl5 | 6.43885574503032e-05 | 0.271687112 | 0.205 | 0.107 | 1 | 28 |
| srsf2a5 | 6.43892106177912e-05 | 0.307064009 | 0.472 | 0.368 | 1 | 28 |
| evla3 | 6.81717603480901e-05 | 0.271933896 | 0.199 | 0.103 | 1 | 28 |
| yrdc6 | 7.09468388004801e-05 | 0.293249973 | 0.224 | 0.124 | 1 | 28 |
| shmt1 | 7.40587135091762e-05 | 0.251257149 | 0.168 | 0.082 | 1 | 28 |
| si:ch211-210c8.73 | 7.4826321321609e-05 | 0.287174669 | 0.23 | 0.128 | 1 | 28 |
| ube2c6 | 7.554042897375e-05 | 0.303259666 | 0.242 | 0.141 | 1 | 28 |
| prr112 | 7.62358338174319e-05 | 0.277227898 | 0.193 | 0.101 | 1 | 28 |
| fbl3 | 8.07867644659926e-05 | 0.329681727 | 0.311 | 0.205 | 1 | 28 |
| rogdi2 | 8.08082928695113e-05 | 0.270484415 | 0.217 | 0.117 | 1 | 28 |
| rbpms2a1 | 9.44115424390665e-05 | 0.254678686 | 0.18 | 0.092 | 1 | 28 |
| si:dkey-102m7.3 | 9.98756527123187e-05 | 0.301802736 | 0.242 | 0.144 | 1 | 28 |
| rbm4.38 | 0.000106195 | 0.28162919 | 0.522 | 0.446 | 1 | 28 |

| | | | | | | |
|---|---|---|---|---|---|---|
| snx10a6 | 0.000113714 | 0.272480123 | 0.23 | 0.127 | 1 | 28 |
| dcun1d11 | 0.000118678 | 0.274258406 | 0.186 | 0.099 | 1 | 28 |
| nudc1 | 0.000119883 | 0.318569089 | 0.248 | 0.152 | 1 | 28 |
| zgc:1738568 | 0.000123385 | 0.299356419 | 0.348 | 0.221 | 1 | 28 |
| nap1l12 | 0.000130027 | 0.294681432 | 0.447 | 0.362 | 1 | 28 |
| cnot81 | 0.000134732 | 0.291586292 | 0.18 | 0.097 | 1 | 28 |
| aurka5 | 0.000140153 | 0.264124453 | 0.217 | 0.12 | 1 | 28 |
| lman11 | 0.000151903 | 0.261388793 | 0.18 | 0.095 | 1 | 28 |
| si:ch73-359m17.23 | 0.000153495 | 0.265375475 | 0.304 | 0.19 | 1 | 28 |
| rbpms2b8 | 0.000171907 | 0.29012077 | 0.342 | 0.222 | 1 | 28 |
| pdia4 | 0.000175271 | 0.303782823 | 0.224 | 0.132 | 1 | 28 |
| cox175 | 0.000176132 | 0.283300349 | 0.478 | 0.378 | 1 | 28 |
| ncl1 | 0.000181422 | 0.26502764 | 0.484 | 0.421 | 1 | 28 |
| zgc:771122 | 0.000192776 | 0.287585342 | 0.211 | 0.121 | 1 | 28 |
| cfl1l4 | 0.000207025 | 0.2828611 | 0.255 | 0.153 | 1 | 28 |
| nol7 | 0.000210578 | 0.268528864 | 0.217 | 0.124 | 1 | 28 |
| sox19b6 | 0.000211712 | 0.251150956 | 0.186 | 0.098 | 1 | 28 |
| eif4e1b3 | 0.000214551 | 0.251155812 | 0.168 | 0.087 | 1 | 28 |
| slc29a22 | 0.000216697 | 0.25643705 | 0.149 | 0.075 | 1 | 28 |
| myl12.21 | 0.000258613 | 0.29325489 | 0.211 | 0.125 | 1 | 28 |
| btg33 | 0.000280928 | 0.318788571 | 0.273 | 0.18 | 1 | 28 |
| rpz24 | 0.000302851 | 0.304960141 | 0.28 | 0.182 | 1 | 28 |
| coil2 | 0.000336236 | 0.26142974 | 0.193 | 0.108 | 1 | 28 |
| pttg1ipb1 | 0.000363607 | 0.277743745 | 0.199 | 0.115 | 1 | 28 |
| txnl4a2 | 0.000363861 | 0.264544663 | 0.199 | 0.114 | 1 | 28 |
| slc37a21 | 0.000388977 | 0.265958774 | 0.168 | 0.092 | 1 | 28 |
| BX957362.14 | 0.000407005 | 0.273961493 | 0.224 | 0.134 | 1 | 28 |
| cnep1r12 | 0.000428825 | 0.271202315 | 0.211 | 0.125 | 1 | 28 |
| mrpl531 | 0.000450064 | 0.250937883 | 0.168 | 0.092 | 1 | 28 |
| pcf113 | 0.000450156 | 0.305448775 | 0.329 | 0.24 | 1 | 28 |
| fdx17 | 0.000477715 | 0.284560683 | 0.323 | 0.222 | 1 | 28 |
| pop43 | 0.00052759 | 0.253607252 | 0.205 | 0.118 | 1 | 28 |
| zgc:1010163 | 0.000536044 | 0.263709458 | 0.174 | 0.098 | 1 | 28 |
| kpna28 | 0.000591857 | 0.261010984 | 0.261 | 0.159 | 1 | 28 |
| tsr2 | 0.000669282 | 0.251208542 | 0.161 | 0.089 | 1 | 28 |

| | | | | | | |
|---|---|---|---|---|---|---|
| naa40 | 0.000853256 | 0.259673305 | 0.18 | 0.104 | 1 | 28 |
| cbx3a5 | 0.000862118 | 0.286243283 | 0.379 | 0.298 | 1 | 28 |
| uck2a | 0.000893141 | 0.288565385 | 0.193 | 0.117 | 1 | 28 |
| anp32b4 | 0.000923266 | 0.283597785 | 0.348 | 0.257 | 1 | 28 |
| tram24 | 0.000984215 | 0.261911318 | 0.217 | 0.134 | 1 | 28 |
| rab5if2 | 0.001103593 | 0.257028637 | 0.211 | 0.131 | 1 | 28 |
| si:ch1073-263o8.28 | 0.001126139 | 0.265032561 | 0.366 | 0.265 | 1 | 28 |
| ckbb7 | 0.001144766 | 0.263715123 | 0.36 | 0.267 | 1 | 28 |
| gtf2a11 | 0.001148544 | 0.267366974 | 0.242 | 0.156 | 1 | 28 |
| wdr83os | 0.001173899 | 0.285300452 | 0.224 | 0.147 | 1 | 28 |
| ndufab1b1 | 0.001238236 | 0.289983668 | 0.236 | 0.159 | 1 | 28 |
| khsrp1 | 0.0014411 | 0.270882144 | 0.286 | 0.2 | 1 | 28 |
| marcksb5 | 0.001563714 | 0.257990555 | 0.267 | 0.177 | 1 | 28 |
| si:ch211-51e12.71 | 0.001616205 | 0.266223922 | 0.261 | 0.178 | 1 | 28 |
| chchd107 | 0.00165827 | 0.25119059 | 0.236 | 0.153 | 1 | 28 |
| sumo3a2 | 0.001663646 | 0.254424948 | 0.429 | 0.371 | 1 | 28 |
| retsatl8 | 0.001877169 | 0.250890695 | 0.298 | 0.205 | 1 | 28 |
| prmt12 | 0.002130994 | 0.284442796 | 0.298 | 0.224 | 1 | 28 |
| g3bp16 | 0.002808243 | 0.250243488 | 0.298 | 0.212 | 1 | 28 |
| STMP11 | 0.002898529 | 0.266264454 | 0.373 | 0.306 | 1 | 28 |
| uqcrfs1 | 0.003405469 | 0.269757278 | 0.261 | 0.19 | 1 | 28 |
| med19a | 0.00367464 | 0.250345773 | 0.199 | 0.13 | 1 | 28 |
| lsm51 | 0.005260322 | 0.265980418 | 0.304 | 0.239 | 1 | 28 |
| cbx52 | 0.006783605 | 0.251629811 | 0.273 | 0.204 | 1 | 28 |
| tpm21 | 4.81132853672661e-93 | 0.666915062 | 0.262 | 0.018 | 1.21096327940872e-88 | 29 |
| acta21 | 2.56949745962695e-61 | 0.491135112 | 0.181 | 0.013 | 6.46716815613506e-57 | 29 |
| zgc:1534999 | 4.26785614285178e-35 | 0.757314518 | 0.953 | 0.51 | 1.07417671259437e-30 | 29 |
| zp2.29 | 1.00338937621708e-34 | 0.738091511 | 0.973 | 0.527 | 2.52543072100076e-30 | 29 |
| si:dkey-240e12.69 | 3.57767265249361e-34 | 0.717502313 | 0.987 | 0.567 | 9.00464429906118e-30 | 29 |
| si:ch73-189n23.110 | 9.24948502212272e-34 | 0.742636973 | 0.973 | 0.522 | 2.32800288521807e-29 | 29 |
| tmsb59 | 1.6806089352414e-33 | 0.750226296 | 0.96 | 0.486 | 4.22992462910908e-29 | 29 |
| si:ch211-125e6.129 | 8.36508033599043e-33 | 0.731759852 | 0.926 | 0.499 | 2.10540706976543e-28 | 29 |
| zp2.39 | 1.7860358387923e-32 | 0.716792769 | 0.926 | 0.517 | 4.49527360265634e-28 | 29 |
| si:dkeyp-98a7.99 | 2.09541069862057e-32 | 0.72228859 | 0.933 | 0.51 | 5.2739391873581e-28 | 29 |
| mt29 | 3.29835723190134e-32 | 0.706260858 | 0.953 | 0.532 | 8.30163531697248e-28 | 29 |

| | | | | | | |
|---|---|---|---|---|---|---|
| si:dkey-241l7.510 | 5.01085455132868e-32 | 0.676845423 | 0.966 | 0.582 | 1.26118198202392e-27 | 29 |
| thy19 | 4.61412050980913e-31 | 0.72006143 | 0.846 | 0.448 | 1.16132799111386e-26 | 29 |
| h2af1al10 | 6.31171629098849e-31 | 0.688866933 | 0.993 | 0.59 | 1.58859587327889e-26 | 29 |
| si:zfos-1505d6.39 | 1.58521115325573e-30 | 0.722127212 | 0.973 | 0.511 | 3.98981795162936e-26 | 29 |
| si:dkey-241l7.29 | 2.53757622684411e-30 | 0.652043421 | 1 | 0.64 | 6.38682560534393e-26 | 29 |
| cldnd10 | 7.31983325166894e-30 | 0.67731151 | 0.993 | 0.602 | 1.84232883111256e-25 | 29 |
| zp310 | 1.08694885375241e-29 | 0.671044093 | 0.966 | 0.582 | 2.73574157000944e-25 | 29 |
| h1m10 | 2.20418104597915e-29 | 0.665877276 | 0.987 | 0.599 | 5.54770327462492e-25 | 29 |
| zp3b9 | 4.54372953030087e-29 | 0.729551871 | 0.799 | 0.386 | 1.14361128548142e-24 | 29 |
| zgc:771189 | 9.52670080270216e-29 | 0.711602787 | 0.893 | 0.455 | 2.39777532503211e-24 | 29 |
| cldng9 | 9.68911370799107e-29 | 0.695589183 | 0.94 | 0.512 | 2.43865302916427e-24 | 29 |
| zp2.510 | 1.03847903580689e-28 | 0.684726902 | 0.966 | 0.561 | 2.61374788522236e-24 | 29 |
| crp29 | 1.81643472506824e-28 | 0.706241291 | 0.913 | 0.487 | 4.57178455952426e-24 | 29 |
| zp3.210 | 2.16008165039502e-28 | 0.649145094 | 0.906 | 0.556 | 5.43670950587922e-24 | 29 |
| zgc:17513510 | 2.6509553774173e-28 | 0.703319792 | 0.899 | 0.478 | 6.67218958942159e-24 | 29 |
| si:ch211-12h2.89 | 4.53298997147594e-28 | 0.66111442 | 0.919 | 0.546 | 1.14090824592078e-23 | 29 |
| si:dkeyp-98a7.89 | 5.02966477390468e-28 | 0.700065025 | 0.852 | 0.456 | 1.26591632694407e-23 | 29 |
| CR790368.110 | 1.44314811854607e-27 | 0.676398575 | 0.926 | 0.518 | 3.6322594995686e-23 | 29 |
| si:dkey-241l7.49 | 2.00872822166943e-27 | 0.619056907 | 0.953 | 0.607 | 5.0557680611198e-23 | 29 |
| rrm29 | 4.49430064650677e-27 | 0.664762143 | 0.893 | 0.496 | 1.13117052971929e-22 | 29 |
| zgc:1362549 | 6.73839085890287e-27 | 0.723783321 | 0.738 | 0.328 | 1.69598559527726e-22 | 29 |
| ccna19 | 8.6740675342469e-27 | 0.678617238 | 0.886 | 0.474 | 2.1831760576946e-22 | 29 |
| zp2.110 | 8.97697941813647e-27 | 0.668862714 | 0.919 | 0.533 | 2.25941594975077e-22 | 29 |
| BX649337.110 | 1.34153354406337e-26 | 0.696874281 | 0.866 | 0.435 | 3.37650577705309e-22 | 29 |
| acp5a10 | 2.74943822672051e-26 | 0.661639083 | 0.946 | 0.538 | 6.92006107283285e-22 | 29 |
| zp2.69 | 2.82529531878686e-26 | 0.680978293 | 0.913 | 0.499 | 7.11098578785465e-22 | 29 |
| si:ch211-226h8.149 | 4.23432490989539e-26 | 0.658481548 | 0.96 | 0.565 | 1.06573723657157e-21 | 29 |
| qdprb29 | 5.78106489644287e-26 | 0.688730364 | 0.819 | 0.431 | 1.4550362237857e-21 | 29 |
| zgc:1717799 | 8.80897246216153e-26 | 0.667602106 | 0.919 | 0.51 | 2.21713027900144e-21 | 29 |
| tuba4l9 | 9.40236333542609e-26 | 0.69691563 | 0.812 | 0.402 | 2.36648082789339e-21 | 29 |
| zar19 | 1.18736577338758e-25 | 0.673367357 | 0.819 | 0.438 | 2.98848091503921e-21 | 29 |
| LOC1005374999 | 1.88906194263279e-25 | 0.704947922 | 0.705 | 0.32 | 4.75458000341248e-21 | 29 |
| si:dkey-229d11.39 | 1.32903515653965e-24 | 0.69511903 | 0.872 | 0.436 | 3.34504858549465e-20 | 29 |
| si:dkeyp-98a7.49 | 3.17546222979143e-24 | 0.666396445 | 0.819 | 0.423 | 7.99232088616204e-20 | 29 |
| si:dkeyp-20e4.89 | 4.78555100190375e-24 | 0.678569493 | 0.779 | 0.406 | 1.20447533166915e-19 | 29 |

| | | | | | | |
|---|---|---|---|---|---|---|
| ccnb29 | 6.32610501609102e-24 | 0.671910448 | 0.765 | 0.382 | 1.59221737149995e-19 | 29 |
| mid1ip1l9 | 1.94642027455754e-23 | 0.602821648 | 0.879 | 0.542 | 4.89894518903389e-19 | 29 |
| zp3a.29 | 6.92901597099166e-23 | 0.6307338 | 0.852 | 0.49 | 1.74396402973889e-18 | 29 |
| zgc:1717769 | 1.46824938089877e-22 | 0.641516086 | 0.886 | 0.494 | 3.69543686678411e-18 | 29 |
| si:dkey-90l23.19 | 1.57179340328447e-22 | 0.64548848 | 0.812 | 0.446 | 3.95604681672669e-18 | 29 |
| si:dkeyp-46h3.69 | 1.94051694433758e-22 | 0.64861398 | 0.779 | 0.409 | 4.88408709720327e-18 | 29 |
| LOC1104378219 | 2.44587888816724e-22 | 0.656834295 | 0.779 | 0.414 | 6.15603257362813e-18 | 29 |
| lamp29 | 4.65419950828655e-22 | 0.594523317 | 0.893 | 0.553 | 1.17141547424064e-17 | 29 |
| zar1l9 | 4.95626458556953e-22 | 0.648783239 | 0.772 | 0.396 | 1.24744223354199e-17 | 29 |
| siva19 | 5.51906430360592e-22 | 0.636427446 | 0.799 | 0.427 | 1.38909329457457e-17 | 29 |
| epcam10 | 7.88089303073732e-22 | 0.622438522 | 0.893 | 0.503 | 1.98354196690628e-17 | 29 |
| CU571163.19 | 9.31639467271394e-22 | 0.660967764 | 0.812 | 0.399 | 2.34484337517537e-17 | 29 |
| zgc:1952459 | 1.0923441406924e-21 | 0.645227207 | 0.752 | 0.387 | 2.74932096770869e-17 | 29 |
| si:dkey-241l7.69 | 4.75713025436985e-21 | 0.587970248 | 0.832 | 0.526 | 1.19732211372235e-16 | 29 |
| gapdh9 | 7.70987448525262e-21 | 0.623758596 | 0.752 | 0.419 | 1.94049830919323e-16 | 29 |
| AL929032.19 | 3.96582537660988e-20 | 0.651754467 | 0.651 | 0.316 | 9.98158589038941e-16 | 29 |
| slbp29 | 7.10357616023501e-20 | 0.623381956 | 0.779 | 0.395 | 1.78789908376955e-15 | 29 |
| zpcx9 | 8.67197466095631e-20 | 0.635585027 | 0.685 | 0.338 | 2.18264930241609e-15 | 29 |
| zgc:1628799 | 1.24132138363376e-19 | 0.612083258 | 0.47 | 0.18 | 3.1242817904678e-15 | 29 |
| gmnn9 | 1.40546849650681e-19 | 0.63489656 | 0.631 | 0.301 | 3.53742365885799e-15 | 29 |
| si:dkeyp-46h3.89 | 2.72970706863498e-19 | 0.624785998 | 0.698 | 0.35 | 6.87039972104738e-15 | 29 |
| gtf3ab10 | 4.04423870239848e-19 | 0.61656657 | 0.671 | 0.319 | 1.01789443900667e-14 | 29 |
| si:dkeyp-50f7.29 | 9.5073958915807e-19 | 0.607500202 | 0.765 | 0.401 | 2.39291647195195e-14 | 29 |
| zgc:1715179 | 1.70066396602136e-18 | 0.60959166 | 0.725 | 0.387 | 4.28040113607916e-14 | 29 |
| si:ch211-226h8.49 | 1.99127007757808e-18 | 0.590352576 | 0.772 | 0.441 | 5.01182765825628e-14 | 29 |
| uchl19 | 2.62087602849493e-18 | 0.598641104 | 0.678 | 0.343 | 6.59648287611888e-14 | 29 |
| zgc:562319 | 2.9066616605562e-18 | 0.604752706 | 0.53 | 0.227 | 7.31577673345389e-14 | 29 |
| si:ch211-12h2.69 | 3.34426985536261e-18 | 0.573940424 | 0.785 | 0.473 | 8.41719279896215e-14 | 29 |
| elovl7a9 | 3.36960219323627e-18 | 0.600499991 | 0.456 | 0.182 | 8.48095176015637e-14 | 29 |
| npm2b9 | 5.03895763524927e-18 | 0.592861392 | 0.705 | 0.395 | 1.26825524721589e-13 | 29 |
| LOC1005351629 | 8.47655670508061e-18 | 0.608203829 | 0.651 | 0.307 | 2.13346455710174e-13 | 29 |
| si:ch211-14a17.79 | 1.55640042921968e-17 | 0.593072619 | 0.758 | 0.407 | 3.91730424030302e-13 | 29 |
| pigrl2.39 | 2.0587148851775e-17 | 0.578358121 | 0.49 | 0.202 | 5.18157949450325e-13 | 29 |
| si:dkeyp-46h3.29 | 2.63468276177922e-17 | 0.577695233 | 0.718 | 0.403 | 6.63123304312211e-13 | 29 |
| lsm14b9 | 6.90437584102358e-17 | 0.58537708 | 0.604 | 0.299 | 1.73776235542723e-12 | 29 |

| | | | | | | |
|---|---|---|---|---|---|---|
| tac3b8 | 7.69139878347471e-17 | 0.544315437 | 0.342 | 0.119 | 1.93584815981275e-12 | 29 |
| btg49 | 8.13748403441925e-17 | 0.587577565 | 0.685 | 0.368 | 2.04812335662298e-12 | 29 |
| nanog9 | 2.14098838095983e-16 | 0.572089155 | 0.678 | 0.359 | 5.3886536560378e-12 | 29 |
| si:dkey-229d11.59 | 5.77225503217887e-16 | 0.562649121 | 0.691 | 0.389 | 1.4528188690491e-11 | 29 |
| hnrnpabb10 | 1.008310328425e-15 | 0.46458991 | 0.805 | 0.596 | 2.53781626561288e-11 | 29 |
| zgc:1734439 | 4.68444201817749e-15 | 0.564288771 | 0.624 | 0.329 | 1.17902721155509e-10 | 29 |
| ctss2.110 | 6.05401746352405e-15 | 0.489809476 | 0.812 | 0.534 | 1.52373565539437e-10 | 29 |
| BX537276.19 | 6.82839176881694e-15 | 0.499667189 | 0.336 | 0.121 | 1.71863792429354e-10 | 29 |
| nasp9 | 7.53263317907606e-15 | 0.559110322 | 0.617 | 0.339 | 1.89588844484165e-10 | 29 |
| snu13b10 | 8.51486450342201e-15 | 0.537849046 | 0.638 | 0.392 | 2.14310624686629e-10 | 29 |
| kpna79 | 9.70506078847158e-15 | 0.553810755 | 0.617 | 0.328 | 2.44266674985041e-10 | 29 |
| AL929192.19 | 1.07213525296117e-14 | 0.523842333 | 0.409 | 0.163 | 2.69845721817797e-10 | 29 |
| si:ch1073-263o8.29 | 1.07400200332909e-14 | 0.553401419 | 0.53 | 0.263 | 2.70315564217898e-10 | 29 |
| LOC1005368609 | 1.07629732796396e-14 | 0.542790481 | 0.624 | 0.35 | 2.70893274475249e-10 | 29 |
| mibp9 | 2.30936191099635e-14 | 0.555936337 | 0.47 | 0.222 | 5.81243299378672e-10 | 29 |
| si:ch73-14h10.29 | 3.35066838448419e-14 | 0.542614919 | 0.544 | 0.271 | 8.43329725690825e-10 | 29 |
| zgc:5546110 | 4.70813965149962e-14 | 0.543749425 | 0.564 | 0.288 | 1.18499166888594e-09 | 29 |
| zgc:1714749 | 6.36778082820496e-14 | 0.548902497 | 0.584 | 0.303 | 1.60270675665091e-09 | 29 |
| si:dkey-188i13.1110 | 6.7984397375245e-14 | 0.537740465 | 0.57 | 0.316 | 1.71109929753754e-09 | 29 |
| s100a19 | 9.73532633271091e-14 | 0.534026866 | 0.678 | 0.363 | 2.45028428468001e-09 | 29 |
| elavl210 | 1.28814115970604e-13 | 0.550143106 | 0.463 | 0.223 | 3.24212248486412e-09 | 29 |
| cpeb1b9 | 1.49008624683403e-13 | 0.545936654 | 0.544 | 0.27 | 3.75039807465657e-09 | 29 |
| mad2l18 | 1.68263694607542e-13 | 0.517577798 | 0.416 | 0.185 | 4.23502892957722e-09 | 29 |
| CABZ01075618.11 | 1.84300700130173e-13 | 0.406427828 | 0.195 | 0.055 | 4.63866432157632e-09 | 29 |
| wu:fi42e039 | 2.30994928340516e-13 | 0.527045571 | 0.624 | 0.349 | 5.81391135140243e-09 | 29 |
| si:dkey-208k4.29 | 3.01055754770524e-13 | 0.519900889 | 0.564 | 0.295 | 7.57727229181933e-09 | 29 |
| rbpms2b9 | 5.47082518234573e-13 | 0.517772152 | 0.463 | 0.221 | 1.3769519901446e-08 | 29 |
| gbgt1l48 | 6.63896949622487e-13 | 0.49122873 | 0.362 | 0.15 | 1.67096223250484e-08 | 29 |
| LOC1018854779 | 6.64108877411227e-13 | 0.546289379 | 0.523 | 0.279 | 1.67149563355632e-08 | 29 |
| cdca88 | 7.26041269780322e-13 | 0.50184843 | 0.383 | 0.164 | 1.82737327191009e-08 | 29 |
| si:dkey-37o8.19 | 9.07441899334513e-13 | 0.518761358 | 0.57 | 0.307 | 2.28394051643504e-08 | 29 |
| si:ch211-250e5.169 | 1.19577478118135e-12 | 0.514703833 | 0.591 | 0.346 | 3.00964554675534e-08 | 29 |
| si:dkeyp-98a7.79 | 1.45230437031748e-12 | 0.502915379 | 0.436 | 0.195 | 3.65530486965207e-08 | 29 |
| zgc:925919 | 1.4723732781534e-12 | 0.518175175 | 0.436 | 0.205 | 3.7058163037843e-08 | 29 |
| si:dkey-19b23.119 | 2.18236855702356e-12 | 0.50399707 | 0.53 | 0.278 | 5.49280342117259e-08 | 29 |

| | | | | | | |
|---|---|---|---|---|---|---|
| snu13a4 | 2.20518454885741e-12 | 0.443528928 | 0.275 | 0.1 | 5.55022899101922e-08 | 29 |
| cth19 | 2.40554274304113e-12 | 0.500529717 | 0.644 | 0.379 | 6.05451052996023e-08 | 29 |
| h3f3a10 | 3.71011172323918e-12 | 0.45581749 | 0.711 | 0.505 | 9.33798019622068e-08 | 29 |
| chia.49 | 3.75308175428568e-12 | 0.481507415 | 0.349 | 0.147 | 9.44613146736162e-08 | 29 |
| elovl7b9 | 4.27033539109827e-12 | 0.505023 | 0.483 | 0.248 | 1.07480071458552e-07 | 29 |
| si:ch211-146l10.89 | 6.52964440613954e-12 | 0.503957876 | 0.564 | 0.306 | 1.64344620058126e-07 | 29 |
| si:ch211-145c1.19 | 6.7448697595179e-12 | 0.505473705 | 0.477 | 0.233 | 1.69761626977306e-07 | 29 |
| gdf39 | 7.50485461214586e-12 | 0.485521218 | 0.463 | 0.223 | 1.88889685733099e-07 | 29 |
| lin28a1 | 9.0056144123001e-12 | 0.396578167 | 0.215 | 0.07 | 2.26662309143181e-07 | 29 |
| zgc:5669910 | 1.01333505314105e-11 | 0.492417219 | 0.497 | 0.251 | 2.55046299525071e-07 | 29 |
| si:dkey-46g23.28 | 1.01832751780486e-11 | 0.461279808 | 0.342 | 0.145 | 2.56302852956305e-07 | 29 |
| haus68 | 1.17761708049218e-11 | 0.491313799 | 0.436 | 0.209 | 2.96394442989078e-07 | 29 |
| zgc:1141236 | 2.2764090488581e-11 | 0.41858833 | 0.275 | 0.103 | 5.72949393507096e-07 | 29 |
| si:dkey-152b24.69 | 2.99480426119836e-11 | 0.489891619 | 0.544 | 0.298 | 7.53762284501016e-07 | 29 |
| zswim7 | 3.00613940802691e-11 | 0.353073529 | 0.161 | 0.046 | 7.56615227606294e-07 | 29 |
| ccnb19 | 3.12270301779185e-11 | 0.485128907 | 0.678 | 0.393 | 7.85953122548031e-07 | 29 |
| si:dkeyp-46h3.59 | 4.55474173577787e-11 | 0.493417048 | 0.537 | 0.289 | 1.14638294747793e-06 | 29 |
| si:ch73-90k17.19 | 6.26762445730776e-11 | 0.483999652 | 0.477 | 0.249 | 1.57749839965979e-06 | 29 |
| si:ch1073-190k2.19 | 7.05937845495361e-11 | 0.469624666 | 0.403 | 0.188 | 1.77677496332727e-06 | 29 |
| gstp17 | 8.70351156927518e-11 | 0.464160502 | 0.584 | 0.365 | 2.19058682687087e-06 | 29 |
| zgc:1532842 | 1.04542756116381e-10 | 0.399130493 | 0.262 | 0.1 | 2.63123662869319e-06 | 29 |
| degs27 | 1.12847821888016e-10 | 0.404593496 | 0.275 | 0.106 | 2.84026682909946e-06 | 29 |
| si:ch211-132b12.89 | 1.25846086642695e-10 | 0.475767378 | 0.577 | 0.342 | 3.16742015471e-06 | 29 |
| si:dkey-90l23.29 | 1.27758159230011e-10 | 0.476567231 | 0.611 | 0.344 | 3.21554510966014e-06 | 29 |
| slbp10 | 1.32345337937125e-10 | 0.483754666 | 0.443 | 0.234 | 3.33099981053951e-06 | 29 |
| ddx39ab8 | 1.4172856804087e-10 | 0.473808434 | 0.503 | 0.289 | 3.56716632902067e-06 | 29 |
| zp3a.19 | 1.68096405375635e-10 | 0.464230682 | 0.617 | 0.381 | 4.23081842689935e-06 | 29 |
| si:dkey-179j5.5 | 2.27467185666687e-10 | 0.271833969 | 0.141 | 0.038 | 5.72512159604485e-06 | 29 |
| birc5b9 | 2.36142267427414e-10 | 0.465464361 | 0.497 | 0.259 | 5.94346472888059e-06 | 29 |
| tmem144b4 | 2.72906696666774e-10 | 0.386318256 | 0.248 | 0.094 | 6.86878864840603e-06 | 29 |
| crp39 | 2.79488628568918e-10 | 0.473838668 | 0.423 | 0.218 | 7.0344492924511e-06 | 29 |
| rnf47 | 2.95735972917773e-10 | 0.433609593 | 0.309 | 0.134 | 7.44337870236742e-06 | 29 |
| spink2.110 | 2.97666491309168e-10 | 0.436429988 | 0.664 | 0.437 | 7.49196791976046e-06 | 29 |
| buc9 | 3.37034938957748e-10 | 0.456706593 | 0.376 | 0.179 | 8.48283237862757e-06 | 29 |
| slc16a39 | 4.67794639314834e-10 | 0.466886895 | 0.47 | 0.249 | 1.1773923276915e-05 | 29 |

| | | | | | | |
|---|---|---|---|---|---|---|
| myl9a4 | 4.90423401549451e-10 | 0.298396792 | 0.121 | 0.031 | 1.23434665935981e-05 | 29 |
| CR847984.19 | 7.01820295074083e-10 | 0.435343571 | 0.356 | 0.164 | 1.76641150067196e-05 | 29 |
| ccna29 | 7.05972849536371e-10 | 0.458164364 | 0.51 | 0.284 | 1.77686306499809e-05 | 29 |
| avd9 | 7.23763181381157e-10 | 0.460604853 | 0.497 | 0.265 | 1.82163955121824e-05 | 29 |
| rogdi3 | 9.01825247294171e-10 | 0.420512585 | 0.275 | 0.116 | 2.2698039649147e-05 | 29 |
| si:ch211-146l10.79 | 1.08701953137779e-09 | 0.425384217 | 0.396 | 0.189 | 2.73591945852475e-05 | 29 |
| zp2l19 | 1.11917858337956e-09 | 0.434015781 | 0.349 | 0.164 | 2.81686057650803e-05 | 29 |
| zgc:923134 | 1.30197562689372e-09 | 0.396849989 | 0.295 | 0.126 | 3.27694245532879e-05 | 29 |
| zgc:1719778 | 1.85987624491167e-09 | 0.442417941 | 0.436 | 0.23 | 4.68112252081817e-05 | 29 |
| phex4 | 2.37819594505043e-09 | 0.348192111 | 0.228 | 0.087 | 5.98568137409744e-05 | 29 |
| arhgef10la9 | 2.4139623215166e-09 | 0.448712963 | 0.483 | 0.275 | 6.07570176702513e-05 | 29 |
| hbba19 | 2.66311006570118e-09 | 0.464964585 | 0.577 | 0.356 | 6.70278172436329e-05 | 29 |
| patl29 | 3.13384847684055e-09 | 0.429511161 | 0.356 | 0.173 | 7.88758323135999e-05 | 29 |
| snx10a7 | 3.67739489277271e-09 | 0.403979214 | 0.289 | 0.126 | 9.25563520561963e-05 | 29 |
| si:ch211-251f6.69 | 3.89916213191727e-09 | 0.441862764 | 0.53 | 0.332 | 9.81380116982257e-05 | 29 |
| slc35f21 | 4.62463378404677e-09 | 0.369071546 | 0.201 | 0.076 | 0.000116397 | 29 |
| zp3c9 | 4.65655321772214e-09 | 0.423572588 | 0.383 | 0.189 | 0.000117201 | 29 |
| si:ch211-286b5.44 | 5.44985064195592e-09 | 0.331647491 | 0.195 | 0.07 | 0.000137167 | 29 |
| zgc:1138868 | 5.64195024531078e-09 | 0.414115144 | 0.309 | 0.142 | 0.000142002 | 29 |
| sox19b7 | 5.8725817341935e-09 | 0.370536586 | 0.242 | 0.098 | 0.000147807 | 29 |
| eif4a1a7 | 5.94520140717071e-09 | 0.40204015 | 0.604 | 0.425 | 0.000149635 | 29 |
| CU019646.11 | 6.67954504268967e-09 | 0.305136956 | 0.161 | 0.053 | 0.000168117 | 29 |
| kpna29 | 6.99931562035124e-09 | 0.404534562 | 0.336 | 0.158 | 0.000176166 | 29 |
| mkrn48 | 7.24252282077524e-09 | 0.437943037 | 0.423 | 0.229 | 0.000182287 | 29 |
| spryd44 | 8.1451654752489e-09 | 0.330680479 | 0.208 | 0.078 | 0.000205006 | 29 |
| dap1b2 | 1.21366439095719e-08 | 0.451104517 | 0.416 | 0.251 | 0.000305467 | 29 |
| hmgb3b3 | 1.28427019015726e-08 | 0.435880717 | 0.342 | 0.178 | 0.000323238 | 29 |
| si:ch211-226l4.610 | 1.30785291271744e-08 | 0.429906336 | 0.51 | 0.293 | 0.000329173 | 29 |
| setb2 | 1.35594307429694e-08 | 0.420037198 | 0.523 | 0.361 | 0.000341277 | 29 |
| lpcat45 | 1.41330488064386e-08 | 0.356844828 | 0.255 | 0.107 | 0.000355715 | 29 |
| pabpc1l9 | 1.68492346708465e-08 | 0.383990158 | 0.309 | 0.142 | 0.000424078 | 29 |
| LOC1104377794 | 1.77139175681924e-08 | 0.339702954 | 0.215 | 0.084 | 0.000445842 | 29 |
| ca15b9 | 2.1196495496314e-08 | 0.420966301 | 0.45 | 0.25 | 0.000533495 | 29 |
| tmem54a7 | 2.22654489763614e-08 | 0.383090831 | 0.309 | 0.141 | 0.000560399 | 29 |
| mrpl9 | 2.39773900518225e-08 | 0.333568276 | 0.201 | 0.078 | 0.000603487 | 29 |

| | | | | | | |
|---|---|---|---|---|---|---|
| cdca94 | 2.45033430517846e-08 | 0.356925482 | 0.215 | 0.086 | 0.000616725 | 29 |
| sinup9 | 2.4731439539839e-08 | 0.42722947 | 0.383 | 0.204 | 0.000622466 | 29 |
| zgc:1017446 | 2.55166161071108e-08 | 0.388140691 | 0.289 | 0.134 | 0.000642228 | 29 |
| rpain5 | 2.6612648594761e-08 | 0.380773026 | 0.268 | 0.12 | 0.000669814 | 29 |
| npm2a1 | 2.85512527111724e-08 | 0.353680532 | 0.201 | 0.079 | 0.000718606 | 29 |
| eif4e1b4 | 3.12486467862098e-08 | 0.348273024 | 0.215 | 0.086 | 0.000786497 | 29 |
| CDPF19 | 3.80724366319807e-08 | 0.423971364 | 0.456 | 0.269 | 0.000958245 | 29 |
| dctpp19 | 5.83260658837375e-08 | 0.3903191 | 0.349 | 0.175 | 0.001468009 | 29 |
| zgc:660249 | 5.89797004097146e-08 | 0.384561608 | 0.302 | 0.142 | 0.00148446 | 29 |
| rcc19 | 6.24536020748224e-08 | 0.400751784 | 0.383 | 0.203 | 0.001571895 | 29 |
| si:dkey-121a9.3 | 6.33896309604474e-08 | 0.267549504 | 0.141 | 0.046 | 0.001595454 | 29 |
| ctsla9 | 6.77124761180248e-08 | 0.391217952 | 0.564 | 0.387 | 0.001704255 | 29 |
| si:ch211-154a22.8 | 7.41306043978784e-08 | 0.281139597 | 0.154 | 0.053 | 0.001865793 | 29 |
| si:ch211-102c2.7 | 7.4909818074066e-08 | 0.285717234 | 0.148 | 0.05 | 0.001885405 | 29 |
| si:ch211-5k11.89 | 7.96091689333165e-08 | 0.332145586 | 0.604 | 0.358 | 0.002003683 | 29 |
| plk18 | 8.54888302795633e-08 | 0.387871953 | 0.349 | 0.177 | 0.002151668 | 29 |
| aurka6 | 8.97633320628788e-08 | 0.338613941 | 0.268 | 0.12 | 0.002259253 | 29 |
| LOC1039099724 | 9.17502087003109e-08 | 0.397515625 | 0.53 | 0.367 | 0.002309261 | 29 |
| zgc:1121461 | 1.03081959966566e-07 | 0.257720496 | 0.128 | 0.04 | 0.00259447 | 29 |
| zgc:868395 | 1.09665006310744e-07 | 0.342894169 | 0.248 | 0.11 | 0.002760159 | 29 |
| zglp19 | 1.10306990389006e-07 | 0.398853865 | 0.376 | 0.199 | 0.002776317 | 29 |
| inka1b9 | 1.18372949047473e-07 | 0.401339426 | 0.349 | 0.185 | 0.002979329 | 29 |
| vps26a | 1.23884607682908e-07 | 0.310143088 | 0.174 | 0.067 | 0.003118052 | 29 |
| sybu9 | 1.42977223921184e-07 | 0.385753644 | 0.329 | 0.167 | 0.003598594 | 29 |
| oclnb | 1.67429602302175e-07 | 0.255572566 | 0.128 | 0.041 | 0.004214036 | 29 |
| si:ch211-113a14.273 | 1.69062332441863e-07 | 0.316782113 | 0.195 | 0.078 | 0.00425513 | 29 |
| zcchc8 | 1.80898572994381e-07 | 0.255184542 | 0.121 | 0.038 | 0.004553036 | 29 |
| ctsba9 | 2.01991927113699e-07 | 0.380334032 | 0.557 | 0.377 | 0.005083935 | 29 |
| zgc:1629459 | 2.24592335553454e-07 | 0.376821971 | 0.322 | 0.163 | 0.005652764 | 29 |
| CU655961.32 | 2.32378789020489e-07 | 0.332746636 | 0.228 | 0.099 | 0.005848742 | 29 |
| si:ch73-78o10.17 | 2.41661721833096e-07 | 0.358949243 | 0.282 | 0.133 | 0.006082384 | 29 |
| cnih12 | 2.42818345009852e-07 | 0.371496318 | 0.248 | 0.117 | 0.006111495 | 29 |
| LOC1018838979 | 2.44996741137284e-07 | 0.378858979 | 0.322 | 0.165 | 0.006166323 | 29 |
| pabpn1l9 | 2.88352414909545e-07 | 0.387525653 | 0.477 | 0.284 | 0.007257542 | 29 |
| zgc:1738569 | 3.24297072278939e-07 | 0.392162694 | 0.389 | 0.221 | 0.008162233 | 29 |

| | | | | | | |
|---|---|---|---|---|---|---|
| si:ch211-14a17.64 | 3.42625240533053e-07 | 0.307950786 | 0.208 | 0.086 | 0.008623535 | 29 |
| zgc:1129804 | 3.61791992704473e-07 | 0.329981883 | 0.215 | 0.093 | 0.009105943 | 29 |
| pttg19 | 3.63297856005372e-07 | 0.359501762 | 0.315 | 0.159 | 0.009143844 | 29 |
| larp6b9 | 3.73946591975366e-07 | 0.369550726 | 0.369 | 0.196 | 0.009411862 | 29 |
| bzw1b6 | 4.33813850414806e-07 | 0.371976732 | 0.544 | 0.403 | 0.010918661 | 29 |
| ppp4ca | 4.80863737862953e-07 | 0.290974199 | 0.181 | 0.072 | 0.012102859 | 29 |
| gpr1604 | 4.810215000968e-07 | 0.322385569 | 0.195 | 0.081 | 0.01210683 | 29 |
| si:dkey-42i9.89 | 5.13118691835219e-07 | 0.346559685 | 0.268 | 0.128 | 0.012914684 | 29 |
| CABZ01059627.29 | 5.29751897453152e-07 | 0.355191515 | 0.275 | 0.134 | 0.013333326 | 29 |
| prr113 | 5.48057964258279e-07 | 0.31507284 | 0.228 | 0.1 | 0.013794071 | 29 |
| si:ch211-276i12.91 | 6.53867083556236e-07 | 0.302763687 | 0.161 | 0.063 | 0.016457181 | 29 |
| dnajc1 | 6.81100818587621e-07 | 0.392223176 | 0.275 | 0.145 | 0.017142627 | 29 |
| retsatl9 | 6.98640726486844e-07 | 0.37462756 | 0.376 | 0.204 | 0.017584088 | 29 |
| yrdc7 | 8.82455972680459e-07 | 0.355638307 | 0.255 | 0.123 | 0.022210534 | 29 |
| dynll2a9 | 9.16441130320956e-07 | 0.328925857 | 0.282 | 0.137 | 0.023065907 | 29 |
| org9 | 1.0613601085937e-06 | 0.378569545 | 0.409 | 0.241 | 0.026713373 | 29 |
| fbxo325 | 1.23132001811607e-06 | 0.313318449 | 0.228 | 0.105 | 0.030991094 | 29 |
| cpm5 | 1.27760150188153e-06 | 0.316818037 | 0.235 | 0.109 | 0.032155952 | 29 |
| si:dkey-216e24.37 | 1.32226391169715e-06 | 0.335607997 | 0.255 | 0.124 | 0.03328006 | 29 |
| BX005461.32 | 1.37467068685217e-06 | 0.289955293 | 0.168 | 0.068 | 0.034599087 | 29 |
| CT009596.15 | 1.38642564650093e-06 | 0.298939719 | 0.208 | 0.091 | 0.034894947 | 29 |
| micall2a | 1.40178573054106e-06 | 0.255312846 | 0.148 | 0.055 | 0.035281545 | 29 |
| btr041 | 1.41611326416128e-06 | 0.276765957 | 0.161 | 0.064 | 0.035642155 | 29 |
| zc3h11a8 | 1.54105622863307e-06 | 0.379325987 | 0.369 | 0.218 | 0.038786844 | 29 |
| brf24 | 1.56063684944469e-06 | 0.300320119 | 0.215 | 0.095 | 0.039279669 | 29 |
| asf1ba2 | 1.60786692583548e-06 | 0.322795655 | 0.215 | 0.098 | 0.040468403 | 29 |
| CU467646.27 | 1.64107137136827e-06 | 0.313895035 | 0.262 | 0.126 | 0.041304125 | 29 |
| sdf2 | 1.69534656831397e-06 | 0.262068107 | 0.141 | 0.053 | 0.042670178 | 29 |
| nol11 | 1.78522210131089e-06 | 0.303378431 | 0.168 | 0.069 | 0.044932255 | 29 |
| f11r.13 | 1.80769546252705e-06 | 0.358646472 | 0.322 | 0.178 | 0.045497887 | 29 |
| ap1s3b | 1.88067018259771e-06 | 0.303057827 | 0.181 | 0.077 | 0.047334588 | 29 |
| rdh10b8 | 1.99045135625913e-06 | 0.372981428 | 0.322 | 0.179 | 0.05009767 | 29 |
| rasl11b7 | 1.99447578956866e-06 | 0.340603141 | 0.268 | 0.135 | 0.050198961 | 29 |
| rbm4.39 | 2.04328293895877e-06 | 0.340496334 | 0.584 | 0.445 | 0.051427388 | 29 |
| mtfr2 | 2.07767373551969e-06 | 0.259793921 | 0.128 | 0.046 | 0.05229297 | 29 |

| | | | | | | |
|---|---|---|---|---|---|---|
| h2afva8 | 2.14950211094707e-06 | 0.380511165 | 0.463 | 0.324 | 0.054100819 | 29 |
| c1qbp1 | 2.31280373914364e-06 | 0.381476774 | 0.302 | 0.172 | 0.058210957 | 29 |
| chrac12 | 2.43444711441625e-06 | 0.345400712 | 0.255 | 0.129 | 0.061272599 | 29 |
| cat1 | 2.69228475953269e-06 | 0.278934232 | 0.181 | 0.078 | 0.067762115 | 29 |
| zgc:1132932 | 2.85090010202144e-06 | 0.269047604 | 0.141 | 0.054 | 0.071754305 | 29 |
| parn7 | 2.93780264553623e-06 | 0.348279919 | 0.282 | 0.148 | 0.073941555 | 29 |
| cdc209 | 3.20979411441187e-06 | 0.342389997 | 0.289 | 0.153 | 0.080787308 | 29 |
| nsun2 | 3.2591141526948e-06 | 0.277993828 | 0.154 | 0.062 | 0.082028644 | 29 |
| ndrg1a | 3.26189462340433e-06 | 0.295300289 | 0.174 | 0.075 | 0.082098626 | 29 |
| tram25 | 3.28608449595274e-06 | 0.329750866 | 0.262 | 0.133 | 0.082707461 | 29 |
| zgc:771123 | 3.44621980159097e-06 | 0.333272628 | 0.242 | 0.121 | 0.086737906 | 29 |
| cdc14b | 3.70780944261424e-06 | 0.360320651 | 0.255 | 0.135 | 0.093321856 | 29 |
| si:ch73-15b2.54 | 3.82220236102737e-06 | 0.301049204 | 0.208 | 0.095 | 0.096201011 | 29 |
| BX571780.16 | 4.1604038492417e-06 | 0.279172546 | 0.188 | 0.082 | 0.104713204 | 29 |
| asz12 | 4.26390600783923e-06 | 0.268289833 | 0.174 | 0.073 | 0.10731825 | 29 |
| ppp1r3b3 | 4.46734580163513e-06 | 0.320525745 | 0.228 | 0.11 | 0.112438626 | 29 |
| dynll110 | 4.46872691442591e-06 | 0.317688915 | 0.591 | 0.455 | 0.112473388 | 29 |
| ccdc581 | 4.47383256608043e-06 | 0.314865833 | 0.228 | 0.111 | 0.112601892 | 29 |
| pcgf1 | 4.49190531555921e-06 | 0.289734733 | 0.154 | 0.064 | 0.113056765 | 29 |
| metap1d1 | 4.82054653903121e-06 | 0.32040322 | 0.195 | 0.09 | 0.121328336 | 29 |
| cfl1l5 | 5.10247564128881e-06 | 0.340936405 | 0.282 | 0.153 | 0.128424209 | 29 |
| map1lc3c9 | 5.13312735903019e-06 | 0.335376367 | 0.362 | 0.202 | 0.129195682 | 29 |
| vps9d14 | 5.38542163683987e-06 | 0.297917356 | 0.208 | 0.097 | 0.135545677 | 29 |
| kif221 | 5.47337318572281e-06 | 0.297494487 | 0.195 | 0.088 | 0.13775933 | 29 |
| tph1a9 | 5.51044268899144e-06 | 0.315376043 | 0.275 | 0.142 | 0.138692332 | 29 |
| si:ch73-160p18.39 | 5.57046787471934e-06 | 0.350643039 | 0.396 | 0.241 | 0.140203106 | 29 |
| hbaa19 | 5.74160785989838e-06 | 0.341784617 | 0.456 | 0.282 | 0.144510528 | 29 |
| si:ch211-226h8.62 | 5.81759665987938e-06 | 0.27498865 | 0.181 | 0.079 | 0.14642309 | 29 |
| st8sia7.12 | 5.85402470451864e-06 | 0.282578118 | 0.168 | 0.072 | 0.147339948 | 29 |
| cdca7a9 | 5.90625104269379e-06 | 0.35272268 | 0.376 | 0.223 | 0.148654432 | 29 |
| CR383662.25 | 6.06981670214154e-06 | 0.281857422 | 0.195 | 0.087 | 0.152771217 | 29 |
| zgc:558883 | 6.28735441240487e-06 | 0.268391448 | 0.195 | 0.086 | 0.158246423 | 29 |
| thop13 | 6.30028070771589e-06 | 0.320929777 | 0.235 | 0.118 | 0.158571765 | 29 |
| tmem703 | 6.87143428728414e-06 | 0.29050977 | 0.221 | 0.106 | 0.17294713 | 29 |
| si:ch211-226h8.83 | 7.12468196528112e-06 | 0.299639293 | 0.208 | 0.098 | 0.17932112 | 29 |

| | | | | | | |
|---|---|---|---|---|---|---|
| zgc:1102395 | 7.19212931353502e-06 | 0.359441446 | 0.349 | 0.215 | 0.181018703 | 29 |
| si:dkey-52l18.4 | 7.43146266709123e-06 | 0.293130963 | 0.134 | 0.054 | 0.187042484 | 29 |
| pdap1a | 7.60021072337267e-06 | 0.318823406 | 0.235 | 0.12 | 0.191289704 | 29 |
| prdx19 | 8.46967967845127e-06 | 0.335927555 | 0.302 | 0.167 | 0.213173368 | 29 |
| zgc:922878 | 8.92876671094031e-06 | 0.319745925 | 0.255 | 0.133 | 0.224728129 | 29 |
| fdx18 | 1.09535198475792e-05 | 0.356793059 | 0.356 | 0.222 | 0.275689141 | 29 |
| si:dkey-241l7.39 | 1.13429445640308e-05 | 0.342782978 | 0.517 | 0.366 | 0.285490572 | 29 |
| ociad1 | 1.14567726943076e-05 | 0.331895476 | 0.208 | 0.104 | 0.288355512 | 29 |
| pdzd11 | 1.16862564425244e-05 | 0.291347236 | 0.201 | 0.096 | 0.294131388 | 29 |
| fbxo5 | 1.25271509805065e-05 | 0.284919081 | 0.161 | 0.071 | 0.315295863 | 29 |
| uba2 | 1.27284661826347e-05 | 0.310938475 | 0.195 | 0.094 | 0.320362765 | 29 |
| g3bp17 | 1.44045659402027e-05 | 0.347073945 | 0.349 | 0.211 | 0.36254852 | 29 |
| nek22 | 1.4476300435221e-05 | 0.255269667 | 0.161 | 0.069 | 0.364354006 | 29 |
| sumo3b2 | 1.52798550952889e-05 | 0.314489308 | 0.242 | 0.126 | 0.384578673 | 29 |
| cx43.45 | 1.54911725264011e-05 | 0.299807192 | 0.242 | 0.124 | 0.389897321 | 29 |
| dnd15 | 1.56146835147906e-05 | 0.292888208 | 0.221 | 0.11 | 0.393005969 | 29 |
| imp37 | 1.6580644607856e-05 | 0.310392918 | 0.255 | 0.135 | 0.417318244 | 29 |
| setd41 | 1.69409406756623e-05 | 0.2635123 | 0.161 | 0.07 | 0.426386536 | 29 |
| cmc2 | 1.85508040081776e-05 | 0.281848654 | 0.141 | 0.06 | 0.466905186 | 29 |
| lman12 | 2.03085511194097e-05 | 0.295679684 | 0.195 | 0.095 | 0.511145923 | 29 |
| zgc:1118686 | 2.03483170840766e-05 | 0.303409894 | 0.275 | 0.148 | 0.512146793 | 29 |
| BX284697.17 | 2.07937073615921e-05 | 0.290364026 | 0.228 | 0.115 | 0.523356821 | 29 |
| cdc62 | 2.1062130155088e-05 | 0.291315128 | 0.208 | 0.103 | 0.530112754 | 29 |
| lsm12 | 2.34707263063391e-05 | 0.302183532 | 0.255 | 0.137 | 0.59073471 | 29 |
| hsd17b12a9 | 2.36257708220854e-05 | 0.307973441 | 0.302 | 0.166 | 0.594637026 | 29 |
| lbh3 | 2.38325097141827e-05 | 0.324333476 | 0.295 | 0.173 | 0.599840437 | 29 |
| cldn7b8 | 2.44460481810109e-05 | 0.31302969 | 0.289 | 0.161 | 0.615282587 | 29 |
| ccne17 | 2.52527278662855e-05 | 0.292516859 | 0.221 | 0.112 | 0.635585908 | 29 |
| cnep1r13 | 2.53391065654476e-05 | 0.317539708 | 0.235 | 0.124 | 0.637759973 | 29 |
| zgc:1737429 | 2.5832623784321e-05 | 0.299240307 | 0.289 | 0.156 | 0.650181308 | 29 |
| zgc:10348210 | 2.60023858288368e-05 | 0.323683955 | 0.329 | 0.19 | 0.654454049 | 29 |
| exosc61 | 2.64537905381398e-05 | 0.304451651 | 0.181 | 0.087 | 0.665815454 | 29 |
| zgc:112185 | 2.76814559058337e-05 | 0.27769062 | 0.174 | 0.081 | 0.696714564 | 29 |
| srp9 | 2.89390191095434e-05 | 0.288307993 | 0.208 | 0.105 | 0.728366172 | 29 |
| top1l2 | 3.03159442223645e-05 | 0.353618295 | 0.349 | 0.229 | 0.763022 | 29 |

| | | | | | | |
|---|---|---|---|---|---|---|
| surf2 | 3.14559448222241e-05 | 0.280661051 | 0.174 | 0.082 | 0.791714675 | 29 |
| zgc:1526527 | 3.19917602623247e-05 | 0.272146043 | 0.228 | 0.115 | 0.805200614 | 29 |
| polr1d | 3.39662907346563e-05 | 0.271186436 | 0.188 | 0.091 | 0.854897572 | 29 |
| rpl7l1 | 3.86708294448343e-05 | 0.317144727 | 0.242 | 0.134 | 0.973306106 | 29 |
| hspa58 | 3.87722332308797e-05 | 0.321031313 | 0.463 | 0.315 | 0.975858338 | 29 |
| ccng1 | 3.96828589818569e-05 | 0.292469148 | 0.55 | 0.448 | 0.998777878 | 29 |
| si:ch211-69g19.27 | 4.06932637048549e-05 | 0.302473408 | 0.228 | 0.123 | 1 | 29 |
| pprc11 | 4.3036976559459e-05 | 0.278865507 | 0.188 | 0.092 | 1 | 29 |
| zgc:171566 | 5.02656857045884e-05 | 0.258356223 | 0.154 | 0.07 | 1 | 29 |
| zgc:1139846 | 5.03060444729573e-05 | 0.253516231 | 0.208 | 0.102 | 1 | 29 |
| ube2c7 | 5.18158981772235e-05 | 0.303001765 | 0.255 | 0.141 | 1 | 29 |
| mylipa4 | 5.99254453965925e-05 | 0.292092507 | 0.221 | 0.116 | 1 | 29 |
| mos8 | 6.06325350584227e-05 | 0.304461776 | 0.248 | 0.137 | 1 | 29 |
| CU655961.44 | 6.6511924943384e-05 | 0.28296276 | 0.181 | 0.089 | 1 | 29 |
| itm2cb5 | 6.75753893219279e-05 | 0.302453804 | 0.248 | 0.139 | 1 | 29 |
| nfr7 | 6.91617344062752e-05 | 0.275371378 | 0.215 | 0.112 | 1 | 29 |
| polr2k2 | 8.90329087846931e-05 | 0.314950897 | 0.309 | 0.191 | 1 | 29 |
| ttc323 | 9.6669988115815e-05 | 0.270651923 | 0.201 | 0.104 | 1 | 29 |
| adipor1a1 | 0.000101221 | 0.272412285 | 0.195 | 0.1 | 1 | 29 |
| emc9 | 0.000108642 | 0.251359132 | 0.148 | 0.068 | 1 | 29 |
| marcksb6 | 0.000109519 | 0.296174564 | 0.295 | 0.177 | 1 | 29 |
| nol71 | 0.000112861 | 0.276860533 | 0.228 | 0.124 | 1 | 29 |
| ypel37 | 0.000114457 | 0.273864675 | 0.235 | 0.128 | 1 | 29 |
| nap1l13 | 0.000121127 | 0.296972743 | 0.47 | 0.362 | 1 | 29 |
| jtb | 0.000121543 | 0.251691437 | 0.134 | 0.06 | 1 | 29 |
| gyg1a10 | 0.000132356 | 0.308088196 | 0.503 | 0.371 | 1 | 29 |
| mrps26 | 0.000133656 | 0.268053703 | 0.161 | 0.079 | 1 | 29 |
| si:dkey-68o6.59 | 0.000137987 | 0.277966195 | 0.228 | 0.123 | 1 | 29 |
| h2afx5 | 0.000146764 | 0.26960495 | 0.248 | 0.139 | 1 | 29 |
| uqcr102 | 0.000163133 | 0.296621307 | 0.47 | 0.358 | 1 | 29 |
| spc242 | 0.000168391 | 0.256906855 | 0.148 | 0.07 | 1 | 29 |
| il13ra23 | 0.000184678 | 0.265248996 | 0.208 | 0.111 | 1 | 29 |
| psmd10 | 0.000185793 | 0.271250889 | 0.154 | 0.077 | 1 | 29 |
| zgc:1630308 | 0.000189013 | 0.26916084 | 0.228 | 0.125 | 1 | 29 |
| larp6a9 | 0.000213844 | 0.311421698 | 0.376 | 0.251 | 1 | 29 |

| | | | | | | |
|---|---|---|---|---|---|---|
| si:dkey-92i17.22 | 0.000220687 | 0.25173476 | 0.181 | 0.093 | 1 | 29 |
| ckbb8 | 0.000304642 | 0.286570748 | 0.396 | 0.267 | 1 | 29 |
| BX001030.17 | 0.000306545 | 0.256830331 | 0.221 | 0.123 | 1 | 29 |
| chchd108 | 0.000347836 | 0.27520748 | 0.255 | 0.152 | 1 | 29 |
| sinhcafl | 0.000352035 | 0.273327506 | 0.228 | 0.133 | 1 | 29 |
| lrrc591 | 0.000369804 | 0.282011439 | 0.242 | 0.144 | 1 | 29 |
| ccnh6 | 0.000373373 | 0.267259038 | 0.221 | 0.126 | 1 | 29 |
| c1d2 | 0.000375912 | 0.278856267 | 0.221 | 0.129 | 1 | 29 |
| cnot82 | 0.000386496 | 0.25126057 | 0.181 | 0.097 | 1 | 29 |
| npc23 | 0.000412836 | 0.27397226 | 0.463 | 0.36 | 1 | 29 |
| akirin11 | 0.000416883 | 0.28532195 | 0.235 | 0.141 | 1 | 29 |
| dazl6 | 0.000419075 | 0.252794198 | 0.195 | 0.107 | 1 | 29 |
| si:ch211-210c8.74 | 0.000421844 | 0.269623948 | 0.221 | 0.129 | 1 | 29 |
| mcfd22 | 0.000427139 | 0.2719083 | 0.201 | 0.114 | 1 | 29 |
| usf1l | 0.000466322 | 0.258389414 | 0.228 | 0.134 | 1 | 29 |
| hsp90b11 | 0.000476477 | 0.283312106 | 0.329 | 0.221 | 1 | 29 |
| si:ch211-262i1.6 | 0.000479048 | 0.250920668 | 0.174 | 0.094 | 1 | 29 |
| ncbp23 | 0.00050953 | 0.285087239 | 0.268 | 0.171 | 1 | 29 |
| alyref2 | 0.000537737 | 0.289022771 | 0.383 | 0.278 | 1 | 29 |
| gtf2a2 | 0.00055972 | 0.2633928 | 0.161 | 0.086 | 1 | 29 |
| chtopa6 | 0.000584105 | 0.31569007 | 0.315 | 0.223 | 1 | 29 |
| STMP12 | 0.00060259 | 0.298256423 | 0.396 | 0.306 | 1 | 29 |
| gar12 | 0.000635659 | 0.250442511 | 0.208 | 0.12 | 1 | 29 |
| faub7 | 0.000638517 | 0.270640454 | 0.309 | 0.202 | 1 | 29 |
| BX957362.15 | 0.000688084 | 0.256983476 | 0.228 | 0.134 | 1 | 29 |
| rpa34 | 0.000714303 | 0.27107453 | 0.262 | 0.164 | 1 | 29 |
| pttg1ipb2 | 0.000772313 | 0.251277957 | 0.201 | 0.115 | 1 | 29 |
| hspd13 | 0.000850331 | 0.28753408 | 0.255 | 0.166 | 1 | 29 |
| lsm12a2 | 0.000954427 | 0.252482671 | 0.215 | 0.127 | 1 | 29 |
| rpz25 | 0.000969258 | 0.286193847 | 0.275 | 0.183 | 1 | 29 |
| sap182 | 0.000977727 | 0.27699657 | 0.315 | 0.219 | 1 | 29 |
| psme31 | 0.000994893 | 0.253212716 | 0.221 | 0.133 | 1 | 29 |
| ilf22 | 0.001032235 | 0.293292228 | 0.336 | 0.246 | 1 | 29 |
| selenoh5 | 0.001067816 | 0.276831904 | 0.349 | 0.245 | 1 | 29 |
| txnl4a3 | 0.001100413 | 0.259187167 | 0.195 | 0.114 | 1 | 29 |

| | | | | | | |
|---|---|---|---|---|---|---|
| arl6ip13 | 0.001132969 | 0.254464668 | 0.45 | 0.371 | 1 | 29 |
| snrpg3 | 0.001134855 | 0.285671223 | 0.362 | 0.269 | 1 | 29 |
| eno32 | 0.001213031 | 0.256374266 | 0.463 | 0.384 | 1 | 29 |
| manf | 0.001277968 | 0.251944197 | 0.174 | 0.1 | 1 | 29 |
| slc25a25a2 | 0.001373535 | 0.261228981 | 0.282 | 0.187 | 1 | 29 |
| rac1a3 | 0.001388918 | 0.266921017 | 0.235 | 0.149 | 1 | 29 |
| thoc72 | 0.00145856 | 0.26975818 | 0.195 | 0.117 | 1 | 29 |
| mrpl532 | 0.001573513 | 0.254455077 | 0.161 | 0.092 | 1 | 29 |
| MED9 | 0.001645835 | 0.27371037 | 0.181 | 0.108 | 1 | 29 |
| si:ch211-51e12.72 | 0.001819018 | 0.27547353 | 0.262 | 0.178 | 1 | 29 |
| cbx3a6 | 0.002239094 | 0.25869617 | 0.383 | 0.298 | 1 | 29 |
| rbm4.21 | 0.002291211 | 0.257865269 | 0.255 | 0.17 | 1 | 29 |
| uqcrfs11 | 0.002338046 | 0.278926594 | 0.268 | 0.19 | 1 | 29 |
| srsf2a6 | 0.00284086 | 0.257356807 | 0.45 | 0.368 | 1 | 29 |
| ube2ia1 | 0.002989428 | 0.267542045 | 0.235 | 0.158 | 1 | 29 |
| zfand5b1 | 0.003365724 | 0.267811753 | 0.302 | 0.223 | 1 | 29 |
| dkc12 | 0.004168031 | 0.253202667 | 0.255 | 0.176 | 1 | 29 |
| lsm71 | 0.004669292 | 0.256658802 | 0.275 | 0.199 | 1 | 29 |
| si:dkey-102m7.31 | 0.005047984 | 0.254569697 | 0.215 | 0.144 | 1 | 29 |
| mrpl36 | 0.006005154 | 0.26332687 | 0.268 | 0.197 | 1 | 29 |
| tomm7 | 0.007604404 | 0.25430453 | 0.282 | 0.215 | 1 | 29 |
| cox7a2l1 | 0.009209314 | 0.251752102 | 0.255 | 0.189 | 1 | 29 |
| lyz | 0 | 2.638397707 | 0.927 | 0.052 | 0 | 30 |
| lect2l1 | 0 | 2.55980028 | 0.968 | 0.024 | 0 | 30 |
| npsn1 | 0 | 2.385495557 | 0.976 | 0.026 | 0 | 30 |
| BX908782.21 | 0 | 2.305793121 | 0.952 | 0.017 | 0 | 30 |
| mpx | 0 | 2.044150276 | 0.952 | 0.006 | 0 | 30 |
| mmp13a | 0 | 1.763033771 | 0.766 | 0.002 | 0 | 30 |
| ch25hl2 | 0 | 1.688672031 | 0.758 | 0.002 | 0 | 30 |
| si:ch1073-429i10.1 | 0 | 1.6886408 | 0.661 | 0.004 | 0 | 30 |
| BX908782.31 | 0 | 1.657595205 | 0.766 | 0.029 | 0 | 30 |
| scpp8 | 0 | 1.625174568 | 0.75 | 0.003 | 0 | 30 |
| il1b2 | 0 | 1.569624727 | 0.669 | 0.018 | 0 | 30 |
| sult2st1 | 0 | 1.370811024 | 0.669 | 0.021 | 0 | 30 |
| il6r | 0 | 1.351451165 | 0.637 | 0.006 | 0 | 30 |

| | | | | | | |
|---|---|---|---|---|---|---|
| cd7al1 | 0 | 1.307955274 | 0.677 | 0.019 | 0 | 30 |
| si:ch73-343l4.82 | 0 | 1.112800434 | 0.556 | 0.018 | 0 | 30 |
| si:ch211-286b5.8 | 0 | 1.026369232 | 0.508 | 0.004 | 0 | 30 |
| CR848841.11 | 0 | 0.986412183 | 0.476 | 0.013 | 0 | 30 |
| ly6m51 | 0 | 0.957919173 | 0.476 | 0.007 | 0 | 30 |
| si:ch211-264f5.21 | 0 | 0.929001074 | 0.427 | 0.009 | 0 | 30 |
| illr41 | 0 | 0.905052194 | 0.435 | 0.006 | 0 | 30 |
| si:ch211-193e13.51 | 0 | 0.831059202 | 0.387 | 0.008 | 0 | 30 |
| si:ch211-284o19.8 | 0 | 0.807923319 | 0.387 | 0.003 | 0 | 30 |
| si:ch211-9d9.81 | 0 | 0.804030456 | 0.379 | 0.008 | 0 | 30 |
| si:ch211-260e23.8 | 0 | 0.747064034 | 0.379 | 0.005 | 0 | 30 |
| chia.6 | 0 | 0.730534124 | 0.331 | 0.003 | 0 | 30 |
| abcb9 | 0 | 0.685321456 | 0.331 | 0.001 | 0 | 30 |
| si:dkey-19b23.14 | 0 | 0.655923389 | 0.315 | 0.002 | 0 | 30 |
| si:ch211-9d9.1 | 0 | 0.647414164 | 0.29 | 0.001 | 0 | 30 |
| si:dkey-238m4.4 | 0 | 0.50577065 | 0.194 | 0 | 0 | 30 |
| CR354540.1 | 5.20310722071918e-284 | 0.335484577 | 0.161 | 0.001 | 1.30957005638281e-279 | 30 |
| adam8a2 | 5.89442364006867e-278 | 1.39992046 | 0.758 | 0.045 | 1.48356748596888e-273 | 30 |
| cpa51 | 3.59739151670495e-271 | 1.452880303 | 0.645 | 0.032 | 9.05427470839468e-267 | 30 |
| ponzr62 | 2.34777237105945e-266 | 0.997947533 | 0.508 | 0.019 | 5.90910828071954e-262 | 30 |
| hpgd | 1.46671430989437e-258 | 0.579228353 | 0.258 | 0.004 | 3.69157324657313e-254 | 30 |
| cnksr2b | 1.32215306756775e-246 | 0.371495242 | 0.177 | 0.001 | 3.32772705576128e-242 | 30 |
| CT573383.1 | 4.13256945047822e-241 | 0.627413883 | 0.315 | 0.007 | 1.04012640499086e-236 | 30 |
| mmp91 | 2.45389476323257e-218 | 1.17163402 | 0.419 | 0.016 | 6.17620772958006e-214 | 30 |
| CT030188.13 | 1.81822831539441e-208 | 1.192943165 | 0.661 | 0.046 | 4.57629884701619e-204 | 30 |
| papss2b | 3.03693834123678e-205 | 0.755730039 | 0.395 | 0.015 | 7.64367011105886e-201 | 30 |
| hbxb | 2.06414504229257e-201 | 0.30493425 | 0.169 | 0.002 | 5.19524665694618e-197 | 30 |
| sms1 | 2.87564171466796e-200 | 1.170313602 | 0.548 | 0.032 | 7.23770263164779e-196 | 30 |
| lta4h4 | 3.30663644173402e-199 | 1.408843631 | 0.75 | 0.066 | 8.32247326020037e-195 | 30 |
| si:ch211-8c17.2 | 5.99121994667941e-199 | 0.572603197 | 0.242 | 0.005 | 1.50793014837974e-194 | 30 |
| ltb4r3 | 3.67567684938536e-195 | 0.906201745 | 0.476 | 0.024 | 9.25131106221801e-191 | 30 |
| abcc13 | 1.35908419559072e-191 | 0.278730878 | 0.145 | 0.001 | 3.42067901188228e-187 | 30 |
| BX908782.1 | 4.06656767086257e-168 | 0.39788664 | 0.129 | 0.001 | 1.0235144170794e-163 | 30 |
| ncf12 | 1.19388981859543e-161 | 0.929392587 | 0.476 | 0.029 | 3.00490128442283e-157 | 30 |
| BX548046.2 | 8.43398855778081e-160 | 0.39923773 | 0.194 | 0.004 | 2.12275058010785e-155 | 30 |

| | | | | | | |
|---|---|---|---|---|---|---|
| alox5ap1 | 1.54196122176524e-159 | 0.784346754 | 0.379 | 0.019 | 3.88096219906093e-155 | 30 |
| ryr3 | 5.48281732865624e-158 | 0.333321391 | 0.161 | 0.002 | 1.37997029344949e-153 | 30 |
| si:ch73-359m17.5 | 5.29675000896196e-157 | 0.482563063 | 0.194 | 0.004 | 1.33313900975564e-152 | 30 |
| CU499336.23 | 1.396146584355e-149 | 0.878271213 | 0.476 | 0.032 | 3.5139613381631e-145 | 30 |
| si:ch1073-443f11.22 | 1.35908570648525e-147 | 0.895859894 | 0.468 | 0.032 | 3.42068281465273e-143 | 30 |
| HACD41 | 5.2261555023921e-147 | 1.04934559 | 0.548 | 0.045 | 1.31537107839707e-142 | 30 |
| ppdpfa4 | 7.91619761525992e-145 | 1.2770955 | 0.669 | 0.071 | 1.99242777778477e-140 | 30 |
| si:ch73-248e21.73 | 5.0088084242671e-144 | 0.966424488 | 0.516 | 0.04 | 1.26066699230379e-139 | 30 |
| triobpa | 5.81212109503891e-139 | 0.528273546 | 0.274 | 0.011 | 1.46285275841034e-134 | 30 |
| mmp25b1 | 3.40376776575138e-137 | 0.512210799 | 0.242 | 0.008 | 8.56694308961966e-133 | 30 |
| adora2b | 4.02749515203423e-135 | 0.462715011 | 0.226 | 0.007 | 1.0136802548155e-130 | 30 |
| slc7a3a3 | 5.33580133878957e-134 | 0.958387149 | 0.492 | 0.039 | 1.34296783895995e-129 | 30 |
| elovl5 | 2.55743953574736e-133 | 0.607947357 | 0.282 | 0.012 | 6.43681956752252e-129 | 30 |
| srgn9 | 7.37570382239852e-133 | 1.813979429 | 0.976 | 0.231 | 1.85639089505948e-128 | 30 |
| slc2a3b2 | 1.59197499962062e-130 | 0.768884358 | 0.403 | 0.026 | 4.00684187654515e-126 | 30 |
| slc7a112 | 3.19633638240384e-127 | 0.557549104 | 0.234 | 0.008 | 8.04485904087223e-123 | 30 |
| si:ch211-137j23.6 | 2.04279553270409e-124 | 0.313296284 | 0.105 | 0.001 | 5.14151207626293e-120 | 30 |
| lrrc15 | 3.54860743504044e-124 | 0.512715696 | 0.25 | 0.01 | 8.93149005325328e-120 | 30 |
| jakmip1 | 3.85776873138053e-124 | 0.428580152 | 0.226 | 0.008 | 9.70961812001166e-120 | 30 |
| vwa11 | 9.02658923524207e-124 | 0.269433372 | 0.113 | 0.001 | 2.27190224461808e-119 | 30 |
| si:ch211-276a23.51 | 6.2790329935291e-123 | 0.601981099 | 0.234 | 0.009 | 1.58036981414134e-118 | 30 |
| alox5a4 | 7.50392133648256e-123 | 0.75491944 | 0.419 | 0.03 | 1.88866196117929e-118 | 30 |
| LOC1018842675 | 3.56082776220519e-122 | 1.126841007 | 0.548 | 0.054 | 8.96224739469425e-118 | 30 |
| agtrap | 3.44516950188709e-121 | 0.497195001 | 0.234 | 0.009 | 8.67114711929962e-117 | 30 |
| rnaset2 | 5.37437650534868e-121 | 0.837262109 | 0.468 | 0.039 | 1.35267682263121e-116 | 30 |
| noctb | 2.12989720613088e-118 | 0.653164626 | 0.29 | 0.014 | 5.3607382781108e-114 | 30 |
| gpr183b | 2.61858479154342e-116 | 0.406532814 | 0.185 | 0.006 | 6.59071606183562e-112 | 30 |
| timp2b2 | 3.13798272014262e-116 | 0.890707641 | 0.427 | 0.034 | 7.89798870832696e-112 | 30 |
| cxcr4b8 | 2.81515886238749e-108 | 1.37596044 | 0.903 | 0.177 | 7.08547334074308e-104 | 30 |
| tcnbb1 | 4.0305628730346e-108 | 0.798494749 | 0.274 | 0.014 | 1.01445236951408e-103 | 30 |
| si:dkey-102g19.33 | 9.45291883252588e-108 | 0.856158771 | 0.476 | 0.044 | 2.37920514095844e-103 | 30 |
| fcer1gl7 | 2.02610604175025e-107 | 1.364638557 | 0.887 | 0.163 | 5.0995062964812e-103 | 30 |
| s1pr4 | 3.22390474058368e-106 | 0.512019179 | 0.202 | 0.008 | 8.11424584157507e-102 | 30 |
| ctss1 | 7.87172423053269e-106 | 0.485221266 | 0.258 | 0.013 | 1.98123427158277e-101 | 30 |
| glipr1a5 | 8.44591327952004e-103 | 0.927744631 | 0.532 | 0.06 | 2.1257519133224e-98 | 30 |

| | | | | | | |
|---|---|---|---|---|---|---|
| si:dkey-27h10.21 | 6.93984342433266e-101 | 0.712045469 | 0.347 | 0.025 | 1.74668919147029e-96 | 30 |
| FAR2 | 1.968592198982e-92 | 0.585165637 | 0.306 | 0.021 | 4.95474970561781e-88 | 30 |
| spi1b4 | 1.13146194387234e-89 | 0.895445518 | 0.492 | 0.058 | 2.84777656653229e-85 | 30 |
| cfbl2 | 4.42069700598243e-89 | 0.944115721 | 0.524 | 0.063 | 1.11264522943572e-84 | 30 |
| hyal31 | 6.38009171603973e-89 | 0.278201925 | 0.137 | 0.004 | 1.60580528401004e-84 | 30 |
| cebpa6 | 2.31187000316215e-88 | 0.830433663 | 0.476 | 0.054 | 5.81874561095883e-84 | 30 |
| LOC101886073 | 1.50345061949304e-87 | 0.272780156 | 0.153 | 0.005 | 3.78403486420203e-83 | 30 |
| CABZ01111959.12 | 2.35648546377492e-87 | 0.620723081 | 0.315 | 0.024 | 5.93103826377509e-83 | 30 |
| si:ch211-66e2.35 | 5.06702416587046e-87 | 0.865504736 | 0.476 | 0.056 | 1.27531931230794e-82 | 30 |
| hlx11 | 1.79721386838012e-86 | 0.393327178 | 0.218 | 0.011 | 4.52340758532593e-82 | 30 |
| mir2233 | 3.24819538406262e-85 | 0.637693858 | 0.339 | 0.028 | 8.17538296214721e-81 | 30 |
| XLOC-004556 | 2.94329793149734e-84 | 0.403893074 | 0.218 | 0.012 | 7.40798656378566e-80 | 30 |
| gadd45aa | 6.11153453072012e-82 | 0.62345891 | 0.258 | 0.017 | 1.53821212603695e-77 | 30 |
| vamp8 | 6.94824833172061e-81 | 0.853617531 | 0.524 | 0.07 | 1.74880462261076e-76 | 30 |
| pnp5a4 | 3.86904966166692e-80 | 1.006688519 | 0.524 | 0.074 | 9.73801109344947e-76 | 30 |
| odc13 | 4.5495537379273e-80 | 1.088136015 | 0.661 | 0.118 | 1.14507718029892e-75 | 30 |
| nccrp13 | 5.48494912764045e-80 | 1.265898749 | 0.766 | 0.19 | 1.38050684593582e-75 | 30 |
| hopx4 | 5.72787340774111e-77 | 0.843931345 | 0.476 | 0.062 | 1.44164845799436e-72 | 30 |
| si:ch211-243a20.42 | 1.73778981109653e-76 | 0.492711026 | 0.242 | 0.016 | 4.37384317554887e-72 | 30 |
| pigrl4.2 | 3.88162199006165e-76 | 0.311520315 | 0.121 | 0.004 | 9.76965438678617e-72 | 30 |
| aldh8a1 | 4.60164016600316e-76 | 0.302911501 | 0.161 | 0.007 | 1.15818681338134e-71 | 30 |
| si:ch73-248e21.53 | 7.0141307704021e-76 | 0.774772585 | 0.387 | 0.043 | 1.7653865736025e-71 | 30 |
| BX323087.1 | 2.69380133101245e-75 | 0.349716708 | 0.161 | 0.007 | 6.78002857002524e-71 | 30 |
| zgc:175264 | 2.97068321486434e-75 | 0.608654851 | 0.347 | 0.033 | 7.47691258349207e-71 | 30 |
| spns31 | 1.42614864526355e-74 | 0.367844288 | 0.21 | 0.012 | 3.58947352526383e-70 | 30 |
| zgc:165573 | 9.79382713374064e-74 | 0.747845174 | 0.452 | 0.056 | 2.46500835129118e-69 | 30 |
| itgb26 | 2.62051478731903e-72 | 0.944255731 | 0.573 | 0.095 | 6.59557366820327e-68 | 30 |
| scinlb6 | 3.465465452256e-72 | 0.871187843 | 0.524 | 0.08 | 8.72222999678313e-68 | 30 |
| si:ch211-171h4.5 | 2.42316896684639e-69 | 0.269547883 | 0.105 | 0.003 | 6.09887397265567e-65 | 30 |
| ank3b2 | 3.36661131197091e-69 | 0.587389981 | 0.347 | 0.036 | 8.47342401109959e-65 | 30 |
| si:dkey-222f2.73 | 4.31293401004056e-69 | 0.59085315 | 0.306 | 0.028 | 1.08552236098711e-64 | 30 |
| rab446 | 4.4952103330233e-69 | 0.81574866 | 0.435 | 0.058 | 1.13139948871863e-64 | 30 |
| itgb3a1 | 9.52264172200796e-69 | 0.460713708 | 0.21 | 0.013 | 2.39675369501218e-64 | 30 |
| cd6313 | 1.20463857320913e-67 | 1.111519463 | 0.903 | 0.337 | 3.03195482491007e-63 | 30 |
| lpcat2 | 6.70197313836249e-67 | 0.544392749 | 0.25 | 0.02 | 1.68681961919446e-62 | 30 |

| | | | | | | |
|---|---|---|---|---|---|---|
| tmsb16 | 2.1574780107058e-66 | 1.141962435 | 0.685 | 0.152 | 5.43015640514543e-62 | 30 |
| mrps156 | 3.92374761162892e-65 | 0.954756365 | 0.621 | 0.121 | 9.87568036370884e-61 | 30 |
| cebpb12 | 1.51148182757374e-64 | 1.171863849 | 0.935 | 0.349 | 3.80424861182036e-60 | 30 |
| alox12 | 5.54432172815195e-64 | 0.329822347 | 0.185 | 0.011 | 1.39545033575856e-59 | 30 |
| fam107b2 | 1.95258068218158e-63 | 0.839388992 | 0.565 | 0.1 | 4.91445031898283e-59 | 30 |
| actb210 | 8.80775301102244e-63 | 0.661892655 | 1 | 0.867 | 2.21682335534424e-58 | 30 |
| b3gnt3.41 | 1.77311629045822e-62 | 0.656975705 | 0.323 | 0.035 | 4.4627563914543e-58 | 30 |
| gsr1 | 3.79206164156054e-62 | 0.537698471 | 0.315 | 0.033 | 9.54423994564372e-58 | 30 |
| capgb6 | 1.65702075914462e-59 | 0.88745817 | 0.573 | 0.111 | 4.17055554869109e-55 | 30 |
| LOC1018850651 | 6.56554707336786e-59 | 0.368709308 | 0.194 | 0.013 | 1.65248254289596e-54 | 30 |
| dhrs92 | 1.5152597817654e-57 | 0.54017639 | 0.298 | 0.032 | 3.81375734472534e-53 | 30 |
| ca61 | 1.64089033955865e-57 | 0.521504628 | 0.226 | 0.019 | 4.12995689563517e-53 | 30 |
| slc6a6b5 | 2.6481432273398e-57 | 0.892364271 | 0.532 | 0.1 | 6.66511168889155e-53 | 30 |
| plek1 | 8.25372159419601e-57 | 0.437664342 | 0.25 | 0.023 | 2.07737918804319e-52 | 30 |
| apol1 | 1.46899393109117e-56 | 0.35731847 | 0.177 | 0.012 | 3.69731082516337e-52 | 30 |
| pgp | 2.5480932583248e-56 | 0.669709724 | 0.419 | 0.061 | 6.41329592187769e-52 | 30 |
| ifngr1 | 4.60604002588649e-56 | 0.263161293 | 0.113 | 0.005 | 1.15929421411537e-51 | 30 |
| cnksr1 | 5.48693493386082e-56 | 0.347068627 | 0.153 | 0.009 | 1.38100665350343e-51 | 30 |
| mapk112 | 1.59272802314986e-55 | 0.41034218 | 0.21 | 0.017 | 4.00873716146588e-51 | 30 |
| bdh1 | 2.54055549727113e-55 | 0.308829327 | 0.177 | 0.012 | 6.39432413108171e-51 | 30 |
| dub5 | 3.80272111713105e-55 | 0.668799601 | 0.435 | 0.069 | 9.57106877970714e-51 | 30 |
| dachd2 | 7.42597131671027e-55 | 0.307820752 | 0.169 | 0.011 | 1.86904272070281e-50 | 30 |
| zmp:00000010824 | 2.13264001966949e-54 | 0.724978758 | 0.379 | 0.055 | 5.36764166550614e-50 | 30 |
| cyp4f31 | 1.07147329873197e-53 | 0.371511563 | 0.194 | 0.015 | 2.69679114557849e-49 | 30 |
| si:ch211-136m16.83 | 1.39027600438373e-53 | 0.589020361 | 0.266 | 0.028 | 3.49918567543342e-49 | 30 |
| dgkzb | 1.39143731614332e-53 | 0.32262815 | 0.145 | 0.008 | 3.50210858100112e-49 | 30 |
| pfn110 | 3.71575044608167e-53 | 0.993970425 | 0.992 | 0.522 | 9.35217229774296e-49 | 30 |
| si:ch211-197k17.3 | 4.51631983539534e-53 | 0.5714487 | 0.298 | 0.035 | 1.13671253937065e-48 | 30 |
| vsir8 | 1.13140610962596e-52 | 0.811123619 | 0.46 | 0.083 | 2.84763603731758e-48 | 30 |
| arpc1b10 | 9.27719629876534e-51 | 1.020820915 | 0.887 | 0.305 | 2.33497753643625e-46 | 30 |
| antxr2a8 | 1.38219096436073e-49 | 0.897787462 | 0.613 | 0.151 | 3.47883643819953e-45 | 30 |
| si:dkey-163m14.7 | 4.56131352989019e-49 | 0.34977531 | 0.121 | 0.006 | 1.14803700233806e-44 | 30 |
| cotl110 | 4.67150827689931e-48 | 0.987154941 | 0.823 | 0.303 | 1.17577191821279e-43 | 30 |
| CABZ01073834.14 | 7.50088645483713e-48 | 0.551803375 | 0.306 | 0.04 | 1.88789811181796e-43 | 30 |
| il164 | 3.10246890930929e-47 | 0.599786021 | 0.331 | 0.047 | 7.80860399784056e-43 | 30 |

| | | | | | | |
|---|---|---|---|---|---|---|
| atp1a3a2 | 3.78009857440004e-47 | 0.411594842 | 0.21 | 0.02 | 9.51413010190746e-43 | 30 |
| rhogb6 | 2.4929456645873e-46 | 0.771941568 | 0.468 | 0.095 | 6.27449494319979e-42 | 30 |
| icn9 | 3.58140068279291e-46 | 1.101286845 | 0.782 | 0.271 | 9.01402737852148e-42 | 30 |
| rac210 | 3.75825856997459e-46 | 0.909268594 | 0.718 | 0.217 | 9.45916099476905e-42 | 30 |
| atf35 | 5.25770906133433e-46 | 0.771765693 | 0.46 | 0.089 | 1.32331279364724e-41 | 30 |
| pim18 | 2.23426761175629e-45 | 0.91573569 | 0.629 | 0.168 | 5.6234281520294e-41 | 30 |
| parm1 | 2.56933973729707e-45 | 0.251586752 | 0.129 | 0.008 | 6.46677118480301e-41 | 30 |
| si:ch211-67e16.31 | 2.9586445332018e-45 | 0.540925772 | 0.185 | 0.016 | 7.44661242561561e-41 | 30 |
| rgs193 | 3.26126354228689e-45 | 0.502143414 | 0.266 | 0.033 | 8.20827420958187e-41 | 30 |
| gpr842 | 4.3348597812879e-45 | 0.398050193 | 0.153 | 0.011 | 1.09104085835235e-40 | 30 |
| lcp17 | 5.95924512216793e-45 | 0.709475311 | 0.444 | 0.086 | 1.49988240479845e-40 | 30 |
| trib33 | 1.31394507384014e-43 | 0.877096759 | 0.468 | 0.103 | 3.30706835634824e-39 | 30 |
| coro1a10 | 2.7623679807909e-43 | 0.889640396 | 0.75 | 0.228 | 6.95260397085262e-39 | 30 |
| nrros3 | 5.43589002217585e-43 | 0.564392567 | 0.315 | 0.046 | 1.36815915968144e-38 | 30 |
| plpp1a2 | 1.04017325765505e-42 | 0.583069055 | 0.387 | 0.066 | 2.61801207219199e-38 | 30 |
| ccdc88b1 | 6.63696587000971e-42 | 0.472410877 | 0.25 | 0.031 | 1.67045793982274e-37 | 30 |
| si:cabz01036022.11 | 8.1421369768583e-42 | 0.301973942 | 0.145 | 0.011 | 2.04929445570547e-37 | 30 |
| adam19b2 | 2.12069421261086e-40 | 0.282418769 | 0.161 | 0.014 | 5.33757526372027e-36 | 30 |
| lxn8 | 2.4162382378832e-40 | 0.787409636 | 0.565 | 0.145 | 6.08143002092824e-36 | 30 |
| XLOC-039817 | 6.03636720364631e-40 | 0.296132354 | 0.137 | 0.01 | 1.51929326148574e-35 | 30 |
| skap25 | 2.08641186682521e-39 | 0.596551599 | 0.387 | 0.073 | 5.25129002761238e-35 | 30 |
| selenow15 | 2.55176980195118e-39 | 0.876799459 | 0.798 | 0.383 | 6.42254941453093e-35 | 30 |
| FP015862.22 | 2.97577965728632e-39 | 0.354430655 | 0.169 | 0.016 | 7.48973981942394e-35 | 30 |
| si:dkey-53k12.24 | 6.60669495350129e-39 | 0.504522922 | 0.29 | 0.043 | 1.66283905284674e-34 | 30 |
| wasb9 | 1.52056894527521e-38 | 0.82327663 | 0.613 | 0.175 | 3.82711997836317e-34 | 30 |
| prkcda | 3.99776393056464e-38 | 0.54262771 | 0.355 | 0.061 | 1.00619720368381e-33 | 30 |
| il344 | 6.0176202704638e-38 | 0.481317371 | 0.306 | 0.048 | 1.51457484587303e-33 | 30 |
| smox4 | 1.08083884223667e-37 | 0.672892336 | 0.403 | 0.083 | 2.72036328202546e-33 | 30 |
| cfl1l6 | 1.70354812477339e-37 | 0.83914675 | 0.556 | 0.15 | 4.28766027524214e-33 | 30 |
| si:dkey-262k9.47 | 2.63619007237314e-37 | 0.63736681 | 0.419 | 0.089 | 6.63502679315595e-33 | 30 |
| COX316 | 4.97033709187484e-37 | 0.655403851 | 0.992 | 0.699 | 1.25098414265398e-32 | 30 |
| sh3bgrl6 | 7.94264937625835e-37 | 0.692579807 | 0.444 | 0.102 | 1.99908542151046e-32 | 30 |
| itgae.23 | 1.04110606119365e-36 | 0.445281163 | 0.25 | 0.034 | 2.62035984541829e-32 | 30 |
| parvg4 | 1.74968462268642e-36 | 0.568814494 | 0.355 | 0.066 | 4.40378122683946e-32 | 30 |
| tmsb4x17 | 2.57340420191218e-36 | 0.681060645 | 1 | 0.685 | 6.47700103579277e-32 | 30 |

| | | | | | | |
|---|---|---|---|---|---|---|
| myh9a2 | 3.62428420956345e-36 | 0.554608228 | 0.315 | 0.053 | 9.12196092705025e-32 | 30 |
| raraa1 | 2.25661856577831e-35 | 0.511984793 | 0.226 | 0.03 | 5.67968326820743e-31 | 30 |
| derl22 | 1.04171174480372e-34 | 0.615370478 | 0.452 | 0.103 | 2.62188429049649e-30 | 30 |
| mbnl17 | 1.35631748779199e-34 | 0.758624138 | 0.597 | 0.181 | 3.41371548502365e-30 | 30 |
| mt-co215 | 2.71072363307522e-34 | 0.621539111 | 0.976 | 0.719 | 6.82262031208701e-30 | 30 |
| xbp15 | 4.98235695403082e-34 | 0.833468487 | 0.79 | 0.341 | 1.25400942176002e-29 | 30 |
| CABZ01001434.12 | 7.16512345454094e-34 | 0.53910478 | 0.137 | 0.012 | 1.80338992227341e-29 | 30 |
| pde4ba9 | 1.00958490054748e-33 | 0.723231569 | 0.508 | 0.138 | 2.54102423618796e-29 | 30 |
| tpd52 | 1.15074729438231e-33 | 0.325308306 | 0.234 | 0.032 | 2.89631586523083e-29 | 30 |
| zdhhc20a | 1.52004650955263e-33 | 0.487830089 | 0.266 | 0.042 | 3.82580505989302e-29 | 30 |
| g6pd | 2.11723198105095e-33 | 0.400126595 | 0.21 | 0.027 | 5.32886117310714e-29 | 30 |
| dusp26 | 2.74457765226259e-33 | 0.804158255 | 0.589 | 0.177 | 6.90782749297972e-29 | 30 |
| rps6ka15 | 5.6366616629118e-33 | 0.571929425 | 0.355 | 0.072 | 1.41869137393827e-28 | 30 |
| gapdhs13 | 1.78276723196598e-32 | 0.809591271 | 0.694 | 0.286 | 4.48704684613518e-28 | 30 |
| plxnc14 | 1.52468113731536e-31 | 0.513681857 | 0.258 | 0.042 | 3.83746995450904e-27 | 30 |
| fabp7a1 | 1.58854234839288e-31 | 0.256120679 | 0.202 | 0.026 | 3.99820223667004e-27 | 30 |
| b3gnt2b | 2.48979660374706e-31 | 0.342397431 | 0.177 | 0.021 | 6.26656907197096e-27 | 30 |
| mrtfab3 | 3.43308117717059e-31 | 0.463827762 | 0.29 | 0.052 | 8.64072201482065e-27 | 30 |
| unm-sa16142 | 7.33170561859209e-31 | 0.37445193 | 0.169 | 0.02 | 1.84531698714344e-26 | 30 |
| ncf42 | 7.68172874632714e-31 | 0.264437953 | 0.161 | 0.018 | 1.93341430816308e-26 | 30 |
| mt-cyb12 | 1.98166076319569e-30 | 0.651036784 | 0.984 | 0.622 | 4.98764197488722e-26 | 30 |
| cyba4 | 2.09868558609409e-30 | 0.500666033 | 0.347 | 0.072 | 5.28218175164022e-26 | 30 |
| CR381673.16 | 2.60921871999969e-30 | 0.496039231 | 0.298 | 0.056 | 6.56714259636721e-26 | 30 |
| prdx56 | 3.37461784176342e-30 | 0.711575113 | 0.508 | 0.157 | 8.49357564593435e-26 | 30 |
| farsa | 3.51112273638444e-30 | 0.351226997 | 0.202 | 0.027 | 8.83714481520601e-26 | 30 |
| si:zfos-741a10.34 | 5.81743829907315e-30 | 0.400682915 | 0.194 | 0.026 | 1.46419104549372e-25 | 30 |
| grasp4 | 7.22518429677684e-30 | 0.598086288 | 0.347 | 0.075 | 1.81850663565576e-25 | 30 |
| arpc5b6 | 8.69299899677707e-30 | 0.622688906 | 0.435 | 0.112 | 2.18794091749882e-25 | 30 |
| crip110 | 9.17916415182111e-30 | 0.781482065 | 0.685 | 0.268 | 2.31030382537185e-25 | 30 |
| itga12 | 1.16130120998454e-29 | 0.384852694 | 0.242 | 0.038 | 2.9228790154101e-25 | 30 |
| spag1a | 1.19627324363098e-29 | 0.278316645 | 0.137 | 0.014 | 3.0109001268948e-25 | 30 |
| mapre1a6 | 1.69379579236177e-29 | 0.695415998 | 0.605 | 0.195 | 4.26311462979533e-25 | 30 |
| mt-atp615 | 2.24428486596404e-29 | 0.662782123 | 0.968 | 0.589 | 5.6486405791449e-25 | 30 |
| arg21 | 3.97506215803659e-29 | 0.422309247 | 0.169 | 0.021 | 1.00048339455623e-24 | 30 |
| sdr16c5b1 | 4.94320060482818e-29 | 0.389078095 | 0.242 | 0.04 | 1.2441541602292e-24 | 30 |

| | | | | | | |
|---|---|---|---|---|---|---|
| cst14b.13 | 4.99162069431566e-29 | 0.469241914 | 0.274 | 0.05 | 1.25634101255231e-24 | 30 |
| sri2 | 7.10390340161474e-29 | 0.616549665 | 0.508 | 0.149 | 1.78798144715241e-24 | 30 |
| LOC1104382371 | 1.91518273316083e-28 | 0.448173373 | 0.282 | 0.053 | 4.8203234210925e-24 | 30 |
| ptprr | 3.12515962878124e-28 | 0.255894255 | 0.145 | 0.016 | 7.86571426967951e-24 | 30 |
| ppp1r14aa | 3.81947288696184e-28 | 0.311274722 | 0.242 | 0.039 | 9.61323130919424e-24 | 30 |
| stk17b4 | 6.92784732183419e-28 | 0.609144888 | 0.419 | 0.109 | 1.74366989243245e-23 | 30 |
| syngr2a3 | 9.32953845328328e-28 | 0.479560598 | 0.282 | 0.055 | 2.34815153330687e-23 | 30 |
| grk6 | 1.21236117522345e-27 | 0.331892154 | 0.177 | 0.023 | 3.05139184191989e-23 | 30 |
| zgc:640519 | 1.34455767437046e-27 | 0.667029389 | 0.524 | 0.16 | 3.384117210623e-23 | 30 |
| mt-co116 | 1.55723610563297e-27 | 0.598272804 | 1 | 0.656 | 3.91940755426763e-23 | 30 |
| si:ch73-208g10.14 | 1.9488713019009e-27 | 0.439011127 | 0.242 | 0.042 | 4.90511417975438e-23 | 30 |
| amd1 | 3.92726403613732e-27 | 0.406282087 | 0.274 | 0.051 | 9.88453085255401e-23 | 30 |
| zgc:91976 | 4.52962913254379e-27 | 0.311611233 | 0.169 | 0.022 | 1.14006235636995e-22 | 30 |
| si:dkey-24p1.72 | 1.17579791156702e-26 | 0.284909244 | 0.129 | 0.013 | 2.95936576362303e-22 | 30 |
| cox6a23 | 1.4423735683214e-26 | 0.374723926 | 0.218 | 0.035 | 3.63031003410814e-22 | 30 |
| arhgdig9 | 1.50406703224592e-26 | 0.698652287 | 0.645 | 0.241 | 3.78558631345976e-22 | 30 |
| wbp2nl2 | 1.52636971359549e-26 | 0.541253145 | 0.355 | 0.086 | 3.84171993214848e-22 | 30 |
| sptlc2a1 | 1.67902330204155e-26 | 0.380408398 | 0.234 | 0.04 | 4.22593374890839e-22 | 30 |
| nfkbiab4 | 1.79864931123041e-26 | 0.743378292 | 0.524 | 0.18 | 4.52702045143582e-22 | 30 |
| zgc:638314 | 2.07199172082665e-26 | 0.557702422 | 0.339 | 0.08 | 5.21499596214861e-22 | 30 |
| rab11fip1b3 | 2.30601043950325e-26 | 0.469171203 | 0.282 | 0.057 | 5.80399767518572e-22 | 30 |
| tpd52l2b | 2.51122557843483e-26 | 0.527101779 | 0.387 | 0.097 | 6.32050365836263e-22 | 30 |
| pgd3 | 2.54896068472802e-26 | 0.467880284 | 0.298 | 0.062 | 6.41547914739196e-22 | 30 |
| itm2bb7 | 3.31022276941769e-26 | 0.68844563 | 0.605 | 0.22 | 8.33149968834737e-22 | 30 |
| kctd12.23 | 5.6829983443618e-26 | 0.504913893 | 0.339 | 0.078 | 1.43035385329242e-21 | 30 |
| cited4a2 | 5.88424088451948e-26 | 0.422704879 | 0.234 | 0.042 | 1.48100458822471e-21 | 30 |
| laptm510 | 8.30749043873943e-26 | 0.625233615 | 0.54 | 0.168 | 2.09091226852633e-21 | 30 |
| cmtm72 | 1.08080426898684e-25 | 0.391459673 | 0.25 | 0.046 | 2.72027626461298e-21 | 30 |
| mt-nd412 | 2.36379156419587e-25 | 0.665134177 | 0.887 | 0.454 | 5.94942698792459e-21 | 30 |
| atg16l1 | 3.08763610847768e-25 | 0.347145244 | 0.185 | 0.027 | 7.77127132142747e-21 | 30 |
| actb14 | 5.17431014022682e-25 | 0.41140558 | 0.96 | 0.784 | 1.30232211919369e-20 | 30 |
| gsto2 | 6.04910201258851e-25 | 0.542809637 | 0.371 | 0.093 | 1.5224984855484e-20 | 30 |
| osgn1 | 6.16019768480793e-25 | 0.406836257 | 0.226 | 0.04 | 1.55046015528931e-20 | 30 |
| LOC1039091156 | 6.94859252862385e-25 | 0.605238924 | 0.508 | 0.164 | 1.74889125352934e-20 | 30 |
| selp2 | 7.74496825671262e-25 | 0.253150458 | 0.113 | 0.011 | 1.949331060532e-20 | 30 |

| | | | | | | |
|---|---|---|---|---|---|---|
| cap110 | 2.00948205425232e-24 | 0.718702095 | 0.589 | 0.232 | 5.05766538234766e-20 | 30 |
| arhgef3l3 | 2.62482368489913e-24 | 0.361590284 | 0.194 | 0.031 | 6.60641873252262e-20 | 30 |
| mcl1b9 | 3.20915038210407e-24 | 0.679189481 | 0.621 | 0.24 | 8.07711059671772e-20 | 30 |
| arpc29 | 6.07274851270668e-24 | 0.671555726 | 0.637 | 0.266 | 1.52845007316314e-19 | 30 |
| tpt18 | 1.655660435935e-23 | 0.369817796 | 0.976 | 0.8 | 4.16713175120481e-19 | 30 |
| ets23 | 2.40452778257888e-23 | 0.519133197 | 0.347 | 0.089 | 6.05195597597279e-19 | 30 |
| msra | 1.32137826349046e-22 | 0.261377409 | 0.153 | 0.021 | 3.32577695137915e-18 | 30 |
| wdr15 | 2.11149993651009e-22 | 0.528179996 | 0.452 | 0.14 | 5.31443419020225e-18 | 30 |
| hbegfa1 | 2.33064036824396e-22 | 0.540419004 | 0.395 | 0.11 | 5.86598874283321e-18 | 30 |
| arpc4l8 | 2.52700671124984e-22 | 0.613374463 | 0.532 | 0.195 | 6.36022319154472e-18 | 30 |
| calm1b7 | 8.94329010395463e-22 | 0.47281234 | 0.355 | 0.094 | 2.25093668626434e-17 | 30 |
| fybb5 | 9.0019282715615e-22 | 0.353445975 | 0.218 | 0.041 | 2.26569532666931e-17 | 30 |
| cited4b2 | 9.64558661036161e-22 | 0.551979182 | 0.435 | 0.13 | 2.42769769396191e-17 | 30 |
| pxk | 1.57174543495282e-21 | 0.350804133 | 0.202 | 0.036 | 3.95592608523275e-17 | 30 |
| si:ch211-105j21.94 | 1.62422906207102e-21 | 0.371526784 | 0.194 | 0.034 | 4.08802212632654e-17 | 30 |
| CR383676.115 | 1.68691329256528e-21 | 0.667846244 | 0.984 | 0.725 | 4.24579206605756e-17 | 30 |
| vaspa7 | 1.95175128410769e-21 | 0.443244611 | 0.306 | 0.074 | 4.91236280697064e-17 | 30 |
| zmp:0000009362 | 2.24771139487985e-21 | 0.271066806 | 0.137 | 0.018 | 5.65726480977309e-17 | 30 |
| fermt3b1 | 2.63976502109213e-21 | 0.263877457 | 0.137 | 0.018 | 6.64402458158677e-17 | 30 |
| rab11ba | 2.75031413899924e-21 | 0.496064536 | 0.387 | 0.112 | 6.92226565644719e-17 | 30 |
| gpia1 | 3.98337604580831e-21 | 0.521048846 | 0.403 | 0.121 | 1.00257591696949e-16 | 30 |
| plekho1b2 | 7.58130505165256e-21 | 0.393664075 | 0.21 | 0.041 | 1.90813866845043e-16 | 30 |
| rab27a1 | 7.60204970965472e-21 | 0.420072872 | 0.226 | 0.046 | 1.913359891423e-16 | 30 |
| hsd3b71 | 8.56467767168616e-21 | 0.41119807 | 0.339 | 0.086 | 2.15564372318669e-16 | 30 |
| h1fx7 | 1.02718815954628e-20 | 0.64791925 | 0.5 | 0.182 | 2.58532987876204e-16 | 30 |
| mt-nd312 | 1.60971573531288e-20 | 0.628835237 | 0.774 | 0.394 | 4.05149353420899e-16 | 30 |
| si:ch211-197h24.93 | 2.04442204509888e-20 | 0.398382994 | 0.282 | 0.068 | 5.14560584530936e-16 | 30 |
| ipcef11 | 6.45543463116913e-20 | 0.269376086 | 0.153 | 0.024 | 1.62476834231896e-15 | 30 |
| rffl | 6.68611563470273e-20 | 0.284691265 | 0.161 | 0.026 | 1.68282844409833e-15 | 30 |
| rps6kb1b | 7.99633170162035e-20 | 0.39337097 | 0.274 | 0.065 | 2.01259672598083e-15 | 30 |
| gapdh10 | 1.45802563220037e-19 | 0.637208642 | 0.766 | 0.419 | 3.66970471368512e-15 | 30 |
| nme2b.2 | 3.33860143456089e-19 | 0.257071162 | 0.129 | 0.018 | 8.4029259506463e-15 | 30 |
| tnfaip8l2a | 4.3151552067509e-19 | 0.287048329 | 0.145 | 0.022 | 1.08608141398713e-14 | 30 |
| fmnl1a2 | 4.95460617301662e-19 | 0.369915253 | 0.194 | 0.038 | 1.24702482768655e-14 | 30 |
| hcar1-43 | 8.07640232035995e-19 | 0.341518761 | 0.185 | 0.036 | 2.0327497000114e-14 | 30 |

| | | | | | | |
|---|---|---|---|---|---|---|
| elovl1b5 | 9.17203862282053e-19 | 0.488900085 | 0.371 | 0.115 | 2.3085104009777e-14 | 30 |
| tagln29 | 9.79084486677607e-19 | 0.527587618 | 0.508 | 0.187 | 2.46425774451887e-14 | 30 |
| arpc39 | 1.12837309563424e-18 | 0.607112708 | 0.645 | 0.309 | 2.84000224440183e-14 | 30 |
| elf18 | 1.42596572776766e-18 | 0.4632592 | 0.347 | 0.103 | 3.58901314021842e-14 | 30 |
| si:ch211-102c2.47 | 1.48077957626836e-18 | 0.501419342 | 0.435 | 0.141 | 3.72697411550984e-14 | 30 |
| fbp1a1 | 3.06954686750904e-18 | 0.307980714 | 0.194 | 0.039 | 7.7257425108335e-14 | 30 |
| s100a10b9 | 5.95869363248489e-18 | 0.614827981 | 0.621 | 0.254 | 1.49974360036012e-13 | 30 |
| ppt15 | 7.28670731739181e-18 | 0.577880411 | 0.435 | 0.162 | 1.83399136471435e-13 | 30 |
| gpx1a5 | 7.33796973542639e-18 | 0.516100422 | 0.573 | 0.223 | 1.84689360270947e-13 | 30 |
| chchd3a | 8.72972518985924e-18 | 0.288195914 | 0.21 | 0.044 | 2.19718453303567e-13 | 30 |
| si:ch73-234b20.54 | 1.06031182907139e-17 | 0.45950896 | 0.331 | 0.1 | 2.66869884258978e-13 | 30 |
| myo1f5 | 1.14788912227061e-17 | 0.341757683 | 0.21 | 0.045 | 2.8891221318429e-13 | 30 |
| zgc:771124 | 1.42937309229492e-17 | 0.46401386 | 0.387 | 0.119 | 3.59758913599707e-13 | 30 |
| capza1b5 | 2.9582468251253e-17 | 0.576356895 | 0.5 | 0.212 | 7.44561143415786e-13 | 30 |
| srd5a2a | 4.09129742991515e-17 | 0.38977242 | 0.315 | 0.088 | 1.02973865013534e-12 | 30 |
| samsn1a3 | 6.17232187466405e-17 | 0.347538981 | 0.177 | 0.036 | 1.5535116926342e-12 | 30 |
| CR392341.15 | 6.38873020451831e-17 | 0.351794921 | 0.242 | 0.059 | 1.60797950517521e-12 | 30 |
| pdhb | 8.57252173666144e-17 | 0.273191349 | 0.194 | 0.041 | 2.15761799590032e-12 | 30 |
| lpar5a5 | 1.16349856701419e-16 | 0.316589363 | 0.194 | 0.041 | 2.928409543318e-12 | 30 |
| sh3kbp13 | 1.22170458077781e-16 | 0.361951123 | 0.266 | 0.071 | 3.07490825935968e-12 | 30 |
| dbi6 | 1.57088945141719e-16 | 0.597817257 | 0.476 | 0.212 | 3.95377166027193e-12 | 30 |
| ptprc9 | 1.87564066085459e-16 | 0.482210284 | 0.379 | 0.127 | 4.72079997930493e-12 | 30 |
| pygb1 | 1.95944356478601e-16 | 0.304838635 | 0.202 | 0.044 | 4.93172350820991e-12 | 30 |
| zgc:1749045 | 2.57100098477013e-16 | 0.367868107 | 0.21 | 0.048 | 6.47095237856794e-12 | 30 |
| tmem30ab | 3.33430962335103e-16 | 0.341208617 | 0.218 | 0.051 | 8.39212389101221e-12 | 30 |
| iqgap12 | 4.85524143021493e-16 | 0.348597223 | 0.258 | 0.068 | 1.2220157155708e-11 | 30 |
| lrrfip1a3 | 5.00209392403401e-16 | 0.301287395 | 0.234 | 0.057 | 1.25897701974012e-11 | 30 |
| ptmaa13 | 5.55681743184594e-16 | 0.5618099 | 0.758 | 0.394 | 1.3985953794213e-11 | 30 |
| abca2 | 6.11500993046534e-16 | 0.289433825 | 0.145 | 0.026 | 1.53908684939882e-11 | 30 |
| actr2a8 | 6.31303143692629e-16 | 0.445319999 | 0.379 | 0.129 | 1.58892688235998e-11 | 30 |
| ak12 | 1.00564724449676e-15 | 0.387495009 | 0.266 | 0.075 | 2.5311135496739e-11 | 30 |
| zgc:664272 | 1.32074746494392e-15 | 0.403757676 | 0.282 | 0.082 | 3.32418929451735e-11 | 30 |
| fam49ba5 | 1.32151571333708e-15 | 0.400940255 | 0.331 | 0.103 | 3.32612289889809e-11 | 30 |
| zgc:77650 | 1.42120136099007e-15 | 0.414271306 | 0.323 | 0.1 | 3.57702170547591e-11 | 30 |
| crema7 | 1.91778331451593e-15 | 0.54454651 | 0.444 | 0.178 | 4.82686882430513e-11 | 30 |

| | | | | | | |
|---|---|---|---|---|---|---|
| dusp17 | 3.21958013353331e-15 | 0.517376997 | 0.419 | 0.159 | 8.10336123808999e-11 | 30 |
| clic17 | 3.80991683799424e-15 | 0.484640326 | 0.331 | 0.113 | 9.5891796895477e-11 | 30 |
| si:ch73-1a9.311 | 4.16534157368154e-15 | 0.558384759 | 0.556 | 0.265 | 1.04837482067991e-10 | 30 |
| aldoaa2 | 4.58078139161177e-15 | 0.517539202 | 0.548 | 0.249 | 1.15293686845477e-10 | 30 |
| add3a8 | 7.83937009429842e-15 | 0.411966132 | 0.363 | 0.123 | 1.97309105903397e-10 | 30 |
| rca2.1 | 8.81031427660284e-15 | 0.409658312 | 0.274 | 0.081 | 2.21746800027817e-10 | 30 |
| tpm35 | 1.13151291901023e-14 | 0.492361837 | 0.556 | 0.253 | 2.84790486585684e-10 | 30 |
| lamtor4 | 1.13660355002124e-14 | 0.357712977 | 0.306 | 0.095 | 2.86071747504846e-10 | 30 |
| si:ch211-132p1.36 | 1.2022944833381e-14 | 0.438358323 | 0.282 | 0.088 | 3.02605498511367e-10 | 30 |
| cox8a7 | 1.27899303736425e-14 | 0.481310781 | 0.516 | 0.226 | 3.21909757574209e-10 | 30 |
| zgc:1528632 | 1.33697627758945e-14 | 0.409474128 | 0.185 | 0.044 | 3.36503559306489e-10 | 30 |
| mtpn3 | 1.63161089124611e-14 | 0.441723277 | 0.387 | 0.146 | 4.10660145217734e-10 | 30 |
| plbd12 | 2.45784752616093e-14 | 0.259631166 | 0.137 | 0.026 | 6.18615643859445e-10 | 30 |
| serpine21 | 2.63473685521107e-14 | 0.322528713 | 0.234 | 0.063 | 6.63136919088074e-10 | 30 |
| arrb2a | 2.71569623121146e-14 | 0.251769821 | 0.169 | 0.036 | 6.83513584433613e-10 | 30 |
| mt-nd113 | 3.5126104117173e-14 | 0.502046768 | 0.742 | 0.41 | 8.84088914525127e-10 | 30 |
| rgs136 | 4.25613150644931e-14 | 0.450275178 | 0.371 | 0.132 | 1.07122573885823e-09 | 30 |
| pkma5 | 4.27283148031708e-14 | 0.454220907 | 0.444 | 0.177 | 1.07542895528101e-09 | 30 |
| CABZ01088428.1 | 6.68195974032379e-14 | 0.28863886 | 0.137 | 0.027 | 1.68178244704209e-09 | 30 |
| msrb1b | 9.13297836948959e-14 | 0.265189871 | 0.202 | 0.05 | 2.29867932581683e-09 | 30 |
| itpkcb4 | 1.48480855686162e-13 | 0.358090356 | 0.258 | 0.077 | 3.73711465676502e-09 | 30 |
| mt-nd211 | 3.28146707302846e-13 | 0.523175193 | 0.694 | 0.399 | 8.25912447610534e-09 | 30 |
| selplg5 | 3.59876629640397e-13 | 0.323239276 | 0.185 | 0.047 | 9.05773489141915e-09 | 30 |
| slc38a22 | 3.90465226425502e-13 | 0.317588234 | 0.185 | 0.047 | 9.82761928390346e-09 | 30 |
| tsc22d37 | 4.01193788082529e-13 | 0.444167566 | 0.371 | 0.143 | 1.00976464522492e-08 | 30 |
| cbx7a4 | 6.58056860983112e-13 | 0.282785878 | 0.202 | 0.053 | 1.65626331340839e-08 | 30 |
| capzb8 | 7.66831221448368e-13 | 0.466550063 | 0.532 | 0.253 | 1.9300375012634e-08 | 30 |
| sptbn21 | 7.86819476290047e-13 | 0.250843327 | 0.169 | 0.04 | 1.98034593987442e-08 | 30 |
| cnp | 8.82708459545825e-13 | 0.446599242 | 0.516 | 0.227 | 2.22168892183089e-08 | 30 |
| pde7a | 9.42377385980962e-13 | 0.252845447 | 0.21 | 0.056 | 2.37186964277548e-08 | 30 |
| lasp17 | 9.75363643602913e-13 | 0.469317718 | 0.508 | 0.236 | 2.45489275458417e-08 | 30 |
| si:rp71-68n21.93 | 1.19335815937127e-12 | 0.272393006 | 0.177 | 0.044 | 3.00356315132155e-08 | 30 |
| hmgn28 | 1.29191935775286e-12 | 0.504367758 | 0.726 | 0.442 | 3.25163183152818e-08 | 30 |
| scamp21 | 1.85747818286647e-12 | 0.332111749 | 0.234 | 0.07 | 4.67508683845663e-08 | 30 |
| LOC5665871 | 2.24484161246553e-12 | 0.324312771 | 0.185 | 0.049 | 5.6500418544145e-08 | 30 |

| | | | | | | |
|---|---|---|---|---|---|---|
| arpc48 | 3.07344779543399e-12 | 0.459422002 | 0.492 | 0.225 | 7.7355607563278e-08 | 30 |
| acsl4b2 | 3.3335749851956e-12 | 0.326152383 | 0.202 | 0.056 | 8.3902748802388e-08 | 30 |
| limd23 | 3.58460918306994e-12 | 0.302851055 | 0.234 | 0.069 | 9.02210285286873e-08 | 30 |
| socs3b8 | 3.60294900435737e-12 | 0.428622375 | 0.387 | 0.155 | 9.06826234906707e-08 | 30 |
| scarb2a2 | 4.61000982927741e-12 | 0.328037683 | 0.218 | 0.064 | 1.16029337393083e-07 | 30 |
| svild6 | 5.75425695743496e-12 | 0.390640954 | 0.25 | 0.081 | 1.4482889336168e-07 | 30 |
| cox6b18 | 8.88305546284424e-12 | 0.323233101 | 0.331 | 0.12 | 2.23577622944327e-07 | 30 |
| ccni10 | 1.2117199799061e-11 | 0.456601397 | 0.565 | 0.294 | 3.04977801742566e-07 | 30 |
| nr4a18 | 1.74271493636803e-11 | 0.454742468 | 0.395 | 0.167 | 4.38623922334471e-07 | 30 |
| myl12.19 | 1.97985749572646e-11 | 0.467836327 | 0.645 | 0.391 | 4.98310333099393e-07 | 30 |
| tpi1b7 | 2.28785472871916e-11 | 0.440419007 | 0.387 | 0.167 | 5.75830156671326e-07 | 30 |
| exoc6 | 2.54837220520643e-11 | 0.281695865 | 0.145 | 0.034 | 6.41399800328406e-07 | 30 |
| LOC1039114302 | 2.967487975734e-11 | 0.28442226 | 0.153 | 0.038 | 7.4688704861249e-07 | 30 |
| rab11fip4b2 | 2.99766256237734e-11 | 0.271927245 | 0.145 | 0.034 | 7.54481690324752e-07 | 30 |
| si:dkey-25e12.34 | 3.80020983660885e-11 | 0.266129436 | 0.185 | 0.051 | 9.56474813776082e-07 | 30 |
| dock82 | 4.28375485187502e-11 | 0.262033174 | 0.153 | 0.038 | 1.07817825866842e-06 | 30 |
| arrb2b6 | 4.88597050528802e-11 | 0.322404244 | 0.266 | 0.091 | 1.22974991647594e-06 | 30 |
| birc24 | 1.15792389768814e-10 | 0.33588917 | 0.29 | 0.107 | 2.91437865809129e-06 | 30 |
| wasa6 | 1.51937184802731e-10 | 0.322148322 | 0.266 | 0.093 | 3.82410700429992e-06 | 30 |
| thy110 | 2.09888062969977e-10 | 0.467792828 | 0.815 | 0.449 | 5.28267265689135e-06 | 30 |
| degs1 | 2.17586613894639e-10 | 0.309155772 | 0.177 | 0.051 | 5.47643748511418e-06 | 30 |
| fam49bb | 5.4968201049287e-10 | 0.261175321 | 0.177 | 0.051 | 1.38349465220951e-05 | 30 |
| si:ch211-226m16.24 | 5.97514996655681e-10 | 0.328769468 | 0.21 | 0.069 | 1.50388549508268e-05 | 30 |
| magt11 | 8.83435258921453e-10 | 0.348004598 | 0.29 | 0.115 | 2.22351820317941e-05 | 30 |
| aldh9a1a.17 | 1.005983529442e-09 | 0.337150708 | 0.379 | 0.164 | 2.53195994525258e-05 | 30 |
| ccdc28a | 1.01693639754555e-09 | 0.29844642 | 0.282 | 0.106 | 2.5595272189824e-05 | 30 |
| cmip3 | 1.14553164247606e-09 | 0.279064046 | 0.202 | 0.065 | 2.883188590948e-05 | 30 |
| flna7 | 1.38488454131202e-09 | 0.282098805 | 0.298 | 0.114 | 3.48561590202823e-05 | 30 |
| tktb2 | 1.57320880619994e-09 | 0.347292558 | 0.395 | 0.173 | 3.95960924432463e-05 | 30 |
| tmem592 | 1.70546515433202e-09 | 0.343204693 | 0.274 | 0.107 | 4.29248524693827e-05 | 30 |
| gpx4b3 | 2.20153047451567e-09 | 0.392542121 | 0.419 | 0.203 | 5.54103205130849e-05 | 30 |
| adgre5b.32 | 2.34707415067517e-09 | 0.280390572 | 0.145 | 0.04 | 5.90735092983432e-05 | 30 |
| chac16 | 2.3862864073931e-09 | 0.417728462 | 0.645 | 0.398 | 6.00604425876769e-05 | 30 |
| grap2b7 | 2.4784337435516e-09 | 0.310822912 | 0.29 | 0.113 | 6.23796988914503e-05 | 30 |
| hmha1b5 | 2.91953359128238e-09 | 0.292349178 | 0.218 | 0.075 | 7.34817409589861e-05 | 30 |

| | | | | | | |
|---|---|---|---|---|---|---|
| eno33 | 2.99993360954137e-09 | 0.429707995 | 0.629 | 0.382 | 7.55053290185468e-05 | 30 |
| pnrc215 | 3.54050023565787e-09 | 0.400939509 | 0.855 | 0.567 | 8.91108504312729e-05 | 30 |
| tln15 | 3.75012752683337e-09 | 0.326284489 | 0.306 | 0.128 | 9.43869597228692e-05 | 30 |
| si:dkey-27i16.24 | 4.21601890445583e-09 | 0.319961272 | 0.25 | 0.091 | 0.000106113 | 30 |
| cdc424 | 4.25186059937759e-09 | 0.382546557 | 0.427 | 0.212 | 0.000107015 | 30 |
| sh3bgrl35 | 4.42567553781698e-09 | 0.344802631 | 0.339 | 0.147 | 0.00011139 | 30 |
| msna10 | 5.02567244169837e-09 | 0.397972612 | 0.468 | 0.242 | 0.000126491 | 30 |
| LO018102.1 | 5.09101470454814e-09 | 0.279926345 | 0.242 | 0.088 | 0.000128136 | 30 |
| npl2 | 6.58561839317844e-09 | 0.252123884 | 0.137 | 0.037 | 0.000165753 | 30 |
| sgk1 | 6.63895386761656e-09 | 0.256935412 | 0.194 | 0.063 | 0.000167096 | 30 |
| mcl1a7 | 7.00628997227019e-09 | 0.367619394 | 0.863 | 0.623 | 0.000176341 | 30 |
| rap1b6 | 7.72671747441317e-09 | 0.349736653 | 0.419 | 0.202 | 0.000194474 | 30 |
| mid1ip1b1 | 8.65131283089135e-09 | 0.260453162 | 0.169 | 0.052 | 0.000217745 | 30 |
| si:dkey-112e7.27 | 1.09626914463686e-08 | 0.327991078 | 0.427 | 0.206 | 0.00027592 | 30 |
| si:ch1073-392o20.22 | 1.22463198092502e-08 | 0.281741577 | 0.169 | 0.054 | 0.000308228 | 30 |
| cd1641 | 1.51151174111475e-08 | 0.309951117 | 0.202 | 0.072 | 0.000380432 | 30 |
| actn42 | 1.86474040371064e-08 | 0.311756246 | 0.242 | 0.094 | 0.000469337 | 30 |
| rab1ab2 | 2.13547746763408e-08 | 0.297440623 | 0.29 | 0.122 | 0.000537478 | 30 |
| capza1a7 | 2.5766517227041e-08 | 0.329799089 | 0.411 | 0.2 | 0.000648517 | 30 |
| hif1ab4 | 3.31352196542874e-08 | 0.291373038 | 0.274 | 0.113 | 0.00083398 | 30 |
| btg111 | 3.42158905634199e-08 | 0.38454788 | 0.589 | 0.332 | 0.00086118 | 30 |
| tuba8l1 | 4.04233320570744e-08 | 0.287625417 | 0.403 | 0.189 | 0.001017415 | 30 |
| cfl18 | 4.61863711844159e-08 | 0.34260226 | 0.734 | 0.577 | 0.001162465 | 30 |
| eif4ebp24 | 6.31108055152165e-08 | 0.293420575 | 0.371 | 0.174 | 0.001588436 | 30 |
| serf210 | 6.37044878658805e-08 | 0.371907573 | 0.565 | 0.343 | 0.001603378 | 30 |
| tnfb4 | 7.28693566687048e-08 | 0.379379046 | 0.161 | 0.053 | 0.001834049 | 30 |
| gadd45ab2 | 7.80396380731004e-08 | 0.250695139 | 0.202 | 0.073 | 0.00196418 | 30 |
| arhgdia8 | 1.07770361553151e-07 | 0.329376552 | 0.444 | 0.229 | 0.002712472 | 30 |
| zgc:16273013 | 1.08010791249256e-07 | 0.396879971 | 0.653 | 0.391 | 0.002718524 | 30 |
| anxa11b4 | 1.53448593180975e-07 | 0.297100994 | 0.323 | 0.15 | 0.003862148 | 30 |
| gnai2a4 | 1.62407867540201e-07 | 0.345191351 | 0.306 | 0.144 | 0.004087644 | 30 |
| myh11a6 | 1.72833304862996e-07 | 0.252895424 | 0.185 | 0.066 | 0.004350041 | 30 |
| f11r.14 | 2.04388947867832e-07 | 0.361142555 | 0.363 | 0.178 | 0.005144265 | 30 |
| nfil3-69 | 2.2595375907736e-07 | 0.311990604 | 0.29 | 0.131 | 0.00568703 | 30 |
| ucp28 | 2.27415904657001e-07 | 0.360950304 | 0.653 | 0.416 | 0.005723831 | 30 |

| | | | | | | |
|---|---|---|---|---|---|---|
| si:ch211-212k18.73 | 3.0441087778093e-07 | 0.293975167 | 0.306 | 0.139 | 0.007661717 | 30 |
| stk17al1 | 3.69007046156875e-07 | 0.253934853 | 0.282 | 0.121 | 0.009287538 | 30 |
| mvb12a1 | 3.79169760384972e-07 | 0.255068662 | 0.202 | 0.078 | 0.009543324 | 30 |
| si:ch211-196l7.412 | 4.50567021922872e-07 | 0.344991229 | 0.524 | 0.314 | 0.011340321 | 30 |
| zgc:564937 | 5.24232761780943e-07 | 0.289215858 | 0.734 | 0.573 | 0.013194414 | 30 |
| sumo2b8 | 1.20446374412482e-06 | 0.318775486 | 0.468 | 0.269 | 0.030315148 | 30 |
| pim25 | 1.58016274525358e-06 | 0.34319201 | 0.21 | 0.09 | 0.039771116 | 30 |
| idh26 | 3.67161104332841e-06 | 0.293399007 | 0.371 | 0.199 | 0.092410778 | 30 |
| tmed71 | 3.98892422738677e-06 | 0.273218405 | 0.387 | 0.207 | 0.100397234 | 30 |
| abracl4 | 5.97236164327063e-06 | 0.279002242 | 0.411 | 0.222 | 0.15031837 | 30 |
| sarnp | 1.00055232759758e-05 | 0.252015774 | 0.339 | 0.178 | 0.251829015 | 30 |
| GCA5 | 1.19720357842295e-05 | 0.261486769 | 0.339 | 0.182 | 0.301324169 | 30 |
| tubb4b2 | 3.73011377561189e-05 | 0.262365218 | 0.742 | 0.585 | 0.938832336 | 30 |
| prelid3b2 | 3.83565267288365e-05 | 0.258478037 | 0.411 | 0.239 | 0.965395421 | 30 |
| cnn28 | 6.45081091429521e-05 | 0.254497931 | 0.258 | 0.135 | 1 | 30 |
| socs3a11 | 8.05168274305986e-05 | 0.329231227 | 0.468 | 0.299 | 1 | 30 |
| romo111 | 8.46765456005805e-05 | 0.287954079 | 0.548 | 0.387 | 1 | 30 |
| eif3f10 | 8.8152719681567e-05 | 0.27764225 | 0.605 | 0.432 | 1 | 30 |
| tomm20b6 | 9.20327972907237e-05 | 0.26975074 | 0.355 | 0.215 | 1 | 30 |
| rsl24d112 | 0.000104746 | 0.293103094 | 0.645 | 0.444 | 1 | 30 |
| nsa211 | 0.000133162 | 0.263939503 | 0.556 | 0.363 | 1 | 30 |
| NC-002333.414 | 0.000150642 | 0.270991152 | 0.879 | 0.622 | 1 | 30 |
| zgc:1538675 | 0.000158382 | 0.26558669 | 0.613 | 0.437 | 1 | 30 |
| LOC103908723 | 0 | 2.061778594 | 0.966 | 0.003 | 0 | 31 |
| zgc:194275 | 0 | 1.974720418 | 0.915 | 0.001 | 0 | 31 |
| LOC1018867661 | 0 | 1.97265098 | 0.949 | 0.005 | 0 | 31 |
| igl4v81 | 0 | 1.883291614 | 0.932 | 0.023 | 0 | 31 |
| cd37 | 0 | 1.833121034 | 0.881 | 0.002 | 0 | 31 |
| CU896602.1 | 0 | 1.677670376 | 0.847 | 0.001 | 0 | 31 |
| igic1s1 | 0 | 1.577561622 | 0.805 | 0.001 | 0 | 31 |
| igl1c3 | 0 | 1.560366202 | 0.754 | 0.001 | 0 | 31 |
| XLOC-026705 | 0 | 1.490755128 | 0.729 | 0.001 | 0 | 31 |
| igl3v5 | 0 | 1.476885238 | 0.737 | 0.001 | 0 | 31 |
| cxcr4a1 | 0 | 1.416762859 | 0.636 | 0.011 | 0 | 31 |
| BX571825.4 | 0 | 1.224411143 | 0.5 | 0.002 | 0 | 31 |

| | | | | | | |
|---|---|---|---|---|---|---|
| LOC101882264 | 0 | 1.043981675 | 0.458 | 0.006 | 0 | 31 |
| pax5 | 0 | 1.009943513 | 0.449 | 0.002 | 0 | 31 |
| BX571825.5 | 0 | 0.965416476 | 0.449 | 0.001 | 0 | 31 |
| epb41a | 0 | 0.953212935 | 0.475 | 0.013 | 0 | 31 |
| CT573342.2 | 0 | 0.951106713 | 0.458 | 0 | 0 | 31 |
| CU571382.4 | 0 | 0.906740169 | 0.39 | 0.001 | 0 | 31 |
| LOC108179343 | 0 | 0.905576248 | 0.398 | 0 | 0 | 31 |
| ly6m2 | 0 | 0.823786815 | 0.314 | 0.001 | 0 | 31 |
| BX571825.3 | 0 | 0.782884777 | 0.331 | 0 | 0 | 31 |
| LOC101886979 | 0 | 0.655070476 | 0.246 | 0.001 | 0 | 31 |
| ighv1-4 | 0 | 0.650998423 | 0.263 | 0.003 | 0 | 31 |
| FP016018.1 | 0 | 0.599008158 | 0.246 | 0 | 0 | 31 |
| rtn4rl2b | 0 | 0.530200246 | 0.22 | 0 | 0 | 31 |
| capga | 0 | 0.487319171 | 0.203 | 0 | 0 | 31 |
| CU571382.1 | 0 | 0.390776051 | 0.153 | 0 | 0 | 31 |
| CT573231.12 | 7.55737785301321e-304 | 1.193925427 | 0.568 | 0.02 | 1.90211643182489e-299 | 31 |
| ncf13 | 1.45232216572554e-278 | 1.206722317 | 0.627 | 0.028 | 3.65534965891461e-274 | 31 |
| CR944667.2 | 3.20058013948879e-277 | 0.359860888 | 0.136 | 0 | 8.05554015307934e-273 | 31 |
| swap70a | 2.6913994131397e-268 | 0.609976353 | 0.271 | 0.004 | 6.77398318293131e-264 | 31 |
| si:ch73-7i4.3 | 2.2404202249284e-261 | 0.629884299 | 0.271 | 0.004 | 5.63891366412228e-257 | 31 |
| XLOC-0183561 | 4.68628203325496e-248 | 1.175424668 | 0.542 | 0.023 | 1.17949032494994e-243 | 31 |
| si:ch211-145b13.6 | 6.36967554437638e-232 | 0.585303244 | 0.254 | 0.004 | 1.60318363776409e-227 | 31 |
| pou2f2a1 | 1.43769147239574e-206 | 0.559515391 | 0.263 | 0.005 | 3.61852566687284e-202 | 31 |
| sftpba2 | 4.27146726949981e-194 | 1.032835301 | 0.483 | 0.024 | 1.07508559706041e-189 | 31 |
| ccr9a6 | 1.08695725972955e-160 | 1.428055956 | 0.763 | 0.082 | 2.73576272701332e-156 | 31 |
| cxcr4b9 | 8.39710705380276e-145 | 1.697497311 | 0.975 | 0.177 | 2.11346787437162e-140 | 31 |
| si:ch73-380l3.21 | 7.8176718355459e-144 | 0.855593223 | 0.39 | 0.021 | 1.96762982428855e-139 | 31 |
| illr11 | 8.09329341354771e-143 | 0.767692197 | 0.314 | 0.013 | 2.03700101925582e-138 | 31 |
| LOC110438240 | 7.59587653390898e-141 | 0.350994859 | 0.153 | 0.002 | 1.91180616481955e-136 | 31 |
| grap2b8 | 5.01570614701164e-140 | 1.358040125 | 0.805 | 0.107 | 1.26240308014136e-135 | 31 |
| btbd11b | 4.69268568264935e-135 | 0.50009929 | 0.229 | 0.007 | 1.18110205946601e-130 | 31 |
| CT027980.1 | 9.28363491391238e-133 | 0.308413446 | 0.136 | 0.002 | 2.33659807148261e-128 | 31 |
| sp6 | 3.98221466083926e-130 | 0.253659238 | 0.119 | 0.001 | 1.00228360798663e-125 | 31 |
| si:dkeyp-117h8.2 | 1.14730991528708e-122 | 0.367191682 | 0.144 | 0.003 | 2.88766432578605e-118 | 31 |
| cobll1a | 8.17669824573604e-113 | 0.363849695 | 0.178 | 0.005 | 2.0579931814693e-108 | 31 |

| | | | | | | |
|---|---|---|---|---|---|---|
| tlr19 | 1.13239143752518e-104 | 0.289353653 | 0.127 | 0.002 | 2.85011600910712e-100 | 31 |
| rbbp8l1 | 2.29975586617911e-100 | 0.467551933 | 0.212 | 0.009 | 5.78825553958621e-96 | 31 |
| pacsin2 | 4.22173680510436e-99 | 0.864749396 | 0.458 | 0.043 | 1.06256893647672e-94 | 31 |
| atp11a1 | 7.29344110620466e-98 | 0.503017938 | 0.229 | 0.01 | 1.83568619202065e-93 | 31 |
| sh2d4a | 9.12306922203668e-96 | 0.478391086 | 0.212 | 0.009 | 2.29618529249441e-91 | 31 |
| lmo2 | 4.23512911562257e-93 | 0.613297079 | 0.288 | 0.018 | 1.06593964711104e-88 | 31 |
| si:ch1073-220m6.12 | 6.73713373896104e-89 | 0.61164583 | 0.288 | 0.019 | 1.6956691907591e-84 | 31 |
| blnk1 | 1.22661809479676e-88 | 0.633629968 | 0.314 | 0.022 | 3.08727508279397e-84 | 31 |
| csrnp1a1 | 1.5954624958044e-88 | 0.849456844 | 0.458 | 0.049 | 4.01561955569008e-84 | 31 |
| si:ch211-119e14.15 | 2.4764052066496e-85 | 1.00725135 | 0.534 | 0.07 | 6.23286426461638e-81 | 31 |
| si:dkey-24p1.1 | 7.33811956999098e-81 | 0.427939515 | 0.178 | 0.008 | 1.84693131457103e-76 | 31 |
| irf4b2 | 6.9499099566794e-80 | 0.718528111 | 0.297 | 0.023 | 1.74922283699664e-75 | 31 |
| XLOC-0425957 | 2.250191902736e-79 | 1.244459288 | 0.805 | 0.16 | 5.66350799999623e-75 | 31 |
| ebf1a | 1.19774395313751e-78 | 0.369215385 | 0.161 | 0.006 | 3.01460175565179e-74 | 31 |
| cd74a7 | 5.37363309324129e-78 | 1.272465169 | 0.915 | 0.214 | 1.3524897132379e-73 | 31 |
| si:busm1-266f07.25 | 1.19876361342768e-77 | 1.117188519 | 0.771 | 0.142 | 3.01716813863613e-73 | 31 |
| mpeg1.12 | 5.65034498622217e-77 | 0.52334935 | 0.237 | 0.015 | 1.42213532958226e-72 | 31 |
| si:dkey-82o10.4 | 2.0218999448501e-74 | 0.407897763 | 0.169 | 0.008 | 5.08891997119322e-70 | 31 |
| arhgap153 | 1.07448335718725e-70 | 0.609311273 | 0.305 | 0.027 | 2.7043671617046e-66 | 31 |
| cd74b8 | 1.35738128055783e-70 | 1.204247017 | 0.881 | 0.219 | 3.41639294503599e-66 | 31 |
| dapp11 | 6.57667339523077e-67 | 0.370727079 | 0.161 | 0.008 | 1.65528292684563e-62 | 31 |
| gng2 | 1.00289207226105e-65 | 0.668057141 | 0.364 | 0.041 | 2.52417905667384e-61 | 31 |
| derl23 | 4.32014524928234e-64 | 0.914999494 | 0.576 | 0.102 | 1.08733735779187e-59 | 31 |
| syk5 | 1.49217337300363e-63 | 0.844104607 | 0.492 | 0.076 | 3.75565116251285e-59 | 31 |
| XLOC-0004319 | 1.16350419313325e-62 | 1.304324085 | 0.915 | 0.274 | 2.92842370369707e-58 | 31 |
| gpr55a1 | 4.92633628157122e-59 | 0.333647996 | 0.144 | 0.007 | 1.23990957870866e-54 | 31 |
| cd7al2 | 5.23158643140878e-57 | 0.522078913 | 0.263 | 0.024 | 1.31673798892128e-52 | 31 |
| cd835 | 2.15699827941611e-56 | 1.082343355 | 0.534 | 0.101 | 5.4289489694624e-52 | 31 |
| kctd5a5 | 8.6389493929122e-56 | 0.954754735 | 0.695 | 0.172 | 2.17433717270207e-51 | 31 |
| bcl11aa | 3.51117111425414e-55 | 0.34196146 | 0.153 | 0.008 | 8.83726657746623e-51 | 31 |
| bpifcl2 | 9.39380483894488e-53 | 0.4138363 | 0.178 | 0.012 | 2.36432673991404e-48 | 31 |
| dtx4a | 5.39269001589573e-52 | 0.30890536 | 0.153 | 0.009 | 1.3572861501008e-47 | 31 |
| XLOC-0139646 | 1.01102393127974e-50 | 1.082823976 | 0.686 | 0.171 | 2.54464613263799e-46 | 31 |
| rgs137 | 5.16874656353324e-50 | 0.902658483 | 0.593 | 0.129 | 1.30092182257568e-45 | 31 |
| mhc2dab4 | 2.16776717458799e-49 | 0.735030084 | 0.517 | 0.095 | 5.4560532017205e-45 | 31 |

| | | | | | | |
|---|---|---|---|---|---|---|
| CABZ01066694.1 | 2.31258481218225e-49 | 0.380011286 | 0.195 | 0.016 | 5.82054471378151e-45 | 31 |
| fam53b | 7.79832009049534e-48 | 0.676569279 | 0.407 | 0.067 | 1.96275918357677e-43 | 31 |
| LOC1039108715 | 2.22544490750981e-46 | 0.933091771 | 0.678 | 0.168 | 5.60122228771144e-42 | 31 |
| ptmaa14 | 4.3893382199711e-46 | 0.965001978 | 0.898 | 0.392 | 1.10475253658453e-41 | 31 |
| rac211 | 7.05192065813176e-46 | 0.905765457 | 0.746 | 0.217 | 1.77489791044518e-41 | 31 |
| tspan2a | 9.80253780073771e-45 | 0.303534924 | 0.144 | 0.009 | 2.46720073906767e-40 | 31 |
| spi1b5 | 1.58184239676705e-44 | 0.629573487 | 0.373 | 0.06 | 3.98133912842299e-40 | 31 |
| s1pr41 | 1.71050866147791e-44 | 0.323435917 | 0.136 | 0.008 | 4.30517925007375e-40 | 31 |
| si:ch73-234b20.55 | 2.40525724381397e-43 | 0.727661674 | 0.475 | 0.098 | 6.05379195695537e-39 | 31 |
| sox121 | 5.60354694829641e-43 | 0.340742783 | 0.153 | 0.011 | 1.41035673141672e-38 | 31 |
| gadd45ga2 | 2.04287007389132e-42 | 0.738155932 | 0.331 | 0.051 | 5.14169968897707e-38 | 31 |
| LOC1039094521 | 2.18507301059584e-41 | 0.413466909 | 0.178 | 0.015 | 5.49961026036866e-37 | 31 |
| gpx1a6 | 1.36505845361141e-39 | 0.895354924 | 0.703 | 0.221 | 3.43571562189456e-35 | 31 |
| rps208 | 8.19314596756871e-39 | 0.348507625 | 1 | 0.916 | 2.06213290857737e-34 | 31 |
| rps2118 | 1.14170518195453e-38 | 0.527459232 | 1 | 0.749 | 2.87355777246136e-34 | 31 |
| si:ch211-197h24.94 | 2.21244407911534e-38 | 0.649409819 | 0.364 | 0.067 | 5.5685005027254e-34 | 31 |
| ccni11 | 3.27470105550317e-38 | 0.863171098 | 0.763 | 0.291 | 8.24209508659592e-34 | 31 |
| rplp19 | 8.52981064249118e-38 | 0.402302291 | 1 | 0.877 | 2.1468680406086e-33 | 31 |
| rps27.112 | 1.74483550688027e-37 | 0.593002388 | 1 | 0.678 | 4.39157648726694e-33 | 31 |
| cited4a3 | 1.87400184939298e-37 | 0.533707427 | 0.28 | 0.041 | 4.7166752547372e-33 | 31 |
| si:dkey-78k11.9 | 2.93680840563761e-37 | 0.277320877 | 0.119 | 0.008 | 7.39165307614929e-33 | 31 |
| XLOC-0108998 | 1.32328549630888e-36 | 0.985317622 | 0.729 | 0.256 | 3.33057726565981e-32 | 31 |
| rplp215 | 1.90898683631643e-36 | 0.520186253 | 1 | 0.757 | 4.80472896832483e-32 | 31 |
| ncf22 | 4.25312638702553e-36 | 0.402346464 | 0.195 | 0.021 | 1.07046938035046e-31 | 31 |
| rpl1112 | 1.9695931862591e-35 | 0.447265738 | 1 | 0.813 | 4.95726909049553e-31 | 31 |
| marcksl1a7 | 4.85789228074558e-35 | 0.822483482 | 0.669 | 0.212 | 1.22268290814086e-30 | 31 |
| crip111 | 6.60454604651391e-35 | 0.815749977 | 0.771 | 0.268 | 1.66229819444709e-30 | 31 |
| LOC1005362135 | 6.65091939786033e-35 | 0.752231011 | 0.449 | 0.104 | 1.67396990324747e-30 | 31 |
| rps15a11 | 8.67680366986819e-34 | 0.494611707 | 1 | 0.779 | 2.18386471566912e-29 | 31 |
| cad1 | 1.06573221878833e-33 | 0.480617766 | 0.212 | 0.027 | 2.68234142146836e-29 | 31 |
| rpl3211 | 1.0442521584141e-32 | 0.43861392 | 1 | 0.812 | 2.62827825751244e-28 | 31 |
| rpl3615 | 1.36107474527428e-32 | 0.464527933 | 1 | 0.789 | 3.42568902638084e-28 | 31 |
| rpl1010 | 1.75130631544816e-32 | 0.448438151 | 0.983 | 0.771 | 4.40786286535147e-28 | 31 |
| rsl24d113 | 1.96684902073252e-32 | 0.75352761 | 0.898 | 0.441 | 4.95036230028167e-28 | 31 |
| pacsin3 | 2.0478974697866e-32 | 0.370937324 | 0.195 | 0.023 | 5.15435314170588e-28 | 31 |

| | | | | | | |
|---|---|---|---|---|---|---|
| rbm38 | 6.74449185347598e-32 | 0.307876278 | 0.161 | 0.016 | 1.69752115460137e-27 | 31 |
| rpl811 | 1.03974195202883e-31 | 0.451826297 | 0.992 | 0.779 | 2.61692651906135e-27 | 31 |
| CR936442.17 | 1.07213138410938e-31 | 0.810270881 | 0.576 | 0.167 | 2.6984474806649e-27 | 31 |
| pfn111 | 2.58348334395484e-31 | 0.823684687 | 1 | 0.522 | 6.50236922839994e-27 | 31 |
| rpl196 | 4.6945909695055e-31 | 0.376402099 | 1 | 0.851 | 1.18158160111484e-26 | 31 |
| rplp06 | 8.3967643235436e-31 | 0.374589957 | 0.992 | 0.851 | 2.11338161259269e-26 | 31 |
| ramp22 | 1.55826017988002e-30 | 0.53463507 | 0.254 | 0.041 | 3.92198504674002e-26 | 31 |
| rpl18a11 | 3.41773320257981e-30 | 0.455163105 | 1 | 0.784 | 8.60209269757313e-26 | 31 |
| parvg5 | 4.30954398947924e-30 | 0.560936598 | 0.331 | 0.066 | 1.08466912671203e-25 | 31 |
| ikzf23 | 5.21414077264774e-30 | 0.404202754 | 0.212 | 0.029 | 1.31234709106771e-25 | 31 |
| h1fx8 | 5.90313952747365e-30 | 0.926662076 | 0.551 | 0.182 | 1.48576118766984e-25 | 31 |
| jam2a2 | 1.32397544463876e-29 | 0.458776043 | 0.237 | 0.036 | 3.33231379661129e-25 | 31 |
| tsc22d38 | 3.17098941436246e-29 | 0.73097895 | 0.483 | 0.142 | 7.98106325700889e-25 | 31 |
| rps511 | 7.57678800068966e-29 | 0.518971328 | 1 | 0.7 | 1.90700177189358e-24 | 31 |
| rpsa16 | 9.71448241082668e-29 | 0.464379155 | 0.992 | 0.751 | 2.44503807798097e-24 | 31 |
| ptmab11 | 1.91466481688895e-28 | 0.676129195 | 0.898 | 0.539 | 4.81901987762779e-24 | 31 |
| rpl1511 | 2.1729925403955e-28 | 0.474934453 | 0.992 | 0.743 | 5.46920492492143e-24 | 31 |
| klf134 | 2.70813274267339e-28 | 0.644659098 | 0.407 | 0.104 | 6.81609930003464e-24 | 31 |
| hmbsa | 6.76970858461813e-28 | 0.374092695 | 0.22 | 0.033 | 1.70386795366254e-23 | 31 |
| rpl379 | 1.26939518616867e-27 | 0.387889166 | 1 | 0.849 | 3.19494074406791e-23 | 31 |
| ralgps21 | 1.50098395326699e-27 | 0.372708302 | 0.169 | 0.021 | 3.77782651197769e-23 | 31 |
| coro1a11 | 1.6352366635443e-27 | 0.704678851 | 0.695 | 0.229 | 4.11572715847464e-23 | 31 |
| rplp2l15 | 1.84874123796577e-27 | 0.457367828 | 1 | 0.773 | 4.65309682183605e-23 | 31 |
| cybb2 | 1.90888416233422e-27 | 0.354236284 | 0.161 | 0.019 | 4.80447054817899e-23 | 31 |
| rps225 | 2.43263629523068e-27 | 0.424376115 | 1 | 0.828 | 6.1227022914661e-23 | 31 |
| naca11 | 3.21319433773883e-27 | 0.575810204 | 0.975 | 0.617 | 8.08728882865487e-23 | 31 |
| rpl10a8 | 3.61717040146607e-27 | 0.439391673 | 0.992 | 0.764 | 9.10405618344995e-23 | 31 |
| rps4x10 | 6.65088019594547e-27 | 0.481895649 | 0.975 | 0.709 | 1.67396003651752e-22 | 31 |
| rps26l13 | 7.37437636133138e-27 | 0.532942752 | 0.975 | 0.663 | 1.85605678638349e-22 | 31 |
| rps2911 | 1.16864239092439e-26 | 0.374047742 | 0.992 | 0.831 | 2.94135603371759e-22 | 31 |
| thrap3a | 1.33214539261431e-26 | 0.289372884 | 0.136 | 0.014 | 3.35287673867096e-22 | 31 |
| TPGS11 | 3.28114044079759e-26 | 0.446671381 | 0.229 | 0.038 | 8.25830237544347e-22 | 31 |
| rpl3413 | 1.27476116058636e-25 | 0.430642375 | 1 | 0.783 | 3.20844636507982e-21 | 31 |
| cyba5 | 2.75442846370938e-25 | 0.521989769 | 0.322 | 0.072 | 6.93262100031014e-21 | 31 |
| rpl76 | 2.77605540146318e-25 | 0.384553046 | 0.983 | 0.804 | 6.98705383994267e-21 | 31 |

| | | | | | | |
|---|---|---|---|---|---|---|
| rpl2812 | 3.90633769053659e-25 | 0.434439587 | 0.983 | 0.748 | 9.83186133331155e-21 | 31 |
| ppiaa10 | 4.57241778172669e-25 | 0.36735103 | 0.966 | 0.803 | 1.15083183148279e-20 | 31 |
| histh1l6 | 6.46910349278884e-25 | 0.649375154 | 0.398 | 0.108 | 1.62820865810002e-20 | 31 |
| rps1313 | 7.87198025271868e-25 | 0.459681888 | 0.983 | 0.723 | 1.98129870980676e-20 | 31 |
| rps68 | 8.14928182034287e-25 | 0.416323032 | 1 | 0.788 | 2.0510927413621e-20 | 31 |
| rpl2110 | 1.24390600145976e-24 | 0.423893183 | 0.983 | 0.755 | 3.13078701507407e-20 | 31 |
| rpl359 | 1.53092669239763e-24 | 0.387798944 | 0.992 | 0.804 | 3.85318939209559e-20 | 31 |
| rpl3013 | 1.70446418991668e-24 | 0.477581234 | 0.992 | 0.707 | 4.2899659196013e-20 | 31 |
| hmgn29 | 1.96202267804763e-24 | 0.919253463 | 0.788 | 0.441 | 4.93821487837808e-20 | 31 |
| pik3r15 | 3.10891200162017e-24 | 0.535034603 | 0.356 | 0.089 | 7.82482061687781e-20 | 31 |
| cnpy31 | 1.34142110049944e-23 | 0.408858966 | 0.254 | 0.049 | 3.37622276784705e-19 | 31 |
| rpl2412 | 2.7252510007906e-23 | 0.408386102 | 0.992 | 0.775 | 6.85918424388987e-19 | 31 |
| b3gnt3.42 | 2.82902440028601e-23 | 0.419601086 | 0.212 | 0.037 | 7.12037151307986e-19 | 31 |
| btg112 | 3.2565101520237e-23 | 0.708112766 | 0.712 | 0.331 | 8.19631040162846e-19 | 31 |
| tfdp2 | 4.62389718663597e-23 | 0.346658973 | 0.186 | 0.029 | 1.16378868290441e-18 | 31 |
| ascc1 | 8.24820771195389e-23 | 0.465240446 | 0.288 | 0.062 | 2.07599139902167e-18 | 31 |
| pde4ba10 | 1.02514178274355e-22 | 0.602281527 | 0.449 | 0.139 | 2.58017935298725e-18 | 31 |
| rpl1812 | 1.21149100757707e-22 | 0.420073252 | 1 | 0.773 | 3.04920171697073e-18 | 31 |
| wu:fb55g092 | 1.99023223094982e-22 | 0.664911573 | 0.636 | 0.249 | 5.00921550207761e-18 | 31 |
| rpl269 | 2.0741535211341e-22 | 0.352028576 | 1 | 0.84 | 5.22043699734243e-18 | 31 |
| rpl36a10 | 3.17329447184286e-22 | 0.372368432 | 1 | 0.809 | 7.9868648561813e-18 | 31 |
| arhgdig10 | 7.88889117561895e-22 | 0.651178937 | 0.619 | 0.241 | 1.98555501999153e-17 | 31 |
| prkcbb | 1.10678646793272e-21 | 0.358422181 | 0.186 | 0.031 | 2.78567086113985e-17 | 31 |
| rpl916 | 1.46238824097022e-21 | 0.544680685 | 0.992 | 0.627 | 3.68068496369794e-17 | 31 |
| LOC1039091157 | 1.65723069499843e-21 | 0.603998863 | 0.483 | 0.164 | 4.17108393624155e-17 | 31 |
| rps1412 | 1.74198165153736e-21 | 0.416559413 | 0.992 | 0.758 | 4.38439361875438e-17 | 31 |
| hsp90ab19 | 2.74779561264569e-21 | 0.367011946 | 0.975 | 0.79 | 6.91592677746794e-17 | 31 |
| rps1614 | 4.71806006373943e-21 | 0.436118751 | 0.983 | 0.737 | 1.18748853744258e-16 | 31 |
| rps127 | 5.48826022605297e-21 | 0.345663507 | 1 | 0.839 | 1.38134021629527e-16 | 31 |
| rpl7a10 | 7.43756122008835e-21 | 0.364011685 | 0.992 | 0.809 | 1.87195978348404e-16 | 31 |
| dap1b3 | 9.28796593995264e-21 | 0.633688792 | 0.636 | 0.249 | 2.33768814742668e-16 | 31 |
| celf213 | 1.5218458879538e-20 | 0.64561501 | 0.585 | 0.241 | 3.83033391539092e-16 | 31 |
| rack17 | 1.58953910881199e-20 | 0.384424505 | 0.949 | 0.744 | 4.00071098296889e-16 | 31 |
| ccnd2a1 | 1.70524378217097e-20 | 0.531594071 | 0.322 | 0.085 | 4.29192807534612e-16 | 31 |
| rps2412 | 2.1391700163309e-20 | 0.402348711 | 0.983 | 0.758 | 5.38407701410324e-16 | 31 |

| | | | | | | |
|---|---|---|---|---|---|---|
| mhc2b2 | 2.96517648868723e-20 | 0.341315188 | 0.178 | 0.029 | 7.4630527043769e-16 | 31 |
| si:dkey-183p4.104 | 3.13112916623581e-20 | 0.329488915 | 0.195 | 0.034 | 7.8807389984989e-16 | 31 |
| rps1813 | 3.54467696431677e-20 | 0.444820047 | 0.992 | 0.723 | 8.92159745148888e-16 | 31 |
| glo13 | 5.03713277714466e-20 | 0.614433718 | 0.508 | 0.195 | 1.26779594867954e-15 | 31 |
| rps178 | 5.19298989083898e-20 | 0.373795217 | 0.958 | 0.766 | 1.30702362562526e-15 | 31 |
| rpl23a15 | 6.91541684883266e-20 | 0.504788335 | 0.966 | 0.633 | 1.74054126668269e-15 | 31 |
| actb211 | 8.3703286281595e-20 | 0.371144628 | 1 | 0.867 | 2.10672801242147e-15 | 31 |
| rpl35a12 | 1.16330964632185e-19 | 0.405896716 | 0.992 | 0.758 | 2.92793404882747e-15 | 31 |
| rpl279 | 1.26401043451726e-19 | 0.401102774 | 0.992 | 0.762 | 3.1813878626365e-15 | 31 |
| hopx5 | 1.4278735010146e-19 | 0.475738356 | 0.271 | 0.064 | 3.59381481470364e-15 | 31 |
| rpl236 | 1.66399744211323e-19 | 0.287329557 | 1 | 0.872 | 4.1881151620548e-15 | 31 |
| rps239 | 1.6752495039154e-19 | 0.417095281 | 0.966 | 0.722 | 4.21643547640468e-15 | 31 |
| faua10 | 3.27825738356528e-19 | 0.460309244 | 0.983 | 0.688 | 8.25104600869544e-15 | 31 |
| ada | 3.36048820580112e-19 | 0.361013564 | 0.203 | 0.04 | 8.45801276518084e-15 | 31 |
| rps1015 | 6.94042442927426e-19 | 0.441736109 | 0.966 | 0.693 | 1.74683542460404e-14 | 31 |
| si:ch211-214p16.17 | 7.48927555130592e-19 | 0.702703131 | 0.644 | 0.256 | 1.88497576350819e-14 | 31 |
| tmc84 | 1.03165075820603e-18 | 0.347688534 | 0.186 | 0.035 | 2.59656179332874e-14 | 31 |
| serf211 | 1.23243934011778e-18 | 0.589871764 | 0.703 | 0.341 | 3.10192657514245e-14 | 31 |
| eif3f11 | 1.42651380366298e-18 | 0.582949354 | 0.78 | 0.43 | 3.59039259243935e-14 | 31 |
| rps198 | 3.31975628831826e-18 | 0.30334003 | 0.992 | 0.863 | 8.35549460206822e-14 | 31 |
| tnfaip8l2a1 | 4.60139629553433e-18 | 0.293596736 | 0.144 | 0.023 | 1.15812543362304e-13 | 31 |
| nr4a19 | 4.79707371597684e-18 | 0.584774167 | 0.466 | 0.166 | 1.20737548357421e-13 | 31 |
| mknk2b11 | 5.34652797494988e-18 | 0.594738599 | 0.5 | 0.196 | 1.34566762601514e-13 | 31 |
| mbnl18 | 6.79284038774338e-18 | 0.54279587 | 0.492 | 0.183 | 1.70968999719113e-13 | 31 |
| rps310 | 1.19857037306759e-17 | 0.412794793 | 0.949 | 0.699 | 3.01668177197382e-13 | 31 |
| rpl614 | 1.67210729636115e-17 | 0.444098946 | 0.949 | 0.666 | 4.20852685421137e-13 | 31 |
| rpl3114 | 1.88651189641936e-17 | 0.45385341 | 0.941 | 0.653 | 4.74816179209788e-13 | 31 |
| tpt19 | 2.29984973105275e-17 | 0.331777089 | 0.975 | 0.8 | 5.78849178808666e-13 | 31 |
| ptprc10 | 2.69973530695795e-17 | 0.515207752 | 0.39 | 0.127 | 6.79496379408246e-13 | 31 |
| rps99 | 3.13425897272921e-17 | 0.36347956 | 0.975 | 0.776 | 7.88861640846214e-13 | 31 |
| nsa212 | 3.64440639764589e-17 | 0.581935051 | 0.72 | 0.361 | 9.17260646223494e-13 | 31 |
| egr2b | 3.64916358378789e-17 | 0.273730345 | 0.102 | 0.012 | 9.18457982403574e-13 | 31 |
| si:ch1073-429i10.3.1 | 6.0253413068804e-17 | 0.576998981 | 0.492 | 0.201 | 1.51651815352873e-12 | 31 |
| plk32 | 7.65729847047296e-17 | 0.484318711 | 0.381 | 0.12 | 1.92726545203334e-12 | 31 |
| rpl3813 | 7.95013095055022e-17 | 0.411620518 | 0.975 | 0.727 | 2.00096845894398e-12 | 31 |

| | | | | | | |
|---|---|---|---|---|---|---|
| rps2813 | 9.6335017856706e-17 | 0.343712761 | 1 | 0.826 | 2.42465606443543e-12 | 31 |
| rpl137 | 9.92601383389012e-17 | 0.323992743 | 0.975 | 0.804 | 2.49827842185181e-12 | 31 |
| btk1 | 1.49915786436387e-16 | 0.257983741 | 0.144 | 0.024 | 3.77323042881742e-12 | 31 |
| ptp4a32 | 2.2134500839447e-16 | 0.38242443 | 0.22 | 0.051 | 5.57103251628041e-12 | 31 |
| rpl399 | 2.31020491128278e-16 | 0.296220736 | 1 | 0.873 | 5.81455474120762e-12 | 31 |
| rpl2212 | 2.49074190988298e-16 | 0.436167359 | 0.975 | 0.693 | 6.26894831298447e-12 | 31 |
| oaz1b11 | 3.3572105230224e-16 | 0.574568524 | 0.636 | 0.322 | 8.44976316539509e-12 | 31 |
| dusp27 | 3.38517752393357e-16 | 0.599588374 | 0.466 | 0.178 | 8.5201533099884e-12 | 31 |
| unc93b12 | 3.92834178345745e-16 | 0.280560385 | 0.136 | 0.022 | 9.88724343478407e-12 | 31 |
| btf311 | 3.94486593007476e-16 | 0.482082851 | 0.907 | 0.586 | 9.92883305940517e-12 | 31 |
| eif3m7 | 4.43926095337493e-16 | 0.552546033 | 0.678 | 0.357 | 1.11731758935494e-11 | 31 |
| rps1513 | 6.79636633260539e-16 | 0.443581212 | 0.975 | 0.683 | 1.71057744225345e-11 | 31 |
| si:ch211-195b13.1 | 9.12913681481572e-16 | 0.34605761 | 0.153 | 0.028 | 2.29771244492097e-11 | 31 |
| rpl5a13 | 1.62476119059103e-15 | 0.514998402 | 0.822 | 0.491 | 4.08936144059856e-11 | 31 |
| rps77 | 3.35600708649919e-15 | 0.314766071 | 0.975 | 0.82 | 8.44673423600981e-11 | 31 |
| zgc:16273014 | 3.59045797789689e-15 | 0.584070968 | 0.805 | 0.39 | 9.03682368456869e-11 | 31 |
| tmc6b | 4.63213269046526e-15 | 0.342505228 | 0.169 | 0.035 | 1.1658614768632e-10 | 31 |
| wasa7 | 4.74130694640603e-15 | 0.426020851 | 0.305 | 0.093 | 1.19333954534093e-10 | 31 |
| slc25a55a1 | 4.81849942756832e-15 | 0.393081587 | 0.246 | 0.066 | 1.21276812092467e-10 | 31 |
| ccl35.14 | 5.55586870878629e-15 | 0.638291641 | 0.28 | 0.082 | 1.39835659531442e-10 | 31 |
| ucp29 | 6.71685439412403e-15 | 0.523343263 | 0.788 | 0.415 | 1.69056508245708e-10 | 31 |
| mbpb1 | 1.72654636182225e-14 | 0.339077473 | 0.229 | 0.059 | 4.34554453807042e-10 | 31 |
| wasb10 | 1.86875040490504e-14 | 0.521547676 | 0.449 | 0.177 | 4.70345789410549e-10 | 31 |
| rpl27a14 | 1.98918587667282e-14 | 0.518937964 | 0.864 | 0.494 | 5.00658193299781e-10 | 31 |
| rps8a6 | 2.14185583325447e-14 | 0.323231469 | 0.966 | 0.787 | 5.39083694671817e-10 | 31 |
| sesn17 | 3.03749756343e-14 | 0.530419646 | 0.424 | 0.167 | 7.64507761739698e-10 | 31 |
| rpl2912 | 4.04261546905872e-14 | 0.332200837 | 0.992 | 0.808 | 1.01748588740739e-09 | 31 |
| rps118 | 4.51048974723446e-14 | 0.335031942 | 0.975 | 0.765 | 1.13524516448144e-09 | 31 |
| xpc | 6.47930933784337e-14 | 0.281790123 | 0.153 | 0.031 | 1.6307773672418e-09 | 31 |
| itpka | 8.97448646155612e-14 | 0.250476089 | 0.127 | 0.023 | 2.25878849750906e-09 | 31 |
| nme2b.13 | 9.135154891284e-14 | 0.265316426 | 1 | 0.857 | 2.29922713458727e-09 | 31 |
| egr35 | 9.47679346321655e-14 | 0.383640516 | 0.22 | 0.058 | 2.38521414675697e-09 | 31 |
| serbp1a12 | 1.15531631432895e-13 | 0.497641618 | 0.814 | 0.481 | 2.90781563153453e-09 | 31 |
| arpc1b11 | 1.22168801658255e-13 | 0.520553556 | 0.686 | 0.307 | 3.07486656893662e-09 | 31 |
| si:dkey-27i16.25 | 1.61878069315372e-13 | 0.436159614 | 0.288 | 0.091 | 4.07430912659859e-09 | 31 |

| | | | | | | |
|---|---|---|---|---|---|---|
| lmnb21 | 2.97236689698072e-13 | 0.357654575 | 0.22 | 0.06 | 7.48115024301077e-09 | 31 |
| rps27a11 | 3.10881809378571e-13 | 0.349034423 | 0.975 | 0.751 | 7.82458426024925e-09 | 31 |
| CABZ01079427.1 | 3.35470211365388e-13 | 0.275899062 | 0.169 | 0.037 | 8.44344974985544e-09 | 31 |
| tmsb4x18 | 3.7089230067691e-13 | 0.515208295 | 1 | 0.685 | 9.33498831573715e-09 | 31 |
| rps3a7 | 4.84947993137969e-13 | 0.317172764 | 0.966 | 0.778 | 1.22056560392895e-08 | 31 |
| rgcc4 | 5.03427459162867e-13 | 0.358039644 | 0.246 | 0.07 | 1.26707657196702e-08 | 31 |
| si:ch211-150o23.21 | 6.53102718955245e-13 | 0.361692536 | 0.246 | 0.071 | 1.64379423333846e-08 | 31 |
| rpl125 | 9.85908983691105e-13 | 0.29377492 | 0.966 | 0.798 | 2.48143432105214e-08 | 31 |
| zgc:11418816 | 9.95220629683229e-13 | 0.451528044 | 0.949 | 0.626 | 2.50487080284972e-08 | 31 |
| rps259 | 1.00284136434767e-12 | 0.367435722 | 0.966 | 0.724 | 2.52405142992666e-08 | 31 |
| rel3 | 1.10883634440416e-12 | 0.284953356 | 0.161 | 0.036 | 2.79083019523082e-08 | 31 |
| herpud1 | 1.27395456716915e-12 | 0.388413398 | 0.246 | 0.074 | 3.20641625010804e-08 | 31 |
| inpp5d2 | 1.5591430989458e-12 | 0.317436852 | 0.153 | 0.034 | 3.92420726573669e-08 | 31 |
| gpr183a5 | 2.19743335160003e-12 | 0.326952466 | 0.229 | 0.065 | 5.53072000264211e-08 | 31 |
| gdi22 | 2.33181358843202e-12 | 0.424913895 | 0.441 | 0.184 | 5.86894162072454e-08 | 31 |
| plcg22 | 2.40782120922439e-12 | 0.314617734 | 0.195 | 0.051 | 6.06024520149687e-08 | 31 |
| gm2a4 | 2.48208355225639e-12 | 0.413693898 | 0.339 | 0.121 | 6.24715609267412e-08 | 31 |
| rps269 | 3.83270122818058e-12 | 0.34010273 | 0.949 | 0.733 | 9.64652572120769e-08 | 31 |
| pnrc216 | 3.93565982614035e-12 | 0.482312464 | 0.924 | 0.566 | 9.90566221641265e-08 | 31 |
| h2afx6 | 4.36235841167723e-12 | 0.498335596 | 0.364 | 0.138 | 1.09796198863504e-07 | 31 |
| nop536 | 4.40253100806339e-12 | 0.495969935 | 0.508 | 0.251 | 1.10807302941947e-07 | 31 |
| si:ch211-198c19.36 | 6.3248470944476e-12 | 0.562825193 | 0.212 | 0.064 | 1.59190076520152e-07 | 31 |
| tob1a2 | 7.93635322480249e-12 | 0.426531882 | 0.339 | 0.129 | 1.99750074315054e-07 | 31 |
| lyn2 | 8.21049230201309e-12 | 0.28144703 | 0.178 | 0.045 | 2.06649880749367e-07 | 31 |
| eef1g10 | 8.28261875181541e-12 | 0.408962573 | 0.898 | 0.619 | 2.08465231364442e-07 | 31 |
| cct26 | 1.34906106691351e-11 | 0.468103439 | 0.475 | 0.227 | 3.39545179931462e-07 | 31 |
| uba5212 | 1.5524610022078e-11 | 0.405005054 | 0.932 | 0.629 | 3.90738909645682e-07 | 31 |
| stk352 | 1.68154500787037e-11 | 0.357649879 | 0.195 | 0.055 | 4.23228063030894e-07 | 31 |
| rhogb7 | 1.78771313235287e-11 | 0.393052596 | 0.28 | 0.097 | 4.49949518281895e-07 | 31 |
| tagapa7 | 1.87684511174214e-11 | 0.441792217 | 0.297 | 0.105 | 4.72383146174378e-07 | 31 |
| myl12.110 | 2.4000823341624e-11 | 0.468862794 | 0.703 | 0.391 | 6.04076722685334e-07 | 31 |
| foxp1b | 2.9815958211002e-11 | 0.264268262 | 0.153 | 0.036 | 7.50437852212709e-07 | 31 |
| nfkbiab5 | 3.01995243842711e-11 | 0.432769514 | 0.424 | 0.181 | 7.60091829227718e-07 | 31 |
| sept9b2 | 3.50239928276059e-11 | 0.31481159 | 0.178 | 0.048 | 8.81518875478012e-07 | 31 |
| wipf1b4 | 3.98299986871668e-11 | 0.310340294 | 0.212 | 0.063 | 1.0024812369573e-06 | 31 |

| | | | | | | |
|---|---|---|---|---|---|---|
| slc25a58 | 4.34038002882801e-11 | 0.286699266 | 0.983 | 0.806 | 1.09243024945572e-06 | 31 |
| irf2bp2b5 | 6.24851903392815e-11 | 0.365227993 | 0.246 | 0.082 | 1.57268975564938e-06 | 31 |
| cirbpb1 | 6.49493094130009e-11 | 0.305977208 | 0.915 | 0.758 | 1.63470916861582e-06 | 31 |
| capzb9 | 8.23754654059894e-11 | 0.457659789 | 0.5 | 0.253 | 2.07330808880335e-06 | 31 |
| mfsd4aa | 9.01447652898479e-11 | 0.261215629 | 0.153 | 0.036 | 2.26885359758018e-06 | 31 |
| fmnl1a3 | 1.04577916950301e-10 | 0.307856317 | 0.153 | 0.039 | 2.63212159172213e-06 | 31 |
| sept122 | 1.49807711222694e-10 | 0.316023442 | 0.203 | 0.061 | 3.77051028376397e-06 | 31 |
| romo112 | 1.53977175753415e-10 | 0.447494302 | 0.661 | 0.385 | 3.87545153653771e-06 | 31 |
| eif5a7 | 1.5993740813502e-10 | 0.423124385 | 0.737 | 0.478 | 4.02546462535031e-06 | 31 |
| mcl1a8 | 2.50802977756615e-10 | 0.394302111 | 0.898 | 0.623 | 6.31246014715623e-06 | 31 |
| CR383676.116 | 2.51021403412657e-10 | 0.574624223 | 0.992 | 0.725 | 6.31795770249316e-06 | 31 |
| ran3 | 2.64398493941694e-10 | 0.339541496 | 0.831 | 0.644 | 6.65464569401849e-06 | 31 |
| rbm39a5 | 3.39686414221112e-10 | 0.410629734 | 0.339 | 0.143 | 8.54956735953117e-06 | 31 |
| capza1a8 | 3.55790985325789e-10 | 0.423313548 | 0.424 | 0.2 | 8.95490330966478e-06 | 31 |
| gnb1a4 | 4.03396651677576e-10 | 0.436457953 | 0.407 | 0.192 | 1.01530903260729e-05 | 31 |
| arf6b5 | 4.21314547272752e-10 | 0.376832145 | 0.271 | 0.101 | 1.06040658403079e-05 | 31 |
| hmha1b6 | 5.31255989503305e-10 | 0.295302724 | 0.229 | 0.075 | 1.33711819998087e-05 | 31 |
| tefa7 | 7.36113141047426e-10 | 0.392378368 | 0.331 | 0.138 | 1.85272316470227e-05 | 31 |
| kdm6bb2 | 8.28639406235187e-10 | 0.281805504 | 0.178 | 0.052 | 2.08560252155334e-05 | 31 |
| COX7A2 (1 of many)5 | 9.52046958518126e-10 | 0.41324484 | 0.508 | 0.26 | 2.39620698989427e-05 | 31 |
| arl8bb1 | 1.05350113139263e-09 | 0.274475903 | 0.153 | 0.041 | 2.6515569976021e-05 | 31 |
| sarnp1 | 1.07801331707992e-09 | 0.405689647 | 0.39 | 0.177 | 2.71325171775846e-05 | 31 |
| arrdc3b1 | 1.140751525849e-09 | 0.258314219 | 0.144 | 0.037 | 2.87115751540935e-05 | 31 |
| fgfr1op2 | 1.55498590847614e-09 | 0.310084959 | 0.195 | 0.061 | 3.91374403304359e-05 | 31 |
| FERMT3 (1 of many)4 | 1.5717981352582e-09 | 0.262870908 | 0.153 | 0.041 | 3.95605872663138e-05 | 31 |
| mibp210 | 1.6967489369921e-09 | 0.405593841 | 0.746 | 0.492 | 4.27054739951541e-05 | 31 |
| zgc:6405110 | 1.98036330985527e-09 | 0.391573874 | 0.373 | 0.162 | 4.98437641457474e-05 | 31 |
| naa101 | 2.43895473727574e-09 | 0.393850677 | 0.458 | 0.222 | 6.1386051782493e-05 | 31 |
| ss18 | 2.7848076126333e-09 | 0.274119972 | 0.212 | 0.069 | 7.00908228023675e-05 | 31 |
| dub6 | 4.38712316700121e-09 | 0.293249109 | 0.212 | 0.071 | 0.00011042 | 31 |
| pycard6 | 4.58454500349829e-09 | 0.322968193 | 0.212 | 0.072 | 0.000115388 | 31 |
| ero1b1 | 4.70280746637131e-09 | 0.284148237 | 0.144 | 0.039 | 0.000118365 | 31 |
| eif3k3 | 4.90819934794325e-09 | 0.376846689 | 0.458 | 0.229 | 0.000123534 | 31 |
| ppp3ccb | 4.97415163498686e-09 | 0.384292596 | 0.297 | 0.123 | 0.000125194 | 31 |
| rpl22l19 | 5.05035812507241e-09 | 0.418144324 | 0.669 | 0.391 | 0.000127112 | 31 |

| | | | | | | |
|---|---|---|---|---|---|---|
| si:dkey-112e7.28 | 7.72000446843107e-09 | 0.396169049 | 0.415 | 0.206 | 0.000194305 | 31 |
| skap26 | 9.49723597579227e-09 | 0.33902043 | 0.212 | 0.075 | 0.000239036 | 31 |
| smox5 | 9.61277640736375e-09 | 0.275404589 | 0.237 | 0.085 | 0.000241944 | 31 |
| acin1a6 | 1.06057972321656e-08 | 0.394067279 | 0.432 | 0.216 | 0.000266937 | 31 |
| nrros4 | 1.10262069848067e-08 | 0.282964967 | 0.161 | 0.048 | 0.000277519 | 31 |
| bcl2l104 | 1.25918396740969e-08 | 0.430206367 | 0.432 | 0.215 | 0.000316924 | 31 |
| arpc4l9 | 1.25991171901045e-08 | 0.39181496 | 0.398 | 0.197 | 0.000317107 | 31 |
| XLOC-0283708 | 1.33553141991791e-08 | 0.382840075 | 0.331 | 0.141 | 0.00033614 | 31 |
| eif3ea8 | 1.45216533872398e-08 | 0.39806842 | 0.5 | 0.277 | 0.000365495 | 31 |
| plekho1b3 | 1.62504303454675e-08 | 0.290437681 | 0.144 | 0.041 | 0.000409007 | 31 |
| lbr | 1.73958690773505e-08 | 0.299543535 | 0.22 | 0.078 | 0.000437837 | 31 |
| tubb2b2 | 2.27628617874947e-08 | 0.450997827 | 0.475 | 0.249 | 0.000572918 | 31 |
| arpc5a7 | 2.3353495205409e-08 | 0.32150834 | 0.28 | 0.115 | 0.000587784 | 31 |
| calm1b8 | 2.42070455154547e-08 | 0.265731149 | 0.254 | 0.095 | 0.000609267 | 31 |
| crlf31 | 2.95223049107055e-08 | 0.261384171 | 0.169 | 0.054 | 0.000743047 | 31 |
| ldha4 | 3.93197612986662e-08 | 0.328375768 | 0.203 | 0.073 | 0.000989639 | 31 |
| CABZ01090890.11 | 4.41050254844912e-08 | 0.273622638 | 0.169 | 0.055 | 0.001110079 | 31 |
| ahcy1 | 5.25251662272379e-08 | 0.366716263 | 0.356 | 0.168 | 0.001322006 | 31 |
| XLOC-0425965 | 5.43825550225128e-08 | 0.250534346 | 0.144 | 0.042 | 0.001368755 | 31 |
| ier2a11 | 6.72987674584225e-08 | 0.409846022 | 0.483 | 0.262 | 0.001693843 | 31 |
| sec61g4 | 6.79737731727017e-08 | 0.324055875 | 0.907 | 0.678 | 0.001710832 | 31 |
| mki671 | 7.52948686288078e-08 | 0.25893366 | 0.144 | 0.042 | 0.001895097 | 31 |
| mych3 | 7.96080949309219e-08 | 0.330896731 | 0.203 | 0.075 | 0.002003656 | 31 |
| si:dkey-262k9.48 | 8.39521528317926e-08 | 0.349430181 | 0.229 | 0.091 | 0.002112992 | 31 |
| jmjd1cb | 8.61851993367485e-08 | 0.272042541 | 0.144 | 0.044 | 0.002169195 | 31 |
| CR381673.17 | 8.89258961660383e-08 | 0.318143123 | 0.169 | 0.057 | 0.002238176 | 31 |
| bhlhe4010 | 1.10247426961437e-07 | 0.456938611 | 0.322 | 0.161 | 0.002774817 | 31 |
| capza1b6 | 1.18955469442316e-07 | 0.34898442 | 0.415 | 0.213 | 0.00299399 | 31 |
| ddit310 | 1.290335367714e-07 | 0.434557123 | 0.347 | 0.177 | 0.003247645 | 31 |
| anapc16 | 1.98926393383685e-07 | 0.289373959 | 0.246 | 0.101 | 0.005006778 | 31 |
| grasp5 | 2.25096109901239e-07 | 0.315450429 | 0.203 | 0.077 | 0.005665444 | 31 |
| eef1da9 | 2.4548361159566e-07 | 0.37631299 | 0.797 | 0.491 | 0.006178577 | 31 |
| rpl5b9 | 2.66347770422587e-07 | 0.333013929 | 0.856 | 0.63 | 0.006703707 | 31 |
| ywhaba5 | 3.29450232454543e-07 | 0.36933469 | 0.534 | 0.321 | 0.008291933 | 31 |
| lasp18 | 4.99353860159627e-07 | 0.360689222 | 0.424 | 0.237 | 0.012568237 | 31 |

| | | | | | | |
|---|---|---|---|---|---|---|
| wasf21 | 5.55991412096265e-07 | 0.270493797 | 0.195 | 0.074 | 0.013993748 | 31 |
| zgc:926062 | 5.89301147518156e-07 | 0.317037502 | 0.212 | 0.087 | 0.014832121 | 31 |
| pabpc1a11 | 6.88269719430111e-07 | 0.347993397 | 0.78 | 0.54 | 0.017323061 | 31 |
| eif3g6 | 8.17078564649816e-07 | 0.364506016 | 0.475 | 0.282 | 0.02056505 | 31 |
| smc1al1 | 8.75414647779526e-07 | 0.262316271 | 0.229 | 0.095 | 0.022033311 | 31 |
| msna11 | 9.79303056371037e-07 | 0.345332926 | 0.432 | 0.242 | 0.024648079 | 31 |
| arf6a1 | 9.99203561769535e-07 | 0.314738078 | 0.339 | 0.164 | 0.025148954 | 31 |
| zgc:564938 | 1.12767292851864e-06 | 0.326439344 | 0.788 | 0.572 | 0.0283824 | 31 |
| si:ch211-214j24.102 | 1.46327120873486e-06 | 0.279901573 | 0.331 | 0.159 | 0.036829073 | 31 |
| zfp36l1a9 | 1.52349606133943e-06 | 0.340528764 | 0.449 | 0.248 | 0.038344872 | 31 |
| rpl48 | 1.5351005823755e-06 | 0.309857662 | 0.864 | 0.627 | 0.038636947 | 31 |
| XLOC-0254239 | 1.81589584377819e-06 | 0.332756083 | 0.347 | 0.17 | 0.045704282 | 31 |
| btg28 | 3.17906935632489e-06 | 0.331407136 | 0.636 | 0.373 | 0.080013997 | 31 |
| ankrd111 | 3.56300378435377e-06 | 0.264419528 | 0.212 | 0.091 | 0.089677242 | 31 |
| erbin1 | 3.95428090381483e-06 | 0.25809284 | 0.153 | 0.056 | 0.099525296 | 31 |
| magt12 | 4.08459842562612e-06 | 0.311074672 | 0.246 | 0.115 | 0.102805258 | 31 |
| arpc310 | 4.51361994254444e-06 | 0.328658453 | 0.517 | 0.311 | 0.1136033 | 31 |
| si:ch211-199g17.22 | 5.32023905918632e-06 | 0.270299269 | 0.169 | 0.066 | 0.133905097 | 31 |
| irf2bp2a5 | 5.54324137163094e-06 | 0.355118541 | 0.322 | 0.177 | 0.139517842 | 31 |
| eif3c6 | 7.23435272422007e-06 | 0.288044786 | 0.322 | 0.169 | 0.182081424 | 31 |
| sumo2b9 | 7.48894521551645e-06 | 0.32758574 | 0.441 | 0.27 | 0.188489262 | 31 |
| ifi304 | 1.15152049387491e-05 | 0.275243053 | 0.339 | 0.175 | 0.289826193 | 31 |
| sat1a.25 | 1.30902120802741e-05 | 0.316057769 | 0.492 | 0.299 | 0.329467548 | 31 |
| itm2bb8 | 1.44506736194282e-05 | 0.289560859 | 0.398 | 0.223 | 0.363709004 | 31 |
| pgam1a5 | 1.47144377279133e-05 | 0.320252238 | 0.297 | 0.158 | 0.370347683 | 31 |
| hmgb1b7 | 1.7753302915246e-05 | 0.300575701 | 0.254 | 0.127 | 0.446832881 | 31 |
| arpc210 | 1.84714607498899e-05 | 0.298311731 | 0.449 | 0.268 | 0.464908196 | 31 |
| mob3a1 | 1.87369320959359e-05 | 0.259668081 | 0.305 | 0.155 | 0.471589844 | 31 |
| arhgdia9 | 1.97509637117424e-05 | 0.308187196 | 0.39 | 0.23 | 0.497112006 | 31 |
| b2m11 | 1.99849356422122e-05 | 0.310130244 | 0.483 | 0.293 | 0.503000845 | 31 |
| rsl1d13 | 2.11978642844874e-05 | 0.300325575 | 0.331 | 0.182 | 0.533529046 | 31 |
| rad21a1 | 2.20622251350229e-05 | 0.292224295 | 0.254 | 0.128 | 0.555284144 | 31 |
| NC-002333.415 | 2.34486639507955e-05 | 0.283781103 | 0.856 | 0.622 | 0.590179423 | 31 |
| sh3kbp14 | 2.52709549678892e-05 | 0.257178536 | 0.169 | 0.072 | 0.636044666 | 31 |
| h3f3c9 | 3.43088599177569e-05 | 0.309480719 | 0.415 | 0.257 | 0.863519695 | 31 |

| | | | | | | |
|---|---|---|---|---|---|---|
| crebzf3 | 3.44704960261523e-05 | 0.292341043 | 0.212 | 0.103 | 0.867587914 | 31 |
| aldoaa3 | 3.57602547217701e-05 | 0.268045248 | 0.432 | 0.251 | 0.900049851 | 31 |
| hmga1a3 | 3.67740326855222e-05 | 0.331768377 | 0.619 | 0.451 | 0.925565629 | 31 |
| atf4b10 | 4.23451957744968e-05 | 0.291468973 | 0.432 | 0.265 | 1 | 31 |
| arl5c6 | 4.79960418487146e-05 | 0.275600849 | 0.229 | 0.113 | 1 | 31 |
| prelid3b3 | 4.81084285169309e-05 | 0.293579165 | 0.398 | 0.239 | 1 | 31 |
| selenok3 | 4.88319712862704e-05 | 0.281743215 | 0.246 | 0.128 | 1 | 31 |
| eif1b8 | 5.81125269363905e-05 | 0.292786125 | 0.576 | 0.383 | 1 | 31 |
| cox6b24 | 7.03349716442334e-05 | 0.294013363 | 0.78 | 0.533 | 1 | 31 |
| hnrnpa0b5 | 8.36552408596704e-05 | 0.304665905 | 0.466 | 0.312 | 1 | 31 |
| cfl19 | 9.57157360564373e-05 | 0.277516956 | 0.797 | 0.577 | 1 | 31 |
| eif4a1b7 | 0.000101029 | 0.26964273 | 0.449 | 0.281 | 1 | 31 |
| si:ch73-1a9.312 | 0.000148654 | 0.325405408 | 0.398 | 0.267 | 1 | 31 |
| eif3ha3 | 0.000154463 | 0.26386622 | 0.347 | 0.209 | 1 | 31 |
| cdc42l6 | 0.000157755 | 0.273525421 | 0.356 | 0.216 | 1 | 31 |
| hmgn611 | 0.000162131 | 0.301042818 | 0.398 | 0.266 | 1 | 31 |
| gng55 | 0.00019165 | 0.254368935 | 0.432 | 0.272 | 1 | 31 |
| ola13 | 0.000226607 | 0.270623129 | 0.364 | 0.23 | 1 | 31 |
| rps27.212 | 0.000300497 | 0.291835926 | 0.763 | 0.517 | 1 | 31 |
| cox7b6 | 0.000325722 | 0.258823399 | 0.619 | 0.417 | 1 | 31 |
| mt-atp616 | 0.000350374 | 0.25983866 | 0.856 | 0.59 | 1 | 31 |
| litaf6 | 0.000677046 | 0.251531862 | 0.237 | 0.136 | 1 | 31 |
| si:ch211-222l21.112 | 0.000710128 | 0.297537661 | 0.398 | 0.277 | 1 | 31 |
| hmgn78 | 0.001048409 | 0.252215891 | 0.407 | 0.281 | 1 | 31 |
| mt-co216 | 0.003026413 | 0.277300421 | 0.975 | 0.719 | 1 | 31 |
| hmgb2a3 | 0.007681781 | 0.27888677 | 0.686 | 0.623 | 1 | 31 |
| lyve1a | 0 | 1.487025921 | 0.5 | 0.002 | 0 | 32 |
| kdr | 0 | 1.049350558 | 0.491 | 0.001 | 0 | 32 |
| she | 0 | 0.948012732 | 0.389 | 0.001 | 0 | 32 |
| mrc1a | 0 | 0.923984763 | 0.37 | 0.004 | 0 | 32 |
| lyve1b | 0 | 0.899992593 | 0.352 | 0 | 0 | 32 |
| si:dkeyp-97a10.2 | 0 | 0.865199458 | 0.361 | 0.003 | 0 | 32 |
| flt4 | 0 | 0.860025422 | 0.37 | 0.002 | 0 | 32 |
| tmem88a | 0 | 0.854415508 | 0.38 | 0.003 | 0 | 32 |
| pcdh12 | 0 | 0.752364431 | 0.315 | 0.001 | 0 | 32 |

| | | | | | | |
|---|---|---|---|---|---|---|
| CT033796.1 | 0 | 0.727997112 | 0.296 | 0.001 | 0 | 32 |
| myct1a | 0 | 0.721923959 | 0.306 | 0.002 | 0 | 32 |
| akap12a | 0 | 0.719177264 | 0.287 | 0.002 | 0 | 32 |
| lgi2a | 0 | 0.706367577 | 0.278 | 0.001 | 0 | 32 |
| si:ch211-152c2.3 | 0 | 0.679258346 | 0.25 | 0.002 | 0 | 32 |
| rca2.2 | 0 | 0.674620621 | 0.25 | 0 | 0 | 32 |
| egfl7 | 0 | 0.646175485 | 0.269 | 0.002 | 0 | 32 |
| clec14a | 0 | 0.631155247 | 0.269 | 0.001 | 0 | 32 |
| rasip1 | 0 | 0.623613441 | 0.269 | 0.001 | 0 | 32 |
| gpr182 | 1.09069502984266e-307 | 0.441524802 | 0.157 | 0 | 2.745170320611e-303 | 32 |
| cldn5b | 2.83307080531059e-279 | 0.891924172 | 0.352 | 0.007 | 7.13055590988622e-275 | 32 |
| parvab | 9.34508350008496e-278 | 0.640877827 | 0.278 | 0.004 | 2.35206406613638e-273 | 32 |
| si:dkey-237i9.8 | 2.58347165987995e-273 | 0.474120206 | 0.176 | 0.001 | 6.50233982075184e-269 | 32 |
| ldb2a | 1.24251123188831e-268 | 0.568969976 | 0.231 | 0.002 | 3.12727651953968e-264 | 32 |
| ehd3 | 4.57369283106205e-266 | 0.970758248 | 0.417 | 0.011 | 1.15115274865001e-261 | 32 |
| stab1 | 2.0620907289557e-265 | 0.677179445 | 0.259 | 0.003 | 5.19007615570859e-261 | 32 |
| si:dkey-126g1.9 | 3.13529473048782e-262 | 0.517211909 | 0.204 | 0.002 | 7.8912233071648e-258 | 32 |
| sele | 1.40523969248231e-257 | 0.597253169 | 0.222 | 0.002 | 3.53684778200872e-253 | 32 |
| sox18 | 5.95558437067461e-250 | 0.82514726 | 0.343 | 0.007 | 1.49896103025509e-245 | 32 |
| kdrl | 9.12697082994801e-249 | 0.567676072 | 0.185 | 0.001 | 2.29716728818961e-244 | 32 |
| igfbp72 | 2.01118228578729e-245 | 1.238546617 | 0.546 | 0.023 | 5.06194469509802e-241 | 32 |
| cdh5 | 3.78101438299736e-221 | 0.513248824 | 0.167 | 0.001 | 9.51643510056605e-217 | 32 |
| si:ch211-248e11.2 | 2.15422381968243e-219 | 0.658482152 | 0.278 | 0.005 | 5.42196593175871e-215 | 32 |
| etv2 | 6.39021279413664e-217 | 0.551900746 | 0.25 | 0.004 | 1.60835265815625e-212 | 32 |
| fabp11a5 | 1.52491132114206e-214 | 1.833676002 | 0.889 | 0.077 | 3.83804930418244e-210 | 32 |
| tie1 | 4.94823670607627e-206 | 0.489215158 | 0.194 | 0.002 | 1.24542169655234e-201 | 32 |
| igf2a | 2.38693436854713e-204 | 0.689491101 | 0.259 | 0.005 | 6.00767511219627e-200 | 32 |
| tm4sf18 | 4.11010067816445e-193 | 0.632496705 | 0.241 | 0.004 | 1.03447123968721e-188 | 32 |
| rab11bb | 6.00490835084183e-190 | 0.88177667 | 0.389 | 0.014 | 1.51137538282338e-185 | 32 |
| b3gnt7 | 1.32732695541237e-189 | 0.418048549 | 0.167 | 0.002 | 3.34074921407738e-185 | 32 |
| plk2b | 3.43772374953382e-186 | 0.80324014 | 0.324 | 0.01 | 8.65240690520166e-182 | 32 |
| fgd5a | 4.26250254193364e-186 | 0.720040444 | 0.333 | 0.01 | 1.07282926477928e-181 | 32 |
| pecam1 | 4.57391069061114e-183 | 0.373506602 | 0.139 | 0.001 | 1.15120758171992e-178 | 32 |
| cdh62 | 1.9202779574343e-181 | 0.990811914 | 0.407 | 0.017 | 4.83314759106639e-177 | 32 |
| exoc3l2a | 1.04450472596861e-180 | 0.44224756 | 0.13 | 0.001 | 2.6289139447904e-176 | 32 |

| | | | | | | |
|---|---|---|---|---|---|---|
| ppp1r13ba | 1.22296649122127e-174 | 0.481909607 | 0.185 | 0.003 | 3.07808436175482e-170 | 32 |
| nrarpb | 9.32736586669064e-174 | 0.467798902 | 0.194 | 0.003 | 2.34760471498737e-169 | 32 |
| shank3b | 9.13334446431691e-173 | 0.28757371 | 0.13 | 0.001 | 2.29877146822392e-168 | 32 |
| podn | 5.58025489738912e-171 | 0.449519293 | 0.185 | 0.003 | 1.40449435512387e-166 | 32 |
| si:ch211-33e4.2 | 1.90439493181435e-168 | 0.366922964 | 0.157 | 0.002 | 4.79317160388353e-164 | 32 |
| ramp23 | 5.92521281062918e-162 | 1.238684555 | 0.556 | 0.038 | 1.49131681230726e-157 | 32 |
| si:ch211-14k19.8 | 2.58428956873006e-159 | 0.313347412 | 0.13 | 0.001 | 6.5043984155367e-155 | 32 |
| nr5a2 | 8.5396964251942e-158 | 0.429945001 | 0.167 | 0.002 | 2.14935619325713e-153 | 32 |
| loxl12 | 3.55489784487416e-157 | 1.036654772 | 0.407 | 0.02 | 8.94732238576378e-153 | 32 |
| fam174b | 5.17677575685131e-157 | 0.7419593 | 0.333 | 0.012 | 1.30294269024191e-152 | 32 |
| CU929150.13 | 7.33396205628756e-149 | 1.125008636 | 0.417 | 0.022 | 1.84588490994702e-144 | 32 |
| erg | 9.18057640058536e-149 | 0.697211416 | 0.296 | 0.01 | 2.31065927426333e-144 | 32 |
| plk2a | 4.14579295294051e-146 | 0.450922654 | 0.185 | 0.004 | 1.0434546283256e-141 | 32 |
| s1pr1 | 1.79927314803274e-143 | 0.578858038 | 0.231 | 0.006 | 4.52859058628361e-139 | 32 |
| arhgef9b | 3.39396899961145e-139 | 0.352997482 | 0.139 | 0.002 | 8.54228057512206e-135 | 32 |
| notchl | 1.11061302761158e-138 | 0.271494695 | 0.111 | 0.001 | 2.7953019291956e-134 | 32 |
| creld1a | 1.14461734755324e-138 | 0.273835631 | 0.111 | 0.001 | 2.88088740205674e-134 | 32 |
| si:dkey-28n18.9 | 1.46589405711659e-137 | 0.364429197 | 0.148 | 0.002 | 3.68950875235675e-133 | 32 |
| dll4 | 3.74383655124043e-136 | 0.442304877 | 0.185 | 0.004 | 9.42286221581705e-132 | 32 |
| slc9a3r1b | 8.59483298436196e-136 | 0.535411465 | 0.204 | 0.005 | 2.16323351383406e-131 | 32 |
| hapln31 | 9.37296474542962e-135 | 0.898193766 | 0.333 | 0.015 | 2.35908149677718e-130 | 32 |
| pgm5 | 9.41815783087133e-130 | 0.512544166 | 0.204 | 0.005 | 2.370456144452e-125 | 32 |
| gpm6ab1 | 1.30129803773901e-126 | 0.750915659 | 0.324 | 0.015 | 3.27523703118531e-122 | 32 |
| piezo2a.2 | 3.05077032484902e-126 | 0.598238945 | 0.25 | 0.009 | 7.6784838306125e-122 | 32 |
| ecscr | 1.05830649293674e-125 | 0.406382462 | 0.157 | 0.003 | 2.66365161207248e-121 | 32 |
| cyp26b11 | 7.55150400807472e-120 | 0.536608878 | 0.185 | 0.005 | 1.90063804379233e-115 | 32 |
| reps2 | 7.94841267683581e-119 | 0.384433572 | 0.148 | 0.003 | 2.0005359866328e-114 | 32 |
| rassf9 | 4.30528660082413e-118 | 0.421469747 | 0.167 | 0.004 | 1.08359758456142e-113 | 32 |
| ednraa1 | 6.52961203027467e-113 | 0.679450259 | 0.296 | 0.014 | 1.64343805189983e-108 | 32 |
| sox7 | 1.03420327325525e-106 | 0.346870114 | 0.139 | 0.003 | 2.60298621845614e-102 | 32 |
| anks1b1 | 6.78389180470241e-105 | 0.651002341 | 0.296 | 0.015 | 1.70743772832555e-100 | 32 |
| cyth3a | 1.19393105624018e-104 | 0.374581929 | 0.148 | 0.003 | 3.0050050754509e-100 | 32 |
| cx39.4 | 1.39726191789105e-104 | 0.273969683 | 0.111 | 0.002 | 3.51676852113998e-100 | 32 |
| si:ch211-145b13.61 | 2.65505696959215e-100 | 0.456576656 | 0.176 | 0.005 | 6.68251288676648e-96 | 32 |
| tmem108 | 1.11942416622205e-99 | 0.407645056 | 0.167 | 0.005 | 2.81747868396428e-95 | 32 |

| | | | | | | |
|---|---|---|---|---|---|---|
| LOC1018822641 | 1.98252505574826e-98 | 0.560085972 | 0.222 | 0.009 | 4.98981731281281e-94 | 32 |
| macf1a1 | 7.04678253819997e-95 | 0.963458142 | 0.491 | 0.05 | 1.77360469703955e-90 | 32 |
| fzd4 | 9.27583935929416e-94 | 0.475748698 | 0.204 | 0.008 | 2.33463600834075e-89 | 32 |
| c1qtnf6a2 | 2.14788844541509e-93 | 0.709119632 | 0.296 | 0.018 | 5.40602042826523e-89 | 32 |
| afap1l1b | 5.10237272280164e-93 | 0.523574301 | 0.204 | 0.008 | 1.28421619060195e-88 | 32 |
| hhex | 4.68594632280799e-92 | 0.428138301 | 0.185 | 0.007 | 1.17940582998754e-87 | 32 |
| calcrlb | 2.10243068632052e-90 | 0.578187379 | 0.25 | 0.013 | 5.29160779440011e-86 | 32 |
| fli1a | 2.8293766711717e-90 | 0.436710301 | 0.176 | 0.006 | 7.12125814367206e-86 | 32 |
| pcdh17 | 8.65165142274977e-90 | 0.37333809 | 0.148 | 0.004 | 2.17753414659189e-85 | 32 |
| aqp8a.1 | 1.79396012427189e-88 | 0.455784901 | 0.13 | 0.003 | 4.51521823677993e-84 | 32 |
| si:ch211-207d6.21 | 3.82127285537179e-88 | 0.551322616 | 0.241 | 0.012 | 9.61776164968525e-84 | 32 |
| si:ch73-334d15.2 | 2.16966207669335e-87 | 0.398259595 | 0.157 | 0.005 | 5.46082248082949e-83 | 32 |
| gas62 | 9.23543002072501e-85 | 0.729500811 | 0.315 | 0.022 | 2.32446538191628e-80 | 32 |
| ebf3a1 | 2.5924496576117e-84 | 0.430227616 | 0.185 | 0.007 | 6.52493654324289e-80 | 32 |
| nova22 | 4.39071098892893e-83 | 0.80423347 | 0.352 | 0.029 | 1.10509804880352e-78 | 32 |
| rapgef5b | 3.52795126965011e-81 | 0.28027373 | 0.111 | 0.002 | 8.87950055058235e-77 | 32 |
| cldn11b | 3.54237038208051e-80 | 0.498199888 | 0.204 | 0.01 | 8.91579201465845e-76 | 32 |
| limch1b | 1.38316552177162e-78 | 0.593568732 | 0.259 | 0.016 | 3.48128930174698e-74 | 32 |
| akap12b5 | 1.27974620251676e-77 | 0.949143797 | 0.481 | 0.057 | 3.22099321711442e-73 | 32 |
| hyal2b | 3.43108574729766e-77 | 0.521123986 | 0.204 | 0.01 | 8.63569971737347e-73 | 32 |
| tnfaip2b | 1.19007258076323e-76 | 0.391231043 | 0.13 | 0.004 | 2.99529367852298e-72 | 32 |
| selenop1 | 4.13685822229768e-73 | 0.648796798 | 0.259 | 0.018 | 1.0412058459701e-68 | 32 |
| marcksl1b8 | 7.52722274255554e-73 | 1.190896591 | 0.639 | 0.113 | 1.8945266920738e-68 | 32 |
| laynb | 8.01451195953617e-70 | 0.295555006 | 0.111 | 0.003 | 2.01717251509566e-65 | 32 |
| cpn14 | 1.15437354138659e-69 | 0.880477568 | 0.435 | 0.053 | 2.90544276631591e-65 | 32 |
| lmo21 | 1.25337843112064e-68 | 0.593778639 | 0.259 | 0.019 | 3.15462817328754e-64 | 32 |
| reep1 | 3.55001267482279e-68 | 0.368381677 | 0.157 | 0.007 | 8.93502690126148e-64 | 32 |
| XLOC-015238 | 5.68396629681163e-68 | 0.279229582 | 0.111 | 0.003 | 1.43059747724452e-63 | 32 |
| cxcl12a4 | 2.6350593034832e-67 | 1.230574568 | 0.528 | 0.08 | 6.63218076093686e-63 | 32 |
| ccl38.11 | 5.01100605137328e-66 | 0.4591878 | 0.157 | 0.007 | 1.26122011307014e-61 | 32 |
| grb10a3 | 1.09588306550671e-63 | 0.802318792 | 0.352 | 0.037 | 2.75822808757383e-59 | 32 |
| tspan4b1 | 1.33931905065264e-63 | 0.876833468 | 0.463 | 0.064 | 3.37093211858763e-59 | 32 |
| adam8a3 | 7.30360730435344e-63 | 0.857568836 | 0.407 | 0.05 | 1.83824492243272e-58 | 32 |
| tpm4a6 | 8.06655301571746e-62 | 0.958889408 | 0.528 | 0.086 | 2.03027072852593e-57 | 32 |
| yrk5 | 2.11850176093509e-60 | 0.83419659 | 0.444 | 0.061 | 5.33205708209754e-56 | 32 |

| | | | | | | |
|---|---|---|---|---|---|---|
| prox1a1 | 5.10598673622991e-59 | 0.547864684 | 0.231 | 0.017 | 1.28512580164171e-54 | 32 |
| klf2a6 | 4.87698782628982e-58 | 1.031551335 | 0.5 | 0.083 | 1.22748906599889e-53 | 32 |
| SEMA4F | 7.7494637515646e-58 | 0.327550949 | 0.12 | 0.004 | 1.95046253163129e-53 | 32 |
| plvapb1 | 2.93516749170719e-57 | 0.637509113 | 0.204 | 0.014 | 7.38752305987783e-53 | 32 |
| tcima5 | 3.99474760723873e-57 | 1.05182339 | 0.444 | 0.066 | 1.00543802526592e-52 | 32 |
| arl4ab | 2.82806467546813e-56 | 0.462275578 | 0.204 | 0.014 | 7.11795598168574e-52 | 32 |
| dab2ipb | 3.65948643426194e-56 | 0.474896851 | 0.194 | 0.013 | 9.21056140639388e-52 | 32 |
| lfng1 | 5.85072231654697e-56 | 0.374355083 | 0.148 | 0.007 | 1.47256829985171e-51 | 32 |
| cdc42ep5 | 2.27337159477311e-55 | 0.687991995 | 0.296 | 0.03 | 5.72184896688443e-51 | 32 |
| oaz2b | 8.0636262062156e-55 | 0.341397888 | 0.139 | 0.006 | 2.0295340798424e-50 | 32 |
| sema3ga | 9.97504733152014e-55 | 0.34512631 | 0.13 | 0.006 | 2.5106196628703e-50 | 32 |
| si:ch211-1a19.35 | 1.15790201251484e-54 | 0.736844952 | 0.389 | 0.052 | 2.91432357529861e-50 | 32 |
| vat17 | 2.59322093396426e-53 | 1.005509368 | 0.611 | 0.132 | 6.52687776869466e-49 | 32 |
| si:ch211-66e2.36 | 4.05597862486636e-52 | 0.788762467 | 0.398 | 0.057 | 1.02084926009261e-47 | 32 |
| fam198b | 1.69853145053267e-50 | 0.272445967 | 0.12 | 0.005 | 4.27503380784569e-46 | 32 |
| exoc61 | 1.89897588380796e-50 | 0.660518327 | 0.296 | 0.033 | 4.77953240195626e-46 | 32 |
| acap3a | 5.77156788370425e-50 | 0.253682282 | 0.111 | 0.004 | 1.45264592064952e-45 | 32 |
| cd81b | 7.72335497048035e-50 | 0.360767436 | 0.157 | 0.009 | 1.9438912125202e-45 | 32 |
| ccdc85b7 | 1.45227995272746e-49 | 0.69220516 | 0.352 | 0.047 | 3.65524341301976e-45 | 32 |
| il13ra11 | 6.0198055094933e-49 | 0.559696762 | 0.278 | 0.03 | 1.51512484868437e-44 | 32 |
| krt810 | 1.81252038402486e-48 | 1.059021758 | 0.88 | 0.273 | 4.56193255455218e-44 | 32 |
| atp1b1a5 | 4.12500501729463e-48 | 1.061324027 | 0.704 | 0.211 | 1.03822251280289e-43 | 32 |
| marcksl1a8 | 5.09919820182338e-48 | 1.127794621 | 0.713 | 0.212 | 1.28341719541693e-43 | 32 |
| myo1ca3 | 1.69432357178646e-47 | 0.543383432 | 0.241 | 0.023 | 4.26444299782934e-43 | 32 |
| epas1b | 3.40710654002397e-46 | 0.434753213 | 0.157 | 0.01 | 8.57534645058634e-42 | 32 |
| gcnt4a1 | 3.61388096679663e-46 | 0.395454768 | 0.185 | 0.014 | 9.09577700533044e-42 | 32 |
| col15a1a3 | 1.18748098312332e-45 | 0.54136732 | 0.222 | 0.021 | 2.98877088642309e-41 | 32 |
| notch1b | 7.45523673619505e-45 | 0.314511079 | 0.139 | 0.008 | 1.87640853413293e-40 | 32 |
| lrch1 | 8.61951105187565e-45 | 0.446584612 | 0.185 | 0.015 | 2.16944473664658e-40 | 32 |
| palm21 | 6.99597692927774e-44 | 0.452188874 | 0.194 | 0.016 | 1.76081743332991e-39 | 32 |
| pdgfd | 7.1586741360886e-44 | 0.31495097 | 0.139 | 0.008 | 1.80176669331214e-39 | 32 |
| ralgapa2 | 7.45861199663452e-44 | 0.396392469 | 0.167 | 0.012 | 1.87725805343294e-39 | 32 |
| crip2 | 2.0042629869899e-43 | 0.430773445 | 0.185 | 0.015 | 5.04452951195487e-39 | 32 |
| CABZ01084131.1 | 2.26407507695753e-43 | 0.441584985 | 0.194 | 0.016 | 5.69845056119441e-39 | 32 |
| lzts2a1 | 1.48754894275526e-42 | 0.489873558 | 0.213 | 0.02 | 3.7440119340207e-38 | 32 |

| | | | | | | |
|---|---|---|---|---|---|---|
| spns2 | 1.68994433134808e-41 | 0.322609872 | 0.12 | 0.007 | 4.25342088756997e-37 | 32 |
| arhgap311 | 2.5647701873902e-41 | 0.524781876 | 0.25 | 0.029 | 6.45527008464239e-37 | 32 |
| npdc1a | 2.83270746355316e-41 | 0.360480398 | 0.176 | 0.014 | 7.12964141501695e-37 | 32 |
| foxp1b1 | 3.98840122148618e-41 | 0.592502566 | 0.278 | 0.035 | 1.00384070343586e-36 | 32 |
| stox2a | 6.68746540511215e-40 | 0.286159113 | 0.12 | 0.007 | 1.68316816781268e-35 | 32 |
| mafba6 | 2.49842279068162e-39 | 0.747724595 | 0.389 | 0.071 | 6.28828032186656e-35 | 32 |
| cttnbp2nlb | 4.37143132065347e-39 | 0.360968134 | 0.13 | 0.008 | 1.10024554909527e-34 | 32 |
| scarf1 | 4.55103916048417e-39 | 0.261722069 | 0.102 | 0.005 | 1.14545104630226e-34 | 32 |
| kank31 | 5.24033870398433e-39 | 0.40616578 | 0.185 | 0.017 | 1.31894084840581e-34 | 32 |
| adgrl3.11 | 9.28121123819421e-39 | 0.524686831 | 0.222 | 0.024 | 2.3359880565411e-34 | 32 |
| fam43a | 1.0962835471121e-38 | 0.368090473 | 0.139 | 0.01 | 2.75923605972645e-34 | 32 |
| krt18a.16 | 1.41995267762019e-37 | 0.838904671 | 0.546 | 0.125 | 3.57387889430225e-33 | 32 |
| spry45 | 2.24916589941099e-37 | 0.631927188 | 0.306 | 0.046 | 5.66092565222751e-33 | 32 |
| grapa2 | 2.2721714745013e-37 | 0.520093414 | 0.231 | 0.027 | 5.71882838417231e-33 | 32 |
| plekhg5a | 1.55940585202662e-36 | 0.314572012 | 0.13 | 0.009 | 3.92486858896579e-32 | 32 |
| fkbp1ab1 | 1.96422040782528e-36 | 0.736651396 | 0.361 | 0.063 | 4.94374634445546e-32 | 32 |
| CU984579.11 | 2.99797603318092e-36 | 0.399401679 | 0.176 | 0.016 | 7.54560587791306e-32 | 32 |
| heg1 | 7.18764707835054e-36 | 0.257951501 | 0.102 | 0.005 | 1.80905889315005e-31 | 32 |
| abcg2a | 7.70827001753362e-36 | 0.352154913 | 0.139 | 0.01 | 1.94009448071304e-31 | 32 |
| klf2b1 | 1.23570078842403e-35 | 0.49024075 | 0.194 | 0.02 | 3.11013531438444e-31 | 32 |
| dlc12 | 1.66150608305697e-35 | 0.510222592 | 0.241 | 0.03 | 4.18184466044608e-31 | 32 |
| XLOC-0160946 | 1.01801340107427e-34 | 0.712832325 | 0.398 | 0.078 | 2.56223792916382e-30 | 32 |
| arl4aa | 1.13078089612889e-34 | 0.644369565 | 0.343 | 0.058 | 2.84606243746681e-30 | 32 |
| sema4c | 2.52578527849592e-34 | 0.373728657 | 0.157 | 0.014 | 6.35714896744638e-30 | 32 |
| oaz2a | 5.7750527588418e-34 | 0.353893622 | 0.157 | 0.014 | 1.45352302887289e-29 | 32 |
| lgmn1 | 1.07444021422291e-33 | 0.686973932 | 0.333 | 0.059 | 2.70425857517764e-29 | 32 |
| cyp3c14 | 1.34812170429902e-33 | 0.657627474 | 0.333 | 0.059 | 3.3930875175502e-29 | 32 |
| c1s | 1.95774089956205e-33 | 0.309671168 | 0.139 | 0.011 | 4.92743807010773e-29 | 32 |
| wasf3b | 3.26410902048115e-33 | 0.446217821 | 0.194 | 0.021 | 8.21543599364901e-29 | 32 |
| rnd3a1 | 5.85945065185386e-33 | 0.535823356 | 0.231 | 0.03 | 1.4747651345651e-28 | 32 |
| pros1 | 1.0011702039674e-32 | 0.554479137 | 0.296 | 0.047 | 2.51984528636555e-28 | 32 |
| mrc1b3 | 1.17385932677941e-32 | 0.563398983 | 0.278 | 0.042 | 2.95448653957111e-28 | 32 |
| acsbg21 | 3.43472571191075e-32 | 0.511339229 | 0.259 | 0.038 | 8.64486114430817e-28 | 32 |
| si:ch211-156j16.12 | 4.83369663177961e-32 | 0.446107264 | 0.185 | 0.02 | 1.21659310525261e-27 | 32 |
| efna1a | 6.77197432710167e-32 | 0.373750127 | 0.148 | 0.013 | 1.70443821838822e-27 | 32 |

| | | | | | | |
|---|---|---|---|---|---|---|
| gstt1b5 | 1.05799199808406e-31 | 0.638870114 | 0.361 | 0.071 | 2.66286005997778e-27 | 32 |
| pmp22b3 | 3.21783252256978e-31 | 0.594267432 | 0.278 | 0.044 | 8.09896267605587e-27 | 32 |
| chka2 | 1.34927626871188e-30 | 0.436271643 | 0.204 | 0.025 | 3.39599344072093e-26 | 32 |
| cavin2b6 | 5.44923471632858e-30 | 0.65342495 | 0.333 | 0.065 | 1.37151788575274e-25 | 32 |
| pdk2b8 | 1.54784052250364e-29 | 0.783857107 | 0.407 | 0.098 | 3.8957598110894e-25 | 32 |
| tcf124 | 3.75392033676331e-29 | 0.605495886 | 0.306 | 0.057 | 9.44824209559958e-25 | 32 |
| LOC103911404 | 5.3483901127907e-28 | 0.285183685 | 0.102 | 0.007 | 1.34613630748829e-23 | 32 |
| si:ch211-71n6.4 | 5.35776335190628e-28 | 0.40347314 | 0.194 | 0.025 | 1.34849545804129e-23 | 32 |
| rps6ka3b | 8.24153770299526e-28 | 0.362847562 | 0.139 | 0.013 | 2.07431262446688e-23 | 32 |
| tead1b1 | 1.12249854170735e-27 | 0.445306091 | 0.204 | 0.028 | 2.82521657962322e-23 | 32 |
| clic25 | 1.12295889379913e-27 | 0.627729013 | 0.306 | 0.059 | 2.82637523980304e-23 | 32 |
| zgc:1533173 | 1.47150211416159e-27 | 0.544809642 | 0.25 | 0.041 | 3.70362367113332e-23 | 32 |
| carhsp11 | 1.51584792966534e-27 | 0.637843295 | 0.333 | 0.069 | 3.81523765417469e-23 | 32 |
| emilin1b1 | 2.34108886257615e-27 | 0.32520293 | 0.139 | 0.013 | 5.89228655821792e-23 | 32 |
| tagln210 | 2.51694609612739e-27 | 0.837090744 | 0.546 | 0.187 | 6.33490162934304e-23 | 32 |
| phlda22 | 3.20843125787839e-26 | 0.597838625 | 0.278 | 0.051 | 8.07530063295412e-22 | 32 |
| srgap1b | 4.3180695071427e-26 | 0.3454467 | 0.139 | 0.014 | 1.08681491425275e-21 | 32 |
| npc11 | 1.15919067517509e-25 | 0.423545838 | 0.194 | 0.027 | 2.91756701034819e-21 | 32 |
| add3a9 | 3.5398759693962e-25 | 0.737004734 | 0.435 | 0.123 | 8.9095138273733e-21 | 32 |
| CR383676.117 | 4.62789086695382e-25 | 0.667820951 | 0.935 | 0.726 | 1.16479385230361e-20 | 32 |
| aplp24 | 5.06312343713522e-25 | 0.611789979 | 0.324 | 0.072 | 1.27433753789256e-20 | 32 |
| aatka1 | 5.97858386330503e-25 | 0.270764638 | 0.12 | 0.011 | 1.50474977255524e-20 | 32 |
| erfl3 | 7.96117743160712e-25 | 0.261126036 | 0.102 | 0.008 | 2.0037487477612e-20 | 32 |
| cbx7a5 | 1.48021512646983e-24 | 0.635716946 | 0.269 | 0.052 | 3.7255534518119e-20 | 32 |
| gng103 | 1.62373507756208e-24 | 0.540631716 | 0.278 | 0.054 | 4.086778816716e-20 | 32 |
| tbc1d41 | 2.27831910848828e-24 | 0.355703553 | 0.157 | 0.019 | 5.73430136415416e-20 | 32 |
| lima1a | 8.21996445525081e-24 | 0.5038264 | 0.269 | 0.05 | 2.06888285374208e-19 | 32 |
| si:ch73-138n13.12 | 2.48448919190123e-23 | 0.467526095 | 0.222 | 0.038 | 6.2532108470962e-19 | 32 |
| ponzr19 | 2.59325247738397e-23 | 0.77965553 | 0.639 | 0.272 | 6.52695716032771e-19 | 32 |
| tmem88b4 | 2.60199548029213e-23 | 0.385398627 | 0.222 | 0.036 | 6.54896242434725e-19 | 32 |
| dhrs3a3 | 3.6137874549506e-23 | 0.394965433 | 0.194 | 0.03 | 9.09554164536515e-19 | 32 |
| rhoca | 5.45664406609203e-23 | 0.501299615 | 0.278 | 0.057 | 1.3733827449947e-18 | 32 |
| tshz23 | 7.21810228493258e-23 | 0.421655889 | 0.185 | 0.027 | 1.81672416409468e-18 | 32 |
| eva1ba5 | 1.09158786408259e-22 | 0.562986614 | 0.296 | 0.065 | 2.74741749510947e-18 | 32 |
| si:dkey-56i24.1 | 1.6313390911485e-22 | 0.323334115 | 0.148 | 0.018 | 4.10591735851167e-18 | 32 |

| | | | | | | |
|---|---|---|---|---|---|---|
| sash1a | 2.49323804965844e-22 | 0.445970391 | 0.213 | 0.036 | 6.27523084718533e-18 | 32 |
| socs3a12 | 2.70776542490553e-22 | 0.815203556 | 0.694 | 0.297 | 6.81517479794473e-18 | 32 |
| si:ch211-284f22.33 | 3.20642497585609e-22 | 0.400803655 | 0.204 | 0.033 | 8.07025102173218e-18 | 32 |
| si:dkey-9i23.165 | 3.77613005203689e-22 | 0.457310389 | 0.222 | 0.04 | 9.50414172797165e-18 | 32 |
| zgc:1951737 | 4.09490574439004e-22 | 0.609477582 | 0.352 | 0.09 | 1.03064682680553e-17 | 32 |
| palm1a1 | 4.24787322227821e-22 | 0.311796007 | 0.139 | 0.016 | 1.0691472113152e-17 | 32 |
| anxa135 | 8.10489159538341e-22 | 0.461754368 | 0.25 | 0.049 | 2.03992016564205e-17 | 32 |
| tuba8l34 | 8.74507011481219e-22 | 0.471858377 | 0.241 | 0.046 | 2.20104669719708e-17 | 32 |
| asap1b | 8.85721458488862e-22 | 0.452726107 | 0.194 | 0.031 | 2.22927233887062e-17 | 32 |
| tpte | 1.27029494401498e-21 | 0.371793725 | 0.176 | 0.026 | 3.1972053445913e-17 | 32 |
| rgcc5 | 1.55678419749107e-21 | 0.739198581 | 0.296 | 0.07 | 3.91827014666527e-17 | 32 |
| jun8 | 2.31225555014909e-21 | 0.789116669 | 0.593 | 0.227 | 5.81971599417024e-17 | 32 |
| pdlim12 | 3.08930241996664e-21 | 0.499871489 | 0.259 | 0.054 | 7.77546526081403e-17 | 32 |
| palm3 | 5.06076382696773e-21 | 0.260801753 | 0.12 | 0.013 | 1.27374364760951e-16 | 32 |
| ckba2 | 5.29403801448738e-21 | 0.53011747 | 0.287 | 0.065 | 1.33245642786633e-16 | 32 |
| ms4a17a.95 | 5.93945053501853e-21 | 0.526012079 | 0.259 | 0.056 | 1.49490030515881e-16 | 32 |
| slc12a21 | 1.08113455375731e-20 | 0.380232527 | 0.167 | 0.025 | 2.72110755835177e-16 | 32 |
| gadd45ba6 | 1.25001574284596e-20 | 0.742067265 | 0.519 | 0.19 | 3.14616462316898e-16 | 32 |
| dab2 | 1.46353109054127e-20 | 0.363652022 | 0.204 | 0.034 | 3.68356140178332e-16 | 32 |
| zfp36l1b7 | 1.71157615959146e-20 | 0.651923899 | 0.426 | 0.129 | 4.30786603607575e-16 | 32 |
| hspg22 | 1.94232147732321e-20 | 0.339050979 | 0.167 | 0.025 | 4.88862892627478e-16 | 32 |
| ier2a12 | 3.27706192098846e-20 | 0.750016568 | 0.602 | 0.261 | 8.24803714893585e-16 | 32 |
| fkbp1aa7 | 4.63794924776881e-20 | 0.666102817 | 0.713 | 0.387 | 1.16732544617093e-15 | 32 |
| mef2cb2 | 5.47913315272563e-20 | 0.36357356 | 0.167 | 0.025 | 1.37904302320951e-15 | 32 |
| mbnl2 | 8.99962422000128e-20 | 0.357319053 | 0.157 | 0.023 | 2.26511541993212e-15 | 32 |
| si:dkey-192g7.3 | 1.47743188731616e-19 | 0.303177643 | 0.13 | 0.016 | 3.71854831718604e-15 | 32 |
| smad11 | 1.5862742307604e-19 | 0.316434641 | 0.139 | 0.018 | 3.99249361140084e-15 | 32 |
| cd276 | 1.58631993657739e-19 | 0.317359409 | 0.139 | 0.019 | 3.99260864837163e-15 | 32 |
| si:dkey-163m14.22 | 1.62005050034323e-19 | 0.261495366 | 0.139 | 0.018 | 4.07750510431389e-15 | 32 |
| dusp54 | 1.78976256390002e-19 | 0.533368799 | 0.296 | 0.072 | 4.50465339707996e-15 | 32 |
| stab2 | 1.80900597652904e-19 | 0.276690706 | 0.13 | 0.016 | 4.55308714232594e-15 | 32 |
| bmpr2a1 | 1.91435481762919e-19 | 0.291491621 | 0.139 | 0.018 | 4.8182396404909e-15 | 32 |
| cav14 | 1.96663437894788e-19 | 0.497797774 | 0.222 | 0.044 | 4.94982206837391e-15 | 32 |
| ctsla10 | 3.20152737969867e-19 | 0.712726968 | 0.704 | 0.386 | 8.0579242619636e-15 | 32 |
| id15 | 3.35632858198244e-19 | 0.683796173 | 0.361 | 0.105 | 8.44754340799161e-15 | 32 |

| | | | | | | |
|---|---|---|---|---|---|---|
| piezo1 | 3.63431030016847e-19 | 0.337702593 | 0.157 | 0.024 | 9.14719559449402e-15 | 32 |
| vamp5 | 1.18179083253247e-18 | 0.341666084 | 0.167 | 0.027 | 2.97444934640098e-14 | 32 |
| acvrl11 | 1.24945416219437e-18 | 0.282621833 | 0.12 | 0.015 | 3.144751180827e-14 | 32 |
| myh103 | 3.21599981145437e-18 | 0.371176588 | 0.194 | 0.036 | 8.09434992544951e-14 | 32 |
| rbp2a2 | 3.44788713401999e-18 | 0.553097561 | 0.111 | 0.013 | 8.67798712761492e-14 | 32 |
| cald1a7 | 5.07551799266777e-18 | 0.563509431 | 0.343 | 0.098 | 1.27745712357455e-13 | 32 |
| appa6 | 5.79035703979638e-18 | 0.50462247 | 0.296 | 0.077 | 1.45737496334635e-13 | 32 |
| tjp1a4 | 6.68618752465931e-18 | 0.463286826 | 0.222 | 0.047 | 1.6828465380815e-13 | 32 |
| lpar15 | 6.87066246822768e-18 | 0.470495053 | 0.269 | 0.064 | 1.72927703662822e-13 | 32 |
| jund7 | 1.00509412763917e-17 | 0.560060428 | 0.324 | 0.092 | 2.52972140985502e-13 | 32 |
| btg29 | 1.10824813711375e-17 | 0.678959365 | 0.694 | 0.372 | 2.78934973630161e-13 | 32 |
| junba9 | 1.14442100896397e-17 | 0.712300638 | 0.667 | 0.308 | 2.88039323746141e-13 | 32 |
| stard8 | 1.54132316211236e-17 | 0.331686187 | 0.157 | 0.025 | 3.87935626672061e-13 | 32 |
| nos1apa1 | 2.2115040289349e-17 | 0.309565808 | 0.139 | 0.021 | 5.56613449042624e-13 | 32 |
| myo1cb2 | 2.85766870001143e-17 | 0.426503733 | 0.185 | 0.035 | 7.19246635105877e-13 | 32 |
| map3k20 | 3.26919458774418e-17 | 0.306715665 | 0.12 | 0.016 | 8.22823585789333e-13 | 32 |
| klf6a7 | 3.48525345297111e-17 | 0.679643068 | 0.444 | 0.167 | 8.772034415783e-13 | 32 |
| prss59.11 | 3.59239985993201e-17 | 0.292896267 | 0.102 | 0.012 | 9.04171120746287e-13 | 32 |
| rapgef5a | 3.99847340849341e-17 | 0.336807444 | 0.157 | 0.025 | 1.00637577218371e-12 | 32 |
| RALGDS | 6.731853481598e-17 | 0.293192897 | 0.13 | 0.019 | 1.6943402027834e-12 | 32 |
| reep3a3 | 7.35938759887823e-17 | 0.335198359 | 0.157 | 0.026 | 1.85228426476166e-12 | 32 |
| soga1 | 2.21214486351897e-16 | 0.289011245 | 0.13 | 0.019 | 5.56774740699089e-12 | 32 |
| nr2f24 | 4.31208938582582e-16 | 0.445802528 | 0.204 | 0.044 | 1.0853097775185e-11 | 32 |
| fmnl31 | 5.3657744320261e-16 | 0.303487802 | 0.13 | 0.019 | 1.35051176679665e-11 | 32 |
| pbx1b3 | 5.41891536716663e-16 | 0.318138714 | 0.157 | 0.028 | 1.36388680876217e-11 | 32 |
| wwtr12 | 5.76812531632624e-16 | 0.298279109 | 0.148 | 0.025 | 1.45177946086615e-11 | 32 |
| ubl3a1 | 6.36558962451695e-16 | 0.415429544 | 0.204 | 0.044 | 1.60215525259467e-11 | 32 |
| qkia6 | 9.58465257432204e-16 | 0.588275089 | 0.435 | 0.168 | 2.41236120643112e-11 | 32 |
| snx9b1 | 1.04723487248034e-15 | 0.445720344 | 0.241 | 0.059 | 2.63578545054577e-11 | 32 |
| cnn3a6 | 1.19127659761179e-15 | 0.449436237 | 0.231 | 0.056 | 2.99832406852911e-11 | 32 |
| krt947 | 1.29065032219019e-15 | 0.631797591 | 0.389 | 0.129 | 3.24843779592049e-11 | 32 |
| tmsb4x19 | 1.84694867211403e-15 | 0.539877165 | 0.991 | 0.685 | 4.64858511284379e-11 | 32 |
| XLOC-0247504 | 1.92866594555693e-15 | 0.525755502 | 0.269 | 0.075 | 4.85425931837224e-11 | 32 |
| ch25h2 | 2.00708809749129e-15 | 0.352097555 | 0.139 | 0.023 | 5.05164003257582e-11 | 32 |
| sypl2a1 | 2.98795057085638e-15 | 0.409851838 | 0.222 | 0.052 | 7.52037279178843e-11 | 32 |

| | | | | | | |
|---|---|---|---|---|---|---|
| fosab10 | 3.48676724632691e-15 | 0.74028167 | 0.639 | 0.349 | 8.7758444822802e-11 | 32 |
| slc25a43 | 4.81719884901023e-15 | 0.349369487 | 0.167 | 0.032 | 1.21244077830738e-10 | 32 |
| ctdspla3 | 4.99106162631561e-15 | 0.37088701 | 0.176 | 0.035 | 1.25620030072738e-10 | 32 |
| cx433 | 7.78616925214548e-15 | 0.483617628 | 0.222 | 0.054 | 1.9597009390725e-10 | 32 |
| sept26 | 9.4043225975985e-15 | 0.565255207 | 0.426 | 0.165 | 2.36697395458957e-10 | 32 |
| fosb6 | 1.10075950100975e-14 | 0.662036338 | 0.361 | 0.125 | 2.77050158809145e-10 | 32 |
| clasp1a | 1.2329563566352e-14 | 0.262303954 | 0.111 | 0.016 | 3.10322785401514e-10 | 32 |
| cd81a7 | 1.7028763698448e-14 | 0.588108197 | 0.5 | 0.206 | 4.28596953526237e-10 | 32 |
| pim19 | 2.07852155142886e-14 | 0.585316815 | 0.435 | 0.171 | 5.2314308927913e-10 | 32 |
| col4a3bpb | 2.16687922184579e-14 | 0.339874061 | 0.157 | 0.03 | 5.45381831346366e-10 | 32 |
| scarb2a3 | 2.91874958977358e-14 | 0.455493748 | 0.241 | 0.064 | 7.34620084250111e-10 | 32 |
| mprip1 | 3.21858987429462e-14 | 0.337244047 | 0.139 | 0.025 | 8.10086885461213e-10 | 32 |
| gstr2 | 3.58754367797363e-14 | 0.524224783 | 0.343 | 0.118 | 9.02948868309183e-10 | 32 |
| ap1b11 | 4.1937264316533e-14 | 0.393411772 | 0.213 | 0.051 | 1.05551900558282e-09 | 32 |
| hspb15 | 6.17388355276852e-14 | 0.438230721 | 0.241 | 0.064 | 1.55390475139631e-09 | 32 |
| mmp14b4 | 7.85722586227745e-14 | 0.424496556 | 0.222 | 0.057 | 1.97758517727661e-09 | 32 |
| pdk4 | 8.76669621590166e-14 | 0.312998046 | 0.148 | 0.028 | 2.20648977058029e-09 | 32 |
| ier2b12 | 1.1615667020018e-13 | 0.610729156 | 0.491 | 0.223 | 2.92354723226832e-09 | 32 |
| cd6314 | 1.70699227330614e-13 | 0.562969962 | 0.63 | 0.34 | 4.29632885268422e-09 | 32 |
| myl67 | 2.04371448571498e-13 | 0.450266307 | 0.315 | 0.101 | 5.14382498909604e-09 | 32 |
| itm2ba10 | 2.5868858156187e-13 | 0.578679997 | 0.537 | 0.273 | 6.51093290933071e-09 | 32 |
| ehd42 | 2.85927220417947e-13 | 0.342410315 | 0.148 | 0.029 | 7.1965022106993e-09 | 32 |
| degs11 | 3.69547969335502e-13 | 0.386306142 | 0.204 | 0.051 | 9.30115284020525e-09 | 32 |
| ahnak9 | 4.24850170275327e-13 | 0.516156692 | 0.352 | 0.126 | 1.06930539356597e-08 | 32 |
| ftr822 | 4.58473518100475e-13 | 0.275420471 | 0.13 | 0.023 | 1.15393199770709e-08 | 32 |
| fblim12 | 4.84078388636599e-13 | 0.283102759 | 0.139 | 0.026 | 1.21837689635946e-08 | 32 |
| cavin1b4 | 1.00571881952575e-12 | 0.408674403 | 0.176 | 0.04 | 2.53129369686435e-08 | 32 |
| trip10a | 1.00578691964375e-12 | 0.251125422 | 0.111 | 0.018 | 2.53146509805135e-08 | 32 |
| tspan18b1 | 1.00901267270162e-12 | 0.333334419 | 0.167 | 0.036 | 2.5395839959227e-08 | 32 |
| antxr1c3 | 1.28301841478652e-12 | 0.356907923 | 0.167 | 0.037 | 3.22922904817619e-08 | 32 |
| tns2a5 | 1.43206554511401e-12 | 0.332381489 | 0.167 | 0.037 | 3.60436577049744e-08 | 32 |
| sgce2 | 1.46933171004728e-12 | 0.288780003 | 0.139 | 0.027 | 3.69816098101799e-08 | 32 |
| si:ch73-375g18.1 | 2.68123297741747e-12 | 0.255240979 | 0.111 | 0.019 | 6.74839528086202e-08 | 32 |
| hecw2b1 | 2.85911770198522e-12 | 0.252405102 | 0.111 | 0.019 | 7.1961133441266e-08 | 32 |
| rgs3a | 3.16972178837477e-12 | 0.268842977 | 0.13 | 0.024 | 7.97787276916047e-08 | 32 |

| | | | | | | |
|---|---|---|---|---|---|---|
| itga92 | 3.4356066587189e-12 | 0.316069943 | 0.148 | 0.031 | 8.64707839932961e-08 | 32 |
| ms4a17a.112 | 4.87384893724755e-12 | 0.328743712 | 0.167 | 0.039 | 1.22669903901584e-07 | 32 |
| cdc425 | 5.10537762622129e-12 | 0.540553095 | 0.454 | 0.212 | 1.28497249474364e-07 | 32 |
| sesn31 | 6.71389208156808e-12 | 0.258659427 | 0.13 | 0.025 | 1.68981949800987e-07 | 32 |
| tspan9a1 | 7.72560078651031e-12 | 0.265695046 | 0.12 | 0.022 | 1.94445646195678e-07 | 32 |
| gpt2 | 9.70806945393711e-12 | 0.254977088 | 0.13 | 0.025 | 2.44342400086143e-07 | 32 |
| klf32 | 1.0499389700316e-11 | 0.330584057 | 0.157 | 0.036 | 2.64259139367254e-07 | 32 |
| rasgef1bb1 | 1.06363415215889e-11 | 0.285439233 | 0.148 | 0.032 | 2.67706079756871e-07 | 32 |
| afdna3 | 1.17590273502003e-11 | 0.323989488 | 0.12 | 0.023 | 2.95962959377191e-07 | 32 |
| ccdc801 | 1.42822408261343e-11 | 0.297863974 | 0.13 | 0.026 | 3.59469719352974e-07 | 32 |
| rplp216 | 1.436221279987e-11 | 0.348676915 | 0.944 | 0.757 | 3.61482533959928e-07 | 32 |
| zfp36l21 | 1.4520663157718e-11 | 0.425602338 | 0.324 | 0.11 | 3.65470571016605e-07 | 32 |
| cox6b19 | 1.85288631895121e-11 | 0.506947618 | 0.315 | 0.12 | 4.6635295761683e-07 | 32 |
| tmem176l.17 | 1.88775281943371e-11 | 0.500847044 | 0.417 | 0.181 | 4.75128507123272e-07 | 32 |
| rgl11 | 1.93482163566602e-11 | 0.298552222 | 0.157 | 0.036 | 4.86975257480781e-07 | 32 |
| foxo1a2 | 2.3099385699324e-11 | 0.344521049 | 0.167 | 0.04 | 5.81388438666286e-07 | 32 |
| pde4ba11 | 2.32676896815309e-11 | 0.497756027 | 0.352 | 0.14 | 5.85624481594452e-07 | 32 |
| rabgap1l1 | 3.56628399580113e-11 | 0.286553459 | 0.157 | 0.037 | 8.97598018903187e-07 | 32 |
| pik3ca | 3.64801373498501e-11 | 0.311869738 | 0.13 | 0.027 | 9.18168576958376e-07 | 32 |
| spred2b4 | 3.66489208814703e-11 | 0.4112369 | 0.222 | 0.067 | 9.22416689665726e-07 | 32 |
| rplp2l16 | 3.90578369051821e-11 | 0.326020052 | 0.944 | 0.774 | 9.83046697066528e-07 | 32 |
| CU459056.11 | 4.06270024140101e-11 | 0.2519865 | 0.12 | 0.023 | 1.02254102375822e-06 | 32 |
| fstl1b6 | 4.18588493673367e-11 | 0.467837854 | 0.333 | 0.125 | 1.0535453797265e-06 | 32 |
| snx13 | 4.41754655516719e-11 | 0.255344421 | 0.12 | 0.023 | 1.11185229247003e-06 | 32 |
| btg113 | 4.61921295624722e-11 | 0.52315739 | 0.593 | 0.333 | 1.16260970895786e-06 | 32 |
| atf4b11 | 6.99065762364557e-11 | 0.519109333 | 0.509 | 0.264 | 1.75947861729535e-06 | 32 |
| si:ch211-196l7.413 | 8.65265340249747e-11 | 0.512260838 | 0.556 | 0.314 | 2.17778633487459e-06 | 32 |
| ralba | 9.37428915691805e-11 | 0.265051274 | 0.12 | 0.024 | 2.3594148379047e-06 | 32 |
| fosl1a5 | 1.12695562656786e-10 | 0.432212136 | 0.269 | 0.092 | 2.83643461650864e-06 | 32 |
| mef2aa3 | 1.13009415445226e-10 | 0.407511259 | 0.222 | 0.069 | 2.8443339773409e-06 | 32 |
| plcg1 | 1.37808152882987e-10 | 0.324074194 | 0.157 | 0.038 | 3.4684933999119e-06 | 32 |
| im:71514493 | 1.75704013992912e-10 | 0.29295707 | 0.13 | 0.027 | 4.42229432818759e-06 | 32 |
| pfn24 | 4.96426992930379e-10 | 0.506693858 | 0.454 | 0.226 | 1.24945709850647e-05 | 32 |
| si:dkeyp-19e1.34 | 7.06871749767426e-10 | 0.411076898 | 0.213 | 0.068 | 1.77912550698964e-05 | 32 |
| mt-co217 | 7.74546796604754e-10 | 0.380893062 | 0.926 | 0.72 | 1.94945683237451e-05 | 32 |

| | | | | | | |
|---|---|---|---|---|---|---|
| entpd15 | 8.68347696838264e-10 | 0.348018091 | 0.222 | 0.071 | 2.18554431817223e-05 | 32 |
| ctrb11 | 9.04686146775579e-10 | 0.29012424 | 0.102 | 0.019 | 2.27700456281946e-05 | 32 |
| itga52 | 9.76898998786831e-10 | 0.305571575 | 0.13 | 0.03 | 2.45875709004657e-05 | 32 |
| txnipa11 | 1.06602233767004e-09 | 0.501195793 | 0.426 | 0.211 | 2.68307162168172e-05 | 32 |
| pnrc217 | 1.16753362406418e-09 | 0.391616221 | 0.769 | 0.568 | 2.93856537840714e-05 | 32 |
| zgc:11418817 | 1.19475003014666e-09 | 0.367862049 | 0.843 | 0.628 | 3.00706635087612e-05 | 32 |
| ehd1b2 | 1.31937061333288e-09 | 0.297917297 | 0.176 | 0.049 | 3.32072389669752e-05 | 32 |
| krt154 | 1.32704012931465e-09 | 0.45012776 | 0.25 | 0.085 | 3.34002730147204e-05 | 32 |
| cdc42se1 | 1.48022082803432e-09 | 0.36677748 | 0.25 | 0.085 | 3.72556780207957e-05 | 32 |
| rhocb | 1.48645066880563e-09 | 0.384959875 | 0.25 | 0.088 | 3.74124768831689e-05 | 32 |
| rpl2913 | 1.80712729018015e-09 | 0.262495281 | 0.935 | 0.809 | 4.54835867665442e-05 | 32 |
| rab5c3 | 1.89744489539262e-09 | 0.381645336 | 0.306 | 0.121 | 4.77567905721368e-05 | 32 |
| rbms1b | 1.95181794711972e-09 | 0.2876646 | 0.139 | 0.034 | 4.91253059110562e-05 | 32 |
| msna12 | 2.01641182853404e-09 | 0.503870645 | 0.454 | 0.242 | 5.07510693123732e-05 | 32 |
| rpl917 | 3.0434291416261e-09 | 0.387927269 | 0.88 | 0.629 | 7.66000680655874e-05 | 32 |
| tns1b4 | 3.91681511362692e-09 | 0.270707128 | 0.139 | 0.034 | 9.85823195948759e-05 | 32 |
| cnn29 | 4.09057227230291e-09 | 0.442206171 | 0.315 | 0.135 | 0.000102956 | 32 |
| il4r.13 | 4.1526335796211e-09 | 0.334740657 | 0.157 | 0.044 | 0.000104518 | 32 |
| tmem176l.3a1 | 4.58028992068831e-09 | 0.307047813 | 0.148 | 0.039 | 0.000115281 | 32 |
| CT030188.14 | 5.24983374214586e-09 | 0.341230747 | 0.176 | 0.052 | 0.000132133 | 32 |
| si:ch211-240l19.81 | 6.34923836620325e-09 | 0.277224991 | 0.102 | 0.021 | 0.000159804 | 32 |
| smad5 | 6.4820315019932e-09 | 0.355317105 | 0.241 | 0.083 | 0.000163146 | 32 |
| col4a15 | 6.97398139569199e-09 | 0.456829746 | 0.185 | 0.059 | 0.000175528 | 32 |
| rps2814 | 9.14998935403245e-09 | 0.255269058 | 0.954 | 0.826 | 0.000230296 | 32 |
| ginm1 | 1.10603859132556e-08 | 0.355611315 | 0.259 | 0.097 | 0.000278379 | 32 |
| parvaa2 | 1.85076974211632e-08 | 0.253366514 | 0.12 | 0.029 | 0.00046582 | 32 |
| rab8b3 | 1.89960858071465e-08 | 0.353190731 | 0.222 | 0.08 | 0.000478112 | 32 |
| ephb35 | 2.11130428049061e-08 | 0.325123961 | 0.148 | 0.042 | 0.000531394 | 32 |
| rpl3616 | 2.23548768800188e-08 | 0.28736305 | 0.944 | 0.789 | 0.00056265 | 32 |
| rap1b7 | 2.99523907614692e-08 | 0.434096619 | 0.398 | 0.203 | 0.000753872 | 32 |
| gabarapb8 | 3.06757865788261e-08 | 0.450147727 | 0.472 | 0.269 | 0.000772079 | 32 |
| mt-nd114 | 4.48175369283758e-08 | 0.442551374 | 0.611 | 0.412 | 0.001128013 | 32 |
| COX317 | 4.49071810637426e-08 | 0.34615274 | 0.889 | 0.7 | 0.001130269 | 32 |
| rhoaa9 | 4.83193247207778e-08 | 0.380925253 | 0.333 | 0.154 | 0.001216149 | 32 |
| si:zfos-464b6.21 | 5.26497588659462e-08 | 0.272084768 | 0.157 | 0.046 | 0.001325142 | 32 |

| | | | | | | |
|---|---|---|---|---|---|---|
| sept123 | 8.87042431649011e-08 | 0.288425114 | 0.185 | 0.062 | 0.002232597 | 32 |
| socs3b9 | 9.45389808998411e-08 | 0.436633258 | 0.333 | 0.156 | 0.002379452 | 32 |
| tspan143 | 9.95270772911941e-08 | 0.398244953 | 0.287 | 0.125 | 0.002504997 | 32 |
| h3f3c10 | 1.43281233163059e-07 | 0.430497803 | 0.444 | 0.257 | 0.003606245 | 32 |
| mt-co117 | 1.45814292065423e-07 | 0.34227937 | 0.861 | 0.657 | 0.00367 | 32 |
| rab1ba | 1.4888890425504e-07 | 0.294378528 | 0.148 | 0.045 | 0.003747385 | 32 |
| abca1a4 | 1.48989350606092e-07 | 0.338823572 | 0.194 | 0.069 | 0.003749913 | 32 |
| mcl1b10 | 1.50550430940328e-07 | 0.435886099 | 0.444 | 0.243 | 0.003789204 | 32 |
| mt-cyb13 | 1.79229749587942e-07 | 0.353549953 | 0.824 | 0.624 | 0.004511034 | 32 |
| shoc2 | 1.90043342031329e-07 | 0.318785123 | 0.185 | 0.065 | 0.004783201 | 32 |
| rhoab8 | 1.98415231524117e-07 | 0.441410986 | 0.361 | 0.19 | 0.004993913 | 32 |
| tacc11 | 2.29914764816582e-07 | 0.255004329 | 0.13 | 0.036 | 0.005786725 | 32 |
| si:ch211-204c21.16 | 2.32608084828286e-07 | 0.30390708 | 0.204 | 0.074 | 0.005854513 | 32 |
| rps2413 | 2.58428828976949e-07 | 0.252522208 | 0.898 | 0.759 | 0.006504395 | 32 |
| sik17 | 2.62036423312916e-07 | 0.391131641 | 0.269 | 0.117 | 0.006595195 | 32 |
| mt-nd212 | 2.63820453279162e-07 | 0.415853207 | 0.583 | 0.401 | 0.006640097 | 32 |
| DDX171 | 2.88964469830434e-07 | 0.306373422 | 0.176 | 0.061 | 0.007272947 | 32 |
| plpp1a3 | 3.09075445559826e-07 | 0.299871212 | 0.194 | 0.069 | 0.00777912 | 32 |
| zgc:1583438 | 3.26779548684242e-07 | 0.438019674 | 0.343 | 0.166 | 0.008224714 | 32 |
| perp3 | 3.36099414061967e-07 | 0.286571446 | 0.167 | 0.055 | 0.008459286 | 32 |
| CR318588.33 | 3.52915320045841e-07 | 0.326339442 | 0.148 | 0.046 | 0.008882526 | 32 |
| si:ch73-335l21.46 | 3.56202405863712e-07 | 0.500249888 | 0.315 | 0.149 | 0.008965258 | 32 |
| rpz58 | 3.59425153469718e-07 | 0.33744939 | 0.25 | 0.103 | 0.009046372 | 32 |
| ier54 | 5.24903630736162e-07 | 0.42206947 | 0.287 | 0.137 | 0.013211299 | 32 |
| sptlc2b1 | 5.57066619653088e-07 | 0.256479951 | 0.148 | 0.047 | 0.01402081 | 32 |
| cst14b.14 | 5.59308849629198e-07 | 0.28696786 | 0.157 | 0.052 | 0.014077244 | 32 |
| mt-atp617 | 6.50329581062297e-07 | 0.361464338 | 0.796 | 0.591 | 0.016368145 | 32 |
| dedd16 | 6.72027858090735e-07 | 0.380416847 | 0.306 | 0.146 | 0.016914269 | 32 |
| ccnl1a2 | 6.8686699312566e-07 | 0.282464307 | 0.176 | 0.063 | 0.017287755 | 32 |
| cox8b3 | 7.05315160376981e-07 | 0.340566664 | 0.231 | 0.096 | 0.017752077 | 32 |
| ptbp3 | 8.20421547545232e-07 | 0.294757203 | 0.176 | 0.063 | 0.02064919 | 32 |
| snx1a1 | 8.29175603643836e-07 | 0.37289245 | 0.259 | 0.115 | 0.020869521 | 32 |
| si:ch211-74m13.34 | 8.56763819167341e-07 | 0.319266187 | 0.185 | 0.069 | 0.021563889 | 32 |
| stau27 | 9.21095765342512e-07 | 0.315895388 | 0.213 | 0.086 | 0.023183059 | 32 |
| per34 | 1.04632337923664e-06 | 0.309116925 | 0.204 | 0.08 | 0.026334913 | 32 |

| | | | | | | |
|---|---|---|---|---|---|---|
| rpsa17 | 1.05012424121798e-06 | 0.275151629 | 0.926 | 0.752 | 0.026430577 | 32 |
| arrb2b7 | 1.09218367921249e-06 | 0.336242504 | 0.222 | 0.092 | 0.027489171 | 32 |
| gnb1a5 | 1.32010248478054e-06 | 0.379163907 | 0.361 | 0.193 | 0.033225659 | 32 |
| LOC1039092846 | 1.42261864019552e-06 | 0.304806411 | 0.176 | 0.065 | 0.035805889 | 32 |
| svild7 | 2.03917573941977e-06 | 0.332469279 | 0.204 | 0.082 | 0.051324014 | 32 |
| hopx6 | 2.48426587175678e-06 | 0.313856879 | 0.176 | 0.066 | 0.062526488 | 32 |
| csrnp1b8 | 2.62254000471368e-06 | 0.421613478 | 0.333 | 0.18 | 0.066006709 | 32 |
| gadd45bb6 | 2.63040224445137e-06 | 0.341780133 | 0.213 | 0.088 | 0.066204594 | 32 |
| ddit311 | 2.95432870893841e-06 | 0.412642512 | 0.333 | 0.177 | 0.074357499 | 32 |
| adipor22 | 3.30774262980326e-06 | 0.30562238 | 0.213 | 0.087 | 0.083252574 | 32 |
| sptan13 | 3.76910647606113e-06 | 0.270528379 | 0.148 | 0.051 | 0.094864641 | 32 |
| il6st3 | 4.48590824092878e-06 | 0.261061725 | 0.148 | 0.052 | 0.112905825 | 32 |
| klf7b | 4.75676752138641e-06 | 0.251485888 | 0.111 | 0.033 | 0.119723082 | 32 |
| ptprga8 | 4.87657091128985e-06 | 0.259230084 | 0.176 | 0.067 | 0.122738413 | 32 |
| srgn10 | 5.08703487269821e-06 | 0.367775157 | 0.426 | 0.238 | 0.128035581 | 32 |
| grinab5 | 5.92844972557323e-06 | 0.292997785 | 0.25 | 0.115 | 0.149213151 | 32 |
| kctd5a6 | 6.52605667172023e-06 | 0.340757753 | 0.333 | 0.176 | 0.16425432 | 32 |
| cyth1a2 | 6.92980752867435e-06 | 0.273981478 | 0.176 | 0.068 | 0.174416326 | 32 |
| b2m12 | 7.2915869743229e-06 | 0.35724197 | 0.481 | 0.293 | 0.183521953 | 32 |
| si:dkey-79d12.55 | 7.97074656925154e-06 | 0.369937015 | 0.185 | 0.077 | 0.20061572 | 32 |
| rps1016 | 8.23368261147143e-06 | 0.267023566 | 0.861 | 0.694 | 0.207233558 | 32 |
| zmiz1a1 | 9.03970442887101e-06 | 0.260834754 | 0.12 | 0.039 | 0.227520321 | 32 |
| NC-002333.416 | 9.51439375439342e-06 | 0.330334251 | 0.815 | 0.623 | 0.239467776 | 32 |
| uba5213 | 1.17428804965587e-05 | 0.262127644 | 0.787 | 0.631 | 0.295556559 | 32 |
| crip112 | 1.5957815485583e-05 | 0.37481839 | 0.444 | 0.271 | 0.401642258 | 32 |
| si:ch73-234b20.56 | 1.64068114338151e-05 | 0.2847206 | 0.222 | 0.101 | 0.412943037 | 32 |
| midn8 | 1.77193971434255e-05 | 0.393785202 | 0.269 | 0.141 | 0.445979507 | 32 |
| rps27.213 | 1.77589539474747e-05 | 0.29888821 | 0.685 | 0.518 | 0.446975112 | 32 |
| elmo16 | 1.78871310098405e-05 | 0.261208912 | 0.176 | 0.071 | 0.4502012 | 32 |
| ucp210 | 1.80622097872637e-05 | 0.335307154 | 0.583 | 0.417 | 0.454607758 | 32 |
| sh2b34 | 2.56308214744012e-05 | 0.263636524 | 0.157 | 0.062 | 0.645102146 | 32 |
| stat1a6 | 2.60696932080811e-05 | 0.346041641 | 0.241 | 0.121 | 0.656148108 | 32 |
| sumo2b10 | 3.28218342979424e-05 | 0.336191155 | 0.426 | 0.27 | 0.826092747 | 32 |
| pkig | 3.85670993223766e-05 | 0.260113212 | 0.213 | 0.095 | 0.970695323 | 32 |
| rps2119 | 4.21433688842531e-05 | 0.2694242 | 0.935 | 0.75 | 1 | 32 |

| | | | | | | |
|---|---|---|---|---|---|---|
| selenot1b | 5.46290493547614e-05 | 0.27629215 | 0.176 | 0.077 | 1 | 32 |
| podxl3 | 6.87841121782303e-05 | 0.260581903 | 0.13 | 0.048 | 1 | 32 |
| sdcbp28 | 6.95490105441548e-05 | 0.324467379 | 0.352 | 0.209 | 1 | 32 |
| aoc27 | 8.50412145690716e-05 | 0.371059611 | 0.157 | 0.067 | 1 | 32 |
| chmp1b2 | 9.4606280721662e-05 | 0.294129863 | 0.241 | 0.122 | 1 | 32 |
| nsa213 | 0.000136863 | 0.311735677 | 0.509 | 0.363 | 1 | 32 |
| fkbp56 | 0.000170805 | 0.311869312 | 0.231 | 0.123 | 1 | 32 |
| mgst3b4 | 0.000216517 | 0.259037526 | 0.306 | 0.173 | 1 | 32 |
| antxr2a9 | 0.000227717 | 0.281229043 | 0.278 | 0.155 | 1 | 32 |
| abcc57 | 0.000228303 | 0.270908834 | 0.194 | 0.095 | 1 | 32 |
| si:dkey-7j14.69 | 0.000259101 | 0.306652912 | 0.296 | 0.176 | 1 | 32 |
| fosl29 | 0.000274688 | 0.289822093 | 0.231 | 0.122 | 1 | 32 |
| nr1d2a3 | 0.000342107 | 0.277294895 | 0.167 | 0.08 | 1 | 32 |
| cltca | 0.000365372 | 0.284846518 | 0.176 | 0.086 | 1 | 32 |
| slc43a2a6 | 0.000403513 | 0.257666118 | 0.204 | 0.103 | 1 | 32 |
| mt-nd413 | 0.000411736 | 0.285303723 | 0.593 | 0.457 | 1 | 32 |
| higd1a8 | 0.000565397 | 0.325857418 | 0.417 | 0.311 | 1 | 32 |
| h2afx111 | 0.000680378 | 0.274799993 | 0.324 | 0.2 | 1 | 32 |
| capns1b5 | 0.00068224 | 0.261280898 | 0.222 | 0.123 | 1 | 32 |
| dusp18 | 0.000704182 | 0.315830154 | 0.269 | 0.161 | 1 | 32 |
| CABZ01072614.18 | 0.000708606 | 0.256522111 | 0.167 | 0.082 | 1 | 32 |
| cdkn1bb4 | 0.000738417 | 0.257311182 | 0.259 | 0.149 | 1 | 32 |
| egr110 | 0.000808365 | 0.276657958 | 0.315 | 0.191 | 1 | 32 |
| maff6 | 0.000809619 | 0.27598139 | 0.167 | 0.084 | 1 | 32 |
| cfd4 | 0.000986972 | 0.284576254 | 0.148 | 0.069 | 1 | 32 |
| dynlt32 | 0.001058175 | 0.26333241 | 0.25 | 0.147 | 1 | 32 |
| rpl22l110 | 0.001372874 | 0.270927193 | 0.509 | 0.393 | 1 | 32 |
| mt-nd313 | 0.002363595 | 0.263675133 | 0.509 | 0.397 | 1 | 32 |
| chd37 | 0.002647548 | 0.275537353 | 0.185 | 0.105 | 1 | 32 |
| ankrd112 | 0.003051542 | 0.252532736 | 0.167 | 0.091 | 1 | 32 |
| epdl1 | 0 | 2.024752447 | 0.957 | 0.029 | 0 | 33 |
| spock3 | 0 | 1.045478579 | 0.505 | 0.01 | 0 | 33 |
| CU928222.3 | 0 | 1.030773945 | 0.495 | 0.001 | 0 | 33 |
| syngr3b | 0 | 0.956985512 | 0.495 | 0.002 | 0 | 33 |
| FP085399.2 | 0 | 0.879943646 | 0.452 | 0.001 | 0 | 33 |

| | | | | | | |
|---|---|---|---|---|---|---|
| zgc:101810 | 0 | 0.87373546 | 0.452 | 0.005 | 0 | 33 |
| AL929237.1 | 0 | 0.730409924 | 0.376 | 0.005 | 0 | 33 |
| ptprh | 0 | 0.338635705 | 0.194 | 0 | 0 | 33 |
| BX323596.11 | 1.91684016350652e-286 | 1.029645591 | 0.398 | 0.007 | 4.82449500752957e-282 | 33 |
| cxcl11.6 | 1.24046297274226e-283 | 0.770177797 | 0.312 | 0.004 | 3.12212125609501e-279 | 33 |
| mxf | 9.95634326680501e-283 | 0.55759239 | 0.237 | 0.002 | 2.50591203682215e-278 | 33 |
| BX323596.2 | 1.20597050926344e-273 | 0.815174553 | 0.366 | 0.006 | 3.03530717476515e-269 | 33 |
| mpeg1.13 | 2.96603232169518e-254 | 0.904567069 | 0.473 | 0.013 | 7.46520675047459e-250 | 33 |
| cxcl11.5 | 6.19109258896303e-253 | 0.72870909 | 0.28 | 0.004 | 1.5582360937161e-248 | 33 |
| si:ch211-1o7.33 | 8.57357133844806e-243 | 0.968410843 | 0.516 | 0.017 | 2.15788217017399e-238 | 33 |
| si:dkey-33i11.91 | 2.07751464024849e-234 | 1.14701076 | 0.527 | 0.019 | 5.22889659804143e-230 | 33 |
| hepacam2 | 6.04631672240908e-208 | 0.476696986 | 0.258 | 0.004 | 1.52179745586314e-203 | 33 |
| si:dkey-148a17.5 | 3.85838007685994e-203 | 0.38641311 | 0.194 | 0.002 | 9.71115681544877e-199 | 33 |
| grn13 | 2.19138149241794e-192 | 1.936102672 | 0.925 | 0.082 | 5.5154880782667e-188 | 33 |
| sftpbb4 | 1.4879817194415e-181 | 1.613105281 | 0.785 | 0.062 | 3.74510118966232e-177 | 33 |
| spi1a3 | 1.56770359969875e-170 | 1.063752864 | 0.581 | 0.033 | 3.9457531900818e-166 | 33 |
| CU657980.2 | 3.791874095179e-166 | 0.27654271 | 0.14 | 0.001 | 9.54376791015602e-162 | 33 |
| tnfa | 6.17565143099499e-164 | 0.449693586 | 0.226 | 0.004 | 1.55434970866713e-159 | 33 |
| arap1a | 8.51646531476362e-164 | 0.374463981 | 0.194 | 0.003 | 2.14350915507286e-159 | 33 |
| ccl35.15 | 1.90394501105066e-162 | 1.760563233 | 0.839 | 0.078 | 4.79203919831341e-158 | 33 |
| si:ch211-214p13.3 | 2.94113831063986e-156 | 0.379900911 | 0.183 | 0.003 | 7.40255101404946e-152 | 33 |
| arhgef15 | 1.11919327002272e-153 | 0.404066619 | 0.204 | 0.004 | 2.81689754132019e-149 | 33 |
| ccl35.2 | 4.27882999321612e-151 | 0.482119714 | 0.204 | 0.004 | 1.07693872099257e-146 | 33 |
| smcr8b | 7.15244385112427e-151 | 0.338623368 | 0.194 | 0.003 | 1.80019859288947e-146 | 33 |
| si:ch211-79k12.1 | 8.87815950173068e-143 | 0.887473242 | 0.419 | 0.02 | 2.2345439649906e-138 | 33 |
| mhc2dab5 | 6.18498949958546e-135 | 1.564979052 | 0.828 | 0.093 | 1.55670000715066e-130 | 33 |
| si:dkey-102c8.3 | 6.18222488075732e-133 | 0.265077366 | 0.151 | 0.002 | 1.55600418023781e-128 | 33 |
| CR385050.11 | 2.00020279891072e-132 | 0.583731508 | 0.323 | 0.012 | 5.0343104245784e-128 | 33 |
| si:busm1-266f07.26 | 2.36383153130481e-131 | 1.777749629 | 0.968 | 0.142 | 5.94952758114107e-127 | 33 |
| gna15.1 | 2.32654198486745e-129 | 0.326552613 | 0.172 | 0.003 | 5.85567352171287e-125 | 33 |
| CR786582.21 | 2.69002117089716e-125 | 0.574941795 | 0.29 | 0.01 | 6.77051428503106e-121 | 33 |
| zgc:1231071 | 2.07081028528797e-121 | 0.540008235 | 0.29 | 0.011 | 5.21202240704129e-117 | 33 |
| CABZ01078737.11 | 8.41892361848867e-120 | 0.491200871 | 0.269 | 0.009 | 2.11895888553741e-115 | 33 |
| abi3a4 | 3.99991726418919e-119 | 0.92654409 | 0.538 | 0.04 | 1.00673917622378e-114 | 33 |
| si:dkey-1h24.6 | 2.13889402912534e-118 | 0.502476987 | 0.247 | 0.008 | 5.38338238190557e-114 | 33 |

| | | | | | | |
|---|---|---|---|---|---|---|
| si:ch73-158p21.31 | 1.00327051952768e-116 | 0.767519728 | 0.419 | 0.025 | 2.52513157059921e-112 | 33 |
| cd74a8 | 3.38719561211893e-106 | 1.912285217 | 1 | 0.215 | 8.52523263614215e-102 | 33 |
| mxc | 3.86102283456101e-105 | 0.376386315 | 0.183 | 0.005 | 9.71780837230662e-101 | 33 |
| si:ch73-236c18.71 | 2.47987044606121e-102 | 0.388507697 | 0.204 | 0.006 | 6.24158592569145e-98 | 33 |
| si:dkey-192g7.31 | 2.77905108124951e-98 | 0.548750968 | 0.301 | 0.015 | 6.99459366639688e-94 | 33 |
| CU459094.3 | 2.31300435417154e-97 | 0.361391277 | 0.129 | 0.002 | 5.82160065901434e-93 | 33 |
| si:ch211-214p16.24 | 5.38684281832397e-97 | 1.108870669 | 0.634 | 0.069 | 1.35581446894396e-92 | 33 |
| kctd12.24 | 1.85836545577273e-95 | 1.052900649 | 0.656 | 0.076 | 4.67732001563438e-91 | 33 |
| arpc5b7 | 2.34668330822652e-91 | 1.189493836 | 0.753 | 0.11 | 5.90636721847533e-87 | 33 |
| rnaset2l4 | 3.99106918270704e-91 | 0.862969867 | 0.473 | 0.041 | 1.00451220259554e-86 | 33 |
| tnfb5 | 2.11108381816761e-90 | 1.153084346 | 0.516 | 0.05 | 5.31338686194605e-86 | 33 |
| tiam1a1 | 9.18360048403759e-88 | 0.438822815 | 0.237 | 0.01 | 2.31142040582742e-83 | 33 |
| cd74b9 | 2.76141021414122e-87 | 1.620046268 | 0.946 | 0.22 | 6.95019336797205e-83 | 33 |
| si:dkey-78o7.32 | 3.27291682435217e-85 | 0.801513781 | 0.344 | 0.023 | 8.23760435521198e-81 | 33 |
| ptgs2b | 9.2365365118925e-85 | 0.395987251 | 0.226 | 0.009 | 2.32474387467822e-80 | 33 |
| ciita | 3.75176546008109e-83 | 0.37858945 | 0.204 | 0.008 | 9.44281848647808e-79 | 33 |
| plekhf1 | 8.63908220000424e-83 | 0.363404741 | 0.151 | 0.004 | 2.17437059891907e-78 | 33 |
| BX004816.2 | 5.26184548694447e-78 | 0.348654021 | 0.172 | 0.006 | 1.32435389060905e-73 | 33 |
| LOC1018854451 | 1.98410538390589e-76 | 0.417793805 | 0.237 | 0.012 | 4.99379484075274e-72 | 33 |
| ccl39.11 | 1.01571861466764e-75 | 0.688401838 | 0.204 | 0.009 | 2.55646218125697e-71 | 33 |
| zgc:1037003 | 3.56872864424968e-73 | 0.922839713 | 0.409 | 0.038 | 8.98213312471202e-69 | 33 |
| si:dkey-7i4.243 | 1.19608727561704e-72 | 0.963344514 | 0.301 | 0.021 | 3.01043206400053e-68 | 33 |
| si:busm1-194e12.111 | 3.63066935088955e-72 | 0.635145216 | 0.28 | 0.018 | 9.1380316892539e-68 | 33 |
| lygl13 | 4.76956300752639e-71 | 1.036286687 | 0.656 | 0.094 | 1.20045131336432e-66 | 33 |
| arpc1b12 | 5.47320214829623e-71 | 1.376908023 | 0.957 | 0.306 | 1.37755024870468e-66 | 33 |
| tmsb17 | 6.88203314531444e-71 | 1.166417823 | 0.828 | 0.152 | 1.73213892234419e-66 | 33 |
| ctss2.23 | 3.21317201776443e-68 | 0.749344369 | 0.409 | 0.04 | 8.0872326515113e-64 | 33 |
| si:dkey-27i16.26 | 3.37578356063691e-68 | 1.043805789 | 0.602 | 0.088 | 8.49650964376705e-64 | 33 |
| gpx1a7 | 4.13195457245658e-68 | 1.324897748 | 0.871 | 0.221 | 1.0399716463416e-63 | 33 |
| grna3 | 9.16922998568695e-68 | 0.823079473 | 0.473 | 0.053 | 2.30780349509755e-63 | 33 |
| spi1b6 | 2.01425829608356e-65 | 0.759274315 | 0.495 | 0.059 | 5.06968670541271e-61 | 33 |
| ccl19a.11 | 3.21642008112599e-65 | 0.657567838 | 0.204 | 0.01 | 8.09540770218601e-61 | 33 |
| nr4a2b | 3.65868700954707e-65 | 0.452104063 | 0.237 | 0.014 | 9.20854933432903e-61 | 33 |
| BX548046.21 | 6.76437053750484e-62 | 0.28050044 | 0.14 | 0.005 | 1.70252442058459e-57 | 33 |
| mhc2b3 | 8.20226122812253e-62 | 0.851068332 | 0.323 | 0.028 | 2.06442712850616e-57 | 33 |

| | | | | | | |
|---|---|---|---|---|---|---|
| LOC1018850652 | 2.63742238681154e-61 | 0.410366 | 0.226 | 0.014 | 6.63812840536595e-57 | 33 |
| si:ch211-8c17.32 | 1.16721115355352e-60 | 0.58254794 | 0.333 | 0.03 | 2.93775375237885e-56 | 33 |
| tmsb4x20 | 8.50908387057348e-60 | 1.0198474 | 1 | 0.685 | 2.14165131938464e-55 | 33 |
| pfn112 | 1.38091555265753e-58 | 1.166058509 | 1 | 0.523 | 3.47562635448375e-54 | 33 |
| XLOC-02542310 | 2.72997982375745e-58 | 1.270327248 | 0.763 | 0.166 | 6.87108621841513e-54 | 33 |
| fgd4b | 5.27763087448439e-58 | 0.594361043 | 0.333 | 0.031 | 1.32832691479898e-53 | 33 |
| lcp18 | 1.09474288750375e-57 | 0.921642135 | 0.548 | 0.086 | 2.75535837355819e-53 | 33 |
| il21r.12 | 7.60775560037263e-57 | 0.487026227 | 0.28 | 0.022 | 1.91479600705779e-52 | 33 |
| si:dkey-53k12.25 | 1.46750692105442e-55 | 0.622566696 | 0.387 | 0.043 | 3.69356816960188e-51 | 33 |
| ccl39.31 | 1.47265055862786e-55 | 0.403294958 | 0.14 | 0.006 | 3.70651419101045e-51 | 33 |
| LO018493.12 | 1.29251540955646e-54 | 0.489207305 | 0.28 | 0.023 | 3.25313203431266e-50 | 33 |
| ptmaa15 | 2.28032708523377e-54 | 1.119749114 | 0.946 | 0.393 | 5.73935524082488e-50 | 33 |
| actb212 | 2.87276456584002e-54 | 0.741316268 | 1 | 0.868 | 7.23046113576276e-50 | 33 |
| TPGS12 | 1.19444097965589e-53 | 0.571771971 | 0.355 | 0.038 | 3.0062885016959e-49 | 33 |
| irak3 | 4.05976748841183e-53 | 0.420755988 | 0.226 | 0.016 | 1.02180287915837e-48 | 33 |
| wasb11 | 9.05596294658083e-51 | 1.020339195 | 0.753 | 0.175 | 2.27929531402493e-46 | 33 |
| CU499330.13 | 1.23972419542841e-50 | 0.817853995 | 0.484 | 0.069 | 3.12026182747375e-46 | 33 |
| cd836 | 1.67600558144415e-50 | 1.090382944 | 0.559 | 0.102 | 4.21833844793677e-46 | 33 |
| cotl111 | 5.25860883495161e-50 | 1.107351684 | 0.925 | 0.304 | 1.32353925766897e-45 | 33 |
| si:ch211-152p11.42 | 2.22868435553115e-49 | 0.553874789 | 0.301 | 0.03 | 5.60937565443635e-45 | 33 |
| actb15 | 7.56515839789416e-49 | 0.712783391 | 1 | 0.784 | 1.90407471716598e-44 | 33 |
| tcima6 | 8.79560649880494e-49 | 0.760594839 | 0.462 | 0.067 | 2.21376619968422e-44 | 33 |
| BX901920.16 | 9.51495889970718e-49 | 0.951921396 | 0.462 | 0.072 | 2.3948200054673e-44 | 33 |
| BX005183.37 | 1.34161993841333e-48 | 0.818296374 | 0.495 | 0.079 | 3.3767232229925e-44 | 33 |
| gstt1a2 | 1.64023288333024e-48 | 0.594872912 | 0.333 | 0.036 | 4.12830214405388e-44 | 33 |
| tfe3a3 | 2.34172423884778e-48 | 0.582191323 | 0.333 | 0.037 | 5.89388573675598e-44 | 33 |
| psap5 | 6.56069023977518e-48 | 1.127790954 | 0.839 | 0.288 | 1.65126012644901e-43 | 33 |
| si:ch211-198c19.1 | 4.63981753485672e-47 | 0.28605199 | 0.161 | 0.009 | 1.16779567534809e-42 | 33 |
| parm11 | 7.1642140475832e-47 | 0.257246655 | 0.151 | 0.008 | 1.80316103363622e-42 | 33 |
| p2rx71 | 7.29345443198071e-47 | 0.336253099 | 0.194 | 0.013 | 1.83568954598522e-42 | 33 |
| aif1l3 | 1.74301976537997e-46 | 1.014681454 | 0.753 | 0.185 | 4.38700644748484e-42 | 33 |
| XLOC-0234927 | 2.89281282280092e-46 | 0.990952899 | 0.516 | 0.093 | 7.28092059370764e-42 | 33 |
| XLOC-0054495 | 6.18535279785997e-46 | 0.745641495 | 0.387 | 0.052 | 1.55679144569337e-41 | 33 |
| sgk3 | 7.0147974389669e-46 | 0.423605682 | 0.269 | 0.025 | 1.76555436741358e-41 | 33 |
| malt3 | 1.02276225913589e-43 | 0.327022263 | 0.194 | 0.014 | 2.57419033001911e-39 | 33 |

| | | | | | | |
|---|---|---|---|---|---|---|
| ccl39.21 | 2.0015765103478e-43 | 0.343852142 | 0.118 | 0.005 | 5.03776791889437e-39 | 33 |
| mbnl33 | 1.3308687550866e-42 | 0.597769097 | 0.387 | 0.054 | 3.34966356967747e-38 | 33 |
| sh3bgrl7 | 4.05960174100719e-42 | 0.790198874 | 0.527 | 0.103 | 1.0217611621941e-37 | 33 |
| krt18b9 | 7.43505828931377e-42 | 0.946812697 | 0.645 | 0.148 | 1.87132982083738e-37 | 33 |
| gabarapb9 | 1.61708978672584e-40 | 0.940208176 | 0.817 | 0.266 | 4.07005328421027e-36 | 33 |
| zgc:1368702 | 3.70432724065721e-40 | 0.381320742 | 0.204 | 0.017 | 9.32342123201014e-36 | 33 |
| gm2a5 | 5.68518402148353e-40 | 0.832732173 | 0.57 | 0.119 | 1.43090396636719e-35 | 33 |
| id2a1 | 7.15554948984501e-40 | 0.983409973 | 0.656 | 0.168 | 1.80098025109909e-35 | 33 |
| arpc211 | 3.4200856348306e-38 | 0.915153611 | 0.806 | 0.265 | 8.60801353430514e-34 | 33 |
| pycard7 | 4.92508085072189e-38 | 0.658252103 | 0.419 | 0.071 | 1.23959359931819e-33 | 33 |
| ppiab5 | 3.5501568935509e-37 | 0.656042014 | 0.989 | 0.656 | 8.93538988537827e-33 | 33 |
| irf23 | 8.24693839898252e-37 | 0.628932022 | 0.398 | 0.067 | 2.07567192563991e-32 | 33 |
| hcls16 | 1.507448117137e-36 | 0.649419173 | 0.409 | 0.069 | 3.79409616602211e-32 | 33 |
| si:ch211-214p16.18 | 1.77226454384128e-36 | 1.015945458 | 0.871 | 0.255 | 4.46061263039413e-32 | 33 |
| CR753876.15 | 8.98177462234573e-36 | 0.871352945 | 0.527 | 0.113 | 2.2606228546982e-31 | 33 |
| tent5ba | 2.50404810803727e-35 | 0.27869056 | 0.151 | 0.011 | 6.30243868311901e-31 | 33 |
| ifi305 | 1.75683203869635e-34 | 0.898410825 | 0.634 | 0.173 | 4.42177055819485e-30 | 33 |
| arpc311 | 2.22166693150093e-34 | 0.886946277 | 0.828 | 0.308 | 5.5917134998947e-30 | 33 |
| unc93b13 | 4.24606724510058e-34 | 0.337856483 | 0.215 | 0.022 | 1.06869266491937e-29 | 33 |
| zgc:638315 | 6.10031358973388e-32 | 0.586103504 | 0.419 | 0.08 | 1.53538792740012e-27 | 33 |
| si:cabz01074946.15 | 6.83668696945336e-32 | 0.521765722 | 0.355 | 0.058 | 1.72072574334172e-27 | 33 |
| pak2b | 1.01866918402541e-31 | 0.550792706 | 0.333 | 0.054 | 2.56388846927356e-27 | 33 |
| mapk113 | 1.8639121576614e-31 | 0.353285257 | 0.183 | 0.018 | 4.69128050961798e-27 | 33 |
| map3k14a1 | 7.4753089975699e-31 | 0.514017974 | 0.247 | 0.032 | 1.88146052159837e-26 | 33 |
| arhgdig11 | 3.63947952768394e-30 | 0.831227075 | 0.731 | 0.241 | 9.1602060232277e-26 | 33 |
| efhd22 | 1.31676335654635e-29 | 0.518734922 | 0.344 | 0.059 | 3.3141616920915e-25 | 33 |
| npc24 | 1.49909694604503e-29 | 0.873664029 | 0.806 | 0.357 | 3.77307710350074e-25 | 33 |
| cox6b110 | 1.81128803180795e-29 | 0.71026759 | 0.495 | 0.119 | 4.55883084725743e-25 | 33 |
| fgl2a6 | 2.24937467750907e-29 | 0.527590215 | 0.366 | 0.065 | 5.66145112582258e-25 | 33 |
| zgc:153733 | 3.75225392405771e-29 | 0.27159292 | 0.183 | 0.018 | 9.44404790146086e-25 | 33 |
| si:ch1073-214b20.2 | 6.65206023251541e-29 | 0.320628149 | 0.237 | 0.03 | 1.6742570399218e-24 | 33 |
| LOC1039101363 | 7.01155663555383e-29 | 0.45432913 | 0.237 | 0.031 | 1.76473868960254e-24 | 33 |
| rgs183 | 8.4289509508745e-29 | 0.414566457 | 0.247 | 0.033 | 2.1214826648256e-24 | 33 |
| triqk1 | 2.55134329135911e-28 | 0.513092203 | 0.301 | 0.05 | 6.42147593002173e-24 | 33 |
| myo1f6 | 2.71691556030738e-28 | 0.455611854 | 0.29 | 0.045 | 6.83820477373765e-24 | 33 |

| | | | | | | |
|---|---|---|---|---|---|---|
| cyb561a3a | 3.4338145536107e-28 | 0.347543004 | 0.204 | 0.024 | 8.64256784998276e-24 | 33 |
| ackr3b3 | 3.77645245397693e-28 | 0.336090901 | 0.204 | 0.023 | 9.50495318141454e-24 | 33 |
| hdac9b2 | 4.03716614591749e-28 | 0.352246849 | 0.204 | 0.024 | 1.01611434726597e-23 | 33 |
| wipf1a6 | 4.7234280996539e-28 | 0.511339765 | 0.355 | 0.064 | 1.18883961840189e-23 | 33 |
| GNAZ | 1.79065620153596e-27 | 0.335974918 | 0.204 | 0.024 | 4.50690259364585e-23 | 33 |
| mrtfab4 | 1.89751275167957e-27 | 0.446400732 | 0.312 | 0.052 | 4.77584984470232e-23 | 33 |
| plekhj12 | 1.90606324365109e-27 | 0.469548283 | 0.312 | 0.052 | 4.79737057794542e-23 | 33 |
| anxa3b3 | 1.91489821543602e-27 | 0.463562628 | 0.237 | 0.032 | 4.81960731843093e-23 | 33 |
| dbn15 | 5.39457107457174e-27 | 0.510031906 | 0.312 | 0.055 | 1.35775959375896e-22 | 33 |
| sept124 | 7.76286132519936e-27 | 0.497961912 | 0.333 | 0.061 | 1.95383456693943e-22 | 33 |
| parvg6 | 1.26835972734327e-26 | 0.485324741 | 0.355 | 0.067 | 3.19233459775028e-22 | 33 |
| cxcr4b10 | 2.02192882068112e-26 | 0.979427815 | 0.559 | 0.182 | 5.08899264877231e-22 | 33 |
| zgc:16273015 | 4.92759208640221e-26 | 0.862127191 | 0.871 | 0.39 | 1.24022565222657e-21 | 33 |
| il12rb2l2 | 1.81173600886471e-25 | 0.427577384 | 0.269 | 0.043 | 4.55995836071159e-21 | 33 |
| ppt16 | 1.83842015878501e-25 | 0.691096756 | 0.559 | 0.162 | 4.627119697646e-21 | 33 |
| capza1b7 | 2.08955776390507e-25 | 0.738037111 | 0.634 | 0.211 | 5.25920793597266e-21 | 33 |
| selplg6 | 2.77070622805074e-25 | 0.414838293 | 0.28 | 0.046 | 6.9735905053809e-21 | 33 |
| coro1a12 | 5.95652635019475e-25 | 0.766170677 | 0.71 | 0.23 | 1.49919811708052e-20 | 33 |
| XLOC-0283709 | 7.10815957125144e-25 | 0.695113766 | 0.527 | 0.14 | 1.78905268248827e-20 | 33 |
| arpc4l10 | 7.12488726116436e-25 | 0.726377392 | 0.602 | 0.196 | 1.79326287476246e-20 | 33 |
| marcksl1a9 | 1.03679161322466e-24 | 0.789046627 | 0.634 | 0.214 | 2.60950081132515e-20 | 33 |
| eif3jb1 | 1.29740736790764e-24 | 0.586092074 | 0.441 | 0.107 | 3.26544460428674e-20 | 33 |
| cyba6 | 3.26631764382653e-24 | 0.489142722 | 0.355 | 0.073 | 8.220994877747e-20 | 33 |
| rac212 | 4.61418522184606e-24 | 0.735477601 | 0.656 | 0.219 | 1.16134427848644e-19 | 33 |
| itpka1 | 9.82894448348925e-24 | 0.265907923 | 0.183 | 0.022 | 2.47384703704941e-19 | 33 |
| dennd2da | 1.36867660156364e-23 | 0.295590288 | 0.172 | 0.02 | 3.44482213847551e-19 | 33 |
| fcer1g4 | 2.65259909474236e-23 | 0.393572373 | 0.258 | 0.042 | 6.67632666155705e-19 | 33 |
| cnn210 | 3.41385686710037e-23 | 0.624282214 | 0.484 | 0.134 | 8.59233634880493e-19 | 33 |
| mao1 | 3.87783024284566e-23 | 0.273474407 | 0.14 | 0.014 | 9.76011093821824e-19 | 33 |
| gng21 | 4.5058892871239e-23 | 0.378496199 | 0.258 | 0.043 | 1.13408727467621e-18 | 33 |
| vsir9 | 6.08943438519684e-23 | 0.529047401 | 0.376 | 0.085 | 1.53264974041019e-18 | 33 |
| arpc49 | 8.82049695796262e-23 | 0.707980811 | 0.645 | 0.225 | 2.22003087934961e-18 | 33 |
| capzb10 | 1.32206157404556e-22 | 0.721492759 | 0.667 | 0.252 | 3.32749677571526e-18 | 33 |
| myl12.111 | 1.97013864562295e-22 | 0.720173543 | 0.828 | 0.391 | 4.95864195716839e-18 | 33 |
| itgb27 | 3.37876587511163e-22 | 0.530563437 | 0.409 | 0.098 | 8.50401583106845e-18 | 33 |

| | | | | | | |
|---|---|---|---|---|---|---|
| ckba3 | 4.16193127798843e-22 | 0.401477067 | 0.323 | 0.065 | 1.04751648335691e-17 | 33 |
| litaf7 | 1.19518565611658e-21 | 0.596947987 | 0.473 | 0.134 | 3.00816277787982e-17 | 33 |
| XLOC-0089933 | 1.66339079526064e-21 | 0.340687795 | 0.215 | 0.033 | 4.1865882925915e-17 | 33 |
| xaf12 | 2.18530671384341e-21 | 0.44195702 | 0.301 | 0.06 | 5.50019846807249e-17 | 33 |
| irf84 | 4.04564511041602e-21 | 0.39584809 | 0.258 | 0.046 | 1.01824841784061e-16 | 33 |
| CR318588.110 | 6.47646440820623e-21 | 0.93095448 | 0.581 | 0.207 | 1.63006132690143e-16 | 33 |
| plbd13 | 1.88236351318909e-20 | 0.289951085 | 0.183 | 0.026 | 4.73772072634561e-16 | 33 |
| csrp1a2 | 2.94571185840991e-20 | 0.446481456 | 0.312 | 0.067 | 7.4140621764319e-16 | 33 |
| abracl5 | 4.07276692359346e-20 | 0.671420115 | 0.624 | 0.22 | 1.02507470699924e-15 | 33 |
| lgals9l11 | 9.07052885569016e-20 | 0.28642209 | 0.172 | 0.024 | 2.28296140768866e-15 | 33 |
| rplp217 | 9.98146558649337e-20 | 0.45373101 | 0.989 | 0.757 | 2.51223507346452e-15 | 33 |
| twf1a4 | 1.33814907756142e-19 | 0.536851423 | 0.505 | 0.15 | 3.36798741331433e-15 | 33 |
| b2m13 | 1.8437648341979e-19 | 0.693013998 | 0.688 | 0.291 | 4.64057171119269e-15 | 33 |
| zgc:6405111 | 6.19727350849882e-19 | 0.599231718 | 0.516 | 0.162 | 1.55979176935407e-14 | 33 |
| XLOC-032112 | 1.69094706273583e-18 | 0.261648854 | 0.151 | 0.02 | 4.25594466219981e-14 | 33 |
| BX908750.13 | 2.13911096775024e-18 | 0.412456003 | 0.194 | 0.032 | 5.38392839473057e-14 | 33 |
| timm101 | 2.39804912516231e-18 | 0.389290872 | 0.28 | 0.058 | 6.03564984312103e-14 | 33 |
| BX322618.14 | 2.93843500575803e-18 | 0.408719974 | 0.183 | 0.029 | 7.39574706599239e-14 | 33 |
| hcst3 | 3.44819408245916e-18 | 0.36327022 | 0.258 | 0.051 | 8.67875968614146e-14 | 33 |
| ufc11 | 6.32000466404544e-18 | 0.454398942 | 0.344 | 0.086 | 1.5906819738936e-13 | 33 |
| ccr9a7 | 6.42959675697634e-18 | 0.53729014 | 0.344 | 0.087 | 1.61826520776337e-13 | 33 |
| arf6b6 | 1.40306951955401e-17 | 0.443964151 | 0.376 | 0.101 | 3.53138567376548e-13 | 33 |
| irf71 | 1.42147462033949e-17 | 0.309612881 | 0.194 | 0.032 | 3.57770947193247e-13 | 33 |
| pak2a1 | 1.71930037556872e-17 | 0.381081421 | 0.28 | 0.061 | 4.3273071152689e-13 | 33 |
| stk17b5 | 1.97576011909457e-17 | 0.493860004 | 0.398 | 0.11 | 4.97279064374912e-13 | 33 |
| prdx57 | 2.41921122722423e-17 | 0.5470619 | 0.484 | 0.158 | 6.08891273780066e-13 | 33 |
| gabarapa1 | 3.19138275935353e-17 | 0.563135619 | 0.484 | 0.16 | 8.03239126701689e-13 | 33 |
| batf31 | 1.18832987178418e-16 | 0.281398631 | 0.226 | 0.043 | 2.99090745429361e-12 | 33 |
| bach1b2 | 1.20306353836497e-16 | 0.297883329 | 0.161 | 0.025 | 3.0279906197108e-12 | 33 |
| cdc42l7 | 1.28461829637393e-16 | 0.581565952 | 0.57 | 0.215 | 3.23325579014353e-12 | 33 |
| wdr16 | 1.66477812473282e-16 | 0.54234386 | 0.441 | 0.141 | 4.19008006214004e-12 | 33 |
| skap27 | 3.45081726939001e-16 | 0.435238063 | 0.301 | 0.074 | 8.68536198532771e-12 | 33 |
| slc15a41 | 4.00870880126529e-16 | 0.255962528 | 0.151 | 0.022 | 1.00895191819046e-11 | 33 |
| gstm.35 | 5.25022011352765e-16 | 0.494163747 | 0.366 | 0.101 | 1.32142790037377e-11 | 33 |
| fam129bb | 5.28517380184155e-16 | 0.271903048 | 0.172 | 0.028 | 1.3302253941855e-11 | 33 |

| | | | | | | |
|---|---|---|---|---|---|---|
| fcer1gl8 | 5.39684373135504e-16 | 0.484830868 | 0.548 | 0.168 | 1.35833159874475e-11 | 33 |
| BX120005.14 | 8.60479138587898e-16 | 0.35706836 | 0.237 | 0.049 | 2.16573994391188e-11 | 33 |
| rab5c4 | 9.59938859291125e-16 | 0.459006617 | 0.398 | 0.12 | 2.41607011494983e-11 | 33 |
| asah1b2 | 1.65374755613199e-15 | 0.293475486 | 0.247 | 0.054 | 4.16231722402861e-11 | 33 |
| XLOC-0331373 | 1.91868716889186e-15 | 0.27263622 | 0.194 | 0.035 | 4.82914373538393e-11 | 33 |
| si:dkey-262k9.49 | 2.34726493925548e-15 | 0.418431675 | 0.333 | 0.09 | 5.90783112561211e-11 | 33 |
| atp6v1g14 | 2.9085143218538e-15 | 0.49879412 | 0.419 | 0.132 | 7.32043969667384e-11 | 33 |
| sema4ab | 4.01338490843692e-15 | 0.268930107 | 0.172 | 0.03 | 1.01012884760449e-10 | 33 |
| wee21 | 4.29439027328418e-15 | 0.41287946 | 0.344 | 0.093 | 1.0808550878829e-10 | 33 |
| itm2bb9 | 4.30885667312321e-15 | 0.560606527 | 0.57 | 0.221 | 1.08449613605838e-10 | 33 |
| cfl110 | 4.70941779690043e-15 | 0.535145087 | 0.882 | 0.576 | 1.18531336530187e-10 | 33 |
| si:ch211-222l21.113 | 4.89829100003568e-15 | 0.596308912 | 0.656 | 0.275 | 1.23285086179898e-10 | 33 |
| ecm1b4 | 7.47064869959935e-15 | 0.536218624 | 0.237 | 0.054 | 1.88028757120216e-10 | 33 |
| plekho1b4 | 1.09243963127331e-14 | 0.296439746 | 0.204 | 0.041 | 2.7495613079518e-10 | 33 |
| ostf13 | 1.31404144507634e-14 | 0.338254247 | 0.237 | 0.053 | 3.30731091311265e-10 | 33 |
| si:dkey-206d17.123 | 1.66575214848178e-14 | 0.306747873 | 0.237 | 0.052 | 4.1925315825138e-10 | 33 |
| rps15a12 | 2.20972407282218e-14 | 0.406447517 | 0.989 | 0.779 | 5.56165451888613e-10 | 33 |
| h1fx9 | 3.0186508764453e-14 | 0.611122321 | 0.484 | 0.183 | 7.59764239092519e-10 | 33 |
| mbd22 | 3.06654380787754e-14 | 0.264269592 | 0.183 | 0.035 | 7.71818411004697e-10 | 33 |
| COX7A2 (1 of many)6 | 3.60906253462546e-14 | 0.557619405 | 0.602 | 0.26 | 9.08364949339881e-10 | 33 |
| ncaldb5 | 5.77003833770269e-14 | 0.347133823 | 0.29 | 0.074 | 1.45226094921639e-09 | 33 |
| si:dkey-112e7.29 | 6.01836153359748e-14 | 0.539786837 | 0.516 | 0.205 | 1.51476141439115e-09 | 33 |
| cmtm3 | 6.36663919359796e-14 | 0.299311367 | 0.204 | 0.043 | 1.60241941863667e-09 | 33 |
| histh1l7 | 6.6853545527969e-14 | 0.498727658 | 0.355 | 0.11 | 1.68263688739345e-09 | 33 |
| itgae.24 | 1.16500095125807e-13 | 0.275669805 | 0.183 | 0.036 | 2.93219089422145e-09 | 33 |
| glrx6 | 1.36920219305505e-13 | 0.565252242 | 0.667 | 0.311 | 3.44614499970024e-09 | 33 |
| si:ch211-153b23.71 | 1.40407451919894e-13 | 0.257814915 | 0.14 | 0.023 | 3.53391515737182e-09 | 33 |
| plekho24 | 1.40534825999376e-13 | 0.387374725 | 0.28 | 0.074 | 3.5371210355783e-09 | 33 |
| arl5c7 | 1.6953019144103e-13 | 0.475208122 | 0.355 | 0.112 | 4.26690538837927e-09 | 33 |
| si:ch73-1a9.313 | 1.88001259493808e-13 | 0.559580517 | 0.613 | 0.265 | 4.73180370019965e-09 | 33 |
| chchd7 | 2.43862841341682e-13 | 0.29436069 | 0.183 | 0.037 | 6.1377838537288e-09 | 33 |
| ctsh3 | 5.19449611771075e-13 | 0.32711665 | 0.226 | 0.054 | 1.30740272786662e-08 | 33 |
| BX248082.16 | 6.78980507011355e-13 | 0.454833022 | 0.194 | 0.042 | 1.70892603809688e-08 | 33 |
| cap111 | 8.39833287332081e-13 | 0.544052257 | 0.548 | 0.233 | 2.11377640088611e-08 | 33 |
| pmaip12 | 8.81765925801522e-13 | 0.488180973 | 0.333 | 0.104 | 2.21931665864985e-08 | 33 |

| | | | | | | |
|---|---|---|---|---|---|---|
| GCA6 | 9.76966081054149e-13 | 0.438680896 | 0.484 | 0.181 | 2.45892592940519e-08 | 33 |
| ak13 | 1.47340203245326e-12 | 0.395378914 | 0.269 | 0.075 | 3.70840557548162e-08 | 33 |
| rpl1813 | 1.52664774775563e-12 | 0.370255912 | 0.989 | 0.773 | 3.84241971632615e-08 | 33 |
| nr4a110 | 1.5289572637058e-12 | 0.546777247 | 0.441 | 0.167 | 3.84823253702114e-08 | 33 |
| gdi23 | 2.01887757662755e-12 | 0.47480029 | 0.473 | 0.185 | 5.08131297261388e-08 | 33 |
| rgs138 | 2.11224480390423e-12 | 0.442105249 | 0.398 | 0.132 | 5.31630894694655e-08 | 33 |
| XLOC-00043110 | 2.22361072178314e-12 | 0.589376619 | 0.634 | 0.278 | 5.59660582565598e-08 | 33 |
| rab8b4 | 3.13595172095265e-12 | 0.353598892 | 0.28 | 0.08 | 7.89287688646572e-08 | 33 |
| XLOC-0054483 | 5.67339189223519e-12 | 0.272845974 | 0.151 | 0.029 | 1.42793600535667e-07 | 33 |
| ptmab12 | 7.33077191552195e-12 | 0.509980603 | 0.86 | 0.54 | 1.84508198341772e-07 | 33 |
| mir21-15 | 7.89975262204862e-12 | 0.392511027 | 0.333 | 0.108 | 1.98828873744342e-07 | 33 |
| stat5a1 | 9.40841985352237e-12 | 0.295257426 | 0.194 | 0.045 | 2.36800519293305e-07 | 33 |
| actr2a9 | 1.59777352538413e-11 | 0.392051072 | 0.376 | 0.13 | 4.02143618603932e-07 | 33 |
| si:dkeyp-68b7.126 | 2.79444122426324e-11 | 0.30005645 | 0.237 | 0.063 | 7.03332911734816e-07 | 33 |
| LOC1039091158 | 3.08267320030285e-11 | 0.44145827 | 0.43 | 0.165 | 7.75878017784224e-07 | 33 |
| rps27.113 | 3.42426324036253e-11 | 0.427662628 | 0.957 | 0.679 | 8.61852814966844e-07 | 33 |
| psme27 | 3.97808941496141e-11 | 0.382210544 | 0.301 | 0.097 | 1.00124532485164e-06 | 33 |
| sdcbp29 | 5.54550546044244e-11 | 0.478132851 | 0.484 | 0.208 | 1.39574826933876e-06 | 33 |
| ccng24 | 6.60419103520792e-11 | 0.342834824 | 0.28 | 0.086 | 1.66220884165148e-06 | 33 |
| ubl3b2 | 7.61366565390431e-11 | 0.276912412 | 0.258 | 0.073 | 1.91628350843117e-06 | 33 |
| map2k62 | 7.63113295463494e-11 | 0.306960184 | 0.29 | 0.089 | 1.92067985335207e-06 | 33 |
| CR392341.16 | 7.74762921287322e-11 | 0.293566343 | 0.226 | 0.06 | 1.95000079658806e-06 | 33 |
| rps226 | 8.47293684669888e-11 | 0.336359868 | 1 | 0.829 | 2.13255347494564e-06 | 33 |
| lgals2a5 | 1.0553993940249e-10 | 0.513440633 | 0.355 | 0.126 | 2.65633473482126e-06 | 33 |
| bloc1s62 | 1.32503292508304e-10 | 0.300555563 | 0.269 | 0.081 | 3.33497536914149e-06 | 33 |
| tpi1b8 | 1.75608742519234e-10 | 0.427399264 | 0.419 | 0.167 | 4.41989644046661e-06 | 33 |
| tagapb6 | 2.63195820361808e-10 | 0.323516148 | 0.237 | 0.068 | 6.62437560268635e-06 | 33 |
| rpl3910 | 2.9231135228963e-10 | 0.258062483 | 0.978 | 0.874 | 7.35718442577769e-06 | 33 |
| ms4a17a.65 | 3.57725444157574e-10 | 0.268730512 | 0.194 | 0.049 | 9.00359170400197e-06 | 33 |
| rps512 | 4.18606035518726e-10 | 0.38389023 | 0.957 | 0.701 | 1.05358953079708e-05 | 33 |
| RASSF55 | 5.00025124266219e-10 | 0.273159977 | 0.215 | 0.058 | 1.25851323526565e-05 | 33 |
| vdac36 | 7.23808683832494e-10 | 0.377013331 | 0.398 | 0.154 | 1.821754076338e-05 | 33 |
| cox8a8 | 8.45514723713693e-10 | 0.457604497 | 0.495 | 0.227 | 2.12807600811499e-05 | 33 |
| naga3 | 8.82363520287552e-10 | 0.256845494 | 0.194 | 0.051 | 2.22082074421174e-05 | 33 |
| tagapa8 | 1.6386531499905e-09 | 0.326558184 | 0.312 | 0.106 | 4.12432611321109e-05 | 33 |

| | | | | | | |
|---|---|---|---|---|---|---|
| sri3 | 1.86661916350732e-09 | 0.412313538 | 0.376 | 0.151 | 4.69809377263158e-05 | 33 |
| tsc22d39 | 1.89223463958734e-09 | 0.402818487 | 0.366 | 0.144 | 4.76256536437737e-05 | 33 |
| glula5 | 2.65999194926608e-09 | 0.364224546 | 0.366 | 0.136 | 6.69493373710779e-05 | 33 |
| rps2120 | 2.76803339136967e-09 | 0.365164162 | 0.968 | 0.75 | 6.96686324273833e-05 | 33 |
| laptm511 | 3.10438437406401e-09 | 0.378887169 | 0.43 | 0.17 | 7.81342503108171e-05 | 33 |
| rps179 | 3.33929937479508e-09 | 0.315554152 | 0.968 | 0.767 | 8.40468259642173e-05 | 33 |
| rps26l14 | 3.50429056045062e-09 | 0.416405361 | 0.968 | 0.664 | 8.81994891159817e-05 | 33 |
| nptna4 | 3.61809134905768e-09 | 0.315753436 | 0.237 | 0.072 | 9.10637411644327e-05 | 33 |
| ctsk3 | 4.0826413568491e-09 | 0.263422503 | 0.215 | 0.063 | 0.000102756 | 33 |
| rpl812 | 5.40642673416892e-09 | 0.31385262 | 0.968 | 0.78 | 0.000136074 | 33 |
| gstm.21 | 5.77175782105149e-09 | 0.281034595 | 0.183 | 0.05 | 0.000145269 | 33 |
| ckbb9 | 5.83447717366177e-09 | 0.436331907 | 0.581 | 0.266 | 0.000146848 | 33 |
| oaz1b12 | 5.94315020338549e-09 | 0.42478339 | 0.634 | 0.322 | 0.000149583 | 33 |
| ppiaa11 | 6.01142815868872e-09 | 0.282830194 | 0.968 | 0.804 | 0.000151302 | 33 |
| tagln211 | 6.02908959201298e-09 | 0.424691017 | 0.43 | 0.188 | 0.000151746 | 33 |
| eif3g7 | 6.18791294364278e-09 | 0.425620421 | 0.57 | 0.282 | 0.000155744 | 33 |
| rhoab9 | 9.26143311466864e-09 | 0.40325028 | 0.43 | 0.189 | 0.000233101 | 33 |
| fuca22 | 9.47068401557718e-09 | 0.26253189 | 0.215 | 0.064 | 0.000238368 | 33 |
| ndufb33 | 9.89726714397828e-09 | 0.403187612 | 0.473 | 0.218 | 0.000249104 | 33 |
| ybx18 | 1.26964320955203e-08 | 0.370702653 | 0.946 | 0.669 | 0.000319556 | 33 |
| grn23 | 1.3100665747446e-08 | 0.299815026 | 0.108 | 0.021 | 0.000329731 | 33 |
| rps1514 | 1.40846881138603e-08 | 0.397757736 | 0.968 | 0.684 | 0.000354498 | 33 |
| zgc:11418818 | 1.56342837168981e-08 | 0.416819196 | 0.935 | 0.627 | 0.000393499 | 33 |
| rps910 | 1.58585440515789e-08 | 0.307449373 | 0.968 | 0.776 | 0.000399144 | 33 |
| arhgdia10 | 1.60620719668703e-08 | 0.405284652 | 0.484 | 0.23 | 0.000404266 | 33 |
| rpl3814 | 1.62222024215419e-08 | 0.350525533 | 0.957 | 0.728 | 0.000408297 | 33 |
| rpl35a13 | 3.40471641583725e-08 | 0.335500375 | 0.989 | 0.759 | 0.000856933 | 33 |
| rpl2111 | 4.14142108532571e-08 | 0.307073705 | 0.957 | 0.756 | 0.001042354 | 33 |
| rpl18a12 | 4.45227760545496e-08 | 0.301198489 | 0.957 | 0.785 | 0.001120594 | 33 |
| si:ch211-212k18.74 | 5.65364948200948e-08 | 0.287895605 | 0.355 | 0.139 | 0.001422967 | 33 |
| atp6ap22 | 6.71976770367485e-08 | 0.277102072 | 0.28 | 0.101 | 0.001691298 | 33 |
| rpsa18 | 7.62289948190362e-08 | 0.32591462 | 0.957 | 0.752 | 0.001918608 | 33 |
| rps1413 | 8.57998296114209e-08 | 0.337501523 | 1 | 0.758 | 0.002159496 | 33 |
| rps311 | 9.6596042971569e-08 | 0.368894092 | 0.978 | 0.699 | 0.002431226 | 33 |
| rpl3617 | 1.11389739720615e-07 | 0.325777929 | 0.989 | 0.789 | 0.002803568 | 33 |

| | | | | | | |
|---|---|---|---|---|---|---|
| sh3bgrl36 | 1.71503548617676e-07 | 0.358583739 | 0.344 | 0.148 | 0.004316573 | 33 |
| clic18 | 1.79879969637566e-07 | 0.305640547 | 0.29 | 0.114 | 0.004527399 | 33 |
| cox4i18 | 1.93448686287306e-07 | 0.400058242 | 0.677 | 0.392 | 0.00486891 | 33 |
| crip113 | 2.05960353637161e-07 | 0.40047051 | 0.548 | 0.271 | 0.005183816 | 33 |
| romo113 | 2.12601351087293e-07 | 0.403715641 | 0.656 | 0.386 | 0.005350963 | 33 |
| fosl210 | 2.16641242026291e-07 | 0.339585528 | 0.301 | 0.122 | 0.005452643 | 33 |
| rpl1512 | 2.20995838890113e-07 | 0.335440263 | 0.978 | 0.744 | 0.005562244 | 33 |
| sod11 | 2.33994267780785e-07 | 0.409094 | 0.581 | 0.328 | 0.005889402 | 33 |
| rpl3212 | 2.61947527896875e-07 | 0.289601403 | 0.978 | 0.813 | 0.006592957 | 33 |
| eif3f12 | 2.81652709091085e-07 | 0.412059949 | 0.699 | 0.432 | 0.007088917 | 33 |
| derl24 | 3.1232358013615e-07 | 0.266310805 | 0.28 | 0.106 | 0.007860872 | 33 |
| rps2815 | 3.1776764606946e-07 | 0.270054875 | 0.978 | 0.826 | 0.007997894 | 33 |
| XLOC-0418703 | 3.46721096814168e-07 | 0.253829785 | 0.118 | 0.029 | 0.008726623 | 33 |
| tln16 | 3.49225626748419e-07 | 0.293703337 | 0.312 | 0.128 | 0.00878966 | 33 |
| fam53b1 | 5.28877514304779e-07 | 0.261291267 | 0.204 | 0.069 | 0.013311318 | 33 |
| rps1615 | 5.36125588848951e-07 | 0.332666849 | 0.968 | 0.737 | 0.013493745 | 33 |
| rpl918 | 6.61075153529975e-07 | 0.385586735 | 0.914 | 0.629 | 0.016638601 | 33 |
| serf212 | 6.71916488173521e-07 | 0.373210654 | 0.624 | 0.343 | 0.016911466 | 33 |
| rpl3710 | 9.03050254315751e-07 | 0.261061915 | 0.989 | 0.849 | 0.022728872 | 33 |
| rps2414 | 9.11812620597895e-07 | 0.298576139 | 0.968 | 0.758 | 0.022949412 | 33 |
| bhlhe4011 | 1.04530421189101e-06 | 0.348708964 | 0.355 | 0.161 | 0.026309262 | 33 |
| mob1a6 | 1.18825297162076e-06 | 0.306601633 | 0.366 | 0.167 | 0.029907139 | 33 |
| sept62 | 1.35495045655279e-06 | 0.316436072 | 0.43 | 0.206 | 0.034102748 | 33 |
| ywhabl4 | 1.44016838281764e-06 | 0.370738496 | 0.505 | 0.279 | 0.036247598 | 33 |
| rpl5a14 | 1.7022905129402e-06 | 0.381905217 | 0.785 | 0.492 | 0.04284495 | 33 |
| mbnl19 | 1.74625100931871e-06 | 0.286993733 | 0.398 | 0.184 | 0.043951392 | 33 |
| rpl2914 | 2.30517488530236e-06 | 0.268264121 | 0.978 | 0.808 | 0.058018947 | 33 |
| fam49ba6 | 2.52234359250934e-06 | 0.290793327 | 0.258 | 0.104 | 0.063484866 | 33 |
| capza1a9 | 2.77433136094482e-06 | 0.334224558 | 0.398 | 0.2 | 0.069827146 | 33 |
| psma22 | 3.01510424516946e-06 | 0.301597477 | 0.333 | 0.153 | 0.075887159 | 33 |
| cdc426 | 3.02620699661499e-06 | 0.301897393 | 0.43 | 0.213 | 0.076166604 | 33 |
| COA41 | 3.03290711338191e-06 | 0.258745509 | 0.226 | 0.085 | 0.076335239 | 33 |
| atf4a3 | 3.19672954324065e-06 | 0.285712883 | 0.387 | 0.183 | 0.080458486 | 33 |
| ube2e36 | 3.21097970864681e-06 | 0.273388442 | 0.344 | 0.156 | 0.080817148 | 33 |
| XLOC-0203015 | 3.69592927453925e-06 | 0.293826498 | 0.226 | 0.086 | 0.093022844 | 33 |

| | | | | | | |
|---|---|---|---|---|---|---|
| rplp2l17 | 4.09683988268455e-06 | 0.297011711 | 0.978 | 0.774 | 0.103113363 | 33 |
| XLOC-0108999 | 4.62470881537891e-06 | 0.278190976 | 0.505 | 0.259 | 0.116399296 | 33 |
| magt13 | 4.76016118299482e-06 | 0.284904473 | 0.269 | 0.115 | 0.119808497 | 33 |
| psme15 | 5.92826782332579e-06 | 0.264601047 | 0.226 | 0.091 | 0.149208573 | 33 |
| atp6v0ca3 | 7.26201621609585e-06 | 0.295438029 | 0.409 | 0.199 | 0.182777686 | 33 |
| mapre1a7 | 8.48437640205683e-06 | 0.259058365 | 0.419 | 0.198 | 0.21354327 | 33 |
| rpl1113 | 8.59055088294678e-06 | 0.262430584 | 0.968 | 0.814 | 0.216215575 | 33 |
| zgc:564939 | 1.08205146631263e-05 | 0.330266651 | 0.817 | 0.573 | 0.272341534 | 33 |
| zgc:1537134 | 1.10567543847849e-05 | 0.285942909 | 0.323 | 0.153 | 0.278287451 | 33 |
| rpl2213 | 1.12431956665764e-05 | 0.305849732 | 0.914 | 0.694 | 0.282979992 | 33 |
| slc6a6b6 | 1.15708296310018e-05 | 0.271027511 | 0.247 | 0.104 | 0.291226211 | 33 |
| naca12 | 1.80287522758685e-05 | 0.326930157 | 0.86 | 0.619 | 0.453765666 | 33 |
| rps4x11 | 1.87792849993706e-05 | 0.301867143 | 0.935 | 0.71 | 0.472655824 | 33 |
| klf6a8 | 2.03965792783808e-05 | 0.295368324 | 0.344 | 0.168 | 0.513361504 | 33 |
| CR318588.34 | 2.08984472154533e-05 | 0.254737532 | 0.14 | 0.046 | 0.525993018 | 33 |
| rpl615 | 2.16446943544236e-05 | 0.32097579 | 0.914 | 0.667 | 0.544775312 | 33 |
| mtpn4 | 2.18256767943068e-05 | 0.256147095 | 0.312 | 0.147 | 0.549330459 | 33 |
| nccrp14 | 2.19999526735315e-05 | 0.261489494 | 0.398 | 0.195 | 0.553716809 | 33 |
| cox5aa3 | 2.80525639821562e-05 | 0.313783325 | 0.602 | 0.352 | 0.706054983 | 33 |
| rps2310 | 3.03002540604553e-05 | 0.27999093 | 0.925 | 0.723 | 0.762627094 | 33 |
| rpl2710 | 3.20874774761618e-05 | 0.265917952 | 0.946 | 0.763 | 0.807609721 | 33 |
| uqcrq4 | 3.42061853069978e-05 | 0.315168707 | 0.473 | 0.272 | 0.860935478 | 33 |
| rpl36a11 | 4.02761249089496e-05 | 0.253884004 | 0.989 | 0.809 | 1 | 33 |
| atp5meb3 | 4.04946145235581e-05 | 0.296419054 | 0.548 | 0.323 | 1 | 33 |
| rpl2413 | 4.08798809751138e-05 | 0.275721145 | 0.978 | 0.776 | 1 | 33 |
| rps1017 | 4.42154523438287e-05 | 0.313044714 | 0.935 | 0.694 | 1 | 33 |
| rpl3414 | 4.6723094258825e-05 | 0.259496282 | 0.957 | 0.784 | 1 | 33 |
| ssr33 | 4.89565362140998e-05 | 0.27820212 | 0.495 | 0.273 | 1 | 33 |
| hmgn210 | 5.55654633590082e-05 | 0.403285726 | 0.645 | 0.443 | 1 | 33 |
| b2ml13 | 5.56560918760965e-05 | 0.293173861 | 0.473 | 0.268 | 1 | 33 |
| tpm36 | 5.83382185245557e-05 | 0.292622567 | 0.452 | 0.255 | 1 | 33 |
| mcl1a9 | 6.45494362631082e-05 | 0.285010786 | 0.806 | 0.624 | 1 | 33 |
| eef1db10 | 6.8469757447133e-05 | 0.284944155 | 0.484 | 0.275 | 1 | 33 |
| sat1a.26 | 7.35449886452687e-05 | 0.275849617 | 0.527 | 0.299 | 1 | 33 |
| cst14a.2 | 7.5096358871167e-05 | 0.271463673 | 0.419 | 0.225 | 1 | 33 |

| | | | | | | |
|---|---|---|---|---|---|---|
| ap2m1a2 | 8.03893538152216e-05 | 0.255050308 | 0.29 | 0.143 | 1 | 33 |
| capgb7 | 8.72075375293643e-05 | 0.27308099 | 0.247 | 0.115 | 1 | 33 |
| rpl27a15 | 9.57620212747993e-05 | 0.349565127 | 0.796 | 0.496 | 1 | 33 |
| ccni12 | 0.000114268 | 0.286773675 | 0.495 | 0.295 | 1 | 33 |
| COX5B4 | 0.000134809 | 0.289423166 | 0.591 | 0.358 | 1 | 33 |
| rps27a12 | 0.000137429 | 0.27899442 | 0.978 | 0.751 | 1 | 33 |
| hmgn79 | 0.000143153 | 0.301106493 | 0.462 | 0.28 | 1 | 33 |
| atp5pb3 | 0.000153275 | 0.262764747 | 0.516 | 0.301 | 1 | 33 |
| rpl178 | 0.000176962 | 0.267106385 | 0.957 | 0.746 | 1 | 33 |
| pomp5 | 0.000189008 | 0.258744456 | 0.409 | 0.232 | 1 | 33 |
| rpl3014 | 0.000195218 | 0.270954775 | 0.925 | 0.708 | 1 | 33 |
| rpl1413 | 0.000276421 | 0.288782654 | 0.882 | 0.635 | 1 | 33 |
| faua11 | 0.000278252 | 0.292879565 | 0.935 | 0.689 | 1 | 33 |
| zgc:1538676 | 0.000424498 | 0.287753621 | 0.667 | 0.437 | 1 | 33 |
| rps27l10 | 0.00049808 | 0.259708746 | 0.355 | 0.204 | 1 | 33 |
| ywhaba6 | 0.000520095 | 0.250838964 | 0.516 | 0.322 | 1 | 33 |
| rps27.214 | 0.000537374 | 0.311988765 | 0.785 | 0.518 | 1 | 33 |
| celf214 | 0.000643516 | 0.255760137 | 0.409 | 0.243 | 1 | 33 |
| mibp211 | 0.000871169 | 0.285582543 | 0.72 | 0.493 | 1 | 33 |
| rps1814 | 0.001033734 | 0.257167215 | 0.935 | 0.725 | 1 | 33 |
| rpl3115 | 0.002074629 | 0.264550661 | 0.871 | 0.655 | 1 | 33 |
| eef1g11 | 0.003072054 | 0.262907487 | 0.849 | 0.621 | 1 | 33 |
| rpl23a16 | 0.00595888 | 0.279157017 | 0.892 | 0.634 | 1 | 33 |
| uba5214 | 0.006533658 | 0.255554056 | 0.882 | 0.63 | 1 | 33 |
| mpx1 | 4.92131939957729e-72 | 0.682059718 | 0.261 | 0.015 | 1.23864687967961e-67 | 34 |
| zgc:17513511 | 2.72210634765991e-23 | 0.787334465 | 0.943 | 0.48 | 6.85126946642523e-19 | 34 |
| cldnd11 | 8.11893085547275e-22 | 0.705666252 | 1 | 0.605 | 2.04345370701394e-17 | 34 |
| zp2.511 | 1.93282158191282e-21 | 0.724425203 | 0.989 | 0.564 | 4.86471863951638e-17 | 34 |
| si:zfos-1505d6.310 | 2.56689862416592e-21 | 0.748186102 | 0.966 | 0.513 | 6.46062714716319e-17 | 34 |
| si:dkeyp-98a7.910 | 3.12054683127388e-21 | 0.733179221 | 0.943 | 0.512 | 7.85410431963324e-17 | 34 |
| si:dkey-240e12.610 | 3.68652688718146e-21 | 0.722009589 | 0.989 | 0.569 | 9.27861952234702e-17 | 34 |
| crp210 | 5.40822850084551e-21 | 0.758522347 | 0.955 | 0.489 | 1.36119703137781e-16 | 34 |
| si:dkey-241l7.210 | 2.58406615722652e-20 | 0.665133844 | 1 | 0.642 | 6.50383611112342e-16 | 34 |
| h2af1al11 | 3.64198564191103e-20 | 0.687540032 | 0.989 | 0.593 | 9.16651366212588e-16 | 34 |
| CR790368.111 | 3.70792431966794e-20 | 0.718321345 | 0.943 | 0.52 | 9.33247472017223e-16 | 34 |

| | | | | | | |
|---|---|---|---|---|---|---|
| zp2.111 | 3.07884927938296e-19 | 0.710960849 | 0.955 | 0.535 | 7.74915575127898e-15 | 34 |
| si:dkey-241l7.511 | 5.87036305806238e-19 | 0.680472475 | 0.989 | 0.584 | 1.47751167808372e-14 | 34 |
| tmsb510 | 1.07776563674033e-18 | 0.722423775 | 0.943 | 0.489 | 2.71262833111173e-14 | 34 |
| si:dkey-241l7.410 | 1.77673695406417e-18 | 0.647870996 | 0.977 | 0.609 | 4.47186923968412e-14 | 34 |
| slbp210 | 2.76450458448618e-18 | 0.731983078 | 0.807 | 0.397 | 6.95798158869327e-14 | 34 |
| si:dkey-241l7.610 | 3.83607145156597e-18 | 0.674512101 | 0.909 | 0.527 | 9.65500823644638e-14 | 34 |
| si:dkeyp-20e4.810 | 5.89945436680304e-18 | 0.73819663 | 0.841 | 0.408 | 1.48483366958066e-13 | 34 |
| LOC10053749910 | 9.56074497828086e-18 | 0.746107509 | 0.727 | 0.322 | 2.40634390358351e-13 | 34 |
| si:ch211-226h8.1410 | 1.11575395151936e-17 | 0.673585611 | 0.977 | 0.567 | 2.80824112057907e-13 | 34 |
| ccna110 | 1.36031286176807e-17 | 0.692269106 | 0.886 | 0.476 | 3.42377144178407e-13 | 34 |
| si:ch211-12h2.810 | 1.43101325139248e-17 | 0.668275751 | 0.955 | 0.548 | 3.60171725242974e-13 | 34 |
| zp2.310 | 1.49479033641471e-17 | 0.696435949 | 0.943 | 0.519 | 3.76223779772219e-13 | 34 |
| zp311 | 1.92421521143811e-17 | 0.648417218 | 0.955 | 0.585 | 4.84305726566858e-13 | 34 |
| si:dkey-229d11.310 | 1.9399162580345e-17 | 0.731082098 | 0.898 | 0.438 | 4.88257522984704e-13 | 34 |
| zgc:15349910 | 2.32586272091932e-17 | 0.712121231 | 0.955 | 0.513 | 5.85396388228185e-13 | 34 |
| cldng10 | 3.3303625285206e-17 | 0.677304691 | 0.909 | 0.514 | 8.3821894480335e-13 | 34 |
| tuba4l10 | 1.04112477827806e-16 | 0.706056023 | 0.818 | 0.404 | 2.62040695444804e-12 | 34 |
| si:ch211-125e6.1210 | 2.30680799590282e-16 | 0.676009398 | 0.898 | 0.501 | 5.80600504488781e-12 | 34 |
| zp2.210 | 2.97424983943258e-16 | 0.661365562 | 0.932 | 0.53 | 7.48588942086785e-12 | 34 |
| zar1l10 | 4.73928245259108e-16 | 0.688125813 | 0.784 | 0.398 | 1.19283000049265e-11 | 34 |
| acp5a11 | 2.34141686969284e-15 | 0.62436296 | 0.886 | 0.541 | 5.8931121193299e-11 | 34 |
| siva110 | 3.28047082999166e-15 | 0.662720287 | 0.773 | 0.429 | 8.25661703200601e-11 | 34 |
| zp2.610 | 3.55231767620087e-15 | 0.649957501 | 0.864 | 0.502 | 8.94082835922998e-11 | 34 |
| zgc:17177610 | 4.71701717314629e-15 | 0.65262882 | 0.886 | 0.496 | 1.18722605230919e-10 | 34 |
| mt210 | 4.96308264890659e-15 | 0.638349684 | 0.92 | 0.535 | 1.2491582719033e-10 | 34 |
| lyz1 | 5.10025907558437e-15 | 0.485509991 | 0.261 | 0.061 | 1.28368420673383e-10 | 34 |
| zp3.211 | 6.26793657493921e-15 | 0.635617848 | 0.932 | 0.558 | 1.57757695654645e-10 | 34 |
| si:dkeyp-98a7.810 | 6.63967715149737e-15 | 0.659246869 | 0.841 | 0.458 | 1.67114034226037e-10 | 34 |
| zgc:5546111 | 8.16850009614649e-15 | 0.690130093 | 0.636 | 0.289 | 2.05592978919911e-10 | 34 |
| thy111 | 1.69595802400203e-14 | 0.643218108 | 0.795 | 0.451 | 4.2685567506107e-10 | 34 |
| si:dkeyp-98a7.410 | 2.13035055581979e-14 | 0.651313928 | 0.784 | 0.426 | 5.36187931394284e-10 | 34 |
| h1m11 | 2.85363539892635e-14 | 0.602545916 | 0.943 | 0.602 | 7.18231493555774e-10 | 34 |
| si:ch73-189n23.111 | 3.400160689929e-14 | 0.642741651 | 0.92 | 0.525 | 8.55786444048231e-10 | 34 |
| slc16a310 | 4.00792092824219e-14 | 0.683062994 | 0.568 | 0.249 | 1.00875361842928e-09 | 34 |
| zgc:7711810 | 4.564868270933e-14 | 0.640341035 | 0.83 | 0.458 | 1.14893169511113e-09 | 34 |

| | | | | | | |
|---|---|---|---|---|---|---|
| ccnb110 | 5.22581428588425e-14 | 0.654817115 | 0.761 | 0.394 | 1.31528519761421e-09 | 34 |
| BX649337.111 | 5.28335513577057e-14 | 0.655043812 | 0.841 | 0.438 | 1.32976765412209e-09 | 34 |
| LOC10188389710 | 5.36830498174011e-14 | 0.642926688 | 0.466 | 0.164 | 1.35114868085417e-09 | 34 |
| zpcx10 | 5.71085771084535e-14 | 0.6744235 | 0.705 | 0.34 | 1.43736577724267e-09 | 34 |
| crp310 | 6.49925179878324e-14 | 0.669333317 | 0.534 | 0.219 | 1.63579668523575e-09 | 34 |
| zgc:17151710 | 8.56513160049656e-14 | 0.65536198 | 0.75 | 0.389 | 2.15575797252898e-09 | 34 |
| ctss2.111 | 1.39397564195187e-13 | 0.576564389 | 0.864 | 0.535 | 3.50849729322867e-09 | 34 |
| si:ch211-226h8.410 | 1.90597996116598e-13 | 0.646497336 | 0.841 | 0.442 | 4.79716096425867e-09 | 34 |
| lamp210 | 2.47212992376312e-13 | 0.5825978 | 0.864 | 0.555 | 6.22210380511941e-09 | 34 |
| qdprb210 | 2.59663436282348e-13 | 0.631901225 | 0.773 | 0.434 | 6.53546902779042e-09 | 34 |
| zp3b10 | 3.47325311676768e-13 | 0.648463995 | 0.761 | 0.389 | 8.74183076959257e-09 | 34 |
| gapdh11 | 3.58140686382471e-13 | 0.621876569 | 0.75 | 0.421 | 9.01404293556041e-09 | 34 |
| CU571163.110 | 4.27811312861357e-13 | 0.640168719 | 0.761 | 0.402 | 1.07675829334075e-08 | 34 |
| si:ch211-250e5.1610 | 4.61580123690859e-13 | 0.653417315 | 0.648 | 0.347 | 1.16175101331752e-08 | 34 |
| zgc:17344310 | 5.424168164223e-13 | 0.641649331 | 0.693 | 0.331 | 1.36520888525329e-08 | 34 |
| ca15b10 | 6.25266006180284e-13 | 0.644624483 | 0.58 | 0.25 | 1.57373201095516e-08 | 34 |
| s100a110 | 6.77654876085588e-13 | 0.627886508 | 0.693 | 0.364 | 1.70558955761982e-08 | 34 |
| si:dkey-90l23.110 | 8.24296761791562e-13 | 0.618823777 | 0.795 | 0.448 | 2.07467251975318e-08 | 34 |
| nasp10 | 9.34724056756248e-13 | 0.643032138 | 0.682 | 0.34 | 2.3526069784498e-08 | 34 |
| gtf3ab11 | 9.41316615046169e-13 | 0.636978467 | 0.682 | 0.321 | 2.3691997884097e-08 | 34 |
| zp3a.110 | 1.26061871966868e-12 | 0.630648494 | 0.705 | 0.382 | 3.17285125553411e-08 | 34 |
| si:dkeyp-50f7.210 | 1.85314750106598e-12 | 0.612206902 | 0.716 | 0.403 | 4.66418694543297e-08 | 34 |
| ccnb210 | 1.89818479141953e-12 | 0.627264284 | 0.727 | 0.385 | 4.77754130152382e-08 | 34 |
| zgc:17177910 | 2.68454368372764e-12 | 0.577214828 | 0.818 | 0.513 | 6.75672799757409e-08 | 34 |
| nanog10 | 3.04642033708413e-12 | 0.627057444 | 0.693 | 0.36 | 7.66753534640706e-08 | 34 |
| mid1ip1l10 | 1.00446431326948e-11 | 0.551336528 | 0.852 | 0.544 | 2.52813623006796e-07 | 34 |
| hnrnpabb11 | 1.04058186906481e-11 | 0.495788295 | 0.818 | 0.598 | 2.61904050624923e-07 | 34 |
| si:dkey-19b23.1110 | 1.19998521230956e-11 | 0.62841336 | 0.568 | 0.28 | 3.02024278086194e-07 | 34 |
| hbba110 | 1.35940640166404e-11 | 0.635366747 | 0.693 | 0.357 | 3.42148997234823e-07 | 34 |
| elovl7a10 | 1.56502176027665e-11 | 0.621930868 | 0.443 | 0.183 | 3.93900326844029e-07 | 34 |
| zp3a.210 | 1.74938409820269e-11 | 0.574158522 | 0.818 | 0.492 | 4.40302483676636e-07 | 34 |
| rrm210 | 4.06228241092722e-11 | 0.557106426 | 0.83 | 0.499 | 1.02243586000627e-06 | 34 |
| AL929032.110 | 4.71204501851937e-11 | 0.611342192 | 0.625 | 0.318 | 1.18597461071114e-06 | 34 |
| zgc:66455 | 5.50674862500012e-11 | 0.396375103 | 0.17 | 0.037 | 1.38599356142628e-06 | 34 |
| wu:fi42e0310 | 5.62008408993882e-11 | 0.606062545 | 0.659 | 0.351 | 1.4145189645967e-06 | 34 |

| | | | | | | |
|---|---|---|---|---|---|---|
| si:dkeyp-46h3.610 | 6.65778450748587e-11 | 0.577352468 | 0.705 | 0.412 | 1.67569778268912e-06 | 34 |
| si:dkeyp-46h3.810 | 1.02455312054335e-10 | 0.590689662 | 0.67 | 0.352 | 2.57869774909555e-06 | 34 |
| gmnn10 | 1.32496626557161e-10 | 0.594537302 | 0.591 | 0.304 | 3.33480759381719e-06 | 34 |
| uchl110 | 1.34933209898261e-10 | 0.582715896 | 0.636 | 0.345 | 3.39613395992932e-06 | 34 |
| cth110 | 1.41242735865268e-10 | 0.57736466 | 0.693 | 0.38 | 3.55493841899293e-06 | 34 |
| si:dkey-90l23.210 | 1.79609291933669e-10 | 0.588222445 | 0.648 | 0.346 | 4.52058626867851e-06 | 34 |
| epcam11 | 1.97796733135352e-10 | 0.545444395 | 0.818 | 0.506 | 4.97834597628367e-06 | 34 |
| zgc:19524510 | 3.23354239558008e-10 | 0.572374455 | 0.693 | 0.389 | 8.1385028554355e-06 | 34 |
| tmem54a8 | 3.47695464993822e-10 | 0.581347281 | 0.364 | 0.142 | 8.75114715842951e-06 | 34 |
| lsm14b10 | 5.82657446931645e-10 | 0.577050641 | 0.58 | 0.301 | 1.46649052818226e-05 | 34 |
| si:ch211-14a17.710 | 8.3932294382595e-10 | 0.554048298 | 0.705 | 0.409 | 2.11249191731553e-05 | 34 |
| kpna710 | 1.09895393906162e-09 | 0.56608322 | 0.614 | 0.329 | 2.76595716922419e-05 | 34 |
| LOC11043782110 | 1.42935592262992e-09 | 0.558140972 | 0.705 | 0.417 | 3.59754592166725e-05 | 34 |
| hnrnpa0l2 | 1.47845172956692e-09 | 0.323400388 | 0.898 | 0.757 | 3.72111515814698e-05 | 34 |
| si:ch211-207i20.3 | 3.04152758277131e-09 | 0.391724125 | 0.182 | 0.047 | 7.6552207730771e-05 | 34 |
| cax11 | 7.83857454229145e-09 | 0.411276157 | 0.205 | 0.059 | 0.000197289 | 34 |
| birc5b10 | 8.19061805129169e-09 | 0.541581841 | 0.523 | 0.26 | 0.00020615 | 34 |
| si:dkeyp-46h3.210 | 8.3494827236273e-09 | 0.530591838 | 0.67 | 0.406 | 0.000210148 | 34 |
| zgc:5623110 | 8.43788824822078e-09 | 0.554460503 | 0.466 | 0.229 | 0.000212373 | 34 |
| ccna210 | 9.46255951927298e-09 | 0.548110758 | 0.534 | 0.285 | 0.000238163 | 34 |
| LOC10053516210 | 1.43095163001408e-08 | 0.540748392 | 0.591 | 0.31 | 0.000360156 | 34 |
| LOC10188547710 | 1.51038826597105e-08 | 0.54232158 | 0.545 | 0.281 | 0.00038015 | 34 |
| si:ch211-132b12.810 | 2.44280303662389e-08 | 0.527293266 | 0.591 | 0.343 | 0.000614829 | 34 |
| zar110 | 2.98189403934301e-08 | 0.507098911 | 0.705 | 0.441 | 0.000750513 | 34 |
| si:ch1073-263o8.210 | 3.15832284282932e-08 | 0.526107204 | 0.511 | 0.265 | 0.000794918 | 34 |
| ttc9c | 4.12813072349133e-08 | 0.368329641 | 0.159 | 0.042 | 0.001039009 | 34 |
| zp3c10 | 4.13289585681392e-08 | 0.528679268 | 0.409 | 0.19 | 0.001040209 | 34 |
| haus69 | 6.77958485395695e-08 | 0.534080485 | 0.432 | 0.21 | 0.001706354 | 34 |
| si:dkey-208k4.210 | 7.06508154899432e-08 | 0.520109127 | 0.523 | 0.297 | 0.00177821 | 34 |
| npm2b10 | 8.56693767175858e-08 | 0.497825284 | 0.659 | 0.397 | 0.002156213 | 34 |
| zgc:13625410 | 1.05874195474251e-07 | 0.502156589 | 0.58 | 0.332 | 0.002664748 | 34 |
| CDPF110 | 1.07895438020287e-07 | 0.532937237 | 0.477 | 0.27 | 0.00271562 | 34 |
| larp6b10 | 1.11751234330931e-07 | 0.505214037 | 0.42 | 0.197 | 0.002812667 | 34 |
| pabpn1l10 | 1.13409880642509e-07 | 0.512784209 | 0.523 | 0.285 | 0.002854413 | 34 |
| si:ch211-12h2.610 | 1.20579636642623e-07 | 0.475044426 | 0.727 | 0.475 | 0.003034869 | 34 |

| | | | | | | |
|---|---|---|---|---|---|---|
| zgc:17385610 | 1.24458146687401e-07 | 0.512654593 | 0.443 | 0.221 | 0.003132487 | 34 |
| zgc:16287910 | 2.395290698194e-07 | 0.475910874 | 0.398 | 0.182 | 0.006028707 | 34 |
| arhgef10la10 | 2.6453045000832e-07 | 0.506855412 | 0.511 | 0.276 | 0.006657967 | 34 |
| btg410 | 2.85253253290223e-07 | 0.483667009 | 0.625 | 0.37 | 0.007179539 | 34 |
| zgc:171717 | 3.14714953897637e-07 | 0.335487202 | 0.159 | 0.045 | 0.007921061 | 34 |
| si:ch211-5k11.810 | 3.81141623579446e-07 | 0.385085806 | 0.682 | 0.359 | 0.009592954 | 34 |
| si:dkey-152b24.610 | 4.19310722254605e-07 | 0.492314683 | 0.534 | 0.299 | 0.010553632 | 34 |
| spink2.111 | 4.20800743947554e-07 | 0.451997041 | 0.67 | 0.438 | 0.010591134 | 34 |
| AL929192.110 | 4.21832377348865e-07 | 0.47415881 | 0.364 | 0.165 | 0.010617099 | 34 |
| hnrnpaba1 | 4.46622161933137e-07 | 0.413668146 | 0.614 | 0.463 | 0.011241033 | 34 |
| si:ch211-145c1.110 | 4.52874720682316e-07 | 0.494080676 | 0.455 | 0.235 | 0.011398404 | 34 |
| si:ch73-109d9.2 | 4.65733736567272e-07 | 0.283113982 | 0.136 | 0.035 | 0.011722052 | 34 |
| si:ch211-251f6.610 | 5.28730421852934e-07 | 0.477943619 | 0.557 | 0.333 | 0.013307616 | 34 |
| osbpl7 | 8.92546476003081e-07 | 0.30709793 | 0.136 | 0.037 | 0.022464502 | 34 |
| sms2 | 9.35566584307503e-07 | 0.326298362 | 0.136 | 0.038 | 0.023547275 | 34 |
| si:dkey-229d11.510 | 1.04217411351787e-06 | 0.476186991 | 0.659 | 0.391 | 0.02623048 | 34 |
| rogdi4 | 1.61706849623815e-06 | 0.442558387 | 0.273 | 0.117 | 0.040699997 | 34 |
| hbaa110 | 1.82041615310971e-06 | 0.462674802 | 0.523 | 0.283 | 0.045818054 | 34 |
| map1lc3c10 | 2.00100528888279e-06 | 0.45562409 | 0.398 | 0.203 | 0.050363302 | 34 |
| tipin | 2.00379531216014e-06 | 0.378647785 | 0.182 | 0.062 | 0.050433524 | 34 |
| CR847984.110 | 2.07284665499577e-06 | 0.465820465 | 0.341 | 0.165 | 0.052171477 | 34 |
| kif222 | 2.09323689901621e-06 | 0.412036689 | 0.227 | 0.088 | 0.05268468 | 34 |
| avd10 | 2.26945058952561e-06 | 0.471287188 | 0.489 | 0.267 | 0.057119802 | 34 |
| gdf310 | 2.49894796410673e-06 | 0.454576965 | 0.432 | 0.225 | 0.062896021 | 34 |
| prdx110 | 2.63440536145289e-06 | 0.44799829 | 0.352 | 0.168 | 0.066305349 | 34 |
| pif16 | 3.37901018207295e-06 | 0.404435587 | 0.261 | 0.108 | 0.085046307 | 34 |
| nanos37 | 3.4216143669324e-06 | 0.422267251 | 0.25 | 0.104 | 0.086118612 | 34 |
| zgc:1134249 | 3.48909016505545e-06 | 0.448633459 | 0.341 | 0.163 | 0.08781691 | 34 |
| inka1b10 | 3.5674648596874e-06 | 0.46030565 | 0.364 | 0.186 | 0.089789523 | 34 |
| si:ch73-15b2.55 | 3.69463660690557e-06 | 0.398329043 | 0.239 | 0.096 | 0.092990309 | 34 |
| zgc:17147410 | 3.7680993534441e-06 | 0.449850516 | 0.523 | 0.306 | 0.094839293 | 34 |
| CABZ01059627.17 | 4.34805139298533e-06 | 0.416122669 | 0.307 | 0.137 | 0.109436106 | 34 |
| zgc:17374210 | 5.37179268624183e-06 | 0.433304688 | 0.33 | 0.157 | 0.13520265 | 34 |
| il13ra24 | 5.8875926579236e-06 | 0.404610897 | 0.261 | 0.111 | 0.14818482 | 34 |
| arid3b | 6.16597197767989e-06 | 0.299479795 | 0.148 | 0.046 | 0.155191349 | 34 |

| | | | | | | |
|---|---|---|---|---|---|---|
| eloca | 6.17619443256632e-06 | 0.420729558 | 0.239 | 0.101 | 0.155448638 | 34 |
| zfand63 | 6.84630343985783e-06 | 0.412200524 | 0.284 | 0.127 | 0.172314611 | 34 |
| timm131 | 6.91366966374059e-06 | 0.402134296 | 0.239 | 0.1 | 0.174010152 | 34 |
| si:ch211-202f5.34 | 8.76806285604268e-06 | 0.367499138 | 0.227 | 0.092 | 0.220683374 | 34 |
| mkrn49 | 9.14230326091813e-06 | 0.439522731 | 0.42 | 0.23 | 0.230102631 | 34 |
| si:ch73-14h10.210 | 1.00087447283963e-05 | 0.443779216 | 0.477 | 0.273 | 0.251910096 | 34 |
| cpeb1b10 | 1.04905565370246e-05 | 0.446344339 | 0.477 | 0.272 | 0.264036817 | 34 |
| si:ch1073-190k2.110 | 1.20179635918658e-05 | 0.434973082 | 0.364 | 0.19 | 0.302480126 | 34 |
| BX537276.110 | 1.24118787936402e-05 | 0.396922174 | 0.273 | 0.123 | 0.312394577 | 34 |
| ddx195 | 1.25979020330183e-05 | 0.435131825 | 0.307 | 0.153 | 0.317076596 | 34 |
| si:dkey-241l7.310 | 1.28929021679283e-05 | 0.441582678 | 0.545 | 0.366 | 0.324501455 | 34 |
| si:ch211-226h8.63 | 1.31807773423222e-05 | 0.351814891 | 0.205 | 0.079 | 0.331746985 | 34 |
| dynll111 | 1.33208421767749e-05 | 0.396564988 | 0.625 | 0.456 | 0.335272277 | 34 |
| srsf2a7 | 1.42378311608207e-05 | 0.426036593 | 0.534 | 0.368 | 0.358351972 | 34 |
| org10 | 1.46435489720149e-05 | 0.412190643 | 0.443 | 0.242 | 0.368563484 | 34 |
| tacc33 | 1.59361562767472e-05 | 0.380942086 | 0.227 | 0.096 | 0.401097117 | 34 |
| zgc:162509 | 1.59915507846084e-05 | 0.267879013 | 0.102 | 0.027 | 0.402491342 | 34 |
| ubc2 | 1.6027583422391e-05 | 0.272088795 | 0.727 | 0.659 | 0.403398247 | 34 |
| med22 | 1.65236810530859e-05 | 0.336099296 | 0.17 | 0.062 | 0.415884528 | 34 |
| rchy11 | 1.68703284730786e-05 | 0.372714058 | 0.193 | 0.077 | 0.424609297 | 34 |
| zgc:1630309 | 1.7074450169968e-05 | 0.40572696 | 0.273 | 0.125 | 0.429746836 | 34 |
| si:dkey-37o8.110 | 1.8608743424518e-05 | 0.433809213 | 0.5 | 0.309 | 0.468363463 | 34 |
| tmco13 | 2.18938143808226e-05 | 0.402122761 | 0.261 | 0.121 | 0.551045414 | 34 |
| si:dkey-42i9.810 | 2.27344135228791e-05 | 0.410648404 | 0.273 | 0.129 | 0.572202454 | 34 |
| sinup10 | 2.49942793077784e-05 | 0.420272011 | 0.375 | 0.205 | 0.629081016 | 34 |
| si:dkeyp-98a7.710 | 2.55918479960455e-05 | 0.403662966 | 0.375 | 0.197 | 0.644121222 | 34 |
| CU467646.28 | 2.61888742721797e-05 | 0.38571136 | 0.273 | 0.127 | 0.659147777 | 34 |
| kmt5ab3 | 2.96072480159366e-05 | 0.362077943 | 0.216 | 0.09 | 0.745184825 | 34 |
| si:ch211-137a8.2 | 3.22519160004097e-05 | 0.321548394 | 0.159 | 0.057 | 0.811748474 | 34 |
| lsm111 | 3.237060163345e-05 | 0.293444532 | 0.136 | 0.044 | 0.814735673 | 34 |
| gyg1a11 | 3.27407831318212e-05 | 0.404679565 | 0.534 | 0.372 | 0.824052771 | 34 |
| si:ch211-69g19.28 | 3.45666675154782e-05 | 0.39587227 | 0.261 | 0.123 | 0.870008455 | 34 |
| zgc:923135 | 3.62279364923052e-05 | 0.372999098 | 0.273 | 0.127 | 0.911820934 | 34 |
| scamp2l | 4.11180128403657e-05 | 0.299409871 | 0.136 | 0.045 | 1 | 34 |
| dazl7 | 4.63703453600824e-05 | 0.367288277 | 0.239 | 0.107 | 1 | 34 |

| | | | | | | |
|---|---|---|---|---|---|---|
| si:ch73-382f3.1 | 4.8169397846596e-05 | 0.277469784 | 0.136 | 0.045 | 1 | 34 |
| coil3 | 4.96441058554595e-05 | 0.373208114 | 0.239 | 0.108 | 1 | 34 |
| fbxo43 | 5.22019730472429e-05 | 0.313290114 | 0.182 | 0.071 | 1 | 34 |
| zglp110 | 5.31392889453575e-05 | 0.398778088 | 0.364 | 0.2 | 1 | 34 |
| alg3 | 6.13510387659368e-05 | 0.258075317 | 0.114 | 0.035 | 1 | 34 |
| hells5 | 6.19828790301035e-05 | 0.386291237 | 0.273 | 0.134 | 1 | 34 |
| prr114 | 6.26370126802307e-05 | 0.367958095 | 0.227 | 0.101 | 1 | 34 |
| retsatl10 | 6.34655341715926e-05 | 0.415474845 | 0.364 | 0.205 | 1 | 34 |
| elovl7b10 | 6.37663762793494e-05 | 0.41196054 | 0.432 | 0.25 | 1 | 34 |
| zgc:9259110 | 6.53574558871246e-05 | 0.402350128 | 0.364 | 0.207 | 1 | 34 |
| zgc:1118687 | 6.71089482175244e-05 | 0.388282591 | 0.295 | 0.149 | 1 | 34 |
| orc65 | 6.81602189299854e-05 | 0.364973583 | 0.239 | 0.108 | 1 | 34 |
| ctsba10 | 6.81868729900178e-05 | 0.376443308 | 0.545 | 0.378 | 1 | 34 |
| slc25a25a3 | 7.89097914075345e-05 | 0.410629311 | 0.33 | 0.188 | 1 | 34 |
| cdca89 | 7.96921788065432e-05 | 0.386689803 | 0.318 | 0.166 | 1 | 34 |
| cdca7a10 | 8.53916018629718e-05 | 0.400573212 | 0.386 | 0.224 | 1 | 34 |
| si:ch211-226l4.611 | 8.7418340129423e-05 | 0.392382218 | 0.477 | 0.294 | 1 | 34 |
| h3f3a11 | 8.88701500796651e-05 | 0.348015497 | 0.636 | 0.507 | 1 | 34 |
| sybu10 | 9.39040803660012e-05 | 0.360808578 | 0.33 | 0.168 | 1 | 34 |
| cdc2010 | 0.000102046 | 0.378915694 | 0.295 | 0.154 | 1 | 34 |
| brf25 | 0.000109435 | 0.346293509 | 0.216 | 0.096 | 1 | 34 |
| zgc:10348211 | 0.000119706 | 0.383175403 | 0.352 | 0.19 | 1 | 34 |
| zgc:1735451 | 0.000121117 | 0.283048089 | 0.136 | 0.049 | 1 | 34 |
| aph1b | 0.000125195 | 0.285264827 | 0.136 | 0.049 | 1 | 34 |
| snu13b11 | 0.000127265 | 0.366265085 | 0.568 | 0.394 | 1 | 34 |
| cdk11 | 0.000128722 | 0.348950583 | 0.205 | 0.092 | 1 | 34 |
| mcm3l4 | 0.000136486 | 0.320789208 | 0.205 | 0.089 | 1 | 34 |
| si:ch211-146l10.810 | 0.000136882 | 0.391539503 | 0.466 | 0.308 | 1 | 34 |
| rbm4.310 | 0.0001384 | 0.353087044 | 0.568 | 0.446 | 1 | 34 |
| sumo2a | 0.000172814 | 0.274288432 | 0.114 | 0.038 | 1 | 34 |
| mcmbp | 0.000175585 | 0.267905278 | 0.148 | 0.055 | 1 | 34 |
| si:dkeyp-46h3.510 | 0.000179057 | 0.380993652 | 0.455 | 0.291 | 1 | 34 |
| ddx39ab9 | 0.000191171 | 0.394835698 | 0.432 | 0.291 | 1 | 34 |
| casd11 | 0.000192859 | 0.321299718 | 0.182 | 0.078 | 1 | 34 |
| si:ch73-90k17.110 | 0.000192864 | 0.379228649 | 0.409 | 0.251 | 1 | 34 |

| | | | | | | |
|---|---|---|---|---|---|---|
| cse1l1 | 0.000197041 | 0.324333821 | 0.182 | 0.078 | 1 | 34 |
| thap11 | 0.000197396 | 0.25469462 | 0.125 | 0.044 | 1 | 34 |
| marcksb7 | 0.000204521 | 0.373155579 | 0.318 | 0.177 | 1 | 34 |
| rbpms2b10 | 0.00021418 | 0.37913272 | 0.375 | 0.223 | 1 | 34 |
| LOC1005380233 | 0.000224175 | 0.315903466 | 0.182 | 0.078 | 1 | 34 |
| zgc:922879 | 0.000231337 | 0.359667184 | 0.261 | 0.134 | 1 | 34 |
| dkc13 | 0.000236348 | 0.387050347 | 0.307 | 0.176 | 1 | 34 |
| si:ch73-160p18.310 | 0.000242133 | 0.373626801 | 0.398 | 0.242 | 1 | 34 |
| ninl | 0.000260273 | 0.274611922 | 0.136 | 0.052 | 1 | 34 |
| si:dkey-188i13.1111 | 0.000260502 | 0.393136694 | 0.466 | 0.318 | 1 | 34 |
| tpx25 | 0.000268633 | 0.356532983 | 0.25 | 0.127 | 1 | 34 |
| lin28a2 | 0.000274388 | 0.296200507 | 0.17 | 0.071 | 1 | 34 |
| zgc:5669911 | 0.000278762 | 0.377736757 | 0.398 | 0.253 | 1 | 34 |
| st8sia7.13 | 0.000280366 | 0.309374189 | 0.17 | 0.072 | 1 | 34 |
| plk19 | 0.000288849 | 0.371845872 | 0.318 | 0.179 | 1 | 34 |
| eloal | 0.000296761 | 0.268616254 | 0.148 | 0.058 | 1 | 34 |
| h2afva9 | 0.00030505 | 0.367255415 | 0.477 | 0.325 | 1 | 34 |
| zc3h11a9 | 0.000316068 | 0.398617548 | 0.352 | 0.219 | 1 | 34 |
| elavl211 | 0.000318618 | 0.370145842 | 0.375 | 0.226 | 1 | 34 |
| cldn7b9 | 0.000329989 | 0.357365883 | 0.295 | 0.161 | 1 | 34 |
| si:dkey-46g23.29 | 0.000343926 | 0.331755692 | 0.284 | 0.146 | 1 | 34 |
| fam173b | 0.00035102 | 0.274639848 | 0.159 | 0.065 | 1 | 34 |
| mcfd23 | 0.000381972 | 0.348515894 | 0.227 | 0.115 | 1 | 34 |
| snu13a5 | 0.000388541 | 0.321759416 | 0.216 | 0.102 | 1 | 34 |
| uqcr103 | 0.000392452 | 0.358377703 | 0.477 | 0.358 | 1 | 34 |
| dnaja2a | 0.000479069 | 0.365457216 | 0.386 | 0.262 | 1 | 34 |
| tmem17 | 0.000481453 | 0.271339087 | 0.148 | 0.06 | 1 | 34 |
| eif4g2a1 | 0.00048711 | 0.350801209 | 0.25 | 0.134 | 1 | 34 |
| LOC10053686010 | 0.000492781 | 0.349998193 | 0.5 | 0.353 | 1 | 34 |
| glipr2l | 0.000511855 | 0.291531048 | 0.159 | 0.067 | 1 | 34 |
| kif7 | 0.000530147 | 0.266639993 | 0.136 | 0.054 | 1 | 34 |
| bmb5 | 0.000572622 | 0.306541041 | 0.193 | 0.089 | 1 | 34 |
| ivns1abpb3 | 0.000577866 | 0.340106581 | 0.227 | 0.118 | 1 | 34 |
| zgc:1717509 | 0.00062731 | 0.355973353 | 0.307 | 0.176 | 1 | 34 |
| dynll2a10 | 0.000643287 | 0.319219753 | 0.261 | 0.138 | 1 | 34 |

| | | | | | | | |
|---|---|---|---|---|---|---|---|
| ap2m1b | 0.000643433 | 0.328798141 | 0.17 | 0.079 | | 1 | 34 |
| BX571780.17 | 0.000647754 | 0.298960612 | 0.182 | 0.082 | | 1 | 34 |
| ankrd10a1 | 0.000653087 | 0.26792914 | 0.136 | 0.055 | | 1 | 34 |
| LOC110440049 | 0.000664433 | 0.25119604 | 0.125 | 0.048 | | 1 | 34 |
| snrpb26 | 0.000686434 | 0.34913059 | 0.273 | 0.151 | | 1 | 34 |
| cyp11a11 | 0.000688615 | 0.291368857 | 0.159 | 0.069 | | 1 | 34 |
| buc2l4 | 0.000723409 | 0.274197177 | 0.182 | 0.081 | | 1 | 34 |
| metap1d2 | 0.000783928 | 0.299621627 | 0.193 | 0.091 | | 1 | 34 |
| exosc81 | 0.000875427 | 0.319633459 | 0.159 | 0.072 | | 1 | 34 |
| CU655961.33 | 0.00089853 | 0.302210464 | 0.205 | 0.1 | | 1 | 34 |
| st3gal82 | 0.00090416 | 0.288867135 | 0.17 | 0.078 | | 1 | 34 |
| lmnl32 | 0.000924315 | 0.273036693 | 0.17 | 0.077 | | 1 | 34 |
| lrp131 | 0.000930811 | 0.280918369 | 0.159 | 0.071 | | 1 | 34 |
| fam122b | 0.00097748 | 0.266956749 | 0.159 | 0.07 | | 1 | 34 |
| cbx3a7 | 0.001015936 | 0.34565046 | 0.42 | 0.298 | | 1 | 34 |
| si:dkey-43p13.53 | 0.001051612 | 0.277740355 | 0.182 | 0.085 | | 1 | 34 |
| tph1a10 | 0.001099907 | 0.333849412 | 0.261 | 0.143 | | 1 | 34 |
| paip2b2 | 0.001138153 | 0.352362093 | 0.42 | 0.306 | | 1 | 34 |
| rac1a4 | 0.001189819 | 0.342437812 | 0.261 | 0.15 | | 1 | 34 |
| zgc:6602410 | 0.001205247 | 0.308040434 | 0.261 | 0.143 | | 1 | 34 |
| BX284697.18 | 0.001212648 | 0.296819074 | 0.227 | 0.116 | | 1 | 34 |
| dnd16 | 0.001252602 | 0.301395372 | 0.216 | 0.111 | | 1 | 34 |
| pttg1ipb3 | 0.001261267 | 0.340560141 | 0.216 | 0.115 | | 1 | 34 |
| ypel38 | 0.001264627 | 0.319639559 | 0.239 | 0.129 | | 1 | 34 |
| adipor1a2 | 0.0013223 | 0.336148473 | 0.193 | 0.101 | | 1 | 34 |
| asz13 | 0.001420961 | 0.298500917 | 0.159 | 0.074 | | 1 | 34 |
| nap1l14 | 0.001422093 | 0.323420673 | 0.466 | 0.363 | | 1 | 34 |
| fdx19 | 0.001618828 | 0.332914649 | 0.352 | 0.223 | | 1 | 34 |
| anp32b5 | 0.001625489 | 0.340023207 | 0.375 | 0.257 | | 1 | 34 |
| si:ch73-78o10.18 | 0.001636951 | 0.292517946 | 0.25 | 0.135 | | 1 | 34 |
| exd21 | 0.001645622 | 0.273402739 | 0.193 | 0.094 | | 1 | 34 |
| lpcat46 | 0.001663771 | 0.284647722 | 0.216 | 0.108 | | 1 | 34 |
| zgc:1719779 | 0.001712752 | 0.334404433 | 0.352 | 0.232 | | 1 | 34 |
| rpain6 | 0.001773871 | 0.301385275 | 0.227 | 0.121 | | 1 | 34 |
| vstm2l6 | 0.001829819 | 0.286548947 | 0.193 | 0.097 | | 1 | 34 |

| | | | | | | |
|---|---|---|---|---|---|---|
| si:ch73-303b9.14 | 0.001858199 | 0.259434103 | 0.17 | 0.08 | 1 | 34 |
| lyar1 | 0.001909897 | 0.265367146 | 0.125 | 0.054 | 1 | 34 |
| polr2k3 | 0.001924025 | 0.345172991 | 0.307 | 0.192 | 1 | 34 |
| gbgt1l49 | 0.002018033 | 0.29164111 | 0.273 | 0.152 | 1 | 34 |
| h2afvb3 | 0.002077387 | 0.275326817 | 0.58 | 0.472 | 1 | 34 |
| rbbp45 | 0.00209704 | 0.324262363 | 0.284 | 0.172 | 1 | 34 |
| rrs11 | 0.002122161 | 0.297492467 | 0.17 | 0.085 | 1 | 34 |
| si:ch73-281n10.25 | 0.002229084 | 0.306436412 | 0.489 | 0.394 | 1 | 34 |
| pigrl2.310 | 0.00229774 | 0.30608863 | 0.341 | 0.205 | 1 | 34 |
| mycla | 0.002342934 | 0.266196228 | 0.159 | 0.075 | 1 | 34 |
| ankrd229 | 0.002424238 | 0.315563179 | 0.25 | 0.145 | 1 | 34 |
| yrdc8 | 0.002488592 | 0.304581218 | 0.227 | 0.124 | 1 | 34 |
| uba21 | 0.002521904 | 0.299187268 | 0.182 | 0.094 | 1 | 34 |
| aurka7 | 0.002619643 | 0.271128884 | 0.227 | 0.121 | 1 | 34 |
| zgc:1526528 | 0.002675783 | 0.281114914 | 0.216 | 0.115 | 1 | 34 |
| tspan362 | 0.002691068 | 0.323880962 | 0.261 | 0.159 | 1 | 34 |
| si:dkey-16l2.16 | 0.002724149 | 0.304235022 | 0.17 | 0.087 | 1 | 34 |
| krcp1 | 0.002749557 | 0.267711532 | 0.159 | 0.076 | 1 | 34 |
| UBL4A2 | 0.002862464 | 0.293189889 | 0.227 | 0.126 | 1 | 34 |
| rcc110 | 0.002925827 | 0.33108997 | 0.318 | 0.204 | 1 | 34 |
| uck2a1 | 0.00293335 | 0.28461654 | 0.216 | 0.118 | 1 | 34 |
| psmd101 | 0.003014876 | 0.264143367 | 0.159 | 0.077 | 1 | 34 |
| mrps18b3 | 0.003159874 | 0.294268932 | 0.227 | 0.127 | 1 | 34 |
| nup58 | 0.003181072 | 0.254179249 | 0.136 | 0.063 | 1 | 34 |
| CT009596.16 | 0.00322459 | 0.262135236 | 0.182 | 0.091 | 1 | 34 |
| buc10 | 0.003501859 | 0.297335488 | 0.295 | 0.18 | 1 | 34 |
| faub8 | 0.003528535 | 0.310629659 | 0.318 | 0.202 | 1 | 34 |
| ccne18 | 0.003561506 | 0.308143804 | 0.205 | 0.113 | 1 | 34 |
| mos9 | 0.003660422 | 0.302642835 | 0.239 | 0.138 | 1 | 34 |
| CABZ01059627.210 | 0.003696462 | 0.29393871 | 0.239 | 0.135 | 1 | 34 |
| pim32 | 0.003996416 | 0.290032718 | 0.227 | 0.13 | 1 | 34 |
| mcm73 | 0.004158955 | 0.301924999 | 0.261 | 0.159 | 1 | 34 |
| pou5f39 | 0.004570333 | 0.293340637 | 0.25 | 0.148 | 1 | 34 |
| cdc14b1 | 0.005138609 | 0.304266792 | 0.227 | 0.136 | 1 | 34 |
| rbm73 | 0.005257421 | 0.272237661 | 0.216 | 0.121 | 1 | 34 |

| | | | | | | |
|---|---|---|---|---|---|---|
| bzw1b7 | 0.005276687 | 0.279994201 | 0.477 | 0.404 | 1 | 34 |
| hsd17b12a10 | 0.005299263 | 0.292383975 | 0.273 | 0.167 | 1 | 34 |
| STMP13 | 0.005393439 | 0.319417169 | 0.398 | 0.307 | 1 | 34 |
| LOC1039099725 | 0.005525101 | 0.298480453 | 0.455 | 0.369 | 1 | 34 |
| zgc:566769 | 0.00572597 | 0.303838836 | 0.398 | 0.297 | 1 | 34 |
| ppp1r3b4 | 0.005936417 | 0.250303816 | 0.205 | 0.111 | 1 | 34 |
| larp6a10 | 0.005960664 | 0.302222554 | 0.364 | 0.252 | 1 | 34 |
| mrpl181 | 0.005975112 | 0.288357982 | 0.193 | 0.11 | 1 | 34 |
| rdh10b9 | 0.006055013 | 0.302436821 | 0.284 | 0.18 | 1 | 34 |
| rcor1 | 0.006126466 | 0.277231161 | 0.193 | 0.107 | 1 | 34 |
| slbp11 | 0.006192211 | 0.293804088 | 0.352 | 0.236 | 1 | 34 |
| rpp211 | 0.006354018 | 0.266174722 | 0.159 | 0.083 | 1 | 34 |
| asf1ba3 | 0.006686027 | 0.259385515 | 0.182 | 0.099 | 1 | 34 |
| lsm12a3 | 0.006813141 | 0.292457905 | 0.216 | 0.128 | 1 | 34 |
| igf2bp3 | 0.006989659 | 0.273054563 | 0.205 | 0.117 | 1 | 34 |
| pttg110 | 0.007089322 | 0.27932089 | 0.261 | 0.16 | 1 | 34 |
| si:dkeyp-75b4.106 | 0.007193092 | 0.250218875 | 0.534 | 0.439 | 1 | 34 |
| ccnh7 | 0.007813904 | 0.271263712 | 0.216 | 0.127 | 1 | 34 |
| thop14 | 0.007902931 | 0.276990706 | 0.205 | 0.118 | 1 | 34 |
| aktip4 | 0.00825817 | 0.29028884 | 0.227 | 0.141 | 1 | 34 |
| zgc:1532843 | 0.00844754 | 0.268639213 | 0.182 | 0.102 | 1 | 34 |
| aurkb4 | 0.008542743 | 0.274972868 | 0.216 | 0.13 | 1 | 34 |
| gemin74 | 0.008686028 | 0.27784157 | 0.25 | 0.156 | 1 | 34 |
| dnmt1 | 0.009398611 | 0.263090755 | 0.182 | 0.103 | 1 | 34 |
| lbh4 | 0.009504219 | 0.297662367 | 0.261 | 0.173 | 1 | 34 |
| itm2cb6 | 0.009508138 | 0.285893998 | 0.227 | 0.139 | 1 | 34 |
| col10a1a | 0 | 2.81215742 | 0.909 | 0.008 | 0 | 35 |
| scpp71 | 0 | 2.641422933 | 0.898 | 0.022 | 0 | 35 |
| ifitm5 | 0 | 2.336795777 | 0.989 | 0.002 | 0 | 35 |
| cd81b1 | 0 | 2.21845384 | 0.898 | 0.003 | 0 | 35 |
| enam1 | 0 | 2.104897526 | 0.795 | 0.007 | 0 | 35 |
| tmem119a | 0 | 1.841292914 | 0.898 | 0.002 | 0 | 35 |
| panx3 | 0 | 1.763704937 | 0.875 | 0.001 | 0 | 35 |
| si:ch211-106n13.3 | 0 | 1.749754623 | 0.875 | 0.001 | 0 | 35 |
| sgms2a1 | 0 | 1.738686036 | 0.875 | 0.005 | 0 | 35 |

| | | | | | | |
|---|---|---|---|---|---|---|
| si:dkey-12l12.11 | 0 | 1.593958876 | 0.852 | 0.017 | 0 | 35 |
| sod3b1 | 0 | 1.565738838 | 0.932 | 0.005 | 0 | 35 |
| cthrc1a | 0 | 1.563951777 | 0.886 | 0.004 | 0 | 35 |
| phospho11 | 0 | 1.548548277 | 0.841 | 0.009 | 0 | 35 |
| bgna1 | 0 | 1.542935806 | 0.841 | 0.01 | 0 | 35 |
| col10a1b | 0 | 1.44895775 | 0.534 | 0.008 | 0 | 35 |
| col11a1a | 0 | 1.444433469 | 0.818 | 0.005 | 0 | 35 |
| loxl4 | 0 | 1.422512634 | 0.807 | 0.002 | 0 | 35 |
| col11a2 | 0 | 1.387895476 | 0.716 | 0.001 | 0 | 35 |
| fgfbp2a | 0 | 1.356997921 | 0.409 | 0.001 | 0 | 35 |
| cdh112 | 0 | 1.266646798 | 0.841 | 0.021 | 0 | 35 |
| ndnf1 | 0 | 1.259686891 | 0.773 | 0.026 | 0 | 35 |
| slc8a4b1 | 0 | 1.20106038 | 0.739 | 0.012 | 0 | 35 |
| spp1 | 0 | 1.158738458 | 0.625 | 0.002 | 0 | 35 |
| zgc:92162 | 0 | 1.154236105 | 0.614 | 0.006 | 0 | 35 |
| chst11 | 0 | 1.123347921 | 0.682 | 0.004 | 0 | 35 |
| smpd3 | 0 | 1.114798198 | 0.682 | 0.006 | 0 | 35 |
| itga10 | 0 | 1.055464972 | 0.705 | 0.002 | 0 | 35 |
| LO018513.1 | 0 | 1.036714033 | 0.545 | 0.001 | 0 | 35 |
| bambib | 0 | 1.009015977 | 0.693 | 0.01 | 0 | 35 |
| ano5a | 0 | 0.985385505 | 0.659 | 0.006 | 0 | 35 |
| bmp8a | 0 | 0.981915161 | 0.625 | 0.005 | 0 | 35 |
| runx31 | 0 | 0.980684062 | 0.67 | 0.016 | 0 | 35 |
| col5a2b | 0 | 0.973617841 | 0.659 | 0.001 | 0 | 35 |
| qpct | 0 | 0.970154216 | 0.591 | 0.005 | 0 | 35 |
| klf2b2 | 0 | 0.964115778 | 0.636 | 0.017 | 0 | 35 |
| entpd5a1 | 0 | 0.941578249 | 0.386 | 0.003 | 0 | 35 |
| pcolce2b | 0 | 0.94062779 | 0.636 | 0.012 | 0 | 35 |
| kazald3 | 0 | 0.863585154 | 0.591 | 0.002 | 0 | 35 |
| dlx3b1 | 0 | 0.861835607 | 0.545 | 0.006 | 0 | 35 |
| sp7 | 0 | 0.815139548 | 0.568 | 0.002 | 0 | 35 |
| si:ch211-170d8.81 | 0 | 0.809819419 | 0.591 | 0.009 | 0 | 35 |
| bmp31 | 0 | 0.789734557 | 0.523 | 0.009 | 0 | 35 |
| COLGALT1 (1 of many) | 0 | 0.754694577 | 0.443 | 0.006 | 0 | 35 |
| myadml2 | 0 | 0.712451198 | 0.466 | 0 | 0 | 35 |

| | | | | | | |
|---|---|---|---|---|---|---|
| thrap3a1 | 0 | 0.708681345 | 0.568 | 0.011 | 0 | 35 |
| nrp2b | 0 | 0.698923664 | 0.455 | 0.009 | 0 | 35 |
| prp | 0 | 0.6729502 | 0.443 | 0.001 | 0 | 35 |
| CABZ01059119.1 | 0 | 0.665168753 | 0.523 | 0.01 | 0 | 35 |
| satb2 | 0 | 0.641176846 | 0.455 | 0.002 | 0 | 35 |
| bmp2a | 0 | 0.639202823 | 0.5 | 0.002 | 0 | 35 |
| ism11 | 0 | 0.622953474 | 0.455 | 0.007 | 0 | 35 |
| dlx5a1 | 0 | 0.570727329 | 0.375 | 0.003 | 0 | 35 |
| pltp | 0 | 0.560249116 | 0.352 | 0.002 | 0 | 35 |
| phlda3 | 0 | 0.552632314 | 0.398 | 0.006 | 0 | 35 |
| plcd3b | 0 | 0.530798629 | 0.386 | 0.001 | 0 | 35 |
| si:ch211-106h11.3 | 0 | 0.514673956 | 0.375 | 0.002 | 0 | 35 |
| ihha | 0 | 0.497580413 | 0.216 | 0 | 0 | 35 |
| dlx2b | 0 | 0.471857372 | 0.341 | 0.001 | 0 | 35 |
| wnt10a | 0 | 0.464900919 | 0.25 | 0.001 | 0 | 35 |
| cspg41 | 0 | 0.459058077 | 0.364 | 0.004 | 0 | 35 |
| zgc:162171 | 0 | 0.434176143 | 0.364 | 0.002 | 0 | 35 |
| scpp1 | 0 | 0.416478904 | 0.33 | 0 | 0 | 35 |
| megf10 | 0 | 0.349200182 | 0.284 | 0 | 0 | 35 |
| ppfibp2a | 0 | 0.304243326 | 0.193 | 0 | 0 | 35 |
| chodl | 0 | 0.285135163 | 0.193 | 0 | 0 | 35 |
| rcn1 | 1.38417896607036e-303 | 0.859735156 | 0.58 | 0.016 | 3.48384003970248e-299 | 35 |
| serpinh1b1 | 5.04457130009371e-300 | 0.989409748 | 0.625 | 0.019 | 1.26966815052059e-295 | 35 |
| fkbp91 | 5.91177377411887e-298 | 0.939933816 | 0.659 | 0.021 | 1.48793434120798e-293 | 35 |
| serpinh1a | 3.74717358436471e-296 | 0.673246856 | 0.5 | 0.011 | 9.43126119448754e-292 | 35 |
| tnfrsf11a | 6.08443925451537e-288 | 0.715204313 | 0.477 | 0.011 | 1.53139251596897e-283 | 35 |
| zgc:1626121 | 7.74994591937158e-288 | 0.558539429 | 0.33 | 0.004 | 1.95058388844663e-283 | 35 |
| mxra5a | 3.91841430673944e-282 | 0.410663248 | 0.284 | 0.003 | 9.8622569686325e-278 | 35 |
| XLOC-005794 | 1.94510585807038e-278 | 0.352606208 | 0.193 | 0.001 | 4.89563693417734e-274 | 35 |
| si:ch211-110e21.3 | 1.93701306631251e-270 | 0.57145285 | 0.352 | 0.006 | 4.87526818660195e-266 | 35 |
| nav2a | 1.04811562696777e-268 | 0.311934034 | 0.193 | 0.001 | 2.63800222151517e-264 | 35 |
| crip21 | 3.45069222660752e-265 | 0.896582735 | 0.489 | 0.013 | 8.68504726514846e-261 | 35 |
| myh11b | 4.75811361448308e-263 | 0.402468011 | 0.284 | 0.003 | 1.19756961562925e-258 | 35 |
| nid2b | 1.38426471380214e-261 | 0.359353867 | 0.261 | 0.003 | 3.48405585816859e-257 | 35 |
| zbtb7c | 3.25007119844007e-260 | 0.390353159 | 0.25 | 0.002 | 8.1801041993538e-256 | 35 |

| | | | | | | |
|---|---|---|---|---|---|---|
| mfsd2b | 5.46009271494785e-250 | 0.281114675 | 0.193 | 0.001 | 1.37425073542523e-245 | 35 |
| olfml3b5 | 8.45931618031367e-246 | 1.172844993 | 0.795 | 0.041 | 2.12912528942315e-241 | 35 |
| xylt1 | 3.23156283997347e-240 | 0.465600878 | 0.33 | 0.006 | 8.13352051192923e-236 | 35 |
| fgfr32 | 6.56455984454805e-234 | 0.7146317 | 0.477 | 0.014 | 1.6522340672743e-229 | 35 |
| ccdc802 | 7.81414161139791e-234 | 0.786967509 | 0.591 | 0.022 | 1.96674130217274e-229 | 35 |
| plod1a | 2.26458525444854e-232 | 0.775151342 | 0.557 | 0.02 | 5.69973462692154e-228 | 35 |
| lmo7a | 1.97151660216687e-231 | 0.480975549 | 0.352 | 0.007 | 4.9621101359938e-227 | 35 |
| sox11a1 | 1.15304567152499e-230 | 0.812992478 | 0.523 | 0.017 | 2.90210065066125e-226 | 35 |
| golim4b | 3.5013718053121e-230 | 0.324416088 | 0.25 | 0.003 | 8.81260269679002e-226 | 35 |
| pcdh171 | 4.4615396294723e-230 | 0.402278893 | 0.261 | 0.003 | 1.12292490934188e-225 | 35 |
| igfbp1a1 | 1.57845990980832e-229 | 1.119205583 | 0.489 | 0.015 | 3.97282574699657e-225 | 35 |
| sema3e1 | 5.8753708341064e-228 | 0.488376375 | 0.375 | 0.008 | 1.47877208523624e-223 | 35 |
| p4ha2 | 7.80021674142176e-226 | 0.5465631 | 0.42 | 0.011 | 1.96323655164844e-221 | 35 |
| heyl1 | 1.34010194753177e-220 | 0.408308523 | 0.205 | 0.002 | 3.37290259174271e-216 | 35 |
| fpr12 | 1.0210758780573e-214 | 0.768799297 | 0.5 | 0.017 | 2.56994587748241e-210 | 35 |
| fam20a | 4.57688835490833e-213 | 0.333035432 | 0.273 | 0.004 | 1.15195703004688e-208 | 35 |
| si:dkey-234i14.3 | 4.11188675968285e-209 | 0.307328194 | 0.136 | 0.001 | 1.03492077854458e-204 | 35 |
| rnd2 | 4.43504492677507e-209 | 0.35215892 | 0.136 | 0.001 | 1.11625645762002e-204 | 35 |
| si:cabz01100188.1 | 6.87326576508774e-207 | 0.337508661 | 0.216 | 0.002 | 1.72993226041493e-202 | 35 |
| tfap2c | 1.9202799725154e-206 | 0.460842689 | 0.239 | 0.003 | 4.83315266282401e-202 | 35 |
| s1pr11 | 3.1789510382774e-205 | 0.479727485 | 0.307 | 0.006 | 8.00110186824039e-201 | 35 |
| p3h1 | 1.68335601211986e-203 | 0.447282766 | 0.386 | 0.01 | 4.23683874690446e-199 | 35 |
| LOC1018861142 | 3.10477724049561e-202 | 0.612269396 | 0.42 | 0.013 | 7.81441383660339e-198 | 35 |
| btbd6a | 8.35880611107339e-201 | 0.499676848 | 0.295 | 0.006 | 2.10382791009606e-196 | 35 |
| her62 | 1.47184310100479e-200 | 0.831173775 | 0.511 | 0.019 | 3.70448190091895e-196 | 35 |
| mef2cb3 | 8.51057970901602e-200 | 0.819667111 | 0.545 | 0.022 | 2.14202780696224e-195 | 35 |
| cdc42ep3 | 5.67341143952637e-197 | 0.500647445 | 0.375 | 0.01 | 1.42794092521439e-192 | 35 |
| si:dkey-193p11.2 | 3.82545690057524e-196 | 1.298116872 | 0.398 | 0.012 | 9.62829247305783e-192 | 35 |
| col11a1b1 | 4.38079075930869e-195 | 0.628965201 | 0.398 | 0.012 | 1.1026012262104e-190 | 35 |
| pcolceb | 7.17267976658761e-194 | 0.301657941 | 0.205 | 0.002 | 1.80529177045244e-189 | 35 |
| pdgfbb | 5.7081685998663e-192 | 0.455158216 | 0.239 | 0.004 | 1.43668895490035e-187 | 35 |
| acvrl12 | 2.67960266427066e-190 | 0.539399079 | 0.409 | 0.013 | 6.74429194570282e-186 | 35 |
| nupr1b4 | 7.02791169245739e-190 | 1.068822203 | 0.727 | 0.044 | 1.7688550938746e-185 | 35 |
| thbs1b1 | 5.53425613854611e-189 | 0.512974125 | 0.352 | 0.009 | 1.39291692751067e-184 | 35 |
| BX322661.2 | 8.5470215867008e-188 | 0.260969344 | 0.136 | 0.001 | 2.15119986315672e-183 | 35 |

| | | | | | | |
|---|---|---|---|---|---|---|
| col14a1b | 2.39873610297268e-186 | 0.505331137 | 0.273 | 0.005 | 6.03737889757194e-182 | 35 |
| ccdc1871 | 3.30729924051028e-180 | 0.747924136 | 0.568 | 0.027 | 8.32414145844033e-176 | 35 |
| twist21 | 7.02693120009892e-179 | 0.629935167 | 0.352 | 0.01 | 1.7686083137529e-174 | 35 |
| cplx2 | 2.79200766570278e-176 | 0.36711303 | 0.25 | 0.005 | 7.02720409380732e-172 | 35 |
| rcn3 | 9.14921678242259e-174 | 0.363595302 | 0.273 | 0.006 | 2.30276637196794e-169 | 35 |
| plekhh1 | 3.31599708922706e-173 | 0.358075392 | 0.25 | 0.005 | 8.34603307387559e-169 | 35 |
| cd82b3 | 3.46812362602796e-172 | 1.248812388 | 0.864 | 0.07 | 8.72892035434977e-168 | 35 |
| serpine22 | 1.91034076517461e-170 | 0.99156437 | 0.807 | 0.059 | 4.80813667186799e-166 | 35 |
| msx2b | 4.09607448014649e-169 | 0.292702083 | 0.182 | 0.002 | 1.03094098590807e-164 | 35 |
| plxna3 | 5.32149785594093e-168 | 0.548077209 | 0.398 | 0.014 | 1.33936779536177e-163 | 35 |
| vkorc1 | 7.20121336965768e-162 | 0.5004693 | 0.364 | 0.012 | 1.81247339300914e-157 | 35 |
| tox3 | 1.16950086002567e-161 | 0.578981093 | 0.443 | 0.018 | 2.94351671459861e-157 | 35 |
| tcf72 | 1.81892994311443e-161 | 0.794766173 | 0.477 | 0.021 | 4.57806477382472e-157 | 35 |
| crtap2 | 8.18993265635728e-156 | 0.69319604 | 0.534 | 0.028 | 2.06132415027857e-151 | 35 |
| fkbp141 | 1.75482471459391e-151 | 0.76628278 | 0.614 | 0.038 | 4.4167183241614e-147 | 35 |
| clec3bb | 2.36789907291608e-150 | 0.630493417 | 0.227 | 0.005 | 5.95976517662249e-146 | 35 |
| zgc:110796 | 9.13768895030351e-150 | 0.369417917 | 0.261 | 0.006 | 2.29986493190189e-145 | 35 |
| tbx15 | 2.17280680612112e-148 | 0.252664345 | 0.17 | 0.002 | 5.46873745032625e-144 | 35 |
| nradd1 | 2.3439073772469e-147 | 0.435998928 | 0.295 | 0.008 | 5.89938047779271e-143 | 35 |
| alcama1 | 3.30812397930059e-146 | 0.74466824 | 0.489 | 0.025 | 8.32621724350167e-142 | 35 |
| rarga2 | 2.56446122651702e-144 | 0.866653702 | 0.705 | 0.053 | 6.45449246102068e-140 | 35 |
| cpe | 5.99381345593842e-137 | 0.391385056 | 0.25 | 0.006 | 1.50858290872514e-132 | 35 |
| bambia2 | 4.35937702191322e-136 | 0.482699658 | 0.409 | 0.018 | 1.09721160264534e-131 | 35 |
| mef2aa4 | 1.51199801378812e-135 | 1.104486232 | 0.739 | 0.065 | 3.80554780090332e-131 | 35 |
| tbx2b1 | 9.36095435987439e-134 | 0.520129393 | 0.341 | 0.013 | 2.35605860283678e-129 | 35 |
| stc2a | 1.53095767689731e-133 | 0.288697622 | 0.182 | 0.003 | 3.85326737698284e-129 | 35 |
| MDFI1 | 1.69626862005503e-133 | 0.524452775 | 0.386 | 0.017 | 4.26933848981651e-129 | 35 |
| farp2 | 2.09331821349666e-133 | 0.489466835 | 0.409 | 0.019 | 5.26867261154975e-129 | 35 |
| twist1a1 | 6.91036015006298e-133 | 0.522172576 | 0.341 | 0.013 | 1.73926854616935e-128 | 35 |
| rbfox12 | 2.1646206764375e-129 | 0.563675383 | 0.443 | 0.023 | 5.44813378052554e-125 | 35 |
| col1a1a6 | 2.61976854947654e-128 | 1.723943695 | 0.989 | 0.135 | 6.5936954621775e-124 | 35 |
| serpinf15 | 5.80297399975161e-128 | 0.812337292 | 0.58 | 0.04 | 1.46055052599748e-123 | 35 |
| tnfaip8l3 | 4.69037941719787e-125 | 0.315257504 | 0.193 | 0.004 | 1.18052159551453e-120 | 35 |
| tmem45a | 3.58282199379128e-124 | 0.480215083 | 0.432 | 0.022 | 9.01760467617327e-120 | 35 |
| htra3b2 | 6.98620954693787e-124 | 0.462572409 | 0.33 | 0.013 | 1.75835908086879e-119 | 35 |

| | | | | | | |
|---|---|---|---|---|---|---|
| sparc7 | 2.41656113322695e-123 | 1.683273547 | 1 | 0.154 | 6.08224271621892e-119 | 35 |
| p4ha1b | 5.2905303886936e-123 | 0.791614832 | 0.727 | 0.062 | 1.33157359353029e-118 | 35 |
| LOC1081790361 | 7.06679265808152e-123 | 0.666610037 | 0.511 | 0.032 | 1.77864104411254e-118 | 35 |
| creb3l1 | 1.6072253270946e-122 | 0.270687162 | 0.227 | 0.006 | 4.04522542576439e-118 | 35 |
| fam20cb | 3.37757004887856e-119 | 0.346367348 | 0.273 | 0.009 | 8.50100605602244e-115 | 35 |
| col1a26 | 1.28299088018188e-118 | 1.737036966 | 1 | 0.151 | 3.22915974632977e-114 | 35 |
| fkbp73 | 2.8924450305079e-118 | 0.748682849 | 0.5 | 0.033 | 7.27999489728533e-114 | 35 |
| LOC100535130 | 1.2541950116504e-117 | 0.386859883 | 0.102 | 0.001 | 3.15668342482289e-113 | 35 |
| FAM163A | 4.53382168649721e-117 | 0.293071584 | 0.159 | 0.003 | 1.14111758027448e-112 | 35 |
| zgc:1533174 | 3.44299864584266e-116 | 0.730058159 | 0.545 | 0.039 | 8.66568329172139e-112 | 35 |
| cdh7a | 2.47959073196446e-114 | 0.38059354 | 0.216 | 0.006 | 6.24088191328135e-110 | 35 |
| cx434 | 8.49078249615986e-113 | 0.892754285 | 0.614 | 0.051 | 2.13704504645847e-108 | 35 |
| znf385a5 | 9.48687626810932e-113 | 0.735256981 | 0.568 | 0.043 | 2.38775188792043e-108 | 35 |
| bcl11aa1 | 1.34212130840499e-112 | 0.345338587 | 0.25 | 0.008 | 3.37798512112451e-108 | 35 |
| zeb2a3 | 1.23653902760185e-111 | 0.68873396 | 0.557 | 0.042 | 3.1122450785711e-107 | 35 |
| kremen1 | 1.56176334039102e-111 | 0.42372061 | 0.261 | 0.009 | 3.93080215143016e-107 | 35 |
| foxd1 | 1.57445089489122e-110 | 0.298386614 | 0.193 | 0.005 | 3.96273545735172e-106 | 35 |
| frmd61 | 2.46441815447209e-110 | 0.399090227 | 0.352 | 0.017 | 6.20269405299081e-106 | 35 |
| XLOC-035413 | 3.97864926868394e-110 | 0.483478932 | 0.432 | 0.026 | 1.00138623443506e-105 | 35 |
| nrp2a2 | 7.89350989262983e-110 | 0.489957809 | 0.341 | 0.016 | 1.986717504876e-105 | 35 |
| mef2ca1 | 5.59426149617654e-109 | 0.381487472 | 0.261 | 0.009 | 1.40801967597267e-104 | 35 |
| marveld1 | 1.20348155175396e-108 | 0.364363099 | 0.273 | 0.01 | 3.02904271760954e-104 | 35 |
| col1a1b6 | 4.19163803387333e-107 | 1.526059632 | 0.966 | 0.143 | 1.05499337674558e-102 | 35 |
| tmem198b | 1.30710492312018e-105 | 0.848793822 | 0.648 | 0.059 | 3.28985238100119e-101 | 35 |
| tspan4a3 | 6.37805626598958e-105 | 0.645882846 | 0.455 | 0.03 | 1.60529298158692e-100 | 35 |
| timp2a6 | 6.39467780532427e-105 | 0.753028777 | 0.614 | 0.054 | 1.60947645682206e-100 | 35 |
| tgfb2l | 5.01403870851877e-102 | 0.350678338 | 0.205 | 0.006 | 1.26198340254709e-97 | 35 |
| ak5 | 1.35874158188282e-101 | 0.292627161 | 0.227 | 0.007 | 3.41981668744087e-97 | 35 |
| pleca4 | 2.5598499916224e-101 | 0.760977776 | 0.614 | 0.056 | 6.44288644391442e-97 | 35 |
| fkbp11 | 1.01634382372997e-100 | 0.550312069 | 0.409 | 0.026 | 2.55803576994596e-96 | 35 |
| bmp61 | 1.13548994010294e-99 | 0.407958876 | 0.273 | 0.011 | 2.8579146302451e-95 | 35 |
| tmeff1b2 | 1.28184889101553e-97 | 0.533500029 | 0.42 | 0.028 | 3.226285473797e-93 | 35 |
| zgc:162939 | 2.67634475451614e-97 | 0.413231139 | 0.284 | 0.013 | 6.73609211264168e-93 | 35 |
| reck2 | 6.70355686238257e-97 | 0.482120166 | 0.398 | 0.025 | 1.68721822669307e-92 | 35 |
| asip11 | 1.28581315632074e-96 | 0.435962821 | 0.341 | 0.018 | 3.23626313314366e-92 | 35 |

| | | | | | | |
|---|---|---|---|---|---|---|
| rtn1a | 2.11007307427978e-95 | 0.411231904 | 0.364 | 0.021 | 5.31084292065479e-91 | 35 |
| FP102018.11 | 2.7990799166994e-94 | 0.460132752 | 0.318 | 0.016 | 7.04500424234072e-90 | 35 |
| cgref1 | 2.97662214863868e-94 | 0.616222867 | 0.375 | 0.023 | 7.49186028590869e-90 | 35 |
| krt18a.17 | 2.1527430674807e-93 | 1.170259319 | 0.886 | 0.123 | 5.41823902654217e-89 | 35 |
| sox63 | 5.22220611604076e-93 | 0.619406583 | 0.398 | 0.026 | 1.3143770573463e-88 | 35 |
| plod31 | 1.43656382514507e-92 | 0.495320085 | 0.443 | 0.032 | 3.61568749150763e-88 | 35 |
| fabp7a2 | 2.74059081637134e-92 | 0.575168054 | 0.386 | 0.025 | 6.89779302572503e-88 | 35 |
| fhl1b1 | 3.7716683545576e-92 | 0.410478896 | 0.307 | 0.015 | 9.49291208158603e-88 | 35 |
| kdelr3 | 1.73532652498121e-91 | 0.386698411 | 0.307 | 0.016 | 4.3676433307252e-87 | 35 |
| pdlim4 | 5.82766700320403e-91 | 0.262024388 | 0.182 | 0.005 | 1.46676550803642e-86 | 35 |
| atp1b3a | 1.28094911837483e-90 | 0.297868434 | 0.227 | 0.008 | 3.2240208360376e-86 | 35 |
| col5a16 | 1.99727595726666e-90 | 1.145021852 | 0.807 | 0.109 | 5.02694385684445e-86 | 35 |
| hoxb7a1 | 6.29169723360242e-90 | 0.362076682 | 0.284 | 0.014 | 1.58355727672539e-85 | 35 |
| jdp2b7 | 9.36753283049425e-90 | 1.380087036 | 0.943 | 0.166 | 2.3577143381071e-85 | 35 |
| ncam1a2 | 2.46046752348617e-89 | 0.404565826 | 0.318 | 0.017 | 6.19275070986234e-85 | 35 |
| mmp16a5 | 4.00477788670053e-88 | 0.801718975 | 0.636 | 0.067 | 1.00796254630366e-83 | 35 |
| pard6gb | 1.18636089026669e-87 | 0.314380715 | 0.307 | 0.016 | 2.98595172471222e-83 | 35 |
| fxyd17 | 1.22636363368389e-87 | 0.779950463 | 0.682 | 0.078 | 3.08663462961898e-83 | 35 |
| cav15 | 1.50808020878901e-87 | 0.793339931 | 0.489 | 0.042 | 3.79568707750107e-83 | 35 |
| srgap3 | 5.34852416948375e-87 | 0.456203973 | 0.341 | 0.02 | 1.34617004821736e-82 | 35 |
| adgra2 | 5.57110650200838e-87 | 0.261854447 | 0.182 | 0.006 | 1.40219179549049e-82 | 35 |
| lhfpl62 | 1.02781424450992e-86 | 0.492729487 | 0.33 | 0.019 | 2.58690567200702e-82 | 35 |
| ckap43 | 1.48798758316098e-86 | 0.564788177 | 0.489 | 0.041 | 3.74511594805788e-82 | 35 |
| CR376783.1 | 2.11802853934081e-86 | 0.296352395 | 0.227 | 0.009 | 5.33086603066688e-82 | 35 |
| VIT1 | 3.37622749589828e-85 | 0.348960849 | 0.216 | 0.008 | 8.49762698442639e-81 | 35 |
| pmp22a6 | 3.4528767278836e-85 | 1.016005535 | 0.92 | 0.135 | 8.69054543641023e-81 | 35 |
| notch31 | 6.85634938733477e-85 | 0.589807462 | 0.341 | 0.021 | 1.72567457729829e-80 | 35 |
| nrgna | 1.31015230515967e-84 | 0.675393417 | 0.216 | 0.008 | 3.29752233685638e-80 | 35 |
| fscn1a3 | 1.70229004213242e-84 | 0.605412032 | 0.568 | 0.055 | 4.28449380704308e-80 | 35 |
| cbfb2 | 2.98393689639891e-84 | 0.557064788 | 0.466 | 0.039 | 7.51027077454642e-80 | 35 |
| id2b2 | 1.57279610237338e-83 | 0.668021485 | 0.42 | 0.032 | 3.95857051006356e-79 | 35 |
| si:ch211-133n4.45 | 4.61289522939186e-80 | 0.581774307 | 0.477 | 0.043 | 1.16101960028564e-75 | 35 |
| fosb7 | 2.15939607136257e-79 | 1.061246841 | 0.818 | 0.122 | 5.43498397201244e-75 | 35 |
| aqp3a2 | 4.7393811281211e-79 | 0.616581361 | 0.375 | 0.027 | 1.1928548361368e-74 | 35 |
| sgcd | 8.5394603359556e-79 | 0.256743726 | 0.193 | 0.007 | 2.14929677195666e-74 | 35 |

| | | | | | | |
|---|---|---|---|---|---|---|
| robo12 | 2.71922978081337e-78 | 0.43604492 | 0.33 | 0.021 | 6.84402943532918e-74 | 35 |
| ier5l2 | 3.9500792539053e-78 | 0.424485181 | 0.307 | 0.018 | 9.94195447415424e-74 | 35 |
| hoxb8a1 | 4.3904409144437e-78 | 0.305324124 | 0.193 | 0.007 | 1.10503007375634e-73 | 35 |
| copz2 | 7.98038564476421e-77 | 0.275735156 | 0.227 | 0.01 | 2.0085832629307e-72 | 35 |
| hsp70l | 8.62210159791232e-77 | 0.2661273 | 0.205 | 0.008 | 2.17009675117855e-72 | 35 |
| efna1a1 | 9.63022032030148e-76 | 0.36225597 | 0.25 | 0.013 | 2.42383015241668e-71 | 35 |
| itgb1a5 | 1.50706370341524e-75 | 0.574334681 | 0.477 | 0.045 | 3.79312863512583e-71 | 35 |
| surf4l | 3.76553644284867e-75 | 0.574480375 | 0.489 | 0.047 | 9.47747867300581e-71 | 35 |
| anxa136 | 4.61982827122263e-73 | 0.49962379 | 0.489 | 0.048 | 1.16276457758402e-68 | 35 |
| paplnb | 5.22141345645182e-72 | 0.506269256 | 0.136 | 0.004 | 1.31417755285436e-67 | 35 |
| rpz59 | 7.94062091440173e-72 | 0.778480757 | 0.716 | 0.099 | 1.99857487794577e-67 | 35 |
| eva1ba6 | 3.58958573532085e-71 | 0.686403861 | 0.557 | 0.063 | 9.03462833722905e-67 | 35 |
| cpamd8 | 3.63899911790943e-71 | 0.252730921 | 0.148 | 0.004 | 9.15899687986624e-67 | 35 |
| adgrl3.12 | 6.91488322637088e-71 | 0.512667838 | 0.33 | 0.024 | 1.74040695924529e-66 | 35 |
| phex5 | 1.11665090231259e-70 | 0.773410615 | 0.659 | 0.084 | 2.81049865603055e-66 | 35 |
| id16 | 1.276801629534e-70 | 0.981605355 | 0.705 | 0.102 | 3.21358202137411e-66 | 35 |
| bckdhbl1 | 1.76106484962592e-70 | 0.374358535 | 0.364 | 0.028 | 4.43242412002348e-66 | 35 |
| serinc54 | 1.86244675424181e-70 | 0.747004722 | 0.625 | 0.08 | 4.6875922357512e-66 | 35 |
| prkd2 | 6.77599757387256e-70 | 0.453546633 | 0.386 | 0.032 | 1.70545082936799e-65 | 35 |
| sox181 | 1.90567149229514e-69 | 0.29973528 | 0.205 | 0.009 | 4.79638457895764e-65 | 35 |
| mmp14a5 | 2.3908931472583e-68 | 0.520023357 | 0.489 | 0.051 | 6.01763896233443e-64 | 35 |
| tuba8l35 | 2.85172961565001e-68 | 0.554578726 | 0.455 | 0.045 | 7.17751826962952e-64 | 35 |
| fgfr1a1 | 6.09436925546569e-68 | 0.395586295 | 0.295 | 0.02 | 1.53389179790816e-63 | 35 |
| slc39a7 | 5.92874343157219e-67 | 0.425706542 | 0.375 | 0.031 | 1.4922054342924e-62 | 35 |
| ctnnb15 | 6.45498707225021e-67 | 0.934363776 | 0.83 | 0.146 | 1.62465569621466e-62 | 35 |
| CU929237.12 | 1.14913317116033e-64 | 0.31086021 | 0.25 | 0.015 | 2.89225327849344e-60 | 35 |
| cntfr2 | 1.19615217474953e-64 | 0.460530393 | 0.25 | 0.015 | 3.01059540862708e-60 | 35 |
| hsp70.32 | 4.62281689171082e-64 | 0.498168813 | 0.295 | 0.021 | 1.1635167834747e-59 | 35 |
| fstl32 | 1.46907453237805e-63 | 0.557078143 | 0.307 | 0.023 | 3.69751369054231e-59 | 35 |
| SCARF22 | 3.79598471356346e-63 | 0.406710785 | 0.307 | 0.023 | 9.55411392556788e-59 | 35 |
| si:ch211-156j16.13 | 6.31953234327225e-63 | 0.429848083 | 0.284 | 0.02 | 1.59056309547819e-58 | 35 |
| icn10 | 1.01873608417068e-61 | 1.399531611 | 0.955 | 0.271 | 2.56405685024919e-57 | 35 |
| sh3pxd2aa2 | 1.50834187727831e-61 | 0.467679443 | 0.352 | 0.031 | 3.79634567092178e-57 | 35 |
| nuak1b | 1.56806737727973e-61 | 0.302946792 | 0.205 | 0.01 | 3.94666878187536e-57 | 35 |
| pcolcea4 | 2.62245700059459e-61 | 0.665426846 | 0.455 | 0.049 | 6.60046202479653e-57 | 35 |

| | | | | | | |
|---|---|---|---|---|---|---|
| fn1a2 | 9.79363753153373e-61 | 0.70180262 | 0.511 | 0.062 | 2.46496063031172e-56 | 35 |
| calua1 | 1.0055755063034e-60 | 0.357531958 | 0.307 | 0.024 | 2.53093299181502e-56 | 35 |
| bhmt3 | 3.18985359212825e-60 | 0.546144198 | 0.42 | 0.043 | 8.02854250602759e-56 | 35 |
| cnn3a7 | 3.79430663690086e-60 | 0.53183835 | 0.477 | 0.055 | 9.54989037441578e-56 | 35 |
| hmgb1b8 | 1.05235378986702e-59 | 0.777809835 | 0.739 | 0.124 | 2.64866925371631e-55 | 35 |
| prkab1b2 | 1.05335773203679e-59 | 0.738338951 | 0.716 | 0.119 | 2.65119607576341e-55 | 35 |
| b3glcta | 2.86633173236481e-59 | 0.294572484 | 0.239 | 0.015 | 7.21427033718899e-55 | 35 |
| angptl45 | 3.64211811939739e-59 | 0.50478701 | 0.443 | 0.048 | 9.16684709471129e-55 | 35 |
| schip11 | 4.83309501868771e-59 | 0.329628495 | 0.273 | 0.019 | 1.21644168525351e-54 | 35 |
| si:ch211-286o17.13 | 9.26933354933026e-59 | 0.522538899 | 0.375 | 0.036 | 2.33299856103093e-54 | 35 |
| mrc23 | 1.47132988476199e-58 | 0.34917123 | 0.307 | 0.024 | 3.70319018695745e-54 | 35 |
| cercam | 1.03510918491643e-57 | 0.25472277 | 0.239 | 0.015 | 2.60526630751616e-53 | 35 |
| dpt1 | 1.63817153387429e-57 | 0.558457557 | 0.227 | 0.014 | 4.1231139336082e-53 | 35 |
| fkbp10b | 1.56135875496714e-56 | 0.343478622 | 0.307 | 0.025 | 3.92978385037679e-52 | 35 |
| perp4 | 1.63708320533263e-56 | 0.516863903 | 0.455 | 0.053 | 4.12037471950171e-52 | 35 |
| yap15 | 1.72281137309859e-56 | 0.509834843 | 0.432 | 0.048 | 4.33614394495185e-52 | 35 |
| cavin1b5 | 2.09434972059791e-56 | 0.507239312 | 0.386 | 0.039 | 5.27126881177289e-52 | 35 |
| sox4b | 3.11641929437164e-56 | 0.362594605 | 0.193 | 0.01 | 7.84371572200397e-52 | 35 |
| emid11 | 6.11449267952443e-56 | 0.435253474 | 0.273 | 0.02 | 1.5389566625095e-51 | 35 |
| ecrg4a3 | 1.87164712411742e-55 | 0.557603054 | 0.33 | 0.03 | 4.71074864669113e-51 | 35 |
| s100v27 | 2.50385789652801e-54 | 0.73219371 | 0.705 | 0.119 | 6.30195993977134e-50 | 35 |
| jun9 | 5.57410011301769e-54 | 1.171985787 | 0.898 | 0.225 | 1.40294525744542e-49 | 35 |
| fosab11 | 9.87177557731332e-54 | 1.271818886 | 1 | 0.346 | 2.48462719505399e-49 | 35 |
| loxa | 1.67713462391903e-53 | 0.375797027 | 0.159 | 0.007 | 4.22118013494181e-49 | 35 |
| zgc:153675 | 2.07931818503869e-53 | 0.562612934 | 0.511 | 0.067 | 5.23343593992388e-49 | 35 |
| fat1a3 | 2.79784490784924e-53 | 0.405311297 | 0.307 | 0.027 | 7.04189584856575e-49 | 35 |
| endouc4 | 3.21785540839271e-53 | 0.647534225 | 0.477 | 0.061 | 8.0990202773836e-49 | 35 |
| nucb2a1 | 6.11498735167765e-53 | 0.473284603 | 0.409 | 0.046 | 1.53908116654375e-48 | 35 |
| ccnd11 | 1.56533764319238e-52 | 0.402701745 | 0.33 | 0.031 | 3.9397983141509e-48 | 35 |
| klf2a7 | 3.11191769124124e-52 | 0.718973185 | 0.557 | 0.083 | 7.83238563708507e-48 | 35 |
| fzd24 | 1.314612555763e-51 | 0.359033738 | 0.307 | 0.027 | 3.3087483415999e-47 | 35 |
| lhfpl2b | 7.78934381977915e-51 | 0.319420272 | 0.227 | 0.016 | 1.96049994600021e-46 | 35 |
| egr111 | 9.2877015879769e-51 | 0.910698843 | 0.852 | 0.187 | 2.33762161267791e-46 | 35 |
| ppfibp1b | 1.75438662656922e-50 | 0.286870949 | 0.25 | 0.019 | 4.41561570041207e-46 | 35 |
| meis1b6 | 1.79229840399259e-50 | 0.486870446 | 0.443 | 0.055 | 4.51103585300895e-46 | 35 |

| | | | | | | |
|---|---|---|---|---|---|---|
| ier2b13 | 2.34892467613658e-50 | 0.973380927 | 0.886 | 0.221 | 5.91200851736815e-46 | 35 |
| igfbp31 | 2.51434078271369e-50 | 0.289627801 | 0.205 | 0.013 | 6.32834431601209e-46 | 35 |
| elovl1b6 | 7.11746958361684e-50 | 0.767961434 | 0.636 | 0.114 | 1.79139591950052e-45 | 35 |
| meis1a1 | 1.32794552321968e-49 | 0.272770646 | 0.216 | 0.014 | 3.34230608739161e-45 | 35 |
| cd992 | 1.48912884884133e-49 | 0.52060627 | 0.42 | 0.052 | 3.74798839964874e-45 | 35 |
| mych4 | 1.54714106373654e-49 | 0.59358106 | 0.511 | 0.073 | 3.8939993433185e-45 | 35 |
| impdh1b | 3.45793066176954e-49 | 0.258268349 | 0.25 | 0.019 | 8.70326568260775e-45 | 35 |
| irf2bpl2 | 3.94182329422928e-48 | 0.449161917 | 0.375 | 0.043 | 9.92117504924567e-44 | 35 |
| ptprfb1 | 9.70218568300769e-48 | 0.297330313 | 0.205 | 0.014 | 2.44194311455621e-43 | 35 |
| lrrc174 | 1.46988425549526e-47 | 0.40799391 | 0.33 | 0.034 | 3.69955168265603e-43 | 35 |
| s100a10b10 | 2.48522359782927e-47 | 1.023092469 | 0.989 | 0.252 | 6.25505927337649e-43 | 35 |
| lgals3b4 | 5.00240006405403e-47 | 0.524707802 | 0.511 | 0.077 | 1.25905407212176e-42 | 35 |
| slc31a1 | 1.08870374946627e-46 | 0.275444227 | 0.25 | 0.02 | 2.74015846703166e-42 | 35 |
| anxa2a6 | 1.28029341047395e-46 | 0.646764027 | 0.614 | 0.104 | 3.22237048482189e-42 | 35 |
| fam168a7 | 2.96926408237194e-46 | 0.462550716 | 0.432 | 0.057 | 7.47334076892193e-42 | 35 |
| atf36 | 3.76882096505047e-46 | 0.726931955 | 0.545 | 0.089 | 9.48574548693553e-42 | 35 |
| mgst3a | 4.05953677356703e-46 | 0.446320262 | 0.443 | 0.059 | 1.02174481053909e-41 | 35 |
| fstl1b7 | 4.24801850838591e-46 | 0.652190657 | 0.682 | 0.122 | 1.06918377837565e-41 | 35 |
| prrx1a1 | 9.1639377693662e-46 | 0.262912921 | 0.17 | 0.01 | 2.30647149717178e-41 | 35 |
| myo1b | 9.72536853407102e-46 | 0.253652374 | 0.182 | 0.011 | 2.44777800634033e-41 | 35 |
| creb3l2 | 4.10398160334024e-45 | 0.414659135 | 0.341 | 0.038 | 1.03293112974471e-40 | 35 |
| junba10 | 4.44310105104573e-45 | 1.099589995 | 0.955 | 0.306 | 1.1182841035377e-40 | 35 |
| plod23 | 5.94546061891336e-45 | 0.355277507 | 0.261 | 0.023 | 1.4964129831743e-40 | 35 |
| cdkn1a2 | 1.84984008259965e-44 | 0.379869283 | 0.307 | 0.031 | 4.65586250389506e-40 | 35 |
| asph1 | 3.12600018682706e-43 | 0.386770279 | 0.318 | 0.035 | 7.86782987022503e-39 | 35 |
| tpm4a7 | 4.54044536629703e-43 | 0.548970935 | 0.534 | 0.087 | 1.1427846942433e-38 | 35 |
| mob2b | 8.04467966272725e-43 | 0.352444504 | 0.284 | 0.028 | 2.02476542431182e-38 | 35 |
| si:ch73-46j18.58 | 1.02794835096671e-42 | 0.70160829 | 0.818 | 0.172 | 2.58724320454812e-38 | 35 |
| midn9 | 1.29948029461678e-42 | 0.705024656 | 0.682 | 0.138 | 3.27066195352099e-38 | 35 |
| h1f07 | 1.56097439589929e-42 | 0.517485397 | 0.432 | 0.061 | 3.92881645703891e-38 | 35 |
| tsc22d310 | 2.46296868329465e-42 | 0.753425484 | 0.67 | 0.142 | 6.1990458789843e-38 | 35 |
| zdhhc8b | 6.90398600044217e-42 | 0.250211997 | 0.193 | 0.014 | 1.73766423645129e-37 | 35 |
| prdx58 | 1.05122784065853e-41 | 0.723408338 | 0.716 | 0.157 | 2.64583535215346e-37 | 35 |
| ptbp31 | 4.30094662273922e-41 | 0.517585608 | 0.42 | 0.061 | 1.08250525547723e-36 | 35 |
| h3f3c11 | 4.71065852630638e-41 | 0.8912735 | 0.875 | 0.254 | 1.18562564448605e-36 | 35 |

| | | | | | | |
|---|---|---|---|---|---|---|
| anxa5b6 | 5.02213420831229e-41 | 0.582227851 | 0.534 | 0.092 | 1.26402095889012e-36 | 35 |
| net11 | 9.06890921941173e-41 | 0.312582443 | 0.239 | 0.021 | 2.28255376143374e-36 | 35 |
| kdelr2a | 2.76321892754036e-39 | 0.476562062 | 0.455 | 0.07 | 6.95474571872635e-35 | 35 |
| gadd45ga3 | 3.35620885270369e-39 | 0.535627914 | 0.375 | 0.052 | 8.44724206136992e-35 | 35 |
| tns3.2 | 1.48786971603302e-38 | 0.31315238 | 0.227 | 0.02 | 3.74481928828351e-34 | 35 |
| txndc5 | 7.0689773154739e-38 | 0.403976716 | 0.443 | 0.068 | 1.77919090053163e-33 | 35 |
| chsy1 | 7.18409307994738e-38 | 0.291873891 | 0.239 | 0.023 | 1.80816438729195e-33 | 35 |
| irf2bp2b6 | 1.1038939023131e-37 | 0.503967159 | 0.477 | 0.081 | 2.77839056273183e-33 | 35 |
| efna1b1 | 1.42603776770209e-37 | 0.314560839 | 0.205 | 0.017 | 3.58919445752939e-33 | 35 |
| tent5ba1 | 1.74907311390595e-37 | 0.278518042 | 0.159 | 0.011 | 4.40224212038988e-33 | 35 |
| mdka7 | 2.03958031469314e-37 | 0.737682415 | 0.5 | 0.092 | 5.13341969405116e-33 | 35 |
| slc38a5b7 | 2.04264505670059e-37 | 0.593607272 | 0.58 | 0.114 | 5.14113334320973e-33 | 35 |
| capns1b6 | 2.22066753611947e-37 | 0.552460752 | 0.602 | 0.12 | 5.58919812165909e-33 | 35 |
| tsc22d1 | 3.87522367225912e-37 | 0.372631782 | 0.318 | 0.039 | 9.75355046070899e-33 | 35 |
| vgll4b | 6.26797036548868e-37 | 0.32706236 | 0.239 | 0.024 | 1.57758546128985e-32 | 35 |
| metrnlb | 8.93432912048857e-37 | 0.316708212 | 0.261 | 0.027 | 2.24868129633577e-32 | 35 |
| LOC1018835071 | 1.48240894538372e-36 | 0.285384066 | 0.205 | 0.018 | 3.73107507463629e-32 | 35 |
| gadd45ba7 | 1.59247128518074e-36 | 0.813197486 | 0.75 | 0.188 | 4.00809097767142e-32 | 35 |
| si:ch73-335l21.47 | 1.78402711507079e-36 | 0.909373373 | 0.636 | 0.147 | 4.49021784592167e-32 | 35 |
| map1lc3b5 | 2.13224631016289e-36 | 0.488955111 | 0.557 | 0.105 | 5.36665073804898e-32 | 35 |
| mmp26 | 9.58939767524994e-36 | 0.444628937 | 0.511 | 0.091 | 2.41355550088366e-31 | 35 |
| socs3a13 | 1.12560492319521e-35 | 1.005298527 | 0.909 | 0.296 | 2.83303503119002e-31 | 35 |
| ier2a13 | 1.42711217322153e-35 | 0.855746483 | 0.886 | 0.26 | 3.59189862878126e-31 | 35 |
| sgce3 | 2.21571692564896e-35 | 0.31893423 | 0.25 | 0.027 | 5.57673793016587e-31 | 35 |
| ppib10 | 5.50630532763294e-35 | 0.889574439 | 0.932 | 0.33 | 1.38588198791193e-30 | 35 |
| dap | 6.33434422173128e-35 | 0.493413367 | 0.545 | 0.104 | 1.59429109716755e-30 | 35 |
| col5a2a6 | 6.72530701987735e-35 | 0.55853134 | 0.477 | 0.085 | 1.69269252383293e-30 | 35 |
| atp2b1a2 | 1.04551607752536e-34 | 0.41900459 | 0.443 | 0.073 | 2.63145941552358e-30 | 35 |
| h2afy27 | 2.47165624554191e-34 | 0.476061215 | 0.511 | 0.096 | 6.22091160440445e-30 | 35 |
| ptprga9 | 2.72043818527562e-34 | 0.414221845 | 0.409 | 0.065 | 6.84707086852021e-30 | 35 |
| selenok4 | 3.03141386357786e-34 | 0.514969434 | 0.602 | 0.125 | 7.62976555323912e-30 | 35 |
| calr1 | 3.35179407113352e-34 | 0.383868452 | 0.352 | 0.051 | 8.43613049763596e-30 | 35 |
| arg22 | 4.03134013327147e-34 | 0.301249671 | 0.216 | 0.021 | 1.0146479981431e-29 | 35 |
| kank1a | 4.50545106534949e-34 | 0.250568884 | 0.193 | 0.017 | 1.13397697863781e-29 | 35 |
| supt3h1 | 5.04871614408161e-34 | 0.323569083 | 0.273 | 0.032 | 1.2707113663039e-29 | 35 |

| | | | | | | |
|---|---|---|---|---|---|---|
| si:busm1-57f23.11 | 1.47128994639033e-33 | 0.3873608 | 0.261 | 0.03 | 3.70308966606981e-29 | 35 |
| klf11a | 2.02551749784195e-33 | 0.381233908 | 0.307 | 0.041 | 5.09802499031841e-29 | 35 |
| snai1a2 | 2.25271334305143e-33 | 0.33274485 | 0.375 | 0.056 | 5.66985421312615e-29 | 35 |
| klf33 | 2.32708483035926e-33 | 0.29844381 | 0.284 | 0.035 | 5.85703980953123e-29 | 35 |
| bcar11 | 1.13106735414285e-32 | 0.254827861 | 0.205 | 0.02 | 2.84678342364213e-28 | 35 |
| triqk2 | 1.34973316298481e-32 | 0.340496752 | 0.341 | 0.05 | 3.39714339791646e-28 | 35 |
| sdc23 | 2.02909033771982e-32 | 0.431140255 | 0.432 | 0.075 | 5.10701747100702e-28 | 35 |
| frzb3 | 3.07125007716916e-32 | 0.425848315 | 0.193 | 0.018 | 7.73002931922705e-28 | 35 |
| sumo1 | 4.04393685024457e-32 | 0.445827021 | 0.557 | 0.114 | 1.01781846583806e-27 | 35 |
| hmgn612 | 4.05551506751689e-32 | 0.752481867 | 0.875 | 0.263 | 1.02073258734333e-27 | 35 |
| ier55 | 4.2744216844444e-32 | 0.563740059 | 0.602 | 0.135 | 1.07582919375781e-27 | 35 |
| si:dkeyp-113d7.1 | 4.29946565546666e-32 | 0.27447283 | 0.284 | 0.036 | 1.0821325108244e-27 | 35 |
| anxa1a7 | 5.10874989454754e-32 | 0.591156503 | 0.636 | 0.141 | 1.28582126095867e-27 | 35 |
| rrbp1a | 5.87320244489639e-32 | 0.353396675 | 0.341 | 0.05 | 1.47822632335597e-27 | 35 |
| hmgn710 | 1.44941254381232e-31 | 0.76265106 | 0.875 | 0.277 | 3.64802643152124e-27 | 35 |
| sar1b1 | 1.56723328604682e-31 | 0.391385716 | 0.443 | 0.079 | 3.94456945765123e-27 | 35 |
| neo1a3 | 2.24464286785442e-31 | 0.304201937 | 0.239 | 0.027 | 5.64954163410279e-27 | 35 |
| calm2a1 | 6.81669821519549e-31 | 0.629937893 | 0.773 | 0.201 | 1.71569477378255e-26 | 35 |
| ssr34 | 1.4867466903694e-30 | 0.729852032 | 0.886 | 0.27 | 3.74199274499075e-26 | 35 |
| klf135 | 1.76096513495182e-30 | 0.462131824 | 0.511 | 0.104 | 4.43217314816023e-26 | 35 |
| tspan72 | 2.48448696667444e-30 | 0.408836376 | 0.432 | 0.077 | 6.2532052464229e-26 | 35 |
| sept15 | 6.8654909505194e-30 | 0.29332356 | 0.295 | 0.041 | 1.72797541733623e-25 | 35 |
| f13a1b4 | 1.10373184976077e-29 | 0.293334166 | 0.25 | 0.031 | 2.77798269266288e-25 | 35 |
| hdlbpa7 | 1.48724176376487e-29 | 0.566439386 | 0.648 | 0.157 | 3.74323879521979e-25 | 35 |
| antxr2a10 | 1.85184960450118e-29 | 0.566359898 | 0.636 | 0.152 | 4.66092026956902e-25 | 35 |
| tspan9a2 | 2.03060329910548e-29 | 0.284328416 | 0.205 | 0.022 | 5.11082544351859e-25 | 35 |
| si:dkey-28b4.78 | 2.63710400973655e-29 | 0.39619181 | 0.466 | 0.091 | 6.63732708210592e-25 | 35 |
| pkig1 | 3.23780715236136e-29 | 0.396725962 | 0.477 | 0.093 | 8.14923682177831e-25 | 35 |
| vdac37 | 3.53365677608756e-29 | 0.539923787 | 0.636 | 0.152 | 8.89386073973479e-25 | 35 |
| zdhhc20a1 | 5.01742835053028e-29 | 0.36203913 | 0.295 | 0.043 | 1.26283654154497e-24 | 35 |
| pfkfb4b4 | 5.28273414404754e-29 | 0.42297981 | 0.477 | 0.095 | 1.32961135671532e-24 | 35 |
| prss352 | 7.54350479762753e-29 | 0.428139759 | 0.193 | 0.02 | 1.89862472251487e-24 | 35 |
| CABZ01080702.17 | 1.51632759922171e-28 | 0.5744241 | 0.42 | 0.084 | 3.81644493448112e-24 | 35 |
| serf213 | 2.61497993093362e-28 | 0.770201084 | 0.92 | 0.341 | 6.58164298816683e-24 | 35 |
| spred2b5 | 3.05838370991304e-28 | 0.406286893 | 0.375 | 0.066 | 7.69764595948014e-24 | 35 |

| | | | | | | |
|---|---|---|---|---|---|---|
| gng22 | 3.76688242252067e-28 | 0.28125359 | 0.295 | 0.043 | 9.48086636924227e-24 | 35 |
| sdc48 | 4.75717064768873e-28 | 0.736127262 | 0.841 | 0.255 | 1.19733228031678e-23 | 35 |
| arl4d1 | 5.5852752388355e-28 | 0.355418995 | 0.318 | 0.049 | 1.40575792486251e-23 | 35 |
| znf704 | 7.45299338928718e-28 | 0.301049432 | 0.239 | 0.03 | 1.87584390614969e-23 | 35 |
| sdf4 | 2.28546120658825e-27 | 0.32402225 | 0.307 | 0.047 | 5.75227731086197e-23 | 35 |
| zgc:16273016 | 2.34754932516111e-27 | 0.876650822 | 0.932 | 0.39 | 5.90854689649801e-23 | 35 |
| twist31 | 2.61486427414084e-27 | 0.281400984 | 0.239 | 0.03 | 6.58135189158509e-23 | 35 |
| nme35 | 5.22296454160831e-27 | 0.391430004 | 0.398 | 0.075 | 1.31456794547739e-22 | 35 |
| pmp22b4 | 7.13609562721055e-27 | 0.30754584 | 0.295 | 0.045 | 1.79608390841262e-22 | 35 |
| spry24 | 3.38129645014306e-26 | 0.385301353 | 0.352 | 0.063 | 8.51038503536507e-22 | 35 |
| sec61a13 | 3.80558397132557e-26 | 0.576548464 | 0.682 | 0.187 | 9.57827429742933e-22 | 35 |
| ip6k2b8 | 7.87785838435033e-26 | 0.538689164 | 0.511 | 0.121 | 1.98277817675714e-21 | 35 |
| enpp12 | 8.75209932506553e-26 | 0.472735713 | 0.261 | 0.039 | 2.20281587912574e-21 | 35 |
| gng76 | 1.31559930562845e-25 | 0.291237665 | 0.318 | 0.053 | 3.31123189233623e-21 | 35 |
| atf4b12 | 1.84617764730796e-25 | 0.648993475 | 0.83 | 0.262 | 4.64664452050942e-21 | 35 |
| nr1d2a4 | 2.24039743754302e-25 | 0.366868879 | 0.398 | 0.078 | 5.63885631055203e-21 | 35 |
| fosl1a6 | 2.72367553388722e-25 | 0.471042276 | 0.432 | 0.091 | 6.85521895124075e-21 | 35 |
| metrnla5 | 3.39138733019699e-25 | 0.486433599 | 0.432 | 0.092 | 8.53578277137281e-21 | 35 |
| tacc12 | 4.47731704412774e-25 | 0.30733107 | 0.25 | 0.036 | 1.12689592683651e-20 | 35 |
| chd6 | 6.2746633318845e-25 | 0.289681986 | 0.261 | 0.039 | 1.57927001400201e-20 | 35 |
| cald1a8 | 1.42486638898148e-24 | 0.410254382 | 0.455 | 0.098 | 3.58624621442748e-20 | 35 |
| emp25 | 3.38827044614689e-24 | 0.300301226 | 0.295 | 0.049 | 8.52793788590712e-20 | 35 |
| lrpap11 | 6.8803963189643e-24 | 0.271755976 | 0.318 | 0.055 | 1.73172694952012e-19 | 35 |
| map1lc3a7 | 1.13517271226987e-23 | 0.307052369 | 0.364 | 0.07 | 2.85711619951204e-19 | 35 |
| klf11b8 | 1.16638710643177e-23 | 0.385851473 | 0.386 | 0.079 | 2.93567970817812e-19 | 35 |
| her9 | 3.25842289240657e-23 | 0.282703614 | 0.17 | 0.019 | 8.2011245778981e-19 | 35 |
| si:ch211-194k22.8 | 4.43726129895877e-23 | 0.287886828 | 0.284 | 0.047 | 1.11681429633493e-18 | 35 |
| ephb36 | 4.50685010344543e-23 | 0.289215061 | 0.261 | 0.041 | 1.13432910253618e-18 | 35 |
| atp2b24 | 5.09282268909949e-23 | 0.297954251 | 0.364 | 0.071 | 1.28181254261945e-18 | 35 |
| kdelr2b | 5.22426638885156e-23 | 0.422840587 | 0.489 | 0.116 | 1.31489560741005e-18 | 35 |
| selenof3 | 5.82924826295822e-23 | 0.582027005 | 0.75 | 0.235 | 1.46716349530395e-18 | 35 |
| arl1 | 8.82895845386682e-23 | 0.297573886 | 0.352 | 0.068 | 2.22216055325374e-18 | 35 |
| arf5 | 9.00460022212522e-23 | 0.509193915 | 0.693 | 0.201 | 2.2663678299067e-18 | 35 |
| myl9b7 | 9.68860528153506e-23 | 0.347204388 | 0.386 | 0.079 | 2.43852506330956e-18 | 35 |
| myca1 | 1.40993604832941e-22 | 0.294950761 | 0.239 | 0.036 | 3.54866804004029e-18 | 35 |

| | | | | | | |
|---|---|---|---|---|---|---|
| flot2a5 | 2.32595717021598e-22 | 0.299822592 | 0.318 | 0.059 | 5.85420160171661e-18 | 35 |
| mgst3b5 | 2.58483494656595e-22 | 0.481787489 | 0.614 | 0.171 | 6.50577107701184e-18 | 35 |
| hnrnpa0a2 | 4.64377766583464e-22 | 0.279244867 | 0.352 | 0.069 | 1.16879240071392e-17 | 35 |
| rhoab10 | 5.62458571774179e-22 | 0.481779841 | 0.659 | 0.188 | 1.41565197929843e-17 | 35 |
| pgrmc16 | 8.92284223416964e-22 | 0.341230738 | 0.557 | 0.14 | 2.24579016191816e-17 | 35 |
| tdh | 1.40246624875196e-21 | 0.341025914 | 0.42 | 0.094 | 3.5298673014838e-17 | 35 |
| btg210 | 1.45478252503914e-21 | 0.696271437 | 0.943 | 0.371 | 3.66154213727102e-17 | 35 |
| mmp14b5 | 1.54497104468395e-21 | 0.301199867 | 0.307 | 0.057 | 3.88853762236504e-17 | 35 |
| klf6a9 | 2.3073874386425e-21 | 0.517780507 | 0.591 | 0.166 | 5.8074634443193e-17 | 35 |
| mafba7 | 2.40217556481501e-21 | 0.335204954 | 0.352 | 0.072 | 6.0460356790829e-17 | 35 |
| cd81a8 | 2.47785560978825e-21 | 0.449919393 | 0.716 | 0.205 | 6.23651478427605e-17 | 35 |
| krt811 | 2.50587268752807e-21 | 0.615068173 | 0.841 | 0.274 | 6.30703096723941e-17 | 35 |
| CABZ01112656.11 | 6.28318713606993e-21 | 0.254540354 | 0.307 | 0.058 | 1.58141537027744e-16 | 35 |
| vamp3 | 9.14915898512015e-21 | 0.42233968 | 0.659 | 0.19 | 2.30275182496489e-16 | 35 |
| ciarta5 | 9.76869663706436e-21 | 0.399436014 | 0.432 | 0.104 | 2.45868325658273e-16 | 35 |
| col6a16 | 1.34064669085876e-20 | 0.371302893 | 0.398 | 0.089 | 3.37427365622241e-16 | 35 |
| appb4 | 3.94435615518456e-20 | 0.279847542 | 0.239 | 0.039 | 9.92755000698403e-16 | 35 |
| osr25 | 4.2488245238369e-20 | 0.563728938 | 0.25 | 0.046 | 1.06938664440451e-15 | 35 |
| histh1l8 | 4.32967746369202e-20 | 0.510213491 | 0.432 | 0.109 | 1.08973652083664e-15 | 35 |
| ywhaqb6 | 4.57536179510895e-20 | 0.39075242 | 0.5 | 0.131 | 1.15157281021097e-15 | 35 |
| ndrg24 | 4.87298486326425e-20 | 0.274927946 | 0.239 | 0.04 | 1.22648156023498e-15 | 35 |
| nfia7 | 5.29499800424095e-20 | 0.33763968 | 0.364 | 0.079 | 1.3326980476874e-15 | 35 |
| pbx1a4 | 6.08102982638699e-20 | 0.25489292 | 0.25 | 0.042 | 1.53053439700334e-15 | 35 |
| gadd45ab3 | 6.46001874997337e-20 | 0.343228912 | 0.341 | 0.072 | 1.6259221191808e-15 | 35 |
| cish5 | 6.58634588125451e-20 | 0.37359783 | 0.284 | 0.054 | 1.65771739485295e-15 | 35 |
| spcs11 | 6.95063267303455e-20 | 0.481543689 | 0.67 | 0.208 | 1.74940473747607e-15 | 35 |
| tgif12 | 7.29458348412738e-20 | 0.29456517 | 0.398 | 0.09 | 1.83597371712002e-15 | 35 |
| capn2a3 | 7.45936041572545e-20 | 0.251051108 | 0.284 | 0.053 | 1.87744642303394e-15 | 35 |
| snd12 | 1.01214981944387e-19 | 0.33444783 | 0.432 | 0.105 | 2.54747988055828e-15 | 35 |
| plecb3 | 1.23595211193565e-19 | 0.286876879 | 0.273 | 0.05 | 3.11076787053085e-15 | 35 |
| rab2a1 | 2.1439012968513e-19 | 0.419285434 | 0.591 | 0.171 | 5.39598517404504e-15 | 35 |
| tfg | 2.44607823028056e-19 | 0.317488947 | 0.398 | 0.093 | 6.15653429779313e-15 | 35 |
| tmem167a | 2.91013911926867e-19 | 0.295225845 | 0.409 | 0.096 | 7.32452914928731e-15 | 35 |
| grpel23 | 2.93762297932579e-19 | 0.284567141 | 0.42 | 0.1 | 7.39370327666509e-15 | 35 |
| tomm20b7 | 3.07022498992855e-19 | 0.460565399 | 0.682 | 0.213 | 7.72744927715117e-15 | 35 |

| | | | | | | |
|---|---|---|---|---|---|---|
| ssr42 | 3.84017058599628e-19 | 0.639873762 | 0.784 | 0.305 | 9.66532534789405e-15 | 35 |
| qkia7 | 3.93194278416095e-19 | 0.422043124 | 0.58 | 0.167 | 9.89630679345469e-15 | 35 |
| ldlrap1a | 4.01285742499026e-19 | 0.327188413 | 0.33 | 0.069 | 1.0099960852958e-14 | 35 |
| svild8 | 5.91202851155075e-19 | 0.303266937 | 0.364 | 0.081 | 1.48799845607221e-14 | 35 |
| chd4b1 | 6.33550253266804e-19 | 0.302454937 | 0.42 | 0.102 | 1.59458263244722e-14 | 35 |
| itga53 | 8.42164231590575e-19 | 0.329095863 | 0.193 | 0.029 | 2.11964315449032e-14 | 35 |
| tpm14 | 9.98487417498506e-19 | 0.490802393 | 0.659 | 0.206 | 2.51309298110199e-14 | 35 |
| sept7a | 1.36911854831573e-18 | 0.30180328 | 0.33 | 0.071 | 3.44593447425586e-14 | 35 |
| atrx7 | 1.41916243510458e-18 | 0.322620162 | 0.42 | 0.104 | 3.57188993291473e-14 | 35 |
| nck1b1 | 1.93759023171722e-18 | 0.253274171 | 0.273 | 0.052 | 4.87672085420907e-14 | 35 |
| cited4a4 | 1.98945521002708e-18 | 0.268696494 | 0.239 | 0.042 | 5.00725981811716e-14 | 35 |
| cd1642 | 2.13920818834576e-18 | 0.271472388 | 0.33 | 0.071 | 5.38417308924745e-14 | 35 |
| id35 | 2.22074931725576e-18 | 0.526331836 | 0.659 | 0.209 | 5.58940395660103e-14 | 35 |
| wls5 | 2.85640286307417e-18 | 0.35611549 | 0.466 | 0.122 | 7.18928036607138e-14 | 35 |
| gsna1 | 3.46986646100589e-18 | 0.258761938 | 0.193 | 0.03 | 8.73330689570572e-14 | 35 |
| ptmaa16 | 6.22287850039154e-18 | 0.625423719 | 0.955 | 0.393 | 1.56623628976355e-13 | 35 |
| mfap26 | 6.74268520063217e-18 | 0.32459132 | 0.398 | 0.096 | 1.69706643814711e-13 | 35 |
| tmem2584 | 7.45564617017816e-18 | 0.484032348 | 0.807 | 0.28 | 1.87651158457214e-13 | 35 |
| si:ch211-286b5.56 | 8.28101696468611e-18 | 0.312532609 | 0.5 | 0.136 | 2.08424915984185e-13 | 35 |
| tpt110 | 8.93408265356415e-18 | 0.367054652 | 0.989 | 0.801 | 2.24861926307556e-13 | 35 |
| ostc1 | 9.40195241498627e-18 | 0.403618415 | 0.591 | 0.179 | 2.36637740332789e-13 | 35 |
| si:zfos-169g10.32 | 1.05200484366297e-17 | 0.296343931 | 0.386 | 0.094 | 2.64779099101532e-13 | 35 |
| eif4a1b8 | 1.10985469036431e-17 | 0.536944973 | 0.773 | 0.279 | 2.79339327017792e-13 | 35 |
| ssr11 | 1.20484317582027e-17 | 0.460442203 | 0.591 | 0.187 | 3.03246978922203e-13 | 35 |
| ywhabb3 | 1.29045958902316e-17 | 0.449009393 | 0.682 | 0.225 | 3.2479577396124e-13 | 35 |
| slc3a2b7 | 1.54306243064469e-17 | 0.352233481 | 0.443 | 0.118 | 3.88373383168962e-13 | 35 |
| krtcap21 | 1.86756206231299e-17 | 0.387646864 | 0.58 | 0.175 | 4.70046695463557e-13 | 35 |
| si:ch73-1a9.314 | 2.90976192292342e-17 | 0.524552916 | 0.75 | 0.264 | 7.32357978380595e-13 | 35 |
| cops9 | 3.04652583171001e-17 | 0.289047699 | 0.409 | 0.104 | 7.66780086583092e-13 | 35 |
| marcksl1b9 | 3.22972408721741e-17 | 0.400448223 | 0.432 | 0.116 | 8.12889255511749e-13 | 35 |
| evla4 | 4.10657902035164e-17 | 0.299211135 | 0.409 | 0.102 | 1.0335848736323e-12 | 35 |
| glud1a3 | 4.1318050638341e-17 | 0.2847974 | 0.375 | 0.09 | 1.0399340165164e-12 | 35 |
| si:ch1073-325m22.2 | 4.26877152939479e-17 | 0.447571764 | 0.682 | 0.228 | 1.07440710623338e-12 | 35 |
| junbb6 | 7.0857596582153e-17 | 0.301182915 | 0.318 | 0.072 | 1.78341484837621e-12 | 35 |
| ier3ip12 | 7.52186848572669e-17 | 0.339619895 | 0.523 | 0.152 | 1.89317907917255e-12 | 35 |

| | | | | | | |
|---|---|---|---|---|---|---|
| LOC1018844515 | 7.75338986142292e-17 | 0.317738521 | 0.352 | 0.085 | 1.95145069422153e-12 | 35 |
| rgcc6 | 1.06798321763913e-16 | 0.406605108 | 0.307 | 0.07 | 2.68800696047593e-12 | 35 |
| elocb2 | 1.34649527832659e-16 | 0.379777016 | 0.557 | 0.17 | 3.38899396602019e-12 | 35 |
| cyr616 | 1.4625277588107e-16 | 0.313615405 | 0.307 | 0.068 | 3.68103611615065e-12 | 35 |
| notch28 | 2.26760944518045e-16 | 0.358020452 | 0.432 | 0.119 | 5.70734621257467e-12 | 35 |
| ahnak10 | 2.7912392347812e-16 | 0.343206231 | 0.455 | 0.126 | 7.02527003002081e-12 | 35 |
| si:dkey-56m19.54 | 3.48299461123862e-16 | 0.291562825 | 0.239 | 0.047 | 8.76634913702649e-12 | 35 |
| idh27 | 4.94589987067322e-16 | 0.361979074 | 0.625 | 0.198 | 1.24483353844974e-11 | 35 |
| kctd12.25 | 6.89461108524674e-16 | 0.311068287 | 0.33 | 0.079 | 1.73530466404575e-11 | 35 |
| jund8 | 1.1833843474058e-15 | 0.28368356 | 0.364 | 0.092 | 2.97846006398567e-11 | 35 |
| nfkbiab6 | 1.40052141041044e-15 | 0.370333799 | 0.568 | 0.18 | 3.52497233786202e-11 | 35 |
| tmem593 | 1.50507406857098e-15 | 0.284001994 | 0.398 | 0.106 | 3.7881209231863e-11 | 35 |
| eef1a1l14 | 1.56913652293545e-15 | 0.256055697 | 1 | 0.915 | 3.94935971457625e-11 | 35 |
| eif1b9 | 1.91627362358826e-15 | 0.568261716 | 0.886 | 0.381 | 4.8230690832093e-11 | 35 |
| rpl22l111 | 2.23276635702882e-15 | 0.604624265 | 0.875 | 0.39 | 5.61964964400582e-11 | 35 |
| ccnd2a2 | 2.29653591046124e-15 | 0.318207874 | 0.341 | 0.086 | 5.78015123303991e-11 | 35 |
| fkbp1aa8 | 2.49642263084386e-15 | 0.568534571 | 0.898 | 0.386 | 6.28324611957091e-11 | 35 |
| ssr21 | 2.73301666976045e-15 | 0.511085551 | 0.83 | 0.32 | 6.87872965612007e-11 | 35 |
| CR383676.118 | 2.76813565665193e-15 | 0.677629587 | 1 | 0.726 | 6.96712063422723e-11 | 35 |
| ptp4a14 | 3.01704870455356e-15 | 0.291052252 | 0.42 | 0.117 | 7.59360988449087e-11 | 35 |
| mtdha1 | 4.15668092751906e-15 | 0.266873638 | 0.477 | 0.138 | 1.04619502264727e-10 | 35 |
| serinc14 | 5.01212207393173e-15 | 0.259484235 | 0.33 | 0.082 | 1.26150100478788e-10 | 35 |
| calm3b2 | 5.85328702531914e-15 | 0.445830988 | 0.705 | 0.255 | 1.47321381140258e-10 | 35 |
| CABZ01045617.1 | 6.28012598520835e-15 | 0.269427028 | 0.227 | 0.046 | 1.58064490921709e-10 | 35 |
| atf4a4 | 7.61970684397543e-15 | 0.359966421 | 0.557 | 0.181 | 1.91780401556018e-10 | 35 |
| ost43 | 8.34066476627931e-15 | 0.443815291 | 0.705 | 0.256 | 2.09926191502484e-10 | 35 |
| reep54 | 8.49237036038892e-15 | 0.324620327 | 0.511 | 0.158 | 2.13744469600629e-10 | 35 |
| zgc:926063 | 9.34597917595417e-15 | 0.251624386 | 0.341 | 0.087 | 2.35228949879591e-10 | 35 |
| cnn211 | 1.05447714635803e-14 | 0.319941796 | 0.455 | 0.134 | 2.65401352966852e-10 | 35 |
| pim26 | 2.99728320190823e-14 | 0.266374119 | 0.341 | 0.089 | 7.54386209088282e-10 | 35 |
| pfn2l5 | 3.14888415037441e-14 | 0.352229414 | 0.625 | 0.208 | 7.92542651807735e-10 | 35 |
| chd4a2 | 3.42711118924532e-14 | 0.302226782 | 0.341 | 0.09 | 8.62569615221156e-10 | 35 |
| cnbpb6 | 5.21574732720674e-14 | 0.371927285 | 0.545 | 0.184 | 1.31275144478466e-09 | 35 |
| tnrc6c19 | 5.30397534105923e-14 | 0.300353726 | 0.443 | 0.133 | 1.3349575535912e-09 | 35 |
| smim151 | 8.32152194475709e-14 | 0.28176843 | 0.466 | 0.143 | 2.09444385827591e-09 | 35 |

| | | | | | | |
|---|---|---|---|---|---|---|
| zgc:77849 | 8.84283997457785e-14 | 0.307339669 | 0.545 | 0.178 | 2.2256543932015e-09 | 35 |
| mt-nd213 | 1.3666972098078e-13 | 0.525800581 | 0.909 | 0.399 | 3.43984020736525e-09 | 35 |
| mcl1b11 | 1.37520310765898e-13 | 0.416245841 | 0.67 | 0.241 | 3.46124870166688e-09 | 35 |
| ybx19 | 1.37526456757173e-13 | 0.463174653 | 1 | 0.669 | 3.46140339012128e-09 | 35 |
| rpl3911 | 1.41654519832582e-13 | 0.303146419 | 1 | 0.874 | 3.56530260966625e-09 | 35 |
| tcf125 | 1.45115148884687e-13 | 0.292269412 | 0.25 | 0.058 | 3.65240318227868e-09 | 35 |
| CU694197.1 | 1.67827164141152e-13 | 0.272436316 | 0.432 | 0.13 | 4.22404189426866e-09 | 35 |
| hp1bp34 | 2.06422904768515e-13 | 0.262875982 | 0.352 | 0.098 | 5.19545809011876e-09 | 35 |
| si:ch211-39i22.14 | 5.2601656699359e-13 | 0.284872754 | 0.284 | 0.072 | 1.32393109746617e-08 | 35 |
| ptmab13 | 6.62253252399122e-13 | 0.525056813 | 0.989 | 0.539 | 1.66682521096335e-08 | 35 |
| jpt1b4 | 6.87212584080577e-13 | 0.394087017 | 0.727 | 0.27 | 1.7296453528724e-08 | 35 |
| mtpn5 | 7.16342796096832e-13 | 0.261411131 | 0.466 | 0.146 | 1.80296318349612e-08 | 35 |
| sem11 | 7.58199910197236e-13 | 0.351072189 | 0.682 | 0.251 | 1.90831335397542e-08 | 35 |
| p4hb2 | 7.6939476018591e-13 | 0.449899831 | 0.568 | 0.215 | 1.93648967191192e-08 | 35 |
| nsa214 | 8.8843018126636e-13 | 0.521696576 | 0.852 | 0.361 | 2.2360899232293e-08 | 35 |
| eif3ea9 | 1.08736436618298e-12 | 0.400690441 | 0.727 | 0.276 | 2.73678737324594e-08 | 35 |
| rhoaa10 | 1.11807472685274e-12 | 0.323583422 | 0.466 | 0.153 | 2.81408228001566e-08 | 35 |
| socs3b10 | 2.85998499256529e-12 | 0.29540293 | 0.477 | 0.155 | 7.19829622778757e-08 | 35 |
| ddit312 | 4.04533075028122e-12 | 0.312160673 | 0.511 | 0.176 | 1.01816929653828e-07 | 35 |
| si:dkey-42i9.44 | 4.40073060474217e-12 | 0.306976724 | 0.443 | 0.147 | 1.10761988590756e-07 | 35 |
| atox12 | 5.33202274464666e-12 | 0.265673164 | 0.364 | 0.11 | 1.34201680460012e-07 | 35 |
| tpm37 | 5.3775482713837e-12 | 0.336257128 | 0.682 | 0.253 | 1.35347512442456e-07 | 35 |
| serp1 | 5.38990374257266e-12 | 0.348146067 | 0.568 | 0.207 | 1.35658487296811e-07 | 35 |
| dusp28 | 6.17623925514214e-12 | 0.364907677 | 0.511 | 0.179 | 1.55449765812673e-07 | 35 |
| mibp212 | 7.73510061672875e-12 | 0.57453561 | 0.898 | 0.491 | 1.94684747422446e-07 | 35 |
| si:ch1073-429i10.3.1 | 8.53031067002058e-12 | 0.292699215 | 0.568 | 0.201 | 2.14699389253748e-07 | 35 |
| eif4ebp25 | 8.65581441236943e-12 | 0.254242081 | 0.511 | 0.174 | 2.17858192944926e-07 | 35 |
| tmed101 | 1.52555025128279e-11 | 0.308310523 | 0.489 | 0.171 | 3.83965742745365e-07 | 35 |
| cdc427 | 2.61485640099321e-11 | 0.339227329 | 0.568 | 0.212 | 6.58133207565982e-07 | 35 |
| ndufa14 | 2.65106982887079e-11 | 0.311751019 | 0.614 | 0.23 | 6.6724776522849e-07 | 35 |
| mycb6 | 2.70792660835747e-11 | 0.390488496 | 0.557 | 0.211 | 6.81558048057491e-07 | 35 |
| gnb1a6 | 3.46785535975365e-11 | 0.340534971 | 0.523 | 0.192 | 8.72824515496396e-07 | 35 |
| ndufa4l9 | 4.1425164819834e-11 | 0.468434925 | 0.898 | 0.393 | 1.0426299733504e-06 | 35 |
| rab105 | 4.93443950328069e-11 | 0.294522179 | 0.534 | 0.194 | 1.24194907858072e-06 | 35 |
| cox6a13 | 5.05605434577854e-11 | 0.450504402 | 0.841 | 0.372 | 1.272558318289e-06 | 35 |

| | | | | | | |
|---|---|---|---|---|---|---|
| itm2ba11 | 6.16630184365032e-11 | 0.338549656 | 0.705 | 0.272 | 1.55199651102835e-06 | 35 |
| si:dkey-79d12.56 | 6.61851359641556e-11 | 0.250892333 | 0.273 | 0.076 | 1.66581368708183e-06 | 35 |
| pno13 | 6.65857697334382e-11 | 0.293392357 | 0.477 | 0.171 | 1.67589723842091e-06 | 35 |
| gtpbp44 | 6.77353184021333e-11 | 0.319322939 | 0.625 | 0.236 | 1.70483022886329e-06 | 35 |
| psmb71 | 7.47725944711581e-11 | 0.277638434 | 0.432 | 0.148 | 1.88195143024458e-06 | 35 |
| mt-nd314 | 7.52772697728402e-11 | 0.428601569 | 0.875 | 0.394 | 1.89465360291262e-06 | 35 |
| mt-nd115 | 9.58809783005988e-11 | 0.451362792 | 0.875 | 0.41 | 2.41322834284777e-06 | 35 |
| si:dkey-7j14.610 | 1.23685676012442e-10 | 0.290455686 | 0.489 | 0.175 | 3.11304477955715e-06 | 35 |
| gabarapb10 | 1.48602117728004e-10 | 0.348257662 | 0.67 | 0.268 | 3.74016670109614e-06 | 35 |
| rps227 | 2.05443811570856e-10 | 0.344268859 | 1 | 0.829 | 5.17081529342689e-06 | 35 |
| tubb2b3 | 2.63514449289443e-10 | 0.259150147 | 0.659 | 0.248 | 6.63239517416599e-06 | 35 |
| cox8a9 | 3.56982796109809e-10 | 0.295308379 | 0.591 | 0.227 | 8.98489999528778e-06 | 35 |
| rpl5a15 | 3.78026564314991e-10 | 0.53322937 | 0.955 | 0.491 | 9.51455059724401e-06 | 35 |
| gng56 | 4.20964249015478e-10 | 0.292211442 | 0.693 | 0.27 | 1.05952491834706e-05 | 35 |
| tegt8 | 4.54869910632905e-10 | 0.271475505 | 0.557 | 0.211 | 1.14486207807196e-05 | 35 |
| tma73 | 4.61260366522901e-10 | 0.359942608 | 0.727 | 0.304 | 1.16094621650149e-05 | 35 |
| cebpb13 | 5.10558678501225e-10 | 0.481395327 | 0.727 | 0.353 | 1.28502513791973e-05 | 35 |
| cox7a2a3 | 5.98128221061397e-10 | 0.287500547 | 0.614 | 0.241 | 1.50542891958943e-05 | 35 |
| rpl2915 | 6.74170620395049e-10 | 0.330761174 | 1 | 0.808 | 1.6968200344723e-05 | 35 |
| rps27.215 | 7.91115250427797e-10 | 0.517516794 | 0.989 | 0.516 | 1.99115797380172e-05 | 35 |
| XLOC-0234928 | 9.3413940197082e-10 | 0.255362054 | 0.307 | 0.095 | 2.35113546082036e-05 | 35 |
| higd1a9 | 1.02133958374535e-09 | 0.370178754 | 0.716 | 0.309 | 2.57060959832868e-05 | 35 |
| eef1g12 | 1.56147767261319e-09 | 0.464435601 | 1 | 0.619 | 3.93008315420014e-05 | 35 |
| sec61b3 | 2.62968785438205e-09 | 0.458259964 | 0.886 | 0.454 | 6.61866136069419e-05 | 35 |
| rpl1814 | 2.80520050889631e-09 | 0.345343747 | 0.989 | 0.774 | 7.06040916084112e-05 | 35 |
| CABZ01072614.19 | 3.27168833279325e-09 | 0.273344001 | 0.261 | 0.081 | 8.23451236480734e-05 | 35 |
| oaz1b13 | 3.46557169484234e-09 | 0.322350986 | 0.773 | 0.321 | 8.72249739874868e-05 | 35 |
| rpl77 | 3.94351817431714e-09 | 0.30098717 | 0.989 | 0.805 | 9.92544089293882e-05 | 35 |
| tob1b4 | 4.79882967852724e-09 | 0.270161556 | 0.455 | 0.174 | 0.000120782 | 35 |
| sumo2b11 | 6.38863209146569e-09 | 0.310296394 | 0.636 | 0.269 | 0.000160795 | 35 |
| rps27.114 | 7.16192748531282e-09 | 0.425868602 | 0.989 | 0.679 | 0.000180259 | 35 |
| mt-nd414 | 7.18909556741152e-09 | 0.4302631 | 0.92 | 0.455 | 0.000180942 | 35 |
| skp12 | 7.96796955571778e-09 | 0.254806639 | 0.614 | 0.251 | 0.000200546 | 35 |
| ubb3 | 8.73996582064223e-09 | 0.354506978 | 0.705 | 0.315 | 0.000219976 | 35 |
| COX318 | 8.93535011634084e-09 | 0.436576406 | 1 | 0.699 | 0.000224894 | 35 |

| | | | | | | |
|---|---|---|---|---|---|---|
| rpl1011 | 9.38289531412126e-09 | 0.328198322 | 0.989 | 0.772 | 0.000236158 | 35 |
| hnrnpa0b6 | 1.00054585823358e-08 | 0.340827318 | 0.716 | 0.311 | 0.000251827 | 35 |
| rpl3213 | 1.17858872821993e-08 | 0.321417371 | 1 | 0.813 | 0.000296639 | 35 |
| rpl919 | 1.53654188777955e-08 | 0.459391697 | 1 | 0.628 | 0.000386732 | 35 |
| lum2 | 2.17390110739559e-08 | 0.287304717 | 0.102 | 0.019 | 0.000547149 | 35 |
| si:ch211-196l7.414 | 2.39910070943835e-08 | 0.312783511 | 0.705 | 0.314 | 0.00060383 | 35 |
| eif2s34 | 3.15130702678529e-08 | 0.288423126 | 0.659 | 0.284 | 0.000793152 | 35 |
| mt-cyb14 | 3.65562063917816e-08 | 0.425511539 | 0.989 | 0.623 | 0.000920083 | 35 |
| rplp110 | 5.86201452359922e-08 | 0.262951967 | 1 | 0.877 | 0.00147541 | 35 |
| mt-atp618 | 7.53713433074091e-08 | 0.437203467 | 0.977 | 0.59 | 0.001897021 | 35 |
| eif3i8 | 7.69207461032585e-08 | 0.271782746 | 0.625 | 0.272 | 0.001936018 | 35 |
| dad12 | 8.05605256053948e-08 | 0.32116672 | 0.773 | 0.35 | 0.002027628 | 35 |
| rpl18a13 | 8.53903327096575e-08 | 0.325262578 | 0.989 | 0.785 | 0.002149189 | 35 |
| ddx52 | 8.72402144849903e-08 | 0.337339862 | 0.773 | 0.356 | 0.002195749 | 35 |
| rpl10a9 | 8.84144339425526e-08 | 0.333021245 | 1 | 0.764 | 0.002225303 | 35 |
| rps1815 | 1.19149577309605e-07 | 0.368232347 | 0.989 | 0.724 | 0.002998876 | 35 |
| rps199 | 1.24523533902912e-07 | 0.259197647 | 1 | 0.863 | 0.003134133 | 35 |
| rpl27a16 | 1.26813992453784e-07 | 0.491686847 | 0.977 | 0.494 | 0.003191781 | 35 |
| rps78 | 1.36708547217209e-07 | 0.289935531 | 1 | 0.82 | 0.003440817 | 35 |
| xbp16 | 1.82533807984521e-07 | 0.322021533 | 0.75 | 0.343 | 0.004594193 | 35 |
| pdia38 | 2.10905079682352e-07 | 0.267470743 | 0.648 | 0.283 | 0.00530827 | 35 |
| rps128 | 2.25139553675927e-07 | 0.269302704 | 1 | 0.84 | 0.005666537 | 35 |
| rps4x12 | 3.84662363498296e-07 | 0.364892073 | 1 | 0.71 | 0.009681567 | 35 |
| sec61g5 | 3.93941694848119e-07 | 0.373483901 | 1 | 0.678 | 0.009915119 | 35 |
| mt-co118 | 4.42331272774432e-07 | 0.410317788 | 1 | 0.657 | 0.011133036 | 35 |
| rps1018 | 5.67683088386881e-07 | 0.391086106 | 1 | 0.693 | 0.014288016 | 35 |
| rps8a7 | 6.19170044086984e-07 | 0.297086921 | 0.989 | 0.788 | 0.015583891 | 35 |
| rpl1114 | 6.24821712569273e-07 | 0.299238651 | 0.989 | 0.814 | 0.015726138 | 35 |
| faua12 | 7.95563923223502e-07 | 0.371175746 | 0.989 | 0.689 | 0.020023548 | 35 |
| zgc:11418819 | 7.98378646908302e-07 | 0.424610295 | 0.977 | 0.627 | 0.020094392 | 35 |
| rps1515 | 8.36276327835707e-07 | 0.391957711 | 0.989 | 0.684 | 0.021048239 | 35 |
| rpl3618 | 9.76164961580501e-07 | 0.322666618 | 0.989 | 0.789 | 0.024569096 | 35 |
| rpl813 | 1.47384317308082e-06 | 0.310369641 | 1 | 0.78 | 0.037095159 | 35 |
| zgc:1935413 | 1.61735909787558e-06 | 0.277289712 | 0.784 | 0.367 | 0.040707311 | 35 |
| rps15a13 | 1.74481783465206e-06 | 0.338314869 | 1 | 0.779 | 0.04391532 | 35 |

| | | | | | | |
|---|---|---|---|---|---|---|
| rpl3711 | 2.26869392765199e-06 | 0.272016964 | 1 | 0.849 | 0.057100757 | 35 |
| rps1616 | 2.46696789133642e-06 | 0.354471124 | 1 | 0.737 | 0.062091115 | 35 |
| rpl1513 | 2.94261748979858e-06 | 0.326852477 | 0.989 | 0.744 | 0.07406274 | 35 |
| rps69 | 2.99646174612756e-06 | 0.287738523 | 0.989 | 0.789 | 0.075417946 | 35 |
| eef2b7 | 3.13381124788194e-06 | 0.398199435 | 0.977 | 0.613 | 0.078874895 | 35 |
| COX5B5 | 3.41189044792416e-06 | 0.256297985 | 0.773 | 0.357 | 0.085873871 | 35 |
| mt-co218 | 3.72574600035466e-06 | 0.392819874 | 1 | 0.72 | 0.093773301 | 35 |
| rpl35a14 | 3.88751227681405e-06 | 0.327001046 | 1 | 0.759 | 0.097844796 | 35 |
| rpsa19 | 8.49960695352463e-06 | 0.322041844 | 0.989 | 0.752 | 0.213926607 | 35 |
| rpl2610 | 9.03679248344776e-06 | 0.256107597 | 1 | 0.841 | 0.22744703 | 35 |
| rps2121 | 1.08772498449257e-05 | 0.337949329 | 0.989 | 0.749 | 0.273769501 | 35 |
| rps1314 | 1.15402451825114e-05 | 0.333917553 | 0.989 | 0.724 | 0.290456431 | 35 |
| rps312 | 1.23408806366871e-05 | 0.349398043 | 1 | 0.699 | 0.310607625 | 35 |
| rpl3415 | 1.26306786788232e-05 | 0.299184842 | 0.989 | 0.783 | 0.317901552 | 35 |
| rpl2414 | 1.78708715251141e-05 | 0.298105885 | 1 | 0.776 | 0.449791965 | 35 |
| rplp2l18 | 2.62915379295989e-05 | 0.304527586 | 1 | 0.774 | 0.661731718 | 35 |
| eef1da10 | 2.77014893426154e-05 | 0.37485931 | 0.886 | 0.491 | 0.697218785 | 35 |
| dynll112 | 3.06234784162465e-05 | 0.288665562 | 0.864 | 0.454 | 0.770762328 | 35 |
| rps2610 | 3.09216592525757e-05 | 0.311705259 | 0.989 | 0.733 | 0.778267242 | 35 |
| rps513 | 3.82603549151087e-05 | 0.344170216 | 0.989 | 0.701 | 0.962974873 | 35 |
| rps3a8 | 4.82285901908969e-05 | 0.273715887 | 0.989 | 0.778 | 1 | 35 |
| ucp211 | 6.46996981898938e-05 | 0.251608396 | 0.841 | 0.415 | 1 | 35 |
| h3f3d | 7.05244527506881e-05 | 0.306174195 | 0.966 | 0.676 | 1 | 35 |
| rpl616 | 8.46830864272561e-05 | 0.363910881 | 1 | 0.667 | 1 | 35 |
| rpl2711 | 0.000100152 | 0.297705645 | 1 | 0.762 | 1 | 35 |
| rack18 | 0.000134641 | 0.293404624 | 0.989 | 0.744 | 1 | 35 |
| eif3f13 | 0.000151728 | 0.307123528 | 0.864 | 0.43 | 1 | 35 |
| rps27a13 | 0.000161371 | 0.294872414 | 0.989 | 0.751 | 1 | 35 |
| eif5a8 | 0.000223896 | 0.318764737 | 0.886 | 0.478 | 1 | 35 |
| rpl3015 | 0.000233335 | 0.327215251 | 0.989 | 0.708 | 1 | 35 |
| eef1b22 | 0.000247593 | 0.279575091 | 0.977 | 0.734 | 1 | 35 |
| rps2415 | 0.000283003 | 0.290415548 | 0.989 | 0.758 | 1 | 35 |
| rpl7a11 | 0.000305436 | 0.254020399 | 1 | 0.81 | 1 | 35 |
| slc25a3b6 | 0.000354734 | 0.286337755 | 0.83 | 0.436 | 1 | 35 |
| rpl23a17 | 0.000406826 | 0.362177487 | 0.977 | 0.633 | 1 | 35 |

| | | | | | | | |
|---|---|---|---|---|---|---|---|
| NC-002333.417 | 0.000541458 | 0.306587459 | 0.977 | 0.622 | | 1 | 35 |
| atp5f1e3 | 0.000603484 | 0.266017825 | 0.92 | 0.482 | | 1 | 35 |
| rps1414 | 0.00066184 | 0.284299728 | 0.989 | 0.759 | | 1 | 35 |
| rpl179 | 0.0009792 | 0.271283105 | 0.989 | 0.746 | | 1 | 35 |
| rps2510 | 0.000981077 | 0.295729944 | 0.989 | 0.724 | | 1 | 35 |
| rps26l15 | 0.001055493 | 0.333049904 | 0.977 | 0.664 | | 1 | 35 |
| rps911 | 0.001315884 | 0.255467865 | 0.989 | 0.776 | | 1 | 35 |
| rpl3116 | 0.001474695 | 0.350694539 | 1 | 0.654 | | 1 | 35 |
| rpl3815 | 0.001740553 | 0.302942657 | 0.989 | 0.727 | | 1 | 35 |
| rpl2214 | 0.006990113 | 0.304488946 | 1 | 0.693 | | 1 | 35 |
| rpl2112 | 0.007816451 | 0.253718766 | 0.989 | 0.756 | | 1 | 35 |
| si:ch211-161c3.6 | 0 | 1.222663348 | 0.588 | 0.009 | | 0 | 36 |
| myb | 0 | 0.80910137 | 0.462 | 0.005 | | 0 | 36 |
| sid1 | 0 | 0.643243836 | 0.212 | 0.001 | | 0 | 36 |
| maptb | 0 | 0.562611907 | 0.338 | 0.003 | | 0 | 36 |
| XLOC-014336 | 2.40679044640105e-106 | 0.607186204 | 0.425 | 0.024 | 6.05765087454681e-102 | | 36 |
| sb:cb812 | 2.67221880910698e-103 | 0.653658879 | 0.425 | 0.025 | 6.72570752064135e-99 | | 36 |
| rag1 | 2.08471859693215e-101 | 0.477607627 | 0.138 | 0.002 | 5.24702823661854e-97 | | 36 |
| lmo22 | 2.05742589605491e-100 | 0.577814108 | 0.362 | 0.018 | 5.1783352377806e-96 | | 36 |
| cd7al3 | 2.09466867611806e-80 | 0.483445482 | 0.375 | 0.024 | 5.27207159092155e-76 | | 36 |
| CU499330.14 | 3.28885859826994e-80 | 1.006271418 | 0.637 | 0.068 | 8.27772820598561e-76 | | 36 |
| kdm6bb3 | 2.67524562506462e-79 | 0.794393289 | 0.525 | 0.05 | 6.73332571372515e-75 | | 36 |
| csf1rb | 4.139952737659e-77 | 0.315138624 | 0.175 | 0.005 | 1.04198470454139e-72 | | 36 |
| si:dkey-217l24.1 | 7.00960277470279e-76 | 0.370825552 | 0.212 | 0.008 | 1.76424692236494e-71 | | 36 |
| cxcr4b11 | 2.15490513222417e-75 | 1.333984204 | 0.963 | 0.18 | 5.423680727295e-71 | | 36 |
| si:dkey-261h17.14 | 1.38930127839751e-74 | 1.011628669 | 0.7 | 0.088 | 3.49673238759869e-70 | | 36 |
| si:dkey-22o12.2 | 5.84349497913914e-66 | 0.319987304 | 0.175 | 0.006 | 1.47074925129953e-61 | | 36 |
| si:ch211-256a21.41 | 2.79714358227895e-62 | 0.337359778 | 0.212 | 0.01 | 7.04013068223789e-58 | | 36 |
| CABZ01066694.11 | 7.29593135562834e-62 | 0.35771448 | 0.262 | 0.016 | 1.8363129628981e-57 | | 36 |
| itgae.25 | 1.66625851624208e-60 | 0.512028088 | 0.388 | 0.034 | 4.19380605952969e-56 | | 36 |
| pdxka | 5.05905777415241e-60 | 0.482005307 | 0.362 | 0.03 | 1.27331425117642e-55 | | 36 |
| cables2b1 | 4.03558074617623e-59 | 0.388508134 | 0.238 | 0.014 | 1.01571531800509e-54 | | 36 |
| msi2a2 | 5.95822832008482e-55 | 0.562520887 | 0.425 | 0.044 | 1.49962648588215e-50 | | 36 |
| tmsb4 | 3.89226039304535e-51 | 0.574066946 | 0.312 | 0.027 | 9.79643018325584e-47 | | 36 |
| s1pr5b1 | 5.25631867090273e-51 | 0.253323973 | 0.162 | 0.007 | 1.32296284627951e-46 | | 36 |

| | | | | | | |
|---|---|---|---|---|---|---|
| si:ch211-222l21.114 | 1.17132746756482e-48 | 1.171186369 | 0.912 | 0.273 | 2.9481141031139e-44 | 36 |
| si:ch211-197h24.95 | 6.98355886034385e-48 | 0.672620371 | 0.488 | 0.067 | 1.75769192955994e-43 | 36 |
| cbfa2t3 | 1.88249665995102e-47 | 0.387061702 | 0.238 | 0.017 | 4.73805584343071e-43 | 36 |
| bahcc1b | 1.03921811219922e-46 | 0.48302569 | 0.3 | 0.027 | 2.61560806659421e-42 | 36 |
| ncf23 | 2.67036822673926e-45 | 0.356693346 | 0.262 | 0.021 | 6.72104978988003e-41 | 36 |
| nr4a111 | 5.12524959773441e-45 | 0.98727715 | 0.762 | 0.165 | 1.28997407125377e-40 | 36 |
| ccr9a8 | 3.62678983127309e-39 | 0.741572515 | 0.512 | 0.086 | 9.12826732633125e-35 | 36 |
| ikzf14 | 3.03905062886035e-38 | 0.417129972 | 0.3 | 0.032 | 7.64898652777861e-34 | 36 |
| tox21 | 3.65242456470643e-38 | 0.283395385 | 0.188 | 0.013 | 9.19278738690961e-34 | 36 |
| rplp111 | 5.66791650762221e-37 | 0.45548567 | 1 | 0.877 | 1.42655790580343e-32 | 36 |
| hsp90ab110 | 3.27741312220393e-35 | 0.509513546 | 0.975 | 0.79 | 8.24892108727508e-31 | 36 |
| fmnl1a4 | 1.29967050562575e-34 | 0.38930273 | 0.312 | 0.038 | 3.27114069560946e-30 | 36 |
| rpl1115 | 1.33260216736528e-34 | 0.494605163 | 0.988 | 0.814 | 3.35402639504167e-30 | 36 |
| rpl1012 | 5.418902875388e-34 | 0.514101022 | 0.975 | 0.772 | 1.36388366470641e-29 | 36 |
| rplp2l19 | 1.06192587298731e-33 | 0.556113423 | 0.988 | 0.774 | 2.67276122972176e-29 | 36 |
| slc16a101 | 1.70399718561268e-33 | 0.28002111 | 0.2 | 0.017 | 4.28879051646855e-29 | 36 |
| eef1a1l15 | 2.4971641441506e-33 | 0.371415624 | 1 | 0.915 | 6.28511243441265e-29 | 36 |
| MCMDC2 | 4.70275332084085e-33 | 0.483765258 | 0.462 | 0.078 | 1.18363598332243e-28 | 36 |
| si:ch211-214p16.25 | 4.84100692564314e-33 | 0.573788018 | 0.438 | 0.072 | 1.21843303311512e-28 | 36 |
| rpl814 | 7.64319443536062e-33 | 0.522226309 | 0.988 | 0.78 | 1.92371560743591e-28 | 36 |
| rps912 | 8.99239883844703e-33 | 0.52221117 | 0.988 | 0.776 | 2.26329686364873e-28 | 36 |
| mybbp1a1 | 1.32384850143695e-32 | 0.30685748 | 0.25 | 0.026 | 3.33199429326667e-28 | 36 |
| rpl2813 | 1.42851684184059e-32 | 0.524397059 | 0.988 | 0.749 | 3.59543403922858e-28 | 36 |
| rps209 | 1.45137917313131e-32 | 0.369425439 | 0.988 | 0.916 | 3.65297624085419e-28 | 36 |
| ebf1a1 | 3.23011257041078e-32 | 0.32261505 | 0.125 | 0.007 | 8.12987032846689e-28 | 36 |
| rplp07 | 4.31148128812602e-32 | 0.437606171 | 0.975 | 0.851 | 1.08515672540844e-27 | 36 |
| rpl138 | 4.92989493489821e-32 | 0.453887657 | 0.975 | 0.805 | 1.24080525616453e-27 | 36 |
| tnrc181 | 9.74416198988977e-32 | 0.409165395 | 0.312 | 0.041 | 2.45250813123536e-27 | 36 |
| tagapb7 | 1.0173942389071e-31 | 0.497737242 | 0.412 | 0.067 | 2.56067955990527e-27 | 36 |
| dub7 | 1.23567518384636e-31 | 0.488983764 | 0.425 | 0.07 | 3.11007087022291e-27 | 36 |
| rpl3712 | 1.53343963495438e-31 | 0.439043826 | 0.975 | 0.85 | 3.85951421721667e-27 | 36 |
| rps15a14 | 1.97560502535065e-31 | 0.527748633 | 0.988 | 0.78 | 4.97240028830505e-27 | 36 |
| rgs139 | 2.96500098465909e-31 | 0.72081037 | 0.6 | 0.131 | 7.46261097828846e-27 | 36 |
| rpl1815 | 4.55235055259151e-31 | 0.514174877 | 0.975 | 0.774 | 1.14578111058176e-26 | 36 |
| faua13 | 5.08767489981352e-31 | 0.581768591 | 0.963 | 0.689 | 1.28051689553407e-26 | 36 |

| | | | | | | |
|---|---|---|---|---|---|---|
| rpl2415 | 6.77370479909977e-31 | 0.501207023 | 0.988 | 0.776 | 1.70487376088542e-26 | 36 |
| rpl617 | 7.46772560382051e-31 | 0.608247527 | 0.975 | 0.667 | 1.87955185722558e-26 | 36 |
| si:dkey-27i16.27 | 1.15308393311887e-30 | 0.653528065 | 0.475 | 0.09 | 2.90219695126688e-26 | 36 |
| gatad2b1 | 1.16103883955886e-30 | 0.444164119 | 0.375 | 0.058 | 2.9222186552857e-26 | 36 |
| rpsa20 | 1.97814001994495e-30 | 0.515161509 | 0.975 | 0.752 | 4.97878061619945e-26 | 36 |
| rps610 | 2.70849772770229e-30 | 0.505258602 | 1 | 0.789 | 6.8170179308539e-26 | 36 |
| rps129 | 4.78708580509856e-30 | 0.44861004 | 0.988 | 0.84 | 1.20486162628526e-25 | 36 |
| rps2122 | 7.04804058696188e-30 | 0.539371533 | 1 | 0.75 | 1.77392133533243e-25 | 36 |
| rps1315 | 1.00470629186528e-29 | 0.538817799 | 0.975 | 0.724 | 2.52874526599572e-25 | 36 |
| rpl1514 | 2.20144842945275e-29 | 0.542589296 | 0.975 | 0.744 | 5.54082555208963e-25 | 36 |
| rpl920 | 3.19719254998185e-29 | 0.644415165 | 0.975 | 0.629 | 8.04701392904932e-25 | 36 |
| rps26l16 | 4.21149621028147e-29 | 0.617062822 | 0.988 | 0.664 | 1.05999148116574e-24 | 36 |
| rpl7a12 | 4.2407729683155e-29 | 0.453056597 | 0.975 | 0.81 | 1.06736014839533e-24 | 36 |
| rpl320 | 4.44601440392626e-29 | 0.361710657 | 0.975 | 0.875 | 1.1190173653242e-24 | 36 |
| rps4x13 | 5.37930649910648e-29 | 0.558231744 | 0.963 | 0.71 | 1.35391765276011e-24 | 36 |
| rps3a9 | 6.71306040356961e-29 | 0.478915853 | 0.963 | 0.778 | 1.68961017297444e-24 | 36 |
| rpl197 | 8.33858150734744e-29 | 0.406015302 | 0.988 | 0.852 | 2.09873757958428e-24 | 36 |
| myca2 | 1.9278760819551e-28 | 0.376994394 | 0.275 | 0.036 | 4.85227131067279e-24 | 36 |
| rps27.115 | 1.97932610350672e-28 | 0.589882543 | 0.963 | 0.679 | 4.98176586991607e-24 | 36 |
| rpl3214 | 2.0567157721701e-28 | 0.457741685 | 0.975 | 0.813 | 5.17654792697491e-24 | 36 |
| rpl18a14 | 4.19357997441827e-28 | 0.486012412 | 0.975 | 0.785 | 1.05548214376133e-23 | 36 |
| rps514 | 4.34454484303613e-28 | 0.567132176 | 0.975 | 0.701 | 1.09347849154376e-23 | 36 |
| rnf445 | 7.27831507490598e-28 | 0.556630298 | 0.45 | 0.089 | 1.83187912120309e-23 | 36 |
| rpl3619 | 7.73408773303358e-28 | 0.472296382 | 0.975 | 0.79 | 1.94659254152722e-23 | 36 |
| wu:fc75a091 | 8.448610394197e-28 | 0.336224266 | 0.212 | 0.023 | 2.12643075011544e-23 | 36 |
| rps1019 | 1.06160453044946e-27 | 0.547706141 | 0.975 | 0.694 | 2.67195244268824e-23 | 36 |
| ahcy2 | 1.40463266510235e-27 | 0.634781406 | 0.662 | 0.166 | 3.53531995479611e-23 | 36 |
| rps119 | 1.9064505974463e-27 | 0.465423179 | 0.988 | 0.766 | 4.7983455087126e-23 | 36 |
| rpl237 | 2.3699551243606e-27 | 0.363222042 | 0.988 | 0.873 | 5.9649400525032e-23 | 36 |
| rpl3912 | 2.42455278399264e-27 | 0.382682617 | 0.988 | 0.874 | 6.10235690203107e-23 | 36 |
| eif3s105 | 2.615940861504e-27 | 0.639498351 | 0.637 | 0.164 | 6.58406155431943e-23 | 36 |
| zfhx3 | 3.89271127175386e-27 | 0.252925545 | 0.225 | 0.025 | 9.7975649998773e-23 | 36 |
| rps79 | 4.18679122328319e-27 | 0.443104191 | 0.988 | 0.82 | 1.05377348298814e-22 | 36 |
| prr5a1 | 5.01872850022153e-27 | 0.331542098 | 0.212 | 0.023 | 1.26316377622076e-22 | 36 |
| rpl22l112 | 5.02768287038895e-27 | 0.807393756 | 0.85 | 0.391 | 1.2654175016482e-22 | 36 |

| | | | | | | |
|---|---|---|---|---|---|---|
| cbfb3 | 5.3780702496291e-27 | 0.371971695 | 0.288 | 0.041 | 1.35360650112915e-22 | 36 |
| znf296 | 5.80743947404804e-27 | 0.280026835 | 0.15 | 0.012 | 1.46167444122315e-22 | 36 |
| rpl10a10 | 7.54783518430027e-27 | 0.484216019 | 0.963 | 0.765 | 1.89971463753654e-22 | 36 |
| eif3c7 | 7.69106536960956e-27 | 0.617240089 | 0.65 | 0.167 | 1.93576424287703e-22 | 36 |
| rps1415 | 8.03431569778806e-27 | 0.488508202 | 0.988 | 0.759 | 2.02215691797628e-22 | 36 |
| sox4a3 | 8.16094545358322e-27 | 0.430138548 | 0.3 | 0.044 | 2.05402836121236e-22 | 36 |
| rpl3510 | 9.58945042840597e-27 | 0.442038233 | 0.988 | 0.804 | 2.4135687783255e-22 | 36 |
| rps1617 | 9.63012313833379e-27 | 0.513679718 | 0.975 | 0.738 | 2.42380569268723e-22 | 36 |
| rpl36a12 | 1.14514899147491e-26 | 0.428741291 | 0.975 | 0.81 | 2.88222549664321e-22 | 36 |
| rps2511 | 1.46613849482376e-26 | 0.485954115 | 0.95 | 0.725 | 3.69012397762191e-22 | 36 |
| rpl2611 | 1.57180347188514e-26 | 0.406601327 | 0.988 | 0.841 | 3.95607215838771e-22 | 36 |
| nop21 | 1.63935124081075e-26 | 0.30870567 | 0.312 | 0.047 | 4.12608313799657e-22 | 36 |
| si:ch211-214p16.19 | 1.82354448630518e-26 | 0.879803248 | 0.85 | 0.256 | 4.58967911758151e-22 | 36 |
| rpl2113 | 1.82562180559265e-26 | 0.470136186 | 0.95 | 0.756 | 4.59490752249613e-22 | 36 |
| rps313 | 2.79741511430437e-26 | 0.536022103 | 0.975 | 0.7 | 7.04081410119267e-22 | 36 |
| pik3c2b | 6.01296605511424e-26 | 0.283633596 | 0.188 | 0.019 | 1.5134034264117e-21 | 36 |
| rbm281 | 7.97989629062169e-26 | 0.486542275 | 0.438 | 0.087 | 2.00846009738657e-21 | 36 |
| si:ch73-248e21.54 | 1.03895592526382e-25 | 0.324310314 | 0.3 | 0.045 | 2.6149481682965e-21 | 36 |
| rpl3016 | 1.48270226425429e-25 | 0.535132396 | 1 | 0.708 | 3.73181332890161e-21 | 36 |
| rps1910 | 1.49100275783561e-25 | 0.378128144 | 0.988 | 0.863 | 3.75270484119645e-21 | 36 |
| gtpbp45 | 1.98770637840703e-25 | 0.756207203 | 0.725 | 0.236 | 5.00285818381266e-21 | 36 |
| rps228 | 2.12377164242515e-25 | 0.461094871 | 1 | 0.829 | 5.34532084681987e-21 | 36 |
| rpl27a17 | 2.21834262876458e-25 | 0.720249813 | 0.925 | 0.495 | 5.58334656233758e-21 | 36 |
| foxp45 | 2.34732506115865e-25 | 0.524095203 | 0.312 | 0.051 | 5.9079824464302e-21 | 36 |
| si:ch73-208g10.15 | 2.69632323646861e-25 | 0.361633661 | 0.288 | 0.042 | 6.78637595386785e-21 | 36 |
| crema8 | 5.92108530996898e-25 | 0.634496138 | 0.662 | 0.177 | 1.49027796166609e-20 | 36 |
| rps2311 | 6.6637637301302e-25 | 0.505323574 | 0.975 | 0.723 | 1.67720269323647e-20 | 36 |
| rack19 | 1.17044053413453e-24 | 0.457617402 | 0.95 | 0.744 | 2.9458817803632e-20 | 36 |
| rps2416 | 1.30439540301675e-24 | 0.477385325 | 1 | 0.758 | 3.28303278985287e-20 | 36 |
| rpl2712 | 1.42748577635672e-24 | 0.453903842 | 0.95 | 0.763 | 3.59283895051223e-20 | 36 |
| rps1816 | 2.22699617823552e-24 | 0.517366636 | 0.988 | 0.724 | 5.60512668100098e-20 | 36 |
| rps8a8 | 2.2370326904637e-24 | 0.437841712 | 0.975 | 0.788 | 5.6303875786281e-20 | 36 |
| rpl3416 | 2.79446940788634e-24 | 0.45437981 | 0.975 | 0.784 | 7.03340005270913e-20 | 36 |
| si:ch211-121a2.22 | 4.21684893755201e-24 | 0.306279497 | 0.25 | 0.034 | 1.06133870909247e-19 | 36 |
| eef2b8 | 7.94197331475066e-24 | 0.578298958 | 0.938 | 0.613 | 1.99891526358959e-19 | 36 |

| | | | | | | |
|---|---|---|---|---|---|---|
| eef1g13 | 1.09824308782488e-23 | 0.587129747 | 0.912 | 0.62 | 2.76416802774644e-19 | 36 |
| rps1516 | 1.71215664818008e-23 | 0.534970655 | 0.963 | 0.684 | 4.30932706780444e-19 | 36 |
| rpl23a18 | 2.64299448016038e-23 | 0.570000986 | 0.938 | 0.634 | 6.65215280711566e-19 | 36 |
| naca13 | 3.08587322259721e-23 | 0.596625084 | 0.963 | 0.618 | 7.76683431395491e-19 | 36 |
| nsa215 | 3.28808552409514e-23 | 0.756937584 | 0.863 | 0.361 | 8.27578245559507e-19 | 36 |
| rpl3117 | 3.34887189499935e-23 | 0.549908751 | 0.963 | 0.654 | 8.42877567252386e-19 | 36 |
| tfr1b | 3.96131996203211e-23 | 0.354412215 | 0.312 | 0.052 | 9.97024621243861e-19 | 36 |
| rps2912 | 5.6874788600509e-23 | 0.386320145 | 0.975 | 0.832 | 1.43148155428621e-18 | 36 |
| rpl2916 | 6.08850727227464e-23 | 0.397423267 | 0.95 | 0.809 | 1.5324163953588e-18 | 36 |
| rsl24d114 | 9.4845332405681e-23 | 0.734614261 | 0.887 | 0.443 | 2.38716217131858e-18 | 36 |
| rpl78 | 1.13871178094256e-22 | 0.417836391 | 0.975 | 0.805 | 2.86602368145433e-18 | 36 |
| BX284696.1 | 1.83665402630522e-22 | 0.312240599 | 0.262 | 0.04 | 4.6226745188076e-18 | 36 |
| epb41a1 | 1.86251027515737e-22 | 0.331797102 | 0.162 | 0.017 | 4.68775211154359e-18 | 36 |
| pho1 | 1.86369533127139e-22 | 0.445011605 | 0.425 | 0.09 | 4.69073477927697e-18 | 36 |
| si:ch211-284f22.34 | 2.40572296241373e-22 | 0.31366825 | 0.238 | 0.034 | 6.05496412409913e-18 | 36 |
| rpl3816 | 2.85830800660625e-22 | 0.494086791 | 0.975 | 0.728 | 7.19407542182726e-18 | 36 |
| b3gnt3.43 | 5.20081654564077e-22 | 0.307772053 | 0.25 | 0.037 | 1.30899351637233e-17 | 36 |
| si:ch73-1a9.315 | 7.54604112997378e-22 | 0.740872928 | 0.75 | 0.264 | 1.8992630920031e-17 | 36 |
| rpl35a15 | 7.83746569089672e-22 | 0.455115233 | 0.975 | 0.759 | 1.97261173974179e-17 | 36 |
| rps27a14 | 1.76060204901604e-21 | 0.43993804 | 0.975 | 0.751 | 4.43125929716846e-17 | 36 |
| rps2816 | 1.78112045883641e-21 | 0.403557986 | 1 | 0.826 | 4.48290208284536e-17 | 36 |
| si:dkey-185m8.23 | 2.04333190732819e-21 | 0.301948681 | 0.288 | 0.048 | 5.14286207755431e-17 | 36 |
| eif3s6ip1 | 2.98224033821721e-21 | 0.48273161 | 0.562 | 0.148 | 7.5060007072589e-17 | 36 |
| zgc:11418820 | 4.1832181768808e-21 | 0.571242893 | 0.95 | 0.627 | 1.05287418293913e-16 | 36 |
| si:ch211-226m16.25 | 7.22972584989417e-21 | 0.382192698 | 0.35 | 0.068 | 1.81964969915986e-16 | 36 |
| hmgn613 | 1.01399205033973e-20 | 0.64665361 | 0.762 | 0.264 | 2.55211659150007e-16 | 36 |
| nop537 | 1.09144013877903e-20 | 0.681269057 | 0.713 | 0.25 | 2.74704568529294e-16 | 36 |
| rpl126 | 1.6513771528545e-20 | 0.367633149 | 0.938 | 0.799 | 4.15635115601949e-16 | 36 |
| rps1710 | 1.69699109399594e-20 | 0.425462836 | 0.963 | 0.767 | 4.27115688447839e-16 | 36 |
| dera | 2.10433544679371e-20 | 0.291808493 | 0.175 | 0.021 | 5.29640188603509e-16 | 36 |
| rpl13a7 | 3.00176073126384e-20 | 0.402243702 | 0.95 | 0.764 | 7.55513158451796e-16 | 36 |
| zbtb2b | 3.10503843471145e-20 | 0.254949423 | 0.25 | 0.039 | 7.81507123632525e-16 | 36 |
| rps2611 | 4.29748482568107e-20 | 0.442158396 | 0.963 | 0.734 | 1.08163395577567e-15 | 36 |
| metap11 | 4.68371830463794e-20 | 0.380116327 | 0.375 | 0.08 | 1.17884506009432e-15 | 36 |
| GNAZ1 | 7.48386377771974e-20 | 0.303547715 | 0.188 | 0.025 | 1.88361367421428e-15 | 36 |

| | | | | | | |
|---|---|---|---|---|---|---|
| si:ch211-262e15.1 | 8.08672273436645e-20 | 0.301788942 | 0.375 | 0.078 | 2.03534724501269e-15 | 36 |
| nop564 | 1.45798676186213e-19 | 0.563651347 | 0.575 | 0.17 | 3.66960688093081e-15 | 36 |
| parvg7 | 1.66879294423308e-19 | 0.354700641 | 0.338 | 0.067 | 4.20018496134023e-15 | 36 |
| calm2b6 | 2.06671483730383e-19 | 0.59141697 | 0.662 | 0.209 | 5.20171457401002e-15 | 36 |
| eif3ha4 | 2.39069365063797e-19 | 0.573729195 | 0.65 | 0.207 | 6.01713684929069e-15 | 36 |
| eef1b23 | 2.41025752231463e-19 | 0.425284212 | 0.938 | 0.735 | 6.0663771579137e-15 | 36 |
| zbtb207 | 3.2984464247856e-19 | 0.540047645 | 0.4 | 0.098 | 8.30185980654287e-15 | 36 |
| h1fx10 | 4.21164692821531e-19 | 0.632389027 | 0.6 | 0.182 | 1.06002941536251e-14 | 36 |
| rpl49 | 5.87103258678974e-19 | 0.521756195 | 0.925 | 0.627 | 1.47768019176911e-14 | 36 |
| uba5215 | 5.99100338874321e-19 | 0.521258765 | 0.925 | 0.63 | 1.50787564291278e-14 | 36 |
| serbp1a13 | 6.26670362984312e-19 | 0.642496859 | 0.875 | 0.482 | 1.57726663659521e-14 | 36 |
| XLOC-00043111 | 6.51611337043154e-19 | 0.885348811 | 0.713 | 0.278 | 1.64004057420391e-14 | 36 |
| cct42 | 6.91987046575698e-19 | 0.594352016 | 0.675 | 0.225 | 1.74166219752637e-14 | 36 |
| adh57 | 6.97759192464458e-19 | 0.577092624 | 0.675 | 0.221 | 1.75619011151379e-14 | 36 |
| cct27 | 8.43248603114803e-19 | 0.60624314 | 0.662 | 0.226 | 2.12237240917965e-14 | 36 |
| tcp14 | 1.02564029185245e-18 | 0.584375149 | 0.675 | 0.225 | 2.58143405056343e-14 | 36 |
| rpl5a16 | 1.0311054137068e-18 | 0.635270711 | 0.9 | 0.491 | 2.59518921575866e-14 | 36 |
| FP074902.21 | 2.14678512649009e-18 | 0.334281868 | 0.362 | 0.079 | 5.40324348486292e-14 | 36 |
| uqcrc2a | 3.29793594689851e-18 | 0.31746244 | 0.312 | 0.063 | 8.30057498474887e-14 | 36 |
| XLOC-0425958 | 3.32238039859054e-18 | 0.679797916 | 0.55 | 0.164 | 8.36209922521252e-14 | 36 |
| pde7a1 | 4.82116067628076e-18 | 0.368501591 | 0.288 | 0.056 | 1.2134379306131e-13 | 36 |
| hnrnpa0b7 | 5.71259235297722e-18 | 0.646665753 | 0.8 | 0.311 | 1.43780236932084e-13 | 36 |
| eif3ea10 | 6.03691929099485e-18 | 0.656436067 | 0.7 | 0.276 | 1.51943221635049e-13 | 36 |
| ascc11 | 6.64567668850223e-18 | 0.336320493 | 0.312 | 0.063 | 1.67265036572913e-13 | 36 |
| mcm61 | 6.72014646811001e-18 | 0.273481277 | 0.225 | 0.036 | 1.69139366455861e-13 | 36 |
| elf19 | 6.88654064415185e-18 | 0.383536472 | 0.425 | 0.104 | 1.73327341472658e-13 | 36 |
| cct86 | 8.89513688487565e-18 | 0.532495403 | 0.625 | 0.201 | 2.23881700255435e-13 | 36 |
| ddx24 | 1.5853192419268e-17 | 0.275331291 | 0.25 | 0.044 | 3.99009000000556e-13 | 36 |
| grap2b9 | 1.64330267669855e-17 | 0.431995969 | 0.438 | 0.112 | 4.13602850698258e-13 | 36 |
| ddx214 | 2.29003518808086e-17 | 0.588458368 | 0.613 | 0.207 | 5.76378956488071e-13 | 36 |
| eef1da11 | 3.18491141568773e-17 | 0.624487054 | 0.912 | 0.491 | 8.01610354214446e-13 | 36 |
| gpr183a6 | 3.87097025433878e-17 | 0.319646803 | 0.312 | 0.065 | 9.74284503314528e-13 | 36 |
| ptmaa17 | 4.99982429211224e-17 | 0.711365492 | 0.85 | 0.394 | 1.25840577608173e-12 | 36 |
| sept125 | 5.11424142105131e-17 | 0.312765465 | 0.3 | 0.061 | 1.2872034232644e-12 | 36 |
| rpl1414 | 6.26340432138371e-17 | 0.51716373 | 0.963 | 0.635 | 1.57643623364907e-12 | 36 |

| | | | | | | |
|---|---|---|---|---|---|---|
| ppiaa12 | 6.38345152566313e-17 | 0.37783 | 1 | 0.804 | 1.60665091449415e-12 | 36 |
| slc25a59 | 6.69336664970534e-17 | 0.351440444 | 0.975 | 0.807 | 1.68465345206434e-12 | 36 |
| rrp1 | 6.94431913453476e-17 | 0.331432106 | 0.388 | 0.093 | 1.74781568297105e-12 | 36 |
| rpl2215 | 8.1799508823819e-17 | 0.471515192 | 0.963 | 0.694 | 2.0588118375867e-12 | 36 |
| CU929418.11 | 8.30666759101846e-17 | 0.25311879 | 0.175 | 0.025 | 2.09070516598344e-12 | 36 |
| tpt111 | 8.40797418479751e-17 | 0.362287133 | 0.975 | 0.801 | 2.11620302257168e-12 | 36 |
| si:dkey-262k9.410 | 9.37986875264576e-17 | 0.37733596 | 0.375 | 0.09 | 2.36081916635341e-12 | 36 |
| eif2s35 | 1.33209227751005e-16 | 0.588963205 | 0.738 | 0.284 | 3.35274305326504e-12 | 36 |
| si:busm1-266f07.27 | 1.3984563966679e-16 | 0.530639596 | 0.512 | 0.146 | 3.51977490477345e-12 | 36 |
| histh1l9 | 1.44866191190683e-16 | 0.592828965 | 0.4 | 0.109 | 3.6461371660783e-12 | 36 |
| rsl1d14 | 1.96700316315266e-16 | 0.484644444 | 0.575 | 0.18 | 4.95075026133892e-12 | 36 |
| egr2a | 2.31735182987333e-16 | 0.274575786 | 0.175 | 0.026 | 5.83254282060818e-12 | 36 |
| caprin1b2 | 4.12127998852222e-16 | 0.491251075 | 0.713 | 0.246 | 1.03728496031116e-11 | 36 |
| si:dkey-102m7.32 | 5.64258894249363e-16 | 0.384327391 | 0.5 | 0.143 | 1.42018321093622e-11 | 36 |
| usp362 | 5.92662312578647e-16 | 0.339313688 | 0.325 | 0.075 | 1.4916717745292e-11 | 36 |
| si:dkey-239h2.31 | 9.68351105975232e-16 | 0.338541703 | 0.362 | 0.089 | 2.43724289862906e-11 | 36 |
| eif3ba5 | 1.03173768815399e-15 | 0.511879703 | 0.613 | 0.209 | 2.59678058731477e-11 | 36 |
| atad2b | 1.25215021805636e-15 | 0.429454048 | 0.188 | 0.031 | 3.15153688382605e-11 | 36 |
| txndc92 | 1.29704340561905e-15 | 0.388410405 | 0.425 | 0.117 | 3.26452854760258e-11 | 36 |
| kdm6ba | 1.38852294550762e-15 | 0.300073407 | 0.25 | 0.049 | 3.49477340154813e-11 | 36 |
| ucp212 | 1.47680284699633e-15 | 0.622531026 | 0.912 | 0.415 | 3.71696508560507e-11 | 36 |
| MRPS121 | 1.58961515493299e-15 | 0.296133028 | 0.35 | 0.084 | 4.00090238345084e-11 | 36 |
| eif3i9 | 1.6888989059932e-15 | 0.539227951 | 0.725 | 0.272 | 4.25078965649428e-11 | 36 |
| cct73 | 1.7183002792904e-15 | 0.553865745 | 0.588 | 0.208 | 4.32478997294602e-11 | 36 |
| btf312 | 1.82508062201826e-15 | 0.530215735 | 0.938 | 0.587 | 4.59354541755775e-11 | 36 |
| RPL37A | 1.89405868157013e-15 | 0.26492892 | 0.3 | 0.066 | 4.76715629564386e-11 | 36 |
| cd837 | 2.6240885780083e-15 | 0.62806694 | 0.375 | 0.104 | 6.60456854198908e-11 | 36 |
| wasb12 | 2.77527513969803e-15 | 0.452059928 | 0.575 | 0.177 | 6.98508999910598e-11 | 36 |
| wu:fb55g093 | 2.91651448565826e-15 | 0.56651919 | 0.675 | 0.25 | 7.34057530895328e-11 | 36 |
| rpl5b10 | 2.94697747173195e-15 | 0.484799873 | 0.925 | 0.63 | 7.41724759860214e-11 | 36 |
| pfdn42 | 3.08387751864283e-15 | 0.401979636 | 0.55 | 0.17 | 7.76181132667214e-11 | 36 |
| rpl1710 | 4.34129299792779e-15 | 0.377930738 | 0.938 | 0.746 | 1.09266003464845e-10 | 36 |
| rplp218 | 5.02372587657394e-15 | 0.437956862 | 0.988 | 0.758 | 1.2644215658749e-10 | 36 |
| CR855337.12 | 5.78460564003862e-15 | 0.250680633 | 0.2 | 0.034 | 1.45592739354132e-10 | 36 |
| cops2 | 6.7150434987462e-15 | 0.268223572 | 0.275 | 0.059 | 1.69010929819943e-10 | 36 |

| | | | | | | |
|---|---|---|---|---|---|---|
| pa2g4a1 | 7.59889333231486e-15 | 0.370748957 | 0.438 | 0.122 | 1.91256546281033e-10 | 36 |
| eif1ad1 | 8.75695278548298e-15 | 0.318967525 | 0.338 | 0.082 | 2.20403744657821e-10 | 36 |
| cebpb14 | 1.384275596971e-14 | 0.669168932 | 0.787 | 0.353 | 3.48408325001632e-10 | 36 |
| tnrc6c110 | 1.38758377135164e-14 | 0.415165593 | 0.45 | 0.133 | 3.49240959411495e-10 | 36 |
| celf215 | 1.4029241071309e-14 | 0.507883421 | 0.675 | 0.242 | 3.53101968523776e-10 | 36 |
| thrap3b1 | 1.53368188823332e-14 | 0.314406066 | 0.35 | 0.088 | 3.86012394449444e-10 | 36 |
| mibp213 | 1.56387211393111e-14 | 0.573874433 | 0.825 | 0.492 | 3.93610972355322e-10 | 36 |
| myg11 | 1.62165461272814e-14 | 0.28838031 | 0.288 | 0.065 | 4.08154249477546e-10 | 36 |
| gtf2f11 | 2.12926013797224e-14 | 0.270954583 | 0.288 | 0.064 | 5.35913484126234e-10 | 36 |
| rac213 | 2.6464858787492e-14 | 0.505468721 | 0.637 | 0.22 | 6.66094030822385e-10 | 36 |
| abce1 | 2.69018260047856e-14 | 0.261492239 | 0.312 | 0.073 | 6.77092058714448e-10 | 36 |
| eif5a9 | 2.95452492703635e-14 | 0.564306304 | 0.85 | 0.478 | 7.43624378885778e-10 | 36 |
| tob1a3 | 2.96612305808172e-14 | 0.41698239 | 0.438 | 0.129 | 7.46543512488588e-10 | 36 |
| LOC5665872 | 4.40829937852223e-14 | 0.291953912 | 0.238 | 0.049 | 1.10952487058026e-09 | 36 |
| hmgn711 | 4.71551236607389e-14 | 0.526959762 | 0.713 | 0.279 | 1.18684730741714e-09 | 36 |
| hmgn211 | 4.82720674860489e-14 | 0.685105959 | 0.838 | 0.442 | 1.21495966655637e-09 | 36 |
| osgn11 | 5.49141221036766e-14 | 0.290323235 | 0.212 | 0.04 | 1.38213353922744e-09 | 36 |
| eif3g8 | 5.52294070752349e-14 | 0.525643804 | 0.713 | 0.281 | 1.39006894667659e-09 | 36 |
| RASSF56 | 6.47540202188067e-14 | 0.341923501 | 0.262 | 0.058 | 1.62979393488715e-09 | 36 |
| si:dkey-250d21.1 | 9.6292479707348e-14 | 0.311523604 | 0.312 | 0.077 | 2.42358542175424e-09 | 36 |
| krt18b10 | 1.03909773171815e-13 | 0.475097517 | 0.475 | 0.15 | 2.6153050809614e-09 | 36 |
| eif4a1b9 | 1.06176579293377e-13 | 0.543103058 | 0.688 | 0.28 | 2.672358324235e-09 | 36 |
| cct32 | 1.11273088375218e-13 | 0.452463133 | 0.588 | 0.206 | 2.80063236131587e-09 | 36 |
| eif3k4 | 1.32866245644677e-13 | 0.467117826 | 0.625 | 0.229 | 3.34411053663088e-09 | 36 |
| tuba8l5 | 1.38352958549134e-13 | 0.54951609 | 0.525 | 0.189 | 3.48220561372316e-09 | 36 |
| gnai2b3 | 1.41106489969415e-13 | 0.277217254 | 0.288 | 0.067 | 3.55150924604021e-09 | 36 |
| aldoaa4 | 1.512594666693e-13 | 0.490305695 | 0.662 | 0.25 | 3.80704951659961e-09 | 36 |
| XLOC-0139647 | 1.8770715617789e-13 | 0.569804673 | 0.512 | 0.174 | 4.72440141384132e-09 | 36 |
| psme28 | 2.25079857917385e-13 | 0.389817073 | 0.35 | 0.097 | 5.66503494392266e-09 | 36 |
| ipo72 | 2.27040584230088e-13 | 0.343605881 | 0.388 | 0.109 | 5.71438446448709e-09 | 36 |
| eif3m8 | 2.67728925914135e-13 | 0.56976328 | 0.787 | 0.357 | 6.73846933633287e-09 | 36 |
| eif3f14 | 3.16495047218466e-13 | 0.588665395 | 0.863 | 0.431 | 7.96586384344157e-09 | 36 |
| si:dkey-11f4.147 | 3.24302488919684e-13 | 0.420494283 | 0.375 | 0.109 | 8.16236934361952e-09 | 36 |
| elavl1a | 3.78431877711424e-13 | 0.32022486 | 0.412 | 0.12 | 9.52475193011883e-09 | 36 |
| hmgn33 | 4.48807368998142e-13 | 0.323029891 | 0.262 | 0.061 | 1.12960326703142e-08 | 36 |

| | | | | | | |
|---|---|---|---|---|---|---|
| si:dkey-112e7.210 | 4.81996246263634e-13 | 0.438279323 | 0.575 | 0.205 | 1.21313635222094e-08 | 36 |
| elmo17 | 5.5018264249531e-13 | 0.340328443 | 0.288 | 0.071 | 1.38475469289644e-08 | 36 |
| gnl32 | 6.38908913570595e-13 | 0.253063082 | 0.262 | 0.06 | 1.60806984456583e-08 | 36 |
| fam120c5 | 8.16881759170404e-13 | 0.272832303 | 0.362 | 0.099 | 2.05600969965599e-08 | 36 |
| eif2s1b4 | 1.13925244883824e-12 | 0.421394011 | 0.5 | 0.173 | 2.86738448848096e-08 | 36 |
| si:dkey-42i9.45 | 1.59342639457805e-12 | 0.334893821 | 0.462 | 0.147 | 4.01049489251348e-08 | 36 |
| sik18 | 1.9824238107796e-12 | 0.389896784 | 0.388 | 0.116 | 4.98956248935118e-08 | 36 |
| egr36 | 2.54317636835022e-12 | 0.313132278 | 0.25 | 0.059 | 6.40092060150067e-08 | 36 |
| b2m14 | 2.66976394658367e-12 | 0.504882706 | 0.713 | 0.291 | 6.71952887715645e-08 | 36 |
| atrx8 | 2.78159778503758e-12 | 0.327558221 | 0.362 | 0.105 | 7.0010034651611e-08 | 36 |
| map1lc3b6 | 2.8546987430709e-12 | 0.37360032 | 0.362 | 0.107 | 7.18499126643514e-08 | 36 |
| jdp2b8 | 3.18053804945996e-12 | 0.415521524 | 0.5 | 0.17 | 8.00509621668578e-08 | 36 |
| cnbpb7 | 3.45998407273408e-12 | 0.389203334 | 0.525 | 0.184 | 8.70843391266441e-08 | 36 |
| igf2bp2a | 3.49087989854922e-12 | 0.271725931 | 0.175 | 0.033 | 8.78619561665853e-08 | 36 |
| mt-nd56 | 4.03739513397724e-12 | 0.306712685 | 0.4 | 0.12 | 1.01617198127073e-07 | 36 |
| ccdc251 | 4.79367639461361e-12 | 0.276148649 | 0.325 | 0.088 | 1.2065204117603e-07 | 36 |
| arhgdig12 | 6.65763561768534e-12 | 0.476173455 | 0.637 | 0.243 | 1.67566030861522e-07 | 36 |
| pinx1 | 6.72246370611266e-12 | 0.324052747 | 0.462 | 0.15 | 1.69197689019149e-07 | 36 |
| lmnb22 | 6.91158304784676e-12 | 0.295312428 | 0.25 | 0.06 | 1.73957633731255e-07 | 36 |
| spi1b7 | 1.00501721422618e-11 | 0.346045104 | 0.25 | 0.062 | 2.52952782648587e-07 | 36 |
| chd4a3 | 1.12154939556377e-11 | 0.29610889 | 0.325 | 0.091 | 2.82282767369445e-07 | 36 |
| hnrnpul1l | 1.48466639999833e-11 | 0.279485832 | 0.225 | 0.052 | 3.73675686215581e-07 | 36 |
| mob1a7 | 1.50843803147649e-11 | 0.403365005 | 0.475 | 0.166 | 3.79658768142317e-07 | 36 |
| btf3l4 | 1.82222661327547e-11 | 0.330252428 | 0.425 | 0.137 | 4.58636216295302e-07 | 36 |
| ola14 | 2.26846357812436e-11 | 0.38129097 | 0.613 | 0.229 | 5.70949597978121e-07 | 36 |
| sept9b3 | 2.29688805687301e-11 | 0.275579231 | 0.212 | 0.048 | 5.78103755034367e-07 | 36 |
| msna13 | 4.33490859635854e-11 | 0.41529096 | 0.625 | 0.241 | 1.09105314461748e-06 | 36 |
| qkia8 | 4.69596810558663e-11 | 0.375272964 | 0.475 | 0.168 | 1.1819282124951e-06 | 36 |
| wasa8 | 5.04557278941512e-11 | 0.293563925 | 0.325 | 0.093 | 1.26992021536789e-06 | 36 |
| ptmab14 | 5.31619342933842e-11 | 0.525055303 | 0.912 | 0.54 | 1.33803272423019e-06 | 36 |
| pbdc11 | 5.5739329285559e-11 | 0.280452236 | 0.4 | 0.127 | 1.40290317878823e-06 | 36 |
| kars | 5.84267055672267e-11 | 0.279789193 | 0.375 | 0.115 | 1.47054175242153e-06 | 36 |
| fabp35 | 5.89489879505856e-11 | 0.471546365 | 0.588 | 0.242 | 1.48368707772829e-06 | 36 |
| nr4a35 | 6.46334640134606e-11 | 0.329913152 | 0.225 | 0.055 | 1.62675965575479e-06 | 36 |
| cd74b10 | 7.58680111464772e-11 | 0.33832841 | 0.613 | 0.223 | 1.90952197254569e-06 | 36 |

| | | | | | | |
|---|---|---|---|---|---|---|
| pgam1a6 | 8.64757780097839e-11 | 0.320787715 | 0.462 | 0.157 | 2.17650885672825e-06 | 36 |
| dusp55 | 8.71986960177758e-11 | 0.254426372 | 0.275 | 0.073 | 2.1947039800714e-06 | 36 |
| hmga1a4 | 1.10162278453096e-10 | 0.534923505 | 0.9 | 0.449 | 2.77267438638597e-06 | 36 |
| pmp22b5 | 1.16499444448562e-10 | 0.299735372 | 0.2 | 0.045 | 2.93217451732585e-06 | 36 |
| mir21-16 | 1.2175535496245e-10 | 0.299995054 | 0.35 | 0.108 | 3.0644605290499e-06 | 36 |
| bccip1 | 1.29628150031736e-10 | 0.327428502 | 0.4 | 0.133 | 3.26261090814878e-06 | 36 |
| si:ch1073-429i10.3.1 | 1.56198016759991e-10 | 0.368048226 | 0.537 | 0.202 | 3.93134788383222e-06 | 36 |
| ran4 | 2.028282240952e-10 | 0.415773057 | 0.925 | 0.644 | 5.10498357225208e-06 | 36 |
| si:ch211-195b13.11 | 2.08144393204292e-10 | 0.25454005 | 0.15 | 0.029 | 5.23878623255882e-06 | 36 |
| nop581 | 2.1402425246762e-10 | 0.351315949 | 0.488 | 0.175 | 5.38677641035752e-06 | 36 |
| romo114 | 2.20334128797488e-10 | 0.498050682 | 0.825 | 0.385 | 5.54558968770398e-06 | 36 |
| anp32a2 | 2.25983056630116e-10 | 0.387923049 | 0.462 | 0.17 | 5.68776755232338e-06 | 36 |
| zgc:6405112 | 3.22167565652626e-10 | 0.290821288 | 0.475 | 0.162 | 8.10863545991095e-06 | 36 |
| atp5pf6 | 3.22886295273337e-10 | 0.406677501 | 0.713 | 0.298 | 8.12672516573461e-06 | 36 |
| pabpc1a12 | 3.33501616832569e-10 | 0.486364908 | 0.912 | 0.539 | 8.39390219405893e-06 | 36 |
| cited4a5 | 3.66658537937371e-10 | 0.26589153 | 0.188 | 0.043 | 9.22842874134568e-06 | 36 |
| slc25a55a2 | 3.92645473233655e-10 | 0.259859179 | 0.25 | 0.067 | 9.88249391581787e-06 | 36 |
| thrap3a2 | 4.80853031685501e-10 | 0.284232028 | 0.1 | 0.015 | 1.21025899544924e-05 | 36 |
| ak23 | 5.01940058725401e-10 | 0.388003199 | 0.575 | 0.232 | 1.26333293380596e-05 | 36 |
| si:dkeyp-68b7.127 | 6.69500119727436e-10 | 0.282595313 | 0.238 | 0.063 | 1.68506485134198e-05 | 36 |
| si:ch211-132p1.37 | 6.9014704143046e-10 | 0.274038372 | 0.3 | 0.089 | 1.73703108857632e-05 | 36 |
| si:dkey-23i12.5 | 7.20593758829628e-10 | 0.283765035 | 0.462 | 0.164 | 1.81366243159829e-05 | 36 |
| hist2h2l3 | 7.26492690668075e-10 | 0.262613162 | 0.275 | 0.079 | 1.82850945314248e-05 | 36 |
| atp1b1a6 | 9.49003409333332e-10 | 0.320100376 | 0.562 | 0.214 | 2.38854668095106e-05 | 36 |
| tefa8 | 9.79664481519044e-10 | 0.254020002 | 0.412 | 0.138 | 2.46571753353528e-05 | 36 |
| ier2b14 | 1.00725985330654e-09 | 0.412671326 | 0.55 | 0.224 | 2.53517232478724e-05 | 36 |
| psma31 | 1.02359875449097e-09 | 0.28223934 | 0.488 | 0.176 | 2.57629570517832e-05 | 36 |
| u2surp2 | 1.12140851217469e-09 | 0.254976953 | 0.312 | 0.095 | 2.82247308429247e-05 | 36 |
| LOC1039108716 | 1.17537393421547e-09 | 0.394151406 | 0.462 | 0.171 | 2.95829865502691e-05 | 36 |
| vdac25 | 1.33503911474711e-09 | 0.411582361 | 0.675 | 0.295 | 3.36015994790701e-05 | 36 |
| crip114 | 1.95653297471473e-09 | 0.420351924 | 0.65 | 0.27 | 4.92439784405952e-05 | 36 |
| b2ml14 | 1.97178188686501e-09 | 0.414776671 | 0.613 | 0.267 | 4.96277783105056e-05 | 36 |
| eif4ebp16 | 2.39998190155187e-09 | 0.355739152 | 0.613 | 0.246 | 6.04051444801591e-05 | 36 |
| larp4ab2 | 2.54522967880256e-09 | 0.256743072 | 0.25 | 0.071 | 6.40608857857816e-05 | 36 |
| psma23 | 2.95006787891993e-09 | 0.287806483 | 0.425 | 0.152 | 7.42502584445358e-05 | 36 |

| | | | | | | |
|---|---|---|---|---|---|---|
| ptena1 | 3.40696562539208e-09 | 0.299439218 | 0.2 | 0.052 | 8.57499178254934e-05 | 36 |
| sumo2b12 | 3.50020407872397e-09 | 0.396475578 | 0.625 | 0.269 | 8.80966364574035e-05 | 36 |
| sarnp2 | 4.42203313324409e-09 | 0.275514178 | 0.475 | 0.177 | 0.000111298 | 36 |
| ywhaqa1 | 5.31003741542775e-09 | 0.341688578 | 0.512 | 0.203 | 0.000133648 | 36 |
| arpc1b13 | 6.23593540379098e-09 | 0.392364314 | 0.75 | 0.308 | 0.000156952 | 36 |
| idh28 | 6.64463385652385e-09 | 0.337212768 | 0.5 | 0.199 | 0.000167239 | 36 |
| ybx110 | 8.64349365560532e-09 | 0.393631595 | 0.95 | 0.669 | 0.000217548 | 36 |
| ddit313 | 9.73041006292763e-09 | 0.421440591 | 0.438 | 0.177 | 0.000244905 | 36 |
| psmb61 | 1.01093629978286e-08 | 0.256721637 | 0.45 | 0.167 | 0.000254443 | 36 |
| rps6ka16 | 1.06567098817599e-08 | 0.263734753 | 0.25 | 0.074 | 0.000268219 | 36 |
| XLOC-01089910 | 1.06740389454584e-08 | 0.430403859 | 0.575 | 0.258 | 0.000268655 | 36 |
| prdx59 | 1.07889305590514e-08 | 0.30772226 | 0.425 | 0.159 | 0.000271547 | 36 |
| ddx611 | 1.15114421958942e-08 | 0.334026255 | 0.525 | 0.215 | 0.000289731 | 36 |
| cd74a9 | 1.15230578026418e-08 | 0.304739689 | 0.55 | 0.219 | 0.000290024 | 36 |
| laptm512 | 1.55768457607483e-08 | 0.298171067 | 0.462 | 0.17 | 0.000392054 | 36 |
| si:ch211-196l7.415 | 1.60463845330867e-08 | 0.383792371 | 0.688 | 0.314 | 0.000403871 | 36 |
| cdk2ap2 | 1.91411495905003e-08 | 0.258660361 | 0.35 | 0.121 | 0.000481764 | 36 |
| plk33 | 1.93966505074433e-08 | 0.363241893 | 0.338 | 0.121 | 0.000488194 | 36 |
| gnb1a7 | 1.98401007086729e-08 | 0.324366861 | 0.475 | 0.193 | 0.000499355 | 36 |
| brd2a | 2.21722151135356e-08 | 0.262552649 | 0.45 | 0.171 | 0.000558052 | 36 |
| cct53 | 2.27060926217236e-08 | 0.37998699 | 0.562 | 0.251 | 0.00057149 | 36 |
| midn10 | 3.0796867233238e-08 | 0.319298605 | 0.375 | 0.141 | 0.000775126 | 36 |
| arpc212 | 4.23318600255407e-08 | 0.346033562 | 0.613 | 0.267 | 0.001065451 | 36 |
| sod12 | 4.72747224904837e-08 | 0.358076192 | 0.725 | 0.327 | 0.001189857 | 36 |
| sept63 | 7.4163683754083e-08 | 0.287866261 | 0.5 | 0.206 | 0.001866626 | 36 |
| fbxl6 | 8.06260549772298e-08 | 0.271388271 | 0.188 | 0.052 | 0.002029277 | 36 |
| oaz1b14 | 1.20378291209857e-07 | 0.36708703 | 0.675 | 0.322 | 0.003029801 | 36 |
| pfn113 | 1.27025833206387e-07 | 0.478437481 | 0.925 | 0.524 | 0.003197113 | 36 |
| mknk2b12 | 1.65453723944886e-07 | 0.373626799 | 0.45 | 0.198 | 0.004164305 | 36 |
| NC-002333.418 | 1.8138838167114e-07 | 0.439434126 | 0.95 | 0.622 | 0.004565364 | 36 |
| pno14 | 2.35688787190063e-07 | 0.256162223 | 0.425 | 0.172 | 0.005932051 | 36 |
| ier2a14 | 2.44143731460829e-07 | 0.351000083 | 0.575 | 0.262 | 0.006144854 | 36 |
| sub1a3 | 2.66250882536063e-07 | 0.290797364 | 0.537 | 0.233 | 0.006701268 | 36 |
| ccnd2a3 | 2.78626971158471e-07 | 0.344370379 | 0.25 | 0.087 | 0.007012762 | 36 |
| nfyba | 2.87122971022936e-07 | 0.251246381 | 0.425 | 0.173 | 0.007226598 | 36 |

| | | | | | | |
|---|---|---|---|---|---|---|
| stmn1a2 | 3.02783290217532e-07 | 0.302149835 | 0.488 | 0.206 | 0.007620753 | 36 |
| gnb1b4 | 3.17955869819679e-07 | 0.288839845 | 0.575 | 0.254 | 0.008002631 | 36 |
| psma11 | 3.42671857770972e-07 | 0.265997387 | 0.575 | 0.25 | 0.008624708 | 36 |
| irf2bp2a6 | 3.60720173775393e-07 | 0.292087223 | 0.425 | 0.177 | 0.009078966 | 36 |
| eif3d7 | 3.6533189103795e-07 | 0.334075631 | 0.55 | 0.253 | 0.009195038 | 36 |
| rps27.216 | 3.66302313390028e-07 | 0.420838694 | 0.838 | 0.518 | 0.009219463 | 36 |
| itm2bb10 | 4.73979635037688e-07 | 0.292160391 | 0.512 | 0.222 | 0.011929593 | 36 |
| ywhaqb7 | 5.94876984163409e-07 | 0.270068114 | 0.338 | 0.133 | 0.014972459 | 36 |
| h2afx7 | 6.44584600137827e-07 | 0.399503326 | 0.338 | 0.139 | 0.01622355 | 36 |
| pdia39 | 6.55333926544828e-07 | 0.300364636 | 0.625 | 0.284 | 0.0164941 | 36 |
| npm1a2 | 7.45410906618914e-07 | 0.381149612 | 0.6 | 0.303 | 0.018761247 | 36 |
| atp5f1d4 | 7.95140602722213e-07 | 0.297655831 | 0.562 | 0.258 | 0.020012894 | 36 |
| atp5mf3 | 8.90394124851286e-07 | 0.285650331 | 0.625 | 0.287 | 0.02241033 | 36 |
| mdh1aa4 | 9.22844978050339e-07 | 0.274468356 | 0.475 | 0.206 | 0.023227085 | 36 |
| txnipa12 | 9.77565875571333e-07 | 0.285468848 | 0.475 | 0.211 | 0.024604356 | 36 |
| kctd5a7 | 1.00211627814918e-06 | 0.281920365 | 0.412 | 0.176 | 0.025222265 | 36 |
| CR383676.119 | 1.0557347438169e-06 | 0.543758191 | 0.988 | 0.726 | 0.026571788 | 36 |
| cox5ab3 | 1.12180461129514e-06 | 0.268351455 | 0.462 | 0.202 | 0.0282347 | 36 |
| chd4b2 | 1.25168743746754e-06 | 0.274879453 | 0.275 | 0.103 | 0.031503721 | 36 |
| arpc5a8 | 1.52416520021612e-06 | 0.255525002 | 0.3 | 0.116 | 0.038361714 | 36 |
| cox6b25 | 1.74064577151715e-06 | 0.410935195 | 0.912 | 0.533 | 0.043810313 | 36 |
| nfkbiab7 | 1.99051342263983e-06 | 0.26084676 | 0.425 | 0.182 | 0.050099232 | 36 |
| zgc:1537135 | 2.05206111031215e-06 | 0.274494527 | 0.362 | 0.153 | 0.051648326 | 36 |
| pomp6 | 2.18805276762635e-06 | 0.28508772 | 0.5 | 0.231 | 0.0550711 | 36 |
| btg114 | 2.6155776821224e-06 | 0.352903902 | 0.65 | 0.333 | 0.065831475 | 36 |
| cnbpa1 | 2.6664395124033e-06 | 0.272656692 | 0.475 | 0.212 | 0.067111616 | 36 |
| ldhba2 | 3.13571652394548e-06 | 0.261053261 | 0.537 | 0.247 | 0.078922849 | 36 |
| hmgb2b3 | 3.47521897080913e-06 | 0.381369956 | 0.6 | 0.32 | 0.087467786 | 36 |
| zgc:1119865 | 4.16435714409295e-06 | 0.276708829 | 0.4 | 0.177 | 0.104812705 | 36 |
| id17 | 4.20275415989043e-06 | 0.396388611 | 0.262 | 0.106 | 0.105779119 | 36 |
| atp5po2 | 4.77856703322127e-06 | 0.250775127 | 0.575 | 0.269 | 0.120271754 | 36 |
| csrnp1b9 | 7.1148266483837e-06 | 0.253736712 | 0.4 | 0.18 | 0.179073072 | 36 |
| COX5B6 | 7.51642748326703e-06 | 0.320368195 | 0.7 | 0.358 | 0.189180963 | 36 |
| syncrip2 | 9.07630289806297e-06 | 0.264359801 | 0.625 | 0.301 | 0.228441468 | 36 |
| hint11 | 9.49595716659353e-06 | 0.260315338 | 0.462 | 0.218 | 0.239003746 | 36 |

| | | | | | | |
|---|---|---|---|---|---|---|
| uqcrh4 | 1.10111604960828e-05 | 0.274807429 | 0.512 | 0.251 | 0.277139899 | 36 |
| cirbpb2 | 1.21382245576748e-05 | 0.296779351 | 0.963 | 0.758 | 0.305506974 | 36 |
| h3f3c12 | 1.4442354362219e-05 | 0.283553207 | 0.525 | 0.257 | 0.363499617 | 36 |
| cap112 | 1.51444233369419e-05 | 0.254555736 | 0.488 | 0.234 | 0.381169991 | 36 |
| mt-atp619 | 1.71142487196146e-05 | 0.36534474 | 0.9 | 0.591 | 0.430748526 | 36 |
| eef1db11 | 2.39667505033009e-05 | 0.261746613 | 0.562 | 0.274 | 0.603219143 | 36 |
| gabarapb11 | 3.65900618251989e-05 | 0.261698746 | 0.525 | 0.269 | 0.920935266 | 36 |
| cox7b7 | 4.00286948920111e-05 | 0.331961946 | 0.762 | 0.417 | 1 | 36 |
| mt-nd214 | 4.02523080793076e-05 | 0.295666047 | 0.738 | 0.4 | 1 | 36 |
| atp5pb4 | 4.86180989343967e-05 | 0.251933447 | 0.588 | 0.3 | 1 | 36 |
| cfl111 | 4.98116587533908e-05 | 0.351038158 | 0.9 | 0.577 | 1 | 36 |
| slc25a3b7 | 6.22389601507606e-05 | 0.338252466 | 0.787 | 0.437 | 1 | 36 |
| prdx24 | 8.01647631601327e-05 | 0.296942356 | 0.75 | 0.401 | 1 | 36 |
| zgc:5649310 | 9.16006633966187e-05 | 0.338419764 | 0.863 | 0.573 | 1 | 36 |
| egr112 | 0.000219099 | 0.253044353 | 0.375 | 0.191 | 1 | 36 |
| mt-nd116 | 0.000266329 | 0.258880987 | 0.713 | 0.411 | 1 | 36 |
| ppiab6 | 0.000322272 | 0.28932678 | 0.9 | 0.657 | 1 | 36 |
| pnrc218 | 0.000363696 | 0.346044531 | 0.925 | 0.567 | 1 | 36 |
| jun10 | 0.000443063 | 0.299929836 | 0.412 | 0.229 | 1 | 36 |
| fkbp1aa9 | 0.000477991 | 0.26065068 | 0.688 | 0.388 | 1 | 36 |
| atp5mc3b2 | 0.000619396 | 0.290626928 | 0.838 | 0.604 | 1 | 36 |
| mt-co219 | 0.001114471 | 0.329802199 | 0.975 | 0.72 | 1 | 36 |
| mcl1a10 | 0.001939952 | 0.298048559 | 0.95 | 0.624 | 1 | 36 |
| tmsb4x21 | 0.006820309 | 0.338108134 | 1 | 0.686 | 1 | 36 |
| si:dkey-21e2.13 | 0 | 2.496485264 | 0.961 | 0.003 | 0 | 37 |
| si:dkey-21e2.15 | 0 | 2.256593584 | 0.857 | 0.004 | 0 | 37 |
| si:dkey-21e2.16 | 0 | 2.155384038 | 0.74 | 0.003 | 0 | 37 |
| si:ch211-212d10.11 | 0 | 1.895536618 | 0.87 | 0.018 | 0 | 37 |
| si:dkey-21e2.14 | 0 | 1.760456851 | 0.74 | 0.002 | 0 | 37 |
| il11b | 0 | 1.349871309 | 0.558 | 0.005 | 0 | 37 |
| si:dkey-78l4.2 | 0 | 1.337692598 | 0.429 | 0.005 | 0 | 37 |
| csf3a | 0 | 1.332932069 | 0.545 | 0.003 | 0 | 37 |
| il13 | 0 | 1.328168537 | 0.506 | 0.001 | 0 | 37 |
| il4 | 0 | 1.318035045 | 0.545 | 0.004 | 0 | 37 |
| si:dkey-21e2.10 | 0 | 1.231962188 | 0.312 | 0.002 | 0 | 37 |

| | | | | | | |
|---|---|---|---|---|---|---|
| si:dkey-78l4.1 | 0 | 0.937253913 | 0.26 | 0.001 | 0 | 37 |
| CR847844.1 | 0 | 0.858161855 | 0.39 | 0 | 0 | 37 |
| si:dkey-21e2.8 | 0 | 0.853653083 | 0.221 | 0.001 | 0 | 37 |
| si:dkey-12f6.5 | 0 | 0.819143629 | 0.403 | 0.002 | 0 | 37 |
| LOC103911745 | 0 | 0.80591714 | 0.416 | 0 | 0 | 37 |
| si:dkey-8e10.2 | 0 | 0.553513917 | 0.299 | 0.001 | 0 | 37 |
| BX571675.1 | 0 | 0.40185694 | 0.195 | 0.001 | 0 | 37 |
| kcnc1b | 0 | 0.339849198 | 0.182 | 0 | 0 | 37 |
| gata3 | 1.48421032382065e-117 | 0.467927689 | 0.234 | 0.006 | 3.7356089640242e-113 | 37 |
| cldni | 2.12344027229342e-116 | 0.63385141 | 0.312 | 0.011 | 5.34448682133531e-112 | 37 |
| si:ch211-67e16.32 | 2.4622308668475e-113 | 0.704012986 | 0.364 | 0.016 | 6.19718886876847e-109 | 37 |
| sox13 | 1.52154508678267e-111 | 0.600823173 | 0.299 | 0.011 | 3.82957682892329e-107 | 37 |
| LOC1001482543 | 1.61593159164312e-109 | 0.762585994 | 0.416 | 0.022 | 4.06713822300656e-105 | 37 |
| si:dkey-33i11.92 | 1.17323769418982e-108 | 0.772773596 | 0.403 | 0.02 | 2.95292195250637e-104 | 37 |
| si:ch211-112c15.8 | 2.60713725375877e-99 | 0.320749243 | 0.143 | 0.002 | 6.56190375398544e-95 | 37 |
| XLOC-024867 | 5.93689101590274e-95 | 0.511907795 | 0.273 | 0.011 | 1.49425609979256e-90 | 37 |
| faslg1 | 2.65210708577561e-94 | 0.614521861 | 0.325 | 0.015 | 6.67508832418862e-90 | 37 |
| slc35f4 | 2.12946233527557e-82 | 0.316196924 | 0.156 | 0.004 | 5.35964375165509e-78 | 37 |
| si:ch211-147k10.5 | 4.71745945478118e-75 | 0.340243691 | 0.156 | 0.004 | 1.18733737017388e-70 | 37 |
| ccl34b.43 | 2.78912224727167e-69 | 0.953323729 | 0.429 | 0.037 | 7.01994178415807e-65 | 37 |
| laptm513 | 4.3014140121006e-68 | 1.439283089 | 0.818 | 0.168 | 1.0826228927056e-63 | 37 |
| CABZ01073834.15 | 3.4786866173974e-67 | 0.778274289 | 0.442 | 0.04 | 8.75550634732751e-63 | 37 |
| tnfrsf9a5 | 2.88722054673266e-65 | 1.110590615 | 0.481 | 0.051 | 7.26684539407142e-61 | 37 |
| ca23 | 2.30722790934381e-60 | 0.919351981 | 0.494 | 0.055 | 5.80706192502744e-56 | 37 |
| scgn3 | 4.74659001329385e-60 | 0.588273783 | 0.312 | 0.022 | 1.19466924044593e-55 | 37 |
| si:cabz01036022.12 | 2.59444819572659e-55 | 0.412391132 | 0.208 | 0.011 | 6.52996666382425e-51 | 37 |
| tfa | 5.67494622022128e-54 | 0.624606944 | 0.195 | 0.01 | 1.42832721416749e-49 | 37 |
| samsn1a4 | 1.85810359196941e-52 | 0.658755433 | 0.364 | 0.035 | 4.6766609306278e-48 | 37 |
| blk | 8.47437698782565e-52 | 0.375893014 | 0.182 | 0.009 | 2.13291594406584e-47 | 37 |
| si:dkey-276j7.2 | 1.66013813143556e-44 | 0.333195558 | 0.156 | 0.008 | 4.17840166301016e-40 | 37 |
| samsn1b2 | 2.30261148605869e-40 | 0.416122468 | 0.26 | 0.023 | 5.79544284926112e-36 | 37 |
| si:ch1073-394i4.1 | 5.05278157033495e-40 | 0.32717989 | 0.13 | 0.006 | 1.2717345934376e-35 | 37 |
| nbeal21 | 8.15749735991269e-40 | 0.419231844 | 0.26 | 0.023 | 2.05316051051642e-35 | 37 |
| abca21 | 9.98537547754804e-36 | 0.46733247 | 0.26 | 0.026 | 2.51321915394407e-31 | 37 |
| pfn114 | 8.00042139402611e-34 | 0.98729247 | 1 | 0.524 | 2.01362606066243e-29 | 37 |

| | | | | | | |
|---|---|---|---|---|---|---|
| zbtb16a2 | 1.60743411621888e-33 | 0.559279784 | 0.351 | 0.048 | 4.04575092711131e-29 | 37 |
| si:dkey-151g10.62 | 6.6550295368471e-33 | 0.271210955 | 1 | 0.992 | 1.67500438412905e-28 | 37 |
| raraa2 | 1.11567417819193e-32 | 0.451522719 | 0.273 | 0.031 | 2.80804033909127e-28 | 37 |
| rgs1310 | 2.33280388378296e-32 | 0.881146931 | 0.584 | 0.131 | 5.87143409509332e-28 | 37 |
| wasb13 | 2.4004148724259e-32 | 0.928526765 | 0.675 | 0.176 | 6.04160419240874e-28 | 37 |
| MFAP4 (1 of many)1 | 1.52499081340903e-31 | 0.707298532 | 0.403 | 0.064 | 3.83824937826918e-27 | 37 |
| zmp:0000013232 | 1.54156355696785e-31 | 1.046505014 | 0.597 | 0.144 | 3.87996131653237e-27 | 37 |
| rplp112 | 5.06151456036125e-31 | 0.438748803 | 1 | 0.877 | 1.27393259969732e-26 | 37 |
| rplp2l20 | 5.43614042669098e-31 | 0.541587793 | 1 | 0.774 | 1.36822218399385e-26 | 37 |
| rplp219 | 7.52101875541788e-31 | 0.563487973 | 1 | 0.758 | 1.89296521055113e-26 | 37 |
| si:ch1073-110a20.31 | 1.21550380554135e-30 | 0.616375322 | 0.351 | 0.05 | 3.05930152816703e-26 | 37 |
| CR392341.17 | 3.14296663788411e-30 | 0.542030938 | 0.377 | 0.059 | 7.91053273089052e-26 | 37 |
| si:dkey-27h10.22 | 7.86895048140224e-30 | 0.415510614 | 0.247 | 0.027 | 1.98053614666413e-25 | 37 |
| si:dkey-262k9.22 | 4.0222678651319e-29 | 0.479849033 | 0.273 | 0.034 | 1.01236459897505e-24 | 37 |
| rps2010 | 1.6277435499688e-28 | 0.36879859 | 1 | 0.916 | 4.09686774091646e-24 | 37 |
| rps1210 | 2.78380585449048e-28 | 0.437686822 | 0.987 | 0.84 | 7.00656095516709e-24 | 37 |
| gpr183a7 | 7.84881917965305e-28 | 0.601967324 | 0.377 | 0.065 | 1.97546929932688e-23 | 37 |
| tnfb6 | 7.95056235474515e-28 | 0.628338034 | 0.338 | 0.052 | 2.00107703906581e-23 | 37 |
| osmr1 | 2.34638109537243e-27 | 0.273203832 | 0.13 | 0.009 | 5.90560657894287e-23 | 37 |
| si:ch211-119e14.16 | 2.35463719103634e-27 | 0.641591608 | 0.403 | 0.073 | 5.92638634611936e-23 | 37 |
| rpl3913 | 3.47981661920433e-27 | 0.390046947 | 0.974 | 0.874 | 8.75835044887538e-23 | 37 |
| rpl3417 | 4.98726497949753e-27 | 0.492512243 | 0.974 | 0.784 | 1.25524472268973e-22 | 37 |
| h1fx11 | 1.10212699450525e-26 | 0.835762731 | 0.662 | 0.182 | 2.77394343247026e-22 | 37 |
| rpl35a16 | 6.67983259904728e-26 | 0.498382899 | 0.987 | 0.759 | 1.68124706685421e-21 | 37 |
| rps229 | 7.00084525301707e-26 | 0.472460356 | 1 | 0.829 | 1.76204274173187e-21 | 37 |
| rps1711 | 9.29850837267222e-26 | 0.506114911 | 1 | 0.767 | 2.34034157231787e-21 | 37 |
| fosl211 | 1.07914892499511e-25 | 0.805466849 | 0.494 | 0.121 | 2.7161099293202e-21 | 37 |
| rpl3817 | 1.41182075561015e-25 | 0.544546093 | 0.987 | 0.728 | 3.5534116597952e-21 | 37 |
| rpl3713 | 2.8238771325278e-25 | 0.429888582 | 1 | 0.85 | 7.10741635485923e-21 | 37 |
| rps1817 | 4.66626789867614e-25 | 0.533547238 | 0.974 | 0.725 | 1.1744529674178e-20 | 37 |
| rps2913 | 7.93436751320116e-25 | 0.430953461 | 1 | 0.832 | 1.9970009593976e-20 | 37 |
| CU855821.12 | 8.57330557157715e-25 | 0.305907976 | 0.156 | 0.014 | 2.15781527931025e-20 | 37 |
| rps2817 | 1.19007554430312e-24 | 0.437702789 | 1 | 0.826 | 2.99530113745653e-20 | 37 |
| srgn11 | 4.44364034656101e-24 | 0.87284414 | 0.688 | 0.236 | 1.11841983882594e-19 | 37 |
| rps1618 | 5.14743970549841e-24 | 0.506730998 | 0.974 | 0.738 | 1.29555909947689e-19 | 37 |

| | | | | | | |
|---|---|---|---|---|---|---|
| ndufa4 | 6.289932298091e-24 | 0.309236729 | 0.156 | 0.014 | 1.58311306010652e-19 | 37 |
| arhgdig13 | 7.14611017498252e-24 | 0.802473173 | 0.727 | 0.242 | 1.79860446994135e-19 | 37 |
| rpl3215 | 1.38824778748061e-23 | 0.448414543 | 0.987 | 0.813 | 3.49408085630995e-19 | 37 |
| rpl2612 | 1.67170713284611e-23 | 0.400729072 | 0.987 | 0.841 | 4.20751968266036e-19 | 37 |
| ctss11 | 2.92457974356898e-23 | 0.293439387 | 0.156 | 0.015 | 7.36087475658876e-19 | 37 |
| si:ch211-207n23.21 | 3.7740158754641e-23 | 0.406001984 | 0.221 | 0.028 | 9.49882055695559e-19 | 37 |
| grap2b10 | 5.15274308403667e-23 | 0.624289694 | 0.481 | 0.112 | 1.29689390682119e-18 | 37 |
| unc119b2 | 1.13791813579928e-22 | 0.425786005 | 0.273 | 0.041 | 2.8640261559932e-18 | 37 |
| rpl1816 | 2.24742444438779e-22 | 0.463964316 | 0.987 | 0.774 | 5.65654258407964e-18 | 37 |
| ada1 | 3.48960355478618e-22 | 0.466192512 | 0.26 | 0.04 | 8.78298318704134e-18 | 37 |
| rpl2713 | 8.9729406285899e-22 | 0.469298878 | 0.987 | 0.763 | 2.25839942680979e-17 | 37 |
| gab21 | 9.64985402668011e-22 | 0.264954524 | 0.156 | 0.016 | 2.42877175997512e-17 | 37 |
| rps1416 | 1.01110712200344e-21 | 0.467519044 | 0.987 | 0.759 | 2.54485551537047e-17 | 37 |
| rps1316 | 1.32962144209702e-21 | 0.497519248 | 0.974 | 0.724 | 3.34652420761399e-17 | 37 |
| fa2h1 | 1.62929747833614e-21 | 0.271195141 | 0.13 | 0.011 | 4.10077882322423e-17 | 37 |
| rpl7a13 | 2.78038730091409e-21 | 0.409785825 | 0.974 | 0.81 | 6.99795679767066e-17 | 37 |
| rpl36a13 | 4.41676384046289e-21 | 0.404714271 | 0.987 | 0.81 | 1.1116552910061e-16 | 37 |
| rpsa21 | 5.86410935137809e-21 | 0.480307428 | 1 | 0.752 | 1.47593768264835e-16 | 37 |
| tnip15 | 6.24241893261216e-21 | 0.576093673 | 0.377 | 0.081 | 1.57115442114915e-16 | 37 |
| rps26l17 | 7.76096468120463e-21 | 0.550941985 | 0.948 | 0.664 | 1.95335720061239e-16 | 37 |
| arpc1b14 | 7.79382309287103e-21 | 0.842058268 | 0.792 | 0.308 | 1.96162733424471e-16 | 37 |
| si:ch211-239m17.1 | 8.02410238109557e-21 | 0.277668431 | 0.156 | 0.016 | 2.01958632829794e-16 | 37 |
| rps15a15 | 9.54167231900602e-21 | 0.468719579 | 0.987 | 0.78 | 2.40154350597063e-16 | 37 |
| rps515 | 1.04477157426756e-20 | 0.521649039 | 0.974 | 0.701 | 2.62958557527402e-16 | 37 |
| rpl1415 | 1.65004103836114e-20 | 0.565919266 | 0.974 | 0.635 | 4.15298828945116e-16 | 37 |
| rps2417 | 2.8731737722994e-20 | 0.45750104 | 1 | 0.758 | 7.23149106750035e-16 | 37 |
| rpl1515 | 2.96921631916856e-20 | 0.481332738 | 0.987 | 0.744 | 7.47322055371534e-16 | 37 |
| rps611 | 3.22968705964154e-20 | 0.407119544 | 0.948 | 0.789 | 8.12879936041178e-16 | 37 |
| rpl3620 | 3.75952180707274e-20 | 0.444831043 | 0.974 | 0.79 | 9.46234043622138e-16 | 37 |
| si:dkey-25e12.35 | 5.32361854092666e-20 | 0.450720474 | 0.286 | 0.051 | 1.33990155056583e-15 | 37 |
| rpl2114 | 6.45506154609199e-20 | 0.436378556 | 0.974 | 0.756 | 1.62467444053589e-15 | 37 |
| sla25 | 6.6433500195047e-20 | 0.603246869 | 0.338 | 0.069 | 1.67206476640914e-15 | 37 |
| rpl815 | 7.14870261278485e-20 | 0.431041049 | 0.974 | 0.78 | 1.79925696061182e-15 | 37 |
| rps27.116 | 1.06209789022238e-19 | 0.523154802 | 0.935 | 0.679 | 2.6731941799007e-15 | 37 |
| rpl1116 | 1.20830156447109e-19 | 0.397214552 | 0.961 | 0.814 | 3.0411742076173e-15 | 37 |

| | | | | | | |
|---|---|---|---|---|---|---|
| rpl238 | 1.47738461184518e-19 | 0.329842841 | 1 | 0.873 | 3.71842932955313e-15 | 37 |
| rpl18a15 | 4.04085452904333e-19 | 0.409705327 | 0.935 | 0.785 | 1.01704267641491e-14 | 37 |
| rpl1711 | 5.67761497691455e-19 | 0.433898672 | 0.974 | 0.746 | 1.42899891353962e-14 | 37 |
| tnfaip8l2b | 6.46745297920353e-19 | 0.256437423 | 0.13 | 0.013 | 1.62779324033574e-14 | 37 |
| zgc:11418821 | 7.93037109698653e-19 | 0.573649074 | 0.974 | 0.627 | 1.99599510140054e-14 | 37 |
| rps1911 | 1.021989677473e-18 | 0.342639783 | 0.987 | 0.863 | 2.57224581923179e-14 | 37 |
| rps710 | 1.14737517910655e-18 | 0.38342694 | 0.974 | 0.821 | 2.88782858829328e-14 | 37 |
| rps2312 | 1.50504558647908e-18 | 0.458859389 | 0.961 | 0.723 | 3.7880492366092e-14 | 37 |
| adma2 | 1.53532232996311e-18 | 0.298628331 | 0.156 | 0.018 | 3.86425277228414e-14 | 37 |
| rpl2917 | 2.0219885747639e-18 | 0.402814621 | 0.987 | 0.809 | 5.08914304382326e-14 | 37 |
| btk2 | 2.21248585880823e-18 | 0.303847424 | 0.182 | 0.024 | 5.56860565803442e-14 | 37 |
| rps2123 | 2.98067983269653e-18 | 0.459193374 | 0.961 | 0.75 | 7.50207307091391e-14 | 37 |
| rplp08 | 3.06862577232582e-18 | 0.363241996 | 1 | 0.851 | 7.72342420636685e-14 | 37 |
| rpl3017 | 3.202032350415e-18 | 0.464331641 | 0.935 | 0.708 | 8.05919522275952e-14 | 37 |
| tnfaip35 | 3.76532401462544e-18 | 0.502823769 | 0.312 | 0.065 | 9.47694401241078e-14 | 37 |
| cebpb15 | 6.04242801932851e-18 | 0.807958296 | 0.766 | 0.353 | 1.52081870818479e-13 | 37 |
| nr4a112 | 7.12797139720671e-18 | 0.691667833 | 0.532 | 0.167 | 1.79403912096296e-13 | 37 |
| glipr1a6 | 7.36433576356352e-18 | 0.422754418 | 0.312 | 0.064 | 1.8535296683313e-13 | 37 |
| rab447 | 1.50606627214975e-17 | 0.457038549 | 0.299 | 0.061 | 3.79061820037371e-13 | 37 |
| rpl2416 | 1.57678855542881e-17 | 0.416465438 | 0.987 | 0.776 | 3.96861911515878e-13 | 37 |
| bhlhe4012 | 2.74358116579573e-17 | 0.678748691 | 0.506 | 0.161 | 6.90531943619128e-13 | 37 |
| rps1517 | 2.89943553006087e-17 | 0.502893429 | 0.961 | 0.684 | 7.29758928561021e-13 | 37 |
| rps4x14 | 3.13899775121097e-17 | 0.464065768 | 0.961 | 0.71 | 7.9005434400229e-13 | 37 |
| rps27a15 | 3.33312408726688e-17 | 0.410600805 | 0.961 | 0.752 | 8.38914001524201e-13 | 37 |
| capgb8 | 3.38463472028318e-17 | 0.577617729 | 0.429 | 0.114 | 8.51878712748075e-13 | 37 |
| rpl3511 | 3.50309813077329e-17 | 0.386349462 | 0.987 | 0.805 | 8.81694768534328e-13 | 37 |
| rpl23a19 | 5.08401244354868e-17 | 0.515227141 | 0.909 | 0.634 | 1.27959509191677e-12 | 37 |
| tagln212 | 7.21589242431172e-17 | 0.637964972 | 0.558 | 0.188 | 1.81616796427502e-12 | 37 |
| n4bp12 | 7.62330028308047e-17 | 0.414592809 | 0.247 | 0.045 | 1.91870844824852e-12 | 37 |
| cox8b4 | 1.13579842247814e-16 | 0.509428611 | 0.377 | 0.095 | 2.85869104953523e-12 | 37 |
| rpl127 | 1.22475006565894e-16 | 0.362987245 | 0.961 | 0.799 | 3.082573440257e-12 | 37 |
| rpl139 | 1.52267510654897e-16 | 0.37590055 | 1 | 0.805 | 3.83242097567309e-12 | 37 |
| si:ch211-66e2.37 | 1.73913885487031e-16 | 0.441748042 | 0.286 | 0.059 | 4.3772385838231e-12 | 37 |
| FP236356.14 | 1.81391843837697e-16 | 0.419367525 | 0.26 | 0.049 | 4.565451317551e-12 | 37 |
| rpl921 | 1.93813287262343e-16 | 0.535108966 | 0.948 | 0.629 | 4.87808662710590e-12 | 37 |

| | | | | | | |
|---|---|---|---|---|---|---|
| nme2b.14 | 2.95786827653383e-16 | 0.311264145 | 0.974 | 0.857 | 7.44465866520799e-12 | 37 |
| rpl198 | 3.15985929471803e-16 | 0.326215404 | 0.974 | 0.852 | 7.9530498588758e-12 | 37 |
| si:zfos-2330d3.11 | 3.25409466243864e-16 | 0.304157084 | 0.182 | 0.026 | 8.19023085589182e-12 | 37 |
| rpl79 | 5.29151206601162e-16 | 0.379220254 | 0.987 | 0.805 | 1.33182067189446e-11 | 37 |
| si:dkey-1c7.11 | 5.95333567643445e-16 | 0.297608565 | 0.169 | 0.024 | 1.49839505640179e-11 | 37 |
| rps2612 | 7.41495629712698e-16 | 0.428526639 | 0.974 | 0.734 | 1.86627035042389e-11 | 37 |
| nfkbiaa6 | 7.53527905024841e-16 | 0.667957032 | 0.481 | 0.152 | 1.89655438415702e-11 | 37 |
| wipf1a7 | 8.43227856746095e-16 | 0.427318036 | 0.299 | 0.065 | 2.12232019264425e-11 | 37 |
| irf4b3 | 1.65216746206422e-15 | 0.323369271 | 0.169 | 0.025 | 4.15834028526942e-11 | 37 |
| coro1a13 | 2.2429567121784e-15 | 0.660337086 | 0.636 | 0.231 | 5.64529774888182e-11 | 37 |
| si:ch211-222l21.115 | 2.26332769182944e-15 | 0.662087472 | 0.688 | 0.275 | 5.69656946756551e-11 | 37 |
| rpl3118 | 2.57857740627484e-15 | 0.481996762 | 0.935 | 0.655 | 6.49002147385315e-11 | 37 |
| rpl10a11 | 4.19292451079288e-15 | 0.408948472 | 0.987 | 0.765 | 1.05531717012146e-10 | 37 |
| ms4a17a.66 | 4.24973429814393e-15 | 0.379176684 | 0.247 | 0.049 | 1.06961562549984e-10 | 37 |
| rps3a10 | 4.71203816376544e-15 | 0.368271638 | 0.961 | 0.778 | 1.18597288543812e-10 | 37 |
| CU499336.24 | 5.05011508427371e-15 | 0.332481407 | 0.208 | 0.036 | 1.27106346556085e-10 | 37 |
| rps2512 | 8.76775236574668e-15 | 0.433299652 | 0.974 | 0.725 | 2.20675559293478e-10 | 37 |
| rps1020 | 1.55868919707866e-14 | 0.453526739 | 0.961 | 0.694 | 3.92306484012728e-10 | 37 |
| rpl2216 | 1.76136908120073e-14 | 0.438508242 | 0.922 | 0.694 | 4.43318984047411e-10 | 37 |
| grap2a5 | 1.78088447482418e-14 | 0.413859581 | 0.234 | 0.047 | 4.48230813468499e-10 | 37 |
| faua14 | 1.84378114243653e-14 | 0.449552979 | 0.935 | 0.69 | 4.64061275739849e-10 | 37 |
| hmha1b7 | 2.19685430750097e-14 | 0.407377853 | 0.312 | 0.075 | 5.5292626065492e-10 | 37 |
| adcy71 | 2.23859518600394e-14 | 0.353906848 | 0.169 | 0.027 | 5.63432022365331e-10 | 37 |
| slc2a3b3 | 2.30447360197404e-14 | 0.259271285 | 0.182 | 0.03 | 5.80012960880846e-10 | 37 |
| ccl34b.13 | 2.9862410476838e-14 | 0.457730293 | 0.182 | 0.03 | 7.51607009291535e-10 | 37 |
| rpl1013 | 3.55432528579018e-14 | 0.373123717 | 0.961 | 0.772 | 8.94588131180531e-10 | 37 |
| rbp56 | 4.76945057484841e-14 | 0.507879072 | 0.351 | 0.095 | 1.2004230151836e-09 | 37 |
| crip115 | 6.69078901405607e-14 | 0.62016206 | 0.675 | 0.27 | 1.68400468694777e-09 | 37 |
| AL954142.12 | 8.99279181097548e-14 | 0.254497096 | 0.156 | 0.023 | 2.26339577090442e-09 | 37 |
| tnfrsf9b1 | 1.16108588678692e-13 | 0.269181578 | 0.13 | 0.017 | 2.922337068454e-09 | 37 |
| CABZ01084575.14 | 1.20348473240728e-13 | 0.339957224 | 0.221 | 0.043 | 3.02905072299588e-09 | 37 |
| rps913 | 2.02874528672627e-13 | 0.376404342 | 0.974 | 0.777 | 5.10614901216135e-09 | 37 |
| mrps157 | 2.5291654284255e-13 | 0.478814117 | 0.416 | 0.125 | 6.36565646680415e-09 | 37 |
| ppp1r186 | 2.94178255996612e-13 | 0.353171544 | 0.247 | 0.054 | 7.40417252517872e-09 | 37 |
| arl5c8 | 3.80798169281885e-13 | 0.505479352 | 0.377 | 0.113 | 9.58430912265575e-09 | 37 |

| | | | | | | |
|---|---|---|---|---|---|---|
| cd247l3 | 4.50181209633044e-13 | 0.257576679 | 0.143 | 0.021 | 1.13306108652541e-08 | 37 |
| ehd1a1 | 5.40513079032167e-13 | 0.270458937 | 0.182 | 0.032 | 1.36041736861606e-08 | 37 |
| uba5216 | 6.26995374231866e-13 | 0.490657748 | 0.974 | 0.63 | 1.57808465740418e-08 | 37 |
| rps8a9 | 7.9881255123071e-13 | 0.361943453 | 0.987 | 0.788 | 2.01053131019257e-08 | 37 |
| rpl618 | 1.37824307788801e-12 | 0.423905315 | 0.909 | 0.668 | 3.46890000273633e-08 | 37 |
| lpar5a6 | 1.41643005548812e-12 | 0.327307529 | 0.208 | 0.042 | 3.56501280665804e-08 | 37 |
| selplg7 | 1.87325480984487e-12 | 0.325340955 | 0.221 | 0.047 | 4.71479503089855e-08 | 37 |
| calm1b9 | 1.98311878653023e-12 | 0.435036353 | 0.338 | 0.095 | 4.99131167381792e-08 | 37 |
| ldha5 | 2.14562655759197e-12 | 0.412643852 | 0.286 | 0.073 | 5.40032748280322e-08 | 37 |
| ets12 | 2.58617455420549e-12 | 0.253258118 | 0.195 | 0.038 | 6.5091427354798e-08 | 37 |
| cfbl3 | 2.98833496438379e-12 | 0.453737214 | 0.273 | 0.067 | 7.52134027185757e-08 | 37 |
| btg115 | 3.8561721980163e-12 | 0.582488123 | 0.714 | 0.332 | 9.70559980518723e-08 | 37 |
| mpp11 | 5.10352427508944e-12 | 0.269322546 | 0.169 | 0.03 | 1.28450602479726e-07 | 37 |
| fli1b3 | 7.19532944645234e-12 | 0.263472385 | 0.182 | 0.035 | 1.81099246837759e-07 | 37 |
| znf7041 | 7.75436515319375e-12 | 0.252194832 | 0.169 | 0.031 | 1.95169616540733e-07 | 37 |
| ptpn66 | 8.03897534337639e-12 | 0.402612061 | 0.325 | 0.091 | 2.0233297041744e-07 | 37 |
| si:ch211-191c10.16 | 8.06986020962287e-12 | 0.36652564 | 0.234 | 0.054 | 2.03110311615998e-07 | 37 |
| si:ch211-226m16.26 | 8.21221164941563e-12 | 0.362362246 | 0.273 | 0.069 | 2.06693155004142e-07 | 37 |
| gnai12 | 9.3255242429192e-12 | 0.360736081 | 0.247 | 0.06 | 2.34714119670033e-07 | 37 |
| mbnl110 | 1.10375239653934e-11 | 0.503031982 | 0.494 | 0.184 | 2.77803440684985e-07 | 37 |
| si:ch73-234b20.57 | 1.12690440322254e-11 | 0.423922106 | 0.338 | 0.101 | 2.83630569247081e-07 | 37 |
| cd6315 | 1.6658208387708e-11 | 0.573211901 | 0.714 | 0.34 | 4.19270446910224e-07 | 37 |
| rpl2814 | 1.91382720945978e-11 | 0.360036614 | 0.948 | 0.75 | 4.81691170348932e-07 | 37 |
| rpl27a18 | 1.98003542368979e-11 | 0.523989065 | 0.818 | 0.496 | 4.98355115788483e-07 | 37 |
| sept126 | 2.51991340997547e-11 | 0.386659852 | 0.247 | 0.062 | 6.34237006156726e-07 | 37 |
| naca14 | 4.94445467182308e-11 | 0.454787381 | 0.909 | 0.619 | 1.24446979635115e-06 | 37 |
| rps1110 | 6.63021863241003e-11 | 0.327465854 | 0.935 | 0.766 | 1.66875972759128e-06 | 37 |
| rpl22l113 | 7.85338588364142e-11 | 0.551686262 | 0.727 | 0.392 | 1.97661869305371e-06 | 37 |
| eif3f15 | 8.53372561003529e-11 | 0.545456127 | 0.818 | 0.431 | 2.14785339878978e-06 | 37 |
| rsl24d115 | 9.25669385327735e-11 | 0.536553848 | 0.779 | 0.444 | 2.32981727593138e-06 | 37 |
| crema9 | 9.47999815026453e-11 | 0.546967664 | 0.455 | 0.179 | 2.38602073444008e-06 | 37 |
| gstk1 | 1.12626908886979e-10 | 0.274710171 | 0.208 | 0.047 | 2.83470666977638e-06 | 37 |
| rpl5a17 | 1.20370239897008e-10 | 0.507243995 | 0.805 | 0.492 | 3.0295985679678e-06 | 37 |
| ets24 | 1.57760926783264e-10 | 0.447783624 | 0.299 | 0.091 | 3.97068476620796e-06 | 37 |
| eno1a6 | 1.74084079440943e-10 | 0.432841371 | 0.39 | 0.135 | 4.38152219544909e-06 | 37 |

| | | | | | | |
|---|---|---|---|---|---|---|
| eef1g14 | 2.50725707283687e-10 | 0.443052524 | 0.909 | 0.62 | 6.31051532662313e-06 | 37 |
| wipf1b5 | 2.64696444385931e-10 | 0.266452071 | 0.247 | 0.063 | 6.66214480874951e-06 | 37 |
| XLOC-0430434 | 2.82304292344623e-10 | 0.303356848 | 0.195 | 0.043 | 7.10531673402183e-06 | 37 |
| ptmaa18 | 2.85179633268411e-10 | 0.561801875 | 0.766 | 0.395 | 7.17768618973263e-06 | 37 |
| tmsb4x22 | 3.49582018091257e-10 | 0.529302426 | 1 | 0.686 | 8.79862981333885e-06 | 37 |
| itga46 | 3.54559726241216e-10 | 0.349460789 | 0.247 | 0.065 | 8.92391374976517e-06 | 37 |
| zgc:1528633 | 3.84012787039895e-10 | 0.301046427 | 0.195 | 0.044 | 9.66521783700712e-06 | 37 |
| dusp19 | 4.76400079699917e-10 | 0.478116959 | 0.429 | 0.16 | 1.19905136059672e-05 | 37 |
| actb213 | 5.5895654684417e-10 | 0.325361011 | 1 | 0.868 | 1.40683773275209e-05 | 37 |
| si:ch1073-443f11.23 | 7.18584058211302e-10 | 0.280159764 | 0.169 | 0.036 | 1.80860421611203e-05 | 37 |
| si:dkeyp-68b7.128 | 1.10608695543533e-09 | 0.356746979 | 0.234 | 0.063 | 2.78391025813518e-05 | 37 |
| si:dkey-206d17.124 | 2.18839690312187e-09 | 0.334129729 | 0.208 | 0.053 | 5.50797616546743e-05 | 37 |
| CR381673.18 | 2.34023094217831e-09 | 0.297714626 | 0.221 | 0.057 | 5.89012725836859e-05 | 37 |
| ltb4r4 | 2.94862556083529e-09 | 0.265817926 | 0.143 | 0.029 | 7.42139567406634e-05 | 37 |
| dbnlb1 | 2.97703723443176e-09 | 0.258359399 | 0.182 | 0.042 | 7.4929050153413e-05 | 37 |
| gapdhs14 | 4.03827507795573e-09 | 0.491164246 | 0.597 | 0.289 | 0.000101639 | 37 |
| CR855311.17 | 4.25530689225079e-09 | 0.538522741 | 0.273 | 0.086 | 0.000107102 | 37 |
| rack110 | 4.81901056595143e-09 | 0.320315756 | 0.935 | 0.744 | 0.00012129 | 37 |
| sec61g6 | 5.64505350730046e-09 | 0.376725158 | 0.909 | 0.679 | 0.00014208 | 37 |
| rac214 | 6.15206047086418e-09 | 0.474867005 | 0.519 | 0.221 | 0.000154841 | 37 |
| rps314 | 6.77170762134396e-09 | 0.354289442 | 0.922 | 0.7 | 0.000170437 | 37 |
| LOC5665873 | 6.96948051588523e-09 | 0.288080502 | 0.195 | 0.049 | 0.000175415 | 37 |
| unc13d2 | 7.30913503576385e-09 | 0.251057563 | 0.143 | 0.03 | 0.000183964 | 37 |
| kdm6ba1 | 7.31082236606767e-09 | 0.289697569 | 0.195 | 0.049 | 0.000184006 | 37 |
| ndfip21 | 8.28032202379492e-09 | 0.324687601 | 0.208 | 0.056 | 0.000208407 | 37 |
| upf3b | 8.35203077919919e-09 | 0.321956763 | 0.195 | 0.05 | 0.000210212 | 37 |
| cox6b111 | 1.07379751046083e-08 | 0.392291833 | 0.338 | 0.121 | 0.000270264 | 37 |
| si:ch211-102c2.48 | 1.3548534936008e-08 | 0.381299305 | 0.39 | 0.143 | 0.000341003 | 37 |
| sh2b35 | 1.41949420966011e-08 | 0.299780566 | 0.221 | 0.062 | 0.000357272 | 37 |
| cotl112 | 2.72301492315857e-08 | 0.508468964 | 0.623 | 0.307 | 0.000685356 | 37 |
| rnf1301 | 3.49290891896996e-08 | 0.282196937 | 0.182 | 0.047 | 0.00087913 | 37 |
| sik19 | 3.78603082356205e-08 | 0.382081578 | 0.325 | 0.117 | 0.000952906 | 37 |
| rps27.217 | 5.75381436335501e-08 | 0.457461873 | 0.883 | 0.517 | 0.001448178 | 37 |
| zfp36l1a10 | 6.60386611708955e-08 | 0.443837386 | 0.532 | 0.248 | 0.001662127 | 37 |
| chac17 | 6.62036022628876e-08 | 0.448929484 | 0.714 | 0.399 | 0.001666278 | 37 |

| | | | | | | |
|---|---|---|---|---|---|---|
| zgc:6405113 | 6.72612373575659e-08 | 0.36973507 | 0.416 | 0.163 | 0.001692898 | 37 |
| LOC1039091159 | 7.6763734941326e-08 | 0.344947755 | 0.416 | 0.166 | 0.001932066 | 37 |
| b3gnt3.44 | 8.33348703135229e-08 | 0.262517368 | 0.156 | 0.038 | 0.002097455 | 37 |
| nsa216 | 8.58184606720724e-08 | 0.464765972 | 0.662 | 0.363 | 0.002159965 | 37 |
| cd28l4 | 1.07823732410946e-07 | 0.25875651 | 0.156 | 0.038 | 0.002713816 | 37 |
| cd74b11 | 1.14299611762802e-07 | 0.351324658 | 0.532 | 0.224 | 0.002876807 | 37 |
| irf2bp2a7 | 1.17598199553434e-07 | 0.353072091 | 0.429 | 0.177 | 0.002959829 | 37 |
| rpl5b11 | 1.33646625654761e-07 | 0.403552444 | 0.909 | 0.63 | 0.003363752 | 37 |
| id36 | 1.47528344636271e-07 | 0.468838136 | 0.468 | 0.211 | 0.003713141 | 37 |
| msna14 | 1.70957944070328e-07 | 0.443820783 | 0.494 | 0.242 | 0.00430284 | 37 |
| arrb2b8 | 1.75659490066858e-07 | 0.27517768 | 0.273 | 0.092 | 0.004421174 | 37 |
| ppiaa13 | 1.90241253431594e-07 | 0.273245359 | 0.961 | 0.804 | 0.004788182 | 37 |
| si:ch211-241f5.34 | 2.22384977231561e-07 | 0.305768682 | 0.221 | 0.068 | 0.005597207 | 37 |
| cap113 | 2.24232487470854e-07 | 0.404627826 | 0.506 | 0.234 | 0.005643707 | 37 |
| tagapb8 | 2.71300016647271e-07 | 0.27216365 | 0.221 | 0.068 | 0.00682835 | 37 |
| BX120005.15 | 2.78681359928115e-07 | 0.251104254 | 0.182 | 0.05 | 0.007014131 | 37 |
| slc38a23 | 4.11018301513612e-07 | 0.31269862 | 0.169 | 0.047 | 0.01034492 | 37 |
| btf313 | 5.75373073411737e-07 | 0.383205602 | 0.857 | 0.588 | 0.014481565 | 37 |
| tagapa9 | 7.33887782295935e-07 | 0.363851046 | 0.286 | 0.106 | 0.018471222 | 37 |
| aldocb4 | 1.25067393377419e-06 | 0.289138972 | 0.208 | 0.068 | 0.031478212 | 37 |
| si:ch73-359m17.24 | 1.47142492269732e-06 | 0.366456635 | 0.429 | 0.19 | 0.037034294 | 37 |
| pim27 | 2.19912826978709e-06 | 0.306620592 | 0.247 | 0.09 | 0.055349859 | 37 |
| myl12.112 | 2.51697794244819e-06 | 0.418114678 | 0.636 | 0.393 | 0.063349818 | 37 |
| tefa9 | 3.79961341933016e-06 | 0.322454276 | 0.325 | 0.139 | 0.09563247 | 37 |
| hic14 | 5.16221449243053e-06 | 0.280303785 | 0.182 | 0.059 | 0.129927777 | 37 |
| ddx215 | 5.42874131093258e-06 | 0.353771206 | 0.429 | 0.208 | 0.13663599 | 37 |
| tnrc6c111 | 5.74283687765329e-06 | 0.331674747 | 0.312 | 0.134 | 0.144541461 | 37 |
| txnipa13 | 8.05832898115006e-06 | 0.36184115 | 0.429 | 0.211 | 0.202820082 | 37 |
| arpc5a9 | 1.03136014070293e-05 | 0.261688413 | 0.286 | 0.116 | 0.259583034 | 37 |
| klf136 | 1.04582702033903e-05 | 0.322742568 | 0.26 | 0.106 | 0.263224203 | 37 |
| rpl13a8 | 1.07242122969038e-05 | 0.262747819 | 0.935 | 0.765 | 0.269917699 | 37 |
| capza1a10 | 1.0935703455787e-05 | 0.353242632 | 0.403 | 0.201 | 0.27524072 | 37 |
| pnrc219 | 1.16316536727561e-05 | 0.374479243 | 0.818 | 0.568 | 0.292757091 | 37 |
| rpl410 | 1.20187226359031e-05 | 0.307883783 | 0.831 | 0.628 | 0.30249923 | 37 |
| slc3a2b8 | 1.36075398681759e-05 | 0.270381663 | 0.286 | 0.119 | 0.342488171 | 37 |

| | | | | | | |
|---|---|---|---|---|---|---|
| serf214 | 1.48298162800659e-05 | 0.360259988 | 0.597 | 0.344 | 0.373251646 | 37 |
| caprin1b3 | 1.63151527823679e-05 | 0.351344113 | 0.468 | 0.248 | 0.41063608 | 37 |
| arpc312 | 1.79261940785919e-05 | 0.384320003 | 0.532 | 0.311 | 0.451184379 | 37 |
| scinlb7 | 1.83366149291721e-05 | 0.293555629 | 0.221 | 0.084 | 0.461514261 | 37 |
| csrnp1b10 | 1.89637956794936e-05 | 0.314960819 | 0.377 | 0.18 | 0.477299773 | 37 |
| tsc22d311 | 2.16945472993495e-05 | 0.343796813 | 0.312 | 0.145 | 0.546030061 | 37 |
| idh29 | 4.02552929745466e-05 | 0.293355963 | 0.403 | 0.2 | 1 | 37 |
| notch29 | 4.6920489117241e-05 | 0.299959681 | 0.273 | 0.12 | 1 | 37 |
| cox8a10 | 5.48950273306351e-05 | 0.321049946 | 0.429 | 0.228 | 1 | 37 |
| hmgn212 | 5.53455446340709e-05 | 0.411756056 | 0.662 | 0.444 | 1 | 37 |
| mob1a8 | 5.84168836929441e-05 | 0.279482601 | 0.351 | 0.167 | 1 | 37 |
| eif3ea11 | 0.000103529 | 0.323002393 | 0.481 | 0.278 | 1 | 37 |
| ppp3ccb1 | 0.000118979 | 0.269310674 | 0.273 | 0.124 | 1 | 37 |
| arpc213 | 0.000158883 | 0.299710519 | 0.481 | 0.268 | 1 | 37 |
| fabp36 | 0.000160164 | 0.336690705 | 0.429 | 0.243 | 1 | 37 |
| dbi7 | 0.000188549 | 0.310377414 | 0.39 | 0.214 | 1 | 37 |
| sesn18 | 0.000194089 | 0.282494093 | 0.338 | 0.169 | 1 | 37 |
| pabpc1a13 | 0.000212026 | 0.316497475 | 0.766 | 0.541 | 1 | 37 |
| sat1a.27 | 0.000235838 | 0.302015744 | 0.519 | 0.299 | 1 | 37 |
| ptprc11 | 0.000247709 | 0.272618015 | 0.273 | 0.128 | 1 | 37 |
| ssr43 | 0.00025598 | 0.303153363 | 0.532 | 0.307 | 1 | 37 |
| eif3s106 | 0.000279032 | 0.274213067 | 0.325 | 0.166 | 1 | 37 |
| zgc:5649311 | 0.000534474 | 0.294894805 | 0.792 | 0.573 | 1 | 37 |
| ssr35 | 0.000534839 | 0.267606345 | 0.481 | 0.273 | 1 | 37 |
| oaz1b15 | 0.000553618 | 0.268467222 | 0.545 | 0.324 | 1 | 37 |
| arpc4l11 | 0.000642166 | 0.254570828 | 0.364 | 0.198 | 1 | 37 |
| mibp214 | 0.000729658 | 0.303432142 | 0.727 | 0.493 | 1 | 37 |
| si:ch211-196l7.416 | 0.001595919 | 0.250305608 | 0.519 | 0.315 | 1 | 37 |
| cox4i19 | 0.00161336 | 0.26994955 | 0.61 | 0.393 | 1 | 37 |
| eef1da12 | 0.001846561 | 0.287981664 | 0.714 | 0.493 | 1 | 37 |
| mcl1a11 | 0.002288658 | 0.267810345 | 0.831 | 0.624 | 1 | 37 |
| romo115 | 0.002340204 | 0.275967797 | 0.571 | 0.387 | 1 | 37 |
| xbp17 | 0.007355828 | 0.256889069 | 0.494 | 0.345 | 1 | 37 |
| scg3 | 6.73833985693582e-155 | 0.86886421 | 0.348 | 0.009 | 1.69597275859218e-150 | 38 |
| dmrt11 | 4.8311346246468e-23 | 0.521489454 | 0.188 | 0.02 | 1.21594827367735e-18 | 38 |

| | | | | | | |
|---|---|---|---|---|---|---|
| zp2.512 | 1.25478593060375e-19 | 0.742895252 | 0.986 | 0.564 | 3.15817070873658e-15 | 38 |
| si:dkey-241l7.211 | 9.62268904542576e-18 | 0.679148941 | 1 | 0.642 | 2.42193460584321e-13 | 38 |
| si:dkeyp-20e4.811 | 3.52007841190704e-17 | 0.797956509 | 0.855 | 0.409 | 8.85968535492884e-13 | 38 |
| mid1ip1l11 | 1.5985044203157e-16 | 0.697401921 | 0.942 | 0.544 | 4.02327577549259e-12 | 38 |
| si:ch211-226h8.411 | 2.01325853924491e-16 | 0.752609826 | 0.884 | 0.442 | 5.06717041742552e-12 | 38 |
| h1m12 | 2.21839702282791e-16 | 0.680511954 | 0.986 | 0.602 | 5.58348346675556e-12 | 38 |
| acp5a12 | 2.73053580038467e-16 | 0.704148032 | 0.942 | 0.541 | 6.87248555598817e-12 | 38 |
| h2af1al12 | 2.87496496584937e-16 | 0.693379106 | 0.986 | 0.593 | 7.23599932254628e-12 | 38 |
| si:dkey-240e12.611 | 3.49428355977301e-16 | 0.707137063 | 0.971 | 0.57 | 8.79476229159268e-12 | 38 |
| si:dkey-229d11.311 | 6.94617707710505e-16 | 0.755698735 | 0.899 | 0.439 | 1.74828330853657e-11 | 38 |
| si:ch73-189n23.112 | 8.67433869500899e-16 | 0.702622548 | 0.913 | 0.526 | 2.18324430614681e-11 | 38 |
| zp2.211 | 1.2131325194465e-15 | 0.697244133 | 0.942 | 0.531 | 3.0533332381949e-11 | 38 |
| cldnd12 | 1.47274063390388e-15 | 0.67629954 | 0.986 | 0.605 | 3.70674090147267e-11 | 38 |
| zp2.611 | 1.73500512351366e-15 | 0.714650484 | 0.899 | 0.503 | 4.36683439537152e-11 | 38 |
| CR790368.112 | 1.92515960555314e-15 | 0.696242091 | 0.942 | 0.521 | 4.84543421121669e-11 | 38 |
| kpna711 | 3.56853208941601e-15 | 0.761020132 | 0.739 | 0.329 | 8.98163841585115e-11 | 38 |
| epcam12 | 4.41445680652868e-15 | 0.709594427 | 0.942 | 0.506 | 1.1110746336352e-10 | 38 |
| si:ch211-226h8.1411 | 4.49725282893822e-15 | 0.68397279 | 0.957 | 0.568 | 1.13191356451546e-10 | 38 |
| zp2.311 | 7.1956973811261e-15 | 0.703105929 | 0.928 | 0.52 | 1.81108507385563e-10 | 38 |
| si:ch211-125e6.1211 | 8.43944402534131e-15 | 0.704678476 | 0.913 | 0.502 | 2.12412366673815e-10 | 38 |
| siva111 | 1.08665390460359e-14 | 0.723756227 | 0.855 | 0.429 | 2.73499921249677e-10 | 38 |
| zp3b11 | 1.17519354734903e-14 | 0.731585697 | 0.768 | 0.389 | 2.95784463932277e-10 | 38 |
| si:dkey-241l7.411 | 1.665409972911e-14 | 0.631373387 | 0.971 | 0.61 | 4.19167036081969e-10 | 38 |
| npm2b11 | 4.26933604358197e-14 | 0.722780963 | 0.783 | 0.397 | 1.07454918880915e-09 | 38 |
| zp3.212 | 4.85241808395495e-14 | 0.664675069 | 0.942 | 0.559 | 1.22130510755062e-09 | 38 |
| LOC11043782111 | 5.97675542350204e-14 | 0.731771646 | 0.855 | 0.416 | 1.50428957254123e-09 | 38 |
| mt211 | 9.24936702902037e-14 | 0.667167817 | 0.942 | 0.536 | 2.32797318753414e-09 | 38 |
| zp312 | 9.4765622997943e-14 | 0.642923681 | 0.957 | 0.585 | 2.38515596523523e-09 | 38 |
| si:ch211-14a17.711 | 1.03642041668292e-13 | 0.711959608 | 0.826 | 0.409 | 2.60856654674924e-09 | 38 |
| ccna111 | 1.12296558951274e-13 | 0.678193792 | 0.855 | 0.477 | 2.82639209224462e-09 | 38 |
| zp2.112 | 1.21948869152027e-13 | 0.675600744 | 0.942 | 0.536 | 3.06933108768737e-09 | 38 |
| tmsb511 | 2.09915487870648e-13 | 0.682490179 | 0.899 | 0.49 | 5.28336291421633e-09 | 38 |
| si:dkeyp-98a7.911 | 2.37352563990909e-13 | 0.675333911 | 0.928 | 0.513 | 5.97392668308719e-09 | 38 |
| zgc:15349911 | 3.50727870197636e-13 | 0.693004105 | 0.942 | 0.514 | 8.8274697650043e-09 | 38 |
| zp3a.211 | 7.56217961887956e-13 | 0.662949024 | 0.87 | 0.492 | 1.9033249882758e-08 | 38 |

| | | | | | | |
|---|---|---|---|---|---|---|
| pabpn1l11 | 2.0112424295117e-12 | 0.692490451 | 0.652 | 0.285 | 5.062096070838e-08 | 38 |
| crp211 | 4.81217094190321e-12 | 0.666061191 | 0.913 | 0.49 | 1.21117530436762e-07 | 38 |
| cldng11 | 4.93579355752696e-12 | 0.645418042 | 0.913 | 0.515 | 1.24228988049396e-07 | 38 |
| zglp111 | 6.49873331915982e-12 | 0.674312222 | 0.522 | 0.199 | 1.63566618909934e-07 | 38 |
| si:ch211-146l10.811 | 6.50442501692558e-12 | 0.688268863 | 0.652 | 0.308 | 1.63709873251e-07 | 38 |
| si:ch211-12h2.811 | 7.02669386062862e-12 | 0.621491146 | 0.913 | 0.549 | 1.76854857778162e-07 | 38 |
| LOC10053516211 | 1.03491881350227e-11 | 0.691432839 | 0.667 | 0.31 | 2.60478716170386e-07 | 38 |
| si:dkey-241l7.512 | 1.61336695495733e-11 | 0.590529764 | 0.913 | 0.585 | 4.0606832889321e-07 | 38 |
| si:ch73-90k17.111 | 2.59549710109271e-11 | 0.679612381 | 0.594 | 0.25 | 6.53260665374024e-07 | 38 |
| zgc:17513512 | 2.62619655118453e-11 | 0.629836217 | 0.826 | 0.481 | 6.60987409967635e-07 | 38 |
| si:dkey-152b24.611 | 3.26098898077605e-11 | 0.674821241 | 0.652 | 0.299 | 8.20758316571523e-07 | 38 |
| lamp211 | 3.42380072078014e-11 | 0.58617982 | 0.87 | 0.556 | 8.61736403413154e-07 | 38 |
| cdca810 | 3.68693814064086e-11 | 0.63839996 | 0.449 | 0.166 | 9.27965460617898e-07 | 38 |
| p2rx2 | 5.95395372720909e-11 | 0.295190355 | 0.101 | 0.012 | 1.49855061360126e-06 | 38 |
| si:zfos-1505d6.311 | 7.57077577046235e-11 | 0.631701238 | 0.913 | 0.514 | 1.90548855366767e-06 | 38 |
| qdprb211 | 9.2170542300269e-11 | 0.630019287 | 0.783 | 0.434 | 2.31984037915547e-06 | 38 |
| zgc:17177611 | 1.6799364592463e-10 | 0.608800016 | 0.855 | 0.497 | 4.22823207427701e-06 | 38 |
| BX649337.112 | 2.51462600178697e-10 | 0.628173124 | 0.826 | 0.439 | 6.32906218389761e-06 | 38 |
| zgc:17177911 | 2.67948023092214e-10 | 0.603238478 | 0.87 | 0.514 | 6.74398379320794e-06 | 38 |
| s100a111 | 2.91523290779623e-10 | 0.62628542 | 0.71 | 0.365 | 7.33734970563233e-06 | 38 |
| si:dkeyp-98a7.411 | 5.21294769433349e-10 | 0.615097052 | 0.797 | 0.426 | 1.3120468051868e-05 | 38 |
| map1lc3c11 | 6.86535341242777e-10 | 0.626590567 | 0.493 | 0.203 | 1.72794080037395e-05 | 38 |
| gtf3ab12 | 1.18176540908841e-09 | 0.623925541 | 0.667 | 0.322 | 2.97438535813461e-05 | 38 |
| cpeb1b11 | 1.65125492597172e-09 | 0.619147754 | 0.565 | 0.272 | 4.15604352317823e-05 | 38 |
| si:dkey-241l7.611 | 1.89731086987092e-09 | 0.560600716 | 0.826 | 0.528 | 4.77534172837811e-05 | 38 |
| rasl11b8 | 1.89745924158162e-09 | 0.58433657 | 0.377 | 0.135 | 4.77571516513679e-05 | 38 |
| zgc:5669912 | 3.00565874092106e-09 | 0.614858817 | 0.551 | 0.252 | 7.5649424850242e-05 | 38 |
| tuba4l11 | 3.47722706023071e-09 | 0.592136452 | 0.739 | 0.406 | 8.75183278789467e-05 | 38 |
| si:dkey-229d11.511 | 3.97376271167536e-09 | 0.587620948 | 0.696 | 0.391 | 0.000100016 | 38 |
| cth | 4.87179670561531e-09 | 0.464852738 | 0.188 | 0.045 | 0.000122618 | 38 |
| zgc:5667610 | 5.58223848369384e-09 | 0.605418591 | 0.565 | 0.296 | 0.000140499 | 38 |
| LOC10188547711 | 5.97501470323866e-09 | 0.610744663 | 0.58 | 0.281 | 0.000150385 | 38 |
| slc35f22 | 7.31851513358153e-09 | 0.483676493 | 0.261 | 0.076 | 0.0001842 | 38 |
| zgc:11342410 | 1.04824450731375e-08 | 0.580709999 | 0.406 | 0.163 | 0.000263833 | 38 |
| AL929032.111 | 1.12356616946991e-08 | 0.596147895 | 0.623 | 0.319 | 0.00028279 | 38 |

| | | | | | | |
|---|---|---|---|---|---|---|
| zgc:5546112 | 1.15579502984766e-08 | 0.595569213 | 0.594 | 0.29 | 0.000290902 | 38 |
| si:dkey-90l23.111 | 1.25730945359676e-08 | 0.56918368 | 0.768 | 0.449 | 0.000316452 | 38 |
| zgc:17344311 | 1.30788918769638e-08 | 0.588187178 | 0.638 | 0.332 | 0.000329183 | 38 |
| zgc:17151711 | 1.4279400371477e-08 | 0.573983486 | 0.71 | 0.39 | 0.000359398 | 38 |
| dnd17 | 1.51290862941616e-08 | 0.54119815 | 0.319 | 0.111 | 0.000380784 | 38 |
| ccnb211 | 1.73306803535981e-08 | 0.575795843 | 0.667 | 0.386 | 0.000436196 | 38 |
| zgc:17147411 | 1.75314282721365e-08 | 0.584688319 | 0.594 | 0.305 | 0.000441249 | 38 |
| si:dkey-90l23.211 | 1.94411067997897e-08 | 0.578319925 | 0.638 | 0.346 | 0.000489313 | 38 |
| si:ch211-132b12.811 | 2.16014239791821e-08 | 0.580435725 | 0.623 | 0.344 | 0.000543686 | 38 |
| slbp211 | 2.67788416189789e-08 | 0.569538546 | 0.696 | 0.399 | 0.000673997 | 38 |
| si:ch211-5k11.811 | 2.68703893351921e-08 | 0.466949494 | 0.725 | 0.359 | 0.000676301 | 38 |
| si:dkeyp-46h3.611 | 2.76069770296226e-08 | 0.564170803 | 0.696 | 0.412 | 0.00069484 | 38 |
| gapdh12 | 2.78746777554917e-08 | 0.548761695 | 0.681 | 0.422 | 0.000701578 | 38 |
| zar111 | 3.36334112675624e-08 | 0.549075687 | 0.739 | 0.441 | 0.000846519 | 38 |
| si:dkeyp-98a7.811 | 4.00318096674746e-08 | 0.554514192 | 0.783 | 0.459 | 0.001007561 | 38 |
| CT009596.17 | 4.05805111332095e-08 | 0.512555465 | 0.275 | 0.091 | 0.001021371 | 38 |
| si:ch211-12h2.611 | 4.70875332873028e-08 | 0.538252644 | 0.768 | 0.476 | 0.001185146 | 38 |
| si:ch211-145c1.111 | 7.00387006801205e-08 | 0.568407944 | 0.507 | 0.235 | 0.001762804 | 38 |
| zgc:13625411 | 8.92094288177681e-08 | 0.553234206 | 0.638 | 0.332 | 0.002245312 | 38 |
| nasp11 | 9.76279955808013e-08 | 0.554978132 | 0.623 | 0.341 | 0.002457199 | 38 |
| fabp7a3 | 1.12786312162049e-07 | 0.331358741 | 0.13 | 0.027 | 0.002838719 | 38 |
| zgc:7711811 | 1.1374349082596e-07 | 0.535636715 | 0.754 | 0.459 | 0.00286281 | 38 |
| si:dkeyp-46h3.811 | 1.33174097417099e-07 | 0.545823831 | 0.638 | 0.353 | 0.003351859 | 38 |
| lto1 | 1.33512717314292e-07 | 0.397675156 | 0.174 | 0.044 | 0.003360382 | 38 |
| ctss2.112 | 1.37302981065718e-07 | 0.477043484 | 0.768 | 0.536 | 0.003455779 | 38 |
| uchl111 | 1.39728302052579e-07 | 0.557633128 | 0.609 | 0.346 | 0.003516822 | 38 |
| grip2a | 1.65089890357164e-07 | 0.381522791 | 0.159 | 0.038 | 0.004155147 | 38 |
| avd11 | 1.78011335612595e-07 | 0.557649111 | 0.536 | 0.267 | 0.004480367 | 38 |
| npm2a2 | 2.35639992915186e-07 | 0.464131065 | 0.246 | 0.08 | 0.005930823 | 38 |
| btg411 | 2.65281688371742e-07 | 0.542513345 | 0.681 | 0.37 | 0.006676875 | 38 |
| asb3 | 3.43893427283765e-07 | 0.314951907 | 0.145 | 0.033 | 0.008655454 | 38 |
| hspa59 | 3.46735022604645e-07 | 0.53853454 | 0.551 | 0.316 | 0.008726974 | 38 |
| birc5b11 | 3.73361231415104e-07 | 0.552072494 | 0.507 | 0.261 | 0.009397129 | 38 |
| si:ch211-250e5.1611 | 5.88678423909659e-07 | 0.521115612 | 0.609 | 0.348 | 0.014816447 | 38 |
| ccng11 | 7.17784559252484e-07 | 0.471429556 | 0.652 | 0.449 | 0.01806592 | 38 |

| | | | | | | |
|---|---|---|---|---|---|---|
| wu:fi42e0311 | 9.74282704755349e-07 | 0.520768824 | 0.58 | 0.352 | 0.024521721 | 38 |
| si:dkeyp-46h3.511 | 1.03124549181595e-06 | 0.527091485 | 0.536 | 0.291 | 0.025955418 | 38 |
| lsm14b11 | 1.22963928371985e-06 | 0.516781091 | 0.565 | 0.301 | 0.030948791 | 38 |
| rcc111 | 1.2324401002209e-06 | 0.531023457 | 0.42 | 0.204 | 0.031019285 | 38 |
| sept4a | 1.73947068898911e-06 | 0.254200971 | 0.101 | 0.02 | 0.043780738 | 38 |
| prdx111 | 1.92690322024244e-06 | 0.498486235 | 0.377 | 0.168 | 0.048498227 | 38 |
| LOC10053749911 | 2.41445456268435e-06 | 0.503480092 | 0.565 | 0.324 | 0.060769407 | 38 |
| hbba111 | 2.68642058079782e-06 | 0.504927124 | 0.623 | 0.358 | 0.06761452 | 38 |
| zp2l110 | 2.69138330777745e-06 | 0.473702217 | 0.377 | 0.165 | 0.067739426 | 38 |
| racgap1 | 2.69836678833651e-06 | 0.370564333 | 0.159 | 0.044 | 0.067915194 | 38 |
| zpcx11 | 2.75297403629343e-06 | 0.500956388 | 0.594 | 0.341 | 0.069289604 | 38 |
| si:ch73-14h10.211 | 2.93708125851171e-06 | 0.498812997 | 0.522 | 0.273 | 0.073923398 | 38 |
| BX537276.111 | 2.95441398290176e-06 | 0.454680156 | 0.304 | 0.123 | 0.074359646 | 38 |
| si:dkey-37o8.111 | 3.31936914449001e-06 | 0.501840982 | 0.551 | 0.309 | 0.083545202 | 38 |
| timm132 | 3.41584126002199e-06 | 0.471442888 | 0.261 | 0.1 | 0.085973309 | 38 |
| pdp2 | 3.47533655479495e-06 | 0.28840893 | 0.116 | 0.026 | 0.087470746 | 38 |
| orc3 | 3.57928363368786e-06 | 0.374456308 | 0.174 | 0.052 | 0.09008699 | 38 |
| zgc:17197710 | 3.82151920281685e-06 | 0.506716819 | 0.449 | 0.232 | 0.096183817 | 38 |
| spink2.112 | 3.87020419660208e-06 | 0.456819638 | 0.667 | 0.438 | 0.097409169 | 38 |
| mibp10 | 4.19013239663065e-06 | 0.497302699 | 0.435 | 0.224 | 0.105461442 | 38 |
| arhgef10la11 | 4.41814386798278e-06 | 0.4905484 | 0.536 | 0.276 | 0.111200263 | 38 |
| pabpc1l10 | 4.93290344990369e-06 | 0.474311912 | 0.333 | 0.143 | 0.124156247 | 38 |
| cth111 | 4.94473155061521e-06 | 0.482962603 | 0.638 | 0.381 | 0.124453948 | 38 |
| cax12 | 5.10541998606178e-06 | 0.370232714 | 0.188 | 0.059 | 0.128498316 | 38 |
| mgme1 | 5.72505333132099e-06 | 0.333622765 | 0.145 | 0.039 | 0.144093867 | 38 |
| LOC10188389711 | 5.91065088185605e-06 | 0.472773803 | 0.362 | 0.166 | 0.148765172 | 38 |
| CU929143.11 | 6.21384223472705e-06 | 0.426549409 | 0.246 | 0.092 | 0.156396195 | 38 |
| zgc:1625091 | 6.91431167356169e-06 | 0.295389972 | 0.116 | 0.027 | 0.174026311 | 38 |
| larp6b11 | 6.92821162893845e-06 | 0.487475044 | 0.406 | 0.198 | 0.174376158 | 38 |
| mrps18b4 | 7.20702680917955e-06 | 0.436446274 | 0.304 | 0.127 | 0.181393658 | 38 |
| org11 | 7.91639108829596e-06 | 0.486328713 | 0.464 | 0.242 | 0.199247647 | 38 |
| elovl7b11 | 8.92200639940548e-06 | 0.490356583 | 0.464 | 0.25 | 0.224557979 | 38 |
| CU855930.1 | 9.2492758474396e-06 | 0.317961095 | 0.145 | 0.04 | 0.232795024 | 38 |
| prr115 | 9.57777195171851e-06 | 0.413238331 | 0.261 | 0.101 | 0.241062942 | 38 |
| zar1l11 | 9.76236615501983e-06 | 0.475317204 | 0.667 | 0.4 | 0.245708994 | 38 |

| | | | | | | |
|---|---|---|---|---|---|---|
| mos10 | 9.77610054880926e-06 | 0.456223527 | 0.319 | 0.138 | 0.246054675 | 38 |
| rbm4.311 | 1.02416165778964e-05 | 0.4402397 | 0.623 | 0.446 | 0.257771248 | 38 |
| si:dkey-241l7.311 | 1.12016209731165e-05 | 0.473379186 | 0.594 | 0.366 | 0.281933598 | 38 |
| CU571163.111 | 1.12315318747569e-05 | 0.476208981 | 0.667 | 0.403 | 0.282686426 | 38 |
| rbpms2b11 | 1.14299394533497e-05 | 0.474118627 | 0.435 | 0.223 | 0.287680146 | 38 |
| zgc:19524511 | 1.14816396140375e-05 | 0.471611692 | 0.667 | 0.39 | 0.288981387 | 38 |
| si:ch211-276i12.92 | 1.21827517900241e-05 | 0.395875744 | 0.188 | 0.064 | 0.30662768 | 38 |
| si:ch73-303b9.15 | 1.52519314203107e-05 | 0.422506873 | 0.217 | 0.08 | 0.383875862 | 38 |
| cdca7a11 | 1.64879416555627e-05 | 0.472622529 | 0.42 | 0.224 | 0.414985004 | 38 |
| zp3a.111 | 1.69992265778795e-05 | 0.461579154 | 0.609 | 0.383 | 0.427853534 | 38 |
| si:dkeyp-46h3.211 | 1.70623777210095e-05 | 0.451430831 | 0.623 | 0.406 | 0.429442985 | 38 |
| gdf311 | 1.76378057649638e-05 | 0.479981408 | 0.435 | 0.225 | 0.443925933 | 38 |
| tyms1 | 1.78960556456128e-05 | 0.393945562 | 0.203 | 0.073 | 0.450425825 | 38 |
| ca15b11 | 1.91731045150579e-05 | 0.465254711 | 0.478 | 0.251 | 0.482567868 | 38 |
| thy112 | 1.97944589371645e-05 | 0.450442699 | 0.71 | 0.452 | 0.498206737 | 38 |
| abcf2a1 | 2.23550806974424e-05 | 0.462242839 | 0.319 | 0.152 | 0.562655026 | 38 |
| ctsla11 | 2.39983417304529e-05 | 0.441102504 | 0.565 | 0.388 | 0.604014263 | 38 |
| gstp18 | 2.59356115338383e-05 | 0.456352801 | 0.565 | 0.367 | 0.652773407 | 38 |
| pde6d | 2.65387939366483e-05 | 0.341339093 | 0.145 | 0.043 | 0.667954905 | 38 |
| fbxo431 | 2.76707149611681e-05 | 0.351039363 | 0.203 | 0.071 | 0.696444225 | 38 |
| si:ch211-226l4.612 | 2.80775343674818e-05 | 0.461816298 | 0.507 | 0.295 | 0.706683462 | 38 |
| CDPF111 | 2.94268519066037e-05 | 0.470355251 | 0.464 | 0.27 | 0.740644436 | 38 |
| anp32b6 | 2.98181633879741e-05 | 0.485528726 | 0.435 | 0.257 | 0.750493354 | 38 |
| tmem266 | 3.00504004302028e-05 | 0.269386871 | 0.116 | 0.03 | 0.756338528 | 38 |
| si:ch1073-190k2.111 | 3.12454183472518e-05 | 0.460941941 | 0.377 | 0.19 | 0.786415934 | 38 |
| ccnb111 | 3.13991813791997e-05 | 0.444299609 | 0.609 | 0.395 | 0.790285996 | 38 |
| dhrs13b | 3.20161436989951e-05 | 0.332951068 | 0.145 | 0.043 | 0.805814321 | 38 |
| rrm211 | 3.30524835771531e-05 | 0.420753966 | 0.739 | 0.5 | 0.831897959 | 38 |
| zgc:17175010 | 3.81700924687033e-05 | 0.447960972 | 0.348 | 0.176 | 0.960703057 | 38 |
| zgc:1532844 | 3.84691990150098e-05 | 0.417276541 | 0.246 | 0.101 | 0.96823127 | 38 |
| srsf2a8 | 4.00644881940402e-05 | 0.433515023 | 0.551 | 0.368 | 1 | 38 |
| cep55l | 4.16527995617925e-05 | 0.281345603 | 0.101 | 0.025 | 1 | 38 |
| degs28 | 4.3212898047827e-05 | 0.386603847 | 0.261 | 0.108 | 1 | 38 |
| CU655961.34 | 4.32562491004115e-05 | 0.392070541 | 0.246 | 0.1 | 1 | 38 |
| snu13b12 | 4.6198525656131e-05 | 0.432484261 | 0.58 | 0.394 | 1 | 38 |

| | | | | | | |
|---|---|---|---|---|---|---|
| CU655961.45 | 4.71940028090936e-05 | 0.366518326 | 0.232 | 0.09 | 1 | 38 |
| chac2 | 5.24261229427858e-05 | 0.258714482 | 0.101 | 0.025 | 1 | 38 |
| tmem54a9 | 5.68721378797788e-05 | 0.433662638 | 0.304 | 0.143 | 1 | 38 |
| patl210 | 5.75302119530839e-05 | 0.452962367 | 0.348 | 0.175 | 1 | 38 |
| zgc:16287911 | 5.83275843430291e-05 | 0.432208635 | 0.362 | 0.183 | 1 | 38 |
| ppp2r2ab | 6.27136954418172e-05 | 0.256033677 | 0.116 | 0.031 | 1 | 38 |
| zgc:1134252 | 6.77093819198848e-05 | 0.373324234 | 0.188 | 0.07 | 1 | 38 |
| si:dkey-208k4.211 | 6.85865131932872e-05 | 0.434347487 | 0.507 | 0.298 | 1 | 38 |
| FP074874.2 | 7.08569362075628e-05 | 0.265658126 | 0.13 | 0.038 | 1 | 38 |
| larp6a11 | 7.97739800306973e-05 | 0.450796728 | 0.435 | 0.251 | 1 | 38 |
| mrpl221 | 8.85854710089709e-05 | 0.369815399 | 0.188 | 0.071 | 1 | 38 |
| thop15 | 9.02687846840937e-05 | 0.427235907 | 0.261 | 0.118 | 1 | 38 |
| zgc:923136 | 9.48538020536883e-05 | 0.427358651 | 0.275 | 0.128 | 1 | 38 |
| LOC1005380234 | 9.78485068684403e-05 | 0.359678774 | 0.203 | 0.078 | 1 | 38 |
| ubc3 | 0.000101578 | 0.271782063 | 0.754 | 0.659 | 1 | 38 |
| ccna211 | 0.000104864 | 0.433945864 | 0.493 | 0.286 | 1 | 38 |
| aurkb5 | 0.000105411 | 0.434284201 | 0.275 | 0.129 | 1 | 38 |
| LOC1104377795 | 0.000108654 | 0.336042441 | 0.217 | 0.085 | 1 | 38 |
| si:ch211-14a17.65 | 0.000109961 | 0.363266404 | 0.217 | 0.087 | 1 | 38 |
| zp3c11 | 0.000112465 | 0.437565905 | 0.362 | 0.191 | 1 | 38 |
| dnajc5ga | 0.000130757 | 0.336602232 | 0.188 | 0.071 | 1 | 38 |
| tmem144b5 | 0.000136369 | 0.355257999 | 0.232 | 0.095 | 1 | 38 |
| si:ch73-160p18.311 | 0.000137732 | 0.421997007 | 0.435 | 0.242 | 1 | 38 |
| ccne19 | 0.000138452 | 0.355252622 | 0.261 | 0.112 | 1 | 38 |
| zgc:1714263 | 0.000142284 | 0.387805122 | 0.246 | 0.109 | 1 | 38 |
| tmem442 | 0.000144049 | 0.30091246 | 0.174 | 0.062 | 1 | 38 |
| hbaa111 | 0.000149841 | 0.360877104 | 0.507 | 0.283 | 1 | 38 |
| si:ch73-204p21.2 | 0.000157277 | 0.336399696 | 0.145 | 0.05 | 1 | 38 |
| si:dkey-19b23.1111 | 0.000157706 | 0.413023432 | 0.478 | 0.281 | 1 | 38 |
| zdhhc18a | 0.000158551 | 0.314288746 | 0.159 | 0.056 | 1 | 38 |
| pigrl2.311 | 0.000159366 | 0.427339902 | 0.391 | 0.205 | 1 | 38 |
| kmt5ab4 | 0.00016251 | 0.378943537 | 0.217 | 0.091 | 1 | 38 |
| tpx26 | 0.000163065 | 0.393459756 | 0.275 | 0.127 | 1 | 38 |
| lrrc47 | 0.00016692 | 0.318291678 | 0.159 | 0.057 | 1 | 38 |
| kif223 | 0.00017016 | 0.341914921 | 0.217 | 0.089 | 1 | 38 |

| | | | | | | |
|---|---|---|---|---|---|---|
| gnmt | 0.000177896 | 0.256445864 | 0.101 | 0.028 | 1 | 38 |
| akirin2 | 0.000178383 | 0.389962081 | 0.232 | 0.103 | 1 | 38 |
| CABZ01075618.12 | 0.000190205 | 0.310474912 | 0.159 | 0.057 | 1 | 38 |
| sept151 | 0.00019561 | 0.314454485 | 0.13 | 0.042 | 1 | 38 |
| cldn7b10 | 0.000205024 | 0.390399738 | 0.333 | 0.161 | 1 | 38 |
| polr1d1 | 0.000217694 | 0.34846156 | 0.217 | 0.092 | 1 | 38 |
| LOC103909742 | 0.00025753 | 0.276334726 | 0.13 | 0.042 | 1 | 38 |
| ppp1r3b5 | 0.000259557 | 0.374301075 | 0.246 | 0.111 | 1 | 38 |
| mcm3l5 | 0.000260203 | 0.31163286 | 0.217 | 0.089 | 1 | 38 |
| sox19b8 | 0.000270904 | 0.328526966 | 0.232 | 0.099 | 1 | 38 |
| hnrnpabb12 | 0.000292369 | 0.326333447 | 0.739 | 0.598 | 1 | 38 |
| si:dkeyp-50f7.211 | 0.000299725 | 0.394331048 | 0.594 | 0.405 | 1 | 38 |
| plk110 | 0.000331127 | 0.414715371 | 0.333 | 0.179 | 1 | 38 |
| tram26 | 0.000358313 | 0.380674535 | 0.275 | 0.134 | 1 | 38 |
| si:ch211-156b7.41 | 0.000362139 | 0.253872609 | 0.116 | 0.036 | 1 | 38 |
| mcfd24 | 0.000369484 | 0.36418264 | 0.246 | 0.115 | 1 | 38 |
| sybu11 | 0.000387996 | 0.370319121 | 0.333 | 0.168 | 1 | 38 |
| CR847984.111 | 0.000391274 | 0.410161204 | 0.319 | 0.166 | 1 | 38 |
| mkrn410 | 0.000395546 | 0.415560826 | 0.391 | 0.23 | 1 | 38 |
| zgc:17374211 | 0.000403002 | 0.397723166 | 0.304 | 0.157 | 1 | 38 |
| gins3 | 0.000406321 | 0.267059586 | 0.13 | 0.044 | 1 | 38 |
| zgc:174935 | 0.000414925 | 0.259224286 | 0.101 | 0.03 | 1 | 38 |
| mpzl2b2 | 0.000425033 | 0.342488158 | 0.217 | 0.096 | 1 | 38 |
| zgc:5623111 | 0.000442934 | 0.388010514 | 0.406 | 0.23 | 1 | 38 |
| BX005461.33 | 0.000496685 | 0.302560837 | 0.174 | 0.068 | 1 | 38 |
| rpp25l | 0.000529002 | 0.319762746 | 0.174 | 0.07 | 1 | 38 |
| si:ch211-251f6.611 | 0.000534621 | 0.401436287 | 0.493 | 0.334 | 1 | 38 |
| cdca95 | 0.000567452 | 0.321721174 | 0.203 | 0.087 | 1 | 38 |
| jtb1 | 0.000579566 | 0.292653546 | 0.159 | 0.061 | 1 | 38 |
| umps2 | 0.000597997 | 0.310716408 | 0.188 | 0.078 | 1 | 38 |
| CR383662.26 | 0.000600097 | 0.336558704 | 0.203 | 0.088 | 1 | 38 |
| h3f3a12 | 0.000606952 | 0.337489686 | 0.638 | 0.507 | 1 | 38 |
| asna1 | 0.000613837 | 0.317249638 | 0.174 | 0.07 | 1 | 38 |
| st14a | 0.000621055 | 0.268819351 | 0.116 | 0.038 | 1 | 38 |
| gmnn11 | 0.000622061 | 0.388421702 | 0.464 | 0.305 | 1 | 38 |

| | | | | | | |
|---|---|---|---|---|---|---|
| CABZ01069022.22 | 0.000629341 | 0.311605204 | 0.159 | 0.062 | 1 | 38 |
| akap1a | 0.000629999 | 0.308052725 | 0.159 | 0.062 | 1 | 38 |
| tubb2b4 | 0.000676016 | 0.381234803 | 0.406 | 0.25 | 1 | 38 |
| si:ch73-340n8.44 | 0.000683538 | 0.363513122 | 0.261 | 0.129 | 1 | 38 |
| hibadhb | 0.000684051 | 0.333931834 | 0.145 | 0.056 | 1 | 38 |
| pcf114 | 0.000713012 | 0.387555744 | 0.391 | 0.241 | 1 | 38 |
| hsd3b21 | 0.000718266 | 0.283088762 | 0.159 | 0.061 | 1 | 38 |
| stard14 | 0.000766175 | 0.269459622 | 0.145 | 0.053 | 1 | 38 |
| ndc13 | 0.000776961 | 0.327687516 | 0.217 | 0.098 | 1 | 38 |
| zgc:1526529 | 0.000806441 | 0.331488916 | 0.246 | 0.115 | 1 | 38 |
| zgc:16303010 | 0.000865317 | 0.337669346 | 0.261 | 0.126 | 1 | 38 |
| gtf2a12 | 0.000897468 | 0.383545992 | 0.29 | 0.157 | 1 | 38 |
| coil4 | 0.000904262 | 0.324659015 | 0.232 | 0.109 | 1 | 38 |
| ube2c8 | 0.000920142 | 0.351274976 | 0.275 | 0.142 | 1 | 38 |
| zgc:165514 | 0.000941512 | 0.281947953 | 0.145 | 0.055 | 1 | 38 |
| zgc:66160 | 0.000946501 | 0.339163684 | 0.174 | 0.074 | 1 | 38 |
| fbxw11b4 | 0.000951278 | 0.378889363 | 0.275 | 0.147 | 1 | 38 |
| mrps6 | 0.000996939 | 0.329499563 | 0.203 | 0.093 | 1 | 38 |
| top1l3 | 0.00103878 | 0.398494432 | 0.362 | 0.23 | 1 | 38 |
| si:dkey-188i13.1112 | 0.001068872 | 0.394555922 | 0.449 | 0.318 | 1 | 38 |
| yrdc9 | 0.00110428 | 0.370009417 | 0.246 | 0.124 | 1 | 38 |
| si:dkey-92i17.23 | 0.00113184 | 0.346017609 | 0.203 | 0.093 | 1 | 38 |
| gfer4 | 0.001154315 | 0.314539579 | 0.217 | 0.1 | 1 | 38 |
| zgc:1118688 | 0.001182071 | 0.378008539 | 0.275 | 0.149 | 1 | 38 |
| si:rp71-1g18.1 | 0.001252725 | 0.261111782 | 0.116 | 0.04 | 1 | 38 |
| retsatl11 | 0.001309067 | 0.383939353 | 0.348 | 0.205 | 1 | 38 |
| haus610 | 0.001324759 | 0.356481094 | 0.377 | 0.211 | 1 | 38 |
| LOC1039099726 | 0.001325498 | 0.361876383 | 0.493 | 0.369 | 1 | 38 |
| BX571780.18 | 0.001349944 | 0.310433261 | 0.188 | 0.082 | 1 | 38 |
| zgc:1742681 | 0.001352518 | 0.321136624 | 0.188 | 0.084 | 1 | 38 |
| gpx81 | 0.001379152 | 0.322835412 | 0.188 | 0.085 | 1 | 38 |
| zgc:6602411 | 0.001413668 | 0.342156806 | 0.275 | 0.143 | 1 | 38 |
| wu:fb18f06 | 0.001467923 | 0.281905897 | 0.145 | 0.057 | 1 | 38 |
| imp38 | 0.001476889 | 0.34335422 | 0.261 | 0.136 | 1 | 38 |
| gyg1a12 | 0.001496656 | 0.360425938 | 0.522 | 0.372 | 1 | 38 |

| | | | | | | |
|---|---|---|---|---|---|---|
| fam89a | 0.001537878 | 0.264793621 | 0.13 | 0.049 | 1 | 38 |
| zgc:1139847 | 0.001581639 | 0.326154048 | 0.217 | 0.103 | 1 | 38 |
| zgc:9259111 | 0.001608548 | 0.346929418 | 0.362 | 0.208 | 1 | 38 |
| nup371 | 0.001621042 | 0.259256005 | 0.116 | 0.042 | 1 | 38 |
| krcp2 | 0.001669076 | 0.303601156 | 0.174 | 0.076 | 1 | 38 |
| pithd1 | 0.001745101 | 0.314626366 | 0.174 | 0.077 | 1 | 38 |
| si:ch73-78o10.19 | 0.001749239 | 0.33601664 | 0.261 | 0.135 | 1 | 38 |
| gpatch4 | 0.001858013 | 0.312774805 | 0.145 | 0.06 | 1 | 38 |
| pou5f310 | 0.001873491 | 0.353789442 | 0.275 | 0.148 | 1 | 38 |
| tom1 | 0.001997791 | 0.256577573 | 0.116 | 0.042 | 1 | 38 |
| apopt1 | 0.002030321 | 0.268006741 | 0.13 | 0.051 | 1 | 38 |
| uck2a2 | 0.002039263 | 0.337680741 | 0.232 | 0.118 | 1 | 38 |
| wdr821 | 0.00208719 | 0.3007573 | 0.188 | 0.087 | 1 | 38 |
| phex6 | 0.002095101 | 0.325607855 | 0.188 | 0.088 | 1 | 38 |
| ndufab1b2 | 0.002221179 | 0.371682655 | 0.275 | 0.16 | 1 | 38 |
| cx43.46 | 0.002289288 | 0.308544179 | 0.246 | 0.125 | 1 | 38 |
| crp311 | 0.002397694 | 0.352679492 | 0.362 | 0.22 | 1 | 38 |
| si:ch73-15b2.56 | 0.002465349 | 0.299585424 | 0.203 | 0.096 | 1 | 38 |
| triap1 | 0.002507165 | 0.265396831 | 0.116 | 0.044 | 1 | 38 |
| adat31 | 0.002507324 | 0.270695313 | 0.116 | 0.044 | 1 | 38 |
| si:dkeyp-114g9.15 | 0.002524254 | 0.290757586 | 0.217 | 0.106 | 1 | 38 |
| ddx414 | 0.002550586 | 0.292962139 | 0.159 | 0.07 | 1 | 38 |
| zgc:1536751 | 0.002557291 | 0.30076882 | 0.159 | 0.07 | 1 | 38 |
| CABZ01059627.211 | 0.002624983 | 0.298259632 | 0.261 | 0.135 | 1 | 38 |
| tcea2 | 0.002640635 | 0.287805062 | 0.13 | 0.053 | 1 | 38 |
| nmt1a | 0.002650726 | 0.313290193 | 0.145 | 0.062 | 1 | 38 |
| BX284697.19 | 0.002746192 | 0.299731311 | 0.232 | 0.116 | 1 | 38 |
| zgc:558884 | 0.002773743 | 0.283038425 | 0.188 | 0.087 | 1 | 38 |
| nanog11 | 0.003050806 | 0.330563372 | 0.522 | 0.362 | 1 | 38 |
| bola31 | 0.003297954 | 0.29183985 | 0.159 | 0.072 | 1 | 38 |
| snu13a6 | 0.00331718 | 0.317141565 | 0.203 | 0.102 | 1 | 38 |
| cdc63 | 0.003385277 | 0.336864513 | 0.203 | 0.104 | 1 | 38 |
| lyrm41 | 0.00342463 | 0.31944759 | 0.203 | 0.103 | 1 | 38 |
| BX957362.16 | 0.003483886 | 0.351062797 | 0.246 | 0.135 | 1 | 38 |
| tph1a11 | 0.003491473 | 0.32369024 | 0.261 | 0.143 | 1 | 38 |

| | | | | | | |
|---|---|---|---|---|---|---|
| zgc:771125 | 0.003594375 | 0.31394164 | 0.232 | 0.121 | 1 | 38 |
| nol72 | 0.003633155 | 0.334455337 | 0.232 | 0.125 | 1 | 38 |
| dynll2a11 | 0.003637542 | 0.307478465 | 0.261 | 0.138 | 1 | 38 |
| ap1s3b1 | 0.003640485 | 0.258725561 | 0.174 | 0.078 | 1 | 38 |
| elovl7a11 | 0.003785222 | 0.332836044 | 0.319 | 0.185 | 1 | 38 |
| orc66 | 0.003868806 | 0.28974867 | 0.217 | 0.109 | 1 | 38 |
| akip11 | 0.00387857 | 0.266388673 | 0.145 | 0.062 | 1 | 38 |
| selenoh6 | 0.003883222 | 0.356083331 | 0.362 | 0.246 | 1 | 38 |
| si:ch1073-263o8.211 | 0.004081451 | 0.325275073 | 0.42 | 0.266 | 1 | 38 |
| wwp2 | 0.004146252 | 0.303339808 | 0.145 | 0.065 | 1 | 38 |
| slbp12 | 0.004188859 | 0.332423228 | 0.377 | 0.236 | 1 | 38 |
| si:ch211-157c3.45 | 0.00438432 | 0.282411024 | 0.188 | 0.091 | 1 | 38 |
| tpst1l | 0.004533824 | 0.275999323 | 0.145 | 0.064 | 1 | 38 |
| efcab11 | 0.004570467 | 0.253563733 | 0.145 | 0.063 | 1 | 38 |
| sde2 | 0.004652128 | 0.281090959 | 0.145 | 0.065 | 1 | 38 |
| vdac11 | 0.004733151 | 0.380785877 | 0.275 | 0.172 | 1 | 38 |
| ppp4ca1 | 0.004866077 | 0.277884647 | 0.159 | 0.073 | 1 | 38 |
| si:dkey-87o1.22 | 0.004936374 | 0.285319613 | 0.174 | 0.083 | 1 | 38 |
| rpain7 | 0.005141465 | 0.288462494 | 0.232 | 0.121 | 1 | 38 |
| si:ch211-218c6.8 | 0.005362591 | 0.296646775 | 0.159 | 0.076 | 1 | 38 |
| gstt1a3 | 0.005388993 | 0.25787304 | 0.101 | 0.039 | 1 | 38 |
| buc2l5 | 0.005477801 | 0.259901567 | 0.174 | 0.082 | 1 | 38 |
| itprip | 0.005527883 | 0.255947047 | 0.116 | 0.047 | 1 | 38 |
| gadd45gip1 | 0.005660763 | 0.30243816 | 0.188 | 0.096 | 1 | 38 |
| elavl212 | 0.005714788 | 0.326824613 | 0.362 | 0.226 | 1 | 38 |
| tatdn24 | 0.005719801 | 0.262485299 | 0.159 | 0.074 | 1 | 38 |
| ppp4r2b1 | 0.006048801 | 0.275214335 | 0.159 | 0.076 | 1 | 38 |
| rhogc | 0.006465706 | 0.261514029 | 0.188 | 0.094 | 1 | 38 |
| alyref3 | 0.006757171 | 0.3269964 | 0.391 | 0.278 | 1 | 38 |
| itm2cb7 | 0.006762549 | 0.295684772 | 0.246 | 0.139 | 1 | 38 |
| diablob | 0.007004112 | 0.264868485 | 0.13 | 0.058 | 1 | 38 |
| snrpd22 | 0.007219242 | 0.352265638 | 0.362 | 0.263 | 1 | 38 |
| hsd17b10 | 0.007300419 | 0.293117185 | 0.159 | 0.078 | 1 | 38 |
| tm2d2 | 0.007306465 | 0.254003995 | 0.159 | 0.076 | 1 | 38 |
| hspe14 | 0.007461133 | 0.309037257 | 0.42 | 0.296 | 1 | 38 |

| | | | | | | |
|---|---|---|---|---|---|---|
| ell2 | 0.007475186 | 0.256922762 | 0.145 | 0.067 | 1 | 38 |
| COQ10A3 | 0.007483645 | 0.261153077 | 0.174 | 0.084 | 1 | 38 |
| AL929192.111 | 0.007608001 | 0.321618712 | 0.275 | 0.166 | 1 | 38 |
| commd4 | 0.007764881 | 0.251456794 | 0.13 | 0.058 | 1 | 38 |
| ccnh8 | 0.007788759 | 0.282213732 | 0.232 | 0.127 | 1 | 38 |
| hsd17b12a11 | 0.007863448 | 0.281258319 | 0.29 | 0.167 | 1 | 38 |
| ivns1abpb4 | 0.007949091 | 0.277889847 | 0.217 | 0.118 | 1 | 38 |
| polr2k4 | 0.007961902 | 0.314125373 | 0.304 | 0.192 | 1 | 38 |
| abhd17c | 0.008056489 | 0.254593257 | 0.13 | 0.058 | 1 | 38 |
| cse1l2 | 0.008918042 | 0.267903208 | 0.159 | 0.079 | 1 | 38 |
| snupn | 0.008954846 | 0.255690618 | 0.159 | 0.077 | 1 | 38 |
| rac1a5 | 0.009082899 | 0.320520924 | 0.246 | 0.15 | 1 | 38 |
| inka1b11 | 0.009389954 | 0.305195968 | 0.304 | 0.187 | 1 | 38 |
| zgc:1138869 | 0.009515339 | 0.288841325 | 0.246 | 0.143 | 1 | 38 |
| faub9 | 0.009625487 | 0.305373 | 0.319 | 0.202 | 1 | 38 |
| buc11 | 0.009642292 | 0.287282595 | 0.304 | 0.181 | 1 | 38 |
| BX005461.15 | 0.009788614 | 0.287041239 | 0.159 | 0.081 | 1 | 38 |
| krt5 | 0 | 2.568460793 | 1 | 0.009 | 0 | 39 |
| si:dkey-248g15.3 | 0 | 2.163154819 | 0.892 | 0.004 | 0 | 39 |
| si:dkey-193p11.21 | 0 | 1.792868911 | 0.723 | 0.011 | 0 | 39 |
| icn21 | 0 | 1.761013907 | 0.677 | 0.005 | 0 | 39 |
| krt97 | 0 | 1.759929592 | 0.662 | 0.005 | 0 | 39 |
| cldnb | 0 | 1.578166038 | 0.538 | 0.008 | 0 | 39 |
| CT027832.1 | 0 | 1.354181271 | 0.415 | 0 | 0 | 39 |
| spaca4l | 0 | 1.313510877 | 0.692 | 0.008 | 0 | 39 |
| apnl | 0 | 1.300988198 | 0.554 | 0.001 | 0 | 39 |
| LO017965.1 | 0 | 1.240344092 | 0.538 | 0.001 | 0 | 39 |
| krt17 | 0 | 1.231032681 | 0.554 | 0.006 | 0 | 39 |
| pkp3a | 0 | 1.166539101 | 0.677 | 0.007 | 0 | 39 |
| cldna | 0 | 1.1292061 | 0.462 | 0.001 | 0 | 39 |
| zgc:1018101 | 0 | 1.084729853 | 0.615 | 0.005 | 0 | 39 |
| si:dkey-33c14.3 | 0 | 1.046524591 | 0.385 | 0.001 | 0 | 39 |
| cldn1 | 0 | 0.90880603 | 0.385 | 0 | 0 | 39 |
| zgc:193505 | 0 | 0.90827362 | 0.446 | 0.005 | 0 | 39 |
| zgc:92380 | 0 | 0.841123016 | 0.508 | 0.008 | 0 | 39 |

| | | | | | | |
|---|---|---|---|---|---|---|
| ms4a17a.5 | 0 | 0.823011528 | 0.477 | 0.004 | 0 | 39 |
| CU682640.1 | 0 | 0.800112591 | 0.369 | 0 | 0 | 39 |
| krtt1c19e | 0 | 0.777413777 | 0.308 | 0.001 | 0 | 39 |
| cxl34b.11 | 0 | 0.722778143 | 0.385 | 0 | 0 | 39 |
| gpa33a | 0 | 0.699297436 | 0.446 | 0.002 | 0 | 39 |
| caspb | 0 | 0.684977156 | 0.415 | 0.001 | 0 | 39 |
| lamc2 | 0 | 0.675969143 | 0.277 | 0.002 | 0 | 39 |
| zgc:162941 | 0 | 0.67459174 | 0.308 | 0 | 0 | 39 |
| si:ch211-95j8.3 | 0 | 0.648336717 | 0.246 | 0 | 0 | 39 |
| RGS13 (1 of many) | 0 | 0.609513156 | 0.292 | 0.001 | 0 | 39 |
| cd44a | 0 | 0.525428158 | 0.292 | 0.002 | 0 | 39 |
| si:ch211-213d14.2 | 0 | 0.476770576 | 0.292 | 0.002 | 0 | 39 |
| ccl25a | 0 | 0.454373474 | 0.169 | 0 | 0 | 39 |
| ms4a17a.4 | 0 | 0.439316529 | 0.323 | 0.002 | 0 | 39 |
| rhbg | 0 | 0.424478687 | 0.246 | 0.001 | 0 | 39 |
| ms4a17a.2 | 0 | 0.414237789 | 0.231 | 0 | 0 | 39 |
| epgn | 0 | 0.372112983 | 0.185 | 0 | 0 | 39 |
| plek2 | 0 | 0.364742503 | 0.185 | 0 | 0 | 39 |
| ccl27b | 0 | 0.360976077 | 0.169 | 0 | 0 | 39 |
| si:rp71-1i20.2 | 0 | 0.343692693 | 0.215 | 0 | 0 | 39 |
| gpr31 | 0 | 0.329963085 | 0.2 | 0 | 0 | 39 |
| upk3b | 0 | 0.295549289 | 0.185 | 0 | 0 | 39 |
| c6ast1 | 1.68897966784777e-304 | 1.042485543 | 0.338 | 0.003 | 4.25099292600605e-300 | 39 |
| si:busm1-71b9.3 | 1.49305068653623e-297 | 0.709735396 | 0.369 | 0.004 | 3.75785927294304e-293 | 39 |
| tp63 | 2.30108685504997e-295 | 0.797779659 | 0.492 | 0.008 | 5.79160550547528e-291 | 39 |
| ptgdsb.1 | 2.75216769869467e-294 | 1.686175149 | 0.523 | 0.01 | 6.9269308808446e-290 | 39 |
| col17a1b | 1.98330001951197e-291 | 0.612230301 | 0.246 | 0.001 | 4.99176781910968e-287 | 39 |
| mep1a.1 | 3.35913285535198e-286 | 0.306679638 | 0.169 | 0 | 8.4546014836354e-282 | 39 |
| mmp20a | 6.41032111732747e-285 | 0.38883219 | 0.123 | 0 | 1.61341372202015e-280 | 39 |
| si:ch211-241e1.3 | 1.26534901773661e-284 | 0.882793126 | 0.385 | 0.005 | 3.18475694274127e-280 | 39 |
| LOC100330348 | 3.51481197414331e-273 | 0.281617911 | 0.154 | 0 | 8.8464302577213e-269 | 39 |
| fabp1b.2 | 1.24248200580247e-271 | 0.580401216 | 0.2 | 0.001 | 3.12720296040423e-267 | 39 |
| ms4a17a.1 | 1.46296433971959e-271 | 0.325708304 | 0.2 | 0.001 | 3.68213494664023e-267 | 39 |
| zgc:77439 | 6.13131787863958e-269 | 1.004257542 | 0.569 | 0.013 | 1.5431913968748e-264 | 39 |
| si:ch211-264f5.61 | 7.11089117559605e-255 | 0.777956465 | 0.508 | 0.011 | 1.78974019998577e-250 | 39 |

| | | | | | | |
|---|---|---|---|---|---|---|
| si:ch1073-406l10.2 | 3.30082000716756e-250 | 0.960845128 | 0.477 | 0.01 | 8.30783387604003e-246 | 39 |
| fam83b | 7.91679644129635e-247 | 0.335125839 | 0.215 | 0.001 | 1.99257849630988e-242 | 39 |
| pof1b | 6.16532811156992e-245 | 0.741328145 | 0.431 | 0.008 | 1.55175143240103e-240 | 39 |
| fermt1 | 4.68806224553851e-238 | 0.40417781 | 0.215 | 0.001 | 1.17993838657959e-233 | 39 |
| aqp3a3 | 9.90297287880641e-228 | 1.316597319 | 0.708 | 0.026 | 2.49247924386679e-223 | 39 |
| cldni1 | 1.41225272199189e-224 | 1.368692994 | 0.462 | 0.011 | 3.55449887598138e-220 | 39 |
| dsg2.1 | 9.99801087239166e-217 | 1.240206073 | 0.646 | 0.023 | 2.51639935647226e-212 | 39 |
| CR936378.1 | 5.0766382462555e-214 | 0.373131626 | 0.169 | 0.001 | 1.27773908020005e-209 | 39 |
| si:dkey-19b23.12 | 6.0102197726585e-214 | 0.256900503 | 0.169 | 0.001 | 1.51271221458042e-209 | 39 |
| krt915 | 2.2177831637302e-213 | 2.323966156 | 1 | 0.064 | 5.58193844479255e-209 | 39 |
| si:ch211-155m12.1 | 2.68183667832431e-212 | 0.831819849 | 0.185 | 0.001 | 6.74991473567446e-208 | 39 |
| scpp72 | 3.14915163828939e-210 | 2.139059414 | 0.662 | 0.025 | 7.92609975841058e-206 | 39 |
| LOC110438215 | 6.3381154227092e-209 | 0.370159175 | 0.215 | 0.002 | 1.59524027074168e-204 | 39 |
| gsnb | 1.34616353982845e-208 | 0.331744565 | 0.215 | 0.002 | 3.38815901339423e-204 | 39 |
| eevs | 1.9164192303407e-208 | 0.27520673 | 0.154 | 0.001 | 4.82343556084451e-204 | 39 |
| dsc2l1 | 7.02205534818483e-207 | 1.095944113 | 0.631 | 0.022 | 1.76738111058464e-202 | 39 |
| dspa | 1.0668462303458e-202 | 1.036704502 | 0.631 | 0.023 | 2.68514527715733e-198 | 39 |
| upk1a | 9.4716891780633e-197 | 0.295068642 | 0.154 | 0.001 | 2.38392944922675e-192 | 39 |
| chl1a | 2.64524621976815e-195 | 0.316149143 | 0.185 | 0.001 | 6.65782021053445e-191 | 39 |
| zgc:1539111 | 6.03078367701523e-192 | 1.221222241 | 0.538 | 0.018 | 1.51788794366796e-187 | 39 |
| slc16a12b | 9.52361014561925e-188 | 0.509476761 | 0.354 | 0.007 | 2.39699743755091e-183 | 39 |
| aldob3 | 7.11759497030485e-176 | 1.350317354 | 0.708 | 0.035 | 1.79142747807603e-171 | 39 |
| mocs2 | 4.62060073269207e-175 | 0.498957957 | 0.308 | 0.006 | 1.16295899841127e-170 | 39 |
| apoeb | 1.68861017327047e-174 | 1.124016527 | 0.446 | 0.013 | 4.25006294510444e-170 | 39 |
| dnase1l4.1 | 4.01183372662487e-173 | 0.410959238 | 0.277 | 0.004 | 1.00973843065421e-168 | 39 |
| gpd1c | 1.38815263539388e-172 | 0.608927055 | 0.431 | 0.012 | 3.49384136802286e-168 | 39 |
| anxa1b | 1.6218858564634e-171 | 0.683780756 | 0.323 | 0.006 | 4.08212451213274e-167 | 39 |
| limk2 | 5.9021817568747e-169 | 0.73715034 | 0.477 | 0.015 | 1.48552012638779e-164 | 39 |
| perp5 | 2.47515186989659e-159 | 1.40856687 | 0.8 | 0.052 | 6.22970974134273e-155 | 39 |
| wu:fb18f061 | 4.19520734128621e-158 | 1.602602812 | 0.815 | 0.053 | 1.05589173572833e-153 | 39 |
| cfd5 | 4.54372730635244e-158 | 1.833978 | 0.877 | 0.064 | 1.14361072573585e-153 | 39 |
| gdpd3a | 9.70709763144939e-156 | 0.320488396 | 0.2 | 0.002 | 2.4431794028595e-151 | 39 |
| evpla | 1.92250708092512e-151 | 0.453547102 | 0.262 | 0.005 | 4.83875807198044e-147 | 39 |
| tmem238a | 3.79381664251664e-149 | 0.588410093 | 0.415 | 0.013 | 9.54865710755014e-145 | 39 |
| esrp1 | 3.03835338648135e-144 | 0.293456215 | 0.185 | 0.002 | 7.64723163843491e-140 | 39 |

| | | | | | | |
|---|---|---|---|---|---|---|
| FP017215.1 | 3.45885193016266e-144 | 0.705061008 | 0.415 | 0.014 | 8.70558442302639e-140 | 39 |
| s100w2 | 8.00400025964193e-137 | 0.706237563 | 0.446 | 0.017 | 2.01452682534928e-132 | 39 |
| si:ch211-12e13.10 | 1.10927761966296e-135 | 0.508026191 | 0.154 | 0.002 | 2.7919408409297e-131 | 39 |
| si:ch211-69b7.6 | 1.42954203045063e-132 | 0.7149657 | 0.277 | 0.006 | 3.59801433644119e-128 | 39 |
| bnipl | 6.97299020257146e-128 | 0.395346675 | 0.262 | 0.006 | 1.75503190408521e-123 | 39 |
| itgb42 | 1.69951518487079e-125 | 1.015034331 | 0.4 | 0.015 | 4.27750976880129e-121 | 39 |
| LOC799914 | 2.94757314447591e-124 | 0.268547317 | 0.138 | 0.001 | 7.41874684733142e-120 | 39 |
| si:dkey-157l19.2 | 2.11920863591669e-122 | 0.441081382 | 0.308 | 0.009 | 5.33383621573873e-118 | 39 |
| jupa | 5.08789085007167e-120 | 1.099848864 | 0.708 | 0.051 | 1.28057124805454e-115 | 39 |
| zgc:194948 | 2.24620383409703e-117 | 0.391374481 | 0.2 | 0.004 | 5.65347043003882e-113 | 39 |
| itgb6 | 1.57507898535102e-116 | 0.261018345 | 0.185 | 0.003 | 3.96431629822999e-112 | 39 |
| f3a | 8.53176314548436e-116 | 0.317562675 | 0.231 | 0.005 | 2.14735946608696e-111 | 39 |
| f2rl1.2 | 1.31488730406318e-115 | 0.722609372 | 0.477 | 0.023 | 3.30943985559661e-111 | 39 |
| cavin2a | 2.45340158589069e-115 | 0.563162127 | 0.354 | 0.012 | 6.17496645152827e-111 | 39 |
| ptgs2a1 | 1.75254712967851e-113 | 0.596051725 | 0.323 | 0.011 | 4.41098587068785e-109 | 39 |
| ppl | 1.11881088675309e-110 | 0.481832738 | 0.308 | 0.01 | 2.81593512086886e-106 | 39 |
| cxcl18a.1 | 1.76868168591219e-108 | 0.281072479 | 0.169 | 0.003 | 4.45159493527239e-104 | 39 |
| scinla | 1.688431882054e-106 | 0.504791394 | 0.308 | 0.01 | 4.24961420394171e-102 | 39 |
| si:ch211-166a6.52 | 4.57321025205482e-105 | 0.805268214 | 0.523 | 0.031 | 1.15103128833968e-100 | 39 |
| tmem176l.4 | 2.67674913502232e-103 | 0.474670426 | 0.277 | 0.008 | 6.73710989793768e-99 | 39 |
| foxq1a | 4.75433355525078e-103 | 0.27031106 | 0.154 | 0.002 | 1.19661821252107e-98 | 39 |
| cfl1l7 | 1.28534044225524e-102 | 1.78948298 | 1 | 0.15 | 3.23507335911221e-98 | 39 |
| tmsb18 | 7.42228204009578e-101 | 1.984574455 | 0.985 | 0.153 | 1.86811416667171e-96 | 39 |
| pkp2 | 2.71822418516812e-99 | 0.341918006 | 0.215 | 0.005 | 6.84149845164965e-95 | 39 |
| serpinb14 | 3.42632132979284e-99 | 0.525210087 | 0.292 | 0.01 | 8.6237081549556e-95 | 39 |
| arhgef54 | 1.77331774851739e-98 | 0.696758588 | 0.477 | 0.027 | 4.46326344124343e-94 | 39 |
| mid1ip1a | 1.33578185308082e-97 | 0.589795866 | 0.323 | 0.012 | 3.36202934601912e-93 | 39 |
| si:dkey-16l2.201 | 1.53472105403831e-97 | 0.572874831 | 0.385 | 0.018 | 3.86273942090903e-93 | 39 |
| serpinb1l4 | 1.00362515043755e-94 | 0.34865381 | 0.215 | 0.005 | 2.52602414113628e-90 | 39 |
| itga6b | 9.67474532392284e-92 | 0.810769821 | 0.369 | 0.018 | 2.43503665057814e-87 | 39 |
| fat2 | 7.39370290389667e-91 | 0.327221806 | 0.215 | 0.006 | 1.86092108388175e-86 | 39 |
| tfap2c1 | 1.10383815734715e-90 | 0.283792223 | 0.185 | 0.004 | 2.77825025822703e-86 | 39 |
| ptk2bb | 6.77992532781116e-88 | 0.341758719 | 0.231 | 0.007 | 1.70643940575679e-83 | 39 |
| CABZ01048956.1 | 1.43195127086689e-87 | 0.413382302 | 0.169 | 0.004 | 3.60407815364487e-83 | 39 |
| pon1 | 1.67186924879174e-87 | 0.522725648 | 0.277 | 0.01 | 4.20792771228392e-83 | 39 |

| | | | | | | |
|---|---|---|---|---|---|---|
| si:ch211-39i22.15 | 1.16459799958887e-86 | 1.105813966 | 0.708 | 0.07 | 2.93117670516523e-82 | 39 |
| krt43 | 8.62941932513582e-85 | 1.176911691 | 0.523 | 0.039 | 2.17193854994343e-80 | 39 |
| phlda23 | 1.19211668015732e-84 | 1.189950236 | 0.585 | 0.05 | 3.00043847228795e-80 | 39 |
| cdh12 | 1.79245206509131e-84 | 0.955263281 | 0.646 | 0.056 | 4.51142260262831e-80 | 39 |
| XLOC-041832 | 8.17864491679236e-84 | 0.421567332 | 0.262 | 0.009 | 2.05848313910747e-79 | 39 |
| ak14 | 1.32137414020679e-81 | 1.084862362 | 0.692 | 0.073 | 3.32576657348647e-77 | 39 |
| gcshb1 | 2.49565738964661e-81 | 0.506277508 | 0.308 | 0.014 | 6.28132008400156e-77 | 39 |
| mapk12a | 1.31968408773202e-80 | 0.345395493 | 0.292 | 0.012 | 3.32151288041272e-76 | 39 |
| efna1b2 | 4.04629030780897e-80 | 0.52728465 | 0.338 | 0.017 | 1.01841080757244e-75 | 39 |
| selenow2b | 4.37735598089631e-79 | 0.322949376 | 0.169 | 0.004 | 1.10173672683179e-74 | 39 |
| tdh1 | 7.30221225091823e-78 | 1.146007347 | 0.769 | 0.093 | 1.83789380143361e-73 | 39 |
| zgc:1533175 | 7.01731983615317e-76 | 0.78728853 | 0.508 | 0.04 | 1.76618922956139e-71 | 39 |
| s100v28 | 5.66827713442206e-75 | 1.298146031 | 0.831 | 0.119 | 1.42664867196269e-70 | 39 |
| lgals3b5 | 2.64265782490815e-74 | 0.995709073 | 0.677 | 0.077 | 6.65130547951132e-70 | 39 |
| zgc:171572 | 2.86891083832869e-74 | 0.330021842 | 0.2 | 0.006 | 7.22076168898949e-70 | 39 |
| elf3 | 4.13651057918445e-74 | 0.412474277 | 0.2 | 0.006 | 1.04111834767493e-69 | 39 |
| amotl2b | 9.31652518024872e-72 | 0.466962154 | 0.292 | 0.014 | 2.3448762226168e-67 | 39 |
| ptgs2b1 | 2.38835308856665e-71 | 0.448420357 | 0.246 | 0.01 | 6.01124588861339e-67 | 39 |
| spint1a1 | 5.39959655379425e-71 | 0.830107588 | 0.585 | 0.054 | 1.35902445662448e-66 | 39 |
| cx35.4 | 1.26595291403287e-69 | 0.29305299 | 0.169 | 0.005 | 3.18627688932932e-65 | 39 |
| si:dkey-16p21.8 | 1.32681903484674e-69 | 1.034200201 | 0.662 | 0.073 | 3.33947082880575e-65 | 39 |
| arg23 | 2.01431606668931e-66 | 0.609190613 | 0.338 | 0.02 | 5.06983210825032e-62 | 39 |
| icn11 | 6.86099134316302e-66 | 1.757037076 | 1 | 0.273 | 1.7268429111607e-61 | 39 |
| anxa2a7 | 8.72240151717187e-66 | 1.194561657 | 0.754 | 0.104 | 2.19534123785699e-61 | 39 |
| ahnak11 | 1.50141582668445e-65 | 1.261143477 | 0.8 | 0.125 | 3.7789134941821e-61 | 39 |
| pnp5a5 | 3.89122964318096e-65 | 1.14948065 | 0.631 | 0.076 | 9.79383588892215e-61 | 39 |
| ptgis1 | 1.02091105694991e-64 | 0.49710353 | 0.277 | 0.014 | 2.56953103923723e-60 | 39 |
| lmna1 | 5.02348884982661e-64 | 0.460605403 | 0.277 | 0.014 | 1.26436190861286e-59 | 39 |
| FAM83G | 7.11168046353199e-64 | 0.359515833 | 0.246 | 0.011 | 1.78993885586637e-59 | 39 |
| anxa1a8 | 3.17767334710968e-63 | 1.377847154 | 0.831 | 0.141 | 7.99788604734035e-59 | 39 |
| klf34 | 6.7666229452226e-63 | 0.618295973 | 0.431 | 0.035 | 1.70309132908308e-58 | 39 |
| crybg1a | 5.66478612637937e-60 | 0.409272161 | 0.292 | 0.017 | 1.42577002014842e-55 | 39 |
| bik | 1.39251139653862e-59 | 0.324889735 | 0.2 | 0.008 | 3.50481193394806e-55 | 39 |
| capn2b1 | 3.59190203550227e-58 | 0.513631532 | 0.338 | 0.023 | 9.04045823315565e-54 | 39 |
| rhoub | 1.53823928933714e-57 | 0.289639094 | 0.169 | 0.006 | 3.87159446733266e-53 | 39 |

| | | | | | | |
|---|---|---|---|---|---|---|
| rnd3a2 | 8.18901131773335e-57 | 0.528713338 | 0.385 | 0.03 | 2.06109225856031e-52 | 39 |
| wu:fi04e121 | 2.17063689783844e-55 | 0.398121176 | 0.308 | 0.02 | 5.46327600816957e-51 | 39 |
| cdo11 | 2.02995964396197e-54 | 0.380077999 | 0.2 | 0.009 | 5.10920542788788e-50 | 39 |
| ripk4 | 2.68833199625907e-53 | 0.259827286 | 0.185 | 0.007 | 6.76626280138445e-49 | 39 |
| krt155 | 9.59718774106312e-53 | 1.350266457 | 0.6 | 0.083 | 2.41551618254818e-48 | 39 |
| gpc1a1 | 2.90077597094304e-52 | 0.281107854 | 0.185 | 0.008 | 7.30096304126655e-48 | 39 |
| plecb4 | 8.80395665602492e-51 | 0.762881885 | 0.462 | 0.05 | 2.21586785075491e-46 | 39 |
| srd5a2a1 | 1.16076160341647e-49 | 0.860966409 | 0.631 | 0.088 | 2.92152087963892e-45 | 39 |
| s100a10b11 | 1.63620905719855e-49 | 1.376953972 | 0.954 | 0.254 | 4.11817457606302e-45 | 39 |
| fosl1a7 | 4.35354036702476e-49 | 0.992417339 | 0.615 | 0.091 | 1.09574257497646e-44 | 39 |
| scel | 1.33533801464499e-48 | 0.251543498 | 0.169 | 0.007 | 3.36091224905996e-44 | 39 |
| mat2ab | 1.49684668812904e-48 | 0.571192283 | 0.354 | 0.03 | 3.76741342935199e-44 | 39 |
| kremen11 | 4.04303692461222e-48 | 0.355363664 | 0.2 | 0.01 | 1.01759196355565e-43 | 39 |
| fam69c | 4.71855784964164e-48 | 0.309284742 | 0.215 | 0.011 | 1.1876138251763e-43 | 39 |
| tuft1a3 | 2.90102875398064e-47 | 0.464174821 | 0.369 | 0.033 | 7.30159927089386e-43 | 39 |
| si:dkey-262k9.23 | 3.67422510469261e-47 | 0.59545934 | 0.369 | 0.034 | 9.24765716600082e-43 | 39 |
| flna8 | 3.83305470058072e-46 | 0.925509634 | 0.677 | 0.113 | 9.6474153758916e-42 | 39 |
| sult2st11 | 6.19879213753999e-46 | 0.48432872 | 0.323 | 0.027 | 1.56017399309744e-41 | 39 |
| cd9b3 | 3.3819356963722e-45 | 0.67748451 | 0.523 | 0.064 | 8.51199395419918e-41 | 39 |
| zgc:92161 | 5.4377785922276e-45 | 0.276901163 | 0.169 | 0.008 | 1.36863449387776e-40 | 39 |
| enam2 | 2.54855058038927e-44 | 0.947889872 | 0.215 | 0.013 | 6.41444695578176e-40 | 39 |
| dlx3b2 | 2.84147609826612e-44 | 0.294049025 | 0.185 | 0.009 | 7.15171119172601e-40 | 39 |
| wsb2 | 9.50216113346068e-44 | 0.346767931 | 0.246 | 0.016 | 2.39159893568072e-39 | 39 |
| zgc:92242 | 9.52759936053506e-44 | 0.405278472 | 0.246 | 0.017 | 2.39800148305307e-39 | 39 |
| exosc1 | 5.55826448686548e-43 | 0.581355571 | 0.385 | 0.04 | 1.39895958869917e-38 | 39 |
| eppk14 | 1.87373185264683e-42 | 0.610550203 | 0.385 | 0.04 | 4.7159956999268e-38 | 39 |
| rab42b | 2.71378783369882e-42 | 0.3285419 | 0.231 | 0.015 | 6.83033259863657e-38 | 39 |
| si:ch211-105c13.34 | 4.29252940613665e-42 | 0.745286793 | 0.477 | 0.06 | 1.08038672623053e-37 | 39 |
| tuba8l24 | 4.53140892813714e-42 | 0.638817768 | 0.477 | 0.06 | 1.14051031312284e-37 | 39 |
| cldne | 2.95643675067818e-41 | 0.316850735 | 0.138 | 0.005 | 7.44105565778192e-37 | 39 |
| cavin2b7 | 6.47491507439614e-41 | 0.657516141 | 0.492 | 0.065 | 1.62967137507476e-36 | 39 |
| CABZ01086029.1 | 1.10059522093639e-39 | 0.335609653 | 0.246 | 0.018 | 2.77008811157481e-35 | 39 |
| ptprfb2 | 1.61631158406997e-39 | 0.281313095 | 0.215 | 0.014 | 4.0680946259457e-35 | 39 |
| ptgdsb.21 | 2.48893491744471e-39 | 1.056896152 | 0.369 | 0.042 | 6.26440029371659e-35 | 39 |
| XLOC-0398171 | 5.04891570001381e-39 | 0.257203855 | 0.185 | 0.01 | 1.27076159253648e-34 | 39 |

| | | | | | | |
|---|---|---|---|---|---|---|
| pdlim13 | 2.23931441608419e-38 | 0.573217484 | 0.431 | 0.054 | 5.63613045384229e-34 | 39 |
| aldh1l1 | 3.24875397189587e-38 | 0.340641929 | 0.262 | 0.021 | 8.1767888718647e-34 | 39 |
| si:ch211-195b11.3 | 4.24648986001382e-38 | 0.650951117 | 0.354 | 0.037 | 1.06879903286688e-33 | 39 |
| vwa12 | 4.96809048588288e-38 | 0.413490705 | 0.262 | 0.021 | 1.25041869439186e-33 | 39 |
| xkr91 | 1.28870192225963e-37 | 0.290164288 | 0.231 | 0.017 | 3.24353386813525e-33 | 39 |
| myh9a3 | 1.74024181573023e-37 | 0.561551374 | 0.431 | 0.054 | 4.38001462601143e-33 | 39 |
| casz11 | 2.29411863788173e-37 | 0.357429943 | 0.262 | 0.021 | 5.77406719968453e-33 | 39 |
| calm2b7 | 8.07509927962161e-37 | 0.96564126 | 0.846 | 0.209 | 2.03242173768796e-32 | 39 |
| capgb9 | 1.7602414899088e-36 | 0.813000182 | 0.631 | 0.113 | 4.43035180595146e-32 | 39 |
| midn11 | 2.60520774750016e-36 | 1.060768845 | 0.646 | 0.139 | 6.55704737968316e-32 | 39 |
| kitlga1 | 4.13113072082812e-36 | 0.326242238 | 0.215 | 0.015 | 1.03976429112523e-31 | 39 |
| jag1b2 | 6.64824033231961e-36 | 0.405378644 | 0.277 | 0.025 | 1.67329560924152e-31 | 39 |
| cast3 | 1.73451677424094e-35 | 0.6284951 | 0.477 | 0.069 | 4.36560526908702e-31 | 39 |
| slc38a5b8 | 7.81458374987647e-35 | 0.750075598 | 0.615 | 0.114 | 1.96685258400641e-30 | 39 |
| lygl14 | 8.94353243743654e-35 | 0.651584287 | 0.585 | 0.096 | 2.2509976791784e-30 | 39 |
| pycard8 | 1.54221612610343e-34 | 0.645126661 | 0.477 | 0.071 | 3.88160376778972e-30 | 39 |
| pmp22b6 | 5.27455513917026e-34 | 0.530147493 | 0.369 | 0.045 | 1.32755278297776e-29 | 39 |
| sat1a.1 | 1.67863436570001e-32 | 0.28804235 | 0.154 | 0.009 | 4.22495483503036e-28 | 39 |
| ywhaz6 | 4.43709267581957e-32 | 0.966336168 | 0.877 | 0.28 | 1.11677185557703e-27 | 39 |
| fxyd18 | 7.17096243410082e-32 | 0.611424439 | 0.492 | 0.08 | 1.80485953503883e-27 | 39 |
| trib34 | 4.0996975873035e-31 | 0.75655473 | 0.554 | 0.105 | 1.03185288574842e-26 | 39 |
| rarga3 | 6.71622884697911e-31 | 0.501061724 | 0.4 | 0.056 | 1.69040763849617e-26 | 39 |
| prdx510 | 1.05895942901422e-30 | 0.830573685 | 0.677 | 0.158 | 2.66529498688589e-26 | 39 |
| cd993 | 1.3647031165554e-29 | 0.401169653 | 0.385 | 0.053 | 3.43482127405829e-25 | 39 |
| lsr | 1.76683836716079e-29 | 0.293230914 | 0.262 | 0.026 | 4.44695548630699e-25 | 39 |
| nfil33 | 2.2551140347492e-29 | 0.398907817 | 0.292 | 0.033 | 5.67589651406026e-25 | 39 |
| capns1a4 | 1.01290228204816e-28 | 0.785218529 | 0.631 | 0.142 | 2.54937375368702e-24 | 39 |
| zgc:1532845 | 1.20051863458659e-28 | 0.608146534 | 0.554 | 0.1 | 3.02158535139098e-24 | 39 |
| si:ch211-134c9.2 | 1.9643057199718e-28 | 0.301945931 | 0.215 | 0.019 | 4.94396106659703e-24 | 39 |
| aldh9a1a.18 | 3.72264613025017e-28 | 0.778184759 | 0.677 | 0.163 | 9.36952804522664e-24 | 39 |
| myof | 4.465488857299e-28 | 0.270008013 | 0.246 | 0.025 | 1.12391889049358e-23 | 39 |
| si:ch73-335l21.1 | 4.88396982897715e-28 | 0.279735223 | 0.185 | 0.015 | 1.22924636625526e-23 | 39 |
| zfand5a1 | 5.09445115260475e-28 | 0.5958063 | 0.446 | 0.076 | 1.28222241059909e-23 | 39 |
| vaspb1 | 1.08547844005332e-27 | 0.438107805 | 0.354 | 0.049 | 2.7320406857702e-23 | 39 |
| cst14b.15 | 1.50509413799275e-27 | 0.543691522 | 0.354 | 0.051 | 3.78817143591396e-23 | 39 |

| | | | | | | |
|---|---|---|---|---|---|---|
| nectin1b3 | 4.41090132208131e-27 | 0.406821183 | 0.323 | 0.042 | 1.11017975375465e-22 | 39 |
| runx32 | 1.07523080862103e-26 | 0.315845257 | 0.215 | 0.02 | 2.70624842221827e-22 | 39 |
| jund9 | 1.16763660417378e-26 | 0.604860986 | 0.492 | 0.092 | 2.93882456904499e-22 | 39 |
| slc20a1b4 | 3.56716931486955e-26 | 0.675710668 | 0.508 | 0.101 | 8.97820844859516e-22 | 39 |
| tagln213 | 6.45275482967892e-26 | 0.833155059 | 0.692 | 0.187 | 1.62409386308189e-21 | 39 |
| itgav4 | 1.46534462194368e-25 | 0.503751358 | 0.369 | 0.058 | 3.68812587897005e-21 | 39 |
| tspan33a1 | 4.28842085805288e-25 | 0.322226593 | 0.185 | 0.016 | 1.07935264576333e-20 | 39 |
| zmp:0000000991 | 4.59358686632524e-25 | 0.261235307 | 0.169 | 0.014 | 1.1561598783854e-20 | 39 |
| gipc2 | 5.77189350853732e-25 | 0.358179376 | 0.277 | 0.034 | 1.45272787716376e-20 | 39 |
| tcima7 | 6.35015160151493e-25 | 0.647573746 | 0.4 | 0.068 | 1.59826965658529e-20 | 39 |
| prr13 | 2.47436700714527e-24 | 0.279465854 | 0.215 | 0.022 | 6.22773432028393e-20 | 39 |
| si:ch211-13k12.2 | 3.44138364417377e-24 | 0.262543574 | 0.185 | 0.017 | 8.66161849402097e-20 | 39 |
| tmc6b1 | 3.74675964526901e-24 | 0.349172273 | 0.277 | 0.035 | 9.43021935117758e-20 | 39 |
| hcar1-44 | 4.11233905657359e-24 | 0.364000836 | 0.277 | 0.036 | 1.03503461714901e-19 | 39 |
| itgb3a2 | 2.39993738141804e-23 | 0.287371086 | 0.169 | 0.015 | 6.04040239529105e-19 | 39 |
| fosb8 | 7.01728388403714e-23 | 0.891191914 | 0.523 | 0.125 | 1.76618018077331e-18 | 39 |
| selenow16 | 1.04920884830278e-22 | 0.840517045 | 0.877 | 0.385 | 2.64075375029328e-18 | 39 |
| zgc:774862 | 1.47061048728871e-22 | 0.403180266 | 0.4 | 0.07 | 3.70137953545696e-18 | 39 |
| dnmbp | 1.48592561760246e-22 | 0.252157943 | 0.169 | 0.015 | 3.73992618694363e-18 | 39 |
| ccl25b5 | 2.15193179118358e-22 | 0.593872651 | 0.415 | 0.076 | 5.41619712522994e-18 | 39 |
| atf37 | 3.91609756759635e-22 | 0.644349449 | 0.446 | 0.091 | 9.85642596788325e-18 | 39 |
| foxp46 | 5.07357265628677e-22 | 0.453587352 | 0.323 | 0.051 | 1.27696750186082e-17 | 39 |
| zbtb16b1 | 2.09890760385381e-21 | 0.325755283 | 0.215 | 0.025 | 5.28274054813966e-17 | 39 |
| si:dkey-180p18.94 | 4.20292445432586e-21 | 0.553930913 | 0.462 | 0.099 | 1.05783405590927e-16 | 39 |
| cavin1b6 | 5.35328008625877e-21 | 0.459254829 | 0.277 | 0.04 | 1.34736706491047e-16 | 39 |
| cav16 | 6.06745689158335e-21 | 0.420798228 | 0.292 | 0.044 | 1.52711822504261e-16 | 39 |
| her91 | 1.28253965918992e-20 | 0.283703807 | 0.185 | 0.019 | 3.22802406821512e-16 | 39 |
| ms4a17a.103 | 1.67212929691819e-20 | 0.370343494 | 0.323 | 0.052 | 4.2085822274134e-16 | 39 |
| fblim13 | 3.12900401563854e-20 | 0.296781062 | 0.215 | 0.026 | 7.87539020696065e-16 | 39 |
| rmdn1 | 4.55852109172094e-20 | 0.422702955 | 0.323 | 0.055 | 1.14733417357524e-15 | 39 |
| ubxn6 | 5.70337341023181e-20 | 0.264721282 | 0.154 | 0.014 | 1.43548205362124e-15 | 39 |
| ets25 | 6.40827562038832e-20 | 0.50509234 | 0.431 | 0.09 | 1.61289889089554e-15 | 39 |
| sult6b1 | 1.02642259356326e-19 | 0.268477296 | 0.169 | 0.017 | 2.58340302573936e-15 | 39 |
| sptlc2b2 | 1.15538777638458e-19 | 0.360596385 | 0.292 | 0.047 | 2.90799549438235e-15 | 39 |
| lama53 | 1.57001867302574e-19 | 0.354224293 | 0.2 | 0.024 | 3.95157999813849e-15 | 39 |

| | | | | | | |
|---|---|---|---|---|---|---|
| anxa47 | 7.88411892946945e-19 | 0.533405628 | 0.538 | 0.136 | 1.98435389335817e-14 | 39 |
| cbsb | 1.2705419014207e-18 | 0.320377217 | 0.246 | 0.035 | 3.19782691168575e-14 | 39 |
| ctnnd11 | 1.38205592962536e-18 | 0.299448572 | 0.338 | 0.06 | 3.47849656927406e-14 | 39 |
| ccng25 | 2.07316675395429e-18 | 0.482838803 | 0.4 | 0.085 | 5.21795340302756e-14 | 39 |
| oclna2 | 3.60428317248846e-18 | 0.330025094 | 0.4 | 0.079 | 9.07162031683621e-14 | 39 |
| zbtb16a3 | 4.51910401258222e-18 | 0.297949923 | 0.292 | 0.049 | 1.13741328892682e-13 | 39 |
| pfn115 | 5.09835928576309e-18 | 0.816813794 | 0.985 | 0.524 | 1.28320604863371e-13 | 39 |
| net12 | 7.23739733399214e-18 | 0.257807919 | 0.185 | 0.022 | 1.82158053499248e-13 | 39 |
| mylipb | 7.67245656509012e-18 | 0.253001619 | 0.185 | 0.022 | 1.93108059286753e-13 | 39 |
| tpm38 | 1.48929286428466e-17 | 0.695923933 | 0.723 | 0.254 | 3.74840121011805e-13 | 39 |
| mcl1b12 | 1.85476027083346e-17 | 0.673936072 | 0.738 | 0.242 | 4.66824612566074e-13 | 39 |
| ybx111 | 2.86809337674026e-17 | 0.527183067 | 0.954 | 0.67 | 7.21870421991755e-13 | 39 |
| gsna2 | 2.92797533864725e-17 | 0.267319912 | 0.215 | 0.03 | 7.36942112984127e-13 | 39 |
| si:ch211-39i2.22 | 4.48284178404866e-17 | 0.418749107 | 0.338 | 0.066 | 1.12828644862721e-12 | 39 |
| si:ch73-335l21.48 | 5.32969694206528e-17 | 0.774054555 | 0.523 | 0.148 | 1.34143142334841e-12 | 39 |
| anxa5b7 | 5.64777772611561e-17 | 0.452451591 | 0.415 | 0.094 | 1.42148917588604e-12 | 39 |
| fbp1a2 | 5.83473633809842e-17 | 0.352128931 | 0.246 | 0.039 | 1.46854478893599e-12 | 39 |
| mpzl2b3 | 6.43685407336583e-17 | 0.404421735 | 0.431 | 0.094 | 1.62009180172545e-12 | 39 |
| myofl2 | 6.57145085356657e-17 | 0.292724961 | 0.169 | 0.02 | 1.65396846533417e-12 | 39 |
| fgfr33 | 9.6947955173701e-17 | 0.252733469 | 0.154 | 0.017 | 2.44008308376688e-12 | 39 |
| si:ch73-234b20.58 | 1.19237904891151e-16 | 0.43312831 | 0.431 | 0.1 | 3.00109882820539e-12 | 39 |
| hdac43 | 1.31969835656855e-16 | 0.260343497 | 0.215 | 0.031 | 3.32154879364737e-12 | 39 |
| tax1bp33 | 1.67458284568946e-16 | 0.393700769 | 0.4 | 0.089 | 4.2147575643158e-12 | 39 |
| itm2bb11 | 3.4996816982789e-16 | 0.58812805 | 0.692 | 0.222 | 8.80834886639816e-12 | 39 |
| tmsb4x23 | 3.69899793307632e-16 | 0.631701738 | 1 | 0.686 | 9.3100078977598e-12 | 39 |
| si:ch73-1a9.316 | 4.11442569251163e-16 | 0.667094954 | 0.738 | 0.265 | 1.03555980254825e-11 | 39 |
| slc1a3a1 | 1.307882061529e-15 | 0.264903162 | 0.154 | 0.018 | 3.29180836066235e-11 | 39 |
| ccnl1a3 | 2.01537236455464e-15 | 0.375839503 | 0.308 | 0.062 | 5.07249070434757e-11 | 39 |
| smox6 | 3.62848273518706e-15 | 0.431870003 | 0.369 | 0.085 | 9.13252819619232e-11 | 39 |
| ywhaqb8 | 4.85759619775224e-15 | 0.4183221 | 0.492 | 0.132 | 1.22260838701226e-10 | 39 |
| jdp2b9 | 5.8785572784115e-15 | 0.705508636 | 0.538 | 0.17 | 1.47957408140339e-10 | 39 |
| dhrs93 | 9.69590966418303e-15 | 0.275900671 | 0.215 | 0.034 | 2.44036350337823e-10 | 39 |
| mdka8 | 1.69231401678157e-14 | 0.65720456 | 0.369 | 0.094 | 4.25938514883754e-10 | 39 |
| postnb5 | 3.28362320230295e-14 | 0.570347101 | 0.338 | 0.079 | 8.2645512378763e-10 | 39 |
| fabp37 | 3.61238049571964e-14 | 0.622080837 | 0.662 | 0.242 | 9.09200046967677e-10 | 39 |

| | | | | | | |
|---|---|---|---|---|---|---|
| nt5c3a | 4.38783749131925e-14 | 0.262574184 | 0.231 | 0.04 | 1.10437481819014e-09 | 39 |
| cebpd8 | 5.61926290530667e-14 | 0.517601779 | 0.462 | 0.129 | 1.41431228063663e-09 | 39 |
| rbfox22 | 5.65350701901843e-14 | 0.28354203 | 0.215 | 0.036 | 1.42293118161675e-09 | 39 |
| pak2a2 | 6.32700016774209e-14 | 0.407513883 | 0.292 | 0.062 | 1.59244267221901e-09 | 39 |
| junba11 | 6.5761758852377e-14 | 0.812038314 | 0.677 | 0.31 | 1.65515770855548e-09 | 39 |
| chmp1b3 | 1.10974981082076e-13 | 0.408473016 | 0.446 | 0.122 | 2.79312929885476e-09 | 39 |
| lamc16 | 1.2258757200449e-13 | 0.278845842 | 0.277 | 0.055 | 3.08540659978101e-09 | 39 |
| smdt1b4 | 1.99049525065387e-13 | 0.443257914 | 0.462 | 0.134 | 5.00987749637072e-09 | 39 |
| fosl212 | 2.63178879374803e-13 | 0.491185117 | 0.431 | 0.122 | 6.62394921498441e-09 | 39 |
| snphb | 2.95689400350529e-13 | 0.285441485 | 0.2 | 0.034 | 7.44220651742246e-09 | 39 |
| eef1a1l16 | 3.0820156141731e-13 | 0.26654626 | 1 | 0.915 | 7.75712509931227e-09 | 39 |
| hbegfa2 | 5.29451368842585e-13 | 0.460043986 | 0.415 | 0.112 | 1.3325761502399e-08 | 39 |
| pim110 | 5.56241359056898e-13 | 0.523337134 | 0.538 | 0.171 | 1.40000387661031e-08 | 39 |
| atp5if1b3 | 6.80059444055814e-13 | 0.46650872 | 0.585 | 0.191 | 1.71164161474408e-08 | 39 |
| csde13 | 7.9729202172065e-13 | 0.476375068 | 0.615 | 0.208 | 2.0067042894687e-08 | 39 |
| znf7061 | 8.42584578989922e-13 | 0.466442012 | 0.569 | 0.182 | 2.12070112685973e-08 | 39 |
| arpc4l12 | 1.02010986357152e-12 | 0.454062267 | 0.6 | 0.197 | 2.56751451562315e-08 | 39 |
| adh58 | 1.24437014536077e-12 | 0.536624488 | 0.615 | 0.222 | 3.13195521885853e-08 | 39 |
| ip6k2b9 | 1.52360744760706e-12 | 0.421267855 | 0.431 | 0.123 | 3.8347675848822e-08 | 39 |
| kdm6ba2 | 2.22022454564039e-12 | 0.270258895 | 0.246 | 0.049 | 5.5880831589223e-08 | 39 |
| wdr17 | 2.89966278505661e-12 | 0.396540957 | 0.477 | 0.142 | 7.29816126370897e-08 | 39 |
| ak24 | 4.8980817712259e-12 | 0.536702618 | 0.615 | 0.233 | 1.23279820099985e-07 | 39 |
| b3gnt5a1 | 5.33345208573562e-12 | 0.278440043 | 0.185 | 0.032 | 1.3423765554588e-07 | 39 |
| itpkcb5 | 7.29953661953019e-12 | 0.271637627 | 0.323 | 0.078 | 1.83722037176955e-07 | 39 |
| ccni13 | 8.22686861463257e-12 | 0.557113006 | 0.723 | 0.294 | 2.07062056161687e-07 | 39 |
| higd1a10 | 8.47813005525093e-12 | 0.622012347 | 0.692 | 0.31 | 2.13386055360611e-07 | 39 |
| ctnna1 | 9.39286708036136e-12 | 0.265606483 | 0.308 | 0.072 | 2.36409071545615e-07 | 39 |
| eif4a1b10 | 1.59464511988717e-11 | 0.561696252 | 0.677 | 0.28 | 4.01356230224401e-07 | 39 |
| sptssa | 1.61817003981598e-11 | 0.288139616 | 0.323 | 0.08 | 4.07277217321283e-07 | 39 |
| tuba8l6 | 1.6509810065806e-11 | 0.451241336 | 0.554 | 0.189 | 4.15535409546271e-07 | 39 |
| rhoca1 | 1.91617053239794e-11 | 0.272069965 | 0.262 | 0.058 | 4.82280961299237e-07 | 39 |
| entpd5b | 2.03974110415599e-11 | 0.328540937 | 0.108 | 0.013 | 5.13382438505022e-07 | 39 |
| myl12.113 | 2.31194761565667e-11 | 0.591332904 | 0.831 | 0.392 | 5.81894095384627e-07 | 39 |
| cotl113 | 2.32638538505696e-11 | 0.584817497 | 0.754 | 0.306 | 5.85527937564986e-07 | 39 |
| zgc:16273017 | 2.40316236148566e-11 | 0.667509965 | 0.815 | 0.392 | 6.04851934762325e-07 | 39 |

| | | | | | | |
|---|---|---|---|---|---|---|
| mvp4 | 2.44328850360079e-11 | 0.301841656 | 0.246 | 0.053 | 6.14951283471282e-07 | 39 |
| cd151l | 3.67936772532846e-11 | 0.255552656 | 0.262 | 0.058 | 9.2606006278792e-07 | 39 |
| baiap2a | 3.72017002901701e-11 | 0.252849286 | 0.2 | 0.038 | 9.36329594603292e-07 | 39 |
| ahcy3 | 4.91855751279297e-11 | 0.420951482 | 0.508 | 0.168 | 1.23795174039486e-06 | 39 |
| maff7 | 4.99400712973701e-11 | 0.30847741 | 0.323 | 0.084 | 1.25694165448351e-06 | 39 |
| hmgn34 | 6.53278190085486e-11 | 0.339337302 | 0.262 | 0.061 | 1.64423587662616e-06 | 39 |
| ddit314 | 8.13617431117556e-11 | 0.549530592 | 0.492 | 0.177 | 2.04779371237978e-06 | 39 |
| slco2b15 | 9.40362915398306e-11 | 0.286114513 | 0.277 | 0.065 | 2.366799421766e-06 | 39 |
| mibp215 | 1.07159083781952e-10 | 0.580956872 | 0.877 | 0.492 | 2.69708697970794e-06 | 39 |
| si:ch1073-429i10.3.1 | 1.18742864883715e-10 | 0.420831966 | 0.569 | 0.202 | 2.98863916625823e-06 | 39 |
| stk353 | 1.26043228733649e-10 | 0.265993263 | 0.246 | 0.056 | 3.17238202399722e-06 | 39 |
| emd3 | 1.55220085410987e-10 | 0.256574935 | 0.246 | 0.056 | 3.90673432970913e-06 | 39 |
| csrnp1b11 | 1.97935877839765e-10 | 0.504340326 | 0.492 | 0.18 | 4.98184810934904e-06 | 39 |
| cdc42se11 | 2.09587439504735e-10 | 0.300227015 | 0.323 | 0.085 | 5.27510626489466e-06 | 39 |
| nsa217 | 2.10600449588958e-10 | 0.565307279 | 0.769 | 0.362 | 5.30060271570448e-06 | 39 |
| si:dkey-79d12.57 | 2.40124025381181e-10 | 0.356857798 | 0.292 | 0.077 | 6.04368159481893e-06 | 39 |
| ppiab7 | 2.73691866216453e-10 | 0.461052017 | 0.985 | 0.657 | 6.8885505808019e-06 | 39 |
| nfkbiaa7 | 2.78617679295239e-10 | 0.4950241 | 0.446 | 0.153 | 7.01252837018186e-06 | 39 |
| csnk1a1 | 2.8612846252211e-10 | 0.473157019 | 0.6 | 0.234 | 7.20156727321898e-06 | 39 |
| pxna2 | 3.60509392976635e-10 | 0.29299531 | 0.338 | 0.092 | 9.07366091182892e-06 | 39 |
| ppib11 | 3.85597750449027e-10 | 0.509267828 | 0.8 | 0.332 | 9.70510978105156e-06 | 39 |
| wls6 | 5.19208528825873e-10 | 0.34963946 | 0.4 | 0.124 | 1.30679594620184e-05 | 39 |
| rpl5a18 | 5.95035493910286e-10 | 0.545916666 | 0.877 | 0.492 | 1.4976448346228e-05 | 39 |
| ier2b15 | 6.05685687951171e-10 | 0.547318582 | 0.554 | 0.224 | 1.5244503080043e-05 | 39 |
| capza1a11 | 7.02646951605303e-10 | 0.434041016 | 0.538 | 0.2 | 1.76849211249539e-05 | 39 |
| arpc214 | 7.1682308973523e-10 | 0.513005934 | 0.631 | 0.268 | 1.8041720345546e-05 | 39 |
| tnfaip36 | 7.22168439864133e-10 | 0.320319625 | 0.262 | 0.066 | 1.81762574629404e-05 | 39 |
| rbp46 | 7.22551846090522e-10 | 0.523163236 | 0.4 | 0.128 | 1.81859074142523e-05 | 39 |
| naca15 | 7.37680022867302e-10 | 0.458851837 | 0.908 | 0.619 | 1.85666684955471e-05 | 39 |
| dusp56 | 7.60244877116221e-10 | 0.384963376 | 0.277 | 0.073 | 1.91346033121382e-05 | 39 |
| si:ch211-15b10.65 | 8.66953995698451e-10 | 0.310166039 | 0.277 | 0.072 | 2.18203651177343e-05 | 39 |
| glo14 | 1.04976719201655e-09 | 0.407195903 | 0.538 | 0.197 | 2.64215904558645e-05 | 39 |
| lxn9 | 1.32667612794642e-09 | 0.364056842 | 0.446 | 0.148 | 3.33911114642835e-05 | 39 |
| mbpb2 | 1.60584684920611e-09 | 0.268719239 | 0.246 | 0.06 | 4.04175593476687e-05 | 39 |
| atf4b13 | 1.71721020682767e-09 | 0.511532448 | 0.615 | 0.265 | 4.32204636956455e-05 | 39 |

| | | | | | | |
|---|---|---|---|---|---|---|
| XLOC-043857 | 1.8867309506444e-09 | 0.275294417 | 0.262 | 0.067 | 4.74871312967688e-05 | 39 |
| pno15 | 2.00340782342171e-09 | 0.346386331 | 0.492 | 0.172 | 5.04237715077009e-05 | 39 |
| rps230 | 2.02275475874199e-09 | 0.350602583 | 1 | 0.829 | 5.09107145227772e-05 | 39 |
| eif3c8 | 2.06695603455555e-09 | 0.36847963 | 0.477 | 0.169 | 5.20232164337287e-05 | 39 |
| cyr617 | 2.17401201478496e-09 | 0.417625982 | 0.262 | 0.069 | 5.47177084001227e-05 | 39 |
| h3f3c13 | 2.25954313562811e-09 | 0.47008354 | 0.631 | 0.257 | 5.68704411806238e-05 | 39 |
| actb214 | 2.4352093897437e-09 | 0.324024239 | 0.985 | 0.868 | 6.12917851304593e-05 | 39 |
| junbb7 | 2.64191560647957e-09 | 0.251644342 | 0.277 | 0.073 | 6.64943738994842e-05 | 39 |
| rpl1117 | 3.02877760648432e-09 | 0.337984622 | 0.985 | 0.814 | 7.62313035776039e-05 | 39 |
| irf2bp2a8 | 3.6254454285155e-09 | 0.428684227 | 0.477 | 0.177 | 9.12488359903066e-05 | 39 |
| rpsa22 | 4.135275295077e-09 | 0.384932871 | 0.985 | 0.752 | 0.000104081 | 39 |
| agrn3 | 4.2178573948308e-09 | 0.274923409 | 0.215 | 0.05 | 0.000106159 | 39 |
| rpl22l114 | 4.33891526210555e-09 | 0.529361201 | 0.8 | 0.392 | 0.000109206 | 39 |
| rab1ab3 | 4.43261364619031e-09 | 0.286953978 | 0.385 | 0.122 | 0.000111564 | 39 |
| chac18 | 4.58101329242079e-09 | 0.531136859 | 0.769 | 0.399 | 0.0001153 | 39 |
| cox8a11 | 5.82589041734767e-09 | 0.444221287 | 0.554 | 0.228 | 0.000146632 | 39 |
| nfe2l2a3 | 6.23288831267367e-09 | 0.250866024 | 0.262 | 0.069 | 0.000156876 | 39 |
| epcam13 | 7.40729139713713e-09 | 0.541060517 | 0.769 | 0.507 | 0.000186434 | 39 |
| tob1a4 | 7.53223759960697e-09 | 0.385251527 | 0.385 | 0.13 | 0.000189579 | 39 |
| btf314 | 9.08525709431838e-09 | 0.454647095 | 0.923 | 0.587 | 0.000228667 | 39 |
| rpl3216 | 1.04517527312664e-08 | 0.340162281 | 1 | 0.813 | 0.00026306 | 39 |
| ier56 | 1.05528986229142e-08 | 0.424745228 | 0.385 | 0.137 | 0.000265606 | 39 |
| ankrd2210 | 1.13018163160726e-08 | 0.340098285 | 0.431 | 0.144 | 0.000284455 | 39 |
| vdac26 | 1.3225296986892e-08 | 0.481000035 | 0.646 | 0.296 | 0.000332867 | 39 |
| gstr3 | 1.38294952682847e-08 | 0.296934426 | 0.369 | 0.119 | 0.000348075 | 39 |
| tgif13 | 1.48509788568037e-08 | 0.26881474 | 0.308 | 0.092 | 0.000373784 | 39 |
| wbp2nl3 | 1.62441327590972e-08 | 0.323367872 | 0.292 | 0.088 | 0.000408849 | 39 |
| reep55 | 1.62598675773138e-08 | 0.328962415 | 0.446 | 0.159 | 0.000409245 | 39 |
| sparc8 | 2.20884621653806e-08 | 0.395070818 | 0.446 | 0.159 | 0.000555945 | 39 |
| rpl3418 | 2.22952326313645e-08 | 0.346908786 | 0.985 | 0.784 | 0.000561149 | 39 |
| cct87 | 2.30411189225215e-08 | 0.363472616 | 0.523 | 0.202 | 0.000579922 | 39 |
| zgc:11418822 | 2.53602907851093e-08 | 0.461326673 | 0.938 | 0.628 | 0.000638293 | 39 |
| rpl27a19 | 2.62635374603617e-08 | 0.519419093 | 0.923 | 0.496 | 0.000661027 | 39 |
| rplp220 | 2.80945347499893e-08 | 0.375290629 | 0.969 | 0.758 | 0.000707111 | 39 |
| eef1g15 | 3.13624829661399e-08 | 0.451027754 | 0.938 | 0.621 | 0.000789362 | 39 |

| | | | | | | |
|---|---|---|---|---|---|---|
| rpl23a20 | 4.4432359837581e-08 | 0.453828049 | 0.969 | 0.634 | 0.001118318 | 39 |
| capns1b7 | 4.62900057781274e-08 | 0.262017678 | 0.369 | 0.122 | 0.001165073 | 39 |
| ppiaa14 | 6.92496402730496e-08 | 0.29730714 | 0.969 | 0.804 | 0.001742944 | 39 |
| capzb11 | 7.0907575300918e-08 | 0.392088688 | 0.6 | 0.254 | 0.001784673 | 39 |
| rplp09 | 8.33137035977519e-08 | 0.279441124 | 1 | 0.851 | 0.002096923 | 39 |
| hopx7 | 1.03241525762896e-07 | 0.374476788 | 0.231 | 0.066 | 0.002598486 | 39 |
| romo116 | 1.09664724979949e-07 | 0.467085806 | 0.754 | 0.386 | 0.002760151 | 39 |
| si:dkey-51e6.15 | 1.40389659653451e-07 | 0.311169909 | 0.338 | 0.117 | 0.003533467 | 39 |
| eef1db12 | 1.69759859049994e-07 | 0.428165542 | 0.6 | 0.274 | 0.004272686 | 39 |
| stau28 | 1.76308224332757e-07 | 0.265188557 | 0.277 | 0.086 | 0.004437502 | 39 |
| rpl3621 | 1.81649022981496e-07 | 0.339295039 | 0.985 | 0.79 | 0.004571924 | 39 |
| rpl2918 | 1.84783060910341e-07 | 0.323177374 | 1 | 0.809 | 0.004650805 | 39 |
| cct28 | 2.03869552555297e-07 | 0.338533956 | 0.554 | 0.228 | 0.005131193 | 39 |
| cap114 | 2.24766303425631e-07 | 0.350785548 | 0.569 | 0.234 | 0.005657143 | 39 |
| ctnnb16 | 2.7225792967622e-07 | 0.25465526 | 0.415 | 0.15 | 0.00685246 | 39 |
| rpl3018 | 2.98357382332459e-07 | 0.386518086 | 0.985 | 0.708 | 0.007509357 | 39 |
| prelid3b4 | 3.0392295143367e-07 | 0.380350163 | 0.554 | 0.239 | 0.007649437 | 39 |
| hdlbpa8 | 3.19453908872769e-07 | 0.33409064 | 0.415 | 0.16 | 0.008040335 | 39 |
| rab106 | 3.26304743063219e-07 | 0.302571497 | 0.492 | 0.195 | 0.008212764 | 39 |
| sdc49 | 3.56952226525944e-07 | 0.468457473 | 0.569 | 0.258 | 0.008984131 | 39 |
| rps1619 | 4.1287295153411e-07 | 0.361885923 | 0.969 | 0.738 | 0.010391599 | 39 |
| rsl24d116 | 4.84229410334176e-07 | 0.468396752 | 0.846 | 0.444 | 0.01218757 | 39 |
| si:ch73-46j18.59 | 5.22975080487887e-07 | 0.298627324 | 0.462 | 0.176 | 0.01316276 | 39 |
| cebpb16 | 6.82493263844803e-07 | 0.530396155 | 0.662 | 0.354 | 0.017177673 | 39 |
| eif1b10 | 7.0057844419915e-07 | 0.460897731 | 0.723 | 0.383 | 0.017632859 | 39 |
| btg211 | 7.10732487076354e-07 | 0.504058724 | 0.631 | 0.374 | 0.017888426 | 39 |
| thy113 | 7.20307960057663e-07 | 0.510516354 | 0.815 | 0.451 | 0.018129431 | 39 |
| rps4x15 | 8.10247576043006e-07 | 0.360089017 | 0.954 | 0.711 | 0.020393121 | 39 |
| rpl18a16 | 8.10307276884957e-07 | 0.31555752 | 0.985 | 0.785 | 0.020394624 | 39 |
| rps1211 | 8.7585056986999e-07 | 0.276265995 | 1 | 0.84 | 0.022044283 | 39 |
| nr4a113 | 9.18025735749782e-07 | 0.327574092 | 0.431 | 0.168 | 0.02310579 | 39 |
| pmaip13 | 9.44043466002419e-07 | 0.259542577 | 0.308 | 0.105 | 0.02376063 | 39 |
| zgc:1537136 | 1.00159609421291e-06 | 0.268895937 | 0.4 | 0.153 | 0.025209172 | 39 |
| rpl3818 | 1.00164994586152e-06 | 0.375374232 | 1 | 0.728 | 0.025210527 | 39 |
| pon3.21 | 1.23764932820352e-06 | 0.250871287 | 0.292 | 0.097 | 0.031150396 | 39 |

| | | | | | | |
|---|---|---|---|---|---|---|
| gapdhs15 | 1.35712224576925e-06 | 0.374044775 | 0.615 | 0.289 | 0.03415741 | 39 |
| mtpn6 | 1.46514869576388e-06 | 0.263280033 | 0.385 | 0.147 | 0.036876328 | 39 |
| eif3f16 | 1.49729419028008e-06 | 0.463930067 | 0.846 | 0.432 | 0.037685397 | 39 |
| zgc:1583439 | 1.51390831400161e-06 | 0.416914724 | 0.4 | 0.167 | 0.038103558 | 39 |
| klf6a10 | 1.66126350245223e-06 | 0.255655239 | 0.431 | 0.168 | 0.041812341 | 39 |
| rpl922 | 1.87633732332111e-06 | 0.396787015 | 0.892 | 0.63 | 0.047225534 | 39 |
| rps1518 | 2.32629397188986e-06 | 0.371994452 | 0.938 | 0.685 | 0.058550493 | 39 |
| si:ch1073-325m22.2! | 2.7595474463205e-06 | 0.311554059 | 0.523 | 0.23 | 0.06945505 | 39 |
| eif3k5 | 3.07374055610458e-06 | 0.34031108 | 0.508 | 0.23 | 0.077362976 | 39 |
| fosab12 | 3.23752665212202e-06 | 0.515075043 | 0.631 | 0.35 | 0.081485308 | 39 |
| chmp2a4 | 3.2898968892482e-06 | 0.251128947 | 0.354 | 0.135 | 0.082803415 | 39 |
| rps1912 | 3.35889804614041e-06 | 0.257430228 | 1 | 0.863 | 0.084540105 | 39 |
| rps27a16 | 3.68787218414045e-06 | 0.298939989 | 0.954 | 0.752 | 0.092820055 | 39 |
| f11r.15 | 4.20620019621911e-06 | 0.259978974 | 0.446 | 0.179 | 0.105865853 | 39 |
| jun11 | 4.70814601619669e-06 | 0.409980885 | 0.492 | 0.229 | 0.118499327 | 39 |
| h2afx112 | 4.7354238384367e-06 | 0.264151199 | 0.477 | 0.199 | 0.119185883 | 39 |
| rpl619 | 4.98170777458215e-06 | 0.383927554 | 0.969 | 0.668 | 0.125384603 | 39 |
| rps1021 | 5.01437519858305e-06 | 0.377659776 | 0.969 | 0.694 | 0.126206809 | 39 |
| rpl2417 | 5.04676652796078e-06 | 0.301500584 | 0.969 | 0.777 | 0.127022067 | 39 |
| fkbp1aa10 | 5.50577158299336e-06 | 0.38613518 | 0.738 | 0.388 | 0.138574765 | 39 |
| rpl3714 | 5.5522240479373e-06 | 0.269577003 | 1 | 0.85 | 0.139743927 | 39 |
| pfn25 | 5.60341946950011e-06 | 0.316003432 | 0.508 | 0.227 | 0.141032465 | 39 |
| rps26l18 | 5.77862397950081e-06 | 0.377000244 | 0.938 | 0.665 | 0.145442187 | 39 |
| eif3d8 | 6.47659800073987e-06 | 0.270376055 | 0.569 | 0.254 | 0.163009495 | 39 |
| rpl1712 | 7.065961345627e-06 | 0.323393918 | 0.985 | 0.746 | 0.177843181 | 39 |
| nop538 | 7.68689997622084e-06 | 0.298814761 | 0.554 | 0.252 | 0.193471586 | 39 |
| si:dkey-112e7.211 | 9.00750758431627e-06 | 0.297417924 | 0.462 | 0.207 | 0.226709958 | 39 |
| rps2513 | 9.24446925438364e-06 | 0.31653592 | 0.954 | 0.725 | 0.232674047 | 39 |
| rpl1516 | 1.06969582007376e-05 | 0.316792866 | 0.969 | 0.745 | 0.269231741 | 39 |
| abracl6 | 1.36589089315574e-05 | 0.270635899 | 0.508 | 0.222 | 0.343781079 | 39 |
| rps1317 | 1.36885345906941e-05 | 0.348354205 | 0.985 | 0.724 | 0.344526727 | 39 |
| ddx216 | 1.53186416496782e-05 | 0.308976404 | 0.462 | 0.208 | 0.385554892 | 39 |
| eef1da13 | 1.68103283302568e-05 | 0.415330407 | 0.831 | 0.493 | 0.423099154 | 39 |
| rps2818 | 1.82499063753648e-05 | 0.267343378 | 0.985 | 0.827 | 0.459331894 | 39 |
| rps2124 | 1.92771258159295e-05 | 0.340057702 | 0.985 | 0.75 | 0.48518598 | 39 |

| | | | | | | |
|---|---|---|---|---|---|---|
| itm2ba12 | 2.00824428438438e-05 | 0.342896559 | 0.554 | 0.274 | 0.505455004 | 39 |
| rps1111 | 2.21494139360451e-05 | 0.280552787 | 0.969 | 0.766 | 0.557478599 | 39 |
| rpl36a14 | 2.26068372090735e-05 | 0.275952811 | 1 | 0.81 | 0.568991486 | 39 |
| rpl2217 | 2.39149159242495e-05 | 0.344615139 | 0.954 | 0.694 | 0.601914519 | 39 |
| rps1417 | 2.59193933981912e-05 | 0.28560387 | 0.954 | 0.759 | 0.652365212 | 39 |
| btg116 | 3.12921433591481e-05 | 0.349829417 | 0.646 | 0.333 | 0.787591956 | 39 |
| ddx53 | 3.96021230051603e-05 | 0.382547458 | 0.631 | 0.358 | 0.996745834 | 39 |
| rplp2l21 | 4.03759524940153e-05 | 0.306457345 | 0.985 | 0.774 | 1 | 39 |
| rpl1310 | 4.22538350969349e-05 | 0.262888231 | 0.985 | 0.805 | 1 | 39 |
| mt-co119 | 4.2654831769694e-05 | 0.3486306 | 0.923 | 0.658 | 1 | 39 |
| rps2418 | 4.57281996459558e-05 | 0.299696697 | 0.969 | 0.759 | 1 | 39 |
| eef2b9 | 5.76378229375568e-05 | 0.377053234 | 0.923 | 0.614 | 1 | 39 |
| rack111 | 6.26514029910109e-05 | 0.281989577 | 0.954 | 0.745 | 1 | 39 |
| rpl1817 | 6.37878071618368e-05 | 0.286198615 | 0.969 | 0.774 | 1 | 39 |
| ywhabl5 | 6.76256853087577e-05 | 0.29587074 | 0.554 | 0.279 | 1 | 39 |
| hsp90ab111 | 7.5698023180289e-05 | 0.269323292 | 0.985 | 0.791 | 1 | 39 |
| rps27.117 | 7.76894398870051e-05 | 0.36099862 | 0.954 | 0.68 | 1 | 39 |
| rpl10a12 | 7.92328113630885e-05 | 0.279903399 | 0.954 | 0.765 | 1 | 39 |
| rpl2815 | 0.000107353 | 0.316823712 | 0.985 | 0.75 | 1 | 39 |
| rpl3119 | 0.000107703 | 0.337623548 | 0.923 | 0.655 | 1 | 39 |
| eif2s36 | 0.000118485 | 0.275686101 | 0.569 | 0.285 | 1 | 39 |
| uba5217 | 0.000124584 | 0.345963585 | 0.908 | 0.631 | 1 | 39 |
| nccrp15 | 0.000127427 | 0.266966633 | 0.415 | 0.195 | 1 | 39 |
| atp5pf7 | 0.000133092 | 0.274265281 | 0.585 | 0.299 | 1 | 39 |
| rpl816 | 0.000152153 | 0.250548662 | 0.938 | 0.781 | 1 | 39 |
| arpc313 | 0.000159396 | 0.278377334 | 0.615 | 0.311 | 1 | 39 |
| eif3ea12 | 0.000174002 | 0.26178384 | 0.554 | 0.278 | 1 | 39 |
| caprin1b4 | 0.000183327 | 0.252867671 | 0.492 | 0.248 | 1 | 39 |
| rps1818 | 0.000191674 | 0.314231539 | 0.969 | 0.725 | 1 | 39 |
| eif4ebp26 | 0.000192193 | 0.264583812 | 0.369 | 0.175 | 1 | 39 |
| pabpc1a14 | 0.000202157 | 0.33713679 | 0.815 | 0.541 | 1 | 39 |
| ptmaa19 | 0.000202391 | 0.343520009 | 0.708 | 0.396 | 1 | 39 |
| hmgn614 | 0.000216427 | 0.263648555 | 0.538 | 0.266 | 1 | 39 |
| fth1a | 0.000228191 | 0.309973457 | 0.846 | 0.637 | 1 | 39 |
| serf215 | 0.000237409 | 0.303648264 | 0.631 | 0.344 | 1 | 39 |

| | | | | | | |
|---|---|---|---|---|---|---|
| ier2a15 | 0.000278757 | 0.293059767 | 0.508 | 0.263 | 1 | 39 |
| rps914 | 0.000313207 | 0.267356671 | 0.985 | 0.777 | 1 | 39 |
| zgc:1538677 | 0.000316438 | 0.322139049 | 0.738 | 0.437 | 1 | 39 |
| serbp1a14 | 0.000429177 | 0.342411949 | 0.785 | 0.483 | 1 | 39 |
| rps516 | 0.000631145 | 0.301411426 | 0.954 | 0.702 | 1 | 39 |
| actb16 | 0.000696427 | 0.260208806 | 0.954 | 0.785 | 1 | 39 |
| eif5a10 | 0.00078582 | 0.313381952 | 0.754 | 0.479 | 1 | 39 |
| pnrc220 | 0.000848051 | 0.293972744 | 0.738 | 0.569 | 1 | 39 |
| faua15 | 0.000982034 | 0.288065375 | 0.908 | 0.69 | 1 | 39 |
| rps2313 | 0.001008413 | 0.278764818 | 0.969 | 0.723 | 1 | 39 |
| si:ch211-157c3.46 | 0.001064315 | 0.253672899 | 0.215 | 0.091 | 1 | 39 |
| rps15a16 | 0.001170198 | 0.261365851 | 0.954 | 0.78 | 1 | 39 |
| rpl2115 | 0.001966772 | 0.257469261 | 0.954 | 0.757 | 1 | 39 |
| ndufa4l10 | 0.003870382 | 0.250311242 | 0.692 | 0.396 | 1 | 39 |
| COX319 | 0.005368284 | 0.276716513 | 0.908 | 0.701 | 1 | 39 |
| rps315 | 0.005817924 | 0.26197991 | 0.908 | 0.7 | 1 | 39 |
| CR383676.120 | 0.009691853 | 0.284556497 | 0.785 | 0.727 | 1 | 39 |
| dmrt12 | 4.27647719131215e-283 | 1.52750704 | 0.661 | 0.017 | 1.07634654428136e-278 | 40 |
| hspb82 | 5.81929210978332e-259 | 1.566559996 | 0.694 | 0.021 | 1.46465763111136e-254 | 40 |
| gsdf1 | 3.26825638781736e-258 | 2.392538762 | 0.984 | 0.047 | 8.22587450249752e-254 | 40 |
| gstt1a4 | 7.30663731613217e-258 | 1.973615575 | 0.855 | 0.034 | 1.83900754609731e-253 | 40 |
| gpx3 | 2.05790031768341e-255 | 1.157539356 | 0.387 | 0.006 | 5.17952930957736e-251 | 40 |
| igf31 | 1.52828581079546e-199 | 1.290779522 | 0.5 | 0.014 | 3.8465425571911e-195 | 40 |
| amh | 2.367774728281e-195 | 0.971036922 | 0.306 | 0.005 | 5.95945221361045e-191 | 40 |
| si:dkeyp-80c12.8 | 1.75628872215186e-154 | 0.499629444 | 0.177 | 0.002 | 4.42040308478401e-150 | 40 |
| cyb5d1 | 5.1046329272358e-152 | 0.566831143 | 0.21 | 0.003 | 1.28478506145598e-147 | 40 |
| zgc:921621 | 4.85512610525394e-125 | 0.843160992 | 0.323 | 0.009 | 1.22198668943137e-120 | 40 |
| scn1ba | 3.78419943072714e-111 | 0.701312018 | 0.258 | 0.006 | 9.52445154719713e-107 | 40 |
| ak41 | 2.15618070838057e-102 | 1.226452762 | 0.5 | 0.03 | 5.42689122492306e-98 | 40 |
| kcnj1a.1 | 1.21024797870317e-93 | 0.448926618 | 0.145 | 0.002 | 3.046073137598e-89 | 40 |
| zgc:1527915 | 1.28287950875486e-88 | 1.594945534 | 0.677 | 0.067 | 3.22887943558511e-84 | 40 |
| nrgna1 | 6.26026854511599e-76 | 0.810021866 | 0.242 | 0.009 | 1.57564699012024e-71 | 40 |
| ppp1r1b | 3.6917980620272e-75 | 0.563709869 | 0.194 | 0.005 | 9.29188654231626e-71 | 40 |
| nkx3.3 | 8.39123691150899e-73 | 0.374452467 | 0.113 | 0.002 | 2.1119904182577e-68 | 40 |
| ftr823 | 4.8849170506538e-67 | 0.835593786 | 0.355 | 0.022 | 1.22948477247905e-62 | 40 |

| | | | | | | |
|---|---|---|---|---|---|---|
| si:ch211-117c9.51 | 1.46064526174317e-64 | 0.830871331 | 0.306 | 0.017 | 3.6762980592814e-60 | 40 |
| cth2 | 1.85053771227814e-64 | 1.139868437 | 0.484 | 0.043 | 4.65761836803286e-60 | 40 |
| isg155 | 2.86118481640452e-61 | 1.311195921 | 0.581 | 0.067 | 7.20131606440854e-57 | 40 |
| ctnnd2b | 1.33991113900571e-58 | 0.357343722 | 0.113 | 0.002 | 3.37242234576347e-54 | 40 |
| sox9a | 3.61384777555871e-58 | 0.589452278 | 0.226 | 0.01 | 9.09569346630372e-54 | 40 |
| fabp7a4 | 5.718967813785e-58 | 0.973458584 | 0.355 | 0.026 | 1.43940700905155e-53 | 40 |
| CABZ01021532.11 | 5.01225800979639e-57 | 0.825979207 | 0.274 | 0.016 | 1.26153521848565e-52 | 40 |
| add2 | 3.33393286885167e-55 | 0.480140669 | 0.161 | 0.005 | 8.39117563761276e-51 | 40 |
| adm2a | 9.76534729115173e-55 | 0.350685644 | 0.129 | 0.003 | 2.45784025970998e-50 | 40 |
| jam2a3 | 8.87520897489005e-53 | 0.979621493 | 0.403 | 0.036 | 2.23380134689008e-48 | 40 |
| ndrg3a | 4.62773183544363e-51 | 0.709097495 | 0.306 | 0.021 | 1.16475382566281e-46 | 40 |
| inha1 | 6.26544453021919e-49 | 0.520497623 | 0.177 | 0.007 | 1.57694973381087e-44 | 40 |
| zgc:194990 | 6.42885123837223e-49 | 0.317804404 | 0.113 | 0.003 | 1.61807756818591e-44 | 40 |
| msx1b | 2.06720967757851e-48 | 0.445568803 | 0.145 | 0.005 | 5.20296003749736e-44 | 40 |
| si:dkey-188i13.91 | 1.50730305902444e-47 | 0.874744441 | 0.306 | 0.023 | 3.79373106925862e-43 | 40 |
| ifi467 | 1.38396816949949e-46 | 1.252494349 | 0.661 | 0.12 | 3.48330948581326e-42 | 40 |
| mmel1 | 6.5965976479999e-46 | 0.362293739 | 0.129 | 0.004 | 1.66029766202509e-41 | 40 |
| ifi458 | 1.12977328519587e-45 | 1.453092177 | 0.677 | 0.128 | 2.84352638150948e-41 | 40 |
| plxnb3 | 3.56332710081638e-45 | 0.419343785 | 0.161 | 0.007 | 8.96853798004474e-41 | 40 |
| krt812 | 2.20405251997334e-43 | 1.41046742 | 0.903 | 0.275 | 5.5473797875209e-39 | 40 |
| scg31 | 2.85204989819303e-42 | 0.579407187 | 0.194 | 0.01 | 7.17832438876205e-38 | 40 |
| cahz2 | 2.69992757443551e-40 | 0.861150335 | 0.371 | 0.039 | 6.79544771209673e-36 | 40 |
| cldnc3 | 7.40997914767021e-35 | 0.736100874 | 0.29 | 0.028 | 1.86501765167712e-30 | 40 |
| hpn1 | 1.3854951011444e-33 | 0.463992573 | 0.145 | 0.007 | 3.48715262007033e-29 | 40 |
| XLOC-042229 | 4.27881322859094e-33 | 0.343431572 | 0.129 | 0.006 | 1.07693450150405e-28 | 40 |
| wfdc2 | 1.388815563038e-32 | 0.34046552 | 0.129 | 0.006 | 3.49550989061034e-28 | 40 |
| rhoub1 | 3.50554856641239e-32 | 0.354998183 | 0.129 | 0.006 | 8.82311518680334e-28 | 40 |
| krt948 | 1.63663952209876e-31 | 1.08325852 | 0.613 | 0.129 | 4.11925801317037e-27 | 40 |
| krt18a.18 | 7.02658822839991e-31 | 1.098370768 | 0.597 | 0.126 | 1.76852199120597e-26 | 40 |
| cyp26a11 | 7.23244858781737e-31 | 0.486971785 | 0.161 | 0.01 | 1.82033498506775e-26 | 40 |
| si:dkey-188i13.106 | 9.08006936583452e-31 | 1.011012072 | 0.516 | 0.104 | 2.28536265868689e-26 | 40 |
| lgals174 | 1.18244022848283e-30 | 0.865295823 | 0.371 | 0.052 | 2.97608381106844e-26 | 40 |
| lgals9l33 | 4.34638678849567e-30 | 0.902380061 | 0.403 | 0.063 | 1.09394209079647e-25 | 40 |
| ckba4 | 1.01998781305937e-28 | 0.897947151 | 0.403 | 0.065 | 2.56720732668913e-24 | 40 |
| calm1a2 | 4.72075331653884e-28 | 1.095930557 | 0.613 | 0.164 | 1.18816640223966e-23 | 40 |

| | | | | | | |
|---|---|---|---|---|---|---|
| aldh1a24 | 2.60784620899022e-27 | 0.991303272 | 0.435 | 0.078 | 6.56368812340747e-23 | 40 |
| cldn11a6 | 5.73216654832455e-27 | 0.860102538 | 0.387 | 0.062 | 1.44272899854781e-22 | 40 |
| fkbp1b1 | 1.39357705679276e-26 | 0.443119391 | 0.177 | 0.014 | 3.5074940942417e-22 | 40 |
| lnx2a | 1.52934105526057e-26 | 0.321355556 | 0.113 | 0.006 | 3.84919850198533e-22 | 40 |
| fyna | 4.87410016523603e-26 | 0.422956193 | 0.161 | 0.012 | 1.22676227058826e-21 | 40 |
| sparc9 | 6.13724666467503e-26 | 0.997770881 | 0.629 | 0.158 | 1.54468361303206e-21 | 40 |
| fam13b1 | 5.09608910439924e-24 | 0.624625091 | 0.258 | 0.032 | 1.28263466668624e-19 | 40 |
| si:ch73-1a9.317 | 6.83243579010854e-23 | 1.014162725 | 0.694 | 0.266 | 1.71965576401242e-18 | 40 |
| ptmaa20 | 3.75123396880092e-22 | 0.933985711 | 0.823 | 0.395 | 9.44148077607503e-18 | 40 |
| marcksl1b10 | 5.22554515950679e-22 | 0.938192105 | 0.484 | 0.116 | 1.31521746119626e-17 | 40 |
| rpl1311 | 1.01226420917262e-21 | 0.437844699 | 0.952 | 0.805 | 2.54776778806656e-17 | 40 |
| cx435 | 3.28824562462533e-21 | 0.717877207 | 0.323 | 0.054 | 8.27618541261949e-17 | 40 |
| alcamb1 | 3.37029284580494e-21 | 0.434265379 | 0.161 | 0.015 | 8.48269006360644e-17 | 40 |
| prss353 | 4.6449421712716e-21 | 0.483144655 | 0.194 | 0.021 | 1.16908549508735e-16 | 40 |
| zgc:927447 | 8.5674948460792e-21 | 0.931126093 | 0.79 | 0.353 | 2.15635277780967e-16 | 40 |
| gstm.36 | 1.10722869060593e-20 | 0.856326371 | 0.452 | 0.101 | 2.78678389138607e-16 | 40 |
| cox5b22 | 1.4126041894288e-20 | 0.405497539 | 0.145 | 0.012 | 3.55538348437336e-16 | 40 |
| selenbp14 | 2.03205041702483e-20 | 0.764032321 | 0.355 | 0.069 | 5.1144676946098e-16 | 40 |
| cmbl | 3.51412548425092e-20 | 0.558057145 | 0.226 | 0.029 | 8.84470243131114e-16 | 40 |
| rbm471 | 1.32744993546645e-19 | 0.482460811 | 0.161 | 0.016 | 3.34105874257551e-15 | 40 |
| hmgn615 | 3.18089709288296e-19 | 0.871188065 | 0.661 | 0.265 | 8.00599989307713e-15 | 40 |
| soul5 | 4.25759173923738e-19 | 0.460297526 | 0.161 | 0.016 | 1.07159326484866e-14 | 40 |
| rpsa23 | 4.35061064976519e-19 | 0.48923298 | 0.984 | 0.752 | 1.0950051944394e-14 | 40 |
| ponzr110 | 4.0975708078142e-18 | 0.921805496 | 0.661 | 0.273 | 1.03131759661876e-13 | 40 |
| si:dkey-204f11.64 | 7.62028261262506e-18 | 0.525369375 | 0.21 | 0.028 | 1.9179489307716e-13 | 40 |
| fthl272 | 1.28065506462709e-17 | 0.576949131 | 0.952 | 0.695 | 3.22328073215993e-13 | 40 |
| gpm6ab2 | 1.79454483337586e-17 | 0.448278815 | 0.161 | 0.018 | 4.51668989112371e-13 | 40 |
| gstt1b6 | 3.92442887583497e-17 | 0.732512299 | 0.339 | 0.073 | 9.87739503758903e-13 | 40 |
| tmsb4x24 | 4.29485735211257e-17 | 0.655365226 | 1 | 0.686 | 1.08097264695321e-12 | 40 |
| rps240 | 4.31234159410198e-17 | 0.431013651 | 0.984 | 0.829 | 1.08537325581953e-12 | 40 |
| nr0b12 | 5.75217150452395e-17 | 0.33132606 | 0.129 | 0.012 | 1.44776404597363e-12 | 40 |
| LOC1104380241 | 1.83565312343639e-16 | 0.357795354 | 0.113 | 0.009 | 4.62015534637706e-12 | 40 |
| cbsb1 | 4.4350467764449e-16 | 0.572911942 | 0.226 | 0.036 | 1.11625692316342e-11 | 40 |
| zgc:11418823 | 8.65051748352629e-16 | 0.579406673 | 0.952 | 0.628 | 2.17724874542873e-11 | 40 |
| znf1104 | 8.97557805473492e-16 | 0.404231263 | 0.145 | 0.016 | 2.25906324059623e-11 | 40 |

| | | | | | | |
|---|---|---|---|---|---|---|
| fkbp1aa11 | 2.77710737418701e-15 | 0.731248513 | 0.726 | 0.388 | 6.98970155009128e-11 | 40 |
| fxyd6l4 | 3.00449593241628e-15 | 0.722624352 | 0.419 | 0.114 | 7.56201581229853e-11 | 40 |
| eef1a1l17 | 3.33961722694979e-15 | 0.270354055 | 0.968 | 0.916 | 8.40548259850993e-11 | 40 |
| ybx112 | 4.67107400454948e-15 | 0.506252298 | 0.887 | 0.67 | 1.17566261620506e-10 | 40 |
| rpl3914 | 1.43985628766172e-14 | 0.345438599 | 1 | 0.874 | 3.62397429041577e-10 | 40 |
| rps1620 | 1.73172953595238e-14 | 0.450640415 | 0.952 | 0.738 | 4.35859006903854e-10 | 40 |
| rpl10a13 | 1.0865062095613e-13 | 0.422392396 | 0.984 | 0.765 | 2.73462747884484e-09 | 40 |
| esr2b2 | 1.16542177937996e-13 | 0.496619089 | 0.21 | 0.036 | 2.93325007652143e-09 | 40 |
| calm2b8 | 1.32367995044145e-13 | 0.800674808 | 0.532 | 0.211 | 3.33157006726608e-09 | 40 |
| rps2419 | 1.61499502242828e-13 | 0.42399074 | 0.968 | 0.759 | 4.06478097194973e-09 | 40 |
| cox6a24 | 1.78465625472754e-13 | 0.523829613 | 0.21 | 0.036 | 4.49180132752376e-09 | 40 |
| hspe15 | 3.16392148142693e-13 | 0.784825999 | 0.613 | 0.295 | 7.96327397660344e-09 | 40 |
| ap2m1a3 | 3.29764091886918e-13 | 0.721142376 | 0.435 | 0.143 | 8.29983242870185e-09 | 40 |
| rpl923 | 4.14684871081293e-13 | 0.49772422 | 0.871 | 0.63 | 1.04372035202451e-08 | 40 |
| rpl2613 | 5.22108644954417e-13 | 0.35015377 | 1 | 0.841 | 1.31409524848577e-08 | 40 |
| rpl3217 | 5.710269523382e-13 | 0.384205577 | 0.984 | 0.814 | 1.43721773634001e-08 | 40 |
| si:ch211-197h24.96 | 5.76369495760919e-13 | 0.617578444 | 0.29 | 0.069 | 1.45066438388066e-08 | 40 |
| phc2a2 | 7.9674598153857e-13 | 0.64853452 | 0.355 | 0.094 | 2.00532996093443e-08 | 40 |
| rpl2116 | 8.3428843351253e-13 | 0.403064695 | 0.952 | 0.757 | 2.09982055830769e-08 | 40 |
| cox6b26 | 1.23062375462776e-12 | 0.584504772 | 0.774 | 0.534 | 3.09735692802261e-08 | 40 |
| rpl18a17 | 1.3006426043006e-12 | 0.391609703 | 0.952 | 0.786 | 3.27358737076419e-08 | 40 |
| rpl620 | 1.77155277865082e-12 | 0.485084821 | 0.935 | 0.668 | 4.45882118858625e-08 | 40 |
| appa7 | 1.82952885692552e-12 | 0.630548547 | 0.306 | 0.078 | 4.60474117999583e-08 | 40 |
| dag14 | 1.97411704896506e-12 | 0.518664924 | 0.242 | 0.05 | 4.96865520054016e-08 | 40 |
| aplp11 | 2.92552447486011e-12 | 0.358337922 | 0.129 | 0.016 | 7.36325255077542e-08 | 40 |
| rps1913 | 5.08718157360609e-12 | 0.327081406 | 1 | 0.863 | 1.28039273026092e-07 | 40 |
| rps1519 | 6.40794266777972e-12 | 0.453023135 | 0.919 | 0.685 | 1.61281509005348e-07 | 40 |
| rpl3120 | 7.19830280363768e-12 | 0.458256175 | 0.871 | 0.655 | 1.81174083264757e-07 | 40 |
| tmem45a1 | 8.1970029739648e-12 | 0.399328408 | 0.161 | 0.025 | 2.0631036785172e-07 | 40 |
| rps27a17 | 1.06233735695695e-11 | 0.367935258 | 0.919 | 0.752 | 2.67379689372495e-07 | 40 |
| naca16 | 1.32159332095797e-11 | 0.511947317 | 0.903 | 0.619 | 3.32631822951912e-07 | 40 |
| rpl2418 | 1.43523176666591e-11 | 0.364650594 | 0.935 | 0.777 | 3.61233483352142e-07 | 40 |
| rpl199 | 1.44566015638606e-11 | 0.327866568 | 1 | 0.852 | 3.63858204760808e-07 | 40 |
| zgc:92140 | 2.20365957886282e-11 | 0.357210783 | 0.161 | 0.026 | 5.54639079403984e-07 | 40 |
| rps2125 | 3.64392436921326e-11 | 0.426714607 | 0.984 | 0.75 | 9.17139324487286e-07 | 40 |

| | | | | | | |
|---|---|---|---|---|---|---|
| rpl3819 | 4.40796773574172e-11 | 0.427728964 | 0.968 | 0.728 | 1.10944139940883e-06 | 40 |
| rpl7a14 | 5.22670976415018e-11 | 0.305623176 | 0.903 | 0.811 | 1.31551058053896e-06 | 40 |
| rpl3419 | 7.55748252529738e-11 | 0.379101682 | 0.968 | 0.784 | 1.9021427767921e-06 | 40 |
| rps4x16 | 7.78047649417396e-11 | 0.437185596 | 0.968 | 0.711 | 1.95826812881864e-06 | 40 |
| slc7a3a4 | 9.26859310986292e-11 | 0.461150669 | 0.21 | 0.043 | 2.3328121998214e-06 | 40 |
| rps711 | 1.02141808680529e-10 | 0.324302885 | 0.952 | 0.821 | 2.57080718268025e-06 | 40 |
| rpl36a15 | 1.03064475446566e-10 | 0.329884103 | 0.952 | 0.81 | 2.59402978251462e-06 | 40 |
| rpl2714 | 1.13244541798059e-10 | 0.370245884 | 0.935 | 0.763 | 2.85025187251535e-06 | 40 |
| rpl2218 | 1.66182275108328e-10 | 0.417009332 | 0.887 | 0.695 | 4.18264168220151e-06 | 40 |
| rpl817 | 1.73315353916024e-10 | 0.370961763 | 0.968 | 0.781 | 4.3621741427124e-06 | 40 |
| oip5-as12 | 1.80890617254537e-10 | 0.370015387 | 0.161 | 0.028 | 4.55283594567945e-06 | 40 |
| eif4ebp17 | 1.97793370372937e-10 | 0.689287113 | 0.532 | 0.247 | 4.97826133891646e-06 | 40 |
| znf326 | 2.22702793685715e-10 | 0.46409829 | 0.226 | 0.05 | 5.60520661427575e-06 | 40 |
| rps1819 | 2.89567877355169e-10 | 0.417713087 | 0.952 | 0.725 | 7.28813390515224e-06 | 40 |
| gstm.15 | 3.05302665210501e-10 | 0.68806088 | 0.548 | 0.261 | 7.68416278068311e-06 | 40 |
| rps1418 | 3.93154732705177e-10 | 0.357020575 | 0.919 | 0.76 | 9.89531146745661e-06 | 40 |
| rpl3512 | 3.94592604709214e-10 | 0.334888709 | 0.952 | 0.805 | 9.9315012679262e-06 | 40 |
| rpl23a21 | 5.44867588878234e-10 | 0.441697138 | 0.871 | 0.635 | 1.37137723444763e-05 | 40 |
| rpl1818 | 5.69924706635336e-10 | 0.335337679 | 0.903 | 0.775 | 1.43444349413048e-05 | 40 |
| ugt1b52 | 5.87480150648714e-10 | 0.397863325 | 0.161 | 0.029 | 1.47862879116775e-05 | 40 |
| rps8a10 | 6.89687036473937e-10 | 0.34886948 | 0.952 | 0.788 | 1.73587330210125e-05 | 40 |
| stat26 | 7.21518843974566e-10 | 0.55979509 | 0.242 | 0.062 | 1.81599077839959e-05 | 40 |
| rplp010 | 8.45943056563254e-10 | 0.292994761 | 0.968 | 0.852 | 2.12915407906405e-05 | 40 |
| ccdc32 | 8.96923018890047e-10 | 0.445315927 | 0.21 | 0.047 | 2.25746554624436e-05 | 40 |
| rps2314 | 1.24581204554623e-09 | 0.382606206 | 0.935 | 0.723 | 3.13558433743529e-05 | 40 |
| rps1212 | 1.49545957453837e-09 | 0.31666531 | 0.984 | 0.84 | 3.76392220315561e-05 | 40 |
| bcam2 | 1.63423153096225e-09 | 0.442514548 | 0.161 | 0.03 | 4.11319734027888e-05 | 40 |
| rps3a11 | 1.69362824031788e-09 | 0.325692417 | 0.935 | 0.779 | 4.26269291805608e-05 | 40 |
| ap1m1 | 1.96935479030365e-09 | 0.550169773 | 0.258 | 0.072 | 4.95666907171526e-05 | 40 |
| hmgb1b9 | 2.30035531603986e-09 | 0.620880452 | 0.355 | 0.127 | 5.78976429494072e-05 | 40 |
| enah | 2.31871167918341e-09 | 0.355312711 | 0.145 | 0.025 | 5.83596542533673e-05 | 40 |
| zbtb10 | 2.58243353066663e-09 | 0.360099565 | 0.161 | 0.031 | 6.49972695333484e-05 | 40 |
| rps2819 | 3.3581318047669e-09 | 0.316238549 | 0.968 | 0.827 | 8.45208193941782e-05 | 40 |
| rpl2919 | 3.41980450162669e-09 | 0.31516244 | 0.952 | 0.809 | 8.60730595014422e-05 | 40 |
| gstm.22 | 3.53092130996597e-09 | 0.44582324 | 0.21 | 0.05 | 8.88697584505334e-05 | 40 |

| | | | | | | |
|---|---|---|---|---|---|---|
| mafba8 | 3.63949667194815e-09 | 0.52436022 | 0.258 | 0.073 | 9.16024917362631e-05 | 40 |
| senp3a | 3.79828212995347e-09 | 0.365785389 | 0.145 | 0.026 | 9.5598962928799e-05 | 40 |
| rpl3715 | 3.88368488923027e-09 | 0.311821894 | 0.984 | 0.85 | 9.77484649770366e-05 | 40 |
| si:ch211-286b5.57 | 4.42195630177727e-09 | 0.60833247 | 0.371 | 0.137 | 0.000111296 | 40 |
| psme16 | 5.16201574117011e-09 | 0.554197557 | 0.29 | 0.091 | 0.000129923 | 40 |
| si:ch211-39k3.2 | 6.0680749678669e-09 | 0.351235715 | 0.145 | 0.027 | 0.000152727 | 40 |
| ahcyl2 | 6.31919333161532e-09 | 0.276170353 | 0.113 | 0.017 | 0.000159048 | 40 |
| gig2e | 6.69914858380048e-09 | 0.272998759 | 0.113 | 0.017 | 0.000168611 | 40 |
| cav17 | 7.87487811887553e-09 | 0.48173641 | 0.194 | 0.045 | 0.000198203 | 40 |
| ppiab8 | 1.01692952026752e-08 | 0.402705473 | 0.839 | 0.657 | 0.000255951 | 40 |
| rps15a17 | 1.06494914420982e-08 | 0.354935701 | 0.952 | 0.78 | 0.000268037 | 40 |
| CR318588.111 | 1.27601927259023e-08 | 0.896330321 | 0.452 | 0.209 | 0.000321161 | 40 |
| ap1s2 | 1.29382881188948e-08 | 0.629033538 | 0.355 | 0.132 | 0.000325644 | 40 |
| si:ch211-270g19.52 | 1.30899580034933e-08 | 0.335918042 | 0.129 | 0.022 | 0.000329461 | 40 |
| rplp2l22 | 1.35189389599749e-08 | 0.360965263 | 0.968 | 0.775 | 0.000340258 | 40 |
| rps612 | 1.64762106599694e-08 | 0.316508226 | 0.935 | 0.789 | 0.00041469 | 40 |
| mtpn7 | 1.68992704407617e-08 | 0.601577118 | 0.371 | 0.147 | 0.000425338 | 40 |
| rpl2816 | 1.90314453412928e-08 | 0.329151909 | 0.903 | 0.75 | 0.000479002 | 40 |
| ywhabl6 | 2.52917992585175e-08 | 0.61742661 | 0.516 | 0.279 | 0.000636569 | 40 |
| faua16 | 2.55216760699682e-08 | 0.375147861 | 0.887 | 0.69 | 0.000642355 | 40 |
| si:ch211-105c13.35 | 3.00080691234105e-08 | 0.52785687 | 0.226 | 0.061 | 0.000755273 | 40 |
| rpl3019 | 3.30868668395663e-08 | 0.36328415 | 0.903 | 0.709 | 0.000832763 | 40 |
| jupa1 | 3.49031372442551e-08 | 0.426480365 | 0.21 | 0.054 | 0.000878477 | 40 |
| rpl3622 | 3.87557422133979e-08 | 0.323062502 | 0.935 | 0.79 | 0.000975443 | 40 |
| rpl1118 | 3.89946933787018e-08 | 0.319999948 | 0.968 | 0.814 | 0.000981457 | 40 |
| rps915 | 3.98869933444089e-08 | 0.342558779 | 0.968 | 0.777 | 0.001003916 | 40 |
| zgc:175088 | 5.48608181354809e-08 | 0.440752811 | 0.21 | 0.056 | 0.001380792 | 40 |
| edf13 | 7.08019541806478e-08 | 0.619769574 | 0.5 | 0.27 | 0.001782014 | 40 |
| atox13 | 8.53609879415978e-08 | 0.548122664 | 0.306 | 0.111 | 0.002148451 | 40 |
| ldlrap1a1 | 8.70317095175757e-08 | 0.466686583 | 0.242 | 0.07 | 0.002190501 | 40 |
| ppiaa15 | 9.17171996595133e-08 | 0.276182872 | 0.919 | 0.805 | 0.00230843 | 40 |
| arpc1a | 1.18591565664995e-07 | 0.599046053 | 0.355 | 0.142 | 0.002984831 | 40 |
| rps2613 | 1.38882097045219e-07 | 0.331405423 | 0.903 | 0.734 | 0.003495524 | 40 |
| rpl1713 | 1.60372357608099e-07 | 0.293020528 | 0.871 | 0.747 | 0.004036412 | 40 |
| rps27.218 | 1.68288678745075e-07 | 0.440293507 | 0.694 | 0.519 | 0.004235658 | 40 |

| | | | | | | |
|---|---|---|---|---|---|---|
| b2m15 | 1.68491325298299e-07 | 0.58516616 | 0.532 | 0.293 | 0.004240758 | 40 |
| rpl1517 | 1.97216406377216e-07 | 0.333068751 | 0.919 | 0.745 | 0.00496374 | 40 |
| gmfb1 | 2.13851200260247e-07 | 0.528071548 | 0.29 | 0.102 | 0.005382421 | 40 |
| got1 | 2.47547720953322e-07 | 0.296784249 | 0.129 | 0.025 | 0.006230529 | 40 |
| rpl35a17 | 2.62907888750468e-07 | 0.308930623 | 0.903 | 0.76 | 0.006617129 | 40 |
| rps26l19 | 2.63418095324863e-07 | 0.38456064 | 0.887 | 0.665 | 0.00662997 | 40 |
| ppdpfb7 | 2.93595712236009e-07 | 0.467014577 | 0.677 | 0.52 | 0.00738951 | 40 |
| rps517 | 3.33606768664289e-07 | 0.351638636 | 0.887 | 0.702 | 0.008396549 | 40 |
| cox8b5 | 3.3498044505285e-07 | 0.49690514 | 0.274 | 0.096 | 0.008431123 | 40 |
| CR318588.35 | 3.45062046062911e-07 | 0.529330799 | 0.177 | 0.046 | 0.008684867 | 40 |
| pik3r24 | 3.49209689908539e-07 | 0.449859499 | 0.21 | 0.061 | 0.008789259 | 40 |
| cox7c5 | 3.50178056075804e-07 | 0.473403048 | 0.677 | 0.482 | 0.008813631 | 40 |
| myl9a5 | 4.9908533699323e-07 | 0.297837526 | 0.145 | 0.032 | 0.012561479 | 40 |
| slc25a510 | 6.16413504852449e-07 | 0.285727302 | 0.968 | 0.807 | 0.015514512 | 40 |
| tuba8l36 | 7.25926891839721e-07 | 0.403500239 | 0.177 | 0.047 | 0.018270854 | 40 |
| rplp221 | 8.26340486088277e-07 | 0.336874587 | 0.935 | 0.758 | 0.020798164 | 40 |
| sub1a4 | 9.28933846923114e-07 | 0.549416794 | 0.452 | 0.234 | 0.023380336 | 40 |
| hmgn712 | 1.08027838807501e-06 | 0.548868104 | 0.484 | 0.281 | 0.027189527 | 40 |
| rpl710 | 1.13521507325714e-06 | 0.251631771 | 0.903 | 0.806 | 0.028572228 | 40 |
| eef1db13 | 1.17276665633863e-06 | 0.5395743 | 0.484 | 0.275 | 0.029517364 | 40 |
| atp6v1e1a1 | 1.17424358179511e-06 | 0.413460848 | 0.177 | 0.049 | 0.029554537 | 40 |
| marcksb8 | 1.42445759976459e-06 | 0.529638659 | 0.403 | 0.177 | 0.035852173 | 40 |
| hmgb3a3 | 1.57158398680023e-06 | 0.393856679 | 0.177 | 0.05 | 0.039555197 | 40 |
| gabarapb12 | 2.37282744709686e-06 | 0.553447636 | 0.468 | 0.27 | 0.059721694 | 40 |
| rps2514 | 2.63834966191428e-06 | 0.28980818 | 0.871 | 0.726 | 0.066404623 | 40 |
| angptl46 | 2.73580187231652e-06 | 0.404138976 | 0.177 | 0.05 | 0.068857397 | 40 |
| rps1022 | 2.7958830911404e-06 | 0.352251134 | 0.919 | 0.695 | 0.070369582 | 40 |
| glrx51 | 3.11882797985284e-06 | 0.497275507 | 0.258 | 0.095 | 0.078497781 | 40 |
| gpx4b4 | 3.18962235892579e-06 | 0.54386332 | 0.403 | 0.205 | 0.080279605 | 40 |
| rps1112 | 3.27225935054353e-06 | 0.267422228 | 0.887 | 0.767 | 0.082359496 | 40 |
| fth1a1 | 4.10086243256649e-06 | 0.345601364 | 0.79 | 0.637 | 0.103214607 | 40 |
| sprypd7a | 4.93180851324964e-06 | 0.268120273 | 0.113 | 0.024 | 0.124128688 | 40 |
| epdl11 | 5.21986072950877e-06 | 0.308707131 | 0.145 | 0.037 | 0.131378675 | 40 |
| vat18 | 5.23522949707127e-06 | 0.479672082 | 0.323 | 0.136 | 0.131765491 | 40 |
| gsna3 | 5.50297737146094e-06 | 0.255862003 | 0.129 | 0.03 | 0.138504437 | 40 |

| | | | | | | |
|---|---|---|---|---|---|---|
| oaz1b16 | 5.57134183874618e-06 | 0.480852405 | 0.532 | 0.324 | 0.140225103 | 40 |
| plekhb2 | 5.95388917169932e-06 | 0.265846415 | 0.113 | 0.024 | 0.149853437 | 40 |
| cgnl13 | 6.49095690896236e-06 | 0.268274166 | 0.113 | 0.025 | 0.163370894 | 40 |
| efhd23 | 6.70284353467049e-06 | 0.384686508 | 0.194 | 0.061 | 0.168703869 | 40 |
| rpl27a20 | 7.02278954405315e-06 | 0.421575416 | 0.726 | 0.497 | 0.17675659 | 40 |
| rps2914 | 7.31286390472696e-06 | 0.278406 | 0.984 | 0.832 | 0.184057472 | 40 |
| zgc:5649312 | 8.97743749050224e-06 | 0.348271273 | 0.726 | 0.574 | 0.225953124 | 40 |
| ywhaz7 | 1.13024194015177e-05 | 0.511423351 | 0.468 | 0.283 | 0.284470594 | 40 |
| rps1318 | 1.16292296320384e-05 | 0.302205317 | 0.887 | 0.725 | 0.292696081 | 40 |
| btf315 | 1.31274451627553e-05 | 0.35382624 | 0.774 | 0.588 | 0.330404667 | 40 |
| eif5a11 | 1.32030717190615e-05 | 0.398877917 | 0.645 | 0.48 | 0.332308112 | 40 |
| tma74 | 1.70341372334894e-05 | 0.5120168 | 0.484 | 0.306 | 0.4287322 | 40 |
| s100v29 | 1.82610620912911e-05 | 0.4441351 | 0.29 | 0.123 | 0.459612672 | 40 |
| calm3b3 | 2.03096598370381e-05 | 0.527973508 | 0.435 | 0.258 | 0.511173828 | 40 |
| mpped2a | 2.22450205700594e-05 | 0.316696713 | 0.129 | 0.033 | 0.559884923 | 40 |
| psmb8a | 2.33081866440621e-05 | 0.310589292 | 0.113 | 0.027 | 0.58664375 | 40 |
| zgc:123105 | 2.45455523852125e-05 | 0.34707797 | 0.161 | 0.048 | 0.617787008 | 40 |
| iscub2 | 2.71415419979054e-05 | 0.402931891 | 0.194 | 0.067 | 0.683125471 | 40 |
| f105 | 2.72244070729415e-05 | 0.359753041 | 0.161 | 0.05 | 0.685211102 | 40 |
| rps316 | 2.77035875069988e-05 | 0.313339209 | 0.887 | 0.701 | 0.697271594 | 40 |
| ndufa4l11 | 2.83856127686559e-05 | 0.40532286 | 0.565 | 0.397 | 0.714437488 | 40 |
| pcyt2 | 3.45019498559891e-05 | 0.284099909 | 0.145 | 0.041 | 0.868379576 | 40 |
| eif4a1b11 | 3.54009722393246e-05 | 0.487724509 | 0.452 | 0.282 | 0.89100707 | 40 |
| uqcrh5 | 3.78701737001354e-05 | 0.472209208 | 0.435 | 0.251 | 0.953154402 | 40 |
| rpl1014 | 4.42758824249303e-05 | 0.293121028 | 0.968 | 0.773 | 1 | 40 |
| eif4ebp27 | 4.53002876057115e-05 | 0.494827652 | 0.339 | 0.176 | 1 | 40 |
| idh11 | 5.08137075938929e-05 | 0.435329145 | 0.29 | 0.127 | 1 | 40 |
| cirbpa1 | 5.11332968174146e-05 | 0.31576827 | 0.726 | 0.619 | 1 | 40 |
| tpm4a8 | 6.54815877218501e-05 | 0.403729904 | 0.226 | 0.09 | 1 | 40 |
| ppp3ccb2 | 6.76296513784979e-05 | 0.4597155 | 0.274 | 0.125 | 1 | 40 |
| itm2bb12 | 7.09865773685646e-05 | 0.508085968 | 0.387 | 0.224 | 1 | 40 |
| arl4d2 | 7.47152713830742e-05 | 0.289908735 | 0.161 | 0.051 | 1 | 40 |
| zgc:1008291 | 7.69393258480399e-05 | 0.305271157 | 0.129 | 0.037 | 1 | 40 |
| s100a10b12 | 8.9681750334353e-05 | 0.465950838 | 0.468 | 0.257 | 1 | 40 |
| si:dkey-28b4.79 | 9.46148638540325e-05 | 0.411805047 | 0.226 | 0.093 | 1 | 40 |

| | | | | | | |
|---|---|---|---|---|---|---|
| ndufb93 | 0.000101806 | 0.450480479 | 0.339 | 0.178 | 1 | 40 |
| mibp216 | 0.000105368 | 0.343194102 | 0.629 | 0.494 | 1 | 40 |
| phlda24 | 0.000107597 | 0.32488743 | 0.161 | 0.052 | 1 | 40 |
| irf2bp2b7 | 0.000128748 | 0.364786795 | 0.21 | 0.083 | 1 | 40 |
| eef1g16 | 0.000139501 | 0.266390287 | 0.726 | 0.622 | 1 | 40 |
| nudcd2 | 0.000146576 | 0.336491545 | 0.177 | 0.063 | 1 | 40 |
| txndc171 | 0.000148298 | 0.336814205 | 0.161 | 0.055 | 1 | 40 |
| psma6l | 0.000149896 | 0.250756952 | 0.129 | 0.038 | 1 | 40 |
| psme29 | 0.000175189 | 0.38946214 | 0.226 | 0.098 | 1 | 40 |
| stat1a7 | 0.000215468 | 0.416566618 | 0.258 | 0.121 | 1 | 40 |
| gng12a | 0.000219782 | 0.285169718 | 0.113 | 0.032 | 1 | 40 |
| si:ch73-46j18.510 | 0.000227468 | 0.419420258 | 0.339 | 0.177 | 1 | 40 |
| itm2ba13 | 0.000230104 | 0.413480341 | 0.435 | 0.275 | 1 | 40 |
| rpl1416 | 0.000234456 | 0.265485353 | 0.758 | 0.636 | 1 | 40 |
| pnrc221 | 0.000238911 | 0.310899463 | 0.677 | 0.569 | 1 | 40 |
| eif1b11 | 0.000245115 | 0.415007359 | 0.516 | 0.384 | 1 | 40 |
| psmf1 | 0.000261609 | 0.31336093 | 0.129 | 0.04 | 1 | 40 |
| cavin2b8 | 0.000282471 | 0.341173052 | 0.177 | 0.067 | 1 | 40 |
| hp1bp35 | 0.000327248 | 0.364545121 | 0.226 | 0.099 | 1 | 40 |
| chmp6b | 0.000334892 | 0.312513112 | 0.145 | 0.05 | 1 | 40 |
| ppp2r1ba | 0.000397405 | 0.341962246 | 0.21 | 0.088 | 1 | 40 |
| eef2b10 | 0.000452626 | 0.263811509 | 0.726 | 0.615 | 1 | 40 |
| atf5b | 0.000453474 | 0.293448926 | 0.113 | 0.034 | 1 | 40 |
| myl68 | 0.000459328 | 0.401812892 | 0.226 | 0.102 | 1 | 40 |
| glod51 | 0.000505438 | 0.285935595 | 0.113 | 0.034 | 1 | 40 |
| si:ch73-42p12.22 | 0.000572846 | 0.380483383 | 0.226 | 0.103 | 1 | 40 |
| map1lc3a8 | 0.000613946 | 0.32147206 | 0.177 | 0.072 | 1 | 40 |
| xaf13 | 0.000614962 | 0.326731042 | 0.161 | 0.062 | 1 | 40 |
| sdcbp210 | 0.000664666 | 0.392000466 | 0.355 | 0.21 | 1 | 40 |
| prdx32 | 0.000700353 | 0.373361281 | 0.274 | 0.136 | 1 | 40 |
| rab11a | 0.000881925 | 0.383378394 | 0.242 | 0.117 | 1 | 40 |
| hsd11b27 | 0.000889333 | 0.319565341 | 0.274 | 0.125 | 1 | 40 |
| ifi35 | 0.000982946 | 0.266250495 | 0.113 | 0.037 | 1 | 40 |
| bin3 | 0.00099147 | 0.251834863 | 0.113 | 0.036 | 1 | 40 |
| capns1b8 | 0.00100337 | 0.418429417 | 0.242 | 0.123 | 1 | 40 |

| | | | | | | | |
|---|---|---|---|---|---|---|---|
| rpl411 | 0.001028865 | 0.262966249 | 0.774 | 0.629 | | 1 | 40 |
| vps25 | 0.001129442 | 0.275283649 | 0.113 | 0.037 | | 1 | 40 |
| atp1b1a7 | 0.001173655 | 0.39255313 | 0.355 | 0.215 | | 1 | 40 |
| cd994 | 0.001186511 | 0.270576751 | 0.145 | 0.054 | | 1 | 40 |
| eif3f17 | 0.001254379 | 0.321835953 | 0.532 | 0.433 | | 1 | 40 |
| kank25 | 0.001265049 | 0.276013238 | 0.145 | 0.055 | | 1 | 40 |
| rps27l11 | 0.001323494 | 0.390819988 | 0.339 | 0.205 | | 1 | 40 |
| cfl112 | 0.001445667 | 0.270144352 | 0.677 | 0.579 | | 1 | 40 |
| LOC1039092847 | 0.001501916 | 0.272370113 | 0.161 | 0.065 | | 1 | 40 |
| ywhah | 0.001583954 | 0.28753712 | 0.129 | 0.047 | | 1 | 40 |
| etfa2 | 0.001687021 | 0.338000452 | 0.194 | 0.089 | | 1 | 40 |
| gabarapa2 | 0.001693809 | 0.366422524 | 0.29 | 0.162 | | 1 | 40 |
| dap1 | 0.001705126 | 0.318747314 | 0.226 | 0.107 | | 1 | 40 |
| uba5218 | 0.00174459 | 0.271113507 | 0.79 | 0.631 | | 1 | 40 |
| sgta | 0.001945287 | 0.293684926 | 0.177 | 0.076 | | 1 | 40 |
| ppib12 | 0.002051033 | 0.344762613 | 0.468 | 0.335 | | 1 | 40 |
| akt1s1 | 0.00214811 | 0.303663147 | 0.145 | 0.058 | | 1 | 40 |
| ctnnb17 | 0.00215368 | 0.372298038 | 0.274 | 0.151 | | 1 | 40 |
| eif3m9 | 0.002448463 | 0.337923265 | 0.468 | 0.36 | | 1 | 40 |
| h2afx113 | 0.002546131 | 0.379956313 | 0.323 | 0.2 | | 1 | 40 |
| prdx511 | 0.003232275 | 0.351505766 | 0.274 | 0.161 | | 1 | 40 |
| psmc1a1 | 0.003291399 | 0.300664173 | 0.161 | 0.07 | | 1 | 40 |
| gatad2b2 | 0.003393775 | 0.263238485 | 0.145 | 0.06 | | 1 | 40 |
| si:dkey-222f8.3 | 0.003399725 | 0.288518429 | 0.145 | 0.061 | | 1 | 40 |
| anxa137 | 0.003467615 | 0.261917454 | 0.129 | 0.051 | | 1 | 40 |
| h3f3c14 | 0.003522944 | 0.372275843 | 0.371 | 0.258 | | 1 | 40 |
| ppp2cb | 0.003596567 | 0.354020731 | 0.29 | 0.169 | | 1 | 40 |
| pfn2l6 | 0.003685679 | 0.362478521 | 0.339 | 0.211 | | 1 | 40 |
| churc1 | 0.003729511 | 0.293649733 | 0.145 | 0.062 | | 1 | 40 |
| myl12.114 | 0.003919136 | 0.315399693 | 0.5 | 0.394 | | 1 | 40 |
| cd1643 | 0.004062143 | 0.303680476 | 0.161 | 0.073 | | 1 | 40 |
| cox8a12 | 0.004363533 | 0.370352353 | 0.339 | 0.229 | | 1 | 40 |
| DHDH3 | 0.004555725 | 0.262192097 | 0.145 | 0.062 | | 1 | 40 |
| prkab1b3 | 0.004755502 | 0.345647525 | 0.226 | 0.123 | | 1 | 40 |
| CU694197.11 | 0.004820689 | 0.320285364 | 0.242 | 0.132 | | 1 | 40 |

| | | | | | | |
|---|---|---|---|---|---|---|
| atp1a1a.11 | 0.005238144 | 0.356431329 | 0.258 | 0.15 | 1 | 40 |
| h1f08 | 0.005385668 | 0.295640319 | 0.145 | 0.064 | 1 | 40 |
| anxa48 | 0.005762715 | 0.335451896 | 0.242 | 0.138 | 1 | 40 |
| csde14 | 0.006318613 | 0.345630981 | 0.323 | 0.209 | 1 | 40 |
| mif2 | 0.006774849 | 0.331134115 | 0.306 | 0.189 | 1 | 40 |
| cnn212 | 0.007357065 | 0.290147281 | 0.242 | 0.136 | 1 | 40 |
| anapc161 | 0.008164625 | 0.313331731 | 0.194 | 0.102 | 1 | 40 |
| sap30l1 | 0.008410559 | 0.316086811 | 0.242 | 0.141 | 1 | 40 |
| lgals3b6 | 0.008895165 | 0.301370619 | 0.161 | 0.08 | 1 | 40 |
| si:dkey-51e6.16 | 0.00919436 | 0.339318695 | 0.21 | 0.118 | 1 | 40 |
| pfn26 | 0.009234398 | 0.392825413 | 0.323 | 0.228 | 1 | 40 |
| rhoac2 | 0.009521675 | 0.325112074 | 0.258 | 0.157 | 1 | 40 |
| sumo11 | 0.009846817 | 0.324180727 | 0.21 | 0.117 | 1 | 40 |
| sod13 | 0.009917861 | 0.306018457 | 0.435 | 0.329 | 1 | 40 |
| si:dkey-78l4.6 | 0 | 2.684086197 | 0.934 | 0.007 | 0 | 41 |
| si:dkey-9i23.41 | 0 | 2.62403675 | 0.967 | 0.014 | 0 | 41 |
| si:dkey-78l4.7 | 0 | 1.909773545 | 0.656 | 0.001 | 0 | 41 |
| LOC1001508821 | 0 | 1.811361716 | 0.639 | 0.008 | 0 | 41 |
| spink4 | 0 | 1.334562507 | 0.59 | 0.002 | 0 | 41 |
| gata31 | 0 | 0.875171268 | 0.426 | 0.005 | 0 | 41 |
| si:dkey-78l4.8 | 0 | 0.496171176 | 0.213 | 0.001 | 0 | 41 |
| si:dkey-78l4.43 | 3.08970864859484e-306 | 2.011200844 | 0.885 | 0.03 | 7.77648769764836e-302 | 41 |
| si:dkey-21e2.31 | 1.73830910916251e-279 | 0.862311938 | 0.393 | 0.005 | 4.37515019685112e-275 | 41 |
| cpa52 | 3.56709017119163e-219 | 1.637019055 | 0.82 | 0.035 | 8.97800925187221e-215 | 41 |
| il11b1 | 6.9065260260087e-217 | 0.765837249 | 0.393 | 0.007 | 1.73830353548613e-212 | 41 |
| CR847936.10 | 3.62116657209852e-208 | 0.619217818 | 0.295 | 0.004 | 9.11411414531477e-204 | 41 |
| adgre131 | 1.09521112818971e-174 | 0.678507013 | 0.344 | 0.007 | 2.75653688854067e-170 | 41 |
| cfbl4 | 2.80517270942189e-164 | 1.846452042 | 0.918 | 0.063 | 7.06033919234395e-160 | 41 |
| tnfrsf9a6 | 5.24525719793326e-156 | 1.588865719 | 0.803 | 0.049 | 1.32017878414782e-151 | 41 |
| acp5b | 3.14033036031711e-155 | 0.780447962 | 0.443 | 0.014 | 7.90389748388214e-151 | 41 |
| AL954142.13 | 1.09442931592056e-152 | 0.983571862 | 0.541 | 0.021 | 2.75456914524045e-148 | 41 |
| ptprjb.1 | 1.29951717691985e-149 | 0.389983022 | 0.197 | 0.002 | 3.27075478258957e-145 | 41 |
| cox6a25 | 4.0243865902173e-147 | 1.133256671 | 0.656 | 0.034 | 1.01289786089179e-142 | 41 |
| CR396595.1 | 1.40759921968801e-144 | 0.345563568 | 0.164 | 0.002 | 3.54278647603276e-140 | 41 |
| eomesa2 | 3.62609066259813e-141 | 1.194694287 | 0.689 | 0.038 | 9.12650758869324e-137 | 41 |

| | | | | | | |
|---|---|---|---|---|---|---|
| kctd7 | 9.02827963687423e-140 | 0.383920893 | 0.213 | 0.003 | 2.27232770180487e-135 | 41 |
| LOC101886102 | 4.5587110260738e-134 | 0.491464304 | 0.246 | 0.004 | 1.14738197815252e-129 | 41 |
| flt3 | 1.36994645502122e-132 | 0.267857689 | 0.115 | 0.001 | 3.4480182326429e-128 | 41 |
| nansb | 9.94257862780324e-124 | 0.281345188 | 0.115 | 0.001 | 2.5024476148318e-119 | 41 |
| prodha | 8.92484757637303e-120 | 0.74621533 | 0.377 | 0.013 | 2.24629488649733e-115 | 41 |
| si:dkey-78l4.5 | 4.7081273438781e-117 | 0.315542957 | 0.148 | 0.002 | 1.18498857118068e-112 | 41 |
| CABZ01073834.16 | 4.84714629018652e-111 | 1.026488554 | 0.623 | 0.04 | 1.21997824977705e-106 | 41 |
| ogfrl1 | 3.94331160661072e-108 | 0.52335458 | 0.262 | 0.007 | 9.92492098267851e-104 | 41 |
| rab36 | 1.0559896876157e-104 | 0.266769429 | 0.148 | 0.002 | 2.65782044475996e-100 | 41 |
| dusp57 | 4.9925769235376e-102 | 1.335644846 | 0.77 | 0.071 | 1.25658168588518e-97 | 41 |
| si:dkey-78l4.21 | 1.87687701778656e-100 | 0.572105517 | 0.246 | 0.006 | 4.72391176606699e-96 | 41 |
| parvb | 7.4612701888169e-99 | 0.502809421 | 0.262 | 0.007 | 1.87792709382333e-94 | 41 |
| ctss12 | 2.18088890476663e-96 | 0.705987113 | 0.344 | 0.014 | 5.48907928440712e-92 | 41 |
| capgb10 | 3.80437056802011e-96 | 1.532762818 | 0.902 | 0.112 | 9.5752202826498e-92 | 41 |
| tnfrsf9b2 | 1.95313911841258e-90 | 0.646867819 | 0.361 | 0.016 | 4.91585584713261e-86 | 41 |
| cremb1 | 1.87072529414509e-89 | 0.707163731 | 0.393 | 0.019 | 4.70842849283378e-85 | 41 |
| yrk6 | 1.72397918192359e-79 | 1.059673209 | 0.656 | 0.062 | 4.33908320298348e-75 | 41 |
| XLOC-0183562 | 1.52577633749521e-75 | 0.782497349 | 0.426 | 0.027 | 3.8402264638417e-71 | 41 |
| rnaset2l5 | 2.01268913789144e-74 | 0.887480341 | 0.525 | 0.042 | 5.06573729115895e-70 | 41 |
| laptm514 | 1.33468918476872e-70 | 1.497518867 | 0.918 | 0.168 | 3.35927920914439e-66 | 41 |
| vaspa8 | 4.11834447892116e-69 | 1.062565523 | 0.672 | 0.074 | 1.03654612189967e-64 | 41 |
| ccr9a9 | 7.18701839107192e-68 | 1.124318868 | 0.721 | 0.086 | 1.80890065884889e-63 | 41 |
| grap2b11 | 9.76440440286658e-67 | 1.251798909 | 0.787 | 0.111 | 2.45760294415749e-62 | 41 |
| CU855821.13 | 2.29754429982507e-65 | 0.499256252 | 0.279 | 0.013 | 5.78268924822972e-61 | 41 |
| ptprc12 | 3.42211309517259e-65 | 1.286008899 | 0.82 | 0.126 | 8.6131164492399e-61 | 41 |
| scgn4 | 2.63726930079997e-64 | 0.613442146 | 0.361 | 0.023 | 6.63774310318345e-60 | 41 |
| fcer1gl9 | 3.23140230518534e-64 | 1.485862752 | 0.918 | 0.167 | 8.13311646192097e-60 | 41 |
| dnajb9b1 | 5.67592150908354e-64 | 0.636460155 | 0.361 | 0.023 | 1.42857268462124e-59 | 41 |
| ndufa41 | 3.60670539661119e-63 | 0.501969481 | 0.279 | 0.014 | 9.07771681273071e-59 | 41 |
| si:zfos-741a10.35 | 2.92781721843117e-62 | 0.647291483 | 0.377 | 0.026 | 7.36902315706942e-58 | 41 |
| si:dkey-33i11.93 | 6.0715771477647e-62 | 0.633608619 | 0.344 | 0.021 | 1.5281552523209e-57 | 41 |
| nptna5 | 4.2974907247155e-60 | 0.963241068 | 0.623 | 0.07 | 1.08163544050364e-55 | 41 |
| hhex1 | 4.3154001571274e-59 | 0.37985944 | 0.197 | 0.007 | 1.08614306554739e-54 | 41 |
| itgb28 | 2.90019508817123e-57 | 1.063794149 | 0.705 | 0.098 | 7.29950101741816e-53 | 41 |
| ca24 | 3.66303696997743e-57 | 0.882237345 | 0.541 | 0.055 | 9.2194977497362e-53 | 41 |

| | | | | | | |
|---|---|---|---|---|---|---|
| SLA (1 of many) | 6.84715294436354e-57 | 0.296202903 | 0.131 | 0.003 | 1.72335992456686e-52 | 41 |
| cyba7 | 4.98069444346907e-53 | 0.866298148 | 0.59 | 0.072 | 1.25359098447673e-48 | 41 |
| mir2234 | 5.56477395575705e-53 | 0.656339986 | 0.377 | 0.03 | 1.40059795692449e-48 | 41 |
| alox5ap2 | 9.74282203507033e-52 | 0.548834173 | 0.311 | 0.021 | 2.45217087800685e-47 | 41 |
| osmr2 | 1.23446355523856e-50 | 0.358534581 | 0.197 | 0.009 | 3.10702132217994e-46 | 41 |
| bhlhe4013 | 8.46155466149236e-49 | 1.221375692 | 0.803 | 0.159 | 2.12968869275101e-44 | 41 |
| fermt3b2 | 1.20231308969582e-48 | 0.531151024 | 0.279 | 0.018 | 3.02610181545542e-44 | 41 |
| ccl35.16 | 1.72299608462571e-48 | 1.140523828 | 0.607 | 0.081 | 4.33660884539445e-44 | 41 |
| zgc:6405114 | 2.10665374112891e-48 | 1.199716042 | 0.803 | 0.161 | 5.30223680104736e-44 | 41 |
| si:ch211-198c19.11 | 5.8616823760931e-47 | 0.333953453 | 0.197 | 0.009 | 1.47532683723887e-42 | 41 |
| si:dkey-1c7.12 | 1.36505138600336e-46 | 0.543994136 | 0.311 | 0.023 | 3.43569783343186e-42 | 41 |
| ccl33.32 | 5.71693831236677e-46 | 1.117374594 | 0.475 | 0.054 | 1.43889620383959e-41 | 41 |
| si:ch211-241f5.35 | 5.80168965109421e-46 | 0.905974361 | 0.525 | 0.067 | 1.4602272682839e-41 | 41 |
| si:ch211-102c2.49 | 5.98921393318487e-46 | 1.1153349 | 0.77 | 0.141 | 1.5074252548433e-41 | 41 |
| hexa | 6.16269448133206e-46 | 0.462013239 | 0.262 | 0.017 | 1.55108857400647e-41 | 41 |
| wasa9 | 1.20908311450868e-45 | 1.023529914 | 0.607 | 0.092 | 3.04314129090689e-41 | 41 |
| il15l1 | 7.01572395862474e-45 | 0.683875175 | 0.344 | 0.029 | 1.76578756314626e-40 | 41 |
| ppp1r187 | 2.24980673606946e-44 | 0.790514206 | 0.459 | 0.053 | 5.66253857401322e-40 | 41 |
| slc41a12 | 3.33989209383373e-44 | 0.397916727 | 0.23 | 0.013 | 8.40617441097011e-40 | 41 |
| mylk5 | 1.2591677631128e-43 | 0.453452265 | 0.262 | 0.018 | 3.16919934297861e-39 | 41 |
| sik110 | 1.77970756361652e-43 | 1.02999259 | 0.672 | 0.115 | 4.47934596686643e-39 | 41 |
| parvg8 | 6.63363552906113e-42 | 0.792711124 | 0.508 | 0.067 | 1.6696197263094e-37 | 41 |
| dgkzb1 | 7.04265408633116e-42 | 0.374657092 | 0.18 | 0.009 | 1.77256560698869e-37 | 41 |
| lpcat21 | 7.76638546041569e-42 | 0.526359657 | 0.279 | 0.021 | 1.95472155653202e-37 | 41 |
| arap3 | 8.24136982869547e-42 | 0.340692465 | 0.18 | 0.009 | 2.07427037218436e-37 | 41 |
| si:ch211-136m16.84 | 5.15194527757165e-41 | 0.5706622 | 0.328 | 0.029 | 1.29669310691201e-36 | 41 |
| hck3 | 6.59478333860702e-41 | 0.644799395 | 0.344 | 0.032 | 1.659841018494e-36 | 41 |
| spi1b8 | 1.76886917572237e-40 | 0.770295176 | 0.475 | 0.061 | 4.45206682837565e-36 | 41 |
| skap28 | 1.04159586441863e-39 | 0.759119139 | 0.525 | 0.074 | 2.62159263115526e-35 | 41 |
| si:dkey-21o19.21 | 1.69548423364067e-39 | 0.425880244 | 0.23 | 0.015 | 4.2673642676502e-35 | 41 |
| il10ra4 | 2.4574415816957e-39 | 0.645175724 | 0.377 | 0.04 | 6.18513471696991e-35 | 41 |
| sort1b3 | 4.06160212975236e-39 | 0.648398297 | 0.328 | 0.031 | 1.02226464003737e-34 | 41 |
| rgs1311 | 1.06185822599797e-38 | 1.033922646 | 0.689 | 0.131 | 2.6725909690143e-34 | 41 |
| viml1 | 1.0993437121985e-38 | 0.308240191 | 0.164 | 0.008 | 2.7669381892324e-34 | 41 |
| cyb5r2 | 1.88618361485165e-38 | 0.338133562 | 0.18 | 0.01 | 4.74733554022013e-34 | 41 |

| | | | | | | |
|---|---|---|---|---|---|---|
| def6c1 | 1.98317472843725e-38 | 0.377357263 | 0.197 | 0.011 | 4.99145247400371e-34 | 41 |
| CU207241.11 | 3.19467461015403e-38 | 0.40937655 | 0.23 | 0.016 | 8.04067652629668e-34 | 41 |
| coro1a14 | 7.75537918411948e-38 | 1.142670171 | 0.869 | 0.23 | 1.95195138685103e-33 | 41 |
| ubash3ba1 | 2.65176077925307e-37 | 0.544512293 | 0.279 | 0.024 | 6.67421670530205e-33 | 41 |
| il165 | 3.87763759184337e-37 | 0.618317923 | 0.41 | 0.049 | 9.75962605491058e-33 | 41 |
| sox51 | 4.31345194281142e-37 | 0.324466646 | 0.197 | 0.012 | 1.08565271948621e-32 | 41 |
| srgn12 | 4.80451595753535e-37 | 1.104596212 | 0.885 | 0.236 | 1.20924862135207e-32 | 41 |
| pla2g4aa1 | 8.95053558825614e-37 | 0.393304303 | 0.23 | 0.016 | 2.25276030220819e-32 | 41 |
| ccr26 | 1.40045104555098e-36 | 0.683019169 | 0.393 | 0.046 | 3.52479523654727e-32 | 41 |
| arpc1b15 | 2.06453288916671e-36 | 1.176680419 | 0.951 | 0.308 | 5.19622282874368e-32 | 41 |
| lcp2b1 | 4.79724158185675e-36 | 0.379963348 | 0.197 | 0.012 | 1.20741773373752e-31 | 41 |
| b3gnt3.45 | 1.13148331703071e-35 | 0.607653418 | 0.344 | 0.037 | 2.8478303606346e-31 | 41 |
| kctd12.26 | 2.31036009167956e-35 | 0.827374743 | 0.508 | 0.079 | 5.81494531474829e-31 | 41 |
| sema7a | 5.40748078533248e-35 | 0.294452205 | 0.148 | 0.007 | 1.36100883886033e-30 | 41 |
| nitr5 | 2.50054605026615e-34 | 0.328276339 | 0.164 | 0.009 | 6.29362435391488e-30 | 41 |
| si:ch211-191c10.17 | 3.75443503195236e-34 | 0.718724054 | 0.41 | 0.054 | 9.44953753192089e-30 | 41 |
| pfn116 | 4.49142005271248e-34 | 1.079409437 | 1 | 0.525 | 1.1304455130672e-29 | 41 |
| tln17 | 1.55414764993822e-33 | 0.891754166 | 0.623 | 0.127 | 3.91163422012952e-29 | 41 |
| csrnp1a2 | 9.46078292033009e-33 | 0.688464275 | 0.393 | 0.051 | 2.38118445321788e-28 | 41 |
| si:dkey-237j10.2 | 7.38224186055759e-32 | 0.275727229 | 0.164 | 0.01 | 1.85803645388374e-27 | 41 |
| si:dkey-172h23.21 | 1.37666457216255e-31 | 0.562780362 | 0.344 | 0.041 | 3.46492706167592e-27 | 41 |
| cd74b12 | 2.35561320852743e-31 | 1.056217373 | 0.852 | 0.223 | 5.9288428845427e-27 | 41 |
| fybb6 | 2.62624728511556e-31 | 0.586447179 | 0.344 | 0.041 | 6.61000179190736e-27 | 41 |
| ptpn67 | 6.35406772190517e-30 | 0.813592849 | 0.508 | 0.091 | 1.59925530492631e-25 | 41 |
| sla26 | 9.12867744925242e-30 | 0.782288758 | 0.443 | 0.069 | 2.29759682720234e-25 | 41 |
| jak2a | 1.71672923223147e-29 | 0.388478581 | 0.246 | 0.023 | 4.32083580460339e-25 | 41 |
| si:ch211-243a20.43 | 3.19829745471873e-29 | 0.368298938 | 0.213 | 0.018 | 8.04979486378157e-25 | 41 |
| atr | 4.14435209621054e-29 | 0.265487149 | 0.115 | 0.005 | 1.04309197909523e-24 | 41 |
| cd6316 | 7.21248341346303e-29 | 0.995240485 | 0.918 | 0.34 | 1.81530995033451e-24 | 41 |
| arhgap20b1 | 2.09519677187892e-28 | 0.440128741 | 0.23 | 0.021 | 5.27340075514205e-24 | 41 |
| fam49ba7 | 2.37888793966228e-28 | 0.76589656 | 0.541 | 0.103 | 5.987423055336e-24 | 41 |
| si:ch73-264p11.15 | 2.71515499373689e-28 | 0.456584326 | 0.262 | 0.027 | 6.83377360373637e-24 | 41 |
| rgs3a1 | 5.46693279217123e-28 | 0.439911972 | 0.246 | 0.024 | 1.37597231446158e-23 | 41 |
| edaradd3 | 6.78255520096162e-28 | 0.424168737 | 0.246 | 0.024 | 1.70710131853003e-23 | 41 |
| glipr1a7 | 1.00919703146561e-27 | 0.655155551 | 0.41 | 0.063 | 2.54004800849578e-23 | 41 |

| | | | | | | |
|---|---|---|---|---|---|---|
| gnsb3 | 1.26142183236544e-27 | 0.567174798 | 0.311 | 0.038 | 3.17487260988057e-23 | 41 |
| LOC1018842676 | 1.43787030795103e-27 | 0.631003541 | 0.393 | 0.058 | 3.61897577808195e-23 | 41 |
| crema10 | 1.6215142992738e-27 | 0.976744518 | 0.672 | 0.178 | 4.08118933984222e-23 | 41 |
| si:ch211-215e19.33 | 2.73392897946119e-27 | 0.413729117 | 0.23 | 0.022 | 6.88102584840586e-23 | 41 |
| myl12.115 | 3.53461367223111e-27 | 0.920850764 | 0.934 | 0.391 | 8.89626915163848e-23 | 41 |
| akt3a | 6.17951543770105e-27 | 0.257162708 | 0.115 | 0.006 | 1.55532224051498e-22 | 41 |
| rhogb8 | 1.65465361434733e-26 | 0.787736328 | 0.492 | 0.097 | 4.1645976819508e-22 | 41 |
| tnikb | 1.81023521872992e-26 | 0.449903448 | 0.23 | 0.022 | 4.55618102202135e-22 | 41 |
| rorab | 3.16252769622935e-26 | 0.295089286 | 0.148 | 0.01 | 7.95976595863966e-22 | 41 |
| itm2bb13 | 5.78223125016303e-26 | 0.920898494 | 0.754 | 0.221 | 1.45532978335353e-21 | 41 |
| rasgrp42 | 7.62590666616004e-26 | 0.47905791 | 0.262 | 0.03 | 1.91936444880582e-21 | 41 |
| aldocb5 | 7.71180485994726e-26 | 0.617222679 | 0.41 | 0.067 | 1.94098416520012e-21 | 41 |
| si:dkey-21e2.151 | 1.25735862855942e-25 | 0.556614396 | 0.148 | 0.01 | 3.16464593222122e-21 | 41 |
| tagapb9 | 1.29578618690822e-25 | 0.614506117 | 0.41 | 0.067 | 3.26136425382931e-21 | 41 |
| mbnl111 | 4.25911438962412e-25 | 0.849364432 | 0.689 | 0.183 | 1.07197650072449e-20 | 41 |
| limd24 | 8.5739187807655e-25 | 0.651160571 | 0.41 | 0.069 | 2.15796961793087e-20 | 41 |
| fam210b | 1.28760777549915e-24 | 0.605341414 | 0.361 | 0.055 | 3.24078001015381e-20 | 41 |
| si:ch211-132b12.71 | 2.46921042801867e-24 | 0.53096894 | 0.344 | 0.051 | 6.2147557262802e-20 | 41 |
| mrps158 | 2.50810775831165e-24 | 0.787312712 | 0.557 | 0.124 | 6.3126564168946e-20 | 41 |
| rab27a2 | 2.84803353664015e-24 | 0.519800534 | 0.328 | 0.047 | 7.16821560836958e-20 | 41 |
| raraa3 | 3.4204243972673e-24 | 0.452115043 | 0.262 | 0.031 | 8.60886616548206e-20 | 41 |
| csf3a1 | 3.98941631179215e-24 | 0.252742638 | 0.115 | 0.006 | 1.00409619151497e-19 | 41 |
| gpr132b2 | 4.92179926116029e-24 | 0.514960199 | 0.279 | 0.035 | 1.23876765604143e-19 | 41 |
| csf2rb4 | 1.21708139498096e-23 | 0.506961977 | 0.295 | 0.039 | 3.06327216302758e-19 | 41 |
| actr2a10 | 1.23075063003164e-23 | 0.764108531 | 0.557 | 0.129 | 3.09767626072663e-19 | 41 |
| cd74a10 | 1.49048589356331e-23 | 0.883249768 | 0.77 | 0.218 | 3.7514039455095e-19 | 41 |
| actb215 | 1.85489815021488e-23 | 0.539284609 | 1 | 0.868 | 4.66859315427584e-19 | 41 |
| si:dkey-262k9.411 | 1.93213437687123e-23 | 0.667217708 | 0.459 | 0.09 | 4.8629890131472e-19 | 41 |
| si:ch211-222l21.116 | 2.55822852164624e-23 | 0.927363808 | 0.803 | 0.275 | 6.43880536613143e-19 | 41 |
| vsir10 | 3.50365840754313e-23 | 0.664220906 | 0.443 | 0.085 | 8.81835784594531e-19 | 41 |
| atp2a31 | 6.33332288591037e-23 | 0.42350736 | 0.262 | 0.032 | 1.59403403715478e-18 | 41 |
| crip116 | 7.99846199971904e-23 | 0.90224336 | 0.82 | 0.27 | 2.01313290070929e-18 | 41 |
| rac215 | 8.57165396446087e-23 | 0.841416046 | 0.754 | 0.22 | 2.15739958631516e-18 | 41 |
| spns32 | 1.01675090139375e-22 | 0.312000332 | 0.164 | 0.014 | 2.55906034371792e-18 | 41 |
| sh2d3ca1 | 1.36565921610545e-22 | 0.325441435 | 0.213 | 0.022 | 3.43722768101581e-18 | 41 |

| | | | | | | |
|---|---|---|---|---|---|---|
| il345 | 1.39029239322962e-22 | 0.529423813 | 0.328 | 0.049 | 3.49922692451964e-18 | 41 |
| sfmbt2 | 1.61375741245734e-22 | 0.273438088 | 0.148 | 0.011 | 4.06166603141388e-18 | 41 |
| rps6ka17 | 2.21087909190986e-22 | 0.554049578 | 0.41 | 0.073 | 5.56456158642793e-18 | 41 |
| rnf1302 | 2.40962374540239e-22 | 0.545104472 | 0.311 | 0.047 | 6.06478200480328e-18 | 41 |
| il41 | 2.45351692877309e-22 | 0.349994577 | 0.115 | 0.007 | 6.17525675802899e-18 | 41 |
| elf110 | 9.28371511050197e-22 | 0.703833543 | 0.475 | 0.104 | 2.33661825616224e-17 | 41 |
| dicp3.13 | 1.03108853506507e-21 | 0.33329184 | 0.18 | 0.017 | 2.59514673390526e-17 | 41 |
| grasp6 | 2.43903070672139e-21 | 0.566203528 | 0.41 | 0.076 | 6.13879638574706e-17 | 41 |
| si:dkey-222f2.74 | 2.79818648955936e-21 | 0.389950056 | 0.246 | 0.03 | 7.04275557557195e-17 | 41 |
| anks1b2 | 4.67860483087374e-21 | 0.314684887 | 0.18 | 0.017 | 1.17755804988261e-16 | 41 |
| BX120005.16 | 5.2143302946943e-21 | 0.653507955 | 0.311 | 0.049 | 1.31239479187161e-16 | 41 |
| ldha6 | 5.44179865499279e-21 | 0.566953358 | 0.393 | 0.073 | 1.36964630347514e-16 | 41 |
| arpc314 | 9.74523629838282e-21 | 0.836894477 | 0.803 | 0.31 | 2.45277852393997e-16 | 41 |
| btg117 | 1.16372823223867e-20 | 0.856661654 | 0.836 | 0.332 | 2.9289875877215e-16 | 41 |
| RASSF57 | 2.24730033946702e-20 | 0.489089382 | 0.344 | 0.058 | 5.65623022440454e-16 | 41 |
| arhgdia11 | 2.51405588890492e-20 | 0.795454615 | 0.705 | 0.229 | 6.32762726678478e-16 | 41 |
| CU499336.25 | 2.90984957249306e-20 | 0.43990302 | 0.262 | 0.036 | 7.32380038900779e-16 | 41 |
| calm1b10 | 4.34034551729738e-20 | 0.656458313 | 0.443 | 0.095 | 1.09242156324858e-15 | 41 |
| atp5if1b4 | 4.55518589657787e-20 | 0.780875315 | 0.639 | 0.19 | 1.14649473830968e-15 | 41 |
| sla12 | 5.73645019262582e-20 | 0.303202392 | 0.197 | 0.021 | 1.44380714898199e-15 | 41 |
| selplg8 | 1.00511986349269e-19 | 0.493930571 | 0.295 | 0.047 | 2.52978618442475e-15 | 41 |
| pik3cd | 1.29885053848087e-19 | 0.365626284 | 0.18 | 0.019 | 3.26907692030251e-15 | 41 |
| sqstm11 | 1.69618758800502e-19 | 0.548456363 | 0.393 | 0.077 | 4.26913454024983e-15 | 41 |
| hif1ab5 | 3.48631070104736e-19 | 0.653242757 | 0.475 | 0.113 | 8.7746954034661e-15 | 41 |
| si:ch211-66e2.38 | 8.53865366005188e-19 | 0.588507588 | 0.328 | 0.059 | 2.14909373969846e-14 | 41 |
| zgc:1538678 | 1.08001400153191e-18 | 0.752288312 | 0.934 | 0.436 | 2.71828724045566e-14 | 41 |
| CABZ01084575.15 | 1.32693825340456e-18 | 0.469717434 | 0.279 | 0.043 | 3.33977088999393e-14 | 41 |
| pdxdc1 | 1.52036780302641e-18 | 0.336977299 | 0.18 | 0.02 | 3.82661372343717e-14 | 41 |
| wasb14 | 1.79747194170585e-18 | 0.736083412 | 0.623 | 0.177 | 4.52405713007946e-14 | 41 |
| ptpn222 | 2.632609341518e-18 | 0.36438191 | 0.197 | 0.023 | 6.62601445166667e-14 | 41 |
| cmah3 | 5.42452956273056e-18 | 0.328929856 | 0.164 | 0.017 | 1.36529984564365e-13 | 41 |
| sh3bgrl8 | 5.70573712646043e-18 | 0.605597369 | 0.443 | 0.104 | 1.43607697735883e-13 | 41 |
| csk6 | 8.94990394974225e-18 | 0.591405232 | 0.443 | 0.101 | 2.25260132511063e-13 | 41 |
| rnf19b1 | 9.26251952839697e-18 | 0.30989075 | 0.197 | 0.024 | 2.33128354010223e-13 | 41 |
| capzb12 | 2.25141327290662e-17 | 0.747175048 | 0.705 | 0.253 | 5.66658206657867e-13 | 41 |

| | | | | | | |
|---|---|---|---|---|---|---|
| syk6 | 2.6582140960606e-17 | 0.563808424 | 0.377 | 0.079 | 6.69045905837491e-13 | 41 |
| cfl113 | 2.99147763272356e-17 | 0.671593663 | 0.918 | 0.577 | 7.52925005380193e-13 | 41 |
| zmp:00000010825 | 3.79086517377677e-17 | 0.500990471 | 0.311 | 0.057 | 9.54122855587876e-13 | 41 |
| gapdhs16 | 4.63513427109018e-17 | 0.753994758 | 0.754 | 0.288 | 1.16661694469069e-12 | 41 |
| pde4ba12 | 7.37281499299583e-17 | 0.67804495 | 0.508 | 0.14 | 1.85566380558712e-12 | 41 |
| hic15 | 9.60921306735867e-17 | 0.492627263 | 0.311 | 0.058 | 2.4185428369235e-12 | 41 |
| arl114 | 1.2629062732366e-16 | 0.363361306 | 0.213 | 0.029 | 3.17860879910919e-12 | 41 |
| gng23 | 2.31955194048298e-16 | 0.401885921 | 0.262 | 0.044 | 5.83808027900162e-12 | 41 |
| LOC10390911510 | 2.40029591900499e-16 | 0.675422123 | 0.557 | 0.165 | 6.04130479854367e-12 | 41 |
| nfkbie4 | 2.45538627423627e-16 | 0.527034557 | 0.41 | 0.094 | 6.17996171362526e-12 | 41 |
| oaz1b17 | 2.65296018764797e-16 | 0.709150822 | 0.803 | 0.322 | 6.67723549629117e-12 | 41 |
| pik3ap1 | 2.70230700785255e-16 | 0.321629616 | 0.164 | 0.019 | 6.80143650806408e-12 | 41 |
| XLOC-0430435 | 4.25782781396594e-16 | 0.419586551 | 0.262 | 0.043 | 1.07165268249709e-11 | 41 |
| ucp213 | 4.49619119491991e-16 | 0.73792266 | 0.82 | 0.417 | 1.13164636184939e-11 | 41 |
| wipf1a8 | 4.92802884098495e-16 | 0.503342466 | 0.328 | 0.065 | 1.2403355789875e-11 | 41 |
| pak1 | 5.44126240035558e-16 | 0.262283625 | 0.164 | 0.019 | 1.3695113335455e-11 | 41 |
| tmc85 | 9.16640680170744e-16 | 0.380492805 | 0.23 | 0.035 | 2.30709292792175e-11 | 41 |
| FP236356.15 | 1.51516687151761e-15 | 0.446426232 | 0.279 | 0.05 | 3.81352349892268e-11 | 41 |
| arpc5a10 | 1.53667487979939e-15 | 0.58321783 | 0.443 | 0.115 | 3.8676570049671e-11 | 41 |
| si:dkey-53k12.26 | 1.6773303197883e-15 | 0.376943212 | 0.262 | 0.045 | 4.22167268187517e-11 | 41 |
| si:ch1073-443f11.24 | 1.70636644929688e-15 | 0.358839116 | 0.23 | 0.036 | 4.29475371623532e-11 | 41 |
| mef2d3 | 2.22075464299324e-15 | 0.449170618 | 0.344 | 0.073 | 5.5894173609497e-11 | 41 |
| mtpn8 | 3.62595089196988e-15 | 0.651525129 | 0.492 | 0.146 | 9.12615579999899e-11 | 41 |
| tspan33a2 | 4.84294551746337e-15 | 0.284697718 | 0.148 | 0.017 | 1.21892095729036e-10 | 41 |
| nme2b.15 | 5.70450790855258e-15 | 0.329184683 | 1 | 0.857 | 1.4357675955036e-10 | 41 |
| zfp36l1a11 | 5.87799484554798e-15 | 0.725727865 | 0.672 | 0.248 | 1.47943252267597e-10 | 41 |
| rasal34 | 6.28102633510406e-15 | 0.303114441 | 0.197 | 0.028 | 1.58087151828234e-10 | 41 |
| fscn1a4 | 6.45906097103166e-15 | 0.477192479 | 0.295 | 0.058 | 1.62568105579896e-10 | 41 |
| dtnbp1a5 | 7.39675786465034e-15 | 0.490043015 | 0.377 | 0.087 | 1.86168998695384e-10 | 41 |
| rplp222 | 7.86884884863329e-15 | 0.473821229 | 1 | 0.758 | 1.98051056671251e-10 | 41 |
| lrrfip1a4 | 1.32471605082715e-14 | 0.413509059 | 0.295 | 0.058 | 3.33417782832686e-10 | 41 |
| pptc7b1 | 1.64288816935818e-14 | 0.340718386 | 0.213 | 0.033 | 4.13498523345759e-10 | 41 |
| si:ch73-208g10.16 | 1.94160351181638e-14 | 0.39476631 | 0.246 | 0.043 | 4.88682187889065e-10 | 41 |
| MAP3K11 | 2.3594383534231e-14 | 0.250555385 | 0.148 | 0.017 | 5.9384703917306e-10 | 41 |
| mrtfaa2 | 2.36911299799251e-14 | 0.278380027 | 0.18 | 0.025 | 5.96282050464734e-10 | 41 |

| | | | | | | |
|---|---|---|---|---|---|---|
| tmsb4x25 | 2.49657261332083e-14 | 0.62449842 | 1 | 0.686 | 6.2836236104672e-10 | 41 |
| rhoab11 | 2.81374675443847e-14 | 0.702771548 | 0.541 | 0.19 | 7.08191920624619e-10 | 41 |
| lasp19 | 3.31326373174234e-14 | 0.652221609 | 0.656 | 0.237 | 8.3391534864223e-10 | 41 |
| arrb2b9 | 3.45373087698054e-14 | 0.497314018 | 0.377 | 0.092 | 8.69269524427231e-10 | 41 |
| BX649485.12 | 3.80168031763752e-14 | 0.324002451 | 0.18 | 0.025 | 9.56844919146188e-10 | 41 |
| mir21-17 | 3.82049945360213e-14 | 0.561236192 | 0.41 | 0.108 | 9.6158150747712e-10 | 41 |
| LOC1104382372 | 7.10824934644426e-14 | 0.365716754 | 0.279 | 0.054 | 1.78907527800656e-09 | 41 |
| si:rp71-36a1.3 | 7.12323529550862e-14 | 0.266011498 | 0.131 | 0.014 | 1.79284709152656e-09 | 41 |
| fli1b4 | 8.22693280322922e-14 | 0.346895487 | 0.213 | 0.035 | 2.07063671724476e-09 | 41 |
| myh11a7 | 9.72031633661474e-14 | 0.434433812 | 0.311 | 0.066 | 2.44650641876256e-09 | 41 |
| pi4k2b | 1.51596035050619e-13 | 0.290984991 | 0.246 | 0.043 | 3.81552060618903e-09 | 41 |
| ncalda | 1.558415971498e-13 | 0.377711192 | 0.262 | 0.05 | 3.92237715866331e-09 | 41 |
| cdkn1bb5 | 2.23977156081601e-13 | 0.608194121 | 0.475 | 0.149 | 5.63728104141781e-09 | 41 |
| ches1 | 2.9254896902443e-13 | 0.287260412 | 0.18 | 0.026 | 7.36316500137587e-09 | 41 |
| cotl114 | 3.0514808645407e-13 | 0.705061855 | 0.754 | 0.307 | 7.68027218796248e-09 | 41 |
| rel4 | 4.78201739565107e-13 | 0.379752931 | 0.213 | 0.037 | 1.20358595831142e-08 | 41 |
| ccni14 | 4.79584502477795e-13 | 0.634213943 | 0.721 | 0.294 | 1.20706623428636e-08 | 41 |
| si:ch211-8c17.33 | 5.98931222418069e-13 | 0.311952029 | 0.197 | 0.032 | 1.50744999370404e-08 | 41 |
| prkcbb1 | 7.45941576563453e-13 | 0.281924171 | 0.197 | 0.032 | 1.87746035405255e-08 | 41 |
| elmo18 | 7.77382062313477e-13 | 0.446388998 | 0.311 | 0.071 | 1.95659291263679e-08 | 41 |
| hcls17 | 7.98119395365269e-13 | 0.429562383 | 0.311 | 0.071 | 2.00878670619485e-08 | 41 |
| snap23.1 | 8.08996504769101e-13 | 0.457844038 | 0.311 | 0.072 | 2.03616330285335e-08 | 41 |
| arfgef2 | 8.12912887499632e-13 | 0.262108416 | 0.148 | 0.019 | 2.04602044654782e-08 | 41 |
| arpc215 | 8.48789715717306e-13 | 0.643141797 | 0.672 | 0.268 | 2.13631883548889e-08 | 41 |
| gstk11 | 9.81201668834247e-13 | 0.343926786 | 0.246 | 0.047 | 2.46958648028892e-08 | 41 |
| gsk3b | 1.02578420038227e-12 | 0.265511758 | 0.18 | 0.028 | 2.58179625394214e-08 | 41 |
| arhgdig14 | 1.09323890418163e-12 | 0.655801326 | 0.639 | 0.243 | 2.75157299793475e-08 | 41 |
| actb17 | 1.16526347321634e-12 | 0.416349865 | 0.934 | 0.785 | 2.93285163573822e-08 | 41 |
| eef1a1l18 | 1.27154518517496e-12 | 0.269985539 | 1 | 0.915 | 3.20035207656686e-08 | 41 |
| tbc1d10b | 1.52647759869688e-12 | 0.287188853 | 0.148 | 0.02 | 3.84199146816018e-08 | 41 |
| myd88 | 1.60473150250282e-12 | 0.407887458 | 0.262 | 0.055 | 4.03894871864935e-08 | 41 |
| rpl3420 | 1.77171984935242e-12 | 0.401334977 | 0.984 | 0.784 | 4.45924168883511e-08 | 41 |
| cebpb17 | 1.79964764043773e-12 | 0.726082269 | 0.787 | 0.353 | 4.52953314621772e-08 | 41 |
| ccdc28a1 | 2.01264878870225e-12 | 0.476122261 | 0.393 | 0.106 | 5.06563573628469e-08 | 41 |
| sh3kbp15 | 2.04137326942588e-12 | 0.382779548 | 0.311 | 0.071 | 5.13793238181801e-08 | 41 |

| | | | | | | |
|---|---|---|---|---|---|---|
| sat1a.28 | 2.26212970483401e-12 | 0.659350773 | 0.689 | 0.299 | 5.69355425409673e-08 | 41 |
| cox6b112 | 2.60016592194099e-12 | 0.560734548 | 0.41 | 0.121 | 6.54435760893328e-08 | 41 |
| rab448 | 2.61446748359571e-12 | 0.453077568 | 0.279 | 0.062 | 6.58035320946205e-08 | 41 |
| plxnc15 | 2.72877321554871e-12 | 0.392148095 | 0.23 | 0.044 | 6.86804930621454e-08 | 41 |
| gabarapb13 | 3.03634640535597e-12 | 0.627803155 | 0.656 | 0.269 | 7.64218026764043e-08 | 41 |
| RASA21 | 3.63158972621359e-12 | 0.350489888 | 0.23 | 0.044 | 9.14034818190698e-08 | 41 |
| parp8 | 4.25215279130988e-12 | 0.307505915 | 0.213 | 0.038 | 1.07022433604478e-07 | 41 |
| cdkn1ba | 4.4788914076984e-12 | 0.28175191 | 0.115 | 0.013 | 1.12729217840361e-07 | 41 |
| slc25a331 | 4.73923279947842e-12 | 0.449751907 | 0.377 | 0.1 | 1.19281750330072e-07 | 41 |
| btk3 | 4.87861510918589e-12 | 0.304677607 | 0.164 | 0.025 | 1.227898636831e-07 | 41 |
| si:dkeyp-19e1.35 | 4.88299421287762e-12 | 0.423483492 | 0.295 | 0.068 | 1.22900081343917e-07 | 41 |
| gng57 | 5.85719103343902e-12 | 0.623769725 | 0.656 | 0.272 | 1.47419641120627e-07 | 41 |
| mhc2dab6 | 6.3052594687236e-12 | 0.44519141 | 0.377 | 0.098 | 1.58697075568304e-07 | 41 |
| tsc22d312 | 8.12927734754967e-12 | 0.588080318 | 0.443 | 0.144 | 2.04605781560478e-07 | 41 |
| dusp110 | 8.201448748622e-12 | 0.561959184 | 0.492 | 0.16 | 2.06422263554067e-07 | 41 |
| mapre1a8 | 9.50482949120325e-12 | 0.569715196 | 0.574 | 0.198 | 2.39227053464095e-07 | 41 |
| fam107b3 | 9.55033751484486e-12 | 0.50235204 | 0.377 | 0.104 | 2.4037244491113e-07 | 41 |
| arpc4l13 | 1.19811127672069e-11 | 0.584928014 | 0.541 | 0.197 | 3.01552627237831e-07 | 41 |
| ets26 | 1.45760449597484e-11 | 0.451110309 | 0.344 | 0.091 | 3.66864475591908e-07 | 41 |
| ndel1a | 1.91278476775328e-11 | 0.260214472 | 0.131 | 0.017 | 4.81428798195823e-07 | 41 |
| CR753876.16 | 1.91898174330264e-11 | 0.491033015 | 0.41 | 0.115 | 4.82988514971842e-07 | 41 |
| nek6 | 2.01743239086094e-11 | 0.263680242 | 0.164 | 0.026 | 5.07767558455791e-07 | 41 |
| cap115 | 2.292963488218e-11 | 0.568479424 | 0.623 | 0.234 | 5.77115980349587e-07 | 41 |
| arpc5b8 | 2.49826622842119e-11 | 0.483756324 | 0.393 | 0.114 | 6.28788627031329e-07 | 41 |
| FP101882.12 | 2.61736401748251e-11 | 0.304365767 | 0.197 | 0.035 | 6.58764349560173e-07 | 41 |
| pnrc222 | 2.71057818013064e-11 | 0.560897955 | 0.918 | 0.568 | 6.82225422157082e-07 | 41 |
| larp4ab3 | 2.8316691127656e-11 | 0.385224692 | 0.295 | 0.071 | 7.12702798991973e-07 | 41 |
| mob1a9 | 2.85653274619101e-11 | 0.476772759 | 0.508 | 0.166 | 7.18960726888814e-07 | 41 |
| mcl1a12 | 4.7388152735563e-11 | 0.513645373 | 0.934 | 0.624 | 1.19271241620138e-06 | 41 |
| far12 | 5.44782381362257e-11 | 0.298272762 | 0.197 | 0.036 | 1.37116277565066e-06 | 41 |
| camk2g21 | 5.81268977027961e-11 | 0.313140381 | 0.197 | 0.037 | 1.46299588828168e-06 | 41 |
| rps2915 | 5.92142222432004e-11 | 0.344753753 | 1 | 0.832 | 1.49036275963911e-06 | 41 |
| ostf14 | 5.9581557083631e-11 | 0.357723504 | 0.246 | 0.054 | 1.49960821023791e-06 | 41 |
| CR392341.18 | 6.15323020984923e-11 | 0.402790713 | 0.262 | 0.06 | 1.54870651151695e-06 | 41 |
| lcp19 | 7.08176349499662e-11 | 0.475415071 | 0.328 | 0.089 | 1.7824090540557e-06 | 41 |

| | | | | | | |
|---|---|---|---|---|---|---|
| cmtm73 | 7.53378338996717e-11 | 0.323592123 | 0.23 | 0.048 | 1.89617794142084e-06 | 41 |
| dbnlb2 | 9.76975047816054e-11 | 0.309798756 | 0.213 | 0.042 | 2.45894849784823e-06 | 41 |
| pdxka1 | 9.94416121892419e-11 | 0.309640152 | 0.18 | 0.032 | 2.50284593719103e-06 | 41 |
| per2 | 1.11554743717723e-10 | 0.299295769 | 0.197 | 0.037 | 2.80772134463136e-06 | 41 |
| stk17al2 | 1.31063888760877e-10 | 0.451624332 | 0.41 | 0.121 | 3.29874701622251e-06 | 41 |
| ccdc88b2 | 1.40555598553449e-10 | 0.30565001 | 0.18 | 0.033 | 3.53764385999176e-06 | 41 |
| arl4aa1 | 1.43675011748076e-10 | 0.378618234 | 0.262 | 0.06 | 3.61615637068731e-06 | 41 |
| rps15a18 | 1.89681933592633e-10 | 0.415609627 | 1 | 0.78 | 4.77410458659299e-06 | 41 |
| bcl10 | 2.00811552131213e-10 | 0.291146735 | 0.148 | 0.023 | 5.05422595559049e-06 | 41 |
| plekho25 | 2.15007450171169e-10 | 0.382644348 | 0.295 | 0.075 | 5.41152251335815e-06 | 41 |
| tyk21 | 2.3536988986649e-10 | 0.251827869 | 0.164 | 0.028 | 5.9240247580497e-06 | 41 |
| cxcr4b12 | 2.63912875377544e-10 | 0.565977411 | 0.525 | 0.184 | 6.6424231603774e-06 | 41 |
| abat | 2.71680555187822e-10 | 0.300846359 | 0.18 | 0.033 | 6.83792789352229e-06 | 41 |
| si:ch211-226m16.27 | 2.79490946834154e-10 | 0.394262964 | 0.279 | 0.069 | 7.03450764086882e-06 | 41 |
| tmcc3 | 2.88821796659493e-10 | 0.261894363 | 0.131 | 0.019 | 7.26935580012279e-06 | 41 |
| clic19 | 4.10531608658629e-10 | 0.425693977 | 0.377 | 0.114 | 1.0332670058329e-05 | 41 |
| mt-co120 | 4.24308415512116e-10 | 0.490541782 | 0.918 | 0.658 | 1.06794185100244e-05 | 41 |
| cox8a13 | 4.72852274766562e-10 | 0.54264075 | 0.557 | 0.228 | 1.19012189035996e-05 | 41 |
| amfra | 6.24152900213904e-10 | 0.285417731 | 0.164 | 0.029 | 1.57093043454838e-05 | 41 |
| CU467110.1 | 6.31882415180601e-10 | 0.359186003 | 0.328 | 0.088 | 1.59038485076805e-05 | 41 |
| CR855337.13 | 6.49638593319087e-10 | 0.351518367 | 0.18 | 0.035 | 1.63507537552481e-05 | 41 |
| tpi1b9 | 8.43976180299052e-10 | 0.523744223 | 0.459 | 0.168 | 2.12420364819468e-05 | 41 |
| CABZ01079663.15 | 8.88353585225531e-10 | 0.308687799 | 0.197 | 0.04 | 2.23589713865414e-05 | 41 |
| cox8b6 | 9.09392266848544e-10 | 0.445426446 | 0.328 | 0.096 | 2.2888493964311e-05 | 41 |
| sec61g7 | 1.19569819323296e-09 | 0.422313761 | 0.918 | 0.679 | 3.00945278254805e-05 | 41 |
| si:busm1-266f07.28 | 1.33943140694101e-09 | 0.434964109 | 0.459 | 0.147 | 3.37121490812982e-05 | 41 |
| dub8 | 1.35243206351678e-09 | 0.349896598 | 0.279 | 0.072 | 3.40393626066539e-05 | 41 |
| rpl1518 | 1.35857972925626e-09 | 0.391066512 | 0.967 | 0.745 | 3.41940932056508e-05 | 41 |
| plek3 | 1.52850760565681e-09 | 0.25040396 | 0.148 | 0.025 | 3.84710079267762e-05 | 41 |
| si:ch211-105j21.95 | 1.6679807940144e-09 | 0.286342939 | 0.18 | 0.035 | 4.19814086045484e-05 | 41 |
| sept7b1 | 2.02513886148975e-09 | 0.392563001 | 0.344 | 0.103 | 5.09707200048355e-05 | 41 |
| rpl2614 | 2.05461802326449e-09 | 0.310636303 | 0.984 | 0.841 | 5.17126810275439e-05 | 41 |
| rps2126 | 2.19141168599951e-09 | 0.424660889 | 1 | 0.75 | 5.51556407249218e-05 | 41 |
| sec11a2 | 2.21214363280187e-09 | 0.446293768 | 0.492 | 0.178 | 5.56774430939903e-05 | 41 |
| COX320 | 2.21509319303612e-09 | 0.483472341 | 0.984 | 0.7 | 5.57516805755262e-05 | 41 |

| | | | | | | |
|---|---|---|---|---|---|---|
| nr4a36 | 2.62756311680575e-09 | 0.404465463 | 0.23 | 0.055 | 6.61331360868838e-05 | 41 |
| rpl3218 | 3.01171551904269e-09 | 0.358130921 | 1 | 0.814 | 7.58018678987854e-05 | 41 |
| cers12 | 3.14504581512725e-09 | 0.262587239 | 0.131 | 0.021 | 7.91576581209378e-05 | 41 |
| rpl5a19 | 3.85586923230592e-09 | 0.531011539 | 0.869 | 0.492 | 9.70483727079076e-05 | 41 |
| zgc:1629445 | 3.95631666241066e-09 | 0.528201389 | 0.59 | 0.253 | 9.95765340762139e-05 | 41 |
| BX901920.17 | 4.30894303555595e-09 | 0.370991298 | 0.279 | 0.074 | 0.000108452 | 41 |
| bcl2l105 | 4.71462229843333e-09 | 0.552890677 | 0.525 | 0.215 | 0.000118662 | 41 |
| CABZ01090890.12 | 5.09228791168863e-09 | 0.316011606 | 0.23 | 0.055 | 0.000128168 | 41 |
| anxa49 | 6.13450417924997e-09 | 0.452449658 | 0.393 | 0.137 | 0.000154399 | 41 |
| iqgap13 | 6.37983358392393e-09 | 0.351410471 | 0.262 | 0.07 | 0.000160574 | 41 |
| capza1a12 | 7.12589855860095e-09 | 0.514570618 | 0.492 | 0.2 | 0.000179352 | 41 |
| rhoaa11 | 7.3711224267037e-09 | 0.439325002 | 0.426 | 0.155 | 0.000185524 | 41 |
| tes1 | 7.62383992024509e-09 | 0.356791495 | 0.246 | 0.063 | 0.000191884 | 41 |
| serf216 | 7.92675322136784e-09 | 0.526876671 | 0.705 | 0.343 | 0.000199508 | 41 |
| si:zfos-2330d3.12 | 8.65564210595097e-09 | 0.361852353 | 0.148 | 0.027 | 0.000217854 | 41 |
| pdcd10a | 9.49559949264122e-09 | 0.28040791 | 0.262 | 0.069 | 0.000238995 | 41 |
| eif3f18 | 1.17408339314515e-08 | 0.523985085 | 0.77 | 0.432 | 0.000295505 | 41 |
| zgc:1537137 | 1.33820867419401e-08 | 0.415980027 | 0.426 | 0.153 | 0.000336814 | 41 |
| tnfaip37 | 1.53607799685097e-08 | 0.368122081 | 0.246 | 0.066 | 0.000386615 | 41 |
| tiparp1 | 1.59662011151964e-08 | 0.317008262 | 0.197 | 0.045 | 0.000401853 | 41 |
| metrnla6 | 1.87199641682705e-08 | 0.381424767 | 0.311 | 0.093 | 0.000471163 | 41 |
| rps250 | 2.09757478720249e-08 | 0.349646587 | 1 | 0.829 | 0.000527939 | 41 |
| rpl18a18 | 2.22752785052089e-08 | 0.354155903 | 0.984 | 0.785 | 0.000560646 | 41 |
| slc3a2b9 | 2.29088241775275e-08 | 0.467230152 | 0.344 | 0.119 | 0.000576592 | 41 |
| si:rp71-36a1.1 | 2.29092188642828e-08 | 0.264687746 | 0.164 | 0.034 | 0.000576602 | 41 |
| ak15 | 2.71847641891234e-08 | 0.404275129 | 0.262 | 0.076 | 0.000684213 | 41 |
| rps2820 | 3.00645244458217e-08 | 0.327484283 | 1 | 0.827 | 0.000756694 | 41 |
| myo1f7 | 3.68602025301179e-08 | 0.29828823 | 0.197 | 0.047 | 0.000927734 | 41 |
| ube2d1b1 | 4.28653502552455e-08 | 0.39642267 | 0.377 | 0.131 | 0.001078878 | 41 |
| rca2.11 | 5.06441589841295e-08 | 0.356236939 | 0.279 | 0.082 | 0.001274663 | 41 |
| sptbn22 | 5.36114083410363e-08 | 0.269334516 | 0.18 | 0.041 | 0.001349346 | 41 |
| prkar2aa3 | 5.89875577688358e-08 | 0.388582193 | 0.311 | 0.099 | 0.001484658 | 41 |
| LOC1001511963 | 7.20802472104042e-08 | 0.282498553 | 0.18 | 0.041 | 0.001814188 | 41 |
| rps27a18 | 8.35756412335322e-08 | 0.345362843 | 0.967 | 0.752 | 0.002103515 | 41 |
| rpl3915 | 8.45860069068e-08 | 0.274687961 | 1 | 0.874 | 0.002128945 | 41 |

| | | | | | | |
|---|---|---|---|---|---|---|
| ubr4 | 8.63648422512583e-08 | 0.278885675 | 0.18 | 0.042 | 0.002173717 | 41 |
| LOC1005362136 | 9.00301467345086e-08 | 0.392474484 | 0.328 | 0.107 | 0.002265969 | 41 |
| rpl2219 | 9.08151817983292e-08 | 0.41231102 | 0.967 | 0.694 | 0.002285727 | 41 |
| lamp1a4 | 9.44412448109786e-08 | 0.360521227 | 0.295 | 0.092 | 0.002376992 | 41 |
| rpl1417 | 1.05630061078172e-07 | 0.415946731 | 0.918 | 0.635 | 0.002658603 | 41 |
| pgam1a7 | 1.28729685123544e-07 | 0.447992613 | 0.41 | 0.158 | 0.003239997 | 41 |
| rpl1910 | 1.32983908150134e-07 | 0.2758603 | 0.984 | 0.852 | 0.003347072 | 41 |
| DHDH4 | 1.36417534238382e-07 | 0.269170593 | 0.23 | 0.062 | 0.003433493 | 41 |
| si:ch211-196l7.417 | 1.47686064778368e-07 | 0.480780227 | 0.656 | 0.315 | 0.003717111 | 41 |
| ptmab15 | 1.56286936006155e-07 | 0.47164033 | 0.852 | 0.541 | 0.003933586 | 41 |
| rps26l20 | 1.65224476330028e-07 | 0.423157046 | 0.951 | 0.665 | 0.004158535 | 41 |
| twf1a5 | 1.68070735115078e-07 | 0.378962577 | 0.41 | 0.152 | 0.004230172 | 41 |
| sept127 | 1.908810803952e-07 | 0.265969837 | 0.23 | 0.062 | 0.004804286 | 41 |
| prelid3b5 | 2.57660346729057e-07 | 0.440677205 | 0.541 | 0.239 | 0.006485053 | 41 |
| psma6l1 | 2.68482297901759e-07 | 0.273380315 | 0.164 | 0.038 | 0.006757431 | 41 |
| rnf446 | 3.19481174742928e-07 | 0.362858913 | 0.279 | 0.09 | 0.008041022 | 41 |
| cdc42l8 | 3.32363075915008e-07 | 0.442294363 | 0.492 | 0.216 | 0.008365246 | 41 |
| rpl3623 | 3.72915823127153e-07 | 0.356600539 | 1 | 0.79 | 0.009385918 | 41 |
| tagapa10 | 3.98578558933229e-07 | 0.418033279 | 0.311 | 0.106 | 0.010031824 | 41 |
| coa6 | 3.99320038488166e-07 | 0.305653251 | 0.279 | 0.088 | 0.010050486 | 41 |
| tec1 | 4.83383407922684e-07 | 0.255996364 | 0.148 | 0.032 | 0.012166277 | 41 |
| rpl924 | 5.62160566773863e-07 | 0.444970205 | 0.951 | 0.63 | 0.014149019 | 41 |
| dok26 | 5.71987503672657e-07 | 0.263460392 | 0.164 | 0.039 | 0.014396353 | 41 |
| selenok5 | 6.36572126740339e-07 | 0.362864584 | 0.344 | 0.128 | 0.016021884 | 41 |
| gm2a6 | 6.56419103903995e-07 | 0.356941736 | 0.344 | 0.122 | 0.016521412 | 41 |
| rps1520 | 6.61886241070421e-07 | 0.401941158 | 0.967 | 0.685 | 0.016659015 | 41 |
| XLOC-0268365 | 6.73216743492431e-07 | 0.309634979 | 0.262 | 0.079 | 0.016944192 | 41 |
| slc6a6b7 | 8.13054100472783e-07 | 0.295316157 | 0.311 | 0.104 | 0.020463759 | 41 |
| rps1820 | 8.19162611720369e-07 | 0.353293899 | 0.934 | 0.725 | 0.020617504 | 41 |
| egr37 | 8.382722750246e-07 | 0.264018046 | 0.213 | 0.059 | 0.021098475 | 41 |
| nek7 | 9.13985770698586e-07 | 0.259941497 | 0.18 | 0.046 | 0.023004108 | 41 |
| GCA7 | 9.68000755194804e-07 | 0.412176147 | 0.426 | 0.182 | 0.024363611 | 41 |
| vamp81 | 1.00195006351914e-06 | 0.29821664 | 0.246 | 0.075 | 0.025218081 | 41 |
| si:dkey-118j18.22 | 1.13375544992626e-06 | 0.294072689 | 0.197 | 0.054 | 0.028535491 | 41 |
| tspan372 | 1.1756511222307e-06 | 0.258613173 | 0.131 | 0.028 | 0.029589963 | 41 |

| | | | | | | |
|---|---|---|---|---|---|---|
| rpl7a15 | 1.19399533067097e-06 | 0.311070484 | 1 | 0.81 | 0.030051668 | 41 |
| rap1b8 | 1.61780180643679e-06 | 0.413097253 | 0.459 | 0.203 | 0.040718454 | 41 |
| si:dkeyp-68b7.129 | 1.72875228788996e-06 | 0.316818254 | 0.213 | 0.064 | 0.043510966 | 41 |
| rpl3820 | 1.75455941450152e-06 | 0.380985345 | 1 | 0.728 | 0.044160506 | 41 |
| creg1 | 1.87310999110132e-06 | 0.273893306 | 0.23 | 0.069 | 0.047144305 | 41 |
| ywhabl7 | 1.9514563884876e-06 | 0.440079349 | 0.574 | 0.279 | 0.049116206 | 41 |
| rpl3716 | 1.96960719734449e-06 | 0.295353358 | 1 | 0.85 | 0.049573044 | 41 |
| rpl3020 | 2.10256127634589e-06 | 0.348104353 | 0.934 | 0.709 | 0.052919365 | 41 |
| rps518 | 2.12636513687086e-06 | 0.357729237 | 0.934 | 0.702 | 0.053518484 | 41 |
| sh2b36 | 2.13573020784544e-06 | 0.26143593 | 0.213 | 0.062 | 0.053754194 | 41 |
| rpl35a18 | 2.30926522530828e-06 | 0.336976576 | 0.984 | 0.76 | 0.058121896 | 41 |
| rarga4 | 2.35724826573948e-06 | 0.33411281 | 0.197 | 0.058 | 0.059329582 | 41 |
| ap2m1a4 | 2.43062931617756e-06 | 0.35597399 | 0.361 | 0.143 | 0.061176509 | 41 |
| anxa11a2 | 2.55757855362218e-06 | 0.336120609 | 0.295 | 0.107 | 0.064371695 | 41 |
| pnp5a6 | 2.57630391779653e-06 | 0.291423479 | 0.246 | 0.079 | 0.064842993 | 41 |
| prkar1ab1 | 2.66235176813386e-06 | 0.276293776 | 0.23 | 0.072 | 0.067008732 | 41 |
| eif4ebp18 | 2.69731970091767e-06 | 0.393616482 | 0.541 | 0.247 | 0.06788884 | 41 |
| ppt17 | 2.79977783070894e-06 | 0.382275976 | 0.393 | 0.164 | 0.070467608 | 41 |
| higd1a11 | 2.84450898719418e-06 | 0.444341427 | 0.607 | 0.311 | 0.071593447 | 41 |
| tpt112 | 3.15312330742581e-06 | 0.28586856 | 0.984 | 0.801 | 0.079360961 | 41 |
| cry23 | 3.44280682954629e-06 | 0.322801882 | 0.246 | 0.082 | 0.086652005 | 41 |
| mbnl34 | 3.47215330407718e-06 | 0.268699225 | 0.197 | 0.057 | 0.087390627 | 41 |
| mt-atp620 | 3.47541015821481e-06 | 0.437148819 | 0.951 | 0.591 | 0.087472598 | 41 |
| rplp113 | 3.47844981499978e-06 | 0.261264737 | 1 | 0.877 | 0.087549103 | 41 |
| si:dkey-112e7.212 | 3.63697529479652e-06 | 0.382069821 | 0.459 | 0.207 | 0.091539031 | 41 |
| cmip4 | 3.91245723300347e-06 | 0.297081831 | 0.213 | 0.065 | 0.098472636 | 41 |
| mt-co220 | 3.98060868670199e-06 | 0.376542227 | 0.934 | 0.721 | 0.10018794 | 41 |
| rab8b5 | 4.02080503752197e-06 | 0.265912723 | 0.246 | 0.08 | 0.101199642 | 41 |
| eif4ebp28 | 4.45522709353572e-06 | 0.363151381 | 0.41 | 0.175 | 0.112133611 | 41 |
| commd31 | 4.66987403293972e-06 | 0.355146089 | 0.295 | 0.109 | 0.11753606 | 41 |
| btf316 | 4.72743671993605e-06 | 0.390473409 | 0.852 | 0.588 | 0.118984855 | 41 |
| rps1419 | 4.82074320800975e-06 | 0.349604862 | 1 | 0.759 | 0.121333286 | 41 |
| mt-nd315 | 4.90609259564229e-06 | 0.419026417 | 0.721 | 0.396 | 0.123481445 | 41 |
| rpl2920 | 5.34254125805823e-06 | 0.305192983 | 1 | 0.809 | 0.134466421 | 41 |
| rpl2817 | 5.73464570013565e-06 | 0.32951726 | 0.967 | 0.75 | 0.144335298 | 41 |

| | | | | | | |
|---|---|---|---|---|---|---|
| rpl23a22 | 6.07392367317107e-06 | 0.41342481 | 0.951 | 0.634 | 0.152874585 | 41 |
| fkbp57 | 6.60623783542973e-06 | 0.368707217 | 0.311 | 0.123 | 0.1662724 | 41 |
| serpinb1l3 | 7.34463350879861e-06 | 0.252694109 | 0.23 | 0.074 | 0.184857081 | 41 |
| nfe2l2a4 | 7.75635732521373e-06 | 0.313960166 | 0.213 | 0.07 | 0.195219758 | 41 |
| eef1db14 | 8.0694388608567e-06 | 0.41510321 | 0.541 | 0.275 | 0.203099707 | 41 |
| eif3s107 | 8.99670111154424e-06 | 0.308296545 | 0.393 | 0.166 | 0.22643797 | 41 |
| uqcrq5 | 9.56352456034362e-06 | 0.399126463 | 0.541 | 0.272 | 0.24070435 | 41 |
| rpsa24 | 9.87767193815415e-06 | 0.343724384 | 0.984 | 0.752 | 0.248611125 | 41 |
| zgc:11418824 | 1.0066449735988e-05 | 0.404195417 | 0.918 | 0.628 | 0.253362473 | 41 |
| h1fx12 | 1.03439880126193e-05 | 0.370950127 | 0.426 | 0.184 | 0.260347834 | 41 |
| efhd24 | 1.0420624576964e-05 | 0.289410152 | 0.197 | 0.061 | 0.2622767 | 41 |
| stk17b6 | 1.06247294144747e-05 | 0.333155779 | 0.295 | 0.112 | 0.267413815 | 41 |
| capza1b8 | 1.11321342470043e-05 | 0.409844025 | 0.443 | 0.214 | 0.280184687 | 41 |
| rpl1119 | 1.1508933723009e-05 | 0.309046181 | 1 | 0.814 | 0.289668353 | 41 |
| ndufb94 | 1.17954419071919e-05 | 0.311755442 | 0.41 | 0.177 | 0.296879477 | 41 |
| mibp217 | 1.26274619011657e-05 | 0.396739168 | 0.738 | 0.493 | 0.317820589 | 41 |
| gnai2a5 | 1.37265988228761e-05 | 0.343941995 | 0.344 | 0.144 | 0.345484766 | 41 |
| litaf8 | 1.48787011638124e-05 | 0.345092542 | 0.328 | 0.136 | 0.37448203 | 41 |
| rpl36a16 | 1.50577707443389e-05 | 0.277118302 | 0.984 | 0.81 | 0.378989032 | 41 |
| rpl818 | 1.71083452823446e-05 | 0.32110299 | 1 | 0.78 | 0.430599942 | 41 |
| rps27.118 | 1.74410084278629e-05 | 0.368787158 | 0.934 | 0.68 | 0.438972741 | 41 |
| rps1621 | 1.75950477782445e-05 | 0.360680819 | 1 | 0.738 | 0.442849758 | 41 |
| zgc:776501 | 1.83455922486728e-05 | 0.26324951 | 0.279 | 0.102 | 0.461740211 | 41 |
| ndufa61 | 1.86628531723553e-05 | 0.370395619 | 0.508 | 0.25 | 0.469725351 | 41 |
| CR383676.121 | 1.87361925118415e-05 | 0.466212493 | 0.902 | 0.727 | 0.471571229 | 41 |
| usp9 | 1.91908472205889e-05 | 0.275427209 | 0.213 | 0.071 | 0.483014434 | 41 |
| tbcb1 | 1.93245307481891e-05 | 0.26539548 | 0.213 | 0.07 | 0.486379114 | 41 |
| bcl2l11 | 2.09362620453289e-05 | 0.304268426 | 0.23 | 0.08 | 0.526944779 | 41 |
| pdk2b9 | 2.15205701759236e-05 | 0.347950146 | 0.262 | 0.101 | 0.541651231 | 41 |
| mt-cyb15 | 2.1886493345349e-05 | 0.379319439 | 0.885 | 0.625 | 0.550861151 | 41 |
| si:ch1073-325m22.2( | 2.4056432496049e-05 | 0.345053712 | 0.475 | 0.231 | 0.605476349 | 41 |
| ndufa81 | 2.60367124972197e-05 | 0.287848431 | 0.328 | 0.134 | 0.655318017 | 41 |
| per35 | 2.67930453174805e-05 | 0.26073512 | 0.23 | 0.08 | 0.674354158 | 41 |
| arl6ip5b5 | 2.74031826944193e-05 | 0.25832111 | 0.262 | 0.097 | 0.689710705 | 41 |
| msna15 | 2.96325890647254e-05 | 0.387025065 | 0.475 | 0.243 | 0.745822634 | 41 |

| | | | | | | |
|---|---|---|---|---|---|---|
| hmgn616 | 3.59040782184312e-05 | 0.369271364 | 0.525 | 0.266 | 0.903669745 | 41 |
| cry-dash | 3.70548957948902e-05 | 0.256926504 | 0.213 | 0.072 | 0.932634672 | 41 |
| cct54 | 3.81568432240902e-05 | 0.336979772 | 0.508 | 0.252 | 0.960369587 | 41 |
| si:ch73-234b20.59 | 4.10199745263607e-05 | 0.321644365 | 0.262 | 0.101 | 1 | 41 |
| eef1da14 | 4.74344075834834e-05 | 0.381582722 | 0.721 | 0.493 | 1 | 41 |
| wdr18 | 4.88250757661849e-05 | 0.344016565 | 0.328 | 0.143 | 1 | 41 |
| gosr2 | 5.09153606259867e-05 | 0.256333303 | 0.197 | 0.066 | 1 | 41 |
| zgc:5649313 | 5.27842001163055e-05 | 0.360439442 | 0.82 | 0.573 | 1 | 41 |
| eno1a7 | 5.57941769890324e-05 | 0.256747067 | 0.328 | 0.135 | 1 | 41 |
| uba5219 | 5.60096003557391e-05 | 0.327915447 | 0.852 | 0.631 | 1 | 41 |
| wu:fi37e092 | 6.01362203271668e-05 | 0.265433766 | 0.295 | 0.119 | 1 | 41 |
| hsp90ab112 | 6.20990218473738e-05 | 0.274101538 | 0.984 | 0.791 | 1 | 41 |
| rps27.219 | 6.23688042780449e-05 | 0.415554157 | 0.869 | 0.518 | 1 | 41 |
| rsl24d117 | 6.42700699859657e-05 | 0.398781512 | 0.738 | 0.445 | 1 | 41 |
| ipo73 | 6.45287014712547e-05 | 0.257810229 | 0.279 | 0.111 | 1 | 41 |
| fosl213 | 7.00813168249364e-05 | 0.297150594 | 0.295 | 0.123 | 1 | 41 |
| cdc428 | 8.42515820880085e-05 | 0.3389123 | 0.426 | 0.213 | 1 | 41 |
| arpc410 | 8.45762333907725e-05 | 0.340205046 | 0.459 | 0.227 | 1 | 41 |
| rpl2715 | 8.47332476339416e-05 | 0.275021653 | 0.934 | 0.763 | 1 | 41 |
| idh210 | 8.64820567290754e-05 | 0.335933294 | 0.41 | 0.2 | 1 | 41 |
| ncaldb6 | 8.95465061829621e-05 | 0.251589875 | 0.213 | 0.075 | 1 | 41 |
| eif4bb | 8.97104218269252e-05 | 0.254552495 | 0.262 | 0.103 | 1 | 41 |
| nsa218 | 9.79547832458803e-05 | 0.371179063 | 0.672 | 0.363 | 1 | 41 |
| jak15 | 0.000106434 | 0.279103621 | 0.246 | 0.097 | 1 | 41 |
| rpl3513 | 0.00010685 | 0.270206198 | 0.984 | 0.805 | 1 | 41 |
| magt14 | 0.000106858 | 0.266130748 | 0.279 | 0.116 | 1 | 41 |
| arf6a2 | 0.000107313 | 0.318202493 | 0.361 | 0.165 | 1 | 41 |
| rplp2l23 | 0.000108717 | 0.301041655 | 0.967 | 0.775 | 1 | 41 |
| romo117 | 0.000112127 | 0.371210454 | 0.639 | 0.387 | 1 | 41 |
| rps613 | 0.000140167 | 0.25088812 | 0.934 | 0.789 | 1 | 41 |
| nr4a114 | 0.000163842 | 0.343155452 | 0.361 | 0.168 | 1 | 41 |
| naca17 | 0.000195856 | 0.340032041 | 0.869 | 0.619 | 1 | 41 |
| notch210 | 0.000207759 | 0.284310117 | 0.279 | 0.121 | 1 | 41 |
| rps1319 | 0.000208749 | 0.318007367 | 0.967 | 0.725 | 1 | 41 |
| rps1023 | 0.000209883 | 0.319716106 | 0.934 | 0.695 | 1 | 41 |

| | | | | | | |
|---|---|---|---|---|---|---|
| spcs12 | 0.000213675 | 0.283038194 | 0.426 | 0.211 | 1 | 41 |
| rps4x17 | 0.000215815 | 0.328120836 | 0.967 | 0.711 | 1 | 41 |
| b2ml15 | 0.000218673 | 0.339051005 | 0.492 | 0.268 | 1 | 41 |
| psenen | 0.000275443 | 0.250564566 | 0.23 | 0.092 | 1 | 41 |
| chmp1b4 | 0.000299993 | 0.2657412 | 0.279 | 0.123 | 1 | 41 |
| dbi8 | 0.000345924 | 0.280394055 | 0.426 | 0.214 | 1 | 41 |
| ube2v24 | 0.000385321 | 0.315271884 | 0.492 | 0.271 | 1 | 41 |
| bccip2 | 0.000386526 | 0.259909601 | 0.295 | 0.134 | 1 | 41 |
| rps8a11 | 0.000402037 | 0.257218436 | 0.967 | 0.788 | 1 | 41 |
| slc25a3b8 | 0.000456332 | 0.344648949 | 0.738 | 0.438 | 1 | 41 |
| pabpc1a15 | 0.000457778 | 0.350759645 | 0.82 | 0.541 | 1 | 41 |
| rpl1015 | 0.000464273 | 0.263054113 | 0.951 | 0.773 | 1 | 41 |
| rps2315 | 0.000497402 | 0.28799459 | 0.951 | 0.723 | 1 | 41 |
| rpl2419 | 0.000501514 | 0.26489729 | 0.951 | 0.777 | 1 | 41 |
| ndufa4l12 | 0.000522129 | 0.327769833 | 0.672 | 0.396 | 1 | 41 |
| rhoac3 | 0.000523683 | 0.256448634 | 0.328 | 0.157 | 1 | 41 |
| rpl621 | 0.000585459 | 0.333182719 | 0.951 | 0.668 | 1 | 41 |
| reep56 | 0.000593553 | 0.265429584 | 0.328 | 0.16 | 1 | 41 |
| atp5pd5 | 0.000626749 | 0.273596561 | 0.492 | 0.263 | 1 | 41 |
| glo15 | 0.000667733 | 0.253434097 | 0.393 | 0.198 | 1 | 41 |
| cox6c3 | 0.000687934 | 0.334228792 | 0.689 | 0.418 | 1 | 41 |
| itm2ba14 | 0.000694382 | 0.281391059 | 0.508 | 0.274 | 1 | 41 |
| ak25 | 0.000825528 | 0.291234679 | 0.426 | 0.234 | 1 | 41 |
| ddx217 | 0.000905201 | 0.283752826 | 0.393 | 0.209 | 1 | 41 |
| rpl1714 | 0.001019007 | 0.279337407 | 0.967 | 0.747 | 1 | 41 |
| rps317 | 0.001289181 | 0.300324819 | 0.934 | 0.7 | 1 | 41 |
| rpl2117 | 0.0013135 | 0.278089222 | 0.984 | 0.757 | 1 | 41 |
| tuba8l7 | 0.001332887 | 0.2815263 | 0.361 | 0.19 | 1 | 41 |
| ppp3ccb3 | 0.001337859 | 0.267876432 | 0.262 | 0.125 | 1 | 41 |
| rpl1819 | 0.00241137 | 0.251813998 | 0.967 | 0.774 | 1 | 41 |
| xbp18 | 0.002752655 | 0.293089621 | 0.557 | 0.345 | 1 | 41 |
| eif2s24 | 0.003117642 | 0.274075292 | 0.492 | 0.296 | 1 | 41 |
| ckbb10 | 0.003405097 | 0.280467759 | 0.459 | 0.267 | 1 | 41 |
| rps916 | 0.003865546 | 0.259583567 | 0.984 | 0.777 | 1 | 41 |
| rpl27a21 | 0.004104798 | 0.315819393 | 0.754 | 0.497 | 1 | 41 |

| | | | | | | | |
|---|---|---|---|---|---|---|---|
| mt-nd415 | 0.004113532 | 0.278321825 | 0.689 | 0.457 | | 1 | 41 |
| sumo2b13 | 0.004541778 | 0.278821468 | 0.426 | 0.271 | | 1 | 41 |
| rps2515 | 0.004860854 | 0.280814575 | 0.967 | 0.725 | | 1 | 41 |
| celf216 | 0.005168701 | 0.285534226 | 0.41 | 0.244 | | 1 | 41 |
| faua17 | 0.007338108 | 0.278940922 | 0.918 | 0.69 | | 1 | 41 |
| NC-002333.419 | 0.00845988 | 0.271970373 | 0.918 | 0.623 | | 1 | 41 |
| atp5f1e4 | 0.009868737 | 0.257841904 | 0.689 | 0.484 | | 1 | 41 |
| si:ch211-212d10.12 | 1.61084132418262e-59 | 0.904322094 | 0.357 | 0.023 | 4.05432652883524e-55 | | 42 |
| si:dkey-12f6.51 | 4.7200110836301e-50 | 0.444762205 | 0.143 | 0.004 | 1.18797958963886e-45 | | 42 |
| si:dkey-21e2.131 | 3.85826136404183e-18 | 0.341719706 | 0.125 | 0.009 | 9.71085802715688e-14 | | 42 |
| si:dkey-21e2.152 | 7.56180894487623e-17 | 0.328475203 | 0.125 | 0.01 | 1.9032316933359e-12 | | 42 |
| h2af1al13 | 8.17971437467291e-16 | 0.727635536 | 1 | 0.594 | 2.05875231096143e-11 | | 42 |
| cldng12 | 4.32037743215079e-14 | 0.720323717 | 0.911 | 0.516 | 1.08739579589803e-09 | | 42 |
| zp313 | 7.84604001942877e-14 | 0.703922532 | 1 | 0.586 | 1.97476981249003e-09 | | 42 |
| acp5a13 | 3.85952281764989e-13 | 0.698647485 | 0.964 | 0.541 | 9.71403297974302e-09 | | 42 |
| zp2.212 | 8.79519906382819e-13 | 0.699408596 | 0.929 | 0.531 | 2.21366365237492e-08 | | 42 |
| si:zfos-1505d6.312 | 1.40069102389521e-12 | 0.719565836 | 0.964 | 0.515 | 3.52539923804185e-08 | | 42 |
| cldnd13 | 1.56659328988625e-12 | 0.67389458 | 1 | 0.606 | 3.94295865131469e-08 | | 42 |
| zgc:17177612 | 1.66589689871512e-12 | 0.716616787 | 0.929 | 0.497 | 4.19289590437609e-08 | | 42 |
| zgc:17177912 | 2.39025421677715e-12 | 0.694411422 | 0.911 | 0.514 | 6.01603083820642e-08 | | 42 |
| zgc:15349912 | 2.56621763201187e-12 | 0.723380954 | 0.946 | 0.514 | 6.45891315801067e-08 | | 42 |
| zp2.312 | 8.90310792509644e-12 | 0.691483878 | 0.946 | 0.52 | 2.24082323366752e-07 | | 42 |
| si:dkey-241l7.212 | 1.24226351089391e-11 | 0.638827107 | 1 | 0.643 | 3.12665303056889e-07 | | 42 |
| epcam14 | 1.52212107988547e-11 | 0.669807209 | 0.893 | 0.507 | 3.83102654596373e-07 | | 42 |
| zgc:7711812 | 2.94921758053753e-11 | 0.685799357 | 0.839 | 0.459 | 7.4228857284549e-07 | | 42 |
| CR790368.113 | 3.02543369484556e-11 | 0.658547766 | 0.893 | 0.522 | 7.61471406655678e-07 | | 42 |
| si:dkey-241l7.513 | 3.8498761145673e-11 | 0.617949282 | 0.911 | 0.586 | 9.68975319275445e-07 | | 42 |
| h1m13 | 3.87810511761149e-11 | 0.638679577 | 0.982 | 0.603 | 9.76080277051637e-07 | | 42 |
| si:ch211-226h8.1412 | 4.17587414580829e-11 | 0.661346125 | 0.964 | 0.568 | 1.05102576375849e-06 | | 42 |
| lamp212 | 4.31616469431125e-11 | 0.629109904 | 0.857 | 0.556 | 1.0863354919112e-06 | | 42 |
| zp3.213 | 5.13601612261256e-11 | 0.658564481 | 0.964 | 0.559 | 1.29268389790036e-06 | | 42 |
| BX649337.113 | 5.37048628120925e-11 | 0.69818483 | 0.857 | 0.439 | 1.35169769211756e-06 | | 42 |
| si:dkey-229d11.312 | 7.53162714515503e-11 | 0.6984596 | 0.857 | 0.44 | 1.89563523616407e-06 | | 42 |
| zp2.513 | 9.9963089414089e-11 | 0.642475471 | 0.929 | 0.565 | 2.51597099746321e-06 | | 42 |
| tmsb512 | 1.0882938173385e-10 | 0.665934221 | 0.893 | 0.49 | 2.73912670885926e-06 | | 42 |

| | | | | | | |
|---|---|---|---|---|---|---|
| zar112 | 1.24120085268948e-10 | 0.673940915 | 0.821 | 0.441 | 3.12397842613415e-06 | 42 |
| zp2.113 | 1.508944035991e-10 | 0.648389197 | 0.911 | 0.537 | 3.79786124418574e-06 | 42 |
| si:ch211-12h2.812 | 1.630158200853e-10 | 0.636392884 | 0.929 | 0.549 | 4.10294517572692e-06 | 42 |
| si:dkey-90l23.112 | 1.67859377237971e-10 | 0.673165307 | 0.821 | 0.449 | 4.22485266570249e-06 | 42 |
| mt212 | 2.10912670503614e-10 | 0.623289886 | 0.893 | 0.536 | 5.30846100390547e-06 | 42 |
| crp212 | 2.70037316463917e-10 | 0.674868296 | 0.929 | 0.49 | 6.79656921808032e-06 | 42 |
| zgc:19524512 | 4.27625496487119e-10 | 0.682012963 | 0.768 | 0.39 | 1.07629061210843e-05 | 42 |
| hbba112 | 4.60650734933425e-10 | 0.693666464 | 0.768 | 0.357 | 1.15941183475394e-05 | 42 |
| si:dkeyp-98a7.812 | 5.95191035201411e-10 | 0.652980529 | 0.839 | 0.459 | 1.49803631649843e-05 | 42 |
| si:ch211-125e6.1212 | 7.58249354884633e-10 | 0.652969418 | 0.911 | 0.502 | 1.90843780130913e-05 | 42 |
| si:ch73-189n23.113 | 7.67716000544369e-10 | 0.639195377 | 0.911 | 0.526 | 1.93226440177012e-05 | 42 |
| si:dkey-241l7.412 | 1.02370044818864e-09 | 0.581169368 | 0.946 | 0.61 | 2.576551658046e-05 | 42 |
| ccna112 | 1.10251409598825e-09 | 0.643938874 | 0.857 | 0.478 | 2.77491772819283e-05 | 42 |
| gtf3ab13 | 1.17595574859792e-09 | 0.679601662 | 0.661 | 0.322 | 2.95976302364611e-05 | 42 |
| zp2.612 | 1.26336976067878e-09 | 0.647191051 | 0.893 | 0.503 | 3.17977535065242e-05 | 42 |
| CU571163.112 | 1.62075235978544e-09 | 0.659922678 | 0.768 | 0.403 | 4.07927161434398e-05 | 42 |
| zgc:17151712 | 2.10565883492761e-09 | 0.653315137 | 0.732 | 0.39 | 5.2997327216293e-05 | 42 |
| chchd6a | 2.61053364007571e-09 | 0.41497273 | 0.143 | 0.023 | 6.57045211870655e-05 | 42 |
| si:dkeyp-20e4.812 | 4.34893377115249e-09 | 0.65605975 | 0.768 | 0.41 | 0.000109458 | 42 |
| si:ch211-12h2.612 | 4.91443748401724e-09 | 0.607143708 | 0.804 | 0.476 | 0.000123691 | 42 |
| zgc:17344312 | 5.0320729772188e-09 | 0.662845047 | 0.679 | 0.332 | 0.000126652 | 42 |
| tuba4l12 | 8.0480488884425e-09 | 0.631418373 | 0.75 | 0.406 | 0.000202561 | 42 |
| buc12 | 8.88351429344754e-09 | 0.640076543 | 0.464 | 0.18 | 0.000223589 | 42 |
| si:dkeyp-98a7.912 | 1.10209570063995e-08 | 0.603214992 | 0.857 | 0.514 | 0.000277386 | 42 |
| hbaa112 | 1.35328518679101e-08 | 0.646981787 | 0.643 | 0.283 | 0.000340608 | 42 |
| si:dkey-241l7.612 | 1.41110523055822e-08 | 0.586393129 | 0.857 | 0.529 | 0.000355161 | 42 |
| si:dkeyp-98a7.412 | 2.08191449254301e-08 | 0.612483209 | 0.768 | 0.427 | 0.000523997 | 42 |
| zgc:17513513 | 2.12081726579672e-08 | 0.600668294 | 0.821 | 0.482 | 0.000533788 | 42 |
| prtfdc1 | 2.82380017583723e-08 | 0.5327068 | 0.232 | 0.06 | 0.000710722 | 42 |
| s100a112 | 3.24577714738621e-08 | 0.612569819 | 0.679 | 0.366 | 0.00081693 | 42 |
| spink2.113 | 5.72526617674122e-08 | 0.58213667 | 0.732 | 0.438 | 0.001440992 | 42 |
| btg412 | 1.34168022987864e-07 | 0.599679837 | 0.679 | 0.371 | 0.003376875 | 42 |
| si:dkey-240e12.612 | 1.4850508628685e-07 | 0.573514994 | 0.911 | 0.571 | 0.003737725 | 42 |
| si:dkeyp-46h3.612 | 1.83501753681871e-07 | 0.577119081 | 0.696 | 0.413 | 0.004618556 | 42 |
| npm2b12 | 1.99156328059679e-07 | 0.585335359 | 0.679 | 0.398 | 0.005012566 | 42 |

| | | | | | | |
|---|---|---|---|---|---|---|
| gapdh13 | 2.08371875110867e-07 | 0.573564599 | 0.714 | 0.422 | 0.005244512 | 42 |
| si:ch211-226h8.412 | 2.28737154086643e-07 | 0.579181687 | 0.786 | 0.444 | 0.005757085 | 42 |
| zp3b12 | 2.39320861791507e-07 | 0.57880453 | 0.679 | 0.39 | 0.006023467 | 42 |
| thy114 | 2.76690409279025e-07 | 0.569767612 | 0.786 | 0.452 | 0.006964021 | 42 |
| zgc:1139848 | 3.65237726742814e-07 | 0.546119445 | 0.304 | 0.102 | 0.009192668 | 42 |
| si:dkey-208k4.212 | 4.06164843607819e-07 | 0.590072586 | 0.571 | 0.298 | 0.010222763 | 42 |
| si:ch211-132b12.812 | 4.42263823607479e-07 | 0.585988612 | 0.643 | 0.344 | 0.011131338 | 42 |
| LOC10053516212 | 4.74824897266851e-07 | 0.585913495 | 0.607 | 0.311 | 0.011950868 | 42 |
| zgc:5546113 | 4.96135137861419e-07 | 0.589166176 | 0.571 | 0.29 | 0.012487225 | 42 |
| cldn7b11 | 4.99997687437003e-07 | 0.579970023 | 0.393 | 0.161 | 0.012584442 | 42 |
| si:dkeyp-46h3.212 | 7.05859155780179e-07 | 0.550692338 | 0.679 | 0.406 | 0.017765769 | 42 |
| uchl112 | 8.83554119569747e-07 | 0.573481988 | 0.643 | 0.346 | 0.022238174 | 42 |
| casd12 | 8.88253286575657e-07 | 0.492100913 | 0.25 | 0.078 | 0.022356447 | 42 |
| slbp212 | 9.31228515094165e-07 | 0.565362434 | 0.714 | 0.399 | 0.02343809 | 42 |
| h3f3d1 | 1.10793394085862e-06 | 0.364096757 | 0.839 | 0.677 | 0.027885589 | 42 |
| LOC10053686011 | 1.33133489312891e-06 | 0.550063763 | 0.571 | 0.353 | 0.033508368 | 42 |
| rrm212 | 1.44444451335855e-06 | 0.511968709 | 0.786 | 0.5 | 0.036355224 | 42 |
| kpna712 | 1.73144059842218e-06 | 0.567885873 | 0.607 | 0.33 | 0.043578628 | 42 |
| zgc:9228710 | 1.79863501321791e-06 | 0.544937776 | 0.339 | 0.134 | 0.045269845 | 42 |
| si:dkeyp-46h3.812 | 1.8198429115064e-06 | 0.551372625 | 0.643 | 0.354 | 0.045803626 | 42 |
| hnrnpabb13 | 1.9970078721352e-06 | 0.451081441 | 0.786 | 0.598 | 0.050262691 | 42 |
| zpcx12 | 2.24961131582623e-06 | 0.555912356 | 0.607 | 0.341 | 0.056620467 | 42 |
| nanog12 | 2.46859730317488e-06 | 0.551926723 | 0.643 | 0.362 | 0.062132126 | 42 |
| cth112 | 2.54724555519507e-06 | 0.538338637 | 0.661 | 0.382 | 0.064111623 | 42 |
| AL929032.112 | 2.95927941511187e-06 | 0.559414838 | 0.607 | 0.319 | 0.074482104 | 42 |
| zar1l12 | 3.33325991739095e-06 | 0.543359434 | 0.696 | 0.4 | 0.083894819 | 42 |
| postna1 | 3.43580522097421e-06 | 0.328118688 | 0.125 | 0.026 | 0.086475782 | 42 |
| siva112 | 3.52994791294854e-06 | 0.521929868 | 0.696 | 0.43 | 0.088845259 | 42 |
| ctss2.113 | 4.15566458743688e-06 | 0.465503081 | 0.768 | 0.536 | 0.104593922 | 42 |
| ccnb212 | 5.41468792008226e-06 | 0.527467135 | 0.643 | 0.386 | 0.13628228 | 42 |
| si:ch211-14a17.712 | 6.64395453641519e-06 | 0.515371074 | 0.679 | 0.41 | 0.167221692 | 42 |
| ap1s3b2 | 9.92916122983479e-06 | 0.448785809 | 0.232 | 0.078 | 0.249907059 | 42 |
| CABZ01059627.3 | 1.032484445522e-05 | 0.393929395 | 0.179 | 0.051 | 0.25986601 | 42 |
| nabp1b | 1.13996595568016e-05 | 0.306091817 | 0.107 | 0.022 | 0.286918031 | 42 |
| ccnb112 | 1.18396636019956e-05 | 0.49759113 | 0.625 | 0.396 | 0.297992493 | 42 |

| | | | | | | |
|---|---|---|---|---|---|---|
| CABZ01059627.212 | 1.21661584719214e-05 | 0.510785465 | 0.321 | 0.135 | 0.306210043 | 42 |
| zgc:1134253 | 1.29445938410005e-05 | 0.438337078 | 0.214 | 0.07 | 0.325802482 | 42 |
| henmt11 | 1.3983383736592e-05 | 0.29021525 | 0.107 | 0.022 | 0.351947785 | 42 |
| dus1l | 1.58352864404318e-05 | 0.374293438 | 0.161 | 0.044 | 0.398558324 | 42 |
| proser3 | 1.59990773092076e-05 | 0.306615242 | 0.107 | 0.022 | 0.402680777 | 42 |
| zp3a.212 | 1.93642065398105e-05 | 0.469509885 | 0.732 | 0.494 | 0.487377714 | 42 |
| ttc324 | 2.01975074578142e-05 | 0.486745297 | 0.268 | 0.105 | 0.508351065 | 42 |
| si:dkeyp-50f7.212 | 2.39422839111449e-05 | 0.495090568 | 0.661 | 0.404 | 0.602603344 | 42 |
| zgc:1141237 | 2.43893251786999e-05 | 0.486700257 | 0.268 | 0.105 | 0.613854925 | 42 |
| plpp2b | 2.58092174823213e-05 | 0.363685197 | 0.143 | 0.038 | 0.649592195 | 42 |
| si:dkey-42i9.811 | 2.79767419543383e-05 | 0.494801926 | 0.304 | 0.129 | 0.704146618 | 42 |
| gdf312 | 2.8773211903842e-05 | 0.529749924 | 0.429 | 0.226 | 0.72419297 | 42 |
| ube2c9 | 2.95021197448118e-05 | 0.498066812 | 0.321 | 0.141 | 0.742538852 | 42 |
| rpl7l11 | 2.97017757681179e-05 | 0.523924627 | 0.304 | 0.135 | 0.747563994 | 42 |
| zgc:9259112 | 3.3604210483652e-05 | 0.517329795 | 0.411 | 0.207 | 0.845784374 | 42 |
| cdc64 | 3.59095724234326e-05 | 0.449504998 | 0.268 | 0.103 | 0.903808028 | 42 |
| si:dkey-152b24.612 | 3.65544896122046e-05 | 0.502080548 | 0.554 | 0.3 | 0.920039949 | 42 |
| tomm341 | 4.12960966519209e-05 | 0.345325889 | 0.143 | 0.039 | 1 | 42 |
| mdh1ab | 4.41161075056045e-05 | 0.308428393 | 0.125 | 0.031 | 1 | 42 |
| vps9d15 | 4.45704233704481e-05 | 0.475325104 | 0.25 | 0.098 | 1 | 42 |
| si:ch73-90k17.112 | 4.47569339150916e-05 | 0.491447515 | 0.482 | 0.251 | 1 | 42 |
| AL772362.1 | 5.10915081926642e-05 | 0.345803744 | 0.161 | 0.046 | 1 | 42 |
| h3f3a13 | 5.2306849803813e-05 | 0.418258315 | 0.643 | 0.507 | 1 | 42 |
| si:ch211-250e5.1612 | 5.66509926791415e-05 | 0.469996065 | 0.571 | 0.348 | 1 | 42 |
| zgc:771126 | 6.40498514765472e-05 | 0.467534565 | 0.286 | 0.121 | 1 | 42 |
| f11r.16 | 6.91527810671956e-05 | 0.501901292 | 0.357 | 0.179 | 1 | 42 |
| ddx196 | 8.11863543932157e-05 | 0.487440735 | 0.321 | 0.153 | 1 | 42 |
| ttc1 | 8.38620043421434e-05 | 0.355273616 | 0.143 | 0.041 | 1 | 42 |
| ptgesl | 9.10677963355553e-05 | 0.392314016 | 0.161 | 0.05 | 1 | 42 |
| slbp13 | 0.000113193 | 0.477776205 | 0.429 | 0.236 | 1 | 42 |
| mid1ip1l12 | 0.000114259 | 0.405323532 | 0.714 | 0.546 | 1 | 42 |
| cfl1l8 | 0.000128531 | 0.475486444 | 0.321 | 0.154 | 1 | 42 |
| asb31 | 0.000130285 | 0.305686262 | 0.125 | 0.033 | 1 | 42 |
| nupl2 | 0.000132392 | 0.33574919 | 0.125 | 0.034 | 1 | 42 |
| avd12 | 0.000138768 | 0.463481762 | 0.482 | 0.267 | 1 | 42 |

| | | | | | | |
|---|---|---|---|---|---|---|
| tomm20a3 | 0.00014402 | 0.500432088 | 0.411 | 0.233 | 1 | 42 |
| zgc:171445 | 0.000151654 | 0.312423482 | 0.125 | 0.034 | 1 | 42 |
| nasp12 | 0.0001539 | 0.463574499 | 0.536 | 0.342 | 1 | 42 |
| spsb12 | 0.000160428 | 0.435372251 | 0.232 | 0.094 | 1 | 42 |
| rnf48 | 0.000165427 | 0.435015069 | 0.304 | 0.136 | 1 | 42 |
| si:dkey-92i17.24 | 0.000167334 | 0.431652821 | 0.232 | 0.093 | 1 | 42 |
| zc3h11a10 | 0.000181813 | 0.48638262 | 0.393 | 0.219 | 1 | 42 |
| COQ10A4 | 0.000202747 | 0.435064341 | 0.214 | 0.084 | 1 | 42 |
| ctsba11 | 0.000205288 | 0.434103134 | 0.571 | 0.378 | 1 | 42 |
| wu:fi42e0312 | 0.000209378 | 0.449332732 | 0.571 | 0.352 | 1 | 42 |
| qdprb212 | 0.000223392 | 0.433024679 | 0.625 | 0.435 | 1 | 42 |
| mettl6 | 0.000233684 | 0.37761138 | 0.161 | 0.054 | 1 | 42 |
| zgc:17147412 | 0.000282796 | 0.440236275 | 0.518 | 0.306 | 1 | 42 |
| trappc6b | 0.000343505 | 0.333410114 | 0.143 | 0.045 | 1 | 42 |
| rad1 | 0.000361483 | 0.334371293 | 0.143 | 0.045 | 1 | 42 |
| zgc:10348212 | 0.000377043 | 0.455132568 | 0.357 | 0.191 | 1 | 42 |
| arid3b1 | 0.000415031 | 0.332360485 | 0.143 | 0.046 | 1 | 42 |
| eif4a1a8 | 0.000421671 | 0.402935988 | 0.554 | 0.426 | 1 | 42 |
| zgc:558885 | 0.000430839 | 0.385420827 | 0.214 | 0.087 | 1 | 42 |
| zgc:1717171 | 0.000488402 | 0.294846675 | 0.143 | 0.045 | 1 | 42 |
| CR847984.112 | 0.000507147 | 0.4423421 | 0.321 | 0.166 | 1 | 42 |
| zgc:16294510 | 0.000515876 | 0.431127564 | 0.321 | 0.164 | 1 | 42 |
| si:dkey-229d11.512 | 0.000551023 | 0.424588739 | 0.607 | 0.392 | 1 | 42 |
| f2rl1.1 | 0.000558677 | 0.299630149 | 0.143 | 0.046 | 1 | 42 |
| ca15b12 | 0.000578459 | 0.436987985 | 0.429 | 0.251 | 1 | 42 |
| tmem183a | 0.000642965 | 0.336070961 | 0.143 | 0.048 | 1 | 42 |
| sybu12 | 0.000659629 | 0.43785024 | 0.321 | 0.168 | 1 | 42 |
| cxxc1b | 0.000690875 | 0.300760298 | 0.125 | 0.039 | 1 | 42 |
| mcm3l6 | 0.000728802 | 0.366102543 | 0.214 | 0.089 | 1 | 42 |
| gbgt1l410 | 0.000733854 | 0.406355729 | 0.304 | 0.152 | 1 | 42 |
| gucd1 | 0.000749522 | 0.285923282 | 0.125 | 0.039 | 1 | 42 |
| si:ch211-226h8.64 | 0.000854448 | 0.370333522 | 0.196 | 0.08 | 1 | 42 |
| spcs31 | 0.000860007 | 0.438083489 | 0.411 | 0.268 | 1 | 42 |
| si:ch211-157c3.47 | 0.000902617 | 0.368662314 | 0.214 | 0.091 | 1 | 42 |
| cdca7a12 | 0.000945796 | 0.425349166 | 0.393 | 0.224 | 1 | 42 |

| | | | | | | |
|---|---|---|---|---|---|---|
| LOC10053749912 | 0.000972297 | 0.41532749 | 0.518 | 0.324 | 1 | 42 |
| zgc:16287912 | 0.000974074 | 0.413845055 | 0.339 | 0.183 | 1 | 42 |
| ccz1 | 0.000983737 | 0.349860562 | 0.143 | 0.05 | 1 | 42 |
| mkrn411 | 0.000995229 | 0.425295524 | 0.393 | 0.23 | 1 | 42 |
| ca7 | 0.001002702 | 0.275729719 | 0.125 | 0.04 | 1 | 42 |
| snrpe2 | 0.001043621 | 0.442391067 | 0.429 | 0.291 | 1 | 42 |
| hmga1a5 | 0.001065345 | 0.373073807 | 0.571 | 0.452 | 1 | 42 |
| gpr1605 | 0.001084108 | 0.375728668 | 0.196 | 0.082 | 1 | 42 |
| coil5 | 0.001100313 | 0.418883624 | 0.232 | 0.109 | 1 | 42 |
| cpeb1b12 | 0.001101719 | 0.411143614 | 0.446 | 0.273 | 1 | 42 |
| brf26 | 0.001105577 | 0.393399766 | 0.214 | 0.096 | 1 | 42 |
| si:ch211-241b2.4 | 0.0011457 | 0.281486195 | 0.125 | 0.04 | 1 | 42 |
| ccnt2b | 0.00115032 | 0.405217625 | 0.214 | 0.098 | 1 | 42 |
| mier1a | 0.001273419 | 0.264483179 | 0.107 | 0.032 | 1 | 42 |
| arhgef10la12 | 0.001276264 | 0.407400554 | 0.446 | 0.277 | 1 | 42 |
| snu13b13 | 0.001278644 | 0.390656826 | 0.536 | 0.394 | 1 | 42 |
| sinup11 | 0.001339306 | 0.41145151 | 0.357 | 0.205 | 1 | 42 |
| ccna212 | 0.001453426 | 0.423312592 | 0.446 | 0.286 | 1 | 42 |
| zgc:5669913 | 0.001570683 | 0.412367447 | 0.411 | 0.254 | 1 | 42 |
| tipin1 | 0.00161074 | 0.334029973 | 0.161 | 0.062 | 1 | 42 |
| zgc:152968 | 0.001781797 | 0.269806886 | 0.125 | 0.042 | 1 | 42 |
| tmem443 | 0.001805642 | 0.320678678 | 0.161 | 0.062 | 1 | 42 |
| LOC1039097421 | 0.001877905 | 0.278372641 | 0.125 | 0.042 | 1 | 42 |
| ints6 | 0.001896551 | 0.261260028 | 0.107 | 0.033 | 1 | 42 |
| slc16a311 | 0.001899791 | 0.401455917 | 0.411 | 0.251 | 1 | 42 |
| orc67 | 0.002011017 | 0.359118742 | 0.232 | 0.109 | 1 | 42 |
| atic | 0.002057805 | 0.329830171 | 0.179 | 0.074 | 1 | 42 |
| LOC11043782112 | 0.002078615 | 0.388370127 | 0.607 | 0.418 | 1 | 42 |
| si:dkeyp-46h3.512 | 0.002201383 | 0.390627951 | 0.464 | 0.292 | 1 | 42 |
| faub10 | 0.002209703 | 0.417493088 | 0.339 | 0.202 | 1 | 42 |
| zglp112 | 0.002238368 | 0.415106074 | 0.339 | 0.2 | 1 | 42 |
| kcnk5b | 0.002488073 | 0.313625909 | 0.125 | 0.045 | 1 | 42 |
| ddx39ab10 | 0.002554105 | 0.412975939 | 0.411 | 0.291 | 1 | 42 |
| ndufs8b | 0.002676244 | 0.275319726 | 0.125 | 0.044 | 1 | 42 |
| CU655961.35 | 0.002728547 | 0.354753124 | 0.214 | 0.1 | 1 | 42 |

| | | | | | | |
|---|---|---|---|---|---|---|
| ankrd10a2 | 0.003049404 | 0.30406012 | 0.143 | 0.055 | 1 | 42 |
| larp6b12 | 0.003137971 | 0.38393854 | 0.339 | 0.198 | 1 | 42 |
| zgc:1655141 | 0.003159689 | 0.287266703 | 0.143 | 0.055 | 1 | 42 |
| si:ch211-5k11.812 | 0.003177581 | 0.331502804 | 0.571 | 0.361 | 1 | 42 |
| pim33 | 0.003304717 | 0.378778676 | 0.25 | 0.13 | 1 | 42 |
| ect2 | 0.003492386 | 0.253579428 | 0.107 | 0.036 | 1 | 42 |
| prdx112 | 0.003632336 | 0.370037887 | 0.304 | 0.169 | 1 | 42 |
| mtrex | 0.003737833 | 0.264890684 | 0.107 | 0.036 | 1 | 42 |
| si:ch211-226l4.613 | 0.003841702 | 0.372647584 | 0.446 | 0.295 | 1 | 42 |
| rrp361 | 0.003849893 | 0.314012916 | 0.125 | 0.047 | 1 | 42 |
| si:dkey-90l23.212 | 0.004014322 | 0.368331104 | 0.5 | 0.347 | 1 | 42 |
| scamp2l1 | 0.004385224 | 0.255453009 | 0.125 | 0.046 | 1 | 42 |
| si:ch211-145c1.112 | 0.004667236 | 0.365475047 | 0.393 | 0.236 | 1 | 42 |
| mfsd4aa1 | 0.004766877 | 0.256257913 | 0.107 | 0.037 | 1 | 42 |
| nol73 | 0.004818604 | 0.402299516 | 0.232 | 0.125 | 1 | 42 |
| trmt61a1 | 0.004923763 | 0.340967794 | 0.179 | 0.082 | 1 | 42 |
| map1lc3c12 | 0.004954815 | 0.363798914 | 0.339 | 0.204 | 1 | 42 |
| cdc2011 | 0.005074023 | 0.399709524 | 0.268 | 0.155 | 1 | 42 |
| gtf3c6 | 0.005271743 | 0.303624994 | 0.143 | 0.059 | 1 | 42 |
| mrgbp1 | 0.005452929 | 0.345932974 | 0.196 | 0.096 | 1 | 42 |
| rfc3 | 0.005649188 | 0.288177759 | 0.125 | 0.049 | 1 | 42 |
| mibp11 | 0.005788903 | 0.377496069 | 0.357 | 0.225 | 1 | 42 |
| cdh13 | 0.006146106 | 0.287792775 | 0.143 | 0.06 | 1 | 42 |
| nusap1 | 0.006183548 | 0.296518324 | 0.143 | 0.06 | 1 | 42 |
| lsm14b12 | 0.006392051 | 0.354449635 | 0.446 | 0.302 | 1 | 42 |
| zp3a.112 | 0.00649866 | 0.34265735 | 0.518 | 0.384 | 1 | 42 |
| nsun5 | 0.006555154 | 0.258093957 | 0.107 | 0.039 | 1 | 42 |
| zgc:1742682 | 0.006663998 | 0.330516025 | 0.179 | 0.085 | 1 | 42 |
| dctpp110 | 0.006736889 | 0.354211971 | 0.304 | 0.177 | 1 | 42 |
| si:ch211-146l10.710 | 0.006751221 | 0.35066177 | 0.321 | 0.191 | 1 | 42 |
| h2afx8 | 0.007060212 | 0.366832321 | 0.25 | 0.14 | 1 | 42 |
| zgc:5667611 | 0.007157214 | 0.37713251 | 0.411 | 0.298 | 1 | 42 |
| fdx110 | 0.007174865 | 0.374234653 | 0.339 | 0.223 | 1 | 42 |
| hsd17b12a12 | 0.007290616 | 0.351765282 | 0.286 | 0.167 | 1 | 42 |
| BX001030.18 | 0.007325487 | 0.343986936 | 0.232 | 0.124 | 1 | 42 |

| | | | | | | | |
|---|---|---|---|---|---|---|---|
| LOC1104377796 | 0.007478073 | 0.313706476 | 0.179 | 0.085 | | 1 | 42 |
| zgc:5623112 | 0.007512674 | 0.343605355 | 0.375 | 0.23 | | 1 | 42 |
| pou5f311 | 0.007692285 | 0.335326326 | 0.268 | 0.149 | | 1 | 42 |
| pcf115 | 0.007858525 | 0.359824371 | 0.357 | 0.241 | | 1 | 42 |
| znf395a3 | 0.007940345 | 0.332831379 | 0.196 | 0.099 | | 1 | 42 |
| sumo3b3 | 0.008317495 | 0.35388287 | 0.232 | 0.127 | | 1 | 42 |
| ppp4ca2 | 0.0084198 | 0.298504468 | 0.161 | 0.073 | | 1 | 42 |
| uchl31 | 0.008620102 | 0.303434782 | 0.143 | 0.063 | | 1 | 42 |
| pigrl2.312 | 0.009054293 | 0.352505081 | 0.339 | 0.205 | | 1 | 42 |
| si:dkey-241l7.312 | 0.009075167 | 0.328428537 | 0.5 | 0.367 | | 1 | 42 |
| cx43.47 | 0.009334184 | 0.324781091 | 0.232 | 0.125 | | 1 | 42 |
| os9 | 0.009451179 | 0.306680963 | 0.143 | 0.064 | | 1 | 42 |
| eif4e1b5 | 0.009503397 | 0.30911817 | 0.179 | 0.087 | | 1 | 42 |
| rbpms2b12 | 0.009528566 | 0.344186761 | 0.357 | 0.223 | | 1 | 42 |
| BX537276.112 | 0.009701799 | 0.31142519 | 0.232 | 0.123 | | 1 | 42 |
| mbd1b | 0.009904817 | 0.264843932 | 0.107 | 0.041 | | 1 | 42 |
| nkl.41 | 1.62649909455929e-133 | 1.201823324 | 0.536 | 0.022 | 4.09373557109628e-129 | | 43 |
| nkl.31 | 9.78808271655712e-33 | 0.477857999 | 0.161 | 0.008 | 2.46356253893026e-28 | | 43 |
| si:zfos-1505d6.313 | 1.66668722744239e-14 | 0.726079201 | 0.911 | 0.515 | 4.19488508274975e-10 | | 43 |
| nanog13 | 2.25752352738545e-14 | 0.792752161 | 0.804 | 0.361 | 5.68196096607644e-10 | | 43 |
| si:dkey-90l23.213 | 6.08111082517344e-14 | 0.78688661 | 0.75 | 0.346 | 1.5305547835879e-09 | | 43 |
| zgc:15349913 | 6.58017784545239e-14 | 0.760449802 | 1 | 0.514 | 1.65616496192191e-09 | | 43 |
| h1m14 | 1.14682474752793e-13 | 0.684394573 | 1 | 0.603 | 2.88644320705304e-09 | | 43 |
| cldng13 | 2.64112234584531e-13 | 0.725060356 | 0.946 | 0.515 | 6.64744083225805e-09 | | 43 |
| si:ch211-226h8.1413 | 7.40587224205623e-13 | 0.686896376 | 0.964 | 0.568 | 1.86398398460313e-08 | | 43 |
| slbp213 | 1.93908231671437e-12 | 0.731212021 | 0.786 | 0.399 | 4.88047628293841e-08 | | 43 |
| nkl.24 | 2.20930266668984e-12 | 0.808730269 | 0.482 | 0.16 | 5.56059388179165e-08 | | 43 |
| si:dkey-241l7.213 | 4.71581854022676e-12 | 0.648785763 | 1 | 0.643 | 1.18692436838967e-07 | | 43 |
| zp314 | 6.47888512591629e-12 | 0.680866526 | 0.982 | 0.586 | 1.63067059734187e-07 | | 43 |
| h2af1al14 | 6.77886180887262e-12 | 0.655727136 | 0.964 | 0.594 | 1.70617172867515e-07 | | 43 |
| si:ch211-125e6.1213 | 8.79054778311486e-12 | 0.694360264 | 0.911 | 0.502 | 2.21249297153218e-07 | | 43 |
| zgc:7711813 | 9.08252985295957e-12 | 0.701395847 | 0.857 | 0.459 | 2.28598193869139e-07 | | 43 |
| hbba113 | 1.09988197471222e-11 | 0.731954292 | 0.768 | 0.357 | 2.76829294215319e-07 | | 43 |
| hbaa113 | 1.34501084935522e-11 | 0.784674349 | 0.679 | 0.283 | 3.38525780674215e-07 | | 43 |
| mt213 | 1.51143914763795e-11 | 0.688696243 | 0.982 | 0.536 | 3.80414119068996e-07 | | 43 |

| | | | | | | |
|---|---|---|---|---|---|---|
| qdprb213 | 1.60941368375532e-11 | 0.6956714 | 0.804 | 0.434 | 4.05073330064377e-07 | 43 |
| zp2.114 | 2.17988598990462e-11 | 0.658120423 | 0.893 | 0.537 | 5.48655504799094e-07 | 43 |
| crp213 | 2.67460919855073e-11 | 0.689500297 | 0.911 | 0.491 | 6.73172389183233e-07 | 43 |
| zgc:17513514 | 5.49552972947576e-11 | 0.674423145 | 0.857 | 0.482 | 1.38316987761175e-06 | 43 |
| zp2.313 | 6.98400006118803e-11 | 0.667630376 | 0.911 | 0.52 | 1.75780297540042e-06 | 43 |
| btg413 | 9.78554807799325e-11 | 0.707027052 | 0.696 | 0.371 | 2.46292459575012e-06 | 43 |
| CU571163.113 | 1.0433281548635e-10 | 0.697900722 | 0.786 | 0.403 | 2.62595263297596e-06 | 43 |
| tmsb513 | 1.10935233247955e-10 | 0.672387562 | 0.911 | 0.49 | 2.79212888561779e-06 | 43 |
| CR790368.114 | 1.58156803095563e-10 | 0.641003743 | 0.875 | 0.522 | 3.98064857711222e-06 | 43 |
| zgc:17177613 | 1.61576298518307e-10 | 0.650796348 | 0.857 | 0.497 | 4.06671385740726e-06 | 43 |
| si:dkeyp-98a7.913 | 2.18848521859368e-10 | 0.651114856 | 0.893 | 0.514 | 5.50819844667843e-06 | 43 |
| cldnd14 | 4.02039406477529e-10 | 0.610129943 | 0.946 | 0.606 | 1.01189298216329e-05 | 43 |
| ccna113 | 4.67035923059247e-10 | 0.653065795 | 0.857 | 0.478 | 1.17548271474782e-05 | 43 |
| zar113 | 5.8468075031053e-10 | 0.65489197 | 0.786 | 0.441 | 1.47158298045657e-05 | 43 |
| si:dkey-208k4.213 | 1.00458637829835e-09 | 0.674828426 | 0.625 | 0.297 | 2.52844345553912e-05 | 43 |
| zp3a.213 | 1.02207094470135e-09 | 0.64246234 | 0.875 | 0.493 | 2.57245036071883e-05 | 43 |
| rrm213 | 1.22052391269247e-09 | 0.622508239 | 0.875 | 0.5 | 3.07193663585567e-05 | 43 |
| BX649337.114 | 1.76178196234541e-09 | 0.650015824 | 0.821 | 0.439 | 4.43422902102716e-05 | 43 |
| lamp213 | 1.76727353986635e-09 | 0.589959532 | 0.857 | 0.556 | 4.44805077248961e-05 | 43 |
| zgc:17177913 | 2.0553039201584e-09 | 0.627905855 | 0.893 | 0.514 | 5.17299443664667e-05 | 43 |
| zp2.213 | 2.8897147346977e-09 | 0.603436786 | 0.875 | 0.531 | 7.27312301576065e-05 | 43 |
| si:dkey-240e12.613 | 3.50320905299102e-09 | 0.622885383 | 0.964 | 0.57 | 8.81722686547309e-05 | 43 |
| epcam15 | 3.59315034889319e-09 | 0.619693088 | 0.893 | 0.507 | 9.04360011312927e-05 | 43 |
| zp2.514 | 4.3701560321562e-09 | 0.581696037 | 0.875 | 0.565 | 0.000109992 | 43 |
| si:ch211-226h8.413 | 5.9796994202038e-09 | 0.627058293 | 0.786 | 0.444 | 0.000150503 | 43 |
| acp5a14 | 6.00226526933168e-09 | 0.599600353 | 0.893 | 0.542 | 0.000151071 | 43 |
| hhla2a.1 | 6.45907433328229e-09 | 0.443195751 | 0.161 | 0.03 | 0.000162568 | 43 |
| cpeb1b13 | 8.70603930489559e-09 | 0.657845395 | 0.554 | 0.272 | 0.000219122 | 43 |
| zp3.214 | 9.0298010565903e-09 | 0.603987485 | 0.929 | 0.559 | 0.000227271 | 43 |
| si:dkey-241l7.514 | 9.97491692291752e-09 | 0.570099987 | 0.911 | 0.586 | 0.000251059 | 43 |
| si:dkey-229d11.513 | 1.36612195512568e-08 | 0.63384428 | 0.714 | 0.391 | 0.000343839 | 43 |
| vrk3 | 1.87524723169247e-08 | 0.496464798 | 0.196 | 0.045 | 0.000471981 | 43 |
| prf1.21 | 1.98152506668789e-08 | 0.346348085 | 0.125 | 0.02 | 0.00049873 | 43 |
| si:ch73-189n23.114 | 2.06805716806884e-08 | 0.59757452 | 0.875 | 0.526 | 0.000520509 | 43 |
| rasl11b9 | 2.10127430961962e-08 | 0.642844497 | 0.375 | 0.135 | 0.00052887 | 43 |

| | | | | | | |
|---|---|---|---|---|---|---|
| si:ch211-145c1.113 | 2.16595587134224e-08 | 0.663503824 | 0.518 | 0.235 | 0.000545149 | 43 |
| si:dkey-241l7.413 | 3.08605617218068e-08 | 0.563666348 | 0.964 | 0.61 | 0.000776729 | 43 |
| si:dkeyp-46h3.613 | 3.39088747074685e-08 | 0.608115111 | 0.714 | 0.412 | 0.000853452 | 43 |
| LOC10053749913 | 4.23178992042038e-08 | 0.624654623 | 0.643 | 0.324 | 0.001065099 | 43 |
| si:dkeyp-46h3.213 | 9.47075158098088e-08 | 0.589486795 | 0.696 | 0.406 | 0.002383693 | 43 |
| si:dkeyp-98a7.813 | 1.09434120096051e-07 | 0.571874838 | 0.75 | 0.46 | 0.002754347 | 43 |
| si:ch211-5k11.813 | 1.21964295356315e-07 | 0.560732342 | 0.714 | 0.36 | 0.003069719 | 43 |
| si:ch211-12h2.613 | 1.37059645505728e-07 | 0.554631692 | 0.75 | 0.476 | 0.003449654 | 43 |
| gapdh14 | 1.42736049330577e-07 | 0.584618514 | 0.714 | 0.422 | 0.003592524 | 43 |
| LOC11043782113 | 1.51507275082643e-07 | 0.60151131 | 0.732 | 0.418 | 0.003813287 | 43 |
| polr2k5 | 1.60698938246967e-07 | 0.626540013 | 0.429 | 0.192 | 0.004044632 | 43 |
| cth113 | 1.83425035678254e-07 | 0.586195125 | 0.625 | 0.382 | 0.004616625 | 43 |
| tuba4l13 | 2.72891679272106e-07 | 0.579536142 | 0.714 | 0.406 | 0.006868411 | 43 |
| si:ch211-12h2.813 | 3.74541080429862e-07 | 0.523559995 | 0.821 | 0.55 | 0.009426824 | 43 |
| zgc:19524513 | 3.99350686237594e-07 | 0.570508275 | 0.643 | 0.391 | 0.010051257 | 43 |
| si:ch211-14a17.713 | 5.05757531060123e-07 | 0.564370786 | 0.696 | 0.41 | 0.012729411 | 43 |
| thy115 | 6.41075744265833e-07 | 0.536157672 | 0.714 | 0.452 | 0.016135235 | 43 |
| zgc:9259113 | 6.73903678355672e-07 | 0.591664511 | 0.446 | 0.207 | 0.016961482 | 43 |
| gtf3ab14 | 7.25633594393776e-07 | 0.572011851 | 0.571 | 0.322 | 0.018263472 | 43 |
| si:ch211-250e5.1613 | 8.57025826024669e-07 | 0.56103813 | 0.607 | 0.348 | 0.021570483 | 43 |
| zp2.613 | 1.49759034035041e-06 | 0.553187374 | 0.839 | 0.503 | 0.037692851 | 43 |
| si:dkeyp-46h3.813 | 1.57356655420071e-06 | 0.554285427 | 0.643 | 0.354 | 0.039605097 | 43 |
| si:dkey-85k15.4 | 1.70943047626503e-06 | 0.415906958 | 0.161 | 0.039 | 0.043024656 | 43 |
| si:ch73-14h10.212 | 1.71519123028933e-06 | 0.569205487 | 0.518 | 0.273 | 0.043169648 | 43 |
| si:ch211-146l10.812 | 2.16537559416049e-06 | 0.569895702 | 0.571 | 0.308 | 0.054500338 | 43 |
| si:dkey-229d11.313 | 2.28419673505085e-06 | 0.53681452 | 0.732 | 0.441 | 0.057490948 | 43 |
| si:dkey-152b24.613 | 2.75839657878719e-06 | 0.546142801 | 0.536 | 0.3 | 0.069426083 | 43 |
| ccna213 | 3.54991910021312e-06 | 0.563341802 | 0.536 | 0.286 | 0.089347914 | 43 |
| mpzl2b4 | 4.04198519180197e-06 | 0.501281527 | 0.268 | 0.096 | 0.101732725 | 43 |
| ccnb113 | 4.67895514779723e-06 | 0.518648374 | 0.625 | 0.396 | 0.117764622 | 43 |
| si:dkey-90l23.113 | 5.68230962870063e-06 | 0.516516454 | 0.696 | 0.45 | 0.143018051 | 43 |
| trmt9b1 | 6.02596566505752e-06 | 0.38948884 | 0.161 | 0.041 | 0.15166753 | 43 |
| polr3h1 | 6.54693467507333e-06 | 0.406943281 | 0.143 | 0.035 | 0.164779799 | 43 |
| si:ch211-132b12.813 | 6.97122842535644e-06 | 0.525593548 | 0.589 | 0.344 | 0.175458848 | 43 |
| zgc:113210 | 7.22049606760259e-06 | 0.347404409 | 0.125 | 0.028 | 0.181732666 | 43 |

| | | | | | | |
|---|---|---|---|---|---|---|
| si:dkey-241l7.613 | 8.11408312538626e-06 | 0.486568466 | 0.786 | 0.529 | 0.204223358 | 43 |
| ctss2.114 | 8.34578673184512e-06 | 0.448708998 | 0.75 | 0.536 | 0.210055106 | 43 |
| zar1l13 | 8.37934930243666e-06 | 0.507255723 | 0.661 | 0.4 | 0.210899843 | 43 |
| zgc:17344313 | 1.06121774249369e-05 | 0.525594946 | 0.589 | 0.332 | 0.267097894 | 43 |
| sinup12 | 1.08768416655132e-05 | 0.535726242 | 0.411 | 0.205 | 0.273759228 | 43 |
| zgc:13625412 | 1.0989103158175e-05 | 0.521169766 | 0.571 | 0.333 | 0.276584737 | 43 |
| htatip2 | 1.12589888440595e-05 | 0.381243005 | 0.143 | 0.036 | 0.28337749 | 43 |
| bphl | 1.2605180606766e-05 | 0.332438729 | 0.125 | 0.028 | 0.317259791 | 43 |
| lsm14b13 | 1.27456461133856e-05 | 0.519389044 | 0.518 | 0.302 | 0.320795167 | 43 |
| npc25 | 1.31480931974125e-05 | 0.504249074 | 0.554 | 0.36 | 0.330924358 | 43 |
| si:dkey-152b24.7 | 1.40444501736328e-05 | 0.398650648 | 0.161 | 0.044 | 0.353484766 | 43 |
| si:ch1073-190k2.112 | 1.6351004905472e-05 | 0.531139117 | 0.393 | 0.19 | 0.411538442 | 43 |
| ca15b13 | 1.69324589176401e-05 | 0.51283306 | 0.482 | 0.251 | 0.426173058 | 43 |
| hnrnpabb14 | 1.90817613457803e-05 | 0.402827001 | 0.75 | 0.599 | 0.480268851 | 43 |
| npm2b13 | 2.16803344115508e-05 | 0.490417418 | 0.607 | 0.398 | 0.545672337 | 43 |
| si:ch211-226l4.614 | 3.23696907275068e-05 | 0.519496316 | 0.518 | 0.295 | 0.814712746 | 43 |
| foxa | 3.86764468736206e-05 | 0.293412506 | 0.107 | 0.024 | 0.973447491 | 43 |
| zgc:11342411 | 3.87598746898183e-05 | 0.538508491 | 0.339 | 0.163 | 0.975547286 | 43 |
| ssna1 | 4.31373284061654e-05 | 0.396911237 | 0.161 | 0.048 | 1 | 43 |
| hnrnpa0l3 | 4.70405202336029e-05 | 0.262032685 | 0.839 | 0.758 | 1 | 43 |
| XLOC-030786 | 4.80822620267677e-05 | 0.328994809 | 0.125 | 0.031 | 1 | 43 |
| vstm2l7 | 4.88928108455314e-05 | 0.461729388 | 0.25 | 0.097 | 1 | 43 |
| zpcx13 | 5.86703446313432e-05 | 0.474273517 | 0.554 | 0.341 | 1 | 43 |
| si:dkeyp-20e4.813 | 5.94742114815879e-05 | 0.482576643 | 0.661 | 0.41 | 1 | 43 |
| si:dkeyp-50f7.213 | 6.60803684277298e-05 | 0.462878409 | 0.625 | 0.405 | 1 | 43 |
| zp3c12 | 6.91710815317063e-05 | 0.510883619 | 0.375 | 0.191 | 1 | 43 |
| zgc:17151713 | 7.09941900621201e-05 | 0.477265247 | 0.643 | 0.39 | 1 | 43 |
| elavl213 | 8.02606152753719e-05 | 0.492051968 | 0.411 | 0.226 | 1 | 43 |
| si:dkeyp-98a7.413 | 8.14687686020688e-05 | 0.453485857 | 0.643 | 0.428 | 1 | 43 |
| cldn7b12 | 8.4680020396958e-05 | 0.480865445 | 0.339 | 0.162 | 1 | 43 |
| siva113 | 8.66211969249627e-05 | 0.443736848 | 0.625 | 0.431 | 1 | 43 |
| npm2a3 | 9.03463145483348e-05 | 0.432320726 | 0.214 | 0.08 | 1 | 43 |
| spink2.22 | 9.71491694481057e-05 | 0.426108968 | 0.304 | 0.133 | 1 | 43 |
| kpna713 | 0.000113955 | 0.471286261 | 0.536 | 0.331 | 1 | 43 |
| mid1ip1l13 | 0.000132732 | 0.406231514 | 0.732 | 0.546 | 1 | 43 |

| | | | | | | | |
|---|---|---|---|---|---|---|---|
| rac1a6 | 0.000134463 | 0.490847751 | 0.304 | 0.15 | | 1 | 43 |
| zp3b13 | 0.00015844 | 0.450865712 | 0.607 | 0.39 | | 1 | 43 |
| zgc:6602412 | 0.000172947 | 0.451356134 | 0.304 | 0.143 | | 1 | 43 |
| AL929192.112 | 0.000180065 | 0.463860328 | 0.339 | 0.166 | | 1 | 43 |
| zgc:17197711 | 0.000189168 | 0.460201227 | 0.411 | 0.232 | | 1 | 43 |
| mibp12 | 0.000211624 | 0.476822801 | 0.411 | 0.225 | | 1 | 43 |
| zglp113 | 0.000215681 | 0.455029638 | 0.375 | 0.2 | | 1 | 43 |
| mad2l19 | 0.000218624 | 0.47240292 | 0.357 | 0.188 | | 1 | 43 |
| zgc:1017447 | 0.000291649 | 0.449251415 | 0.286 | 0.135 | | 1 | 43 |
| tmem9b | 0.000318859 | 0.410942443 | 0.196 | 0.077 | | 1 | 43 |
| ccnb213 | 0.000340515 | 0.426417148 | 0.589 | 0.387 | | 1 | 43 |
| si:ch73-78o10.110 | 0.000354787 | 0.455230379 | 0.286 | 0.135 | | 1 | 43 |
| mfsd4aa2 | 0.000355464 | 0.320105375 | 0.125 | 0.037 | | 1 | 43 |
| LOC10188547712 | 0.000356564 | 0.441082481 | 0.464 | 0.282 | | 1 | 43 |
| s100a113 | 0.000437842 | 0.415971288 | 0.536 | 0.366 | | 1 | 43 |
| eif4e1b6 | 0.000448952 | 0.38681152 | 0.214 | 0.087 | | 1 | 43 |
| hccsa.1 | 0.000450597 | 0.304967309 | 0.107 | 0.03 | | 1 | 43 |
| si:dkeyp-2e4.2 | 0.000452922 | 0.309557465 | 0.125 | 0.038 | | 1 | 43 |
| mkrn412 | 0.000459644 | 0.449298748 | 0.393 | 0.23 | | 1 | 43 |
| glulb2 | 0.000478414 | 0.443573923 | 0.304 | 0.153 | | 1 | 43 |
| AL929032.113 | 0.000478849 | 0.448967146 | 0.518 | 0.32 | | 1 | 43 |
| manbal | 0.000515843 | 0.331612277 | 0.125 | 0.039 | | 1 | 43 |
| nfr8 | 0.000536586 | 0.402614072 | 0.25 | 0.112 | | 1 | 43 |
| elovl7a12 | 0.000546028 | 0.431571894 | 0.339 | 0.185 | | 1 | 43 |
| zcchc81 | 0.000572066 | 0.318724164 | 0.125 | 0.039 | | 1 | 43 |
| pabpn1l12 | 0.000667865 | 0.425152528 | 0.464 | 0.286 | | 1 | 43 |
| sc:d02021 | 0.000674613 | 0.334107627 | 0.143 | 0.049 | | 1 | 43 |
| mob1bb | 0.000735753 | 0.382115692 | 0.179 | 0.07 | | 1 | 43 |
| si:dkeyp-98a7.711 | 0.000750872 | 0.432312441 | 0.357 | 0.198 | | 1 | 43 |
| zgc:17385611 | 0.000799609 | 0.447772695 | 0.375 | 0.222 | | 1 | 43 |
| uros | 0.000801112 | 0.304258225 | 0.125 | 0.04 | | 1 | 43 |
| shmt11 | 0.000854741 | 0.38839601 | 0.196 | 0.083 | | 1 | 43 |
| tarbp2 | 0.000910908 | 0.362342309 | 0.161 | 0.06 | | 1 | 43 |
| si:ch211-146l10.711 | 0.000982577 | 0.431653243 | 0.339 | 0.191 | | 1 | 43 |
| fbxo432 | 0.00100953 | 0.378157443 | 0.179 | 0.071 | | 1 | 43 |

| | | | | | | | |
|---|---|---|---|---|---|---|---|
| gbgt1l1 | 0.00103995 | 0.334776736 | 0.143 | 0.051 | | 1 | 43 |
| gmnn12 | 0.001211618 | 0.400864394 | 0.464 | 0.305 | | 1 | 43 |
| marcksb9 | 0.001223428 | 0.423940066 | 0.321 | 0.178 | | 1 | 43 |
| tdp2b | 0.001224035 | 0.27063112 | 0.107 | 0.032 | | 1 | 43 |
| LOC1104377797 | 0.001253878 | 0.379155879 | 0.196 | 0.085 | | 1 | 43 |
| si:dkey-188i13.1113 | 0.001323329 | 0.437298122 | 0.446 | 0.318 | | 1 | 43 |
| si:ch211-251f6.612 | 0.001363948 | 0.403942489 | 0.464 | 0.334 | | 1 | 43 |
| uchl113 | 0.001432428 | 0.389676984 | 0.5 | 0.347 | | 1 | 43 |
| rnf175 | 0.001458366 | 0.276831931 | 0.107 | 0.033 | | 1 | 43 |
| LOC1039099727 | 0.001462265 | 0.395341123 | 0.482 | 0.369 | | 1 | 43 |
| rnf49 | 0.001467463 | 0.401255238 | 0.268 | 0.136 | | 1 | 43 |
| si:ch211-59d15.9 | 0.001559066 | 0.285671996 | 0.125 | 0.042 | | 1 | 43 |
| cnep1r14 | 0.001712396 | 0.409262413 | 0.25 | 0.125 | | 1 | 43 |
| eif4a1a9 | 0.001734951 | 0.374220265 | 0.518 | 0.427 | | 1 | 43 |
| wu:fi42e0313 | 0.001740025 | 0.397137336 | 0.5 | 0.353 | | 1 | 43 |
| fdx111 | 0.001910377 | 0.418873119 | 0.357 | 0.223 | | 1 | 43 |
| zgc:10348213 | 0.002108466 | 0.392103493 | 0.339 | 0.191 | | 1 | 43 |
| ess2 | 0.002138793 | 0.294709161 | 0.125 | 0.044 | | 1 | 43 |
| LOC101885376 | 0.002173194 | 0.317350894 | 0.143 | 0.054 | | 1 | 43 |
| prr116 | 0.002302864 | 0.39342543 | 0.214 | 0.102 | | 1 | 43 |
| mcm3l7 | 0.00247846 | 0.359361234 | 0.196 | 0.089 | | 1 | 43 |
| hddc21 | 0.00248024 | 0.310042628 | 0.143 | 0.054 | | 1 | 43 |
| ankrd2211 | 0.002522099 | 0.389302133 | 0.268 | 0.145 | | 1 | 43 |
| si:dkeyp-2e4.3 | 0.00257419 | 0.278690758 | 0.125 | 0.044 | | 1 | 43 |
| sybu13 | 0.002577675 | 0.410642836 | 0.304 | 0.168 | | 1 | 43 |
| mad2l2 | 0.002628269 | 0.279593951 | 0.107 | 0.035 | | 1 | 43 |
| dhx161 | 0.002724588 | 0.326318755 | 0.143 | 0.055 | | 1 | 43 |
| ckbb11 | 0.002789038 | 0.38584331 | 0.393 | 0.268 | | 1 | 43 |
| arhgef10la13 | 0.002836643 | 0.384735793 | 0.429 | 0.277 | | 1 | 43 |
| thoc73 | 0.002848965 | 0.392697047 | 0.232 | 0.117 | | 1 | 43 |
| CR383662.27 | 0.002962288 | 0.328938662 | 0.196 | 0.088 | | 1 | 43 |
| ell21 | 0.003054513 | 0.336510035 | 0.161 | 0.067 | | 1 | 43 |
| zgc:1735443 | 0.003148542 | 0.329053033 | 0.179 | 0.078 | | 1 | 43 |
| pdia41 | 0.003165679 | 0.389883715 | 0.25 | 0.132 | | 1 | 43 |
| st6galnac1.21 | 0.003386279 | 0.286297987 | 0.143 | 0.055 | | 1 | 43 |

| | | | | | | |
|---|---|---|---|---|---|---|
| rnaseh2b | 0.003392698 | 0.301284549 | 0.143 | 0.056 | 1 | 43 |
| pgam51 | 0.003396846 | 0.396245649 | 0.214 | 0.109 | 1 | 43 |
| metap1d3 | 0.003427578 | 0.354186341 | 0.196 | 0.091 | 1 | 43 |
| kmt5ab5 | 0.003461065 | 0.342119673 | 0.196 | 0.091 | 1 | 43 |
| tp53rk | 0.003708729 | 0.310859752 | 0.143 | 0.057 | 1 | 43 |
| si:ch73-281n10.26 | 0.003859957 | 0.354745153 | 0.5 | 0.394 | 1 | 43 |
| retsatl12 | 0.003863829 | 0.383937332 | 0.339 | 0.206 | 1 | 43 |
| CU655961.46 | 0.003927687 | 0.339822247 | 0.196 | 0.09 | 1 | 43 |
| spink2.114 | 0.003934461 | 0.341791572 | 0.589 | 0.439 | 1 | 43 |
| si:ch211-137a8.21 | 0.004167609 | 0.302741855 | 0.143 | 0.057 | 1 | 43 |
| tph1a12 | 0.004242333 | 0.355804423 | 0.268 | 0.143 | 1 | 43 |
| sh2d5 | 0.00442018 | 0.299590965 | 0.143 | 0.057 | 1 | 43 |
| BX005461.34 | 0.004473165 | 0.309246096 | 0.161 | 0.068 | 1 | 43 |
| rnf128a | 0.004691759 | 0.328329156 | 0.161 | 0.07 | 1 | 43 |
| slbp14 | 0.004951565 | 0.40908989 | 0.357 | 0.236 | 1 | 43 |
| nifk3 | 0.005111531 | 0.373022745 | 0.232 | 0.124 | 1 | 43 |
| baiap2a1 | 0.005261662 | 0.272534363 | 0.107 | 0.038 | 1 | 43 |
| BX571701.3 | 0.005276368 | 0.261278481 | 0.107 | 0.038 | 1 | 43 |
| snrnp271 | 0.005332398 | 0.398285969 | 0.232 | 0.128 | 1 | 43 |
| spsb13 | 0.005454705 | 0.330291513 | 0.196 | 0.094 | 1 | 43 |
| wu:fa19b122 | 0.005551538 | 0.308449526 | 0.161 | 0.07 | 1 | 43 |
| zgc:17147413 | 0.005590328 | 0.353583322 | 0.446 | 0.307 | 1 | 43 |
| cnot83 | 0.00573241 | 0.366141276 | 0.196 | 0.098 | 1 | 43 |
| selenou1a1 | 0.0059129 | 0.272504758 | 0.107 | 0.039 | 1 | 43 |
| dkc14 | 0.005950126 | 0.416294871 | 0.286 | 0.176 | 1 | 43 |
| LOC10188389712 | 0.0060644 | 0.356133341 | 0.286 | 0.166 | 1 | 43 |
| cenps1 | 0.00608107 | 0.251466268 | 0.107 | 0.038 | 1 | 43 |
| si:dkeyp-114g9.16 | 0.00616998 | 0.326635837 | 0.214 | 0.106 | 1 | 43 |
| zmp:00000013233 | 0.006281033 | 0.33914418 | 0.268 | 0.146 | 1 | 43 |
| cax13 | 0.00635755 | 0.299459786 | 0.143 | 0.06 | 1 | 43 |
| ints12 | 0.006562627 | 0.284951663 | 0.125 | 0.05 | 1 | 43 |
| heatr3 | 0.006716442 | 0.262990212 | 0.107 | 0.039 | 1 | 43 |
| si:dkey-241l7.313 | 0.006789576 | 0.355015025 | 0.482 | 0.367 | 1 | 43 |
| drg1 | 0.007077813 | 0.328412055 | 0.143 | 0.062 | 1 | 43 |
| BX001030.19 | 0.007209736 | 0.343789982 | 0.232 | 0.124 | 1 | 43 |

| | | | | | | |
|---|---|---|---|---|---|---|
| pigrl2.313 | 0.007259777 | 0.351130176 | 0.339 | 0.205 | 1 | 43 |
| nasp13 | 0.007285835 | 0.356761842 | 0.464 | 0.342 | 1 | 43 |
| rcc112 | 0.007389434 | 0.373410738 | 0.321 | 0.205 | 1 | 43 |
| ap1g1 | 0.007530205 | 0.321043606 | 0.161 | 0.074 | 1 | 43 |
| wdr43 | 0.007764387 | 0.353975947 | 0.232 | 0.127 | 1 | 43 |
| dynll113 | 0.007851194 | 0.312300031 | 0.536 | 0.457 | 1 | 43 |
| il13ra25 | 0.007935829 | 0.338817755 | 0.214 | 0.112 | 1 | 43 |
| LOC10053516213 | 0.007979183 | 0.3446824 | 0.464 | 0.311 | 1 | 43 |
| ccz11 | 0.008047674 | 0.276028911 | 0.125 | 0.05 | 1 | 43 |
| hexim11 | 0.008090564 | 0.327383754 | 0.179 | 0.087 | 1 | 43 |
| eif1axb | 0.008184572 | 0.386903416 | 0.393 | 0.303 | 1 | 43 |
| lin28a3 | 0.008575963 | 0.276707826 | 0.161 | 0.071 | 1 | 43 |
| haus611 | 0.008608626 | 0.364267812 | 0.321 | 0.212 | 1 | 43 |
| snu13a7 | 0.008970523 | 0.332279987 | 0.196 | 0.102 | 1 | 43 |
| zgc:5546114 | 0.009004161 | 0.343801867 | 0.429 | 0.291 | 1 | 43 |
| ccdc47 | 0.009110841 | 0.283575334 | 0.125 | 0.052 | 1 | 43 |
| chrac13 | 0.009164646 | 0.352012303 | 0.232 | 0.13 | 1 | 43 |
| btg34 | 0.009399021 | 0.372714149 | 0.286 | 0.181 | 1 | 43 |
| zgc:868396 | 0.009847282 | 0.320099927 | 0.214 | 0.111 | 1 | 43 |
| cldnb1 | 0 | 2.573363982 | 1 | 0.007 | 0 | 44 |
| icn22 | 0 | 2.476579204 | 1 | 0.005 | 0 | 44 |
| zgc:1935051 | 0 | 2.456110929 | 0.979 | 0.003 | 0 | 44 |
| cldne1 | 0 | 2.40672388 | 1 | 0.002 | 0 | 44 |
| zgc:111983 | 0 | 2.333759761 | 1 | 0.001 | 0 | 44 |
| si:dkey-7c18.24 | 0 | 2.326374537 | 0.957 | 0.001 | 0 | 44 |
| si:dkey-193p11.22 | 0 | 2.323029779 | 0.957 | 0.011 | 0 | 44 |
| si:ch211-69b7.61 | 0 | 2.260526971 | 0.979 | 0.004 | 0 | 44 |
| krt51 | 0 | 2.213615188 | 0.979 | 0.011 | 0 | 44 |
| mid1ip1a1 | 0 | 2.170912989 | 0.957 | 0.01 | 0 | 44 |
| si:dkey-247k7.2 | 0 | 2.149774051 | 0.957 | 0 | 0 | 44 |
| krt171 | 0 | 2.115315264 | 0.936 | 0.005 | 0 | 44 |
| si:ch211-217k17.9 | 0 | 2.101758399 | 0.957 | 0.001 | 0 | 44 |
| lye | 0 | 2.000009402 | 0.915 | 0.001 | 0 | 44 |
| si:ch211-207n23.22 | 0 | 1.965595698 | 0.936 | 0.025 | 0 | 44 |
| zgc:1539112 | 0 | 1.935085067 | 0.872 | 0.017 | 0 | 44 |

| | | | | | | |
|---|---|---|---|---|---|---|
| dsg2.11 | 0 | 1.887393117 | 0.957 | 0.022 | 0 | 44 |
| ppl1 | 0 | 1.886986358 | 0.957 | 0.007 | 0 | 44 |
| dsc2l2 | 0 | 1.86563054 | 0.957 | 0.022 | 0 | 44 |
| pkp3a1 | 0 | 1.755535428 | 0.915 | 0.007 | 0 | 44 |
| evpla1 | 0 | 1.704976276 | 0.915 | 0.002 | 0 | 44 |
| cldn23a | 0 | 1.68284027 | 0.851 | 0.002 | 0 | 44 |
| si:dkey-248g15.31 | 0 | 1.663865106 | 0.723 | 0.007 | 0 | 44 |
| tnks1bp1 | 0 | 1.64510537 | 0.872 | 0.004 | 0 | 44 |
| soul2 | 0 | 1.64217815 | 0.83 | 0 | 0 | 44 |
| zgc:153665 | 0 | 1.639131127 | 0.851 | 0.019 | 0 | 44 |
| CABZ01048956.11 | 0 | 1.638671253 | 0.766 | 0.001 | 0 | 44 |
| anxa1c | 0 | 1.633444346 | 0.787 | 0.001 | 0 | 44 |
| LOC108190590 | 0 | 1.62822474 | 0.872 | 0.001 | 0 | 44 |
| krt971 | 0 | 1.615214322 | 0.681 | 0.006 | 0 | 44 |
| scel1 | 0 | 1.613505987 | 0.851 | 0.004 | 0 | 44 |
| krt1-19d | 0 | 1.609300672 | 0.745 | 0.001 | 0 | 44 |
| si:ch211-125o16.4 | 0 | 1.548130804 | 0.83 | 0 | 0 | 44 |
| zmp:0000000906 | 0 | 1.457051367 | 0.745 | 0.002 | 0 | 44 |
| si:ch211-98n17.5 | 0 | 1.449743443 | 0.787 | 0.001 | 0 | 44 |
| tgm5l | 0 | 1.421858742 | 0.702 | 0.002 | 0 | 44 |
| znf185 | 0 | 1.420922733 | 0.766 | 0.002 | 0 | 44 |
| anxa1b1 | 0 | 1.372447514 | 0.723 | 0.005 | 0 | 44 |
| rhcgl1 | 0 | 1.319934726 | 0.617 | 0.001 | 0 | 44 |
| lmo7a1 | 0 | 1.30151463 | 0.702 | 0.007 | 0 | 44 |
| ftr83 | 0 | 1.276105054 | 0.766 | 0.004 | 0 | 44 |
| zgc:110333 | 0 | 1.272498725 | 0.745 | 0.001 | 0 | 44 |
| ptgs2b2 | 0 | 1.241016795 | 0.617 | 0.009 | 0 | 44 |
| LOC100330250 | 0 | 1.23381921 | 0.596 | 0.003 | 0 | 44 |
| hrc | 0 | 1.228348794 | 0.638 | 0.006 | 0 | 44 |
| AL929237.11 | 0 | 1.219346604 | 0.596 | 0.006 | 0 | 44 |
| sts | 0 | 1.211203828 | 0.702 | 0.008 | 0 | 44 |
| chst6 | 0 | 1.186029947 | 0.638 | 0.005 | 0 | 44 |
| bicdl2l | 0 | 1.175267902 | 0.617 | 0.001 | 0 | 44 |
| mir203a | 0 | 1.175179816 | 0.617 | 0.001 | 0 | 44 |
| agr1 | 0 | 1.170227048 | 0.574 | 0.001 | 0 | 44 |

| | | | | | | |
|---|---|---|---|---|---|---|
| slc16a12b1 | 0 | 1.161483109 | 0.66 | 0.006 | 0 | 44 |
| CR854839.1 | 0 | 1.147922974 | 0.638 | 0.002 | 0 | 44 |
| evplb | 0 | 1.126721782 | 0.66 | 0 | 0 | 44 |
| apodb | 0 | 1.109444768 | 0.596 | 0.001 | 0 | 44 |
| sox3 | 0 | 1.101921966 | 0.596 | 0.004 | 0 | 44 |
| LOC101882729 | 0 | 1.096931165 | 0.596 | 0.001 | 0 | 44 |
| stx11b.1 | 0 | 1.080961914 | 0.574 | 0.001 | 0 | 44 |
| myzap | 0 | 1.065097045 | 0.553 | 0.005 | 0 | 44 |
| pkp21 | 0 | 1.029281902 | 0.553 | 0.004 | 0 | 44 |
| fam129aa | 0 | 1.02154943 | 0.553 | 0 | 0 | 44 |
| CABZ01067657.1 | 0 | 1.001549569 | 0.553 | 0.001 | 0 | 44 |
| cyldb | 0 | 1.000553613 | 0.532 | 0.001 | 0 | 44 |
| bik1 | 0 | 0.993731043 | 0.553 | 0.007 | 0 | 44 |
| FO681323.1 | 0 | 0.975473041 | 0.553 | 0 | 0 | 44 |
| CABZ01084230.1 | 0 | 0.944578093 | 0.511 | 0.001 | 0 | 44 |
| cldnh | 0 | 0.931722758 | 0.34 | 0.001 | 0 | 44 |
| si:ch211-182p11.1 | 0 | 0.921189268 | 0.489 | 0.002 | 0 | 44 |
| si:dkeyp-51b9.3 | 0 | 0.907643708 | 0.489 | 0 | 0 | 44 |
| capn9 | 0 | 0.907471249 | 0.532 | 0.002 | 0 | 44 |
| si:dkeyp-86h10.3 | 0 | 0.907218065 | 0.426 | 0.001 | 0 | 44 |
| si:ch73-52f15.5 | 0 | 0.868267031 | 0.468 | 0.001 | 0 | 44 |
| zgc:100868 | 0 | 0.855231894 | 0.489 | 0 | 0 | 44 |
| ANXA1 (1 of many) | 0 | 0.845862552 | 0.426 | 0.001 | 0 | 44 |
| cx35.41 | 0 | 0.841597484 | 0.468 | 0.004 | 0 | 44 |
| tmem176l.3b | 0 | 0.838462771 | 0.447 | 0.001 | 0 | 44 |
| CR388148.2 | 0 | 0.834522672 | 0.447 | 0 | 0 | 44 |
| ponzr10 | 0 | 0.814819483 | 0.383 | 0 | 0 | 44 |
| btr12 | 0 | 0.810545359 | 0.447 | 0.003 | 0 | 44 |
| rnf222 | 0 | 0.800613962 | 0.447 | 0 | 0 | 44 |
| si:dkeyp-1h4.6 | 0 | 0.800437731 | 0.426 | 0.003 | 0 | 44 |
| LOC100149234 | 0 | 0.799550484 | 0.468 | 0 | 0 | 44 |
| eps8l1a | 0 | 0.797660845 | 0.426 | 0.001 | 0 | 44 |
| wu:fc34e06 | 0 | 0.796245785 | 0.447 | 0 | 0 | 44 |
| si:dkey-175d9.2 | 0 | 0.783226248 | 0.447 | 0 | 0 | 44 |
| si:ch211-197e7.1 | 0 | 0.782839369 | 0.447 | 0 | 0 | 44 |

| | | | | | | |
|---|---|---|---|---|---|---|
| rassf7b | 0 | 0.78266861 | 0.404 | 0.002 | 0 | 44 |
| CABZ01015815.1 | 0 | 0.781369211 | 0.447 | 0.002 | 0 | 44 |
| CABZ01068469.1 | 0 | 0.738780644 | 0.404 | 0.001 | 0 | 44 |
| BX005027.2 | 0 | 0.733279307 | 0.383 | 0 | 0 | 44 |
| CABZ01068248.1 | 0 | 0.724577574 | 0.404 | 0.002 | 0 | 44 |
| LOC100333235 | 0 | 0.721093806 | 0.362 | 0.001 | 0 | 44 |
| cldnf | 0 | 0.714333645 | 0.404 | 0.001 | 0 | 44 |
| st14b | 0 | 0.713435906 | 0.404 | 0.002 | 0 | 44 |
| CR589947.2 | 0 | 0.706784318 | 0.383 | 0 | 0 | 44 |
| si:ch73-184c24.1 | 0 | 0.702143753 | 0.404 | 0.001 | 0 | 44 |
| CU914622.2 | 0 | 0.697695899 | 0.34 | 0 | 0 | 44 |
| erbb3a | 0 | 0.68997761 | 0.383 | 0.001 | 0 | 44 |
| ccl20b | 0 | 0.68571268 | 0.404 | 0.002 | 0 | 44 |
| paqr5b | 0 | 0.67795531 | 0.362 | 0 | 0 | 44 |
| si:rp71-1h20.9 | 0 | 0.67722882 | 0.383 | 0.002 | 0 | 44 |
| XLOC-041945 | 0 | 0.67363614 | 0.383 | 0.001 | 0 | 44 |
| esrp11 | 0 | 0.659748188 | 0.34 | 0.002 | 0 | 44 |
| ponzr2 | 0 | 0.647175567 | 0.362 | 0 | 0 | 44 |
| foxq1a1 | 0 | 0.643087443 | 0.319 | 0.002 | 0 | 44 |
| adgrf6 | 0 | 0.629734588 | 0.362 | 0.001 | 0 | 44 |
| zgc:136892 | 0 | 0.621911966 | 0.34 | 0 | 0 | 44 |
| LOC1104382151 | 0 | 0.614586225 | 0.34 | 0.002 | 0 | 44 |
| si:dkey-73p2.1 | 0 | 0.606168198 | 0.298 | 0 | 0 | 44 |
| si:ch73-347e22.8 | 0 | 0.604531441 | 0.319 | 0 | 0 | 44 |
| noxa1 | 0 | 0.602206207 | 0.277 | 0.001 | 0 | 44 |
| si:dkey-165e24.1 | 0 | 0.600188748 | 0.319 | 0 | 0 | 44 |
| XLOC-027129 | 0 | 0.596767734 | 0.34 | 0 | 0 | 44 |
| nectin4a | 0 | 0.590395088 | 0.34 | 0.001 | 0 | 44 |
| LOC103909616 | 0 | 0.588673122 | 0.319 | 0 | 0 | 44 |
| CU062645.1 | 0 | 0.584760693 | 0.319 | 0 | 0 | 44 |
| si:ch211-226h7.5 | 0 | 0.576951828 | 0.255 | 0 | 0 | 44 |
| si:busm1-194e12.8 | 0 | 0.572403205 | 0.319 | 0 | 0 | 44 |
| si:ch73-196i15.3 | 0 | 0.563111864 | 0.298 | 0.001 | 0 | 44 |
| adgrg11 | 0 | 0.559836721 | 0.255 | 0.001 | 0 | 44 |
| zgc:113229 | 0 | 0.556057048 | 0.298 | 0 | 0 | 44 |

| | | | | | | | |
|---|---|---|---|---|---|---|---|
| si:dkey-221j11.3 | 0 | 0.547844206 | 0.319 | 0 | | 0 | 44 |
| LOC407643 | 0 | 0.541624134 | 0.277 | 0.001 | | 0 | 44 |
| si:dkey-27n6.4 | 0 | 0.53758266 | 0.319 | 0 | | 0 | 44 |
| pkp1b | 0 | 0.521162609 | 0.319 | 0.001 | | 0 | 44 |
| zgc:194839 | 0 | 0.511083708 | 0.255 | 0.001 | | 0 | 44 |
| LOC100149931 | 0 | 0.503685853 | 0.255 | 0 | | 0 | 44 |
| anxa3a | 0 | 0.502574298 | 0.298 | 0.001 | | 0 | 44 |
| LOC100001846 | 0 | 0.495490163 | 0.277 | 0.001 | | 0 | 44 |
| il17rc | 0 | 0.494900548 | 0.298 | 0 | | 0 | 44 |
| vsig8b | 0 | 0.492177307 | 0.298 | 0 | | 0 | 44 |
| tgm1 | 0 | 0.486554297 | 0.255 | 0 | | 0 | 44 |
| gna14 | 0 | 0.472842511 | 0.234 | 0 | | 0 | 44 |
| ttc22 | 0 | 0.45295315 | 0.213 | 0 | | 0 | 44 |
| BX004785.3 | 0 | 0.452247696 | 0.255 | 0.001 | | 0 | 44 |
| il20ra | 0 | 0.4280559 | 0.255 | 0.001 | | 0 | 44 |
| card14 | 0 | 0.427891376 | 0.255 | 0.001 | | 0 | 44 |
| zgc:165423 | 0 | 0.416090263 | 0.149 | 0 | | 0 | 44 |
| si:dkey-11o15.7 | 0 | 0.409188289 | 0.234 | 0 | | 0 | 44 |
| CABZ01049361.1 | 0 | 0.404845274 | 0.191 | 0 | | 0 | 44 |
| adgrg2a | 0 | 0.404482322 | 0.213 | 0 | | 0 | 44 |
| CABZ01046954.1 | 0 | 0.397872864 | 0.234 | 0 | | 0 | 44 |
| zgc:123295 | 0 | 0.386641688 | 0.17 | 0 | | 0 | 44 |
| mala | 0 | 0.38626183 | 0.191 | 0 | | 0 | 44 |
| XLOC-011066 | 0 | 0.37552423 | 0.191 | 0 | | 0 | 44 |
| LOC101882034 | 0 | 0.370389431 | 0.191 | 0 | | 0 | 44 |
| ltc4s | 0 | 0.36854438 | 0.17 | 0 | | 0 | 44 |
| LOC100537138 | 0 | 0.36572884 | 0.213 | 0 | | 0 | 44 |
| si:ch73-208h1.1 | 0 | 0.35412718 | 0.17 | 0 | | 0 | 44 |
| muc13a | 0 | 0.352436815 | 0.149 | 0 | | 0 | 44 |
| si:ch211-232i5.1 | 0 | 0.332087789 | 0.17 | 0 | | 0 | 44 |
| si:ch211-149a19.3 | 0 | 0.330753447 | 0.191 | 0 | | 0 | 44 |
| CR926130.1 | 0 | 0.323735403 | 0.149 | 0 | | 0 | 44 |
| sp8a | 0 | 0.305655563 | 0.17 | 0 | | 0 | 44 |
| BX901957.1 | 0 | 0.29644349 | 0.17 | 0 | | 0 | 44 |
| ankk1 | 0 | 0.256148171 | 0.17 | 0 | | 0 | 44 |

| | | | | | | |
|---|---|---|---|---|---|---|
| tmem45b | 6.80655797402151e-307 | 0.435149588 | 0.213 | 0.001 | 1.71314257648147e-302 | 44 |
| si:ch1073-340i21.3 | 4.79107306943404e-306 | 0.354035825 | 0.191 | 0 | 1.20586518084585e-301 | 44 |
| si:dkey-61f9.2 | 1.1182384361691e-301 | 0.328977795 | 0.149 | 0 | 2.814494319994e-297 | 44 |
| papss2a | 1.9525397089691e-301 | 0.871962636 | 0.447 | 0.005 | 4.91434719350433e-297 | 44 |
| elf31 | 6.39547850323231e-300 | 0.86377115 | 0.468 | 0.005 | 1.60967798447854e-295 | 44 |
| LOC110438610 | 3.02775538231762e-299 | 0.545079902 | 0.298 | 0.002 | 7.62055752175523e-295 | 44 |
| gpa33a1 | 1.10383890055212e-297 | 0.662670364 | 0.383 | 0.003 | 2.77825212879963e-293 | 44 |
| si:ch211-79k12.11 | 1.91123011991187e-294 | 1.391101955 | 0.83 | 0.02 | 4.81037508880619e-290 | 44 |
| tmem176l.41 | 2.9149668029749e-283 | 0.967673935 | 0.532 | 0.008 | 7.33667994640752e-279 | 44 |
| zgc:91849 | 1.451016954174e-281 | 0.476707121 | 0.298 | 0.002 | 3.65206457196053e-277 | 44 |
| CABZ01039863.1 | 2.58298174615895e-281 | 0.511966257 | 0.298 | 0.002 | 6.50110675690747e-277 | 44 |
| sec23b | 1.19562003639999e-277 | 1.730855855 | 0.979 | 0.031 | 3.00925606961514e-273 | 44 |
| dbnla | 2.38196845513411e-275 | 0.633347496 | 0.362 | 0.003 | 5.99517640472705e-271 | 44 |
| pof1b1 | 4.13697014962907e-273 | 0.969798094 | 0.532 | 0.008 | 1.04123401696014e-268 | 44 |
| hmgcra | 4.59169800646203e-271 | 0.623129485 | 0.34 | 0.003 | 1.15568447124643e-266 | 44 |
| f2rl1.21 | 8.95670532056576e-271 | 1.4898565 | 0.83 | 0.022 | 2.2543131621332e-266 | 44 |
| CR677617.1 | 1.29668489484005e-267 | 0.581028634 | 0.319 | 0.002 | 3.26362621182293e-263 | 44 |
| selenow2b1 | 1.70842654458803e-264 | 0.625895294 | 0.362 | 0.003 | 4.29993877007361e-260 | 44 |
| dspa1 | 1.06687693345676e-263 | 1.487236541 | 0.83 | 0.023 | 2.68522255381733e-259 | 44 |
| plekhg6 | 3.40527953834361e-262 | 0.267731245 | 0.17 | 0 | 8.57074807005703e-258 | 44 |
| ftr39p | 4.27229622405203e-262 | 0.268412246 | 0.17 | 0 | 1.07529423663166e-257 | 44 |
| klf5l | 1.36118239570894e-257 | 0.404488947 | 0.234 | 0.001 | 3.42595997175983e-253 | 44 |
| si:dkeyp-82b4.2 | 1.35471626833155e-253 | 0.305254545 | 0.128 | 0 | 3.40968537576369e-249 | 44 |
| frmd4ba | 2.58754343791095e-250 | 1.106624939 | 0.617 | 0.013 | 6.51258807887808e-246 | 44 |
| zgc:172075 | 6.56919699563368e-247 | 0.28005712 | 0.106 | 0 | 1.65340119183104e-242 | 44 |
| adgrf7 | 6.56919699563368e-247 | 0.257445753 | 0.106 | 0 | 1.65340119183104e-242 | 44 |
| zgc:922421 | 3.31395671613322e-245 | 1.170311414 | 0.66 | 0.015 | 8.34089765883571e-241 | 44 |
| ttc39a | 8.22538254783279e-243 | 0.480570558 | 0.255 | 0.002 | 2.07024653346403e-238 | 44 |
| si:dkeyp-75b4.8 | 8.26627519266873e-242 | 0.275001273 | 0.17 | 0 | 2.08053880324279e-237 | 44 |
| LOC100005723 | 3.81458297876976e-241 | 0.292369641 | 0.149 | 0 | 9.60092389926561e-237 | 44 |
| ccl20a.3 | 5.0204492927846e-234 | 0.461534996 | 0.255 | 0.002 | 1.26359688250096e-229 | 44 |
| XLOC-003147 | 5.79066767764479e-234 | 0.31624003 | 0.191 | 0.001 | 1.45745314778642e-229 | 44 |
| ano1 | 8.86246522357397e-232 | 0.756592205 | 0.468 | 0.007 | 2.23059387212133e-227 | 44 |
| krt44 | 7.29899296377441e-230 | 2.272252734 | 0.957 | 0.038 | 1.83708353905238e-225 | 44 |
| abcb5 | 2.8440720137744e-229 | 1.737540923 | 0.83 | 0.027 | 7.15824485146879e-225 | 44 |

| | | | | | | |
|---|---|---|---|---|---|---|
| CU693379.1 | 1.22490552516514e-222 | 0.38664766 | 0.213 | 0.001 | 3.08296471628813e-218 | 44 |
| si:ch211-282j17.8 | 8.17198352970912e-222 | 0.265073467 | 0.128 | 0 | 2.05680653459249e-217 | 44 |
| sparta | 8.43701667543709e-222 | 0.262059589 | 0.128 | 0 | 2.12351272704076e-217 | 44 |
| si:ch211-195b11.31 | 8.69763633657796e-220 | 1.798533284 | 0.915 | 0.035 | 2.18910808955331e-215 | 44 |
| nkx3-1 | 1.71669885984327e-218 | 0.945851903 | 0.404 | 0.006 | 4.32075936033954e-214 | 44 |
| si:dkey-228d14.5 | 3.14309245237817e-217 | 0.610089946 | 0.34 | 0.004 | 7.91084939339062e-213 | 44 |
| LO017816.1 | 1.66490824650266e-212 | 0.346371434 | 0.213 | 0.001 | 4.19040756562255e-208 | 44 |
| zgc:194398 | 3.23333053302192e-211 | 0.394367886 | 0.234 | 0.002 | 8.13796961856286e-207 | 44 |
| erbb3b | 1.26892192196516e-209 | 0.333618061 | 0.17 | 0.001 | 3.19374958539412e-205 | 44 |
| abca12 | 1.70357805439376e-209 | 0.340199044 | 0.17 | 0.001 | 4.28773560510365e-205 | 44 |
| LOC110438361 | 2.23570083819226e-209 | 0.263703435 | 0.17 | 0.001 | 5.62703543964609e-205 | 44 |
| styk1b | 1.08304679917728e-203 | 0.36828921 | 0.213 | 0.001 | 2.7259204888493e-199 | 44 |
| si:dkey-160o24.3 | 7.75808079672304e-201 | 0.318079711 | 0.149 | 0 | 1.95263135572722e-196 | 44 |
| oat | 9.21997537523774e-200 | 0.645558827 | 0.362 | 0.005 | 2.32057560219359e-195 | 44 |
| si:dkey-19b23.121 | 1.10249066065464e-198 | 0.354600366 | 0.191 | 0.001 | 2.77485874380166e-194 | 44 |
| xkr92 | 9.36490761776189e-198 | 1.110786189 | 0.596 | 0.015 | 2.35705359831449e-193 | 44 |
| macc1 | 2.00954638694636e-197 | 0.80736006 | 0.468 | 0.009 | 5.05782730130528e-193 | 44 |
| CABZ01070852.1 | 3.01335879231766e-196 | 0.542282092 | 0.277 | 0.003 | 7.58432274438432e-192 | 44 |
| s100w3 | 7.89620357994628e-196 | 1.09090132 | 0.617 | 0.017 | 1.98739547903668e-191 | 44 |
| aldob4 | 1.50403273705456e-191 | 1.709575217 | 0.851 | 0.036 | 3.78549999589261e-187 | 44 |
| si:ch73-22a13.3 | 7.04433475800808e-188 | 0.431833095 | 0.213 | 0.002 | 1.77298861524305e-183 | 44 |
| LOC110438236 | 8.52587091462959e-185 | 0.501006984 | 0.255 | 0.002 | 2.14587645050312e-180 | 44 |
| CABZ01056052.1 | 1.341353974506e-184 | 0.324950977 | 0.17 | 0.001 | 3.37605381843414e-180 | 44 |
| dhrs13a.21 | 2.13211108155212e-184 | 1.998767456 | 0.957 | 0.048 | 5.36631038115854e-180 | 44 |
| si:dkey-242g16.2 | 2.14458749870907e-184 | 0.433719823 | 0.255 | 0.002 | 5.39771227550085e-180 | 44 |
| wsb21 | 2.43729823331368e-184 | 0.964580746 | 0.574 | 0.015 | 6.1344359234272e-180 | 44 |
| si:busm1-104n07.3 | 5.20229476628067e-181 | 0.3844133 | 0.213 | 0.002 | 1.30936556972518e-176 | 44 |
| limk21 | 1.09124053926998e-180 | 0.951323514 | 0.574 | 0.016 | 2.74654331328861e-176 | 44 |
| nlrp16 | 2.10784904739979e-177 | 0.259708085 | 0.128 | 0 | 5.30524526740052e-173 | 44 |
| zgc:162184 | 2.53634208963557e-175 | 0.475786106 | 0.255 | 0.003 | 6.38371940540377e-171 | 44 |
| fam83b1 | 8.25464021449556e-175 | 0.423986351 | 0.213 | 0.002 | 2.07761039558639e-170 | 44 |
| tmprss13b | 2.26209189927579e-174 | 0.340898755 | 0.17 | 0.001 | 5.69345910128724e-170 | 44 |
| stard141 | 4.62514345335566e-174 | 1.75313576 | 0.957 | 0.05 | 1.16410235577509e-169 | 44 |
| phlda25 | 1.69860629656882e-172 | 1.83227682 | 0.936 | 0.049 | 4.27522218783407e-168 | 44 |
| si:ch211-161h7.5 | 4.3673392602381e-172 | 0.351308654 | 0.149 | 0.001 | 1.09921561840933e-167 | 44 |

| | | | | | | |
|---|---|---|---|---|---|---|
| zgc:112433 | 2.30465898933997e-168 | 0.407258021 | 0.213 | 0.002 | 5.80059621026977e-164 | 44 |
| tuft1a4 | 1.49672995581924e-167 | 1.293122681 | 0.766 | 0.032 | 3.76711962580143e-163 | 44 |
| BX548046.22 | 4.1964708807855e-167 | 0.545528278 | 0.319 | 0.005 | 1.0562097559849e-162 | 44 |
| slc9a2 | 6.51858906059442e-165 | 0.316501486 | 0.191 | 0.001 | 1.64066368066101e-160 | 44 |
| wu:fb18f062 | 1.71505408472206e-163 | 2.041903026 | 0.936 | 0.054 | 4.31661962583696e-159 | 44 |
| mpeg1.2 | 1.59274946519643e-160 | 0.25357604 | 0.149 | 0.001 | 4.0087911289529e-156 | 44 |
| si:ch211-153b23.4 | 1.62461197239889e-160 | 0.369359204 | 0.149 | 0.001 | 4.08898587333077e-156 | 44 |
| perp6 | 4.13562021758386e-158 | 1.783456526 | 0.915 | 0.052 | 1.04089425256368e-153 | 44 |
| si:dkey-223p19.1 | 5.91005513991577e-157 | 0.435018379 | 0.17 | 0.001 | 1.4875017781654e-152 | 44 |
| cyp8b2 | 3.09173205023935e-152 | 0.442663798 | 0.213 | 0.002 | 7.78158039724743e-148 | 44 |
| fam169aa | 4.72850651504235e-148 | 0.414903087 | 0.213 | 0.002 | 1.19011780477101e-143 | 44 |
| tlcd2 | 7.03465038834534e-148 | 0.677511545 | 0.404 | 0.009 | 1.77055115624264e-143 | 44 |
| rnh1 | 4.84442121795194e-145 | 0.435716018 | 0.255 | 0.003 | 1.21929237634632e-140 | 44 |
| plekhf11 | 8.57319014910136e-144 | 0.601887895 | 0.277 | 0.004 | 2.15778622862732e-139 | 44 |
| rarres3 | 1.58124594469504e-143 | 0.43787553 | 0.234 | 0.003 | 3.97983791820294e-139 | 44 |
| plekhn1 | 2.09571442601662e-143 | 0.382571975 | 0.213 | 0.002 | 5.27470363884122e-139 | 44 |
| si:ch1073-385f13.3 | 2.47596886005422e-143 | 0.621356553 | 0.319 | 0.006 | 6.23176602387046e-139 | 44 |
| LO018605.2 | 1.50599988484785e-142 | 0.348606109 | 0.17 | 0.001 | 3.79045111017355e-138 | 44 |
| ugt8 | 1.60173239997242e-142 | 0.341280712 | 0.17 | 0.001 | 4.03140027749058e-138 | 44 |
| bbox1 | 2.35271634442142e-142 | 0.74888581 | 0.468 | 0.013 | 5.92155176727427e-138 | 44 |
| XLOC-019059 | 1.33729000738813e-141 | 0.253386946 | 0.149 | 0.001 | 3.36582521959519e-137 | 44 |
| rhpn22 | 2.94520300243096e-140 | 1.211394537 | 0.723 | 0.033 | 7.41278143681848e-136 | 44 |
| tmem30b | 1.59960938221517e-137 | 0.820180463 | 0.489 | 0.015 | 4.02605685409735e-133 | 44 |
| elovl1a | 1.85107969368297e-137 | 1.495010385 | 0.851 | 0.048 | 4.65898248103065e-133 | 44 |
| rhcga | 3.64793578685638e-137 | 0.284010889 | 0.106 | 0 | 9.18148958193882e-133 | 44 |
| frk | 3.91568424042511e-137 | 0.270888309 | 0.106 | 0 | 9.85538566472597e-133 | 44 |
| BX005421.3 | 3.79746500261726e-136 | 0.303100665 | 0.17 | 0.001 | 9.55783966508739e-132 | 44 |
| si:dkey-262k9.24 | 4.33734783099738e-135 | 1.155164014 | 0.702 | 0.033 | 1.09166707558373e-130 | 44 |
| tnfrsf11b | 3.14109570148279e-134 | 0.756091391 | 0.277 | 0.005 | 7.90582377106204e-130 | 44 |
| si:ch211-158d24.4 | 7.99044332591598e-134 | 0.542817076 | 0.234 | 0.003 | 2.01111468069979e-129 | 44 |
| oclnb1 | 6.75781269914266e-133 | 1.309977181 | 0.766 | 0.039 | 1.70087387824722e-128 | 44 |
| ildr1a | 9.56995052931401e-133 | 1.158317862 | 0.702 | 0.032 | 2.40866084872304e-128 | 44 |
| klf35 | 2.16973007073399e-132 | 1.153223879 | 0.702 | 0.034 | 5.46099361503038e-128 | 44 |
| rgs23 | 4.42729096257223e-132 | 1.248075818 | 0.617 | 0.026 | 1.1143048623698e-127 | 44 |
| krt916 | 7.69017612514183e-131 | 2.032760181 | 0.936 | 0.066 | 1.93554042893695e-126 | 44 |

| | | | | | | |
|---|---|---|---|---|---|---|
| trpm4b.2 | 1.46579547399752e-130 | 0.319286699 | 0.17 | 0.002 | 3.68926062850436e-126 | 44 |
| gdpd3a1 | 4.78474072765043e-128 | 0.373914007 | 0.213 | 0.003 | 1.20427139374234e-123 | 44 |
| tlcd1 | 2.97919907771219e-127 | 0.298126071 | 0.191 | 0.002 | 7.49834615869381e-123 | 44 |
| ch25hl1.1 | 3.49697780215071e-127 | 0.33505113 | 0.234 | 0.003 | 8.80154343023313e-123 | 44 |
| tfap2c2 | 5.27021220336264e-127 | 0.503282702 | 0.255 | 0.004 | 1.32645970946434e-122 | 44 |
| barx2 | 1.41933955605874e-126 | 0.261315373 | 0.128 | 0.001 | 3.57233572864423e-122 | 44 |
| si:ch211-217k17.10 | 6.65577907868287e-125 | 0.572192672 | 0.234 | 0.003 | 1.67519303631369e-120 | 44 |
| dnmbp1 | 7.20732025073168e-122 | 0.761338065 | 0.447 | 0.014 | 1.81401043390666e-117 | 44 |
| LOC108180802 | 4.31813215366989e-118 | 0.26746664 | 0.128 | 0.001 | 1.08683068175717e-113 | 44 |
| si:dkey-33i11.94 | 5.74113308585316e-118 | 0.899436915 | 0.532 | 0.021 | 1.44498578637838e-113 | 44 |
| si:dkey-28d5.11 | 5.9592449271058e-118 | 0.251852119 | 0.128 | 0.001 | 1.49988235570326e-113 | 44 |
| wu:fi04e122 | 6.57054986963174e-118 | 0.912329614 | 0.511 | 0.02 | 1.65374169668761e-113 | 44 |
| si:ch211-264f5.62 | 3.08472341493359e-116 | 0.669460404 | 0.404 | 0.012 | 7.76394036304634e-112 | 44 |
| SEMA4F1 | 1.01645112400916e-115 | 0.42772061 | 0.255 | 0.005 | 2.55830583401866e-111 | 44 |
| nqo1 | 1.30254794663853e-115 | 0.695309563 | 0.426 | 0.014 | 3.27838292689452e-111 | 44 |
| arhgef55 | 2.58302691145353e-115 | 0.952216324 | 0.596 | 0.027 | 6.50122043343738e-111 | 44 |
| zgc:173737 | 2.15193768307103e-114 | 0.268929072 | 0.149 | 0.001 | 5.41621195452146e-110 | 44 |
| iqsec2b | 8.39074224681764e-113 | 0.368620537 | 0.213 | 0.003 | 2.11186591610153e-108 | 44 |
| jupa2 | 2.10402539794996e-112 | 1.336263118 | 0.787 | 0.052 | 5.29562152410024e-108 | 44 |
| trib1 | 5.4151517114771e-112 | 0.469115168 | 0.255 | 0.005 | 1.36293953426167e-107 | 44 |
| casz12 | 1.68699203448099e-111 | 0.858124851 | 0.511 | 0.021 | 4.24599025158521e-107 | 44 |
| zgc:153932 | 1.77498426734001e-111 | 0.318510169 | 0.17 | 0.002 | 4.46745790246808e-107 | 44 |
| FO704779.1 | 1.28007064122452e-110 | 0.269552317 | 0.128 | 0.001 | 3.22180979689798e-106 | 44 |
| cd9b4 | 4.85869621200505e-110 | 1.629517891 | 0.851 | 0.063 | 1.22288524959955e-105 | 44 |
| ptpn13 | 2.37342204902486e-109 | 0.687469188 | 0.447 | 0.016 | 5.97366595519066e-105 | 44 |
| zgc:158263 | 2.88502992905814e-109 | 0.30955897 | 0.191 | 0.003 | 7.26133182844643e-105 | 44 |
| CT027980.11 | 1.80787299943069e-106 | 0.345194771 | 0.191 | 0.003 | 4.55023555226712e-102 | 44 |
| slc2a1b1 | 2.24188079575638e-106 | 0.833117672 | 0.468 | 0.018 | 5.64258977483922e-102 | 44 |
| si:ch73-359m17.51 | 1.98236647778642e-105 | 0.708147786 | 0.255 | 0.005 | 4.98941818794063e-101 | 44 |
| epdl12 | 1.24721031148554e-104 | 1.041992811 | 0.638 | 0.034 | 3.13910363297795e-100 | 44 |
| si:ch211-157c3.48 | 2.06754111562304e-104 | 1.829452012 | 0.936 | 0.088 | 5.20379423391162e-100 | 44 |
| kremen12 | 2.44163221994086e-103 | 0.57977655 | 0.34 | 0.01 | 6.14534413436914e-99 | 44 |
| gcnt4a2 | 3.32001999885005e-102 | 0.75523854 | 0.404 | 0.014 | 8.3561583351057e-98 | 44 |
| net13 | 5.32127749101344e-102 | 0.863710747 | 0.489 | 0.021 | 1.33931233171317e-97 | 44 |
| nox1 | 9.34061946346142e-101 | 0.357579026 | 0.191 | 0.003 | 2.3509405127586e-96 | 44 |

| | | | | | | |
|---|---|---|---|---|---|---|
| krt222 | 1.0747801307086e-100 | 0.292600427 | 0.17 | 0.002 | 2.70511411098047e-96 | 44 |
| LOC108179621 | 3.66515905767471e-100 | 0.314357687 | 0.149 | 0.002 | 9.22483883226148e-96 | 44 |
| elovl6l | 8.26959026876737e-100 | 0.762046521 | 0.447 | 0.017 | 2.08137317474606e-95 | 44 |
| si:dkey-30c15.2 | 1.92938634611333e-96 | 0.411838578 | 0.213 | 0.004 | 4.85607249453264e-92 | 44 |
| homer2 | 1.5723906692302e-93 | 0.642221651 | 0.362 | 0.012 | 3.9575500753855e-89 | 44 |
| ehf | 3.91527115092903e-93 | 0.270465009 | 0.128 | 0.001 | 9.85434595977328e-89 | 44 |
| arrdc1a | 1.17095363575041e-92 | 0.814344214 | 0.532 | 0.026 | 2.9471732058202e-88 | 44 |
| kras | 1.74003832932384e-92 | 1.009254405 | 0.638 | 0.04 | 4.37950247107517e-88 | 44 |
| si:dkey-204f11.641 | 2.26742719446338e-92 | 0.813038201 | 0.532 | 0.027 | 5.70688750574487e-88 | 44 |
| ms4a17a.51 | 1.1480737794484e-90 | 0.410407274 | 0.255 | 0.006 | 2.88958689549369e-86 | 44 |
| spint21 | 7.68720730301887e-89 | 1.480281329 | 0.851 | 0.076 | 1.93479320609682e-84 | 44 |
| FP017215.11 | 1.44679251590493e-88 | 0.658716614 | 0.383 | 0.014 | 3.64143208328111e-84 | 44 |
| cast4 | 7.59934897471434e-88 | 1.229117765 | 0.809 | 0.068 | 1.91268014344585e-83 | 44 |
| eppk15 | 2.77441587653445e-86 | 1.022926434 | 0.617 | 0.039 | 6.98292731964956e-82 | 44 |
| tead3b | 4.66479887127333e-86 | 0.798293611 | 0.426 | 0.019 | 1.17408322791078e-81 | 44 |
| mpp6a | 5.82196301103569e-86 | 0.627164521 | 0.383 | 0.015 | 1.46532987024757e-81 | 44 |
| myo5b | 6.18990948879775e-86 | 0.36339571 | 0.213 | 0.004 | 1.5579383192355e-81 | 44 |
| ftr36 | 9.70840887658267e-85 | 0.499997853 | 0.234 | 0.005 | 2.44350943014709e-80 | 44 |
| cx34.4 | 1.39563516394239e-84 | 0.311036581 | 0.213 | 0.004 | 3.5126741441266e-80 | 44 |
| rhov | 2.28652771139393e-84 | 0.518460476 | 0.34 | 0.012 | 5.75496159680738e-80 | 44 |
| myo1ea | 8.41110089283048e-84 | 0.42875606 | 0.191 | 0.004 | 2.1169899837165e-79 | 44 |
| si:dkey-157l19.21 | 1.06205857567726e-83 | 0.521873489 | 0.298 | 0.009 | 2.67309522912209e-79 | 44 |
| gna15.11 | 6.38702094224571e-82 | 0.395293201 | 0.191 | 0.004 | 1.60754930095382e-77 | 44 |
| zbtb16b2 | 2.4375083185736e-81 | 0.83916586 | 0.468 | 0.024 | 6.13496468701789e-77 | 44 |
| ptgs2a2 | 6.79197107135934e-81 | 0.647743581 | 0.319 | 0.011 | 1.70947119895043e-76 | 44 |
| usp2b | 1.54382520026093e-80 | 0.411853325 | 0.213 | 0.005 | 3.88565364653674e-76 | 44 |
| si:ch211-285f17.1 | 5.04371493539042e-80 | 0.339903088 | 0.191 | 0.004 | 1.26945261208841e-75 | 44 |
| atp8b1 | 4.39354956632821e-79 | 0.384073559 | 0.234 | 0.006 | 1.10581249034915e-74 | 44 |
| samd10a | 1.00074017556323e-78 | 0.4283994 | 0.255 | 0.007 | 2.51876294787508e-74 | 44 |
| apnl1 | 4.35190208145677e-78 | 0.355171852 | 0.191 | 0.004 | 1.09533023488186e-73 | 44 |
| fam160a1a | 1.88418198287294e-77 | 0.370752219 | 0.213 | 0.005 | 4.74229763269291e-73 | 44 |
| cdh14 | 2.51225517578639e-76 | 1.153806958 | 0.702 | 0.057 | 6.32309505193676e-72 | 44 |
| ssh2a | 1.69057853138274e-75 | 0.841738138 | 0.511 | 0.031 | 4.25501710563722e-71 | 44 |
| alox121 | 2.8732573660956e-75 | 0.503836522 | 0.319 | 0.012 | 7.23170146472601e-71 | 44 |
| paqr6 | 6.33962457278331e-75 | 0.64466819 | 0.362 | 0.015 | 1.59562010872383e-70 | 44 |

| | | | | | | |
|---|---|---|---|---|---|---|
| hopx8 | 6.49903357677381e-75 | 1.183731438 | 0.723 | 0.064 | 1.6357417609382e-70 | 44 |
| pard6b | 1.15491549717224e-73 | 0.428753738 | 0.255 | 0.008 | 2.9068068148328e-69 | 44 |
| gsnb1 | 1.78439878749546e-72 | 0.290342139 | 0.149 | 0.002 | 4.49115330824733e-68 | 44 |
| tiam1a2 | 7.07665082828453e-72 | 0.524373707 | 0.298 | 0.011 | 1.78112224697093e-67 | 44 |
| si:dkey-16l2.202 | 6.75107459998414e-71 | 0.649452262 | 0.383 | 0.018 | 1.69917796607001e-66 | 44 |
| pld2 | 1.67206070943087e-69 | 0.349482505 | 0.213 | 0.006 | 4.20840959956656e-65 | 44 |
| zgc:92481 | 4.76188355963209e-69 | 0.674113404 | 0.404 | 0.021 | 1.1985184731238e-64 | 44 |
| xiap | 5.50055500563354e-69 | 0.938947836 | 0.553 | 0.04 | 1.38443468936791e-64 | 44 |
| chmp4c | 1.02093167250731e-68 | 0.753451773 | 0.489 | 0.03 | 2.56958292653365e-64 | 44 |
| lmbr1l1 | 2.59319310058832e-68 | 0.587263761 | 0.34 | 0.015 | 6.52680771487074e-64 | 44 |
| mgst1.26 | 4.45588918568516e-68 | 1.331451521 | 0.851 | 0.101 | 1.1215027491451e-63 | 44 |
| si:ch73-204p21.21 | 5.23965149945943e-68 | 0.949505728 | 0.617 | 0.048 | 1.31876788589895e-63 | 44 |
| osbpl71 | 8.46089074513148e-68 | 0.837700012 | 0.532 | 0.036 | 2.12952159164214e-63 | 44 |
| plcd1a | 3.57627962283811e-65 | 0.387628516 | 0.191 | 0.005 | 9.00113818272125e-61 | 44 |
| ccng26 | 6.64347212663385e-65 | 1.175727011 | 0.766 | 0.084 | 1.67209549955247e-60 | 44 |
| cd44a1 | 1.39979331739585e-64 | 0.297798631 | 0.149 | 0.003 | 3.52313980055361e-60 | 44 |
| dusp58 | 8.20645550444826e-64 | 1.225666659 | 0.702 | 0.072 | 2.06548278591458e-59 | 44 |
| ptgis2 | 1.23265371516683e-63 | 0.641317001 | 0.319 | 0.014 | 3.1024661357034e-59 | 44 |
| zgc:162193 | 3.3678683618429e-63 | 0.292365452 | 0.17 | 0.004 | 8.47658787992239e-59 | 44 |
| spint1a2 | 3.70411649206061e-63 | 0.930608164 | 0.638 | 0.055 | 9.32289079886734e-59 | 44 |
| CABZ01039859.1 | 6.01237211941362e-63 | 0.252163417 | 0.128 | 0.002 | 1.51325393873521e-58 | 44 |
| LO017965.11 | 8.35732776084772e-62 | 0.325476606 | 0.17 | 0.004 | 2.10345582412776e-57 | 44 |
| igf2bp2b | 1.96386557311763e-61 | 0.505115632 | 0.34 | 0.017 | 4.94285326097976e-57 | 44 |
| si:ch73-141c7.1 | 3.27370308899292e-61 | 0.300108777 | 0.213 | 0.006 | 8.23958330468629e-57 | 44 |
| foxc1b | 1.5619262638891e-60 | 0.289493128 | 0.17 | 0.004 | 3.93121221358247e-56 | 44 |
| cfl1l9 | 6.47144026921876e-60 | 1.515801004 | 0.936 | 0.151 | 1.62879680135967e-55 | 44 |
| slc7a3b1 | 1.06484341096867e-59 | 0.760545141 | 0.447 | 0.03 | 2.68010438106704e-55 | 44 |
| b4galnt1a | 1.62242046617019e-59 | 0.387924339 | 0.213 | 0.007 | 4.08347007130376e-55 | 44 |
| FAM83G1 | 1.91220006239902e-59 | 0.492879806 | 0.277 | 0.011 | 4.8128163370521e-55 | 44 |
| ptpn212 | 2.14048826400185e-59 | 0.807205756 | 0.468 | 0.033 | 5.38739491166625e-55 | 44 |
| cavin2a1 | 5.84242804965443e-59 | 0.482502888 | 0.298 | 0.013 | 1.47048071581752e-54 | 44 |
| XLOC-0418321 | 9.70858346889602e-59 | 0.520113772 | 0.255 | 0.01 | 2.44355337328644e-54 | 44 |
| ahnak12 | 1.40077375400491e-58 | 1.340028125 | 0.872 | 0.125 | 3.52560746145495e-54 | 44 |
| si:dkey-87o1.23 | 5.99841886157104e-58 | 1.119880923 | 0.745 | 0.08 | 1.50974204326881e-53 | 44 |
| rab11al | 6.00596829887961e-58 | 0.407882636 | 0.255 | 0.01 | 1.51164216114501e-53 | 44 |

| | | | | | | |
|---|---|---|---|---|---|---|
| ptprfb3 | 6.39241256268015e-57 | 0.520433845 | 0.298 | 0.014 | 1.60890631790097e-52 | 44 |
| cdc42ep51 | 1.2201898896406e-56 | 0.715700855 | 0.447 | 0.031 | 3.07109593323643e-52 | 44 |
| anxa1a9 | 1.39338561943637e-56 | 1.503715284 | 0.894 | 0.142 | 3.5070122655594e-52 | 44 |
| tcnbb2 | 1.75022272759104e-56 | 0.747338531 | 0.319 | 0.016 | 4.40513558307388e-52 | 44 |
| pdlim2 | 8.06508068134858e-56 | 0.433001226 | 0.255 | 0.01 | 2.02990015668862e-51 | 44 |
| cdaa3 | 8.74047226029711e-56 | 0.755936483 | 0.468 | 0.034 | 2.19988946319418e-51 | 44 |
| c1galt1b | 3.07716481013683e-55 | 0.783252638 | 0.489 | 0.038 | 7.74491611063338e-51 | 44 |
| zgc:774863 | 3.42192411794731e-55 | 0.99004574 | 0.66 | 0.07 | 8.61264081246158e-51 | 44 |
| si:ch211-166a6.53 | 5.66215244357687e-55 | 0.711724167 | 0.447 | 0.032 | 1.42510714852386e-50 | 44 |
| serpinb141 | 6.29652069940826e-55 | 0.479078552 | 0.255 | 0.011 | 1.58477129483407e-50 | 44 |
| XLOC-044191 | 5.58245236174034e-54 | 0.523431768 | 0.319 | 0.017 | 1.40504743492643e-49 | 44 |
| nipa2 | 1.05968671298714e-53 | 0.879178522 | 0.553 | 0.049 | 2.66712548791733e-49 | 44 |
| nr1i2 | 2.09339151783433e-53 | 0.255372731 | 0.128 | 0.003 | 5.26885711123722e-49 | 44 |
| mboat1 | 5.18975022828453e-53 | 0.37278284 | 0.191 | 0.006 | 1.30620823495693e-48 | 44 |
| kdm6ba3 | 8.46536702094629e-53 | 0.910139826 | 0.532 | 0.048 | 2.13064822550197e-48 | 44 |
| osbpl3a | 2.53230597092953e-52 | 0.630574656 | 0.383 | 0.025 | 6.37356089823254e-48 | 44 |
| rnd3a3 | 3.4583322019332e-52 | 0.663768468 | 0.426 | 0.031 | 8.70427631904566e-48 | 44 |
| efna1b3 | 7.6886277805038e-52 | 0.530019218 | 0.319 | 0.017 | 1.935150726075e-47 | 44 |
| midn12 | 8.75423201482085e-52 | 1.46581128 | 0.83 | 0.139 | 2.20335265581026e-47 | 44 |
| galnt7 | 1.04582852745063e-51 | 0.735083641 | 0.447 | 0.034 | 2.63224582074048e-47 | 44 |
| ephb6 | 1.50886787050178e-51 | 0.256547403 | 0.149 | 0.004 | 3.79766954326593e-47 | 44 |
| ldlrb | 2.03001827533514e-51 | 0.330568502 | 0.191 | 0.006 | 5.10935299719102e-47 | 44 |
| cxadr | 2.05379258308989e-51 | 0.745570344 | 0.447 | 0.034 | 5.16919055237895e-47 | 44 |
| nfkbiz | 2.17124528473664e-51 | 0.58056971 | 0.404 | 0.027 | 5.46480725715365e-47 | 44 |
| mhc2dab7 | 3.4365310793105e-51 | 1.122512156 | 0.766 | 0.096 | 8.64940507351659e-47 | 44 |
| tob1b5 | 8.83768228707254e-51 | 1.411739614 | 0.915 | 0.173 | 2.22435625483329e-46 | 44 |
| cpeb4b | 1.27494118115413e-50 | 0.749373477 | 0.511 | 0.045 | 3.20889945884682e-46 | 44 |
| si:dkey-261p22.1 | 2.31337004688334e-50 | 0.290180762 | 0.128 | 0.003 | 5.82252107100067e-46 | 44 |
| grhl12 | 3.81475166292414e-50 | 0.697441787 | 0.383 | 0.026 | 9.60134846041376e-46 | 44 |
| bicdl2 | 3.86813806650615e-50 | 0.497029674 | 0.298 | 0.015 | 9.73571669958932e-46 | 44 |
| ifngr11 | 9.88808408125528e-50 | 0.294534436 | 0.17 | 0.005 | 2.48873188241114e-45 | 44 |
| cst14b.16 | 9.93571100272868e-50 | 0.865743992 | 0.532 | 0.051 | 2.50071910227678e-45 | 44 |
| epha2a | 1.86934785414469e-49 | 0.453504632 | 0.213 | 0.008 | 4.70496161409676e-45 | 44 |
| snphb1 | 2.60764898403152e-49 | 0.696541219 | 0.426 | 0.033 | 6.56319172790894e-45 | 44 |
| s100v210 | 3.03701683364307e-49 | 1.28218964 | 0.787 | 0.121 | 7.64386766859625e-45 | 44 |

| | | | | | | |
|---|---|---|---|---|---|---|
| fosb9 | 3.06705377096481e-49 | 1.387527021 | 0.809 | 0.125 | 7.71946763614134e-45 | 44 |
| adrb2a | 3.16401858578406e-49 | 0.463708064 | 0.298 | 0.016 | 7.96351837855991e-45 | 44 |
| tnfaip38 | 4.79231887097045e-49 | 0.950802407 | 0.596 | 0.064 | 1.20617873663455e-44 | 44 |
| si:ch211-152p11.43 | 1.16627975729377e-47 | 0.715934223 | 0.404 | 0.031 | 2.93540952113268e-43 | 44 |
| CT027702.1 | 1.38448883411846e-47 | 0.27683459 | 0.128 | 0.003 | 3.48461994659274e-43 | 44 |
| si:dkey-30g5.1 | 4.30214672272245e-47 | 0.337264753 | 0.191 | 0.007 | 1.08280730864201e-42 | 44 |
| lmna2 | 1.04982059830163e-46 | 0.481225948 | 0.277 | 0.015 | 2.64229346386537e-42 | 44 |
| chmp1b5 | 1.14269960232428e-46 | 1.103758682 | 0.787 | 0.121 | 2.87606062908997e-42 | 44 |
| ngef | 2.34493753224707e-46 | 0.269403113 | 0.149 | 0.004 | 5.90197327491265e-42 | 44 |
| pleca5 | 2.40954468401089e-46 | 0.927354326 | 0.553 | 0.058 | 6.06458301518701e-42 | 44 |
| CABZ01072077.1 | 3.01475221146717e-46 | 0.26986818 | 0.128 | 0.003 | 7.58782984104171e-42 | 44 |
| cpeb2 | 4.8535860088674e-46 | 0.363066632 | 0.17 | 0.006 | 1.22159906257184e-41 | 44 |
| btr25 | 7.82536561085174e-46 | 0.347051279 | 0.191 | 0.007 | 1.96956627059527e-41 | 44 |
| errfi1a | 1.58139586875125e-45 | 0.513327592 | 0.319 | 0.02 | 3.98021526206003e-41 | 44 |
| bcl6ab | 4.54234352695223e-45 | 0.527349745 | 0.277 | 0.015 | 1.14326244229861e-40 | 44 |
| tent5ba2 | 7.05306486627901e-45 | 0.342586253 | 0.234 | 0.011 | 1.77518589619376e-40 | 44 |
| zfand5a2 | 2.11215363870646e-44 | 0.930933857 | 0.617 | 0.076 | 5.3160794932603e-40 | 44 |
| cldna1 | 7.18575743055586e-44 | 0.331464544 | 0.128 | 0.003 | 1.80858328769661e-39 | 44 |
| akap9 | 8.91539882190773e-44 | 0.408285522 | 0.213 | 0.009 | 2.24391672948596e-39 | 44 |
| sqstm12 | 9.43889856660677e-44 | 0.971048751 | 0.617 | 0.076 | 2.37567638022926e-39 | 44 |
| prkcdb | 1.24520986457972e-43 | 0.481554706 | 0.255 | 0.013 | 3.13406870816069e-39 | 44 |
| nt5c2b2 | 1.28106870035142e-43 | 0.778946679 | 0.468 | 0.045 | 3.22432181191448e-39 | 44 |
| si:ch211-39i22.16 | 1.83020311198099e-43 | 0.952839544 | 0.596 | 0.071 | 4.60643821254495e-39 | 44 |
| dlx3b3 | 2.01640844838441e-43 | 0.426137399 | 0.213 | 0.009 | 5.07509842373872e-39 | 44 |
| fam83hb | 2.5813288053603e-43 | 0.320942447 | 0.191 | 0.008 | 6.49694647021133e-39 | 44 |
| s100v1 | 2.91502697125616e-43 | 0.524675253 | 0.255 | 0.013 | 7.33683138395462e-39 | 44 |
| pnp5a7 | 3.24591498138956e-43 | 1.000251457 | 0.617 | 0.077 | 8.16964341665938e-39 | 44 |
| stk10 | 6.67140314081655e-43 | 0.70954292 | 0.489 | 0.047 | 1.67912545651212e-38 | 44 |
| ier2b16 | 9.6076938508933e-43 | 1.439585118 | 0.915 | 0.223 | 2.41816046533134e-38 | 44 |
| si:busm1-57f23.12 | 6.30382140413083e-42 | 0.640426561 | 0.383 | 0.03 | 1.58660880920569e-37 | 44 |
| zgc:154054 | 6.9309165205429e-42 | 0.287180978 | 0.191 | 0.008 | 1.74444237905544e-37 | 44 |
| trib35 | 8.44976098240122e-42 | 1.043072406 | 0.702 | 0.105 | 2.12672034166056e-37 | 44 |
| gprc5c | 8.81952542489281e-42 | 0.285199802 | 0.128 | 0.003 | 2.21978635419127e-37 | 44 |
| fosl1a8 | 1.40638462218904e-41 | 1.024398909 | 0.66 | 0.091 | 3.5397294555876e-37 | 44 |
| zgc:1532846 | 1.6410992808893e-41 | 1.106070058 | 0.702 | 0.1 | 4.13048278007029e-37 | 44 |

| | | | | | | |
|---|---|---|---|---|---|---|
| scpp73 | 1.73598933592304e-41 | 0.565277565 | 0.362 | 0.027 | 4.36931155958469e-37 | 44 |
| jdp2b10 | 2.27115081375713e-41 | 1.472794068 | 0.83 | 0.17 | 5.71625948314531e-37 | 44 |
| b3gnt5a2 | 3.14539964824733e-41 | 0.580721771 | 0.383 | 0.031 | 7.91665637467372e-37 | 44 |
| prodha1 | 4.16766965210816e-41 | 0.386066856 | 0.255 | 0.014 | 1.0489607747391e-36 | 44 |
| jag1b3 | 4.47980466590065e-41 | 0.559627929 | 0.34 | 0.025 | 1.12752203636054e-36 | 44 |
| tmsb19 | 5.52622092550227e-41 | 1.409900101 | 0.809 | 0.155 | 1.39089454473967e-36 | 44 |
| BX664618.1 | 1.13665779117334e-40 | 0.307543534 | 0.17 | 0.006 | 2.86085399460417e-36 | 44 |
| sorl1 | 1.24806058342834e-40 | 0.522845612 | 0.319 | 0.022 | 3.1412436824308e-36 | 44 |
| zgc:153759 | 2.23537320912669e-40 | 0.32916335 | 0.17 | 0.006 | 5.62621083005097e-36 | 44 |
| slc38a5a | 3.58573285608151e-40 | 0.529887778 | 0.277 | 0.017 | 9.02493102547154e-36 | 44 |
| si:dkey-105i14.1 | 4.36856176703373e-40 | 0.304199338 | 0.149 | 0.005 | 1.09952331114472e-35 | 44 |
| icn12 | 6.71281853433287e-40 | 1.495337786 | 1 | 0.274 | 1.68954929690624e-35 | 44 |
| anxa2a8 | 7.99461690390598e-40 | 1.064499165 | 0.702 | 0.105 | 2.0121651285441e-35 | 44 |
| bnipl1 | 2.01155633097185e-39 | 0.322775134 | 0.17 | 0.007 | 5.06288612942305e-35 | 44 |
| efnb2a1 | 7.61132986056241e-39 | 0.401186614 | 0.213 | 0.01 | 1.91569561260495e-34 | 44 |
| si:ch211-202f5.35 | 9.97656114451585e-39 | 0.964091622 | 0.66 | 0.09 | 2.51100067446319e-34 | 44 |
| chac19 | 1.17044593812061e-38 | 1.423630083 | 1 | 0.398 | 2.94589538165577e-34 | 44 |
| b2m16 | 3.02312059330094e-38 | 1.37880647 | 0.957 | 0.292 | 7.60889222127913e-34 | 44 |
| ezrb | 3.86329302558261e-38 | 0.863267419 | 0.574 | 0.073 | 9.72352221608887e-34 | 44 |
| vsig10 | 2.17181367932005e-37 | 0.505361772 | 0.319 | 0.024 | 5.46623784948063e-33 | 44 |
| spns21 | 2.33284241204632e-37 | 0.369373489 | 0.17 | 0.007 | 5.87153106687938e-33 | 44 |
| klc1b | 2.0981360730265e-36 | 0.384853619 | 0.255 | 0.016 | 5.28079868220041e-32 | 44 |
| fuca1.1 | 2.20816069556209e-36 | 0.667221695 | 0.404 | 0.039 | 5.55771965466022e-32 | 44 |
| si:ch211-207d6.22 | 2.43429187597107e-36 | 0.379674136 | 0.234 | 0.013 | 6.12686922263158e-32 | 44 |
| ckap44 | 6.85206454575767e-36 | 0.593719173 | 0.426 | 0.043 | 1.72459612552175e-31 | 44 |
| sh3d21 | 7.11163730951526e-36 | 0.649248236 | 0.404 | 0.039 | 1.7899279944319e-31 | 44 |
| cd995 | 1.16219973171509e-35 | 0.730803541 | 0.468 | 0.053 | 2.92514050475371e-31 | 44 |
| selenow17 | 1.60314850315772e-35 | 1.26496815 | 0.979 | 0.385 | 4.03496446759766e-31 | 44 |
| uba7 | 1.66674648074575e-35 | 0.298048828 | 0.17 | 0.007 | 4.19503421738898e-31 | 44 |
| zbtb16a4 | 2.34367382747939e-35 | 0.68016774 | 0.447 | 0.048 | 5.89879265638288e-31 | 44 |
| oclna3 | 2.55662830939979e-35 | 0.844922732 | 0.596 | 0.078 | 6.43477779192833e-31 | 44 |
| chmp2bb | 4.96277270823062e-35 | 0.689838931 | 0.511 | 0.062 | 1.24908026293456e-30 | 44 |
| erbb21 | 8.16110817251432e-35 | 0.610755021 | 0.34 | 0.03 | 2.05406931594013e-30 | 44 |
| mgst1.15 | 1.06806938609466e-34 | 0.630714543 | 0.404 | 0.041 | 2.68822383786165e-30 | 44 |
| si:ch73-22o12.11 | 1.35281488125482e-34 | 0.454344384 | 0.319 | 0.026 | 3.40489977463025e-30 | 44 |

| | | | | | | |
|---|---|---|---|---|---|---|
| sipa1l3 | 2.39771170142985e-34 | 0.433692433 | 0.255 | 0.017 | 6.03480058132878e-30 | 44 |
| nfkbiaa8 | 2.85762559549381e-34 | 1.199569885 | 0.766 | 0.152 | 7.19235786129836e-30 | 44 |
| cishb | 2.99838870094461e-34 | 0.319291505 | 0.17 | 0.008 | 7.54664452140749e-30 | 44 |
| epcam16 | 4.82160824003463e-34 | 1.140037971 | 1 | 0.507 | 1.21355057793432e-29 | 44 |
| si:ch1073-429i10.3.1 | 4.92796013074709e-34 | 1.086155856 | 0.872 | 0.201 | 1.24031828530773e-29 | 44 |
| si:dkey-79d12.58 | 5.67621737475705e-34 | 0.812657729 | 0.553 | 0.076 | 1.4286471510526e-29 | 44 |
| si:ch73-335l21.11 | 5.81448495388422e-34 | 0.455482327 | 0.234 | 0.015 | 1.46344771804312e-29 | 44 |
| zgc:1533176 | 5.90461027724394e-34 | 0.629904858 | 0.404 | 0.042 | 1.48613136067953e-29 | 44 |
| rpz4 | 5.99674923509149e-34 | 0.426103715 | 0.298 | 0.023 | 1.50932181498018e-29 | 44 |
| nrg1 | 3.91212048620386e-33 | 0.435062283 | 0.277 | 0.02 | 9.84641605172649e-29 | 44 |
| b2ml16 | 4.05947988671809e-33 | 1.209888939 | 0.915 | 0.267 | 1.02173049268808e-28 | 44 |
| ier57 | 5.53384660381571e-33 | 1.039347669 | 0.702 | 0.136 | 1.39281385171437e-28 | 44 |
| cx30.9 | 7.14784604708804e-33 | 0.281906107 | 0.149 | 0.006 | 1.79904137159159e-28 | 44 |
| CABZ01029938.1 | 1.34926064660898e-32 | 0.547277416 | 0.362 | 0.035 | 3.39595412145014e-28 | 44 |
| rhoca2 | 1.68239324050619e-32 | 0.69430185 | 0.468 | 0.057 | 4.23441554703002e-28 | 44 |
| myo6a | 1.69721804031489e-32 | 0.278099705 | 0.17 | 0.008 | 4.27172808566855e-28 | 44 |
| reep3b | 1.90533735385971e-32 | 0.699285536 | 0.468 | 0.057 | 4.79554358592951e-28 | 44 |
| tjp3 | 2.66354266156057e-32 | 0.462234577 | 0.298 | 0.024 | 6.70387052488179e-28 | 44 |
| zgc:923801 | 2.82326338188521e-32 | 0.343458662 | 0.191 | 0.01 | 7.10587160586689e-28 | 44 |
| relt | 2.88701379954254e-32 | 0.310615598 | 0.149 | 0.006 | 7.26632503206863e-28 | 44 |
| sh3d19 | 3.82841157857008e-32 | 0.403254891 | 0.234 | 0.015 | 9.63572910210302e-28 | 44 |
| CR847566.15 | 4.65207997991669e-32 | 0.573913874 | 0.362 | 0.035 | 1.17088201014523e-27 | 44 |
| tiparp2 | 5.32022507488852e-32 | 0.650479647 | 0.404 | 0.044 | 1.33904744909869e-27 | 44 |
| b4galnt3b | 5.49780387709188e-32 | 0.271847623 | 0.17 | 0.008 | 1.38374225782526e-27 | 44 |
| gpatch81 | 6.12605197399122e-32 | 0.676264809 | 0.447 | 0.054 | 1.54186602133385e-27 | 44 |
| s100a10b13 | 7.64614309040801e-32 | 1.240492697 | 0.957 | 0.255 | 1.92445775442479e-27 | 44 |
| ddit315 | 1.489494964524e-31 | 1.176099212 | 0.766 | 0.176 | 3.74890987621046e-27 | 44 |
| adipor23 | 2.68815505033281e-31 | 0.804509629 | 0.574 | 0.086 | 6.76581744618264e-27 | 44 |
| s100u3 | 4.01861948659266e-31 | 0.479540809 | 0.362 | 0.036 | 1.01144633858051e-26 | 44 |
| si:ch73-335l21.49 | 5.41701727736743e-31 | 1.236092312 | 0.723 | 0.148 | 1.36340907854061e-26 | 44 |
| tdh2 | 5.74974729059857e-31 | 0.843397529 | 0.596 | 0.095 | 1.44715389557076e-26 | 44 |
| acaca | 1.08161639785442e-30 | 0.32661589 | 0.191 | 0.011 | 2.72232031175979e-26 | 44 |
| gadd45aa1 | 1.39852454064215e-30 | 0.415884224 | 0.255 | 0.019 | 3.51994641634223e-26 | 44 |
| ccdc50 | 1.40649556155432e-30 | 0.527013319 | 0.319 | 0.029 | 3.54000867887606e-26 | 44 |
| cldn7b13 | 1.6354072950378e-30 | 1.064383664 | 0.787 | 0.16 | 4.11615662088063e-26 | 44 |

| | | | | | | |
|---|---|---|---|---|---|---|
| tmem594 | 3.4535452559892e-30 | 0.844727761 | 0.617 | 0.106 | 8.69222805479921e-26 | 44 |
| ubb4 | 3.71015208396866e-30 | 1.24577442 | 0.915 | 0.316 | 9.33808178014072e-26 | 44 |
| mhc1zba | 3.83044051064631e-30 | 0.284385717 | 0.149 | 0.007 | 9.64083572124571e-26 | 44 |
| rab11fip1a | 5.31904092540813e-30 | 0.266317604 | 0.106 | 0.003 | 1.33874941051597e-25 | 44 |
| dhrs13a.11 | 6.10923150605225e-30 | 0.436326333 | 0.277 | 0.023 | 1.53763247775829e-25 | 44 |
| znfx11 | 8.69469105945582e-30 | 0.478526813 | 0.319 | 0.03 | 2.18836679275444e-25 | 44 |
| cavin2b9 | 9.49043004526218e-30 | 0.664418439 | 0.489 | 0.066 | 2.38864633809204e-25 | 44 |
| rnf128a1 | 1.07459014785661e-29 | 0.662695892 | 0.511 | 0.068 | 2.7046359431403e-25 | 44 |
| porb1 | 1.24659680600898e-29 | 0.614897659 | 0.383 | 0.043 | 3.137559501044e-25 | 44 |
| etnk11 | 1.91059563242249e-29 | 0.570596045 | 0.34 | 0.034 | 4.80877814724416e-25 | 44 |
| cd74b13 | 3.99296754194344e-29 | 1.17519772 | 0.894 | 0.223 | 1.00499000063174e-24 | 44 |
| plce1 | 5.86823541051645e-29 | 0.325321093 | 0.17 | 0.009 | 1.47697617047288e-24 | 44 |
| slc20a1b5 | 6.62939126549369e-29 | 0.853681434 | 0.596 | 0.102 | 1.66855148761211e-24 | 44 |
| daam1a1 | 8.45790797433477e-29 | 0.456504291 | 0.319 | 0.031 | 2.12877085806032e-24 | 44 |
| zgc:92107 | 1.16804995131355e-28 | 0.378746063 | 0.213 | 0.014 | 2.93986492246108e-24 | 44 |
| fxyd19 | 1.25453148024739e-28 | 0.720940549 | 0.532 | 0.081 | 3.15753028263466e-24 | 44 |
| cd74a11 | 2.81434597123202e-28 | 1.063508412 | 0.915 | 0.219 | 7.08342737499388e-24 | 44 |
| CR318588.36 | 2.97662302358862e-28 | 0.861416531 | 0.383 | 0.045 | 7.49186248807021e-24 | 44 |
| tjp1b | 3.61217427540337e-28 | 0.509592965 | 0.298 | 0.028 | 9.09148143376275e-24 | 44 |
| jun12 | 4.86652261677408e-28 | 1.20724069 | 0.851 | 0.228 | 1.22485507741587e-23 | 44 |
| evi5b | 5.85157912890287e-28 | 0.540182812 | 0.298 | 0.028 | 1.47278395095356e-23 | 44 |
| tax1bp34 | 5.92745900091152e-28 | 0.744507726 | 0.553 | 0.089 | 1.49188215593942e-23 | 44 |
| si:busm1-266f07.29 | 8.36657841592061e-28 | 0.970597882 | 0.745 | 0.146 | 2.10578412150306e-23 | 44 |
| itsn2b | 1.18026074543588e-27 | 0.472661382 | 0.34 | 0.036 | 2.97059827018757e-23 | 44 |
| si:ch1073-345a8.1 | 1.50763853794031e-27 | 0.270810739 | 0.191 | 0.012 | 3.79457543614196e-23 | 44 |
| arid3a | 1.61578525276108e-27 | 0.280722482 | 0.17 | 0.01 | 4.06676990267435e-23 | 44 |
| abca3b | 2.86677930265036e-27 | 0.361910881 | 0.234 | 0.018 | 7.21539682684069e-23 | 44 |
| tuba8l25 | 4.71932042579871e-27 | 0.63127033 | 0.447 | 0.061 | 1.18780575796928e-22 | 44 |
| btbd10b | 5.5916020606218e-27 | 0.677137017 | 0.489 | 0.071 | 1.4073503226379e-22 | 44 |
| esyt2a | 1.18989827980426e-26 | 0.314679715 | 0.191 | 0.012 | 2.99485498043934e-22 | 44 |
| itprid2 | 1.53777343613234e-26 | 0.349863595 | 0.213 | 0.015 | 3.87042196140149e-22 | 44 |
| traf4a | 1.56598881427932e-26 | 0.268738598 | 0.149 | 0.008 | 3.94143724665963e-22 | 44 |
| chmp5b1 | 2.4421436287492e-26 | 0.771684887 | 0.617 | 0.114 | 6.14663129919887e-22 | 44 |
| fgd1 | 3.15761436771354e-26 | 0.309770407 | 0.191 | 0.013 | 7.9473996020982e-22 | 44 |
| csrp2 | 5.03649991978175e-26 | 0.315570112 | 0.213 | 0.015 | 1.26763666480987e-21 | 44 |

| | | | | | | |
|---|---|---|---|---|---|---|
| gnb1a8 | 5.79471521370408e-26 | 0.95976883 | 0.766 | 0.192 | 1.45847187213718e-21 | 44 |
| cbx3b | 1.18280119076973e-25 | 0.50543975 | 0.319 | 0.034 | 2.97699231704832e-21 | 44 |
| pim111 | 1.21471424161758e-25 | 0.943553785 | 0.745 | 0.171 | 3.0573142747273e-21 | 44 |
| mcl1b13 | 1.26846799338552e-25 | 1.035457858 | 0.851 | 0.242 | 3.19260709255203e-21 | 44 |
| dcun1d2b | 1.94558198193105e-25 | 0.340348515 | 0.213 | 0.016 | 4.89683529032227e-21 | 44 |
| gipc21 | 3.88314123686811e-25 | 0.479987382 | 0.319 | 0.034 | 9.77347817907333e-21 | 44 |
| aste1a1 | 4.03224855619977e-25 | 0.338637518 | 0.255 | 0.023 | 1.01487663910992e-20 | 44 |
| XLOC-018182 | 6.59681828150105e-25 | 0.392862938 | 0.191 | 0.013 | 1.660353193271e-20 | 44 |
| fosab13 | 1.0160956196697e-24 | 1.190511212 | 0.957 | 0.349 | 2.55741106514667e-20 | 44 |
| atf4b14 | 1.07677785618547e-24 | 0.988808525 | 0.851 | 0.264 | 2.71014218623322e-20 | 44 |
| jund10 | 1.4504178636587e-24 | 0.741833476 | 0.532 | 0.093 | 3.65055672104257e-20 | 44 |
| chmp4bb1 | 1.6670710452304e-24 | 0.634043048 | 0.511 | 0.083 | 4.1958511137404e-20 | 44 |
| esrp2 | 2.64536553841663e-24 | 0.259425855 | 0.149 | 0.008 | 6.65812052364081e-20 | 44 |
| zmp:0000001114 | 2.70443002774761e-24 | 0.312263113 | 0.17 | 0.011 | 6.80677993683797e-20 | 44 |
| zdhhc16a | 2.75131989535313e-24 | 0.409281952 | 0.255 | 0.023 | 6.92479704461429e-20 | 44 |
| nfe2l2a5 | 5.73106354330894e-24 | 0.622664566 | 0.447 | 0.069 | 1.44245138321543e-19 | 44 |
| zmp:00000009911 | 6.44860809256209e-24 | 0.344763761 | 0.191 | 0.014 | 1.62305017081695e-19 | 44 |
| efna1a2 | 6.6867173902395e-24 | 0.265213197 | 0.191 | 0.014 | 1.68297989994938e-19 | 44 |
| arl4d3 | 7.29444255441674e-24 | 0.533374325 | 0.383 | 0.05 | 1.83593824652115e-19 | 44 |
| fam129ba | 1.15981159273669e-23 | 0.430876773 | 0.277 | 0.028 | 2.91912979775898e-19 | 44 |
| f11r.17 | 1.59866085648917e-23 | 0.962464625 | 0.745 | 0.178 | 4.02366950969759e-19 | 44 |
| zgc:113333 | 1.87603473429327e-23 | 0.273436814 | 0.149 | 0.009 | 4.72179182274274e-19 | 44 |
| crybg1a1 | 5.63170410756035e-23 | 0.337329145 | 0.213 | 0.018 | 1.41744360683187e-18 | 44 |
| mcl1a13 | 6.87647608876147e-23 | 0.811389903 | 0.957 | 0.624 | 1.73074026678038e-18 | 44 |
| slc10a3 | 8.98477226091169e-23 | 0.339128078 | 0.213 | 0.018 | 2.26137733034886e-18 | 44 |
| plcb3 | 9.60585715612612e-23 | 0.369991576 | 0.234 | 0.021 | 2.41769818762538e-18 | 44 |
| pvrl2l | 9.87468523950421e-23 | 0.270825264 | 0.17 | 0.012 | 2.48535952793081e-18 | 44 |
| stk354 | 1.49193644578369e-22 | 0.625847523 | 0.383 | 0.055 | 3.75505484039298e-18 | 44 |
| baiap2a2 | 1.63384675011085e-22 | 0.424364771 | 0.319 | 0.037 | 4.11222888535401e-18 | 44 |
| lipia | 2.20237884545529e-22 | 0.361379025 | 0.234 | 0.021 | 5.54316731612643e-18 | 44 |
| uacab | 2.58620543294249e-22 | 0.283712276 | 0.149 | 0.009 | 6.50922045417294e-18 | 44 |
| st14a1 | 3.28562263806463e-22 | 0.418721801 | 0.319 | 0.038 | 8.26958361774486e-18 | 44 |
| gmds | 8.25588635955972e-22 | 0.496324091 | 0.362 | 0.049 | 2.07792403783758e-17 | 44 |
| zyg11 | 8.79005719171618e-22 | 0.422961264 | 0.277 | 0.03 | 2.21236949458304e-17 | 44 |
| mpc11 | 8.83497066688843e-22 | 0.627836998 | 0.447 | 0.075 | 2.22367376714915e-17 | 44 |

| | | | | | | |
|---|---|---|---|---|---|---|
| ankrd2212 | 1.20738757498748e-21 | 0.772012607 | 0.681 | 0.143 | 3.03887378748599e-17 | 44 |
| smox7 | 1.53978752174634e-21 | 0.592968056 | 0.489 | 0.085 | 3.87549121348336e-17 | 44 |
| slc37a1 | 1.80165765903189e-21 | 0.326499532 | 0.191 | 0.015 | 4.53459216201737e-17 | 44 |
| bada | 2.07653433064179e-21 | 0.474128411 | 0.277 | 0.031 | 5.22642925679233e-17 | 44 |
| crema11 | 2.99276091229693e-21 | 0.865449952 | 0.702 | 0.178 | 7.53247994016015e-17 | 44 |
| tmem79b | 4.12346461805646e-21 | 0.332588613 | 0.17 | 0.012 | 1.03783480971863e-16 | 44 |
| afdna4 | 4.48295945760797e-21 | 0.323034777 | 0.234 | 0.023 | 1.12831606588535e-16 | 44 |
| zgc:1748635 | 5.10830842704695e-21 | 0.750153627 | 0.511 | 0.102 | 1.28571014800345e-16 | 44 |
| cx30.3 | 5.40198263973283e-21 | 0.415249919 | 0.234 | 0.023 | 1.35962501059436e-16 | 44 |
| kitlga2 | 5.63462940931469e-21 | 0.31087006 | 0.191 | 0.016 | 1.41817987603041e-16 | 44 |
| capns1a5 | 6.17317729229523e-21 | 0.821022485 | 0.617 | 0.143 | 1.55372699269779e-16 | 44 |
| lsr1 | 6.75555516421883e-21 | 0.355408287 | 0.255 | 0.027 | 1.70030567928224e-16 | 44 |
| rhoab12 | 7.01324497011238e-21 | 0.819542655 | 0.723 | 0.189 | 1.76516362652759e-16 | 44 |
| lgals3b7 | 7.49829285475184e-21 | 0.617364667 | 0.447 | 0.079 | 1.88724532861249e-16 | 44 |
| higd1a12 | 9.01343740130849e-21 | 0.947084501 | 0.83 | 0.31 | 2.26859205953533e-16 | 44 |
| ftr824 | 1.4495247338443e-20 | 0.324041492 | 0.234 | 0.023 | 3.64830880261272e-16 | 44 |
| ifi459 | 1.99638253725036e-20 | 0.678670252 | 0.617 | 0.129 | 5.02469520800543e-16 | 44 |
| micu2 | 2.32575172361441e-20 | 0.476134242 | 0.277 | 0.033 | 5.85368451316511e-16 | 44 |
| si:ch73-103b11.2 | 3.12506251676385e-20 | 0.316165384 | 0.213 | 0.02 | 7.86546984844294e-16 | 44 |
| mhc1uba | 4.20352855185246e-20 | 0.30519713 | 0.213 | 0.02 | 1.05798610121575e-15 | 44 |
| CABZ01021599.1 | 4.27267760616539e-20 | 0.283263906 | 0.17 | 0.013 | 1.07539022669577e-15 | 44 |
| zgc:1527916 | 4.58249595993606e-20 | 0.526184829 | 0.426 | 0.069 | 1.15336840815631e-15 | 44 |
| ciartb | 5.36911684568966e-20 | 0.460630386 | 0.319 | 0.043 | 1.35135301889163e-15 | 44 |
| krt156 | 5.70910726127534e-20 | 1.066433839 | 0.447 | 0.085 | 1.43692520659039e-15 | 44 |
| tjp1a5 | 6.1123175227961e-20 | 0.467520522 | 0.34 | 0.047 | 1.53840919731255e-15 | 44 |
| btg212 | 6.36930079711277e-20 | 0.943362432 | 0.894 | 0.373 | 1.60308931762531e-15 | 44 |
| kalrnb1 | 6.4382443294359e-20 | 0.253736743 | 0.149 | 0.01 | 1.62044171527572e-15 | 44 |
| rasgef1ba3 | 1.26762254391655e-19 | 0.579027888 | 0.404 | 0.066 | 3.19047918078356e-15 | 44 |
| si:ch211-286b5.58 | 1.27398510165124e-19 | 0.737617286 | 0.596 | 0.137 | 3.206493102346e-15 | 44 |
| slc25a55a3 | 1.32166203871095e-19 | 0.536530089 | 0.404 | 0.067 | 3.32649118523159e-15 | 44 |
| cds11 | 1.3614514458646e-19 | 0.275353225 | 0.128 | 0.008 | 3.42663714409662e-15 | 44 |
| jmjd1cb1 | 1.43535729350746e-19 | 0.492238302 | 0.319 | 0.044 | 3.61265077202894e-15 | 44 |
| tnfrsfa1 | 1.63936120970021e-19 | 0.369727876 | 0.255 | 0.029 | 4.12610822869446e-15 | 44 |
| ucp214 | 1.76454248766752e-19 | 0.886635372 | 0.936 | 0.417 | 4.44117698721037e-15 | 44 |
| pls3 | 2.38982816505927e-19 | 0.28033306 | 0.149 | 0.011 | 6.01495850863767e-15 | 44 |

| | | | | | | |
|---|---|---|---|---|---|---|
| baiap2l1a | 2.45042019418515e-19 | 0.353708914 | 0.319 | 0.042 | 6.16746258674462e-15 | 44 |
| si:ch73-138n13.13 | 3.02602939415712e-19 | 0.407620037 | 0.298 | 0.039 | 7.61621338215405e-15 | 44 |
| vamp82 | 3.3736403760681e-19 | 0.62493383 | 0.426 | 0.074 | 8.4911154625258e-15 | 44 |
| ccdc1711 | 4.56506879575437e-19 | 0.338997924 | 0.213 | 0.021 | 1.14898216520342e-14 | 44 |
| mboat2a2 | 4.86127815999768e-19 | 0.389984397 | 0.234 | 0.025 | 1.22353510008982e-14 | 44 |
| rab5ab | 5.01395019342792e-19 | 0.585316586 | 0.447 | 0.082 | 1.26196112418387e-14 | 44 |
| gabarapl21 | 5.29464499429622e-19 | 0.524345351 | 0.426 | 0.074 | 1.33260919861442e-14 | 44 |
| arhgef42 | 5.76469956957985e-19 | 0.441343801 | 0.277 | 0.035 | 1.45091723466755e-14 | 44 |
| si:ch211-191a24.4 | 5.89673988531677e-19 | 0.502924617 | 0.34 | 0.049 | 1.48415046173538e-14 | 44 |
| sh3yl1 | 6.9225101823798e-19 | 0.429555902 | 0.277 | 0.034 | 1.74232658780317e-14 | 44 |
| rsrp12 | 6.95617777330626e-19 | 0.696430251 | 0.553 | 0.12 | 1.75080038376345e-14 | 44 |
| plecb5 | 7.14724013850486e-19 | 0.496937791 | 0.34 | 0.051 | 1.79888887046029e-14 | 44 |
| irf2bp2a9 | 7.59546783532977e-19 | 0.814092989 | 0.66 | 0.176 | 1.91170329947415e-14 | 44 |
| cd82b4 | 9.17924568859946e-19 | 0.570225568 | 0.426 | 0.075 | 2.3103243473636e-14 | 44 |
| atp1a1b | 9.82538612428056e-19 | 0.318861851 | 0.17 | 0.014 | 2.47295143362017e-14 | 44 |
| ampd3b | 1.13781950221078e-18 | 0.311290813 | 0.17 | 0.014 | 2.86377790511432e-14 | 44 |
| reep57 | 1.30082741385669e-18 | 0.7777957 | 0.617 | 0.159 | 3.27405251793591e-14 | 44 |
| rap2b1 | 1.66657282755859e-18 | 0.487095236 | 0.34 | 0.051 | 4.19459714968221e-14 | 44 |
| arg24 | 2.22045285583737e-18 | 0.319240315 | 0.213 | 0.022 | 5.58865779285709e-14 | 44 |
| junba12 | 2.64294569754926e-18 | 0.966397286 | 0.872 | 0.309 | 6.65203002616172e-14 | 44 |
| taok1a | 3.53571569971594e-18 | 0.335640711 | 0.191 | 0.018 | 8.89904284461504e-14 | 44 |
| calm2b9 | 3.84899596242971e-18 | 0.823952794 | 0.723 | 0.211 | 9.68753793783934e-14 | 44 |
| dusp61 | 4.49366968236529e-18 | 0.516524585 | 0.319 | 0.047 | 1.13101172235452e-13 | 44 |
| si:ch1073-443f11.25 | 5.33314973918778e-18 | 0.395857338 | 0.277 | 0.036 | 1.34230045785617e-13 | 44 |
| ms4a17a.104 | 6.17408467055322e-18 | 0.573295592 | 0.34 | 0.052 | 1.55395537073154e-13 | 44 |
| cebpa7 | 6.72917527304734e-18 | 0.491631081 | 0.362 | 0.058 | 1.69366612447328e-13 | 44 |
| pycard9 | 7.58690848868371e-18 | 0.580061216 | 0.404 | 0.072 | 1.9095489975168e-13 | 44 |
| abcc32 | 8.46777373500179e-18 | 0.409012066 | 0.213 | 0.023 | 2.1312539713626e-13 | 44 |
| sdcbp211 | 1.02708551513627e-17 | 0.799482591 | 0.702 | 0.209 | 2.58507153304647e-13 | 44 |
| strbp2 | 1.22636377248075e-17 | 0.300488092 | 0.191 | 0.019 | 3.08663497895679e-13 | 44 |
| fkbp58 | 1.229293045503e-17 | 0.693295466 | 0.532 | 0.123 | 3.09400766622651e-13 | 44 |
| per21 | 1.25875104133239e-17 | 0.456294419 | 0.277 | 0.037 | 3.1681504959295e-13 | 44 |
| mt-co121 | 1.40419048177979e-17 | 0.699136777 | 1 | 0.658 | 3.53420702359156e-13 | 44 |
| si:ch211-107o10.3 | 1.45482587982804e-17 | 0.4238449 | 0.277 | 0.037 | 3.66165125693919e-13 | 44 |
| plekha62 | 1.54365111200402e-17 | 0.253849781 | 0.191 | 0.019 | 3.88521548380291e-13 | 44 |

| | | | | | | |
|---|---|---|---|---|---|---|
| fam199x | 1.59041343366432e-17 | 0.309274168 | 0.191 | 0.019 | 4.00291157118973e-13 | 44 |
| irs2b1 | 1.81542864175681e-17 | 0.314422852 | 0.191 | 0.019 | 4.56925234843772e-13 | 44 |
| snrkb1 | 2.02510174565219e-17 | 0.261105052 | 0.191 | 0.019 | 5.09697858363201e-13 | 44 |
| zgc:919761 | 2.23527597164951e-17 | 0.275185829 | 0.213 | 0.023 | 5.62596609304465e-13 | 44 |
| CR383676.122 | 2.53366679869385e-17 | 0.801034964 | 1 | 0.727 | 6.37698596563255e-13 | 44 |
| si:dkey-71l4.4 | 2.82754590103495e-17 | 0.255671355 | 0.149 | 0.012 | 7.11665027831488e-13 | 44 |
| rin21 | 3.68953269032484e-17 | 0.27530366 | 0.213 | 0.023 | 9.28618482827859e-13 | 44 |
| prdx512 | 4.06258567062261e-17 | 0.692421008 | 0.617 | 0.159 | 1.02251218743901e-12 | 44 |
| dhrs13l13 | 4.21497509939562e-17 | 0.40924059 | 0.255 | 0.033 | 1.06086708276688e-12 | 44 |
| ovol1a | 4.4153428516367e-17 | 0.270028133 | 0.106 | 0.006 | 1.11129764232844e-12 | 44 |
| zswim51 | 4.8499704335834e-17 | 0.435920641 | 0.298 | 0.043 | 1.22068905842861e-12 | 44 |
| si:ch211-212k18.75 | 5.58583728761694e-17 | 0.711147236 | 0.574 | 0.139 | 1.40589938692031e-12 | 44 |
| mark2b | 5.71976039411012e-17 | 0.326914903 | 0.213 | 0.023 | 1.43960649359358e-12 | 44 |
| zgc:15834310 | 5.86128513938688e-17 | 0.774724586 | 0.638 | 0.166 | 1.47522685673228e-12 | 44 |
| rabac1 | 8.03850419351097e-17 | 0.423629168 | 0.298 | 0.044 | 2.02321112046478e-12 | 44 |
| plpp2b1 | 8.18283888112523e-17 | 0.387208484 | 0.277 | 0.037 | 2.05953871799041e-12 | 44 |
| trim16 | 8.93220498479108e-17 | 0.251137265 | 0.191 | 0.019 | 2.24814667262207e-12 | 44 |
| sox131 | 9.32746957620345e-17 | 0.269709756 | 0.149 | 0.012 | 2.34763081763465e-12 | 44 |
| plk2b1 | 1.11247929274892e-16 | 0.251636643 | 0.149 | 0.012 | 2.79999913191975e-12 | 44 |
| alox5a5 | 1.12917876668671e-16 | 0.469458014 | 0.255 | 0.033 | 2.84203003787378e-12 | 44 |
| skia1 | 1.13554528876705e-16 | 0.414558382 | 0.298 | 0.044 | 2.85805393729779e-12 | 44 |
| atxn1a | 1.32030244882595e-16 | 0.30634969 | 0.213 | 0.024 | 3.32306923345004e-12 | 44 |
| ptbp32 | 1.45626471496807e-16 | 0.492308438 | 0.362 | 0.063 | 3.66527266110314e-12 | 44 |
| fosl214 | 1.91348170204418e-16 | 0.61025903 | 0.532 | 0.122 | 4.816042095875e-12 | 44 |
| fam102ab1 | 2.00679129957677e-16 | 0.420226421 | 0.255 | 0.034 | 5.05089302190476e-12 | 44 |
| zgc:774391 | 2.06500032226156e-16 | 0.268921177 | 0.17 | 0.016 | 5.19739931110012e-12 | 44 |
| adam28 | 2.19219460716153e-16 | 0.276126189 | 0.128 | 0.009 | 5.51753460676485e-12 | 44 |
| arfgap3 | 2.55065305944753e-16 | 0.345011656 | 0.213 | 0.024 | 6.4197386853235e-12 | 44 |
| waslb | 2.76240096454637e-16 | 0.396416762 | 0.255 | 0.033 | 6.95268698766675e-12 | 44 |
| rnf213a7 | 3.94690849473589e-16 | 0.645891175 | 0.426 | 0.089 | 9.93397399040076e-12 | 44 |
| vps4b1 | 4.21990990906864e-16 | 0.557031584 | 0.468 | 0.1 | 1.06210912501349e-11 | 44 |
| LOC1039092848 | 4.85373537665414e-16 | 0.508371967 | 0.362 | 0.065 | 1.22163665695008e-11 | 44 |
| fuca23 | 5.41734774920225e-16 | 0.515738375 | 0.362 | 0.064 | 1.36349225499671e-11 | 44 |
| degs29 | 5.63003883147972e-16 | 0.576407288 | 0.511 | 0.107 | 1.41702447349513e-11 | 44 |
| CU633745.1 | 8.88222544749087e-16 | 0.273366576 | 0.191 | 0.021 | 2.23556732287898e-11 | 44 |

| | | | | | | |
|---|---|---|---|---|---|---|
| tmbim1a | 1.1718781888181e-15 | 0.334327926 | 0.213 | 0.025 | 2.94950021343627e-11 | 44 |
| tmod22 | 1.22086481825836e-15 | 0.480863428 | 0.298 | 0.047 | 3.07279466107446e-11 | 44 |
| abhd2a | 1.3026589711154e-15 | 0.295175834 | 0.234 | 0.03 | 3.27866236440035e-11 | 44 |
| cnn213 | 1.30765799897919e-15 | 0.68902342 | 0.532 | 0.135 | 3.29124441763072e-11 | 44 |
| far13 | 1.61762012695776e-15 | 0.485278957 | 0.255 | 0.036 | 4.07138809753998e-11 | 44 |
| mef2d4 | 1.68012909801796e-15 | 0.522483648 | 0.383 | 0.073 | 4.22871692680141e-11 | 44 |
| si:dkey-177p2.61 | 1.81320334803291e-15 | 0.566832289 | 0.383 | 0.075 | 4.56365150666404e-11 | 44 |
| namptb | 1.89200957712981e-15 | 0.305363976 | 0.17 | 0.017 | 4.76199890467803e-11 | 44 |
| itpkcb6 | 1.95601021072759e-15 | 0.46110635 | 0.404 | 0.078 | 4.92308209938027e-11 | 44 |
| klf2b3 | 2.24657957471453e-15 | 0.328825988 | 0.191 | 0.021 | 5.65441613159901e-11 | 44 |
| si:ch211-213a13.2 | 2.45483652043421e-15 | 0.346704749 | 0.213 | 0.026 | 6.17857803828086e-11 | 44 |
| ist11 | 2.55745399158526e-15 | 0.589317724 | 0.447 | 0.097 | 6.43685595142094e-11 | 44 |
| fgfr34 | 2.57365173451052e-15 | 0.283250505 | 0.17 | 0.017 | 6.47762405058952e-11 | 44 |
| klf8 | 2.82950988322249e-15 | 0.381254795 | 0.234 | 0.031 | 7.12159342508269e-11 | 44 |
| furina1 | 4.17606767384946e-15 | 0.532505362 | 0.34 | 0.062 | 1.05107447283117e-10 | 44 |
| rnasekb | 4.22430351327385e-15 | 0.300039485 | 0.213 | 0.026 | 1.06321495125589e-10 | 44 |
| prelid3b6 | 5.23878731214162e-15 | 0.714089041 | 0.745 | 0.238 | 1.31855037859293e-10 | 44 |
| CU929259.1 | 5.88040389965217e-15 | 0.279574098 | 0.191 | 0.022 | 1.48003885750346e-10 | 44 |
| slc38a5b9 | 6.23785749745095e-15 | 0.590116202 | 0.489 | 0.116 | 1.57000635353343e-10 | 44 |
| atf38 | 7.09143030655676e-15 | 0.638337342 | 0.426 | 0.091 | 1.78484209385727e-10 | 44 |
| COX321 | 7.30590073501687e-15 | 0.639304601 | 1 | 0.701 | 1.8388221559964e-10 | 44 |
| rps6kb1a | 9.55378354965639e-15 | 0.267689396 | 0.191 | 0.022 | 2.40459178161302e-10 | 44 |
| stap2b | 1.00941695876306e-14 | 0.279208939 | 0.149 | 0.014 | 2.54060154351075e-10 | 44 |
| fxyd6l5 | 1.70480037449885e-14 | 0.576772577 | 0.489 | 0.114 | 4.29081206257615e-10 | 44 |
| ehbp1l1a | 1.89406207730369e-14 | 0.265589803 | 0.17 | 0.018 | 4.76716484236567e-10 | 44 |
| csnk1da | 2.47910527693258e-14 | 0.594917412 | 0.489 | 0.117 | 6.23966007151162e-10 | 44 |
| per36 | 2.50350799905049e-14 | 0.560916092 | 0.383 | 0.08 | 6.30107928281017e-10 | 44 |
| map1lc3a9 | 2.79798670229731e-14 | 0.4799514 | 0.362 | 0.071 | 7.0422527310121e-10 | 44 |
| gadd45ba8 | 2.86370970608785e-14 | 0.690127861 | 0.66 | 0.191 | 7.20767095925251e-10 | 44 |
| cebpd9 | 2.87310584431309e-14 | 0.584598453 | 0.532 | 0.13 | 7.23132009955161e-10 | 44 |
| camsap1b | 3.10202163650402e-14 | 0.259925249 | 0.149 | 0.014 | 7.80747825691698e-10 | 44 |
| eif1b12 | 3.24861460930863e-14 | 0.744029638 | 0.872 | 0.383 | 8.1764381101689e-10 | 44 |
| elovl7a13 | 4.03637746642487e-14 | 0.653541164 | 0.681 | 0.183 | 1.01591584452447e-09 | 44 |
| btg118 | 4.92721060956206e-14 | 0.765104142 | 0.851 | 0.333 | 1.24012963832068e-09 | 44 |
| dusp7 | 5.42765396911436e-14 | 0.253816829 | 0.191 | 0.023 | 1.36608622748639e-09 | 44 |

| | | | | | | |
|---|---|---|---|---|---|---|
| dusp111 | 5.71396396088756e-14 | 0.608590331 | 0.596 | 0.16 | 1.43814758931579e-09 | 44 |
| prox1a2 | 7.6742249801556e-14 | 0.267408942 | 0.17 | 0.019 | 1.93152568525536e-09 | 44 |
| fut9b | 9.21394633957385e-14 | 0.31632777 | 0.191 | 0.023 | 2.31905815420734e-09 | 44 |
| foxp47 | 9.30587443266806e-14 | 0.436742716 | 0.298 | 0.052 | 2.34219553595823e-09 | 44 |
| LOC103910015 | 1.05590220637244e-13 | 0.31722777 | 0.17 | 0.019 | 2.65760026321879e-09 | 44 |
| SBNO22 | 1.14619248275424e-13 | 0.343720341 | 0.234 | 0.034 | 2.88485185984414e-09 | 44 |
| zgc:123258 | 1.20307166715498e-13 | 0.394316404 | 0.277 | 0.046 | 3.02801107906236e-09 | 44 |
| zgc:1017151 | 1.2366692091391e-13 | 0.324918626 | 0.191 | 0.024 | 3.1125727324822e-09 | 44 |
| si:ch211-132b12.72 | 1.30427822940015e-13 | 0.404199588 | 0.298 | 0.052 | 3.28273787557723e-09 | 44 |
| ube2d1b2 | 1.35866120522704e-13 | 0.57171 | 0.511 | 0.131 | 3.41961438743593e-09 | 44 |
| tmem176l.3a2 | 1.41185696404925e-13 | 0.360571727 | 0.255 | 0.04 | 3.55350279281555e-09 | 44 |
| slmapa | 3.54129887437033e-13 | 0.265297549 | 0.234 | 0.034 | 8.91309513690268e-09 | 44 |
| uckl1b | 4.17942933519035e-13 | 0.274201144 | 0.17 | 0.02 | 1.05192056937406e-08 | 44 |
| exosc11 | 4.45776433239386e-13 | 0.341729868 | 0.255 | 0.041 | 1.12197470482021e-08 | 44 |
| si:ch211-251p5.5 | 4.8181150874112e-13 | 0.329984397 | 0.213 | 0.029 | 1.21267138635052e-08 | 44 |
| ier2a16 | 5.66438188163125e-13 | 0.767715651 | 0.723 | 0.263 | 1.42566827578777e-08 | 44 |
| egr113 | 5.83298389158724e-13 | 0.706831451 | 0.617 | 0.19 | 1.46810371567359e-08 | 44 |
| ppp2r2d1 | 6.13848526488726e-13 | 0.381942702 | 0.298 | 0.054 | 1.54499535631947e-08 | 44 |
| h3f3c15 | 6.20883982466267e-13 | 0.672965577 | 0.745 | 0.257 | 1.56270289546935e-08 | 44 |
| pcdh111 | 1.01141827034792e-12 | 0.293259808 | 0.149 | 0.016 | 2.54563864463867e-08 | 44 |
| wbp1lb1 | 1.43245702957377e-12 | 0.360478486 | 0.277 | 0.049 | 3.60535109773422e-08 | 44 |
| nr4a37 | 1.65219804347962e-12 | 0.381153227 | 0.298 | 0.055 | 4.15841725563386e-08 | 44 |
| kmt2ca | 1.697403279702e-12 | 0.253388106 | 0.213 | 0.03 | 4.27219431468196e-08 | 44 |
| galnt12 | 1.84542657771963e-12 | 0.329580435 | 0.191 | 0.026 | 4.64475415346254e-08 | 44 |
| actn43 | 2.09738359475061e-12 | 0.488730273 | 0.404 | 0.094 | 5.27890476962781e-08 | 44 |
| egr2a1 | 2.20249614558394e-12 | 0.34471996 | 0.191 | 0.026 | 5.54346254882023e-08 | 44 |
| emd4 | 2.42104370663071e-12 | 0.353118311 | 0.298 | 0.056 | 6.09352490521883e-08 | 44 |
| pdcd6ip1 | 4.04109318037644e-12 | 0.332498198 | 0.277 | 0.049 | 1.01710274256895e-07 | 44 |
| hsd17b12a13 | 4.2289140616772e-12 | 0.627718157 | 0.596 | 0.166 | 1.06437538018354e-07 | 44 |
| selenot2 | 4.4133110165479e-12 | 0.417453718 | 0.298 | 0.058 | 1.11078624975494e-07 | 44 |
| naga4 | 4.57193505380531e-12 | 0.409802681 | 0.277 | 0.051 | 1.15071033369226e-07 | 44 |
| csrnp1b12 | 4.76964126804936e-12 | 0.642367024 | 0.574 | 0.18 | 1.20047101075534e-07 | 44 |
| tax1bp1b | 4.94868677311601e-12 | 0.428245487 | 0.319 | 0.066 | 1.24553497392557e-07 | 44 |
| golgb1 | 5.91808997704988e-12 | 0.341047155 | 0.255 | 0.044 | 1.48952406632369e-07 | 44 |
| CABZ01055347.11 | 5.95303763674539e-12 | 0.350212676 | 0.213 | 0.032 | 1.49832004279245e-07 | 44 |

| | | | | | | |
|---|---|---|---|---|---|---|
| itm2bb14 | 5.96079065148959e-12 | 0.60178737 | 0.681 | 0.222 | 1.50027139907341e-07 | 44 |
| nmi | 6.1124223443062e-12 | 0.42160008 | 0.362 | 0.08 | 1.53843557983843e-07 | 44 |
| pde4ba13 | 7.09624721146549e-12 | 0.54599825 | 0.511 | 0.14 | 1.78605446065375e-07 | 44 |
| wbp2nl4 | 8.00935156968292e-12 | 0.404042553 | 0.383 | 0.087 | 2.01587369657349e-07 | 44 |
| rffl1 | 8.78689798676387e-12 | 0.290108326 | 0.191 | 0.027 | 2.2115743542886e-07 | 44 |
| ar5 | 9.82945701958989e-12 | 0.476801637 | 0.298 | 0.06 | 2.47397603726058e-07 | 44 |
| si:ch211-196l7.418 | 9.9671397019078e-12 | 0.702156359 | 0.745 | 0.315 | 2.50862939157317e-07 | 44 |
| pmp22b7 | 1.0905405404339e-11 | 0.44221211 | 0.255 | 0.046 | 2.74478148621809e-07 | 44 |
| tprg1 | 1.54226922354425e-11 | 0.31616989 | 0.255 | 0.044 | 3.88173740873852e-07 | 44 |
| herpud2 | 1.55612656270237e-11 | 0.350525707 | 0.213 | 0.034 | 3.9166149456656e-07 | 44 |
| si:ch1073-190k2.113 | 1.55682282176145e-11 | 0.627951089 | 0.638 | 0.189 | 3.91836736009138e-07 | 44 |
| pitpnb | 1.64262902424938e-11 | 0.369783852 | 0.255 | 0.046 | 4.13433299113326e-07 | 44 |
| atp2a32 | 1.8997553995957e-11 | 0.286088485 | 0.213 | 0.033 | 4.78149436524242e-07 | 44 |
| cdc42bpb1 | 2.21040855050764e-11 | 0.328073427 | 0.191 | 0.028 | 5.56337728077269e-07 | 44 |
| sult2st12 | 2.37496987303001e-11 | 0.271743947 | 0.191 | 0.028 | 5.97756167342922e-07 | 44 |
| mycbp2 | 2.45362760703251e-11 | 0.396338793 | 0.234 | 0.041 | 6.17553532414013e-07 | 44 |
| chmp1a | 2.51482422247621e-11 | 0.373517847 | 0.277 | 0.053 | 6.32956108555038e-07 | 44 |
| hmgn35 | 2.63423224518428e-11 | 0.447800069 | 0.298 | 0.061 | 6.63009913790431e-07 | 44 |
| npc12 | 3.74476431707189e-11 | 0.276925648 | 0.191 | 0.028 | 9.42519730963824e-07 | 44 |
| clint1a | 3.87042824132885e-11 | 0.378000747 | 0.255 | 0.048 | 9.74148084060059e-07 | 44 |
| tpm39 | 3.98176143597561e-11 | 0.654762004 | 0.681 | 0.255 | 1.0021695358207e-06 | 44 |
| mvp5 | 4.11050858770351e-11 | 0.363962571 | 0.277 | 0.054 | 1.0345739064391e-06 | 44 |
| ptdss2 | 5.10023932024323e-11 | 0.255455327 | 0.128 | 0.014 | 1.28367923451202e-06 | 44 |
| hexb2 | 5.50954078495465e-11 | 0.289936385 | 0.213 | 0.035 | 1.38669632016524e-06 | 44 |
| capns1b9 | 7.02597979191383e-11 | 0.46702196 | 0.447 | 0.122 | 1.76836885382679e-06 | 44 |
| zgc:16273018 | 7.60383782406977e-11 | 0.737784457 | 0.894 | 0.392 | 1.91380994194012e-06 | 44 |
| tnfrsf11a1 | 7.9850468676556e-11 | 0.267936399 | 0.128 | 0.014 | 2.00975644612024e-06 | 44 |
| plpp1a4 | 8.20600166799788e-11 | 0.416467775 | 0.319 | 0.069 | 2.06536855981839e-06 | 44 |
| elf111 | 8.32691723690075e-11 | 0.494976211 | 0.404 | 0.105 | 2.09580179935555e-06 | 44 |
| scaf1 | 8.33368246476844e-11 | 0.358228615 | 0.213 | 0.035 | 2.09750453955757e-06 | 44 |
| gnal | 8.60962341582477e-11 | 0.380390455 | 0.213 | 0.036 | 2.16695611752894e-06 | 44 |
| cflara3 | 9.17353827792745e-11 | 0.412294239 | 0.298 | 0.064 | 2.30888784917156e-06 | 44 |
| CABZ01086029.11 | 1.10056248706896e-10 | 0.266685675 | 0.149 | 0.019 | 2.77000572370386e-06 | 44 |
| usp24 | 1.10929183178825e-10 | 0.261379307 | 0.149 | 0.019 | 2.79197661142784e-06 | 44 |
| klf6a11 | 1.24063747359101e-10 | 0.582423775 | 0.532 | 0.168 | 3.12256045728121e-06 | 44 |

| | | | | | | |
|---|---|---|---|---|---|---|
| rtn32 | 1.31166098334521e-10 | 0.446059972 | 0.34 | 0.079 | 3.30131952898155e-06 | 44 |
| gstt1b7 | 1.73828303174698e-10 | 0.480987355 | 0.319 | 0.073 | 4.37508456260398e-06 | 44 |
| ponzr63 | 2.02445569614914e-10 | 0.303414576 | 0.17 | 0.024 | 5.09535254163776e-06 | 44 |
| prrg25 | 2.47364865252655e-10 | 0.347485863 | 0.383 | 0.09 | 6.22592629354406e-06 | 44 |
| si:ch73-233k15.2 | 2.58497603376721e-10 | 0.301277883 | 0.17 | 0.024 | 6.50612617938869e-06 | 44 |
| nfil3-610 | 2.82049175295571e-10 | 0.582388303 | 0.447 | 0.131 | 7.09889569301423e-06 | 44 |
| taok1b1 | 2.96112730434354e-10 | 0.251934509 | 0.17 | 0.025 | 7.45286131230224e-06 | 44 |
| rbfox23 | 3.47215803334337e-10 | 0.273599944 | 0.213 | 0.037 | 8.73907455412192e-06 | 44 |
| cnn3a8 | 3.83661219841557e-10 | 0.323898519 | 0.277 | 0.057 | 9.65636924219214e-06 | 44 |
| nktr | 4.11776560466659e-10 | 0.298774675 | 0.213 | 0.037 | 1.03640042503853e-05 | 44 |
| mgst2 | 5.18917594235205e-10 | 0.26135613 | 0.149 | 0.02 | 1.30606369293059e-05 | 44 |
| qkia9 | 5.77460274325856e-10 | 0.503322909 | 0.532 | 0.169 | 1.45340976445075e-05 | 44 |
| nfixb5 | 8.25412871173877e-10 | 0.351830572 | 0.298 | 0.066 | 2.07748165545753e-05 | 44 |
| mid1ip1b2 | 1.02342194514321e-09 | 0.390887484 | 0.255 | 0.053 | 2.57585069373093e-05 | 44 |
| atl3 | 1.14987617814463e-09 | 0.401310458 | 0.298 | 0.068 | 2.89412335277223e-05 | 44 |
| nudt9 | 1.17416451878739e-09 | 0.250350773 | 0.17 | 0.026 | 2.95525467733598e-05 | 44 |
| stk40 | 1.18177631050018e-09 | 0.258082748 | 0.191 | 0.032 | 2.97441279589791e-05 | 44 |
| fam126a | 1.36734158826442e-09 | 0.251928489 | 0.213 | 0.038 | 3.44146204350273e-05 | 44 |
| psph | 1.45750072731585e-09 | 0.268890936 | 0.17 | 0.026 | 3.66838358058127e-05 | 44 |
| ccnl1b | 1.61977793769463e-09 | 0.436433251 | 0.383 | 0.102 | 4.07681909138361e-05 | 44 |
| arl5c9 | 1.65785466616652e-09 | 0.449327448 | 0.404 | 0.113 | 4.17265440927451e-05 | 44 |
| atp8a11 | 1.73328650463262e-09 | 0.293660954 | 0.213 | 0.039 | 4.36250880350983e-05 | 44 |
| CABZ01079427.11 | 1.74286103235288e-09 | 0.25195535 | 0.213 | 0.038 | 4.38660693232896e-05 | 44 |
| CR318588.112 | 1.9711157792002e-09 | 0.640086627 | 0.596 | 0.208 | 4.961101304669e-05 | 44 |
| u2surp3 | 2.63190379660513e-09 | 0.393393327 | 0.362 | 0.096 | 6.62423866567544e-05 | 44 |
| tcima8 | 2.63726886520494e-09 | 0.437595127 | 0.298 | 0.069 | 6.63774200683432e-05 | 44 |
| psmb8a1 | 2.80762695305775e-09 | 0.252486665 | 0.17 | 0.027 | 7.06651627815104e-05 | 44 |
| arpc1a1 | 3.02797090354139e-09 | 0.453835689 | 0.468 | 0.142 | 7.62109996712333e-05 | 44 |
| gnpnat1 | 3.06653789703126e-09 | 0.257866205 | 0.191 | 0.033 | 7.71816923303798e-05 | 44 |
| utrn | 3.88848930884075e-09 | 0.261733848 | 0.191 | 0.033 | 9.78693874142128e-05 | 44 |
| alas12 | 4.97135928496827e-09 | 0.412527991 | 0.468 | 0.14 | 0.000125124 | 44 |
| sdf41 | 4.99602825181812e-09 | 0.375076168 | 0.234 | 0.048 | 0.000125745 | 44 |
| ddx3xa | 5.1452613543465e-09 | 0.390246424 | 0.362 | 0.097 | 0.000129501 | 44 |
| csnk1a11 | 6.58422469103381e-09 | 0.533584462 | 0.617 | 0.234 | 0.000165718 | 44 |
| junbb8 | 8.07645270317425e-09 | 0.378988026 | 0.298 | 0.074 | 0.000203276 | 44 |

| | | | | | | |
|---|---|---|---|---|---|---|
| ftr86 | 8.85408868271567e-09 | 0.33386877 | 0.234 | 0.047 | 0.000222849 | 44 |
| stk17al3 | 9.38159047101726e-09 | 0.401216743 | 0.426 | 0.121 | 0.000236125 | 44 |
| mt-co221 | 1.03040371019857e-08 | 0.514091193 | 1 | 0.721 | 0.000259342 | 44 |
| mat2aa1 | 1.05615280999325e-08 | 0.541632384 | 0.532 | 0.184 | 0.000265823 | 44 |
| id2b3 | 1.13576425992097e-08 | 0.284998104 | 0.191 | 0.035 | 0.000285861 | 44 |
| meis1b7 | 1.46039723619002e-08 | 0.329303318 | 0.255 | 0.058 | 0.000367567 | 44 |
| yap16 | 1.49048585757362e-08 | 0.372423749 | 0.234 | 0.051 | 0.00037514 | 44 |
| sec24a | 1.57435290230142e-08 | 0.299461216 | 0.213 | 0.043 | 0.000396249 | 44 |
| rarga5 | 1.65568633111856e-08 | 0.29711163 | 0.255 | 0.057 | 0.00041672 | 44 |
| il154 | 1.68179419903333e-08 | 0.327851182 | 0.255 | 0.057 | 0.000423291 | 44 |
| vaspb2 | 1.81661691215701e-08 | 0.294894362 | 0.234 | 0.05 | 0.000457224 | 44 |
| ak16 | 1.82276188850249e-08 | 0.365001135 | 0.298 | 0.076 | 0.000458771 | 44 |
| egfra3 | 1.86803388089056e-08 | 0.284075694 | 0.17 | 0.029 | 0.000470165 | 44 |
| cmip5 | 1.96381855552672e-08 | 0.2958998 | 0.277 | 0.065 | 0.000494273 | 44 |
| letm1 | 2.04693064200147e-08 | 0.263773286 | 0.17 | 0.03 | 0.000515192 | 44 |
| limch1b1 | 2.50296279246743e-08 | 0.260021702 | 0.128 | 0.018 | 0.000629971 | 44 |
| tsg101b | 2.81958452561977e-08 | 0.256257962 | 0.17 | 0.03 | 0.000709661 | 44 |
| zgc:1008292 | 2.9305770575042e-08 | 0.262380976 | 0.191 | 0.037 | 0.000737597 | 44 |
| lygl15 | 3.12702255205985e-08 | 0.30129691 | 0.362 | 0.098 | 0.00078704 | 44 |
| ca25 | 3.13371991779796e-08 | 0.287571918 | 0.255 | 0.057 | 0.000788726 | 44 |
| zgc:923137 | 3.15478691481757e-08 | 0.448287537 | 0.426 | 0.127 | 0.000794028 | 44 |
| si:ch211-39i2.23 | 3.43225251436526e-08 | 0.343935018 | 0.277 | 0.067 | 0.000863864 | 44 |
| si:ch211-114c12.2 | 3.50371830587312e-08 | 0.304291273 | 0.213 | 0.044 | 0.000881851 | 44 |
| elovl7b12 | 3.55001296158677e-08 | 0.622186243 | 0.617 | 0.25 | 0.000893503 | 44 |
| ppp4cb | 3.55888460621705e-08 | 0.367292917 | 0.404 | 0.12 | 0.000895736 | 44 |
| susd61 | 3.70525041034659e-08 | 0.27716444 | 0.255 | 0.059 | 0.000932574 | 44 |
| gbgt1l411 | 3.80873815594103e-08 | 0.453128256 | 0.489 | 0.151 | 0.000958621 | 44 |
| mt-cyb16 | 3.96277604841445e-08 | 0.530722708 | 0.957 | 0.625 | 0.000997391 | 44 |
| sptssa1 | 4.33446291712543e-08 | 0.448819872 | 0.298 | 0.081 | 0.001090941 | 44 |
| creb3l3l1 | 4.73184354466519e-08 | 0.350804643 | 0.234 | 0.053 | 0.001190958 | 44 |
| zbtb101 | 5.52235924790768e-08 | 0.298729724 | 0.17 | 0.031 | 0.001389923 | 44 |
| tpd52l2a | 6.10538847021317e-08 | 0.271461149 | 0.234 | 0.052 | 0.001536665 | 44 |
| tegt9 | 6.86575896290428e-08 | 0.479632503 | 0.553 | 0.212 | 0.001728043 | 44 |
| ppm1aa | 7.35547575961217e-08 | 0.323936855 | 0.255 | 0.062 | 0.0018513 | 44 |
| kcnk5b1 | 8.4285511613687e-08 | 0.297456777 | 0.213 | 0.045 | 0.002121382 | 44 |

| | | | | | | |
|---|---|---|---|---|---|---|
| ank3b3 | 1.37085873796292e-07 | 0.255947902 | 0.191 | 0.039 | 0.003450314 | 44 |
| cd6317 | 1.48616893931069e-07 | 0.511223025 | 0.787 | 0.341 | 0.003740539 | 44 |
| rab107 | 1.51937677452816e-07 | 0.435206277 | 0.532 | 0.196 | 0.003824119 | 44 |
| ctsk4 | 1.57503142327602e-07 | 0.294418439 | 0.255 | 0.063 | 0.003964197 | 44 |
| notch211 | 1.64915629748314e-07 | 0.375145125 | 0.383 | 0.12 | 0.004150761 | 44 |
| chmp4ba1 | 2.65866725405778e-07 | 0.319794165 | 0.234 | 0.057 | 0.0066916 | 44 |
| CABZ01090890.13 | 2.74060438958202e-07 | 0.252620626 | 0.234 | 0.055 | 0.006897827 | 44 |
| h1f09 | 2.74749705217918e-07 | 0.272843778 | 0.255 | 0.063 | 0.006915175 | 44 |
| ddb21 | 4.45558115061371e-07 | 0.323122173 | 0.298 | 0.084 | 0.011214252 | 44 |
| si:ch211-191j22.32 | 4.47066433680053e-07 | 0.348916359 | 0.362 | 0.115 | 0.011252215 | 44 |
| emp26 | 4.60116626048913e-07 | 0.313306234 | 0.213 | 0.05 | 0.011580675 | 44 |
| NC-002333.420 | 4.63199283193458e-07 | 0.510597153 | 1 | 0.623 | 0.011658263 | 44 |
| sdc410 | 4.78211727437601e-07 | 0.537605865 | 0.596 | 0.258 | 0.012036111 | 44 |
| psme210 | 4.89254755207754e-07 | 0.372149144 | 0.319 | 0.098 | 0.012314053 | 44 |
| mt-nd416 | 5.59383456683315e-07 | 0.519073128 | 0.894 | 0.457 | 0.014079122 | 44 |
| cd838 | 6.13286127029233e-07 | 0.419062513 | 0.34 | 0.105 | 0.015435799 | 44 |
| vps28 | 6.21500130042379e-07 | 0.375509109 | 0.319 | 0.097 | 0.015642537 | 44 |
| stag2b | 6.64756150004068e-07 | 0.252425331 | 0.234 | 0.058 | 0.016731248 | 44 |
| ddx54 | 6.69457478171404e-07 | 0.521344916 | 0.745 | 0.358 | 0.016849575 | 44 |
| brd4 | 7.34659072618223e-07 | 0.325873667 | 0.298 | 0.087 | 0.018490634 | 44 |
| klf2a8 | 7.51077920997863e-07 | 0.377117243 | 0.298 | 0.086 | 0.01890388 | 44 |
| pmaip14 | 7.51313232477149e-07 | 0.361607385 | 0.34 | 0.105 | 0.018909803 | 44 |
| syap1 | 8.29414367898219e-07 | 0.336788671 | 0.255 | 0.069 | 0.02087553 | 44 |
| ubald2 | 9.65407187330034e-07 | 0.276766173 | 0.191 | 0.044 | 0.024298333 | 44 |
| ptp4a2b2 | 1.0210816960154e-06 | 0.399458426 | 0.426 | 0.151 | 0.025699605 | 44 |
| tnrc6c112 | 1.06422416503539e-06 | 0.405546364 | 0.383 | 0.134 | 0.026785458 | 44 |
| coro1ca | 1.26470375821226e-06 | 0.289040999 | 0.255 | 0.069 | 0.031831329 | 44 |
| sar1b2 | 1.28796493798804e-06 | 0.353031595 | 0.277 | 0.081 | 0.03241679 | 44 |
| si:ch211-270g19.53 | 1.30541487909077e-06 | 0.287058707 | 0.128 | 0.022 | 0.032855987 | 44 |
| adipor1b | 1.32887788866228e-06 | 0.258620881 | 0.213 | 0.051 | 0.033446528 | 44 |
| calm1b11 | 1.43181138184407e-06 | 0.31081784 | 0.319 | 0.096 | 0.036037261 | 44 |
| hmgcl | 1.7258990022893e-06 | 0.267172487 | 0.277 | 0.077 | 0.043439152 | 44 |
| ankrd113 | 1.83555391307691e-06 | 0.33599043 | 0.298 | 0.091 | 0.046199056 | 44 |
| dazap2 | 2.51860426972224e-06 | 0.353465959 | 0.298 | 0.093 | 0.063390751 | 44 |
| ets27 | 3.02996905986701e-06 | 0.310957967 | 0.298 | 0.091 | 0.076261291 | 44 |

| | | | | | | |
|---|---|---|---|---|---|---|
| ndel1b | 3.1088852243295e-06 | 0.258025652 | 0.213 | 0.054 | 0.078247532 | 44 |
| ankrd121 | 3.16261930773079e-06 | 0.261094339 | 0.255 | 0.072 | 0.079599965 | 44 |
| sf3b1 | 4.35758190758111e-06 | 0.296086209 | 0.319 | 0.103 | 0.109675979 | 44 |
| txnrd3 | 4.46410424703992e-06 | 0.27740853 | 0.319 | 0.101 | 0.11235704 | 44 |
| kmt2bb2 | 4.50050625458666e-06 | 0.339005294 | 0.277 | 0.087 | 0.113273242 | 44 |
| pef1 | 5.83245147999395e-06 | 0.279142865 | 0.234 | 0.066 | 0.146796971 | 44 |
| birc25 | 6.11901954543098e-06 | 0.354046718 | 0.319 | 0.108 | 0.154009603 | 44 |
| bcap31 | 6.46922772750896e-06 | 0.263144203 | 0.234 | 0.065 | 0.162823993 | 44 |
| slc3a2b10 | 6.75161760107418e-06 | 0.348362677 | 0.34 | 0.119 | 0.169931463 | 44 |
| zgc:1584634 | 6.97375201740211e-06 | 0.259290895 | 0.255 | 0.075 | 0.175522365 | 44 |
| lonrf1l | 7.44114077552308e-06 | 0.266259015 | 0.234 | 0.066 | 0.187286072 | 44 |
| itm2ba15 | 7.54466563001146e-06 | 0.403153046 | 0.617 | 0.274 | 0.189891689 | 44 |
| cdc42l9 | 7.91151576094799e-06 | 0.402559428 | 0.511 | 0.217 | 0.19912494 | 44 |
| sik111 | 8.98209284640664e-06 | 0.329583387 | 0.34 | 0.117 | 0.226070295 | 44 |
| ncor12 | 9.02367678313767e-06 | 0.257948236 | 0.213 | 0.058 | 0.227116921 | 44 |
| smdt1b5 | 1.04859324222953e-05 | 0.348019916 | 0.362 | 0.135 | 0.263920433 | 44 |
| per1b2 | 1.32985461555722e-05 | 0.258262727 | 0.234 | 0.068 | 0.334711108 | 44 |
| cfd6 | 1.70144406940018e-05 | 0.293021392 | 0.234 | 0.069 | 0.428236458 | 44 |
| zgc:163080 | 1.70889062592904e-05 | 0.300509365 | 0.255 | 0.08 | 0.430110682 | 44 |
| pik3r3b1 | 1.77064304982496e-05 | 0.258765795 | 0.191 | 0.051 | 0.445653149 | 44 |
| map2k63 | 2.10585687029082e-05 | 0.272099919 | 0.277 | 0.09 | 0.530023116 | 44 |
| ralbp1 | 2.1732117030245e-05 | 0.261385967 | 0.191 | 0.052 | 0.546975654 | 44 |
| rnf447 | 2.22081333564206e-05 | 0.290843992 | 0.277 | 0.091 | 0.558956508 | 44 |
| dedd17 | 2.69729579465123e-05 | 0.322160554 | 0.383 | 0.147 | 0.678882379 | 44 |
| glo16 | 2.86683268439652e-05 | 0.348281073 | 0.468 | 0.198 | 0.721553118 | 44 |
| si:dkey-112a7.41 | 3.18080557950145e-05 | 0.263910053 | 0.34 | 0.122 | 0.800576956 | 44 |
| plk34 | 3.23550993771829e-05 | 0.30287881 | 0.34 | 0.122 | 0.814345496 | 44 |
| cnot11 | 3.32277413847761e-05 | 0.302605311 | 0.213 | 0.063 | 0.836309023 | 44 |
| dnajb1b | 3.8911855183566e-05 | 0.295235463 | 0.191 | 0.054 | 0.979372483 | 44 |
| gng58 | 4.24534539480485e-05 | 0.358631993 | 0.596 | 0.273 | 1 | 44 |
| si:ch73-1a9.318 | 4.49933924530094e-05 | 0.416199668 | 0.553 | 0.267 | 1 | 44 |
| litaf9 | 5.07366245340342e-05 | 0.253596194 | 0.362 | 0.136 | 1 | 44 |
| h2afx114 | 6.20813882972704e-05 | 0.273510145 | 0.489 | 0.2 | 1 | 44 |
| hbegfa3 | 6.27305874770118e-05 | 0.256174732 | 0.319 | 0.113 | 1 | 44 |
| cish6 | 6.56572666443084e-05 | 0.264725239 | 0.191 | 0.056 | 1 | 44 |

| | | | | | | |
|---|---|---|---|---|---|---|
| brd2a1 | 7.72846175691689e-05 | 0.271059505 | 0.426 | 0.172 | 1 | 44 |
| arhgdig15 | 7.80493039831468e-05 | 0.374372964 | 0.532 | 0.244 | 1 | 44 |
| sptlc2b3 | 7.85174810896579e-05 | 0.259772531 | 0.17 | 0.048 | 1 | 44 |
| nfkbiab8 | 0.000112176 | 0.488471626 | 0.383 | 0.183 | 1 | 44 |
| lrrfip1a5 | 0.000125332 | 0.276292963 | 0.191 | 0.059 | 1 | 44 |
| cdkn1bb6 | 0.000143469 | 0.30416515 | 0.362 | 0.15 | 1 | 44 |
| ifi306 | 0.000146495 | 0.345172479 | 0.404 | 0.176 | 1 | 44 |
| psme17 | 0.000157776 | 0.261501102 | 0.255 | 0.091 | 1 | 44 |
| cnbpb8 | 0.000171309 | 0.293178273 | 0.426 | 0.185 | 1 | 44 |
| atf4a5 | 0.00020093 | 0.344687604 | 0.404 | 0.183 | 1 | 44 |
| znf7062 | 0.000203416 | 0.280172436 | 0.426 | 0.184 | 1 | 44 |
| mt-nd117 | 0.000222838 | 0.363955542 | 0.745 | 0.412 | 1 | 44 |
| pik3r25 | 0.000224572 | 0.264157024 | 0.191 | 0.061 | 1 | 44 |
| mt-nd316 | 0.000275375 | 0.403995114 | 0.681 | 0.397 | 1 | 44 |
| mt-atp621 | 0.000275492 | 0.404852327 | 0.936 | 0.591 | 1 | 44 |
| LOC1039104381 | 0.000316345 | 0.282344237 | 0.383 | 0.166 | 1 | 44 |
| ppp3ccb4 | 0.000351169 | 0.330809361 | 0.298 | 0.125 | 1 | 44 |
| ndufa53 | 0.000358689 | 0.264132055 | 0.34 | 0.144 | 1 | 44 |
| actb18 | 0.000360751 | 0.27644222 | 0.936 | 0.786 | 1 | 44 |
| ccni15 | 0.000385003 | 0.337000662 | 0.574 | 0.295 | 1 | 44 |
| sumo2b14 | 0.000433735 | 0.308712162 | 0.553 | 0.27 | 1 | 44 |
| ywhabl8 | 0.000466443 | 0.32582787 | 0.553 | 0.279 | 1 | 44 |
| zgc:778491 | 0.000486438 | 0.253731753 | 0.404 | 0.18 | 1 | 44 |
| hmgn617 | 0.000530104 | 0.309083506 | 0.553 | 0.266 | 1 | 44 |
| pfn27 | 0.000565935 | 0.315433803 | 0.468 | 0.227 | 1 | 44 |
| selenok6 | 0.000840501 | 0.270335096 | 0.298 | 0.129 | 1 | 44 |
| pomp7 | 0.000945406 | 0.284103556 | 0.468 | 0.232 | 1 | 44 |
| pnrc223 | 0.001044202 | 0.373621947 | 0.851 | 0.569 | 1 | 44 |
| cox8a14 | 0.001309759 | 0.253313697 | 0.468 | 0.229 | 1 | 44 |
| cst14a.21 | 0.001361522 | 0.300173456 | 0.447 | 0.226 | 1 | 44 |
| nop539 | 0.002116052 | 0.26501212 | 0.489 | 0.252 | 1 | 44 |
| rplp223 | 0.002771484 | 0.279518425 | 0.936 | 0.758 | 1 | 44 |
| mt-nd215 | 0.002873743 | 0.285821215 | 0.723 | 0.401 | 1 | 44 |
| oaz1b18 | 0.003032094 | 0.278437235 | 0.596 | 0.324 | 1 | 44 |
| ybx113 | 0.003818804 | 0.268733452 | 0.851 | 0.671 | 1 | 44 |

| | | | | | | | |
|---|---|---|---|---|---|---|---|
| gnb1b5 | 0.003970337 | 0.261826732 | 0.468 | 0.256 | | 1 | 44 |
| ywhaz8 | 0.004696133 | 0.264742521 | 0.511 | 0.283 | | 1 | 44 |
| chmp2a5 | 0.004936016 | 0.251185022 | 0.277 | 0.135 | | 1 | 44 |
| xbp19 | 0.004960808 | 0.281621144 | 0.617 | 0.345 | | 1 | 44 |
| myl12.116 | 0.005078905 | 0.314164267 | 0.638 | 0.393 | | 1 | 44 |
| bzw1b8 | 0.005465401 | 0.283474359 | 0.681 | 0.404 | | 1 | 44 |
| lum3 | 0 | 2.108508247 | 1 | 0.016 | | 0 | 45 |
| ogna1 | 0 | 1.790071375 | 0.953 | 0.006 | | 0 | 45 |
| cgref11 | 0 | 1.585599599 | 0.953 | 0.022 | | 0 | 45 |
| fgl2b2 | 0 | 1.457348962 | 0.93 | 0.012 | | 0 | 45 |
| tmem119b2 | 0 | 1.41206035 | 0.907 | 0.014 | | 0 | 45 |
| mettl24 | 0 | 1.388733715 | 0.884 | 0.009 | | 0 | 45 |
| erg1 | 0 | 1.383903145 | 0.884 | 0.01 | | 0 | 45 |
| htra3b3 | 0 | 1.348352475 | 0.93 | 0.012 | | 0 | 45 |
| igfbp32 | 0 | 1.297605598 | 0.767 | 0.011 | | 0 | 45 |
| mmp23bb | 0 | 1.280767207 | 0.814 | 0.006 | | 0 | 45 |
| si:ch211-243a20.31 | 0 | 1.277752514 | 0.884 | 0.011 | | 0 | 45 |
| ak51 | 0 | 1.148407709 | 0.814 | 0.006 | | 0 | 45 |
| olfml11 | 0 | 1.109108291 | 0.744 | 0.004 | | 0 | 45 |
| col5a3a1 | 0 | 0.984262648 | 0.698 | 0.009 | | 0 | 45 |
| creb3l11 | 0 | 0.874718891 | 0.628 | 0.005 | | 0 | 45 |
| apcdd1l1 | 0 | 0.872101545 | 0.628 | 0.006 | | 0 | 45 |
| omd | 0 | 0.850318565 | 0.395 | 0.002 | | 0 | 45 |
| npnt | 0 | 0.797544852 | 0.581 | 0.004 | | 0 | 45 |
| wisp1b | 0 | 0.764640689 | 0.512 | 0.001 | | 0 | 45 |
| fgl1 | 0 | 0.711779945 | 0.419 | 0.001 | | 0 | 45 |
| si:ch211-37e10.2 | 0 | 0.550912253 | 0.419 | 0.002 | | 0 | 45 |
| c1qtnf6b | 0 | 0.514822496 | 0.372 | 0.001 | | 0 | 45 |
| wnt2 | 4.66030961673119e-307 | 0.427453002 | 0.326 | 0.002 | 1.17295332743507e-302 | | 45 |
| scinla1 | 1.30091419590768e-305 | 1.245876932 | 0.628 | 0.009 | 3.27427093968004e-301 | | 45 |
| XLOC-008935 | 1.29048984142132e-297 | 0.310893747 | 0.233 | 0.001 | 3.24803388187331e-293 | | 45 |
| col16a13 | 5.70223101308357e-280 | 1.97714969 | 1 | 0.031 | 1.435194523683e-275 | | 45 |
| twist32 | 2.19657298953619e-278 | 1.541328953 | 0.977 | 0.028 | 5.52855455736364e-274 | | 45 |
| cspg5a1 | 4.43840332591416e-266 | 0.771375786 | 0.512 | 0.007 | 1.11710173309934e-261 | | 45 |
| twist22 | 8.21273355619511e-264 | 0.969512749 | 0.605 | 0.01 | 2.06706290875875e-259 | | 45 |

| | | | | | | |
|---|---|---|---|---|---|---|
| col7a1 | 5.05497940699089e-255 | 0.345383614 | 0.233 | 0.001 | 1.27228776694554e-250 | 45 |
| CABZ01074363.14 | 8.74937418121803e-254 | 1.453918971 | 0.907 | 0.026 | 2.20212998767077e-249 | 45 |
| fam19a1a | 1.16769128467257e-247 | 0.401141347 | 0.279 | 0.002 | 2.93896219439239e-243 | 45 |
| dnm3a | 7.56092648219532e-247 | 0.496497819 | 0.372 | 0.004 | 1.90300958630374e-242 | 45 |
| irx1b | 1.71533948778847e-243 | 0.466168291 | 0.302 | 0.002 | 4.31733795681481e-239 | 45 |
| olfml2bb | 4.03387656462534e-229 | 0.291724253 | 0.186 | 0.001 | 1.01528639255055e-224 | 45 |
| rcn31 | 1.81319030706108e-226 | 0.572547136 | 0.442 | 0.006 | 4.56361868384203e-222 | 45 |
| srpx21 | 3.67945058135558e-224 | 0.734496995 | 0.465 | 0.007 | 9.26080916821385e-220 | 45 |
| cdo12 | 1.75297208556112e-221 | 0.817057208 | 0.488 | 0.008 | 4.41205544214877e-217 | 45 |
| scara33 | 1.97568381638083e-214 | 1.226711142 | 0.884 | 0.029 | 4.9725985974489e-210 | 45 |
| cd248b | 8.34406189480173e-214 | 0.482048317 | 0.302 | 0.003 | 2.10011693830265e-209 | 45 |
| wnt10b | 1.11678584841757e-207 | 0.428804087 | 0.326 | 0.003 | 2.81083830188218e-203 | 45 |
| serpinf16 | 4.63697834067817e-200 | 1.556624837 | 0.977 | 0.041 | 1.16708107856529e-195 | 45 |
| col15a1a4 | 4.6146457663889e-194 | 1.055649927 | 0.698 | 0.02 | 1.16146019294242e-189 | 45 |
| vwa13 | 5.7565899854299e-193 | 0.96215848 | 0.698 | 0.02 | 1.44887613343285e-188 | 45 |
| megf6a | 1.42833400175833e-189 | 0.538437764 | 0.395 | 0.006 | 3.59497384902553e-185 | 45 |
| pigr1 | 1.13424129395692e-180 | 0.831250061 | 0.651 | 0.019 | 2.85477191276017e-176 | 45 |
| fkbp111 | 2.36901319576263e-176 | 1.087813316 | 0.744 | 0.026 | 5.96256931241497e-172 | 45 |
| pdgfrl4 | 4.80172394822496e-175 | 1.14885933 | 0.744 | 0.026 | 1.20854590052874e-170 | 45 |
| fkbp74 | 1.37501834304326e-172 | 1.182749857 | 0.837 | 0.034 | 3.46078366760557e-168 | 45 |
| clu3 | 2.5146952128579e-169 | 1.164112976 | 0.721 | 0.025 | 6.32923638124205e-165 | 45 |
| kdelr31 | 2.55069875677682e-169 | 0.803508025 | 0.581 | 0.016 | 6.41985370093158e-165 | 45 |
| hhipl21 | 9.98390149305407e-165 | 0.358059191 | 0.256 | 0.003 | 2.51284816678678e-160 | 45 |
| copz21 | 1.44666942166837e-164 | 0.648329716 | 0.465 | 0.01 | 3.64112226739712e-160 | 45 |
| c1qtnf22 | 3.45096764956324e-162 | 0.841185397 | 0.651 | 0.021 | 8.68574047718571e-158 | 45 |
| si:dkeyp-59c12.1 | 1.72571843139719e-161 | 0.356397529 | 0.233 | 0.002 | 4.3434607199836e-157 | 45 |
| rgs5a4 | 1.42592804927048e-159 | 0.997908362 | 0.721 | 0.026 | 3.58891830720887e-155 | 45 |
| CMKLR1 (1 of many) | 4.45284419264161e-155 | 0.489907974 | 0.349 | 0.006 | 1.12073635484597e-150 | 45 |
| si:dkey-12l12.12 | 8.33834995917234e-155 | 0.859899399 | 0.651 | 0.021 | 2.09867930122409e-150 | 45 |
| bhmt4 | 3.81328303486697e-154 | 1.440117574 | 0.884 | 0.043 | 9.59765207045667e-150 | 45 |
| fkbp92 | 1.73006857238434e-151 | 0.89638126 | 0.674 | 0.024 | 4.35440958983414e-147 | 45 |
| ccdc803 | 2.94229035832109e-151 | 0.902192282 | 0.674 | 0.024 | 7.40545060285834e-147 | 45 |
| atoh83 | 8.12466879023351e-151 | 0.731898717 | 0.581 | 0.018 | 2.04489788781387e-146 | 45 |
| twist1a2 | 2.70870062022292e-150 | 0.782314821 | 0.512 | 0.014 | 6.81752859103907e-146 | 45 |
| rgma1 | 5.19364743125829e-149 | 0.504785843 | 0.395 | 0.008 | 1.3071891219734e-144 | 45 |

| | | | | | | |
|---|---|---|---|---|---|---|
| TENT5A | 1.69007346367977e-147 | 0.482601958 | 0.326 | 0.005 | 4.25374590073562e-143 | 45 |
| gtdc1 | 2.6983947403201e-146 | 0.516747283 | 0.419 | 0.009 | 6.79158972191166e-142 | 45 |
| timp2a7 | 1.45036115587525e-144 | 1.451988799 | 0.953 | 0.055 | 3.65041399322242e-140 | 45 |
| asip12 | 2.45826892718123e-144 | 0.818617804 | 0.581 | 0.019 | 6.18721706282245e-140 | 45 |
| si:ch211-51c14.1 | 1.21477232223859e-140 | 0.343944885 | 0.279 | 0.004 | 3.05746045784231e-136 | 45 |
| hpdb1 | 7.65475837029772e-139 | 0.69218051 | 0.488 | 0.013 | 1.92662613422023e-134 | 45 |
| ccdc3b3 | 1.05320274331202e-136 | 0.856822531 | 0.628 | 0.023 | 2.65080598464203e-132 | 45 |
| abi3bpa2 | 8.19438829666713e-135 | 0.585143363 | 0.465 | 0.012 | 2.06244559038815e-130 | 45 |
| hkdc1 | 1.04691648347149e-134 | 0.283931796 | 0.209 | 0.002 | 2.63498409724939e-130 | 45 |
| ecrg4a4 | 2.13619106645041e-134 | 1.179279388 | 0.698 | 0.029 | 5.37657929514903e-130 | 45 |
| casz13 | 7.19417554599593e-131 | 0.785236789 | 0.581 | 0.021 | 1.81070204317171e-126 | 45 |
| nradd2 | 2.33010781465581e-130 | 0.494440699 | 0.395 | 0.009 | 5.8646483587072e-126 | 45 |
| ckap45 | 9.16739937400525e-130 | 1.029152693 | 0.814 | 0.042 | 2.30734274844338e-125 | 45 |
| serpinh1b2 | 3.22263333510988e-129 | 1.149351785 | 0.581 | 0.022 | 8.11104584113806e-125 | 45 |
| srpx1 | 4.23278721864775e-128 | 1.004077088 | 0.674 | 0.029 | 1.06535021506145e-123 | 45 |
| si:ch211-133n4.46 | 9.29314132454906e-124 | 1.22730798 | 0.791 | 0.043 | 2.33899073997575e-119 | 45 |
| p4ha21 | 3.10696525352961e-123 | 0.620010817 | 0.442 | 0.013 | 7.81992084660867e-119 | 45 |
| nkd2a | 7.64952416854137e-122 | 0.457957869 | 0.372 | 0.009 | 1.92530873798018e-117 | 45 |
| cygb15 | 5.99917076092377e-119 | 1.394184715 | 0.86 | 0.053 | 1.5099312888169e-114 | 45 |
| FQ378018.1 | 1.64676184744851e-118 | 0.420888032 | 0.256 | 0.004 | 4.14473489384316e-114 | 45 |
| vkorc11 | 2.17273376656076e-118 | 0.571137254 | 0.442 | 0.013 | 5.46855361705678e-114 | 45 |
| krt917 | 5.65831991869344e-117 | 1.710985559 | 0.953 | 0.066 | 1.42414254033595e-112 | 45 |
| col5a2a7 | 5.99423888891638e-116 | 1.913306191 | 1 | 0.084 | 1.50868998595136e-111 | 45 |
| reck3 | 9.32342548805977e-116 | 0.797108391 | 0.605 | 0.026 | 2.34661296108976e-111 | 45 |
| si:ch211-106h4.126 | 1.57309090438046e-115 | 1.588400513 | 0.884 | 0.057 | 3.95931249723517e-111 | 45 |
| fkbp142 | 2.50150283131792e-115 | 1.059278583 | 0.744 | 0.04 | 6.29603247614407e-111 | 45 |
| cd82a3 | 2.15202220299824e-114 | 1.221753866 | 0.86 | 0.055 | 5.41642468272627e-110 | 45 |
| col21a12 | 5.18031266782833e-114 | 0.740749108 | 0.488 | 0.017 | 1.30383289536571e-109 | 45 |
| postnb6 | 7.93887543722743e-111 | 1.800007277 | 0.977 | 0.077 | 1.99813555879577e-106 | 45 |
| zgc:113307 | 3.79691209608214e-109 | 0.265368915 | 0.163 | 0.002 | 9.55644805462914e-105 | 45 |
| ms4a17a.113 | 1.76447333941091e-106 | 0.905026226 | 0.698 | 0.037 | 4.44100294796332e-102 | 45 |
| cxcl8a2 | 1.80877392506255e-106 | 0.753078444 | 0.442 | 0.015 | 4.55250309198994e-102 | 45 |
| nupr1b5 | 1.63260862139781e-103 | 1.106286817 | 0.767 | 0.047 | 4.10911263919614e-99 | 45 |
| nucb2a2 | 5.77540446005151e-103 | 0.915007407 | 0.767 | 0.047 | 1.45361154855037e-98 | 45 |
| irx1a1 | 7.23643494002471e-103 | 0.444944399 | 0.256 | 0.005 | 1.82133831005482e-98 | 45 |

| | | | | | | |
|---|---|---|---|---|---|---|
| olfml3b6 | 6.27113598642649e-102 | 0.971681632 | 0.744 | 0.044 | 1.57838221642368e-97 | 45 |
| klf12a | 3.39652982825231e-101 | 0.572176887 | 0.465 | 0.017 | 8.54872592472825e-97 | 45 |
| plxdc1 | 1.01570655727696e-100 | 0.267294228 | 0.163 | 0.002 | 2.55643183401039e-96 | 45 |
| nfixb6 | 2.83772319600725e-99 | 1.116593418 | 0.884 | 0.064 | 7.14226551203066e-95 | 45 |
| wnt11r1 | 6.03512160251927e-99 | 0.446906581 | 0.302 | 0.007 | 1.51897975613807e-94 | 45 |
| dpt2 | 6.73017286499703e-99 | 0.700088015 | 0.419 | 0.014 | 1.6939172083911e-94 | 45 |
| ehd2b2 | 8.46972710666823e-99 | 0.783176851 | 0.605 | 0.03 | 2.13174561547733e-94 | 45 |
| fam198a1 | 1.53319155578526e-98 | 0.528275805 | 0.419 | 0.014 | 3.85888982675593e-94 | 45 |
| calua2 | 6.90905991230845e-97 | 0.670444624 | 0.535 | 0.024 | 1.73894128932891e-92 | 45 |
| pvalb6 | 1.25506356178951e-96 | 0.284125275 | 0.14 | 0.001 | 3.15886947866801e-92 | 45 |
| pabpc1b6 | 4.19785219756034e-94 | 1.257719899 | 0.86 | 0.067 | 1.05655741960396e-89 | 45 |
| colec125 | 4.82294417305852e-94 | 0.946682357 | 0.791 | 0.052 | 1.2138868189171e-89 | 45 |
| prelp2 | 6.1751231546434e-94 | 0.624553635 | 0.349 | 0.01 | 1.5542167467922e-89 | 45 |
| ecm21 | 1.34418826636077e-93 | 0.425952556 | 0.302 | 0.008 | 3.38318744760342e-89 | 45 |
| thbs4b | 2.0722424285552e-93 | 0.269147987 | 0.163 | 0.002 | 5.21562696843058e-89 | 45 |
| cercam1 | 2.79992416817641e-92 | 0.572749936 | 0.419 | 0.015 | 7.04712913888321e-88 | 45 |
| aebp15 | 7.3197064274546e-92 | 0.98134223 | 0.651 | 0.038 | 1.84229691072605e-87 | 45 |
| epha4b1 | 2.19483369623025e-91 | 0.519190401 | 0.326 | 0.009 | 5.52417693004191e-87 | 45 |
| col5a17 | 1.333530089374e-90 | 1.865374755 | 1 | 0.111 | 3.35636188194542e-86 | 45 |
| si:ch1073-459j12.16 | 7.28102997230867e-90 | 1.229432363 | 0.837 | 0.065 | 1.83256243373037e-85 | 45 |
| tgfbi3 | 5.9853136276309e-86 | 0.849722228 | 0.512 | 0.025 | 1.50644358693842e-81 | 45 |
| CABZ01092746.14 | 6.43036268514443e-86 | 1.211452522 | 0.837 | 0.066 | 1.618457984224e-81 | 45 |
| fam69c1 | 9.63120400736792e-86 | 0.46686833 | 0.349 | 0.011 | 2.42407773661443e-81 | 45 |
| traf4a1 | 5.01122107627832e-85 | 0.355705751 | 0.279 | 0.007 | 1.26127423268849e-80 | 45 |
| fkbp10b1 | 6.1245990898994e-82 | 0.582304857 | 0.512 | 0.025 | 1.54150034493678e-77 | 45 |
| pdgfra4 | 7.22434746634322e-82 | 0.893747222 | 0.698 | 0.047 | 1.81829601380392e-77 | 45 |
| XLOC-041451 | 1.12881350948438e-81 | 0.257226775 | 0.163 | 0.002 | 2.84111072202125e-77 | 45 |
| dcn6 | 1.18729106876265e-81 | 1.744286733 | 1 | 0.118 | 2.98829289096871e-77 | 45 |
| glis2a2 | 3.61448429713198e-81 | 0.640913064 | 0.512 | 0.026 | 9.09729552745148e-77 | 45 |
| tmeff1b3 | 1.14866757140789e-78 | 0.59661316 | 0.535 | 0.029 | 2.89108141047652e-74 | 45 |
| col6a36 | 1.44335418737363e-78 | 1.333355605 | 0.977 | 0.099 | 3.6327781542007e-74 | 45 |
| ctgfb1 | 2.88826931382323e-77 | 0.504602278 | 0.326 | 0.011 | 7.26948503596169e-73 | 45 |
| col1a1a7 | 3.63567037521443e-77 | 2.341844201 | 1 | 0.139 | 9.1506187673772e-73 | 45 |
| CABZ01039304.12 | 3.88344281431507e-76 | 0.474261811 | 0.372 | 0.015 | 9.7742372193496e-72 | 45 |
| atf5b1 | 6.3784748981353e-76 | 0.651961689 | 0.558 | 0.033 | 1.60539834711167e-71 | 45 |

| | | | | | | |
|---|---|---|---|---|---|---|
| mmp27 | 1.58428915423425e-74 | 1.203968275 | 0.907 | 0.091 | 3.98749737229219e-70 | 45 |
| col1a1b7 | 2.1531143563889e-74 | 2.351903159 | 1 | 0.146 | 5.41917352359523e-70 | 45 |
| ier5l3 | 7.04313843964589e-74 | 0.6040903 | 0.419 | 0.019 | 1.77268751387447e-69 | 45 |
| htra1a2 | 2.88785175517639e-73 | 0.63257532 | 0.419 | 0.019 | 7.26843408260346e-69 | 45 |
| sdc24 | 3.66824453966945e-73 | 0.955583427 | 0.837 | 0.075 | 9.23260468189405e-69 | 45 |
| plpp1a5 | 1.73188162753801e-72 | 0.987996938 | 0.791 | 0.067 | 4.35897286835042e-68 | 45 |
| tns2b1 | 1.00406950888243e-71 | 0.43050988 | 0.326 | 0.012 | 2.52714254690618e-67 | 45 |
| col1a27 | 5.08047319120114e-71 | 2.399624778 | 1 | 0.155 | 1.27870429749342e-66 | 45 |
| zgc:565851 | 1.48935808279168e-70 | 0.421453282 | 0.302 | 0.01 | 3.74856535857839e-66 | 45 |
| crtap3 | 1.85018041692404e-70 | 0.686828005 | 0.512 | 0.03 | 4.65671909135612e-66 | 45 |
| sparc10 | 3.56840955634632e-70 | 2.293359164 | 1 | 0.157 | 8.98133001236805e-66 | 45 |
| olfml2ba | 8.59071018572416e-70 | 0.310528002 | 0.209 | 0.005 | 2.16219584664491e-65 | 45 |
| sgcd1 | 3.73277312153178e-69 | 0.362899993 | 0.256 | 0.008 | 9.39501666958335e-65 | 45 |
| p3h4 | 2.01701021757937e-68 | 0.412463879 | 0.326 | 0.012 | 5.07661301662552e-64 | 45 |
| selenop2 | 3.86209032955251e-68 | 0.516038513 | 0.395 | 0.019 | 9.7204951504507e-64 | 45 |
| crhbp1 | 6.20812041819711e-68 | 0.686538689 | 0.372 | 0.016 | 1.56252182805603e-63 | 45 |
| rcn11 | 2.23904683009448e-67 | 0.540141252 | 0.395 | 0.019 | 5.63545696666478e-63 | 45 |
| apbb2b | 9.8221425490471e-67 | 0.511421557 | 0.372 | 0.017 | 2.47213505816966e-62 | 45 |
| tcf42 | 8.42567254678299e-66 | 0.504336352 | 0.419 | 0.021 | 2.12065752329981e-61 | 45 |
| si:ch73-138n13.14 | 4.548955787279e-65 | 0.633350788 | 0.558 | 0.038 | 1.14492668210025e-60 | 45 |
| CU929237.13 | 1.20216004758103e-64 | 0.5783824 | 0.349 | 0.015 | 3.02571662375669e-60 | 45 |
| irx3a | 2.17453054518925e-64 | 0.352168247 | 0.256 | 0.008 | 5.47307592918682e-60 | 45 |
| col11a1b2 | 1.4265441541432e-63 | 0.494533908 | 0.326 | 0.014 | 3.59046898156303e-59 | 45 |
| mfap27 | 2.09444154455442e-63 | 1.107178121 | 0.884 | 0.095 | 5.27149992348901e-59 | 45 |
| smoc22 | 2.40758824568095e-63 | 0.471133852 | 0.302 | 0.012 | 6.05965885555437e-59 | 45 |
| nfia8 | 1.08568747985083e-61 | 0.933297789 | 0.791 | 0.079 | 2.73256681803655e-57 | 45 |
| pmp22a7 | 1.68897005281261e-61 | 1.433534372 | 1 | 0.138 | 4.25096872592405e-57 | 45 |
| irx5a | 2.70578981541384e-60 | 0.256185748 | 0.186 | 0.005 | 6.8102023864151e-56 | 45 |
| antxr2a11 | 3.10182145519848e-60 | 1.463169375 | 0.977 | 0.153 | 7.80697442058907e-56 | 45 |
| ephb37 | 3.80075554593695e-60 | 0.671318495 | 0.558 | 0.041 | 9.56612163356871e-56 | 45 |
| antxr1b | 3.84862583491904e-60 | 0.250287494 | 0.186 | 0.005 | 9.68660636390774e-56 | 45 |
| FAT41 | 5.81114388114348e-60 | 0.517195211 | 0.349 | 0.017 | 1.462606803445e-55 | 45 |
| lamb1b3 | 5.95582300997819e-60 | 0.600614337 | 0.442 | 0.026 | 1.49902109338141e-55 | 45 |
| BX957346.1 | 1.93645280576013e-59 | 0.272670545 | 0.233 | 0.007 | 4.87385806681767e-55 | 45 |
| plpp34 | 4.72939034232873e-59 | 0.745812192 | 0.651 | 0.054 | 1.19034025526072e-54 | 45 |

| | | | | | | |
|---|---|---|---|---|---|---|
| c1qtnf6a3 | 3.60101159122634e-58 | 0.491178536 | 0.372 | 0.019 | 9.06338607395758e-54 | 45 |
| slc7a10a | 5.24862926448283e-58 | 0.386780312 | 0.302 | 0.013 | 1.32102749957768e-53 | 45 |
| p3h3 | 5.69310964810363e-58 | 0.349578043 | 0.256 | 0.009 | 1.4328987673312e-53 | 45 |
| appb5 | 1.31667469257716e-57 | 0.614339883 | 0.535 | 0.039 | 3.31393853374747e-53 | 45 |
| glis2b | 5.03706326164652e-56 | 0.338715944 | 0.302 | 0.013 | 1.26777845232381e-51 | 45 |
| bcl11ba1 | 5.10431596104708e-56 | 0.404735968 | 0.326 | 0.015 | 1.28470528423594e-51 | 45 |
| angptl47 | 2.12969200290416e-55 | 0.866984293 | 0.581 | 0.049 | 5.36022180210947e-51 | 45 |
| fndc3ba1 | 5.72989776283434e-55 | 0.434773528 | 0.326 | 0.016 | 1.44215796792778e-50 | 45 |
| wipi1 | 5.78413952611663e-55 | 0.337367786 | 0.256 | 0.01 | 1.45581007732829e-50 | 45 |
| fxyd110 | 1.90059410722456e-54 | 0.929728197 | 0.744 | 0.08 | 4.78360530847348e-50 | 45 |
| CR847566.16 | 4.62132500793303e-54 | 0.621898557 | 0.488 | 0.035 | 1.16314129124667e-49 | 45 |
| klf7a | 6.77407669671531e-54 | 0.317788831 | 0.233 | 0.008 | 1.70496736379628e-49 | 45 |
| dag15 | 1.06672541347362e-53 | 0.70613815 | 0.581 | 0.049 | 2.68484119317175e-49 | 45 |
| XLOC-0418322 | 2.78058516416284e-53 | 0.363059923 | 0.256 | 0.01 | 6.99845479968145e-49 | 45 |
| rrbp1a1 | 7.15955718596306e-53 | 0.739614689 | 0.581 | 0.05 | 1.80198894813504e-48 | 45 |
| si:ch73-46j18.511 | 8.90849018857545e-53 | 1.425037265 | 1 | 0.174 | 2.24217789556256e-48 | 45 |
| p3h11 | 6.94732064251596e-52 | 0.366129891 | 0.279 | 0.012 | 1.74857113251484e-47 | 45 |
| yaf2 | 2.32663631652263e-51 | 0.612305723 | 0.512 | 0.04 | 5.85591094505581e-47 | 45 |
| tuba1b3 | 1.00877393665317e-50 | 0.542430276 | 0.442 | 0.031 | 2.53898312116236e-46 | 45 |
| zgc:1101581 | 1.33634790471161e-50 | 0.54160835 | 0.442 | 0.031 | 3.36345404136865e-46 | 45 |
| wfdc11 | 3.70485218698141e-50 | 0.294291956 | 0.209 | 0.007 | 9.32474246941352e-46 | 45 |
| cyr618 | 3.0039139836553e-49 | 1.028691276 | 0.651 | 0.068 | 7.56055110546203e-45 | 45 |
| nt5e | 1.76049038105438e-48 | 0.41175949 | 0.279 | 0.013 | 4.43097824007577e-44 | 45 |
| klf6a12 | 1.51579533111163e-46 | 1.28808192 | 0.953 | 0.167 | 3.81510526887487e-42 | 45 |
| zgc:194981 | 1.91360490185333e-46 | 0.758819594 | 0.698 | 0.077 | 4.81635217747465e-42 | 45 |
| ehd1b3 | 2.46923713948556e-45 | 0.596893661 | 0.535 | 0.048 | 6.2148229563712e-41 | 45 |
| dnase1l13 | 2.65295775553084e-45 | 0.563931375 | 0.512 | 0.045 | 6.67722937489558e-41 | 45 |
| zgc:1533177 | 3.36256674027209e-45 | 0.628395606 | 0.488 | 0.041 | 8.46324422859083e-41 | 45 |
| col2a1b2 | 4.91534826239477e-45 | 0.510686717 | 0.279 | 0.014 | 1.23714400416214e-40 | 45 |
| f3b6 | 1.36389722185184e-44 | 1.056960858 | 0.767 | 0.101 | 3.4327929176789e-40 | 45 |
| marcksl1a10 | 1.84221618056134e-44 | 1.408852154 | 1 | 0.214 | 4.63667390485484e-40 | 45 |
| smoc11 | 2.02925353228177e-44 | 0.267711653 | 0.209 | 0.008 | 5.10742821539998e-40 | 45 |
| col6a17 | 3.0565118715464e-44 | 0.958665421 | 0.721 | 0.089 | 7.69293472949513e-40 | 45 |
| mrc24 | 3.1364499634343e-44 | 0.485779478 | 0.372 | 0.025 | 7.8941309129678e-40 | 45 |
| sh3pxd2aa3 | 5.7877887055443e-44 | 0.525361051 | 0.419 | 0.032 | 1.45672853929845e-39 | 45 |

| | | | | | | |
|---|---|---|---|---|---|---|
| tuba8l37 | 9.61245184267969e-44 | 0.609175807 | 0.512 | 0.046 | 2.41935800428405e-39 | 45 |
| ca8 | 1.0288275201033e-43 | 0.315823991 | 0.256 | 0.012 | 2.589455985348e-39 | 45 |
| sertad41 | 2.86618550421624e-43 | 0.347767975 | 0.279 | 0.015 | 7.21390229556187e-39 | 45 |
| mych5 | 2.9675636983051e-43 | 0.803863185 | 0.651 | 0.074 | 7.46906107226411e-39 | 45 |
| prrx1a2 | 6.11161192288771e-43 | 0.315812713 | 0.233 | 0.01 | 1.53823160487161e-38 | 45 |
| mmp14b6 | 8.95880046818929e-43 | 0.729757084 | 0.558 | 0.057 | 2.25484048983856e-38 | 45 |
| cebpd10 | 3.88029067210385e-42 | 1.031472703 | 0.86 | 0.128 | 9.76630359261818e-38 | 45 |
| zgc:1536752 | 1.1179325338464e-41 | 0.648941571 | 0.628 | 0.069 | 2.813724394438e-37 | 45 |
| znf7042 | 3.54634203159392e-41 | 0.514870365 | 0.395 | 0.03 | 8.92578825931875e-37 | 45 |
| fam102ab2 | 6.36655906735551e-41 | 0.465544005 | 0.419 | 0.033 | 1.60239925166271e-36 | 45 |
| cd28l5 | 1.87876204406389e-40 | 0.531490217 | 0.442 | 0.037 | 4.72865618870441e-36 | 45 |
| tcf126 | 2.29702325776241e-40 | 0.593434174 | 0.558 | 0.058 | 5.7813778374622e-36 | 45 |
| ak42 | 2.57284457114823e-40 | 0.47617704 | 0.395 | 0.031 | 6.47559250112297e-36 | 45 |
| dnmt3ab4 | 4.44845964120179e-40 | 0.703326406 | 0.651 | 0.079 | 1.11963280709408e-35 | 45 |
| cd2761 | 5.66230204137768e-40 | 0.400998641 | 0.302 | 0.019 | 1.42514480079435e-35 | 45 |
| LOC101886030 | 7.1168975625338e-40 | 0.330540029 | 0.256 | 0.013 | 1.79125194751413e-35 | 45 |
| calhm2 | 1.13999391413648e-39 | 0.26377635 | 0.186 | 0.007 | 2.86925068249011e-35 | 45 |
| gas1a5 | 1.71239054420019e-39 | 0.567948148 | 0.465 | 0.043 | 4.30991576069747e-35 | 45 |
| LOC1018823342 | 2.41946626806087e-39 | 0.412252335 | 0.279 | 0.016 | 6.0895546500824e-35 | 45 |
| adrb2a1 | 3.37290672046262e-39 | 0.373186035 | 0.279 | 0.016 | 8.48926892473236e-35 | 45 |
| fosl1a9 | 4.16181597869231e-39 | 0.812334488 | 0.698 | 0.092 | 1.04748746367707e-34 | 45 |
| sept103 | 4.45191010985324e-39 | 0.561629036 | 0.488 | 0.047 | 1.12050125554896e-34 | 45 |
| flrt2 | 5.2907817128652e-39 | 0.287853924 | 0.233 | 0.011 | 1.33163684931104e-34 | 45 |
| fosb10 | 6.1424721735782e-39 | 1.03396884 | 0.814 | 0.125 | 1.5459988213679e-34 | 45 |
| lrrc58b | 9.93390015498412e-39 | 0.374039319 | 0.302 | 0.019 | 2.50026333000795e-34 | 45 |
| si:ch211-39i22.17 | 1.6447976939215e-38 | 0.771488184 | 0.605 | 0.071 | 4.13979131583103e-34 | 45 |
| foxo1a3 | 3.45646413586431e-38 | 0.552690388 | 0.442 | 0.04 | 8.69957458355689e-34 | 45 |
| p4ha1b1 | 7.79494127085051e-38 | 0.683534291 | 0.581 | 0.066 | 1.96190876846037e-33 | 45 |
| loxl13 | 1.59136918657416e-37 | 0.390936332 | 0.326 | 0.022 | 4.00531710568849e-33 | 45 |
| s100v211 | 2.31520387921355e-37 | 0.812752659 | 0.814 | 0.121 | 5.82713664359258e-33 | 45 |
| arhgef1b2 | 7.54787877050871e-37 | 0.776188915 | 0.581 | 0.071 | 1.89972560774934e-32 | 45 |
| si:dkey-101k6.52 | 1.46196499289233e-36 | 0.350587578 | 0.302 | 0.02 | 3.6796196906107e-32 | 45 |
| si:ch211-284f22.35 | 1.66305964128369e-36 | 0.49682247 | 0.395 | 0.034 | 4.18575481114691e-32 | 45 |
| ms4a17a.96 | 3.68585589809353e-36 | 0.63551626 | 0.512 | 0.056 | 9.27693070991161e-32 | 45 |
| vat19 | 3.94761164803716e-36 | 0.901116428 | 0.837 | 0.134 | 9.93574375694473e-32 | 45 |

| | | | | | | |
|---|---|---|---|---|---|---|
| sema6e | 4.10750666818791e-36 | 0.299578054 | 0.186 | 0.008 | 1.03381835331621e-31 | 45 |
| cbfa2t31 | 5.70510704936124e-36 | 0.352594122 | 0.279 | 0.018 | 1.43591839325373e-31 | 45 |
| znf385a6 | 7.38666947470529e-36 | 0.550969948 | 0.465 | 0.046 | 1.85915084008858e-31 | 45 |
| rgcc7 | 1.08068601341086e-35 | 0.725844905 | 0.581 | 0.07 | 2.71997862715379e-31 | 45 |
| klf151 | 1.27355324134105e-35 | 0.252403688 | 0.209 | 0.01 | 3.2054061531313e-31 | 45 |
| lrp1aa5 | 1.62585377655788e-35 | 0.576689286 | 0.512 | 0.055 | 4.09211137021853e-31 | 45 |
| meis1a2 | 2.80682418026208e-35 | 0.323802734 | 0.256 | 0.015 | 7.06449577930164e-31 | 45 |
| LOC1018835072 | 6.46416009660731e-35 | 0.365138614 | 0.279 | 0.018 | 1.62696445471509e-30 | 45 |
| fam114a1 | 1.82418055514068e-34 | 0.263185285 | 0.209 | 0.01 | 4.59128003923359e-30 | 45 |
| fstl1b8 | 1.89150751003925e-34 | 0.865078392 | 0.791 | 0.124 | 4.7607352520178e-30 | 45 |
| ppm1la | 2.50058105490012e-34 | 0.46582127 | 0.395 | 0.035 | 6.29371245707812e-30 | 45 |
| crip22 | 2.63216462918764e-34 | 0.395491634 | 0.256 | 0.016 | 6.62489515520238e-30 | 45 |
| tmem45a2 | 2.94327581360679e-34 | 0.381639589 | 0.326 | 0.025 | 7.40793089526692e-30 | 45 |
| rasgef1bb2 | 2.9869499785243e-34 | 0.413996963 | 0.372 | 0.032 | 7.51785440094781e-30 | 45 |
| efnb2a2 | 3.30121759883082e-34 | 0.302023563 | 0.209 | 0.011 | 8.30883457449729e-30 | 45 |
| slc38a5b10 | 4.50589850938743e-34 | 0.782839589 | 0.744 | 0.115 | 1.13408959582772e-29 | 45 |
| serf217 | 5.6124549915331e-34 | 1.205332381 | 0.977 | 0.343 | 1.41259879681897e-29 | 45 |
| marcksl1b11 | 6.72346874692606e-34 | 0.730999954 | 0.767 | 0.116 | 1.69222984891382e-29 | 45 |
| jdp2b11 | 8.60124141923442e-34 | 1.355530688 | 0.814 | 0.17 | 2.16484645280711e-29 | 45 |
| adh8a5 | 1.93698939835017e-33 | 0.613235831 | 0.442 | 0.044 | 4.87520861670754e-29 | 45 |
| egr114 | 1.96977929940537e-33 | 1.051745719 | 0.93 | 0.189 | 4.95773751867338e-29 | 45 |
| soul51 | 5.73642176241754e-33 | 0.295603627 | 0.256 | 0.016 | 1.44379999338287e-28 | 45 |
| col27a1b2 | 8.05549800257481e-33 | 0.424958658 | 0.279 | 0.019 | 2.02748829226805e-28 | 45 |
| id18 | 1.1971883345657e-32 | 0.803655791 | 0.698 | 0.105 | 3.01320331926841e-28 | 45 |
| nanos13 | 4.01167506849989e-32 | 0.392201541 | 0.279 | 0.02 | 1.00969849799074e-27 | 45 |
| loxl2b3 | 5.1888741999954e-32 | 0.4644444 | 0.419 | 0.042 | 1.30598774739684e-27 | 45 |
| plod1a1 | 7.72173048918836e-32 | 0.372754918 | 0.302 | 0.023 | 1.94348234682382e-27 | 45 |
| mafba9 | 1.09430025254615e-31 | 0.632101683 | 0.558 | 0.072 | 2.75424430563341e-27 | 45 |
| gstcd | 1.16608438163622e-31 | 0.306232799 | 0.186 | 0.009 | 2.93491778014021e-27 | 45 |
| si:dkey-56m19.55 | 1.21542151754357e-31 | 0.520839165 | 0.442 | 0.047 | 3.05909441750541e-27 | 45 |
| fpr13 | 1.36635192730624e-31 | 0.408463899 | 0.279 | 0.02 | 3.43897116583708e-27 | 45 |
| vdac38 | 1.53883140462599e-31 | 0.862266474 | 0.837 | 0.154 | 3.87308476230315e-27 | 45 |
| marveld11 | 1.97882932564655e-31 | 0.25977004 | 0.209 | 0.012 | 4.98051552971981e-27 | 45 |
| zgc:922422 | 2.76847267769192e-31 | 0.310376691 | 0.256 | 0.017 | 6.96796888248279e-27 | 45 |
| ppib13 | 2.79879005125915e-31 | 1.132598537 | 1 | 0.333 | 7.04427468001415e-27 | 45 |

| | | | | | | |
|---|---|---|---|---|---|---|
| tgfbr33 | 6.34037679889681e-31 | 0.463667152 | 0.419 | 0.043 | 1.59580943651434e-26 | 45 |
| XLOC-0354131 | 7.12232300537316e-31 | 0.438822595 | 0.326 | 0.028 | 1.79261747722237e-26 | 45 |
| zgc:1716792 | 9.4023591613515e-31 | 0.39214702 | 0.326 | 0.027 | 2.36647977732056e-26 | 45 |
| sulf2b2 | 9.98505440014727e-31 | 0.429780692 | 0.395 | 0.039 | 2.51313834197307e-26 | 45 |
| SCARF23 | 1.25854420401978e-30 | 0.389835141 | 0.302 | 0.024 | 3.16762990709738e-26 | 45 |
| eef1da15 | 2.63165464427813e-30 | 1.028174785 | 1 | 0.493 | 6.62361157418362e-26 | 45 |
| sept7a1 | 2.88894104679657e-30 | 0.673910954 | 0.535 | 0.071 | 7.27117572068228e-26 | 45 |
| emp27 | 3.24450123575451e-30 | 0.520217902 | 0.442 | 0.049 | 8.16608516027052e-26 | 45 |
| ift20 | 3.69596703531952e-30 | 0.338470376 | 0.279 | 0.021 | 9.3023794311957e-26 | 45 |
| map7d1b | 7.37523755090426e-30 | 0.323514575 | 0.256 | 0.018 | 1.85627353918709e-25 | 45 |
| hdlbpa9 | 8.72619776912517e-30 | 0.831874763 | 0.837 | 0.159 | 2.19629671651111e-25 | 45 |
| gpc43 | 9.36099471936033e-30 | 0.339700949 | 0.326 | 0.028 | 2.3560687609158e-25 | 45 |
| ecm1b5 | 1.22417104701299e-29 | 0.523967492 | 0.465 | 0.054 | 3.081116108227e-25 | 45 |
| fam110b | 1.34669595992385e-29 | 0.337914763 | 0.256 | 0.018 | 3.38949906153235e-25 | 45 |
| lzts2a2 | 2.44794107499468e-29 | 0.312330456 | 0.279 | 0.021 | 6.16122289165411e-25 | 45 |
| sdc411 | 4.08616804039525e-29 | 1.159502358 | 0.93 | 0.257 | 1.02844763408708e-24 | 45 |
| rbfox13 | 7.6592543847772e-29 | 0.369552736 | 0.302 | 0.025 | 1.92775773610457e-24 | 45 |
| calm3a | 8.87456908558241e-29 | 0.549361673 | 0.535 | 0.071 | 2.23364029315024e-24 | 45 |
| mycb7 | 1.07424235812892e-28 | 1.067646855 | 0.884 | 0.211 | 2.70376059117467e-24 | 45 |
| cyp2n135 | 1.09752101668147e-28 | 0.476673457 | 0.372 | 0.037 | 2.76235064688559e-24 | 45 |
| jun13 | 1.56617772646505e-28 | 1.122233754 | 0.953 | 0.228 | 3.94191271973987e-24 | 45 |
| si:ch211-286o17.14 | 1.61931298591988e-28 | 0.446484138 | 0.372 | 0.037 | 4.07564885426175e-24 | 45 |
| her63 | 1.96499541758786e-28 | 0.501553497 | 0.279 | 0.022 | 4.94569696652689e-24 | 45 |
| zgc:15834311 | 2.96211615383968e-28 | 0.911741987 | 0.837 | 0.165 | 7.45535014759908e-24 | 45 |
| cxcl18b7 | 3.4696601200467e-28 | 1.109482293 | 0.628 | 0.107 | 8.73278755614555e-24 | 45 |
| ddit316 | 3.60204731803346e-28 | 0.940793751 | 0.837 | 0.176 | 9.0659928947584e-24 | 45 |
| fosab14 | 5.72619302053939e-28 | 1.306620025 | 0.977 | 0.349 | 1.44122552133956e-23 | 45 |
| tspan73 | 5.73758437762046e-28 | 0.590409013 | 0.558 | 0.078 | 1.44409261200329e-23 | 45 |
| ier2a17 | 6.00502636369505e-28 | 1.161564938 | 0.93 | 0.262 | 1.51140508547841e-23 | 45 |
| st7 | 6.87149978981036e-28 | 0.401536071 | 0.326 | 0.03 | 1.72948778209737e-23 | 45 |
| sec61a14 | 8.99152820977347e-28 | 0.904910764 | 0.86 | 0.189 | 2.26307773511788e-23 | 45 |
| stau29 | 1.00402408839706e-27 | 0.585250507 | 0.581 | 0.085 | 2.52702822808656e-23 | 45 |
| wbp1lb2 | 1.06200981322885e-27 | 0.492286514 | 0.419 | 0.048 | 2.67297249891569e-23 | 45 |
| anxa410 | 1.36032075391823e-27 | 0.743169023 | 0.744 | 0.136 | 3.4237913055368e-23 | 45 |
| reep58 | 1.80072102891695e-27 | 0.756312526 | 0.814 | 0.158 | 4.53223475768108e-23 | 45 |

| | | | | | | |
|---|---|---|---|---|---|---|
| ip6k2b10 | 1.85155673546619e-27 | 0.749817295 | 0.698 | 0.122 | 4.66018314749486e-23 | 45 |
| krt813 | 1.90030397696115e-27 | 1.156639107 | 1 | 0.276 | 4.78287507961353e-23 | 45 |
| ghra5 | 3.56612423523145e-27 | 0.386144226 | 0.326 | 0.031 | 8.97557808765405e-23 | 45 |
| tnfsf123 | 4.08462121030603e-27 | 0.335002572 | 0.279 | 0.023 | 1.02805831242193e-22 | 45 |
| atf39 | 4.14734952504046e-27 | 0.804054615 | 0.581 | 0.091 | 1.04384640195743e-22 | 45 |
| tob1a5 | 4.14831050252381e-27 | 0.731453442 | 0.721 | 0.129 | 1.04408827038022e-22 | 45 |
| tmem39a | 1.48149961904966e-26 | 0.364969927 | 0.349 | 0.035 | 3.7287863911861e-22 | 45 |
| map1lc3a10 | 1.5182464612696e-26 | 0.511947673 | 0.512 | 0.07 | 3.82127451836945e-22 | 45 |
| dip2ca | 1.92396770092086e-26 | 0.270606576 | 0.233 | 0.017 | 4.84243430644771e-22 | 45 |
| hcar1-45 | 2.03151122843107e-26 | 0.432673025 | 0.349 | 0.036 | 5.11311061083816e-22 | 45 |
| pdk2b10 | 4.32372425502849e-26 | 0.700224299 | 0.605 | 0.1 | 1.08823815774812e-21 | 45 |
| ptn1 | 4.52584101397358e-26 | 0.264680086 | 0.163 | 0.009 | 1.13910892480701e-21 | 45 |
| triqk3 | 5.2167795629914e-26 | 0.441442259 | 0.419 | 0.05 | 1.3130112482093e-21 | 45 |
| BX640522.24 | 5.54246724659259e-26 | 0.297085307 | 0.302 | 0.027 | 1.39498358129489e-21 | 45 |
| lysmd3 | 9.93376483199796e-26 | 0.383104313 | 0.372 | 0.041 | 2.50022927056557e-21 | 45 |
| ptk2aa1 | 1.55987883477154e-25 | 0.261007793 | 0.233 | 0.017 | 3.92605903923648e-21 | 45 |
| dusp62 | 3.30692490178187e-25 | 0.480383259 | 0.395 | 0.047 | 8.32319928529479e-21 | 45 |
| dhrs13l14 | 6.90118111992804e-25 | 0.348036549 | 0.326 | 0.033 | 1.73695827607469e-20 | 45 |
| si:dkey-100n23.5 | 7.57763504377665e-25 | 0.367984274 | 0.372 | 0.042 | 1.90721496416815e-20 | 45 |
| tmem595 | 1.45134568792367e-24 | 0.572393322 | 0.628 | 0.107 | 3.65289196193509e-20 | 45 |
| susd62 | 1.66209630714103e-24 | 0.458728748 | 0.442 | 0.058 | 4.18333019544325e-20 | 45 |
| nme36 | 2.2068032383395e-24 | 0.525717711 | 0.512 | 0.076 | 5.55430307057668e-20 | 45 |
| kdelr2b1 | 2.57233796382521e-24 | 0.680603419 | 0.651 | 0.117 | 6.47431742115166e-20 | 45 |
| zgc:1533721 | 2.73847286771846e-24 | 0.376076951 | 0.302 | 0.03 | 6.89246236076059e-20 | 45 |
| ANKRD50 | 3.05304088595004e-24 | 0.319760933 | 0.256 | 0.022 | 7.68419860584764e-20 | 45 |
| junba13 | 5.68340993889434e-24 | 1.144873425 | 0.93 | 0.309 | 1.43045744752032e-19 | 45 |
| fbp1a3 | 9.39752378066151e-24 | 0.398691052 | 0.349 | 0.039 | 2.3652627603547e-19 | 45 |
| ugp2b | 1.0171225926337e-23 | 0.493072894 | 0.488 | 0.07 | 2.55999585339976e-19 | 45 |
| capns1b10 | 1.17202166946471e-23 | 0.5708724 | 0.674 | 0.122 | 2.94986133987574e-19 | 45 |
| smco4 | 1.38557889789459e-23 | 0.305001471 | 0.256 | 0.022 | 3.4873635281109e-19 | 45 |
| snd13 | 4.28058308664957e-23 | 0.568566218 | 0.605 | 0.105 | 1.07737995707883e-18 | 45 |
| zgc:16273019 | 4.91744942227751e-23 | 1.123023526 | 0.953 | 0.392 | 1.23767284509303e-18 | 45 |
| foxo3b5 | 5.35094506073463e-23 | 0.456513611 | 0.465 | 0.066 | 1.3467793623363e-18 | 45 |
| laptm4a2 | 7.19904487919578e-23 | 0.614363202 | 0.605 | 0.109 | 1.81192760564478e-18 | 45 |
| nova23 | 7.50070024447974e-23 | 0.313364063 | 0.302 | 0.031 | 1.88785124453311e-18 | 45 |

| | | | | | | |
|---|---|---|---|---|---|---|
| pdgfrb6 | 8.8863237725921e-23 | 0.465950834 | 0.395 | 0.051 | 2.23659883032371e-18 | 45 |
| spred2b6 | 9.03580535187783e-23 | 0.461907347 | 0.465 | 0.067 | 2.27422184901413e-18 | 45 |
| rab2a2 | 1.02401718719129e-22 | 0.725075057 | 0.791 | 0.172 | 2.57734885844176e-18 | 45 |
| eif4a1b12 | 1.77513396979389e-22 | 0.920185944 | 0.93 | 0.28 | 4.46783468857424e-18 | 45 |
| myl9b8 | 2.09639030135018e-22 | 0.542029019 | 0.512 | 0.08 | 5.27640474946827e-18 | 45 |
| ctsk5 | 2.76982900836018e-22 | 0.446591591 | 0.442 | 0.063 | 6.97138263114173e-18 | 45 |
| spry46 | 3.83472904042251e-22 | 0.48275666 | 0.372 | 0.047 | 9.65162952183942e-18 | 45 |
| dusp59 | 5.47002959586244e-22 | 0.677521636 | 0.465 | 0.073 | 1.37675174898262e-17 | 45 |
| ccnd12 | 8.77882143697648e-22 | 0.309532734 | 0.302 | 0.032 | 2.20954156747261e-17 | 45 |
| ablim1b4 | 1.01497265773868e-21 | 0.331003093 | 0.326 | 0.037 | 2.55458468226249e-17 | 45 |
| prdx4 | 1.04128104025351e-21 | 0.664254629 | 0.744 | 0.156 | 2.62080025021407e-17 | 45 |
| fxyd6l6 | 1.12779607367155e-21 | 0.551483918 | 0.628 | 0.114 | 2.83854993782393e-17 | 45 |
| nfic7 | 1.20734234864186e-21 | 0.503667838 | 0.488 | 0.077 | 3.0387599572967e-17 | 45 |
| ssr36 | 1.7065086826114e-21 | 0.864191957 | 0.977 | 0.272 | 4.29511170326464e-17 | 45 |
| bnip3lb1 | 2.15281372605946e-21 | 0.411152083 | 0.442 | 0.064 | 5.41841686711906e-17 | 45 |
| arf51 | 2.60608229255066e-21 | 0.719987966 | 0.86 | 0.203 | 6.55924852212076e-17 | 45 |
| fzd25 | 2.70366419441365e-21 | 0.321180365 | 0.279 | 0.029 | 6.80485241091973e-17 | 45 |
| slc39a71 | 2.74866565840515e-21 | 0.331616723 | 0.302 | 0.033 | 6.91811659563992e-17 | 45 |
| CABZ01058650.1 | 4.1229913862331e-21 | 0.342348975 | 0.326 | 0.038 | 1.03771570200101e-16 | 45 |
| zfp36l1b8 | 4.99080050615668e-21 | 0.705037472 | 0.651 | 0.13 | 1.25613457939457e-16 | 45 |
| rabac11 | 6.72019281216517e-21 | 0.348290845 | 0.349 | 0.043 | 1.69140532889385e-16 | 45 |
| ssr44 | 1.0456930235487e-20 | 0.913549681 | 0.93 | 0.307 | 2.63190477096972e-16 | 45 |
| XLOC-0441911 | 1.07792378065533e-20 | 0.276254352 | 0.209 | 0.017 | 2.71302636353139e-16 | 45 |
| boc6 | 1.08757263513759e-20 | 0.400558413 | 0.372 | 0.049 | 2.7373115653778e-16 | 45 |
| selenom2 | 1.26315986108267e-20 | 0.49708488 | 0.628 | 0.116 | 3.17924705435898e-16 | 45 |
| pgrmc17 | 1.52184996656823e-20 | 0.577791612 | 0.698 | 0.141 | 3.83034418085558e-16 | 45 |
| emd5 | 2.66029025792787e-20 | 0.391321096 | 0.395 | 0.056 | 6.69568455017866e-16 | 45 |
| anxa138 | 2.98166836062516e-20 | 0.381046979 | 0.372 | 0.05 | 7.50456109685746e-16 | 45 |
| eef1a1l19 | 3.42169646764474e-20 | 0.366876926 | 1 | 0.916 | 8.61206783941504e-16 | 45 |
| grb10b4 | 4.41781244456595e-20 | 0.500092021 | 0.302 | 0.036 | 1.1119192141728e-15 | 45 |
| adgre5b.33 | 5.48349665086188e-20 | 0.378050578 | 0.326 | 0.04 | 1.38014127205543e-15 | 45 |
| selenow18 | 6.49498635422358e-20 | 0.897516722 | 0.977 | 0.385 | 1.63472311549453e-15 | 45 |
| rassf1 | 7.62439480705212e-20 | 0.286510322 | 0.256 | 0.026 | 1.91898392898695e-15 | 45 |
| atf4b15 | 9.49911307983422e-20 | 0.841676375 | 0.907 | 0.264 | 2.39083177106348e-15 | 45 |
| fgl2a7 | 1.15094675415475e-19 | 0.644104911 | 0.419 | 0.066 | 2.8968178855321e-15 | 45 |

| | | | | | | |
|---|---|---|---|---|---|---|
| aldh9a1a.19 | 1.15342718004402e-19 | 0.686897924 | 0.721 | 0.164 | 2.90306086945279e-15 | 45 |
| vasna4 | 1.17329754668446e-19 | 0.325942268 | 0.302 | 0.035 | 2.95307259525011e-15 | 45 |
| mylipb1 | 1.22649273409851e-19 | 0.334451447 | 0.233 | 0.022 | 3.08695956245253e-15 | 45 |
| h3f3c16 | 1.55120485222778e-19 | 0.817231713 | 0.907 | 0.256 | 3.90422749257211e-15 | 45 |
| cdon5 | 1.87355003945284e-19 | 0.422296136 | 0.395 | 0.058 | 4.71553809429886e-15 | 45 |
| sdr16c5b2 | 2.03339457980785e-19 | 0.318294261 | 0.326 | 0.041 | 5.11785081791838e-15 | 45 |
| sh3gl1b2 | 2.15797514212961e-19 | 0.382359957 | 0.302 | 0.037 | 5.43140763522602e-15 | 45 |
| clstn11 | 2.33312957007269e-19 | 0.259749685 | 0.233 | 0.022 | 5.87225381491595e-15 | 45 |
| lhfpl2a | 2.68477408949859e-19 | 0.268824862 | 0.256 | 0.027 | 6.757307905859e-15 | 45 |
| rbfox24 | 3.07751519258853e-19 | 0.323706231 | 0.302 | 0.036 | 7.74579798822606e-15 | 45 |
| yif1a | 3.39225220026376e-19 | 0.349635949 | 0.419 | 0.063 | 8.53795956284385e-15 | 45 |
| ier2b17 | 5.9687292112893e-19 | 0.834967252 | 0.814 | 0.224 | 1.5022694551894e-14 | 45 |
| epdl13 | 6.50691510641325e-19 | 0.262448338 | 0.302 | 0.036 | 1.63772546313315e-14 | 45 |
| gulp1a5 | 6.87906955907445e-19 | 0.311364078 | 0.279 | 0.032 | 1.73139301732345e-14 | 45 |
| slc3a2b11 | 7.18309519260024e-19 | 0.605652691 | 0.581 | 0.118 | 1.80791322902555e-14 | 45 |
| gars | 7.84380756944977e-19 | 0.384311047 | 0.419 | 0.065 | 1.97420792715481e-14 | 45 |
| sgce4 | 1.0697218322969e-18 | 0.287717477 | 0.256 | 0.027 | 2.69238287970807e-14 | 45 |
| socs3b11 | 1.63232223856089e-18 | 0.646109048 | 0.698 | 0.155 | 4.1083918422339e-14 | 45 |
| rrbp1b | 2.03458656979215e-18 | 0.310597284 | 0.326 | 0.042 | 5.12085093750985e-14 | 45 |
| anxa2a9 | 2.43743723390731e-18 | 0.510507271 | 0.558 | 0.106 | 6.13478577402131e-14 | 45 |
| atox14 | 2.71322279651767e-18 | 0.519988485 | 0.558 | 0.11 | 6.82891045655532e-14 | 45 |
| midn13 | 2.89709907765175e-18 | 0.674689945 | 0.628 | 0.14 | 7.29170866854168e-14 | 45 |
| tpd52l2b1 | 3.19303668509176e-18 | 0.4146763 | 0.535 | 0.099 | 8.03655403270746e-14 | 45 |
| f106 | 3.88959985524801e-18 | 0.407809112 | 0.349 | 0.049 | 9.78973387567371e-14 | 45 |
| inka1a2 | 5.19149090491876e-18 | 0.274718111 | 0.209 | 0.02 | 1.306646345859e-13 | 45 |
| klf11b9 | 5.54780911715423e-18 | 0.422408398 | 0.465 | 0.08 | 1.39632807669655e-13 | 45 |
| wls7 | 6.54817439703777e-18 | 0.509576752 | 0.605 | 0.123 | 1.64811001399044e-13 | 45 |
| rarga6 | 7.16956131513581e-18 | 0.45203171 | 0.372 | 0.057 | 1.80450688740653e-13 | 45 |
| si:dkey-238c7.162 | 7.45823430693589e-18 | 0.483264516 | 0.395 | 0.063 | 1.87716299271269e-13 | 45 |
| s100a10b14 | 8.13495985120792e-18 | 0.778784069 | 0.953 | 0.255 | 2.04748804495052e-13 | 45 |
| fstl33 | 9.59252394201765e-18 | 0.349582444 | 0.233 | 0.025 | 2.41434235096642e-13 | 45 |
| ier58 | 1.10081981054564e-17 | 0.566594669 | 0.628 | 0.136 | 2.77065338116232e-13 | 45 |
| ATP11B | 1.20331460113236e-17 | 0.299242441 | 0.233 | 0.024 | 3.02862251959003e-13 | 45 |
| zgc:1132086 | 1.49802937207352e-17 | 0.396750735 | 0.326 | 0.045 | 3.77039012657184e-13 | 45 |
| LO018493.13 | 1.67431431047974e-17 | 0.322304412 | 0.233 | 0.025 | 4.21408168804646e-13 | 45 |

| | | | | | | |
|---|---|---|---|---|---|---|
| mgst3b6 | 1.73078436419903e-17 | 0.60693944 | 0.721 | 0.172 | 4.35621116625255e-13 | 45 |
| tnfrsfa2 | 1.79663808746064e-17 | 0.256620071 | 0.256 | 0.029 | 4.52195840232968e-13 | 45 |
| si:dkey-9i23.166 | 3.0825305127826e-17 | 0.387851896 | 0.302 | 0.04 | 7.75842104762253e-13 | 45 |
| spry25 | 3.85611703251942e-17 | 0.412586215 | 0.395 | 0.064 | 9.70546095914813e-13 | 45 |
| cers2a | 3.87734875267768e-17 | 0.322471821 | 0.279 | 0.035 | 9.75889907561445e-13 | 45 |
| asph2 | 7.03295144969788e-17 | 0.391894557 | 0.279 | 0.036 | 1.77012355037446e-12 | 45 |
| mfap54 | 7.33233002235074e-17 | 0.316657471 | 0.326 | 0.045 | 1.84547414332546e-12 | 45 |
| jag1b4 | 7.39713822856618e-17 | 0.315757493 | 0.233 | 0.026 | 1.86178572074782e-12 | 45 |
| hspg23 | 7.91382582589537e-17 | 0.275363797 | 0.233 | 0.025 | 1.99183082211961e-12 | 45 |
| pleca6 | 8.23560320065078e-17 | 0.433134473 | 0.372 | 0.059 | 2.0728189695718e-12 | 45 |
| fam114a2 | 9.02922315363691e-17 | 0.276690867 | 0.279 | 0.035 | 2.27256517553887e-12 | 45 |
| rimkla | 9.31410009305474e-17 | 0.283084193 | 0.302 | 0.04 | 2.34426585242095e-12 | 45 |
| gadd45ga4 | 1.08894451411826e-16 | 0.429925657 | 0.349 | 0.053 | 2.74076444758425e-12 | 45 |
| arl6ip5b6 | 1.12861660995467e-16 | 0.535495265 | 0.488 | 0.097 | 2.84061514559491e-12 | 45 |
| fhl1b2 | 1.33693377092469e-16 | 0.25193176 | 0.186 | 0.017 | 3.36492860804035e-12 | 45 |
| myo1cb3 | 1.71165627345826e-16 | 0.273313656 | 0.279 | 0.036 | 4.3080676746671e-12 | 45 |
| zgc:1008293 | 1.79982375662576e-16 | 0.336739901 | 0.279 | 0.036 | 4.52997641305138e-12 | 45 |
| cybrd11 | 2.09459299654959e-16 | 0.352084712 | 0.349 | 0.053 | 5.27188111301565e-12 | 45 |
| sept9a | 2.31842840377003e-16 | 0.301192249 | 0.326 | 0.046 | 5.83525244944879e-12 | 45 |
| klf11a1 | 2.50045399604624e-16 | 0.335914175 | 0.302 | 0.042 | 6.29339266264878e-12 | 45 |
| sar1b3 | 2.61723470210126e-16 | 0.435752825 | 0.442 | 0.08 | 6.58731802171865e-12 | 45 |
| socs3a14 | 2.91687265264223e-16 | 0.913355622 | 0.907 | 0.298 | 7.34147677943524e-12 | 45 |
| ctdspla4 | 3.04982238662505e-16 | 0.276589147 | 0.279 | 0.036 | 7.67609796489658e-12 | 45 |
| gpatch82 | 3.85546805240353e-16 | 0.391873261 | 0.349 | 0.054 | 9.70382754109446e-12 | 45 |
| hmbox1a1 | 4.44385325805645e-16 | 0.300943322 | 0.256 | 0.032 | 1.11847342652023e-11 | 45 |
| mibp218 | 7.08340456139506e-16 | 0.762951515 | 0.977 | 0.493 | 1.78282209405752e-11 | 45 |
| ostc2 | 8.146002058704e-16 | 0.568029984 | 0.721 | 0.18 | 2.05026725815521e-11 | 45 |
| si:dkey-7j14.611 | 8.32801042641322e-16 | 0.608880226 | 0.698 | 0.175 | 2.09607694422394e-11 | 45 |
| alg5 | 8.44428750639407e-16 | 0.329870613 | 0.395 | 0.067 | 2.12534272248432e-11 | 45 |
| tmem167a1 | 9.85829742914898e-16 | 0.439658278 | 0.488 | 0.097 | 2.48123487994251e-11 | 45 |
| gng24 | 1.82299826877364e-15 | 0.347865642 | 0.302 | 0.044 | 4.58830434267636e-11 | 45 |
| skia2 | 1.90124327032184e-15 | 0.33220373 | 0.302 | 0.044 | 4.78523918707304e-11 | 45 |
| si:dkey-28b4.710 | 2.090715628974e-15 | 0.456027025 | 0.465 | 0.092 | 5.26212216656466e-11 | 45 |
| cd82b5 | 2.11055802012332e-15 | 0.363603546 | 0.419 | 0.075 | 5.31206348084838e-11 | 45 |
| CABZ01075242.1 | 2.77144567532325e-15 | 0.257265808 | 0.186 | 0.019 | 6.97545162022109e-11 | 45 |

| | | | | | | |
|---|---|---|---|---|---|---|
| nid1b4 | 3.34542816456151e-15 | 0.357522663 | 0.349 | 0.056 | 8.42010814738486e-11 | 45 |
| ube2e37 | 3.76864761943806e-15 | 0.504324193 | 0.651 | 0.156 | 9.48530919336365e-11 | 45 |
| rpl3916 | 4.02090226311186e-15 | 0.387870992 | 1 | 0.874 | 1.01202089060262e-10 | 45 |
| yap17 | 5.52881409723286e-15 | 0.300541821 | 0.326 | 0.05 | 1.39154722013254e-10 | 45 |
| sec62 | 5.85043005768562e-15 | 0.337403301 | 0.395 | 0.07 | 1.47249474121889e-10 | 45 |
| calm1a3 | 6.0267865512767e-15 | 0.50880025 | 0.674 | 0.165 | 1.51688190709083e-10 | 45 |
| tnrc6c113 | 7.17711662949054e-15 | 0.499571463 | 0.581 | 0.133 | 1.80640848447647e-10 | 45 |
| rpl1016 | 7.86022394560287e-15 | 0.469903596 | 1 | 0.773 | 1.97833976486879e-10 | 45 |
| cox4i23 | 8.48163863444651e-15 | 0.272143286 | 0.256 | 0.034 | 2.13474362790384e-10 | 45 |
| dap2 | 8.69501580902826e-15 | 0.375604289 | 0.512 | 0.106 | 2.18844852897432e-10 | 45 |
| nsa219 | 8.70959880914781e-15 | 0.779239431 | 0.977 | 0.362 | 2.19211892427441e-10 | 45 |
| sox4a4 | 9.22703900079891e-15 | 0.304125417 | 0.302 | 0.045 | 2.32235344611108e-10 | 45 |
| slc38a10 | 9.36896033347276e-15 | 0.277245799 | 0.233 | 0.029 | 2.35807362633176e-10 | 45 |
| tead1b2 | 9.4831197124122e-15 | 0.299098385 | 0.233 | 0.029 | 2.38680640041703e-10 | 45 |
| cald1a9 | 1.01057183385606e-14 | 0.409957107 | 0.488 | 0.099 | 2.54350824863231e-10 | 45 |
| kdelr2a1 | 1.18726886146235e-14 | 0.409194445 | 0.395 | 0.072 | 2.98823699741459e-10 | 45 |
| ifi307 | 1.33682232516645e-14 | 0.518442043 | 0.698 | 0.175 | 3.36464811021144e-10 | 45 |
| gpx82 | 1.38315894345553e-14 | 0.408924631 | 0.442 | 0.084 | 3.48127274478323e-10 | 45 |
| trib36 | 1.686378430821e-14 | 0.532088496 | 0.488 | 0.106 | 4.24444587253338e-10 | 45 |
| rps260 | 1.73924229532804e-14 | 0.460079714 | 1 | 0.83 | 4.37749893311115e-10 | 45 |
| si:ch211-140m22.74 | 1.8662129654056e-14 | 0.408024023 | 0.372 | 0.067 | 4.69707141262935e-10 | 45 |
| ybx114 | 1.90523426704817e-14 | 0.563863671 | 1 | 0.67 | 4.79528412673354e-10 | 45 |
| tsc22d313 | 2.09600538009414e-14 | 0.508204001 | 0.605 | 0.144 | 5.27543594115894e-10 | 45 |
| serp11 | 2.52253823741113e-14 | 0.557712626 | 0.767 | 0.207 | 6.34897648974008e-10 | 45 |
| pno16 | 2.77654909514259e-14 | 0.507651842 | 0.674 | 0.171 | 6.98829641756439e-10 | 45 |
| sec61g8 | 2.82059009918779e-14 | 0.552809677 | 1 | 0.679 | 7.09914322064576e-10 | 45 |
| chd38 | 2.90786856352021e-14 | 0.450526813 | 0.488 | 0.105 | 7.31881438752402e-10 | 45 |
| rgs3a2 | 3.54465112562154e-14 | 0.275853672 | 0.209 | 0.025 | 8.92153241807685e-10 | 45 |
| cdca4 | 4.01996984801864e-14 | 0.331797985 | 0.349 | 0.06 | 1.01178621104781e-09 | 45 |
| eef2b11 | 4.56647580144476e-14 | 0.592247215 | 0.977 | 0.614 | 1.14933629446563e-09 | 45 |
| cx436 | 4.94082519635173e-14 | 0.435973351 | 0.326 | 0.055 | 1.24355629366977e-09 | 45 |
| arhgap312 | 5.67886644002756e-14 | 0.260062564 | 0.233 | 0.03 | 1.42931389429054e-09 | 45 |
| FO082781.16 | 7.38166014572017e-14 | 0.332757625 | 0.302 | 0.048 | 1.85789004207631e-09 | 45 |
| sec13 | 9.16962750047805e-14 | 0.310015831 | 0.326 | 0.054 | 2.30790354559532e-09 | 45 |
| ramp24 | 1.52044510629505e-13 | 0.288155464 | 0.279 | 0.042 | 3.82680828803402e-09 | 45 |

| | | | | | | |
|---|---|---|---|---|---|---|
| bhlhe4014 | 1.54796054083312e-13 | 0.624439333 | 0.605 | 0.161 | 3.89606188522288e-09 | 45 |
| yipf5 | 1.69480925141163e-13 | 0.296916185 | 0.395 | 0.075 | 4.26566540487793e-09 | 45 |
| selenof4 | 2.32302833174283e-13 | 0.604944648 | 0.791 | 0.237 | 5.84683000816352e-09 | 45 |
| nfkbib | 2.48740077889606e-13 | 0.265322659 | 0.279 | 0.043 | 6.26053902040349e-09 | 45 |
| unc50 | 2.65639557894611e-13 | 0.250250083 | 0.279 | 0.042 | 6.68588203264945e-09 | 45 |
| txndc51 | 2.88454658401479e-13 | 0.362772123 | 0.372 | 0.07 | 7.26011529730683e-09 | 45 |
| pbx1a5 | 4.03016722928636e-13 | 0.270978482 | 0.279 | 0.043 | 1.01435278993908e-08 | 45 |
| sec23a | 4.18394533971676e-13 | 0.264181192 | 0.186 | 0.022 | 1.05305720255331e-08 | 45 |
| rpl128 | 5.45679071852162e-13 | 0.395029008 | 0.977 | 0.799 | 1.37341965594471e-08 | 45 |
| anxa5b8 | 6.41599568110864e-13 | 0.400830834 | 0.442 | 0.094 | 1.61484195297823e-08 | 45 |
| LOC1039092849 | 7.13816385117872e-13 | 0.358149781 | 0.349 | 0.065 | 1.79660445970317e-08 | 45 |
| zfyve211 | 7.15172145034861e-13 | 0.369476879 | 0.326 | 0.058 | 1.80001677183824e-08 | 45 |
| zfand3 | 7.22029070142764e-13 | 0.300764148 | 0.326 | 0.057 | 1.81727496664232e-08 | 45 |
| eif4ebp3l6 | 7.68872407342401e-13 | 0.667011342 | 0.698 | 0.207 | 1.93517496204009e-08 | 45 |
| si:dkey-239h2.32 | 9.15038473778113e-13 | 0.426866517 | 0.419 | 0.089 | 2.30306033465213e-08 | 45 |
| gstz12 | 9.51534751882786e-13 | 0.272424418 | 0.233 | 0.033 | 2.39491781701378e-08 | 45 |
| spcs13 | 9.79432266051455e-13 | 0.5973556 | 0.698 | 0.21 | 2.46513307042491e-08 | 45 |
| mmp16a6 | 1.15022889491433e-12 | 0.280635709 | 0.372 | 0.07 | 2.89501110560989e-08 | 45 |
| map1lc3b7 | 1.22093214813975e-12 | 0.449741876 | 0.465 | 0.107 | 3.07296412365293e-08 | 45 |
| fbxo9 | 1.27098460871607e-12 | 0.293511832 | 0.279 | 0.045 | 3.19894116167749e-08 | 45 |
| ssr22 | 1.36589028015589e-12 | 0.650109957 | 0.953 | 0.322 | 3.43780924612437e-08 | 45 |
| si:dkey-199f5.8 | 1.53195237050107e-12 | 0.291931679 | 0.233 | 0.033 | 3.85577092131415e-08 | 45 |
| plecb6 | 1.80911723310999e-12 | 0.253958855 | 0.302 | 0.051 | 4.55336716401453e-08 | 45 |
| angptl1a4 | 1.89300788406863e-12 | 0.303860952 | 0.209 | 0.028 | 4.76451154341235e-08 | 45 |
| flot2a6 | 1.98278184883532e-12 | 0.341590438 | 0.326 | 0.06 | 4.99046363533362e-08 | 45 |
| nupr1a | 2.14911717123036e-12 | 0.484988481 | 0.605 | 0.161 | 5.40911300826969e-08 | 45 |
| rhoaa12 | 2.81038129564292e-12 | 0.488712223 | 0.581 | 0.154 | 7.07344868300366e-08 | 45 |
| rps1521 | 3.73380264271583e-12 | 0.536439954 | 1 | 0.685 | 9.39760787145147e-08 | 45 |
| anxa11b5 | 4.63460788522184e-12 | 0.428995039 | 0.581 | 0.15 | 1.16648445863148e-07 | 45 |
| ddx218 | 4.90046151352364e-12 | 0.549731389 | 0.698 | 0.208 | 1.23339715833877e-07 | 45 |
| mxra8a4 | 4.92700333648281e-12 | 0.28881255 | 0.279 | 0.046 | 1.24007746975936e-07 | 45 |
| sec61b4 | 9.96909599404654e-12 | 0.691826666 | 0.953 | 0.455 | 2.50912177074157e-07 | 45 |
| ciarta6 | 1.03929027611027e-11 | 0.304155829 | 0.465 | 0.105 | 2.61578969594194e-07 | 45 |
| csrnp1b13 | 1.2848848777144e-11 | 0.519439806 | 0.628 | 0.18 | 3.23392674871937e-07 | 45 |
| pik3r26 | 1.31815942646093e-11 | 0.261030521 | 0.326 | 0.061 | 3.31767546045953e-07 | 45 |

| | | | | | | |
|---|---|---|---|---|---|---|
| rnd3b5 | 1.70561805040539e-11 | 0.274138274 | 0.256 | 0.042 | 4.29287007106534e-07 | 45 |
| tpt113 | 1.92133027394986e-11 | 0.397127084 | 1 | 0.801 | 4.83579616650439e-07 | 45 |
| eif2s37 | 2.01314712752013e-11 | 0.578644228 | 0.86 | 0.285 | 5.06689000525541e-07 | 45 |
| arl12 | 2.25083739049588e-11 | 0.265961298 | 0.349 | 0.069 | 5.66513262813909e-07 | 45 |
| lgals2a6 | 2.54809509077907e-11 | 0.43346204 | 0.512 | 0.126 | 6.41330053398183e-07 | 45 |
| ssr12 | 2.69696548510264e-11 | 0.462488652 | 0.651 | 0.188 | 6.78799242945482e-07 | 45 |
| rab1ab4 | 2.77794304998669e-11 | 0.393138427 | 0.488 | 0.123 | 6.99180486251151e-07 | 45 |
| ghrb5 | 3.41396526682045e-11 | 0.29435404 | 0.256 | 0.043 | 8.59260918006039e-07 | 45 |
| dcaf7 | 3.7767154993773e-11 | 0.304688001 | 0.465 | 0.11 | 9.50561524038272e-07 | 45 |
| cpn15 | 4.18748667303264e-11 | 0.261438104 | 0.302 | 0.056 | 1.05394852073558e-06 | 45 |
| serpine23 | 4.37420712750003e-11 | 0.288569279 | 0.326 | 0.064 | 1.10094419192048e-06 | 45 |
| rps27l12 | 4.40585256605769e-11 | 0.44628884 | 0.698 | 0.204 | 1.10890903235106e-06 | 45 |
| h2afy28 | 4.65673463649447e-11 | 0.377961031 | 0.419 | 0.098 | 1.17205354065929e-06 | 45 |
| prelid3b7 | 4.72031795086246e-11 | 0.576677349 | 0.721 | 0.239 | 1.18805682505257e-06 | 45 |
| zgc:926064 | 4.90202817737088e-11 | 0.318174034 | 0.395 | 0.087 | 1.23379147196248e-06 | 45 |
| col6a26 | 4.95470406971648e-11 | 0.392491473 | 0.326 | 0.066 | 1.24704946730694e-06 | 45 |
| LOC1081802691 | 5.03818555455357e-11 | 0.268620837 | 0.302 | 0.056 | 1.26806092222559e-06 | 45 |
| her92 | 5.42180562448114e-11 | 0.251190726 | 0.163 | 0.02 | 1.36461425762566e-06 | 45 |
| glo17 | 5.80490851058878e-11 | 0.439708589 | 0.674 | 0.197 | 1.46103742303009e-06 | 45 |
| btg213 | 7.34164400012277e-11 | 0.692223134 | 0.907 | 0.373 | 1.8478183783909e-06 | 45 |
| rps1213 | 9.11682869275866e-11 | 0.367561211 | 1 | 0.84 | 2.29461461368043e-06 | 45 |
| serpine12 | 1.16057556113424e-10 | 0.377247916 | 0.256 | 0.045 | 2.92105262981876e-06 | 45 |
| gstr4 | 1.69118327205273e-10 | 0.383784007 | 0.465 | 0.119 | 4.2565391774295e-06 | 45 |
| add3a10 | 1.90514607567916e-10 | 0.362655818 | 0.488 | 0.125 | 4.79506215787687e-06 | 45 |
| cyb5r3 | 2.19638815511685e-10 | 0.274630967 | 0.233 | 0.039 | 5.5280893476136e-06 | 45 |
| ap2m1a5 | 2.22632894263486e-10 | 0.343026963 | 0.535 | 0.143 | 5.60344731571767e-06 | 45 |
| ldlrad4a4 | 2.25218592475839e-10 | 0.263465416 | 0.209 | 0.032 | 5.66852675402439e-06 | 45 |
| si:ch211-66k16.27 | 2.93868614571724e-10 | 0.304046449 | 0.349 | 0.076 | 7.39637916015573e-06 | 45 |
| rpl622 | 3.07870513377684e-10 | 0.513135484 | 1 | 0.668 | 7.74879295120292e-06 | 45 |
| si:ch73-335l21.410 | 3.22484054493868e-10 | 0.616572916 | 0.512 | 0.149 | 8.11660116755616e-06 | 45 |
| vapb | 3.22487789449612e-10 | 0.315433033 | 0.372 | 0.084 | 8.11669517265727e-06 | 45 |
| zgc:1232581 | 3.3064858375288e-10 | 0.250665978 | 0.256 | 0.046 | 8.32209420447625e-06 | 45 |
| tmem2585 | 3.39132377689189e-10 | 0.530345954 | 0.814 | 0.283 | 8.53562281405919e-06 | 45 |
| nr4a115 | 3.40833255994737e-10 | 0.456760862 | 0.581 | 0.168 | 8.57843222013154e-06 | 45 |
| crip117 | 3.74780426892653e-10 | 0.586179812 | 0.791 | 0.271 | 9.43284856446118e-06 | 45 |

| | | | | | | |
|---|---|---|---|---|---|---|
| zgc:11418825 | 3.86392527175472e-10 | 0.566967359 | 1 | 0.628 | 9.72511351647946e-06 | 45 |
| mtdha2 | 4.03626010373525e-10 | 0.37502387 | 0.512 | 0.139 | 1.01588630550912e-05 | 45 |
| cd81a9 | 4.40751539614369e-10 | 0.417113498 | 0.698 | 0.207 | 1.10932755005541e-05 | 45 |
| rpl2220 | 4.46266130764266e-10 | 0.495976663 | 1 | 0.695 | 1.12320722452058e-05 | 45 |
| srprb | 4.50513064371635e-10 | 0.371605948 | 0.558 | 0.157 | 1.13389633171697e-05 | 45 |
| rps614 | 4.75440188038546e-10 | 0.407739178 | 1 | 0.789 | 1.19663540927422e-05 | 45 |
| rps3a12 | 4.9261073032049e-10 | 0.402347805 | 1 | 0.779 | 1.23985194714364e-05 | 45 |
| rps1320 | 4.98964868930924e-10 | 0.463610335 | 1 | 0.725 | 1.25584467861224e-05 | 45 |
| cnbpb9 | 5.20586361709478e-10 | 0.398135225 | 0.628 | 0.185 | 1.31026381378659e-05 | 45 |
| h2afx115 | 6.75806297144246e-10 | 0.451245303 | 0.651 | 0.199 | 1.70093686928235e-05 | 45 |
| tram13 | 7.04593075354701e-10 | 0.326133378 | 0.419 | 0.103 | 1.77339031136025e-05 | 45 |
| rap1b9 | 7.36054881177063e-10 | 0.398866157 | 0.674 | 0.203 | 1.85257653043455e-05 | 45 |
| rpl1715 | 7.86056098975265e-10 | 0.428022751 | 1 | 0.747 | 1.97842459551084e-05 | 45 |
| b2ml17 | 7.90024249570488e-10 | 0.48498696 | 0.814 | 0.267 | 1.98841203374396e-05 | 45 |
| hbegfa4 | 1.22711955078372e-09 | 0.352713318 | 0.442 | 0.112 | 3.08853719736754e-05 | 45 |
| rpl18a19 | 1.29327987291108e-09 | 0.419432622 | 1 | 0.786 | 3.25505611212991e-05 | 45 |
| krt18a.19 | 1.49198288741304e-09 | 0.278261911 | 0.488 | 0.128 | 3.75517172932987e-05 | 45 |
| eef1g17 | 1.55041036714867e-09 | 0.531119703 | 0.977 | 0.621 | 3.9022278530765e-05 | 45 |
| rps1024 | 1.62954367135787e-09 | 0.484787192 | 1 | 0.695 | 4.10139846644063e-05 | 45 |
| selenok7 | 1.64587777491186e-09 | 0.387823098 | 0.465 | 0.128 | 4.14250977167566e-05 | 45 |
| tdh3 | 1.81730994007122e-09 | 0.261502761 | 0.395 | 0.096 | 4.57398738816525e-05 | 45 |
| rpl1120 | 1.83726962851312e-09 | 0.395508908 | 1 | 0.814 | 4.62422392800467e-05 | 45 |
| prdx513 | 1.96127090798872e-09 | 0.346212669 | 0.558 | 0.16 | 4.9363227483168e-05 | 45 |
| pim112 | 2.06145916812709e-09 | 0.395982528 | 0.581 | 0.172 | 5.18848658025907e-05 | 45 |
| surf4l1 | 2.09993218494869e-09 | 0.298086472 | 0.256 | 0.05 | 5.28531931629737e-05 | 45 |
| rpl35a19 | 2.54824466567252e-09 | 0.426034962 | 1 | 0.76 | 6.41367699903117e-05 | 45 |
| rps2011 | 2.74461369169709e-09 | 0.279033628 | 1 | 0.917 | 6.90791820063242e-05 | 45 |
| faua18 | 2.97511938579948e-09 | 0.471927499 | 0.977 | 0.69 | 7.48807798211872e-05 | 45 |
| s100u4 | 3.40610329790784e-09 | 0.28351761 | 0.209 | 0.037 | 8.57282139050424e-05 | 45 |
| ndfip1 | 3.55962001602018e-09 | 0.345006468 | 0.558 | 0.163 | 8.95920761832118e-05 | 45 |
| ipo74 | 3.70951263112357e-09 | 0.34711554 | 0.419 | 0.11 | 9.33647234127491e-05 | 45 |
| rps27a19 | 3.86286777794199e-09 | 0.415428143 | 1 | 0.752 | 9.72245191030221e-05 | 45 |
| arpc1a2 | 4.12061503621093e-09 | 0.284911954 | 0.512 | 0.142 | 0.000103712 | 45 |
| cavin2b10 | 5.77640083189866e-09 | 0.298563605 | 0.302 | 0.067 | 0.000145386 | 45 |
| si:ch211-204c21.17 | 5.97822081781602e-09 | 0.264543443 | 0.326 | 0.074 | 0.000150466 | 45 |

| | | | | | | |
|---|---|---|---|---|---|---|
| ndufa4l13 | 6.13059457140374e-09 | 0.586634958 | 0.953 | 0.395 | 0.000154301 | 45 |
| gabarapb14 | 6.35740814142444e-09 | 0.478183774 | 0.767 | 0.269 | 0.00016001 | 45 |
| rhoab13 | 6.36555253639722e-09 | 0.410810551 | 0.605 | 0.19 | 0.000160215 | 45 |
| rpl925 | 7.1928880229003e-09 | 0.546730433 | 1 | 0.63 | 0.000181038 | 45 |
| hic16 | 7.55397616362431e-09 | 0.258915695 | 0.279 | 0.059 | 0.000190126 | 45 |
| eif3ba6 | 8.41578848962032e-09 | 0.407290879 | 0.651 | 0.211 | 0.000211817 | 45 |
| usp363 | 8.48310228655178e-09 | 0.25645139 | 0.326 | 0.075 | 0.000213511 | 45 |
| romo118 | 8.61704637303974e-09 | 0.587874561 | 0.93 | 0.386 | 0.000216882 | 45 |
| si:ch1073-325m22.2 | 1.00212596371778e-08 | 0.402422993 | 0.698 | 0.23 | 0.000252225 | 45 |
| tob1b6 | 1.14344605400708e-08 | 0.412399401 | 0.558 | 0.174 | 0.000287794 | 45 |
| nck1b2 | 1.17792286232438e-08 | 0.28192207 | 0.256 | 0.053 | 0.000296471 | 45 |
| ccl25b6 | 1.21168611142187e-08 | 0.370683173 | 0.326 | 0.077 | 0.000304969 | 45 |
| dbi9 | 1.31866205962482e-08 | 0.453771745 | 0.628 | 0.214 | 0.000331894 | 45 |
| rps1622 | 1.35951686110665e-08 | 0.447328718 | 1 | 0.738 | 0.000342177 | 45 |
| ier3ip13 | 1.42894833064114e-08 | 0.340351312 | 0.512 | 0.154 | 0.000359652 | 45 |
| tmem176l.18 | 1.69430738670986e-08 | 0.38021719 | 0.581 | 0.181 | 0.00042644 | 45 |
| rps1821 | 1.70084437882104e-08 | 0.450834236 | 1 | 0.725 | 0.000428086 | 45 |
| rpl711 | 1.9641929706854e-08 | 0.343885766 | 0.977 | 0.806 | 0.000494368 | 45 |
| yipf4 | 1.97159569786685e-08 | 0.260996318 | 0.349 | 0.086 | 0.000496231 | 45 |
| rpl36a17 | 2.08407035844974e-08 | 0.361793188 | 1 | 0.81 | 0.00052454 | 45 |
| rpl3021 | 2.18624957441558e-08 | 0.454666763 | 1 | 0.709 | 0.000550257 | 45 |
| rps27.220 | 2.20401054478966e-08 | 0.605204322 | 1 | 0.518 | 0.000554727 | 45 |
| celf217 | 2.29203430249495e-08 | 0.409711119 | 0.721 | 0.243 | 0.000576882 | 45 |
| CR383676.123 | 2.42332477306479e-08 | 0.678488501 | 1 | 0.727 | 0.000609927 | 45 |
| chmp5b2 | 2.62590292727549e-08 | 0.291832057 | 0.419 | 0.115 | 0.000660914 | 45 |
| hmgn618 | 2.75538986494321e-08 | 0.474633661 | 0.744 | 0.266 | 0.000693504 | 45 |
| rpl2921 | 2.76320380927825e-08 | 0.37085757 | 1 | 0.809 | 0.000695471 | 45 |
| eif2s1b5 | 2.91592352609099e-08 | 0.332246942 | 0.558 | 0.174 | 0.000733909 | 45 |
| rpl5a20 | 3.06802669613739e-08 | 0.619450927 | 1 | 0.492 | 0.000772192 | 45 |
| ptp4a15 | 3.14975194327103e-08 | 0.33927606 | 0.419 | 0.118 | 0.000792761 | 45 |
| cnn214 | 3.78264344846834e-08 | 0.319235884 | 0.465 | 0.135 | 0.000952054 | 45 |
| cd1644 | 3.9320381921372e-08 | 0.272543892 | 0.302 | 0.072 | 0.000989655 | 45 |
| rpl2818 | 4.79437925815373e-08 | 0.409894319 | 1 | 0.75 | 0.001206697 | 45 |
| rplp114 | 5.01273215760548e-08 | 0.320178926 | 1 | 0.877 | 0.001261655 | 45 |
| rpsa25 | 5.03696103923904e-08 | 0.417486786 | 1 | 0.753 | 0.001267753 | 45 |

| | | | | | | |
|---|---|---|---|---|---|---|
| junbb9 | 5.07110336419324e-08 | 0.291581219 | 0.302 | 0.074 | 0.001276346 | 45 |
| syf2 | 6.11351997824525e-08 | 0.289609548 | 0.488 | 0.147 | 0.001538712 | 45 |
| rps4x18 | 6.12665238231296e-08 | 0.443588126 | 1 | 0.711 | 0.001542017 | 45 |
| zgc:927448 | 6.92305909140002e-08 | 0.525879259 | 0.884 | 0.354 | 0.001742465 | 45 |
| ahnak13 | 7.11508341505712e-08 | 0.291591402 | 0.442 | 0.128 | 0.001790795 | 45 |
| slc25a332 | 7.64241183910059e-08 | 0.273675239 | 0.372 | 0.1 | 0.001923519 | 45 |
| rpl3421 | 9.40685602071813e-08 | 0.39068634 | 1 | 0.784 | 0.002367612 | 45 |
| myl69 | 9.92042553215609e-08 | 0.296339744 | 0.372 | 0.102 | 0.002496872 | 45 |
| rpl27a22 | 1.02968982387684e-07 | 0.591815328 | 1 | 0.496 | 0.002591626 | 45 |
| maff8 | 1.25011249997789e-07 | 0.264459219 | 0.326 | 0.084 | 0.003146408 | 45 |
| rps2127 | 1.25461672741781e-07 | 0.439493163 | 1 | 0.75 | 0.003157745 | 45 |
| cops91 | 1.33618563374727e-07 | 0.318743347 | 0.372 | 0.105 | 0.003363046 | 45 |
| rps2516 | 1.36120786647303e-07 | 0.415000566 | 1 | 0.726 | 0.003426024 | 45 |
| aimp21 | 1.37412609852121e-07 | 0.2977906 | 0.419 | 0.124 | 0.003458538 | 45 |
| ptprga10 | 1.41605609034815e-07 | 0.269178486 | 0.279 | 0.067 | 0.003564072 | 45 |
| itm2ba16 | 1.43599302069767e-07 | 0.401704818 | 0.767 | 0.274 | 0.003614251 | 45 |
| xbp110 | 1.47869398663259e-07 | 0.517700001 | 0.837 | 0.344 | 0.003721725 | 45 |
| pim34 | 1.48587928847879e-07 | 0.296423822 | 0.442 | 0.129 | 0.00373981 | 45 |
| rpl7a16 | 1.64866244016909e-07 | 0.352973239 | 1 | 0.81 | 0.004149518 | 45 |
| rpl1820 | 1.84250672360339e-07 | 0.392435975 | 1 | 0.774 | 0.004637405 | 45 |
| rpl3121 | 1.88163796576665e-07 | 0.475166068 | 1 | 0.655 | 0.004735895 | 45 |
| rps712 | 1.94035012385965e-07 | 0.345616349 | 1 | 0.821 | 0.004883667 | 45 |
| vdac27 | 1.94369044638523e-07 | 0.402595416 | 0.814 | 0.296 | 0.004892074 | 45 |
| rpl819 | 2.0608570104644e-07 | 0.384033731 | 1 | 0.781 | 0.005186971 | 45 |
| rpl3219 | 2.12749297528546e-07 | 0.366858982 | 1 | 0.814 | 0.005354687 | 45 |
| rpl330 | 2.16174981863201e-07 | 0.27033228 | 1 | 0.876 | 0.005440908 | 45 |
| rack112 | 2.35688011030935e-07 | 0.387209567 | 1 | 0.745 | 0.005932032 | 45 |
| p4hb3 | 2.4352760012574e-07 | 0.384907919 | 0.605 | 0.216 | 0.006129346 | 45 |
| rab5aa3 | 2.6742476962435e-07 | 0.269265908 | 0.302 | 0.078 | 0.006730814 | 45 |
| histh1l10 | 2.84887849994893e-07 | 0.398996335 | 0.372 | 0.111 | 0.007170342 | 45 |
| rps1914 | 2.90038913305128e-07 | 0.305005566 | 1 | 0.864 | 0.007299989 | 45 |
| irf2bp2a10 | 2.95262588841762e-07 | 0.335921167 | 0.535 | 0.177 | 0.007431464 | 45 |
| rplp2l24 | 2.98528971480999e-07 | 0.401602574 | 1 | 0.775 | 0.007513676 | 45 |
| rpl1519 | 3.61191086513952e-07 | 0.41176791 | 1 | 0.745 | 0.009090818 | 45 |
| si:ch73-1a9.319 | 3.62084902415298e-07 | 0.426492408 | 0.698 | 0.266 | 0.009113315 | 45 |

| | | | | | | |
|---|---|---|---|---|---|---|
| eif1b13 | 3.63464244864218e-07 | 0.498189443 | 0.93 | 0.383 | 0.009148032 | 45 |
| mdh1aa5 | 3.93622691948579e-07 | 0.31645131 | 0.605 | 0.207 | 0.00990709 | 45 |
| rpl2615 | 4.12063733661003e-07 | 0.321834047 | 1 | 0.841 | 0.010371232 | 45 |
| rpl3717 | 4.16170857352121e-07 | 0.337785901 | 1 | 0.85 | 0.010474604 | 45 |
| rpl23a23 | 4.17190865147306e-07 | 0.48250083 | 0.977 | 0.635 | 0.010500277 | 45 |
| ginm11 | 4.28520451386852e-07 | 0.265071673 | 0.349 | 0.098 | 0.010785431 | 45 |
| nr1d2a5 | 4.36326514548344e-07 | 0.26518792 | 0.302 | 0.08 | 0.010981902 | 45 |
| mcl1b14 | 4.77800373689275e-07 | 0.37301965 | 0.674 | 0.243 | 0.012025758 | 45 |
| XLOC-00043112 | 4.85541663741364e-07 | 0.314669177 | 0.767 | 0.279 | 0.012220598 | 45 |
| rpl3514 | 5.39688225348697e-07 | 0.347194742 | 1 | 0.805 | 0.013583413 | 45 |
| rpl1312 | 5.51963061738149e-07 | 0.335425643 | 1 | 0.805 | 0.013892358 | 45 |
| nop565 | 6.33952184100707e-07 | 0.318661093 | 0.512 | 0.172 | 0.015955943 | 45 |
| nme2b.16 | 7.44897198858437e-07 | 0.267529131 | 1 | 0.858 | 0.018748318 | 45 |
| gnb1a9 | 7.86048901731096e-07 | 0.325894966 | 0.558 | 0.193 | 0.019784065 | 45 |
| rpl2716 | 8.64235325416714e-07 | 0.385844824 | 1 | 0.763 | 0.021751939 | 45 |
| rplp011 | 8.78358287650966e-07 | 0.300735053 | 1 | 0.852 | 0.0221074 | 45 |
| adh59 | 9.24321196574e-07 | 0.316356169 | 0.628 | 0.222 | 0.02326424 | 45 |
| eef1b24 | 9.24951693988071e-07 | 0.375140926 | 0.977 | 0.735 | 0.023280109 | 45 |
| gadd45ba9 | 9.27674907774373e-07 | 0.386676835 | 0.535 | 0.191 | 0.02334865 | 45 |
| sumo2b15 | 9.41284500552375e-07 | 0.398070781 | 0.698 | 0.27 | 0.02369119 | 45 |
| pfn28 | 1.01241979855407e-06 | 0.316797822 | 0.628 | 0.227 | 0.025481594 | 45 |
| rps2420 | 1.06325134360383e-06 | 0.38496292 | 1 | 0.759 | 0.026760973 | 45 |
| gtpbp46 | 1.16280742405451e-06 | 0.350551133 | 0.651 | 0.238 | 0.0292667 | 45 |
| zgc:778492 | 1.25421723808514e-06 | 0.331761871 | 0.512 | 0.18 | 0.031567394 | 45 |
| btg119 | 1.59742050658134e-06 | 0.420280976 | 0.837 | 0.333 | 0.040205477 | 45 |
| rpl1911 | 1.83925760709004e-06 | 0.298959655 | 1 | 0.852 | 0.046292275 | 45 |
| rpl10a14 | 2.14149724163441e-06 | 0.373314758 | 1 | 0.765 | 0.053899344 | 45 |
| rps2316 | 2.29939466201084e-06 | 0.400870559 | 1 | 0.723 | 0.057873464 | 45 |
| eif3c9 | 2.43012532643156e-06 | 0.300730606 | 0.488 | 0.17 | 0.061163824 | 45 |
| txndc93 | 2.50442338100436e-06 | 0.285037544 | 0.372 | 0.118 | 0.063033832 | 45 |
| pdia310 | 2.67746727757173e-06 | 0.363662727 | 0.744 | 0.284 | 0.067389174 | 45 |
| cd6318 | 2.68175967957472e-06 | 0.410869437 | 0.86 | 0.341 | 0.067497209 | 45 |
| rps15a19 | 2.85544502716453e-06 | 0.39619376 | 1 | 0.78 | 0.071868696 | 45 |
| rpl1418 | 3.04594240494143e-06 | 0.461386331 | 1 | 0.635 | 0.076663324 | 45 |
| rps519 | 3.09353108697957e-06 | 0.427097355 | 1 | 0.702 | 0.077861084 | 45 |

| | | | | | | |
|---|---|---|---|---|---|---|
| cdc429 | 3.68611789217088e-06 | 0.311416311 | 0.581 | 0.213 | 0.092775901 | 45 |
| si:dkey-11f4.148 | 3.72130518251822e-06 | 0.31747545 | 0.349 | 0.11 | 0.09366153 | 45 |
| rps1113 | 3.83955766045601e-06 | 0.351125448 | 1 | 0.766 | 0.096637827 | 45 |
| ak26 | 4.60441050217626e-06 | 0.335802905 | 0.605 | 0.233 | 0.115888408 | 45 |
| edf14 | 5.81693655779917e-06 | 0.30628547 | 0.698 | 0.269 | 0.146406476 | 45 |
| rps8a12 | 5.93814533772056e-06 | 0.325850758 | 0.977 | 0.788 | 0.14945718 | 45 |
| krtcap22 | 7.71879971949672e-06 | 0.282613813 | 0.488 | 0.178 | 0.19427447 | 45 |
| tomm55 | 7.97887660813152e-06 | 0.2602983 | 0.581 | 0.215 | 0.200820345 | 45 |
| zgc:1102396 | 8.391307666675e-06 | 0.273318863 | 0.581 | 0.215 | 0.211200823 | 45 |
| mknk2b13 | 1.0459190103176e-05 | 0.351917385 | 0.512 | 0.198 | 0.263247356 | 45 |
| rps1420 | 1.06672444835443e-05 | 0.368601078 | 1 | 0.76 | 0.268483876 | 45 |
| eif5a12 | 1.07154088487013e-05 | 0.513579117 | 0.977 | 0.479 | 0.269696125 | 45 |
| jpt1b5 | 1.57990678314193e-05 | 0.296500502 | 0.698 | 0.272 | 0.397646738 | 45 |
| rpl2118 | 1.62231785449382e-05 | 0.344510763 | 0.977 | 0.757 | 0.408321181 | 45 |
| arpc216 | 1.76235706923714e-05 | 0.27599245 | 0.698 | 0.268 | 0.443567651 | 45 |
| si:ch211-196l7.419 | 2.01733855125491e-05 | 0.368289828 | 0.721 | 0.315 | 0.50774394 | 45 |
| rps27.119 | 2.0891779888644e-05 | 0.440922276 | 1 | 0.68 | 0.525825208 | 45 |
| eif3ea13 | 2.12809320401455e-05 | 0.312053104 | 0.698 | 0.278 | 0.535619779 | 45 |
| zfp36l1a12 | 2.15828559522013e-05 | 0.260354199 | 0.651 | 0.249 | 0.543218901 | 45 |
| rab108 | 2.41840453812393e-05 | 0.269608992 | 0.512 | 0.196 | 0.608688238 | 45 |
| eif3i10 | 2.63610495650479e-05 | 0.315229465 | 0.674 | 0.274 | 0.663481257 | 45 |
| rpl2420 | 3.2066615650848e-05 | 0.347800539 | 1 | 0.777 | 0.807084649 | 45 |
| rps318 | 3.87954364063504e-05 | 0.401878352 | 1 | 0.7 | 0.976442339 | 45 |
| rps26l21 | 4.67847374856915e-05 | 0.442153632 | 1 | 0.665 | 1 | 45 |
| calm2b10 | 5.47328400400989e-05 | 0.254633928 | 0.535 | 0.212 | 1 | 45 |
| serbp1a15 | 6.00940292929144e-05 | 0.49372607 | 0.977 | 0.482 | 1 | 45 |
| rpl3624 | 7.13459115817189e-05 | 0.353533714 | 1 | 0.79 | 1 | 45 |
| rps2916 | 8.03891378829459e-05 | 0.297178941 | 1 | 0.833 | 1 | 45 |
| ppiaa16 | 8.7982938339103e-05 | 0.300138356 | 1 | 0.804 | 1 | 45 |
| rps917 | 9.82109384111757e-05 | 0.329585358 | 1 | 0.777 | 1 | 45 |
| eif3f19 | 0.000100874 | 0.457146141 | 0.907 | 0.432 | 1 | 45 |
| wu:fb55g094 | 0.000109504 | 0.276365418 | 0.605 | 0.251 | 1 | 45 |
| eif3m10 | 0.000114416 | 0.369553153 | 0.791 | 0.359 | 1 | 45 |
| nfil3-611 | 0.000129278 | 0.288756264 | 0.349 | 0.132 | 1 | 45 |
| ctsla12 | 0.000140752 | 0.349655777 | 0.884 | 0.387 | 1 | 45 |

| | | | | | | |
|---|---|---|---|---|---|---|
| uba5220 | 0.00016415 | 0.435479832 | 1 | 0.631 | 1 | 45 |
| hmgn713 | 0.000330318 | 0.255601043 | 0.651 | 0.28 | 1 | 45 |
| ccng12 | 0.000430213 | 0.338991602 | 0.977 | 0.448 | 1 | 45 |
| rps2614 | 0.000477784 | 0.342771433 | 1 | 0.734 | 1 | 45 |
| dap1b4 | 0.000533882 | 0.275188 | 0.558 | 0.252 | 1 | 45 |
| rps2821 | 0.000623234 | 0.29263263 | 1 | 0.827 | 1 | 45 |
| higd1a13 | 0.000650367 | 0.30972033 | 0.651 | 0.311 | 1 | 45 |
| rpl3821 | 0.001165501 | 0.356222725 | 1 | 0.729 | 1 | 45 |
| rsl24d118 | 0.001176521 | 0.400449295 | 0.953 | 0.444 | 1 | 45 |
| ptmaa21 | 0.00130287 | 0.281738316 | 0.86 | 0.396 | 1 | 45 |
| rplp224 | 0.001462642 | 0.345324972 | 1 | 0.758 | 1 | 45 |
| ppdpfb8 | 0.00195831 | 0.382130044 | 0.93 | 0.52 | 1 | 45 |
| ddx55 | 0.002475089 | 0.251440501 | 0.744 | 0.358 | 1 | 45 |
| rpl412 | 0.003452473 | 0.370957751 | 0.977 | 0.628 | 1 | 45 |
| rpl13a9 | 0.003666331 | 0.29104989 | 1 | 0.765 | 1 | 45 |
| fkbp1aa12 | 0.005763408 | 0.254884976 | 0.791 | 0.388 | 1 | 45 |
| rpl22l115 | 0.008990592 | 0.254160104 | 0.791 | 0.393 | 1 | 45 |
| LOC1104380242 | 0 | 2.11776984 | 0.976 | 0.006 | 0 | 46 |
| igf32 | 0 | 1.967562106 | 0.951 | 0.013 | 0 | 46 |
| LOC1039109031 | 0 | 1.839956182 | 0.902 | 0.01 | 0 | 46 |
| znf11041 | 0 | 1.727434233 | 0.878 | 0.013 | 0 | 46 |
| dmrt13 | 0 | 1.720373167 | 0.927 | 0.017 | 0 | 46 |
| gpx31 | 0 | 1.719961297 | 0.854 | 0.005 | 0 | 46 |
| LOC110437926 | 0 | 1.576840391 | 0.829 | 0.002 | 0 | 46 |
| cyp26a12 | 0 | 1.483502805 | 0.78 | 0.008 | 0 | 46 |
| si:ch211-117c9.52 | 0 | 1.445505723 | 0.829 | 0.016 | 0 | 46 |
| scg32 | 0 | 1.39116123 | 0.829 | 0.008 | 0 | 46 |
| amh1 | 0 | 1.389431204 | 0.707 | 0.004 | 0 | 46 |
| inha2 | 0 | 1.362316261 | 0.756 | 0.005 | 0 | 46 |
| zgc:921622 | 0 | 1.324643995 | 0.732 | 0.008 | 0 | 46 |
| ugt5g11 | 0 | 1.31907184 | 0.78 | 0.009 | 0 | 46 |
| alcamb2 | 0 | 1.268191712 | 0.756 | 0.013 | 0 | 46 |
| mmel11 | 0 | 1.233014569 | 0.707 | 0.002 | 0 | 46 |
| inhbb | 0 | 1.226856614 | 0.683 | 0.003 | 0 | 46 |
| nrgna2 | 0 | 1.211561805 | 0.61 | 0.008 | 0 | 46 |

| | | | | | | |
|---|---|---|---|---|---|---|
| XLOC-0422291 | 0 | 1.203999954 | 0.707 | 0.004 | 0 | 46 |
| scn1ba1 | 0 | 1.187421319 | 0.634 | 0.005 | 0 | 46 |
| sv2bb | 0 | 1.149565958 | 0.683 | 0.006 | 0 | 46 |
| kirrel3l | 0 | 1.103655888 | 0.634 | 0.006 | 0 | 46 |
| LOC103908925 | 0 | 1.074969584 | 0.585 | 0.002 | 0 | 46 |
| col28a2a | 0 | 1.060169479 | 0.61 | 0.002 | 0 | 46 |
| cnga1a | 0 | 0.949467358 | 0.537 | 0.002 | 0 | 46 |
| kcnj1a.11 | 0 | 0.938215982 | 0.512 | 0.001 | 0 | 46 |
| ppp1r1b1 | 0 | 0.872282362 | 0.512 | 0.005 | 0 | 46 |
| LOC110438394 | 0 | 0.793640917 | 0.366 | 0.001 | 0 | 46 |
| nts | 0 | 0.765671464 | 0.317 | 0.001 | 0 | 46 |
| CT573799.2 | 0 | 0.745961709 | 0.439 | 0.001 | 0 | 46 |
| si:dkeyp-80c12.81 | 0 | 0.738594131 | 0.415 | 0.001 | 0 | 46 |
| slc4a2a | 0 | 0.722096941 | 0.463 | 0.003 | 0 | 46 |
| adm2a1 | 0 | 0.671123061 | 0.39 | 0.003 | 0 | 46 |
| XLOC-042806 | 0 | 0.662195372 | 0.341 | 0.001 | 0 | 46 |
| dlb | 0 | 0.492747314 | 0.244 | 0.001 | 0 | 46 |
| LOC103910515 | 0 | 0.474232601 | 0.244 | 0 | 0 | 46 |
| sstr5 | 0 | 0.451907471 | 0.268 | 0 | 0 | 46 |
| flrt1b | 0 | 0.422454192 | 0.22 | 0 | 0 | 46 |
| si:dkey-165n16.5 | 0 | 0.369252094 | 0.195 | 0 | 0 | 46 |
| plxnb31 | 1.5854613666252e-307 | 0.907108838 | 0.512 | 0.006 | 3.99044771365895e-303 | 46 |
| tyro3 | 1.70062145385729e-305 | 0.551372704 | 0.317 | 0.002 | 4.28029413721341e-301 | 46 |
| hpn2 | 2.38781909190095e-281 | 0.889315794 | 0.512 | 0.006 | 6.00990187240549e-277 | 46 |
| dmrt3a | 3.5968637869875e-254 | 0.260133548 | 0.146 | 0 | 9.05294646546884e-250 | 46 |
| add21 | 4.01020108010979e-244 | 0.784624228 | 0.415 | 0.005 | 1.00932750985283e-239 | 46 |
| sox9a1 | 1.1924017820834e-242 | 0.944315923 | 0.561 | 0.009 | 3.00115604532572e-238 | 46 |
| sema7a1 | 2.01979667865554e-239 | 0.856393043 | 0.463 | 0.006 | 5.08362626050813e-235 | 46 |
| cx34.51 | 1.74973599433798e-237 | 0.83217283 | 0.39 | 0.004 | 4.40391052414927e-233 | 46 |
| CABZ01058222.1 | 8.44248586889453e-236 | 0.307266744 | 0.122 | 0 | 2.12488926834206e-231 | 46 |
| msx1b1 | 2.03669078075747e-233 | 0.69679156 | 0.39 | 0.004 | 5.12614702608847e-229 | 46 |
| XLOC-040516 | 6.14494340962416e-233 | 0.657507918 | 0.341 | 0.003 | 1.5466208067683e-228 | 46 |
| cyb5d11 | 9.19333026294943e-230 | 0.527568398 | 0.317 | 0.003 | 2.31386929388174e-225 | 46 |
| nkx3.31 | 2.8563975097166e-224 | 0.450244982 | 0.244 | 0.001 | 7.1892668922057e-220 | 46 |
| gpm6ab3 | 1.46990665700845e-221 | 1.149959691 | 0.683 | 0.016 | 3.69960806502457e-217 | 46 |

| | | | | | | |
|---|---|---|---|---|---|---|
| ctnnd2b1 | 1.42077309525902e-218 | 0.464027588 | 0.268 | 0.002 | 3.57594380345744e-214 | 46 |
| slc2a1b2 | 6.49421990143968e-217 | 1.203170945 | 0.707 | 0.018 | 1.63453020699335e-212 | 46 |
| XLOC-033950 | 5.51138487241644e-212 | 0.438934426 | 0.195 | 0.001 | 1.3871604585385e-207 | 46 |
| soul52 | 1.09122193109151e-207 | 1.172152586 | 0.634 | 0.015 | 2.74649647836423e-203 | 46 |
| rxfp3.2a | 2.29710213199449e-203 | 0.32790882 | 0.146 | 0 | 5.78157635601694e-199 | 46 |
| si:ch211-212k18.6 | 2.81900702340835e-203 | 0.287196102 | 0.146 | 0 | 7.09515877721647e-199 | 46 |
| cldnc4 | 3.71092381316102e-203 | 1.46290988 | 0.829 | 0.026 | 9.34002414534496e-199 | 46 |
| large21 | 1.12280644806763e-200 | 0.762869075 | 0.415 | 0.006 | 2.82599154914142e-196 | 46 |
| scube11 | 1.08983691371938e-199 | 1.051620376 | 0.683 | 0.018 | 2.74301052814031e-195 | 46 |
| gstt1a5 | 4.4225297178714e-196 | 1.701644021 | 0.927 | 0.036 | 1.11310650469105e-191 | 46 |
| tmem229b | 1.36179278863335e-195 | 0.68007401 | 0.366 | 0.005 | 3.42749626971127e-191 | 46 |
| jam2a4 | 1.47318513515483e-193 | 1.898425585 | 0.902 | 0.035 | 3.70785966667119e-189 | 46 |
| fat21 | 1.40398873748617e-192 | 0.730160467 | 0.39 | 0.005 | 3.53369925337893e-188 | 46 |
| si:dkey-6n6.11 | 1.03265525308811e-191 | 0.744241251 | 0.341 | 0.004 | 2.59909000649746e-187 | 46 |
| LOC100149700 | 2.20432620911138e-189 | 0.375844797 | 0.195 | 0.001 | 5.54806863571244e-185 | 46 |
| map1ab | 9.81396678714454e-187 | 0.70489138 | 0.366 | 0.005 | 2.47007730065641e-182 | 46 |
| zgc:1949901 | 5.9849398950651e-183 | 0.495953049 | 0.268 | 0.002 | 1.50634952218894e-178 | 46 |
| fhod3b | 3.06754465474482e-182 | 0.726206515 | 0.341 | 0.004 | 7.72070314152725e-178 | 46 |
| gsdf2 | 1.41280806826657e-179 | 2.524706226 | 1 | 0.048 | 3.55589662702013e-175 | 46 |
| CDK18 | 2.79246592960937e-178 | 0.526911968 | 0.293 | 0.003 | 7.02835749823381e-174 | 46 |
| gh1 | 6.24003981767252e-178 | 0.622377855 | 0.268 | 0.003 | 1.57055562171e-173 | 46 |
| bcam3 | 7.17860515683974e-170 | 1.371796804 | 0.78 | 0.028 | 1.80678313192499e-165 | 46 |
| XLOC-040108 | 3.60938097698655e-169 | 0.5024819 | 0.268 | 0.003 | 9.08445098097745e-165 | 46 |
| slc16a7 | 3.759186916219e-169 | 0.591656825 | 0.341 | 0.005 | 9.4614975494316e-165 | 46 |
| plpp7 | 3.04502477144977e-168 | 0.365834445 | 0.22 | 0.002 | 7.66402284726192e-164 | 46 |
| atp11a2 | 1.7969870869882e-160 | 0.799235867 | 0.488 | 0.011 | 4.52283679924061e-156 | 46 |
| lpar3 | 1.8173700329939e-160 | 0.494343105 | 0.293 | 0.004 | 4.57413863604234e-156 | 46 |
| si:ch211-198m17.1 | 7.52393336825771e-160 | 0.420071869 | 0.244 | 0.002 | 1.89369878945678e-155 | 46 |
| CU929418.2 | 4.11205378716761e-157 | 0.28590365 | 0.122 | 0 | 1.03496281769221e-152 | 46 |
| ftr825 | 5.98734058167071e-156 | 1.070872798 | 0.659 | 0.022 | 1.5069537510007e-151 | 46 |
| mylka | 1.52974140089204e-152 | 0.814216479 | 0.488 | 0.012 | 3.85020613190517e-148 | 46 |
| rhoub2 | 3.34637726953292e-151 | 0.656287494 | 0.341 | 0.005 | 8.42249694968742e-147 | 46 |
| hspb83 | 2.53434745420952e-150 | 1.239119387 | 0.659 | 0.023 | 6.37869910749993e-146 | 46 |
| fyna1 | 9.58934893242299e-147 | 0.75223299 | 0.463 | 0.011 | 2.41354323280154e-142 | 46 |
| si:ch73-156l6.2 | 3.70233671353946e-145 | 0.257285514 | 0.146 | 0.001 | 9.31841127430746e-141 | 46 |

| | | | | | | |
|---|---|---|---|---|---|---|
| dag16 | 8.36533856521772e-144 | 1.560039606 | 0.902 | 0.048 | 2.10547206347965e-139 | 46 |
| slc7a3a5 | 1.73165510028232e-143 | 1.387020998 | 0.854 | 0.041 | 4.35840272190058e-139 | 46 |
| nat8l | 9.30930122600683e-140 | 0.411999831 | 0.244 | 0.003 | 2.34305802557366e-135 | 46 |
| lnx2a1 | 3.80021729427691e-139 | 0.522450829 | 0.317 | 0.005 | 9.56476690796557e-135 | 46 |
| lcat | 4.14147073100887e-138 | 0.381293667 | 0.195 | 0.002 | 1.04236676828762e-133 | 46 |
| prr15lb | 1.44751378776219e-137 | 0.283641794 | 0.171 | 0.001 | 3.64324745241865e-133 | 46 |
| fndc5b | 2.2684231716773e-137 | 0.507005695 | 0.268 | 0.004 | 5.70939428079459e-133 | 46 |
| nhsl1a | 8.52213698098456e-136 | 0.612243365 | 0.341 | 0.006 | 2.14493665674401e-131 | 46 |
| cgnl14 | 6.96827908810224e-128 | 0.990987945 | 0.61 | 0.023 | 1.75384616368445e-123 | 46 |
| abtb2a | 9.68499817257898e-124 | 0.364324881 | 0.195 | 0.002 | 2.4376171900564e-119 | 46 |
| itga3b1 | 1.11582256228064e-123 | 0.420804381 | 0.244 | 0.003 | 2.80841380700415e-119 | 46 |
| cldn11a7 | 8.04162793623994e-123 | 1.772523596 | 0.927 | 0.061 | 2.02399733527223e-118 | 46 |
| rnf24 | 3.97830555170496e-122 | 1.052989729 | 0.634 | 0.026 | 1.00129972430862e-117 | 46 |
| cx437 | 1.08211810557177e-121 | 1.534983236 | 0.878 | 0.052 | 2.72358305991359e-117 | 46 |
| stard9 | 1.98553107797842e-119 | 0.642723403 | 0.366 | 0.008 | 4.99738317016388e-115 | 46 |
| rims2b | 7.37665736417318e-118 | 0.541618504 | 0.317 | 0.006 | 1.85663089198875e-113 | 46 |
| sobpa | 4.05023900312234e-116 | 0.687049653 | 0.39 | 0.01 | 1.01940465469586e-111 | 46 |
| trim3b | 2.412991953754e-115 | 0.467771998 | 0.293 | 0.005 | 6.07325944840344e-111 | 46 |
| cox6a26 | 3.13131330621249e-115 | 1.123058928 | 0.707 | 0.035 | 7.88120246040621e-111 | 46 |
| ncam1a3 | 2.93807106482442e-114 | 0.804158077 | 0.512 | 0.018 | 7.39483106305658e-110 | 46 |
| appa8 | 6.58870443119345e-114 | 1.539675088 | 1 | 0.075 | 1.65831101828708e-109 | 46 |
| nos1apa2 | 1.12625769287683e-113 | 0.865939934 | 0.537 | 0.02 | 2.8346779872017e-109 | 46 |
| phlda26 | 2.82751196643706e-113 | 1.57321057 | 0.829 | 0.05 | 7.11656486832544e-109 | 46 |
| nfasca | 2.71277631844605e-111 | 0.418714725 | 0.244 | 0.004 | 6.82778671589687e-107 | 46 |
| rbm472 | 4.24300458361527e-111 | 0.771989691 | 0.463 | 0.015 | 1.06792182365013e-106 | 46 |
| map4l | 3.50937504936832e-110 | 0.305747684 | 0.171 | 0.002 | 8.83274606175513e-106 | 46 |
| pyroxd2 | 4.92764780867735e-110 | 0.294311286 | 0.171 | 0.002 | 1.240239676966e-105 | 46 |
| si:ch211-225b11.11 | 1.78809349653909e-107 | 1.121870269 | 0.659 | 0.032 | 4.50045252143923e-103 | 46 |
| adgrl2a | 6.34732045126497e-107 | 0.731044161 | 0.439 | 0.014 | 1.59755708437888e-102 | 46 |
| csgalnact1a | 2.06463063013446e-106 | 0.813161626 | 0.488 | 0.017 | 5.19646883298542e-102 | 46 |
| epdl14 | 3.55862123093322e-106 | 1.166589084 | 0.683 | 0.035 | 8.95669377613582e-102 | 46 |
| LOC110439817 | 9.43403779209807e-102 | 0.33592019 | 0.171 | 0.002 | 2.37445297189316e-97 | 46 |
| ak43 | 6.93266320431735e-100 | 1.116679916 | 0.61 | 0.03 | 1.74488200189463e-95 | 46 |
| igsf31 | 2.54412242043844e-98 | 0.44870515 | 0.268 | 0.005 | 6.40330172000152e-94 | 46 |
| metrn1 | 1.06177809290771e-97 | 0.736592588 | 0.39 | 0.012 | 2.67238928203941e-93 | 46 |

| | | | | | | |
|---|---|---|---|---|---|---|
| asb5b | 1.60510207340971e-96 | 0.302919785 | 0.195 | 0.003 | 4.0398814085649e-92 | 46 |
| mmp11a | 7.98399157579268e-95 | 0.395535698 | 0.171 | 0.002 | 2.00949083971126e-90 | 46 |
| bag33 | 3.85509284385929e-92 | 1.106797113 | 0.634 | 0.035 | 9.70288317870946e-88 | 46 |
| rhcgb | 4.25634366853305e-92 | 0.34716548 | 0.146 | 0.002 | 1.07127913793308e-87 | 46 |
| prss121 | 7.37436217990605e-92 | 0.421284407 | 0.22 | 0.004 | 1.85605321706055e-87 | 46 |
| cth3 | 2.25695114461955e-90 | 1.163230177 | 0.707 | 0.043 | 5.68052033589295e-86 | 46 |
| myh104 | 2.00363846871789e-89 | 0.933866043 | 0.634 | 0.035 | 5.04295766191607e-85 | 46 |
| XLOC-031367 | 5.49940285572481e-88 | 0.435431337 | 0.244 | 0.005 | 1.38414470475738e-83 | 46 |
| pkdcca | 2.0550745570478e-87 | 0.384080837 | 0.195 | 0.003 | 5.17241715263361e-83 | 46 |
| fam13b2 | 1.35190459401682e-86 | 0.929271234 | 0.585 | 0.031 | 3.40260867268094e-82 | 46 |
| elovl51 | 2.28233758711798e-86 | 0.691354858 | 0.39 | 0.014 | 5.74441547301725e-82 | 46 |
| hkdc11 | 6.87273980391961e-86 | 0.322826416 | 0.171 | 0.002 | 1.72979988124853e-81 | 46 |
| znfx12 | 1.0417940435636e-85 | 0.885487727 | 0.561 | 0.029 | 2.62209142824524e-81 | 46 |
| cnnm2b | 5.52777936492796e-85 | 0.33701092 | 0.195 | 0.003 | 1.39128678835872e-80 | 46 |
| XLOC-031372 | 1.89026161093069e-84 | 0.256102445 | 0.146 | 0.002 | 4.75759944855144e-80 | 46 |
| RTKN | 4.01408152433651e-81 | 0.476998142 | 0.268 | 0.007 | 1.01030417886026e-76 | 46 |
| si:dkey-194e6.11 | 1.24687424562275e-80 | 0.342378574 | 0.171 | 0.003 | 3.13825778880789e-76 | 46 |
| lpcat22 | 1.75374444444462e-80 | 0.712032282 | 0.463 | 0.021 | 4.41399939222266e-76 | 46 |
| lgr4 | 3.81532253232287e-80 | 0.353060284 | 0.22 | 0.004 | 9.60278528160343e-76 | 46 |
| mfge8b1 | 6.06419439697281e-80 | 0.924948888 | 0.585 | 0.034 | 1.52629708777409e-75 | 46 |
| cahz3 | 6.19984836033305e-79 | 0.971789905 | 0.634 | 0.039 | 1.56043983381223e-74 | 46 |
| si:dkeyp-80c12.7 | 1.10862467992577e-78 | 0.284710408 | 0.122 | 0.001 | 2.79029745690516e-74 | 46 |
| lamb1a5 | 3.73766639960688e-78 | 0.945345392 | 0.61 | 0.038 | 9.40733256117055e-74 | 46 |
| slc31a2 | 8.65238602447492e-78 | 0.545695259 | 0.366 | 0.013 | 2.17771903850009e-73 | 46 |
| prss354 | 3.82254463465132e-76 | 0.852913667 | 0.439 | 0.02 | 9.6209625909539e-72 | 46 |
| gdpd4a1 | 2.55963040534495e-75 | 0.546715142 | 0.317 | 0.01 | 6.44233376721271e-71 | 46 |
| shank3a | 6.05726752854018e-75 | 0.292997047 | 0.146 | 0.002 | 1.52455366425828e-70 | 46 |
| itgb8 | 6.89069572462721e-75 | 0.25968248 | 0.146 | 0.002 | 1.73431920693142e-70 | 46 |
| gadd45aa2 | 1.67398328838457e-73 | 0.796893628 | 0.415 | 0.019 | 4.21324853853512e-69 | 46 |
| nlgn4xa1 | 5.21516476507843e-73 | 0.498945595 | 0.317 | 0.011 | 1.31260481972259e-68 | 46 |
| fabp7a5 | 1.07136286159747e-72 | 0.961724427 | 0.488 | 0.026 | 2.69651318635468e-68 | 46 |
| ctsf1 | 1.94419794468868e-72 | 0.784164849 | 0.488 | 0.026 | 4.89335180698694e-68 | 46 |
| FAT42 | 1.99740963126567e-72 | 0.643392025 | 0.39 | 0.017 | 5.02728030093256e-68 | 46 |
| aplp25 | 6.1669603539677e-72 | 1.130236332 | 0.805 | 0.071 | 1.55216225149013e-67 | 46 |
| gata63 | 9.91909297463918e-72 | 0.620634741 | 0.415 | 0.019 | 2.49653651078694e-67 | 46 |

| | | | | | | |
|---|---|---|---|---|---|---|
| podxl4 | 2.3492813293092e-70 | 0.979006668 | 0.659 | 0.046 | 5.91290617773832e-66 | 46 |
| esr2b3 | 2.47233656745394e-70 | 0.818417348 | 0.561 | 0.035 | 6.22262390662483e-66 | 46 |
| blf | 1.31778697188069e-69 | 0.35523646 | 0.195 | 0.004 | 3.31673802952651e-65 | 46 |
| kank26 | 1.93156562025886e-69 | 1.020085035 | 0.683 | 0.053 | 4.86155750962952e-65 | 46 |
| entpd8 | 2.29918430137847e-69 | 0.445624557 | 0.244 | 0.007 | 5.78681696813948e-65 | 46 |
| FO704858.1 | 2.3876145479582e-68 | 0.291678035 | 0.171 | 0.003 | 6.009387055756e-64 | 46 |
| osbpl3a1 | 5.34608610106336e-68 | 0.745738126 | 0.463 | 0.025 | 1.34555641077664e-63 | 46 |
| pard3ab | 1.09352935560521e-67 | 0.459108552 | 0.268 | 0.008 | 2.75230403512275e-63 | 46 |
| DOCK9 | 1.1406170851006e-67 | 0.419999961 | 0.244 | 0.007 | 2.87081914148971e-63 | 46 |
| CABZ01021532.12 | 1.13525823850752e-66 | 0.661081209 | 0.366 | 0.016 | 2.85733146049958e-62 | 46 |
| ap1m11 | 1.4967828831539e-65 | 1.104867301 | 0.756 | 0.07 | 3.76725283861005e-61 | 46 |
| ahcyl21 | 3.51987197607326e-65 | 0.613669509 | 0.366 | 0.016 | 8.85916577657879e-61 | 46 |
| mgll | 1.5095121913435e-64 | 0.392001592 | 0.195 | 0.004 | 3.79929123439244e-60 | 46 |
| zmp:00000011141 | 2.18848230283062e-63 | 0.466992013 | 0.293 | 0.01 | 5.5081911079944e-59 | 46 |
| atf5b2 | 3.44501631928426e-63 | 0.811703292 | 0.512 | 0.033 | 8.67076157400657e-59 | 46 |
| tmem45a3 | 6.13218030258786e-63 | 0.785178193 | 0.439 | 0.024 | 1.54340846035834e-58 | 46 |
| wfdc21 | 9.03055033618306e-63 | 0.369294501 | 0.22 | 0.006 | 2.27289921411391e-58 | 46 |
| emilin1a3 | 1.315490211671e-62 | 0.627821383 | 0.39 | 0.019 | 3.31095731375473e-58 | 46 |
| rassf91 | 3.49454210020141e-62 | 0.360395851 | 0.195 | 0.005 | 8.79541301199693e-58 | 46 |
| inhbab1 | 4.1093647406402e-62 | 0.497142471 | 0.268 | 0.009 | 1.03428601157173e-57 | 46 |
| tjp1a6 | 2.11235568259277e-61 | 0.856539008 | 0.61 | 0.047 | 5.31658801751773e-57 | 46 |
| lmbrd2b | 2.56410878164305e-61 | 0.400762535 | 0.22 | 0.006 | 6.4536053925174e-57 | 46 |
| wu:fc27b112 | 8.55435931504775e-61 | 0.538240966 | 0.317 | 0.013 | 2.15304669600437e-56 | 46 |
| ugt1b53 | 2.13312116639759e-60 | 0.834957872 | 0.463 | 0.028 | 5.3688526637061e-56 | 46 |
| mmp15b | 3.41027640383263e-60 | 0.42335419 | 0.244 | 0.008 | 8.58332468080635e-56 | 46 |
| slc7a61 | 4.39041642064066e-60 | 0.580207199 | 0.366 | 0.017 | 1.10502390891105e-55 | 46 |
| senp3a1 | 8.13186793252054e-60 | 0.758078388 | 0.439 | 0.025 | 2.04670983993609e-55 | 46 |
| zgc:1527917 | 1.68118904900148e-59 | 1.2644642 | 0.707 | 0.068 | 4.23138471743183e-55 | 46 |
| aplp12 | 6.17073442654029e-59 | 0.610261264 | 0.341 | 0.015 | 1.55311214781593e-54 | 46 |
| angptl48 | 1.1204498881848e-58 | 0.919219705 | 0.61 | 0.049 | 2.82006032357232e-54 | 46 |
| ube2ql1 | 1.38074478717873e-58 | 0.407754399 | 0.22 | 0.006 | 3.47519655485015e-54 | 46 |
| lrp2a | 6.83765439473706e-58 | 0.321948174 | 0.195 | 0.005 | 1.72096923461137e-53 | 46 |
| lgals9l34 | 8.05520826359226e-58 | 0.948938357 | 0.683 | 0.063 | 2.02741536786354e-53 | 46 |
| tle2b1 | 1.17814610285958e-57 | 0.547989791 | 0.317 | 0.014 | 2.96527592628728e-53 | 46 |
| bmpr2a2 | 1.47658938565089e-57 | 0.639650476 | 0.366 | 0.018 | 3.71642782474472e-53 | 46 |

| | | | | | | |
|---|---|---|---|---|---|---|
| ndrg25 | 1.51194820474736e-57 | 0.793444717 | 0.537 | 0.039 | 3.80542243652864e-53 | 46 |
| actn14 | 3.22863033359267e-57 | 0.909664982 | 0.561 | 0.043 | 8.1261396866194e-53 | 46 |
| ca71 | 3.09072987146096e-56 | 0.784509745 | 0.537 | 0.038 | 7.77905801348008e-52 | 46 |
| ptpn131 | 1.06940488688635e-55 | 0.615524175 | 0.341 | 0.017 | 2.69158515980425e-51 | 46 |
| sparc11 | 5.47908466454069e-55 | 1.508789909 | 1 | 0.158 | 1.37903081921825e-50 | 46 |
| sema3b1 | 2.15933409522566e-54 | 0.620163462 | 0.39 | 0.022 | 5.43482798427347e-50 | 46 |
| marcksl1b12 | 1.47263041461857e-53 | 1.439356043 | 0.854 | 0.115 | 3.70646349055347e-49 | 46 |
| fosl1a10 | 7.32041916224263e-53 | 1.152601737 | 0.78 | 0.091 | 1.84247629894485e-48 | 46 |
| phc2a3 | 8.47043626200397e-53 | 1.171112262 | 0.805 | 0.092 | 2.13192410278378e-48 | 46 |
| slc4a4a2 | 1.13627700261185e-52 | 0.484306531 | 0.268 | 0.011 | 2.85989558787378e-48 | 46 |
| pon11 | 2.96638522950406e-52 | 0.499060739 | 0.268 | 0.011 | 7.46609498413877e-48 | 46 |
| plat1 | 6.04876380516773e-52 | 0.498850949 | 0.268 | 0.011 | 1.52241336212267e-47 | 46 |
| si:dkey-188i13.92 | 9.84650830628228e-51 | 0.73344178 | 0.39 | 0.024 | 2.47826767560819e-46 | 46 |
| fam184b | 2.01303388400446e-50 | 0.303400681 | 0.146 | 0.003 | 5.06660498265083e-46 | 46 |
| p2rx72 | 1.34957304488238e-49 | 0.502179676 | 0.293 | 0.014 | 3.39674039666446e-45 | 46 |
| agrn4 | 2.26273579409801e-49 | 0.887078968 | 0.561 | 0.049 | 5.69507972016529e-45 | 46 |
| shisa9b | 2.37820510187612e-49 | 0.296414719 | 0.171 | 0.005 | 5.98570442091201e-45 | 46 |
| sema4ba6 | 7.60042847967087e-49 | 0.865440502 | 0.537 | 0.046 | 1.91295184404836e-44 | 46 |
| tmem245 | 1.79403747050281e-48 | 0.45170203 | 0.268 | 0.012 | 4.51541290950852e-44 | 46 |
| ifi468 | 2.30032596778984e-48 | 1.311848384 | 0.829 | 0.121 | 5.78969042833025e-44 | 46 |
| arhgef18a1 | 3.57432224563756e-48 | 0.542621914 | 0.341 | 0.019 | 8.99621166004517e-44 | 46 |
| znf5361 | 3.85976606117904e-48 | 0.387595906 | 0.22 | 0.008 | 9.71464519938152e-44 | 46 |
| CR376783.11 | 6.0447805111051e-48 | 0.476535977 | 0.244 | 0.01 | 1.52141080684004e-43 | 46 |
| clstn12 | 7.75846760652624e-48 | 0.568300892 | 0.366 | 0.022 | 1.95272871188659e-43 | 46 |
| hmcn1 | 1.37102039521536e-47 | 0.377168444 | 0.195 | 0.006 | 3.45072123271754e-43 | 46 |
| plxna42 | 1.39544892292233e-47 | 0.645043389 | 0.366 | 0.022 | 3.51220539410322e-43 | 46 |
| f107 | 1.07358446681885e-46 | 0.874295861 | 0.537 | 0.048 | 2.70210474453637e-42 | 46 |
| rhoj1 | 1.2796850962405e-46 | 0.54830839 | 0.293 | 0.015 | 3.22083941872771e-42 | 46 |
| prnprs3 | 2.0546847721353e-46 | 0.488615779 | 0.293 | 0.015 | 5.17143610298734e-42 | 46 |
| hmgb3a4 | 2.05741790608835e-46 | 0.794366362 | 0.537 | 0.049 | 5.17831512783377e-42 | 46 |
| si:ch211-151g22.1 | 5.74564160556793e-46 | 0.510808038 | 0.317 | 0.017 | 1.44612053570539e-41 | 46 |
| aldh1a25 | 1.38787815233601e-45 | 1.108520369 | 0.683 | 0.078 | 3.49315052161451e-41 | 46 |
| jupa3 | 1.49502297939408e-45 | 0.829786075 | 0.561 | 0.053 | 3.76282333683695e-41 | 46 |
| erbb22 | 2.14381300010032e-45 | 0.621662112 | 0.415 | 0.03 | 5.3957629399525e-41 | 46 |
| LOC108179567 | 1.28320290734074e-44 | 0.374236692 | 0.195 | 0.007 | 3.2296933974859e-40 | 46 |

| | | | | | | |
|---|---|---|---|---|---|---|
| LOC110438541 | 1.30168225759416e-44 | 0.541419881 | 0.366 | 0.023 | 3.27620407413875e-40 | 46 |
| CR318588.37 | 2.07038106928078e-44 | 0.886258824 | 0.512 | 0.045 | 5.2109421132728e-40 | 46 |
| plekha63 | 3.2401033766574e-44 | 0.588266867 | 0.317 | 0.018 | 8.155016188709e-40 | 46 |
| fosb11 | 1.1481554495158e-43 | 1.313497894 | 0.829 | 0.125 | 2.88979245088631e-39 | 46 |
| igfbp5b4 | 1.56224391097507e-43 | 0.917063564 | 0.659 | 0.072 | 3.93201169953316e-39 | 46 |
| pgr3 | 2.34590914848368e-43 | 0.490215481 | 0.317 | 0.018 | 5.90441873581858e-39 | 46 |
| clmn | 8.08799978202014e-43 | 0.292542633 | 0.146 | 0.004 | 2.03566866513665e-38 | 46 |
| tapbp.2 | 1.83846262993738e-42 | 0.32006361 | 0.195 | 0.007 | 4.62722659328938e-38 | 46 |
| FP245456.1 | 1.97498164626044e-42 | 0.384215983 | 0.22 | 0.009 | 4.97083130547291e-38 | 46 |
| atp1a1a.12 | 2.19468017633554e-42 | 1.172367636 | 0.902 | 0.148 | 5.52379053581893e-38 | 46 |
| dsc2l3 | 6.72634323882245e-42 | 0.589154837 | 0.366 | 0.025 | 1.69295332977922e-37 | 46 |
| tspan9b4 | 1.86780422144259e-41 | 0.680862231 | 0.488 | 0.044 | 4.70107644494886e-37 | 46 |
| prr131 | 2.31976389319663e-41 | 0.54307324 | 0.341 | 0.022 | 5.83861374278659e-37 | 46 |
| dspa2 | 6.43796340537141e-41 | 0.542099563 | 0.366 | 0.025 | 1.62037100949793e-36 | 46 |
| afdna5 | 6.47161879444597e-41 | 0.569839067 | 0.341 | 0.022 | 1.62884173437411e-36 | 46 |
| nr0b13 | 1.54448213320503e-40 | 0.382334756 | 0.244 | 0.012 | 3.88730708106374e-36 | 46 |
| CABZ01068366.14 | 3.6164331177582e-40 | 0.615815507 | 0.39 | 0.03 | 9.10220051408562e-36 | 46 |
| fam102ab3 | 8.60810633563831e-40 | 0.654653495 | 0.415 | 0.034 | 2.16657428361681e-35 | 46 |
| mov10b.1 | 2.13550554740407e-39 | 0.527499636 | 0.317 | 0.02 | 5.37485391226132e-35 | 46 |
| CABZ01076275.11 | 4.01884119701256e-39 | 0.516851391 | 0.317 | 0.02 | 1.01150214087609e-34 | 46 |
| met1 | 5.81145008994473e-39 | 0.369868722 | 0.195 | 0.008 | 1.46268387313819e-34 | 46 |
| bambia3 | 6.46415719240017e-39 | 0.583432284 | 0.317 | 0.02 | 1.6269637237552e-34 | 46 |
| ap2m1a6 | 7.68601195823102e-39 | 1.028200867 | 0.854 | 0.142 | 1.93449234976717e-34 | 46 |
| krt18a.110 | 1.09852689663777e-38 | 1.24098633 | 0.805 | 0.126 | 2.7648823461476e-34 | 46 |
| gstt1b8 | 1.86383734061989e-38 | 0.83590079 | 0.61 | 0.072 | 4.69109220260621e-34 | 46 |
| stat1a8 | 2.05063924758453e-38 | 1.004829956 | 0.78 | 0.12 | 5.1612539222455e-34 | 46 |
| gnav1 | 2.76443206191976e-38 | 0.442333755 | 0.317 | 0.02 | 6.95779905664584e-34 | 46 |
| si:ch211-276k2.1 | 3.73831911430305e-38 | 0.28131905 | 0.122 | 0.003 | 9.40897537878934e-34 | 46 |
| fam69aa | 4.36201802332688e-38 | 0.483785102 | 0.268 | 0.015 | 1.09787631629114e-33 | 46 |
| coro2ba | 5.66440801772575e-38 | 0.321854091 | 0.195 | 0.008 | 1.42567485398139e-33 | 46 |
| lfng2 | 6.35653726041579e-38 | 0.348347323 | 0.195 | 0.008 | 1.59987686307405e-33 | 46 |
| phospho12 | 3.58941178722927e-37 | 0.401139173 | 0.268 | 0.015 | 9.03419052727735e-33 | 46 |
| atp1b1a8 | 3.64129002774488e-37 | 1.225277478 | 0.951 | 0.213 | 9.1647628708311e-33 | 46 |
| map1aa1 | 3.9988587548562e-37 | 0.372946054 | 0.171 | 0.006 | 1.00647276000976e-32 | 46 |
| slc20a2 | 4.26830254620353e-37 | 0.523205845 | 0.341 | 0.025 | 1.07428906785397e-32 | 46 |

| | | | | | | |
|---|---|---|---|---|---|---|
| hs3st3b1b | 6.62724219788216e-37 | 0.337642406 | 0.195 | 0.008 | 1.66801058878496e-32 | 46 |
| mafba10 | 1.57337364489368e-36 | 0.879430697 | 0.585 | 0.072 | 3.96002412683291e-32 | 46 |
| ar6 | 3.00152494175905e-36 | 0.706359572 | 0.537 | 0.059 | 7.55453812591336e-32 | 46 |
| csrnp2 | 4.15138219100453e-36 | 0.39445394 | 0.22 | 0.011 | 1.04486138365393e-31 | 46 |
| st3gal41 | 4.36595025620956e-36 | 0.511295217 | 0.317 | 0.022 | 1.09886601998538e-31 | 46 |
| pcdh7b | 4.58380909135099e-36 | 0.344112294 | 0.171 | 0.006 | 1.15369891020213e-31 | 46 |
| ndrg3a1 | 1.28584028173608e-35 | 0.474625804 | 0.317 | 0.022 | 3.23633140510154e-31 | 46 |
| lgals175 | 1.56548755430165e-35 | 0.885322747 | 0.488 | 0.052 | 3.94017562542183e-31 | 46 |
| palm31 | 1.59761927561618e-35 | 0.435895238 | 0.244 | 0.013 | 4.02104795479836e-31 | 46 |
| rcan1a | 2.01934414537918e-35 | 0.683486473 | 0.439 | 0.042 | 5.08248727950487e-31 | 46 |
| itm2bb15 | 3.24570400286543e-35 | 1.336822862 | 0.902 | 0.222 | 8.169112404812e-31 | 46 |
| vat110 | 3.37376527987147e-35 | 1.050315996 | 0.805 | 0.134 | 8.4914298329085e-31 | 46 |
| tdp2a | 5.44193215146833e-35 | 0.391633613 | 0.195 | 0.009 | 1.36967990320307e-30 | 46 |
| cttnbp2nlb1 | 5.60934642355028e-35 | 0.359024956 | 0.195 | 0.009 | 1.41181640134337e-30 | 46 |
| tent5ba3 | 7.67972809138622e-35 | 0.39794071 | 0.22 | 0.011 | 1.932910763321e-30 | 46 |
| si:dkeyp-77h1.4 | 1.38414741439861e-34 | 0.365830344 | 0.195 | 0.009 | 3.48376062729985e-30 | 46 |
| pi15a3 | 1.8910827422506e-34 | 0.723216664 | 0.463 | 0.046 | 4.75966615397053e-30 | 46 |
| numbl | 2.17168707553594e-34 | 0.253926841 | 0.122 | 0.003 | 5.46591920041641e-30 | 46 |
| krt814 | 2.7955910671345e-34 | 1.396385853 | 1 | 0.276 | 7.03622315687081e-30 | 46 |
| wbp1lb3 | 2.91020056496955e-34 | 0.660791698 | 0.463 | 0.048 | 7.32468380197185e-30 | 46 |
| pcnx1 | 3.64034694907957e-34 | 0.369861704 | 0.195 | 0.009 | 9.16238923613836e-30 | 46 |
| ckba5 | 3.72736423188986e-34 | 0.905172065 | 0.537 | 0.065 | 9.38140303524358e-30 | 46 |
| fxyd6l7 | 1.03166785113998e-33 | 1.065418824 | 0.707 | 0.114 | 2.59660481453421e-29 | 46 |
| tbc1d1 | 1.11577668639385e-33 | 0.428611089 | 0.268 | 0.017 | 2.80829834198468e-29 | 46 |
| hsd11b28 | 1.27272838281277e-33 | 1.099051879 | 0.78 | 0.123 | 3.20333006670145e-29 | 46 |
| rnf11b | 1.40344739299164e-33 | 0.494356317 | 0.293 | 0.02 | 3.53233674342065e-29 | 46 |
| ptprfa | 2.7028250667405e-33 | 0.257927673 | 0.146 | 0.005 | 6.80274041047916e-29 | 46 |
| jun14 | 3.15222687268172e-33 | 1.366974134 | 0.927 | 0.228 | 7.93383981585263e-29 | 46 |
| si:ch73-61d6.3 | 3.74007718865767e-33 | 0.330213928 | 0.195 | 0.009 | 9.41340027613248e-29 | 46 |
| ptprfb4 | 4.04970043254992e-33 | 0.396840002 | 0.244 | 0.014 | 1.01926910186849e-28 | 46 |
| si:cabz01069012.2 | 8.30563695109509e-33 | 0.257666571 | 0.122 | 0.004 | 2.09044576422112e-28 | 46 |
| fuk | 8.53568717553018e-33 | 0.268707891 | 0.171 | 0.007 | 2.14834710520919e-28 | 46 |
| cbsb2 | 8.99682780055326e-33 | 0.643882663 | 0.39 | 0.035 | 2.26441158912125e-28 | 46 |
| nit2 | 1.32276647846627e-32 | 0.449020421 | 0.268 | 0.017 | 3.32927094965176e-28 | 46 |
| st51 | 1.53785623032207e-32 | 0.386559985 | 0.22 | 0.012 | 3.87063034609762e-28 | 46 |

| | | | | | | |
|---|---|---|---|---|---|---|
| pxnb | 2.78343510004732e-32 | 0.457031306 | 0.244 | 0.015 | 7.00562780330909e-28 | 46 |
| mbnl35 | 4.31630857673738e-32 | 0.707572857 | 0.488 | 0.056 | 1.08637170567903e-27 | 46 |
| col4a61 | 4.8703522430707e-32 | 0.317842127 | 0.195 | 0.009 | 1.22581895605847e-27 | 46 |
| ifi4510 | 5.63232563733632e-32 | 1.233496131 | 0.732 | 0.128 | 1.41760003966118e-27 | 46 |
| FP102018.12 | 6.05581945367256e-32 | 0.462244834 | 0.268 | 0.018 | 1.52418919829485e-27 | 46 |
| b2m17 | 6.83078344266699e-32 | 1.228902958 | 0.976 | 0.292 | 1.71923988468485e-27 | 46 |
| foxp48 | 1.08964292124327e-31 | 0.684571644 | 0.463 | 0.051 | 2.74252226847719e-27 | 46 |
| scarf11 | 1.1051839618246e-31 | 0.277800181 | 0.146 | 0.005 | 2.78163751351634e-27 | 46 |
| gab1 | 1.79626652083119e-31 | 0.394943464 | 0.244 | 0.015 | 4.52102320628002e-27 | 46 |
| cemip21 | 2.6139895592517e-31 | 0.460056024 | 0.293 | 0.021 | 6.57915032168061e-27 | 46 |
| bmp1a | 5.05891214368033e-31 | 0.403025026 | 0.244 | 0.015 | 1.2732775974429e-26 | 46 |
| clip2 | 9.38828491277688e-31 | 0.263887835 | 0.146 | 0.006 | 2.36293742969681e-26 | 46 |
| si:ch73-335l21.411 | 1.11159972065977e-30 | 1.295112568 | 0.756 | 0.148 | 2.79778533692857e-26 | 46 |
| pdgfaa | 1.49914970216477e-30 | 0.353309292 | 0.195 | 0.01 | 3.7732098853785e-26 | 46 |
| si:dkey-188i13.107 | 1.76983468714947e-30 | 0.99208804 | 0.634 | 0.104 | 4.45449692408651e-26 | 46 |
| dchs1a1 | 2.23577293757884e-30 | 0.318861405 | 0.195 | 0.01 | 5.62721690659218e-26 | 46 |
| hexb3 | 2.30152343448562e-30 | 0.529326535 | 0.366 | 0.034 | 5.79270433225686e-26 | 46 |
| slc22a23 | 2.66955273087363e-30 | 0.264988052 | 0.146 | 0.006 | 6.71899726833584e-26 | 46 |
| si:ch211-195e19.1 | 2.78876280529536e-30 | 0.436424906 | 0.268 | 0.019 | 7.0190371046479e-26 | 46 |
| gadd45ba10 | 3.15579052925425e-30 | 1.185014508 | 0.854 | 0.19 | 7.94280918308001e-26 | 46 |
| arhgef43 | 5.7204877072358e-30 | 0.556541939 | 0.366 | 0.034 | 1.43978955103418e-25 | 46 |
| ccdc1872 | 7.17789275567083e-30 | 0.561947384 | 0.341 | 0.03 | 1.80660382767479e-25 | 46 |
| sox4a5 | 1.30057851617292e-29 | 0.707538994 | 0.415 | 0.044 | 3.27342606735561e-25 | 46 |
| cebpd11 | 1.36719130936811e-29 | 0.927858091 | 0.756 | 0.129 | 3.44108380654859e-25 | 46 |
| isg156 | 1.85363822591559e-29 | 0.978859284 | 0.512 | 0.068 | 4.66542205080695e-25 | 46 |
| ildr2 | 1.91942630800357e-29 | 0.302345229 | 0.195 | 0.01 | 4.83100407461418e-25 | 46 |
| cxadr1 | 3.48161059699975e-29 | 0.540613094 | 0.366 | 0.034 | 8.76286571158867e-25 | 46 |
| socs3a15 | 6.93143811065172e-29 | 1.340214494 | 0.951 | 0.298 | 1.74457365806993e-24 | 46 |
| ano5a1 | 1.17751467961994e-28 | 0.40995938 | 0.195 | 0.011 | 2.96368669713544e-24 | 46 |
| gig2e1 | 1.95782645865148e-28 | 0.434525484 | 0.244 | 0.017 | 4.92765341377991e-24 | 46 |
| sntb1 | 2.07798259556346e-28 | 0.3621021 | 0.195 | 0.011 | 5.23007439477368e-24 | 46 |
| itm2ba17 | 4.23007699008555e-28 | 1.061938773 | 0.976 | 0.273 | 1.06466807763463e-23 | 46 |
| bmpr1bb1 | 7.69870570362593e-28 | 0.30779773 | 0.195 | 0.011 | 1.93768723854561e-23 | 46 |
| rpz510 | 7.71470173275841e-28 | 0.828212148 | 0.634 | 0.102 | 1.94171327911796e-23 | 46 |
| calm1a4 | 7.92136274447774e-28 | 1.034485719 | 0.78 | 0.164 | 1.9937277891576e-23 | 46 |

| | | | | | | |
|---|---|---|---|---|---|---|
| fgfr35 | 7.95969697669045e-28 | 0.378764602 | 0.244 | 0.017 | 2.00337613206322e-23 | 46 |
| man1b1b1 | 1.02567953386214e-27 | 0.487640622 | 0.317 | 0.028 | 2.58153281877761e-23 | 46 |
| dclk2a | 1.06290214679674e-27 | 0.285951115 | 0.146 | 0.006 | 2.67521841327272e-23 | 46 |
| XLOC-015291 | 1.27076530431802e-27 | 0.304723235 | 0.171 | 0.008 | 3.19838919443802e-23 | 46 |
| gadd45bb7 | 1.5536577031319e-27 | 0.934459172 | 0.561 | 0.088 | 3.91040107301267e-23 | 46 |
| pcdh1b1 | 2.24170264611385e-27 | 0.329010069 | 0.171 | 0.009 | 5.64214139000395e-23 | 46 |
| pik3r27 | 3.41437757887593e-27 | 0.745062677 | 0.463 | 0.06 | 8.59364692827284e-23 | 46 |
| sipa1l31 | 3.57838873281594e-27 | 0.345587009 | 0.244 | 0.017 | 9.00644660162445e-23 | 46 |
| akap7 | 3.78139771418099e-27 | 0.330451314 | 0.195 | 0.011 | 9.51739990682214e-23 | 46 |
| metrnla7 | 4.50830874208759e-27 | 0.701548669 | 0.61 | 0.093 | 1.13469622729602e-22 | 46 |
| ccdc1251 | 6.67283193054928e-27 | 0.384953676 | 0.22 | 0.014 | 1.67948506859995e-22 | 46 |
| notch1b1 | 7.15573021220401e-27 | 0.341216355 | 0.171 | 0.009 | 1.80102573710963e-22 | 46 |
| wwtr13 | 7.70507408448776e-27 | 0.461279995 | 0.293 | 0.025 | 1.93929009632472e-22 | 46 |
| crybg1a2 | 8.70916337250775e-27 | 0.382252288 | 0.244 | 0.018 | 2.19200932922647e-22 | 46 |
| bmpr2b | 1.53347060973626e-26 | 0.250482708 | 0.146 | 0.007 | 3.8595921776452e-22 | 46 |
| jdp2b12 | 2.1127245653374e-26 | 1.031427315 | 0.805 | 0.17 | 5.31751645849769e-22 | 46 |
| plod32 | 2.26119441842206e-26 | 0.528972467 | 0.341 | 0.034 | 5.69120023172648e-22 | 46 |
| efhd25 | 2.35251075106148e-26 | 0.668542779 | 0.463 | 0.06 | 5.92103430934665e-22 | 46 |
| nqo11 | 2.38227852246729e-26 | 0.368886844 | 0.22 | 0.015 | 5.99595681319791e-22 | 46 |
| pfkfb4b5 | 2.97359133777081e-26 | 0.824704222 | 0.585 | 0.096 | 7.48423203803535e-22 | 46 |
| id19 | 3.3524548284137e-26 | 1.048480099 | 0.61 | 0.105 | 8.43779355763444e-22 | 46 |
| rgrb | 4.23740309637566e-26 | 0.329356523 | 0.171 | 0.009 | 1.06651198532679e-21 | 46 |
| nol4lb1 | 5.30749141662308e-26 | 0.478212395 | 0.293 | 0.026 | 1.33584251464986e-21 | 46 |
| ptprga11 | 5.35548108679139e-26 | 0.643387543 | 0.488 | 0.066 | 1.34792103473453e-21 | 46 |
| marcksb10 | 5.73208880771397e-26 | 0.925055866 | 0.878 | 0.176 | 1.44270943201353e-21 | 46 |
| mdka9 | 7.60611246975762e-26 | 0.84506423 | 0.585 | 0.094 | 1.91438244751329e-21 | 46 |
| efhd1 | 1.1195185426257e-25 | 0.375609252 | 0.244 | 0.018 | 2.81771621993463e-21 | 46 |
| CABZ01086041.11 | 1.20425388679896e-25 | 0.357262887 | 0.195 | 0.012 | 3.03098660768431e-21 | 46 |
| mkln11 | 1.25715031831622e-25 | 0.564658107 | 0.366 | 0.039 | 3.16412163617008e-21 | 46 |
| apoeb1 | 1.27469247003969e-25 | 0.431409149 | 0.22 | 0.015 | 3.20827347784289e-21 | 46 |
| magi1b | 1.29695927591479e-25 | 0.318118334 | 0.195 | 0.012 | 3.26431680154993e-21 | 46 |
| grk41 | 1.45315559833039e-25 | 0.519631285 | 0.317 | 0.031 | 3.65744732543777e-21 | 46 |
| LOC100333572 | 1.78084616157752e-25 | 0.338024867 | 0.195 | 0.012 | 4.48221170407446e-21 | 46 |
| fdxr1 | 1.81850686996105e-25 | 0.553242359 | 0.293 | 0.026 | 4.57699994100497e-21 | 46 |
| ankrd46a | 2.17611068361473e-25 | 0.405010176 | 0.244 | 0.019 | 5.47705297958991e-21 | 46 |

| | | | | | | |
|---|---|---|---|---|---|---|
| CABZ01086029.12 | 3.03572984345474e-25 | 0.370157246 | 0.244 | 0.019 | 7.64062844299124e-21 | 46 |
| pelp1 | 3.28456039963833e-25 | 0.378277128 | 0.293 | 0.026 | 8.26691006984972e-21 | 46 |
| smad12 | 3.53219059485284e-25 | 0.44630246 | 0.244 | 0.019 | 8.89017050818512e-21 | 46 |
| arl4d4 | 3.92433852172228e-25 | 0.615488163 | 0.415 | 0.05 | 9.87716762532281e-21 | 46 |
| serinc15 | 4.76882479079535e-25 | 0.689958318 | 0.537 | 0.082 | 1.20026551159528e-20 | 46 |
| papss2b1 | 4.93559622714586e-25 | 0.340086157 | 0.244 | 0.019 | 1.24224021441034e-20 | 46 |
| slc44a5b1 | 5.34300381019396e-25 | 0.309906017 | 0.195 | 0.012 | 1.34478062898772e-20 | 46 |
| rasl11a1 | 6.00556495722666e-25 | 0.288338719 | 0.195 | 0.012 | 1.51154064408438e-20 | 46 |
| ywhae2 | 8.15538533678632e-25 | 0.515825361 | 0.293 | 0.027 | 2.05262893541575e-20 | 46 |
| CABZ01092156.1 | 8.94615524383783e-25 | 0.401254862 | 0.317 | 0.03 | 2.25165781332154e-20 | 46 |
| znf3261 | 9.2298692192022e-25 | 0.585085799 | 0.415 | 0.049 | 2.323065783781e-20 | 46 |
| selenbp15 | 1.01951864611272e-24 | 0.66954428 | 0.488 | 0.069 | 2.56602648040111e-20 | 46 |
| si:rp71-36a1.2 | 1.37786354649877e-24 | 0.270608725 | 0.146 | 0.007 | 3.46794476018276e-20 | 46 |
| vasnb4 | 2.54982312628669e-24 | 0.493553407 | 0.317 | 0.032 | 6.41764982655098e-20 | 46 |
| bmp62 | 2.56246448594404e-24 | 0.390366351 | 0.195 | 0.013 | 6.44946686467255e-20 | 46 |
| ap1s21 | 3.10729845676947e-24 | 0.838505804 | 0.683 | 0.131 | 7.82075948584309e-20 | 46 |
| wu:fb16f035 | 3.56720233651998e-24 | 0.785187484 | 0.537 | 0.087 | 8.97829156078714e-20 | 46 |
| myl9a6 | 3.91577482286486e-24 | 0.480793481 | 0.317 | 0.032 | 9.85561365166857e-20 | 46 |
| fosab15 | 4.72946831188937e-24 | 1.246902083 | 0.951 | 0.349 | 1.19035987941944e-19 | 46 |
| alcama2 | 6.12756501590046e-24 | 0.499899033 | 0.293 | 0.028 | 1.54224683885199e-19 | 46 |
| trim2a | 6.48932473853061e-24 | 0.298929379 | 0.171 | 0.01 | 1.63329814344077e-19 | 46 |
| cited4a6 | 7.79777960908012e-24 | 0.547555395 | 0.366 | 0.042 | 1.96262314980938e-19 | 46 |
| cav21 | 8.59422920444858e-24 | 0.308745159 | 0.195 | 0.013 | 2.16308154846766e-19 | 46 |
| stat27 | 1.10323989163263e-23 | 0.701904925 | 0.439 | 0.061 | 2.77674448325016e-19 | 46 |
| nectin1b4 | 1.13807657366373e-23 | 0.603360176 | 0.366 | 0.042 | 2.86442492825425e-19 | 46 |
| egr115 | 1.32124310260331e-23 | 0.982572303 | 0.805 | 0.19 | 3.32543676494227e-19 | 46 |
| cmtm31 | 1.82319277478881e-23 | 0.538258966 | 0.366 | 0.043 | 4.58879389486596e-19 | 46 |
| bahcc1b1 | 2.1323973384928e-23 | 0.431234544 | 0.293 | 0.028 | 5.36703086125252e-19 | 46 |
| abl12 | 2.27358058755532e-23 | 0.489870978 | 0.317 | 0.033 | 5.72237498081799e-19 | 46 |
| cadm2a2 | 2.55125614487171e-23 | 0.419788747 | 0.22 | 0.017 | 6.42125659102761e-19 | 46 |
| b3gnt2l1 | 2.6237595468228e-23 | 0.401050073 | 0.244 | 0.02 | 6.60374040339831e-19 | 46 |
| mpped2a1 | 2.85484235071556e-23 | 0.502033634 | 0.317 | 0.033 | 7.185352712516e-19 | 46 |
| piezo11 | 3.53058490237561e-23 | 0.390264147 | 0.268 | 0.024 | 8.88612914078917e-19 | 46 |
| slc12a7a | 3.89863385190918e-23 | 0.303578976 | 0.146 | 0.008 | 9.81247154187021e-19 | 46 |
| kalrnb2 | 5.68805356439849e-23 | 0.276334084 | 0.171 | 0.01 | 1.43162620162345e-18 | 46 |

| | | | | | | |
|---|---|---|---|---|---|---|
| cpn16 | 6.11533580402203e-23 | 0.63243299 | 0.415 | 0.055 | 1.5391688685143e-18 | 46 |
| dbn16 | 7.85997542029217e-23 | 0.574539349 | 0.415 | 0.056 | 1.97827721353334e-18 | 46 |
| lhfpl2b1 | 8.57650107174345e-23 | 0.298922094 | 0.22 | 0.017 | 2.15861955474711e-18 | 46 |
| tegt10 | 9.67977079173864e-23 | 0.889888119 | 0.829 | 0.211 | 2.4363015105727e-18 | 46 |
| cdh15 | 9.81178073383793e-23 | 0.571450975 | 0.439 | 0.059 | 2.46952709289967e-18 | 46 |
| mogat21 | 1.15563727413943e-22 | 0.358584888 | 0.22 | 0.017 | 2.90862345528153e-18 | 46 |
| foxl2b1 | 1.27076062164329e-22 | 0.31364527 | 0.171 | 0.01 | 3.19837740861399e-18 | 46 |
| si:dkey-42i9.46 | 1.33492119447103e-22 | 0.787544509 | 0.707 | 0.147 | 3.35986315436414e-18 | 46 |
| cdv32 | 1.60324823191185e-22 | 0.772608179 | 0.683 | 0.137 | 4.03521547489893e-18 | 46 |
| gamt3 | 1.70354744556741e-22 | 0.491828073 | 0.366 | 0.044 | 4.28765856574862e-18 | 46 |
| ctnnb18 | 2.19658724014396e-22 | 0.914368267 | 0.683 | 0.15 | 5.52859042471833e-18 | 46 |
| zfp36l1b9 | 2.63085664943604e-22 | 0.85855148 | 0.659 | 0.13 | 6.62160310096556e-18 | 46 |
| got11 | 3.49391709558631e-22 | 0.430622989 | 0.268 | 0.025 | 8.79383993788119e-18 | 46 |
| tmem205 | 4.13638627471566e-22 | 0.286075311 | 0.171 | 0.011 | 1.04108706148318e-17 | 46 |
| si:ch211-214p13.9 | 4.15789831443149e-22 | 0.281494391 | 0.195 | 0.014 | 1.04650142675926e-17 | 46 |
| appb6 | 4.33006793267969e-22 | 0.500197942 | 0.341 | 0.04 | 1.08983479797615e-17 | 46 |
| ppme1 | 4.35021097380559e-22 | 0.50444322 | 0.366 | 0.045 | 1.09490459999713e-17 | 46 |
| CU984600.21 | 5.40937713765067e-22 | 0.45991824 | 0.268 | 0.026 | 1.3614861317753e-17 | 46 |
| dab2ipb1 | 5.84318418213063e-22 | 0.322306678 | 0.195 | 0.014 | 1.47067102680046e-17 | 46 |
| ldb1a | 6.90934308410539e-22 | 0.387657064 | 0.268 | 0.025 | 1.73901256083848e-17 | 46 |
| tspan5a | 7.81333199868765e-22 | 0.3822101 | 0.195 | 0.014 | 1.96653753074969e-17 | 46 |
| cd1645 | 8.60997299481655e-22 | 0.65435412 | 0.463 | 0.072 | 2.16704410306538e-17 | 46 |
| pgrmc18 | 9.21632474477532e-22 | 0.789666052 | 0.683 | 0.141 | 2.3196567750125e-17 | 46 |
| ier59 | 1.53582088644199e-21 | 0.875716607 | 0.634 | 0.136 | 3.86550758908585e-17 | 46 |
| hmgn619 | 1.71636056561972e-21 | 0.929649379 | 0.902 | 0.265 | 4.31990790760826e-17 | 46 |
| slc38a24 | 1.90466722240919e-21 | 0.551666153 | 0.366 | 0.047 | 4.7938569320817e-17 | 46 |
| sptlc2b4 | 2.24579435578951e-21 | 0.515527768 | 0.366 | 0.047 | 5.65243981408661e-17 | 46 |
| hist2h2l4 | 2.45403415634234e-21 | 0.628712794 | 0.488 | 0.078 | 6.17655856809805e-17 | 46 |
| mgst1.16 | 2.52273118718156e-21 | 0.524356293 | 0.341 | 0.042 | 6.34946212501726e-17 | 46 |
| pnrc224 | 2.75403137900795e-21 | 0.866107365 | 1 | 0.568 | 6.9316215778251e-17 | 46 |
| tcima9 | 3.1083908749186e-21 | 0.907405535 | 0.439 | 0.069 | 7.82350899308262e-17 | 46 |
| chka3 | 3.47655684167699e-21 | 0.380316339 | 0.268 | 0.026 | 8.75014591481682e-17 | 46 |
| hsp70.33 | 4.22271564562128e-21 | 0.547174621 | 0.244 | 0.022 | 1.06281530084642e-16 | 46 |
| ctnnd2a1 | 4.32798120935536e-21 | 0.286353031 | 0.171 | 0.011 | 1.08930959058265e-16 | 46 |
| b2ml18 | 7.18070873993955e-21 | 0.944081595 | 0.878 | 0.267 | 1.80731258275539e-16 | 46 |

| | | | | | | |
|---|---|---|---|---|---|---|
| wsb1 | 7.22866978640852e-21 | 0.604758809 | 0.488 | 0.079 | 1.81938389854116e-16 | 46 |
| mt-nd57 | 9.26153376053583e-21 | 0.708188654 | 0.61 | 0.12 | 2.33103543218926e-16 | 46 |
| si:dkey-23i12.74 | 9.38630298895523e-21 | 0.569371854 | 0.293 | 0.032 | 2.36243859929014e-16 | 46 |
| gng77 | 1.05484974959182e-20 | 0.531321166 | 0.39 | 0.054 | 2.65495133474766e-16 | 46 |
| csrnp1b14 | 1.22470178919317e-20 | 0.884770709 | 0.732 | 0.18 | 3.0824519332203e-16 | 46 |
| ponzr111 | 1.46778903305576e-20 | 0.95612593 | 0.902 | 0.273 | 3.69427821729803e-16 | 46 |
| junba14 | 1.61287675287758e-20 | 1.077537792 | 0.927 | 0.309 | 4.05944949931757e-16 | 46 |
| her64 | 1.67793027459923e-20 | 0.406905074 | 0.244 | 0.023 | 4.22318270813879e-16 | 46 |
| zgc:114200 | 1.79105814977815e-20 | 0.44404806 | 0.293 | 0.032 | 4.50791425717664e-16 | 46 |
| tjap1 | 2.08863778587732e-20 | 0.28962687 | 0.146 | 0.009 | 5.25689244327464e-16 | 46 |
| btg214 | 2.37193935745735e-20 | 1.018097306 | 0.902 | 0.374 | 5.96993416878439e-16 | 46 |
| mt-nd417 | 2.84441768507191e-20 | 0.947222543 | 1 | 0.457 | 7.1591148715575e-16 | 46 |
| XLOC-012680 | 3.37900963503711e-20 | 0.327464872 | 0.195 | 0.015 | 8.50462935042491e-16 | 46 |
| zgc:1230101 | 3.64690116963045e-20 | 0.448293754 | 0.317 | 0.037 | 9.17888555384289e-16 | 46 |
| edem11 | 4.43853293841303e-20 | 0.467349067 | 0.293 | 0.033 | 1.11713435526918e-15 | 46 |
| sdc412 | 4.4566741583411e-20 | 0.927899286 | 0.927 | 0.257 | 1.12170031891287e-15 | 46 |
| nomo | 4.52347649815425e-20 | 0.307770077 | 0.22 | 0.019 | 1.13851379982044e-15 | 46 |
| cd99l22 | 4.53385559771409e-20 | 0.448327002 | 0.244 | 0.024 | 1.14112611538866e-15 | 46 |
| sorcs2 | 5.58788796447625e-20 | 0.255379244 | 0.146 | 0.009 | 1.40641552177903e-15 | 46 |
| zgc:1750881 | 6.43543460733253e-20 | 0.532122069 | 0.39 | 0.055 | 1.61973453631952e-15 | 46 |
| ctsk6 | 7.76992277240984e-20 | 0.583682128 | 0.415 | 0.063 | 1.95561186258783e-15 | 46 |
| prkar2aa4 | 8.83320228704908e-20 | 0.660325768 | 0.537 | 0.099 | 2.22322868362738e-15 | 46 |
| midn14 | 1.01062262893442e-19 | 0.737323729 | 0.659 | 0.14 | 2.54363609476505e-15 | 46 |
| LOC10390928410 | 1.03560884823252e-19 | 0.650217391 | 0.415 | 0.065 | 2.60652391011644e-15 | 46 |
| kalrna | 1.05858443581686e-19 | 0.282142688 | 0.171 | 0.012 | 2.66435116650745e-15 | 46 |
| pak2a3 | 1.09674782897656e-19 | 0.531556625 | 0.415 | 0.062 | 2.76040461075111e-15 | 46 |
| rab11a1 | 1.32016742565346e-19 | 0.706172059 | 0.585 | 0.116 | 3.32272939362721e-15 | 46 |
| ackr4b1 | 1.53288011446945e-19 | 0.303085324 | 0.146 | 0.009 | 3.85810596010817e-15 | 46 |
| cald1a10 | 1.83115429421667e-19 | 0.69149696 | 0.537 | 0.099 | 4.60883224311394e-15 | 46 |
| cav18 | 1.87463576156356e-19 | 0.57607583 | 0.341 | 0.044 | 4.71827074827933e-15 | 46 |
| si:ch211-114c12.21 | 1.88316169698177e-19 | 0.433274377 | 0.341 | 0.044 | 4.73972967513342e-15 | 46 |
| ralba1 | 1.97568792052679e-19 | 0.412047091 | 0.244 | 0.024 | 4.97260892717388e-15 | 46 |
| col1a1b8 | 2.07484744326294e-19 | 0.702727826 | 0.707 | 0.147 | 5.22218352994849e-15 | 46 |
| rgs5b6 | 2.30298289001388e-19 | 0.695501789 | 0.463 | 0.076 | 5.79637763587595e-15 | 46 |
| zc2hc1a | 2.78893835882807e-19 | 0.292719798 | 0.171 | 0.012 | 7.01947895533438e-15 | 46 |

| | | | | | | |
|---|---|---|---|---|---|---|
| rab11fip1b4 | 4.11442211668781e-19 | 0.584755705 | 0.39 | 0.058 | 1.03555890254916e-14 | 46 |
| cdkn1ca | 5.26088209742763e-19 | 0.254830789 | 0.146 | 0.009 | 1.32411141510156e-14 | 46 |
| mt-co122 | 5.71140274275566e-19 | 0.757744703 | 1 | 0.658 | 1.43750295632417e-14 | 46 |
| arl5c10 | 6.35404330436023e-19 | 0.725586563 | 0.561 | 0.113 | 1.59924915927443e-14 | 46 |
| tenm34 | 9.76333558104822e-19 | 0.513695585 | 0.341 | 0.046 | 2.45733393239403e-14 | 46 |
| trim8b | 1.0301370359646e-18 | 0.406101585 | 0.244 | 0.025 | 2.59275190581931e-14 | 46 |
| mmp28 | 1.25379060300331e-18 | 0.596609823 | 0.512 | 0.092 | 3.15566556869902e-14 | 46 |
| cmbl1 | 1.47926603153213e-18 | 0.351291572 | 0.268 | 0.029 | 3.72316467476321e-14 | 46 |
| myl610 | 1.6260361597049e-18 | 0.629302434 | 0.537 | 0.101 | 4.09257041036127e-14 | 46 |
| tcf7l21 | 1.64681238833716e-18 | 0.304519766 | 0.195 | 0.016 | 4.14486210020579e-14 | 46 |
| tpbgb | 1.69041903736545e-18 | 0.330237276 | 0.22 | 0.02 | 4.2546156751451e-14 | 46 |
| si:dkeyp-115e12.6 | 1.74855588699856e-18 | 0.278593672 | 0.171 | 0.013 | 4.40094031198668e-14 | 46 |
| man2c1 | 1.75927296827358e-18 | 0.332230472 | 0.171 | 0.013 | 4.42791413384776e-14 | 46 |
| ppp3ccb5 | 1.82433358127968e-18 | 0.714316431 | 0.585 | 0.124 | 4.59166519072284e-14 | 46 |
| fgd5a1 | 2.77953683759878e-18 | 0.293114474 | 0.171 | 0.013 | 6.99581626655238e-14 | 46 |
| arhgap11 | 2.78102290107584e-18 | 0.371701883 | 0.268 | 0.03 | 6.99955653971777e-14 | 46 |
| COX322 | 2.81098525583678e-18 | 0.737040995 | 1 | 0.701 | 7.0749687904156e-14 | 46 |
| cmtm4 | 3.38258928445063e-18 | 0.289033421 | 0.171 | 0.013 | 8.5136389700338e-14 | 46 |
| foxo3b6 | 3.54768319023203e-18 | 0.537431461 | 0.415 | 0.066 | 8.929163821495e-14 | 46 |
| auts2a | 3.9004520208844e-18 | 0.292681271 | 0.171 | 0.013 | 9.81704769136396e-14 | 46 |
| ier2a18 | 5.70240495823486e-18 | 0.970765278 | 0.805 | 0.263 | 1.43523830393813e-13 | 46 |
| zbtb102 | 5.82467035251842e-18 | 0.422464703 | 0.268 | 0.031 | 1.46601128102536e-13 | 46 |
| stat1b | 5.86842121990597e-18 | 0.295110919 | 0.171 | 0.013 | 1.47702293683813e-13 | 46 |
| ppp1r14aa1 | 6.62368263720951e-18 | 0.393773555 | 0.317 | 0.04 | 1.66711468295926e-13 | 46 |
| pcyt21 | 7.54460365143592e-18 | 0.383892967 | 0.317 | 0.041 | 1.89890129302991e-13 | 46 |
| si:ch211-197h24.97 | 7.65912736220633e-18 | 0.579049603 | 0.415 | 0.069 | 1.92772576579371e-13 | 46 |
| zgc:161969 | 7.85038424938885e-18 | 0.326681943 | 0.195 | 0.017 | 1.97586321172868e-13 | 46 |
| thrab | 8.5316444844966e-18 | 0.297062238 | 0.171 | 0.013 | 2.14732960030295e-13 | 46 |
| zgc:153916 | 1.02339640423159e-17 | 0.276301731 | 0.171 | 0.013 | 2.57578640981049e-13 | 46 |
| pde4ba14 | 1.83767266920236e-17 | 0.762996438 | 0.61 | 0.14 | 4.62523834111541e-13 | 46 |
| ier2b18 | 1.90587349381892e-17 | 0.85881877 | 0.78 | 0.224 | 4.79689299659285e-13 | 46 |
| pitpnc1b | 2.49380924411147e-17 | 0.31048653 | 0.195 | 0.018 | 6.27666848650417e-13 | 46 |
| mt-nd118 | 2.56174672747027e-17 | 0.891857514 | 0.976 | 0.411 | 6.44766033836992e-13 | 46 |
| phactr4a4 | 2.89172900920632e-17 | 0.442347045 | 0.293 | 0.037 | 7.27819274327139e-13 | 46 |
| gmfb2 | 3.55629514637805e-17 | 0.646985116 | 0.512 | 0.102 | 8.95083925391891e-13 | 46 |

| | | | | | | |
|---|---|---|---|---|---|---|
| zgc:1951738 | 3.70195335398294e-17 | 0.605199339 | 0.488 | 0.092 | 9.31744639663965e-13 | 46 |
| CR318588.113 | 4.49595427527405e-17 | 1.154880987 | 0.707 | 0.208 | 1.13158673154373e-12 | 46 |
| qsox11 | 4.74093564439362e-17 | 0.353362854 | 0.268 | 0.031 | 1.19324609233743e-12 | 46 |
| purab1 | 6.01617579158637e-17 | 0.460782433 | 0.341 | 0.05 | 1.51421128498437e-12 | 46 |
| ccdc85b8 | 6.8164366514469e-17 | 0.416895353 | 0.341 | 0.049 | 1.71562894080267e-12 | 46 |
| h1f010 | 6.89844353615078e-17 | 0.539472598 | 0.39 | 0.063 | 1.73626925361379e-12 | 46 |
| limch1b2 | 8.17565119912824e-17 | 0.273401182 | 0.195 | 0.018 | 2.05772965030859e-12 | 46 |
| lrrc8da1 | 9.61535221568136e-17 | 0.371191894 | 0.22 | 0.023 | 2.42008799916484e-12 | 46 |
| taok1a1 | 9.66541107620214e-17 | 0.348207481 | 0.195 | 0.018 | 2.43268731376932e-12 | 46 |
| si:ch1073-440b2.1 | 1.07159425969414e-16 | 0.345363791 | 0.22 | 0.023 | 2.69709559222419e-12 | 46 |
| cxxc5a | 1.16683922454576e-16 | 0.265511258 | 0.171 | 0.014 | 2.93681764425922e-12 | 46 |
| oip5-as13 | 1.30723980414192e-16 | 0.406851279 | 0.244 | 0.028 | 3.29019186304479e-12 | 46 |
| CR383676.124 | 1.34478867636865e-16 | 0.818344163 | 0.976 | 0.727 | 3.38469861955227e-12 | 46 |
| LOC5533161 | 1.64887059719839e-16 | 0.337028409 | 0.171 | 0.014 | 4.15004240608863e-12 | 46 |
| si:dkey-199f5.81 | 2.27147593910725e-16 | 0.38429034 | 0.268 | 0.033 | 5.71707779113903e-12 | 46 |
| zgc:927449 | 2.3489958541256e-16 | 0.883654542 | 0.927 | 0.354 | 5.91218766524873e-12 | 46 |
| ctgfa3 | 2.39428201322403e-16 | 0.520208477 | 0.244 | 0.028 | 6.02616839908357e-12 | 46 |
| txnipa14 | 2.93655349120034e-16 | 0.775331967 | 0.756 | 0.211 | 7.39101148200214e-12 | 46 |
| cadm41 | 4.0582561352425e-16 | 0.367183311 | 0.22 | 0.023 | 1.02142248667918e-11 | 46 |
| tnfrsf9a7 | 4.20780699562713e-16 | 0.654024887 | 0.341 | 0.053 | 1.05906294272939e-11 | 46 |
| tjp1b1 | 4.54253815496927e-16 | 0.339257297 | 0.244 | 0.028 | 1.14331142822421e-11 | 46 |
| tgfb3 | 4.90982811865432e-16 | 0.363166627 | 0.22 | 0.023 | 1.23575463918411e-11 | 46 |
| rdh10a4 | 7.16549814551405e-16 | 0.575818412 | 0.293 | 0.041 | 1.80348422824443e-11 | 46 |
| wt1a3 | 7.27903216851039e-16 | 0.434808497 | 0.268 | 0.034 | 1.83205960649238e-11 | 46 |
| optn | 8.109125388704e-16 | 0.363029748 | 0.22 | 0.024 | 2.04098576908291e-11 | 46 |
| tgfb21 | 8.14599835719691e-16 | 0.281686584 | 0.171 | 0.015 | 2.05026632652289e-11 | 46 |
| si:ch211-195b13.12 | 8.29298712422845e-16 | 0.388169934 | 0.244 | 0.029 | 2.08726192929706e-11 | 46 |
| swap70b | 9.77489782575738e-16 | 0.337425422 | 0.195 | 0.019 | 2.46024403376488e-11 | 46 |
| ap2a1 | 1.11642318914547e-15 | 0.47979921 | 0.366 | 0.059 | 2.80992552476023e-11 | 46 |
| gusb | 1.55700410331776e-15 | 0.282658117 | 0.171 | 0.015 | 3.91882362764047e-11 | 46 |
| ptmaa22 | 1.60207290031874e-15 | 0.906275381 | 0.927 | 0.396 | 4.03225728281222e-11 | 46 |
| LO017872.1 | 1.61669021933085e-15 | 0.390967196 | 0.293 | 0.04 | 4.06904761303381e-11 | 46 |
| ywhaqb9 | 1.6312724844285e-15 | 0.658067546 | 0.561 | 0.132 | 4.10574971605808e-11 | 46 |
| tuba8l38 | 1.73417474106664e-15 | 0.444413579 | 0.317 | 0.047 | 4.36474440579061e-11 | 46 |
| wu:fj29h11 | 1.81353787862329e-15 | 0.40396563 | 0.22 | 0.024 | 4.56449348670696e-11 | 46 |

| | | | | | | |
|---|---|---|---|---|---|---|
| tns1b5 | 1.97933221308669e-15 | 0.3741467 | 0.268 | 0.035 | 4.98178124711789e-11 | 46 |
| mt-nd216 | 2.07796989026991e-15 | 0.84802066 | 0.927 | 0.401 | 5.23004241682033e-11 | 46 |
| CABZ01068367.13 | 2.11116373700041e-15 | 0.341587401 | 0.195 | 0.02 | 5.31358800965634e-11 | 46 |
| msi2a3 | 2.55909643090356e-15 | 0.375352208 | 0.317 | 0.046 | 6.44098980694118e-11 | 46 |
| neu1 | 2.60313791336788e-15 | 0.287467893 | 0.195 | 0.02 | 6.55183781415562e-11 | 46 |
| si:ch211-244b2.4 | 2.61139396028695e-15 | 0.305059981 | 0.171 | 0.015 | 6.57261745864621e-11 | 46 |
| tsg101b1 | 2.98040969942087e-15 | 0.333295865 | 0.244 | 0.03 | 7.50139317247239e-11 | 46 |
| loxl2b4 | 3.00468728689784e-15 | 0.484361109 | 0.293 | 0.042 | 7.56249743239318e-11 | 46 |
| si:dkey-56m19.56 | 3.426518470671e-15 | 0.415899249 | 0.317 | 0.047 | 8.62420433883185e-11 | 46 |
| si:dkey-222b8.1 | 3.56789139355711e-15 | 0.256758244 | 0.171 | 0.015 | 8.9800258484439e-11 | 46 |
| zgc:664331 | 3.82124594793697e-15 | 0.321756875 | 0.244 | 0.029 | 9.61769392636257e-11 | 46 |
| hmgb1a11 | 4.02391904453477e-15 | 0.777464055 | 0.659 | 0.189 | 1.01278018431896e-10 | 46 |
| CABZ01026681.1 | 4.56082983591348e-15 | 0.4304273 | 0.317 | 0.048 | 1.14791526140106e-10 | 46 |
| prkab1b4 | 4.76605853005192e-15 | 0.585917546 | 0.537 | 0.122 | 1.19956927142877e-10 | 46 |
| atp10a6 | 4.90496061742054e-15 | 0.556297673 | 0.341 | 0.057 | 1.23452953779858e-10 | 46 |
| cpne12 | 5.20330522038759e-15 | 0.63001163 | 0.537 | 0.124 | 1.30961989091935e-10 | 46 |
| ecm1b6 | 5.29460703230883e-15 | 0.461230808 | 0.341 | 0.055 | 1.33259964396181e-10 | 46 |
| mhc1uba1 | 6.01942947889531e-15 | 0.29016794 | 0.195 | 0.02 | 1.51503020554316e-10 | 46 |
| idh12 | 6.24020723253823e-15 | 0.661383369 | 0.561 | 0.127 | 1.57059775835755e-10 | 46 |
| cflara4 | 6.39672324032268e-15 | 0.498058249 | 0.366 | 0.063 | 1.60999127235681e-10 | 46 |
| mt-cyb17 | 6.71921284262629e-15 | 0.731746444 | 1 | 0.625 | 1.69115868036061e-10 | 46 |
| adamts21 | 7.2263861573516e-15 | 0.314603861 | 0.22 | 0.025 | 1.81880913194382e-10 | 46 |
| lgals3b8 | 7.24441829351456e-15 | 0.501708973 | 0.415 | 0.079 | 1.82334764029468e-10 | 46 |
| zbtb208 | 8.01003405661279e-15 | 0.619672814 | 0.463 | 0.099 | 2.01604547170887e-10 | 46 |
| oga | 8.21752338714523e-15 | 0.331296511 | 0.244 | 0.03 | 2.06826846131058e-10 | 46 |
| rnf213a8 | 8.50868614237091e-15 | 0.598106744 | 0.439 | 0.089 | 2.14155121517333e-10 | 46 |
| cgref12 | 8.69291354734311e-15 | 0.315680455 | 0.22 | 0.025 | 2.18791941073079e-10 | 46 |
| prrc2c | 9.34226816173472e-15 | 0.392358613 | 0.293 | 0.042 | 2.35135547362701e-10 | 46 |
| mt-atp622 | 1.03647477381956e-14 | 0.760153937 | 1 | 0.591 | 2.60870335822644e-10 | 46 |
| crim14 | 1.10739151155318e-14 | 0.388005064 | 0.268 | 0.037 | 2.7871936954282e-10 | 46 |
| h2afx116 | 1.14487414470649e-14 | 0.763149523 | 0.683 | 0.199 | 2.88153373481176e-10 | 46 |
| znf516 | 1.20841800091077e-14 | 0.271183213 | 0.171 | 0.016 | 3.04146726649233e-10 | 46 |
| gab22 | 1.26429700026266e-14 | 0.262886307 | 0.171 | 0.016 | 3.1821091199611e-10 | 46 |
| enah1 | 1.26726005619827e-14 | 0.337889165 | 0.22 | 0.025 | 3.18956683544543e-10 | 46 |
| si:ch211-178n15.13 | 1.28982109176132e-14 | 0.399722337 | 0.244 | 0.031 | 3.24635070585407e-10 | 46 |

| | | | | | | |
|---|---|---|---|---|---|---|
| vgll4a | 1.49660935269311e-14 | 0.259186243 | 0.146 | 0.012 | 3.7668160797933e-10 | 46 |
| dusp4 | 1.67013149672185e-14 | 0.356650005 | 0.22 | 0.025 | 4.20355396409922e-10 | 46 |
| erbin2 | 1.67262009845794e-14 | 0.425664578 | 0.341 | 0.056 | 4.20981752580879e-10 | 46 |
| psme18 | 1.69061669169225e-14 | 0.553239955 | 0.439 | 0.09 | 4.25511315132023e-10 | 46 |
| arpp19a | 1.889618649657e-14 | 0.277271029 | 0.195 | 0.021 | 4.7559811793217e-10 | 46 |
| clpxa | 1.89037290782218e-14 | 0.314877727 | 0.244 | 0.031 | 4.75787957169764e-10 | 46 |
| tsc2 | 1.94814605175132e-14 | 0.259107826 | 0.146 | 0.012 | 4.90328879765291e-10 | 46 |
| rgs161 | 2.00345133632168e-14 | 0.350785084 | 0.22 | 0.025 | 5.04248666838804e-10 | 46 |
| fam3c | 2.14080119961905e-14 | 0.358523223 | 0.293 | 0.043 | 5.38818253932119e-10 | 46 |
| mef2cb4 | 2.17051620804866e-14 | 0.372536202 | 0.22 | 0.026 | 5.46297224403767e-10 | 46 |
| ncor13 | 2.28989148363036e-14 | 0.467236171 | 0.341 | 0.057 | 5.76342787514925e-10 | 46 |
| tpcn1 | 2.34127643858139e-14 | 0.258383193 | 0.146 | 0.012 | 5.8927586682655e-10 | 46 |
| zfhx31 | 2.35740434751728e-14 | 0.35747009 | 0.22 | 0.026 | 5.93335100226625e-10 | 46 |
| puraa | 2.58071556277844e-14 | 0.305719497 | 0.244 | 0.031 | 6.49540299995705e-10 | 46 |
| gadd45ab4 | 2.87279108706706e-14 | 0.547059683 | 0.39 | 0.073 | 7.23052788703909e-10 | 46 |
| cntfr3 | 3.13211569888786e-14 | 0.274654347 | 0.171 | 0.016 | 7.88322200253084e-10 | 46 |
| edf15 | 3.28552535517723e-14 | 0.75544972 | 0.805 | 0.269 | 8.26933876644558e-10 | 46 |
| cpda | 3.41185001770869e-14 | 0.250663492 | 0.171 | 0.016 | 8.587285309571e-10 | 46 |
| zgc:921401 | 3.42433690122533e-14 | 0.28489214 | 0.22 | 0.026 | 8.61871354669402e-10 | 46 |
| mt-co222 | 3.9502562688576e-14 | 0.651334753 | 1 | 0.721 | 9.94240000308769e-10 | 46 |
| LO018340.12 | 4.2161038655557e-14 | 0.324288328 | 0.22 | 0.026 | 1.06115118192171e-09 | 46 |
| cox5b23 | 4.38849260638656e-14 | 0.279905449 | 0.146 | 0.012 | 1.10453970410143e-09 | 46 |
| hgs | 5.09463756957907e-14 | 0.428903786 | 0.293 | 0.045 | 1.28226932988736e-09 | 46 |
| atf310 | 5.6509639790017e-14 | 0.633110626 | 0.439 | 0.092 | 1.42229112387494e-09 | 46 |
| msrb1b1 | 7.8870670535284e-14 | 0.361736096 | 0.317 | 0.051 | 1.98509590670256e-09 | 46 |
| pcolce2b1 | 8.00312406331821e-14 | 0.252177409 | 0.171 | 0.017 | 2.01430629549656e-09 | 46 |
| cd1511 | 8.01319999502548e-14 | 0.315383309 | 0.171 | 0.017 | 2.01684230674796e-09 | 46 |
| NC-002333.421 | 1.01313720557499e-13 | 0.715805634 | 0.976 | 0.623 | 2.5499650327117e-09 | 46 |
| lsr2 | 1.30771376328925e-13 | 0.343861244 | 0.22 | 0.027 | 3.2913847708227e-09 | 46 |
| mtpn9 | 1.35760434212688e-13 | 0.560201077 | 0.585 | 0.147 | 3.41695436869915e-09 | 46 |
| zgc:922423 | 1.51819638458766e-13 | 0.297205794 | 0.171 | 0.017 | 3.82114848036869e-09 | 46 |
| rps6kb1a1 | 1.54927690736011e-13 | 0.267549269 | 0.195 | 0.022 | 3.89937504813467e-09 | 46 |
| macf1a2 | 1.70396552164092e-13 | 0.455748707 | 0.317 | 0.053 | 4.28871082141803e-09 | 46 |
| aldh2.11 | 1.7260105415465e-13 | 0.309829319 | 0.244 | 0.033 | 4.34419593201839e-09 | 46 |
| gpatch83 | 1.9534806106756e-13 | 0.498874628 | 0.317 | 0.055 | 4.91671534900941e-09 | 46 |

| | | | | | | |
|---|---|---|---|---|---|---|
| araf | 2.10805559231155e-13 | 0.331234009 | 0.244 | 0.033 | 5.30576512028895e-09 | 46 |
| lgmn2 | 2.16720369892339e-13 | 0.470181148 | 0.341 | 0.061 | 5.45463498982027e-09 | 46 |
| si:ch211-74m13.35 | 2.20153711320792e-13 | 0.499086994 | 0.366 | 0.069 | 5.54104876023301e-09 | 46 |
| ucp215 | 2.38424421319739e-13 | 0.775426427 | 0.927 | 0.417 | 6.00090426019652e-09 | 46 |
| si:dkey-28b4.711 | 2.46289585623891e-13 | 0.478742227 | 0.439 | 0.092 | 6.19886258056772e-09 | 46 |
| rab4b | 2.54187342589021e-13 | 0.375196555 | 0.293 | 0.046 | 6.39764122562307e-09 | 46 |
| zgc:1124373 | 2.98260557445741e-13 | 0.296933296 | 0.171 | 0.018 | 7.50691997035185e-09 | 46 |
| krt949 | 3.2044880773263e-13 | 0.70132306 | 0.537 | 0.13 | 8.06537604182256e-09 | 46 |
| pon3.22 | 4.1637372076349e-13 | 0.460568146 | 0.463 | 0.097 | 1.04797101778963e-08 | 46 |
| zgc:110783 | 4.42232348344258e-13 | 0.296989527 | 0.244 | 0.033 | 1.11305459754766e-08 | 46 |
| diaph22 | 4.55154016966776e-13 | 0.386726533 | 0.244 | 0.035 | 1.14557714530368e-08 | 46 |
| XLOC-0242481 | 5.11249154695275e-13 | 0.29899717 | 0.195 | 0.023 | 1.28676299745254e-08 | 46 |
| kctd12.27 | 5.42708216052549e-13 | 0.573264876 | 0.39 | 0.08 | 1.36594230898266e-08 | 46 |
| pum2 | 6.16961291009865e-13 | 0.289361926 | 0.195 | 0.023 | 1.55282987334273e-08 | 46 |
| pim113 | 6.30652331430994e-13 | 0.705528074 | 0.61 | 0.172 | 1.58728885297867e-08 | 46 |
| map1lc3b8 | 6.74825215234609e-13 | 0.546604228 | 0.463 | 0.107 | 1.69846758422399e-08 | 46 |
| hipk3b1 | 7.06725512563596e-13 | 0.395713263 | 0.268 | 0.041 | 1.77875744257131e-08 | 46 |
| arl8a | 7.40468317551106e-13 | 0.378515156 | 0.317 | 0.054 | 1.86368470844438e-08 | 46 |
| myclb | 7.4393736279949e-13 | 0.604293586 | 0.415 | 0.088 | 1.87241594843004e-08 | 46 |
| lamc17 | 7.55311486905406e-13 | 0.473821118 | 0.317 | 0.056 | 1.90104348139222e-08 | 46 |
| htra1b4 | 7.81963176118933e-13 | 0.363073162 | 0.268 | 0.04 | 1.96812311797374e-08 | 46 |
| cnn215 | 7.84317705373901e-13 | 0.601252495 | 0.537 | 0.135 | 1.97404923265557e-08 | 46 |
| si:dkeyp-69e1.8 | 8.63330987942672e-13 | 0.27343487 | 0.195 | 0.023 | 2.17291776355291e-08 | 46 |
| rp2 | 9.40947520530658e-13 | 0.393297178 | 0.317 | 0.054 | 2.36827081442361e-08 | 46 |
| si:dkey-204f11.642 | 9.77437964171567e-13 | 0.282993823 | 0.22 | 0.029 | 2.46011361202342e-08 | 46 |
| tead1b3 | 1.01733586626012e-12 | 0.319004982 | 0.22 | 0.029 | 2.56053264179009e-08 | 46 |
| ppp1r10 | 1.03435703768072e-12 | 0.432766648 | 0.317 | 0.056 | 2.60337322813861e-08 | 46 |
| aqp3a4 | 1.24171074440376e-12 | 0.450474125 | 0.22 | 0.029 | 3.12526177258983e-08 | 46 |
| eif4ebp19 | 1.26480719202069e-12 | 0.737782512 | 0.732 | 0.246 | 3.18339322159687e-08 | 46 |
| prrc2a | 1.27893030796829e-12 | 0.356151777 | 0.268 | 0.041 | 3.21893969212539e-08 | 46 |
| CABZ01113812.1 | 1.69733264217688e-12 | 0.327622558 | 0.244 | 0.035 | 4.27201652709499e-08 | 46 |
| irf2bp2b8 | 1.70439337341522e-12 | 0.514322204 | 0.39 | 0.083 | 4.28978768154877e-08 | 46 |
| calm2b11 | 1.80332649477819e-12 | 0.730035517 | 0.659 | 0.211 | 4.53879245470722e-08 | 46 |
| ppib14 | 1.87237851807778e-12 | 0.762360459 | 0.854 | 0.333 | 4.71258949214997e-08 | 46 |
| vps37b | 1.87729791351636e-12 | 0.348581179 | 0.244 | 0.035 | 4.72497111852931e-08 | 46 |

| | | | | | | |
|---|---|---|---|---|---|---|
| map1sa | 1.89264101574094e-12 | 0.312640769 | 0.195 | 0.024 | 4.76358817251838e-08 | 46 |
| ankrd114 | 2.09777996034938e-12 | 0.503323546 | 0.415 | 0.091 | 5.27990238220335e-08 | 46 |
| si:dkey-6n21.122 | 2.1190860674917e-12 | 0.31431088 | 0.244 | 0.035 | 5.33352772326986e-08 | 46 |
| pkp4 | 2.24630187772258e-12 | 0.260629306 | 0.146 | 0.014 | 5.65371719603995e-08 | 46 |
| tbc1d5 | 2.45728644256245e-12 | 0.338991221 | 0.22 | 0.03 | 6.18474424728543e-08 | 46 |
| selenof5 | 2.63250965779838e-12 | 0.671435267 | 0.732 | 0.237 | 6.62576355771275e-08 | 46 |
| fndc3bb | 2.88707548694017e-12 | 0.273784884 | 0.146 | 0.014 | 7.26648029307971e-08 | 46 |
| crema12 | 3.05898024214414e-12 | 0.608039154 | 0.634 | 0.179 | 7.69914737145259e-08 | 46 |
| spag9a | 3.23928249471265e-12 | 0.290815633 | 0.171 | 0.019 | 8.15295011094228e-08 | 46 |
| rnf213b | 3.37736118177862e-12 | 0.288648249 | 0.146 | 0.014 | 8.50048035841861e-08 | 46 |
| si:dkeyp-44a8.21 | 3.42820877731357e-12 | 0.390882291 | 0.317 | 0.057 | 8.62845867162054e-08 | 46 |
| gstm.23 | 3.44492479021369e-12 | 0.409473984 | 0.293 | 0.05 | 8.67053120448884e-08 | 46 |
| si:ch211-286b5.59 | 3.597505385295e-12 | 0.636657437 | 0.512 | 0.137 | 9.05456130424899e-08 | 46 |
| cyfip12 | 3.73413981569871e-12 | 0.407755948 | 0.317 | 0.057 | 9.39845650213208e-08 | 46 |
| ptprsa2 | 3.93017739423202e-12 | 0.358008208 | 0.22 | 0.031 | 9.89186348354258e-08 | 46 |
| ppt18 | 4.20036527601819e-12 | 0.60937232 | 0.585 | 0.164 | 1.05718993632102e-07 | 46 |
| tax1bp1b1 | 4.32235391319221e-12 | 0.408867714 | 0.341 | 0.066 | 1.08789325641135e-07 | 46 |
| psme211 | 4.326372319274e-12 | 0.441014434 | 0.439 | 0.098 | 1.08890464903807e-07 | 46 |
| tgif14 | 4.3494543506473e-12 | 0.497655984 | 0.415 | 0.092 | 1.09471416551442e-07 | 46 |
| chd4b3 | 4.50972841536476e-12 | 0.551918184 | 0.439 | 0.103 | 1.13505354486316e-07 | 46 |
| copb2 | 4.70952005352239e-12 | 0.408668101 | 0.317 | 0.058 | 1.18533910227105e-07 | 46 |
| gstm.37 | 4.75809490762e-12 | 0.627538972 | 0.439 | 0.102 | 1.19756490729888e-07 | 46 |
| ppp1cb1 | 4.96972037509322e-12 | 0.396780301 | 0.341 | 0.065 | 1.25082892120721e-07 | 46 |
| pcyox12 | 5.23169767663118e-12 | 0.410029635 | 0.293 | 0.051 | 1.3167659882313e-07 | 46 |
| si:ch73-1a9.320 | 5.34425707597793e-12 | 0.733809442 | 0.756 | 0.266 | 1.34509606345288e-07 | 46 |
| si:ch211-232b12.51 | 5.644927347021e-12 | 0.257340813 | 0.171 | 0.019 | 1.42077176397172e-07 | 46 |
| sat1a.29 | 5.90943550632797e-12 | 0.748717505 | 0.78 | 0.299 | 1.48734582258769e-07 | 46 |
| timp2a8 | 5.94410104885859e-12 | 0.388082301 | 0.317 | 0.058 | 1.49607079298722e-07 | 46 |
| snx12 | 6.27911226000199e-12 | 0.496717654 | 0.366 | 0.076 | 1.5803897647199e-07 | 46 |
| ptk7a | 6.28887088273083e-12 | 0.270774251 | 0.195 | 0.025 | 1.58284591247452e-07 | 46 |
| gatad2b3 | 6.62312083136153e-12 | 0.442520958 | 0.317 | 0.06 | 1.66697328204538e-07 | 46 |
| yap18 | 6.65869775380329e-12 | 0.361323723 | 0.293 | 0.051 | 1.67592763765475e-07 | 46 |
| lpar16 | 7.2869791262275e-12 | 0.411003873 | 0.341 | 0.065 | 1.8340597762802e-07 | 46 |
| tob1a6 | 7.34359740711474e-12 | 0.535430178 | 0.512 | 0.13 | 1.84831003139671e-07 | 46 |
| hnrpkl | 7.48188468477095e-12 | 0.407209466 | 0.366 | 0.074 | 1.88311555631e-07 | 46 |

| | | | | | | |
|---|---|---|---|---|---|---|
| lin7c | 8.15060528164218e-12 | 0.33422389 | 0.22 | 0.031 | 2.05142584333652e-07 | 46 |
| crtap4 | 8.28454009292019e-12 | 0.336228326 | 0.22 | 0.031 | 2.08513589598708e-07 | 46 |
| btr09 | 9.08174833010254e-12 | 0.251595053 | 0.146 | 0.015 | 2.28578523720351e-07 | 46 |
| pfn2l7 | 1.03363034097965e-11 | 0.592513235 | 0.707 | 0.21 | 2.60154420521168e-07 | 46 |
| h3f3c17 | 1.04894622201027e-11 | 0.676004787 | 0.756 | 0.257 | 2.64009274617764e-07 | 46 |
| adam10a5 | 1.06918197906875e-11 | 0.503687831 | 0.415 | 0.095 | 2.69102412311813e-07 | 46 |
| fkbp1aa13 | 1.07728429482152e-11 | 0.719290323 | 0.878 | 0.388 | 2.71141684163628e-07 | 46 |
| LOC1018844516 | 1.20654659573011e-11 | 0.510854126 | 0.39 | 0.086 | 3.03675712679312e-07 | 46 |
| unm-hu7910 | 1.32523372899975e-11 | 0.33142782 | 0.22 | 0.031 | 3.33548077251946e-07 | 46 |
| cox7a13 | 1.34559537145784e-11 | 0.330903767 | 0.22 | 0.031 | 3.38672899042224e-07 | 46 |
| antxr1c4 | 1.44998535330621e-11 | 0.330423565 | 0.244 | 0.038 | 3.6494681357364e-07 | 46 |
| atf4b16 | 1.49840250413205e-11 | 0.677095911 | 0.756 | 0.265 | 3.77132926264995e-07 | 46 |
| CABZ01110881.1 | 1.63225014503188e-11 | 0.31826239 | 0.268 | 0.044 | 4.10821039003073e-07 | 46 |
| acox1 | 1.68582400962437e-11 | 0.291262138 | 0.195 | 0.025 | 4.24305044982357e-07 | 46 |
| ctsd4 | 1.96858505971919e-11 | 0.499473477 | 0.537 | 0.137 | 4.95473173680723e-07 | 46 |
| smox8 | 2.2181180491088e-11 | 0.45936797 | 0.39 | 0.085 | 5.58278131780194e-07 | 46 |
| dsg2.12 | 2.31410312154397e-11 | 0.314005485 | 0.195 | 0.026 | 5.82436614661403e-07 | 46 |
| crebrf | 2.48453202417399e-11 | 0.306360388 | 0.244 | 0.038 | 6.25331865164352e-07 | 46 |
| rsad2 | 2.49336399698187e-11 | 0.30725995 | 0.122 | 0.011 | 6.27554784400367e-07 | 46 |
| CABZ01080702.18 | 2.51669145710942e-11 | 0.44830197 | 0.39 | 0.085 | 6.33426072839869e-07 | 46 |
| ppp2cb1 | 2.68134245657888e-11 | 0.571944756 | 0.585 | 0.168 | 6.74867082896338e-07 | 46 |
| gnpda13 | 2.89661764632736e-11 | 0.293091591 | 0.268 | 0.045 | 7.29049695404134e-07 | 46 |
| cox8a15 | 3.1264760837076e-11 | 0.601900955 | 0.707 | 0.228 | 7.86902765508365e-07 | 46 |
| tmem596 | 3.22741472555021e-11 | 0.487095 | 0.439 | 0.107 | 8.12308012273734e-07 | 46 |
| psmc51 | 3.37045949464539e-11 | 0.443464633 | 0.39 | 0.087 | 8.48310950207299e-07 | 46 |
| si:ch211-216l23.2 | 3.45398640603627e-11 | 0.339164366 | 0.195 | 0.026 | 8.69333838535269e-07 | 46 |
| hivep1 | 3.6864075431954e-11 | 0.31818577 | 0.244 | 0.039 | 9.27831914546851e-07 | 46 |
| irf1b4 | 3.9960272792066e-11 | 0.364836736 | 0.244 | 0.039 | 1.00576010590351e-06 | 46 |
| ndufa4l14 | 4.19595746218457e-11 | 0.689755994 | 0.927 | 0.396 | 1.05608053365723e-06 | 46 |
| si:ch73-22o12.12 | 4.50998138930467e-11 | 0.263503184 | 0.195 | 0.026 | 1.13511721587409e-06 | 46 |
| kmt2cb2 | 5.7047224944175e-11 | 0.378001429 | 0.293 | 0.054 | 1.43582160461994e-06 | 46 |
| dynlt33 | 8.04104439426053e-11 | 0.479165636 | 0.537 | 0.147 | 2.02385046359143e-06 | 46 |
| ube2e38 | 8.25558682273329e-11 | 0.56328453 | 0.537 | 0.156 | 2.07784864741374e-06 | 46 |
| cox6b27 | 8.29270044406796e-11 | 0.648144872 | 0.951 | 0.534 | 2.08718977476747e-06 | 46 |
| ndufa15 | 8.82067941087711e-11 | 0.579443491 | 0.707 | 0.231 | 2.22007680092366e-06 | 46 |

| | | | | | | |
|---|---|---|---|---|---|---|
| creb3l21 | 1.00196042813642e-10 | 0.271378653 | 0.244 | 0.04 | 2.52183420157655e-06 | 46 |
| clint1a1 | 1.05672260549983e-10 | 0.333816624 | 0.268 | 0.048 | 2.65966512578251e-06 | 46 |
| CABZ01072614.110 | 1.05766862433163e-10 | 0.455756702 | 0.366 | 0.081 | 2.66204616058028e-06 | 46 |
| prkd21 | 1.18474328631227e-10 | 0.366555489 | 0.22 | 0.034 | 2.98188037731934e-06 | 46 |
| si:ch73-46j18.512 | 1.21804665323944e-10 | 0.573607208 | 0.61 | 0.176 | 3.06570162153835e-06 | 46 |
| NC-002333.171 | 1.25960270132883e-10 | 0.389673707 | 0.293 | 0.056 | 3.17029403897453e-06 | 46 |
| ldlrap1a2 | 1.29937421852562e-10 | 0.394567558 | 0.341 | 0.07 | 3.27039497060713e-06 | 46 |
| tmub2 | 1.35532244811753e-10 | 0.318515596 | 0.195 | 0.028 | 3.41121106966701e-06 | 46 |
| serinc22 | 1.50649510550722e-10 | 0.501694203 | 0.439 | 0.113 | 3.79169753105112e-06 | 46 |
| homeza1 | 1.53352111300539e-10 | 0.389841609 | 0.317 | 0.063 | 3.85971928932328e-06 | 46 |
| junbb10 | 1.55696084308797e-10 | 0.40829922 | 0.341 | 0.073 | 3.91871474596812e-06 | 46 |
| sptbn23 | 1.65336999752187e-10 | 0.3013342 | 0.244 | 0.041 | 4.16136694676279e-06 | 46 |
| gsk3b1 | 1.7083571582675e-10 | 0.278679126 | 0.195 | 0.028 | 4.29976413164346e-06 | 46 |
| pgd4 | 1.80847888138462e-10 | 0.365613675 | 0.317 | 0.064 | 4.55176049655696e-06 | 46 |
| CABZ01068358.17 | 1.85355364055399e-10 | 0.429745144 | 0.317 | 0.065 | 4.66520915791034e-06 | 46 |
| elavl1a1 | 2.40444316277012e-10 | 0.473217303 | 0.463 | 0.121 | 6.05174299637611e-06 | 46 |
| fat1a4 | 2.52980294171909e-10 | 0.302777539 | 0.195 | 0.029 | 6.36726102401277e-06 | 46 |
| zfand31 | 2.71731578497398e-10 | 0.365873019 | 0.293 | 0.057 | 6.83921209920101e-06 | 46 |
| rnf102 | 2.75560154177172e-10 | 0.502151568 | 0.415 | 0.107 | 6.93557352048523e-06 | 46 |
| si:dkey-79d12.59 | 2.77842460509954e-10 | 0.479632316 | 0.341 | 0.077 | 6.99301688857503e-06 | 46 |
| hephl1a | 2.79773930385788e-10 | 0.265609824 | 0.171 | 0.022 | 7.0416300538799e-06 | 46 |
| ifrd1 | 3.13176468495155e-10 | 0.486720561 | 0.415 | 0.104 | 7.88233853555454e-06 | 46 |
| pank2 | 3.17813514420164e-10 | 0.301831917 | 0.22 | 0.035 | 7.9990483444411e-06 | 46 |
| lrpap12 | 3.19016073394757e-10 | 0.319471335 | 0.293 | 0.057 | 8.02931555127265e-06 | 46 |
| tmsb6 | 3.20257274914936e-10 | 0.352223629 | 0.195 | 0.029 | 8.06055535233402e-06 | 46 |
| tnrc6a1 | 3.27427720500335e-10 | 0.388043803 | 0.341 | 0.074 | 8.24102829727294e-06 | 46 |
| fstl1b9 | 3.57941894754144e-10 | 0.444171134 | 0.488 | 0.126 | 9.00903954906705e-06 | 46 |
| epsti1 | 3.59792334179665e-10 | 0.252384744 | 0.146 | 0.017 | 9.05561325896799e-06 | 46 |
| sdcbp212 | 3.60718002127015e-10 | 0.576396836 | 0.634 | 0.209 | 9.07891139553485e-06 | 46 |
| ganab | 3.70486689707459e-10 | 0.308149413 | 0.22 | 0.035 | 9.32477949324703e-06 | 46 |
| rab109 | 3.96720221453626e-10 | 0.56145186 | 0.61 | 0.196 | 9.9850512537663e-06 | 46 |
| si:dkey-42i9.73 | 4.15604519419228e-10 | 0.297633229 | 0.171 | 0.023 | 1.04603501492625e-05 | 46 |
| exoc62 | 4.3935925132115e-10 | 0.280790326 | 0.22 | 0.035 | 1.1058232996502e-05 | 46 |
| PDPR | 4.54854365809384e-10 | 0.372782138 | 0.22 | 0.036 | 1.14482295330564e-05 | 46 |
| ewsr1a1 | 4.59823013501168e-10 | 0.294920982 | 0.244 | 0.042 | 1.15732854268109e-05 | 46 |

| | | | | | | |
|---|---|---|---|---|---|---|
| stau210 | 4.71940318891986e-10 | 0.44880727 | 0.366 | 0.086 | 1.18782658861924e-05 | 46 |
| mphosph85 | 5.080134759114e-10 | 0.53641822 | 0.439 | 0.119 | 1.2786191175214e-05 | 46 |
| CABZ01029938.11 | 5.13934225842393e-10 | 0.302172782 | 0.22 | 0.036 | 1.29352105302272e-05 | 46 |
| selenok8 | 5.17637803232946e-10 | 0.48914257 | 0.463 | 0.128 | 1.302842586957e-05 | 46 |
| aste1a2 | 5.18522523685668e-10 | 0.300729789 | 0.171 | 0.023 | 1.30506933986446e-05 | 46 |
| bbip1 | 5.37422358042564e-10 | 0.38821701 | 0.317 | 0.066 | 1.35263833295733e-05 | 46 |
| strip1 | 5.59326831533357e-10 | 0.255226361 | 0.195 | 0.029 | 1.40776970228631e-05 | 46 |
| degs12 | 6.20831960550652e-10 | 0.395909588 | 0.268 | 0.052 | 1.56257196150994e-05 | 46 |
| bin31 | 6.31283787354479e-10 | 0.326560254 | 0.22 | 0.036 | 1.58887816439249e-05 | 46 |
| serpinf17 | 6.3755805437244e-10 | 0.406673473 | 0.244 | 0.044 | 1.60466986704999e-05 | 46 |
| u2surp4 | 6.79117672400339e-10 | 0.428697276 | 0.39 | 0.096 | 1.70927126966441e-05 | 46 |
| tcf127 | 7.48192324480359e-10 | 0.350548539 | 0.293 | 0.059 | 1.88312526148462e-05 | 46 |
| si:ch211-212d10.21 | 7.51792344746497e-10 | 0.255365212 | 0.146 | 0.018 | 1.89218615249246e-05 | 46 |
| parvaa3 | 7.62849825267733e-10 | 0.261724881 | 0.195 | 0.029 | 1.92001672521636e-05 | 46 |
| klf36 | 8.49701563553214e-10 | 0.296932773 | 0.22 | 0.036 | 2.13861386530709e-05 | 46 |
| wbp2nl5 | 8.77155688334983e-10 | 0.426709426 | 0.366 | 0.088 | 2.20771315197032e-05 | 46 |
| ifi351 | 8.82925780191589e-10 | 0.305597556 | 0.22 | 0.036 | 2.22223589616421e-05 | 46 |
| igf2b6 | 8.86339756632727e-10 | 0.421770308 | 0.293 | 0.06 | 2.23082853346891e-05 | 46 |
| dctn3 | 9.2549199069593e-10 | 0.405689781 | 0.366 | 0.087 | 2.32937079138259e-05 | 46 |
| stxbp62 | 9.52251324772002e-10 | 0.372827858 | 0.244 | 0.044 | 2.39672135931865e-05 | 46 |
| zgc:1008294 | 9.57360962207249e-10 | 0.310165074 | 0.22 | 0.037 | 2.40958180577942e-05 | 46 |
| sptan14 | 1.09321059699025e-09 | 0.327942617 | 0.268 | 0.051 | 2.75150175156475e-05 | 46 |
| CABZ01078261.1 | 1.10135277551697e-09 | 0.270493545 | 0.195 | 0.03 | 2.77199480069865e-05 | 46 |
| atg13 | 1.10267721199324e-09 | 0.3357948 | 0.244 | 0.044 | 2.77532827486578e-05 | 46 |
| golgb11 | 1.13370977048153e-09 | 0.328926108 | 0.244 | 0.044 | 2.85343412132496e-05 | 46 |
| hspe16 | 1.13752375858306e-09 | 0.65488674 | 0.78 | 0.295 | 2.86303354797771e-05 | 46 |
| mt-nd317 | 1.23329912680069e-09 | 0.631779731 | 0.927 | 0.396 | 3.10409057224467e-05 | 46 |
| nedd81 | 1.3270288120363e-09 | 0.387777174 | 0.341 | 0.078 | 3.33999881701415e-05 | 46 |
| arf2a2 | 1.48565543219784e-09 | 0.462227093 | 0.463 | 0.128 | 3.73924615729873e-05 | 46 |
| nr1d2a6 | 1.57851289143199e-09 | 0.41572008 | 0.341 | 0.079 | 3.97295909644519e-05 | 46 |
| scrib | 1.61129136315329e-09 | 0.250200836 | 0.171 | 0.024 | 4.05545923192052e-05 | 46 |
| eif1b14 | 1.62292951931321e-09 | 0.648355422 | 0.878 | 0.383 | 4.08475130715941e-05 | 46 |
| kdm6bb4 | 1.80589786304135e-09 | 0.326820954 | 0.268 | 0.053 | 4.54526433148876e-05 | 46 |
| si:dkey-187e18.12 | 2.05652531359793e-09 | 0.37535605 | 0.317 | 0.07 | 5.17606856179463e-05 | 46 |
| si:ch211-108d22.2 | 2.20349670125215e-09 | 0.254703366 | 0.195 | 0.03 | 5.54598084738155e-05 | 46 |

| | | | | | | |
|---|---|---|---|---|---|---|
| map1lc3a11 | 2.48195161023968e-09 | 0.380196021 | 0.317 | 0.071 | 6.24682400781224e-05 | 46 |
| tmem176l.19 | 2.7220419098666e-09 | 0.501782221 | 0.585 | 0.181 | 6.85110728294326e-05 | 46 |
| rhoaa13 | 2.77094562088296e-09 | 0.474168961 | 0.512 | 0.155 | 6.97419303320032e-05 | 46 |
| rhocb1 | 2.78782543113667e-09 | 0.377493883 | 0.366 | 0.089 | 7.01667782762789e-05 | 46 |
| parp12b1 | 2.85618799198546e-09 | 0.276366623 | 0.195 | 0.031 | 7.18873955702819e-05 | 46 |
| hdac44 | 2.86688288822352e-09 | 0.324734849 | 0.195 | 0.031 | 7.21565754136977e-05 | 46 |
| CU694197.12 | 3.14274589332427e-09 | 0.449667353 | 0.463 | 0.131 | 7.90997713890786e-05 | 46 |
| pi4kb2 | 3.15930643549217e-09 | 0.265100847 | 0.22 | 0.038 | 7.95165836749024e-05 | 46 |
| nt5c2b3 | 3.18074162544646e-09 | 0.311015375 | 0.244 | 0.046 | 8.0056085970862e-05 | 46 |
| ctnnd12 | 3.20955555683217e-09 | 0.335370746 | 0.293 | 0.061 | 8.07813038099089e-05 | 46 |
| rbm39a6 | 3.25012853082773e-09 | 0.476909991 | 0.488 | 0.143 | 8.1802484992403e-05 | 46 |
| pmp22b8 | 3.31462879478724e-09 | 0.363594252 | 0.244 | 0.046 | 8.34258921360002e-05 | 46 |
| si:ch73-233k15.21 | 3.42033269607198e-09 | 0.286510107 | 0.171 | 0.025 | 8.60863536274357e-05 | 46 |
| ylpm1 | 3.54575660602993e-09 | 0.338731923 | 0.268 | 0.053 | 8.92431480171674e-05 | 46 |
| ivns1abpa1 | 3.83229283575049e-09 | 0.275798619 | 0.195 | 0.031 | 9.6454978383004e-05 | 46 |
| si:ch211-105c13.36 | 4.15962997000916e-09 | 0.373825701 | 0.293 | 0.061 | 0.000104694 | 46 |
| chd4a4 | 4.20559526842205e-09 | 0.421638709 | 0.366 | 0.092 | 0.000105851 | 46 |
| thtpa | 4.68037795052357e-09 | 0.33914864 | 0.317 | 0.07 | 0.0001178 | 46 |
| adssl1 | 4.70902387388832e-09 | 0.414013023 | 0.415 | 0.108 | 0.000118521 | 46 |
| sesn32 | 4.86303492814851e-09 | 0.316989925 | 0.171 | 0.025 | 0.000122398 | 46 |
| si:ch211-87m7.2 | 5.05409175062408e-09 | 0.2921481 | 0.195 | 0.032 | 0.000127206 | 46 |
| fzd7b3 | 5.060928834457e-09 | 0.329015649 | 0.22 | 0.039 | 0.000127379 | 46 |
| matr3l1.11 | 5.07701959612729e-09 | 0.354628907 | 0.341 | 0.08 | 0.000127784 | 46 |
| ndfip22 | 5.20580240433583e-09 | 0.389827619 | 0.268 | 0.056 | 0.000131025 | 46 |
| gstm.16 | 5.32072818418308e-09 | 0.606117 | 0.707 | 0.261 | 0.000133917 | 46 |
| atp1b3b1 | 5.39690585630842e-09 | 0.41936065 | 0.293 | 0.063 | 0.000135835 | 46 |
| litaf10 | 5.61211394517972e-09 | 0.469853837 | 0.463 | 0.136 | 0.000141251 | 46 |
| slc38a5b11 | 5.62577194537995e-09 | 0.481385877 | 0.415 | 0.116 | 0.000141595 | 46 |
| flot1b3 | 5.74557862326581e-09 | 0.388027947 | 0.341 | 0.083 | 0.00014461 | 46 |
| sh3bp5b | 5.85113768485216e-09 | 0.272242774 | 0.195 | 0.031 | 0.000147267 | 46 |
| si:ch211-39i22.18 | 5.97200665789517e-09 | 0.406007029 | 0.317 | 0.073 | 0.000150309 | 46 |
| ywhaz9 | 6.09070685793235e-09 | 0.613365041 | 0.707 | 0.282 | 0.000153297 | 46 |
| st14a2 | 6.46966502822204e-09 | 0.257129564 | 0.22 | 0.038 | 0.000162835 | 46 |
| asah1b3 | 6.86031881803988e-09 | 0.285555301 | 0.268 | 0.054 | 0.000172667 | 46 |
| calcoco1a1 | 7.09803221964571e-09 | 0.282884294 | 0.195 | 0.032 | 0.00017865 | 46 |

| | | | | | | |
|---|---|---|---|---|---|---|
| ralab | 7.34139117229174e-09 | 0.281379183 | 0.171 | 0.026 | 0.000184775 | 46 |
| polr2b | 7.34837236014566e-09 | 0.259191277 | 0.22 | 0.039 | 0.000184951 | 46 |
| mpzl2b5 | 7.51466129286908e-09 | 0.340624483 | 0.39 | 0.095 | 0.000189137 | 46 |
| ywhah1 | 8.13694819159606e-09 | 0.295253841 | 0.244 | 0.047 | 0.000204799 | 46 |
| rn7sk1 | 8.756626779534e-09 | 0.41831776 | 0.22 | 0.041 | 0.000220396 | 46 |
| sh3pxd2aa4 | 9.18563882168559e-09 | 0.284594169 | 0.195 | 0.033 | 0.000231193 | 46 |
| perp7 | 9.5917807909056e-09 | 0.287524729 | 0.268 | 0.055 | 0.000241416 | 46 |
| gabarapb15 | 1.08039780426416e-08 | 0.583839453 | 0.683 | 0.27 | 0.000271925 | 46 |
| ndufb95 | 1.10347505019489e-08 | 0.511721156 | 0.537 | 0.177 | 0.000277734 | 46 |
| eif4a1b13 | 1.36995126724324e-08 | 0.540346663 | 0.756 | 0.281 | 0.000344803 | 46 |
| tfcp2 | 1.52129287906824e-08 | 0.289235811 | 0.195 | 0.033 | 0.000382894 | 46 |
| tmed102 | 1.56690920077885e-08 | 0.465444505 | 0.537 | 0.172 | 0.000394375 | 46 |
| wls8 | 1.58124804481564e-08 | 0.409968228 | 0.439 | 0.124 | 0.000397984 | 46 |
| plbd14 | 1.584956344672e-08 | 0.262571381 | 0.171 | 0.027 | 0.000398918 | 46 |
| ndfip11 | 1.59697032863845e-08 | 0.565822411 | 0.488 | 0.164 | 0.000401941 | 46 |
| glod52 | 1.702785075732e-08 | 0.328525082 | 0.195 | 0.034 | 0.000428574 | 46 |
| inhbaa4 | 1.8500730128e-08 | 0.348490921 | 0.22 | 0.041 | 0.000465645 | 46 |
| pros11 | 1.90174176887678e-08 | 0.334986877 | 0.244 | 0.049 | 0.000478649 | 46 |
| csde15 | 1.93266578119761e-08 | 0.478635739 | 0.61 | 0.209 | 0.000486433 | 46 |
| msi2b1 | 1.93554879745421e-08 | 0.405550216 | 0.366 | 0.094 | 0.000487158 | 46 |
| si:ch211-140m22.75 | 1.98034046942321e-08 | 0.361323697 | 0.293 | 0.067 | 0.000498432 | 46 |
| b4galt62 | 2.04351112349243e-08 | 0.369891776 | 0.293 | 0.068 | 0.000514331 | 46 |
| hmgb1b10 | 2.12206208535915e-08 | 0.438653958 | 0.439 | 0.127 | 0.000534102 | 46 |
| zfand5a3 | 2.46272444574811e-08 | 0.37276684 | 0.317 | 0.077 | 0.000619843 | 46 |
| arhgap35a | 2.54392421212251e-08 | 0.252206112 | 0.171 | 0.027 | 0.00064028 | 46 |
| tspan3a | 2.54512186179706e-08 | 0.305972285 | 0.293 | 0.066 | 0.000640582 | 46 |
| cox8b7 | 2.55003930584908e-08 | 0.395416416 | 0.366 | 0.096 | 0.000641819 | 46 |
| ctsla13 | 2.56508483912969e-08 | 0.632803544 | 0.854 | 0.387 | 0.000645606 | 46 |
| abca22 | 2.57556396099486e-08 | 0.253606204 | 0.171 | 0.027 | 0.000648244 | 46 |
| arf2b1 | 2.64136679199418e-08 | 0.48554938 | 0.512 | 0.163 | 0.000664806 | 46 |
| jak16 | 2.69409799453656e-08 | 0.389280824 | 0.366 | 0.097 | 0.000678078 | 46 |
| canx1 | 2.72408643323587e-08 | 0.421201346 | 0.341 | 0.089 | 0.000685625 | 46 |
| nfil3-612 | 2.86686584982605e-08 | 0.485456662 | 0.439 | 0.132 | 0.000721561 | 46 |
| arid1ab | 2.91930565784295e-08 | 0.258390907 | 0.171 | 0.027 | 0.00073476 | 46 |
| s100v212 | 2.94054213596931e-08 | 0.373157693 | 0.439 | 0.123 | 0.000740105 | 46 |

| | | | | | | |
|---|---|---|---|---|---|---|
| cpvl | 3.12124388761395e-08 | 0.258947587 | 0.244 | 0.049 | 0.000785586 | 46 |
| clcn7 | 3.16086072940244e-08 | 0.275358713 | 0.171 | 0.027 | 0.000795557 | 46 |
| bcl7a1 | 3.17651427955155e-08 | 0.263611518 | 0.195 | 0.034 | 0.000799497 | 46 |
| CABZ01112656.12 | 3.19849605714678e-08 | 0.298722503 | 0.268 | 0.059 | 0.000805029 | 46 |
| birc26 | 3.35812719179841e-08 | 0.40079045 | 0.39 | 0.108 | 0.000845207 | 46 |
| foxo4 | 3.71530918355434e-08 | 0.27233653 | 0.22 | 0.042 | 0.000935106 | 46 |
| ddx3xa1 | 3.85809845560746e-08 | 0.361978868 | 0.366 | 0.097 | 0.000971045 | 46 |
| pfn29 | 3.99520115016116e-08 | 0.514624053 | 0.634 | 0.227 | 0.001005552 | 46 |
| abhd17ab | 4.64871840549351e-08 | 0.252379755 | 0.171 | 0.028 | 0.001170036 | 46 |
| oaz1b19 | 4.70931761001068e-08 | 0.555316007 | 0.78 | 0.323 | 0.001185288 | 46 |
| trip12 | 4.73640286677041e-08 | 0.330111759 | 0.22 | 0.043 | 0.001192105 | 46 |
| tmem50a | 4.7447912843308e-08 | 0.429947865 | 0.463 | 0.142 | 0.001194217 | 46 |
| laptm4a3 | 4.8656011471581e-08 | 0.387719512 | 0.39 | 0.11 | 0.001224623 | 46 |
| dbnlb3 | 4.94812570567921e-08 | 0.27886691 | 0.22 | 0.043 | 0.001245394 | 46 |
| appbp22 | 5.18656654832849e-08 | 0.277408445 | 0.22 | 0.043 | 0.001305407 | 46 |
| bsdc1 | 5.39754829795256e-08 | 0.346246719 | 0.268 | 0.061 | 0.001358509 | 46 |
| rnaseka2 | 6.37245051150527e-08 | 0.413419925 | 0.439 | 0.13 | 0.001603882 | 46 |
| chd39 | 6.37977857536187e-08 | 0.451889392 | 0.366 | 0.105 | 0.001605726 | 46 |
| hif1ab6 | 6.64048561793121e-08 | 0.425111089 | 0.39 | 0.114 | 0.001671344 | 46 |
| cst31 | 6.83778104149033e-08 | 0.339592224 | 0.22 | 0.045 | 0.001721001 | 46 |
| ralbp11 | 7.23428040089363e-08 | 0.287329218 | 0.244 | 0.052 | 0.001820796 | 46 |
| katnb1 | 7.28506592517561e-08 | 0.261764656 | 0.171 | 0.028 | 0.001833578 | 46 |
| vdac39 | 7.51177375167676e-08 | 0.427485469 | 0.488 | 0.155 | 0.001890638 | 46 |
| ciao2b | 8.02066244864109e-08 | 0.281235034 | 0.244 | 0.052 | 0.002018721 | 46 |
| caprin1a1 | 8.32474440378414e-08 | 0.336841976 | 0.244 | 0.053 | 0.002095255 | 46 |
| hcfc1b | 8.72518022173338e-08 | 0.303131448 | 0.22 | 0.044 | 0.002196041 | 46 |
| fstb3 | 9.57496457900447e-08 | 0.260462171 | 0.146 | 0.022 | 0.002409923 | 46 |
| XLOC-0009461 | 9.6755621732608e-08 | 0.314330382 | 0.22 | 0.045 | 0.002435242 | 46 |
| rnf71 | 1.0243066036152e-07 | 0.341331008 | 0.366 | 0.1 | 0.002578077 | 46 |
| tpm4a9 | 1.02646386191277e-07 | 0.350004743 | 0.341 | 0.089 | 0.002583507 | 46 |
| pik3ip1 | 1.05526064102857e-07 | 0.349883532 | 0.341 | 0.088 | 0.002655986 | 46 |
| tomm61 | 1.06723217631148e-07 | 0.361629783 | 0.39 | 0.111 | 0.002686117 | 46 |
| tpm310 | 1.07602293609625e-07 | 0.484323895 | 0.683 | 0.255 | 0.002708242 | 46 |
| si:ch73-42p12.23 | 1.15248158511401e-07 | 0.373353467 | 0.366 | 0.103 | 0.002900681 | 46 |
| chchd21 | 1.17945750378388e-07 | 0.358938316 | 0.293 | 0.072 | 0.002968577 | 46 |

| | | | | | | |
|---|---|---|---|---|---|---|
| grinab6 | 1.28884630274799e-07 | 0.403505862 | 0.39 | 0.115 | 0.003243897 | 46 |
| fthl273 | 1.3094436086406e-07 | 0.472410983 | 1 | 0.696 | 0.003295739 | 46 |
| taf3 | 1.36509302946465e-07 | 0.315168721 | 0.244 | 0.054 | 0.003435803 | 46 |
| hp1bp36 | 1.39380208410849e-07 | 0.458387426 | 0.341 | 0.099 | 0.00350806 | 46 |
| MYADM1 | 1.39550169223224e-07 | 0.266533426 | 0.195 | 0.037 | 0.003512338 | 46 |
| ccnl1a4 | 1.51814107373357e-07 | 0.299154786 | 0.268 | 0.063 | 0.003821009 | 46 |
| myd881 | 1.54292608898401e-07 | 0.349216796 | 0.244 | 0.055 | 0.003883391 | 46 |
| srgn13 | 1.56068635490642e-07 | 0.492470547 | 0.659 | 0.238 | 0.003928091 | 46 |
| arrdc3a5 | 1.58959342188399e-07 | 0.395165939 | 0.293 | 0.073 | 0.004000848 | 46 |
| nfkbiaa9 | 1.6887162110078e-07 | 0.497758774 | 0.463 | 0.153 | 0.00425033 | 46 |
| psmb11 | 1.8017052958334e-07 | 0.419721445 | 0.512 | 0.171 | 0.004534712 | 46 |
| dync1i2a1 | 1.98229169031191e-07 | 0.319613428 | 0.244 | 0.055 | 0.00498923 | 46 |
| rbms2a1 | 2.03781122031065e-07 | 0.251874206 | 0.171 | 0.03 | 0.005128967 | 46 |
| bcl2l106 | 2.0559329601116e-07 | 0.510110673 | 0.585 | 0.216 | 0.005174578 | 46 |
| jund11 | 2.11506015528114e-07 | 0.364657318 | 0.341 | 0.094 | 0.005323395 | 46 |
| arhgap313 | 2.1349686449856e-07 | 0.271675153 | 0.171 | 0.03 | 0.005373503 | 46 |
| zgc:1232582 | 2.18278324846892e-07 | 0.288665302 | 0.22 | 0.046 | 0.005493847 | 46 |
| XLOC-0160947 | 2.25717785890393e-07 | 0.301523177 | 0.317 | 0.08 | 0.005681091 | 46 |
| cux1a4 | 2.27159290616367e-07 | 0.287721652 | 0.22 | 0.046 | 0.005717372 | 46 |
| morc21 | 2.47909926073679e-07 | 0.280961851 | 0.22 | 0.046 | 0.006239645 | 46 |
| si:ch211-191j22.33 | 2.54754159869213e-07 | 0.344883211 | 0.39 | 0.115 | 0.006411907 | 46 |
| ctbp1 | 2.55325661269778e-07 | 0.390455417 | 0.439 | 0.136 | 0.006426292 | 46 |
| slc3a2b12 | 2.65075164363004e-07 | 0.41119051 | 0.39 | 0.119 | 0.006671677 | 46 |
| rps27l13 | 2.68446819872779e-07 | 0.515555507 | 0.537 | 0.204 | 0.006756538 | 46 |
| bnip3lb2 | 2.93703344558146e-07 | 0.303147842 | 0.268 | 0.065 | 0.007392219 | 46 |
| gchfr2 | 3.10698628326256e-07 | 0.268721715 | 0.195 | 0.038 | 0.007819974 | 46 |
| pdlim14 | 3.28978201370143e-07 | 0.28527774 | 0.244 | 0.055 | 0.008280052 | 46 |
| baiap2a3 | 3.49797697730538e-07 | 0.260186885 | 0.195 | 0.038 | 0.008804058 | 46 |
| ndufa33 | 3.53210869427856e-07 | 0.399904472 | 0.537 | 0.185 | 0.008889964 | 46 |
| lsm14aa | 3.59589776156283e-07 | 0.262998708 | 0.244 | 0.055 | 0.009050515 | 46 |
| cdh25 | 3.67051950796195e-07 | 0.340264033 | 0.244 | 0.056 | 0.009238331 | 46 |
| aif1l4 | 3.72429302389748e-07 | 0.4635051 | 0.561 | 0.189 | 0.009373673 | 46 |
| kmt2bb3 | 3.84486618636627e-07 | 0.348613063 | 0.317 | 0.087 | 0.009677144 | 46 |
| mbnl21 | 3.86041880139608e-07 | 0.263281518 | 0.146 | 0.024 | 0.009716288 | 46 |
| atp6ap1b | 3.86726289341179e-07 | 0.256688316 | 0.195 | 0.039 | 0.009733514 | 46 |

| | | | | | | |
|---|---|---|---|---|---|---|
| atox15 | 3.9176386846273e-07 | 0.402958603 | 0.366 | 0.111 | 0.009860305 | 46 |
| si:ch211-196l7.420 | 3.94525237653333e-07 | 0.526206262 | 0.732 | 0.315 | 0.009929806 | 46 |
| elovl1b7 | 4.13575851856849e-07 | 0.373112699 | 0.39 | 0.117 | 0.010409291 | 46 |
| elocb3 | 4.23966823280844e-07 | 0.44773491 | 0.488 | 0.172 | 0.010670821 | 46 |
| smdt1b6 | 4.39569474822392e-07 | 0.400225012 | 0.415 | 0.134 | 0.011063524 | 46 |
| tbca1 | 4.52801972457721e-07 | 0.44844987 | 0.537 | 0.196 | 0.011396573 | 46 |
| ergic2 | 4.6294423725348e-07 | 0.257334818 | 0.22 | 0.047 | 0.011651844 | 46 |
| si:dkey-7j14.612 | 4.76383592435179e-07 | 0.489203183 | 0.488 | 0.176 | 0.011990099 | 46 |
| csnk1a12 | 4.86737018179672e-07 | 0.465661469 | 0.61 | 0.234 | 0.012250684 | 46 |
| enc14 | 4.93956086668218e-07 | 0.256793522 | 0.171 | 0.031 | 0.012432381 | 46 |
| tma75 | 5.06050491001129e-07 | 0.501754107 | 0.732 | 0.306 | 0.012736785 | 46 |
| si:dkeyp-75h12.52 | 5.27766182275897e-07 | 0.268128658 | 0.195 | 0.039 | 0.013283347 | 46 |
| bcap311 | 5.65412235883802e-07 | 0.254842775 | 0.268 | 0.065 | 0.014230861 | 46 |
| gng12a1 | 5.81616772723973e-07 | 0.259210677 | 0.171 | 0.032 | 0.014638713 | 46 |
| ahnak14 | 5.8894455357048e-07 | 0.366752129 | 0.415 | 0.128 | 0.014823145 | 46 |
| si:dkey-250d21.11 | 5.90842428580483e-07 | 0.349301072 | 0.293 | 0.078 | 0.014870913 | 46 |
| notch212 | 6.06823292455351e-07 | 0.376925354 | 0.39 | 0.121 | 0.015273135 | 46 |
| dpp73 | 6.40738956863708e-07 | 0.307111277 | 0.22 | 0.049 | 0.016126759 | 46 |
| psmb51 | 6.4739888491233e-07 | 0.338634694 | 0.268 | 0.068 | 0.016294383 | 46 |
| vps281 | 6.67599156318316e-07 | 0.324030995 | 0.341 | 0.097 | 0.016802803 | 46 |
| ece2b2 | 6.84275766622412e-07 | 0.287301643 | 0.22 | 0.048 | 0.017222537 | 46 |
| rab5c5 | 6.93613428381034e-07 | 0.379677715 | 0.39 | 0.121 | 0.017457556 | 46 |
| yy1a | 7.17124329624963e-07 | 0.302094197 | 0.22 | 0.049 | 0.018049302 | 46 |
| casp3a | 7.34984956323919e-07 | 0.261081424 | 0.22 | 0.048 | 0.018498836 | 46 |
| mtdha3 | 7.50300960992881e-07 | 0.322564484 | 0.439 | 0.14 | 0.018884325 | 46 |
| vps35 | 7.98636154602615e-07 | 0.346145271 | 0.244 | 0.059 | 0.020100873 | 46 |
| si:rp71-36a1.52 | 8.00087962689255e-07 | 0.253824563 | 0.22 | 0.048 | 0.020137414 | 46 |
| fam168a8 | 8.49220190821165e-07 | 0.30443314 | 0.244 | 0.059 | 0.021374023 | 46 |
| f11r.18 | 8.63646385081278e-07 | 0.363731222 | 0.537 | 0.179 | 0.021737116 | 46 |
| srrm11 | 8.6844683471787e-07 | 0.350498089 | 0.39 | 0.12 | 0.021857938 | 46 |
| higd1a14 | 9.08639806205906e-07 | 0.50855115 | 0.732 | 0.311 | 0.022869555 | 46 |
| ddt1 | 9.58173807911771e-07 | 0.335139216 | 0.341 | 0.1 | 0.024116277 | 46 |
| baz1a1 | 9.60714026191436e-07 | 0.295940916 | 0.293 | 0.077 | 0.024180211 | 46 |
| fxr2 | 9.80501776747209e-07 | 0.274330903 | 0.244 | 0.058 | 0.024678249 | 46 |
| fundc22 | 9.81140904549843e-07 | 0.348135104 | 0.317 | 0.09 | 0.024694335 | 46 |

| | | | | | | |
|---|---|---|---|---|---|---|
| dldh1 | 1.18045571258538e-06 | 0.287326235 | 0.317 | 0.089 | 0.02971089 | 46 |
| itgav5 | 1.19624343595823e-06 | 0.274860338 | 0.244 | 0.059 | 0.030108251 | 46 |
| clic43 | 1.39195824231367e-06 | 0.261248655 | 0.22 | 0.05 | 0.035034197 | 46 |
| mvb12a2 | 1.42940034036036e-06 | 0.282686431 | 0.293 | 0.078 | 0.035976577 | 46 |
| pomp8 | 1.43058796693869e-06 | 0.455204316 | 0.585 | 0.232 | 0.036006469 | 46 |
| gnai13 | 1.44472428225111e-06 | 0.289834029 | 0.244 | 0.06 | 0.036362265 | 46 |
| zgc:162698 | 1.44842988648711e-06 | 0.284663313 | 0.22 | 0.051 | 0.036455532 | 46 |
| anxa411 | 1.48048756382752e-06 | 0.375316762 | 0.415 | 0.138 | 0.037262391 | 46 |
| naa38 | 1.52774036808634e-06 | 0.282671098 | 0.244 | 0.059 | 0.038451697 | 46 |
| flot2a7 | 1.56254502894841e-06 | 0.270820841 | 0.244 | 0.06 | 0.039327696 | 46 |
| kdm5c | 1.6268045769274e-06 | 0.274475626 | 0.22 | 0.051 | 0.040945044 | 46 |
| si:dkey-222f8.31 | 1.65525383715533e-06 | 0.27730383 | 0.244 | 0.06 | 0.041661084 | 46 |
| si:ch211-212k18.76 | 1.67016595892848e-06 | 0.430508523 | 0.415 | 0.14 | 0.042036407 | 46 |
| lrrc8aa | 1.67125146393749e-06 | 0.267960791 | 0.22 | 0.051 | 0.042063728 | 46 |
| chmp2a6 | 1.73360492962413e-06 | 0.341077116 | 0.415 | 0.135 | 0.043633102 | 46 |
| mcl1a14 | 1.77886964245656e-06 | 0.467551661 | 0.951 | 0.625 | 0.04477237 | 46 |
| elmo19 | 1.83211924809548e-06 | 0.354458736 | 0.268 | 0.072 | 0.046112609 | 46 |
| rxrba | 1.83872524481358e-06 | 0.295686005 | 0.22 | 0.052 | 0.046278876 | 46 |
| etfb1 | 1.85924158608021e-06 | 0.323065293 | 0.317 | 0.092 | 0.046795251 | 46 |
| ash1l1 | 1.94315411129629e-06 | 0.294885544 | 0.195 | 0.043 | 0.048907246 | 46 |
| chmp5b3 | 1.9493501132077e-06 | 0.359801893 | 0.366 | 0.116 | 0.049063193 | 46 |
| si:ch211-241f5.36 | 1.98600630196e-06 | 0.27038805 | 0.268 | 0.069 | 0.049985793 | 46 |
| tmem30aa2 | 2.00864336438705e-06 | 0.311829174 | 0.244 | 0.062 | 0.050555545 | 46 |
| syncripl | 2.05348689649753e-06 | 0.261979691 | 0.293 | 0.08 | 0.051684212 | 46 |
| smad2 | 2.19882861232597e-06 | 0.308376387 | 0.317 | 0.091 | 0.055342317 | 46 |
| glud1a4 | 2.24876991645675e-06 | 0.324970486 | 0.317 | 0.092 | 0.05659929 | 46 |
| zgc:16273020 | 2.26272207592476e-06 | 0.608839874 | 0.756 | 0.393 | 0.056950452 | 46 |
| xaf14 | 2.28117576401893e-06 | 0.305165142 | 0.244 | 0.062 | 0.057414913 | 46 |
| dbi10 | 2.36414865510242e-06 | 0.417538795 | 0.561 | 0.214 | 0.059503258 | 46 |
| ghrb6 | 2.61033350698735e-06 | 0.273458794 | 0.195 | 0.043 | 0.065699484 | 46 |
| zgc:56095 | 2.63469971482788e-06 | 0.312349619 | 0.341 | 0.102 | 0.066312757 | 46 |
| trappc12 | 2.94670271298857e-06 | 0.254926284 | 0.341 | 0.101 | 0.074165561 | 46 |
| lrp1ab5 | 2.94805285535479e-06 | 0.286081629 | 0.268 | 0.071 | 0.074199542 | 46 |
| ddx56 | 3.03933454411415e-06 | 0.492935695 | 0.78 | 0.358 | 0.076497011 | 46 |
| ptp4a2a | 3.05486936960808e-06 | 0.289972048 | 0.268 | 0.072 | 0.076888007 | 46 |

| | | | | | | |
|---|---|---|---|---|---|---|
| golga7 | 3.14118764401077e-06 | 0.351938473 | 0.463 | 0.162 | 0.079060552 | 46 |
| akap12b6 | 3.23557996156446e-06 | 0.269575929 | 0.244 | 0.061 | 0.081436312 | 46 |
| emc61 | 3.32395749527196e-06 | 0.289265648 | 0.415 | 0.133 | 0.083660686 | 46 |
| ddx612 | 3.449250247824e-06 | 0.403390772 | 0.561 | 0.216 | 0.086814179 | 46 |
| mgst3b7 | 3.47053082263973e-06 | 0.416983411 | 0.463 | 0.174 | 0.08734979 | 46 |
| dazap21 | 3.61437819662773e-06 | 0.29734206 | 0.317 | 0.093 | 0.090970285 | 46 |
| mef2d5 | 3.84586432319524e-06 | 0.317487062 | 0.268 | 0.074 | 0.096796559 | 46 |
| kpna4 | 3.87252720160254e-06 | 0.256197023 | 0.244 | 0.062 | 0.097467637 | 46 |
| cd6319 | 3.93864251576876e-06 | 0.485068094 | 0.78 | 0.341 | 0.099131693 | 46 |
| acadvl | 3.9452276343971e-06 | 0.33060866 | 0.244 | 0.064 | 0.099297434 | 46 |
| si:zfos-169g10.33 | 4.2506034133796e-06 | 0.313415892 | 0.317 | 0.096 | 0.106983437 | 46 |
| purba | 4.34036982351324e-06 | 0.286677464 | 0.244 | 0.064 | 0.109242768 | 46 |
| atp2b25 | 4.43734928852292e-06 | 0.252778299 | 0.268 | 0.073 | 0.111683644 | 46 |
| wu:fu71h07 | 4.44635017646239e-06 | 0.27188623 | 0.171 | 0.036 | 0.111910188 | 46 |
| elf2b | 4.78896599610772e-06 | 0.253264596 | 0.22 | 0.053 | 0.120533485 | 46 |
| eif4a22 | 5.20397672260523e-06 | 0.323512728 | 0.293 | 0.086 | 0.13097889 | 46 |
| mycb8 | 5.34719635653292e-06 | 0.468383365 | 0.537 | 0.213 | 0.134583585 | 46 |
| si:dkey-42i9.62 | 5.5777916182094e-06 | 0.320003424 | 0.268 | 0.076 | 0.140387437 | 46 |
| emc10 | 5.6609389390195e-06 | 0.282591054 | 0.293 | 0.084 | 0.142480172 | 46 |
| nocta2 | 5.81521828529954e-06 | 0.314130839 | 0.195 | 0.046 | 0.146363229 | 46 |
| mcl1b15 | 6.11663000974086e-06 | 0.426618728 | 0.61 | 0.243 | 0.153949461 | 46 |
| sub1a5 | 6.12895566054032e-06 | 0.408019377 | 0.585 | 0.234 | 0.154259685 | 46 |
| eif3c10 | 6.5786884201606e-06 | 0.35379846 | 0.463 | 0.17 | 0.165579009 | 46 |
| nme37 | 6.93267363260057e-06 | 0.308674785 | 0.268 | 0.077 | 0.174488463 | 46 |
| mknk2b14 | 7.75183399314967e-06 | 0.399637979 | 0.512 | 0.198 | 0.19510591 | 46 |
| gnb1b6 | 8.82506650104909e-06 | 0.411061784 | 0.61 | 0.255 | 0.222118099 | 46 |
| shoc21 | 8.98808845284769e-06 | 0.257944432 | 0.244 | 0.066 | 0.226221198 | 46 |
| gtf2f12 | 9.19207772648499e-06 | 0.257369389 | 0.244 | 0.065 | 0.231355404 | 46 |
| acin1a7 | 9.4075415083588e-06 | 0.416495095 | 0.537 | 0.218 | 0.236778412 | 46 |
| calm3b4 | 9.48315162919864e-06 | 0.425058578 | 0.61 | 0.257 | 0.238681443 | 46 |
| nenf | 1.01034243382432e-05 | 0.286968881 | 0.244 | 0.066 | 0.254293087 | 46 |
| fkbp59 | 1.07154681707536e-05 | 0.330473372 | 0.366 | 0.124 | 0.269697618 | 46 |
| msna16 | 1.12056562943226e-05 | 0.422726006 | 0.585 | 0.243 | 0.282035163 | 46 |
| uba11 | 1.14223306372638e-05 | 0.293529576 | 0.341 | 0.11 | 0.28748864 | 46 |
| nmi1 | 1.18387838982339e-05 | 0.344984107 | 0.268 | 0.081 | 0.297970352 | 46 |

| | | | | | | |
|---|---|---|---|---|---|---|
| tax1bp35 | 1.32096798745395e-05 | 0.302107945 | 0.293 | 0.09 | 0.332474433 | 46 |
| fosl215 | 1.4331813349687e-05 | 0.305848609 | 0.366 | 0.123 | 0.36071741 | 46 |
| eif5b1 | 1.46308902035057e-05 | 0.391142715 | 0.39 | 0.144 | 0.368244876 | 46 |
| eef1db15 | 1.5272811346052e-05 | 0.449782132 | 0.61 | 0.275 | 0.384401389 | 46 |
| spred2b7 | 1.57192119343717e-05 | 0.26065655 | 0.244 | 0.068 | 0.395636845 | 46 |
| tln18 | 1.60350430823597e-05 | 0.343105972 | 0.366 | 0.129 | 0.403585999 | 46 |
| reep3b1 | 1.63767087165619e-05 | 0.278980477 | 0.22 | 0.058 | 0.412185382 | 46 |
| akt1s11 | 1.63781256739324e-05 | 0.256170225 | 0.22 | 0.058 | 0.412221045 | 46 |
| gng59 | 1.64844412393146e-05 | 0.407009351 | 0.634 | 0.273 | 0.414896902 | 46 |
| eif4g1a | 1.75357928731084e-05 | 0.344562162 | 0.341 | 0.117 | 0.441358371 | 46 |
| trim25 | 1.80614873905347e-05 | 0.266493059 | 0.195 | 0.049 | 0.454589576 | 46 |
| zgc:100918 | 1.88330557454067e-05 | 0.283234596 | 0.22 | 0.059 | 0.47400918 | 46 |
| myl9b9 | 1.91300335341071e-05 | 0.323330206 | 0.268 | 0.081 | 0.481483814 | 46 |
| atp5meb4 | 2.039517970586e-05 | 0.420898699 | 0.707 | 0.323 | 0.513326278 | 46 |
| ubb5 | 2.09471040975727e-05 | 0.447417664 | 0.683 | 0.317 | 0.527217663 | 46 |
| rras | 2.13367506996434e-05 | 0.26088185 | 0.341 | 0.11 | 0.537024678 | 46 |
| LEPROTL1 | 2.21470588080254e-05 | 0.302459493 | 0.293 | 0.092 | 0.557419323 | 46 |
| cd151l1 | 2.49033660728666e-05 | 0.260008663 | 0.22 | 0.059 | 0.626792821 | 46 |
| capns1b11 | 2.68025593942467e-05 | 0.366101458 | 0.341 | 0.123 | 0.674593617 | 46 |
| psmf11 | 2.80997643077731e-05 | 0.25072743 | 0.171 | 0.04 | 0.707242968 | 46 |
| ogt.1 | 2.97544403698292e-05 | 0.280157796 | 0.244 | 0.071 | 0.74888951 | 46 |
| glula6 | 3.02099299309874e-05 | 0.319707765 | 0.39 | 0.137 | 0.760353726 | 46 |
| ccng27 | 3.27802680267606e-05 | 0.367936603 | 0.268 | 0.087 | 0.825046566 | 46 |
| ube4b | 3.31313236471896e-05 | 0.258951565 | 0.195 | 0.05 | 0.833882285 | 46 |
| ndufa111 | 3.32145820852757e-05 | 0.315121703 | 0.415 | 0.155 | 0.835977817 | 46 |
| sltm1 | 3.59747324141956e-05 | 0.26576417 | 0.22 | 0.061 | 0.90544804 | 46 |
| ost44 | 3.90548838030663e-05 | 0.356472115 | 0.61 | 0.258 | 0.98297237 | 46 |
| trib37 | 3.9381380265728e-05 | 0.308331529 | 0.317 | 0.107 | 0.99118996 | 46 |
| cox7c6 | 4.11293016176777e-05 | 0.459752161 | 0.829 | 0.482 | 1 | 46 |
| klf6a13 | 4.19081692355254e-05 | 0.452217912 | 0.415 | 0.169 | 1 | 46 |
| zgc:1119866 | 4.21222868723649e-05 | 0.373364454 | 0.439 | 0.178 | 1 | 46 |
| eppk16 | 4.21600451508682e-05 | 0.281793289 | 0.171 | 0.042 | 1 | 46 |
| vamp31 | 4.33114452208933e-05 | 0.371073304 | 0.463 | 0.193 | 1 | 46 |
| smc1al2 | 4.64386465745408e-05 | 0.278022224 | 0.293 | 0.095 | 1 | 46 |
| tbl1xr1a | 4.81696056554485e-05 | 0.265866733 | 0.171 | 0.043 | 1 | 46 |

| | | | | | | |
|---|---|---|---|---|---|---|
| usp7 | 5.09058876996265e-05 | 0.257860071 | 0.22 | 0.062 | 1 | 46 |
| cct29 | 5.41166542678674e-05 | 0.362384989 | 0.537 | 0.229 | 1 | 46 |
| socs3b12 | 5.45036006862931e-05 | 0.339632761 | 0.415 | 0.157 | 1 | 46 |
| sec11a3 | 5.51783138588856e-05 | 0.280768835 | 0.463 | 0.179 | 1 | 46 |
| fam120c6 | 5.75121711375356e-05 | 0.310926019 | 0.293 | 0.1 | 1 | 46 |
| ndufc23 | 6.00857973279769e-05 | 0.312969922 | 0.366 | 0.137 | 1 | 46 |
| gabarapa3 | 6.18298133293172e-05 | 0.323829123 | 0.415 | 0.162 | 1 | 46 |
| sfpq1 | 6.48855622106755e-05 | 0.275335235 | 0.317 | 0.109 | 1 | 46 |
| relb1 | 6.8008162376795e-05 | 0.26755583 | 0.171 | 0.044 | 1 | 46 |
| cltca1 | 7.00310551767605e-05 | 0.258707287 | 0.268 | 0.086 | 1 | 46 |
| calr3b2 | 7.5943995690788e-05 | 0.344859364 | 0.244 | 0.078 | 1 | 46 |
| akirin12 | 7.86948427759694e-05 | 0.255274765 | 0.39 | 0.141 | 1 | 46 |
| srsf5a6 | 8.01916968329657e-05 | 0.379703451 | 0.634 | 0.289 | 1 | 46 |
| fkbp21 | 8.77544859315308e-05 | 0.298595154 | 0.366 | 0.136 | 1 | 46 |
| hsbp1b | 8.98464649674178e-05 | 0.316825815 | 0.463 | 0.191 | 1 | 46 |
| ywhae12 | 9.81235749714811e-05 | 0.392932976 | 0.61 | 0.284 | 1 | 46 |
| chtopa7 | 0.000113034 | 0.334617063 | 0.537 | 0.223 | 1 | 46 |
| sri4 | 0.000113535 | 0.301890679 | 0.39 | 0.152 | 1 | 46 |
| slc20a1b6 | 0.00011786 | 0.316494283 | 0.293 | 0.103 | 1 | 46 |
| sh3bgrl37 | 0.00013135 | 0.264833097 | 0.39 | 0.149 | 1 | 46 |
| emd6 | 0.000135715 | 0.255942408 | 0.195 | 0.056 | 1 | 46 |
| selenow19 | 0.000139631 | 0.424327566 | 0.756 | 0.386 | 1 | 46 |
| atp5pf8 | 0.000149127 | 0.397818419 | 0.61 | 0.3 | 1 | 46 |
| map2k64 | 0.000150451 | 0.255627243 | 0.268 | 0.09 | 1 | 46 |
| ptp4a2b3 | 0.000155957 | 0.281362025 | 0.39 | 0.151 | 1 | 46 |
| dnmt3ab5 | 0.000156029 | 0.27286694 | 0.244 | 0.08 | 1 | 46 |
| zgc:1748636 | 0.000169276 | 0.257574602 | 0.293 | 0.103 | 1 | 46 |
| gpx4b5 | 0.000171012 | 0.284068734 | 0.488 | 0.205 | 1 | 46 |
| stk35l1 | 0.000173903 | 0.272720784 | 0.146 | 0.037 | 1 | 46 |
| ap2s1 | 0.000197794 | 0.253733038 | 0.22 | 0.069 | 1 | 46 |
| zgc:778493 | 0.000203595 | 0.276475117 | 0.439 | 0.18 | 1 | 46 |
| atrx9 | 0.000210397 | 0.273040131 | 0.293 | 0.106 | 1 | 46 |
| ptp4a16 | 0.00021148 | 0.280032182 | 0.317 | 0.119 | 1 | 46 |
| tmsb4x26 | 0.000217524 | 0.477445082 | 1 | 0.687 | 1 | 46 |
| hint12 | 0.000220092 | 0.285168086 | 0.512 | 0.219 | 1 | 46 |

| | | | | | | |
|---|---|---|---|---|---|---|
| nfia9 | 0.000222691 | 0.265222449 | 0.244 | 0.081 | 1 | 46 |
| s100a10b15 | 0.000261437 | 0.278517046 | 0.61 | 0.257 | 1 | 46 |
| prdx41 | 0.000264713 | 0.280295335 | 0.39 | 0.158 | 1 | 46 |
| hmgn714 | 0.000276716 | 0.395225688 | 0.561 | 0.281 | 1 | 46 |
| pdia61 | 0.000278659 | 0.29814567 | 0.317 | 0.12 | 1 | 46 |
| ube2d1b3 | 0.000302732 | 0.254359092 | 0.341 | 0.132 | 1 | 46 |
| tomm56 | 0.000336006 | 0.290992108 | 0.488 | 0.215 | 1 | 46 |
| pdia311 | 0.000338775 | 0.339980731 | 0.61 | 0.285 | 1 | 46 |
| ndufb101 | 0.000342773 | 0.274579853 | 0.366 | 0.149 | 1 | 46 |
| tagln214 | 0.000356939 | 0.320224647 | 0.439 | 0.19 | 1 | 46 |
| maff9 | 0.000361457 | 0.254883483 | 0.244 | 0.084 | 1 | 46 |
| rbx12 | 0.000374914 | 0.336147899 | 0.561 | 0.267 | 1 | 46 |
| aimp22 | 0.000396735 | 0.267047381 | 0.317 | 0.124 | 1 | 46 |
| nars1 | 0.00039867 | 0.25362874 | 0.22 | 0.073 | 1 | 46 |
| ndufb34 | 0.000416383 | 0.297990775 | 0.488 | 0.219 | 1 | 46 |
| zgc:92818 | 0.000437063 | 0.25888227 | 0.268 | 0.095 | 1 | 46 |
| ppdpfb9 | 0.000438355 | 0.415027652 | 0.829 | 0.52 | 1 | 46 |
| ddit317 | 0.000458389 | 0.309521721 | 0.415 | 0.178 | 1 | 46 |
| cdc42l10 | 0.000460879 | 0.286576723 | 0.488 | 0.217 | 1 | 46 |
| skp13 | 0.000484932 | 0.35808173 | 0.512 | 0.253 | 1 | 46 |
| hnrnpa0b8 | 0.000499242 | 0.332332019 | 0.659 | 0.313 | 1 | 46 |
| psmb22 | 0.000503777 | 0.270581026 | 0.39 | 0.164 | 1 | 46 |
| sf3b61 | 0.000505165 | 0.291618565 | 0.537 | 0.247 | 1 | 46 |
| prdx25 | 0.00050523 | 0.395046762 | 0.756 | 0.402 | 1 | 46 |
| csnk2b2 | 0.000506807 | 0.272020938 | 0.463 | 0.204 | 1 | 46 |
| pabpn12 | 0.00054302 | 0.312755614 | 0.585 | 0.276 | 1 | 46 |
| si:dkey-188i13.1114 | 0.000555225 | 0.27750772 | 0.707 | 0.318 | 1 | 46 |
| atp6v0ca4 | 0.000609792 | 0.328700917 | 0.439 | 0.2 | 1 | 46 |
| myl12.117 | 0.000615726 | 0.384923874 | 0.756 | 0.393 | 1 | 46 |
| mif3 | 0.000659319 | 0.323971988 | 0.415 | 0.189 | 1 | 46 |
| XLOC-0247505 | 0.000663217 | 0.268417654 | 0.22 | 0.077 | 1 | 46 |
| gnb1a10 | 0.000675113 | 0.275249108 | 0.439 | 0.194 | 1 | 46 |
| zfp36l1a13 | 0.000695133 | 0.317446085 | 0.537 | 0.249 | 1 | 46 |
| ube2nb | 0.000885223 | 0.258893174 | 0.341 | 0.142 | 1 | 46 |
| atp5pb5 | 0.00092854 | 0.31337678 | 0.61 | 0.301 | 1 | 46 |

| | | | | | | |
|---|---|---|---|---|---|---|
| tmed72 | 0.001015227 | 0.304346186 | 0.439 | 0.208 | 1 | 46 |
| tsc22d314 | 0.001044272 | 0.268619348 | 0.341 | 0.145 | 1 | 46 |
| cox5ab4 | 0.001054141 | 0.282494975 | 0.439 | 0.203 | 1 | 46 |
| rbbp61 | 0.001118367 | 0.275288985 | 0.268 | 0.106 | 1 | 46 |
| f3b7 | 0.001129792 | 0.255951234 | 0.268 | 0.103 | 1 | 46 |
| atp5f1d5 | 0.001144929 | 0.327917021 | 0.512 | 0.26 | 1 | 46 |
| abcf11 | 0.001368201 | 0.265449601 | 0.244 | 0.094 | 1 | 46 |
| si:dkey-11f4.149 | 0.001386405 | 0.292583219 | 0.268 | 0.11 | 1 | 46 |
| hmgb2b4 | 0.00157134 | 0.291180187 | 0.659 | 0.321 | 1 | 46 |
| capzb13 | 0.001679295 | 0.254115609 | 0.537 | 0.255 | 1 | 46 |
| uqcrh6 | 0.001769463 | 0.271863213 | 0.512 | 0.251 | 1 | 46 |
| eif3s108 | 0.001796549 | 0.252551914 | 0.366 | 0.166 | 1 | 46 |
| aldh9a1a.110 | 0.001967097 | 0.255606907 | 0.366 | 0.166 | 1 | 46 |
| ndufb61 | 0.001981708 | 0.255594586 | 0.268 | 0.112 | 1 | 46 |
| zgc:1935414 | 0.002009431 | 0.343195599 | 0.659 | 0.369 | 1 | 46 |
| taf13 | 0.002016421 | 0.2518876 | 0.488 | 0.233 | 1 | 46 |
| COX7A2 (1 of many)7 | 0.002039057 | 0.260593411 | 0.537 | 0.262 | 1 | 46 |
| GCA8 | 0.002054267 | 0.263631181 | 0.39 | 0.183 | 1 | 46 |
| atp5f1e5 | 0.002237182 | 0.359835617 | 0.78 | 0.484 | 1 | 46 |
| cdc4210 | 0.002267142 | 0.320581358 | 0.415 | 0.214 | 1 | 46 |
| histh1l11 | 0.002298896 | 0.264302915 | 0.268 | 0.111 | 1 | 46 |
| zgc:1629446 | 0.002557671 | 0.270343944 | 0.512 | 0.254 | 1 | 46 |
| ywhabl9 | 0.003128037 | 0.283687119 | 0.537 | 0.28 | 1 | 46 |
| oaz1a2 | 0.003220478 | 0.34815389 | 0.732 | 0.482 | 1 | 46 |
| serf218 | 0.00328696 | 0.305176208 | 0.659 | 0.344 | 1 | 46 |
| prdx514 | 0.003354003 | 0.259403647 | 0.341 | 0.161 | 1 | 46 |
| psap6 | 0.003494522 | 0.255837289 | 0.585 | 0.292 | 1 | 46 |
| tnrc6c114 | 0.004099467 | 0.268205485 | 0.293 | 0.134 | 1 | 46 |
| dad13 | 0.005047894 | 0.292938398 | 0.634 | 0.353 | 1 | 46 |
| rpl1313 | 0.005316769 | 0.258931085 | 1 | 0.805 | 1 | 46 |
| rps27.221 | 0.005417053 | 0.332274811 | 0.78 | 0.519 | 1 | 46 |
| sumo3a3 | 0.006248346 | 0.282934343 | 0.683 | 0.371 | 1 | 46 |
| serbp1a16 | 0.006565249 | 0.354714903 | 0.854 | 0.483 | 1 | 46 |
| sod14 | 0.009537138 | 0.259956933 | 0.585 | 0.329 | 1 | 46 |
| chac110 | 0.009541976 | 0.267482261 | 0.707 | 0.4 | 1 | 46 |

| | | | | | | |
|---|---|---|---|---|---|---|
| gp1bb | 0 | 1.839351536 | 0.861 | 0.002 | 0 | 47 |
| thbs1b2 | 0 | 1.572020256 | 0.694 | 0.01 | 0 | 47 |
| f5 | 0 | 1.076548771 | 0.417 | 0.002 | 0 | 47 |
| rac3a | 0 | 1.050725802 | 0.444 | 0.003 | 0 | 47 |
| si:ch211-33e4.21 | 0 | 1.020642843 | 0.417 | 0.002 | 0 | 47 |
| itga2b | 0 | 0.760455857 | 0.306 | 0.001 | 0 | 47 |
| hbegfb | 1.67114470098831e-305 | 1.573468516 | 0.639 | 0.008 | 4.20610409791747e-301 | 47 |
| apln1 | 1.19908883043403e-280 | 1.285956592 | 0.472 | 0.005 | 3.01798667731942e-276 | 47 |
| sele1 | 3.2082949207812e-266 | 0.998194573 | 0.389 | 0.003 | 8.0749574861142e-262 | 47 |
| tubb1 | 5.11133009615418e-236 | 0.662346402 | 0.25 | 0.001 | 1.28647067190105e-231 | 47 |
| rpgra | 2.30141403306334e-228 | 0.726789933 | 0.222 | 0.001 | 5.79242897981711e-224 | 47 |
| mpl | 5.40927607052075e-225 | 0.564371157 | 0.194 | 0.001 | 1.36146069418937e-220 | 47 |
| itgb3b1 | 1.21506347795164e-209 | 1.274528465 | 0.556 | 0.01 | 3.05819326765648e-205 | 47 |
| rprma | 2.31793453153255e-201 | 0.449440379 | 0.139 | 0 | 5.83400942241426e-197 | 47 |
| rprml | 2.95546493343291e-201 | 0.309250481 | 0.139 | 0 | 7.43860969095729e-197 | 47 |
| vegfc | 6.72180262253221e-173 | 0.682319891 | 0.25 | 0.002 | 1.69181050206513e-168 | 47 |
| tnr | 2.02128420059113e-167 | 0.690164848 | 0.25 | 0.002 | 5.0873702044678e-163 | 47 |
| tal1 | 8.79192324643906e-150 | 0.456062046 | 0.194 | 0.001 | 2.21283916189625e-145 | 47 |
| ppp2r2ba | 1.17058054165698e-147 | 0.360427056 | 0.111 | 0 | 2.94623416529645e-143 | 47 |
| zgc:171686 | 4.57198420564668e-133 | 0.783181792 | 0.306 | 0.004 | 1.15072270471921e-128 | 47 |
| si:ch211-195b11.32 | 3.48644831623242e-132 | 1.859080334 | 0.806 | 0.037 | 8.77504176712538e-128 | 47 |
| parvb1 | 2.67802704636339e-131 | 0.875998484 | 0.389 | 0.008 | 6.74032627299202e-127 | 47 |
| bmp161 | 1.36680893941142e-108 | 0.855758787 | 0.306 | 0.006 | 3.44012141960461e-104 | 47 |
| gata1a | 4.54407333012994e-105 | 0.365283207 | 0.167 | 0.002 | 1.1436978164604e-100 | 47 |
| blf1 | 3.3888063542795e-101 | 0.674919777 | 0.25 | 0.004 | 8.52928671308608e-97 | 47 |
| zgc:162989 | 3.44616946892103e-94 | 0.364571032 | 0.111 | 0.001 | 8.67366393632735e-90 | 47 |
| fgl2a8 | 2.9323848356281e-88 | 1.728204222 | 0.861 | 0.065 | 7.38051939279238e-84 | 47 |
| admb | 3.67451515922142e-85 | 0.771254965 | 0.222 | 0.004 | 9.24838720424438e-81 | 47 |
| zmat4b | 1.61148616840194e-80 | 0.367263785 | 0.139 | 0.001 | 4.05594953725085e-76 | 47 |
| si:ch211-153b23.5 | 1.17589855878155e-74 | 0.434621471 | 0.167 | 0.002 | 2.95961908259729e-70 | 47 |
| myocd1 | 3.39652960611267e-71 | 0.689627691 | 0.278 | 0.007 | 8.54872536562497e-67 | 47 |
| lpp | 5.62635343509905e-71 | 1.316899782 | 0.556 | 0.032 | 1.41609689608008e-66 | 47 |
| reps21 | 1.70817032991993e-69 | 0.462935817 | 0.194 | 0.004 | 4.29929390337547e-65 | 47 |
| fhl1a | 2.78121580984004e-69 | 0.690243195 | 0.222 | 0.005 | 7.00004207178641e-65 | 47 |
| arhgef9b1 | 6.01095990217665e-68 | 0.43127485 | 0.167 | 0.003 | 1.51289849777884e-63 | 47 |

| | | | | | | |
|---|---|---|---|---|---|---|
| adra2a | 7.23399308820429e-65 | 0.332645646 | 0.111 | 0.001 | 1.82072372037014e-60 | 47 |
| p2ry13 | 2.22588232671528e-64 | 0.322333187 | 0.139 | 0.002 | 5.60232322810968e-60 | 47 |
| hacd31 | 7.94499875463133e-64 | 1.18408782 | 0.556 | 0.035 | 1.99967673655316e-59 | 47 |
| tpm4a10 | 3.94242241116815e-63 | 1.600539909 | 0.833 | 0.088 | 9.92268296666912e-59 | 47 |
| spns22 | 8.95071899554833e-63 | 0.675197399 | 0.25 | 0.007 | 2.25280646398956e-58 | 47 |
| cald1a11 | 2.23671930819391e-62 | 1.709225156 | 0.861 | 0.098 | 5.62959882679325e-58 | 47 |
| lmo23 | 4.65195229662599e-55 | 0.911850915 | 0.389 | 0.02 | 1.17084987353779e-50 | 47 |
| mir2235 | 9.76673313009269e-54 | 1.163237212 | 0.472 | 0.03 | 2.45818906151303e-49 | 47 |
| XLOC-0203016 | 1.11714688586541e-50 | 1.566803632 | 0.75 | 0.085 | 2.81174699703464e-46 | 47 |
| si:ch211-214p13.91 | 4.78215769428553e-50 | 0.869187039 | 0.306 | 0.013 | 1.20362127007472e-45 | 47 |
| fhl2a1 | 1.6136077071219e-47 | 0.95146057 | 0.333 | 0.017 | 4.0612892380551e-43 | 47 |
| wnt21 | 1.75240792265351e-47 | 0.474513039 | 0.139 | 0.003 | 4.41063550052661e-43 | 47 |
| gp9 | 3.06249039263927e-47 | 0.250519003 | 0.111 | 0.002 | 7.70798206923377e-43 | 47 |
| myh11a8 | 2.49214342757724e-46 | 1.26174398 | 0.639 | 0.066 | 6.27247579286916e-42 | 47 |
| LOC1081907175 | 3.82784620758952e-46 | 1.417669106 | 0.778 | 0.096 | 9.63430611988205e-42 | 47 |
| susd1 | 1.23365411286841e-45 | 0.589768407 | 0.222 | 0.008 | 3.10498403667851e-41 | 47 |
| hpca | 3.9873711549804e-45 | 0.788581105 | 0.333 | 0.017 | 1.00358144599702e-40 | 47 |
| plek4 | 5.57225032140907e-45 | 0.935870468 | 0.389 | 0.024 | 1.40247968339545e-40 | 47 |
| arhgap20b2 | 8.50526662894121e-45 | 0.876400419 | 0.361 | 0.021 | 2.14069055783821e-40 | 47 |
| plac8l1 | 1.83756482267197e-42 | 0.467696191 | 0.139 | 0.003 | 4.62496690218309e-38 | 47 |
| sept128 | 4.34969132687437e-41 | 1.12263246 | 0.583 | 0.061 | 1.09477381006101e-36 | 47 |
| timd4 | 5.16111005523182e-40 | 0.328646381 | 0.111 | 0.002 | 1.2989997898013e-35 | 47 |
| pdgfd1 | 1.17698650050516e-39 | 0.543528877 | 0.222 | 0.009 | 2.96235732312144e-35 | 47 |
| ctgfa4 | 2.90704216346616e-39 | 1.073546402 | 0.389 | 0.028 | 7.31673442122798e-35 | 47 |
| XLOC-02837010 | 5.68966120482609e-39 | 1.561398526 | 0.833 | 0.141 | 1.43203082864268e-34 | 47 |
| fli1a1 | 5.94562838907492e-38 | 0.503363042 | 0.194 | 0.007 | 1.49645520924627e-33 | 47 |
| dera1 | 8.20417487418229e-38 | 0.782028632 | 0.333 | 0.021 | 2.06490877408294e-33 | 47 |
| celf41 | 3.81685194402886e-37 | 0.436111744 | 0.167 | 0.005 | 9.60663465792623e-33 | 47 |
| cldn5b1 | 4.82526863304839e-37 | 0.539568036 | 0.222 | 0.009 | 1.21447186225195e-32 | 47 |
| bmp63 | 3.50054655618729e-36 | 0.787672881 | 0.25 | 0.012 | 8.81052562726778e-32 | 47 |
| ptger2b | 5.25096677502298e-36 | 0.431798921 | 0.167 | 0.005 | 1.32161582760554e-31 | 47 |
| smtna | 7.38746097712992e-36 | 0.305982486 | 0.111 | 0.002 | 1.85935005333383e-31 | 47 |
| tln19 | 1.8756145907058e-35 | 1.357415606 | 0.75 | 0.128 | 4.72073436334743e-31 | 47 |
| tpte1 | 4.54323966953398e-35 | 0.747016448 | 0.361 | 0.026 | 1.14348799242501e-30 | 47 |
| CR753876.17 | 1.23481257314709e-34 | 1.269437339 | 0.75 | 0.115 | 3.10789976535392e-30 | 47 |

| | | | | | | |
|---|---|---|---|---|---|---|
| myl9b10 | 2.15093626507315e-34 | 1.130632777 | 0.611 | 0.08 | 5.41369148556262e-30 | 47 |
| b3gnt3.46 | 5.04039316035754e-34 | 0.973063306 | 0.417 | 0.037 | 1.26861655453039e-29 | 47 |
| derl25 | 8.25205341929643e-34 | 1.249766394 | 0.694 | 0.105 | 2.07695932510272e-29 | 47 |
| cavin2b11 | 1.38801046021459e-33 | 1.045043527 | 0.556 | 0.066 | 3.4934835273141e-29 | 47 |
| fn1b5 | 3.97947924320092e-32 | 1.189761872 | 0.639 | 0.088 | 1.00159513072124e-27 | 47 |
| UNC13B | 1.73992843021875e-30 | 0.285516344 | 0.111 | 0.003 | 4.37922586601758e-26 | 47 |
| tmem35 | 3.30170535867719e-30 | 0.438027008 | 0.167 | 0.007 | 8.31006221725463e-26 | 47 |
| lrrc8da2 | 4.42719326950178e-30 | 0.719061291 | 0.306 | 0.022 | 1.1142802740009e-25 | 47 |
| XLOC-0268366 | 4.95309257522369e-30 | 1.152501027 | 0.583 | 0.079 | 1.24664387025805e-25 | 47 |
| si:dkey-217l24.11 | 1.00636586919713e-29 | 0.493033684 | 0.194 | 0.009 | 2.53292225618225e-25 | 47 |
| si:ch211-222l21.117 | 1.55365340928891e-29 | 1.380240836 | 0.944 | 0.276 | 3.91039026583925e-25 | 47 |
| mocs21 | 4.28966476312924e-29 | 0.4908295 | 0.167 | 0.007 | 1.079665724232e-24 | 47 |
| boka | 1.70176883234333e-28 | 0.577860286 | 0.222 | 0.012 | 4.28318197412492e-24 | 47 |
| nuak1a | 2.56705675844391e-28 | 0.435273056 | 0.139 | 0.005 | 6.46102515532747e-24 | 47 |
| mylka1 | 6.70031071970229e-28 | 0.574673911 | 0.222 | 0.013 | 1.68640120504187e-23 | 47 |
| rap1b10 | 8.65727007258523e-28 | 1.403142137 | 0.806 | 0.203 | 2.17894830456898e-23 | 47 |
| nfe21 | 7.38613951500375e-27 | 0.576527353 | 0.222 | 0.013 | 1.85901745453129e-22 | 47 |
| XLOC-019062 | 2.06808666172953e-26 | 0.439591783 | 0.139 | 0.005 | 5.20516731890705e-22 | 47 |
| blvrb | 1.55492745720069e-25 | 0.884453393 | 0.389 | 0.042 | 3.91359691702842e-21 | 47 |
| rhag1 | 4.02081163671006e-25 | 0.480849205 | 0.194 | 0.011 | 1.01199808084355e-20 | 47 |
| f13a1b5 | 5.35741555695091e-25 | 0.704860995 | 0.333 | 0.031 | 1.34840792152897e-20 | 47 |
| actn15 | 1.23972761588324e-24 | 0.875959529 | 0.389 | 0.044 | 3.12027043641652e-20 | 47 |
| aatka2 | 1.07135445492825e-23 | 0.51403441 | 0.194 | 0.012 | 2.69649202760892e-19 | 47 |
| AL954142.14 | 1.55032450298957e-23 | 0.672329102 | 0.278 | 0.024 | 3.90201174157445e-19 | 47 |
| pim28 | 2.28022980719894e-23 | 1.25529488 | 0.528 | 0.09 | 5.739110401739e-19 | 47 |
| zgc:171506 | 3.83003835136089e-23 | 0.433600961 | 0.167 | 0.009 | 9.63982352654023e-19 | 47 |
| itprid21 | 5.30865244749214e-23 | 0.52410853 | 0.222 | 0.015 | 1.3361347345093e-18 | 47 |
| slc5a8l | 6.78712208950871e-23 | 0.36267899 | 0.139 | 0.006 | 1.70825075870845e-18 | 47 |
| si:ch211-226m16.28 | 9.03310152840164e-23 | 0.902263347 | 0.472 | 0.069 | 2.27354132368341e-18 | 47 |
| ano5b | 2.61907368058204e-22 | 0.64030937 | 0.25 | 0.02 | 6.59194654665694e-18 | 47 |
| zgc:921402 | 1.62677261867114e-21 | 0.635748133 | 0.278 | 0.026 | 4.09442400393339e-17 | 47 |
| rap1gap2a | 2.37484891918024e-21 | 0.450480836 | 0.167 | 0.01 | 5.97725724468474e-17 | 47 |
| iqgap2 | 3.45794578075944e-21 | 0.398679244 | 0.139 | 0.007 | 8.70330373559344e-17 | 47 |
| si:ch211-214p16.26 | 5.06077422014543e-21 | 0.952144431 | 0.472 | 0.073 | 1.2737462634684e-16 | 47 |
| itpr3 | 6.71547595989799e-21 | 0.448809813 | 0.167 | 0.01 | 1.69021814434673e-16 | 47 |

| | | | | | | |
|---|---|---|---|---|---|---|
| srgn14 | 1.19831437302655e-20 | 1.142274246 | 0.833 | 0.238 | 3.01603744547052e-16 | 47 |
| ptmaa23 | 1.3289677543536e-20 | 1.137673376 | 0.917 | 0.396 | 3.34487894093257e-16 | 47 |
| mef2aa5 | 2.3451641661695e-20 | 0.949068503 | 0.444 | 0.069 | 5.90254368983201e-16 | 47 |
| pmp22b9 | 3.37302805168295e-20 | 0.852216455 | 0.361 | 0.046 | 8.48957430328083e-16 | 47 |
| tmsb4x27 | 4.14648802528648e-20 | 0.858471878 | 1 | 0.687 | 1.04362957108435e-15 | 47 |
| limd25 | 4.71772025939299e-20 | 0.945851815 | 0.444 | 0.07 | 1.18740301208662e-15 | 47 |
| p2rx73 | 1.14361296617528e-19 | 0.501978509 | 0.194 | 0.014 | 2.87835947456656e-15 | 47 |
| hif1aa | 1.22813760219895e-19 | 0.319549278 | 0.111 | 0.005 | 3.09109953097453e-15 | 47 |
| btg120 | 1.2990022054255e-19 | 1.107916833 | 0.861 | 0.333 | 3.26945865083545e-15 | 47 |
| si:ch211-284f22.36 | 1.49630093241003e-19 | 0.677152749 | 0.306 | 0.034 | 3.7660398167828e-15 | 47 |
| svila1 | 3.53880591931401e-19 | 0.655910924 | 0.278 | 0.029 | 8.90682061832143e-15 | 47 |
| rpz511 | 4.53122610106157e-19 | 0.996419673 | 0.528 | 0.103 | 1.14046429737619e-14 | 47 |
| fermt3b3 | 5.26451861852014e-19 | 0.59193978 | 0.222 | 0.019 | 1.32502669109533e-14 | 47 |
| ophn1 | 5.61450019295667e-19 | 0.686831447 | 0.306 | 0.035 | 1.41311355356526e-14 | 47 |
| flna9 | 8.95175304622663e-19 | 0.973787262 | 0.556 | 0.115 | 2.25306672420478e-14 | 47 |
| mgst1.27 | 1.17596988726306e-18 | 0.961227684 | 0.528 | 0.103 | 2.95979860925239e-14 | 47 |
| meis1b8 | 1.79353916481422e-18 | 0.786270907 | 0.389 | 0.057 | 4.5141587239209e-14 | 47 |
| gcnt4a3 | 2.06615606525697e-18 | 0.555853582 | 0.194 | 0.015 | 5.20030820064527e-14 | 47 |
| btk4 | 4.58252553588151e-18 | 0.514528738 | 0.25 | 0.025 | 1.15337585212602e-13 | 47 |
| asap1b1 | 4.35686446435808e-17 | 0.710217952 | 0.278 | 0.032 | 1.09657921703428e-12 | 47 |
| ilk | 5.22006458288734e-17 | 0.868316754 | 0.444 | 0.081 | 1.31383805486691e-12 | 47 |
| tagln215 | 5.60654102862348e-17 | 1.017205447 | 0.667 | 0.189 | 1.41111031149424e-12 | 47 |
| XLOC-02542311 | 5.72117208518186e-17 | 1.087450088 | 0.667 | 0.17 | 1.43996180211942e-12 | 47 |
| rbpms | 1.59888858409386e-16 | 0.552400829 | 0.222 | 0.021 | 4.02424267730584e-12 | 47 |
| si:dkey-238c7.163 | 3.16049314861071e-16 | 0.752520245 | 0.389 | 0.063 | 7.95464520573828e-12 | 47 |
| pfn117 | 3.18092163980299e-16 | 0.970959019 | 1 | 0.526 | 8.00606167522014e-12 | 47 |
| fosab16 | 8.23354566175996e-16 | 1.160512438 | 0.833 | 0.35 | 2.07230110760836e-11 | 47 |
| phldb1b | 8.65650681056476e-16 | 0.26303779 | 0.111 | 0.006 | 2.17875619915104e-11 | 47 |
| si:ch73-234b20.510 | 1.26646615353551e-15 | 0.903904634 | 0.472 | 0.101 | 3.18756866183353e-11 | 47 |
| zgc:1538679 | 2.4345857396615e-15 | 0.909877413 | 0.889 | 0.437 | 6.12760884815402e-11 | 47 |
| ppiab9 | 3.39652059457034e-15 | 0.66654216 | 0.972 | 0.657 | 8.54870268447408e-11 | 47 |
| dennd2c | 4.35669751264815e-15 | 0.41042424 | 0.167 | 0.013 | 1.09653719695841e-10 | 47 |
| creg11 | 7.61924104008449e-15 | 0.794253966 | 0.389 | 0.069 | 1.91768677737886e-10 | 47 |
| actb216 | 1.29055875295953e-14 | 0.555403383 | 1 | 0.868 | 3.24820732532383e-10 | 47 |
| bcl2l107 | 1.61592495066717e-14 | 0.93942085 | 0.694 | 0.215 | 4.0671215083342e-10 | 47 |

| | | | | | | |
|---|---|---|---|---|---|---|
| tspan5a1 | 2.10381206134284e-14 | 0.461375234 | 0.167 | 0.014 | 5.29508457719379e-10 | 47 |
| CR735102.1 | 3.78933222500047e-14 | 0.305305503 | 0.111 | 0.007 | 9.53737027710370e-10 | 47 |
| ccnd2a4 | 8.23954136470058e-14 | 0.866878786 | 0.417 | 0.087 | 2.07381016608149e-09 | 47 |
| gapdhs17 | 1.1837211382558e-13 | 0.92923514 | 0.722 | 0.29 | 2.97930773287603e-09 | 47 |
| bambib1 | 1.25267244920893e-13 | 0.362787624 | 0.167 | 0.015 | 3.15285128741396e-09 | 47 |
| XLOC-0320023 | 2.77322951570217e-13 | 0.641941195 | 0.306 | 0.047 | 6.97994136807079e-09 | 47 |
| si:dkey-51e6.17 | 6.78199751518837e-13 | 0.859761698 | 0.472 | 0.117 | 1.70696095459776e-08 | 47 |
| lzts2a3 | 1.02300800897083e-12 | 0.494786914 | 0.194 | 0.022 | 2.57480885777869e-08 | 47 |
| BX005183.38 | 1.15256400569311e-12 | 0.796031734 | 0.389 | 0.082 | 2.90088834592899e-08 | 47 |
| adgrg1 | 2.3735464782665e-12 | 0.327347298 | 0.139 | 0.012 | 5.97397913114897e-08 | 47 |
| CR847543.1 | 5.89690885465823e-12 | 0.372676781 | 0.139 | 0.012 | 1.48419298962893e-07 | 47 |
| ptpn11b1 | 7.67849432573313e-12 | 0.498427353 | 0.194 | 0.023 | 1.93260023684377e-07 | 47 |
| zgc:664332 | 9.47367171513043e-12 | 0.503369439 | 0.222 | 0.03 | 2.38442843398118e-07 | 47 |
| CR383676.125 | 1.12372595726614e-11 | 0.744077006 | 0.944 | 0.727 | 2.82830586184314e-07 | 47 |
| lpar17 | 1.16731888747147e-11 | 0.733868277 | 0.333 | 0.065 | 2.93802490787695e-07 | 47 |
| hdr | 1.82485435838731e-11 | 0.652097092 | 0.306 | 0.053 | 4.59297593462501e-07 | 47 |
| ap1s11 | 2.10200196636815e-11 | 0.757847561 | 0.417 | 0.099 | 5.29052874915199e-07 | 47 |
| gng104 | 2.22828057437578e-11 | 0.635295477 | 0.306 | 0.055 | 5.60835937764641e-07 | 47 |
| ing5a | 2.28610974183417e-11 | 0.451520877 | 0.194 | 0.024 | 5.75390960922243e-07 | 47 |
| zgc:153409 | 2.35077854332677e-11 | 0.336621574 | 0.139 | 0.013 | 5.91667451569915e-07 | 47 |
| myl12.118 | 3.3404250785851e-11 | 0.811552327 | 0.778 | 0.393 | 8.40751588029085e-07 | 47 |
| si:ch211-102c2.410 | 4.59888117257504e-11 | 0.849745135 | 0.5 | 0.144 | 1.15749240232541e-06 | 47 |
| arpc1b16 | 4.9141032811859e-11 | 0.890966805 | 0.75 | 0.31 | 1.23683065484168e-06 | 47 |
| vash2 | 8.66819587427326e-11 | 0.316546075 | 0.111 | 0.009 | 2.18169821959584e-06 | 47 |
| si:ch211-218g4.2 | 9.67094849642365e-11 | 0.283150408 | 0.111 | 0.009 | 2.43408102706487e-06 | 47 |
| emilin1a4 | 2.99858379782074e-10 | 0.435650995 | 0.167 | 0.02 | 7.54713556073502e-06 | 47 |
| cyr619 | 3.58342639100038e-10 | 0.656358763 | 0.333 | 0.069 | 9.01912588350885e-06 | 47 |
| bmp2k | 3.76915593091939e-10 | 0.3528956 | 0.167 | 0.02 | 9.48658856253102e-06 | 47 |
| nup88 | 4.61317416241592e-10 | 0.32416322 | 0.111 | 0.009 | 1.16108980493846e-05 | 47 |
| pbx1a6 | 4.81436530209342e-10 | 0.602601616 | 0.25 | 0.043 | 1.21172760288389e-05 | 47 |
| sb:cb813 | 5.13791025568512e-10 | 0.473093705 | 0.194 | 0.027 | 1.29316063225339e-05 | 47 |
| gna11b | 6.56356476672686e-10 | 0.677704753 | 0.306 | 0.063 | 1.65198361613748e-05 | 47 |
| dusp510 | 6.74274241253572e-10 | 0.665930255 | 0.333 | 0.074 | 1.69708083781111e-05 | 47 |
| fkbp1b2 | 6.80231010633825e-10 | 0.302821289 | 0.139 | 0.015 | 1.71207343066427e-05 | 47 |
| mcl1a15 | 9.04937125536339e-10 | 0.590760376 | 0.833 | 0.625 | 2.27763625126241e-05 | 47 |

| | | | | | | |
|---|---|---|---|---|---|---|
| macf1a3 | 1.18390543225531e-09 | 0.549925109 | 0.278 | 0.054 | 2.97977158244339e-05 | 47 |
| pbx1b4 | 1.3967011028512e-09 | 0.451345804 | 0.194 | 0.028 | 3.51535700576619e-05 | 47 |
| cotl115 | 1.43083992730398e-09 | 0.826080781 | 0.694 | 0.308 | 3.60128101303138e-05 | 47 |
| zgc:1625092 | 1.82752579597134e-09 | 0.381273676 | 0.194 | 0.027 | 4.59969967588026e-05 | 47 |
| stk26 | 2.53027624418327e-09 | 0.409572922 | 0.167 | 0.022 | 6.36845227898486e-05 | 47 |
| si:zfos-464b6.22 | 3.25560990039876e-09 | 0.612598262 | 0.25 | 0.047 | 8.19404455831364e-05 | 47 |
| zgc:6405115 | 3.64159662479291e-09 | 0.756082757 | 0.5 | 0.164 | 9.16553454494127e-05 | 47 |
| gsna4 | 5.77233574281339e-09 | 0.491216022 | 0.194 | 0.03 | 0.000145284 | 47 |
| pnrc225 | 6.91422075778702e-09 | 0.602589997 | 0.861 | 0.569 | 0.000174024 | 47 |
| actb19 | 8.96305950768901e-09 | 0.415479919 | 0.917 | 0.786 | 0.000225591 | 47 |
| snx5 | 9.45613510173026e-09 | 0.612992945 | 0.278 | 0.059 | 0.000238001 | 47 |
| cyth1b | 1.2197659353325e-08 | 0.434020893 | 0.194 | 0.031 | 0.000307003 | 47 |
| mtus1b1 | 1.92794046580332e-08 | 0.373665868 | 0.167 | 0.024 | 0.000485243 | 47 |
| si:cabz01074946.16 | 2.28154211242641e-08 | 0.552190789 | 0.278 | 0.06 | 0.000574241 | 47 |
| akap71 | 2.96731044987552e-08 | 0.317120686 | 0.111 | 0.012 | 0.000746842 | 47 |
| fhl1b3 | 3.23350233718625e-08 | 0.331435947 | 0.139 | 0.017 | 0.00081384 | 47 |
| vaspb3 | 3.36875434642051e-08 | 0.463755218 | 0.25 | 0.05 | 0.000847882 | 47 |
| prss13 | 4.00647963394148e-08 | 0.332955397 | 0.167 | 0.024 | 0.001008391 | 47 |
| atp2a33 | 4.27570438293029e-08 | 0.467385061 | 0.194 | 0.033 | 0.001076152 | 47 |
| kdm6bb5 | 4.72373637128803e-08 | 0.524049402 | 0.25 | 0.053 | 0.001188917 | 47 |
| XLOC-0437605 | 5.21820373567699e-08 | 0.551170702 | 0.25 | 0.052 | 0.00131337 | 47 |
| si:ch211-196l7.421 | 6.15019582281e-08 | 0.690341542 | 0.667 | 0.316 | 0.001547943 | 47 |
| ehbp1l1a1 | 6.25886538852372e-08 | 0.364582441 | 0.139 | 0.018 | 0.001575294 | 47 |
| yrk7 | 6.59711879497347e-08 | 0.601235475 | 0.278 | 0.064 | 0.001660429 | 47 |
| crema13 | 6.87851177568941e-08 | 0.775207353 | 0.472 | 0.18 | 0.001731253 | 47 |
| alox5a6 | 7.70510566595468e-08 | 0.480405505 | 0.194 | 0.034 | 0.001939298 | 47 |
| cbfa2t32 | 7.77746075331027e-08 | 0.298746819 | 0.139 | 0.018 | 0.001957509 | 47 |
| trib38 | 1.59032944085563e-07 | 0.635753398 | 0.361 | 0.107 | 0.0040027 | 47 |
| si:ch211-105j21.96 | 1.63599783939089e-07 | 0.439402517 | 0.194 | 0.035 | 0.004117643 | 47 |
| zeb2a4 | 1.67241698186567e-07 | 0.516251739 | 0.222 | 0.046 | 0.004209306 | 47 |
| pdlim15 | 1.70799056186406e-07 | 0.505545599 | 0.25 | 0.055 | 0.004298841 | 47 |
| prkcaa1 | 2.13528375738459e-07 | 0.358004854 | 0.139 | 0.02 | 0.005374296 | 47 |
| mt-cyb18 | 2.27884946479386e-07 | 0.558197655 | 1 | 0.625 | 0.005735636 | 47 |
| mtss1 | 3.00406960412801e-07 | 0.372012293 | 0.167 | 0.027 | 0.007560943 | 47 |
| rassf2b | 3.02701401407341e-07 | 0.301345928 | 0.139 | 0.02 | 0.007618692 | 47 |

| | | | | | | |
|---|---|---|---|---|---|---|
| ier2a19 | 3.13795923113901e-07 | 0.725000224 | 0.583 | 0.264 | 0.00789793 | 47 |
| LOC10390911511 | 4.88492760049155e-07 | 0.665937723 | 0.444 | 0.167 | 0.012294874 | 47 |
| glcci1a | 5.19135919659575e-07 | 0.256346514 | 0.111 | 0.013 | 0.013066132 | 47 |
| rab11fip1b5 | 5.57490648010416e-07 | 0.49857578 | 0.25 | 0.059 | 0.014031482 | 47 |
| sgms1 | 6.51000976471847e-07 | 0.396296479 | 0.139 | 0.021 | 0.016385044 | 47 |
| kcne41 | 7.38146685227603e-07 | 0.367710769 | 0.111 | 0.014 | 0.018578414 | 47 |
| jdp2b13 | 7.45001703721776e-07 | 0.808845786 | 0.444 | 0.172 | 0.018750948 | 47 |
| dusp63 | 8.14862980548617e-07 | 0.397472223 | 0.222 | 0.048 | 0.020509286 | 47 |
| nfatc3a | 1.16072167914575e-06 | 0.262434234 | 0.111 | 0.014 | 0.029214204 | 47 |
| selenow110 | 1.22823566277646e-06 | 0.628597414 | 0.667 | 0.387 | 0.030913463 | 47 |
| mibp219 | 1.30807914611591e-06 | 0.568515342 | 0.75 | 0.494 | 0.032923044 | 47 |
| LOC101886015 | 1.71584494871264e-06 | 0.364349691 | 0.139 | 0.022 | 0.043186102 | 47 |
| XLOC-0437612 | 1.78295095535716e-06 | 0.365665283 | 0.139 | 0.022 | 0.044875093 | 47 |
| mylipb2 | 2.57385725506417e-06 | 0.356220871 | 0.139 | 0.023 | 0.064781413 | 47 |
| phlda27 | 2.60619548620139e-06 | 0.579792449 | 0.222 | 0.052 | 0.065595334 | 47 |
| itgb3a3 | 2.98052413582499e-06 | 0.318645832 | 0.111 | 0.015 | 0.075016812 | 47 |
| msna17 | 3.03822741181305e-06 | 0.624957981 | 0.528 | 0.243 | 0.076469146 | 47 |
| ehd1a2 | 4.64971279770352e-06 | 0.429708157 | 0.167 | 0.033 | 0.117028621 | 47 |
| cdc26 | 4.8272971944466e-06 | 0.536595627 | 0.278 | 0.081 | 0.121498243 | 47 |
| inpp5ka | 5.46262492390642e-06 | 0.252016417 | 0.111 | 0.016 | 0.137488807 | 47 |
| si:dkey-52l18.41 | 5.90959213463384e-06 | 0.490069955 | 0.222 | 0.054 | 0.148738524 | 47 |
| tspan74 | 6.9359367293506e-06 | 0.529559224 | 0.278 | 0.08 | 0.174570592 | 47 |
| fosb12 | 7.24555075347535e-06 | 0.66747598 | 0.361 | 0.127 | 0.182363267 | 47 |
| cfl114 | 8.75990531377386e-06 | 0.517005519 | 0.806 | 0.578 | 0.220478057 | 47 |
| ubl3a2 | 9.90613838987851e-06 | 0.422413545 | 0.194 | 0.046 | 0.249327597 | 47 |
| rhoab14 | 1.24860753958684e-05 | 0.585206611 | 0.444 | 0.191 | 0.314262032 | 47 |
| orai1a2 | 1.28781228913063e-05 | 0.307522931 | 0.139 | 0.025 | 0.324129475 | 47 |
| tpm311 | 1.42514772490492e-05 | 0.642808696 | 0.5 | 0.256 | 0.358695431 | 47 |
| foxn3 | 1.42661386259918e-05 | 0.294101668 | 0.111 | 0.017 | 0.359064443 | 47 |
| RASSF58 | 1.82736069589975e-05 | 0.442528902 | 0.222 | 0.059 | 0.459928414 | 47 |
| dub9 | 1.98559679294615e-05 | 0.467899317 | 0.25 | 0.072 | 0.499754857 | 47 |
| elmo110 | 2.04653821919195e-05 | 0.450417155 | 0.25 | 0.072 | 0.515093204 | 47 |
| lrrfip1a6 | 2.18672276362637e-05 | 0.410767675 | 0.222 | 0.059 | 0.550376252 | 47 |
| sept9a1 | 2.36534285399036e-05 | 0.425128761 | 0.194 | 0.047 | 0.595333143 | 47 |
| arpc217 | 2.38865610393927e-05 | 0.622275036 | 0.5 | 0.269 | 0.601200855 | 47 |

| | | | | | | |
|---|---|---|---|---|---|---|
| btg215 | 2.51837544942869e-05 | 0.575035282 | 0.667 | 0.375 | 0.633849917 | 47 |
| zgc:560951 | 2.74431271653106e-05 | 0.5393298 | 0.306 | 0.102 | 0.690716068 | 47 |
| atp2b1a3 | 2.78196811012814e-05 | 0.524396489 | 0.25 | 0.075 | 0.700193554 | 47 |
| cap116 | 2.89884703715688e-05 | 0.667249686 | 0.472 | 0.235 | 0.729610811 | 47 |
| nup931 | 3.49164996332125e-05 | 0.386750049 | 0.167 | 0.037 | 0.878813379 | 47 |
| dynll2b2 | 3.55618890188027e-05 | 0.540496578 | 0.361 | 0.134 | 0.895057185 | 47 |
| abhd3 | 5.52880857490679e-05 | 0.290500484 | 0.111 | 0.019 | 1 | 47 |
| hmgn620 | 5.81155434208447e-05 | 0.54973456 | 0.5 | 0.267 | 1 | 47 |
| rgs24 | 6.21412189888726e-05 | 0.321883829 | 0.139 | 0.028 | 1 | 47 |
| mt-co223 | 6.21496391208256e-05 | 0.439913129 | 0.972 | 0.721 | 1 | 47 |
| actn44 | 6.9264079651705e-05 | 0.488807288 | 0.278 | 0.095 | 1 | 47 |
| rfx5 | 8.15058554307344e-05 | 0.370978681 | 0.139 | 0.03 | 1 | 47 |
| ralgapa1 | 8.98675362526974e-05 | 0.382101865 | 0.139 | 0.03 | 1 | 47 |
| arpc5a11 | 9.42719960526925e-05 | 0.53473088 | 0.306 | 0.116 | 1 | 47 |
| cmtm6 | 0.000100054 | 0.428875383 | 0.194 | 0.054 | 1 | 47 |
| sh3bgrl38 | 0.000101906 | 0.510204171 | 0.361 | 0.149 | 1 | 47 |
| slc25a55a4 | 0.00012447 | 0.422203604 | 0.222 | 0.068 | 1 | 47 |
| cpn17 | 0.000134799 | 0.484248014 | 0.194 | 0.056 | 1 | 47 |
| map3k3 | 0.000139673 | 0.271322315 | 0.111 | 0.021 | 1 | 47 |
| mt-co123 | 0.0001399 | 0.421470756 | 0.972 | 0.658 | 1 | 47 |
| sdc413 | 0.000147682 | 0.523143237 | 0.528 | 0.259 | 1 | 47 |
| jupa4 | 0.00014952 | 0.390486114 | 0.194 | 0.054 | 1 | 47 |
| COX323 | 0.000150252 | 0.41412182 | 0.972 | 0.701 | 1 | 47 |
| leo1 | 0.000156712 | 0.313382336 | 0.139 | 0.031 | 1 | 47 |
| h3f3c18 | 0.000171668 | 0.548426032 | 0.472 | 0.258 | 1 | 47 |
| sh3kbp16 | 0.000212961 | 0.45816406 | 0.222 | 0.072 | 1 | 47 |
| sh3bgrl9 | 0.00021644 | 0.50250423 | 0.278 | 0.106 | 1 | 47 |
| myl9a7 | 0.000220293 | 0.423069345 | 0.139 | 0.032 | 1 | 47 |
| rhof | 0.000226279 | 0.309146395 | 0.111 | 0.022 | 1 | 47 |
| myh9a4 | 0.000226831 | 0.40684062 | 0.194 | 0.056 | 1 | 47 |
| plcb31 | 0.000233045 | 0.270885778 | 0.111 | 0.022 | 1 | 47 |
| erbin3 | 0.000251999 | 0.363092387 | 0.194 | 0.057 | 1 | 47 |
| srsf3a1 | 0.000276337 | 0.299331813 | 0.167 | 0.044 | 1 | 47 |
| anapc15 | 0.000313769 | 0.294348589 | 0.167 | 0.044 | 1 | 47 |
| ddx57 | 0.000328857 | 0.482839297 | 0.583 | 0.359 | 1 | 47 |

| | | | | | | |
|---|---|---|---|---|---|---|
| TCIM (1 of many)3 | 0.000354961 | 0.409421839 | 0.167 | 0.045 | 1 | 47 |
| jun15 | 0.000355078 | 0.578072707 | 0.472 | 0.23 | 1 | 47 |
| capza1b9 | 0.000403985 | 0.516378817 | 0.417 | 0.214 | 1 | 47 |
| si:ch211-225b11.12 | 0.000448029 | 0.320109814 | 0.139 | 0.034 | 1 | 47 |
| oaz1b20 | 0.000454826 | 0.4863477 | 0.528 | 0.324 | 1 | 47 |
| tuba8l39 | 0.000505261 | 0.368015151 | 0.167 | 0.048 | 1 | 47 |
| ppm1ba | 0.00050596 | 0.279279403 | 0.111 | 0.024 | 1 | 47 |
| mxd4 | 0.000573293 | 0.313298655 | 0.111 | 0.024 | 1 | 47 |
| sept9b4 | 0.000616648 | 0.361592174 | 0.167 | 0.049 | 1 | 47 |
| ier2b19 | 0.000671698 | 0.548636437 | 0.417 | 0.226 | 1 | 47 |
| ppp3r1a | 0.00074907 | 0.303300245 | 0.167 | 0.048 | 1 | 47 |
| purab2 | 0.00079191 | 0.379565312 | 0.167 | 0.05 | 1 | 47 |
| sec23b1 | 0.000814805 | 0.250142635 | 0.139 | 0.035 | 1 | 47 |
| sft2d1 | 0.000822622 | 0.264967925 | 0.139 | 0.036 | 1 | 47 |
| chmp2a7 | 0.000867577 | 0.478761486 | 0.306 | 0.135 | 1 | 47 |
| rgl12 | 0.000889331 | 0.311549454 | 0.139 | 0.037 | 1 | 47 |
| dyrk1b | 0.000889822 | 0.276728504 | 0.111 | 0.025 | 1 | 47 |
| rhogb9 | 0.000896075 | 0.453004236 | 0.25 | 0.098 | 1 | 47 |
| rabgap1l2 | 0.000937953 | 0.342574557 | 0.139 | 0.037 | 1 | 47 |
| GNAZ2 | 0.000939301 | 0.286417855 | 0.111 | 0.026 | 1 | 47 |
| tes2 | 0.001076286 | 0.359552747 | 0.194 | 0.064 | 1 | 47 |
| nxpe3 | 0.001101918 | 0.287843294 | 0.111 | 0.026 | 1 | 47 |
| crebrf1 | 0.001166394 | 0.358567706 | 0.139 | 0.039 | 1 | 47 |
| ppp1cbl1 | 0.001309787 | 0.434000753 | 0.194 | 0.069 | 1 | 47 |
| kctd5a8 | 0.001326781 | 0.430520921 | 0.361 | 0.177 | 1 | 47 |
| taf71 | 0.001366925 | 0.358404414 | 0.167 | 0.052 | 1 | 47 |
| ggh | 0.001395305 | 0.297567756 | 0.139 | 0.038 | 1 | 47 |
| ppt19 | 0.001411746 | 0.524258889 | 0.333 | 0.165 | 1 | 47 |
| anp32e | 0.001428122 | 0.395841945 | 0.306 | 0.13 | 1 | 47 |
| rplp225 | 0.001532609 | 0.293279866 | 0.917 | 0.759 | 1 | 47 |
| rap2c | 0.001615651 | 0.424393199 | 0.278 | 0.115 | 1 | 47 |
| selenok9 | 0.001700784 | 0.492650762 | 0.278 | 0.129 | 1 | 47 |
| pim114 | 0.001888111 | 0.509156842 | 0.333 | 0.173 | 1 | 47 |
| si:dkeyp-75h12.53 | 0.001934068 | 0.285529043 | 0.139 | 0.04 | 1 | 47 |
| hmgb1a12 | 0.001962136 | 0.461398243 | 0.361 | 0.191 | 1 | 47 |

| | | | | | | |
|---|---|---|---|---|---|---|
| klf6a14 | 0.002100431 | 0.509192919 | 0.333 | 0.169 | 1 | 47 |
| erh1 | 0.00218814 | 0.41921942 | 0.556 | 0.344 | 1 | 47 |
| mbnl112 | 0.002207729 | 0.44521636 | 0.361 | 0.185 | 1 | 47 |
| trappc13 | 0.002360613 | 0.340631044 | 0.25 | 0.102 | 1 | 47 |
| tuba8l8 | 0.002366335 | 0.462715954 | 0.361 | 0.191 | 1 | 47 |
| sept27 | 0.00243088 | 0.428311285 | 0.333 | 0.167 | 1 | 47 |
| tubb4b3 | 0.002689587 | 0.3371953 | 0.722 | 0.586 | 1 | 47 |
| mt-atp623 | 0.002753447 | 0.355419297 | 0.778 | 0.592 | 1 | 47 |
| si:ch211-214p16.110 | 0.002880517 | 0.467037571 | 0.5 | 0.26 | 1 | 47 |
| fam53b2 | 0.002955305 | 0.302452615 | 0.194 | 0.07 | 1 | 47 |
| cd6320 | 0.003192936 | 0.445461195 | 0.528 | 0.343 | 1 | 47 |
| ndfip23 | 0.003421312 | 0.300881302 | 0.167 | 0.057 | 1 | 47 |
| ramp25 | 0.003445409 | 0.316295454 | 0.139 | 0.043 | 1 | 47 |
| qkia10 | 0.003527866 | 0.417249096 | 0.333 | 0.17 | 1 | 47 |
| parp16 | 0.003738916 | 0.264271396 | 0.111 | 0.03 | 1 | 47 |
| inka1b12 | 0.003744526 | 0.388856135 | 0.389 | 0.187 | 1 | 47 |
| mfng5 | 0.003955992 | 0.277050379 | 0.139 | 0.043 | 1 | 47 |
| skap29 | 0.004048751 | 0.372820587 | 0.194 | 0.076 | 1 | 47 |
| histh1l12 | 0.004062537 | 0.40636579 | 0.25 | 0.111 | 1 | 47 |
| lrrfip2 | 0.004373547 | 0.267369995 | 0.139 | 0.043 | 1 | 47 |
| susd63 | 0.004527155 | 0.308017266 | 0.167 | 0.059 | 1 | 47 |
| taf101 | 0.005270174 | 0.261211644 | 0.194 | 0.072 | 1 | 47 |
| sik112 | 0.005367393 | 0.449812562 | 0.25 | 0.118 | 1 | 47 |
| gnai14 | 0.005554688 | 0.297547153 | 0.167 | 0.061 | 1 | 47 |
| calm2a2 | 0.005956963 | 0.447502835 | 0.361 | 0.205 | 1 | 47 |
| capza1a13 | 0.006913453 | 0.381815377 | 0.361 | 0.202 | 1 | 47 |
| gabarapa4 | 0.006942299 | 0.41621134 | 0.306 | 0.163 | 1 | 47 |
| lyn3 | 0.006946768 | 0.275059945 | 0.139 | 0.046 | 1 | 47 |
| fkbp510 | 0.007512543 | 0.469324522 | 0.25 | 0.124 | 1 | 47 |
| cops7a | 0.00769869 | 0.259846636 | 0.167 | 0.061 | 1 | 47 |
| wdr110 | 0.007710033 | 0.415183525 | 0.278 | 0.143 | 1 | 47 |
| klf7b1 | 0.008029091 | 0.26260949 | 0.111 | 0.034 | 1 | 47 |
| shq1 | 0.00806129 | 0.262289388 | 0.111 | 0.033 | 1 | 47 |
| suv39h1b1 | 0.008130239 | 0.299122765 | 0.139 | 0.047 | 1 | 47 |
| serf219 | 0.008412227 | 0.366226449 | 0.5 | 0.345 | 1 | 47 |

| | | | | | | | |
|---|---|---|---|---|---|---|---|
| zgc:1585641 | 0.008492173 | 0.32623934 | 0.194 | 0.08 | | 1 | 47 |
| fosl216 | 0.008707116 | 0.410349766 | 0.25 | 0.123 | | 1 | 47 |
| herpud21 | 0.008787518 | 0.257179393 | 0.111 | 0.034 | | 1 | 47 |
| si:dkey-261h17.15 | 0.008968835 | 0.285864269 | 0.222 | 0.093 | | 1 | 47 |
| psma41 | 0.008991793 | 0.350875178 | 0.333 | 0.176 | | 1 | 47 |
| nr4a116 | 0.00918522 | 0.447412501 | 0.306 | 0.169 | | 1 | 47 |
| kif5ba | 0.009297687 | 0.274804314 | 0.139 | 0.048 | | 1 | 47 |
| gadd45ba11 | 0.009847103 | 0.34973861 | 0.361 | 0.192 | | 1 | 47 |

| gene |
|---|
| zgc:153499 |
| cldnd |
| h1m |
| h2af1al |
| si:dkey-241l7.2 |
| zp3 |
| mt2 |
| si:dkey-240e12.6 |
| zp2.5 |
| zp3.2 |
| acp5a |
| si:ch73-189n23.1 |
| zp2.3 |
| si:ch211-226h8.14 |
| zp2.6 |
| tmsb5 |
| si:dkeyp-98a7.9 |
| ccna1 |
| zp2.2 |
| si:ch211-125e6.12 |
| cldng |
| si:dkey-90l23.1 |
| si:dkey-241l7.5 |
| BX649337.1 |
| zgc:171776 |
| si:zfos-1505d6.3 |
| crp2 |
| zp2.1 |
| zgc:77118 |
| zgc:175135 |
| zgc:171779 |
| si:dkey-241l7.4 |
| si:ch211-226h8.4 |
| CR790368.1 |

si:dkey-241l7.6
si:dkey-229d11.3
rrm2
lamp2
epcam
si:ch211-12h2.8
CU571163.1
ccnb2
si:dkeyp-98a7.8
LOC110437821
zar1l
qdprb2
thy1
mid1ip1l
si:dkeyp-20e4.8
si:ch211-12h2.6
si:dkeyp-46h3.6
zp3a.2
hbba1
si:dkeyp-98a7.4
si:ch211-14a17.7
zp3b
zar1
zgc:195245
si:dkey-229d11.5
tuba4l
ctss2.1
LOC100537499
nanog
si:dkeyp-50f7.2
slbp2
LOC100535162
zgc:173443
ccnb1
gtf3ab

siva1
btg4
cth1
zpcx
si:ch211-5k11.8
si:dkeyp-46h3.8
AL929032.1
si:ch211-132b12.8
hnrnpabb
kpna7
s100a1
zgc:171517
npm2b
hbaa1
zgc:136254
si:dkey-208k4.2
si:dkeyp-46h3.2
cpeb1b
si:dkey-90l23.2
gapdh
wu:fi42e03
uchl1
si:dkeyp-46h3.5
lsm14b
nasp
ca15b
si:ch211-146l10.8
si:ch211-226l4.6
avd
si:ch73-90k17.1
si:dkeyp-98a7.7
arhgef10la
zgc:56699
si:dkey-37o8.1
si:ch211-251f6.6

si:dkey-152b24.6
si:ch73-14h10.2
zp3a.1
zgc:171474
pigrl2.3
si:ch211-250e5.16
LOC101885477
si:ch211-145c1.1
zgc:55461
pabpn1l
birc5b
zgc:56231
sinup
elovl7b
gmnn
si:ch1073-263o8.2
haus6
ccna2
mkrn4
zgc:113424
rcc1
zp3c
buc
slc16a3
gdf3
map1lc3c
zgc:92591
rbpms2b
si:ch1073-190k2.1
si:dkey-19b23.11
ctsba
spink2.1
sybu
zgc:173856
zgc:113886

si:dkey-188i13.11
rbm4.3
AL929192.1
snu13b
slbp
gyg1a
h3f3a
cdca7a
larp6a
LOC100536860
crp3
zgc:103482
elavl2
zgc:162945
larp6b
nfr
inka1b
LOC101883897
CR383662.2
zglp1
tram2
gbgt1l4
tph1a
si:dkey-42i9.8
mibp
zgc:162879
org
tac3b
CU467646.2
patl2
dynll1
pttg1
hsd17b12a
cdc20
retsatl

plk1
elovl7a
gstp1
dctpp1
si:ch73-78o10.1
hspa5
ankrd22
ddx19
si:ch211-146l10.7
si:dkey-46g23.2
tmem54a
CR847984.1
zp2l1
tmprss4b
si:dkey-241l7.3
lpcat4
LOC103909972
cldn7b
snu13a
BX571780.1
ppp1r3b
h2afva
cox17
CABZ01059627.2
sox19b
cx44.2
zgc:173742
degs2
eif4a1a
zgc:165539
BX001030.1
zgc:152986
zgc:56676
pou5f3
orc6

CDPF1
BX537276.1
zc3h11a
nanos3
gpr160
si:dkeyp-114g9.1
si:ch211-14a17.6
si:ch73-160p18.3
rnf4
zgc:113984
ccne1
cdca8
zgc:171750
CU655961.4
zgc:114123
ctsla
si:dkey-68o6.5
tmem144b
srsf2a
zgc:66024
chia.4
ccnh
CABZ01059627.1
prrg2
aurka
il13ra2
rpz2
itm2cb
CT009596.1
parn
si:ch211-69g19.2
vstm2l
mad2l1
g3bp1
BX284697.1

brf2
pabpc1l
prr11
prdx1
figla
rasl11b
cpm
ddx39ab
chtopa
fdx1
hells
tpx2
kpna2
imp3
faub
tent5c
rdh10b
nifk
nutf2l
anp32b
rab5if
si:dkey-241l7.2
crp2
si:dkey-229d11.3
zp3
zp2.2
si:ch73-189n23.1
si:dkey-240e12.6
zp3.2
si:ch211-125e6.12
tmsb5
zp2.5
zp2.1
zp2.6
si:dkeyp-98a7.9

si:ch211-226h8.14
cldnd
si:zfos-1505d6.3
zgc:153499
si:dkey-241l7.5
BX649337.1
mt2
h2af1al
h1m
zp2.3
zp3a.2
si:ch211-12h2.8
LOC110437821
cldng
zgc:175135
si:ch211-226h8.4
zgc:171779
acp5a
si:dkeyp-20e4.8
rrm2
zgc:171776
si:dkey-241l7.4
si:dkey-90l23.1
CR790368.1
zgc:77118
si:dkeyp-98a7.8
epcam
si:dkeyp-98a7.4
si:ch211-12h2.6
LOC100535162
si:dkey-229d11.5
CU571163.1
si:dkeyp-46h3.8
si:dkey-241l7.6
si:dkeyp-50f7.2

ccnb2
si:ch211-14a17.7
kpna7
zp3b
qdprb2
zar1
zgc:136254
zgc:171517
zgc:173443
zgc:195245
slbp2
siva1
si:dkey-152b24.6
zar1l
AL929032.1
ddx4
mid1ip1l
ccna1
thy1
s100a1
ccnb1
si:dkeyp-46h3.6
nasp
hnrnpabb
tuba4l
npm2b
nanog
lamp2
si:dkeyp-46h3.2
zpcx
avd
uchl1
si:ch211-132b12.8
si:ch211-226l4.6
cth1

si:ch211-145c1.1
stmn1a
wu:fi42e03
ccna2
LOC100537499
si:dkey-90l23.2
zgc:171474
si:dkeyp-46h3.5
gmnn
gtf3ab
zgc:56699
si:dkey-19b23.11
pigrl2.3
si:ch211-146l10.8
ctss2.1
si:ch73-90k17.1
zp3a.1
si:ch73-14h10.2
btg4
si:ch211-251f6.6
gapdh
arhgef10la
ca15b
si:dkey-208k4.2
AL929192.1
si:dkey-37o8.1
LOC101885477
zgc:56231
zgc:55461
mkrn4
CU467646.2
elovl7b
si:dkeyp-98a7.7
gdf3
CABZ01059627.1

tmem54a
cpeb1b
lsm14b
org
zgc:173742
si:ch211-250e5.16
larp6b
zgc:66024
crp3
LOC101883897
chia.4
patl2
si:ch1073-263o8.2
pabpn1l
birc5b
sinup
rbpms2b
zgc:173856
zgc:103482
elovl7a
zgc:162879
zp3c
map1lc3c
snu13b
slc16a3
zgc:111868
zgc:162945
zgc:92591
retsatl
hsp90aa1.2
larp6a
si:dkey-188i13.11
dctpp1
haus6
hsd17b12a

top2a
pttg1
rcc1
zgc:114130
gbgt1l4
zgc:113424
sybu
zgc:171977
buc
si:ch211-146l10.7
mad2l1
inka1b
tac3b
cdca7a
zglp1
CR847984.1
zgc:101744
zgc:113886
cdc6
zgc:171750
sox19b
slbp
gyg1a
spink2.1
si:ch211-14a17.6
LOC100536860
si:ch73-78o10.1
si:ch1073-190k2.1
CDPF1
pabpc1l
cdc20
zgc:171426
CU655961.3
zgc:92287
mcm3l

elavl2
h3f3a
aurka
zgc:113984
si:dkey-216e24.3
rpain
pou5f3
BX548000.1
BX537276.1
zp2l1
si:dkey-46g23.2
BX284697.1
buc2l
zgc:56676
gstp1
hmgb2a
rasl11b
rdh10b
ubc
plk1
parn
si:ch73-160p18.3
ypel3
prr11
LOC110437779
snx10a
kpna2
BX001030.1
zgc:152652
evla
si:ch73-15b2.5
eif4e1b
rbm4.3
zgc:114123
CABZ01059627.2

brf2
CR383662.2
si:dkey-241l7.3
zgc:86839
zgc:165539
tph1a
pif1
cx44.2
dazl
degs2
cpm
si:dkeyp-114g9.1
tpx2
thop1
vstm2l
il13ra2
exd2
zgc:152986
yrdc
tmem144b
orc6
BX571780.1
srsf2a
st8sia7.1
cdca8
dnd1
ankrd22
nfr
krcp
piwil1
cldn7b
si:ch73-303b9.1
mos
aurkb
si:ch211-286b5.4

ddx19
rogdi
si:ch211-69g19.2
LOC103909972
coil
CU326366.2
tram2
hells
zfand6
depdc7b
BX005461.1
si:dkey-43p13.5
si:ch211-113a14.27
ndc1
CT009596.1
zgc:113425
znf395a
zgc:163030
rpa3
tatdn2
znf1173
prdx1
tmem44
gpx8
lpcat4
umps
zgc:55888
zgc:173544
gpr160
prc1a
cdca9
mibp
zgc:101016
nek2
si:dkey-42i9.8

ddx39ab
h2afva
ctsba
tspan13a
lmnl3
rbbp4
metap1d
ttc32
dynll2a
selenoh
si:dkey-189g17.2
cx43.4
ccnh
mrps18b
zgc:92313
fbxo32
hspa5
bmb
zc3h11a
ccne1
fdx1
npm2a
snu13a
BX957362.1
cited1
lrp13
pcna
dynll1
si:dkey-87o1.2
BX005461.3
LOC100538023
itm2cb
lrrc59
exd1
wu:fa19b12

zgc:175136
cox17
dcun1d1
ddx41
bzw1b
CU655961.4
si:ch211-157c3.4
pias4b
snrpb2
g3bp1
si:ch73-340n8.4
mylipa
cdt1
sec61a1l
chrac1
gemin7
ccdc58
rpp30
aif1l
ctsla
si:ch73-281n10.2
bokb
lman1
eif4a1a
timm13
cfl1l
zgc:153284
marcksb
mrpl22
si:dkey-68o6.5
ube2c
hmga1a
hmgb3b
CABZ01069022.2
slc29a2

nfs1
si:ch211-210c8.7
spsb1
chchd10
tmprss4b
gar1
vps9d1
kmt5ab
golt1bb
lipg
ncbp2
ppp1r3b
spryd4
mcfd2
nanos3
mcm7
trub2
rpz2
dnajc9
COQ10A
mrpl48
faub
figla
fen1
rbm7
casd1
tmem70
h2afx
polr2k
pop4
cnih1
gtf2a1
srsf7a
zgc:110239
zgc:112980

asf1ba
snrpg
arpp19b
tent5c
eif5a2
fbxw11b
ckbb
rbm4.2
cnep1r1
si:ch211-226h8.8
akirin1
atp5mc1
hsp90b1
rac1a
tomm20a
nutf2l
ppm1g
eno3
snrpa1
gfer
anp32b
imp3
tacc3
dkc1
top1l
cbx3a
cstf1
mpzl2b
pon3.2
nap1l1
glulb
txnl4a
mrps18c
ptgr1
fbl

mat2aa
ebna1bp2
ivns1abpb
exosc6
nifk
selenom
hprt1
ncl
paip2b
btg3
pim3
zgc:77112
emc6
f11r.1
dut
khsrp
spcs3
sumo3b
ube2al
UBL4A
si:dkey-33c12.4
uqcr10
hnrnpaba
ube2g1b
lsm1
thoc7
cbx5
lsm12a
snrpd2
STMP1
sub1b
npm1a
dap1b
rtn4a
aktip

tuba8l4
snrpe
tubb2b
lsm6
pcf11
slc25a25a
snrpc
cycsb
rpf1
dad1
snrpb
chtopa
hspd1
setb
prmt1
zfand5b
si:dkey-240e12.6
zgc:153499
h2af1al
zp2.2
cldnd
si:dkey-241l7.2
si:zfos-1505d6.3
zp3
zp2.6
h1m
si:dkey-241l7.5
zp2.1
zp2.5
si:ch211-226h8.14
cldng
si:dkey-229d11.3
si:ch73-189n23.1
crp2
tmsb5

zp3.2
si:dkeyp-98a7.9
si:ch211-125e6.12
zp2.3
acp5a
mt2
zgc:175135
zgc:171776
si:dkey-241l7.4
si:dkey-90l23.1
BX649337.1
ccna1
CR790368.1
si:ch211-12h2.8
si:ch211-226h8.4
zgc:171779
epcam
si:dkey-241l7.6
si:dkeyp-98a7.8
rrm2
mid1ip1l
si:dkeyp-98a7.4
LOC110437821
qdprb2
ccnb2
CU571163.1
zgc:77118
si:dkeyp-20e4.8
zp3b
zar1
si:dkeyp-46h3.6
zp3a.2
zgc:195245
npm2b
si:dkey-90l23.2

si:dkeyp-50f7.2
si:ch211-12h2.6
tuba4l
LOC100537499
ccnb1
zar1l
lamp2
LOC100535162
thy1
slbp2
s100a1
siva1
gapdh
si:ch211-14a17.7
si:dkey-229d11.5
ctss2.1
AL929032.1
nanog
si:dkeyp-46h3.8
zpcx
cth1
zgc:171517
zgc:136254
btg4
zgc:173443
si:dkeyp-46h3.2
si:ch211-132b12.8
hnrnpabb
gtf3ab
uchl1
si:ch211-146l10.8
si:ch73-90k17.1
zgc:171474
nasp
wu:fi42e03

avd
zgc:55461
si:ch73-14h10.2
kpna7
arhgef10la
si:dkey-37o8.1
si:ch211-145c1.1
birc5b
si:dkey-152b24.6
LOC101885477
si:ch211-250e5.16
spink2.1
si:dkeyp-46h3.5
si:dkey-208k4.2
cd9b
si:dkeyp-98a7.7
si:ch211-226l4.6
hbba1
gmnn
slc16a3
si:dkey-19b23.11
cpeb1b
mkrn4
pabpn1l
zp3a.1
si:ch1073-263o8.2
zgc:173856
elovl7b
si:ch211-5k11.8
haus6
h3f3a
elovl7a
si:ch211-251f6.6
lsm14b
ccna2

ca15b
larp6b
si:ch211-157c3.4
zp3c
crp3
zgc:171977
dhrs13a.2
map1lc3c
AL929192.1
zgc:113424
hbaa1
org
zglp1
zgc:56231
zp2l1
gdf3
pou5f3
zgc:92591
si:ch1073-190k2.1
sinup
pigrl2.3
patl2
rcc1
gbgt1l4
si:ch73-160p18.3
sybu
zgc:56699
rbm4.3
slbp
rbpms2b
CU467646.2
LOC101883897
zgc:162879
LOC100536860
snu13b

chia.4
si:dkey-188i13.11
rasl11b
CR847984.1
gyg1a
h2afva
inka1b
plk1
zgc:113886
cdc20
cldn7b
si:dkey-241l7.3
ubc
mibp
si:dkey-46g23.2
tph1a
ddx39ab
pabpc1l
CABZ01059627.1
buc
cdca7a
larp6a
CABZ01059627.2
dctpp1
vstm2l
zgc:103482
hspa5
hsd17b12a
zgc:152652
retsatl
rdh10b
zgc:163030
si:dkey-43p13.5
mad2l1
si:ch73-15b2.5

BX537276.1
tmem54a
zgc:165539
trim35-29
zgc:92313
faub
zgc:162945
si:ch211-146l10.7
zgc:101744
tmem70
zc3h11a
gstp1
degs2
prrg2
zgc:66024
tac3b
CDPF1
lpcat4
ctsla
CR383662.2
si:ch73-78o10.1
dynll1
chchd10
BX001030.1
pif1
pttg1
CU655961.3
mos
zgc:173742
ctsba
zgc:111868
srsf2a
cx43.4
BX649350.1
zgc:171750

kpna2
BX571780.1
nanos3
aurka
buc2l
zgc:113293
cx44.2
dph3
zgc:92287
sox19b
cdca8
ccnh
il13ra2
yrdc
ttc32
ypel3
zgc:113984
BX284697.1
hsd3b2
mcm3l
dynll2a
zgc:171426
tmprss4b
abcf2a
rpain
e2f5
si:dkey-216e24.3
tatdn2
phex
si:dkey-68o6.5
rnf4
eif4a1a
ube2c
si:dkeyp-114g9.1
snx10a

gpr160
spint2
zgc:55888
elavl2
nfr
dck
si:dkey-42i9.8
itm2cb
prkrip1
prdx1
h2afx
evla
zgc:112980
znf395a
si:ch211-69g19.2
umps
bzw1b
lmnl3
cdca9
selenoh
si:ch211-226h8.6
rogdi
ppp1r3b
si:ch73-303b9.1
lipg
spryd4
thop1
lsm12a
nifk
h2afvb
cox17
ccne1
zgc:173544
mrps18b
bmb

ddx19
tram2
rpz2
tmem144b
BX957362.1
snu13a
psme3
ncbp2
ankrd22
brf2
tpx2
fdx1
mylipa
cfl1l
marcksb
dnd1
gfer
parn
vps9d1
cdt1
cnot8
si:ch73-281n10.2
rbbp4
zgc:114123
ckbb
hmga1a
zgc:56676
g3bp1
hells
idh1
btg3
tuba8l4
zgc:77112
gemin8
zgc:86839

zfand6
snrpb2
ube2g1a
ebna1bp2
chtopa
alyref
mphosph6
dap1b
imp3
gemin7
zgc:110239
f11r.1
selenom
atp5mc1
tmco1
fbxw11b
cbx5
sap18
top1l
ilf2
ctsd
rtn4a
cbx3a
etf1b
snrpg
h2af1al
h1m
cldnd
tmsb5
zp2.5
zgc:153499
si:dkey-241l7.2
si:ch211-226h8.14
si:zfos-1505d6.3
si:dkey-241l7.4

si:dkey-240e12.6
acp5a
zp2.6
si:ch73-189n23.1
zp2.1
zp3
zp2.3
mt2
si:dkeyp-98a7.9
zp3.2
zgc:175135
si:ch211-125e6.12
zgc:171776
hbaa1
crp2
CR790368.1
cldng
si:dkey-241l7.6
ccna1
hbba1
zgc:171779
zgc:77118
si:dkey-241l7.5
zp3a.2
si:ch211-12h2.8
si:ch211-226h8.4
si:dkeyp-75b4.10
lamp2
zp2.2
si:dkey-229d11.3
BX649337.1
si:ch211-5k11.8
mid1ip1l
epcam
si:dkeyp-98a7.8

qdprb2
si:dkeyp-98a7.4
rrm2
si:dkeyp-20e4.8
CU571163.1
LOC110437821
si:dkeyp-46h3.8
si:ch211-14a17.7
zar1
si:dkeyp-50f7.2
si:ch211-12h2.6
zgc:195245
zar1l
si:dkey-229d11.5
slbp2
siva1
si:dkey-90l23.1
npm2b
nanog
btg4
s100a1
zpcx
AL929032.1
ctss2.1
si:dkeyp-46h3.2
spink2.1
LOC100535162
zgc:136254
ccnb1
gapdh
si:dkey-203a12.2
zp3b
tuba4l
zgc:173443
kpna7

LOC100537499
thy1
cth1
gtf3ab
si:ch211-132b12.8
ccnb2
si:dkeyp-46h3.6
si:dkey-90l23.2
uchl1
zgc:171517
zp3a.1
avd
si:ch73-14h10.2
wu:fi42e03
spink2.2
hnrnpabb
si:ch211-226l4.6
si:ch211-146l10.8
ccna2
lsm14b
si:dkey-152b24.6
si:dkeyp-98a7.7
zgc:171474
si:dkey-37o8.1
birc5b
slbp
si:ch211-250e5.16
cpeb1b
zgc:55461
pabpn1l
gmnn
LOC101885477
LOC100536860
si:dkey-241l7.3
nasp

zmp:0000001323
BX322787.1
h3f3a
si:ch1073-263o8.2
si:ch211-14a17.10
crp3
ca15b
elovl7b
arhgef10la
pigrl2.3
hsd17b12a
map1lc3c
si:ch211-251f6.6
slc16a3
buc
si:dkey-208k4.2
si:ch73-359m17.2
zgc:92591
si:dkey-203a12.9
spink2.4
si:dkey-19b23.11
haus6
si:ch211-145c1.1
si:dkeyp-46h3.5
mibp
rbpms2b
elavl2
si:ch73-90k17.1
zgc:56231
retsatl
gyg1a
gdf3
org
zgc:171977
zglp1

zgc:113424
ypel3
BX001030.1
sybu
zgc:56699
bmb
gbgt1l4
zgc:113886
CDPF1
CR847984.1
cldn7b
si:ch73-160p18.3
snu13b
si:ch211-146l10.7
zgc:171750
zgc:92287
zgc:173856
zgc:162945
elovl7a
larp6a
tph1a
zgc:111868
kpna2
AL929192.1
si:ch211-226h8.6
nfr
CR383662.2
gstp1
mkrn4
si:ch73-78o10.1
rasl11b
cdca7a
sox19b
rdh10b
pou5f3

si:dkey-46g23.2
zp3c
mad2l1
CT009596.1
ctsla
zp2l1
cdca8
ctsba
BX284697.1
si:ch1073-190k2.1
zgc:162879
h2afva
si:dkey-68o6.5
cx43.4
inka1b
dynll1
zgc:103482
pabpc1l
rcc1
tent5c
yrdc
cdc20
CABZ01059627.2
sinup
fdx1
zgc:173742
degs2
pttg1ipb
ddx39ab
si:dkey-216e24.3
mcm3l
CU467646.2
znf395a
dynll2a
larp6b

BX537276.1
zgc:153284
zgc:86839
BX005461.1
patl2
prdx1
LOC101883897
dazl
zgc:165539
tacc3
tac3b
dctpp1
si:dkeyp-114g9.1
pttg1
plk1
pif1
zgc:114123
prrg2
hspa5
faub
si:ch211-226h8.8
mos
nanos3
rbm4.3
zgc:152652
fbxo32
zgc:66024
si:ch211-69g19.2
chia.4
si:dkey-188i13.11
eif4a1a
parn
cox17
si:ch73-340n8.4
si:dkey-42i9.8

zgc:163030
g3bp1
cx44.2
si:ch211-210c8.7
rnf4
tram2
cstf1
aurkb
f11r.1
ankrd22
zc3h11a
marcksb
chchd10
BX957362.1
snrpb2
hprt1
ddx19
cbx3a
zgc:110239
selenoh
zgc:56676
drap1
hbba1
si:ch211-5k11.8
hbaa1
si:dkey-240e12.6
zgc:153499
h1m
h2af1al
cldnd
si:dkey-241l7.2
mt2
si:dkey-241l7.5
tmsb5
zp3

zgc:171776
zp2.2
si:ch211-226h8.14
si:ch73-189n23.1
crp2
cldng
si:dkeyp-98a7.9
zp2.5
zp3.2
zp2.3
zp2.1
acp5a
zgc:175135
si:ch211-12h2.8
zgc:77118
zgc:171779
CR790368.1
si:dkey-241l7.4
si:dkey-229d11.3
si:zfos-1505d6.3
BX649337.1
zp2.6
epcam
rrm2
zar1l
si:ch211-125e6.12
ccna1
CU571163.1
lamp2
si:dkeyp-98a7.4
zp3b
si:dkeyp-98a7.8
zar1
si:ch211-226h8.4
si:dkey-241l7.6

mid1ip1l
tuba4l
LOC110437821
npm2b
zp3a.2
zgc:195245
si:ch211-12h2.6
si:dkeyp-20e4.8
spink2.1
slbp2
si:dkey-90l23.1
thy1
si:ch211-14a17.7
qdprb2
siva1
cth1
AL929032.1
btg4
gapdh
si:dkeyp-46h3.8
si:dkey-229d11.5
nanog
zgc:136254
zgc:171517
zgc:173443
ccnb2
si:dkeyp-50f7.2
LOC100537499
LOC100535162
gtf3ab
si:dkeyp-46h3.2
si:dkeyp-46h3.6
ctss2.1
si:ch211-132b12.8
kpna7

si:dkeyp-75b4.10
si:dkey-152b24.6
haus6
s100a1
si:dkey-90l23.2
wu:fi42e03
zpcx
si:dkey-208k4.2
ccnb1
hnrnpabb
ca15b
zgc:171977
si:ch211-145c1.1
gmnn
avd
ccna2
zgc:171474
pigrl2.3
larp6b
cpeb1b
gdf3
lsm14b
zgc:92591
arhgef10la
zglp1
si:ch211-250e5.16
si:ch211-251f6.6
nasp
si:dkey-37o8.1
si:ch73-14h10.2
uchl1
birc5b
slbp
pabpn1l
h3f3a

elovl7b
si:ch211-146l10.8
zp3a.1
si:dkeyp-98a7.7
zgc:55461
LOC101885477
si:dkey-241l7.3
mkrn4
si:ch73-90k17.1
si:dkeyp-46h3.5
si:dkey-19b23.11
si:ch211-146l10.7
si:ch1073-263o8.2
si:ch211-226l4.6
AL929192.1
org
zgc:56231
inka1b
slc16a3
LOC101883897
elavl2
zgc:165539
rbpms2b
sinup
mibp
zgc:56699
cldn7b
tph1a
zgc:152652
zgc:173856
rasl11b
si:dkey-203a12.2
zgc:171750
zgc:162879
BX537276.1

si:dkey-68o6.5
pabpc1l
zp3c
rbm4.3
rcc1
crp3
si:ch73-15b2.5
ddx39ab
snu13b
zgc:103482
zgc:173742
buc
chia.4
pou5f3
zgc:66024
CR847984.1
si:ch73-78o10.1
mos
si:ch73-359m17.2
CABZ01075618.1
zgc:114130
dynll2a
ankrd22
LOC100536860
kmt5ab
retsatl
zgc:56676
patl2
CDPF1
CU655961.4
si:ch73-160p18.3
cdt1
lpcat4
cdca7a
si:ch73-281n10.2

tac3b
nfr
tmem54a
si:ch211-202f5.3
hsd17b12a
dynll1
map1lc3c
snx10a
elovl7a
dctpp1
gyg1a
ypel3
zgc:113424
oclna
vstm2l
si:ch1073-190k2.1
si:ch211-157c3.4
zgc:113886
cx44.2
sybu
ctsla
sox19b
phex
mad2l1
h2afx
rnf4
dnd1
zgc:114123
zgc:101744
cdc20
cox17
zgc:92287
snrpb2
aif1l
bzw1b

si:dkey-42i9.8
zgc:112980
zgc:163030
si:dkey-46g23.2
cpm
si:dkey-92i17.2
fbxo32
pttg1
CABZ01059627.2
smn1
larp6a
pif1
si:dkey-203a12.9
zp2l1
si:dkey-188i13.11
CABZ01059627.1
tent5c
zgc:162945
zgc:86839
BX284697.1
cnih1
aurkb
pcf11
aktip
fdx1
anp32b
ccl36.1
XLOC-042595
XLOC-000431
ccl38.6
XLOC-013964
XLOC-010899
LOC103910871
si:ch211-214p16.1
CR936442.1

dusp2
XLOC-028370
XLOC-025423
XLOC-020301
XLOC-026836
ccl34b.4
LOC100536213
CR753876.1
ccl32b.3
LOC108190717
BX005105.1
XLOC-032002
si:dkey-183i3.6
si:ch73-223p23.2
LOC101882070
si:dkey-109a10.2
CR848749.2
LOC100004978
si:dkey-183p4.10
LOC108185170
si:dkey-260g12.1
FP016106.1
cxcr3.1
si:ch211-93g21.1
XLOC-043760
si:ch73-285p12.4
zgc:136605
si:rp71-81e14.2
tnfsf14
faslg
XLOC-023492
c3a.5
gzm3
wasb
si:ch211-119e14.1

si:ch211-167j9.5
gpr183a
FP236356.1
si:ch73-226l13.2
ptpn6
wasa
BX005183.3
LOC103910136
si:ch211-132g1.1
gzmk
sla2
LOC103909107
id3
LOC100535806
XLOC-008993
si:ch73-266m14.1
lpar6b
ccl38a.5
si:dkey-27i16.2
mrps15
kctd5a
arpc1b
cabz01093075.1
btg1
cd247l
XLOC-043761
il2rb
si:ch73-46n24.1
ncaldb
XLOC-033137
si:dkeyp-85e10.3
ccl38a.4
LOC110439931
coro1a
cyth4a

rgs13
pfn1
rplp2
LOC101885675
entpd3
si:ch211-196f2.6
CU499336.2
XLOC-042596
hcst
ptprc
nitr2b
nitr10a
si:ch211-226m16.2
rplp1
FP101882.1
rac2
rpl18a
rplp2l
rpl37
si:dkey-171c9.3
CR855337.1
themis
rps20
si:ch211-25d12.7
h1fx
rps27.1
rps29
laptm5
rps2
tagapb
rps6
sh2d1ab
itk
wdr19
sept12

rps17
capgb
arhgdig
slc6a15
rpl11
rps15a
rps26l
rpl36
wu:fb55g09
rps5
zgc:64051
calm1b
rpl19
XLOC-011085
si:dkey-78o7.3
rpl34
rps21
adgre5b.3
samsn1b
rpl15
rpl32
cxcr4b
rps16
BX511079.2
cd40lg
skap1
ptmaa
grap2a
rsl24d1
rpl7a
zgc:174945
csk
hmgn2
rpl35
rpl8

rps7
tox
rpl26
si:dkey-185m8.2
srgn
XLOC-005449
tspan33a
si:ch211-132g1.4
rpl27
syk
cebpb
rpl18
pik3r1
BX901920.1
sh2b3
rps28
tox2
si:ch73-264p11.1
rps14
rpl23
tagapa
rps9
cytip
rpl10
il12rb2l
rpl10a
rpsa
rps12
rps19
nitr1m
slc25a5
si:dkey-211g8.4
cmah
LOC103909115
tmsb4x

rpl22
rpl38
rplp0
si:ch211-105j21.9
rpl28
rpl21
rps3
rpl7
rps15
ptger4b
CR786582.2
sesn1
CABZ01079663.1
zfp36l1a
faua
rpl39
rps13
si:ch211-284f22.3
rps24
naca
gzm3.3
cotl1
rpl9
rpl29
rpl35a
b2m
rps4x
rps26
LOC100151196
mbnl1
rpl30
rps18
CR385050.1
scinlb
si:ch211-191c10.1

rpl36a
hsp90ab1
CR381673.1
pip5k1ba
rps10
ms4a17a.6
mfng
gnai2b
actb2
lcp1
rps25
rpl24
tpt1
map2k6
si:dkeyp-68b7.12
rasal3
arhgef3l
ccni
il34
si:ch1073-220m6.1
lbh
crip1
nsa2
si:ch211-121a2.2
XLOC-014079
rps3a
si:ch211-102c2.4
si:dkey-262k9.2
ikzf1
unc119b
rps23
rpl13
eif3f
rpl23a
arrdc3b

cd74a
gpr132b
si:ch211-230g15.5
prkacba
zbtb32
sept9b
rps8a
dok2
phf11
RASSF5
prf1.9
lgals17
fli1b
ptpn4b
nme2b.1
msna
XLOC-041870
wipf1a
rpl27a
itga4
gfi1ab
rps11
myl12.1
nkl.2
slc44a2
CR318588.1
fam49a
ccl33.3
glipr1a
XLOC-043043
p2ry10
FERMT3 (1 of many)
tmc8
zgc:152791
arpc3

si:ch211-132p1.3
tefa
rpl3
arhgap27
il21r.1
ifi45
rpl31
ccr9a
rpl5a
uba52
rpl6
zgc:114188
CR855389.1
si:ch211-196l7.4
fcer1g
lpar5a
ppiaa
histh1l
arpc4
sik1
rack1
rpl13a
bcl2l10
sat1a.2
pycard
mcl1a
hmgn3
ikzf2
FQ323156.1
notch2
eef1g
vaspa
btf3
lasp1
si:ch73-234b20.5

vsir
tnip1
hmgn6
si:ch211-241f5.3
tnrc6c1
pnrc2
zgc:162944
cd74b
si:rp71-68n21.9
nfkbie
eef1da
rpl12
celf2
elf1
rps27a
sept2
zgc:152863
spry2
serbp1a
nptna
abi3a
ube2e3
rps27.2
si:dkey-165a24.10
arl11
skia
ptmab
ccr2
cfl1
mknk2b
cox6b1
si:ch73-248e21.7
cish
clic1
arpc2

mob1a
il16
rpl17
si:dkey-112e7.2
litaf
pgam1a
BX248082.1
acap1
ercc1
fscn1a
rbpjb
zgc:162730
socs1a
tmem131l
rps6ka1
wipf1b
cap1
cd63
gapdhs
sumo2b
si:dkey-187e18.1
ponzr1
isg15
ppp1r18
rnf213a
fam49ba
CR383676.1
rnf44
rpl4
arpc5a
GCA
hmha1b
stat2
psme2
pabpc1a

cdkn1bb
romo1
pam16
eif3m
psme1
ier2b
nr4a1
dnmt3ab
tnfrsf9a
nrros
irf2bp2a
eif3ba
hmgb1a
txnipa
actr2a
tpi1b
si:dkey-262k9.4
rpl14
elmo1
fabp3
dtnbp1a
plk3
cox7b
ran
arhgdia
ucp2
mt-co1
serf2
rpl22l1
si:ch211-222l21.1
eif3ea
mt-co2
gtpbp4
eno1a
mibp2

slc25a3b
rpl5b
gng5
nop53
cdc42l
sh3bgrl3
arpc4l
zgc:56493
ddx21
gnai2a
capzb
b2ml
COX3
rhoab
arl5c
tomm20b
oaz1b
zgc:153713
cox4i1
si:dkey-188i13.10
stat1a
cct2
eif3g
eif4ebp2
ywhabl
capza1a
glrx
crema
eif3s10
si:ch73-1a9.3
cox8a
eif3c
pde4ba
mt-atp6
eif5a

eif3i
atp5l
h2afx1
NC-002333.4
grn1
cd74a
lgals2a
c1qb
c1qc
si:dkey-5n18.1
lgals3bpb
ccl34b.1
si:busm1-266f07.2
lygl1
cd74b
zgc:153118
fcer1gl
NPC2 (1 of many)
c1qa
CU499330.1
marco
si:ch211-147m6.1
tmsb4x
LOC100537504
mhc2dab
actb2
ms4a17a.10
pfn1
si:dkey-7i4.24
CU929150.1
si:ch211-283g2.2
slc3a2a
grna
slc40a1
zgc:103700

tmsb1
havcr1
psap
krt18b
sftpbb
ccl35.1
grn2
si:ch211-194m7.3
zgc:174904
cotl1
si:ch211-165i18.2
BX649485.2
rnaset2l
ccl34b.8
LOC100536946
ifi30
CR318588.1
rbp2a
rplp2
txn
rps20
cmklr1
rps15a
rpl39
cfp
spi1a
gpx1a
mhc2b
itgae.1
lta4h
gng10
ppiab
rps2
rps29
ctss2.2

fabp11a
scpep1
rps27.1
arpc1b
mfap4
rpl35a
rpl18a
cndp2
rpl22
zgc:64065
rpl36
rpl29
rps21
slc25a5
rpl35
rpl11
rps28
rpl18
dpp7
rpl24
rps24
rpl32
si:ch211-1o7.3
rpl37
rps9
rpl17
rps5
rpl15
rpl8
f13a1b
rpl36a
arpc5b
atp6v1g1
faua
actb1

rpl38
rps16
mrc1b
anxa3b
si:ch211-15b10.6
rps15
zgc:152830
tuba8l2
sec61g
ckbb
rpsa
rpl21
atp6v0ca
LOC110437731
naga
rps14
rpl34
tpp1
npc2
laptm5
rpl9
rpl28
ybx1
kctd12.2
itm2bb
zgc:114188
ctsba
arl6ip1
rplp2l
spi1b
mdh1aa
naca
rpl30
cxcr3.2
gm2a

wasb
si:ch211-212k18.7
gstt1a
atp6v1e1b
tcima
rpl6
rpl23a
rpl27a
crip1
coro1a
CR855311.1
mapre1a
pepd
XLOC-023492
rps26l
ctsd
pycard
pdia3
rac2
fgl2a
zgc:153317
ppdpfb
gabarapb
arpc2
gapdhs
BX248082.1
cst14b.1
cd63
eef1db
ctsa
zgc:162730
clic2
cebpa
cap1
arpc4l

btg1
xaf1
rap1b
si:ch211-66e2.3
ptmaa
ms4a17a.6
eif4ebp1
ctsh
cox7b
ctsk
prdx5
smox
COX7A2 (1 of many)
arpc3
suclg1
si:dkey-180p18.9
nccrp1
zgc:64051
glrx
ctsz
vdac3
slc6a6b
si:dkey-112e7.2
igfbp5b
si:dkey-261h17.1
pmp22a
rbp4
krt94
endouc
col1a1b
col1a2
dcn
pcolcea
si:ch211-105c13.3
plpp3

fstl1b
fn1a
CABZ01092746.1
neu3.5
spon2b
htra1b
AL732488.1
thbs3a
mfap5
mgp
emilin2a
si:ch73-173h19.3
CABZ01039096.1
im:7151449
slit3
BX663503.3
osr1
efnb1
crhbp
adamts15b
flnb
mustn1a
plvapb
crabp2b
dock9b
hapln3
itih5
tspan35
tgfb2
crabp2a
slc6a4a
adgrg6
tll1
col1a1a
emid1

si:ch211-227m13.1
plpp1a
fbln2
abca1a
mfge8b
LOC108180269
fbln5
gfpt2
htra1a
lrp1aa
XLOC-016094
ablim1b
nid1a
tgfbr3
CR855311.1
si:ch211-67e16.11
vasna
kirrel1a
crabp1b
si:ch211-198c19.3
kcne4
dpysl3
fstb
tgfbr2b
si:ch211-39i22.1
gpc6a
id2b
slc6a2
neu3.4
ca10b
nptx1l
ccl25b
nova2
tmem100a
add3a

col5a2a
col5a1
marcksl1a
icn
cbln2b
hapln1b
tgm1l1
tsku
eva1ba
june
bag2
pamr1
akap12b
LOC100537342
aplp1
ebf3a
zgc:195173
s100a10b
gpr146
nfixb
emp2
has2
cyp2n13
zcchc24
cx43
CR383676.1
colec12
angptl2a
tnfsf10l
anxa1a
xkr9
fgl2b
peli1a
mdka
ddr2l

krt8
ppap2d
itga5
pmp22b
c7a
galr1b
c6
ackr3b
angptl1a
crabp1a
arrdc3a
angptl4
BX640522.2
mast2
sema3d
lpar1
znf385a
bicc1b
kank2
adamts15a
klf2a
mfap2
c1qtnf2
ramp2
clec19a
col6a3
rnd3b
rgs5a
cyp2ad3
ifitm1
tnfaip2a
tns2b
fkbp1ab
adrb3a
antxr2a

cyr61
ahnak
cyp2p6
slc7a1
eda
sparc
wscd2
dio2
si:dkey-240h12.3
jund
ca5a
pdgfra
bgnb
abhd15a
ank3b
arhgap21a
sema4ba
dhrs3b
ip6k2b
itpkcb
rgcc
sost
socs3a
ptx3a
hspb1
abi3bpa
si:ch73-46j18.5
synm
itgb1a
nectin1b
fam107b
fgfr2
rem1
cebpd
tmsb1

ackr4b
cxcl12a
junba
ugt5g1
loxl1
anxa5b
tns1b
cyp2ad6
agtr2
si:ch1073-459j12.1
fbn2b
scara3
NC-002333.4
si:ch211-212k18.8
fabp11a
prelp
s100v2
vim
nfia
klf3
emilin1b
lhfpl6
lrp1ab
hspb8
klf6a
cd81a
tshz2
LOC103909284
vat1
grb10a
fosab
cyp2p7
tmem176
iqsec1b
prss59.2

zfp36l1b
rpz5
plod2
clu
lix1l
tril
s100w
pdgfrb
pabpc1b
lama4
si:ch211-106h4.12
tlr5b
COX3
tuft1a
prss59.1
slc38a5b
col6a2
lxn
klf13
col4a1
mmp14a
mt-atp6
magi3a
XLOC-040020
htra3b
sesn3
XLOC-042917
zgc:114181
sult1st6
ddit3
LOC566587
dub
dedd1
hic1
LO018340.1

zgc:162730
neu3.2
gadd45ba
rras2
cald1a
dhrs3a
si:ch1073-291c23.2
myl6
CABZ01039304.1
adh8a
rbp5
sostdc1a
il15
cxcl8b.1
ccl19a.1
ptges
mt-cyb
lpin1
capn2a
mpzl1l
col6a1
si:ch211-1a19.3
mmp14b
kitlga
myo1ca
igf2b
mt-co2
boc
tagln2
cldn5a
btg2
stom
celf2
egfra
uap1

fgfr1a
loxl2a
CABZ01072614.1
il11ra
sema3b
ctdspla
sc:d0202
tcf12
sptan1
lpl
cxcl18b
cdon
dlc1
hdac4
cdh6
ldlrad4a
rbms2a
krt4
paplna
si:ch211-212k18.9
diaph2
fxyd1
jun
mgst1.2
zgc:158463
lamb1b
rdh10a
ghra
tns2a
ndfip1l
nfil3-6
crispld2
jam3b
tuba8l3
crip1

mych
llgl1
XLOC-000946
ehd1b
anxa13
sik2b
anxa2a
plekho2
qki2
krt15
fbxo34
dusp8a
ulk2
gulp1a
mir199-3
hsd11b2
ccdc85b
pim1
glis2a
b2ml
CR847566.1
dcbld2
mef2aa
timp2a
mt-nd4
appa
flna
mt-co1
pdlim1
svila
TCIM (1 of many)
glula
spon1b
BX322530.1
rbms3

wu:fj16a03
spry4
CABZ01074363.1
appb
reep3a
arl6ip5b
cast
svild
tspan14
fzd2
si:ch73-237c6.1
hopx
krt18b
foxp4
cst3
lrrc17
cd28l
pnpla6
mt-nd2
chd3
abcc5
mknk2b
nrp1a
lamc1
lamb1a
ckap4
zmiz2
cish
si:ch73-335l21.4
tshz3b
foxo3b
si:ch211-133n4.4
sdc4
adam10a
ubl3a

sntb2
c4
grk4
tmod2
gli3
krt91
etnk1
XLOC-024750
sesn1
eif4ebp3l
prss1
mt-nd1
fkbp5
ptp4a3
mcl1b
perp
eppk1
rtn3
si:dkey-238c7.16
FO082781.1
si:dkey-9i23.16
cebpa
tjp1a
pbx1a
junbb
nfic
hdlbpa
slc43a2a
f3b
ash1l
rbpjb
rab5aa
si:dkeyp-19e1.3
mt-nd3
egr1

fosl1a
itm2ba
dbn1
zgc:113208
stau2
mrtfab
zfp36l2
rerea
gpx4a
pdk2b
cyp3c1
myl9b
ptprga
sulf2b
sdcbp2
pdcd6
calm2b
fxyd6l
plecb
csrnp1b
cd63
rab10
ier2b
sdc2
clic2
aoc2
ppib
itgav
si:ch211-196l7.4
tmem176l.1
atp2b2
ywhaqb
h2afx1
fam168a
rbm39a

tmsb4x
per3
txnipa
ciarta
marcksl1b
pnrc2
klf11b
rhoaa
socs3b
aldh9a1a.1
wls
slc3a2b
tob1b
notch2
acin1a
arf2a
higd1a
zfp36l1a
si:dkey-7j14.6
tcima
h3f3c
ier2a
krt94
icn
epcam
krt8
si:ch211-39i22.1
BX649337.1
gtf3ab
cldnd
acp5a
si:ch211-5k11.8
hbba1
CR790368.1
h1m

si:dkey-241l7.5
zp3.2
zp2.5
zp3
h2af1al
hbaa1
zgc:175135
zp2.1
si:ch73-189n23.1
cxcl18b
tagln
rbp4
hsd11b2
mmp16a
ccl25b
fn1b
wu:fj16a03
nid2a
col6a1
aoc2
col4a1
cald1a
mfap2
si:ch1073-291c23.2
cpn1
cd82a
tmem176
inhbaa
si:dkey-23i12.7
adamts15a
dkk3b
rnd1b
cd248a
mdkb
cyp1c1

fhl2a
si:ch211-270g19.5
fank1
ccdc3b
si:ch211-106j24.1
mmrn2a
myocd
rxfp3.3b
rbp5
myl9b
angptl2b
loxl2a
selenbp1
cyp2ad3
dcn
slco2b1
col6a4a
nrp1a
krt18a.1
csrp1b
acta2
asic1b
c6
lrrc17
col1a2
zfp36l1b
prss23
atp1a3b
aebp1
pdgfra
si:ch1073-459j12.1
col18a1a
col6a3
ntn1a
cyp1c2

cygb1
col1a1a
igf2b
nid1b
mylkb
tenm3
zgc:158343
col6a2
col5a1
lama4
si:ch73-46j18.5
lamc1
pmp22a
cdon
cd81a
socs3b
col1a1b
spon1b
myl9a
cyp3c1
col12a1b
sfrp1b
irf8
zgc:195173
vcam1b
mdka
rgs5b
cxcl12a
gstm.3
snx17
socs3a
fosl1a
vegfaa
cfd
si:ch211-160j14.3

tpm4a
robo2
fstl1b
TCIM (1 of many)
CABZ01068367.1
sparc
CU459094.2
cdh2
si:dkey-9i23.16
jun
col4a2
olfml3b
si:ch211-212k18.9
cnn1b
bag3
mxra8a
eva1ba
im:7152348
spry4
gadd45ba
si:dkey-226m8.10
col5a2a
mgst1.1
BX088707.3
cyp27c1
rdh10a
krt8
si:dkey-261h17.1
cebpd
nrp2a
zgc:162730
c1qtnf5
sdc4
pdgfrb
zgc:152791

kif26ba
si:ch211-212k18.8
f3b
lxn
junba
pcdh18b
cbx7a
svild
LOC103909099
fosab
CABZ01068358.1
pbx1a
lamb1a
tmem47
pim1
fosb
gadd45ab
nme3
cyp2p6
si:ch73-86n18.1
f10
si:ch211-178n15.1
cldn11a
igfbp7
cplx2l
rasl11a
si:ch73-335l21.4
MMP23B
anos1a
gchfr
gstt1b
si:ch211-191i18.2
cpox
bhmt
sema4ba

il13ra1
chd3
csrnp1b
lpar1
cpxm1a
mafba
nog1
esr2a
lrp1ab
s100v2
aldh9a1a.1
ms4a17a.11
boc
caln2
pbx1b
aldh1a2
col16a1
tpm4b
ephb3
mmp2
cnn3a
si:dkey-188i13.10
ier2a
gsta.1
adh8a
si:dkey-7j14.6
zgc:112437
jdp2b
CABZ01092746.1
mmp14a
h2afy2
nr2f1a
flna
cox4i2
c1qtnf6a

pik3r2
c4b
tshz3b
nfia
nfil3-6
s100a10b
zeb2a
col15a1a
gpc1b
CABZ01068366.1
cldn5a
marcksl1a
xirp2a
dclk1a
atf3
myl6
emilin1a
colec12
tnfsf12
mtss1la
crim1
gas1a
il15
ifi45
mrc2
rgl1
lgals9l3
tns2a
fstl3
anxa1a
spry2
tmem176l.1
nfic
sh3gl1b
slc7a1

ar
ms4a17a.9
LOC103909284
postnb
h3f3c
CABZ01080702.1
grb10a
znf385a
zgc:154058
si:ch211-106h4.12
meis2b
zgc:92744
slc38a5b
palm1a
col27a1b
ece1
meis1b
inka1a
cyp2n13
nr2f2
fkbp7
rbp7b
p3h2
gamt
junbb
klf11b
cebpb
s1pr2
cyp2p7
lrp1aa
BX322530.1
tmem88b
itga9
b2ml
aldob

socs2
plekha6
FP102052.2
si:ch211-286o17.1
clec11a
spred2b
deptor
tmeff1b
nfil3
CR855311.1
ier5
osr2
hmgn6
nav3
kank2
gng7
rhoua
sfrp1a
ptk2aa
lgals17
LOC553316
ghrb
prss35
hmgb1b
c4
CR383676.1
ppib
adh5
actn1
maff
fblim1
cox8b
fhl3b
atf4b
irf2bp2b

egr1
hic1
cdh11
ptges
agtr2
thbs1a
krt91
ppp1cbl
zgc:113208
yap1
gadd45bb
b2m
rpz5
ppap2d
gstm.1
srpx
si:dkeyp-75h12.5
mgst1.2
slc43a2a
rnd3b
fam168a
si:dkey-163m14.2
fat1a
loxl2b
irf1b
asip2b
si:dkey-51e6.1
appbp2
zgc:114188
crispld2
ckap4
pdgfrl
selenow1
notch3
ybx1

hmgb3a
si:ch73-237c6.1
tshz2
CU459056.1
tmem119b
tmem100a
si:dkey-58f10.11
qkia
rpl9
gas6
krt94
mvp
zeb1b
snai2
cebpa
mgst3b
pgrmc1
clic2
grinab
higd1a
sh3pxd2aa
sult1st1
ctdsp2
fzd7b
mxra8b
tagln2
celf2
pabpc1b
ifi46
dhrs13l1
si:ch211-74m13.3
ptbp1b
rps27.2
hoxc3a
si:dkey-239h2.3

vdac3
grb10b
fxyd1
si:ch211-133n4.4
timp2a
ca10a
CABZ01074363.1
ppdpfa
fgfr2
CR318588.1
hsp70.3
ptenb
vim
CR847953.1
fermt2
arrdc3a
cyr61
angptl4
ghra
arhgap31
tspan4a
ptgs2a
creg2
btg2
ctdspla
antxr2a
si:dkey-79d12.5
mycb
eif4a2
si:ch211-204c21.1
ier5l
rps2
klf6a
map1lc3b
si:ch73-27e22.2

BX569787.1
serinc2
edil3a
tsku
cux1a
rpl27a
tuba8l3
tpm1
rps15
foxo3b
ctgfa
rps9
tns1b
tnfrsf1a
vasnb
arhgap5
c7a
cnbpb
itm2ba
tspan9b
si:ch211-197h24.9
abl1
rpl29
sgce
rps28
homeza
rps18
pi4kb
cxcl8a
ccdc85b
ldlrad4a
mprip
jpt1b
raraa
ywhaz

zcchc24
rpl36
SBNO2
mob2a
si:ch73-1a9.3
atp10a
XLOC-041269
adh8b
eif1b
fbn2b
itgb1a
si:ch211-1a19.3
tgfb1a
rps16
rpl6
wu:fb16f03
dhrs3a
gstz1
rplp1
hmgn7
plpp3
nsa2
ckba
rps13
pcolcea
rpl3
tnfsf10
crtap
rbms3
si:ch211-222l21.1
pnrc2
rps19
clic4
sumo2b
rpsa

sptlc2a
rpl39
cavin2b
rpl34
ptprga
mych
isg15
rps8a
rhoab
rpl32
FO082781.1
tax1bp3
stxbp6
rpl7a
sept2
mcl1b
her6
myo1ca
aste1a
pdk2a
mmp14b
stat1a
angptl1a
ablim1b
slc3a2b
CU984600.2
eef1db
il6st
igfbp5b
si:ch211-286b5.5
rbfox2
proza
ets1
rps7
ndufa4l

rpl26
rpl13
rpl37
myh11a
foxp4
akap12b
stau2
gli3
si:ch211-198c19.3
rpl24
rgs5a
ndrg2
ptmaa
enc1
rpl36a
rpl23a
rpl10a
rpl11
rps24
lims1
ccnl1a
ponzr1
rps27a
anxa13
sulf2b
rpl28
h2afx1
s100u
si:dkey-42i9.6
tpm2
neo1a
hspb1
zgc:174863
aldh2.2
rplp2l

pdia3
sh3pxd2b
col12a1a
rps14
ctdsp1
rps15a
ptena
rpl18
flot2a
bgnb
rps10
eppk1
ddit3
rpl8
rps4x
si:dkey-28b4.7
rpl35a
appb
capns1a
rps6
ptx3a
nfixb
rpl22l1
faua
sept10
gnl3
dedd1
rps11
rpl38
rpl17
b4galt6
abcc5
gpx4a
eef1g
rps21

emp2
tnrc6c1
rpl5a
tpm3
si:ch211-15b10.6
ahnak
rps26l
rpl30
rpl7
rps3a
capns1b
eif4a1b
rps26
calm2a
add3a
anxa4
rps29
pno1
mt-nd2
rpl18a
fosl2
gapdhs
snai1a
dbi
cldnc
tgif1
tmsb4x
hdlbpa
CABZ01072614.1
pleca
pdk2b
uba52
eif3g
serf2
serpinf1

cflara
rpl22
romo1
rpl15
serinc5
rpl27
dnmt3ab
jund
rps5
tob1b
arl6ip5b
krt18b
rplp0
ywhaqb
flot1b
h1f0
baz2ba
hmgb1a
ube2e3
lgals3b
tomm5
rpl35
rpl12
tspan7
txnipa
map1lc3a
prkab1b
dbn1
pim2
rps12
rpl21
naca
ctnnb1
pfn2l
rps27l

rps3
emd
bhlhe40
si:ch211-140m22.7
rps25
XLOC-016094
cry2
ddt
serbp1a
atrx
nop53
entpd1
foxo1a
icn
zbtb20
cct2
nop56
btg1
fabp3
zgc:153867
stat2
appa
rpl10
rpl31
eif4ebp3l
crebzf
pin1
metap1
eif3m
msna
ndfip1l
slc20a1b
cdc42
mt-nd1
wls

LOC101884451
zgc:174904
usp36
cxcl8b.1
gstr
ppiaa
zgc:110339
zfp36l1a
marcksl1b
tpt1
hnrnpa0b
si:ch211-132p1.3
skp1
larp4ab
rhoaa
rpl13a
rplp2
zfand5a
vdac1
fn1a
anxa11b
sdcbp2
rps27.1
cd82b
eif3ha
midn
fxyd6l
rbm28
ier2b
cd63
ciarta
lgals2a
eif3i
mt-nd4
sesn1

NDUFB1
pfn2
rack1
cnn2
rps23
tcp1
prdx5
acin1a
tegt
eif3f
rpl4
mtpn
COX3
tomm20b
cct8
ppdpfb
rsl1d1
irf2bp2a
laptm4a
eif3s10
srsf5a
pabpc1a
mt-atp6
si:dkey-42i9.4
eif2s3
eif2s2
COX7A2 (1 of many)
gabarapb
crema
mibp2
sh3bgrl
rpl5b
caprin1b
btf3
mphosph8

ip6k2b
ak2
hsp90ab1
rsl24d1
eif3c
uqcrh
rpl14
prdx2
ddx21
rab10
eif3ea
mt-nd3
eif3d
oaz1b
ccni
tmem258
ddx5
mknk2b
syncrip
si:ch211-196l7.4
NC-002333.4
eif5a
ubb
mcl1a
eef1da
eef2b
fkbp1aa
mt-cyb
sec61b
mt-co1
zgc:56493
ptmab
mt-co2
si:ch211-14a17.10
spink2.2

spink2.3
spink2.4
si:dkey-203a12.2
zmp:0000001323
si:ch211-194m7.8
ccl34b.9
LOC110437886
itln1
si:dkey-77g12.4
BX322787.1
MFAP4 (1 of many)
si:ch73-359m17.2
si:ch1073-110a20.3
clca1
si:dkey-203a12.5
cfbl
LOC110438970
LOC110437731
CABZ01112049.1
FO203432.1
cfp
cfi
si:ch1073-376c22.1
c3a.5
cfhl2
BX908782.3
agr2
spink2.5
si:zfos-2330d3.1
c3a.4
si:dkey-203a12.4
si:dkey-203a12.7
ca6
LOC108179162
LOC101883382

cfh
timp4.2
LOC103909452
cd59
pdia2
nfe2
LOC110438722
osmr
LOC110437920
BX323555.3
cfhl1
si:dkey-102g19.3
viml
itgb3a
si:ch211-212d10.1
si:dkey-27h10.2
cdaa
si:dkey-203a12.9
h6pd
si:ch211-214p16.2
si:dkeyp-66d1.7
cxcr2
si:dkeyp-75b4.10
CR848841.1
nfkbiaa
si:ch211-194m7.5
adgre13
clstn1
si:ch73-160i9.2
cnr2
si:ch211-214p16.1
spink2.1
si:ch211-270n8.1
si:ch211-276a23.5
myofl

cxcl8b.1
gata2a
adam8a
BX936393.1
ank3b
ltb4r
qsox1
cyp4f3
sult5a1
si:ch211-74f19.2
srgn
myh11a
itgb2
cd81a
mmp25b
fcer1gl
lrrc8da
alox5a
cd63
cd7al
furina
selp
si:cabz01036022.1
slc7a3a
atp1a3a
mctp2b
dhrs13a.1
alox5ap
illr4
pfn1
slc6a6b
lpar5b
LOC100148254
spns3
si:ch73-248e21.7

rgcc
si:ch73-237c6.1
mt-co1
mt-atp6
def6c
coro1a
mt-co2
ehd1a
limd2
si:ch211-136m16.8
si:dkey-206d17.12
samsn1a
capgb
sema4aa
CT030188.1
glipr1a
LOC108190702
anks1b
pygb
mt-nd4
tagapa
zmp:0000001082
actb2
mt-cyb
eno1a
cyba
pla2g4aa
zgc:158343
MYADM
pim2
COX3
calm1b
grap2b
nr4a1
dicp3.1

ncf1
cahz
xbp1
m17
si:ch1073-443f11.2
grasp
moxd1
myh9a
hacd3
CR383676.1
mir223
vsir
hcar1-4
mbnl1
rgs19
unc13d
camk1b
si:dkey-24p1.7
vaspa
ssr3
syngr2a
lta4h
cebpb
ikzf2
si:zfos-741a10.3
si:dkey-73n10.1
slc2a3b
pnp5a
si:ch211-215e19.3
relb
camk2g2
cxcr4b
p4hb
gsr
ptpn6

msna
arl5c
hck
XLOC-016094
CABZ01073834.1
rab11fip4b
mt-nd1
LOC101884267
smox
tnfaip3
mt-nd2
plxnc1
atp2a3
cap1
mrtfab
cotl1
rasal3
antxr2a
itga4
marcksl1b
far1
LOC103911430
birc2
ccr9a
yrk
sort1b
actn1
si:ch73-208g10.1
gapdhs
si:dkey-118j18.2
rab44
cd74b
wdr1
fybb
itgae.2

rac2
cox6a2
rhogb
mt-nd3
arpc1b
pnrc2
ccng2
arpc5a
ptprc
zbtb20
itgav
pgd
cmtm7
ahr2
arrb2b
nfkbiab
laptm5
tram1
n4bp1
dbnlb
zgc:66427
parvg
cyth1a
fam107b
sec11a
arhgdig
stk17b
tmsb1
sh3kbp1
actr2a
tagln2
maff
tmsb4x
cox6b1
gyg1a

socs3a
capzb
fosl2
ddit3
rps27l
cyp17a1
cyp11c1
rgs16
star
hsd3b1
pi15a
XLOC-018328
ptx3a
cyp11a2
prkag2a
ece2b
pik3r3b
fdx1b
osr2
fam184a
esr2b
nr2f1a
robo2
fdxr
nell2a
islr2
mctp1b
CABZ01074994.1
nr5a1b
meis3
tspan9a
nol4lb
CABZ01079281.1
arhgap20b
CABZ01038709.1

prox1a
nlgn4xa
camkvl
inhbab
si:dkey-183j2.10
stmn2b
pcdh8
zgc:153395
XLOC-001115
unc5db
cyp4t8
pklr
rprm3
chrnb3a
celf4
sypl1
BX901962.2
nrxn1a
cyp17a2
lrrtm1
hmgb3a
insl3
cox7a1
hsd11b2
oprd1a
pex5lb
ar
hs6st1b
ift172
col18a1a
ppdpfa
brinp2
tmsb2
dachd
jag1b

dab1a
LOC101882765
nme3
sipa1l2.1
atf3
skib
CR318588.3
dclk1a
atp6ap1lb
mtbl
fam222a
cox6a2
vamp2
zgc:92744
ctnnd2a
atp1a3b
cited4a
nr2f2
fundc2
hmgb1b
inhbaa
b4galt6
socs2
atp10a
ckba
irs2b
bag3
acsbg2
cxcl18b
tbx2b
cldn11a
cdon
pgr
meis1b
pcdh11

pgrmc1
acss1
dhcr7
pde4ba
robo1
gpr27
socs3a
jun
tmsb
hspe1
gramd1a
zgc:92606
fam89b
tp53inp1
edil3a
si:dkey-151g10.6
cux1a
nt5c2b
sox12
hist2h2l
asph
FO082781.1
ghrb
dnmt3ab
socs3b
pde8a
ptprda
si:ch73-46j18.5
syngr1a
slc4a4a
msi2a
hipk2
irf2bpl
slc51a
junba

mtus1b
tpst1
gfra1a
marcksl1b
nfic
ifi45
klf11b
aldh2.1
cplx2l
dusp5
atf4b
ifi46
fosl1a
CABZ01086041.1
rps28
cbx7a
lifrb
rpl34
rpl9
gbp
rpl32
mdka
cadm2a
tgfbr3
gadd45ga
enc1
slc38a2
rpl37
h1f0
rps7
prkar2aa
crim1
zgc:114188
si:dkey-188i13.10
nt5dc2

rps26
mafba
rpl18
fmnl3
si:dkey-44g23.5
NC-002333.4
fosab
znf503
ptp4a3
msi2b
hmgn6
rpl36
zbtb4
rpl22l1
rps16
rps9
bnip3lb
rps27.1
rpl27a
rpl29
b2m
tubb5
nlk2
rplp0
adh5
rplp2l
rps19
tbc1d4
cox8a
nupr1b
plxna4
rpl11
rps18
CR318588.1
faua

rpl28
rpl24
aoc2
rps8a
rps25
rps12
fndc3a
znfx1
fat1a
rpl6
rps15
maff
lgals17
rpl36a
cx43
rpl7a
rpl38
rps10
rpl26
rps27.2
rps13
si:dkey-28b4.7
rps26l
si:ch211-286b5.5
rps21
rpl35
rpl35a
cd99l2
rplp1
cyb5a
rps3a
CR383676.1
rpl27
si:ch211-178n15.1
ptprga

rps4x
wu:fc75a09
rps24
slc25a5
XLOC-041269
stat1a
eif1b
rpl31
elmsan1b
mt-nd1
eef1g
rpl30
rps11
rps14
h3f3c
rpl18a
kif5bb
rgs4
c4
si:dkey-23i12.7
atp1b1a
fam168a
anxa13
nipsnap2
mvp
hspd1
mt-nd2
rps29
antxr1c
diaph2
rps6
rpl23a
eef1a1l1
si:ch73-1a9.3
zgc:152791

gas1a
rps15a
rpl10a
rpl23
rpl8
ier2a
zgc:113208
mt-co2
nfia
flot2a
sept10
mgst1.1
rpl10
isg15
rpl13
rpl12
rpl15
fam102ab
zbtb20
junbb
rps27a
higd1a
ewsr1a
echs1
eif4ebp1
rps2
pdia3
rpl5b
COX3
rpl13a
il4r.1
egr1
rack1
rpl39
rpl3

rpl17
hnrnpa0a
hmgb1a
ywhaqb
uba52
sfpq
rsl24d1
serinc1
mt-co1
nop53
sema4ba
porb
pkma
h2afx1
zgc:195173
ybx1
naca
irf1b
rps27l
arhgef4
phactr4a
si:dkey-42i9.4
tmem243b
gabarapl2
il6st
hp1bp3
mt-nd3
jam2a
rpl19
rps20
rpl14
mt-nd4
b2ml
parp12b
rps23

stat2
rps3
DHDH
eif3f
hnrnpa0b
oaz1b
hipk3b
irs1
mibp2
map1lc3a
aldh7a1
tjp2b
bcl2l1
rpl21
mknk2a
rpl5a
CABZ01080702.1
fabp3
rps5
rpsa
rpl22
eef1db
si:dkey-51e6.1
aldh6a1
chd3
eif2s1b
ddx5
cish
nsa2
rbbp6
brd3b
hsp90ab1
ccnl1a
si:ch211-222l21.1
CABZ01072614.1

mphosph8
NDUFB1
mid1ip1b
rpl7
txndc9
ptp4a1
ube2e3
uqcrc1
ccnd1
rab11fip1b
atrx
rnf213a
tomm5
laptm4a
agrn
mt-nd5
si:ch211-196l7.4
ndufc2
si:dkeyp-44a8.2
eef2b
gtpbp4
serpine1
cct2
gadd45ab
btf3
gng7
dldh
anxa4
atp5l
zgc:158343
stau2
cbx1b
appa
cct7
rnf7

pno1
elocb
eif3m
grinab
cnbpb
sesn1
ppib
irf2bp2b
h2afy2
mt-atp6
srsf5a
add3a
rnf10
eif3i
eif4a1b
cct8
eif3c
ppiaa
MRPS12
rpl4
pim2
etfa
ndufa5
eif3ea
pfn2
grpel2
romo1
ola1
atp5f1e
zgc:111986
cox6b2
tcp1
idh2
smc1al
eif3ba

mt-cyb
u2surp
tktb
atp5mf
esd
ndufb8
cyc1
si:dkey-79d12.5
prdx3
rhoaa
gadd45bb
eif3g
atp5mc3a
cebpd
map1lc3b
sumo2b
caprin1b
ostc
eef1b2
hmgn7
rpz5
selenok
tegt
rnf44
cox6b1
cirbpb
mycb
eif2s3
prdx5
si:dkey-7j14.6
qkia
prdx2
pfn2l
hnrnpa1a
cct4

atf4a
ndufa4l
eif3k
dynlrb1
tomm20b
ndufb9
ptmab
atp5pf
jpt1b
itm2ba
eif3ha
ldhba
atp5pd
rplp2
hmgb2b
gabarapb
tnrc6c1
rbm39a
ywhaz
rgcc
eif3d
tma7
tpt1
cct5
sdc4
midn
atp5f1d
nfil3-6
gnb1b
eif2s2
pnrc2
vdac2
fkbp1aa
serbp1a
serf2

eif5a
slc25a3b
eef1da
cox4i1
cox7c
si:ch211-132g1.4
si:dkey-109a10.2
zgc:136605
si:ch211-196f2.6
foxp3a
si:ch73-361p23.3
tcf7
XLOC-042595
zap70
si:ch211-67e16.3
XLOC-013964
sla2
LOC100535806
LOC103910871
CR936442.1
dusp2
si:dkey-27i16.2
tnfrsf9b
XLOC-000431
XLOC-010899
cd28
gpr184
si:dkey-260g12.1
si:dkey-266f7.4
FP236356.1
gzmk
tagapa
rgs13
si:ch211-119e14.1
XLOC-042596

cxcr4a
il7r
si:ch1073-220m6.1
tnfrsf18
LOC101885675
b3gnt5a
si:ch211-214p16.1
si:ch211-93g21.1
FP016106.1
CR848749.2
zfp36l1a
nab1a
tnfrsf9a
si:dkey-183i3.6
sh2d1ab
skap1
cxcr4b
c3a.5
si:ch73-40i7.2
si:ch73-223p23.2
si:dkey-262k9.2
cebpb
s1pr5b
irf4b
ncaldb
samsn1b
LOC100536213
si:dkeyp-85e10.3
ccr9a
wasb
adgre5b.3
ptpn4b
si:ch211-132g1.1
zgc:64051
cd4-1

ptpn11b
si:dkey-183p4.10
itk
arhgdig
gpr55a
pfn1
hcst
btg1
myca
wu:fb55g09
abo
coro1a
srgn
rplp1
rps27.1
sik1
rps5
rpl18a
cxcr3.1
rplp2l
si:ch211-25d12.7
si:ch211-105j21.9
cytip
rps17
rplp2
si:dkeyp-68b7.12
rac2
rps6
tagapb
pde4ba
rpl32
rps12
rpl37
cremb
rpl7a

mxd
rps20
rpl11
rps2
si:ch73-285p12.4
wasa
rps21
rpl15
rpl8
arpc1b
rpl36
tpt1
si:ch211-226m16.2
rpl7
ccl38.6
ptprc
rsl24d1
sept9b
LOC101882375
rps26l
rpl27
rpl34
cyth4a
rps7
rpl18
rps16
ikzf2
il12rb2l
rps15a
h1fx
LOC103909115
XLOC-011085
LOC110439931
rps29
rpl19

themis
rpl10a
rpl10
laptm5
rps19
rps4x
si:dkey-42i9.7
BX005105.1
rps9
sept12
si:ch73-208g10.1
rplp0
il21r.1
nr4a1
rpl35
slc25a5
rpl38
LOC103909107
rpl26
calm1b
rpsa
rel
dgkaa
rps14
faua
BX511079.2
rps18
csrnp1a
rpl23
rpl9
hsp90ab1
rps13
ptmaa
agap2
rpl35a

rps3
CU207241.1
ms4a17a.6
rps23
vsir
si:dkey-262k9.4
wipf1a
rpl28
CR855337.1
tnip1
rps15
rpl13
lpar5b
RASA2
capgb
rpl24
rpl3
cabz01093075.1
rpl30
prkacba
litaf
rpl39
rps28
nfkbie
rps24
LOC100151196
irf2bp2a
rpl21
tnfaip3
rpl36a
mcl1a
rps10
si:dkey-53k12.2
si:dkey-171c9.3
nr4a3

fam49a
eef1a1l1
wipf1b
naca
ets2
hmha1b
sat1a.2
rps3a
acap1
zgc:195282
rps8a
cd28l
b2m
rpl29
si:ch211-256a21.4
si:rp71-81e14.2
ccr2
pnrc2
ptpn6
ldha
p2ry10
gpr183a
rack1
sh2b3
traf1
runx3
arhgef3l
si:ch73-226l13.2
rps25
si:ch211-167j9.5
rpl12
ccni
rps26
abi3a
zgc:152774

kctd5a
rpl22
ucp2
btf3
rasal3
tnip2
rpl23a
ikzf1
XLOC-008993
mbnl1
eef1g
rnf44
fmnl1a
ppp1r18
ets1
id3
lbh
nsa2
cd83
CU499336.2
chmp2a
CR392341.1
si:dkey-185m8.2
AL954142.1
tnfsf14
rps11
rpl31
lcp2a
gtpbp4
mob1a
si:ch211-196l7.4
hmgn2
tefa
selplg
uba52

scinlb
rpl13a
CR381673.1
si:ch73-248e21.5
CU929418.1
rpl6
stk17al
crema
vaspa
il2rb
rpl27a
si:ch211-191c10.1
rpl5a
CABZ01079663.1
msna
si:ch211-132p1.3
gnai2b
celf2
igfbp1a
sept2
actb2
arhgap15
tox
lpar5a
mrps15
mknk2b
lcp1
spint1a
eif3f
cd247l
myl12.1
si:ch211-284f22.3
grap2a
rps27a
ddb2

ppiaa
LOC101882070
dnajb9b
dtnbp1a
arhgap4b
tes
eef1da
cd74a
slc44a2
tnfb
prex1
rpl22l1
si:ch211-150o23.2
cotl1
csk
serbp1a
n4bp1
tmsb4x
XLOC-025423
si:dkey-25e12.3
si:ch73-234b20.5
si:dkey-187e18.1
arl5c
si:cabz01074946.1
batf3
tiparp
si:ch211-121a2.2
FERMT3 (1 of many)
crip1
pgam1a
CR383676.1
ddx21
sept6
mfng
elf1

kmt2bb
XLOC-024750
aldocb
zgc:174945
FP101882.1
tmc8
fli1b
myh11a
cdkn1bb
hmgb1a
rpl17
cd74b
si:dkey-118j18.2
psme1
ccl36.1
rbp5
cd63
CABZ01084575.1
ier2b
arg2
rpl5b
il15l
pabpc1a
hmgn3
nop53
serpinb1l1
tpi1b
eif3m
gapdhs
arpc2
unc119b
zgc:162944
dub
pik3r1
rpl14

ccnd2a
rpl4
fam49ba
ptmab
eno1a
eif3ea
si:dkey-112e7.2
grap2b
tnrc6c1
zgc:162730
sumo2b
arpc5a
BX005183.3
slc25a33
dusp1
elmo1
ponzr1
si:rp71-68n21.9
si:dkey-172h23.2
si:dkey-112a7.4
b2ml
rgs5b
bcl2l10
nfkbiab
nfkbiaa
clic1
mbpb
psme2
eif5a
eif3g
arpc3
cap1
prkar1ab
birc2
romo1

ccl32b.3
oaz1b
cdc42l
higd1a
eif3ba
cfl1
mibp2
XLOC-023492
zgc:114188
hcls1
XLOC-005449
prelid3b
tsc22d3
si:busm1-266f07.2
kdm6bb
dok2
rhoab
capza1a
ccl34b.4
midn
serf2
lasp1
gng5
sesn1
rnf213a
pycard
si:dkey-11f4.14
histh1l
rps27.2
rhogb
arf6b
NC-002333.4
sh3bgrl
egr3
anxa11a

fosl2
ankrd11
slc25a3b
itm2bb
cct8
chac1
atf4b
csrnp1b
ddit3
eif3i
eif3s10
GCA
idh2
XLOC-028370
gnb1a
cnbpb
notch2
nop56
tob1a
si:ch211-222l21.1
eif4ebp2
ppt1
arpc4l
eif3c
eef2b
hnrnpa0b
arhgdia
srsf5a
eif3d
ywhaba
arpc4
ier2a
hmgn6
zgc:56493
txnipa

ran
CR318588.1
mt-co2
mt-co1
COX3
si:ch211-226h8.14
si:dkey-240e12.6
CU571163.1
h2af1al
si:ch211-125e6.12
cldnd
zp3
si:dkey-241l7.2
mt2
acp5a
zgc:153499
zgc:175135
si:dkey-241l7.5
tmsb5
zgc:171779
si:zfos-1505d6.3
si:dkey-229d11.3
crp2
si:ch73-189n23.1
zp2.1
h1m
zp2.5
zp2.3
zp2.2
cldng
si:dkeyp-98a7.9
si:dkey-241l7.4
BX649337.1
zgc:171776
si:ch211-226h8.4

si:dkeyp-46h3.6
zp3.2
slbp2
lamp2
rrm2
si:dkey-90l23.1
CR790368.1
si:ch211-12h2.8
zp2.6
qdprb2
si:dkeyp-20e4.8
si:dkeyp-98a7.8
ccna1
si:ch211-12h2.6
zp3a.2
LOC110437821
siva1
zgc:171517
zar1l
mid1ip1l
epcam
zp3b
si:ch211-14a17.7
uchl1
gtf3ab
zgc:195245
wu:fi42e03
si:dkeyp-46h3.8
zgc:77118
si:dkeyp-50f7.2
thy1
si:dkey-229d11.5
AL929032.1
si:dkey-241l7.6
btg4

s100a1
ccnb2
si:dkeyp-98a7.4
tuba4l
hbba1
zgc:55461
npm2b
ccnb1
si:dkey-152b24.6
ctss2.1
si:ch211-132b12.8
zar1
bambia
nanog
zp3a.1
zgc:56231
zgc:173443
avd
zpcx
cth1
LOC100537499
LOC100535162
zglp1
si:ch1073-263o8.2
si:ch73-14h10.2
si:ch211-5k11.8
si:ch211-226l4.6
cpeb1b
si:dkeyp-46h3.2
zgc:136254
zgc:171474
LOC101885477
birc5b
si:dkeyp-46h3.5
arhgef10la

si:ch211-146l10.8
ttll12
CU467646.2
si:ch211-145c1.1
elovl7b
gapdh
hnrnpabb
kpna7
pigrl2.3
pabpn1l
LOC100536860
si:dkey-90l23.2
si:ch211-250e5.16
AL929192.1
si:dkey-19b23.11
gdf3
zgc:92591
sybu
si:dkey-208k4.2
hbaa1
zgc:56699
zgc:173545
si:dkey-46g23.2
nasp
cdca7a
lsm14b
gmnn
chchd10
snu13b
org
ccna2
si:dkeyp-98a7.7
si:ch211-146l10.7
si:dkey-37o8.1
zgc:113886

imp3
crp3
si:ch73-90k17.1
slbp
elavl2
yrdc
ccne1
cyp11a1
ca15b
spink2.1
slc16a3
sinup
adat3
kif23
h3f3a
patl2
larp6a
rcc1
LOC101883897
pou5f3
pabpc1l
trmt9b
depdc7b
cax1
si:ch1073-190k2.1
si:ch211-69g19.2
si:dkey-216e24.3
cx44.2
COQ10A
buc
zgc:112980
haus6
si:ch73-78o10.1
dctpp1
tmem54a

zgc:173856
zgc:113293
BX571780.1
nek2
ypel3
zgc:163030
vstm2l
ddx39ab
rbm4.3
ccnh
tph1b
zgc:103482
BX537276.1
zgc:113424
zgc:111868
zgc:162879
tph1a
itm2cb
si:ch211-251f6.6
larp6b
zgc:101744
snx10a
gyg1a
CABZ01059627.1
kpna2
si:ch211-286b5.4
si:dkey-42i9.8
tatdn2
zgc:162945
elovl7a
dynll1
retsatl
si:ch211-157c3.4
si:ch211-14a17.6
mkrn4

CT009596.1
CR847984.1
ankrd22
pop4
gbgt1l4
lipg
rasl11b
ctsba
hsd17b12a
map1lc3c
rpain
pif1
zgc:173742
oclna
si:dkey-68o6.5
si:ch73-281n10.2
zgc:55888
zp3c
nfr
gstp1
rbpms2b
anp32b
cdc20
tbcc
rdh10b
cdca9
kif22
si:ch211-202f5.3
pttg1
orc6
snrpc
si:ch211-241j12.3
cpm
zgc:92287
zgc:113984

akip1
si:dkey-87o1.2
prrg2
zgc:171750
BX005461.1
LOC100538023
zgc:56676
si:ch73-160p18.3
chia.4
tac3b
BX001030.1
LOC110437779
cdca8
kmt5ab
LOC103909972
si:ch211-113a14.27
tent5c
si:ch73-15b2.5
cx43.4
zgc:66024
dazl
id2a
ttc32
zgc:171977
slc48a1b
mibp
fdx1
nanos3
plk1
vps9d1
ube2c
zgc:92313
rbbp4
zgc:101016
mrps18b

figla
sumo3a
eno3
aurkb
degs2
prdx1
chtopa
parn
rab5if
fbxo32
dck
rnf4
faub
CABZ01059627.2
si:dkey-241l7.3
zgc:152652
zc3h11a
ctsla
dynll2a
cldn7b
snrpb2
mos
cnep1r1
evla
hspa5
si:dkey-188i13.11
tpx2
ckbb
mrgbp
CDPF1
bzw1b
si:ch73-340n8.4
inka1b
rpa3
adipor1a

h2afva
zp2l1
ivns1abpb
BX957362.1
gemin7
g3bp1
aktip
sub1b
tia1l
tspan36
btg3
ccl36.1
si:dkey-21e2.12
si:dkey-21e2.11
ncaldb
zbtb32
ca2
si:ch211-277c7.6
il34
si:ch211-207n23.2
si:dkey-78l4.4
ccr2
si:rp71-81e14.2
blnk
dicp1.5-6
BX000438.2
fam167b
XLOC-026336
scinlb
CABZ01084575.1
si:ch211-102c2.4
ccl38.6
ponzr6
LOC108190717
znrf1

obsl1a
si:ch211-241f5.3
XLOC-013964
LOC103910871
mctp1a
si:ch73-40a2.1
si:ch211-214p16.1
CR936442.1
LOC103910169
XLOC-042595
tnxba
itpr1b
sla2
nitr1m
XLOC-020301
si:ch211-191c10.1
CT573231.1
creb5b
tnfb
XLOC-042596
XLOC-026836
XLOC-010899
XLOC-028370
dusp2
XLOC-032002
XLOC-000431
CR392341.1
XLOC-025423
syk
arpc1b
stim1a
si:dkey-183i3.6
LOC100536213
zgc:64051
CU914617.1

prkcba
gramd2aa
BX901920.1
orai1a
mbnl1
XLOC-043043
rgs13
laptm5
il2rb
XLOC-043760
ttyh2l
si:ch211-214p16.2
gfi1ab
wdr19
gpx1a
calm1b
CABZ01117603.1
CABZ01044746.1
XLOC-023492
CR753876.1
srgn
slc41a1
CU972454.4
cd44b
mef2d
id3
camk1db
bhlhe40
gzm3.4
atp2b1a
sik1
arhgdig
fcer1g
adam19b
FP236356.1

dock10
LOC103909107
XLOC-041870
irf4b
zgc:153018
kdm2ba
BX322618.1
ptprc
tagapa
wasa
si:ch211-167j9.5
XLOC-014079
tmsb4x
plcg2
capgb
hcls1
plp2
XLOC-039757
CABZ01111959.1
pik3r1
ptmaa
BX005183.3
caspa
XLOC-043761
atp11c
gsdmea
cd247l
grapa
coro1a
tmc6a
pxna
cd40lg
BX908750.1
si:dkey-183p4.10
sla1

lck
atp5if1b
tmc8
aldocb
cotl1
si:ch73-208g10.1
ptmab
rplp2
XLOC-005449
si:dkey-1c7.1
tagapb
LOC110439931
fscn1a
si:ch211-93g21.1
si:dkey-53k12.2
arpc5a
lasp1
tnip1
zgc:174945
cox6b1
mef2ca
vaspb
CR381673.1
myl12.1
si:rp71-68n21.9
ponzr1
elf1
rac2
si:ch211-119e14.1
hmgn2
ikzf1
unc13d
CABZ01054965.1
fpr1
CR392341.4

si:dkey-151g10.3
casz1
dok2
si:ch211-222l21.1
crema
LOC110438754
XLOC-011085
RASSF5
yrk
arpc2
arpc3
XLOC-033137
nptna
vaspa
cap1
wasb
fybb
si:ch211-132g1.1
pycard
si:cabz01074946.1
entpd3
rps29
fmnl1a
si:dkeyp-19e1.3
cyth4b
si:dkey-260g12.1
gcnt4a
itga4
h1fx
rplp1
wipf1a
si:ch211-215e19.3
strbp
samd11
CABZ01088229.1

si:dkeyp-68b7.12
arpc4
cers1
LOC103910136
bcl2l10
csf2rb
rps6ka1
pip5k1ba
dtnbp1a
sox5
XLOC-008993
zgc:195175
pfn1
rin2
rps21
LOC101882375
rpl11
rps27.1
rpl37
msna
arhgdia
actb2
arl11
clic1
pim2
cmah
ldha
megf8l
arhgef3l
rpl19
gapdhs
cebpb
cyfip2
rpl27
inpp5d

si:dkey-262k9.4
cd63
dub
rps17
LOC100151196
ptpn22
rps28
celf2
rpl34
CU571074.1
rps26l
prkcaa
plxnc1
mctp1b
rps15
si:ch211-226m16.2
tefa
egr3
mt-co1
rps2
myo1f
arf6b
nfkbiaa
rps15a
skap2
pgam1a
BX120005.1
lrrfip1a
rplp2l
kif21b
syne1b
mrps15
rhoab
cox8a
dusp1

tagln2
rps20
tln1
ankrd44
selplg
grap2a
irf2bp2a
si:dkey-171c9.3
capzb
nkl.2
rpl18a
pnrc2
crip1
sort1b
rpl35
rpl36
fli1b
btg1
CR855389.1
sept2
rpl38
si:ch73-223p23.2
edaradd
rpl7a
cfl1
rpl23
gng5
si:ch73-264p11.1
pik3cg
si:ch211-196l7.4
nr4a1
rps6
sh2b3
rps10
si:dkey-185m8.2

rps16
prelid1b
rpl26
rpl39
kctd5a
rpl8
serf2
rpsa
carhsp1
rpl18
dock8
krt18b
rpl10a
rpl15
sdcbp2
rps19
il16
rpl22
dusp5
ankrd12
cd83
bcar1
histh1l
rps18
BX248082.1
ccdc88b
actr2a
elovl1b
grap2b
capza1a
rps7
cmtm7
unm-sa1614
zgc:56493
reep5

rps24
rps13
rps23
rps14
zgc:114188
CABZ01079663.1
vsir
rpl21
rps3
rpl24
rpl5a
si:dkey-78o7.3
rpl32
ywhabl
rpl10
rps9
rps5
si:dkey-177p2.6
pabpc1a
erbin
lcp1
mt-atp6
capza1b
hcst
eno1a
nfkbie
rpl35a
LOC103909115
zgc:153713
psme2
bida
rasal3
fam49ba
cdc42l
arl6ip5b

nab1a
si:ch211-105j21.9
mob1a
rpl14
fam13b
rps12
hmgn6
ccni
XLOC-005448
lpar5a
actn4
rpl36a
gnai2a
rpl9
actb1
glrx
per3
rel
zgc:162944
tpi1b
rpl31
grasp
sh3kbp1
rpl30
rpl29
slc25a5
hsp90ab1
eif3f
plekho2
iqgap1
chmp2a
tmem30aa
litaf
rpl28
arrb2b

gnai2b
rab8b
mt-co2
si:dkey-25e12.3
LOC101884451
faua
mcl1a
cmip
rsl24d1
glipr1a
rps11
zdhhc18b
uba52
si:ch73-234b20.5
mob3a
rps25
c3a.5
nfkbiab
tgif1
cpne1
ccr9a
slc9a3r1a
rin3
fosl2
CR383676.1
zgc:152863
rps4x
rps3a
cox7b
cox7a2a
anxa11a
tnfaip3
rps8a
rab44
cib1

tnrc6c1
lbh
oaz1b
zgc:63831
rpl7
mef2aa
scamp2
kdm6bb
arpc5b
COX3
zmynd11
sh3bgrl3
arpc4l
sh3bgrl
rps26
zgc:162730
si:dkey-112e7.2
ndufa4l
nr4a3
pde4ba
lmnb2
ywhaba
gnb1a
mtpn
ier2b
mt-cyb
sub1a
cry1aa
slc25a55a
zgc:153867
csk
rnf44
abracl
ccl39.7
gnb1b

uqcrh
pkma
naca
hmgb1a
ppiab
tpt1
si:ch211-132p1.3
GCA
sept7b
atp5f1e
atp5pf
arf6a
jak1
h2afx1
ppiaa
spcs2
mrpl33
cox4i1
rps27a
gadd45bb
ap1s1
rpl27a
rpl23a
cox6a1
ywhabb
atp5pd
ip6k2b
rpl13a
mvb12a
elmo1
brk1
twf1a
ptpn6
edf1
rps27.2

NC-002333.4
psme1
ucp2
COX5B
tpm1
rhoac
hmgn7
hnrnpa0b
cdkn1bb
itgb2
notch2
rack1
sumo2b
mibp2
NDUFB1
b2m
atp5f1d
ppt1
cox7c
eif4ebp2
selenof
cxcr4b
mt-nd4
btg2
rhoaa
hmgb2a
atp5meb
mt-nd1
serbp1a
si:ch1073-325m22.2
ube2v2
txnipa
romo1
tomm20b
pomp

atrx
rpl6
uqcrq
tmed7
sri
aldoaa
eif3i
ywhae1
hnrnpa1a
egr1
b2ml
atp5l
fkbp1aa
rap1b
rpl4
btf3
tegt
rpl5b
xbp1
eif3ea
nsa2
mt-nd3
eef1g
anp32a
calm3b
ybx1
chac1
vdac2
cox5aa
slc25a3b
ctrb1
cldnd
hbba1
zgc:175135
mt2

h2af1al
h1m
rrm2
si:dkey-241l7.4
si:dkey-240e12.6
zp3
CR790368.1
zp2.5
prss59.2
si:zfos-1505d6.3
zp2.3
acp5a
si:dkeyp-98a7.9
zp2.2
si:dkey-241l7.5
si:dkey-241l7.2
zp2.1
hbaa1
zp2.6
si:ch211-226h8.14
ccna1
crp2
si:ch211-5k11.8
si:dkey-229d11.3
si:ch73-189n23.1
prss1
si:ch211-125e6.12
zgc:171776
zgc:77118
zgc:153499
tmsb5
tuba4l
zar1
BX649337.1
cldng

ela2l
zgc:171779
si:ch211-12h2.6
si:ch211-12h2.8
si:dkeyp-98a7.8
cth1
epcam
si:ch211-226h8.4
si:dkeyp-50f7.2
zp3a.2
zp3b
mid1ip1l
hnrnpabb
lamp2
gapdh
ccnb1
siva1
si:dkeyp-98a7.4
zp3.2
si:dkey-241l7.3
si:dkey-241l7.6
slbp2
btg4
zgc:195245
si:dkey-90l23.1
s100a1
CU571163.1
si:dkeyp-46h3.6
wu:fi42e03
si:dkeyp-46h3.8
zpcx
si:ch211-14a17.7
zp3a.1
zar1l
npm2b

ccnb2
si:dkey-152b24.6
qdprb2
si:dkey-19b23.11
birc5b
si:ch211-132b12.8
nanog
si:ch211-240l19.8
zgc:171517
ca15b
gmnn
spink2.1
zgc:136254
lsm14b
kpna7
zgc:173443
gtf3ab
si:ch73-90k17.1
LOC110437821
hnrnpa0l
zgc:55461
si:dkeyp-20e4.8
thy1
avd
si:dkey-90l23.2
si:ch73-14h10.2
zgc:113424
pabpn1l
zgc:56699
h3f3a
AL929192.1
gbgt1l4
si:dkey-229d11.5
si:ch211-146l10.7
si:ch211-145c1.1

haus6
zgc:162945
elovl7b
snu13b
gdf3
si:ch211-251f6.6
si:ch211-146l10.8
AL929032.1
uchl1
LOC100535162
ddx39ab
si:dkey-208k4.2
si:dkey-37o8.1
LOC100537499
si:dkeyp-46h3.2
ctss2.1
slc16a3
si:ch73-160p18.3
ccna2
rcc1
slbp
si:dkeyp-75b4.10
crp3
cpeb1b
cdc20
LOC101885477
larp6b
arhgef10la
si:ch73-78o10.1
si:ch211-250e5.16
cx44.2
inka1b
zgc:56231
org
zgc:171474

si:dkeyp-46h3.5
zgc:173856
zgc:92591
nasp
CDPF1
LOC100536860
rdh10b
zgc:162879
larp6a
LOC101883897
srsf2a
nanos3
st6galnac1.2
si:dkeyp-98a7.7
si:ch211-226l4.6
dynll1
lin28a
mos
si:ch211-226h8.8
si:dkey-42i9.8
LOC110437779
cbx3a
eif4a1a
zgc:103482
pabpc1l
si:dkey-46g23.2
zgc:66024
lhx8a
orc6
patl2
zp2l1
plk1
elovl7a
si:dkey-216e24.3
cdca8

zgc:163030
rbm4.3
sybu
yrdc
map1lc3c
dazl
BX005461.1
rbpms2b
buc
retsatl
ankrd22
ddx41
buc2l
snx10a
si:ch211-202f5.3
hspa5
kpna2
aurka
hsd17b12a
si:ch73-132k15.2
BX537276.1
snu13a
rbbp4
dynll2a
tent5c
si:dkey-68o6.5
faub
hmgb3b
ctsla
bmb
zgc:165539
si:ch73-303b9.1
pou5f3
rpain
LOC100538023

zfand6
rpf1
CR847984.1
ccne1
CABZ01059627.2
gyg1a
zgc:175136
prdx1
elavl2
alyref
mibp
marcksb
pttg1
ube2c
nabp1a
gfer
zgc:171977
si:ch1073-190k2.1
h2afx
CU467646.2
si:ch1073-263o8.2
zgc:101016
imp3
sinup
lsm1
mylipa
cdca7a
thop1
cdc6
tph1a
snrpc
chchd10
zp3c
parn
zgc:171750

dctpp1
chia.4
pigrl2.3
tomm20a
zgc:56676
zglp1
rpa3
mcm7
chtopa
si:dkey-188i13.11
zc3h11a
ebna1bp2
gemin7
ncbp2
ddx3xb
CU499330.1
LOC110437731
grn1
lygl1
BX649485.2
c1qb
lgals2a
mrc1b
c1qc
cfp
CR855311.1
si:ch211-165i18.2
c1qa
si:dkey-5n18.1
ccl34b.8
cx32.2
ms4a17a.10
ccl34a.4
si:dkey-102g19.3
LOC100536946

spi1a
itgae.1
grna
timp2b
si:ch73-343l4.8
cxcr3.2
CABZ01001434.1
cdaa
samsn1a
dhrs9
moxd1
ccr12b.2
LOC564660
jac3
wu:fb64b08
si:ch73-343l4.2
si:ch211-264f5.2
cybb
sftpbb
cd22
zgc:153118
selp
clic2
cst14b.1
si:ch211-147m6.1
NPC2 (1 of many)
si:ch211-194m7.3
si:busm1-266f07.2
grn2
fcer1gl
spi1b
krt18b
mfap4
slc3a2a
ms4a17a.6

si:ch211-283g2.2
jac9
zgc:152830
lgals3bpb
BX649485.1
marco
mir21-1
calhm6
cxcl8b.1
smox
gpr84
mhc2dab
fgl2a
LO018493.1
ccl39.1
zgc:136870
ctss2.2
si:dkey-56f14.7
sb:cb81
si:ch211-9d9.8
cyba
ltb4r
ncf2
havcr1
tcnbb
tmsb1
enpp1
mapk11
abca1b
LOC100536983
zgc:64065
ppp1r14ab
hlx1
FP015862.2
pnp5a

f13a1b
lta4h
nrros
cd74b
m17
si:ch211-152p11.4
itgb2
plekho1b
gpx1a
cmklr1
cd83
slc16a9a
hdac9b
asah1b
cd74a
fosl2
ccl35.1
ms4a17a.9
si:dkey-262k9.4
atp6v1e1b
AL928650.1
ccl39.3
il1b
xpnpep2
slc6a6b
slc2a6
si:ch211-193e13.5
BX324132.2
lect2l
ly6m5
anxa5b
pmaip1
naga
prdx5
ncf4

txn
glb1l
zmp:0000000936
FP102158.1
atp1a3a
snx10b
si:dkey-7i4.24
si:ch211-74m13.3
XLOC-005448
LOC100537504
tcima
plbd1
stat3
myo1f
tgfbi
il4r.1
si:dkey-11f4.14
adgrg3
il10ra
rnaset2l
laptm5
vsir
parvg
LOC100150882
trib3
LOC101884267
si:ch211-15b10.6
cebpa
cxcr4b
atp6v1ba
nr4a1
rasgrp4
s100a10b
ccl39.2
entpd1

scpep1
ctsh
pgd
atp6v1g1
si:cabz01074946.1
dpp7
prcp
hexb
slc7a11
zgc:63831
icn
dnase2
klf2b
zgc:174904
unc93b1
crema
il4r.2
plin2
tcirg1b
BX248082.1
ets2
zgc:64051
slc7a7
ctsa
ccng2
acsl4b
atp6v0ca
si:ch73-248e21.5
bach1b
LOC100148254
cndp2
ugt5b2
XLOC-024516
gnpda1
rab44

ptgis
cebpb
fuca2
aplp2
sec61a1
si:dkeyp-23e4.3
tpp1
cd63
lxn
glipr1a
rhogb
si:ch211-66e2.3
dicp3.1
ssr3
arl5c
XLOC-023492
csf2rb
XLOC-005449
gpr183a
lpar5a
si:dkey-206d17.12
zmp:0000001082
nccrp1
cotl1
pdia3
fermt3b
tp53i11b
zgc:101715
si:ch211-1o7.3
ubl3b
myofl
gnsb
anxa2a
actb2
si:ch1073-429i10.3.1

si:zfos-1069f5.1
BX901920.1
snd1
atp7a
mef2cb
tmsb4x
rgs2
si:ch211-204c21.1
XLOC-000431
gm2a
atp6v1aa
arpc5b
cap1
antxr2a
plxnc1
echdc2
nfkbiaa
gapdhs
junba
derl2
pepd
ppiaa
psap
nfil3-6
tram1
tpi1b
ifngr1l
rgs19
dram2b
coro1a
itm2bb
slc2a3b
si:ch211-102c2.4
rps15a
gsdmea

ctsd
tspan37
arhgef5
stx11a
mboat2a
nfe2l2a
zgc:162730
tuba1b
srgn
arpc2
dbi
CT030188.1
lyn
si:zfos-741a10.3
tuba8l2
itgae.2
slc43a2a
rab11fip4b
canx
si:ch1073-443f11.2
crip1
anxa4
abcc3
ehd4
stat5a
tfe3a
gnao1a
si:ch73-248e21.7
bhlhe40
dbn1
klf13
wdr1
si:dkey-112e7.2
nme2b.1
si:dkey-25e12.3

csrnp1b
birc2
si:ch211-39i2.2
nocta
irf8
triqk
rpl36a
si:ch211-212k18.7
BX908782.2
zgc:103700
xbp1
atp6ap2
rassf2a
cflarb
atp6v1d
ssr4
rpl24
lcp1
rap1b
rps14
arpc1b
wasb
gadd45bb
wipf1b
rabgap1l
si:ch73-237c6.1
grap2b
suclg1
BX323596.1
rpl15
elovl1b
rps9
sec61g
rps21
gng10

rplp2l
rpl39
rnf144b
pfn1
ifi30
ahr2
flna
atf3
FP074902.2
rps20
cpox
vma21
si:ch1073-392o20.2
btg1
rps19
rpl35a
ppiab
rpl36
hcls1
gdi2
pkma
rpl29
pim1
rac2
celf2
osbp
zgc:114188
ccr9a
smdt1b
dnase1l1
wbp2nl
ostf1
rpl37
ehbp1l1b
cfbl

rpl18
eef1a1l1
rpl23
tagln2
XLOC-025423
rpl38
prdx6
metrnla
pycard
arpc4l
si:dkey-180p18.9
rps27.1
cox6b1
CABZ01073834.1
rgs18
rpl27a
rps26l
LOC103909115
eef1db
znf827
rplp1
rps23
clic1
hmha1b
rps16
ckbb
rps12
DHDH
ppp1r18
rps15
rab8b
bri3
sdcbp2
rps10
ppt1

rpl18a
grinab
skap2
mafbb
rpl30
pdk2b
rps29
arhgap15
mbd2
rpl28
rps5
jak1
CR936442.1
rps28
rpl26
mrtfab
rpl11
rpl23a
mdh1aa
zgc:110591
rpl9
rpl32
atp6v1h
fybb
si:ch211-74f19.2
ddit3
kdm7aa
rpl17
dusp1
npl
rpl10
dennd4a
oaz1b
si:dkey-27i16.2
pak2a

rplp2
ufm1
XLOC-042595
ssr1
krtcap2
rps25
CR392341.1
rps3
rpl19
aldoaa
rps8a
rpl10a
slc25a5
myh9a
rpl27
rps6
mob1a
abi2b
psme2
arpc5a
ncf1
arhgdia
FERMT3 (1 of many)
CABZ01068358.1
gnai1
rps7
rpl6
rplp0
rpl8
m6pr
krt8
cox8a
rpn1
sri
bloc1s6

rab1ab
slc7a3a
COX5B
rps27a
rps24
rpl31
vdac3
cfl1
rpl34
rpsa
si:dkey-53k12.2
slc20a1b
smim15
rpl22
syngr2a
rps4x
si:ch1073-325m22.2
tln1
rpl7a
capzb
egr1
gstm.2
rpl21
mapre1a
gnai2a
rps3a
rpl12
rps13
hsp90ab1
lgals3b
plp2
faua
rps2
csrp1a
cited4a

sucla2
ppib
gpia
atp6v0b
txnipa
dub
mt-nd5
zgc:162944
vdac2
capza1b
csde1
zgc:66427
si:dkey-238c7.16
mt-atp6
ssr2
arpc3
rack1
rnaseka
tsc22d3
XLOC-010897
lamp1a
cited4b
mfng
arhgdig
calr3b
COX7A2 (1 of many)
pcyox1
atp2b2
eef1g
eno1a
ldhba
zgc:153713
ncor2
ddost
rpl35

ier3ip1
msna
ap2m1a
uqcrq
mbnl3
ahnak
socs3b
atp5pd
mt-nd3
hdlbpa
ptprc
spcs1
rpl13
pde4ba
aoc2
COX3
spg21
adipor2
BX322618.1
adh5
etfb
rps18
gnb1a
capza1a
ccl39.7
rps11
gabarapb
ifi46
uba52
si:ch73-42p12.2
GCA
arrb2b
anxa1a
mt-co2
rpl5a

arf6b
si:dkey-79d12.5
zgc:158463
mrps15
rpl7
jdp2b
eif4ebp1
naca
actr2a
zbtb16a
atp1b1a
tmem258
mrpl33
ctsc
eif3k
fkbp5
capgb
tpt1
zeb2a
NDUFB1
nt5dc2
mt-nd1
ier2b
fkbp2
irf2
eif3m
commd7
eif3s6ip
cox4i1
eif3c
psme1
ier2a
XLOC-010899
midn
eif2s1b

selenof
eif3s10
chmp2a
ak2
rasgef1ba
ccni
zgc:158343
lasp1
rab5c
rpl22l1
cbr1l
atp5pf
snx1a
rhoab
rsl24d1
litaf
arpc4
twf1a
zfp36l1a
egr3
ndufa2
rps26
lgals9l3
ybx1
b2ml
ptmaa
zgc:77486
cox7b
CU929150.1
mibp2
XLOC-028370
ywhaba
ctsz
cox5ab
glo1

cyfip1
ddx21
mcl1b
tnfrsf1a
si:ch73-1a9.3
socs3a
mt-cyb
atp5f1d
BX005183.3
ccl34b.1
cyth1a
lamtor5
aimp1
calr3a
eef2b
pgam1a
rpl14
ola1
sumo2b
cct2
eif3ha
chmp1b
eif3f
cmip
reep5
mt-co1
cnn2
rpl4
btf3
rpl3
fosab
sec61b
gpx4b
selenok
atp5mf

atp5pb
eif3i
atf4b
si:dkey-42i9.6
mt-nd2
mknk2b
cox6c
stk17b
aldh9a1a.1
dad1
mgst1.2
cox6b2
ube2v2
sec11a
mtdha
eif3ba
magt1
sh3bgrl3
nsa2
tomm5
myl12.1
nr4a3
elf1
ndufb3
psmb2
alas1
ost4
eif3ea
glrx
atp5l
vps29
rps17
rnf10
aimp2
cox5aa

tmed2
pfdn4
cct7
cox6a1
pnrc2
pomp
pgk1
anxa11b
atf4a
itm2ba
cct8
elmo1
hnrnpa1a
metap2b
rnf213a
mt-nd4
tnfb
uqcrh
mcl1a
cox7a2l
tomm20b
ndufa4l
anxa3b
odc1
ponzr1
dusp5
ipo7
gtpbp4
fam120c
ndufa3
rpl13a
pdia6
b2m
si:dkey-7j14.6
si:ch211-196l7.4

p4hb
psma2
sat1a.2
prdx2
rsl1d1
rps27.2
zgc:56493
zgc:193541
eif3ja
nop56
rps27l
cct5
map3k14a
psma8
csnk2b
tmed10
arl6ip1
serbp1a
eif3g
eif4a1b
atp5mc3a
arf2a
atp5po
abracl
cox7a2a
eif3d
idh2
cd9b
eif2s3
cct3
ywhabb
serf2
atp5md
rpl5b
nop53

gng5
cox7c
NC-002333.4
atp5f1c
tcp1
slc25a3b
tegt
romo1
eef1b2
CR383676.1
hspa8
atp5mc3b
pabpc1a
ywhaz
eif2s2
atp5fa1
atp5meb
npc2
histh1l
fkbp1aa
ecm1b
eif1b
higd1a
ppdpfb
chac1
eif5a
CR318588.1
eef1da
ucp2
ptmab
dcn
col6a4a
asip2b
c6
hsd11b2

col1a1a
col1a2
rbp4
si:ch211-106j24.1
col6a3
col5a1
fn1b
postnb
c1qtnf5
mmp16a
col1a1b
meis2b
zgc:113531
crispld2
si:ch1073-291c23.2
mfap2
si:ch1073-459j12.1
pbx1a
col6a1
pmp22a
pdgfra
si:ch73-46j18.5
fstl1b
si:ch211-270g19.5
tshz3b
col6a2
wfdc1
XLOC-041269
loxl2a
BX640522.2
col4a1
si:ch211-106h4.12
nrp1a
mxra8a
nid1b

lrp1aa
igf2b
nid2a
aebp1
sparc
ccl25b
col5a2a
prss23
adamts10
pdgfrb
gas1a
cebpd
slco2b1
tmem119b
cyp2ad3
cxcl18b
cd81a
si:dkey-261h17.1
CABZ01092746.1
ptges
abi3bpb
tmem176
col16a1
mdka
cldn11a
cpn1
fosab
vim
adh8a
zgc:195173
cald1a
rhoua
tagln
BX569787.1
socs3a

col12a1a
zgc:112437
jun
ece1
lrp1ab
mmp2
bhmt
serpinf1
nfixb
mrc2
dhrs3a
angptl1a
CR383676.1
mfap5
lxn
nfia
pcdh18b
igfbp5b
nr2f2
tenm3
s100a10b
inhbaa
lrrc17
foxo3b
colec12
adamts15a
col12a1b
slc38a5b
pcolcea
cdon
junba
p3h2
marcksl1a
ms4a17a.8
mmrn2a

mir199-3
si:dkey-163m14.2
sult1st6
c1qtnf6a
boc
si:dkey-9i23.16
ier2a
rbp5
mmp14a
cfd
BX322530.1
fibinb
inka1a
slc7a1
mmp14b
chd3
lpar1
cd82a
si:ch211-198c19.3
FP102783.1
anxa1a
eva1ba
ptprga
fblim1
cyp3c1
gpr27
c4
cyp2p7
cyp2n13
col15a1a
tsku
gadd45ba
tmem176l.1
ephb3
cldn5a

cyp2p6
svild
cyp27c1
icn
hoxc9a
clec11a
loxl2b
wu:fj16a03
emp2
tshz2
dkk3b
tnfsf10
nupr1b
lama4
zfp36l1b
zgc:162730
tril
tnfrsf19
dedd1
fosb
col2a1b
LOC103909284
rgs7bpb
meis1b
b2ml
pabpc1b
timp2a
ccbe1
krt8
f3b
myl9b
fam198a
ip6k2b
efnb1
dhrs3b

hic1
cnn3a
endouc
sox4a
si:ch211-212k18.9
fzd7b
ldlrad4a
foxp4
lgals2a
pbx1b
ahnak
mgst1.1
id1
tns2a
olfml3b
sema3d
col18a1a
yap1
ntn1a
krt91
ppib
aoc2
mmp15a
tnfsf12
nfil3-6
vcam1b
cygb1
map1lc3a
s100v2
vdrb
lamb1a
pdk2a
si:dkey-7j14.6
bgnb
ar

aqp3a
pdgfrl
rgs5a
f10
frzb
cd248a
XLOC-016094
rerea
sostdc1a
atoh8
mdkb
spry2
itgb1a
spred2b
rbms3
krt94
aldh9a1a.1
hoxc3a
antxr2a
cox4i2
spon1b
si:ch211-212k18.8
deptor
tpm4a
smo
fzd2
gstt1b
CU929237.1
pid1
si:ch211-133n4.4
slit3
ciarta
ghra
BX663516.2
grb10b

ctdsp2
pik3r2
ch25h
egr1
hmgn6
LOC101882334
pfkfb4b
grb10a
edil3a
tcf12
nr2f1a
cxcl12a
grhl1
scube1
zcchc24
ccdc3b
stau2
cdh2
celf2
kirrel1a
egfra
tns1b
h1f0
arrdc3a
vasna
FO744854.1
myl6
txnipa
rbfox1
ier5
SCARF2
qkia
ackr3b
nfic
ulk2

ptenb
zgc:114188
zgc:171679
h3f3c
zgc:158343
plpp3
krt18b
selenow1
si:ch73-335l21.4
btg2
mcl1b
entpd1
mgst1.2
bnc2
fxyd1
lamc1
robo2
socs3b
ddit3
si:ch211-39i22.1
ip6k2a
junbb
znf385a
osr2
myl9a
rnd3b
ppap2d
CABZ01074363.1
itm2ba
cyr61
sesn1
CR847566.1
fkbp5
rps27.2
FO082781.1

zeb1b
mt-nd4
appb
si:ch73-1a9.3
sema4ba
klf2a
si:rp71-79p20.2
jdp2b
gstm.3
creg2
TCIM (1 of many)
gpx4a
rpl9
crim1
c7a
CABZ01080702.1
adh5
nme3
cavin2b
serinc5
COX3
add3a
ptprk
dbpb
pnrc2
vasnb
ccdc85b
rpl36
cux1a
si:ch211-222l21.1
flot2a
rpl29
s1pr2
si:ch211-132b12.7
hspb1

gng7
mknk2b
mt-nd2
reck
cd28l
map1lc3b
selenbp1
klf11b
ablim1b
rpsa
mt-atp6
BX663503.3
rps16
rplp1
mgst3b
h2afx1
htra1b
il15
zgc:92744
tnrc6c1
rbm39a
h2afy2
rps28
klf6a
zgc:113208
tgfbi
mafba
rpl34
ybx1
NC-002333.4
mt-cyb
rpl27a
hmgb1b
mt-nd1
flna

abcc5
mt-co2
lum
CABZ01068358.1
nsa2
slc43a2a
si:dkey-51e6.1
sh3pxd2b
tsc22d3
rpl24
wls
si:ch211-204c21.1
hmgb1a
rpl6
rps15
rps13
anxa4
ghrb
zgc:174863
cnbpb
itpkcb
spry4
si:dkey-28b4.7
b2m
eif4ebp3
si:ch211-196l7.4
rps2
pik3r3a
pgrmc1
rps21
rps18
ms4a17a.9
rpl7a
jpt1b
rpl28

eif4ebp3l
arl6ip5b
si:ch211-105c13.3
rsrp1
sdc4
krt15
krt18a.1
fam168a
tomm5
rplp2
hmgn7
gstm.1
angptl4
per1b
hist2h2l
hdlbpa
eif4a1b
rps27l
rpz5
eef1g
wu:fb16f03
rgs5b
rpl11
XLOC-000431
atf4b
si:ch211-132p1.3
rps10
cd82b
rps24
rps26
sdc2
rps4x
gbp
akap12b
zbtb20

sfrp1a
rps27a
si:ch211-140m22.7
rps14
cry2
slc3a2b
si:dkey-11f4.14
rpl23a
CABZ01072614.1
mt-co1
adam10a
rplp2l
rpl18
ier2b
rpl17
CR855311.1
rpl22l1
rpl30
si:dkey-188i13.10
fxyd6l
serinc1
capns1b
rpl14
si:dkeyp-19e1.3
uba52
rpl31
rpl38
vdac3
btg1
irf2bp2a
marcksl1b
rps3
zgc:111986
romo1
bhlhe40

pim1
serf2
cebpb
ppdpfb
eif1b
jund
tmsb4x
vat1
mycb
pdk2b
tob1b
pabpc1a
rps26l
mir21-1
si:ch73-376l24.3
rps25
tpm1
rpl5a
pno1
mt-nd3
pdia3
klf13
nop53
eif3d
nfkbiab
eef1db
naca
serbp1a
ccni
rpl5b
glula
ywhaz
eef2b
acin1a
csrnp1b

eef1da
tcima
btf3
rsl24d1
ccl33.3
sftpba
npsn
si:ch211-220f21.3
FO393424.2
si:dkey-222f2.7
XLOC-018356
fcer1g
CABZ01046427.2
vaspa
dicp1.1
CT573231.1
si:ch73-380l3.4
ccl35.1
si:ch211-102c2.4
fcer1gl
rnaset2l
si:ch211-214k5.3
arpc1b
CABZ01046432.1
coro1a
pfn1
si:dkey-172o19.2
pou2f2a
LOC101882805
lcp2b
gpr132b
ephx2
spi1b
kctd12.2
si:dkey-33i11.9

CU972454.4
actb2
mef2b
cotl1
bpifcl
LOC100536213
wipf1a
si:ch211-214p16.1
cd74a
cd74b
XLOC-028370
tmsb4x
LOC100535743
arpc5b
abi3a
itga1
zdhhc23a
XLOC-020301
ptmaa
si:busm1-266f07.2
CU855821.1
yrk
grap2b
si:ch73-264p11.1
fam49ba
LOC108190717
rac2
si:ch73-380l3.2
arpc3
lcp1
zgc:64051
wasa
arpc2
cnpy3
BX120005.1

hcls1
ptpn6
sftpbb
ppp1r18
arpc4l
ptprc
arpc4
actb1
il7r
mhc2dab
si:dkey-9i23.4
arl11
dtnbp1a
CR381673.1
fgd4a
XLOC-026836
plekho2
arhgdig
rplp2
rhogb
fam49al
myl12.1
CR753876.1
capzb
rgs13
capza1b
XLOC-025423
cfl1
si:ch211-243a20.4
washc2c
si:ch211-214p16.2
b3gnt2l
LOC101885445
ipcef1
AL954142.1

XLOC-032002
tln1
wasb
apbb1ip
cnn2
zgc:162730
eomesa
syk
ppiab
mir223
rab44
scgn
mpeg1.1
laptm5
si:ch211-66e2.3
hck
cers1
LOC103909115
itm2bb
si:ch211-153b23.7
ccni
ccr2
mrtfaa
ptmab
naca
ckbb
cyba
gm2a
si:ch211-277c7.6
eif3f
grasp
rpl26
stk17b
RASSF5
il10ra

mob1a
selplg
mcl1a
rps29
dok2
XLOC-043760
gabarapb
tagapa
rgs18
zgc:174945
mapre1a
si:dkey-262k9.4
myh11a
ucp2
srgn
rps27.1
tagapb
cd83
ptprea
rps2
lasp1
skap2
si:ch211-119e14.1
ppiaa
pnrc2
h1fx
crip1
itgb2
abracl
rpl22
si:ch211-105j21.9
hsp90ab1
alox5a
LOC101884267
nr4a1

fybb
myo1f
hmgn2
gapdhs
rpl32
CABZ01109481.1
rps18
rpl38
calm1b
rps10
btg1
tpi1b
rpl15
rap1b
rpl36
elf1
rpl34
csk
BX005183.3
si:dkey-112e7.2
mbnl1
rpl35a
arhgap15
rsl24d1
mfng
rpl8
cap1
rps21
rps15a
cdkn1bb
zgc:153867
nptna
si:ch211-213a13.1
rps17
rps6ka1

rpsa
cox8a
idh2
arl5c
cox8b
rps16
rpl14
rpl10
XLOC-023492
rps5
rpl11
vsir
romo1
limd2
parvg
actr2a
cd63
lrrfip1a
wdr1
fgl2a
arhgdia
gng5
plekhj1
twf1a
zgc:152791
rps26l
rpl9
zgc:92744
prkar2aa
rps15
rpl18a
clic1
eif4ebp1
rps14
litaf

arrb2b
arf6b
rpl6
rpl30
cxcr4b
si:ch1073-429i10.3.1
sik1
ak1
scinlb
rplp2l
pde4ba
ifi45
rpl31
cox6b1
sumo2b
sept6
pabpc1a
msna
cdc42l
zgc:114188
capza1a
btf3
rps27.2
serf2
hif1ab
atp6v1g1
cox4i1
mtpn
zfp36l1a
oaz1b
nsa2
cdc42
glo1
CR318588.1
c1qb

c1qc
si:dkey-5n18.1
CU499330.1
grn1
lygl1
c1qa
ccl35.1
mhc2dab
ms4a17a.10
cmklr1
lgals3bpb
zgc:174904
havcr1
mrc1b
sftpbb
itgae.1
grna
marco
si:ch211-165i18.2
LOC100537504
NPC2 (1 of many)
cx32.2
si:ch211-147m6.1
ccl34b.1
slc3a2a
ctss2.2
ccl34a.4
CU929150.1
zgc:64065
si:ch211-194m7.3
mafbb
si:ch211-283g2.2
slc7a7
abca1b
CABZ01076275.1

zgc:153118
mfap4
lgals2a
si:ch73-158p21.3
si:busm1-266f07.2
adgrg3
cfp
si:zfos-2330d3.3
zgc:103700
si:zfos-1069f5.1
rnaset2l
naga
spi1a
cd83
tcima
zgc:136870
LO018493.1
si:ch211-74m13.3
aatka
cxcr3.2
il10ra
fgl2a
asah1b
fcer1gl
cd74a
zmp:0000000936
zgc:152830
slc6a6b
cd74b
stoml3b
plin2
BX649485.1
plekho1b
slc40a1
tmem163a

pmaip1
ctsh
mhc2b
tpp1
slc16a9a
dnase2
CR759843.2
csf2rb
grn2
si:busm1-194e12.11
smox
FP015862.2
plaua
si:cabz01074946.1
rbp2a
mpeg1.1
gng10
si:dkey-56f14.7
krt18b
nrros
psap
si:dkey-27i16.2
atp7a
cndp2
si:ch211-39i2.2
fuca2
cxcr4b
clic2
LOC560447
ifi30
si:dkeyp-23e4.3
gnsb
ms4a17a.6
slc2a6
nr4a1

p2rx7
robo4
calhm6
scpep1
AL928650.1
si:ch211-66e2.3
spi1b
slc43a2a
lipf
gm2a
npl
tmsb1
gpr84
mir21-1
ctsa
enpp1
mpp1
zgc:153317
ccng2
si:dkey-11f4.14
sb:cb81
CABZ01056054.1
ctsk
si:dkey-23a13.11
BX324132.2
crema
nfil3-6
sowahd
tnfb
abi2b
rassf2a
laptm5
CR855311.1
arl5c
plbd1

fosl2
si:ch73-236c18.7
ctsd
ccr9a
cybb
dpp7
ncf2
lta4h
si:ch211-212k18.7
LOC108190616
rab44
anxa3b
cebpb
pepd
XLOC-024516
cd22
atp6v1e1b
si:ch211-1o7.3
ccl38.1
samsn1a
dusp1
rab8b
mertka
atp6v0ca
mbnl3
slc43a2b
itgae.2
timp4.3
snx27a
wdfy4
tmsb4x
si:ch211-8c17.3
glb1l
si:ch211-204c21.1
CABZ01078737.1

itm2bb
LOC101885706
bach1b
ctsc
zgc:162730
f13a1b
slc43a3b
pdk2b
slc36a1
cebpa
si:ch73-343l4.8
kctd12.2
LOC101885065
trpv1
XLOC-023492
mapk11
tnfrsf21
LOC100148254
tuba8l2
rgs19
gpx1a
dbn1
atp6v1d
cadm4
FERMT3 (1 of many)
tnfrsf18
zgc:64051
XLOC-005449
slc7a8a
cyba
tnfrsf1b
lgmn
rxrga
BX511021.2
timp2b

gstt1a
atp6v1g1
actb2
si:dkey-24p1.7
si:zfos-741a10.3
ets2
grapa
wasb
ehbp1l1b
ncf4
gnpda1
tmem86a
slc30a1a
sdr16c5b
tspan37
zgc:63831
bri3
si:dkey-262k9.4
cflarb
mir22a
zfp36l1a
entpd2a.1
ctsz
ehd4
arpc5b
dennd4a
si:ch211-15b10.6
zgc:123107
BX649485.2
ifngr1l
rnaseka
si:ch211-152p11.4
csrnp1b
cd63
FP102158.1

tspan36
m17
dicp3.1
git2b
wipf1b
mgat4a
pcyox1
CABZ01068358.1
arhgef12b
mfsd1
vsir
XLOC-000431
xaf1
junba
rac2
nck1a
cds1
nocta
txnipa
si:dkey-206d17.12
BX248082.1
rgs2
lyn
mt-co1
aplp2
metrnla
egr3
pfn1
klf13
irf8
rbm47
dhrs9
flvcr2b
LOC101886114
nfe2l2a

si:ch73-248e21.5
cotl1
acsl4b
slc1a3a
atp6v1ba
COX3
CABZ01001434.1
klf6a
ddit3
tfe3a
me2
si:dkey-25e12.3
ccl34b.8
coro1a
nfkbiaa
abcc5
rnf130
mcl1b
gabarapb
btg1
si:ch211-102c2.4
abca1a
LOC101884267
LO018340.1
si:ch1073-392o20.2
fybb
LOC100536946
mfng
rnf144b
cap1
grap2b
bpifcl
itgb2
il4r.1
sdcbp2

arrb2b
slc20a1b
cst14b.1
b2ml
txn
mboat2a
stat3
irf2
atp2b2
slc39a1
pparg
si:ch211-243a20.4
gapdhs
tcirg1b
crip1
si:dkey-222f2.7
CABZ01073834.1
mt-co2
rgs18
sat1a.2
si:dkey-112e7.2
cd2ap
me1
zbtb16a
si:ch211-212g7.6
tacc1
jdp2b
rasgrp4
mt-cyb
abcc3
FBXO48
arpc1b
il1b
lcp1
celf2

CABZ01020840.1
selplg
dnase1l1
arpc2
b3gnt3.4
kdm7aa
plek
cul5a
ltb4r
adipor2
hexb
anxa4
rhogb
clic1
egr1
slc15a4
lamp1a
klf2a
pfkfb4b
si:dkey-53k12.2
xpr1b
sdc4
LOC110437731
ndfip1l
CR318588.1
CR936442.1
ctsba
plxnc1
elovl1b
si:dkey-79d12.5
b2m
unc93b1
tnfaip3
fabp11a
tsc22d3

marcksl1a
unm-sa1614
chmp4ba
ch25h
rasgef1ba
myo1f
npc2
skap2
ugt5b2
rplp2
arpc4l
midn
zgc:77486
si:dkey-69o16.5
sh3pxd2b
jund
BX120005.1
nr4a3
BX901920.1
CR383676.1
gnao1a
elmo1
gadd45bb
arhgef5
atp6ap2
si:ch73-237c6.1
TPGS1
prss16
nt5dc2
ptprc
gpr183a
ypel1
mgrn1b
pnrc2
mt-atp6

slc12a2
mcl1a
rap1b
LOC103910136
ucp2
mknk2b
mef2cb
hdac9b
eif3jb
bhlhe40
ms4a17a.9
lgals9l1
osbp
litaf
lpar5b
snx9b
eif4ebp1
si:dkey-7i4.24
zmp:0000001082
mdh1aa
dusp5
fosab
si:ch211-136m16.8
trib3
arl6ip1
CABZ01055347.1
LOC103911430
mt-nd4
mt-nd3
parvg
efhd2
ppt1
npc1
rab5aa
rps15a

pycard
CR392341.1
atp6v0b
ubl3b
si:ch1073-429i10.3.1
si:dkeyp-68b7.12
rnf213a
bloc1s6
actr2a
sik1
zgc:158343
tmem131
si:ch211-106a19.1
gnai2a
NC-002333.4
chmp2a
cpox
dusp2
pnp5a
tram1
si:dkey-102g19.3
XLOC-010897
arhgdig
si:ch211-215e19.3
wdr1
CABZ01080702.1
pim1
hcls1
lxn
BX005183.3
atp1b1a
prdx5
chmp4bb
mgst1.2
grasp

histh1l
lpar5a
XLOC-010899
abi3a
mapre1a
rhoab
smdt1b
rps27.1
wbp2nl
tln1
ppiaa
mbd2
cfl1
glmp
rpl39
arhgdia
zgc:174863
rps20
itm2ba
oaz1b
mob1a
itga4
cited4b
nfkbie
mtss1la
pkma
BX322618.1
rpl36
capza1a
mt-nd1
wasf2
ier2b
XLOC-025423
m6pr
fosb

znf827
eef1db
rpl37
ifi46
gdi2
rpl38
arl8bb
XLOC-033137
BX908750.1
tpi1b
si:ch211-198c19.3
gnai1
ube2v2
zgc:110591
pdcd6ip
psme2
ostf1
rpl22
ahnak
slc35f6
zfyve21
itpkcb
rpl24
syngr2a
COX7A2 (1 of many)
tagln2
igf2b
dusp6
krt8
si:ch211-196l7.4
chmp1b
XLOC-028370
derl2
gabarapa
ptenb

elf1
sec61a1
LOC103909115
rap2b
zgc:114188
si:dkey-56m19.5
rps21
rpl35a
irf7
arpc3
si:rp71-36a1.5
nck1b
sqstm1
rarga
xbp1
stk35
hif1ab
pdia3
si:ch211-260e23.9
cyfip1
ppiab
cdc42l
rps29
rps2
sec61g
rpl18a
myl12.1
glula
mt-nd5
rab1ab
jak1
fxyd1
rpl11
sptlc2b
cox8a

lamtor5
uqcrq
si:ch1073-325m22.2
hmha1b
GCA
XLOC-042595
anxa5b
DHDH
rps9
nme2b.1
tmem59
glud1b
rps26l
fkbp5
srgn
grb2b
twf1a
slc37a2
si:ch211-191c10.1
msna
cd9b
arpc4
vat1
rpl29
rps23
faua
si:dkey-180p18.9
dync1i2a
rps14
ktn1
rpl36a
stat2
rpl35
rps15
tegt

rpl18
rps5
tagapb
rpl27a
commd3
coq10b
rab5c
rpl15
kctd5a
vdac3
ptmaa
ier2a
hif1al
rgs5b
pgd
rpl32
si:ch73-1a9.3
ppdpfb
maff
rps28
rps10
tob1a
ip6k2b
gnb1a
wee2
capzb
slc25a5
romo1
zgc:162944
zgc:153713
zgc:153911
higd1a
rpl30
grinab
snd1

cnn2
ptp4a2b
rpl9
notch2
sh3bgrl3
rps18
capza1b
atf4b
csde1
wu:fi37e09
antxr2a
dbi
mt-nd2
rplp2l
s100a10b
rpl23a
cox7b
rpl28
atp5pf
icn
COX5B
rpl31
rpl7a
vma21
ccni
slc3a2b
rps19
klf11b
glrx
ndufb3
rps27a
arf6b
rps16
cox4i1
nccrp1

rhoaa
eif3f
eif3m
tnfrsf9a
btg2
bsg
alas1
vmp1
rpl34
rps27l
mafba
rps27.2
socs3a
naca
rps25
ckbb
nsa2
ssr3
rpl21
chac1
rps4x
ndufa4l
rpsa
rpl8
ywhaba
rpl27
rps17
rpl6
fthl27
atp5l
tpm3
rps26
rpl17
uba52
rpl10

NDUFB1
zgc:193541
rps24
cox6c
zgc:56493
zgc:92744
ppib
eif1b
rps11
rps13
rps3
btf3
rpl10a
rpl5a
h1fx
pabpc1a
rsl24d1
rpl14
cox6b2
mibp2
ctss2.1
serbp1a
eef1g
cxcl12a
podxl
krt15
nid1b
aldh1a2
c6
tmem88b
postnb
slco2b1
XLOC-016094
il15
akap12b

hspb1
ndnf
ace
jam2b
slc29a1a
col12a1b
c4
gpr17
wt1a
ptprk
grhl1
tmsb
si:ch211-191i18.2
cd151
gria3b
vegfd
foxl2a
phc2a
ctgfa
arxa
CR786577.1
steap4
cyp19a1a
krt18a.1
csf1a
entpd1
CR751602.2
s1pr3a
fn1b
tspan9b
lpar1
FO744854.1
bnc2
cldn5a
spon1b

c7a
wt1b
anxa2a
cebpd
sh3pxd2aa
si:ch211-204c21.1
itga9
rbms3
mao
vasnb
lrp1ab
col6a3
BX322530.1
tmem98
phactr4a
adma
cdh2
eppk1
krt94
s1pr2
si:ch211-197h24.9
CABZ01080702.1
gabra5
si:ch211-106h4.12
vcam1b
ugt1b5
pdgfc
gstm.3
met
pbx1b
si:dkey-56m19.5
hsd11b2
tmem176
cd81a
lama5

BX569787.1
kirrel1b
ghrb
mmp16a
CR749775.2
krt8
cygb1
cxcl14
paplna
vat1
nid2a
mmp2
pleca
nr2f1a
wnt11r
cavin2b
itm2bb
sox4a
abl2
si:ch211-198c19.3
c4b
fstl1b
cd248a
AL954191.1
fabp11a
zeb2b
ptprga
bmp3
adamts2
rgs7bpb
ftr82
pabpc1b
gdpd4a
col6a1
ahnak

clu
fhl3b
col5a1
tfpia
zfp36l1b
add3a
gna12a
tns2a
arhgap42b
myl9b
gas6
col6a2
dkk3b
si:ch211-136a13.1
tmem176l.1
anxa1a
rasgef1bb
jcada
clic4
slc43a3b
cfd
ext1a
bmp4
f3b
zbtb16b
mtss1la
gpc4
aadac
pard6a
ywhag1
antxr1c
slc7a6
f10
sorbs1
mxra8a

acvrl1
aebp1
sparc
eva1ba
nr2f2
pgr
wu:fc27b11
adamts8a
tagln
CU539058.1
hdac4
slc43a2a
si:ch73-46j18.5
pdgfrb
hspg2
rbp4
cald1a
col1a1a
tnfaip6
atp1a3b
lxn
col27a1b
ppdpfa
jun
svild
dag1
dchs1a
cxcl18b
grb10b
rhobtb4
CABZ01068366.1
ptprua
junba
greb1l
dcn

nr0b1
socs3a
gstt1b
pmp22a
col1a1b
clec11a
fosab
ms4a17a.8
si:ch73-22o12.1
scarb2a
xirp2a
col18a1a
id1
stard13b
tspan4a
eif4ebp3l
col1a2
s100a10b
foxo3b
si:dkey-7j14.6
rab6ba
FP016056.1
mcl1b
eif4ebp3
cav1
desi1a
mir199-3
si:dkeyp-9d4.2
s100v2
mfap2
arhgef5
colec12
si:dkey-28b4.7
si:ch211-204a13.2
sema4ba

agrn
gata6
selenbp1
enc1
grip1
zgc:110182
loxl2a
col21a1
ugt5b2
XLOC-024248
lipf
tcf12
dachd
tmem204
mknk2b
si:ch211-250c4.4
plod2
CR383676.1
slc38a5b
cnn2
robo2
dlc1
ptprsa
CABZ01068367.1
pdgfrl
ywhaz
si:ch211-222l21.1
pfkfb4b
nudt4a
gsta.1
tnfsf12
znf385a
phldb2a
neo1a
col5a2a

CR848833.1
lrp1aa
hdlbpa
frzb
cavin1b
zgc:195175
mgst1.2
rhpn2
fam171a2a
epb41l5
CR318588.3
XLOC-034538
anos1a
nrg2a
tle2b
rasgef1ba
nrxn1a
egfra
il6st
gulp1a
zgc:162730
chd3
si:ch73-86n18.1
cntfr
lamb1a
gadd45ba
olfml3b
stxbp6
robo1
deptor
si:ch211-1a19.3
rhoj
si:ch211-74m13.3
ksr1b
phactr4b

marcksl1b
pi15a
wbp1lb
appa
hipk2
tjp2b
ppp1r14bb
mir21-1
itga1
ephb3
kank3
cdc42bpb
si:dkey-9i23.16
usp43b
ar
mdka
hbp1
pik3r2
mxra8b
LOC101884451
si:dkey-58f10.11
ccdc85b
tiam1a
b2ml
gamt
si:dkeyp-75h12.5
prss35
nr1d2a
NAV1 (1 of many)
hspb8
ifi46
txnipa
mogat2
mob2a
COX3

pgrmc1
pkp3b
kank2
calcoco1a
itgb5
zgc:195173
mt-nd1
sox6
dpp3
fkbp5
ptpn21
adam10a
mt-co2
si:ch211-212k18.8
mmp14a
rpz5
si:dkey-11f4.14
cybrd1
egr1
FO082781.1
nfil3-6
mpp5a
mt-nd4
tcf4
plxnb2a
fzd2
actn1
ctdsp2
tnfrsfa
calm2b
mt-atp6
mt-nd2
nt5c2b
si:ch211-140m22.7
lrrc1

pdk2a
creg2
dpyda.1
fkbp7
BX663516.2
tspan14
mt-cyb
sptan1
sik2b
itgav
serinc1
fbn2b
zbtb46
si:dkey-6n21.12
rbpms2a
tpm4a
plpp1a
sh2d3cb
si:dkey-101k6.5
myo1cb
nrp2a
atoh8
kmt2cb
cyp2p7
cgnl1
rab11fip1b
reep3a
si:dkey-188i13.10
erbb2
cyp2ad3
mt-nd3
grinab
ralgps2
b2m
cbx7a

wls
cyp3c1
palm2
hmgn6
junbb
mt-co1
cyr61
itm2ba
atp10a
serinc5
tjp1a
mphosph8
ngdn
si:rp71-79p20.2
ier2a
si:ch211-74f19.2
si:ch211-212k18.9
btg2
pard3bb
nid1a
lamc1
sypl2a
afdna
si:ch73-237c6.1
pdlim1
pbx4
si:ch73-1a9.3
si:ch211-196l7.4
gng7
ccl25b
znf609b
pabpn1
itgb1a
celf2
zbtb4

h1f0
net1
NC-002333.4
jak1
zgc:66433
hcar1-4
si:dkey-42i9.7
si:ch211-199g17.2
si:ch211-166a6.5
ctbp2a
si:ch211-132p1.3
zgc:174863
serinc2
si:ch211-15b10.6
rsrp1
tfe3a
marcksl1a
ier2b
map1lc3a
crebzf
h3f3c
rsu1
si:zfos-464b6.2
mgst1.1
arhgef1b
ddit3
tspan18b
igfbp7
klf6a
mfap5
ifi45
cast
fermt2
lama4
aldh9a1a.1

fam168a
stat1a
selenow1
sh3pxd2b
anxa4
oip5-as1
si:ch1073-291c23.2
ppfibp1a
arhgef4
wwtr1
tp53inp1
tmsb4x
flna
si:ch73-335l21.4
ulk2
hmgb1b
si:dkey-23i12.7
krt18b
nfia
stk35l
ecm1b
atf4b
dhrs13l1
map1lc3b
socs3b
si:ch211-286b5.5
sdcbp2
SBNO2
pi4kb
tefa
fbln5
rnd3b
CR847566.1
meis1b
irf2bp2b

cflara
dedd1
CR318588.1
timp2a
gpx1a
fundc2
qkia
ca5a
anxa5b
klf11b
cebpb
ppap2d
zmiz1a
map2k6
si:ch211-133n4.4
scara3
tmem176l.3a
boc
esr2b
mvp
nupr1b
pnrc2
si:dkey-17m8.1
appbp2
cnn3a
pdk2b
zbtb20
rnf19a
emd
si:ch1073-459j12.1
gstm.1
nr3c1
wu:fj16a03
cebpg
arl4d

hspe1
serpinf1
anxa13
osr2
jdp2b
cpn1
wu:fi37e09
crim1
stk35
XLOC-043861
ccng2
ier5
ndfip1l
klf2a
tnfrsf1a
zgc:113208
ppp1cb
vdrb
ak1
nckap1
tax1bp3
ube2e3
CABZ01072614.1
rhoab
ponzr1
macf1a
hopx
si:dkey-51e6.1
CABZ01068358.1
ppdpfb
sdc4
wu:fb16f03
flot2a
ecrg4a
capns1b

yap1
csrp1a
rhoaa
ctnnb1
pik3r3a
slc7a1
gmfb
mafba
adamts15a
rbm39a
si:ch211-213a13.1
nt5dc2
rnf213a
tagln2
si:ch211-214p16.2
si:ch73-376l24.2
zgc:114188
ppib
capn2a
irf2
fxyd1
mt-nd5
psme2
tsc22d3
tnrc6a
rerea
si:ch211-260e23.9
acin1a
abca1a
stau2
XLOC-024750
lrpap1
slc3a2b
aplp2
notch2

mycb
ptp4a1
capns1a
jpt1b
gpatch8
rplp2l
hif1al
stat2
srsf5a
dynlt3
rps28
mgst3b
sumo2b
ubb
prkab1b
atrx
snx17
jund
rps13
rab5aa
fkbp1aa
smdt1b
tpm1
tomm5
gapdhs
si:dkey-42i9.4
fam120c
plk3
pim1
ywhaqb
eif1b
pfn2l
sh3bgrl
btg1
si:ch211-191j22.3

atp1b1a
hmgb1a
rplp2
zgc:111986
rps27l
si:ch211-214j24.10
antxr2a
glula
rps24
cldn11a
dbi
calm1a
hnrnpa0a
ciarta
nfkbiaa
tpm3
rpl36
chac1
higd1a
pomp
rab10
sept2
anxa11b
capza1a
rap1b
fabp3
bhlhe40
tegt
ndufa4l
fosb
tmem258
ywhabl
tnrc6c1
zgc:153867
eef1db

ptmab
cd63
rpl9
hmgn7
zgc:158463
hmgn2
rps21
LOC103910438
zgc:92744
rps18
glrx
gnb1b
rps10
rpl14
oaz1b
rps27.2
ptmaa
rpsa
serbp1a
ucp2
rpl22l1
eef1da
rpl23a
rpl31
rpl27a
pabpc1a
uba52
rpl5a
si:ch73-167f10.1
sycp1
stmn1b
si:dkey-40g16.5
tuba7l
smc1b
LOC100149647

CU929391.1
hsp90aa1.2
hormad1
sycp3
ddx4
hnrnpa3
si:ch211-176l24.4
rnf212b
pcp4l1
stag3
LOC101886261
parp1
CCNB1IP1
si:ch211-241j8.2
ddx39b
cox5b2
zgc:193690
zte38
spata22
tdrd1
chtopb
piwil1
CU927934.3
zgc:194246
gcshb
hmgb2b
adad1
mrto4
mgmt
CABZ01080371.2
tdrd9
dpy30
dmrt1
si:ch211-277c7.7
dut

stmn1a
si:dkeyp-93d12.1
syce2
dnph1
si:ch211-173m16.2
tuba1c
calm2b
iqck
fanci
si:dkey-276j7.1
ddx43
terb1
top2a
pld6
piwil2
rnf17
CABZ01083945.1
si:ch211-222l21.1
mycbp
chd4a
lsm6
lsm7
h2afx1
mcm3
gpd1b
iqcb1
sub1b
nsmce1
llph
zgc:161973
odf3l2a
snrpd1
adad2
morc3a
trmt112

isg15
srsf2b
si:dkey-4p15.3
zgc:162025
esco2
acot13
chaf1a
minos1
hmgb1b
rbx1
snrpf
stk31
CR382383.2
snrpg
sms
gnl3
zgc:153681
srsf7b
tdrkh
rpa2
hirip3
pole4
nub1
parp3
smfn
snrpd3l
cmss1
ssrp1a
ppih
si:dkey-119f1.1
auh
ccnd3
hells
sart1
lsm4

cpsf2
tdrd6
odc1
dlgap5
psmb7
ndufb6
dcun1d5
lrrc20
hpf1
npm1a
gra
timm10
hint1
pcna
supt3h
ppa1b
dynll2b
fkbp3
rabif
h2afvb
lin52
tubb2b
mcm6
wdr82
cenpx
ube2t
hmgn2
polr2l
henmt1
setb
si:ch211-212d10.2
dnajc9
mki67
ppid
efcab1

atp5mc3b
nars
eif4e1b
cirbpa
ascl1b
txndc17
med28
tomm20a
wdr76
sub1a
cenps
ubtf
ss18l2
psmb3
hibadha
msh6
lsm5
ndufa2
ccdc25
nutf2l
nup107
tubb4b
rbm8a
gstr
psmc5
pfdn6
chek1
nudc
cks2
snrnp27
atp5mea
snrnp200
edc4
alkbh3
snrpe

zgc:56106
erh
dnd1
mybbp1a
atp5pd
zcchc2
pomp
si:dkey-148h10.5
mif
rad21l1
eftud2
vrk1
nelfcd
dazl
seta
cct7
hnrnpul1
smc3
si:ch211-156b7.4
baz1a
ap3s2
dhfr
nuf2
exd1
mbd3b
rad51
nup93
emd
zgc:66472
ift46
aaas
sf3b5
sf3a2
nudt21
psmc1a

srsf1b
txlna
lsm3
birc5a
ndufc2
eif4a3
tomm6
spata6l
psma5
ndufs5
mphosph8
mrps25
ndufa8
zgc:158564
dusp22b
cenph
tfdp1a
nop16
phb2b
hddc2
hdhd2
nsmce2
gtf2f2a
zgc:110540
ndufs7
rbmx2
mcm7
calm3b
si:ch73-41e3.7
dgcr6
tcp1
ppie
hddc3
cox5aa
ilf2

exosc2
gtf2h5
eif2s2
ccdc137
tbcb
mtrf1
myl6
nedd8
ubxn1
yeats4
eif4e1c
cops3
xrcc5
ube2c
rbb4l
ndufab1b
fip1l1b
pin1
ebna1bp2
crnkl1
taf15
isoc2
pbdc1
ppig
ndnl2
psmd8
tuba8l
ndufab1a
rfc4
ptgr1
ruvbl1
denr
hmgb1a
luc7l3
psmd4b

gtf2e2
cdk11b
COX7A2 (1 of many)
ifi45
metap2b
psma8
psmc3ip
snrpa1
hnrnpd
elavl2
timm23a
CR762420.1
ddx18
fundc1
dmap1
eif4eb
utp11l
srrm2
si:dkey-6i22.5
nup50
otub1b
rcc2
aifm1
srsf3b
si:ch211-235e9.6
psmd7
mtx2
psmb1
ndufa12
snip1
tyms
bcl7a
etfa
CABZ01112656.1
unga

si:dkey-170l10.1
si:dkey-57a22.11
psmd1
polr2f
asf1bb
hnrnpa0b
gcsha
psmd11b
ptges3b
usp1
zpr1
ndufa6
gemin2
psmb5
si:ch211-22i13.2
prpf6
atp6v0cb
mrpl38
ciapin1
spc24
drg2
snu13b
ufc1
ttf1
eif1ad
pqbp1
cops5
ruvbl2
sf3b4
tsn
magoh
ankrd10a
hp1bp3
rpa3
pcmt

ik
apex1
pfdn4
ppil1
esf1
cox7a2a
fabp7a
dhps
gkap1
cnot1
dhx16
snrpd2
pcm1
top1mt
rpa1
phf20b
zgc:112496
cox8b
mcts1
rsl1d1
larp7
prdx3
dus3l
cct2
psmb6
tomm34
g3bp2
si:ch211-68a17.7
prpf8
phb
nolc1
hspa4a
ralaa
eif5a2
ube2f

mrps33
supt16h
prmt1
nip7
selenou1a
cox7b
atp5f1d
si:ch211-226l4.6
krr1
si:ch211-51e12.7
cbx3a
mrps14
hypk
psmc6
zswim5
cox7c
rplp0
LOC110438491
pole3
nap1l1
caprin1a
rpsa
CABZ01068251.1
mrpl21
ndufb5
gins1
zgc:173742
csnk2b
ddx23
sf3b6
trip13
ndufa9a
alkbh5
mkln1
ccdc106b

msh2
zgc:92287
zgc:101851
si:ch211-286b5.5
cks1b
psma3
cope
polr2d
zgc:63470
cdk2
sdhc
dynlrb1
zgc:103697
dnajc8
cdk1
mrpl30
atp5mf
eif5b
ilf3b
phax
uchl3
swt1
tuba1b
prpf39
pfdn1
uqcc2
fam162a
chchd2
sephs2
skp1
pcgf6
cse1l
coil
mphosph10
gtf2b

chmp5b
ndufb7
fen1
pes
npm3
cript
ISCU (1 of many)
mrpl46
dynll1
polr1c
idh3g
si:dkey-67c22.2
flot1b
pop5
rchy1
sod1
dpp3
fam133b
eny2
bud31
mrps21
maco1a
si:ch211-284b7.3
selenoh
gins2
nsfl1c
nup37
pane1
polr3h
trnau1apa
psma6a
baz1b
smim8
rfc2
tma7

trappc1
si:dkey-92i17.2
pdrg1
atp5md
tuba8l2
UTP14C
rrp36
ing5b
si:dkeyp-55f12.3
cenpw
hnrnpa0l
gnpda1
mtap
pho
eme1
trir
med20
cct3
c1qbp
atp5f1c
abcf1
ndufs4
ddx61
suclg1
si:dkey-189g17.2
nkap
atp5l
bida
ube2d1b
psmd14
bola3
ndufa1
parga
ndufa3
snw1

mrpl13
safb
cops4
si:zfos-169g10.3
hspe1
mrpl14
csnk2a1
sap18
polr2h
orc4
rpf1
atp5mc3a
myg1
atp5pf
rps19bp1
fam168b
taf11
rbm25b
mcm2
rpl23a
si:ch211-276i12.9
h3f3c
sem1
cstf2
prpf31
mrpl11
prdx1
u2af2a
fbl
cct5
dynll2a
srsf3a
s100u
hdac1
naa20

gstm.1
ndufb10
zgc:56699
alg13
cox5ab
hmgb2a
idh3b
uba1
lyar
slirp
ncaph
zgc:56304
mfap1
rbbp4
zgc:103482
ptp4a1
prpf3
gpn2
srpk1a
gtf2f1
mrps11
COX5B
fhit
rexo4
uqcrc1
si:ch73-1a9.3
ythdf2
lsm14ab
hars
exosc8
ndufb2
atp5mc1
atp5fa1
tufm
hmgn3

btr04
morc2
eif3ja
clns1a
txndc9
cnot9
prmt5
khdrbs1a
si:dkey-180p18.9
atp5meb
pfn2l
uqcrh
chrac1
pcnp
brk1
pgam5
srsf5a
psmd3
cox14
rnaseh2a
rad23b
ndufv2
psma2
nop2
esd
cct6a
smndc1
kpna2
cox6c
tomm22
htatsf1
mrpl53
cfap20
ndufb9
thrap3b

lyrm9
ndufa11
taf7
dctpp1
zgc:55461
lig1
mrps18c
rrs1
smu1a
pabpc1b
polr2eb
tnrc6b
si:dkey-188i13.11
txn2
phf5a
mpc1
psip1b
rrp8
psma1
ndufs2
timm17a
hexim1
rbm19
rrp9
hspd1
ubap2b
psmd13
ndufa10
rplp2l
strap
znf593
vcp
seh1l
iscub
ywhabb

pa2g4a
mad2l1
imp3
bccip
vps4b
u2af2b
mrpl52
park7
pfdn2
sf3b2
rwdd1
ppp2r2d
rpia
pdcd4a
psmd12
aimp1
dkc1
ddx27
cct4
rtca
atp5pb
ndufb3
ran
psma4
taf10
atox1
sod2
copa
e2f5
ndufb8
psmd6
nhp2
nop10
ranbp1
arl14ep

cnbpa
smn1
ube2a
txnl4a
atrx
matr3l1.1
ndufa5
sumo3b
srrm1
si:ch211-282b22.1
h2afy2
ddx39aa
lsm8
hnrnpl2
mrpl32
mcm5
slc25a3b
wdr46
ube2e1
UBL4A
psmc3
adh5
eif3k
si:zfos-1192g2.3
nxt2
uqcrq
snrpb
ost4
hmgn7
nfr
psmb4
pprc1
mrpl57
prdx2
myl12.2

COA4
naa10
lyrm4
ube2ia
polr2k
cox4i1
tktb
nol12
grpel2
api5
si:dkey-46a10.3
adi1
mapre1a
eif6
zmat2
ube2v2
ppp1cab
h2afva
ywhae1
si:rp71-45k5.4
lamtor5
gpx4b
supt4h1
si:dkey-28b4.7
gapdhs
rps10
zgc:86598
tuba8l4
si:ch211-217k17.7
sumo3a
ewsr1b
atp5f1b
ndufa4l
eif4h
chchd10

mrpl12
med10
tbca
sap30l
fubp1
zgc:66388
ube2d2
cdv3
st13
rps21
hnrnpabb
pgk1
nop58
znf706
ifi46
zgc:193541
elocb
kpnb1
paip2b
meaf6
eif3d
syncrip
rad21a
adssl
dohh
eif4a2
psmb2
ahcy
serbp1b
oaz1a
hnrnpa1a
u2surp
si:ch1073-325m22.2
mdh1aa
eif3g

atp5if1a
cct8
eef1da
si:ch211-214j24.10
atp5po
cox6a1
h3f3a
zgc:111986
pabpn1
ube2v1
tmem258
idh2
fam32a
nop56
ywhaqa
rps27l
ak2
eif2s1b
uqcr10
vdac2
u2af1
si:dkey-12l12.1
lum
ecrg4a
hpdb
asip1
tgfbi
scara3
zmp:0000001268
ctgfb
si:ch211-243a20.3
rhag
si:dkey-282h22.5
tat
hoxb8a

ttc36
prrx1b
tnfrsf19
vcana
hgd
aqp3b
hyal4
FQ323119.1
tpbga
sp9
col5a2a
clu
mfap2
fgfr3
bhmt
aspn
smoc2
CU929237.1
dpt
twist1a
cxcl8a
ccdc80
twist2
igfbp3
sparc
ehd2b
col5a1
cyr61
col1a2
mmp14b
col1a1b
mmp11b
si:dkey-85k7.7
nradd
CABZ01092746.1

col6a3
filip1l
col6a1
si:ch211-39i22.1
fmoda
lhfpl6
col1a1a
hoxb9a
pmp22a
twist3
cldn19
htra1a
ccl44
lpl
sod3b
si:dkey-178e17.3
CABZ01074363.1
ms4a17a.7
hhipl2
col2a1b
cdh11
mmp14a
zgc:162612
postna
fah
epha4b
sertad4
krt91
crabp2b
ier5l
fstl1b
LO018188.1
mmp2
s100v2
fxyd1

si:ch211-286o17.1
c1qtnf2
jun
rarga
ms4a17a.9
atoh8
prrx1a
f3b
fosl1a
si:ch73-335l21.4
alx4a
zgc:56585
si:ch211-170d8.8
heyl
fgl2b
sema3aa
thbs1b
rnd3b
serpine1
enpp2
bcl11ba
gpm6bb
olfml1
XLOC-040020
adh8a
fosab
ms4a17a.11
loxl1
dcn
sema3e
s100a10b
fosb
cntfr
pigr
anxa1a

gstz1
pcdh18b
sox11a
capn2b
si:ch73-46j18.5
apln
sdc4
zgc:154093
postnb
ier2a
ctsk
vmo1a
tcf12
junba
fstb
egr1
lix1l
bgnb
ecm2
spon2b
rgma
ier2b
serpinf1
vwa1
junbb
lrrc17
klf6a
socs3a
marcksl1a
dlx5a
col6a2
qdpra
mxra8b
mamdc2b
col5a3a

pcolcea
jdp2b
colec12
myo1ca
cd99
cspg4
serpinh1b
nfixb
cyp26b1
zeb2a
si:ch211-133n4.4
spock2
klf15
osr2
zgc:162730
icn
apcdd1l
hspg2
tcf7
ccdc3b
MDFI
endouc
krt8
ckap4
fap
srpx2
dlx3b
vim
anxa5b
plpp3
jag1b
pleca
antxr2a
atf3
cdo1

FAT4
rgs5b
tcf4
h3f3c
rbp4
emilin2a
cxcl18b
cd81a
gpc1a
slc16a10
rbfox1
fstl3
sfrp1b
ism1
si:dkey-261h17.1
grb10b
pmp22b
akap12b
im:7142702
efna5b
timp2a
irx1a
slc25a48
cspg5a
scara5
c1qtnf5
ddit3
si:busm1-57f23.1
zcchc24
bmp16
ablim1b
plxdc2
fkbp7
angptl1a
cox4i2

smad1
klf2a
btg2
zfp36l1b
meis1a
ecm1b
tnfaip2a
tgfbr3
si:ch211-178n15.1
nupr1b
id1
spred2b
zgc:153317
ppap2d
cnn3a
entpd5a
ramp2
lfng
f13a1b
mex3b
gas1a
fbn2b
itgb4
hoxb7a
tuba8l3
lmna
atf4b
BX640522.2
vstm4a
pcsk6
cd82a
tenm3
tpm4a
aldob
ghra

ier5
her6
XLOC-042917
arhgap21a
rgcc
CR383676.1
socs3b
abi3bpa
col6a4a
cebpd
wnt5b
lsp1
fgl2a
gng7
fkbp14
mtmr11
ahnak
sertad2a
BX663503.3
nova2
im:7152348
efnb2a
rgs5a
tuft1a
LOC101883507
s100w
csrnp1b
cd248a
htra3b
col11a1b
hmbox1a
fpr1
chd3
SCARF2
si:ch73-1a9.3

edil3a
vat1
zgc:171679
si:ch211-251b21.1
dpysl3
tnfsf10l
marcksl1b
kank2
pdgfrl
anxa2a
krt18b
gadd45ba
fkbp9
angptl4
si:ch73-138n13.1
pdgfrb
nectin1b
inhbaa
ppib
hbegfa
tmem176
ednraa
si:ch211-212k18.8
fxyd6l
loxl2b
sfrp1a
kirrel1a
pdgfab
si:dkey-101k6.5
FP102018.1
s1pr5a
gpc4
sept10
sh3pxd2b
fhl1b

si:dkey-9i23.16
tmeff1b
kalrnb
LOC101886114
pdgfra
rnd3a
si:ch1073-459j12.1
jund
capns1b
capn2a
ip6k2b
gli3
meis1b
CR847953.1
mycb
plecb
sdc2
serpine2
lamb1b
nkd1
tagln2
prelp
si:dkey-226m8.10
tmem100a
mcl1b
VIT
cygb1
sox6
CR847566.1
cxcl14
pcbd1
frmd6
ptprga
rhoua
cavin2b

eva1ba
selenop
ogna
bckdhbl
s100u
mych
anxa4
gulp1a
midn
ccl25b
CABZ01039096.1
cd28l
wls
ccdc85b
krt4
si:ch211-284f22.3
reck
dhrs13l1
sh3gl1b
itgb1a
sulf1
bicc1b
hoxc3a
lrp1aa
myl9b
si:ch211-105c13.3
calua
myl6
olfml3b
crtap
nfia
phactr4b
wu:fi04e12
itga5
aoc2

spry4
mrc2
CABZ01039304.1
mafba
XLOC-000431
foxo1a
hcar1-4
slc16a1b
rbfox2
zgc:110158
eif1b
pard3bb
col12a1a
nfil3
ciarta
osr1
lzts2a
htra1b
tcf7l2
ctnnb1
dedd1
hic1
vasna
mgp
znf503
hmgn6
wwtr1
spry2
higd1a
tspan4b
mfap5
gsna
enam
zgc:153372
si:dkey-6n21.12

antxr1c
pdk2b
nanos1
ntn1a
pim1
flna
aqp3a
nsa2
si:dkey-26m3.3
ndrg2
LOC103909284
proza
nme3
adgrl3.1
fbln2
yap1
cav1
si:ch211-212k18.9
col16a1
adamts15a
nid1b
sgce
cyp2n13
phlda2
id2b
dpysl2b
atp2b2
tspan14
appb
ubtd2
si:dkey-7j14.6
agtr2
h1f0
glud1a
cnn2

ak4
AL732488.1
si:ch211-227m13.1
calm2b
klhl24b
scpp7
glis2a
fbp1a
zgc:113208
tead1b
col21a1
eif4a1b
parvaa
histh1l
ehd1b
im:7151449
neo1a
ybx1
irf2bp2b
serf2
igf2b
boc
acsl1a
ptbp1b
notch2
rem1
vdac3
cdkn1a
cbfb
tuba1b
hmgn7
adma
svild
add3a
ctdsp2

rgs4
nfil3-6
myh10
cavin1b
pbx1a
mef2aa
perp
col15a1a
zgc:174904
gadd45ga
calr
cish
fam168a
ackr3b
dnase1l1
DDX17
tmem176l.1
dusp8a
ephb3
flot2a
klf3
mgst1.1
prdx5
slc3a2b
mgst3b
slc38a5b
plod3
fzd2
palm1b
sema4ba
slc7a1
emp2
zgc:92606
anxa13
nr1d2a

celf2
dag1
daam1a
FO082781.1
CT030188.1
nfe2l2a
rhoab
rps2
dbn1
hsp70.3
eef1da
si:ch1073-291c23.2
anxa11b
selenow1
tpt1
arl6ip5b
rpz5
rpl32
reep5
pnrc2
tns1b
lgals3b
XLOC-010899
si:dkey-28b4.7
zgc:114188
HACD4
CABZ01068358.1
col4a1
si:ch211-196l7.4
LOC108179036
tnrc6c1
mdka
nfic
slit3
klf11b

pdcd6
maff
cnbpb
krt18a.1
cald1a
si:ch211-106h4.12
tspan7
creb3l3l
trib3
june
tnrc18
rpl9
tmod2
itm2ba
aldh9a1a.1
irf2bpl
adam10a
hdlbpa
dnmt3ab
map1lc3a
hopx
hmgb1b
h2afy2
ube2e3
smdt1b
atox1
snai1a
rps16
si:dkey-56m19.5
cebpb
cd82b
cdc42
pdia3
stau2
sec61a1

rps18
mmp9
COX3
mmp16a
cd63
lxn
si:dkey-79d12.5
faua
rps15
pgrmc1
tob1b
sar1b
rpl22
rpl28
fosl2
ptp4a1
rpl36
ppiaa
glo1
col12a1b
fkbp1aa
hp1bp3
mxra8a
ptgdsb.2
abcc5
irf1b
pim2
rpl18
rps27.2
rpl6
gabarapb
rpl18a
tsc22d3
gatad2b
rpl5a

qkia
grpel2
btg1
slco2b1
eef2b
sdcbp2
h2afx1
ssr4
rps4x
cebpg
rpl10
map1lc3b
CABZ01072614.1
si:ch211-286b5.5
zgc:158343
metrnla
rab2a
rpl23a
aebp1
sltm
rhoaa
mknk2b
rpl11
taf12
tpm3
rpl24
ppdpfb
rps15a
zfp36l1a
serinc5
capns1a
XLOC-013964
adh5
cpne1
rpl34

mibp2
sec61g
rab10
glula
rps6
anp32a
rps21
arhgef1b
rpl8
eef1g
rps10
rps24
romo1
eif4ebp3l
tax1bp3
mt-co1
rpl27a
hmgb1a
prelid3b
b2ml
si:dkey-11f4.14
tegt
rpl27
mt-atp6
dusp1
rps13
eif2s3
ppt1
naca
eif3i
gbp
rpl30
NC-002333.4
eif3ea
eif3m

acin1a
rps27a
ubb
rps27.1
mt-nd2
serbp1a
rpl35a
rsl24d1
bsg
mt-nd1
eif3f
rpl22l1
rpsa
eif5a
fabp3
sec61b
myl12.1
tubb4b
rplp2
mt-cyb
bhlhe40
rpl38
tmsb4x
rps26l
CR855311.1
mt-co2
prf1.9
ccl36.1
ccl38.6
si:ch211-214p16.1
FP236356.1
prf1.7
LOC103910871
si:dkey-151g10.6
CR936442.1

rps29
zbtb32
rplp2
rplp2l
il34
rps28
arpc1b
nkl.2
XLOC-000431
XLOC-010899
rpl35
si:dkey-183i3.6
rps20
rps21
rps15a
CR753876.1
rplp1
scgn
LOC108190717
coro1a
syk
rpl11
ncaldb
rpl38
rpl37
BX005105.1
rpl39
XLOC-013964
rps27.1
tmsb4x
rpl34
rps26l
pfn1
rpl27
rpl18

rpl29
rps18
CABZ01084575.1
CU855821.1
rps17
si:ch211-241f5.3
ca2
si:dkey-78l4.4
sla2
rpl23
rpl22
uba52
rps19
rpl36
rpl19
alox5a
rpl10a
dicp1.5-6
rpl36a
rps12
rps14
ela3l
cotl1
actb2
fscn1a
rps24
ptpn6
mbnl1
rps2
cmah
si:ch211-277c7.6
rpl26
id3
XLOC-042595
rps11

dock10
rps13
ptmaa
rps15
rpl15
rps25
rpl30
rgs13
rpl18a
dusp2
hsp90ab1
rpl21
rpl31
rps26
rps23
rpl24
rpl7a
entpd3
rpl13
grap2a
rps9
rpl10
rps5
si:rp71-81e14.2
rpsa
rps10
arhgdig
si:ch211-119e14.1
rac2
zgc:114188
rpl35a
rpl23a
srgn
rpl14
ppiaa

XLOC-025423
il16
capgb
ptprc
faua
rps3a
rps27a
rpl5b
rpl32
rps3
si:ch211-102c2.4
creb5b
si:ch211-191c10.1
rpl6
ptmab
ldha
ccl34b.4
rps7
rsl24d1
rpl13a
rpl8
eif3f
rps16
yrk
rpl5a
itga4
wasb
edaradd
myl12.1
tagapb
scinlb
si:ch73-264p11.1
arpc3
si:dkeyp-68b7.12
elf1

wipf1a
arpc5a
rps6
cfl1
wdr19
tagapa
wasa
XLOC-042596
rpl9
hmgn2
pabpc1a
rpl28
tefa
arhgdia
ponzr1
kctd5a
si:dkey-183p4.10
XLOC-043043
sik1
rpl4
CR381673.1
calm1b
XLOC-043760
dusp1
zgc:56493
lasp1
CU499336.2
RASSF5
mrps15
msna
laptm5
h1fx
LOC100536213
XLOC-028370
aldocb

mt-co1
si:ch211-93g21.1
dtnbp1a
cap1
ccr2
BX901920.1
atp5if1b
zgc:64051
fcer1g
rpl22l1
si:ch211-222l21.1
gapdhs
XLOC-020301
CABZ01079663.1
gpr183a
dok2
csk
fcer1gl
stim1a
pde4ba
ppp1r18
lamp1a
cebpb
vaspa
rps6ka1
cox6b1
XLOC-026836
COX3
mef2d
si:ch73-234b20.5
pik3r1
mt-atp6
arpc4
actr2a
tnip1

rpl27a
sept2
pxna
crip1
arpc2
ywhaba
ccni
capzb
celf2
hmgn6
bcl2l10
bhlhe40
gsdf
bcam
alcama
igf3
LOC103910903
lamb2l
smoc2
map1aa
foxl2b
LOC110438024
bmp6
si:dkey-194e6.1
hsd17b1
cdh4
emx2
nectin1b
metrn
slc4a4a
col4a5
jam2a
cldn11a
cx34.5
lhx9

col4a6
sema3ab
ccdc3a
ecrg4a
znf536
foxl2a
inha
aldh1a2
agrn
cemip2
itga3b
si:ch211-207d6.2
rpz5
lama5
si:dkey-6n6.1
bmpr2a
si:dkey-42i9.7
itm2bb
thbs3b
pi15a
chka
trim35-33
f10
nr0b1
podxl
XLOC-005605
plxna4
wu:fc27b11
igsf3
scn4bb
slc44a5b
ivns1abpa
nog1
emilin1a
negr1

CABZ01021532.1
cldnc
sparc
rab11fip1b
XLOC-009815
cahz
ccdc187
gstt1b
cadm2a
ptn
appa
ace
si:ch211-117c9.5
st5
large2
hpn
ar
nos1apa
CABZ01068358.1
slc7a3a
F7 (1 of many)
pgr
sdk2b
si:dkey-188i13.9
colec11
ifi45
bambia
ncam1a
hsbp1a
pcdh1b
cx43
fgfr3
htra1b
phlda2
hdac4

cdh1
nanos1
atp1b3b
ttll12
cav1
zgc:123010
gata6
cxcl14
b2m
wt1a
ccdc125
jam3b
vasnb
mpzl2b
CR318588.3
krt18a.1
zgc:114181
isg15
ifi46
ponzr1
gpx1b
schip1
alcamb
parp12a
si:ch211-156j16.1
bmpr1bb
mmp2
cyp26a1
gstm.3
hey2
s100v2
fndc3ba
mbnl3
atp11a
phactr4a

cox5b2
dag1
aplp2
zgc:92744
ece2b
cd99l2
tfr1a
CABZ01068366.1
myl9a
tp53inp1
tspan9b
fkbp1b
ptprfb
wu:fj16a03
myh10
h1f0
lamb1a
atp1b1a
slc38a2
lamc1
si:ch73-1a9.3
PDE3B
si:ch211-264f5.6
arhgef18a
id1
foxp4
parvaa
arhgef5
enc1
rbp5
man1b1b
chpfa
pwwp2b
fkbp1aa
COX3

spred2b
si:dkey-178e17.3
tjp1a
CR318588.1
si:dkey-73n10.1
si:ch211-225b11.1
svild
cavin1b
mt-co2
ecm1b
si:ch211-286b5.5
rdh10a
mt-co1
hsd11b2
c7a
uxs1
cox6a2
prkar2aa
sptbn2
s100a10b
edem1
cgnl1
oaz1b
cebpd
krt8
oip5-as1
abhd15a
mt-cyb
stat1a
adam10a
glulb
zgc:100829
pnp5a
mphosph8
slc2a1b

CABZ01068367.1
ap2m1a
afdna
tmem176l.1
mt-atp6
lgals9l3
prss1
yap1
pde4ba
txnipa
serpinf1
cdh2
glod5
anxa2a
msi2a
NC-002333.4
gulp1a
ahnak
si:ch211-222l21.1
si:dkey-7j14.6
atp1a1a.1
myl9b
si:dkey-188i13.10
nfic
ptprga
zgc:195173
tuba8l3
cd74b
hmgb3a
cald1a
atp10a
arf2b
zbtb20
gli3
selenof

zgc:174863
mt-nd2
h2afx1
rnf213a
cox7a1
ndrg2
rn7sk
si:ch211-199g17.2
CABZ01080702.1
mknk2b
map1lc3a
cd164
cux1a
mt-nd3
fam168a
itm2ba
gng7
kmt2cb
ccdc85b
rpia
abl1
mt-nd4
atp5mea
si:dkey-28b4.7
ctnnb1
phc2a
tmem88b
gpx4a
hmgb1a
calm2b
mt-nd1
atp8a1
LOC103909284
trappc1
hbp1

zfp36l1b
ywhaqb
cad
rgs2
sptan1
cavin2b
LOC101884451
hspe1
b2ml
ywhaz
purab
zgc:110182
sox4a
wu:fb16f03
hist2h2l
hmgn6
robo2
lgals17
pgrmc1
pfn2
hmgb1b
ncor1
pfn2l
hsd3b7
smarca2
srpra
socs3a
XLOC-016094
ppib
stat2
nucb2a
angptl4
serinc1
tpm4a
ctnnd1

psme1
chchd6b
si:ch211-197h24.9
gabarapb
slc38a5b
fn1b
tnfrsf9a
flot1b
eif4ebp1
fam120c
si:ch211-1a19.3
krt94
hp1bp3
oaz1a
atrx
fthl27
atp6v1e1a
atp5pb
add3a
si:ch73-376l24.3
dynlt3
marcksl1b
ndufb9
cd74a
hmgb2b
dynll2b
nucks1a
mvp
mcl1b
ndufa4l
cnn3a
glud1a
arrdc3a
iscub
calm1a

ucp2
h2afy2
metrnla
ndufa5
cdv3
zgc:152791
cdc42
hnrnpa1a
tp53
ndufab1a
sdcbp2
rhoaa
si:ch73-376l24.2
tnrc6c1
chd4a
myl6
chd4b
ier3ip1
ndufc2
hif1ab
si:zfos-169g10.3
ndufa1
psme2
ptmaa
notch2
rab10
gnb1b
eef1db
rab5c
h3f3c
jun
ifi30
ndufa3
gpx4b
si:ch73-46j18.5

abcc5
tmem59
dbi
mt-nd5
cnn2
rpl27a
atf4b
selenok
atp5pf
dpy30
cox6b2
midn
edf1
si:ch73-42p12.2
si:ch211-191j22.3
tegt
CR383676.1
cox7c
sub1a
atp5mc3a
fosab
odc1
eif1b
cfl1l
fam133b
cox5ab
rps27l
zgc:114188
dynlrb1
hmgn2
mif
tma7
vat1
atp5pd
psap

atp5f1c
ier2a
ost4
si:dkey-42i9.4
ier2b
atp5f1e
rbx1
pomp
tmsb4x
rpl9
nkl.4
prf1.2
LOC101884267
cpa5
prf1.7
XLOC-043043
XLOC-039757
tspan2b
nkl.3
si:dkey-21e2.3
CABZ01084575.1
nkl.2
XLOC-014079
BX248410.3
si:ch211-173d10.4
csf2rb
BX120005.1
LOC108190717
si:ch211-277c7.6
mef2b
CABZ01088229.1
ca2
si:dkey-78l4.4
hck
alox5a

ptk2ba
stim1a
CR753876.1
CABZ01073834.1
zbtb32
creb5b
sort1b
gcnt7
edaradd
prr5a
gzm3.3
srgn
efhd2
ponzr6
arl11
illr1
BX005105.1
sftpba
slc41a1
XLOC-043760
arpc1b
LOC110440216
si:ch211-102c2.4
plcg2
rps6ka1
cyfip2
mrtfaa
si:dkey-21o19.2
rbbp8l
XLOC-005448
ephx2
LOC110437825
znrf1
coro1a
adam19b

tbx21
pfn1
ccl39.9
il12rb2
myh11a
syk
wipf1a
fa2h
fcer1gl
strbp
zgc:64051
si:ch211-241f5.3
gramd2aa
zgc:152891
mbnl1
nr4a3
atp11c
BX901920.1
vaspa
ly75
gfi1ab
hyal3
bhlhe40
stat4
pstpip2
si:ch73-208g10.1
actb2
ptprja
plat
egr3
arhgdig
elf1
sh3kbp1
BX322618.1
laptm5

sh3bp2
dapp1
actr2a
dicp1.1
uba52
gab2
pla2g15
tent5ab
wasb
sla1
tagapa
eef1a1l1
itga4
si:ch211-214p16.1
dok2
ptprc
cables2b
sik1
ndfip2
BX908750.1
XLOC-028370
pabpc1a
dicp1.5-6
camk1b
dock10
stk17b
tec
rpl4
dock8
LOC103910871
wdr19
yrk
aldocb
XLOC-025423
sla2

rps10
hsp90ab1
irak1
prkcha
eef2b
cotl1
rps18
rpl31
si:zfos-741a10.3
tmsb4x
tyk2
rpl6
si:dkey-222f2.7
eomesa
CABZ01079663.1
CR392341.1
si:ch211-226m16.2
rps26l
actn4
hcls1
si:rp71-36a1.5
gnsb
slc25a5
cap1
sh2d3ca
XLOC-010899
scgn
rpl14
atp2b1a
rpl21
cfl1
rps4x
zgc:114188
btk
ponzr1

rpl5a
msna
st3gal4
wdr1
rnf19b
lamp1a
ubash3ba
rpl36a
il16
CR936442.1
slc7a3b
si:ch73-264p11.1
rps23
abo
per3
il34
samd11
ptpn22
plp2
si:dkey-10h3.2
capzb
far1
tnfrsf9a
csk
pptc7b
tmc8
wdfy2
RASSF5
zgc:153018
adcy7
edf1
megf8l
mir223
dock11
XLOC-020301

rpsa
grap2b
arhgdia
XLOC-019150
aif1l
si:dkey-151g10.3
ikzf1
limd2
orai1a
rpl9
CABZ01111959.1
cmip
tmem30aa
rgs13
rpl24
si:cabz01074946.1
lcp1
zeb1b
ldha
slc1a5
rps14
ap1b1
nptna
dtnba
fryl
scinlb
ptmaa
susd6
pnp5b
icn
ccr2
si:dkey-262k9.4
prf1.9
rps13
slc6a6b

atp5if1b
rac2
btf3
rab27a
ptp4a2b
rpl15
aars
myo1f
si:ch211-136m16.8
ccdc171
rpl13
lcp2a
rplp1
itgb2
rpl7a
acap1
ccni
adam17b
cox8a
CU972454.4
mthfd2
arpc5a
agpat3
BX005183.3
mt-atp6
mibp2
adam8a
rps28
rps29
XLOC-000431
arhgap1
jak1
hspa8
glipr1a
kif21b

arpc5b
arhgef6
rps3
zmp:0000000521
cdaa
rpl22
actb1
si:ch211-200p22.4
XLOC-013964
mt-co1
zyx
cd63
rel
zmp:0000001082
elmo3
arpc2
rpl26
XLOC-041870
rps16
ccl36.1
rps21
rpl5b
rpl38
taok3a
si:ch211-8c17.3
si:dkeyp-68b7.12
rpl29
nbeal2
pde4ba
fam120c
dusp1
lrrfip1a
tpt1
pik3cg
mef2d

nsa2
arpc3
ppp1r18
kdm2ba
rpl18
CR381673.1
rack1
si:dkeyp-110e4.6
ipo7
CABZ01044746.1
id3
serbp1a
inpp5d
CABZ01072614.1
stxbp2
cxcr4b
vdac2
oaz1b
sat1a.2
ankrd44
il10ra
COX3
myo18ab
abracl
plekhj1
ostf1
wasa
rps27.1
fam49ba
reep5
dnajc5gb
casc4
iqgap1
tomm20b
rplp2

selplg
NC-002333.4
prkar1aa
snx27a
metrnla
XLOC-026836
eef1db
rps5
arpc4
ncaldb
calr3b
rps15
rsl24d1
rps27a
cct8
calm1b
skap2
cbfb
zmynd11
sh3bgrl
NC-002333.17
hmha1b
grb2b
ero1b
rpl35a
nr4a1
eif4ebp2
celf2
plekho2
cox6b1
myl12.1
clic1
dub
XLOC-042596
crlf3

rpl34
tpi1b
LOC110438237
eno1a
per1b
elmo1
mt-co2
larp4ab
rps17
tnfaip3
vsir
abca1a
atp10a
tnip1
pum1
birc2
arpc4l
eef1g
elovl1b
rpl23a
rpl36
rplp2l
tefb
hnrnpa1a
eif3d
si:ch211-222l21.1
si:dkeyp-19e1.3
arrb2b
XLOC-042595
chac1
rpl28
capza1b
zgc:63831
pdia3
fosl2

dusp2
xbp1
eif3c
hif1ab
grap2a
pik3r1
tefa
si:ch211-191c10.1
cct6a
zgc:153867
tmem131
CABZ01090890.1
pgam1a
rab44
sh2b3
rhogb
lasp1
rhoac
pnrc2
mapre1a
tln1
mt-nd4
eif3s10
sh3bgrl3
crema
eif2s1b
mrps15
psap
pkma
ptmab
cox4i1
eif3i
eif4a1b
ist1
gapdhs

BX248082.1
slc3a2b
ip6k2b
ola1
eif3ba
atf4a
rnf44
cct2
cdc42l
idh2
eif3ea
mt-nd1
ywhabl
csde1
serf2
mt-nd3
h1fx
ccl39.7
CR383676.1
cavin1b
podxl
cldnc
s100a10a
gcga
krt15
krt4
gstm.3
ucmaa
scn4ba
si:ch211-156j16.1
cyp2ad2
fgf24
slc29a1b
fn1b
itgb4

fsta
rap1gap
pdlim3b
eppk1
frya
sccpdha
cav1
aldh1a2
LOC110438169
sox6
postnb
bnc2
krt18a.1
krt94
wt1a
cygb1
si:ch211-105c13.3
si:ch211-137i24.10
col6a4a
cavin2b
krt8
sfrp1a
krt91
FP016056.1
kcnq5a
anxa1a
plod2
cfd
si:ch211-250c4.4
usp43b
frzb
tmsb1
hspb1
si:ch211-106h4.12
CU984579.1

dkk3b
paplna
si:ch211-63p21.1
anxa2a
c1qtnf12
tmem98
slc43a3b
s100a10b
mmp2
lurap1
cldn5a
col1a1b
slc43a1b
col1a2
si:ch73-46j18.5
jam2b
col1a1a
cebpd
ehd2b
c6
im:7151449
zfp36l1b
ptprq
ppdpfa
adh8a
col6a3
col4a1
endouc
icn
gata6
hsd11b2
efna1b
bmp2b
nfia
cdh2

zgc:112437
smoc1
sparc
emilin1a
pleca
sema3d
f3b
daam1b
myo1cb
col6a1
ncam1a
bag3
XLOC-041269
junba
cyp1c2
AL954191.1
socs3a
cxcl18b
fosb
cdh6
CABZ01080702.1
fosab
itgb3b
sost
f10
mical2a
nrp1a
nanos1
snrkb
thbs1a
jun
dcn
tjp1a
col6a2
flnb

LOC101882334
vim
id1
LO018188.1
cyp27c1
edil3a
rnd1b
plekha6
sh3bgrl2
gadd45ba
rhpn2
zgc:154058
znf385a
colec11
eif4ebp3l
cyp2n13
s100v2
tpm4a
si:ch211-198c19.3
loxl2a
hdlbpa
LOC103909284
mob2a
ctgfa
tgm1l1
si:ch211-1a19.3
wt1b
fap
cyp2ad3
ghra
reep3a
tagln
dsc2l
igf2b
col5a1

lamb1b
cd81a
ahnak
lrp1ab
gpm6ab
tnfsf10l
esr2a
si:ch1073-459j12.1
si:dkey-16l2.20
prss23
pik3r2
ftr82
slco2b1
fstl1b
igfbp5b
LOC101885758
pbx3b
phactr4a
fosl1a
zgc:162730
BX322530.1
nfil3-6
nfic
ptprsa
cdon
deptor
zgc:158463
ptges
c7a
cyp2p6
jdp2b
ugt1b5
slc12a7b
cav2
h1f0

sult1st6
pim1
aoc2
boc
fstb
xirp2a
pmp22a
cldn11a
appa
pdk2a
meis1b
btg2
epb41l5
cyr61
ptpn21
epha3
chka
rpz5
tspan4a
slc7a1
zgc:110182
stau2
aldh9a1a.1
BX663516.2
CR383676.1
gchfr
si:ch211-166a6.5
LOC110439416
ctsf
tspan9b
gamt
lmbr1l
lama5
rhoua
klf2a

vasna
scarb2a
pnrc2
sostdc1a
si:ch211-66e2.3
lxn
ier2a
pkp3b
dedd1
foxp4
rtn3
cebpb
bcam
mknk2b
pawr
clic4
egr1
gas1a
cald1a
col5a2a
cgnl1
dag1
cdkn1a
csrnp1b
gpr146
tmem176
hecw2b
plecb
cast
pabpc1b
tmem88b
myl9a
selenbp1
afdna
si:ch73-138n13.1

rsu1
slc38a5b
itgav
rdh10a
olfml3b
gstt1b
pfkfb4b
si:ch211-232b12.5
cyp2p7
ulk2
ptprga
si:dkey-51e6.1
CABZ01072614.1
LOC101884451
irf2bp2b
map1lc3a
tns2a
gulp1a
chd3
sdc4
txnipa
actn1
nfixb
aldob
perp
si:ch211-222l21.1
tshz3b
irf8
ephb3
gli3
rpsa
FO744854.1
zbtb20
midn
gpc4

pdk2b
mycb
fkbp5
selenow1
fxyd1
foxo3b
myh10
per3
col12a1a
socs3b
zgc:158343
ier2b
NC-002333.4
abcc5
nfil3
myl9b
CABZ01068366.1
taok1b
cpn1
ctdspla
XLOC-024750
klf11b
atp10a
gstm.1
si:dkey-28b4.7
tenm3
fgl2a
lgals3b
dbpb
ghrb
b2ml
entpd1
dclk1a
si:dkey-23i12.7
itpkcb

rgs5b
ccdc85b
sort1a
pmp22b
stat1a
si:ch211-204c21.1
hmgb1a
mfap2
cd99
si:dkey-56m19.5
pdgfrb
fam120c
tuft1a
lpar1
si:ch73-1a9.3
fat1a
timp2a
ldlrad4a
fzd7b
cbx7a
calm2b
ptmaa
spry4
ndrg2
adamts15a
fosl2
mt-nd2
cnn3a
cox7a1
si:ch211-286o17.1
CR847566.1
qkia
ifi45
atf3
mir21-1

mt-nd1
rbp5
atf4b
rbp4
flna
si:ch1073-291c23.2
ncor1
emp2
h3f3c
tagln2
myl6
ywhaqb
rps28
tmsb4x
zgc:195173
myh11a
adh5
kank2
cry2
aebp1
mcl1b
ube2e3
gadd45bb
cebpa
sesn1
si:ch211-140m22.7
rpl29
ctdsp2
capns1b
glud1a
irf1b
rbm5
rpl6
rpl9
zgc:114188

wu:fb16f03
cish
cflara
rps10
mt-atp6
rpl5a
tefa
kmt2bb
rpl36
ppib
cnn2
hmgn6
nr1d2a
rps27.2
jund
eif4ebp3
si:ch211-196l7.4
rplp2
maff
BX640522.2
slc20a1b
COX3
ddit3
lamc1
nme3
si:ch73-335l21.4
mt-cyb
capns1a
ndufa4l
elovl1b
hmgn7
mt-nd4
ctnnb1
tmem176l.1
rbm39a

vat1
pfn2
rpl30
rpl14
anxa5b
ybx1
tsc22d3
usp36
add3a
h2afx1
ip6k2b
h2afy2
acin1a
hopx
klf6a
rplp2l
rps16
mmp16a
si:dkey-7j14.6
eef1db
b2m
zgc:111986
tpm3
crebzf
atrx
rps13
mt-nd5
mt-co2
fabp11a
arrdc3a
bhlhe40
uba52
marcksl1a
dbi
rpl27a

itm2ba
rhoaa
zfp36l1a
rps27l
mdka
btf3
si:dkey-11f4.14
crip1
celf2
serbp1a
rpl23a
srsf5a
romo1
cnbpb
mt-co1
XLOC-025423
ptmab
eif4a1b
mt-nd3
igl4v8
LOC101886766
zp3
cldnd
si:dkey-241l7.2
zgc:153499
si:ch211-125e6.12
si:dkeyp-98a7.9
mt2
zp2.3
zp2.5
zp3.2
h2af1al
si:ch73-189n23.1
zp2.6
tmsb5

si:zfos-1505d6.3
h1m
si:dkey-241l7.4
si:ch211-12h2.8
si:ch211-226h8.14
zp3a.2
cldng
si:dkey-240e12.6
si:dkey-229d11.3
zgc:175135
si:dkey-241l7.5
acp5a
zp2.1
si:ch211-14a17.7
zgc:171779
ccna1
zp2.2
crp2
si:dkeyp-46h3.6
CR790368.1
si:dkeyp-98a7.8
si:ch211-226h8.4
zgc:77118
zgc:171776
mid1ip1l
si:dkeyp-98a7.4
epcam
nanog
cth1
si:dkey-229d11.5
si:dkey-19b23.11
si:dkeyp-20e4.8
LOC110437821
zgc:173443
si:dkeyp-50f7.2

BX649337.1
LOC100537499
zgc:171517
btg4
lamp2
zar1l
npm2b
AL929032.1
slbp2
zp3b
si:dkey-90l23.1
rrm2
zgc:195245
wu:fi42e03
si:dkey-37o8.1
si:dkey-241l7.6
ccnb2
si:ch1073-263o8.2
si:ch211-132b12.8
ccnb1
qdprb2
zgc:163030
LOC101885477
kpna7
CU571163.1
LOC100535162
s100a1
tuba4l
hnrnpabb
siva1
si:ch211-5k11.8
zar1
si:ch211-12h2.6
zgc:152652
hbba1

arhgef10la
si:dkey-208k4.2
si:ch211-226l4.6
gdf3
si:dkey-152b24.6
zgc:136254
ca15b
zgc:103482
gbgt1l4
cpeb1b
si:dkey-90l23.2
LOC101883897
si:dkeyp-46h3.8
hbaa1
si:dkeyp-46h3.2
si:ch211-146l10.8
zgc:171474
uchl1
zpcx
zgc:55461
ctss2.1
spink2.1
zp3a.1
gapdh
avd
mos
gmnn
si:dkeyp-98a7.7
gtf3ab
si:ch211-286b5.4
plk1
thy1
si:ch211-251f6.6
zgc:173742
bzw1b

zgc:113984
AL929192.1
si:ch211-146l10.7
LOC100536860
tmem70
eif4e1b
pigrl2.3
nasp
srsf2a
sybu
mkrn4
si:ch73-14h10.2
zglp1
zgc:56699
pabpn1l
si:dkeyp-46h3.5
si:ch211-250e5.16
sox19b
cdca8
mibp
setd4
b4galnt2.2
crp3
fdx1
st8sia7.1
birc5b
bach2a
CDPF1
elovl7a
cldn7b
zgc:56231
zgc:113424
yrdc
ppfibp1a
zgc:173856

si:dkey-188i13.11
si:ch73-160p18.3
suv39h1b
si:dkey-42i9.8
zp2l1
gstp1
cdca9
zgc:113886
LOC110437779
larp6a
lsm14b
rcc1
snx10a
elavl2
dynll1
rbm4.3
spsb1
pou5f3
rbpms2b
zgc:162879
slbp
tac3b
snu13b
zgc:92591
si:dkeyp-75b4.10
h2afva
ddx41
ccna2
slc16a3
LOC100334439
hspa5
zgc:162945
zgc:86839
h3f3a
dctpp1

zgc:66024
BX005461.3
si:ch1073-190k2.1
CABZ01059627.1
g3bp1
degs2
rpz2
si:ch73-90k17.1
zp3c
map1lc3c
cat
si:dkey-241l7.3
sinup
zgc:165539
tmem54a
hsd17b12a
org
zgc:171426
si:ch211-210c8.7
si:ch211-202f5.3
si:ch211-113a14.27
larp6b
mad2l1
aktip
zgc:171750
CR847984.1
mcm3l
si:ch211-157c3.4
BX571780.1
si:dkey-81e3.2
cdc20
tmem44
retsatl
si:dkey-189g17.2
unga

zgc:171977
asz1
asf1ba
CU467646.2
ppp4r2b
BX284697.1
hmga1a
tmco1
slc29a2
eif4a1a
chia.4
gpr160
CABZ01059627.2
si:dkey-46g23.2
si:dkey-203a12.2
pttg1
cx44.2
cpm
patl2
phex
pabpc1l
tuba8l4
rnf4
si:dkey-216e24.3
tmem144b
zgc:56676
zgc:92287
ccne1
zgc:111868
rdh10b
rpf1
h2afvb
ube2c
orc6
ruvbl1

sprydd4
ndc1
ccnh
thoc7
ankrd22
CT573476.1
cdca7a
slc35f2
st3gal8
si:ch211-145c1.1
elovl7b
zgc:101744
spc24
si:ch211-69g19.2
fbxo32
ivns1abpb
aurka
ypel3
si:dkey-68o6.5
rangrf
faub
gyg1a
lipg
tph1a
BX537276.1
c1d
tomm40
pop4
chtopa
fbxw11b
kpna2
snrpb2
hells
mcm5
rpain

ctsla
rac1a
prdx1
rps19bp1
sec61a1l
ctsba
si:ch73-281n10.2
gar1
chchd10
dynll2a
cfl1l
buc
BX001030.1
mrpl18
rbm7
pcf11
nop16
mcfd2
prps1a
zgc:110239
marcksb
mrps16
inka1b
zc3h11a
snrpc
bud31
hmgb3b
slc25a25a
eif4g2a
pcbp2
fbl
phospho1
slc8a4b
si:ch73-189n23.1
sgms2a

h1m
zgc:153499
si:dkey-240e12.6
zp2.3
zp2.5
tmsb5
zp2.2
CR790368.1
zgc:171776
h2af1al
si:ch211-226h8.4
si:dkey-241l7.2
acp5a
crp2
cldnd
cldng
si:zfos-1505d6.3
mt2
si:ch211-125e6.12
si:ch211-12h2.8
si:dkeyp-20e4.8
si:dkey-241l7.4
BX649337.1
zp3.2
si:ch211-226h8.14
si:ch211-12h2.6
zp3
si:dkeyp-98a7.9
epcam
rrm2
ccna1
zgc:171474
si:dkey-241l7.5
zp2.1
zar1l

si:dkeyp-98a7.4
si:dkey-241l7.6
zp3a.2
si:ch211-14a17.7
zp2.6
si:dkey-229d11.3
zgc:171779
si:dkey-90l23.1
ccnb2
si:dkeyp-46h3.6
CU571163.1
zgc:175135
zgc:171517
LOC110437821
si:dkeyp-98a7.8
si:dkey-208k4.2
zar1
slbp2
si:ch211-250e5.16
thy1
zpcx
qdprb2
si:dkey-90l23.2
LOC100537499
LOC100535162
hnrnpabb
nasp
btg4
lamp2
zp3b
s100a1
avd
nanog
mid1ip1l
cpeb1b

zgc:77118
si:ch211-132b12.8
npm2b
si:dkeyp-50f7.2
gapdh
si:dkeyp-46h3.2
elovl7b
cth1
bgna
si:dkeyp-98a7.7
AL929032.1
zgc:173443
si:dkeyp-46h3.8
gtf3ab
si:ch211-146l10.8
siva1
ccna2
tuba4l
LOC101885477
si:ch211-145c1.1
uchl1
si:dkey-229d11.5
zgc:195245
kpna7
hbba1
chia.4
si:ch73-90k17.1
ccne1
CDPF1
zgc:66024
arhgef10la
wu:fi42e03
zgc:136254
si:dkey-19b23.11
ctss2.1

ddx41
cdca8
rcc1
BX284697.1
si:dkey-241l7.3
si:dkeyp-46h3.5
h3f3a
plk1
buc
LOC101883897
si:dkey-152b24.6
si:ch211-59o9.10
spink2.1
ccnb1
larp6b
larp6a
si:dkey-37o8.1
LOC103909972
crp3
elovl7a
zgc:55461
zgc:92591
mkrn4
birc5b
si:ch1073-190k2.1
pabpn1l
LOC100536860
si:ch73-14h10.2
gmnn
rbm7
zgc:114130
si:ch211-5k11.8
gfer
zgc:162945
gdf3

si:ch211-226l4.6
haus6
BX571780.1
zgc:56699
CR383662.2
COQ10A
elavl2
zp3c
CR847984.1
tph1a
skp2
wu:fa19b12
tpx2
si:dkey-42i9.8
zgc:114123
zp3a.1
CABZ01069022.2
zgc:56231
si:ch211-251f6.6
nanos3
cdca7a
hbaa1
pif1
dynll1
ctsla
zgc:152986
pabpc1l
pou5f3
sinup
slc16a3
CU856358.1
lpcat4
BX571701.1
zgc:113886
ypel3

vps9d1
ddx19
sybu
zgc:162879
rasl11b
zglp1
si:ch211-69g19.2
gstp1
zgc:112146
h2afx
imp3
si:dkey-81e3.2
dnd1
prdx1
prrg2
zgc:171977
buc2l
org
si:dkey-43p13.5
inka1b
rpp21
mylipa
dynll2a
use1
h2afva
tent5c
map1lc3c
bmb
zgc:103482
tacc3
tmem33
selenoh
ankrd22
BX537276.1
rdh10b

BX649350.1
slbp
birc5a
si:ch73-340n8.4
CU929143.1
si:dkey-68o6.5
zgc:113424
si:ch73-132k15.2
si:dkey-216e24.3
CU019646.1
ca15b
zgc:92313
lsm14b
cldn7b
hspa5
CT009596.1
pttg1
tac3b
snu13b
zgc:56676
mos
gbgt1l4
BX001030.1
si:ch211-266i6.3
ccnh
si:dkey-188i13.11
si:dkeyp-75b4.10
zc3h11a
zgc:165539
degs2
zgc:174268
zgc:110239
patl2
ctsba
si:ch211-286b5.4

CABZ01059627.2
si:ch73-303b9.1
pim3
AL929192.1
zgc:113425
cx43.4
si:ch211-14a17.6
zgc:171750
asz1
st3gal8
zp2l1
zgc:163030
BX005461.1
si:ch211-11k18.4
spryd4
cdc20
ctnnbip1
vstm2l
zgc:92287
brf2
cdt1
fbxw11b
CU655961.4
si:ch211-146l10.7
si:dkeyp-114g9.1
parn
zgc:113984
BX323557.1
rac1a
ndc1
pigrl2.3
eif5a2
c1d
eif4a1a
zgc:152652

tatdn2
zgc:111868
zgc:101744
gyg1a
hsd17b12a
ube2l3a
itm2cb
bzw1b
trmt61a
phex
si:ch73-160p18.3
ddx39ab
tmco1
gemin8
lipg
CABZ01059627.1
mibp
zgc:173742
e2f5
tmem54a
mad2l1
zgc:173544
dazl
srsf2a
evla
yrdc
shmt1
si:ch211-210c8.7
ube2c
prr11
fbl
rogdi
rbpms2a
si:dkey-102m7.3
rbm4.3

snx10a
dcun1d1
nudc
zgc:173856
nap1l1
cnot8
aurka
lman1
si:ch73-359m17.2
rbpms2b
pdia4
cox17
ncl
zgc:77112
cfl1l
nol7
sox19b
eif4e1b
slc29a2
myl12.2
btg3
rpz2
coil
pttg1ipb
txnl4a
slc37a2
BX957362.1
cnep1r1
mrpl53
pcf11
fdx1
pop4
zgc:101016
kpna2
tsr2

naa40
cbx3a
uck2a
anp32b
tram2
rab5if
si:ch1073-263o8.2
ckbb
gtf2a1
wdr83os
ndufab1b
khsrp
marcksb
si:ch211-51e12.7
chchd10
sumo3a
retsatl
prmt1
g3bp1
STMP1
uqcrfs1
med19a
lsm5
cbx5
tpm2
acta2
zgc:153499
zp2.2
si:dkey-240e12.6
si:ch73-189n23.1
tmsb5
si:ch211-125e6.12
zp2.3
si:dkeyp-98a7.9
mt2

si:dkey-241l7.5
thy1
h2af1al
si:zfos-1505d6.3
si:dkey-241l7.2
cldnd
zp3
h1m
zp3b
zgc:77118
cldng
zp2.5
crp2
zp3.2
zgc:175135
si:ch211-12h2.8
si:dkeyp-98a7.8
CR790368.1
si:dkey-241l7.4
rrm2
zgc:136254
ccna1
zp2.1
BX649337.1
acp5a
zp2.6
si:ch211-226h8.14
qdprb2
zgc:171779
tuba4l
zar1
LOC100537499
si:dkey-229d11.3
si:dkeyp-98a7.4
si:dkeyp-20e4.8

ccnb2
mid1ip1l
zp3a.2
zgc:171776
si:dkey-90l23.1
si:dkeyp-46h3.6
LOC110437821
lamp2
zar1l
siva1
epcam
CU571163.1
zgc:195245
si:dkey-241l7.6
gapdh
AL929032.1
slbp2
zpcx
zgc:162879
gmnn
si:dkeyp-46h3.8
gtf3ab
si:dkeyp-50f7.2
zgc:171517
si:ch211-226h8.4
uchl1
zgc:56231
si:ch211-12h2.6
elovl7a
npm2b
LOC100535162
si:ch211-14a17.7
pigrl2.3
si:dkeyp-46h3.2
lsm14b

tac3b
btg4
nanog
si:dkey-229d11.5
hnrnpabb
zgc:173443
ctss2.1
BX537276.1
nasp
snu13b
kpna7
AL929192.1
si:ch1073-263o8.2
LOC100536860
mibp
si:ch73-14h10.2
zgc:55461
zgc:171474
si:dkey-188i13.11
s100a1
elavl2
cpeb1b
mad2l1
CABZ01075618.1
wu:fi42e03
si:dkey-208k4.2
rbpms2b
gbgt1l4
LOC101885477
cdca8
si:dkey-37o8.1
si:ch211-250e5.16
si:dkeyp-98a7.7
zgc:92591
si:dkey-19b23.11

snu13a
cth1
h3f3a
chia.4
elovl7b
si:ch211-146l10.8
si:ch211-145c1.1
gdf3
lin28a
zgc:56699
si:dkey-46g23.2
haus6
zgc:114123
si:dkey-152b24.6
zswim7
ccnb1
si:dkeyp-46h3.5
si:ch73-90k17.1
si:ch1073-190k2.1
gstp1
zgc:153284
degs2
si:ch211-132b12.8
si:dkey-90l23.2
slbp
ddx39ab
zp3a.1
si:dkey-179j5.5
birc5b
tmem144b
crp3
rnf4
spink2.1
buc
slc16a3

myl9a
CR847984.1
ccna2
avd
rogdi
si:ch211-146l10.7
zp2l1
zgc:92313
zgc:171977
phex
arhgef10la
hbba1
patl2
snx10a
si:ch211-251f6.6
slc35f2
zp3c
si:ch211-286b5.4
zgc:113886
sox19b
eif4a1a
CU019646.1
kpna2
mkrn4
spryd4
dap1b
hmgb3b
si:ch211-226l4.6
setb
lpcat4
pabpc1l
LOC110437779
ca15b
tmem54a
mrpl9

cdca9
sinup
zgc:101744
rpain
npm2a
eif4e1b
CDPF1
dctpp1
zgc:66024
rcc1
si:dkey-121a9.3
ctsla
si:ch211-154a22.8
si:ch211-102c2.7
si:ch211-5k11.8
plk1
aurka
LOC103909972
zgc:112146
zgc:86839
zglp1
inka1b
vps26a
sybu
oclnb
si:ch211-113a14.27
zcchc8
ctsba
zgc:162945
CU655961.3
si:ch73-78o10.1
cnih1
LOC101883897
pabpn1l
zgc:173856

si:ch211-14a17.6
zgc:112980
pttg1
larp6b
bzw1b
ppp4ca
gpr160
si:dkey-42i9.8
CABZ01059627.2
prr11
si:ch211-276i12.9
dnajc1
retsatl
yrdc
dynll2a
org
fbxo32
cpm
si:dkey-216e24.3
BX005461.3
CT009596.1
micall2a
btr04
zc3h11a
brf2
asf1ba
CU467646.2
sdf2
nol11
f11r.1
ap1s3b
rdh10b
rasl11b
rbm4.3
mtfr2

h2afva
c1qbp
chrac1
cat
zgc:113293
parn
cdc20
nsun2
ndrg1a
tram2
zgc:77112
cdc14b
si:ch73-15b2.5
BX571780.1
asz1
ppp1r3b
dynll1
ccdc58
pcgf1
metap1d
cfl1l
map1lc3c
vps9d1
kif22
tph1a
si:ch73-160p18.3
hbaa1
si:ch211-226h8.6
st8sia7.1
cdca7a
CR383662.2
zgc:55888
thop1
tmem70
si:ch211-226h8.8

zgc:110239
si:dkey-52l18.4
pdap1a
prdx1
zgc:92287
fdx1
si:dkey-241l7.3
ociad1
pdzd11
fbxo5
uba2
g3bp1
nek2
sumo3b
cx43.4
dnd1
imp3
setd4
cmc2
lman1
zgc:111868
BX284697.1
cdc6
lsm1
hsd17b12a
lbh
cldn7b
ccne1
cnep1r1
zgc:173742
zgc:103482
exosc6
zgc:112185
srp9
top1l

surf2
zgc:152652
polr1d
rpl7l1
hspa5
ccng1
si:ch211-69g19.2
pprc1
zgc:171566
zgc:113984
ube2c
mylipa
mos
CU655961.4
itm2cb
nfr
polr2k
ttc32
adipor1a
emc9
marcksb
nol7
ypel3
nap1l1
jtb
gyg1a
mrps26
si:dkey-68o6.5
h2afx
uqcr10
spc24
il13ra2
psmd10
zgc:163030
larp6a

si:dkey-92i17.2
ckbb
BX001030.1
chchd10
sinhcafl
lrrc59
ccnh
c1d
cnot8
npc2
akirin1
dazl
si:ch211-210c8.7
mcfd2
usf1l
hsp90b1
si:ch211-262i1.6
ncbp2
alyref
gtf2a2
chtopa
STMP1
gar1
faub
BX957362.1
rpa3
pttg1ipb
hspd1
lsm12a
rpz2
sap18
psme3
ilf2
selenoh
txnl4a

arl6ip1
snrpg
eno3
manf
slc25a25a
rac1a
thoc7
mrpl53
MED9
si:ch211-51e12.7
cbx3a
rbm4.2
uqcrfs1
srsf2a
ube2ia
zfand5b
dkc1
lsm7
si:dkey-102m7.3
mrpl36
tomm7
cox7a2l
lyz
lect2l
npsn
BX908782.2
mpx
mmp13a
ch25hl2
si:ch1073-429i10.1
BX908782.3
scpp8
il1b
sult2st1
il6r

cd7al
si:ch73-343l4.8
si:ch211-286b5.8
CR848841.1
ly6m5
si:ch211-264f5.2
illr4
si:ch211-193e13.5
si:ch211-284o19.8
si:ch211-9d9.8
si:ch211-260e23.8
chia.6
abcb9
si:dkey-19b23.14
si:ch211-9d9.1
si:dkey-238m4.4
CR354540.1
adam8a
cpa5
ponzr6
hpgd
cnksr2b
CT573383.1
mmp9
CT030188.1
papss2b
hbxb
sms
lta4h
si:ch211-8c17.2
ltb4r
abcc13
BX908782.1
ncf1
BX548046.2

alox5ap
ryr3
si:ch73-359m17.5
CU499336.2
si:ch1073-443f11.2
HACD4
ppdpfa
si:ch73-248e21.7
triobpa
mmp25b
adora2b
slc7a3a
elovl5
srgn
slc2a3b
slc7a11
si:ch211-137j23.6
lrrc15
jakmip1
vwa11
si:ch211-276a23.5
alox5a
LOC101884267
agtrap
rnaset2
noctb
gpr183b
timp2b
cxcr4b
tcnbb
si:dkey-102g19.3
fcer1gl
s1pr4
ctss1
glipr1a

si:dkey-27h10.2
FAR2
spi1b
cfbl
hyal3
cebpa
LOC101886073
CABZ01111959.1
si:ch211-66e2.3
hlx1
mir223
XLOC-004556
gadd45aa
vamp8
pnp5a
odc1
nccrp1
hopx
si:ch211-243a20.4
pigrl4.2
aldh8a1
si:ch73-248e21.5
BX323087.1
zgc:175264
spns3
zgc:165573
itgb2
scinlb
si:ch211-171h4.5
ank3b
si:dkey-222f2.7
rab44
itgb3a
cd63
lpcat2

tmsb1
mrps15
cebpb
alox12
fam107b
actb2
b3gnt3.4
gsr
capgb
LOC101885065
dhrs9
ca6
slc6a6b
plek
apol1
pgp
ifngr1
cnksr1
mapk11
bdh1
dub
dachd
zmp:0000001082
cyp4f3
si:ch211-136m16.8
dgkzb
pfn1
si:ch211-197k17.3
vsir
arpc1b
antxr2a
si:dkey-163m14.7
cotl1
CABZ01073834.1
il16

atp1a3a
rhogb
icn
rac2
atf3
pim1
parm1
si:ch211-67e16.3
rgs19
gpr84
lcp1
trib3
coro1a
nrros
plpp1a
ccdc88b
si:cabz01036022.1
adam19b
lxn
XLOC-039817
skap2
selenow1
FP015862.2
si:dkey-53k12.2
wasb
prkcda
il34
smox
cfl1l
si:dkey-262k9.4
COX3
sh3bgrl
itgae.2
parvg
tmsb4x

myh9a
raraa
derl2
mbnl1
mt-co2
xbp1
CABZ01001434.1
pde4ba
tpd52
zdhhc20a
g6pd
dusp2
rps6ka1
gapdhs
plxnc1
fabp7a
b3gnt2b
mrtfab
unm-sa1614
ncf4
mt-cyb
cyba
CR381673.1
prdx5
farsa
si:zfos-741a10.3
grasp
arpc5b
crip1
itga1
spag1a
mapre1a
mt-atp6
arg2
sdr16c5b

cst14b.1
sri
LOC110438237
ptprr
ppp1r14aa
stk17b
syngr2a
grk6
zgc:64051
mt-co1
si:ch73-208g10.1
amd1
zgc:91976
si:dkey-24p1.7
cox6a2
arhgdig
wbp2nl
sptlc2a
nfkbiab
zgc:63831
rab11fip1b
tpd52l2b
pgd
itm2bb
kctd12.2
cited4a
laptm5
cmtm7
mt-nd4
atg16l1
actb1
gsto2
osgn1
LOC103909115
selp

cap1
arhgef3l
mcl1b
arpc2
tpt1
ets2
msra
wdr1
hbegfa
arpc4l
calm1b
fybb
cited4b
pxk
si:ch211-105j21.9
CR383676.1
vaspa
zmp:0000000936
fermt3b
rab11ba
gpia
plekho1b
rab27a
hsd3b7
h1fx
mt-nd3
si:ch211-197h24.9
ipcef1
rffl
rps6kb1b
gapdh
nme2b.2
tnfaip8l2a
fmnl1a
hcar1-4

elovl1b
tagln2
arpc3
elf1
si:ch211-102c2.4
fbp1a
s100a10b
ppt1
gpx1a
chchd3a
si:ch73-234b20.5
myo1f
zgc:77112
capza1b
srd5a2a
samsn1a
CR392341.1
pdhb
lpar5a
sh3kbp1
dbi
ptprc
pygb
zgc:174904
tmem30ab
iqgap1
lrrfip1a
ptmaa
abca2
actr2a
ak1
zgc:66427
fam49ba
zgc:77650
crema

dusp1
clic1
si:ch73-1a9.3
aldoaa
add3a
rca2.1
tpm3
lamtor4
si:ch211-132p1.3
cox8a
zgc:152863
mtpn
plbd1
serpine2
arrb2a
mt-nd1
rgs13
pkma
CABZ01088428.1
msrb1b
itpkcb
mt-nd2
selplg
slc38a2
tsc22d3
cbx7a
capzb
sptbn2
cnp
pde7a
lasp1
si:rp71-68n21.9
hmgn2
scamp2
LOC566587

arpc4
acsl4b
limd2
socs3b
scarb2a
svild
cox6b1
ccni
nr4a1
myl12.1
tpi1b
exoc6
LOC103911430
rab11fip4b
si:dkey-25e12.3
dock8
arrb2b
birc2
wasa
thy1
degs1
fam49bb
si:ch211-226m16.2
magt1
aldh9a1a.1
ccdc28a
cmip
flna
tktb
tmem59
gpx4b
adgre5b.3
chac1
grap2b
hmha1b

eno3
pnrc2
tln1
si:dkey-27i16.2
cdc42
sh3bgrl3
msna
LO018102.1
npl
sgk1
mcl1a
rap1b
mid1ip1b
si:dkey-112e7.2
si:ch1073-392o20.2
cd164
actn4
rab1ab
capza1a
hif1ab
btg1
tuba8l
cfl1
eif4ebp2
serf2
tnfb
gadd45ab
arhgdia
zgc:162730
anxa11b
gnai2a
myh11a
f11r.1
nfil3-6
ucp2

si:ch211-212k18.7
stk17al
mvb12a
si:ch211-196l7.4
zgc:56493
sumo2b
pim2
idh2
tmed7
abracl
sarnp
GCA
tubb4b
prelid3b
cnn2
socs3a
romo1
eif3f
tomm20b
rsl24d1
nsa2
NC-002333.4
zgc:153867
LOC103908723
zgc:194275
LOC101886766
igl4v8
cd37
CU896602.1
igic1s1
igl1c3
XLOC-026705
igl3v5
cxcr4a
BX571825.4

LOC101882264
pax5
BX571825.5
epb41a
CT573342.2
CU571382.4
LOC108179343
ly6m2
BX571825.3
LOC101886979
ighv1-4
FP016018.1
rtn4rl2b
capga
CU571382.1
CT573231.1
ncf1
CR944667.2
swap70a
si:ch73-7i4.3
XLOC-018356
si:ch211-145b13.6
pou2f2a
sftpba
ccr9a
cxcr4b
si:ch73-380l3.2
illr1
LOC110438240
grap2b
btbd11b
CT027980.1
sp6
si:dkeyp-117h8.2
cobll1a

tlr19
rbbp8l
pacsin2
atp11a
sh2d4a
lmo2
si:ch1073-220m6.1
blnk
csrnp1a
si:ch211-119e14.1
si:dkey-24p1.1
irf4b
XLOC-042595
ebf1a
cd74a
si:busm1-266f07.2
mpeg1.1
si:dkey-82o10.4
arhgap15
cd74b
dapp1
gng2
derl2
syk
XLOC-000431
gpr55a
cd7al
cd83
kctd5a
bcl11aa
bpifcl
dtx4a
XLOC-013964
rgs13
mhc2dab

CABZ01066694.1
fam53b
LOC103910871
ptmaa
rac2
tspan2a
spi1b
s1pr4
si:ch73-234b20.5
sox12
gadd45ga
LOC103909452
gpx1a
rps20
rps21
si:ch211-197h24.9
ccni
rplp1
rps27.1
cited4a
si:dkey-78k11.9
XLOC-010899
rplp2
ncf2
rpl11
marcksl1a
crip1
LOC100536213
rps15a
cad
rpl32
rpl36
rpl10
rsl24d1
pacsin3

rbm38
rpl8
CR936442.1
pfn1
rpl19
rplp0
ramp2
rpl18a
parvg
ikzf2
h1fx
jam2a
tsc22d3
rps5
rpsa
ptmab
rpl15
klf13
hmbsa
rpl37
ralgps2
coro1a
rplp2l
cybb
rps2
naca
rpl10a
rps4x
rps26l
rps29
thrap3a
TPGS1
rpl34
cyba
rpl7

rpl28
ppiaa
histh1l
rps13
rps6
rpl21
rpl35
rpl30
hmgn2
pik3r1
cnpy3
rpl24
b3gnt3.4
btg1
tfdp2
ascc1
pde4ba
rpl18
wu:fb55g09
rpl26
rpl36a
arhgdig
prkcbb
rpl9
LOC103909115
rps14
hsp90ab1
rps16
rps12
rpl7a
dap1b
celf2
rack1
ccnd2a
rps24

mhc2b
si:dkey-183p4.10
rps18
glo1
rps17
rpl23a
actb2
rpl35a
rpl27
hopx
rpl23
rps23
faua
ada
rps10
si:ch211-214p16.1
tmc8
serf2
eif3f
rps19
tnfaip8l2a
nr4a1
mknk2b
mbnl1
rps3
rpl6
rpl31
tpt1
ptprc
rps9
nsa2
egr2b
si:ch1073-429i10.3.1
plk3
rpl38

rps28
rpl13
btk
ptp4a3
rpl39
rpl22
oaz1b
dusp2
unc93b1
btf3
eif3m
rps15
si:ch211-195b13.1
rpl5a
rps7
zgc:162730
tmc6b
wasa
slc25a55a
ccl35.1
ucp2
mbpb
wasb
rpl27a
rps8a
sesn1
rpl29
rps11
xpc
itpka
nme2b.1
egr3
serbp1a
arpc1b
si:dkey-27i16.2

lmnb2
rps27a
CABZ01079427.1
tmsb4x
rps3a
rgcc
si:ch211-150o23.2
rpl12
zgc:114188
rps25
rel
herpud1
inpp5d
gpr183a
gdi2
plcg2
gm2a
rps26
pnrc2
h2afx
nop53
si:ch211-198c19.3
tob1a
lyn
eef1g
cct2
uba52
stk35
rhogb
tagapa
myl12.1
foxp1b
nfkbiab
sept9b
wipf1b

slc25a5
irf2bp2b
cirbpb
capzb
mfsd4aa
fmnl1a
sept12
romo1
eif5a
mcl1a
CR383676.1
ran
rbm39a
capza1a
gnb1a
arf6b
hmha1b
tefa
kdm6bb
COX7A2 (1 of many)
arl8bb
sarnp
arrdc3b
fgfr1op2
FERMT3 (1 of many)
mibp2
zgc:64051
naa10
ss18
dub
pycard
ero1b
eif3k
ppp3ccb
rpl22l1

si:dkey-112e7.2
skap2
smox
acin1a
nrros
bcl2l10
arpc4l
XLOC-028370
eif3ea
plekho1b
lbr
tubb2b
arpc5a
calm1b
crlf3
ldha
CABZ01090890.1
ahcy
XLOC-042596
ier2a
sec61g
mki67
mych
si:dkey-262k9.4
jmjd1cb
CR381673.1
bhlhe40
capza1b
ddit3
anapc16
grasp
eef1da
rpl5b
ywhaba
lasp1

wasf2
zgc:92606
pabpc1a
eif3g
smc1al
msna
arf6a
zgc:56493
si:ch211-214j24.10
zfp36l1a
rpl4
XLOC-025423
btg2
ankrd11
erbin
magt1
arpc3
si:ch211-199g17.2
irf2bp2a
eif3c
sumo2b
ifi30
sat1a.2
itm2bb
pgam1a
hmgb1b
arpc2
mob3a
arhgdia
b2m
rsl1d1
rad21a
NC-002333.4
sh3kbp1
h3f3c

crebzf
aldoaa
hmga1a
atf4b
arl5c
prelid3b
selenok
eif1b
cox6b2
hnrnpa0b
cfl1
eif4a1b
si:ch73-1a9.3
eif3ha
cdc42l
hmgn6
gng5
ola1
rps27.2
cox7b
mt-atp6
litaf
si:ch211-222l21.1
hmgn7
mt-co2
hmgb2a
lyve1a
kdr
she
mrc1a
lyve1b
si:dkeyp-97a10.2
flt4
tmem88a
pcdh12

CT033796.1
myct1a
akap12a
lgi2a
si:ch211-152c2.3
rca2.2
egfl7
clec14a
rasip1
gpr182
cldn5b
parvab
si:dkey-237i9.8
ldb2a
ehd3
stab1
si:dkey-126g1.9
sele
sox18
kdrl
igfbp7
cdh5
si:ch211-248e11.2
etv2
fabp11a
tie1
igf2a
tm4sf18
rab11bb
b3gnt7
plk2b
fgd5a
pecam1
cdh6
exoc3l2a

ppp1r13ba
nrarpb
shank3b
podn
si:ch211-33e4.2
ramp2
si:ch211-14k19.8
nr5a2
loxl1
fam174b
CU929150.1
erg
plk2a
s1pr1
arhgef9b
notchl
creld1a
si:dkey-28n18.9
dll4
slc9a3r1b
hapln3
pgm5
gpm6ab
piezo2a.2
ecscr
cyp26b1
reps2
rassf9
ednraa
sox7
anks1b
cyth3a
cx39.4
si:ch211-145b13.6
tmem108

LOC101882264
macf1a
fzd4
c1qtnf6a
afap1l1b
hhex
calcrlb
fli1a
pcdh17
aqp8a.1
si:ch211-207d6.2
si:ch73-334d15.2
gas6
ebf3a
nova2
rapgef5b
cldn11b
limch1b
akap12b
hyal2b
tnfaip2b
selenop
marcksl1b
laynb
cpn1
lmo2
reep1
XLOC-015238
cxcl12a
ccl38.1
grb10a
tspan4b
adam8a
tpm4a
yrk

prox1a
klf2a
SEMA4F
plvapb
tcima
arl4ab
dab2ipb
lfng
cdc42ep5
oaz2b
sema3ga
si:ch211-1a19.3
vat1
si:ch211-66e2.3
fam198b
exoc6
acap3a
cd81b
ccdc85b
il13ra1
krt8
atp1b1a
marcksl1a
myo1ca
epas1b
gcnt4a
col15a1a
notch1b
lrch1
palm2
pdgfd
ralgapa2
crip2
CABZ01084131.1
lzts2a

spns2
arhgap31
npdc1a
foxp1b
stox2a
mafba
cttnbp2nlb
scarf1
kank3
adgrl3.1
fam43a
krt18a.1
spry4
grapa
plekhg5a
fkbp1ab
CU984579.1
heg1
abcg2a
klf2b
dlc1
XLOC-016094
arl4aa
sema4c
oaz2a
lgmn
cyp3c1
c1s
wasf3b
rnd3a
pros1
mrc1b
acsbg2
si:ch211-156j16.1
efna1a

gstt1b
pmp22b
chka
cavin2b
pdk2b
tcf12
LOC103911404
si:ch211-71n6.4
rps6ka3b
tead1b
clic2
zgc:153317
carhsp1
emilin1b
tagln2
phlda2
srgap1b
npc1
add3a
CR383676.1
aplp2
aatka
erfl3
cbx7a
gng10
tbc1d4
lima1a
si:ch73-138n13.1
ponzr1
tmem88b
dhrs3a
rhoca
tshz2
eva1ba
si:dkey-56i24.1

sash1a
socs3a
si:ch211-284f22.3
si:dkey-9i23.16
zgc:195173
palm1a
anxa13
tuba8l3
asap1b
tpte
rgcc
jun
pdlim1
palm3
ckba
ms4a17a.9
slc12a2
gadd45ba
dab2
zfp36l1b
hspg2
ier2a
fkbp1aa
mef2cb
mbnl2
si:dkey-192g7.3
smad1
cd276
si:dkey-163m14.2
dusp5
stab2
bmpr2a
cav1
ctsla
id1

piezo1
vamp5
acvrl1
myh10
rbp2a
cald1a
appa
tjp1a
lpar1
jund
btg2
junba
stard8
nos1apa
myo1cb
map3k20
klf6a
prss59.1
rapgef5a
RALGDS
reep3a
soga1
nr2f2
fmnl3
pbx1b
wwtr1
ubl3a
qkia
snx9b
cnn3a
krt94
tmsb4x
XLOC-024750
ch25h
sypl2a

fosab
slc25a43
ctdspla
cx43
sept2
fosb
clasp1a
cd81a
pim1
col4a3bpb
scarb2a
mprip
gstr
ap1b1
hspb1
mmp14b
pdk4
ier2b
cd63
myl6
itm2ba
ehd4
degs1
ahnak
ftr82
fblim1
cavin1b
trip10a
tspan18b
antxr1c
tns2a
sgce
si:ch73-375g18.1
hecw2b
rgs3a

itga9
ms4a17a.11
cdc42
sesn3
tspan9a
gpt2
klf3
rasgef1bb
afdna
ccdc80
rplp2
zfp36l2
cox6b1
tmem176l.1
rgl1
foxo1a
pde4ba
rabgap1l
pik3ca
spred2b
rplp2l
CU459056.1
fstl1b
snx13
btg1
atf4b
si:ch211-196l7.4
ralba
fosl1a
mef2aa
plcg1
im:7151449
pfn2
si:dkeyp-19e1.3
mt-co2

entpd1
ctrb1
itga5
txnipa
pnrc2
zgc:114188
ehd1b
krt15
cdc42se1
rhocb
rpl29
rab5c
rbms1b
msna
rpl9
tns1b
cnn2
il4r.1
tmem176l.3a
CT030188.1
si:ch211-240l19.8
smad5
col4a1
rps28
ginm1
parvaa
rab8b
ephb3
rpl36
rap1b
gabarapb
mt-nd1
COX3
rhoaa
si:zfos-464b6.2

sept12
socs3b
tspan14
h3f3c
mt-co1
rab1ba
abca1a
mcl1b
mt-cyb
shoc2
rhoab
tacc1
si:ch211-204c21.1
rps24
sik1
mt-nd2
DDX17
plpp1a
zgc:158343
perp
CR318588.3
si:ch73-335l21.4
rpz5
ier5
sptlc2b
cst14b.1
mt-atp6
dedd1
ccnl1a
cox8b
ptbp3
snx1a
si:ch211-74m13.3
stau2
per3

rpsa
arrb2b
gnb1a
LOC103909284
svild
hopx
csrnp1b
gadd45bb
ddit3
adipor2
sptan1
il6st
klf7b
ptprga
srgn
grinab
kctd5a
cyth1a
b2m
si:dkey-79d12.5
rps10
zmiz1a
NC-002333.4
uba52
crip1
si:ch73-234b20.5
midn
rps27.2
elmo1
ucp2
sh2b3
stat1a
sumo2b
pkig
rps21

selenot1b
podxl
sdcbp2
aoc2
chmp1b
nsa2
fkbp5
mgst3b
antxr2a
abcc5
si:dkey-7j14.6
fosl2
nr1d2a
cltca
slc43a2a
mt-nd4
higd1a
h2afx1
capns1b
dusp1
CABZ01072614.1
cdkn1bb
egr1
maff
cfd
dynlt3
rpl22l1
mt-nd3
chd3
ankrd11
epdl1
spock3
CU928222.3
syngr3b
FP085399.2

zgc:101810
AL929237.1
ptprh
BX323596.1
cxcl11.6
mxf
BX323596.2
mpeg1.1
cxcl11.5
si:ch211-1o7.3
si:dkey-33i11.9
hepacam2
si:dkey-148a17.5
grn1
sftpbb
spi1a
CU657980.2
tnfa
arap1a
ccl35.1
si:ch211-214p13.3
arhgef15
ccl35.2
smcr8b
si:ch211-79k12.1
mhc2dab
si:dkey-102c8.3
CR385050.1
si:busm1-266f07.2
gna15.1
CR786582.2
zgc:123107
CABZ01078737.1
abi3a
si:dkey-1h24.6

si:ch73-158p21.3
cd74a
mxc
si:ch73-236c18.7
si:dkey-192g7.3
CU459094.3
si:ch211-214p16.2
kctd12.2
arpc5b
rnaset2l
tnfb
tiam1a
cd74b
si:dkey-78o7.3
ptgs2b
ciita
plekhf1
BX004816.2
LOC101885445
ccl39.1
zgc:103700
si:dkey-7i4.24
si:busm1-194e12.11
lygl1
arpc1b
tmsb1
ctss2.2
si:dkey-27i16.2
gpx1a
grna
spi1b
ccl19a.1
nr4a2b
BX548046.2
mhc2b

LOC101885065
si:ch211-8c17.3
tmsb4x
pfn1
XLOC-025423
fgd4b
lcp1
il21r.1
si:dkey-53k12.2
ccl39.3
LO018493.1
ptmaa
actb2
TPGS1
irak3
wasb
CU499330.1
cd83
cotl1
si:ch211-152p11.4
actb1
tcima
BX901920.1
BX005183.3
gstt1a
tfe3a
psap
si:ch211-198c19.1
parm1
p2rx7
aif1l
XLOC-023492
XLOC-005449
sgk3
malt3

ccl39.2
mbnl3
sh3bgrl
krt18b
gabarapb
zgc:136870
gm2a
id2a
arpc2
pycard
ppiab
irf2
hcls1
si:ch211-214p16.1
CR753876.1
tent5ba
ifi30
arpc3
unc93b1
zgc:63831
si:cabz01074946.1
pak2b
mapk11
map3k14a
arhgdig
efhd2
npc2
cox6b1
fgl2a
zgc:153733
si:ch1073-214b20.2
LOC103910136
rgs18
triqk
myo1f

cyb561a3a
ackr3b
hdac9b
wipf1a
GNAZ
mrtfab
plekhj1
anxa3b
dbn1
sept12
parvg
cxcr4b
zgc:162730
il12rb2l
ppt1
capza1b
selplg
coro1a
XLOC-028370
arpc4l
marcksl1a
eif3jb
cyba
rac2
itpka
dennd2da
fcer1g
cnn2
mao
gng2
vsir
arpc4
capzb
myl12.1
itgb2

ckba
litaf
XLOC-008993
xaf1
irf8
CR318588.1
plbd1
csrp1a
abracl
lgals9l1
rplp2
twf1a
b2m
zgc:64051
XLOC-032112
BX908750.1
timm10
BX322618.1
hcst
ufc1
ccr9a
arf6b
irf7
pak2a
stk17b
prdx5
gabarapa
batf3
bach1b
cdc42l
wdr1
skap2
slc15a4
gstm.3
fam129bb

fcer1gl
BX120005.1
rab5c
asah1b
XLOC-033137
si:dkey-262k9.4
atp6v1g1
sema4ab
wee2
itm2bb
cfl1
si:ch211-222l21.1
ecm1b
plekho1b
ostf1
si:dkey-206d17.12
rps15a
h1fx
mbd2
COX7A2 (1 of many)
ncaldb
si:dkey-112e7.2
cmtm3
histh1l
itgae.2
glrx
si:ch211-153b23.7
plekho2
arl5c
si:ch73-1a9.3
chchd7
ctsh
BX248082.1
cap1
pmaip1

GCA
ak1
rpl18
nr4a1
gdi2
rgs13
XLOC-000431
rab8b
XLOC-005448
ptmab
mir21-1
stat5a
actr2a
si:dkeyp-68b7.12
LOC103909115
rps27.1
psme2
sdcbp2
ccng2
ubl3b
map2k6
CR392341.1
rps2
lgals2a
bloc1s6
tpi1b
tagapb
rpl39
ms4a17a.6
rps5
RASSF5
vdac3
cox8a
naga
tagapa

sri
tsc22d3
glula
rps21
laptm5
rps17
rps26l
nptna
ctsk
rpl8
gstm.2
ckbb
oaz1b
ppiaa
tagln2
eif3g
rhoab
fuca2
ndufb3
ybx1
grn2
rps15
zgc:114188
rps9
arhgdia
rpl38
rpl35a
rpl21
rpl18a
si:ch211-212k18.7
atp6ap2
rpsa
rps14
rps3
rpl36

sh3bgrl3
clic1
cox4i1
crip1
romo1
fosl2
rpl15
sod1
rpl32
eif3f
derl2
rps28
XLOC-041870
tln1
fam53b
rps16
rpl9
serf2
rpl37
rps24
bhlhe40
mob1a
sept6
ywhabl
rpl5a
mbnl1
rpl29
fam49ba
capza1a
psma2
cdc42
COA4
atf4a
ube2e3
XLOC-020301

rplp2l
XLOC-010899
magt1
psme1
atp6v0ca
mapre1a
rpl11
zgc:56493
zgc:153713
rpl22
slc6a6b
naca
rps4x
klf6a
CR318588.3
rpl6
mtpn
nccrp1
cox5aa
rps23
rpl27
uqcrq
rpl36a
atp5meb
rpl24
rps10
rpl34
ssr3
hmgn2
b2ml
tpm3
mcl1a
eef1db
sat1a.2
cst14a.2

ap2m1a
capgb
rpl27a
ccni
COX5B
rps27a
hmgn7
atp5pb
rpl17
pomp
rpl30
rpl14
faua
zgc:153867
rps27l
ywhaba
rps27.2
celf2
mibp2
rps18
rpl31
eef1g
rpl23a
uba52
mpx
zgc:175135
cldnd
zp2.5
si:zfos-1505d6.3
si:dkeyp-98a7.9
si:dkey-240e12.6
crp2
si:dkey-241l7.2
h2af1al
CR790368.1

zp2.1
si:dkey-241l7.5
tmsb5
si:dkey-241l7.4
slbp2
si:dkey-241l7.6
si:dkeyp-20e4.8
LOC100537499
si:ch211-226h8.14
ccna1
si:ch211-12h2.8
zp2.3
zp3
si:dkey-229d11.3
zgc:153499
cldng
tuba4l
si:ch211-125e6.12
zp2.2
zar1l
acp5a
siva1
zp2.6
zgc:171776
mt2
lyz
zp3.2
si:dkeyp-98a7.8
zgc:55461
thy1
si:dkeyp-98a7.4
h1m
si:ch73-189n23.1
slc16a3
zgc:77118

ccnb1
BX649337.1
LOC101883897
zpcx
crp3
zgc:171517
ctss2.1
si:ch211-226h8.4
lamp2
qdprb2
zp3b
gapdh
CU571163.1
si:ch211-250e5.16
zgc:173443
ca15b
s100a1
si:dkey-90l23.1
nasp
gtf3ab
zp3a.1
si:dkeyp-50f7.2
ccnb2
zgc:171779
nanog
mid1ip1l
hnrnpabb
si:dkey-19b23.11
hbba1
elovl7a
zp3a.2
rrm2
AL929032.1
zgc:66455
wu:fi42e03

si:dkeyp-46h3.6
si:dkeyp-46h3.8
gmnn
uchl1
cth1
si:dkey-90l23.2
epcam
zgc:195245
tmem54a
lsm14b
si:ch211-14a17.7
kpna7
LOC110437821
hnrnpa0l
si:ch211-207i20.3
cax1
birc5b
si:dkeyp-46h3.2
zgc:56231
ccna2
LOC100535162
LOC101885477
si:ch211-132b12.8
zar1
si:ch1073-263o8.2
ttc9c
zp3c
haus6
si:dkey-208k4.2
npm2b
zgc:136254
CDPF1
larp6b
pabpn1l
si:ch211-12h2.6

zgc:173856
zgc:162879
arhgef10la
btg4
zgc:171717
si:ch211-5k11.8
si:dkey-152b24.6
spink2.1
AL929192.1
hnrnpaba
si:ch211-145c1.1
si:ch73-109d9.2
si:ch211-251f6.6
osbpl7
sms
si:dkey-229d11.5
rogdi
hbaa1
map1lc3c
tipin
CR847984.1
kif22
avd
gdf3
prdx1
pif1
nanos3
zgc:113424
inka1b
si:ch73-15b2.5
zgc:171474
CABZ01059627.1
zgc:173742
il13ra2
arid3b

eloca
zfand6
timm13
si:ch211-202f5.3
mkrn4
si:ch73-14h10.2
cpeb1b
si:ch1073-190k2.1
BX537276.1
ddx19
si:dkey-241l7.3
si:ch211-226h8.6
dynll1
srsf2a
org
tacc3
zgc:162509
ubc
med22
rchy1
zgc:163030
si:dkey-37o8.1
tmco1
si:dkey-42i9.8
sinup
si:dkeyp-98a7.7
CU467646.2
kmt5ab
si:ch211-137a8.2
lsm11
gyg1a
si:ch211-69g19.2
zgc:92313
scamp2l
dazl

si:ch73-382f3.1
coil
fbxo43
zglp1
alg3
hells
prr11
retsatl
elovl7b
zgc:92591
zgc:111868
orc6
ctsba
slc25a25a
cdca8
cdca7a
si:ch211-226l4.6
h3f3a
sybu
cdc20
brf2
zgc:103482
zgc:173545
aph1b
snu13b
cdk1
mcm3l
si:ch211-146l10.8
rbm4.3
sumo2a
mcmbp
si:dkeyp-46h3.5
ddx39ab
casd1
si:ch73-90k17.1

cse1l
thap11
marcksb
rbpms2b
LOC100538023
zgc:92287
dkc1
si:ch73-160p18.3
ninl
si:dkey-188i13.11
tpx2
lin28a
zgc:56699
st8sia7.1
plk1
eloal
h2afva
zc3h11a
elavl2
cldn7b
si:dkey-46g23.2
fam173b
mcfd2
snu13a
uqcr10
dnaja2a
tmem17
eif4g2a
LOC100536860
glipr2l
kif7
bmb
ivns1abpb
zgc:171750
dynll2a

ap2m1b
BX571780.1
ankrd10a
LOC110440049
snrpb2
cyp11a1
buc2l
metap1d
exosc8
CU655961.3
st3gal8
lmnl3
lrp13
fam122b
cbx3a
si:dkey-43p13.5
tph1a
paip2b
rac1a
zgc:66024
BX284697.1
dnd1
pttg1ipb
ypel3
adipor1a
asz1
nap1l1
fdx1
anp32b
si:ch73-78o10.1
exd2
lpcat4
zgc:171977
rpain
vstm2l

si:ch73-303b9.1
lyar
polr2k
gbgt1l4
h2afvb
rbbp4
rrs1
si:ch73-281n10.2
pigrl2.3
mycla
ankrd22
yrdc
uba2
aurka
zgc:152652
tspan36
si:dkey-16l2.16
krcp
UBL4A
rcc1
uck2a
psmd10
mrps18b
nup58
CT009596.1
buc
faub
ccne1
mos
CABZ01059627.2
pim3
mcm7
pou5f3
cdc14b
rbm7

bzw1b
hsd17b12a
STMP1
LOC103909972
zgc:56676
ppp1r3b
larp6a
mrpl18
rdh10b
rcor1
slbp
rpp21
asf1ba
lsm12a
igf2bp3
pttg1
si:dkeyp-75b4.10
ccnh
thop1
aktip
zgc:153284
aurkb
gemin7
dnmt1
lbh
itm2cb
col10a1a
scpp7
ifitm5
cd81b
enam
tmem119a
panx3
si:ch211-106n13.3
sgms2a

si:dkey-12l12.1
sod3b
cthrc1a
phospho1
bgna
col10a1b
col11a1a
loxl4
col11a2
fgfbp2a
cdh11
ndnf
slc8a4b
spp1
zgc:92162
chst11
smpd3
itga10
LO018513.1
bambib
ano5a
bmp8a
runx3
col5a2b
qpct
klf2b
entpd5a
pcolce2b
kazald3
dlx3b
sp7
si:ch211-170d8.8
bmp3
COLGALT1 (1 of many)
myadml2

thrap3a
nrp2b
prp
CABZ01059119.1
satb2
bmp2a
ism1
dlx5a
pltp
phlda3
plcd3b
si:ch211-106h11.3
ihha
dlx2b
wnt10a
cspg4
zgc:162171
scpp1
megf10
ppfibp2a
chodl
rcn1
serpinh1b
fkbp9
serpinh1a
tnfrsf11a
zgc:162612
mxra5a
XLOC-005794
si:ch211-110e21.3
nav2a
crip2
myh11b
nid2b
zbtb7c

mfsd2b
olfml3b
xylt1
fgfr3
ccdc80
plod1a
lmo7a
sox11a
golim4b
pcdh17
igfbp1a
sema3e
p4ha2
heyl
fpr1
fam20a
si:dkey-234i14.3
rnd2
si:cabz01100188.1
tfap2c
s1pr1
p3h1
LOC101886114
btbd6a
her6
mef2cb
cdc42ep3
si:dkey-193p11.2
col11a1b
pcolceb
pdgfbb
acvrl1
nupr1b
thbs1b
BX322661.2

col14a1b
ccdc187
twist2
cplx2
rcn3
plekhh1
cd82b
serpine2
msx2b
plxna3
vkorc1
tox
tcf7
crtap
fkbp14
clec3bb
zgc:110796
tbx15
nradd
alcama
rarga
cpe
bambia
mef2aa
tbx2b
stc2a
MDFI
farp2
twist1a
rbfox1
col1a1a
serpinf1
tnfaip8l3
tmem45a
htra3b

sparc
p4ha1b
LOC108179036
creb3l1
fam20cb
col1a2
fkbp7
LOC100535130
FAM163A
zgc:153317
cdh7a
cx43
znf385a
bcl11aa
zeb2a
kremen1
foxd1
frmd6
XLOC-035413
nrp2a
mef2ca
marveld1
col1a1b
tmem198b
tspan4a
timp2a
tgfb2l
ak5
pleca
fkbp11
bmp6
tmeff1b
zgc:162939
reck
asip1

rtn1a
FP102018.1
cgref1
krt18a.1
sox6
plod3
fabp7a
fhl1b
kdelr3
pdlim4
atp1b3a
col5a1
hoxb7a
jdp2b
ncam1a
mmp16a
pard6gb
fxyd1
cav1
srgap3
adgra2
lhfpl6
ckap4
CR376783.1
VIT
pmp22a
notch3
nrgna
fscn1a
cbfb
id2b
si:ch211-133n4.4
fosb
aqp3a
sgcd

robo1
ier5l
hoxb8a
copz2
hsp70l
efna1a
itgb1a
surf4l
anxa13
paplnb
rpz5
eva1ba
cpamd8
adgrl3.1
phex
id1
bckdhbl
serinc5
prkd2
sox18
mmp14a
tuba8l3
fgfr1a
slc39a7
ctnnb1
CU929237.1
cntfr
hsp70.3
fstl3
SCARF2
si:ch211-156j16.1
icn
sh3pxd2aa
nuak1b
pcolcea

fn1a
calua
bhmt
cnn3a
hmgb1b
prkab1b
b3glcta
angptl4
schip1
si:ch211-286o17.1
mrc2
cercam
dpt
fkbp10b
perp
yap1
cavin1b
sox4b
emid1
ecrg4a
s100v2
jun
fosab
loxa
zgc:153675
fat1a
endouc
nucb2a
ccnd1
klf2a
fzd2
lhfpl2b
egr1
ppfibp1b
meis1b

ier2b
igfbp3
elovl1b
meis1a
cd99
mych
impdh1b
irf2bpl
ptprfb
lrrc17
s100a10b
lgals3b
slc31a1
anxa2a
fam168a
atf3
mgst3a
fstl1b
prrx1a
myo1b
creb3l2
junba
plod2
cdkn1a
asph
tpm4a
mob2b
si:ch73-46j18.5
midn
h1f0
tsc22d3
zdhhc8b
prdx5
ptbp3
h3f3c

anxa5b
net1
kdelr2a
gadd45ga
tns3.2
txndc5
chsy1
irf2bp2b
efna1b
tent5ba
mdka
slc38a5b
capns1b
tsc22d1
vgll4b
metrnlb
LOC101883507
gadd45ba
si:ch73-335l21.4
map1lc3b
mmp2
socs3a
ier2a
sgce
ppib
dap
col5a2a
atp2b1a
h2afy2
ptprga
selenok
calr
arg2
kank1a
supt3h

si:busm1-57f23.1
klf11a
snai1a
klf3
bcar1
triqk
sdc2
frzb
sumo1
hmgn6
ier5
si:dkeyp-113d7.1
anxa1a
rrbp1a
hmgn7
sar1b
neo1a
calm2a
ssr3
klf13
tspan7
sept15
f13a1b
hdlbpa
antxr2a
tspan9a
si:dkey-28b4.7
pkig
vdac3
zdhhc20a
pfkfb4b
prss35
CABZ01080702.1
serf2
spred2b

gng2
sdc4
arl4d
znf704
sdf4
zgc:162730
twist3
nme3
pmp22b
spry2
sec61a1
ip6k2b
enpp1
gng7
atf4b
nr1d2a
fosl1a
metrnla
tacc1
chd6
cald1a
emp2
lrpap1
map1lc3a
klf11b
her9
si:ch211-194k22.8
ephb3
atp2b2
kdelr2b
selenof
arl1
arf5
myl9b
myca

flot2a
mgst3b
hnrnpa0a
rhoab
pgrmc1
tdh
btg2
mmp14b
klf6a
mafba
cd81a
krt8
CABZ01112656.1
vamp3
ciarta
col6a1
appb
osr2
histh1l
ywhaqb
ndrg2
nfia
pbx1a
gadd45ab
cish
spcs1
tgif1
capn2a
snd1
plecb
rab2a
tfg
tmem167a
grpel2
tomm20b

ssr4
qkia
ldlrap1a
svild
chd4b
itga5
tpm1
sept7a
atrx
nck1b
cited4a
cd164
id3
wls
gsna
ptmaa
mfap2
tmem258
si:ch211-286b5.5
tpt1
ostc
si:zfos-169g10.3
eif4a1b
ssr1
ywhabb
slc3a2b
krtcap2
si:ch73-1a9.3
cops9
marcksl1b
evla
glud1a
si:ch1073-325m22.2
junbb
ier3ip1

LOC101884451
rgcc
elocb
cyr61
notch2
ahnak
si:dkey-56m19.5
idh2
kctd12.2
jund
nfkbiab
tmem59
eef1a1l1
eif1b
rpl22l1
ccnd2a
fkbp1aa
ssr2
CR383676.1
ptp4a1
mtdha
serinc1
calm3b
CABZ01045617.1
atf4a
ost4
reep5
zgc:92606
cnn2
pim2
pfn2l
chd4a
cnbpb
tnrc6c1
smim15

zgc:77849
mt-nd2
mcl1b
ybx1
rpl39
tcf12
CU694197.1
hp1bp3
si:ch211-39i22.1
ptmab
jpt1b
mtpn
sem1
p4hb
nsa2
eif3ea
rhoaa
socs3b
ddit3
si:dkey-42i9.4
atox1
tpm3
serp1
dusp2
mibp2
si:ch1073-429i10.3.1
eif4ebp2
tmed10
cdc42
ndufa1
mycb
gnb1a
ndufa4l
rab10
cox6a1

itm2ba
si:dkey-79d12.5
pno1
gtpbp4
psmb7
mt-nd3
mt-nd1
si:dkey-7j14.6
gabarapb
rps2
tubb2b
cox8a
rpl5a
gng5
tegt
tma7
cebpb
cox7a2a
rpl29
rps27.2
XLOC-023492
higd1a
eef1g
sec61b
rpl18
CABZ01072614.1
oaz1b
rpl7
tob1b
sumo2b
rps27.1
mt-nd4
skp1
ubb
COX3

rpl10
hnrnpa0b
rpl32
rpl9
lum
si:ch211-196l7.4
eif2s3
mt-cyb
rplp1
mt-atp6
eif3i
dad1
rpl18a
ddx5
rpl10a
rps18
rps19
rpl27a
rps7
xbp1
pdia3
rps12
rps4x
sec61g
mt-co1
rps10
rps8a
rpl11
faua
zgc:114188
rps15
rpl36
rpl8
zgc:193541
rps15a

rpl37
rps16
rpl15
rps6
eef2b
COX5B
mt-co2
rpl35a
rpsa
rpl26
rps21
rps13
rps3
rpl34
rpl24
rplp2l
eef1da
dynll1
rps26
rps5
rps3a
ucp2
h3f3d
rpl6
rpl27
rack1
eif3f
rps27a
eif5a
rpl30
eef1b2
rps24
rpl7a
slc25a3b
rpl23a

NC-002333.4
atp5f1e
rps14
rpl17
rps25
rps26l
rps9
rpl31
rpl38
rpl22
rpl21
si:ch211-161c3.6
myb
sid1
maptb
XLOC-014336
sb:cb81
rag1
lmo2
cd7al
CU499330.1
kdm6bb
csf1rb
si:dkey-217l24.1
cxcr4b
si:dkey-261h17.1
si:dkey-22o12.2
si:ch211-256a21.4
CABZ01066694.1
itgae.2
pdxka
cables2b
msi2a
tmsb
s1pr5b

si:ch211-222l21.1
si:ch211-197h24.9
cbfa2t3
bahcc1b
ncf2
nr4a1
ccr9a
ikzf1
tox2
rplp1
hsp90ab1
fmnl1a
rpl11
rpl10
rplp2l
slc16a10
eef1a1l1
MCMDC2
si:ch211-214p16.2
rpl8
rps9
mybbp1a
rpl28
rps20
ebf1a
rplp0
rpl13
tnrc18
tagapb
dub
rpl37
rps15a
rgs13
rpl18
faua

rpl24
rpl6
si:dkey-27i16.2
gatad2b
rpsa
rps6
rps12
rps21
rps13
rpl15
rpl9
rps26l
rpl7a
rpl3
rps4x
rps3a
rpl19
myca
rps27.1
rpl32
rpl18a
rps5
rnf44
rpl36
wu:fc75a09
rps10
ahcy
rps11
rpl23
rpl39
eif3s10
zfhx3
rps7
prr5a
rpl22l1

cbfb
znf296
rpl10a
eif3c
rps14
sox4a
rpl35
rps16
rpl36a
rps25
rpl26
nop2
si:ch211-214p16.1
rpl21
rps3
pik3c2b
rbm28
si:ch73-248e21.5
rpl30
rps19
gtpbp4
rps2
rpl27a
foxp4
si:ch73-208g10.1
crema
rps23
rack1
rps24
rpl27
rps18
rps8a
rpl34
si:ch211-121a2.2
eef2b

eef1g
rps15
rpl23a
naca
nsa2
rpl31
tfr1b
rps29
rpl29
rsl24d1
rpl7
BX284696.1
epb41a
pho
si:ch211-284f22.3
rpl38
b3gnt3.4
si:ch73-1a9.3
rpl35a
rps27a
rps28
si:dkey-185m8.2
eif3s6ip
zgc:114188
si:ch211-226m16.2
hmgn6
nop53
rpl12
rps17
dera
rpl13a
zbtb2b
rps26
metap1
GNAZ

si:ch211-262e15.1
nop56
parvg
calm2b
eif3ha
eef1b2
zbtb20
h1fx
rpl4
uba52
serbp1a
XLOC-000431
cct4
adh5
cct2
tcp1
rpl5a
FP074902.2
uqcrc2a
XLOC-042595
pde7a
hnrnpa0b
eif3ea
ascc1
mcm6
elf1
cct8
ddx24
grap2b
ddx21
eef1da
gpr183a
ptmaa
sept12
rpl14

ppiaa
slc25a5
rrp1
rpl22
CU929418.1
tpt1
si:dkey-262k9.4
eif2s3
si:busm1-266f07.2
histh1l
rsl1d1
egr2a
caprin1b
si:dkey-102m7.3
usp36
si:dkey-239h2.3
eif3ba
atad2b
txndc9
kdm6ba
ucp2
MRPS12
eif3i
cct7
btf3
RPL37A
cd83
wasb
wu:fb55g09
rpl5b
pfdn4
rpl17
rplp2
CR855337.1
cops2

pa2g4a
eif1ad
cebpb
tnrc6c1
celf2
thrap3b
mibp2
myg1
gtf2f1
rac2
abce1
eif5a
tob1a
LOC566587
hmgn7
hmgn2
osgn1
eif3g
RASSF5
si:dkey-250d21.1
krt18b
eif4a1b
cct3
eif3k
tuba8l
gnai2b
aldoaa
XLOC-013964
psme2
ipo7
eif3m
eif3f
si:dkey-11f4.14
elavl1a
hmgn3

si:dkey-112e7.2
elmo1
gnl3
fam120c
eif2s1b
si:dkey-42i9.4
sik1
egr3
b2m
atrx
map1lc3b
jdp2b
cnbpb
igf2bp2a
mt-nd5
ccdc25
arhgdig
pinx1
lmnb2
spi1b
chd4a
hnrnpul1l
mob1a
btf3l4
ola1
sept9b
msna
qkia
wasa
ptmab
pbdc1
kars
fabp3
nr4a3
cd74b

pgam1a
dusp5
hmga1a
pmp22b
mir21-1
bccip
si:ch1073-429i10.3.1
ran
si:ch211-195b13.1
nop58
romo1
anp32a
zgc:64051
atp5pf
pabpc1a
cited4a
slc25a55a
thrap3a
ak2
si:dkeyp-68b7.12
si:ch211-132p1.3
si:dkey-23i12.5
hist2h2l
atp1b1a
tefa
ier2b
psma3
u2surp
LOC103910871
vdac2
crip1
b2ml
eif4ebp1
larp4ab
psma2

ptena
sumo2b
sarnp
ywhaqa
arpc1b
idh2
ybx1
ddit3
psmb6
rps6ka1
XLOC-010899
prdx5
ddx61
cd74a
laptm5
si:ch211-196l7.4
cdk2ap2
plk3
gnb1a
brd2a
cct5
midn
arpc2
sod1
sept6
fbxl6
oaz1b
pfn1
mknk2b
NC-002333.4
pno1
ier2a
sub1a
ccnd2a
nfyba

stmn1a
gnb1b
psma1
irf2bp2a
eif3d
rps27.2
itm2bb
ywhaqb
h2afx
pdia3
npm1a
atp5f1d
atp5mf
mdh1aa
txnipa
kctd5a
CR383676.1
cox5ab
chd4b
arpc5a
cox6b2
nfkbiab
zgc:153713
pomp
btg1
cnbpa
ldhba
hmgb2b
zgc:111986
id1
atp5po
csrnp1b
COX5B
syncrip
hint1

uqcrh
cirbpb
h3f3c
cap1
mt-atp6
eef1db
gabarapb
cox7b
mt-nd2
atp5pb
cfl1
slc25a3b
prdx2
zgc:56493
egr1
mt-nd1
ppiab
pnrc2
jun
fkbp1aa
atp5mc3b
mt-co2
mcl1a
tmsb4x
si:dkey-21e2.13
si:dkey-21e2.15
si:dkey-21e2.16
si:ch211-212d10.1
si:dkey-21e2.14
il11b
si:dkey-78l4.2
csf3a
il13
il4
si:dkey-21e2.10

si:dkey-78l4.1
CR847844.1
si:dkey-21e2.8
si:dkey-12f6.5
LOC103911745
si:dkey-8e10.2
BX571675.1
kcnc1b
gata3
cldni
si:ch211-67e16.3
sox13
LOC100148254
si:dkey-33i11.9
si:ch211-112c15.8
XLOC-024867
faslg
slc35f4
si:ch211-147k10.5
ccl34b.4
laptm5
CABZ01073834.1
tnfrsf9a
ca2
scgn
si:cabz01036022.1
tfa
samsn1a
blk
si:dkey-276j7.2
samsn1b
si:ch1073-394i4.1
nbeal2
abca2
pfn1

zbtb16a
si:dkey-151g10.6
raraa
rgs13
wasb
MFAP4 (1 of many)
zmp:0000001323
rplp1
rplp2l
rplp2
si:ch1073-110a20.3
CR392341.1
si:dkey-27h10.2
si:dkey-262k9.2
rps20
rps12
gpr183a
tnfb
osmr
si:ch211-119e14.1
rpl39
rpl34
h1fx
rpl35a
rps2
rps17
fosl2
rpl38
rpl37
rps18
rps29
CU855821.1
rps28
srgn
rps16

ndufa4
arhgdig
rpl32
rpl26
ctss1
si:ch211-207n23.2
grap2b
unc119b
rpl18
ada
rpl27
gab2
rps14
rps13
fa2h
rpl7a
rpl36a
rpsa
tnip1
rps26l
arpc1b
si:ch211-239m17.1
rps15a
rps5
rpl14
rps24
rpl15
rps6
rpl36
si:dkey-25e12.3
rpl21
sla2
rpl8
rps27.1
rpl11

rpl23
rpl18a
rpl17
tnfaip8l2b
zgc:114188
rps19
rps7
rps23
adma
rpl29
btk
rps21
rplp0
rpl30
tnfaip3
cebpb
nr4a1
glipr1a
rab44
rpl24
bhlhe40
rps15
rps4x
rps27a
capgb
rpl35
rpl23a
tagln2
n4bp1
cox8b
rpl12
rpl13
si:ch211-66e2.3
FP236356.1
rpl9

nme2b.1
rpl19
si:zfos-2330d3.1
rpl7
si:dkey-1c7.1
rps26
nfkbiaa
wipf1a
irf4b
coro1a
si:ch211-222l21.1
rpl31
rpl10a
ms4a17a.6
rps3a
CU499336.2
rps25
rps10
rpl22
grap2a
faua
hmha1b
adcy7
slc2a3b
ccl34b.1
rpl10
rbp5
crip1
AL954142.1
tnfrsf9b
CABZ01084575.1
rps9
mrps15
ppp1r18
arl5c

cd247l
ehd1a
uba52
rps8a
rpl6
lpar5a
selplg
calm1b
ldha
ets1
cfbl
btg1
mpp1
fli1b
znf704
ptpn6
si:ch211-191c10.1
si:ch211-226m16.2
gnai1
mbnl1
si:ch73-234b20.5
cd63
rpl28
rpl27a
sept12
naca
rps11
rpl22l1
eif3f
rsl24d1
crema
gstk1
rpl5a
ets2
eno1a

eef1g
wipf1b
XLOC-043043
ptmaa
tmsb4x
itga4
zgc:152863
dusp1
actb2
si:ch1073-443f11.2
si:dkeyp-68b7.12
si:dkey-206d17.12
CR381673.1
ltb4r
dbnlb
gapdhs
CR855311.1
rack1
sec61g
rac2
rps3
LOC566587
unc13d
kdm6ba
ndfip2
upf3b
cox6b1
si:ch211-102c2.4
sh2b3
cotl1
rnf130
sik1
rps27.2
zfp36l1a
chac1

zgc:64051
LOC103909115
b3gnt3.4
nsa2
cd28l
cd74b
irf2bp2a
rpl5b
id3
msna
arrb2b
ppiaa
si:ch211-241f5.3
cap1
tagapb
BX120005.1
slc38a2
btf3
tagapa
aldocb
si:ch73-359m17.2
pim2
myl12.1
tefa
hic1
ddx21
tnrc6c1
txnipa
arpc5a
klf13
rpl13a
capza1a
pnrc2
rpl4
slc3a2b

serf2
caprin1b
arpc3
scinlb
csrnp1b
tsc22d3
idh2
notch2
cox8a
hmgn2
mob1a
eif3ea
ppp3ccb
arpc2
fabp3
dbi
sesn1
pabpc1a
sat1a.2
ptprc
ssr4
eif3s10
zgc:56493
ssr3
oaz1b
arpc4l
mibp2
si:ch211-196l7.4
cox4i1
eef1da
mcl1a
romo1
xbp1
scg3
dmrt1

zp2.5
si:dkey-241l7.2
si:dkeyp-20e4.8
mid1ip1l
si:ch211-226h8.4
h1m
acp5a
h2af1al
si:dkey-240e12.6
si:dkey-229d11.3
si:ch73-189n23.1
zp2.2
cldnd
zp2.6
CR790368.1
kpna7
epcam
si:ch211-226h8.14
zp2.3
si:ch211-125e6.12
siva1
zp3b
si:dkey-241l7.4
npm2b
zp3.2
LOC110437821
mt2
zp3
si:ch211-14a17.7
ccna1
zp2.1
tmsb5
si:dkeyp-98a7.9
zgc:153499
zp3a.2

pabpn1l
crp2
cldng
zglp1
si:ch211-146l10.8
si:ch211-12h2.8
LOC100535162
si:dkey-241l7.5
si:ch73-90k17.1
zgc:175135
si:dkey-152b24.6
lamp2
cdca8
p2rx2
si:zfos-1505d6.3
qdprb2
zgc:171776
BX649337.1
zgc:171779
s100a1
si:dkeyp-98a7.4
map1lc3c
gtf3ab
cpeb1b
si:dkey-241l7.6
rasl11b
zgc:56699
tuba4l
si:dkey-229d11.5
cth
zgc:56676
LOC101885477
slc35f2
zgc:113424
AL929032.1

zgc:55461
si:dkey-90l23.1
zgc:173443
zgc:171517
dnd1
ccnb2
zgc:171474
si:dkey-90l23.2
si:ch211-132b12.8
slbp2
si:ch211-5k11.8
si:dkeyp-46h3.6
gapdh
zar1
si:dkeyp-98a7.8
CT009596.1
si:ch211-12h2.6
si:ch211-145c1.1
zgc:136254
nasp
fabp7a
zgc:77118
si:dkeyp-46h3.8
lto1
ctss2.1
uchl1
grip2a
avd
npm2a
btg4
asb3
hspa5
birc5b
si:ch211-250e5.16
ccng1

wu:fi42e03
si:dkeyp-46h3.5
lsm14b
rcc1
sept4a
prdx1
LOC100537499
hbba1
zp2l1
racgap1
zpcx
si:ch73-14h10.2
BX537276.1
si:dkey-37o8.1
timm13
pdp2
orc3
zgc:171977
spink2.1
mibp
arhgef10la
pabpc1l
cth1
cax1
mgme1
LOC101883897
CU929143.1
zgc:162509
larp6b
mrps18b
org
elovl7b
CU855930.1
prr11
zar1l

mos
rbm4.3
si:dkey-241l7.3
CU571163.1
rbpms2b
zgc:195245
si:ch211-276i12.9
si:ch73-303b9.1
cdca7a
zp3a.1
si:dkeyp-46h3.2
gdf3
tyms
ca15b
thy1
abcf2a
ctsla
gstp1
pde6d
fbxo43
si:ch211-226l4.6
CDPF1
anp32b
tmem266
si:ch1073-190k2.1
ccnb1
dhrs13b
rrm2
zgc:171750
zgc:153284
srsf2a
cep55l
degs2
CU655961.3
snu13b

CU655961.4
chac2
tmem54a
patl2
zgc:162879
ppp2r2ab
zgc:113425
si:dkey-208k4.2
FP074874.2
larp6a
mrpl22
thop1
zgc:92313
LOC100538023
ubc
ccna2
aurkb
LOC110437779
si:ch211-14a17.6
zp3c
dnajc5ga
tmem144b
si:ch73-160p18.3
ccne1
zgc:171426
tmem44
hbaa1
si:ch73-204p21.2
si:dkey-19b23.11
zdhhc18a
pigrl2.3
kmt5ab
tpx2
lrrc47
kif22

gnmt
akirin2
CABZ01075618.1
sept15
cldn7b
polr1d
LOC103909742
ppp1r3b
mcm3l
sox19b
hnrnpabb
si:dkeyp-50f7.2
plk1
tram2
si:ch211-156b7.4
mcfd2
sybu
CR847984.1
mkrn4
zgc:173742
gins3
zgc:174935
mpzl2b
zgc:56231
BX005461.3
rpp25l
si:ch211-251f6.6
cdca9
jtb
umps
CR383662.2
h3f3a
asna1
st14a
gmnn

CABZ01069022.2
akap1a
tubb2b
si:ch73-340n8.4
hibadhb
pcf11
hsd3b2
stard14
ndc1
zgc:152652
zgc:163030
gtf2a1
coil
ube2c
zgc:165514
zgc:66160
fbxw11b
mrps6
top1l
si:dkey-188i13.11
yrdc
si:dkey-92i17.2
gfer
zgc:111868
si:rp71-1g18.1
retsatl
haus6
LOC103909972
BX571780.1
zgc:174268
gpx8
zgc:66024
wu:fb18f06
imp3
gyg1a

fam89a
zgc:113984
zgc:92591
nup37
krcp
pithd1
si:ch73-78o10.1
gpatch4
pou5f3
tom1
apopt1
uck2a
wdr82
phex
ndufab1b
cx43.4
crp3
si:ch73-15b2.5
triap1
adat3
si:dkeyp-114g9.1
ddx41
zgc:153675
CABZ01059627.2
tcea2
nmt1a
BX284697.1
zgc:55888
nanog
bola3
snu13a
cdc6
lyrm4
BX957362.1
tph1a

zgc:77112
nol7
dynll2a
ap1s3b
elovl7a
orc6
akip1
selenoh
si:ch1073-263o8.2
wwp2
slbp
si:ch211-157c3.4
tpst1l
efcab11
sde2
vdac1
ppp4ca
si:dkey-87o1.2
rpain
si:ch211-218c6.8
gstt1a
buc2l
itprip
gadd45gip1
elavl2
tatdn2
ppp4r2b
rhogc
alyref
itm2cb
diablob
snrpd2
hsd17b10
tm2d2
hspe1

ell2
COQ10A
AL929192.1
commd4
ccnh
hsd17b12a
ivns1abpb
polr2k
abhd17c
cse1l
snupn
rac1a
inka1b
zgc:113886
faub
buc
BX005461.1
krt5
si:dkey-248g15.3
si:dkey-193p11.2
icn2
krt97
cldnb
CT027832.1
spaca4l
apnl
LO017965.1
krt17
pkp3a
cldna
zgc:101810
si:dkey-33c14.3
cldn1
zgc:193505
zgc:92380

ms4a17a.5
CU682640.1
krtt1c19e
cxl34b.11
gpa33a
caspb
lamc2
zgc:162941
si:ch211-95j8.3
RGS13 (1 of many)
cd44a
si:ch211-213d14.2
ccl25a
ms4a17a.4
rhbg
ms4a17a.2
epgn
plek2
ccl27b
si:rp71-1i20.2
gpr31
upk3b
c6ast1
si:busm1-71b9.3
tp63
ptgdsb.1
col17a1b
mep1a.1
mmp20a
si:ch211-241e1.3
LOC100330348
fabp1b.2
ms4a17a.1
zgc:77439
si:ch211-264f5.6

si:ch1073-406l10.2
fam83b
pof1b
fermt1
aqp3a
cldni
dsg2.1
CR936378.1
si:dkey-19b23.12
krt91
si:ch211-155m12.1
scpp7
LOC110438215
gsnb
eevs
dsc2l
dspa
upk1a
chl1a
zgc:153911
slc16a12b
aldob
mocs2
apoeb
dnase1l4.1
gpd1c
anxa1b
limk2
perp
wu:fb18f06
cfd
gdpd3a
evpla
tmem238a
esrp1

FP017215.1
s100w
si:ch211-12e13.10
si:ch211-69b7.6
bnipl
itgb4
LOC799914
si:dkey-157l19.2
jupa
zgc:194948
itgb6
f3a
f2rl1.2
cavin2a
ptgs2a
ppl
cxcl18a.1
scinla
si:ch211-166a6.5
tmem176l.4
foxq1a
cfl1l
tmsb1
pkp2
serpinb14
arhgef5
mid1ip1a
si:dkey-16l2.20
serpinb1l4
itga6b
fat2
tfap2c
ptk2bb
CABZ01048956.1
pon1

si:ch211-39i22.1
krt4
phlda2
cdh1
XLOC-041832
ak1
gcshb
mapk12a
efna1b
selenow2b
tdh
zgc:153317
s100v2
lgals3b
zgc:171572
elf3
amotl2b
ptgs2b
spint1a
cx35.4
si:dkey-16p21.8
arg2
icn
anxa2a
ahnak
pnp5a
ptgis
lmna
FAM83G
anxa1a
klf3
crybg1a
bik
capn2b
rhoub

rnd3a
wu:fi04e12
cdo1
ripk4
krt15
gpc1a
plecb
srd5a2a
s100a10b
fosl1a
scel
mat2ab
kremen1
fam69c
tuft1a
si:dkey-262k9.2
flna
sult2st1
cd9b
zgc:92161
enam
dlx3b
wsb2
zgc:92242
exosc1
eppk1
rab42b
si:ch211-105c13.3
tuba8l2
cldne
cavin2b
CABZ01086029.1
ptprfb
ptgdsb.2
XLOC-039817

pdlim1
aldh1l1
si:ch211-195b11.3
vwa1
xkr9
myh9a
casz1
calm2b
capgb
midn
kitlga
jag1b
cast
slc38a5b
lygl1
pycard
pmp22b
sat1a.1
ywhaz
fxyd1
trib3
rarga
prdx5
cd99
lsr
nfil3
capns1a
zgc:153284
si:ch211-134c9.2
aldh9a1a.1
myof
si:ch73-335l21.1
zfand5a
vaspb
cst14b.1

nectin1b
runx3
jund
slc20a1b
tagln2
itgav
tspan33a
zmp:0000000991
gipc2
tcima
prr13
si:ch211-13k12.2
tmc6b
hcar1-4
itgb3a
fosb
selenow1
zgc:77486
dnmbp
ccl25b
atf3
foxp4
zbtb16b
si:dkey-180p18.9
cavin1b
cav1
her9
ms4a17a.10
fblim1
rmdn1
ubxn6
ets2
sult6b1
sptlc2b
lama5

anxa4
cbsb
ctnnd1
ccng2
oclna
zbtb16a
pfn1
net1
mylipb
tpm3
mcl1b
ybx1
gsna
si:ch211-39i2.2
si:ch73-335l21.4
anxa5b
fbp1a
mpzl2b
myofl
fgfr3
si:ch73-234b20.5
hdac4
tax1bp3
itm2bb
tmsb4x
si:ch73-1a9.3
slc1a3a
ccnl1a
smox
ywhaqb
jdp2b
dhrs9
mdka
postnb
fabp3

nt5c3a
cebpd
rbfox2
pak2a
junba
chmp1b
lamc1
smdt1b
fosl2
snphb
eef1a1l1
hbegfa
pim1
atp5if1b
csde1
znf706
arpc4l
adh5
ip6k2b
kdm6ba
wdr1
ak2
b3gnt5a
itpkcb
ccni
higd1a
ctnna1
eif4a1b
sptssa
tuba8l
rhoca
entpd5b
myl12.1
cotl1
zgc:162730

mvp
cd151l
baiap2a
ahcy
maff
hmgn3
ddit3
slco2b1
mibp2
si:ch1073-429i10.3.1
stk35
emd
csrnp1b
cdc42se1
nsa2
si:dkey-79d12.5
ppiab
nfkbiaa
csnk1a1
pxna
ppib
wls
rpl5a
ier2b
capza1a
arpc2
tnfaip3
rbp4
naca
dusp5
si:ch211-15b10.6
glo1
lxn
mbpb
atf4b

XLOC-043857
pno1
rps2
eif3c
cyr61
h3f3c
actb2
junbb
rpl11
irf2bp2a
rpsa
agrn
rpl22l1
rab1ab
chac1
cox8a
nfe2l2a
epcam
tob1a
btf3
rpl32
ier5
ankrd22
vdac2
gstr
tgif1
wbp2nl
reep5
sparc
rpl34
cct8
zgc:114188
rpl27a
rplp2
eef1g

rpl23a
capns1b
ppiaa
capzb
rplp0
hopx
romo1
si:dkey-51e6.1
eef1db
stau2
rpl36
rpl29
cct2
cap1
ctnnb1
rpl30
prelid3b
hdlbpa
rab10
sdc4
rps16
rsl24d1
si:ch73-46j18.5
cebpb
eif1b
btg2
thy1
rps4x
rpl18a
rps12
nr4a1
pmaip1
zgc:153713
rpl38
pon3.2

gapdhs
mtpn
eif3f
zgc:158343
klf6a
rpl9
rps15
si:ch1073-325m22.2
eif3k
fosab
chmp2a
rps19
rps27a
f11r.1
jun
h2afx1
rpl6
rps10
rpl24
fkbp1aa
rpl37
pfn2
rps26l
eif3d
rpl17
nop53
si:dkey-112e7.2
rps25
rpl15
abracl
rps13
ddx21
eef1da
rps28
rps21

itm2ba
rps11
rpl36a
rpl22
rps14
btg1
ddx5
rplp2l
rpl13
mt-co1
rps24
eef2b
rack1
rpl18
ywhabl
hsp90ab1
rps27.1
rpl10a
rpl28
rpl31
eif2s3
uba52
nccrp1
atp5pf
rpl8
arpc3
eif3ea
caprin1b
rps18
eif4ebp2
pabpc1a
ptmaa
hmgn6
fth1a
serf2

ier2a
rps9
zgc:153867
serbp1a
rps5
actb1
eif5a
pnrc2
faua
rps23
si:ch211-157c3.4
rps15a
rpl21
ndufa4l
COX3
rps3
CR383676.1
dmrt1
hspb8
gsdf
gstt1a
gpx3
igf3
amh
si:dkeyp-80c12.8
cyb5d1
zgc:92162
scn1ba
ak4
kcnj1a.1
zgc:152791
nrgna
ppp1r1b
nkx3.3
ftr82

si:ch211-117c9.5
cth
isg15
ctnnd2b
sox9a
fabp7a
CABZ01021532.1
add2
adm2a
jam2a
ndrg3a
inha
zgc:194990
msx1b
si:dkey-188i13.9
ifi46
mmel1
ifi45
plxnb3
krt8
scg3
cahz
cldnc
hpn
XLOC-042229
wfdc2
rhoub
krt94
krt18a.1
cyp26a1
si:dkey-188i13.10
lgals17
lgals9l3
ckba
calm1a

aldh1a2
cldn11a
fkbp1b
lnx2a
fyna
sparc
fam13b
si:ch73-1a9.3
ptmaa
marcksl1b
rpl13
cx43
alcamb
prss35
zgc:92744
gstm.3
cox5b2
selenbp1
cmbl
rbm47
hmgn6
soul5
rpsa
ponzr1
si:dkey-204f11.64
fthl27
gpm6ab
gstt1b
tmsb4x
rps2
nr0b1
LOC110438024
cbsb
zgc:114188
znf1104

fkbp1aa
fxyd6l
eef1a1l1
ybx1
rpl39
rps16
rpl10a
esr2b
calm2b
rps24
cox6a2
hspe1
ap2m1a
rpl9
rpl26
rpl32
si:ch211-197h24.9
phc2a
rpl21
cox6b2
rpl18a
rpl6
appa
dag1
aplp1
rps19
rps15
rpl31
tmem45a
rps27a
naca
rpl24
rpl19
zgc:92140
rps21

rpl38
rpl7a
rpl34
rps4x
slc7a3a
rps7
rpl36a
rpl27
rpl22
rpl8
oip5-as1
eif4ebp1
znf326
rps18
gstm.1
rps14
rpl35
rpl23a
rpl18
ugt1b5
rps8a
stat2
rplp0
ccdc32
rps23
rps12
bcam
rps3a
ap1m1
hmgb1b
enah
zbtb10
rps28
rpl29
gstm.2

mafba
senp3a
rpl37
si:ch211-286b5.5
psme1
si:ch211-39k3.2
ahcyl2
gig2e
cav1
ppiab
rps15a
CR318588.1
ap1s2
si:ch211-270g19.5
rplp2l
rps6
mtpn
rpl28
ywhabl
faua
si:ch211-105c13.3
rpl30
jupa
rpl36
rpl11
rps9
zgc:175088
edf1
atox1
ldlrap1a
ppiaa
arpc1a
rps26
rpl17
rps27.2

b2m
rpl15
gmfb
got1
rpl35a
rps26l
ppdpfb
rps5
cox8b
CR318588.3
pik3r2
cox7c
myl9a
slc25a5
tuba8l3
rplp2
sub1a
hmgn7
rpl7
eef1db
atp6v1e1a
marcksb
hmgb3a
gabarapb
rps25
angptl4
rps10
glrx5
gpx4b
rps11
fth1a
sprydd7a
epdl1
vat1
gsna

oaz1b
plekhb2
cgnl1
efhd2
rpl27a
rps29
zgc:56493
ywhaz
rps13
btf3
eif5a
tma7
s100v2
calm3b
mpped2a
psmb8a
zgc:123105
iscub
f10
rps3
ndufa4l
pcyt2
eif4a1b
uqcrh
rpl10
eif4ebp2
idh1
cirbpa
tpm4a
ppp3ccb
itm2bb
arl4d
zgc:100829
s100a10b
si:dkey-28b4.7

ndufb9
mibp2
phlda2
irf2bp2b
eef1g
nudcd2
txndc17
psma6l
psme2
stat1a
gng12a
si:ch73-46j18.5
itm2ba
rpl14
pnrc2
eif1b
psmf1
cavin2b
hp1bp3
chmp6b
ppp2r1ba
eef2b
atf5b
myl6
glod5
si:ch73-42p12.2
map1lc3a
xaf1
sdcbp2
prdx3
rab11a
hsd11b2
ifi35
bin3
capns1b

rpl4
vps25
atp1b1a
cd99
eif3f
kank2
rps27l
cfl1
LOC103909284
ywhah
etfa
gabarapa
dap
uba52
sgta
ppib
akt1s1
ctnnb1
eif3m
h2afx1
prdx5
psmc1a
gatad2b
si:dkey-222f8.3
anxa13
h3f3c
ppp2cb
pfn2l
churc1
myl12.1
cd164
cox8a
DHDH
prkab1b
CU694197.1

atp1a1a.1
h1f0
anxa4
csde1
mif
cnn2
anapc16
sap30l
lgals3b
si:dkey-51e6.1
pfn2
rhoac
sumo1
sod1
si:dkey-78l4.6
si:dkey-9i23.4
si:dkey-78l4.7
LOC100150882
spink4
gata3
si:dkey-78l4.8
si:dkey-78l4.4
si:dkey-21e2.3
cpa5
il11b
CR847936.10
adgre13
cfbl
tnfrsf9a
acp5b
AL954142.1
ptprjb.1
cox6a2
CR396595.1
eomesa

kctd7
LOC101886102
flt3
nansb
prodha
si:dkey-78l4.5
CABZ01073834.1
ogfrl1
rab36
dusp5
si:dkey-78l4.2
parvb
ctss1
capgb
tnfrsf9b
cremb
yrk
XLOC-018356
rnaset2l
laptm5
vaspa
ccr9a
grap2b
CU855821.1
ptprc
scgn
fcer1gl
dnajb9b
ndufa4
si:zfos-741a10.3
si:dkey-33i11.9
nptna
hhex
itgb2
ca2

SLA (1 of many)
cyba
mir223
alox5ap
osmr
bhlhe40
fermt3b
ccl35.1
zgc:64051
si:ch211-198c19.1
si:dkey-1c7.1
ccl33.3
si:ch211-241f5.3
si:ch211-102c2.4
hexa
wasa
il15l
ppp1r18
slc41a1
mylk5
sik1
parvg
dgkzb
lpcat2
arap3
si:ch211-136m16.8
hck
spi1b
skap2
si:dkey-21o19.2
il10ra
sort1b
rgs13
viml
cyb5r2

def6c
CU207241.1
coro1a
ubash3ba
il16
sox5
srgn
pla2g4aa
ccr2
arpc1b
lcp2b
b3gnt3.4
kctd12.2
sema7a
nitr5
si:ch211-191c10.1
pfn1
tln1
csrnp1a
si:dkey-237j10.2
si:dkey-172h23.2
cd74b
fybb
ptpn6
sla2
jak2a
si:ch211-243a20.4
atr
cd63
arhgap20b
fam49ba
si:ch73-264p11.1
rgs3a
edaradd
glipr1a

gnsb
LOC101884267
crema
si:ch211-215e19.3
myl12.1
akt3a
rhogb
tnikb
rorab
itm2bb
rasgrp4
aldocb
si:dkey-21e2.15
tagapb
mbnl1
limd2
fam210b
si:ch211-132b12.7
mrps15
rab27a
raraa
csf3a
gpr132b
csf2rb
actr2a
cd74a
actb2
si:dkey-262k9.4
si:ch211-222l21.1
vsir
atp2a3
crip1
rac2
spns3
sh2d3ca

il34
sfmbt2
rps6ka1
rnf130
il4
elf1
dicp3.1
grasp
si:dkey-222f2.7
anks1b
BX120005.1
ldha
arpc3
btg1
RASSF5
arhgdia
CU499336.2
calm1b
atp5if1b
sla1
selplg
pik3cd
sqstm1
hif1ab
si:ch211-66e2.3
zgc:153867
CABZ01084575.1
pdxdc1
wasb
ptpn22
cmah
sh3bgrl
csk
rnf19b
capzb

syk
cfl1
zmp:0000001082
gapdhs
pde4ba
hic1
arl11
gng2
LOC103909115
nfkbie
oaz1b
pik3ap1
XLOC-043043
ucp2
wipf1a
pak1
tmc8
FP236356.1
arpc5a
si:dkey-53k12.2
si:ch1073-443f11.2
mef2d
mtpn
tspan33a
nme2b.1
zfp36l1a
rasal3
fscn1a
dtnbp1a
rplp2
lrrfip1a
pptc7b
si:ch73-208g10.1
MAP3K11
mrtfaa

tmsb4x
rhoab
lasp1
arrb2b
BX649485.1
mir21-1
LOC110438237
si:rp71-36a1.3
fli1b
myh11a
pi4k2b
ncalda
cdkn1bb
ches1
cotl1
rel
ccni
si:ch211-8c17.3
prkcbb
elmo1
hcls1
snap23.1
arfgef2
arpc2
gstk1
gsk3b
arhgdig
actb1
eef1a1l1
tbc1d10b
myd88
rpl34
cebpb
ccdc28a
sh3kbp1

sat1a.2
cox6b1
rab44
plxnc1
gabarapb
RASA2
parp8
cdkn1ba
slc25a33
btk
si:dkeyp-19e1.3
gng5
mhc2dab
tsc22d3
dusp1
mapre1a
fam107b
arpc4l
ets2
ndel1a
CR753876.1
nek6
cap1
arpc5b
FP101882.1
pnrc2
larp4ab
mob1a
mcl1a
far1
camk2g2
rps29
ostf1
CR392341.1
lcp1

cmtm7
dbnlb
pdxka
per2
stk17al
ccdc88b
arl4aa
rps15a
bcl10
plekho2
tyk2
cxcr4b
abat
si:ch211-226m16.2
tmcc3
clic1
mt-co1
cox8a
amfra
CU467110.1
CR855337.1
tpi1b
CABZ01079663.1
cox8b
sec61g
si:busm1-266f07.2
dub
rpl15
plek
si:ch211-105j21.9
sept7b
rpl26
rps21
sec11a
COX3

nr4a3
rpl32
cers1
rpl5a
zgc:162944
BX901920.1
bcl2l10
CABZ01090890.1
anxa4
iqgap1
capza1a
rhoaa
tes
serf2
si:zfos-2330d3.1
pdcd10a
eif3f
zgc:153713
tnfaip3
tiparp
metrnla
rps2
rpl18a
slc3a2b
si:rp71-36a1.1
ak1
rps28
myo1f
ube2d1b
rca2.1
sptbn2
prkar2aa
LOC100151196
rps27a
rpl39

ubr4
LOC100536213
rpl22
lamp1a
rpl14
pgam1a
rpl19
DHDH
si:ch211-196l7.4
ptmab
rps26l
twf1a
sept12
prelid3b
psma6l
rnf44
cdc42l
rpl36
tagapa
coa6
tec
rpl9
dok2
selenok
gm2a
rps15
XLOC-026836
slc6a6b
rps18
egr3
nek7
GCA
vamp8
si:dkey-118j18.2
tspan37

rpl7a
rap1b
si:dkeyp-68b7.12
rpl38
creg1
ywhabl
rpl37
rpl30
rps5
sh2b3
rpl35a
rarga
ap2m1a
anxa11a
pnp5a
prkar1ab
eif4ebp1
ppt1
higd1a
tpt1
cry2
mbnl3
mt-atp6
rplp1
si:dkey-112e7.2
cmip
mt-co2
rab8b
eif4ebp2
commd3
btf3
rps14
mt-nd3
rpl29
rpl28

rpl23a
fkbp5
serpinb1l3
nfe2l2a
eef1db
eif3s10
uqcrq
rpsa
zgc:114188
h1fx
efhd2
stk17b
capza1b
rpl11
ndufb9
mibp2
gnai2a
litaf
rpl36a
rpl8
rps27.1
rps16
zgc:77650
ndufa6
CR383676.1
usp9
tbcb
bcl2l1
pdk2b
mt-cyb
si:ch1073-325m22.2
ndufa8
per3
arl6ip5b
msna

hmgn6
cry-dash
cct5
si:ch73-234b20.5
eef1da
wdr1
gosr2
zgc:56493
eno1a
uba52
wu:fi37e09
hsp90ab1
rps27.2
rsl24d1
ipo7
fosl2
cdc42
arpc4
rpl27
idh2
ncaldb
eif4bb
nsa2
jak1
rpl35
magt1
arf6a
rplp2l
romo1
rps6
nr4a1
naca
notch2
rps13
rps10

spcs1
rps4x
b2ml
psenen
chmp1b
dbi
ube2v2
bccip
rps8a
slc25a3b
pabpc1a
rpl10
rps23
rpl24
ndufa4l
rhoac
rpl6
reep5
atp5pd
glo1
cox6c
itm2ba
ak2
ddx21
rpl17
rps3
rpl21
tuba8l
ppp3ccb
rpl18
xbp1
eif2s2
ckbb
rps9
rpl27a

mt-nd4
sumo2b
rps25
celf2
faua
NC-002333.4
atp5f1e
si:ch211-212d10.1
si:dkey-12f6.5
si:dkey-21e2.13
si:dkey-21e2.15
h2af1al
cldng
zp3
acp5a
zp2.2
si:zfos-1505d6.3
cldnd
zgc:171776
zgc:171779
zgc:153499
zp2.3
si:dkey-241l7.2
epcam
zgc:77118
CR790368.1
si:dkey-241l7.5
h1m
si:ch211-226h8.14
lamp2
zp3.2
BX649337.1
si:dkey-229d11.3
zp2.5
tmsb5

zar1
zp2.1
si:ch211-12h2.8
si:dkey-90l23.1
mt2
crp2
zgc:195245
hbba1
si:dkeyp-98a7.8
si:ch211-125e6.12
si:ch73-189n23.1
si:dkey-241l7.4
ccna1
gtf3ab
zp2.6
CU571163.1
zgc:171517
chchd6a
si:dkeyp-20e4.8
si:ch211-12h2.6
zgc:173443
tuba4l
buc
si:dkeyp-98a7.9
hbaa1
si:dkey-241l7.6
si:dkeyp-98a7.4
zgc:175135
prtfdc1
s100a1
spink2.1
btg4
si:dkey-240e12.6
si:dkeyp-46h3.6
npm2b

gapdh
si:ch211-226h8.4
zp3b
thy1
zgc:113984
si:dkey-208k4.2
si:ch211-132b12.8
LOC100535162
zgc:55461
cldn7b
si:dkeyp-46h3.2
uchl1
casd1
slbp2
h3f3d
LOC100536860
rrm2
kpna7
zgc:92287
si:dkeyp-46h3.8
hnrnpabb
zpcx
nanog
cth1
AL929032.1
zar1l
postna
siva1
ctss2.1
ccnb2
si:ch211-14a17.7
ap1s3b
CABZ01059627.3
nabp1b
ccnb1

CABZ01059627.2
zgc:113425
henmt1
dus1l
proser3
zp3a.2
ttc32
si:dkeyp-50f7.2
zgc:114123
plpp2b
si:dkey-42i9.8
gdf3
ube2c
rpl7l1
zgc:92591
cdc6
si:dkey-152b24.6
tomm34
mdh1ab
vps9d1
si:ch73-90k17.1
AL772362.1
h3f3a
si:ch211-250e5.16
zgc:77112
f11r.1
ddx19
ttc1
ptgesl
slbp
mid1ip1l
cfl1l
asb3
nupl2
avd

tomm20a
zgc:171445
nasp
spsb1
rnf4
si:dkey-92i17.2
zc3h11a
COQ10A
ctsba
wu:fi42e03
qdprb2
mettl6
zgc:171474
trappc6b
rad1
zgc:103482
arid3b
eif4a1a
zgc:55888
zgc:171717
CR847984.1
zgc:162945
si:dkey-229d11.5
f2rl1.1
ca15b
tmem183a
sybu
cxxc1b
mcm3l
gbgt1l4
gucd1
si:ch211-226h8.6
spcs3
si:ch211-157c3.4
cdca7a

LOC100537499
zgc:162879
ccz1
mkrn4
ca7
snrpe
hmga1a
gpr160
coil
cpeb1b
brf2
si:ch211-241b2.4
ccnt2b
mier1a
arhgef10la
snu13b
sinup
ccna2
zgc:56699
tipin
zgc:152968
tmem44
LOC103909742
ints6
slc16a3
orc6
atic
LOC110437821
si:dkeyp-46h3.5
faub
zglp1
kcnk5b
ddx39ab
ndufs8b
CU655961.3

ankrd10a
larp6b
zgc:165514
si:ch211-5k11.8
pim3
ect2
prdx1
mtrex
si:ch211-226l4.6
rrp36
si:dkey-90l23.2
scamp2l
si:ch211-145c1.1
mfsd4aa
nol7
trmt61a
map1lc3c
cdc20
gtf3c6
mrgbp
rfc3
mibp
cdh1
nusap1
lsm14b
zp3a.1
nsun5
zgc:174268
dctpp1
si:ch211-146l10.7
h2afx
zgc:56676
fdx1
hsd17b12a
BX001030.1

LOC110437779
zgc:56231
pou5f3
pcf11
znf395a
sumo3b
ppp4ca
uchl3
pigrl2.3
si:dkey-241l7.3
cx43.4
os9
eif4e1b
rbpms2b
BX537276.1
mbd1b
nkl.4
nkl.3
si:zfos-1505d6.3
nanog
si:dkey-90l23.2
zgc:153499
h1m
cldng
si:ch211-226h8.14
slbp2
nkl.2
si:dkey-241l7.2
zp3
h2af1al
si:ch211-125e6.12
zgc:77118
hbba1
hbaa1
mt2

qdprb2
zp2.1
crp2
zgc:175135
zp2.3
btg4
CU571163.1
tmsb5
CR790368.1
zgc:171776
si:dkeyp-98a7.9
cldnd
ccna1
zar1
si:dkey-208k4.2
zp3a.2
rrm2
BX649337.1
lamp2
zgc:171779
zp2.2
si:dkey-240e12.6
epcam
zp2.5
si:ch211-226h8.4
acp5a
hhla2a.1
cpeb1b
zp3.2
si:dkey-241l7.5
si:dkey-229d11.5
vrk3
prf1.2
si:ch73-189n23.1
rasl11b

si:ch211-145c1.1
si:dkey-241l7.4
si:dkeyp-46h3.6
LOC100537499
si:dkeyp-46h3.2
si:dkeyp-98a7.8
si:ch211-5k11.8
si:ch211-12h2.6
gapdh
LOC110437821
polr2k
cth1
tuba4l
si:ch211-12h2.8
zgc:195245
si:ch211-14a17.7
thy1
zgc:92591
gtf3ab
si:ch211-250e5.16
zp2.6
si:dkeyp-46h3.8
si:dkey-85k15.4
si:ch73-14h10.2
si:ch211-146l10.8
si:dkey-229d11.3
si:dkey-152b24.6
ccna2
mpzl2b
ccnb1
si:dkey-90l23.1
trmt9b
polr3h
si:ch211-132b12.8
zgc:113210

si:dkey-241l7.6
ctss2.1
zar1l
zgc:173443
sinup
zgc:136254
htatip2
bphl
lsm14b
npc2
si:dkey-152b24.7
si:ch1073-190k2.1
ca15b
hnrnpabb
npm2b
si:ch211-226l4.6
foxa
zgc:113424
ssna1
hnrnpa0l
XLOC-030786
vstm2l
zpcx
si:dkeyp-20e4.8
si:dkeyp-50f7.2
zp3c
zgc:171517
elavl2
si:dkeyp-98a7.4
cldn7b
siva1
npm2a
spink2.2
kpna7
mid1ip1l

rac1a
zp3b
zgc:66024
AL929192.1
zgc:171977
mibp
zglp1
mad2l1
zgc:101744
tmem9b
ccnb2
si:ch73-78o10.1
mfsd4aa
LOC101885477
s100a1
eif4e1b
hccsa.1
si:dkeyp-2e4.2
mkrn4
glulb
AL929032.1
manbal
nfr
elovl7a
zcchc8
pabpn1l
sc:d0202
mob1bb
si:dkeyp-98a7.7
zgc:173856
uros
shmt1
tarbp2
si:ch211-146l10.7
fbxo43

gbgt1l1
gmnn
marcksb
tdp2b
LOC110437779
si:dkey-188i13.11
si:ch211-251f6.6
uchl1
rnf175
LOC103909972
rnf4
si:ch211-59d15.9
cnep1r1
eif4a1a
wu:fi42e03
fdx1
zgc:103482
ess2
LOC101885376
prr11
mcm3l
hddc2
ankrd22
si:dkeyp-2e4.3
sybu
mad2l2
dhx16
ckbb
arhgef10la
thoc7
CR383662.2
ell2
zgc:173544
pdia4
st6galnac1.2

rnaseh2b
pgam5
metap1d
kmt5ab
tp53rk
si:ch73-281n10.2
retsatl
CU655961.4
spink2.1
si:ch211-137a8.2
tph1a
sh2d5
BX005461.3
rnf128a
slbp
nifk
baiap2a
BX571701.3
snrnp27
spsb1
wu:fa19b12
zgc:171474
cnot8
selenou1a
dkc1
LOC101883897
cenps
si:dkeyp-114g9.1
zmp:0000001323
cax1
ints12
heatr3
si:dkey-241l7.3
drg1
BX001030.1

pigrl2.3
nasp
rcc1
ap1g1
wdr43
dynll1
il13ra2
LOC100535162
ccz1
hexim1
eif1axb
lin28a
haus6
snu13a
zgc:55461
ccdc47
chrac1
btg3
zgc:86839
cldnb
icn2
zgc:193505
cldne
zgc:111983
si:dkey-7c18.24
si:dkey-193p11.2
si:ch211-69b7.6
krt5
mid1ip1a
si:dkey-247k7.2
krt17
si:ch211-217k17.9
lye
si:ch211-207n23.2
zgc:153911

dsg2.1
ppl
dsc2l
pkp3a
evpla
cldn23a
si:dkey-248g15.3
tnks1bp1
soul2
zgc:153665
CABZ01048956.1
anxa1c
LOC108190590
krt97
scel
krt1-19d
si:ch211-125o16.4
zmp:0000000906
si:ch211-98n17.5
tgm5l
znf185
anxa1b
rhcgl1
lmo7a
ftr83
zgc:110333
ptgs2b
LOC100330250
hrc
AL929237.1
sts
chst6
bicdl2l
mir203a
agr1

slc16a12b
CR854839.1
evplb
apodb
sox3
LOC101882729
stx11b.1
myzap
pkp2
fam129aa
CABZ01067657.1
cyldb
bik
FO681323.1
CABZ01084230.1
cldnh
si:ch211-182p11.1
si:dkeyp-51b9.3
capn9
si:dkeyp-86h10.3
si:ch73-52f15.5
zgc:100868
ANXA1 (1 of many)
cx35.4
tmem176l.3b
CR388148.2
ponzr10
btr12
rnf222
si:dkeyp-1h4.6
LOC100149234
eps8l1a
wu:fc34e06
si:dkey-175d9.2
si:ch211-197e7.1

rassf7b
CABZ01015815.1
CABZ01068469.1
BX005027.2
CABZ01068248.1
LOC100333235
cldnf
st14b
CR589947.2
si:ch73-184c24.1
CU914622.2
erbb3a
ccl20b
paqr5b
si:rp71-1h20.9
XLOC-041945
esrp1
ponzr2
foxq1a
adgrf6
zgc:136892
LOC110438215
si:dkey-73p2.1
si:ch73-347e22.8
noxa1
si:dkey-165e24.1
XLOC-027129
nectin4a
LOC103909616
CU062645.1
si:ch211-226h7.5
si:busm1-194e12.8
si:ch73-196i15.3
adgrg11
zgc:113229

si:dkey-221j11.3
LOC407643
si:dkey-27n6.4
pkp1b
zgc:194839
LOC100149931
anxa3a
LOC100001846
il17rc
vsig8b
tgm1
gna14
ttc22
BX004785.3
il20ra
card14
zgc:165423
si:dkey-11o15.7
CABZ01049361.1
adgrg2a
CABZ01046954.1
zgc:123295
mala
XLOC-011066
LOC101882034
ltc4s
LOC100537138
si:ch73-208h1.1
muc13a
si:ch211-232i5.1
si:ch211-149a19.3
CR926130.1
sp8a
BX901957.1
ankk1

tmem45b
si:ch1073-340i21.3
si:dkey-61f9.2
papss2a
elf3
LOC110438610
gpa33a
si:ch211-79k12.1
tmem176l.4
zgc:91849
CABZ01039863.1
sec23b
dbnla
pof1b
hmgcra
f2rl1.2
CR677617.1
selenow2b
dspa
plekhg6
ftr39p
klf5l
si:dkeyp-82b4.2
frmd4ba
zgc:172075
adgrf7
zgc:92242
ttc39a
si:dkeyp-75b4.8
LOC100005723
ccl20a.3
XLOC-003147
ano1
krt4
abcb5

CU693379.1
si:ch211-282j17.8
sparta
si:ch211-195b11.3
nkx3-1
si:dkey-228d14.5
LO017816.1
zgc:194398
erbb3b
abca12
LOC110438361
styk1b
si:dkey-160o24.3
oat
si:dkey-19b23.12
xkr9
macc1
CABZ01070852.1
s100w
aldob
si:ch73-22a13.3
LOC110438236
CABZ01056052.1
dhrs13a.2
si:dkey-242g16.2
wsb2
si:busm1-104n07.3
limk2
nlrp16
zgc:162184
fam83b
tmprss13b
stard14
phlda2
si:ch211-161h7.5

zgc:112433
tuft1a
BX548046.2
slc9a2
wu:fb18f06
mpeg1.2
si:ch211-153b23.4
perp
si:dkey-223p19.1
cyp8b2
fam169aa
tlcd2
rnh1
plekhf1
rarres3
plekhn1
si:ch1073-385f13.3
LO018605.2
ugt8
bbox1
XLOC-019059
rhpn2
tmem30b
elovl1a
rhcga
frk
BX005421.3
si:dkey-262k9.2
tnfrsf11b
si:ch211-158d24.4
oclnb
ildr1a
klf3
rgs2
krt91

trpm4b.2
gdpd3a
tlcd1
ch25hl1.1
tfap2c
barx2
si:ch211-217k17.10
dnmbp
LOC108180802
si:dkey-33i11.9
si:dkey-28d5.11
wu:fi04e12
si:ch211-264f5.6
SEMA4F
nqo1
arhgef5
zgc:173737
iqsec2b
jupa
trib1
casz1
zgc:153932
FO704779.1
cd9b
ptpn13
zgc:158263
CT027980.1
slc2a1b
si:ch73-359m17.5
epdl1
si:ch211-157c3.4
kremen1
gcnt4a
net1
nox1

krt222
LOC108179621
elovl6l
si:dkey-30c15.2
homer2
ehf
arrdc1a
kras
si:dkey-204f11.64
ms4a17a.5
spint2
FP017215.1
cast
eppk1
tead3b
mpp6a
myo5b
ftr36
cx34.4
rhov
myo1ea
si:dkey-157l19.2
gna15.1
zbtb16b
ptgs2a
usp2b
si:ch211-285f17.1
atp8b1
samd10a
apnl
fam160a1a
cdh1
ssh2a
alox12
paqr6

hopx
pard6b
gsnb
tiam1a
si:dkey-16l2.20
pld2
zgc:92481
xiap
chmp4c
lmbr1l
mgst1.2
si:ch73-204p21.2
osbpl7
plcd1a
ccng2
cd44a
dusp5
ptgis
zgc:162193
spint1a
CABZ01039859.1
LO017965.1
igf2bp2b
si:ch73-141c7.1
foxc1b
cfl1l
slc7a3b
b4galnt1a
FAM83G
ptpn21
cavin2a
XLOC-041832
ahnak
si:dkey-87o1.2
rab11al

ptprfb
cdc42ep5
anxa1a
tcnbb
pdlim2
cdaa
c1galt1b
zgc:77486
si:ch211-166a6.5
serpinb14
XLOC-044191
nipa2
nr1i2
mboat1
kdm6ba
osbpl3a
rnd3a
efna1b
midn
galnt7
ephb6
ldlrb
cxadr
nfkbiz
mhc2dab
tob1b
cpeb4b
si:dkey-261p22.1
grhl1
bicdl2
ifngr1
cst14b.1
epha2a
snphb
s100v2

fosb
adrb2a
tnfaip3
si:ch211-152p11.4
CT027702.1
si:dkey-30g5.1
lmna
chmp1b
ngef
pleca
CABZ01072077.1
cpeb2
btr25
errfi1a
bcl6ab
tent5ba
zfand5a
cldna
akap9
sqstm1
prkcdb
nt5c2b
si:ch211-39i22.1
dlx3b
fam83hb
s100v1
pnp5a
stk10
ier2b
si:busm1-57f23.1
zgc:154054
trib3
gprc5c
fosl1a
zgc:153284

scpp7
jdp2b
b3gnt5a
prodha
jag1b
tmsb1
BX664618.1
sorl1
zgc:153759
slc38a5a
si:dkey-105i14.1
icn
anxa2a
bnipl
efnb2a
si:ch211-202f5.3
chac1
b2m
ezrb
vsig10
spns2
klc1b
fuca1.1
si:ch211-207d6.2
ckap4
sh3d21
cd99
selenow1
uba7
zbtb16a
oclna
chmp2bb
erbb2
mgst1.1
si:ch73-22o12.1

sipa1l3
nfkbiaa
cishb
epcam
si:ch1073-429i10.3.1
si:dkey-79d12.5
si:ch73-335l21.1
zgc:153317
rpz4
nrg1
b2ml
ier5
cx30.9
CABZ01029938.1
rhoca
myo6a
reep3b
tjp3
zgc:92380
relt
sh3d19
CR847566.1
tiparp
b4galnt3b
gpatch8
s100a10b
ddit3
adipor2
s100u
si:ch73-335l21.4
tdh
acaca
gadd45aa
ccdc50
cldn7b

tmem59
ubb
mhc1zba
rab11fip1a
dhrs13a.1
znfx1
cavin2b
rnf128a
porb
etnk1
cd74b
plce1
slc20a1b
daam1a
zgc:92107
fxyd1
cd74a
CR318588.3
tjp1b
jun
evi5b
tax1bp3
si:busm1-266f07.2
itsn2b
si:ch1073-345a8.1
arid3a
abca3b
tuba8l2
btbd10b
esyt2a
itprid2
traf4a
chmp5b
fgd1
csrp2

gnb1a
cbx3b
pim1
mcl1b
dcun1d2b
gipc2
aste1a
XLOC-018182
fosab
atf4b
jund
chmp4bb
esrp2
zmp:0000001114
zdhhc16a
nfe2l2a
zmp:0000000991
efna1a
arl4d
fam129ba
f11r.1
zgc:113333
crybg1a
mcl1a
slc10a3
plcb3
pvrl2l
stk35
baiap2a
lipia
uacab
st14a
gmds
zyg11
mpc1

ankrd22
smox
slc37a1
bada
crema
tmem79b
afdna
zgc:174863
cx30.3
kitlga
capns1a
lsr
rhoab
lgals3b
higd1a
ftr82
ifi45
micu2
si:ch73-103b11.2
mhc1uba
CABZ01021599.1
zgc:152791
ciartb
krt15
tjp1a
btg2
kalrnb
rasgef1ba
si:ch211-286b5.5
slc25a55a
cds1
jmjd1cb
tnfrsfa
ucp2
pls3

baiap2l1a
si:ch73-138n13.1
vamp8
ccdc171
mboat2a
rab5ab
gabarapl2
arhgef4
si:ch211-191a24.4
sh3yl1
rsrp1
plecb
irf2bp2a
cd82b
atp1a1b
ampd3b
reep5
rap2b
arg2
junba
taok1a
calm2b
dusp6
si:ch1073-443f11.2
ms4a17a.10
cebpa
pycard
abcc3
sdcbp2
strbp
fkbp5
per2
mt-co1
si:ch211-107o10.3
plekha6

fam199x
irs2b
snrkb
zgc:91976
CR383676.1
si:dkey-71l4.4
rin2
prdx5
dhrs13l1
ovol1a
zswim5
si:ch211-212k18.7
mark2b
zgc:158343
rabac1
plpp2b
trim16
sox13
plk2b
alox5a
skia
atxn1a
ptbp3
fosl2
fam102ab
zgc:77439
adam28
arfgap3
waslb
rnf213a
vps4b
LOC103909284
fuca2
degs2
CU633745.1

tmbim1a
tmod2
abhd2a
cnn2
far1
mef2d
si:dkey-177p2.6
namptb
itpkcb
klf2b
si:ch211-213a13.2
ist1
fgfr3
klf8
furina
rnasekb
prelid3b
CU929259.1
slc38a5b
atf3
COX3
rps6kb1a
stap2b
fxyd6l
ehbp1l1a
csnk1da
per3
map1lc3a
gadd45ba
cebpd
camsap1b
eif1b
elovl7a
btg1
dusp7

dusp1
prox1a
fut9b
foxp4
LOC103910015
SBNO2
zgc:123258
zgc:101715
si:ch211-132b12.7
ube2d1b
tmem176l.3a
slmapa
uckl1b
exosc1
si:ch211-251p5.5
ier2a
egr1
ppp2r2d
h3f3c
pcdh11
wbp1lb
nr4a3
kmt2ca
galnt12
actn4
egr2a
emd
pdcd6ip
hsd17b12a
selenot2
naga
csrnp1b
tax1bp1b
golgb1
CABZ01055347.1

itm2bb
nmi
pde4ba
wbp2nl
rffl
ar
si:ch211-196l7.4
pmp22b
tprg1
herpud2
si:ch1073-190k2.1
pitpnb
atp2a3
cdc42bpb
sult2st1
mycbp2
chmp1a
hmgn3
npc1
clint1a
tpm3
mvp
ptdss2
hexb
capns1b
zgc:162730
tnfrsf11a
plpp1a
elf1
scaf1
gnal
cflara
CABZ01086029.1
usp24
klf6a

rtn3
gstt1b
ponzr6
prrg2
si:ch73-233k15.2
nfil3-6
taok1b
rbfox2
cnn3a
nktr
mgst2
qkia
nfixb
mid1ip1b
atl3
nudt9
stk40
fam126a
psph
ccnl1b
arl5c
atp8a1
CABZ01079427.1
CR318588.1
u2surp
tcima
psmb8a
arpc1a
gnpnat1
utrn
alas1
sdf4
ddx3xa
csnk1a1
junbb

ftr86
stk17al
mt-co2
mat2aa
id2b
meis1b
yap1
sec24a
rarga
il15
vaspb
ak1
egfra
cmip
letm1
limch1b
tsg101b
zgc:100829
lygl1
ca2
zgc:92313
si:ch211-39i2.2
si:ch211-114c12.2
elovl7b
ppp4cb
susd6
gbgt1l4
mt-cyb
sptssa
creb3l3l
zbtb10
tpd52l2a
tegt
ppm1aa
kcnk5b

ank3b
cd63
rab10
ctsk
notch2
chmp4ba
CABZ01090890.1
h1f0
ddb2
si:ch211-191j22.3
emp2
NC-002333.4
sdc4
psme2
mt-nd4
cd83
vps28
stag2b
ddx5
brd4
klf2a
pmaip1
syap1
ubald2
ptp4a2b
tnrc6c1
coro1ca
sar1b
si:ch211-270g19.5
adipor1b
calm1b
hmgcl
ankrd11
dazap2
ets2

ndel1b
ankrd12
sf3b1
txnrd3
kmt2bb
pef1
birc2
bcap31
slc3a2b
zgc:158463
lonrf1l
itm2ba
cdc42l
sik1
ncor1
smdt1b
per1b
cfd
zgc:163080
pik3r3b
map2k6
ralbp1
rnf44
dedd1
glo1
si:dkey-112a7.4
plk3
cnot1
dnajb1b
gng5
si:ch73-1a9.3
litaf
h2afx1
hbegfa
cish

brd2a
arhgdig
sptlc2b
nfkbiab
lrrfip1a
cdkn1bb
ifi30
psme1
cnbpb
atf4a
znf706
mt-nd1
pik3r2
mt-nd3
mt-atp6
LOC103910438
ppp3ccb
ndufa5
actb1
ccni
sumo2b
ywhabl
zgc:77849
hmgn6
pfn2
selenok
pomp
pnrc2
cox8a
cst14a.2
nop53
rplp2
mt-nd2
oaz1b
ybx1

gnb1b
ywhaz
chmp2a
xbp1
myl12.1
bzw1b
lum
ogna
cgref1
fgl2b
tmem119b
mettl24
erg
htra3b
igfbp3
mmp23bb
si:ch211-243a20.3
ak5
olfml1
col5a3a
creb3l1
apcdd1l
omd
npnt
wisp1b
fgl1
si:ch211-37e10.2
c1qtnf6b
wnt2
scinla
XLOC-008935
col16a1
twist3
cspg5a
twist2

col7a1
CABZ01074363.1
fam19a1a
dnm3a
irx1b
olfml2bb
rcn3
srpx2
cdo1
scara3
cd248b
wnt10b
serpinf1
col15a1a
vwa1
megf6a
pigr
fkbp11
pdgfrl
fkbp7
clu
kdelr3
hhipl2
copz2
c1qtnf2
si:dkeyp-59c12.1
rgs5a
CMKLR1 (1 of many)
si:dkey-12l12.1
bhmt
fkbp9
ccdc80
atoh8
twist1a
rgma

TENT5A
gtdc1
timp2a
asip1
si:ch211-51c14.1
hpdb
ccdc3b
abi3bpa
hkdc1
ecrg4a
casz1
nradd
ckap4
serpinh1b
srpx
si:ch211-133n4.4
p4ha2
nkd2a
cygb1
FQ378018.1
vkorc1
krt91
col5a2a
reck
si:ch211-106h4.12
fkbp14
cd82a
col21a1
postnb
zgc:113307
ms4a17a.11
cxcl8a
nupr1b
nucb2a
irx1a

olfml3b
klf12a
plxdc1
nfixb
wnt11r
dpt
ehd2b
fam198a
calua
pvalb6
pabpc1b
colec12
prelp
ecm2
thbs4b
cercam
aebp1
epha4b
col5a1
si:ch1073-459j12.1
tgfbi
CABZ01092746.1
fam69c
traf4a
fkbp10b
pdgfra
XLOC-041451
dcn
glis2a
tmeff1b
col6a3
ctgfb
col1a1a
CABZ01039304.1
atf5b

mmp2
col1a1b
ier5l
htra1a
sdc2
plpp1a
tns2b
col1a2
zgc:56585
crtap
sparc
olfml2ba
sgcd
p3h4
selenop
crhbp
rcn1
apbb2b
tcf4
si:ch73-138n13.1
CU929237.1
irx3a
col11a1b
mfap2
smoc2
nfia
pmp22a
irx5a
antxr2a
ephb3
antxr1b
FAT4
lamb1b
BX957346.1
plpp3

c1qtnf6a
slc7a10a
p3h3
appb
glis2b
bcl11ba
angptl4
fndc3ba
wipi1
fxyd1
CR847566.1
klf7a
dag1
XLOC-041832
rrbp1a
si:ch73-46j18.5
p3h1
yaf2
tuba1b
zgc:110158
wfdc1
cyr61
nt5e
klf6a
zgc:194981
ehd1b
dnase1l1
zgc:153317
col2a1b
f3b
marcksl1a
smoc1
col6a1
mrc2
sh3pxd2aa

tuba8l3
ca8
sertad4
mych
prrx1a
mmp14b
cebpd
zgc:153675
znf704
fam102ab
cd28l
tcf12
ak4
dnmt3ab
cd276
LOC101886030
calhm2
gas1a
LOC101882334
adrb2a
fosl1a
sept10
flrt2
fosb
lrrc58b
si:ch211-39i22.1
foxo1a
p4ha1b
loxl1
s100v2
arhgef1b
si:dkey-101k6.5
si:ch211-284f22.3
ms4a17a.9
vat1

sema6e
cbfa2t3
znf385a
rgcc
klf15
lrp1aa
meis1a
LOC101883507
fam114a1
fstl1b
ppm1la
crip2
tmem45a
rasgef1bb
efnb2a
slc38a5b
serf2
marcksl1b
jdp2b
adh8a
egr1
soul5
col27a1b
id1
nanos1
loxl2b
plod1a
mafba
gstcd
si:dkey-56m19.5
fpr1
vdac3
marveld1
zgc:92242
ppib

tgfbr3
XLOC-035413
zgc:171679
sulf2b
SCARF2
eef1da
sept7a
emp2
ift20
map7d1b
hdlbpa
gpc4
ecm1b
fam110b
lzts2a
sdc4
rbfox1
calm3a
mycb
cyp2n13
jun
si:ch211-286o17.1
her6
zgc:158343
cxcl18b
ddit3
fosab
tspan7
ier2a
st7
sec61a1
stau2
wbp1lb
anxa4
reep5

ip6k2b
krt8
ghra
tnfsf12
atf3
tob1a
tmem39a
map1lc3a
dip2ca
hcar1-4
pdk2b
ptn
triqk
BX640522.2
lysmd3
ptk2aa
dusp6
dhrs13l1
si:dkey-100n23.5
tmem59
susd6
nme3
kdelr2b
zgc:153372
ANKRD50
junba
fbp1a
ugp2b
capns1b
smco4
snd1
zgc:162730
foxo3b
laptm4a
nova2

pdgfrb
spred2b
rab2a
eif4a1b
myl9b
ctsk
spry4
dusp5
ccnd1
ablim1b
prdx4
fxyd6l
nfic
ssr3
bnip3lb
arf5
fzd2
slc39a7
CABZ01058650.1
zfp36l1b
rabac1
ssr4
XLOC-044191
boc
selenom
pgrmc1
emd
anxa13
eef1a1l1
grb10b
adgre5b.3
selenow1
rassf1
atf4b
fgl2a

aldh9a1a.1
vasna
mylipb
h3f3c
cdon
sdr16c5b
sh3gl1b
clstn1
lhfpl2a
rbfox2
yif1a
ier2b
epdl1
gulp1a
slc3a2b
gars
sgce
socs3b
rrbp1b
anxa2a
atox1
midn
tpd52l2b
f10
inka1a
klf11b
wls
rarga
si:dkey-238c7.16
s100a10b
fstl3
ier5
ATP11B
zgc:113208
LO018493.1

mgst3b
tnfrsfa
si:dkey-9i23.16
spry2
cers2a
asph
mfap5
jag1b
hspg2
pleca
fam114a2
rimkla
gadd45ga
arl6ip5b
fhl1b
myo1cb
zgc:100829
cybrd1
sept9a
klf11a
sar1b
socs3a
ctdspla
gpatch8
hmbox1a
mibp2
ostc
si:dkey-7j14.6
alg5
tmem167a
gng2
skia
si:dkey-28b4.7
cd82b
CABZ01075242.1

nid1b
ube2e3
rpl39
yap1
sec62
calm1a
tnrc6c1
rpl10
cox4i2
dap
nsa2
sox4a
slc38a10
tead1b
cald1a
kdelr2a
ifi30
gpx8
trib3
rps2
si:ch211-140m22.7
ybx1
tsc22d3
serp1
pno1
sec61g
chd3
rgs3a
cdca4
eef2b
cx43
arhgap31
FO082781.1
sec13
ramp2

bhlhe40
yipf5
selenof
nfkbib
unc50
txndc5
pbx1a
sec23a
rpl12
anxa5b
LOC103909284
zfyve21
zfand3
eif4ebp3l
si:dkey-239h2.3
gstz1
spcs1
mmp16a
map1lc3b
fbxo9
ssr2
si:dkey-199f5.8
plecb
angptl1a
flot2a
nupr1a
rhoaa
rps15
anxa11b
ddx21
mxra8a
sec61b
ciarta
csrnp1b
pik3r2

rnd3b
tpt1
eif2s3
arl1
lgals2a
ssr1
rab1ab
ghrb
dcaf7
cpn1
serpine2
rps27l
h2afy2
prelid3b
zgc:92606
col6a2
LOC108180269
her9
glo1
btg2
rps12
serpine1
gstr
add3a
cyb5r3
ap2m1a
ldlrad4a
si:ch211-66k16.27
rpl6
si:ch73-335l21.4
vapb
zgc:123258
tmem258
nr4a1
crip1

zgc:114188
mtdha
cd81a
rpl22
srprb
rps6
rps3a
rps13
cnbpb
h2afx1
tram1
rap1b
rpl17
b2ml
hbegfa
rpl18a
krt18a.1
eef1g
rps10
selenok
tdh
rpl11
prdx5
pim1
surf4l
rpl35a
rps20
faua
s100u
ndfip1
ipo7
rps27a
arpc1a
cavin2b
si:ch211-204c21.1

ndufa4l
gabarapb
rhoab
rpl9
hic1
eif3ba
usp36
romo1
si:ch1073-325m22.2
tob1b
nck1b
ccl25b
dbi
rps16
ier3ip1
tmem176l.1
rps18
rpl7
yipf4
rpl36a
rpl30
rps27.2
celf2
CR383676.1
chmp5b
hmgn6
rpl29
eif2s1b
rpl5a
ptp4a1
cnn2
cd164
rpl28
rplp1
rpsa

junbb
syf2
rps4x
zgc:92744
ahnak
slc25a33
rpl34
myl6
rpl27a
maff
rps21
cops9
rps25
aimp2
ptprga
itm2ba
xbp1
pim3
rpl7a
rpl18
rpl31
rps7
vdac2
rpl8
rpl32
rpl3
rack1
p4hb
rab5aa
histh1l
rps19
irf2bp2a
rplp2l
rpl15
si:ch73-1a9.3

eif1b
mdh1aa
rpl26
rpl37
rpl23a
ginm1
nr1d2a
mcl1b
XLOC-000431
rpl35
rpl13
nop56
nme2b.1
gnb1a
rpl27
rplp0
adh5
eef1b2
gadd45ba
sumo2b
pfn2
rps24
gtpbp4
zgc:77849
btg1
rpl19
rpl10a
rps23
eif3c
txndc9
pdia3
cd63
rps15a
rpl14
rps5

cdc42
si:dkey-11f4.14
rps11
ak2
edf1
rps8a
krtcap2
tomm5
zgc:110239
mknk2b
rps14
eif5a
jpt1b
rpl21
arpc2
si:ch211-196l7.4
rps27.1
eif3ea
zfp36l1a
rab10
eif3i
rpl24
rps3
rps26l
calm2b
serbp1a
rpl36
rps29
ppiaa
rps9
eif3f
wu:fb55g09
eif3m
nfil3-6
ctsla

uba52
hmgn7
ccng1
rps26
dap1b
rps28
higd1a
rpl38
rsl24d1
ptmaa
rplp2
ppdpfb
ddx5
rpl4
rpl13a
fkbp1aa
rpl22l1
LOC110438024
igf3
LOC103910903
znf1104
dmrt1
gpx3
LOC110437926
cyp26a1
si:ch211-117c9.5
scg3
amh
inha
zgc:92162
ugt5g1
alcamb
mmel1
inhbb
nrgna

XLOC-042229
scn1ba
sv2bb
kirrel3l
LOC103908925
col28a2a
cnga1a
kcnj1a.1
ppp1r1b
LOC110438394
nts
CT573799.2
si:dkeyp-80c12.8
slc4a2a
adm2a
XLOC-042806
dlb
LOC103910515
sstr5
flrt1b
si:dkey-165n16.5
plxnb3
tyro3
hpn
dmrt3a
add2
sox9a
sema7a
cx34.5
CABZ01058222.1
msx1b
XLOC-040516
cyb5d1
nkx3.3
gpm6ab

ctnnd2b
slc2a1b
XLOC-033950
soul5
rxfp3.2a
si:ch211-212k18.6
cldnc
large2
scube1
gstt1a
tmem229b
jam2a
fat2
si:dkey-6n6.1
LOC100149700
map1ab
zgc:194990
fhod3b
gsdf
CDK18
gh1
bcam
XLOC-040108
slc16a7
plpp7
atp11a
lpar3
si:ch211-198m17.1
CU929418.2
ftr82
mylka
rhoub
hspb8
fyna
si:ch73-156l6.2

dag1
slc7a3a
nat8l
lnx2a
lcat
prr15lb
fndc5b
nhsl1a
cgnl1
abtb2a
itga3b
cldn11a
rnf24
cx43
stard9
rims2b
sobpa
trim3b
cox6a2
ncam1a
appa
nos1apa
phlda2
nfasca
rbm47
map4l
pyroxd2
si:ch211-225b11.1
adgrl2a
csgalnact1a
epdl1
LOC110439817
ak4
igsf3
metrn

asb5b
mmp11a
bag3
rhcgb
prss12
cth
myh10
XLOC-031367
pkdcca
fam13b
elovl5
hkdc1
znfx1
cnnm2b
XLOC-031372
RTKN
si:dkey-194e6.1
lpcat2
lgr4
mfge8b
cahz
si:dkeyp-80c12.7
lamb1a
slc31a2
prss35
gdpd4a
shank3a
itgb8
gadd45aa
nlgn4xa
fabp7a
ctsf
FAT4
aplp2
gata6

podxl
esr2b
blf
kank2
entpd8
FO704858.1
osbpl3a
pard3ab
DOCK9
CABZ01021532.1
ap1m1
ahcyl2
mgll
zmp:0000001114
atf5b
tmem45a
wfdc2
emilin1a
rassf9
inhbab
tjp1a
lmbrd2b
wu:fc27b11
ugt1b5
mmp15b
slc7a6
senp3a
zgc:152791
aplp1
angptl4
ube2ql1
lrp2a
lgals9l3
tle2b
bmpr2a

ndrg2
actn1
ca7
ptpn13
sparc
sema3b
marcksl1b
fosl1a
phc2a
slc4a4a
pon1
plat
si:dkey-188i13.9
fam184b
p2rx7
agrn
shisa9b
sema4ba
tmem245
ifi46
arhgef18a
znf536
CR376783.1
clstn1
hmcn1
plxna4
f10
rhoj
prnprs3
hmgb3a
si:ch211-151g22.1
aldh1a2
jupa
erbb2
LOC108179567

LOC110438541
CR318588.3
plekha6
fosb
igfbp5b
pgr
clmn
tapbp.2
FP245456.1
atp1a1a.1
dsc2l
tspan9b
prr13
dspa
afdna
nr0b1
CABZ01068366.1
fam102ab
mov10b.1
CABZ01076275.1
met
bambia
ap2m1a
krt18a.1
gstt1b
stat1a
gnav1
si:ch211-276k2.1
fam69aa
coro2ba
lfng
phospho1
atp1b1a
map1aa
slc20a2

hs3st3b1b
mafba
ar
csrnp2
st3gal4
pcdh7b
ndrg3a
lgals17
palm3
rcan1a
itm2bb
vat1
tdp2a
cttnbp2nlb
tent5ba
si:dkeyp-77h1.4
pi15a
numbl
krt8
wbp1lb
pcnx1
ckba
fxyd6l
tbc1d1
hsd11b2
rnf11b
ptprfa
jun
si:ch73-61d6.3
ptprfb
si:cabz01069012.2
fuk
cbsb
nit2
st5

pxnb
mbnl3
col4a6
ifi45
FP102018.1
b2m
foxp4
scarf1
gab1
cemip2
bmp1a
clip2
si:ch73-335l21.4
pdgfaa
si:dkey-188i13.10
dchs1a
hexb
slc22a23
si:ch211-195e19.1
gadd45ba
arhgef4
ccdc187
sox4a
cebpd
isg15
ildr2
cxadr
socs3a
ano5a
gig2e
sntb1
itm2ba
bmpr1bb
rpz5
calm1a

fgfr3
man1b1b
dclk2a
XLOC-015291
gadd45bb
pcdh1b
pik3r2
sipa1l3
akap7
metrnla
ccdc125
notch1b
wwtr1
crybg1a
bmpr2b
jdp2b
plod3
efhd2
nqo1
pfkfb4b
id1
rgrb
nol4lb
ptprga
marcksb
mdka
efhd1
CABZ01086041.1
mkln1
apoeb
magi1b
grk4
LOC100333572
fdxr
ankrd46a

CABZ01086029.1
pelp1
smad1
arl4d
serinc1
papss2b
slc44a5b
rasl11a
ywhae2
CABZ01092156.1
znf326
selenbp1
si:rp71-36a1.2
vasnb
bmp6
ap1s2
wu:fb16f03
myl9a
fosab
alcama
trim2a
cited4a
cav2
stat2
nectin1b
egr1
cmtm3
bahcc1b
abl1
cadm2a
b3gnt2l
mpped2a
piezo1
slc12a7a
kalrnb

cpn1
dbn1
lhfpl2b
tegt
cdh1
mogat2
foxl2b
si:dkey-42i9.4
cdv3
gamt
ctnnb1
zfp36l1b
got1
tmem205
si:ch211-214p13.9
appb
ppme1
CU984600.2
dab2ipb
ldb1a
tspan5a
cd164
pgrmc1
ier5
hmgn6
slc38a2
sptlc2b
hist2h2l
mgst1.1
pnrc2
tcima
chka
hsp70.3
ctnnd2a
b2ml

wsb1
mt-nd5
si:dkey-23i12.7
gng7
csrnp1b
ponzr1
junba
her6
zgc:114200
tjap1
btg2
mt-nd4
XLOC-012680
zgc:123010
edem1
sdc4
nomo
cd99l2
sorcs2
zgc:175088
ctsk
prkar2aa
midn
LOC103909284
kalrna
pak2a
rab11a
ackr4b
cald1a
cav1
si:ch211-114c12.2
ralba
col1a1b
rgs5b
zc2hc1a

rab11fip1b
cdkn1ca
mt-co1
arl5c
tenm3
trim8b
mmp2
cmbl
myl6
tcf7l2
tpbgb
si:dkeyp-115e12.6
man2c1
ppp3ccb
fgd5a
arhgap1
COX3
cmtm4
foxo3b
auts2a
ier2a
zbtb10
stat1b
ppp1r14aa
pcyt2
si:ch211-197h24.9
zgc:161969
thrab
zgc:153916
pde4ba
ier2b
pitpnc1b
mt-nd1
phactr4a
gmfb

zgc:195173
CR318588.1
qsox1
purab
ccdc85b
h1f0
limch1b
lrrc8da
taok1a
si:ch1073-440b2.1
cxxc5a
oip5-as1
CR383676.1
LOC553316
si:dkey-199f5.8
zgc:92744
ctgfa
txnipa
cadm4
tnfrsf9a
tjp1b
tgfb3
rdh10a
wt1a
optn
tgfb2
si:ch211-195b13.1
swap70b
ap2a1
gusb
ptmaa
LO017872.1
ywhaqb
tuba8l3
wu:fj29h11

tns1b
mt-nd2
CABZ01068367.1
msi2a
neu1
si:ch211-244b2.4
tsg101b
loxl2b
si:dkey-56m19.5
si:dkey-222b8.1
zgc:66433
hmgb1a
CABZ01026681.1
prkab1b
atp10a
cpne1
ecm1b
mhc1uba
idh1
cflara
mt-cyb
adamts2
lgals3b
zbtb20
oga
rnf213a
cgref1
prrc2c
mt-atp6
crim1
h2afx1
znf516
gab2
enah
si:ch211-178n15.1

vgll4a
dusp4
erbin
psme1
arpp19a
clpxa
tsc2
rgs16
fam3c
mef2cb
ncor1
tpcn1
zfhx3
puraa
gadd45ab
cntfr
edf1
cpda
zgc:92140
mt-co2
LO018340.1
cox5b2
hgs
atf3
msrb1b
pcolce2b
cd151
NC-002333.4
lsr
mtpn
zgc:92242
rps6kb1a
macf1a
aldh2.1
gpatch8

araf
lgmn
si:ch211-74m13.3
ucp2
si:dkey-28b4.7
rab4b
zgc:112437
krt94
pon3.2
zgc:110783
diaph2
XLOC-024248
kctd12.2
pum2
pim1
map1lc3b
hipk3b
arl8a
myclb
lamc1
htra1b
cnn2
si:dkeyp-69e1.8
rp2
si:dkey-204f11.64
tead1b
ppp1r10
aqp3a
eif4ebp1
prrc2a
CABZ01113812.1
irf2bp2b
calm2b
ppib
vps37b

map1sa
ankrd11
si:dkey-6n21.12
pkp4
tbc1d5
selenof
fndc3bb
crema
spag9a
rnf213b
si:dkeyp-44a8.2
gstm.2
si:ch211-286b5.5
cyfip1
ptprsa
ppt1
tax1bp1b
psme2
tgif1
chd4b
copb2
gstm.3
ppp1cb
pcyox1
si:ch73-1a9.3
si:ch211-232b12.5
sat1a.2
timp2a
snx12
ptk7a
gatad2b
yap1
lpar1
tob1a
hnrpkl

lin7c
crtap
btr09
pfn2l
h3f3c
adam10a
fkbp1aa
LOC101884451
unm-hu7910
cox7a1
antxr1c
atf4b
CABZ01110881.1
acox1
ctsd
smox
dsg2.1
crebrf
rsad2
CABZ01080702.1
ppp2cb
gnpda1
cox8a
tmem59
psmc5
si:ch211-216l23.2
hivep1
irf1b
ndufa4l
si:ch73-22o12.1
kmt2cb
dynlt3
ube2e3
cox6b2
ndufa1

creb3l2
clint1a
CABZ01072614.1
prkd2
si:ch73-46j18.5
NC-002333.17
ldlrap1a
tmub2
serinc2
homeza
junbb
sptbn2
gsk3b
pgd
CABZ01068358.1
elavl1a
fat1a
zfand3
rnf10
si:dkey-79d12.5
hephl1a
ifrd1
pank2
lrpap1
tmsb
tnrc6a
fstl1b
epsti1
sdcbp2
ganab
rab10
si:dkey-42i9.7
exoc6
PDPR
ewsr1a

stau2
mphosph8
CABZ01029938.1
selenok
aste1a
bbip1
strip1
degs1
bin3
serpinf1
u2surp
tcf12
si:ch211-212d10.2
parvaa
klf3
wbp2nl
ifi35
igf2b
dctn3
stxbp6
zgc:100829
sptan1
CABZ01078261.1
atg13
golgb1
hspe1
mt-nd3
nedd8
arf2a
nr1d2a
scrib
eif1b
kdm6bb
si:dkey-187e18.1
si:ch211-108d22.2

map1lc3a
tmem176l.1
rhoaa
rhocb
parp12b
hdac4
CU694197.1
pi4kb
nt5c2b
ctnnd1
rbm39a
pmp22b
si:ch73-233k15.2
ylpm1
ivns1abpa
si:ch211-105c13.3
chd4a
thtpa
adssl
sesn3
si:ch211-87m7.2
fzd7b
matr3l1.1
ndfip2
gstm.1
atp1b3b
litaf
slc38a5b
flot1b
sh3bp5b
si:ch211-39i22.1
ywhaz
st14a
asah1b
calcoco1a

ralab
polr2b
mpzl2b
ywhah
rn7sk
sh3pxd2aa
perp
gabarapb
ndufb9
eif4a1b
tfcp2
tmed10
wls
plbd1
ndfip1
glod5
inhbaa
pros1
csde1
msi2b
si:ch211-140m22.7
b4galt6
hmgb1b
zfand5a
arhgap35a
tspan3a
cox8b
ctsla
abca2
arf2b
jak1
canx
nfil3-6
arid1ab
s100v2

cpvl
clcn7
bcl7a
CABZ01112656.1
birc2
foxo4
ddx3xa
pfn2
abhd17ab
oaz1b
trip12
tmem50a
laptm4a
dbnlb
appbp2
bsdc1
rnaseka
chd3
hif1ab
cst3
ralbp1
katnb1
vdac3
ciao2b
caprin1a
hcfc1b
fstb
XLOC-000946
rnf7
tpm4a
pik3ip1
tomm6
tpm3
si:ch73-42p12.2
chchd2

grinab
fthl27
taf3
hp1bp3
MYADM
ccnl1a
myd88
srgn
arrdc3a
nfkbiaa
psmb1
dync1i2a
rbms2a
bcl2l10
jund
arhgap31
zgc:123258
XLOC-016094
cux1a
morc2
si:ch211-191j22.3
ctbp1
slc3a2b
rps27l
bnip3lb
gchfr
pdlim1
baiap2a
ndufa3
lsm14aa
cdh2
aif1l
kmt2bb
mbnl2
atp6ap1b

atox1
si:ch211-196l7.4
elovl1b
elocb
smdt1b
tbca
ergic2
si:dkey-7j14.6
csnk1a1
enc1
tma7
si:dkeyp-75h12.5
bcap31
gng12a
ahnak
si:dkey-250d21.1
notch2
dpp7
psmb5
vps28
ece2b
rab5c
yy1a
casp3a
mtdha
vps35
si:rp71-36a1.5
fam168a
f11r.1
srrm1
higd1a
ddt
baz1a
fxr2
fundc2

dldh
itgav
clic4
mvb12a
pomp
gnai1
zgc:162698
anxa4
naa38
flot2a
kdm5c
si:dkey-222f8.3
si:ch211-212k18.7
lrrc8aa
chmp2a
mcl1a
elmo1
rxrba
etfb
ash1l
chmp5b
si:ch211-241f5.3
tmem30aa
syncripl
smad2
glud1a
zgc:162730
xaf1
dbi
ghrb
zgc:56095
trappc1
lrp1ab
ddx5
ptp4a2a

golga7
akap12b
emc6
ddx61
mgst3b
dazap2
mef2d
kpna4
cd63
acadvl
si:zfos-169g10.3
purba
atp2b2
wu:fu71h07
elf2b
eif4a2
mycb
si:dkey-42i9.6
emc10
nocta
mcl1b
sub1a
eif3c
nme3
mknk2b
gnb1b
shoc2
gtf2f1
acin1a
calm3b
nenf
fkbp5
msna
uba1
nmi

tax1bp3
fosl2
eif5b
eef1db
spred2b
tln1
reep3b
akt1s1
gng5
eif4g1a
trim25
zgc:100918
myl9b
atp5meb
ubb
rras
LEPROTL1
cd151l
capns1b
psmf1
ogt.1
glula
ccng2
ube4b
ndufa11
sltm
ost4
trib3
cox7c
klf6a
zgc:111986
eppk1
vamp3
smc1al
tbl1xr1a

usp7
cct2
socs3b
sec11a
fam120c
ndufc2
gabarapa
sfpq
relb
cltca
calr3b
akirin1
srsf5a
fkbp2
hsbp1b
ywhae1
chtopa
sri
slc20a1b
sh3bgrl3
emd
selenow1
atp5pf
map2k6
ptp4a2b
dnmt3ab
zgc:174863
gpx4b
stk35l
ap2s1
zgc:77849
atrx
ptp4a1
tmsb4x
hint1

nfia
s100a10b
prdx4
hmgn7
pdia6
ube2d1b
tomm5
pdia3
ndufb10
tagln2
maff
rbx1
aimp2
nars
ndufb3
zgc:92818
ppdpfb
ddit3
cdc42l
skp1
hnrnpa0b
psmb2
sf3b6
prdx2
csnk2b
pabpn1
si:dkey-188i13.11
atp6v0ca
myl12.1
mif
XLOC-024750
gnb1a
zfp36l1a
ube2nb
atp5pb

tmed7
tsc22d3
cox5ab
rbbp6
f3b
atp5f1d
abcf1
si:dkey-11f4.14
hmgb2b
capzb
uqcrh
eif3s10
aldh9a1a.1
ndufb6
zgc:193541
taf13
COX7A2 (1 of many)
GCA
atp5f1e
cdc42
histh1l
zgc:162944
ywhabl
oaz1a
serf2
prdx5
psap
tnrc6c1
dad1
rpl13
rps27.2
sumo3a
serbp1a
sod1
chac1

gp1bb
thbs1b
f5
rac3a
si:ch211-33e4.2
itga2b
hbegfb
apln
sele
tubb1
rpgra
mpl
itgb3b
rprma
rprml
vegfc
tnr
tal1
ppp2r2ba
zgc:171686
si:ch211-195b11.3
parvb
bmp16
gata1a
blf
zgc:162989
fgl2a
admb
zmat4b
si:ch211-153b23.5
myocd
lpp
reps2
fhl1a
arhgef9b

adra2a
p2ry13
hacd3
tpm4a
spns2
cald1a
lmo2
mir223
XLOC-020301
si:ch211-214p13.9
fhl2a
wnt2
gp9
myh11a
LOC108190717
susd1
hpca
plek
arhgap20b
plac8l1
sept12
timd4
pdgfd
ctgfa
XLOC-028370
fli1a
dera
celf4
cldn5b
bmp6
ptger2b
smtna
tln1
tpte
CR753876.1

myl9b
b3gnt3.4
derl2
cavin2b
fn1b
UNC13B
tmem35
lrrc8da
XLOC-026836
si:dkey-217l24.1
si:ch211-222l21.1
mocs2
boka
nuak1a
mylka
rap1b
nfe2
XLOC-019062
blvrb
rhag
f13a1b
actn1
aatka
AL954142.1
pim2
zgc:171506
itprid2
slc5a8l
si:ch211-226m16.2
ano5b
zgc:92140
rap1gap2a
iqgap2
si:ch211-214p16.2
itpr3

srgn
ptmaa
mef2aa
pmp22b
tmsb4x
limd2
p2rx7
hif1aa
btg1
si:ch211-284f22.3
svila
rpz5
fermt3b
ophn1
flna
mgst1.2
meis1b
gcnt4a
btk
asap1b
ilk
tagln2
XLOC-025423
rbpms
si:dkey-238c7.16
pfn1
fosab
phldb1b
si:ch73-234b20.5
zgc:153867
ppiab
dennd2c
creg1
actb2
bcl2l10

tspan5a
CR735102.1
ccnd2a
gapdhs
bambib
XLOC-032002
si:dkey-51e6.1
lzts2a
BX005183.3
adgrg1
CR847543.1
ptpn11b
zgc:66433
CR383676.1
lpar1
hdr
ap1s1
gng10
ing5a
zgc:153409
myl12.1
si:ch211-102c2.4
arpc1b
vash2
si:ch211-218g4.2
emilin1a
cyr61
bmp2k
nup88
pbx1a
sb:cb81
gna11b
dusp5
fkbp1b
mcl1a

macf1a
pbx1b
cotl1
zgc:162509
stk26
si:zfos-464b6.2
zgc:64051
gsna
pnrc2
actb1
snx5
cyth1b
mtus1b
si:cabz01074946.1
akap7
fhl1b
vaspb
prss1
atp2a3
kdm6bb
XLOC-043760
si:ch211-196l7.4
ehbp1l1a
yrk
crema
alox5a
cbfa2t3
trib3
si:ch211-105j21.9
zeb2a
pdlim1
prkcaa
mt-cyb
mtss1
rassf2b

ier2a
LOC103909115
glcci1a
rab11fip1b
sgms1
kcne4
jdp2b
dusp6
nfatc3a
selenow1
mibp2
LOC101886015
XLOC-043761
mylipb
phlda2
itgb3a
msna
ehd1a
cdc26
inpp5ka
si:dkey-52l18.4
tspan7
fosb
cfl1
ubl3a
rhoab
orai1a
tpm3
foxn3
RASSF5
dub
elmo1
lrrfip1a
sept9a
arpc2

btg2
zgc:56095
atp2b1a
cap1
nup93
dynll2b
abhd3
hmgn6
rgs2
mt-co2
actn4
rfx5
ralgapa1
arpc5a
cmtm6
sh3bgrl3
slc25a55a
cpn1
map3k3
mt-co1
sdc4
jupa
COX3
leo1
h3f3c
sh3kbp1
sh3bgrl
myl9a
rhof
myh9a
plcb3
erbin
srsf3a
anapc15
ddx5

TCIM (1 of many)
jun
capza1b
si:ch211-225b11.1
oaz1b
tuba8l3
ppm1ba
mxd4
sept9b
ier2b
ppp3r1a
purab
sec23b
sft2d1
chmp2a
rgl1
dyrk1b
rhogb
rabgap1l
GNAZ
tes
nxpe3
crebrf
ppp1cbl
kctd5a
taf7
ggh
ppt1
anp32e
rplp2
rap2c
selenok
pim1
si:dkeyp-75h12.5
hmgb1a

klf6a
erh
mbnl1
trappc1
tuba8l
sept2
tubb4b
mt-atp6
si:ch211-214p16.1
fam53b
cd63
ndfip2
ramp2
qkia
parp16
inka1b
mfng
skap2
histh1l
lrrfip2
susd6
taf10
sik1
gnai1
calm2a
capza1a
gabarapa
lyn
fkbp5
cops7a
wdr1
klf7b
shq1
suv39h1b
serf2

zgc:158564
fosl2
herpud2
si:dkey-261h17.1
psma4
nr4a1
kif5ba
gadd45ba
