## Supplementary material for "Direct Male Development in Chromosomally ZZ Zebrafish": SuppTable S6 Merged 19dpf 30dpf Cell Counts per Cluster

| Cluster | ZW_19dpf cell number | ZW_30dpf cell number | ZZ_19dpf cell number | ZZ_30dpf cell number |
|---|---|---|---|---|
| 0 | 0 | 1442 | 0 | 0 |
| 1 | 2 | 934 | 0 | 0 |
| 2 | 23 | 582 | 52 | 276 |
| 3 | 8 | 875 | 2 | 7 |
| 4 | 117 | 261 | 214 | 261 |
| 5 | 0 | 756 | 0 | 0 |
| 6 | 90 | 131 | 111 | 285 |
| 7 | 44 | 261 | 99 | 162 |
| 8 | 141 | 154 | 186 | 67 |
| 9 | 26 | 287 | 67 | 96 |
| 10 | 88 | 131 | 91 | 69 |
| 11 | 17 | 174 | 68 | 113 |
| 12 | 1 | 14 | 3 | 269 |
| 13 | 6 | 169 | 13 | 82 |
| 14 | 34 | 57 | 73 | 100 |
| 15 | 0 | 253 | 1 | 0 |
| 16 | 35 | 107 | 17 | 91 |
| 17 | 2 | 216 | 0 | 0 |
| 18 | 11 | 194 | 6 | 1 |
| 19 | 0 | 81 | 4 | 103 |
| 20 | 0 | 0 | 0 | 177 |
| 21 | 35 | 66 | 73 | 2 |
| 22 | 28 | 55 | 33 | 57 |
| 23 | 59 | 15 | 89 | 8 |
| 24 | 0 | 167 | 0 | 0 |
| 25 | 4 | 55 | 30 | 69 |
| 26 | 3 | 91 | 21 | 27 |
| 27 | 8 | 60 | 32 | 36 |
| 28 | 3 | 91 | 13 | 20 |
| 29 | 12 | 10 | 14 | 74 |
| 30 | 0 | 1 | 0 | 101 |
| 31 | 8 | 46 | 12 | 27 |
| 32 | 0 | 2 | 0 | 87 |
| 33 | 8 | 29 | 14 | 36 |
| 34 | 0 | 3 | 52 | 21 |
| 35 | 3 | 0 | 11 | 47 |
| 36 | 19 | 3 | 24 | 9 |
| 37 | 7 | 27 | 8 | 9 |
| 38 | 0 | 0 | 0 | 42 |
| 39 | 4 | 0 | 4 | 12 |
