## Supplementary material for "Direct Male Development in Chromosomally ZZ Zebrafish": SuppTable S7 Merged19and30dpf_AllMarkers_minpct10

| | p_val | avg_log2FC | pct.1 | pct.2 | p_val_adj | cluster | gene |
|---|---|---|---|---|---|---|---|
| zgc:153499 | 0 | 1.071272133 | 0.968 | 0.356 | 0 | 0 | zgc:153499 |
| si:dkey-240e12.6 | 0 | 1.039205277 | 0.982 | 0.406 | 0 | 0 | si:dkey-240e12.6 |
| si:ch73-189n23.1 | 0 | 1.020939141 | 0.946 | 0.371 | 0 | 0 | si:ch73-189n23.1 |
| cldnd | 0 | 1.016529552 | 0.991 | 0.439 | 0 | 0 | cldnd |
| h2af1al | 0 | 1.01568497 | 0.992 | 0.43 | 0 | 0 | h2af1al |
| tmsb5 | 0 | 1.006442166 | 0.915 | 0.341 | 0 | 0 | tmsb5 |
| zp2.2 | 0 | 1.00295896 | 0.939 | 0.375 | 0 | 0 | zp2.2 |
| si:ch211-226h8.14 | 0 | 1.000927387 | 0.958 | 0.408 | 0 | 0 | si:ch211-226h8.14 |
| cldng | 0 | 0.999104041 | 0.921 | 0.365 | 0 | 0 | cldng |
| zp3 | 0 | 0.998247951 | 0.972 | 0.422 | 0 | 0 | zp3 |
| zgc:175135 | 0 | 0.997445766 | 0.887 | 0.336 | 0 | 0 | zgc:175135 |
| crp2 | 0 | 0.99727291 | 0.904 | 0.343 | 0 | 0 | crp2 |
| mt2 | 0 | 0.997010892 | 0.95 | 0.38 | 0 | 0 | mt2 |
| zp2.5 | 0 | 0.994693954 | 0.958 | 0.405 | 0 | 0 | zp2.5 |
| si:zfos-1505d6.3 | 0 | 0.994042971 | 0.92 | 0.363 | 0 | 0 | si:zfos-1505d6.3 |
| si:dkeyp-98a7.9 | 0 | 0.992705944 | 0.915 | 0.363 | 0 | 0 | si:dkeyp-98a7.9 |
| si:dkey-229d11.3 | 0 | 0.992582411 | 0.85 | 0.302 | 0 | 0 | si:dkey-229d11.3 |
| si:dkey-241l7.2 | 0 | 0.992232059 | 0.997 | 0.473 | 0 | 0 | si:dkey-241l7.2 |
| zp2.6 | 0 | 0.989133733 | 0.902 | 0.354 | 0 | 0 | zp2.6 |
| si:ch211-125e6.12 | 0 | 0.987490174 | 0.904 | 0.354 | 0 | 0 | si:ch211-125e6.12 |
| h1m | 0 | 0.986786812 | 0.982 | 0.439 | 0 | 0 | h1m |
| zp2.1 | 0 | 0.985854001 | 0.928 | 0.382 | 0 | 0 | zp2.1 |
| acp5a | 0 | 0.983984306 | 0.939 | 0.387 | 0 | 0 | acp5a |
| zp2.3 | 0 | 0.981817249 | 0.915 | 0.368 | 0 | 0 | zp2.3 |
| BX649337.1 | 0 | 0.976371884 | 0.844 | 0.303 | 0 | 0 | BX649337.1 |
| zgc:171776 | 0 | 0.973259996 | 0.89 | 0.35 | 0 | 0 | zgc:171776 |
| si:dkey-241l7.5 | 0 | 0.969481672 | 0.96 | 0.424 | 0 | 0 | si:dkey-241l7.5 |
| ccna1 | 0 | 0.964361142 | 0.863 | 0.336 | 0 | 0 | ccna1 |
| zp3.2 | 0 | 0.963745595 | 0.935 | 0.403 | 0 | 0 | zp3.2 |
| si:ch211-226h8.4 | 0 | 0.956465171 | 0.83 | 0.308 | 0 | 0 | si:ch211-226h8.4 |
| LOC110437821 | 0 | 0.952564333 | 0.785 | 0.289 | 0 | 0 | LOC110437821 |
| si:dkeyp-98a7.8 | 0 | 0.949718072 | 0.829 | 0.323 | 0 | 0 | si:dkeyp-98a7.8 |
| CR790368.1 | 0 | 0.944986073 | 0.896 | 0.373 | 0 | 0 | CR790368.1 |
| hbba1 | 0 | 0.944519099 | 0.731 | 0.243 | 0 | 0 | hbba1 |

| | | | | | | | |
|---|---|---|---|---|---|---|---|
| si:dkeyp-20e4.8 | 0 | 0.942894246 | 0.775 | 0.283 | 0 | 0 | si:dkeyp-20e4.8 |
| CU571163.1 | 0 | 0.935508655 | 0.773 | 0.278 | 0 | 0 | CU571163.1 |
| zgc:171779 | 0 | 0.934526886 | 0.878 | 0.367 | 0 | 0 | zgc:171779 |
| zp3b | 0 | 0.931276255 | 0.755 | 0.268 | 0 | 0 | zp3b |
| si:dkey-241l7.4 | 0 | 0.923686624 | 0.958 | 0.447 | 0 | 0 | si:dkey-241l7.4 |
| si:dkey-90l23.1 | 0 | 0.921886993 | 0.811 | 0.316 | 0 | 0 | si:dkey-90l23.1 |
| si:dkeyp-98a7.4 | 0 | 0.920804038 | 0.788 | 0.297 | 0 | 0 | si:dkeyp-98a7.4 |
| si:ch211-12h2.8 | 0 | 0.917658635 | 0.899 | 0.398 | 0 | 0 | si:ch211-12h2.8 |
| epcam | 0 | 0.914738261 | 0.87 | 0.367 | 0 | 0 | epcam |
| qdprb2 | 0 | 0.913612293 | 0.782 | 0.306 | 0 | 0 | qdprb2 |
| zgc:77118 | 0 | 0.910853134 | 0.817 | 0.325 | 0 | 0 | zgc:77118 |
| rrm2 | 0 | 0.89404823 | 0.868 | 0.37 | 0 | 0 | rrm2 |
| si:dkey-241l7.6 | 0 | 0.877821132 | 0.847 | 0.385 | 0 | 0 | si:dkey-241l7.6 |
| zp3a.2 | 0 | 0.872927463 | 0.825 | 0.355 | 0 | 0 | zp3a.2 |
| mid1ip1l | 0 | 0.851375521 | 0.852 | 0.414 | 0 | 0 | mid1ip1l |
| lamp2 | 0 | 0.848605057 | 0.868 | 0.439 | 0 | 0 | lamp2 |
| zar1l | 3.690332 | 0.893047157 | 0.739 | 0.279 | 9.3752901 | 0 | zar1l |
| si:dkeyp-46h3.6 | 1.829545 | 0.89473812 | 0.755 | 0.288 | 4.6479602 | 0 | si:dkeyp-46h3.6 |
| ccnb2 | 2.038497 | 0.891473904 | 0.725 | 0.269 | 5.1788029 | 0 | ccnb2 |
| si:ch211-5k11.8 | 2.444848 | 0.837633325 | 0.719 | 0.247 | 6.2111375 | 0 | si:ch211-5k11.8 |
| slbp2 | 4.680522 | 0.880892489 | 0.738 | 0.279 | 1.1890868 | 0 | slbp2 |
| hbaa1 | 5.992874 | 0.889544316 | 0.618 | 0.188 | 1.5224896 | 0 | hbaa1 |
| zar1 | 5.107356 | 0.861476215 | 0.763 | 0.316 | 1.2975238 | 0 | zar1 |
| si:dkeyp-50f7.2 | 1.436045 | 0.861858266 | 0.729 | 0.285 | 3.6482734 | 0 | si:dkeyp-50f7.2 |
| si:ch211-12h2.6 | 2.844876 | 0.849152798 | 0.787 | 0.343 | 7.2274082 | 0 | si:ch211-12h2.6 |
| tuba4l | 2.056531 | 0.861649044 | 0.732 | 0.287 | 5.2246179 | 0 | tuba4l |
| zgc:195245 | 3.678201 | 0.862493946 | 0.714 | 0.273 | 9.3444697 | 0 | zgc:195245 |
| npm2b | 3.363085 | 0.856948399 | 0.718 | 0.281 | 8.5439197 | 0 | npm2b |
| si:dkeyp-46h3.8 | 4.034345 | 0.857811982 | 0.671 | 0.244 | 1.0249254 | 0 | si:dkeyp-46h3.8 |
| siva1 | 7.096863 | 0.835643026 | 0.741 | 0.317 | 1.8029581 | 0 | siva1 |
| thy1 | 1.167437 | 0.831267733 | 0.766 | 0.333 | 2.9658748 | 0 | thy1 |
| zgc:171517 | 8.466646 | 0.846306721 | 0.695 | 0.275 | 2.1509515 | 0 | zgc:171517 |
| cth1 | 1.443385 | 0.836606624 | 0.686 | 0.268 | 3.6669204 | 0 | cth1 |
| LOC100535162 | 7.420299 | 0.843471582 | 0.623 | 0.211 | 1.8851270 | 0 | LOC100535162 |
| zpcx | 1.010220 | 0.837198351 | 0.641 | 0.236 | 2.5664650 | 0 | zpcx |

| | | | | | | | |
|---|---|---|---|---|---|---|---|
| si:ch211-14a17.7 | 1.965613 | 0.832887845 | 0.718 | 0.292 | 4.9936402 | 0 | si:ch211-14a17.7 |
| si:dkey-229d11.5 | 9.494159 | 0.833068291 | 0.698 | 0.276 | 2.4119912 | 0 | si:dkey-229d11.5 |
| LOC100537499 | 2.522341 | 0.834824059 | 0.632 | 0.222 | 6.4080089 | 0 | LOC100537499 |
| gapdh | 1.278400 | 0.815823052 | 0.717 | 0.304 | 3.2477754 | 0 | gapdh |
| s100a1 | 9.575492 | 0.819672296 | 0.675 | 0.256 | 2.4326538 | 0 | s100a1 |
| AL929032.1 | 2.467409 | 0.836154236 | 0.619 | 0.219 | 6.2684535 | 0 | AL929032.1 |
| ctss2.1 | 1.203995 | 0.759554621 | 0.806 | 0.41 | 3.0587498 | 0 | ctss2.1 |
| ccnb1 | 2.610960 | 0.810173723 | 0.691 | 0.284 | 6.6331458 | 0 | ccnb1 |
| si:dkey-90l23.2 | 6.437594 | 0.818383354 | 0.64 | 0.242 | 1.6354709 | 0 | si:dkey-90l23.2 |
| nanog | 2.537588 | 0.809514005 | 0.66 | 0.254 | 6.4467440 | 0 | nanog |
| btg4 | 2.796782 | 0.808892696 | 0.666 | 0.261 | 7.1052269 | 0 | btg4 |
| zgc:173443 | 3.425358 | 0.806782074 | 0.628 | 0.23 | 8.7021243 | 0 | zgc:173443 |
| gtf3ab | 2.588751 | 0.794393478 | 0.614 | 0.223 | 6.5767240 | 0 | gtf3ab |
| spink2.1 | 7.090815 | 0.76027375 | 0.729 | 0.332 | 1.8014217 | 0 | spink2.1 |
| zgc:136254 | 3.989982 | 0.794356561 | 0.615 | 0.232 | 1.0136549 | 0 | zgc:136254 |
| si:ch211-132b12.8 | 9.515734 | 0.789314599 | 0.628 | 0.242 | 2.4174724 | 0 | si:ch211-132b12.8 |
| wu:fi42e03 | 4.845855 | 0.781045793 | 0.625 | 0.249 | 1.2310895 | 0 | wu:fi42e03 |
| kpna7 | 4.456223 | 0.7829622 | 0.607 | 0.232 | 1.1321035 | 0 | kpna7 |
| si:dkeyp-46h3.2 | 4.136050 | 0.76648932 | 0.665 | 0.294 | 1.0507637 | 0 | si:dkeyp-46h3.2 |
| hnrnpabb | 2.798795 | 0.621997951 | 0.797 | 0.531 | 7.1103402 | 0 | hnrnpabb |
| uchl1 | 4.442726 | 0.75234419 | 0.612 | 0.253 | 1.1286747 | 0 | uchl1 |
| zgc:171977 | 7.947209 | 0.747612946 | 0.471 | 0.156 | 2.0189885 | 0 | zgc:171977 |
| cpeb1b | 2.277348 | 0.743732955 | 0.525 | 0.188 | 5.7856037 | 0 | cpeb1b |
| zgc:171474 | 1.298121 | 0.744826684 | 0.556 | 0.215 | 3.2978787 | 0 | zgc:171474 |
| si:dkey-152b24.6 | 1.539321 | 0.731430942 | 0.55 | 0.209 | 3.9106475 | 0 | si:dkey-152b24.6 |
| si:ch73-90k17.1 | 5.839808 | 0.722955504 | 0.492 | 0.171 | 1.4836033 | 0 | si:ch73-90k17.1 |
| zp3a.1 | 4.573688 | 0.725046113 | 0.615 | 0.279 | 1.1619456 | 0 | zp3a.1 |
| LOC101885477 | 3.823788 | 0.726078469 | 0.519 | 0.197 | 9.7143349 | 0 | LOC101885477 |
| avd | 2.829384 | 0.716844276 | 0.507 | 0.185 | 7.1880522 | 0 | avd |
| si:dkey-208k4.2 | 2.310735 | 0.717649189 | 0.539 | 0.21 | 5.8704232 | 0 | si:dkey-208k4.2 |
| si:dkey-37o8.1 | 1.387862 | 0.706574032 | 0.544 | 0.219 | 3.5258640 | 0 | si:dkey-37o8.1 |
| gmnn | 2.228368 | 0.704566012 | 0.54 | 0.222 | 5.6611699 | 0 | gmnn |
| si:ch211-146l10.8 | 5.877175 | 0.708982067 | 0.538 | 0.219 | 1.4930964 | 0 | si:ch211-146l10.8 |
| ca15b | 4.272648 | 0.687192388 | 0.476 | 0.174 | 1.0854663 | 0 | ca15b |
| arhgef10la | 4.014019 | 0.697822626 | 0.504 | 0.195 | 1.0197617 | 0 | arhgef10la |

| | | | | | | | |
|---|---|---|---|---|---|---|---|
| lsm14b | 1.639053 | 0.687978842 | 0.524 | 0.215 | 4.1640163 | 0 | lsm14b |
| si:ch211-145c1.1 | 3.639198 | 0.683997713 | 0.457 | 0.162 | 9.2453827 | 0 | si:ch211-145c1.1 |
| si:ch73-14h10.2 | 4.885845 | 0.686358441 | 0.497 | 0.192 | 1.2412490 | 0 | si:ch73-14h10.2 |
| nasp | 1.720593 | 0.694799019 | 0.572 | 0.262 | 4.3711689 | 0 | nasp |
| zgc:55461 | 3.707428 | 0.687355291 | 0.517 | 0.21 | 9.4187229 | 0 | zgc:55461 |
| elovl7b | 6.419514 | 0.668772271 | 0.467 | 0.175 | 1.6308776 | 0 | elovl7b |
| haus6 | 8.749558 | 0.660615316 | 0.42 | 0.145 | 2.2228252 | 0 | haus6 |
| si:dkeyp-46h3.5 | 3.617548 | 0.676641973 | 0.508 | 0.207 | 9.1903811 | 0 | si:dkeyp-46h3.5 |
| si:dkeyp-98a7.7 | 3.711334 | 0.647397416 | 0.404 | 0.133 | 9.4286455 | 0 | si:dkeyp-98a7.7 |
| si:dkey-19b23.11 | 7.903951 | 0.674118172 | 0.492 | 0.199 | 2.0079989 | 0 | si:dkey-19b23.11 |
| ccna2 | 8.563447 | 0.668047967 | 0.505 | 0.209 | 2.1755439 | 0 | ccna2 |
| birc5b | 8.906856 | 0.66407522 | 0.475 | 0.184 | 2.2627868 | 0 | birc5b |
| si:ch211-250e5.16 | 6.597618 | 0.668684869 | 0.559 | 0.254 | 1.6761249 | 0 | si:ch211-250e5.16 |
| zgc:56231 | 2.789439 | 0.644119746 | 0.434 | 0.159 | 7.0865715 | 0 | zgc:56231 |
| pigrl2.3 | 1.380367 | 0.635297033 | 0.406 | 0.141 | 3.5068239 | 0 | pigrl2.3 |
| mkrn4 | 5.992689 | 0.652996234 | 0.433 | 0.166 | 1.5224428 | 0 | mkrn4 |
| zgc:92591 | 7.697325 | 0.635664325 | 0.404 | 0.143 | 1.9555056 | 0 | zgc:92591 |
| crp3 | 3.825901 | 0.638356099 | 0.418 | 0.152 | 9.7197032 | 0 | crp3 |
| si:ch211-251f6.6 | 2.618390 | 0.65506864 | 0.531 | 0.243 | 6.6520208 | 0 | si:ch211-251f6.6 |
| larp6b | 1.354757 | 0.625998261 | 0.392 | 0.135 | 3.4417615 | 0 | larp6b |
| pabpn1l | 2.982345 | 0.649504786 | 0.493 | 0.206 | 7.5766477 | 0 | pabpn1l |
| gdf3 | 2.385318 | 0.625884273 | 0.423 | 0.157 | 6.0599014 | 0 | gdf3 |
| si:ch1073-263o8.2 | 2.393803 | 0.645908078 | 0.46 | 0.189 | 6.0814578 | 0 | si:ch1073-263o8.2 |
| si:ch211-226l4.6 | 1.440481 | 0.633056472 | 0.5 | 0.214 | 3.6595429 | 0 | si:ch211-226l4.6 |
| zgc:56699 | 2.867680 | 0.62797948 | 0.447 | 0.183 | 7.2853411 | 0 | zgc:56699 |
| slc16a3 | 8.513826 | 0.621639237 | 0.438 | 0.18 | 2.1629376 | 0 | slc16a3 |
| si:ch211-157c3.4 | 1.076621 | 0.499778661 | 0.232 | 0.057 | 2.7351574 | 0 | si:ch211-157c3.4 |
| zglp1 | 1.439434 | 0.593006554 | 0.38 | 0.139 | 3.6568836 | 0 | zglp1 |
| LOC100536860 | 2.252491 | 0.614880251 | 0.527 | 0.261 | 5.7224557 | 0 | LOC100536860 |
| sinup | 8.950023 | 0.594826202 | 0.384 | 0.143 | 2.2737534 | 0 | sinup |
| si:ch73-160p18.3 | 9.529021 | 0.615250031 | 0.419 | 0.173 | 2.4208478 | 0 | si:ch73-160p18.3 |
| LOC101883897 | 2.799379 | 0.561542722 | 0.334 | 0.113 | 7.1118240 | 0 | LOC101883897 |
| zgc:173856 | 6.090790 | 0.588215684 | 0.394 | 0.157 | 1.5473653 | 0 | zgc:173856 |
| org | 1.119340 | 0.605257426 | 0.416 | 0.175 | 2.8436832 | 0 | org |
| map1lc3c | 7.710330 | 0.583307783 | 0.37 | 0.143 | 1.9588094 | 0 | map1lc3c |

| | | | | | | | |
|---|---|---|---|---|---|---|---|
| zgc:162879 | 4.26086( | 0.55932236 | 0.35 | 0.127 | 1.0824715 | 0 | zgc:162879 |
| h3f3a | 5.013694 | 0.526970652 | 0.665 | 0.462 | 1.2737290 | 0 | h3f3a |
| slbp | 7.611218 | 0.588676989 | 0.413 | 0.175 | 1.9336300 | 0 | slbp |
| si:dkey-241l7.3 | 4.422506 | 0.594128091 | 0.526 | 0.275 | 1.1235377 | 0 | si:dkey-241l7.3 |
| AL929192.1 | 4.532554 | 0.541874221 | 0.326 | 0.113 | 1.1514953 | 0 | AL929192.1 |
| zp3c | 2.841665 | 0.550926986 | 0.353 | 0.133 | 7.2192513 | 0 | zp3c |
| zgc:113424 | 2.68316( | 0.528880047 | 0.319 | 0.112 | 6.8165691 | 0 | zgc:113424 |
| cdca7a | 2.694799 | 0.570236228 | 0.393 | 0.17 | 6.8461390 | 0 | cdca7a |
| hnrnpa0l | 1.017331 | 0.301338949 | 0.816 | 0.732 | 2.5845297 | 0 | hnrnpa0l |
| si:ch1073-190k2.1 | 1.618661 | 0.541455069 | 0.346 | 0.133 | 4.1122091 | 0 | si:ch1073-190k2.1 |
| rcc1 | 2.922173 | 0.553577599 | 0.37 | 0.151 | 7.4237820 | 0 | rcc1 |
| elovl7a | 8.077789 | 0.537230231 | 0.338 | 0.132 | 2.0521625 | 0 | elovl7a |
| si:ch211-146l10.7 | 5.943072 | 0.535376229 | 0.342 | 0.134 | 1.5098375 | 0 | si:ch211-146l10.7 |
| tph1a | 1.206196 | 0.497638664 | 0.285 | 0.098 | 3.0643410 | 0 | tph1a |
| patl2 | 4.654587 | 0.525932562 | 0.322 | 0.123 | 1.1824980 | 0 | patl2 |
| buc | 2.345075 | 0.526791383 | 0.329 | 0.127 | 5.9576650 | 0 | buc |
| CR847984.1 | 8.916192 | 0.520994841 | 0.316 | 0.119 | 2.2651587 | 0 | CR847984.1 |
| si:dkeyp-75b4.10 | 1.116164 | 0.483567357 | 0.603 | 0.339 | 2.8356164 | 0 | si:dkeyp-75b4.10 |
| retsatl | 2.158396 | 0.536892079 | 0.353 | 0.147 | 5.4834054 | 0 | retsatl |
| rbpms2b | 1.480307 | 0.539701568 | 0.381 | 0.169 | 3.7607202 | 0 | rbpms2b |
| pou5f3 | 2.46632( | 0.492911471 | 0.288 | 0.102 | 6.2656870 | 0 | pou5f3 |
| BX537276.1 | 2.426948 | 0.469966827 | 0.253 | 0.083 | 6.1656638 | 0 | BX537276.1 |
| tmem54a | 3.977211 | 0.488192252 | 0.279 | 0.099 | 1.0104106 | 0 | tmem54a |
| tac3b | 4.590564 | 0.470503649 | 0.25 | 0.083 | 1.1662330 | 0 | tac3b |
| hsd17b12a | 2.820416 | 0.500303324 | 0.307 | 0.118 | 7.1652678 | 0 | hsd17b12a |
| zgc:113886 | 3.119478 | 0.485246514 | 0.277 | 0.098 | 7.9250347 | 0 | zgc:113886 |
| inka1b | 2.516172 | 0.509404684 | 0.333 | 0.136 | 6.3923362 | 0 | inka1b |
| zgc:171750 | 4.550425 | 0.506063496 | 0.314 | 0.125 | 1.1560356 | 0 | zgc:171750 |
| CU467646.2 | 5.047139 | 0.453576183 | 0.257 | 0.086 | 1.2822257 | 0 | CU467646.2 |
| snu13b | 5.989255 | 0.519130316 | 0.549 | 0.351 | 1.5215704 | 0 | snu13b |
| gyg1a | 1.224489 | 0.540338677 | 0.504 | 0.298 | 3.1108153 | 0 | gyg1a |
| rasl11b | 4.060661 | 0.474739811 | 0.268 | 0.096 | 1.0316110 | 0 | rasl11b |
| sybu | 9.158853 | 0.490403782 | 0.304 | 0.118 | 2.3268068 | 0 | sybu |
| chia.4 | 1.214938 | 0.480217477 | 0.279 | 0.103 | 3.0865511 | 0 | chia.4 |
| mibp | 1.372315 | 0.516700577 | 0.383 | 0.179 | 3.4863686 | 0 | mibp |

| | | | | | | | |
|---|---|---|---|---|---|---|---|
| phex | 4.165085 | 0.416097881 | 0.198 | 0.058 | 1.0581400 | 0 | phex |
| zgc:165539 | 1.821100 | 0.448468174 | 0.244 | 0.082 | 4.6265063 | 0 | zgc:165539 |
| larp6a | 3.182068 | 0.529679708 | 0.397 | 0.193 | 8.0840459 | 0 | larp6a |
| si:dkey-46g23.2 | 1.424828 | 0.467297744 | 0.274 | 0.102 | 3.6197767 | 0 | si:dkey-46g23.2 |
| vstm2l | 1.832207 | 0.418368366 | 0.209 | 0.064 | 4.6547242 | 0 | vstm2l |
| zgc:103482 | 2.218552 | 0.492739308 | 0.328 | 0.138 | 5.6362317 | 0 | zgc:103482 |
| rbm4.3 | 3.223982 | 0.474771006 | 0.568 | 0.405 | 8.1905279 | 0 | rbm4.3 |
| plk1 | 8.972351 | 0.486585701 | 0.313 | 0.13 | 2.2794258 | 0 | plk1 |
| rdh10b | 1.061716 | 0.500060204 | 0.313 | 0.133 | 2.6972897 | 0 | rdh10b |
| CU655961.3 | 3.149061 | 0.409217428 | 0.211 | 0.067 | 8.0001917 | 0 | CU655961.3 |
| pabpc1l | 3.517490 | 0.452012384 | 0.268 | 0.101 | 8.9361837 | 0 | pabpc1l |
| CABZ01059627.1 | 4.929711 | 0.448663932 | 0.259 | 0.096 | 1.2523931 | 0 | CABZ01059627.1 |
| gbgt1l4 | 7.667068 | 0.454297875 | 0.279 | 0.107 | 1.9478186 | 0 | gbgt1l4 |
| CDPF1 | 5.372271 | 0.524253357 | 0.408 | 0.216 | 1.3648254 | 0 | CDPF1 |
| CABZ01059627.2 | 3.487380 | 0.440415605 | 0.252 | 0.094 | 8.8596903 | 0 | CABZ01059627.2 |
| si:ch73-78o10.1 | 2.066045 | 0.43375565 | 0.253 | 0.094 | 5.2487873 | 0 | si:ch73-78o10.1 |
| zgc:173742 | 1.087761 | 0.45747729 | 0.281 | 0.113 | 2.7634574 | 0 | zgc:173742 |
| zp2l1 | 6.271539 | 0.458791589 | 0.286 | 0.119 | 1.5932845 | 0 | zp2l1 |
| elavl2 | 1.090403 | 0.485167109 | 0.355 | 0.168 | 2.7701709 | 0 | elavl2 |
| zgc:111868 | 1.537928 | 0.444889563 | 0.268 | 0.106 | 3.9071061 | 0 | zgc:111868 |
| zgc:66024 | 1.002578 | 0.432123571 | 0.261 | 0.101 | 2.5470498 | 0 | zgc:66024 |
| BX001030.1 | 1.596420 | 0.416751069 | 0.234 | 0.086 | 4.0557053 | 0 | BX001030.1 |
| si:dkey-42i9.8 | 6.765053 | 0.423717468 | 0.241 | 0.091 | 1.7186618 | 0 | si:dkey-42i9.8 |
| zgc:101744 | 2.454378 | 0.433361099 | 0.249 | 0.097 | 6.2353485 | 0 | zgc:101744 |
| lpcat4 | 6.595003 | 0.404522481 | 0.216 | 0.077 | 1.6754605 | 0 | lpcat4 |
| degs2 | 7.452195 | 0.401879757 | 0.214 | 0.075 | 1.8932303 | 0 | degs2 |
| zgc:152652 | 3.491150 | 0.408824323 | 0.221 | 0.08 | 8.8692666 | 0 | zgc:152652 |
| cldn7b | 8.349070 | 0.443941275 | 0.279 | 0.117 | 2.1210812 | 0 | cldn7b |
| mad2l1 | 2.812406 | 0.454706334 | 0.311 | 0.142 | 7.1449181 | 0 | mad2l1 |
| BX284697.1 | 3.449124 | 0.404274776 | 0.22 | 0.08 | 8.7625014 | 0 | BX284697.1 |
| CU929143.1 | 1.247662 | 0.39456402 | 0.186 | 0.063 | 3.1696871 | 0 | CU929143.1 |
| pif1 | 3.683209 | 0.395575788 | 0.209 | 0.075 | 9.3571932 | 0 | pif1 |
| zc3h11a | 9.276330 | 0.472659815 | 0.338 | 0.173 | 2.3566516 | 0 | zc3h11a |
| zgc:162945 | 1.166541 | 0.428162761 | 0.279 | 0.121 | 2.9635988 | 0 | zgc:162945 |
| spint2 | 1.646041 | 0.370003219 | 0.175 | 0.057 | 4.1817676 | 0 | spint2 |

| | | | | | | | |
|---|---|---|---|---|---|---|---|
| si:dkey-188i13.11 | 6.182172 | 0.475250283 | 0.447 | 0.274 | 1.5705808 | 0 | si:dkey-188i13.11 |
| rpain | 7.512322 | 0.406501259 | 0.225 | 0.087 | 1.9085054 | 0 | rpain |
| ubc | 1.027851 | 0.279295641 | 0.703 | 0.649 | 2.6112572 | 0 | ubc |
| mos | 1.795742 | 0.415159436 | 0.245 | 0.1 | 4.5620845 | 0 | mos |
| h2afva | 4.802249 | 0.469608501 | 0.449 | 0.291 | 1.2200114 | 0 | h2afva |
| sox19b | 5.815531 | 0.360472157 | 0.196 | 0.068 | 1.4774358 | 0 | sox19b |
| ddx39ab | 1.416558 | 0.475249167 | 0.418 | 0.254 | 3.5987663 | 0 | ddx39ab |
| il13ra2 | 5.572926 | 0.38737237 | 0.212 | 0.08 | 1.4158019 | 0 | il13ra2 |
| si:ch73-15b2.5 | 1.050804 | 0.366725745 | 0.189 | 0.066 | 2.6695696 | 0 | si:ch73-15b2.5 |
| dynll1 | 1.525805 | 0.408667539 | 0.566 | 0.428 | 3.8763096 | 0 | dynll1 |
| zgc:163030 | 2.173433 | 0.40145773 | 0.232 | 0.093 | 5.5216079 | 0 | zgc:163030 |
| gstp1 | 3.043944 | 0.445203596 | 0.491 | 0.332 | 7.7331413 | 0 | gstp1 |
| CT009596.1 | 8.184161 | 0.35873023 | 0.182 | 0.062 | 2.0791862 | 0 | CT009596.1 |
| CR383662.2 | 1.452599 | 0.347961407 | 0.178 | 0.06 | 3.6903284 | 0 | CR383662.2 |
| cdc20 | 5.909411 | 0.413323833 | 0.264 | 0.115 | 1.5012858 | 0 | cdc20 |
| zgc:113984 | 6.555694 | 0.371348525 | 0.196 | 0.071 | 1.6654740 | 0 | zgc:113984 |
| dctpp1 | 2.305395 | 0.430828417 | 0.288 | 0.134 | 5.8568561 | 0 | dctpp1 |
| cdca9 | 3.697395 | 0.350798935 | 0.175 | 0.06 | 9.3932322 | 0 | cdca9 |
| si:dkey-216e24.3 | 5.897812 | 0.387802906 | 0.223 | 0.089 | 1.4983392 | 0 | si:dkey-216e24.3 |
| BX571780.1 | 8.719383 | 0.338761148 | 0.169 | 0.056 | 2.2151593 | 0 | BX571780.1 |
| dynll2a | 3.116447 | 0.400845418 | 0.244 | 0.104 | 7.9173344 | 0 | dynll2a |
| si:dkeyp-114g9.1 | 3.700407 | 0.368146792 | 0.198 | 0.074 | 9.4008857 | 0 | si:dkeyp-114g9.1 |
| nanos3 | 5.274340 | 0.371757419 | 0.196 | 0.073 | 1.3399462 | 0 | nanos3 |
| ccne1 | 5.365675 | 0.387715726 | 0.209 | 0.081 | 1.3631497 | 0 | ccne1 |
| buc2l | 5.522440 | 0.335440515 | 0.166 | 0.055 | 1.4029758 | 0 | buc2l |
| nfr | 6.333816 | 0.373366117 | 0.207 | 0.079 | 1.6091060 | 0 | nfr |
| pttg1 | 1.690367 | 0.408523191 | 0.27 | 0.122 | 4.2943794 | 0 | pttg1 |
| ctsba | 2.257481 | 0.43238981 | 0.497 | 0.337 | 5.7351328 | 0 | ctsba |
| kpna2 | 1.962846 | 0.404131581 | 0.268 | 0.122 | 4.9866115 | 0 | kpna2 |
| bmb | 3.250518 | 0.349322777 | 0.175 | 0.062 | 8.2579433 | 0 | bmb |
| tmem144b | 1.339969 | 0.346195712 | 0.183 | 0.066 | 3.4041914 | 0 | tmem144b |
| cx44.2 | 2.857225 | 0.369086276 | 0.212 | 0.084 | 7.2587808 | 0 | cx44.2 |
| ddx41 | 3.684125 | 0.301556611 | 0.148 | 0.047 | 9.3595206 | 0 | ddx41 |
| prdx1 | 4.825504 | 0.420048519 | 0.28 | 0.134 | 1.2259193 | 0 | prdx1 |
| gpr160 | 6.023875 | 0.331352332 | 0.166 | 0.057 | 1.5303656 | 0 | gpr160 |

| | | | | | | | |
|---|---|---|---|---|---|---|---|
| cox17 | 9.040495 | 0.428584288 | 0.467 | 0.328 | 2.2967379 | 0 | cox17 |
| si:ch211-113a14.27 | 1.849954 | 0.331445142 | 0.161 | 0.055 | 4.6998098 | 0 | si:ch211-113a14.27 |
| aurka | 2.083987 | 0.370158379 | 0.215 | 0.088 | 5.2943709 | 0 | aurka |
| zgc:92287 | 2.830059 | 0.374022482 | 0.233 | 0.098 | 7.1897666 | 0 | zgc:92287 |
| zgc:112980 | 3.830781 | 0.350096645 | 0.181 | 0.067 | 9.7320995 | 0 | zgc:112980 |
| dazl | 6.031918 | 0.358790715 | 0.199 | 0.078 | 1.5324089 | 0 | dazl |
| si:dkey-43p13.5 | 7.531217 | 0.325794814 | 0.166 | 0.058 | 1.9133057 | 0 | si:dkey-43p13.5 |
| mcm3l | 1.379730 | 0.324660697 | 0.173 | 0.062 | 3.5052065 | 0 | mcm3l |
| yrdc | 2.187842 | 0.379301797 | 0.217 | 0.091 | 5.5582131 | 0 | yrdc |
| cdca8 | 2.944685 | 0.400437755 | 0.268 | 0.127 | 7.4809738 | 0 | cdca8 |
| zgc:162592 | 3.901516 | 0.266718399 | 0.121 | 0.035 | 9.9118024 | 0 | zgc:162592 |
| si:dkey-68o6.5 | 4.213532 | 0.36902839 | 0.214 | 0.089 | 1.0704478 | 0 | si:dkey-68o6.5 |
| st8sia7.1 | 7.715974 | 0.303696709 | 0.148 | 0.049 | 1.9602434 | 0 | st8sia7.1 |
| ankrd22 | 2.979770 | 0.381839582 | 0.24 | 0.109 | 7.5701074 | 0 | ankrd22 |
| faub | 8.836668 | 0.428022886 | 0.309 | 0.168 | 2.2449556 | 0 | faub |
| COQ10A | 9.129376 | 0.328953677 | 0.164 | 0.06 | 2.3193181 | 0 | COQ10A |
| snx10a | 9.335598 | 0.365671754 | 0.217 | 0.093 | 2.3717087 | 0 | snx10a |
| zgc:114123 | 1.500621 | 0.342886598 | 0.189 | 0.075 | 3.8123293 | 0 | zgc:114123 |
| zgc:173544 | 2.648292 | 0.313837994 | 0.153 | 0.053 | 6.7279873 | 0 | zgc:173544 |
| si:ch211-69g19.2 | 3.524395 | 0.364527874 | 0.214 | 0.092 | 8.9537275 | 0 | si:ch211-69g19.2 |
| alox5b.3 | 4.091092 | 0.257596429 | 0.115 | 0.033 | 1.0393420 | 0 | alox5b.3 |
| tmprss4b | 7.271239 | 0.296659345 | 0.144 | 0.048 | 1.8472584 | 0 | tmprss4b |
| npm2a | 8.461868 | 0.309457364 | 0.156 | 0.055 | 2.1497377 | 0 | npm2a |
| h2afx | 9.660605 | 0.389166106 | 0.237 | 0.11 | 2.4542768 | 0 | h2afx |
| prrg2 | 1.165968 | 0.335351093 | 0.175 | 0.067 | 2.9621429 | 0 | prrg2 |
| zgc:171426 | 1.174768 | 0.339986671 | 0.192 | 0.078 | 2.9845004 | 0 | zgc:171426 |
| dhrs13a.2 | 2.184802 | 0.273483653 | 0.119 | 0.035 | 5.5504896 | 0 | dhrs13a.2 |
| BX957362.1 | 2.221240 | 0.378135213 | 0.225 | 0.102 | 5.6430602 | 0 | BX957362.1 |
| LOC103909972 | 3.419403 | 0.413303203 | 0.443 | 0.31 | 8.6869936 | 0 | LOC103909972 |
| zgc:86839 | 8.322099 | 0.356109299 | 0.197 | 0.083 | 2.1142292 | 0 | zgc:86839 |
| srsf2a | 1.300666 | 0.409530337 | 0.458 | 0.337 | 3.3043420 | 0 | srsf2a |
| lipg | 2.289480 | 0.311540906 | 0.163 | 0.061 | 5.8164255 | 0 | lipg |
| hspa5 | 2.362319 | 0.40770267 | 0.422 | 0.282 | 6.0014715 | 0 | hspa5 |
| BX005461.1 | 3.774109 | 0.30259768 | 0.155 | 0.056 | 9.5881262 | 0 | BX005461.1 |
| stard14 | 4.360960 | 0.275551752 | 0.121 | 0.037 | 1.1079019 | 0 | stard14 |

| | | | | | | | |
|---|---|---|---|---|---|---|---|
| tatdn2 | 6.260737 | 0.294448247 | 0.146 | 0.051 | 1.5905404 | 0 | tatdn2 |
| zgc:92313 | 9.783890 | 0.359252455 | 0.216 | 0.096 | 2.4855974 | 0 | zgc:92313 |
| ccnh | 1.566760 | 0.357709729 | 0.214 | 0.095 | 3.9803547 | 0 | ccnh |
| zgc:56676 | 4.059057 | 0.422459963 | 0.397 | 0.261 | 1.0312035 | 0 | zgc:56676 |
| hsd3b2 | 4.443494 | 0.269513979 | 0.128 | 0.041 | 1.1288698 | 0 | hsd3b2 |
| cpm | 8.503448 | 0.33409189 | 0.19 | 0.079 | 2.1603010 | 0 | cpm |
| si:dkey-81e3.2 | 1.580734 | 0.289278251 | 0.132 | 0.044 | 4.0158560 | 0 | si:dkey-81e3.2 |
| orc6 | 1.722453 | 0.346241257 | 0.194 | 0.082 | 4.3758939 | 0 | orc6 |
| g3bp1 | 1.221344 | 0.408861414 | 0.313 | 0.181 | 3.1028250 | 0 | g3bp1 |
| ypel3 | 1.234212 | 0.357812234 | 0.216 | 0.099 | 3.1355163 | 0 | ypel3 |
| ube2c | 1.530051 | 0.36640659 | 0.234 | 0.112 | 3.8870949 | 0 | ube2c |
| ctsla | 2.181468 | 0.377834862 | 0.488 | 0.362 | 5.5420218 | 0 | ctsla |
| si:ch73-303b9.1 | 2.567472 | 0.296774592 | 0.151 | 0.056 | 6.5226646 | 0 | si:ch73-303b9.1 |
| dph3 | 6.921428 | 0.306080948 | 0.152 | 0.057 | 1.7583887 | 0 | dph3 |
| vps9d1 | 7.054353 | 0.326279083 | 0.175 | 0.072 | 1.7921584 | 0 | vps9d1 |
| rnf4 | 1.453842 | 0.360330183 | 0.223 | 0.105 | 3.6934857 | 0 | rnf4 |
| si:ch211-286b5.4 | 2.493388 | 0.276799658 | 0.139 | 0.049 | 6.3344537 | 0 | si:ch211-286b5.4 |
| fbxo32 | 3.043891 | 0.322545459 | 0.184 | 0.077 | 7.7330058 | 0 | fbxo32 |
| cd9b | 3.367675 | 0.27990435 | 0.144 | 0.052 | 8.5555785 | 0 | cd9b |
| umps | 4.071935 | 0.303871843 | 0.153 | 0.058 | 1.0344751 | 0 | umps |
| zgc:113293 | 9.774103 | 0.254124734 | 0.115 | 0.036 | 2.4831109 | 0 | zgc:113293 |
| exd2 | 1.141160 | 0.314569999 | 0.171 | 0.07 | 2.8991175 | 0 | exd2 |
| ppp1r3b | 5.868816 | 0.333687985 | 0.188 | 0.082 | 1.4909728 | 0 | ppp1r3b |
| zgc:152986 | 2.213267 | 0.298449136 | 0.162 | 0.064 | 5.6228061 | 0 | zgc:152986 |
| aurkb | 2.576190 | 0.34613638 | 0.213 | 0.1 | 6.5448116 | 0 | aurkb |
| asf1ba | 3.819757 | 0.313474889 | 0.18 | 0.077 | 9.7040934 | 0 | asf1ba |
| dnd1 | 6.255245 | 0.319109606 | 0.188 | 0.082 | 1.5891452 | 0 | dnd1 |
| tpx2 | 6.541836 | 0.336446626 | 0.208 | 0.096 | 1.6619536 | 0 | tpx2 |
| bzw1b | 7.549196 | 0.373567424 | 0.469 | 0.369 | 1.9178732 | 0 | bzw1b |
| ttc32 | 1.185624 | 0.318658579 | 0.182 | 0.079 | 3.0120786 | 0 | ttc32 |
| zgc:55888 | 1.293709 | 0.28865887 | 0.157 | 0.061 | 3.2866687 | 0 | zgc:55888 |
| si:ch211-226h8.6 | 2.848951 | 0.287797848 | 0.147 | 0.056 | 7.2377611 | 0 | si:ch211-226h8.6 |
| parn | 3.024640 | 0.359059481 | 0.236 | 0.119 | 7.6841001 | 0 | parn |
| eif4a1a | 5.998972 | 0.354162299 | 0.49 | 0.397 | 1.5240389 | 0 | eif4a1a |
| tram2 | 1.309015 | 0.352444555 | 0.224 | 0.111 | 3.3255540 | 0 | tram2 |

| | | | | | | | |
|---|---|---|---|---|---|---|---|
| nek2 | 1.374292 | 0.265411461 | 0.133 | 0.048 | 3.4913906 | 0 | nek2 |
| cfl1l | 1.477152 | 0.351376546 | 0.245 | 0.125 | 3.7527053 | 0 | cfl1l |
| CU655961.4 | 3.913624 | 0.287622604 | 0.159 | 0.064 | 9.9425639 | 0 | CU655961.4 |
| spryd4 | 4.766991 | 0.290681712 | 0.151 | 0.06 | 1.2110542 | 0 | spryd4 |
| lin28a | 5.022805 | 0.273248088 | 0.135 | 0.05 | 1.2760438 | 0 | lin28a |
| ap1s3b | 8.534873 | 0.27704913 | 0.148 | 0.058 | 2.1682845 | 0 | ap1s3b |
| cited1 | 9.463440 | 0.257659851 | 0.127 | 0.045 | 2.4041869 | 0 | cited1 |
| ddx19 | 1.374391 | 0.35369634 | 0.239 | 0.124 | 3.4916424 | 0 | ddx19 |
| prr11 | 1.951321 | 0.303171695 | 0.174 | 0.075 | 4.9573313 | 0 | prr11 |
| cx43.4 | 2.350922 | 0.332922558 | 0.209 | 0.101 | 5.9725193 | 0 | cx43.4 |
| rogdi | 2.752047 | 0.330201662 | 0.195 | 0.091 | 6.9915776 | 0 | rogdi |
| bokb | 7.394282 | 0.250159134 | 0.121 | 0.043 | 1.8785175 | 0 | bokb |
| LOC110437779 | 1.350619 | 0.275364065 | 0.15 | 0.06 | 3.4312488 | 0 | LOC110437779 |
| snu13a | 2.187810 | 0.31171904 | 0.179 | 0.08 | 5.5581316 | 0 | snu13a |
| tent5c | 2.681127 | 0.342749722 | 0.223 | 0.113 | 6.8114036 | 0 | tent5c |
| tmem70 | 3.429250 | 0.316567181 | 0.181 | 0.082 | 8.7120101 | 0 | tmem70 |
| chchd10 | 3.791694 | 0.35515957 | 0.236 | 0.123 | 9.6328002 | 0 | chchd10 |
| zgc:175136 | 7.713637 | 0.300975115 | 0.178 | 0.079 | 1.9596495 | 0 | zgc:175136 |
| mylipa | 8.310656 | 0.319700774 | 0.189 | 0.089 | 2.1113223 | 0 | mylipa |
| zgc:114130 | 9.832282 | 0.259207286 | 0.142 | 0.057 | 2.4978913 | 0 | zgc:114130 |
| LOC100538023 | 1.152387 | 0.272831998 | 0.139 | 0.055 | 2.9276393 | 0 | LOC100538023 |
| si:ch211-14a17.6 | 2.326754 | 0.273415975 | 0.151 | 0.063 | 5.9111195 | 0 | si:ch211-14a17.6 |
| cyp11a1 | 5.727790 | 0.250658632 | 0.128 | 0.049 | 1.4551450 | 0 | cyp11a1 |
| fdx1 | 6.545380 | 0.371175066 | 0.304 | 0.187 | 1.6628540 | 0 | fdx1 |
| asz1 | 2.565630 | 0.255783215 | 0.135 | 0.053 | 6.5179853 | 0 | asz1 |
| coil | 3.144217 | 0.323148186 | 0.183 | 0.088 | 7.9878849 | 0 | coil |
| si:dkey-92i17.2 | 3.658008 | 0.294123654 | 0.158 | 0.069 | 9.2931697 | 0 | si:dkey-92i17.2 |
| itm2cb | 4.435173 | 0.331657741 | 0.215 | 0.111 | 1.1267558 | 0 | itm2cb |
| BX005461.3 | 4.699474 | 0.250304686 | 0.127 | 0.049 | 1.1939015 | 0 | BX005461.3 |
| gfer | 5.547626 | 0.294946097 | 0.171 | 0.078 | 1.4093745 | 0 | gfer |
| mrps18b | 5.808831 | 0.330293168 | 0.207 | 0.106 | 1.4757336 | 0 | mrps18b |
| lmnl3 | 6.643932 | 0.257025153 | 0.138 | 0.055 | 1.6878910 | 0 | lmnl3 |
| kmt5ab | 7.873611 | 0.277369482 | 0.156 | 0.067 | 2.0002909 | 0 | kmt5ab |
| eif4e1b | 1.005204 | 0.266102041 | 0.15 | 0.064 | 2.5537228 | 0 | eif4e1b |
| zgc:153284 | 2.462903 | 0.288267536 | 0.168 | 0.076 | 6.2570073 | 0 | zgc:153284 |

| | | | | | | | |
|---|---|---|---|---|---|---|---|
| evla | 2.578893 | 0.300857638 | 0.174 | 0.081 | 6.5516778 | 0 | evla |
| brf2 | 3.234849 | 0.28087659 | 0.162 | 0.072 | 8.2181346 | 0 | brf2 |
| imp3 | 1.843765 | 0.330646776 | 0.216 | 0.114 | 4.6840873 | 0 | imp3 |
| fbxo5 | 3.115698 | 0.257377415 | 0.13 | 0.053 | 7.9154320 | 0 | fbxo5 |
| si:ch211-202f5.3 | 3.322574 | 0.265748034 | 0.155 | 0.068 | 8.4409994 | 0 | si:ch211-202f5.3 |
| thop1 | 8.826257 | 0.310858255 | 0.19 | 0.096 | 2.2423107 | 0 | thop1 |
| hmga1a | 1.101703 | 0.298702903 | 0.516 | 0.435 | 2.7988789 | 0 | hmga1a |
| krcp | 2.093562 | 0.255222533 | 0.135 | 0.057 | 5.3186944 | 0 | krcp |
| btg3 | 2.569689 | 0.355036391 | 0.255 | 0.152 | 6.5282957 | 0 | btg3 |
| zgc:110239 | 2.820263 | 0.357048761 | 0.293 | 0.187 | 7.1648788 | 0 | zgc:110239 |
| snrpb2 | 3.964212 | 0.33966709 | 0.23 | 0.129 | 1.0071082 | 0 | snrpb2 |
| zgc:77112 | 9.852396 | 0.297227289 | 0.184 | 0.092 | 2.5030013 | 0 | zgc:77112 |
| marcksb | 1.675240 | 0.334091461 | 0.255 | 0.151 | 4.2559480 | 0 | marcksb |
| si:dkey-87o1.2 | 2.320777 | 0.257974205 | 0.142 | 0.062 | 5.8959357 | 0 | si:dkey-87o1.2 |
| oclna | 3.549733 | 0.260327101 | 0.137 | 0.059 | 9.0180987 | 0 | oclna |
| pop4 | 7.250593 | 0.306507139 | 0.186 | 0.095 | 1.8420133 | 0 | pop4 |
| ddx4 | 9.062819 | 0.281851345 | 0.184 | 0.091 | 2.3024093 | 0 | ddx4 |
| birc5a | 9.513843 | 0.274684793 | 0.165 | 0.079 | 2.4169918 | 0 | birc5a |
| figla | 2.364982 | 0.268185912 | 0.154 | 0.071 | 6.0082377 | 0 | figla |
| cdt1 | 5.317023 | 0.262413357 | 0.146 | 0.066 | 1.3507897 | 0 | cdt1 |
| pttg1ipb | 5.795231 | 0.296394456 | 0.182 | 0.093 | 1.4722785 | 0 | pttg1ipb |
| ccdc58 | 6.038771 | 0.298041074 | 0.179 | 0.091 | 1.5341499 | 0 | ccdc58 |
| f11r.1 | 8.331779 | 0.331758268 | 0.255 | 0.155 | 2.1166886 | 0 | f11r.1 |
| znf395a | 9.097486 | 0.266815826 | 0.161 | 0.077 | 2.3112164 | 0 | znf395a |
| prkrip1 | 9.148499 | 0.266972875 | 0.145 | 0.066 | 2.3241761 | 0 | prkrip1 |
| cdc6 | 2.008746 | 0.270778647 | 0.165 | 0.08 | 5.1032217 | 0 | cdc6 |
| spsb1 | 7.140193 | 0.25302902 | 0.15 | 0.071 | 1.8139661 | 0 | spsb1 |
| metap1d | 8.008042 | 0.263452621 | 0.148 | 0.071 | 2.0344432 | 0 | metap1d |
| cstf1 | 1.323054 | 0.279429875 | 0.163 | 0.082 | 3.3612209 | 0 | cstf1 |
| exosc6 | 1.517562 | 0.273308731 | 0.151 | 0.074 | 3.8553677 | 0 | exosc6 |
| lrrc59 | 2.074224 | 0.32584963 | 0.216 | 0.126 | 5.2695663 | 0 | lrrc59 |
| zfand6 | 2.156688 | 0.296366298 | 0.191 | 0.104 | 5.4790670 | 0 | zfand6 |
| si:ch73-281n10.2 | 2.179064 | 0.289976483 | 0.47 | 0.389 | 5.5359142 | 0 | si:ch73-281n10.2 |
| ckbb | 2.800339 | 0.302199616 | 0.353 | 0.241 | 7.1142626 | 0 | ckbb |
| rpz2 | 3.921341 | 0.328288692 | 0.249 | 0.156 | 9.9621672 | 0 | rpz2 |

| | | | | | | | |
|---|---|---|---|---|---|---|---|
| selenoh | 4.361867 | 0.334698053 | 0.315 | 0.219 | 1.1081323 | 0 | selenoh |
| polr2k | 5.093915 | 0.326840153 | 0.261 | 0.168 | 1.2941093 | 0 | polr2k |
| dnmt1 | 9.767944 | 0.264690528 | 0.161 | 0.081 | 2.4815463 | 0 | dnmt1 |
| hprt1 | 1.221048 | 0.298493349 | 0.19 | 0.105 | 3.1020729 | 0 | hprt1 |
| si:ch73-340n8.4 | 1.667846 | 0.288567834 | 0.193 | 0.106 | 4.2371631 | 0 | si:ch73-340n8.4 |
| dck | 4.091986 | 0.261760713 | 0.149 | 0.074 | 1.0395690 | 0 | dck |
| tacc3 | 4.878862 | 0.254732311 | 0.151 | 0.075 | 1.2394750 | 0 | tacc3 |
| rbm7 | 6.496776 | 0.287538427 | 0.184 | 0.101 | 1.6505061 | 0 | rbm7 |
| ociad1 | 1.729278 | 0.289200176 | 0.166 | 0.089 | 4.3932328 | 0 | ociad1 |
| cnep1r1 | 2.645577 | 0.291119942 | 0.185 | 0.103 | 6.7210897 | 0 | cnep1r1 |
| adipor1a | 4.803773 | 0.26328463 | 0.16 | 0.083 | 1.2203987 | 0 | adipor1a |
| dcun1d1 | 1.391752 | 0.256056142 | 0.156 | 0.08 | 3.5357462 | 0 | dcun1d1 |
| abcf2a | 1.476718 | 0.302750993 | 0.214 | 0.13 | 3.7516028 | 0 | abcf2a |
| alyref | 3.347956 | 0.327036122 | 0.327 | 0.244 | 8.5054836 | 0 | alyref |
| timm13 | 3.693341 | 0.271415486 | 0.167 | 0.09 | 9.3829335 | 0 | timm13 |
| rpa3 | 5.044594 | 0.301020935 | 0.232 | 0.146 | 1.2815792 | 0 | rpa3 |
| c1d | 3.387802 | 0.289495624 | 0.191 | 0.112 | 8.6067118 | 0 | c1d |
| anp32b | 4.254198 | 0.325519326 | 0.323 | 0.246 | 1.0807792 | 0 | anp32b |
| cdc14b | 6.146257 | 0.282835789 | 0.196 | 0.117 | 1.5614565 | 0 | cdc14b |
| rbbp4 | 9.504908 | 0.293896866 | 0.241 | 0.155 | 2.4147221 | 0 | rbbp4 |
| chtopa | 6.598085 | 0.306121252 | 0.282 | 0.2 | 1.6762436 | 0 | chtopa |
| thoc7 | 1.783634 | 0.271406283 | 0.175 | 0.101 | 4.5313242 | 0 | thoc7 |
| hsp90b1 | 2.622157 | 0.297854976 | 0.284 | 0.203 | 6.6615911 | 0 | hsp90b1 |
| gemin7 | 6.127294 | 0.281857845 | 0.213 | 0.135 | 1.5566392 | 0 | gemin7 |
| fbxw11b | 6.633810 | 0.268406251 | 0.202 | 0.125 | 1.6853194 | 0 | fbxw11b |
| nol7 | 1.633142 | 0.269363805 | 0.179 | 0.106 | 4.1489990 | 0 | nol7 |
| aktip | 2.126875 | 0.272177195 | 0.194 | 0.119 | 5.4033262 | 0 | aktip |
| ncbp2 | 3.116496 | 0.290487671 | 0.232 | 0.155 | 7.9174588 | 0 | ncbp2 |
| gtf2a1 | 3.187759 | 0.274593451 | 0.212 | 0.135 | 8.0985041 | 0 | gtf2a1 |
| pcf11 | 3.391537 | 0.296185216 | 0.286 | 0.21 | 8.6162015 | 0 | pcf11 |
| psme3 | 4.631031 | 0.275145786 | 0.191 | 0.118 | 1.1765135 | 0 | psme3 |
| slc25a25a | 5.336313 | 0.288546038 | 0.233 | 0.156 | 1.3556905 | 0 | slc25a25a |
| si:ch211-210c8.7 | 6.362912 | 0.256604444 | 0.182 | 0.109 | 1.6164979 | 0 | si:ch211-210c8.7 |
| tia1l | 5.300264 | 0.283582855 | 0.207 | 0.136 | 1.3465322 | 0 | tia1l |
| cycsb | 1.306061 | 0.279027665 | 0.365 | 0.31 | 3.3180480 | 0 | cycsb |

| | | | | | | | |
|---|---|---|---|---|---|---|---|
| glulb | 1.347782 | 0.26111757 | 0.206 | 0.133 | 3.4240410 | 0 | glulb |
| nabp1a | 1.474450 | 0.254810082 | 0.17 | 0.103 | 3.7458421 | 0 | nabp1a |
| rac1a | 1.583250 | 0.265117079 | 0.206 | 0.134 | 4.0222469 | 0 | rac1a |
| chrac1 | 1.645742 | 0.255085491 | 0.187 | 0.116 | 4.1810083 | 0 | chrac1 |
| timm8b | 2.152091 | 0.257909401 | 0.168 | 0.102 | 5.4673896 | 0 | timm8b |
| sumo3b | 2.203598 | 0.262630954 | 0.177 | 0.109 | 5.5982431 | 0 | sumo3b |
| snrpc | 2.467428 | 0.284353476 | 0.242 | 0.17 | 6.2685029 | 0 | snrpc |
| STMP1 | 2.624878 | 0.292116866 | 0.347 | 0.289 | 6.6685046 | 0 | STMP1 |
| tmco1 | 5.640903 | 0.252981093 | 0.171 | 0.105 | 1.4330716 | 0 | tmco1 |
| top1l | 1.067985 | 0.292231627 | 0.277 | 0.213 | 2.7132164 | 0 | top1l |
| nutf2l | 1.938290 | 0.281153196 | 0.284 | 0.216 | 4.9242271 | 0 | nutf2l |
| rbm4.2 | 2.175197 | 0.272366154 | 0.22 | 0.151 | 5.5260889 | 0 | rbm4.2 |
| zfand5b | 2.914776 | 0.285603201 | 0.266 | 0.199 | 7.4049903 | 0 | zfand5b |
| lsm1 | 3.508292 | 0.255330514 | 0.187 | 0.12 | 8.9128167 | 0 | lsm1 |
| hmgb3b | 4.836565 | 0.271647183 | 0.234 | 0.165 | 1.2287295 | 0 | hmgb3b |
| aif1l | 6.138537 | 0.250127343 | 0.237 | 0.164 | 1.5594955 | 0 | aif1l |
| snrpg | 4.126325 | 0.26323094 | 0.313 | 0.261 | 1.0482929 | 0 | snrpg |
| dkc1 | 5.512841 | 0.255712207 | 0.227 | 0.17 | 1.4005372 | 0 | dkc1 |
| dap1b | 6.028349 | 0.253712827 | 0.297 | 0.246 | 1.5315021 | 0 | dap1b |
| rtn4a | 3.785837 | 0.261709301 | 0.284 | 0.237 | 9.6179206 | 0 | rtn4a |
| sap18 | 4.681986 | 0.260840723 | 0.267 | 0.217 | 1.1894586 | 0 | sap18 |
| zgc:1534991 | 6.492456 | 1.004346089 | 0.954 | 0.383 | 1.6494086 | 1 | zgc:153499 |
| cldnd1 | 1.602153 | 0.961530183 | 0.993 | 0.462 | 4.0702712 | 1 | cldnd |
| si:dkey-241l7.21 | 8.987257 | 0.935783981 | 0.996 | 0.495 | 2.2832127 | 1 | si:dkey-241l7.2 |
| si:dkey-240e12.61 | 5.998455 | 0.96320473 | 0.973 | 0.431 | 1.5239077 | 1 | si:dkey-240e12.6 |
| mt21 | 6.967066 | 0.956891515 | 0.971 | 0.402 | 1.7699831 | 1 | mt2 |
| h2af1al1 | 3.202943 | 0.942228158 | 0.98 | 0.455 | 8.1370787 | 1 | h2af1al |
| h1m1 | 9.291298 | 0.930833197 | 0.98 | 0.462 | 2.3604544 | 1 | h1m |
| zp2.51 | 2.616975 | 0.936304001 | 0.957 | 0.429 | 6.6484269 | 1 | zp2.5 |
| si:ch73-189n23.11 | 4.225926 | 0.947722845 | 0.934 | 0.396 | 1.0735965 | 1 | si:ch73-189n23.1 |
| zp31 | 1.930110 | 0.926841272 | 0.97 | 0.445 | 4.9034457 | 1 | zp3 |
| zp2.21 | 7.689013 | 0.931482569 | 0.937 | 0.399 | 1.9533939 | 1 | zp2.2 |
| si:ch211-226h8.141 | 1.863704 | 0.919333857 | 0.954 | 0.432 | 4.7347424 | 1 | si:ch211-226h8.14 |
| CR790368.11 | 3.949420 | 0.917952667 | 0.908 | 0.394 | 1.0033503 | 1 | CR790368.1 |
| tmsb51 | 2.425477 | 0.944016869 | 0.908 | 0.366 | 6.1619267 | 1 | tmsb5 |

| | | | | | | | |
|---|---|---|---|---|---|---|---|
| zgc:1751351 | 3.114488 | 0.93855924 | 0.888 | 0.359 | 7.9123590 | 1 | zgc:175135 |
| zp2.31 | 1.485409 | 0.9253429 | 0.919 | 0.391 | 3.7736819 | 1 | zp2.3 |
| zp2.11 | 1.024500 | 0.920618901 | 0.924 | 0.405 | 2.6027445 | 1 | zp2.1 |
| si:dkeyp-98a7.91 | 6.634660 | 0.923235164 | 0.909 | 0.387 | 1.6855355 | 1 | si:dkeyp-98a7.9 |
| si:zfos-1505d6.31 | 9.365762 | 0.922057947 | 0.904 | 0.387 | 2.3793719 | 1 | si:zfos-1505d6.3 |
| zp3.21 | 1.766761 | 0.903773155 | 0.928 | 0.426 | 4.4884574 | 1 | zp3.2 |
| si:dkey-241l7.51 | 9.247602 | 0.890429912 | 0.944 | 0.448 | 2.3493535 | 1 | si:dkey-241l7.5 |
| cldng1 | 9.703797 | 0.912554192 | 0.896 | 0.391 | 2.4652496 | 1 | cldng |
| crp21 | 5.367158 | 0.918547702 | 0.887 | 0.368 | 1.3635266 | 1 | crp2 |
| acp5a1 | 2.258778 | 0.892346263 | 0.913 | 0.413 | 5.7384277 | 1 | acp5a |
| zgc:1717791 | 2.671949 | 0.902767878 | 0.88 | 0.389 | 6.7880886 | 1 | zgc:171779 |
| zgc:1717761 | 3.090801 | 0.910822982 | 0.876 | 0.374 | 7.8521819 | 1 | zgc:171776 |
| zgc:771181 | 1.769675 | 0.912286864 | 0.843 | 0.343 | 4.4958617 | 1 | zgc:77118 |
| zp2.61 | 4.159122 | 0.911422624 | 0.887 | 0.378 | 1.0566249 | 1 | zp2.6 |
| BX649337.11 | 3.960630 | 0.91592561 | 0.837 | 0.326 | 1.0061982 | 1 | BX649337.1 |
| ccna11 | 5.185791 | 0.906460183 | 0.86 | 0.358 | 1.3174504 | 1 | ccna1 |
| si:dkey-229d11.31 | 9.388525 | 0.926941075 | 0.83 | 0.327 | 2.3851549 | 1 | si:dkey-229d11.3 |
| si:dkey-90l23.11 | 2.015533 | 0.910136475 | 0.822 | 0.336 | 5.1204628 | 1 | si:dkey-90l23.1 |
| si:ch211-226h8.41 | 6.631731 | 0.912913068 | 0.824 | 0.33 | 1.6847914 | 1 | si:ch211-226h8.4 |
| si:dkey-241l7.41 | 4.822327 | 0.858422714 | 0.958 | 0.469 | 1.2251121 | 1 | si:dkey-241l7.4 |
| si:ch211-125e6.121 | 3.121434 | 0.89702668 | 0.878 | 0.379 | 7.9300043 | 1 | si:ch211-125e6.12 |
| CU571163.11 | 5.781820 | 0.907189752 | 0.78 | 0.298 | 1.4688715 | 1 | CU571163.1 |
| si:ch211-12h2.81 | 2.912442 | 0.843344657 | 0.875 | 0.421 | 7.3990606 | 1 | si:ch211-12h2.8 |
| si:dkey-241l7.61 | 4.122028 | 0.839121258 | 0.857 | 0.404 | 1.0472012 | 1 | si:dkey-241l7.6 |
| mid1ip1l1 | 4.424629 | 0.798714862 | 0.847 | 0.433 | 1.1240770 | 1 | mid1ip1l |
| si:dkeyp-98a7.81 | 4.356198 | 0.842152155 | 0.803 | 0.346 | 1.1066922 | 1 | si:dkeyp-98a7.8 |
| si:ch211-12h2.61 | 7.944290 | 0.828910114 | 0.79 | 0.362 | 2.0182468 | 1 | si:ch211-12h2.6 |
| zar1l1 | 6.426976 | 0.84789998 | 0.737 | 0.299 | 1.6327733 | 1 | zar1l |
| LOC1104378211 | 6.988850 | 0.857250723 | 0.753 | 0.313 | 1.7755173 | 1 | LOC110437821 |
| zp3a.21 | 1.203848 | 0.818757449 | 0.813 | 0.376 | 3.0583760 | 1 | zp3a.2 |
| epcam1 | 6.015876 | 0.815350843 | 0.842 | 0.391 | 1.5283334 | 1 | epcam |
| rrm21 | 1.021139 | 0.802589929 | 0.832 | 0.394 | 2.5942059 | 1 | rrm2 |
| si:dkeyp-20e4.81 | 1.473686 | 0.848299181 | 0.739 | 0.307 | 3.7439013 | 1 | si:dkeyp-20e4.8 |
| lamp21 | 1.632526 | 0.774919561 | 0.837 | 0.459 | 4.1474327 | 1 | lamp2 |
| slbp21 | 2.568037 | 0.829797378 | 0.725 | 0.299 | 6.5240980 | 1 | slbp2 |

| | | | | | | | |
|---|---|---|---|---|---|---|---|
| qdprb21 | 4.146525 | 0.826890774 | 0.756 | 0.328 | 1.0534249 | 1 | qdprb2 |
| zar11 | 1.415485 | 0.819211697 | 0.765 | 0.334 | 3.5960405 | 1 | zar1 |
| ccnb21 | 1.741671 | 0.828915248 | 0.717 | 0.288 | 4.4247167 | 1 | ccnb2 |
| si:dkeyp-98a7.41 | 2.025821 | 0.817765147 | 0.757 | 0.321 | 5.1465984 | 1 | si:dkeyp-98a7.4 |
| thy11 | 3.118190 | 0.799834479 | 0.763 | 0.351 | 7.9217639 | 1 | thy1 |
| si:ch211-14a17.71 | 2.111774 | 0.813630119 | 0.724 | 0.309 | 5.3649642 | 1 | si:ch211-14a17.7 |
| gtf3ab1 | 7.802566 | 0.814557653 | 0.632 | 0.238 | 1.9822420 | 1 | gtf3ab |
| si:dkeyp-46h3.61 | 6.252905 | 0.806755838 | 0.714 | 0.311 | 1.5885506 | 1 | si:dkeyp-46h3.6 |
| si:dkeyp-50f7.21 | 1.204278 | 0.80203915 | 0.72 | 0.305 | 3.0594689 | 1 | si:dkeyp-50f7.2 |
| tuba4l1 | 2.901935 | 0.794047526 | 0.7 | 0.308 | 7.3723670 | 1 | tuba4l |
| zp3b1 | 1.758528 | 0.795190408 | 0.704 | 0.292 | 4.4675428 | 1 | zp3b |
| si:dkeyp-46h3.81 | 1.322071 | 0.797122634 | 0.659 | 0.263 | 3.3587233 | 1 | si:dkeyp-46h3.8 |
| si:dkey-229d11.51 | 1.163164 | 0.791352046 | 0.691 | 0.294 | 2.9550191 | 1 | si:dkey-229d11.5 |
| LOC1005374991 | 2.727008 | 0.796081444 | 0.624 | 0.24 | 6.9279642 | 1 | LOC100537499 |
| siva11 | 5.873112 | 0.768797333 | 0.721 | 0.336 | 1.4920643 | 1 | siva1 |
| nanog1 | 2.031330 | 0.780747068 | 0.656 | 0.272 | 5.1605944 | 1 | nanog |
| AL929032.11 | 2.350636 | 0.788581072 | 0.616 | 0.236 | 5.9717924 | 1 | AL929032.1 |
| cth11 | 2.965152 | 0.768851748 | 0.67 | 0.287 | 7.5329697 | 1 | cth1 |
| ctss2.11 | 7.417039 | 0.709295313 | 0.794 | 0.428 | 1.8842989 | 1 | ctss2.1 |
| zgc:1952451 | 1.709753 | 0.76709085 | 0.686 | 0.294 | 4.3436275 | 1 | zgc:195245 |
| LOC1005351621 | 3.499618 | 0.777654441 | 0.605 | 0.23 | 8.8907819 | 1 | LOC100535162 |
| npm2b1 | 5.643761 | 0.762925923 | 0.675 | 0.303 | 1.4337975 | 1 | npm2b |
| s100a11 | 2.153357 | 0.759128928 | 0.643 | 0.276 | 5.4706043 | 1 | s100a1 |
| btg41 | 2.876584 | 0.755825115 | 0.643 | 0.28 | 7.3079639 | 1 | btg4 |
| zgc:1362541 | 1.723292 | 0.752250442 | 0.607 | 0.249 | 4.3780237 | 1 | zgc:136254 |
| zpcx1 | 6.609894 | 0.749286038 | 0.619 | 0.255 | 1.6792437 | 1 | zpcx |
| zgc:1715171 | 2.005430 | 0.744971077 | 0.657 | 0.296 | 5.0947969 | 1 | zgc:171517 |
| kpna71 | 1.779271 | 0.747141685 | 0.596 | 0.248 | 4.5202390 | 1 | kpna7 |
| si:dkeyp-46h3.21 | 7.021249 | 0.736157638 | 0.66 | 0.31 | 1.7837483 | 1 | si:dkeyp-46h3.2 |
| ccnb11 | 1.252743 | 0.734642263 | 0.662 | 0.303 | 3.1825938 | 1 | ccnb1 |
| wu:fi42e031 | 8.901615 | 0.736897424 | 0.612 | 0.266 | 2.2614554 | 1 | wu:fi42e03 |
| gapdh1 | 2.761911 | 0.711938268 | 0.679 | 0.324 | 7.0166350 | 1 | gapdh |
| zgc:1734431 | 3.667519 | 0.7175454 | 0.592 | 0.25 | 9.3173322 | 1 | zgc:173443 |
| si:ch211-132b12.81 | 5.150023 | 0.716318428 | 0.594 | 0.261 | 1.3083635 | 1 | si:ch211-132b12.8 |
| si:dkey-90l23.21 | 3.417517 | 0.713307303 | 0.594 | 0.262 | 8.6822021 | 1 | si:dkey-90l23.2 |

| | | | | | | | |
|---|---|---|---|---|---|---|---|
| si:ch211-226l4.61 | 5.818934 | 0.696894957 | 0.537 | 0.223 | 1.4783002 | 1 | si:ch211-226l4.6 |
| hbba11 | 1.598357 | 0.693846905 | 0.641 | 0.271 | 4.0606282 | 1 | hbba1 |
| si:dkeyp-46h3.51 | 1.207406 | 0.69058376 | 0.525 | 0.219 | 3.0674173 | 1 | si:dkeyp-46h3.5 |
| si:ch211-146l10.81 | 1.317860 | 0.699295323 | 0.54 | 0.233 | 3.3480252 | 1 | si:ch211-146l10.8 |
| uchl11 | 1.134003 | 0.686646996 | 0.583 | 0.27 | 2.8809352 | 1 | uchl1 |
| si:dkey-208k4.21 | 7.627989 | 0.681384082 | 0.532 | 0.224 | 1.9378907 | 1 | si:dkey-208k4.2 |
| hnrnpabb1 | 2.219431 | 0.553972723 | 0.776 | 0.544 | 5.6384663 | 1 | hnrnpabb |
| avd1 | 3.935504 | 0.6716781 | 0.494 | 0.199 | 9.9981495 | 1 | avd |
| lsm14b1 | 4.085353 | 0.674593265 | 0.525 | 0.228 | 1.0378840 | 1 | lsm14b |
| si:dkeyp-98a7.71 | 3.972369 | 0.651108086 | 0.409 | 0.144 | 1.0091804 | 1 | si:dkeyp-98a7.7 |
| si:dkey-152b24.61 | 3.239782 | 0.669008487 | 0.527 | 0.226 | 8.2306670 | 1 | si:dkey-152b24.6 |
| ca15b1 | 4.201803 | 0.660048538 | 0.471 | 0.187 | 1.0674681 | 1 | ca15b |
| si:dkey-37o8.11 | 2.363710 | 0.664598339 | 0.526 | 0.235 | 6.0050073 | 1 | si:dkey-37o8.1 |
| nasp1 | 2.258768 | 0.656691339 | 0.571 | 0.275 | 5.7384021 | 1 | nasp |
| si:ch73-14h10.21 | 2.435838 | 0.655399291 | 0.489 | 0.206 | 6.1882488 | 1 | si:ch73-14h10.2 |
| si:ch211-5k11.81 | 3.816671 | 0.541912434 | 0.631 | 0.274 | 9.6962536 | 1 | si:ch211-5k11.8 |
| si:ch211-250e5.161 | 2.660366 | 0.64641111 | 0.557 | 0.267 | 6.7586609 | 1 | si:ch211-250e5.16 |
| si:ch211-145c1.11 | 4.044038 | 0.639763948 | 0.44 | 0.176 | 1.0273879 | 1 | si:ch211-145c1.1 |
| si:ch73-90k17.11 | 6.377132 | 0.639348082 | 0.453 | 0.188 | 1.6201103 | 1 | si:ch73-90k17.1 |
| zgc:1714741 | 2.306709 | 0.637882054 | 0.515 | 0.232 | 5.8601945 | 1 | zgc:171474 |
| arhgef10la1 | 1.399475 | 0.628737679 | 0.474 | 0.21 | 3.5553672 | 1 | arhgef10la |
| zgc:554611 | 2.236539 | 0.624256198 | 0.493 | 0.225 | 5.6819287 | 1 | zgc:55461 |
| zp3a.11 | 1.027017 | 0.613751962 | 0.571 | 0.297 | 2.6091388 | 1 | zp3a.1 |
| birc5b1 | 2.034375 | 0.617970384 | 0.456 | 0.198 | 5.1683305 | 1 | birc5b |
| zgc:566991 | 2.188891 | 0.619182434 | 0.446 | 0.194 | 5.5608789 | 1 | zgc:56699 |
| pabpn1l1 | 7.907611 | 0.610001375 | 0.473 | 0.219 | 2.0089286 | 1 | pabpn1l |
| gmnn1 | 5.726631 | 0.611977439 | 0.495 | 0.239 | 1.4548506 | 1 | gmnn |
| cpeb1b1 | 2.531558 | 0.59870309 | 0.46 | 0.207 | 6.4314241 | 1 | cpeb1b |
| ccna21 | 5.881856 | 0.600853977 | 0.475 | 0.223 | 1.4942855 | 1 | ccna2 |
| elovl7b1 | 4.520245 | 0.591221039 | 0.433 | 0.19 | 1.1483684 | 1 | elovl7b |
| si:ch211-251f6.61 | 9.188229 | 0.597407939 | 0.512 | 0.257 | 2.3342696 | 1 | si:ch211-251f6.6 |
| si:ch1073-263o8.21 | 2.676221 | 0.587369725 | 0.443 | 0.202 | 6.7989409 | 1 | si:ch1073-263o8.2 |
| LOC1018854771 | 2.694156 | 0.593109054 | 0.463 | 0.215 | 6.8445044 | 1 | LOC101885477 |
| zgc:562311 | 1.410661 | 0.580557961 | 0.403 | 0.173 | 3.5837860 | 1 | zgc:56231 |
| pigrl2.31 | 1.115311 | 0.553864194 | 0.372 | 0.155 | 2.8334500 | 1 | pigrl2.3 |

| | | | | | | | |
|---|---|---|---|---|---|---|---|
| slc16a31 | 1.864971 | 0.56577466 | 0.417 | 0.192 | 4.7379591 | 1 | slc16a3 |
| AL929192.11 | 3.460486 | 0.526288586 | 0.324 | 0.123 | 8.7913649 | 1 | AL929192.1 |
| LOC1005368601 | 4.967190 | 0.546256682 | 0.504 | 0.274 | 1.2619148 | 1 | LOC100536860 |
| elovl7a1 | 6.552934 | 0.531656 | 0.343 | 0.14 | 1.6647729 | 1 | elovl7a |
| si:dkey-19b23.111 | 1.053471 | 0.546009359 | 0.447 | 0.215 | 2.6763436 | 1 | si:dkey-19b23.11 |
| gdf31 | 4.703440 | 0.539313055 | 0.382 | 0.171 | 1.1949090 | 1 | gdf3 |
| larp6b1 | 2.868655 | 0.531614847 | 0.35 | 0.149 | 7.2878202 | 1 | larp6b |
| map1lc3c1 | 8.819887 | 0.529363085 | 0.354 | 0.154 | 2.2406923 | 1 | map1lc3c |
| zgc:1134241 | 4.101266 | 0.510963817 | 0.306 | 0.122 | 1.0419266 | 1 | zgc:113424 |
| hbaa11 | 5.234409 | 0.487714906 | 0.472 | 0.218 | 1.3298016 | 1 | hbaa1 |
| sybu1 | 1.076312 | 0.503404213 | 0.312 | 0.126 | 2.7343711 | 1 | sybu |
| zgc:925911 | 2.307874 | 0.521493016 | 0.356 | 0.158 | 5.8631545 | 1 | zgc:92591 |
| buc1 | 4.238866 | 0.504768618 | 0.326 | 0.136 | 1.0768840 | 1 | buc |
| si:ch1073-190k2.11 | 1.073998 | 0.517800153 | 0.332 | 0.143 | 2.7284923 | 1 | si:ch1073-190k2.1 |
| hnrnpa0l1 | 8.546645 | 0.265696217 | 0.792 | 0.738 | 2.1712752 | 1 | hnrnpa0l |
| gyg1a1 | 2.348671 | 0.524291005 | 0.509 | 0.307 | 5.9667997 | 1 | gyg1a |
| crp31 | 7.876809 | 0.51514894 | 0.364 | 0.168 | 2.0011034 | 1 | crp3 |
| haus61 | 1.387294 | 0.516272228 | 0.355 | 0.162 | 3.5244216 | 1 | haus6 |
| zp3c1 | 5.981554 | 0.503169102 | 0.329 | 0.145 | 1.5196139 | 1 | zp3c |
| LOC1018838971 | 1.270471 | 0.485695641 | 0.302 | 0.124 | 3.2276324 | 1 | LOC101883897 |
| elavl21 | 1.410742 | 0.515594331 | 0.369 | 0.174 | 3.5839913 | 1 | elavl2 |
| rcc11 | 3.337611 | 0.51520848 | 0.35 | 0.162 | 8.4792009 | 1 | rcc1 |
| zgc:1628791 | 1.467409 | 0.490318186 | 0.321 | 0.139 | 3.7279529 | 1 | zgc:162879 |
| sinup1 | 1.323425 | 0.499105078 | 0.343 | 0.157 | 3.3621630 | 1 | sinup |
| inka1b1 | 1.794566 | 0.493296753 | 0.323 | 0.146 | 4.5590973 | 1 | inka1b |
| slbp1 | 2.408889 | 0.498981622 | 0.378 | 0.188 | 6.1197832 | 1 | slbp |
| zglp11 | 6.591000 | 0.482972884 | 0.333 | 0.153 | 1.6744437 | 1 | zglp1 |
| zgc:1034821 | 2.587582 | 0.478983654 | 0.323 | 0.147 | 6.5737531 | 1 | zgc:103482 |
| larp6a1 | 8.021322 | 0.505887617 | 0.386 | 0.203 | 2.0378169 | 1 | larp6a |
| CU467646.21 | 6.401401 | 0.433464617 | 0.245 | 0.095 | 1.6262761 | 1 | CU467646.2 |
| zgc:1629451 | 9.282775 | 0.472825401 | 0.29 | 0.127 | 2.3582890 | 1 | zgc:162945 |
| rbpms2b1 | 9.789452 | 0.489529248 | 0.358 | 0.18 | 2.4870104 | 1 | rbpms2b |
| tph1a1 | 1.643956 | 0.449891827 | 0.261 | 0.107 | 4.1764711 | 1 | tph1a |
| zgc:1138861 | 6.903488 | 0.442523964 | 0.262 | 0.107 | 1.7538313 | 1 | zgc:113886 |
| zgc:1738561 | 1.004694 | 0.472604529 | 0.348 | 0.171 | 2.5524253 | 1 | zgc:173856 |

| | | | | | | | |
|---|---|---|---|---|---|---|---|
| mkrn41 | 3.277381 | 0.47747023 | 0.361 | 0.183 | 8.3261874 | 1 | mkrn4 |
| org1 | 8.155693 | 0.469485443 | 0.371 | 0.189 | 2.0719538 | 1 | org |
| snu13b1 | 3.481322 | 0.461391516 | 0.522 | 0.362 | 8.8443007 | 1 | snu13b |
| h3f3a1 | 7.332734 | 0.418426901 | 0.607 | 0.475 | 1.8628810 | 1 | h3f3a |
| si:ch73-78o10.11 | 1.820033 | 0.426197317 | 0.246 | 0.101 | 4.6237939 | 1 | si:ch73-78o10.1 |
| si:dkey-188i13.111 | 2.154711 | 0.483052811 | 0.446 | 0.281 | 5.4740435 | 1 | si:dkey-188i13.11 |
| si:ch211-146l10.71 | 2.537777 | 0.445625886 | 0.309 | 0.146 | 6.4472244 | 1 | si:ch211-146l10.7 |
| cdc201 | 1.663961 | 0.438923556 | 0.271 | 0.121 | 4.2272934 | 1 | cdc20 |
| gbgt1l41 | 2.420462 | 0.425939483 | 0.266 | 0.115 | 6.1491842 | 1 | gbgt1l4 |
| BX537276.11 | 3.252361 | 0.407748743 | 0.23 | 0.092 | 8.2626256 | 1 | BX537276.1 |
| hsd17b12a1 | 2.075242 | 0.43093384 | 0.28 | 0.128 | 5.2721528 | 1 | hsd17b12a |
| cdca7a1 | 2.729520 | 0.460620894 | 0.347 | 0.183 | 6.9343474 | 1 | cdca7a |
| retsatl1 | 2.385999 | 0.442363288 | 0.319 | 0.158 | 6.0616320 | 1 | retsatl |
| pttg11 | 5.395754 | 0.431835597 | 0.277 | 0.128 | 1.3707913 | 1 | pttg1 |
| ubc1 | 1.419910 | 0.260548195 | 0.691 | 0.652 | 3.6072825 | 1 | ubc |
| BX001030.11 | 1.548726 | 0.406496939 | 0.224 | 0.093 | 3.9345385 | 1 | BX001030.1 |
| plk11 | 4.319082 | 0.43575548 | 0.29 | 0.14 | 1.0972628 | 1 | plk1 |
| patl21 | 5.708373 | 0.427937489 | 0.284 | 0.134 | 1.4502122 | 1 | patl2 |
| CDPF11 | 6.106989 | 0.469585938 | 0.385 | 0.226 | 1.5514806 | 1 | CDPF1 |
| si:dkey-241l7.31 | 3.699622 | 0.448236513 | 0.463 | 0.291 | 9.3988917 | 1 | si:dkey-241l7.3 |
| tac3b1 | 3.748310 | 0.391012066 | 0.222 | 0.092 | 9.5225824 | 1 | tac3b |
| nfr1 | 9.198951 | 0.382112453 | 0.209 | 0.084 | 2.3369936 | 1 | nfr |
| mibp1 | 1.565483 | 0.442616584 | 0.349 | 0.19 | 3.9771115 | 1 | mibp |
| tmem54a1 | 1.732556 | 0.407340363 | 0.247 | 0.11 | 4.4015585 | 1 | tmem54a |
| cdca81 | 1.008714 | 0.421440722 | 0.274 | 0.132 | 2.5626395 | 1 | cdca8 |
| chia.41 | 1.260524 | 0.399648055 | 0.251 | 0.113 | 3.2023625 | 1 | chia.4 |
| sox19b1 | 2.181751 | 0.355565549 | 0.191 | 0.074 | 5.5427389 | 1 | sox19b |
| ddx39ab1 | 2.365350 | 0.457945431 | 0.407 | 0.262 | 6.0091721 | 1 | ddx39ab |
| zgc:660241 | 3.165719 | 0.396799744 | 0.244 | 0.109 | 8.0425110 | 1 | zgc:66024 |
| ctsba1 | 4.920346 | 0.421936684 | 0.494 | 0.344 | 1.2500139 | 1 | ctsba |
| CR847984.11 | 1.682907 | 0.410242291 | 0.271 | 0.13 | 4.2754262 | 1 | CR847984.1 |
| BX571780.11 | 3.661086 | 0.342925442 | 0.167 | 0.061 | 9.3009889 | 1 | BX571780.1 |
| cx44.21 | 2.399676 | 0.374563789 | 0.212 | 0.089 | 6.0963780 | 1 | cx44.2 |
| zgc:558881 | 5.050343 | 0.338903995 | 0.171 | 0.064 | 1.2830397 | 1 | zgc:55888 |
| ankrd221 | 7.685109 | 0.397620175 | 0.244 | 0.114 | 1.9524021 | 1 | ankrd22 |

| | | | | | | | |
|---|---|---|---|---|---|---|---|
| pou5f31 | 2.517307 | 0.396800644 | 0.243 | 0.114 | 6.3952208 | 1 | pou5f3 |
| CR383662.21 | 3.424011 | 0.330055041 | 0.172 | 0.065 | 8.6987001 | 1 | CR383662.2 |
| spink2.11 | 5.802507 | 0.394928525 | 0.543 | 0.364 | 1.4741270 | 1 | spink2.1 |
| zgc:922871 | 8.346830 | 0.381512433 | 0.229 | 0.105 | 2.1205121 | 1 | zgc:92287 |
| zp2l11 | 1.080120 | 0.391639343 | 0.262 | 0.128 | 2.7440451 | 1 | zp2l1 |
| si:dkey-42i9.81 | 1.364423 | 0.373852248 | 0.221 | 0.099 | 3.4663179 | 1 | si:dkey-42i9.8 |
| zgc:1737421 | 1.739574 | 0.391901133 | 0.254 | 0.122 | 4.4193879 | 1 | zgc:173742 |
| cldn7b1 | 1.913827 | 0.393198566 | 0.259 | 0.126 | 4.8620780 | 1 | cldn7b |
| LOC1039099721 | 2.616821 | 0.428856792 | 0.439 | 0.316 | 6.6480339 | 1 | LOC103909972 |
| CABZ01059627.11 | 4.225221 | 0.370344274 | 0.231 | 0.105 | 1.0734176 | 1 | CABZ01059627.1 |
| dctpp11 | 5.504905 | 0.40220273 | 0.277 | 0.142 | 1.3985211 | 1 | dctpp1 |
| hspa51 | 1.704605 | 0.413331907 | 0.422 | 0.288 | 4.3305498 | 1 | hspa5 |
| zgc:1139841 | 9.681827 | 0.340792428 | 0.186 | 0.077 | 2.4596683 | 1 | zgc:113984 |
| degs21 | 9.873528 | 0.349956702 | 0.193 | 0.082 | 2.5083700 | 1 | degs2 |
| zc3h11a1 | 1.311288 | 0.423194975 | 0.316 | 0.182 | 3.3313288 | 1 | zc3h11a |
| rbm4.31 | 2.706306 | 0.37133386 | 0.515 | 0.416 | 6.8753722 | 1 | rbm4.3 |
| si:dkey-68o6.51 | 3.199019 | 0.37143463 | 0.21 | 0.095 | 8.1271088 | 1 | si:dkey-68o6.5 |
| CABZ01059627.21 | 4.844175 | 0.364752463 | 0.222 | 0.103 | 1.2306628 | 1 | CABZ01059627.2 |
| pabpc1l1 | 1.590648 | 0.363710021 | 0.233 | 0.11 | 4.0410413 | 1 | pabpc1l |
| si:dkeyp-114g9.11 | 2.397999 | 0.345117674 | 0.187 | 0.08 | 6.0921174 | 1 | si:dkeyp-114g9.1 |
| mad2l11 | 4.242408 | 0.388947139 | 0.282 | 0.152 | 1.0777839 | 1 | mad2l1 |
| dynll11 | 4.268705 | 0.358912173 | 0.532 | 0.436 | 1.0844645 | 1 | dynll1 |
| lpcat41 | 4.596861 | 0.344439296 | 0.193 | 0.084 | 1.1678327 | 1 | lpcat4 |
| CT009596.11 | 4.792606 | 0.32902308 | 0.169 | 0.069 | 1.2175615 | 1 | CT009596.1 |
| fdx11 | 9.401994 | 0.414705971 | 0.321 | 0.191 | 2.3885766 | 1 | fdx1 |
| tpx21 | 1.305727 | 0.360055983 | 0.215 | 0.101 | 3.3172008 | 1 | tpx2 |
| CU655961.41 | 2.863663 | 0.322711766 | 0.166 | 0.067 | 7.2751378 | 1 | CU655961.4 |
| si:dkey-46g23.21 | 4.966839 | 0.356480308 | 0.232 | 0.113 | 1.2618255 | 1 | si:dkey-46g23.2 |
| nanos31 | 6.228009 | 0.333908448 | 0.183 | 0.079 | 1.5822257 | 1 | nanos3 |
| si:ch73-160p18.31 | 7.628222 | 0.391936931 | 0.326 | 0.191 | 1.9379499 | 1 | si:ch73-160p18.3 |
| dynll2a1 | 2.723169 | 0.361626671 | 0.226 | 0.111 | 6.9182111 | 1 | dynll2a |
| cpm1 | 4.428060 | 0.334566331 | 0.188 | 0.084 | 1.1249488 | 1 | cpm |
| rnf41 | 1.194271 | 0.359948866 | 0.223 | 0.11 | 3.0340471 | 1 | rnf4 |
| orc61 | 1.536296 | 0.3408617 | 0.191 | 0.087 | 3.9029600 | 1 | orc6 |
| pif11 | 1.588953 | 0.331778765 | 0.186 | 0.083 | 4.0367359 | 1 | pif1 |

| | | | | | | | |
|---|---|---|---|---|---|---|---|
| BX284697.11 | 2.08147( | 0.332581991 | 0.194 | 0.088 | 5.2879746 | 1 | BX284697.1 |
| si:ch211-14a17.61 | 2.49910( | 0.302984722 | 0.16 | 0.066 | 6.3489645 | 1 | si:ch211-14a17.6 |
| zgc:1526521 | 2.58675 | 0.321626804 | 0.194 | 0.088 | 6.5716488 | 1 | zgc:152652 |
| itm2cb1 | 4.76510 | 0.363673751 | 0.226 | 0.115 | 1.2105748 | 1 | itm2cb |
| zgc:1118681 | 5.42415( | 0.3540306 | 0.23 | 0.115 | 1.3780054 | 1 | zgc:111868 |
| depdc7b | 7.19290 | 0.278181579 | 0.135 | 0.051 | 1.8273586 | 1 | depdc7b |
| snu13a1 | 7.99710 | 0.333802619 | 0.185 | 0.084 | 2.0316653 | 1 | snu13a |
| si:ch211-69g19.21 | 1.444951 | 0.33785747 | 0.205 | 0.098 | 3.6708986 | 1 | si:ch211-69g19.2 |
| kpna21 | 2.63774 | 0.355152338 | 0.248 | 0.13 | 6.7011849 | 1 | kpna2 |
| tmem701 | 3.01170 | 0.336306306 | 0.186 | 0.086 | 7.6512395 | 1 | tmem70 |
| ctsla1 | 4.79590( | 0.361507421 | 0.471 | 0.369 | 1.2183984 | 1 | ctsla |
| zgc:1655391 | 5.09615( | 0.327838766 | 0.198 | 0.093 | 1.2946771 | 1 | zgc:165539 |
| cdca91 | 8.40171 | 0.301246166 | 0.157 | 0.066 | 2.1344565 | 1 | cdca9 |
| mos1 | 1.07602 | 0.342848498 | 0.218 | 0.108 | 2.7336494 | 1 | mos |
| CU655961.31 | 2.75048( | 0.317115407 | 0.171 | 0.076 | 6.9875949 | 1 | CU655961.3 |
| zgc:1529861 | 4.55532 | 0.29930149 | 0.16 | 0.069 | 1.1572815 | 1 | zgc:152986 |
| zgc:1129801 | 5.51322 | 0.303475194 | 0.166 | 0.073 | 1.4006350 | 1 | zgc:112980 |
| yrdc1 | 9.63355 | 0.333738963 | 0.201 | 0.098 | 2.4474052 | 1 | yrdc |
| ppp1r3b1 | 1.37291 | 0.316915094 | 0.185 | 0.086 | 3.4879007 | 1 | ppp1r3b |
| zgc:1630301 | 1.46234 | 0.337450781 | 0.204 | 0.101 | 3.7150998 | 1 | zgc:163030 |
| si:ch211-286b5.41 | 1.76321 | 0.273336324 | 0.135 | 0.053 | 4.4794421 | 1 | si:ch211-286b5.4 |
| brf21 | 2.25693 | 0.306276642 | 0.168 | 0.075 | 5.7337435 | 1 | brf2 |
| si:ch73-303b9.11 | 2.71346 | 0.283381818 | 0.145 | 0.06 | 6.8935656 | 1 | si:ch73-303b9.1 |
| zgc:1141231 | 3.45929 | 0.312851582 | 0.175 | 0.081 | 8.7883366 | 1 | zgc:114123 |
| CABZ01069022.2 | 3.74202 | 0.260378835 | 0.124 | 0.047 | 9.5066106 | 1 | CABZ01069022.2 |
| tram21 | 3.890521 | 0.354437312 | 0.221 | 0.116 | 9.8838693 | 1 | tram2 |
| gstp11 | 4.29354 | 0.355598774 | 0.444 | 0.342 | 1.0907754 | 1 | gstp1 |
| tmem144b1 | 4.53052 | 0.291807503 | 0.165 | 0.073 | 1.1509796 | 1 | tmem144b |
| zgc:1717501 | 5.13432 | 0.347688909 | 0.252 | 0.138 | 1.3043762 | 1 | zgc:171750 |
| si:dkey-92i17.21 | 6.48186 | 0.314018262 | 0.162 | 0.073 | 1.6467179 | 1 | si:dkey-92i17.2 |
| cox171 | 6.75889( | 0.351090904 | 0.44 | 0.336 | 1.7170960 | 1 | cox17 |
| vstm2l1 | 7.65137 | 0.296820025 | 0.166 | 0.074 | 1.9438311 | 1 | vstm2l |
| kmt5ab1 | 8.47139 | 0.297762853 | 0.16 | 0.071 | 2.1521573 | 1 | kmt5ab |
| snx10a1 | 1.07165 | 0.325301262 | 0.201 | 0.1 | 2.7225324 | 1 | snx10a |
| chtopa1 | 1.303241 | 0.384280117 | 0.31 | 0.201 | 3.3108853 | 1 | chtopa |

| | | | | | | | |
|---|---|---|---|---|---|---|---|
| buc2l1 | 1.336221 | 0.273776419 | 0.146 | 0.061 | 3.3946702 | 1 | buc2l |
| si:ch73-15b2.51 | 2.385991 | 0.294794043 | 0.163 | 0.073 | 6.0616110 | 1 | si:ch73-15b2.5 |
| zgc:566761 | 3.11799[illegible] | 0.379972789 | 0.375 | 0.269 | 7.9212552 | 1 | zgc:56676 |
| ddx191 | 3.13217[illegible] | 0.351055184 | 0.234 | 0.129 | 7.9572846 | 1 | ddx19 |
| zgc:1714261 | 3.43055[illegible] | 0.308322615 | 0.177 | 0.084 | 8.7153302 | 1 | zgc:171426 |
| mcm3l1 | 4.72034[illegible] | 0.281929916 | 0.155 | 0.068 | 1.1992043 | 1 | mcm3l |
| ccne11 | 5.38648[illegible] | 0.311809798 | 0.185 | 0.089 | 1.3684365 | 1 | ccne1 |
| rasl11b1 | 6.58042[illegible] | 0.326315662 | 0.21 | 0.108 | 1.6717571 | 1 | rasl11b |
| nek21 | 1.108001 | 0.263310622 | 0.129 | 0.052 | 2.8148767 | 1 | nek2 |
| LOC1104377791 | 1.74726[illegible] | 0.282230105 | 0.147 | 0.064 | 4.4389205 | 1 | LOC110437779 |
| bzw1b1 | 1.884561 | 0.344041565 | 0.458 | 0.374 | 4.7877290 | 1 | bzw1b |
| zgc:1017441 | 3.08729[illegible] | 0.316847232 | 0.207 | 0.107 | 7.8432705 | 1 | zgc:101744 |
| chchd101 | 4.30297[illegible] | 0.34699182 | 0.232 | 0.128 | 1.0931702 | 1 | chchd10 |
| lin28a1 | 4.467451 | 0.263322891 | 0.13 | 0.053 | 1.1349560 | 1 | lin28a |
| rdh10b1 | 5.82409[illegible] | 0.339048301 | 0.254 | 0.145 | 1.4796114 | 1 | rdh10b |
| bmb1 | 1.297831 | 0.276844658 | 0.153 | 0.068 | 3.2971397 | 1 | bmb |
| si:ch211-226h8.61 | 4.38125[illegible] | 0.260524015 | 0.14 | 0.061 | 1.1130586 | 1 | si:ch211-226h8.6 |
| ccnh1 | 5.642251 | 0.313807049 | 0.196 | 0.101 | 1.4334139 | 1 | ccnh |
| tmprss4b1 | 7.31384[illegible] | 0.252423764 | 0.128 | 0.053 | 1.8580823 | 1 | tmprss4b |
| npm2a1 | 9.19151[illegible] | 0.255832354 | 0.14 | 0.061 | 2.3351045 | 1 | npm2a |
| prrg21 | 1.39681[illegible] | 0.279726804 | 0.157 | 0.073 | 3.5486177 | 1 | prrg2 |
| zgc:923131 | 1.70923[illegible] | 0.318570739 | 0.196 | 0.103 | 4.3423097 | 1 | zgc:92313 |
| si:dkey-216e24.31 | 1.95297[illegible] | 0.304811473 | 0.189 | 0.097 | 4.9615344 | 1 | si:dkey-216e24.3 |
| parn1 | 2.97008[illegible] | 0.33543251 | 0.221 | 0.125 | 7.5455020 | 1 | parn |
| si:dkeyp-75b4.101 | 3.07830[illegible] | 0.274901965 | 0.494 | 0.359 | 7.8204222 | 1 | si:dkeyp-75b4.10 |
| prr111 | 4.04717[illegible] | 0.292336562 | 0.165 | 0.08 | 1.0281845 | 1 | prr11 |
| zgc:1532841 | 5.52506[illegible] | 0.28482297 | 0.166 | 0.08 | 1.4036439 | 1 | zgc:153284 |
| pus3 | 5.86010[illegible] | 0.258872858 | 0.132 | 0.058 | 1.4887592 | 1 | pus3 |
| dcun1d11 | 6.54145[illegible] | 0.300597877 | 0.167 | 0.083 | 1.6618573 | 1 | dcun1d1 |
| vps9d11 | 7.87005[illegible] | 0.288294498 | 0.16 | 0.077 | 1.9993877 | 1 | vps9d1 |
| e2f5 | 1.23315[illegible] | 0.258938069 | 0.131 | 0.057 | 3.1328243 | 1 | e2f5 |
| rpain1 | 1.997213 | 0.303446572 | 0.185 | 0.096 | 5.0739203 | 1 | rpain |
| si:ch211-210c8.71 | 2.18450[illegible] | 0.32479248 | 0.201 | 0.11 | 5.5497356 | 1 | si:ch211-210c8.7 |
| zgc:868391 | 2.200071 | 0.293002058 | 0.176 | 0.089 | 5.5892805 | 1 | zgc:86839 |
| cdt11 | 3.33428[illegible] | 0.274251613 | 0.147 | 0.069 | 8.4707621 | 1 | cdt1 |

| | | | | | | | |
|---|---|---|---|---|---|---|---|
| gpr1601 | 3.80929( | 0.254859043 | 0.14 | 0.063 | 9.6775029 | 1 | gpr160 |
| eif4a1a1 | 4.923725 | 0.307902434 | 0.465 | 0.403 | 1.2508724 | 1 | eif4a1a |
| lipg1 | 5.864142 | 0.262960367 | 0.143 | 0.066 | 1.4897853 | 1 | lipg |
| srsf2a1 | 1.815986 | 0.338336381 | 0.417 | 0.346 | 4.6135127 | 1 | srsf2a |
| si:dkey-43p13.51 | 2.599236 | 0.256515234 | 0.14 | 0.065 | 6.6033607 | 1 | si:dkey-43p13.5 |
| rogdi1 | 4.240928 | 0.291522642 | 0.182 | 0.096 | 1.0774079 | 1 | rogdi |
| ttc321 | 4.392704 | 0.282821442 | 0.166 | 0.084 | 1.1159666 | 1 | ttc32 |
| mylipa1 | 5.332643 | 0.293342877 | 0.178 | 0.094 | 1.3547581 | 1 | mylipa |
| aurka1 | 8.219443 | 0.280662807 | 0.181 | 0.096 | 2.0881495 | 1 | aurka |
| cdc61 | 8.595114 | 0.274642087 | 0.165 | 0.084 | 2.1835889 | 1 | cdc6 |
| prdx11 | 9.094767 | 0.321497645 | 0.238 | 0.144 | 2.3105257 | 1 | prdx1 |
| si:ch211-202f5.31 | 1.420356 | 0.269544623 | 0.147 | 0.072 | 3.6084165 | 1 | si:ch211-202f5.3 |
| h2afva1 | 1.818777 | 0.331926283 | 0.386 | 0.302 | 4.6206038 | 1 | h2afva |
| pithd1 | 4.41720( | 0.252115745 | 0.136 | 0.064 | 1.1221898 | 1 | pithd1 |
| zgc:1751361 | 1.04036( | 0.268205153 | 0.163 | 0.085 | 2.6430363 | 1 | zgc:175136 |
| selenoh1 | 1.455356 | 0.339629637 | 0.31 | 0.224 | 3.6973319 | 1 | selenoh |
| gfer1 | 2.013342 | 0.271556897 | 0.159 | 0.083 | 5.1148965 | 1 | gfer |
| dazl1 | 2.201541 | 0.260915559 | 0.165 | 0.086 | 5.5930150 | 1 | dazl |
| tent5c1 | 3.306567 | 0.29528924 | 0.204 | 0.119 | 8.4003337 | 1 | tent5c |
| rpz21 | 5.55108( | 0.319763303 | 0.248 | 0.16 | 1.4102520 | 1 | rpz2 |
| ypel31 | 1.151566 | 0.281369922 | 0.188 | 0.106 | 2.9255548 | 1 | ypel3 |
| metap1d1 | 1.305187 | 0.252412082 | 0.146 | 0.074 | 3.3158290 | 1 | metap1d |
| aurkb1 | 1.363889 | 0.279406647 | 0.188 | 0.107 | 3.4649603 | 1 | aurkb |
| spsb11 | 1.715931 | 0.250406147 | 0.147 | 0.075 | 4.3593250 | 1 | spsb1 |
| il13ra21 | 1.936799 | 0.260511049 | 0.166 | 0.089 | 4.9204390 | 1 | il13ra2 |
| faub1 | 6.207407 | 0.314038132 | 0.264 | 0.178 | 1.5769918 | 1 | faub |
| pop41 | 6.257855 | 0.281450048 | 0.176 | 0.099 | 1.5898082 | 1 | pop4 |
| hells | 9.60927( | 0.289856485 | 0.201 | 0.121 | 2.4412352 | 1 | hells |
| si:ch73-281n10.21 | 1.114625 | 0.270783268 | 0.452 | 0.394 | 2.8317061 | 1 | si:ch73-281n10.2 |
| imp31 | 1.221084 | 0.287675078 | 0.2 | 0.12 | 3.1021642 | 1 | imp3 |
| zfand61 | 2.794491 | 0.288494738 | 0.184 | 0.108 | 7.0994058 | 1 | zfand6 |
| h2afx1 | 3.251058 | 0.279221365 | 0.199 | 0.119 | 8.2593139 | 1 | h2afx |
| pim3 | 2.36534( | 0.280735399 | 0.186 | 0.113 | 6.0091477 | 1 | pim3 |
| pttg1ipb1 | 2.764189 | 0.269979938 | 0.168 | 0.097 | 7.0224223 | 1 | pttg1ipb |
| gtf2a11 | 3.000686 | 0.296103812 | 0.214 | 0.138 | 7.6232442 | 1 | gtf2a1 |

| | | | | | | | |
|---|---|---|---|---|---|---|---|
| si:ch73-340n8.41 | 2.008419 | 0.270692216 | 0.182 | 0.111 | 5.1023888 | 1 | si:ch73-340n8.4 |
| marcksb1 | 2.560756 | 0.283372167 | 0.233 | 0.157 | 6.5056020 | 1 | marcksb |
| nol71 | 2.782074 | 0.275514752 | 0.178 | 0.11 | 7.0678607 | 1 | nol7 |
| thop11 | 3.805549 | 0.256099613 | 0.17 | 0.101 | 9.6679984 | 1 | thop1 |
| BX957362.11 | 4.427134 | 0.260944439 | 0.182 | 0.111 | 1.1247135 | 1 | BX957362.1 |
| snrpb21 | 7.983323 | 0.286908665 | 0.206 | 0.135 | 2.0281633 | 1 | snrpb2 |
| rbm4.21 | 8.332489 | 0.287094266 | 0.225 | 0.154 | 2.1168690 | 1 | rbm4.2 |
| aktip1 | 9.849222 | 0.2775376 | 0.191 | 0.123 | 2.5021949 | 1 | aktip |
| cx43.41 | 1.042845 | 0.250448565 | 0.177 | 0.108 | 2.6493496 | 1 | cx43.4 |
| lsm12a | 1.410018 | 0.276244267 | 0.179 | 0.113 | 3.5821520 | 1 | lsm12a |
| chrac11 | 1.545443 | 0.274251575 | 0.187 | 0.119 | 3.9261981 | 1 | chrac1 |
| g3bp11 | 2.410738 | 0.286436312 | 0.261 | 0.19 | 6.1244804 | 1 | g3bp1 |
| gemin71 | 2.555494 | 0.285225671 | 0.207 | 0.139 | 6.4922338 | 1 | gemin7 |
| btg31 | 4.827784 | 0.29049959 | 0.226 | 0.159 | 1.2264985 | 1 | btg3 |
| fbxw11b1 | 7.571696 | 0.270090221 | 0.194 | 0.129 | 1.9235894 | 1 | fbxw11b |
| rbm71 | 9.242914 | 0.252467265 | 0.17 | 0.105 | 2.3481624 | 1 | rbm7 |
| lsm11 | 1.518689 | 0.265734834 | 0.188 | 0.123 | 3.8582318 | 1 | lsm1 |
| sumo3b1 | 2.088775 | 0.260885734 | 0.176 | 0.112 | 5.3065342 | 1 | sumo3b |
| f11r.11 | 2.257622 | 0.275495102 | 0.229 | 0.161 | 5.7354887 | 1 | f11r.1 |
| abcf2a1 | 2.657331 | 0.270347836 | 0.199 | 0.134 | 6.7509519 | 1 | abcf2a |
| psme31 | 3.437276 | 0.265880688 | 0.185 | 0.122 | 8.7323998 | 1 | psme3 |
| polr2k1 | 4.388476 | 0.279098861 | 0.238 | 0.173 | 1.1148923 | 1 | polr2k |
| nabp1a1 | 6.999812 | 0.257294753 | 0.166 | 0.106 | 1.7783024 | 1 | nabp1a |
| nifk | 7.318582 | 0.260192007 | 0.175 | 0.115 | 1.8592858 | 1 | nifk |
| nutf2l1 | 7.408746 | 0.282118459 | 0.279 | 0.219 | 1.8821921 | 1 | nutf2l |
| rpa31 | 1.255342 | 0.261363339 | 0.215 | 0.151 | 3.1891980 | 1 | rpa3 |
| pcf111 | 1.676596 | 0.277572946 | 0.272 | 0.215 | 4.2593939 | 1 | pcf11 |
| ddx6 | 6.568433 | 0.270245375 | 0.234 | 0.175 | 1.6687105 | 1 | ddx6 |
| slc25a25a1 | 4.424334 | 0.256970288 | 0.217 | 0.161 | 0.000112 | 1 | slc25a25a |
| hmgb3b1 | 1.177478 | 0.262648596 | 0.222 | 0.169 | 0.000299 | 1 | hmgb3b |
| rab5if | 1.456479 | 0.250718247 | 0.177 | 0.126 | 0.00037 | 1 | rab5if |
| cbx5 | 5.850173 | 0.252719961 | 0.25 | 0.203 | 0.001486 | 1 | cbx5 |
| XLOC-000431 | 0 | 1.840542583 | 0.956 | 0.222 | 0 | 2 | XLOC-000431 |
| XLOC-042595 | 0 | 1.785624189 | 0.87 | 0.099 | 0 | 2 | XLOC-042595 |
| XLOC-010899 | 0 | 1.730783909 | 0.924 | 0.189 | 0 | 2 | XLOC-010899 |

| | | | | | | | |
|---|---|---|---|---|---|---|---|
| XLOC-013964 | 0 | 1.722818063 | 0.89 | 0.116 | 0 | 2 | XLOC-013964 |
| ccl36.1 | 0 | 1.662012374 | 0.642 | 0.045 | 0 | 2 | ccl36.1 |
| LOC103910871 | 0 | 1.660630033 | 0.881 | 0.103 | 0 | 2 | LOC103910871 |
| ccl38.6 | 0 | 1.594747008 | 0.687 | 0.05 | 0 | 2 | ccl38.6 |
| si:ch211-214p16.1 | 0 | 1.582281535 | 0.95 | 0.192 | 0 | 2 | si:ch211-214p16.1 |
| CR936442.1 | 0 | 1.568369053 | 0.868 | 0.101 | 0 | 2 | CR936442.1 |
| dusp2 | 0 | 1.394285123 | 0.79 | 0.117 | 0 | 2 | dusp2 |
| arpc1b | 0 | 1.031532231 | 0.885 | 0.272 | 0 | 2 | arpc1b |
| pfn1 | 0 | 1.012821068 | 0.988 | 0.466 | 0 | 2 | pfn1 |
| XLOC-025423 | 0 | 1.012476603 | 0.611 | 0.138 | 0 | 2 | XLOC-025423 |
| LOC100536213 | 0 | 1.010938709 | 0.587 | 0.076 | 0 | 2 | LOC100536213 |
| id3 | 0 | 1.008381955 | 0.675 | 0.163 | 0 | 2 | id3 |
| wasb | 0 | 0.992000359 | 0.683 | 0.135 | 0 | 2 | wasb |
| rgs13 | 0 | 0.9884299 | 0.579 | 0.092 | 0 | 2 | rgs13 |
| coro1a | 0 | 0.986906355 | 0.779 | 0.197 | 0 | 2 | coro1a |
| XLOC-028370 | 0 | 0.967359115 | 0.512 | 0.097 | 0 | 2 | XLOC-028370 |
| si:dkey-183i3.6 | 0 | 0.958449082 | 0.416 | 0.012 | 0 | 2 | si:dkey-183i3.6 |
| sla2 | 0 | 0.930231812 | 0.462 | 0.031 | 0 | 2 | sla2 |
| CR753876.1 | 0 | 0.899471243 | 0.486 | 0.084 | 0 | 2 | CR753876.1 |
| FP236356.1 | 0 | 0.895197749 | 0.421 | 0.018 | 0 | 2 | FP236356.1 |
| ccl34b.4 | 0 | 0.894150101 | 0.338 | 0.012 | 0 | 2 | ccl34b.4 |
| XLOC-020301 | 0 | 0.885869779 | 0.401 | 0.046 | 0 | 2 | XLOC-020301 |
| LOC108190717 | 0 | 0.881345636 | 0.439 | 0.056 | 0 | 2 | LOC108190717 |
| XLOC-026836 | 0 | 0.870271165 | 0.411 | 0.055 | 0 | 2 | XLOC-026836 |
| BX005105.1 | 0 | 0.860347164 | 0.413 | 0.01 | 0 | 2 | BX005105.1 |
| si:ch211-119e14.1 | 0 | 0.857537964 | 0.473 | 0.041 | 0 | 2 | si:ch211-119e14.1 |
| tagapa | 0 | 0.856326888 | 0.463 | 0.071 | 0 | 2 | tagapa |
| si:dkey-27i16.2 | 0 | 0.841256558 | 0.468 | 0.064 | 0 | 2 | si:dkey-27i16.2 |
| si:dkey-109a10.2 | 0 | 0.805085855 | 0.372 | 0.004 | 0 | 2 | si:dkey-109a10.2 |
| ccl32b.3 | 0 | 0.785242662 | 0.324 | 0.007 | 0 | 2 | ccl32b.3 |
| c3a.5 | 0 | 0.766365723 | 0.398 | 0.037 | 0 | 2 | c3a.5 |
| wasa | 0 | 0.752535216 | 0.467 | 0.07 | 0 | 2 | wasa |
| ncaldb | 0 | 0.75150191 | 0.429 | 0.045 | 0 | 2 | ncaldb |
| si:ch73-223p23.2 | 0 | 0.682035875 | 0.325 | 0.006 | 0 | 2 | si:ch73-223p23.2 |
| CR848749.2 | 0 | 0.671365554 | 0.293 | 0.005 | 0 | 2 | CR848749.2 |

| | | | | | | | |
|---|---|---|---|---|---|---|---|
| si:dkey-183p4.10 | 0 | 0.665766088 | 0.325 | 0.011 | 0 | 2 | si:dkey-183p4.10 |
| si:dkey-260g12.1 | 0 | 0.662634323 | 0.328 | 0.009 | 0 | 2 | si:dkey-260g12.1 |
| si:ch211-93g21.1 | 0 | 0.653945722 | 0.312 | 0.012 | 0 | 2 | si:ch211-93g21.1 |
| XLOC-042596 | 0 | 0.645638652 | 0.313 | 0.019 | 0 | 2 | XLOC-042596 |
| zgc:136605 | 0 | 0.622139549 | 0.266 | 0.002 | 0 | 2 | zgc:136605 |
| LOC100535806 | 0 | 0.596552026 | 0.274 | 0.014 | 0 | 2 | LOC100535806 |
| si:ch211-132g1.4 | 0 | 0.593916569 | 0.253 | 0.002 | 0 | 2 | si:ch211-132g1.4 |
| FP016106.1 | 0 | 0.580560398 | 0.269 | 0.004 | 0 | 2 | FP016106.1 |
| si:rp71-81e14.2 | 0 | 0.57819367 | 0.269 | 0.014 | 0 | 2 | si:rp71-81e14.2 |
| LOC101882070 | 0 | 0.573512164 | 0.249 | 0.002 | 0 | 2 | LOC101882070 |
| rplp2 | 0 | 0.544890599 | 0.994 | 0.768 | 0 | 2 | rplp2 |
| si:ch211-196f2.6 | 0 | 0.488565267 | 0.211 | 0.002 | 0 | 2 | si:ch211-196f2.6 |
| LOC103909107 | 0 | 0.48817321 | 0.205 | 0.009 | 0 | 2 | LOC103909107 |
| cxcr3.1 | 0 | 0.47308379 | 0.223 | 0.005 | 0 | 2 | cxcr3.1 |
| si:ch211-132g1.1 | 0 | 0.454319995 | 0.222 | 0.007 | 0 | 2 | si:ch211-132g1.1 |
| si:ch211-167j9.5 | 0 | 0.446821747 | 0.211 | 0.011 | 0 | 2 | si:ch211-167j9.5 |
| LOC100004978 | 0 | 0.4453144 | 0.184 | 0.001 | 0 | 2 | LOC100004978 |
| gzmk | 0 | 0.436291154 | 0.188 | 0.002 | 0 | 2 | gzmk |
| LOC108185170 | 0 | 0.420235457 | 0.177 | 0.001 | 0 | 2 | LOC108185170 |
| si:ch73-285p12.4 | 0 | 0.39540835 | 0.186 | 0.004 | 0 | 2 | si:ch73-285p12.4 |
| tnfsf14 | 0 | 0.390331253 | 0.165 | 0.003 | 0 | 2 | tnfsf14 |
| sh2d1ab | 0 | 0.367814344 | 0.156 | 0.003 | 0 | 2 | sh2d1ab |
| LOC101885675 | 0 | 0.350526585 | 0.158 | 0.002 | 0 | 2 | LOC101885675 |
| itk | 0 | 0.336433225 | 0.152 | 0.003 | 0 | 2 | itk |
| LOC110439931 | 1.671420 | 0.410523051 | 0.197 | 0.009 | 4.2462438 | 2 | LOC110439931 |
| si:dkeyp-85e10.3 | 3.733561 | 0.406411114 | 0.196 | 0.009 | 9.4851130 | 2 | si:dkeyp-85e10.3 |
| skap1 | 4.307549 | 0.410335518 | 0.197 | 0.01 | 1.0943329 | 2 | skap1 |
| ptpn6 | 4.134046 | 0.750769535 | 0.444 | 0.074 | 1.0502545 | 2 | ptpn6 |
| rplp2l | 3.212205 | 0.508495669 | 0.989 | 0.781 | 8.1606072 | 2 | rplp2l |
| ptprc | 7.763222 | 0.750072188 | 0.504 | 0.097 | 1.9722466 | 2 | ptprc |
| gpr183a | 6.052898 | 0.65199019 | 0.343 | 0.045 | 1.5377389 | 2 | gpr183a |
| rps27.1 | 2.519326 | 0.582531892 | 0.981 | 0.691 | 6.4003477 | 2 | rps27.1 |
| si:ch73-226l13.2 | 1.128889 | 0.374543423 | 0.161 | 0.006 | 2.8679437 | 2 | si:ch73-226l13.2 |
| rplp1 | 1.459193 | 0.404452351 | 0.998 | 0.874 | 3.7070802 | 2 | rplp1 |
| rps20 | 2.087395 | 0.356036328 | 0.994 | 0.904 | 5.3030274 | 2 | rps20 |

| | | | | | | | |
|---|---|---|---|---|---|---|---|
| rps29 | 3.176663 | 0.425398227 | 0.994 | 0.834 | 8.0703131 | 2 | rps29 |
| rpl37 | 5.675567 | 0.423585954 | 0.994 | 0.845 | 1.4418780 | 2 | rpl37 |
| hcst | 7.795656 | 0.535756738 | 0.283 | 0.028 | 1.9804864 | 2 | hcst |
| rac2 | 7.405461 | 0.856289388 | 0.699 | 0.19 | 1.8813573 | 2 | rac2 |
| btg1 | 8.359451 | 0.895184141 | 0.815 | 0.322 | 2.1237185 | 2 | btg1 |
| rps17 | 1.710265 | 0.485027909 | 0.97 | 0.766 | 4.3449300 | 2 | rps17 |
| il2rb | 1.333386 | 0.372781198 | 0.17 | 0.008 | 3.3874688 | 2 | il2rb |
| tagapb | 9.671467 | 0.621495468 | 0.349 | 0.049 | 2.4570363 | 2 | tagapb |
| XLOC-032002 | 1.039242 | 0.61559626 | 0.271 | 0.028 | 2.6401958 | 2 | XLOC-032002 |
| XLOC-008993 | 2.174399 | 0.439199144 | 0.208 | 0.014 | 5.5240630 | 2 | XLOC-008993 |
| XLOC-043760 | 2.761492 | 0.597208991 | 0.277 | 0.029 | 7.0155729 | 2 | XLOC-043760 |
| si:ch211-25d12.7 | 1.250786 | 0.328806799 | 0.16 | 0.006 | 3.1776242 | 2 | si:ch211-25d12.7 |
| si:ch1073-220m6.1 | 9.105336 | 0.354725857 | 0.176 | 0.009 | 2.3132108 | 2 | si:ch1073-220m6.1 |
| zap70 | 6.161456 | 0.286017108 | 0.131 | 0.003 | 1.5653180 | 2 | zap70 |
| samsn1b | 1.216393 | 0.386630635 | 0.186 | 0.011 | 3.0902489 | 2 | samsn1b |
| rps21 | 4.932172 | 0.495298202 | 0.987 | 0.761 | 1.2530184 | 2 | rps21 |
| arhgdig | 1.388892 | 0.852068927 | 0.714 | 0.227 | 3.5284821 | 2 | arhgdig |
| rpl11 | 1.230865 | 0.438644793 | 0.991 | 0.818 | 3.1270141 | 2 | rpl11 |
| rpl34 | 2.433979 | 0.461178498 | 0.984 | 0.79 | 6.1835245 | 2 | rpl34 |
| cytip | 5.874848 | 0.305485038 | 0.149 | 0.006 | 1.4925051 | 2 | cytip |
| rpl36 | 5.432456 | 0.45597228 | 0.99 | 0.793 | 1.3801156 | 2 | rpl36 |
| cd247l | 6.014714 | 0.350557936 | 0.17 | 0.009 | 1.5280381 | 2 | cd247l |
| rpl18a | 9.825499 | 0.4623164 | 0.985 | 0.788 | 2.4961682 | 2 | rpl18a |
| mrps15 | 3.136103 | 0.711843246 | 0.448 | 0.089 | 7.9672711 | 2 | mrps15 |
| rps15a | 3.054893 | 0.479059311 | 0.988 | 0.784 | 7.7609569 | 2 | rps15a |
| rps2 | 6.365731 | 0.439561829 | 0.99 | 0.831 | 1.6172141 | 2 | rps2 |
| cyth4a | 1.212526 | 0.337320849 | 0.166 | 0.009 | 3.0804233 | 2 | cyth4a |
| cebpb | 3.221936 | 0.926943203 | 0.804 | 0.306 | 8.1853294 | 2 | cebpb |
| rps26l | 3.722623 | 0.559513434 | 0.98 | 0.68 | 9.4573246 | 2 | rps26l |
| lpar6b | 4.943478 | 0.331710831 | 0.152 | 0.007 | 1.2558906 | 2 | lpar6b |
| cabz01093075.1 | 4.655439 | 0.346026953 | 0.161 | 0.008 | 1.1827143 | 2 | cabz01093075.1 |
| h1fx | 1.283025 | 0.890975469 | 0.582 | 0.162 | 3.2595260 | 2 | h1fx |
| cxcr4b | 5.851762 | 0.850040441 | 0.574 | 0.149 | 1.4866401 | 2 | cxcr4b |
| srgn | 6.123368 | 0.796161344 | 0.689 | 0.202 | 1.5556418 | 2 | srgn |
| faslg | 6.922564 | 0.327293521 | 0.15 | 0.007 | 1.7586775 | 2 | faslg |

| | | | | | | | |
|---|---|---|---|---|---|---|---|
| zgc:64051 | 5.206324 | 0.715118452 | 0.557 | 0.134 | 1.3226667 | 2 | zgc:64051 |
| capgb | 5.886648 | 0.691356182 | 0.445 | 0.091 | 1.4955030 | 2 | capgb |
| rpl19 | 2.667399 | 0.383537166 | 0.984 | 0.845 | 6.7765278 | 2 | rpl19 |
| entpd3 | 1.579531 | 0.329617693 | 0.146 | 0.007 | 4.0127988 | 2 | entpd3 |
| rps6 | 3.314389 | 0.433694867 | 0.971 | 0.792 | 8.4202071 | 2 | rps6 |
| ccl38a.4 | 3.347419 | 0.413539452 | 0.123 | 0.004 | 8.5041189 | 2 | ccl38a.4 |
| rps5 | 1.178651 | 0.515664554 | 0.968 | 0.712 | 2.9943631 | 2 | rps5 |
| rpl35 | 4.245615 | 0.426003088 | 0.988 | 0.802 | 1.0785985 | 2 | rpl35 |
| rsl24d1 | 9.592870 | 0.750178326 | 0.861 | 0.432 | 2.4370686 | 2 | rsl24d1 |
| kctd5a | 5.088124 | 0.763086747 | 0.551 | 0.159 | 1.2926380 | 2 | kctd5a |
| XLOC-023492 | 6.169954 | 0.677341106 | 0.383 | 0.073 | 1.5674769 | 2 | XLOC-023492 |
| prf1.9 | 6.370512 | 0.47276508 | 0.141 | 0.007 | 1.6184287 | 2 | prf1.9 |
| CU499336.2 | 1.855057 | 0.422750987 | 0.213 | 0.02 | 4.7127728 | 2 | CU499336.2 |
| si:ch211-226m16.2 | 6.245193 | 0.568564538 | 0.319 | 0.051 | 1.5865913 | 2 | si:ch211-226m16.2 |
| rpl15 | 7.063326 | 0.477327407 | 0.977 | 0.749 | 1.7944379 | 2 | rpl15 |
| ccl38a.5 | 1.462725 | 0.36905075 | 0.121 | 0.004 | 3.7160529 | 2 | ccl38a.5 |
| rpl7a | 8.157617 | 0.412858123 | 0.99 | 0.812 | 2.0724427 | 2 | rpl7a |
| laptm5 | 1.287360 | 0.707163182 | 0.571 | 0.148 | 3.2705404 | 2 | laptm5 |
| rpl27 | 5.397113 | 0.448649373 | 0.977 | 0.766 | 1.3711366 | 2 | rpl27 |
| rpl18 | 1.000286 | 0.445244183 | 0.985 | 0.779 | 2.5412276 | 2 | rpl18 |
| rps28 | 8.532708 | 0.392906894 | 0.991 | 0.83 | 2.1677344 | 2 | rps28 |
| rpl32 | 4.171353 | 0.414196533 | 0.986 | 0.813 | 1.0597322 | 2 | rpl32 |
| si:dkey-171c9.3 | 1.005803 | 0.397324215 | 0.208 | 0.02 | 2.5552441 | 2 | si:dkey-171c9.3 |
| il7r | 1.174530 | 0.320213828 | 0.149 | 0.009 | 2.9838953 | 2 | il7r |
| XLOC-011085 | 1.502685 | 0.522278152 | 0.283 | 0.042 | 3.8175725 | 2 | XLOC-011085 |
| grap2a | 7.349893 | 0.456503771 | 0.254 | 0.033 | 1.8672405 | 2 | grap2a |
| tcf7 | 2.679867 | 0.354713677 | 0.169 | 0.013 | 6.8082028 | 2 | tcf7 |
| zfp36l1a | 6.473032 | 0.787423148 | 0.647 | 0.226 | 1.6444739 | 2 | zfp36l1a |
| rps12 | 1.331348 | 0.387910119 | 0.986 | 0.831 | 3.3822903 | 2 | rps12 |
| rpl38 | 2.164951 | 0.473694011 | 0.976 | 0.738 | 5.5000588 | 2 | rpl38 |
| calm1b | 2.682990 | 0.5754343 | 0.397 | 0.082 | 6.8161382 | 2 | calm1b |
| adgre5b.3 | 5.777376 | 0.422035204 | 0.218 | 0.025 | 1.4677424 | 2 | adgre5b.3 |
| rpl23 | 7.203301 | 0.328372431 | 0.987 | 0.865 | 1.8299986 | 2 | rpl23 |
| rps19 | 8.124047 | 0.344183808 | 0.99 | 0.86 | 2.0639142 | 2 | rps19 |
| rpl8 | 1.131917 | 0.424980307 | 0.977 | 0.782 | 2.8756361 | 2 | rpl8 |

| | | | | | | | |
|---|---|---|---|---|---|---|---|
| BX005183.3 | 3.31310 | 0.581729682 | 0.325 | 0.059 | 8.4169480 | 2 | BX005183.3 |
| rpl10a | 6.73814 | 0.431964216 | 0.979 | 0.765 | 1.7118253 | 2 | rpl10a |
| sept12 | 1.54759 | 0.502170138 | 0.294 | 0.049 | 3.9316536 | 2 | sept12 |
| wu:fb55g09 | 1.63332 | 0.777460355 | 0.637 | 0.232 | 4.1494718 | 2 | wu:fb55g09 |
| rps16 | 6.67999 | 0.451942975 | 0.976 | 0.751 | 1.6970536 | 2 | rps16 |
| rps7 | 1.42088 | 0.388036101 | 0.981 | 0.818 | 3.6097639 | 2 | rps7 |
| rps14 | 1.45736 | 0.427893359 | 0.98 | 0.767 | 3.7024424 | 2 | rps14 |
| si:ch211-67e16.3 | 3.80790 | 0.325503051 | 0.143 | 0.009 | 9.6739768 | 2 | si:ch211-67e16.3 |
| ptpn4b | 2.14909 | 0.333776041 | 0.165 | 0.013 | 5.4597713 | 2 | ptpn4b |
| si:dkeyp-68b7.12 | 9.31250 | 0.491805533 | 0.279 | 0.045 | 2.3658428 | 2 | si:dkeyp-68b7.12 |
| abo | 1.38585 | 0.301602762 | 0.145 | 0.01 | 3.5207620 | 2 | abo |
| rpl39 | 9.48475 | 0.33924529 | 0.989 | 0.869 | 2.4096016 | 2 | rpl39 |
| ptmaa | 5.64908 | 0.726296258 | 0.856 | 0.402 | 1.4351493 | 2 | ptmaa |
| rpl10 | 6.07318 | 0.412663692 | 0.979 | 0.777 | 1.5428915 | 2 | rpl10 |
| il12rb2l | 1.70335 | 0.416521434 | 0.222 | 0.029 | 4.3273790 | 2 | il12rb2l |
| rps9 | 3.449151 | 0.415450545 | 0.98 | 0.777 | 8.7625704 | 2 | rps9 |
| zgc:174945 | 6.19313 | 0.334299083 | 0.165 | 0.014 | 1.5733662 | 2 | zgc:174945 |
| LOC103910136 | 7.74939 | 0.372955892 | 0.174 | 0.016 | 1.9687346 | 2 | LOC103910136 |
| rpl26 | 4.66034 | 0.351465723 | 0.986 | 0.842 | 1.1839611 | 2 | rpl26 |
| mbnl1 | 2.56036 | 0.670553366 | 0.527 | 0.162 | 6.5046187 | 2 | mbnl1 |
| si:ch211-105j21.9 | 3.40249 | 0.376630507 | 0.2 | 0.023 | 8.6440444 | 2 | si:ch211-105j21.9 |
| CR855337.1 | 3.55636 | 0.375400486 | 0.192 | 0.022 | 9.0349381 | 2 | CR855337.1 |
| rps18 | 4.85780 | 0.448703031 | 0.977 | 0.738 | 1.2341255 | 2 | rps18 |
| slc25a5 | 6.02242 | 0.368050237 | 0.967 | 0.803 | 1.5299975 | 2 | slc25a5 |
| si:dkey-262k9.2 | 4.70017 | 0.362718031 | 0.191 | 0.022 | 1.1940797 | 2 | si:dkey-262k9.2 |
| hsp90ab1 | 5.93444 | 0.389243668 | 0.981 | 0.788 | 1.5076454 | 2 | hsp90ab1 |
| rpl29 | 2.35975 | 0.375697981 | 0.981 | 0.812 | 5.9949659 | 2 | rpl29 |
| rpl7 | 3.06619 | 0.381731236 | 0.972 | 0.798 | 7.7896577 | 2 | rpl7 |
| rps13 | 1.51166 | 0.440208144 | 0.973 | 0.733 | 3.8403834 | 2 | rps13 |
| sik1 | 1.03165 | 0.596747552 | 0.375 | 0.092 | 2.6209249 | 2 | sik1 |
| scinlb | 1.15112 | 0.501861436 | 0.326 | 0.067 | 2.9244296 | 2 | scinlb |
| tpt1 | 3.15435 | 0.372221008 | 0.974 | 0.8 | 8.0136262 | 2 | tpt1 |
| si:dkey-185m8.2 | 6.968281 | 0.417277981 | 0.228 | 0.034 | 1.7702920 | 2 | si:dkey-185m8.2 |
| rpsa | 1.64983 | 0.408390888 | 0.984 | 0.772 | 4.1914147 | 2 | rpsa |
| zbtb32 | 3.76199 | 0.404212816 | 0.214 | 0.029 | 9.5573587 | 2 | zbtb32 |

| | | | | | | | |
|---|---|---|---|---|---|---|---|
| LOC103909115 | 2.400855 | 0.597771896 | 0.487 | 0.146 | 6.0993733 | 2 | LOC103909115 |
| arhgef3l | 4.686693 | 0.335317123 | 0.179 | 0.02 | 1.1906545 | 2 | arhgef3l |
| rps3 | 6.461957 | 0.447488213 | 0.96 | 0.71 | 1.6416603 | 2 | rps3 |
| rpl30 | 2.784453 | 0.454243736 | 0.971 | 0.713 | 7.0739030 | 2 | rpl30 |
| ccr9a | 6.135371 | 0.569701089 | 0.329 | 0.071 | 1.5586911 | 2 | ccr9a |
| rpl21 | 1.194228 | 0.406744693 | 0.977 | 0.763 | 3.0339384 | 2 | rpl21 |
| rpl22 | 4.377068 | 0.465961739 | 0.969 | 0.703 | 1.1119942 | 2 | rpl22 |
| rplp0 | 3.654083 | 0.33070592 | 0.98 | 0.85 | 9.2831996 | 2 | rplp0 |
| si:ch211-191c10.1 | 5.423977 | 0.424124221 | 0.232 | 0.037 | 1.3779614 | 2 | si:ch211-191c10.1 |
| rps24 | 1.087700 | 0.401837099 | 0.975 | 0.765 | 2.7633019 | 2 | rps24 |
| sept9b | 1.521928 | 0.429325348 | 0.226 | 0.035 | 3.8664592 | 2 | sept9b |
| rps15 | 2.769720 | 0.471005939 | 0.972 | 0.699 | 7.0364758 | 2 | rps15 |
| CABZ01079663.1 | 2.824474 | 0.371201668 | 0.197 | 0.026 | 7.1755785 | 2 | CABZ01079663.1 |
| cmah | 2.902333 | 0.25922081 | 0.122 | 0.009 | 7.3733779 | 2 | cmah |
| rps26 | 4.213347 | 0.416683198 | 0.967 | 0.736 | 1.0704010 | 2 | rps26 |
| faua | 1.272232 | 0.458217293 | 0.96 | 0.701 | 3.2321073 | 2 | faua |
| rpl35a | 1.571507 | 0.395878216 | 0.971 | 0.769 | 3.9924146 | 2 | rpl35a |
| csk | 3.217933 | 0.521364544 | 0.359 | 0.086 | 8.1751609 | 2 | csk |
| ikzf1 | 2.276257 | 0.329025298 | 0.186 | 0.024 | 5.7828316 | 2 | ikzf1 |
| bcl11ba | 2.336862 | 0.251341157 | 0.119 | 0.008 | 5.9367998 | 2 | bcl11ba |
| LOC100151196 | 3.264422 | 0.366692937 | 0.203 | 0.029 | 8.2932646 | 2 | LOC100151196 |
| rpl36a | 7.273925 | 0.358128957 | 0.988 | 0.812 | 1.8479407 | 2 | rpl36a |
| tnfrsf9a | 8.099959 | 0.499041795 | 0.225 | 0.036 | 2.0577946 | 2 | tnfrsf9a |
| hmgn2 | 3.936392 | 0.737609782 | 0.825 | 0.442 | 1.0000403 | 2 | hmgn2 |
| rps23 | 1.402784 | 0.425534526 | 0.972 | 0.728 | 3.5637731 | 2 | rps23 |
| syk | 5.522110 | 0.499609873 | 0.296 | 0.063 | 1.4028920 | 2 | syk |
| rpl28 | 1.646593 | 0.396502198 | 0.976 | 0.763 | 4.1831696 | 2 | rpl28 |
| rpl24 | 2.819203 | 0.385815002 | 0.98 | 0.781 | 7.1621867 | 2 | rpl24 |
| sh2b3 | 2.237185 | 0.446192381 | 0.254 | 0.049 | 5.6835686 | 2 | sh2b3 |
| cotl1 | 1.714975 | 0.665874416 | 0.718 | 0.291 | 4.3568948 | 2 | cotl1 |
| wipf1a | 3.421090 | 0.440203887 | 0.27 | 0.054 | 8.6912814 | 2 | wipf1a |
| il21r.1 | 8.174445 | 0.286666817 | 0.144 | 0.015 | 2.0767177 | 2 | il21r.1 |
| CR381673.1 | 1.280511 | 0.42567814 | 0.241 | 0.044 | 3.2531402 | 2 | CR381673.1 |
| pik3r1 | 2.429037 | 0.508678853 | 0.315 | 0.076 | 6.1709704 | 2 | pik3r1 |
| ms4a17a.6 | 8.203918 | 0.387079921 | 0.228 | 0.039 | 2.0842054 | 2 | ms4a17a.6 |

| | | | | | | | |
|---|---|---|---|---|---|---|---|
| lbh | 2.551127 | 0.550488308 | 0.454 | 0.131 | 6.4811399 | 2 | lbh |
| rpl13 | 1.222093 | 0.346205552 | 0.986 | 0.807 | 3.1047277 | 2 | rpl13 |
| rpl9 | 5.233523 | 0.493138105 | 0.955 | 0.648 | 1.3295766 | 2 | rpl9 |
| XLOC-033137 | 5.394055 | 0.366571366 | 0.178 | 0.024 | 1.3703597 | 2 | XLOC-033137 |
| rasal3 | 1.071772 | 0.296886791 | 0.156 | 0.018 | 2.7228390 | 2 | rasal3 |
| rps25 | 9.142437 | 0.407781502 | 0.977 | 0.733 | 2.3226363 | 2 | rps25 |
| tox | 4.069556 | 0.284088863 | 0.136 | 0.014 | 1.0338707 | 2 | tox |
| rps10 | 6.089099 | 0.430341602 | 0.968 | 0.71 | 1.5469357 | 2 | rps10 |
| FP101882.1 | 9.751406 | 0.345536038 | 0.185 | 0.027 | 2.4773447 | 2 | FP101882.1 |
| naca | 1.375725 | 0.477522072 | 0.917 | 0.633 | 3.4950309 | 2 | naca |
| prkacba | 2.886881 | 0.265968886 | 0.133 | 0.013 | 7.3341231 | 2 | prkacba |
| rps3a | 2.163529 | 0.361156618 | 0.967 | 0.779 | 5.4964478 | 2 | rps3a |
| tmsb4x | 2.753433 | 0.544628592 | 0.995 | 0.699 | 6.9950989 | 2 | tmsb4x |
| ptpn11b | 2.693925 | 0.25904068 | 0.136 | 0.014 | 6.8439168 | 2 | ptpn11b |
| scgn | 3.681979 | 0.282478816 | 0.14 | 0.015 | 9.3540677 | 2 | scgn |
| si:dkey-78o7.3 | 8.141117 | 0.296919557 | 0.131 | 0.013 | 2.0682509 | 2 | si:dkey-78o7.3 |
| p2ry10 | 3.638778 | 0.260474064 | 0.133 | 0.014 | 9.2443155 | 2 | p2ry10 |
| rps11 | 2.676628 | 0.357774767 | 0.966 | 0.772 | 6.7999743 | 2 | rps11 |
| ikzf2 | 5.935937 | 0.328059565 | 0.171 | 0.025 | 1.5080248 | 2 | ikzf2 |
| rps4x | 2.561541 | 0.400816218 | 0.937 | 0.723 | 6.5075971 | 2 | rps4x |
| actb2 | 3.601287 | 0.35741415 | 0.992 | 0.858 | 9.1490698 | 2 | actb2 |
| gnai2b | 4.421037 | 0.397558431 | 0.242 | 0.05 | 1.1231646 | 2 | gnai2b |
| b3gnt5a | 5.649159 | 0.342630922 | 0.171 | 0.025 | 1.4351689 | 2 | b3gnt5a |
| ccr2 | 3.693819 | 0.366691737 | 0.202 | 0.034 | 9.3841485 | 2 | ccr2 |
| il34 | 2.124233 | 0.3962211 | 0.2 | 0.035 | 5.3966158 | 2 | il34 |
| uba52 | 7.398262 | 0.460169375 | 0.936 | 0.642 | 1.8795286 | 2 | uba52 |
| si:ch73-208g10.1 | 2.909406 | 0.328368268 | 0.186 | 0.031 | 7.3913461 | 2 | si:ch73-208g10.1 |
| tnfrsf9b | 4.251117 | 0.250055745 | 0.107 | 0.009 | 1.0799963 | 2 | tnfrsf9b |
| XLOC-043761 | 8.730720 | 0.278754022 | 0.12 | 0.012 | 2.2180394 | 2 | XLOC-043761 |
| eif3f | 6.383699 | 0.55577494 | 0.811 | 0.441 | 1.6217788 | 2 | eif3f |
| ccni | 1.125973 | 0.581514195 | 0.63 | 0.297 | 2.8605359 | 2 | ccni |
| irf4b | 6.758539 | 0.275872133 | 0.132 | 0.015 | 1.7170069 | 2 | irf4b |
| si:ch211-241f5.3 | 1.878823 | 0.420052305 | 0.235 | 0.05 | 4.7731508 | 2 | si:ch211-241f5.3 |
| rpl3 | 2.816064 | 0.271408193 | 0.985 | 0.865 | 7.1542118 | 2 | rpl3 |
| tnip1 | 7.685770 | 0.452951547 | 0.271 | 0.067 | 1.9525699 | 2 | tnip1 |

| | | | | | | | |
|---|---|---|---|---|---|---|---|
| rpl23a | 2.992515 | 0.4528841 | 0.936 | 0.651 | 7.6024850 | 2 | rpl23a |
| unc119b | 6.472024 | 0.31830929 | 0.174 | 0.028 | 1.6442177 | 2 | unc119b |
| tmc8 | 8.394256 | 0.299489013 | 0.164 | 0.025 | 2.1325609 | 2 | tmc8 |
| XLOC-043043 | 1.418147 | 0.363180747 | 0.194 | 0.034 | 3.6028044 | 2 | XLOC-043043 |
| tefa | 1.370566 | 0.504136337 | 0.379 | 0.124 | 3.4819235 | 2 | tefa |
| wdr19 | 3.856908 | 0.331871275 | 0.158 | 0.024 | 9.7984757 | 2 | wdr19 |
| lcp1 | 9.403427 | 0.403832996 | 0.282 | 0.071 | 2.3889406 | 2 | lcp1 |
| XLOC-014079 | 8.736295 | 0.298623391 | 0.158 | 0.024 | 2.2194557 | 2 | XLOC-014079 |
| fli1b | 1.082871 | 0.301333067 | 0.163 | 0.026 | 2.7510341 | 2 | fli1b |
| BX901920.1 | 2.173806 | 0.433768643 | 0.257 | 0.061 | 5.5225566 | 2 | BX901920.1 |
| sat1a.2 | 4.327269 | 0.560236898 | 0.614 | 0.28 | 1.0993427 | 2 | sat1a.2 |
| si:ch73-264p11.1 | 1.690046 | 0.299201182 | 0.144 | 0.02 | 4.2935619 | 2 | si:ch73-264p11.1 |
| bcl2l10 | 4.056729 | 0.541764659 | 0.487 | 0.18 | 1.0306122 | 2 | bcl2l10 |
| rpl31 | 8.448468 | 0.421955548 | 0.942 | 0.669 | 2.1463334 | 2 | rpl31 |
| rps8a | 3.916094 | 0.337393779 | 0.972 | 0.789 | 9.9488386 | 2 | rps8a |
| nr4a1 | 3.290417 | 0.538081884 | 0.408 | 0.141 | 8.3593063 | 2 | nr4a1 |
| msna | 1.734249 | 0.554305092 | 0.552 | 0.246 | 4.4058609 | 2 | msna |
| ppiaa | 7.989122 | 0.314737644 | 0.974 | 0.803 | 2.0296365 | 2 | ppiaa |
| nsa2 | 9.930119 | 0.540624981 | 0.707 | 0.366 | 2.5227467 | 2 | nsa2 |
| nab1a | 2.902625 | 0.2748954 | 0.117 | 0.014 | 7.3741200 | 2 | nab1a |
| fam49a | 3.105525 | 0.342911779 | 0.207 | 0.044 | 7.8895877 | 2 | fam49a |
| crip1 | 5.191676 | 0.539484671 | 0.616 | 0.261 | 1.3189454 | 2 | crip1 |
| myl12.1 | 2.522330 | 0.536581977 | 0.76 | 0.402 | 6.4079806 | 2 | myl12.1 |
| rpl27a | 1.863714 | 0.515475383 | 0.855 | 0.499 | 4.7347658 | 2 | rpl27a |
| vsir | 2.675118 | 0.402481038 | 0.279 | 0.075 | 6.7961393 | 2 | vsir |
| rpl5a | 2.801986 | 0.503608189 | 0.857 | 0.514 | 7.1184465 | 2 | rpl5a |
| mcl1a | 2.492583 | 0.464913467 | 0.87 | 0.595 | 6.3324073 | 2 | mcl1a |
| elf1 | 3.072380 | 0.430598124 | 0.303 | 0.089 | 7.8053819 | 2 | elf1 |
| dok2 | 6.130714 | 0.319738428 | 0.169 | 0.031 | 1.5575081 | 2 | dok2 |
| XLOC-005449 | 1.305035 | 0.366135391 | 0.21 | 0.046 | 3.3154419 | 2 | XLOC-005449 |
| nkl.2 | 8.163067 | 0.522127808 | 0.358 | 0.118 | 2.0738273 | 2 | nkl.2 |
| FERMT3 (1 of many) | 1.759881 | 0.307734161 | 0.18 | 0.036 | 4.4709798 | 2 | FERMT3 (1 of many) |
| b2m | 2.904680 | 0.556523476 | 0.582 | 0.279 | 7.3793415 | 2 | b2m |
| pde4ba | 1.359895 | 0.523969252 | 0.358 | 0.127 | 3.4548140 | 2 | pde4ba |
| nfkbie | 2.090357 | 0.418537824 | 0.287 | 0.084 | 5.3105531 | 2 | nfkbie |

| | | | | | | | |
|---|---|---|---|---|---|---|---|
| btf3 | 2.268754 | 0.443551151 | 0.88 | 0.592 | 5.7637705 | 2 | btf3 |
| rpl13a | 6.130590 | 0.322458264 | 0.958 | 0.77 | 1.5574766 | 2 | rpl13a |
| rpl6 | 1.130723 | 0.404534389 | 0.949 | 0.684 | 2.8726028 | 2 | rpl6 |
| acap1 | 2.429329 | 0.285981924 | 0.165 | 0.031 | 6.1717105 | 2 | acap1 |
| pnrc2 | 1.697169 | 0.470460077 | 0.897 | 0.577 | 4.3116581 | 2 | pnrc2 |
| hmha1b | 3.114595 | 0.353413481 | 0.243 | 0.062 | 7.9126294 | 2 | hmha1b |
| rack1 | 5.054232 | 0.333523255 | 0.947 | 0.751 | 1.2840278 | 2 | rack1 |
| si:ch211-284f22.3 | 1.290766 | 0.281436111 | 0.151 | 0.027 | 3.2791933 | 2 | si:ch211-284f22.3 |
| RASSF5 | 6.890522 | 0.339190113 | 0.214 | 0.051 | 1.7505371 | 2 | RASSF5 |
| vaspa | 8.411234 | 0.357373084 | 0.26 | 0.069 | 2.1368740 | 2 | vaspa |
| lpar5a | 2.599689 | 0.303169336 | 0.176 | 0.036 | 6.6045110 | 2 | lpar5a |
| sesn1 | 7.074956 | 0.505738787 | 0.406 | 0.159 | 1.7973926 | 2 | sesn1 |
| rpl12 | 1.358939 | 0.294039526 | 0.962 | 0.802 | 3.4523853 | 2 | rpl12 |
| si:ch211-121a2.2 | 4.269117 | 0.255469293 | 0.14 | 0.024 | 1.0845692 | 2 | si:ch211-121a2.2 |
| mfng | 3.513380 | 0.289873842 | 0.175 | 0.037 | 8.9257435 | 2 | mfng |
| ppp1r18 | 7.403535 | 0.313504546 | 0.198 | 0.046 | 1.8808682 | 2 | ppp1r18 |
| si:ch73-234b20.5 | 9.239225 | 0.388485256 | 0.294 | 0.093 | 2.3472251 | 2 | si:ch73-234b20.5 |
| arpc3 | 1.092685 | 0.494727233 | 0.631 | 0.313 | 2.7759674 | 2 | arpc3 |
| itga4 | 6.869294 | 0.357033472 | 0.214 | 0.054 | 1.7451441 | 2 | itga4 |
| si:ch211-196l7.4 | 2.085670 | 0.49211449 | 0.612 | 0.31 | 5.2986451 | 2 | si:ch211-196l7.4 |
| si:dkey-262k9.4 | 3.103984 | 0.378539722 | 0.265 | 0.078 | 7.8856736 | 2 | si:dkey-262k9.4 |
| eef1g | 4.192654 | 0.402333428 | 0.891 | 0.636 | 1.0651439 | 2 | eef1g |
| sept2 | 1.448744 | 0.447474604 | 0.415 | 0.167 | 3.6805347 | 2 | sept2 |
| lcp2a | 3.089698 | 0.265677162 | 0.153 | 0.03 | 7.8493786 | 2 | lcp2a |
| slc44a2 | 3.932277 | 0.317401372 | 0.193 | 0.046 | 9.9899509 | 2 | slc44a2 |
| fscn1a | 1.569854 | 0.330138318 | 0.19 | 0.044 | 3.9882146 | 2 | fscn1a |
| wipf1b | 2.256576 | 0.326456174 | 0.212 | 0.055 | 5.7328326 | 2 | wipf1b |
| rps27a | 9.784775 | 0.327447414 | 0.962 | 0.762 | 2.4858222 | 2 | rps27a |
| arrdc3b | 2.013335 | 0.288042223 | 0.15 | 0.029 | 5.1148795 | 2 | arrdc3b |
| litaf | 6.501427 | 0.424305597 | 0.35 | 0.132 | 1.6516877 | 2 | litaf |
| si:rp71-68n21.9 | 3.608621 | 0.290118994 | 0.171 | 0.038 | 9.1677041 | 2 | si:rp71-68n21.9 |
| il16 | 6.665667 | 0.292487569 | 0.176 | 0.04 | 1.6934129 | 2 | il16 |
| myca | 9.943215 | 0.295648986 | 0.156 | 0.033 | 2.5260739 | 2 | myca |
| nme2b.1 | 1.244254 | 0.253404441 | 0.982 | 0.848 | 3.1610285 | 2 | nme2b.1 |
| CABZ01084575.1 | 6.381613 | 0.289662707 | 0.156 | 0.032 | 1.6212490 | 2 | CABZ01084575.1 |

| | | | | | | | |
|---|---|---|---|---|---|---|---|
| si:ch211-132p1.3 | 1.031810 | 0.382134779 | 0.267 | 0.086 | 2.6213135 | 2 | si:ch211-132p1.3 |
| dtnbp1a | 4.899593 | 0.361045295 | 0.254 | 0.077 | 1.2447417 | 2 | dtnbp1a |
| si:dkey-53k12.2 | 1.640998 | 0.273170744 | 0.165 | 0.036 | 4.1689570 | 2 | si:dkey-53k12.2 |
| celf2 | 1.772973 | 0.45609268 | 0.513 | 0.24 | 4.5042388 | 2 | celf2 |
| abi3a | 2.019691 | 0.260509742 | 0.171 | 0.038 | 5.1310268 | 2 | abi3a |
| cd74a | 2.332727 | 0.332712518 | 0.505 | 0.196 | 5.9262944 | 2 | cd74a |
| serbp1a | 2.467250 | 0.448904648 | 0.786 | 0.487 | 6.2680493 | 2 | serbp1a |
| pgam1a | 2.985833 | 0.426213014 | 0.363 | 0.141 | 7.5855108 | 2 | pgam1a |
| mob1a | 2.310275 | 0.411624039 | 0.39 | 0.158 | 5.8692540 | 2 | mob1a |
| ldha | 4.445300 | 0.32180456 | 0.212 | 0.06 | 1.1293285 | 2 | ldha |
| CR392341.1 | 4.794728 | 0.293285147 | 0.193 | 0.051 | 1.2181007 | 2 | CR392341.1 |
| ptmab | 2.785862 | 0.43009427 | 0.832 | 0.552 | 7.0774836 | 2 | ptmab |
| FQ323156.1 | 3.438045 | 0.365873959 | 0.367 | 0.138 | 8.7343534 | 2 | FQ323156.1 |
| rpl5b | 4.039522 | 0.369859924 | 0.877 | 0.629 | 1.0262407 | 2 | rpl5b |
| zgc:152863 | 4.234693 | 0.250979177 | 0.148 | 0.032 | 1.0758238 | 2 | zgc:152863 |
| eef1da | 2.115311 | 0.424392864 | 0.762 | 0.488 | 5.3739489 | 2 | eef1da |
| RASA2 | 3.616436 | 0.26921542 | 0.159 | 0.038 | 9.1875566 | 2 | RASA2 |
| rnf44 | 1.219515 | 0.346446338 | 0.255 | 0.086 | 3.0981780 | 2 | rnf44 |
| lasp1 | 5.827257 | 0.420105823 | 0.488 | 0.237 | 1.4804148 | 2 | lasp1 |
| cfl1 | 2.114467 | 0.410125077 | 0.865 | 0.582 | 5.3718047 | 2 | cfl1 |
| cap1 | 3.198365 | 0.442280981 | 0.477 | 0.23 | 8.1254481 | 2 | cap1 |
| cox6b1 | 4.694477 | 0.353222208 | 0.297 | 0.11 | 1.1926319 | 2 | cox6b1 |
| fam49ba | 1.048640 | 0.318024322 | 0.274 | 0.094 | 2.6640701 | 2 | fam49ba |
| arpc5a | 1.493528 | 0.376067203 | 0.298 | 0.114 | 3.7943095 | 2 | arpc5a |
| rps6ka1 | 2.137918 | 0.297090767 | 0.206 | 0.061 | 5.4313812 | 2 | rps6ka1 |
| zgc:162944 | 2.461553 | 0.426008142 | 0.506 | 0.26 | 6.2535778 | 2 | zgc:162944 |
| arpc4 | 2.893928 | 0.406866962 | 0.471 | 0.222 | 7.3520253 | 2 | arpc4 |
| si:ch211-102c2.4 | 4.608630 | 0.395139629 | 0.342 | 0.133 | 1.1708224 | 2 | si:ch211-102c2.4 |
| aldocb | 5.477219 | 0.307550713 | 0.196 | 0.057 | 1.3914876 | 2 | aldocb |
| ponzr1 | 6.408531 | 0.430981175 | 0.547 | 0.272 | 1.6280875 | 2 | ponzr1 |
| zgc:114188 | 9.336805 | 0.401301262 | 0.946 | 0.648 | 2.3720153 | 2 | zgc:114188 |
| histh1l | 1.499848 | 0.403502215 | 0.293 | 0.111 | 3.8103661 | 2 | histh1l |
| pabpc1a | 6.989137 | 0.398262282 | 0.846 | 0.551 | 1.7755903 | 2 | pabpc1a |
| irf2bp2a | 1.090457 | 0.400107174 | 0.385 | 0.172 | 2.7703083 | 2 | irf2bp2a |
| ucp2 | 1.332826 | 0.426896572 | 0.689 | 0.395 | 3.3860453 | 2 | ucp2 |

| | | | | | | | |
|---|---|---|---|---|---|---|---|
| rpl4 | 3.607039 | 0.351218364 | 0.881 | 0.642 | 9.1636846 | 2 | rpl4 |
| mknk2b | 4.511489 | 0.402469178 | 0.404 | 0.186 | 1.1461439 | 2 | mknk2b |
| stk17al | 3.286356 | 0.323166976 | 0.275 | 0.097 | 8.3489884 | 2 | stk17al |
| clic1 | 7.646030 | 0.33574392 | 0.299 | 0.118 | 1.9424739 | 2 | clic1 |
| rpl22l1 | 2.103528 | 0.420323524 | 0.683 | 0.408 | 5.3440138 | 2 | rpl22l1 |
| pycard | 6.108119 | 0.314228736 | 0.224 | 0.075 | 1.5517676 | 2 | pycard |
| rpl14 | 6.276191 | 0.350478949 | 0.906 | 0.653 | 1.5944664 | 2 | rpl14 |
| si:dkey-112e7.2 | 1.197694 | 0.388799121 | 0.417 | 0.201 | 3.0427417 | 2 | si:dkey-112e7.2 |
| dub | 1.337551 | 0.280204538 | 0.206 | 0.065 | 3.3980490 | 2 | dub |
| arpc2 | 2.030602 | 0.396055391 | 0.538 | 0.283 | 5.1587443 | 2 | arpc2 |
| glipr1a | 4.290505 | 0.279503699 | 0.195 | 0.06 | 1.0900028 | 2 | glipr1a |
| gapdhs | 6.721031 | 0.403381684 | 0.558 | 0.299 | 1.7074780 | 2 | gapdhs |
| arl5c | 9.920157 | 0.34699346 | 0.263 | 0.098 | 2.5202160 | 2 | arl5c |
| hmgn3 | 1.157390 | 0.324029752 | 0.209 | 0.069 | 2.9403494 | 2 | hmgn3 |
| tes | 2.518697 | 0.266450045 | 0.188 | 0.057 | 6.3987518 | 2 | tes |
| elmo1 | 8.314468 | 0.287551555 | 0.196 | 0.062 | 2.1122906 | 2 | elmo1 |
| notch2 | 2.824861 | 0.34528021 | 0.278 | 0.112 | 7.1765602 | 2 | notch2 |
| crema | 6.651492 | 0.397299321 | 0.354 | 0.162 | 1.6898116 | 2 | crema |
| tnrc6c1 | 7.760768 | 0.359853918 | 0.317 | 0.139 | 1.9716232 | 2 | tnrc6c1 |
| tnfb | 8.362990 | 0.277771624 | 0.156 | 0.043 | 2.1246177 | 2 | tnfb |
| rpl17 | 1.151630 | 0.284347177 | 0.939 | 0.751 | 2.9257161 | 2 | rpl17 |
| gtpbp4 | 4.060315 | 0.389054506 | 0.447 | 0.233 | 1.0315231 | 2 | gtpbp4 |
| ddx21 | 1.014245 | 0.375002436 | 0.398 | 0.197 | 2.5766898 | 2 | ddx21 |
| cd74b | 1.678298 | 0.289901529 | 0.472 | 0.209 | 4.2637168 | 2 | cd74b |
| dusp1 | 4.856923 | 0.359455972 | 0.316 | 0.138 | 1.2339014 | 2 | dusp1 |
| BX248082.1 | 6.043243 | 0.259314952 | 0.125 | 0.031 | 1.5352859 | 2 | BX248082.1 |
| cdkn1bb | 3.650307 | 0.336421901 | 0.318 | 0.143 | 9.2736055 | 2 | cdkn1bb |
| arhgdia | 4.120913 | 0.377726787 | 0.448 | 0.238 | 1.0469181 | 2 | arhgdia |
| eno1a | 1.308042 | 0.301554926 | 0.3 | 0.131 | 3.3230816 | 2 | eno1a |
| map2k6 | 1.444748 | 0.294265394 | 0.229 | 0.09 | 3.6703839 | 2 | map2k6 |
| si:dkey-187e18.1 | 2.250715 | 0.271949555 | 0.185 | 0.064 | 5.7179417 | 2 | si:dkey-187e18.1 |
| tpi1b | 1.487854 | 0.313035891 | 0.347 | 0.168 | 3.7798941 | 2 | tpi1b |
| psme2 | 1.536897 | 0.291950008 | 0.249 | 0.104 | 3.9044874 | 2 | psme2 |
| cd63 | 5.610933 | 0.348703169 | 0.618 | 0.357 | 1.4254576 | 2 | cd63 |
| grap2b | 1.569316 | 0.282492935 | 0.274 | 0.115 | 3.9868490 | 2 | grap2b |

| | | | | | | | |
|---|---|---|---|---|---|---|---|
| sept6 | 1.613547 | 0.313365706 | 0.384 | 0.19 | 4.0992170 | 2 | sept6 |
| ddb2 | 1.979406 | 0.253348429 | 0.194 | 0.07 | 5.0286820 | 2 | ddb2 |
| zgc:162730 | 1.547007 | 0.364166495 | 0.676 | 0.384 | 3.9301730 | 2 | zgc:162730 |
| ier2b | 1.790255 | 0.342230874 | 0.424 | 0.225 | 4.5481452 | 2 | ier2b |
| plk3 | 4.836108 | 0.269624218 | 0.248 | 0.1 | 1.2286134 | 2 | plk3 |
| eif3m | 8.319326 | 0.341795975 | 0.602 | 0.369 | 2.1135248 | 2 | eif3m |
| zgc:152791 | 2.065325 | 0.266356777 | 0.175 | 0.061 | 5.2469582 | 2 | zgc:152791 |
| mibp2 | 6.652102 | 0.348268669 | 0.768 | 0.504 | 1.6899667 | 2 | mibp2 |
| fabp3 | 3.131587 | 0.343112088 | 0.405 | 0.217 | 7.9557989 | 2 | fabp3 |
| psme1 | 1.889719 | 0.260083311 | 0.24 | 0.104 | 4.8008315 | 2 | psme1 |
| romo1 | 2.023205 | 0.329038649 | 0.631 | 0.394 | 5.1399525 | 2 | romo1 |
| sumo2b | 2.203590 | 0.326781254 | 0.486 | 0.284 | 5.5982207 | 2 | sumo2b |
| rps27.2 | 2.882152 | 0.350183354 | 0.819 | 0.535 | 7.3221071 | 2 | rps27.2 |
| cdc42l | 3.157323 | 0.314540637 | 0.409 | 0.222 | 8.0211803 | 2 | cdc42l |
| chmp2a | 4.825773 | 0.312572394 | 0.281 | 0.135 | 1.2259878 | 2 | chmp2a |
| CR383676.1 | 7.591936 | 0.491718557 | 0.956 | 0.7 | 1.9287315 | 2 | CR383676.1 |
| eif3ea | 8.411153 | 0.314595141 | 0.502 | 0.292 | 2.1368534 | 2 | eif3ea |
| nop53 | 8.373259 | 0.323800676 | 0.442 | 0.255 | 2.1272265 | 2 | nop53 |
| CR318588.1 | 2.226726 | 0.526391024 | 0.363 | 0.206 | 5.6569993 | 2 | CR318588.1 |
| mt-co1 | 1.812548 | 0.334645633 | 0.9 | 0.622 | 4.6047806 | 2 | mt-co1 |
| zgc:56493 | 2.821676 | 0.309061859 | 0.809 | 0.575 | 7.1684698 | 2 | zgc:56493 |
| gng5 | 6.504897 | 0.311821992 | 0.466 | 0.276 | 1.6525693 | 2 | gng5 |
| si:ch211-222l21.1 | 1.281074 | 0.343713631 | 0.507 | 0.302 | 3.2545691 | 2 | si:ch211-222l21.1 |
| cd83 | 8.085993 | 0.302463618 | 0.207 | 0.086 | 2.0542465 | 2 | cd83 |
| rnf213a | 3.689384 | 0.258054645 | 0.205 | 0.089 | 9.3728824 | 2 | rnf213a |
| rhoab | 4.455513 | 0.290722027 | 0.36 | 0.198 | 1.1319231 | 2 | rhoab |
| eif3ba | 5.399707 | 0.286974622 | 0.405 | 0.231 | 1.3717957 | 2 | eif3ba |
| gnai2a | 9.397565 | 0.254733788 | 0.285 | 0.14 | 2.3874513 | 2 | gnai2a |
| serf2 | 2.363310 | 0.298474749 | 0.574 | 0.365 | 6.0039911 | 2 | serf2 |
| ran | 3.273413 | 0.279774099 | 0.839 | 0.626 | 8.3161067 | 2 | ran |
| hmgn6 | 4.101388 | 0.300686354 | 0.499 | 0.299 | 1.0419576 | 2 | hmgn6 |
| actr2a | 4.300863 | 0.259068543 | 0.262 | 0.125 | 1.0926344 | 2 | actr2a |
| eif3g | 1.661227 | 0.289541718 | 0.475 | 0.289 | 4.2203480 | 2 | eif3g |
| slc25a3b | 1.697106 | 0.303982356 | 0.672 | 0.444 | 4.3114998 | 2 | slc25a3b |
| hmgb1a | 1.141749 | 0.2777864 | 0.384 | 0.218 | 2.9006151 | 2 | hmgb1a |

| | | | | | | | |
|---|---|---|---|---|---|---|---|
| GCA | 6.397108 | 0.262082888 | 0.345 | 0.191 | 1.6251854 | 2 | GCA |
| eif3s10 | 9.068545 | 0.266738821 | 0.312 | 0.171 | 2.3038640 | 2 | eif3s10 |
| mt-co2 | 9.659271 | 0.328333302 | 0.898 | 0.655 | 2.4539378 | 2 | mt-co2 |
| ifi45 | 1.223891 | 0.255667685 | 0.269 | 0.131 | 3.1092970 | 2 | ifi45 |
| capzb | 1.297556 | 0.273948958 | 0.432 | 0.261 | 3.2964412 | 2 | capzb |
| capza1a | 2.621640 | 0.255267296 | 0.37 | 0.211 | 6.6602775 | 2 | capza1a |
| eif3c | 3.684818 | 0.262418052 | 0.319 | 0.18 | 9.3612823 | 2 | eif3c |
| oaz1b | 2.637569 | 0.269697477 | 0.542 | 0.348 | 6.7007462 | 2 | oaz1b |
| cox7b | 2.810596 | 0.280149432 | 0.625 | 0.414 | 7.1403208 | 2 | cox7b |
| eif5a | 3.266095 | 0.285508937 | 0.708 | 0.481 | 8.2975162 | 2 | eif5a |
| ywhaba | 1.706764 | 0.2660987 | 0.498 | 0.318 | 4.3360352 | 2 | ywhaba |
| b2ml | 1.445466 | 0.262126537 | 0.454 | 0.284 | 3.6722088 | 2 | b2ml |
| atp5if1b | 2.125005 | 0.253736053 | 0.341 | 0.199 | 5.3985757 | 2 | atp5if1b |
| COX3 | 1.112559 | 0.285562134 | 0.907 | 0.66 | 2.8264577 | 2 | COX3 |
| zp2.22 | 3.217361 | 0.964500751 | 0.952 | 0.4 | 8.1737066 | 3 | zp2.2 |
| crp22 | 3.358906 | 0.968872035 | 0.925 | 0.367 | 8.5333013 | 3 | crp2 |
| si:ch211-125e6.122 | 1.005713 | 0.967020483 | 0.925 | 0.377 | 2.5550159 | 3 | si:ch211-125e6.12 |
| tmsb52 | 3.647188 | 0.97175518 | 0.934 | 0.366 | 9.2656823 | 3 | tmsb5 |
| zp32 | 7.382410 | 0.935725996 | 0.965 | 0.447 | 1.8755014 | 3 | zp3 |
| si:dkey-241l7.22 | 6.080457 | 0.914463308 | 0.98 | 0.498 | 1.5447401 | 3 | si:dkey-241l7.2 |
| zp3.22 | 4.501085 | 0.930101019 | 0.941 | 0.427 | 1.1435007 | 3 | zp3.2 |
| si:dkey-240e12.62 | 5.280553 | 0.939541534 | 0.956 | 0.434 | 1.3415247 | 3 | si:dkey-240e12.6 |
| si:dkeyp-98a7.92 | 8.333876 | 0.94458697 | 0.925 | 0.387 | 2.1172213 | 3 | si:dkeyp-98a7.9 |
| si:ch73-189n23.12 | 9.414603 | 0.945655223 | 0.934 | 0.398 | 2.3917800 | 3 | si:ch73-189n23.1 |
| BX649337.12 | 2.820002 | 0.952323372 | 0.883 | 0.324 | 7.1642151 | 3 | BX649337.1 |
| si:dkey-229d11.32 | 2.126599 | 0.959945458 | 0.87 | 0.326 | 5.4026260 | 3 | si:dkey-229d11.3 |
| si:dkey-241l7.52 | 2.685623 | 0.904877621 | 0.95 | 0.449 | 6.8228259 | 3 | si:dkey-241l7.5 |
| zp2.52 | 9.078061 | 0.92497119 | 0.946 | 0.431 | 2.3062815 | 3 | zp2.5 |
| zp2.32 | 1.569736 | 0.936629404 | 0.929 | 0.392 | 3.9879153 | 3 | zp2.3 |
| si:zfos-1505d6.32 | 9.891146 | 0.937802568 | 0.925 | 0.388 | 2.5128457 | 3 | si:zfos-1505d6.3 |
| zp2.12 | 3.588612 | 0.92795542 | 0.926 | 0.407 | 9.1168694 | 3 | zp2.1 |
| zgc:1534992 | 7.923990 | 0.950927146 | 0.937 | 0.387 | 2.0130898 | 3 | zgc:153499 |
| zp2.62 | 9.302282 | 0.936565141 | 0.908 | 0.379 | 2.3632448 | 3 | zp2.6 |
| si:ch211-226h8.142 | 1.114395 | 0.914055232 | 0.948 | 0.434 | 2.8311216 | 3 | si:ch211-226h8.14 |
| h2af1al2 | 5.096713 | 0.913127956 | 0.966 | 0.458 | 1.2948200 | 3 | h2af1al |

| | | | | | | | |
|---|---|---|---|---|---|---|---|
| cldnd2 | 3.290296 | 0.91731359 | 0.973 | 0.466 | 8.3589972 | 3 | cldnd |
| mt22 | 9.240121 | 0.924131272 | 0.946 | 0.406 | 2.3474528 | 3 | mt2 |
| cldng2 | 2.617938 | 0.916030472 | 0.914 | 0.391 | 6.6508725 | 3 | cldng |
| h1m2 | 5.270189 | 0.890407195 | 0.96 | 0.465 | 1.3388916 | 3 | h1m |
| zgc:1751352 | 1.163027 | 0.922608846 | 0.888 | 0.361 | 2.9546717 | 3 | zgc:175135 |
| zp3a.22 | 8.516492 | 0.898032808 | 0.871 | 0.373 | 2.1636148 | 3 | zp3a.2 |
| si:ch211-226h8.42 | 1.566613 | 0.919755683 | 0.853 | 0.33 | 3.9799821 | 3 | si:ch211-226h8.4 |
| acp5a2 | 7.677569 | 0.89682776 | 0.947 | 0.412 | 1.9504865 | 3 | acp5a |
| si:ch211-12h2.82 | 1.723948 | 0.876256297 | 0.907 | 0.42 | 4.3796914 | 3 | si:ch211-12h2.8 |
| si:dkey-90l23.12 | 1.304345 | 0.90605267 | 0.843 | 0.336 | 3.3136892 | 3 | si:dkey-90l23.1 |
| zgc:1717792 | 1.222259 | 0.893780612 | 0.897 | 0.389 | 3.1051512 | 3 | zgc:171779 |
| si:dkeyp-20e4.82 | 5.875353 | 0.923928095 | 0.799 | 0.304 | 1.4926334 | 3 | si:dkeyp-20e4.8 |
| LOC1104378212 | 3.554346 | 0.917015938 | 0.812 | 0.31 | 9.0298172 | 3 | LOC110437821 |
| epcam2 | 4.035929 | 0.878868595 | 0.893 | 0.389 | 1.0253277 | 3 | epcam |
| zgc:771182 | 1.084919 | 0.890684526 | 0.854 | 0.344 | 2.7562389 | 3 | zgc:77118 |
| zgc:1717762 | 1.840095 | 0.882593795 | 0.892 | 0.374 | 4.6747616 | 3 | zgc:171776 |
| si:dkey-241l7.42 | 4.743805 | 0.837251108 | 0.945 | 0.472 | 1.2051637 | 3 | si:dkey-241l7.4 |
| si:dkey-241l7.62 | 2.039173 | 0.852425044 | 0.865 | 0.405 | 5.1805197 | 3 | si:dkey-241l7.6 |
| CR790368.12 | 3.235811 | 0.861679056 | 0.893 | 0.397 | 8.2205795 | 3 | CR790368.1 |
| rrm22 | 5.487812 | 0.843720121 | 0.878 | 0.392 | 1.3941786 | 3 | rrm2 |
| LOC1005351622 | 1.203174 | 0.873821566 | 0.692 | 0.225 | 3.0566647 | 3 | LOC100535162 |
| zgc:1734432 | 1.307867 | 0.86908085 | 0.714 | 0.242 | 3.3226377 | 3 | zgc:173443 |
| si:dkeyp-98a7.42 | 1.142399 | 0.864662534 | 0.799 | 0.319 | 2.9022646 | 3 | si:dkeyp-98a7.4 |
| si:ch211-14a17.72 | 9.356036 | 0.868866366 | 0.78 | 0.307 | 2.3769010 | 3 | si:ch211-14a17.7 |
| si:dkeyp-98a7.82 | 7.553738 | 0.863330326 | 0.832 | 0.346 | 1.9190273 | 3 | si:dkeyp-98a7.8 |
| CU571163.12 | 2.400747 | 0.878334177 | 0.775 | 0.3 | 6.0990997 | 3 | CU571163.1 |
| zp3b2 | 6.844019 | 0.868221121 | 0.761 | 0.29 | 1.7387232 | 3 | zp3b |
| zgc:1952452 | 2.498106 | 0.858437705 | 0.767 | 0.289 | 6.3464405 | 3 | zgc:195245 |
| ccna12 | 7.188351 | 0.852832166 | 0.857 | 0.36 | 1.8262005 | 3 | ccna1 |
| qdprb22 | 1.336642 | 0.858617819 | 0.796 | 0.327 | 3.3957403 | 3 | qdprb2 |
| si:ch211-12h2.62 | 7.385547 | 0.835149678 | 0.826 | 0.36 | 1.8762982 | 3 | si:ch211-12h2.6 |
| zar12 | 2.454694 | 0.843335947 | 0.803 | 0.333 | 6.2361511 | 3 | zar1 |
| ccnb22 | 2.772135 | 0.851065701 | 0.753 | 0.287 | 7.0426104 | 3 | ccnb2 |
| si:dkey-229d11.52 | 1.430174 | 0.846471339 | 0.743 | 0.291 | 3.6333591 | 3 | si:dkey-229d11.5 |
| zgc:1715172 | 1.990917 | 0.84942801 | 0.738 | 0.291 | 5.0579269 | 3 | zgc:171517 |

| | | | | | | | |
|---|---|---|---|---|---|---|---|
| tuba4l2 | 7.470435 | 0.842139751 | 0.77 | 0.304 | 1.8978642 | 3 | tuba4l |
| AL929032.12 | 1.185388 | 0.847482085 | 0.675 | 0.233 | 3.0114789 | 3 | AL929032.1 |
| si:dkeyp-50f7.22 | 3.669103 | 0.837147008 | 0.76 | 0.303 | 9.3213563 | 3 | si:dkeyp-50f7.2 |
| thy12 | 8.196295 | 0.820122847 | 0.803 | 0.35 | 2.0822689 | 3 | thy1 |
| zar1l2 | 3.117810 | 0.830783429 | 0.753 | 0.299 | 7.9207985 | 3 | zar1l |
| si:dkeyp-46h3.62 | 4.184420 | 0.832102651 | 0.752 | 0.31 | 1.0630520 | 3 | si:dkeyp-46h3.6 |
| ccnb12 | 1.952527 | 0.820392052 | 0.753 | 0.298 | 4.9603951 | 3 | ccnb1 |
| slbp22 | 6.861645 | 0.819654918 | 0.762 | 0.298 | 1.7432009 | 3 | slbp2 |
| npm2b2 | 1.445791 | 0.82138206 | 0.753 | 0.298 | 3.6730332 | 3 | npm2b |
| zgc:1362542 | 1.674688 | 0.822435475 | 0.682 | 0.245 | 4.2545472 | 3 | zgc:136254 |
| mid1ip1l2 | 2.148745 | 0.77624141 | 0.862 | 0.433 | 5.4588877 | 3 | mid1ip1l |
| si:dkeyp-46h3.22 | 1.428123 | 0.806179756 | 0.738 | 0.306 | 3.6281469 | 3 | si:dkeyp-46h3.2 |
| kpna72 | 1.186245 | 0.81164911 | 0.669 | 0.244 | 3.0136569 | 3 | kpna7 |
| s100a12 | 1.573050 | 0.808073075 | 0.712 | 0.273 | 3.9963349 | 3 | s100a1 |
| si:dkeyp-46h3.82 | 5.407017 | 0.811178771 | 0.694 | 0.262 | 1.3736528 | 3 | si:dkeyp-46h3.8 |
| nanog2 | 1.784338 | 0.806804239 | 0.701 | 0.27 | 4.5331114 | 3 | nanog |
| lamp22 | 2.584639 | 0.755098214 | 0.867 | 0.458 | 6.5662754 | 3 | lamp2 |
| si:dkey-152b24.62 | 3.895714 | 0.798950467 | 0.631 | 0.219 | 9.8970636 | 3 | si:dkey-152b24.6 |
| siva12 | 2.987131 | 0.794572688 | 0.77 | 0.334 | 7.5888082 | 3 | siva1 |
| si:dkey-90l23.22 | 9.637191 | 0.801966755 | 0.676 | 0.257 | 2.4483285 | 3 | si:dkey-90l23.2 |
| uchl12 | 1.426277 | 0.788795991 | 0.684 | 0.264 | 3.6234587 | 3 | uchl1 |
| nasp2 | 2.151373 | 0.799418917 | 0.673 | 0.269 | 5.4655642 | 3 | nasp |
| cth12 | 5.972169 | 0.790295318 | 0.719 | 0.285 | 1.5172295 | 3 | cth1 |
| si:ch211-132b12.82 | 3.745250 | 0.788922114 | 0.677 | 0.256 | 9.5148096 | 3 | si:ch211-132b12.8 |
| wu:fi42e032 | 3.909942 | 0.787737017 | 0.675 | 0.262 | 9.9332081 | 3 | wu:fi42e03 |
| si:ch211-146l10.82 | 3.337007 | 0.78709547 | 0.628 | 0.227 | 8.4776676 | 3 | si:ch211-146l10.8 |
| zpcx2 | 9.170920 | 0.786375821 | 0.668 | 0.253 | 2.3298722 | 3 | zpcx |
| ddx42 | 9.444466 | 0.660396738 | 0.363 | 0.082 | 2.3993667 | 3 | ddx4 |
| LOC1005374992 | 1.550326 | 0.781662776 | 0.65 | 0.24 | 3.9386054 | 3 | LOC100537499 |
| gtf3ab2 | 2.657997 | 0.763725631 | 0.658 | 0.237 | 6.7526431 | 3 | gtf3ab |
| ctss2.12 | 2.229165 | 0.719139288 | 0.82 | 0.427 | 5.6631953 | 3 | ctss2.1 |
| btg42 | 1.420941 | 0.770814279 | 0.696 | 0.277 | 3.6099008 | 3 | btg4 |
| zgc:1714742 | 3.194204 | 0.76166923 | 0.617 | 0.226 | 8.1148753 | 3 | zgc:171474 |
| avd2 | 6.766433 | 0.747603251 | 0.573 | 0.195 | 1.7190124 | 3 | avd |
| si:ch211-145c1.12 | 8.247889 | 0.74313526 | 0.527 | 0.17 | 2.0953762 | 3 | si:ch211-145c1.1 |

| | | | | | | | |
|---|---|---|---|---|---|---|---|
| gapdh2 | 2.700366 | 0.748300859 | 0.731 | 0.321 | 6.8602803 | 3 | gapdh |
| hnrnpabb2 | 1.891035 | 0.628106573 | 0.839 | 0.54 | 4.8041767 | 3 | hnrnpabb |
| si:dkeyp-46h3.52 | 1.631515 | 0.741274551 | 0.587 | 0.215 | 4.1448649 | 3 | si:dkeyp-46h3.5 |
| LOC1018854772 | 1.217668 | 0.739337581 | 0.577 | 0.207 | 3.0934875 | 3 | LOC101885477 |
| si:dkey-208k4.22 | 1.358704 | 0.724016696 | 0.599 | 0.22 | 3.4517885 | 3 | si:dkey-208k4.2 |
| si:ch211-250e5.162 | 1.698891 | 0.730905423 | 0.632 | 0.262 | 4.3160342 | 3 | si:ch211-250e5.16 |
| zgc:566992 | 3.537759 | 0.708382185 | 0.544 | 0.188 | 8.9876779 | 3 | zgc:56699 |
| si:ch73-14h10.22 | 4.329432 | 0.726763477 | 0.561 | 0.202 | 1.0998924 | 3 | si:ch73-14h10.2 |
| si:dkey-19b23.112 | 1.011875 | 0.723843336 | 0.567 | 0.207 | 2.5706686 | 3 | si:dkey-19b23.11 |
| lsm14b2 | 2.807212 | 0.714088619 | 0.595 | 0.224 | 7.1317240 | 3 | lsm14b |
| si:ch73-90k17.12 | 2.826929 | 0.71417728 | 0.531 | 0.183 | 7.1818150 | 3 | si:ch73-90k17.1 |
| si:ch211-226l4.62 | 1.104305 | 0.717348448 | 0.587 | 0.22 | 2.8054875 | 3 | si:ch211-226l4.6 |
| si:ch211-251f6.62 | 2.503807 | 0.719523501 | 0.614 | 0.25 | 6.3609242 | 3 | si:ch211-251f6.6 |
| pigrl2.32 | 2.200309 | 0.673734084 | 0.47 | 0.149 | 5.5898856 | 3 | pigrl2.3 |
| zp3a.12 | 7.125072 | 0.71632725 | 0.659 | 0.291 | 1.8101245 | 3 | zp3a.1 |
| gmnn2 | 1.423391 | 0.715218746 | 0.593 | 0.232 | 3.6161254 | 3 | gmnn |
| zgc:554612 | 1.984213 | 0.709069856 | 0.575 | 0.219 | 5.0408939 | 3 | zgc:55461 |
| si:dkeyp-98a7.72 | 1.387138 | 0.665946924 | 0.453 | 0.142 | 3.5240244 | 3 | si:dkeyp-98a7.7 |
| si:dkey-37o8.12 | 1.750117 | 0.690187821 | 0.593 | 0.231 | 4.4461725 | 3 | si:dkey-37o8.1 |
| mkrn42 | 1.256941 | 0.669269281 | 0.499 | 0.173 | 3.1932600 | 3 | mkrn4 |
| arhgef10la2 | 8.730788 | 0.687631879 | 0.548 | 0.206 | 2.2180568 | 3 | arhgef10la |
| birc5b2 | 1.732955 | 0.680388192 | 0.534 | 0.193 | 4.4025744 | 3 | birc5b |
| zgc:562312 | 1.000273 | 0.666962266 | 0.491 | 0.168 | 2.5411943 | 3 | zgc:56231 |
| si:ch1073-263o8.22 | 2.151516 | 0.682794218 | 0.527 | 0.196 | 5.4659277 | 3 | si:ch1073-263o8.2 |
| ccna22 | 6.467309 | 0.683155158 | 0.556 | 0.218 | 1.6430200 | 3 | ccna2 |
| gdf32 | 1.903944 | 0.652185598 | 0.479 | 0.165 | 4.8369708 | 3 | gdf3 |
| ca15b2 | 6.159093 | 0.659997194 | 0.512 | 0.185 | 1.5647175 | 3 | ca15b |
| cpeb1b2 | 1.229247 | 0.668952856 | 0.538 | 0.202 | 3.1229045 | 3 | cpeb1b |
| zgc:1034822 | 3.949731 | 0.62617342 | 0.423 | 0.14 | 1.0034293 | 3 | zgc:103482 |
| CU467646.22 | 1.558481 | 0.559686235 | 0.33 | 0.089 | 3.9593233 | 3 | CU467646.2 |
| pabpn1l2 | 2.475356 | 0.646643319 | 0.547 | 0.215 | 6.2886429 | 3 | pabpn1l |
| elovl7b2 | 1.907280 | 0.640304295 | 0.501 | 0.185 | 4.8454472 | 3 | elovl7b |
| AL929192.12 | 1.247109 | 0.59278038 | 0.386 | 0.119 | 3.1682828 | 3 | AL929192.1 |
| stmn1a | 1.279818 | 0.631290286 | 0.462 | 0.17 | 3.2513796 | 3 | stmn1a |
| sybu2 | 9.058429 | 0.589061359 | 0.385 | 0.121 | 2.3012940 | 3 | sybu |

| | | | | | | | |
|---|---|---|---|---|---|---|---|
| zp3c2 | 2.967412 | 0.593804147 | 0.414 | 0.139 | 7.5387111 | 3 | zp3c |
| rbpms2b2 | 3.076227 | 0.622434441 | 0.463 | 0.173 | 7.8151569 | 3 | rbpms2b |
| org2 | 4.547806 | 0.624464102 | 0.478 | 0.182 | 1.1553703 | 3 | org |
| haus62 | 8.168441 | 0.600587404 | 0.439 | 0.156 | 2.0751924 | 3 | haus6 |
| crp32 | 1.323332 | 0.616864444 | 0.444 | 0.163 | 3.3619258 | 3 | crp3 |
| zgc:925912 | 2.111143 | 0.597062039 | 0.434 | 0.152 | 5.3633605 | 3 | zgc:92591 |
| CABZ01059627.12 | 3.432299 | 0.546412691 | 0.33 | 0.098 | 8.7197578 | 3 | CABZ01059627.1 |
| gbgt1l42 | 4.128784 | 0.560965594 | 0.353 | 0.109 | 1.0489177 | 3 | gbgt1l4 |
| patl22 | 5.038831 | 0.566648197 | 0.388 | 0.127 | 1.2801151 | 3 | patl2 |
| hsp90aa1.2 | 1.767895 | 0.432550612 | 0.174 | 0.032 | 4.4913397 | 3 | hsp90aa1.2 |
| zgc:1737422 | 2.606575 | 0.56853549 | 0.36 | 0.115 | 6.6220047 | 3 | zgc:173742 |
| slc16a32 | 8.918096 | 0.615859469 | 0.481 | 0.188 | 2.2656424 | 3 | slc16a3 |
| dctpp12 | 4.005009 | 0.574357566 | 0.391 | 0.134 | 1.0174726 | 3 | dctpp1 |
| zgc:660242 | 4.760846 | 0.542237684 | 0.337 | 0.103 | 1.2094930 | 3 | zgc:66024 |
| zgc:1628792 | 1.118197 | 0.578375395 | 0.392 | 0.134 | 2.8407804 | 3 | zgc:162879 |
| LOC1018838972 | 1.877337 | 0.551109643 | 0.371 | 0.12 | 4.7693757 | 3 | LOC101883897 |
| elovl7a2 | 2.391029 | 0.569652293 | 0.399 | 0.137 | 6.0744101 | 3 | elovl7a |
| retsatl2 | 5.781198 | 0.592416839 | 0.416 | 0.151 | 1.4687135 | 3 | retsatl |
| zglp12 | 8.012543 | 0.571303652 | 0.418 | 0.147 | 2.0355865 | 3 | zglp1 |
| tmem54a2 | 1.920288 | 0.538887414 | 0.334 | 0.104 | 4.8784926 | 3 | tmem54a |
| sinup2 | 2.376105 | 0.584143973 | 0.418 | 0.152 | 6.0364949 | 3 | sinup |
| si:ch1073-190k2.12 | 2.921072 | 0.575118333 | 0.397 | 0.139 | 7.4209844 | 3 | si:ch1073-190k2.1 |
| spink2.12 | 3.427553 | 0.597385482 | 0.688 | 0.353 | 8.7076996 | 3 | spink2.1 |
| zgc:1629452 | 6.599443 | 0.550893672 | 0.365 | 0.122 | 1.6765887 | 3 | zgc:162945 |
| zgc:1738562 | 2.129565 | 0.592932082 | 0.43 | 0.165 | 5.4101607 | 3 | zgc:173856 |
| hsd17b12a2 | 3.531269 | 0.55028099 | 0.364 | 0.122 | 8.9711892 | 3 | hsd17b12a |
| map1lc3c2 | 4.588882 | 0.564551682 | 0.415 | 0.15 | 1.1658055 | 3 | map1lc3c |
| zgc:1134242 | 1.162569 | 0.532075423 | 0.361 | 0.118 | 2.9535072 | 3 | zgc:113424 |
| buc2 | 3.206413 | 0.553825378 | 0.38 | 0.132 | 8.1458922 | 3 | buc |
| slbp3 | 6.474509 | 0.583223649 | 0.457 | 0.182 | 1.6448491 | 3 | slbp |
| larp6a2 | 5.360323 | 0.59452813 | 0.467 | 0.197 | 1.3617901 | 3 | larp6a |
| elavl22 | 8.952155 | 0.569863449 | 0.437 | 0.17 | 2.2742951 | 3 | elavl2 |
| zgc:1138862 | 1.582262 | 0.516532435 | 0.326 | 0.103 | 4.0197366 | 3 | zgc:113886 |
| pttg12 | 1.865577 | 0.544251525 | 0.357 | 0.123 | 4.7395003 | 3 | pttg1 |
| larp6b2 | 2.570324 | 0.55114223 | 0.402 | 0.146 | 6.5299094 | 3 | larp6b |

| | | | | | | | |
|---|---|---|---|---|---|---|---|
| zgc:1717502 | 2.635685 | 0.546332299 | 0.37 | 0.13 | 6.6959602 | 3 | zgc:171750 |
| mad2l12 | 3.821370 | 0.553288441 | 0.392 | 0.144 | 9.7081917 | 3 | mad2l1 |
| si:dkey-188i13.112 | 2.597129 | 0.599687875 | 0.546 | 0.274 | 6.5980080 | 3 | si:dkey-188i13.11 |
| CDPF12 | 8.079384 | 0.588308026 | 0.489 | 0.219 | 2.0525675 | 3 | CDPF1 |
| si:dkey-46g23.22 | 8.718628 | 0.516639226 | 0.326 | 0.106 | 2.2149675 | 3 | si:dkey-46g23.2 |
| snx10a2 | 2.611836 | 0.500442681 | 0.299 | 0.093 | 6.6353706 | 3 | snx10a |
| aurka2 | 1.526316 | 0.483539902 | 0.29 | 0.088 | 3.8776078 | 3 | aurka |
| chia.42 | 3.946222 | 0.505785532 | 0.328 | 0.108 | 1.0025378 | 3 | chia.4 |
| LOC1005368602 | 1.598305 | 0.577294508 | 0.554 | 0.271 | 4.0604950 | 3 | LOC100536860 |
| zgc:1719771 | 3.591478 | 0.561624406 | 0.43 | 0.174 | 9.1241509 | 3 | zgc:171977 |
| CR847984.12 | 5.703926 | 0.525309268 | 0.355 | 0.125 | 1.4490825 | 3 | CR847984.1 |
| cdc202 | 7.042582 | 0.512743233 | 0.342 | 0.116 | 1.7891681 | 3 | cdc20 |
| si:ch211-146l10.72 | 6.604193 | 0.530466703 | 0.382 | 0.141 | 1.6777953 | 3 | si:ch211-146l10.7 |
| zp2l12 | 1.648769 | 0.516176775 | 0.346 | 0.122 | 4.1886989 | 3 | zp2l1 |
| rcc12 | 6.429105 | 0.550084521 | 0.402 | 0.159 | 1.6333142 | 3 | rcc1 |
| pabpc1l2 | 7.340444 | 0.482264019 | 0.316 | 0.105 | 1.8648399 | 3 | pabpc1l |
| tac3b2 | 1.029471 | 0.471630636 | 0.283 | 0.088 | 2.6153714 | 3 | tac3b |
| cdca7a2 | 1.672693 | 0.54647918 | 0.427 | 0.177 | 4.2494778 | 3 | cdca7a |
| si:ch211-14a17.62 | 2.111569 | 0.421636243 | 0.228 | 0.061 | 5.3644423 | 3 | si:ch211-14a17.6 |
| zgc:1017442 | 3.620730 | 0.479532213 | 0.304 | 0.1 | 9.1984664 | 3 | zgc:101744 |
| BX284697.12 | 7.605725 | 0.455718703 | 0.271 | 0.083 | 1.9322345 | 3 | BX284697.1 |
| si:ch73-78o10.12 | 9.184659 | 0.469765658 | 0.302 | 0.098 | 2.3333627 | 3 | si:ch73-78o10.1 |
| si:ch73-15b2.52 | 6.309054 | 0.4264068 | 0.241 | 0.068 | 1.6028153 | 3 | si:ch73-15b2.5 |
| cdc62 | 1.470643 | 0.43636807 | 0.258 | 0.077 | 3.7361704 | 3 | cdc6 |
| snu13b2 | 5.352987 | 0.54886194 | 0.603 | 0.356 | 1.3599264 | 3 | snu13b |
| zgc:922872 | 5.721070 | 0.468686678 | 0.3 | 0.1 | 1.4534378 | 3 | zgc:92287 |
| zgc:1139842 | 1.117144 | 0.422379957 | 0.25 | 0.073 | 2.8381047 | 3 | zgc:113984 |
| zgc:1118682 | 9.365429 | 0.490071116 | 0.314 | 0.11 | 2.3792872 | 3 | zgc:111868 |
| zgc:1141232 | 9.669781 | 0.435691263 | 0.251 | 0.075 | 2.4566078 | 3 | zgc:114123 |
| plk12 | 1.316005 | 0.496244058 | 0.358 | 0.136 | 3.3433109 | 3 | plk1 |
| gyg1a2 | 1.236146 | 0.546724571 | 0.557 | 0.304 | 3.1404302 | 3 | gyg1a |
| ypel32 | 1.326280 | 0.466390374 | 0.289 | 0.099 | 3.3694158 | 3 | ypel3 |
| LOC1104377792 | 2.076864 | 0.399928942 | 0.215 | 0.059 | 5.2762732 | 3 | LOC110437779 |
| cdca82 | 3.983165 | 0.487263147 | 0.34 | 0.128 | 1.0119233 | 3 | cdca8 |
| si:dkey-241l7.32 | 6.168212 | 0.540938319 | 0.547 | 0.285 | 1.5670344 | 3 | si:dkey-241l7.3 |

| | | | | | | | |
|---|---|---|---|---|---|---|---|
| BX537276.12 | 9.23355( | 0.442643107 | 0.275 | 0.089 | 2.3457834 | 3 | BX537276.1 |
| si:ch73-160p18.32 | 2.64444 | 0.526652348 | 0.422 | 0.184 | 6.7182101 | 3 | si:ch73-160p18.3 |
| parn2 | 4.17535( | 0.485854796 | 0.317 | 0.118 | 1.0607494 | 3 | parn |
| inka1b2 | 8.76163( | 0.495247734 | 0.362 | 0.143 | 2.2258937 | 3 | inka1b |
| h3f3a2 | 6.17706 | 0.483047579 | 0.689 | 0.47 | 1.5692839 | 3 | h3f3a |
| buc2l2 | 1.82107 | 0.380600821 | 0.207 | 0.057 | 4.6264398 | 3 | buc2l |
| rasl11b2 | 2.99895( | 0.453129775 | 0.293 | 0.102 | 7.6188330 | 3 | rasl11b |
| BX001030.12 | 1.39787 | 0.431080323 | 0.27 | 0.09 | 3.5513067 | 3 | BX001030.1 |
| cx44.22 | 1.866291 | 0.42569809 | 0.263 | 0.086 | 4.7413136 | 3 | cx44.2 |
| CR383662.22 | 2.26790 | 0.376778454 | 0.217 | 0.062 | 5.7616089 | 3 | CR383662.2 |
| zgc:868392 | 2.41478( | 0.431046645 | 0.256 | 0.083 | 6.1347660 | 3 | zgc:86839 |
| piwil1 | 4.61579 | 0.33595953 | 0.142 | 0.031 | 1.1726435 | 3 | piwil1 |
| pou5f32 | 9.57352 | 0.451401485 | 0.305 | 0.11 | 2.4321534 | 3 | pou5f3 |
| pif12 | 3.36794( | 0.411574208 | 0.248 | 0.079 | 8.5562533 | 3 | pif1 |
| rbm4.32 | 1.44819( | 0.483249076 | 0.621 | 0.409 | 3.6791427 | 3 | rbm4.3 |
| rdh10b2 | 2.28793( | 0.488155266 | 0.342 | 0.139 | 5.8125030 | 3 | rdh10b |
| tph1a2 | 5.52561 | 0.443293131 | 0.294 | 0.105 | 1.4037837 | 3 | tph1a |
| CABZ01059627.22 | 7.81525 | 0.430040674 | 0.281 | 0.099 | 1.9854664 | 3 | CABZ01059627.2 |
| gstp12 | 8.20840 | 0.504546469 | 0.557 | 0.334 | 2.0853452 | 3 | gstp1 |
| kpna22 | 1.01243 | 0.459911757 | 0.322 | 0.125 | 2.5721002 | 3 | kpna2 |
| zgc:1526522 | 2.31242( | 0.418907739 | 0.252 | 0.084 | 5.8747048 | 3 | zgc:152652 |
| prr112 | 1.10034 | 0.407938742 | 0.233 | 0.075 | 2.7954356 | 3 | prr11 |
| cldn7b2 | 1.60724 | 0.438048911 | 0.32 | 0.122 | 4.0832142 | 3 | cldn7b |
| dnd11 | 3.98693 | 0.407034169 | 0.247 | 0.082 | 1.0128807 | 3 | dnd1 |
| rpain2 | 6.58589( | 0.410622827 | 0.262 | 0.091 | 1.6731454 | 3 | rpain |
| zgc:1655392 | 1.940771 | 0.416114432 | 0.258 | 0.089 | 4.9305300 | 3 | zgc:165539 |
| nfr2 | 4.01008 | 0.392195369 | 0.247 | 0.082 | 1.0187620 | 3 | nfr |
| il13ra22 | 1.43415 | 0.411042709 | 0.244 | 0.083 | 3.6434787 | 3 | il13ra2 |
| ankrd222 | 3.86024 | 0.444302535 | 0.291 | 0.111 | 9.8069611 | 3 | ankrd22 |
| zgc:566762 | 1.08775 | 0.504435775 | 0.479 | 0.261 | 2.7634341 | 3 | zgc:56676 |
| vstm2l2 | 1.26101( | 0.368346883 | 0.222 | 0.07 | 3.2036125 | 3 | vstm2l |
| BX571780.12 | 1.36202 | 0.353232949 | 0.2 | 0.059 | 3.4602225 | 3 | BX571780.1 |
| brf22 | 1.548651 | 0.388923866 | 0.222 | 0.071 | 3.9343499 | 3 | brf2 |
| zgc:923132 | 9.16805 | 0.408131824 | 0.269 | 0.098 | 2.3291445 | 3 | zgc:92313 |
| depdc7b1 | 1.16429( | 0.323973071 | 0.175 | 0.048 | 2.9578790 | 3 | depdc7b |

| | | | | | | | |
|---|---|---|---|---|---|---|---|
| zgc:1714262 | 1.249915 | 0.379420094 | 0.238 | 0.079 | 3.1754105 | 3 | zgc:171426 |
| tatdn21 | 2.269667 | 0.340170025 | 0.184 | 0.053 | 5.7660899 | 3 | tatdn2 |
| sox19b2 | 9.773533 | 0.364436411 | 0.223 | 0.072 | 2.4829661 | 3 | sox19b |
| si:dkey-216e24.32 | 1.392379 | 0.405485876 | 0.257 | 0.092 | 3.5373413 | 3 | si:dkey-216e24.3 |
| zgc:558882 | 6.253890 | 0.357477124 | 0.202 | 0.062 | 1.5888007 | 3 | zgc:55888 |
| hnrnpa0l2 | 7.512075 | 0.270839218 | 0.826 | 0.736 | 1.9084427 | 3 | hnrnpa0l |
| si:dkeyp-114g9.12 | 8.800414 | 0.377219148 | 0.23 | 0.077 | 2.2357453 | 3 | si:dkeyp-114g9.1 |
| krcp1 | 1.769885 | 0.342940733 | 0.188 | 0.056 | 4.4963932 | 3 | krcp |
| eif4e1b1 | 1.871072 | 0.366341857 | 0.203 | 0.064 | 4.7534599 | 3 | eif4e1b |
| ctsba2 | 2.312480 | 0.469174115 | 0.563 | 0.34 | 5.8748571 | 3 | ctsba |
| zgc:1141301 | 2.965929 | 0.318274953 | 0.191 | 0.057 | 7.5349435 | 3 | zgc:114130 |
| yrdc2 | 4.118073 | 0.395308584 | 0.259 | 0.094 | 1.0461964 | 3 | yrdc |
| ddx39ab2 | 7.824546 | 0.485994104 | 0.47 | 0.258 | 1.9878260 | 3 | ddx39ab |
| CU655961.32 | 1.057594 | 0.356698616 | 0.222 | 0.073 | 2.6868198 | 3 | CU655961.3 |
| tmem144b2 | 1.540927 | 0.350736273 | 0.214 | 0.069 | 3.9147257 | 3 | tmem144b |
| LOC1039099722 | 7.596254 | 0.478972096 | 0.512 | 0.311 | 1.9298284 | 3 | LOC103909972 |
| si:dkey-179j5.5 | 8.658988 | 0.253432616 | 0.119 | 0.026 | 2.1998159 | 3 | si:dkey-179j5.5 |
| gbgt1l1 | 9.985106 | 0.285099547 | 0.14 | 0.035 | 2.5367162 | 3 | gbgt1l1 |
| degs22 | 1.052518 | 0.380872654 | 0.231 | 0.08 | 2.6739235 | 3 | degs2 |
| ttc322 | 1.211355 | 0.380537117 | 0.229 | 0.08 | 3.0774486 | 3 | ttc32 |
| zgc:1529862 | 1.942134 | 0.349802662 | 0.205 | 0.066 | 4.9339915 | 3 | zgc:152986 |
| si:dkey-43p13.52 | 2.172861 | 0.342177229 | 0.196 | 0.061 | 5.5201539 | 3 | si:dkey-43p13.5 |
| BX005461.11 | 2.968902 | 0.332354328 | 0.189 | 0.058 | 7.5424970 | 3 | BX005461.1 |
| prdx12 | 3.619652 | 0.439702683 | 0.321 | 0.138 | 9.1957264 | 3 | prdx1 |
| zc3h11a2 | 4.898126 | 0.464849642 | 0.373 | 0.178 | 1.2443689 | 3 | zc3h11a |
| h2afva2 | 7.447903 | 0.473777684 | 0.499 | 0.294 | 1.8921399 | 3 | h2afva |
| zgc:1532842 | 1.199270 | 0.372881692 | 0.222 | 0.076 | 3.0467461 | 3 | zgc:153284 |
| BX548000.1 | 1.309871 | 0.268456314 | 0.124 | 0.029 | 3.3277272 | 3 | BX548000.1 |
| si:dkey-42i9.82 | 1.507141 | 0.386399384 | 0.259 | 0.096 | 3.8288932 | 3 | si:dkey-42i9.8 |
| ppp1r3b2 | 1.513531 | 0.378351239 | 0.234 | 0.083 | 3.8451258 | 3 | ppp1r3b |
| thop12 | 2.276002 | 0.386214027 | 0.253 | 0.095 | 5.7821844 | 3 | thop1 |
| kmt5ab2 | 3.024658 | 0.358040977 | 0.205 | 0.068 | 7.6841451 | 3 | kmt5ab |
| cpm2 | 3.600157 | 0.366980492 | 0.231 | 0.081 | 9.1462005 | 3 | cpm |
| srsf2a2 | 3.638006 | 0.464483692 | 0.531 | 0.337 | 9.2423566 | 3 | srsf2a |
| mcm3l2 | 4.231306 | 0.341486516 | 0.202 | 0.065 | 1.0749634 | 3 | mcm3l |

| | | | | | | | |
|---|---|---|---|---|---|---|---|
| mibp3 | 6.025052 | 0.452380704 | 0.392 | 0.188 | 1.5306645 | 3 | mibp |
| orc62 | 2.927010 | 0.378057002 | 0.234 | 0.085 | 7.4360707 | 3 | orc6 |
| fdx12 | 8.942912 | 0.460541278 | 0.377 | 0.187 | 2.2719470 | 3 | fdx1 |
| st8sia7.11 | 1.188961 | 0.313768716 | 0.174 | 0.052 | 3.0205565 | 3 | st8sia7.1 |
| si:ch211-69g19.22 | 1.581869 | 0.380260756 | 0.251 | 0.095 | 4.0187401 | 3 | si:ch211-69g19.2 |
| cox172 | 2.551374 | 0.453596577 | 0.524 | 0.33 | 6.4817667 | 3 | cox17 |
| si:ch211-113a14.271 | 3.557188 | 0.331498285 | 0.184 | 0.058 | 9.0370364 | 3 | si:ch211-113a14.27 |
| zgc:56628 | 5.506170 | 0.267986717 | 0.129 | 0.032 | 1.3988426 | 3 | zgc:56628 |
| si:dkey-68o6.52 | 6.685350 | 0.388061883 | 0.247 | 0.093 | 1.6984134 | 3 | si:dkey-68o6.5 |
| ddx192 | 8.007338 | 0.415849225 | 0.294 | 0.125 | 2.0342643 | 3 | ddx19 |
| znf395a1 | 1.149157 | 0.368986994 | 0.215 | 0.076 | 2.9194358 | 3 | znf395a |
| CU655961.42 | 2.963227 | 0.344376783 | 0.197 | 0.065 | 7.5280794 | 3 | CU655961.4 |
| si:ch211-286b5.42 | 3.975840 | 0.306951568 | 0.169 | 0.051 | 1.0100623 | 3 | si:ch211-286b5.4 |
| cx43.42 | 5.631658 | 0.398084543 | 0.257 | 0.102 | 1.4307228 | 3 | cx43.4 |
| figla1 | 1.112199 | 0.345401906 | 0.206 | 0.071 | 2.8255437 | 3 | figla |
| tmem44 | 1.222181 | 0.284828991 | 0.154 | 0.044 | 3.1049513 | 3 | tmem44 |
| ccne12 | 3.387628 | 0.358053854 | 0.232 | 0.085 | 8.6062690 | 3 | ccne1 |
| mos2 | 3.414106 | 0.394264298 | 0.263 | 0.105 | 8.6735386 | 3 | mos |
| prc1a | 4.923528 | 0.303716795 | 0.167 | 0.051 | 1.2508224 | 3 | prc1a |
| itm2cb2 | 6.583657 | 0.400472998 | 0.27 | 0.112 | 1.6725782 | 3 | itm2cb |
| ndc1 | 8.625180 | 0.359790392 | 0.217 | 0.08 | 2.1912271 | 3 | ndc1 |
| ccnh2 | 2.759811 | 0.381338313 | 0.248 | 0.097 | 7.0112998 | 3 | ccnh |
| rogdi2 | 3.706537 | 0.379365 | 0.238 | 0.092 | 9.4164584 | 3 | rogdi |
| si:ch211-202f5.32 | 3.991181 | 0.332331488 | 0.2 | 0.069 | 1.0139597 | 3 | si:ch211-202f5.3 |
| BX957362.12 | 4.916114 | 0.392072229 | 0.258 | 0.106 | 1.2489388 | 3 | BX957362.1 |
| hspa52 | 8.253982 | 0.444355635 | 0.482 | 0.284 | 2.0969243 | 3 | hspa5 |
| evla1 | 8.261859 | 0.367947861 | 0.22 | 0.082 | 2.0989254 | 3 | evla |
| gpr1602 | 8.816723 | 0.326554055 | 0.184 | 0.06 | 2.2398885 | 3 | gpr160 |
| ctsla2 | 1.080138 | 0.430175464 | 0.552 | 0.363 | 2.7440917 | 3 | ctsla |
| lpcat42 | 5.431312 | 0.345456761 | 0.223 | 0.083 | 1.3798250 | 3 | lpcat4 |
| bmp15 | 5.998986 | 0.268069729 | 0.13 | 0.035 | 1.5240425 | 3 | bmp15 |
| top2a | 6.122367 | 0.287350141 | 0.126 | 0.034 | 1.5553874 | 3 | top2a |
| ubc2 | 9.628083 | 0.284939515 | 0.738 | 0.649 | 2.4460145 | 3 | ubc |
| bokb1 | 1.122421 | 0.28561301 | 0.15 | 0.044 | 2.8515106 | 3 | bokb |
| dazl2 | 1.189780 | 0.349922702 | 0.221 | 0.082 | 3.0226366 | 3 | dazl |

| | | | | | | | |
|---|---|---|---|---|---|---|---|
| lmnl31 | 1.337557 | 0.307270502 | 0.175 | 0.057 | 3.3980643 | 3 | lmnl3 |
| dynll2a2 | 1.407825 | 0.376552516 | 0.265 | 0.109 | 3.5765799 | 3 | dynll2a |
| umps1 | 1.483884 | 0.320138064 | 0.18 | 0.06 | 3.7698087 | 3 | umps |
| exd21 | 2.146549 | 0.340101806 | 0.202 | 0.072 | 5.4533102 | 3 | exd2 |
| si:ch211-226h8.62 | 2.250867 | 0.301110832 | 0.178 | 0.058 | 5.7183285 | 3 | si:ch211-226h8.6 |
| zgc:1630302 | 4.210529 | 0.375718462 | 0.246 | 0.098 | 1.0696849 | 3 | zgc:163030 |
| si:ch211-137a8.2 | 4.378849 | 0.275238864 | 0.142 | 0.041 | 1.1124467 | 3 | si:ch211-137a8.2 |
| aurkb2 | 4.487424 | 0.375368812 | 0.251 | 0.103 | 1.1400301 | 3 | aurkb |
| si:ch73-303b9.12 | 4.984115 | 0.295464016 | 0.178 | 0.058 | 1.2662145 | 3 | si:ch73-303b9.1 |
| si:dkey-87o1.21 | 8.180722 | 0.312326 | 0.185 | 0.062 | 2.0783124 | 3 | si:dkey-87o1.2 |
| cax1 | 9.402404 | 0.281901549 | 0.145 | 0.042 | 2.3886808 | 3 | cax1 |
| mylipa2 | 1.324716 | 0.3710149 | 0.231 | 0.091 | 3.3654416 | 3 | mylipa |
| si:ch211-5k11.82 | 2.108042 | 0.371232994 | 0.516 | 0.284 | 5.3554826 | 3 | si:ch211-5k11.8 |
| nek22 | 3.447425 | 0.294786867 | 0.16 | 0.05 | 8.7581840 | 3 | nek2 |
| snu13a2 | 3.646612 | 0.353926335 | 0.216 | 0.082 | 9.2642194 | 3 | snu13a |
| tspan13a | 5.432292 | 0.290491244 | 0.151 | 0.046 | 1.3800738 | 3 | tspan13a |
| metap1d2 | 7.252222 | 0.336208361 | 0.196 | 0.071 | 1.8424272 | 3 | metap1d |
| tpx22 | 9.416427 | 0.360403316 | 0.244 | 0.099 | 2.3922434 | 3 | tpx2 |
| lrp13 | 9.564786 | 0.298861143 | 0.161 | 0.052 | 2.4299341 | 3 | lrp13 |
| ube2c1 | 2.138999 | 0.364351895 | 0.27 | 0.115 | 5.4341279 | 3 | ube2c |
| hmgb2a | 2.533396 | 0.317255206 | 0.747 | 0.601 | 6.4360945 | 3 | hmgb2a |
| zfand62 | 3.030483 | 0.375608417 | 0.248 | 0.104 | 7.6989421 | 3 | zfand6 |
| nanos32 | 4.501186 | 0.336592871 | 0.209 | 0.077 | 1.1435265 | 3 | nanos3 |
| casd1 | 7.796106 | 0.305336556 | 0.174 | 0.059 | 1.9806008 | 3 | casd1 |
| hells1 | 1.096190 | 0.382820792 | 0.266 | 0.116 | 2.7848730 | 3 | hells |
| hbba12 | 1.688962 | 0.41320605 | 0.509 | 0.282 | 4.2908094 | 3 | hbba1 |
| rpz22 | 1.920875 | 0.408052718 | 0.318 | 0.155 | 4.8799841 | 3 | rpz2 |
| faub2 | 2.496317 | 0.419383992 | 0.34 | 0.172 | 6.3418955 | 3 | faub |
| tram22 | 3.508335 | 0.373822745 | 0.261 | 0.114 | 8.9129257 | 3 | tram2 |
| BX005461.31 | 3.618700 | 0.284058679 | 0.157 | 0.05 | 9.1933081 | 3 | BX005461.3 |
| rbbp41 | 5.057632 | 0.404975339 | 0.315 | 0.154 | 1.2848916 | 3 | rbbp4 |
| si:ch73-340n8.42 | 6.838357 | 0.367211618 | 0.249 | 0.106 | 1.7372848 | 3 | si:ch73-340n8.4 |
| aif1l1 | 7.696520 | 0.391739396 | 0.328 | 0.161 | 1.9553010 | 3 | aif1l |
| ddx411 | 7.949955 | 0.278828391 | 0.158 | 0.051 | 2.0196861 | 3 | ddx41 |
| tmem702 | 8.021460 | 0.334131144 | 0.216 | 0.084 | 2.0378519 | 3 | tmem70 |

| | | | | | | | |
|---|---|---|---|---|---|---|---|
| npm2a2 | 1.04042( | 0.298916456 | 0.173 | 0.059 | 2.6431873 | 3 | npm2a |
| CT009596.12 | 1.138311 | 0.308562744 | 0.189 | 0.067 | 2.8918807 | 3 | CT009596.1 |
| LOC100334439 | 1.20046¢ | 0.256433953 | 0.136 | 0.04 | 3.0497939 | 3 | LOC100334439 |
| cdca92 | 1.362805 | 0.303209255 | 0.184 | 0.064 | 3.4622065 | 3 | cdca9 |
| fbxo321 | 2.090342 | 0.329486867 | 0.21 | 0.08 | 5.3105140 | 3 | fbxo32 |
| asz11 | 2.42076( | 0.287989296 | 0.165 | 0.055 | 6.1499433 | 3 | asz1 |
| pop42 | 3.040995 | 0.358044519 | 0.232 | 0.096 | 7.7256499 | 3 | pop4 |
| wu:fa19b12 | 3.058864 | 0.289693187 | 0.165 | 0.055 | 7.7710444 | 3 | wu:fa19b12 |
| g3bp12 | 3.901222 | 0.419662953 | 0.348 | 0.184 | 9.9110545 | 3 | g3bp1 |
| si:ch211-210c8.72 | 5.756208 | 0.370341759 | 0.248 | 0.107 | 1.4623646 | 3 | si:ch211-210c8.7 |
| cited11 | 6.996488 | 0.266633683 | 0.15 | 0.047 | 1.7774579 | 3 | cited1 |
| sec61a1l | 7.53505( | 0.372396888 | 0.247 | 0.108 | 1.9142796 | 3 | sec61a1l |
| zgc:113425 | 1.689483 | 0.283556957 | 0.157 | 0.051 | 4.2921331 | 3 | zgc:113425 |
| gpx8 | 3.705155 | 0.314376288 | 0.182 | 0.066 | 9.4129475 | 3 | gpx8 |
| rnf42 | 4.061424 | 0.357717148 | 0.251 | 0.109 | 1.0318047 | 3 | rnf4 |
| exd1 | 4.247005 | 0.281381761 | 0.143 | 0.045 | 1.0789517 | 3 | exd1 |
| tent5c2 | 6.875252 | 0.365319122 | 0.26 | 0.115 | 1.7466578 | 3 | tent5c |
| coil1 | 6.887992 | 0.34441744 | 0.22 | 0.089 | 1.7498945 | 3 | coil |
| si:dkey-189g17.2 | 7.599879 | 0.265680676 | 0.14 | 0.044 | 1.9307494 | 3 | si:dkey-189g17.2 |
| eif4a1a2 | 8.709093 | 0.38743936 | 0.55 | 0.396 | 2.2125451 | 3 | eif4a1a |
| zgc:1102391 | 1.277871 | 0.408410473 | 0.349 | 0.188 | 3.2464315 | 3 | zgc:110239 |
| rpp30 | 2.348978 | 0.288507688 | 0.151 | 0.05 | 5.9675790 | 3 | rpp30 |
| zgc:1735441 | 2.825953 | 0.279661182 | 0.166 | 0.057 | 7.1793351 | 3 | zgc:173544 |
| zgc:101016 | 6.527487 | 0.325378226 | 0.203 | 0.081 | 1.6583082 | 3 | zgc:101016 |
| pias4b | 7.666894 | 0.26761067 | 0.148 | 0.048 | 1.9477746 | 3 | pias4b |
| marcksb2 | 7.687243 | 0.382676461 | 0.307 | 0.152 | 1.9529441 | 3 | marcksb |
| spsb12 | 9.395632 | 0.309576665 | 0.191 | 0.072 | 2.3869603 | 3 | spsb1 |
| LOC1005380231 | 9.476726 | 0.276448182 | 0.166 | 0.057 | 2.4075623 | 3 | LOC100538023 |
| CABZ01075618.1 | 1.011543 | 0.253530821 | 0.133 | 0.041 | 2.5698262 | 3 | CABZ01075618.1 |
| micall2a | 1.174694 | 0.25984089 | 0.132 | 0.04 | 2.9843118 | 3 | micall2a |
| st3gal8 | 1.243799 | 0.296107746 | 0.167 | 0.059 | 3.1598727 | 3 | st3gal8 |
| efcab11 | 2.122629 | 0.254704129 | 0.145 | 0.046 | 5.3925391 | 3 | efcab11 |
| zgc:1751362 | 2.703144 | 0.328653682 | 0.205 | 0.082 | 6.8673388 | 3 | zgc:175136 |
| spryd41 | 3.053779 | 0.300686883 | 0.173 | 0.063 | 7.7581269 | 3 | spryd4 |
| CCDC69 | 4.13447( | 0.273122261 | 0.148 | 0.049 | 1.0503621 | 3 | CCDC69 |

| | | | | | | | |
|---|---|---|---|---|---|---|---|
| bmb2 | 4.358812 | 0.290583917 | 0.182 | 0.067 | 1.1073561 | 3 | bmb |
| bzw1b2 | 1.217395 | 0.391728895 | 0.525 | 0.369 | 3.0927936 | 3 | bzw1b |
| alg13 | 1.431892 | 0.261136792 | 0.148 | 0.049 | 3.6377231 | 3 | alg13 |
| zgc:171566 | 1.515132 | 0.280351106 | 0.157 | 0.054 | 3.8491931 | 3 | zgc:171566 |
| fbxo43 | 5.763108 | 0.272525014 | 0.154 | 0.053 | 1.4641178 | 3 | fbxo43 |
| si:dkey-81e3.21 | 6.027113 | 0.257705951 | 0.145 | 0.048 | 1.5311880 | 3 | si:dkey-81e3.2 |
| si:ch211-226h8.8 | 9.465038 | 0.309080991 | 0.189 | 0.074 | 2.4045930 | 3 | si:ch211-226h8.8 |
| akap1a | 1.077535 | 0.250327097 | 0.139 | 0.045 | 2.7374790 | 3 | akap1a |
| rbm72 | 1.227424 | 0.345155189 | 0.23 | 0.101 | 3.1182713 | 3 | rbm7 |
| ncbp21 | 1.362715 | 0.384676649 | 0.299 | 0.153 | 3.4619793 | 3 | ncbp2 |
| rpa32 | 1.970954 | 0.376412643 | 0.29 | 0.146 | 5.0072109 | 3 | rpa3 |
| COQ10A1 | 2.260699 | 0.294688064 | 0.173 | 0.064 | 5.7433068 | 3 | COQ10A |
| pon3.2 | 8.004745 | 0.315946292 | 0.194 | 0.079 | 2.0336057 | 3 | pon3.2 |
| setd4 | 9.313706 | 0.270152635 | 0.152 | 0.054 | 2.3661472 | 3 | setd4 |
| cfl1l1 | 1.010101 | 0.341892746 | 0.274 | 0.129 | 2.5661638 | 3 | cfl1l |
| dynll12 | 1.589759 | 0.359607741 | 0.584 | 0.433 | 4.0387827 | 3 | dynll1 |
| ccdc581 | 2.558748 | 0.329834644 | 0.214 | 0.093 | 6.5004993 | 3 | ccdc58 |
| e2f51 | 3.697329 | 0.271683976 | 0.156 | 0.056 | 9.3930660 | 3 | e2f5 |
| slc29a2 | 6.673358 | 0.272696394 | 0.158 | 0.058 | 1.6953667 | 3 | slc29a2 |
| mrps18b1 | 7.475059 | 0.34554365 | 0.235 | 0.108 | 1.8990387 | 3 | mrps18b |
| selenoh2 | 1.728334 | 0.387471184 | 0.373 | 0.219 | 4.3908341 | 3 | selenoh |
| pim31 | 4.814056 | 0.328907096 | 0.237 | 0.109 | 1.2230110 | 3 | pim3 |
| eno3 | 5.319908 | 0.368037688 | 0.5 | 0.355 | 1.3515228 | 3 | eno3 |
| zgc:1129802 | 6.129894 | 0.281792161 | 0.182 | 0.072 | 1.5572996 | 3 | zgc:112980 |
| h2afx2 | 6.850670 | 0.322712609 | 0.249 | 0.115 | 1.7404128 | 3 | h2afx |
| lman1 | 9.216955 | 0.311847935 | 0.193 | 0.081 | 2.3415676 | 3 | lman1 |
| chrac12 | 1.071201 | 0.339346971 | 0.243 | 0.115 | 2.7213863 | 3 | chrac1 |
| vps9d12 | 1.248434 | 0.28474691 | 0.187 | 0.076 | 3.1716486 | 3 | vps9d1 |
| chchd102 | 1.625283 | 0.341922247 | 0.262 | 0.126 | 4.1290320 | 3 | chchd10 |
| polr2k2 | 2.315180 | 0.377902735 | 0.309 | 0.168 | 5.8817155 | 3 | polr2k |
| snrpb22 | 3.048869 | 0.356340477 | 0.263 | 0.131 | 7.7456525 | 3 | snrpb2 |
| cdk1 | 3.269778 | 0.293513255 | 0.187 | 0.077 | 8.3068725 | 3 | cdk1 |
| slc35f2 | 4.515641 | 0.272079089 | 0.159 | 0.06 | 1.1471986 | 3 | slc35f2 |
| zgc:771121 | 4.585139 | 0.309530863 | 0.214 | 0.094 | 1.1648546 | 3 | zgc:77112 |
| gemin72 | 4.778299 | 0.353734229 | 0.267 | 0.135 | 1.2139269 | 3 | gemin7 |

| | | | | | | | |
|---|---|---|---|---|---|---|---|
| si:ch211-157c3.41 | 4.896671 | 0.265914697 | 0.177 | 0.069 | 1.2439993 | 3 | si:ch211-157c3.4 |
| gtf2a12 | 4.978733 | 0.353585425 | 0.267 | 0.134 | 1.2648473 | 3 | gtf2a1 |
| fam122b | 5.191096 | 0.270323218 | 0.15 | 0.055 | 1.3187979 | 3 | fam122b |
| cdt12 | 6.759694 | 0.275618416 | 0.173 | 0.068 | 1.7173004 | 3 | cdt1 |
| oclna1 | 1.692208 | 0.272363943 | 0.16 | 0.061 | 4.2990560 | 3 | oclna |
| hsp90b11 | 1.955147 | 0.371296283 | 0.349 | 0.202 | 4.9670521 | 3 | hsp90b1 |
| dph31 | 2.239140 | 0.266118905 | 0.159 | 0.061 | 5.6885357 | 3 | dph3 |
| pus31 | 2.281626 | 0.26521306 | 0.151 | 0.056 | 5.7964731 | 3 | pus3 |
| mcm7 | 2.620198 | 0.348225979 | 0.272 | 0.14 | 6.6566144 | 3 | mcm7 |
| lipg2 | 2.276349 | 0.253085605 | 0.166 | 0.065 | 5.7830656 | 3 | lipg |
| snrpg1 | 3.998227 | 0.376985153 | 0.397 | 0.257 | 1.0157496 | 3 | snrpg |
| fbxw11b2 | 4.244002 | 0.333033464 | 0.25 | 0.125 | 1.0781889 | 3 | fbxw11b |
| khsrp | 5.295287 | 0.372583625 | 0.32 | 0.186 | 1.3452678 | 3 | khsrp |
| pcna | 8.270998 | 0.334040577 | 0.271 | 0.141 | 2.1012471 | 3 | pcna |
| imp32 | 1.277319 | 0.331187039 | 0.239 | 0.117 | 3.2450305 | 3 | imp3 |
| hmgb3b2 | 1.761339 | 0.362086556 | 0.295 | 0.164 | 4.4746836 | 3 | hmgb3b |
| rac1a1 | 1.914974 | 0.341082989 | 0.258 | 0.134 | 4.8649920 | 3 | rac1a |
| prrg22 | 2.011601 | 0.276668055 | 0.174 | 0.072 | 5.1104726 | 3 | prrg2 |
| nutf2l2 | 2.638266 | 0.364549951 | 0.353 | 0.214 | 6.7025157 | 3 | nutf2l |
| cdc34a | 3.346227 | 0.257676468 | 0.148 | 0.057 | 8.5010903 | 3 | cdc34a |
| zgc:112185 | 3.632340 | 0.26739166 | 0.163 | 0.066 | 9.2279609 | 3 | zgc:112185 |
| BX323557.1 | 4.782155 | 0.286205871 | 0.176 | 0.075 | 1.2149064 | 3 | BX323557.1 |
| gfer2 | 5.545363 | 0.276950714 | 0.187 | 0.081 | 1.4087996 | 3 | gfer |
| nap1l1 | 8.832664 | 0.351775717 | 0.474 | 0.35 | 2.2439383 | 3 | nap1l1 |
| cnih1 | 8.978458 | 0.314070614 | 0.22 | 0.106 | 2.2809774 | 3 | cnih1 |
| golt1bb | 1.035091 | 0.286202674 | 0.179 | 0.078 | 2.6296511 | 3 | golt1bb |
| aktip2 | 1.079438 | 0.326157099 | 0.239 | 0.119 | 2.7423129 | 3 | aktip |
| si:dkey-92i17.22 | 1.112255 | 0.263446972 | 0.174 | 0.072 | 2.8256854 | 3 | si:dkey-92i17.2 |
| rcor1 | 1.145935 | 0.299362593 | 0.203 | 0.094 | 2.9112502 | 3 | rcor1 |
| ncl | 5.097591 | 0.325663932 | 0.519 | 0.409 | 1.2950430 | 3 | ncl |
| slc25a25a2 | 5.120296 | 0.346226488 | 0.281 | 0.156 | 1.3008112 | 3 | slc25a25a |
| dnajc9 | 5.829299 | 0.278837483 | 0.184 | 0.082 | 1.4809335 | 3 | dnajc9 |
| hmga1a1 | 6.103977 | 0.319899833 | 0.559 | 0.436 | 1.5507153 | 3 | hmga1a |
| pithd11 | 1.096023 | 0.259126861 | 0.155 | 0.063 | 2.7844484 | 3 | pithd1 |
| cnep1r11 | 4.383607 | 0.307140005 | 0.215 | 0.105 | 1.1136553 | 3 | cnep1r1 |

| | | | | | | | |
|---|---|---|---|---|---|---|---|
| cbx3a | 4.828276 | 0.355599145 | 0.415 | 0.285 | 1.2266235 | 3 | cbx3a |
| nfs1 | 5.389237 | 0.266433907 | 0.169 | 0.074 | 1.3691357 | 3 | nfs1 |
| si:ch73-281n10.22 | 7.847695 | 0.322104631 | 0.527 | 0.389 | 1.9937071 | 3 | si:ch73-281n10.2 |
| lrrc591 | 8.797808 | 0.3147038 | 0.247 | 0.128 | 2.2350833 | 3 | lrrc59 |
| rangrf | 1.083199 | 0.250882549 | 0.152 | 0.062 | 2.7518671 | 3 | rangrf |
| srsf7a | 1.724998 | 0.357897362 | 0.379 | 0.256 | 4.3823587 | 3 | srsf7a |
| mcfd2 | 2.779988 | 0.294842699 | 0.209 | 0.102 | 7.0625612 | 3 | mcfd2 |
| asf1ba1 | 3.286644 | 0.269525975 | 0.18 | 0.081 | 8.3497213 | 3 | asf1ba |
| prpf31 | 3.764035 | 0.264384185 | 0.164 | 0.071 | 9.5625321 | 3 | prpf31 |
| dcun1d12 | 3.976848 | 0.27267426 | 0.182 | 0.082 | 1.0103184 | 3 | dcun1d1 |
| sumo3b2 | 4.404931 | 0.323605247 | 0.216 | 0.109 | 1.1190728 | 3 | sumo3b |
| tipin | 6.367370 | 0.250854611 | 0.13 | 0.051 | 1.6176304 | 3 | tipin |
| STMP11 | 9.620704 | 0.360613724 | 0.405 | 0.287 | 2.4441399 | 3 | STMP1 |
| top1l1 | 1.082482 | 0.341220886 | 0.336 | 0.211 | 2.7500477 | 3 | top1l |
| fen1 | 1.102959 | 0.26025932 | 0.175 | 0.079 | 2.8020676 | 3 | fen1 |
| gar1 | 1.225258 | 0.29760586 | 0.216 | 0.109 | 3.1127698 | 3 | gar1 |
| mcm4 | 1.417533 | 0.262203535 | 0.169 | 0.076 | 3.6012442 | 3 | mcm4 |
| si:dkeyp-75b4.102 | 1.644643 | 0.269733037 | 0.536 | 0.356 | 4.1782158 | 3 | si:dkeyp-75b4.10 |
| tacc31 | 1.871529 | 0.256645357 | 0.173 | 0.077 | 4.7546203 | 3 | tacc3 |
| f11r.12 | 1.882953 | 0.322149605 | 0.28 | 0.158 | 4.7836437 | 3 | f11r.1 |
| spcs3 | 1.948873 | 0.345013508 | 0.377 | 0.254 | 4.9511120 | 3 | spcs3 |
| lsm12a1 | 2.077486 | 0.294393288 | 0.219 | 0.11 | 5.2778557 | 3 | lsm12a |
| mrpl48 | 2.568249 | 0.254970969 | 0.161 | 0.07 | 6.5246373 | 3 | mrpl48 |
| btg32 | 3.068686 | 0.332242551 | 0.274 | 0.155 | 7.7959973 | 3 | btg3 |
| mrgbp | 3.521177 | 0.277705105 | 0.179 | 0.083 | 8.9455523 | 3 | mrgbp |
| snupn | 6.007745 | 0.250901254 | 0.152 | 0.066 | 1.5262678 | 3 | snupn |
| ckbb1 | 1.065108 | 0.322835548 | 0.388 | 0.243 | 2.7059079 | 3 | ckbb |
| tomm20a | 1.111476 | 0.336529301 | 0.345 | 0.221 | 2.8237048 | 3 | tomm20a |
| aldh18a1 | 1.179779 | 0.25660029 | 0.155 | 0.067 | 2.9972297 | 3 | aldh18a1 |
| selenom | 3.269267 | 0.287609139 | 0.209 | 0.106 | 8.3055732 | 3 | selenom |
| UBL4A | 3.269514 | 0.302566374 | 0.216 | 0.113 | 8.3062025 | 3 | UBL4A |
| npm1a | 6.172831 | 0.333517504 | 0.413 | 0.294 | 1.5682077 | 3 | npm1a |
| chtopa2 | 8.838118 | 0.32715467 | 0.322 | 0.2 | 2.2453239 | 3 | chtopa |
| mat2aa | 9.765930 | 0.316134789 | 0.277 | 0.162 | 2.4810345 | 3 | mat2aa |
| eif5a2 | 1.169691 | 0.303649665 | 0.504 | 0.403 | 2.9716024 | 3 | eif5a2 |

| | | | | | | | |
|---|---|---|---|---|---|---|---|
| pttg1ipb2 | 1.185804 | 0.271156025 | 0.195 | 0.096 | 3.0125372 | 3 | pttg1ipb |
| timm131 | 1.527485 | 0.276579586 | 0.188 | 0.092 | 3.8805764 | 3 | timm13 |
| fbl | 1.898169 | 0.330679184 | 0.311 | 0.195 | 4.8222993 | 3 | fbl |
| mpzl2b | 2.761836 | 0.260496805 | 0.176 | 0.083 | 7.0164451 | 3 | mpzl2b |
| c1d1 | 2.814475 | 0.294001665 | 0.216 | 0.114 | 7.1501757 | 3 | c1d |
| ptgr1 | 3.587922 | 0.285039407 | 0.201 | 0.103 | 9.1151177 | 3 | ptgr1 |
| paip2b | 4.195955 | 0.331453361 | 0.408 | 0.303 | 1.0659824 | 3 | paip2b |
| naa40 | 4.335682 | 0.266510252 | 0.188 | 0.093 | 1.1014801 | 3 | naa40 |
| akirin1 | 4.548580 | 0.299450144 | 0.237 | 0.131 | 1.1555668 | 3 | akirin1 |
| hnrnpaba | 5.638658 | 0.26859553 | 0.545 | 0.464 | 1.4325011 | 3 | hnrnpaba |
| arpp19b | 5.876436 | 0.297082929 | 0.233 | 0.128 | 1.4929087 | 3 | arpp19b |
| cycsb1 | 7.621642 | 0.320240127 | 0.425 | 0.308 | 1.9362783 | 3 | cycsb |
| tssc4 | 8.117473 | 0.253046071 | 0.157 | 0.072 | 2.0622440 | 3 | tssc4 |
| nifk1 | 9.020954 | 0.278470632 | 0.214 | 0.112 | 2.2917734 | 3 | nifk |
| pdia4 | 1.161029 | 0.283717506 | 0.221 | 0.118 | 2.9495943 | 3 | pdia4 |
| pprc1 | 1.357149 | 0.254080142 | 0.169 | 0.081 | 3.4478387 | 3 | pprc1 |
| atp5mc1 | 3.258890 | 0.317282217 | 0.435 | 0.324 | 8.2792102 | 3 | atp5mc1 |
| lbh1 | 4.747878 | 0.290116164 | 0.257 | 0.147 | 1.2061985 | 3 | lbh |
| setb | 7.061175 | 0.312265637 | 0.456 | 0.356 | 1.7938917 | 3 | setb |
| ppm1g | 7.977953 | 0.292436691 | 0.232 | 0.13 | 2.0267991 | 3 | ppm1g |
| ivns1abpb | 1.092027 | 0.261093969 | 0.193 | 0.099 | 2.7742966 | 3 | ivns1abpb |
| dap1b1 | 1.109347 | 0.313220682 | 0.358 | 0.244 | 2.8182975 | 3 | dap1b |
| txnl4a | 1.162829 | 0.266118439 | 0.201 | 0.105 | 2.9541692 | 3 | txnl4a |
| rbm4.22 | 1.246360 | 0.304675322 | 0.258 | 0.151 | 3.1663785 | 3 | rbm4.2 |
| sub1b | 1.575282 | 0.316890628 | 0.324 | 0.21 | 4.0020053 | 3 | sub1b |
| ssb | 1.613545 | 0.278132709 | 0.189 | 0.099 | 4.0992112 | 3 | ssb |
| abcf2a2 | 1.729168 | 0.274495408 | 0.237 | 0.132 | 4.3929528 | 3 | abcf2a |
| mrps18c | 1.953237 | 0.288011755 | 0.205 | 0.111 | 4.9621998 | 3 | mrps18c |
| nol72 | 2.568285 | 0.273297497 | 0.204 | 0.108 | 6.5247286 | 3 | nol7 |
| thoc71 | 3.353123 | 0.279157408 | 0.196 | 0.103 | 8.5186112 | 3 | thoc7 |
| ebna1bp2 | 4.042930 | 0.286023277 | 0.23 | 0.129 | 1.0271065 | 3 | ebna1bp2 |
| hprt11 | 4.738022 | 0.267818244 | 0.204 | 0.108 | 1.2036946 | 3 | hprt1 |
| idh1 | 5.945307 | 0.286204963 | 0.207 | 0.113 | 1.5104052 | 3 | idh1 |
| ube2al | 9.449660 | 0.291594518 | 0.246 | 0.144 | 2.4006861 | 3 | ube2al |
| uck2a | 1.322962 | 0.268134551 | 0.196 | 0.104 | 3.3609850 | 3 | uck2a |

| | | | | | | | |
|---|---|---|---|---|---|---|---|
| anp32b1 | 1.533029 | 0.314869024 | 0.358 | 0.247 | 3.8946621 | 3 | anp32b |
| uqcr10 | 1.844685 | 0.302478532 | 0.442 | 0.343 | 4.6864238 | 3 | uqcr10 |
| hbaa12 | 2.812215 | 0.282644757 | 0.361 | 0.227 | 7.1444323 | 3 | hbaa1 |
| eloca | 5.814120 | 0.259276578 | 0.175 | 0.09 | 1.4770772 | 3 | eloca |
| lsm6 | 6.235906 | 0.310172542 | 0.328 | 0.22 | 1.5842321 | 3 | lsm6 |
| tmco11 | 6.309354 | 0.259068245 | 0.198 | 0.106 | 1.6028915 | 3 | tmco1 |
| psme32 | 2.846780 | 0.271640689 | 0.212 | 0.12 | 7.2322467 | 3 | psme3 |
| dkc11 | 2.975539 | 0.292851717 | 0.269 | 0.169 | 7.5593587 | 3 | dkc1 |
| glulb1 | 4.840484 | 0.257715659 | 0.233 | 0.134 | 1.2297250 | 3 | glulb |
| snrpe | 6.296848 | 0.308686243 | 0.378 | 0.279 | 1.5997144 | 3 | snrpe |
| ube2g1b | 7.058485 | 0.285261859 | 0.217 | 0.128 | 1.7932081 | 3 | ube2g1b |
| h2afvb | 8.634962 | 0.253356931 | 0.571 | 0.47 | 2.1937122 | 3 | h2afvb |
| zgc:171480 | 9.010078 | 0.269891266 | 0.185 | 0.101 | 2.2890105 | 3 | zgc:171480 |
| sinhcafl | 1.391997 | 0.265657895 | 0.211 | 0.121 | 3.5363706 | 3 | sinhcafl |
| rpf1 | 1.474514 | 0.280141814 | 0.216 | 0.127 | 3.7460045 | 3 | rpf1 |
| alyref1 | 1.483895 | 0.300062615 | 0.349 | 0.246 | 3.7698377 | 3 | alyref |
| zfand5b1 | 2.413690 | 0.295626143 | 0.299 | 0.2 | 6.1319809 | 3 | zfand5b |
| cdc14b1 | 7.273271 | 0.251621301 | 0.209 | 0.12 | 1.8477746 | 3 | cdc14b |
| tspan36 | 1.109593 | 0.266321653 | 0.248 | 0.152 | 2.8189217 | 3 | tspan36 |
| cbx51 | 1.465832 | 0.292312063 | 0.295 | 0.2 | 3.7239463 | 3 | cbx5 |
| snrpa1 | 1.760912 | 0.255816713 | 0.202 | 0.117 | 4.4735970 | 3 | snrpa1 |
| FQ323156.11 | 2.007293 | 0.256920717 | 0.242 | 0.148 | 5.0995293 | 3 | FQ323156.1 |
| lsm12 | 2.592489 | 0.251921453 | 0.21 | 0.122 | 6.5862184 | 3 | lsm1 |
| lsm8 | 5.381984 | 0.267648974 | 0.237 | 0.148 | 1.3672932 | 3 | lsm8 |
| ddx61 | 8.515376 | 0.277135349 | 0.265 | 0.173 | 2.1633314 | 3 | ddx6 |
| pcf112 | 1.064333 | 0.2745528 | 0.309 | 0.212 | 2.7039399 | 3 | pcf11 |
| hspd1 | 5.566732 | 0.272876436 | 0.244 | 0.16 | 1.4142283 | 3 | hspd1 |
| snrpb | 5.995197 | 0.2652442 | 0.426 | 0.354 | 1.5230798 | 3 | snrpb |
| sap181 | 1.870175 | 0.272638742 | 0.304 | 0.217 | 4.7511806 | 3 | sap18 |
| snrpd2 | 3.416241 | 0.26755782 | 0.342 | 0.255 | 8.6789612 | 3 | snrpd2 |
| snrpc1 | 6.633851 | 0.262663617 | 0.257 | 0.172 | 1.6853300 | 3 | snrpc |
| hnrnpa1b | 1.615696 | 0.266970999 | 0.346 | 0.269 | 4.1046773 | 3 | hnrnpa1b |
| ilf2 | 2.128821 | 0.27336547 | 0.32 | 0.24 | 5.4082717 | 3 | ilf2 |
| ranbp1 | 5.493581 | 0.262707835 | 0.353 | 0.279 | 1.3956444 | 3 | ranbp1 |
| rtn4a1 | 7.486379 | 0.253103632 | 0.317 | 0.236 | 1.9019147 | 3 | rtn4a |

| | | | | | | | |
|---|---|---|---|---|---|---|---|
| mrpl36 | 7.499794 | 0.252685457 | 0.269 | 0.189 | 1.9053226 | 3 | mrpl36 |
| tktb | 9.190869 | 0.251198058 | 0.237 | 0.16 | 2.3349404 | 3 | tktb |
| nop58 | 2.003414 | 0.251248989 | 0.252 | 0.177 | 5.0896757 | 3 | nop58 |
| si:ch211-246m6.4 | 4.946455 | 0.262456875 | 0.244 | 0.171 | 1.2566469 | 3 | si:ch211-246m6.4 |
| zgc:91910 | 7.496134 | 0.251281608 | 0.339 | 0.267 | 1.9043930 | 3 | zgc:91910 |
| igfbp5b | 0 | 1.653669857 | 0.708 | 0.047 | 0 | 4 | igfbp5b |
| si:dkey-261h17.1 | 0 | 1.553160445 | 0.733 | 0.074 | 0 | 4 | si:dkey-261h17.1 |
| col1a2 | 0 | 1.520685394 | 0.875 | 0.135 | 0 | 4 | col1a2 |
| pmp22a | 0 | 1.492772702 | 0.841 | 0.121 | 0 | 4 | pmp22a |
| col1a1a | 0 | 1.412847041 | 0.817 | 0.122 | 0 | 4 | col1a1a |
| col1a1b | 0 | 1.410264625 | 0.809 | 0.127 | 0 | 4 | col1a1b |
| dcn | 0 | 1.407520815 | 0.782 | 0.101 | 0 | 4 | dcn |
| pcolcea | 0 | 1.397730273 | 0.648 | 0.028 | 0 | 4 | pcolcea |
| rbp4 | 0 | 1.373951193 | 0.695 | 0.104 | 0 | 4 | rbp4 |
| ccl25b | 0 | 1.308388116 | 0.557 | 0.063 | 0 | 4 | ccl25b |
| krt94 | 0 | 1.302763558 | 0.728 | 0.129 | 0 | 4 | krt94 |
| fstl1b | 0 | 1.259199913 | 0.725 | 0.109 | 0 | 4 | fstl1b |
| endouc | 0 | 1.195701325 | 0.538 | 0.04 | 0 | 4 | endouc |
| fn1a | 0 | 1.158851104 | 0.562 | 0.051 | 0 | 4 | fn1a |
| plpp3 | 0 | 1.13917688 | 0.536 | 0.034 | 0 | 4 | plpp3 |
| CABZ01092746.1 | 0 | 1.133248981 | 0.562 | 0.057 | 0 | 4 | CABZ01092746.1 |
| marcksl1a | 0 | 1.109761055 | 0.749 | 0.229 | 0 | 4 | marcksl1a |
| mfap5 | 0 | 1.101747531 | 0.523 | 0.033 | 0 | 4 | mfap5 |
| col5a1 | 0 | 1.10039024 | 0.639 | 0.098 | 0 | 4 | col5a1 |
| sparc | 0 | 1.062834802 | 0.698 | 0.162 | 0 | 4 | sparc |
| mfap2 | 0 | 1.043673895 | 0.57 | 0.09 | 0 | 4 | mfap2 |
| si:ch211-105c13.3 | 0 | 0.988782113 | 0.47 | 0.04 | 0 | 4 | si:ch211-105c13.3 |
| nfixb | 0 | 0.965854809 | 0.499 | 0.055 | 0 | 4 | nfixb |
| col6a3 | 0 | 0.948769043 | 0.536 | 0.087 | 0 | 4 | col6a3 |
| BX640522.2 | 0 | 0.947746511 | 0.385 | 0.019 | 0 | 4 | BX640522.2 |
| col5a2a | 0 | 0.922511786 | 0.505 | 0.074 | 0 | 4 | col5a2a |
| osr1 | 0 | 0.877449715 | 0.395 | 0.02 | 0 | 4 | osr1 |
| CR383676.11 | 0 | 0.868479585 | 0.957 | 0.702 | 0 | 4 | CR383676.1 |
| plpp1a | 0 | 0.868071946 | 0.409 | 0.056 | 0 | 4 | plpp1a |
| slit3 | 0 | 0.855850431 | 0.386 | 0.013 | 0 | 4 | slit3 |

| | | | | | | | |
|---|---|---|---|---|---|---|---|
| vasna | 0 | 0.794632704 | 0.366 | 0.024 | 0 | 4 | vasna |
| lrp1aa | 0 | 0.786688929 | 0.395 | 0.044 | 0 | 4 | lrp1aa |
| spon2b | 0 | 0.776513052 | 0.261 | 0.008 | 0 | 4 | spon2b |
| thbs3a | 0 | 0.771250323 | 0.3 | 0.003 | 0 | 4 | thbs3a |
| efnb1 | 0 | 0.762707451 | 0.352 | 0.024 | 0 | 4 | efnb1 |
| neu3.5 | 0 | 0.753287751 | 0.28 | 0.002 | 0 | 4 | neu3.5 |
| mgp | 0 | 0.726996638 | 0.251 | 0.007 | 0 | 4 | mgp |
| htra1b | 0 | 0.725188253 | 0.286 | 0.021 | 0 | 4 | htra1b |
| AL732488.1 | 0 | 0.720908777 | 0.306 | 0.024 | 0 | 4 | AL732488.1 |
| BX663503.3 | 0 | 0.706976711 | 0.304 | 0.016 | 0 | 4 | BX663503.3 |
| emilin2a | 0 | 0.664779685 | 0.268 | 0.007 | 0 | 4 | emilin2a |
| rgs5a | 0 | 0.65784729 | 0.288 | 0.022 | 0 | 4 | rgs5a |
| emid1 | 0 | 0.651638878 | 0.266 | 0.014 | 0 | 4 | emid1 |
| angptl1a | 0 | 0.641507412 | 0.265 | 0.019 | 0 | 4 | angptl1a |
| CABZ01039096.1 | 0 | 0.633534972 | 0.261 | 0.008 | 0 | 4 | CABZ01039096.1 |
| crhbp | 0 | 0.619675605 | 0.219 | 0.006 | 0 | 4 | crhbp |
| htra1a | 0 | 0.610195061 | 0.247 | 0.009 | 0 | 4 | htra1a |
| tsku | 0 | 0.605741228 | 0.29 | 0.021 | 0 | 4 | tsku |
| adamts15b | 0 | 0.604531649 | 0.25 | 0.01 | 0 | 4 | adamts15b |
| si:ch73-173h19.3 | 0 | 0.589659643 | 0.232 | 0.007 | 0 | 4 | si:ch73-173h19.3 |
| sost | 0 | 0.56894414 | 0.225 | 0.006 | 0 | 4 | sost |
| mustn1a | 0 | 0.562255496 | 0.234 | 0.009 | 0 | 4 | mustn1a |
| flnb | 0 | 0.55907202 | 0.229 | 0.013 | 0 | 4 | flnb |
| itih5 | 0 | 0.34482945 | 0.129 | 0.001 | 0 | 4 | itih5 |
| s100a10b | 1.703946 | 1.09110549 | 0.808 | 0.251 | 4.3288751 | 4 | s100a10b |
| kirrel1a | 3.702498 | 0.579218099 | 0.258 | 0.019 | 9.4061981 | 4 | kirrel1a |
| crabp2a | 1.777080 | 0.379331437 | 0.134 | 0.001 | 4.5146717 | 4 | crabp2a |
| colec12 | 3.413952 | 0.716959012 | 0.359 | 0.043 | 8.6731464 | 4 | colec12 |
| crabp2b | 2.052204 | 0.4808085 | 0.199 | 0.009 | 5.2136244 | 4 | crabp2b |
| si:ch1073-459j12.1 | 7.239349 | 0.814186487 | 0.421 | 0.063 | 1.8391567 | 4 | si:ch1073-459j12.1 |
| fbln5 | 9.605425 | 0.681689824 | 0.311 | 0.031 | 2.4402584 | 4 | fbln5 |
| fstb | 9.705687 | 0.560418061 | 0.22 | 0.013 | 2.4657299 | 4 | fstb |
| icn | 4.869033 | 1.125311781 | 0.762 | 0.235 | 1.2369779 | 4 | icn |
| tgfbr3 | 1.075583 | 0.684508274 | 0.298 | 0.03 | 2.7325199 | 4 | tgfbr3 |
| LOC108180269 | 2.442713 | 0.729309998 | 0.34 | 0.04 | 6.2057134 | 4 | LOC108180269 |

| | | | | | | | |
|---|---|---|---|---|---|---|---|
| mdka | 1.330865 | 0.912607485 | 0.484 | 0.089 | 3.3810632 | 4 | mdka |
| si:ch211-39i22.1 | 3.361028 | 0.810887923 | 0.419 | 0.066 | 8.5386933 | 4 | si:ch211-39i22.1 |
| zcchc24 | 5.699579 | 0.575530167 | 0.265 | 0.023 | 1.4479781 | 4 | zcchc24 |
| tnfsf10l | 6.790508 | 0.5760017 | 0.27 | 0.024 | 1.7251287 | 4 | tnfsf10l |
| LOC100537342 | 2.905934 | 0.383116204 | 0.152 | 0.004 | 7.3825278 | 4 | LOC100537342 |
| c6 | 6.816712 | 0.949459058 | 0.472 | 0.09 | 1.7317857 | 4 | c6 |
| dpysl3 | 9.119891 | 0.437754219 | 0.192 | 0.01 | 2.3169083 | 4 | dpysl3 |
| zgc:195173 | 5.637665 | 0.841847806 | 0.471 | 0.09 | 1.4322490 | 4 | zgc:195173 |
| add3a | 1.163057 | 0.880045598 | 0.519 | 0.113 | 2.9547470 | 4 | add3a |
| id2b | 2.112493 | 0.67462744 | 0.271 | 0.026 | 5.3667888 | 4 | id2b |
| akap12b | 1.614315 | 0.727986879 | 0.377 | 0.054 | 4.1011697 | 4 | akap12b |
| hapln3 | 3.395753 | 0.455084869 | 0.178 | 0.008 | 8.6269114 | 4 | hapln3 |
| asip2b | 2.904893 | 0.619964251 | 0.216 | 0.014 | 7.3798827 | 4 | asip2b |
| nova2 | 2.252546 | 0.574427955 | 0.253 | 0.022 | 5.7225953 | 4 | nova2 |
| si:ch211-198c19.3 | 8.359167 | 0.827365768 | 0.399 | 0.062 | 2.1236466 | 4 | si:ch211-198c19.3 |
| fbn2b | 1.845487 | 0.639961902 | 0.301 | 0.034 | 4.6884607 | 4 | fbn2b |
| plvapb | 1.831848 | 0.452782555 | 0.165 | 0.007 | 4.6538118 | 4 | plvapb |
| nid1a | 7.638167 | 0.565367694 | 0.265 | 0.025 | 1.9404765 | 4 | nid1a |
| XLOC-016094 | 3.116825 | 0.771155455 | 0.394 | 0.063 | 7.9182944 | 4 | XLOC-016094 |
| slc6a4a | 8.095904 | 0.380519485 | 0.154 | 0.005 | 2.0567646 | 4 | slc6a4a |
| emp2 | 1.498900 | 0.670220806 | 0.317 | 0.04 | 3.8079557 | 4 | emp2 |
| FP102783.1 | 2.750881 | 0.365046478 | 0.14 | 0.004 | 6.9886148 | 4 | FP102783.1 |
| fbln2 | 3.090088 | 0.490956708 | 0.213 | 0.015 | 7.8503688 | 4 | fbln2 |
| im:7151449 | 1.510808 | 0.564441233 | 0.223 | 0.018 | 3.8382090 | 4 | im:7151449 |
| c1qtnf5 | 3.524359 | 0.568283891 | 0.24 | 0.021 | 8.9536352 | 4 | c1qtnf5 |
| scara3 | 3.108345 | 0.536045744 | 0.242 | 0.022 | 7.8967512 | 4 | scara3 |
| tmem100a | 1.740083 | 0.471585687 | 0.204 | 0.014 | 4.4206832 | 4 | tmem100a |
| si:ch73-46j18.5 | 8.655693 | 0.895405578 | 0.642 | 0.18 | 2.1989790 | 4 | si:ch73-46j18.5 |
| ablim1b | 2.893142 | 0.588270522 | 0.267 | 0.028 | 7.3500284 | 4 | ablim1b |
| nfia | 3.359584 | 0.749081331 | 0.42 | 0.078 | 8.5350239 | 4 | nfia |
| tgfb2 | 6.783271 | 0.371417814 | 0.149 | 0.006 | 1.7232901 | 4 | tgfb2 |
| vim | 6.609458 | 0.568734056 | 0.275 | 0.031 | 1.6791329 | 4 | vim |
| znf385a | 6.364768 | 0.609020411 | 0.299 | 0.038 | 1.6169695 | 4 | znf385a |
| adgrg6 | 2.171238 | 0.358201018 | 0.132 | 0.004 | 5.5160310 | 4 | adgrg6 |
| cdh10a | 9.997024 | 0.276696775 | 0.111 | 0.002 | 2.5397440 | 4 | cdh10a |

| | | | | | | | |
|---|---|---|---|---|---|---|---|
| cdh6 | 1.080038 | 0.497030206 | 0.217 | 0.018 | 2.7438375 | 4 | cdh6 |
| tspan35 | 2.148208 | 0.400416919 | 0.161 | 0.008 | 5.4575225 | 4 | tspan35 |
| si:ch211-227m13.1 | 2.564908 | 0.378814072 | 0.165 | 0.009 | 6.5161502 | 4 | si:ch211-227m13.1 |
| dhrs3b | 1.827035 | 0.44192367 | 0.191 | 0.014 | 4.6415847 | 4 | dhrs3b |
| gpc6a | 7.250926 | 0.335931892 | 0.142 | 0.006 | 1.8420979 | 4 | gpc6a |
| dock9b | 6.305125 | 0.440308341 | 0.183 | 0.012 | 1.6018171 | 4 | dock9b |
| hsd11b2 | 1.214624 | 0.923380828 | 0.53 | 0.124 | 3.0857545 | 4 | hsd11b2 |
| crabp1b | 1.831542 | 0.294691219 | 0.109 | 0.002 | 4.6530348 | 4 | crabp1b |
| abca1a | 9.086649 | 0.720123461 | 0.336 | 0.055 | 2.3084633 | 4 | abca1a |
| klf2a | 2.314760 | 0.788703214 | 0.407 | 0.08 | 5.8806484 | 4 | klf2a |
| eda | 1.553541 | 0.347864022 | 0.135 | 0.005 | 3.9467726 | 4 | eda |
| anxa1a | 1.296089 | 0.865573075 | 0.535 | 0.139 | 3.2927166 | 4 | anxa1a |
| ptx3a | 3.108078 | 0.625160516 | 0.25 | 0.028 | 7.8960723 | 4 | ptx3a |
| igf2b | 2.941608 | 0.694314638 | 0.342 | 0.058 | 7.4731552 | 4 | igf2b |
| rbp5 | 4.592677 | 0.761597296 | 0.423 | 0.09 | 1.1667697 | 4 | rbp5 |
| col6a2 | 3.275887 | 0.633306811 | 0.34 | 0.057 | 8.3223918 | 4 | col6a2 |
| ackr3b | 3.361329 | 0.491992487 | 0.213 | 0.021 | 8.5394584 | 4 | ackr3b |
| si:ch211-251b21.1 | 2.039566 | 0.327826941 | 0.123 | 0.004 | 5.1815191 | 4 | si:ch211-251b21.1 |
| tgfbr2b | 2.884691 | 0.429362114 | 0.178 | 0.013 | 7.3285581 | 4 | tgfbr2b |
| adamts15a | 3.727220 | 0.605535694 | 0.294 | 0.043 | 9.4690045 | 4 | adamts15a |
| loxa | 3.775279 | 0.342017944 | 0.142 | 0.007 | 9.5910964 | 4 | loxa |
| loxl1 | 1.448328 | 0.466393817 | 0.206 | 0.02 | 3.6794788 | 4 | loxl1 |
| CR847566.1 | 5.342203 | 0.57093586 | 0.27 | 0.037 | 1.3571868 | 4 | CR847566.1 |
| cd81a | 1.643397 | 0.836397282 | 0.651 | 0.215 | 4.1750502 | 4 | cd81a |
| eva1ba | 3.071631 | 0.626653238 | 0.314 | 0.051 | 7.8034800 | 4 | eva1ba |
| CR855311.1 | 2.356572 | 0.844244951 | 0.366 | 0.071 | 5.9868723 | 4 | CR855311.1 |
| tll1 | 1.368800 | 0.276222512 | 0.101 | 0.002 | 3.4774370 | 4 | tll1 |
| lrrc17 | 5.480721 | 0.588084411 | 0.29 | 0.043 | 1.3923773 | 4 | lrrc17 |
| mfge8b | 2.335372 | 0.575614528 | 0.218 | 0.024 | 5.9330137 | 4 | mfge8b |
| nptx1l | 7.732889 | 0.282031679 | 0.107 | 0.003 | 1.9645404 | 4 | nptx1l |
| col6a1 | 1.236897 | 0.677856182 | 0.389 | 0.079 | 3.1423388 | 4 | col6a1 |
| crispld2 | 2.420850 | 0.420827317 | 0.159 | 0.011 | 6.1501703 | 4 | crispld2 |
| ppap2d | 3.091824 | 0.473966702 | 0.222 | 0.024 | 7.8547794 | 4 | ppap2d |
| cyp2ad3 | 3.555825 | 0.595458212 | 0.298 | 0.047 | 9.0335736 | 4 | cyp2ad3 |
| cyp2n13 | 4.531691 | 0.51697111 | 0.237 | 0.028 | 1.1512762 | 4 | cyp2n13 |

| | | | | | | | |
|---|---|---|---|---|---|---|---|
| lpar1 | 8.591558 | 0.611850271 | 0.317 | 0.054 | 2.1826855 | 4 | lpar1 |
| LOC103909284 | 2.438849 | 0.6215949 | 0.331 | 0.061 | 6.1958961 | 4 | LOC103909284 |
| ddr2l | 1.107452 | 0.430759825 | 0.19 | 0.018 | 2.8134821 | 4 | ddr2l |
| c1qtnf2 | 8.081047 | 0.386152119 | 0.181 | 0.016 | 2.0529901 | 4 | c1qtnf2 |
| krt8 | 9.142301 | 0.840564567 | 0.746 | 0.284 | 2.3226018 | 4 | krt8 |
| pamr1 | 6.410671 | 0.299462739 | 0.121 | 0.005 | 1.6286312 | 4 | pamr1 |
| cebpd | 4.319348 | 0.761671215 | 0.485 | 0.13 | 1.0973304 | 4 | cebpd |
| crabp1a | 5.140333 | 0.286434882 | 0.108 | 0.004 | 1.3059017 | 4 | crabp1a |
| rnd3b | 2.208873 | 0.495828007 | 0.253 | 0.037 | 5.6116426 | 4 | rnd3b |
| lrp1ab | 4.056673 | 0.600865624 | 0.325 | 0.062 | 1.0305978 | 4 | lrp1ab |
| bicc1b | 1.223008 | 0.387346057 | 0.171 | 0.015 | 3.1070522 | 4 | bicc1b |
| pdgfra | 5.139030 | 0.539463084 | 0.285 | 0.047 | 1.3055707 | 4 | pdgfra |
| fgl2b | 2.084900 | 0.315932624 | 0.13 | 0.008 | 5.2966894 | 4 | fgl2b |
| antxr2a | 1.323882 | 0.720185192 | 0.495 | 0.148 | 3.3633230 | 4 | antxr2a |
| pdgfrb | 1.351481 | 0.491212004 | 0.257 | 0.039 | 3.4334376 | 4 | pdgfrb |
| wscd2 | 1.787540 | 0.301051848 | 0.125 | 0.007 | 4.5412460 | 4 | wscd2 |
| s100v2 | 7.119795 | 0.677207856 | 0.449 | 0.121 | 1.8087841 | 4 | s100v2 |
| c7a | 9.072439 | 0.515346264 | 0.234 | 0.033 | 2.3048532 | 4 | c7a |
| ahnak | 2.083718 | 0.693268977 | 0.437 | 0.117 | 5.2936870 | 4 | ahnak |
| rspo3 | 2.520717 | 0.292055963 | 0.125 | 0.008 | 6.4038816 | 4 | rspo3 |
| bag2 | 1.398100 | 0.320240927 | 0.132 | 0.009 | 3.5518751 | 4 | bag2 |
| boc | 4.214971 | 0.544739227 | 0.292 | 0.053 | 1.0708135 | 4 | boc |
| sema3d | 3.093486 | 0.343878987 | 0.143 | 0.011 | 7.8590035 | 4 | sema3d |
| tril | 4.497845 | 0.386382944 | 0.172 | 0.017 | 1.1426776 | 4 | tril |
| si:ch1073-291c23.2 | 1.299761 | 0.547440227 | 0.304 | 0.058 | 3.3020449 | 4 | si:ch1073-291c23.2 |
| sostdc1a | 3.604278 | 0.466236135 | 0.163 | 0.015 | 9.1566702 | 4 | sostdc1a |
| foxp4 | 7.547700 | 0.57920143 | 0.301 | 0.059 | 1.9174933 | 4 | foxp4 |
| fgfr2 | 2.092277 | 0.412819197 | 0.188 | 0.021 | 5.3154306 | 4 | fgfr2 |
| peli1a | 6.787855 | 0.281174762 | 0.104 | 0.005 | 1.7244546 | 4 | peli1a |
| tgm1l1 | 6.332399 | 0.492058876 | 0.208 | 0.028 | 1.6087460 | 4 | tgm1l1 |
| clu | 4.570565 | 0.432655356 | 0.185 | 0.022 | 1.1611522 | 4 | clu |
| slc7a1 | 1.877967 | 0.436464117 | 0.199 | 0.027 | 4.7709761 | 4 | slc7a1 |
| gpr146 | 3.800350 | 0.338279666 | 0.137 | 0.011 | 9.6547897 | 4 | gpr146 |
| angptl4 | 9.794624 | 0.509551809 | 0.251 | 0.043 | 2.4883242 | 4 | angptl4 |
| tmem176 | 1.839681 | 0.537126373 | 0.311 | 0.066 | 4.6737116 | 4 | tmem176 |

| | | | | | | | |
|---|---|---|---|---|---|---|---|
| CU929237.1 | 5.745084 | 0.359354943 | 0.15 | 0.014 | 1.4595386 | 4 | CU929237.1 |
| ptges | 2.954523 | 0.38338654 | 0.158 | 0.016 | 7.5059675 | 4 | ptges |
| si:ch211-106h4.12 | 6.637489 | 0.645514144 | 0.257 | 0.045 | 1.6862541 | 4 | si:ch211-106h4.12 |
| arrdc3a | 1.446993 | 0.57168381 | 0.291 | 0.059 | 3.6760857 | 4 | arrdc3a |
| kcne4 | 2.259823 | 0.312822777 | 0.11 | 0.007 | 5.7410819 | 4 | kcne4 |
| cyp2p7 | 3.485442 | 0.386597965 | 0.177 | 0.021 | 8.8547663 | 4 | cyp2p7 |
| kank2 | 6.171842 | 0.501557958 | 0.245 | 0.043 | 1.5679566 | 4 | kank2 |
| mmp14a | 7.740350 | 0.459482592 | 0.258 | 0.047 | 1.9664360 | 4 | mmp14a |
| sema3bl | 8.035479 | 0.300287122 | 0.123 | 0.009 | 2.0414134 | 4 | sema3bl |
| hic1 | 2.806990 | 0.508042286 | 0.284 | 0.057 | 7.1311603 | 4 | hic1 |
| pabpc1b | 1.304280 | 0.531961423 | 0.297 | 0.063 | 3.3135240 | 4 | pabpc1b |
| tns1b | 6.366700 | 0.414371608 | 0.192 | 0.026 | 1.6174602 | 4 | tns1b |
| dhrs3a | 6.496935 | 0.420444809 | 0.197 | 0.028 | 1.6505464 | 4 | dhrs3a |
| meis2b | 3.977970 | 0.362768672 | 0.169 | 0.02 | 1.0106033 | 4 | meis2b |
| cyp2p6 | 1.082929 | 0.344216862 | 0.161 | 0.018 | 2.7511832 | 4 | cyp2p6 |
| cyr61 | 1.055219 | 0.590813991 | 0.274 | 0.055 | 2.6807840 | 4 | cyr61 |
| socs3a | 1.457527 | 0.774462941 | 0.674 | 0.292 | 3.7028485 | 4 | socs3a |
| mmp14b | 2.181699 | 0.508834945 | 0.27 | 0.054 | 5.5426077 | 4 | mmp14b |
| june | 2.216846 | 0.396860624 | 0.17 | 0.021 | 5.6318990 | 4 | june |
| abhd15a | 6.239813 | 0.438914674 | 0.206 | 0.031 | 1.5852245 | 4 | abhd15a |
| cxcl12a | 1.129178 | 0.658245377 | 0.311 | 0.069 | 2.8686776 | 4 | cxcl12a |
| itgb1a | 1.155573 | 0.462450719 | 0.233 | 0.041 | 2.9357342 | 4 | itgb1a |
| NC-002333.4 | 1.102020 | 0.599617594 | 0.912 | 0.59 | 2.7996821 | 4 | NC-002333.4 |
| pbx1a | 3.165071 | 0.461540785 | 0.236 | 0.043 | 8.0408640 | 4 | pbx1a |
| junba | 4.645195 | 0.743183104 | 0.669 | 0.298 | 1.1801118 | 4 | junba |
| cald1a | 6.686809 | 0.593975709 | 0.377 | 0.104 | 1.6987838 | 4 | cald1a |
| hspb1 | 1.215309 | 0.53683151 | 0.302 | 0.068 | 3.0874942 | 4 | hspb1 |
| agtr2 | 5.773968 | 0.335801653 | 0.143 | 0.015 | 1.4668768 | 4 | agtr2 |
| bmper | 5.796672 | 0.305048075 | 0.14 | 0.015 | 1.4726445 | 4 | bmper |
| mast2 | 3.015365 | 0.449962587 | 0.206 | 0.035 | 7.6605369 | 4 | mast2 |
| col6a4a | 5.454343 | 0.498615983 | 0.26 | 0.054 | 1.3856760 | 4 | col6a4a |
| jun | 4.004679 | 0.686980598 | 0.587 | 0.229 | 1.0173887 | 4 | jun |
| fosab | 2.171832 | 0.768942431 | 0.679 | 0.342 | 5.5175402 | 4 | fosab |
| cdon | 2.198880 | 0.486390644 | 0.267 | 0.059 | 5.5862566 | 4 | cdon |
| myl6 | 3.349146 | 0.564334891 | 0.36 | 0.101 | 8.5085072 | 4 | myl6 |

| | | | | | | | |
|---|---|---|---|---|---|---|---|
| rem1 | 3.354552 | 0.383824457 | 0.17 | 0.023 | 8.5222418 | 4 | rem1 |
| ip6k2b | 4.411179 | 0.608471122 | 0.373 | 0.113 | 1.1206602 | 4 | ip6k2b |
| itga5 | 1.418320 | 0.385388123 | 0.164 | 0.023 | 3.6032433 | 4 | itga5 |
| slc38a5b | 2.028146 | 0.570133782 | 0.373 | 0.113 | 5.1525051 | 4 | slc38a5b |
| adh8a | 2.764465 | 0.499948218 | 0.213 | 0.038 | 7.0231236 | 4 | adh8a |
| tenm3 | 5.375527 | 0.457336813 | 0.244 | 0.05 | 1.3656527 | 4 | tenm3 |
| col4a1 | 1.422489 | 0.520239001 | 0.264 | 0.058 | 3.6138338 | 4 | col4a1 |
| xkr9 | 4.123865 | 0.286480614 | 0.116 | 0.011 | 1.0476679 | 4 | xkr9 |
| si:ch211-212k18.8 | 1.818002 | 0.382303839 | 0.21 | 0.038 | 4.6186356 | 4 | si:ch211-212k18.8 |
| dedd1 | 3.497669 | 0.613787744 | 0.414 | 0.141 | 8.8858296 | 4 | dedd1 |
| tshz2 | 9.478955 | 0.359311019 | 0.159 | 0.022 | 2.4081285 | 4 | tshz2 |
| ramp2 | 3.994354 | 0.466859889 | 0.197 | 0.035 | 1.0147656 | 4 | ramp2 |
| zfp36l1b | 4.120112 | 0.603790628 | 0.394 | 0.124 | 1.0467146 | 4 | zfp36l1b |
| ca5a | 8.685902 | 0.348315576 | 0.161 | 0.023 | 2.2066534 | 4 | ca5a |
| olfml3a | 1.908015 | 0.263119658 | 0.12 | 0.012 | 4.8473134 | 4 | olfml3a |
| jund | 2.487252 | 0.5246924 | 0.297 | 0.077 | 6.3188643 | 4 | jund |
| klf6a | 2.935135 | 0.612987877 | 0.453 | 0.163 | 7.4567114 | 4 | klf6a |
| bgnb | 2.953046 | 0.337843199 | 0.145 | 0.018 | 7.5022143 | 4 | bgnb |
| pmp22b | 3.513283 | 0.428395312 | 0.203 | 0.037 | 8.9254956 | 4 | pmp22b |
| sult1st6 | 2.248240 | 0.273126592 | 0.121 | 0.013 | 5.7116544 | 4 | sult1st6 |
| chd3 | 5.316978 | 0.534042092 | 0.353 | 0.111 | 1.3507785 | 4 | chd3 |
| lxn | 9.314060 | 0.612550912 | 0.419 | 0.152 | 2.3662371 | 4 | lxn |
| loxl2a | 5.576069 | 0.402605193 | 0.198 | 0.038 | 1.4166005 | 4 | loxl2a |
| fzd2 | 1.947360 | 0.363312484 | 0.185 | 0.034 | 4.9472691 | 4 | fzd2 |
| iqsec1b | 2.438664 | 0.267733082 | 0.114 | 0.012 | 6.1954260 | 4 | iqsec1b |
| lama4 | 6.312384 | 0.384317681 | 0.197 | 0.038 | 1.6036613 | 4 | lama4 |
| si:ch211-133n4.4 | 3.412509 | 0.403225568 | 0.224 | 0.049 | 8.6694799 | 4 | si:ch211-133n4.4 |
| grb10a | 1.274327 | 0.368355212 | 0.186 | 0.034 | 3.2374283 | 4 | grb10a |
| lhfpl6 | 8.539456 | 0.281459929 | 0.132 | 0.017 | 2.1694488 | 4 | lhfpl6 |
| fgfr1a | 6.818608 | 0.274935996 | 0.127 | 0.016 | 1.7322675 | 4 | fgfr1a |
| mir199-3 | 2.328029 | 0.295035189 | 0.128 | 0.016 | 5.9143588 | 4 | mir199-3 |
| lamb1a | 3.365345 | 0.395081467 | 0.212 | 0.045 | 8.5496598 | 4 | lamb1a |
| tuba8l3 | 4.777851 | 0.417955699 | 0.253 | 0.061 | 1.2138131 | 4 | tuba8l3 |
| CABZ01072614.1 | 1.190986 | 0.507505861 | 0.292 | 0.082 | 3.0257014 | 4 | CABZ01072614.1 |
| kitlga | 4.495006 | 0.254964064 | 0.111 | 0.012 | 1.1419564 | 4 | kitlga |

| | | | | | | | |
|---|---|---|---|---|---|---|---|
| vat1 | 1.029941 | 0.558136428 | 0.385 | 0.133 | 2.6165671 | 4 | vat1 |
| cxcl8b.1 | 1.038792 | 0.497149945 | 0.195 | 0.038 | 2.6390531 | 4 | cxcl8b.1 |
| rpz5 | 6.610396 | 0.512391228 | 0.352 | 0.113 | 1.6793712 | 4 | rpz5 |
| fkbp1ab | 1.357823 | 0.437208302 | 0.225 | 0.05 | 3.4495509 | 4 | fkbp1ab |
| atoh8 | 1.566214 | 0.281553078 | 0.135 | 0.019 | 3.9789680 | 4 | atoh8 |
| fblim1 | 9.654334 | 0.30564095 | 0.168 | 0.029 | 2.4526837 | 4 | fblim1 |
| si:ch211-212k18.9 | 2.213160 | 0.349359853 | 0.197 | 0.041 | 5.6225345 | 4 | si:ch211-212k18.9 |
| ank3b | 3.130322 | 0.36184492 | 0.17 | 0.03 | 7.9525831 | 4 | ank3b |
| cxcl18b | 3.997469 | 0.559448324 | 0.324 | 0.097 | 1.0155572 | 4 | cxcl18b |
| fzd7b | 1.463101 | 0.36459019 | 0.193 | 0.04 | 3.7170095 | 4 | fzd7b |
| anxa5b | 2.066154 | 0.495285158 | 0.305 | 0.093 | 5.2490659 | 4 | anxa5b |
| gulp1a | 3.038625 | 0.317731773 | 0.164 | 0.029 | 7.7196279 | 4 | gulp1a |
| XLOC-041269 | 3.369979 | 0.302675167 | 0.145 | 0.023 | 8.5614331 | 4 | XLOC-041269 |
| LO018188.1 | 1.549033 | 0.307611309 | 0.141 | 0.022 | 3.9353208 | 4 | LO018188.1 |
| meis1a | 2.310415 | 0.25377059 | 0.116 | 0.015 | 5.8696112 | 4 | meis1a |
| spon1b | 2.413024 | 0.394836112 | 0.196 | 0.043 | 6.1302896 | 4 | spon1b |
| ddit3 | 5.182879 | 0.577969914 | 0.415 | 0.164 | 1.3167105 | 4 | ddit3 |
| ppib | 3.213344 | 0.559298859 | 0.646 | 0.363 | 8.1635006 | 4 | ppib |
| timp2a | 9.769547 | 0.405695471 | 0.224 | 0.055 | 2.4819534 | 4 | timp2a |
| col16a1 | 2.784770 | 0.330363503 | 0.169 | 0.033 | 7.0747101 | 4 | col16a1 |
| itpkcb | 6.658069 | 0.422396581 | 0.239 | 0.065 | 1.6914826 | 4 | itpkcb |
| klf3 | 8.225323 | 0.319405719 | 0.163 | 0.032 | 2.0896434 | 4 | klf3 |
| ghra | 3.829001 | 0.32708386 | 0.149 | 0.027 | 9.7275792 | 4 | ghra |
| gadd45ba | 1.272906 | 0.575676482 | 0.436 | 0.183 | 3.2338196 | 4 | gadd45ba |
| id1 | 3.413532 | 0.615668634 | 0.334 | 0.119 | 8.6720780 | 4 | id1 |
| myl9b | 3.865558 | 0.420857486 | 0.286 | 0.087 | 9.8204521 | 4 | myl9b |
| sema4ba | 4.776076 | 0.35764881 | 0.168 | 0.034 | 1.2133622 | 4 | sema4ba |
| tnfaip2a | 9.093242 | 0.275673971 | 0.118 | 0.017 | 2.3101382 | 4 | tnfaip2a |
| mmp16a | 9.164229 | 0.445481525 | 0.246 | 0.068 | 2.3281725 | 4 | mmp16a |
| rgcc | 3.235187 | 0.426057023 | 0.232 | 0.062 | 8.2189945 | 4 | rgcc |
| ckap4 | 4.038912 | 0.363020221 | 0.206 | 0.052 | 1.0260857 | 4 | ckap4 |
| cyp1c2 | 7.045251 | 0.254359866 | 0.103 | 0.013 | 1.7898461 | 4 | cyp1c2 |
| stom | 3.390592 | 0.262126804 | 0.116 | 0.017 | 8.6138000 | 4 | stom |
| COX31 | 4.606948 | 0.441308878 | 0.898 | 0.662 | 1.1703953 | 4 | COX3 |
| si:dkey-240h12.3 | 1.506130 | 0.25804224 | 0.11 | 0.016 | 3.8263256 | 4 | si:dkey-240h12.3 |

| | | | | | | | |
|---|---|---|---|---|---|---|---|
| meis1b | 2.91404( | 0.387092993 | 0.229 | 0.064 | 7.4031187 | 4 | meis1b |
| gas1a | 6.421246 | 0.347315598 | 0.199 | 0.05 | 1.6313177 | 4 | gas1a |
| ccdc3b | 6.21555( | 0.278092225 | 0.136 | 0.025 | 1.5790606 | 4 | ccdc3b |
| cldn5a | 6.77328( | 0.407810793 | 0.206 | 0.053 | 1.7207518 | 4 | cldn5a |
| olfml3b | 1.86394( | 0.360506442 | 0.192 | 0.047 | 4.7353420 | 4 | olfml3b |
| plod2 | 2.273975 | 0.283073995 | 0.132 | 0.024 | 5.7770335 | 4 | plod2 |
| mex3b | 2.769561 | 0.251833026 | 0.103 | 0.014 | 7.0360699 | 4 | mex3b |
| mt-atp6 | 1.326893 | 0.480426416 | 0.795 | 0.541 | 3.3709739 | 4 | mt-atp6 |
| egfra | 2.026182 | 0.282426136 | 0.138 | 0.026 | 5.1475174 | 4 | egfra |
| cox4i2 | 4.853141 | 0.264596963 | 0.149 | 0.03 | 1.2329404 | 4 | cox4i2 |
| fkbp9 | 2.989388 | 0.29582043 | 0.147 | 0.03 | 7.5945424 | 4 | fkbp9 |
| fam107b | 4.141643 | 0.453761132 | 0.293 | 0.102 | 1.0521845 | 4 | fam107b |
| nrp1a | 5.02422( | 0.388004539 | 0.206 | 0.056 | 1.2764031 | 4 | nrp1a |
| hoxc3a | 2.00637( | 0.250800579 | 0.128 | 0.023 | 5.0971834 | 4 | hoxc3a |
| BX322530.1 | 8.375795 | 0.340661683 | 0.179 | 0.044 | 2.1278708 | 4 | BX322530.1 |
| prss23 | 2.086561 | 0.281568487 | 0.137 | 0.027 | 5.3009102 | 4 | prss23 |
| pdgfrl | 4.937234 | 0.266305258 | 0.137 | 0.027 | 1.2543044 | 4 | pdgfrl |
| cnn3a | 5.298284 | 0.339521835 | 0.215 | 0.06 | 1.3460291 | 4 | cnn3a |
| b2ml1 | 5.576975 | 0.490952641 | 0.529 | 0.28 | 1.4168306 | 4 | b2ml |
| cx43 | 2.11141( | 0.427352324 | 0.19 | 0.05 | 5.3640374 | 4 | cx43 |
| tead1b | 3.028143 | 0.28274038 | 0.137 | 0.027 | 7.6929973 | 4 | tead1b |
| nectin1b | 3.72904( | 0.324122834 | 0.165 | 0.038 | 9.4736283 | 4 | nectin1b |
| ldlrad4a | 1.000068 | 0.273548897 | 0.138 | 0.028 | 2.5406745 | 4 | ldlrad4a |
| si:dkey-9i23.16 | 1.588734 | 0.303193163 | 0.174 | 0.042 | 4.0361797 | 4 | si:dkey-9i23.16 |
| cd28l | 2.05908( | 0.270451296 | 0.152 | 0.033 | 5.2310929 | 4 | cd28l |
| celf21 | 2.52270( | 0.498432559 | 0.483 | 0.244 | 6.4089207 | 4 | celf2 |
| rbms3 | 6.877449 | 0.272594464 | 0.143 | 0.03 | 1.7472160 | 4 | rbms3 |
| klf13 | 2.494307 | 0.442178146 | 0.267 | 0.094 | 6.3367882 | 4 | klf13 |
| mmp2 | 3.32736( | 0.417405882 | 0.285 | 0.101 | 8.4531599 | 4 | mmp2 |
| tgfb3 | 1.192378 | 0.254814922 | 0.135 | 0.027 | 3.0292382 | 4 | tgfb3 |
| rnd3a | 1.425226 | 0.275575024 | 0.14 | 0.029 | 3.6207873 | 4 | rnd3a |
| anxa2a | 7.082722 | 0.453461485 | 0.295 | 0.107 | 1.7993656 | 4 | anxa2a |
| capn2a | 1.691347 | 0.328424468 | 0.178 | 0.047 | 4.2968685 | 4 | capn2a |
| fxyd6l | 2.311305 | 0.390826997 | 0.297 | 0.107 | 5.8718727 | 4 | fxyd6l |
| tmem176l.1 | 3.647064 | 0.446981276 | 0.42 | 0.189 | 9.2653683 | 4 | tmem176l.1 |

| | | | | | | | |
|---|---|---|---|---|---|---|---|
| tns2a | 4.899875 | 0.268802875 | 0.141 | 0.031 | 1.2448133 | 4 | tns2a |
| pard3bb | 5.365583 | 0.257592047 | 0.13 | 0.027 | 1.3631265 | 4 | pard3bb |
| ece1 | 1.769193 | 0.271027897 | 0.148 | 0.034 | 4.4946360 | 4 | ece1 |
| si:dkey-238c7.16 | 2.675021 | 0.320371548 | 0.21 | 0.062 | 6.7958920 | 4 | si:dkey-238c7.16 |
| fbxo34 | 5.337197 | 0.298274684 | 0.164 | 0.041 | 1.3559150 | 4 | fbxo34 |
| svild | 5.391802 | 0.357667257 | 0.232 | 0.075 | 1.3697873 | 4 | svild |
| sdc2 | 3.985041 | 0.351948774 | 0.227 | 0.074 | 1.0123996 | 4 | sdc2 |
| fabp11a | 5.089209 | 0.401603119 | 0.242 | 0.078 | 1.2929136 | 4 | fabp11a |
| sptan1 | 5.479236 | 0.316826649 | 0.182 | 0.051 | 1.3920000 | 4 | sptan1 |
| gstt1b | 2.614427 | 0.323552456 | 0.242 | 0.081 | 6.6419543 | 4 | gstt1b |
| sntb2 | 5.724766 | 0.278624246 | 0.17 | 0.046 | 1.4543768 | 4 | sntb2 |
| sik2b | 1.251608 | 0.297669696 | 0.152 | 0.038 | 3.1797126 | 4 | sik2b |
| zgc:1627301 | 1.695836 | 0.485085388 | 0.66 | 0.387 | 4.3082727 | 4 | zgc:162730 |
| appb | 4.725148 | 0.284028955 | 0.156 | 0.04 | 1.2004238 | 4 | appb |
| spred2b | 5.694484 | 0.344997805 | 0.227 | 0.076 | 1.4466836 | 4 | spred2b |
| serpinf1 | 6.871912 | 0.278086713 | 0.154 | 0.039 | 1.7458094 | 4 | serpinf1 |
| krt18b | 8.694540 | 0.405271726 | 0.382 | 0.164 | 2.2088480 | 4 | krt18b |
| lgals2a | 1.078892 | 0.409990526 | 0.329 | 0.127 | 2.7409254 | 4 | lgals2a |
| ulk2 | 2.076680 | 0.291808474 | 0.17 | 0.047 | 5.2758078 | 4 | ulk2 |
| cebpa | 2.273944 | 0.371855534 | 0.212 | 0.068 | 5.7769569 | 4 | cebpa |
| zgc:158463 | 2.939634 | 0.381361043 | 0.218 | 0.072 | 7.4681425 | 4 | zgc:158463 |
| mt-nd2 | 4.678889 | 0.438504566 | 0.632 | 0.377 | 1.1886718 | 4 | mt-nd2 |
| fkbp7 | 6.131834 | 0.265780888 | 0.163 | 0.044 | 1.5577924 | 4 | fkbp7 |
| foxo3b | 7.753081 | 0.342131481 | 0.2 | 0.063 | 1.9696702 | 4 | foxo3b |
| mt-nd4 | 8.972129 | 0.427953364 | 0.692 | 0.433 | 2.2793694 | 4 | mt-nd4 |
| stau2 | 1.291656 | 0.347274516 | 0.244 | 0.087 | 3.2814537 | 4 | stau2 |
| spry4 | 2.322740 | 0.289433227 | 0.161 | 0.043 | 5.9009229 | 4 | spry4 |
| sc:d0202 | 5.283295 | 0.292296503 | 0.155 | 0.04 | 1.3422211 | 4 | sc:d0202 |
| tagln2 | 5.407759 | 0.428890477 | 0.413 | 0.198 | 1.3738413 | 4 | tagln2 |
| tcf12 | 1.314657 | 0.307320943 | 0.196 | 0.061 | 3.3398864 | 4 | tcf12 |
| mt-cyb | 4.414037 | 0.401747207 | 0.776 | 0.565 | 1.1213861 | 4 | mt-cyb |
| mgst1.2 | 8.039563 | 0.37611044 | 0.277 | 0.107 | 2.0424509 | 4 | mgst1.2 |
| cpn1 | 9.130613 | 0.315774171 | 0.213 | 0.069 | 2.3196323 | 4 | cpn1 |
| dkk3b | 7.329422 | 0.312197191 | 0.181 | 0.054 | 1.8620397 | 4 | dkk3b |
| flna | 8.605874 | 0.372901151 | 0.283 | 0.112 | 2.1863224 | 4 | flna |

| | | | | | | | |
|---|---|---|---|---|---|---|---|
| hdlbpa | 5.212503 | 0.396310341 | 0.358 | 0.164 | 1.3242365 | 4 | hdlbpa |
| arl6ip5b | 6.695173 | 0.342659874 | 0.251 | 0.095 | 1.7009089 | 4 | arl6ip5b |
| nfil3-6 | 5.179617 | 0.410043704 | 0.292 | 0.122 | 1.3158817 | 4 | nfil3-6 |
| mcl1b | 2.214139 | 0.424009697 | 0.454 | 0.239 | 5.6250205 | 4 | mcl1b |
| rerea | 4.232372 | 0.274184721 | 0.163 | 0.048 | 1.0752341 | 4 | rerea |
| rbfox2 | 9.157258 | 0.257947817 | 0.138 | 0.037 | 2.3264015 | 4 | rbfox2 |
| rdh10a | 2.535042 | 0.27236787 | 0.151 | 0.042 | 6.4402749 | 4 | rdh10a |
| ndfip1l | 2.695993 | 0.335960341 | 0.198 | 0.068 | 6.8491718 | 4 | ndfip1l |
| mt-co21 | 3.567672 | 0.35957415 | 0.839 | 0.661 | 9.0636730 | 4 | mt-co2 |
| fam168a | 1.173704 | 0.303884718 | 0.195 | 0.067 | 2.9817967 | 4 | fam168a |
| ccdc85b | 1.933700 | 0.254742291 | 0.155 | 0.045 | 4.9125664 | 4 | ccdc85b |
| si:ch73-335l21.4 | 1.762458 | 0.465009178 | 0.301 | 0.132 | 4.4775261 | 4 | si:ch73-335l21.4 |
| fxyd1 | 2.345984 | 0.320873068 | 0.211 | 0.077 | 5.9599735 | 4 | fxyd1 |
| tnfsf10 | 7.020045 | 0.25864485 | 0.138 | 0.038 | 1.7834426 | 4 | tnfsf10 |
| pdlim1 | 8.695686 | 0.273988207 | 0.171 | 0.054 | 2.2091391 | 4 | pdlim1 |
| si:dkey-7j14.6 | 2.757127 | 0.378213519 | 0.375 | 0.186 | 7.0044815 | 4 | si:dkey-7j14.6 |
| mych | 4.296908 | 0.325441148 | 0.22 | 0.082 | 1.0916297 | 4 | mych |
| krt15 | 4.934273 | 0.29854169 | 0.24 | 0.088 | 1.2535521 | 4 | krt15 |
| gng7 | 1.478561 | 0.273159555 | 0.171 | 0.056 | 3.7562853 | 4 | gng7 |
| rbpjb | 3.368758 | 0.289986304 | 0.171 | 0.056 | 8.5583305 | 4 | rbpjb |
| aoc2 | 9.046702 | 0.338932801 | 0.196 | 0.07 | 2.2983146 | 4 | aoc2 |
| TCIM (1 of many) | 1.328320 | 0.344635476 | 0.147 | 0.043 | 3.3745984 | 4 | TCIM (1 of many) |
| ephb3 | 1.707094 | 0.253653738 | 0.152 | 0.046 | 4.3368739 | 4 | ephb3 |
| cast | 2.038383 | 0.286940339 | 0.19 | 0.066 | 5.1785124 | 4 | cast |
| h3f3c | 2.058056 | 0.410052295 | 0.485 | 0.294 | 5.2284932 | 4 | h3f3c |
| FO082781.1 | 6.218523 | 0.264765601 | 0.157 | 0.05 | 1.5798159 | 4 | FO082781.1 |
| dbn1 | 2.128137 | 0.272581765 | 0.175 | 0.06 | 5.4065344 | 4 | dbn1 |
| plekho2 | 2.322593 | 0.301841808 | 0.193 | 0.07 | 5.9005490 | 4 | plekho2 |
| f3b | 1.791140 | 0.336070359 | 0.231 | 0.091 | 4.5503927 | 4 | f3b |
| c4 | 3.108559 | 0.259345637 | 0.123 | 0.034 | 7.8972964 | 4 | c4 |
| eif4ebp3l | 6.848382 | 0.372445969 | 0.383 | 0.195 | 1.7398316 | 4 | eif4ebp3l |
| BX284696.1 | 7.759649 | 0.277381443 | 0.161 | 0.054 | 1.9713390 | 4 | BX284696.1 |
| mt-nd1 | 8.208250 | 0.385027152 | 0.594 | 0.385 | 2.0853060 | 4 | mt-nd1 |
| marcksl1b | 1.263334 | 0.356117743 | 0.294 | 0.133 | 3.2095017 | 4 | marcksl1b |
| tspan14 | 1.325345 | 0.342972998 | 0.274 | 0.124 | 3.3670408 | 4 | tspan14 |

| | | | | | | | |
|---|---|---|---|---|---|---|---|
| btg2 | 3.901343 | 0.431701442 | 0.516 | 0.34 | 9.9113642 | 4 | btg2 |
| tmsb1 | 5.898730 | 0.396530634 | 0.315 | 0.151 | 1.4985724 | 4 | tmsb1 |
| ctdsp2 | 1.617595 | 0.289466358 | 0.2 | 0.079 | 4.1095004 | 4 | ctdsp2 |
| mef2aa | 2.342482 | 0.274063506 | 0.183 | 0.067 | 5.9510764 | 4 | mef2aa |
| si:ch211-1a19.3 | 3.795235 | 0.270093005 | 0.155 | 0.052 | 9.6417969 | 4 | si:ch211-1a19.3 |
| purab | 3.837097 | 0.264697089 | 0.157 | 0.054 | 9.7481451 | 4 | purab |
| abcc5 | 7.179821 | 0.306679198 | 0.213 | 0.088 | 1.8240335 | 4 | abcc5 |
| ptprga | 3.108851 | 0.277056575 | 0.177 | 0.066 | 7.8980373 | 4 | ptprga |
| h2afx11 | 6.421916 | 0.35653899 | 0.394 | 0.218 | 1.6314878 | 4 | h2afx1 |
| il15 | 1.175342 | 0.257235027 | 0.157 | 0.054 | 2.9859581 | 4 | il15 |
| nfic | 2.547968 | 0.277680456 | 0.195 | 0.078 | 6.4731134 | 4 | nfic |
| si:ch73-1a9.3 | 7.704647 | 0.360976699 | 0.496 | 0.305 | 1.9573656 | 4 | si:ch73-1a9.3 |
| wls | 2.969951 | 0.296927351 | 0.266 | 0.123 | 7.5451614 | 4 | wls |
| aldh9a1a.1 | 3.725540 | 0.344079457 | 0.342 | 0.185 | 9.4647350 | 4 | aldh9a1a.1 |
| adam10a | 5.629916 | 0.300083529 | 0.225 | 0.099 | 1.4302803 | 4 | adam10a |
| mknk2b1 | 8.569990 | 0.340889118 | 0.354 | 0.191 | 2.1772061 | 4 | mknk2b |
| rarga | 9.899839 | 0.250108143 | 0.155 | 0.055 | 2.5150541 | 4 | rarga |
| rbm39a | 3.656814 | 0.311786215 | 0.29 | 0.146 | 9.2901370 | 4 | rbm39a |
| ciarta | 1.097223 | 0.293369832 | 0.24 | 0.111 | 2.7874953 | 4 | ciarta |
| dnmt3ab | 2.790817 | 0.282065836 | 0.202 | 0.086 | 7.0900708 | 4 | dnmt3ab |
| crip11 | 4.311084 | 0.397353477 | 0.433 | 0.276 | 1.0952310 | 4 | crip1 |
| mt-co11 | 6.677922 | 0.30396672 | 0.841 | 0.628 | 1.6965262 | 4 | mt-co1 |
| tpm4a | 7.987383 | 0.267444142 | 0.238 | 0.107 | 2.0291946 | 4 | tpm4a |
| mafba | 8.573948 | 0.266977504 | 0.19 | 0.079 | 2.1782115 | 4 | mafba |
| hmgb3a | 2.006266 | 0.268532786 | 0.188 | 0.077 | 5.0969197 | 4 | hmgb3a |
| sdc4 | 2.247555 | 0.320626046 | 0.435 | 0.244 | 5.7099144 | 4 | sdc4 |
| appa | 6.472323 | 0.263088358 | 0.195 | 0.082 | 1.6442937 | 4 | appa |
| cyp3c1 | 6.579180 | 0.254557696 | 0.157 | 0.06 | 1.6714407 | 4 | cyp3c1 |
| pim1 | 1.229096 | 0.340852103 | 0.305 | 0.162 | 3.1225199 | 4 | pim1 |
| LOC566587 | 1.493120 | 0.252184148 | 0.121 | 0.04 | 3.7932721 | 4 | LOC566587 |
| fn1b | 1.893419 | 0.290943323 | 0.219 | 0.095 | 4.8102311 | 4 | fn1b |
| sdcbp2 | 4.908564 | 0.328936187 | 0.374 | 0.22 | 1.2470207 | 4 | sdcbp2 |
| qkia | 1.432747 | 0.315080569 | 0.342 | 0.191 | 3.6398943 | 4 | qkia |
| jpt1b | 4.858617 | 0.310849999 | 0.447 | 0.267 | 1.2343317 | 4 | jpt1b |
| fkbp5 | 1.366143 | 0.304544753 | 0.242 | 0.121 | 3.4706865 | 4 | fkbp5 |

| | | | | | | | |
|---|---|---|---|---|---|---|---|
| calm2b | 4.322621 | 0.310394095 | 0.399 | 0.238 | 1.0981618 | 4 | calm2b |
| CABZ01080702.1 | 1.189259 | 0.254109899 | 0.203 | 0.092 | 3.0213135 | 4 | CABZ01080702.1 |
| hmgb1a1 | 1.505067 | 0.310834876 | 0.363 | 0.221 | 3.8236235 | 4 | hmgb1a |
| itm2ba | 1.630676 | 0.303836618 | 0.445 | 0.289 | 4.1427347 | 4 | itm2ba |
| si:ch211-196l7.41 | 3.385025 | 0.290994073 | 0.482 | 0.322 | 8.5996572 | 4 | si:ch211-196l7.4 |
| postnb | 7.708302 | 0.309020623 | 0.195 | 0.09 | 1.9582941 | 4 | postnb |
| slc43a2a | 1.290918 | 0.268025271 | 0.2 | 0.097 | 3.2795785 | 4 | slc43a2a |
| ywhaqb | 1.840626 | 0.271206585 | 0.27 | 0.149 | 4.6761123 | 4 | ywhaqb |
| dub1 | 2.350703 | 0.264995779 | 0.157 | 0.069 | 5.9719614 | 4 | dub |
| glula | 2.655968 | 0.278209339 | 0.236 | 0.119 | 6.7474869 | 4 | glula |
| ier2b1 | 2.787953 | 0.316624126 | 0.367 | 0.23 | 7.0827963 | 4 | ier2b |
| rab10 | 5.096894 | 0.2809308 | 0.342 | 0.207 | 1.2948660 | 4 | rab10 |
| acin1a | 1.056575 | 0.298578434 | 0.356 | 0.222 | 2.6842299 | 4 | acin1a |
| notch21 | 1.382032 | 0.251996711 | 0.223 | 0.117 | 3.5110538 | 4 | notch2 |
| sesn11 | 1.521080 | 0.274934116 | 0.294 | 0.169 | 3.8643048 | 4 | sesn1 |
| mt-nd3 | 2.681307 | 0.27956836 | 0.519 | 0.364 | 6.8118623 | 4 | mt-nd3 |
| hmgn61 | 5.184416 | 0.290516521 | 0.441 | 0.304 | 1.3171009 | 4 | hmgn6 |
| cd631 | 8.817821 | 0.255669066 | 0.54 | 0.364 | 2.2401675 | 4 | cd63 |
| zfp36l1a1 | 4.136114 | 0.278310151 | 0.387 | 0.248 | 1.0507799 | 4 | zfp36l1a |
| ier2a | 6.753460 | 0.320103709 | 0.393 | 0.268 | 1.7157167 | 4 | ier2a |
| selenow1 | 4.712438 | 0.254480768 | 0.576 | 0.406 | 1.1971949 | 4 | selenow1 |
| egr1 | 7.607520 | 0.281656064 | 0.278 | 0.179 | 1.9326905 | 4 | egr1 |
| tmsb4x1 | 8.214817 | 0.25607626 | 0.958 | 0.703 | 0.002087 | 4 | tmsb4x |
| h1m3 | 2.640469 | 0.921166499 | 0.985 | 0.469 | 6.7081132 | 5 | h1m |
| zgc:1534993 | 1.554856 | 0.970721145 | 0.955 | 0.392 | 3.9501127 | 5 | zgc:153499 |
| h2af1al3 | 5.915447 | 0.91912709 | 0.987 | 0.462 | 1.5028193 | 5 | h2af1al |
| cldnd3 | 3.880479 | 0.921837124 | 0.988 | 0.47 | 9.8583583 | 5 | cldnd |
| hbba13 | 3.070727 | 0.988216962 | 0.786 | 0.267 | 7.8011833 | 5 | hbba1 |
| tmsb53 | 1.564040 | 0.937096446 | 0.914 | 0.374 | 3.9734454 | 5 | tmsb5 |
| zp2.53 | 6.244789 | 0.911495177 | 0.956 | 0.436 | 1.5864886 | 5 | zp2.5 |
| hbaa13 | 1.349989 | 0.968074611 | 0.689 | 0.208 | 3.4296491 | 5 | hbaa1 |
| zp33 | 6.559918 | 0.899867278 | 0.967 | 0.453 | 1.6665473 | 5 | zp3 |
| si:dkey-241l7.23 | 1.443995 | 0.884300668 | 0.991 | 0.503 | 3.6684700 | 5 | si:dkey-241l7.2 |
| acp5a3 | 1.239835 | 0.8902828 | 0.925 | 0.419 | 3.1498025 | 5 | acp5a |
| mt23 | 5.576147 | 0.894241131 | 0.93 | 0.413 | 1.4166202 | 5 | mt2 |

| | | | | | | | |
|---|---|---|---|---|---|---|---|
| zgc:1717763 | 1.205200 | 0.900155763 | 0.88 | 0.381 | 3.0618126 | 5 | zgc:171776 |
| si:dkey-240e12.63 | 1.331486 | 0.905966826 | 0.963 | 0.44 | 3.3826418 | 5 | si:dkey-240e12.6 |
| crp23 | 8.624520 | 0.904034564 | 0.899 | 0.375 | 2.1910594 | 5 | crp2 |
| zp2.33 | 1.072287 | 0.899581006 | 0.914 | 0.399 | 2.7241467 | 5 | zp2.3 |
| si:zfos-1505d6.33 | 1.135622 | 0.895212076 | 0.913 | 0.394 | 2.8850492 | 5 | si:zfos-1505d6.3 |
| si:ch73-189n23.13 | 7.372368 | 0.893941061 | 0.917 | 0.405 | 1.8729501 | 5 | si:ch73-189n23.1 |
| zp2.13 | 3.298404 | 0.883636576 | 0.907 | 0.414 | 8.3795959 | 5 | zp2.1 |
| si:ch211-125e6.123 | 6.014790 | 0.894879422 | 0.897 | 0.385 | 1.5280574 | 5 | si:ch211-125e6.12 |
| si:dkey-241l7.43 | 2.821727 | 0.83150665 | 0.934 | 0.478 | 7.1685980 | 5 | si:dkey-241l7.4 |
| si:dkeyp-98a7.93 | 5.246875 | 0.885898619 | 0.901 | 0.395 | 1.3329686 | 5 | si:dkeyp-98a7.9 |
| si:ch211-226h8.143 | 9.619417 | 0.869247737 | 0.929 | 0.441 | 2.4438130 | 5 | si:ch211-226h8.14 |
| zp3.23 | 2.212909 | 0.861255387 | 0.917 | 0.434 | 5.6218967 | 5 | zp3.2 |
| zp2.63 | 4.907045 | 0.878685188 | 0.878 | 0.386 | 1.2466348 | 5 | zp2.6 |
| si:dkey-241l7.53 | 7.896314 | 0.844931536 | 0.931 | 0.456 | 2.0060587 | 5 | si:dkey-241l7.5 |
| si:ch211-5k11.83 | 2.539153 | 0.849928892 | 0.75 | 0.272 | 6.4507193 | 5 | si:ch211-5k11.8 |
| zgc:1751353 | 1.010491 | 0.874379569 | 0.865 | 0.368 | 2.5671544 | 5 | zgc:175135 |
| zgc:1717793 | 5.782932 | 0.854315073 | 0.866 | 0.397 | 1.4691539 | 5 | zgc:171779 |
| zp2.23 | 8.842671 | 0.85195323 | 0.893 | 0.41 | 2.2464808 | 5 | zp2.2 |
| cldng3 | 1.669417 | 0.855147776 | 0.877 | 0.399 | 4.2411541 | 5 | cldng |
| zgc:771183 | 6.720092 | 0.856708754 | 0.823 | 0.352 | 1.7072394 | 5 | zgc:77118 |
| ccna13 | 5.095430 | 0.842148169 | 0.831 | 0.367 | 1.2944941 | 5 | ccna1 |
| BX649337.13 | 1.208141 | 0.845826893 | 0.803 | 0.336 | 3.0692826 | 5 | BX649337.1 |
| CR790368.13 | 3.138996 | 0.823005575 | 0.866 | 0.405 | 7.9746194 | 5 | CR790368.1 |
| si:dkeyp-98a7.83 | 5.255960 | 0.826458385 | 0.803 | 0.353 | 1.3352768 | 5 | si:dkeyp-98a7.8 |
| epcam3 | 1.047994 | 0.807760124 | 0.854 | 0.397 | 2.6624289 | 5 | epcam |
| zp3a.23 | 9.635766 | 0.807225006 | 0.817 | 0.382 | 2.4479665 | 5 | zp3a.2 |
| si:ch211-12h2.83 | 2.595551 | 0.79277 | 0.866 | 0.428 | 6.5939974 | 5 | si:ch211-12h2.8 |
| si:dkey-229d11.33 | 1.899326 | 0.827775435 | 0.79 | 0.337 | 4.8252395 | 5 | si:dkey-229d11.3 |
| qdprb23 | 6.973609 | 0.809168492 | 0.759 | 0.334 | 1.7716456 | 5 | qdprb2 |
| si:ch211-226h8.43 | 6.010342 | 0.810835923 | 0.784 | 0.34 | 1.5269276 | 5 | si:ch211-226h8.4 |
| si:ch211-12h2.63 | 2.689121 | 0.786713116 | 0.787 | 0.368 | 6.8317126 | 5 | si:ch211-12h2.6 |
| si:dkey-241l7.63 | 1.273631 | 0.77711934 | 0.831 | 0.412 | 3.2356614 | 5 | si:dkey-241l7.6 |
| si:ch211-14a17.73 | 6.512579 | 0.799790258 | 0.729 | 0.315 | 1.6545208 | 5 | si:ch211-14a17.7 |
| rrm23 | 9.402346 | 0.764693874 | 0.819 | 0.401 | 2.3886660 | 5 | rrm2 |
| lamp23 | 2.568973 | 0.744306066 | 0.835 | 0.465 | 6.5264784 | 5 | lamp2 |

| | | | | | | | |
|---|---|---|---|---|---|---|---|
| si:dkeyp-98a7.43 | 1.245200 | 0.776104965 | 0.734 | 0.329 | 3.1634321 | 5 | si:dkeyp-98a7.4 |
| slbp23 | 1.053652 | 0.775188625 | 0.698 | 0.307 | 2.6768034 | 5 | slbp2 |
| zar1l3 | 2.067611 | 0.773880254 | 0.706 | 0.307 | 5.2527678 | 5 | zar1l |
| CU571163.13 | 5.825478 | 0.783570395 | 0.714 | 0.309 | 1.4799628 | 5 | CU571163.1 |
| mid1ip1l3 | 2.892191 | 0.72128776 | 0.808 | 0.441 | 7.3476131 | 5 | mid1ip1l |
| zar13 | 5.017058 | 0.763687338 | 0.737 | 0.343 | 1.2745836 | 5 | zar1 |
| btg43 | 4.447818 | 0.770493537 | 0.659 | 0.284 | 1.1299682 | 5 | btg4 |
| nanog3 | 5.149438 | 0.767983932 | 0.657 | 0.277 | 1.3082147 | 5 | nanog |
| si:dkeyp-20e4.83 | 6.683200 | 0.776406395 | 0.717 | 0.315 | 1.6978670 | 5 | si:dkeyp-20e4.8 |
| si:dkey-90l23.13 | 8.638164 | 0.75507667 | 0.739 | 0.348 | 2.1945258 | 5 | si:dkey-90l23.1 |
| siva13 | 1.400012 | 0.747501741 | 0.718 | 0.342 | 3.5567325 | 5 | siva1 |
| zgc:1952453 | 3.184380 | 0.750095571 | 0.68 | 0.3 | 8.0899193 | 5 | zgc:195245 |
| tuba4l3 | 9.761224 | 0.74829648 | 0.692 | 0.315 | 2.4798391 | 5 | tuba4l |
| si:dkey-229d11.53 | 8.563169 | 0.741329816 | 0.665 | 0.301 | 2.1754732 | 5 | si:dkey-229d11.5 |
| ccnb13 | 1.310809 | 0.735612696 | 0.664 | 0.308 | 3.3301115 | 5 | ccnb1 |
| LOC1104378213 | 1.531796 | 0.750428511 | 0.697 | 0.323 | 3.8915285 | 5 | LOC110437821 |
| thy13 | 1.439793 | 0.714841315 | 0.726 | 0.359 | 3.6577942 | 5 | thy1 |
| ccnb23 | 2.370178 | 0.735132987 | 0.66 | 0.298 | 6.0214374 | 5 | ccnb2 |
| si:dkeyp-50f7.23 | 2.716853 | 0.725593911 | 0.683 | 0.313 | 6.9021655 | 5 | si:dkeyp-50f7.2 |
| si:dkeyp-46h3.63 | 3.486747 | 0.721723535 | 0.681 | 0.319 | 8.8580831 | 5 | si:dkeyp-46h3.6 |
| zgc:1734433 | 6.602693 | 0.721079904 | 0.601 | 0.254 | 1.6774143 | 5 | zgc:173443 |
| zpcx3 | 2.796874 | 0.720763544 | 0.604 | 0.261 | 7.1054608 | 5 | zpcx |
| s100a13 | 2.554710 | 0.712151101 | 0.623 | 0.283 | 6.4902428 | 5 | s100a1 |
| npm2b3 | 7.089664 | 0.714561481 | 0.656 | 0.31 | 1.8011293 | 5 | npm2b |
| si:ch211-132b12.83 | 6.162092 | 0.712720797 | 0.603 | 0.265 | 1.5654795 | 5 | si:ch211-132b12.8 |
| ctss2.13 | 3.295307 | 0.648453388 | 0.758 | 0.436 | 8.3717292 | 5 | ctss2.1 |
| si:dkeyp-46h3.83 | 2.609779 | 0.709654699 | 0.614 | 0.272 | 6.6301456 | 5 | si:dkeyp-46h3.8 |
| zgc:1362543 | 2.272325 | 0.70241404 | 0.589 | 0.255 | 5.7728441 | 5 | zgc:136254 |
| AL929032.13 | 1.311215 | 0.707503747 | 0.578 | 0.244 | 3.3311418 | 5 | AL929032.1 |
| zgc:1715173 | 3.013706 | 0.699726111 | 0.635 | 0.303 | 7.6563223 | 5 | zgc:171517 |
| LOC1005374993 | 5.224612 | 0.69760777 | 0.579 | 0.249 | 1.3273128 | 5 | LOC100537499 |
| si:dkeyp-46h3.23 | 7.052193 | 0.687032991 | 0.647 | 0.316 | 1.7916097 | 5 | si:dkeyp-46h3.2 |
| cth13 | 1.332428 | 0.686269265 | 0.62 | 0.296 | 3.3850352 | 5 | cth1 |
| spink2.13 | 1.395757 | 0.658463076 | 0.69 | 0.357 | 3.5459211 | 5 | spink2.1 |
| si:dkeyp-75b4.103 | 4.632287 | 0.638386689 | 0.664 | 0.35 | 1.1768327 | 5 | si:dkeyp-75b4.10 |

| | | | | | | | |
|---|---|---|---|---|---|---|---|
| kpna73 | 2.050815 | 0.685610626 | 0.581 | 0.255 | 5.2100956 | 5 | kpna7 |
| LOC1005351623 | 7.224428 | 0.691277648 | 0.557 | 0.238 | 1.8353659 | 5 | LOC100535162 |
| gtf3ab3 | 8.806958 | 0.677223299 | 0.567 | 0.248 | 2.2374077 | 5 | gtf3ab |
| zp3b3 | 1.468785 | 0.67627718 | 0.638 | 0.303 | 3.7314488 | 5 | zp3b |
| uchl13 | 1.679084 | 0.673339759 | 0.583 | 0.275 | 4.2657147 | 5 | uchl1 |
| cpeb1b3 | 2.649414 | 0.675577318 | 0.5 | 0.208 | 6.7308363 | 5 | cpeb1b |
| si:dkey-152b24.63 | 1.977696 | 0.671876359 | 0.533 | 0.23 | 5.0243369 | 5 | si:dkey-152b24.6 |
| gapdh3 | 7.878491 | 0.657649403 | 0.64 | 0.332 | 2.0015308 | 5 | gapdh |
| si:dkey-90l23.23 | 1.509157 | 0.666880729 | 0.571 | 0.269 | 3.8340154 | 5 | si:dkey-90l23.2 |
| avd3 | 6.139595 | 0.66133841 | 0.489 | 0.204 | 1.5597641 | 5 | avd |
| si:ch73-14h10.23 | 1.121703 | 0.646389015 | 0.496 | 0.21 | 2.8496887 | 5 | si:ch73-14h10.2 |
| ccna23 | 1.732605 | 0.652183841 | 0.503 | 0.225 | 4.4016842 | 5 | ccna2 |
| si:ch211-226l4.63 | 9.817043 | 0.647547044 | 0.511 | 0.229 | 2.4940199 | 5 | si:ch211-226l4.6 |
| wu:fi42e033 | 2.270669 | 0.637080064 | 0.56 | 0.274 | 5.7686346 | 5 | wu:fi42e03 |
| si:ch211-250e5.163 | 2.615707 | 0.629487114 | 0.548 | 0.272 | 6.6452058 | 5 | si:ch211-250e5.16 |
| zgc:554613 | 1.900578 | 0.626311522 | 0.503 | 0.228 | 4.8284188 | 5 | zgc:55461 |
| si:ch211-146l10.83 | 5.975678 | 0.630313674 | 0.516 | 0.239 | 1.5181211 | 5 | si:ch211-146l10.8 |
| gmnn3 | 4.516742 | 0.62583745 | 0.511 | 0.241 | 1.1474785 | 5 | gmnn |
| si:dkey-208k4.23 | 5.416196 | 0.616815278 | 0.499 | 0.231 | 1.3759846 | 5 | si:dkey-208k4.2 |
| zp3a.13 | 2.731219 | 0.615684521 | 0.577 | 0.3 | 6.9386631 | 5 | zp3a.1 |
| si:dkey-241l7.33 | 3.759900 | 0.628181916 | 0.545 | 0.288 | 9.5520264 | 5 | si:dkey-241l7.3 |
| lsm14b3 | 2.148110 | 0.610534705 | 0.495 | 0.234 | 5.4572736 | 5 | lsm14b |
| zgc:1714743 | 1.268141 | 0.608465299 | 0.501 | 0.237 | 3.2217140 | 5 | zgc:171474 |
| pabpn1l3 | 8.869161 | 0.606378372 | 0.479 | 0.223 | 2.2532104 | 5 | pabpn1l |
| birc5b3 | 9.424707 | 0.602035924 | 0.451 | 0.202 | 2.3943468 | 5 | birc5b |
| arhgef10la3 | 3.800393 | 0.604182158 | 0.468 | 0.215 | 9.6548988 | 5 | arhgef10la |
| hnrnpabb3 | 1.580192 | 0.480178131 | 0.734 | 0.55 | 4.0144802 | 5 | hnrnpabb |
| LOC1005368603 | 3.282818 | 0.590210927 | 0.521 | 0.277 | 8.3399995 | 5 | LOC100536860 |
| LOC1018854773 | 3.192109 | 0.583805442 | 0.464 | 0.218 | 8.1095532 | 5 | LOC101885477 |
| slbp4 | 2.057709 | 0.589444387 | 0.413 | 0.188 | 5.2276097 | 5 | slbp |
| si:dkey-203a12.2 | 9.520939 | 0.526485605 | 0.434 | 0.198 | 2.4187947 | 5 | si:dkey-203a12.2 |
| si:dkey-37o8.13 | 2.291203 | 0.573204343 | 0.483 | 0.242 | 5.8208012 | 5 | si:dkey-37o8.1 |
| nasp3 | 3.100551 | 0.568101177 | 0.521 | 0.283 | 7.8769499 | 5 | nasp |
| ca15b3 | 4.776673 | 0.557174859 | 0.422 | 0.194 | 1.2135138 | 5 | ca15b |
| elovl7b3 | 3.068094 | 0.559356257 | 0.414 | 0.194 | 7.7944931 | 5 | elovl7b |

| | | | | | | | |
|---|---|---|---|---|---|---|---|
| si:ch211-251f6.63 | 9.526089 | 0.556611286 | 0.492 | 0.262 | 2.4201029 | 5 | si:ch211-251f6.6 |
| si:dkey-19b23.113 | 7.927646 | 0.554362505 | 0.444 | 0.219 | 2.0140186 | 5 | si:dkey-19b23.11 |
| slc16a33 | 9.524798 | 0.545879633 | 0.414 | 0.196 | 2.4197750 | 5 | slc16a3 |
| haus63 | 3.017865 | 0.533059594 | 0.373 | 0.164 | 7.6668871 | 5 | haus6 |
| elavl23 | 7.219118 | 0.542720323 | 0.382 | 0.176 | 1.8340171 | 5 | elavl2 |
| pigrl2.33 | 1.622096 | 0.532956292 | 0.358 | 0.159 | 4.1209367 | 5 | pigrl2.3 |
| buc3 | 3.960765 | 0.514893767 | 0.329 | 0.138 | 1.0062325 | 5 | buc |
| gdf33 | 6.469101 | 0.531306259 | 0.377 | 0.175 | 1.6434752 | 5 | gdf3 |
| spink2.2 | 7.588126 | 0.441976664 | 0.288 | 0.107 | 1.9277634 | 5 | spink2.2 |
| zgc:1738563 | 8.521371 | 0.532103181 | 0.37 | 0.172 | 2.1648543 | 5 | zgc:173856 |
| si:ch211-145c1.13 | 6.578412 | 0.530570474 | 0.392 | 0.183 | 1.6712456 | 5 | si:ch211-145c1.1 |
| zgc:562313 | 7.016873 | 0.525443689 | 0.381 | 0.178 | 1.7826367 | 5 | zgc:56231 |
| si:ch1073-263o8.23 | 1.255403 | 0.529194536 | 0.418 | 0.207 | 3.1893532 | 5 | si:ch1073-263o8.2 |
| hnrnpa0l3 | 2.167133 | 0.264646745 | 0.79 | 0.739 | 5.5056027 | 5 | hnrnpa0l |
| zgc:925913 | 5.385553 | 0.503982745 | 0.354 | 0.161 | 1.3681998 | 5 | zgc:92591 |
| cldn7b3 | 1.147023 | 0.49801983 | 0.299 | 0.125 | 2.9140131 | 5 | cldn7b |
| map1lc3c3 | 1.152160 | 0.503912606 | 0.345 | 0.157 | 2.9270648 | 5 | map1lc3c |
| si:dkeyp-98a7.73 | 3.206722 | 0.489988072 | 0.345 | 0.152 | 8.1466792 | 5 | si:dkeyp-98a7.7 |
| si:ch73-90k17.13 | 7.115467 | 0.50937411 | 0.397 | 0.195 | 1.8076846 | 5 | si:ch73-90k17.1 |
| si:dkeyp-46h3.53 | 4.268800 | 0.501357339 | 0.427 | 0.229 | 1.0844888 | 5 | si:dkeyp-46h3.5 |
| sybu3 | 4.221650 | 0.476039578 | 0.296 | 0.129 | 1.0725103 | 5 | sybu |
| zp3c3 | 4.557748 | 0.483143282 | 0.321 | 0.148 | 1.1578959 | 5 | zp3c |
| zglp13 | 3.337081 | 0.483128293 | 0.328 | 0.156 | 8.4778544 | 5 | zglp1 |
| AL929192.13 | 8.789075 | 0.44786421 | 0.292 | 0.127 | 2.2328646 | 5 | AL929192.1 |
| sinup3 | 1.110193 | 0.472435673 | 0.333 | 0.16 | 2.8204456 | 5 | sinup |
| crp33 | 1.812382 | 0.468592016 | 0.352 | 0.172 | 4.6043570 | 5 | crp3 |
| org3 | 2.131959 | 0.483502047 | 0.372 | 0.192 | 5.4162420 | 5 | org |
| h3f3a3 | 7.914724 | 0.423280614 | 0.608 | 0.477 | 2.0107357 | 5 | h3f3a |
| rbpms2b3 | 1.136904 | 0.477575043 | 0.357 | 0.183 | 2.8883068 | 5 | rbpms2b |
| zgc:1134243 | 7.764502 | 0.448877607 | 0.283 | 0.126 | 1.9725718 | 5 | zgc:113424 |
| zgc:566993 | 8.585795 | 0.459894936 | 0.384 | 0.201 | 2.1812213 | 5 | zgc:56699 |
| zgc:1719772 | 5.559722 | 0.464289242 | 0.351 | 0.182 | 1.4124475 | 5 | zgc:171977 |
| gyg1a3 | 1.609927 | 0.465829493 | 0.475 | 0.312 | 4.0900196 | 5 | gyg1a |
| pabpc1l3 | 2.012655 | 0.429216376 | 0.254 | 0.111 | 5.1131511 | 5 | pabpc1l |
| hsd17b12a3 | 2.451183 | 0.436952524 | 0.282 | 0.13 | 6.2272322 | 5 | hsd17b12a |

| | | | | | | | |
|---|---|---|---|---|---|---|---|
| pou5f33 | 3.615417 | 0.428986216 | 0.261 | 0.115 | 9.1849678 | 5 | pou5f3 |
| sox19b3 | 6.799101 | 0.384402346 | 0.2 | 0.075 | 1.7273117 | 5 | sox19b |
| zgc:1629453 | 8.649495 | 0.433836232 | 0.28 | 0.13 | 2.1974044 | 5 | zgc:162945 |
| zp2l13 | 3.472395 | 0.434645694 | 0.276 | 0.129 | 8.8216213 | 5 | zp2l1 |
| si:ch211-146l10.73 | 7.635480 | 0.43793902 | 0.303 | 0.149 | 1.9397938 | 5 | si:ch211-146l10.7 |
| si:ch73-359m17.2 | 1.017056 | 0.404166427 | 0.321 | 0.157 | 2.5838310 | 5 | si:ch73-359m17.2 |
| retsatl3 | 1.035039 | 0.443640871 | 0.319 | 0.16 | 2.6295176 | 5 | retsatl |
| elovl7a3 | 1.332528 | 0.429738233 | 0.3 | 0.146 | 3.3852890 | 5 | elovl7a |
| CR383662.23 | 2.923128 | 0.355354176 | 0.183 | 0.066 | 7.4262078 | 5 | CR383662.2 |
| si:ch73-78o10.13 | 4.485429 | 0.406406701 | 0.241 | 0.104 | 1.1395234 | 5 | si:ch73-78o10.1 |
| zgc:1141302 | 1.895754 | 0.352747501 | 0.168 | 0.06 | 4.8161639 | 5 | zgc:114130 |
| gbgt1l43 | 2.185822 | 0.414889626 | 0.259 | 0.118 | 5.5530825 | 5 | gbgt1l4 |
| zgc:1628793 | 1.005382 | 0.42383561 | 0.291 | 0.143 | 2.5541730 | 5 | zgc:162879 |
| mkrn43 | 2.805157 | 0.432707555 | 0.345 | 0.187 | 7.1265034 | 5 | mkrn4 |
| si:dkey-203a12.9 | 2.812320 | 0.395013691 | 0.249 | 0.111 | 7.1447012 | 5 | si:dkey-203a12.9 |
| rasl11b3 | 4.128148 | 0.402697972 | 0.241 | 0.108 | 1.0487560 | 5 | rasl11b |
| CR847984.13 | 9.462095 | 0.420950581 | 0.272 | 0.132 | 2.4038452 | 5 | CR847984.1 |
| tph1a3 | 1.221136 | 0.393440318 | 0.245 | 0.111 | 3.1022965 | 5 | tph1a |
| snu13b3 | 4.745417 | 0.419362905 | 0.499 | 0.366 | 1.2055732 | 5 | snu13b |
| zgc:1118683 | 5.624312 | 0.398155449 | 0.25 | 0.116 | 1.4288565 | 5 | zgc:111868 |
| zmp:0000001323 | 9.959179 | 0.350314441 | 0.262 | 0.12 | 2.5301295 | 5 | zmp:0000001323 |
| rcc13 | 1.038363 | 0.428422151 | 0.315 | 0.167 | 2.6379616 | 5 | rcc1 |
| inka1b3 | 1.733784 | 0.417219482 | 0.294 | 0.15 | 4.4046795 | 5 | inka1b |
| zgc:1034823 | 3.259349 | 0.406708361 | 0.296 | 0.151 | 8.2803765 | 5 | zgc:103482 |
| dctpp13 | 5.575845 | 0.418420719 | 0.28 | 0.143 | 1.4165436 | 5 | dctpp1 |
| CABZ01059627.23 | 5.986250 | 0.386858432 | 0.23 | 0.104 | 1.5208068 | 5 | CABZ01059627.2 |
| BX322787.1 | 1.253176 | 0.360042063 | 0.253 | 0.116 | 3.1836948 | 5 | BX322787.1 |
| si:dkey-46g23.23 | 1.344702 | 0.391083153 | 0.242 | 0.114 | 3.4162168 | 5 | si:dkey-46g23.2 |
| si:dkey-216e24.33 | 1.428535 | 0.381461025 | 0.218 | 0.096 | 3.6291952 | 5 | si:dkey-216e24.3 |
| patl23 | 1.535083 | 0.400096773 | 0.275 | 0.137 | 3.8998798 | 5 | patl2 |
| larp6b3 | 2.830842 | 0.404524634 | 0.299 | 0.156 | 7.1917544 | 5 | larp6b |
| zgc:1717503 | 8.476212 | 0.404342317 | 0.272 | 0.138 | 2.1533818 | 5 | zgc:171750 |
| mos3 | 2.615617 | 0.386310736 | 0.233 | 0.109 | 6.6449753 | 5 | mos |
| plk13 | 1.014122 | 0.394607819 | 0.276 | 0.143 | 2.5763770 | 5 | plk1 |
| si:ch73-160p18.33 | 1.313905 | 0.408084893 | 0.336 | 0.192 | 3.3379759 | 5 | si:ch73-160p18.3 |

| | | | | | | | |
|---|---|---|---|---|---|---|---|
| zgc:1138863 | 3.512610 | 0.369234702 | 0.235 | 0.111 | 8.9237867 | 5 | zgc:113886 |
| zgc:1737423 | 4.741497 | 0.382170177 | 0.253 | 0.124 | 1.2045774 | 5 | zgc:173742 |
| zgc:660243 | 5.814834 | 0.369225063 | 0.234 | 0.112 | 1.4772586 | 5 | zgc:66024 |
| cdca7a3 | 6.171331 | 0.412615875 | 0.324 | 0.187 | 1.5678267 | 5 | cdca7a |
| mcm3l3 | 1.198979 | 0.329810157 | 0.169 | 0.068 | 3.0460070 | 5 | mcm3l |
| chia.43 | 6.077841 | 0.369242341 | 0.237 | 0.116 | 1.5440756 | 5 | chia.4 |
| zgc:922873 | 1.370876 | 0.361777811 | 0.222 | 0.107 | 3.4827114 | 5 | zgc:92287 |
| nfr3 | 1.053907 | 0.341196923 | 0.193 | 0.087 | 2.6774525 | 5 | nfr |
| larp6a3 | 1.106972 | 0.401371244 | 0.34 | 0.208 | 2.8122627 | 5 | larp6a |
| CDPF13 | 1.110537 | 0.41448109 | 0.356 | 0.23 | 2.8213199 | 5 | CDPF1 |
| mad2l13 | 1.731717 | 0.376465232 | 0.28 | 0.154 | 4.3994282 | 5 | mad2l1 |
| si:ch1073-190k2.13 | 1.877539 | 0.378644571 | 0.276 | 0.15 | 4.7698889 | 5 | si:ch1073-190k2.1 |
| LOC1018838973 | 2.361287 | 0.361787693 | 0.253 | 0.13 | 5.9988520 | 5 | LOC101883897 |
| figla2 | 2.500085 | 0.329885818 | 0.173 | 0.075 | 6.3514675 | 5 | figla |
| dynll13 | 6.157746 | 0.345452914 | 0.529 | 0.438 | 1.5643753 | 5 | dynll1 |
| cdc203 | 7.785491 | 0.358967014 | 0.243 | 0.125 | 1.9779042 | 5 | cdc20 |
| tac3b3 | 9.292400 | 0.33534484 | 0.204 | 0.095 | 2.3607342 | 5 | tac3b |
| zgc:1017443 | 1.235248 | 0.354106038 | 0.218 | 0.108 | 3.1381477 | 5 | zgc:101744 |
| cyp11a11 | 1.616945 | 0.288197532 | 0.138 | 0.053 | 4.1078498 | 5 | cyp11a1 |
| mibp4 | 1.734099 | 0.383349414 | 0.323 | 0.194 | 4.4054810 | 5 | mibp |
| gstp13 | 2.596312 | 0.379545553 | 0.45 | 0.343 | 6.5959310 | 5 | gstp1 |
| si:dkey-42i9.83 | 2.690575 | 0.339371683 | 0.21 | 0.101 | 6.8354072 | 5 | si:dkey-42i9.8 |
| ypel33 | 8.534563 | 0.352699007 | 0.213 | 0.106 | 2.1682059 | 5 | ypel3 |
| BX005461.12 | 8.545860 | 0.29847428 | 0.151 | 0.062 | 2.1710758 | 5 | BX005461.1 |
| si:dkey-68o6.53 | 1.159927 | 0.346029821 | 0.202 | 0.097 | 2.9467962 | 5 | si:dkey-68o6.5 |
| si:ch211-226h8.81 | 3.473853 | 0.324805511 | 0.171 | 0.076 | 8.8253241 | 5 | si:ch211-226h8.8 |
| pttg13 | 2.264047 | 0.360417296 | 0.242 | 0.132 | 5.7518121 | 5 | pttg1 |
| ccne13 | 2.571825 | 0.32931514 | 0.188 | 0.09 | 6.5337230 | 5 | ccne1 |
| BX001030.13 | 2.671029 | 0.323362756 | 0.198 | 0.097 | 6.7857492 | 5 | BX001030.1 |
| zgc:1141233 | 3.083078 | 0.307256208 | 0.177 | 0.082 | 7.8325619 | 5 | zgc:114123 |
| ankrd223 | 3.182230 | 0.345354663 | 0.225 | 0.117 | 8.0844571 | 5 | ankrd22 |
| eif4a1a3 | 8.429469 | 0.344488612 | 0.471 | 0.403 | 2.1415066 | 5 | eif4a1a |
| CU467646.23 | 9.383168 | 0.318517848 | 0.201 | 0.1 | 2.3837939 | 5 | CU467646.2 |
| LOC1005380232 | 1.257952 | 0.284148018 | 0.143 | 0.06 | 3.1958276 | 5 | LOC100538023 |
| CT009596.13 | 1.863460 | 0.292494156 | 0.159 | 0.071 | 4.7341205 | 5 | CT009596.1 |

| | | | | | | | |
|---|---|---|---|---|---|---|---|
| ccnh3 | 2.490411 | 0.330505269 | 0.201 | 0.102 | 6.3268910 | 5 | ccnh |
| yrdc3 | 2.593977 | 0.328552402 | 0.198 | 0.1 | 6.5900000 | 5 | yrdc |
| pif13 | 3.052293 | 0.312095892 | 0.179 | 0.085 | 7.7543527 | 5 | pif1 |
| zgc:1655393 | 4.504763 | 0.317351819 | 0.192 | 0.095 | 1.1444352 | 5 | zgc:165539 |
| CABZ01059627.13 | 5.163778 | 0.322023626 | 0.21 | 0.108 | 1.3118580 | 5 | CABZ01059627.1 |
| si:dkey-43p13.53 | 7.110893 | 0.285317589 | 0.149 | 0.065 | 1.8065224 | 5 | si:dkey-43p13.5 |
| kpna23 | 7.912588 | 0.337461945 | 0.239 | 0.132 | 2.0101931 | 5 | kpna2 |
| vstm2l3 | 8.361647 | 0.295531253 | 0.164 | 0.075 | 2.1242765 | 5 | vstm2l |
| h2afva3 | 1.005056 | 0.367300896 | 0.402 | 0.302 | 2.5533461 | 5 | h2afva |
| cx44.23 | 1.096921 | 0.313805592 | 0.188 | 0.093 | 2.7867288 | 5 | cx44.2 |
| dynll2a3 | 1.114786 | 0.328439673 | 0.216 | 0.114 | 2.8321162 | 5 | dynll2a |
| bzw1b3 | 1.442531 | 0.338384013 | 0.456 | 0.375 | 3.6647506 | 5 | bzw1b |
| cdca83 | 2.358472 | 0.336738473 | 0.243 | 0.136 | 5.9916988 | 5 | cdca8 |
| si:ch73-132k15.2 | 2.569608 | 0.266968664 | 0.123 | 0.049 | 6.5280892 | 5 | si:ch73-132k15.2 |
| npm2a3 | 2.661408 | 0.270856004 | 0.143 | 0.062 | 6.7613089 | 5 | npm2a |
| rbm4.33 | 3.444986 | 0.321520646 | 0.489 | 0.419 | 8.7519872 | 5 | rbm4.3 |
| CU655961.43 | 3.711148 | 0.29114789 | 0.153 | 0.07 | 9.4281725 | 5 | CU655961.4 |
| fdx13 | 4.738038 | 0.366241718 | 0.3 | 0.194 | 1.2036985 | 5 | fdx1 |
| lmnl32 | 4.955137 | 0.27621949 | 0.139 | 0.06 | 1.2588527 | 5 | lmnl3 |
| nanos33 | 5.623636 | 0.295151366 | 0.171 | 0.081 | 1.4286848 | 5 | nanos3 |
| zgc:1529863 | 7.408502 | 0.283576499 | 0.155 | 0.07 | 1.8821299 | 5 | zgc:152986 |
| lpcat43 | 7.473271 | 0.296123198 | 0.179 | 0.087 | 1.8985847 | 5 | lpcat4 |
| si:ch211-69g19.23 | 7.975895 | 0.318616727 | 0.194 | 0.1 | 2.0262761 | 5 | si:ch211-69g19.2 |
| si:dkeyp-114g9.13 | 9.883472 | 0.295335969 | 0.172 | 0.082 | 2.5108963 | 5 | si:dkeyp-114g9.1 |
| si:dkey-188i13.113 | 1.636668 | 0.360281442 | 0.39 | 0.287 | 4.1579562 | 5 | si:dkey-188i13.11 |
| degs23 | 1.883372 | 0.292621376 | 0.175 | 0.085 | 4.7847079 | 5 | degs2 |
| cpm3 | 1.898038 | 0.297530498 | 0.176 | 0.086 | 4.8219675 | 5 | cpm |
| si:ch211-202f5.33 | 2.454556 | 0.286781514 | 0.156 | 0.073 | 6.2358003 | 5 | si:ch211-202f5.3 |
| si:ch211-14a17.63 | 3.820877 | 0.271076584 | 0.149 | 0.068 | 9.7069382 | 5 | si:ch211-14a17.6 |
| faub3 | 7.159846 | 0.354794661 | 0.28 | 0.178 | 1.8189589 | 5 | faub |
| BX284697.13 | 1.148998 | 0.29761454 | 0.18 | 0.091 | 2.9190296 | 5 | BX284697.1 |
| rnf43 | 1.357025 | 0.329047916 | 0.205 | 0.113 | 3.4475238 | 5 | rnf4 |
| BX537276.13 | 1.658893 | 0.29046515 | 0.188 | 0.096 | 4.2144186 | 5 | BX537276.1 |
| rdh10b3 | 1.678998 | 0.32969243 | 0.249 | 0.147 | 4.2654946 | 5 | rdh10b |
| tmem144b3 | 2.295190 | 0.280491479 | 0.156 | 0.075 | 5.8309314 | 5 | tmem144b |

| | | | | | | | |
|---|---|---|---|---|---|---|---|
| cdt13 | 2.297979 | 0.278283099 | 0.149 | 0.07 | 5.8380174 | 5 | cdt1 |
| hspa53 | 2.331514 | 0.339584052 | 0.388 | 0.292 | 5.9232125 | 5 | hspa5 |
| ctsba3 | 3.228289 | 0.327136559 | 0.447 | 0.349 | 8.2014693 | 5 | ctsba |
| zgc:1526523 | 3.506458 | 0.289280814 | 0.179 | 0.09 | 8.9081577 | 5 | zgc:152652 |
| marcksb3 | 3.632833 | 0.345042805 | 0.254 | 0.157 | 9.2292122 | 5 | marcksb |
| snx10a3 | 5.559063 | 0.298119223 | 0.192 | 0.102 | 1.4122799 | 5 | snx10a |
| kif22 | 7.777447 | 0.295038059 | 0.153 | 0.075 | 1.9758606 | 5 | kif22 |
| si:ch211-226h8.63 | 1.007274 | 0.262030607 | 0.136 | 0.062 | 2.5589809 | 5 | si:ch211-226h8.6 |
| zgc:1714263 | 1.267758 | 0.28020983 | 0.169 | 0.085 | 3.2207399 | 5 | zgc:171426 |
| zgc:1630303 | 1.567891 | 0.296055023 | 0.193 | 0.103 | 3.9832273 | 5 | zgc:163030 |
| zgc:923133 | 1.632333 | 0.305569495 | 0.193 | 0.105 | 4.1469444 | 5 | zgc:92313 |
| zgc:558883 | 2.152560 | 0.257421098 | 0.144 | 0.068 | 5.4685808 | 5 | zgc:55888 |
| COQ10A2 | 2.430604 | 0.264844451 | 0.143 | 0.067 | 6.1749503 | 5 | COQ10A |
| cdca93 | 2.753838 | 0.252991725 | 0.146 | 0.068 | 6.9961278 | 5 | cdca9 |
| brf23 | 2.771887 | 0.269447994 | 0.157 | 0.077 | 7.0419809 | 5 | brf2 |
| zgc:1735442 | 3.149377 | 0.251955785 | 0.132 | 0.06 | 8.0009939 | 5 | zgc:173544 |
| fbxo322 | 3.507576 | 0.283031025 | 0.167 | 0.085 | 8.9109983 | 5 | fbxo32 |
| bmb3 | 3.521888 | 0.264835788 | 0.147 | 0.07 | 8.9473572 | 5 | bmb |
| zgc:566763 | 3.546725 | 0.34924199 | 0.361 | 0.271 | 9.0104560 | 5 | zgc:56676 |
| cox173 | 5.209787 | 0.322657492 | 0.415 | 0.339 | 1.3235466 | 5 | cox17 |
| CU655961.33 | 6.001357 | 0.264895448 | 0.159 | 0.079 | 1.5246448 | 5 | CU655961.3 |
| si:ch73-340n8.43 | 6.309256 | 0.310749741 | 0.198 | 0.111 | 1.6028665 | 5 | si:ch73-340n8.4 |
| zgc:1129803 | 1.286450 | 0.262306002 | 0.152 | 0.075 | 3.2682264 | 5 | zgc:112980 |
| parn3 | 1.555417 | 0.316576729 | 0.216 | 0.127 | 3.9515391 | 5 | parn |
| tram23 | 1.928930 | 0.314341708 | 0.205 | 0.119 | 4.9004473 | 5 | tram2 |
| dazl3 | 2.045270 | 0.268577244 | 0.168 | 0.087 | 5.1960092 | 5 | dazl |
| pttg1ipb3 | 2.352318 | 0.296914804 | 0.18 | 0.098 | 5.9760648 | 5 | pttg1ipb |
| mylipa3 | 2.719776 | 0.286004253 | 0.179 | 0.095 | 6.9095919 | 5 | mylipa |
| znf395a2 | 3.245980 | 0.276784649 | 0.159 | 0.081 | 8.2464145 | 5 | znf395a |
| hmga1a2 | 4.577680 | 0.266993544 | 0.492 | 0.441 | 1.1629597 | 5 | hmga1a |
| chchd103 | 4.837943 | 0.318770448 | 0.22 | 0.131 | 1.2290795 | 5 | chchd10 |
| gpr1603 | 8.568107 | 0.250667743 | 0.135 | 0.065 | 2.1767277 | 5 | gpr160 |
| zgc:771122 | 9.970493 | 0.290201346 | 0.177 | 0.097 | 2.5330038 | 5 | zgc:77112 |
| prrg23 | 1.364692 | 0.264474482 | 0.148 | 0.075 | 3.4670004 | 5 | prrg2 |
| dnd12 | 1.540005 | 0.266429673 | 0.168 | 0.089 | 3.9123845 | 5 | dnd1 |

| | | | | | | | |
|---|---|---|---|---|---|---|---|
| ppp1r3b3 | 1.944388 | 0.275234706 | 0.167 | 0.089 | 4.9397178 | 5 | ppp1r3b |
| kmt5ab3 | 2.599127 | 0.251740746 | 0.146 | 0.073 | 6.6030842 | 5 | kmt5ab |
| ctsla3 | 3.521660 | 0.29486285 | 0.438 | 0.372 | 8.9467777 | 5 | ctsla |
| gfer3 | 3.974051 | 0.269757249 | 0.159 | 0.084 | 1.0096077 | 5 | gfer |
| tent5c3 | 4.452712 | 0.294426907 | 0.204 | 0.12 | 1.1312114 | 5 | tent5c |
| cdc63 | 4.694262 | 0.269317748 | 0.16 | 0.085 | 1.1925774 | 5 | cdc6 |
| tmem54a3 | 4.876587 | 0.275658277 | 0.2 | 0.115 | 1.2388970 | 5 | tmem54a |
| il13ra23 | 5.063245 | 0.269907287 | 0.167 | 0.09 | 1.2863174 | 5 | il13ra2 |
| LOC1039099723 | 7.206756 | 0.312708407 | 0.393 | 0.32 | 1.8308763 | 5 | LOC103909972 |
| ddx39ab3 | 1.440158 | 0.32794355 | 0.345 | 0.268 | 3.6587220 | 5 | ddx39ab |
| aurka3 | 1.479142 | 0.258095598 | 0.176 | 0.097 | 3.7577610 | 5 | aurka |
| zgc:868393 | 1.594441 | 0.267302297 | 0.167 | 0.091 | 4.0506779 | 5 | zgc:86839 |
| aurkb3 | 2.479330 | 0.279166661 | 0.187 | 0.108 | 6.2987399 | 5 | aurkb |
| nifk2 | 2.924433 | 0.306123909 | 0.19 | 0.115 | 7.4295222 | 5 | nifk |
| ube2c2 | 3.744772 | 0.274166628 | 0.202 | 0.121 | 9.5135934 | 5 | ube2c |
| zgc:1139843 | 3.920951 | 0.250618043 | 0.152 | 0.081 | 9.9611767 | 5 | zgc:113984 |
| cx43.43 | 6.557488 | 0.281793599 | 0.185 | 0.109 | 1.6659298 | 5 | cx43.4 |
| tpx23 | 9.984555 | 0.267823608 | 0.18 | 0.105 | 2.5365762 | 5 | tpx2 |
| rogdi3 | 1.042952 | 0.261524538 | 0.173 | 0.098 | 2.6496197 | 5 | rogdi |
| pop43 | 3.556658 | 0.266766756 | 0.173 | 0.101 | 9.0356910 | 5 | pop4 |
| rpain3 | 4.139425 | 0.262264462 | 0.171 | 0.098 | 1.0516210 | 5 | rpain |
| prdx13 | 7.110127 | 0.283666514 | 0.225 | 0.146 | 1.8063279 | 5 | prdx1 |
| sec61a1l1 | 1.577838 | 0.292272365 | 0.183 | 0.113 | 4.0084996 | 5 | sec61a1l |
| zc3h11a3 | 1.735747 | 0.300249153 | 0.263 | 0.187 | 4.4096672 | 5 | zc3h11a |
| alyref2 | 2.833555 | 0.306709483 | 0.315 | 0.249 | 7.1986476 | 5 | alyref |
| f11r.13 | 4.208912 | 0.294211094 | 0.235 | 0.162 | 1.0692742 | 5 | f11r.1 |
| selenoh3 | 4.408503 | 0.298855096 | 0.296 | 0.226 | 1.1199803 | 5 | selenoh |
| thop13 | 6.024216 | 0.262143197 | 0.169 | 0.102 | 1.5304521 | 5 | thop1 |
| h2afx3 | 6.169078 | 0.263301878 | 0.192 | 0.12 | 1.5672543 | 5 | h2afx |
| si:ch211-210c8.73 | 1.582135 | 0.270641372 | 0.18 | 0.113 | 4.0194149 | 5 | si:ch211-210c8.7 |
| thoc72 | 2.240963 | 0.272039261 | 0.171 | 0.106 | 5.6931670 | 5 | thoc7 |
| rpz23 | 2.293177 | 0.285606037 | 0.233 | 0.162 | 5.8258177 | 5 | rpz2 |
| cbx3a1 | 2.474595 | 0.286423313 | 0.345 | 0.291 | 6.2867089 | 5 | cbx3a |
| snrpb23 | 4.839207 | 0.27985004 | 0.202 | 0.136 | 1.2294006 | 5 | snrpb2 |
| cnep1r12 | 7.067133 | 0.259250976 | 0.172 | 0.109 | 1.7954052 | 5 | cnep1r1 |

| | | | | | | |
|---|---|---|---|---|---|---|
| chtopa3 | 9.621543 | 0.288886967 | 0.269 | 0.205 | 2.4443531 | 5 | chtopa |
| rbm73 | 1.680543 | 0.257122524 | 0.168 | 0.106 | 4.2694213 | 5 | rbm7 |
| nol73 | 2.383460 | 0.257686983 | 0.172 | 0.111 | 6.0551820 | 5 | nol7 |
| ddx193 | 6.177429 | 0.253934896 | 0.196 | 0.133 | 0.000157 | 5 | ddx19 |
| STMP12 | 6.748003 | 0.284680824 | 0.335 | 0.293 | 0.000171 | 5 | STMP1 |
| polr2k3 | 8.052079 | 0.269283617 | 0.237 | 0.174 | 0.000205 | 5 | polr2k |
| FQ323156.12 | 1.264905 | 0.262122395 | 0.21 | 0.151 | 0.000321 | 5 | FQ323156.1 |
| g3bp13 | 2.531862 | 0.266499363 | 0.25 | 0.192 | 0.000643 | 5 | g3bp1 |
| hmgb3b3 | 2.704644 | 0.255384255 | 0.224 | 0.17 | 0.006871 | 5 | hmgb3b |
| zgc:1102392 | 3.250799 | 0.254201574 | 0.246 | 0.196 | 0.008259 | 5 | zgc:110239 |
| ncbp22 | 3.828022 | 0.250533138 | 0.213 | 0.16 | 0.009725 | 5 | ncbp2 |
| anp32b2 | 1.845773 | 0.251960207 | 0.29 | 0.252 | 0.046892 | 5 | anp32b |
| rbp41 | 0 | 1.625642516 | 0.912 | 0.105 | 0 | 6 | rbp4 |
| hsd11b21 | 0 | 1.565883301 | 0.908 | 0.113 | 0 | 6 | hsd11b2 |
| cxcl18b1 | 0 | 1.375322748 | 0.66 | 0.085 | 0 | 6 | cxcl18b |
| fn1b1 | 0 | 1.303216785 | 0.778 | 0.069 | 0 | 6 | fn1b |
| nid2a | 0 | 1.296726783 | 0.75 | 0.032 | 0 | 6 | nid2a |
| tagln | 0 | 1.283734044 | 0.626 | 0.048 | 0 | 6 | tagln |
| col1a21 | 0 | 1.224689327 | 0.916 | 0.148 | 0 | 6 | col1a2 |
| cd81a1 | 0 | 1.211954553 | 0.895 | 0.211 | 0 | 6 | cd81a |
| mmp16a1 | 0 | 1.203215297 | 0.655 | 0.051 | 0 | 6 | mmp16a |
| zgc:158343 | 0 | 1.199870474 | 0.699 | 0.145 | 0 | 6 | zgc:158343 |
| dcn1 | 0 | 1.183791055 | 0.827 | 0.112 | 0 | 6 | dcn |
| c61 | 0 | 1.147332767 | 0.776 | 0.082 | 0 | 6 | c6 |
| col1a1a1 | 0 | 1.143724232 | 0.872 | 0.133 | 0 | 6 | col1a1a |
| inhbaa | 0 | 1.124966426 | 0.582 | 0.028 | 0 | 6 | inhbaa |
| si:ch73-46j18.51 | 0 | 1.114252065 | 0.859 | 0.178 | 0 | 6 | si:ch73-46j18.5 |
| pmp22a1 | 0 | 1.112851657 | 0.823 | 0.136 | 0 | 6 | pmp22a |
| ccl25b1 | 0 | 1.094777509 | 0.562 | 0.072 | 0 | 6 | ccl25b |
| gstm.3 | 0 | 1.077243993 | 0.661 | 0.106 | 0 | 6 | gstm.3 |
| mdka1 | 0 | 1.063900659 | 0.671 | 0.087 | 0 | 6 | mdka |
| col5a11 | 0 | 1.016874961 | 0.746 | 0.104 | 0 | 6 | col5a1 |
| col6a4a1 | 0 | 0.998275419 | 0.58 | 0.042 | 0 | 6 | col6a4a |
| mfap21 | 0 | 0.996552445 | 0.7 | 0.092 | 0 | 6 | mfap2 |
| col1a1b1 | 0 | 0.987817151 | 0.809 | 0.14 | 0 | 6 | col1a1b |

| | | | | | | | |
|---|---|---|---|---|---|---|---|
| col6a11 | 0 | 0.987088374 | 0.629 | 0.072 | 0 | 6 | col6a1 |
| col4a11 | 0 | 0.973471623 | 0.579 | 0.046 | 0 | 6 | col4a1 |
| cald1a1 | 0 | 0.966628232 | 0.674 | 0.094 | 0 | 6 | cald1a |
| pdgfra1 | 0 | 0.965308488 | 0.579 | 0.037 | 0 | 6 | pdgfra |
| nid1b | 0 | 0.960136622 | 0.572 | 0.045 | 0 | 6 | nid1b |
| si:ch1073-291c23.21 | 0 | 0.958386185 | 0.616 | 0.047 | 0 | 6 | si:ch1073-291c23.2 |
| wu:fj16a03 | 0 | 0.953530203 | 0.394 | 0.029 | 0 | 6 | wu:fj16a03 |
| cpn11 | 0 | 0.918432145 | 0.574 | 0.054 | 0 | 6 | cpn1 |
| zgc:1951731 | 0 | 0.90866061 | 0.64 | 0.089 | 0 | 6 | zgc:195173 |
| col18a1a | 0 | 0.893922433 | 0.464 | 0.03 | 0 | 6 | col18a1a |
| tmem1761 | 0 | 0.88488432 | 0.583 | 0.057 | 0 | 6 | tmem176 |
| aoc21 | 0 | 0.882546126 | 0.476 | 0.058 | 0 | 6 | aoc2 |
| cd82a | 0 | 0.882030975 | 0.559 | 0.046 | 0 | 6 | cd82a |
| slco2b1 | 0 | 0.8808556 | 0.517 | 0.046 | 0 | 6 | slco2b1 |
| col6a31 | 0 | 0.869309739 | 0.622 | 0.091 | 0 | 6 | col6a3 |
| postnb1 | 0 | 0.864835873 | 0.562 | 0.074 | 0 | 6 | postnb |
| myl9b1 | 0 | 0.852967451 | 0.566 | 0.077 | 0 | 6 | myl9b |
| cd248a | 0 | 0.820303348 | 0.489 | 0.027 | 0 | 6 | cd248a |
| dkk3b1 | 0 | 0.809290116 | 0.502 | 0.04 | 0 | 6 | dkk3b |
| nrp1a1 | 0 | 0.802933035 | 0.499 | 0.044 | 0 | 6 | nrp1a |
| si:ch1073-459j12.11 | 0 | 0.800786504 | 0.519 | 0.065 | 0 | 6 | si:ch1073-459j12.1 |
| nr2f1a | 0 | 0.771902413 | 0.452 | 0.045 | 0 | 6 | nr2f1a |
| mxra8a | 0 | 0.771125874 | 0.475 | 0.043 | 0 | 6 | mxra8a |
| robo2 | 0 | 0.740777797 | 0.444 | 0.036 | 0 | 6 | robo2 |
| lrrc171 | 0 | 0.736131356 | 0.455 | 0.04 | 0 | 6 | lrrc17 |
| cyp2ad31 | 0 | 0.716966847 | 0.428 | 0.045 | 0 | 6 | cyp2ad3 |
| cyp3c11 | 0 | 0.698079392 | 0.449 | 0.047 | 0 | 6 | cyp3c1 |
| osr2 | 0 | 0.681674313 | 0.391 | 0.037 | 0 | 6 | osr2 |
| loxl2a1 | 0 | 0.669396862 | 0.387 | 0.031 | 0 | 6 | loxl2a |
| aebp1 | 0 | 0.662678613 | 0.413 | 0.031 | 0 | 6 | aebp1 |
| spon1b1 | 0 | 0.647290381 | 0.4 | 0.036 | 0 | 6 | spon1b |
| si:dkey-9i23.161 | 0 | 0.585116232 | 0.382 | 0.034 | 0 | 6 | si:dkey-9i23.16 |
| atp1a3b | 0 | 0.585025119 | 0.332 | 0.02 | 0 | 6 | atp1a3b |
| inka1a | 0 | 0.548557051 | 0.297 | 0.017 | 0 | 6 | inka1a |
| si:ch211-106j24.1 | 0 | 0.529067595 | 0.28 | 0.008 | 0 | 6 | si:ch211-106j24.1 |

| | | | | | | | |
|---|---|---|---|---|---|---|---|
| si:ch211-270g19.5 | 0 | 0.507569662 | 0.243 | 0.009 | 0 | 6 | si:ch211-270g19.5 |
| pcdh18b | 0 | 0.489048843 | 0.267 | 0.014 | 0 | 6 | pcdh18b |
| fhl2a | 0 | 0.460413237 | 0.232 | 0.008 | 0 | 6 | fhl2a |
| rxfp3.3b | 0 | 0.316493349 | 0.149 | 0.001 | 0 | 6 | rxfp3.3b |
| angptl2b | 0 | 0.297433957 | 0.16 | 0.003 | 0 | 6 | angptl2b |
| adamts15a1 | 1.743163 | 0.660384662 | 0.407 | 0.042 | 4.4285065 | 6 | adamts15a |
| ntn1a | 1.153278 | 0.359167658 | 0.19 | 0.006 | 2.9299048 | 6 | ntn1a |
| vegfaa | 1.756702 | 0.430679134 | 0.24 | 0.012 | 4.4629025 | 6 | vegfaa |
| selenbp1 | 2.444613 | 0.734978898 | 0.48 | 0.063 | 6.2105401 | 6 | selenbp1 |
| CU459094.2 | 2.880775 | 0.357005952 | 0.199 | 0.007 | 7.3186105 | 6 | CU459094.2 |
| pbx1a1 | 3.097434 | 0.606743207 | 0.386 | 0.039 | 7.8690325 | 6 | pbx1a |
| fank1 | 1.705687 | 0.409117321 | 0.217 | 0.01 | 4.3332988 | 6 | fank1 |
| igf2b1 | 7.089742 | 0.777666464 | 0.457 | 0.058 | 1.8011491 | 6 | igf2b |
| socs3a1 | 4.875380 | 1.173835937 | 0.906 | 0.288 | 1.2385903 | 6 | socs3a |
| jun1 | 6.075838 | 1.110334756 | 0.848 | 0.222 | 1.5435666 | 6 | jun |
| cebpd1 | 2.896016 | 0.908090157 | 0.676 | 0.128 | 7.3573301 | 6 | cebpd |
| cdon1 | 4.043152 | 0.664533942 | 0.441 | 0.054 | 1.0271628 | 6 | cdon |
| cyp27c1 | 7.425636 | 0.412939964 | 0.23 | 0.012 | 1.8864828 | 6 | cyp27c1 |
| fstl1b1 | 1.236160 | 0.843873186 | 0.677 | 0.123 | 3.1404650 | 6 | fstl1b |
| caln2 | 5.068642 | 0.334624409 | 0.185 | 0.006 | 1.2876886 | 6 | caln2 |
| col12a1b | 5.596349 | 0.553601323 | 0.308 | 0.025 | 1.4217526 | 6 | col12a1b |
| tshz3b | 8.722532 | 0.533640083 | 0.293 | 0.022 | 2.2159593 | 6 | tshz3b |
| col6a21 | 3.410267 | 0.672375329 | 0.451 | 0.057 | 8.6637850 | 6 | col6a2 |
| tenm31 | 9.354385 | 0.622298256 | 0.404 | 0.046 | 2.3764817 | 6 | tenm3 |
| CABZ01092746.11 | 2.013779 | 0.759669493 | 0.499 | 0.07 | 5.1160064 | 6 | CABZ01092746.1 |
| si:ch211-212k18.91 | 2.979581 | 0.537766369 | 0.357 | 0.036 | 7.5696257 | 6 | si:ch211-212k18.9 |
| cldn11a | 8.983787 | 0.759535462 | 0.475 | 0.065 | 2.2823311 | 6 | cldn11a |
| myocd | 2.163075 | 0.280955329 | 0.141 | 0.003 | 5.4952936 | 6 | myocd |
| gas1a1 | 3.820433 | 0.613840476 | 0.379 | 0.043 | 9.7058116 | 6 | gas1a |
| cygb1 | 6.286960 | 0.728772205 | 0.391 | 0.046 | 1.5972022 | 6 | cygb1 |
| fosab1 | 2.641713 | 1.132809543 | 0.925 | 0.336 | 6.7112727 | 6 | fosab |
| col5a2a1 | 3.580380 | 0.687516144 | 0.527 | 0.081 | 9.0959573 | 6 | col5a2a |
| cldn5a1 | 1.716586 | 0.676089699 | 0.392 | 0.047 | 4.3609883 | 6 | cldn5a |
| rnd1b | 1.156687 | 0.474148108 | 0.229 | 0.014 | 2.9385654 | 6 | rnd1b |
| junba1 | 1.024435 | 1.056709012 | 0.909 | 0.293 | 2.6025790 | 6 | junba |

| | | | | | | | |
|---|---|---|---|---|---|---|---|
| BX569787.1 | 4.991855 | 0.512588826 | 0.293 | 0.025 | 1.2681808 | 6 | BX569787.1 |
| lama41 | 2.027686 | 0.566974601 | 0.332 | 0.034 | 5.1513369 | 6 | lama4 |
| mmrn2a | 1.229723 | 0.354042465 | 0.185 | 0.008 | 3.1241117 | 6 | mmrn2a |
| rbp51 | 1.471698 | 0.919283417 | 0.519 | 0.092 | 3.7388507 | 6 | rbp5 |
| sparc1 | 3.734756 | 0.870449343 | 0.771 | 0.169 | 9.4881490 | 6 | sparc |
| nr2f2 | 1.119995 | 0.575001724 | 0.35 | 0.039 | 2.8453479 | 6 | nr2f2 |
| si:dkey-261h17.11 | 1.829487 | 0.760618195 | 0.556 | 0.095 | 4.6478125 | 6 | si:dkey-261h17.1 |
| krt18a.1 | 1.616612 | 0.924150414 | 0.627 | 0.128 | 4.1070039 | 6 | krt18a.1 |
| ccdc3b1 | 4.331214 | 0.43356435 | 0.261 | 0.021 | 1.1003449 | 6 | ccdc3b |
| lamc1 | 3.287536 | 0.582099605 | 0.366 | 0.044 | 8.3519871 | 6 | lamc1 |
| pbx1b | 1.113797 | 0.449959806 | 0.274 | 0.023 | 2.8296020 | 6 | pbx1b |
| col27a1b | 9.253228 | 0.439896222 | 0.245 | 0.018 | 2.3507826 | 6 | col27a1b |
| si:dkey-23i12.7 | 9.874218 | 0.524828893 | 0.243 | 0.018 | 2.5085451 | 6 | si:dkey-23i12.7 |
| mgst1.1 | 1.425191 | 0.551076571 | 0.353 | 0.042 | 3.6206989 | 6 | mgst1.1 |
| ar | 2.820885 | 0.570992004 | 0.394 | 0.052 | 7.1664583 | 6 | ar |
| cxcl12a1 | 4.395429 | 0.766324435 | 0.451 | 0.067 | 1.1166587 | 6 | cxcl12a |
| cdh2 | 6.031232 | 0.590882322 | 0.392 | 0.052 | 1.5322345 | 6 | cdh2 |
| eva1ba1 | 4.398712 | 0.603141817 | 0.394 | 0.052 | 1.1174930 | 6 | eva1ba |
| zfp36l1b1 | 5.199153 | 0.909724079 | 0.574 | 0.12 | 1.3208448 | 6 | zfp36l1b |
| ece11 | 1.509315 | 0.501086638 | 0.295 | 0.029 | 3.8344156 | 6 | ece1 |
| si:dkey-7j14.61 | 3.541225 | 0.832873861 | 0.7 | 0.173 | 8.9964836 | 6 | si:dkey-7j14.6 |
| tpm4a1 | 1.006216 | 0.75258444 | 0.52 | 0.095 | 2.5562922 | 6 | tpm4a |
| mdkb | 2.402068 | 0.451091649 | 0.203 | 0.012 | 6.1024545 | 6 | mdkb |
| asic1b | 4.633324 | 0.336038396 | 0.183 | 0.009 | 1.1770960 | 6 | asic1b |
| socs3b | 5.244582 | 0.865824878 | 0.595 | 0.131 | 1.3323862 | 6 | socs3b |
| tmem176l.11 | 3.087779 | 0.837230547 | 0.708 | 0.179 | 7.8445035 | 6 | tmem176l.1 |
| kif26ba | 1.144188 | 0.28533566 | 0.149 | 0.005 | 2.9068118 | 6 | kif26ba |
| prss231 | 1.298165 | 0.47342244 | 0.259 | 0.023 | 3.2979885 | 6 | prss23 |
| snx17 | 4.387199 | 0.581345067 | 0.378 | 0.051 | 1.1145681 | 6 | snx17 |
| acta2 | 6.305669 | 0.603072897 | 0.167 | 0.008 | 1.6019553 | 6 | acta2 |
| olfml3b1 | 6.149041 | 0.535465856 | 0.344 | 0.043 | 1.5621639 | 6 | olfml3b |
| ier2a1 | 1.784659 | 0.947920418 | 0.786 | 0.251 | 4.5339270 | 6 | ier2a |
| chd31 | 3.628612 | 0.690304353 | 0.538 | 0.106 | 9.2184904 | 6 | chd3 |
| c1qtnf51 | 1.352000 | 0.451824373 | 0.261 | 0.024 | 3.4347575 | 6 | c1qtnf5 |
| tmem119b | 1.169170 | 0.394385454 | 0.203 | 0.014 | 2.9702773 | 6 | tmem119b |

| | | | | | | | |
|---|---|---|---|---|---|---|---|
| tnfsf101 | 1.557952 | 0.502541167 | 0.293 | 0.032 | 3.9579792 | 6 | tnfsf10 |
| csrp1b | 6.390665 | 0.308664015 | 0.104 | 0.002 | 1.6235486 | 6 | csrp1b |
| tmem88b | 6.312206 | 0.49819578 | 0.331 | 0.041 | 1.6036159 | 6 | tmem88b |
| ms4a17a.11 | 3.032993 | 0.48375759 | 0.319 | 0.04 | 7.7053198 | 6 | ms4a17a.11 |
| lxn1 | 4.342147 | 0.799302462 | 0.609 | 0.147 | 1.1031225 | 6 | lxn |
| si:ch211-212k18.81 | 5.001891 | 0.459189396 | 0.306 | 0.037 | 1.2707306 | 6 | si:ch211-212k18.8 |
| si:dkey-226m8.10 | 1.139524 | 0.25942019 | 0.135 | 0.005 | 2.8949621 | 6 | si:dkey-226m8.10 |
| cfd | 5.620248 | 0.764405135 | 0.374 | 0.057 | 1.4278241 | 6 | cfd |
| boc1 | 1.279244 | 0.553605864 | 0.368 | 0.054 | 3.2499202 | 6 | boc |
| f10 | 7.751975 | 0.504935696 | 0.344 | 0.047 | 1.9693894 | 6 | f10 |
| krt81 | 1.243699 | 0.910344845 | 0.862 | 0.287 | 3.1596195 | 6 | krt8 |
| vstm4a | 9.100073 | 0.288370373 | 0.167 | 0.01 | 2.3118735 | 6 | vstm4a |
| cplx2l | 5.493005 | 0.305835942 | 0.185 | 0.013 | 1.3954981 | 6 | cplx2l |
| luzp2 | 1.941781 | 0.253976253 | 0.13 | 0.005 | 4.9330969 | 6 | luzp2 |
| nfic1 | 7.889608 | 0.574145177 | 0.407 | 0.069 | 2.0043550 | 6 | nfic |
| c1qtnf6a | 1.995129 | 0.372093502 | 0.217 | 0.019 | 5.0686276 | 6 | c1qtnf6a |
| mmp21 | 5.970842 | 0.613198789 | 0.488 | 0.094 | 1.5168924 | 6 | mmp2 |
| tmem47 | 4.307980 | 0.412446641 | 0.261 | 0.028 | 1.0944423 | 6 | tmem47 |
| ephb31 | 4.497119 | 0.48502004 | 0.31 | 0.041 | 1.1424932 | 6 | ephb3 |
| nme3 | 2.873914 | 0.549779417 | 0.418 | 0.074 | 7.3011785 | 6 | nme3 |
| lamb1a1 | 7.950426 | 0.477778421 | 0.318 | 0.043 | 2.0198058 | 6 | lamb1a |
| nme2b.2 | 1.114504 | 0.42573627 | 0.211 | 0.018 | 2.8313995 | 6 | nme2b.2 |
| h2afy2 | 4.601912 | 0.631117448 | 0.493 | 0.106 | 1.1691157 | 6 | h2afy2 |
| colec121 | 5.407417 | 0.508648418 | 0.344 | 0.05 | 1.3737544 | 6 | colec12 |
| pdgfrb1 | 8.754507 | 0.520001412 | 0.306 | 0.041 | 2.2240826 | 6 | pdgfrb |
| mrc2 | 1.058399 | 0.388140817 | 0.225 | 0.021 | 2.6888650 | 6 | mrc2 |
| spry2 | 1.102033 | 0.511239279 | 0.37 | 0.059 | 2.7997153 | 6 | spry2 |
| BX088707.3 | 2.410336 | 0.271064301 | 0.113 | 0.004 | 6.1234592 | 6 | BX088707.3 |
| si:dkey-188i13.10 | 8.237264 | 0.620476809 | 0.478 | 0.097 | 2.0926769 | 6 | si:dkey-188i13.10 |
| cyp1c1 | 3.625517 | 0.39512285 | 0.175 | 0.012 | 9.2106266 | 6 | cyp1c1 |
| zgc:1627302 | 1.055993 | 0.935464082 | 0.865 | 0.382 | 2.6827509 | 6 | zgc:162730 |
| aldh9a1a.11 | 2.159874 | 0.72154479 | 0.64 | 0.173 | 5.4871609 | 6 | aldh9a1a.1 |
| si:ch211-286o17.1 | 1.267549 | 0.452884149 | 0.297 | 0.038 | 3.2202099 | 6 | si:ch211-286o17.1 |
| nrp2a | 2.005168 | 0.319806757 | 0.183 | 0.013 | 5.0941293 | 6 | nrp2a |
| rhoua | 1.367221 | 0.388815427 | 0.225 | 0.022 | 3.4734264 | 6 | rhoua |

| | | | | | | | |
|---|---|---|---|---|---|---|---|
| gadd45ba1 | 1.604623 | 0.833244226 | 0.634 | 0.178 | 4.0765447 | 6 | gadd45ba |
| dio2 | 2.026914 | 0.284545919 | 0.144 | 0.008 | 5.1493751 | 6 | dio2 |
| marcksl1a1 | 2.810304 | 0.799725078 | 0.771 | 0.238 | 7.1395777 | 6 | marcksl1a |
| svild1 | 5.241967 | 0.571012646 | 0.399 | 0.07 | 1.3317218 | 6 | svild |
| cpxm1a1 | 8.299937 | 0.315373981 | 0.172 | 0.012 | 2.1085991 | 6 | cpxm1a |
| nfia1 | 2.701788 | 0.584928374 | 0.434 | 0.084 | 6.8638938 | 6 | nfia |
| mtss1la | 1.391892 | 0.349656035 | 0.203 | 0.018 | 3.5361028 | 6 | mtss1la |
| igfbp7 | 4.799696 | 0.384612959 | 0.23 | 0.024 | 1.2193628 | 6 | igfbp7 |
| itga9 | 2.215217 | 0.41558876 | 0.25 | 0.028 | 5.6277608 | 6 | itga9 |
| cnn3a1 | 3.253712 | 0.48661621 | 0.35 | 0.056 | 8.2660562 | 6 | cnn3a |
| cpox | 4.915559 | 0.486584191 | 0.306 | 0.044 | 1.2487979 | 6 | cpox |
| lpar11 | 1.150805 | 0.499086665 | 0.357 | 0.057 | 2.9236215 | 6 | lpar1 |
| CR383676.12 | 3.135412 | 0.777991426 | 0.994 | 0.705 | 7.9655153 | 6 | CR383676.1 |
| rasl11a | 6.061967 | 0.26728914 | 0.157 | 0.01 | 1.5400429 | 6 | rasl11a |
| col15a1a | 1.513070 | 0.327112603 | 0.198 | 0.017 | 3.8439550 | 6 | col15a1a |
| lrp1ab1 | 1.848133 | 0.540500401 | 0.374 | 0.065 | 4.6951835 | 6 | lrp1ab |
| meis2b1 | 6.648280 | 0.364738192 | 0.212 | 0.021 | 1.6889956 | 6 | meis2b |
| si:ch211-197h24.9 | 2.149147 | 0.567195184 | 0.382 | 0.07 | 5.4599081 | 6 | si:ch211-197h24.9 |
| c41 | 2.896703 | 0.435134536 | 0.25 | 0.029 | 7.3590750 | 6 | c4 |
| qkia1 | 2.958664 | 0.714128925 | 0.634 | 0.18 | 7.5164879 | 6 | qkia |
| spry41 | 3.157476 | 0.508126865 | 0.284 | 0.039 | 8.0215701 | 6 | spry4 |
| mmp14a1 | 3.547405 | 0.47768831 | 0.318 | 0.048 | 9.0121848 | 6 | mmp14a |
| lrrtm1 | 7.095783 | 0.293286783 | 0.12 | 0.005 | 1.8026839 | 6 | lrrtm1 |
| irf8 | 2.240112 | 0.583562284 | 0.279 | 0.038 | 5.6910068 | 6 | irf8 |
| hic11 | 1.004764 | 0.521811533 | 0.348 | 0.058 | 2.5526030 | 6 | hic1 |
| cox4i21 | 1.193320 | 0.374982828 | 0.241 | 0.028 | 3.0316304 | 6 | cox4i2 |
| pik3r2 | 3.693824 | 0.497900628 | 0.337 | 0.055 | 9.3841609 | 6 | pik3r2 |
| sh3gl1b | 3.874200 | 0.383781417 | 0.256 | 0.032 | 9.8424067 | 6 | sh3gl1b |
| vcam1b | 6.396035 | 0.376964798 | 0.185 | 0.016 | 1.6249127 | 6 | vcam1b |
| ptprga1 | 2.000275 | 0.538033651 | 0.348 | 0.059 | 5.0816991 | 6 | ptprga |
| esr2a | 5.563058 | 0.368205614 | 0.227 | 0.025 | 1.4132950 | 6 | esr2a |
| nog1 | 6.026065 | 0.26112333 | 0.128 | 0.006 | 1.5309220 | 6 | nog1 |
| CABZ01068367.1 | 7.065278 | 0.304493553 | 0.188 | 0.016 | 1.7949339 | 6 | CABZ01068367.1 |
| ccbe1 | 4.233547 | 0.346007346 | 0.196 | 0.018 | 1.0755327 | 6 | ccbe1 |
| cux1a | 3.697570 | 0.472180021 | 0.298 | 0.045 | 9.3936768 | 6 | cux1a |

| | | | | | | | |
|---|---|---|---|---|---|---|---|
| rgs5b | 5.138079 | 0.764145989 | 0.342 | 0.061 | 1.3053290 | 6 | rgs5b |
| f3b1 | 7.569470 | 0.644283278 | 0.418 | 0.085 | 1.9230240 | 6 | f3b |
| CABZ01068366.1 | 3.523197 | 0.387963227 | 0.256 | 0.032 | 8.9506830 | 6 | CABZ01068366.1 |
| b2ml2 | 1.371576 | 0.728616015 | 0.775 | 0.272 | 3.4844913 | 6 | b2ml |
| xirp2a | 7.145280 | 0.372293106 | 0.229 | 0.026 | 1.8152586 | 6 | xirp2a |
| slc38a5b1 | 9.189994 | 0.568121664 | 0.486 | 0.113 | 2.3347181 | 6 | slc38a5b |
| gsta.1 | 5.238190 | 0.416682511 | 0.282 | 0.041 | 1.3307621 | 6 | gsta.1 |
| lrp1aa1 | 8.604595 | 0.473499012 | 0.329 | 0.054 | 2.1859974 | 6 | lrp1aa |
| hmgn62 | 1.174145 | 0.752334418 | 0.78 | 0.29 | 2.9829174 | 6 | hmgn6 |
| h3f3c1 | 1.101475 | 0.750691931 | 0.763 | 0.284 | 2.7982973 | 6 | h3f3c |
| tpbga | 9.206465 | 0.250454096 | 0.135 | 0.008 | 2.3389024 | 6 | tpbga |
| fosb | 1.085757 | 0.718202892 | 0.46 | 0.106 | 2.7583658 | 6 | fosb |
| fam198a | 4.654166 | 0.287090182 | 0.17 | 0.014 | 1.1823910 | 6 | fam198a |
| zgc:112437 | 1.783966 | 0.340723226 | 0.186 | 0.018 | 4.5321659 | 6 | zgc:112437 |
| gstt1b1 | 2.333651 | 0.492273583 | 0.391 | 0.076 | 5.9286419 | 6 | gstt1b |
| gstm.1 | 3.845731 | 0.732628817 | 0.755 | 0.257 | 9.7700809 | 6 | gstm.1 |
| mafba1 | 5.185242 | 0.511151042 | 0.371 | 0.072 | 1.3173108 | 6 | mafba |
| scube1 | 2.688092 | 0.294792128 | 0.172 | 0.015 | 6.8290988 | 6 | scube1 |
| hmgb1b | 5.609156 | 0.609109964 | 0.532 | 0.138 | 1.4250062 | 6 | hmgb1b |
| snai2 | 2.564508 | 0.320506751 | 0.204 | 0.022 | 6.5151329 | 6 | snai2 |
| ppib1 | 3.529144 | 0.745232857 | 0.851 | 0.358 | 8.9657924 | 6 | ppib |
| spred2b1 | 2.452509 | 0.494082017 | 0.37 | 0.072 | 6.2306008 | 6 | spred2b |
| im:7152348 | 1.077315 | 0.277146209 | 0.139 | 0.01 | 2.7369195 | 6 | im:7152348 |
| sdc41 | 1.093946 | 0.787893092 | 0.676 | 0.236 | 2.7791708 | 6 | sdc4 |
| BX322530.11 | 6.253981 | 0.405529786 | 0.277 | 0.042 | 1.5888239 | 6 | BX322530.1 |
| sema4c | 3.021761 | 0.261758789 | 0.144 | 0.011 | 7.6767853 | 6 | sema4c |
| cbx7a | 4.412059 | 0.47004522 | 0.276 | 0.043 | 1.1208836 | 6 | cbx7a |
| dclk1a | 7.807287 | 0.352744622 | 0.22 | 0.027 | 1.9834415 | 6 | dclk1a |
| anxa1a1 | 1.065559 | 0.638872415 | 0.551 | 0.145 | 2.7070529 | 6 | anxa1a |
| grb10a1 | 5.510606 | 0.377615005 | 0.245 | 0.034 | 1.3999696 | 6 | grb10a |
| hmgb3a1 | 1.840313 | 0.503652615 | 0.357 | 0.071 | 4.6753164 | 6 | hmgb3a |
| loxl2b | 2.157635 | 0.404289282 | 0.253 | 0.037 | 5.4814736 | 6 | loxl2b |
| XLOC-016198 | 3.090904 | 0.25435326 | 0.151 | 0.012 | 7.8524422 | 6 | XLOC-016198 |
| s100v21 | 8.954054 | 0.534883642 | 0.502 | 0.124 | 2.2747775 | 6 | s100v2 |
| il151 | 1.242402 | 0.415256647 | 0.297 | 0.049 | 3.1563223 | 6 | il15 |

| | | | | | | | |
|---|---|---|---|---|---|---|---|
| crim1 | 1.410597 | 0.386714028 | 0.237 | 0.033 | 3.5836228 | 6 | crim1 |
| cyp2p61 | 3.259419 | 0.321253303 | 0.185 | 0.019 | 8.2805559 | 6 | cyp2p6 |
| tmeff1b | 3.280211 | 0.351854205 | 0.24 | 0.033 | 8.3333775 | 6 | tmeff1b |
| si:dkey-163m14.2 | 6.203168 | 0.300540993 | 0.165 | 0.015 | 1.5759150 | 6 | si:dkey-163m14.2 |
| anos1a | 1.194408 | 0.252354352 | 0.143 | 0.011 | 3.0343945 | 6 | anos1a |
| ghrb | 1.048738 | 0.373974178 | 0.245 | 0.035 | 2.6643208 | 6 | ghrb |
| ms4a17a.9 | 1.236385 | 0.425725715 | 0.319 | 0.058 | 3.1410368 | 6 | ms4a17a.9 |
| CABZ01068358.1 | 1.985632 | 0.512104915 | 0.31 | 0.056 | 5.0445005 | 6 | CABZ01068358.1 |
| adh5 | 2.888331 | 0.630243305 | 0.673 | 0.223 | 7.3378067 | 6 | adh5 |
| ndnf | 3.093973 | 0.386940413 | 0.22 | 0.029 | 7.8602390 | 6 | ndnf |
| si:ch73-335l21.41 | 3.871291 | 0.809630173 | 0.467 | 0.127 | 9.8350149 | 6 | si:ch73-335l21.4 |
| yap1 | 8.888629 | 0.389413849 | 0.284 | 0.047 | 2.2581564 | 6 | yap1 |
| zgc:92744 | 1.126903 | 0.760633367 | 0.734 | 0.309 | 2.8628984 | 6 | zgc:92744 |
| pgrmc1 | 1.307004 | 0.543775778 | 0.515 | 0.136 | 3.3204436 | 6 | pgrmc1 |
| s100a10b1 | 3.187807 | 0.713133549 | 0.776 | 0.263 | 8.0986238 | 6 | s100a10b |
| gadd45ab | 3.448888 | 0.486985619 | 0.314 | 0.059 | 8.7619013 | 6 | gadd45ab |
| sema4ba1 | 7.295602 | 0.3780294 | 0.237 | 0.034 | 1.8534476 | 6 | sema4ba |
| fosl1a | 7.847001 | 0.50051361 | 0.352 | 0.072 | 1.9935308 | 6 | fosl1a |
| myl9a | 2.279098 | 0.445459245 | 0.245 | 0.036 | 5.7900500 | 6 | myl9a |
| ifi451 | 2.679719 | 0.732832363 | 0.467 | 0.125 | 6.8078282 | 6 | ifi45 |
| fkbp71 | 9.130002 | 0.361363276 | 0.261 | 0.041 | 2.3194771 | 6 | fkbp7 |
| si:ch211-191i18.2 | 2.645811 | 0.309090615 | 0.167 | 0.017 | 6.7216841 | 6 | si:ch211-191i18.2 |
| ptenb | 4.341692 | 0.477765331 | 0.358 | 0.075 | 1.1030069 | 6 | ptenb |
| jdp2b | 1.148446 | 0.72800802 | 0.511 | 0.149 | 2.9176280 | 6 | jdp2b |
| flna1 | 1.912038 | 0.566444287 | 0.431 | 0.108 | 4.8575326 | 6 | flna |
| tns2a1 | 7.761959 | 0.334370941 | 0.219 | 0.03 | 1.9719258 | 6 | tns2a |
| gpr27 | 7.815925 | 0.255789755 | 0.139 | 0.011 | 1.9856358 | 6 | gpr27 |
| celf22 | 3.048037 | 0.66542815 | 0.674 | 0.239 | 7.7435382 | 6 | celf2 |
| cyp1c21 | 9.741339 | 0.322858528 | 0.146 | 0.013 | 2.4747871 | 6 | cyp1c2 |
| slc7a11 | 5.137771 | 0.350535939 | 0.216 | 0.029 | 1.3052508 | 6 | slc7a1 |
| zgc:1527911 | 5.022224 | 0.525539073 | 0.303 | 0.057 | 1.2758961 | 6 | zgc:152791 |
| aldh1a2 | 1.072024 | 0.479846736 | 0.391 | 0.085 | 2.7234777 | 6 | aldh1a2 |
| zgc:1141881 | 1.421661 | 0.524286921 | 0.961 | 0.655 | 3.6117302 | 6 | zgc:114188 |
| si:ch211-222l21.11 | 1.781805 | 0.695402724 | 0.76 | 0.294 | 4.5266761 | 6 | si:ch211-222l21.1 |
| sptlc2a | 3.139847 | 0.362976082 | 0.243 | 0.038 | 7.9767828 | 6 | sptlc2a |

| | | | | | | | |
|---|---|---|---|---|---|---|---|
| clec11a | 6.826067 | 0.258192953 | 0.146 | 0.013 | 1.7341623 | 6 | clec11a |
| gng71 | 9.114235 | 0.378295029 | 0.287 | 0.052 | 2.3154716 | 6 | gng7 |
| sfrp1b | 2.835174 | 0.255514599 | 0.135 | 0.011 | 7.2027598 | 6 | sfrp1b |
| myl61 | 2.975627 | 0.592296414 | 0.412 | 0.104 | 7.5595808 | 6 | myl6 |
| rpl91 | 1.509379 | 0.52579652 | 0.972 | 0.655 | 3.8345782 | 6 | rpl9 |
| meis1b1 | 5.930806 | 0.433486927 | 0.311 | 0.063 | 1.5067214 | 6 | meis1b |
| lgals9l3 | 1.702995 | 0.42058702 | 0.339 | 0.072 | 4.3264599 | 6 | lgals9l3 |
| zeb2a | 4.672423 | 0.383932951 | 0.238 | 0.038 | 1.1870292 | 6 | zeb2a |
| zgc:113531 | 3.076111 | 0.297699586 | 0.164 | 0.018 | 7.8148625 | 6 | zgc:113531 |
| b2m1 | 1.186636 | 0.648740993 | 0.726 | 0.279 | 3.0146494 | 6 | b2m |
| col16a11 | 2.122566 | 0.340322777 | 0.22 | 0.033 | 5.3923813 | 6 | col16a1 |
| krt941 | 2.180720 | 0.577202227 | 0.532 | 0.15 | 5.5401192 | 6 | krt94 |
| mgst3b | 5.756320 | 0.551251376 | 0.558 | 0.177 | 1.4623932 | 6 | mgst3b |
| si:ch211-133n4.41 | 2.466544 | 0.388489058 | 0.271 | 0.05 | 6.2662561 | 6 | si:ch211-133n4.4 |
| cyp2n131 | 5.835951 | 0.345316068 | 0.22 | 0.033 | 1.4826235 | 6 | cyp2n13 |
| ctdsp21 | 4.921227 | 0.42640871 | 0.335 | 0.075 | 1.2502377 | 6 | ctdsp2 |
| s1pr2 | 5.332389 | 0.314004362 | 0.206 | 0.03 | 1.3546934 | 6 | s1pr2 |
| bhmt | 1.348764 | 0.373062367 | 0.222 | 0.034 | 3.4265367 | 6 | bhmt |
| sult1st1 | 1.715452 | 0.308601734 | 0.194 | 0.027 | 4.3581072 | 6 | sult1st1 |
| rdh10a1 | 2.163347 | 0.382627253 | 0.24 | 0.04 | 5.4959833 | 6 | rdh10a |
| nfil3-61 | 2.918904 | 0.566220595 | 0.431 | 0.118 | 7.4154773 | 6 | nfil3-6 |
| cyp2p71 | 6.005597 | 0.313203193 | 0.183 | 0.024 | 1.5257220 | 6 | cyp2p7 |
| fblim11 | 1.091779 | 0.306594283 | 0.206 | 0.03 | 2.7736648 | 6 | fblim1 |
| tmem100a1 | 1.203048 | 0.272570616 | 0.167 | 0.02 | 3.0563446 | 6 | tmem100a |
| selenow11 | 5.192544 | 0.636249593 | 0.857 | 0.395 | 1.3191659 | 6 | selenow1 |
| enc1 | 2.583024 | 0.335681707 | 0.209 | 0.032 | 6.5621747 | 6 | enc1 |
| esr2b | 6.214395 | 0.299434573 | 0.211 | 0.032 | 1.5787672 | 6 | esr2b |
| smo | 1.102948 | 0.292586677 | 0.175 | 0.023 | 2.8020402 | 6 | smo |
| mxra8b | 3.385120 | 0.277098197 | 0.178 | 0.023 | 8.5998996 | 6 | mxra8b |
| ybx1 | 4.664929 | 0.460902671 | 0.95 | 0.685 | 1.1851254 | 6 | ybx1 |
| zeb1b | 1.098238 | 0.295649624 | 0.18 | 0.024 | 2.7900757 | 6 | zeb1b |
| LOC1039092841 | 2.455143 | 0.415493725 | 0.31 | 0.067 | 6.2372908 | 6 | LOC103909284 |
| tnfrsf19 | 1.426840 | 0.269413204 | 0.146 | 0.016 | 3.6248876 | 6 | tnfrsf19 |
| tuba8l31 | 3.938681 | 0.392863143 | 0.298 | 0.062 | 1.0006220 | 6 | tuba8l3 |
| ctdspla | 4.114516 | 0.307875643 | 0.211 | 0.033 | 1.0452929 | 6 | ctdspla |

| | | | | | | | |
|---|---|---|---|---|---|---|---|
| dhrs13l1 | 1.765103 | 0.30780088 | 0.207 | 0.033 | 4.4842454 | 6 | dhrs13l1 |
| fam168a1 | 1.279908 | 0.38486992 | 0.297 | 0.064 | 3.2516071 | 6 | fam168a |
| foxo3b1 | 4.175884 | 0.404209287 | 0.288 | 0.061 | 1.0608834 | 6 | foxo3b |
| si:dkey-239h2.3 | 6.964952 | 0.413542625 | 0.358 | 0.088 | 1.7694462 | 6 | si:dkey-239h2.3 |
| atf4b | 1.163679 | 0.582139198 | 0.671 | 0.26 | 2.9563271 | 6 | atf4b |
| junbb | 1.247087 | 0.379492644 | 0.287 | 0.061 | 3.1682264 | 6 | junbb |
| rgl1 | 1.321428 | 0.320567052 | 0.203 | 0.032 | 3.3570890 | 6 | rgl1 |
| egr11 | 2.932928 | 0.580595959 | 0.519 | 0.168 | 7.4511037 | 6 | egr1 |
| serinc2 | 3.849356 | 0.431755617 | 0.413 | 0.112 | 9.7792908 | 6 | serinc2 |
| ywhaz | 1.331245 | 0.606911069 | 0.692 | 0.288 | 3.3820295 | 6 | ywhaz |
| cnbpb | 1.545518 | 0.514588686 | 0.566 | 0.194 | 3.9263898 | 6 | cnbpb |
| raraa | 1.798761 | 0.295544292 | 0.199 | 0.031 | 4.5697537 | 6 | raraa |
| CABZ01080702.11 | 4.214787 | 0.456578965 | 0.35 | 0.087 | 1.0707666 | 6 | CABZ01080702.1 |
| ckba | 8.518441 | 0.428746818 | 0.293 | 0.064 | 2.1641099 | 6 | ckba |
| bag3 | 9.305142 | 0.325150368 | 0.186 | 0.028 | 2.3639713 | 6 | bag3 |
| grhl1 | 1.748787 | 0.299262816 | 0.175 | 0.024 | 4.4427946 | 6 | grhl1 |
| ier5 | 7.105389 | 0.49535457 | 0.446 | 0.134 | 1.8051241 | 6 | ier5 |
| sgce | 9.000265 | 0.26160084 | 0.18 | 0.026 | 2.2865175 | 6 | sgce |
| fbn2b1 | 1.132634 | 0.355137876 | 0.235 | 0.043 | 2.8774584 | 6 | fbn2b |
| antxr2a1 | 7.954392 | 0.508579529 | 0.494 | 0.155 | 2.0208135 | 6 | antxr2a |
| fzd7b1 | 2.990185 | 0.326064479 | 0.229 | 0.041 | 7.5965674 | 6 | fzd7b |
| rps131 | 5.180952 | 0.403128244 | 0.956 | 0.74 | 1.3162210 | 6 | rps13 |
| rps181 | 6.209840 | 0.407747116 | 0.963 | 0.745 | 1.5776101 | 6 | rps18 |
| lgals17 | 7.734310 | 0.400486309 | 0.245 | 0.047 | 1.9649015 | 6 | lgals17 |
| sulf2b | 1.108712 | 0.320581389 | 0.207 | 0.035 | 2.8166828 | 6 | sulf2b |
| rps22 | 2.899798 | 0.338821792 | 0.963 | 0.837 | 7.3669386 | 6 | rps2 |
| c4b | 2.961265 | 0.274822942 | 0.135 | 0.015 | 7.5230947 | 6 | c4b |
| cox8b | 5.688561 | 0.387776631 | 0.357 | 0.09 | 1.4451790 | 6 | cox8b |
| si:ch211-178n15.1 | 8.431814 | 0.298744213 | 0.167 | 0.023 | 2.1421025 | 6 | si:ch211-178n15.1 |
| FO082781.11 | 9.659087 | 0.342890971 | 0.243 | 0.047 | 2.4538911 | 6 | FO082781.1 |
| XLOC-0412691 | 2.106747 | 0.320869497 | 0.169 | 0.024 | 5.3521913 | 6 | XLOC-041269 |
| fzd21 | 2.189354 | 0.297486778 | 0.207 | 0.035 | 5.5620553 | 6 | fzd2 |
| asip2b1 | 3.007047 | 0.373184777 | 0.159 | 0.021 | 7.6394053 | 6 | asip2b |
| si:dkeyp-75h12.5 | 1.249134 | 0.288366098 | 0.212 | 0.037 | 3.1734256 | 6 | si:dkeyp-75h12.5 |
| vasnb | 1.251556 | 0.304385114 | 0.188 | 0.03 | 3.1795804 | 6 | vasnb |

| | | | | | | | |
|---|---|---|---|---|---|---|---|
| SCARF2 | 1.499601 | 0.269436554 | 0.16 | 0.022 | 3.8097375 | 6 | SCARF2 |
| tnfsf12 | 2.072155 | 0.257344713 | 0.159 | 0.022 | 5.2643102 | 6 | tnfsf12 |
| ifi46 | 3.285518 | 0.475525315 | 0.412 | 0.12 | 8.3468598 | 6 | ifi46 |
| socs2 | 7.655767 | 0.263658352 | 0.16 | 0.022 | 1.9449477 | 6 | socs2 |
| ablim1b1 | 1.465731 | 0.310372783 | 0.207 | 0.036 | 3.7236920 | 6 | ablim1b |
| ptbp1b | 1.976598 | 0.279982698 | 0.199 | 0.034 | 5.0215491 | 6 | ptbp1b |
| entpd1 | 2.006232 | 0.373208344 | 0.308 | 0.073 | 5.0968340 | 6 | entpd1 |
| si:ch73-1a9.31 | 4.586638 | 0.577330899 | 0.72 | 0.298 | 1.1652353 | 6 | si:ch73-1a9.3 |
| rps27.21 | 9.784670 | 0.538650897 | 0.908 | 0.538 | 2.4857956 | 6 | rps27.2 |
| rps151 | 1.153069 | 0.420784582 | 0.945 | 0.707 | 2.9293724 | 6 | rps15 |
| rps91 | 1.210304 | 0.359716365 | 0.964 | 0.783 | 3.0747773 | 6 | rps9 |
| deptor | 1.908046 | 0.291800521 | 0.185 | 0.03 | 4.8473911 | 6 | deptor |
| rplp11 | 2.393589 | 0.304682998 | 0.99 | 0.878 | 6.0809142 | 6 | rplp1 |
| jpt1b1 | 2.397516 | 0.540873545 | 0.676 | 0.259 | 6.0908895 | 6 | jpt1b |
| arhgap31 | 2.474349 | 0.264985579 | 0.175 | 0.027 | 6.2860836 | 6 | arhgap31 |
| XLOC-005605 | 2.914271 | 0.266111945 | 0.164 | 0.023 | 7.4037069 | 6 | XLOC-005605 |
| rps191 | 3.871064 | 0.294827469 | 0.987 | 0.864 | 9.8344404 | 6 | rps19 |
| vim1 | 3.977229 | 0.334989041 | 0.214 | 0.039 | 1.0104151 | 6 | vim |
| rpl291 | 4.255401 | 0.339423489 | 0.982 | 0.816 | 1.0810847 | 6 | rpl29 |
| tcf121 | 1.334257 | 0.369917668 | 0.269 | 0.06 | 3.3896813 | 6 | tcf12 |
| irf2bp2b | 1.535185 | 0.411184817 | 0.344 | 0.092 | 3.9001375 | 6 | irf2bp2b |
| si:dkey-51e6.1 | 2.154995 | 0.406642858 | 0.381 | 0.107 | 5.4747665 | 6 | si:dkey-51e6.1 |
| TCIM (1 of many)1 | 2.242040 | 0.537531927 | 0.217 | 0.042 | 5.6959036 | 6 | TCIM (1 of many) |
| cdh11 | 5.808988 | 0.273595925 | 0.164 | 0.024 | 1.4757735 | 6 | cdh11 |
| si:dkey-58f10.11 | 1.017672 | 0.254419694 | 0.156 | 0.022 | 2.5853965 | 6 | si:dkey-58f10.11 |
| rpz51 | 1.376781 | 0.424104675 | 0.402 | 0.115 | 3.4977124 | 6 | rpz5 |
| rps281 | 1.431005 | 0.319144429 | 0.979 | 0.835 | 3.6354701 | 6 | rps28 |
| mvp | 1.633964 | 0.333710091 | 0.25 | 0.052 | 4.1510868 | 6 | mvp |
| hmgn7 | 5.367752 | 0.56783036 | 0.699 | 0.301 | 1.3636775 | 6 | hmgn7 |
| ctbp2a | 6.530827 | 0.26794748 | 0.198 | 0.034 | 1.6591567 | 6 | ctbp2a |
| rps8a1 | 9.145404 | 0.338856224 | 0.959 | 0.795 | 2.3233899 | 6 | rps8a |
| rpl361 | 8.010728 | 0.360850656 | 0.977 | 0.799 | 2.0351255 | 6 | rpl36 |
| rps161 | 1.650010 | 0.38909046 | 0.971 | 0.757 | 4.1918517 | 6 | rps16 |
| si:ch211-106h4.121 | 4.777820 | 0.460766093 | 0.241 | 0.05 | 1.2138054 | 6 | si:ch211-106h4.12 |
| emilin1a | 5.742811 | 0.253018738 | 0.149 | 0.02 | 1.4589612 | 6 | emilin1a |

| | | | | | | | |
|---|---|---|---|---|---|---|---|
| foxp41 | 1.452458 | 0.374297126 | 0.277 | 0.065 | 3.6899696 | 6 | foxp4 |
| rpl61 | 2.091963 | 0.433044975 | 0.956 | 0.691 | 5.3146338 | 6 | rpl6 |
| rpl27a1 | 3.613857 | 0.549708383 | 0.885 | 0.507 | 9.1810061 | 6 | rpl27a |
| rbfox21 | 3.833349 | 0.289822038 | 0.198 | 0.036 | 9.7386236 | 6 | rbfox2 |
| sh3pxd2aa | 8.453491 | 0.27374136 | 0.173 | 0.028 | 2.1476094 | 6 | sh3pxd2aa |
| grinab | 1.201846 | 0.399878382 | 0.382 | 0.112 | 3.0532910 | 6 | grinab |
| fstl3 | 1.804367 | 0.252129051 | 0.144 | 0.02 | 4.5839954 | 6 | fstl3 |
| actn1 | 7.004554 | 0.314947014 | 0.212 | 0.041 | 1.7795071 | 6 | actn1 |
| ckap41 | 7.103330 | 0.31713002 | 0.245 | 0.053 | 1.8046011 | 6 | ckap4 |
| rpsa1 | 1.028591 | 0.36342019 | 0.974 | 0.778 | 2.6131379 | 6 | rpsa |
| ch25h | 1.694692 | 0.272926122 | 0.143 | 0.019 | 4.3053663 | 6 | ch25h |
| dedd11 | 2.000718 | 0.469069415 | 0.441 | 0.145 | 5.0828241 | 6 | dedd1 |
| rpl7a1 | 2.204672 | 0.326540113 | 0.977 | 0.817 | 5.6009710 | 6 | rpl7a |
| cebpb1 | 4.061951 | 0.627474223 | 0.712 | 0.323 | 1.0319387 | 6 | cebpb |
| rpl321 | 6.597883 | 0.320765401 | 0.955 | 0.819 | 1.6761922 | 6 | rpl32 |
| pim11 | 1.301629 | 0.563295048 | 0.447 | 0.157 | 3.3067900 | 6 | pim1 |
| stxbp6 | 1.537472 | 0.264611458 | 0.209 | 0.039 | 3.9059500 | 6 | stxbp6 |
| rps101 | 1.539727 | 0.404180994 | 0.959 | 0.717 | 3.9116785 | 6 | rps10 |
| crtap | 2.554015 | 0.261642562 | 0.19 | 0.034 | 6.4884769 | 6 | crtap |
| ets1 | 3.437628 | 0.290651409 | 0.19 | 0.034 | 8.7332962 | 6 | ets1 |
| itm2ba1 | 5.116118 | 0.517372213 | 0.666 | 0.281 | 1.2997498 | 6 | itm2ba |
| si:ch211-132p1.31 | 8.383935 | 0.389612445 | 0.324 | 0.087 | 2.1299389 | 6 | si:ch211-132p1.3 |
| rpl23a1 | 1.399349 | 0.446161307 | 0.935 | 0.658 | 3.5550470 | 6 | rpl23a |
| aldh2.2 | 3.294133 | 0.349500399 | 0.308 | 0.08 | 8.3687469 | 6 | aldh2.2 |
| XLOC-0160941 | 5.623602 | 0.359487568 | 0.298 | 0.074 | 1.4286762 | 6 | XLOC-016094 |
| ponzr11 | 9.342392 | 0.554391909 | 0.652 | 0.274 | 2.3734349 | 6 | ponzr1 |
| csrnp1b | 1.228771 | 0.51979776 | 0.439 | 0.153 | 3.1216928 | 6 | csrnp1b |
| grb10b | 1.495782 | 0.269532156 | 0.188 | 0.034 | 3.8000344 | 6 | grb10b |
| adh8a1 | 1.962532 | 0.412561801 | 0.209 | 0.041 | 4.9858150 | 6 | adh8a |
| atf5b | 2.529739 | 0.261206418 | 0.185 | 0.033 | 6.4268038 | 6 | atf5b |
| rpl281 | 7.330388 | 0.353107761 | 0.959 | 0.769 | 1.8622852 | 6 | rpl28 |
| rpl261 | 1.143582 | 0.298524765 | 0.981 | 0.846 | 2.9052707 | 6 | rpl26 |
| igfbp5b1 | 4.608919 | 0.369800962 | 0.31 | 0.079 | 1.1708958 | 6 | igfbp5b |
| add3a1 | 6.046430 | 0.430132881 | 0.404 | 0.126 | 1.5360956 | 6 | add3a |
| si:ch211-39i22.11 | 1.125438 | 0.365302638 | 0.305 | 0.079 | 2.8591752 | 6 | si:ch211-39i22.1 |

| | | | | | | | |
|---|---|---|---|---|---|---|---|
| rps241 | 2.645677 | 0.350515412 | 0.964 | 0.771 | 6.7213431 | 6 | rps24 |
| mgst1.21 | 3.533895 | 0.41542994 | 0.358 | 0.106 | 8.9778603 | 6 | mgst1.2 |
| rpl10a1 | 1.177396 | 0.344185675 | 0.956 | 0.772 | 2.9911767 | 6 | rpl10a |
| mmp14b1 | 1.287369 | 0.343214819 | 0.251 | 0.059 | 3.2705624 | 6 | mmp14b |
| irf1b | 2.154751 | 0.274098894 | 0.173 | 0.03 | 5.4741451 | 6 | irf1b |
| rpl371 | 2.176320 | 0.307869333 | 0.984 | 0.85 | 5.5289417 | 6 | rpl37 |
| gchfr | 2.935289 | 0.319779038 | 0.18 | 0.033 | 7.4571018 | 6 | gchfr |
| rbms31 | 6.497279 | 0.262007136 | 0.175 | 0.031 | 1.6506338 | 6 | rbms3 |
| rps27a1 | 2.031171 | 0.345358829 | 0.961 | 0.767 | 5.1601924 | 6 | rps27a |
| rpl131 | 4.284959 | 0.307725119 | 0.972 | 0.812 | 1.0885939 | 6 | rpl13 |
| tspan4a | 1.204974 | 0.255052397 | 0.177 | 0.032 | 3.0612377 | 6 | tspan4a |
| pdia3 | 1.558248 | 0.508343709 | 0.687 | 0.297 | 3.9587300 | 6 | pdia3 |
| rps4x1 | 1.737147 | 0.384254766 | 0.955 | 0.727 | 4.4132235 | 6 | rps4x |
| rpl391 | 4.336657 | 0.265136146 | 0.972 | 0.873 | 1.1017277 | 6 | rpl39 |
| appbp2 | 6.223690 | 0.278297358 | 0.191 | 0.037 | 1.5811284 | 6 | appbp2 |
| dhrs3a1 | 1.242009 | 0.265359766 | 0.177 | 0.032 | 3.1553260 | 6 | dhrs3a |
| pabpc1b1 | 2.569119 | 0.320101447 | 0.274 | 0.069 | 6.5268477 | 6 | pabpc1b |
| si:ch211-286b5.5 | 2.735240 | 0.374659282 | 0.425 | 0.139 | 6.9488774 | 6 | si:ch211-286b5.5 |
| flot2a | 2.885920 | 0.320994645 | 0.263 | 0.066 | 7.3316817 | 6 | flot2a |
| eif4a2 | 3.347039 | 0.333209757 | 0.298 | 0.081 | 8.5031544 | 6 | eif4a2 |
| rpl171 | 3.817309 | 0.346981794 | 0.956 | 0.755 | 9.6978741 | 6 | rpl17 |
| rps71 | 6.613403 | 0.29137731 | 0.942 | 0.824 | 1.6801352 | 6 | rps7 |
| h2afx12 | 8.105885 | 0.471066497 | 0.548 | 0.213 | 2.0593003 | 6 | h2afx1 |
| rpl36a1 | 9.031422 | 0.301231254 | 0.964 | 0.818 | 2.2944327 | 6 | rpl36a |
| zgc:174863 | 9.716971 | 0.367232312 | 0.34 | 0.101 | 2.4685966 | 6 | zgc:174863 |
| tax1bp3 | 9.902251 | 0.33273949 | 0.324 | 0.092 | 2.5156670 | 6 | tax1bp3 |
| hspb11 | 1.637594 | 0.317962581 | 0.287 | 0.073 | 4.1603078 | 6 | hspb1 |
| rplp2l1 | 3.585764 | 0.361546518 | 0.981 | 0.787 | 9.1096355 | 6 | rplp2l |
| marcksl1b1 | 4.225371 | 0.461581143 | 0.399 | 0.131 | 1.0734555 | 6 | marcksl1b |
| lama5 | 6.880640 | 0.268949968 | 0.152 | 0.025 | 1.7480267 | 6 | lama5 |
| ccdc85b1 | 8.951739 | 0.297089093 | 0.207 | 0.044 | 2.2741895 | 6 | ccdc85b |
| rpl181 | 1.278537 | 0.343086717 | 0.971 | 0.785 | 3.2481240 | 6 | rpl18 |
| snai1a | 1.635116 | 0.276433103 | 0.224 | 0.049 | 4.1540130 | 6 | snai1a |
| rnd3b1 | 3.531847 | 0.273908505 | 0.206 | 0.043 | 8.9726593 | 6 | rnd3b |
| egfra1 | 4.611742 | 0.252531242 | 0.159 | 0.027 | 1.1716130 | 6 | egfra |

| | | | | | | | |
|---|---|---|---|---|---|---|---|
| nsa21 | 8.021408 | 0.519221675 | 0.757 | 0.373 | 2.0378388 | 6 | nsa2 |
| vdac3 | 8.544983 | 0.387550951 | 0.451 | 0.156 | 2.1708531 | 6 | vdac3 |
| zgc:113208 | 8.545452 | 0.276153071 | 0.203 | 0.043 | 2.1709722 | 6 | zgc:113208 |
| serinc5 | 1.626184 | 0.350952221 | 0.29 | 0.079 | 4.1313207 | 6 | serinc5 |
| hmgb1a2 | 2.602031 | 0.464519072 | 0.536 | 0.215 | 6.6104612 | 6 | hmgb1a |
| CR855311.11 | 1.041448 | 0.459492295 | 0.29 | 0.08 | 2.6457989 | 6 | CR855311.1 |
| rps3a1 | 1.944468 | 0.322137032 | 0.958 | 0.785 | 4.9399212 | 6 | rps3a |
| znf385a1 | 3.697718 | 0.296412563 | 0.214 | 0.048 | 9.3940545 | 6 | znf385a |
| atf3 | 4.125938 | 0.453707036 | 0.271 | 0.074 | 1.0481946 | 6 | atf3 |
| il6st | 4.158575 | 0.313535077 | 0.216 | 0.049 | 1.0564862 | 6 | il6st |
| tspan9b | 4.507634 | 0.26513581 | 0.204 | 0.044 | 1.1451646 | 6 | tspan9b |
| faua1 | 5.364437 | 0.385676535 | 0.943 | 0.708 | 1.3628353 | 6 | faua |
| stau21 | 1.082663 | 0.337827187 | 0.305 | 0.087 | 2.7505071 | 6 | stau2 |
| rps141 | 1.310510 | 0.346695961 | 0.971 | 0.773 | 3.3293506 | 6 | rps14 |
| mycb | 2.096940 | 0.459891731 | 0.499 | 0.188 | 5.3272785 | 6 | mycb |
| klf11b | 2.862060 | 0.354411591 | 0.274 | 0.075 | 7.2710648 | 6 | klf11b |
| map1lc3b | 3.050774 | 0.374442021 | 0.355 | 0.114 | 7.7504928 | 6 | map1lc3b |
| ptena | 4.073247 | 0.287195076 | 0.237 | 0.058 | 1.0348084 | 6 | ptena |
| ppp1cbl | 5.835016 | 0.303281597 | 0.24 | 0.06 | 1.4823859 | 6 | ppp1cbl |
| si:rp71-79p20.2 | 6.568625 | 0.278989197 | 0.175 | 0.034 | 1.6687593 | 6 | si:rp71-79p20.2 |
| tnfrsf1a | 9.056616 | 0.306347059 | 0.224 | 0.053 | 2.3008334 | 6 | tnfrsf1a |
| rpl341 | 3.041046 | 0.330294973 | 0.972 | 0.795 | 7.7257774 | 6 | rpl34 |
| sumo2b1 | 3.054410 | 0.464228633 | 0.639 | 0.282 | 7.7597301 | 6 | sumo2b |
| rpl241 | 3.302555 | 0.325194875 | 0.963 | 0.787 | 8.3901419 | 6 | rpl24 |
| rps261 | 3.515556 | 0.34845148 | 0.951 | 0.742 | 8.9312702 | 6 | rps26 |
| rpl111 | 3.854193 | 0.321826104 | 0.982 | 0.823 | 9.7915780 | 6 | rpl11 |
| rgs5a1 | 6.119365 | 0.27161869 | 0.172 | 0.033 | 1.5546248 | 6 | rgs5a |
| sfrp1a | 1.452128 | 0.263181681 | 0.165 | 0.031 | 3.6891326 | 6 | sfrp1a |
| tpm1 | 4.000980 | 0.485533287 | 0.486 | 0.191 | 1.0164491 | 6 | tpm1 |
| eef1db | 4.378877 | 0.462195442 | 0.621 | 0.27 | 1.1124538 | 6 | eef1db |
| itgb1a1 | 5.275899 | 0.281854264 | 0.206 | 0.046 | 1.3403422 | 6 | itgb1a |
| rpl22l11 | 8.033383 | 0.510591407 | 0.788 | 0.409 | 2.0408810 | 6 | rpl22l1 |
| eif4a1b | 8.219507 | 0.473936771 | 0.653 | 0.297 | 2.0881659 | 6 | eif4a1b |
| rpl121 | 9.537137 | 0.296766146 | 0.968 | 0.806 | 2.4229097 | 6 | rpl12 |
| sept21 | 2.444456 | 0.402427919 | 0.46 | 0.171 | 6.2101406 | 6 | sept2 |

| | | | | | | | |
|---|---|---|---|---|---|---|---|
| wls1 | 3.239301 | 0.375878538 | 0.368 | 0.121 | 8.2294452 | 6 | wls |
| clic4 | 4.571947 | 0.25028177 | 0.22 | 0.052 | 1.1615032 | 6 | clic4 |
| tnrc6c11 | 4.807639 | 0.409419665 | 0.394 | 0.14 | 1.2213808 | 6 | tnrc6c1 |
| CABZ01072614.11 | 5.353957 | 0.364580737 | 0.293 | 0.086 | 1.3601730 | 6 | CABZ01072614.1 |
| rps61 | 7.633892 | 0.314715335 | 0.964 | 0.797 | 1.9393904 | 6 | rps6 |
| rps111 | 1.441405 | 0.317759421 | 0.958 | 0.777 | 3.6618902 | 6 | rps11 |
| timp2a1 | 1.456664 | 0.292158997 | 0.233 | 0.058 | 3.7006572 | 6 | timp2a |
| eef1g1 | 2.742562 | 0.419538032 | 0.922 | 0.641 | 6.9674798 | 6 | eef1g |
| rpl81 | 3.579220 | 0.308331148 | 0.943 | 0.789 | 9.0930095 | 6 | rpl8 |
| rps31 | 4.664553 | 0.3596422 | 0.934 | 0.717 | 1.1850298 | 6 | rps3 |
| vdrb | 5.970425 | 0.276959756 | 0.186 | 0.04 | 1.5167864 | 6 | vdrb |
| slc43a2a1 | 1.099193 | 0.360774182 | 0.308 | 0.093 | 2.7925006 | 6 | slc43a2a |
| si:dkey-28b4.7 | 1.313785 | 0.341128883 | 0.323 | 0.101 | 3.3376722 | 6 | si:dkey-28b4.7 |
| mcl1b1 | 1.455051 | 0.446229108 | 0.574 | 0.237 | 3.6965577 | 6 | mcl1b |
| emd | 1.953255 | 0.284118227 | 0.243 | 0.064 | 4.9622452 | 6 | emd |
| sh3pxd2b | 2.474986 | 0.265716804 | 0.196 | 0.044 | 6.2877042 | 6 | sh3pxd2b |
| appb1 | 1.209579 | 0.263144837 | 0.186 | 0.041 | 3.0729370 | 6 | appb |
| kank21 | 1.295530 | 0.302121753 | 0.206 | 0.049 | 3.2912939 | 6 | kank2 |
| h1f0 | 1.442128 | 0.342865886 | 0.243 | 0.064 | 3.6637263 | 6 | h1f0 |
| tgif1 | 2.294341 | 0.311733578 | 0.305 | 0.091 | 5.8287733 | 6 | tgif1 |
| rps251 | 2.022127 | 0.345974206 | 0.956 | 0.74 | 5.1372141 | 6 | rps25 |
| rps15a1 | 2.959273 | 0.324073338 | 0.942 | 0.792 | 7.5180355 | 6 | rps15a |
| si:dkey-6n21.12 | 4.178045 | 0.251423867 | 0.173 | 0.037 | 1.0614324 | 6 | si:dkey-6n21.12 |
| map1lc3a | 5.353879 | 0.29214973 | 0.261 | 0.073 | 1.3601529 | 6 | map1lc3a |
| tob1b | 1.449810 | 0.365517508 | 0.431 | 0.157 | 3.6832446 | 6 | tob1b |
| si:ch211-1a19.31 | 2.606469 | 0.321264337 | 0.209 | 0.051 | 6.6217352 | 6 | si:ch211-1a19.3 |
| maff | 3.553240 | 0.316220037 | 0.258 | 0.073 | 9.0270065 | 6 | maff |
| rpl35a1 | 3.785611 | 0.316118637 | 0.951 | 0.775 | 9.6173451 | 6 | rpl35a |
| nfixb1 | 5.939981 | 0.306973871 | 0.266 | 0.075 | 1.5090522 | 6 | nfixb |
| ube2e3 | 9.681897 | 0.378761238 | 0.423 | 0.159 | 2.4596860 | 6 | ube2e3 |
| emp21 | 1.161707 | 0.270469381 | 0.207 | 0.05 | 2.9513185 | 6 | emp2 |
| si:ch73-237c6.1 | 1.273634 | 0.256268455 | 0.182 | 0.04 | 3.2356693 | 6 | si:ch73-237c6.1 |
| mych1 | 2.348640 | 0.329182435 | 0.277 | 0.082 | 5.9667218 | 6 | mych |
| rps211 | 2.439022 | 0.350998139 | 0.964 | 0.768 | 6.1963377 | 6 | rps21 |
| agrn | 3.519258 | 0.289677703 | 0.214 | 0.054 | 8.9406762 | 6 | agrn |

| | | | | | | | |
|---|---|---|---|---|---|---|---|
| gamt | 4.183753 | 0.251255629 | 0.175 | 0.038 | 1.0628826 | 6 | gamt |
| rpl301 | 5.805650 | 0.360425514 | 0.951 | 0.721 | 1.4749255 | 6 | rpl30 |
| krt18b1 | 1.343498 | 0.358811137 | 0.451 | 0.164 | 3.4131572 | 6 | krt18b |
| serpinf11 | 1.866073 | 0.260428272 | 0.178 | 0.04 | 4.7407597 | 6 | serpinf1 |
| id11 | 2.075158 | 0.40984753 | 0.355 | 0.122 | 5.2719406 | 6 | id1 |
| atp10a | 4.844702 | 0.289556866 | 0.209 | 0.053 | 1.2307965 | 6 | atp10a |
| sox4a | 5.019762 | 0.318656984 | 0.217 | 0.056 | 1.2752706 | 6 | sox4a |
| klf6a1 | 9.894572 | 0.405289654 | 0.438 | 0.169 | 2.5137161 | 6 | klf6a |
| calm2a | 1.359183 | 0.391911018 | 0.493 | 0.197 | 3.4530059 | 6 | calm2a |
| osr11 | 1.746370 | 0.260601458 | 0.175 | 0.038 | 4.4366541 | 6 | osr1 |
| btg21 | 1.842685 | 0.499266836 | 0.661 | 0.336 | 4.6813422 | 6 | btg2 |
| rps26l1 | 2.170983 | 0.392986072 | 0.948 | 0.689 | 5.5153827 | 6 | rps26l |
| ahnak1 | 2.474670 | 0.346445771 | 0.366 | 0.127 | 6.2868999 | 6 | ahnak |
| ndufa4l | 3.312041 | 0.456643172 | 0.791 | 0.406 | 8.4142409 | 6 | ndufa4l |
| tomm5 | 1.073097 | 0.382492843 | 0.515 | 0.218 | 2.7262051 | 6 | tomm5 |
| flot1b | 3.708168 | 0.287620962 | 0.293 | 0.093 | 9.4206031 | 6 | flot1b |
| anxa4 | 4.516095 | 0.333621406 | 0.386 | 0.141 | 1.1473141 | 6 | anxa4 |
| rpl13a1 | 4.718881 | 0.296657732 | 0.95 | 0.775 | 1.1988317 | 6 | rpl13a |
| rpl221 | 6.612141 | 0.346232781 | 0.921 | 0.712 | 1.6798146 | 6 | rpl22 |
| vdac1 | 1.874135 | 0.309514026 | 0.431 | 0.159 | 4.7612416 | 6 | vdac1 |
| pnrc21 | 1.903423 | 0.45097887 | 0.929 | 0.583 | 4.8356462 | 6 | pnrc2 |
| hist2h2l | 2.491199 | 0.330220575 | 0.287 | 0.091 | 6.3288916 | 6 | hist2h2l |
| rpl18a1 | 3.200443 | 0.297142914 | 0.947 | 0.795 | 8.1307258 | 6 | rpl18a |
| tspan7 | 4.001032 | 0.26052621 | 0.258 | 0.074 | 1.0164624 | 6 | tspan7 |
| rps291 | 4.193954 | 0.279203578 | 0.982 | 0.838 | 1.0654741 | 6 | rps29 |
| cflara | 4.412261 | 0.274021911 | 0.216 | 0.057 | 1.1209349 | 6 | cflara |
| rpl271 | 6.092000 | 0.304321557 | 0.945 | 0.773 | 1.5476727 | 6 | rpl27 |
| dnmt3ab1 | 6.327051 | 0.318157846 | 0.272 | 0.085 | 1.6073875 | 6 | dnmt3ab |
| rpl5a1 | 9.599477 | 0.455454021 | 0.87 | 0.522 | 2.4387471 | 6 | rpl5a |
| rpl151 | 2.676885 | 0.325171186 | 0.948 | 0.756 | 6.8006273 | 6 | rpl15 |
| ciarta1 | 5.913496 | 0.320210247 | 0.321 | 0.109 | 1.5023237 | 6 | ciarta |
| rpl381 | 5.966429 | 0.344003184 | 0.963 | 0.745 | 1.5157714 | 6 | rpl38 |
| uba521 | 6.095320 | 0.393569787 | 0.929 | 0.65 | 1.5485161 | 6 | uba52 |
| isg15 | 1.012774 | 0.340041214 | 0.216 | 0.058 | 2.5729534 | 6 | isg15 |
| rpl71 | 2.251017 | 0.268834932 | 0.943 | 0.804 | 5.7187088 | 6 | rpl7 |

| | | | | | | | |
|---|---|---|---|---|---|---|---|
| eif1b | 5.82680[illegible] | 0.439383013 | 0.755 | 0.392 | 1.4803006 | 6 | eif1b |
| ip6k2a | 7.51523[illegible] | 0.26482719 | 0.224 | 0.063 | 1.9092462 | 6 | ip6k2a |
| sesn12 | 8.24804[illegible] | 0.390702046 | 0.417 | 0.165 | 2.0954162 | 6 | sesn1 |
| eif3g1 | 9.67608[illegible] | 0.395911018 | 0.614 | 0.286 | 2.4582097 | 6 | eif3g |
| rps51 | 1.27128[illegible] | 0.341527974 | 0.93 | 0.72 | 3.2296891 | 6 | rps5 |
| krt91 | 2.34006[illegible] | 0.324679822 | 0.232 | 0.066 | 5.9449296 | 6 | krt91 |
| ywhaqb1 | 3.71565[illegible] | 0.356063384 | 0.378 | 0.146 | 9.4396297 | 6 | ywhaqb |
| rpl351 | 4.37545[illegible] | 0.271382473 | 0.947 | 0.809 | 1.1115831 | 6 | rpl35 |
| ddit31 | 7.24614[illegible] | 0.385370623 | 0.42 | 0.168 | 1.8408829 | 6 | ddit3 |
| si:ch211-15b10.6 | 9.14714[illegible] | 0.272600343 | 0.237 | 0.07 | 2.3238320 | 6 | si:ch211-15b10.6 |
| serbp1a1 | 3.52686[illegible] | 0.451340836 | 0.844 | 0.491 | 8.9600120 | 6 | serbp1a |
| capns1a | 5.43418[illegible] | 0.306217624 | 0.394 | 0.149 | 1.3805554 | 6 | capns1a |
| si:ch211-204c21.1 | 7.11138[illegible] | 0.282549849 | 0.233 | 0.068 | 1.8066470 | 6 | si:ch211-204c21.1 |
| wu:fb16f03 | 8.29273[illegible] | 0.301110089 | 0.261 | 0.083 | 2.1067698 | 6 | wu:fb16f03 |
| nop531 | 2.46918[illegible] | 0.379677217 | 0.559 | 0.254 | 6.2729743 | 6 | nop53 |
| capns1b | 4.90386[illegible] | 0.309895632 | 0.363 | 0.137 | 1.2458263 | 6 | capns1b |
| ppdpfa | 5.79415[illegible] | 0.296286685 | 0.225 | 0.065 | 1.4720052 | 6 | ppdpfa |
| nop56 | 9.75865[illegible] | 0.357616533 | 0.426 | 0.177 | 2.4791868 | 6 | nop56 |
| cavin2b | 1.39954[illegible] | 0.260833049 | 0.241 | 0.072 | 3.5555507 | 6 | cavin2b |
| cct2 | 2.09644[illegible] | 0.360175942 | 0.54 | 0.242 | 5.3260216 | 6 | cct2 |
| rpl211 | 2.60252[illegible] | 0.289994707 | 0.943 | 0.77 | 6.6117132 | 6 | rpl21 |
| anxa11b | 4.11879[illegible] | 0.313782119 | 0.412 | 0.164 | 1.0463795 | 6 | anxa11b |
| higd1a | 6.69768[illegible] | 0.417411759 | 0.595 | 0.291 | 1.7015460 | 6 | higd1a |
| hdlbpa1 | 1.05717[illegible] | 0.316075171 | 0.413 | 0.165 | 2.6857608 | 6 | hdlbpa |
| CR318588.11 | 1.07769[illegible] | 0.564201981 | 0.451 | 0.206 | 2.7378828 | 6 | CR318588.1 |
| khdrbs1b | 1.34498[illegible] | 0.271388094 | 0.29 | 0.098 | 3.4169241 | 6 | khdrbs1b |
| serf21 | 2.11362[illegible] | 0.408247581 | 0.713 | 0.363 | 5.3696746 | 6 | serf2 |
| naca1 | 3.05775[illegible] | 0.375723268 | 0.908 | 0.641 | 7.7682256 | 6 | naca |
| dbi | 4.91951[illegible] | 0.346871565 | 0.519 | 0.231 | 1.2498029 | 6 | dbi |
| pfn2l | 9.58777[illegible] | 0.331269987 | 0.489 | 0.204 | 2.4357753 | 6 | pfn2l |
| hnrnpa0b | 1.37700[illegible] | 0.406499007 | 0.658 | 0.328 | 3.4982893 | 6 | hnrnpa0b |
| rpl311 | 3.08896[illegible] | 0.365024388 | 0.94 | 0.677 | 7.8475057 | 6 | rpl31 |
| clic2 | 4.44045[illegible] | 0.255312995 | 0.22 | 0.065 | 1.1280967 | 6 | clic2 |
| zfp36l1a2 | 1.08140[illegible] | 0.385258673 | 0.533 | 0.243 | 2.7473062 | 6 | zfp36l1a |
| ctnnb1 | 1.23344[illegible] | 0.309964188 | 0.381 | 0.149 | 3.1335628 | 6 | ctnnb1 |

| | | | | | | | |
|---|---|---|---|---|---|---|---|
| tegt | 1.512949 | 0.34635878 | 0.502 | 0.227 | 3.8436471 | 6 | tegt |
| pik3r3a | 1.690801 | 0.252541338 | 0.263 | 0.086 | 4.2954807 | 6 | pik3r3a |
| arrdc3a1 | 1.898818 | 0.307157586 | 0.22 | 0.067 | 4.8239482 | 6 | arrdc3a |
| mt-nd21 | 3.290305 | 0.407457664 | 0.718 | 0.378 | 8.3590202 | 6 | mt-nd2 |
| rpl101 | 6.191058 | 0.278149265 | 0.942 | 0.784 | 1.5728384 | 6 | rpl10 |
| romo11 | 7.742689 | 0.405381874 | 0.741 | 0.394 | 1.9670301 | 6 | romo1 |
| rps27l | 9.876215 | 0.312678596 | 0.475 | 0.207 | 2.5090526 | 6 | rps27l |
| myh11a | 2.110494 | 0.310235225 | 0.216 | 0.066 | 5.3617124 | 6 | myh11a |
| ndfip1l1 | 3.547303 | 0.264231987 | 0.224 | 0.069 | 9.0119257 | 6 | ndfip1l |
| txnipa | 2.636071 | 0.369087156 | 0.504 | 0.233 | 6.6969391 | 6 | txnipa |
| eif3ha | 7.292336 | 0.304987666 | 0.494 | 0.219 | 1.8526181 | 6 | eif3ha |
| pno1 | 8.118284 | 0.295859395 | 0.392 | 0.162 | 2.0624501 | 6 | pno1 |
| hp1bp3 | 9.322979 | 0.273723022 | 0.305 | 0.113 | 2.3685028 | 6 | hp1bp3 |
| stat1a | 2.990228 | 0.290717475 | 0.308 | 0.116 | 7.5966749 | 6 | stat1a |
| rps231 | 4.169521 | 0.303591315 | 0.934 | 0.737 | 1.0592669 | 6 | rps23 |
| atp2b2 | 4.303810 | 0.259975124 | 0.235 | 0.077 | 1.0933830 | 6 | atp2b2 |
| si:ch211-198c19.31 | 8.144609 | 0.273994105 | 0.238 | 0.077 | 2.0691379 | 6 | si:ch211-198c19.3 |
| rhoab1 | 8.888092 | 0.337373739 | 0.441 | 0.198 | 2.2580198 | 6 | rhoab |
| slc3a2b | 2.513626 | 0.324412285 | 0.31 | 0.121 | 6.3858681 | 6 | slc3a2b |
| map2k61 | 3.613670 | 0.265453604 | 0.261 | 0.092 | 9.1805311 | 6 | map2k6 |
| tagln21 | 5.322881 | 0.340473018 | 0.449 | 0.2 | 1.3522779 | 6 | tagln2 |
| laptm4a | 5.458093 | 0.281095466 | 0.301 | 0.115 | 1.3866285 | 6 | laptm4a |
| crebzf | 9.492809 | 0.28179713 | 0.279 | 0.102 | 2.4116481 | 6 | crebzf |
| prdx5 | 1.252283 | 0.280835574 | 0.405 | 0.17 | 3.1814274 | 6 | prdx5 |
| cyr611 | 1.298091 | 0.302125825 | 0.203 | 0.062 | 3.2978024 | 6 | cyr61 |
| mt-nd41 | 3.069474 | 0.386919864 | 0.791 | 0.433 | 7.7979991 | 6 | mt-nd4 |
| rack11 | 3.203092 | 0.2840382 | 0.945 | 0.756 | 8.1374575 | 6 | rack1 |
| skp1 | 8.206244 | 0.317747186 | 0.541 | 0.257 | 2.0847963 | 6 | skp1 |
| pin1 | 1.103046 | 0.258932051 | 0.365 | 0.149 | 2.8022902 | 6 | pin1 |
| si:dkey-11f4.14 | 1.219955 | 0.305145658 | 0.298 | 0.116 | 3.0992963 | 6 | si:dkey-11f4.14 |
| tmem50a | 1.299560 | 0.25869582 | 0.355 | 0.143 | 3.3015322 | 6 | tmem50a |
| ppiaa1 | 3.249070 | 0.25922097 | 0.966 | 0.808 | 8.2542646 | 6 | ppiaa |
| fxyd11 | 3.506641 | 0.280660678 | 0.229 | 0.078 | 8.9086220 | 6 | fxyd1 |
| abcc51 | 7.132676 | 0.261760237 | 0.25 | 0.089 | 1.8120565 | 6 | abcc5 |
| acin1a1 | 7.741330 | 0.331306401 | 0.467 | 0.219 | 1.9666851 | 6 | acin1a |

| | | | | | | | |
|---|---|---|---|---|---|---|---|
| tpm3 | 1.360286 | 0.332672767 | 0.556 | 0.273 | 3.4558075 | 6 | tpm3 |
| NC-002333.41 | 1.523166 | 0.401083512 | 0.924 | 0.595 | 3.8696046 | 6 | NC-002333.4 |
| eif4ebp3l1 | 2.215660 | 0.308843959 | 0.444 | 0.196 | 5.6288843 | 6 | eif4ebp3l |
| atrx | 3.141785 | 0.27919844 | 0.305 | 0.121 | 7.9817060 | 6 | atrx |
| bccip | 5.375261 | 0.252403612 | 0.344 | 0.141 | 1.3655851 | 6 | bccip |
| XLOC-0004311 | 5.433571 | 0.275847404 | 0.561 | 0.261 | 1.3803989 | 6 | XLOC-000431 |
| cct8 | 6.960364 | 0.300652654 | 0.462 | 0.212 | 1.7682807 | 6 | cct8 |
| mt-nd11 | 7.115770 | 0.368667163 | 0.695 | 0.384 | 1.8077615 | 6 | mt-nd1 |
| bhlhe40 | 5.842133 | 0.280719238 | 0.376 | 0.159 | 1.4841939 | 6 | bhlhe40 |
| rpl41 | 1.274982 | 0.341771668 | 0.909 | 0.646 | 3.2390932 | 6 | rpl4 |
| rpl141 | 1.318860 | 0.356057078 | 0.943 | 0.658 | 3.3505643 | 6 | rpl14 |
| cdc42 | 2.793854 | 0.294581074 | 0.475 | 0.224 | 7.0977869 | 6 | cdc42 |
| srsf5a | 3.536649 | 0.31698026 | 0.582 | 0.292 | 8.9848579 | 6 | srsf5a |
| wu:fi37e09 | 5.964982 | 0.260416451 | 0.305 | 0.122 | 1.5154039 | 6 | wu:fi37e09 |
| rhoaa | 6.936577 | 0.289004996 | 0.378 | 0.169 | 1.7622374 | 6 | rhoaa |
| eif3m1 | 1.376832 | 0.357399593 | 0.686 | 0.371 | 3.4978436 | 6 | eif3m |
| rps27.11 | 1.604504 | 0.342683814 | 0.943 | 0.7 | 4.0762439 | 6 | rps27.1 |
| ppdpfb | 1.711562 | 0.372887796 | 0.815 | 0.52 | 4.3482256 | 6 | ppdpfb |
| mtpn | 2.183973 | 0.255477397 | 0.36 | 0.156 | 5.5483856 | 6 | mtpn |
| midn | 3.158255 | 0.286804633 | 0.326 | 0.138 | 8.0235483 | 6 | midn |
| eif2s3 | 9.828228 | 0.317514086 | 0.58 | 0.297 | 2.4968614 | 6 | eif2s3 |
| gadd45bb | 2.075549 | 0.256221212 | 0.217 | 0.077 | 5.2729339 | 6 | gadd45bb |
| btg11 | 7.364079 | 0.321534207 | 0.666 | 0.342 | 1.8708444 | 6 | btg1 |
| pfn2 | 1.052233 | 0.284637423 | 0.48 | 0.229 | 2.6731992 | 6 | pfn2 |
| tsc22d3 | 1.577843 | 0.291153931 | 0.342 | 0.152 | 4.0085112 | 6 | tsc22d3 |
| ier2b2 | 5.665028 | 0.312783766 | 0.464 | 0.228 | 1.4392005 | 6 | ier2b |
| si:ch211-196l7.42 | 9.152664 | 0.319124775 | 0.601 | 0.319 | 2.3252344 | 6 | si:ch211-196l7.4 |
| COX32 | 1.436360 | 0.375071051 | 0.937 | 0.665 | 3.6490748 | 6 | COX3 |
| irf2bp2a1 | 2.564923 | 0.280263363 | 0.384 | 0.178 | 6.5161892 | 6 | irf2bp2a |
| zgc:153867 | 4.052063 | 0.360632573 | 0.739 | 0.453 | 1.0294267 | 6 | zgc:153867 |
| ip6k2b1 | 6.925578 | 0.278319674 | 0.288 | 0.123 | 1.7594432 | 6 | ip6k2b |
| tcp1 | 9.765010 | 0.251807025 | 0.486 | 0.236 | 2.4808009 | 6 | tcp1 |
| zgc:111986 | 1.476614 | 0.259911051 | 0.384 | 0.178 | 3.7513386 | 6 | zgc:111986 |
| rpl5b1 | 1.843200 | 0.33800406 | 0.908 | 0.634 | 4.6826499 | 6 | rpl5b |
| eif3d | 2.491284 | 0.285884705 | 0.528 | 0.271 | 6.3291092 | 6 | eif3d |

| | | | | | | | |
|---|---|---|---|---|---|---|---|
| ptmaa1 | 5.971855 | 0.351375146 | 0.76 | 0.419 | 1.5171498 | 6 | ptmaa |
| rsl24d11 | 2.510943 | 0.360800191 | 0.755 | 0.448 | 6.3790512 | 6 | rsl24d1 |
| rab101 | 8.490822 | 0.254200018 | 0.423 | 0.206 | 2.1570935 | 6 | rab10 |
| tomm20b | 1.776086 | 0.253944182 | 0.442 | 0.22 | 4.5121481 | 6 | tomm20b |
| eif3ea1 | 4.538839 | 0.285243203 | 0.556 | 0.294 | 1.1530921 | 6 | eif3ea |
| btf31 | 5.622032 | 0.350424135 | 0.901 | 0.598 | 1.4282774 | 6 | btf3 |
| gapdhs1 | 6.409158 | 0.28100923 | 0.582 | 0.304 | 1.6282466 | 6 | gapdhs |
| prdx2 | 1.005564 | 0.31829235 | 0.721 | 0.41 | 2.5546366 | 6 | prdx2 |
| ptmab1 | 1.433747 | 0.348821012 | 0.849 | 0.558 | 3.6424364 | 6 | ptmab |
| eif3i | 1.587990 | 0.276968412 | 0.532 | 0.281 | 4.0342892 | 6 | eif3i |
| syncrip | 2.089569 | 0.284170462 | 0.569 | 0.302 | 5.3085507 | 6 | syncrip |
| pabpc1a1 | 2.324257 | 0.345739468 | 0.849 | 0.558 | 5.9047749 | 6 | pabpc1a |
| tmsb4x2 | 1.835100 | 0.391250171 | 0.99 | 0.707 | 4.6620724 | 6 | tmsb4x |
| fkbp1aa | 4.618580 | 0.315205189 | 0.692 | 0.403 | 1.1733504 | 6 | fkbp1aa |
| cd632 | 7.516405 | 0.283186224 | 0.671 | 0.361 | 1.9095427 | 6 | cd63 |
| eif5a1 | 1.082829 | 0.336250819 | 0.78 | 0.483 | 2.7509276 | 6 | eif5a |
| eif3f1 | 2.311305 | 0.342655733 | 0.765 | 0.453 | 5.8718707 | 6 | eif3f |
| gabarapb | 2.440420 | 0.252735561 | 0.556 | 0.295 | 6.1998895 | 6 | gabarapb |
| eif2s2 | 1.035967 | 0.252406033 | 0.556 | 0.313 | 2.6318756 | 6 | eif2s2 |
| mt-atp61 | 3.012641 | 0.314110648 | 0.799 | 0.546 | 7.6536149 | 6 | mt-atp6 |
| rplp21 | 8.611730 | 0.272075748 | 0.969 | 0.775 | 2.1878101 | 6 | rplp2 |
| mibp21 | 4.044288 | 0.294551148 | 0.783 | 0.51 | 1.0274515 | 6 | mibp2 |
| eef2b | 1.248908 | 0.284043547 | 0.883 | 0.626 | 3.1728518 | 6 | eef2b |
| eef1da1 | 5.769333 | 0.272298586 | 0.783 | 0.494 | 1.4656990 | 6 | eef1da |
| c1qb | 0 | 1.841613282 | 0.933 | 0.036 | 0 | 7 | c1qb |
| CU499330.1 | 0 | 1.752583604 | 0.89 | 0.037 | 0 | 7 | CU499330.1 |
| grn1 | 0 | 1.724404412 | 0.894 | 0.055 | 0 | 7 | grn1 |
| c1qc | 0 | 1.684333266 | 0.91 | 0.033 | 0 | 7 | c1qc |
| lygl1 | 0 | 1.680665801 | 0.919 | 0.064 | 0 | 7 | lygl1 |
| si:busm1-266f07.2 | 0 | 1.650576286 | 0.935 | 0.114 | 0 | 7 | si:busm1-266f07.2 |
| si:dkey-5n18.1 | 0 | 1.650413433 | 0.816 | 0.032 | 0 | 7 | si:dkey-5n18.1 |
| cd74a1 | 0 | 1.640716198 | 0.938 | 0.185 | 0 | 7 | cd74a |
| lgals2a1 | 0 | 1.619483406 | 0.94 | 0.103 | 0 | 7 | lgals2a |
| cd74b1 | 0 | 1.575546694 | 0.972 | 0.194 | 0 | 7 | cd74b |
| c1qa | 0 | 1.543558362 | 0.853 | 0.021 | 0 | 7 | c1qa |

| | | | | | | | |
|---|---|---|---|---|---|---|---|
| fcer1gl | 0 | 1.534330974 | 0.97 | 0.148 | 0 | 7 | fcer1gl |
| mrc1b | 0 | 1.476452403 | 0.784 | 0.013 | 0 | 7 | mrc1b |
| si:ch211-165i18.2 | 0 | 1.383455108 | 0.703 | 0.01 | 0 | 7 | si:ch211-165i18.2 |
| ms4a17a.10 | 0 | 1.380293463 | 0.763 | 0.023 | 0 | 7 | ms4a17a.10 |
| zgc:153118 | 0 | 1.376851441 | 0.601 | 0.02 | 0 | 7 | zgc:153118 |
| mhc2dab | 0 | 1.37461337 | 0.765 | 0.066 | 0 | 7 | mhc2dab |
| LOC110437731 | 0 | 1.354048062 | 0.659 | 0.067 | 0 | 7 | LOC110437731 |
| cfp | 0 | 1.327770418 | 0.772 | 0.046 | 0 | 7 | cfp |
| BX649485.2 | 0 | 1.31655645 | 0.56 | 0.013 | 0 | 7 | BX649485.2 |
| ccl35.1 | 0 | 1.282150139 | 0.613 | 0.061 | 0 | 7 | ccl35.1 |
| lgals3bpb | 0 | 1.275996775 | 0.572 | 0.021 | 0 | 7 | lgals3bpb |
| krt18b2 | 0 | 1.262029654 | 0.843 | 0.148 | 0 | 7 | krt18b |
| marco | 0 | 1.219989268 | 0.615 | 0.017 | 0 | 7 | marco |
| NPC2 (1 of many) | 0 | 1.207889446 | 0.613 | 0.016 | 0 | 7 | NPC2 (1 of many) |
| itgae.1 | 0 | 1.163568603 | 0.655 | 0.01 | 0 | 7 | itgae.1 |
| ccl34b.8 | 0 | 1.156855072 | 0.445 | 0.008 | 0 | 7 | ccl34b.8 |
| sftpbb | 0 | 1.155185556 | 0.7 | 0.043 | 0 | 7 | sftpbb |
| gpx1a | 0 | 1.133766987 | 0.866 | 0.209 | 0 | 7 | gpx1a |
| cx32.2 | 0 | 1.123940856 | 0.648 | 0.005 | 0 | 7 | cx32.2 |
| CR855311.12 | 0 | 1.120269236 | 0.64 | 0.065 | 0 | 7 | CR855311.1 |
| grna | 0 | 1.101176199 | 0.684 | 0.03 | 0 | 7 | grna |
| tmsb11 | 0 | 1.09631881 | 0.8 | 0.133 | 0 | 7 | tmsb1 |
| cd831 | 0 | 1.093297224 | 0.599 | 0.072 | 0 | 7 | cd83 |
| ccl34a.4 | 0 | 1.093161724 | 0.519 | 0.002 | 0 | 7 | ccl34a.4 |
| ctss2.2 | 0 | 1.092319047 | 0.608 | 0.027 | 0 | 7 | ctss2.2 |
| havcr1 | 0 | 1.073356849 | 0.572 | 0.013 | 0 | 7 | havcr1 |
| fgl2a | 0 | 1.070634451 | 0.62 | 0.052 | 0 | 7 | fgl2a |
| zgc:174904 | 0 | 1.039836807 | 0.525 | 0.031 | 0 | 7 | zgc:174904 |
| cmklr1 | 0 | 1.018262368 | 0.525 | 0.011 | 0 | 7 | cmklr1 |
| pmaip1 | 0 | 1.002152468 | 0.587 | 0.08 | 0 | 7 | pmaip1 |
| si:ch211-147m6.1 | 0 | 0.974492438 | 0.516 | 0.012 | 0 | 7 | si:ch211-147m6.1 |
| mfap4 | 0 | 0.967151396 | 0.468 | 0.011 | 0 | 7 | mfap4 |
| zgc:103700 | 0 | 0.961584447 | 0.477 | 0.036 | 0 | 7 | zgc:103700 |
| smox | 0 | 0.94888465 | 0.648 | 0.063 | 0 | 7 | smox |
| tcima | 0 | 0.932029178 | 0.542 | 0.051 | 0 | 7 | tcima |

| | | | | | | | |
|---|---|---|---|---|---|---|---|
| clic21 | 0 | 0.913458747 | 0.572 | 0.05 | 0 | 7 | clic2 |
| spi1a | 0 | 0.910602824 | 0.551 | 0.017 | 0 | 7 | spi1a |
| si:dkey-7i4.24 | 0 | 0.902419714 | 0.293 | 0.016 | 0 | 7 | si:dkey-7i4.24 |
| LOC100536946 | 0 | 0.900291434 | 0.371 | 0.007 | 0 | 7 | LOC100536946 |
| scpep1 | 0 | 0.898835034 | 0.664 | 0.097 | 0 | 7 | scpep1 |
| si:ch211-74m13.3 | 0 | 0.886733673 | 0.518 | 0.049 | 0 | 7 | si:ch211-74m13.3 |
| slc6a6b | 0 | 0.886316538 | 0.611 | 0.075 | 0 | 7 | slc6a6b |
| mir21-1 | 0 | 0.884407234 | 0.595 | 0.087 | 0 | 7 | mir21-1 |
| zgc:152830 | 0 | 0.873850362 | 0.58 | 0.046 | 0 | 7 | zgc:152830 |
| slc3a2a | 0 | 0.873067388 | 0.521 | 0.017 | 0 | 7 | slc3a2a |
| txn | 0 | 0.837253507 | 0.41 | 0.037 | 0 | 7 | txn |
| ms4a17a.61 | 0 | 0.829340767 | 0.523 | 0.031 | 0 | 7 | ms4a17a.6 |
| zgc:64065 | 0 | 0.785033131 | 0.396 | 0.008 | 0 | 7 | zgc:64065 |
| naga | 0 | 0.780498869 | 0.504 | 0.037 | 0 | 7 | naga |
| si:ch211-194m7.3 | 0 | 0.77878534 | 0.389 | 0.008 | 0 | 7 | si:ch211-194m7.3 |
| LOC564660 | 0 | 0.77221517 | 0.249 | 0.003 | 0 | 7 | LOC564660 |
| LOC100537504 | 0 | 0.763872415 | 0.327 | 0.009 | 0 | 7 | LOC100537504 |
| tpp1 | 0 | 0.763423011 | 0.514 | 0.058 | 0 | 7 | tpp1 |
| lta4h | 0 | 0.760190759 | 0.534 | 0.056 | 0 | 7 | lta4h |
| jac3 | 0 | 0.755757773 | 0.253 | 0.002 | 0 | 7 | jac3 |
| rnaset2l | 0 | 0.754955142 | 0.472 | 0.031 | 0 | 7 | rnaset2l |
| cxcr3.2 | 0 | 0.75397642 | 0.482 | 0.016 | 0 | 7 | cxcr3.2 |
| ctsh | 0 | 0.747547824 | 0.47 | 0.05 | 0 | 7 | ctsh |
| asah1b | 0 | 0.747427724 | 0.475 | 0.045 | 0 | 7 | asah1b |
| ccl34b.1 | 0 | 0.736336914 | 0.322 | 0.017 | 0 | 7 | ccl34b.1 |
| si:cabz01074946.1 | 0 | 0.732730748 | 0.443 | 0.045 | 0 | 7 | si:cabz01074946.1 |
| si:dkey-102g19.3 | 0 | 0.73199815 | 0.442 | 0.032 | 0 | 7 | si:dkey-102g19.3 |
| cndp2 | 0 | 0.720120556 | 0.489 | 0.055 | 0 | 7 | cndp2 |
| grn2 | 0 | 0.716195966 | 0.244 | 0.008 | 0 | 7 | grn2 |
| gng10 | 0 | 0.713939643 | 0.456 | 0.044 | 0 | 7 | gng10 |
| spi1b | 0 | 0.68566007 | 0.507 | 0.053 | 0 | 7 | spi1b |
| BX649485.1 | 0 | 0.68426961 | 0.357 | 0.012 | 0 | 7 | BX649485.1 |
| mafbb | 0 | 0.683741556 | 0.332 | 0.012 | 0 | 7 | mafbb |
| samsn1a | 0 | 0.674374429 | 0.426 | 0.022 | 0 | 7 | samsn1a |
| timp2b | 0 | 0.668155907 | 0.367 | 0.024 | 0 | 7 | timp2b |

| | | | | | | | |
|---|---|---|---|---|---|---|---|
| plekho1b | 0 | 0.657540298 | 0.422 | 0.027 | 0 | 7 | plekho1b |
| si:ch211-283g2.2 | 0 | 0.633388248 | 0.36 | 0.008 | 0 | 7 | si:ch211-283g2.2 |
| f13a1b | 0 | 0.631256841 | 0.367 | 0.018 | 0 | 7 | f13a1b |
| nrros | 0 | 0.627202796 | 0.406 | 0.033 | 0 | 7 | nrros |
| CABZ01001434.1 | 0 | 0.62468831 | 0.253 | 0.005 | 0 | 7 | CABZ01001434.1 |
| ccl39.1 | 0 | 0.618679738 | 0.198 | 0.005 | 0 | 7 | ccl39.1 |
| LO018493.1 | 0 | 0.61010483 | 0.371 | 0.012 | 0 | 7 | LO018493.1 |
| dhrs9 | 0 | 0.575418255 | 0.373 | 0.022 | 0 | 7 | dhrs9 |
| ccr12b.2 | 0 | 0.574537527 | 0.29 | 0.003 | 0 | 7 | ccr12b.2 |
| il10ra | 0 | 0.574236941 | 0.367 | 0.028 | 0 | 7 | il10ra |
| dnase2 | 0 | 0.568099775 | 0.367 | 0.029 | 0 | 7 | dnase2 |
| si:ch73-343l4.8 | 0 | 0.565813944 | 0.314 | 0.008 | 0 | 7 | si:ch73-343l4.8 |
| csf2rb | 0 | 0.561606321 | 0.346 | 0.026 | 0 | 7 | csf2rb |
| enpp1 | 0 | 0.550648999 | 0.35 | 0.026 | 0 | 7 | enpp1 |
| si:ch73-158p21.3 | 0 | 0.548961001 | 0.288 | 0.016 | 0 | 7 | si:ch73-158p21.3 |
| abca1b | 0 | 0.532088605 | 0.304 | 0.008 | 0 | 7 | abca1b |
| slc7a7 | 0 | 0.520095602 | 0.272 | 0.006 | 0 | 7 | slc7a7 |
| adgrg3 | 0 | 0.514036111 | 0.295 | 0.011 | 0 | 7 | adgrg3 |
| cybb | 0 | 0.474770922 | 0.292 | 0.009 | 0 | 7 | cybb |
| AL928650.1 | 0 | 0.463204295 | 0.205 | 0.004 | 0 | 7 | AL928650.1 |
| sb:cb81 | 0 | 0.46062287 | 0.297 | 0.018 | 0 | 7 | sb:cb81 |
| zgc:136870 | 0 | 0.439735215 | 0.274 | 0.008 | 0 | 7 | zgc:136870 |
| jac9 | 0 | 0.437875382 | 0.171 | 0.001 | 0 | 7 | jac9 |
| calhm6 | 0 | 0.420515835 | 0.26 | 0.008 | 0 | 7 | calhm6 |
| moxd1 | 0 | 0.417337083 | 0.235 | 0.005 | 0 | 7 | moxd1 |
| si:ch211-283g2.3 | 0 | 0.412368758 | 0.161 | 0.002 | 0 | 7 | si:ch211-283g2.3 |
| zmp:0000000936 | 0 | 0.399606627 | 0.24 | 0.008 | 0 | 7 | zmp:0000000936 |
| si:dkey-56f14.7 | 0 | 0.381935204 | 0.235 | 0.005 | 0 | 7 | si:dkey-56f14.7 |
| mapk11 | 0 | 0.36655892 | 0.239 | 0.01 | 0 | 7 | mapk11 |
| slc16a9a | 0 | 0.346350327 | 0.187 | 0.003 | 0 | 7 | slc16a9a |
| si:zfos-1069f5.1 | 0 | 0.343064078 | 0.196 | 0.004 | 0 | 7 | si:zfos-1069f5.1 |
| cd22 | 0 | 0.342572394 | 0.201 | 0.002 | 0 | 7 | cd22 |
| gpr84 | 0 | 0.323583778 | 0.182 | 0.003 | 0 | 7 | gpr84 |
| si:ch73-343l4.2 | 0 | 0.319969031 | 0.2 | 0.002 | 0 | 7 | si:ch73-343l4.2 |
| havcr2 | 0 | 0.284191377 | 0.159 | 0.002 | 0 | 7 | havcr2 |

| | | | | | | | |
|---|---|---|---|---|---|---|---|
| slc2a6 | 0 | 0.275794448 | 0.152 | 0.001 | 0 | 7 | slc2a6 |
| m17 | 3.739326 | 0.403902575 | 0.207 | 0.007 | 9.4997591 | 7 | m17 |
| cxcr4b1 | 4.339153 | 1.079753666 | 0.781 | 0.152 | 1.1023620 | 7 | cxcr4b |
| plin2 | 7.505974 | 0.536429898 | 0.29 | 0.017 | 1.9068928 | 7 | plin2 |
| mhc2b | 1.527415 | 0.762351732 | 0.346 | 0.028 | 3.8803980 | 7 | mhc2b |
| si:zfos-2330d3.3 | 1.894938 | 0.429706358 | 0.173 | 0.004 | 4.8140908 | 7 | si:zfos-2330d3.3 |
| atp6v1e1b | 2.639404 | 0.724273321 | 0.542 | 0.072 | 6.7054081 | 7 | atp6v1e1b |
| CU929150.1 | 4.343792 | 0.668485129 | 0.281 | 0.017 | 1.1035405 | 7 | CU929150.1 |
| tcnbb | 1.149075 | 0.509959348 | 0.235 | 0.01 | 2.9192264 | 7 | tcnbb |
| dpp7 | 6.897848 | 0.633851647 | 0.426 | 0.045 | 1.7523984 | 7 | dpp7 |
| cyba | 2.842761 | 0.711114744 | 0.507 | 0.064 | 7.2220368 | 7 | cyba |
| si:ch211-264f5.2 | 5.352661 | 0.330812201 | 0.196 | 0.006 | 1.3598435 | 7 | si:ch211-264f5.2 |
| glb1l | 6.933129 | 0.469725494 | 0.279 | 0.017 | 1.7613615 | 7 | glb1l |
| si:dkey-262k9.41 | 1.431454 | 0.686266187 | 0.537 | 0.071 | 3.6366109 | 7 | si:dkey-262k9.4 |
| fosl2 | 5.270981 | 0.840975661 | 0.592 | 0.093 | 1.3390927 | 7 | fosl2 |
| laptm51 | 1.388802 | 0.943459727 | 0.776 | 0.151 | 3.5282517 | 7 | laptm5 |
| BX324132.2 | 1.375046 | 0.27012562 | 0.147 | 0.002 | 3.4933057 | 7 | BX324132.2 |
| FP015862.2 | 5.316661 | 0.331828019 | 0.189 | 0.006 | 1.3506978 | 7 | FP015862.2 |
| psap | 6.949207 | 1.10028321 | 0.905 | 0.262 | 1.7654461 | 7 | psap |
| si:busm1-194e12.11 | 1.902584 | 0.532920527 | 0.269 | 0.017 | 4.8335167 | 7 | si:busm1-194e12.11 |
| ncf2 | 1.625582 | 0.393787015 | 0.247 | 0.014 | 4.1297919 | 7 | ncf2 |
| cst14b.1 | 7.669670 | 0.589685411 | 0.415 | 0.047 | 1.9484798 | 7 | cst14b.1 |
| ctsa | 3.222360 | 0.704787134 | 0.532 | 0.078 | 8.1864068 | 7 | ctsa |
| fuca2 | 4.222862 | 0.629048314 | 0.438 | 0.053 | 1.0728181 | 7 | fuca2 |
| si:ch211-152p11.4 | 1.172957 | 0.419543076 | 0.27 | 0.018 | 2.9798979 | 7 | si:ch211-152p11.4 |
| itgb2 | 2.115289 | 0.757896617 | 0.569 | 0.089 | 5.3738920 | 7 | itgb2 |
| vsir1 | 3.461010 | 0.702883266 | 0.504 | 0.071 | 8.7926960 | 7 | vsir |
| ccl39.3 | 4.014764 | 0.449658231 | 0.147 | 0.003 | 1.0199510 | 7 | ccl39.3 |
| ms4a17a.91 | 3.778341 | 0.65083915 | 0.429 | 0.054 | 9.5988776 | 7 | ms4a17a.9 |
| il4r.1 | 3.689242 | 0.583140979 | 0.346 | 0.035 | 9.3725213 | 7 | il4r.1 |
| gnpda1 | 7.110525 | 0.501603731 | 0.348 | 0.033 | 1.8064290 | 7 | gnpda1 |
| lect2l | 6.087006 | 0.585368102 | 0.307 | 0.026 | 1.5464039 | 7 | lect2l |
| ehd4 | 3.078070 | 0.460229262 | 0.295 | 0.024 | 7.8198387 | 7 | ehd4 |
| hdac9b | 1.740188 | 0.394661614 | 0.256 | 0.017 | 4.4209493 | 7 | hdac9b |
| gnsb | 6.184814 | 0.499305285 | 0.313 | 0.028 | 1.5712520 | 7 | gnsb |

| | | | | | | | |
|---|---|---|---|---|---|---|---|
| nr4a11 | 1.982289 | 0.907755383 | 0.666 | 0.137 | 5.0360067 | 7 | nr4a1 |
| gm2a | 1.318706 | 0.740594531 | 0.592 | 0.101 | 3.3501736 | 7 | gm2a |
| cebpb2 | 3.596555 | 1.10847314 | 0.894 | 0.316 | 9.1370496 | 7 | cebpb |
| atp6v0ca | 3.784731 | 0.914687777 | 0.751 | 0.181 | 9.6151106 | 7 | atp6v0ca |
| si:ch211-39i2.2 | 5.755101 | 0.586031697 | 0.399 | 0.047 | 1.4620834 | 7 | si:ch211-39i2.2 |
| ltb4r | 2.251267 | 0.383759094 | 0.256 | 0.018 | 5.7193446 | 7 | ltb4r |
| ifi30 | 3.559283 | 0.993255071 | 0.716 | 0.174 | 9.0423591 | 7 | ifi30 |
| ets2 | 2.624277 | 0.620632826 | 0.481 | 0.071 | 6.6669757 | 7 | ets2 |
| stoml3b | 8.304038 | 0.348075401 | 0.18 | 0.008 | 2.1096409 | 7 | stoml3b |
| si:ch211-1o7.3 | 3.674577 | 0.3546085 | 0.221 | 0.013 | 9.3352642 | 7 | si:ch211-1o7.3 |
| cdaa | 5.369859 | 0.436323846 | 0.283 | 0.024 | 1.3642127 | 7 | cdaa |
| zgc:640511 | 1.135907 | 0.813383089 | 0.686 | 0.141 | 2.8857730 | 7 | zgc:64051 |
| atp6v1g1 | 2.187097 | 0.749275172 | 0.627 | 0.122 | 5.5563217 | 7 | atp6v1g1 |
| crema1 | 2.459908 | 0.839041059 | 0.684 | 0.153 | 6.2493985 | 7 | crema |
| FP102158.1 | 3.317161 | 0.302849494 | 0.198 | 0.011 | 8.4272489 | 7 | FP102158.1 |
| si:dkey-11f4.141 | 5.201089 | 0.810012077 | 0.551 | 0.105 | 1.3213366 | 7 | si:dkey-11f4.14 |
| si:ch211-212k18.7 | 8.131346 | 0.788953758 | 0.624 | 0.126 | 2.0657685 | 7 | si:ch211-212k18.7 |
| ctsd | 2.448056 | 0.85222854 | 0.624 | 0.128 | 6.2192879 | 7 | ctsd |
| parvg | 1.494375 | 0.593500963 | 0.426 | 0.059 | 3.7964608 | 7 | parvg |
| si:dkeyp-23e4.3 | 5.218247 | 0.296391706 | 0.173 | 0.008 | 1.3256957 | 7 | si:dkeyp-23e4.3 |
| tmsb4x3 | 6.503407 | 0.726093475 | 1 | 0.707 | 1.6521906 | 7 | tmsb4x |
| XLOC-0234921 | 6.775025 | 0.722813776 | 0.484 | 0.077 | 1.7211953 | 7 | XLOC-023492 |
| tspan37 | 4.805830 | 0.399415178 | 0.265 | 0.022 | 1.2209212 | 7 | tspan37 |
| plbd1 | 6.416194 | 0.358125393 | 0.226 | 0.015 | 1.6300341 | 7 | plbd1 |
| hlx1 | 4.824148 | 0.326290262 | 0.198 | 0.011 | 1.2255748 | 7 | hlx1 |
| slc40a1 | 8.018209 | 0.549808707 | 0.302 | 0.031 | 2.0370260 | 7 | slc40a1 |
| tgfbi | 8.070886 | 0.419728174 | 0.265 | 0.023 | 2.0504087 | 7 | tgfbi |
| cd633 | 2.881581 | 0.96530487 | 0.905 | 0.352 | 7.3206584 | 7 | cd63 |
| cebpa1 | 2.348745 | 0.644053457 | 0.422 | 0.062 | 5.9669876 | 7 | cebpa |
| rab44 | 1.411118 | 0.524745105 | 0.378 | 0.048 | 3.5849457 | 7 | rab44 |
| snx10b | 4.193982 | 0.270682599 | 0.154 | 0.006 | 1.0654813 | 7 | snx10b |
| XLOC-024516 | 2.737496 | 0.257692196 | 0.157 | 0.007 | 6.9546105 | 7 | XLOC-024516 |
| si:ch73-248e21.5 | 4.213625 | 0.461433649 | 0.316 | 0.034 | 1.0704716 | 7 | si:ch73-248e21.5 |
| pepd | 7.723365 | 0.534278635 | 0.422 | 0.062 | 1.9621210 | 7 | pepd |
| atp6v1ba | 1.464677 | 0.508514178 | 0.355 | 0.044 | 3.7210123 | 7 | atp6v1ba |

| | | | | | | | |
|---|---|---|---|---|---|---|---|
| XLOC-0054491 | 2.052418 | 0.588239339 | 0.357 | 0.045 | 5.2141682 | 7 | XLOC-005449 |
| cotl11 | 2.281831 | 0.9608293 | 0.901 | 0.296 | 5.7969934 | 7 | cotl1 |
| zgc:63831 | 3.457821 | 0.568726776 | 0.452 | 0.07 | 8.7845951 | 7 | zgc:63831 |
| si:ch211-15b10.61 | 1.425676 | 0.560049286 | 0.419 | 0.062 | 3.6219308 | 7 | si:ch211-15b10.6 |
| stat3 | 1.150913 | 0.498959266 | 0.352 | 0.044 | 2.9238948 | 7 | stat3 |
| actb21 | 1.864271 | 0.560261314 | 1 | 0.862 | 4.7361825 | 7 | actb2 |
| ncf4 | 4.869126 | 0.297798185 | 0.196 | 0.012 | 1.2370016 | 7 | ncf4 |
| pnp5a | 1.073718 | 0.626213352 | 0.41 | 0.061 | 2.7277813 | 7 | pnp5a |
| si:ch211-193e13.5 | 5.539357 | 0.256232321 | 0.159 | 0.007 | 1.4072737 | 7 | si:ch211-193e13.5 |
| ccng2 | 2.171455 | 0.611348197 | 0.436 | 0.07 | 5.5165817 | 7 | ccng2 |
| aplp2 | 1.216695 | 0.568695963 | 0.417 | 0.063 | 3.0910155 | 7 | aplp2 |
| si:dkey-27i16.21 | 1.503675 | 0.704246586 | 0.456 | 0.076 | 3.8200870 | 7 | si:dkey-27i16.2 |
| slc43a2a2 | 2.852267 | 0.667595022 | 0.484 | 0.086 | 7.2461852 | 7 | slc43a2a |
| si:ch211-204c21.11 | 6.669673 | 0.617549274 | 0.405 | 0.061 | 1.6944305 | 7 | si:ch211-204c21.1 |
| nfil3-62 | 1.881609 | 0.745347869 | 0.555 | 0.114 | 4.7802292 | 7 | nfil3-6 |
| bach1b | 6.834653 | 0.36725666 | 0.231 | 0.019 | 1.7363437 | 7 | bach1b |
| irf81 | 4.717385 | 0.470067217 | 0.32 | 0.037 | 1.1984517 | 7 | irf8 |
| si:ch211-102c2.41 | 9.016749 | 0.723660034 | 0.613 | 0.127 | 2.2907051 | 7 | si:ch211-102c2.4 |
| atp1a3a | 1.662331 | 0.301379696 | 0.21 | 0.015 | 4.2231537 | 7 | atp1a3a |
| arl5c1 | 2.218408 | 0.654165312 | 0.498 | 0.092 | 5.6358662 | 7 | arl5c |
| itgae.2 | 1.668686 | 0.426437223 | 0.27 | 0.027 | 4.2392976 | 7 | itgae.2 |
| si:ch211-66e2.3 | 2.043522 | 0.524399781 | 0.367 | 0.05 | 5.1915690 | 7 | si:ch211-66e2.3 |
| acsl4b | 1.953463 | 0.464566977 | 0.334 | 0.042 | 4.9627727 | 7 | acsl4b |
| il1b | 3.844572 | 0.353017894 | 0.207 | 0.015 | 9.7671358 | 7 | il1b |
| aatka | 3.852260 | 0.266603247 | 0.143 | 0.006 | 9.7866687 | 7 | aatka |
| cxcl8b.11 | 4.106132 | 0.76292028 | 0.306 | 0.037 | 1.0431629 | 7 | cxcl8b.1 |
| LOC100148254 | 1.314216 | 0.348587331 | 0.217 | 0.017 | 3.3387680 | 7 | LOC100148254 |
| CABZ01076275.1 | 9.714492 | 0.386050585 | 0.193 | 0.013 | 2.4679668 | 7 | CABZ01076275.1 |
| tuba8l2 | 1.109991 | 0.500692347 | 0.38 | 0.056 | 2.8199335 | 7 | tuba8l2 |
| prdx51 | 3.018507 | 0.768692386 | 0.625 | 0.16 | 7.6685186 | 7 | prdx5 |
| zgc:1627303 | 1.010031 | 0.947260774 | 0.926 | 0.381 | 2.5659841 | 7 | zgc:162730 |
| itm2bb | 1.816667 | 0.784323714 | 0.776 | 0.22 | 4.6152435 | 7 | itm2bb |
| lyn | 6.345572 | 0.418557548 | 0.299 | 0.035 | 1.6120925 | 7 | lyn |
| LOC101885706 | 9.170139 | 0.267000303 | 0.148 | 0.007 | 2.3296738 | 7 | LOC101885706 |
| npl | 5.445326 | 0.435315822 | 0.265 | 0.029 | 1.3833850 | 7 | npl |

| | | | | | | | |
|---|---|---|---|---|---|---|---|
| rab8b | 1.262758 | 0.548067311 | 0.415 | 0.071 | 3.2080371 | 7 | rab8b |
| rasgrp4 | 2.906997 | 0.335107429 | 0.224 | 0.02 | 7.3852262 | 7 | rasgrp4 |
| BX901920.11 | 1.138562 | 0.597892014 | 0.382 | 0.061 | 2.8925176 | 7 | BX901920.1 |
| si:zfos-741a10.3 | 1.342756 | 0.353095468 | 0.235 | 0.023 | 3.4112725 | 7 | si:zfos-741a10.3 |
| tcirg1b | 1.135037 | 0.405592071 | 0.309 | 0.04 | 2.8835617 | 7 | tcirg1b |
| pfn11 | 4.796233 | 0.937213911 | 0.998 | 0.481 | 1.2184830 | 7 | pfn1 |
| coro1a1 | 2.010715 | 0.775311732 | 0.797 | 0.213 | 5.1082239 | 7 | coro1a |
| atp7a | 8.963352 | 0.266941246 | 0.168 | 0.011 | 2.2771396 | 7 | atp7a |
| myo1f | 9.101322 | 0.380062098 | 0.297 | 0.038 | 2.3121909 | 7 | myo1f |
| wasb1 | 1.352956 | 0.652381484 | 0.647 | 0.153 | 3.4371859 | 7 | wasb |
| cap11 | 3.945584 | 0.7550315 | 0.754 | 0.225 | 1.0023757 | 7 | cap1 |
| arpc5b | 5.712938 | 0.588401736 | 0.532 | 0.115 | 1.4513720 | 7 | arpc5b |
| zgc:153317 | 1.823787 | 0.403812837 | 0.263 | 0.031 | 4.6333311 | 7 | zgc:153317 |
| LOC101885065 | 7.420579 | 0.26236843 | 0.157 | 0.01 | 1.8851981 | 7 | LOC101885065 |
| sowahd | 2.918842 | 0.258229665 | 0.157 | 0.01 | 7.4153185 | 7 | sowahd |
| rac21 | 2.885062 | 0.743097208 | 0.746 | 0.203 | 7.3295016 | 7 | rac2 |
| XLOC-005448 | 1.574195 | 0.414924387 | 0.214 | 0.021 | 3.9992446 | 7 | XLOC-005448 |
| rgs2 | 3.228897 | 0.409491965 | 0.221 | 0.022 | 8.2030153 | 7 | rgs2 |
| si:ch1073-392o20.2 | 3.656619 | 0.432001893 | 0.316 | 0.046 | 9.2896414 | 7 | si:ch1073-392o20.2 |
| plxnc1 | 2.047172 | 0.4001985 | 0.284 | 0.037 | 5.2008417 | 7 | plxnc1 |
| ctsc | 6.659496 | 0.527510478 | 0.357 | 0.058 | 1.6918451 | 7 | ctsc |
| trpv1 | 2.550689 | 0.253385339 | 0.15 | 0.009 | 6.4800277 | 7 | trpv1 |
| abi2b | 2.696304 | 0.468159898 | 0.307 | 0.044 | 6.8499622 | 7 | abi2b |
| XLOC-0004312 | 2.574542 | 0.775061116 | 0.822 | 0.25 | 6.5406250 | 7 | XLOC-000431 |
| rassf2a | 2.020167 | 0.308736413 | 0.2 | 0.018 | 5.1322343 | 7 | rassf2a |
| rgs19 | 3.062127 | 0.348120294 | 0.24 | 0.027 | 7.7793345 | 7 | rgs19 |
| zgc:123107 | 3.155833 | 0.256213209 | 0.141 | 0.008 | 8.0173943 | 7 | zgc:123107 |
| ccr9a1 | 3.181972 | 0.580975639 | 0.405 | 0.075 | 8.0838010 | 7 | ccr9a |
| nfkbiaa | 3.811501 | 0.719737243 | 0.546 | 0.136 | 9.6831185 | 7 | nfkbiaa |
| dusp11 | 4.404284 | 0.711117452 | 0.537 | 0.133 | 1.1189085 | 7 | dusp1 |
| unc93b1 | 8.878631 | 0.295663638 | 0.189 | 0.017 | 2.2556163 | 7 | unc93b1 |
| BX323596.2 | 4.817453 | 0.326103749 | 0.125 | 0.006 | 1.2238740 | 7 | BX323596.2 |
| gsdmea | 4.952019 | 0.331527153 | 0.189 | 0.017 | 1.2580604 | 7 | gsdmea |
| BX248082.11 | 6.481606 | 0.595567156 | 0.239 | 0.028 | 1.6466521 | 7 | BX248082.1 |
| il4r.2 | 7.264366 | 0.28636257 | 0.141 | 0.009 | 1.8455123 | 7 | il4r.2 |

| | | | | | | | |
|---|---|---|---|---|---|---|---|
| prcp | 1.174240 | 0.258129549 | 0.168 | 0.013 | 2.9831577 | 7 | prcp |
| entpd11 | 1.861756 | 0.527471868 | 0.383 | 0.07 | 4.7297924 | 7 | entpd1 |
| atp6v1d | 4.254233 | 0.311276917 | 0.214 | 0.022 | 1.0807880 | 7 | atp6v1d |
| ifngr1l | 6.733907 | 0.313323356 | 0.221 | 0.024 | 1.7107493 | 7 | ifngr1l |
| pdk2b | 4.649387 | 0.606320228 | 0.435 | 0.092 | 1.1811768 | 7 | pdk2b |
| trib3 | 1.884656 | 0.568759994 | 0.445 | 0.094 | 4.7879702 | 7 | trib3 |
| arpc1b1 | 5.075839 | 0.815747811 | 0.885 | 0.29 | 1.2895170 | 7 | arpc1b |
| CR392341.11 | 5.715251 | 0.461652273 | 0.316 | 0.049 | 1.4519597 | 7 | CR392341.1 |
| rhogb | 5.866649 | 0.510889826 | 0.454 | 0.094 | 1.4904224 | 7 | rhogb |
| gpr183a1 | 1.023449 | 0.420915467 | 0.336 | 0.054 | 2.6000727 | 7 | gpr183a |
| nocta | 1.722626 | 0.378828027 | 0.258 | 0.033 | 4.3763315 | 7 | nocta |
| grapa | 4.145731 | 0.324686873 | 0.205 | 0.021 | 1.0532229 | 7 | grapa |
| crip12 | 6.526779 | 0.76409617 | 0.797 | 0.263 | 1.6581283 | 7 | crip1 |
| ptgis | 4.230279 | 0.257259145 | 0.148 | 0.01 | 1.0747024 | 7 | ptgis |
| CABZ01073834.1 | 5.807500 | 0.366507078 | 0.267 | 0.035 | 1.4753954 | 7 | CABZ01073834.1 |
| si:dkey-206d17.12 | 1.829812 | 0.397519308 | 0.286 | 0.041 | 4.6486381 | 7 | si:dkey-206d17.12 |
| gapdhs2 | 2.113565 | 0.743901631 | 0.797 | 0.295 | 5.3695138 | 7 | gapdhs |
| zgc:101715 | 4.060765 | 0.266857419 | 0.193 | 0.019 | 1.0316375 | 7 | zgc:101715 |
| LOC101884267 | 7.082668 | 0.385962474 | 0.29 | 0.042 | 1.7993519 | 7 | LOC101884267 |
| grap2b1 | 9.169167 | 0.529489441 | 0.495 | 0.109 | 2.3294271 | 7 | grap2b |
| pgd | 2.072614 | 0.443104112 | 0.345 | 0.06 | 5.2654767 | 7 | pgd |
| lpar5a1 | 2.814765 | 0.372913847 | 0.265 | 0.036 | 7.1509119 | 7 | lpar5a |
| mpp1 | 5.958761 | 0.340062569 | 0.217 | 0.025 | 1.5138232 | 7 | mpp1 |
| wipf1b1 | 1.624664 | 0.41795836 | 0.325 | 0.054 | 4.1274598 | 7 | wipf1b |
| FERMT3 (1 of many)1 | 1.776378 | 0.369269454 | 0.263 | 0.036 | 4.5128894 | 7 | FERMT3 (1 of many) |
| zmp:0000001082 | 3.572231 | 0.381770609 | 0.299 | 0.046 | 9.0752539 | 7 | zmp:0000001082 |
| hexb | 5.551790 | 0.349641295 | 0.254 | 0.035 | 1.4104323 | 7 | hexb |
| nccrp1 | 7.183361 | 0.674605517 | 0.622 | 0.179 | 1.8249330 | 7 | nccrp1 |
| gnao1a | 1.870675 | 0.290307931 | 0.186 | 0.018 | 4.7524502 | 7 | gnao1a |
| nfe2l2a | 7.708059 | 0.445944238 | 0.33 | 0.058 | 1.9582325 | 7 | nfe2l2a |
| entpd2a.1 | 1.338263 | 0.333604768 | 0.223 | 0.026 | 3.3998590 | 7 | entpd2a.1 |
| arhgef5 | 2.355615 | 0.323508336 | 0.216 | 0.025 | 5.9844423 | 7 | arhgef5 |
| tfe3a | 4.951042 | 0.341224108 | 0.246 | 0.033 | 1.2578122 | 7 | tfe3a |
| csrnp1b1 | 7.329335 | 0.625957315 | 0.553 | 0.149 | 1.8620175 | 7 | csrnp1b |
| si:dkey-25e12.3 | 1.047022 | 0.398951667 | 0.27 | 0.04 | 2.6599600 | 7 | si:dkey-25e12.3 |

| | | | | | | | |
|---|---|---|---|---|---|---|---|
| klf131 | 2.090318 | 0.509634802 | 0.422 | 0.091 | 5.3104544 | 7 | klf13 |
| snx27a | 2.587556 | 0.346988166 | 0.246 | 0.033 | 6.5736874 | 7 | snx27a |
| pycard1 | 6.602042 | 0.466196651 | 0.375 | 0.072 | 1.6772488 | 7 | pycard |
| junba2 | 1.860715 | 0.780622515 | 0.832 | 0.299 | 4.7271465 | 7 | junba |
| abcc3 | 1.851018 | 0.264150482 | 0.182 | 0.018 | 4.7025114 | 7 | abcc3 |
| si:ch73-237c6.11 | 2.794929 | 0.353509773 | 0.261 | 0.037 | 7.1005181 | 7 | si:ch73-237c6.1 |
| tram1 | 4.258320 | 0.470722309 | 0.433 | 0.093 | 1.0818263 | 7 | tram1 |
| anxa5b1 | 2.001013 | 0.508491955 | 0.422 | 0.092 | 5.0835741 | 7 | anxa5b |
| fermt3b | 2.293104 | 0.251190857 | 0.17 | 0.016 | 5.8256314 | 7 | fermt3b |
| dbn11 | 6.274655 | 0.400698641 | 0.32 | 0.056 | 1.5940762 | 7 | dbn1 |
| derl2 | 1.568201 | 0.48793743 | 0.431 | 0.094 | 3.9840155 | 7 | derl2 |
| ctsz | 2.023119 | 0.559860529 | 0.387 | 0.081 | 5.1397338 | 7 | ctsz |
| si:ch1073-429i10.3.1 | 4.451857 | 0.623301555 | 0.67 | 0.208 | 1.1309943 | 7 | si:ch1073-429i10.3.1 |
| tnfrsf1b | 6.310628 | 0.259435391 | 0.159 | 0.014 | 1.6032152 | 7 | tnfrsf1b |
| bri3 | 2.157599 | 0.386475146 | 0.3 | 0.05 | 5.4813804 | 7 | bri3 |
| srgn1 | 3.741714 | 0.672756733 | 0.696 | 0.216 | 9.5058246 | 7 | srgn |
| bhlhe401 | 6.929481 | 0.62697313 | 0.548 | 0.152 | 1.7604348 | 7 | bhlhe40 |
| git2b | 1.509962 | 0.286354128 | 0.189 | 0.021 | 3.8360600 | 7 | git2b |
| dennd4a | 2.102689 | 0.386439105 | 0.228 | 0.031 | 5.3418836 | 7 | dennd4a |
| mef2cb | 2.163030 | 0.325886928 | 0.2 | 0.023 | 5.4951784 | 7 | mef2cb |
| slc30a1a | 6.319218 | 0.283790014 | 0.182 | 0.019 | 1.6053974 | 7 | slc30a1a |
| ctsk | 1.371572 | 0.517530545 | 0.332 | 0.064 | 3.4844796 | 7 | ctsk |
| ubl3b | 1.830188 | 0.366563574 | 0.325 | 0.058 | 4.6495938 | 7 | ubl3b |
| ugt5b2 | 7.074968 | 0.282075797 | 0.208 | 0.026 | 1.7973957 | 7 | ugt5b2 |
| arpc21 | 7.144606 | 0.66823578 | 0.765 | 0.28 | 1.8150872 | 7 | arpc2 |
| atp6ap2 | 1.060224 | 0.473685641 | 0.415 | 0.094 | 2.6935008 | 7 | atp6ap2 |
| elovl1b | 2.523135 | 0.483430656 | 0.456 | 0.109 | 6.4100265 | 7 | elovl1b |
| dnase1l1 | 3.568254 | 0.373845097 | 0.272 | 0.044 | 9.0651507 | 7 | dnase1l1 |
| si:dkey-53k12.21 | 1.002945 | 0.34198138 | 0.247 | 0.036 | 2.5479837 | 7 | si:dkey-53k12.2 |
| lxn2 | 1.096167 | 0.578186447 | 0.548 | 0.152 | 2.7848129 | 7 | lxn |
| mfng1 | 2.344597 | 0.352752461 | 0.249 | 0.037 | 5.9564497 | 7 | mfng |
| stx11a | 1.333878 | 0.264328786 | 0.191 | 0.022 | 3.3887191 | 7 | stx11a |
| mapre1a | 2.779873 | 0.556747333 | 0.62 | 0.179 | 7.0622687 | 7 | mapre1a |
| rnaseka | 2.971785 | 0.514503192 | 0.465 | 0.116 | 7.5498212 | 7 | rnaseka |
| rnf144b | 3.707854 | 0.284330157 | 0.198 | 0.024 | 9.4198043 | 7 | rnf144b |

| | | | | | | | |
|---|---|---|---|---|---|---|---|
| ppiaa2 | 3.311985 | 0.37785144 | 0.986 | 0.808 | 8.4140981 | 7 | ppiaa |
| CABZ01068358.11 | 1.792965 | 0.432585183 | 0.307 | 0.058 | 4.5550292 | 7 | CABZ01068358.1 |
| si:dkey-112e7.21 | 2.498683 | 0.593225851 | 0.617 | 0.198 | 6.3479051 | 7 | si:dkey-112e7.2 |
| sec61a1 | 9.399158 | 0.573001178 | 0.574 | 0.177 | 2.3878562 | 7 | sec61a1 |
| rps15a2 | 2.060614 | 0.426804519 | 0.988 | 0.791 | 5.2349904 | 7 | rps15a |
| gadd45bb1 | 2.453421 | 0.452012154 | 0.346 | 0.072 | 6.2329181 | 7 | gadd45bb |
| mfsd1 | 4.130580 | 0.277937723 | 0.178 | 0.02 | 1.0493739 | 7 | mfsd1 |
| cpox1 | 9.704677 | 0.348174038 | 0.274 | 0.046 | 2.4654734 | 7 | cpox |
| echdc2 | 1.111286 | 0.314345986 | 0.224 | 0.032 | 2.8232222 | 7 | echdc2 |
| mboat2a | 1.768066 | 0.260885152 | 0.178 | 0.02 | 4.4917734 | 7 | mboat2a |
| hmox1a | 4.476278 | 0.271099926 | 0.124 | 0.009 | 1.1371985 | 7 | hmox1a |
| si:ch73-248e21.7 | 7.304489 | 0.345931088 | 0.258 | 0.042 | 1.8557055 | 7 | si:ch73-248e21.7 |
| me2 | 2.131820 | 0.278579271 | 0.18 | 0.021 | 5.4158898 | 7 | me2 |
| hcls1 | 2.690751 | 0.378489965 | 0.336 | 0.066 | 6.8358544 | 7 | hcls1 |
| atp6v1aa | 5.230920 | 0.311933989 | 0.224 | 0.032 | 1.3289153 | 7 | atp6v1aa |
| arpc4l | 2.493155 | 0.589823804 | 0.618 | 0.203 | 6.3338623 | 7 | arpc4l |
| mbnl3 | 9.495366 | 0.410833285 | 0.288 | 0.053 | 2.4122978 | 7 | mbnl3 |
| rap1b | 1.280313 | 0.583234284 | 0.624 | 0.204 | 3.2526364 | 7 | rap1b |
| tnfb1 | 8.071704 | 0.438371884 | 0.253 | 0.042 | 2.0506165 | 7 | tnfb |
| pcyox1 | 1.130120 | 0.386600096 | 0.281 | 0.051 | 2.8710703 | 7 | pcyox1 |
| sdr16c5b | 1.363215 | 0.338261462 | 0.246 | 0.04 | 3.4632484 | 7 | sdr16c5b |
| sdcbp21 | 2.537682 | 0.59916077 | 0.627 | 0.212 | 6.4469814 | 7 | sdcbp2 |
| lcp11 | 2.689561 | 0.421372333 | 0.348 | 0.074 | 6.8328312 | 7 | lcp1 |
| lgals9l1 | 4.545862 | 0.356070988 | 0.182 | 0.022 | 1.1548763 | 7 | lgals9l1 |
| ckbb2 | 2.499490 | 0.639415826 | 0.701 | 0.233 | 6.3499549 | 7 | ckbb |
| pdia31 | 4.165178 | 0.653856906 | 0.74 | 0.296 | 1.0581636 | 7 | pdia3 |
| ssr3 | 1.530049 | 0.645273612 | 0.707 | 0.271 | 3.8870918 | 7 | ssr3 |
| kdm7aa | 1.552668 | 0.282606077 | 0.187 | 0.024 | 3.9445533 | 7 | kdm7aa |
| tagln22 | 1.911579 | 0.596096649 | 0.599 | 0.194 | 4.8563682 | 7 | tagln2 |
| CR936442.11 | 8.657439 | 0.472184908 | 0.521 | 0.139 | 2.1994224 | 7 | CR936442.1 |
| tec | 2.235626 | 0.269545886 | 0.201 | 0.028 | 5.6796100 | 7 | tec |
| btg12 | 3.225425 | 0.657343676 | 0.797 | 0.338 | 8.1941943 | 7 | btg1 |
| tpi1b1 | 3.505538 | 0.507635074 | 0.544 | 0.164 | 8.9058214 | 7 | tpi1b |
| tspan361 | 3.907711 | 0.582463131 | 0.498 | 0.143 | 9.9275416 | 7 | tspan36 |
| slc20a1b | 9.202204 | 0.430213649 | 0.367 | 0.084 | 2.3378201 | 7 | slc20a1b |

| | | | | | | | |
|---|---|---|---|---|---|---|---|
| wdr1 | 2.176412 | 0.492604685 | 0.5 | 0.142 | 5.5291751 | 7 | wdr1 |
| anxa41 | 3.047369 | 0.491868207 | 0.482 | 0.137 | 7.7418411 | 7 | anxa4 |
| pkma | 1.911985 | 0.510089832 | 0.546 | 0.164 | 4.8573980 | 7 | pkma |
| atp2b21 | 1.057940 | 0.392371863 | 0.336 | 0.073 | 2.6876971 | 7 | atp2b2 |
| rplp22 | 4.478067 | 0.425319081 | 0.989 | 0.775 | 1.1376530 | 7 | rplp2 |
| hmha1b1 | 4.936114 | 0.364656822 | 0.314 | 0.064 | 1.2540199 | 7 | hmha1b |
| XLOC-0254231 | 1.027646 | 0.688896956 | 0.511 | 0.157 | 2.6107370 | 7 | XLOC-025423 |
| LO018340.1 | 1.060374 | 0.259200112 | 0.18 | 0.023 | 2.6938816 | 7 | LO018340.1 |
| lipf | 1.808619 | 0.337299418 | 0.239 | 0.04 | 4.5947968 | 7 | lipf |
| cfl11 | 2.824829 | 0.586408935 | 0.965 | 0.586 | 7.1764787 | 7 | cfl1 |
| ppt1 | 3.667941 | 0.521256422 | 0.541 | 0.167 | 9.3184061 | 7 | ppt1 |
| metrnla | 5.352238 | 0.41475757 | 0.364 | 0.084 | 1.3597362 | 7 | metrnla |
| clic11 | 6.106295 | 0.461720423 | 0.436 | 0.117 | 1.5513043 | 7 | clic1 |
| s100a10b2 | 7.659077 | 0.752783703 | 0.68 | 0.27 | 1.9457887 | 7 | s100a10b |
| antxr2a2 | 1.703024 | 0.590537531 | 0.504 | 0.156 | 4.3265338 | 7 | antxr2a |
| nme2b.11 | 3.395825 | 0.310764929 | 0.993 | 0.851 | 8.6270957 | 7 | nme2b.1 |
| xbp1 | 1.474715 | 0.665968165 | 0.761 | 0.339 | 3.7465146 | 7 | xbp1 |
| anxa3b | 3.854134 | 0.33250262 | 0.18 | 0.024 | 9.7914295 | 7 | anxa3b |
| icn1 | 1.470334 | 0.770700475 | 0.654 | 0.253 | 3.7353844 | 7 | icn |
| zbtb16a | 2.459343 | 0.29666121 | 0.224 | 0.036 | 6.2479619 | 7 | zbtb16a |
| egr12 | 3.551121 | 0.569228672 | 0.537 | 0.169 | 9.0216235 | 7 | egr1 |
| sec61g | 3.986508 | 0.482800853 | 0.949 | 0.671 | 1.0127724 | 7 | sec61g |
| dram2b | 1.346094 | 0.261685965 | 0.189 | 0.027 | 3.4197543 | 7 | dram2b |
| slc7a8a | 3.935233 | 0.255624718 | 0.182 | 0.025 | 9.9974613 | 7 | slc7a8a |
| txnipa1 | 4.087935 | 0.626773709 | 0.615 | 0.23 | 1.0385400 | 7 | txnipa |
| ddit32 | 6.847765 | 0.565052029 | 0.519 | 0.165 | 1.7396748 | 7 | ddit3 |
| nck1a | 4.011725 | 0.296143106 | 0.201 | 0.031 | 1.0191788 | 7 | nck1a |
| pim12 | 9.389398 | 0.52781837 | 0.509 | 0.156 | 2.3853767 | 7 | pim1 |
| lamp1a | 1.281556 | 0.404466509 | 0.353 | 0.083 | 3.2557930 | 7 | lamp1a |
| skap2 | 1.371014 | 0.348100452 | 0.325 | 0.071 | 3.4830625 | 7 | skap2 |
| rpl242 | 1.811619 | 0.385973172 | 0.986 | 0.786 | 4.6024206 | 7 | rpl24 |
| mbd2 | 2.966634 | 0.277470601 | 0.21 | 0.033 | 7.5367361 | 7 | mbd2 |
| rpl392 | 4.907220 | 0.312932717 | 0.991 | 0.873 | 1.2466793 | 7 | rpl39 |
| rps201 | 6.738547 | 0.2814751 | 0.995 | 0.907 | 1.7119279 | 7 | rps20 |
| osbp | 2.017168 | 0.258322647 | 0.205 | 0.032 | 5.1246172 | 7 | osbp |

| | | | | | | | |
|---|---|---|---|---|---|---|---|
| celf23 | 4.450117 | 0.568361851 | 0.655 | 0.241 | 1.1305524 | 7 | celf2 |
| rps212 | 9.315566 | 0.41328272 | 0.991 | 0.768 | 2.3666197 | 7 | rps21 |
| rpl362 | 1.579773 | 0.38688113 | 0.991 | 0.799 | 4.0134153 | 7 | rpl36 |
| rgs18 | 2.154808 | 0.253882737 | 0.189 | 0.028 | 5.4742911 | 7 | rgs18 |
| ppiab | 3.556801 | 0.474041407 | 0.965 | 0.663 | 9.0360553 | 7 | ppiab |
| fybb | 3.564178 | 0.30460317 | 0.221 | 0.037 | 9.0547962 | 7 | fybb |
| ostf1 | 5.156636 | 0.322420936 | 0.246 | 0.046 | 1.3100435 | 7 | ostf1 |
| rps27.12 | 7.355570 | 0.46941551 | 0.982 | 0.7 | 1.8686827 | 7 | rps27.1 |
| atp6v1h | 2.722685 | 0.287168498 | 0.233 | 0.042 | 6.9169831 | 7 | atp6v1h |
| jak1 | 7.525782 | 0.398926341 | 0.357 | 0.09 | 1.9119250 | 7 | jak1 |
| mdh1aa | 1.127413 | 0.510096406 | 0.602 | 0.214 | 2.8641930 | 7 | mdh1aa |
| DHDH | 1.176690 | 0.316444084 | 0.284 | 0.061 | 2.9893816 | 7 | DHDH |
| rpl36a2 | 1.330659 | 0.350737365 | 0.996 | 0.817 | 3.3805415 | 7 | rpl36a |
| zfp36l1a3 | 2.494869 | 0.587792114 | 0.636 | 0.239 | 6.3382169 | 7 | zfp36l1a |
| dbi1 | 7.338784 | 0.532718196 | 0.62 | 0.228 | 1.8644181 | 7 | dbi |
| wbp2nl | 2.000110 | 0.357141404 | 0.339 | 0.083 | 5.0812808 | 7 | wbp2nl |
| suclg1 | 3.893999 | 0.384600493 | 0.403 | 0.111 | 9.8927062 | 7 | suclg1 |
| rps142 | 3.514376 | 0.381406824 | 0.982 | 0.774 | 8.9282740 | 7 | rps14 |
| rps92 | 6.361461 | 0.38420746 | 0.989 | 0.783 | 1.6161292 | 7 | rps9 |
| glipr1a1 | 7.128603 | 0.307864887 | 0.283 | 0.06 | 1.8110216 | 7 | glipr1a |
| si:ch211-8c17.3 | 3.646290 | 0.267486145 | 0.163 | 0.022 | 9.2634018 | 7 | si:ch211-8c17.3 |
| arhgdig1 | 3.862676 | 0.535512073 | 0.67 | 0.243 | 9.8131290 | 7 | arhgdig |
| ptprc1 | 7.806135 | 0.408837928 | 0.413 | 0.113 | 1.9831486 | 7 | ptprc |
| oaz1b1 | 1.674058 | 0.569209066 | 0.77 | 0.344 | 4.2529461 | 7 | oaz1b |
| gabarapb1 | 2.195601 | 0.545733101 | 0.705 | 0.289 | 5.5779266 | 7 | gabarapb |
| sat1a.21 | 2.224476 | 0.562634038 | 0.708 | 0.285 | 5.6512831 | 7 | sat1a.2 |
| rpl382 | 2.244554 | 0.416208925 | 0.989 | 0.745 | 5.7022905 | 7 | rpl38 |
| lgmn | 1.625017 | 0.436091967 | 0.283 | 0.065 | 4.1283562 | 7 | lgmn |
| LOC1039091151 | 1.936823 | 0.433945679 | 0.5 | 0.156 | 4.9205004 | 7 | LOC103909115 |
| ctsba4 | 2.132260 | 0.629171447 | 0.76 | 0.336 | 5.4170081 | 7 | ctsba |
| COX7A2 (1 of many) | 2.201510 | 0.505655187 | 0.657 | 0.251 | 5.5929363 | 7 | COX7A2 (1 of many) |
| snd1 | 2.465773 | 0.393307018 | 0.383 | 0.107 | 6.2642972 | 7 | snd1 |
| gdi2 | 2.474179 | 0.473245967 | 0.548 | 0.189 | 6.2856518 | 7 | gdi2 |
| rpl35a2 | 6.315876 | 0.382055136 | 0.986 | 0.774 | 1.6045485 | 7 | rpl35a |
| XLOC-0425951 | 8.651899 | 0.335208151 | 0.488 | 0.14 | 2.1980150 | 7 | XLOC-042595 |

| | | | | | | | |
|---|---|---|---|---|---|---|---|
| selplg | 1.416696 | 0.294703226 | 0.216 | 0.039 | 3.5991176 | 7 | selplg |
| litaf1 | 2.217265 | 0.435227471 | 0.442 | 0.134 | 5.6329617 | 7 | litaf |
| eef1db1 | 2.263357 | 0.529682104 | 0.677 | 0.269 | 5.7500604 | 7 | eef1db |
| COX33 | 1.461080 | 0.517072175 | 0.917 | 0.667 | 3.7118750 | 7 | COX3 |
| vma21 | 2.911004 | 0.374609586 | 0.373 | 0.101 | 7.3954058 | 7 | vma21 |
| adipor2 | 3.417305 | 0.352153773 | 0.314 | 0.076 | 8.6816640 | 7 | adipor2 |
| rpl372 | 3.507548 | 0.33180302 | 0.993 | 0.85 | 8.9109276 | 7 | rpl37 |
| grinab1 | 4.794349 | 0.396985263 | 0.394 | 0.112 | 1.2180044 | 7 | grinab |
| irf2 | 4.962653 | 0.346839281 | 0.288 | 0.067 | 1.2607622 | 7 | irf2 |
| rpl152 | 6.480717 | 0.401745384 | 0.982 | 0.755 | 1.6464263 | 7 | rpl15 |
| gnai1 | 1.147543 | 0.333413683 | 0.274 | 0.061 | 2.9153345 | 7 | gnai1 |
| m6pr | 2.651276 | 0.346635585 | 0.329 | 0.082 | 6.7355687 | 7 | m6pr |
| arrb2b | 5.662151 | 0.358277068 | 0.337 | 0.087 | 1.4384695 | 7 | arrb2b |
| triqk | 1.215058 | 0.282403072 | 0.239 | 0.048 | 3.0868565 | 7 | triqk |
| arl6ip1 | 3.575276 | 0.562970163 | 0.783 | 0.341 | 9.0829889 | 7 | arl6ip1 |
| si:dkey-180p18.9 | 2.533082 | 0.347826195 | 0.362 | 0.099 | 6.4352952 | 7 | si:dkey-180p18.9 |
| mob1a1 | 3.140772 | 0.404484864 | 0.496 | 0.16 | 7.9791334 | 7 | mob1a |
| b3gnt3.4 | 3.213857 | 0.252608678 | 0.191 | 0.033 | 8.1648055 | 7 | b3gnt3.4 |
| ifi461 | 3.549591 | 0.377676013 | 0.417 | 0.12 | 9.0177361 | 7 | ifi46 |
| mt-co12 | 5.228962 | 0.516028124 | 0.919 | 0.629 | 1.3284178 | 7 | mt-co1 |
| atp6v0b | 7.936648 | 0.353524704 | 0.357 | 0.096 | 2.0163055 | 7 | atp6v0b |
| mcl1b2 | 9.449829 | 0.525053472 | 0.617 | 0.236 | 2.4007292 | 7 | mcl1b |
| rps232 | 1.536626 | 0.395855263 | 0.973 | 0.736 | 3.9037991 | 7 | rps23 |
| arhgdia1 | 1.621582 | 0.479223826 | 0.629 | 0.236 | 4.1196314 | 7 | arhgdia |
| gstt1a | 1.804252 | 0.317935662 | 0.196 | 0.035 | 4.5837028 | 7 | gstt1a |
| canx | 3.104908 | 0.360415743 | 0.345 | 0.093 | 7.8880194 | 7 | canx |
| rps26l2 | 3.474223 | 0.447352561 | 0.968 | 0.689 | 8.8262645 | 7 | rps26l |
| kctd12.2 | 8.207601 | 0.403369817 | 0.339 | 0.091 | 2.0851412 | 7 | kctd12.2 |
| xaf1 | 4.350986 | 0.32517188 | 0.249 | 0.054 | 1.1053680 | 7 | xaf1 |
| stat5a | 4.479077 | 0.26732958 | 0.205 | 0.038 | 1.1379096 | 7 | stat5a |
| XLOC-0283701 | 6.273852 | 0.478036886 | 0.392 | 0.115 | 1.5938722 | 7 | XLOC-028370 |
| zgc:77486 | 6.856469 | 0.305227218 | 0.265 | 0.06 | 1.7418859 | 7 | zgc:77486 |
| gnai2a1 | 9.215076 | 0.406992872 | 0.438 | 0.138 | 2.3410900 | 7 | gnai2a |
| bloc1s6 | 1.101236 | 0.320963819 | 0.33 | 0.087 | 2.7976906 | 7 | bloc1s6 |
| rpl292 | 1.349523 | 0.333720663 | 0.986 | 0.817 | 3.4284656 | 7 | rpl29 |

| | | | | | | | |
|---|---|---|---|---|---|---|---|
| rps192 | 1.978675 | 0.300765338 | 0.998 | 0.864 | 5.0268253 | 7 | rps19 |
| ppp1r181 | 2.571028 | 0.271951054 | 0.237 | 0.049 | 6.5316986 | 7 | ppp1r18 |
| birc2 | 2.789988 | 0.369803875 | 0.353 | 0.098 | 7.0879666 | 7 | birc2 |
| arpc31 | 1.045518 | 0.537125457 | 0.735 | 0.318 | 2.6561392 | 7 | arpc3 |
| rpl182 | 1.406880 | 0.371329157 | 0.988 | 0.785 | 3.5741788 | 7 | rpl18 |
| glmp | 1.410177 | 0.261433538 | 0.203 | 0.038 | 3.5825554 | 7 | glmp |
| jdp2b1 | 1.959377 | 0.523878492 | 0.454 | 0.153 | 4.9777987 | 7 | jdp2b |
| zgc:1141882 | 2.025816 | 0.476782206 | 0.975 | 0.656 | 5.1465863 | 7 | zgc:114188 |
| atp1b1a | 3.046296 | 0.462858969 | 0.581 | 0.216 | 7.7391171 | 7 | atp1b1a |
| rpl302 | 6.317670 | 0.409161218 | 0.981 | 0.72 | 1.6050043 | 7 | rpl30 |
| egr3 | 7.652558 | 0.303896398 | 0.228 | 0.047 | 1.9441323 | 7 | egr3 |
| rpl231 | 2.053087 | 0.269509502 | 0.986 | 0.868 | 5.2158698 | 7 | rpl23 |
| si:dkey-79d12.5 | 1.390732 | 0.333211242 | 0.26 | 0.061 | 3.5331565 | 7 | si:dkey-79d12.5 |
| rps292 | 9.535027 | 0.311314516 | 0.988 | 0.839 | 2.4223738 | 7 | rps29 |
| smdt1b | 1.243791 | 0.376830239 | 0.435 | 0.141 | 3.1598527 | 7 | smdt1b |
| rpl18a2 | 1.381325 | 0.361117968 | 0.986 | 0.794 | 3.5092568 | 7 | rpl18a |
| tsc22d31 | 3.251895 | 0.423871435 | 0.443 | 0.148 | 8.2614399 | 7 | tsc22d3 |
| rpl112 | 4.378702 | 0.340931076 | 0.991 | 0.823 | 1.1124094 | 7 | rpl11 |
| elmo11 | 6.284564 | 0.305148503 | 0.265 | 0.063 | 1.5965935 | 7 | elmo1 |
| rasgef1ba | 7.619967 | 0.345619546 | 0.24 | 0.054 | 1.9358528 | 7 | rasgef1ba |
| syngr2a | 8.021791 | 0.252806644 | 0.233 | 0.05 | 2.0379360 | 7 | syngr2a |
| b2ml3 | 1.064682 | 0.482747211 | 0.678 | 0.278 | 2.7048250 | 7 | b2ml |
| si:ch1073-325m22.2 | 1.888314 | 0.44847673 | 0.597 | 0.236 | 4.7972620 | 7 | si:ch1073-325m22.2 |
| rpl222 | 2.501715 | 0.413422484 | 0.973 | 0.711 | 6.3556072 | 7 | rpl22 |
| mcl1a1 | 3.016379 | 0.494925589 | 0.928 | 0.601 | 7.6631120 | 7 | mcl1a |
| msna1 | 4.376865 | 0.497388571 | 0.622 | 0.252 | 1.1119428 | 7 | msna |
| rpl27a2 | 4.455934 | 0.530455778 | 0.887 | 0.508 | 1.1320301 | 7 | rpl27a |
| tln1 | 6.982992 | 0.38497171 | 0.413 | 0.132 | 1.7740293 | 7 | tln1 |
| npc2 | 1.160282 | 0.529592735 | 0.737 | 0.327 | 2.9476981 | 7 | npc2 |
| alas1 | 2.004161 | 0.378662105 | 0.408 | 0.127 | 5.0915715 | 7 | alas1 |
| cyfip1 | 6.254829 | 0.286880927 | 0.242 | 0.055 | 1.5890394 | 7 | cyfip1 |
| lamtor5 | 2.597347 | 0.346437677 | 0.403 | 0.123 | 6.5985606 | 7 | lamtor5 |
| efhd2 | 3.415657 | 0.288169054 | 0.237 | 0.053 | 8.6774773 | 7 | efhd2 |
| slc25a51 | 4.006198 | 0.320091078 | 0.984 | 0.807 | 1.0177746 | 7 | slc25a5 |
| rab1ab | 4.409144 | 0.368799719 | 0.401 | 0.128 | 1.1201431 | 7 | rab1ab |

| | | | | | | | |
|---|---|---|---|---|---|---|---|
| zgc:110591 | 4.99550( | 0.257410732 | 0.223 | 0.048 | 1.2691069 | 7 | zgc:110591 |
| pak2a | 6.22673‌8 | 0.269045861 | 0.258 | 0.061 | 1.5819028 | 7 | pak2a |
| capza1a1 | 6.99506‌6 | 0.430851627 | 0.551 | 0.208 | 1.7770965 | 7 | capza1a |
| rplp2l2 | 7.32300‌5 | 0.366770161 | 0.988 | 0.787 | 1.8604094 | 7 | rplp2l |
| rps102 | 7.65496‌3 | 0.410110141 | 0.981 | 0.717 | 1.9447435 | 7 | rps10 |
| rpl23a2 | 2.09101( | 0.450866615 | 0.968 | 0.658 | 5.3122119 | 7 | rpl23a |
| ktn1 | 5.32409‌8 | 0.275907098 | 0.24 | 0.056 | 1.3525872 | 7 | ktn1 |
| rps52 | 7.14537‌5 | 0.404111227 | 0.97 | 0.719 | 1.8152825 | 7 | rps5 |
| lgals9l31 | 7.33557‌3 | 0.3611229 | 0.284 | 0.076 | 1.8636024 | 7 | lgals9l3 |
| actr2a1 | 1.03870‌6 | 0.37364719 | 0.392 | 0.123 | 2.6388340 | 7 | actr2a |
| rps282 | 1.13940‌7 | 0.320199015 | 0.995 | 0.835 | 2.8946650 | 7 | rps28 |
| midn1 | 1.15639‌5 | 0.400548142 | 0.41 | 0.135 | 2.9378233 | 7 | midn |
| rps121 | 1.41940‌1 | 0.305330433 | 0.984 | 0.836 | 3.6059903 | 7 | rps12 |
| eif4ebp1 | 1.67623‌8 | 0.453023245 | 0.611 | 0.244 | 4.2584843 | 7 | eif4ebp1 |
| capza1b | 1.79598‌2 | 0.427913869 | 0.576 | 0.221 | 4.5626948 | 7 | capza1b |
| BX005183.31 | 2.53099‌5 | 0.353375026 | 0.272 | 0.069 | 6.4299944 | 7 | BX005183.3 |
| XLOC-0108991 | 3.31460‌8 | 0.356843492 | 0.602 | 0.225 | 8.4207635 | 7 | XLOC-010899 |
| rps152 | 4.58072‌4 | 0.411505674 | 0.966 | 0.707 | 1.1637329 | 7 | rps15 |
| uqcrq | 7.13495‌1 | 0.454046891 | 0.65 | 0.272 | 1.8126344 | 7 | uqcrq |
| BX322618.1 | 1.60723‌7 | 0.305748651 | 0.168 | 0.03 | 4.0831868 | 7 | BX322618.1 |
| atf31 | 9.58555‌4 | 0.336184613 | 0.283 | 0.074 | 2.4352100 | 7 | atf3 |
| ier2b3 | 9.81378‌8 | 0.456757354 | 0.571 | 0.224 | 2.4931930 | 7 | ier2b |
| GCA1 | 1.55955‌4 | 0.407559077 | 0.509 | 0.189 | 3.9620484 | 7 | GCA |
| zgc:1629441 | 1.97795‌5 | 0.447272155 | 0.634 | 0.261 | 5.0249948 | 7 | zgc:162944 |
| rhoab2 | 9.88435‌5 | 0.433649715 | 0.509 | 0.196 | 2.5111204 | 7 | rhoab |
| sdc42 | 2.05677‌6 | 0.524555584 | 0.574 | 0.242 | 5.2252399 | 7 | sdc4 |
| rps162 | 2.72027‌8 | 0.36904692 | 0.975 | 0.758 | 6.9108680 | 7 | rps16 |
| rps27a2 | 3.80463‌9 | 0.344590687 | 0.977 | 0.767 | 9.6656858 | 7 | rps27a |
| nt5dc2 | 1.37077‌1 | 0.28281707 | 0.279 | 0.074 | 3.4824451 | 7 | nt5dc2 |
| zgc:1583431 | 2.01610‌6 | 0.428549709 | 0.447 | 0.159 | 5.1219185 | 7 | zgc:158343 |
| rpl322 | 2.52087‌9 | 0.331451441 | 0.989 | 0.818 | 6.4042943 | 7 | rpl32 |
| rpl92 | 3.22929‌9 | 0.447277186 | 0.968 | 0.657 | 8.2040342 | 7 | rpl9 |
| BX908750.1 | 3.27028‌3 | 0.258825304 | 0.154 | 0.027 | 8.3081548 | 7 | BX908750.1 |
| ssr4 | 1.34256‌3 | 0.483156453 | 0.686 | 0.318 | 3.4107825 | 7 | ssr4 |
| rpl312 | 1.69423‌4 | 0.421063765 | 0.975 | 0.676 | 4.3042032 | 7 | rpl31 |

| | | | | | | | |
|---|---|---|---|---|---|---|---|
| rpl282 | 1.726532 | 0.350709222 | 0.981 | 0.769 | 4.3862559 | 7 | rpl28 |
| rps252 | 2.418291 | 0.368572873 | 0.977 | 0.74 | 6.1436684 | 7 | rps25 |
| dusp5 | 2.677463 | 0.303399712 | 0.246 | 0.061 | 6.8020949 | 7 | dusp5 |
| rpl262 | 4.637336 | 0.288198278 | 0.982 | 0.846 | 1.1781154 | 7 | rpl26 |
| csde1 | 5.573255 | 0.369030117 | 0.518 | 0.195 | 1.4158854 | 7 | csde1 |
| capzb1 | 1.646155 | 0.439969518 | 0.613 | 0.258 | 4.1820580 | 7 | capzb |
| ptpn61 | 2.277267 | 0.289362369 | 0.318 | 0.091 | 5.7853968 | 7 | ptpn6 |
| mt-nd31 | 2.342645 | 0.48493231 | 0.735 | 0.357 | 5.9514917 | 7 | mt-nd3 |
| si:dkeyp-68b7.121 | 2.812615 | 0.25518314 | 0.228 | 0.054 | 7.1454488 | 7 | si:dkeyp-68b7.12 |
| smim15 | 3.246718 | 0.332760094 | 0.417 | 0.143 | 8.2482874 | 7 | smim15 |
| COX5B | 6.036786 | 0.477480467 | 0.747 | 0.362 | 1.5336455 | 7 | COX5B |
| rpl191 | 9.069123 | 0.278763712 | 0.981 | 0.849 | 2.3040107 | 7 | rpl19 |
| rplp12 | 2.219795 | 0.282773145 | 0.993 | 0.878 | 5.6393905 | 7 | rplp1 |
| map3k14a | 3.379390 | 0.304662824 | 0.163 | 0.032 | 8.5853424 | 7 | map3k14a |
| glrx | 4.523579 | 0.46308038 | 0.687 | 0.312 | 1.1492154 | 7 | glrx |
| rpl352 | 9.003316 | 0.312605796 | 0.981 | 0.808 | 2.2872924 | 7 | rpl35 |
| ccl39.7 | 1.547740 | 0.479937709 | 0.221 | 0.056 | 3.9320347 | 7 | ccl39.7 |
| zgc:153713 | 1.722064 | 0.363146038 | 0.443 | 0.162 | 4.3749044 | 7 | zgc:153713 |
| grasp | 2.063755 | 0.262917724 | 0.254 | 0.066 | 5.2429711 | 7 | grasp |
| aldoaa | 2.843355 | 0.423380047 | 0.581 | 0.24 | 7.2235450 | 7 | aldoaa |
| prdx6 | 4.511276 | 0.38111761 | 0.468 | 0.174 | 1.1460898 | 7 | prdx6 |
| arpc41 | 1.443656 | 0.383597729 | 0.571 | 0.225 | 3.6676086 | 7 | arpc4 |
| cox4i1 | 2.329989 | 0.47927157 | 0.804 | 0.402 | 5.9193386 | 7 | cox4i1 |
| rpl172 | 3.277633 | 0.335432123 | 0.961 | 0.756 | 8.3268280 | 7 | rpl17 |
| cox8a | 3.654534 | 0.41871053 | 0.597 | 0.255 | 9.2843452 | 7 | cox8a |
| stk17b | 3.660972 | 0.304279847 | 0.339 | 0.106 | 9.3007010 | 7 | stk17b |
| krt82 | 3.856021 | 0.438858804 | 0.708 | 0.296 | 9.7962224 | 7 | krt8 |
| mt-co22 | 4.452241 | 0.446741196 | 0.86 | 0.664 | 1.1310918 | 7 | mt-co2 |
| rnf213a1 | 9.491325 | 0.338487172 | 0.293 | 0.088 | 2.4112713 | 7 | rnf213a |
| marcksl1a2 | 9.518765 | 0.453978945 | 0.588 | 0.248 | 2.4182424 | 7 | marcksl1a |
| cited4b | 6.148656 | 0.313693906 | 0.353 | 0.112 | 1.5620663 | 7 | cited4b |
| vdac31 | 6.691823 | 0.317552365 | 0.44 | 0.158 | 1.7000577 | 7 | vdac3 |
| fosab2 | 7.429415 | 0.487781877 | 0.749 | 0.346 | 1.8874430 | 7 | fosab |
| rpl102 | 8.006345 | 0.314698416 | 0.958 | 0.784 | 2.0340121 | 7 | rpl10 |
| cox7b1 | 1.095746 | 0.47673306 | 0.793 | 0.413 | 2.7837443 | 7 | cox7b |

| | | | | | | | |
|---|---|---|---|---|---|---|---|
| rps27 | 1.770875 | 0.320615401 | 0.988 | 0.836 | 4.4989092 | 7 | rps2 |
| psme21 | 3.688204 | 0.318163394 | 0.327 | 0.105 | 9.3698830 | 7 | psme2 |
| CR318588.12 | 8.674299 | 0.617042511 | 0.484 | 0.205 | 2.2037058 | 7 | CR318588.1 |
| faua2 | 1.747167 | 0.377372571 | 0.956 | 0.708 | 4.4386785 | 7 | faua |
| mt-atp62 | 2.814073 | 0.490400175 | 0.832 | 0.546 | 7.1491539 | 7 | mt-atp6 |
| tnfrsf1a1 | 4.602765 | 0.259881963 | 0.216 | 0.054 | 1.1693324 | 7 | tnfrsf1a |
| FP074902.2 | 8.060272 | 0.253049557 | 0.254 | 0.071 | 2.0477121 | 7 | FP074902.2 |
| klf6a2 | 2.891437 | 0.41229315 | 0.438 | 0.17 | 7.3456973 | 7 | klf6a |
| arpc5a1 | 3.019125 | 0.307093743 | 0.352 | 0.117 | 7.6700890 | 7 | arpc5a |
| rab5c | 3.317753 | 0.303517059 | 0.36 | 0.123 | 8.4287523 | 7 | rab5c |
| rpl272 | 3.796699 | 0.337663339 | 0.982 | 0.772 | 9.6455149 | 7 | rpl27 |
| fkbp51 | 4.299795 | 0.316070369 | 0.352 | 0.119 | 1.0923630 | 7 | fkbp5 |
| tnfaip3 | 5.254975 | 0.272556301 | 0.214 | 0.054 | 1.3350264 | 7 | tnfaip3 |
| chmp4bb | 6.039451 | 0.275012876 | 0.27 | 0.079 | 1.5343227 | 7 | chmp4bb |
| rpl82 | 1.040462 | 0.326956345 | 0.981 | 0.788 | 2.6432950 | 7 | rpl8 |
| ssr2 | 1.150285 | 0.445814583 | 0.673 | 0.32 | 2.9223001 | 7 | ssr2 |
| ube2v2 | 1.173507 | 0.406977835 | 0.61 | 0.27 | 2.9812955 | 7 | ube2v2 |
| rpl7a2 | 1.531438 | 0.300360131 | 0.984 | 0.818 | 3.8906206 | 7 | rpl7a |
| snx1a | 2.455478 | 0.276290261 | 0.345 | 0.114 | 6.2381421 | 7 | snx1a |
| b2m2 | 3.676432 | 0.419607963 | 0.641 | 0.285 | 9.3399772 | 7 | b2m |
| flna2 | 3.897324 | 0.299541638 | 0.343 | 0.114 | 9.9011527 | 7 | flna |
| rpl342 | 4.769218 | 0.325588229 | 0.988 | 0.795 | 1.2116200 | 7 | rpl34 |
| rpl122 | 5.003232 | 0.292606788 | 0.979 | 0.806 | 1.2710710 | 7 | rpl12 |
| abca1a1 | 7.402015 | 0.282233803 | 0.239 | 0.066 | 1.8804819 | 7 | abca1a |
| chmp2a1 | 9.221133 | 0.31262812 | 0.38 | 0.135 | 2.3426290 | 7 | chmp2a |
| rpl212 | 1.669607 | 0.322593051 | 0.965 | 0.769 | 4.2416386 | 7 | rpl21 |
| myl12.11 | 2.622172 | 0.467834157 | 0.8 | 0.411 | 6.6616303 | 7 | myl12.1 |
| rps242 | 1.640679 | 0.333651437 | 0.986 | 0.771 | 4.1681474 | 7 | rps24 |
| hsp90ab11 | 1.999444 | 0.296531692 | 0.972 | 0.794 | 5.0795899 | 7 | hsp90ab1 |
| mt-cyb1 | 3.939590 | 0.457683388 | 0.823 | 0.567 | 1.0008530 | 7 | mt-cyb |
| fkbp2 | 8.108327 | 0.275952628 | 0.385 | 0.137 | 2.0599205 | 7 | fkbp2 |
| wasf2 | 1.083120 | 0.260084947 | 0.251 | 0.073 | 2.7516687 | 7 | wasf2 |
| rps32 | 1.311597 | 0.356192102 | 0.956 | 0.717 | 3.3321145 | 7 | rps3 |
| rps4x2 | 1.334099 | 0.36583159 | 0.972 | 0.728 | 3.3892798 | 7 | rps4x |
| rps8a2 | 1.462257 | 0.31137179 | 0.988 | 0.794 | 3.7148661 | 7 | rps8a |

| | | | | | | | |
|---|---|---|---|---|---|---|---|
| anxa2a1 | 2.101114 | 0.303610008 | 0.33 | 0.11 | 5.3378811 | 7 | anxa2a |
| ahnak2 | 2.625021 | 0.323286052 | 0.362 | 0.128 | 6.6688665 | 7 | ahnak |
| rps72 | 2.659776 | 0.294198227 | 0.988 | 0.822 | 6.7571627 | 7 | rps7 |
| eno1a1 | 2.695348 | 0.312516256 | 0.371 | 0.133 | 6.8475338 | 7 | eno1a |
| rpl62 | 3.133916 | 0.38641677 | 0.97 | 0.691 | 7.9617141 | 7 | rpl6 |
| gnb1a | 3.545833 | 0.356652909 | 0.472 | 0.192 | 9.0081901 | 7 | gnb1a |
| naca2 | 7.546192 | 0.404540299 | 0.942 | 0.641 | 1.9171102 | 7 | naca |
| pfkfb4b | 7.995631 | 0.32000275 | 0.327 | 0.113 | 2.0312900 | 7 | pfkfb4b |
| twf1a | 9.029802 | 0.290440942 | 0.405 | 0.148 | 2.2940214 | 7 | twf1a |
| abcc52 | 9.252322 | 0.289666077 | 0.279 | 0.088 | 2.3505526 | 7 | abcc5 |
| atp5f1d | 1.973278 | 0.3578044 | 0.59 | 0.261 | 5.0131132 | 7 | atp5f1d |
| sec61b | 2.187738 | 0.456443058 | 0.813 | 0.456 | 5.5579497 | 7 | sec61b |
| pnrc22 | 7.368081 | 0.438133171 | 0.917 | 0.585 | 1.8718609 | 7 | pnrc2 |
| capgb1 | 1.165888 | 0.282048041 | 0.325 | 0.107 | 2.9619396 | 7 | capgb |
| rps3a2 | 1.361937 | 0.309414433 | 0.982 | 0.784 | 3.4600021 | 7 | rps3a |
| itm2ba2 | 2.072639 | 0.386779434 | 0.62 | 0.284 | 5.2655412 | 7 | itm2ba |
| ldhba | 2.884277 | 0.363118619 | 0.565 | 0.249 | 7.3275075 | 7 | ldhba |
| nr4a3 | 5.825978 | 0.276355296 | 0.173 | 0.042 | 1.4800899 | 7 | nr4a3 |
| chmp1b | 1.264826 | 0.282623078 | 0.348 | 0.125 | 3.2132917 | 7 | chmp1b |
| hdlbpa2 | 3.416237 | 0.296736311 | 0.426 | 0.166 | 8.6789500 | 7 | hdlbpa |
| atp5pd | 3.461025 | 0.36162133 | 0.595 | 0.271 | 8.7927356 | 7 | atp5pd |
| eif3m2 | 4.827140 | 0.423369455 | 0.731 | 0.37 | 1.2263350 | 7 | eif3m |
| atp5l | 1.150119 | 0.433569597 | 0.811 | 0.443 | 2.9218782 | 7 | atp5l |
| ier2a2 | 1.338178 | 0.392064298 | 0.581 | 0.263 | 3.3996428 | 7 | ier2a |
| rps62 | 1.962312 | 0.298699354 | 0.979 | 0.797 | 4.9852550 | 7 | rps6 |
| mt-nd5 | 2.061288 | 0.262395004 | 0.339 | 0.122 | 5.2367044 | 7 | mt-nd5 |
| elf11 | 2.202838 | 0.254925991 | 0.293 | 0.096 | 5.5963116 | 7 | elf1 |
| rpn1 | 2.650205 | 0.278677918 | 0.399 | 0.152 | 6.7328481 | 7 | rpn1 |
| rpl10a2 | 2.893346 | 0.314475396 | 0.97 | 0.772 | 7.3505475 | 7 | rpl10a |
| rps182 | 3.020439 | 0.352246881 | 0.981 | 0.745 | 7.6734269 | 7 | rps18 |
| eif3jb | 3.706246 | 0.296007951 | 0.316 | 0.111 | 9.4157196 | 7 | eif3jb |
| ppib2 | 3.900919 | 0.422601208 | 0.76 | 0.364 | 9.9102858 | 7 | ppib |
| CABZ01080702.12 | 4.119244 | 0.292741354 | 0.277 | 0.091 | 1.0464941 | 7 | CABZ01080702.1 |
| jund1 | 1.543441 | 0.292726188 | 0.26 | 0.084 | 3.9211126 | 7 | jund |
| atf4a | 1.624562 | 0.297948157 | 0.44 | 0.176 | 4.1272014 | 7 | atf4a |

| | | | | | | | |
|---|---|---|---|---|---|---|---|
| arf6b | 1.919973 | 0.276045431 | 0.306 | 0.106 | 4.8776934 | 7 | arf6b |
| mknk2b2 | 2.710119 | 0.352679622 | 0.452 | 0.191 | 6.8850587 | 7 | mknk2b |
| anxa11b1 | 2.927477 | 0.297242796 | 0.415 | 0.165 | 7.4372560 | 7 | anxa11b |
| gabarapa | 3.242772 | 0.331029674 | 0.422 | 0.172 | 8.2382638 | 7 | gabarapa |
| socs3a2 | 3.670975 | 0.388321087 | 0.663 | 0.301 | 9.3261139 | 7 | socs3a |
| mgst1.22 | 3.838101 | 0.262827521 | 0.314 | 0.109 | 9.7506974 | 7 | mgst1.2 |
| sri | 9.493326 | 0.298678449 | 0.39 | 0.152 | 2.4117797 | 7 | sri |
| rps171 | 1.318651 | 0.30440298 | 0.972 | 0.772 | 3.3500333 | 7 | rps17 |
| nfkbie1 | 1.601439 | 0.252738049 | 0.276 | 0.09 | 4.0684564 | 7 | nfkbie |
| eif3f2 | 1.882557 | 0.44673594 | 0.825 | 0.451 | 4.7826369 | 7 | eif3f |
| cox6b11 | 2.390584 | 0.260598438 | 0.323 | 0.115 | 6.0732810 | 7 | cox6b1 |
| ap2m1a | 3.623748 | 0.2943805 | 0.406 | 0.162 | 9.2061326 | 7 | ap2m1a |
| mt-nd12 | 4.389801 | 0.419924949 | 0.71 | 0.385 | 1.1152291 | 7 | mt-nd1 |
| cnn2 | 6.006695 | 0.31870219 | 0.378 | 0.149 | 1.5260009 | 7 | cnn2 |
| atp5pf | 1.348433 | 0.376706005 | 0.629 | 0.308 | 3.4256960 | 7 | atp5pf |
| uba522 | 1.783035 | 0.386184602 | 0.947 | 0.65 | 4.5298014 | 7 | uba52 |
| eif3s101 | 1.960636 | 0.298002939 | 0.415 | 0.17 | 4.9809957 | 7 | eif3s10 |
| krtcap2 | 2.026251 | 0.310211494 | 0.443 | 0.186 | 5.1476913 | 7 | krtcap2 |
| spcs1 | 2.585214 | 0.310830542 | 0.507 | 0.221 | 6.5677384 | 7 | spcs1 |
| cox6c | 5.040058 | 0.396468044 | 0.769 | 0.397 | 1.2804267 | 7 | cox6c |
| zgc:1748631 | 5.966136 | 0.260590301 | 0.297 | 0.104 | 1.5156970 | 7 | zgc:174863 |
| cd9b1 | 1.245075 | 0.295826993 | 0.198 | 0.056 | 3.1631136 | 7 | cd9b |
| eif3c1 | 3.140121 | 0.293788202 | 0.429 | 0.179 | 7.9774795 | 7 | eif3c |
| mrpl33 | 3.253776 | 0.263872812 | 0.284 | 0.1 | 8.2662188 | 7 | mrpl33 |
| ost4 | 7.235572 | 0.296950551 | 0.564 | 0.25 | 1.8381971 | 7 | ost4 |
| rps112 | 1.900073 | 0.302469796 | 0.984 | 0.777 | 4.8271378 | 7 | rps11 |
| abracl | 2.190466 | 0.280726971 | 0.505 | 0.211 | 5.5648804 | 7 | abracl |
| ptmaa2 | 2.307931 | 0.429145706 | 0.811 | 0.418 | 5.8633000 | 7 | ptmaa |
| rpsa2 | 2.440405 | 0.308374838 | 0.973 | 0.779 | 6.1998509 | 7 | rpsa |
| cox6b2 | 5.213289 | 0.403252 | 0.859 | 0.538 | 1.3244362 | 7 | cox6b2 |
| tpt11 | 6.434148 | 0.262413209 | 0.965 | 0.806 | 1.6345953 | 7 | tpt1 |
| rps132 | 7.300089 | 0.337266686 | 0.984 | 0.74 | 1.8545877 | 7 | rps13 |
| NDUFB1 | 1.098939 | 0.282259888 | 0.468 | 0.202 | 2.7918560 | 7 | NDUFB1 |
| cdc42l1 | 1.449350 | 0.318957384 | 0.498 | 0.224 | 3.6820748 | 7 | cdc42l |
| cox6a1 | 4.107845 | 0.384918221 | 0.719 | 0.377 | 1.0435982 | 7 | cox6a1 |

| | | | | | | | |
|---|---|---|---|---|---|---|---|
| ndufb3 | 4.168711 | 0.298696369 | 0.505 | 0.228 | 1.0590612 | 7 | ndufb3 |
| mt-nd42 | 5.048895 | 0.409583876 | 0.753 | 0.436 | 1.2826719 | 7 | mt-nd4 |
| zgc:564931 | 8.650238 | 0.403316391 | 0.899 | 0.578 | 2.1975930 | 7 | zgc:56493 |
| vdac2 | 1.898129 | 0.357720575 | 0.62 | 0.315 | 4.8221980 | 7 | vdac2 |
| pgam1a1 | 2.065456 | 0.260415363 | 0.371 | 0.147 | 5.2472924 | 7 | pgam1a |
| magt1 | 3.162411 | 0.253020539 | 0.32 | 0.122 | 8.0341067 | 7 | magt1 |
| rpl132 | 5.941540 | 0.266267679 | 0.982 | 0.812 | 1.5094484 | 7 | rpl13 |
| ccni1 | 8.408472 | 0.339765113 | 0.627 | 0.307 | 2.1361724 | 7 | ccni |
| ywhaba1 | 1.526431 | 0.345331306 | 0.643 | 0.317 | 3.8778988 | 7 | ywhaba |
| ptp4a2b | 1.979052 | 0.26554089 | 0.357 | 0.142 | 5.0277831 | 7 | ptp4a2b |
| reep5 | 2.174280 | 0.25916523 | 0.398 | 0.165 | 5.5237605 | 7 | reep5 |
| lasp11 | 2.689463 | 0.310635219 | 0.527 | 0.243 | 6.8325829 | 7 | lasp1 |
| selenof | 3.327531 | 0.317165386 | 0.544 | 0.256 | 8.4535937 | 7 | selenof |
| eif2s1b | 4.565506 | 0.281589466 | 0.426 | 0.186 | 1.1598668 | 7 | eif2s1b |
| ecm1b | 5.109468 | 0.452139372 | 0.194 | 0.063 | 1.2980604 | 7 | ecm1b |
| rps262 | 1.383781 | 0.317902497 | 0.973 | 0.742 | 3.5154965 | 7 | rps26 |
| btf32 | 1.551952 | 0.377719661 | 0.903 | 0.599 | 3.9427352 | 7 | btf3 |
| rpl5a2 | 3.039483 | 0.411017825 | 0.866 | 0.524 | 7.7218090 | 7 | rpl5a |
| tmem258 | 3.449253 | 0.325342839 | 0.595 | 0.294 | 8.7628289 | 7 | tmem258 |
| actb1 | 4.548091 | 0.31743565 | 0.973 | 0.759 | 1.1554427 | 7 | actb1 |
| tmed2 | 1.037245 | 0.284785542 | 0.527 | 0.245 | 2.6351212 | 7 | tmed2 |
| sh3bgrl3 | 1.093147 | 0.251974205 | 0.362 | 0.148 | 2.7771413 | 7 | sh3bgrl3 |
| rsl24d12 | 1.732730 | 0.403457303 | 0.799 | 0.447 | 4.4020012 | 7 | rsl24d1 |
| rack12 | 3.498013 | 0.295962391 | 0.966 | 0.756 | 8.8867023 | 7 | rack1 |
| uqcrh | 5.383613 | 0.28540104 | 0.527 | 0.25 | 1.3677070 | 7 | uqcrh |
| nsa22 | 7.747011 | 0.375438671 | 0.712 | 0.376 | 1.9681283 | 7 | nsa2 |
| ndufa4l1 | 1.656275 | 0.372175085 | 0.758 | 0.409 | 4.2077674 | 7 | ndufa4l |
| ucp21 | 3.461627 | 0.374506491 | 0.742 | 0.402 | 8.7942651 | 7 | ucp2 |
| slc43a3b | 5.472890 | 0.286054473 | 0.113 | 0.026 | 1.3903877 | 7 | slc43a3b |
| socs3b1 | 5.478090 | 0.270462601 | 0.352 | 0.144 | 1.3917088 | 7 | socs3b |
| atf4b1 | 6.757378 | 0.306470738 | 0.557 | 0.267 | 1.7167120 | 7 | atf4b |
| atp5mf | 8.834138 | 0.268192442 | 0.58 | 0.278 | 2.2443129 | 7 | atp5mf |
| si:ch211-196l7.43 | 2.134987 | 0.337816777 | 0.61 | 0.319 | 5.4239351 | 7 | si:ch211-196l7.4 |
| zgc:193541 | 2.186946 | 0.338847585 | 0.689 | 0.37 | 5.5559374 | 7 | zgc:193541 |
| eef1g2 | 5.309788 | 0.351687265 | 0.926 | 0.642 | 1.3489518 | 7 | eef1g |

| | | | | | | | |
|---|---|---|---|---|---|---|---|
| rps27.22 | 8.274500 | 0.412435258 | 0.91 | 0.539 | 2.1021367 | 7 | rps27.2 |
| rpl142 | 8.428900 | 0.348561562 | 0.947 | 0.659 | 2.1413621 | 7 | rpl14 |
| klf2a1 | 2.446566 | 0.262896309 | 0.253 | 0.094 | 6.2155011 | 7 | klf2a |
| cox5aa | 3.880956 | 0.329083427 | 0.655 | 0.347 | 9.8595699 | 7 | cox5aa |
| si:ch211-198c19.32 | 1.794765 | 0.3255904 | 0.217 | 0.078 | 4.5596019 | 7 | si:ch211-198c19.3 |
| ppdpfb1 | 2.373491 | 0.369746853 | 0.82 | 0.521 | 6.0298544 | 7 | ppdpfb |
| ak2 | 5.044882 | 0.254135843 | 0.505 | 0.245 | 1.2816523 | 7 | ak2 |
| higd1a1 | 2.503277 | 0.299720089 | 0.576 | 0.293 | 6.3595771 | 7 | higd1a |
| NC-002333.42 | 5.751683 | 0.366948081 | 0.866 | 0.599 | 1.4612152 | 7 | NC-002333.4 |
| rps27l1 | 5.778609 | 0.258172806 | 0.44 | 0.21 | 1.4680558 | 7 | rps27l |
| ybx11 | 1.248540 | 0.314852507 | 0.94 | 0.686 | 3.1719166 | 7 | ybx1 |
| rpl22l12 | 1.651318 | 0.361616869 | 0.731 | 0.414 | 4.1951738 | 7 | rpl22l1 |
| romo12 | 2.797737 | 0.335770789 | 0.712 | 0.397 | 7.1076515 | 7 | romo1 |
| cox7c | 3.618134 | 0.351658601 | 0.8 | 0.482 | 9.1918696 | 7 | cox7c |
| si:ch73-1a9.32 | 2.648115 | 0.274682726 | 0.606 | 0.305 | 6.7275382 | 7 | si:ch73-1a9.3 |
| sumo2b2 | 2.679638 | 0.264099729 | 0.565 | 0.286 | 6.8076212 | 7 | sumo2b |
| eef2b1 | 2.796144 | 0.331407075 | 0.898 | 0.626 | 7.1036051 | 7 | eef2b |
| dad1 | 7.695894 | 0.295859508 | 0.652 | 0.347 | 1.9551419 | 7 | dad1 |
| atp5pb | 2.954364 | 0.265848512 | 0.611 | 0.322 | 7.5055636 | 7 | atp5pb |
| atp5md | 5.771441 | 0.271664466 | 0.65 | 0.345 | 1.4662347 | 7 | atp5md |
| CR383676.13 | 7.080489 | 0.454660729 | 0.88 | 0.711 | 1.7987984 | 7 | CR383676.1 |
| mt-nd22 | 7.412819 | 0.321039951 | 0.664 | 0.382 | 1.8832267 | 7 | mt-nd2 |
| mibp22 | 3.144859 | 0.340118574 | 0.807 | 0.51 | 7.9895143 | 7 | mibp2 |
| pabpc1a2 | 3.662936 | 0.355861822 | 0.885 | 0.558 | 9.3056911 | 7 | pabpc1a |
| histh1l1 | 2.366291 | 0.254162313 | 0.269 | 0.117 | 6.0115646 | 7 | histh1l |
| eif3g2 | 2.734260 | 0.25159212 | 0.551 | 0.291 | 6.9463892 | 7 | eif3g |
| ponzr12 | 2.066129 | 0.26024801 | 0.539 | 0.28 | 5.2490012 | 7 | ponzr1 |
| rpl42 | 1.385890 | 0.317013766 | 0.915 | 0.647 | 3.5208559 | 7 | rpl4 |
| slc25a3b1 | 3.297446 | 0.310938784 | 0.744 | 0.447 | 8.3771625 | 7 | slc25a3b |
| serbp1a2 | 3.994626 | 0.340338034 | 0.814 | 0.494 | 1.0148349 | 7 | serbp1a |
| chac1 | 2.323166 | 0.265142957 | 0.643 | 0.368 | 5.9020048 | 7 | chac1 |
| serf22 | 4.836632 | 0.256486631 | 0.654 | 0.367 | 1.2287464 | 7 | serf2 |
| fabp11a1 | 1.933125 | 0.288009162 | 0.196 | 0.084 | 4.9111057 | 7 | fabp11a |
| rpl5b2 | 4.655920 | 0.289792387 | 0.898 | 0.636 | 1.1828367 | 7 | rpl5b |
| atp5mc3b | 1.432171 | 0.25434491 | 0.825 | 0.594 | 3.6384329 | 7 | atp5mc3b |

| | | | | | | | |
|---|---|---|---|---|---|---|---|
| gstm.31 | 7.76694[illegible] | 1.237306928 | 0.515 | 0.116 | 1.9731912 | 8 | gstm.3 |
| krt151 | 5.42361[illegible] | 0.847384391 | 0.33 | 0.088 | 1.3778683 | 8 | krt15 |
| aldh1a21 | 8.84030[illegible] | 0.716371958 | 0.336 | 0.09 | 2.2458783 | 8 | aldh1a2 |
| krt4 | 6.97648[illegible] | 0.655984763 | 0.223 | 0.044 | 1.7723755 | 8 | krt4 |
| gsdf | 6.38806[illegible] | 0.731245115 | 0.228 | 0.05 | 1.6228889 | 8 | gsdf |
| ybx12 | 2.06197[illegible] | 0.407977423 | 0.856 | 0.69 | 5.2384524 | 8 | ybx1 |
| krt83 | 4.52253[illegible] | 0.747366969 | 0.6 | 0.301 | 1.1489507 | 8 | krt8 |
| s100a10b3 | 1.51915[illegible] | 0.793302128 | 0.569 | 0.275 | 3.8594133 | 8 | s100a10b |
| rps122 | 1.12163[illegible] | 0.290943007 | 0.938 | 0.838 | 2.8495013 | 8 | rps12 |
| krt18a.11 | 9.95066[illegible] | 0.706602686 | 0.389 | 0.142 | 2.5279670 | 8 | krt18a.1 |
| zgc:1141883 | 5.46170[illegible] | 0.405934824 | 0.85 | 0.662 | 1.3875458 | 8 | zgc:114188 |
| rps8a3 | 1.88176[illegible] | 0.297874716 | 0.905 | 0.798 | 4.7806193 | 8 | rps8a |
| rplp2l3 | 2.59342[illegible] | 0.344775559 | 0.951 | 0.789 | 6.5885897 | 8 | rplp2l |
| rps163 | 3.45765[illegible] | 0.331540621 | 0.9 | 0.761 | 8.7841664 | 8 | rps16 |
| s100a10a | 4.90699[illegible] | 0.34328641 | 0.1 | 0.011 | 1.2466229 | 8 | s100a10a |
| rpsa3 | 3.51410[illegible] | 0.328560269 | 0.931 | 0.781 | 8.9275943 | 8 | rpsa |
| rps183 | 8.69627[illegible] | 0.345948395 | 0.894 | 0.75 | 2.2092881 | 8 | rps18 |
| rplp13 | 4.83754[illegible] | 0.262531994 | 0.965 | 0.879 | 1.2289776 | 8 | rplp1 |
| krt942 | 5.16633[illegible] | 0.711738195 | 0.403 | 0.158 | 1.3125062 | 8 | krt94 |
| rps210 | 4.16020[illegible] | 0.299540149 | 0.947 | 0.838 | 1.0568999 | 8 | rps2 |
| rps73 | 1.77340[illegible] | 0.28270464 | 0.932 | 0.825 | 4.5053313 | 8 | rps7 |
| rps243 | 3.93631[illegible] | 0.313126453 | 0.903 | 0.775 | 1.0000207 | 8 | rps24 |
| rps193 | 6.62989[illegible] | 0.254537749 | 0.956 | 0.866 | 1.6843247 | 8 | rps19 |
| rpl93 | 7.59852[illegible] | 0.404008995 | 0.87 | 0.661 | 1.9304046 | 8 | rpl9 |
| rps153 | 2.66163[illegible] | 0.352501225 | 0.865 | 0.712 | 6.7618711 | 8 | rps15 |
| rpl113 | 1.12492[illegible] | 0.290148616 | 0.945 | 0.826 | 2.8578689 | 8 | rpl11 |
| rpl323 | 2.84512[illegible] | 0.29264815 | 0.945 | 0.82 | 7.2280403 | 8 | rpl32 |
| rps15a3 | 1.21170[illegible] | 0.32083347 | 0.934 | 0.793 | 3.0783254 | 8 | rps15a |
| rpl36a3 | 1.50170[illegible] | 0.261539299 | 0.914 | 0.821 | 3.8150741 | 8 | rpl36a |
| rpl353 | 4.58712[illegible] | 0.280088378 | 0.92 | 0.811 | 1.1653589 | 8 | rpl35 |
| rpl23a3 | 4.69042[illegible] | 0.371391907 | 0.83 | 0.664 | 1.1916031 | 8 | rpl23a |
| rps93 | 2.16756[illegible] | 0.298452791 | 0.912 | 0.787 | 5.5066955 | 8 | rps9 |
| faua3 | 4.03712[illegible] | 0.344981966 | 0.869 | 0.713 | 1.0256305 | 8 | faua |
| rpl173 | 6.84928[illegible] | 0.303787523 | 0.885 | 0.76 | 1.7400596 | 8 | rpl17 |
| rps133 | 7.79450[illegible] | 0.318464899 | 0.885 | 0.745 | 1.9801943 | 8 | rps13 |

| | | | | | | | |
|---|---|---|---|---|---|---|---|
| rps4x3 | 6.757675 | 0.3142843 | 0.858 | 0.733 | 1.7167874 | 8 | rps4x |
| rpl243 | 6.807668 | 0.300428299 | 0.927 | 0.789 | 1.7294880 | 8 | rpl24 |
| rps27a3 | 9.703227 | 0.286638436 | 0.891 | 0.771 | 2.4651050 | 8 | rps27a |
| rpl35a3 | 1.670851 | 0.299901012 | 0.911 | 0.778 | 4.2447994 | 8 | rpl35a |
| rpl103 | 1.284823 | 0.27991133 | 0.901 | 0.787 | 3.2640951 | 8 | rpl10 |
| rpl123 | 1.475447 | 0.256581971 | 0.901 | 0.81 | 3.7483732 | 8 | rpl12 |
| rpl7a3 | 3.566764 | 0.274464986 | 0.938 | 0.82 | 9.0613645 | 8 | rpl7a |
| rpl18a3 | 5.858049 | 0.292234458 | 0.922 | 0.797 | 1.4882373 | 8 | rpl18a |
| rpl83 | 1.857558 | 0.269866131 | 0.889 | 0.792 | 4.7191263 | 8 | rpl8 |
| rpl183 | 2.517256 | 0.291346995 | 0.912 | 0.789 | 6.3950892 | 8 | rpl18 |
| rpl133 | 4.101019 | 0.262665111 | 0.922 | 0.815 | 1.0418639 | 8 | rpl13 |
| rpl10a3 | 1.522981 | 0.292243177 | 0.903 | 0.775 | 3.8691357 | 8 | rpl10a |
| rpl293 | 1.630104 | 0.267733963 | 0.932 | 0.819 | 4.1412793 | 8 | rpl29 |
| rpl273 | 2.239328 | 0.282814413 | 0.881 | 0.777 | 5.6890130 | 8 | rpl27 |
| rps3a3 | 4.102535 | 0.267370855 | 0.892 | 0.788 | 1.0422492 | 8 | rps3a |
| rpl13a2 | 1.729603 | 0.257151882 | 0.861 | 0.78 | 4.3940564 | 8 | rpl13a |
| rpl213 | 2.135316 | 0.288209606 | 0.9 | 0.773 | 5.4247728 | 8 | rpl21 |
| rps283 | 1.154072 | 0.255325443 | 0.947 | 0.837 | 2.9319216 | 8 | rps28 |
| rpl363 | 2.091637 | 0.279289919 | 0.923 | 0.803 | 5.3138052 | 8 | rpl36 |
| rpl283 | 3.378120 | 0.278923017 | 0.891 | 0.773 | 8.5821143 | 8 | rpl28 |
| rpl303 | 1.113374 | 0.305640181 | 0.865 | 0.726 | 2.8285266 | 8 | rpl30 |
| rpl383 | 4.623576 | 0.298313239 | 0.883 | 0.75 | 1.1746195 | 8 | rpl38 |
| rps63 | 2.604073 | 0.26511395 | 0.905 | 0.8 | 6.6156488 | 8 | rps6 |
| rpl153 | 3.423772 | 0.302576422 | 0.901 | 0.759 | 8.6980933 | 8 | rpl15 |
| rpl63 | 2.501923 | 0.309306538 | 0.818 | 0.698 | 6.3561354 | 8 | rpl6 |
| rps53 | 7.764189 | 0.310255059 | 0.869 | 0.724 | 1.9724923 | 8 | rps5 |
| sparc2 | 2.932654 | 0.537190754 | 0.403 | 0.188 | 7.4504099 | 8 | sparc |
| rps233 | 2.888572 | 0.272594588 | 0.854 | 0.741 | 7.3384189 | 8 | rps23 |
| rps103 | 3.631358 | 0.291420295 | 0.859 | 0.723 | 9.2254658 | 8 | rps10 |
| ppdpfa1 | 2.682480 | 0.436201311 | 0.208 | 0.067 | 6.8148419 | 8 | ppdpfa |
| anxa1a2 | 4.246591 | 0.642798834 | 0.332 | 0.157 | 1.0788465 | 8 | anxa1a |
| rps253 | 1.157387 | 0.253130635 | 0.847 | 0.746 | 2.9403434 | 8 | rps25 |
| rps27.23 | 3.394187 | 0.353585956 | 0.682 | 0.55 | 8.6229334 | 8 | rps27.2 |
| rps213 | 6.979870 | 0.282118072 | 0.916 | 0.771 | 1.7732361 | 8 | rps21 |
| rpl223 | 2.511518 | 0.285836471 | 0.856 | 0.716 | 6.3805117 | 8 | rpl22 |

| | | | | | | | |
|---|---|---|---|---|---|---|---|
| si:ch73-1a9.33 | 2.686558 | 0.529317232 | 0.485 | 0.31 | 6.8252010 | 8 | si:ch73-1a9.3 |
| rpl313 | 8.057519 | 0.27903296 | 0.803 | 0.684 | 2.0470127 | 8 | rpl31 |
| rps33 | 1.355439 | 0.256731315 | 0.821 | 0.724 | 3.4434927 | 8 | rps3 |
| rps27.13 | 1.638831 | 0.288063941 | 0.852 | 0.706 | 4.1634525 | 8 | rps27.1 |
| si:ch73-46j18.52 | 4.288192 | 0.456660177 | 0.387 | 0.203 | 1.0894152 | 8 | si:ch73-46j18.5 |
| eef1g3 | 9.786572 | 0.27980765 | 0.755 | 0.65 | 2.4862786 | 8 | eef1g |
| hmgn63 | 3.369749 | 0.457106085 | 0.458 | 0.307 | 8.5608488 | 8 | hmgn6 |
| anxa2a2 | 4.098925 | 0.479270547 | 0.255 | 0.113 | 1.0413319 | 8 | anxa2a |
| si:ch211-222l21.12 | 2.257313 | 0.482148502 | 0.469 | 0.31 | 5.7347044 | 8 | si:ch211-222l21.1 |
| podxl | 2.394797 | 0.341055886 | 0.164 | 0.054 | 6.0839835 | 8 | podxl |
| rps26l3 | 3.085128 | 0.276108715 | 0.841 | 0.695 | 7.8377688 | 8 | rps26l |
| cavin1b | 1.039983 | 0.300178344 | 0.139 | 0.042 | 2.6420768 | 8 | cavin1b |
| cygb11 | 3.561147 | 0.358114583 | 0.166 | 0.058 | 9.0470960 | 8 | cygb1 |
| rpl143 | 1.406379 | 0.254254779 | 0.768 | 0.667 | 3.5729070 | 8 | rpl14 |
| tuba8l32 | 7.633763 | 0.379923927 | 0.179 | 0.069 | 1.9393576 | 8 | tuba8l3 |
| fkbp1aa1 | 1.076808 | 0.38336775 | 0.538 | 0.411 | 2.7356311 | 8 | fkbp1aa |
| krt911 | 2.464025 | 0.445138057 | 0.181 | 0.069 | 6.2598555 | 8 | krt91 |
| rpl5a3 | 3.832174 | 0.300715528 | 0.633 | 0.535 | 9.7356388 | 8 | rpl5a |
| rpl22l13 | 4.186140 | 0.363960387 | 0.536 | 0.423 | 1.0634889 | 8 | rpl22l1 |
| naca3 | 7.263986 | 0.256276388 | 0.761 | 0.649 | 1.8454156 | 8 | naca |
| tmsb4x4 | 1.984650 | 0.316337455 | 0.92 | 0.711 | 5.0420047 | 8 | tmsb4x |
| gstm.11 | 1.716206 | 0.388527717 | 0.403 | 0.276 | 4.3600230 | 8 | gstm.1 |
| id12 | 5.980948 | 0.396902585 | 0.245 | 0.128 | 1.5194598 | 8 | id1 |
| jam2a | 7.505211 | 0.301190113 | 0.13 | 0.049 | 1.9066990 | 8 | jam2a |
| hspb12 | 8.352719 | 0.322526801 | 0.177 | 0.079 | 2.1220085 | 8 | hspb1 |
| s100v22 | 1.099086 | 0.375484773 | 0.25 | 0.137 | 2.7922298 | 8 | s100v2 |
| bcam | 5.250198 | 0.251260314 | 0.111 | 0.039 | 1.3338129 | 8 | bcam |
| ptmab2 | 2.949815 | 0.260349845 | 0.65 | 0.569 | 7.4940060 | 8 | ptmab |
| rpz52 | 1.138942 | 0.375850194 | 0.226 | 0.124 | 2.8934833 | 8 | rpz5 |
| hmgb1b1 | 1.667854 | 0.377908027 | 0.257 | 0.152 | 4.2371852 | 8 | hmgb1b |
| gstt1b2 | 2.694165 | 0.330452734 | 0.177 | 0.087 | 6.8445279 | 8 | gstt1b |
| hmgn71 | 3.239731 | 0.351191157 | 0.418 | 0.316 | 8.2305374 | 8 | hmgn7 |
| hmgb3a2 | 5.579287 | 0.284103176 | 0.17 | 0.081 | 1.4174178 | 8 | hmgb3a |
| CR318588.13 | 1.428157 | 0.421418605 | 0.336 | 0.212 | 3.6282336 | 8 | CR318588.1 |
| calm2b1 | 9.469126 | 0.366861041 | 0.345 | 0.244 | 2.4056315 | 8 | calm2b |

| | | | | | | | |
|---|---|---|---|---|---|---|---|
| dbi2 | 2.301843 | 0.354407175 | 0.338 | 0.241 | 5.8478339 | 8 | dbi |
| ptmaa3 | 3.263381 | 0.286999524 | 0.535 | 0.431 | 8.2906198 | 8 | ptmaa |
| hspe1 | 3.743134 | 0.350667457 | 0.391 | 0.289 | 9.5094327 | 8 | hspe1 |
| ppib3 | 5.444520 | 0.291276852 | 0.474 | 0.377 | 1.3831804 | 8 | ppib |
| sox4a1 | 9.845865 | 0.251598504 | 0.133 | 0.061 | 2.5013421 | 8 | sox4a |
| marcksl1b2 | 9.819812 | 0.324236055 | 0.232 | 0.14 | 2.4947234 | 8 | marcksl1b |
| hmgn21 | 1.074574 | 0.314291178 | 0.538 | 0.467 | 2.7299556 | 8 | hmgn2 |
| ifi452 | 3.586425 | 0.322070288 | 0.228 | 0.137 | 9.1113142 | 8 | ifi45 |
| myl62 | 5.346351 | 0.279223853 | 0.199 | 0.115 | 1.3582406 | 8 | myl6 |
| prdx21 | 1.506804 | 0.252484336 | 0.484 | 0.422 | 3.8280371 | 8 | prdx2 |
| tagln1 | 3.175841 | 0.307059579 | 0.141 | 0.073 | 8.0682244 | 8 | tagln |
| h3f3c2 | 5.753035 | 0.289433745 | 0.378 | 0.303 | 1.4615586 | 8 | h3f3c |
| icn2 | 8.986821 | 0.356349792 | 0.358 | 0.266 | 2.2831020 | 8 | icn |
| pfn21 | 9.158236 | 0.287775094 | 0.325 | 0.237 | 2.3266499 | 8 | pfn2 |
| tpm4a2 | 9.257828 | 0.271925543 | 0.188 | 0.112 | 2.3519512 | 8 | tpm4a |
| aldh9a1a.12 | 1.804433 | 0.322896368 | 0.266 | 0.192 | 4.5841626 | 8 | aldh9a1a.1 |
| hmgb2b | 1.180510 | 0.288124267 | 0.401 | 0.336 | 0.0003 | 8 | hmgb2b |
| zgc:927441 | 1.874480 | 0.268904848 | 0.396 | 0.326 | 0.000476 | 8 | zgc:92744 |
| ponzr13 | 4.550186 | 0.302917836 | 0.347 | 0.289 | 0.001156 | 8 | ponzr1 |
| si:ch73-281n10.23 | 5.994221 | 0.255289583 | 0.449 | 0.396 | 0.001523 | 8 | si:ch73-281n10.2 |
| eif4ebp3l2 | 6.291116 | 0.26648004 | 0.281 | 0.204 | 0.001598 | 8 | eif4ebp3l |
| tmsb12 | 4.536414 | 0.294584199 | 0.226 | 0.159 | 0.011525 | 8 | tmsb1 |
| hmgb1a3 | 1.168378 | 0.250548993 | 0.277 | 0.228 | 0.296827 | 8 | hmgb1a |
| grn11 | 0 | 1.754524313 | 0.817 | 0.064 | 0 | 9 | grn1 |
| cd74a2 | 0 | 1.680387144 | 0.903 | 0.192 | 0 | 9 | cd74a |
| lgals2a2 | 0 | 1.551275574 | 0.817 | 0.114 | 0 | 9 | lgals2a |
| c1qb1 | 0 | 1.530766999 | 0.71 | 0.051 | 0 | 9 | c1qb |
| c1qc1 | 0 | 1.461332073 | 0.664 | 0.049 | 0 | 9 | c1qc |
| cd74b2 | 0 | 1.446756218 | 0.868 | 0.204 | 0 | 9 | cd74b |
| si:dkey-5n18.11 | 0 | 1.421845421 | 0.63 | 0.044 | 0 | 9 | si:dkey-5n18.1 |
| si:busm1-266f07.21 | 0 | 1.396174729 | 0.737 | 0.127 | 0 | 9 | si:busm1-266f07.2 |
| lygl11 | 0 | 1.294297218 | 0.641 | 0.081 | 0 | 9 | lygl1 |
| lgals3bpb1 | 0 | 1.152558018 | 0.414 | 0.031 | 0 | 9 | lgals3bpb |
| ccl34b.11 | 3.729377 | 0.932246634 | 0.313 | 0.02 | 9.4744823 | 9 | ccl34b.1 |
| zgc:1531181 | 1.623081 | 1.034635847 | 0.384 | 0.032 | 4.1234395 | 9 | zgc:153118 |

| | | | | | | | |
|---|---|---|---|---|---|---|---|
| NPC2 (1 of many)1 | 1.989234 | 0.942057128 | 0.361 | 0.03 | 5.0536493 | 9 | NPC2 (1 of many) |
| fcer1gl1 | 4.087239 | 1.259018214 | 0.788 | 0.161 | 1.0383630 | 9 | fcer1gl |
| c1qa1 | 3.726202 | 0.956530254 | 0.418 | 0.044 | 9.4664179 | 9 | c1qa |
| marco1 | 1.009325 | 0.824098814 | 0.34 | 0.032 | 2.5641901 | 9 | marco |
| CU499330.11 | 1.484063 | 0.974006867 | 0.439 | 0.06 | 3.7702627 | 9 | CU499330.1 |
| tmsb4x5 | 1.072669 | 0.746722528 | 0.998 | 0.71 | 2.7251161 | 9 | tmsb4x |
| CU929150.11 | 9.946053 | 0.717771303 | 0.246 | 0.02 | 2.5267950 | 9 | CU929150.1 |
| slc3a2a1 | 1.276646 | 0.708222886 | 0.277 | 0.03 | 3.2433202 | 9 | slc3a2a |
| actb22 | 4.981117 | 0.572883949 | 0.998 | 0.863 | 1.2654528 | 9 | actb2 |
| si:ch211-147m6.11 | 7.700688 | 0.675012361 | 0.254 | 0.026 | 1.9563598 | 9 | si:ch211-147m6.1 |
| si:dkey-7i4.241 | 6.801392 | 0.709525725 | 0.227 | 0.02 | 1.7278938 | 9 | si:dkey-7i4.24 |
| pfn12 | 1.102387 | 0.956850068 | 0.979 | 0.486 | 2.8006148 | 9 | pfn1 |
| LOC1005375041 | 9.184093 | 0.594659364 | 0.197 | 0.016 | 2.3332189 | 9 | LOC100537504 |
| grna1 | 7.708573 | 0.754432669 | 0.328 | 0.049 | 1.9583631 | 9 | grna |
| mhc2dab1 | 3.058730 | 0.927200198 | 0.429 | 0.084 | 7.7707037 | 9 | mhc2dab |
| ms4a17a.101 | 3.764577 | 0.735562353 | 0.315 | 0.046 | 9.5639080 | 9 | ms4a17a.10 |
| havcr11 | 8.763617 | 0.617836685 | 0.242 | 0.03 | 2.2263969 | 9 | havcr1 |
| si:ch211-283g2.21 | 3.983469 | 0.484587037 | 0.181 | 0.017 | 1.0120004 | 9 | si:ch211-283g2.2 |
| LOC1018857061 | 2.764002 | 0.395226817 | 0.134 | 0.009 | 7.0219488 | 9 | LOC101885706 |
| slc40a11 | 8.776789 | 0.635260514 | 0.248 | 0.035 | 2.2297434 | 9 | slc40a1 |
| zgc:1037001 | 1.032490 | 0.700737278 | 0.29 | 0.047 | 2.6230410 | 9 | zgc:103700 |
| zgc:1749041 | 3.365382 | 0.711530229 | 0.275 | 0.044 | 8.5497548 | 9 | zgc:174904 |
| krt18b3 | 1.939088 | 0.894738693 | 0.525 | 0.165 | 4.9262555 | 9 | krt18b |
| si:ch211-194m7.31 | 1.412782 | 0.492931656 | 0.174 | 0.019 | 3.5891731 | 9 | si:ch211-194m7.3 |
| ccl35.11 | 3.853240 | 0.815204919 | 0.349 | 0.075 | 9.7891586 | 9 | ccl35.1 |
| psap1 | 7.313913 | 0.864569252 | 0.632 | 0.277 | 1.8580996 | 9 | psap |
| cmklr11 | 5.749118 | 0.509995206 | 0.197 | 0.027 | 1.4605634 | 9 | cmklr1 |
| cotl12 | 2.647093 | 0.770599977 | 0.683 | 0.308 | 6.7249407 | 9 | cotl1 |
| si:ch211-165i18.21 | 3.281060 | 0.519072808 | 0.214 | 0.034 | 8.3355341 | 9 | si:ch211-165i18.2 |
| itgae.11 | 1.222489 | 0.507049291 | 0.204 | 0.032 | 3.1057355 | 9 | itgae.1 |
| ifi301 | 2.722155 | 0.811268434 | 0.498 | 0.186 | 6.9156368 | 9 | ifi30 |
| tmsb13 | 3.955901 | 0.708033684 | 0.471 | 0.15 | 1.0049968 | 9 | tmsb1 |
| grn21 | 1.321778 | 0.443171688 | 0.13 | 0.014 | 3.3579773 | 9 | grn2 |
| txn1 | 3.902869 | 0.590519781 | 0.239 | 0.046 | 9.9152393 | 9 | txn |
| ccl34b.81 | 2.703158 | 0.415670708 | 0.164 | 0.022 | 6.8673734 | 9 | ccl34b.8 |

| | | | | | | | |
|---|---|---|---|---|---|---|---|
| BX649485.21 | 5.295637 | 0.54643157 | 0.193 | 0.031 | 1.3453568 | 9 | BX649485.2 |
| rnaset2l1 | 5.454573 | 0.536757108 | 0.229 | 0.044 | 1.3857344 | 9 | rnaset2l |
| ctss2.21 | 5.318331 | 0.559849005 | 0.231 | 0.046 | 1.3511221 | 9 | ctss2.2 |
| spi1a1 | 2.304978 | 0.488054024 | 0.2 | 0.035 | 5.8557973 | 9 | spi1a |
| zgc:640651 | 1.727486 | 0.419093621 | 0.149 | 0.02 | 4.3886803 | 9 | zgc:64065 |
| lgals9l11 | 1.594662 | 0.482964881 | 0.164 | 0.024 | 4.0512404 | 9 | lgals9l1 |
| mfap41 | 1.724311 | 0.456923357 | 0.166 | 0.026 | 4.3806146 | 9 | mfap4 |
| mhc2b1 | 9.368220 | 0.483894419 | 0.195 | 0.036 | 2.3799963 | 9 | mhc2b |
| fabp11a2 | 1.153901 | 0.896371989 | 0.3 | 0.081 | 2.9314858 | 9 | fabp11a |
| sftpbb1 | 4.354691 | 0.593955672 | 0.267 | 0.065 | 1.1063094 | 9 | sftpbb |
| cndp21 | 1.791817 | 0.593332985 | 0.267 | 0.067 | 4.5521134 | 9 | cndp2 |
| gng101 | 2.826334 | 0.544727371 | 0.242 | 0.055 | 7.1803033 | 9 | gng10 |
| CR318588.14 | 3.580037 | 0.852813609 | 0.502 | 0.207 | 9.0950846 | 9 | CR318588.1 |
| rplp23 | 4.274240 | 0.369156525 | 0.952 | 0.778 | 1.0858707 | 9 | rplp2 |
| LOC1005369461 | 4.649989 | 0.382276142 | 0.134 | 0.019 | 1.1813297 | 9 | LOC100536946 |
| lta4h1 | 1.837559 | 0.536465448 | 0.267 | 0.07 | 4.6683188 | 9 | lta4h |
| gpx1a1 | 1.761219 | 0.638881327 | 0.504 | 0.228 | 4.4743789 | 9 | gpx1a |
| rps15a4 | 2.353749 | 0.349268094 | 0.96 | 0.793 | 5.9797005 | 9 | rps15a |
| apoeb | 9.127643 | 0.410161399 | 0.113 | 0.016 | 2.3188777 | 9 | apoeb |
| arpc1b2 | 1.582606 | 0.623909173 | 0.626 | 0.304 | 4.0206116 | 9 | arpc1b |
| cfp1 | 2.806406 | 0.492994568 | 0.256 | 0.071 | 7.1296751 | 9 | cfp |
| mafbb1 | 1.439194 | 0.338153134 | 0.128 | 0.022 | 3.6562740 | 9 | mafbb |
| ppiab1 | 2.206605 | 0.354961804 | 0.824 | 0.671 | 5.6058816 | 9 | ppiab |
| zgc:1528301 | 4.702372 | 0.473106858 | 0.223 | 0.063 | 1.1946378 | 9 | zgc:152830 |
| laptm52 | 2.564560 | 0.548654383 | 0.401 | 0.17 | 6.5152669 | 9 | laptm5 |
| tuba8l21 | 9.098880 | 0.452804513 | 0.223 | 0.064 | 2.3115706 | 9 | tuba8l2 |
| atp6v1g11 | 1.681075 | 0.56938062 | 0.336 | 0.137 | 4.2707710 | 9 | atp6v1g1 |
| mrc1b1 | 2.389534 | 0.386039426 | 0.174 | 0.041 | 6.0706127 | 9 | mrc1b |
| scpep11 | 5.125053 | 0.521659652 | 0.305 | 0.115 | 1.3020197 | 9 | scpep1 |
| actb11 | 6.004623 | 0.309473082 | 0.878 | 0.765 | 1.5254745 | 9 | actb1 |
| rps214 | 1.057417 | 0.289375915 | 0.971 | 0.838 | 2.6863695 | 9 | rps2 |
| rps27.14 | 2.379480 | 0.352603831 | 0.91 | 0.704 | 6.0450705 | 9 | rps27.1 |
| tpp11 | 4.448107 | 0.449352053 | 0.227 | 0.073 | 1.1300417 | 9 | tpp1 |
| gstt1a1 | 1.812098 | 0.353704172 | 0.155 | 0.038 | 4.6036365 | 9 | gstt1a |
| LO018493.11 | 2.481504 | 0.312390243 | 0.12 | 0.024 | 6.3042615 | 9 | LO018493.1 |

| | | | | | | | |
|---|---|---|---|---|---|---|---|
| si:ch211-1o7.31 | 3.559664 | 0.274464491 | 0.105 | 0.019 | 9.0433282 | 9 | si:ch211-1o7.3 |
| spi1b1 | 5.165747 | 0.414014166 | 0.214 | 0.068 | 1.3123580 | 9 | spi1b |
| ctsba5 | 5.326362 | 0.552172447 | 0.563 | 0.347 | 1.3531623 | 9 | ctsba |
| ckbb3 | 1.199056 | 0.524606458 | 0.477 | 0.245 | 3.0462033 | 9 | ckbb |
| ncf21 | 2.288224 | 0.269246369 | 0.109 | 0.021 | 5.8132349 | 9 | ncf2 |
| atp6v0ca1 | 2.645983 | 0.545663506 | 0.393 | 0.198 | 6.7221206 | 9 | atp6v0ca |
| ctsd1 | 2.734748 | 0.57081781 | 0.326 | 0.143 | 6.9476273 | 9 | ctsd |
| rpl18a4 | 7.443746 | 0.275943302 | 0.931 | 0.797 | 1.8910838 | 9 | rpl18a |
| f13a1b1 | 9.066048 | 0.317481682 | 0.13 | 0.029 | 2.3032296 | 9 | f13a1b |
| dpp71 | 1.237644 | 0.410547919 | 0.187 | 0.057 | 3.1442355 | 9 | dpp7 |
| rpl35a4 | 3.708344 | 0.274195543 | 0.916 | 0.778 | 9.4210499 | 9 | rpl35a |
| rpl294 | 2.791440 | 0.260627904 | 0.954 | 0.819 | 7.0916558 | 9 | rpl29 |
| rps293 | 3.369899 | 0.253079082 | 0.973 | 0.84 | 8.5612293 | 9 | rps29 |
| mdh1aa1 | 8.139202 | 0.515895131 | 0.414 | 0.224 | 2.0677644 | 9 | mdh1aa |
| rpl224 | 2.666524 | 0.299531297 | 0.87 | 0.717 | 6.7743061 | 9 | rpl22 |
| cst14b.11 | 4.243030 | 0.424200138 | 0.185 | 0.058 | 1.0779420 | 9 | cst14b.1 |
| atp6v1e1b1 | 5.584850 | 0.439128779 | 0.233 | 0.087 | 1.4188312 | 9 | atp6v1e1b |
| arpc5b1 | 9.222901 | 0.468321078 | 0.294 | 0.127 | 2.3430782 | 9 | arpc5b |
| wasb2 | 1.248133 | 0.485815494 | 0.351 | 0.168 | 3.1708826 | 9 | wasb |
| gm2a1 | 2.312867 | 0.457535411 | 0.279 | 0.117 | 5.8758389 | 9 | gm2a |
| rpl364 | 5.076646 | 0.266038357 | 0.941 | 0.803 | 1.2897220 | 9 | rpl36 |
| npc21 | 5.883014 | 0.510211109 | 0.519 | 0.338 | 1.4945798 | 9 | npc2 |
| rpl184 | 9.296534 | 0.270047547 | 0.935 | 0.789 | 2.3617845 | 9 | rpl18 |
| si:ch211-15b10.62 | 1.190110 | 0.427773285 | 0.204 | 0.073 | 3.0234747 | 9 | si:ch211-15b10.6 |
| rps215 | 1.386215 | 0.275347524 | 0.916 | 0.772 | 3.5216812 | 9 | rps21 |
| kctd12.21 | 2.289867 | 0.441040166 | 0.244 | 0.097 | 5.8174089 | 9 | kctd12.2 |
| cxcr3.21 | 6.300413 | 0.301894541 | 0.128 | 0.033 | 1.6006201 | 9 | cxcr3.2 |
| naga1 | 8.254797 | 0.365420841 | 0.168 | 0.053 | 2.0971312 | 9 | naga |
| rpl154 | 6.665283 | 0.261650896 | 0.895 | 0.76 | 1.6933152 | 9 | rpl15 |
| rps244 | 1.148559 | 0.259233483 | 0.924 | 0.775 | 2.9179155 | 9 | rps24 |
| ctsk1 | 1.168435 | 0.449700547 | 0.193 | 0.071 | 2.9684097 | 9 | ctsk |
| si:ch211-212k18.71 | 1.440590 | 0.447874821 | 0.3 | 0.142 | 3.6598197 | 9 | si:ch211-212k18.7 |
| arl6ip11 | 1.745532 | 0.45714842 | 0.519 | 0.355 | 4.4345253 | 9 | arl6ip1 |
| sec61g1 | 3.838258 | 0.27371031 | 0.784 | 0.679 | 9.7510967 | 9 | sec61g |
| rps54 | 9.340194 | 0.278562132 | 0.878 | 0.725 | 2.3728763 | 9 | rps5 |

| | | | | | | | |
|---|---|---|---|---|---|---|---|
| faua4 | 1.402548 | 0.277396311 | 0.866 | 0.714 | 3.5631733 | 9 | faua |
| si:busm1-194e12.111 | 3.195668 | 0.250670945 | 0.103 | 0.025 | 8.1185954 | 9 | si:busm1-194e12.11 |
| cebpa2 | 2.235827 | 0.3793478 | 0.191 | 0.073 | 5.6801209 | 9 | cebpa |
| tcima1 | 5.585065 | 0.366552669 | 0.185 | 0.069 | 1.4188860 | 9 | tcima |
| rps154 | 1.868161 | 0.278500292 | 0.88 | 0.712 | 4.7460631 | 9 | rps15 |
| CR855311.13 | 2.299097 | 0.365951088 | 0.214 | 0.085 | 5.8408583 | 9 | CR855311.1 |
| LOC1104377311 | 3.188032 | 0.399384492 | 0.218 | 0.088 | 8.0991962 | 9 | LOC110437731 |
| rps164 | 3.478788 | 0.260557242 | 0.916 | 0.762 | 8.8378625 | 9 | rps16 |
| ctsa1 | 8.852850 | 0.386768629 | 0.216 | 0.094 | 2.2490666 | 9 | ctsa |
| pepd1 | 1.783618 | 0.354362379 | 0.185 | 0.073 | 4.5312817 | 9 | pepd |
| coro1a2 | 2.285617 | 0.387629938 | 0.42 | 0.232 | 5.8066106 | 9 | coro1a |
| rac22 | 3.386572 | 0.410543652 | 0.38 | 0.221 | 8.6035882 | 9 | rac2 |
| cd9b2 | 8.410846 | 0.475018364 | 0.158 | 0.058 | 2.1367755 | 9 | cd9b |
| zgc:1627304 | 9.133842 | 0.392520549 | 0.611 | 0.397 | 2.3204527 | 9 | zgc:162730 |
| enpp11 | 2.289619 | 0.266277191 | 0.12 | 0.037 | 5.8167777 | 9 | enpp1 |
| arpc22 | 2.516654 | 0.408410198 | 0.435 | 0.296 | 6.3935598 | 9 | arpc2 |
| gabarapb2 | 2.581883 | 0.423456285 | 0.433 | 0.302 | 6.5592753 | 9 | gabarapb |
| lipf1 | 1.104554 | 0.332759028 | 0.132 | 0.045 | 2.8061207 | 9 | lipf |
| rpl94 | 1.788034 | 0.281756829 | 0.832 | 0.664 | 4.5425020 | 9 | rpl9 |
| itm2bb1 | 1.891575 | 0.391507309 | 0.391 | 0.239 | 4.8055465 | 9 | itm2bb |
| cd634 | 2.618389 | 0.388821269 | 0.502 | 0.371 | 6.6520183 | 9 | cd63 |
| rap1b1 | 8.358169 | 0.42034017 | 0.349 | 0.217 | 2.1233930 | 9 | rap1b |
| crip13 | 2.974878 | 0.379696355 | 0.433 | 0.281 | 7.5576791 | 9 | crip1 |
| zgc:1141884 | 5.741328 | 0.264876762 | 0.815 | 0.664 | 1.4585845 | 9 | zgc:114188 |
| pycard2 | 6.572150 | 0.340003838 | 0.185 | 0.082 | 1.6696547 | 9 | pycard |
| cap12 | 2.744397 | 0.397240516 | 0.376 | 0.243 | 6.9721415 | 9 | cap1 |
| mapre1a1 | 3.180128 | 0.37141075 | 0.332 | 0.193 | 8.0791160 | 9 | mapre1a |
| dnase21 | 4.446151 | 0.255147929 | 0.12 | 0.041 | 1.1295447 | 9 | dnase2 |
| fuca21 | 1.072489 | 0.304638898 | 0.16 | 0.067 | 2.7246594 | 9 | fuca2 |
| zgc:564932 | 1.784567 | 0.251888739 | 0.687 | 0.589 | 4.5336939 | 9 | zgc:56493 |
| tspan362 | 1.977360 | 0.380495395 | 0.282 | 0.154 | 5.0234847 | 9 | tspan36 |
| ctsh1 | 2.396737 | 0.313355319 | 0.155 | 0.065 | 6.0889117 | 9 | ctsh |
| atp6v1ba1 | 4.621630 | 0.289304657 | 0.139 | 0.055 | 1.1741253 | 9 | atp6v1ba |
| asah1b1 | 5.794545 | 0.290637212 | 0.147 | 0.06 | 1.4721044 | 9 | asah1b |
| ms4a17a.62 | 8.800659 | 0.280590368 | 0.13 | 0.05 | 2.2358074 | 9 | ms4a17a.6 |

| | | | | | | | |
|---|---|---|---|---|---|---|---|
| arpc4l1 | 1.350841 | 0.38224124 | 0.336 | 0.217 | 3.4318122 | 9 | arpc4l |
| eif4ebp11 | 1.654347 | 0.365423116 | 0.38 | 0.256 | 4.2028694 | 9 | eif4ebp1 |
| ctsz1 | 3.042212 | 0.360214687 | 0.189 | 0.091 | 7.7287397 | 9 | ctsz |
| fgl2a1 | 7.731263 | 0.300032653 | 0.166 | 0.074 | 1.9641274 | 9 | fgl2a |
| pdia32 | 1.758431 | 0.343328832 | 0.42 | 0.311 | 4.4672958 | 9 | pdia3 |
| gapdhs3 | 3.165708 | 0.354640727 | 0.416 | 0.314 | 8.0424835 | 9 | gapdhs |
| si:cabz01074946.11 | 3.184970 | 0.277735465 | 0.141 | 0.06 | 8.0914172 | 9 | si:cabz01074946.1 |
| ppdpfb2 | 6.267946 | 0.289483279 | 0.576 | 0.532 | 1.5923717 | 9 | ppdpfb |
| si:dkey-27i16.22 | 6.360096 | 0.309012721 | 0.185 | 0.09 | 1.6157824 | 9 | si:dkey-27i16.2 |
| smox1 | 9.338661 | 0.31139138 | 0.176 | 0.085 | 2.3724869 | 9 | smox |
| si:ch211-66e2.31 | 2.004441 | 0.270889638 | 0.141 | 0.061 | 5.0922848 | 9 | si:ch211-66e2.3 |
| mcl1b3 | 6.939764 | 0.351375182 | 0.357 | 0.249 | 1.7630471 | 9 | mcl1b |
| clic22 | 1.115387 | 0.266305451 | 0.151 | 0.07 | 2.8336412 | 9 | clic2 |
| si:dkey-112e7.22 | 7.573800 | 0.334275161 | 0.311 | 0.213 | 1.9241240 | 9 | si:dkey-112e7.2 |
| arpc32 | 8.245487 | 0.308752772 | 0.431 | 0.333 | 2.0947661 | 9 | arpc3 |
| cyba1 | 1.106387 | 0.267079901 | 0.162 | 0.081 | 2.8107770 | 9 | cyba |
| btg13 | 1.121750 | 0.311087409 | 0.447 | 0.354 | 2.8498063 | 9 | btg1 |
| zgc:640512 | 3.688494 | 0.298573833 | 0.263 | 0.161 | 9.3706214 | 9 | zgc:64051 |
| si:ch211-198c19.33 | 7.553112 | 0.29883684 | 0.162 | 0.081 | 1.9188681 | 9 | si:ch211-198c19.3 |
| nccrp11 | 5.636478 | 0.288832327 | 0.296 | 0.195 | 1.4319474 | 9 | nccrp1 |
| glrx1 | 8.640466 | 0.282439676 | 0.41 | 0.325 | 2.1951105 | 9 | glrx |
| slc6a6b1 | 1.057380 | 0.284360887 | 0.172 | 0.096 | 2.6862742 | 9 | slc6a6b |
| XLOC-0234922 | 1.804502 | 0.270585524 | 0.168 | 0.092 | 4.5843395 | 9 | XLOC-023492 |
| suclg11 | 3.517361 | 0.289411647 | 0.2 | 0.121 | 8.9358581 | 9 | suclg1 |
| cebpb3 | 5.313131 | 0.258466958 | 0.452 | 0.337 | 0.00135 | 9 | cebpb |
| lgmn1 | 8.456343 | 0.255816608 | 0.132 | 0.073 | 0.002148 | 9 | lgmn |
| sdcbp22 | 1.030470 | 0.28163217 | 0.298 | 0.227 | 0.002618 | 9 | sdcbp2 |
| eef1db2 | 1.180871 | 0.278830254 | 0.349 | 0.284 | 0.003 | 9 | eef1db |
| prdx52 | 1.083584 | 0.257811067 | 0.244 | 0.178 | 0.027528 | 9 | prdx5 |
| ccl33.3 | 0 | 2.761999806 | 0.953 | 0.035 | 0 | 10 | ccl33.3 |
| fcer1gl2 | 0 | 1.541155789 | 0.902 | 0.163 | 0 | 10 | fcer1gl |
| vaspa1 | 0 | 1.354488182 | 0.681 | 0.065 | 0 | 10 | vaspa |
| sftpba | 0 | 1.34786888 | 0.604 | 0.019 | 0 | 10 | sftpba |
| fcer1g | 0 | 0.953300588 | 0.446 | 0.033 | 0 | 10 | fcer1g |
| si:ch211-220f21.3 | 0 | 0.775455173 | 0.288 | 0.001 | 0 | 10 | si:ch211-220f21.3 |

| | | | | | | | |
|---|---|---|---|---|---|---|---|
| si:ch73-380l3.4 | 0 | 0.63502937 | 0.251 | 0.007 | 0 | 10 | si:ch73-380l3.4 |
| dicp1.1 | 0 | 0.631109262 | 0.259 | 0.008 | 0 | 10 | dicp1.1 |
| FO393424.2 | 0 | 0.470670249 | 0.174 | 0.002 | 0 | 10 | FO393424.2 |
| LOC108190565 | 0 | 0.431448757 | 0.161 | 0.001 | 0 | 10 | LOC108190565 |
| si:dkey-222f2.7 | 2.361301 | 0.868245452 | 0.361 | 0.023 | 5.9988871 | 10 | si:dkey-222f2.7 |
| LOC110438870 | 1.872080 | 0.329035731 | 0.121 | 0.001 | 4.7560208 | 10 | LOC110438870 |
| npsn | 2.931818 | 0.824093369 | 0.369 | 0.026 | 7.4482860 | 10 | npsn |
| coro1a3 | 3.069861 | 1.337045088 | 0.881 | 0.219 | 7.7989840 | 10 | coro1a |
| CABZ01046427.2 | 1.214562 | 0.418229728 | 0.172 | 0.004 | 3.0855967 | 10 | CABZ01046427.2 |
| CT573231.1 | 2.099047 | 0.717338548 | 0.309 | 0.018 | 5.3326304 | 10 | CT573231.1 |
| XLOC-018356 | 3.150531 | 0.859878823 | 0.325 | 0.021 | 8.0039257 | 10 | XLOC-018356 |
| si:ch211-102c2.42 | 3.454460 | 1.274217263 | 0.702 | 0.131 | 8.7760561 | 10 | si:ch211-102c2.4 |
| arpc1b3 | 5.468834 | 1.330074117 | 0.929 | 0.297 | 1.3893574 | 10 | arpc1b |
| pfn13 | 2.152856 | 1.265989651 | 1 | 0.489 | 5.4693326 | 10 | pfn1 |
| si:ch211-215e19.4 | 2.918543 | 0.305851663 | 0.119 | 0.001 | 7.4145598 | 10 | si:ch211-215e19.4 |
| lcp2b | 2.938655 | 0.543415645 | 0.222 | 0.01 | 7.4656551 | 10 | lcp2b |
| LOC101882805 | 1.322524 | 0.341027215 | 0.132 | 0.002 | 3.3598735 | 10 | LOC101882805 |
| LOC100535743 | 3.576779 | 0.410335472 | 0.156 | 0.004 | 9.0868076 | 10 | LOC100535743 |
| arpc5b2 | 1.410288 | 1.150478528 | 0.623 | 0.118 | 3.5828390 | 10 | arpc5b |
| ccl35.12 | 4.461411 | 1.124960267 | 0.52 | 0.072 | 1.1334216 | 10 | ccl35.1 |
| cotl13 | 4.455901 | 1.232423734 | 0.887 | 0.305 | 1.1320217 | 10 | cotl1 |
| kctd12.22 | 2.071742 | 1.029644639 | 0.546 | 0.089 | 5.2632620 | 10 | kctd12.2 |
| mef2b | 2.112483 | 0.482637032 | 0.198 | 0.009 | 5.3667641 | 10 | mef2b |
| si:dkey-172o19.2 | 4.826316 | 0.402580917 | 0.153 | 0.005 | 1.2261256 | 10 | si:dkey-172o19.2 |
| wipf1a1 | 6.171651 | 0.886120035 | 0.443 | 0.058 | 1.5679080 | 10 | wipf1a |
| LOC1005362131 | 3.429475 | 1.033042635 | 0.57 | 0.099 | 8.7125817 | 10 | LOC100536213 |
| si:dkey-33i11.9 | 6.630512 | 0.623624893 | 0.266 | 0.019 | 1.6844816 | 10 | si:dkey-33i11.9 |
| ephx2 | 1.890781 | 0.72268492 | 0.327 | 0.031 | 4.8035300 | 10 | ephx2 |
| spi1b2 | 8.559047 | 0.905633676 | 0.446 | 0.062 | 2.1744259 | 10 | spi1b |
| si:ch211-214k5.3 | 4.960879 | 0.459018551 | 0.19 | 0.01 | 1.2603115 | 10 | si:ch211-214k5.3 |
| actb23 | 1.016552 | 0.677052655 | 0.992 | 0.864 | 2.5825508 | 10 | actb2 |
| abi3a1 | 1.036398 | 0.727285128 | 0.348 | 0.039 | 2.6329696 | 10 | abi3a |
| tmsb4x6 | 1.433747 | 0.776982264 | 0.997 | 0.712 | 3.6424363 | 10 | tmsb4x |
| gpr132b | 2.247158 | 0.689819346 | 0.29 | 0.029 | 5.7089066 | 10 | gpr132b |
| si:ch211-214p16.11 | 1.229418 | 1.1818058 | 0.813 | 0.229 | 3.1233371 | 10 | si:ch211-214p16.1 |

| | | | | | | | |
|---|---|---|---|---|---|---|---|
| pou2f2a | 3.105359 | 0.350782343 | 0.142 | 0.006 | 7.8891647 | 10 | pou2f2a |
| ptpn62 | 6.305248 | 0.95290126 | 0.485 | 0.089 | 1.6018483 | 10 | ptpn6 |
| itga1 | 4.189305 | 0.731335145 | 0.311 | 0.036 | 1.0642930 | 10 | itga1 |
| actb12 | 1.093300 | 0.653540188 | 0.979 | 0.762 | 2.7775299 | 10 | actb1 |
| grap2b2 | 1.148178 | 0.931502771 | 0.536 | 0.114 | 2.9169465 | 10 | grap2b |
| fam49ba1 | 3.573199 | 0.880653477 | 0.485 | 0.095 | 9.0777132 | 10 | fam49ba |
| cd74a3 | 5.894256 | 0.990104901 | 0.741 | 0.203 | 1.4974358 | 10 | cd74a |
| si:ch73-264p11.11 | 5.331794 | 0.526736183 | 0.237 | 0.023 | 1.3545424 | 10 | si:ch73-264p11.1 |
| hcls11 | 2.346617 | 0.777306723 | 0.401 | 0.068 | 5.9615827 | 10 | hcls1 |
| arpc33 | 2.640886 | 0.974854443 | 0.778 | 0.323 | 6.7091711 | 10 | arpc3 |
| eomesa | 2.258366 | 0.631420716 | 0.288 | 0.034 | 5.7373811 | 10 | eomesa |
| cd74b3 | 3.838950 | 0.997692325 | 0.731 | 0.213 | 9.7528543 | 10 | cd74b |
| arpc23 | 2.154220 | 0.962612803 | 0.723 | 0.289 | 5.4727971 | 10 | arpc2 |
| ptmaa4 | 2.165776 | 0.88410412 | 0.852 | 0.423 | 5.5021554 | 10 | ptmaa |
| rnaset2l2 | 9.844478 | 0.609439557 | 0.306 | 0.043 | 2.5009898 | 10 | rnaset2l |
| arpc42 | 1.654696 | 0.934127593 | 0.657 | 0.227 | 4.2037574 | 10 | arpc4 |
| CU972454.4 | 1.007029 | 0.469453387 | 0.193 | 0.018 | 2.5583578 | 10 | CU972454.4 |
| wasa1 | 1.885134 | 0.768911532 | 0.422 | 0.088 | 4.7891846 | 10 | wasa |
| CU855821.1 | 3.175773 | 0.387714645 | 0.145 | 0.01 | 8.0680531 | 10 | CU855821.1 |
| zdhhc23a | 1.104073 | 0.301771745 | 0.124 | 0.007 | 2.8048991 | 10 | zdhhc23a |
| yrk | 2.057326 | 0.668548686 | 0.343 | 0.06 | 5.2266377 | 10 | yrk |
| rac23 | 2.647715 | 0.874738693 | 0.644 | 0.215 | 6.7265211 | 10 | rac2 |
| ckbb4 | 1.415189 | 0.938349284 | 0.668 | 0.241 | 3.5952876 | 10 | ckbb |
| CABZ01046432.1 | 2.138048 | 0.376226564 | 0.161 | 0.013 | 5.4317131 | 10 | CABZ01046432.1 |
| cfl12 | 6.042618 | 0.701942837 | 0.889 | 0.594 | 1.5351272 | 10 | cfl1 |
| CR381673.11 | 7.308879 | 0.632777988 | 0.298 | 0.051 | 1.8568209 | 10 | CR381673.1 |
| washc2c | 1.997043 | 0.529051153 | 0.243 | 0.035 | 5.0734894 | 10 | washc2c |
| myl12.12 | 2.441797 | 0.785074667 | 0.776 | 0.418 | 6.2033854 | 10 | myl12.1 |
| arpc4l2 | 9.769162 | 0.843033817 | 0.586 | 0.21 | 2.4818557 | 10 | arpc4l |
| CR753876.11 | 3.134043 | 0.760769517 | 0.425 | 0.104 | 7.9620367 | 10 | CR753876.1 |
| BX120005.1 | 5.099518 | 0.588507519 | 0.256 | 0.041 | 1.2955326 | 10 | BX120005.1 |
| lcp12 | 2.788987 | 0.654217948 | 0.351 | 0.078 | 7.0854234 | 10 | lcp1 |
| si:ch73-380l3.2 | 5.246684 | 0.449217684 | 0.195 | 0.026 | 1.3329201 | 10 | si:ch73-380l3.2 |
| zgc:640513 | 1.002928 | 0.785227394 | 0.488 | 0.155 | 2.5479397 | 10 | zgc:64051 |
| arhgdig2 | 8.454854 | 0.783991691 | 0.617 | 0.251 | 2.1479557 | 10 | arhgdig |

| | | | | | | | |
|---|---|---|---|---|---|---|---|
| si:ch211-214p16.2 | 2.374894 | 0.641318291 | 0.317 | 0.069 | 6.0334203 | 10 | si:ch211-214p16.2 |
| rhogb1 | 2.503436 | 0.661501973 | 0.383 | 0.102 | 6.3599797 | 10 | rhogb |
| bpifcl | 1.627894 | 0.286313676 | 0.129 | 0.012 | 4.1356666 | 10 | bpifcl |
| rplp24 | 3.001528 | 0.423967641 | 0.979 | 0.778 | 7.6253827 | 10 | rplp2 |
| cnpy3 | 1.404231 | 0.503356807 | 0.253 | 0.051 | 3.5674494 | 10 | cnpy3 |
| XLOC-0268361 | 2.048554 | 0.626592964 | 0.314 | 0.073 | 5.2043517 | 10 | XLOC-026836 |
| itgb21 | 3.085896 | 0.700737749 | 0.364 | 0.103 | 7.8397193 | 10 | itgb2 |
| capza1b1 | 1.018675 | 0.72274888 | 0.549 | 0.227 | 2.5879457 | 10 | capza1b |
| AL954142.1 | 4.124351 | 0.373585697 | 0.161 | 0.022 | 1.0477915 | 10 | AL954142.1 |
| si:busm1-266f07.22 | 5.142061 | 0.676218865 | 0.467 | 0.14 | 1.3063407 | 10 | si:busm1-266f07.2 |
| ppp1r182 | 3.549267 | 0.491820271 | 0.251 | 0.051 | 9.0169146 | 10 | ppp1r18 |
| ptprc2 | 6.325244 | 0.64498496 | 0.383 | 0.119 | 1.6069283 | 10 | ptprc |
| capzb2 | 2.214624 | 0.686609308 | 0.573 | 0.265 | 5.6262531 | 10 | capzb |
| skap21 | 1.336848 | 0.522389729 | 0.298 | 0.076 | 3.3962637 | 10 | skap2 |
| laptm53 | 2.534299 | 0.635010777 | 0.464 | 0.169 | 6.4383870 | 10 | laptm5 |
| sftpbb2 | 3.779155 | 0.499568665 | 0.274 | 0.066 | 9.6009435 | 10 | sftpbb |
| h1fx1 | 1.317734 | 0.737611371 | 0.472 | 0.184 | 3.3477037 | 10 | h1fx |
| il7r1 | 2.808541 | 0.27466993 | 0.124 | 0.016 | 7.1351003 | 10 | il7r |
| gm2a2 | 4.383086 | 0.596837731 | 0.367 | 0.115 | 1.1135231 | 10 | gm2a |
| stk17b1 | 5.159604 | 0.57094509 | 0.354 | 0.109 | 1.3107974 | 10 | stk17b |
| fgd4a | 1.808849 | 0.260013773 | 0.116 | 0.015 | 4.5953830 | 10 | fgd4a |
| rgs131 | 5.600736 | 0.61686193 | 0.367 | 0.12 | 1.4228670 | 10 | rgs13 |
| mapre1a2 | 2.480716 | 0.597993718 | 0.483 | 0.19 | 6.3022608 | 10 | mapre1a |
| ppiab2 | 6.114911 | 0.413442806 | 0.868 | 0.671 | 1.5534933 | 10 | ppiab |
| tln11 | 1.012580 | 0.588474239 | 0.38 | 0.137 | 2.5724600 | 10 | tln1 |
| csk1 | 2.856706 | 0.52993905 | 0.325 | 0.099 | 7.2574628 | 10 | csk |
| hck | 3.040193 | 0.344234922 | 0.169 | 0.031 | 7.7236125 | 10 | hck |
| ptmab3 | 9.611785 | 0.51230957 | 0.842 | 0.564 | 2.4418742 | 10 | ptmab |
| dtnbp1a1 | 1.969661 | 0.523545415 | 0.29 | 0.084 | 5.0039256 | 10 | dtnbp1a |
| cnn21 | 3.825342 | 0.592176115 | 0.401 | 0.152 | 9.7182832 | 10 | cnn2 |
| ipcef1 | 1.136798 | 0.326884106 | 0.142 | 0.023 | 2.8880357 | 10 | ipcef1 |
| zgc:1037002 | 2.876132 | 0.444115541 | 0.222 | 0.051 | 7.3068158 | 10 | zgc:103700 |
| abracl1 | 1.115529 | 0.61262048 | 0.493 | 0.216 | 2.8340036 | 10 | abracl |
| ikzf21 | 2.785557 | 0.335448696 | 0.166 | 0.031 | 7.0767094 | 10 | ikzf2 |
| si:ch211-243a20.4 | 6.859092 | 0.255718527 | 0.121 | 0.018 | 1.7425525 | 10 | si:ch211-243a20.4 |

| | | | | | | | |
|---|---|---|---|---|---|---|---|
| si:zfos-741a10.31 | 1.128760 | 0.337170685 | 0.156 | 0.028 | 2.8676168 | 10 | si:zfos-741a10.3 |
| dok21 | 2.253277 | 0.372380361 | 0.179 | 0.037 | 5.7244507 | 10 | dok2 |
| syk1 | 2.466918 | 0.433891351 | 0.269 | 0.074 | 6.2672067 | 10 | syk |
| zgc:1538671 | 3.592560 | 0.541403773 | 0.704 | 0.46 | 9.1269012 | 10 | zgc:153867 |
| si:ch211-277c7.6 | 3.690284 | 0.402460295 | 0.172 | 0.034 | 9.3751676 | 10 | si:ch211-277c7.6 |
| myo1f1 | 6.449895 | 0.386031938 | 0.193 | 0.045 | 1.6385959 | 10 | myo1f |
| plekho21 | 9.085718 | 0.446096191 | 0.253 | 0.073 | 2.3082266 | 10 | plekho2 |
| cap13 | 4.961343 | 0.599010987 | 0.499 | 0.24 | 1.2604292 | 10 | cap1 |
| ppiaa3 | 6.504473 | 0.285256291 | 0.945 | 0.812 | 1.6524615 | 10 | ppiaa |
| zgc:1627305 | 7.529452 | 0.58251859 | 0.72 | 0.396 | 1.9128574 | 10 | zgc:162730 |
| rgs181 | 1.997251 | 0.333237704 | 0.153 | 0.031 | 5.0740162 | 10 | rgs18 |
| mir223 | 2.732120 | 0.326842045 | 0.156 | 0.032 | 6.9409530 | 10 | mir223 |
| ptprea | 6.145992 | 0.301482828 | 0.135 | 0.025 | 1.5613893 | 10 | ptprea |
| LOC1039091152 | 9.853257 | 0.52399406 | 0.393 | 0.164 | 2.5032200 | 10 | LOC103909115 |
| sla1 | 2.056393 | 0.273923445 | 0.124 | 0.022 | 5.2242670 | 10 | sla1 |
| zgc:1749451 | 2.549080 | 0.274395742 | 0.124 | 0.022 | 6.4759380 | 10 | zgc:174945 |
| mcl1a2 | 7.412532 | 0.412636405 | 0.794 | 0.61 | 1.8831537 | 10 | mcl1a |
| XLOC-0320021 | 4.517863 | 0.37905145 | 0.174 | 0.041 | 1.1477632 | 10 | XLOC-032002 |
| LOC1018842671 | 2.316460 | 0.36411797 | 0.19 | 0.048 | 5.8849670 | 10 | LOC101884267 |
| srgn2 | 2.433989 | 0.514904999 | 0.496 | 0.229 | 6.1835507 | 10 | srgn |
| hmgn22 | 3.463015 | 0.606189989 | 0.715 | 0.463 | 8.7977916 | 10 | hmgn2 |
| rps294 | 3.564081 | 0.271136319 | 0.968 | 0.842 | 9.0545485 | 10 | rps29 |
| tagapb1 | 1.159676 | 0.398562085 | 0.224 | 0.066 | 2.9461587 | 10 | tagapb |
| rpl263 | 1.752448 | 0.260732964 | 0.976 | 0.848 | 4.4520946 | 10 | rpl26 |
| grasp1 | 9.011321 | 0.388631985 | 0.23 | 0.069 | 2.2893263 | 10 | grasp |
| si:ch211-153b23.7 | 3.373895 | 0.252871398 | 0.116 | 0.022 | 8.5713814 | 10 | si:ch211-153b23.7 |
| idh2 | 6.222598 | 0.539789478 | 0.441 | 0.223 | 1.5808510 | 10 | idh2 |
| rab441 | 1.193034 | 0.370764345 | 0.203 | 0.059 | 3.0309047 | 10 | rab44 |
| wdr11 | 1.424037 | 0.50799092 | 0.354 | 0.152 | 3.6177678 | 10 | wdr1 |
| rps172 | 2.376085 | 0.282688293 | 0.918 | 0.776 | 6.0364439 | 10 | rps17 |
| arl11 | 4.917478 | 0.27705354 | 0.119 | 0.023 | 1.2492855 | 10 | arl11 |
| mhc2dab2 | 6.714278 | 0.399490997 | 0.272 | 0.091 | 1.7057624 | 10 | mhc2dab |
| myh11a1 | 1.509464 | 0.365878537 | 0.222 | 0.068 | 3.8347956 | 10 | myh11a |
| nptna | 2.944879 | 0.354256648 | 0.216 | 0.065 | 7.4814651 | 10 | nptna |
| XLOC-0254232 | 3.613176 | 0.495775684 | 0.393 | 0.166 | 9.1792743 | 10 | XLOC-025423 |

| | | | | | | | |
|---|---|---|---|---|---|---|---|
| mob1a2 | 1.822918 | 0.48208383 | 0.369 | 0.169 | 4.6311239 | 10 | mob1a |
| itm2bb2 | 1.910363 | 0.49790048 | 0.464 | 0.238 | 4.8532786 | 10 | itm2bb |
| fybb1 | 9.403090 | 0.307345015 | 0.158 | 0.042 | 2.3888550 | 10 | fybb |
| gapdhs4 | 1.592867 | 0.475559876 | 0.533 | 0.311 | 4.0466802 | 10 | gapdhs |
| pnrc23 | 1.638616 | 0.371436702 | 0.792 | 0.594 | 4.1629054 | 10 | pnrc2 |
| rpl144 | 1.682909 | 0.339874085 | 0.871 | 0.665 | 4.2754317 | 10 | rpl14 |
| il10ra1 | 2.009542 | 0.295931693 | 0.153 | 0.04 | 5.1052435 | 10 | il10ra |
| lasp12 | 3.094349 | 0.48687605 | 0.454 | 0.249 | 7.8611959 | 10 | lasp1 |
| CABZ01109481.1 | 4.735335 | 0.270935862 | 0.121 | 0.027 | 1.2030120 | 10 | CABZ01109481.1 |
| lrrfip1a | 7.286022 | 0.340263109 | 0.19 | 0.058 | 1.8510140 | 10 | lrrfip1a |
| eif3f3 | 2.338372 | 0.410895894 | 0.673 | 0.462 | 5.9406361 | 10 | eif3f |
| rps27.15 | 5.336919 | 0.322713745 | 0.892 | 0.706 | 1.3558444 | 10 | rps27.1 |
| XLOC-0283702 | 9.512878 | 0.455391582 | 0.301 | 0.122 | 2.4167468 | 10 | XLOC-028370 |
| calm1b1 | 1.159416 | 0.39383426 | 0.261 | 0.1 | 2.9454967 | 10 | calm1b |
| mfng2 | 2.305451 | 0.284034191 | 0.153 | 0.043 | 5.8570006 | 10 | mfng |
| nr4a12 | 3.759808 | 0.473934676 | 0.335 | 0.155 | 9.5517922 | 10 | nr4a1 |
| LOC1081907171 | 3.840244 | 0.366442213 | 0.227 | 0.08 | 9.7561399 | 10 | LOC108190717 |
| ccni2 | 1.040258 | 0.453171886 | 0.507 | 0.316 | 2.6427767 | 10 | ccni |
| cfbl | 1.094128 | 0.279347093 | 0.208 | 0.068 | 2.7796324 | 10 | cfbl |
| RASSF51 | 1.112482 | 0.32557851 | 0.182 | 0.059 | 2.8262621 | 10 | RASSF5 |
| si:ch211-66e2.32 | 1.683393 | 0.338202868 | 0.185 | 0.06 | 4.2766609 | 10 | si:ch211-66e2.3 |
| si:ch211-119e14.11 | 3.780791 | 0.322522434 | 0.201 | 0.068 | 9.6051017 | 10 | si:ch211-119e14.1 |
| arhgdia2 | 4.768225 | 0.448799881 | 0.438 | 0.248 | 1.2113676 | 10 | arhgdia |
| cdc42l2 | 1.180880 | 0.440736503 | 0.417 | 0.23 | 3.0000276 | 10 | cdc42l |
| wasb3 | 1.763550 | 0.416269069 | 0.356 | 0.169 | 4.4802996 | 10 | wasb |
| XLOC-0203011 | 2.115314 | 0.356445838 | 0.198 | 0.068 | 5.3739555 | 10 | XLOC-020301 |
| sh3bgrl | 7.526381 | 0.399367858 | 0.269 | 0.118 | 1.9120772 | 10 | sh3bgrl |
| rpl225 | 1.098936 | 0.286032934 | 0.879 | 0.718 | 2.7918473 | 10 | rpl22 |
| crip14 | 1.251939 | 0.406703656 | 0.507 | 0.28 | 3.1805526 | 10 | crip1 |
| cyba2 | 4.217582 | 0.338298984 | 0.211 | 0.08 | 1.0714767 | 10 | cyba |
| ucp22 | 4.511667 | 0.393620103 | 0.602 | 0.411 | 1.1461891 | 10 | ucp2 |
| tagapa1 | 6.448013 | 0.379069398 | 0.237 | 0.095 | 1.6381177 | 10 | tagapa |
| naca4 | 2.183087 | 0.317031032 | 0.847 | 0.648 | 5.5461343 | 10 | naca |
| si:dkey-262k9.42 | 2.590885 | 0.336996877 | 0.222 | 0.088 | 6.5821441 | 10 | si:dkey-262k9.4 |
| si:rp71-68n21.91 | 3.455061 | 0.259036969 | 0.145 | 0.045 | 8.7775845 | 10 | si:rp71-68n21.9 |

| | | | | | | | |
|---|---|---|---|---|---|---|---|
| cdkn1bb1 | 4.757829 | 0.398089447 | 0.309 | 0.151 | 1.2087266 | 10 | cdkn1bb |
| parvg1 | 6.603995 | 0.315470771 | 0.193 | 0.071 | 1.6777451 | 10 | parvg |
| cox8b1 | 8.684785 | 0.367361392 | 0.235 | 0.099 | 2.2063696 | 10 | cox8b |
| pycard3 | 9.876570 | 0.339743014 | 0.208 | 0.082 | 2.5091427 | 10 | pycard |
| mob3a | 2.311828 | 0.400849775 | 0.314 | 0.156 | 5.8732006 | 10 | mob3a |
| cox8a1 | 2.774024 | 0.414101085 | 0.43 | 0.265 | 7.0474086 | 10 | cox8a |
| glipr1a2 | 3.487254 | 0.291282661 | 0.182 | 0.066 | 8.8593689 | 10 | glipr1a |
| ccr21 | 4.854424 | 0.275527644 | 0.14 | 0.044 | 1.2332664 | 10 | ccr2 |
| rps15a5 | 1.651643 | 0.270248539 | 0.947 | 0.795 | 4.1959995 | 10 | rps15a |
| tpi1b2 | 1.080140 | 0.393119935 | 0.327 | 0.176 | 2.7440975 | 10 | tpi1b |
| rps184 | 2.403411 | 0.272421602 | 0.942 | 0.75 | 6.1058658 | 10 | rps18 |
| clic12 | 3.798220 | 0.355421373 | 0.261 | 0.127 | 9.6493799 | 10 | clic1 |
| actr2a2 | 5.716830 | 0.364591969 | 0.269 | 0.131 | 1.4523608 | 10 | actr2a |
| twf1a1 | 5.718229 | 0.360454457 | 0.303 | 0.155 | 1.4527162 | 10 | twf1a |
| gabarapb3 | 7.750953 | 0.39013818 | 0.459 | 0.303 | 1.9691298 | 10 | gabarapb |
| rap1b2 | 3.026826 | 0.366605534 | 0.375 | 0.218 | 7.6896522 | 10 | rap1b |
| rpl314 | 5.498548 | 0.253217716 | 0.839 | 0.685 | 1.3969063 | 10 | rpl31 |
| BX005183.32 | 6.308422 | 0.295043243 | 0.185 | 0.075 | 1.6026548 | 10 | BX005183.3 |
| rps55 | 1.967080 | 0.254721121 | 0.892 | 0.725 | 4.9973684 | 10 | rps5 |
| scinlb1 | 5.822581 | 0.296751312 | 0.193 | 0.083 | 1.4792268 | 10 | scinlb |
| gng51 | 1.603031 | 0.358149132 | 0.43 | 0.285 | 4.0725008 | 10 | gng5 |
| mbnl11 | 3.189934 | 0.349228129 | 0.325 | 0.185 | 8.1040287 | 10 | mbnl1 |
| pxna | 3.275508 | 0.278099808 | 0.201 | 0.088 | 8.3214289 | 10 | pxna |
| rps104 | 8.072874 | 0.252400711 | 0.9 | 0.723 | 2.0509136 | 10 | rps10 |
| arpc5a2 | 1.451852 | 0.31205576 | 0.243 | 0.124 | 3.6884307 | 10 | arpc5a |
| zgc:564933 | 3.569402 | 0.255160917 | 0.718 | 0.589 | 9.0680658 | 10 | zgc:56493 |
| btg14 | 4.411577 | 0.338074191 | 0.491 | 0.354 | 1.1207611 | 10 | btg1 |
| msna2 | 8.061667 | 0.340208814 | 0.401 | 0.264 | 2.0480665 | 10 | msna |
| capza1a2 | 2.639270 | 0.323786869 | 0.348 | 0.219 | 6.7050665 | 10 | capza1a |
| arrb2b1 | 3.001046 | 0.266917204 | 0.198 | 0.095 | 7.6241581 | 10 | arrb2b |
| si:dkey-112e7.23 | 4.418366 | 0.326154035 | 0.338 | 0.213 | 1.1224860 | 10 | si:dkey-112e7.2 |
| sept61 | 1.760316 | 0.312894957 | 0.327 | 0.2 | 4.4720851 | 10 | sept6 |
| rps26l4 | 2.187721 | 0.255574828 | 0.884 | 0.696 | 5.5579060 | 10 | rps26l |
| cd635 | 2.225483 | 0.293170882 | 0.525 | 0.371 | 5.6538407 | 10 | cd63 |
| ccr9a2 | 3.047397 | 0.250545401 | 0.185 | 0.087 | 7.7419127 | 10 | ccr9a |

| | | | | | | | |
|---|---|---|---|---|---|---|---|
| cox4i11 | 3.422745 | 0.288118058 | 0.544 | 0.416 | 8.6954852 | 10 | cox4i1 |
| mibp23 | 3.610198 | 0.250137163 | 0.633 | 0.52 | 9.1717104 | 10 | mibp2 |
| cox6b12 | 9.530834 | 0.261484107 | 0.227 | 0.121 | 2.4213086 | 10 | cox6b1 |
| rpl5a4 | 1.559996 | 0.252386618 | 0.694 | 0.534 | 3.9631707 | 10 | rpl5a |
| si:ch1073-429i10.3.1 | 9.662450 | 0.311997978 | 0.335 | 0.225 | 2.4547454 | 10 | si:ch1073-429i10.3.1 |
| tpm31 | 2.340420 | 0.29738123 | 0.388 | 0.283 | 5.9458373 | 10 | tpm3 |
| si:ch211-213a13.1 | 3.712637 | 0.251105856 | 0.182 | 0.097 | 9.4319557 | 10 | si:ch211-213a13.1 |
| glo1 | 2.206143 | 0.275273441 | 0.298 | 0.209 | 0.005605 | 10 | glo1 |
| GCA2 | 4.707734 | 0.255453817 | 0.288 | 0.2 | 0.01196 | 10 | GCA |
| CR318588.15 | 5.638271 | 0.288128908 | 0.306 | 0.215 | 0.14324 | 10 | CR318588.1 |
| si:ch211-14a17.10 | 0 | 2.454969546 | 0.978 | 0.055 | 0 | 11 | si:ch211-14a17.10 |
| spink2.21 | 0 | 2.304010788 | 0.952 | 0.093 | 0 | 11 | spink2.2 |
| si:dkey-203a12.21 | 0 | 2.265752438 | 0.997 | 0.188 | 0 | 11 | si:dkey-203a12.2 |
| spink2.3 | 0 | 2.201500088 | 0.919 | 0.042 | 0 | 11 | spink2.3 |
| spink2.4 | 0 | 2.199217227 | 0.879 | 0.043 | 0 | 11 | spink2.4 |
| zmp:00000013231 | 0 | 2.180379851 | 0.952 | 0.104 | 0 | 11 | zmp:0000001323 |
| si:ch211-194m7.8 | 0 | 2.136594075 | 0.925 | 0.031 | 0 | 11 | si:ch211-194m7.8 |
| MFAP4 (1 of many) | 0 | 2.127717823 | 0.906 | 0.038 | 0 | 11 | MFAP4 (1 of many) |
| BX322787.11 | 0 | 2.111459969 | 0.944 | 0.099 | 0 | 11 | BX322787.1 |
| itln1 | 0 | 2.086934781 | 0.917 | 0.033 | 0 | 11 | itln1 |
| ccl34b.9 | 0 | 2.068679146 | 0.895 | 0.026 | 0 | 11 | ccl34b.9 |
| si:dkey-77g12.4 | 0 | 2.058943785 | 0.858 | 0.024 | 0 | 11 | si:dkey-77g12.4 |
| si:ch73-359m17.21 | 0 | 2.054353157 | 0.976 | 0.142 | 0 | 11 | si:ch73-359m17.2 |
| LOC110437886 | 0 | 2.020059711 | 0.887 | 0.013 | 0 | 11 | LOC110437886 |
| si:ch1073-110a20.3 | 0 | 1.984818643 | 0.868 | 0.025 | 0 | 11 | si:ch1073-110a20.3 |
| clca1 | 0 | 1.93449763 | 0.852 | 0.01 | 0 | 11 | clca1 |
| si:dkey-203a12.5 | 0 | 1.893695854 | 0.806 | 0.027 | 0 | 11 | si:dkey-203a12.5 |
| cfbl1 | 0 | 1.884756645 | 0.866 | 0.049 | 0 | 11 | cfbl |
| si:dkeyp-75b4.104 | 0 | 1.87043967 | 0.995 | 0.35 | 0 | 11 | si:dkeyp-75b4.10 |
| LOC110438970 | 0 | 1.83802894 | 0.809 | 0.011 | 0 | 11 | LOC110438970 |
| LOC1104377312 | 0 | 1.789991008 | 0.866 | 0.07 | 0 | 11 | LOC110437731 |
| CABZ01112049.1 | 0 | 1.764155492 | 0.772 | 0.007 | 0 | 11 | CABZ01112049.1 |
| FO203432.1 | 0 | 1.691959712 | 0.753 | 0.021 | 0 | 11 | FO203432.1 |
| cfi | 0 | 1.548222889 | 0.72 | 0.005 | 0 | 11 | cfi |
| cfp2 | 0 | 1.491273299 | 0.745 | 0.058 | 0 | 11 | cfp |

| | | | | | | | |
|---|---|---|---|---|---|---|---|
| c3a.51 | 0 | 1.489208743 | 0.675 | 0.045 | 0 | 11 | c3a.5 |
| si:dkey-203a12.7 | 0 | 1.481763348 | 0.527 | 0.007 | 0 | 11 | si:dkey-203a12.7 |
| spink2.5 | 0 | 1.478950584 | 0.465 | 0.01 | 0 | 11 | spink2.5 |
| agr2 | 0 | 1.472858561 | 0.629 | 0.014 | 0 | 11 | agr2 |
| si:ch1073-376c22.1 | 0 | 1.460474313 | 0.659 | 0.011 | 0 | 11 | si:ch1073-376c22.1 |
| si:zfos-2330d3.1 | 0 | 1.448283661 | 0.578 | 0.013 | 0 | 11 | si:zfos-2330d3.1 |
| cfhl2 | 0 | 1.379139894 | 0.634 | 0.006 | 0 | 11 | cfhl2 |
| BX908782.3 | 0 | 1.266277738 | 0.551 | 0.023 | 0 | 11 | BX908782.3 |
| c3a.4 | 0 | 1.250124665 | 0.581 | 0.004 | 0 | 11 | c3a.4 |
| si:dkey-203a12.4 | 0 | 1.22788379 | 0.36 | 0.008 | 0 | 11 | si:dkey-203a12.4 |
| si:ch211-214p16.21 | 0 | 1.224196419 | 0.618 | 0.06 | 0 | 11 | si:ch211-214p16.2 |
| LOC108179162 | 0 | 1.081823051 | 0.473 | 0.005 | 0 | 11 | LOC108179162 |
| si:dkey-102g19.31 | 0 | 1.071170981 | 0.53 | 0.036 | 0 | 11 | si:dkey-102g19.3 |
| ca6 | 0 | 1.011768017 | 0.449 | 0.008 | 0 | 11 | ca6 |
| si:ch211-212d10.1 | 0 | 0.915717375 | 0.387 | 0.015 | 0 | 11 | si:ch211-212d10.1 |
| LOC101883382 | 0 | 0.876501669 | 0.387 | 0.003 | 0 | 11 | LOC101883382 |
| cfh | 0 | 0.874631332 | 0.384 | 0.007 | 0 | 11 | cfh |
| timp4.2 | 0 | 0.872341891 | 0.376 | 0.006 | 0 | 11 | timp4.2 |
| cdaa1 | 0 | 0.849846686 | 0.398 | 0.024 | 0 | 11 | cdaa |
| LOC103909452 | 0 | 0.848086223 | 0.379 | 0.008 | 0 | 11 | LOC103909452 |
| si:dkey-27h10.2 | 0 | 0.818354322 | 0.371 | 0.018 | 0 | 11 | si:dkey-27h10.2 |
| nfe2 | 0 | 0.754115054 | 0.344 | 0.007 | 0 | 11 | nfe2 |
| cd59 | 0 | 0.746055898 | 0.312 | 0.006 | 0 | 11 | cd59 |
| LOC110438722 | 0 | 0.693000512 | 0.28 | 0.001 | 0 | 11 | LOC110438722 |
| itgb3a | 0 | 0.586816888 | 0.25 | 0.007 | 0 | 11 | itgb3a |
| viml | 0 | 0.564453382 | 0.237 | 0.003 | 0 | 11 | viml |
| osmr | 0 | 0.563541827 | 0.223 | 0.003 | 0 | 11 | osmr |
| si:dkeyp-66d1.7 | 0 | 0.557389309 | 0.231 | 0.006 | 0 | 11 | si:dkeyp-66d1.7 |
| LOC110437920 | 0 | 0.541997115 | 0.22 | 0.001 | 0 | 11 | LOC110437920 |
| cfhl1 | 0 | 0.407875624 | 0.172 | 0.001 | 0 | 11 | cfhl1 |
| BX323555.3 | 0 | 0.402761095 | 0.167 | 0.001 | 0 | 11 | BX323555.3 |
| cxcr2 | 1.391877 | 0.379797098 | 0.161 | 0.002 | 3.5360653 | 11 | cxcr2 |
| pdia2 | 1.084437 | 0.620944607 | 0.263 | 0.01 | 2.7550131 | 11 | pdia2 |
| adgre13 | 8.553323 | 0.457502917 | 0.191 | 0.004 | 2.1729717 | 11 | adgre13 |
| si:ch211-194m7.5 | 2.026791 | 0.372627154 | 0.156 | 0.002 | 5.1490629 | 11 | si:ch211-194m7.5 |

| | | | | | | | |
|---|---|---|---|---|---|---|---|
| h6pd | 4.904511 | 0.567474353 | 0.237 | 0.009 | 1.2459912 | 11 | h6pd |
| cst14b.2 | 4.086582 | 0.718173449 | 0.21 | 0.006 | 1.0381962 | 11 | cst14b.2 |
| si:dkey-203a12.91 | 1.579085 | 1.494064792 | 0.656 | 0.103 | 4.0116668 | 11 | si:dkey-203a12.9 |
| si:ch211-214p16.12 | 5.604157 | 1.382515785 | 0.938 | 0.226 | 1.4237361 | 11 | si:ch211-214p16.1 |
| si:ch211-276a23.5 | 1.813074 | 0.470952502 | 0.207 | 0.008 | 4.6061147 | 11 | si:ch211-276a23.5 |
| CR848841.1 | 3.140850 | 0.572743476 | 0.237 | 0.011 | 7.9793309 | 11 | CR848841.1 |
| si:ch73-160i9.2 | 3.935802 | 0.266559197 | 0.11 | 0.001 | 9.9989050 | 11 | si:ch73-160i9.2 |
| spink2.14 | 2.131726 | 1.347858125 | 0.935 | 0.36 | 5.4156504 | 11 | spink2.1 |
| si:ch211-270n8.1 | 2.395355 | 0.346858463 | 0.14 | 0.003 | 6.0853999 | 11 | si:ch211-270n8.1 |
| gata2a | 1.097199 | 0.571919542 | 0.261 | 0.016 | 2.7874364 | 11 | gata2a |
| sult5a1 | 4.124949 | 0.444409166 | 0.18 | 0.006 | 1.0479433 | 11 | sult5a1 |
| adam8a | 8.254707 | 0.768125367 | 0.398 | 0.041 | 2.0971085 | 11 | adam8a |
| nfkbiaa1 | 1.082464 | 1.21848891 | 0.667 | 0.139 | 2.7500016 | 11 | nfkbiaa |
| si:ch1073-67j19.1 | 1.019471 | 0.407394813 | 0.11 | 0.002 | 2.5899684 | 11 | si:ch1073-67j19.1 |
| clstn1 | 1.472966 | 0.581355275 | 0.258 | 0.017 | 3.7420717 | 11 | clstn1 |
| myofl | 5.025177 | 0.540327125 | 0.247 | 0.017 | 1.2766463 | 11 | myofl |
| BX936393.1 | 1.265571 | 0.327965427 | 0.142 | 0.005 | 3.2151842 | 11 | BX936393.1 |
| myh11a2 | 1.171975 | 0.768708232 | 0.409 | 0.063 | 2.9774045 | 11 | myh11a |
| qsox1 | 7.343609 | 0.548440518 | 0.269 | 0.028 | 1.8656439 | 11 | qsox1 |
| si:ch211-74f19.2 | 3.634576 | 0.560562464 | 0.253 | 0.026 | 9.2336413 | 11 | si:ch211-74f19.2 |
| itgb22 | 3.088939 | 0.831429141 | 0.487 | 0.099 | 7.8474502 | 11 | itgb2 |
| cyp4f3 | 4.611993 | 0.417061248 | 0.185 | 0.014 | 1.1716768 | 11 | cyp4f3 |
| srgn3 | 2.060665 | 0.982752033 | 0.712 | 0.223 | 5.2351207 | 11 | srgn |
| ank3b1 | 3.740857 | 0.592626569 | 0.274 | 0.033 | 9.5036481 | 11 | ank3b |
| cd7al | 4.734794 | 0.479664964 | 0.231 | 0.023 | 1.2028744 | 11 | cd7al |
| ltb4r1 | 8.012116 | 0.499282638 | 0.226 | 0.023 | 2.0354782 | 11 | ltb4r |
| mctp2b | 3.697545 | 0.414977017 | 0.177 | 0.014 | 9.3936147 | 11 | mctp2b |
| spns3 | 9.531836 | 0.358222674 | 0.161 | 0.012 | 2.4215630 | 11 | spns3 |
| lrrc8da | 2.042931 | 0.457517272 | 0.218 | 0.023 | 5.1900665 | 11 | lrrc8da |
| atp1a3a1 | 2.877631 | 0.42837951 | 0.196 | 0.018 | 7.3106227 | 11 | atp1a3a |
| cxcl8b.12 | 7.261386 | 0.62616392 | 0.293 | 0.041 | 1.8447551 | 11 | cxcl8b.1 |
| mmp25b | 2.029652 | 0.279622243 | 0.124 | 0.007 | 5.1563326 | 11 | mmp25b |
| fcer1gl3 | 2.945445 | 0.842584651 | 0.661 | 0.17 | 7.4829045 | 11 | fcer1gl |
| illr4 | 1.891398 | 0.308210504 | 0.129 | 0.008 | 4.8050984 | 11 | illr4 |
| alox5a | 1.128477 | 0.516581196 | 0.234 | 0.028 | 2.8668980 | 11 | alox5a |

| | | | | | | | |
|---|---|---|---|---|---|---|---|
| si:ch73-237c6.12 | 2.815114 | 0.520731407 | 0.272 | 0.041 | 7.1517993 | 11 | si:ch73-237c6.1 |
| CT030188.1 | 3.506317 | 0.579224935 | 0.301 | 0.049 | 8.9077991 | 11 | CT030188.1 |
| coro1a4 | 6.983007 | 0.880818835 | 0.656 | 0.226 | 1.7740330 | 11 | coro1a |
| calm1b2 | 1.088505 | 0.717237716 | 0.419 | 0.095 | 2.7653471 | 11 | calm1b |
| pfn14 | 2.701443 | 0.860314841 | 0.957 | 0.49 | 6.8630176 | 11 | pfn1 |
| cd81a2 | 3.862322 | 0.869008717 | 0.653 | 0.231 | 9.8122314 | 11 | cd81a |
| capgb2 | 9.820288 | 0.730019856 | 0.444 | 0.107 | 2.4948442 | 11 | capgb |
| si:cabz01036022.1 | 3.456064 | 0.31005263 | 0.126 | 0.009 | 8.7801312 | 11 | si:cabz01036022.1 |
| alox5ap | 4.467513 | 0.375729247 | 0.188 | 0.02 | 1.1349717 | 11 | alox5ap |
| cd636 | 1.630212 | 0.821537482 | 0.755 | 0.365 | 4.1415560 | 11 | cd63 |
| LOC1001482541 | 7.529535 | 0.409313844 | 0.188 | 0.021 | 1.9128786 | 11 | LOC100148254 |
| limd2 | 1.960055 | 0.609004362 | 0.325 | 0.062 | 4.9795218 | 11 | limd2 |
| rgcc1 | 5.936973 | 0.602156164 | 0.333 | 0.065 | 1.5082880 | 11 | rgcc |
| furina | 6.183815 | 0.582940591 | 0.312 | 0.059 | 1.5709983 | 11 | furina |
| glipr1a3 | 2.445108 | 0.591742381 | 0.317 | 0.062 | 6.2117983 | 11 | glipr1a |
| def6c | 9.389945 | 0.267129327 | 0.129 | 0.01 | 2.3855155 | 11 | def6c |
| dhrs13a.1 | 6.856028 | 0.4073633 | 0.177 | 0.02 | 1.7417740 | 11 | dhrs13a.1 |
| si:ch1073-443f11.2 | 2.628279 | 0.443066973 | 0.239 | 0.036 | 6.6771441 | 11 | si:ch1073-443f11.2 |
| pygb | 2.901765 | 0.476997838 | 0.25 | 0.04 | 7.3719353 | 11 | pygb |
| slc6a6b2 | 3.117032 | 0.675744252 | 0.379 | 0.09 | 7.9188206 | 11 | slc6a6b |
| si:ch73-248e21.71 | 1.191807 | 0.496514394 | 0.261 | 0.045 | 3.0277864 | 11 | si:ch73-248e21.7 |
| ehd1a | 1.888437 | 0.405517521 | 0.207 | 0.029 | 4.7975752 | 11 | ehd1a |
| selp | 6.551142 | 0.28943114 | 0.121 | 0.01 | 1.6643178 | 11 | selp |
| mir2231 | 3.831854 | 0.454845018 | 0.21 | 0.03 | 9.7348252 | 11 | mir223 |
| tspan2b | 3.112680 | 0.257000783 | 0.118 | 0.01 | 7.9077641 | 11 | tspan2b |
| grap2b3 | 1.116303 | 0.697813412 | 0.419 | 0.117 | 2.8359702 | 11 | grap2b |
| eno1a2 | 1.429299 | 0.714856594 | 0.441 | 0.135 | 3.6311347 | 11 | eno1a |
| cyba3 | 1.080251 | 0.565171789 | 0.331 | 0.076 | 2.7443790 | 11 | cyba |
| si:ch211-136m16.8 | 1.323015 | 0.379103355 | 0.191 | 0.027 | 3.3611197 | 11 | si:ch211-136m16.8 |
| pla2g4aa | 4.395458 | 0.309137824 | 0.14 | 0.015 | 1.1166662 | 11 | pla2g4aa |
| hacd3 | 2.698798 | 0.397626502 | 0.212 | 0.033 | 6.8562981 | 11 | hacd3 |
| slc7a3a | 1.172659 | 0.485833768 | 0.237 | 0.043 | 2.9791422 | 11 | slc7a3a |
| zmp:00000010821 | 3.670050 | 0.459168587 | 0.258 | 0.051 | 9.3237621 | 11 | zmp:0000001082 |
| anks1b | 2.646021 | 0.309039476 | 0.14 | 0.016 | 6.7222177 | 11 | anks1b |
| lpar5b | 7.088938 | 0.373247944 | 0.159 | 0.021 | 1.8009449 | 11 | lpar5b |

| | | | | | | | |
|---|---|---|---|---|---|---|---|
| ssr31 | 8.882793 | 0.740613062 | 0.602 | 0.281 | 2.2566735 | 11 | ssr3 |
| unc13d | 2.381259 | 0.38020582 | 0.177 | 0.027 | 6.0495909 | 11 | unc13d |
| MYADM | 1.761146 | 0.407034506 | 0.207 | 0.037 | 4.4741923 | 11 | MYADM |
| xbp11 | 4.534043 | 0.687558511 | 0.672 | 0.348 | 1.1518737 | 11 | xbp1 |
| myh9a | 6.255844 | 0.426083926 | 0.239 | 0.047 | 1.5892974 | 11 | myh9a |
| ikzf22 | 2.168515 | 0.358006156 | 0.188 | 0.031 | 5.5091148 | 11 | ikzf2 |
| nr4a13 | 2.261758 | 0.630686297 | 0.452 | 0.152 | 5.7459984 | 11 | nr4a1 |
| actb24 | 2.358088 | 0.377542901 | 0.984 | 0.865 | 5.9907246 | 11 | actb2 |
| mt-co13 | 3.712126 | 0.536236557 | 0.892 | 0.634 | 9.4306565 | 11 | mt-co1 |
| samsn1a1 | 3.813898 | 0.42601708 | 0.196 | 0.035 | 9.6892093 | 11 | samsn1a |
| si:dkey-24p1.7 | 2.982312 | 0.253898562 | 0.11 | 0.012 | 7.5765638 | 11 | si:dkey-24p1.7 |
| moxd11 | 1.664709 | 0.264594277 | 0.113 | 0.013 | 4.2291939 | 11 | moxd1 |
| mt-atp63 | 4.470721 | 0.588027504 | 0.812 | 0.551 | 1.1357866 | 11 | mt-atp6 |
| vaspa2 | 4.803863 | 0.465680213 | 0.301 | 0.077 | 1.2204214 | 11 | vaspa |
| lta4h2 | 9.697370 | 0.469829106 | 0.285 | 0.071 | 2.4636168 | 11 | lta4h |
| si:dkey-206d17.121 | 1.772618 | 0.468564413 | 0.22 | 0.047 | 4.5033369 | 11 | si:dkey-206d17.12 |
| ncf1 | 2.771208 | 0.391386672 | 0.175 | 0.03 | 7.0402560 | 11 | ncf1 |
| syngr2a1 | 1.186598 | 0.408076895 | 0.237 | 0.053 | 3.0145532 | 11 | syngr2a |
| p4hb | 2.936919 | 0.644897009 | 0.495 | 0.206 | 7.4612432 | 11 | p4hb |
| grasp2 | 3.792824 | 0.498751324 | 0.266 | 0.068 | 9.6356711 | 11 | grasp |
| camk1b | 8.061373 | 0.292029434 | 0.134 | 0.02 | 2.0479918 | 11 | camk1b |
| m171 | 2.657512 | 0.285715735 | 0.108 | 0.013 | 6.7514098 | 11 | m17 |
| atp2a3 | 3.258161 | 0.340864684 | 0.175 | 0.032 | 8.2773598 | 11 | atp2a3 |
| slc2a3b | 2.571524 | 0.324466375 | 0.151 | 0.025 | 6.5329583 | 11 | slc2a3b |
| zgc:1583432 | 3.944335 | 0.655244287 | 0.427 | 0.164 | 1.0020583 | 11 | zgc:158343 |
| mt-co23 | 1.182335 | 0.469684458 | 0.844 | 0.668 | 3.0037239 | 11 | mt-co2 |
| si:zfos-741a10.32 | 2.394976 | 0.316080933 | 0.159 | 0.028 | 6.0844379 | 11 | si:zfos-741a10.3 |
| rgs191 | 6.833845 | 0.343853165 | 0.169 | 0.033 | 1.7361385 | 11 | rgs19 |
| cotl14 | 5.927800 | 0.619156202 | 0.626 | 0.313 | 1.5059578 | 11 | cotl1 |
| mt-nd43 | 1.178680 | 0.567032049 | 0.726 | 0.442 | 2.9944368 | 11 | mt-nd4 |
| cebpb4 | 2.786178 | 0.606472836 | 0.691 | 0.331 | 7.0782859 | 11 | cebpb |
| mbnl12 | 4.652498 | 0.567859469 | 0.444 | 0.181 | 1.1819673 | 11 | mbnl1 |
| gsr | 1.477433 | 0.350446847 | 0.172 | 0.035 | 3.7534196 | 11 | gsr |
| pim2 | 9.814303 | 0.520964857 | 0.277 | 0.084 | 2.4933237 | 11 | pim2 |
| camk2g2 | 3.271723 | 0.343866336 | 0.172 | 0.036 | 8.3118128 | 11 | camk2g2 |

| | | | | | | | |
|---|---|---|---|---|---|---|---|
| si:ch211-215e19.3 | 2.137501 | 0.256119166 | 0.116 | 0.017 | 5.4303225 | 11 | si:ch211-215e19.3 |
| tagapa2 | 3.212534 | 0.525479128 | 0.296 | 0.093 | 8.1614439 | 11 | tagapa |
| COX34 | 6.103240 | 0.445603858 | 0.855 | 0.673 | 1.5505282 | 11 | COX3 |
| mt-cyb2 | 1.478010 | 0.498071705 | 0.777 | 0.573 | 3.7548857 | 11 | mt-cyb |
| ptpn63 | 4.774826 | 0.441600603 | 0.301 | 0.095 | 1.2130445 | 11 | ptpn6 |
| ccr9a3 | 5.454876 | 0.444519522 | 0.28 | 0.084 | 1.3858113 | 11 | ccr9a |
| si:dkey-73n10.1 | 2.363802 | 0.279869418 | 0.126 | 0.022 | 6.0052401 | 11 | si:dkey-73n10.1 |
| rasal31 | 4.690953 | 0.267313867 | 0.137 | 0.025 | 1.1917367 | 11 | rasal3 |
| CABZ01073834.11 | 1.376265 | 0.309093665 | 0.18 | 0.042 | 3.4964029 | 11 | CABZ01073834.1 |
| msna3 | 2.864262 | 0.549978471 | 0.513 | 0.26 | 7.2766577 | 11 | msna |
| si:ch73-208g10.11 | 2.866090 | 0.326005809 | 0.169 | 0.038 | 7.2813027 | 11 | si:ch73-208g10.1 |
| rac24 | 4.385514 | 0.520023192 | 0.495 | 0.219 | 1.1141400 | 11 | rac2 |
| far1 | 5.101139 | 0.318789208 | 0.159 | 0.034 | 1.2959444 | 11 | far1 |
| cxcr4b2 | 1.995425 | 0.490363982 | 0.43 | 0.172 | 5.0693793 | 11 | cxcr4b |
| hck1 | 5.087008 | 0.283301581 | 0.148 | 0.031 | 1.2923544 | 11 | hck |
| fybb2 | 7.180512 | 0.280114895 | 0.175 | 0.042 | 1.8242092 | 11 | fybb |
| plxnc11 | 8.819472 | 0.340034607 | 0.177 | 0.044 | 2.2405870 | 11 | plxnc1 |
| vsir2 | 1.698407 | 0.45121038 | 0.258 | 0.085 | 4.3148049 | 11 | vsir |
| relb | 7.085816 | 0.328779769 | 0.161 | 0.039 | 1.8001516 | 11 | relb |
| yrk1 | 1.070977 | 0.335314994 | 0.22 | 0.064 | 2.7208183 | 11 | yrk |
| LOC103911430 | 4.676318 | 0.292139919 | 0.153 | 0.036 | 1.1880186 | 11 | LOC103911430 |
| CR383676.14 | 5.343890 | 0.511202056 | 0.917 | 0.713 | 1.3576152 | 11 | CR383676.1 |
| arpc1b4 | 3.322764 | 0.511042528 | 0.616 | 0.307 | 8.4414835 | 11 | arpc1b |
| cap14 | 6.377895 | 0.5041451 | 0.473 | 0.241 | 1.6203042 | 11 | cap1 |
| mrtfab | 1.341356 | 0.354605748 | 0.18 | 0.05 | 3.4077154 | 11 | mrtfab |
| scinlb2 | 1.746428 | 0.352007736 | 0.247 | 0.081 | 4.4368018 | 11 | scinlb |
| hcar1-4 | 2.338204 | 0.293582694 | 0.137 | 0.031 | 5.9402088 | 11 | hcar1-4 |
| si:dkey-118j18.2 | 7.661841 | 0.319441839 | 0.188 | 0.054 | 1.9464907 | 11 | si:dkey-118j18.2 |
| rab442 | 1.187218 | 0.329258887 | 0.199 | 0.059 | 3.0161294 | 11 | rab44 |
| prkcbb | 1.959454 | 0.254178299 | 0.132 | 0.029 | 4.9779932 | 11 | prkcbb |
| actn11 | 4.425981 | 0.291171145 | 0.169 | 0.046 | 1.1244205 | 11 | actn1 |
| wdr12 | 1.372022 | 0.437866193 | 0.344 | 0.152 | 3.4856226 | 11 | wdr1 |
| il161 | 1.544297 | 0.289324104 | 0.167 | 0.046 | 3.9232878 | 11 | il16 |
| arpc5a3 | 2.918282 | 0.417635831 | 0.298 | 0.122 | 7.4138965 | 11 | arpc5a |
| rab11fip4b | 5.504558 | 0.284176749 | 0.132 | 0.032 | 1.3984330 | 11 | rab11fip4b |

| | | | | | | | |
|---|---|---|---|---|---|---|---|
| arl5c2 | 2.18430[illegible] | 0.433161331 | 0.266 | 0.105 | 5.5492301 | 11 | arl5c |
| itgae.21 | 3.89022[illegible] | 0.254387442 | 0.137 | 0.034 | 9.8831068 | 11 | itgae.2 |
| tmsb14 | 4.46241[illegible] | 0.403734496 | 0.368 | 0.156 | 1.1336765 | 11 | tmsb1 |
| rhogb2 | 4.84438[illegible] | 0.384554777 | 0.269 | 0.105 | 1.2307161 | 11 | rhogb |
| gapdhs5 | 9.69658[illegible] | 0.448608659 | 0.522 | 0.311 | 2.4634186 | 11 | gapdhs |
| arhgdig3 | 5.75142[illegible] | 0.45029742 | 0.47 | 0.256 | 1.4611497 | 11 | arhgdig |
| pnp5a1 | 1.00761[illegible] | 0.401345117 | 0.207 | 0.072 | 2.5598444 | 11 | pnp5a |
| tram11 | 2.49357[illegible] | 0.357603476 | 0.263 | 0.104 | 6.3349286 | 11 | tram1 |
| antxr2a3 | 3.20236[illegible] | 0.411435879 | 0.352 | 0.166 | 8.1356207 | 11 | antxr2a |
| cox6b13 | 7.06166[illegible] | 0.374466686 | 0.282 | 0.119 | 1.7940166 | 11 | cox6b1 |
| itga41 | 1.00487[illegible] | 0.350259603 | 0.185 | 0.062 | 2.5528943 | 11 | itga4 |
| pycard4 | 1.06041[illegible] | 0.322258053 | 0.223 | 0.081 | 2.6939945 | 11 | pycard |
| actb13 | 2.46644[illegible] | 0.26214527 | 0.884 | 0.765 | 6.2660055 | 11 | actb1 |
| birc21 | 2.46859[illegible] | 0.377596339 | 0.255 | 0.105 | 6.2714627 | 11 | birc2 |
| smox2 | 4.60594[illegible] | 0.343456223 | 0.226 | 0.085 | 1.1701401 | 11 | smox |
| laptm54 | 1.44729[illegible] | 0.383430547 | 0.366 | 0.173 | 3.6768489 | 11 | laptm5 |
| tnfaip31 | 1.67087[illegible] | 0.377827961 | 0.172 | 0.058 | 4.2448594 | 11 | tnfaip3 |
| XLOC-0160942 | 6.66253[illegible] | 0.316042999 | 0.218 | 0.081 | 1.6926177 | 11 | XLOC-016094 |
| ptprc3 | 2.18360[illegible] | 0.339820066 | 0.282 | 0.122 | 5.5474557 | 11 | ptprc |
| actr2a3 | 5.03623[illegible] | 0.352942062 | 0.29 | 0.131 | 1.2794547 | 11 | actr2a |
| mt-nd23 | 5.98794[illegible] | 0.408392677 | 0.578 | 0.389 | 1.5212373 | 11 | mt-nd2 |
| LOC1018842672 | 6.72281[illegible] | 0.281982275 | 0.156 | 0.049 | 1.7079306 | 11 | LOC101884267 |
| zbtb20 | 2.67184[illegible] | 0.381918662 | 0.242 | 0.105 | 6.7878337 | 11 | zbtb20 |
| arrb2b2 | 2.96635[illegible] | 0.336539183 | 0.228 | 0.094 | 7.5360190 | 11 | arrb2b |
| mt-nd13 | 9.40789[illegible] | 0.404861183 | 0.57 | 0.394 | 2.3900764 | 11 | mt-nd1 |
| sec11a | 1.25795[illegible] | 0.374582634 | 0.358 | 0.187 | 3.1958437 | 11 | sec11a |
| pgd1 | 1.56718[illegible] | 0.261468504 | 0.188 | 0.069 | 3.9814363 | 11 | pgd |
| cyth1a | 1.26717[illegible] | 0.279450493 | 0.175 | 0.065 | 3.2192660 | 11 | cyth1a |
| cebpa3 | 2.92923[illegible] | 0.256189815 | 0.194 | 0.074 | 7.4417217 | 11 | cebpa |
| itgav | 4.47608[illegible] | 0.270664421 | 0.167 | 0.061 | 1.1371493 | 11 | itgav |
| parvg2 | 8.15542[illegible] | 0.262447018 | 0.185 | 0.072 | 2.0718858 | 11 | parvg |
| fam107b1 | 1.48527[illegible] | 0.272456896 | 0.25 | 0.11 | 3.7733308 | 11 | fam107b |
| cfl13 | 3.02991[illegible] | 0.28877856 | 0.718 | 0.599 | 7.6974951 | 11 | cfl1 |
| tes1 | 4.66819[illegible] | 0.263408287 | 0.167 | 0.063 | 1.1859542 | 11 | tes |
| sh3kbp1 | 9.40822[illegible] | 0.271688116 | 0.185 | 0.075 | 2.3901602 | 11 | sh3kbp1 |

| | | | | | | | |
|---|---|---|---|---|---|---|---|
| stk17b2 | 2.270358 | 0.305270685 | 0.242 | 0.112 | 5.7678467 | 11 | stk17b |
| nfkbiab | 1.044160 | 0.349423562 | 0.349 | 0.199 | 2.6526888 | 11 | nfkbiab |
| wasa2 | 1.279508 | 0.269527825 | 0.215 | 0.095 | 3.2505910 | 11 | wasa |
| pnrc24 | 2.009549 | 0.266088097 | 0.755 | 0.595 | 5.1052604 | 11 | pnrc2 |
| capzb3 | 4.395712 | 0.318320912 | 0.409 | 0.27 | 1.1167306 | 11 | capzb |
| ccng21 | 1.004397 | 0.28928989 | 0.177 | 0.084 | 2.5516706 | 11 | ccng2 |
| mt-nd32 | 1.305926 | 0.329093059 | 0.5 | 0.37 | 3.3177061 | 11 | mt-nd3 |
| maff1 | 1.393097 | 0.259961555 | 0.172 | 0.079 | 3.5391632 | 11 | maff |
| gyg1a4 | 1.895660 | 0.313465206 | 0.481 | 0.316 | 4.8159253 | 11 | gyg1a |
| tagln23 | 2.337930 | 0.28895005 | 0.344 | 0.208 | 5.9395119 | 11 | tagln2 |
| wasb4 | 3.077560 | 0.278923366 | 0.304 | 0.171 | 7.8185432 | 11 | wasb |
| calr3b | 3.747561 | 0.291639842 | 0.188 | 0.093 | 9.5206799 | 11 | calr3b |
| tmsb4x7 | 7.622613 | 0.305626472 | 0.946 | 0.714 | 1.9365248 | 11 | tmsb4x |
| h1fx2 | 1.050842 | 0.286030161 | 0.323 | 0.189 | 2.6696654 | 11 | h1fx |
| crip15 | 3.098758 | 0.286387943 | 0.433 | 0.282 | 7.8723967 | 11 | crip1 |
| marcksl1b3 | 2.815927 | 0.29926857 | 0.234 | 0.141 | 0.000715 | 11 | marcksl1b |
| pdia33 | 6.562672 | 0.261787515 | 0.417 | 0.312 | 0.001667 | 11 | pdia3 |
| cox8a2 | 1.336842 | 0.264868749 | 0.358 | 0.267 | 0.003396 | 11 | cox8a |
| cyp17a1 | 0 | 2.279176839 | 0.983 | 0.049 | 0 | 12 | cyp17a1 |
| cyp11c1 | 0 | 1.970253402 | 0.892 | 0.004 | 0 | 12 | cyp11c1 |
| star | 0 | 1.842477693 | 0.739 | 0.004 | 0 | 12 | star |
| rgs16 | 0 | 1.836288081 | 0.787 | 0.005 | 0 | 12 | rgs16 |
| hsd3b1 | 0 | 1.694178843 | 0.857 | 0.002 | 0 | 12 | hsd3b1 |
| XLOC-018328 | 0 | 1.551184575 | 0.822 | 0.003 | 0 | 12 | XLOC-018328 |
| pi15a | 0 | 1.521582708 | 0.801 | 0.039 | 0 | 12 | pi15a |
| cyp11a2 | 0 | 1.425648047 | 0.763 | 0.001 | 0 | 12 | cyp11a2 |
| ptx3a1 | 0 | 1.407375376 | 0.7 | 0.028 | 0 | 12 | ptx3a |
| prkag2a | 0 | 1.342358476 | 0.767 | 0.024 | 0 | 12 | prkag2a |
| ece2b | 0 | 1.244356273 | 0.704 | 0.035 | 0 | 12 | ece2b |
| pik3r3b | 0 | 1.212001541 | 0.669 | 0.036 | 0 | 12 | pik3r3b |
| fdx1b | 0 | 1.198183815 | 0.634 | 0.006 | 0 | 12 | fdx1b |
| fam184a | 0 | 1.06892943 | 0.582 | 0.003 | 0 | 12 | fam184a |
| osr21 | 0 | 1.065344823 | 0.627 | 0.041 | 0 | 12 | osr2 |
| esr2b1 | 0 | 1.006238864 | 0.596 | 0.028 | 0 | 12 | esr2b |
| fdxr | 0 | 0.982734541 | 0.554 | 0.017 | 0 | 12 | fdxr |

| | | | | | | | |
|---|---|---|---|---|---|---|---|
| robo21 | 0 | 0.952852412 | 0.613 | 0.043 | 0 | 12 | robo2 |
| nr2f1a1 | 0 | 0.948424209 | 0.652 | 0.051 | 0 | 12 | nr2f1a |
| nell2a | 0 | 0.928429594 | 0.547 | 0.013 | 0 | 12 | nell2a |
| mctp1b | 0 | 0.875917631 | 0.495 | 0.011 | 0 | 12 | mctp1b |
| islr2 | 0 | 0.87264538 | 0.46 | 0.005 | 0 | 12 | islr2 |
| CABZ01074994.1 | 0 | 0.835380425 | 0.425 | 0.003 | 0 | 12 | CABZ01074994.1 |
| nr5a1b | 0 | 0.817909013 | 0.477 | 0.003 | 0 | 12 | nr5a1b |
| BX901962.2 | 0 | 0.746746187 | 0.477 | 0.027 | 0 | 12 | BX901962.2 |
| meis3 | 0 | 0.734280251 | 0.39 | 0.01 | 0 | 12 | meis3 |
| tspan9a | 0 | 0.665865838 | 0.38 | 0.016 | 0 | 12 | tspan9a |
| nol4lb | 0 | 0.653347256 | 0.404 | 0.02 | 0 | 12 | nol4lb |
| CABZ01079281.1 | 0 | 0.610770883 | 0.366 | 0.005 | 0 | 12 | CABZ01079281.1 |
| CABZ01038709.1 | 0 | 0.589241277 | 0.303 | 0.005 | 0 | 12 | CABZ01038709.1 |
| arhgap20b | 0 | 0.579220775 | 0.362 | 0.015 | 0 | 12 | arhgap20b |
| prox1a | 0 | 0.550836297 | 0.328 | 0.011 | 0 | 12 | prox1a |
| camkvl | 0 | 0.44108418 | 0.261 | 0.004 | 0 | 12 | camkvl |
| si:dkey-183j2.10 | 0 | 0.385888702 | 0.202 | 0.002 | 0 | 12 | si:dkey-183j2.10 |
| stmn2b | 0 | 0.370710339 | 0.202 | 0.001 | 0 | 12 | stmn2b |
| pcdh8 | 0 | 0.368897789 | 0.199 | 0 | 0 | 12 | pcdh8 |
| zgc:153395 | 0 | 0.368762889 | 0.209 | 0.001 | 0 | 12 | zgc:153395 |
| XLOC-001115 | 0 | 0.358342723 | 0.188 | 0.001 | 0 | 12 | XLOC-001115 |
| unc5db | 0 | 0.351013736 | 0.195 | 0.001 | 0 | 12 | unc5db |
| insl3 | 0 | 0.348235152 | 0.125 | 0 | 0 | 12 | insl3 |
| cyp4t8 | 0 | 0.345338729 | 0.195 | 0.001 | 0 | 12 | cyp4t8 |
| pklr | 0 | 0.327969522 | 0.171 | 0.002 | 0 | 12 | pklr |
| cyp17a2 | 0 | 0.318620215 | 0.185 | 0.002 | 0 | 12 | cyp17a2 |
| rprm3 | 0 | 0.317463855 | 0.164 | 0.001 | 0 | 12 | rprm3 |
| chrnb3a | 0 | 0.281130304 | 0.157 | 0 | 0 | 12 | chrnb3a |
| nlgn4xa | 5.765813 | 0.444957874 | 0.265 | 0.008 | 1.4648050 | 12 | nlgn4xa |
| pex5lb | 1.950472 | 0.302135565 | 0.157 | 0.002 | 4.9551747 | 12 | pex5lb |
| inhbab | 1.666217 | 0.413707349 | 0.226 | 0.005 | 4.2330254 | 12 | inhbab |
| sypl1 | 2.787022 | 0.573890247 | 0.348 | 0.017 | 7.0804310 | 12 | sypl1 |
| celf4 | 4.482878 | 0.328466487 | 0.171 | 0.003 | 1.1388751 | 12 | celf4 |
| dachd | 9.785712 | 0.383011109 | 0.226 | 0.007 | 2.4860601 | 12 | dachd |
| ppdpfa2 | 1.909779 | 0.899914179 | 0.589 | 0.061 | 4.8517938 | 12 | ppdpfa |

| | | | | | | | |
|---|---|---|---|---|---|---|---|
| atf32 | 4.414699 | 1.133816575 | 0.613 | 0.071 | 1.1215544 | 12 | atf3 |
| hs6st1b | 9.735101 | 0.468251534 | 0.265 | 0.011 | 2.4732025 | 12 | hs6st1b |
| oprd1a | 4.610572 | 0.285019749 | 0.125 | 0.001 | 1.1713159 | 12 | oprd1a |
| ift172 | 1.340077 | 0.343705643 | 0.206 | 0.006 | 3.4044677 | 12 | ift172 |
| nrxn1a | 7.783130 | 0.384881961 | 0.233 | 0.008 | 1.9773043 | 12 | nrxn1a |
| LOC101882765 | 6.043937 | 0.278957527 | 0.167 | 0.004 | 1.5354622 | 12 | LOC101882765 |
| ar1 | 5.880729 | 0.825779311 | 0.537 | 0.058 | 1.4939994 | 12 | ar |
| dab1a | 3.283283 | 0.284218631 | 0.15 | 0.003 | 8.3411813 | 12 | dab1a |
| dclk1a1 | 6.814565 | 0.589943392 | 0.383 | 0.028 | 1.7312403 | 12 | dclk1a |
| ctnnd2a | 1.429420 | 0.340651415 | 0.192 | 0.006 | 3.6314416 | 12 | ctnnd2a |
| cox7a1 | 4.672999 | 0.650874559 | 0.401 | 0.032 | 1.1871754 | 12 | cox7a1 |
| hsd11b22 | 1.354766 | 1.408040121 | 0.763 | 0.137 | 3.4417851 | 12 | hsd11b2 |
| atp6ap1lb | 8.882278 | 0.286298311 | 0.174 | 0.005 | 2.2565428 | 12 | atp6ap1lb |
| sipa1l2.1 | 1.052276 | 0.320673669 | 0.192 | 0.006 | 2.6733094 | 12 | sipa1l2.1 |
| jag1b | 1.469579 | 0.518655239 | 0.331 | 0.023 | 3.7334659 | 12 | jag1b |
| fam222a | 1.944218 | 0.365352965 | 0.233 | 0.011 | 4.9392872 | 12 | fam222a |
| brinp2 | 6.067462 | 0.288899719 | 0.157 | 0.004 | 1.5414389 | 12 | brinp2 |
| col18a1a1 | 5.808546 | 0.65299418 | 0.436 | 0.042 | 1.4756612 | 12 | col18a1a |
| tmsb2 | 1.276963 | 0.300638253 | 0.174 | 0.005 | 3.2441261 | 12 | tmsb2 |
| zgc:927442 | 9.891406 | 1.288902705 | 0.927 | 0.315 | 2.5129119 | 12 | zgc:92744 |
| cited4a | 1.755461 | 0.646448926 | 0.38 | 0.035 | 4.4597511 | 12 | cited4a |
| nme31 | 5.201207 | 0.774616674 | 0.568 | 0.08 | 1.3213666 | 12 | nme3 |
| irs2b | 1.503443 | 0.413574722 | 0.233 | 0.012 | 3.8194992 | 12 | irs2b |
| CR318588.3 | 9.413096 | 0.881492054 | 0.418 | 0.044 | 2.3913972 | 12 | CR318588.3 |
| cox6a2 | 4.377666 | 0.568313493 | 0.359 | 0.032 | 1.1121460 | 12 | cox6a2 |
| vamp2 | 1.653414 | 0.254884072 | 0.15 | 0.005 | 4.2005000 | 12 | vamp2 |
| bag31 | 1.122336 | 0.586120428 | 0.331 | 0.028 | 2.8512967 | 12 | bag3 |
| acsbg2 | 1.075077 | 0.530801863 | 0.348 | 0.032 | 2.7312332 | 12 | acsbg2 |
| b4galt6 | 1.087591 | 0.767260502 | 0.467 | 0.06 | 2.7630271 | 12 | b4galt6 |
| socs21 | 1.248049 | 0.493961484 | 0.289 | 0.023 | 3.1706704 | 12 | socs2 |
| mtbl | 3.215975 | 0.405927004 | 0.237 | 0.015 | 8.1701861 | 12 | mtbl |
| atp10a1 | 4.586921 | 0.641872687 | 0.432 | 0.052 | 1.1653072 | 12 | atp10a |
| fundc2 | 1.011427 | 0.733271519 | 0.557 | 0.088 | 2.5695318 | 12 | fundc2 |
| skib | 3.366704 | 0.413259197 | 0.265 | 0.019 | 8.5531126 | 12 | skib |
| atp1a3b1 | 8.795249 | 0.515993237 | 0.314 | 0.028 | 2.2344332 | 12 | atp1a3b |

| | | | | | | | |
|---|---|---|---|---|---|---|---|
| lrrtm11 | 2.777076 | 0.301497102 | 0.167 | 0.007 | 7.0551632 | 12 | lrrtm1 |
| cxcl18b2 | 5.330199 | 0.874259195 | 0.603 | 0.101 | 1.3541370 | 12 | cxcl18b |
| hmgb3a3 | 6.007082 | 0.661879462 | 0.512 | 0.075 | 1.5260992 | 12 | hmgb3a |
| pcdh11 | 8.153834 | 0.3666045 | 0.209 | 0.012 | 2.0714816 | 12 | pcdh11 |
| nr2f21 | 7.518870 | 0.614661672 | 0.394 | 0.046 | 1.9101691 | 12 | nr2f2 |
| acss1 | 8.252577 | 0.505307586 | 0.321 | 0.031 | 2.0965673 | 12 | acss1 |
| syngr1a | 1.020432 | 0.266697008 | 0.167 | 0.008 | 2.5924079 | 12 | syngr1a |
| slc51a | 4.908514 | 0.272618165 | 0.171 | 0.008 | 1.2470080 | 12 | slc51a |
| pgr | 2.204288 | 0.3555425 | 0.226 | 0.016 | 5.5999959 | 12 | pgr |
| ckba1 | 8.128892 | 0.646334811 | 0.463 | 0.066 | 2.0651450 | 12 | ckba |
| pde4ba1 | 1.129970 | 0.86188092 | 0.652 | 0.132 | 2.8706903 | 12 | pde4ba |
| dhcr7 | 1.432991 | 0.328124036 | 0.213 | 0.014 | 3.6405144 | 12 | dhcr7 |
| hspe11 | 4.938901 | 1.060825471 | 0.878 | 0.28 | 1.2547279 | 12 | hspe1 |
| pgrmc11 | 9.301449 | 0.888952755 | 0.659 | 0.143 | 2.3630332 | 12 | pgrmc1 |
| hmgb1b2 | 1.771468 | 0.80240232 | 0.683 | 0.144 | 4.5004148 | 12 | hmgb1b |
| gramd1a | 4.533569 | 0.303202695 | 0.185 | 0.011 | 1.1517534 | 12 | gramd1a |
| socs3a3 | 7.432439 | 1.128814638 | 0.895 | 0.304 | 1.8882111 | 12 | socs3a |
| pde8a | 1.788594 | 0.265087594 | 0.164 | 0.009 | 4.5439244 | 12 | pde8a |
| gpr271 | 1.412663 | 0.340660912 | 0.199 | 0.013 | 3.5888707 | 12 | gpr27 |
| si:dkey-151g10.6 | 3.283105 | 0.270048774 | 1 | 0.976 | 8.3407300 | 12 | si:dkey-151g10.6 |
| nt5c2b | 1.579982 | 0.526361435 | 0.341 | 0.041 | 4.0139447 | 12 | nt5c2b |
| edil3a | 3.852080 | 0.366480001 | 0.24 | 0.021 | 9.7862099 | 12 | edil3a |
| socs3b2 | 1.771581 | 0.825134993 | 0.645 | 0.142 | 4.5007022 | 12 | socs3b |
| tbx2b | 1.966010 | 0.372839474 | 0.209 | 0.016 | 4.9946497 | 12 | tbx2b |
| jun2 | 1.281858 | 1.011780006 | 0.819 | 0.239 | 3.2565615 | 12 | jun |
| robo1 | 1.444784 | 0.37426649 | 0.251 | 0.023 | 3.6704757 | 12 | robo1 |
| inhbaa1 | 1.142049 | 0.545475155 | 0.362 | 0.047 | 2.9013760 | 12 | inhbaa |
| tp53inp1 | 1.426504 | 0.573621782 | 0.38 | 0.054 | 3.6240352 | 12 | tp53inp1 |
| gfra1a | 3.542942 | 0.272373464 | 0.164 | 0.009 | 9.0008463 | 12 | gfra1a |
| zgc:92606 | 6.731611 | 0.625678694 | 0.491 | 0.088 | 1.7101657 | 12 | zgc:92606 |
| ghrb1 | 3.567323 | 0.463890734 | 0.321 | 0.039 | 9.0627851 | 12 | ghrb |
| msi2a | 4.150457 | 0.462529132 | 0.334 | 0.042 | 1.0544236 | 12 | msi2a |
| meis1b2 | 8.269627 | 0.574913607 | 0.422 | 0.067 | 2.1008988 | 12 | meis1b |
| cdon2 | 5.555458 | 0.575389667 | 0.415 | 0.065 | 1.4113643 | 12 | cdon |
| sox12 | 6.112568 | 0.300039339 | 0.164 | 0.01 | 1.5528981 | 12 | sox12 |

| | | | | | | | |
|---|---|---|---|---|---|---|---|
| fosl1a1 | 2.198058 | 0.637646948 | 0.453 | 0.077 | 5.5841686 | 12 | fosl1a |
| hipk2 | 8.599276 | 0.341444602 | 0.216 | 0.018 | 2.1846460 | 12 | hipk2 |
| tmsb | 1.709805 | 0.480991788 | 0.268 | 0.029 | 4.3437615 | 12 | tmsb |
| slc4a4a | 3.509336 | 0.292442924 | 0.157 | 0.009 | 8.9154687 | 12 | slc4a4a |
| ptprda | 4.655982 | 0.270914719 | 0.164 | 0.01 | 1.1828522 | 12 | ptprda |
| irf2bpl | 1.123941 | 0.473243314 | 0.321 | 0.041 | 2.8553733 | 12 | irf2bpl |
| junba3 | 1.294290 | 0.986683377 | 0.882 | 0.31 | 3.2881452 | 12 | junba |
| CABZ01086041.1 | 6.511227 | 0.257759197 | 0.15 | 0.009 | 1.6541772 | 12 | CABZ01086041.1 |
| fam89b | 6.749374 | 0.338636316 | 0.206 | 0.017 | 1.7146785 | 12 | fam89b |
| mtus1b | 2.978139 | 0.340149109 | 0.226 | 0.021 | 7.5659641 | 12 | mtus1b |
| dusp51 | 4.177596 | 0.676767677 | 0.387 | 0.062 | 1.0613183 | 12 | dusp5 |
| FO082781.12 | 2.823382 | 0.548131927 | 0.341 | 0.05 | 7.1728042 | 12 | FO082781.1 |
| cldn11a1 | 4.232464 | 0.67648407 | 0.436 | 0.077 | 1.0752575 | 12 | cldn11a |
| si:dkey-44g23.5 | 7.652263 | 0.546398715 | 0.383 | 0.06 | 1.9440575 | 12 | si:dkey-44g23.5 |
| cux1a1 | 1.294137 | 0.47477659 | 0.341 | 0.05 | 3.2877575 | 12 | cux1a |
| cbx7a1 | 2.449771 | 0.557851445 | 0.328 | 0.048 | 6.2236449 | 12 | cbx7a |
| dnmt3ab2 | 1.627566 | 0.564954406 | 0.453 | 0.086 | 4.1348318 | 12 | dnmt3ab |
| lifrb | 4.461918 | 0.257617139 | 0.167 | 0.012 | 1.1335502 | 12 | lifrb |
| asph | 1.431041 | 0.430957442 | 0.293 | 0.039 | 3.6355610 | 12 | asph |
| msi2b | 2.976049 | 0.55037816 | 0.439 | 0.08 | 7.5606526 | 12 | msi2b |
| klf11b1 | 4.517359 | 0.558947338 | 0.418 | 0.077 | 1.1476352 | 12 | klf11b |
| tpst1 | 3.808332 | 0.385055322 | 0.279 | 0.036 | 9.6750686 | 12 | tpst1 |
| nfic2 | 8.839764 | 0.544026491 | 0.418 | 0.078 | 2.2457421 | 12 | nfic |
| slc38a2 | 2.571904 | 0.456930013 | 0.296 | 0.042 | 6.5339224 | 12 | slc38a2 |
| hist2h2l1 | 8.340558 | 0.621338452 | 0.453 | 0.093 | 2.1189188 | 12 | hist2h2l |
| rpl343 | 4.720014 | 0.460817329 | 1 | 0.799 | 1.1991195 | 12 | rpl34 |
| rps284 | 7.177862 | 0.412186207 | 1 | 0.838 | 1.8235359 | 12 | rps28 |
| atf4b2 | 1.427169 | 0.791927464 | 0.777 | 0.268 | 3.6257228 | 12 | atf4b |
| gadd45ga | 2.708003 | 0.527950082 | 0.317 | 0.049 | 6.8796840 | 12 | gadd45ga |
| ifi453 | 1.214290 | 0.829975634 | 0.537 | 0.132 | 3.0849057 | 12 | ifi45 |
| rpl373 | 4.288029 | 0.399265974 | 1 | 0.853 | 1.0893737 | 12 | rpl37 |
| rpl324 | 2.471822 | 0.427757147 | 0.99 | 0.822 | 6.2796638 | 12 | rpl32 |
| ptp4a3 | 6.809264 | 0.420870817 | 0.31 | 0.047 | 1.7298937 | 12 | ptp4a3 |
| si:ch73-46j18.53 | 9.176254 | 0.712802102 | 0.721 | 0.199 | 2.3312274 | 12 | si:ch73-46j18.5 |
| ifi462 | 1.669535 | 0.668316008 | 0.519 | 0.125 | 4.2414546 | 12 | ifi46 |

| | | | | | | | |
|---|---|---|---|---|---|---|---|
| tgfbr31 | 1.268887 | 0.401619231 | 0.289 | 0.042 | 3.2236080 | 12 | tgfbr3 |
| NC-002333.43 | 2.019753 | 0.719172945 | 0.909 | 0.604 | 5.1311840 | 12 | NC-002333.4 |
| cplx2l1 | 2.563737 | 0.273268898 | 0.178 | 0.017 | 6.5131762 | 12 | cplx2l |
| fosab3 | 3.386796 | 0.887873604 | 0.871 | 0.352 | 8.6041559 | 12 | fosab |
| rps263 | 5.104026 | 0.464878502 | 0.976 | 0.747 | 1.2966780 | 12 | rps26 |
| rps74 | 1.387680 | 0.402129849 | 0.979 | 0.826 | 3.5254028 | 12 | rps7 |
| nupr1b | 3.522467 | 0.457914808 | 0.275 | 0.041 | 8.9488289 | 12 | nupr1b |
| marcksl1b4 | 3.695039 | 0.635848854 | 0.544 | 0.134 | 9.3872480 | 12 | marcksl1b |
| h1f01 | 2.122511 | 0.557759756 | 0.355 | 0.066 | 5.3922394 | 12 | h1f0 |
| fmnl3 | 3.790454 | 0.25818298 | 0.181 | 0.018 | 9.6296493 | 12 | fmnl3 |
| rpl365 | 8.036362 | 0.434888873 | 1 | 0.803 | 2.0416380 | 12 | rpl36 |
| nt5dc21 | 1.376508 | 0.476048779 | 0.387 | 0.076 | 3.4970206 | 12 | nt5dc2 |
| rpl95 | 2.857903 | 0.573486707 | 0.997 | 0.663 | 7.2605046 | 12 | rpl9 |
| aldh2.1 | 9.979001 | 0.341118781 | 0.251 | 0.035 | 2.5351654 | 12 | aldh2.1 |
| b2m3 | 4.506049 | 0.759126962 | 0.78 | 0.29 | 1.1447619 | 12 | b2m |
| prkar2aa | 1.831735 | 0.52663404 | 0.449 | 0.102 | 4.6535228 | 12 | prkar2aa |
| si:dkey-188i13.101 | 1.242535 | 0.598368616 | 0.453 | 0.108 | 3.1566603 | 12 | si:dkey-188i13.10 |
| rpl185 | 2.826473 | 0.442951198 | 0.997 | 0.789 | 7.1806552 | 12 | rpl18 |
| tbc1d4 | 7.317524 | 0.272113662 | 0.171 | 0.017 | 1.8590171 | 12 | tbc1d4 |
| gbp | 2.154283 | 0.558961236 | 0.474 | 0.116 | 5.4729567 | 12 | gbp |
| rpl27a3 | 2.459918 | 0.663268629 | 0.906 | 0.516 | 6.2494241 | 12 | rpl27a |
| zbtb4 | 3.489237 | 0.289616816 | 0.188 | 0.021 | 8.8644076 | 12 | zbtb4 |
| rpl295 | 1.169532 | 0.400375217 | 0.997 | 0.82 | 2.9711976 | 12 | rpl29 |
| rps27.16 | 1.508369 | 0.517665968 | 0.983 | 0.706 | 3.8320134 | 12 | rps27.1 |
| tubb5 | 2.138614 | 0.42313748 | 0.31 | 0.055 | 5.4331501 | 12 | tubb5 |
| si:dkey-23i12.71 | 4.976430 | 0.370410772 | 0.202 | 0.025 | 1.2642621 | 12 | si:dkey-23i12.7 |
| plxna4 | 9.767462 | 0.278780448 | 0.185 | 0.021 | 2.4814238 | 12 | plxna4 |
| si:ch211-178n15.11 | 1.225614 | 0.317851729 | 0.206 | 0.026 | 3.1136735 | 12 | si:ch211-178n15.1 |
| mafba2 | 1.699652 | 0.533975336 | 0.373 | 0.079 | 4.3179681 | 12 | mafba |
| fndc3a | 1.756656 | 0.250718979 | 0.153 | 0.015 | 4.4627858 | 12 | fndc3a |
| maff2 | 6.522379 | 0.455139687 | 0.366 | 0.075 | 1.6570104 | 12 | maff |
| rpl22l14 | 1.054176 | 0.716005198 | 0.892 | 0.417 | 2.6781353 | 12 | rpl22l1 |
| rps165 | 1.418099 | 0.450681789 | 0.993 | 0.762 | 3.6026813 | 12 | rps16 |
| rplp01 | 1.515458 | 0.364298272 | 0.993 | 0.857 | 3.8500210 | 12 | rplp0 |
| rps94 | 3.623744 | 0.42466373 | 0.99 | 0.787 | 9.2061223 | 12 | rps9 |

| | | | | | | | |
|---|---|---|---|---|---|---|---|
| cadm2a | 8.14262( | 0.26416127 | 0.171 | 0.019 | 2.0686327 | 12 | cadm2a |
| zgc:1141885 | 8.761516 | 0.546507147 | 0.979 | 0.663 | 2.2258633 | 12 | zgc:114188 |
| crim11 | 9.134159 | 0.356806034 | 0.247 | 0.038 | 2.3205331 | 12 | crim1 |
| rgs4 | 1.118160 | 0.437956257 | 0.247 | 0.038 | 2.8406878 | 12 | rgs4 |
| CR318588.16 | 9.036003 | 0.857279053 | 0.638 | 0.208 | 2.2955966 | 12 | CR318588.1 |
| rpl114 | 1.297729 | 0.400563839 | 1 | 0.827 | 3.2968820 | 12 | rpl11 |
| adh51 | 2.025381 | 0.65852816 | 0.679 | 0.234 | 5.1454825 | 12 | adh5 |
| znf503 | 2.182777 | 0.276716954 | 0.188 | 0.023 | 5.5453463 | 12 | znf503 |
| c42 | 9.233803 | 0.345414797 | 0.237 | 0.035 | 2.3458477 | 12 | c4 |
| enc11 | 3.751078 | 0.366087118 | 0.233 | 0.036 | 9.5296146 | 12 | enc1 |
| rps194 | 4.510950 | 0.33393087 | 0.993 | 0.867 | 1.1460070 | 12 | rps19 |
| rpl244 | 8.120173 | 0.412791831 | 0.993 | 0.791 | 2.0629300 | 12 | rpl24 |
| mdka2 | 8.308457 | 0.528193724 | 0.446 | 0.108 | 2.1107637 | 12 | mdka |
| mt-nd14 | 1.246977 | 0.716763171 | 0.829 | 0.389 | 3.1679460 | 12 | mt-nd1 |
| rplp2l4 | 1.509351 | 0.426513721 | 0.99 | 0.792 | 3.8345076 | 12 | rplp2l |
| wu:fc75a09 | 1.974970 | 0.2896981 | 0.174 | 0.021 | 5.0174124 | 12 | wu:fc75a09 |
| fat1a | 2.583667 | 0.272172234 | 0.202 | 0.028 | 6.5638081 | 12 | fat1a |
| rpl384 | 2.481251 | 0.442294968 | 0.979 | 0.75 | 6.3036202 | 12 | rpl38 |
| rps185 | 6.994456 | 0.445999681 | 0.983 | 0.751 | 1.7769417 | 12 | rps18 |
| rps254 | 9.088544 | 0.430782752 | 0.979 | 0.745 | 2.3089447 | 12 | rps25 |
| hspd11 | 1.763358 | 0.581843165 | 0.537 | 0.157 | 4.4798115 | 12 | hspd1 |
| elmsan1b | 2.024973 | 0.356479131 | 0.251 | 0.043 | 5.1444452 | 12 | elmsan1b |
| znfx1 | 2.441021 | 0.340262773 | 0.206 | 0.03 | 6.2014159 | 12 | znfx1 |
| rps123 | 3.306097 | 0.357364388 | 0.983 | 0.839 | 8.3991410 | 12 | rps12 |
| rps216 | 3.421025 | 0.434051424 | 0.993 | 0.772 | 8.6911157 | 12 | rps21 |
| rpl284 | 3.580644 | 0.412135565 | 0.986 | 0.774 | 9.0966283 | 12 | rpl28 |
| cx431 | 3.976636 | 0.372748066 | 0.289 | 0.054 | 1.0102644 | 12 | cx43 |
| rps8a4 | 4.187886 | 0.386179538 | 0.979 | 0.799 | 1.0639326 | 12 | rps8a |
| slc25a52 | 4.270126 | 0.377228245 | 0.983 | 0.811 | 1.0848257 | 12 | slc25a5 |
| mt-nd24 | 1.060409 | 0.689996935 | 0.836 | 0.384 | 2.6939696 | 12 | mt-nd2 |
| stat1a1 | 1.605899 | 0.533397327 | 0.439 | 0.118 | 4.0797880 | 12 | stat1a |
| lgals171 | 1.627025 | 0.456859158 | 0.275 | 0.052 | 4.1334591 | 12 | lgals17 |
| faua5 | 3.422191 | 0.466646177 | 0.965 | 0.714 | 8.6940768 | 12 | faua |
| rpl7a4 | 4.968888 | 0.371412623 | 0.983 | 0.821 | 1.2623462 | 12 | rpl7a |
| CR383676.15 | 2.046487 | 0.642107751 | 0.923 | 0.714 | 5.1991020 | 12 | CR383676.1 |

| | | | | | | | |
|---|---|---|---|---|---|---|---|
| rps26l5 | 3.629034 | 0.480825345 | 0.962 | 0.696 | 9.2195633 | 12 | rps26l |
| ptprga2 | 4.501452 | 0.372833023 | 0.324 | 0.067 | 1.1435938 | 12 | ptprga |
| mt-co24 | 9.481072 | 0.576890986 | 0.923 | 0.667 | 2.4086664 | 12 | mt-co2 |
| eif1b1 | 2.067377 | 0.655806325 | 0.836 | 0.399 | 5.2521714 | 12 | eif1b |
| sema4ba2 | 2.660385 | 0.340228658 | 0.233 | 0.039 | 6.7587104 | 12 | sema4ba |
| rpl354 | 2.978277 | 0.36978063 | 0.983 | 0.811 | 7.5663128 | 12 | rpl35 |
| aoc22 | 3.424366 | 0.410175915 | 0.334 | 0.073 | 8.6996031 | 12 | aoc2 |
| zgc:1527912 | 3.797422 | 0.539786459 | 0.303 | 0.064 | 9.6473516 | 12 | zgc:152791 |
| cyb5a | 6.801444 | 0.369467111 | 0.296 | 0.06 | 1.7279070 | 12 | cyb5a |
| higd1a2 | 9.933199 | 0.626131395 | 0.76 | 0.295 | 2.5235294 | 12 | higd1a |
| isg151 | 1.154136 | 0.556736738 | 0.293 | 0.061 | 2.9320834 | 12 | isg15 |
| hmgn64 | 2.833336 | 0.645574189 | 0.756 | 0.303 | 7.1980906 | 12 | hmgn6 |
| antxr1c | 5.125470 | 0.321678085 | 0.223 | 0.037 | 1.3021258 | 12 | antxr1c |
| nipsnap2 | 5.253765 | 0.389948616 | 0.369 | 0.088 | 1.3347190 | 12 | nipsnap2 |
| egr13 | 1.880807 | 0.641512776 | 0.54 | 0.177 | 4.7781914 | 12 | egr1 |
| rpl36a4 | 2.208176 | 0.356012551 | 0.99 | 0.821 | 5.6098735 | 12 | rpl36a |
| rps27.24 | 3.544017 | 0.574863997 | 0.916 | 0.547 | 9.0035761 | 12 | rps27.2 |
| rps155 | 3.748533 | 0.463559547 | 0.979 | 0.713 | 9.5231481 | 12 | rps15 |
| si:ch211-286b5.51 | 6.508118 | 0.49016632 | 0.495 | 0.145 | 1.6533875 | 12 | si:ch211-286b5.5 |
| rpl64 | 6.998298 | 0.471893734 | 0.979 | 0.697 | 1.7779176 | 12 | rpl6 |
| rps3a4 | 1.146129 | 0.382101894 | 0.979 | 0.789 | 2.9117422 | 12 | rps3a |
| rpl264 | 3.333389 | 0.334440646 | 0.99 | 0.849 | 8.4684762 | 12 | rpl26 |
| rpl35a5 | 3.672676 | 0.400456816 | 0.986 | 0.779 | 9.3304346 | 12 | rpl35a |
| junbb1 | 5.410019 | 0.369188401 | 0.307 | 0.066 | 1.3744153 | 12 | junbb |
| cox8a3 | 6.090745 | 0.586800987 | 0.686 | 0.26 | 1.5473539 | 12 | cox8a |
| rps134 | 7.242300 | 0.432784031 | 0.986 | 0.745 | 1.8399065 | 12 | rps13 |
| bnip3lb | 9.915439 | 0.377932409 | 0.334 | 0.077 | 2.5190173 | 12 | bnip3lb |
| si:dkey-28b4.71 | 2.167451 | 0.418507801 | 0.404 | 0.105 | 5.5064115 | 12 | si:dkey-28b4.7 |
| rpl274 | 6.933656 | 0.385204012 | 0.969 | 0.777 | 1.7614953 | 12 | rpl27 |
| diaph2 | 7.694220 | 0.28283152 | 0.202 | 0.032 | 1.9547165 | 12 | diaph2 |
| rps105 | 1.275745 | 0.444482749 | 0.986 | 0.723 | 3.2410314 | 12 | rps10 |
| rps295 | 5.247876 | 0.334799217 | 0.986 | 0.842 | 1.3332229 | 12 | rps29 |
| ier2a3 | 5.507318 | 0.655345833 | 0.679 | 0.267 | 1.3991342 | 12 | ier2a |
| rpl18a5 | 5.661738 | 0.377703294 | 0.969 | 0.798 | 1.4383647 | 12 | rpl18a |
| rps113 | 6.707362 | 0.370490147 | 0.972 | 0.782 | 1.7040054 | 12 | rps11 |

| | | | | | | | |
|---|---|---|---|---|---|---|---|
| eef1g4 | 8.635672 | 0.498127046 | 0.944 | 0.648 | 2.1938925 | 12 | eef1g |
| rpl304 | 2.044358 | 0.441637572 | 0.979 | 0.726 | 5.1936937 | 12 | rpl30 |
| rps245 | 2.519410 | 0.392153027 | 0.979 | 0.776 | 6.4005636 | 12 | rps24 |
| atp1b1a1 | 3.395976 | 0.592640483 | 0.603 | 0.224 | 8.6274788 | 12 | atp1b1a |
| tmem243b | 3.619547 | 0.273773029 | 0.244 | 0.045 | 9.1954591 | 12 | tmem243b |
| irf1b1 | 4.140226 | 0.326576269 | 0.202 | 0.033 | 1.0518245 | 12 | irf1b |
| rplp14 | 7.008187 | 0.322167602 | 0.997 | 0.88 | 1.7804300 | 12 | rplp1 |
| mt-nd33 | 8.423424 | 0.622896731 | 0.805 | 0.364 | 2.1399710 | 12 | mt-nd3 |
| rps64 | 2.844471 | 0.372298167 | 0.983 | 0.801 | 7.2263800 | 12 | rps6 |
| fam102ab | 3.715331 | 0.284222849 | 0.199 | 0.033 | 9.4387994 | 12 | fam102ab |
| rpl315 | 5.066416 | 0.472360587 | 0.979 | 0.683 | 1.2871229 | 12 | rpl31 |
| COX35 | 5.173510 | 0.535853945 | 0.927 | 0.672 | 1.3143304 | 12 | COX3 |
| mt-co14 | 5.213280 | 0.545759202 | 0.899 | 0.636 | 1.3244339 | 12 | mt-co1 |
| anxa13 | 1.183224 | 0.323398082 | 0.254 | 0.051 | 3.0059811 | 12 | anxa13 |
| porb | 1.616023 | 0.293410721 | 0.223 | 0.04 | 4.1055088 | 12 | porb |
| eef1a1l1 | 1.894078 | 0.259498186 | 1 | 0.914 | 4.8119054 | 12 | eef1a1l1 |
| mvp1 | 4.322622 | 0.36995365 | 0.268 | 0.057 | 1.0981623 | 12 | mvp |
| eif4ebp12 | 4.390607 | 0.569733459 | 0.659 | 0.251 | 1.1154337 | 12 | eif4ebp1 |
| rps4x4 | 6.503021 | 0.426645257 | 0.976 | 0.733 | 1.6520925 | 12 | rps4x |
| rps143 | 6.580828 | 0.391480575 | 0.99 | 0.778 | 1.6718594 | 12 | rps14 |
| rpl5b3 | 9.222894 | 0.476043089 | 0.927 | 0.641 | 2.3430764 | 12 | rpl5b |
| zgc:1132081 | 1.026856 | 0.338992355 | 0.237 | 0.046 | 2.6087295 | 12 | zgc:113208 |
| XLOC-0412692 | 4.437584 | 0.271899168 | 0.178 | 0.027 | 1.1273683 | 12 | XLOC-041269 |
| mknk2a | 7.302930 | 0.286177921 | 0.226 | 0.042 | 1.8553094 | 12 | mknk2a |
| pkma1 | 8.781258 | 0.46491619 | 0.533 | 0.173 | 2.2308787 | 12 | pkma |
| rps15a6 | 9.608690 | 0.390804262 | 0.979 | 0.795 | 2.4410877 | 12 | rps15a |
| rsl24d13 | 1.970361 | 0.604224427 | 0.875 | 0.454 | 5.0057038 | 12 | rsl24d1 |
| rpl232 | 2.111172 | 0.285577252 | 0.993 | 0.871 | 5.3634343 | 12 | rpl23 |
| rpl10a4 | 5.998242 | 0.37647426 | 0.972 | 0.776 | 1.5238536 | 12 | rpl10a |
| sept10 | 6.168126 | 0.315503227 | 0.244 | 0.049 | 1.5670124 | 12 | sept10 |
| zbtb201 | 2.253920 | 0.404144921 | 0.376 | 0.103 | 5.7260852 | 12 | zbtb20 |
| rpl33 | 2.363545 | 0.286640307 | 0.99 | 0.871 | 6.0045872 | 12 | rpl3 |
| rpl84 | 3.174211 | 0.361052295 | 0.962 | 0.793 | 8.0640844 | 12 | rpl8 |
| bcl2l1 | 3.722021 | 0.333001331 | 0.303 | 0.072 | 9.4557966 | 12 | bcl2l1 |
| mt-nd44 | 4.234222 | 0.597720595 | 0.836 | 0.441 | 1.0757041 | 12 | mt-nd4 |

| | | | | | | | |
|---|---|---|---|---|---|---|---|
| rpl134 | 4.606059 | 0.329767003 | 0.983 | 0.816 | 1.1701695 | 12 | rpl13 |
| sfpq | 5.056117 | 0.389484701 | 0.38 | 0.106 | 1.2845066 | 12 | sfpq |
| fabp31 | 5.378488 | 0.509626805 | 0.61 | 0.222 | 1.3664050 | 12 | fabp3 |
| rpl23a4 | 3.272313 | 0.45558609 | 0.937 | 0.665 | 8.3133115 | 12 | rpl23a |
| egr2a | 7.535785 | 0.252527985 | 0.15 | 0.022 | 1.9144663 | 12 | egr2a |
| nop532 | 8.000342 | 0.535376909 | 0.645 | 0.26 | 2.0324870 | 12 | nop53 |
| fam168a2 | 1.589966 | 0.337035312 | 0.289 | 0.07 | 4.0393110 | 12 | fam168a |
| rpl124 | 2.132412 | 0.335678756 | 0.986 | 0.81 | 5.4173939 | 12 | rpl12 |
| rpl155 | 2.773598 | 0.392158878 | 0.976 | 0.761 | 7.0463278 | 12 | rpl15 |
| arhgef4 | 6.596146 | 0.261167641 | 0.188 | 0.034 | 1.6757509 | 12 | arhgef4 |
| rps217 | 6.797231 | 0.345379833 | 0.979 | 0.84 | 1.7268365 | 12 | rps2 |
| rpl393 | 8.148849 | 0.285641786 | 0.983 | 0.876 | 2.0702151 | 12 | rpl39 |
| rpl104 | 2.590133 | 0.355257733 | 0.972 | 0.787 | 6.5802347 | 12 | rpl10 |
| serinc1 | 4.844370 | 0.313669854 | 0.321 | 0.083 | 1.2307123 | 12 | serinc1 |
| uba523 | 5.426376 | 0.44803782 | 0.944 | 0.656 | 1.3785710 | 12 | uba52 |
| il4r.11 | 2.522269 | 0.291673131 | 0.216 | 0.044 | 6.4078267 | 12 | il4r.1 |
| cd99l2 | 3.315512 | 0.253159374 | 0.181 | 0.033 | 8.4230586 | 12 | cd99l2 |
| rack13 | 1.528657 | 0.369424769 | 0.986 | 0.76 | 3.8835542 | 12 | rack1 |
| rps27a4 | 2.274400 | 0.353018853 | 0.965 | 0.772 | 5.7781140 | 12 | rps27a |
| serpine1 | 2.346714 | 0.306523592 | 0.195 | 0.037 | 5.9618285 | 12 | serpine1 |
| echs1 | 3.190577 | 0.316513229 | 0.317 | 0.085 | 8.1056616 | 12 | echs1 |
| ddx5 | 4.962636 | 0.534767148 | 0.774 | 0.347 | 1.2607578 | 12 | ddx5 |
| flot2a1 | 5.236661 | 0.306631383 | 0.282 | 0.071 | 1.3303738 | 12 | flot2a |
| phactr4a | 6.030587 | 0.287861463 | 0.195 | 0.038 | 1.5320707 | 12 | phactr4a |
| ewsr1a | 6.471796 | 0.262872304 | 0.223 | 0.048 | 1.6441599 | 12 | ewsr1a |
| h3f3c3 | 9.198158 | 0.522463443 | 0.704 | 0.297 | 2.3367920 | 12 | h3f3c |
| rpl13a3 | 9.804037 | 0.336540663 | 0.965 | 0.779 | 2.4907157 | 12 | rpl13a |
| rps27l2 | 1.331406 | 0.459063795 | 0.557 | 0.212 | 3.3824387 | 12 | rps27l |
| naca5 | 1.488966 | 0.445758622 | 0.92 | 0.648 | 3.7827188 | 12 | naca |
| pdia34 | 3.313624 | 0.529920834 | 0.7 | 0.307 | 8.4182640 | 12 | pdia3 |
| stat2 | 3.734581 | 0.320940512 | 0.254 | 0.06 | 9.4877038 | 12 | stat2 |
| il6st1 | 5.027144 | 0.267896032 | 0.237 | 0.053 | 1.2771460 | 12 | il6st |
| hnrnpa0a | 1.072857 | 0.325148553 | 0.296 | 0.078 | 2.7255934 | 12 | hnrnpa0a |
| hipk3b | 2.811717 | 0.254733295 | 0.199 | 0.04 | 7.1431683 | 12 | hipk3b |
| si:dkey-42i9.4 | 4.125594 | 0.422580401 | 0.443 | 0.152 | 1.0481072 | 12 | si:dkey-42i9.4 |

| | | | | | | | |
|---|---|---|---|---|---|---|---|
| nfia2 | 8.814457 | 0.344151438 | 0.334 | 0.095 | 2.2393129 | 12 | nfia |
| rpl192 | 1.075557 | 0.284158661 | 0.976 | 0.852 | 2.7324542 | 12 | rpl19 |
| si:ch73-1a9.34 | 1.207686 | 0.530399182 | 0.7 | 0.309 | 3.0681279 | 12 | si:ch73-1a9.3 |
| eef1db3 | 1.221065 | 0.49770461 | 0.648 | 0.278 | 3.1021156 | 12 | eef1db |
| ccnl1a | 1.443985 | 0.294978042 | 0.244 | 0.058 | 3.6684445 | 12 | ccnl1a |
| cish | 1.858149 | 0.273737823 | 0.23 | 0.052 | 4.7206292 | 12 | cish |
| mibp24 | 3.223645 | 0.513001522 | 0.854 | 0.516 | 8.1896716 | 12 | mibp2 |
| rps202 | 9.170134 | 0.255439931 | 1 | 0.909 | 2.3296725 | 12 | rps20 |
| rpl174 | 1.057774 | 0.355579785 | 0.962 | 0.761 | 2.6872753 | 12 | rpl17 |
| ywhaqb2 | 2.793796 | 0.411846297 | 0.432 | 0.15 | 7.0976396 | 12 | ywhaqb |
| eif3f4 | 3.708326 | 0.532075643 | 0.833 | 0.46 | 9.4210039 | 12 | eif3f |
| arih2 | 5.120609 | 0.251141579 | 0.244 | 0.058 | 1.3008909 | 12 | arih2 |
| h2afx13 | 6.239855 | 0.524124069 | 0.537 | 0.222 | 1.5852353 | 12 | h2afx1 |
| aldh7a1 | 6.316985 | 0.289290463 | 0.261 | 0.067 | 1.6048302 | 12 | aldh7a1 |
| mid1ip1b | 9.811460 | 0.27213012 | 0.223 | 0.051 | 2.4926014 | 12 | mid1ip1b |
| DHDH1 | 1.177910 | 0.284204557 | 0.258 | 0.066 | 2.9924816 | 12 | DHDH |
| gas1a2 | 1.508851 | 0.273451891 | 0.233 | 0.056 | 3.8332361 | 12 | gas1a |
| gabarapl2 | 2.139062 | 0.297299991 | 0.293 | 0.081 | 5.4342885 | 12 | gabarapl2 |
| aldh6a1 | 2.548736 | 0.301153533 | 0.31 | 0.089 | 6.4750641 | 12 | aldh6a1 |
| hnrnpa0b1 | 3.059736 | 0.502689083 | 0.725 | 0.335 | 7.7732614 | 12 | hnrnpa0b |
| si:dkey-51e6.11 | 3.297185 | 0.355497974 | 0.362 | 0.115 | 8.3765000 | 12 | si:dkey-51e6.1 |
| ybx13 | 4.589944 | 0.422065355 | 0.972 | 0.691 | 1.1660752 | 12 | ybx1 |
| tjp2b | 6.510770 | 0.268654514 | 0.247 | 0.061 | 1.6540613 | 12 | tjp2b |
| hmgb1a4 | 5.097111 | 0.44679961 | 0.551 | 0.223 | 1.2949211 | 12 | hmgb1a |
| rbbp6 | 6.308991 | 0.333270845 | 0.341 | 0.106 | 1.6027992 | 12 | rbbp6 |
| b2ml4 | 7.006693 | 0.479696571 | 0.645 | 0.288 | 1.7800504 | 12 | b2ml |
| gadd45ab1 | 8.166975 | 0.309719427 | 0.254 | 0.067 | 2.0748200 | 12 | gadd45ab |
| NDUFB11 | 1.058509 | 0.424810359 | 0.516 | 0.207 | 2.6891421 | 12 | NDUFB1 |
| hp1bp31 | 2.249080 | 0.347218647 | 0.359 | 0.117 | 5.7137900 | 12 | hp1bp3 |
| atp5l1 | 2.381342 | 0.506799315 | 0.812 | 0.451 | 6.0498010 | 12 | atp5l |
| rps234 | 3.660784 | 0.346037452 | 0.937 | 0.742 | 9.3002236 | 12 | rps23 |
| rps34 | 9.728497 | 0.367958883 | 0.948 | 0.723 | 2.4715247 | 12 | rps3 |
| rpl145 | 1.827810 | 0.425817103 | 0.972 | 0.664 | 4.6435518 | 12 | rpl14 |
| rpl226 | 2.063233 | 0.382958163 | 0.955 | 0.717 | 5.2416456 | 12 | rpl22 |
| rab11fip1b | 2.538729 | 0.257868943 | 0.23 | 0.058 | 6.4496422 | 12 | rab11fip1b |

| | | | | | | | |
|---|---|---|---|---|---|---|---|
| txndc9 | 3.520239 | 0.319199007 | 0.373 | 0.125 | 8.9431688 | 12 | txndc9 |
| mt-atp64 | 3.755844 | 0.503690895 | 0.875 | 0.551 | 9.5417225 | 12 | mt-atp6 |
| map1lc3a1 | 9.572019 | 0.292873125 | 0.272 | 0.078 | 2.4317715 | 12 | map1lc3a |
| nsa23 | 1.156261 | 0.502413755 | 0.732 | 0.383 | 2.9374819 | 12 | nsa2 |
| oaz1b2 | 2.120017 | 0.4669648 | 0.732 | 0.354 | 5.3859052 | 12 | oaz1b |
| btf33 | 3.026665 | 0.455568927 | 0.934 | 0.605 | 7.6892436 | 12 | btf3 |
| ptp4a1 | 4.813882 | 0.336657625 | 0.373 | 0.128 | 1.2229668 | 12 | ptp4a1 |
| si:ch211-196l7.44 | 5.593104 | 0.453052575 | 0.693 | 0.324 | 1.4209282 | 12 | si:ch211-196l7.4 |
| brd3b | 9.981779 | 0.266132957 | 0.244 | 0.066 | 2.5358711 | 12 | brd3b |
| rpl214 | 1.112092 | 0.328743451 | 0.969 | 0.774 | 2.8252721 | 12 | rpl21 |
| rps56 | 3.301923 | 0.376700147 | 0.965 | 0.725 | 8.3885358 | 12 | rps5 |
| CABZ01080702.13 | 3.372082 | 0.297070215 | 0.307 | 0.095 | 8.5667755 | 12 | CABZ01080702.1 |
| mt-nd51 | 4.190552 | 0.328114575 | 0.362 | 0.126 | 1.0646097 | 12 | mt-nd5 |
| mt-cyb3 | 4.408576 | 0.477050464 | 0.871 | 0.572 | 1.1199988 | 12 | mt-cyb |
| zgc:1951732 | 4.478938 | 0.310996032 | 0.341 | 0.11 | 1.1378742 | 12 | zgc:195173 |
| pno11 | 5.836264 | 0.356276522 | 0.439 | 0.167 | 1.4827029 | 12 | pno1 |
| rpl5a5 | 6.060314 | 0.484149186 | 0.899 | 0.531 | 1.5396229 | 12 | rpl5a |
| tomm51 | 6.455043 | 0.389075528 | 0.537 | 0.226 | 1.6399036 | 12 | tomm5 |
| rpl72 | 6.668061 | 0.296094833 | 0.962 | 0.807 | 1.6940211 | 12 | rpl7 |
| eef2b2 | 6.787804 | 0.427300192 | 0.937 | 0.631 | 1.7244418 | 12 | eef2b |
| hsp90ab12 | 7.418704 | 0.313149929 | 0.99 | 0.798 | 1.8847218 | 12 | hsp90ab1 |
| ube2e31 | 1.215760 | 0.361999648 | 0.436 | 0.166 | 3.0886393 | 12 | ube2e3 |
| gtpbp41 | 1.531148 | 0.412294768 | 0.557 | 0.242 | 3.8898827 | 12 | gtpbp4 |
| zgc:1583433 | 2.083344 | 0.468513949 | 0.422 | 0.166 | 5.2927375 | 12 | zgc:158343 |
| rnf213a2 | 4.414045 | 0.324346915 | 0.293 | 0.093 | 1.1213881 | 12 | rnf213a |
| sesn13 | 4.485369 | 0.418356808 | 0.436 | 0.171 | 1.1395080 | 12 | sesn1 |
| eif2s1b1 | 6.487794 | 0.356031837 | 0.474 | 0.19 | 1.6482242 | 12 | eif2s1b |
| si:dkey-79d12.51 | 2.696086 | 0.278629049 | 0.233 | 0.066 | 6.8494080 | 12 | si:dkey-79d12.5 |
| pim21 | 3.084756 | 0.26137675 | 0.279 | 0.086 | 7.8368250 | 12 | pim2 |
| rnf7 | 3.655988 | 0.300840209 | 0.314 | 0.104 | 9.2880379 | 12 | rnf7 |
| dldh | 4.529903 | 0.293610399 | 0.296 | 0.097 | 1.1508218 | 12 | dldh |
| tktb1 | 5.512279 | 0.329153576 | 0.425 | 0.16 | 1.4003945 | 12 | tktb |
| gadd45bb2 | 5.902294 | 0.317065291 | 0.261 | 0.08 | 1.4994780 | 12 | gadd45bb |
| atp5f1e | 6.528863 | 0.460862393 | 0.819 | 0.485 | 1.6586578 | 12 | atp5f1e |
| chd32 | 7.639720 | 0.320250858 | 0.348 | 0.122 | 1.9408708 | 12 | chd3 |

| | | | | | | | |
|---|---|---|---|---|---|---|---|
| gng72 | 7.658224 | 0.250245588 | 0.22 | 0.06 | 1.9455718 | 12 | gng7 |
| uqcrc1 | 1.020321 | 0.291700665 | 0.314 | 0.105 | 2.5921266 | 12 | uqcrc1 |
| CABZ01072614.12 | 1.720571 | 0.301521307 | 0.286 | 0.092 | 4.3711128 | 12 | CABZ01072614.1 |
| eif3m3 | 3.198200 | 0.449996743 | 0.728 | 0.378 | 8.1250285 | 12 | eif3m |
| grinab2 | 3.995738 | 0.288780292 | 0.341 | 0.12 | 1.0151173 | 12 | grinab |
| anxa42 | 8.873881 | 0.331456141 | 0.387 | 0.147 | 2.2544095 | 12 | anxa4 |
| laptm4a1 | 1.321741 | 0.284305198 | 0.338 | 0.119 | 3.3578832 | 12 | laptm4a |
| rpsa4 | 2.055612 | 0.3353498 | 0.99 | 0.782 | 5.2222829 | 12 | rpsa |
| rtca | 2.074137 | 0.259413782 | 0.286 | 0.091 | 5.2693466 | 12 | rtca |
| srsf5a1 | 2.133119 | 0.401603693 | 0.631 | 0.298 | 5.4191902 | 12 | srsf5a |
| atp5mf1 | 4.153924 | 0.407775831 | 0.596 | 0.284 | 1.0553045 | 12 | atp5mf |
| cct21 | 6.419436 | 0.38991095 | 0.544 | 0.25 | 1.6308579 | 12 | cct2 |
| ndufc2 | 1.311210 | 0.308304545 | 0.387 | 0.15 | 3.3311290 | 12 | ndufc2 |
| stau22 | 1.334988 | 0.280307619 | 0.282 | 0.094 | 3.3915389 | 12 | stau2 |
| mphosph8 | 1.692616 | 0.305536202 | 0.366 | 0.137 | 4.3000911 | 12 | mphosph8 |
| grpel2 | 2.009521 | 0.270989574 | 0.303 | 0.104 | 5.1051898 | 12 | grpel2 |
| eif3i1 | 2.774663 | 0.393415621 | 0.596 | 0.286 | 7.0490322 | 12 | eif3i |
| si:ch211-222l21.13 | 4.380979 | 0.416016009 | 0.662 | 0.309 | 1.1129878 | 12 | si:ch211-222l21.1 |
| appa1 | 1.141905 | 0.254665346 | 0.265 | 0.086 | 2.9010106 | 12 | appa |
| cct7 | 1.482362 | 0.356824334 | 0.505 | 0.224 | 3.7659410 | 12 | cct7 |
| mycb1 | 1.731013 | 0.350687538 | 0.47 | 0.197 | 4.3976403 | 12 | mycb |
| eif3c2 | 2.274838 | 0.334364233 | 0.436 | 0.184 | 5.7792260 | 12 | eif3c |
| cox6b21 | 2.605195 | 0.415093818 | 0.847 | 0.545 | 6.6184984 | 12 | cox6b2 |
| add3a2 | 3.095882 | 0.274993294 | 0.362 | 0.135 | 7.8650897 | 12 | add3a |
| atrx1 | 4.040671 | 0.27933195 | 0.338 | 0.125 | 1.0265327 | 12 | atrx |
| prdx3 | 1.559691 | 0.296467952 | 0.359 | 0.14 | 3.9623952 | 12 | prdx3 |
| caprin1b | 1.761787 | 0.339397577 | 0.533 | 0.242 | 4.4758199 | 12 | caprin1b |
| elocb | 4.415657 | 0.325554403 | 0.443 | 0.192 | 1.1217978 | 12 | elocb |
| ola1 | 6.620257 | 0.323496729 | 0.53 | 0.24 | 1.6818765 | 12 | ola1 |
| romo13 | 7.809824 | 0.404416458 | 0.742 | 0.403 | 1.9840859 | 12 | romo1 |
| cct81 | 1.048601 | 0.319194987 | 0.491 | 0.218 | 2.6639722 | 12 | cct8 |
| pfn22 | 1.294981 | 0.32590082 | 0.523 | 0.235 | 3.2899013 | 12 | pfn2 |
| zgc:1119861 | 1.968132 | 0.293827454 | 0.432 | 0.182 | 5.0000408 | 12 | zgc:111986 |
| eif4a1b1 | 3.116833 | 0.376408255 | 0.61 | 0.307 | 7.9183147 | 12 | eif4a1b |
| cnbpb1 | 5.954701 | 0.301182561 | 0.467 | 0.206 | 1.5127919 | 12 | cnbpb |

| | | | | | | | |
|---|---|---|---|---|---|---|---|
| eef1b2 | 1.661295 | 0.285008664 | 0.909 | 0.739 | 4.2205217 | 12 | eef1b2 |
| ndufb8 | 1.816612 | 0.332058297 | 0.544 | 0.261 | 4.6151039 | 12 | ndufb8 |
| ppiaa4 | 2.434123 | 0.275860736 | 0.986 | 0.812 | 6.1838919 | 12 | ppiaa |
| rpl43 | 2.820979 | 0.380997089 | 0.948 | 0.652 | 7.1666980 | 12 | rpl4 |
| cirbpb | 7.277018 | 0.304934802 | 0.962 | 0.754 | 1.8487264 | 12 | cirbpb |
| ndufa5 | 1.062005 | 0.276332865 | 0.373 | 0.157 | 2.6980249 | 12 | ndufa5 |
| tcp11 | 2.420089 | 0.315522442 | 0.509 | 0.242 | 6.1482383 | 12 | tcp1 |
| eif3g3 | 2.669173 | 0.345920193 | 0.582 | 0.296 | 6.7810351 | 12 | eif3g |
| eif3ea2 | 2.691858 | 0.340938533 | 0.599 | 0.3 | 6.8386656 | 12 | eif3ea |
| atp5mc3a | 3.328021 | 0.312685735 | 0.502 | 0.237 | 8.4548396 | 12 | atp5mc3a |
| idh21 | 3.837464 | 0.306373563 | 0.477 | 0.224 | 9.7490797 | 12 | idh2 |
| cyc1 | 5.707870 | 0.268632377 | 0.355 | 0.149 | 1.4500845 | 12 | cyc1 |
| ostc | 9.230224 | 0.261086235 | 0.425 | 0.186 | 2.3449384 | 12 | ostc |
| atf4a1 | 3.163993 | 0.258842587 | 0.415 | 0.182 | 8.0381242 | 12 | atf4a |
| ppib4 | 6.003941 | 0.37109761 | 0.725 | 0.373 | 1.5253013 | 12 | ppib |
| prdx22 | 6.426346 | 0.386161205 | 0.7 | 0.418 | 1.6326134 | 12 | prdx2 |
| csrnp1b2 | 1.635719 | 0.281687835 | 0.369 | 0.162 | 4.1555445 | 12 | csrnp1b |
| eif3ba1 | 2.772141 | 0.312226604 | 0.481 | 0.238 | 7.0426248 | 12 | eif3ba |
| sdc43 | 2.829195 | 0.313505286 | 0.523 | 0.25 | 7.1875720 | 12 | sdc4 |
| pfn2l1 | 3.107604 | 0.255567998 | 0.467 | 0.212 | 7.8948683 | 12 | pfn2l |
| hnrnpa1a | 4.859878 | 0.27617992 | 0.366 | 0.163 | 1.2346521 | 12 | hnrnpa1a |
| rhoaa1 | 9.989877 | 0.256502757 | 0.387 | 0.174 | 2.5379283 | 12 | rhoaa |
| eif2s31 | 1.609577 | 0.318934136 | 0.578 | 0.304 | 4.0891320 | 12 | eif2s3 |
| jpt1b2 | 2.034091 | 0.285406932 | 0.561 | 0.273 | 5.1676093 | 12 | jpt1b |
| eif3k | 2.489602 | 0.26495358 | 0.495 | 0.241 | 6.3248346 | 12 | eif3k |
| cct4 | 3.084227 | 0.275304977 | 0.491 | 0.241 | 7.8354791 | 12 | cct4 |
| atp5pf1 | 8.086012 | 0.319712761 | 0.582 | 0.316 | 2.0542515 | 12 | atp5pf |
| ldhba1 | 1.019455 | 0.288408876 | 0.505 | 0.258 | 2.5899279 | 12 | ldhba |
| nfil3-63 | 1.233092 | 0.274487755 | 0.3 | 0.129 | 3.1326725 | 12 | nfil3-6 |
| sumo2b3 | 1.885759 | 0.290723798 | 0.564 | 0.293 | 4.7907710 | 12 | sumo2b |
| tomm20b1 | 1.888345 | 0.272195453 | 0.456 | 0.225 | 4.7973429 | 12 | tomm20b |
| alas11 | 3.879935 | 0.251382228 | 0.31 | 0.135 | 9.8569764 | 12 | alas1 |
| tegt1 | 5.069466 | 0.27435052 | 0.463 | 0.235 | 1.2878978 | 12 | tegt |
| ndufa4l2 | 9.122396 | 0.345720907 | 0.693 | 0.419 | 2.3175448 | 12 | ndufa4l |
| hmgb2b1 | 9.914589 | 0.27433107 | 0.645 | 0.332 | 2.5188014 | 12 | hmgb2b |

| | | | | | | | |
|---|---|---|---|---|---|---|---|
| rplp25 | 1.019700 | 0.299202866 | 0.979 | 0.78 | 2.5905496 | 12 | rplp2 |
| cebpd2 | 1.187178 | 0.279816818 | 0.331 | 0.15 | 3.0160266 | 12 | cebpd |
| midn2 | 1.352088 | 0.289410717 | 0.31 | 0.143 | 3.4349820 | 12 | midn |
| atp5pd1 | 2.119019 | 0.287030814 | 0.523 | 0.28 | 5.3833690 | 12 | atp5pd |
| atp5f1d1 | 4.078455 | 0.274524271 | 0.509 | 0.27 | 1.0361316 | 12 | atp5f1d |
| hmgn72 | 5.390821 | 0.277617675 | 0.596 | 0.314 | 1.3695383 | 12 | hmgn7 |
| pnrc25 | 2.271681 | 0.336489998 | 0.878 | 0.593 | 5.7712069 | 12 | pnrc2 |
| ywhaz1 | 2.242002 | 0.255255352 | 0.557 | 0.301 | 5.6958069 | 12 | ywhaz |
| ptmab4 | 5.232730 | 0.323786605 | 0.854 | 0.566 | 1.3293750 | 12 | ptmab |
| gabarapb4 | 9.542976 | 0.26286538 | 0.53 | 0.302 | 2.4243931 | 12 | gabarapb |
| tma7 | 1.170709 | 0.255358211 | 0.578 | 0.328 | 2.9741871 | 12 | tma7 |
| eef1da2 | 2.273593 | 0.315804064 | 0.805 | 0.501 | 5.7760652 | 12 | eef1da |
| serbp1a3 | 5.544743 | 0.313272655 | 0.787 | 0.502 | 1.4086419 | 12 | serbp1a |
| eif5a3 | 6.070151 | 0.301385432 | 0.774 | 0.491 | 1.5421219 | 12 | eif5a |
| fkbp1aa2 | 8.010336 | 0.263607939 | 0.69 | 0.41 | 2.0350260 | 12 | fkbp1aa |
| slc25a3b2 | 1.770500 | 0.277770086 | 0.728 | 0.454 | 4.4979567 | 12 | slc25a3b |
| cox7c1 | 1.727731 | 0.254547176 | 0.739 | 0.49 | 0.000439 | 12 | cox7c |
| ccl36.11 | 0 | 2.069288306 | 0.967 | 0.069 | 0 | 13 | ccl36.1 |
| si:dkey-21e2.12 | 0 | 1.696631014 | 0.752 | 0.004 | 0 | 13 | si:dkey-21e2.12 |
| ccl38.61 | 0 | 1.529092957 | 0.756 | 0.082 | 0 | 13 | ccl38.6 |
| si:dkey-21e2.11 | 0 | 1.408444163 | 0.559 | 0.002 | 0 | 13 | si:dkey-21e2.11 |
| ncaldb1 | 0 | 1.367173359 | 0.77 | 0.057 | 0 | 13 | ncaldb |
| zbtb321 | 0 | 1.361110268 | 0.719 | 0.028 | 0 | 13 | zbtb32 |
| ca2 | 0 | 1.313305098 | 0.667 | 0.041 | 0 | 13 | ca2 |
| si:ch211-277c7.61 | 0 | 1.282132316 | 0.607 | 0.026 | 0 | 13 | si:ch211-277c7.6 |
| scinlb3 | 0 | 1.166754857 | 0.707 | 0.073 | 0 | 13 | scinlb |
| il341 | 0 | 1.147453998 | 0.604 | 0.035 | 0 | 13 | il34 |
| si:ch211-207n23.2 | 0 | 1.138689936 | 0.596 | 0.016 | 0 | 13 | si:ch211-207n23.2 |
| si:dkey-78l4.4 | 0 | 1.125887627 | 0.563 | 0.021 | 0 | 13 | si:dkey-78l4.4 |
| ccr22 | 0 | 1.084892181 | 0.611 | 0.034 | 0 | 13 | ccr2 |
| si:rp71-81e14.21 | 0 | 0.978373053 | 0.478 | 0.022 | 0 | 13 | si:rp71-81e14.2 |
| CABZ01084575.11 | 0 | 0.950736364 | 0.504 | 0.031 | 0 | 13 | CABZ01084575.1 |
| blnk | 0 | 0.820965847 | 0.393 | 0.017 | 0 | 13 | blnk |
| ponzr6 | 0 | 0.744303224 | 0.367 | 0.015 | 0 | 13 | ponzr6 |
| dicp1.5-6 | 0 | 0.676505361 | 0.344 | 0.01 | 0 | 13 | dicp1.5-6 |

| | | | | | | | |
|---|---|---|---|---|---|---|---|
| BX000438.2 | 0 | 0.65808531 | 0.27 | 0.001 | 0 | 13 | BX000438.2 |
| fam167b | 0 | 0.5736612 | 0.278 | 0.006 | 0 | 13 | fam167b |
| XLOC-026336 | 0 | 0.407586956 | 0.185 | 0.002 | 0 | 13 | XLOC-026336 |
| obsl1a | 7.178215 | 0.401308526 | 0.178 | 0.002 | 1.8236257 | 13 | obsl1a |
| LOC1081907172 | 1.547910 | 1.254584915 | 0.685 | 0.071 | 3.9324670 | 13 | LOC108190717 |
| mctp1a | 1.646412 | 0.48613045 | 0.233 | 0.006 | 4.1827103 | 13 | mctp1a |
| si:ch211-102c2.43 | 3.614817 | 1.386944639 | 0.878 | 0.133 | 9.1834434 | 13 | si:ch211-102c2.4 |
| si:ch211-241f5.31 | 6.174687 | 1.030172994 | 0.581 | 0.052 | 1.5686793 | 13 | si:ch211-241f5.3 |
| LOC1039108711 | 6.640239 | 1.495429745 | 0.907 | 0.143 | 1.6869528 | 13 | LOC103910871 |
| XLOC-0139641 | 1.449782 | 1.573948544 | 0.907 | 0.156 | 3.6831727 | 13 | XLOC-013964 |
| LOC103910169 | 3.960867 | 0.338503876 | 0.163 | 0.002 | 1.0062584 | 13 | LOC103910169 |
| znrf1 | 1.015625 | 0.544239837 | 0.274 | 0.01 | 2.5801956 | 13 | znrf1 |
| XLOC-0425952 | 2.150919 | 1.476775471 | 0.885 | 0.139 | 5.4644097 | 13 | XLOC-042595 |
| itpr1b | 7.870874 | 0.575918635 | 0.27 | 0.01 | 1.9995957 | 13 | itpr1b |
| CR936442.12 | 9.724565 | 1.463284156 | 0.852 | 0.141 | 2.4705257 | 13 | CR936442.1 |
| sla21 | 1.902271 | 1.013419336 | 0.556 | 0.052 | 4.8327195 | 13 | sla2 |
| si:ch211-214p16.13 | 1.919265 | 1.650337482 | 0.985 | 0.231 | 4.8758935 | 13 | si:ch211-214p16.1 |
| tnxba | 2.498366 | 0.392883547 | 0.189 | 0.004 | 6.3471009 | 13 | tnxba |
| XLOC-0425961 | 1.311670 | 0.832108686 | 0.433 | 0.032 | 3.3322984 | 13 | XLOC-042596 |
| si:ch73-40a2.1 | 1.161383 | 0.427690566 | 0.215 | 0.006 | 2.9504954 | 13 | si:ch73-40a2.1 |
| XLOC-0203012 | 3.186961 | 1.094597609 | 0.578 | 0.061 | 8.0964756 | 13 | XLOC-020301 |
| creb5b | 2.126031 | 0.586953162 | 0.281 | 0.013 | 5.4011827 | 13 | creb5b |
| tnfb2 | 1.118305 | 1.010487169 | 0.47 | 0.042 | 2.8410545 | 13 | tnfb |
| nitr1m | 3.034728 | 0.495560324 | 0.207 | 0.006 | 7.7097273 | 13 | nitr1m |
| si:ch211-191c10.11 | 3.378578 | 0.845543368 | 0.459 | 0.042 | 8.5832789 | 13 | si:ch211-191c10.1 |
| dusp21 | 5.206145 | 1.265069022 | 0.841 | 0.151 | 1.3226211 | 13 | dusp2 |
| XLOC-0108992 | 6.888442 | 1.473657794 | 0.944 | 0.227 | 1.7500088 | 13 | XLOC-010899 |
| CR392341.12 | 2.496625 | 0.873088454 | 0.496 | 0.052 | 6.3426776 | 13 | CR392341.1 |
| XLOC-0283703 | 2.503003 | 1.259102336 | 0.715 | 0.115 | 6.3588814 | 13 | XLOC-028370 |
| CT573231.11 | 4.637072 | 0.630102574 | 0.322 | 0.021 | 1.1780483 | 13 | CT573231.1 |
| XLOC-0268362 | 1.070651 | 1.115281081 | 0.563 | 0.07 | 2.7199895 | 13 | XLOC-026836 |
| XLOC-0004313 | 1.392476 | 1.566080557 | 0.959 | 0.261 | 3.5375863 | 13 | XLOC-000431 |
| stim1a | 4.256304 | 0.860965747 | 0.504 | 0.057 | 1.0813141 | 13 | stim1a |
| ttyh2l | 8.933754 | 0.391980663 | 0.193 | 0.007 | 2.2696204 | 13 | ttyh2l |
| XLOC-0430431 | 3.825527 | 0.8744615 | 0.415 | 0.038 | 9.7187514 | 13 | XLOC-043043 |

| | | | | | | | |
|---|---|---|---|---|---|---|---|
| XLOC-0254233 | 1.427588 | 1.342249361 | 0.789 | 0.159 | 3.6267885 | 13 | XLOC-025423 |
| XLOC-0320022 | 2.487025 | 0.806070242 | 0.407 | 0.038 | 6.3182895 | 13 | XLOC-032002 |
| si:dkey-183i3.61 | 3.874967 | 0.719428666 | 0.389 | 0.034 | 9.8443544 | 13 | si:dkey-183i3.6 |
| CU914617.1 | 9.236010 | 0.265941981 | 0.104 | 0.001 | 2.3464085 | 13 | CU914617.1 |
| syk2 | 1.263101 | 0.877735961 | 0.522 | 0.07 | 3.2089094 | 13 | syk |
| prkcba | 4.377871 | 0.501632005 | 0.248 | 0.015 | 1.1121983 | 13 | prkcba |
| zgc:640514 | 6.241835 | 1.081973367 | 0.752 | 0.152 | 1.5857383 | 13 | zgc:64051 |
| orai1a | 8.055439 | 0.527508376 | 0.278 | 0.019 | 2.0464844 | 13 | orai1a |
| gramd2aa | 5.513538 | 0.465981132 | 0.237 | 0.014 | 1.4007143 | 13 | gramd2aa |
| gfi1ab | 6.947238 | 0.537860706 | 0.278 | 0.019 | 1.7649459 | 13 | gfi1ab |
| CABZ01044746.1 | 7.333328 | 0.924573938 | 0.57 | 0.083 | 1.8630321 | 13 | CABZ01044746.1 |
| arpc1b5 | 6.075427 | 1.214455996 | 0.967 | 0.302 | 1.5434624 | 13 | arpc1b |
| BX901920.12 | 1.156595 | 0.881075111 | 0.496 | 0.066 | 2.9383313 | 13 | BX901920.1 |
| LOC1005362132 | 1.232761 | 0.928404924 | 0.626 | 0.102 | 3.1318294 | 13 | LOC100536213 |
| mbnl13 | 1.039723 | 1.043299596 | 0.778 | 0.176 | 2.6414168 | 13 | mbnl1 |
| rgs132 | 1.676224 | 0.980713166 | 0.656 | 0.116 | 4.2584471 | 13 | rgs13 |
| il2rb1 | 1.813231 | 0.485660686 | 0.237 | 0.015 | 4.6065156 | 13 | il2rb |
| XLOC-0437601 | 1.498844 | 0.773175175 | 0.378 | 0.04 | 3.8078147 | 13 | XLOC-043760 |
| wdr191 | 3.045565 | 0.611641398 | 0.315 | 0.027 | 7.7372582 | 13 | wdr19 |
| slc41a1 | 3.475557 | 0.395086978 | 0.196 | 0.01 | 8.8296534 | 13 | slc41a1 |
| CABZ01117603.1 | 6.131660 | 0.4492975 | 0.23 | 0.014 | 1.5577484 | 13 | CABZ01117603.1 |
| XLOC-039757 | 1.641780 | 0.581017235 | 0.281 | 0.022 | 4.1709440 | 13 | XLOC-039757 |
| XLOC-0234923 | 2.571613 | 0.943971926 | 0.541 | 0.085 | 6.5331851 | 13 | XLOC-023492 |
| XLOC-0140791 | 5.910542 | 0.617249454 | 0.307 | 0.027 | 1.5015732 | 13 | XLOC-014079 |
| camk1db | 2.740379 | 0.423802284 | 0.222 | 0.014 | 6.9619336 | 13 | camk1db |
| cd44b | 1.214311 | 0.590624438 | 0.307 | 0.028 | 3.0849574 | 13 | cd44b |
| si:ch211-214p16.22 | 1.422829 | 0.821322889 | 0.481 | 0.068 | 3.6146979 | 13 | si:ch211-214p16.2 |
| CU972454.41 | 5.495773 | 0.475044171 | 0.248 | 0.018 | 1.3962011 | 13 | CU972454.4 |
| laptm55 | 1.424733 | 1.020283413 | 0.737 | 0.166 | 3.6195361 | 13 | laptm5 |
| tagapa3 | 1.534878 | 0.915124339 | 0.541 | 0.09 | 3.8993579 | 13 | tagapa |
| gpx1a2 | 8.881854 | 1.264798894 | 0.767 | 0.227 | 2.2564350 | 13 | gpx1a |
| calm1b3 | 8.977613 | 0.835764483 | 0.552 | 0.095 | 2.2807627 | 13 | calm1b |
| irf4b1 | 1.592997 | 0.520295259 | 0.248 | 0.019 | 4.0470096 | 13 | irf4b |
| CR753876.12 | 2.638929 | 0.961407379 | 0.578 | 0.103 | 6.7042013 | 13 | CR753876.1 |
| srgn4 | 6.710024 | 1.023894181 | 0.852 | 0.224 | 1.7046817 | 13 | srgn |

| | | | | | | | |
|---|---|---|---|---|---|---|---|
| atp2b1a | 1.147945 | 0.789011666 | 0.456 | 0.065 | 2.9163547 | 13 | atp2b1a |
| sik11 | 1.362376 | 0.849778419 | 0.57 | 0.102 | 3.4611171 | 13 | sik1 |
| si:ch211-167j9.51 | 3.560877 | 0.464089373 | 0.256 | 0.02 | 9.0464085 | 13 | si:ch211-167j9.5 |
| cd40lg | 6.979467 | 0.334591943 | 0.167 | 0.008 | 1.7731337 | 13 | cd40lg |
| id31 | 1.297645 | 1.047251858 | 0.737 | 0.188 | 3.2966680 | 13 | id3 |
| dock10 | 9.671068 | 0.514825693 | 0.274 | 0.025 | 2.4569348 | 13 | dock10 |
| LOC1039091071 | 2.763291 | 0.515743341 | 0.237 | 0.019 | 7.0201422 | 13 | LOC103909107 |
| fcer1g1 | 5.862297 | 0.59117199 | 0.337 | 0.038 | 1.4893167 | 13 | fcer1g |
| FP236356.11 | 5.464338 | 0.643073199 | 0.344 | 0.041 | 1.3882151 | 13 | FP236356.1 |
| mef2d | 6.568481 | 0.742048952 | 0.441 | 0.069 | 1.6687226 | 13 | mef2d |
| XLOC-0437611 | 2.281647 | 0.476774789 | 0.215 | 0.016 | 5.7965254 | 13 | XLOC-043761 |
| bhlhe402 | 1.302145 | 0.948878584 | 0.663 | 0.159 | 3.3081002 | 13 | bhlhe40 |
| BX005183.33 | 1.197493 | 0.720984294 | 0.444 | 0.07 | 3.0422310 | 13 | BX005183.3 |
| hcls12 | 7.682276 | 0.687084994 | 0.441 | 0.07 | 1.9516824 | 13 | hcls1 |
| arhgdig4 | 2.233643 | 0.959392115 | 0.815 | 0.25 | 5.6745717 | 13 | arhgdig |
| atp11c | 2.637753 | 0.517260196 | 0.27 | 0.027 | 6.7012138 | 13 | atp11c |
| plcg2 | 5.136314 | 0.638965893 | 0.344 | 0.045 | 1.3048807 | 13 | plcg2 |
| adam19b | 6.218478 | 0.376556419 | 0.178 | 0.012 | 1.5798043 | 13 | adam19b |
| gzm3.4 | 4.588553 | 0.298433702 | 0.122 | 0.005 | 1.1657219 | 13 | gzm3.4 |
| aldocb1 | 6.999146 | 0.622513711 | 0.393 | 0.06 | 1.7781330 | 13 | aldocb |
| ptprc4 | 4.791259 | 0.777993187 | 0.559 | 0.117 | 1.2172194 | 13 | ptprc |
| XLOC-0054492 | 1.054611 | 0.666711323 | 0.359 | 0.052 | 2.6792393 | 13 | XLOC-005449 |
| BX322618.11 | 2.119770 | 0.598416513 | 0.278 | 0.031 | 5.3852779 | 13 | BX322618.1 |
| capgb3 | 5.421555 | 0.780890414 | 0.533 | 0.108 | 1.3773462 | 13 | capgb |
| pxna1 | 2.877228 | 0.711418455 | 0.463 | 0.083 | 7.3095991 | 13 | pxna |
| kdm2ba | 1.547167 | 0.521355748 | 0.296 | 0.037 | 3.9305784 | 13 | kdm2ba |
| XLOC-041870 | 1.242086 | 0.563380916 | 0.256 | 0.027 | 3.1555211 | 13 | XLOC-041870 |
| CABZ01111959.1 | 2.887328 | 0.423820815 | 0.233 | 0.023 | 7.3352591 | 13 | CABZ01111959.1 |
| si:dkey-151g10.3 | 5.182122 | 0.349728724 | 0.181 | 0.014 | 1.3165183 | 13 | si:dkey-151g10.3 |
| lck | 2.371192 | 0.2930129 | 0.141 | 0.008 | 6.0240136 | 13 | lck |
| coro1a5 | 7.786820 | 0.895714079 | 0.789 | 0.227 | 1.9782416 | 13 | coro1a |
| tmsb4x8 | 9.099683 | 0.67805869 | 1 | 0.714 | 2.3117746 | 13 | tmsb4x |
| pik3r11 | 3.658631 | 0.67470385 | 0.448 | 0.085 | 9.2947534 | 13 | pik3r1 |
| zgc:153018 | 1.969309 | 0.470287899 | 0.23 | 0.024 | 5.0030314 | 13 | zgc:153018 |
| wasa3 | 2.938013 | 0.661943534 | 0.47 | 0.09 | 7.4640223 | 13 | wasa |

| | | | | | | | |
|---|---|---|---|---|---|---|---|
| plp2 | 3.101437 | 0.555623911 | 0.333 | 0.049 | 7.8792018 | 13 | plp2 |
| BX908750.11 | 3.818658 | 0.466657966 | 0.252 | 0.028 | 9.7013008 | 13 | BX908750.1 |
| cd247l1 | 9.811361 | 0.362641942 | 0.196 | 0.017 | 2.4925764 | 13 | cd247l |
| tmc6a | 2.305742 | 0.328269345 | 0.156 | 0.011 | 5.8577395 | 13 | tmc6a |
| LOC1104399311 | 1.050362 | 0.39293087 | 0.204 | 0.019 | 2.6684469 | 13 | LOC110439931 |
| CR381673.12 | 5.233297 | 0.55269107 | 0.344 | 0.052 | 1.3295192 | 13 | CR381673.1 |
| fscn1a1 | 2.149427 | 0.581143111 | 0.33 | 0.049 | 5.4606194 | 13 | fscn1a |
| sla11 | 1.513870 | 0.378672266 | 0.215 | 0.021 | 3.8459879 | 13 | sla1 |
| vaspb | 1.931311 | 0.574775437 | 0.322 | 0.048 | 4.9064960 | 13 | vaspb |
| si:dkey-53k12.22 | 4.437562 | 0.517059186 | 0.296 | 0.04 | 1.1273626 | 13 | si:dkey-53k12.2 |
| casz1 | 1.106319 | 0.373103151 | 0.193 | 0.017 | 2.8106049 | 13 | casz1 |
| si:ch73-208g10.12 | 3.253111 | 0.512224367 | 0.281 | 0.037 | 8.2645295 | 13 | si:ch73-208g10.1 |
| atp5if1b1 | 5.059593 | 0.816060731 | 0.67 | 0.199 | 1.2853897 | 13 | atp5if1b |
| si:ch211-215e19.31 | 8.331665 | 0.369004839 | 0.189 | 0.017 | 2.1166597 | 13 | si:ch211-215e19.3 |
| tnip11 | 8.355905 | 0.701126019 | 0.4 | 0.075 | 2.1228178 | 13 | tnip1 |
| si:dkey-1c7.1 | 8.677580 | 0.347232565 | 0.193 | 0.017 | 2.2045392 | 13 | si:dkey-1c7.1 |
| mef2ca | 1.251463 | 0.273632325 | 0.13 | 0.008 | 3.1793431 | 13 | mef2ca |
| elf12 | 6.592222 | 0.700222363 | 0.463 | 0.097 | 1.6747540 | 13 | elf1 |
| ptmaa5 | 7.476751 | 0.856780326 | 0.933 | 0.425 | 1.8994687 | 13 | ptmaa |
| CR392341.4 | 1.425594 | 0.277753587 | 0.13 | 0.008 | 3.6217238 | 13 | CR392341.4 |
| caspa | 4.187297 | 0.401131851 | 0.215 | 0.022 | 1.0637828 | 13 | caspa |
| LOC110438754 | 1.705933 | 0.290314058 | 0.159 | 0.012 | 4.3339238 | 13 | LOC110438754 |
| si:ch211-93g21.11 | 1.797335 | 0.424588818 | 0.248 | 0.029 | 4.5661304 | 13 | si:ch211-93g21.1 |
| XLOC-0331371 | 6.342534 | 0.495671417 | 0.252 | 0.031 | 1.6113208 | 13 | XLOC-033137 |
| arpc5a4 | 2.489175 | 0.702383163 | 0.511 | 0.119 | 6.3237507 | 13 | arpc5a |
| rplp26 | 8.473488 | 0.490842203 | 0.993 | 0.78 | 2.1526896 | 13 | rplp2 |
| CABZ01054965.1 | 9.694261 | 0.354027239 | 0.163 | 0.013 | 2.4628270 | 13 | CABZ01054965.1 |
| gsdmea1 | 4.220113 | 0.406617041 | 0.204 | 0.021 | 1.0721198 | 13 | gsdmea |
| tmc81 | 7.383604 | 0.445667472 | 0.248 | 0.03 | 1.8758046 | 13 | tmc8 |
| grapa1 | 1.240446 | 0.441911007 | 0.222 | 0.025 | 3.1513540 | 13 | grapa |
| cotl15 | 6.557745 | 0.870808913 | 0.848 | 0.311 | 1.6659952 | 13 | cotl1 |
| fpr1 | 3.044455 | 0.379417086 | 0.193 | 0.019 | 7.7344396 | 13 | fpr1 |
| si:cabz01074946.12 | 3.303687 | 0.55248567 | 0.341 | 0.057 | 8.3930187 | 13 | si:cabz01074946.1 |
| cox6b14 | 7.561880 | 0.681023427 | 0.496 | 0.116 | 1.9210957 | 13 | cox6b1 |
| tagapb2 | 3.129953 | 0.599173306 | 0.363 | 0.065 | 7.9516456 | 13 | tagapb |

| | | | | | | | |
|---|---|---|---|---|---|---|---|
| rac25 | 6.156798 | 0.792230061 | 0.726 | 0.216 | 1.5641347 | 13 | rac2 |
| zgc:1749452 | 2.616259 | 0.376377313 | 0.204 | 0.021 | 6.6466084 | 13 | zgc:174945 |
| si:ch211-119e14.12 | 3.393731 | 0.603083306 | 0.367 | 0.066 | 8.6217740 | 13 | si:ch211-119e14.1 |
| si:rp71-68n21.92 | 4.807775 | 0.487055336 | 0.289 | 0.043 | 1.2214154 | 13 | si:rp71-68n21.9 |
| si:dkeyp-19e1.3 | 6.120613 | 0.560532088 | 0.341 | 0.058 | 1.5549417 | 13 | si:dkeyp-19e1.3 |
| dok22 | 7.383730 | 0.45687113 | 0.267 | 0.036 | 1.8758366 | 13 | dok2 |
| wasb5 | 1.671077 | 0.771038959 | 0.607 | 0.166 | 4.2453712 | 13 | wasb |
| si:dkey-183p4.101 | 3.051691 | 0.415323103 | 0.237 | 0.029 | 7.7528231 | 13 | si:dkey-183p4.10 |
| lasp13 | 1.634674 | 0.780036944 | 0.715 | 0.246 | 4.1528905 | 13 | lasp1 |
| unc13d1 | 1.704102 | 0.393660803 | 0.226 | 0.027 | 4.3292733 | 13 | unc13d |
| XLOC-0110851 | 2.019468 | 0.530921231 | 0.322 | 0.054 | 5.1304608 | 13 | XLOC-011085 |
| ponzr14 | 2.039373 | 0.837432948 | 0.752 | 0.282 | 5.1810277 | 13 | ponzr1 |
| ikzf11 | 3.352297 | 0.41006472 | 0.241 | 0.031 | 8.5165123 | 13 | ikzf1 |
| ptmab5 | 9.088999 | 0.705267882 | 0.941 | 0.564 | 2.3090603 | 13 | ptmab |
| CABZ01088229.1 | 1.208342 | 0.269447944 | 0.141 | 0.011 | 3.0697945 | 13 | CABZ01088229.1 |
| rps6ka11 | 3.135436 | 0.558843477 | 0.356 | 0.065 | 7.9655764 | 13 | rps6ka1 |
| dtnbp1a2 | 9.421391 | 0.571748004 | 0.407 | 0.083 | 2.3935044 | 13 | dtnbp1a |
| bcl2l101 | 2.654712 | 0.795595899 | 0.633 | 0.193 | 6.7442960 | 13 | bcl2l10 |
| LOC1039101361 | 9.167329 | 0.399268184 | 0.207 | 0.024 | 2.3289600 | 13 | LOC103910136 |
| si:dkey-260g12.11 | 1.549588 | 0.428207102 | 0.226 | 0.028 | 3.9367297 | 13 | si:dkey-260g12.1 |
| csf2rb1 | 1.442619 | 0.477527657 | 0.252 | 0.035 | 3.6649757 | 13 | csf2rb |
| XLOC-0089931 | 2.213601 | 0.401207329 | 0.207 | 0.024 | 5.6236547 | 13 | XLOC-008993 |
| nptna1 | 2.542474 | 0.562640353 | 0.348 | 0.063 | 6.4591568 | 13 | nptna |
| RASSF52 | 3.239893 | 0.545307741 | 0.322 | 0.057 | 8.2309504 | 13 | RASSF5 |
| fybb3 | 6.749311 | 0.458075451 | 0.27 | 0.041 | 1.7146626 | 13 | fybb |
| strbp | 9.914838 | 0.320175703 | 0.167 | 0.016 | 2.5188647 | 13 | strbp |
| si:ch211-132g1.11 | 1.468406 | 0.364855143 | 0.181 | 0.019 | 3.7304875 | 13 | si:ch211-132g1.1 |
| crema2 | 3.235143 | 0.770023395 | 0.563 | 0.168 | 8.2188811 | 13 | crema |
| vaspa3 | 1.957207 | 0.570724586 | 0.381 | 0.077 | 4.9722857 | 13 | vaspa |
| entpd31 | 6.095192 | 0.326394969 | 0.152 | 0.014 | 1.5484837 | 13 | entpd3 |
| si:dkeyp-68b7.122 | 1.332237 | 0.499066897 | 0.315 | 0.056 | 3.3845495 | 13 | si:dkeyp-68b7.12 |
| cap15 | 1.746105 | 0.735183945 | 0.707 | 0.238 | 4.4359821 | 13 | cap1 |
| itga42 | 3.001125 | 0.524511681 | 0.326 | 0.06 | 7.6243604 | 13 | itga4 |
| arpc34 | 5.263127 | 0.765455905 | 0.789 | 0.327 | 1.3370975 | 13 | arpc3 |
| myl12.13 | 6.073080 | 0.73748278 | 0.863 | 0.419 | 1.5428662 | 13 | myl12.1 |

| | | | | | | | |
|---|---|---|---|---|---|---|---|
| rin2 | 7.132933 | 0.385696891 | 0.189 | 0.022 | 1.8121216 | 13 | rin2 |
| yrk2 | 1.149230 | 0.508055069 | 0.337 | 0.063 | 2.9196209 | 13 | yrk |
| wipf1a2 | 1.736404 | 0.525317821 | 0.337 | 0.064 | 4.4113368 | 13 | wipf1a |
| fmnl1a | 1.943035 | 0.455442064 | 0.241 | 0.035 | 4.9362805 | 13 | fmnl1a |
| cers1 | 5.420254 | 0.349901642 | 0.174 | 0.019 | 1.3770157 | 13 | cers1 |
| hmgn23 | 6.654551 | 0.853732493 | 0.881 | 0.461 | 1.6905886 | 13 | hmgn2 |
| cmah1 | 5.834635 | 0.295269373 | 0.148 | 0.014 | 1.4822891 | 13 | cmah |
| pip5k1ba | 7.450839 | 0.2803023 | 0.137 | 0.012 | 1.8928856 | 13 | pip5k1ba |
| samd11 | 3.113967 | 0.331971352 | 0.159 | 0.016 | 7.9110341 | 13 | samd11 |
| arl111 | 5.312279 | 0.36377041 | 0.189 | 0.023 | 1.3495846 | 13 | arl11 |
| h1fx3 | 1.834445 | 0.787875831 | 0.581 | 0.184 | 4.6604090 | 13 | h1fx |
| arpc24 | 2.748245 | 0.723988033 | 0.73 | 0.292 | 6.9819182 | 13 | arpc2 |
| rps296 | 7.651290 | 0.365052226 | 0.985 | 0.842 | 1.9438103 | 13 | rps29 |
| megf8l | 9.660514 | 0.2658417 | 0.133 | 0.012 | 2.4542536 | 13 | megf8l |
| LOC101882375 | 1.205396 | 0.292425452 | 0.133 | 0.012 | 3.0623098 | 13 | LOC101882375 |
| arpc43 | 1.579398 | 0.709465654 | 0.656 | 0.231 | 4.0124629 | 13 | arpc4 |
| pycard5 | 4.624125 | 0.532578899 | 0.363 | 0.079 | 1.1747591 | 13 | pycard |
| ptpn22 | 1.136425 | 0.332834292 | 0.178 | 0.021 | 2.8870879 | 13 | ptpn22 |
| zgc:195175 | 1.481120 | 0.406685101 | 0.226 | 0.034 | 3.7627867 | 13 | zgc:195175 |
| cyth4b | 1.588542 | 0.296635721 | 0.152 | 0.016 | 4.0356931 | 13 | cyth4b |
| pfn15 | 2.062401 | 0.813637031 | 0.985 | 0.494 | 5.2395314 | 13 | pfn1 |
| si:ch211-222l21.14 | 3.366389 | 0.74713861 | 0.774 | 0.307 | 8.5523137 | 13 | si:ch211-222l21.1 |
| egr31 | 3.927158 | 0.504454019 | 0.278 | 0.05 | 9.9769468 | 13 | egr3 |
| mctp1b1 | 1.106557 | 0.318056453 | 0.167 | 0.019 | 2.8112086 | 13 | mctp1b |
| BX120005.11 | 7.521194 | 0.476495456 | 0.256 | 0.043 | 1.9107594 | 13 | BX120005.1 |
| pim22 | 9.399806 | 0.572069659 | 0.363 | 0.084 | 2.3880207 | 13 | pim2 |
| kif21b | 1.484727 | 0.340001029 | 0.174 | 0.022 | 3.7719513 | 13 | kif21b |
| sox5 | 6.866540 | 0.25771256 | 0.122 | 0.011 | 1.7444447 | 13 | sox5 |
| inpp5d | 1.308923 | 0.378502295 | 0.215 | 0.033 | 3.3253193 | 13 | inpp5d |
| clic13 | 3.873505 | 0.59148692 | 0.444 | 0.124 | 9.8406408 | 13 | clic1 |
| ldha1 | 6.104205 | 0.487241272 | 0.315 | 0.066 | 1.5507734 | 13 | ldha |
| nkl.21 | 2.461457 | 0.656731895 | 0.463 | 0.128 | 6.2533334 | 13 | nkl.2 |
| pgam1a2 | 2.469031 | 0.601902869 | 0.5 | 0.15 | 6.2725735 | 13 | pgam1a |
| gcnt4a | 9.606795 | 0.280350348 | 0.133 | 0.013 | 2.4406064 | 13 | gcnt4a |
| dub2 | 2.416103 | 0.466778397 | 0.322 | 0.07 | 6.1381112 | 13 | dub |

| | | | | | | | |
|---|---|---|---|---|---|---|---|
| arhgef3l1 | 2.617098 | 0.326983314 | 0.196 | 0.028 | 6.6487382 | 13 | arhgef3l |
| rplp15 | 1.382263 | 0.334987172 | 0.996 | 0.881 | 3.5116409 | 13 | rplp1 |
| rps218 | 2.501488 | 0.428748289 | 0.985 | 0.773 | 6.3550323 | 13 | rps21 |
| abhd3 | 3.929822 | 0.26743934 | 0.152 | 0.018 | 9.9837149 | 13 | abhd3 |
| si:dkey-262k9.43 | 2.132255 | 0.515008376 | 0.356 | 0.086 | 5.4169941 | 13 | si:dkey-262k9.4 |
| msna4 | 9.107069 | 0.66926275 | 0.652 | 0.26 | 2.3136508 | 13 | msna |
| rpl374 | 1.052143 | 0.345604603 | 0.985 | 0.853 | 2.6729695 | 13 | rpl37 |
| rps27.17 | 1.354202 | 0.475089874 | 0.97 | 0.706 | 3.4403511 | 13 | rps27.1 |
| cebpb5 | 1.840833 | 0.740238244 | 0.781 | 0.332 | 4.6766362 | 13 | cebpb |
| mef2b1 | 6.396228 | 0.263359269 | 0.122 | 0.012 | 1.6249618 | 13 | mef2b |
| cyfip2 | 3.999540 | 0.298368542 | 0.17 | 0.023 | 1.0160831 | 13 | cyfip2 |
| rpl115 | 7.885535 | 0.365735766 | 0.985 | 0.827 | 2.0033202 | 13 | rpl11 |
| actb25 | 8.053920 | 0.398258041 | 1 | 0.865 | 2.0460985 | 13 | actb2 |
| dusp12 | 1.302406 | 0.624911613 | 0.463 | 0.145 | 3.3087643 | 13 | dusp1 |
| arhgdia3 | 2.193804 | 0.629799771 | 0.626 | 0.246 | 5.5733605 | 13 | arhgdia |
| si:dkey-171c9.31 | 4.718724 | 0.33952119 | 0.193 | 0.03 | 1.1987918 | 13 | si:dkey-171c9.3 |
| tefa1 | 6.239290 | 0.558255504 | 0.444 | 0.136 | 1.5850918 | 13 | tefa |
| grap2a1 | 1.522246 | 0.391255394 | 0.237 | 0.045 | 3.8672661 | 13 | grap2a |
| nr4a14 | 1.987770 | 0.613809982 | 0.485 | 0.153 | 5.0499304 | 13 | nr4a1 |
| BX248082.12 | 3.054974 | 0.441582926 | 0.204 | 0.034 | 7.7611637 | 13 | BX248082.1 |
| rps173 | 6.098596 | 0.389553949 | 0.974 | 0.776 | 1.5493484 | 13 | rps17 |
| si:ch73-264p11.12 | 2.103183 | 0.314686788 | 0.174 | 0.026 | 5.3431376 | 13 | si:ch73-264p11.1 |
| lrrfip1a1 | 3.176577 | 0.407533784 | 0.27 | 0.058 | 8.0700957 | 13 | lrrfip1a |
| nfkbiaa2 | 5.407857 | 0.676966384 | 0.452 | 0.148 | 1.3738660 | 13 | nfkbiaa |
| LOC1001511961 | 1.118706 | 0.362301281 | 0.211 | 0.038 | 2.8420744 | 13 | LOC100151196 |
| mrps151 | 2.916616 | 0.525733657 | 0.389 | 0.109 | 7.4096645 | 13 | mrps15 |
| si:ch73-223p23.21 | 3.419114 | 0.286431359 | 0.174 | 0.026 | 8.6862600 | 13 | si:ch73-223p23.2 |
| edaradd | 4.075400 | 0.270484775 | 0.148 | 0.02 | 1.0353555 | 13 | edaradd |
| rpl193 | 5.697796 | 0.322199838 | 0.985 | 0.852 | 1.4475251 | 13 | rpl19 |
| si:dkey-185m8.21 | 6.650277 | 0.363286131 | 0.23 | 0.044 | 1.6895029 | 13 | si:dkey-185m8.2 |
| mt-co15 | 7.584207 | 0.560319821 | 0.981 | 0.635 | 1.9267678 | 13 | mt-co1 |
| skap22 | 1.522438 | 0.442864217 | 0.315 | 0.077 | 3.8677544 | 13 | skap2 |
| cd832 | 2.978646 | 0.56969941 | 0.344 | 0.09 | 7.5672521 | 13 | cd83 |
| plxnc12 | 5.555141 | 0.378291148 | 0.226 | 0.044 | 1.4112835 | 13 | plxnc1 |
| camk1b1 | 6.527834 | 0.262180287 | 0.148 | 0.02 | 1.6583964 | 13 | camk1b |

| | | | | | | | |
|---|---|---|---|---|---|---|---|
| si:ch211-226m16.21 | 2.843902 | 0.447184486 | 0.281 | 0.066 | 7.2249334 | 13 | si:ch211-226m16.2 |
| sort1b | 2.913200 | 0.32253244 | 0.174 | 0.028 | 7.4009850 | 13 | sort1b |
| rps285 | 4.683489 | 0.329179155 | 0.978 | 0.839 | 1.1898406 | 13 | rps28 |
| ankrd44 | 5.423154 | 0.310625745 | 0.17 | 0.027 | 1.3777524 | 13 | ankrd44 |
| gapdhs6 | 1.240035 | 0.603596947 | 0.696 | 0.309 | 3.1503102 | 13 | gapdhs |
| rps26l6 | 5.100831 | 0.452295831 | 0.963 | 0.696 | 1.2958662 | 13 | rps26l |
| rpl344 | 5.294812 | 0.373639961 | 0.993 | 0.8 | 1.3451470 | 13 | rpl34 |
| syne1b | 7.124373 | 0.281068754 | 0.148 | 0.021 | 1.8099470 | 13 | syne1b |
| si:dkey-78o7.31 | 7.376406 | 0.287683355 | 0.141 | 0.019 | 1.8739761 | 13 | si:dkey-78o7.3 |
| dock8 | 1.210986 | 0.318015143 | 0.196 | 0.035 | 3.0765121 | 13 | dock8 |
| sh2b31 | 1.369927 | 0.441511941 | 0.259 | 0.059 | 3.4802999 | 13 | sh2b3 |
| selplg1 | 1.543537 | 0.349272613 | 0.219 | 0.043 | 3.9213576 | 13 | selplg |
| prkcaa | 2.287897 | 0.252515558 | 0.13 | 0.017 | 5.8124023 | 13 | prkcaa |
| CU571074.1 | 2.379246 | 0.311739487 | 0.178 | 0.03 | 6.0444748 | 13 | CU571074.1 |
| arf6b1 | 9.987460 | 0.460061711 | 0.37 | 0.11 | 2.5373142 | 13 | arf6b |
| celf24 | 1.823044 | 0.576388425 | 0.611 | 0.252 | 4.6314434 | 13 | celf2 |
| rpl275 | 2.223820 | 0.378107874 | 0.978 | 0.777 | 5.6496154 | 13 | rpl27 |
| actr2a4 | 2.989238 | 0.493820756 | 0.411 | 0.129 | 7.5941592 | 13 | actr2a |
| prelid1b | 5.554411 | 0.266397562 | 0.148 | 0.022 | 1.4110982 | 13 | prelid1b |
| cd637 | 5.208130 | 0.592004248 | 0.748 | 0.368 | 1.3231255 | 13 | cd63 |
| myo1f2 | 5.832666 | 0.368750071 | 0.219 | 0.046 | 1.4817888 | 13 | myo1f |
| rel | 7.243725 | 0.334307284 | 0.181 | 0.033 | 1.8402685 | 13 | rel |
| dock11 | 1.630894 | 0.259477916 | 0.148 | 0.023 | 4.1432883 | 13 | dock11 |
| il162 | 2.349069 | 0.364743692 | 0.219 | 0.046 | 5.9678106 | 13 | il16 |
| pnrc26 | 2.414773 | 0.522915311 | 0.944 | 0.592 | 6.1347311 | 13 | pnrc2 |
| crip16 | 3.245516 | 0.588880689 | 0.659 | 0.278 | 8.2452345 | 13 | crip1 |
| nab1a1 | 3.697231 | 0.294786333 | 0.133 | 0.019 | 9.3928164 | 13 | nab1a |
| irf2bp2a2 | 4.274711 | 0.540448664 | 0.485 | 0.181 | 1.0859904 | 13 | irf2bp2a |
| btg15 | 5.006819 | 0.595649903 | 0.719 | 0.35 | 1.2719825 | 13 | btg1 |
| rhoab3 | 1.215979 | 0.540191789 | 0.515 | 0.203 | 3.0891956 | 13 | rhoab |
| rps15a7 | 1.248965 | 0.379724278 | 0.978 | 0.795 | 3.1729980 | 13 | rps15a |
| capzb4 | 1.324612 | 0.559054774 | 0.607 | 0.267 | 3.3651790 | 13 | capzb |
| gng52 | 1.459567 | 0.561328811 | 0.63 | 0.282 | 3.7080301 | 13 | gng5 |
| fli1b1 | 4.776442 | 0.300134822 | 0.178 | 0.033 | 1.2134551 | 13 | fli1b |
| CR855389.1 | 1.353934 | 0.277456373 | 0.163 | 0.028 | 3.4396703 | 13 | CR855389.1 |

| | | | | | | | |
|---|---|---|---|---|---|---|---|
| cox8a4 | 2.247505 | 0.557244166 | 0.589 | 0.263 | 5.7097885 | 13 | cox8a |
| pik3cg | 2.311135 | 0.260328079 | 0.144 | 0.023 | 5.8714387 | 13 | pik3cg |
| ccdc88b | 3.353729 | 0.269568564 | 0.17 | 0.031 | 8.5201492 | 13 | ccdc88b |
| rps156 | 3.882064 | 0.41497452 | 0.956 | 0.713 | 9.8623846 | 13 | rps15 |
| CABZ01079663.11 | 4.121925 | 0.296478952 | 0.185 | 0.035 | 1.0471752 | 13 | CABZ01079663.1 |
| cfl14 | 4.199186 | 0.509091832 | 0.919 | 0.595 | 1.0668033 | 13 | cfl1 |
| rps219 | 5.236579 | 0.343605936 | 0.985 | 0.84 | 1.3303529 | 13 | rps2 |
| unm-sa1614 | 7.797853 | 0.263274165 | 0.122 | 0.017 | 1.9810446 | 13 | unm-sa1614 |
| hcst1 | 2.518697 | 0.313812379 | 0.204 | 0.043 | 6.3987500 | 13 | hcst |
| rpl385 | 3.604199 | 0.385597458 | 0.97 | 0.751 | 9.1564678 | 13 | rpl38 |
| kctd5a1 | 4.582297 | 0.510088439 | 0.47 | 0.181 | 1.1641327 | 13 | kctd5a |
| rpl366 | 4.840154 | 0.353630431 | 0.985 | 0.804 | 1.2296411 | 13 | rpl36 |
| rpl18a6 | 1.055717 | 0.339175431 | 0.963 | 0.799 | 2.6820495 | 13 | rpl18a |
| carhsp1 | 1.081590 | 0.371575804 | 0.27 | 0.072 | 2.7477816 | 13 | carhsp1 |
| rplp2l5 | 2.211929 | 0.358725115 | 0.981 | 0.792 | 5.6194065 | 13 | rplp2l |
| elovl1b1 | 3.741531 | 0.442219055 | 0.363 | 0.119 | 9.5053618 | 13 | elovl1b |
| lcp13 | 4.329526 | 0.373022655 | 0.293 | 0.081 | 1.0999160 | 13 | lcp1 |
| si:ch211-196l7.45 | 6.783322 | 0.542208099 | 0.663 | 0.325 | 1.7233031 | 13 | si:ch211-196l7.4 |
| LOC1039091153 | 9.553041 | 0.464006817 | 0.448 | 0.165 | 2.4269501 | 13 | LOC103909115 |
| grap2b4 | 1.254293 | 0.431124557 | 0.374 | 0.121 | 3.1865323 | 13 | grap2b |
| rpl355 | 1.650430 | 0.322659151 | 0.978 | 0.812 | 4.1929183 | 13 | rpl35 |
| sept22 | 1.893990 | 0.472970075 | 0.463 | 0.178 | 4.8116821 | 13 | sept2 |
| tagln24 | 2.053594 | 0.521447254 | 0.504 | 0.206 | 5.2171564 | 13 | tagln2 |
| rpl7a5 | 2.206527 | 0.314405097 | 0.981 | 0.822 | 5.6056830 | 13 | rpl7a |
| rpl233 | 2.450811 | 0.257807461 | 0.985 | 0.871 | 6.2262872 | 13 | rpl23 |
| grasp3 | 6.024973 | 0.359738066 | 0.263 | 0.07 | 1.5306444 | 13 | grasp |
| tln12 | 1.187345 | 0.442265083 | 0.389 | 0.139 | 3.0164515 | 13 | tln1 |
| histh1l2 | 1.324301 | 0.504896609 | 0.352 | 0.119 | 3.3643889 | 13 | histh1l |
| zgc:564934 | 3.346732 | 0.449707278 | 0.87 | 0.586 | 8.5023748 | 13 | zgc:56493 |
| rps65 | 3.614150 | 0.312847643 | 0.952 | 0.802 | 9.1817491 | 13 | rps6 |
| vsir3 | 4.636700 | 0.382122161 | 0.293 | 0.086 | 1.1779536 | 13 | vsir |
| actb14 | 5.359097 | 0.361880393 | 0.933 | 0.765 | 1.3614787 | 13 | actb1 |
| rps166 | 9.324209 | 0.373457755 | 0.985 | 0.763 | 2.3688153 | 13 | rps16 |
| si:ch211-105j21.91 | 1.060374 | 0.258293745 | 0.167 | 0.033 | 2.6938824 | 13 | si:ch211-105j21.9 |
| dusp52 | 1.227680 | 0.401450661 | 0.244 | 0.066 | 3.1189219 | 13 | dusp5 |

| | | | | | | | |
|---|---|---|---|---|---|---|---|
| capza1a3 | 2.034016 | 0.469789398 | 0.515 | 0.217 | 5.1674191 | 13 | capza1a |
| lbh2 | 2.944744 | 0.414692311 | 0.43 | 0.149 | 7.4811226 | 13 | lbh |
| krt18b4 | 7.343792 | 0.510633913 | 0.444 | 0.172 | 1.8656903 | 13 | krt18b |
| rpl85 | 1.020352 | 0.329382698 | 0.97 | 0.793 | 2.5922051 | 13 | rpl8 |
| eno1a3 | 1.579156 | 0.437397407 | 0.381 | 0.139 | 4.0118469 | 13 | eno1a |
| ankrd12 | 1.690631 | 0.341476747 | 0.252 | 0.07 | 4.2950503 | 13 | ankrd12 |
| sh3kbp11 | 1.997409 | 0.364777388 | 0.259 | 0.074 | 5.0744176 | 13 | sh3kbp1 |
| serf23 | 2.312782 | 0.491869544 | 0.722 | 0.373 | 5.8756249 | 13 | serf2 |
| nfkbie2 | 3.548667 | 0.392262296 | 0.3 | 0.094 | 9.0153903 | 13 | nfkbie |
| rpl227 | 3.728629 | 0.384618472 | 0.956 | 0.717 | 9.4725829 | 13 | rpl22 |
| glrx2 | 7.623186 | 0.493609284 | 0.656 | 0.322 | 1.9366705 | 13 | glrx |
| mob1a3 | 8.875082 | 0.421291827 | 0.437 | 0.169 | 2.2547145 | 13 | mob1a |
| mcl1a3 | 1.268695 | 0.444121049 | 0.893 | 0.609 | 3.2231197 | 13 | mcl1a |
| ywhabl | 2.046349 | 0.476758386 | 0.619 | 0.296 | 5.1987500 | 13 | ywhabl |
| rpl265 | 2.191219 | 0.276840187 | 0.978 | 0.849 | 5.5667939 | 13 | rpl26 |
| XLOC-0054481 | 2.285740 | 0.283414948 | 0.141 | 0.027 | 5.8069238 | 13 | XLOC-005448 |
| sdcbp23 | 2.647571 | 0.486238975 | 0.504 | 0.224 | 6.7261566 | 13 | sdcbp2 |
| rpl394 | 6.648561 | 0.261268554 | 0.981 | 0.876 | 1.6890671 | 13 | rpl39 |
| cmtm7 | 1.126509 | 0.311511392 | 0.196 | 0.049 | 2.8618980 | 13 | cmtm7 |
| rpl156 | 1.195687 | 0.350823456 | 0.963 | 0.761 | 3.0376451 | 13 | rpl15 |
| pabpc1a3 | 1.538845 | 0.441238026 | 0.867 | 0.566 | 3.9094364 | 13 | pabpc1a |
| erbin | 1.619782 | 0.323195842 | 0.211 | 0.055 | 4.1150579 | 13 | erbin |
| mt-atp65 | 3.155059 | 0.48797033 | 0.911 | 0.551 | 8.0154276 | 13 | mt-atp6 |
| tpi1b3 | 3.176882 | 0.424895298 | 0.433 | 0.175 | 8.0708693 | 13 | tpi1b |
| gnai2a2 | 4.892857 | 0.409430461 | 0.385 | 0.146 | 1.2430305 | 13 | gnai2a |
| rps75 | 5.050876 | 0.284181074 | 0.959 | 0.827 | 1.2831751 | 13 | rps7 |
| fosl21 | 5.079020 | 0.426262012 | 0.319 | 0.111 | 1.2903252 | 13 | fosl2 |
| zdhhc18b | 5.597512 | 0.317756007 | 0.233 | 0.063 | 1.4220480 | 13 | zdhhc18b |
| lcp2a1 | 5.825386 | 0.257728814 | 0.163 | 0.036 | 1.4799395 | 13 | lcp2a |
| gnai2b1 | 6.108064 | 0.313012506 | 0.222 | 0.06 | 1.5517537 | 13 | gnai2b |
| litaf2 | 6.307675 | 0.408679863 | 0.378 | 0.143 | 1.6024649 | 13 | litaf |
| lpar5a2 | 7.758273 | 0.287767665 | 0.181 | 0.043 | 1.9709893 | 13 | lpar5a |
| nr4a31 | 8.174523 | 0.316966497 | 0.185 | 0.045 | 2.0767375 | 13 | nr4a3 |
| rps106 | 8.261795 | 0.376529507 | 0.974 | 0.723 | 2.0989090 | 13 | rps10 |
| rpl186 | 9.152032 | 0.327856656 | 0.97 | 0.79 | 2.3250737 | 13 | rpl18 |

| | | | | | | | |
|---|---|---|---|---|---|---|---|
| reep51 | 1.430984 | 0.438196374 | 0.415 | 0.17 | 3.6354166 | 13 | reep5 |
| rpl10a5 | 1.490914 | 0.323757912 | 0.963 | 0.777 | 3.7876684 | 13 | rpl10a |
| mob3a1 | 2.155434 | 0.386697687 | 0.407 | 0.155 | 5.4758809 | 13 | mob3a |
| LOC101884451 | 2.197212 | 0.368688549 | 0.278 | 0.089 | 5.5820178 | 13 | LOC101884451 |
| slc25a53 | 2.396394 | 0.289969666 | 0.967 | 0.811 | 6.0880392 | 13 | slc25a5 |
| per3 | 2.751762 | 0.337649844 | 0.252 | 0.076 | 6.9908534 | 13 | per3 |
| fam49ba2 | 4.235112 | 0.355373332 | 0.311 | 0.103 | 1.0759302 | 13 | fam49ba |
| rps195 | 5.877890 | 0.261759722 | 0.985 | 0.867 | 1.4932779 | 13 | rps19 |
| iqgap1 | 7.386364 | 0.317710219 | 0.237 | 0.069 | 1.8765059 | 13 | iqgap1 |
| capza1b2 | 1.211586 | 0.445710107 | 0.507 | 0.231 | 3.0780343 | 13 | capza1b |
| si:dkey-177p2.6 | 1.313403 | 0.353320698 | 0.259 | 0.081 | 3.3367014 | 13 | si:dkey-177p2.6 |
| rps144 | 2.671356 | 0.325895935 | 0.974 | 0.779 | 6.7865809 | 13 | rps14 |
| rpsa5 | 3.067968 | 0.327779898 | 0.974 | 0.783 | 7.7941743 | 13 | rpsa |
| rpl245 | 4.072540 | 0.312829285 | 0.967 | 0.792 | 1.0346289 | 13 | rpl24 |
| glipr1a4 | 4.748694 | 0.312633087 | 0.23 | 0.066 | 1.2064059 | 13 | glipr1a |
| rab8b1 | 5.596798 | 0.35106048 | 0.259 | 0.083 | 1.4218667 | 13 | rab8b |
| rsl24d14 | 7.037981 | 0.47521316 | 0.793 | 0.456 | 1.7879991 | 13 | rsl24d1 |
| rps246 | 7.510742 | 0.32301163 | 0.97 | 0.776 | 1.9081040 | 13 | rps24 |
| plekho22 | 7.564425 | 0.339736004 | 0.244 | 0.075 | 1.9217423 | 13 | plekho2 |
| rpl215 | 8.493277 | 0.309145675 | 0.952 | 0.774 | 2.1577170 | 13 | rpl21 |
| arl6ip5b1 | 8.702138 | 0.363680736 | 0.293 | 0.101 | 2.2107781 | 13 | arl6ip5b |
| slc9a3r1a | 2.282174 | 0.267191416 | 0.189 | 0.049 | 5.7978636 | 13 | slc9a3r1a |
| rpl5a6 | 2.723151 | 0.448084226 | 0.859 | 0.532 | 6.9181654 | 13 | rpl5a |
| si:dkey-25e12.31 | 2.814624 | 0.304266907 | 0.181 | 0.047 | 7.1505539 | 13 | si:dkey-25e12.3 |
| rps186 | 3.644189 | 0.342073971 | 0.959 | 0.751 | 9.2580622 | 13 | rps18 |
| zgc:1537131 | 4.887190 | 0.412707637 | 0.4 | 0.169 | 1.2415907 | 13 | zgc:153713 |
| rps235 | 5.128103 | 0.335135944 | 0.956 | 0.742 | 1.3027948 | 13 | rps23 |
| cdc42l3 | 5.795100 | 0.427572672 | 0.5 | 0.23 | 1.4722452 | 13 | cdc42l |
| ccni3 | 7.568044 | 0.456910763 | 0.604 | 0.315 | 1.9226615 | 13 | ccni |
| rps135 | 1.448090 | 0.354817281 | 0.981 | 0.746 | 3.6788727 | 13 | rps13 |
| bida | 1.689011 | 0.290161278 | 0.233 | 0.071 | 4.2909348 | 13 | bida |
| rps57 | 2.489488 | 0.34599483 | 0.941 | 0.726 | 6.3245450 | 13 | rps5 |
| tnfaip32 | 2.713936 | 0.322899217 | 0.204 | 0.058 | 6.8947568 | 13 | tnfaip3 |
| rpl105 | 3.316165 | 0.307893223 | 0.97 | 0.788 | 8.4247172 | 13 | rpl10 |
| mt-co25 | 4.560369 | 0.44773115 | 0.944 | 0.667 | 1.1585619 | 13 | mt-co2 |

| | | | | | | | |
|---|---|---|---|---|---|---|---|
| zgc:1629442 | 5.657763 | 0.425041147 | 0.552 | 0.271 | 1.4373549 | 13 | zgc:162944 |
| zgc:1141886 | 6.203061 | 0.409763442 | 0.952 | 0.664 | 1.5758877 | 13 | zgc:114188 |
| actn4 | 6.476388 | 0.336121035 | 0.278 | 0.095 | 1.6453266 | 13 | actn4 |
| rps95 | 1.130876 | 0.305691659 | 0.967 | 0.788 | 2.8729911 | 13 | rps9 |
| psme22 | 1.145054 | 0.351955034 | 0.3 | 0.111 | 2.9090101 | 13 | psme2 |
| chmp2a2 | 1.524707 | 0.377875284 | 0.352 | 0.141 | 3.8735195 | 13 | chmp2a |
| rpl296 | 1.589411 | 0.292837282 | 0.985 | 0.821 | 4.0379007 | 13 | rpl29 |
| CU467110.1 | 1.916248 | 0.261717782 | 0.248 | 0.077 | 4.8682290 | 13 | CU467110.1 |
| rpl325 | 2.848213 | 0.291998447 | 0.974 | 0.822 | 7.2358858 | 13 | rpl32 |
| rps124 | 4.767855 | 0.265891528 | 0.978 | 0.84 | 1.2112736 | 13 | rps12 |
| rpl146 | 5.334780 | 0.38260921 | 0.963 | 0.665 | 1.3553010 | 13 | rpl14 |
| zmynd11 | 8.457736 | 0.309793585 | 0.196 | 0.057 | 2.1486879 | 13 | zmynd11 |
| hsp90ab13 | 9.615981 | 0.283897462 | 0.967 | 0.798 | 2.4429399 | 13 | hsp90ab1 |
| rps35 | 1.397292 | 0.347793305 | 0.963 | 0.723 | 3.5498208 | 13 | rps3 |
| arrb2b3 | 1.447151 | 0.31400398 | 0.27 | 0.094 | 3.6764884 | 13 | arrb2b |
| si:ch73-234b20.51 | 1.658716 | 0.331101518 | 0.285 | 0.103 | 4.2139689 | 13 | si:ch73-234b20.5 |
| ccr9a4 | 1.866919 | 0.354214521 | 0.256 | 0.086 | 4.7429097 | 13 | ccr9a |
| eif3f5 | 4.891547 | 0.42989674 | 0.763 | 0.462 | 1.2426977 | 13 | eif3f |
| arpc5b3 | 4.967942 | 0.360038622 | 0.33 | 0.129 | 1.2621056 | 13 | arpc5b |
| zgc:1627306 | 6.904787 | 0.449085866 | 0.774 | 0.397 | 1.7541612 | 13 | zgc:162730 |
| rpl36a5 | 9.576448 | 0.26863329 | 0.967 | 0.822 | 2.4328966 | 13 | rpl36a |
| pkma2 | 1.137350 | 0.386119543 | 0.4 | 0.176 | 2.8894381 | 13 | pkma |
| rpl35a6 | 1.294506 | 0.308165334 | 0.97 | 0.779 | 3.2886940 | 13 | rpl35a |
| c3a.52 | 1.443823 | 0.296068152 | 0.204 | 0.06 | 3.6680340 | 13 | c3a.5 |
| cmip | 1.752432 | 0.296331903 | 0.204 | 0.063 | 4.4520541 | 13 | cmip |
| rpl316 | 2.025695 | 0.358398334 | 0.948 | 0.684 | 5.1462792 | 13 | rpl31 |
| nfkbiab1 | 2.219416 | 0.422891319 | 0.422 | 0.198 | 5.6384265 | 13 | nfkbiab |
| tmem30aa | 2.496240 | 0.311373248 | 0.211 | 0.068 | 6.3416999 | 13 | tmem30aa |
| si:dkey-112e7.24 | 2.774545 | 0.393193458 | 0.444 | 0.212 | 7.0487317 | 13 | si:dkey-112e7.2 |
| cox7a2a | 6.573092 | 0.381920097 | 0.507 | 0.248 | 1.6698942 | 13 | cox7a2a |
| zgc:638311 | 7.037818 | 0.320625466 | 0.241 | 0.084 | 1.7879577 | 13 | zgc:63831 |
| ywhaba2 | 1.092806 | 0.413271426 | 0.593 | 0.325 | 2.7762757 | 13 | ywhaba |
| rpl305 | 1.468644 | 0.330214488 | 0.948 | 0.727 | 3.7310921 | 13 | rpl30 |
| mvb12a | 2.356395 | 0.290593057 | 0.226 | 0.076 | 5.9864225 | 13 | mvb12a |
| cib1 | 2.752063 | 0.267628683 | 0.237 | 0.081 | 6.9916165 | 13 | cib1 |

| | | | | | | | |
|---|---|---|---|---|---|---|---|
| ccl39.71 | 3.024927 | 0.363623672 | 0.193 | 0.06 | 7.6848282 | 13 | ccl39.7 |
| cox7b2 | 4.202198 | 0.410700006 | 0.707 | 0.423 | 1.0675685 | 13 | cox7b |
| rpl96 | 5.136974 | 0.379980799 | 0.937 | 0.665 | 1.3050483 | 13 | rpl9 |
| rps114 | 1.114136 | 0.274586614 | 0.956 | 0.782 | 2.8304643 | 13 | rps11 |
| GCA3 | 1.290535 | 0.34965932 | 0.422 | 0.198 | 3.2786060 | 13 | GCA |
| arf6a | 1.434080 | 0.326735341 | 0.374 | 0.16 | 3.6432821 | 13 | arf6a |
| abracl2 | 1.497063 | 0.378037483 | 0.463 | 0.219 | 3.8032889 | 13 | abracl |
| twf1a2 | 1.570427 | 0.32165442 | 0.363 | 0.155 | 3.9896718 | 13 | twf1a |
| uba524 | 1.727503 | 0.359346373 | 0.926 | 0.657 | 4.3887214 | 13 | uba52 |
| uqcrh1 | 1.892038 | 0.367715608 | 0.504 | 0.257 | 4.8067242 | 13 | uqcrh |
| cpne1 | 2.832293 | 0.322024859 | 0.315 | 0.131 | 7.1954424 | 13 | cpne1 |
| hmgn65 | 2.847847 | 0.39424546 | 0.578 | 0.308 | 7.2349559 | 13 | hmgn6 |
| gadd45bb3 | 5.148611 | 0.323646545 | 0.226 | 0.081 | 1.3080046 | 13 | gadd45bb |
| rpl285 | 6.541200 | 0.294845321 | 0.963 | 0.774 | 1.6617920 | 13 | rpl28 |
| faua6 | 8.387721 | 0.332908499 | 0.944 | 0.715 | 2.1309007 | 13 | faua |
| sept7b | 1.045244 | 0.28391759 | 0.278 | 0.108 | 2.6554430 | 13 | sept7b |
| arpc4l3 | 1.286435 | 0.348692975 | 0.444 | 0.216 | 3.2681901 | 13 | arpc4l |
| anxa11a | 1.568938 | 0.316301751 | 0.281 | 0.115 | 3.9858873 | 13 | anxa11a |
| gnb1a1 | 1.598192 | 0.35710052 | 0.411 | 0.2 | 4.0602082 | 13 | gnb1a |
| cry1aa | 2.071862 | 0.26056489 | 0.181 | 0.058 | 5.2635663 | 13 | cry1aa |
| mt-cyb4 | 2.518337 | 0.395237986 | 0.856 | 0.573 | 6.3978352 | 13 | mt-cyb |
| btg22 | 2.741682 | 0.390888796 | 0.637 | 0.346 | 6.9652449 | 13 | btg2 |
| ier2b4 | 3.007379 | 0.391936943 | 0.459 | 0.234 | 7.6402468 | 13 | ier2b |
| csk2 | 3.158032 | 0.287497829 | 0.267 | 0.102 | 8.0229818 | 13 | csk |
| tgif11 | 3.835432 | 0.291709364 | 0.256 | 0.098 | 9.7439170 | 13 | tgif1 |
| rab443 | 4.557657 | 0.273145547 | 0.185 | 0.06 | 1.1578728 | 13 | rab44 |
| atp5f1e1 | 8.591650 | 0.377858619 | 0.793 | 0.486 | 2.1827088 | 13 | atp5f1e |
| si:ch211-132p1.32 | 9.018506 | 0.288783702 | 0.248 | 0.096 | 2.2911516 | 13 | si:ch211-132p1.3 |
| sh3bgrl31 | 9.777365 | 0.317251532 | 0.344 | 0.154 | 2.4839397 | 13 | sh3bgrl3 |
| COX36 | 2.184784 | 0.381436898 | 0.937 | 0.672 | 5.5504444 | 13 | COX3 |
| CR383676.16 | 2.447074 | 0.530707312 | 0.981 | 0.713 | 6.2167915 | 13 | CR383676.1 |
| rps255 | 2.886054 | 0.295409342 | 0.948 | 0.746 | 7.3320217 | 13 | rps25 |
| jak11 | 2.951399 | 0.29973861 | 0.248 | 0.098 | 7.4980299 | 13 | jak1 |
| ap1s1 | 3.638346 | 0.264277969 | 0.252 | 0.097 | 9.2432191 | 13 | ap1s1 |
| rps4x5 | 4.950940 | 0.296130462 | 0.93 | 0.734 | 1.2577863 | 13 | rps4x |

| | | | | | | | |
|---|---|---|---|---|---|---|---|
| rnf441 | 9.803454 | 0.292909139 | 0.241 | 0.095 | 2.4905675 | 13 | rnf44 |
| rps3a5 | 1.086928 | 0.256236876 | 0.948 | 0.79 | 2.7613414 | 13 | rps3a |
| ndufa4l3 | 1.340861 | 0.366546355 | 0.711 | 0.419 | 3.4064594 | 13 | ndufa4l |
| gnb1b | 1.683837 | 0.345007263 | 0.489 | 0.263 | 4.2777894 | 13 | gnb1b |
| pde4ba2 | 1.902142 | 0.345745767 | 0.311 | 0.14 | 4.8323936 | 13 | pde4ba |
| edf1 | 2.514388 | 0.363426219 | 0.504 | 0.285 | 6.3878037 | 13 | edf1 |
| COX5B1 | 5.765471 | 0.345709349 | 0.637 | 0.373 | 1.4647181 | 13 | COX5B |
| mrpl331 | 6.473909 | 0.259153822 | 0.256 | 0.105 | 1.6446968 | 13 | mrpl33 |
| spcs2 | 6.950915 | 0.30478537 | 0.359 | 0.172 | 1.7658801 | 13 | spcs2 |
| cox4i12 | 1.041050 | 0.350526155 | 0.678 | 0.414 | 2.6447886 | 13 | cox4i1 |
| cxcr4b3 | 1.054658 | 0.328292552 | 0.378 | 0.175 | 2.6793587 | 13 | cxcr4b |
| ip6k2b2 | 1.153568 | 0.292330383 | 0.289 | 0.127 | 2.9306406 | 13 | ip6k2b |
| naca6 | 1.671436 | 0.330625253 | 0.896 | 0.649 | 4.2462852 | 13 | naca |
| FQ323156.13 | 1.939901 | 0.289785908 | 0.337 | 0.15 | 4.9283206 | 13 | FQ323156.1 |
| hmgb2a1 | 1.948495 | 0.361643951 | 0.819 | 0.607 | 4.9501527 | 13 | hmgb2a |
| oaz1b3 | 2.599261 | 0.355370386 | 0.578 | 0.358 | 6.6034236 | 13 | oaz1b |
| ucp23 | 2.774144 | 0.365379601 | 0.656 | 0.411 | 7.0477147 | 13 | ucp2 |
| rps264 | 3.595306 | 0.265334189 | 0.926 | 0.749 | 9.1338759 | 13 | rps26 |
| sh3bgrl1 | 3.605927 | 0.293076783 | 0.27 | 0.12 | 9.1608582 | 13 | sh3bgrl |
| sub1a | 3.805152 | 0.316092822 | 0.47 | 0.25 | 9.6669899 | 13 | sub1a |
| slc25a55a | 4.290065 | 0.251660611 | 0.196 | 0.074 | 1.0898911 | 13 | slc25a55a |
| aldoaa1 | 6.066627 | 0.315383654 | 0.47 | 0.251 | 1.5412266 | 13 | aldoaa |
| mtpn1 | 6.183219 | 0.29518731 | 0.333 | 0.162 | 1.5708468 | 13 | mtpn |
| cdkn1bb2 | 6.571159 | 0.297356432 | 0.322 | 0.152 | 1.6694030 | 13 | cdkn1bb |
| ube2v21 | 9.001701 | 0.318574346 | 0.504 | 0.28 | 2.2868823 | 13 | ube2v2 |
| zgc:1538672 | 1.076901 | 0.347234082 | 0.722 | 0.462 | 2.7358683 | 13 | zgc:153867 |
| atp5pf2 | 1.099431 | 0.320694302 | 0.556 | 0.317 | 2.7931054 | 13 | atp5pf |
| brk1 | 1.493523 | 0.285895397 | 0.378 | 0.19 | 3.7942970 | 13 | brk1 |
| ywhabb | 4.014586 | 0.294219855 | 0.463 | 0.251 | 1.0199056 | 13 | ywhabb |
| tpm11 | 5.053437 | 0.252703823 | 0.411 | 0.2 | 1.2838258 | 13 | tpm1 |
| rpl27a4 | 6.771108 | 0.352585106 | 0.785 | 0.519 | 1.7202001 | 13 | rpl27a |
| cox6a11 | 7.761990 | 0.321418944 | 0.648 | 0.387 | 1.9719336 | 13 | cox6a1 |
| h2afx14 | 9.116623 | 0.316726775 | 0.419 | 0.225 | 2.3160782 | 13 | h2afx1 |
| eif3i2 | 1.080961 | 0.306863582 | 0.504 | 0.288 | 2.7461833 | 13 | eif3i |
| rhoac | 1.570076 | 0.263997406 | 0.341 | 0.17 | 3.9887795 | 13 | rhoac |

| | | | | | | | |
|---|---|---|---|---|---|---|---|
| cox7c2 | 1.959997 | 0.322241525 | 0.726 | 0.491 | 4.9793726 | 13 | cox7c |
| egr14 | 2.218521 | 0.346879346 | 0.348 | 0.182 | 5.6361537 | 13 | egr1 |
| hif1al | 2.226216 | 0.258959335 | 0.263 | 0.119 | 5.6557023 | 13 | hif1al |
| psme11 | 3.046945 | 0.257074411 | 0.244 | 0.111 | 7.7407659 | 13 | psme1 |
| tomm20b2 | 3.649583 | 0.291391928 | 0.411 | 0.226 | 9.2717668 | 13 | tomm20b |
| atp5f1d2 | 7.100588 | 0.289155851 | 0.474 | 0.271 | 1.8039044 | 13 | atp5f1d |
| eif4ebp2 | 7.970745 | 0.273025646 | 0.363 | 0.191 | 2.0249677 | 13 | eif4ebp2 |
| tnrc6c12 | 8.590640 | 0.291292721 | 0.296 | 0.149 | 2.1824521 | 13 | tnrc6c1 |
| mibp25 | 1.141619 | 0.327788745 | 0.8 | 0.517 | 2.9002853 | 13 | mibp2 |
| serbp1a4 | 1.227258 | 0.320085627 | 0.744 | 0.503 | 3.1178496 | 13 | serbp1a |
| si:ch1073-325m22.2: | 1.335040 | 0.285710171 | 0.433 | 0.248 | 3.3916710 | 13 | si:ch1073-325m22.2 |
| ppiab3 | 1.447530 | 0.278574454 | 0.889 | 0.672 | 3.6774506 | 13 | ppiab |
| rack14 | 2.053783 | 0.253158562 | 0.948 | 0.761 | 5.2176369 | 13 | rack1 |
| NC-002333.44 | 2.725658 | 0.321027762 | 0.893 | 0.605 | 6.9245354 | 13 | NC-002333.4 |
| wdr13 | 3.415885 | 0.258254426 | 0.307 | 0.154 | 8.6780580 | 13 | wdr1 |
| NDUFB12 | 3.601986 | 0.273997928 | 0.378 | 0.21 | 9.1508475 | 13 | NDUFB1 |
| mt-nd15 | 4.442492 | 0.288421637 | 0.656 | 0.394 | 1.1286151 | 13 | mt-nd1 |
| hmgb1a5 | 4.553002 | 0.278628471 | 0.404 | 0.226 | 1.1566903 | 13 | hmgb1a |
| uqcrq1 | 6.065088 | 0.282026999 | 0.485 | 0.285 | 1.5408356 | 13 | uqcrq |
| chac11 | 6.346956 | 0.301796471 | 0.607 | 0.375 | 1.6124443 | 13 | chac1 |
| mt-nd45 | 8.568580 | 0.308211411 | 0.696 | 0.445 | 2.1768479 | 13 | mt-nd4 |
| mt-nd34 | 1.099470 | 0.292141136 | 0.596 | 0.369 | 2.7932049 | 13 | mt-nd3 |
| ppt11 | 1.114256 | 0.286368067 | 0.333 | 0.18 | 2.8307685 | 13 | ppt1 |
| ywhae1 | 1.253074 | 0.270614754 | 0.5 | 0.293 | 3.1834365 | 13 | ywhae1 |
| hnrnpa0b2 | 1.406540 | 0.288899038 | 0.559 | 0.339 | 3.5733163 | 13 | hnrnpa0b |
| atp5pd2 | 1.626759 | 0.268632159 | 0.478 | 0.281 | 4.1327835 | 13 | atp5pd |
| tmed7 | 1.935711 | 0.26170061 | 0.389 | 0.219 | 4.9176741 | 13 | tmed7 |
| cox174 | 1.962310 | 0.270806682 | 0.596 | 0.338 | 4.9852485 | 13 | cox17 |
| sumo2b4 | 2.810232 | 0.277219445 | 0.485 | 0.295 | 7.1393958 | 13 | sumo2b |
| rps27.25 | 2.814548 | 0.320568891 | 0.826 | 0.55 | 7.1503594 | 13 | rps27.2 |
| romo14 | 3.595354 | 0.288711534 | 0.63 | 0.406 | 9.1339984 | 13 | romo1 |
| rpl5b4 | 4.323347 | 0.289913406 | 0.881 | 0.642 | 1.0983464 | 13 | rpl5b |
| atp5l2 | 4.737350 | 0.294728095 | 0.674 | 0.455 | 1.2035238 | 13 | atp5l |
| b2m4 | 5.504721 | 0.280601368 | 0.496 | 0.296 | 1.3984743 | 13 | b2m |
| atp5meb | 6.817390 | 0.265808215 | 0.537 | 0.337 | 1.7319581 | 13 | atp5meb |

| | | | | | | | |
|---|---|---|---|---|---|---|---|
| rpl23a5 | 6.954897 | 0.296893518 | 0.911 | 0.666 | 1.7668917 | 13 | rpl23a |
| arl6ip12 | 7.934940 | 0.268487431 | 0.593 | 0.356 | 2.0158715 | 13 | arl6ip1 |
| sat1a.22 | 8.351506 | 0.271142188 | 0.504 | 0.299 | 2.1217003 | 13 | sat1a.2 |
| selenof1 | 1.305760 | 0.25626593 | 0.448 | 0.265 | 3.3172847 | 13 | selenof |
| xbp12 | 6.068484 | 0.277479248 | 0.548 | 0.353 | 0.000154 | 13 | xbp1 |
| nsa24 | 2.190805 | 0.266593907 | 0.585 | 0.387 | 0.000557 | 13 | nsa2 |
| btf34 | 2.235708 | 0.281943648 | 0.856 | 0.607 | 0.000568 | 13 | btf3 |
| rpl44 | 6.016583 | 0.262527936 | 0.878 | 0.654 | 0.001529 | 13 | rpl4 |
| rpl65 | 9.620030 | 0.254296604 | 0.926 | 0.698 | 0.02444 | 13 | rpl6 |
| eef1g5 | 1.365132 | 0.256690269 | 0.878 | 0.65 | 0.034681 | 13 | eef1g |
| krt152 | 0 | 1.57847052 | 0.754 | 0.085 | 0 | 14 | krt15 |
| cldnc | 0 | 1.51604176 | 0.701 | 0.023 | 0 | 14 | cldnc |
| podxl1 | 0 | 1.509595255 | 0.773 | 0.044 | 0 | 14 | podxl |
| cavin1b1 | 0 | 1.469627647 | 0.773 | 0.031 | 0 | 14 | cavin1b |
| fn1b2 | 0 | 1.460113474 | 0.826 | 0.088 | 0 | 14 | fn1b |
| gcga | 0 | 1.032532084 | 0.447 | 0.006 | 0 | 14 | gcga |
| cav1 | 0 | 1.025109868 | 0.557 | 0.04 | 0 | 14 | cav1 |
| wt1a | 0 | 0.925841629 | 0.542 | 0.033 | 0 | 14 | wt1a |
| bnc2 | 0 | 0.925204784 | 0.53 | 0.031 | 0 | 14 | bnc2 |
| si:ch211-156j16.1 | 0 | 0.920583937 | 0.519 | 0.019 | 0 | 14 | si:ch211-156j16.1 |
| s100a10a1 | 0 | 0.845269664 | 0.326 | 0.009 | 0 | 14 | s100a10a |
| jam2b | 0 | 0.693806533 | 0.386 | 0.016 | 0 | 14 | jam2b |
| slc29a1b | 0 | 0.627257525 | 0.284 | 0.008 | 0 | 14 | slc29a1b |
| fgf24 | 1.292033 | 0.463446297 | 0.22 | 0.004 | 3.2824104 | 14 | fgf24 |
| eppk1 | 1.475412 | 0.937771603 | 0.504 | 0.035 | 3.7482857 | 14 | eppk1 |
| sfrp1a1 | 6.444904 | 0.835988703 | 0.458 | 0.029 | 1.6373278 | 14 | sfrp1a |
| sox6 | 2.660551 | 0.767789041 | 0.428 | 0.025 | 6.7591318 | 14 | sox6 |
| pdlim3b | 4.333638 | 0.385228456 | 0.205 | 0.004 | 1.1009609 | 14 | pdlim3b |
| ucmaa | 1.030526 | 0.641702355 | 0.288 | 0.01 | 2.6180531 | 14 | ucmaa |
| cyp2ad2 | 1.239335 | 0.482925401 | 0.208 | 0.005 | 3.1485325 | 14 | cyp2ad2 |
| aldh1a22 | 2.230520 | 1.162319537 | 0.735 | 0.087 | 5.6666377 | 14 | aldh1a2 |
| gstm.32 | 5.130591 | 1.569858219 | 0.803 | 0.119 | 1.3034267 | 14 | gstm.3 |
| frzb | 1.807902 | 0.727122799 | 0.314 | 0.014 | 4.5929754 | 14 | frzb |
| scn4ba | 4.599828 | 0.325560173 | 0.155 | 0.002 | 1.1685864 | 14 | scn4ba |
| emilin1a1 | 3.679144 | 0.612027451 | 0.36 | 0.02 | 9.3468672 | 14 | emilin1a |

| | | | | | | | |
|---|---|---|---|---|---|---|---|
| krt41 | 2.059185 | 1.100874028 | 0.504 | 0.042 | 5.2313612 | 14 | krt4 |
| postnb2 | 1.032780 | 1.39567672 | 0.682 | 0.085 | 2.6237779 | 14 | postnb |
| BNC1 | 1.935792 | 0.28585765 | 0.159 | 0.002 | 4.9178814 | 14 | BNC1 |
| FP016056.1 | 8.217290 | 0.424988002 | 0.223 | 0.006 | 2.0876026 | 14 | FP016056.1 |
| nanos1 | 1.734287 | 0.88626939 | 0.394 | 0.026 | 4.4059581 | 14 | nanos1 |
| lhx9 | 2.543109 | 0.439175721 | 0.17 | 0.003 | 6.4607695 | 14 | lhx9 |
| usp43b | 1.458019 | 0.553445958 | 0.295 | 0.013 | 3.7040987 | 14 | usp43b |
| tmem98 | 2.386169 | 0.479130564 | 0.292 | 0.013 | 6.0620625 | 14 | tmem98 |
| hspb13 | 1.520709 | 1.022676935 | 0.629 | 0.072 | 3.8633614 | 14 | hspb1 |
| cldn5a2 | 5.480112 | 0.970852285 | 0.542 | 0.053 | 1.3922226 | 14 | cldn5a |
| lurap1 | 7.781061 | 0.391951776 | 0.201 | 0.006 | 1.9767786 | 14 | lurap1 |
| plod21 | 5.511066 | 0.63996714 | 0.367 | 0.024 | 1.4000864 | 14 | plod2 |
| paplna | 6.670957 | 0.652775526 | 0.352 | 0.022 | 1.6947566 | 14 | paplna |
| cdh21 | 1.455189 | 0.880790171 | 0.549 | 0.058 | 3.6969099 | 14 | cdh2 |
| fsta | 5.695411 | 0.341795105 | 0.167 | 0.004 | 1.4469193 | 14 | fsta |
| itgb4 | 8.379171 | 0.543574007 | 0.273 | 0.013 | 2.1287285 | 14 | itgb4 |
| pleca | 6.522477 | 0.896066926 | 0.519 | 0.054 | 1.6570355 | 14 | pleca |
| cavin2b1 | 8.838287 | 0.987580508 | 0.587 | 0.07 | 2.2453669 | 14 | cavin2b |
| rap1gap | 2.794202 | 0.435409831 | 0.205 | 0.007 | 7.0986708 | 14 | rap1gap |
| si:ch211-137i24.10 | 3.035100 | 0.432618641 | 0.201 | 0.006 | 7.7106738 | 14 | si:ch211-137i24.10 |
| krt18a.12 | 1.218961 | 1.301266059 | 0.807 | 0.139 | 3.0967708 | 14 | krt18a.1 |
| ncam1a | 1.974804 | 0.573407074 | 0.292 | 0.016 | 5.0169907 | 14 | ncam1a |
| frya | 4.477410 | 0.37587918 | 0.186 | 0.006 | 1.1374862 | 14 | frya |
| hsd11b23 | 1.717555 | 1.244166525 | 0.822 | 0.137 | 4.3634500 | 14 | hsd11b2 |
| kcnq5a | 6.247848 | 0.364836274 | 0.17 | 0.005 | 1.5872659 | 14 | kcnq5a |
| mmp22 | 3.233159 | 0.990572798 | 0.678 | 0.101 | 8.2138406 | 14 | mmp2 |
| zgc:1124371 | 1.120353 | 0.56984862 | 0.311 | 0.02 | 2.8462577 | 14 | zgc:112437 |
| dkk3b2 | 1.165782 | 0.818132433 | 0.496 | 0.053 | 2.9616697 | 14 | dkk3b |
| slc43a3b1 | 1.599971 | 0.614992106 | 0.337 | 0.024 | 4.0647264 | 14 | slc43a3b |
| ehd2b | 4.487895 | 0.608235117 | 0.364 | 0.028 | 1.1401497 | 14 | ehd2b |
| gata6 | 7.006967 | 0.536514504 | 0.314 | 0.021 | 1.7801200 | 14 | gata6 |
| LOC110438169 | 8.815308 | 0.25832856 | 0.125 | 0.002 | 2.2395290 | 14 | LOC110438169 |
| col6a4a2 | 5.630692 | 0.907437556 | 0.508 | 0.058 | 1.4304773 | 14 | col6a4a |
| si:ch211-250c4.4 | 1.531422 | 0.452278459 | 0.261 | 0.014 | 3.8905780 | 14 | si:ch211-250c4.4 |
| si:ch211-63p21.1 | 2.862372 | 0.368211332 | 0.178 | 0.006 | 7.2718566 | 14 | si:ch211-63p21.1 |

| | | | | | | | |
|---|---|---|---|---|---|---|---|
| si:ch211-105c13.31 | 1.260721 | 0.793743644 | 0.515 | 0.059 | 3.2028631 | 14 | si:ch211-105c13.3 |
| pcdh10b | 2.353301 | 0.303019816 | 0.148 | 0.004 | 5.9785628 | 14 | pcdh10b |
| sccpdha | 4.735923 | 0.317121405 | 0.14 | 0.003 | 1.2031612 | 14 | sccpdha |
| krt84 | 1.210347 | 1.341178141 | 0.955 | 0.301 | 3.0748880 | 14 | krt8 |
| c62 | 1.121856 | 0.954507557 | 0.667 | 0.104 | 2.8500771 | 14 | c6 |
| serpine3 | 1.711627 | 0.336935869 | 0.167 | 0.006 | 4.3483891 | 14 | serpine3 |
| cebpd3 | 6.597423 | 1.056445189 | 0.754 | 0.141 | 1.6760754 | 14 | cebpd |
| col4a12 | 2.254296 | 0.847494563 | 0.504 | 0.063 | 5.7270391 | 14 | col4a1 |
| si:ch73-46j18.54 | 1.259079 | 1.166582234 | 0.833 | 0.198 | 3.1986925 | 14 | si:ch73-46j18.5 |
| krt943 | 2.716346 | 1.180947033 | 0.777 | 0.155 | 6.9008792 | 14 | krt94 |
| CU984579.1 | 1.217266 | 0.495599103 | 0.242 | 0.014 | 3.0924659 | 14 | CU984579.1 |
| si:ch211-106h4.122 | 1.869371 | 1.017003941 | 0.443 | 0.051 | 4.7491390 | 14 | si:ch211-106h4.12 |
| im:71514491 | 6.271813 | 0.511601164 | 0.318 | 0.025 | 1.5933542 | 14 | im:7151449 |
| ftr82 | 1.547244 | 0.553432569 | 0.348 | 0.031 | 3.9307735 | 14 | ftr82 |
| col1a1a2 | 2.550438 | 1.169677576 | 0.765 | 0.156 | 6.4793899 | 14 | col1a1a |
| anxa2a3 | 3.648496 | 1.035603498 | 0.629 | 0.109 | 9.2690046 | 14 | anxa2a |
| anxa1a3 | 1.988943 | 1.149450791 | 0.723 | 0.153 | 5.0529099 | 14 | anxa1a |
| col6a32 | 2.289538 | 0.943328828 | 0.621 | 0.106 | 5.8165716 | 14 | col6a3 |
| efna1b | 2.091228 | 0.448618079 | 0.242 | 0.016 | 5.3127650 | 14 | efna1b |
| cyp27c11 | 1.852628 | 0.435638622 | 0.254 | 0.017 | 4.7066020 | 14 | cyp27c1 |
| sparc3 | 1.005373 | 1.089878274 | 0.818 | 0.184 | 2.5541517 | 14 | sparc |
| col1a1b2 | 6.042262 | 1.148816583 | 0.739 | 0.16 | 1.5350367 | 14 | col1a1b |
| cygb12 | 4.099176 | 0.818889852 | 0.436 | 0.055 | 1.0413958 | 14 | cygb1 |
| ugt1b5 | 1.088681 | 0.536263679 | 0.337 | 0.032 | 2.7657955 | 14 | ugt1b5 |
| si:ch211-207d6.2 | 3.297889 | 0.588577785 | 0.25 | 0.018 | 8.3782891 | 14 | si:ch211-207d6.2 |
| c1qtnf12 | 2.377167 | 0.251674054 | 0.133 | 0.004 | 6.0391931 | 14 | c1qtnf12 |
| bmper1 | 1.932668 | 0.487822539 | 0.25 | 0.018 | 4.9099442 | 14 | bmper |
| LOC1039092842 | 4.070557 | 0.735350798 | 0.485 | 0.07 | 1.0341250 | 14 | LOC103909284 |
| col1a22 | 1.594861 | 1.150782114 | 0.758 | 0.172 | 4.0517467 | 14 | col1a2 |
| slc43a1b | 9.537166 | 0.377279649 | 0.189 | 0.01 | 2.4229171 | 14 | slc43a1b |
| tjp1a | 9.972261 | 0.597227704 | 0.398 | 0.047 | 2.5334529 | 14 | tjp1a |
| zgc:110182 | 4.803127 | 0.717424194 | 0.523 | 0.078 | 1.2202345 | 14 | zgc:110182 |
| nfia3 | 3.206151 | 0.77357673 | 0.545 | 0.091 | 8.1452282 | 14 | nfia |
| epha4l | 2.430550 | 0.268909707 | 0.148 | 0.006 | 6.1748127 | 14 | epha4l |
| flnb1 | 4.133267 | 0.487279811 | 0.265 | 0.022 | 1.0500566 | 14 | flnb |

| | | | | | | | |
|---|---|---|---|---|---|---|---|
| wt1b | 7.09645( | 0.514127909 | 0.284 | 0.026 | 1.8028531 | 14 | wt1b |
| s100a10b4 | 1.762271 | 1.214602656 | 0.864 | 0.276 | 4.4770517 | 14 | s100a10b |
| nog11 | 1.478497 | 0.420893527 | 0.17 | 0.009 | 3.7561222 | 14 | nog1 |
| bcam1 | 1.026077 | 0.710344233 | 0.33 | 0.036 | 2.6067502 | 14 | bcam |
| epha3 | 2.409585 | 0.462169967 | 0.235 | 0.018 | 6.1215511 | 14 | epha3 |
| prss35 | 6.351464 | 0.511120427 | 0.273 | 0.025 | 1.6135896 | 14 | prss35 |
| sema3fa | 1.193061 | 0.338194433 | 0.189 | 0.012 | 3.0309727 | 14 | sema3fa |
| si:ch211-1a19.32 | 1.632523 | 0.657059138 | 0.394 | 0.052 | 4.1474262 | 14 | si:ch211-1a19.3 |
| oaz2b | 4.568553 | 0.296222502 | 0.17 | 0.009 | 1.1606411 | 14 | oaz2b |
| itgb3b | 4.624744 | 0.373232142 | 0.182 | 0.011 | 1.1749163 | 14 | itgb3b |
| tpm4a3 | 1.078649 | 0.781039781 | 0.568 | 0.106 | 2.7403078 | 14 | tpm4a |
| epb41l4a | 5.722102 | 0.256443454 | 0.148 | 0.007 | 1.4537002 | 14 | epb41l4a |
| XLOC-0412693 | 1.088446 | 0.514257427 | 0.273 | 0.026 | 2.7651970 | 14 | XLOC-041269 |
| CABZ01080702.14 | 2.047752 | 0.770266096 | 0.511 | 0.091 | 5.2023140 | 14 | CABZ01080702.1 |
| myo1cb | 1.455101 | 0.498922837 | 0.303 | 0.032 | 3.6966841 | 14 | myo1cb |
| ptprsa | 9.964275 | 0.456793997 | 0.284 | 0.028 | 2.5314240 | 14 | ptprsa |
| id13 | 9.263559 | 0.976963263 | 0.591 | 0.123 | 2.3534072 | 14 | id1 |
| mical2a | 2.239617 | 0.298182086 | 0.14 | 0.006 | 5.6897482 | 14 | mical2a |
| socs3a4 | 8.270161 | 1.080601673 | 0.902 | 0.305 | 2.1010346 | 14 | socs3a |
| col5a12 | 9.342664 | 0.845236692 | 0.61 | 0.124 | 2.3735039 | 14 | col5a1 |
| zfp36l1b2 | 1.247446 | 0.862586755 | 0.621 | 0.132 | 3.1691385 | 14 | zfp36l1b |
| endouc1 | 8.591495 | 0.768088363 | 0.428 | 0.065 | 2.1826694 | 14 | endouc |
| ahnak3 | 1.445613 | 0.812342488 | 0.61 | 0.129 | 3.6725798 | 14 | ahnak |
| slc12a7b | 8.978257 | 0.421731131 | 0.239 | 0.021 | 2.2809264 | 14 | slc12a7b |
| hsbp1a | 9.856571 | 0.508664301 | 0.28 | 0.029 | 2.5040620 | 14 | hsbp1a |
| meis1b3 | 1.581357 | 0.655657803 | 0.428 | 0.067 | 4.0174398 | 14 | meis1b |
| krt912 | 2.262197 | 0.894017187 | 0.424 | 0.066 | 5.7471136 | 14 | krt91 |
| boc2 | 5.396510 | 0.665315662 | 0.409 | 0.062 | 1.3709833 | 14 | boc |
| jun3 | 5.987885 | 1.005109148 | 0.83 | 0.24 | 1.5212222 | 14 | jun |
| nrp1a2 | 8.847910 | 0.652408304 | 0.398 | 0.059 | 2.2478116 | 14 | nrp1a |
| fosb1 | 9.795900 | 0.980155033 | 0.553 | 0.114 | 2.4886485 | 14 | fosb |
| cfd1 | 2.153704 | 0.914666586 | 0.417 | 0.065 | 5.4714875 | 14 | cfd |
| hoxb3a | 3.019973 | 0.288854867 | 0.155 | 0.009 | 7.6722429 | 14 | hoxb3a |
| rdh10a2 | 3.615203 | 0.559248583 | 0.341 | 0.043 | 9.1844239 | 14 | rdh10a |
| si:ch211-66e2.33 | 7.610918 | 0.657634444 | 0.39 | 0.057 | 1.9335539 | 14 | si:ch211-66e2.3 |

| | | | | | | | |
|---|---|---|---|---|---|---|---|
| XLOC-015291 | 1.475542 | 0.267890836 | 0.163 | 0.01 | 3.7486163 | 14 | XLOC-015291 |
| ackr3b1 | 1.492848 | 0.49109559 | 0.273 | 0.028 | 3.7925817 | 14 | ackr3b |
| bmp2b | 1.033278 | 0.263434424 | 0.144 | 0.008 | 2.6250445 | 14 | bmp2b |
| edil3a1 | 1.518002 | 0.406787902 | 0.235 | 0.021 | 3.8564843 | 14 | edil3a |
| fosab4 | 4.651916 | 1.087636203 | 0.909 | 0.352 | 1.1818193 | 14 | fosab |
| LO018188.11 | 6.484551 | 0.458603486 | 0.254 | 0.025 | 1.6474004 | 14 | LO018188.1 |
| rhpn2 | 5.760884 | 0.46953858 | 0.284 | 0.031 | 1.4635526 | 14 | rhpn2 |
| esr2a1 | 2.853824 | 0.469517449 | 0.273 | 0.029 | 7.2501418 | 14 | esr2a |
| rpz53 | 8.179366 | 0.754607596 | 0.572 | 0.119 | 2.0779680 | 14 | rpz5 |
| f101 | 2.841147 | 0.571117334 | 0.375 | 0.055 | 7.2179339 | 14 | f10 |
| ctgfa | 8.952234 | 0.567948377 | 0.273 | 0.03 | 2.2743152 | 14 | ctgfa |
| gadd45ba2 | 8.956462 | 0.926349533 | 0.701 | 0.189 | 2.2753893 | 14 | gadd45ba |
| f3b2 | 3.022833 | 0.804875359 | 0.489 | 0.093 | 7.6795088 | 14 | f3b |
| cgnl1 | 4.029062 | 0.413778882 | 0.254 | 0.026 | 1.0235834 | 14 | cgnl1 |
| c7a1 | 3.152597 | 0.557972451 | 0.318 | 0.041 | 8.0091737 | 14 | c7a |
| fstl1b2 | 1.501571 | 0.812438432 | 0.614 | 0.14 | 3.8147424 | 14 | fstl1b |
| cd151 | 2.165489 | 0.339811774 | 0.212 | 0.018 | 5.5014264 | 14 | cd151 |
| cxcl18b3 | 5.923752 | 0.891846919 | 0.515 | 0.104 | 1.5049294 | 14 | cxcl18b |
| cald1a2 | 4.511031 | 0.728297579 | 0.542 | 0.113 | 1.1460276 | 14 | cald1a |
| foxp42 | 6.877102 | 0.620138473 | 0.409 | 0.068 | 1.7471280 | 14 | foxp4 |
| si:ch1073-459j12.12 | 1.189258 | 0.709304961 | 0.439 | 0.079 | 3.0213112 | 14 | si:ch1073-459j12.1 |
| gpm6ab | 2.118373 | 0.357619288 | 0.193 | 0.016 | 5.3817276 | 14 | gpm6ab |
| tnfsf10l1 | 5.453672 | 0.511071518 | 0.288 | 0.035 | 1.3855054 | 14 | tnfsf10l |
| loxl2a2 | 2.598602 | 0.511436774 | 0.318 | 0.043 | 6.6017491 | 14 | loxl2a |
| junba4 | 3.565214 | 0.99545376 | 0.852 | 0.311 | 9.0574275 | 14 | junba |
| tmem88b1 | 7.694663 | 0.554517538 | 0.341 | 0.049 | 1.9548292 | 14 | tmem88b |
| arvcfb | 8.179143 | 0.298399628 | 0.17 | 0.013 | 2.0779113 | 14 | arvcfb |
| cdon3 | 1.609077 | 0.584045185 | 0.39 | 0.066 | 4.0878602 | 14 | cdon |
| cdh61 | 8.502953 | 0.487388262 | 0.242 | 0.027 | 2.1601753 | 14 | cdh6 |
| pkp3b | 1.468697 | 0.481201036 | 0.318 | 0.044 | 3.7312250 | 14 | pkp3b |
| chka | 3.490858 | 0.387351297 | 0.242 | 0.027 | 8.8685255 | 14 | chka |
| itgav1 | 9.957249 | 0.552108882 | 0.36 | 0.058 | 2.5296392 | 14 | itgav |
| pmp22a2 | 1.389972 | 0.737564277 | 0.663 | 0.159 | 3.5312251 | 14 | pmp22a |
| CR383676.17 | 1.919931 | 0.792375916 | 0.973 | 0.713 | 4.8775859 | 14 | CR383676.1 |
| s100v23 | 2.098103 | 0.716859052 | 0.572 | 0.133 | 5.3302315 | 14 | s100v2 |

| | | | | | | | |
|---|---|---|---|---|---|---|---|
| phactr4a1 | 2.582963 | 0.454884474 | 0.284 | 0.037 | 6.5620190 | 14 | phactr4a |
| tspan9b1 | 1.503521 | 0.480985113 | 0.318 | 0.046 | 3.8196965 | 14 | tspan9b |
| si:dkey-28b4.72 | 1.986734 | 0.680541149 | 0.481 | 0.104 | 5.0472978 | 14 | si:dkey-28b4.7 |
| col6a12 | 2.437101 | 0.692872446 | 0.462 | 0.091 | 6.1914559 | 14 | col6a1 |
| bag32 | 2.197829 | 0.499170608 | 0.254 | 0.031 | 5.5835857 | 14 | bag3 |
| daam1b | 3.832453 | 0.307587647 | 0.174 | 0.014 | 9.7363471 | 14 | daam1b |
| tspan4a1 | 4.197541 | 0.421114058 | 0.269 | 0.034 | 1.0663854 | 14 | tspan4a |
| sema3d1 | 5.222791 | 0.340986338 | 0.186 | 0.016 | 1.3268500 | 14 | sema3d |
| col6a22 | 5.611252 | 0.59875529 | 0.394 | 0.069 | 1.4255388 | 14 | col6a2 |
| tmsb15 | 8.752888 | 0.958286829 | 0.58 | 0.154 | 2.2236713 | 14 | tmsb1 |
| h1f02 | 1.745035 | 0.652990612 | 0.379 | 0.066 | 4.4332636 | 14 | h1f0 |
| cyp2ad32 | 6.937452 | 0.572178087 | 0.352 | 0.057 | 1.7624597 | 14 | cyp2ad3 |
| cldn11a2 | 5.454415 | 0.63400622 | 0.42 | 0.078 | 1.3856941 | 14 | cldn11a |
| aplp1 | 1.431301 | 0.306261822 | 0.174 | 0.015 | 3.6362224 | 14 | aplp1 |
| cd81a3 | 2.947592 | 0.853678844 | 0.746 | 0.233 | 7.4883584 | 14 | cd81a |
| myl9a1 | 1.208228 | 0.431437981 | 0.295 | 0.041 | 3.0695053 | 14 | myl9a |
| lama51 | 3.312260 | 0.393741144 | 0.235 | 0.027 | 8.4147973 | 14 | lama5 |
| epb41l5 | 4.539438 | 0.431104616 | 0.288 | 0.039 | 1.1532442 | 14 | epb41l5 |
| adh8a2 | 6.711321 | 0.579150277 | 0.303 | 0.044 | 1.7050112 | 14 | adh8a |
| dsc2l | 8.871114 | 0.376421526 | 0.22 | 0.024 | 2.2537065 | 14 | dsc2l |
| sult1st61 | 8.476813 | 0.309190063 | 0.182 | 0.017 | 2.1535343 | 14 | sult1st6 |
| vim2 | 8.041478 | 0.483132349 | 0.292 | 0.042 | 2.0429376 | 14 | vim |
| fat1a1 | 1.579718 | 0.391876464 | 0.231 | 0.027 | 4.0132743 | 14 | fat1a |
| afdna | 1.656904 | 0.399892169 | 0.212 | 0.023 | 4.2093664 | 14 | afdna |
| slc4a4a1 | 5.841711 | 0.29767466 | 0.14 | 0.01 | 1.4840869 | 14 | slc4a4a |
| lrp1ab2 | 4.046207 | 0.558888466 | 0.39 | 0.073 | 1.0279390 | 14 | lrp1ab |
| rnd1b1 | 1.352957 | 0.371241758 | 0.197 | 0.02 | 3.4371887 | 14 | rnd1b |
| rai14 | 1.366574 | 0.259079667 | 0.152 | 0.012 | 3.4717834 | 14 | rai14 |
| reep3a | 1.474539 | 0.368770451 | 0.227 | 0.027 | 3.7460666 | 14 | reep3a |
| tmem238a | 1.779257 | 0.261559469 | 0.17 | 0.015 | 4.5202040 | 14 | tmem238a |
| col12a1a | 3.209402 | 0.440994799 | 0.231 | 0.028 | 8.1534873 | 14 | col12a1a |
| net1 | 4.590905 | 0.38822336 | 0.235 | 0.029 | 1.1663195 | 14 | net1 |
| chd33 | 5.141892 | 0.637131162 | 0.504 | 0.119 | 1.3062977 | 14 | chd3 |
| ghra1 | 8.900662 | 0.405604174 | 0.242 | 0.031 | 2.2612132 | 14 | ghra |
| clic41 | 1.722666 | 0.495875586 | 0.326 | 0.054 | 4.3764345 | 14 | clic4 |

| | | | | | | | |
|---|---|---|---|---|---|---|---|
| nfic3 | 2.567435 | 0.565670732 | 0.398 | 0.079 | 6.5225692 | 14 | nfic |
| sostdc1a1 | 3.646809 | 0.423848485 | 0.201 | 0.021 | 9.2647193 | 14 | sostdc1a |
| plekha6 | 7.440378 | 0.32895282 | 0.174 | 0.016 | 1.8902282 | 14 | plekha6 |
| tagln3 | 9.545950 | 0.614738511 | 0.375 | 0.069 | 2.4251487 | 14 | tagln |
| CR847566.11 | 2.496558 | 0.512868904 | 0.299 | 0.047 | 6.3425057 | 14 | CR847566.1 |
| zgc:154058 | 4.018790 | 0.283276313 | 0.163 | 0.014 | 1.0209736 | 14 | zgc:154058 |
| pik3r21 | 5.371591 | 0.524596522 | 0.348 | 0.063 | 1.3646528 | 14 | pik3r2 |
| hmga2 | 1.182552 | 0.296972831 | 0.148 | 0.012 | 3.0042750 | 14 | hmga2 |
| lmbr1l | 5.484110 | 0.294161795 | 0.163 | 0.015 | 1.3932383 | 14 | lmbr1l |
| si:ch211-198c19.34 | 6.511151 | 0.658824294 | 0.394 | 0.078 | 1.6541580 | 14 | si:ch211-198c19.3 |
| col5a2a2 | 8.878966 | 0.605018972 | 0.443 | 0.095 | 2.2557015 | 14 | col5a2a |
| icn3 | 1.360135 | 0.958853214 | 0.735 | 0.26 | 3.4554250 | 14 | icn |
| dcn2 | 1.382425 | 0.782734377 | 0.542 | 0.137 | 3.5120516 | 14 | dcn |
| mob2a | 8.320861 | 0.467509788 | 0.258 | 0.037 | 2.1139147 | 14 | mob2a |
| zgc:1584631 | 1.279367 | 0.634851042 | 0.375 | 0.076 | 3.2502342 | 14 | zgc:158463 |
| znf385a2 | 2.892325 | 0.475062733 | 0.307 | 0.05 | 7.3479523 | 14 | znf385a |
| slco2b11 | 9.622320 | 0.528980694 | 0.345 | 0.062 | 2.4445505 | 14 | slco2b1 |
| fam171a2a | 1.505211 | 0.294726311 | 0.182 | 0.019 | 3.8239901 | 14 | fam171a2a |
| si:ch211-222l21.15 | 2.836337 | 0.8123156 | 0.799 | 0.306 | 7.2057165 | 14 | si:ch211-222l21.1 |
| pbx3b | 3.205270 | 0.336099961 | 0.182 | 0.019 | 8.1429902 | 14 | pbx3b |
| BX322530.12 | 3.388891 | 0.467157113 | 0.295 | 0.048 | 8.6094779 | 14 | BX322530.1 |
| fap | 5.656415 | 0.278343624 | 0.148 | 0.013 | 1.4370123 | 14 | fap |
| cav2 | 1.360472 | 0.265866692 | 0.159 | 0.015 | 3.4562791 | 14 | cav2 |
| snrkb | 1.702423 | 0.284561846 | 0.163 | 0.015 | 4.3250063 | 14 | snrkb |
| si:dkey-16l2.20 | 1.612880 | 0.33213093 | 0.186 | 0.02 | 4.0975230 | 14 | si:dkey-16l2.20 |
| ctsf | 3.115019 | 0.351860451 | 0.242 | 0.034 | 7.9137061 | 14 | ctsf |
| igf2b2 | 4.367078 | 0.559148344 | 0.364 | 0.071 | 1.1094564 | 14 | igf2b |
| zbtb202 | 5.247787 | 0.564931929 | 0.443 | 0.102 | 1.3332003 | 14 | zbtb20 |
| cast1 | 3.229538 | 0.484856 | 0.356 | 0.068 | 8.2046418 | 14 | cast |
| ptpn21 | 7.369873 | 0.374101712 | 0.239 | 0.033 | 1.8723164 | 14 | ptpn21 |
| AL954191.1 | 1.688793 | 0.319634239 | 0.159 | 0.015 | 4.2903788 | 14 | AL954191.1 |
| rab3db | 1.985888 | 0.259925623 | 0.152 | 0.014 | 5.0451508 | 14 | rab3db |
| dag1 | 2.919143 | 0.460811798 | 0.299 | 0.051 | 7.4160845 | 14 | dag1 |
| xirp2a1 | 6.183049 | 0.349176995 | 0.231 | 0.032 | 1.5708038 | 14 | xirp2a |
| prss232 | 8.581211 | 0.397155978 | 0.223 | 0.03 | 2.1800568 | 14 | prss23 |

| | | | | | | | |
|---|---|---|---|---|---|---|---|
| tenm32 | 1.590007 | 0.501626904 | 0.314 | 0.058 | 4.0394142 | 14 | tenm3 |
| LOC1018844511 | 2.095327 | 0.569955812 | 0.39 | 0.086 | 5.3231786 | 14 | LOC101884451 |
| hdlbpa3 | 2.401225 | 0.683356777 | 0.561 | 0.169 | 6.1003122 | 14 | hdlbpa |
| appa2 | 2.404377 | 0.537360225 | 0.386 | 0.083 | 6.1083211 | 14 | appa |
| zgc:114181 | 2.409378 | 0.268511939 | 0.159 | 0.016 | 6.1210271 | 14 | zgc:114181 |
| atp10a2 | 2.710748 | 0.461787165 | 0.307 | 0.055 | 6.8866566 | 14 | atp10a |
| entpd12 | 3.260844 | 0.538396381 | 0.371 | 0.078 | 8.2841761 | 14 | entpd1 |
| CABZ01072614.13 | 7.500115 | 0.576706145 | 0.398 | 0.09 | 1.9054044 | 14 | CABZ01072614.1 |
| fzd4 | 3.433542 | 0.314651356 | 0.155 | 0.015 | 8.7229158 | 14 | fzd4 |
| dbn12 | 4.859342 | 0.473267418 | 0.322 | 0.062 | 1.2345159 | 14 | dbn1 |
| vasna1 | 6.726121 | 0.399280211 | 0.265 | 0.042 | 1.7087712 | 14 | vasna |
| fxyd12 | 1.050719 | 0.464140838 | 0.375 | 0.079 | 2.6693531 | 14 | fxyd1 |
| tspan18b | 4.069038 | 0.369735099 | 0.25 | 0.038 | 1.0337391 | 14 | tspan18b |
| fibinb | 8.124900 | 0.318989252 | 0.178 | 0.021 | 2.0641309 | 14 | fibinb |
| LOC101882334 | 8.557375 | 0.286905453 | 0.148 | 0.015 | 2.1740013 | 14 | LOC101882334 |
| si:ch73-138n13.1 | 1.748817 | 0.385154979 | 0.254 | 0.041 | 4.4428717 | 14 | si:ch73-138n13.1 |
| rsu1 | 2.889313 | 0.412470072 | 0.28 | 0.049 | 7.3403001 | 14 | rsu1 |
| slc38a5b2 | 3.003749 | 0.569434643 | 0.47 | 0.123 | 7.6310247 | 14 | slc38a5b |
| zgc:1951733 | 7.033185 | 0.526306141 | 0.447 | 0.109 | 1.7867808 | 14 | zgc:195173 |
| olfml3b2 | 8.859285 | 0.458410262 | 0.288 | 0.052 | 2.2507015 | 14 | olfml3b |
| gpc4 | 1.266404 | 0.328133042 | 0.208 | 0.029 | 3.2173014 | 14 | gpc4 |
| bmp6 | 5.970113 | 0.319221677 | 0.136 | 0.013 | 1.5167074 | 14 | bmp6 |
| NC-002333.45 | 1.785211 | 0.652662813 | 0.943 | 0.604 | 4.5353297 | 14 | NC-002333.4 |
| b2ml5 | 3.884673 | 0.678981146 | 0.739 | 0.287 | 9.8690135 | 14 | b2ml |
| sorbs1 | 5.175569 | 0.305645094 | 0.174 | 0.021 | 1.3148535 | 14 | sorbs1 |
| rtn3 | 1.993683 | 0.461260097 | 0.348 | 0.075 | 5.0649531 | 14 | rtn3 |
| wnt5b | 2.514295 | 0.284239401 | 0.14 | 0.014 | 6.3875673 | 14 | wnt5b |
| tmem1762 | 7.527294 | 0.475810569 | 0.352 | 0.076 | 1.9123091 | 14 | tmem176 |
| sox4a2 | 8.224149 | 0.459750793 | 0.303 | 0.059 | 2.0893452 | 14 | sox4a |
| emx2 | 9.257021 | 0.314911134 | 0.178 | 0.022 | 2.3517463 | 14 | emx2 |
| CU539058.1 | 1.176776 | 0.253983347 | 0.155 | 0.017 | 2.9896005 | 14 | CU539058.1 |
| gulp1a1 | 1.714082 | 0.343108781 | 0.223 | 0.034 | 4.3546255 | 14 | gulp1a |
| thbs1a | 3.694267 | 0.273899062 | 0.129 | 0.012 | 9.3852864 | 14 | thbs1a |
| selenbp11 | 7.700581 | 0.488573306 | 0.348 | 0.077 | 1.9563328 | 14 | selenbp1 |
| cxxc5a | 1.561600 | 0.277096773 | 0.182 | 0.023 | 3.9672469 | 14 | cxxc5a |

| | | | | | | | |
|---|---|---|---|---|---|---|---|
| fstb1 | 1.677680 | 0.317742609 | 0.182 | 0.023 | 4.2621472 | 14 | fstb |
| dpyda.1 | 2.547604 | 0.254061507 | 0.14 | 0.014 | 6.4721883 | 14 | dpyda.1 |
| col27a1b1 | 3.324369 | 0.341631618 | 0.189 | 0.026 | 8.4455602 | 14 | col27a1b |
| lamb1b | 8.224386 | 0.300156195 | 0.178 | 0.023 | 2.0894055 | 14 | lamb1b |
| nfixb2 | 8.949863 | 0.522767515 | 0.348 | 0.078 | 2.2737129 | 14 | nfixb |
| zgc:153372 | 1.890541 | 0.310855352 | 0.208 | 0.031 | 4.8029207 | 14 | zgc:153372 |
| meis31 | 2.168892 | 0.282085941 | 0.148 | 0.016 | 5.5100709 | 14 | meis3 |
| scarb2a | 2.452191 | 0.462702721 | 0.295 | 0.059 | 6.2297920 | 14 | scarb2a |
| schip1 | 3.684178 | 0.300834562 | 0.167 | 0.021 | 9.3596547 | 14 | schip1 |
| aoc23 | 1.072138 | 0.578610173 | 0.33 | 0.073 | 2.7237686 | 14 | aoc2 |
| stau23 | 1.615065 | 0.477172149 | 0.379 | 0.092 | 4.1030730 | 14 | stau2 |
| cyp1c22 | 3.152213 | 0.313572732 | 0.148 | 0.017 | 8.0081991 | 14 | cyp1c2 |
| ier2a4 | 9.558217 | 0.728247039 | 0.674 | 0.268 | 2.4282651 | 14 | ier2a |
| cxadr | 1.340224 | 0.312281014 | 0.208 | 0.031 | 3.4048396 | 14 | cxadr |
| gas1a3 | 1.622858 | 0.417812438 | 0.28 | 0.055 | 4.1228724 | 14 | gas1a |
| spred2b2 | 6.873349 | 0.499145418 | 0.345 | 0.081 | 1.7461743 | 14 | spred2b |
| dedd12 | 7.495805 | 0.612029509 | 0.492 | 0.152 | 1.9043093 | 14 | dedd1 |
| qkia2 | 8.020574 | 0.582726095 | 0.58 | 0.193 | 2.0376269 | 14 | qkia |
| myl9b2 | 9.100333 | 0.476682091 | 0.383 | 0.095 | 2.3119396 | 14 | myl9b |
| fosl1a2 | 1.197946 | 0.552167703 | 0.337 | 0.08 | 3.0433834 | 14 | fosl1a |
| calm2b2 | 1.406368 | 0.605351075 | 0.667 | 0.24 | 3.5728779 | 14 | calm2b |
| itga91 | 2.219361 | 0.35815779 | 0.216 | 0.035 | 5.6382880 | 14 | itga9 |
| jdp2b2 | 1.977260 | 0.729648667 | 0.492 | 0.16 | 5.0232301 | 14 | jdp2b |
| mfap22 | 2.223586 | 0.485762243 | 0.436 | 0.115 | 5.6490215 | 14 | mfap2 |
| pim13 | 3.140387 | 0.646831983 | 0.504 | 0.164 | 7.9781544 | 14 | pim1 |
| plecb | 3.557439 | 0.370598865 | 0.258 | 0.049 | 9.0376744 | 14 | plecb |
| erbb2 | 4.787010 | 0.298058456 | 0.197 | 0.031 | 1.2161399 | 14 | erbb2 |
| myof | 7.146453 | 0.258976978 | 0.163 | 0.021 | 1.8155563 | 14 | myof |
| fstl31 | 8.574924 | 0.29784326 | 0.167 | 0.023 | 2.1784595 | 14 | fstl3 |
| cyp2n132 | 1.167281 | 0.356275722 | 0.223 | 0.038 | 2.9654777 | 14 | cyp2n13 |
| si:dkey-56m19.5 | 1.465772 | 0.396154629 | 0.265 | 0.053 | 3.7237957 | 14 | si:dkey-56m19.5 |
| ulk21 | 1.622129 | 0.348466485 | 0.261 | 0.051 | 4.1210196 | 14 | ulk2 |
| tgm1l11 | 1.710607 | 0.355752363 | 0.216 | 0.036 | 4.3457972 | 14 | tgm1l1 |
| cdkn1a | 1.935758 | 0.364342126 | 0.246 | 0.046 | 4.9177940 | 14 | cdkn1a |
| stat1a2 | 2.217430 | 0.507849202 | 0.42 | 0.119 | 5.6333827 | 14 | stat1a |

| | | | | | | | |
|---|---|---|---|---|---|---|---|
| angptl41 | 4.685432 | 0.389148026 | 0.265 | 0.052 | 1.1903340 | 14 | angptl4 |
| ptprk | 3.369272 | 0.325433289 | 0.205 | 0.034 | 8.5596372 | 14 | ptprk |
| ccdc85b2 | 4.604157 | 0.33320725 | 0.25 | 0.048 | 1.1696863 | 14 | ccdc85b |
| btg23 | 5.170365 | 0.705291267 | 0.735 | 0.344 | 1.3135313 | 14 | btg2 |
| mfap51 | 6.930295 | 0.453395194 | 0.284 | 0.06 | 1.7606414 | 14 | mfap5 |
| gli3 | 8.796260 | 0.347453724 | 0.201 | 0.033 | 2.2346898 | 14 | gli3 |
| tjp2b1 | 1.356679 | 0.345533095 | 0.288 | 0.061 | 3.4466449 | 14 | tjp2b |
| ifi454 | 1.386531 | 0.698212421 | 0.439 | 0.135 | 3.5224830 | 14 | ifi45 |
| klf2a2 | 2.346614 | 0.557035985 | 0.364 | 0.096 | 5.9615740 | 14 | klf2a |
| si:ch73-22o12.1 | 4.624893 | 0.285735994 | 0.186 | 0.029 | 1.1749543 | 14 | si:ch73-22o12.1 |
| meis2b2 | 5.258341 | 0.290828258 | 0.178 | 0.027 | 1.3358817 | 14 | meis2b |
| ptges1 | 6.483537 | 0.324934052 | 0.163 | 0.023 | 1.6471427 | 14 | ptges |
| sost1 | 8.995680 | 0.27645912 | 0.144 | 0.018 | 2.2853525 | 14 | sost |
| fbn2b2 | 1.032084 | 0.357790568 | 0.246 | 0.048 | 2.6220103 | 14 | fbn2b |
| igfbp5b2 | 1.253541 | 0.458904047 | 0.348 | 0.085 | 3.1846225 | 14 | igfbp5b |
| aldh9a1a.13 | 1.258882 | 0.56007965 | 0.542 | 0.188 | 3.1981899 | 14 | aldh9a1a.1 |
| map1lc3a2 | 1.547392 | 0.400576118 | 0.322 | 0.077 | 3.9311495 | 14 | map1lc3a |
| flna3 | 1.553242 | 0.475977319 | 0.417 | 0.118 | 3.9460129 | 14 | flna |
| mdka3 | 2.041563 | 0.608645959 | 0.39 | 0.11 | 5.1865924 | 14 | mdka |
| CABZ01045617.1 | 3.238336 | 0.34480368 | 0.265 | 0.055 | 8.2269932 | 14 | CABZ01045617.1 |
| dhrs13l11 | 3.857712 | 0.303235225 | 0.212 | 0.038 | 9.8005180 | 14 | dhrs13l1 |
| foxo3b2 | 4.775066 | 0.41003218 | 0.295 | 0.067 | 1.2131056 | 14 | foxo3b |
| ar2 | 5.018189 | 0.427603258 | 0.284 | 0.064 | 1.2748711 | 14 | ar |
| egr15 | 5.704107 | 0.62308239 | 0.515 | 0.178 | 1.4491284 | 14 | egr1 |
| txnipa2 | 6.466081 | 0.579164069 | 0.633 | 0.238 | 1.6427081 | 14 | txnipa |
| ephb32 | 9.422119 | 0.385352036 | 0.246 | 0.049 | 2.3936894 | 14 | ephb3 |
| hmgb1a6 | 2.119247 | 0.546924494 | 0.598 | 0.222 | 5.3839488 | 14 | hmgb1a |
| glis2a | 2.713922 | 0.271748068 | 0.17 | 0.025 | 6.8947199 | 14 | glis2a |
| actn12 | 3.290754 | 0.336505961 | 0.235 | 0.045 | 8.3601630 | 14 | actn1 |
| cyr612 | 3.406222 | 0.521742944 | 0.284 | 0.065 | 8.6535090 | 14 | cyr61 |
| klf11b2 | 5.300465 | 0.424579878 | 0.322 | 0.08 | 1.3465833 | 14 | klf11b |
| BX640522.21 | 1.377098 | 0.489357641 | 0.212 | 0.039 | 3.4985185 | 14 | BX640522.2 |
| fzd7b2 | 1.931574 | 0.344032566 | 0.235 | 0.046 | 4.9071648 | 14 | fzd7b |
| rhoua1 | 2.168672 | 0.314842697 | 0.178 | 0.028 | 5.5095118 | 14 | rhoua |
| cux1a2 | 7.803359 | 0.375164821 | 0.25 | 0.053 | 1.9824435 | 14 | cux1a |

| | | | | | | | |
|---|---|---|---|---|---|---|---|
| myl63 | 1.042217 | 0.469242974 | 0.398 | 0.112 | 2.6477532 | 14 | myl6 |
| myh10 | 1.521841 | 0.330800512 | 0.216 | 0.04 | 3.8662379 | 14 | myh10 |
| ctnnb11 | 3.310989 | 0.497253983 | 0.477 | 0.154 | 8.4115686 | 14 | ctnnb1 |
| pdk2a | 3.323626 | 0.289558254 | 0.174 | 0.028 | 8.4436720 | 14 | pdk2a |
| fam168a3 | 3.871150 | 0.371572919 | 0.295 | 0.071 | 9.8346569 | 14 | fam168a |
| cyp2p62 | 3.983712 | 0.279128147 | 0.163 | 0.024 | 1.0120621 | 14 | cyp2p6 |
| pard3bb1 | 5.784701 | 0.268984369 | 0.186 | 0.031 | 1.4696035 | 14 | pard3bb |
| cd99 | 8.104156 | 0.33667 | 0.242 | 0.05 | 2.0588609 | 14 | cd99 |
| irf2bp2b1 | 8.221223 | 0.456334024 | 0.356 | 0.098 | 2.0886018 | 14 | irf2bp2b |
| ptprga3 | 8.473758 | 0.426384003 | 0.288 | 0.069 | 2.1527582 | 14 | ptprga |
| eif4ebp3l3 | 9.140415 | 0.626644662 | 0.538 | 0.201 | 2.3221226 | 14 | eif4ebp3l |
| calcoco1a | 1.994674 | 0.285950874 | 0.193 | 0.034 | 5.0674703 | 14 | calcoco1a |
| ier2b5 | 2.391565 | 0.576797077 | 0.595 | 0.232 | 6.0757716 | 14 | ier2b |
| mt-nd16 | 3.302437 | 0.633323409 | 0.761 | 0.392 | 8.3898435 | 14 | mt-nd1 |
| gstt1b3 | 3.797965 | 0.393645267 | 0.333 | 0.086 | 9.6487316 | 14 | gstt1b |
| si:ch73-1a9.35 | 4.276532 | 0.605484444 | 0.705 | 0.31 | 1.0864530 | 14 | si:ch73-1a9.3 |
| wu:fb16f031 | 5.310280 | 0.416637008 | 0.33 | 0.086 | 1.3490766 | 14 | wu:fb16f03 |
| mphosph81 | 6.038016 | 0.477273429 | 0.436 | 0.136 | 1.5339581 | 14 | mphosph8 |
| ywhaqb3 | 1.282141 | 0.481311506 | 0.458 | 0.15 | 3.2572802 | 14 | ywhaqb |
| map3k1 | 1.505652 | 0.260127823 | 0.152 | 0.022 | 3.8251089 | 14 | map3k1 |
| mknk2b3 | 1.794703 | 0.526442869 | 0.542 | 0.195 | 4.5594435 | 14 | mknk2b |
| lama42 | 1.951423 | 0.322540127 | 0.223 | 0.044 | 4.9575911 | 14 | lama4 |
| cyp2p72 | 2.765662 | 0.274335455 | 0.174 | 0.029 | 7.0261652 | 14 | cyp2p7 |
| dhrs3a2 | 2.768450 | 0.30284936 | 0.197 | 0.036 | 7.0332493 | 14 | dhrs3a |
| tshz3b1 | 4.172917 | 0.287978816 | 0.186 | 0.032 | 1.0601297 | 14 | tshz3b |
| tnfsf102 | 5.316526 | 0.335818141 | 0.212 | 0.041 | 1.3506635 | 14 | tnfsf10 |
| pabpc1b2 | 5.941731 | 0.391425038 | 0.299 | 0.074 | 1.5094968 | 14 | pabpc1b |
| pfkfb4b1 | 6.638846 | 0.432979221 | 0.394 | 0.116 | 1.6865988 | 14 | pfkfb4b |
| fgl2a2 | 7.760193 | 0.38666366 | 0.299 | 0.073 | 1.9714771 | 14 | fgl2a |
| hmgn66 | 8.388401 | 0.603550604 | 0.674 | 0.306 | 2.1310734 | 14 | hmgn6 |
| dbpb | 1.141942 | 0.304366674 | 0.216 | 0.042 | 2.9011053 | 14 | dbpb |
| hecw2b | 1.920632 | 0.250417087 | 0.14 | 0.02 | 4.8793662 | 14 | hecw2b |
| hoxc3a1 | 5.479899 | 0.253514772 | 0.167 | 0.027 | 1.3921684 | 14 | hoxc3a |
| ube2e32 | 1.069051 | 0.484438799 | 0.477 | 0.165 | 2.7159249 | 14 | ube2e3 |
| pbx4 | 1.153750 | 0.345795558 | 0.292 | 0.071 | 2.9311026 | 14 | pbx4 |

| | | | | | | | |
|---|---|---|---|---|---|---|---|
| abcc53 | 1.295918 | 0.442003108 | 0.33 | 0.092 | 3.2922816 | 14 | abcc5 |
| zbtb41 | 2.131929 | 0.258042823 | 0.148 | 0.022 | 5.4161662 | 14 | zbtb4 |
| gstm.12 | 2.212700 | 0.560945083 | 0.659 | 0.273 | 5.6213654 | 14 | gstm.1 |
| si:ch211-166a6.5 | 2.462079 | 0.287886195 | 0.193 | 0.036 | 6.2549130 | 14 | si:ch211-166a6.5 |
| tmem176l.12 | 2.923245 | 0.506048853 | 0.545 | 0.197 | 7.4265058 | 14 | tmem176l.1 |
| slc7a12 | 3.611929 | 0.312601058 | 0.189 | 0.035 | 9.1761068 | 14 | slc7a1 |
| si:ch211-286o17.11 | 4.739847 | 0.344611388 | 0.223 | 0.047 | 1.2041583 | 14 | si:ch211-286o17.1 |
| pdk2b1 | 5.361778 | 0.412945426 | 0.356 | 0.102 | 1.3621599 | 14 | pdk2b |
| fam120c | 6.856685 | 0.409027373 | 0.352 | 0.102 | 1.7419410 | 14 | fam120c |
| add3a3 | 7.302786 | 0.47178928 | 0.42 | 0.134 | 1.8552728 | 14 | add3a |
| sypl2a | 9.844267 | 0.320866396 | 0.254 | 0.058 | 2.5009361 | 14 | sypl2a |
| her9 | 1.870166 | 0.283797333 | 0.148 | 0.023 | 4.7511581 | 14 | her9 |
| mt-nd46 | 1.970331 | 0.586343283 | 0.803 | 0.443 | 5.0056260 | 14 | mt-nd4 |
| tgfb31 | 2.894294 | 0.252674439 | 0.178 | 0.031 | 7.3529549 | 14 | tgfb3 |
| adh8b | 3.306309 | 0.253251524 | 0.22 | 0.045 | 8.3996788 | 14 | adh8b |
| elovl1b2 | 4.580660 | 0.415806573 | 0.39 | 0.119 | 1.1637168 | 14 | elovl1b |
| h2afy21 | 5.204003 | 0.413000268 | 0.386 | 0.119 | 1.3220771 | 14 | h2afy2 |
| h3f3c4 | 1.796579 | 0.554943924 | 0.67 | 0.299 | 4.5642104 | 14 | h3f3c |
| nfil3-64 | 1.997109 | 0.505682129 | 0.394 | 0.128 | 5.0736561 | 14 | nfil3-6 |
| kmt2bb | 2.317524 | 0.398886092 | 0.33 | 0.093 | 5.8876716 | 14 | kmt2bb |
| tmeff1b1 | 2.661879 | 0.356069024 | 0.197 | 0.04 | 6.7625050 | 14 | tmeff1b |
| acin1a2 | 2.811806 | 0.510959882 | 0.568 | 0.224 | 7.1433954 | 14 | acin1a |
| aebp11 | 5.081399 | 0.331506809 | 0.216 | 0.046 | 1.2909295 | 14 | aebp1 |
| mt-nd25 | 1.404017 | 0.590911925 | 0.727 | 0.387 | 3.5669060 | 14 | mt-nd2 |
| si:dkey-188i13.102 | 1.646768 | 0.392784184 | 0.367 | 0.11 | 4.1836144 | 14 | si:dkey-188i13.10 |
| dspa | 1.822238 | 0.257901228 | 0.159 | 0.027 | 4.6293966 | 14 | dspa |
| aldh2.21 | 3.040877 | 0.338465329 | 0.314 | 0.086 | 7.7253484 | 14 | aldh2.2 |
| ppdpfa3 | 3.800571 | 0.438963484 | 0.269 | 0.069 | 9.6553530 | 14 | ppdpfa |
| sesn14 | 4.688020 | 0.545900531 | 0.462 | 0.171 | 1.1909916 | 14 | sesn1 |
| timp2a2 | 6.180646 | 0.325397614 | 0.258 | 0.063 | 1.5701933 | 14 | timp2a |
| pfn23 | 7.256346 | 0.494209891 | 0.587 | 0.234 | 1.8434748 | 14 | pfn2 |
| grhl11 | 8.479791 | 0.271538889 | 0.163 | 0.029 | 2.1542910 | 14 | grhl1 |
| rnd3b2 | 1.107503 | 0.267845353 | 0.22 | 0.047 | 2.8136117 | 14 | rnd3b |
| fkbp52 | 1.441955 | 0.460102567 | 0.379 | 0.124 | 3.6632889 | 14 | fkbp5 |
| deptor1 | 1.680752 | 0.279450459 | 0.178 | 0.034 | 4.2699526 | 14 | deptor |

| | | | | | | | |
|---|---|---|---|---|---|---|---|
| agrn1 | 2.190018 | 0.339206757 | 0.242 | 0.057 | 5.5637431 | 14 | agrn |
| angptl1a1 | 2.207846 | 0.343158629 | 0.174 | 0.033 | 5.6090339 | 14 | angptl1a |
| zgc:1627307 | 2.429635 | 0.60155909 | 0.788 | 0.397 | 6.1724879 | 14 | zgc:162730 |
| cnn22 | 2.616291 | 0.437163974 | 0.443 | 0.153 | 6.6466888 | 14 | cnn2 |
| selenow12 | 2.947941 | 0.55620321 | 0.765 | 0.41 | 7.4892446 | 14 | selenow1 |
| midn3 | 4.125959 | 0.49399182 | 0.409 | 0.141 | 1.0482001 | 14 | midn |
| cpn12 | 8.476012 | 0.345750252 | 0.284 | 0.075 | 2.1533309 | 14 | cpn1 |
| csrnp1b3 | 1.029065 | 0.508160651 | 0.443 | 0.161 | 2.6143396 | 14 | csrnp1b |
| sh3pxd2aa1 | 1.321485 | 0.260066745 | 0.17 | 0.032 | 3.3572344 | 14 | sh3pxd2aa |
| ldlrad4a1 | 2.709131 | 0.272514458 | 0.17 | 0.032 | 6.8825481 | 14 | ldlrad4a |
| rbm39a1 | 7.950814 | 0.425075874 | 0.428 | 0.149 | 2.0199044 | 14 | rbm39a |
| si:ch211-39i22.12 | 8.211403 | 0.383744903 | 0.303 | 0.085 | 2.0861069 | 14 | si:ch211-39i22.1 |
| lxn3 | 1.244734 | 0.465522121 | 0.455 | 0.163 | 3.1622469 | 14 | lxn |
| crim12 | 1.271338 | 0.273368353 | 0.189 | 0.039 | 3.2298355 | 14 | crim1 |
| b2m5 | 1.402591 | 0.533437911 | 0.644 | 0.293 | 3.5632841 | 14 | b2m |
| mir21-11 | 1.442097 | 0.421583279 | 0.337 | 0.105 | 3.6636480 | 14 | mir21-1 |
| tnrc6c13 | 1.526260 | 0.455270359 | 0.413 | 0.146 | 3.8774649 | 14 | tnrc6c1 |
| gchfr1 | 1.690254 | 0.29231351 | 0.182 | 0.037 | 4.2940904 | 14 | gchfr |
| alcama | 1.995890 | 0.26203087 | 0.163 | 0.03 | 5.0705587 | 14 | alcama |
| ckap42 | 2.368768 | 0.291912314 | 0.239 | 0.058 | 6.0178556 | 14 | ckap4 |
| adamts15a2 | 3.480132 | 0.354354681 | 0.231 | 0.056 | 8.8412759 | 14 | adamts15a |
| csrp1a | 4.643973 | 0.327842687 | 0.284 | 0.078 | 1.1798014 | 14 | csrp1a |
| kank22 | 5.932577 | 0.308513499 | 0.223 | 0.053 | 1.5071713 | 14 | kank2 |
| jmjd1cb | 9.321006 | 0.270346008 | 0.201 | 0.044 | 2.3680018 | 14 | jmjd1cb |
| CABZ01068366.11 | 1.325853 | 0.262741331 | 0.189 | 0.04 | 3.3683316 | 14 | CABZ01068366.1 |
| flot2a2 | 1.491283 | 0.327960985 | 0.265 | 0.071 | 3.7886059 | 14 | flot2a |
| kmt2cb | 2.457719 | 0.308572132 | 0.235 | 0.058 | 6.2438372 | 14 | kmt2cb |
| lgals3b | 2.648815 | 0.391521007 | 0.311 | 0.096 | 6.7293150 | 14 | lgals3b |
| arl4d | 2.751075 | 0.277054004 | 0.22 | 0.051 | 6.9891078 | 14 | arl4d |
| COX37 | 3.097979 | 0.468061343 | 0.894 | 0.673 | 7.8704178 | 14 | COX3 |
| kdm6bb | 4.840618 | 0.312774878 | 0.239 | 0.06 | 1.2297590 | 14 | kdm6bb |
| si:dkey-7j14.62 | 5.280268 | 0.453300534 | 0.496 | 0.192 | 1.3414521 | 14 | si:dkey-7j14.6 |
| ehd1b | 5.979272 | 0.27670995 | 0.216 | 0.05 | 1.5190342 | 14 | ehd1b |
| si:ch211-286b5.52 | 6.170655 | 0.404481217 | 0.417 | 0.148 | 1.5676550 | 14 | si:ch211-286b5.5 |
| cebpb6 | 7.312350 | 0.58347352 | 0.689 | 0.334 | 1.8577026 | 14 | cebpb |

| | | | | | | | |
|---|---|---|---|---|---|---|---|
| aldh2.11 | 7.972042 | 0.251784378 | 0.178 | 0.037 | 2.0252975 | 14 | aldh2.1 |
| glud1a | 8.103116 | 0.36886765 | 0.318 | 0.097 | 2.0585966 | 14 | glud1a |
| ncor1 | 8.522098 | 0.316545097 | 0.227 | 0.056 | 2.1650390 | 14 | ncor1 |
| tead1b1 | 1.063512 | 0.268384739 | 0.163 | 0.032 | 2.7018533 | 14 | tead1b |
| coro1ca | 2.113998 | 0.308640814 | 0.254 | 0.067 | 5.3706140 | 14 | coro1ca |
| vat11 | 4.136927 | 0.429899524 | 0.409 | 0.144 | 1.0509864 | 14 | vat1 |
| cry2 | 4.184490 | 0.315612176 | 0.277 | 0.077 | 1.0630697 | 14 | cry2 |
| atp2b22 | 4.362529 | 0.307934645 | 0.284 | 0.081 | 1.1083005 | 14 | atp2b2 |
| hmgn73 | 8.934691 | 0.511553881 | 0.648 | 0.313 | 2.2698584 | 14 | hmgn7 |
| hmgb3a4 | 9.654480 | 0.38385691 | 0.277 | 0.081 | 2.4527208 | 14 | hmgb3a |
| atrx2 | 2.152013 | 0.399306645 | 0.364 | 0.125 | 5.4671897 | 14 | atrx |
| perp | 3.710203 | 0.281774942 | 0.231 | 0.059 | 9.4257723 | 14 | perp |
| mcl1b4 | 4.325616 | 0.475996439 | 0.576 | 0.246 | 1.0989228 | 14 | mcl1b |
| si:dkey-51e6.12 | 4.482301 | 0.335515757 | 0.352 | 0.116 | 1.1387286 | 14 | si:dkey-51e6.1 |
| pnrc27 | 4.886421 | 0.493301013 | 0.92 | 0.593 | 1.2413954 | 14 | pnrc2 |
| mxra8a1 | 7.155193 | 0.265721844 | 0.235 | 0.06 | 1.8177770 | 14 | mxra8a |
| socs3b3 | 7.776219 | 0.453310214 | 0.405 | 0.148 | 1.9755486 | 14 | socs3b |
| vasnb1 | 8.010222 | 0.269218254 | 0.167 | 0.035 | 2.0349970 | 14 | vasnb |
| ctdspla1 | 9.542837 | 0.254705131 | 0.178 | 0.039 | 2.4243577 | 14 | ctdspla |
| akap12b1 | 1.278919 | 0.319170683 | 0.261 | 0.072 | 3.2490949 | 14 | akap12b |
| purab1 | 1.467621 | 0.293316903 | 0.223 | 0.057 | 3.7284933 | 14 | purab |
| egfra2 | 1.811802 | 0.259225852 | 0.155 | 0.031 | 4.6028836 | 14 | egfra |
| slc25a55a1 | 1.919258 | 0.322260671 | 0.258 | 0.073 | 4.8758760 | 14 | slc25a55a |
| mycb2 | 1.970359 | 0.464606727 | 0.489 | 0.197 | 5.0056981 | 14 | mycb |
| atf4b3 | 2.132807 | 0.472367103 | 0.602 | 0.273 | 5.4183972 | 14 | atf4b |
| ncor2 | 2.521067 | 0.319664183 | 0.269 | 0.078 | 6.4047708 | 14 | ncor2 |
| ppib5 | 2.766743 | 0.507198659 | 0.731 | 0.374 | 7.0289106 | 14 | ppib |
| si:ch211-196l7.46 | 5.297906 | 0.473428211 | 0.667 | 0.325 | 1.3459330 | 14 | si:ch211-196l7.4 |
| ciarta2 | 5.359924 | 0.34642793 | 0.345 | 0.115 | 1.3616888 | 14 | ciarta |
| nme32 | 5.880582 | 0.305798151 | 0.288 | 0.086 | 1.4939618 | 14 | nme3 |
| fkbp72 | 8.032785 | 0.250574886 | 0.201 | 0.048 | 2.0407290 | 14 | fkbp7 |
| aplp21 | 9.174581 | 0.316269589 | 0.261 | 0.075 | 2.3308024 | 14 | aplp2 |
| hist2h2l2 | 1.177933 | 0.371664354 | 0.303 | 0.096 | 2.9925407 | 14 | hist2h2l |
| lrpap1 | 1.303052 | 0.286813033 | 0.231 | 0.062 | 3.3104058 | 14 | lrpap1 |
| si:ch1073-291c23.22 | 2.552588 | 0.288732782 | 0.254 | 0.071 | 6.4848513 | 14 | si:ch1073-291c23.2 |

| | | | | | | | |
|---|---|---|---|---|---|---|---|
| mt-atp66 | 3.151585 | 0.500203488 | 0.795 | 0.553 | 8.0066030 | 14 | mt-atp6 |
| nfil3 | 3.557970 | 0.250264218 | 0.155 | 0.032 | 9.0390230 | 14 | nfil3 |
| per31 | 4.666638 | 0.342804748 | 0.258 | 0.076 | 1.1855594 | 14 | per3 |
| rmdn1 | 6.096749 | 0.252301035 | 0.231 | 0.062 | 1.5488791 | 14 | rmdn1 |
| myh11a3 | 1.460439 | 0.307810811 | 0.246 | 0.069 | 3.7102456 | 14 | myh11a |
| cnn3a2 | 2.112960 | 0.295224923 | 0.239 | 0.067 | 5.3679766 | 14 | cnn3a |
| mgst1.11 | 3.508846 | 0.269869537 | 0.208 | 0.054 | 8.9142250 | 14 | mgst1.1 |
| cd151l | 4.715232 | 0.254343008 | 0.227 | 0.061 | 1.1979048 | 14 | cd151l |
| gsta.11 | 5.237826 | 0.250324753 | 0.197 | 0.049 | 1.3306697 | 14 | gsta.1 |
| cebpg | 7.025162 | 0.265199439 | 0.235 | 0.065 | 1.7847426 | 14 | cebpg |
| ctdsp22 | 1.237633 | 0.310754575 | 0.269 | 0.083 | 3.1442082 | 14 | ctdsp2 |
| sdc44 | 1.308920 | 0.497904825 | 0.545 | 0.251 | 3.3253125 | 14 | sdc4 |
| si:ch211-140m22.7 | 1.488746 | 0.28491915 | 0.231 | 0.065 | 3.7821603 | 14 | si:ch211-140m22.7 |
| wls2 | 1.707194 | 0.349640647 | 0.36 | 0.128 | 4.3371278 | 14 | wls |
| tax1bp1b | 1.983925 | 0.256630723 | 0.246 | 0.071 | 5.0401622 | 14 | tax1bp1b |
| ywhah | 2.201002 | 0.274175542 | 0.216 | 0.058 | 5.5916474 | 14 | ywhah |
| id2a | 2.328823 | 0.467073091 | 0.413 | 0.164 | 5.9163766 | 14 | id2a |
| tnrc18 | 2.739167 | 0.256154628 | 0.178 | 0.043 | 6.9588555 | 14 | tnrc18 |
| jupa | 2.944690 | 0.267904269 | 0.216 | 0.058 | 7.4809857 | 14 | jupa |
| irf82 | 3.838163 | 0.365158819 | 0.186 | 0.047 | 9.7508545 | 14 | irf8 |
| mt-nd52 | 5.023050 | 0.373691799 | 0.345 | 0.127 | 1.2761059 | 14 | mt-nd5 |
| serinc11 | 7.486376 | 0.312341178 | 0.269 | 0.084 | 1.9019138 | 14 | serinc1 |
| tagln25 | 1.498913 | 0.412571816 | 0.485 | 0.206 | 3.8079897 | 14 | tagln2 |
| fam3c | 1.679095 | 0.257366258 | 0.178 | 0.044 | 4.2657408 | 14 | fam3c |
| marcksl1a3 | 2.259482 | 0.45126832 | 0.564 | 0.257 | 5.7402165 | 14 | marcksl1a |
| ifi463 | 4.078250 | 0.359101663 | 0.352 | 0.129 | 1.0360796 | 14 | ifi46 |
| capns1b1 | 4.516235 | 0.355415401 | 0.371 | 0.143 | 1.1473495 | 14 | capns1b |
| rbm5 | 7.279555 | 0.285516504 | 0.242 | 0.073 | 1.8493709 | 14 | rbm5 |
| ppp1cb | 7.491707 | 0.273571874 | 0.231 | 0.068 | 1.9032681 | 14 | ppp1cb |
| mt-cyb5 | 8.666801 | 0.457299697 | 0.769 | 0.575 | 2.2018009 | 14 | mt-cyb |
| lamb1a2 | 1.640503 | 0.271171655 | 0.197 | 0.053 | 4.1676983 | 14 | lamb1a |
| serinc21 | 1.947491 | 0.317883861 | 0.337 | 0.122 | 4.9476011 | 14 | serinc2 |
| si:ch211-204c21.12 | 2.192064 | 0.320689398 | 0.239 | 0.073 | 5.5689386 | 14 | si:ch211-204c21.1 |
| brd3b1 | 2.975213 | 0.26780032 | 0.227 | 0.067 | 7.5585290 | 14 | brd3b |
| ndfip1l2 | 3.128663 | 0.313437168 | 0.239 | 0.073 | 7.9483692 | 14 | ndfip1l |

| | | | | | | | |
|---|---|---|---|---|---|---|---|
| klf6a3 | 3.431764 | 0.381868706 | 0.432 | 0.177 | 8.7183966 | 14 | klf6a |
| si:ch211-132b12.7 | 4.180833 | 0.260668264 | 0.193 | 0.052 | 1.0621408 | 14 | si:ch211-132b12.7 |
| mmp14b2 | 4.908815 | 0.263370148 | 0.223 | 0.065 | 1.2470846 | 14 | mmp14b |
| add1 | 5.594990 | 0.278342098 | 0.212 | 0.061 | 1.4214072 | 14 | add1 |
| h2afx15 | 6.978719 | 0.402618433 | 0.504 | 0.224 | 1.7729437 | 14 | h2afx1 |
| rbbp61 | 7.152338 | 0.305351104 | 0.307 | 0.107 | 1.8170515 | 14 | rbbp6 |
| tefa2 | 1.190251 | 0.364357242 | 0.356 | 0.138 | 3.0238351 | 14 | tefa |
| ptmaa6 | 1.293170 | 0.471062949 | 0.773 | 0.428 | 3.2853004 | 14 | ptmaa |
| capns1a1 | 1.418124 | 0.337341424 | 0.394 | 0.156 | 3.6027441 | 14 | capns1a |
| ogt.1 | 1.997809 | 0.273374991 | 0.235 | 0.072 | 5.0754352 | 14 | ogt.1 |
| nr3c1 | 2.122216 | 0.256642605 | 0.182 | 0.048 | 5.3914916 | 14 | nr3c1 |
| adh52 | 2.492336 | 0.387697622 | 0.519 | 0.239 | 6.3317804 | 14 | adh5 |
| zgc:1583434 | 2.501386 | 0.462350698 | 0.398 | 0.167 | 6.3547714 | 14 | zgc:158343 |
| CABZ01092746.12 | 3.507825 | 0.315455619 | 0.269 | 0.087 | 8.9116318 | 14 | CABZ01092746.1 |
| XLOC-024750 | 3.770809 | 0.276540663 | 0.212 | 0.062 | 9.5797407 | 14 | XLOC-024750 |
| fbxl6 | 4.527877 | 0.263440989 | 0.212 | 0.062 | 1.1503073 | 14 | fbxl6 |
| lgals9l32 | 5.182810 | 0.273695564 | 0.254 | 0.081 | 1.3166931 | 14 | lgals9l3 |
| gna11b | 8.141668 | 0.250202929 | 0.216 | 0.063 | 2.0683908 | 14 | gna11b |
| si:ch211-197h24.91 | 8.492411 | 0.254478348 | 0.254 | 0.081 | 2.1574972 | 14 | si:ch211-197h24.9 |
| tpm32 | 9.937246 | 0.393247455 | 0.583 | 0.28 | 2.5245575 | 14 | tpm3 |
| usp36 | 1.072289 | 0.276360994 | 0.246 | 0.08 | 2.7241520 | 14 | usp36 |
| mycbp | 1.556409 | 0.30783293 | 0.47 | 0.197 | 3.9540577 | 14 | mycbp |
| nr1d2a | 1.604853 | 0.293250733 | 0.261 | 0.087 | 4.0771293 | 14 | nr1d2a |
| celf25 | 1.652145 | 0.40455482 | 0.538 | 0.254 | 4.1972755 | 14 | celf2 |
| itm2ba3 | 1.699269 | 0.410570952 | 0.583 | 0.293 | 4.3169939 | 14 | itm2ba |
| dclk1a2 | 1.728045 | 0.255674129 | 0.144 | 0.034 | 4.3900994 | 14 | dclk1a |
| rbp52 | 2.201936 | 0.279252037 | 0.307 | 0.108 | 5.5940203 | 14 | rbp5 |
| jund2 | 2.613520 | 0.301972863 | 0.261 | 0.088 | 6.6396496 | 14 | jund |
| ptmab6 | 3.442512 | 0.413394556 | 0.814 | 0.567 | 8.7457031 | 14 | ptmab |
| rnf213a3 | 4.988019 | 0.319279439 | 0.269 | 0.094 | 1.2672062 | 14 | rnf213a |
| itpkcb1 | 5.884429 | 0.270523748 | 0.231 | 0.073 | 1.4949394 | 14 | itpkcb |
| si:ch73-335l21.42 | 7.041534 | 0.471917965 | 0.345 | 0.139 | 1.7889019 | 14 | si:ch73-335l21.4 |
| hmgb1b3 | 1.309650 | 0.306437583 | 0.379 | 0.152 | 3.3271675 | 14 | hmgb1b |
| zfp36l1a4 | 1.653117 | 0.404612138 | 0.527 | 0.251 | 4.1997459 | 14 | zfp36l1a |
| anxa5b2 | 3.830348 | 0.3033615 | 0.284 | 0.103 | 9.7310012 | 14 | anxa5b |

| | | | | | | | |
|---|---|---|---|---|---|---|---|
| si:dkeyp-44a8.2 | 5.450057 | 0.251645154 | 0.197 | 0.059 | 1.3845871 | 14 | si:dkeyp-44a8.2 |
| rhoaa2 | 5.458418 | 0.335888532 | 0.402 | 0.174 | 1.3867111 | 14 | rhoaa |
| FO082781.13 | 6.047224 | 0.256945114 | 0.186 | 0.054 | 1.5362974 | 14 | FO082781.1 |
| crebzf1 | 1.028316 | 0.313127236 | 0.288 | 0.107 | 2.6124368 | 14 | crebzf |
| lrrc172 | 1.058088 | 0.254327711 | 0.193 | 0.057 | 2.6880729 | 14 | lrrc17 |
| fam133b | 1.242893 | 0.274931896 | 0.379 | 0.153 | 3.1575701 | 14 | fam133b |
| mt-co16 | 1.306449 | 0.388560891 | 0.883 | 0.637 | 3.3190340 | 14 | mt-co1 |
| mmp16a2 | 1.317359 | 0.327611773 | 0.231 | 0.076 | 3.3467525 | 14 | mmp16a |
| snx171 | 3.342366 | 0.27342016 | 0.205 | 0.064 | 8.4912833 | 14 | snx17 |
| cflara1 | 3.583381 | 0.250193308 | 0.201 | 0.062 | 9.1035797 | 14 | cflara |
| grinab3 | 4.030327 | 0.292493863 | 0.311 | 0.121 | 1.0239047 | 14 | grinab |
| rgs5b1 | 5.382830 | 0.377661923 | 0.216 | 0.071 | 1.3675079 | 14 | rgs5b |
| chd4a | 5.841734 | 0.275020352 | 0.284 | 0.105 | 1.4840925 | 14 | chd4a |
| arhgef1b | 6.517403 | 0.265189929 | 0.231 | 0.078 | 1.6557464 | 14 | arhgef1b |
| mt-co26 | 1.161154 | 0.36924755 | 0.818 | 0.67 | 2.9499131 | 14 | mt-co2 |
| u2surp | 1.178659 | 0.299084955 | 0.273 | 0.102 | 2.9943849 | 14 | u2surp |
| srrm1 | 1.209317 | 0.299503114 | 0.307 | 0.119 | 3.0722710 | 14 | srrm1 |
| srsf5a2 | 1.636687 | 0.365091094 | 0.583 | 0.3 | 4.1580055 | 14 | srsf5a |
| cebpa4 | 2.140542 | 0.265766477 | 0.223 | 0.074 | 5.4380478 | 14 | cebpa |
| CABZ01068358.12 | 3.336961 | 0.259840739 | 0.205 | 0.066 | 8.4775505 | 14 | CABZ01068358.1 |
| fosl22 | 4.582169 | 0.315344441 | 0.288 | 0.112 | 1.1641001 | 14 | fosl2 |
| rsrp1 | 5.275786 | 0.290330596 | 0.307 | 0.12 | 1.3403136 | 14 | rsrp1 |
| rps107 | 6.620775 | 0.319317304 | 0.936 | 0.724 | 1.6820080 | 14 | rps10 |
| bhlhe403 | 1.039750 | 0.332603063 | 0.379 | 0.165 | 2.6414856 | 14 | bhlhe40 |
| serbp1a5 | 1.659714 | 0.400035859 | 0.818 | 0.502 | 4.2165058 | 14 | serbp1a |
| adam10a1 | 2.839942 | 0.265282244 | 0.273 | 0.104 | 7.2148740 | 14 | adam10a |
| rps27l3 | 4.114171 | 0.293546997 | 0.458 | 0.215 | 1.0452052 | 14 | rps27l |
| kctd12.23 | 5.079569 | 0.283767837 | 0.261 | 0.099 | 1.2904645 | 14 | kctd12.2 |
| ndufa4l4 | 6.068500 | 0.372769013 | 0.731 | 0.418 | 1.5417025 | 14 | ndufa4l |
| rbp42 | 6.184567 | 0.350524807 | 0.337 | 0.139 | 1.5711893 | 14 | rbp4 |
| zgc:1119862 | 1.065369 | 0.297684108 | 0.402 | 0.183 | 2.7065715 | 14 | zgc:111986 |
| pi15a1 | 1.176192 | 0.270216672 | 0.174 | 0.053 | 2.9881159 | 14 | pi15a |
| rnf10 | 1.344765 | 0.276959245 | 0.303 | 0.126 | 3.4163775 | 14 | rnf10 |
| notch22 | 1.512066 | 0.274165445 | 0.295 | 0.121 | 3.8414048 | 14 | notch2 |
| LOC103910438 | 2.192077 | 0.320945832 | 0.364 | 0.164 | 5.5689727 | 14 | LOC103910438 |

| | | | | | | | |
|---|---|---|---|---|---|---|---|
| hp1bp32 | 2.330706 | 0.260491512 | 0.292 | 0.119 | 5.9211586 | 14 | hp1bp3 |
| spry42 | 2.366004 | 0.266090985 | 0.159 | 0.048 | 6.0108339 | 14 | spry4 |
| fxyd6l1 | 3.557058 | 0.281429457 | 0.288 | 0.116 | 9.0367072 | 14 | fxyd6l |
| pfn2l2 | 6.228864 | 0.281270872 | 0.458 | 0.212 | 1.5824430 | 14 | pfn2l |
| rpl147 | 8.724257 | 0.313737233 | 0.886 | 0.667 | 2.2163977 | 14 | rpl14 |
| pabpn1 | 1.000572 | 0.316429428 | 0.534 | 0.272 | 2.5419535 | 14 | pabpn1 |
| calm1a | 1.901066 | 0.257801845 | 0.383 | 0.172 | 4.8296602 | 14 | calm1a |
| si:dkey-11f4.142 | 9.387085 | 0.282642862 | 0.284 | 0.121 | 2.3847890 | 14 | si:dkey-11f4.14 |
| nop561 | 1.114626 | 0.261109961 | 0.394 | 0.184 | 2.8317076 | 14 | nop56 |
| nucks1a | 1.309803 | 0.278528664 | 0.394 | 0.186 | 3.3275569 | 14 | nucks1a |
| ywhaqa | 1.439934 | 0.265898217 | 0.432 | 0.207 | 3.6581539 | 14 | ywhaqa |
| zgc:1141887 | 2.278405 | 0.33384045 | 0.909 | 0.665 | 5.7882896 | 14 | zgc:114188 |
| rab102 | 7.775696 | 0.264646787 | 0.428 | 0.212 | 1.9754156 | 14 | rab10 |
| ddit33 | 9.098397 | 0.329916629 | 0.36 | 0.177 | 2.3114479 | 14 | ddit3 |
| tegt2 | 1.154765 | 0.279089873 | 0.458 | 0.235 | 2.9336827 | 14 | tegt |
| mt-nd35 | 1.158411 | 0.351962398 | 0.595 | 0.369 | 2.9429439 | 14 | mt-nd3 |
| ubb | 1.173621 | 0.308997605 | 0.583 | 0.322 | 2.9815842 | 14 | ubb |
| tmsb4x9 | 1.237379 | 0.383190293 | 0.985 | 0.715 | 3.1435636 | 14 | tmsb4x |
| arrdc3a2 | 1.835039 | 0.252453672 | 0.193 | 0.072 | 4.6619175 | 14 | arrdc3a |
| ccl25b2 | 2.086398 | 0.309283964 | 0.231 | 0.092 | 5.3004946 | 14 | ccl25b |
| ddx611 | 3.626861 | 0.272578432 | 0.451 | 0.233 | 9.2140423 | 14 | ddx61 |
| rpl97 | 7.545971 | 0.31986971 | 0.905 | 0.665 | 1.9170540 | 14 | rpl9 |
| tuba8l | 9.073375 | 0.256637023 | 0.409 | 0.207 | 2.3050909 | 14 | tuba8l |
| rpl66 | 1.011462 | 0.295955569 | 0.928 | 0.699 | 2.5696199 | 14 | rpl6 |
| rps27.26 | 1.786179 | 0.339560355 | 0.811 | 0.55 | 4.5377891 | 14 | rps27.2 |
| cnbpb2 | 2.377607 | 0.263498827 | 0.402 | 0.208 | 6.0403106 | 14 | cnbpb |
| msna5 | 2.950665 | 0.289095388 | 0.485 | 0.263 | 7.4961661 | 14 | msna |
| rpl5a7 | 3.015165 | 0.342848513 | 0.822 | 0.533 | 7.6600290 | 14 | rpl5a |
| eif1b2 | 3.033068 | 0.310714113 | 0.689 | 0.403 | 7.7055112 | 14 | eif1b |
| uba525 | 3.721703 | 0.298227608 | 0.883 | 0.658 | 9.4549882 | 14 | uba52 |
| pdia35 | 4.096623 | 0.294499429 | 0.553 | 0.311 | 1.0407473 | 14 | pdia3 |
| syncrip1 | 7.743015 | 0.269227147 | 0.561 | 0.31 | 1.9671131 | 14 | syncrip |
| jpt1b3 | 1.219377 | 0.257286811 | 0.519 | 0.274 | 3.0978288 | 14 | jpt1b |
| si:ch211-288g17.3 | 3.307386 | 0.264795234 | 0.42 | 0.224 | 8.4024155 | 14 | si:ch211-288g17.3 |
| ywhaz2 | 3.868584 | 0.26445004 | 0.542 | 0.302 | 9.8281391 | 14 | ywhaz |

| | | | | | | | |
|---|---|---|---|---|---|---|---|
| atp1b1a2 | 9.244597 | 0.281692086 | 0.405 | 0.229 | 2.3485898 | 14 | atp1b1a |
| atf33 | 1.965573 | 0.271045064 | 0.189 | 0.081 | 4.9935401 | 14 | atf3 |
| fkbp1aa3 | 3.995723 | 0.282978106 | 0.655 | 0.412 | 1.0151135 | 14 | fkbp1aa |
| hmgb2b2 | 2.425259 | 0.253028308 | 0.553 | 0.334 | 6.1613723 | 14 | hmgb2b |
| hnrnpa0b3 | 3.083716 | 0.254036566 | 0.561 | 0.339 | 7.8341808 | 14 | hnrnpa0b |
| rpl22l15 | 1.550580 | 0.267274111 | 0.67 | 0.422 | 0.000394 | 14 | rpl22l1 |
| rpl45 | 3.084937 | 0.253121659 | 0.852 | 0.655 | 0.000784 | 14 | rpl4 |
| CR318588.17 | 1.598030 | 0.329736939 | 0.333 | 0.215 | 0.040598 | 14 | CR318588.1 |
| ctrb1 | 2.991522 | 0.469812537 | 0.189 | 0.017 | 7.5999634 | 15 | ctrb1 |
| cldnd4 | 2.565118 | 0.897323037 | 0.992 | 0.491 | 6.5166839 | 15 | cldnd |
| h2af1al4 | 2.756437 | 0.881506196 | 0.98 | 0.483 | 7.0027294 | 15 | h2af1al |
| mt24 | 9.410587 | 0.892811707 | 0.953 | 0.433 | 2.3907598 | 15 | mt2 |
| hbba14 | 8.452153 | 0.935522401 | 0.799 | 0.287 | 2.1472696 | 15 | hbba1 |
| zgc:1751354 | 1.549549 | 0.889755269 | 0.894 | 0.387 | 3.9366295 | 15 | zgc:175135 |
| h1m4 | 1.587021 | 0.848653284 | 0.976 | 0.49 | 4.0318269 | 15 | h1m |
| si:dkey-240e12.64 | 1.311988 | 0.886431129 | 0.976 | 0.46 | 3.3331075 | 15 | si:dkey-240e12.6 |
| zp2.34 | 2.327538 | 0.87971894 | 0.937 | 0.419 | 5.9131124 | 15 | zp2.3 |
| zp34 | 2.715324 | 0.861207578 | 0.98 | 0.473 | 6.8982813 | 15 | zp3 |
| si:dkeyp-98a7.94 | 3.100373 | 0.875542665 | 0.933 | 0.414 | 7.8764987 | 15 | si:dkeyp-98a7.9 |
| CR790368.14 | 6.707415 | 0.854829378 | 0.902 | 0.422 | 1.7040188 | 15 | CR790368.1 |
| rrm24 | 3.538763 | 0.837029497 | 0.894 | 0.416 | 8.9902286 | 15 | rrm2 |
| zp2.54 | 6.489316 | 0.856913949 | 0.949 | 0.457 | 1.6486109 | 15 | zp2.5 |
| si:dkey-241l7.44 | 1.329286 | 0.815862393 | 0.957 | 0.495 | 3.3770521 | 15 | si:dkey-241l7.4 |
| zp2.14 | 1.775251 | 0.845740378 | 0.917 | 0.433 | 4.5100267 | 15 | zp2.1 |
| tmsb54 | 3.024710 | 0.862006856 | 0.917 | 0.395 | 7.6842773 | 15 | tmsb5 |
| zgc:1534994 | 9.034440 | 0.870105367 | 0.937 | 0.414 | 2.2951996 | 15 | zgc:153499 |
| si:dkey-241l7.24 | 1.388780 | 0.831810965 | 0.992 | 0.522 | 3.5281967 | 15 | si:dkey-241l7.2 |
| si:ch211-226h8.144 | 1.833151 | 0.833948288 | 0.949 | 0.46 | 4.6571210 | 15 | si:ch211-226h8.14 |
| si:zfos-1505d6.34 | 1.481325 | 0.843909094 | 0.909 | 0.415 | 3.7633086 | 15 | si:zfos-1505d6.3 |
| si:dkey-241l7.54 | 2.446471 | 0.812531706 | 0.937 | 0.475 | 6.2152604 | 15 | si:dkey-241l7.5 |
| ccna14 | 6.436047 | 0.834407489 | 0.862 | 0.385 | 1.6350778 | 15 | ccna1 |
| zgc:1717764 | 6.925278 | 0.834630727 | 0.866 | 0.401 | 1.7593670 | 15 | zgc:171776 |
| si:ch73-189n23.14 | 8.801218 | 0.841690397 | 0.913 | 0.425 | 2.2359494 | 15 | si:ch73-189n23.1 |
| zp2.64 | 9.790716 | 0.847436685 | 0.902 | 0.405 | 2.4873315 | 15 | zp2.6 |
| prss1 | 2.965525 | 0.486125388 | 0.181 | 0.021 | 7.5339187 | 15 | prss1 |

| | | | | | | | |
|---|---|---|---|---|---|---|---|
| zp2.24 | 5.842241 | 0.825855784 | 0.906 | 0.429 | 1.4842214 | 15 | zp2.2 |
| acp5a4 | 6.786411 | 0.815727159 | 0.906 | 0.44 | 1.7240878 | 15 | acp5a |
| si:dkey-229d11.34 | 1.723365 | 0.853724263 | 0.846 | 0.354 | 4.3782109 | 15 | si:dkey-229d11.3 |
| hbaa14 | 9.310899 | 0.849282174 | 0.65 | 0.228 | 2.3654340 | 15 | hbaa1 |
| si:ch211-125e6.124 | 1.145416 | 0.823976172 | 0.874 | 0.406 | 2.9099302 | 15 | si:ch211-125e6.12 |
| zgc:771184 | 1.297523 | 0.822473409 | 0.827 | 0.371 | 3.2963576 | 15 | zgc:77118 |
| tuba4l4 | 7.120681 | 0.821890069 | 0.752 | 0.328 | 1.8090091 | 15 | tuba4l |
| si:ch211-5k11.84 | 7.730265 | 0.812339179 | 0.736 | 0.291 | 1.9638740 | 15 | si:ch211-5k11.8 |
| crp24 | 1.206106 | 0.823367433 | 0.878 | 0.396 | 3.0641124 | 15 | crp2 |
| si:ch211-12h2.84 | 3.278720 | 0.788496431 | 0.882 | 0.445 | 8.3295886 | 15 | si:ch211-12h2.8 |
| zgc:1717794 | 5.467491 | 0.8093594 | 0.894 | 0.415 | 1.3890161 | 15 | zgc:171779 |
| cldng4 | 6.066849 | 0.811032621 | 0.894 | 0.418 | 1.5412832 | 15 | cldng |
| zar14 | 9.485780 | 0.813517875 | 0.799 | 0.357 | 2.4098624 | 15 | zar1 |
| siva14 | 5.285066 | 0.799910483 | 0.776 | 0.356 | 1.3426712 | 15 | siva1 |
| si:ch211-226h8.44 | 8.008061 | 0.802924068 | 0.803 | 0.357 | 2.0344480 | 15 | si:ch211-226h8.4 |
| si:dkeyp-98a7.84 | 2.826660 | 0.793368162 | 0.811 | 0.371 | 7.1811320 | 15 | si:dkeyp-98a7.8 |
| si:ch211-12h2.64 | 3.184809 | 0.78232368 | 0.815 | 0.384 | 8.0910081 | 15 | si:ch211-12h2.6 |
| BX649337.14 | 4.618518 | 0.797120035 | 0.807 | 0.354 | 1.1733346 | 15 | BX649337.1 |
| epcam4 | 1.251458 | 0.767797152 | 0.846 | 0.415 | 3.1793295 | 15 | epcam |
| btg44 | 1.987076 | 0.783745686 | 0.697 | 0.299 | 5.0481681 | 15 | btg4 |
| prss59.2 | 1.270812 | 0.369798131 | 0.138 | 0.015 | 3.2284996 | 15 | prss59.2 |
| zp3b4 | 5.153598 | 0.776087877 | 0.697 | 0.315 | 1.3092715 | 15 | zp3b |
| zp3.24 | 9.690438 | 0.74742951 | 0.882 | 0.454 | 2.4618558 | 15 | zp3.2 |
| CU571163.14 | 5.956012 | 0.77063662 | 0.736 | 0.325 | 1.5131248 | 15 | CU571163.1 |
| zp3a.24 | 3.143162 | 0.73267917 | 0.791 | 0.4 | 7.9852031 | 15 | zp3a.2 |
| lamp24 | 3.744478 | 0.696090972 | 0.823 | 0.48 | 9.5128478 | 15 | lamp2 |
| mid1ip1l4 | 1.255372 | 0.696602452 | 0.811 | 0.456 | 3.1892728 | 15 | mid1ip1l |
| si:dkeyp-50f7.24 | 2.118414 | 0.729809794 | 0.705 | 0.327 | 5.3818312 | 15 | si:dkeyp-50f7.2 |
| si:dkeyp-98a7.44 | 2.221850 | 0.730213979 | 0.744 | 0.344 | 5.6446100 | 15 | si:dkeyp-98a7.4 |
| cth14 | 3.881679 | 0.728479233 | 0.681 | 0.308 | 9.8614063 | 15 | cth1 |
| si:dkey-241l7.64 | 1.302712 | 0.705029989 | 0.803 | 0.429 | 3.3095403 | 15 | si:dkey-241l7.6 |
| si:ch211-14a17.74 | 5.971564 | 0.7249801 | 0.713 | 0.332 | 1.5170759 | 15 | si:ch211-14a17.7 |
| gapdh4 | 1.546417 | 0.715352439 | 0.713 | 0.342 | 3.9286730 | 15 | gapdh |
| si:dkey-90l23.14 | 2.055795 | 0.712091028 | 0.752 | 0.364 | 5.2227489 | 15 | si:dkey-90l23.1 |
| ccnb14 | 7.165712 | 0.712525483 | 0.669 | 0.322 | 1.8204493 | 15 | ccnb1 |

| | | | | | | | |
|---|---|---|---|---|---|---|---|
| si:dkeyp-46h3.84 | 9.132342 | 0.719128858 | 0.638 | 0.285 | 2.3200716 | 15 | si:dkeyp-46h3.8 |
| zgc:1952454 | 7.945450 | 0.699948491 | 0.677 | 0.315 | 2.0185415 | 15 | zgc:195245 |
| qdprb24 | 1.789315 | 0.700717624 | 0.724 | 0.352 | 4.5457556 | 15 | qdprb2 |
| s100a14 | 1.598672 | 0.696321008 | 0.638 | 0.296 | 4.0614264 | 15 | s100a1 |
| npm2b4 | 1.873133 | 0.692932989 | 0.685 | 0.323 | 4.7586964 | 15 | npm2b |
| slbp24 | 6.368200 | 0.688688666 | 0.669 | 0.323 | 1.6178412 | 15 | slbp2 |
| zar1l4 | 7.583193 | 0.687478616 | 0.681 | 0.324 | 1.9265103 | 15 | zar1l |
| si:dkeyp-46h3.64 | 9.321341 | 0.691159475 | 0.681 | 0.333 | 2.3680868 | 15 | si:dkeyp-46h3.6 |
| si:ch211-132b12.84 | 4.967847 | 0.676938119 | 0.594 | 0.279 | 1.2620816 | 15 | si:ch211-132b12.8 |
| si:dkey-241l7.34 | 7.466481 | 0.686042889 | 0.594 | 0.297 | 1.8968595 | 15 | si:dkey-241l7.3 |
| LOC1104378214 | 2.423731 | 0.67200701 | 0.673 | 0.338 | 6.1574892 | 15 | LOC110437821 |
| nanog4 | 3.222486 | 0.658171766 | 0.626 | 0.293 | 8.1867274 | 15 | nanog |
| zgc:1734434 | 5.746684 | 0.661271685 | 0.591 | 0.268 | 1.4599451 | 15 | zgc:173443 |
| si:dkey-152b24.64 | 6.006845 | 0.666484345 | 0.543 | 0.241 | 1.5260391 | 15 | si:dkey-152b24.6 |
| zgc:1362544 | 1.269259 | 0.653043037 | 0.591 | 0.269 | 3.2245525 | 15 | zgc:136254 |
| hnrnpabb4 | 1.657432 | 0.536951594 | 0.78 | 0.557 | 4.2107077 | 15 | hnrnpabb |
| ca15b4 | 8.083865 | 0.656434249 | 0.48 | 0.202 | 2.0537061 | 15 | ca15b |
| zp3a.14 | 1.167770 | 0.65043991 | 0.606 | 0.311 | 2.9667197 | 15 | zp3a.1 |
| ccnb24 | 1.391448 | 0.651220297 | 0.634 | 0.313 | 3.5349753 | 15 | ccnb2 |
| kpna74 | 8.098361 | 0.647085141 | 0.571 | 0.268 | 2.0573886 | 15 | kpna7 |
| gtf3ab4 | 1.983222 | 0.638564527 | 0.555 | 0.261 | 5.0383763 | 15 | gtf3ab |
| zpcx4 | 1.031339 | 0.633081813 | 0.571 | 0.276 | 2.6201169 | 15 | zpcx |
| wu:fi42e034 | 1.152380 | 0.639539899 | 0.575 | 0.285 | 2.9276216 | 15 | wu:fi42e03 |
| lsm14b4 | 2.677918 | 0.6334575 | 0.524 | 0.244 | 6.8032514 | 15 | lsm14b |
| zgc:554614 | 4.358306 | 0.62573692 | 0.524 | 0.238 | 1.1072278 | 15 | zgc:55461 |
| si:dkeyp-20e4.84 | 6.100484 | 0.635322271 | 0.646 | 0.332 | 1.5498281 | 15 | si:dkeyp-20e4.8 |
| thy14 | 1.656106 | 0.605578503 | 0.665 | 0.375 | 4.2073386 | 15 | thy1 |
| zgc:1715174 | 1.743046 | 0.621771444 | 0.626 | 0.316 | 4.4282099 | 15 | zgc:171517 |
| pabpn1l4 | 1.884178 | 0.619390168 | 0.512 | 0.232 | 4.7867547 | 15 | pabpn1l |
| si:dkey-19b23.114 | 6.013581 | 0.629878946 | 0.496 | 0.227 | 1.5277502 | 15 | si:dkey-19b23.11 |
| avd4 | 8.067075 | 0.610385024 | 0.488 | 0.215 | 2.0494405 | 15 | avd |
| si:dkey-90l23.24 | 9.171121 | 0.616800993 | 0.567 | 0.281 | 2.3299235 | 15 | si:dkey-90l23.2 |
| si:dkey-229d11.54 | 9.213310 | 0.612911097 | 0.618 | 0.317 | 2.3406416 | 15 | si:dkey-229d11.5 |
| gmnn4 | 4.440296 | 0.605674484 | 0.524 | 0.252 | 1.1280573 | 15 | gmnn |
| si:ch211-145c1.14 | 7.030702 | 0.608483099 | 0.441 | 0.19 | 1.7861498 | 15 | si:ch211-145c1.1 |

| | | | | | | | |
|---|---|---|---|---|---|---|---|
| birc5b4 | 1.008103 | 0.61063377 | 0.465 | 0.211 | 2.5610862 | 15 | birc5b |
| si:dkeyp-75b4.105 | 3.357369 | 0.548937377 | 0.646 | 0.363 | 8.5293972 | 15 | si:dkeyp-75b4.10 |
| si:dkey-208k4.24 | 5.375190 | 0.599486474 | 0.5 | 0.241 | 1.3655672 | 15 | si:dkey-208k4.2 |
| si:ch211-251f6.64 | 5.642430 | 0.602855068 | 0.528 | 0.27 | 1.4334595 | 15 | si:ch211-251f6.6 |
| gdf34 | 6.059719 | 0.599160005 | 0.425 | 0.182 | 1.5394717 | 15 | gdf3 |
| spink2.15 | 1.641129 | 0.567131555 | 0.646 | 0.371 | 4.1692890 | 15 | spink2.1 |
| elovl7b4 | 1.742413 | 0.584690189 | 0.457 | 0.202 | 4.4266020 | 15 | elovl7b |
| gbgt1l44 | 3.036850 | 0.542008757 | 0.335 | 0.122 | 7.7151193 | 15 | gbgt1l4 |
| si:dkeyp-46h3.24 | 6.062747 | 0.584820342 | 0.602 | 0.33 | 1.5402409 | 15 | si:dkeyp-46h3.2 |
| si:ch73-14h10.24 | 8.171403 | 0.595367999 | 0.469 | 0.222 | 2.0759450 | 15 | si:ch73-14h10.2 |
| si:dkey-37o8.14 | 2.703841 | 0.591116439 | 0.5 | 0.251 | 6.8691104 | 15 | si:dkey-37o8.1 |
| LOC1005351624 | 6.415035 | 0.574879315 | 0.528 | 0.251 | 1.6297396 | 15 | LOC100535162 |
| hnrnpa0l4 | 2.031934 | 0.314265706 | 0.85 | 0.74 | 5.1621307 | 15 | hnrnpa0l |
| AL929032.14 | 3.546795 | 0.567831187 | 0.528 | 0.258 | 9.0106338 | 15 | AL929032.1 |
| zgc:1134244 | 5.471271 | 0.5466367 | 0.331 | 0.131 | 1.3899765 | 15 | zgc:113424 |
| ctss2.14 | 1.087618 | 0.51838019 | 0.701 | 0.449 | 2.7630949 | 15 | ctss2.1 |
| LOC1005374994 | 1.162837 | 0.56311614 | 0.528 | 0.263 | 2.9541898 | 15 | LOC100537499 |
| si:ch73-90k17.14 | 1.306034 | 0.560760426 | 0.437 | 0.203 | 3.3179806 | 15 | si:ch73-90k17.1 |
| si:ch73-160p18.34 | 1.461991 | 0.563752301 | 0.421 | 0.196 | 3.7141896 | 15 | si:ch73-160p18.3 |
| org4 | 2.461758 | 0.566576525 | 0.421 | 0.198 | 6.2540964 | 15 | org |
| uchl14 | 2.724089 | 0.556133666 | 0.539 | 0.288 | 6.9205498 | 15 | uchl1 |
| cpeb1b4 | 2.939747 | 0.547203363 | 0.469 | 0.221 | 7.4684284 | 15 | cpeb1b |
| si:ch211-146l10.74 | 5.270003 | 0.537034906 | 0.362 | 0.153 | 1.3388444 | 15 | si:ch211-146l10.7 |
| si:ch211-250e5.164 | 1.067767 | 0.557563178 | 0.524 | 0.283 | 2.7126637 | 15 | si:ch211-250e5.16 |
| LOC1018854774 | 2.813368 | 0.552802609 | 0.465 | 0.228 | 7.1473621 | 15 | LOC101885477 |
| AL929192.14 | 7.100083 | 0.498501255 | 0.335 | 0.133 | 1.8037761 | 15 | AL929192.1 |
| zgc:925914 | 2.338278 | 0.521143169 | 0.378 | 0.168 | 5.9403953 | 15 | zgc:92591 |
| si:dkeyp-46h3.54 | 3.130041 | 0.535716939 | 0.465 | 0.236 | 7.9518695 | 15 | si:dkeyp-46h3.5 |
| ccna24 | 3.481470 | 0.539102494 | 0.461 | 0.237 | 8.8446751 | 15 | ccna2 |
| zgc:566994 | 9.951922 | 0.533586937 | 0.421 | 0.208 | 2.5282859 | 15 | zgc:56699 |
| si:ch211-146l10.84 | 3.319369 | 0.527174665 | 0.476 | 0.25 | 8.4328588 | 15 | si:ch211-146l10.8 |
| h3f3a4 | 4.397485 | 0.463678721 | 0.65 | 0.482 | 1.1171811 | 15 | h3f3a |
| larp6b4 | 5.358889 | 0.507214737 | 0.358 | 0.16 | 1.3614257 | 15 | larp6b |
| zgc:1714744 | 5.674224 | 0.520405013 | 0.48 | 0.248 | 1.4415366 | 15 | zgc:171474 |
| slc16a34 | 9.147429 | 0.522167222 | 0.413 | 0.204 | 2.3239044 | 15 | slc16a3 |

| | | | | | | | |
|---|---|---|---|---|---|---|---|
| si:dkeyp-98a7.74 | 1.048146 | 0.498331718 | 0.358 | 0.159 | 2.6628154 | 15 | si:dkeyp-98a7.7 |
| arhgef10la4 | 1.231253 | 0.521914003 | 0.433 | 0.225 | 3.1279992 | 15 | arhgef10la |
| inka1b4 | 1.472199 | 0.501594898 | 0.343 | 0.155 | 3.7401239 | 15 | inka1b |
| crp34 | 1.809622 | 0.512308996 | 0.374 | 0.178 | 4.5973448 | 15 | crp3 |
| LOC1005368604 | 2.093792 | 0.511947019 | 0.492 | 0.287 | 5.3192797 | 15 | LOC100536860 |
| nasp4 | 4.858789 | 0.517361446 | 0.5 | 0.293 | 1.2343753 | 15 | nasp |
| haus64 | 1.214094 | 0.492882565 | 0.366 | 0.172 | 3.0844059 | 15 | haus6 |
| si:ch211-226l4.64 | 4.276998 | 0.497828637 | 0.445 | 0.242 | 1.0865714 | 15 | si:ch211-226l4.6 |
| snu13b4 | 5.659851 | 0.48842826 | 0.539 | 0.37 | 1.4378853 | 15 | snu13b |
| slbp5 | 1.009719 | 0.498885367 | 0.39 | 0.198 | 2.5651923 | 15 | slbp |
| zgc:562314 | 1.259363 | 0.487618794 | 0.378 | 0.186 | 3.1994138 | 15 | zgc:56231 |
| LOC1018838974 | 2.432323 | 0.467097357 | 0.303 | 0.134 | 6.1793179 | 15 | LOC101883897 |
| zgc:1738564 | 3.189104 | 0.47139567 | 0.37 | 0.18 | 8.1019191 | 15 | zgc:173856 |
| cx44.24 | 1.143112 | 0.426194921 | 0.24 | 0.095 | 2.9040773 | 15 | cx44.2 |
| zgc:1629454 | 1.956463 | 0.452444857 | 0.299 | 0.135 | 4.9703948 | 15 | zgc:162945 |
| ddx39ab4 | 2.085870 | 0.492357439 | 0.441 | 0.269 | 5.2991550 | 15 | ddx39ab |
| larp6a4 | 4.839695 | 0.486028094 | 0.39 | 0.213 | 1.2295245 | 15 | larp6a |
| cdc204 | 4.921958 | 0.450324391 | 0.283 | 0.129 | 1.2504234 | 15 | cdc20 |
| rcc14 | 5.485026 | 0.485598031 | 0.339 | 0.172 | 1.3934709 | 15 | rcc1 |
| patl24 | 6.209226 | 0.454345314 | 0.303 | 0.142 | 1.5774540 | 15 | patl2 |
| si:ch73-78o10.14 | 1.261484 | 0.424775243 | 0.256 | 0.109 | 3.2048015 | 15 | si:ch73-78o10.1 |
| zp2l14 | 2.583768 | 0.438663015 | 0.287 | 0.134 | 6.5640643 | 15 | zp2l1 |
| hsd17b12a4 | 5.840848 | 0.426783412 | 0.291 | 0.136 | 1.4838675 | 15 | hsd17b12a |
| rbpms2b4 | 9.789474 | 0.445144269 | 0.362 | 0.19 | 2.4870159 | 15 | rbpms2b |
| map1lc3c4 | 1.250517 | 0.442807616 | 0.327 | 0.165 | 3.1769388 | 15 | map1lc3c |
| cdca84 | 1.815402 | 0.428213021 | 0.291 | 0.139 | 4.6120298 | 15 | cdca8 |
| si:ch1073-263o8.24 | 1.827484 | 0.452828061 | 0.394 | 0.216 | 4.6427239 | 15 | si:ch1073-263o8.2 |
| si:ch211-226h8.82 | 2.184945 | 0.405341965 | 0.201 | 0.079 | 5.5508552 | 15 | si:ch211-226h8.8 |
| retsatl4 | 2.566077 | 0.441950225 | 0.327 | 0.167 | 6.5191207 | 15 | retsatl |
| elavl24 | 2.814271 | 0.442099103 | 0.354 | 0.185 | 7.1496571 | 15 | elavl2 |
| plk14 | 4.960153 | 0.424832714 | 0.299 | 0.148 | 1.2601269 | 15 | plk1 |
| eif4a1a4 | 5.203836 | 0.412166387 | 0.531 | 0.405 | 1.3220347 | 15 | eif4a1a |
| mos4 | 7.751895 | 0.40785894 | 0.252 | 0.113 | 1.9693689 | 15 | mos |
| si:dkey-216e24.34 | 9.050981 | 0.38934109 | 0.232 | 0.101 | 2.2994018 | 15 | si:dkey-216e24.3 |
| si:dkey-46g23.24 | 1.151139 | 0.394887923 | 0.26 | 0.118 | 2.9244705 | 15 | si:dkey-46g23.2 |

| | | | | | | | |
|---|---|---|---|---|---|---|---|
| zgc:1628794 | 1.351175 | 0.414769937 | 0.299 | 0.149 | 3.4326611 | 15 | zgc:162879 |
| zgc:660244 | 2.217270 | 0.394460244 | 0.252 | 0.116 | 5.6329755 | 15 | zgc:66024 |
| orc63 | 2.735982 | 0.378832521 | 0.217 | 0.092 | 6.9507647 | 15 | orc6 |
| rdh10b4 | 4.167513 | 0.434322478 | 0.291 | 0.15 | 1.0587567 | 15 | rdh10b |
| pou5f34 | 5.066822 | 0.397535428 | 0.256 | 0.12 | 1.2872263 | 15 | pou5f3 |
| buc2l3 | 5.335391 | 0.325173456 | 0.173 | 0.065 | 1.3554561 | 15 | buc2l |
| LOC1104377793 | 6.657172 | 0.33406732 | 0.177 | 0.068 | 1.6912546 | 15 | LOC110437779 |
| nanos34 | 6.721487 | 0.371843138 | 0.201 | 0.084 | 1.7075938 | 15 | nanos3 |
| BX537276.14 | 9.279029 | 0.373382691 | 0.224 | 0.099 | 2.3573375 | 15 | BX537276.1 |
| ankrd224 | 1.325873 | 0.402588704 | 0.252 | 0.121 | 3.3683817 | 15 | ankrd22 |
| chia.44 | 1.417310 | 0.390603912 | 0.252 | 0.12 | 3.6006763 | 15 | chia.4 |
| dynll14 | 1.851018 | 0.378413552 | 0.547 | 0.441 | 4.7025130 | 15 | dynll1 |
| zglp14 | 2.104804 | 0.404162689 | 0.311 | 0.163 | 5.3472559 | 15 | zglp1 |
| si:dkey-42i9.84 | 2.765458 | 0.3795056 | 0.228 | 0.105 | 7.0256471 | 15 | si:dkey-42i9.8 |
| pigrl2.34 | 3.403323 | 0.404560807 | 0.315 | 0.168 | 8.6461421 | 15 | pigrl2.3 |
| st6galnac1.2 | 3.532700 | 0.287902051 | 0.13 | 0.044 | 8.9748267 | 15 | st6galnac1.2 |
| buc4 | 3.759679 | 0.40504484 | 0.283 | 0.147 | 9.5514665 | 15 | buc |
| zgc:1034824 | 4.459939 | 0.397168911 | 0.295 | 0.157 | 1.1330475 | 15 | zgc:103482 |
| sinup4 | 4.699695 | 0.403771354 | 0.311 | 0.167 | 1.1939577 | 15 | sinup |
| rbm4.34 | 8.446130 | 0.371365227 | 0.524 | 0.421 | 2.1457394 | 15 | rbm4.3 |
| dctpp14 | 1.225500 | 0.397670902 | 0.28 | 0.149 | 3.1133836 | 15 | dctpp1 |
| CU655961.44 | 1.273973 | 0.322868435 | 0.177 | 0.072 | 3.2365300 | 15 | CU655961.4 |
| zgc:1630304 | 1.735681 | 0.365207159 | 0.224 | 0.106 | 4.4094995 | 15 | zgc:163030 |
| pttg14 | 1.825806 | 0.387555009 | 0.264 | 0.136 | 4.6384614 | 15 | pttg1 |
| sybu4 | 2.703366 | 0.37120432 | 0.268 | 0.136 | 6.8679023 | 15 | sybu |
| CDPF14 | 2.804908 | 0.407181263 | 0.378 | 0.234 | 7.1258710 | 15 | CDPF1 |
| lin28a2 | 3.264564 | 0.29696056 | 0.15 | 0.057 | 8.2936257 | 15 | lin28a |
| CABZ01059627.24 | 3.665518 | 0.35342072 | 0.228 | 0.109 | 9.3122489 | 15 | CABZ01059627.2 |
| mibp5 | 4.068367 | 0.397908246 | 0.339 | 0.199 | 1.0335686 | 15 | mibp |
| pabpc1l4 | 4.331755 | 0.365305571 | 0.236 | 0.117 | 1.1004824 | 15 | pabpc1l |
| si:dkey-203a12.22 | 4.420960 | 0.32138313 | 0.362 | 0.209 | 1.1231448 | 15 | si:dkey-203a12.2 |
| si:ch73-132k15.21 | 5.233156 | 0.306937471 | 0.138 | 0.052 | 1.3294833 | 15 | si:ch73-132k15.2 |
| elovl7a4 | 6.519446 | 0.368937986 | 0.287 | 0.152 | 1.6562655 | 15 | elovl7a |
| f2rl1.1 | 7.523586 | 0.259078102 | 0.11 | 0.037 | 1.9113672 | 15 | f2rl1.1 |
| ddx412 | 7.551751 | 0.283287131 | 0.146 | 0.056 | 1.9185223 | 15 | ddx41 |

| | | | | | | | |
|---|---|---|---|---|---|---|---|
| zgc:1138864 | 1.028801 | 0.349332249 | 0.236 | 0.116 | 2.6136710 | 15 | zgc:113886 |
| LOC110440049 | 1.182224 | 0.259847529 | 0.114 | 0.039 | 3.0034410 | 15 | LOC110440049 |
| si:dkey-68o6.54 | 1.209068 | 0.354664873 | 0.213 | 0.101 | 3.0716384 | 15 | si:dkey-68o6.5 |
| si:ch211-154a22.8 | 1.341726 | 0.257267756 | 0.122 | 0.043 | 3.4086558 | 15 | si:ch211-154a22.8 |
| CR847984.14 | 1.452940 | 0.356557104 | 0.264 | 0.138 | 3.6911951 | 15 | CR847984.1 |
| srsf2a3 | 1.486992 | 0.387570456 | 0.457 | 0.349 | 3.7777055 | 15 | srsf2a |
| snx10a4 | 1.547568 | 0.342279484 | 0.217 | 0.105 | 3.9315969 | 15 | snx10a |
| aurka4 | 1.611200 | 0.343122827 | 0.209 | 0.1 | 4.0932542 | 15 | aurka |
| rasl11b4 | 1.629915 | 0.336344571 | 0.232 | 0.113 | 4.1408007 | 15 | rasl11b |
| CU467646.24 | 1.914239 | 0.326145772 | 0.217 | 0.103 | 4.8631248 | 15 | CU467646.2 |
| BX571780.13 | 2.062903 | 0.286342312 | 0.161 | 0.066 | 5.2408059 | 15 | BX571780.1 |
| dazl4 | 2.399134 | 0.3305239 | 0.193 | 0.089 | 6.0950004 | 15 | dazl |
| zp3c4 | 2.560596 | 0.359804688 | 0.283 | 0.155 | 6.5051964 | 15 | zp3c |
| prdx14 | 2.713474 | 0.378677077 | 0.268 | 0.148 | 6.8935828 | 15 | prdx1 |
| wu:fa19b121 | 3.161605 | 0.29342387 | 0.15 | 0.061 | 8.0320595 | 15 | wu:fa19b12 |
| zgc:1655394 | 3.180986 | 0.325834518 | 0.209 | 0.098 | 8.0812965 | 15 | zgc:165539 |
| faub4 | 3.401638 | 0.389848859 | 0.303 | 0.182 | 8.6418630 | 15 | faub |
| lhx8a | 3.640008 | 0.285089292 | 0.126 | 0.048 | 9.2474410 | 15 | lhx8a |
| CR383662.24 | 4.126015 | 0.295327546 | 0.165 | 0.071 | 0.000105 | 15 | CR383662.2 |
| zgc:1717504 | 5.308224 | 0.357423894 | 0.264 | 0.144 | 0.000135 | 15 | zgc:171750 |
| zc3h11a4 | 5.457728 | 0.392380419 | 0.311 | 0.189 | 0.000139 | 15 | zc3h11a |
| zgc:558884 | 8.150991 | 0.296462566 | 0.161 | 0.07 | 0.000207 | 15 | zgc:55888 |
| cdc64 | 8.385125 | 0.332472603 | 0.185 | 0.088 | 0.000213 | 15 | cdc6 |
| ing5b | 8.632324 | 0.270617704 | 0.118 | 0.044 | 0.000219 | 15 | ing5b |
| tbpl2 | 8.903956 | 0.252622361 | 0.122 | 0.046 | 0.000226 | 15 | tbpl2 |
| dynll2a4 | 1.036785 | 0.357549691 | 0.224 | 0.117 | 0.000263 | 15 | dynll2a |
| CABZ01069022.21 | 1.186103 | 0.269819838 | 0.13 | 0.051 | 0.000301 | 15 | CABZ01069022.2 |
| brf24 | 1.354888 | 0.314469489 | 0.173 | 0.08 | 0.000344 | 15 | brf2 |
| pias4b1 | 1.362947 | 0.26370271 | 0.134 | 0.053 | 0.000346 | 15 | pias4b |
| gyg1a5 | 1.537870 | 0.366846626 | 0.445 | 0.319 | 0.000391 | 15 | gyg1a |
| zgc:1529864 | 1.762865 | 0.291244473 | 0.165 | 0.074 | 0.000448 | 15 | zgc:152986 |
| cbx3a2 | 1.926730 | 0.3740924 | 0.398 | 0.292 | 0.000489 | 15 | cbx3a |
| mad2l14 | 2.088030 | 0.355144162 | 0.276 | 0.159 | 0.00053 | 15 | mad2l1 |
| h2afx4 | 2.110282 | 0.334695884 | 0.232 | 0.122 | 0.000536 | 15 | h2afx |
| marcksb4 | 3.707862 | 0.359369674 | 0.272 | 0.16 | 0.000942 | 15 | marcksb |

| | | | | | | | |
|---|---|---|---|---|---|---|---|
| zgc:1719773 | 3.937777 | 0.355664894 | 0.311 | 0.189 | 0.001 | 15 | zgc:171977 |
| zgc:1526524 | 4.056072 | 0.29570439 | 0.193 | 0.094 | 0.00103 | 15 | zgc:152652 |
| snu13a3 | 4.109192 | 0.306785144 | 0.185 | 0.089 | 0.001044 | 15 | snu13a |
| si:ch211-286b5.43 | 4.171860 | 0.259837736 | 0.138 | 0.058 | 0.00106 | 15 | si:ch211-286b5.4 |
| LOC1005380233 | 4.247885 | 0.279801533 | 0.146 | 0.063 | 0.001079 | 15 | LOC100538023 |
| kpna24 | 4.408300 | 0.331495608 | 0.248 | 0.136 | 0.00112 | 15 | kpna2 |
| zgc:1751363 | 5.120290 | 0.317939501 | 0.181 | 0.089 | 0.001301 | 15 | zgc:175136 |
| zgc:1735443 | 5.762929 | 0.26408313 | 0.146 | 0.063 | 0.001464 | 15 | zgc:173544 |
| metap1d3 | 6.520254 | 0.300054745 | 0.165 | 0.078 | 0.001656 | 15 | metap1d |
| rpain4 | 6.615713 | 0.320876181 | 0.197 | 0.101 | 0.001681 | 15 | rpain |
| ube2c3 | 7.706231 | 0.323975697 | 0.228 | 0.123 | 0.001958 | 15 | ube2c |
| hspa54 | 8.146392 | 0.351374171 | 0.402 | 0.295 | 0.00207 | 15 | hspa5 |
| tent5c4 | 8.686012 | 0.340825091 | 0.224 | 0.123 | 0.002207 | 15 | tent5c |
| st8sia7.12 | 9.028791 | 0.258879946 | 0.138 | 0.059 | 0.002294 | 15 | st8sia7.1 |
| mkrn44 | 9.338052 | 0.342675745 | 0.315 | 0.193 | 0.002372 | 15 | mkrn4 |
| BX005461.13 | 1.215573 | 0.276832616 | 0.146 | 0.065 | 0.003088 | 15 | BX005461.1 |
| si:ch211-202f5.34 | 1.222079 | 0.29106736 | 0.161 | 0.076 | 0.003105 | 15 | si:ch211-202f5.3 |
| bmb4 | 1.332533 | 0.278262924 | 0.157 | 0.073 | 0.003385 | 15 | bmb |
| tmem144b4 | 1.341450 | 0.280566312 | 0.165 | 0.078 | 0.003408 | 15 | tmem144b |
| zgc:922874 | 1.354327 | 0.319235324 | 0.209 | 0.112 | 0.003441 | 15 | zgc:92287 |
| yrdc4 | 1.400104 | 0.329007365 | 0.197 | 0.103 | 0.003557 | 15 | yrdc |
| zgc:1118684 | 1.432356 | 0.315238958 | 0.224 | 0.122 | 0.003639 | 15 | zgc:111868 |
| BX001030.14 | 1.555662 | 0.298996546 | 0.197 | 0.101 | 0.003952 | 15 | BX001030.1 |
| alyref3 | 1.616864 | 0.361624221 | 0.354 | 0.251 | 0.004108 | 15 | alyref |
| si:ch73-303b9.13 | 1.765005 | 0.260535587 | 0.146 | 0.065 | 0.004484 | 15 | si:ch73-303b9.1 |
| chchd104 | 2.217813 | 0.330431227 | 0.236 | 0.134 | 0.005634 | 15 | chchd10 |
| cdca7a4 | 2.240141 | 0.346748115 | 0.303 | 0.192 | 0.005691 | 15 | cdca7a |
| mylipa4 | 2.308721 | 0.317721228 | 0.189 | 0.099 | 0.005865 | 15 | mylipa |
| ccnh4 | 2.343591 | 0.305101946 | 0.201 | 0.106 | 0.005954 | 15 | ccnh |
| ccne14 | 2.392544 | 0.298585029 | 0.185 | 0.094 | 0.006078 | 15 | ccne1 |
| zfand63 | 2.650273 | 0.328322996 | 0.205 | 0.112 | 0.006733 | 15 | zfand6 |
| cldn7b4 | 2.966074 | 0.311829315 | 0.236 | 0.133 | 0.007535 | 15 | cldn7b |
| LOC1039099724 | 3.320794 | 0.34115124 | 0.417 | 0.323 | 0.008436 | 15 | LOC103909972 |
| lsm13 | 3.887333 | 0.339585097 | 0.22 | 0.126 | 0.009876 | 15 | lsm1 |
| tph1a4 | 5.692365 | 0.29669396 | 0.213 | 0.117 | 0.014461 | 15 | tph1a |

| | | | | | | | |
|---|---|---|---|---|---|---|---|
| rogdi4 | 7.380677 | 0.296647171 | 0.189 | 0.1 | 0.018751 | 15 | rogdi |
| asf1ba2 | 7.653221 | 0.284389567 | 0.169 | 0.086 | 0.019443 | 15 | asf1ba |
| si:ch211-14a17.64 | 8.195703 | 0.256880621 | 0.15 | 0.071 | 0.020821 | 15 | si:ch211-14a17.6 |
| coil2 | 9.479135 | 0.317061795 | 0.181 | 0.097 | 0.024082 | 15 | coil |
| zgc:566764 | 9.481885 | 0.354122444 | 0.366 | 0.275 | 0.024089 | 15 | zgc:56676 |
| zgc:1141234 | 1.271980 | 0.271600114 | 0.169 | 0.086 | 0.032315 | 15 | zgc:114123 |
| si:dkey-188i13.114 | 1.308413 | 0.338182917 | 0.386 | 0.291 | 0.03324 | 15 | si:dkey-188i13.11 |
| zgc:1134251 | 1.351524 | 0.251712442 | 0.126 | 0.057 | 0.034335 | 15 | zgc:113425 |
| si:ch1073-190k2.14 | 1.561341 | 0.30673636 | 0.256 | 0.155 | 0.039666 | 15 | si:ch1073-190k2.1 |
| gfer4 | 2.250550 | 0.283022556 | 0.165 | 0.087 | 0.057175 | 15 | gfer |
| lmnl33 | 2.296395 | 0.255669924 | 0.134 | 0.063 | 0.05834 | 15 | lmnl3 |
| dnd13 | 2.906953 | 0.273721419 | 0.173 | 0.092 | 0.073851 | 15 | dnd1 |
| CU655961.34 | 2.972002 | 0.251646542 | 0.161 | 0.082 | 0.075504 | 15 | CU655961.3 |
| zgc:1010161 | 3.148484 | 0.285670638 | 0.165 | 0.088 | 0.079987 | 15 | zgc:101016 |
| ctsla4 | 4.075415 | 0.302694233 | 0.437 | 0.375 | 0.103536 | 15 | ctsla |
| hmgb3b4 | 4.428904 | 0.331752117 | 0.26 | 0.171 | 0.112516 | 15 | hmgb3b |
| mcm3l4 | 4.442719 | 0.252338707 | 0.146 | 0.073 | 0.112867 | 15 | mcm3l |
| zgc:923134 | 5.009778 | 0.274518878 | 0.193 | 0.108 | 0.127273 | 15 | zgc:92313 |
| lpcat44 | 5.064693 | 0.275488841 | 0.169 | 0.091 | 0.128669 | 15 | lpcat4 |
| snrpc2 | 6.102920 | 0.340360957 | 0.26 | 0.177 | 0.155045 | 15 | snrpc |
| ncbp23 | 6.577265 | 0.325866551 | 0.248 | 0.162 | 0.167095 | 15 | ncbp2 |
| snrpb24 | 7.634966 | 0.305689704 | 0.224 | 0.138 | 0.193966 | 15 | snrpb2 |
| rnf128a | 8.024554 | 0.253566747 | 0.126 | 0.062 | 0.203864 | 15 | rnf128a |
| aktip3 | 8.138102 | 0.303710075 | 0.209 | 0.126 | 0.206748 | 15 | aktip |
| BX284697.14 | 8.776045 | 0.259501857 | 0.173 | 0.094 | 0.222955 | 15 | BX284697.1 |
| hells2 | 8.897670 | 0.289931604 | 0.209 | 0.125 | 0.226045 | 15 | hells |
| si:ch211-69g19.24 | 8.936113 | 0.263662978 | 0.185 | 0.104 | 0.227022 | 15 | si:ch211-69g19.2 |
| si:ch73-281n10.24 | 8.989537 | 0.282522537 | 0.457 | 0.397 | 0.228379 | 15 | si:ch73-281n10.2 |
| chtopa4 | 9.003033 | 0.332414582 | 0.287 | 0.207 | 0.228722 | 15 | chtopa |
| pop44 | 1.093574 | 0.285351837 | 0.181 | 0.103 | 0.277823 | 15 | pop4 |
| znf395a3 | 1.125560 | 0.257281911 | 0.157 | 0.085 | 0.285949 | 15 | znf395a |
| ivns1abpb1 | 1.142864 | 0.277541484 | 0.181 | 0.104 | 0.290345 | 15 | ivns1abpb |
| fbxo323 | 1.193896 | 0.263276571 | 0.161 | 0.088 | 0.303309 | 15 | fbxo32 |
| fdx14 | 1.209380 | 0.310320126 | 0.287 | 0.199 | 0.307243 | 15 | fdx1 |
| imp33 | 1.260554 | 0.292676815 | 0.205 | 0.124 | 0.320244 | 15 | imp3 |

| | | | | | | | |
|---|---|---|---|---|---|---|---|
| rpf11 | 1.362192 | 0.319981383 | 0.209 | 0.131 | 0.346065 | 15 | rpf1 |
| tpx24 | 1.362677 | 0.260455552 | 0.189 | 0.107 | 0.346188 | 15 | tpx2 |
| rnf45 | 1.444258 | 0.264197388 | 0.201 | 0.117 | 0.366914 | 15 | rnf4 |
| zgc:1737424 | 1.591477 | 0.26858544 | 0.217 | 0.13 | 0.404315 | 15 | zgc:173742 |
| ddx194 | 1.623933 | 0.295016548 | 0.217 | 0.135 | 0.41256 | 15 | ddx19 |
| ebna1bp21 | 1.695341 | 0.312969499 | 0.213 | 0.135 | 0.430702 | 15 | ebna1bp2 |
| pim32 | 2.290604 | 0.28478939 | 0.193 | 0.116 | 0.581928 | 15 | pim3 |
| thop14 | 2.302400 | 0.263620898 | 0.181 | 0.105 | 0.584925 | 15 | thop1 |
| evla2 | 2.411945 | 0.258679943 | 0.161 | 0.09 | 0.612755 | 15 | evla |
| aif1l2 | 2.534364 | 0.284222714 | 0.256 | 0.17 | 0.643855 | 15 | aif1l |
| prmt1 | 2.561713 | 0.330397847 | 0.295 | 0.222 | 0.650803 | 15 | prmt1 |
| ccng1 | 3.197461 | 0.251117816 | 0.469 | 0.445 | 0.812315 | 15 | ccng1 |
| tmem54a4 | 3.229150 | 0.261669689 | 0.197 | 0.118 | 0.820366 | 15 | tmem54a |
| ctsba6 | 3.436239 | 0.285020192 | 0.421 | 0.354 | 0.872977 | 15 | ctsba |
| selenoh4 | 4.302227 | 0.301276776 | 0.307 | 0.228 | 1 | 15 | selenoh |
| nabp1a2 | 4.318324 | 0.275840678 | 0.181 | 0.109 | 1 | 15 | nabp1a |
| parn4 | 4.464951 | 0.272637024 | 0.209 | 0.13 | 1 | 15 | parn |
| gemin73 | 4.770112 | 0.301148278 | 0.217 | 0.142 | 1 | 15 | gemin7 |
| cox175 | 4.852807 | 0.278450618 | 0.402 | 0.342 | 1 | 15 | cox17 |
| cstf11 | 5.124036 | 0.257470184 | 0.157 | 0.09 | 1 | 15 | cstf1 |
| tomm20a1 | 6.430349 | 0.322972957 | 0.295 | 0.228 | 1 | 15 | tomm20a |
| si:ch73-340n8.44 | 9.405207 | 0.261710734 | 0.185 | 0.114 | 1 | 15 | si:ch73-340n8.4 |
| rbm4.23 | 0.0001 | 0.291547503 | 0.228 | 0.157 | 1 | 15 | rbm4.2 |
| sec61a1l2 | 0.0002 | 0.25873853 | 0.181 | 0.116 | 1 | 15 | sec61a1l |
| lrrc592 | 0.0002 | 0.285484847 | 0.201 | 0.135 | 1 | 15 | lrrc59 |
| ddx3xb | 0.0003 | 0.287870718 | 0.201 | 0.137 | 1 | 15 | ddx3xb |
| rpz24 | 0.0003 | 0.278000574 | 0.232 | 0.165 | 1 | 15 | rpz2 |
| nifk3 | 0.0003 | 0.256081202 | 0.181 | 0.118 | 1 | 15 | nifk |
| mcm71 | 0.0004 | 0.259233439 | 0.213 | 0.147 | 1 | 15 | mcm7 |
| tia1l1 | 0.0006 | 0.265139714 | 0.205 | 0.143 | 1 | 15 | tia1l |
| rbbp42 | 0.0007 | 0.253859935 | 0.228 | 0.164 | 1 | 15 | rbbp4 |
| rtn4a2 | 0.0007 | 0.277068584 | 0.295 | 0.241 | 1 | 15 | rtn4a |
| lsm61 | 0.0007 | 0.282895103 | 0.283 | 0.226 | 1 | 15 | lsm6 |
| pdap1a | 0.0009 | 0.25065338 | 0.177 | 0.12 | 1 | 15 | pdap1a |
| lsm12a2 | 0.001 | 0.251503178 | 0.173 | 0.116 | 1 | 15 | lsm12a |

| | | | | | | | |
|---|---|---|---|---|---|---|---|
| nutf2l3 | 0.0016 | 0.259330206 | 0.276 | 0.222 | 1 | 15 | nutf2l |
| zgc:1102393 | 0.0016 | 0.258359986 | 0.252 | 0.198 | 1 | 15 | zgc:110239 |
| rbm8a | 0.0017 | 0.256815233 | 0.346 | 0.321 | 1 | 15 | rbm8a |
| serp1 | 0.0022 | 0.273012045 | 0.248 | 0.199 | 1 | 15 | serp1 |
| c1qbp | 0.0051 | 0.261159129 | 0.213 | 0.168 | 1 | 15 | c1qbp |
| ddx44 | 0 | 1.532493288 | 0.804 | 0.087 | 0 | 16 | ddx4 |
| hsp90aa1.21 | 0 | 1.235093906 | 0.596 | 0.031 | 0 | 16 | hsp90aa1.2 |
| si:ch73-167f10.1 | 0 | 1.178892605 | 0.54 | 0.006 | 0 | 16 | si:ch73-167f10.1 |
| sycp1 | 0 | 0.931254275 | 0.368 | 0.005 | 0 | 16 | sycp1 |
| stmn1b | 0 | 0.81848417 | 0.36 | 0.007 | 0 | 16 | stmn1b |
| si:dkey-40g16.5 | 0 | 0.781630641 | 0.316 | 0.003 | 0 | 16 | si:dkey-40g16.5 |
| hormad1 | 0 | 0.671174811 | 0.296 | 0.005 | 0 | 16 | hormad1 |
| sycp3 | 0 | 0.636088267 | 0.248 | 0.004 | 0 | 16 | sycp3 |
| tuba7l | 0 | 0.627964196 | 0.252 | 0.002 | 0 | 16 | tuba7l |
| smc1b | 0 | 0.552628399 | 0.228 | 0.002 | 0 | 16 | smc1b |
| LOC100149647 | 0 | 0.414277172 | 0.152 | 0 | 0 | 16 | LOC100149647 |
| CU929391.1 | 6.213032 | 0.357191909 | 0.144 | 0.001 | 1.5784209 | 16 | CU929391.1 |
| pcp4l1 | 4.080813 | 0.501949061 | 0.208 | 0.004 | 1.0367307 | 16 | pcp4l1 |
| si:ch211-176l24.4 | 5.677044 | 0.574451873 | 0.212 | 0.004 | 1.4422532 | 16 | si:ch211-176l24.4 |
| hnrnpa3 | 7.479505 | 0.502278605 | 0.208 | 0.004 | 1.9001682 | 16 | hnrnpa3 |
| stag3 | 1.015857 | 0.548029171 | 0.236 | 0.007 | 2.5807857 | 16 | stag3 |
| rnf212b | 4.535765 | 0.278939688 | 0.108 | 0.001 | 1.1523113 | 16 | rnf212b |
| zgc:193690 | 1.288444 | 0.778978565 | 0.384 | 0.025 | 3.2732926 | 16 | zgc:193690 |
| LOC101886261 | 3.926075 | 0.315194658 | 0.116 | 0.001 | 9.9741947 | 16 | LOC101886261 |
| si:ch211-241j8.2 | 1.299491 | 0.356970254 | 0.136 | 0.002 | 3.3013585 | 16 | si:ch211-241j8.2 |
| chtopb | 8.407508 | 0.77083393 | 0.38 | 0.029 | 2.1359274 | 16 | chtopb |
| nsg2 | 5.779087 | 0.308986127 | 0.12 | 0.002 | 1.4681771 | 16 | nsg2 |
| spata22 | 1.051183 | 0.389053789 | 0.168 | 0.005 | 2.6705316 | 16 | spata22 |
| ddx39b | 2.269968 | 0.696582835 | 0.324 | 0.024 | 5.7668543 | 16 | ddx39b |
| LOC101883959 | 7.677028 | 0.287437085 | 0.112 | 0.002 | 1.9503491 | 16 | LOC101883959 |
| cox5b2 | 4.861264 | 0.490262752 | 0.208 | 0.009 | 1.2350041 | 16 | cox5b2 |
| tdrd1 | 2.283589 | 0.706461054 | 0.336 | 0.026 | 5.8014596 | 16 | tdrd1 |
| CCNB1IP1 | 6.230934 | 0.322115619 | 0.128 | 0.003 | 1.5829690 | 16 | CCNB1IP1 |
| parp1 | 5.183317 | 0.736966439 | 0.336 | 0.029 | 1.3168218 | 16 | parp1 |
| piwil11 | 8.111599 | 0.803475153 | 0.356 | 0.032 | 2.0607517 | 16 | piwil1 |

| | | | | | | | |
|---|---|---|---|---|---|---|---|
| arl15a | 1.526824 | 0.27193579 | 0.108 | 0.002 | 3.8788981 | 16 | arl15a |
| adad1 | 3.355343 | 0.574234375 | 0.264 | 0.018 | 8.5242509 | 16 | adad1 |
| tdrd9 | 1.188790 | 0.59284826 | 0.284 | 0.021 | 3.0201230 | 16 | tdrd9 |
| syce2 | 1.619236 | 0.572614942 | 0.24 | 0.015 | 4.1136707 | 16 | syce2 |
| CABZ01083945.1 | 1.367275 | 0.47526105 | 0.204 | 0.011 | 3.4735633 | 16 | CABZ01083945.1 |
| dut | 4.560215 | 1.022210073 | 0.612 | 0.111 | 1.1585226 | 16 | dut |
| hmgb2b3 | 8.287175 | 1.175683726 | 0.864 | 0.328 | 2.1053568 | 16 | hmgb2b |
| gcshb | 2.245279 | 0.457230531 | 0.204 | 0.013 | 5.7041337 | 16 | gcshb |
| piwil2 | 3.161724 | 0.586699163 | 0.292 | 0.026 | 8.0323604 | 16 | piwil2 |
| zgc:194246 | 3.498273 | 0.743669099 | 0.34 | 0.037 | 8.8873639 | 16 | zgc:194246 |
| mgmt | 9.114714 | 0.90819502 | 0.516 | 0.087 | 2.3155931 | 16 | mgmt |
| pld6 | 1.620271 | 0.678211936 | 0.34 | 0.038 | 4.1162999 | 16 | pld6 |
| stmn1a1 | 2.228155 | 1.071228049 | 0.716 | 0.18 | 5.6606278 | 16 | stmn1a |
| si:ch211-173m16.2 | 2.524893 | 0.282374818 | 0.12 | 0.004 | 6.4144919 | 16 | si:ch211-173m16.2 |
| CU927934.3 | 2.469000 | 0.467885463 | 0.132 | 0.005 | 6.2724947 | 16 | CU927934.3 |
| zgc:161973 | 1.974120 | 0.375566492 | 0.168 | 0.009 | 5.0152533 | 16 | zgc:161973 |
| mrto4 | 1.006020 | 0.697047779 | 0.344 | 0.044 | 2.5557957 | 16 | mrto4 |
| dpy30 | 1.552265 | 1.001053427 | 0.628 | 0.156 | 3.9435310 | 16 | dpy30 |
| si:dkeyp-93d12.1 | 2.341310 | 0.473447307 | 0.224 | 0.018 | 5.9480981 | 16 | si:dkeyp-93d12.1 |
| si:ch211-277c7.7 | 3.427932 | 0.270023587 | 0.1 | 0.003 | 8.7086614 | 16 | si:ch211-277c7.7 |
| terb1 | 1.546312 | 0.352091893 | 0.156 | 0.009 | 3.9284066 | 16 | terb1 |
| tuba1c | 6.819801 | 0.341995151 | 0.132 | 0.007 | 1.7325704 | 16 | tuba1c |
| dmrt1 | 5.094628 | 0.42912086 | 0.192 | 0.015 | 1.2942902 | 16 | dmrt1 |
| odf3l2a | 1.762989 | 0.385457205 | 0.168 | 0.011 | 4.4788753 | 16 | odf3l2a |
| calm2b3 | 4.276375 | 1.029511373 | 0.712 | 0.24 | 1.0864131 | 16 | calm2b |
| esco2 | 6.239969 | 0.616227262 | 0.284 | 0.034 | 1.5852643 | 16 | esco2 |
| ccnd3 | 2.767472 | 0.414129313 | 0.184 | 0.014 | 7.0307642 | 16 | ccnd3 |
| dnph1 | 8.489477 | 0.519675752 | 0.248 | 0.027 | 2.1567518 | 16 | dnph1 |
| nsmce1 | 2.374347 | 0.520552626 | 0.24 | 0.025 | 6.0320307 | 16 | nsmce1 |
| top2a1 | 3.571048 | 0.592905184 | 0.284 | 0.035 | 9.0722484 | 16 | top2a |
| ddx43 | 3.173608 | 0.297288168 | 0.128 | 0.007 | 8.0625516 | 16 | ddx43 |
| lsm62 | 8.138871 | 0.975032197 | 0.688 | 0.219 | 2.0676802 | 16 | lsm6 |
| sub1b1 | 1.630291 | 0.960145586 | 0.676 | 0.209 | 4.1417563 | 16 | sub1b |
| CABZ01080371.2 | 7.246921 | 0.314533181 | 0.124 | 0.007 | 1.8410802 | 16 | CABZ01080371.2 |
| fanci | 9.481189 | 0.305787358 | 0.116 | 0.006 | 2.4086962 | 16 | fanci |

| | | | | | | | |
|---|---|---|---|---|---|---|---|
| lsm7 | 2.422813 | 0.946753944 | 0.644 | 0.195 | 6.1551566 | 16 | lsm7 |
| rnf17 | 8.352475 | 0.472093129 | 0.22 | 0.022 | 2.1219463 | 16 | rnf17 |
| tdrkh | 4.251371 | 0.545121379 | 0.264 | 0.032 | 1.0800609 | 16 | tdrkh |
| morc3a | 1.071512 | 0.467363467 | 0.192 | 0.018 | 2.7221772 | 16 | morc3a |
| mycbp1 | 1.736194 | 0.949610689 | 0.628 | 0.194 | 4.4108031 | 16 | mycbp |
| si:ch211-222l21.16 | 3.349840 | 0.956022829 | 0.776 | 0.307 | 8.5102698 | 16 | si:ch211-222l21.1 |
| stk31 | 2.547021 | 0.352162798 | 0.16 | 0.012 | 6.4707069 | 16 | stk31 |
| srsf2b | 6.325608 | 0.518479238 | 0.26 | 0.033 | 1.6070209 | 16 | srsf2b |
| si:dkey-276j7.1 | 5.938343 | 0.571723192 | 0.276 | 0.038 | 1.5086360 | 16 | si:dkey-276j7.1 |
| snrpd1 | 2.249010 | 0.934605662 | 0.672 | 0.233 | 5.7136115 | 16 | snrpd1 |
| chaf1a | 2.518654 | 0.633586329 | 0.36 | 0.063 | 6.3986423 | 16 | chaf1a |
| trmt112 | 4.443358 | 0.784979124 | 0.464 | 0.109 | 1.1288351 | 16 | trmt112 |
| adad2 | 2.300179 | 0.330450379 | 0.164 | 0.014 | 5.8436071 | 16 | adad2 |
| hmgb1b4 | 2.002133 | 0.909244871 | 0.524 | 0.149 | 5.0864201 | 16 | hmgb1b |
| iqcb1 | 2.207668 | 0.255105522 | 0.1 | 0.005 | 5.6085819 | 16 | iqcb1 |
| sms | 2.220245 | 0.526974989 | 0.264 | 0.037 | 5.6405338 | 16 | sms |
| iqck | 3.354551 | 0.330041001 | 0.144 | 0.011 | 8.5222388 | 16 | iqck |
| hells3 | 4.999782 | 0.767613976 | 0.504 | 0.119 | 1.2701947 | 16 | hells |
| CR382383.2 | 5.000496 | 0.461192735 | 0.24 | 0.03 | 1.2703760 | 16 | CR382383.2 |
| hirip3 | 1.157402 | 0.719147211 | 0.392 | 0.079 | 2.9403805 | 16 | hirip3 |
| hpf1 | 1.261563 | 0.664310382 | 0.336 | 0.059 | 3.2050010 | 16 | hpf1 |
| mcm3 | 3.686529 | 0.516193952 | 0.248 | 0.034 | 9.3656274 | 16 | mcm3 |
| gpd1b | 1.157546 | 0.294175803 | 0.14 | 0.011 | 2.9407471 | 16 | gpd1b |
| h2afx16 | 1.357890 | 0.884458654 | 0.628 | 0.221 | 3.4497195 | 16 | h2afx1 |
| snrpg2 | 1.198133 | 0.874826077 | 0.688 | 0.259 | 3.0438578 | 16 | snrpg |
| smfn | 2.517238 | 0.657119912 | 0.372 | 0.074 | 6.3950436 | 16 | smfn |
| minos1 | 3.166112 | 0.79941966 | 0.544 | 0.16 | 8.0435078 | 16 | minos1 |
| snrpf | 5.521564 | 0.854339489 | 0.68 | 0.264 | 1.4027535 | 16 | snrpf |
| zgc:162025 | 7.321916 | 0.624568802 | 0.316 | 0.057 | 1.8601329 | 16 | zgc:162025 |
| rad21l1 | 3.562187 | 0.337155811 | 0.144 | 0.012 | 9.0497385 | 16 | rad21l1 |
| cmss1 | 6.600107 | 0.511572315 | 0.28 | 0.045 | 1.6767572 | 16 | cmss1 |
| snrpd3l | 8.119629 | 0.801265206 | 0.548 | 0.163 | 2.0627917 | 16 | snrpd3l |
| si:ch211-212d10.2 | 9.480265 | 0.374881345 | 0.16 | 0.015 | 2.4084613 | 16 | si:ch211-212d10.2 |
| ssrp1a | 3.263420 | 0.617174413 | 0.328 | 0.061 | 8.2907209 | 16 | ssrp1a |
| dcun1d5 | 1.155691 | 0.638992362 | 0.376 | 0.079 | 2.9360347 | 16 | dcun1d5 |

| | | | | | | | |
|---|---|---|---|---|---|---|---|
| parp3 | 3.264039 | 0.43814217 | 0.188 | 0.023 | 8.2922918 | 16 | parp3 |
| si:dkey-119f1.1 | 5.862048 | 0.280215492 | 0.12 | 0.009 | 1.4892534 | 16 | si:dkey-119f1.1 |
| rpa2 | 2.461941 | 0.640446798 | 0.376 | 0.083 | 6.2545625 | 16 | rpa2 |
| llph | 3.674728 | 0.600313295 | 0.308 | 0.06 | 9.3356485 | 16 | llph |
| srsf7b | 6.137572 | 0.287117558 | 0.136 | 0.012 | 1.5592502 | 16 | srsf7b |
| zgc:153681 | 7.242692 | 0.3423509 | 0.152 | 0.015 | 1.8400061 | 16 | zgc:153681 |
| cpsf2 | 1.150359 | 0.495441152 | 0.256 | 0.042 | 2.9224892 | 16 | cpsf2 |
| tomm20a2 | 1.812301 | 0.794509641 | 0.62 | 0.222 | 4.6041518 | 16 | tomm20a |
| eif4e1b2 | 2.609605 | 0.575436908 | 0.344 | 0.068 | 6.6297033 | 16 | eif4e1b |
| sart1 | 4.201089 | 0.65233239 | 0.388 | 0.093 | 1.0672868 | 16 | sart1 |
| ppid | 7.800964 | 0.654731499 | 0.38 | 0.089 | 1.9818350 | 16 | ppid |
| rbx1 | 1.576464 | 0.795256601 | 0.664 | 0.276 | 4.0050086 | 16 | rbx1 |
| dynll2b | 9.214505 | 0.732355757 | 0.456 | 0.126 | 2.3409451 | 16 | dynll2b |
| auh | 1.093107 | 0.434968044 | 0.208 | 0.029 | 2.7770401 | 16 | auh |
| rabif | 1.846582 | 0.54385112 | 0.276 | 0.05 | 4.6912423 | 16 | rabif |
| gnl3 | 1.960182 | 0.619537996 | 0.316 | 0.066 | 4.9798424 | 16 | gnl3 |
| npm1a1 | 2.373537 | 0.826903018 | 0.668 | 0.295 | 6.0299726 | 16 | npm1a |
| lsm4 | 4.852209 | 0.780642775 | 0.588 | 0.212 | 1.2327038 | 16 | lsm4 |
| gra | 2.871989 | 0.47325276 | 0.252 | 0.041 | 7.2962885 | 16 | gra |
| hint1 | 1.129734 | 0.781668819 | 0.592 | 0.225 | 2.8700894 | 16 | hint1 |
| h2afvb1 | 1.568634 | 0.716415047 | 0.82 | 0.47 | 3.9851166 | 16 | h2afvb |
| dlgap5 | 2.354816 | 0.42093295 | 0.228 | 0.035 | 5.9824123 | 16 | dlgap5 |
| pcna1 | 2.784351 | 0.721485095 | 0.488 | 0.143 | 7.0736449 | 16 | pcna |
| wdr76 | 3.195366 | 0.354822378 | 0.172 | 0.021 | 8.1178296 | 16 | wdr76 |
| ube2t | 6.114211 | 0.478582109 | 0.252 | 0.042 | 1.5533155 | 16 | ube2t |
| chek1 | 2.437325 | 0.332214286 | 0.16 | 0.018 | 6.1920248 | 16 | chek1 |
| tubb2b | 6.500731 | 0.800642476 | 0.636 | 0.254 | 1.6515107 | 16 | tubb2b |
| rad51 | 9.645928 | 0.436272553 | 0.228 | 0.037 | 2.4505480 | 16 | rad51 |
| ppih | 1.271249 | 0.568551891 | 0.328 | 0.073 | 3.2296085 | 16 | ppih |
| chd4a1 | 2.773950 | 0.636903281 | 0.388 | 0.104 | 7.0472201 | 16 | chd4a |
| tdrd6 | 3.318049 | 0.347254949 | 0.152 | 0.017 | 8.4295039 | 16 | tdrd6 |
| acot13 | 5.662533 | 0.367649138 | 0.172 | 0.022 | 1.4385666 | 16 | acot13 |
| nutf2l4 | 6.110208 | 0.73177457 | 0.596 | 0.216 | 1.5522984 | 16 | nutf2l |
| ppa1b | 6.731437 | 0.628030956 | 0.348 | 0.083 | 1.7101217 | 16 | ppa1b |
| nub1 | 1.228067 | 0.359309879 | 0.16 | 0.02 | 3.1199044 | 16 | nub1 |

| | | | | | | | |
|---|---|---|---|---|---|---|---|
| lsm5 | 5.343058 | 0.751957632 | 0.592 | 0.23 | 1.3574041 | 16 | lsm5 |
| setb1 | 1.493668 | 0.740320827 | 0.712 | 0.356 | 3.7946654 | 16 | setb |
| spata6l | 3.047127 | 0.425990723 | 0.208 | 0.032 | 7.7412266 | 16 | spata6l |
| rbm8a1 | 4.021049 | 0.787135453 | 0.652 | 0.315 | 1.0215477 | 16 | rbm8a |
| cenps | 4.393146 | 0.398244437 | 0.208 | 0.032 | 1.1160788 | 16 | cenps |
| atp5mc3b1 | 4.753980 | 0.557114259 | 0.856 | 0.599 | 1.2077488 | 16 | atp5mc3b |
| supt3h | 7.551403 | 0.446630227 | 0.216 | 0.036 | 1.9184339 | 16 | supt3h |
| efcab1 | 9.444733 | 0.272743074 | 0.124 | 0.012 | 2.3994344 | 16 | efcab1 |
| nudc | 9.574087 | 0.698491708 | 0.476 | 0.152 | 2.4322968 | 16 | nudc |
| odc1 | 1.625226 | 0.652371739 | 0.416 | 0.117 | 4.1288886 | 16 | odc1 |
| lin52 | 1.808211 | 0.501659424 | 0.264 | 0.052 | 4.5937606 | 16 | lin52 |
| cks2 | 1.890400 | 0.473988863 | 0.228 | 0.039 | 4.8025626 | 16 | cks2 |
| wdr82 | 3.828912 | 0.575466278 | 0.348 | 0.085 | 9.7273532 | 16 | wdr82 |
| dnajc91 | 4.808122 | 0.585019899 | 0.348 | 0.083 | 1.2215035 | 16 | dnajc9 |
| dazl5 | 7.429439 | 0.59169305 | 0.36 | 0.086 | 1.8874490 | 16 | dazl |
| psmb7 | 8.154689 | 0.701755666 | 0.496 | 0.169 | 2.0716989 | 16 | psmb7 |
| snrnp27 | 3.507337 | 0.634764851 | 0.42 | 0.122 | 8.9103897 | 16 | snrnp27 |
| hibadha | 5.005692 | 0.284112811 | 0.124 | 0.013 | 1.2716962 | 16 | hibadha |
| ss18l2 | 5.397035 | 0.484314593 | 0.276 | 0.058 | 1.3711167 | 16 | ss18l2 |
| dnd14 | 1.208430 | 0.597489006 | 0.36 | 0.088 | 3.0700184 | 16 | dnd1 |
| snrpe1 | 2.681721 | 0.732310707 | 0.64 | 0.279 | 6.8129141 | 16 | snrpe |
| pole4 | 4.112249 | 0.52920793 | 0.32 | 0.075 | 1.0447170 | 16 | pole4 |
| polr2l | 5.423861 | 0.703848529 | 0.54 | 0.199 | 1.3779319 | 16 | polr2l |
| isg152 | 1.087870 | 0.627218262 | 0.28 | 0.062 | 2.7637359 | 16 | isg15 |
| rbm46 | 1.344375 | 0.261437444 | 0.112 | 0.011 | 3.4153870 | 16 | rbm46 |
| timm10 | 1.585672 | 0.510113132 | 0.284 | 0.062 | 4.0284008 | 16 | timm10 |
| ndufb6 | 2.025236 | 0.644155718 | 0.4 | 0.12 | 5.1451145 | 16 | ndufb6 |
| cenpx | 2.640902 | 0.560786254 | 0.328 | 0.08 | 6.7092120 | 16 | cenpx |
| nsmce2 | 4.129494 | 0.417406281 | 0.204 | 0.034 | 1.0490979 | 16 | nsmce2 |
| fkbp3 | 5.723140 | 0.703534352 | 0.484 | 0.174 | 1.4539638 | 16 | fkbp3 |
| pfdn6 | 6.265845 | 0.620397795 | 0.392 | 0.114 | 1.5918380 | 16 | pfdn6 |
| phb2b | 7.657642 | 0.455975764 | 0.256 | 0.051 | 1.9454240 | 16 | phb2b |
| med28 | 1.375094 | 0.576867034 | 0.34 | 0.087 | 3.4934267 | 16 | med28 |
| exd11 | 2.018679 | 0.420482556 | 0.252 | 0.048 | 5.1284552 | 16 | exd1 |
| seta | 3.627244 | 0.717460243 | 0.628 | 0.289 | 9.2150153 | 16 | seta |

| | | | | | | | |
|---|---|---|---|---|---|---|---|
| cirbpa | 4.970061 | 0.522658489 | 0.84 | 0.621 | 1.2626441 | 16 | cirbpa |
| mcm6 | 5.302490 | 0.454170064 | 0.224 | 0.042 | 1.3470976 | 16 | mcm6 |
| erh | 1.414302 | 0.70349847 | 0.668 | 0.342 | 3.5930360 | 16 | erh |
| alkbh3 | 1.945083 | 0.307918806 | 0.136 | 0.016 | 4.9414853 | 16 | alkbh3 |
| zcchc2 | 2.014722 | 0.302551299 | 0.124 | 0.014 | 5.1184035 | 16 | zcchc2 |
| birc5a1 | 3.337771 | 0.558145525 | 0.336 | 0.083 | 8.4796086 | 16 | birc5a |
| tubb4b | 4.543217 | 0.567342393 | 0.832 | 0.589 | 1.1542043 | 16 | tubb4b |
| ube2c4 | 5.750493 | 0.589480143 | 0.42 | 0.12 | 1.4609129 | 16 | ube2c |
| mki67 | 1.306326 | 0.454342188 | 0.212 | 0.038 | 3.3187232 | 16 | mki67 |
| hddc3 | 1.937910 | 0.406783507 | 0.196 | 0.033 | 4.9232603 | 16 | hddc3 |
| nuf2 | 2.390016 | 0.350317968 | 0.172 | 0.026 | 6.0718379 | 16 | nuf2 |
| si:dkey-148h10.5 | 2.957442 | 0.25328665 | 0.116 | 0.012 | 7.5133830 | 16 | si:dkey-148h10.5 |
| si:dkey-189g17.21 | 3.789208 | 0.429986411 | 0.24 | 0.047 | 9.6264839 | 16 | si:dkey-189g17.2 |
| hmgn24 | 5.079159 | 0.666415939 | 0.796 | 0.463 | 1.2903604 | 16 | hmgn2 |
| nars | 5.431472 | 0.572168477 | 0.308 | 0.079 | 1.3798656 | 16 | nars |
| cct71 | 2.702589 | 0.68367023 | 0.536 | 0.225 | 6.8659274 | 16 | cct7 |
| vrk1 | 6.217543 | 0.452915682 | 0.24 | 0.049 | 1.5795669 | 16 | vrk1 |
| aaas | 8.775759 | 0.290423742 | 0.136 | 0.018 | 2.2294817 | 16 | aaas |
| atp5mea | 9.614265 | 0.656162396 | 0.436 | 0.149 | 2.4425040 | 16 | atp5mea |
| gtf2f2a | 1.178161 | 0.543589004 | 0.284 | 0.07 | 2.9931185 | 16 | gtf2f2a |
| eif4eb | 6.462713 | 0.455535903 | 0.236 | 0.049 | 1.6418524 | 16 | eif4eb |
| elavl25 | 1.275982 | 0.628445931 | 0.528 | 0.182 | 3.2416335 | 16 | elavl2 |
| nelfcd | 1.348768 | 0.288275624 | 0.156 | 0.023 | 3.4265467 | 16 | nelfcd |
| edc4 | 1.880632 | 0.365759711 | 0.18 | 0.03 | 4.7777472 | 16 | edc4 |
| lsm3 | 2.231757 | 0.635306538 | 0.46 | 0.161 | 5.6697810 | 16 | lsm3 |
| hddc2 | 3.636517 | 0.413212814 | 0.236 | 0.049 | 9.2385720 | 16 | hddc2 |
| snrpa11 | 7.647298 | 0.596271752 | 0.384 | 0.117 | 1.9427961 | 16 | snrpa1 |
| timm23a | 1.478618 | 0.589585386 | 0.38 | 0.119 | 3.7564308 | 16 | timm23a |
| mybbp1a | 3.490987 | 0.356029156 | 0.168 | 0.028 | 8.8688531 | 16 | mybbp1a |
| eif4a3 | 5.224800 | 0.61518096 | 0.44 | 0.156 | 1.3273605 | 16 | eif4a3 |
| mcm72 | 6.968523 | 0.601692567 | 0.432 | 0.143 | 1.7703535 | 16 | mcm7 |
| zgc:56106 | 9.165717 | 0.415795718 | 0.224 | 0.047 | 2.3285505 | 16 | zgc:56106 |
| srrm2 | 9.675300 | 0.529046385 | 0.308 | 0.083 | 2.4580101 | 16 | srrm2 |
| lrrc20 | 1.246033 | 0.272935014 | 0.132 | 0.018 | 3.1655474 | 16 | lrrc20 |
| psmc5 | 1.465763 | 0.538980824 | 0.34 | 0.1 | 3.7237723 | 16 | psmc5 |

| | | | | | | | |
|---|---|---|---|---|---|---|---|
| henmt1 | 2.175284 | 0.267128618 | 0.128 | 0.017 | 5.5263107 | 16 | henmt1 |
| zgc:66472 | 4.583736 | 0.343221709 | 0.188 | 0.034 | 1.1644982 | 16 | zgc:66472 |
| ndufa8 | 6.718341 | 0.59789357 | 0.412 | 0.144 | 1.7067946 | 16 | ndufa8 |
| baz1a | 1.010022 | 0.473950045 | 0.288 | 0.074 | 2.5659610 | 16 | baz1a |
| gstr | 1.066631 | 0.594909081 | 0.376 | 0.124 | 2.7097767 | 16 | gstr |
| sub1a1 | 1.094172 | 0.684661455 | 0.544 | 0.249 | 2.7797461 | 16 | sub1a |
| ap3s2 | 1.375225 | 0.394700476 | 0.228 | 0.049 | 3.4937614 | 16 | ap3s2 |
| msh6 | 1.595960 | 0.317318641 | 0.164 | 0.027 | 4.0545366 | 16 | msh6 |
| mbd3b | 2.291568 | 0.441581196 | 0.24 | 0.054 | 5.8217298 | 16 | mbd3b |
| tuba8l1 | 2.741119 | 0.6505617 | 0.5 | 0.205 | 6.9638142 | 16 | tuba8l |
| psma5 | 3.037356 | 0.625512137 | 0.452 | 0.171 | 7.7164053 | 16 | psma5 |
| rcc2 | 7.292471 | 0.588275153 | 0.412 | 0.142 | 1.8526524 | 16 | rcc2 |
| nedd8 | 1.029038 | 0.521739741 | 0.3 | 0.084 | 2.6142715 | 16 | nedd8 |
| ruvbl1 | 1.106377 | 0.473690889 | 0.308 | 0.082 | 2.8107519 | 16 | ruvbl1 |
| txlna | 1.497937 | 0.418984046 | 0.22 | 0.047 | 3.8055103 | 16 | txlna |
| calm3b | 1.726648 | 0.660206965 | 0.568 | 0.275 | 4.3865514 | 16 | calm3b |
| tfdp1a | 2.694035 | 0.362839633 | 0.188 | 0.036 | 6.8441968 | 16 | tfdp1a |
| ptgr11 | 3.077893 | 0.520670498 | 0.352 | 0.105 | 7.8193892 | 16 | ptgr1 |
| ddx18 | 4.075515 | 0.543642179 | 0.34 | 0.102 | 1.0353846 | 16 | ddx18 |
| ilf21 | 6.539418 | 0.627399647 | 0.556 | 0.239 | 1.6613393 | 16 | ilf2 |
| ubtf | 9.576532 | 0.437813689 | 0.236 | 0.056 | 2.4329179 | 16 | ubtf |
| ndufab1b | 9.746745 | 0.597955554 | 0.436 | 0.161 | 2.4761607 | 16 | ndufab1b |
| ndufs7 | 3.462412 | 0.559892858 | 0.368 | 0.124 | 8.7962584 | 16 | ndufs7 |
| nup50 | 3.729845 | 0.490629431 | 0.328 | 0.096 | 9.4756713 | 16 | nup50 |
| zgc:922875 | 4.016113 | 0.488977195 | 0.372 | 0.108 | 1.0202936 | 16 | zgc:92287 |
| ankrd10a | 4.939753 | 0.355974118 | 0.22 | 0.047 | 1.2549443 | 16 | ankrd10a |
| gtf2b | 5.968185 | 0.498457768 | 0.284 | 0.078 | 1.5162175 | 16 | gtf2b |
| si:ch211-156b7.4 | 6.649608 | 0.368859129 | 0.188 | 0.037 | 1.6893330 | 16 | si:ch211-156b7.4 |
| zgc:1737425 | 7.011404 | 0.509477216 | 0.408 | 0.126 | 1.7812473 | 16 | zgc:173742 |
| ralaa | 7.964613 | 0.304027849 | 0.144 | 0.023 | 2.0234101 | 16 | ralaa |
| tcp12 | 8.153853 | 0.626165826 | 0.528 | 0.242 | 2.0714864 | 16 | tcp1 |
| nup93 | 1.190318 | 0.363371736 | 0.184 | 0.036 | 3.0240041 | 16 | nup93 |
| psmb3 | 1.237076 | 0.593388358 | 0.412 | 0.152 | 3.1427915 | 16 | psmb3 |
| sf3b5 | 1.328622 | 0.549327169 | 0.424 | 0.152 | 3.3753650 | 16 | sf3b5 |
| nudt21 | 1.617258 | 0.476394791 | 0.284 | 0.078 | 4.1086459 | 16 | nudt21 |

| | | | | | | | |
|---|---|---|---|---|---|---|---|
| nup107 | 1.903207 | 0.329235552 | 0.164 | 0.029 | 4.8350993 | 16 | nup107 |
| psmc3ip | 2.101443 | 0.325935133 | 0.152 | 0.025 | 5.3387178 | 16 | psmc3ip |
| snu13b5 | 2.758719 | 0.616265378 | 0.704 | 0.367 | 7.0085276 | 16 | snu13b |
| dhx16 | 3.364736 | 0.392950549 | 0.228 | 0.052 | 8.5481122 | 16 | dhx16 |
| acaa1 | 3.485287 | 0.268293992 | 0.128 | 0.019 | 8.8543731 | 16 | acaa1 |
| COX7A2 (1 of many)1 | 4.238750 | 0.636565124 | 0.548 | 0.263 | 1.0768544 | 16 | COX7A2 (1 of many) |
| nme4 | 5.216600 | 0.27351676 | 0.144 | 0.023 | 1.3252774 | 16 | nme4 |
| eftud2 | 5.694847 | 0.305506692 | 0.156 | 0.027 | 1.4467759 | 16 | eftud2 |
| hnrnpul1 | 7.358754 | 0.325632302 | 0.168 | 0.032 | 1.8694914 | 16 | hnrnpul1 |
| ndufa2 | 7.580924 | 0.587277745 | 0.408 | 0.152 | 1.9259339 | 16 | ndufa2 |
| rbb4l | 1.102369 | 0.475688027 | 0.292 | 0.082 | 2.8005696 | 16 | rbb4l |
| txndc17 | 1.211489 | 0.401400952 | 0.232 | 0.055 | 3.0777879 | 16 | txndc17 |
| eif4e1c | 1.767750 | 0.491883772 | 0.328 | 0.098 | 4.4909712 | 16 | eif4e1c |
| polr2f | 1.966069 | 0.564350872 | 0.38 | 0.134 | 4.9948002 | 16 | polr2f |
| unga | 2.072011 | 0.429632966 | 0.264 | 0.067 | 5.2639441 | 16 | unga |
| cenph | 2.079602 | 0.276956156 | 0.132 | 0.02 | 5.2832309 | 16 | cenph |
| xrcc5 | 2.994185 | 0.258736836 | 0.12 | 0.017 | 7.6067289 | 16 | xrcc5 |
| LOC110438491 | 3.309333 | 0.292117829 | 0.152 | 0.026 | 8.4073605 | 16 | LOC110438491 |
| eif2s21 | 3.409307 | 0.59604894 | 0.616 | 0.319 | 8.6613446 | 16 | eif2s2 |
| psmc1a | 5.110722 | 0.473638829 | 0.284 | 0.081 | 1.2983790 | 16 | psmc1a |
| larp7 | 6.136859 | 0.412666022 | 0.244 | 0.06 | 1.5590692 | 16 | larp7 |
| dhfr | 6.528191 | 0.367174888 | 0.212 | 0.047 | 1.6584870 | 16 | dhfr |
| snrnp200 | 8.396774 | 0.364540732 | 0.192 | 0.041 | 2.1332006 | 16 | snrnp200 |
| sf3a2 | 1.074341 | 0.549910732 | 0.38 | 0.134 | 2.7293648 | 16 | sf3a2 |
| tyms | 2.064275 | 0.40486619 | 0.26 | 0.066 | 5.2442913 | 16 | tyms |
| gcsha | 2.264791 | 0.433665812 | 0.256 | 0.066 | 5.7537040 | 16 | gcsha |
| fen11 | 3.306014 | 0.485778997 | 0.288 | 0.081 | 8.3989295 | 16 | fen1 |
| si:ch211-226l4.65 | 3.404388 | 0.585120036 | 0.592 | 0.239 | 8.6488487 | 16 | si:ch211-226l4.6 |
| cops5 | 3.759019 | 0.407719259 | 0.22 | 0.052 | 9.5497887 | 16 | cops5 |
| ppie | 4.197937 | 0.337298796 | 0.184 | 0.038 | 1.0664859 | 16 | ppie |
| ccdc137 | 5.198120 | 0.323830246 | 0.156 | 0.028 | 1.3205825 | 16 | ccdc137 |
| cops3 | 5.413184 | 0.429518361 | 0.236 | 0.059 | 1.3752194 | 16 | cops3 |
| smc3 | 7.993914 | 0.407297295 | 0.212 | 0.05 | 2.0308539 | 16 | smc3 |
| ndufab1a | 9.142329 | 0.567239247 | 0.38 | 0.139 | 2.3226087 | 16 | ndufab1a |
| dgcr6 | 1.003468 | 0.324918504 | 0.164 | 0.031 | 2.5493113 | 16 | dgcr6 |

| | | | | | | | |
|---|---|---|---|---|---|---|---|
| ccdc25 | 1.096436 | 0.508411931 | 0.312 | 0.099 | 2.7854972 | 16 | ccdc25 |
| dnajc18 | 1.991704 | 0.262655068 | 0.124 | 0.019 | 5.0599242 | 16 | dnajc18 |
| ptges3b | 2.052516 | 0.590849876 | 0.444 | 0.178 | 5.2144183 | 16 | ptges3b |
| cox5aa1 | 2.347830 | 0.590308075 | 0.628 | 0.355 | 5.9646630 | 16 | cox5aa |
| trip13 | 3.598873 | 0.310796231 | 0.16 | 0.03 | 9.1429384 | 16 | trip13 |
| mif | 3.872864 | 0.611231921 | 0.464 | 0.204 | 9.8390133 | 16 | mif |
| nap1l11 | 4.447470 | 0.579297806 | 0.652 | 0.353 | 1.1298798 | 16 | nap1l1 |
| rfc4 | 4.912594 | 0.439017833 | 0.264 | 0.072 | 1.2480446 | 16 | rfc4 |
| mphosph82 | 5.150803 | 0.532931131 | 0.376 | 0.138 | 1.3085615 | 16 | mphosph8 |
| nop16 | 8.117563 | 0.499442788 | 0.328 | 0.106 | 2.0622670 | 16 | nop16 |
| si:ch211-51e12.7 | 8.123859 | 0.554551662 | 0.444 | 0.174 | 2.0638665 | 16 | si:ch211-51e12.7 |
| mrps25 | 1.029167 | 0.468023591 | 0.284 | 0.085 | 2.6145987 | 16 | mrps25 |
| taf15 | 1.226333 | 0.557450904 | 0.448 | 0.18 | 3.1154999 | 16 | taf15 |
| ndufs5 | 1.531106 | 0.579049472 | 0.404 | 0.162 | 3.8897758 | 16 | ndufs5 |
| tomm6 | 1.855154 | 0.540038872 | 0.352 | 0.126 | 4.7130203 | 16 | tomm6 |
| atp5pd3 | 2.913391 | 0.596146136 | 0.544 | 0.28 | 7.4014699 | 16 | atp5pd |
| pomp | 3.124030 | 0.581797115 | 0.524 | 0.257 | 7.9365992 | 16 | pomp |
| pane1 | 3.206209 | 0.311671052 | 0.156 | 0.03 | 8.1453746 | 16 | pane1 |
| srsf3b | 3.653403 | 0.57482783 | 0.532 | 0.25 | 9.2814717 | 16 | srsf3b |
| drg2 | 4.522119 | 0.43214637 | 0.268 | 0.077 | 1.1488444 | 16 | drg2 |
| cdk11 | 4.952017 | 0.4335982 | 0.284 | 0.08 | 1.2580599 | 16 | cdk1 |
| cbx3a3 | 5.430071 | 0.580405445 | 0.584 | 0.288 | 1.3795096 | 16 | cbx3a |
| psma8 | 1.285082 | 0.549747811 | 0.492 | 0.22 | 3.2647517 | 16 | psma8 |
| chtf18 | 1.465980 | 0.256560498 | 0.128 | 0.021 | 3.7243231 | 16 | chtf18 |
| si:dkey-6i22.5 | 1.788250 | 0.342095675 | 0.168 | 0.034 | 4.5430514 | 16 | si:dkey-6i22.5 |
| tsn | 2.067068 | 0.354254265 | 0.2 | 0.046 | 5.2513886 | 16 | tsn |
| srsf1b | 2.368269 | 0.416770018 | 0.24 | 0.065 | 6.0165882 | 16 | srsf1b |
| phax | 3.332390 | 0.396902013 | 0.232 | 0.06 | 8.4659379 | 16 | phax |
| idh3g | 3.851001 | 0.381274329 | 0.18 | 0.04 | 9.7834700 | 16 | idh3g |
| yeats4 | 4.011939 | 0.400518944 | 0.22 | 0.057 | 1.0192332 | 16 | yeats4 |
| spc24 | 5.166782 | 0.373283249 | 0.232 | 0.059 | 1.3126210 | 16 | spc24 |
| gtf2h5 | 5.628465 | 0.484266083 | 0.328 | 0.109 | 1.4299116 | 16 | gtf2h5 |
| dusp22b | 6.371757 | 0.331361241 | 0.156 | 0.031 | 1.6187450 | 16 | dusp22b |
| gtf2e2 | 7.052833 | 0.42355102 | 0.256 | 0.073 | 1.7917723 | 16 | gtf2e2 |
| snrpd21 | 9.073083 | 0.578081815 | 0.532 | 0.256 | 2.3050169 | 16 | snrpd2 |

| | | | | | | | |
|---|---|---|---|---|---|---|---|
| ciapin1 | 9.626673 | 0.355168911 | 0.192 | 0.045 | 2.4456564 | 16 | ciapin1 |
| psmd7 | 1.916009 | 0.490925295 | 0.332 | 0.116 | 4.8676213 | 16 | psmd7 |
| psmd8 | 1.962573 | 0.481652042 | 0.34 | 0.118 | 4.9859175 | 16 | psmd8 |
| rpa1 | 1.973098 | 0.432359524 | 0.264 | 0.077 | 5.0126565 | 16 | rpa1 |
| cenpw | 2.768472 | 0.325268718 | 0.168 | 0.036 | 7.0333056 | 16 | cenpw |
| ttf1 | 2.854055 | 0.376967179 | 0.224 | 0.058 | 7.2507285 | 16 | ttf1 |
| cdk2 | 3.163673 | 0.333435256 | 0.18 | 0.04 | 8.0373116 | 16 | cdk2 |
| cct22 | 3.499271 | 0.564802198 | 0.508 | 0.251 | 8.8898993 | 16 | cct2 |
| rsl1d1 | 6.387424 | 0.541444945 | 0.432 | 0.185 | 1.6227252 | 16 | rsl1d1 |
| pbdc1 | 7.413993 | 0.502018654 | 0.36 | 0.137 | 1.8835251 | 16 | pbdc1 |
| rpa33 | 1.005060 | 0.490337783 | 0.404 | 0.151 | 2.5533568 | 16 | rpa3 |
| utp11l | 1.109352 | 0.366418138 | 0.228 | 0.061 | 2.8183088 | 16 | utp11l |
| selenou1a | 1.357272 | 0.354316734 | 0.176 | 0.04 | 3.4481501 | 16 | selenou1a |
| ubxn1 | 1.522406 | 0.360079706 | 0.2 | 0.05 | 3.8676744 | 16 | ubxn1 |
| mtrf1 | 1.587577 | 0.250557055 | 0.12 | 0.02 | 4.0332405 | 16 | mtrf1 |
| magoh | 1.891446 | 0.505291313 | 0.336 | 0.12 | 4.8052190 | 16 | magoh |
| mtx2 | 2.162990 | 0.362025657 | 0.188 | 0.045 | 5.4950766 | 16 | mtx2 |
| zpr1 | 2.289877 | 0.295995455 | 0.148 | 0.029 | 5.8174336 | 16 | zpr1 |
| sf3b6 | 2.374749 | 0.557718586 | 0.512 | 0.255 | 6.0330501 | 16 | sf3b6 |
| gpat2 | 2.605298 | 0.253906156 | 0.12 | 0.02 | 6.6187619 | 16 | gpat2 |
| prpf6 | 2.750370 | 0.30900504 | 0.16 | 0.034 | 6.9873175 | 16 | prpf6 |
| ppig | 2.856962 | 0.386844487 | 0.232 | 0.064 | 7.2581119 | 16 | ppig |
| ccdc106b | 5.670089 | 0.328426215 | 0.168 | 0.036 | 1.4404862 | 16 | ccdc106b |
| smim8 | 6.969157 | 0.336388524 | 0.176 | 0.041 | 1.7705144 | 16 | smim8 |
| hspa4a | 7.928735 | 0.338909864 | 0.192 | 0.047 | 2.0142951 | 16 | hspa4a |
| rbmx2 | 7.977143 | 0.299736225 | 0.152 | 0.032 | 2.0265933 | 16 | rbmx2 |
| alkbh5 | 8.095520 | 0.329439821 | 0.18 | 0.042 | 2.0566670 | 16 | alkbh5 |
| exosc2 | 1.280904 | 0.346621637 | 0.188 | 0.045 | 3.2541367 | 16 | exosc2 |
| nolc1 | 1.381049 | 0.423422427 | 0.276 | 0.086 | 3.5085566 | 16 | nolc1 |
| ndufc21 | 1.394602 | 0.513451663 | 0.372 | 0.151 | 3.5429876 | 16 | ndufc2 |
| ebna1bp22 | 1.422258 | 0.499056063 | 0.356 | 0.132 | 3.6132480 | 16 | ebna1bp2 |
| etfa | 1.548474 | 0.431048937 | 0.288 | 0.096 | 3.9338990 | 16 | etfa |
| ufc1 | 2.137042 | 0.440015026 | 0.272 | 0.087 | 5.4291564 | 16 | ufc1 |
| psmb5 | 2.653700 | 0.401640479 | 0.252 | 0.076 | 6.7417260 | 16 | psmb5 |
| cse1l | 3.230437 | 0.405235956 | 0.24 | 0.068 | 8.2069269 | 16 | cse1l |

| | | | | | | | |
|---|---|---|---|---|---|---|---|
| ik | 3.594012 | 0.344770984 | 0.2 | 0.052 | 9.1305885 | 16 | ik |
| cox7a2a1 | 4.429757 | 0.551602742 | 0.5 | 0.249 | 1.1253798 | 16 | cox7a2a |
| g3bp2 | 4.449237 | 0.316871071 | 0.164 | 0.037 | 1.1303286 | 16 | g3bp2 |
| zgc:158564 | 4.480196 | 0.420282704 | 0.256 | 0.079 | 1.1381938 | 16 | zgc:158564 |
| polr3f | 6.015016 | 0.288960769 | 0.18 | 0.042 | 1.5281148 | 16 | polr3f |
| fip1l1b | 7.119541 | 0.325034299 | 0.192 | 0.048 | 1.8087195 | 16 | fip1l1b |
| usp1 | 9.341532 | 0.340179685 | 0.184 | 0.045 | 2.3732162 | 16 | usp1 |
| pcmt | 1.071226 | 0.319549179 | 0.168 | 0.039 | 2.7214514 | 16 | pcmt |
| tomm34 | 1.271139 | 0.292751503 | 0.16 | 0.035 | 3.2293294 | 16 | tomm34 |
| top1mt | 1.464621 | 0.283641256 | 0.172 | 0.039 | 3.7208720 | 16 | top1mt |
| asf1bb | 1.636285 | 0.366457114 | 0.22 | 0.061 | 4.1569833 | 16 | asf1bb |
| rpf12 | 1.696026 | 0.473328276 | 0.348 | 0.129 | 4.3087552 | 16 | rpf1 |
| skp11 | 1.855843 | 0.547580297 | 0.512 | 0.266 | 4.7147705 | 16 | skp1 |
| hnrnpa0b4 | 2.026830 | 0.542368907 | 0.576 | 0.339 | 5.1491627 | 16 | hnrnpa0b |
| aifm1 | 2.246351 | 0.269177391 | 0.132 | 0.026 | 5.7068552 | 16 | aifm1 |
| prdx31 | 2.522377 | 0.498809366 | 0.356 | 0.14 | 6.4080991 | 16 | prdx3 |
| ndufb5 | 2.617333 | 0.492748054 | 0.376 | 0.153 | 6.6493351 | 16 | ndufb5 |
| tbcb | 2.722399 | 0.429195117 | 0.244 | 0.076 | 6.9162570 | 16 | tbcb |
| zgc:566995 | 9.294723 | 0.498746986 | 0.5 | 0.206 | 2.3613244 | 16 | zgc:56699 |
| ndufa6 | 1.028425 | 0.531945354 | 0.504 | 0.256 | 2.6127143 | 16 | ndufa6 |
| gins1 | 1.755684 | 0.263259043 | 0.132 | 0.026 | 4.4603154 | 16 | gins1 |
| si:dkey-170l10.1 | 1.989903 | 0.250829713 | 0.124 | 0.023 | 5.0553498 | 16 | si:dkey-170l10.1 |
| ruvbl2 | 2.309872 | 0.29977411 | 0.184 | 0.046 | 5.8682321 | 16 | ruvbl2 |
| hp1bp33 | 2.409235 | 0.463454505 | 0.312 | 0.119 | 6.1206628 | 16 | hp1bp3 |
| atp6v0cb | 2.778014 | 0.278079771 | 0.12 | 0.023 | 7.0575462 | 16 | atp6v0cb |
| cdk11b | 3.030229 | 0.331811665 | 0.184 | 0.047 | 7.6982979 | 16 | cdk11b |
| pfdn4 | 3.044949 | 0.503802484 | 0.412 | 0.185 | 7.7356929 | 16 | pfdn4 |
| snip1 | 4.224027 | 0.302257726 | 0.156 | 0.036 | 1.0731141 | 16 | snip1 |
| metap2b | 4.327257 | 0.525034191 | 0.452 | 0.219 | 1.0993397 | 16 | metap2b |
| fundc1 | 4.398759 | 0.410879456 | 0.248 | 0.078 | 1.1175048 | 16 | fundc1 |
| zgc:554615 | 4.904532 | 0.508642851 | 0.54 | 0.238 | 1.2459965 | 16 | zgc:55461 |
| CABZ01044053.1 | 4.992961 | 0.281231661 | 0.172 | 0.042 | 1.2684617 | 16 | CABZ01044053.1 |
| gemin2 | 5.645624 | 0.340010991 | 0.196 | 0.053 | 1.4342709 | 16 | gemin2 |
| krr1 | 5.909129 | 0.281673629 | 0.148 | 0.033 | 1.5012142 | 16 | krr1 |
| mrpl21 | 6.304958 | 0.307924353 | 0.196 | 0.051 | 1.6017747 | 16 | mrpl21 |

| | | | | | | | |
|---|---|---|---|---|---|---|---|
| csnk2b | 8.570898 | 0.514032477 | 0.448 | 0.218 | 2.1774367 | 16 | csnk2b |
| CABZ01112656.1 | 9.222765 | 0.380506062 | 0.208 | 0.06 | 2.3430434 | 16 | CABZ01112656.1 |
| emd1 | 9.928736 | 0.402079303 | 0.224 | 0.069 | 2.5223954 | 16 | emd |
| luc7l3 | 1.047902 | 0.335611822 | 0.204 | 0.057 | 2.6621963 | 16 | luc7l3 |
| fabp7a | 1.218623 | 0.275515049 | 0.116 | 0.022 | 3.0959124 | 16 | fabp7a |
| apex1 | 1.515715 | 0.38037238 | 0.236 | 0.072 | 3.8506746 | 16 | apex1 |
| zgc:1034825 | 1.869401 | 0.438370914 | 0.408 | 0.154 | 4.7492156 | 16 | zgc:103482 |
| pin11 | 1.942997 | 0.477890359 | 0.368 | 0.155 | 4.9361852 | 16 | pin1 |
| prmt11 | 2.108244 | 0.50247994 | 0.464 | 0.218 | 5.3559944 | 16 | prmt1 |
| denr | 2.334849 | 0.467490136 | 0.34 | 0.136 | 5.9316849 | 16 | denr |
| dynlrb1 | 2.337991 | 0.508415002 | 0.444 | 0.21 | 5.9396667 | 16 | dynlrb1 |
| eif1ad | 3.133489 | 0.429581493 | 0.264 | 0.091 | 7.9606289 | 16 | eif1ad |
| bcl7a | 3.289402 | 0.270535223 | 0.152 | 0.035 | 8.3567279 | 16 | bcl7a |
| mrpl38 | 3.796591 | 0.318407653 | 0.164 | 0.04 | 9.6452413 | 16 | mrpl38 |
| ddx23 | 4.153533 | 0.336885346 | 0.2 | 0.056 | 1.0552051 | 16 | ddx23 |
| sbds | 4.476596 | 0.334496711 | 0.192 | 0.052 | 1.1372793 | 16 | sbds |
| pop5 | 4.579016 | 0.28775102 | 0.18 | 0.046 | 1.1632990 | 16 | pop5 |
| si:dkey-92i17.23 | 4.622257 | 0.369723451 | 0.248 | 0.076 | 1.1742844 | 16 | si:dkey-92i17.2 |
| pcgf6 | 5.067705 | 0.280082313 | 0.156 | 0.036 | 1.2874504 | 16 | pcgf6 |
| psmb1 | 5.122841 | 0.504932233 | 0.416 | 0.195 | 1.3014579 | 16 | psmb1 |
| prdx15 | 6.160919 | 0.472030502 | 0.372 | 0.146 | 1.5651815 | 16 | prdx1 |
| zgc:101851 | 6.171565 | 0.373040931 | 0.256 | 0.082 | 1.5678861 | 16 | zgc:101851 |
| coil3 | 6.825364 | 0.386137496 | 0.284 | 0.095 | 1.7339837 | 16 | coil |
| pcm1 | 7.096434 | 0.258002071 | 0.116 | 0.022 | 1.8028492 | 16 | pcm1 |
| hnrnpa0l5 | 7.325821 | 0.318355422 | 0.88 | 0.739 | 1.8611249 | 16 | hnrnpa0l |
| selenoh5 | 7.679787 | 0.474514326 | 0.488 | 0.225 | 1.9510500 | 16 | selenoh |
| ube2f | 8.738021 | 0.298524152 | 0.144 | 0.033 | 2.2198944 | 16 | ube2f |
| si:ch211-235e9.6 | 9.354394 | 0.493024597 | 0.364 | 0.158 | 2.3764839 | 16 | si:ch211-235e9.6 |
| atp5md1 | 1.076932 | 0.518694286 | 0.576 | 0.354 | 2.7359464 | 16 | atp5md |
| hmgb1a7 | 1.387271 | 0.589159369 | 0.44 | 0.226 | 3.5243642 | 16 | hmgb1a |
| psmd11b | 1.685609 | 0.333165802 | 0.184 | 0.05 | 4.2822904 | 16 | psmd11b |
| gkap1 | 2.006201 | 0.429339006 | 0.308 | 0.116 | 5.0967556 | 16 | gkap1 |
| ndufa12 | 2.066508 | 0.438138079 | 0.324 | 0.128 | 5.2499649 | 16 | ndufa12 |
| pdrg1 | 2.122503 | 0.282746733 | 0.16 | 0.039 | 5.3922207 | 16 | pdrg1 |
| cript | 2.195345 | 0.279688648 | 0.16 | 0.039 | 5.5772758 | 16 | cript |

| | | | | | | | |
|---|---|---|---|---|---|---|---|
| mrps14 | 2.279202 | 0.406749358 | 0.272 | 0.096 | 5.7903129 | 16 | mrps14 |
| CABZ01068251.1 | 2.980796 | 0.268685459 | 0.172 | 0.044 | 7.5727122 | 16 | CABZ01068251.1 |
| caprin1a | 3.361567 | 0.349099462 | 0.192 | 0.055 | 8.5400625 | 16 | caprin1a |
| eif5a21 | 3.888820 | 0.484802517 | 0.636 | 0.405 | 9.8795476 | 16 | eif5a2 |
| pole3 | 4.173184 | 0.307013722 | 0.18 | 0.049 | 1.0601976 | 16 | pole3 |
| clns1a | 4.408770 | 0.368237855 | 0.232 | 0.073 | 1.1200481 | 16 | clns1a |
| orc4 | 4.707733 | 0.295856581 | 0.16 | 0.04 | 1.1959997 | 16 | orc4 |
| ndnl2 | 5.052901 | 0.283107884 | 0.16 | 0.04 | 1.2836895 | 16 | ndnl2 |
| isoc2 | 5.338928 | 0.292385454 | 0.16 | 0.041 | 1.3563547 | 16 | isoc2 |
| bud31 | 5.488517 | 0.497572852 | 0.46 | 0.227 | 1.3943578 | 16 | bud31 |
| psmd1 | 6.220285 | 0.370364981 | 0.256 | 0.087 | 1.5802635 | 16 | psmd1 |
| pgam5 | 7.223380 | 0.400379421 | 0.288 | 0.103 | 1.8350996 | 16 | pgam5 |
| cox7b3 | 7.233218 | 0.466188931 | 0.636 | 0.425 | 1.8375991 | 16 | cox7b |
| msh2 | 8.456286 | 0.260156474 | 0.14 | 0.032 | 2.1483195 | 16 | msh2 |
| dhps | 9.001682 | 0.280590951 | 0.136 | 0.031 | 2.2868774 | 16 | dhps |
| sod1 | 9.330092 | 0.486665669 | 0.572 | 0.343 | 2.3703101 | 16 | sod1 |
| si:ch211-22i13.2 | 1.298521 | 0.392501888 | 0.208 | 0.064 | 3.2988946 | 16 | si:ch211-22i13.2 |
| phb | 1.304054 | 0.356596843 | 0.216 | 0.068 | 3.3129503 | 16 | phb |
| psmd4b | 1.353947 | 0.368963391 | 0.212 | 0.066 | 3.4397037 | 16 | psmd4b |
| cct5 | 1.424663 | 0.510279059 | 0.496 | 0.272 | 3.6193569 | 16 | cct5 |
| eif5b | 1.741436 | 0.444641393 | 0.352 | 0.15 | 4.4241187 | 16 | eif5b |
| cnot1 | 1.977748 | 0.341719622 | 0.204 | 0.061 | 5.0244694 | 16 | cnot1 |
| cops4 | 2.493919 | 0.386228299 | 0.24 | 0.081 | 6.3358035 | 16 | cops4 |
| kpna25 | 2.863814 | 0.42273904 | 0.348 | 0.134 | 7.2755216 | 16 | kpna2 |
| mrps33 | 3.606995 | 0.356062497 | 0.244 | 0.081 | 9.1635725 | 16 | mrps33 |
| atp5f1d3 | 4.306778 | 0.496127956 | 0.496 | 0.271 | 1.0941371 | 16 | atp5f1d |
| pfdn1 | 4.368389 | 0.482526756 | 0.352 | 0.157 | 1.1097892 | 16 | pfdn1 |
| psmc6 | 4.822567 | 0.363458749 | 0.264 | 0.094 | 1.2251731 | 16 | psmc6 |
| mfap1 | 5.728725 | 0.327572512 | 0.192 | 0.056 | 1.4553826 | 16 | mfap1 |
| strap | 6.052511 | 0.40980558 | 0.268 | 0.097 | 1.5376406 | 16 | strap |
| uchl3 | 6.651119 | 0.335651694 | 0.2 | 0.06 | 1.6897168 | 16 | uchl3 |
| zgc:63470 | 6.786114 | 0.320231143 | 0.164 | 0.044 | 1.7240123 | 16 | zgc:63470 |
| zgc:112496 | 7.910886 | 0.252402965 | 0.14 | 0.033 | 2.0097605 | 16 | zgc:112496 |
| ppil1 | 7.911783 | 0.322784798 | 0.228 | 0.072 | 2.0099886 | 16 | ppil1 |
| mcm2 | 8.049707 | 0.374153041 | 0.244 | 0.082 | 2.0450282 | 16 | mcm2 |

| | | | | | | | |
|---|---|---|---|---|---|---|---|
| otub1b | 9.079125 | 0.281130607 | 0.144 | 0.035 | 2.3065518 | 16 | otub1b |
| tma71 | 1.482735 | 0.486343953 | 0.552 | 0.329 | 3.7668892 | 16 | tma7 |
| sf3b4 | 1.644302 | 0.329105209 | 0.228 | 0.074 | 4.1773512 | 16 | sf3b4 |
| nsfl1c | 1.692292 | 0.394957383 | 0.232 | 0.08 | 4.2992689 | 16 | nsfl1c |
| cox7c3 | 1.843827 | 0.425762171 | 0.668 | 0.492 | 4.6842448 | 16 | cox7c |
| rfc5 | 2.040379 | 0.282201852 | 0.156 | 0.04 | 5.1835841 | 16 | rfc5 |
| ubc3 | 2.409191 | 0.354926662 | 0.808 | 0.652 | 6.1205506 | 16 | ubc |
| prpf311 | 2.660591 | 0.325841044 | 0.232 | 0.074 | 6.7592320 | 16 | prpf31 |
| apooa | 4.037393 | 0.287804553 | 0.148 | 0.037 | 1.0256996 | 16 | apooa |
| hypk | 4.053153 | 0.463872366 | 0.436 | 0.218 | 1.0297036 | 16 | hypk |
| dus3l | 4.291086 | 0.356514326 | 0.208 | 0.066 | 1.0901504 | 16 | dus3l |
| nip7 | 5.499422 | 0.356452261 | 0.2 | 0.063 | 1.3971282 | 16 | nip7 |
| ndufa9a | 5.814590 | 0.398104356 | 0.236 | 0.084 | 1.4771966 | 16 | ndufa9a |
| ifi455 | 6.898107 | 0.444671094 | 0.332 | 0.137 | 1.7524642 | 16 | ifi45 |
| psmd14 | 7.394527 | 0.386891479 | 0.28 | 0.107 | 1.8785798 | 16 | psmd14 |
| esf1 | 7.735832 | 0.340037792 | 0.172 | 0.05 | 1.9652883 | 16 | esf1 |
| supt16h | 7.778578 | 0.346403093 | 0.224 | 0.074 | 1.9761479 | 16 | supt16h |
| smu1a | 8.408999 | 0.329354671 | 0.216 | 0.07 | 2.1363063 | 16 | smu1a |
| ing5b1 | 8.412494 | 0.261907314 | 0.164 | 0.043 | 2.1371942 | 16 | ing5b |
| hnrnpd | 8.912640 | 0.336429769 | 0.196 | 0.061 | 2.2642563 | 16 | hnrnpd |
| mtap | 9.443119 | 0.38479745 | 0.244 | 0.087 | 2.3990245 | 16 | mtap |
| CABZ01005379.1 | 1.047112 | 0.277893921 | 0.16 | 0.043 | 2.6601894 | 16 | CABZ01005379.1 |
| atp5mc11 | 1.111780 | 0.465348291 | 0.568 | 0.327 | 2.8244779 | 16 | atp5mc1 |
| swt1 | 1.231603 | 0.297262793 | 0.152 | 0.04 | 3.1288874 | 16 | swt1 |
| atp5fa1 | 1.233870 | 0.466029367 | 0.588 | 0.368 | 3.1346487 | 16 | atp5fa1 |
| uqcc2 | 1.315688 | 0.355824368 | 0.208 | 0.068 | 3.3425069 | 16 | uqcc2 |
| cstf2 | 1.400039 | 0.340605738 | 0.22 | 0.073 | 3.5568011 | 16 | cstf2 |
| uqcrc11 | 1.420257 | 0.414624502 | 0.272 | 0.107 | 3.6081637 | 16 | uqcrc1 |
| psmb6 | 1.495198 | 0.436621981 | 0.392 | 0.186 | 3.7985518 | 16 | psmb6 |
| dctpp15 | 1.544588 | 0.391697736 | 0.364 | 0.147 | 3.9240264 | 16 | dctpp1 |
| si:ch211-284b7.3 | 1.579948 | 0.344070662 | 0.18 | 0.054 | 4.0138595 | 16 | si:ch211-284b7.3 |
| atp5f1c | 1.704137 | 0.488162879 | 0.476 | 0.267 | 4.3293608 | 16 | atp5f1c |
| myl64 | 1.829212 | 0.398617907 | 0.292 | 0.115 | 4.6471137 | 16 | myl6 |
| cct3 | 1.969588 | 0.451221194 | 0.436 | 0.223 | 5.0037395 | 16 | cct3 |
| polr2d | 2.047473 | 0.428918153 | 0.324 | 0.14 | 5.2016074 | 16 | polr2d |

| | | | | | | | |
|---|---|---|---|---|---|---|---|
| polr1c | 2.073333 | 0.267141322 | 0.148 | 0.038 | 5.2673047 | 16 | polr1c |
| ddx612 | 2.107952 | 0.490334606 | 0.436 | 0.233 | 5.3552541 | 16 | ddx61 |
| rpsa6 | 2.245651 | 0.285584427 | 0.948 | 0.784 | 5.7050777 | 16 | rpsa |
| fam168b | 2.406060 | 0.340882805 | 0.232 | 0.079 | 6.1125973 | 16 | fam168b |
| flot1b1 | 2.490308 | 0.388673535 | 0.26 | 0.099 | 6.3266281 | 16 | flot1b |
| hmgb2a2 | 3.348347 | 0.385184172 | 0.8 | 0.608 | 8.5064769 | 16 | hmgb2a |
| psma6a | 3.429054 | 0.356508485 | 0.284 | 0.109 | 8.7115121 | 16 | psma6a |
| sap182 | 3.533131 | 0.468178091 | 0.432 | 0.219 | 8.9759216 | 16 | sap18 |
| ndufb7 | 3.822668 | 0.469336502 | 0.388 | 0.193 | 9.7114895 | 16 | ndufb7 |
| alg131 | 3.857938 | 0.303442298 | 0.184 | 0.053 | 9.8010931 | 16 | alg13 |
| rchy1 | 3.925197 | 0.325110539 | 0.216 | 0.07 | 9.9719652 | 16 | rchy1 |
| mcts1 | 4.045035 | 0.405075554 | 0.284 | 0.113 | 1.0276413 | 16 | mcts1 |
| prmt5 | 4.261764 | 0.297762495 | 0.164 | 0.046 | 1.0827013 | 16 | prmt5 |
| mrpl14 | 4.348435 | 0.357439281 | 0.264 | 0.097 | 1.1047200 | 16 | mrpl14 |
| chrac13 | 6.325977 | 0.399410944 | 0.304 | 0.121 | 1.6071146 | 16 | chrac1 |
| si:ch211-68a17.7 | 6.368787 | 0.285653433 | 0.172 | 0.05 | 1.6179903 | 16 | si:ch211-68a17.7 |
| rtca1 | 7.031804 | 0.371584115 | 0.252 | 0.092 | 1.7864299 | 16 | rtca |
| lin37 | 7.120707 | 0.269681876 | 0.144 | 0.037 | 1.8090157 | 16 | lin37 |
| si:ch211-286b5.53 | 7.389478 | 0.429058007 | 0.336 | 0.149 | 1.8772970 | 16 | si:ch211-286b5.5 |
| ilf3b | 7.542334 | 0.374670765 | 0.288 | 0.113 | 1.9161301 | 16 | ilf3b |
| rps19bp1 | 8.138617 | 0.361896842 | 0.248 | 0.09 | 2.0676158 | 16 | rps19bp1 |
| polr2h | 9.214935 | 0.312228256 | 0.208 | 0.067 | 2.3410544 | 16 | polr2h |
| trappc1 | 1.016488 | 0.422794242 | 0.28 | 0.115 | 2.5823892 | 16 | trappc1 |
| sympk | 1.217836 | 0.253906335 | 0.128 | 0.031 | 3.0939127 | 16 | sympk |
| gnpda11 | 1.249472 | 0.284892712 | 0.16 | 0.045 | 3.1742847 | 16 | gnpda1 |
| lyrm9 | 1.377629 | 0.285522837 | 0.192 | 0.059 | 3.4998668 | 16 | lyrm9 |
| maco1a | 1.391991 | 0.303440778 | 0.156 | 0.044 | 3.5363544 | 16 | maco1a |
| dynll15 | 1.622283 | 0.443715487 | 0.652 | 0.439 | 4.1214117 | 16 | dynll1 |
| cope | 1.874372 | 0.413421966 | 0.328 | 0.144 | 4.7618443 | 16 | cope |
| psmd10 | 1.985335 | 0.306656566 | 0.216 | 0.071 | 5.0437445 | 16 | psmd10 |
| polr3h | 2.128695 | 0.278353961 | 0.148 | 0.04 | 5.4079496 | 16 | polr3h |
| sdhc | 2.239213 | 0.398062503 | 0.332 | 0.146 | 5.6887228 | 16 | sdhc |
| dynll2a5 | 2.458547 | 0.368902957 | 0.3 | 0.116 | 6.2459406 | 16 | dynll2a |
| ndufv2 | 2.647645 | 0.341256584 | 0.224 | 0.079 | 6.7263441 | 16 | ndufv2 |
| ndufb2 | 2.792049 | 0.458130016 | 0.388 | 0.193 | 7.0932022 | 16 | ndufb2 |

| | | | | | | | |
|---|---|---|---|---|---|---|---|
| lyar | 3.383690 | 0.309098479 | 0.18 | 0.056 | 8.5962666 | 16 | lyar |
| ncln | 3.531570 | 0.29771229 | 0.164 | 0.048 | 8.9719544 | 16 | ncln |
| e2f52 | 3.583278 | 0.327501413 | 0.192 | 0.06 | 9.1033193 | 16 | e2f5 |
| si:ch211-276i12.11 | 3.611435 | 0.29325041 | 0.2 | 0.065 | 9.1748511 | 16 | si:ch211-276i12.11 |
| gstm.13 | 3.822917 | 0.469234233 | 0.488 | 0.277 | 9.7121224 | 16 | gstm.1 |
| chchd2 | 3.890221 | 0.350217652 | 0.22 | 0.078 | 9.8831079 | 16 | chchd2 |
| rbbp43 | 4.485233 | 0.412731754 | 0.364 | 0.161 | 1.1394736 | 16 | rbbp4 |
| prpf39 | 4.543398 | 0.344882879 | 0.204 | 0.07 | 1.1542502 | 16 | prpf39 |
| atp5mf2 | 5.857863 | 0.453641457 | 0.496 | 0.287 | 1.4881901 | 16 | atp5mf |
| mob1bb | 5.884928 | 0.305079174 | 0.196 | 0.063 | 1.4950661 | 16 | mob1bb |
| usp5 | 6.090882 | 0.274729534 | 0.152 | 0.043 | 1.5473886 | 16 | usp5 |
| ndufa3 | 6.092407 | 0.434306357 | 0.384 | 0.19 | 1.5477761 | 16 | ndufa3 |
| lmo4a | 9.943631 | 0.259570072 | 0.136 | 0.035 | 2.5261796 | 16 | lmo4a |
| rrp36 | 1.091251 | 0.284563736 | 0.16 | 0.047 | 2.7723233 | 16 | rrp36 |
| brk11 | 1.096510 | 0.456623851 | 0.376 | 0.19 | 2.7856858 | 16 | brk1 |
| cnot9 | 1.113122 | 0.326749482 | 0.192 | 0.064 | 2.8278881 | 16 | cnot9 |
| uba1 | 1.165878 | 0.388114314 | 0.28 | 0.117 | 2.9619141 | 16 | uba1 |
| sod2 | 1.236972 | 0.393358987 | 0.312 | 0.135 | 3.1425288 | 16 | sod2 |
| tuba8l22 | 1.472365 | 0.320280871 | 0.2 | 0.068 | 3.7405445 | 16 | tuba8l2 |
| rfc2 | 1.492913 | 0.268912812 | 0.152 | 0.043 | 3.7927465 | 16 | rfc2 |
| ndufs4 | 1.498743 | 0.378658066 | 0.268 | 0.109 | 3.8075567 | 16 | ndufs4 |
| cks1b | 1.580309 | 0.275620113 | 0.192 | 0.061 | 4.0147757 | 16 | cks1b |
| baz1b | 1.623332 | 0.342038243 | 0.204 | 0.071 | 4.1240756 | 16 | baz1b |
| COX5B2 | 1.772996 | 0.43540595 | 0.568 | 0.375 | 4.5042981 | 16 | COX5B |
| gins2 | 1.783644 | 0.309960077 | 0.192 | 0.061 | 4.5313480 | 16 | gins2 |
| zswim5 | 1.807254 | 0.278195004 | 0.148 | 0.042 | 4.5913301 | 16 | zswim5 |
| si:ch211-276i12.9 | 2.074515 | 0.292093461 | 0.188 | 0.06 | 5.2703068 | 16 | si:ch211-276i12.9 |
| zgc:158803 | 2.082352 | 0.304784231 | 0.172 | 0.054 | 5.2902159 | 16 | zgc:158803 |
| atp5mc3a1 | 2.172133 | 0.458084396 | 0.432 | 0.239 | 5.5183045 | 16 | atp5mc3a |
| hdac1 | 2.380912 | 0.437500209 | 0.424 | 0.225 | 6.0487089 | 16 | hdac1 |
| atp5f1b | 2.545828 | 0.396498593 | 0.672 | 0.491 | 6.4676782 | 16 | atp5f1b |
| nxt2 | 2.659561 | 0.343416929 | 0.232 | 0.084 | 6.7566155 | 16 | nxt2 |
| hspd12 | 3.210532 | 0.404685457 | 0.352 | 0.162 | 8.1563569 | 16 | hspd1 |
| khdrbs1a | 3.235797 | 0.364470965 | 0.728 | 0.561 | 8.2205446 | 16 | khdrbs1a |
| txn2 | 3.903636 | 0.371969101 | 0.24 | 0.093 | 9.9171879 | 16 | txn2 |

| | | | | | | | |
|---|---|---|---|---|---|---|---|
| bida1 | 3.936811 | 0.339265635 | 0.204 | 0.071 | 1.0001470 | 16 | bida |
| slirp | 4.054889 | 0.387536117 | 0.296 | 0.127 | 1.0301446 | 16 | slirp |
| zgc:103697 | 4.717536 | 0.314233217 | 0.18 | 0.058 | 1.1984901 | 16 | zgc:103697 |
| ndufa1 | 4.880030 | 0.449567812 | 0.432 | 0.24 | 1.2397717 | 16 | ndufa1 |
| orc3 | 5.127079 | 0.255033246 | 0.156 | 0.045 | 1.3025345 | 16 | orc3 |
| pho | 5.593390 | 0.395623985 | 0.24 | 0.094 | 1.4210009 | 16 | pho |
| ing1 | 5.690833 | 0.250027233 | 0.12 | 0.03 | 1.4457562 | 16 | ing1 |
| sephs2 | 6.739062 | 0.266834869 | 0.164 | 0.05 | 1.7120587 | 16 | sephs2 |
| rexo4 | 6.955554 | 0.305583732 | 0.2 | 0.068 | 1.7670586 | 16 | rexo4 |
| bola3 | 7.511868 | 0.264627268 | 0.196 | 0.065 | 1.9083902 | 16 | bola3 |
| med20 | 7.632166 | 0.250735601 | 0.144 | 0.041 | 1.9389520 | 16 | med20 |
| prpf8 | 8.371321 | 0.299136333 | 0.176 | 0.057 | 2.1267343 | 16 | prpf8 |
| ndufb10 | 8.861572 | 0.427320813 | 0.336 | 0.162 | 2.2512824 | 16 | ndufb10 |
| hspe12 | 9.272370 | 0.449570451 | 0.508 | 0.289 | 2.3556458 | 16 | hspe1 |
| ppme1 | 9.641559 | 0.275301519 | 0.16 | 0.049 | 2.4494382 | 16 | ppme1 |
| c1qbp1 | 1.006826 | 0.423750741 | 0.348 | 0.166 | 2.5578426 | 16 | c1qbp |
| psmd13 | 1.076573 | 0.345524372 | 0.244 | 0.095 | 2.7350355 | 16 | psmd13 |
| snw1 | 1.316258 | 0.343477757 | 0.216 | 0.08 | 3.3439539 | 16 | snw1 |
| nkap | 1.362066 | 0.373968848 | 0.28 | 0.119 | 3.4603299 | 16 | nkap |
| fbl1 | 1.793436 | 0.430089711 | 0.392 | 0.2 | 4.5562250 | 16 | fbl |
| rpia | 1.884005 | 0.318199978 | 0.192 | 0.066 | 4.7863169 | 16 | rpia |
| pes | 1.933023 | 0.301375896 | 0.16 | 0.05 | 4.9108465 | 16 | pes |
| mrps21 | 1.946231 | 0.330846196 | 0.232 | 0.088 | 4.9444003 | 16 | mrps21 |
| eny2 | 2.027189 | 0.364311906 | 0.268 | 0.111 | 5.1500744 | 16 | eny2 |
| vcp | 2.182234 | 0.40562588 | 0.308 | 0.14 | 5.5439664 | 16 | vcp |
| mrpl46 | 2.238775 | 0.277828418 | 0.14 | 0.04 | 5.6876087 | 16 | mrpl46 |
| rbm19 | 2.312988 | 0.328841532 | 0.168 | 0.055 | 5.8761483 | 16 | rbm19 |
| phf5a | 3.063235 | 0.387085344 | 0.344 | 0.163 | 7.7821492 | 16 | phf5a |
| csnk2a1 | 3.090524 | 0.438461602 | 0.364 | 0.185 | 7.8514776 | 16 | csnk2a1 |
| cox8b2 | 3.114215 | 0.394812442 | 0.244 | 0.1 | 7.9116635 | 16 | cox8b |
| exosc8 | 3.533631 | 0.292574284 | 0.196 | 0.067 | 8.9771917 | 16 | exosc8 |
| si:dkey-180p18.91 | 3.942473 | 0.369447844 | 0.256 | 0.107 | 1.0015854 | 16 | si:dkey-180p18.9 |
| pfdn2 | 4.703570 | 0.380680236 | 0.292 | 0.129 | 1.1949421 | 16 | pfdn2 |
| psma3 | 5.110387 | 0.429695847 | 0.368 | 0.191 | 1.2982938 | 16 | psma3 |
| trnau1apa | 5.292543 | 0.255176706 | 0.16 | 0.049 | 1.3445707 | 16 | trnau1apa |

| | | | | | | | |
|---|---|---|---|---|---|---|---|
| rad23b | 5.343231 | 0.411664705 | 0.356 | 0.179 | 1.3574479 | 16 | rad23b |
| mrpl30 | 6.025412 | 0.353901307 | 0.264 | 0.111 | 1.5307561 | 16 | mrpl30 |
| ube2d1b | 6.088906 | 0.382809996 | 0.308 | 0.142 | 1.5468866 | 16 | ube2d1b |
| pcnp | 6.092486 | 0.271300743 | 0.16 | 0.05 | 1.5477961 | 16 | pcnp |
| taf11 | 7.149695 | 0.263023539 | 0.156 | 0.049 | 1.8163800 | 16 | taf11 |
| gtf2f1 | 7.226613 | 0.305775902 | 0.196 | 0.07 | 1.8359212 | 16 | gtf2f1 |
| cct6a | 7.231217 | 0.422631713 | 0.4 | 0.219 | 1.8370907 | 16 | cct6a |
| imp34 | 7.685241 | 0.350612864 | 0.292 | 0.122 | 1.9524357 | 16 | imp3 |
| mad2l15 | 8.107237 | 0.337316952 | 0.36 | 0.157 | 2.0596437 | 16 | mad2l1 |
| rnaseh2b | 9.561038 | 0.26780303 | 0.164 | 0.052 | 2.4289818 | 16 | rnaseh2b |
| api5 | 1.043147 | 0.324081297 | 0.228 | 0.087 | 2.6501172 | 16 | api5 |
| srsf5a3 | 1.052582 | 0.413100941 | 0.492 | 0.302 | 2.6740859 | 16 | srsf5a |
| cox5ab | 1.071738 | 0.408759907 | 0.4 | 0.216 | 2.7227512 | 16 | cox5ab |
| phf20b | 1.101602 | 0.270395118 | 0.16 | 0.051 | 2.7986217 | 16 | phf20b |
| FQ976913.1 | 1.171947 | 0.250122424 | 0.136 | 0.039 | 2.9773315 | 16 | FQ976913.1 |
| atp5pb1 | 1.194008 | 0.429789921 | 0.512 | 0.331 | 3.0333789 | 16 | atp5pb |
| ran1 | 1.555092 | 0.29059868 | 0.764 | 0.639 | 3.9507124 | 16 | ran |
| hnrnpabb5 | 1.618718 | 0.371350094 | 0.748 | 0.557 | 4.1123555 | 16 | hnrnpabb |
| btr04 | 1.659960 | 0.257699157 | 0.168 | 0.053 | 4.2171288 | 16 | btr04 |
| ndufs2 | 1.741345 | 0.331278145 | 0.236 | 0.095 | 4.4238878 | 16 | ndufs2 |
| aldh7a11 | 1.812636 | 0.294166169 | 0.192 | 0.069 | 4.6050041 | 16 | aldh7a1 |
| mcm5 | 2.014555 | 0.323827981 | 0.236 | 0.091 | 5.1179791 | 16 | mcm5 |
| tcea2 | 2.266267 | 0.252529208 | 0.152 | 0.047 | 5.7574534 | 16 | tcea2 |
| mrpl53 | 2.270410 | 0.318417346 | 0.224 | 0.086 | 5.7679774 | 16 | mrpl53 |
| glrx3 | 2.624311 | 0.29223741 | 0.188 | 0.067 | 6.6670639 | 16 | glrx3 |
| polr2c | 2.710895 | 0.261521459 | 0.172 | 0.057 | 6.8870293 | 16 | polr2c |
| tuba1b | 3.131988 | 0.250420838 | 0.132 | 0.038 | 7.9568173 | 16 | tuba1b |
| dnajc8 | 3.526683 | 0.317195903 | 0.2 | 0.074 | 8.9595395 | 16 | dnajc8 |
| trappc3 | 3.599681 | 0.253576522 | 0.148 | 0.046 | 9.1449914 | 16 | trappc3 |
| ndufa11 | 4.154681 | 0.389767964 | 0.328 | 0.162 | 1.0554968 | 16 | ndufa11 |
| sem1 | 4.268641 | 0.418486673 | 0.456 | 0.272 | 1.0844484 | 16 | sem1 |
| myg1 | 4.922731 | 0.306122146 | 0.192 | 0.071 | 1.2506199 | 16 | myg1 |
| u2af2a | 5.157847 | 0.257518325 | 0.172 | 0.058 | 1.3103511 | 16 | u2af2a |
| esd | 5.228557 | 0.352822828 | 0.256 | 0.11 | 1.3283150 | 16 | esd |
| fam133b1 | 5.488486 | 0.380810333 | 0.32 | 0.155 | 1.3943500 | 16 | fam133b |

| | | | | | | | |
|---|---|---|---|---|---|---|---|
| atp5meb1 | 6.312726 | 0.424636995 | 0.508 | 0.338 | 1.6037482 | 16 | atp5meb |
| rpp14 | 6.613339 | 0.250975978 | 0.16 | 0.052 | 1.6801188 | 16 | rpp14 |
| smndc1 | 8.556208 | 0.367802189 | 0.268 | 0.12 | 2.1737047 | 16 | smndc1 |
| tuba8l4 | 8.833796 | 0.407512995 | 0.612 | 0.413 | 2.2442259 | 16 | tuba8l4 |
| srpk1a | 9.061891 | 0.28050659 | 0.184 | 0.065 | 2.3021736 | 16 | srpk1a |
| si:dkeyp-55f12.3 | 9.235354 | 0.254389261 | 0.168 | 0.057 | 2.3462418 | 16 | si:dkeyp-55f12.3 |
| ing2 | 9.510981 | 0.273476422 | 0.196 | 0.07 | 2.4162649 | 16 | ing2 |
| nudcd2 | 9.657104 | 0.284054568 | 0.184 | 0.066 | 2.4533873 | 16 | nudcd2 |
| pabpc1b3 | 9.943802 | 0.320787374 | 0.2 | 0.076 | 2.5262229 | 16 | pabpc1b |
| mrpl13 | 1.019249 | 0.285565827 | 0.168 | 0.058 | 2.5894022 | 16 | mrpl13 |
| mrpl11 | 1.026519 | 0.307067216 | 0.192 | 0.072 | 2.6078739 | 16 | mrpl11 |
| pa2g4a | 1.066020 | 0.312161033 | 0.284 | 0.124 | 2.7082245 | 16 | pa2g4a |
| rnaseh2a | 1.173976 | 0.25831294 | 0.152 | 0.049 | 2.9824860 | 16 | rnaseh2a |
| safb | 1.357717 | 0.279734503 | 0.168 | 0.058 | 3.4492819 | 16 | safb |
| akap1a1 | 1.412931 | 0.258901964 | 0.156 | 0.049 | 3.5895529 | 16 | akap1a |
| ubap2b | 1.672627 | 0.306859687 | 0.216 | 0.085 | 4.2493090 | 16 | ubap2b |
| si:dkey-188i13.115 | 1.963902 | 0.404189212 | 0.508 | 0.289 | 4.9892937 | 16 | si:dkey-188i13.11 |
| zgc:123105 | 2.093623 | 0.250455548 | 0.148 | 0.047 | 5.3188496 | 16 | zgc:123105 |
| psma2 | 2.197494 | 0.377292267 | 0.324 | 0.163 | 5.5827341 | 16 | psma2 |
| atp5l3 | 2.228371 | 0.358503196 | 0.624 | 0.456 | 5.6611786 | 16 | atp5l |
| fhit | 2.251066 | 0.262184741 | 0.152 | 0.049 | 5.7188337 | 16 | fhit |
| zgc:56304 | 3.053263 | 0.336983586 | 0.268 | 0.121 | 7.7568163 | 16 | zgc:56304 |
| ISCU (1 of many) | 3.085262 | 0.344808335 | 0.272 | 0.123 | 7.8381099 | 16 | ISCU (1 of many) |
| rpl23a6 | 3.186308 | 0.286558946 | 0.832 | 0.668 | 8.0948172 | 16 | rpl23a |
| ptp4a11 | 3.309859 | 0.338850282 | 0.28 | 0.131 | 8.4086968 | 16 | ptp4a1 |
| u2af2b | 3.545979 | 0.347399062 | 0.276 | 0.126 | 9.0085612 | 16 | u2af2b |
| ndufb81 | 3.803433 | 0.387108155 | 0.448 | 0.264 | 9.6626230 | 16 | ndufb8 |
| gapdhs7 | 4.204378 | 0.412203726 | 0.48 | 0.314 | 1.0681224 | 16 | gapdhs |
| lsm12a3 | 4.834914 | 0.300987252 | 0.268 | 0.115 | 1.2283100 | 16 | lsm12a |
| aurkb4 | 4.855469 | 0.271845871 | 0.268 | 0.11 | 1.2335320 | 16 | aurkb |
| abcf1 | 4.999864 | 0.329277917 | 0.248 | 0.108 | 1.2702154 | 16 | abcf1 |
| h3f3a5 | 5.186005 | 0.376681873 | 0.688 | 0.481 | 1.3175046 | 16 | h3f3a |
| mrps18c1 | 5.454015 | 0.314559365 | 0.264 | 0.115 | 1.3855925 | 16 | mrps18c |
| polr1d | 5.612026 | 0.276891532 | 0.216 | 0.084 | 1.4257352 | 16 | polr1d |
| actl6a | 6.134711 | 0.265486937 | 0.18 | 0.065 | 1.5585235 | 16 | actl6a |

| | | | | | | | |
|---|---|---|---|---|---|---|---|
| lig1 | 6.519617 | 0.290904895 | 0.168 | 0.059 | 1.6563089 | 16 | lig1 |
| ube2ia | 6.612444 | 0.37132125 | 0.32 | 0.159 | 1.6798914 | 16 | ube2ia |
| snrpb1 | 7.221950 | 0.399281778 | 0.532 | 0.355 | 1.8347365 | 16 | snrpb |
| aprt | 7.380356 | 0.297685996 | 0.212 | 0.083 | 1.8749795 | 16 | aprt |
| nol12 | 8.029724 | 0.282083118 | 0.196 | 0.074 | 2.0399515 | 16 | nol12 |
| fam162a | 8.954240 | 0.296399772 | 0.216 | 0.088 | 2.2748246 | 16 | fam162a |
| cct41 | 9.699490 | 0.393451159 | 0.416 | 0.244 | 2.4641556 | 16 | cct4 |
| lsm14ab | 9.952711 | 0.286657557 | 0.2 | 0.079 | 2.5284862 | 16 | lsm14ab |
| sf3b2 | 1.048198 | 0.313141696 | 0.208 | 0.084 | 2.6629471 | 16 | sf3b2 |
| npm3 | 1.067123 | 0.34945453 | 0.26 | 0.119 | 2.7110283 | 16 | npm3 |
| uqcrh2 | 1.220361 | 0.390888407 | 0.432 | 0.259 | 3.1003276 | 16 | uqcrh |
| gpn2 | 1.284862 | 0.258836316 | 0.156 | 0.053 | 3.2641924 | 16 | gpn2 |
| si:dkey-67c22.2 | 1.343006 | 0.310047551 | 0.228 | 0.097 | 3.4119082 | 16 | si:dkey-67c22.2 |
| eif3ja | 1.410380 | 0.400768468 | 0.356 | 0.196 | 3.5830713 | 16 | eif3ja |
| si:zfos-169g10.3 | 1.456113 | 0.35979661 | 0.248 | 0.114 | 3.6992561 | 16 | si:zfos-169g10.3 |
| dcaf13 | 1.494865 | 0.252079333 | 0.152 | 0.051 | 3.7977067 | 16 | dcaf13 |
| ufd1l | 1.615299 | 0.28285649 | 0.176 | 0.066 | 4.1036693 | 16 | ufd1l |
| sae1 | 1.696386 | 0.250771556 | 0.148 | 0.049 | 4.3096695 | 16 | sae1 |
| emc9 | 1.733471 | 0.258736099 | 0.176 | 0.063 | 4.4038834 | 16 | emc9 |
| psip1b | 1.960526 | 0.311854517 | 0.184 | 0.071 | 4.9807176 | 16 | psip1b |
| nfr4 | 2.216248 | 0.253395263 | 0.232 | 0.091 | 5.6303788 | 16 | nfr |
| trir | 2.239523 | 0.287076591 | 0.164 | 0.059 | 5.6895095 | 16 | trir |
| zgc:86598 | 2.414584 | 0.320080973 | 0.252 | 0.114 | 6.1342525 | 16 | zgc:86598 |
| arl14ep | 2.720073 | 0.265182611 | 0.196 | 0.075 | 6.9103464 | 16 | arl14ep |
| chmp5b | 2.803362 | 0.352415963 | 0.268 | 0.127 | 7.1219426 | 16 | chmp5b |
| tsr2 | 2.887075 | 0.262669123 | 0.216 | 0.086 | 7.3346154 | 16 | tsr2 |
| rrp8 | 2.941005 | 0.294452565 | 0.18 | 0.068 | 7.4716250 | 16 | rrp8 |
| ywhabb1 | 3.123528 | 0.384213656 | 0.42 | 0.252 | 7.9353232 | 16 | ywhabb |
| suclg12 | 4.346466 | 0.327713216 | 0.26 | 0.121 | 1.1042197 | 16 | suclg1 |
| nop2 | 4.491175 | 0.264894875 | 0.152 | 0.054 | 1.1409831 | 16 | nop2 |
| mapre1a3 | 4.587402 | 0.355157348 | 0.372 | 0.195 | 1.1654296 | 16 | mapre1a |
| pdcd4a | 4.788033 | 0.263277012 | 0.16 | 0.058 | 1.2163998 | 16 | pdcd4a |
| bccip1 | 5.113566 | 0.358183435 | 0.292 | 0.148 | 1.2991014 | 16 | bccip |
| cox6c1 | 5.130588 | 0.370977075 | 0.572 | 0.41 | 1.3034259 | 16 | cox6c |
| ss18 | 5.335548 | 0.286913796 | 0.184 | 0.072 | 1.3554959 | 16 | ss18 |

| | | | | | | | |
|---|---|---|---|---|---|---|---|
| polr2k4 | 6.324987 | 0.356624141 | 0.344 | 0.175 | 1.6068630 | 16 | polr2k |
| eloca1 | 6.568470 | 0.294689877 | 0.224 | 0.094 | 1.6687198 | 16 | eloca |
| h2afva4 | 6.662061 | 0.376025378 | 0.5 | 0.304 | 1.6924967 | 16 | h2afva |
| tufm | 6.803505 | 0.274100276 | 0.212 | 0.088 | 1.7284306 | 16 | tufm |
| rbm25b | 7.311070 | 0.251349548 | 0.156 | 0.056 | 1.8573773 | 16 | rbm25b |
| mrps11 | 7.607331 | 0.299508169 | 0.184 | 0.073 | 1.9326426 | 16 | mrps11 |
| ccdc12 | 8.089309 | 0.264510114 | 0.184 | 0.072 | 2.0550891 | 16 | ccdc12 |
| tomm22 | 8.308077 | 0.29252787 | 0.22 | 0.094 | 2.1106670 | 16 | tomm22 |
| hnrnpl2 | 8.570642 | 0.32340692 | 0.264 | 0.123 | 2.1773717 | 16 | hnrnpl2 |
| kdm5c | 9.655027 | 0.250320124 | 0.148 | 0.052 | 2.4528596 | 16 | kdm5c |
| tnrc6b | 9.852379 | 0.299464246 | 0.204 | 0.085 | 2.5029971 | 16 | tnrc6b |
| smn1 | 1.005561 | 0.272969681 | 0.212 | 0.086 | 2.5546277 | 16 | smn1 |
| rrs1 | 1.045157 | 0.294235141 | 0.204 | 0.084 | 2.6552226 | 16 | rrs1 |
| polr2i | 1.095805 | 0.25990152 | 0.18 | 0.069 | 2.7838945 | 16 | polr2i |
| cops7a | 1.223378 | 0.265697992 | 0.168 | 0.063 | 3.1079933 | 16 | cops7a |
| pfn2l3 | 1.285590 | 0.348465746 | 0.388 | 0.214 | 3.2660433 | 16 | pfn2l |
| znf593 | 1.300810 | 0.306752944 | 0.256 | 0.117 | 3.3047089 | 16 | znf593 |
| med10 | 1.518327 | 0.32128306 | 0.308 | 0.155 | 3.8573104 | 16 | med10 |
| mpc1 | 1.618443 | 0.26464937 | 0.184 | 0.073 | 4.1116557 | 16 | mpc1 |
| atp5pf3 | 1.800362 | 0.374732831 | 0.476 | 0.319 | 4.5738207 | 16 | atp5pf |
| naa20 | 1.822806 | 0.26781546 | 0.176 | 0.07 | 4.6308386 | 16 | naa20 |
| mrpl17 | 1.901506 | 0.264690161 | 0.188 | 0.074 | 4.8307761 | 16 | mrpl17 |
| ranbp11 | 1.917739 | 0.384415559 | 0.448 | 0.281 | 4.8720160 | 16 | ranbp1 |
| dkc12 | 2.184835 | 0.355284652 | 0.332 | 0.173 | 5.5505738 | 16 | dkc1 |
| ndufb9 | 2.269384 | 0.347277987 | 0.356 | 0.199 | 5.7653719 | 16 | ndufb9 |
| rbmx | 2.660450 | 0.318049801 | 0.288 | 0.141 | 6.7588751 | 16 | rbmx |
| si:dkey-46a10.3 | 3.070149 | 0.295664053 | 0.188 | 0.077 | 7.7997158 | 16 | si:dkey-46a10.3 |
| cox4i13 | 3.101885 | 0.335247146 | 0.58 | 0.416 | 7.8803397 | 16 | cox4i1 |
| tktb2 | 3.272073 | 0.320269277 | 0.32 | 0.163 | 8.3127034 | 16 | tktb |
| cdc65 | 4.601848 | 0.260789807 | 0.212 | 0.087 | 1.1690995 | 16 | cdc6 |
| psmd12 | 4.871775 | 0.262616999 | 0.172 | 0.067 | 1.2376746 | 16 | psmd12 |
| nop10 | 5.549597 | 0.361884672 | 0.4 | 0.238 | 1.4098752 | 16 | nop10 |
| mcm41 | 5.978624 | 0.265985053 | 0.196 | 0.08 | 1.5188696 | 16 | mcm4 |
| nap1l4a | 5.981083 | 0.252467281 | 0.172 | 0.067 | 1.5194943 | 16 | nap1l4a |
| timm132 | 6.187106 | 0.276152899 | 0.224 | 0.096 | 1.5718343 | 16 | timm13 |

| | | | | | | | |
|---|---|---|---|---|---|---|---|
| mrpl57 | 7.150845 | 0.325514407 | 0.252 | 0.121 | 1.8166721 | 16 | mrpl57 |
| wdr121 | 7.489676 | 0.259585588 | 0.156 | 0.059 | 1.9027523 | 16 | wdr12 |
| sumo3a | 8.292775 | 0.359927322 | 0.532 | 0.37 | 2.1067797 | 16 | sumo3a |
| cox14 | 8.473995 | 0.285842409 | 0.236 | 0.109 | 2.1528186 | 16 | cox14 |
| si:ch211-282b22.1 | 1.051857 | 0.293840093 | 0.224 | 0.1 | 2.6722449 | 16 | si:ch211-282b22.1 |
| brdt | 1.140165 | 0.288466629 | 0.204 | 0.088 | 2.8965914 | 16 | brdt |
| naa10 | 1.234269 | 0.380033828 | 0.376 | 0.229 | 3.1356620 | 16 | naa10 |
| h2afy22 | 1.300969 | 0.341759727 | 0.244 | 0.122 | 3.3051130 | 16 | h2afy2 |
| ppp2r2d | 1.440484 | 0.256018517 | 0.152 | 0.058 | 3.6595512 | 16 | ppp2r2d |
| uqcrq2 | 1.482977 | 0.346602522 | 0.448 | 0.286 | 3.7675053 | 16 | uqcrq |
| hmgn31 | 1.604228 | 0.262520397 | 0.184 | 0.077 | 4.0755416 | 16 | hmgn3 |
| si:ch211-214j24.10 | 1.654235 | 0.338475149 | 0.296 | 0.157 | 4.2025863 | 16 | si:ch211-214j24.10 |
| slc25a3b3 | 1.967102 | 0.310815099 | 0.604 | 0.457 | 4.9974245 | 16 | slc25a3b |
| ube2v22 | 1.989533 | 0.356078624 | 0.44 | 0.281 | 5.0544090 | 16 | ube2v2 |
| ube2a | 2.010454 | 0.259866743 | 0.188 | 0.079 | 5.1075584 | 16 | ube2a |
| ube2e1 | 2.193519 | 0.329770222 | 0.352 | 0.198 | 5.5726363 | 16 | ube2e1 |
| srrm11 | 2.282921 | 0.303567786 | 0.248 | 0.121 | 5.7997632 | 16 | srrm1 |
| psmc3 | 2.356062 | 0.269610356 | 0.236 | 0.109 | 5.9855755 | 16 | psmc3 |
| ddx39aa | 2.372134 | 0.32710257 | 0.312 | 0.168 | 6.0264069 | 16 | ddx39aa |
| gtf2a2 | 2.450665 | 0.276861887 | 0.196 | 0.084 | 6.2259164 | 16 | gtf2a2 |
| ythdf2 | 2.494889 | 0.265742918 | 0.188 | 0.079 | 6.3382655 | 16 | ythdf2 |
| ahcy | 2.740188 | 0.322126266 | 0.3 | 0.157 | 6.9614484 | 16 | ahcy |
| srsf4 | 2.868025 | 0.274075262 | 0.22 | 0.099 | 7.2862199 | 16 | srsf4 |
| supt4h1 | 2.963156 | 0.297450517 | 0.208 | 0.095 | 7.5278992 | 16 | supt4h1 |
| polr2eb | 3.137617 | 0.281992585 | 0.2 | 0.089 | 7.9711171 | 16 | polr2eb |
| taf7 | 3.180027 | 0.25518413 | 0.152 | 0.059 | 8.0788600 | 16 | taf7 |
| cnbpa | 3.578630 | 0.348445721 | 0.368 | 0.216 | 9.0915117 | 16 | cnbpa |
| prdx23 | 3.654471 | 0.331177828 | 0.584 | 0.421 | 9.2841840 | 16 | prdx2 |
| h3f3c5 | 3.664946 | 0.343938736 | 0.464 | 0.303 | 9.3107972 | 16 | h3f3c |
| txnl4a1 | 4.444083 | 0.266811763 | 0.236 | 0.109 | 1.1290195 | 16 | txnl4a |
| vps4b | 4.939752 | 0.303738712 | 0.22 | 0.105 | 1.2549440 | 16 | vps4b |
| aimp1 | 5.026633 | 0.302648155 | 0.22 | 0.104 | 1.2770161 | 16 | aimp1 |
| timm17a | 5.264183 | 0.264029296 | 0.176 | 0.074 | 1.3373658 | 16 | timm17a |
| bxdc2 | 6.253046 | 0.267301599 | 0.212 | 0.096 | 1.5885865 | 16 | bxdc2 |
| mrpl47 | 6.786926 | 0.269756648 | 0.18 | 0.076 | 1.7242186 | 16 | mrpl47 |

| | | | | | | | |
|---|---|---|---|---|---|---|---|
| mrpl18 | 6.989733 | 0.272307364 | 0.228 | 0.107 | 1.7757418 | 16 | mrpl18 |
| paip2b1 | 8.089063 | 0.341331705 | 0.472 | 0.307 | 2.0550266 | 16 | paip2b |
| myl12.2 | 8.970235 | 0.302774565 | 0.248 | 0.122 | 2.2788883 | 16 | myl12.2 |
| ewsr1b | 9.104623 | 0.327158878 | 0.348 | 0.198 | 2.3130295 | 16 | ewsr1b |
| ddx27 | 9.461029 | 0.252142347 | 0.192 | 0.084 | 2.4035746 | 16 | ddx27 |
| park7 | 9.771760 | 0.267177659 | 0.188 | 0.082 | 2.4825158 | 16 | park7 |
| cct82 | 1.135518 | 0.355365508 | 0.36 | 0.222 | 2.8847854 | 16 | cct8 |
| txndc91 | 1.149084 | 0.318115957 | 0.248 | 0.128 | 2.9192495 | 16 | txndc9 |
| taf10 | 1.205289 | 0.253262035 | 0.172 | 0.072 | 3.0620386 | 16 | taf10 |
| oaz1a | 1.220606 | 0.296065459 | 0.628 | 0.496 | 3.1009518 | 16 | oaz1a |
| matr3l1.1 | 1.234689 | 0.2639121 | 0.188 | 0.083 | 3.1367294 | 16 | matr3l1.1 |
| pdap1a1 | 1.382412 | 0.272118178 | 0.244 | 0.118 | 3.5120178 | 16 | pdap1a |
| puf60a | 1.693173 | 0.27236184 | 0.228 | 0.109 | 4.3015078 | 16 | puf60a |
| wrb | 1.772214 | 0.250971413 | 0.168 | 0.07 | 4.5023104 | 16 | wrb |
| nhp2 | 1.983369 | 0.306802916 | 0.348 | 0.197 | 5.0387503 | 16 | nhp2 |
| ube2d2 | 2.197997 | 0.323025134 | 0.34 | 0.2 | 5.5840134 | 16 | ube2d2 |
| stip1 | 2.325264 | 0.270442555 | 0.228 | 0.11 | 5.9073346 | 16 | stip1 |
| chchd105 | 2.375314 | 0.287556294 | 0.268 | 0.133 | 6.0344863 | 16 | chchd10 |
| org5 | 2.419959 | 0.29856485 | 0.372 | 0.199 | 6.1479081 | 16 | org |
| adi1 | 2.497719 | 0.25412742 | 0.188 | 0.083 | 6.3454556 | 16 | adi1 |
| rwdd1 | 2.568908 | 0.306162514 | 0.264 | 0.139 | 6.5263129 | 16 | rwdd1 |
| wdr46 | 2.797682 | 0.262652555 | 0.168 | 0.071 | 7.1075115 | 16 | wdr46 |
| adh53 | 2.901307 | 0.358837069 | 0.376 | 0.242 | 7.3707726 | 16 | adh5 |
| si:rp71-45k5.4 | 3.049552 | 0.302773292 | 0.244 | 0.126 | 7.7473892 | 16 | si:rp71-45k5.4 |
| ndufa4l5 | 3.204383 | 0.308547544 | 0.556 | 0.422 | 8.1407371 | 16 | ndufa4l |
| sumo3b3 | 4.089752 | 0.278156349 | 0.236 | 0.114 | 0.000104 | 16 | sumo3b |
| hspa9 | 4.612985 | 0.259453275 | 0.22 | 0.105 | 0.000117 | 16 | hspa9 |
| eif6 | 4.919308 | 0.302373524 | 0.26 | 0.136 | 0.000125 | 16 | eif6 |
| tbca | 5.023478 | 0.323471705 | 0.348 | 0.211 | 0.000128 | 16 | tbca |
| rpl7l1 | 5.680496 | 0.273409211 | 0.256 | 0.13 | 0.000144 | 16 | rpl7l1 |
| ndufa51 | 7.564070 | 0.332340354 | 0.28 | 0.16 | 0.000192 | 16 | ndufa5 |
| grpel21 | 7.996401 | 0.269946584 | 0.216 | 0.107 | 0.000203 | 16 | grpel2 |
| fis1 | 8.525051 | 0.251356513 | 0.148 | 0.062 | 0.000217 | 16 | fis1 |
| elocb1 | 9.118797 | 0.320217606 | 0.324 | 0.195 | 0.000232 | 16 | elocb |
| psma1 | 1.069744 | 0.307147902 | 0.416 | 0.272 | 0.000272 | 16 | psma1 |

| | | | | | | | |
|---|---|---|---|---|---|---|---|
| UBL4A1 | 1.074122 | 0.274989708 | 0.236 | 0.117 | 0.000273 | 16 | UBL4A |
| zgc:1935411 | 1.091328 | 0.315270814 | 0.516 | 0.381 | 0.000277 | 16 | zgc:193541 |
| si:zfos-1192g2.3 | 1.164431 | 0.26225403 | 0.2 | 0.096 | 0.000296 | 16 | si:zfos-1192g2.3 |
| psma4 | 1.205814 | 0.317480035 | 0.32 | 0.191 | 0.000306 | 16 | psma4 |
| ndufb31 | 1.322551 | 0.334297842 | 0.372 | 0.238 | 0.000336 | 16 | ndufb3 |
| atp5if1a | 1.579811 | 0.31562052 | 0.496 | 0.344 | 0.000401 | 16 | atp5if1a |
| psmc4 | 1.665428 | 0.271562253 | 0.212 | 0.105 | 0.000423 | 16 | psmc4 |
| ost41 | 1.916388 | 0.298884133 | 0.412 | 0.261 | 0.000487 | 16 | ost4 |
| ywhaqa1 | 1.941481 | 0.307218494 | 0.344 | 0.209 | 0.000493 | 16 | ywhaqa |
| lamtor51 | 2.086114 | 0.283281161 | 0.252 | 0.133 | 0.00053 | 16 | lamtor5 |
| eif3k1 | 2.352979 | 0.323381428 | 0.372 | 0.244 | 0.000598 | 16 | eif3k |
| si:ch73-1a9.36 | 2.369470 | 0.333209577 | 0.448 | 0.315 | 0.000602 | 16 | si:ch73-1a9.3 |
| ywhae11 | 2.484528 | 0.309025716 | 0.432 | 0.294 | 0.000631 | 16 | ywhae1 |
| psmb4 | 3.436957 | 0.31509826 | 0.316 | 0.192 | 0.000873 | 16 | psmb4 |
| zmat2 | 5.113556 | 0.278409538 | 0.26 | 0.145 | 0.001299 | 16 | zmat2 |
| btf3l4 | 5.317715 | 0.271715064 | 0.264 | 0.146 | 0.001351 | 16 | btf3l4 |
| mrpl32 | 6.103196 | 0.258810847 | 0.208 | 0.105 | 0.001551 | 16 | mrpl32 |
| lyrm4 | 6.193124 | 0.25754499 | 0.208 | 0.104 | 0.001573 | 16 | lyrm4 |
| pabpn1l5 | 6.958027 | 0.300213577 | 0.388 | 0.235 | 0.001768 | 16 | pabpn1l |
| pgk1 | 8.090987 | 0.252619551 | 0.232 | 0.122 | 0.002056 | 16 | pgk1 |
| lsm81 | 8.735841 | 0.254303668 | 0.276 | 0.151 | 0.002219 | 16 | lsm8 |
| gpia | 1.072572 | 0.260092642 | 0.232 | 0.123 | 0.002725 | 16 | gpia |
| nop581 | 1.110089 | 0.290305029 | 0.304 | 0.18 | 0.00282 | 16 | nop58 |
| eif3g4 | 1.234150 | 0.300135268 | 0.42 | 0.3 | 0.003135 | 16 | eif3g |
| sinhcafl1 | 1.352332 | 0.262319782 | 0.236 | 0.125 | 0.003436 | 16 | sinhcafl |
| cycsb2 | 1.453486 | 0.283202712 | 0.468 | 0.313 | 0.003693 | 16 | cycsb |
| ndufs6 | 1.760978 | 0.273015924 | 0.28 | 0.162 | 0.004474 | 16 | ndufs6 |
| vdac21 | 1.823155 | 0.29254051 | 0.452 | 0.326 | 0.004632 | 16 | vdac2 |
| si:dkey-33c12.4 | 1.909568 | 0.270843968 | 0.312 | 0.184 | 0.004851 | 16 | si:dkey-33c12.4 |
| si:dkey-28b4.73 | 2.108844 | 0.257254446 | 0.208 | 0.109 | 0.005358 | 16 | si:dkey-28b4.7 |
| znf706 | 2.251885 | 0.264051968 | 0.308 | 0.183 | 0.005721 | 16 | znf706 |
| mdh1aa2 | 2.576454 | 0.301878391 | 0.348 | 0.229 | 0.006545 | 16 | mdh1aa |
| ube2l3a | 3.823759 | 0.265671317 | 0.276 | 0.16 | 0.009714 | 16 | ube2l3a |
| syncrip2 | 4.568763 | 0.283156029 | 0.44 | 0.312 | 0.011607 | 16 | syncrip |
| pabpn11 | 4.742836 | 0.282945894 | 0.404 | 0.274 | 0.012049 | 16 | pabpn1 |

| | | | | | | | |
|---|---|---|---|---|---|---|---|
| anp32b3 | 4.984217 | 0.280195342 | 0.388 | 0.252 | 0.012662 | 16 | anp32b |
| cox6a12 | 5.727902 | 0.272131781 | 0.516 | 0.39 | 0.014552 | 16 | cox6a1 |
| atp5po | 5.961849 | 0.278248241 | 0.404 | 0.276 | 0.015146 | 16 | atp5po |
| taf13 | 7.871748 | 0.270324735 | 0.364 | 0.238 | 0.019998 | 16 | taf13 |
| meaf6 | 1.266425 | 0.258318595 | 0.248 | 0.145 | 0.032174 | 16 | meaf6 |
| rad21a | 1.720478 | 0.255284079 | 0.248 | 0.147 | 0.043709 | 16 | rad21a |
| atrx3 | 1.796715 | 0.266231194 | 0.22 | 0.128 | 0.045646 | 16 | atrx |
| nasp5 | 1.820102 | 0.265476538 | 0.468 | 0.293 | 0.04624 | 16 | nasp |
| u2af1 | 2.364989 | 0.285593871 | 0.296 | 0.192 | 0.060083 | 16 | u2af1 |
| si:ch1073-325m22.2 | 7.064336 | 0.297300606 | 0.348 | 0.25 | 0.179469 | 16 | si:ch1073-325m22.2 |
| fam32a | 1.304321 | 0.264637268 | 0.396 | 0.291 | 0.331363 | 16 | fam32a |
| gpx4b | 2.915429 | 0.251017081 | 0.316 | 0.22 | 0.740665 | 16 | gpx4b |
| rps27l4 | 3.023415 | 0.272333524 | 0.308 | 0.218 | 0.768099 | 16 | rps27l |
| clta | 3.844975 | 0.252487807 | 0.288 | 0.195 | 0.976816 | 16 | clta |
| phospho1 | 3.717532 | 0.550528359 | 0.197 | 0.011 | 9.4443924 | 17 | phospho1 |
| zgc:1534995 | 2.071841 | 0.927291976 | 0.954 | 0.416 | 5.2635138 | 17 | zgc:153499 |
| h1m5 | 5.556438 | 0.86910725 | 0.972 | 0.491 | 1.4116132 | 17 | h1m |
| zp2.35 | 3.986854 | 0.896014084 | 0.936 | 0.421 | 1.0128605 | 17 | zp2.3 |
| zp2.55 | 2.946925 | 0.882371654 | 0.963 | 0.458 | 7.4866643 | 17 | zp2.5 |
| tmsb55 | 3.237357 | 0.905594742 | 0.931 | 0.396 | 8.2245079 | 17 | tmsb5 |
| h2af1al5 | 5.921774 | 0.873806374 | 0.977 | 0.485 | 1.5044267 | 17 | h2af1al |
| si:dkey-240e12.65 | 1.993298 | 0.878910396 | 0.959 | 0.462 | 5.0639739 | 17 | si:dkey-240e12.6 |
| si:ch211-125e6.125 | 5.555257 | 0.889175299 | 0.927 | 0.406 | 1.4113132 | 17 | si:ch211-125e6.12 |
| si:zfos-1505d6.35 | 3.518572 | 0.881950062 | 0.931 | 0.416 | 8.9389331 | 17 | si:zfos-1505d6.3 |
| zp2.25 | 3.757268 | 0.87640084 | 0.95 | 0.43 | 9.5453411 | 17 | zp2.2 |
| acp5a5 | 5.317968 | 0.858671608 | 0.94 | 0.44 | 1.3510299 | 17 | acp5a |
| zgc:1717765 | 6.212076 | 0.879857173 | 0.904 | 0.402 | 1.5781781 | 17 | zgc:171776 |
| scg3 | 2.066335 | 0.329205297 | 0.119 | 0.008 | 5.2495258 | 17 | scg3 |
| si:ch211-226h8.145 | 2.662343 | 0.858208903 | 0.963 | 0.461 | 6.7636845 | 17 | si:ch211-226h8.14 |
| CR790368.15 | 2.595821 | 0.852297725 | 0.917 | 0.423 | 6.5946841 | 17 | CR790368.1 |
| cldnd5 | 3.251619 | 0.861626651 | 0.982 | 0.492 | 8.2607382 | 17 | cldnd |
| zp35 | 1.084113 | 0.852754321 | 0.963 | 0.475 | 2.7541910 | 17 | zp3 |
| si:ch73-189n23.15 | 1.096385 | 0.862231272 | 0.913 | 0.427 | 2.7853663 | 17 | si:ch73-189n23.1 |
| si:dkey-241l7.25 | 1.631137 | 0.840547263 | 0.986 | 0.524 | 4.1439037 | 17 | si:dkey-241l7.2 |
| si:dkey-241l7.45 | 2.184193 | 0.819276077 | 0.959 | 0.497 | 5.5489433 | 17 | si:dkey-241l7.4 |

| | | | | | | | |
|---|---|---|---|---|---|---|---|
| mt25 | 3.857956 | 0.851414636 | 0.94 | 0.435 | 9.8011382 | 17 | mt2 |
| BX649337.15 | 1.885768 | 0.866361686 | 0.853 | 0.355 | 4.7907946 | 17 | BX649337.1 |
| si:dkey-229d11.35 | 1.944585 | 0.876450885 | 0.862 | 0.355 | 4.9402184 | 17 | si:dkey-229d11.3 |
| zp3.25 | 3.189175 | 0.836321025 | 0.927 | 0.454 | 8.1021006 | 17 | zp3.2 |
| si:ch211-12h2.85 | 4.983108 | 0.831365959 | 0.922 | 0.445 | 1.2659588 | 17 | si:ch211-12h2.8 |
| crp25 | 6.687376 | 0.859405415 | 0.908 | 0.397 | 1.6989280 | 17 | crp2 |
| ccna15 | 2.497034 | 0.848031836 | 0.872 | 0.386 | 6.3437173 | 17 | ccna1 |
| si:dkeyp-20e4.85 | 1.687035 | 0.871915862 | 0.812 | 0.33 | 4.2859129 | 17 | si:dkeyp-20e4.8 |
| zp3b5 | 3.184950 | 0.860246959 | 0.784 | 0.314 | 8.0913666 | 17 | zp3b |
| si:dkeyp-98a7.95 | 3.345402 | 0.846794421 | 0.908 | 0.416 | 8.4989940 | 17 | si:dkeyp-98a7.9 |
| cldng5 | 1.267956 | 0.83691146 | 0.904 | 0.419 | 3.2212435 | 17 | cldng |
| zgc:1717795 | 1.642900 | 0.836757337 | 0.904 | 0.416 | 4.1737882 | 17 | zgc:171779 |
| epcam5 | 5.317794 | 0.827453648 | 0.885 | 0.416 | 1.3509857 | 17 | epcam |
| zp2.15 | 5.420570 | 0.835552776 | 0.913 | 0.435 | 1.3770960 | 17 | zp2.1 |
| si:dkey-241l7.65 | 1.289008 | 0.817644218 | 0.89 | 0.429 | 3.2747252 | 17 | si:dkey-241l7.6 |
| zp2.65 | 1.096129 | 0.837639395 | 0.899 | 0.407 | 2.7847165 | 17 | zp2.6 |
| zgc:1751355 | 5.768110 | 0.829353772 | 0.862 | 0.389 | 1.4653884 | 17 | zgc:175135 |
| si:dkey-241l7.55 | 1.341123 | 0.791070188 | 0.922 | 0.476 | 3.4071239 | 17 | si:dkey-241l7.5 |
| mid1ip1l5 | 1.423831 | 0.769393028 | 0.872 | 0.456 | 3.6172437 | 17 | mid1ip1l |
| si:ch211-226h8.45 | 2.241744 | 0.807611621 | 0.803 | 0.359 | 5.6951509 | 17 | si:ch211-226h8.4 |
| si:dkeyp-46h3.65 | 3.066716 | 0.810552727 | 0.78 | 0.333 | 7.7909927 | 17 | si:dkeyp-46h3.6 |
| LOC1104378215 | 8.036601 | 0.820872325 | 0.794 | 0.337 | 2.0416985 | 17 | LOC110437821 |
| zar1l5 | 1.264176 | 0.802533137 | 0.771 | 0.323 | 3.2116398 | 17 | zar1l |
| s100a15 | 1.564852 | 0.806095933 | 0.739 | 0.296 | 3.9755069 | 17 | s100a1 |
| CU571163.15 | 2.026643 | 0.819891676 | 0.766 | 0.326 | 5.1486876 | 17 | CU571163.1 |
| rrm25 | 3.916698 | 0.773708599 | 0.885 | 0.418 | 9.9503718 | 17 | rrm2 |
| slc8a4b | 4.001020 | 0.404516682 | 0.138 | 0.014 | 1.0164591 | 17 | slc8a4b |
| si:dkeyp-98a7.45 | 4.941041 | 0.799994513 | 0.789 | 0.345 | 1.2552714 | 17 | si:dkeyp-98a7.4 |
| zp3a.25 | 8.911085 | 0.786730329 | 0.844 | 0.4 | 2.2638611 | 17 | zp3a.2 |
| gapdh5 | 1.143513 | 0.788126901 | 0.775 | 0.342 | 2.9050971 | 17 | gapdh |
| si:dkeyp-98a7.85 | 2.968979 | 0.795446453 | 0.821 | 0.372 | 7.5426920 | 17 | si:dkeyp-98a7.8 |
| si:dkey-90l23.15 | 3.212456 | 0.796146457 | 0.803 | 0.364 | 8.1612446 | 17 | si:dkey-90l23.1 |
| npm2b5 | 1.836943 | 0.79314364 | 0.743 | 0.323 | 4.6667560 | 17 | npm2b |
| LOC1005374995 | 7.934559 | 0.788557398 | 0.674 | 0.261 | 2.0157748 | 17 | LOC100537499 |
| si:ch211-12h2.65 | 1.065021 | 0.770430135 | 0.794 | 0.386 | 2.7056862 | 17 | si:ch211-12h2.6 |

| | | | | | | | |
|---|---|---|---|---|---|---|---|
| si:ch211-14a17.75 | 8.035661 | 0.782876125 | 0.761 | 0.332 | 2.0414599 | 17 | si:ch211-14a17.7 |
| avd5 | 1.115026 | 0.776623902 | 0.606 | 0.214 | 2.8327242 | 17 | avd |
| zgc:1714745 | 8.944593 | 0.774788714 | 0.628 | 0.246 | 2.2723739 | 17 | zgc:171474 |
| siva15 | 1.571352 | 0.751550033 | 0.775 | 0.357 | 3.9920207 | 17 | siva1 |
| zpcx5 | 1.852736 | 0.768912548 | 0.665 | 0.275 | 4.7068771 | 17 | zpcx |
| zgc:771185 | 2.493240 | 0.75700923 | 0.798 | 0.372 | 6.3340763 | 17 | zgc:77118 |
| gtf3ab5 | 4.205561 | 0.758029989 | 0.656 | 0.26 | 1.0684229 | 17 | gtf3ab |
| nanog5 | 5.967898 | 0.752514506 | 0.711 | 0.292 | 1.5161446 | 17 | nanog |
| si:ch211-250e5.165 | 8.270746 | 0.750538449 | 0.665 | 0.281 | 2.1011831 | 17 | si:ch211-250e5.16 |
| qdprb25 | 1.804996 | 0.754683683 | 0.761 | 0.352 | 4.5855928 | 17 | qdprb2 |
| si:dkeyp-46h3.85 | 3.195664 | 0.764953473 | 0.679 | 0.285 | 8.1185861 | 17 | si:dkeyp-46h3.8 |
| zar15 | 4.974191 | 0.741521568 | 0.757 | 0.359 | 1.2636934 | 17 | zar1 |
| LOC1005351625 | 5.532208 | 0.761779042 | 0.647 | 0.25 | 1.4054575 | 17 | LOC100535162 |
| zgc:1715175 | 1.323707 | 0.747827093 | 0.716 | 0.315 | 3.3628784 | 17 | zgc:171517 |
| zgc:1734435 | 5.993661 | 0.739981716 | 0.665 | 0.268 | 1.5226897 | 17 | zgc:173443 |
| si:ch73-90k17.15 | 1.015477 | 0.73194726 | 0.56 | 0.201 | 2.5798210 | 17 | si:ch73-90k17.1 |
| si:ch211-146l10.85 | 1.915480 | 0.743306088 | 0.619 | 0.249 | 4.8662782 | 17 | si:ch211-146l10.8 |
| si:dkeyp-46h3.25 | 2.636973 | 0.736665747 | 0.711 | 0.329 | 6.6992307 | 17 | si:dkeyp-46h3.2 |
| btg45 | 4.506974 | 0.737479881 | 0.697 | 0.3 | 1.1449967 | 17 | btg4 |
| slbp25 | 8.310876 | 0.731626805 | 0.72 | 0.323 | 2.1113782 | 17 | slbp2 |
| thy15 | 8.966858 | 0.717024509 | 0.775 | 0.374 | 2.2780303 | 17 | thy1 |
| si:ch211-145c1.15 | 1.274762 | 0.73278588 | 0.528 | 0.189 | 3.2385345 | 17 | si:ch211-145c1.1 |
| si:ch211-132b12.85 | 3.660451 | 0.732405517 | 0.661 | 0.279 | 9.2993780 | 17 | si:ch211-132b12.8 |
| lamp25 | 2.781933 | 0.673858512 | 0.83 | 0.481 | 7.0675029 | 17 | lamp2 |
| si:dkey-208k4.25 | 6.423253 | 0.715355467 | 0.596 | 0.24 | 1.6318276 | 17 | si:dkey-208k4.2 |
| si:dkey-229d11.55 | 9.067497 | 0.714638239 | 0.688 | 0.316 | 2.3035976 | 17 | si:dkey-229d11.5 |
| hnrnpabb6 | 1.634146 | 0.585386114 | 0.835 | 0.556 | 4.1515481 | 17 | hnrnpabb |
| ccnb25 | 3.512192 | 0.706821556 | 0.702 | 0.313 | 8.9227252 | 17 | ccnb2 |
| nasp6 | 5.552567 | 0.700174274 | 0.651 | 0.291 | 1.4106297 | 17 | nasp |
| ctss2.15 | 6.150655 | 0.647105043 | 0.798 | 0.449 | 1.5625739 | 17 | ctss2.1 |
| uchl15 | 8.898803 | 0.699060039 | 0.638 | 0.287 | 2.2607410 | 17 | uchl1 |
| zgc:1952455 | 9.750021 | 0.69749755 | 0.665 | 0.316 | 2.4769930 | 17 | zgc:195245 |
| kpna75 | 1.002749 | 0.699256384 | 0.619 | 0.268 | 2.5474859 | 17 | kpna7 |
| LOC1018854775 | 4.238446 | 0.702100171 | 0.555 | 0.227 | 1.0767772 | 17 | LOC101885477 |
| si:dkeyp-50f7.25 | 4.799779 | 0.690153468 | 0.688 | 0.329 | 1.2193840 | 17 | si:dkeyp-50f7.2 |

| | | | | | | | |
|---|---|---|---|---|---|---|---|
| cth15 | 1.197221 | 0.688269571 | 0.67 | 0.309 | 3.0415418 | 17 | cth1 |
| cpeb1b5 | 7.032567 | 0.677401528 | 0.541 | 0.22 | 1.7866238 | 17 | cpeb1b |
| arhgef10la5 | 8.151470 | 0.673530921 | 0.555 | 0.224 | 2.0708810 | 17 | arhgef10la |
| hbba15 | 8.405587 | 0.699748059 | 0.656 | 0.291 | 2.1354395 | 17 | hbba1 |
| si:dkey-90l23.25 | 1.218553 | 0.681470273 | 0.633 | 0.28 | 3.0957363 | 17 | si:dkey-90l23.2 |
| elovl7b5 | 1.605232 | 0.674849465 | 0.514 | 0.202 | 4.0780925 | 17 | elovl7b |
| wu:fi42e035 | 2.467546 | 0.685736638 | 0.624 | 0.285 | 6.2688027 | 17 | wu:fi42e03 |
| AL929032.15 | 3.366024 | 0.685380598 | 0.606 | 0.258 | 8.5513850 | 17 | AL929032.1 |
| pabpn1l6 | 2.062328 | 0.664192929 | 0.555 | 0.232 | 5.2393462 | 17 | pabpn1l |
| zgc:554616 | 3.398671 | 0.660196006 | 0.569 | 0.238 | 8.6343248 | 17 | zgc:55461 |
| zgc:1362545 | 2.423941 | 0.655116947 | 0.615 | 0.269 | 6.1580233 | 17 | zgc:136254 |
| si:dkey-152b24.65 | 4.664836 | 0.663306921 | 0.56 | 0.242 | 1.1851017 | 17 | si:dkey-152b24.6 |
| crp35 | 4.155900 | 0.638324237 | 0.454 | 0.177 | 1.0558064 | 17 | crp3 |
| si:dkey-19b23.115 | 4.334331 | 0.650273108 | 0.528 | 0.227 | 1.1011368 | 17 | si:dkey-19b23.11 |
| cdca85 | 6.235388 | 0.589374456 | 0.394 | 0.138 | 1.5841003 | 17 | cdca8 |
| tuba4l5 | 8.084846 | 0.649106044 | 0.665 | 0.331 | 2.0539552 | 17 | tuba4l |
| si:ch211-5k11.85 | 3.506077 | 0.606970369 | 0.628 | 0.294 | 8.9071906 | 17 | si:ch211-5k11.8 |
| si:dkeyp-98a7.75 | 5.418387 | 0.607855854 | 0.431 | 0.158 | 1.3765414 | 17 | si:dkeyp-98a7.7 |
| rcc15 | 8.064320 | 0.616918753 | 0.44 | 0.171 | 2.0487406 | 17 | rcc1 |
| ccnb15 | 8.298275 | 0.635679087 | 0.642 | 0.324 | 2.1081769 | 17 | ccnb1 |
| si:dkey-241l7.35 | 2.102310 | 0.635651455 | 0.606 | 0.298 | 5.3409190 | 17 | si:dkey-241l7.3 |
| chia.45 | 5.102457 | 0.557904111 | 0.353 | 0.119 | 1.2962793 | 17 | chia.4 |
| hbaa15 | 9.213823 | 0.593951965 | 0.55 | 0.231 | 2.3407717 | 17 | hbaa1 |
| birc5b5 | 5.537330 | 0.618227465 | 0.486 | 0.212 | 1.4067588 | 17 | birc5b |
| rasl11b5 | 7.299907 | 0.54423996 | 0.335 | 0.112 | 1.8545414 | 17 | rasl11b |
| zp3a.15 | 1.559381 | 0.612308985 | 0.606 | 0.312 | 3.9616097 | 17 | zp3a.1 |
| zgc:660245 | 1.808244 | 0.543684171 | 0.339 | 0.115 | 4.5938448 | 17 | zgc:66024 |
| ccna25 | 3.640914 | 0.599785507 | 0.523 | 0.237 | 9.2497436 | 17 | ccna2 |
| mkrn45 | 9.170621 | 0.604410928 | 0.445 | 0.192 | 2.3297963 | 17 | mkrn4 |
| si:ch1073-190k2.15 | 1.366359 | 0.581710843 | 0.394 | 0.153 | 3.4712366 | 17 | si:ch1073-190k2.1 |
| LOC1018838975 | 1.880261 | 0.544008919 | 0.367 | 0.133 | 4.7768034 | 17 | LOC101883897 |
| si:dkey-37o8.15 | 2.188962 | 0.591342644 | 0.537 | 0.251 | 5.5610603 | 17 | si:dkey-37o8.1 |
| spink2.16 | 2.597519 | 0.575765407 | 0.674 | 0.371 | 6.5989981 | 17 | spink2.1 |
| larp6b5 | 4.153966 | 0.569152813 | 0.408 | 0.16 | 1.0553152 | 17 | larp6b |
| org6 | 4.348668 | 0.596587823 | 0.459 | 0.198 | 1.1047792 | 17 | org |

| | | | | | | | |
|---|---|---|---|---|---|---|---|
| dmrt11 | 1.204402 | 0.261394955 | 0.106 | 0.017 | 3.0597844 | 17 | dmrt1 |
| si:ch211-226l4.66 | 1.673909 | 0.576051296 | 0.523 | 0.241 | 4.2525670 | 17 | si:ch211-226l4.6 |
| zgc:925915 | 1.864910 | 0.560511243 | 0.417 | 0.168 | 4.7378048 | 17 | zgc:92591 |
| gdf35 | 2.514553 | 0.577815089 | 0.436 | 0.182 | 6.3882223 | 17 | gdf3 |
| haus65 | 3.515935 | 0.561546864 | 0.417 | 0.172 | 8.9322347 | 17 | haus6 |
| zgc:562315 | 6.116278 | 0.566019309 | 0.44 | 0.186 | 1.5538404 | 17 | zgc:56231 |
| buc5 | 7.504567 | 0.55449422 | 0.372 | 0.145 | 1.9065354 | 17 | buc |
| ca15b5 | 8.357676 | 0.57243416 | 0.463 | 0.203 | 2.1232678 | 17 | ca15b |
| si:ch73-14h10.25 | 1.087950 | 0.571211186 | 0.491 | 0.222 | 2.7639385 | 17 | si:ch73-14h10.2 |
| zgc:566996 | 1.134883 | 0.573132247 | 0.468 | 0.208 | 2.8831717 | 17 | zgc:56699 |
| gmnn5 | 2.385181 | 0.572608019 | 0.523 | 0.253 | 6.0595542 | 17 | gmnn |
| si:dkeyp-46h3.55 | 7.375770 | 0.571010589 | 0.5 | 0.236 | 1.8738145 | 17 | si:dkeyp-46h3.5 |
| CDPF15 | 7.663438 | 0.582956448 | 0.477 | 0.233 | 1.9468966 | 17 | CDPF1 |
| ccne15 | 4.521959 | 0.468251075 | 0.275 | 0.092 | 1.1488038 | 17 | ccne1 |
| zglp15 | 5.792685 | 0.513371875 | 0.394 | 0.162 | 1.4716318 | 17 | zglp1 |
| patl25 | 3.161816 | 0.509114562 | 0.353 | 0.141 | 8.0325956 | 17 | patl2 |
| si:ch211-251f6.65 | 4.424458 | 0.545191761 | 0.523 | 0.271 | 1.1240335 | 17 | si:ch211-251f6.6 |
| AL929192.15 | 4.917415 | 0.512821883 | 0.339 | 0.134 | 1.2492694 | 17 | AL929192.1 |
| zgc:1719774 | 4.961140 | 0.532403897 | 0.417 | 0.188 | 1.2603777 | 17 | zgc:171977 |
| ddx413 | 5.401522 | 0.389525248 | 0.197 | 0.056 | 1.3722568 | 17 | ddx41 |
| gbgt1l45 | 5.892996 | 0.496240944 | 0.321 | 0.123 | 1.4971157 | 17 | gbgt1l4 |
| BX537276.15 | 7.785599 | 0.46741636 | 0.28 | 0.099 | 1.9779316 | 17 | BX537276.1 |
| larp6a5 | 1.664469 | 0.539306327 | 0.445 | 0.212 | 4.2285857 | 17 | larp6a |
| sybu5 | 1.902268 | 0.487732518 | 0.344 | 0.136 | 4.8327120 | 17 | sybu |
| h3f3a6 | 2.208932 | 0.475077269 | 0.679 | 0.482 | 5.6117931 | 17 | h3f3a |
| slc16a35 | 7.127985 | 0.523940548 | 0.436 | 0.204 | 1.8108646 | 17 | slc16a3 |
| lsm14b5 | 8.407284 | 0.5311646 | 0.495 | 0.245 | 2.1358705 | 17 | lsm14b |
| pigrl2.35 | 1.043195 | 0.510223429 | 0.385 | 0.167 | 2.6502373 | 17 | pigrl2.3 |
| zgc:1526525 | 1.102666 | 0.431289291 | 0.266 | 0.093 | 2.8013229 | 17 | zgc:152652 |
| si:ch211-146l10.75 | 1.283726 | 0.499155554 | 0.362 | 0.154 | 3.2613077 | 17 | si:ch211-146l10.7 |
| zgc:1628795 | 3.068855 | 0.487860505 | 0.353 | 0.148 | 7.7964285 | 17 | zgc:162879 |
| prr113 | 3.445498 | 0.427661079 | 0.243 | 0.083 | 8.7532889 | 17 | prr11 |
| elovl7a5 | 8.186128 | 0.495777073 | 0.353 | 0.151 | 2.0796860 | 17 | elovl7a |
| zgc:1714264 | 8.389792 | 0.43819731 | 0.248 | 0.088 | 2.1314267 | 17 | zgc:171426 |
| sinup5 | 1.075976 | 0.495412943 | 0.376 | 0.167 | 2.7335173 | 17 | sinup |

| | | | | | | | |
|---|---|---|---|---|---|---|---|
| tpx25 | 1.127957 | 0.44867262 | 0.28 | 0.106 | 2.8655759 | 17 | tpx2 |
| dnd15 | 1.152722 | 0.415291632 | 0.257 | 0.091 | 2.9284921 | 17 | dnd1 |
| elavl26 | 1.208527 | 0.504342127 | 0.399 | 0.185 | 3.0702631 | 17 | elavl2 |
| zgc:1629455 | 2.098180 | 0.469882338 | 0.326 | 0.135 | 5.3304281 | 17 | zgc:162945 |
| zgc:1138865 | 2.130438 | 0.456610205 | 0.294 | 0.115 | 5.4123786 | 17 | zgc:113886 |
| prdx16 | 2.375339 | 0.481714567 | 0.339 | 0.147 | 6.0345508 | 17 | prdx1 |
| zgc:112146 | 3.069785 | 0.302853786 | 0.133 | 0.032 | 7.7987895 | 17 | zgc:112146 |
| si:dkey-42i9.85 | 3.112374 | 0.440973395 | 0.275 | 0.105 | 7.9069879 | 17 | si:dkey-42i9.8 |
| ctsla5 | 3.242418 | 0.486968034 | 0.583 | 0.373 | 8.2373654 | 17 | ctsla |
| ypel34 | 3.571141 | 0.451540523 | 0.28 | 0.109 | 9.0724849 | 17 | ypel3 |
| zgc:1139844 | 3.709009 | 0.406927563 | 0.239 | 0.082 | 9.4227374 | 17 | zgc:113984 |
| BX571701.1 | 3.874674 | 0.331776779 | 0.156 | 0.042 | 9.8436118 | 17 | BX571701.1 |
| mibp6 | 4.937309 | 0.493536259 | 0.408 | 0.198 | 1.2543235 | 17 | mibp |
| pcf113 | 6.329157 | 0.517009245 | 0.408 | 0.216 | 1.6079224 | 17 | pcf11 |
| cldn7b5 | 7.590946 | 0.455326221 | 0.321 | 0.132 | 1.9284798 | 17 | cldn7b |
| degs24 | 8.419839 | 0.424407471 | 0.243 | 0.088 | 2.1390601 | 17 | degs2 |
| map1lc3c5 | 1.237850 | 0.4698856 | 0.372 | 0.165 | 3.1447581 | 17 | map1lc3c |
| cdca7a5 | 1.463474 | 0.49010806 | 0.399 | 0.191 | 3.7179577 | 17 | cdca7a |
| zp3c5 | 2.076160 | 0.466683735 | 0.349 | 0.155 | 5.2744854 | 17 | zp3c |
| LOC1104377794 | 2.786578 | 0.368889277 | 0.206 | 0.068 | 7.0793015 | 17 | LOC110437779 |
| BX284697.15 | 3.139179 | 0.409725274 | 0.252 | 0.093 | 7.9750858 | 17 | BX284697.1 |
| zgc:566765 | 3.421329 | 0.495172895 | 0.477 | 0.273 | 8.6918887 | 17 | zgc:56676 |
| mos5 | 4.624184 | 0.451317138 | 0.28 | 0.113 | 1.1747740 | 17 | mos |
| CR383662.25 | 4.954321 | 0.373252829 | 0.211 | 0.071 | 1.2586454 | 17 | CR383662.2 |
| zgc:174268 | 8.011184 | 0.39515365 | 0.216 | 0.076 | 2.0352413 | 17 | zgc:174268 |
| snu13b6 | 9.438755 | 0.470621389 | 0.578 | 0.37 | 2.3979159 | 17 | snu13b |
| use1 | 1.007476 | 0.338509599 | 0.147 | 0.041 | 2.5594938 | 17 | use1 |
| si:ch211-14a17.65 | 1.363527 | 0.365997671 | 0.206 | 0.07 | 3.4640426 | 17 | si:ch211-14a17.6 |
| gyg1a6 | 1.378646 | 0.485442104 | 0.523 | 0.318 | 3.5024521 | 17 | gyg1a |
| ddx195 | 1.721742 | 0.46231309 | 0.303 | 0.134 | 4.3740867 | 17 | ddx19 |
| zgc:1134245 | 2.093731 | 0.434156722 | 0.312 | 0.132 | 5.3191248 | 17 | zgc:113424 |
| CU929143.11 | 2.184650 | 0.384853008 | 0.211 | 0.074 | 5.5501035 | 17 | CU929143.1 |
| CU856358.1 | 2.202494 | 0.284829712 | 0.124 | 0.031 | 5.5954373 | 17 | CU856358.1 |
| pttg15 | 2.343194 | 0.442581364 | 0.312 | 0.136 | 5.9528866 | 17 | pttg1 |
| gfer5 | 3.663791 | 0.401491562 | 0.229 | 0.086 | 9.3078620 | 17 | gfer |

| | | | | | | | |
|---|---|---|---|---|---|---|---|
| CU655961.35 | 3.831658 | 0.378621795 | 0.225 | 0.081 | 9.7343285 | 17 | CU655961.3 |
| si:dkeyp-75b4.106 | 3.854465 | 0.41293488 | 0.601 | 0.365 | 9.7922707 | 17 | si:dkeyp-75b4.10 |
| si:ch73-303b9.14 | 4.105819 | 0.354406472 | 0.193 | 0.064 | 1.0430833 | 17 | si:ch73-303b9.1 |
| rbm4.35 | 5.758264 | 0.439526268 | 0.583 | 0.421 | 1.4628870 | 17 | rbm4.3 |
| cdc205 | 8.693976 | 0.42538087 | 0.298 | 0.129 | 2.2087046 | 17 | cdc20 |
| si:ch73-132k15.22 | 1.012066 | 0.348194018 | 0.165 | 0.052 | 2.5711546 | 17 | si:ch73-132k15.2 |
| gpr1604 | 1.019403 | 0.340978816 | 0.197 | 0.067 | 2.5897942 | 17 | gpr160 |
| si:dkey-68o6.55 | 1.130929 | 0.418987926 | 0.252 | 0.101 | 2.8731264 | 17 | si:dkey-68o6.5 |
| hdr | 1.480888 | 0.316472578 | 0.151 | 0.045 | 3.7621975 | 17 | hdr |
| dazl6 | 1.682647 | 0.376033459 | 0.234 | 0.089 | 4.2747669 | 17 | dazl |
| LOC1005368605 | 1.958434 | 0.460401828 | 0.491 | 0.288 | 4.9754021 | 17 | LOC100536860 |
| LOC1039099725 | 1.982894 | 0.471667969 | 0.5 | 0.322 | 5.0375442 | 17 | LOC103909972 |
| CR847984.15 | 2.063799 | 0.428034851 | 0.312 | 0.138 | 5.2430820 | 17 | CR847984.1 |
| si:ch211-226h8.83 | 2.067926 | 0.378604271 | 0.216 | 0.079 | 5.2535674 | 17 | si:ch211-226h8.8 |
| zgc:1717505 | 2.204056 | 0.442450054 | 0.312 | 0.143 | 5.5994051 | 17 | zgc:171750 |
| zgc:1141303 | 2.731159 | 0.321151428 | 0.188 | 0.064 | 6.9385099 | 17 | zgc:114130 |
| plk15 | 2.781503 | 0.444278791 | 0.321 | 0.148 | 7.0664089 | 17 | plk1 |
| si:ch1073-263o8.25 | 2.905321 | 0.45908069 | 0.413 | 0.216 | 7.3809690 | 17 | si:ch1073-263o8.2 |
| si:dkey-81e3.22 | 3.022157 | 0.316033693 | 0.165 | 0.052 | 7.6777917 | 17 | si:dkey-81e3.2 |
| CABZ01059627.25 | 4.881621 | 0.399491976 | 0.261 | 0.109 | 1.2401758 | 17 | CABZ01059627.2 |
| bmb5 | 5.214475 | 0.34754869 | 0.202 | 0.072 | 1.3247376 | 17 | bmb |
| figla3 | 5.554062 | 0.369205546 | 0.211 | 0.078 | 1.4110094 | 17 | figla |
| aurkb5 | 8.152307 | 0.405065637 | 0.261 | 0.11 | 2.0710938 | 17 | aurkb |
| eif4e1b3 | 1.014000 | 0.356139078 | 0.197 | 0.071 | 2.5760687 | 17 | eif4e1b |
| sox19b4 | 1.027610 | 0.350485617 | 0.216 | 0.08 | 2.6106447 | 17 | sox19b |
| fbxo51 | 1.059311 | 0.342057531 | 0.174 | 0.059 | 2.6911809 | 17 | fbxo5 |
| zp2l15 | 1.114336 | 0.41729334 | 0.298 | 0.134 | 2.8309706 | 17 | zp2l1 |
| baiap2l1a | 1.629047 | 0.281896361 | 0.124 | 0.035 | 4.1385961 | 17 | baiap2l1a |
| imp35 | 1.631490 | 0.416201592 | 0.275 | 0.123 | 4.1448008 | 17 | imp3 |
| zgc:1738565 | 1.723927 | 0.431931699 | 0.367 | 0.18 | 4.3796387 | 17 | zgc:173856 |
| CT009596.14 | 1.838445 | 0.332132674 | 0.202 | 0.074 | 4.6705713 | 17 | CT009596.1 |
| tent5c5 | 2.264561 | 0.418187795 | 0.275 | 0.122 | 5.7531190 | 17 | tent5c |
| selenoh6 | 2.265450 | 0.451605548 | 0.404 | 0.227 | 5.7553772 | 17 | selenoh |
| lto1 | 3.002157 | 0.299478632 | 0.133 | 0.04 | 7.6269803 | 17 | lto1 |
| CU655961.45 | 3.097528 | 0.339387806 | 0.197 | 0.072 | 7.8692704 | 17 | CU655961.4 |

| | | | | | | | |
|---|---|---|---|---|---|---|---|
| BX001030.15 | 3.14778( | 0.37858998 | 0.243 | 0.1 | 7.9969373 | 17 | BX001030.1 |
| zgc:923135 | 3.29855( | 0.389065721 | 0.252 | 0.107 | 8.3799903 | 17 | zgc:92313 |
| phex1 | 3.74473( | 0.355412718 | 0.193 | 0.071 | 9.5134869 | 17 | phex |
| LOC1005380234 | 5.00023! | 0.329482968 | 0.179 | 0.063 | 1.2703098 | 17 | LOC100538023 |
| zgc:1630305 | 6.15192( | 0.368038218 | 0.252 | 0.106 | 1.5628976 | 17 | zgc:163030 |
| slbp6 | 6.36029; | 0.433908793 | 0.381 | 0.198 | 1.6158322 | 17 | slbp |
| retsatl5 | 7.08715{ | 0.439540639 | 0.33 | 0.167 | 1.8004925 | 17 | retsatl |
| brf25 | 7.57231( | 0.357186746 | 0.206 | 0.08 | 1.9237469 | 17 | brf2 |
| cdca94 | 8.19581{ | 0.327557076 | 0.193 | 0.071 | 2.0821477 | 17 | cdca9 |
| fbxw11b3 | 9.33992: | 0.406515469 | 0.28 | 0.131 | 2.3728074 | 17 | fbxw11b |
| COQ10A3 | 9.48026: | 0.349370677 | 0.188 | 0.07 | 2.4084609 | 17 | COQ10A |
| si:ch211-69g19.25 | 1.07392: | 0.380131841 | 0.243 | 0.103 | 2.7282969 | 17 | si:ch211-69g19.2 |
| tmem54a5 | 1.45035: | 0.399786661 | 0.261 | 0.117 | 3.6846189 | 17 | tmem54a |
| zgc:1034826 | 1.51459; | 0.407697737 | 0.321 | 0.157 | 3.8478222 | 17 | zgc:103482 |
| aurka5 | 1.60325⁄ | 0.357465838 | 0.239 | 0.099 | 4.0730680 | 17 | aurka |
| cfl1l2 | 1.73530( | 0.405496965 | 0.289 | 0.136 | 4.4085531 | 17 | cfl1l |
| BX649350.1 | 1.92788( | 0.268035487 | 0.142 | 0.045 | 4.8977808 | 17 | BX649350.1 |
| si:ch211-286b5.44 | 1.93782⁄ | 0.303430361 | 0.165 | 0.057 | 4.9230434 | 17 | si:ch211-286b5.4 |
| rbpms2b5 | 2.09089! | 0.42077542 | 0.362 | 0.19 | 5.3119203 | 17 | rbpms2b |
| zgc:1529865 | 2.11538; | 0.338447835 | 0.193 | 0.073 | 5.3741420 | 17 | zgc:152986 |
| zgc:771123 | 2.29731( | 0.382695185 | 0.234 | 0.1 | 5.8363398 | 17 | zgc:77112 |
| si:dkey-188i13.116 | 2.49173! | 0.459156841 | 0.45 | 0.29 | 6.3302531 | 17 | si:dkey-188i13.11 |
| cax11 | 2.49361{ | 0.28008512 | 0.147 | 0.048 | 6.3350371 | 17 | cax1 |
| rdh10b5 | 2.59964; | 0.413442246 | 0.307 | 0.15 | 6.6044050 | 17 | rdh10b |
| yrdc5 | 2.62961: | 0.37897832 | 0.239 | 0.103 | 6.6805319 | 17 | yrdc |
| pabpc1l5 | 2.67381⁄ | 0.387743711 | 0.261 | 0.117 | 6.7928250 | 17 | pabpc1l |
| inka1b5 | 3.21703{ | 0.4048481 | 0.317 | 0.156 | 8.1728857 | 17 | inka1b |
| si:dkey-92i17.24 | 3.90922: | 0.355068997 | 0.197 | 0.077 | 9.9313811 | 17 | si:dkey-92i17.2 |
| cth | 4.91856( | 0.283897933 | 0.124 | 0.038 | 1.2495604 | 17 | cth |
| lipg3 | 5.19016{ | 0.339235476 | 0.183 | 0.07 | 1.3185623 | 17 | lipg |
| zgc:1737426 | 9.20941⁄ | 0.398261206 | 0.275 | 0.129 | 2.3396516 | 17 | zgc:173742 |
| si:ch211-157c3.42 | 9.95850: | 0.31604151 | 0.193 | 0.074 | 2.5299572 | 17 | si:ch211-157c3.4 |
| lrp131 | 1.06685; | 0.292408733 | 0.161 | 0.057 | 2.7103511 | 17 | lrp13 |
| cdt14 | 1.10906; | 0.327385958 | 0.188 | 0.073 | 2.8175722 | 17 | cdt1 |
| cited12 | 1.13595⁄ | 0.30754754 | 0.151 | 0.053 | 2.8858922 | 17 | cited1 |

| | | | | | | | |
|---|---|---|---|---|---|---|---|
| apopt1 | 1.136954 | 0.283244664 | 0.138 | 0.046 | 2.8884340 | 17 | apopt1 |
| rbm74 | 1.278635 | 0.384417904 | 0.239 | 0.108 | 3.2483742 | 17 | rbm7 |
| tatdn22 | 1.287867 | 0.304479205 | 0.165 | 0.06 | 3.2718268 | 17 | tatdn2 |
| fbxo431 | 1.341734 | 0.307300962 | 0.161 | 0.058 | 3.4086772 | 17 | fbxo43 |
| tmem441 | 1.365558 | 0.27362178 | 0.147 | 0.05 | 3.4692005 | 17 | tmem44 |
| si:ch73-160p18.35 | 1.398576 | 0.411204873 | 0.367 | 0.198 | 3.5530840 | 17 | si:ch73-160p18.3 |
| zc3h11a5 | 1.400884 | 0.413287633 | 0.349 | 0.189 | 3.5589460 | 17 | zc3h11a |
| ndc11 | 1.575388 | 0.357032652 | 0.206 | 0.087 | 4.0022747 | 17 | ndc1 |
| CU019646.1 | 1.712255 | 0.263131532 | 0.133 | 0.043 | 4.3499851 | 17 | CU019646.1 |
| wu:fa19b122 | 1.774531 | 0.32180103 | 0.165 | 0.061 | 4.5081985 | 17 | wu:fa19b12 |
| hsd17b12a5 | 1.785573 | 0.36863916 | 0.284 | 0.136 | 4.5362504 | 17 | hsd17b12a |
| hmga1a3 | 2.605457 | 0.364825564 | 0.587 | 0.442 | 6.6191656 | 17 | hmga1a |
| mcm3l5 | 2.611248 | 0.280488442 | 0.188 | 0.072 | 6.6338771 | 17 | mcm3l |
| si:ch211-167j6.3 | 2.850722 | 0.2533052 | 0.115 | 0.035 | 7.2422599 | 17 | si:ch211-167j6.3 |
| gstp14 | 2.859000 | 0.401416174 | 0.5 | 0.347 | 7.2632915 | 17 | gstp1 |
| fbxo324 | 3.041945 | 0.360023636 | 0.206 | 0.087 | 7.7280633 | 17 | fbxo32 |
| BX571780.14 | 3.245540 | 0.31072346 | 0.174 | 0.066 | 8.2452950 | 17 | BX571780.1 |
| si:ch211-266i6.3 | 3.503529 | 0.294122782 | 0.156 | 0.057 | 8.9007166 | 17 | si:ch211-266i6.3 |
| tacc32 | 3.508741 | 0.330271486 | 0.197 | 0.081 | 8.9139566 | 17 | tacc3 |
| snx10a5 | 5.485083 | 0.352954032 | 0.234 | 0.105 | 1.3934855 | 17 | snx10a |
| slc48a1b | 6.225857 | 0.325053249 | 0.188 | 0.077 | 1.5816792 | 17 | slc48a1b |
| npm2a4 | 6.357783 | 0.295046166 | 0.17 | 0.065 | 1.6151950 | 17 | npm2a |
| hsd3b21 | 7.193679 | 0.269983966 | 0.142 | 0.05 | 1.8275542 | 17 | hsd3b2 |
| hspa55 | 7.359060 | 0.397115559 | 0.454 | 0.295 | 1.8695692 | 17 | hspa5 |
| st3gal81 | 7.792766 | 0.292817104 | 0.17 | 0.065 | 1.9797522 | 17 | st3gal8 |
| eif4a1a5 | 8.760529 | 0.375831771 | 0.532 | 0.405 | 2.2256125 | 17 | eif4a1a |
| chchd106 | 9.085642 | 0.377214193 | 0.271 | 0.134 | 2.3082073 | 17 | chchd10 |
| buc2l4 | 9.408836 | 0.30568232 | 0.17 | 0.066 | 2.3903150 | 17 | buc2l |
| tph1a5 | 9.446217 | 0.345316953 | 0.252 | 0.116 | 2.3998115 | 17 | tph1a |
| si:dkey-46g23.25 | 9.934414 | 0.340139251 | 0.257 | 0.119 | 2.5238379 | 17 | si:dkey-46g23.2 |
| BX005461.14 | 1.030432 | 0.291622131 | 0.17 | 0.065 | 2.6178144 | 17 | BX005461.1 |
| hmgb2a3 | 1.056885 | 0.283099717 | 0.725 | 0.609 | 2.6850174 | 17 | hmgb2a |
| zgc:1102394 | 1.081380 | 0.40910512 | 0.339 | 0.196 | 2.7472483 | 17 | zgc:110239 |
| zgc:1017444 | 1.136913 | 0.3431379 | 0.243 | 0.112 | 2.8883296 | 17 | zgc:101744 |
| vps9d13 | 1.434027 | 0.333841535 | 0.193 | 0.081 | 3.6431461 | 17 | vps9d1 |

| | | | | | | | |
|---|---|---|---|---|---|---|---|
| ctsba7 | 1.461445 | 0.383179801 | 0.509 | 0.352 | 3.7128015 | 17 | ctsba |
| h2afva5 | 1.579629 | 0.40413853 | 0.454 | 0.306 | 4.0130495 | 17 | h2afva |
| zgc:1118685 | 1.720555 | 0.367778868 | 0.252 | 0.121 | 4.3710708 | 17 | zgc:111868 |
| thop15 | 1.992792 | 0.355579828 | 0.225 | 0.104 | 5.0626897 | 17 | thop1 |
| zgc:1134252 | 2.390858 | 0.293864031 | 0.151 | 0.057 | 6.0739769 | 17 | zgc:113425 |
| nanos35 | 2.701764 | 0.32192826 | 0.197 | 0.084 | 6.8638327 | 17 | nanos3 |
| si:ch73-340n8.45 | 2.782197 | 0.355377993 | 0.239 | 0.114 | 7.0681727 | 17 | si:ch73-340n8.4 |
| lpcat45 | 2.848900 | 0.321359108 | 0.206 | 0.09 | 7.2376306 | 17 | lpcat4 |
| si:dkeyp-114g9.14 | 3.140688 | 0.299329803 | 0.202 | 0.086 | 7.9789203 | 17 | si:dkeyp-114g9.1 |
| spryd42 | 3.186345 | 0.309854143 | 0.17 | 0.069 | 8.0949097 | 17 | spryd4 |
| ankrd225 | 3.206851 | 0.347449024 | 0.252 | 0.121 | 8.1470051 | 17 | ankrd22 |
| tmem33 | 3.443556 | 0.326865905 | 0.188 | 0.081 | 8.7483552 | 17 | tmem33 |
| si:ch211-113a14.272 | 3.847964 | 0.285998184 | 0.165 | 0.065 | 9.7757543 | 17 | si:ch211-113a14.27 |
| metap1d4 | 3.922556 | 0.325717115 | 0.183 | 0.078 | 9.9652541 | 17 | metap1d |
| cnep1r13 | 5.368859 | 0.352603754 | 0.229 | 0.11 | 0.000136 | 17 | cnep1r1 |
| birc5a2 | 6.028498 | 0.315100551 | 0.197 | 0.086 | 0.000153 | 17 | birc5a |
| pif14 | 6.143565 | 0.313552669 | 0.202 | 0.088 | 0.000156 | 17 | pif1 |
| CU467646.25 | 7.186419 | 0.317716939 | 0.225 | 0.103 | 0.000183 | 17 | CU467646.2 |
| zgc:1655395 | 7.227029 | 0.327760052 | 0.216 | 0.099 | 0.000184 | 17 | zgc:165539 |
| g3bp14 | 7.318762 | 0.377249438 | 0.335 | 0.193 | 0.000186 | 17 | g3bp1 |
| bzw1b4 | 7.679359 | 0.373205613 | 0.5 | 0.378 | 0.000195 | 17 | bzw1b |
| lhx8a1 | 7.743293 | 0.259155872 | 0.133 | 0.048 | 0.000197 | 17 | lhx8a |
| dynll2a6 | 8.061837 | 0.329043722 | 0.243 | 0.117 | 0.000205 | 17 | dynll2a |
| ap1s3b1 | 8.673443 | 0.283675559 | 0.165 | 0.067 | 0.00022 | 17 | ap1s3b |
| ivns1abpb2 | 9.383693 | 0.314785259 | 0.22 | 0.103 | 0.000238 | 17 | ivns1abpb |
| si:ch211-210c8.74 | 9.859802 | 0.347328423 | 0.234 | 0.115 | 0.00025 | 17 | si:ch211-210c8.7 |
| zgc:1129804 | 1.010391 | 0.289029894 | 0.183 | 0.078 | 0.000257 | 17 | zgc:112980 |
| cpm4 | 1.080249 | 0.303260678 | 0.202 | 0.09 | 0.000274 | 17 | cpm |
| atic | 1.216518 | 0.278363251 | 0.161 | 0.065 | 0.000309 | 17 | atic |
| zgc:1751364 | 1.505103 | 0.313868145 | 0.197 | 0.089 | 0.000382 | 17 | zgc:175136 |
| kti12 | 1.514233 | 0.2702918 | 0.142 | 0.055 | 0.000385 | 17 | kti12 |
| tmem144b5 | 1.526031 | 0.278047024 | 0.183 | 0.078 | 0.000388 | 17 | tmem144b |
| ccnh5 | 1.627268 | 0.314480775 | 0.225 | 0.106 | 0.000413 | 17 | ccnh |
| pou5f35 | 1.651065 | 0.331590055 | 0.248 | 0.121 | 0.000419 | 17 | pou5f3 |
| marcksb5 | 1.743464 | 0.35837975 | 0.294 | 0.16 | 0.000443 | 17 | marcksb |

| | | | | | | | |
|---|---|---|---|---|---|---|---|
| efcab111 | 1.795021 | 0.251427936 | 0.138 | 0.052 | 0.000456 | 17 | efcab11 |
| rpp21 | 1.940558 | 0.311241735 | 0.174 | 0.076 | 0.000493 | 17 | rpp21 |
| scarb1 | 2.423398 | 0.261239149 | 0.138 | 0.053 | 0.000616 | 17 | scarb1 |
| zgc:558885 | 2.491588 | 0.272345594 | 0.17 | 0.07 | 0.000633 | 17 | zgc:55888 |
| h2afx5 | 2.645025 | 0.338922973 | 0.243 | 0.122 | 0.000672 | 17 | h2afx |
| CABZ01059627.14 | 3.284234 | 0.321542404 | 0.229 | 0.112 | 0.000834 | 17 | CABZ01059627.1 |
| tmem703 | 4.516427 | 0.304058452 | 0.197 | 0.091 | 0.001147 | 17 | tmem70 |
| cdc66 | 4.809825 | 0.299242968 | 0.193 | 0.088 | 0.001222 | 17 | cdc6 |
| nol74 | 5.235411 | 0.328922962 | 0.225 | 0.113 | 0.00133 | 17 | nol7 |
| snu13a4 | 6.081400 | 0.311582651 | 0.193 | 0.089 | 0.001545 | 17 | snu13a |
| cx43.44 | 7.012511 | 0.311361881 | 0.225 | 0.111 | 0.001782 | 17 | cx43.4 |
| btr041 | 7.252507 | 0.258511343 | 0.138 | 0.054 | 0.001842 | 17 | btr04 |
| atpaf2 | 7.548925 | 0.260119129 | 0.133 | 0.052 | 0.001918 | 17 | atpaf2 |
| zgc:1141235 | 7.930770 | 0.288604692 | 0.188 | 0.086 | 0.002015 | 17 | zgc:114123 |
| si:dkey-216e24.35 | 8.338093 | 0.283073579 | 0.216 | 0.102 | 0.002118 | 17 | si:dkey-216e24.3 |
| kpna26 | 8.410915 | 0.325762345 | 0.261 | 0.136 | 0.002137 | 17 | kpna2 |
| zgc:1735444 | 9.932612 | 0.263432712 | 0.151 | 0.063 | 0.002523 | 17 | zgc:173544 |
| polr1d1 | 1.020650 | 0.292866701 | 0.183 | 0.085 | 0.002593 | 17 | polr1d |
| BX957362.13 | 1.027851 | 0.329398652 | 0.225 | 0.114 | 0.002611 | 17 | BX957362.1 |
| gpx81 | 1.137233 | 0.284229377 | 0.165 | 0.073 | 0.002889 | 17 | gpx8 |
| asz12 | 1.291369 | 0.26558374 | 0.147 | 0.061 | 0.003281 | 17 | asz1 |
| ppp1r3b4 | 1.413631 | 0.282869026 | 0.197 | 0.092 | 0.003591 | 17 | ppp1r3b |
| dctpp16 | 1.759639 | 0.324318848 | 0.275 | 0.149 | 0.00447 | 17 | dctpp1 |
| ccdc582 | 1.866581 | 0.310244933 | 0.202 | 0.099 | 0.004742 | 17 | ccdc58 |
| BX323557.11 | 2.284863 | 0.284308821 | 0.174 | 0.08 | 0.005805 | 17 | BX323557.1 |
| fbl2 | 2.550401 | 0.369699167 | 0.317 | 0.201 | 0.006479 | 17 | fbl |
| zgc:922876 | 2.617126 | 0.284038901 | 0.225 | 0.112 | 0.006649 | 17 | zgc:92287 |
| cbx3a4 | 3.315486 | 0.351614844 | 0.408 | 0.292 | 0.008423 | 17 | cbx3a |
| nfr5 | 3.602451 | 0.274627025 | 0.193 | 0.092 | 0.009152 | 17 | nfr |
| coil4 | 3.789715 | 0.29189872 | 0.197 | 0.097 | 0.009628 | 17 | coil |
| mylipa5 | 4.337506 | 0.283898635 | 0.202 | 0.099 | 0.011019 | 17 | mylipa |
| pttg1ipb4 | 4.819390 | 0.296023311 | 0.202 | 0.101 | 0.012244 | 17 | pttg1ipb |
| mad2l16 | 5.180324 | 0.316244749 | 0.28 | 0.159 | 0.013161 | 17 | mad2l1 |
| gtf2a13 | 5.400300 | 0.331641344 | 0.252 | 0.142 | 0.013719 | 17 | gtf2a1 |
| tmco12 | 6.388159 | 0.321371052 | 0.211 | 0.111 | 0.016229 | 17 | tmco1 |

| | | | | | | | |
|---|---|---|---|---|---|---|---|
| cx44.25 | 7.590337 | 0.277085648 | 0.197 | 0.096 | 0.019283 | 17 | cx44.2 |
| nsun2 | 8.224872 | 0.255784803 | 0.133 | 0.057 | 0.020895 | 17 | nsun2 |
| exd22 | 8.742439 | 0.265070466 | 0.17 | 0.08 | 0.02221 | 17 | exd2 |
| pim33 | 8.966576 | 0.291884791 | 0.22 | 0.116 | 0.02278 | 17 | pim3 |
| faub5 | 9.384428 | 0.320791389 | 0.303 | 0.182 | 0.023841 | 17 | faub |
| txnl4a2 | 9.954943 | 0.315075996 | 0.206 | 0.11 | 0.025291 | 17 | txnl4a |
| si:ch73-78o10.15 | 1.024779 | 0.274934021 | 0.216 | 0.11 | 0.026035 | 17 | si:ch73-78o10.1 |
| slc29a21 | 1.101356 | 0.265290448 | 0.142 | 0.063 | 0.02798 | 17 | slc29a2 |
| vstm2l4 | 1.196611 | 0.251659226 | 0.17 | 0.079 | 0.0304 | 17 | vstm2l |
| fdx15 | 1.300368 | 0.332355718 | 0.317 | 0.198 | 0.033036 | 17 | fdx1 |
| pprc11 | 1.346521 | 0.274062675 | 0.174 | 0.085 | 0.034208 | 17 | pprc1 |
| aif1l3 | 1.492269 | 0.318428678 | 0.284 | 0.17 | 0.037911 | 17 | aif1l |
| mrpl531 | 1.661520 | 0.281303411 | 0.174 | 0.087 | 0.042211 | 17 | mrpl53 |
| mrpl9 | 1.789845 | 0.27035628 | 0.151 | 0.071 | 0.045471 | 17 | mrpl9 |
| golt1bb1 | 1.934112 | 0.281486491 | 0.17 | 0.083 | 0.049136 | 17 | golt1bb |
| parn5 | 2.500371 | 0.295441607 | 0.234 | 0.13 | 0.063522 | 17 | parn |
| timm133 | 2.730402 | 0.282543442 | 0.188 | 0.097 | 0.069366 | 17 | timm13 |
| tram24 | 2.945126 | 0.302454541 | 0.22 | 0.122 | 0.074821 | 17 | tram2 |
| cnih11 | 2.981392 | 0.297740652 | 0.206 | 0.112 | 0.075742 | 17 | cnih1 |
| rnf46 | 3.196720 | 0.285430944 | 0.216 | 0.117 | 0.081213 | 17 | rnf4 |
| lrrc593 | 3.370042 | 0.312952606 | 0.234 | 0.135 | 0.085616 | 17 | lrrc59 |
| c1d2 | 3.388805 | 0.302997772 | 0.216 | 0.119 | 0.086093 | 17 | c1d |
| dcun1d13 | 3.439904 | 0.270741236 | 0.174 | 0.087 | 0.087391 | 17 | dcun1d1 |
| gemin8 | 3.578316 | 0.295117925 | 0.188 | 0.1 | 0.090907 | 17 | gemin8 |
| srsf2a4 | 3.613327 | 0.318013294 | 0.45 | 0.349 | 0.091797 | 17 | srsf2a |
| rpz25 | 3.911261 | 0.31820593 | 0.271 | 0.164 | 0.099366 | 17 | rpz2 |
| hprt12 | 5.451425 | 0.285911832 | 0.206 | 0.113 | 0.138493 | 17 | hprt1 |
| mrps18b2 | 5.587391 | 0.278018717 | 0.211 | 0.116 | 0.141948 | 17 | mrps18b |
| emc91 | 6.129228 | 0.252264458 | 0.138 | 0.064 | 0.155713 | 17 | emc9 |
| nap1l12 | 7.059736 | 0.300573937 | 0.445 | 0.357 | 0.179353 | 17 | nap1l1 |
| ddx39ab5 | 7.101686 | 0.32702368 | 0.376 | 0.27 | 0.180418 | 17 | ddx39ab |
| ddx45 | 7.903993 | 0.251181443 | 0.193 | 0.1 | 0.200801 | 17 | ddx4 |
| adipor1a1 | 8.465605 | 0.259768877 | 0.174 | 0.09 | 0.215069 | 17 | adipor1a |
| znf395a4 | 1.278030 | 0.254908462 | 0.165 | 0.085 | 0.324684 | 17 | znf395a |
| pop45 | 1.314942 | 0.260199182 | 0.193 | 0.103 | 0.334061 | 17 | pop4 |

| | | | | | | | |
|---|---|---|---|---|---|---|---|
| arpp19b1 | 1.447059 | 0.299495036 | 0.225 | 0.134 | 0.367626 | 17 | arpp19b |
| uck2a1 | 1.979687 | 0.292918953 | 0.193 | 0.109 | 0.50294 | 17 | uck2a |
| cox176 | 2.047400 | 0.293415591 | 0.436 | 0.342 | 0.520142 | 17 | cox17 |
| etf1b | 2.052820 | 0.310518628 | 0.298 | 0.205 | 0.521519 | 17 | etf1b |
| hspd13 | 2.229131 | 0.306411418 | 0.257 | 0.164 | 0.566311 | 17 | hspd1 |
| itm2cb3 | 2.575813 | 0.267323944 | 0.211 | 0.121 | 0.654385 | 17 | itm2cb |
| ccng11 | 2.578022 | 0.258243884 | 0.514 | 0.444 | 0.654947 | 17 | ccng1 |
| anp32b4 | 2.600426 | 0.311418793 | 0.349 | 0.253 | 0.660638 | 17 | anp32b |
| pgrmc2 | 2.813167 | 0.260573006 | 0.165 | 0.089 | 0.714685 | 17 | pgrmc2 |
| dynll16 | 3.377510 | 0.265469951 | 0.518 | 0.442 | 0.858057 | 17 | dynll1 |
| nutf2l5 | 3.444479 | 0.304280005 | 0.317 | 0.222 | 0.87507 | 17 | nutf2l |
| UBL4A2 | 4.618424 | 0.263284601 | 0.202 | 0.118 | 1 | 17 | UBL4A |
| dap1b2 | 6.044190 | 0.293457668 | 0.339 | 0.25 | 1 | 17 | dap1b |
| mphosph6 | 6.094482 | 0.2549032 | 0.174 | 0.098 | 1 | 17 | mphosph6 |
| slc37a2 | 6.181116 | 0.250608699 | 0.156 | 0.085 | 1 | 17 | slc37a2 |
| ube2c5 | 7.625632 | 0.259774087 | 0.206 | 0.124 | 1 | 17 | ube2c |
| glulb2 | 7.676226 | 0.27657249 | 0.225 | 0.14 | 1 | 17 | glulb |
| khsrp1 | 7.978798 | 0.299154949 | 0.28 | 0.194 | 1 | 17 | khsrp |
| hmgb3b5 | 8.867386 | 0.294547578 | 0.257 | 0.171 | 1 | 17 | hmgb3b |
| mat2aa1 | 8.868266 | 0.295652819 | 0.252 | 0.168 | 1 | 17 | mat2aa |
| eif5a22 | 9.148103 | 0.258513142 | 0.463 | 0.409 | 1 | 17 | eif5a2 |
| med19a | 9.330646 | 0.269858765 | 0.206 | 0.127 | 1 | 17 | med19a |
| rbm4.24 | 9.850953 | 0.278879101 | 0.243 | 0.157 | 1 | 17 | rbm4.2 |
| stmn1a2 | 9.896658 | 0.259854261 | 0.289 | 0.189 | 1 | 17 | stmn1a |
| btg33 | 0.0001 | 0.285193183 | 0.248 | 0.162 | 1 | 17 | btg3 |
| snrpa | 0.0001 | 0.27503752 | 0.216 | 0.136 | 1 | 17 | snrpa |
| lbh3 | 0.0001 | 0.262288564 | 0.239 | 0.153 | 1 | 17 | lbh |
| atp5if1a1 | 0.0001 | 0.277997934 | 0.408 | 0.346 | 1 | 17 | atp5if1a |
| rbbp44 | 0.0001 | 0.277320411 | 0.248 | 0.164 | 1 | 17 | rbbp4 |
| pdia41 | 0.0001 | 0.271223926 | 0.202 | 0.124 | 1 | 17 | pdia4 |
| hnrnpub | 0.0002 | 0.28045324 | 0.229 | 0.154 | 1 | 17 | hnrnpub |
| srsf1a | 0.0002 | 0.256791329 | 0.17 | 0.101 | 1 | 17 | srsf1a |
| alyref4 | 0.0002 | 0.270251321 | 0.335 | 0.251 | 1 | 17 | alyref |
| snrpd22 | 0.0002 | 0.295496767 | 0.33 | 0.26 | 1 | 17 | snrpd2 |
| serp11 | 0.0003 | 0.276390576 | 0.28 | 0.199 | 1 | 17 | serp1 |

| | | | | | | | |
|---|---|---|---|---|---|---|---|
| STMP13 | 0.0005 | 0.27347648 | 0.362 | 0.294 | 1 | 17 | STMP1 |
| sumo3a1 | 0.0007 | 0.250136248 | 0.417 | 0.373 | 1 | 17 | sumo3a |
| lsm63 | 0.0007 | 0.27931432 | 0.298 | 0.226 | 1 | 17 | lsm6 |
| pcbp2 | 0.0008 | 0.281509362 | 0.284 | 0.218 | 1 | 17 | pcbp2 |
| cycsb3 | 0.0008 | 0.255656742 | 0.376 | 0.315 | 1 | 17 | cycsb |
| lsm82 | 0.0008 | 0.267461765 | 0.22 | 0.153 | 1 | 17 | lsm8 |
| cbx52 | 0.001 | 0.262931552 | 0.275 | 0.205 | 1 | 17 | cbx5 |
| ube2al1 | 0.001 | 0.263301251 | 0.216 | 0.15 | 1 | 17 | ube2al |
| drap1 | 0.0012 | 0.269094631 | 0.28 | 0.213 | 1 | 17 | drap1 |
| ndufab1b1 | 0.0022 | 0.266768873 | 0.225 | 0.166 | 1 | 17 | ndufab1b |
| cxcl12a2 | 0 | 1.917960815 | 0.925 | 0.071 | 0 | 18 | cxcl12a |
| podxl2 | 0 | 1.624614352 | 0.877 | 0.045 | 0 | 18 | podxl |
| nid1b1 | 0 | 1.590439352 | 0.868 | 0.057 | 0 | 18 | nid1b |
| krt153 | 0 | 1.58549599 | 0.854 | 0.086 | 0 | 18 | krt15 |
| aldh1a23 | 0 | 1.530004331 | 0.882 | 0.087 | 0 | 18 | aldh1a2 |
| c63 | 0 | 1.506501127 | 0.906 | 0.102 | 0 | 18 | c6 |
| tmem88b2 | 0 | 1.504619521 | 0.807 | 0.042 | 0 | 18 | tmem88b |
| phc2a | 0 | 1.474567584 | 0.821 | 0.082 | 0 | 18 | phc2a |
| slco2b12 | 0 | 1.442213189 | 0.802 | 0.056 | 0 | 18 | slco2b1 |
| il152 | 0 | 1.338750957 | 0.75 | 0.05 | 0 | 18 | il15 |
| XLOC-0160943 | 0 | 1.305272469 | 0.792 | 0.073 | 0 | 18 | XLOC-016094 |
| ndnf1 | 0 | 1.236448729 | 0.675 | 0.027 | 0 | 18 | ndnf |
| jam2b1 | 0 | 1.123111442 | 0.594 | 0.014 | 0 | 18 | jam2b |
| ace | 0 | 1.102977229 | 0.566 | 0.022 | 0 | 18 | ace |
| slc29a1a | 0 | 1.094542803 | 0.613 | 0.031 | 0 | 18 | slc29a1a |
| col12a1b1 | 0 | 1.070809531 | 0.594 | 0.029 | 0 | 18 | col12a1b |
| c43 | 0 | 1.037707074 | 0.575 | 0.031 | 0 | 18 | c4 |
| gpr17 | 0 | 1.033826298 | 0.547 | 0.006 | 0 | 18 | gpr17 |
| si:ch211-191i18.21 | 0 | 0.945654639 | 0.453 | 0.017 | 0 | 18 | si:ch211-191i18.2 |
| grhl12 | 0 | 0.939056625 | 0.528 | 0.023 | 0 | 18 | grhl1 |
| ptprk1 | 0 | 0.938117009 | 0.542 | 0.029 | 0 | 18 | ptprk |
| cd1511 | 0 | 0.745648996 | 0.42 | 0.016 | 0 | 18 | cd151 |
| gria3b | 0 | 0.596828552 | 0.292 | 0.002 | 0 | 18 | gria3b |
| foxl2a | 0 | 0.593331184 | 0.311 | 0.008 | 0 | 18 | foxl2a |
| vegfd | 0 | 0.521221578 | 0.278 | 0.005 | 0 | 18 | vegfd |

| | | | | | | | |
|---|---|---|---|---|---|---|---|
| CR786577.1 | 0 | 0.357102857 | 0.165 | 0.001 | 0 | 18 | CR786577.1 |
| s1pr3a | 0 | 0.346047745 | 0.156 | 0.001 | 0 | 18 | s1pr3a |
| akap12b2 | 2.999039 | 1.282279491 | 0.731 | 0.065 | 7.6190588 | 18 | akap12b |
| arxa | 4.641856 | 0.378307263 | 0.193 | 0.003 | 1.1792637 | 18 | arxa |
| wt1a1 | 4.394791 | 0.958654646 | 0.552 | 0.035 | 1.1164968 | 18 | wt1a |
| csf1a | 8.909469 | 0.457086468 | 0.231 | 0.004 | 2.2634506 | 18 | csf1a |
| postnb3 | 5.275105 | 1.425371468 | 0.821 | 0.085 | 1.3401406 | 18 | postnb |
| steap4 | 2.949380 | 0.633901495 | 0.307 | 0.01 | 7.4929013 | 18 | steap4 |
| hspb14 | 1.759553 | 1.212289874 | 0.75 | 0.072 | 4.4701448 | 18 | hspb1 |
| tmsb3 | 1.893397 | 0.913138593 | 0.472 | 0.027 | 4.8101751 | 18 | tmsb |
| cyp19a1a | 4.342293 | 0.580052576 | 0.203 | 0.003 | 1.1031596 | 18 | cyp19a1a |
| krt18a.13 | 4.286464 | 1.686851715 | 0.915 | 0.139 | 1.0889762 | 18 | krt18a.1 |
| fn1b3 | 4.658069 | 1.361791762 | 0.792 | 0.092 | 1.1833824 | 18 | fn1b |
| entpd13 | 2.052489 | 1.220962091 | 0.693 | 0.074 | 5.2143496 | 18 | entpd1 |
| CR751602.2 | 1.367829 | 0.567050551 | 0.278 | 0.009 | 3.4749711 | 18 | CR751602.2 |
| lpar12 | 3.339283 | 1.082539294 | 0.651 | 0.062 | 8.4834496 | 18 | lpar1 |
| ctgfa1 | 2.658709 | 0.938657134 | 0.448 | 0.028 | 6.7544524 | 18 | ctgfa |
| tspan9b2 | 6.371608 | 0.84817387 | 0.528 | 0.044 | 1.6187071 | 18 | tspan9b |
| anxa2a4 | 6.733494 | 1.416455941 | 0.783 | 0.108 | 1.7106442 | 18 | anxa2a |
| c7a2 | 1.149969 | 0.916241178 | 0.5 | 0.039 | 2.9214965 | 18 | c7a |
| FO744854.1 | 4.183278 | 0.597095206 | 0.321 | 0.015 | 1.0627618 | 18 | FO744854.1 |
| bnc21 | 3.056444 | 0.799257652 | 0.462 | 0.034 | 7.7648983 | 18 | bnc2 |
| si:ch211-204c21.13 | 3.809560 | 1.037963128 | 0.627 | 0.067 | 9.6781875 | 18 | si:ch211-204c21.1 |
| gabra5 | 2.246281 | 0.310360963 | 0.151 | 0.002 | 5.7066777 | 18 | gabra5 |
| sh3pxd2aa2 | 3.099404 | 0.78111932 | 0.41 | 0.028 | 7.8740377 | 18 | sh3pxd2aa |
| cldn5a3 | 1.701339 | 0.954704152 | 0.566 | 0.055 | 4.3222528 | 18 | cldn5a |
| adma | 9.068220 | 0.661685399 | 0.292 | 0.014 | 2.3037815 | 18 | adma |
| phactr4a2 | 2.213631 | 0.804393448 | 0.443 | 0.035 | 5.6237310 | 18 | phactr4a |
| spon1b2 | 7.415615 | 0.840318717 | 0.505 | 0.046 | 1.8839370 | 18 | spon1b |
| eppk11 | 4.287228 | 0.850336756 | 0.458 | 0.037 | 1.0891704 | 18 | eppk1 |
| mao | 3.311780 | 0.519587161 | 0.264 | 0.011 | 8.4135780 | 18 | mao |
| rbms32 | 6.939748 | 0.729644722 | 0.42 | 0.031 | 1.7630431 | 18 | rbms3 |
| wt1b1 | 4.876936 | 0.672039886 | 0.377 | 0.025 | 1.2389858 | 18 | wt1b |
| cebpd4 | 1.149714 | 1.263442535 | 0.849 | 0.142 | 2.9208509 | 18 | cebpd |
| vasnb2 | 9.627705 | 0.715615062 | 0.415 | 0.031 | 2.4459184 | 18 | vasnb |

| | | | | | | | |
|---|---|---|---|---|---|---|---|
| vcam1b1 | 1.015037 | 0.672582502 | 0.325 | 0.019 | 2.5787033 | 18 | vcam1b |
| itga92 | 4.427868 | 0.762382598 | 0.415 | 0.033 | 1.1249001 | 18 | itga9 |
| BX322530.13 | 4.359453 | 0.801775698 | 0.486 | 0.046 | 1.1075192 | 18 | BX322530.1 |
| col6a33 | 7.860647 | 1.081275069 | 0.741 | 0.106 | 1.9969974 | 18 | col6a3 |
| si:ch211-106h4.123 | 4.626625 | 1.044907696 | 0.519 | 0.052 | 1.1753941 | 18 | si:ch211-106h4.12 |
| s1pr21 | 8.679575 | 0.686978518 | 0.406 | 0.032 | 2.2050460 | 18 | s1pr2 |
| pdgfc | 5.895113 | 0.34778657 | 0.198 | 0.007 | 1.4976536 | 18 | pdgfc |
| lrp1ab3 | 6.499425 | 1.016886617 | 0.585 | 0.071 | 1.6511791 | 18 | lrp1ab |
| CABZ01080702.15 | 5.757029 | 0.977955043 | 0.66 | 0.09 | 1.4625733 | 18 | CABZ01080702.1 |
| cdh22 | 7.149390 | 0.864466287 | 0.542 | 0.06 | 1.8163026 | 18 | cdh2 |
| tmem981 | 6.875972 | 0.479148226 | 0.269 | 0.015 | 1.7468407 | 18 | tmem98 |
| met | 1.381107 | 0.400608355 | 0.193 | 0.007 | 3.5087046 | 18 | met |
| si:ch211-197h24.92 | 1.103853 | 0.949406643 | 0.575 | 0.077 | 2.8043386 | 18 | si:ch211-197h24.9 |
| ghrb2 | 7.980671 | 0.693862776 | 0.415 | 0.039 | 2.0274895 | 18 | ghrb |
| kirrel1b | 1.286283 | 0.354763383 | 0.165 | 0.005 | 3.2678034 | 18 | kirrel1b |
| si:dkey-56m19.51 | 1.385734 | 0.783294198 | 0.467 | 0.05 | 3.5204581 | 18 | si:dkey-56m19.5 |
| cxcl14 | 6.298030 | 0.422862077 | 0.217 | 0.01 | 1.6000145 | 18 | cxcl14 |
| pbx1b1 | 1.254667 | 0.62539469 | 0.358 | 0.03 | 3.1874824 | 18 | pbx1b |
| krt944 | 2.891846 | 1.226203151 | 0.816 | 0.157 | 7.3467357 | 18 | krt94 |
| krt85 | 2.851089 | 1.360059879 | 0.967 | 0.303 | 7.2431921 | 18 | krt8 |
| pleca1 | 4.330464 | 0.818004529 | 0.486 | 0.057 | 1.1001545 | 18 | pleca |
| lama52 | 2.141307 | 0.622334445 | 0.335 | 0.026 | 5.4399925 | 18 | lama5 |
| cd81a4 | 3.222885 | 1.242158752 | 0.901 | 0.232 | 8.1877402 | 18 | cd81a |
| mmp16a3 | 2.107760 | 0.892725639 | 0.552 | 0.072 | 5.3547662 | 18 | mmp16a |
| zeb2b | 1.756913 | 0.38300703 | 0.193 | 0.008 | 4.4634377 | 18 | zeb2b |
| ugt1b51 | 4.445939 | 0.650857758 | 0.368 | 0.033 | 1.1294908 | 18 | ugt1b5 |
| gstm.33 | 7.895868 | 1.19665635 | 0.703 | 0.123 | 2.0059453 | 18 | gstm.3 |
| CR749775.2 | 1.340289 | 0.391360062 | 0.175 | 0.007 | 3.4050049 | 18 | CR749775.2 |
| tmem1763 | 2.133952 | 0.811677907 | 0.547 | 0.074 | 5.4213075 | 18 | tmem176 |
| abl2 | 9.029095 | 0.4762824 | 0.245 | 0.015 | 2.2938417 | 18 | abl2 |
| itm2bb3 | 1.240044 | 1.147114868 | 0.887 | 0.234 | 3.1503324 | 18 | itm2bb |
| cygb13 | 1.156904 | 0.750486815 | 0.472 | 0.056 | 2.9391163 | 18 | cygb1 |
| vat12 | 4.654408 | 1.055866476 | 0.722 | 0.14 | 1.1824525 | 18 | vat1 |
| gdpd4a | 2.914109 | 0.427691293 | 0.193 | 0.009 | 7.4032948 | 18 | gdpd4a |
| nid2a1 | 1.158185 | 0.751470045 | 0.476 | 0.059 | 2.9423713 | 18 | nid2a |

| | | | | | | | |
|---|---|---|---|---|---|---|---|
| cavin2b2 | 3.069146 | 0.859281593 | 0.519 | 0.073 | 7.7971669 | 18 | cavin2b |
| adamts2 | 2.997643 | 0.553313764 | 0.297 | 0.024 | 7.6155136 | 18 | adamts2 |
| c4b1 | 1.854362 | 0.508212606 | 0.25 | 0.017 | 4.7110081 | 18 | c4b |
| mmp23 | 4.502275 | 0.904113144 | 0.623 | 0.105 | 1.1438029 | 18 | mmp2 |
| ptprga4 | 8.014250 | 0.79875317 | 0.486 | 0.066 | 2.0360203 | 18 | ptprga |
| fhl3b | 8.573152 | 0.379488075 | 0.208 | 0.012 | 2.1780095 | 18 | fhl3b |
| fabp11a3 | 1.258714 | 1.038412467 | 0.542 | 0.082 | 3.1977636 | 18 | fabp11a |
| zfp36l1b3 | 1.655644 | 0.992884406 | 0.693 | 0.133 | 4.2061645 | 18 | zfp36l1b |
| nr2f1a2 | 1.675651 | 0.749308822 | 0.453 | 0.057 | 4.2569930 | 18 | nr2f1a |
| hsd11b24 | 4.293203 | 1.098284738 | 0.736 | 0.141 | 1.0906884 | 18 | hsd11b2 |
| bmp3 | 3.867218 | 0.42541146 | 0.198 | 0.011 | 9.8246674 | 18 | bmp3 |
| zbtb16b | 4.428536 | 0.478457264 | 0.255 | 0.019 | 1.1250697 | 18 | zbtb16b |
| BX569787.11 | 3.411191 | 0.608174502 | 0.335 | 0.033 | 8.6661322 | 18 | BX569787.1 |
| paplna1 | 2.636215 | 0.497604657 | 0.288 | 0.024 | 6.6973045 | 18 | paplna |
| bmp4 | 1.721219 | 0.387342784 | 0.217 | 0.014 | 4.3727572 | 18 | bmp4 |
| pard6a | 1.748994 | 0.260057626 | 0.142 | 0.005 | 4.4433211 | 18 | pard6a |
| si:ch211-136a13.1 | 5.089081 | 0.367519893 | 0.175 | 0.009 | 1.2928810 | 18 | si:ch211-136a13.1 |
| fstl1b3 | 5.158796 | 0.916839445 | 0.703 | 0.14 | 1.3105921 | 18 | fstl1b |
| tfpia | 1.872452 | 0.470099084 | 0.245 | 0.018 | 4.7569654 | 18 | tfpia |
| slc7a6 | 2.024139 | 0.410360136 | 0.212 | 0.014 | 5.1423264 | 18 | slc7a6 |
| lrp2a | 8.744640 | 0.250949368 | 0.118 | 0.004 | 2.2215758 | 18 | lrp2a |
| col6a13 | 3.134507 | 0.809612327 | 0.557 | 0.091 | 7.9632160 | 18 | col6a1 |
| slc43a3b2 | 4.323264 | 0.515457871 | 0.288 | 0.026 | 1.0983252 | 18 | slc43a3b |
| tns2a2 | 4.779419 | 0.578587465 | 0.325 | 0.034 | 1.2142115 | 18 | tns2a |
| ahnak4 | 5.608075 | 0.912992624 | 0.642 | 0.13 | 1.4247314 | 18 | ahnak |
| rgs7bpb | 1.054395 | 0.420953332 | 0.236 | 0.018 | 2.6786927 | 18 | rgs7bpb |
| AL954191.11 | 6.343019 | 0.433515886 | 0.217 | 0.015 | 1.6114440 | 18 | AL954191.1 |
| eva1ba2 | 1.035563 | 0.701006085 | 0.443 | 0.062 | 2.6308484 | 18 | eva1ba |
| f3b3 | 2.185666 | 0.844224335 | 0.547 | 0.093 | 5.5526857 | 18 | f3b |
| rasgef1bb | 1.085475 | 0.561536487 | 0.302 | 0.03 | 2.7576515 | 18 | rasgef1bb |
| sox4a3 | 2.699645 | 0.749436479 | 0.42 | 0.058 | 6.8584505 | 18 | sox4a |
| cd248a1 | 2.466853 | 0.632077711 | 0.363 | 0.044 | 6.2670414 | 18 | cd248a |
| antxr1c1 | 2.495106 | 0.534113929 | 0.33 | 0.036 | 6.3388176 | 18 | antxr1c |
| tmem176l.13 | 4.369344 | 0.962702356 | 0.764 | 0.195 | 1.1100319 | 18 | tmem176l.1 |
| wnt11r | 7.485032 | 0.310829794 | 0.16 | 0.008 | 1.9015724 | 18 | wnt11r |

| | | | | | | | |
|---|---|---|---|---|---|---|---|
| pabpc1b4 | 1.316365 | 0.745768993 | 0.467 | 0.072 | 3.3442258 | 18 | pabpc1b |
| atp1a3b2 | 1.157213 | 0.538113455 | 0.297 | 0.03 | 2.9399006 | 18 | atp1a3b |
| cfd2 | 2.387358 | 0.840434251 | 0.443 | 0.066 | 6.0650832 | 18 | cfd |
| col5a13 | 5.981234 | 0.826689427 | 0.642 | 0.126 | 1.5195326 | 18 | col5a1 |
| cxcl18b4 | 1.211534 | 0.867544902 | 0.575 | 0.104 | 3.0779034 | 18 | cxcl18b |
| add3a4 | 4.705265 | 0.889752269 | 0.618 | 0.132 | 1.1953726 | 18 | add3a |
| aebp12 | 5.773883 | 0.578041321 | 0.358 | 0.044 | 1.4668552 | 18 | aebp1 |
| ftr821 | 1.163571 | 0.539052154 | 0.307 | 0.033 | 2.9560533 | 18 | ftr82 |
| svild2 | 3.671817 | 0.741381775 | 0.481 | 0.079 | 9.3282527 | 18 | svild |
| mtss1la1 | 3.683387 | 0.466525128 | 0.255 | 0.023 | 9.3576466 | 18 | mtss1la |
| junba5 | 4.080184 | 1.081131852 | 0.906 | 0.312 | 1.0365709 | 18 | junba |
| gpc41 | 4.618649 | 0.460943006 | 0.283 | 0.028 | 1.1733678 | 18 | gpc4 |
| dag11 | 1.658844 | 0.632617378 | 0.382 | 0.051 | 4.2142937 | 18 | dag1 |
| si:ch211-198c19.35 | 1.689237 | 0.806866432 | 0.481 | 0.078 | 4.2915084 | 18 | si:ch211-198c19.3 |
| ext1a | 1.894333 | 0.386006772 | 0.212 | 0.016 | 4.8125547 | 18 | ext1a |
| arhgap42b | 3.344726 | 0.297875002 | 0.16 | 0.009 | 8.4972785 | 18 | arhgap42b |
| slc43a2a3 | 5.812104 | 0.815388322 | 0.524 | 0.096 | 1.4765652 | 18 | slc43a2a |
| scarb2a1 | 2.652796 | 0.634234925 | 0.406 | 0.058 | 6.7394306 | 18 | scarb2a |
| pdgfrb2 | 3.383959 | 0.609319533 | 0.368 | 0.048 | 8.5969485 | 18 | pdgfrb |
| f102 | 2.543229 | 0.597228477 | 0.396 | 0.056 | 6.4610733 | 18 | f10 |
| jun4 | 2.860514 | 1.047169142 | 0.825 | 0.243 | 7.2671366 | 18 | jun |
| acvrl1 | 4.232423 | 0.385893362 | 0.208 | 0.016 | 1.0752472 | 18 | acvrl1 |
| hdac4 | 1.445637 | 0.511981453 | 0.283 | 0.03 | 3.6726425 | 18 | hdac4 |
| CU539058.11 | 5.167450 | 0.383757631 | 0.212 | 0.017 | 1.3127909 | 18 | CU539058.1 |
| col6a23 | 6.756943 | 0.666091524 | 0.443 | 0.07 | 1.7166015 | 18 | col6a2 |
| ppdpfa4 | 3.932204 | 0.661316771 | 0.434 | 0.067 | 9.9897667 | 18 | ppdpfa |
| socs3a5 | 4.571662 | 1.106484153 | 0.868 | 0.308 | 1.1614308 | 18 | socs3a |
| dachd1 | 7.653150 | 0.299285951 | 0.16 | 0.009 | 1.9442829 | 18 | dachd |
| fosab5 | 4.122286 | 1.095665486 | 0.92 | 0.354 | 1.0472669 | 18 | fosab |
| ptprua | 7.423585 | 0.255748897 | 0.137 | 0.007 | 1.8859618 | 18 | ptprua |
| anxa1a4 | 1.309322 | 0.931913455 | 0.67 | 0.156 | 3.3263335 | 18 | anxa1a |
| nr2f22 | 5.373488 | 0.603741865 | 0.358 | 0.049 | 1.3651347 | 18 | nr2f2 |
| gna12a | 1.063738 | 0.469967225 | 0.259 | 0.026 | 2.7024267 | 18 | gna12a |
| gas6 | 1.084671 | 0.478478654 | 0.269 | 0.028 | 2.7556070 | 18 | gas6 |
| mxra8a2 | 3.628723 | 0.614155559 | 0.392 | 0.058 | 9.2187721 | 18 | mxra8a |

| | | | | | | | |
|---|---|---|---|---|---|---|---|
| clu1 | 3.910266 | 0.498673803 | 0.274 | 0.029 | 9.9340327 | 18 | clu |
| dchs1a | 1.154100 | 0.2882395 | 0.156 | 0.009 | 2.9319926 | 18 | dchs1a |
| clic42 | 1.679983 | 0.592410447 | 0.373 | 0.055 | 4.2679984 | 18 | clic4 |
| zgc:1101821 | 3.102266 | 0.661998104 | 0.476 | 0.081 | 7.8813083 | 18 | zgc:110182 |
| sorbs11 | 8.047843 | 0.404863462 | 0.226 | 0.021 | 2.0445547 | 18 | sorbs1 |
| myl9b3 | 8.953459 | 0.729617373 | 0.495 | 0.094 | 2.2746264 | 18 | myl9b |
| hspg2 | 1.202000 | 0.421543304 | 0.255 | 0.026 | 3.0536832 | 18 | hspg2 |
| wu:fc27b11 | 3.397484 | 0.36932541 | 0.193 | 0.015 | 8.6313089 | 18 | wu:fc27b11 |
| clec11a1 | 8.182755 | 0.364306536 | 0.203 | 0.017 | 2.0788291 | 18 | clec11a |
| aadac | 1.473427 | 0.26620586 | 0.132 | 0.007 | 3.7432414 | 18 | aadac |
| greb1l | 1.002631 | 0.255355679 | 0.123 | 0.006 | 2.5471846 | 18 | greb1l |
| eif4ebp3 | 1.543048 | 0.91910044 | 0.552 | 0.124 | 3.9201138 | 18 | eif4ebp3 |
| grip1 | 1.708588 | 0.286610767 | 0.146 | 0.009 | 4.3406689 | 18 | grip1 |
| rbp43 | 4.342550 | 0.782441308 | 0.642 | 0.135 | 1.1032248 | 18 | rbp4 |
| si:ch73-86n18.1 | 4.407959 | 0.352485764 | 0.179 | 0.013 | 1.1198421 | 18 | si:ch73-86n18.1 |
| sema4ba3 | 5.589190 | 0.552402442 | 0.307 | 0.039 | 1.4199337 | 18 | sema4ba |
| si:dkeyp-9d4.2 | 6.837047 | 0.311407279 | 0.17 | 0.012 | 1.7369518 | 18 | si:dkeyp-9d4.2 |
| pgr1 | 1.158374 | 0.35615936 | 0.203 | 0.017 | 2.9428511 | 18 | pgr |
| sparc4 | 1.935382 | 0.876534565 | 0.741 | 0.188 | 4.9168385 | 18 | sparc |
| cald1a3 | 5.486537 | 0.751890524 | 0.542 | 0.115 | 1.3938548 | 18 | cald1a |
| eif4ebp3l4 | 9.507061 | 0.875259113 | 0.722 | 0.199 | 2.4152688 | 18 | eif4ebp3l |
| mcl1b5 | 3.194191 | 0.901978515 | 0.783 | 0.244 | 8.1148436 | 18 | mcl1b |
| jcada | 3.927036 | 0.36397842 | 0.208 | 0.019 | 9.9766373 | 18 | jcada |
| rhobtb4 | 3.499485 | 0.38572263 | 0.208 | 0.019 | 8.8904424 | 18 | rhobtb4 |
| tnfaip6 | 7.124913 | 0.359504833 | 0.193 | 0.016 | 1.8100842 | 18 | tnfaip6 |
| CR848833.1 | 1.050487 | 0.256020256 | 0.113 | 0.005 | 2.6687637 | 18 | CR848833.1 |
| col18a1a2 | 1.650468 | 0.623512355 | 0.325 | 0.046 | 4.1930139 | 18 | col18a1a |
| si:dkey-7j14.63 | 8.528402 | 0.844618896 | 0.675 | 0.19 | 2.1666407 | 18 | si:dkey-7j14.6 |
| desi1a | 8.886025 | 0.415031519 | 0.231 | 0.024 | 2.2574947 | 18 | desi1a |
| si:ch73-46j18.55 | 3.111601 | 0.899826214 | 0.708 | 0.202 | 7.9050235 | 18 | si:ch73-46j18.5 |
| ywhag1 | 9.304544 | 0.253739312 | 0.156 | 0.011 | 2.3638194 | 18 | ywhag1 |
| grb10b1 | 1.084036 | 0.481216062 | 0.288 | 0.037 | 2.7539946 | 18 | grb10b |
| dkk3b3 | 2.363389 | 0.525440291 | 0.368 | 0.057 | 6.0041899 | 18 | dkk3b |
| CABZ01068366.12 | 1.401762 | 0.509325638 | 0.292 | 0.039 | 3.5611784 | 18 | CABZ01068366.1 |
| col27a1b2 | 5.384573 | 0.427260836 | 0.236 | 0.026 | 1.3679508 | 18 | col27a1b |

| | | | | | | | |
|---|---|---|---|---|---|---|---|
| gstt1b4 | 5.658023 | 0.612978574 | 0.448 | 0.085 | 1.4374209 | 18 | gstt1b |
| lxn4 | 1.572511 | 0.850436937 | 0.608 | 0.162 | 3.9949663 | 18 | lxn |
| col1a1a3 | 1.224819 | 0.748353126 | 0.67 | 0.16 | 3.1116545 | 18 | col1a1a |
| XLOC-024248 | 1.443761 | 0.357710846 | 0.208 | 0.02 | 3.6678750 | 18 | XLOC-024248 |
| si:ch73-22o12.11 | 1.605001 | 0.458597211 | 0.245 | 0.028 | 4.0775075 | 18 | si:ch73-22o12.1 |
| mknk2b4 | 5.279756 | 0.813529868 | 0.675 | 0.194 | 1.3413220 | 18 | mknk2b |
| FP016056.11 | 9.688132 | 0.270604161 | 0.137 | 0.009 | 2.4612700 | 18 | FP016056.1 |
| id14 | 9.305888 | 0.856191904 | 0.538 | 0.126 | 2.3641610 | 18 | id1 |
| tagln4 | 1.299238 | 0.611775297 | 0.401 | 0.07 | 3.3007156 | 18 | tagln |
| nr0b1 | 1.522081 | 0.318160649 | 0.175 | 0.015 | 3.8668471 | 18 | nr0b1 |
| dlc1 | 1.709301 | 0.458691096 | 0.25 | 0.03 | 4.3424812 | 18 | dlc1 |
| tspan4a2 | 5.485768 | 0.441101477 | 0.269 | 0.035 | 1.3936594 | 18 | tspan4a |
| plod22 | 6.338730 | 0.411468526 | 0.236 | 0.027 | 1.6103545 | 18 | plod2 |
| arhgef51 | 1.088411 | 0.438580404 | 0.245 | 0.03 | 2.7651082 | 18 | arhgef5 |
| s100a10b5 | 1.548782 | 0.92527222 | 0.825 | 0.279 | 3.9346822 | 18 | s100a10b |
| CABZ01068367.11 | 3.045267 | 0.360377849 | 0.208 | 0.022 | 7.7365029 | 18 | CABZ01068367.1 |
| col1a1b3 | 6.525048 | 0.726520576 | 0.656 | 0.164 | 1.6576884 | 18 | col1a1b |
| CR383676.18 | 1.038500 | 0.779248149 | 1 | 0.714 | 2.6383103 | 18 | CR383676.1 |
| si:dkey-28b4.74 | 1.050636 | 0.636589055 | 0.476 | 0.105 | 2.6691429 | 18 | si:dkey-28b4.7 |
| pmp22a3 | 1.302962 | 0.755016587 | 0.642 | 0.161 | 3.3101759 | 18 | pmp22a |
| selenbp12 | 1.392405 | 0.590522521 | 0.406 | 0.077 | 3.5374069 | 18 | selenbp1 |
| CR318588.31 | 3.175783 | 0.583129179 | 0.316 | 0.048 | 8.0680776 | 18 | CR318588.3 |
| enc12 | 3.964243 | 0.48547666 | 0.269 | 0.036 | 1.0071159 | 18 | enc1 |
| epb41l51 | 1.210090 | 0.453233801 | 0.288 | 0.04 | 3.0742350 | 18 | epb41l5 |
| ywhaz3 | 2.852012 | 0.845554037 | 0.764 | 0.299 | 7.2455382 | 18 | ywhaz |
| ugt5b21 | 7.750601 | 0.427332337 | 0.241 | 0.03 | 1.9690403 | 18 | ugt5b2 |
| col1a23 | 1.140488 | 0.751164374 | 0.689 | 0.176 | 2.8974102 | 18 | col1a2 |
| stard13b | 1.156059 | 0.258882555 | 0.142 | 0.01 | 2.9369687 | 18 | stard13b |
| cav11 | 1.496775 | 0.499816618 | 0.307 | 0.046 | 3.8025587 | 18 | cav1 |
| rab6ba | 1.694547 | 0.298866591 | 0.156 | 0.013 | 4.3049986 | 18 | rab6ba |
| foxo3b3 | 2.134217 | 0.579282827 | 0.368 | 0.067 | 5.4219795 | 18 | foxo3b |
| s100v24 | 4.301797 | 0.710606179 | 0.542 | 0.136 | 1.0928715 | 18 | s100v2 |
| tnfsf121 | 5.165579 | 0.361020863 | 0.217 | 0.025 | 1.3123155 | 18 | tnfsf12 |
| tcf122 | 2.869922 | 0.529878134 | 0.358 | 0.065 | 7.2910372 | 18 | tcf12 |
| neo1a | 2.908989 | 0.40764861 | 0.236 | 0.03 | 7.3902878 | 18 | neo1a |

| | | | | | | | |
|---|---|---|---|---|---|---|---|
| tjp2b2 | 5.208774 | 0.501418393 | 0.349 | 0.061 | 1.3232890 | 18 | tjp2b |
| slc38a5b3 | 1.037483 | 0.647135551 | 0.509 | 0.124 | 2.6357278 | 18 | slc38a5b |
| col21a1 | 1.138710 | 0.36916164 | 0.184 | 0.019 | 2.8928927 | 18 | col21a1 |
| tmem204 | 1.826933 | 0.256851586 | 0.127 | 0.009 | 4.6413243 | 18 | tmem204 |
| ptprsa1 | 3.770639 | 0.385294456 | 0.236 | 0.03 | 9.5793103 | 18 | ptprsa |
| rasgef1ba1 | 8.371715 | 0.519113 | 0.335 | 0.058 | 2.1268341 | 18 | rasgef1ba |
| loxl2a3 | 1.837092 | 0.50502969 | 0.288 | 0.044 | 4.6671336 | 18 | loxl2a |
| lipf2 | 2.503142 | 0.486755279 | 0.288 | 0.045 | 6.3592337 | 18 | lipf |
| dcn3 | 9.200741 | 0.650523359 | 0.58 | 0.139 | 2.3374484 | 18 | dcn |
| zgc:1627308 | 4.150021 | 0.860696491 | 0.896 | 0.397 | 1.0543128 | 18 | zgc:162730 |
| phactr4b | 3.017914 | 0.333255114 | 0.189 | 0.021 | 7.6670127 | 18 | phactr4b |
| si:ch211-74m13.31 | 3.553633 | 0.531235747 | 0.349 | 0.065 | 9.0280049 | 18 | si:ch211-74m13.3 |
| agrn2 | 4.088101 | 0.499433969 | 0.325 | 0.057 | 1.0385822 | 18 | agrn |
| gadd45ba3 | 6.222887 | 0.762013711 | 0.637 | 0.192 | 1.5809246 | 18 | gadd45ba |
| il6st2 | 6.612916 | 0.467392367 | 0.311 | 0.053 | 1.6800113 | 18 | il6st |
| stxbp61 | 6.830253 | 0.415398315 | 0.283 | 0.043 | 1.7352260 | 18 | stxbp6 |
| mir199-31 | 8.768242 | 0.373828008 | 0.189 | 0.021 | 2.2275720 | 18 | mir199-3 |
| gata61 | 1.067681 | 0.364002129 | 0.203 | 0.024 | 2.7124454 | 18 | gata6 |
| mir21-12 | 1.251808 | 0.622986685 | 0.448 | 0.104 | 3.1802197 | 18 | mir21-1 |
| hipk21 | 2.216215 | 0.349531915 | 0.184 | 0.02 | 5.6302962 | 18 | hipk2 |
| robo11 | 5.109410 | 0.354573215 | 0.208 | 0.025 | 1.2980458 | 18 | robo1 |
| robo22 | 5.312504 | 0.455586826 | 0.307 | 0.052 | 1.3496417 | 18 | robo2 |
| deptor2 | 9.477865 | 0.409076688 | 0.241 | 0.034 | 2.4078517 | 18 | deptor |
| rhpn21 | 1.049122 | 0.389278314 | 0.241 | 0.033 | 2.6652957 | 18 | rhpn2 |
| XLOC-034538 | 1.320613 | 0.298743172 | 0.156 | 0.015 | 3.3550192 | 18 | XLOC-034538 |
| colec122 | 1.470148 | 0.508928962 | 0.33 | 0.06 | 3.7349125 | 18 | colec12 |
| zgc:1951751 | 1.646539 | 0.397090675 | 0.245 | 0.035 | 4.1830341 | 18 | zgc:195175 |
| xirp2a2 | 2.709225 | 0.381572694 | 0.236 | 0.032 | 6.8827878 | 18 | xirp2a |
| dpp3 | 3.223109 | 0.460079488 | 0.283 | 0.045 | 8.1883091 | 18 | dpp3 |
| lamb1a3 | 3.621740 | 0.508433151 | 0.302 | 0.052 | 9.2010322 | 18 | lamb1a |
| usp43b1 | 4.461703 | 0.32431707 | 0.165 | 0.016 | 1.1334956 | 18 | usp43b |
| fam171a2a1 | 1.155343 | 0.315509987 | 0.179 | 0.02 | 2.9351506 | 18 | fam171a2a |
| kank3 | 1.209108 | 0.273351509 | 0.16 | 0.016 | 3.0717409 | 18 | kank3 |
| si:ch211-222l21.17 | 1.322562 | 0.796652224 | 0.802 | 0.308 | 3.3599697 | 18 | si:ch211-222l21.1 |
| lrp1aa2 | 3.275592 | 0.529421918 | 0.335 | 0.063 | 8.3216423 | 18 | lrp1aa |

| | | | | | | | |
|---|---|---|---|---|---|---|---|
| nrg2a | 4.429048 | 0.26622012 | 0.142 | 0.012 | 1.1251998 | 18 | nrg2a |
| wbp1lb | 6.921254 | 0.454954111 | 0.283 | 0.047 | 1.7583445 | 18 | wbp1lb |
| anos1a1 | 1.073649 | 0.280934915 | 0.156 | 0.015 | 2.7276072 | 18 | anos1a |
| si:ch211-1a19.33 | 1.199528 | 0.47295585 | 0.311 | 0.055 | 3.0474033 | 18 | si:ch211-1a19.3 |
| col5a2a3 | 1.311630 | 0.567311388 | 0.434 | 0.097 | 3.3321962 | 18 | col5a2a |
| itgb5 | 1.839196 | 0.264374942 | 0.151 | 0.014 | 4.6724794 | 18 | itgb5 |
| hspb8 | 1.907264 | 0.321481865 | 0.189 | 0.022 | 4.8454064 | 18 | hspb8 |
| si:ch211-250c4.41 | 2.312075 | 0.283885805 | 0.165 | 0.017 | 5.8738270 | 18 | si:ch211-250c4.4 |
| nudt4a | 2.788301 | 0.325989837 | 0.17 | 0.018 | 7.0836799 | 18 | nudt4a |
| pdgfrl1 | 3.644920 | 0.367756601 | 0.226 | 0.031 | 9.2599207 | 18 | pdgfrl |
| cnn23 | 3.672103 | 0.653230016 | 0.542 | 0.153 | 9.3289797 | 18 | cnn2 |
| gsta.12 | 4.897208 | 0.420772987 | 0.288 | 0.048 | 1.2441357 | 18 | gsta.1 |
| frzb1 | 1.100501 | 0.345880119 | 0.17 | 0.018 | 2.7958229 | 18 | frzb |
| mfap23 | 1.167152 | 0.597413023 | 0.481 | 0.115 | 2.9651502 | 18 | mfap2 |
| hdlbpa4 | 1.196005 | 0.668527638 | 0.575 | 0.17 | 3.0384507 | 18 | hdlbpa |
| mgst1.23 | 2.473458 | 0.613482366 | 0.458 | 0.113 | 6.2838225 | 18 | mgst1.2 |
| ms4a17a.8 | 2.740748 | 0.254011838 | 0.165 | 0.017 | 6.9628709 | 18 | ms4a17a.8 |
| pi15a2 | 3.266798 | 0.49927808 | 0.302 | 0.052 | 8.2993008 | 18 | pi15a |
| tle2b | 6.056822 | 0.287490695 | 0.156 | 0.016 | 1.5387357 | 18 | tle2b |
| znf385a3 | 9.038468 | 0.464211996 | 0.292 | 0.052 | 2.2962230 | 18 | znf385a |
| si:dkey-58f10.111 | 1.047996 | 0.332380397 | 0.198 | 0.025 | 2.6624351 | 18 | si:dkey-58f10.11 |
| pgrmc12 | 1.294911 | 0.639413883 | 0.528 | 0.148 | 3.2897223 | 18 | pgrmc1 |
| pik3r22 | 3.689956 | 0.508640512 | 0.325 | 0.064 | 9.3743355 | 18 | pik3r2 |
| ar3 | 3.718629 | 0.508868899 | 0.325 | 0.064 | 9.4471782 | 18 | ar |
| pfkfb4b2 | 1.442422 | 0.60498034 | 0.453 | 0.117 | 3.6644748 | 18 | pfkfb4b |
| appa3 | 7.512539 | 0.50722949 | 0.382 | 0.085 | 1.9085605 | 18 | appa |
| COX38 | 1.468139 | 0.620580334 | 0.991 | 0.673 | 3.7298087 | 18 | COX3 |
| mob2a1 | 5.431244 | 0.399736483 | 0.241 | 0.038 | 1.3798076 | 18 | mob2a |
| cavin1b2 | 5.447041 | 0.436813985 | 0.259 | 0.043 | 1.3838208 | 18 | cavin1b |
| rhoj | 8.153758 | 0.272230339 | 0.146 | 0.015 | 2.0714623 | 18 | rhoj |
| junbb2 | 1.292791 | 0.489639401 | 0.325 | 0.067 | 3.2843358 | 18 | junbb |
| mxra8b1 | 1.299247 | 0.330986812 | 0.203 | 0.028 | 3.3007383 | 18 | mxra8b |
| rbpms2a | 5.531269 | 0.461745401 | 0.373 | 0.079 | 1.4052189 | 18 | rbpms2a |
| mt-co27 | 6.477357 | 0.633630101 | 0.986 | 0.668 | 1.6455725 | 18 | mt-co2 |
| ppp1r14bb | 8.515302 | 0.381512616 | 0.245 | 0.04 | 2.1633127 | 18 | ppp1r14bb |

| | | | | | | | |
|---|---|---|---|---|---|---|---|
| gamt1 | 1.356582 | 0.37082161 | 0.25 | 0.041 | 3.4463978 | 18 | gamt |
| olfml3b3 | 1.369535 | 0.426071155 | 0.288 | 0.053 | 3.4793059 | 18 | olfml3b |
| egfra3 | 1.676923 | 0.338301131 | 0.212 | 0.031 | 4.2602249 | 18 | egfra |
| mt-nd47 | 1.694010 | 0.706308936 | 0.906 | 0.443 | 4.3036325 | 18 | mt-nd4 |
| ifi464 | 3.694414 | 0.576517839 | 0.476 | 0.128 | 9.3856602 | 18 | ifi46 |
| LOC1018844512 | 7.101429 | 0.556329666 | 0.373 | 0.088 | 1.8041182 | 18 | LOC101884451 |
| chd34 | 1.014548 | 0.567008642 | 0.453 | 0.122 | 2.5774614 | 18 | chd3 |
| ksr1b | 1.742449 | 0.261952686 | 0.137 | 0.014 | 4.4266917 | 18 | ksr1b |
| mt-nd17 | 4.221473 | 0.690974902 | 0.863 | 0.391 | 1.0724654 | 18 | mt-nd1 |
| mogat2 | 9.378088 | 0.29617213 | 0.142 | 0.015 | 2.3825034 | 18 | mogat2 |
| ephb33 | 1.580407 | 0.441237104 | 0.269 | 0.05 | 4.0150250 | 18 | ephb3 |
| hbp1 | 4.486319 | 0.44243324 | 0.288 | 0.056 | 1.1397495 | 18 | hbp1 |
| egr16 | 1.139260 | 0.655075585 | 0.561 | 0.179 | 2.8942918 | 18 | egr1 |
| nrxn1a1 | 1.674333 | 0.25243411 | 0.123 | 0.012 | 4.2536441 | 18 | nrxn1a |
| efna1b1 | 2.256370 | 0.256565626 | 0.156 | 0.018 | 5.7323100 | 18 | efna1b |
| nr1d2a1 | 3.044521 | 0.511491706 | 0.363 | 0.086 | 7.7346067 | 18 | nr1d2a |
| ccdc85b3 | 2.019469 | 0.42929751 | 0.259 | 0.049 | 5.1304625 | 18 | ccdc85b |
| mmp14a2 | 2.321009 | 0.407129161 | 0.288 | 0.057 | 5.8965235 | 18 | mmp14a |
| ptpn211 | 3.050676 | 0.333025531 | 0.217 | 0.035 | 7.7502448 | 18 | ptpn21 |
| mt-nd26 | 6.370370 | 0.679950938 | 0.844 | 0.387 | 1.6183925 | 18 | mt-nd2 |
| tcf4 | 7.234551 | 0.293392991 | 0.17 | 0.022 | 1.8379378 | 18 | tcf4 |
| gulp1a2 | 8.850357 | 0.342074872 | 0.217 | 0.035 | 2.2484332 | 18 | gulp1a |
| pkp3b1 | 9.175333 | 0.404210561 | 0.255 | 0.046 | 2.3309934 | 18 | pkp3b |
| cybrd1 | 1.013748 | 0.387446629 | 0.264 | 0.05 | 2.5754289 | 18 | cybrd1 |
| kank23 | 1.107089 | 0.382722654 | 0.274 | 0.052 | 2.8125609 | 18 | kank2 |
| cntfr | 1.715589 | 0.271333517 | 0.156 | 0.019 | 4.3584561 | 18 | cntfr |
| tnfrsfa | 2.021525 | 0.319808772 | 0.198 | 0.03 | 5.1356859 | 18 | tnfrsfa |
| fkbp53 | 2.333370 | 0.549350947 | 0.439 | 0.124 | 5.9279274 | 18 | fkbp5 |
| mt-atp67 | 2.532254 | 0.653960317 | 0.948 | 0.552 | 6.4331924 | 18 | mt-atp6 |
| mt-cyb6 | 2.742137 | 0.642126306 | 0.976 | 0.572 | 6.9664015 | 18 | mt-cyb |
| lrrc1 | 3.952967 | 0.294460774 | 0.179 | 0.025 | 1.0042514 | 18 | lrrc1 |
| b2ml6 | 4.932746 | 0.654074202 | 0.708 | 0.289 | 1.2531642 | 18 | b2ml |
| marcksl1b5 | 4.951868 | 0.62306956 | 0.472 | 0.138 | 1.2580221 | 18 | marcksl1b |
| cdc42bpb | 5.751127 | 0.297240124 | 0.189 | 0.027 | 1.4610739 | 18 | cdc42bpb |
| mt-nd36 | 6.838555 | 0.669586206 | 0.821 | 0.366 | 1.7373349 | 18 | mt-nd3 |

| | | | | | | | |
|---|---|---|---|---|---|---|---|
| si:dkey-11f4.143 | 1.161695 | 0.604009172 | 0.42 | 0.119 | 2.9512865 | 18 | si:dkey-11f4.14 |
| itga11 | 3.301951 | 0.389044188 | 0.231 | 0.041 | 8.3886082 | 18 | itga1 |
| nfil3-65 | 4.383080 | 0.581775592 | 0.443 | 0.128 | 1.1135216 | 18 | nfil3-6 |
| actn13 | 4.564325 | 0.433957843 | 0.245 | 0.046 | 1.1595667 | 18 | actn1 |
| itgav2 | 4.806813 | 0.424802467 | 0.288 | 0.06 | 1.2211709 | 18 | itgav |
| grinab4 | 6.178171 | 0.521910411 | 0.425 | 0.119 | 1.5695643 | 18 | grinab |
| mt-co17 | 2.158085 | 0.587352043 | 0.991 | 0.636 | 5.4826166 | 18 | mt-co1 |
| b2m6 | 2.359415 | 0.662478101 | 0.708 | 0.294 | 5.9940963 | 18 | b2m |
| mdka4 | 2.370037 | 0.588038146 | 0.41 | 0.11 | 6.0210811 | 18 | mdka |
| adam10a2 | 2.741013 | 0.473126616 | 0.392 | 0.103 | 6.9635446 | 18 | adam10a |
| wls3 | 3.170545 | 0.537621759 | 0.443 | 0.128 | 8.0547699 | 18 | wls |
| zgc:1951734 | 3.846656 | 0.523907773 | 0.41 | 0.111 | 9.7724320 | 18 | zgc:195173 |
| calcoco1a1 | 4.269332 | 0.340343052 | 0.208 | 0.034 | 1.0846238 | 18 | calcoco1a |
| mpp5a | 5.485001 | 0.250364977 | 0.132 | 0.015 | 1.3934647 | 18 | mpp5a |
| serinc51 | 6.932513 | 0.455420162 | 0.349 | 0.085 | 1.7612050 | 18 | serinc5 |
| si:dkey-42i9.7 | 7.317953 | 0.333733261 | 0.175 | 0.025 | 1.8591261 | 18 | si:dkey-42i9.7 |
| nrp2a1 | 2.977511 | 0.285183286 | 0.151 | 0.019 | 7.5643676 | 18 | nrp2a |
| txnipa3 | 3.580760 | 0.648004086 | 0.623 | 0.24 | 9.0969212 | 18 | txnipa |
| si:dkey-188i13.103 | 5.271995 | 0.483026808 | 0.406 | 0.111 | 1.3393503 | 18 | si:dkey-188i13.10 |
| si:dkeyp-75h12.51 | 5.531703 | 0.361919606 | 0.231 | 0.042 | 1.4053293 | 18 | si:dkeyp-75h12.5 |
| cbx7a2 | 6.323234 | 0.405090454 | 0.255 | 0.05 | 1.6064176 | 18 | cbx7a |
| pdk2a1 | 9.278459 | 0.316116972 | 0.184 | 0.028 | 2.3571927 | 18 | pdk2a |
| btg24 | 1.155315 | 0.682324001 | 0.75 | 0.345 | 2.9350800 | 18 | btg2 |
| fstl32 | 2.370605 | 0.276491653 | 0.165 | 0.023 | 6.0225243 | 18 | fstl3 |
| NAV1 (1 of many) | 2.495743 | 0.311072077 | 0.175 | 0.026 | 6.3404356 | 18 | NAV1 (1 of many) |
| si:ch211-212k18.82 | 3.821169 | 0.364968474 | 0.241 | 0.046 | 9.7076820 | 18 | si:ch211-212k18.8 |
| ralgps2 | 6.448197 | 0.264962934 | 0.16 | 0.022 | 1.6381645 | 18 | ralgps2 |
| si:ch211-140m22.71 | 1.229059 | 0.428602608 | 0.288 | 0.065 | 3.1224257 | 18 | si:ch211-140m22.7 |
| ier2a5 | 1.568290 | 0.69445561 | 0.642 | 0.27 | 3.9842413 | 18 | ier2a |
| si:dkey-101k6.5 | 1.997755 | 0.307580739 | 0.142 | 0.018 | 5.0752989 | 18 | si:dkey-101k6.5 |
| sox61 | 2.036534 | 0.313000252 | 0.189 | 0.03 | 5.1738169 | 18 | sox6 |
| si:dkey-23i12.72 | 5.738105 | 0.339003083 | 0.175 | 0.027 | 1.4577656 | 18 | si:dkey-23i12.7 |
| sptan11 | 9.559135 | 0.387988714 | 0.264 | 0.056 | 2.4284984 | 18 | sptan1 |
| plpp1a1 | 2.250891 | 0.430535255 | 0.316 | 0.075 | 5.7183909 | 18 | plpp1a |
| serinc12 | 4.531338 | 0.445499923 | 0.33 | 0.084 | 1.1511866 | 18 | serinc1 |

| | | | | | | | |
|---|---|---|---|---|---|---|---|
| si:dkey-6n21.121 | 5.007927 | 0.328771787 | 0.217 | 0.04 | 1.2722641 | 18 | si:dkey-6n21.12 |
| plxnb2a | 5.583211 | 0.311970173 | 0.17 | 0.026 | 1.4184147 | 18 | plxnb2a |
| reep3a1 | 7.895440 | 0.275338056 | 0.179 | 0.028 | 2.0058367 | 18 | reep3a |
| zbtb46 | 1.051202 | 0.303356156 | 0.17 | 0.026 | 2.6705802 | 18 | zbtb46 |
| prss351 | 1.833026 | 0.304843066 | 0.175 | 0.027 | 4.6568041 | 18 | prss35 |
| rsrp11 | 2.020442 | 0.48597495 | 0.41 | 0.119 | 5.1329350 | 18 | rsrp1 |
| si:dkey-9i23.162 | 2.784138 | 0.379288759 | 0.236 | 0.048 | 7.0731043 | 18 | si:dkey-9i23.16 |
| pbx41 | 3.038065 | 0.415090339 | 0.302 | 0.072 | 7.7182042 | 18 | pbx4 |
| nt5c2b1 | 3.142403 | 0.389285437 | 0.226 | 0.045 | 7.9832749 | 18 | nt5c2b |
| sik2b1 | 3.538178 | 0.363619774 | 0.222 | 0.043 | 8.9887424 | 18 | sik2b |
| tspan141 | 3.920754 | 0.496964804 | 0.425 | 0.129 | 9.9606759 | 18 | tspan14 |
| atp10a3 | 8.058362 | 0.392378975 | 0.259 | 0.057 | 2.0472270 | 18 | atp10a |
| cyr613 | 1.568361 | 0.472656291 | 0.283 | 0.065 | 3.9844235 | 18 | cyr61 |
| pabpn12 | 2.498418 | 0.580156952 | 0.665 | 0.27 | 6.3472322 | 18 | pabpn1 |
| rpz54 | 3.417134 | 0.513497849 | 0.41 | 0.124 | 8.6812302 | 18 | rpz5 |
| ppfibp1a | 3.515951 | 0.313456314 | 0.222 | 0.042 | 8.9322735 | 18 | ppfibp1a |
| si:ch211-15b10.63 | 1.188242 | 0.422924273 | 0.297 | 0.074 | 3.0187294 | 18 | si:ch211-15b10.6 |
| NC-002333.46 | 1.282576 | 0.569581038 | 0.976 | 0.605 | 3.2583852 | 18 | NC-002333.4 |
| FO082781.14 | 1.459430 | 0.401063914 | 0.245 | 0.053 | 3.7076820 | 18 | FO082781.1 |
| rab11fip1b1 | 1.760546 | 0.401903927 | 0.259 | 0.058 | 4.4726692 | 18 | rab11fip1b |
| zgc:66433 | 1.946550 | 0.300857694 | 0.189 | 0.033 | 4.9452112 | 18 | zgc:66433 |
| kmt2cb1 | 2.400907 | 0.384892358 | 0.259 | 0.059 | 6.0995046 | 18 | kmt2cb |
| myo1cb1 | 2.414732 | 0.321867461 | 0.193 | 0.035 | 6.1346289 | 18 | myo1cb |
| tjp1a1 | 3.316226 | 0.371255518 | 0.241 | 0.051 | 8.4248741 | 18 | tjp1a |
| ctdsp23 | 3.499907 | 0.418421154 | 0.316 | 0.083 | 8.8915160 | 18 | ctdsp2 |
| cyp3c12 | 1.111169 | 0.382787934 | 0.269 | 0.063 | 2.8229249 | 18 | cyp3c1 |
| zbtb42 | 1.149060 | 0.273856831 | 0.151 | 0.023 | 2.9191886 | 18 | zbtb4 |
| cgnl11 | 1.802710 | 0.2633683 | 0.17 | 0.028 | 4.5797860 | 18 | cgnl1 |
| si:ch211-196l7.47 | 1.833408 | 0.587375933 | 0.703 | 0.326 | 4.6577752 | 18 | si:ch211-196l7.4 |
| calm2b4 | 1.891265 | 0.563735606 | 0.608 | 0.243 | 4.8047589 | 18 | calm2b |
| fzd22 | 3.949011 | 0.310061533 | 0.208 | 0.041 | 1.0032463 | 18 | fzd2 |
| zgc:1748632 | 6.943130 | 0.431860104 | 0.368 | 0.109 | 1.7639022 | 18 | zgc:174863 |
| ngdn | 9.228988 | 0.455865884 | 0.335 | 0.094 | 2.3446245 | 18 | ngdn |
| si:ch211-74f19.21 | 1.299302 | 0.308631456 | 0.175 | 0.03 | 3.3008773 | 18 | si:ch211-74f19.2 |
| hcar1-41 | 1.432141 | 0.290268188 | 0.179 | 0.032 | 3.6383565 | 18 | hcar1-4 |

| | | | | | | | |
|---|---|---|---|---|---|---|---|
| cyp2ad33 | 1.529963 | 0.380704171 | 0.259 | 0.06 | 3.8868729 | 18 | cyp2ad3 |
| arhgef1b1 | 1.557294 | 0.424826772 | 0.297 | 0.077 | 3.9563065 | 18 | arhgef1b |
| wwtr1 | 1.922851 | 0.27283333 | 0.165 | 0.028 | 4.8850030 | 18 | wwtr1 |
| si:ch211-212k18.92 | 1.986631 | 0.32063387 | 0.226 | 0.048 | 5.0470363 | 18 | si:ch211-212k18.9 |
| si:ch211-166a6.51 | 2.208445 | 0.300591444 | 0.193 | 0.037 | 5.6105559 | 18 | si:ch211-166a6.5 |
| stat1a3 | 3.761603 | 0.461945559 | 0.387 | 0.121 | 9.5563527 | 18 | stat1a |
| cyp2p73 | 4.883399 | 0.294001719 | 0.17 | 0.029 | 1.2406276 | 18 | cyp2p7 |
| lamc11 | 6.299804 | 0.376783209 | 0.245 | 0.056 | 1.6004652 | 18 | lamc1 |
| h1f03 | 9.847024 | 0.483906308 | 0.274 | 0.069 | 2.5016366 | 18 | h1f0 |
| nid1a1 | 1.693043 | 0.297272091 | 0.198 | 0.038 | 4.3011772 | 18 | nid1a |
| mphosph83 | 3.156955 | 0.476178164 | 0.415 | 0.138 | 8.0202456 | 18 | mphosph8 |
| erbb21 | 3.478281 | 0.273630734 | 0.175 | 0.032 | 8.8365737 | 18 | erbb2 |
| si:ch211-199g17.2 | 3.895780 | 0.408450867 | 0.269 | 0.067 | 9.8972313 | 18 | si:ch211-199g17.2 |
| si:rp71-79p20.21 | 4.926562 | 0.351438318 | 0.193 | 0.039 | 1.2515932 | 18 | si:rp71-79p20.2 |
| ctbp2a1 | 5.460587 | 0.285155087 | 0.198 | 0.039 | 1.3872621 | 18 | ctbp2a |
| itm2ba4 | 8.227205 | 0.550645664 | 0.646 | 0.293 | 2.0901215 | 18 | itm2ba |
| si:ch73-335l21.43 | 8.331276 | 0.656612782 | 0.406 | 0.139 | 2.1165606 | 18 | si:ch73-335l21.4 |
| si:ch73-237c6.13 | 8.426377 | 0.298988981 | 0.212 | 0.044 | 2.1407213 | 18 | si:ch73-237c6.1 |
| tp53inp11 | 8.563411 | 0.37324013 | 0.245 | 0.058 | 2.1755347 | 18 | tp53inp1 |
| zgc:92140 | 8.745683 | 0.254690932 | 0.17 | 0.03 | 2.2218410 | 18 | zgc:92140 |
| socs3b4 | 8.803924 | 0.552574645 | 0.429 | 0.149 | 2.2366369 | 18 | socs3b |
| celf26 | 1.688521 | 0.547461989 | 0.604 | 0.254 | 4.2896892 | 18 | celf2 |
| jak12 | 1.782275 | 0.451159811 | 0.33 | 0.098 | 4.5278697 | 18 | jak1 |
| si:zfos-464b6.2 | 1.884027 | 0.310880008 | 0.217 | 0.046 | 4.7863730 | 18 | si:zfos-464b6.2 |
| gng73 | 5.887874 | 0.334154274 | 0.25 | 0.06 | 1.4958144 | 18 | gng7 |
| tfe3a1 | 8.980249 | 0.302732259 | 0.193 | 0.039 | 2.2814325 | 18 | tfe3a |
| ifi456 | 1.320756 | 0.534475266 | 0.41 | 0.136 | 3.3553822 | 18 | ifi45 |
| sypl2a1 | 2.232972 | 0.321192336 | 0.245 | 0.059 | 5.6728671 | 18 | sypl2a |
| crebzf2 | 2.565943 | 0.445671034 | 0.344 | 0.107 | 6.5187794 | 18 | crebzf |
| atf4b4 | 3.867745 | 0.52892451 | 0.623 | 0.274 | 9.8260079 | 18 | atf4b |
| pdlim11 | 3.998252 | 0.315828354 | 0.245 | 0.059 | 1.0157559 | 18 | pdlim1 |
| tpm4a4 | 4.508407 | 0.410167953 | 0.363 | 0.111 | 1.1453609 | 18 | tpm4a |
| ddit34 | 4.906386 | 0.545426229 | 0.462 | 0.176 | 1.2464675 | 18 | ddit3 |
| tspan18b1 | 8.810458 | 0.316673466 | 0.193 | 0.04 | 2.2382969 | 18 | tspan18b |
| ier2b6 | 1.084439 | 0.541781915 | 0.557 | 0.234 | 2.7550195 | 18 | ier2b |

| | | | | | | | |
|---|---|---|---|---|---|---|---|
| itgb1a2 | 1.760026 | 0.327092241 | 0.222 | 0.051 | 4.4713485 | 18 | itgb1a |
| cast2 | 1.795834 | 0.387579266 | 0.269 | 0.071 | 4.5623167 | 18 | cast |
| CABZ01068358.13 | 2.352888 | 0.386337414 | 0.255 | 0.065 | 5.9775124 | 18 | CABZ01068358.1 |
| serinc22 | 2.443268 | 0.431563352 | 0.373 | 0.122 | 6.2071238 | 18 | serinc2 |
| cebpb7 | 2.472167 | 0.592018019 | 0.736 | 0.335 | 6.2805403 | 18 | cebpb |
| creg2 | 3.052300 | 0.272809272 | 0.175 | 0.034 | 7.7543701 | 18 | creg2 |
| afdna1 | 3.534293 | 0.259190679 | 0.146 | 0.025 | 8.9788731 | 18 | afdna |
| hmgn67 | 3.912206 | 0.532177036 | 0.67 | 0.307 | 9.9389602 | 18 | hmgn6 |
| CR318588.18 | 6.679047 | 0.703301732 | 0.514 | 0.213 | 1.6968120 | 18 | CR318588.1 |
| fundc21 | 9.600459 | 0.364186956 | 0.316 | 0.094 | 2.4389966 | 18 | fundc2 |
| selenow13 | 9.884967 | 0.542998444 | 0.764 | 0.412 | 2.5112760 | 18 | selenow1 |
| map2k62 | 1.172837 | 0.381713026 | 0.321 | 0.096 | 2.9795944 | 18 | map2k6 |
| pi4kb | 1.734130 | 0.280005249 | 0.179 | 0.037 | 4.4055576 | 18 | pi4kb |
| fkbp73 | 1.895838 | 0.290391978 | 0.212 | 0.049 | 4.8163767 | 18 | fkbp7 |
| ecm1b1 | 2.104807 | 0.33517507 | 0.255 | 0.065 | 5.3472641 | 18 | ecm1b |
| klf11b3 | 4.644785 | 0.39013355 | 0.283 | 0.081 | 1.1800078 | 18 | klf11b |
| flna4 | 6.927121 | 0.418001366 | 0.363 | 0.12 | 1.7598352 | 18 | flna |
| meis1b4 | 8.484658 | 0.345613045 | 0.264 | 0.071 | 2.1555274 | 18 | meis1b |
| ulk22 | 8.627224 | 0.339147113 | 0.217 | 0.052 | 2.1917465 | 18 | ulk2 |
| si:ch211-132p1.33 | 9.000415 | 0.405755957 | 0.311 | 0.095 | 2.2865556 | 18 | si:ch211-132p1.3 |
| gstm.14 | 1.602730 | 0.505966314 | 0.632 | 0.275 | 4.0717357 | 18 | gstm.1 |
| ccl25b3 | 1.664104 | 0.467645335 | 0.311 | 0.092 | 4.2276586 | 18 | ccl25b |
| nupr1b1 | 2.671197 | 0.363266915 | 0.193 | 0.044 | 6.7861773 | 18 | nupr1b |
| esr2b2 | 2.806611 | 0.286177982 | 0.179 | 0.038 | 7.1301970 | 18 | esr2b |
| si:ch73-1a9.37 | 3.135754 | 0.533321009 | 0.656 | 0.312 | 7.9663839 | 18 | si:ch73-1a9.3 |
| ecrg4a | 3.735313 | 0.315747821 | 0.175 | 0.036 | 9.4895632 | 18 | ecrg4a |
| znf609b | 4.506774 | 0.2505217 | 0.146 | 0.027 | 1.1449461 | 18 | znf609b |
| hspe13 | 5.627961 | 0.520145072 | 0.646 | 0.287 | 1.4297836 | 18 | hspe1 |
| arhgef41 | 7.573520 | 0.256452811 | 0.17 | 0.035 | 1.9240528 | 18 | arhgef4 |
| map1lc3a3 | 8.394627 | 0.36237431 | 0.274 | 0.079 | 2.1326550 | 18 | map1lc3a |
| pard3bb2 | 1.364160 | 0.259772732 | 0.16 | 0.032 | 3.4656507 | 18 | pard3bb |
| si:dkey-17m8.1 | 2.655071 | 0.280408248 | 0.193 | 0.044 | 6.7452086 | 18 | si:dkey-17m8.1 |
| XLOC-043861 | 2.893002 | 0.292316481 | 0.198 | 0.046 | 7.3496723 | 18 | XLOC-043861 |
| sh3pxd2b1 | 2.956049 | 0.327262058 | 0.203 | 0.049 | 7.5098448 | 18 | sh3pxd2b |
| jdp2b3 | 3.185055 | 0.528521799 | 0.425 | 0.162 | 8.0916328 | 18 | jdp2b |

| | | | | | | | |
|---|---|---|---|---|---|---|---|
| tefa3 | 3.759788 | 0.423271868 | 0.387 | 0.139 | 9.5517423 | 18 | tefa |
| stk35l | 3.874571 | 0.280401319 | 0.179 | 0.039 | 9.8433485 | 18 | stk35l |
| net11 | 6.626601 | 0.261835984 | 0.156 | 0.031 | 1.6834881 | 18 | net1 |
| mgst1.12 | 6.806493 | 0.297804512 | 0.217 | 0.054 | 1.7291897 | 18 | mgst1.1 |
| map1lc3b1 | 7.374069 | 0.387589719 | 0.354 | 0.122 | 1.8733823 | 18 | map1lc3b |
| cflara2 | 1.024586 | 0.296057495 | 0.236 | 0.062 | 2.6029624 | 18 | cflara |
| rsu11 | 1.029981 | 0.320055696 | 0.208 | 0.051 | 2.6166688 | 18 | rsu1 |
| crim13 | 1.150415 | 0.303327709 | 0.179 | 0.04 | 2.9226314 | 18 | crim1 |
| arl4d1 | 2.221683 | 0.27305654 | 0.212 | 0.052 | 5.6441878 | 18 | arl4d |
| fermt2 | 3.733380 | 0.251705744 | 0.156 | 0.032 | 9.4846532 | 18 | fermt2 |
| klf6a4 | 4.498522 | 0.480010403 | 0.443 | 0.178 | 1.1428496 | 18 | klf6a |
| marcksl1a4 | 7.747112 | 0.459502967 | 0.599 | 0.257 | 1.9681539 | 18 | marcksl1a |
| krt18b5 | 8.738055 | 0.408218247 | 0.462 | 0.173 | 2.2199031 | 18 | krt18b |
| ppap2d1 | 1.150832 | 0.251649932 | 0.165 | 0.035 | 2.9236896 | 18 | ppap2d |
| anxa43 | 1.305051 | 0.392669505 | 0.396 | 0.148 | 3.3154837 | 18 | anxa4 |
| wu:fj16a031 | 1.696819 | 0.316341595 | 0.189 | 0.044 | 4.3107707 | 18 | wu:fj16a03 |
| fam168a4 | 1.926485 | 0.315699148 | 0.25 | 0.072 | 4.8942369 | 18 | fam168a |
| hmgb1b5 | 3.992521 | 0.415385872 | 0.401 | 0.152 | 1.0143000 | 18 | hmgb1b |
| SBNO2 | 5.950572 | 0.273487777 | 0.156 | 0.034 | 1.5117429 | 18 | SBNO2 |
| lama43 | 6.343251 | 0.291610119 | 0.189 | 0.046 | 1.6115029 | 18 | lama4 |
| si:ch1073-291c23.23 | 9.249969 | 0.308268681 | 0.25 | 0.071 | 2.3499547 | 18 | si:ch1073-291c23.2 |
| si:ch211-286b5.54 | 9.281543 | 0.383140375 | 0.396 | 0.149 | 2.3579761 | 18 | si:ch211-286b5.5 |
| timp2a3 | 1.007505 | 0.299690944 | 0.231 | 0.064 | 2.5595677 | 18 | timp2a |
| sdcbp24 | 1.366647 | 0.407134809 | 0.528 | 0.225 | 3.4719669 | 18 | sdcbp2 |
| aldh9a1a.14 | 1.900597 | 0.441960356 | 0.453 | 0.191 | 4.8284673 | 18 | aldh9a1a.1 |
| irf2bp2b2 | 1.903294 | 0.355370582 | 0.302 | 0.1 | 4.8353203 | 18 | irf2bp2b |
| ccng22 | 1.918331 | 0.340866773 | 0.269 | 0.083 | 4.8735217 | 18 | ccng2 |
| sdc45 | 2.967325 | 0.462909785 | 0.571 | 0.251 | 7.5384902 | 18 | sdc4 |
| dedd13 | 3.079027 | 0.44551667 | 0.392 | 0.155 | 7.8222701 | 18 | dedd1 |
| ier51 | 3.586602 | 0.411581535 | 0.377 | 0.145 | 9.1117624 | 18 | ier5 |
| wu:fb16f032 | 3.956080 | 0.339865119 | 0.278 | 0.088 | 1.0050421 | 18 | wu:fb16f03 |
| stk35 | 4.979906 | 0.31526721 | 0.217 | 0.06 | 1.2651452 | 18 | stk35 |
| mvp2 | 6.037693 | 0.28102589 | 0.217 | 0.059 | 1.5338759 | 18 | mvp |
| emd2 | 1.360761 | 0.298490564 | 0.236 | 0.07 | 3.4570137 | 18 | emd |
| pnrc28 | 1.963890 | 0.440869014 | 0.892 | 0.595 | 4.9892632 | 18 | pnrc2 |

| | | | | | | | |
|---|---|---|---|---|---|---|---|
| serpinf12 | 2.119238 | 0.268572096 | 0.179 | 0.044 | 5.3839256 | 18 | serpinf1 |
| boc3 | 2.139219 | 0.305347908 | 0.231 | 0.066 | 5.4346871 | 18 | boc |
| fbn2b3 | 3.338730 | 0.253866256 | 0.193 | 0.049 | 8.4820454 | 18 | fbn2b |
| qkia3 | 4.724898 | 0.415231672 | 0.462 | 0.197 | 1.2003603 | 18 | qkia |
| pdk2b2 | 5.743394 | 0.356147423 | 0.302 | 0.104 | 1.4591093 | 18 | pdk2b |
| tmsb4x10 | 6.253689 | 0.459656647 | 0.991 | 0.716 | 1.5887496 | 18 | tmsb4x |
| h3f3c6 | 8.066027 | 0.446289892 | 0.618 | 0.301 | 2.0491743 | 18 | h3f3c |
| anxa5b3 | 1.159698 | 0.336893784 | 0.302 | 0.103 | 2.9462148 | 18 | anxa5b |
| mycb3 | 1.803282 | 0.41034964 | 0.472 | 0.198 | 4.5812393 | 18 | mycb |
| CABZ01072614.14 | 1.814667 | 0.351634356 | 0.278 | 0.093 | 4.6101620 | 18 | CABZ01072614.1 |
| stat21 | 1.819811 | 0.289400723 | 0.217 | 0.062 | 4.6232313 | 18 | stat2 |
| ube2e33 | 2.186620 | 0.395252579 | 0.406 | 0.168 | 5.5551104 | 18 | ube2e3 |
| tnfrsf1a2 | 2.582460 | 0.288772268 | 0.208 | 0.059 | 6.5607398 | 18 | tnfrsf1a |
| zgc:1132082 | 3.284330 | 0.263964216 | 0.184 | 0.048 | 8.3438405 | 18 | zgc:113208 |
| gmfb | 4.367847 | 0.30613852 | 0.316 | 0.11 | 1.1096517 | 18 | gmfb |
| cnn3a3 | 4.674572 | 0.298628856 | 0.226 | 0.068 | 1.1875752 | 18 | cnn3a |
| wu:fi37e091 | 4.939662 | 0.334884369 | 0.344 | 0.127 | 1.2549213 | 18 | wu:fi37e09 |
| gpx1a3 | 5.071582 | 0.381060506 | 0.538 | 0.233 | 1.2884356 | 18 | gpx1a |
| tmem176l.3a | 5.385326 | 0.255246854 | 0.179 | 0.047 | 1.3681422 | 18 | tmem176l.3a |
| rnf213a4 | 5.440557 | 0.333740006 | 0.278 | 0.094 | 1.3821737 | 18 | rnf213a |
| mafba3 | 8.486860 | 0.331308514 | 0.255 | 0.083 | 2.1560867 | 18 | mafba |
| cebpg1 | 9.241302 | 0.29065321 | 0.222 | 0.066 | 2.3477528 | 18 | cebpg |
| anxa131 | 1.022106 | 0.256358912 | 0.193 | 0.053 | 2.5966628 | 18 | anxa13 |
| ndfip1l3 | 1.342498 | 0.319809254 | 0.236 | 0.074 | 3.4106165 | 18 | ndfip1l |
| osr22 | 1.879205 | 0.295126726 | 0.189 | 0.051 | 4.7741223 | 18 | osr2 |
| flot2a3 | 1.880036 | 0.304293907 | 0.231 | 0.073 | 4.7762323 | 18 | flot2a |
| nfia4 | 2.314270 | 0.366222038 | 0.278 | 0.097 | 5.8794038 | 18 | nfia |
| tax1bp31 | 3.882670 | 0.293508471 | 0.288 | 0.1 | 9.8639253 | 18 | tax1bp3 |
| ppdpfb3 | 4.734622 | 0.430292262 | 0.807 | 0.529 | 1.2028309 | 18 | ppdpfb |
| appbp21 | 8.661931 | 0.255117249 | 0.165 | 0.043 | 2.2005637 | 18 | appbp2 |
| ponzr15 | 1.122557 | 0.413757123 | 0.604 | 0.287 | 2.8518581 | 18 | ponzr1 |
| ctnnb12 | 1.370770 | 0.342961882 | 0.387 | 0.157 | 3.4824435 | 18 | ctnnb1 |
| nt5dc22 | 1.747183 | 0.294836994 | 0.245 | 0.08 | 4.4387187 | 18 | nt5dc2 |
| vdrb1 | 1.785575 | 0.259554452 | 0.17 | 0.045 | 4.5362545 | 18 | vdrb |
| rerea1 | 2.221169 | 0.256326343 | 0.189 | 0.053 | 5.6428822 | 18 | rerea |

| | | | | | | | |
|---|---|---|---|---|---|---|---|
| XLOC-0247501 | 3.095147 | 0.284432454 | 0.208 | 0.063 | 7.8632215 | 18 | XLOC-024750 |
| zbtb203 | 3.684781 | 0.348354968 | 0.288 | 0.106 | 9.3611866 | 18 | zbtb20 |
| si:ch1073-459j12.13 | 5.353060 | 0.277216781 | 0.255 | 0.084 | 1.3599448 | 18 | si:ch1073-459j12.1 |
| macf1a | 6.039397 | 0.270828377 | 0.198 | 0.059 | 1.5343089 | 18 | macf1a |
| si:dkey-51e6.13 | 7.575862 | 0.327523946 | 0.307 | 0.117 | 1.9246478 | 18 | si:dkey-51e6.1 |
| ppp1cb1 | 1.103649 | 0.270724147 | 0.217 | 0.069 | 2.8038222 | 18 | ppp1cb |
| ubb1 | 1.106958 | 0.405822143 | 0.618 | 0.322 | 2.8122276 | 18 | ubb |
| csrp1a1 | 1.191709 | 0.259826093 | 0.241 | 0.08 | 3.0275384 | 18 | csrp1a |
| rbm39a2 | 1.246305 | 0.370983713 | 0.358 | 0.152 | 3.1662379 | 18 | rbm39a |
| adamts15a3 | 1.487852 | 0.29059338 | 0.193 | 0.057 | 3.7798893 | 18 | adamts15a |
| lrpap11 | 2.221403 | 0.269539671 | 0.203 | 0.063 | 5.6434766 | 18 | lrpap1 |
| si:dkey-42i9.41 | 3.282238 | 0.358449412 | 0.363 | 0.155 | 8.3385267 | 18 | si:dkey-42i9.4 |
| hopx | 4.242288 | 0.254787369 | 0.208 | 0.064 | 1.0777534 | 18 | hopx |
| jpt1b4 | 5.926155 | 0.364150558 | 0.575 | 0.274 | 1.5055398 | 18 | jpt1b |
| rhoab4 | 8.043456 | 0.373067251 | 0.439 | 0.205 | 2.0434400 | 18 | rhoab |
| aplp22 | 2.693543 | 0.260770089 | 0.226 | 0.076 | 6.8429481 | 18 | aplp2 |
| notch23 | 3.247116 | 0.320720851 | 0.302 | 0.121 | 8.2493001 | 18 | notch2 |
| pik3r3a1 | 3.451543 | 0.273178739 | 0.255 | 0.091 | 8.7686456 | 18 | pik3r3a |
| pim14 | 4.759052 | 0.374308492 | 0.377 | 0.168 | 1.2090372 | 18 | pim1 |
| psme23 | 6.577295 | 0.281475965 | 0.288 | 0.112 | 1.6709618 | 18 | psme2 |
| acin1a3 | 6.630334 | 0.3523403 | 0.472 | 0.227 | 1.6844364 | 18 | acin1a |
| hif1al1 | 7.047829 | 0.30089816 | 0.302 | 0.119 | 1.7905011 | 18 | hif1al |
| si:ch211-213a13.11 | 8.260954 | 0.275879649 | 0.259 | 0.097 | 2.0986954 | 18 | si:ch211-213a13.1 |
| si:ch73-376l24.2 | 1.239138 | 0.324163889 | 0.302 | 0.122 | 3.1480306 | 18 | si:ch73-376l24.2 |
| plk31 | 1.270769 | 0.268075623 | 0.288 | 0.107 | 3.2283886 | 18 | plk3 |
| abca1a2 | 1.636548 | 0.250537547 | 0.212 | 0.071 | 4.1576517 | 18 | abca1a |
| ptp4a12 | 1.867469 | 0.295221712 | 0.316 | 0.131 | 4.7443073 | 18 | ptp4a1 |
| tnrc6a | 2.491496 | 0.256199526 | 0.226 | 0.079 | 6.3296460 | 18 | tnrc6a |
| brd3b2 | 3.125169 | 0.250526216 | 0.203 | 0.068 | 7.9394936 | 18 | brd3b |
| ppib6 | 3.544151 | 0.400551827 | 0.665 | 0.377 | 9.0039164 | 18 | ppib |
| irf21 | 3.565003 | 0.285073893 | 0.212 | 0.075 | 9.0568920 | 18 | irf2 |
| cpn13 | 4.390619 | 0.253911497 | 0.222 | 0.076 | 1.1154367 | 18 | cpn1 |
| jund3 | 5.204983 | 0.280176367 | 0.241 | 0.089 | 1.3223259 | 18 | jund |
| tsc22d32 | 5.934921 | 0.342090503 | 0.349 | 0.158 | 1.5077668 | 18 | tsc22d3 |
| capns1a2 | 6.221201 | 0.30113176 | 0.358 | 0.157 | 1.5804962 | 18 | capns1a |

| | | | | | | | |
|---|---|---|---|---|---|---|---|
| rhoaa3 | 9.186192 | 0.326731157 | 0.377 | 0.175 | 2.3337521 | 18 | rhoaa |
| stau24 | 1.052982 | 0.270236613 | 0.25 | 0.095 | 2.6751011 | 18 | stau2 |
| fosb2 | 1.496777 | 0.352456678 | 0.292 | 0.12 | 3.8025626 | 18 | fosb |
| slc3a2b1 | 1.535885 | 0.304372147 | 0.302 | 0.127 | 3.9019182 | 18 | slc3a2b |
| cldn11a3 | 2.194909 | 0.31728074 | 0.226 | 0.083 | 5.5761671 | 18 | cldn11a |
| mt-nd53 | 5.965598 | 0.298483603 | 0.297 | 0.129 | 1.5155603 | 18 | mt-nd5 |
| si:ch211-191j22.3 | 6.189758 | 0.2614644 | 0.292 | 0.122 | 1.5725081 | 18 | si:ch211-191j22.3 |
| chac12 | 7.329671 | 0.358725865 | 0.66 | 0.375 | 1.8621030 | 18 | chac1 |
| glula1 | 8.121233 | 0.274280025 | 0.302 | 0.124 | 2.0631993 | 18 | glula |
| prkab1b1 | 1.516517 | 0.270001078 | 0.307 | 0.135 | 3.8527117 | 18 | prkab1b |
| capns1b2 | 1.955338 | 0.286813136 | 0.321 | 0.144 | 4.9675378 | 18 | capns1b |
| srsf5a4 | 1.982582 | 0.327843099 | 0.557 | 0.301 | 5.0367517 | 18 | srsf5a |
| klf2a3 | 2.010443 | 0.283177569 | 0.25 | 0.099 | 5.1075309 | 18 | klf2a |
| fkbp1aa4 | 2.584261 | 0.359575663 | 0.693 | 0.412 | 6.5653160 | 18 | fkbp1aa |
| mgst3b1 | 3.255669 | 0.305354254 | 0.392 | 0.192 | 8.2710276 | 18 | mgst3b |
| pfn2l4 | 3.339252 | 0.277579288 | 0.448 | 0.214 | 8.4833715 | 18 | pfn2l |
| tpm12 | 3.573765 | 0.332138699 | 0.41 | 0.201 | 9.0791502 | 18 | tpm1 |
| tagln26 | 5.020839 | 0.293607588 | 0.429 | 0.208 | 1.2755442 | 18 | tagln2 |
| atp1b1a3 | 1.696221 | 0.328670521 | 0.434 | 0.229 | 4.3092501 | 18 | atp1b1a |
| sumo2b5 | 2.356835 | 0.310209273 | 0.538 | 0.295 | 5.9875394 | 18 | sumo2b |
| atrx4 | 3.929437 | 0.268973705 | 0.283 | 0.127 | 9.9827350 | 18 | atrx |
| zgc:1141888 | 4.060152 | 0.324930117 | 0.896 | 0.666 | 1.0314817 | 18 | zgc:114188 |
| tnrc6c14 | 8.864656 | 0.268917048 | 0.316 | 0.149 | 2.2520660 | 18 | tnrc6c1 |
| calm1a1 | 8.946752 | 0.262532997 | 0.358 | 0.173 | 2.2729224 | 18 | calm1a |
| tgif12 | 9.133386 | 0.254683196 | 0.236 | 0.099 | 2.3203368 | 18 | tgif1 |
| zgc:1119863 | 9.834917 | 0.28353861 | 0.368 | 0.185 | 2.4985607 | 18 | zgc:111986 |
| dynlt3 | 1.788734 | 0.250594735 | 0.325 | 0.157 | 4.5442795 | 18 | dynlt3 |
| higd1a3 | 1.848138 | 0.305363899 | 0.542 | 0.302 | 4.6951961 | 18 | higd1a |
| fabp32 | 1.871450 | 0.258169751 | 0.448 | 0.227 | 4.7544204 | 18 | fabp3 |
| btg16 | 1.943184 | 0.306054309 | 0.618 | 0.353 | 4.9366604 | 18 | btg1 |
| eif1b3 | 2.244850 | 0.322220032 | 0.66 | 0.405 | 5.7030426 | 18 | eif1b |
| bhlhe404 | 2.392279 | 0.286397156 | 0.34 | 0.167 | 6.0775850 | 18 | bhlhe40 |
| pfn24 | 2.771047 | 0.261808499 | 0.458 | 0.237 | 7.0398465 | 18 | pfn2 |
| zgc:927443 | 2.777750 | 0.299853345 | 0.585 | 0.325 | 7.0568762 | 18 | zgc:92744 |
| tomm52 | 3.127014 | 0.269624688 | 0.429 | 0.229 | 7.9441791 | 18 | tomm5 |

| | | | | | | | |
|---|---|---|---|---|---|---|---|
| rab103 | 4.298393 | 0.281310726 | 0.401 | 0.213 | 1.0920067 | 18 | rab10 |
| rps27l5 | 4.775800 | 0.262196786 | 0.41 | 0.217 | 1.2132921 | 18 | rps27l |
| rplp2l6 | 5.498527 | 0.256370852 | 0.953 | 0.793 | 1.3969010 | 18 | rplp2l |
| ywhaqb4 | 1.165980 | 0.301323488 | 0.302 | 0.154 | 2.9621725 | 18 | ywhaqb |
| nfkbiaa3 | 1.185748 | 0.273588269 | 0.311 | 0.151 | 3.0123948 | 18 | nfkbiaa |
| eef1db4 | 3.057744 | 0.280920832 | 0.495 | 0.283 | 7.7681994 | 18 | eef1db |
| brd2a | 3.864163 | 0.258605846 | 0.325 | 0.164 | 9.8169069 | 18 | brd2a |
| gapdhs8 | 4.626106 | 0.276709901 | 0.552 | 0.313 | 0.000118 | 18 | gapdhs |
| dbi3 | 4.699164 | 0.265049677 | 0.434 | 0.242 | 0.000119 | 18 | dbi |
| ywhabl1 | 5.169570 | 0.266835759 | 0.524 | 0.299 | 0.000131 | 18 | ywhabl |
| tmem2581 | 6.178944 | 0.275392637 | 0.514 | 0.304 | 0.000157 | 18 | tmem258 |
| pomp1 | 7.239726 | 0.25733299 | 0.458 | 0.259 | 0.000184 | 18 | pomp |
| ndufa4l6 | 7.807407 | 0.295044645 | 0.66 | 0.421 | 0.000198 | 18 | ndufa4l |
| tob1b1 | 7.891447 | 0.256664606 | 0.325 | 0.167 | 0.0002 | 18 | tob1b |
| ucp24 | 1.181048 | 0.303562826 | 0.642 | 0.413 | 0.0003 | 18 | ucp2 |
| cd638 | 1.494467 | 0.284471979 | 0.608 | 0.372 | 0.00038 | 18 | cd63 |
| zgc:1584632 | 2.552330 | 0.253556746 | 0.184 | 0.08 | 0.000648 | 18 | zgc:158463 |
| LOC1039104381 | 2.815190 | 0.265611366 | 0.311 | 0.165 | 0.000715 | 18 | LOC103910438 |
| glrx4 | 3.121641 | 0.254749898 | 0.552 | 0.325 | 0.000793 | 18 | glrx |
| zgc:1538673 | 3.533521 | 0.290257314 | 0.717 | 0.463 | 0.000898 | 18 | zgc:153867 |
| tpm33 | 4.431141 | 0.259158418 | 0.476 | 0.283 | 0.001126 | 18 | tpm3 |
| gnb1b1 | 5.560473 | 0.255249851 | 0.448 | 0.265 | 0.001413 | 18 | gnb1b |
| ptmab7 | 1.773888 | 0.287536236 | 0.816 | 0.568 | 0.004507 | 18 | ptmab |
| hmgb1a8 | 2.411163 | 0.253535526 | 0.382 | 0.227 | 0.006126 | 18 | hmgb1a |
| eef1da3 | 2.136242 | 0.2626673 | 0.708 | 0.505 | 0.054271 | 18 | eef1da |
| ptmaa7 | 2.723461 | 0.26065126 | 0.67 | 0.431 | 0.06919 | 18 | ptmaa |
| rpl27a5 | 3.684844 | 0.262098482 | 0.717 | 0.522 | 0.093613 | 18 | rpl27a |
| rpl98 | 1.616705 | 0.27565055 | 0.901 | 0.667 | 0.410724 | 18 | rpl9 |
| serbp1a6 | 3.341809 | 0.255077124 | 0.741 | 0.504 | 0.848987 | 18 | serbp1a |
| nkl.22 | 0 | 2.380588264 | 0.989 | 0.122 | 0 | 19 | nkl.2 |
| nkl.4 | 0 | 1.754532942 | 0.766 | 0.01 | 0 | 19 | nkl.4 |
| prf1.2 | 0 | 1.695024238 | 0.755 | 0.006 | 0 | 19 | prf1.2 |
| LOC1018842673 | 0 | 1.674812043 | 0.75 | 0.042 | 0 | 19 | LOC101884267 |
| prf1.7 | 0 | 1.486035419 | 0.649 | 0.007 | 0 | 19 | prf1.7 |
| cpa5 | 0 | 1.46605211 | 0.622 | 0.028 | 0 | 19 | cpa5 |

| | | | | | | | |
|---|---|---|---|---|---|---|---|
| XLOC-0430432 | 0 | 1.410535711 | 0.638 | 0.037 | 0 | 19 | XLOC-043043 |
| CABZ01084575.12 | 0 | 1.139872604 | 0.574 | 0.033 | 0 | 19 | CABZ01084575.1 |
| XLOC-0397571 | 0 | 1.129104211 | 0.505 | 0.02 | 0 | 19 | XLOC-039757 |
| tspan2b1 | 0 | 0.832921089 | 0.367 | 0.008 | 0 | 19 | tspan2b |
| nkl.3 | 0 | 0.823371087 | 0.314 | 0.004 | 0 | 19 | nkl.3 |
| BX248410.3 | 0 | 0.364518097 | 0.154 | 0.001 | 0 | 19 | BX248410.3 |
| si:dkey-21e2.3 | 5.766872 | 0.511425034 | 0.234 | 0.004 | 1.4650740 | 19 | si:dkey-21e2.3 |
| XLOC-0140792 | 1.614837 | 1.13005861 | 0.5 | 0.026 | 4.1024953 | 19 | XLOC-014079 |
| si:ch211-173d10.4 | 1.789018 | 0.458286504 | 0.202 | 0.003 | 4.5450023 | 19 | si:ch211-173d10.4 |
| LOC1081907173 | 4.891703 | 1.39663574 | 0.75 | 0.074 | 1.2427373 | 19 | LOC108190717 |
| BX120005.12 | 5.401416 | 1.199923031 | 0.537 | 0.04 | 1.3722298 | 19 | BX120005.1 |
| si:ch211-277c7.62 | 3.101835 | 1.069816688 | 0.479 | 0.031 | 7.8802139 | 19 | si:ch211-277c7.6 |
| csf2rb2 | 6.343400 | 1.00765714 | 0.479 | 0.034 | 1.6115409 | 19 | csf2rb |
| si:dkey-78l4.41 | 1.153572 | 0.86180485 | 0.42 | 0.026 | 2.9306514 | 19 | si:dkey-78l4.4 |
| ca21 | 1.053831 | 1.142195668 | 0.543 | 0.047 | 2.6772596 | 19 | ca2 |
| zbtb322 | 7.232861 | 0.927204281 | 0.463 | 0.036 | 1.8375085 | 19 | zbtb32 |
| CABZ01088229.11 | 2.340064 | 0.586035322 | 0.245 | 0.01 | 5.9449347 | 19 | CABZ01088229.1 |
| mef2b2 | 6.202120 | 0.572118281 | 0.255 | 0.011 | 1.5756486 | 19 | mef2b |
| alox5a1 | 1.160898 | 0.830408748 | 0.394 | 0.029 | 2.9492634 | 19 | alox5a |
| stim1a1 | 6.221346 | 1.046897891 | 0.548 | 0.059 | 1.5805331 | 19 | stim1a |
| CR753876.13 | 4.206083 | 1.248509563 | 0.723 | 0.104 | 1.0685555 | 19 | CR753876.1 |
| ptk2ba | 8.391569 | 0.697815174 | 0.319 | 0.019 | 2.1318781 | 19 | ptk2ba |
| hck2 | 4.038574 | 0.791223582 | 0.388 | 0.029 | 1.0259998 | 19 | hck |
| ponzr61 | 9.004359 | 0.678700496 | 0.298 | 0.018 | 2.2875575 | 19 | ponzr6 |
| creb5b1 | 1.039825 | 0.612814713 | 0.271 | 0.015 | 2.6416772 | 19 | creb5b |
| sort1b1 | 6.357582 | 0.731674178 | 0.346 | 0.026 | 1.6151438 | 19 | sort1b |
| gcnt7 | 1.140004 | 0.310629352 | 0.128 | 0.003 | 2.8961819 | 19 | gcnt7 |
| edaradd1 | 5.729068 | 0.679519607 | 0.287 | 0.018 | 1.4554699 | 19 | edaradd |
| gzm3.3 | 1.675278 | 0.490165122 | 0.207 | 0.009 | 4.2560456 | 19 | gzm3.3 |
| slc41a11 | 3.695517 | 0.484634612 | 0.223 | 0.011 | 9.3884625 | 19 | slc41a1 |
| CABZ01073834.12 | 1.151213 | 0.828443341 | 0.415 | 0.04 | 2.9246577 | 19 | CABZ01073834.1 |
| illr1 | 8.838798 | 0.519411065 | 0.229 | 0.012 | 2.2454967 | 19 | illr1 |
| arl112 | 9.119821 | 0.71987749 | 0.303 | 0.022 | 2.3168906 | 19 | arl11 |
| BX005105.11 | 1.221838 | 0.762444221 | 0.378 | 0.034 | 3.1040816 | 19 | BX005105.1 |
| srgn5 | 8.799081 | 1.304230211 | 0.872 | 0.228 | 2.2354067 | 19 | srgn |

| | | | | | | | |
|---|---|---|---|---|---|---|---|
| efhd21 | 1.330609 | 0.946635581 | 0.463 | 0.055 | 3.3804129 | 19 | efhd2 |
| XLOC-0437602 | 4.095731 | 0.888569889 | 0.399 | 0.042 | 1.0405205 | 19 | XLOC-043760 |
| prr5a | 3.096837 | 0.635045132 | 0.282 | 0.021 | 7.8675166 | 19 | prr5a |
| ccl39.9 | 7.088368 | 0.400290651 | 0.117 | 0.003 | 1.8008000 | 19 | ccl39.9 |
| mrtfaa | 2.993068 | 0.561962485 | 0.266 | 0.02 | 7.6038905 | 19 | mrtfaa |
| rps6ka12 | 8.844702 | 0.875415792 | 0.468 | 0.065 | 2.2469966 | 19 | rps6ka1 |
| tbx21 | 1.605871 | 0.365208298 | 0.144 | 0.006 | 4.0797158 | 19 | tbx21 |
| znrf11 | 5.501024 | 0.4904586 | 0.207 | 0.013 | 1.3975353 | 19 | znrf1 |
| arpc1b6 | 1.150097 | 1.201542967 | 0.899 | 0.307 | 2.9218229 | 19 | arpc1b |
| si:ch211-102c2.44 | 5.435242 | 1.047839466 | 0.665 | 0.14 | 1.3808233 | 19 | si:ch211-102c2.4 |
| si:dkey-21o19.2 | 1.050938 | 0.440912489 | 0.207 | 0.014 | 2.6699092 | 19 | si:dkey-21o19.2 |
| cyfip21 | 1.557951 | 0.561744407 | 0.266 | 0.023 | 3.9579750 | 19 | cyfip2 |
| plcg21 | 4.792614 | 0.745644796 | 0.378 | 0.046 | 1.2175637 | 19 | plcg2 |
| XLOC-0054482 | 2.208736 | 0.672598038 | 0.271 | 0.026 | 5.6112942 | 19 | XLOC-005448 |
| LOC110440216 | 2.809700 | 0.346414844 | 0.149 | 0.007 | 7.1380428 | 19 | LOC110440216 |
| syk3 | 3.997621 | 0.843398626 | 0.463 | 0.074 | 1.0155958 | 19 | syk |
| fa2h | 1.043641 | 0.345964689 | 0.16 | 0.009 | 2.6513705 | 19 | fa2h |
| rbbp8l | 3.140536 | 0.347779148 | 0.16 | 0.009 | 7.9785323 | 19 | rbbp8l |
| nr4a32 | 8.354384 | 0.769229422 | 0.346 | 0.043 | 2.1224312 | 19 | nr4a3 |
| gramd2aa1 | 1.047964 | 0.430466029 | 0.207 | 0.015 | 2.6623543 | 19 | gramd2aa |
| si:ch211-241f5.32 | 1.798223 | 0.77545847 | 0.399 | 0.058 | 4.5683862 | 19 | si:ch211-241f5.3 |
| adam19b1 | 1.867064 | 0.383752658 | 0.186 | 0.012 | 4.7432776 | 19 | adam19b |
| coro1a6 | 2.434351 | 1.047814741 | 0.782 | 0.231 | 6.1844697 | 19 | coro1a |
| LOC110437825 | 4.278103 | 0.338293175 | 0.138 | 0.007 | 1.0868521 | 19 | LOC110437825 |
| pfn16 | 4.763529 | 1.027719699 | 0.989 | 0.497 | 1.2101747 | 19 | pfn1 |
| wipf1a3 | 1.606954 | 0.721357246 | 0.415 | 0.064 | 4.0824684 | 19 | wipf1a |
| il12rb2 | 2.677659 | 0.309532722 | 0.138 | 0.007 | 6.8025947 | 19 | il12rb2 |
| egr32 | 3.454435 | 0.737057895 | 0.362 | 0.051 | 8.7759945 | 19 | egr3 |
| sftpba1 | 1.766003 | 0.60029699 | 0.293 | 0.032 | 4.4865316 | 19 | sftpba |
| atp11c1 | 9.948537 | 0.566867877 | 0.271 | 0.029 | 2.5274259 | 19 | atp11c |
| ptprja | 1.123286 | 0.353317345 | 0.154 | 0.009 | 2.8537101 | 19 | ptprja |
| ephx21 | 1.730565 | 0.592611059 | 0.303 | 0.036 | 4.3965013 | 19 | ephx2 |
| strbp1 | 3.763491 | 0.454804512 | 0.202 | 0.016 | 9.5611497 | 19 | strbp |
| mbnl14 | 8.847595 | 1.021515049 | 0.654 | 0.182 | 2.2477316 | 19 | mbnl1 |
| zgc:640515 | 2.205077 | 0.922369261 | 0.638 | 0.158 | 5.6019984 | 19 | zgc:64051 |

| | | | | | | | |
|---|---|---|---|---|---|---|---|
| myh11a4 | 4.475802 | 0.756042331 | 0.41 | 0.068 | 1.1370775 | 19 | myh11a |
| gfi1ab1 | 6.579045 | 0.486088015 | 0.229 | 0.022 | 1.6714065 | 19 | gfi1ab |
| BX901920.13 | 4.056240 | 0.806038445 | 0.415 | 0.07 | 1.0304878 | 19 | BX901920.1 |
| pstpip2 | 1.205940 | 0.481754021 | 0.213 | 0.02 | 3.0636921 | 19 | pstpip2 |
| ly75 | 3.754071 | 0.496267303 | 0.234 | 0.024 | 9.5372189 | 19 | ly75 |
| fcer1gl4 | 1.122820 | 0.935705475 | 0.723 | 0.176 | 2.8525264 | 19 | fcer1gl |
| elf13 | 6.546820 | 0.802380602 | 0.479 | 0.099 | 1.6632196 | 19 | elf1 |
| zgc:152891 | 3.262494 | 0.38760699 | 0.17 | 0.013 | 8.2883682 | 19 | zgc:152891 |
| stat4 | 3.498058 | 0.303202959 | 0.128 | 0.007 | 8.8868187 | 19 | stat4 |
| bhlhe405 | 1.338835 | 0.958324634 | 0.601 | 0.163 | 3.4013118 | 19 | bhlhe40 |
| si:ch73-208g10.13 | 1.048806 | 0.566407353 | 0.287 | 0.038 | 2.6644937 | 19 | si:ch73-208g10.1 |
| XLOC-0283704 | 1.324734 | 0.861977719 | 0.537 | 0.121 | 3.3654884 | 19 | XLOC-028370 |
| vaspa4 | 9.324206 | 0.668485083 | 0.426 | 0.078 | 2.3688147 | 19 | vaspa |
| arhgdig5 | 9.978030 | 0.968503005 | 0.718 | 0.255 | 2.5349187 | 19 | arhgdig |
| si:ch211-214p16.14 | 4.952111 | 1.027503413 | 0.755 | 0.239 | 1.2580839 | 19 | si:ch211-214p16.1 |
| actb26 | 2.827532 | 0.52795478 | 1 | 0.866 | 7.1833461 | 19 | actb2 |
| tagapa4 | 6.507581 | 0.805894382 | 0.441 | 0.094 | 1.6532511 | 19 | tagapa |
| sh3bp2 | 1.835095 | 0.36230495 | 0.176 | 0.016 | 4.6620592 | 19 | sh3bp2 |
| dock101 | 6.779258 | 0.488286023 | 0.229 | 0.027 | 1.7222706 | 19 | dock10 |
| sh3kbp12 | 6.994117 | 0.668628154 | 0.383 | 0.074 | 1.7768555 | 19 | sh3kbp1 |
| wasb6 | 8.982166 | 0.849633547 | 0.601 | 0.169 | 2.2819194 | 19 | wasb |
| LOC1039108712 | 7.873410 | 0.862862441 | 0.585 | 0.153 | 2.0002399 | 19 | LOC103910871 |
| laptm56 | 1.645954 | 0.828627822 | 0.612 | 0.172 | 4.1815470 | 19 | laptm5 |
| BX908750.12 | 3.003219 | 0.484786142 | 0.234 | 0.03 | 7.6296793 | 19 | BX908750.1 |
| ptprc5 | 4.305041 | 0.782750913 | 0.489 | 0.121 | 1.0936957 | 19 | ptprc |
| camk1b2 | 3.057596 | 0.387851997 | 0.191 | 0.02 | 7.7678233 | 19 | camk1b |
| sik12 | 3.241504 | 0.855477821 | 0.447 | 0.107 | 8.2350420 | 19 | sik1 |
| wdr192 | 6.465859 | 0.523400523 | 0.234 | 0.03 | 1.6426515 | 19 | wdr19 |
| tent5ab | 6.960593 | 0.310314389 | 0.128 | 0.009 | 1.7683386 | 19 | tent5ab |
| pla2g15 | 9.027478 | 0.566151079 | 0.293 | 0.046 | 2.2934308 | 19 | pla2g15 |
| uba526 | 1.007724 | 0.61109454 | 0.915 | 0.659 | 2.5601235 | 19 | uba52 |
| dicp1.5-61 | 1.954779 | 0.373115526 | 0.16 | 0.014 | 4.9661174 | 19 | dicp1.5-6 |
| aldocb2 | 6.474377 | 0.632157921 | 0.34 | 0.063 | 1.6448156 | 19 | aldocb |
| sla22 | 7.429167 | 0.654271131 | 0.33 | 0.058 | 1.8873799 | 19 | sla2 |
| actr2a5 | 2.160308 | 0.767820337 | 0.495 | 0.13 | 5.4882628 | 19 | actr2a |

| | | | | | | | |
|---|---|---|---|---|---|---|---|
| eef1a1l11 | 2.417787 | 0.312502726 | 0.989 | 0.915 | 6.1423896 | 19 | eef1a1l1 |
| plat | 8.142296 | 0.294564359 | 0.138 | 0.011 | 2.0685503 | 19 | plat |
| dapp1 | 1.592095 | 0.27661153 | 0.117 | 0.008 | 4.0447183 | 19 | dapp1 |
| BX322618.12 | 4.546296 | 0.583930557 | 0.239 | 0.033 | 1.1549866 | 19 | BX322618.1 |
| itga43 | 5.285727 | 0.649416303 | 0.33 | 0.062 | 1.3428390 | 19 | itga4 |
| XLOC-0254234 | 7.026619 | 0.830038325 | 0.59 | 0.166 | 1.7851127 | 19 | XLOC-025423 |
| prkcha | 2.128553 | 0.380514015 | 0.17 | 0.017 | 5.4075908 | 19 | prkcha |
| dok23 | 1.376689 | 0.471768153 | 0.255 | 0.038 | 3.4974800 | 19 | dok2 |
| dock81 | 3.373896 | 0.47133505 | 0.245 | 0.036 | 8.5713838 | 19 | dock8 |
| cables2b | 6.288547 | 0.348494252 | 0.149 | 0.014 | 1.5976054 | 19 | cables2b |
| gab2 | 1.171268 | 0.342761703 | 0.154 | 0.015 | 2.9756084 | 19 | gab2 |
| sla12 | 2.912904 | 0.392439105 | 0.191 | 0.023 | 7.4002335 | 19 | sla1 |
| pabpc1a4 | 9.320577 | 0.64877109 | 0.867 | 0.568 | 2.3678926 | 19 | pabpc1a |
| ndfip2 | 4.641008 | 0.551723632 | 0.309 | 0.058 | 1.1790483 | 19 | ndfip2 |
| irak1 | 5.703543 | 0.41483472 | 0.191 | 0.024 | 1.4489852 | 19 | irak1 |
| stk17b3 | 1.793529 | 0.688614362 | 0.436 | 0.111 | 4.5564626 | 19 | stk17b |
| rpl46 | 7.547502 | 0.54189375 | 0.883 | 0.656 | 1.9174430 | 19 | rpl4 |
| XLOC-0108993 | 1.412258 | 0.85426524 | 0.66 | 0.236 | 3.5878428 | 19 | XLOC-010899 |
| tyk2 | 6.340136 | 0.403147385 | 0.191 | 0.025 | 1.6107117 | 19 | tyk2 |
| eomesa1 | 3.760890 | 0.500754533 | 0.245 | 0.039 | 9.5545430 | 19 | eomesa |
| dicp1.11 | 5.024572 | 0.337131307 | 0.138 | 0.013 | 1.2764925 | 19 | dicp1.1 |
| tec1 | 1.549728 | 0.4415946 | 0.213 | 0.033 | 3.9370845 | 19 | tec |
| hsp90ab14 | 2.726397 | 0.392581122 | 0.968 | 0.799 | 6.9264123 | 19 | hsp90ab1 |
| yrk3 | 3.543079 | 0.527099812 | 0.314 | 0.065 | 9.0011937 | 19 | yrk |
| XLOC-0203013 | 8.305735 | 0.565874453 | 0.324 | 0.068 | 2.1100720 | 19 | XLOC-020301 |
| CR936442.13 | 1.059148 | 0.743942928 | 0.505 | 0.151 | 2.6907661 | 19 | CR936442.1 |
| CABZ01079663.12 | 1.916322 | 0.423656268 | 0.223 | 0.036 | 4.8684173 | 19 | CABZ01079663.1 |
| si:ch211-226m16.22 | 3.079800 | 0.580242876 | 0.309 | 0.067 | 7.8242322 | 19 | si:ch211-226m16.2 |
| sh2d3ca | 3.952145 | 0.372247182 | 0.165 | 0.021 | 1.0040426 | 19 | sh2d3ca |
| si:rp71-36a1.5 | 6.921679 | 0.464625156 | 0.245 | 0.043 | 1.7584527 | 19 | si:rp71-36a1.5 |
| scgn1 | 3.073975 | 0.337535667 | 0.17 | 0.022 | 7.8094338 | 19 | scgn |
| rpl317 | 4.835463 | 0.465100315 | 0.894 | 0.686 | 1.2284494 | 19 | rpl31 |
| eef2b3 | 5.569373 | 0.539751242 | 0.883 | 0.634 | 1.4148993 | 19 | eef2b |
| CR392341.13 | 6.135889 | 0.529725772 | 0.282 | 0.058 | 1.5588226 | 19 | CR392341.1 |
| rnf19b | 7.842995 | 0.341510479 | 0.165 | 0.021 | 1.9925130 | 19 | rnf19b |

| | | | | | | | |
|---|---|---|---|---|---|---|---|
| cotl16 | 1.127070 | 0.757470025 | 0.729 | 0.316 | 2.8633228 | 19 | cotl1 |
| rps187 | 1.771780 | 0.417275531 | 0.931 | 0.753 | 4.5012079 | 19 | rps18 |
| rps108 | 1.983285 | 0.444390731 | 0.957 | 0.725 | 5.0385373 | 19 | rps10 |
| st3gal4 | 7.391793 | 0.34021682 | 0.154 | 0.019 | 1.8778851 | 19 | st3gal4 |
| atp2b1a1 | 1.636460 | 0.542785961 | 0.314 | 0.07 | 4.1574267 | 19 | atp2b1a |
| rps26l7 | 1.782817 | 0.456670617 | 0.91 | 0.698 | 4.5292472 | 19 | rps26l |
| gnsb1 | 7.609501 | 0.459282981 | 0.218 | 0.038 | 1.9331937 | 19 | gnsb |
| tmsb4x11 | 7.853267 | 0.553274154 | 0.989 | 0.716 | 1.9951226 | 19 | tmsb4x |
| rpl67 | 9.305715 | 0.453211847 | 0.926 | 0.7 | 2.3641170 | 19 | rpl6 |
| slc7a3b | 1.396635 | 0.37049681 | 0.181 | 0.027 | 3.5481520 | 19 | slc7a3b |
| slc25a54 | 1.476847 | 0.343632923 | 0.941 | 0.813 | 3.7519319 | 19 | slc25a5 |
| si:dkey-10h3.2 | 2.009363 | 0.296330014 | 0.133 | 0.015 | 5.1047868 | 19 | si:dkey-10h3.2 |
| tnfrsf9a1 | 3.951944 | 0.46719505 | 0.245 | 0.046 | 1.0039913 | 19 | tnfrsf9a |
| cap16 | 7.786342 | 0.713589063 | 0.59 | 0.243 | 1.9781201 | 19 | cap1 |
| il342 | 1.007862 | 0.465337264 | 0.234 | 0.044 | 2.5604751 | 19 | il34 |
| nuggc.3 | 2.006512 | 0.261540208 | 0.112 | 0.011 | 5.0975438 | 19 | nuggc.3 |
| actn41 | 2.121640 | 0.551529178 | 0.356 | 0.095 | 5.3900269 | 19 | actn4 |
| ikzf12 | 3.002989 | 0.380893673 | 0.197 | 0.033 | 7.6290946 | 19 | ikzf1 |
| si:zfos-741a10.33 | 3.982120 | 0.38762834 | 0.186 | 0.03 | 1.0116576 | 19 | si:zfos-741a10.3 |
| ubash3ba | 4.373415 | 0.345175245 | 0.16 | 0.022 | 1.1110662 | 19 | ubash3ba |
| aif1l4 | 5.350738 | 0.62392699 | 0.521 | 0.167 | 1.3593550 | 19 | aif1l |
| si:dkey-222f2.71 | 6.960259 | 0.354670027 | 0.191 | 0.031 | 1.7682538 | 19 | si:dkey-222f2.7 |
| prf1.91 | 1.107232 | 0.334161267 | 0.128 | 0.015 | 2.8129240 | 19 | prf1.9 |
| rpl216 | 1.309632 | 0.361375038 | 0.926 | 0.776 | 3.3271203 | 19 | rpl21 |
| rpl148 | 5.141457 | 0.461096754 | 0.867 | 0.668 | 1.3061872 | 19 | rpl14 |
| il163 | 5.317646 | 0.462706753 | 0.234 | 0.047 | 1.3509480 | 19 | il16 |
| abo1 | 5.454373 | 0.326329585 | 0.138 | 0.018 | 1.3856835 | 19 | abo |
| samd111 | 6.770892 | 0.26838041 | 0.138 | 0.018 | 1.7201451 | 19 | samd11 |
| megf8l1 | 8.007280 | 0.254956761 | 0.117 | 0.013 | 2.0342495 | 19 | megf8l |
| hcls13 | 2.673391 | 0.520837806 | 0.303 | 0.075 | 6.7917508 | 19 | hcls1 |
| ponzr16 | 4.126293 | 0.682680713 | 0.644 | 0.287 | 1.0482848 | 19 | ponzr1 |
| btk | 7.938805 | 0.339737524 | 0.17 | 0.027 | 2.0168534 | 19 | btk |
| nptna2 | 8.040237 | 0.464553004 | 0.287 | 0.066 | 2.0426222 | 19 | nptna |
| orai1a1 | 1.501596 | 0.326523481 | 0.154 | 0.023 | 3.8148050 | 19 | orai1a |
| msna6 | 2.448752 | 0.675383556 | 0.585 | 0.263 | 6.2210568 | 19 | msna |

| | | | | | | | |
|---|---|---|---|---|---|---|---|
| si:ch73-264p11.13 | 2.502402 | 0.334306242 | 0.17 | 0.027 | 6.3573527 | 19 | si:ch73-264p11.1 |
| wdr14 | 3.004475 | 0.612724677 | 0.447 | 0.153 | 7.6328694 | 19 | wdr1 |
| ptpn221 | 3.126565 | 0.319431535 | 0.154 | 0.023 | 7.9430388 | 19 | ptpn22 |
| tmc82 | 9.793082 | 0.391111511 | 0.186 | 0.033 | 2.4879327 | 19 | tmc8 |
| cfl15 | 1.301631 | 0.529818373 | 0.835 | 0.599 | 3.3067938 | 19 | cfl1 |
| per32 | 1.720020 | 0.490677787 | 0.298 | 0.076 | 4.3697122 | 19 | per3 |
| lamp1a1 | 2.149317 | 0.571582474 | 0.33 | 0.092 | 5.4603413 | 19 | lamp1a |
| limd21 | 3.664203 | 0.484130295 | 0.277 | 0.066 | 9.3089096 | 19 | limd2 |
| XLOC-019150 | 3.874194 | 0.30900414 | 0.149 | 0.022 | 9.8423899 | 19 | XLOC-019150 |
| capzb5 | 4.761856 | 0.644558843 | 0.59 | 0.269 | 1.2097497 | 19 | capzb |
| rpl36a6 | 5.430376 | 0.329607503 | 0.973 | 0.822 | 1.3795870 | 19 | rpl36a |
| jak3 | 6.631072 | 0.256230386 | 0.101 | 0.011 | 1.6846240 | 19 | jak3 |
| slc1a5 | 1.086884 | 0.261119594 | 0.128 | 0.017 | 2.7612311 | 19 | slc1a5 |
| rps4x6 | 1.089443 | 0.405054317 | 0.947 | 0.735 | 2.7677321 | 19 | rps4x |
| rps236 | 1.127924 | 0.376236647 | 0.915 | 0.744 | 2.8654913 | 19 | rps23 |
| ccr23 | 1.130926 | 0.410924757 | 0.218 | 0.044 | 2.8731177 | 19 | ccr2 |
| CABZ01111959.11 | 1.133387 | 0.313709962 | 0.16 | 0.025 | 2.8793712 | 19 | CABZ01111959.1 |
| fryl | 1.196654 | 0.377291271 | 0.191 | 0.035 | 3.0401017 | 19 | fryl |
| rpl5a8 | 1.412981 | 0.538968183 | 0.835 | 0.534 | 3.5896800 | 19 | rpl5a |
| RASSF53 | 1.413232 | 0.485377082 | 0.255 | 0.06 | 3.5903183 | 19 | RASSF5 |
| zgc:1141889 | 1.690239 | 0.461960366 | 0.926 | 0.666 | 4.2940523 | 19 | zgc:114188 |
| wdfy2 | 2.953565 | 0.269873889 | 0.122 | 0.016 | 7.5035337 | 19 | wdfy2 |
| cmip1 | 5.709237 | 0.493825574 | 0.261 | 0.063 | 1.4504316 | 19 | cmip |
| pptc7b | 7.559268 | 0.34511463 | 0.176 | 0.031 | 1.9204322 | 19 | pptc7b |
| csk3 | 1.152638 | 0.546409511 | 0.346 | 0.102 | 2.9282788 | 19 | csk |
| rgs133 | 1.436444 | 0.599352099 | 0.388 | 0.123 | 3.6492871 | 19 | rgs13 |
| plp21 | 2.183146 | 0.409706039 | 0.234 | 0.053 | 5.5462846 | 19 | plp2 |
| BX005183.34 | 4.028742 | 0.505516592 | 0.287 | 0.075 | 1.0235020 | 19 | BX005183.3 |
| ccl36.12 | 5.966730 | 0.559488071 | 0.324 | 0.084 | 1.5158480 | 19 | ccl36.1 |
| dock111 | 6.490975 | 0.308188649 | 0.149 | 0.023 | 1.6490323 | 19 | dock11 |
| slc6a6b3 | 1.087183 | 0.512426483 | 0.33 | 0.095 | 2.7619899 | 19 | slc6a6b |
| far11 | 1.256471 | 0.381408085 | 0.186 | 0.036 | 3.1920649 | 19 | far1 |
| pnp5b | 1.282600 | 0.295599595 | 0.144 | 0.022 | 3.2584468 | 19 | pnp5b |
| adcy7 | 1.832637 | 0.330347865 | 0.16 | 0.027 | 4.6558148 | 19 | adcy7 |
| susd6 | 2.190740 | 0.425152729 | 0.239 | 0.056 | 5.5655752 | 19 | susd6 |

| | | | | | | | |
|---|---|---|---|---|---|---|---|
| lcp14 | 3.186969 | 0.506022896 | 0.298 | 0.083 | 8.0964956 | 19 | lcp1 |
| CU972454.42 | 1.170873 | 0.27373086 | 0.138 | 0.021 | 2.9746037 | 19 | CU972454.4 |
| si:dkey-151g10.31 | 1.645042 | 0.288940952 | 0.117 | 0.016 | 4.1792308 | 19 | si:dkey-151g10.3 |
| ptp4a2b1 | 1.693120 | 0.572901975 | 0.415 | 0.148 | 4.3013733 | 19 | ptp4a2b |
| edf11 | 2.173695 | 0.627263816 | 0.58 | 0.285 | 5.5222726 | 19 | edf1 |
| mir2232 | 2.624669 | 0.398877818 | 0.176 | 0.033 | 6.6679737 | 19 | mir223 |
| XLOC-0004314 | 5.632235 | 0.576418003 | 0.654 | 0.27 | 1.4308695 | 19 | XLOC-000431 |
| ldha2 | 7.326939 | 0.473712289 | 0.261 | 0.068 | 1.8614089 | 19 | ldha |
| scinlb4 | 1.224220 | 0.439148357 | 0.298 | 0.083 | 3.1101329 | 19 | scinlb |
| acap11 | 5.180107 | 0.33310631 | 0.186 | 0.038 | 1.3160064 | 19 | acap1 |
| arhgdia4 | 6.809691 | 0.601156627 | 0.537 | 0.249 | 1.7300021 | 19 | arhgdia |
| ap1b1 | 6.973394 | 0.386671556 | 0.218 | 0.05 | 1.7715907 | 19 | ap1b1 |
| si:cabz01074946.13 | 7.035336 | 0.445863189 | 0.239 | 0.06 | 1.7873271 | 19 | si:cabz01074946.1 |
| grap2b5 | 1.182233 | 0.530298568 | 0.367 | 0.123 | 3.0034640 | 19 | grap2b |
| btf35 | 1.501875 | 0.451121652 | 0.846 | 0.609 | 3.8155136 | 19 | btf3 |
| si:ch211-136m16.81 | 1.765364 | 0.335825457 | 0.16 | 0.03 | 4.4849075 | 19 | si:ch211-136m16.8 |
| rac26 | 1.863690 | 0.553770333 | 0.548 | 0.222 | 4.7347067 | 19 | rac2 |
| icn4 | 2.241966 | 0.687365752 | 0.59 | 0.265 | 5.6957168 | 19 | icn |
| rab27a | 3.158041 | 0.344171936 | 0.207 | 0.047 | 8.0230047 | 19 | rab27a |
| mt-atp68 | 3.492384 | 0.540274575 | 0.856 | 0.554 | 8.8724016 | 19 | mt-atp6 |
| ccdc171 | 5.263523 | 0.263180085 | 0.128 | 0.02 | 1.3371980 | 19 | ccdc171 |
| rpl246 | 6.402678 | 0.331634971 | 0.963 | 0.793 | 1.6266004 | 19 | rpl24 |
| adam8a1 | 7.125801 | 0.361361211 | 0.213 | 0.049 | 1.8103099 | 19 | adam8a |
| rpsa7 | 1.338258 | 0.325600178 | 0.936 | 0.785 | 3.3998462 | 19 | rpsa |
| ptmaa8 | 2.792736 | 0.570550941 | 0.739 | 0.431 | 7.0949474 | 19 | ptmaa |
| myo1f3 | 4.195558 | 0.363924429 | 0.202 | 0.047 | 1.0658817 | 19 | myo1f |
| dtnba | 4.858588 | 0.332648159 | 0.17 | 0.035 | 1.2343243 | 19 | dtnba |
| taok3a | 5.231744 | 0.317079866 | 0.149 | 0.028 | 1.3291247 | 19 | taok3a |
| tmem30aa1 | 6.342686 | 0.39153053 | 0.25 | 0.068 | 1.6113594 | 19 | tmem30aa |
| XLOC-0139642 | 7.403426 | 0.540267785 | 0.452 | 0.168 | 1.8808404 | 19 | XLOC-013964 |
| zgc:1530181 | 1.054962 | 0.292886239 | 0.144 | 0.026 | 2.6801311 | 19 | zgc:153018 |
| mt-co18 | 1.373794 | 0.475933115 | 0.904 | 0.638 | 3.4901238 | 19 | mt-co1 |
| si:dkeyp-68b7.123 | 1.942781 | 0.39731493 | 0.229 | 0.059 | 4.9356364 | 19 | si:dkeyp-68b7.12 |
| rps297 | 5.005784 | 0.26897121 | 0.947 | 0.844 | 1.2717195 | 19 | rps29 |
| si:dkey-262k9.44 | 5.007296 | 0.460909468 | 0.287 | 0.089 | 1.2721037 | 19 | si:dkey-262k9.4 |

| | | | | | | | |
|---|---|---|---|---|---|---|---|
| rps145 | 5.355416 | 0.321425195 | 0.936 | 0.78 | 1.3605436 | 19 | rps14 |
| XLOC-0418701 | 6.898528 | 0.348435102 | 0.154 | 0.03 | 1.7525711 | 19 | XLOC-041870 |
| si:ch211-8c17.31 | 1.102374 | 0.268160386 | 0.144 | 0.027 | 2.8005830 | 19 | si:ch211-8c17.3 |
| rps136 | 1.177291 | 0.345334259 | 0.926 | 0.748 | 2.9909090 | 19 | rps13 |
| lcp2a2 | 1.217055 | 0.316050025 | 0.17 | 0.037 | 3.0919302 | 19 | lcp2a |
| rpl135 | 1.758775 | 0.270364124 | 0.931 | 0.818 | 4.4681696 | 19 | rpl13 |
| hspa8 | 1.764666 | 0.275398997 | 0.926 | 0.821 | 4.4831343 | 19 | hspa8 |
| zmp:00000010822 | 2.304286 | 0.389210814 | 0.213 | 0.055 | 5.8540392 | 19 | zmp:0000001082 |
| arpc5a5 | 2.547588 | 0.476293803 | 0.346 | 0.124 | 6.4721475 | 19 | arpc5a |
| atp5if1b2 | 2.547842 | 0.545614477 | 0.463 | 0.205 | 6.4727945 | 19 | atp5if1b |
| rps286 | 3.107162 | 0.280591509 | 0.963 | 0.84 | 7.8937458 | 19 | rps28 |
| zeb1b1 | 6.943091 | 0.285721526 | 0.149 | 0.03 | 1.7638924 | 19 | zeb1b |
| CABZ01044746.11 | 7.256676 | 0.442283596 | 0.293 | 0.09 | 1.8435585 | 19 | CABZ01044746.1 |
| rpl99 | 9.818407 | 0.400835819 | 0.899 | 0.667 | 2.4943664 | 19 | rpl9 |
| rel1 | 1.005166 | 0.345704308 | 0.16 | 0.034 | 2.5536256 | 19 | rel |
| glipr1a5 | 1.023136 | 0.382188032 | 0.239 | 0.067 | 2.5992794 | 19 | glipr1a |
| zmp:0000000521 | 1.461121 | 0.288913928 | 0.122 | 0.022 | 3.7119791 | 19 | zmp:0000000521 |
| rpl157 | 1.462357 | 0.329429322 | 0.926 | 0.763 | 3.7151202 | 19 | rpl15 |
| id32 | 2.328881 | 0.562100215 | 0.457 | 0.196 | 5.9165240 | 19 | id3 |
| adam17b | 2.616898 | 0.270915617 | 0.128 | 0.023 | 6.6482297 | 19 | adam17b |
| kif21b1 | 2.626145 | 0.280407335 | 0.128 | 0.023 | 6.6717230 | 19 | kif21b |
| mibp26 | 6.157074 | 0.458532455 | 0.766 | 0.52 | 1.5642047 | 19 | mibp2 |
| nbeal2 | 6.607548 | 0.268320144 | 0.122 | 0.022 | 1.6786477 | 19 | nbeal2 |
| arhgap1 | 6.611062 | 0.280526663 | 0.144 | 0.029 | 1.6795403 | 19 | arhgap1 |
| rplp16 | 1.148124 | 0.260898631 | 0.979 | 0.882 | 2.9168115 | 19 | rplp1 |
| cdaa2 | 1.536243 | 0.323309412 | 0.154 | 0.033 | 3.9028262 | 19 | cdaa |
| rpl386 | 1.751457 | 0.321439395 | 0.92 | 0.753 | 4.4495767 | 19 | rpl38 |
| aars | 1.887378 | 0.388295059 | 0.25 | 0.076 | 4.7948847 | 19 | aars |
| arhgef6 | 2.115252 | 0.258851925 | 0.138 | 0.028 | 5.3737999 | 19 | arhgef6 |
| rpl5b5 | 2.181886 | 0.38466053 | 0.851 | 0.644 | 5.5430817 | 19 | rpl5b |
| mthfd2 | 2.793584 | 0.275462739 | 0.149 | 0.032 | 7.0971017 | 19 | mthfd2 |
| rpl228 | 3.395302 | 0.339763644 | 0.894 | 0.72 | 8.6257652 | 19 | rpl22 |
| elmo3 | 3.758656 | 0.296855212 | 0.144 | 0.03 | 9.5488663 | 19 | elmo3 |
| LOC101884253 | 4.829172 | 0.262833504 | 0.138 | 0.027 | 1.2268512 | 19 | LOC101884253 |
| rpl7a6 | 5.389337 | 0.265173247 | 0.931 | 0.823 | 1.3691612 | 19 | rpl7a |

| | | | | | | | |
|---|---|---|---|---|---|---|---|
| si:dkeyp-110e4.6 | 6.501187 | 0.254954689 | 0.122 | 0.023 | 1.6516266 | 19 | si:dkeyp-110e4.6 |
| dusp13 | 8.014972 | 0.538917677 | 0.367 | 0.148 | 2.0362037 | 19 | dusp1 |
| si:ch211-200p22.4 | 9.301863 | 0.263003539 | 0.128 | 0.025 | 2.3631385 | 19 | si:ch211-200p22.4 |
| ccni4 | 9.386289 | 0.525040121 | 0.569 | 0.318 | 2.3845867 | 19 | ccni |
| rps2110 | 9.511326 | 0.303367752 | 0.915 | 0.775 | 2.4163525 | 19 | rps21 |
| actb15 | 1.685308 | 0.316459557 | 0.91 | 0.767 | 4.2815251 | 19 | actb1 |
| agpat3 | 1.863054 | 0.301825395 | 0.16 | 0.037 | 4.7330887 | 19 | agpat3 |
| ppp1r183 | 2.656011 | 0.333666524 | 0.202 | 0.055 | 6.7475977 | 19 | ppp1r18 |
| rpl297 | 4.196011 | 0.274812324 | 0.957 | 0.822 | 1.0659968 | 19 | rpl29 |
| jak13 | 4.499526 | 0.436351918 | 0.282 | 0.099 | 1.1431047 | 19 | jak1 |
| wasa4 | 1.078838 | 0.394080554 | 0.282 | 0.096 | 2.7407893 | 19 | wasa |
| zyx | 1.352178 | 0.419078014 | 0.234 | 0.074 | 3.4352103 | 19 | zyx |
| cxcr4b4 | 2.063834 | 0.45782578 | 0.431 | 0.176 | 5.2431726 | 19 | cxcr4b |
| NC-002333.17 | 3.589510 | 0.367014765 | 0.191 | 0.054 | 9.1191521 | 19 | NC-002333.17 |
| itgb23 | 4.384261 | 0.426637329 | 0.298 | 0.107 | 1.1138215 | 19 | itgb2 |
| fam120c1 | 5.496783 | 0.394962283 | 0.287 | 0.105 | 1.3964578 | 19 | fam120c |
| pde4ba3 | 6.148596 | 0.463557505 | 0.346 | 0.141 | 1.5620510 | 19 | pde4ba |
| fam49ba3 | 6.179125 | 0.405265285 | 0.293 | 0.104 | 1.5698068 | 19 | fam49ba |
| COX39 | 1.172613 | 0.417484427 | 0.936 | 0.674 | 2.9790245 | 19 | COX3 |
| myo18ab | 2.696613 | 0.258956981 | 0.149 | 0.036 | 6.8507475 | 19 | myo18ab |
| arpc5b4 | 3.655194 | 0.420655794 | 0.33 | 0.13 | 9.2860213 | 19 | arpc5b |
| lrrfip1a2 | 3.834306 | 0.33524506 | 0.202 | 0.06 | 9.7410548 | 19 | lrrfip1a |
| rps36 | 4.179277 | 0.315699058 | 0.904 | 0.725 | 1.0617454 | 19 | rps3 |
| selplg2 | 5.290167 | 0.268219255 | 0.17 | 0.045 | 1.3439669 | 19 | selplg |
| dnajc5gb | 6.341965 | 0.257942896 | 0.138 | 0.032 | 1.6111763 | 19 | dnajc5gb |
| cox8a5 | 6.846249 | 0.485260845 | 0.489 | 0.266 | 1.7392896 | 19 | cox8a |
| serbp1a7 | 8.164792 | 0.417014961 | 0.761 | 0.505 | 2.0742656 | 19 | serbp1a |
| rps167 | 9.230117 | 0.28861701 | 0.91 | 0.765 | 2.3449114 | 19 | rps16 |
| XLOC-0268363 | 9.580572 | 0.396283943 | 0.239 | 0.078 | 2.4339444 | 19 | XLOC-026836 |
| nsa25 | 1.352183 | 0.466550661 | 0.617 | 0.388 | 3.4352228 | 19 | nsa2 |
| casc4 | 1.431245 | 0.282206282 | 0.154 | 0.039 | 3.6360794 | 19 | casc4 |
| cd639 | 1.752952 | 0.464607177 | 0.633 | 0.372 | 4.4533768 | 19 | cd63 |
| iqgap11 | 2.190484 | 0.337974329 | 0.218 | 0.07 | 5.5649252 | 19 | iqgap1 |
| CR381673.13 | 2.365346 | 0.31051768 | 0.191 | 0.056 | 6.0091624 | 19 | CR381673.1 |
| ipo7 | 3.728022 | 0.408260287 | 0.293 | 0.115 | 9.4710420 | 19 | ipo7 |

| | | | | | | | |
|---|---|---|---|---|---|---|---|
| abracl3 | 4.383534 | 0.451320266 | 0.457 | 0.22 | 1.1136369 | 19 | abracl |
| rplp27 | 6.426643 | 0.308145828 | 0.957 | 0.782 | 1.6326888 | 19 | rplp2 |
| rpl187 | 7.110902 | 0.270446891 | 0.926 | 0.792 | 1.8065248 | 19 | rpl18 |
| ncaldb2 | 7.420433 | 0.319155331 | 0.218 | 0.07 | 1.8851611 | 19 | ncaldb |
| mef2d1 | 8.041353 | 0.339940403 | 0.223 | 0.075 | 2.0429059 | 19 | mef2d |
| sat1a.23 | 8.141411 | 0.474178777 | 0.527 | 0.3 | 2.0683256 | 19 | sat1a.2 |
| arpc25 | 8.296061 | 0.457975567 | 0.532 | 0.298 | 2.1076143 | 19 | arpc2 |
| stxbp2 | 8.892765 | 0.314985071 | 0.191 | 0.058 | 2.2592071 | 19 | stxbp2 |
| il10ra2 | 1.261443 | 0.301467927 | 0.154 | 0.042 | 3.2046981 | 19 | il10ra |
| arpc35 | 1.857990 | 0.446553569 | 0.58 | 0.333 | 4.7202236 | 19 | arpc3 |
| tomm20b3 | 1.858048 | 0.437457395 | 0.441 | 0.227 | 4.7203714 | 19 | tomm20b |
| nr4a15 | 1.903258 | 0.427030684 | 0.362 | 0.157 | 4.8352285 | 19 | nr4a1 |
| dusp22 | 1.989213 | 0.479242963 | 0.367 | 0.163 | 5.0535973 | 19 | dusp2 |
| plekhj1 | 2.396928 | 0.29772836 | 0.176 | 0.051 | 6.0893978 | 19 | plekhj1 |
| XLOC-0425962 | 2.405052 | 0.26564259 | 0.149 | 0.039 | 6.1100356 | 19 | XLOC-042596 |
| inpp5d1 | 2.805211 | 0.257539209 | 0.138 | 0.035 | 7.1266393 | 19 | inpp5d |
| hmha1b2 | 4.130923 | 0.326095985 | 0.218 | 0.073 | 1.0494610 | 19 | hmha1b |
| NC-002333.47 | 6.172540 | 0.393307591 | 0.862 | 0.607 | 1.5681340 | 19 | NC-002333.4 |
| prkar1aa | 6.604984 | 0.320037312 | 0.186 | 0.058 | 1.6779963 | 19 | prkar1aa |
| kdm2ba1 | 9.134523 | 0.269461717 | 0.149 | 0.041 | 2.3206255 | 19 | kdm2ba |
| snx27a1 | 1.062002 | 0.251303464 | 0.149 | 0.04 | 2.6980184 | 19 | snx27a |
| oaz1b4 | 1.286663 | 0.431901466 | 0.569 | 0.359 | 3.2687685 | 19 | oaz1b |
| rps27.18 | 1.481522 | 0.325994727 | 0.904 | 0.709 | 3.7638075 | 19 | rps27.1 |
| ostf11 | 1.678217 | 0.268649377 | 0.176 | 0.053 | 4.2635120 | 19 | ostf1 |
| si:dkeyp-19e1.31 | 2.268398 | 0.323684908 | 0.191 | 0.062 | 5.7628676 | 19 | si:dkeyp-19e1.3 |
| mt-co28 | 2.382613 | 0.401522895 | 0.926 | 0.669 | 6.0530291 | 19 | mt-co2 |
| rsl24d15 | 8.594096 | 0.39825774 | 0.67 | 0.46 | 2.1833303 | 19 | rsl24d1 |
| ero1b | 9.561627 | 0.265723845 | 0.138 | 0.038 | 2.4291315 | 19 | ero1b |
| chac13 | 1.113340 | 0.447935876 | 0.569 | 0.377 | 2.8284410 | 19 | chac1 |
| metrnla1 | 1.443764 | 0.291546843 | 0.25 | 0.094 | 3.6678845 | 19 | metrnla |
| si:ch211-191c10.12 | 2.850861 | 0.288004563 | 0.16 | 0.049 | 7.2426146 | 19 | si:ch211-191c10.1 |
| skap23 | 3.152466 | 0.31855997 | 0.218 | 0.08 | 8.0088413 | 19 | skap2 |
| CABZ01072614.15 | 3.750836 | 0.362853173 | 0.239 | 0.094 | 9.5289992 | 19 | CABZ01072614.1 |
| vdac22 | 4.510604 | 0.417471142 | 0.527 | 0.325 | 1.1459191 | 19 | vdac2 |
| arpc44 | 4.658919 | 0.402269763 | 0.447 | 0.237 | 1.1835985 | 19 | arpc4 |

| | | | | | | | |
|---|---|---|---|---|---|---|---|
| cox6b15 | 4.969285 | 0.363644942 | 0.282 | 0.122 | 1.2624470 | 19 | cox6b1 |
| grap2a2 | 4.990776 | 0.288384305 | 0.154 | 0.048 | 1.2679067 | 19 | grap2a |
| cbfb | 5.296374 | 0.25927016 | 0.144 | 0.042 | 1.3455439 | 19 | cbfb |
| LOC110438237 | 5.613323 | 0.267005522 | 0.17 | 0.054 | 1.4260648 | 19 | LOC110438237 |
| eef1db5 | 6.865537 | 0.432496973 | 0.473 | 0.284 | 1.7441898 | 19 | eef1db |
| reep52 | 8.039136 | 0.373172635 | 0.351 | 0.173 | 2.0423426 | 19 | reep5 |
| fosl23 | 9.286943 | 0.369803867 | 0.266 | 0.113 | 2.3593479 | 19 | fosl2 |
| plekho23 | 9.324396 | 0.310619964 | 0.207 | 0.076 | 2.3688629 | 19 | plekho2 |
| zmynd111 | 9.658866 | 0.279950593 | 0.176 | 0.058 | 2.4538350 | 19 | zmynd11 |
| XLOC-0425953 | 9.752618 | 0.315075245 | 0.351 | 0.152 | 2.4776527 | 19 | XLOC-042595 |
| dub3 | 1.010338 | 0.298382815 | 0.202 | 0.073 | 2.5667659 | 19 | dub |
| tnfaip33 | 1.619757 | 0.288833377 | 0.176 | 0.06 | 4.1149944 | 19 | tnfaip3 |
| mrps152 | 3.623125 | 0.342685078 | 0.266 | 0.113 | 9.2045495 | 19 | mrps15 |
| grb2b | 3.651898 | 0.342948701 | 0.202 | 0.077 | 9.2776488 | 19 | grb2b |
| calm1b4 | 4.449604 | 0.31519785 | 0.25 | 0.102 | 1.1304220 | 19 | calm1b |
| larp4ab | 5.287519 | 0.291913673 | 0.202 | 0.076 | 1.3432942 | 19 | larp4ab |
| tnip12 | 6.364313 | 0.305795061 | 0.207 | 0.08 | 1.6168538 | 19 | tnip1 |
| eno1a4 | 1.235248 | 0.327605573 | 0.303 | 0.141 | 3.1381486 | 19 | eno1a |
| pum1 | 2.025410 | 0.252070491 | 0.176 | 0.062 | 5.1455544 | 19 | pum1 |
| vsir4 | 2.038125 | 0.317727084 | 0.218 | 0.088 | 5.1778569 | 19 | vsir |
| eif4ebp21 | 2.118657 | 0.374415142 | 0.362 | 0.192 | 5.3824499 | 19 | eif4ebp2 |
| cct83 | 2.266834 | 0.370264327 | 0.399 | 0.222 | 5.7588942 | 19 | cct8 |
| tefb | 2.270026 | 0.267494926 | 0.181 | 0.066 | 5.7670035 | 19 | tefb |
| rps58 | 2.508400 | 0.268972415 | 0.888 | 0.728 | 6.3725915 | 19 | rps5 |
| elmo12 | 2.895083 | 0.286726345 | 0.186 | 0.07 | 7.3549585 | 19 | elmo1 |
| calr3b1 | 4.059976 | 0.348132932 | 0.223 | 0.094 | 1.0314370 | 19 | calr3b |
| atp10a4 | 5.227771 | 0.258144723 | 0.165 | 0.059 | 1.3281153 | 19 | atp10a |
| per1b | 5.718055 | 0.281942758 | 0.186 | 0.072 | 1.4526720 | 19 | per1b |
| celf27 | 6.388740 | 0.37783821 | 0.436 | 0.257 | 1.6230593 | 19 | celf2 |
| tefa4 | 1.074668 | 0.379926831 | 0.282 | 0.141 | 2.7301941 | 19 | tefa |
| tmem131 | 1.388864 | 0.256632948 | 0.165 | 0.06 | 3.5284100 | 19 | tmem131 |
| tpi1b4 | 1.598054 | 0.351743201 | 0.335 | 0.178 | 4.0598569 | 19 | tpi1b |
| birc22 | 1.616262 | 0.314757853 | 0.239 | 0.107 | 4.1061144 | 19 | birc2 |
| elovl1b3 | 1.622606 | 0.322395722 | 0.261 | 0.122 | 4.1222318 | 19 | elovl1b |
| xbp13 | 2.019784 | 0.370913382 | 0.543 | 0.355 | 5.1312614 | 19 | xbp1 |

| | | | | | | | |
|---|---|---|---|---|---|---|---|
| pgam1a3 | 2.112391 | 0.33704692 | 0.309 | 0.155 | 5.3665295 | 19 | pgam1a |
| hnrnpa1a1 | 2.44762( | 0.364711202 | 0.314 | 0.166 | 6.2181796 | 19 | hnrnpa1a |
| mapre1a4 | 2.465715 | 0.32450139 | 0.378 | 0.196 | 6.2641491 | 19 | mapre1a |
| abca1a3 | 4.014212 | 0.287756857 | 0.181 | 0.072 | 0.000102 | 19 | abca1a |
| clic14 | 5.452014 | 0.318528021 | 0.266 | 0.129 | 0.000139 | 19 | clic1 |
| myl12.14 | 7.080690 | 0.339220211 | 0.596 | 0.426 | 0.00018 | 19 | myl12.1 |
| arrb2b4 | 7.288340 | 0.290440777 | 0.218 | 0.096 | 0.000185 | 19 | arrb2b |
| sh3bgrl2 | 8.504299 | 0.328174367 | 0.25 | 0.121 | 0.000216 | 19 | sh3bgrl |
| mt-nd48 | 1.119598 | 0.332346411 | 0.66 | 0.447 | 0.000284 | 19 | mt-nd4 |
| rps157 | 1.213419 | 0.259770201 | 0.878 | 0.716 | 0.000308 | 19 | rps15 |
| rhogb3 | 1.423894 | 0.289333111 | 0.234 | 0.108 | 0.000362 | 19 | rhogb |
| rab444 | 1.874728 | 0.254169354 | 0.16 | 0.061 | 0.000476 | 19 | rab44 |
| arpc4l4 | 1.897669 | 0.350016933 | 0.372 | 0.219 | 0.000482 | 19 | arpc4l |
| eif3c3 | 2.022044 | 0.323934943 | 0.335 | 0.188 | 0.000514 | 19 | eif3c |
| eif3d1 | 2.497474 | 0.342154364 | 0.441 | 0.281 | 0.000634 | 19 | eif3d |
| lasp14 | 2.502645 | 0.323923732 | 0.42 | 0.253 | 0.000636 | 19 | lasp1 |
| crema3 | 4.592989 | 0.332344202 | 0.319 | 0.174 | 0.001167 | 19 | crema |
| BX248082.13 | 4.640569 | 0.273028595 | 0.112 | 0.037 | 0.001179 | 19 | BX248082.1 |
| eef1g6 | 6.741488 | 0.270642064 | 0.83 | 0.652 | 0.001713 | 19 | eef1g |
| pik3r12 | 7.794150 | 0.265070379 | 0.202 | 0.091 | 0.00198 | 19 | pik3r1 |
| zgc:638312 | 7.936829 | 0.279678756 | 0.191 | 0.085 | 0.002016 | 19 | zgc:63831 |
| capza1b3 | 9.376444 | 0.32020623 | 0.388 | 0.234 | 0.002382 | 19 | capza1b |
| cct6a1 | 1.012659 | 0.30844949 | 0.372 | 0.22 | 0.002573 | 19 | cct6a |
| pkma3 | 1.259615 | 0.327450142 | 0.314 | 0.179 | 0.0032 | 19 | pkma |
| mt-cyb7 | 1.458870 | 0.302416766 | 0.814 | 0.575 | 0.003706 | 19 | mt-cyb |
| rhoac1 | 2.162502 | 0.31328226 | 0.303 | 0.172 | 0.005494 | 19 | rhoac |
| pdia36 | 3.086919 | 0.323551329 | 0.468 | 0.313 | 0.007842 | 19 | pdia3 |
| psap2 | 3.292735 | 0.305351025 | 0.463 | 0.287 | 0.008365 | 19 | psap |
| pnrc29 | 3.768482 | 0.28295548 | 0.809 | 0.597 | 0.009574 | 19 | pnrc2 |
| ist1 | 4.229491 | 0.274600901 | 0.202 | 0.096 | 0.010745 | 19 | ist1 |
| si:ch211-222l21.18 | 4.258906 | 0.307809368 | 0.495 | 0.314 | 0.01082 | 19 | si:ch211-222l21.1 |
| hif1ab | 5.807638 | 0.294823932 | 0.234 | 0.122 | 0.014754 | 19 | hif1ab |
| zgc:1538674 | 8.617993 | 0.280981552 | 0.606 | 0.465 | 0.021894 | 19 | zgc:153867 |
| eif3s102 | 1.042655 | 0.29865971 | 0.303 | 0.179 | 0.026489 | 19 | eif3s10 |
| sh3bgrl32 | 1.161222 | 0.265198729 | 0.282 | 0.156 | 0.029501 | 19 | sh3bgrl3 |

| | | | | | | | |
|---|---|---|---|---|---|---|---|
| mt-nd37 | 1.342886 | 0.299600933 | 0.532 | 0.372 | 0.034116 | 19 | mt-nd3 |
| tln13 | 1.611645 | 0.254454006 | 0.261 | 0.142 | 0.040944 | 19 | tln1 |
| csde11 | 2.951394 | 0.3036824 | 0.33 | 0.207 | 0.07498 | 19 | csde1 |
| cox4i14 | 3.708978 | 0.268713879 | 0.569 | 0.417 | 0.094227 | 19 | cox4i1 |
| eif2s1b2 | 3.949443 | 0.261229762 | 0.319 | 0.195 | 0.100336 | 19 | eif2s1b |
| mt-nd18 | 4.875262 | 0.280574579 | 0.564 | 0.397 | 0.123856 | 19 | mt-nd1 |
| eif3i3 | 4.906678 | 0.297534107 | 0.415 | 0.291 | 0.124654 | 19 | eif3i |
| atf4a2 | 7.064044 | 0.262781501 | 0.309 | 0.186 | 0.179462 | 19 | atf4a |
| h1fx4 | 5.204002 | 0.258547961 | 0.309 | 0.191 | 1 | 19 | h1fx |
| ccl39.72 | 0.0003 | 0.341202746 | 0.122 | 0.062 | 1 | 19 | ccl39.7 |
| CR383676.19 | 0.0088 | 0.265803324 | 0.835 | 0.717 | 1 | 19 | CR383676.1 |
| si:dkey-12l12.1 | 0 | 1.391712204 | 0.61 | 0.015 | 0 | 20 | si:dkey-12l12.1 |
| lum | 0 | 1.247580427 | 0.559 | 0.015 | 0 | 20 | lum |
| hpdb | 0 | 1.157821095 | 0.537 | 0.007 | 0 | 20 | hpdb |
| asip1 | 0 | 1.143012227 | 0.571 | 0.013 | 0 | 20 | asip1 |
| zmp:0000001268 | 0 | 0.96914272 | 0.22 | 0.001 | 0 | 20 | zmp:0000001268 |
| ctgfb | 0 | 0.793619444 | 0.333 | 0.007 | 0 | 20 | ctgfb |
| si:dkey-282h22.5 | 0 | 0.689569898 | 0.271 | 0.002 | 0 | 20 | si:dkey-282h22.5 |
| hgd | 0 | 0.65280703 | 0.288 | 0.006 | 0 | 20 | hgd |
| tat | 0 | 0.634419728 | 0.237 | 0.001 | 0 | 20 | tat |
| vcana | 0 | 0.622870214 | 0.226 | 0.002 | 0 | 20 | vcana |
| hyal4 | 0 | 0.57056739 | 0.203 | 0.002 | 0 | 20 | hyal4 |
| ttc36 | 0 | 0.520690887 | 0.22 | 0.001 | 0 | 20 | ttc36 |
| aqp3b | 0 | 0.373750463 | 0.13 | 0 | 0 | 20 | aqp3b |
| sp9 | 0 | 0.317982707 | 0.124 | 0 | 0 | 20 | sp9 |
| rhag | 3.407340 | 0.751454346 | 0.333 | 0.009 | 8.6563476 | 20 | rhag |
| scara31 | 2.517995 | 0.988117012 | 0.548 | 0.029 | 6.3969677 | 20 | scara3 |
| si:ch211-243a20.3 | 1.694116 | 0.751528407 | 0.35 | 0.01 | 4.3039031 | 20 | si:ch211-243a20.3 |
| tgfbi1 | 7.187522 | 1.052319053 | 0.503 | 0.027 | 1.8259901 | 20 | tgfbi |
| ecrg4a1 | 1.315620 | 1.171927701 | 0.537 | 0.031 | 3.3423343 | 20 | ecrg4a |
| cxcl8a | 6.905317 | 0.764016636 | 0.316 | 0.01 | 1.7542958 | 20 | cxcl8a |
| bhmt1 | 4.381565 | 1.141982546 | 0.548 | 0.036 | 1.1131367 | 20 | bhmt |
| twist2 | 2.974925 | 0.608190783 | 0.282 | 0.008 | 7.5577980 | 20 | twist2 |
| smoc2 | 9.692262 | 0.566791967 | 0.294 | 0.009 | 2.4623192 | 20 | smoc2 |
| FQ323119.1 | 2.603723 | 0.448473533 | 0.226 | 0.005 | 6.6147603 | 20 | FQ323119.1 |

| | | | | | | | |
|---|---|---|---|---|---|---|---|
| prrx1b | 3.877078 | 0.497076002 | 0.237 | 0.006 | 9.8497174 | 20 | prrx1b |
| hoxb8a | 5.645080 | 0.512614944 | 0.266 | 0.008 | 1.4341327 | 20 | hoxb8a |
| nradd | 6.470992 | 0.490085676 | 0.243 | 0.006 | 1.6439556 | 20 | nradd |
| igfbp3 | 3.402398 | 0.669236004 | 0.288 | 0.01 | 8.6437939 | 20 | igfbp3 |
| fmoda | 1.385690 | 0.346421304 | 0.141 | 0.001 | 3.5203461 | 20 | fmoda |
| sod3b | 7.797671 | 0.459413084 | 0.249 | 0.007 | 1.9809983 | 20 | sod3b |
| col5a2a4 | 1.373007 | 1.303724353 | 0.78 | 0.093 | 3.4881254 | 20 | col5a2a |
| cldn19 | 2.304936 | 0.458947477 | 0.198 | 0.004 | 5.8556914 | 20 | cldn19 |
| tnfrsf191 | 4.133680 | 0.636737036 | 0.362 | 0.017 | 1.0501614 | 20 | tnfrsf19 |
| cyr614 | 5.212999 | 1.227237341 | 0.638 | 0.061 | 1.3243624 | 20 | cyr61 |
| clu2 | 2.074393 | 0.989222284 | 0.435 | 0.027 | 5.2699962 | 20 | clu |
| postna | 2.426772 | 1.046101175 | 0.367 | 0.019 | 6.1652144 | 20 | postna |
| hoxb9a | 3.032979 | 0.40601526 | 0.198 | 0.005 | 7.7052851 | 20 | hoxb9a |
| dpt | 5.193761 | 0.811078781 | 0.311 | 0.014 | 1.3194750 | 20 | dpt |
| twist3 | 1.562433 | 0.745497672 | 0.418 | 0.026 | 3.9693624 | 20 | twist3 |
| aspn | 5.509299 | 0.422945348 | 0.13 | 0.002 | 1.3996375 | 20 | aspn |
| ms4a17a.7 | 1.455737 | 0.436493101 | 0.203 | 0.005 | 3.6983008 | 20 | ms4a17a.7 |
| tpbga1 | 4.024577 | 0.569427928 | 0.271 | 0.011 | 1.0224438 | 20 | tpbga |
| CABZ01074363.1 | 1.441909 | 0.796905954 | 0.384 | 0.024 | 3.6631712 | 20 | CABZ01074363.1 |
| fah | 3.599962 | 0.386090314 | 0.198 | 0.005 | 9.1457035 | 20 | fah |
| mfap24 | 1.729555 | 1.259197791 | 0.797 | 0.112 | 4.3939352 | 20 | mfap2 |
| ehd2b1 | 7.331687 | 0.781244282 | 0.424 | 0.029 | 1.8626151 | 20 | ehd2b |
| fgfr3 | 9.261869 | 0.684250895 | 0.328 | 0.017 | 2.3529780 | 20 | fgfr3 |
| col6a14 | 4.595559 | 1.210871689 | 0.701 | 0.091 | 1.1675018 | 20 | col6a1 |
| zgc:162612 | 4.922207 | 0.397076559 | 0.181 | 0.005 | 1.2504867 | 20 | zgc:162612 |
| col1a1b4 | 6.372184 | 1.513871755 | 0.904 | 0.161 | 1.6188533 | 20 | col1a1b |
| col5a14 | 2.155055 | 1.323052661 | 0.808 | 0.125 | 5.4749186 | 20 | col5a1 |
| si:dkey-178e17.3 | 2.396217 | 0.567507949 | 0.288 | 0.014 | 6.0875902 | 20 | si:dkey-178e17.3 |
| ccdc80 | 1.839336 | 0.731601812 | 0.407 | 0.029 | 4.6728354 | 20 | ccdc80 |
| col6a34 | 6.798040 | 1.207133513 | 0.751 | 0.107 | 1.7270420 | 20 | col6a3 |
| col1a24 | 6.900325 | 1.578632033 | 0.915 | 0.174 | 1.7530277 | 20 | col1a2 |
| lpl | 5.747094 | 0.67461947 | 0.322 | 0.018 | 1.4600492 | 20 | lpl |
| heyl | 2.259463 | 0.274411094 | 0.113 | 0.001 | 5.7401669 | 20 | heyl |
| nog2 | 5.453458 | 0.253642693 | 0.102 | 0.001 | 1.3854511 | 20 | nog2 |
| si:dkey-85k7.7 | 2.811876 | 0.442721613 | 0.203 | 0.007 | 7.1435731 | 20 | si:dkey-85k7.7 |

| | | | | | | | |
|---|---|---|---|---|---|---|---|
| sparc5 | 1.091971 | 1.454608991 | 0.927 | 0.187 | 2.7741539 | 20 | sparc |
| twist1a | 5.178131 | 0.537797002 | 0.299 | 0.016 | 1.3155042 | 20 | twist1a |
| mmp14b3 | 3.209378 | 0.99550846 | 0.559 | 0.061 | 8.1534250 | 20 | mmp14b |
| fosl1a3 | 7.028716 | 1.135871732 | 0.616 | 0.078 | 1.7856454 | 20 | fosl1a |
| krt913 | 1.473481 | 1.236781219 | 0.582 | 0.066 | 3.7433806 | 20 | krt91 |
| col1a1a4 | 1.607321 | 1.442456191 | 0.87 | 0.159 | 4.0833992 | 20 | col1a1a |
| si:ch211-170d8.8 | 2.444994 | 0.508190676 | 0.237 | 0.01 | 6.2115091 | 20 | si:ch211-170d8.8 |
| serpine11 | 9.826965 | 0.966963785 | 0.424 | 0.036 | 2.4965405 | 20 | serpine1 |
| cdh111 | 7.780196 | 0.753066535 | 0.362 | 0.026 | 1.9765589 | 20 | cdh11 |
| mmp11b | 1.069372 | 0.504794944 | 0.164 | 0.004 | 2.7167399 | 20 | mmp11b |
| hhipl2 | 1.402351 | 0.277714416 | 0.13 | 0.003 | 3.5626751 | 20 | hhipl2 |
| lhfpl61 | 5.095149 | 0.616377402 | 0.322 | 0.021 | 1.2944228 | 20 | lhfpl6 |
| fxyd13 | 1.627983 | 0.964161099 | 0.61 | 0.078 | 4.1358929 | 20 | fxyd1 |
| sema3e | 2.554159 | 0.44903443 | 0.203 | 0.008 | 6.4888421 | 20 | sema3e |
| CABZ01092746.13 | 3.540000 | 1.148044002 | 0.627 | 0.083 | 8.9933708 | 20 | CABZ01092746.1 |
| CU929237.11 | 5.553364 | 0.621440584 | 0.311 | 0.019 | 1.4108322 | 20 | CU929237.1 |
| epha4b | 1.016708 | 0.465371941 | 0.226 | 0.01 | 2.5829483 | 20 | epha4b |
| si:ch211-39i22.13 | 2.345755 | 1.080196348 | 0.621 | 0.082 | 5.9593910 | 20 | si:ch211-39i22.1 |
| pmp22a4 | 1.436099 | 1.253571605 | 0.87 | 0.159 | 3.6484117 | 20 | pmp22a |
| ccl44 | 2.706746 | 0.399613262 | 0.141 | 0.003 | 6.8764885 | 20 | ccl44 |
| f3b4 | 4.462266 | 1.278860075 | 0.638 | 0.093 | 1.1336387 | 20 | f3b |
| XLOC-040020 | 4.713729 | 0.385966965 | 0.203 | 0.008 | 1.1975230 | 20 | XLOC-040020 |
| col2a1b | 6.078210 | 0.533122109 | 0.266 | 0.014 | 1.5441692 | 20 | col2a1b |
| si:ch73-335l21.44 | 1.936865 | 1.474892066 | 0.751 | 0.135 | 4.9206064 | 20 | si:ch73-335l21.4 |
| vmo1a | 1.164830 | 0.352340496 | 0.153 | 0.004 | 2.9592511 | 20 | vmo1a |
| sertad4 | 4.463884 | 0.477208544 | 0.266 | 0.014 | 1.1340499 | 20 | sertad4 |
| mmp14a3 | 8.285875 | 0.823462142 | 0.503 | 0.055 | 2.1050267 | 20 | mmp14a |
| crabp2b1 | 9.963702 | 0.598663836 | 0.294 | 0.018 | 2.5312785 | 20 | crabp2b |
| c1qtnf21 | 1.084270 | 0.618692977 | 0.328 | 0.023 | 2.7545896 | 20 | c1qtnf2 |
| fgl2b1 | 1.146918 | 0.555968411 | 0.249 | 0.013 | 2.9137465 | 20 | fgl2b |
| olfml1 | 2.353432 | 0.354333926 | 0.164 | 0.005 | 5.9788955 | 20 | olfml1 |
| mamdc2b | 1.361824 | 0.300848295 | 0.119 | 0.002 | 3.4597145 | 20 | mamdc2b |
| jun5 | 5.642674 | 1.419809499 | 0.938 | 0.243 | 1.4335215 | 20 | jun |
| htra1a1 | 2.610416 | 0.647396101 | 0.311 | 0.021 | 6.6317638 | 20 | htra1a |
| thbs1b | 4.444125 | 0.493312324 | 0.209 | 0.009 | 1.1290300 | 20 | thbs1b |

| | | | | | | | |
|---|---|---|---|---|---|---|---|
| dlx5a | 2.258010 | 0.293861157 | 0.13 | 0.003 | 5.7364747 | 20 | dlx5a |
| rarga1 | 3.624257 | 0.829017058 | 0.492 | 0.056 | 9.2074268 | 20 | rarga |
| s100v25 | 2.582241 | 1.103157301 | 0.751 | 0.134 | 6.5601856 | 20 | s100v2 |
| bcl11ba1 | 3.600971 | 0.524078121 | 0.243 | 0.013 | 9.1482689 | 20 | bcl11ba |
| filip1l | 5.163972 | 0.582616239 | 0.266 | 0.016 | 1.3119071 | 20 | filip1l |
| sema3aa | 8.219232 | 0.274652459 | 0.136 | 0.004 | 2.0880958 | 20 | sema3aa |
| mmp24 | 1.361330 | 1.035821255 | 0.667 | 0.105 | 3.4584593 | 20 | mmp2 |
| fstl1b4 | 7.155139 | 1.105568598 | 0.774 | 0.141 | 1.8177630 | 20 | fstl1b |
| qdpra | 3.506776 | 0.361406777 | 0.175 | 0.007 | 8.9089647 | 20 | qdpra |
| fosb3 | 4.087198 | 1.186504598 | 0.689 | 0.115 | 1.0383528 | 20 | fosb |
| bgnb1 | 7.177834 | 0.619596709 | 0.311 | 0.023 | 1.8235289 | 20 | bgnb |
| ier5l | 2.691436 | 0.604011162 | 0.311 | 0.023 | 6.8375948 | 20 | ier5l |
| fosab6 | 3.444577 | 1.402929382 | 0.966 | 0.355 | 8.7509494 | 20 | fosab |
| zgc:56585 | 3.228831 | 0.412759896 | 0.203 | 0.01 | 8.2028470 | 20 | zgc:56585 |
| sdc46 | 4.235489 | 1.243511395 | 0.91 | 0.247 | 1.0760261 | 20 | sdc4 |
| gstz1 | 4.055059 | 0.63019272 | 0.345 | 0.031 | 1.0301879 | 20 | gstz1 |
| apln | 2.387142 | 0.304950083 | 0.141 | 0.004 | 6.0645360 | 20 | apln |
| egr17 | 1.381229 | 1.15531386 | 0.797 | 0.177 | 3.5090124 | 20 | egr1 |
| alx4a | 3.451133 | 0.288074778 | 0.136 | 0.004 | 8.7676042 | 20 | alx4a |
| ms4a17a.92 | 3.638353 | 0.739426057 | 0.497 | 0.065 | 9.2432377 | 20 | ms4a17a.9 |
| gpm6bb | 4.179990 | 0.515152086 | 0.294 | 0.023 | 1.0619264 | 20 | gpm6bb |
| prrx1a | 6.686115 | 0.383937911 | 0.209 | 0.011 | 1.6986077 | 20 | prrx1a |
| adh8a3 | 1.131989 | 1.051128362 | 0.401 | 0.044 | 2.8758204 | 20 | adh8a |
| capn2b | 1.793257 | 0.525420673 | 0.294 | 0.023 | 4.5557709 | 20 | capn2b |
| junba6 | 3.521071 | 1.262441449 | 0.955 | 0.313 | 8.9452827 | 20 | junba |
| si:ch211-286o17.12 | 7.493613 | 0.74607772 | 0.407 | 0.045 | 1.9037524 | 20 | si:ch211-286o17.1 |
| ier2a6 | 1.148203 | 1.23824413 | 0.898 | 0.268 | 2.9170105 | 20 | ier2a |
| junbb3 | 3.372342 | 0.739793654 | 0.486 | 0.066 | 8.5674361 | 20 | junbb |
| serpinf13 | 3.613651 | 0.70041182 | 0.384 | 0.041 | 9.1804810 | 20 | serpinf1 |
| enpp2 | 6.027631 | 0.269948751 | 0.147 | 0.005 | 1.5313198 | 20 | enpp2 |
| col5a3a | 2.193261 | 0.370542345 | 0.175 | 0.008 | 5.5719801 | 20 | col5a3a |
| pigr | 3.153715 | 0.469456439 | 0.271 | 0.021 | 8.0120135 | 20 | pigr |
| vwa1 | 1.576306 | 0.509180753 | 0.26 | 0.019 | 4.0046067 | 20 | vwa1 |
| zgc:154093 | 1.880451 | 0.4111951 | 0.226 | 0.014 | 4.7772869 | 20 | zgc:154093 |
| spon2b1 | 2.908618 | 0.653595862 | 0.277 | 0.021 | 7.3893442 | 20 | spon2b |

| | | | | | | | |
|---|---|---|---|---|---|---|---|
| s100a10b6 | 1.349485 | 1.265707621 | 0.932 | 0.279 | 3.4283685 | 20 | s100a10b |
| spock2 | 9.425345 | 0.334598426 | 0.124 | 0.004 | 2.3945091 | 20 | spock2 |
| atoh81 | 2.331651 | 0.516777072 | 0.282 | 0.023 | 5.9235608 | 20 | atoh8 |
| tcf71 | 4.507419 | 0.573438758 | 0.266 | 0.021 | 1.1451098 | 20 | tcf7 |
| rnd3b3 | 1.008562 | 0.717918173 | 0.395 | 0.046 | 2.5622518 | 20 | rnd3b |
| jdp2b4 | 5.203951 | 1.27910548 | 0.701 | 0.159 | 1.3220639 | 20 | jdp2b |
| cdo1 | 6.443054 | 0.313687136 | 0.153 | 0.006 | 1.6368580 | 20 | cdo1 |
| lix1l | 1.160920 | 0.436510829 | 0.226 | 0.015 | 2.9493193 | 20 | lix1l |
| socs3a6 | 1.820515 | 1.234422397 | 0.938 | 0.308 | 4.6250190 | 20 | socs3a |
| dlx3b | 9.000738 | 0.378443885 | 0.158 | 0.007 | 2.2866375 | 20 | dlx3b |
| klf6a5 | 3.154594 | 1.132603409 | 0.723 | 0.175 | 8.0142478 | 20 | klf6a |
| dcn4 | 5.965305 | 1.033592902 | 0.695 | 0.138 | 1.5154859 | 20 | dcn |
| cd991 | 6.493008 | 0.691559207 | 0.401 | 0.049 | 1.6495487 | 20 | cd99 |
| ier2b7 | 1.230511 | 1.115260438 | 0.831 | 0.231 | 3.1261150 | 20 | ier2b |
| anxa1a5 | 2.899450 | 1.092191062 | 0.718 | 0.157 | 7.3660531 | 20 | anxa1a |
| atf34 | 3.508701 | 0.948341355 | 0.497 | 0.077 | 8.9138565 | 20 | atf3 |
| tcf123 | 4.568985 | 0.707387601 | 0.458 | 0.064 | 1.1607507 | 20 | tcf12 |
| fstb2 | 7.516615 | 0.628995561 | 0.271 | 0.023 | 1.9095961 | 20 | fstb |
| si:ch73-46j18.56 | 7.482172 | 1.034628892 | 0.819 | 0.202 | 1.9008458 | 20 | si:ch73-46j18.5 |
| zgc:1627309 | 3.131590 | 1.187794915 | 0.949 | 0.398 | 7.9558058 | 20 | zgc:162730 |
| cntfr1 | 5.207514 | 0.556193805 | 0.237 | 0.018 | 1.3229691 | 20 | cntfr |
| slc25a48 | 1.825148 | 0.291178241 | 0.147 | 0.007 | 4.6367908 | 20 | slc25a48 |
| icn5 | 1.956835 | 1.293343458 | 0.853 | 0.262 | 4.9713418 | 20 | icn |
| ctsk2 | 2.460953 | 0.675606187 | 0.475 | 0.07 | 6.2520529 | 20 | ctsk |
| zeb2a1 | 2.627652 | 0.738874102 | 0.362 | 0.043 | 6.6755516 | 20 | zeb2a |
| cspg4 | 3.457455 | 0.283215527 | 0.136 | 0.006 | 8.7836665 | 20 | cspg4 |
| ms4a17a.111 | 6.286531 | 0.62271439 | 0.384 | 0.049 | 1.5970933 | 20 | ms4a17a.11 |
| ecm2 | 9.301587 | 0.33492485 | 0.164 | 0.009 | 2.3630681 | 20 | ecm2 |
| col6a24 | 4.367511 | 0.775985321 | 0.463 | 0.071 | 1.1095663 | 20 | col6a2 |
| entpd5a | 1.784687 | 0.518238253 | 0.107 | 0.004 | 4.5339979 | 20 | entpd5a |
| fstl33 | 8.731532 | 0.50907449 | 0.254 | 0.022 | 2.2182459 | 20 | fstl3 |
| irx1a | 1.263119 | 0.258530963 | 0.113 | 0.004 | 3.2089545 | 20 | irx1a |
| postnb4 | 3.925459 | 1.069431778 | 0.514 | 0.091 | 9.9726295 | 20 | postnb |
| colec123 | 4.005831 | 0.693050888 | 0.418 | 0.059 | 1.0176816 | 20 | colec12 |
| emilin2a1 | 5.155320 | 0.515078089 | 0.249 | 0.021 | 1.3097091 | 20 | emilin2a |

| | | | | | | | |
|---|---|---|---|---|---|---|---|
| rgma | 6.31010[illegible] | 0.33161715 | 0.169 | 0.01 | 1.6030832 | 20 | rgma |
| mxra8b2 | 5.24223[illegible] | 0.485774959 | 0.277 | 0.027 | 1.3317909 | 20 | mxra8b |
| rgs5b2 | 2.36016[illegible] | 1.032030443 | 0.435 | 0.069 | 5.9960002 | 20 | rgs5b |
| LO018188.12 | 4.83192[illegible] | 0.491718556 | 0.271 | 0.026 | 1.2275504 | 20 | LO018188.1 |
| loxl11 | 1.02093[illegible] | 0.515149809 | 0.282 | 0.028 | 2.5936978 | 20 | loxl1 |
| cxcl18b5 | 1.19885[illegible] | 1.065807175 | 0.548 | 0.106 | 3.0456895 | 20 | cxcl18b |
| btg25 | 8.32866[illegible] | 1.042675617 | 0.91 | 0.344 | 2.1158972 | 20 | btg2 |
| klf15 | 2.20746[illegible] | 0.296704695 | 0.175 | 0.011 | 5.6080699 | 20 | klf15 |
| myo1ca | 1.26600[illegible] | 0.47081305 | 0.266 | 0.026 | 3.2162979 | 20 | myo1ca |
| pmp22b1 | 4.84239[illegible] | 0.660486535 | 0.345 | 0.044 | 1.2302116 | 20 | pmp22b |
| tnfaip2a1 | 5.95340[illegible] | 0.461747329 | 0.237 | 0.021 | 1.5124636 | 20 | tnfaip2a |
| marcksl1a5 | 9.74361[illegible] | 1.066383157 | 0.825 | 0.255 | 2.4753649 | 20 | marcksl1a |
| im:7142702 | 2.05096[illegible] | 0.264787001 | 0.141 | 0.007 | 5.2104647 | 20 | im:7142702 |
| srpx2 | 2.26529[illegible] | 0.30716648 | 0.147 | 0.008 | 5.7549905 | 20 | srpx2 |
| hspg21 | 2.36008[illegible] | 0.476846622 | 0.266 | 0.027 | 5.9957843 | 20 | hspg2 |
| nupr1b2 | 5.54495[illegible] | 0.712337289 | 0.333 | 0.042 | 1.4086953 | 20 | nupr1b |
| osr23 | 9.97828[illegible] | 0.661194226 | 0.362 | 0.049 | 2.5349830 | 20 | osr2 |
| pleca2 | 1.86389[illegible] | 0.672313247 | 0.395 | 0.059 | 4.7352234 | 20 | pleca |
| ddit35 | 2.02730[illegible] | 0.970346093 | 0.684 | 0.173 | 5.1503753 | 20 | ddit3 |
| pcdh18b1 | 1.54516[illegible] | 0.42709121 | 0.243 | 0.023 | 3.9254798 | 20 | pcdh18b |
| sox11a | 2.69123[illegible] | 0.585226514 | 0.26 | 0.027 | 6.8370903 | 20 | sox11a |
| si:busm1-57f23.1 | 4.16571[illegible] | 0.554835355 | 0.266 | 0.028 | 1.0583010 | 20 | si:busm1-57f23.1 |
| anxa5b4 | 1.01929[illegible] | 0.796026669 | 0.514 | 0.101 | 2.5895208 | 20 | anxa5b |
| si:ch211-178n15.12 | 1.39091[illegible] | 0.483960791 | 0.26 | 0.027 | 3.5336233 | 20 | si:ch211-178n15.1 |
| fap1 | 1.52468[illegible] | 0.418304467 | 0.181 | 0.013 | 3.8734638 | 20 | fap |
| zgc:1533171 | 3.63252[illegible] | 0.565234348 | 0.305 | 0.037 | 9.2284376 | 20 | zgc:153317 |
| antxr2a4 | 5.86077[illegible] | 0.885283309 | 0.655 | 0.164 | 1.4889308 | 20 | antxr2a |
| jag1b1 | 6.42334[illegible] | 0.493701878 | 0.254 | 0.026 | 1.6318513 | 20 | jag1b |
| plpp31 | 1.01131[illegible] | 0.644343176 | 0.407 | 0.062 | 2.5692569 | 20 | plpp3 |
| tcf41 | 1.56154[illegible] | 0.418113317 | 0.232 | 0.022 | 3.9670937 | 20 | tcf4 |
| krt86 | 2.08434[illegible] | 1.028632525 | 0.893 | 0.306 | 5.2952728 | 20 | krt8 |
| nfixb3 | 7.18666[illegible] | 0.700596695 | 0.452 | 0.079 | 1.8257726 | 20 | nfixb |
| aldob | 7.38614[illegible] | 0.449555241 | 0.288 | 0.033 | 1.8764492 | 20 | aldob |
| tuft1a | 1.45249[illegible] | 0.455328119 | 0.26 | 0.028 | 3.6900548 | 20 | tuft1a |
| grb10b2 | 4.64119[illegible] | 0.530113792 | 0.299 | 0.037 | 1.1790951 | 20 | grb10b |

| | | | | | | | |
|---|---|---|---|---|---|---|---|
| gpc1a | 5.21122[illegible] | 0.296480866 | 0.13 | 0.007 | 1.3239106 | 20 | gpc1a |
| rbfox1 | 1.69842[illegible] | 0.453191448 | 0.254 | 0.027 | 4.3148422 | 20 | rbfox1 |
| apcdd1l | 3.08949[illegible] | 0.285778209 | 0.141 | 0.008 | 7.8488502 | 20 | apcdd1l |
| abi3bpa | 4.42540[illegible] | 0.312885047 | 0.164 | 0.011 | 1.1242730 | 20 | abi3bpa |
| rbp44 | 5.44579[illegible] | 1.086082261 | 0.582 | 0.137 | 1.3835040 | 20 | rbp4 |
| ppap2d2 | 6.28023[illegible] | 0.539175926 | 0.282 | 0.034 | 1.5954929 | 20 | ppap2d |
| socs3b5 | 7.56065[illegible] | 0.885705607 | 0.605 | 0.147 | 1.9207847 | 20 | socs3b |
| endouc2 | 8.56842[illegible] | 0.718086391 | 0.412 | 0.068 | 2.1768083 | 20 | endouc |
| lmna | 1.27671[illegible] | 0.340737674 | 0.169 | 0.012 | 3.2434983 | 20 | lmna |
| MDFI | 1.49211[illegible] | 0.394800796 | 0.226 | 0.022 | 3.7907261 | 20 | MDFI |
| zfp36l1b4 | 4.82902[illegible] | 0.953431946 | 0.565 | 0.136 | 1.2268125 | 20 | zfp36l1b |
| pcolcea1 | 5.96433[illegible] | 0.706339369 | 0.401 | 0.064 | 1.5152387 | 20 | pcolcea |
| im:71523481 | 3.35742[illegible] | 0.36313468 | 0.175 | 0.014 | 8.5295324 | 20 | im:7152348 |
| csrnp1b4 | 4.22071[illegible] | 0.826311309 | 0.633 | 0.16 | 1.0722735 | 20 | csrnp1b |
| timp2a4 | 4.98346[illegible] | 0.71652405 | 0.379 | 0.062 | 1.2660482 | 20 | timp2a |
| tgfbr32 | 1.55011[illegible] | 0.513965135 | 0.322 | 0.044 | 3.9380674 | 20 | tgfbr3 |
| FAT4 | 2.51298[illegible] | 0.340854145 | 0.209 | 0.019 | 6.3842402 | 20 | FAT4 |
| si:ch211-133n4.42 | 3.97304[illegible] | 0.554053538 | 0.362 | 0.056 | 1.0093509 | 20 | si:ch211-133n4.4 |
| htra3b | 1.51075[illegible] | 0.361292669 | 0.169 | 0.013 | 3.8380637 | 20 | htra3b |
| ccdc3b2 | 2.07709[illegible] | 0.44405393 | 0.254 | 0.029 | 5.2768520 | 20 | ccdc3b |
| sfrp1b1 | 4.34284[illegible] | 0.402887985 | 0.181 | 0.015 | 1.1033008 | 20 | sfrp1b |
| f13a1b2 | 1.29299[illegible] | 0.433461192 | 0.26 | 0.03 | 3.2848639 | 20 | f13a1b |
| ism1 | 4.93838[illegible] | 0.344285599 | 0.141 | 0.009 | 1.2545962 | 20 | ism1 |
| cspg5a | 7.38875[illegible] | 0.281493994 | 0.147 | 0.01 | 1.8771139 | 20 | cspg5a |
| ramp21 | 2.40143[illegible] | 0.542473762 | 0.305 | 0.042 | 6.1008410 | 20 | ramp2 |
| serpinh1b | 2.67984[illegible] | 0.489940172 | 0.249 | 0.029 | 6.8081378 | 20 | serpinh1b |
| hoxb7a | 5.16826[illegible] | 0.341886273 | 0.175 | 0.015 | 1.3129966 | 20 | hoxb7a |
| itgb41 | 6.14807[illegible] | 0.402009211 | 0.181 | 0.016 | 1.5619184 | 20 | itgb4 |
| ablim1b2 | 1.04804[illegible] | 0.577185227 | 0.294 | 0.041 | 2.6625644 | 20 | ablim1b |
| fkbp14 | 1.19774[illegible] | 0.473871321 | 0.294 | 0.04 | 3.0428739 | 20 | fkbp14 |
| cox4i22 | 1.97442[illegible] | 0.449600813 | 0.271 | 0.035 | 5.0160216 | 20 | cox4i2 |
| zcchc241 | 3.99459[illegible] | 0.476680884 | 0.277 | 0.036 | 1.0148262 | 20 | zcchc24 |
| sertad2a | 1.07990[illegible] | 0.381971534 | 0.215 | 0.022 | 2.7435040 | 20 | sertad2a |
| rgcc2 | 2.88934[illegible] | 0.735320183 | 0.384 | 0.068 | 7.3403827 | 20 | rgcc |
| klf2a4 | 3.93012[illegible] | 0.8222533 | 0.458 | 0.096 | 9.9844699 | 20 | klf2a |

| | | | | | | | |
|---|---|---|---|---|---|---|---|
| h3f3c7 | 4.502559 | 0.83048727 | 0.831 | 0.299 | 1.1438752 | 20 | h3f3c |
| atf4b5 | 1.703835 | 0.858522356 | 0.757 | 0.273 | 4.3285946 | 20 | atf4b |
| cd81a5 | 2.014444 | 0.831067635 | 0.78 | 0.236 | 5.1176967 | 20 | cd81a |
| arhgap21a | 2.168706 | 0.309475851 | 0.153 | 0.012 | 5.5095992 | 20 | arhgap21a |
| XLOC-042917 | 2.822436 | 0.316550948 | 0.158 | 0.012 | 7.1703990 | 20 | XLOC-042917 |
| smad1 | 3.138804 | 0.351584116 | 0.198 | 0.019 | 7.9741322 | 20 | smad1 |
| si:dkey-261h17.12 | 2.470870 | 0.815055063 | 0.503 | 0.112 | 6.2772462 | 20 | si:dkey-261h17.1 |
| LOC101883507 | 3.261382 | 0.33685985 | 0.186 | 0.018 | 8.2855426 | 20 | LOC101883507 |
| efna5b | 5.770732 | 0.25436709 | 0.136 | 0.009 | 1.4660544 | 20 | efna5b |
| lfng | 1.049548 | 0.350583288 | 0.13 | 0.009 | 2.6663768 | 20 | lfng |
| her6 | 1.286582 | 0.463235368 | 0.215 | 0.024 | 3.2685628 | 20 | her6 |
| zgc:171679 | 6.325931 | 0.395088874 | 0.226 | 0.026 | 1.6071029 | 20 | zgc:171679 |
| CR383676.110 | 7.079836 | 0.772222136 | 0.989 | 0.715 | 1.7986323 | 20 | CR383676.1 |
| akap12b3 | 1.563415 | 0.670477984 | 0.384 | 0.071 | 3.9718565 | 20 | akap12b |
| bmp16 | 2.529305 | 0.291503212 | 0.119 | 0.007 | 6.4256999 | 20 | bmp16 |
| ier52 | 3.565557 | 0.779541862 | 0.542 | 0.143 | 9.0582985 | 20 | ier5 |
| s100w | 4.050817 | 0.35945422 | 0.186 | 0.018 | 1.0291102 | 20 | s100w |
| plxdc2 | 6.591808 | 0.305397923 | 0.147 | 0.012 | 1.6746488 | 20 | plxdc2 |
| vim3 | 7.712804 | 0.51687556 | 0.294 | 0.044 | 1.9594380 | 20 | vim |
| hbegfa | 1.327941 | 0.768869498 | 0.452 | 0.099 | 3.3736365 | 20 | hbegfa |
| ghra2 | 1.914529 | 0.435005557 | 0.249 | 0.032 | 4.8638631 | 20 | ghra |
| pcsk6 | 3.840963 | 0.266367129 | 0.136 | 0.01 | 9.7579685 | 20 | pcsk6 |
| ckap43 | 5.787138 | 0.507060875 | 0.339 | 0.058 | 1.4702226 | 20 | ckap4 |
| cyp26b1 | 6.623904 | 0.283301594 | 0.113 | 0.007 | 1.6828028 | 20 | cyp26b1 |
| angptl1a2 | 8.654299 | 0.544209302 | 0.249 | 0.033 | 2.1986246 | 20 | angptl1a |
| ecm1b2 | 1.018095 | 0.550719932 | 0.362 | 0.064 | 2.5864712 | 20 | ecm1b |
| prelp | 1.826337 | 0.348746318 | 0.124 | 0.008 | 4.6398111 | 20 | prelp |
| ahnak5 | 2.013058 | 0.725654796 | 0.531 | 0.133 | 5.1141746 | 20 | ahnak |
| slc16a10 | 2.525301 | 0.366839441 | 0.209 | 0.024 | 6.4155281 | 20 | slc16a10 |
| mtmr11 | 3.724154 | 0.27637615 | 0.147 | 0.012 | 9.4612135 | 20 | mtmr11 |
| fpr11 | 4.835921 | 0.360125792 | 0.192 | 0.02 | 1.2285658 | 20 | fpr1 |
| si:dkey-101k6.51 | 5.033205 | 0.32625796 | 0.181 | 0.018 | 1.2786859 | 20 | si:dkey-101k6.5 |
| kank24 | 1.314161 | 0.554701993 | 0.316 | 0.052 | 3.3386271 | 20 | kank2 |
| c1qtnf52 | 3.019075 | 0.609405168 | 0.243 | 0.033 | 7.6699608 | 20 | c1qtnf5 |
| VIT | 3.403876 | 0.285452134 | 0.113 | 0.007 | 8.6475472 | 20 | VIT |

| | | | | | | | |
|---|---|---|---|---|---|---|---|
| id15 | 5.346033 | 0.869044451 | 0.508 | 0.128 | 1.3581597 | 20 | id1 |
| fxyd6l2 | 7.167282 | 0.650649101 | 0.492 | 0.114 | 1.8208482 | 20 | fxyd6l |
| cd82a1 | 1.165437 | 0.522591783 | 0.362 | 0.067 | 2.9607937 | 20 | cd82a |
| lrrc173 | 1.169216 | 0.539632341 | 0.328 | 0.056 | 2.9703935 | 20 | lrrc17 |
| spred2b3 | 1.385794 | 0.580799743 | 0.401 | 0.082 | 3.5206113 | 20 | spred2b |
| cxcl141 | 7.018215 | 0.283793522 | 0.141 | 0.011 | 1.7829777 | 20 | cxcl14 |
| mycb4 | 7.769866 | 0.804557357 | 0.638 | 0.197 | 1.9739345 | 20 | mycb |
| gadd45ba4 | 1.186456 | 0.871536125 | 0.621 | 0.194 | 3.0141938 | 20 | gadd45ba |
| BX663503.31 | 1.326167 | 0.422360052 | 0.243 | 0.032 | 3.3691296 | 20 | BX663503.3 |
| nova21 | 4.269168 | 0.450377029 | 0.249 | 0.034 | 1.0845822 | 20 | nova2 |
| cnn3a4 | 6.173487 | 0.493213156 | 0.356 | 0.067 | 1.5683744 | 20 | cnn3a |
| CR847953.1 | 7.207149 | 0.365645582 | 0.186 | 0.02 | 1.8309762 | 20 | CR847953.1 |
| gng74 | 1.498664 | 0.473782867 | 0.333 | 0.06 | 3.8073579 | 20 | gng7 |
| pcbd1 | 6.138539 | 0.528195966 | 0.322 | 0.056 | 1.5594958 | 20 | pcbd1 |
| col11a1a | 1.931124 | 0.25372689 | 0.119 | 0.008 | 4.9060226 | 20 | col11a1a |
| meis1a1 | 1.956156 | 0.332275274 | 0.181 | 0.019 | 4.9696159 | 20 | meis1a |
| fgl2a3 | 2.088426 | 0.636500536 | 0.367 | 0.073 | 5.3056487 | 20 | fgl2a |
| jund4 | 2.361526 | 0.62366018 | 0.401 | 0.087 | 5.9994581 | 20 | jund |
| pdgfrb3 | 2.419275 | 0.481086184 | 0.299 | 0.05 | 6.1461682 | 20 | pdgfrb |
| cebpd5 | 7.638701 | 0.776379105 | 0.537 | 0.148 | 1.9406120 | 20 | cebpd |
| fkbp74 | 1.150786 | 0.480660479 | 0.288 | 0.048 | 2.9235732 | 20 | fkbp7 |
| serpine2 | 3.023386 | 0.547236854 | 0.322 | 0.059 | 7.6809127 | 20 | serpine2 |
| si:dkey-226m8.101 | 4.677380 | 0.261921344 | 0.124 | 0.01 | 1.1882884 | 20 | si:dkey-226m8.10 |
| pdgfab | 6.365139 | 0.317783212 | 0.158 | 0.016 | 1.6170636 | 20 | pdgfab |
| wnt5b1 | 1.510704 | 0.298866128 | 0.153 | 0.015 | 3.8379458 | 20 | wnt5b |
| si:ch73-138n13.11 | 1.699250 | 0.459471033 | 0.266 | 0.042 | 4.3169461 | 20 | si:ch73-138n13.1 |
| gpc42 | 7.456538 | 0.347740669 | 0.22 | 0.03 | 1.8943335 | 20 | gpc4 |
| edil3a2 | 7.766427 | 0.397996931 | 0.192 | 0.023 | 1.9730609 | 20 | edil3a |
| frmd6 | 8.873807 | 0.285103657 | 0.164 | 0.017 | 2.2543908 | 20 | frmd6 |
| lsp1 | 1.401186 | 0.337396104 | 0.169 | 0.018 | 3.5597151 | 20 | lsp1 |
| kalrnb | 1.568132 | 0.271915174 | 0.119 | 0.009 | 3.9838415 | 20 | kalrnb |
| SCARF21 | 3.231801 | 0.378985514 | 0.203 | 0.026 | 8.2103926 | 20 | SCARF2 |
| col11a1b | 4.010957 | 0.34885519 | 0.158 | 0.016 | 1.0189837 | 20 | col11a1b |
| ogna | 4.156695 | 0.336415374 | 0.113 | 0.008 | 1.0560085 | 20 | ogna |
| plecb1 | 5.101751 | 0.482789858 | 0.288 | 0.05 | 1.2960998 | 20 | plecb |

| | | | | | | | |
|---|---|---|---|---|---|---|---|
| si:ch211-212k18.83 | 8.312949 | 0.431044701 | 0.277 | 0.046 | 2.1119047 | 20 | si:ch211-212k18.8 |
| tenm33 | 2.201263 | 0.512218788 | 0.316 | 0.059 | 5.5923104 | 20 | tenm3 |
| vstm4a1 | 2.923278 | 0.281448808 | 0.153 | 0.015 | 7.4265881 | 20 | vstm4a |
| lamb1b1 | 3.727933 | 0.351851769 | 0.192 | 0.024 | 9.4708149 | 20 | lamb1b |
| ip6k2b3 | 5.859721 | 0.650592277 | 0.475 | 0.126 | 1.4886622 | 20 | ip6k2b |
| sept101 | 5.898237 | 0.431261256 | 0.288 | 0.05 | 1.4984472 | 20 | sept10 |
| angptl42 | 5.940937 | 0.565637579 | 0.294 | 0.053 | 1.5092950 | 20 | angptl4 |
| LOC101886114 | 7.936931 | 0.318921999 | 0.153 | 0.016 | 2.0163773 | 20 | LOC101886114 |
| nectin1b1 | 9.231482 | 0.456094716 | 0.266 | 0.044 | 2.3452582 | 20 | nectin1b |
| sh3pxd2b2 | 1.354694 | 0.477451167 | 0.277 | 0.048 | 3.4416007 | 20 | sh3pxd2b |
| loxl2b1 | 1.777267 | 0.428710909 | 0.266 | 0.044 | 4.5151471 | 20 | loxl2b |
| vat13 | 3.885035 | 0.655922945 | 0.52 | 0.144 | 9.8699324 | 20 | vat1 |
| hcar1-42 | 4.260744 | 0.388964443 | 0.22 | 0.032 | 1.0824422 | 20 | hcar1-4 |
| mcl1b6 | 6.866693 | 0.746369371 | 0.695 | 0.247 | 1.7444834 | 20 | mcl1b |
| rgs5a2 | 1.162754 | 0.527080284 | 0.237 | 0.037 | 2.9539779 | 20 | rgs5a |
| enam | 1.260830 | 0.496209148 | 0.119 | 0.01 | 3.2031398 | 20 | enam |
| rnd3a1 | 1.272335 | 0.407880314 | 0.226 | 0.034 | 3.2323685 | 20 | rnd3a |
| phactr4b1 | 2.201956 | 0.311722924 | 0.175 | 0.021 | 5.5940713 | 20 | phactr4b |
| hmbox1a | 2.212474 | 0.405814836 | 0.237 | 0.038 | 5.6207905 | 20 | hmbox1a |
| tpm4a5 | 2.917830 | 0.587131901 | 0.441 | 0.111 | 7.4127490 | 20 | tpm4a |
| XLOC-0004315 | 3.255908 | 0.694276295 | 0.785 | 0.268 | 8.2716343 | 20 | XLOC-000431 |
| chd35 | 3.989175 | 0.565065516 | 0.463 | 0.122 | 1.0134499 | 20 | chd3 |
| dpysl31 | 4.556997 | 0.315179213 | 0.169 | 0.02 | 1.1577052 | 20 | dpysl3 |
| sdc21 | 6.653587 | 0.514057849 | 0.362 | 0.08 | 1.6903438 | 20 | sdc2 |
| bckdhbl | 7.071567 | 0.346598712 | 0.209 | 0.03 | 1.7965317 | 20 | bckdhbl |
| si:ch211-284f22.31 | 8.665156 | 0.36870168 | 0.22 | 0.033 | 2.2013829 | 20 | si:ch211-284f22.3 |
| pdgfrl2 | 1.035480 | 0.37313064 | 0.215 | 0.032 | 2.6306390 | 20 | pdgfrl |
| LOC101886030 | 1.064718 | 0.250362488 | 0.141 | 0.014 | 2.7049185 | 20 | LOC101886030 |
| gas1a4 | 1.664601 | 0.460304695 | 0.294 | 0.056 | 4.2289210 | 20 | gas1a |
| capn2a1 | 1.897227 | 0.44583267 | 0.282 | 0.052 | 4.8199057 | 20 | capn2a |
| eva1ba3 | 2.084462 | 0.481912745 | 0.322 | 0.065 | 5.2955775 | 20 | eva1ba |
| midn4 | 2.235489 | 0.650553396 | 0.492 | 0.142 | 5.6792604 | 20 | midn |
| ccdc85b4 | 2.424789 | 0.442289537 | 0.271 | 0.049 | 6.1601775 | 20 | ccdc85b |
| anxa2a5 | 5.561704 | 0.62371067 | 0.446 | 0.115 | 1.4129511 | 20 | anxa2a |
| col6a4a3 | 6.058303 | 0.537527278 | 0.316 | 0.064 | 1.5391121 | 20 | col6a4a |

| | | | | | | | |
|---|---|---|---|---|---|---|---|
| efnb2a | 6.473782 | 0.291339981 | 0.141 | 0.015 | 1.6446645 | 20 | efnb2a |
| krt18b6 | 1.106702 | 0.683251557 | 0.559 | 0.173 | 2.8115783 | 20 | krt18b |
| wls4 | 1.359078 | 0.582062404 | 0.469 | 0.128 | 3.4527376 | 20 | wls |
| cygb14 | 2.030755 | 0.468175736 | 0.305 | 0.06 | 5.1591339 | 20 | cygb1 |
| FP102018.1 | 3.321405 | 0.298019228 | 0.164 | 0.02 | 8.4380297 | 20 | FP102018.1 |
| wu:fi04e12 | 3.855291 | 0.279122829 | 0.169 | 0.021 | 9.7943692 | 20 | wu:fi04e12 |
| tmem1764 | 7.041117 | 0.497465396 | 0.35 | 0.078 | 1.7887958 | 20 | tmem176 |
| sox62 | 2.611618 | 0.433063006 | 0.203 | 0.031 | 6.6348169 | 20 | sox6 |
| cd28l1 | 4.355057 | 0.393367327 | 0.232 | 0.038 | 1.1064024 | 20 | cd28l |
| fhl1b | 4.920972 | 0.298059207 | 0.158 | 0.019 | 1.2501730 | 20 | fhl1b |
| foxo1a | 5.553676 | 0.371672212 | 0.237 | 0.04 | 1.4109115 | 20 | foxo1a |
| si:dkey-9i23.163 | 8.422782 | 0.428207556 | 0.26 | 0.048 | 2.1398077 | 20 | si:dkey-9i23.16 |
| htra1b1 | 1.223216 | 0.38841396 | 0.226 | 0.036 | 3.1075808 | 20 | htra1b |
| anxa44 | 1.271721 | 0.606830695 | 0.486 | 0.148 | 3.2308076 | 20 | anxa4 |
| marcksl1b6 | 2.134710 | 0.737489646 | 0.469 | 0.139 | 5.4232310 | 20 | marcksl1b |
| ednraa | 3.439976 | 0.300955209 | 0.169 | 0.022 | 8.7392597 | 20 | ednraa |
| wnt11r1 | 6.965116 | 0.279179058 | 0.107 | 0.009 | 1.7694879 | 20 | wnt11r |
| ppib7 | 7.879415 | 0.683113243 | 0.825 | 0.375 | 2.0017655 | 20 | ppib |
| cd248a2 | 9.153635 | 0.39493136 | 0.254 | 0.046 | 2.3254810 | 20 | cd248a |
| gli31 | 1.429455 | 0.326224957 | 0.215 | 0.034 | 3.6315309 | 20 | gli3 |
| tagln27 | 1.764510 | 0.726984077 | 0.576 | 0.207 | 4.4827388 | 20 | tagln2 |
| tuba8l33 | 2.557978 | 0.458627854 | 0.322 | 0.07 | 6.4985448 | 20 | tuba8l3 |
| reck | 2.664779 | 0.33055173 | 0.192 | 0.028 | 6.7698725 | 20 | reck |
| si:ch73-1a9.38 | 2.851333 | 0.676682197 | 0.763 | 0.312 | 7.2438139 | 20 | si:ch73-1a9.3 |
| sfrp1a2 | 3.228176 | 0.451882296 | 0.215 | 0.035 | 8.2011821 | 20 | sfrp1a |
| sulf1 | 3.594707 | 0.311335401 | 0.153 | 0.018 | 9.1323549 | 20 | sulf1 |
| higd1a4 | 5.496512 | 0.699626821 | 0.706 | 0.3 | 1.3963890 | 20 | higd1a |
| CABZ01039096.11 | 6.646852 | 0.346051416 | 0.169 | 0.023 | 1.6886329 | 20 | CABZ01039096.1 |
| spry43 | 8.348517 | 0.427688154 | 0.254 | 0.048 | 2.1209407 | 20 | spry4 |
| pdgfra2 | 1.083195 | 0.499992504 | 0.288 | 0.06 | 2.7518571 | 20 | pdgfra |
| si:ch1073-459j12.14 | 1.289983 | 0.522276253 | 0.35 | 0.083 | 3.2772028 | 20 | si:ch1073-459j12.1 |
| s100u | 3.697393 | 0.34951698 | 0.226 | 0.039 | 9.3932292 | 20 | s100u |
| mych2 | 4.286742 | 0.528322917 | 0.356 | 0.088 | 1.0890468 | 20 | mych |
| tmem100a2 | 5.311499 | 0.339158455 | 0.175 | 0.025 | 1.3493863 | 20 | tmem100a |
| inhbaa2 | 5.722027 | 0.552524097 | 0.26 | 0.052 | 1.4536811 | 20 | inhbaa |

| | | | | | | | |
|---|---|---|---|---|---|---|---|
| sh3gl1b1 | 7.682125 | 0.375203236 | 0.226 | 0.04 | 1.9516439 | 20 | sh3gl1b |
| si:ch211-251b21.11 | 1.431897 | 0.262332304 | 0.113 | 0.011 | 3.6377344 | 20 | si:ch211-251b21.1 |
| tnfsf10l2 | 1.439533 | 0.402898401 | 0.22 | 0.038 | 3.6571356 | 20 | tnfsf10l |
| kirrel1a1 | 1.689590 | 0.344152555 | 0.203 | 0.032 | 4.2924049 | 20 | kirrel1a |
| itga51 | 2.711380 | 0.336787175 | 0.192 | 0.03 | 6.8882621 | 20 | itga5 |
| slc16a1b | 3.947900 | 0.275134491 | 0.153 | 0.02 | 1.0029642 | 20 | slc16a1b |
| eif1b4 | 4.973089 | 0.641347126 | 0.814 | 0.403 | 1.2634133 | 20 | eif1b |
| bicc1b1 | 6.948612 | 0.302520236 | 0.169 | 0.024 | 1.7652951 | 20 | bicc1b |
| fkbp91 | 6.958135 | 0.357722147 | 0.209 | 0.035 | 1.7677144 | 20 | fkbp9 |
| cavin2b3 | 1.094602 | 0.474896298 | 0.328 | 0.077 | 2.7808363 | 20 | cavin2b |
| s1pr5a | 1.539131 | 0.26527412 | 0.124 | 0.014 | 3.9101636 | 20 | s1pr5a |
| capns1b3 | 1.653171 | 0.543168594 | 0.463 | 0.143 | 4.1998834 | 20 | capns1b |
| zgc:110158 | 2.707960 | 0.36183943 | 0.203 | 0.034 | 6.8795738 | 20 | zgc:110158 |
| nfil31 | 2.937061 | 0.379216651 | 0.198 | 0.033 | 7.4616057 | 20 | nfil3 |
| lzts2a | 4.669032 | 0.276105494 | 0.158 | 0.021 | 1.1861676 | 20 | lzts2a |
| ptprga5 | 5.634153 | 0.454122254 | 0.305 | 0.07 | 1.4313568 | 20 | ptprga |
| nkd1 | 5.942290 | 0.277301719 | 0.147 | 0.019 | 1.5096389 | 20 | nkd1 |
| CABZ01039304.1 | 1.686911 | 0.252964365 | 0.136 | 0.016 | 4.2855977 | 20 | CABZ01039304.1 |
| BX640522.22 | 2.321804 | 0.328586548 | 0.226 | 0.04 | 5.8985431 | 20 | BX640522.2 |
| pim15 | 4.205404 | 0.625451578 | 0.497 | 0.166 | 1.0683829 | 20 | pim1 |
| pdk2b3 | 5.425819 | 0.504507815 | 0.379 | 0.103 | 1.3784293 | 20 | pdk2b |
| tmeff1b2 | 8.845188 | 0.341248185 | 0.22 | 0.041 | 2.2471202 | 20 | tmeff1b |
| ctnnb13 | 1.413489 | 0.518211035 | 0.492 | 0.156 | 3.5909699 | 20 | ctnnb1 |
| si:ch211-105c13.32 | 1.662436 | 0.441789771 | 0.294 | 0.065 | 4.2234209 | 20 | si:ch211-105c13.3 |
| itgb1a3 | 1.935610 | 0.41367888 | 0.249 | 0.051 | 4.9174193 | 20 | itgb1a |
| krt42 | 1.986034 | 0.405846457 | 0.249 | 0.049 | 5.0455202 | 20 | krt4 |
| gulp1a3 | 3.476621 | 0.334013962 | 0.203 | 0.036 | 8.8323559 | 20 | gulp1a |
| olfml3b4 | 3.655354 | 0.369286778 | 0.26 | 0.054 | 9.2864291 | 20 | olfml3b |
| ccl25b4 | 5.599788 | 0.868468081 | 0.339 | 0.092 | 1.4226261 | 20 | ccl25b |
| mrc21 | 1.010191 | 0.2709638 | 0.181 | 0.029 | 2.5663925 | 20 | mrc2 |
| mex3b1 | 1.049833 | 0.251859719 | 0.141 | 0.019 | 2.6671024 | 20 | mex3b |
| wwtr11 | 1.191900 | 0.316690441 | 0.175 | 0.028 | 3.0280242 | 20 | wwtr1 |
| antxr1c2 | 1.248007 | 0.37325085 | 0.209 | 0.039 | 3.1705632 | 20 | antxr1c |
| tspan4b | 1.772745 | 0.427175392 | 0.299 | 0.07 | 4.5036612 | 20 | tspan4b |
| rhoua2 | 1.850585 | 0.308798432 | 0.181 | 0.029 | 4.7014118 | 20 | rhoua |

| | | | | | | | |
|---|---|---|---|---|---|---|---|
| meis1b5 | 2.175445 | 0.425641572 | 0.299 | 0.071 | 5.5267198 | 20 | meis1b |
| mafba4 | 3.225176 | 0.466711093 | 0.322 | 0.083 | 8.1935621 | 20 | mafba |
| tcf7l2 | 6.111801 | 0.263935072 | 0.136 | 0.018 | 1.5527032 | 20 | tcf7l2 |
| lrp1aa3 | 8.164894 | 0.39634183 | 0.282 | 0.064 | 2.0742915 | 20 | lrp1aa |
| aqp3a | 8.631013 | 0.361853353 | 0.175 | 0.028 | 2.1927089 | 20 | aqp3a |
| scpp7 | 2.040300 | 0.286524397 | 0.153 | 0.022 | 5.1833829 | 20 | scpp7 |
| fbn2b4 | 3.489063 | 0.398106529 | 0.237 | 0.049 | 8.8639646 | 20 | fbn2b |
| zbtb16b1 | 6.501111 | 0.252399945 | 0.147 | 0.021 | 1.6516073 | 20 | zbtb16b |
| myl65 | 7.716622 | 0.509221471 | 0.39 | 0.115 | 1.9604078 | 20 | myl6 |
| ciarta3 | 9.036258 | 0.505237505 | 0.384 | 0.116 | 2.2956614 | 20 | ciarta |
| klhl24b | 1.091206 | 0.269739456 | 0.124 | 0.016 | 2.7722100 | 20 | klhl24b |
| nsa26 | 1.602348 | 0.607474402 | 0.774 | 0.386 | 4.0707674 | 20 | nsa2 |
| spry21 | 2.011029 | 0.411595283 | 0.288 | 0.071 | 5.1090217 | 20 | spry2 |
| dhrs13l12 | 2.396818 | 0.324114736 | 0.203 | 0.039 | 6.0891168 | 20 | dhrs13l1 |
| selenop | 2.516399 | 0.27969332 | 0.158 | 0.025 | 6.3929119 | 20 | selenop |
| aoc24 | 3.147248 | 0.471216045 | 0.299 | 0.075 | 7.9955855 | 20 | aoc2 |
| mgp1 | 3.615561 | 0.324819828 | 0.147 | 0.021 | 9.1853351 | 20 | mgp |
| ntn1a1 | 4.194077 | 0.256774371 | 0.113 | 0.013 | 1.0655052 | 20 | ntn1a |
| nfia5 | 4.416053 | 0.45003411 | 0.35 | 0.097 | 1.1218983 | 20 | nfia |
| rbfox22 | 5.719193 | 0.334790268 | 0.209 | 0.041 | 1.4529611 | 20 | rbfox2 |
| phlda2 | 6.814037 | 0.501715119 | 0.237 | 0.052 | 1.7311061 | 20 | phlda2 |
| crtap1 | 8.088329 | 0.298612628 | 0.203 | 0.039 | 2.0548401 | 20 | crtap |
| znf5031 | 8.162601 | 0.255926215 | 0.158 | 0.025 | 2.0737088 | 20 | znf503 |
| rem11 | 8.986446 | 0.315194084 | 0.181 | 0.031 | 2.2830066 | 20 | rem1 |
| hoxc3a2 | 1.135502 | 0.251675223 | 0.169 | 0.028 | 2.8847429 | 20 | hoxc3a |
| col12a1a1 | 1.186574 | 0.36458469 | 0.175 | 0.03 | 3.0144915 | 20 | col12a1a |
| fbp1a | 2.147750 | 0.328098243 | 0.209 | 0.042 | 5.4563605 | 20 | fbp1a |
| dedd14 | 3.875695 | 0.526271385 | 0.452 | 0.155 | 9.8462046 | 20 | dedd1 |
| flna5 | 3.951526 | 0.483542322 | 0.39 | 0.12 | 1.0038853 | 20 | flna |
| tspan142 | 4.627142 | 0.503102886 | 0.407 | 0.13 | 1.1755255 | 20 | tspan14 |
| ehd1b1 | 5.160992 | 0.331732134 | 0.237 | 0.051 | 1.3111500 | 20 | ehd1b |
| cav12 | 6.860603 | 0.40325399 | 0.226 | 0.048 | 1.7429362 | 20 | cav1 |
| AL732488.11 | 8.199777 | 0.378708424 | 0.203 | 0.04 | 2.0831535 | 20 | AL732488.1 |
| gsna | 8.727765 | 0.301083694 | 0.181 | 0.033 | 2.2172887 | 20 | gsna |
| ak4 | 9.380671 | 0.274646154 | 0.181 | 0.033 | 2.3831596 | 20 | ak4 |

| | | | | | | | |
|---|---|---|---|---|---|---|---|
| zgc:1533721 | 1.025603 | 0.267625626 | 0.181 | 0.032 | 2.6055457 | 20 | zgc:153372 |
| proza | 1.423085 | 0.290027205 | 0.192 | 0.036 | 3.6153481 | 20 | proza |
| col21a11 | 2.464759 | 0.267101028 | 0.136 | 0.02 | 6.2617213 | 20 | col21a1 |
| myl9b4 | 2.759505 | 0.423648307 | 0.345 | 0.097 | 7.0105224 | 20 | myl9b |
| tmem119b1 | 4.273676 | 0.250686909 | 0.141 | 0.021 | 1.0857274 | 20 | tmem119b |
| calua | 5.074290 | 0.279389552 | 0.175 | 0.032 | 1.2891236 | 20 | calua |
| yap11 | 7.353836 | 0.347494306 | 0.243 | 0.056 | 1.8682421 | 20 | yap1 |
| si:dkey-26m3.3 | 1.354979 | 0.251713095 | 0.13 | 0.019 | 3.4423265 | 20 | si:dkey-26m3.3 |
| glis2a1 | 1.387224 | 0.275782872 | 0.158 | 0.027 | 3.5242427 | 20 | glis2a |
| zgc:1132083 | 1.834415 | 0.328011935 | 0.22 | 0.048 | 4.6603313 | 20 | zgc:113208 |
| gadd45ga1 | 2.015144 | 0.369824502 | 0.232 | 0.052 | 5.1194754 | 20 | gadd45ga |
| pard3bb3 | 2.459492 | 0.276609563 | 0.175 | 0.032 | 6.2483401 | 20 | pard3bb |
| fbln21 | 2.748606 | 0.268417334 | 0.158 | 0.027 | 6.9828339 | 20 | fbln2 |
| cyp2n133 | 2.956894 | 0.347109603 | 0.198 | 0.04 | 7.5119912 | 20 | cyp2n13 |
| glud1a1 | 3.048791 | 0.405854379 | 0.339 | 0.098 | 7.7454555 | 20 | glud1a |
| adgrl3.1 | 8.178437 | 0.321247903 | 0.164 | 0.029 | 2.0777320 | 20 | adgrl3.1 |
| rgs41 | 9.255939 | 0.371605463 | 0.198 | 0.04 | 2.3514714 | 20 | rgs4 |
| nfil3-66 | 1.155643 | 0.491172512 | 0.395 | 0.129 | 2.9359124 | 20 | nfil3-6 |
| nme33 | 1.227972 | 0.383702263 | 0.311 | 0.087 | 3.1196639 | 20 | nme3 |
| cbfb1 | 1.333025 | 0.351249956 | 0.198 | 0.042 | 3.3865507 | 20 | cbfb |
| slc20a2 | 1.459618 | 0.25073632 | 0.153 | 0.026 | 3.7081606 | 20 | slc20a2 |
| si:dkey-6n21.122 | 1.911604 | 0.298351591 | 0.198 | 0.041 | 4.8564303 | 20 | si:dkey-6n21.12 |
| CR847566.12 | 1.974574 | 0.38569213 | 0.22 | 0.05 | 5.0164060 | 20 | CR847566.1 |
| notch24 | 2.249279 | 0.456900239 | 0.373 | 0.121 | 5.7142953 | 20 | notch2 |
| atp2b23 | 3.422835 | 0.411627129 | 0.294 | 0.082 | 8.6957135 | 20 | atp2b2 |
| adamts15a4 | 4.491988 | 0.409460169 | 0.237 | 0.057 | 1.1411897 | 20 | adamts15a |
| histh1l3 | 5.263896 | 0.5405575 | 0.367 | 0.121 | 1.3372928 | 20 | histh1l |
| adma1 | 5.536244 | 0.313875981 | 0.119 | 0.017 | 1.4064828 | 20 | adma |
| si:dkey-7j14.64 | 5.645245 | 0.470896061 | 0.52 | 0.194 | 1.4341745 | 20 | si:dkey-7j14.6 |
| bmp31 | 6.898222 | 0.250278778 | 0.102 | 0.013 | 1.7524934 | 20 | bmp3 |
| hic12 | 7.016885 | 0.39599601 | 0.266 | 0.069 | 1.7826398 | 20 | hic1 |
| col16a12 | 7.041291 | 0.344137242 | 0.192 | 0.04 | 1.7888401 | 20 | col16a1 |
| cish1 | 7.082709 | 0.415164635 | 0.226 | 0.054 | 1.7993622 | 20 | cish |
| vdac32 | 9.890479 | 0.470947525 | 0.458 | 0.166 | 2.5126762 | 20 | vdac3 |
| acsl1a | 1.133332 | 0.261616565 | 0.136 | 0.022 | 2.8792320 | 20 | acsl1a |

| | | | | | | | |
|---|---|---|---|---|---|---|---|
| h1f04 | 2.983904 | 0.417624432 | 0.266 | 0.07 | 7.5806083 | 20 | h1f0 |
| svild3 | 3.369711 | 0.412824523 | 0.294 | 0.083 | 8.5607531 | 20 | svild |
| agtr21 | 4.172197 | 0.277521912 | 0.136 | 0.022 | 1.0599466 | 20 | agtr2 |
| tead1b2 | 4.479918 | 0.279399894 | 0.169 | 0.033 | 1.1381233 | 20 | tead1b |
| sgce1 | 4.523153 | 0.289264606 | 0.164 | 0.031 | 1.1491071 | 20 | sgce |
| XLOC-0108994 | 4.597886 | 0.384977097 | 0.616 | 0.237 | 1.1680931 | 20 | XLOC-010899 |
| im:71514492 | 6.196506 | 0.34772513 | 0.158 | 0.029 | 1.5742225 | 20 | im:7151449 |
| sema4ba4 | 9.967692 | 0.31908894 | 0.192 | 0.041 | 2.5322923 | 20 | sema4ba |
| col15a1a1 | 1.614021 | 0.270294778 | 0.141 | 0.024 | 4.1004219 | 20 | col15a1a |
| appb2 | 1.616962 | 0.292688733 | 0.203 | 0.046 | 4.1078943 | 20 | appb |
| neo1a1 | 1.775531 | 0.263565637 | 0.164 | 0.031 | 4.5107377 | 20 | neo1a |
| serf24 | 2.077062 | 0.523333968 | 0.734 | 0.375 | 5.2767769 | 20 | serf2 |
| id2b1 | 2.768048 | 0.393906446 | 0.186 | 0.04 | 7.0322272 | 20 | id2b |
| hsp70.3 | 2.786654 | 0.252270866 | 0.124 | 0.019 | 7.0794956 | 20 | hsp70.3 |
| DDX17 | 3.304146 | 0.312866764 | 0.243 | 0.062 | 8.3941839 | 20 | DDX17 |
| perp1 | 3.924985 | 0.342080201 | 0.237 | 0.06 | 9.9714262 | 20 | perp |
| nfe2l2a1 | 7.078962 | 0.320254386 | 0.254 | 0.067 | 1.7984104 | 20 | nfe2l2a |
| palm1b | 1.506007 | 0.299055349 | 0.232 | 0.057 | 3.8260128 | 20 | palm1b |
| slc7a13 | 2.125519 | 0.256526168 | 0.175 | 0.036 | 5.3998823 | 20 | slc7a1 |
| eif4a1b2 | 2.452545 | 0.514466877 | 0.638 | 0.31 | 6.2306921 | 20 | eif4a1b |
| ubtd2 | 2.727492 | 0.260387954 | 0.158 | 0.031 | 6.9291937 | 20 | ubtd2 |
| nid1b2 | 3.181547 | 0.361488987 | 0.254 | 0.068 | 8.0827208 | 20 | nid1b |
| eef1da4 | 3.271317 | 0.49580846 | 0.808 | 0.504 | 8.3107812 | 20 | eef1da |
| anxa132 | 3.417522 | 0.335440975 | 0.215 | 0.053 | 8.6822159 | 20 | anxa13 |
| vasna2 | 5.265032 | 0.267582893 | 0.198 | 0.045 | 1.3375815 | 20 | vasna |
| cnn24 | 6.348169 | 0.451600143 | 0.418 | 0.155 | 1.6127524 | 20 | cnn2 |
| slc3a2b2 | 6.871475 | 0.41701718 | 0.367 | 0.127 | 1.7456982 | 20 | slc3a2b |
| maff3 | 8.976511 | 0.37111498 | 0.271 | 0.079 | 2.2804828 | 20 | maff |
| HACD4 | 1.034184 | 0.270168865 | 0.203 | 0.048 | 2.6273456 | 20 | HACD4 |
| zgc:1749042 | 1.096023 | 0.388008167 | 0.209 | 0.051 | 2.7844479 | 20 | zgc:174904 |
| ybx14 | 1.320378 | 0.394890053 | 0.944 | 0.694 | 3.3544215 | 20 | ybx1 |
| add3a5 | 1.517551 | 0.440382669 | 0.384 | 0.136 | 3.8553402 | 20 | add3a |
| LOC1039092843 | 2.292773 | 0.349488031 | 0.266 | 0.076 | 5.8247909 | 20 | LOC103909284 |
| si:ch211-106h4.124 | 2.351932 | 0.548941647 | 0.22 | 0.057 | 5.9750849 | 20 | si:ch211-106h4.12 |
| hmgn68 | 2.527814 | 0.486427268 | 0.672 | 0.308 | 6.4219130 | 20 | hmgn6 |

| | | | | | | | |
|---|---|---|---|---|---|---|---|
| si:ch211-227m13.11 | 3.496160 | 0.264277447 | 0.113 | 0.018 | 8.8819946 | 20 | si:ch211-227m13.1 |
| cavin1b3 | 3.916410 | 0.319503125 | 0.192 | 0.044 | 9.9496398 | 20 | cavin1b |
| mef2aa1 | 4.652330 | 0.37916413 | 0.254 | 0.073 | 1.1819245 | 20 | mef2aa |
| arl6ip5b2 | 7.914179 | 0.397481607 | 0.311 | 0.102 | 2.0105973 | 20 | arl6ip5b |
| tpt12 | 1.286330 | 0.287185716 | 0.949 | 0.811 | 3.2679220 | 20 | tpt1 |
| CT030188.11 | 1.322611 | 0.314065123 | 0.215 | 0.055 | 3.3600957 | 20 | CT030188.1 |
| ndrg2 | 1.369455 | 0.302531979 | 0.203 | 0.051 | 3.4791010 | 20 | ndrg2 |
| daam1a | 1.387493 | 0.25695331 | 0.153 | 0.031 | 3.5249261 | 20 | daam1a |
| mmp9 | 1.658270 | 0.329699711 | 0.107 | 0.017 | 4.2128372 | 20 | mmp9 |
| klf11b4 | 1.961438 | 0.356082529 | 0.271 | 0.082 | 4.9830349 | 20 | klf11b |
| plod3 | 2.600538 | 0.267061154 | 0.169 | 0.038 | 6.6066683 | 20 | plod3 |
| si:dkey-79d12.52 | 2.814133 | 0.372862314 | 0.237 | 0.067 | 7.1493057 | 20 | si:dkey-79d12.5 |
| ptbp1b1 | 3.131688 | 0.252359333 | 0.175 | 0.04 | 7.9560556 | 20 | ptbp1b |
| zgc:926061 | 3.288995 | 0.368790451 | 0.294 | 0.095 | 8.3556921 | 20 | zgc:92606 |
| irf2bp2b3 | 4.246243 | 0.387006669 | 0.305 | 0.101 | 1.0787580 | 20 | irf2bp2b |
| tns1b1 | 5.569542 | 0.257097383 | 0.164 | 0.036 | 1.4149421 | 20 | tns1b |
| tuba1b1 | 5.584170 | 0.302685207 | 0.169 | 0.038 | 1.4186586 | 20 | tuba1b |
| myh101 | 6.755299 | 0.263816042 | 0.181 | 0.042 | 1.7161838 | 20 | myh10 |
| mgst3b2 | 6.881752 | 0.44374612 | 0.458 | 0.192 | 1.7483091 | 20 | mgst3b |
| calm2b5 | 9.070344 | 0.497995269 | 0.531 | 0.245 | 2.3043209 | 20 | calm2b |
| prdx53 | 9.099737 | 0.45697727 | 0.429 | 0.177 | 2.3117884 | 20 | prdx5 |
| snai1a1 | 9.457774 | 0.280300088 | 0.215 | 0.055 | 2.4027476 | 20 | snai1a |
| tmem176l.14 | 9.848152 | 0.425760041 | 0.492 | 0.2 | 2.5019230 | 20 | tmem176l.1 |
| mfap52 | 9.953176 | 0.325323401 | 0.232 | 0.063 | 2.5286045 | 20 | mfap5 |
| cebpb8 | 1.019500 | 0.567167833 | 0.65 | 0.337 | 2.5900408 | 20 | cebpb |
| klf31 | 1.318747 | 0.280405492 | 0.169 | 0.039 | 3.3502769 | 20 | klf3 |
| parvaa | 1.395575 | 0.25602833 | 0.158 | 0.034 | 3.5454583 | 20 | parvaa |
| celf28 | 1.526447 | 0.510803162 | 0.537 | 0.255 | 3.8779393 | 20 | celf2 |
| si:ch211-196l7.48 | 1.668367 | 0.487317884 | 0.644 | 0.328 | 4.2384869 | 20 | si:ch211-196l7.4 |
| trib31 | 2.787261 | 0.415036252 | 0.311 | 0.107 | 7.0810372 | 20 | trib3 |
| nr1d2a2 | 4.013430 | 0.348084425 | 0.277 | 0.088 | 1.0196121 | 20 | nr1d2a |
| pnrc210 | 4.481287 | 0.445808607 | 0.887 | 0.596 | 1.1384711 | 20 | pnrc2 |
| igf2b3 | 5.316150 | 0.318868225 | 0.254 | 0.075 | 1.3505679 | 20 | igf2b |
| si:ch211-212k18.93 | 6.139656 | 0.277148066 | 0.192 | 0.049 | 1.5597796 | 20 | si:ch211-212k18.9 |
| june1 | 7.574646 | 0.255817596 | 0.141 | 0.029 | 1.9243390 | 20 | june |

| | | | | | | | |
|---|---|---|---|---|---|---|---|
| tob1b2 | 9.17865 | 0.421440619 | 0.418 | 0.166 | 2.3318375 | 20 | tob1b |
| pdcd6 | 9.543097 | 0.348379332 | 0.26 | 0.082 | 2.4244240 | 20 | pdcd6 |
| rhoab5 | 1.195302 | 0.459596013 | 0.463 | 0.206 | 3.0366653 | 20 | rhoab |
| FO082781.15 | 1.269206 | 0.302596438 | 0.203 | 0.055 | 3.2244182 | 20 | FO082781.1 |
| nanos11 | 1.41955 | 0.263021135 | 0.147 | 0.031 | 3.6063914 | 20 | nanos1 |
| rps220 | 1.85013 | 0.295098969 | 0.972 | 0.841 | 4.7002661 | 20 | rps2 |
| slc38a5b4 | 3.534048 | 0.375333837 | 0.345 | 0.128 | 8.9782497 | 20 | slc38a5b |
| osr12 | 4.52870 | 0.310778336 | 0.175 | 0.043 | 1.1505173 | 20 | osr1 |
| tnrc181 | 5.61324 | 0.262106698 | 0.175 | 0.044 | 1.4260436 | 20 | tnrc18 |
| LOC5665871 | 7.52253 | 0.269472376 | 0.175 | 0.044 | 1.9110997 | 20 | LOC566587 |
| ephb34 | 7.90196 | 0.311623848 | 0.192 | 0.051 | 2.0074953 | 20 | ephb3 |
| emp22 | 9.22943 | 0.289132774 | 0.203 | 0.056 | 2.3447380 | 20 | emp2 |
| ptgdsb.2 | 1.01477 | 0.325189906 | 0.158 | 0.037 | 2.5780233 | 20 | ptgdsb.2 |
| hmgn74 | 1.043792 | 0.467313307 | 0.61 | 0.316 | 2.6517537 | 20 | hmgn7 |
| CABZ01068358.14 | 1.05529 | 0.308840156 | 0.226 | 0.066 | 2.6809742 | 20 | CABZ01068358.1 |
| dbn13 | 1.11641 | 0.319600886 | 0.22 | 0.065 | 2.8362636 | 20 | dbn1 |
| selenow14 | 1.25825 | 0.461057077 | 0.74 | 0.413 | 3.1965971 | 20 | selenow1 |
| hopx1 | 1.838198 | 0.335431224 | 0.22 | 0.064 | 4.6699443 | 20 | hopx |
| irf1b2 | 1.934677 | 0.266059241 | 0.153 | 0.036 | 4.9150487 | 20 | irf1b |
| dnase1l11 | 2.14194 | 0.272474316 | 0.192 | 0.052 | 5.4416005 | 20 | dnase1l1 |
| ctdsp24 | 2.69495 | 0.314433888 | 0.26 | 0.085 | 6.8465446 | 20 | ctdsp2 |
| nfic4 | 2.910362 | 0.342613333 | 0.254 | 0.083 | 7.3937764 | 20 | nfic |
| si:ch1073-291c23.24 | 3.38888 | 0.30499872 | 0.237 | 0.072 | 8.6094634 | 20 | si:ch1073-291c23.2 |
| ackr3b2 | 5.23688 | 0.285798035 | 0.141 | 0.032 | 1.3304308 | 20 | ackr3b |
| irf2bpl1 | 5.922646 | 0.25906562 | 0.175 | 0.045 | 1.5046483 | 20 | irf2bpl |
| itm2ba5 | 6.241962 | 0.427124165 | 0.593 | 0.295 | 1.5857706 | 20 | itm2ba |
| tspan71 | 6.423452 | 0.289477729 | 0.254 | 0.081 | 1.6318781 | 20 | tspan7 |
| tnrc6c15 | 6.770876 | 0.360185386 | 0.373 | 0.149 | 1.7201411 | 20 | tnrc6c1 |
| anxa11b2 | 7.62551 | 0.383570517 | 0.407 | 0.173 | 1.9372622 | 20 | anxa11b |
| COX310 | 8.08827 | 0.44816395 | 0.994 | 0.673 | 2.0548262 | 20 | COX3 |
| tmod2 | 9.47593 | 0.273120311 | 0.181 | 0.048 | 2.4073613 | 20 | tmod2 |
| pim23 | 9.694048 | 0.283075564 | 0.266 | 0.087 | 2.4627730 | 20 | pim2 |
| fam168a5 | 1.19632 | 0.290966214 | 0.232 | 0.073 | 3.0392642 | 20 | fam168a |
| reep53 | 1.725528 | 0.364153161 | 0.407 | 0.172 | 4.3837040 | 20 | reep5 |
| sec61a11 | 1.941766 | 0.41048158 | 0.429 | 0.191 | 4.9330582 | 20 | sec61a1 |

| | | | | | | | |
|---|---|---|---|---|---|---|---|
| pbx1a2 | 2.010600 | 0.251960078 | 0.192 | 0.053 | 5.1079307 | 20 | pbx1a |
| dag12 | 2.534378 | 0.299700216 | 0.192 | 0.055 | 6.4385878 | 20 | dag1 |
| fosl24 | 2.780526 | 0.360355855 | 0.305 | 0.112 | 7.0639273 | 20 | fosl2 |
| ube2e34 | 3.318532 | 0.339547888 | 0.407 | 0.168 | 8.4307316 | 20 | ube2e3 |
| calr | 3.435184 | 0.307451483 | 0.215 | 0.066 | 8.7270863 | 20 | calr |
| rpl326 | 1.318080 | 0.285888876 | 0.977 | 0.823 | 3.3485845 | 20 | rpl32 |
| lgals3b1 | 1.372691 | 0.325001453 | 0.271 | 0.098 | 3.4873225 | 20 | lgals3b |
| cdkn1a1 | 1.380696 | 0.273123037 | 0.175 | 0.048 | 3.5076599 | 20 | cdkn1a |
| flot2a4 | 1.522577 | 0.282046585 | 0.226 | 0.073 | 3.8681070 | 20 | flot2a |
| cd82b | 1.740812 | 0.31115129 | 0.232 | 0.075 | 4.4225343 | 20 | cd82b |
| cnbpb3 | 2.318312 | 0.371246113 | 0.452 | 0.208 | 5.8896718 | 20 | cnbpb |
| boc4 | 5.445231 | 0.257590616 | 0.215 | 0.067 | 1.3833611 | 20 | boc |
| si:dkey-28b4.75 | 6.491208 | 0.316855827 | 0.288 | 0.109 | 1.6490915 | 20 | si:dkey-28b4.7 |
| adam10a3 | 6.810493 | 0.305570675 | 0.282 | 0.105 | 1.7302059 | 20 | adam10a |
| hdlbpa5 | 7.257511 | 0.344083818 | 0.401 | 0.174 | 1.8437709 | 20 | hdlbpa |
| zfyve21 | 7.286199 | 0.260373652 | 0.198 | 0.06 | 1.8510590 | 20 | zfyve21 |
| creb3l3l | 7.882967 | 0.252365241 | 0.198 | 0.06 | 2.0026679 | 20 | creb3l3l |
| rpz55 | 1.417752 | 0.441204669 | 0.305 | 0.126 | 3.6017992 | 20 | rpz5 |
| pdia37 | 1.520487 | 0.404303285 | 0.593 | 0.312 | 3.8627978 | 20 | pdia3 |
| ptp4a13 | 2.086856 | 0.32277786 | 0.322 | 0.131 | 5.3016584 | 20 | ptp4a1 |
| krt18a.14 | 2.171450 | 0.421032344 | 0.356 | 0.149 | 5.5165694 | 20 | krt18a.1 |
| col4a13 | 3.152884 | 0.296305191 | 0.215 | 0.07 | 8.0099026 | 20 | col4a1 |
| cdc421 | 3.393029 | 0.354779782 | 0.48 | 0.232 | 8.6199915 | 20 | cdc42 |
| mdka5 | 3.481306 | 0.416183284 | 0.288 | 0.113 | 8.8442599 | 20 | mdka |
| atox1 | 4.012564 | 0.276504728 | 0.311 | 0.123 | 1.0193920 | 20 | atox1 |
| zgc:11418810 | 4.125989 | 0.378740841 | 0.944 | 0.666 | 1.0482075 | 20 | zgc:114188 |
| map1lc3a4 | 4.335294 | 0.268431207 | 0.232 | 0.08 | 1.1013815 | 20 | map1lc3a |
| aldh9a1a.15 | 5.234596 | 0.368937917 | 0.412 | 0.192 | 1.3298492 | 20 | aldh9a1a.1 |
| dnmt3ab3 | 5.682829 | 0.31335891 | 0.249 | 0.092 | 1.4437227 | 20 | dnmt3ab |
| glula2 | 5.739505 | 0.370581468 | 0.311 | 0.124 | 1.4581213 | 20 | glula |
| mknk2b5 | 7.749422 | 0.373919085 | 0.424 | 0.199 | 1.9687407 | 20 | mknk2b |
| smdt1b1 | 8.184513 | 0.342956534 | 0.345 | 0.152 | 2.0792756 | 20 | smdt1b |
| prelid3b | 8.200783 | 0.351054693 | 0.469 | 0.225 | 2.0834090 | 20 | prelid3b |
| rab2a | 9.735101 | 0.315831641 | 0.395 | 0.176 | 2.4732025 | 20 | rab2a |
| slco2b13 | 1.096951 | 0.288751274 | 0.203 | 0.066 | 2.7868060 | 20 | slco2b1 |

| | | | | | | | |
|---|---|---|---|---|---|---|---|
| cd6310 | 1.215899 | 0.383743257 | 0.701 | 0.371 | 3.0889923 | 20 | cd63 |
| rpl910 | 1.313760 | 0.371231461 | 0.932 | 0.667 | 3.3376096 | 20 | rpl9 |
| glo11 | 1.441471 | 0.314751466 | 0.446 | 0.208 | 3.6620577 | 20 | glo1 |
| pgrmc13 | 1.579145 | 0.314103663 | 0.356 | 0.151 | 4.0118202 | 20 | pgrmc1 |
| sar1b | 1.913502 | 0.280619908 | 0.243 | 0.089 | 4.8612543 | 20 | sar1b |
| lxn5 | 1.978068 | 0.314746622 | 0.384 | 0.166 | 5.0252829 | 20 | lxn |
| abcc54 | 2.644338 | 0.264101311 | 0.254 | 0.094 | 6.7179425 | 20 | abcc5 |
| XLOC-0139643 | 2.750997 | 0.291519547 | 0.395 | 0.169 | 6.9889101 | 20 | XLOC-013964 |
| col12a1b2 | 3.371942 | 0.33239516 | 0.136 | 0.037 | 8.5664196 | 20 | col12a1b |
| eef2b4 | 3.485540 | 0.349259466 | 0.887 | 0.634 | 8.8550159 | 20 | eef2b |
| metrnla2 | 3.542555 | 0.274512953 | 0.254 | 0.094 | 8.9998617 | 20 | metrnla |
| mt-co19 | 4.290279 | 0.399377324 | 0.994 | 0.637 | 1.0899455 | 20 | mt-co1 |
| sec61g2 | 5.545316 | 0.328863998 | 0.915 | 0.68 | 1.4087876 | 20 | sec61g |
| btg17 | 5.736666 | 0.371846863 | 0.644 | 0.354 | 1.4574001 | 20 | btg1 |
| rpl229 | 6.801087 | 0.329345883 | 0.949 | 0.719 | 1.7278162 | 20 | rpl22 |
| cald1a4 | 8.041706 | 0.312035064 | 0.294 | 0.119 | 2.0429956 | 20 | cald1a |
| rpl367 | 9.413130 | 0.291642464 | 0.977 | 0.805 | 2.3914057 | 20 | rpl36 |
| mmp16a4 | 9.987321 | 0.283244787 | 0.22 | 0.078 | 2.5372789 | 20 | mmp16a |
| zgc:1583435 | 1.042831 | 0.426303786 | 0.362 | 0.169 | 2.6493138 | 20 | zgc:158343 |
| sdcbp25 | 1.087322 | 0.325909318 | 0.463 | 0.227 | 2.7623435 | 20 | sdcbp2 |
| eif4ebp3l5 | 1.105091 | 0.32614159 | 0.441 | 0.204 | 2.8074860 | 20 | eif4ebp3l |
| tsc22d33 | 1.818420 | 0.333872664 | 0.345 | 0.158 | 4.6196970 | 20 | tsc22d3 |
| ppdpfb4 | 2.328425 | 0.377167625 | 0.825 | 0.53 | 5.9153655 | 20 | ppdpfb |
| rps168 | 2.507122 | 0.301892411 | 0.96 | 0.765 | 6.3693444 | 20 | rps16 |
| dusp14 | 3.845333 | 0.332922635 | 0.333 | 0.149 | 9.7690690 | 20 | dusp1 |
| zfp36l1a5 | 4.219918 | 0.350070416 | 0.497 | 0.254 | 1.0720703 | 20 | zfp36l1a |
| stau25 | 4.608986 | 0.278189761 | 0.243 | 0.096 | 1.1709131 | 20 | stau2 |
| rps188 | 4.712715 | 0.290257313 | 0.944 | 0.753 | 1.1972654 | 20 | rps18 |
| cebpg2 | 6.100337 | 0.251342128 | 0.192 | 0.067 | 1.5497907 | 20 | cebpg |
| grpel22 | 7.820544 | 0.276771003 | 0.26 | 0.107 | 1.9868093 | 20 | grpel2 |
| serinc52 | 1.098401 | 0.284091289 | 0.226 | 0.087 | 2.7904899 | 20 | serinc5 |
| fkbp1aa5 | 1.148773 | 0.359372854 | 0.723 | 0.412 | 2.9184591 | 20 | fkbp1aa |
| ssr41 | 1.373570 | 0.351739354 | 0.588 | 0.33 | 3.4895555 | 20 | ssr4 |
| faua7 | 1.578726 | 0.320289084 | 0.949 | 0.716 | 4.0107557 | 20 | faua |
| rpl188 | 1.959781 | 0.268685992 | 0.96 | 0.792 | 4.9788241 | 20 | rpl18 |

| | | | | | | | |
|---|---|---|---|---|---|---|---|
| mt-atp69 | 2.369198 | 0.355881585 | 0.898 | 0.553 | 6.0189488 | 20 | mt-atp6 |
| hmgb1b6 | 3.136082 | 0.271024286 | 0.339 | 0.154 | 7.9672173 | 20 | hmgb1b |
| gabarapb5 | 4.225466 | 0.320174629 | 0.554 | 0.304 | 1.0734798 | 20 | gabarapb |
| hp1bp34 | 4.616141 | 0.263262 | 0.277 | 0.12 | 1.1727308 | 20 | hp1bp3 |
| cpne11 | 5.863421 | 0.283979781 | 0.294 | 0.133 | 1.4896022 | 20 | cpne1 |
| h2afy23 | 7.331321 | 0.265668953 | 0.277 | 0.122 | 1.8625223 | 20 | h2afy2 |
| rpl18a7 | 8.945451 | 0.258346643 | 0.949 | 0.8 | 2.2725918 | 20 | rpl18a |
| si:ch211-286b5.55 | 1.213899 | 0.271663303 | 0.322 | 0.151 | 3.0839105 | 20 | si:ch211-286b5.5 |
| sltm | 1.827270 | 0.251892404 | 0.186 | 0.07 | 4.6421815 | 20 | sltm |
| rps158 | 1.922676 | 0.313050395 | 0.938 | 0.716 | 4.8845604 | 20 | rps15 |
| tpm34 | 1.928050 | 0.317158779 | 0.508 | 0.283 | 4.8982114 | 20 | tpm3 |
| rpl345 | 2.325692 | 0.262354318 | 0.966 | 0.801 | 5.9084212 | 20 | rpl34 |
| h2afx17 | 2.354360 | 0.330783211 | 0.424 | 0.227 | 5.9812541 | 20 | h2afx1 |
| rpl27a6 | 2.820443 | 0.36282706 | 0.802 | 0.521 | 7.1653368 | 20 | rpl27a |
| rab104 | 3.509092 | 0.307652097 | 0.401 | 0.213 | 8.9148498 | 20 | rab10 |
| arhgef1b2 | 4.009465 | 0.277581654 | 0.198 | 0.079 | 0.000102 | 20 | arhgef1b |
| adh54 | 4.467519 | 0.286731082 | 0.452 | 0.241 | 0.000113 | 20 | adh5 |
| mibp27 | 8.406592 | 0.380294129 | 0.768 | 0.52 | 0.000214 | 20 | mibp2 |
| rps27.27 | 1.169393 | 0.346348257 | 0.847 | 0.551 | 0.000297 | 20 | rps27.2 |
| romo15 | 1.323983 | 0.312084966 | 0.678 | 0.407 | 0.000336 | 20 | romo1 |
| rhoaa4 | 1.690773 | 0.263803707 | 0.345 | 0.176 | 0.00043 | 20 | rhoaa |
| qkia4 | 1.773921 | 0.264530456 | 0.384 | 0.199 | 0.000451 | 20 | qkia |
| rpl68 | 2.515199 | 0.287968313 | 0.921 | 0.7 | 0.000639 | 20 | rpl6 |
| capns1a3 | 2.573375 | 0.268443423 | 0.316 | 0.159 | 0.000654 | 20 | capns1a |
| rpl5a9 | 3.168643 | 0.349605864 | 0.842 | 0.535 | 0.000805 | 20 | rpl5a |
| CABZ01072614.16 | 3.990350 | 0.294233536 | 0.215 | 0.095 | 0.001014 | 20 | CABZ01072614.1 |
| si:dkey-11f4.144 | 5.242770 | 0.261019733 | 0.26 | 0.122 | 0.001332 | 20 | si:dkey-11f4.14 |
| bsg | 9.207883 | 0.281506034 | 0.644 | 0.375 | 0.002339 | 20 | bsg |
| mt-cyb8 | 1.254179 | 0.311450817 | 0.91 | 0.574 | 0.003186 | 20 | mt-cyb |
| rps4x7 | 1.487682 | 0.263435428 | 0.927 | 0.736 | 0.003779 | 20 | rps4x |
| rpl23a7 | 1.544035 | 0.301654591 | 0.921 | 0.668 | 0.003923 | 20 | rpl23a |
| eef1g7 | 1.566207 | 0.290366977 | 0.87 | 0.651 | 0.003979 | 20 | eef1g |
| rps2111 | 1.647299 | 0.265318846 | 0.949 | 0.775 | 0.004185 | 20 | rps21 |
| mt-nd19 | 1.870285 | 0.279374298 | 0.65 | 0.396 | 0.004751 | 20 | mt-nd1 |
| rps15a8 | 1.880496 | 0.268162718 | 0.977 | 0.797 | 0.004777 | 20 | rps15a |

| | | | | | | | |
|---|---|---|---|---|---|---|---|
| NC-002333.48 | 1.984167 | 0.309509468 | 0.898 | 0.607 | 0.005041 | 20 | NC-002333.4 |
| mt-nd27 | 2.085132 | 0.278285769 | 0.655 | 0.39 | 0.005297 | 20 | mt-nd2 |
| eif3i4 | 3.343771 | 0.262089606 | 0.492 | 0.29 | 0.008495 | 20 | eif3i |
| eif2s32 | 3.756988 | 0.255056094 | 0.52 | 0.307 | 0.009545 | 20 | eif2s3 |
| rsl24d16 | 5.191287 | 0.311105006 | 0.729 | 0.459 | 0.013188 | 20 | rsl24d1 |
| fabp33 | 5.273453 | 0.268612104 | 0.401 | 0.228 | 0.013397 | 20 | fabp3 |
| eif3m4 | 6.105777 | 0.273239006 | 0.621 | 0.383 | 0.015512 | 20 | eif3m |
| mt-nd49 | 1.270254 | 0.257837358 | 0.729 | 0.446 | 0.032271 | 20 | mt-nd4 |
| rpl306 | 1.659989 | 0.26408709 | 0.944 | 0.729 | 0.042172 | 20 | rpl30 |
| b2ml7 | 1.741196 | 0.251951199 | 0.48 | 0.293 | 0.044235 | 20 | b2ml |
| naca7 | 3.124532 | 0.264534223 | 0.859 | 0.651 | 0.079379 | 20 | naca |
| serbp1a8 | 3.766671 | 0.284455643 | 0.751 | 0.505 | 0.095692 | 20 | serbp1a |
| rps109 | 6.964969 | 0.275179085 | 0.955 | 0.725 | 0.176945 | 20 | rps10 |
| eif5a4 | 7.788906 | 0.273921842 | 0.74 | 0.494 | 0.197877 | 20 | eif5a |
| sec61b1 | 1.068341 | 0.258364013 | 0.689 | 0.468 | 0.271412 | 20 | sec61b |
| eif3f6 | 1.189549 | 0.277341369 | 0.74 | 0.464 | 0.302205 | 20 | eif3f |
| tubb4b1 | 1.837001 | 0.261623375 | 0.825 | 0.591 | 0.46669 | 20 | tubb4b |
| mt-co29 | 2.608664 | 0.313576562 | 0.972 | 0.669 | 0.662731 | 20 | mt-co2 |
| rps27.19 | 2.950037 | 0.268212037 | 0.932 | 0.709 | 0.749457 | 20 | rps27.1 |
| bhlhe406 | 6.779144 | 0.253448447 | 0.277 | 0.168 | 1 | 20 | bhlhe40 |
| rpl5b6 | 9.568606 | 0.250632744 | 0.864 | 0.644 | 1 | 20 | rpl5b |
| CR855311.14 | 0.0004 | 0.263002348 | 0.164 | 0.089 | 1 | 20 | CR855311.1 |
| gsdf1 | 0 | 1.882664035 | 0.881 | 0.046 | 0 | 21 | gsdf |
| cldn11a4 | 0 | 1.795098493 | 0.983 | 0.073 | 0 | 21 | cldn11a |
| bcam2 | 0 | 1.697285212 | 0.898 | 0.03 | 0 | 21 | bcam |
| jam2a1 | 0 | 1.667537547 | 0.898 | 0.041 | 0 | 21 | jam2a |
| igf3 | 0 | 1.606996421 | 0.801 | 0.011 | 0 | 21 | igf3 |
| LOC103910903 | 0 | 1.52728479 | 0.722 | 0.008 | 0 | 21 | LOC103910903 |
| pi15a3 | 0 | 1.519370515 | 0.812 | 0.045 | 0 | 21 | pi15a |
| ecrg4a2 | 0 | 1.451090142 | 0.676 | 0.029 | 0 | 21 | ecrg4a |
| cx432 | 0 | 1.378921238 | 0.733 | 0.05 | 0 | 21 | cx43 |
| alcama1 | 0 | 1.373201663 | 0.75 | 0.023 | 0 | 21 | alcama |
| foxl2b | 0 | 1.181355963 | 0.636 | 0.01 | 0 | 21 | foxl2b |
| XLOC-0056051 | 0 | 1.104003235 | 0.602 | 0.022 | 0 | 21 | XLOC-005605 |
| bambia | 0 | 1.025491603 | 0.54 | 0.021 | 0 | 21 | bambia |

| | | | | | | | |
|---|---|---|---|---|---|---|---|
| cx34.5 | 0 | 0.996834546 | 0.466 | 0.003 | 0 | 21 | cx34.5 |
| fgfr31 | 0 | 0.985142269 | 0.483 | 0.015 | 0 | 21 | fgfr3 |
| LOC110438024 | 0 | 0.884253704 | 0.432 | 0.006 | 0 | 21 | LOC110438024 |
| si:dkey-6n6.1 | 0 | 0.879720697 | 0.358 | 0.004 | 0 | 21 | si:dkey-6n6.1 |
| alcamb | 0 | 0.853979336 | 0.472 | 0.015 | 0 | 21 | alcamb |
| sema3ab | 0 | 0.821071962 | 0.375 | 0.009 | 0 | 21 | sema3ab |
| smoc21 | 0 | 0.815160305 | 0.358 | 0.008 | 0 | 21 | smoc2 |
| cdh4 | 0 | 0.799901138 | 0.398 | 0.007 | 0 | 21 | cdh4 |
| metrn | 0 | 0.785178029 | 0.409 | 0.011 | 0 | 21 | metrn |
| XLOC-009815 | 0 | 0.772074594 | 0.403 | 0.014 | 0 | 21 | XLOC-009815 |
| col4a6 | 0 | 0.742147313 | 0.403 | 0.01 | 0 | 21 | col4a6 |
| ackr4b | 0 | 0.699362655 | 0.352 | 0.01 | 0 | 21 | ackr4b |
| fez1 | 0 | 0.68688633 | 0.347 | 0.005 | 0 | 21 | fez1 |
| inha | 0 | 0.650337037 | 0.318 | 0.005 | 0 | 21 | inha |
| foxl2a1 | 0 | 0.650042884 | 0.352 | 0.009 | 0 | 21 | foxl2a |
| col4a5 | 0 | 0.633778388 | 0.364 | 0.007 | 0 | 21 | col4a5 |
| XLOC-042229 | 0 | 0.62612504 | 0.352 | 0.005 | 0 | 21 | XLOC-042229 |
| znf536 | 0 | 0.621061994 | 0.312 | 0.006 | 0 | 21 | znf536 |
| scn4bb | 0 | 0.545299059 | 0.295 | 0.006 | 0 | 21 | scn4bb |
| lamb2l | 0 | 0.527243048 | 0.273 | 0.002 | 0 | 21 | lamb2l |
| asb11 | 0 | 0.512012358 | 0.256 | 0.002 | 0 | 21 | asb11 |
| phlda1 | 0 | 0.488088365 | 0.239 | 0.003 | 0 | 21 | phlda1 |
| itga3b | 0 | 0.486563477 | 0.273 | 0.003 | 0 | 21 | itga3b |
| hsd17b1 | 0 | 0.39617487 | 0.182 | 0.001 | 0 | 21 | hsd17b1 |
| si:ch211-117c9.5 | 2.615598 | 0.741107544 | 0.426 | 0.016 | 6.6449288 | 21 | si:ch211-117c9.5 |
| slc7a3a1 | 3.194695 | 1.044004995 | 0.636 | 0.041 | 8.1161242 | 21 | slc7a3a |
| CABZ01068366.13 | 1.380353 | 0.954754093 | 0.585 | 0.036 | 3.5067872 | 21 | CABZ01068366.1 |
| nectin1b2 | 8.240347 | 1.04915711 | 0.608 | 0.039 | 2.0934603 | 21 | nectin1b |
| bmpr2a | 5.015680 | 0.766271968 | 0.42 | 0.018 | 1.2742337 | 21 | bmpr2a |
| aldh1a24 | 1.025277 | 1.497943374 | 0.841 | 0.09 | 2.6047177 | 21 | aldh1a2 |
| f103 | 7.876760 | 1.080143942 | 0.665 | 0.053 | 2.0010910 | 21 | f10 |
| cadm2a1 | 4.158489 | 0.731903703 | 0.392 | 0.017 | 1.0564643 | 21 | cadm2a |
| cemip2 | 3.621613 | 0.789586562 | 0.415 | 0.019 | 9.2007079 | 21 | cemip2 |
| nr0b11 | 1.270437 | 0.617476608 | 0.347 | 0.013 | 3.2275455 | 21 | nr0b1 |
| bmp61 | 1.176042 | 0.673271488 | 0.324 | 0.011 | 2.9877352 | 21 | bmp6 |

| | | | | | | | |
|---|---|---|---|---|---|---|---|
| pcdh1b | 1.111398 | 0.622796653 | 0.307 | 0.01 | 2.8235067 | 21 | pcdh1b |
| plxna41 | 3.908875 | 0.749303555 | 0.409 | 0.019 | 9.9304983 | 21 | plxna4 |
| ccdc187 | 6.724975 | 0.895190851 | 0.517 | 0.033 | 1.7084800 | 21 | ccdc187 |
| thbs3b | 7.562836 | 0.441656829 | 0.205 | 0.004 | 1.9213387 | 21 | thbs3b |
| nos1apa | 1.165088 | 0.712394967 | 0.449 | 0.024 | 2.9599060 | 21 | nos1apa |
| emx21 | 1.533584 | 0.766848248 | 0.415 | 0.02 | 3.8960721 | 21 | emx2 |
| rpz56 | 4.451808 | 1.539255123 | 0.892 | 0.118 | 1.1309818 | 21 | rpz5 |
| si:dkey-194e6.1 | 1.527147 | 0.351777115 | 0.142 | 0.001 | 3.8797190 | 21 | si:dkey-194e6.1 |
| ccdc3a | 1.506281 | 0.304448217 | 0.136 | 0.001 | 3.8267073 | 21 | ccdc3a |
| si:dkey-42i9.71 | 4.690918 | 0.837916497 | 0.426 | 0.022 | 1.1917277 | 21 | si:dkey-42i9.7 |
| agrn3 | 1.330026 | 1.050478051 | 0.636 | 0.053 | 3.3789334 | 21 | agrn |
| map1aa | 1.050063 | 0.414959522 | 0.205 | 0.004 | 2.6676861 | 21 | map1aa |
| XLOC-004335 | 7.182216 | 0.304368471 | 0.142 | 0.001 | 1.8246420 | 21 | XLOC-004335 |
| corin | 1.131994 | 0.446578377 | 0.21 | 0.005 | 2.8758314 | 21 | corin |
| atp1b3b | 1.421000 | 1.023092518 | 0.642 | 0.055 | 3.6100528 | 21 | atp1b3b |
| appa4 | 3.518808 | 1.175712879 | 0.744 | 0.08 | 8.9395317 | 21 | appa |
| tgfb2l | 5.500241 | 0.429673245 | 0.239 | 0.006 | 1.3973363 | 21 | tgfb2l |
| gata62 | 1.937974 | 0.698183776 | 0.403 | 0.022 | 4.9234245 | 21 | gata6 |
| spock1 | 3.771803 | 0.370301437 | 0.188 | 0.004 | 9.5822657 | 21 | spock1 |
| hpn | 1.488745 | 0.442094034 | 0.233 | 0.006 | 3.7821574 | 21 | hpn |
| chka1 | 5.546895 | 0.720098327 | 0.432 | 0.025 | 1.4091887 | 21 | chka |
| trim35-33 | 1.903680 | 0.35159338 | 0.188 | 0.004 | 4.8362994 | 21 | trim35-33 |
| si:ch211-207d6.21 | 4.690942 | 0.791657297 | 0.364 | 0.018 | 1.1917339 | 21 | si:ch211-207d6.2 |
| sema7a | 6.830799 | 0.589843289 | 0.267 | 0.009 | 1.7353645 | 21 | sema7a |
| si:ch211-225b11.1 | 4.530702 | 0.83691792 | 0.477 | 0.032 | 1.1510249 | 21 | si:ch211-225b11.1 |
| lama53 | 1.429863 | 0.764757709 | 0.426 | 0.026 | 3.6325675 | 21 | lama5 |
| islr21 | 6.360373 | 0.555056312 | 0.295 | 0.012 | 1.6158529 | 21 | islr2 |
| podxl3 | 2.227952 | 1.016009526 | 0.602 | 0.051 | 5.6601124 | 21 | podxl |
| ndp | 4.237424 | 0.349211568 | 0.182 | 0.004 | 1.0765177 | 21 | ndp |
| hkdc1 | 6.540748 | 0.31217772 | 0.159 | 0.003 | 1.6616772 | 21 | hkdc1 |
| wu:fc27b111 | 2.482653 | 0.561765257 | 0.318 | 0.014 | 6.3071805 | 21 | wu:fc27b11 |
| si:ch73-61d6.3 | 3.698085 | 0.495324367 | 0.278 | 0.011 | 9.3949862 | 21 | si:ch73-61d6.3 |
| zgc:92275 | 3.002132 | 0.266387246 | 0.119 | 0.001 | 7.6269186 | 21 | zgc:92275 |
| igsf3 | 3.814410 | 0.393048908 | 0.205 | 0.005 | 9.6905110 | 21 | igsf3 |
| itm2bb4 | 1.246747 | 1.635603754 | 0.994 | 0.234 | 3.1673613 | 21 | itm2bb |

| | | | | | | | |
|---|---|---|---|---|---|---|---|
| mpzl2b1 | 6.518231 | 1.112518472 | 0.716 | 0.081 | 1.6559567 | 21 | mpzl2b |
| fzd10 | 6.059742 | 0.29615794 | 0.165 | 0.003 | 1.5394775 | 21 | fzd10 |
| LOC553316 | 8.801269 | 0.616809204 | 0.347 | 0.019 | 2.2359626 | 21 | LOC553316 |
| rasd1 | 4.444058 | 0.406799519 | 0.17 | 0.004 | 1.1290129 | 21 | rasd1 |
| ece2b1 | 6.051343 | 0.847613505 | 0.511 | 0.043 | 1.5373437 | 21 | ece2b |
| mbnl31 | 1.796233 | 0.865918627 | 0.58 | 0.056 | 4.5633321 | 21 | mbnl3 |
| ifi465 | 5.800009 | 1.186988396 | 0.812 | 0.124 | 1.4734923 | 21 | ifi46 |
| CABZ01049847.1 | 4.206966 | 0.395602803 | 0.199 | 0.006 | 1.0687797 | 21 | CABZ01049847.1 |
| slc44a5b | 3.295507 | 0.496475221 | 0.273 | 0.012 | 8.3722365 | 21 | slc44a5b |
| XLOC-0152911 | 4.890417 | 0.436056955 | 0.244 | 0.01 | 1.2424105 | 21 | XLOC-015291 |
| large2 | 3.860601 | 0.342101306 | 0.193 | 0.006 | 9.8078579 | 21 | large2 |
| rn7sk | 6.252975 | 0.983056648 | 0.494 | 0.045 | 1.5885683 | 21 | rn7sk |
| slc4a4a2 | 4.512532 | 0.487234399 | 0.239 | 0.01 | 1.1464088 | 21 | slc4a4a |
| greb1 | 3.083444 | 0.258707177 | 0.142 | 0.003 | 7.8334902 | 21 | greb1 |
| jam3b | 6.176688 | 0.632294613 | 0.398 | 0.029 | 1.5691877 | 21 | jam3b |
| parp12a | 2.423093 | 0.609770633 | 0.358 | 0.024 | 6.1558682 | 21 | parp12a |
| tmem88b3 | 5.272810 | 0.871927091 | 0.5 | 0.049 | 1.3395574 | 21 | tmem88b |
| si:dkey-188i13.9 | 9.498518 | 0.637650898 | 0.352 | 0.024 | 2.4130987 | 21 | si:dkey-188i13.9 |
| vasnb3 | 9.503307 | 0.684184214 | 0.409 | 0.032 | 2.4143152 | 21 | vasnb |
| sv2bb | 1.834374 | 0.374925704 | 0.205 | 0.007 | 4.6602271 | 21 | sv2bb |
| CABZ01068358.15 | 5.533841 | 0.906187249 | 0.557 | 0.062 | 1.4058724 | 21 | CABZ01068358.1 |
| ptprfb | 6.549651 | 0.495459828 | 0.273 | 0.014 | 1.6639389 | 21 | ptprfb |
| robo12 | 9.215485 | 0.611493319 | 0.347 | 0.024 | 2.3411940 | 21 | robo1 |
| fgfr4 | 1.462633 | 0.512172558 | 0.256 | 0.012 | 3.7158203 | 21 | fgfr4 |
| XLOC-040516 | 3.605368 | 0.298009209 | 0.153 | 0.004 | 9.1594395 | 21 | XLOC-040516 |
| cd99l21 | 8.427791 | 0.641767935 | 0.392 | 0.031 | 2.1410804 | 21 | cd99l2 |
| si:dkey-26c10.5 | 4.268301 | 0.472069878 | 0.222 | 0.009 | 1.0843621 | 21 | si:dkey-26c10.5 |
| lamb1a4 | 1.265458 | 0.776664972 | 0.494 | 0.05 | 3.2148980 | 21 | lamb1a |
| CABZ01068367.12 | 2.881790 | 0.586804686 | 0.318 | 0.021 | 7.3211900 | 21 | CABZ01068367.1 |
| pcdh10a | 2.155211 | 0.268275748 | 0.148 | 0.004 | 5.4753139 | 21 | pcdh10a |
| nav3 | 4.534249 | 0.528166692 | 0.301 | 0.019 | 1.1519261 | 21 | nav3 |
| ptn | 7.601743 | 0.370625948 | 0.188 | 0.007 | 1.9312229 | 21 | ptn |
| sparc6 | 4.315438 | 1.226387457 | 0.938 | 0.187 | 1.0963372 | 21 | sparc |
| emilin1a2 | 1.238579 | 0.64700885 | 0.324 | 0.023 | 3.1466114 | 21 | emilin1a |
| gpx1b | 3.065898 | 0.794665839 | 0.341 | 0.025 | 7.7889145 | 21 | gpx1b |

| | | | | | | | |
|---|---|---|---|---|---|---|---|
| si:ch211-276k2.1 | 7.786040 | 0.297250934 | 0.136 | 0.003 | 1.9780435 | 21 | si:ch211-276k2.1 |
| dnmt3bb.1 | 2.032381 | 0.433911921 | 0.244 | 0.012 | 5.1632652 | 21 | dnmt3bb.1 |
| ecm1b3 | 4.014467 | 1.001213519 | 0.528 | 0.062 | 1.0198754 | 21 | ecm1b |
| ace1 | 3.722751 | 0.646319793 | 0.347 | 0.026 | 9.4576490 | 21 | ace |
| CR318588.32 | 3.806494 | 0.804068222 | 0.466 | 0.047 | 9.6703999 | 21 | CR318588.3 |
| fgfrl1a | 4.730893 | 0.503347805 | 0.25 | 0.013 | 1.2018834 | 21 | fgfrl1a |
| ar4 | 6.455871 | 0.76725749 | 0.528 | 0.062 | 1.6401141 | 21 | ar |
| dag13 | 1.856508 | 0.79227824 | 0.472 | 0.051 | 4.7164600 | 21 | dag1 |
| gstm.34 | 2.866508 | 1.300466958 | 0.733 | 0.125 | 7.2823661 | 21 | gstm.3 |
| CABZ01021532.1 | 5.118588 | 0.549907732 | 0.261 | 0.015 | 1.3003775 | 21 | CABZ01021532.1 |
| XLOC-002129 | 7.187152 | 0.290522082 | 0.142 | 0.004 | 1.8258962 | 21 | XLOC-002129 |
| hs3st3b1b | 2.504190 | 0.361027704 | 0.199 | 0.009 | 6.3618957 | 21 | hs3st3b1b |
| edem1 | 1.572391 | 0.610753809 | 0.375 | 0.033 | 3.9946606 | 21 | edem1 |
| mmp25 | 1.870795 | 0.926711983 | 0.682 | 0.105 | 4.7527562 | 21 | mmp2 |
| hmgb3a5 | 4.518789 | 0.892121411 | 0.568 | 0.078 | 1.1479985 | 21 | hmgb3a |
| atp11a | 2.578429 | 0.42161293 | 0.21 | 0.01 | 6.5505006 | 21 | atp11a |
| phactr4a3 | 1.669274 | 0.649327655 | 0.392 | 0.037 | 4.2407930 | 21 | phactr4a |
| sox9a | 1.257105 | 0.430619327 | 0.21 | 0.01 | 3.1936771 | 21 | sox9a |
| tent5ba | 3.257573 | 0.391618279 | 0.216 | 0.011 | 8.2758660 | 21 | tent5ba |
| hsd3b7 | 5.941368 | 0.919750207 | 0.58 | 0.079 | 1.5094047 | 21 | hsd3b7 |
| man2c1 | 4.833049 | 0.507895748 | 0.239 | 0.014 | 1.2278360 | 21 | man2c1 |
| prkar2aa1 | 3.031678 | 0.880267329 | 0.648 | 0.102 | 7.7019804 | 21 | prkar2aa |
| st5 | 3.199591 | 0.435567915 | 0.244 | 0.015 | 8.1285615 | 21 | st5 |
| myh102 | 4.631715 | 0.636174909 | 0.398 | 0.039 | 1.1766874 | 21 | myh10 |
| mxra8a3 | 7.820084 | 0.747736993 | 0.483 | 0.058 | 1.9866925 | 21 | mxra8a |
| slc38a21 | 2.198740 | 0.684366894 | 0.409 | 0.042 | 5.5859006 | 21 | slc38a2 |
| rdh10a3 | 4.039592 | 0.864522703 | 0.415 | 0.044 | 1.0262585 | 21 | rdh10a |
| apoeb1 | 1.405542 | 0.455343722 | 0.256 | 0.016 | 3.5707797 | 21 | apoeb |
| ifi457 | 1.600275 | 1.176725306 | 0.71 | 0.133 | 4.0655009 | 21 | ifi45 |
| tuba8l34 | 1.889224 | 0.879183434 | 0.506 | 0.068 | 4.7995754 | 21 | tuba8l3 |
| zgc:100829 | 1.769166 | 0.62221209 | 0.386 | 0.039 | 4.4945677 | 21 | zgc:100829 |
| gstt1b5 | 5.148415 | 0.839454048 | 0.568 | 0.085 | 1.3079549 | 21 | gstt1b |
| CABZ01084501.3 | 7.053676 | 0.292289355 | 0.119 | 0.003 | 1.7919865 | 21 | CABZ01084501.3 |
| lamc12 | 2.207231 | 0.723391546 | 0.449 | 0.054 | 5.6074715 | 21 | lamc1 |
| ponzr17 | 7.713723 | 1.177496071 | 0.909 | 0.283 | 1.9596713 | 21 | ponzr1 |

| | | | | | | | |
|---|---|---|---|---|---|---|---|
| arhgef18a | 2.347219 | 0.425243309 | 0.25 | 0.017 | 5.9631117 | 21 | arhgef18a |
| msi2a1 | 5.018808 | 0.586742452 | 0.398 | 0.044 | 1.2750282 | 21 | msi2a |
| fgd5a | 8.856572 | 0.373904105 | 0.205 | 0.012 | 2.2500122 | 21 | fgd5a |
| lgals9l33 | 2.336246 | 0.741627988 | 0.528 | 0.079 | 5.9352340 | 21 | lgals9l3 |
| nat8l | 5.168026 | 0.320062137 | 0.142 | 0.005 | 1.3129370 | 21 | nat8l |
| ccdc125 | 1.723426 | 0.382106189 | 0.21 | 0.013 | 4.3783640 | 21 | ccdc125 |
| wnt4a | 1.813899 | 0.310765916 | 0.159 | 0.007 | 4.6082117 | 21 | wnt4a |
| si:dkey-7j14.65 | 1.162704 | 0.961201007 | 0.818 | 0.19 | 2.9538509 | 21 | si:dkey-7j14.6 |
| lsr | 1.614569 | 0.456974351 | 0.301 | 0.026 | 4.1018141 | 21 | lsr |
| si:dkey-188i13.104 | 4.026778 | 0.831914632 | 0.614 | 0.109 | 1.0230030 | 21 | si:dkey-188i13.10 |
| znf1104 | 1.263988 | 0.454350419 | 0.222 | 0.015 | 3.2111621 | 21 | znf1104 |
| enc13 | 3.627333 | 0.55904324 | 0.347 | 0.036 | 9.2152411 | 21 | enc1 |
| ttll12 | 7.459395 | 0.618028219 | 0.284 | 0.024 | 1.8950594 | 21 | ttll12 |
| adam10a4 | 1.568863 | 0.827876114 | 0.58 | 0.101 | 3.9856977 | 21 | adam10a |
| slit2 | 1.941994 | 0.341900082 | 0.188 | 0.01 | 4.9336380 | 21 | slit2 |
| notch1b | 6.356182 | 0.323169372 | 0.17 | 0.009 | 1.6147881 | 21 | notch1b |
| cxxc5a1 | 8.164459 | 0.483452826 | 0.273 | 0.023 | 2.0741808 | 21 | cxxc5a |
| pgrmc14 | 1.262269 | 0.912507039 | 0.693 | 0.147 | 3.2067964 | 21 | pgrmc1 |
| etv4 | 4.302949 | 0.40951954 | 0.216 | 0.015 | 1.0931643 | 21 | etv4 |
| b2m7 | 8.786749 | 1.085440756 | 0.892 | 0.292 | 2.2322737 | 21 | b2m |
| tgfb32 | 1.234630 | 0.487869543 | 0.312 | 0.031 | 3.1365793 | 21 | tgfb3 |
| slc16a101 | 2.005661 | 0.460185369 | 0.267 | 0.023 | 5.0953831 | 21 | slc16a10 |
| zgc:123010 | 9.174696 | 0.558204738 | 0.324 | 0.034 | 2.3308315 | 21 | zgc:123010 |
| tmem176l.15 | 1.033191 | 0.94273992 | 0.795 | 0.196 | 2.6248222 | 21 | tmem176l.1 |
| gig2e | 1.610411 | 0.400762766 | 0.222 | 0.016 | 4.0912491 | 21 | gig2e |
| si:ch73-1a9.39 | 2.033117 | 1.065339058 | 0.938 | 0.309 | 5.1651341 | 21 | si:ch73-1a9.3 |
| CR318588.19 | 1.934430 | 1.242181669 | 0.801 | 0.21 | 4.9144197 | 21 | CR318588.1 |
| si:ch211-199g17.21 | 8.062481 | 0.620672897 | 0.455 | 0.065 | 2.0482735 | 21 | si:ch211-199g17.2 |
| cdh1 | 5.097628 | 0.663715319 | 0.403 | 0.052 | 1.2950524 | 21 | cdh1 |
| atp1b1a4 | 8.930643 | 0.948003575 | 0.818 | 0.224 | 2.2688298 | 21 | atp1b1a |
| rpia1 | 5.834871 | 0.671689261 | 0.438 | 0.063 | 1.4823490 | 21 | rpia |
| rab11fip1b2 | 1.239897 | 0.669904123 | 0.409 | 0.057 | 3.1499587 | 21 | rab11fip1b |
| svild4 | 1.217105 | 0.733264962 | 0.489 | 0.08 | 3.0920571 | 21 | svild |
| slc2a1b | 1.817107 | 0.427354634 | 0.222 | 0.017 | 4.6163615 | 21 | slc2a1b |
| hspe14 | 1.966879 | 1.034358672 | 0.892 | 0.285 | 4.9968572 | 21 | hspe1 |

| | | | | | | | |
|---|---|---|---|---|---|---|---|
| phlda21 | 3.672031 | 0.69359776 | 0.381 | 0.05 | 9.3287954 | 21 | phlda2 |
| cox7a11 | 2.753797 | 0.456609863 | 0.324 | 0.037 | 6.9960235 | 21 | cox7a1 |
| plcl1 | 6.132610 | 0.332686285 | 0.176 | 0.011 | 1.5579897 | 21 | plcl1 |
| abhd15a1 | 7.070580 | 0.529468974 | 0.335 | 0.039 | 1.7962810 | 21 | abhd15a |
| si:dkey-22o12.2 | 1.691545 | 0.303770686 | 0.159 | 0.009 | 4.2973705 | 21 | si:dkey-22o12.2 |
| dpp31 | 3.582183 | 0.570731508 | 0.358 | 0.045 | 9.1005375 | 21 | dpp3 |
| krt18a.15 | 5.366668 | 0.910404275 | 0.676 | 0.145 | 1.3634021 | 21 | krt18a.1 |
| adgrl3.11 | 7.040276 | 0.47646115 | 0.278 | 0.028 | 1.7885821 | 21 | adgrl3.1 |
| ivns1abpa | 9.688379 | 0.53714316 | 0.278 | 0.028 | 2.4613329 | 21 | ivns1abpa |
| sox4a4 | 1.160163 | 0.687758115 | 0.409 | 0.059 | 2.9473949 | 21 | sox4a |
| pdgfaa | 1.761852 | 0.386078628 | 0.21 | 0.016 | 4.4759874 | 21 | pdgfaa |
| si:dkey-73n10.11 | 6.338252 | 0.386094293 | 0.244 | 0.021 | 1.6102331 | 21 | si:dkey-73n10.1 |
| nacc2 | 8.285853 | 0.381310684 | 0.239 | 0.021 | 2.1050209 | 21 | nacc2 |
| hist2h2l3 | 1.195171 | 0.804645735 | 0.511 | 0.095 | 3.0363336 | 21 | hist2h2l |
| h1f05 | 1.205121 | 0.715273394 | 0.438 | 0.068 | 3.0616103 | 21 | h1f0 |
| lgals172 | 1.580866 | 0.668436295 | 0.381 | 0.052 | 4.0161914 | 21 | lgals17 |
| ap2m1a1 | 1.775577 | 0.861123404 | 0.688 | 0.166 | 4.5108542 | 21 | ap2m1a |
| h2afx18 | 4.200507 | 0.949247455 | 0.79 | 0.221 | 1.0671389 | 21 | h2afx1 |
| COLEC10 | 1.255500 | 0.27625263 | 0.136 | 0.007 | 3.1895990 | 21 | COLEC10 |
| mphosph84 | 3.767994 | 0.788343391 | 0.625 | 0.136 | 9.5725892 | 21 | mphosph8 |
| cldnc1 | 9.247398 | 0.541865548 | 0.301 | 0.033 | 2.3493015 | 21 | cldnc |
| uxs1 | 1.791870 | 0.419285068 | 0.284 | 0.03 | 4.5522479 | 21 | uxs1 |
| tspan9b3 | 2.771126 | 0.54904126 | 0.358 | 0.047 | 7.0400471 | 21 | tspan9b |
| robo23 | 5.526023 | 0.55795328 | 0.375 | 0.051 | 1.4038863 | 21 | robo2 |
| ndrg21 | 6.111993 | 0.520946291 | 0.364 | 0.049 | 1.5527518 | 21 | ndrg2 |
| ptprga6 | 1.103422 | 0.655369392 | 0.432 | 0.068 | 2.8032439 | 21 | ptprga |
| aplp23 | 2.420621 | 0.653842099 | 0.449 | 0.073 | 6.1495899 | 21 | aplp2 |
| sptbn2 | 4.673941 | 0.510402126 | 0.33 | 0.041 | 1.1874147 | 21 | sptbn2 |
| rgrb | 7.474828 | 0.282983355 | 0.176 | 0.012 | 1.8989801 | 21 | rgrb |
| arf2b | 1.235800 | 0.7942185 | 0.699 | 0.163 | 3.1395503 | 21 | arf2b |
| rhoj1 | 7.480841 | 0.326683424 | 0.193 | 0.015 | 1.9005078 | 21 | rhoj |
| pde4ba4 | 8.611963 | 0.818556387 | 0.619 | 0.137 | 2.1878692 | 21 | pde4ba |
| znf609b1 | 1.371114 | 0.427124378 | 0.256 | 0.026 | 3.4833166 | 21 | znf609b |
| sat1a.1 | 2.967415 | 0.320686348 | 0.148 | 0.008 | 7.5387191 | 21 | sat1a.1 |
| pgr2 | 4.975164 | 0.392400368 | 0.21 | 0.018 | 1.2639405 | 21 | pgr |

| | | | | | | | |
|---|---|---|---|---|---|---|---|
| colec11 | 5.427827 | 0.304948734 | 0.142 | 0.008 | 1.3789395 | 21 | colec11 |
| hdac41 | 7.057376 | 0.475725462 | 0.278 | 0.03 | 1.7929264 | 21 | hdac4 |
| cebpd6 | 7.242175 | 0.823772321 | 0.659 | 0.147 | 1.8398747 | 21 | cebpd |
| tnfaip61 | 2.085219 | 0.40108643 | 0.205 | 0.017 | 5.2975008 | 21 | tnfaip6 |
| bmpr1bb | 1.033012 | 0.304155553 | 0.153 | 0.009 | 2.6243692 | 21 | bmpr1bb |
| nanos12 | 2.505429 | 0.499249066 | 0.273 | 0.03 | 6.3650428 | 21 | nanos1 |
| tnfrsfa1 | 8.573135 | 0.403900655 | 0.267 | 0.03 | 2.1780050 | 21 | tnfrsfa |
| si:ch211-264f5.6 | 1.016863 | 0.355142033 | 0.17 | 0.012 | 2.5833420 | 21 | si:ch211-264f5.6 |
| hey2 | 1.222708 | 0.280129446 | 0.148 | 0.009 | 3.1062901 | 21 | hey2 |
| gas61 | 5.777205 | 0.437448729 | 0.261 | 0.029 | 1.4676991 | 21 | gas6 |
| hmgb1b7 | 1.865325 | 0.74537844 | 0.636 | 0.15 | 4.7388604 | 21 | hmgb1b |
| zgc:921401 | 1.265369 | 0.427214987 | 0.261 | 0.029 | 3.2146720 | 21 | zgc:92140 |
| gdpd4a1 | 9.270043 | 0.31446168 | 0.153 | 0.01 | 2.3550546 | 21 | gdpd4a |
| zgc:927444 | 3.752240 | 0.928767355 | 0.864 | 0.322 | 9.5325665 | 21 | zgc:92744 |
| hmgb1a9 | 3.981430 | 0.831382124 | 0.75 | 0.223 | 1.0114823 | 21 | hmgb1a |
| selenof2 | 4.115807 | 0.845523092 | 0.812 | 0.262 | 1.0456209 | 21 | selenof |
| prr13 | 4.406568 | 0.397386123 | 0.227 | 0.023 | 1.1194887 | 21 | prr13 |
| tusc3 | 4.713985 | 0.269918522 | 0.165 | 0.012 | 1.1975880 | 21 | tusc3 |
| crybg1a | 4.888448 | 0.327758849 | 0.199 | 0.018 | 1.2419103 | 21 | crybg1a |
| csrnp2 | 4.943600 | 0.281970497 | 0.165 | 0.012 | 1.2559217 | 21 | csrnp2 |
| errfi1a | 1.497364 | 0.400135708 | 0.205 | 0.019 | 3.8040555 | 21 | errfi1a |
| angptl43 | 2.724835 | 0.558615994 | 0.352 | 0.053 | 6.9224443 | 21 | angptl4 |
| txnipa4 | 1.000423 | 0.852376215 | 0.778 | 0.239 | 2.5415765 | 21 | txnipa |
| XLOC-035202 | 1.045701 | 0.286355817 | 0.148 | 0.01 | 2.6566053 | 21 | XLOC-035202 |
| igflr1 | 2.138443 | 0.357051138 | 0.193 | 0.017 | 5.4327159 | 21 | igflr1 |
| fn1a1 | 2.471313 | 0.567309239 | 0.449 | 0.079 | 6.2783723 | 21 | fn1a |
| man1b1b | 2.769041 | 0.439402448 | 0.256 | 0.03 | 7.0347507 | 21 | man1b1b |
| ppib8 | 5.813693 | 0.855331007 | 0.915 | 0.374 | 1.4769688 | 21 | ppib |
| phc2a1 | 5.859320 | 0.633956035 | 0.466 | 0.089 | 1.4885604 | 21 | phc2a |
| aff2 | 1.779590 | 0.263175536 | 0.148 | 0.01 | 4.5210506 | 21 | aff2 |
| isg153 | 8.729625 | 0.646719362 | 0.369 | 0.062 | 2.2177612 | 21 | isg15 |
| fam234a | 1.841004 | 0.299079567 | 0.176 | 0.015 | 4.6770720 | 21 | fam234a |
| jcada1 | 1.947294 | 0.353206702 | 0.199 | 0.019 | 4.9471017 | 21 | jcada |
| zfp36l1b5 | 5.242535 | 0.713128172 | 0.58 | 0.136 | 1.3318660 | 21 | zfp36l1b |
| si:dkey-178e17.31 | 1.768729 | 0.303904195 | 0.176 | 0.015 | 4.4934574 | 21 | si:dkey-178e17.3 |

| | | | | | | | |
|---|---|---|---|---|---|---|---|
| psme12 | 8.629008 | 0.612462165 | 0.494 | 0.109 | 2.1921997 | 21 | psme1 |
| wu:fc75a091 | 8.780801 | 0.358416904 | 0.21 | 0.022 | 2.2307626 | 21 | wu:fc75a09 |
| fkbp1aa6 | 9.510414 | 0.823572115 | 0.909 | 0.41 | 2.4161207 | 21 | fkbp1aa |
| marcksl1b7 | 1.215634 | 0.822150212 | 0.557 | 0.138 | 3.0883188 | 21 | marcksl1b |
| ifi302 | 1.757032 | 0.785730222 | 0.682 | 0.191 | 4.4637410 | 21 | ifi30 |
| wt1a2 | 1.876550 | 0.45940558 | 0.29 | 0.04 | 4.7673754 | 21 | wt1a |
| ywhaz4 | 2.473646 | 0.807358325 | 0.835 | 0.3 | 6.2842998 | 21 | ywhaz |
| tp53inp12 | 4.054509 | 0.636024152 | 0.341 | 0.057 | 1.0300481 | 21 | tp53inp1 |
| hspg22 | 4.932803 | 0.374912497 | 0.233 | 0.027 | 1.2531786 | 21 | hspg2 |
| adgrl2a | 5.260405 | 0.344794766 | 0.188 | 0.018 | 1.3364059 | 21 | adgrl2a |
| htra1b2 | 2.393294 | 0.562390583 | 0.267 | 0.036 | 6.0801652 | 21 | htra1b |
| tle2b1 | 2.653498 | 0.296728727 | 0.176 | 0.016 | 6.7412122 | 21 | tle2b |
| nme34 | 2.818850 | 0.561225112 | 0.432 | 0.086 | 7.1612900 | 21 | nme3 |
| decr1 | 3.162221 | 0.434628374 | 0.301 | 0.044 | 8.0336241 | 21 | decr1 |
| fam168a6 | 3.518284 | 0.527459444 | 0.386 | 0.071 | 8.9382026 | 21 | fam168a |
| arrdc3a3 | 3.545939 | 0.601040977 | 0.386 | 0.07 | 9.0084584 | 21 | arrdc3a |
| zfpm1 | 1.419837 | 0.271003912 | 0.159 | 0.013 | 3.6070963 | 21 | zfpm1 |
| si:ch211-214p13.9 | 4.744540 | 0.297576537 | 0.17 | 0.015 | 1.2053506 | 21 | si:ch211-214p13.9 |
| tegt3 | 6.207616 | 0.7851958 | 0.71 | 0.233 | 1.5770450 | 21 | tegt |
| zgc:1951735 | 1.588811 | 0.65608223 | 0.494 | 0.11 | 4.0363760 | 21 | zgc:195173 |
| gdi1 | 2.112075 | 0.313985706 | 0.182 | 0.018 | 5.3657274 | 21 | gdi1 |
| spred2b4 | 2.425999 | 0.615747764 | 0.409 | 0.082 | 6.1632511 | 21 | spred2b |
| id2b2 | 4.749173 | 0.423043074 | 0.278 | 0.039 | 1.2065275 | 21 | id2b |
| irf2bp2b4 | 6.944566 | 0.609704492 | 0.455 | 0.099 | 1.7642671 | 21 | irf2bp2b |
| si:ch211-286b5.56 | 1.193882 | 0.635698615 | 0.58 | 0.147 | 3.0330576 | 21 | si:ch211-286b5.5 |
| atp10a5 | 3.128875 | 0.498040481 | 0.335 | 0.057 | 7.9489076 | 21 | atp10a |
| kremen1 | 5.419559 | 0.268808631 | 0.142 | 0.011 | 1.3768390 | 21 | kremen1 |
| wbp1la | 8.238024 | 0.289539828 | 0.17 | 0.016 | 2.0928700 | 21 | wbp1la |
| inka1a1 | 9.063538 | 0.375838928 | 0.227 | 0.028 | 2.3025918 | 21 | inka1a |
| rhoq | 1.706480 | 0.278254278 | 0.165 | 0.015 | 4.3353133 | 21 | rhoq |
| csgalnact1a | 3.666409 | 0.325165862 | 0.193 | 0.02 | 9.3145132 | 21 | csgalnact1a |
| odc11 | 8.355234 | 0.713033324 | 0.494 | 0.118 | 2.1226474 | 21 | odc1 |
| hp1bp35 | 1.882479 | 0.589461896 | 0.494 | 0.117 | 4.7824395 | 21 | hp1bp3 |
| pwwp2b | 2.502935 | 0.384233743 | 0.261 | 0.037 | 6.3587067 | 21 | pwwp2b |
| mydgf | 5.492059 | 0.414544263 | 0.273 | 0.04 | 1.3952575 | 21 | mydgf |

| | | | | | | | |
|---|---|---|---|---|---|---|---|
| serpinf14 | 7.284211 | 0.439146259 | 0.284 | 0.043 | 1.8505539 | 21 | serpinf1 |
| sema3b | 9.042291 | 0.335746592 | 0.193 | 0.021 | 2.2971942 | 21 | sema3b |
| nid1b3 | 9.099185 | 0.52868674 | 0.364 | 0.066 | 2.3116481 | 21 | nid1b |
| zgc:1141811 | 1.214624 | 0.327919583 | 0.17 | 0.017 | 3.0857542 | 21 | zgc:114181 |
| s100v26 | 1.414432 | 0.628540745 | 0.551 | 0.137 | 3.5933650 | 21 | s100v2 |
| cox6a21 | 2.073328 | 0.408306947 | 0.256 | 0.036 | 5.2672914 | 21 | cox6a2 |
| hsbp1a1 | 2.290875 | 0.360451208 | 0.239 | 0.031 | 5.8199696 | 21 | hsbp1a |
| swap70b | 3.430931 | 0.329894597 | 0.188 | 0.02 | 8.7162825 | 21 | swap70b |
| pcyox11 | 5.666759 | 0.454519267 | 0.33 | 0.057 | 1.4396402 | 21 | pcyox1 |
| cpn14 | 5.959647 | 0.575302723 | 0.381 | 0.075 | 1.5140484 | 21 | cpn1 |
| si:ch211-195o20.7 | 7.350786 | 0.262552138 | 0.159 | 0.015 | 1.8674672 | 21 | si:ch211-195o20.7 |
| zgc:1748633 | 9.969515 | 0.611941109 | 0.46 | 0.108 | 2.5327553 | 21 | zgc:174863 |
| pde7a | 1.091713 | 0.472840555 | 0.347 | 0.063 | 2.7734968 | 21 | pde7a |
| prnprs3 | 1.615263 | 0.26844158 | 0.153 | 0.014 | 4.1035769 | 21 | prnprs3 |
| hmgn69 | 1.754849 | 0.768163315 | 0.807 | 0.307 | 4.4581958 | 21 | hmgn6 |
| ywhaqb5 | 2.179550 | 0.648604624 | 0.557 | 0.151 | 5.5371476 | 21 | ywhaqb |
| cdh23 | 2.960880 | 0.523890421 | 0.347 | 0.064 | 7.5221172 | 21 | cdh2 |
| kmt2cb2 | 3.156421 | 0.46115748 | 0.33 | 0.058 | 8.0188893 | 21 | kmt2cb |
| znfx11 | 3.780680 | 0.392293702 | 0.233 | 0.031 | 9.6048185 | 21 | znfx1 |
| ghrb3 | 7.486006 | 0.441669285 | 0.273 | 0.042 | 1.9018200 | 21 | ghrb |
| oaz1b5 | 8.525620 | 0.736423945 | 0.869 | 0.355 | 2.1659338 | 21 | oaz1b |
| itm2ba6 | 1.810800 | 0.742204985 | 0.784 | 0.292 | 4.6003394 | 21 | itm2ba |
| si:dkey-187e18.11 | 1.949938 | 0.510274216 | 0.358 | 0.069 | 4.9538184 | 21 | si:dkey-187e18.1 |
| atrx5 | 3.349758 | 0.613117988 | 0.494 | 0.125 | 8.5100605 | 21 | atrx |
| glulb3 | 3.606559 | 0.767739246 | 0.517 | 0.136 | 9.1624643 | 21 | glulb |
| flot1b2 | 9.629756 | 0.505501695 | 0.438 | 0.098 | 2.4464395 | 21 | flot1b |
| cpxm1a1 | 9.656435 | 0.308873999 | 0.17 | 0.018 | 2.4532174 | 21 | cpxm1a |
| si:ch211-222l21.19 | 9.799524 | 0.783059662 | 0.835 | 0.309 | 2.4895692 | 21 | si:ch211-222l21.1 |
| ehd3 | 1.064503 | 0.299270029 | 0.159 | 0.015 | 2.7043719 | 21 | ehd3 |
| tcf42 | 1.149763 | 0.334801126 | 0.193 | 0.022 | 2.9209733 | 21 | tcf4 |
| crim14 | 1.200154 | 0.407240919 | 0.261 | 0.039 | 3.0489922 | 21 | crim1 |
| si:ch211-197h24.93 | 1.242439 | 0.551737859 | 0.386 | 0.081 | 3.1564170 | 21 | si:ch211-197h24.9 |
| rnf213b | 1.559645 | 0.280263374 | 0.153 | 0.014 | 3.9622786 | 21 | rnf213b |
| psme24 | 1.877713 | 0.573225068 | 0.46 | 0.11 | 4.7703318 | 21 | psme2 |
| zgc:175088 | 1.878698 | 0.453654293 | 0.341 | 0.063 | 4.7728336 | 21 | zgc:175088 |

| | | | | | | | |
|---|---|---|---|---|---|---|---|
| zgc:114200 | 2.080338 | 0.383246361 | 0.239 | 0.033 | 5.2850999 | 21 | zgc:114200 |
| schip11 | 4.823756 | 0.304814077 | 0.188 | 0.021 | 1.2254752 | 21 | schip1 |
| pik3c2a | 7.195880 | 0.319167959 | 0.205 | 0.025 | 1.8281135 | 21 | pik3c2a |
| map1lc3a5 | 7.543786 | 0.484768728 | 0.381 | 0.078 | 1.9164990 | 21 | map1lc3a |
| tjp1a2 | 1.118564 | 0.430100485 | 0.301 | 0.051 | 2.8417141 | 21 | tjp1a |
| rnf11b | 1.303282 | 0.296071148 | 0.182 | 0.02 | 3.3109884 | 21 | rnf11b |
| iscub | 1.656858 | 0.450993639 | 0.375 | 0.075 | 4.2092499 | 21 | iscub |
| txndc12 | 1.950570 | 0.425670116 | 0.301 | 0.052 | 4.9554240 | 21 | txndc12 |
| nucb2a | 2.163886 | 0.466700622 | 0.318 | 0.058 | 5.4973532 | 21 | nucb2a |
| metrnla3 | 2.758898 | 0.530764117 | 0.42 | 0.092 | 7.0089825 | 21 | metrnla |
| tfr1a | 6.218323 | 0.388784237 | 0.176 | 0.019 | 1.5797650 | 21 | tfr1a |
| cux1a3 | 8.129108 | 0.42467812 | 0.301 | 0.053 | 2.0651999 | 21 | cux1a |
| esd1 | 2.575923 | 0.499394476 | 0.455 | 0.109 | 6.5441332 | 21 | esd |
| tmem50a1 | 2.747428 | 0.562795936 | 0.545 | 0.147 | 6.9798419 | 21 | tmem50a |
| cgnl12 | 3.709500 | 0.338373816 | 0.21 | 0.028 | 9.4239858 | 21 | cgnl1 |
| krt945 | 5.142053 | 0.621649722 | 0.602 | 0.162 | 1.3063386 | 21 | krt94 |
| ndufc22 | 5.738076 | 0.55366289 | 0.545 | 0.149 | 1.4577582 | 21 | ndufc2 |
| lama44 | 6.401181 | 0.41424164 | 0.273 | 0.045 | 1.6262200 | 21 | lama4 |
| CABZ01086029.1 | 1.003290 | 0.297727179 | 0.176 | 0.02 | 2.5488603 | 21 | CABZ01086029.1 |
| dhrs3a3 | 1.459957 | 0.438085844 | 0.239 | 0.036 | 3.7090209 | 21 | dhrs3a |
| ckap44 | 1.520802 | 0.438988619 | 0.312 | 0.058 | 3.8635987 | 21 | ckap4 |
| chd6 | 2.335798 | 0.392386072 | 0.273 | 0.046 | 5.9340951 | 21 | chd6 |
| si:ch73-46j18.57 | 2.470796 | 0.669627793 | 0.682 | 0.204 | 6.2770572 | 21 | si:ch73-46j18.5 |
| appb3 | 2.587419 | 0.422844169 | 0.267 | 0.045 | 6.5733387 | 21 | appb |
| arhgap5 | 2.695093 | 0.324282948 | 0.222 | 0.031 | 6.8468851 | 21 | arhgap5 |
| chpfa | 9.874501 | 0.265396517 | 0.136 | 0.013 | 2.5086171 | 21 | chpfa |
| irs1 | 4.520796 | 0.304325634 | 0.205 | 0.028 | 1.1485084 | 21 | irs1 |
| srpra | 1.328587 | 0.462481371 | 0.312 | 0.06 | 3.3752759 | 21 | srpra |
| znf292a | 1.833593 | 0.29036656 | 0.188 | 0.024 | 4.6582448 | 21 | znf292a |
| foxp43 | 5.324570 | 0.490384982 | 0.341 | 0.071 | 1.3527070 | 21 | foxp4 |
| ppp1r14bb1 | 5.466382 | 0.341739154 | 0.25 | 0.041 | 1.3887345 | 21 | ppp1r14bb |
| zgc:1527913 | 5.693154 | 0.481046028 | 0.33 | 0.066 | 1.4463459 | 21 | zgc:152791 |
| lhfpl2a | 6.172299 | 0.363708755 | 0.227 | 0.035 | 1.5680727 | 21 | lhfpl2a |
| cd82a2 | 7.450550 | 0.470745844 | 0.33 | 0.067 | 1.8928122 | 21 | cd82a |
| stat1a4 | 8.057195 | 0.542517774 | 0.46 | 0.12 | 2.0469305 | 21 | stat1a |

| | | | | | | | |
|---|---|---|---|---|---|---|---|
| mfge8b1 | 9.242997 | 0.327611201 | 0.227 | 0.034 | 2.3481834 | 21 | mfge8b |
| spry22 | 1.060100 | 0.466938819 | 0.335 | 0.07 | 2.6931862 | 21 | spry2 |
| tspan3a | 2.111848 | 0.426448278 | 0.341 | 0.071 | 5.3651500 | 21 | tspan3a |
| dynll2b1 | 4.443914 | 0.49799405 | 0.494 | 0.127 | 1.1289765 | 21 | dynll2b |
| nab1b | 5.629357 | 0.267289755 | 0.153 | 0.017 | 1.4301382 | 21 | nab1b |
| NC-002333.49 | 6.079592 | 0.636453312 | 0.955 | 0.606 | 1.5445205 | 21 | NC-002333.4 |
| trappc11 | 1.514159 | 0.500739308 | 0.443 | 0.114 | 3.8467220 | 21 | trappc1 |
| b2ml8 | 1.925203 | 0.660959258 | 0.756 | 0.29 | 4.8909784 | 21 | b2ml |
| chd4a2 | 1.957416 | 0.501858785 | 0.42 | 0.105 | 4.9728161 | 21 | chd4a |
| eif4ebp13 | 2.480893 | 0.629516633 | 0.722 | 0.254 | 6.3027107 | 21 | eif4ebp1 |
| cxcl142 | 2.824476 | 0.304002772 | 0.125 | 0.012 | 7.1755825 | 21 | cxcl14 |
| reep3b | 3.653986 | 0.417324953 | 0.318 | 0.065 | 9.2829529 | 21 | reep3b |
| rnf130 | 7.967971 | 0.377434792 | 0.278 | 0.051 | 2.0242632 | 21 | rnf130 |
| canx1 | 1.578009 | 0.489433261 | 0.403 | 0.099 | 4.0089320 | 21 | canx |
| zbtb204 | 3.132852 | 0.503556227 | 0.415 | 0.105 | 7.9590106 | 21 | zbtb20 |
| slc29a1a1 | 3.528451 | 0.349947072 | 0.233 | 0.037 | 8.9640314 | 21 | slc29a1a |
| tmsb4 | 4.112266 | 0.370760678 | 0.21 | 0.032 | 1.0447213 | 21 | tmsb |
| emc10 | 4.230431 | 0.45321642 | 0.375 | 0.087 | 1.0747410 | 21 | emc10 |
| PDE3B | 6.953146 | 0.265374223 | 0.159 | 0.019 | 1.7664469 | 21 | PDE3B |
| CR383676.111 | 7.400890 | 0.684346099 | 0.972 | 0.715 | 1.8801961 | 21 | CR383676.1 |
| sdcbp26 | 7.401792 | 0.600579183 | 0.653 | 0.224 | 1.8804253 | 21 | sdcbp2 |
| lrrc58b | 7.777621 | 0.261777098 | 0.165 | 0.02 | 1.9759048 | 21 | lrrc58b |
| zgc:664331 | 7.863493 | 0.326492794 | 0.216 | 0.033 | 1.9977205 | 21 | zgc:66433 |
| mknk2b6 | 1.321666 | 0.667479223 | 0.58 | 0.197 | 3.3576928 | 21 | mknk2b |
| adamts21 | 1.556027 | 0.308051904 | 0.188 | 0.026 | 3.9530876 | 21 | adamts2 |
| oip5-as1 | 1.595667 | 0.330393388 | 0.205 | 0.031 | 4.0537944 | 21 | oip5-as1 |
| fam13b | 1.937058 | 0.319376018 | 0.222 | 0.035 | 4.9210964 | 21 | fam13b |
| calm2b6 | 1.997170 | 0.642580265 | 0.682 | 0.243 | 5.0738109 | 21 | calm2b |
| prdx4 | 1.999962 | 0.453998892 | 0.562 | 0.165 | 5.0809048 | 21 | prdx4 |
| atp1a1a.1 | 2.289040 | 0.571810681 | 0.494 | 0.142 | 5.8153064 | 21 | atp1a1a.1 |
| rnf213a5 | 4.397192 | 0.489719634 | 0.381 | 0.093 | 1.1171066 | 21 | rnf213a |
| nr2f1a3 | 7.048467 | 0.488891551 | 0.295 | 0.061 | 1.7906632 | 21 | nr2f1a |
| neo1a2 | 1.098145 | 0.290130019 | 0.205 | 0.031 | 2.7898378 | 21 | neo1a |
| glod5 | 1.759693 | 0.329979241 | 0.222 | 0.036 | 4.4705012 | 21 | glod5 |
| asph1 | 1.926988 | 0.368585049 | 0.239 | 0.042 | 4.8955133 | 21 | asph |

| | | | | | | | |
|---|---|---|---|---|---|---|---|
| ctnnb14 | 2.049145 | 0.540474028 | 0.523 | 0.156 | 5.2058540 | 21 | ctnnb1 |
| h3f3c8 | 2.143873 | 0.645807694 | 0.761 | 0.3 | 5.4465109 | 21 | h3f3c |
| ewsr1a1 | 2.147630 | 0.353542817 | 0.261 | 0.049 | 5.4560552 | 21 | ewsr1a |
| socs3a7 | 2.173902 | 0.720990399 | 0.79 | 0.311 | 5.5228001 | 21 | socs3a |
| erbb22 | 2.513457 | 0.317797896 | 0.205 | 0.032 | 6.3854395 | 21 | erbb2 |
| serinc13 | 2.546657 | 0.456115631 | 0.358 | 0.084 | 6.4697841 | 21 | serinc1 |
| slc38a5b5 | 2.732690 | 0.48875195 | 0.46 | 0.126 | 6.9423997 | 21 | slc38a5b |
| emd3 | 3.003525 | 0.398894335 | 0.318 | 0.069 | 7.6304564 | 21 | emd |
| ndufa4l7 | 5.759407 | 0.640760961 | 0.869 | 0.419 | 1.4631774 | 21 | ndufa4l |
| pard3bb4 | 7.191259 | 0.312652088 | 0.205 | 0.032 | 1.8269394 | 21 | pard3bb |
| psmb11 | 7.365491 | 0.54602173 | 0.597 | 0.194 | 1.8712030 | 21 | psmb1 |
| atp6v0cb1 | 8.326570 | 0.263502753 | 0.17 | 0.023 | 2.1153651 | 21 | atp6v0cb |
| ncam1a1 | 1.388778 | 0.276986864 | 0.159 | 0.02 | 3.5281920 | 21 | ncam1a |
| cyp17a11 | 1.697106 | 0.456706848 | 0.318 | 0.067 | 4.3114978 | 21 | cyp17a1 |
| pnp5a2 | 2.812294 | 0.490685786 | 0.324 | 0.073 | 7.1446353 | 21 | pnp5a |
| fkbp21 | 3.816663 | 0.489797554 | 0.494 | 0.143 | 9.6962344 | 21 | fkbp2 |
| h2afy24 | 6.461984 | 0.484485415 | 0.432 | 0.12 | 1.6416671 | 21 | h2afy2 |
| epb41l52 | 6.600443 | 0.328496164 | 0.239 | 0.042 | 1.6768426 | 21 | epb41l5 |
| gstm.15 | 7.229615 | 0.632523501 | 0.727 | 0.275 | 1.8366837 | 21 | gstm.1 |
| krt87 | 9.065661 | 0.649434447 | 0.812 | 0.307 | 2.3031313 | 21 | krt8 |
| dspa1 | 9.370937 | 0.278425612 | 0.188 | 0.028 | 2.3806866 | 21 | dspa |
| gng75 | 9.468827 | 0.385808338 | 0.29 | 0.06 | 2.4055555 | 21 | gng7 |
| tnrc6c16 | 1.009460 | 0.528807022 | 0.489 | 0.147 | 2.5645333 | 21 | tnrc6c1 |
| pdia6 | 1.522247 | 0.472848177 | 0.449 | 0.123 | 3.8672693 | 21 | pdia6 |
| ndfip1 | 1.751069 | 0.540389537 | 0.528 | 0.168 | 4.4485922 | 21 | ndfip1 |
| selenok | 2.342656 | 0.496983319 | 0.466 | 0.137 | 5.9515189 | 21 | selenok |
| clstn11 | 3.419992 | 0.268118726 | 0.165 | 0.022 | 8.6884903 | 21 | clstn1 |
| hbp11 | 3.499141 | 0.393983213 | 0.278 | 0.057 | 8.8895693 | 21 | hbp1 |
| pik3ip1 | 4.045403 | 0.468660205 | 0.341 | 0.079 | 1.0277347 | 21 | pik3ip1 |
| tmed10 | 4.557028 | 0.509920909 | 0.557 | 0.178 | 1.1577132 | 21 | tmed10 |
| nrp2a2 | 4.594372 | 0.292169525 | 0.153 | 0.02 | 1.1672002 | 21 | nrp2a |
| slc20a21 | 5.293474 | 0.301616575 | 0.176 | 0.025 | 1.3448073 | 21 | slc20a2 |
| calr3a | 7.057596 | 0.556405729 | 0.506 | 0.155 | 1.7929824 | 21 | calr3a |
| psmc51 | 1.195401 | 0.441519937 | 0.386 | 0.101 | 3.0369174 | 21 | psmc5 |
| si:dkey-17m8.11 | 1.442947 | 0.34163066 | 0.239 | 0.044 | 3.6658068 | 21 | si:dkey-17m8.1 |

| | | | | | | | |
|---|---|---|---|---|---|---|---|
| ldlrap1a | 1.716173 | 0.352510208 | 0.295 | 0.061 | 4.3599387 | 21 | ldlrap1a |
| parvaa1 | 1.814676 | 0.313678576 | 0.205 | 0.034 | 4.6101845 | 21 | parvaa |
| purab2 | 1.957790 | 0.371748427 | 0.278 | 0.058 | 4.9737670 | 21 | purab |
| prkd2 | 2.002994 | 0.303857596 | 0.205 | 0.034 | 5.0886079 | 21 | prkd2 |
| gpx4b1 | 2.135605 | 0.543596912 | 0.619 | 0.217 | 5.4255069 | 21 | gpx4b |
| cox8a6 | 2.206038 | 0.588335699 | 0.688 | 0.264 | 5.6044399 | 21 | cox8a |
| ifi35 | 2.357466 | 0.301419045 | 0.216 | 0.037 | 5.9891444 | 21 | ifi35 |
| tma72 | 3.054570 | 0.594385196 | 0.801 | 0.327 | 7.7601362 | 21 | tma7 |
| ctsf1 | 4.692489 | 0.299480902 | 0.21 | 0.036 | 1.1921269 | 21 | ctsf |
| zgc:162509 | 5.826304 | 0.266919989 | 0.193 | 0.03 | 1.4801727 | 21 | zgc:162509 |
| FAT41 | 6.242063 | 0.255029194 | 0.153 | 0.02 | 1.5857963 | 21 | FAT4 |
| blmh | 9.313729 | 0.280356248 | 0.244 | 0.046 | 2.3661529 | 21 | blmh |
| COX311 | 1.133318 | 0.554948434 | 0.955 | 0.674 | 2.8791948 | 21 | COX3 |
| fndc3ba | 1.165831 | 0.250387457 | 0.136 | 0.016 | 2.9617941 | 21 | fndc3ba |
| tcf124 | 1.243172 | 0.431725571 | 0.295 | 0.066 | 3.1582795 | 21 | tcf12 |
| hnrnpa0b5 | 1.562283 | 0.625916002 | 0.773 | 0.338 | 3.9689799 | 21 | hnrnpa0b |
| smdt1b2 | 1.635675 | 0.463414628 | 0.489 | 0.15 | 4.1554334 | 21 | smdt1b |
| ankrd11 | 1.786955 | 0.4332458 | 0.375 | 0.098 | 4.5397616 | 21 | ankrd11 |
| grinab5 | 2.513048 | 0.486290697 | 0.42 | 0.12 | 6.3844007 | 21 | grinab |
| chd4b | 2.785056 | 0.461648584 | 0.392 | 0.106 | 7.0754352 | 21 | chd4b |
| id16 | 2.989712 | 0.618372153 | 0.438 | 0.129 | 7.5953650 | 21 | id1 |
| ctbp2a2 | 3.025618 | 0.298499946 | 0.222 | 0.039 | 7.6865834 | 21 | ctbp2a |
| ext2 | 3.036761 | 0.261035948 | 0.176 | 0.026 | 7.7148931 | 21 | ext2 |
| yap12 | 6.188328 | 0.361418949 | 0.267 | 0.056 | 1.5721449 | 21 | yap1 |
| tpm4a6 | 6.666517 | 0.499222065 | 0.403 | 0.111 | 1.6936288 | 21 | tpm4a |
| cd164 | 7.053588 | 0.410930395 | 0.312 | 0.074 | 1.7919640 | 21 | cd164 |
| nsfl1c1 | 1.216144 | 0.370803837 | 0.33 | 0.079 | 3.0896148 | 21 | nsfl1c |
| nktr | 1.252734 | 0.308981864 | 0.216 | 0.039 | 3.1825720 | 21 | nktr |
| zc3h7ba | 1.268848 | 0.273349032 | 0.182 | 0.028 | 3.2235089 | 21 | zc3h7ba |
| calr3b2 | 1.320791 | 0.437982432 | 0.358 | 0.092 | 3.3554712 | 21 | calr3b |
| tmem59 | 1.691494 | 0.465233029 | 0.426 | 0.124 | 4.2972423 | 21 | tmem59 |
| kif5bb | 1.719859 | 0.275196858 | 0.182 | 0.029 | 4.3693039 | 21 | kif5bb |
| pfn25 | 1.874488 | 0.593864164 | 0.631 | 0.236 | 4.7621370 | 21 | pfn2 |
| tomm61 | 1.964730 | 0.431659197 | 0.438 | 0.126 | 4.9913986 | 21 | tomm6 |
| ndrg3a | 2.503617 | 0.251344476 | 0.159 | 0.022 | 6.3604406 | 21 | ndrg3a |

| | | | | | | | |
|---|---|---|---|---|---|---|---|
| si:ch73-42p12.2 | 4.334450 | 0.433576904 | 0.398 | 0.111 | 1.1011671 | 21 | si:ch73-42p12.2 |
| zgc:1951752 | 4.625778 | 0.323758369 | 0.205 | 0.036 | 1.1751789 | 21 | zgc:195175 |
| arl4d2 | 4.894835 | 0.348140208 | 0.256 | 0.052 | 1.2435329 | 21 | arl4d |
| il153 | 5.255964 | 0.371150861 | 0.273 | 0.058 | 1.3352778 | 21 | il15 |
| si:zfos-464b6.21 | 7.254363 | 0.334601076 | 0.239 | 0.047 | 1.8429710 | 21 | si:zfos-464b6.2 |
| pfn2l5 | 7.814057 | 0.514917177 | 0.619 | 0.212 | 1.9851613 | 21 | pfn2l |
| rrbp1a | 8.420815 | 0.365037556 | 0.267 | 0.057 | 2.1393082 | 21 | rrbp1a |
| arhgef52 | 1.209342 | 0.331409746 | 0.188 | 0.031 | 3.0723350 | 21 | arhgef5 |
| CABZ01072614.17 | 1.834962 | 0.389107704 | 0.358 | 0.093 | 4.6617225 | 21 | CABZ01072614.1 |
| med4 | 1.877084 | 0.33009848 | 0.227 | 0.043 | 4.7687324 | 21 | med4 |
| LEPROTL1 | 2.033313 | 0.440362346 | 0.358 | 0.095 | 5.1656336 | 21 | LEPROTL1 |
| itga93 | 2.468416 | 0.366112199 | 0.205 | 0.037 | 6.2710115 | 21 | itga9 |
| gabarapb6 | 2.503621 | 0.567036599 | 0.727 | 0.301 | 6.3604516 | 21 | gabarapb |
| pfkfb4b3 | 2.666589 | 0.543812338 | 0.398 | 0.118 | 6.7744713 | 21 | pfkfb4b |
| cdc422 | 3.009352 | 0.526909524 | 0.625 | 0.23 | 7.6452593 | 21 | cdc42 |
| CU984600.2 | 4.229681 | 0.261121125 | 0.165 | 0.025 | 1.0745505 | 21 | CU984600.2 |
| cited4a1 | 4.579616 | 0.3157009 | 0.216 | 0.04 | 1.1634515 | 21 | cited4a |
| ier3ip1 | 5.004490 | 0.498893483 | 0.489 | 0.161 | 1.2713906 | 21 | ier3ip1 |
| ostc1 | 6.087264 | 0.51430583 | 0.545 | 0.187 | 1.5464694 | 21 | ostc |
| LOC1018844513 | 7.717292 | 0.442513623 | 0.341 | 0.089 | 1.9605782 | 21 | LOC101884451 |
| tmem30aa2 | 7.957849 | 0.385186273 | 0.29 | 0.068 | 2.0216917 | 21 | tmem30aa |
| tp53 | 8.294769 | 0.446053238 | 0.398 | 0.113 | 2.1072862 | 21 | tp53 |
| krtcap21 | 1.036622 | 0.49824903 | 0.557 | 0.192 | 2.6335385 | 21 | krtcap2 |
| mgst3b3 | 1.051646 | 0.519567532 | 0.54 | 0.191 | 2.6717074 | 21 | mgst3b |
| fstl34 | 1.441088 | 0.278763793 | 0.159 | 0.024 | 3.6610848 | 21 | fstl3 |
| ndufs71 | 1.493550 | 0.398280439 | 0.426 | 0.124 | 3.7943658 | 21 | ndufs7 |
| oaz1a1 | 1.628270 | 0.573379064 | 0.881 | 0.493 | 4.1366214 | 21 | oaz1a |
| bcap31 | 1.727134 | 0.384449391 | 0.29 | 0.068 | 4.3877841 | 21 | bcap31 |
| tgif13 | 2.397195 | 0.406982118 | 0.364 | 0.098 | 6.0900758 | 21 | tgif1 |
| atp1a3b3 | 3.551057 | 0.330779686 | 0.188 | 0.032 | 9.0214615 | 21 | atp1a3b |
| puraa | 4.245003 | 0.304296456 | 0.216 | 0.041 | 1.0784430 | 21 | puraa |
| stk35l1 | 4.600024 | 0.298307226 | 0.21 | 0.039 | 1.1686362 | 21 | stk35l |
| fam120c2 | 5.429078 | 0.42808804 | 0.369 | 0.104 | 1.3792575 | 21 | fam120c |
| ralbp1 | 5.546893 | 0.330072102 | 0.267 | 0.06 | 1.4091883 | 21 | ralbp1 |
| psmd4b1 | 5.815807 | 0.345637172 | 0.284 | 0.066 | 1.4775058 | 21 | psmd4b |

| | | | | | | | |
|---|---|---|---|---|---|---|---|
| abcf11 | 5.894021 | 0.428700902 | 0.381 | 0.107 | 1.4973761 | 21 | abcf1 |
| nucks1a1 | 7.513980 | 0.51890283 | 0.534 | 0.185 | 1.9089268 | 21 | nucks1a |
| cdv3 | 7.533164 | 0.421011725 | 0.466 | 0.145 | 1.9138003 | 21 | cdv3 |
| atf4b6 | 7.860057 | 0.549932544 | 0.688 | 0.274 | 1.9968476 | 21 | atf4b |
| ing4 | 8.380182 | 0.285958771 | 0.222 | 0.043 | 2.1289854 | 21 | ing4 |
| psmb71 | 9.137760 | 0.456174301 | 0.517 | 0.17 | 2.3214479 | 21 | psmb7 |
| si:ch211-132p1.34 | 1.164113 | 0.457172044 | 0.347 | 0.095 | 2.9574309 | 21 | si:ch211-132p1.3 |
| hvcn1 | 1.256380 | 0.265507133 | 0.188 | 0.032 | 3.1918340 | 21 | hvcn1 |
| akap12b4 | 1.474264 | 0.418869668 | 0.301 | 0.072 | 3.7453676 | 21 | akap12b |
| tob1a | 1.843007 | 0.470380142 | 0.449 | 0.142 | 4.6821599 | 21 | tob1a |
| pomp2 | 2.365845 | 0.551738757 | 0.636 | 0.257 | 6.0104300 | 21 | pomp |
| ccnd1 | 3.328604 | 0.324477451 | 0.216 | 0.043 | 8.4563198 | 21 | ccnd1 |
| glula3 | 3.333481 | 0.484279136 | 0.415 | 0.123 | 8.4687099 | 21 | glula |
| etnk1 | 3.813199 | 0.293254701 | 0.199 | 0.037 | 9.6874322 | 21 | etnk1 |
| zgc:1101822 | 8.149871 | 0.36494986 | 0.335 | 0.084 | 2.0704749 | 21 | zgc:110182 |
| egfra4 | 9.886069 | 0.275397336 | 0.182 | 0.032 | 2.5115559 | 21 | egfra |
| mvp3 | 1.094060 | 0.334148165 | 0.261 | 0.059 | 2.7794598 | 21 | mvp |
| hspd14 | 1.183684 | 0.466689332 | 0.494 | 0.161 | 3.0071508 | 21 | hspd1 |
| uqcrc12 | 1.200498 | 0.373243616 | 0.375 | 0.106 | 3.0498662 | 21 | uqcrc1 |
| si:dkey-6n21.123 | 1.657940 | 0.289368692 | 0.21 | 0.041 | 4.2119985 | 21 | si:dkey-6n21.12 |
| ndufb4 | 1.710654 | 0.46556312 | 0.608 | 0.225 | 4.3459175 | 21 | ndufb4 |
| fbxl61 | 1.889406 | 0.359762093 | 0.267 | 0.062 | 4.8000367 | 21 | fbxl6 |
| CABZ01080702.16 | 1.995309 | 0.430458383 | 0.347 | 0.096 | 5.0690843 | 21 | CABZ01080702.1 |
| nt5c2b2 | 2.358516 | 0.328885362 | 0.222 | 0.046 | 5.9918108 | 21 | nt5c2b |
| atp6v1e1a | 2.673741 | 0.345435741 | 0.25 | 0.056 | 6.7926408 | 21 | atp6v1e1a |
| si:ch211-114c12.2 | 2.763164 | 0.329501221 | 0.216 | 0.044 | 7.0198187 | 21 | si:ch211-114c12.2 |
| si:ch73-376l24.21 | 2.992632 | 0.40172573 | 0.409 | 0.121 | 7.6027834 | 21 | si:ch73-376l24.2 |
| nars1 | 3.386471 | 0.358204173 | 0.312 | 0.081 | 8.6033309 | 21 | nars |
| tcf3b | 4.748068 | 0.257327208 | 0.176 | 0.03 | 1.2062468 | 21 | tcf3b |
| zgc:926062 | 5.145754 | 0.382477365 | 0.341 | 0.094 | 1.3072789 | 21 | zgc:92606 |
| mafba5 | 6.533247 | 0.41503094 | 0.312 | 0.083 | 1.6597714 | 21 | mafba |
| chd7 | 7.731845 | 0.324974858 | 0.244 | 0.053 | 1.9642752 | 21 | chd7 |
| gli32 | 7.987688 | 0.29803025 | 0.188 | 0.035 | 2.0292722 | 21 | gli3 |
| fdxr1 | 9.062646 | 0.266077529 | 0.165 | 0.027 | 2.3023653 | 21 | fdxr |
| mif1 | 1.286975 | 0.473652783 | 0.557 | 0.204 | 3.2695620 | 21 | mif |

| | | | | | | | |
|---|---|---|---|---|---|---|---|
| si:ch211-191j22.31 | 1.309563 | 0.411220166 | 0.398 | 0.121 | 3.3269453 | 21 | si:ch211-191j22.3 |
| wls5 | 1.357729 | 0.411019861 | 0.42 | 0.129 | 3.4493117 | 21 | wls |
| tmed71 | 1.388242 | 0.479553434 | 0.58 | 0.218 | 3.5268301 | 21 | tmed7 |
| sltm1 | 1.688767 | 0.377245465 | 0.278 | 0.069 | 4.2903146 | 21 | sltm |
| pam | 2.012366 | 0.270372627 | 0.153 | 0.024 | 5.1124179 | 21 | pam |
| stat22 | 2.725421 | 0.352800122 | 0.261 | 0.062 | 6.9239324 | 21 | stat2 |
| anxa133 | 2.994637 | 0.310254759 | 0.239 | 0.053 | 7.6078757 | 21 | anxa13 |
| flot2a5 | 3.875698 | 0.3677013 | 0.284 | 0.072 | 9.8462132 | 21 | flot2a |
| igf2b4 | 3.923862 | 0.373695777 | 0.295 | 0.074 | 9.9685728 | 21 | igf2b |
| parp12b | 4.831032 | 0.287292358 | 0.193 | 0.037 | 1.2273238 | 21 | parp12b |
| atp5mea1 | 5.062907 | 0.438541697 | 0.455 | 0.151 | 1.2862315 | 21 | atp5mea |
| nfic5 | 5.365071 | 0.402785783 | 0.307 | 0.082 | 1.3629964 | 21 | nfic |
| rab105 | 5.447745 | 0.454288777 | 0.574 | 0.211 | 1.3839996 | 21 | rab10 |
| slc38a10 | 6.333046 | 0.271809711 | 0.176 | 0.032 | 1.6089104 | 21 | slc38a10 |
| si:ch211-212k18.72 | 6.362583 | 0.452399976 | 0.443 | 0.144 | 1.6164143 | 21 | si:ch211-212k18.7 |
| ndufs51 | 7.248034 | 0.394706953 | 0.483 | 0.162 | 1.8413632 | 21 | ndufs5 |
| prdx32 | 7.552400 | 0.40030147 | 0.438 | 0.14 | 1.9186872 | 21 | prdx3 |
| edf12 | 8.730175 | 0.490483648 | 0.688 | 0.284 | 2.2179010 | 21 | edf1 |
| gnb1b2 | 9.851832 | 0.511587335 | 0.636 | 0.263 | 2.5028580 | 21 | gnb1b |
| qkia5 | 1.200757 | 0.490586792 | 0.534 | 0.197 | 3.0505240 | 21 | qkia |
| cmtm6 | 1.205758 | 0.342421298 | 0.244 | 0.056 | 3.0632307 | 21 | cmtm6 |
| rhoaa5 | 1.325419 | 0.473543331 | 0.483 | 0.174 | 3.3672281 | 21 | rhoaa |
| ptbp1a | 1.350031 | 0.38944315 | 0.318 | 0.088 | 3.4297543 | 21 | ptbp1a |
| cald1a5 | 1.485904 | 0.415854908 | 0.392 | 0.118 | 3.7749414 | 21 | cald1a |
| ube2e35 | 1.514810 | 0.461581483 | 0.477 | 0.168 | 3.8483763 | 21 | ube2e3 |
| atp5f1c1 | 1.839094 | 0.46312783 | 0.67 | 0.266 | 4.6722204 | 21 | atp5f1c |
| zbtb461 | 1.882435 | 0.252025507 | 0.159 | 0.027 | 4.7823263 | 21 | zbtb46 |
| ndufb91 | 2.094442 | 0.438280941 | 0.54 | 0.198 | 5.3209318 | 21 | ndufb9 |
| dido1 | 2.386601 | 0.376567357 | 0.341 | 0.097 | 6.0631618 | 21 | dido1 |
| ap1m1 | 2.522220 | 0.360433414 | 0.29 | 0.075 | 6.4077006 | 21 | ap1m1 |
| si:dkey-28b4.76 | 3.119869 | 0.387609132 | 0.364 | 0.108 | 7.9260274 | 21 | si:dkey-28b4.7 |
| zgc:1629443 | 3.402801 | 0.498533637 | 0.665 | 0.272 | 8.6448164 | 21 | zgc:162944 |
| si:dkey-56m19.52 | 3.926006 | 0.338936615 | 0.239 | 0.055 | 9.9740186 | 21 | si:dkey-56m19.5 |
| zgc:77086 | 3.940943 | 0.336158243 | 0.256 | 0.061 | 1.0011967 | 21 | zgc:77086 |
| vdac33 | 4.382777 | 0.423565723 | 0.483 | 0.166 | 1.1134446 | 21 | vdac3 |

| | | | | | | | |
|---|---|---|---|---|---|---|---|
| irf9 | 4.594835 | 0.253918887 | 0.148 | 0.024 | 1.1673178 | 21 | irf9 |
| pkp3b2 | 4.696047 | 0.303043108 | 0.222 | 0.048 | 1.1930307 | 21 | pkp3b |
| rab5c1 | 5.690896 | 0.418488493 | 0.403 | 0.129 | 1.4457722 | 21 | rab5c |
| ucp25 | 6.140090 | 0.544297808 | 0.847 | 0.411 | 1.5598900 | 21 | ucp2 |
| sptan12 | 6.404333 | 0.324437321 | 0.244 | 0.057 | 1.6270208 | 21 | sptan1 |
| ndufa31 | 8.108274 | 0.447246521 | 0.517 | 0.189 | 2.0599072 | 21 | ndufa3 |
| zfand3 | 8.592458 | 0.299896709 | 0.267 | 0.066 | 2.1829140 | 21 | zfand3 |
| hif1ab1 | 1.017623 | 0.424223157 | 0.381 | 0.12 | 2.5852719 | 21 | hif1ab |
| ndufs21 | 1.212730 | 0.329524921 | 0.335 | 0.095 | 3.0809418 | 21 | ndufs2 |
| ch25hl3 | 1.290645 | 0.292814832 | 0.233 | 0.052 | 3.2788853 | 21 | ch25hl3 |
| sypl2a2 | 1.387818 | 0.307887767 | 0.25 | 0.059 | 3.5257531 | 21 | sypl2a |
| rbx11 | 1.659102 | 0.48116988 | 0.67 | 0.278 | 4.2149500 | 21 | rbx1 |
| myl9a2 | 1.766480 | 0.281992979 | 0.21 | 0.044 | 4.4877430 | 21 | myl9a |
| jun6 | 1.921193 | 0.570713386 | 0.614 | 0.247 | 4.8807911 | 21 | jun |
| ccl19a.1 | 1.997705 | 0.280554784 | 0.108 | 0.014 | 5.0751696 | 21 | ccl19a.1 |
| mtdha | 2.043102 | 0.395152246 | 0.438 | 0.145 | 5.1905030 | 21 | mtdha |
| elocb2 | 2.100201 | 0.452457255 | 0.517 | 0.193 | 5.3355608 | 21 | elocb |
| agpat31 | 2.107509 | 0.263497298 | 0.188 | 0.037 | 5.3541281 | 21 | agpat3 |
| spcs21 | 2.320066 | 0.441309452 | 0.483 | 0.171 | 5.8941282 | 21 | spcs2 |
| rbp53 | 2.339062 | 0.462204422 | 0.358 | 0.109 | 5.9423888 | 21 | rbp5 |
| hsd11b25 | 2.718479 | 0.49392965 | 0.455 | 0.147 | 6.9062975 | 21 | hsd11b2 |
| zgc:1584633 | 2.862622 | 0.331169509 | 0.295 | 0.079 | 7.2724912 | 21 | zgc:158463 |
| ndufs41 | 2.975111 | 0.341414431 | 0.364 | 0.109 | 7.5582700 | 21 | ndufs4 |
| crtap2 | 3.229099 | 0.270924244 | 0.193 | 0.039 | 8.2035261 | 21 | crtap |
| psmb31 | 3.548825 | 0.409762238 | 0.449 | 0.153 | 9.0157906 | 21 | psmb3 |
| spcs11 | 3.570197 | 0.473191971 | 0.574 | 0.229 | 9.0700865 | 21 | spcs1 |
| her61 | 3.887461 | 0.280125975 | 0.148 | 0.025 | 9.8760965 | 21 | her6 |
| si:zfos-169g10.31 | 3.919824 | 0.395550297 | 0.364 | 0.113 | 9.9583150 | 21 | si:zfos-169g10.3 |
| fmr1 | 4.017173 | 0.284776126 | 0.216 | 0.047 | 1.0205628 | 21 | fmr1 |
| midn5 | 4.063735 | 0.463613678 | 0.42 | 0.143 | 1.0323921 | 21 | midn |
| perp2 | 4.707279 | 0.288064525 | 0.25 | 0.06 | 1.1958842 | 21 | perp |
| psmd3 | 5.029803 | 0.342160041 | 0.307 | 0.084 | 1.2778216 | 21 | psmd3 |
| atp5pb2 | 8.739596 | 0.486095143 | 0.739 | 0.329 | 2.2202944 | 21 | atp5pb |
| crebzf3 | 9.233874 | 0.396371312 | 0.352 | 0.108 | 2.3458657 | 21 | crebzf |
| sumo3a2 | 1.170366 | 0.525867489 | 0.795 | 0.368 | 2.9733153 | 21 | sumo3a |

| | | | | | | | |
|---|---|---|---|---|---|---|---|
| zgc:1231051 | 1.212069 | 0.250247717 | 0.216 | 0.047 | 3.0792626 | 21 | zgc:123105 |
| si:ch73-376l24.3 | 1.9570966 | 0.402989911 | 0.381 | 0.121 | 4.9720025 | 21 | si:ch73-376l24.3 |
| erp44 | 1.995625 | 0.35488937 | 0.312 | 0.089 | 5.0698854 | 21 | erp44 |
| ppt12 | 2.0110899 | 0.431297919 | 0.494 | 0.179 | 5.1091740 | 21 | ppt1 |
| tmem30ab | 2.1302188 | 0.304816317 | 0.239 | 0.057 | 5.4118207 | 21 | tmem30ab |
| eef1db6 | 3.296871 | 0.490790999 | 0.659 | 0.281 | 8.3757018 | 21 | eef1db |
| fundc22 | 3.433305 | 0.344369031 | 0.324 | 0.095 | 8.7223127 | 21 | fundc2 |
| ctsd2 | 3.963345 | 0.321302027 | 0.449 | 0.145 | 1.0068878 | 21 | ctsd |
| top2b | 4.085631 | 0.279035258 | 0.188 | 0.039 | 1.0379547 | 21 | top2b |
| marcksl1a6 | 4.548515 | 0.466566404 | 0.648 | 0.258 | 1.1555504 | 21 | marcksl1a |
| selenot2 | 4.642291 | 0.316661721 | 0.256 | 0.065 | 1.1793742 | 21 | selenot2 |
| tnrc6a1 | 5.113682 | 0.324919453 | 0.29 | 0.079 | 1.2991309 | 21 | tnrc6a |
| sf3b51 | 5.670072 | 0.411637841 | 0.443 | 0.153 | 1.4404819 | 21 | sf3b5 |
| ctsh2 | 6.583237 | 0.345302215 | 0.256 | 0.066 | 1.6724715 | 21 | ctsh |
| si:ch211-156j16.11 | 7.159702 | 0.26711834 | 0.153 | 0.027 | 1.8189225 | 21 | si:ch211-156j16.1 |
| arhgef42 | 9.066118 | 0.275991807 | 0.176 | 0.035 | 2.3032473 | 21 | arhgef4 |
| ndufab1a1 | 9.596700 | 0.389641604 | 0.415 | 0.14 | 2.4380416 | 21 | ndufab1a |
| smarca4a | 1.072284 | 0.307027362 | 0.25 | 0.063 | 2.7241397 | 21 | smarca4a |
| rbm39a3 | 1.211137 | 0.414485105 | 0.432 | 0.151 | 3.0768944 | 21 | rbm39a |
| mt-nd54 | 1.267523 | 0.346961245 | 0.392 | 0.128 | 3.2201423 | 21 | mt-nd5 |
| ppp2cb | 1.353460 | 0.407921466 | 0.483 | 0.176 | 3.4384654 | 21 | ppp2cb |
| ccdc85b5 | 1.358628 | 0.290454748 | 0.216 | 0.05 | 3.4515957 | 21 | ccdc85b |
| mt-nd410 | 1.502677 | 0.535370153 | 0.807 | 0.445 | 3.8175527 | 21 | mt-nd4 |
| matr3l1.11 | 1.555519 | 0.322761993 | 0.295 | 0.082 | 3.9517976 | 21 | matr3l1.1 |
| golgb1 | 1.943785 | 0.276705861 | 0.216 | 0.05 | 4.9381863 | 21 | golgb1 |
| pgd2 | 2.147026 | 0.299324432 | 0.267 | 0.07 | 5.4545205 | 21 | pgd |
| actr1 | 2.252258 | 0.282456799 | 0.244 | 0.06 | 5.7218620 | 21 | actr1 |
| tmed9 | 2.420779 | 0.378706212 | 0.443 | 0.152 | 6.1499893 | 21 | tmed9 |
| gstm.2 | 2.611846 | 0.300809313 | 0.239 | 0.06 | 6.6353949 | 21 | gstm.2 |
| mt-nd28 | 2.777520 | 0.532829 | 0.756 | 0.389 | 7.0562910 | 21 | mt-nd2 |
| sdc47 | 2.993526 | 0.46321201 | 0.636 | 0.251 | 7.6050542 | 21 | sdc4 |
| srp54 | 3.139429 | 0.292293693 | 0.233 | 0.057 | 7.9757213 | 21 | srp54 |
| ddx613 | 3.268584 | 0.45157305 | 0.574 | 0.232 | 8.3038377 | 21 | ddx61 |
| glrx5 | 3.433032 | 0.320146994 | 0.324 | 0.095 | 8.7216202 | 21 | glrx5 |
| mtm1 | 3.784051 | 0.25570354 | 0.17 | 0.034 | 9.6133826 | 21 | mtm1 |

| | | | | | | | |
|---|---|---|---|---|---|---|---|
| rpn11 | 4.495583 | 0.414630301 | 0.443 | 0.159 | 1.1421028 | 21 | rpn1 |
| mycbp2 | 4.656635 | 0.266256425 | 0.193 | 0.042 | 1.1830181 | 21 | mycbp2 |
| cpne12 | 4.838393 | 0.381985854 | 0.392 | 0.132 | 1.2291939 | 21 | cpne1 |
| tmem14ca | 4.846403 | 0.302055458 | 0.324 | 0.096 | 1.2312287 | 21 | tmem14ca |
| atf5b1 | 5.109792 | 0.273595772 | 0.182 | 0.038 | 1.2981428 | 21 | atf5b |
| hmgb2b4 | 5.726094 | 0.522132854 | 0.722 | 0.333 | 1.4547142 | 21 | hmgb2b |
| tbca1 | 6.410747 | 0.433123746 | 0.528 | 0.209 | 1.6286504 | 21 | tbca |
| rras | 6.905115 | 0.346389629 | 0.352 | 0.108 | 1.7542445 | 21 | rras |
| si:rp71-45k5.41 | 7.825479 | 0.356340357 | 0.381 | 0.125 | 1.9880629 | 21 | si:rp71-45k5.4 |
| kif5ba | 7.978484 | 0.270180508 | 0.222 | 0.053 | 2.0269339 | 21 | kif5ba |
| psma11 | 8.650900 | 0.465758892 | 0.625 | 0.269 | 2.1977611 | 21 | psma1 |
| snx12 | 8.789361 | 0.316024194 | 0.29 | 0.082 | 2.2329371 | 21 | snx12 |
| rsf1b.1 | 9.800853 | 0.313642685 | 0.233 | 0.059 | 2.4899069 | 21 | rsf1b.1 |
| tm2d3 | 1.036484 | 0.306338351 | 0.267 | 0.072 | 2.6331883 | 21 | tm2d3 |
| chchd6b | 1.050828 | 0.332564263 | 0.301 | 0.087 | 2.6696306 | 21 | chchd6b |
| ogdha | 1.297099 | 0.321141392 | 0.267 | 0.073 | 3.2952817 | 21 | ogdha |
| cd151l1 | 1.694021 | 0.290863186 | 0.244 | 0.062 | 4.3036607 | 21 | cd151l |
| tsc22d34 | 1.732121 | 0.405814671 | 0.438 | 0.157 | 4.4004553 | 21 | tsc22d3 |
| psmb10 | 1.919141 | 0.265236364 | 0.21 | 0.049 | 4.8755782 | 21 | psmb10 |
| abl1 | 1.942402 | 0.253559861 | 0.176 | 0.037 | 4.9346723 | 21 | abl1 |
| rbm4.1 | 2.097435 | 0.299196226 | 0.278 | 0.076 | 5.3285358 | 21 | rbm4.1 |
| myl9b5 | 2.117322 | 0.39899684 | 0.318 | 0.097 | 5.3790573 | 21 | myl9b |
| dpy301 | 2.307928 | 0.343621092 | 0.455 | 0.161 | 5.8632927 | 21 | dpy30 |
| antxr1c3 | 2.500369 | 0.267680828 | 0.182 | 0.039 | 6.3521878 | 21 | antxr1c |
| gatad2b | 2.708135 | 0.333302671 | 0.25 | 0.067 | 6.8800180 | 21 | gatad2b |
| pik3r23 | 3.233553 | 0.313959339 | 0.25 | 0.066 | 8.2148436 | 21 | pik3r2 |
| dnajc81 | 3.302366 | 0.26986649 | 0.273 | 0.074 | 8.3896615 | 21 | dnajc8 |
| kmt2bb1 | 3.498523 | 0.351119403 | 0.312 | 0.095 | 8.8879983 | 21 | kmt2bb |
| brd9 | 4.176290 | 0.28546869 | 0.205 | 0.048 | 1.0609865 | 21 | brd9 |
| aldh9a1a.16 | 5.457173 | 0.472706123 | 0.477 | 0.191 | 1.3863948 | 21 | aldh9a1a.1 |
| srsf5a5 | 5.460207 | 0.459687231 | 0.67 | 0.301 | 1.3871657 | 21 | srsf5a |
| mesd | 5.779177 | 0.279557646 | 0.256 | 0.068 | 1.4682000 | 21 | mesd |
| cox5ab1 | 6.123107 | 0.364324544 | 0.551 | 0.215 | 1.5555754 | 21 | cox5ab |
| tjp2b3 | 6.431353 | 0.280988065 | 0.244 | 0.063 | 1.6338853 | 21 | tjp2b |
| wu:fb16f033 | 6.634990 | 0.359667329 | 0.295 | 0.088 | 1.6856192 | 21 | wu:fb16f03 |

| | | | | | | | |
|---|---|---|---|---|---|---|---|
| ndufb71 | 6.954687 | 0.364406373 | 0.506 | 0.192 | 1.7668384 | 21 | ndufb7 |
| tpp12 | 8.554582 | 0.320700897 | 0.273 | 0.076 | 2.1732917 | 21 | tpp1 |
| bcl2a | 9.821986 | 0.269974373 | 0.159 | 0.032 | 2.4952755 | 21 | bcl2a |
| sar1b1 | 1.043758 | 0.341845728 | 0.295 | 0.088 | 2.6516676 | 21 | sar1b |
| si:ch211-106e7.2 | 1.186248 | 0.255870833 | 0.188 | 0.041 | 3.0136638 | 21 | si:ch211-106e7.2 |
| stau26 | 1.347614 | 0.355212805 | 0.307 | 0.095 | 3.4236157 | 21 | stau2 |
| sap30l | 1.396361 | 0.345050329 | 0.42 | 0.149 | 3.5474566 | 21 | sap30l |
| calm3b1 | 1.504484 | 0.469663837 | 0.619 | 0.276 | 3.8221427 | 21 | calm3b |
| si:dkey-42i9.42 | 1.683092 | 0.432276222 | 0.415 | 0.155 | 4.2758961 | 21 | si:dkey-42i9.4 |
| hmgn75 | 1.877326 | 0.467974696 | 0.693 | 0.315 | 4.7693490 | 21 | hmgn7 |
| dynlt31 | 1.952623 | 0.377156018 | 0.426 | 0.156 | 4.9606408 | 21 | dynlt3 |
| pkig | 1.977973 | 0.334521428 | 0.318 | 0.097 | 5.0250406 | 21 | pkig |
| txndc15 | 2.006673 | 0.305031678 | 0.267 | 0.075 | 5.0979543 | 21 | txndc15 |
| diaph21 | 2.253132 | 0.258562502 | 0.165 | 0.034 | 5.7240821 | 21 | diaph2 |
| si:dkey-44g23.51 | 2.300062 | 0.25628293 | 0.25 | 0.065 | 5.8433076 | 21 | si:dkey-44g23.5 |
| ms4a17a.112 | 2.314865 | 0.26150867 | 0.21 | 0.051 | 5.8809166 | 21 | ms4a17a.11 |
| atp5meb2 | 2.362137 | 0.423025581 | 0.739 | 0.336 | 6.0010104 | 21 | atp5meb |
| atp8a1 | 2.419794 | 0.295327947 | 0.17 | 0.036 | 6.1474871 | 21 | atp8a1 |
| slka | 3.080999 | 0.256692754 | 0.176 | 0.038 | 7.8272804 | 21 | slka |
| ap2s1 | 3.175374 | 0.264134994 | 0.273 | 0.077 | 8.0670397 | 21 | ap2s1 |
| FO082781.16 | 3.227593 | 0.318986336 | 0.216 | 0.054 | 8.1997002 | 21 | FO082781.1 |
| trappc6b | 3.712044 | 0.255756613 | 0.193 | 0.044 | 9.4304483 | 21 | trappc6b |
| BX284696.11 | 3.742104 | 0.281916623 | 0.227 | 0.059 | 9.5068171 | 21 | BX284696.1 |
| rnf101 | 4.826528 | 0.350698604 | 0.369 | 0.127 | 1.2261796 | 21 | rnf10 |
| ptmaa9 | 5.884646 | 0.508114947 | 0.83 | 0.43 | 1.4949944 | 21 | ptmaa |
| capns1b4 | 6.520251 | 0.380107618 | 0.398 | 0.144 | 1.6564699 | 21 | capns1b |
| ap1s2 | 6.905359 | 0.355542097 | 0.386 | 0.133 | 1.7543065 | 21 | ap1s2 |
| csnk2b1 | 7.364610 | 0.415349329 | 0.528 | 0.218 | 1.8709793 | 21 | csnk2b |
| ndufa7 | 8.043291 | 0.333407585 | 0.335 | 0.108 | 2.0433981 | 21 | ndufa7 |
| eny21 | 8.384874 | 0.34474696 | 0.341 | 0.111 | 2.1301772 | 21 | eny2 |
| dad11 | 8.942835 | 0.469996351 | 0.739 | 0.355 | 2.2719274 | 21 | dad1 |
| calm1a2 | 9.826130 | 0.397390136 | 0.455 | 0.172 | 2.4963284 | 21 | calm1a |
| serinc23 | 1.026030 | 0.371027417 | 0.358 | 0.123 | 2.6066316 | 21 | serinc2 |
| sec11a1 | 1.105026 | 0.387209243 | 0.483 | 0.188 | 2.8073194 | 21 | sec11a |
| mt-co110 | 1.158637 | 0.463801927 | 0.909 | 0.638 | 2.9435175 | 21 | mt-co1 |

| | | | | | | | |
|---|---|---|---|---|---|---|---|
| rps1010 | 1.166492 | 0.383580385 | 0.983 | 0.725 | 2.9634740 | 21 | rps10 |
| mtpn2 | 1.187022 | 0.392024634 | 0.426 | 0.162 | 3.0156296 | 21 | mtpn |
| si:ch211-213a13.12 | 1.196906 | 0.335464871 | 0.307 | 0.096 | 3.0407419 | 21 | si:ch211-213a13.1 |
| eci1 | 1.435302 | 0.270523111 | 0.227 | 0.059 | 3.6463866 | 21 | eci1 |
| snx6 | 1.443450 | 0.250987969 | 0.267 | 0.075 | 3.6670853 | 21 | snx6 |
| ogt.11 | 1.841366 | 0.314356605 | 0.256 | 0.073 | 4.6779926 | 21 | ogt.1 |
| zgc:56095 | 1.881912 | 0.258073667 | 0.324 | 0.1 | 4.7809990 | 21 | zgc:56095 |
| ncor11 | 2.182457 | 0.275284497 | 0.222 | 0.058 | 5.5445339 | 21 | ncor1 |
| wdr83os | 2.504528 | 0.348505906 | 0.415 | 0.149 | 6.3627539 | 21 | wdr83os |
| ctbp1 | 4.226432 | 0.351698248 | 0.392 | 0.138 | 1.0737251 | 21 | ctbp1 |
| lgmn2 | 4.570086 | 0.268934601 | 0.256 | 0.072 | 1.1610305 | 21 | lgmn |
| ganab | 5.170289 | 0.253110941 | 0.159 | 0.034 | 1.3135121 | 21 | ganab |
| echs11 | 5.171220 | 0.31176854 | 0.284 | 0.087 | 1.3137486 | 21 | echs1 |
| atp2b24 | 5.441746 | 0.320750842 | 0.273 | 0.082 | 1.3824757 | 21 | atp2b2 |
| slc39a7 | 5.780582 | 0.253368991 | 0.182 | 0.043 | 1.4685570 | 21 | slc39a7 |
| rmdn11 | 5.859475 | 0.26143065 | 0.233 | 0.063 | 1.4885997 | 21 | rmdn1 |
| ctdsp25 | 7.304051 | 0.300626557 | 0.278 | 0.084 | 1.8555942 | 21 | ctdsp2 |
| rab2a1 | 8.659520 | 0.365507518 | 0.455 | 0.175 | 2.1999512 | 21 | rab2a |
| pdia5 | 8.984109 | 0.268559342 | 0.199 | 0.049 | 2.2824131 | 21 | pdia5 |
| fthl27 | 9.226706 | 0.401666375 | 0.943 | 0.68 | 2.3440448 | 21 | fthl27 |
| atp5pd4 | 9.292654 | 0.392457546 | 0.631 | 0.28 | 2.3607988 | 21 | atp5pd |
| selenot1b | 1.014938 | 0.286386885 | 0.273 | 0.082 | 2.5784501 | 21 | selenot1b |
| dctn3 | 1.085075 | 0.313819156 | 0.307 | 0.099 | 2.7566350 | 21 | dctn3 |
| psmb61 | 1.118761 | 0.368214288 | 0.472 | 0.186 | 2.8422141 | 21 | psmb6 |
| sik13 | 1.120943 | 0.358807105 | 0.33 | 0.109 | 2.8477558 | 21 | sik1 |
| ndufb101 | 1.181881 | 0.327255701 | 0.432 | 0.162 | 3.0025697 | 21 | ndufb10 |
| laptm4a2 | 1.336972 | 0.314304176 | 0.352 | 0.121 | 3.3965780 | 21 | laptm4a |
| rab7a | 1.480030 | 0.29637712 | 0.244 | 0.07 | 3.7600164 | 21 | rab7a |
| nudcd21 | 1.842211 | 0.256289309 | 0.239 | 0.066 | 4.6801384 | 21 | nudcd2 |
| rab11a | 1.877838 | 0.312642252 | 0.364 | 0.126 | 4.7706498 | 21 | rab11a |
| trappc2l | 1.942027 | 0.251732493 | 0.21 | 0.055 | 4.9337205 | 21 | trappc2l |
| eif5b1 | 2.085735 | 0.361311663 | 0.403 | 0.15 | 5.2988114 | 21 | eif5b |
| reep54 | 2.695224 | 0.415940637 | 0.426 | 0.172 | 6.8472177 | 21 | reep5 |
| si:ch73-281n10.25 | 3.226129 | 0.490060311 | 0.784 | 0.393 | 8.1959831 | 21 | si:ch73-281n10.2 |
| gpatch8 | 3.330670 | 0.273743994 | 0.227 | 0.063 | 8.4615685 | 21 | gpatch8 |

| | | | | | | | |
|---|---|---|---|---|---|---|---|
| smarca2 | 3.896263 | 0.267584893 | 0.193 | 0.049 | 9.8984563 | 21 | smarca2 |
| chd36 | 4.325990 | 0.346316262 | 0.352 | 0.124 | 1.0990178 | 21 | chd3 |
| sik2b2 | 4.426871 | 0.273972278 | 0.182 | 0.044 | 1.1246468 | 21 | sik2b |
| ywhabl2 | 4.622324 | 0.422934501 | 0.648 | 0.298 | 1.1743014 | 21 | ywhabl |
| ptp4a2b2 | 4.684660 | 0.399880767 | 0.392 | 0.148 | 1.1901381 | 21 | ptp4a2b |
| smim13 | 5.096147 | 0.271456979 | 0.222 | 0.06 | 1.2946762 | 21 | smim13 |
| mgst3a | 5.269193 | 0.281731865 | 0.25 | 0.073 | 1.3386386 | 21 | mgst3a |
| tax1bp32 | 5.378845 | 0.304769242 | 0.307 | 0.1 | 1.3664955 | 21 | tax1bp3 |
| XLOC-0160944 | 5.744136 | 0.309496472 | 0.273 | 0.082 | 1.4592978 | 21 | XLOC-016094 |
| eif4g1a | 6.518382 | 0.331166887 | 0.341 | 0.118 | 1.6559950 | 21 | eif4g1a |
| psma51 | 1.030869 | 0.342363417 | 0.443 | 0.173 | 2.6189237 | 21 | psma5 |
| chmp5b1 | 1.087643 | 0.345003919 | 0.352 | 0.127 | 2.7631584 | 21 | chmp5b |
| cnbpb4 | 1.164945 | 0.395402457 | 0.489 | 0.208 | 2.9595431 | 21 | cnbpb |
| zgc:1119864 | 1.172630 | 0.360596342 | 0.46 | 0.184 | 2.9790675 | 21 | zgc:111986 |
| top1l2 | 1.214021 | 0.374253105 | 0.528 | 0.216 | 3.0842223 | 21 | top1l |
| tspan31 | 1.264479 | 0.267878338 | 0.199 | 0.051 | 3.2124096 | 21 | tspan31 |
| ptgr12 | 1.422277 | 0.305794206 | 0.324 | 0.107 | 3.6132971 | 21 | ptgr1 |
| rad21a1 | 1.461357 | 0.345785868 | 0.392 | 0.146 | 3.7125785 | 21 | rad21a |
| si:dkey-250d21.1 | 1.792033 | 0.307870896 | 0.267 | 0.084 | 4.5526611 | 21 | si:dkey-250d21.1 |
| hnrnpl21 | 2.178901 | 0.300961706 | 0.352 | 0.123 | 5.5354993 | 21 | hnrnpl2 |
| hnrnpa0a1 | 2.195606 | 0.295461103 | 0.261 | 0.081 | 5.5779381 | 21 | hnrnpa0a |
| ndufa81 | 2.243492 | 0.300343097 | 0.392 | 0.146 | 5.6995922 | 21 | ndufa8 |
| fam107b2 | 2.275531 | 0.392799668 | 0.318 | 0.111 | 5.7809884 | 21 | fam107b |
| psma41 | 2.554765 | 0.364313438 | 0.466 | 0.19 | 6.4903808 | 21 | psma4 |
| golga7 | 2.744497 | 0.334712175 | 0.42 | 0.161 | 6.9723948 | 21 | golga7 |
| eif1b5 | 2.825872 | 0.448314255 | 0.784 | 0.404 | 7.1791282 | 21 | eif1b |
| si:ch211-235e9.61 | 2.880391 | 0.298583131 | 0.42 | 0.159 | 7.3176357 | 21 | si:ch211-235e9.6 |
| mmgt1 | 2.982738 | 0.295492432 | 0.375 | 0.135 | 7.5776467 | 21 | mmgt1 |
| sem11 | 3.153419 | 0.396896614 | 0.597 | 0.271 | 8.0112611 | 21 | sem1 |
| lman2 | 3.585306 | 0.320345091 | 0.386 | 0.143 | 9.1084719 | 21 | lman2 |
| emc7 | 3.666103 | 0.280779321 | 0.278 | 0.088 | 9.3137365 | 21 | emc7 |
| ubb2 | 3.690677 | 0.437623494 | 0.665 | 0.323 | 9.3761674 | 21 | ubb |
| psenen | 4.401382 | 0.306301858 | 0.295 | 0.098 | 1.1181710 | 21 | psenen |
| rps27.28 | 4.673721 | 0.449287181 | 0.892 | 0.551 | 1.1873590 | 21 | rps27.2 |
| ube2v1 | 4.754973 | 0.389588049 | 0.5 | 0.217 | 1.2080010 | 21 | ube2v1 |

| | | | | | | | |
|---|---|---|---|---|---|---|---|
| arhgef1b3 | 5.218987 | 0.295834133 | 0.256 | 0.078 | 1.3258838 | 21 | arhgef1b |
| ppp2r1ba | 5.382205 | 0.272614466 | 0.29 | 0.094 | 1.3673492 | 21 | ppp2r1ba |
| eif4a1b3 | 5.437282 | 0.411089512 | 0.653 | 0.309 | 1.3813415 | 21 | eif4a1b |
| stat31 | 5.772453 | 0.251891504 | 0.205 | 0.055 | 1.4664919 | 21 | stat3 |
| txndc92 | 5.861326 | 0.33713947 | 0.347 | 0.128 | 1.4890700 | 21 | txndc9 |
| ywhae12 | 6.491398 | 0.4123347 | 0.625 | 0.292 | 1.6491396 | 21 | ywhae1 |
| etfa1 | 6.582836 | 0.265742691 | 0.295 | 0.097 | 1.6723695 | 21 | etfa |
| sub1a2 | 7.937306 | 0.37365347 | 0.568 | 0.251 | 2.0164726 | 21 | sub1a |
| psma31 | 8.085709 | 0.372827136 | 0.46 | 0.191 | 2.0541744 | 21 | psma3 |
| eif2s22 | 8.106964 | 0.402865141 | 0.67 | 0.32 | 2.0595744 | 21 | eif2s2 |
| arf2a | 8.783491 | 0.317199685 | 0.358 | 0.13 | 2.2314460 | 21 | arf2a |
| atp5mc3a2 | 9.330987 | 0.392763011 | 0.534 | 0.239 | 2.3705372 | 21 | atp5mc3a |
| psmc61 | 1.033951 | 0.267053241 | 0.29 | 0.094 | 2.6267540 | 21 | psmc6 |
| prkab1b1 | 1.072661 | 0.33261823 | 0.358 | 0.135 | 2.7250954 | 21 | prkab1b |
| mt-nd110 | 1.121014 | 0.464194258 | 0.705 | 0.395 | 2.8479366 | 21 | mt-nd1 |
| dlst | 1.177150 | 0.302719256 | 0.324 | 0.114 | 2.9905508 | 21 | dlst |
| atox11 | 1.383567 | 0.328407708 | 0.335 | 0.122 | 3.5149529 | 21 | atox1 |
| btf3l41 | 1.436706 | 0.298705359 | 0.386 | 0.145 | 3.6499520 | 21 | btf3l4 |
| lsm41 | 1.527336 | 0.332796108 | 0.517 | 0.215 | 3.8801989 | 21 | lsm4 |
| ndufb21 | 1.589110 | 0.333010537 | 0.472 | 0.193 | 4.0371355 | 21 | ndufb2 |
| ndufb32 | 1.593322 | 0.355087295 | 0.54 | 0.236 | 4.0478368 | 21 | ndufb3 |
| psmd131 | 1.672148 | 0.277195165 | 0.29 | 0.095 | 4.2480940 | 21 | psmd13 |
| sdhb | 1.776375 | 0.305545442 | 0.307 | 0.106 | 4.5128816 | 21 | sdhb |
| acin1a4 | 1.826283 | 0.371123878 | 0.523 | 0.227 | 4.6396730 | 21 | acin1a |
| ctsla6 | 1.950712 | 0.410903407 | 0.79 | 0.37 | 4.9557851 | 21 | ctsla |
| ube2v23 | 2.209013 | 0.3829741 | 0.608 | 0.28 | 5.6119996 | 21 | ube2v2 |
| cav13 | 2.306244 | 0.263359606 | 0.188 | 0.049 | 5.8590152 | 21 | cav1 |
| gmfb1 | 2.736537 | 0.298867375 | 0.318 | 0.111 | 6.9521740 | 21 | gmfb |
| atp5pf4 | 2.965887 | 0.381698276 | 0.659 | 0.317 | 7.5348373 | 21 | atp5pf |
| emc6 | 3.443471 | 0.282769432 | 0.358 | 0.128 | 8.7481405 | 21 | emc6 |
| ndufa13 | 3.623300 | 0.32852201 | 0.545 | 0.24 | 9.2049953 | 21 | ndufa1 |
| mrps6 | 3.749760 | 0.265184376 | 0.284 | 0.093 | 9.5262671 | 21 | mrps6 |
| hsbp1b | 4.655700 | 0.345069472 | 0.466 | 0.197 | 1.1827805 | 21 | hsbp1b |
| ndufa52 | 5.086715 | 0.308853608 | 0.403 | 0.159 | 1.2922800 | 21 | ndufa5 |
| si:ch211-66k16.27 | 6.399768 | 0.291233768 | 0.273 | 0.091 | 1.6258610 | 21 | si:ch211-66k16.27 |

| | | | | | | | |
|---|---|---|---|---|---|---|---|
| calr1 | 6.643134 | 0.281178251 | 0.222 | 0.066 | 1.6876882 | 21 | calr |
| add3a6 | 7.750360 | 0.328672102 | 0.364 | 0.137 | 1.9689789 | 21 | add3a |
| psmg1 | 7.793564 | 0.260115793 | 0.222 | 0.065 | 1.9799551 | 21 | psmg1 |
| chchd21 | 9.461118 | 0.259006269 | 0.25 | 0.079 | 2.4035971 | 21 | chchd2 |
| ddt | 1.015710 | 0.260632334 | 0.301 | 0.104 | 2.5804123 | 21 | ddt |
| s100a10b7 | 1.025856 | 0.308556907 | 0.653 | 0.283 | 2.6061882 | 21 | s100a10b |
| pbx42 | 1.115640 | 0.270070062 | 0.239 | 0.073 | 2.8342840 | 21 | pbx4 |
| rer1 | 1.210799 | 0.294893595 | 0.335 | 0.123 | 3.0760364 | 21 | rer1 |
| zgc:11418811 | 1.342313 | 0.40801862 | 0.966 | 0.666 | 3.4101464 | 21 | zgc:114188 |
| anapc16 | 1.597712 | 0.279068943 | 0.312 | 0.111 | 4.0589884 | 21 | anapc16 |
| rtn4a3 | 1.613189 | 0.335905673 | 0.545 | 0.238 | 4.0983080 | 21 | rtn4a |
| cox6b22 | 1.675433 | 0.422080567 | 0.892 | 0.547 | 4.2564380 | 21 | cox6b2 |
| vapb | 1.675755 | 0.256906759 | 0.261 | 0.084 | 4.2572560 | 21 | vapb |
| hectd1 | 1.688135 | 0.261206812 | 0.205 | 0.059 | 4.2887083 | 21 | hectd1 |
| cox6a13 | 1.843799 | 0.410246555 | 0.75 | 0.387 | 4.6841718 | 21 | cox6a1 |
| ctsa2 | 2.027759 | 0.263313382 | 0.284 | 0.096 | 5.1515217 | 21 | ctsa |
| ube2ib | 2.031566 | 0.29701649 | 0.386 | 0.151 | 5.1611950 | 21 | ube2ib |
| ssr42 | 2.129923 | 0.379168173 | 0.67 | 0.329 | 5.4110703 | 21 | ssr4 |
| zgc:1935412 | 2.426676 | 0.383354373 | 0.75 | 0.379 | 6.1649728 | 21 | zgc:193541 |
| psmb51 | 2.912606 | 0.257696982 | 0.244 | 0.077 | 7.3994763 | 21 | psmb5 |
| pin4 | 2.916234 | 0.257922833 | 0.273 | 0.091 | 7.4086936 | 21 | pin4 |
| rbbp62 | 3.625882 | 0.310697251 | 0.301 | 0.108 | 9.2115543 | 21 | rbbp6 |
| zc4h2 | 3.982356 | 0.261024561 | 0.278 | 0.093 | 1.0117177 | 21 | zc4h2 |
| rps137 | 4.082767 | 0.333714116 | 0.966 | 0.748 | 1.0372271 | 21 | rps13 |
| tmed21 | 4.119378 | 0.359968801 | 0.551 | 0.253 | 1.0465282 | 21 | tmed2 |
| ahnak6 | 4.453011 | 0.339825025 | 0.352 | 0.135 | 1.1312874 | 21 | ahnak |
| atp5mc12 | 4.647065 | 0.352670951 | 0.699 | 0.326 | 1.1805870 | 21 | atp5mc1 |
| ier2a7 | 5.035488 | 0.460927737 | 0.545 | 0.273 | 1.2792658 | 21 | ier2a |
| rps27l6 | 5.370616 | 0.349161506 | 0.483 | 0.216 | 1.3644050 | 21 | rps27l |
| si:ch1073-429i10.3.1 | 5.387448 | 0.336418149 | 0.511 | 0.225 | 1.3686813 | 21 | si:ch1073-429i10.3.1 |
| eif3ba2 | 5.477860 | 0.344967888 | 0.528 | 0.239 | 1.3916504 | 21 | eif3ba |
| ndufa21 | 5.581211 | 0.297818317 | 0.386 | 0.154 | 1.4179068 | 21 | ndufa2 |
| psmd81 | 6.169956 | 0.252046268 | 0.33 | 0.119 | 1.5674774 | 21 | psmd8 |
| sec62 | 6.900752 | 0.25861368 | 0.239 | 0.076 | 1.7531362 | 21 | sec62 |
| myclb | 7.506110 | 0.29696854 | 0.261 | 0.086 | 1.9069273 | 21 | myclb |

| | | | | | | | |
|---|---|---|---|---|---|---|---|
| ssr1 | 7.634609 | 0.311035581 | 0.46 | 0.199 | 1.9395725 | 21 | ssr1 |
| ndufa121 | 9.587837 | 0.276302461 | 0.341 | 0.129 | 2.4357900 | 21 | ndufa12 |
| atp6ap21 | 9.789066 | 0.282091 | 0.295 | 0.105 | 2.4869123 | 21 | atp6ap2 |
| snrpd11 | 1.047612 | 0.32609963 | 0.54 | 0.237 | 2.6614604 | 21 | snrpd1 |
| ptges3b1 | 1.133898 | 0.304786302 | 0.438 | 0.18 | 2.8806686 | 21 | ptges3b |
| smndc11 | 1.798343 | 0.263931048 | 0.324 | 0.12 | 4.5686921 | 21 | smndc1 |
| gabarapa1 | 1.865945 | 0.31621943 | 0.426 | 0.18 | 4.7404351 | 21 | gabarapa |
| atp5f1d4 | 1.875461 | 0.308199026 | 0.585 | 0.271 | 4.7646099 | 21 | atp5f1d |
| rpl327 | 2.206848 | 0.264108999 | 0.949 | 0.824 | 5.6064993 | 21 | rpl32 |
| fkbp54 | 2.251759 | 0.331302321 | 0.324 | 0.127 | 5.7205962 | 21 | fkbp5 |
| slc3a2b3 | 2.254567 | 0.317051491 | 0.33 | 0.128 | 5.7277278 | 21 | slc3a2b |
| cnn3a5 | 2.612497 | 0.289072779 | 0.216 | 0.068 | 6.6370498 | 21 | cnn3a |
| ptmab8 | 2.823047 | 0.412267338 | 0.881 | 0.568 | 7.1719512 | 21 | ptmab |
| map1lc3b2 | 3.403262 | 0.311359683 | 0.318 | 0.123 | 8.6459879 | 21 | map1lc3b |
| rab1ab1 | 3.404345 | 0.26621698 | 0.352 | 0.137 | 8.6487396 | 21 | rab1ab |
| wu:fb55g091 | 3.414809 | 0.359059152 | 0.557 | 0.257 | 8.6753225 | 21 | wu:fb55g09 |
| chmp2a3 | 3.861347 | 0.292390491 | 0.358 | 0.143 | 9.8097526 | 21 | chmp2a |
| vamp3 | 4.029539 | 0.306004025 | 0.449 | 0.194 | 1.0237044 | 21 | vamp3 |
| brd3b3 | 4.355312 | 0.251849037 | 0.216 | 0.068 | 1.1064671 | 21 | brd3b |
| lsm71 | 4.896101 | 0.305885512 | 0.466 | 0.2 | 1.2438544 | 21 | lsm7 |
| notch25 | 5.089591 | 0.330806456 | 0.312 | 0.122 | 1.2930107 | 21 | notch2 |
| ap2b1 | 5.533128 | 0.252614584 | 0.188 | 0.055 | 1.4056913 | 21 | ap2b1 |
| dnmt3ab4 | 6.897820 | 0.279280622 | 0.261 | 0.092 | 1.7523913 | 21 | dnmt3ab |
| sumo1 | 7.299677 | 0.286208051 | 0.324 | 0.126 | 1.8544831 | 21 | sumo1 |
| rhoca | 8.630834 | 0.258484679 | 0.21 | 0.066 | 2.1926634 | 21 | rhoca |
| etfb | 9.830950 | 0.252269579 | 0.278 | 0.101 | 2.4975528 | 21 | etfb |
| si:dkey-177p2.61 | 1.132580 | 0.254772707 | 0.244 | 0.083 | 2.8773203 | 21 | si:dkey-177p2.6 |
| u2surp1 | 1.233060 | 0.295616047 | 0.278 | 0.103 | 3.1325912 | 21 | u2surp |
| serinc53 | 1.256516 | 0.300031373 | 0.25 | 0.087 | 3.1921799 | 21 | serinc5 |
| CABZ01115113.1 | 1.420193 | 0.263547662 | 0.261 | 0.093 | 3.6080009 | 21 | CABZ01115113.1 |
| inhbaa3 | 1.436267 | 0.264925866 | 0.182 | 0.053 | 3.6488380 | 21 | inhbaa |
| foxo3b4 | 1.652168 | 0.263214343 | 0.216 | 0.07 | 4.1973340 | 21 | foxo3b |
| brk12 | 1.726094 | 0.314957983 | 0.432 | 0.19 | 4.3851430 | 21 | brk1 |
| fosab7 | 1.824740 | 0.451548127 | 0.659 | 0.36 | 4.6357526 | 21 | fosab |
| rps159 | 1.892630 | 0.328355956 | 0.938 | 0.716 | 4.8082282 | 21 | rps15 |

| | | | | | | | |
|---|---|---|---|---|---|---|---|
| med101 | 1.911287 | 0.264788889 | 0.381 | 0.155 | 4.8556266 | 21 | med10 |
| fam133b2 | 2.047224 | 0.289852473 | 0.375 | 0.155 | 5.2009743 | 21 | fam133b |
| rpl23a8 | 2.094332 | 0.370346652 | 0.949 | 0.668 | 5.3206521 | 21 | rpl23a |
| psmb41 | 2.096580 | 0.29343617 | 0.438 | 0.191 | 5.3263635 | 21 | psmb4 |
| arpc1a | 2.107077 | 0.290441823 | 0.352 | 0.141 | 5.3530313 | 21 | arpc1a |
| eif4ebp22 | 2.184400 | 0.297186113 | 0.438 | 0.191 | 5.5494692 | 21 | eif4ebp2 |
| cox7c4 | 2.186324 | 0.388743356 | 0.841 | 0.491 | 5.5543583 | 21 | cox7c |
| wu:fj16a032 | 2.369368 | 0.431617915 | 0.159 | 0.045 | 6.0193804 | 21 | wu:fj16a03 |
| ywhah1 | 2.436561 | 0.259052113 | 0.193 | 0.06 | 6.1900857 | 21 | ywhah |
| CU694197.1 | 2.660701 | 0.258026136 | 0.358 | 0.144 | 6.7595109 | 21 | CU694197.1 |
| si:dkey-11f4.145 | 2.976060 | 0.301033752 | 0.312 | 0.122 | 7.5606821 | 21 | si:dkey-11f4.14 |
| si:ch211-1a19.34 | 3.208131 | 0.288627108 | 0.188 | 0.057 | 8.1502585 | 21 | si:ch211-1a19.3 |
| anp32b5 | 3.211652 | 0.302773832 | 0.551 | 0.251 | 8.1592036 | 21 | anp32b |
| pin12 | 3.434675 | 0.284539913 | 0.375 | 0.156 | 8.7257940 | 21 | pin1 |
| hint11 | 3.588985 | 0.302191561 | 0.5 | 0.228 | 9.1178172 | 21 | hint1 |
| rab11ba | 4.138474 | 0.261354571 | 0.312 | 0.121 | 1.0513794 | 21 | rab11ba |
| myl66 | 4.381698 | 0.307310939 | 0.301 | 0.116 | 1.1131705 | 21 | myl6 |
| psap3 | 4.554079 | 0.344558848 | 0.591 | 0.286 | 1.1569638 | 21 | psap |
| slc43a2a4 | 5.129583 | 0.303681691 | 0.273 | 0.101 | 1.3031707 | 21 | slc43a2a |
| zmat21 | 5.164380 | 0.271309175 | 0.352 | 0.144 | 1.3120108 | 21 | zmat2 |
| ilf3b1 | 6.772860 | 0.259111527 | 0.301 | 0.114 | 1.7206452 | 21 | ilf3b |
| dynlrb11 | 6.929726 | 0.297665776 | 0.466 | 0.211 | 1.7604971 | 21 | dynlrb1 |
| adh55 | 7.943460 | 0.310881465 | 0.511 | 0.241 | 2.0180360 | 21 | adh5 |
| skp12 | 1.003404 | 0.307776602 | 0.557 | 0.267 | 2.5491501 | 21 | skp1 |
| srsf3b1 | 1.027829 | 0.336975722 | 0.523 | 0.252 | 2.6112018 | 21 | srsf3b |
| cd81a6 | 1.200000 | 0.30512432 | 0.534 | 0.239 | 3.0486007 | 21 | cd81a |
| csnk1a1 | 1.281701 | 0.313401132 | 0.5 | 0.236 | 3.2561621 | 21 | csnk1a1 |
| rbm8a2 | 1.473069 | 0.300973283 | 0.653 | 0.317 | 3.7423342 | 21 | rbm8a |
| anxa45 | 1.752591 | 0.267985342 | 0.358 | 0.15 | 4.4524586 | 21 | anxa4 |
| sap183 | 1.848454 | 0.262646698 | 0.489 | 0.219 | 4.6959989 | 21 | sap18 |
| eif3jb1 | 1.900124 | 0.261088525 | 0.301 | 0.118 | 4.8272666 | 21 | eif3jb |
| COX5B3 | 2.327109 | 0.3313211 | 0.716 | 0.374 | 5.9120225 | 21 | COX5B |
| chmp1b1 | 3.012036 | 0.25715332 | 0.324 | 0.132 | 7.6520783 | 21 | chmp1b |
| rpl276 | 3.357995 | 0.281900962 | 0.96 | 0.779 | 8.5309878 | 21 | rpl27 |
| hnrnpa1a2 | 3.415038 | 0.278709719 | 0.381 | 0.165 | 8.6759047 | 21 | hnrnpa1a |

| | | | | | | | |
|---|---|---|---|---|---|---|---|
| ndufb82 | 3.44666 | 0.313397833 | 0.534 | 0.264 | 8.7562645 | 21 | ndufb8 |
| crema4 | 3.45257 | 0.316056351 | 0.392 | 0.173 | 8.7712792 | 21 | crema |
| wu:fi37e092 | 3.53966 | 0.263783796 | 0.318 | 0.128 | 8.9925268 | 21 | wu:fi37e09 |
| lsm64 | 6.50579 | 0.250510555 | 0.494 | 0.224 | 1.6527962 | 21 | lsm6 |
| sdhc1 | 8.05508 | 0.260887608 | 0.347 | 0.147 | 2.0463935 | 21 | sdhc |
| psmb2 | 9.31274 | 0.2746041 | 0.398 | 0.179 | 2.3659025 | 21 | psmb2 |
| pdia38 | 9.86056 | 0.345489158 | 0.602 | 0.312 | 2.5050760 | 21 | pdia3 |
| dbi4 | 1.03098 | 0.300030234 | 0.494 | 0.242 | 2.6192104 | 21 | dbi |
| sumo2b6 | 1.30655 | 0.326070891 | 0.58 | 0.295 | 3.3193086 | 21 | sumo2b |
| psmd11 | 1.31606 | 0.255780861 | 0.239 | 0.089 | 3.3434756 | 21 | psmd1 |
| smarce1 | 1.37722 | 0.268213695 | 0.398 | 0.178 | 3.4988431 | 21 | smarce1 |
| cops9 | 1.38361 | 0.250876322 | 0.284 | 0.113 | 3.5150825 | 21 | cops9 |
| cst14a.2 | 1.38689 | 0.301320969 | 0.489 | 0.234 | 3.5234144 | 21 | cst14a.2 |
| sf3b61 | 1.84814 | 0.291190631 | 0.523 | 0.257 | 4.6952017 | 21 | sf3b6 |
| anxa11b3 | 1.87367 | 0.280417514 | 0.381 | 0.173 | 4.7600619 | 21 | anxa11b |
| rpl911 | 1.88806 | 0.37609932 | 0.972 | 0.666 | 4.7966192 | 21 | rpl9 |
| eif4a31 | 1.98303 | 0.250870856 | 0.364 | 0.159 | 5.0379089 | 21 | eif4a3 |
| cdc42l4 | 2.00839 | 0.284200194 | 0.489 | 0.232 | 5.1023276 | 21 | cdc42l |
| mt-co210 | 2.74532 | 0.344522298 | 0.812 | 0.671 | 6.9744946 | 21 | mt-co2 |
| snrpf1 | 2.78580 | 0.26967266 | 0.557 | 0.268 | 7.0773249 | 21 | snrpf |
| hypk1 | 3.04124 | 0.277848935 | 0.46 | 0.219 | 7.7262816 | 21 | hypk |
| slc25a3b4 | 3.80669 | 0.357279559 | 0.801 | 0.456 | 9.6708964 | 21 | slc25a3b |
| hdac11 | 5.11661 | 0.288439585 | 0.466 | 0.225 | 1.2998762 | 21 | hdac1 |
| cd74b4 | 6.75354 | 0.396993836 | 0.449 | 0.225 | 1.7157369 | 21 | cd74b |
| ost42 | 6.92934 | 0.275414024 | 0.528 | 0.26 | 1.7603991 | 21 | ost4 |
| uqcrq3 | 7.05800 | 0.250341933 | 0.568 | 0.285 | 1.7930852 | 21 | uqcrq |
| h3f3d | 7.84974 | 0.327152836 | 0.938 | 0.675 | 1.9942268 | 21 | h3f3d |
| ybx15 | 7.88885 | 0.332106312 | 0.96 | 0.694 | 2.0041646 | 21 | ybx1 |
| vat14 | 8.58657 | 0.261151898 | 0.341 | 0.147 | 2.1814197 | 21 | vat1 |
| sod11 | 8.84431 | 0.310889137 | 0.631 | 0.343 | 2.2468986 | 21 | sod1 |
| rpl10a6 | 9.70020 | 0.29192302 | 0.989 | 0.778 | 2.4643380 | 21 | rpl10a |
| junba7 | 1.03006 | 0.376131267 | 0.591 | 0.319 | 2.6168680 | 21 | junba |
| ddx51 | 1.04682 | 0.328378278 | 0.659 | 0.352 | 2.6594641 | 21 | ddx5 |
| psma81 | 1.06057 | 0.269103089 | 0.46 | 0.222 | 2.6943966 | 21 | psma8 |
| cct23 | 1.06887 | 0.280119212 | 0.506 | 0.253 | 2.7154866 | 21 | cct2 |

| | | | | | | | |
|---|---|---|---|---|---|---|---|
| rpl27a7 | 1.198498 | 0.383496717 | 0.881 | 0.52 | 3.0447844 | 21 | rpl27a |
| eif3k2 | 1.372505 | 0.290516778 | 0.483 | 0.243 | 3.4868514 | 21 | eif3k |
| arf5 | 1.391092 | 0.286516732 | 0.432 | 0.211 | 3.5340715 | 21 | arf5 |
| eif3s103 | 1.479148 | 0.264371995 | 0.381 | 0.178 | 3.7577776 | 21 | eif3s10 |
| vcp1 | 1.482250 | 0.252960378 | 0.324 | 0.141 | 3.7656578 | 21 | vcp |
| myl12.15 | 1.564440 | 0.342981281 | 0.761 | 0.424 | 3.9744619 | 21 | myl12.1 |
| tomm53 | 1.955518 | 0.26413907 | 0.466 | 0.229 | 4.9679950 | 21 | tomm5 |
| ier2b8 | 2.176159 | 0.328415331 | 0.46 | 0.236 | 5.5285326 | 21 | ier2b |
| atp6v0ca2 | 2.211284 | 0.252508222 | 0.432 | 0.202 | 5.6177693 | 21 | atp6v0ca |
| gng53 | 2.673566 | 0.296958793 | 0.551 | 0.286 | 6.7921954 | 21 | gng5 |
| eif1axb | 2.809320 | 0.277335968 | 0.591 | 0.305 | 7.1370788 | 21 | eif1axb |
| cdc26 | 3.087360 | 0.25100106 | 0.227 | 0.089 | 7.8434398 | 21 | cdc26 |
| jpt1b5 | 3.718680 | 0.298462347 | 0.545 | 0.275 | 9.4473079 | 21 | jpt1b |
| paip2b2 | 4.314617 | 0.273714127 | 0.597 | 0.306 | 1.0961285 | 21 | paip2b |
| rps169 | 4.394363 | 0.302080148 | 0.977 | 0.764 | 1.1163881 | 21 | rps16 |
| sat1a.24 | 4.905595 | 0.342991848 | 0.54 | 0.3 | 1.2462664 | 21 | sat1a.2 |
| si:ch211-288g17.31 | 5.258396 | 0.301254221 | 0.449 | 0.225 | 1.3358956 | 21 | si:ch211-288g17.3 |
| si:ch211-196l7.49 | 6.475817 | 0.297061358 | 0.614 | 0.328 | 1.6451813 | 21 | si:ch211-196l7.4 |
| NDUFB13 | 7.140148 | 0.261596879 | 0.426 | 0.211 | 1.8139548 | 21 | NDUFB1 |
| dap1b3 | 8.395555 | 0.266161237 | 0.5 | 0.248 | 2.1328908 | 21 | dap1b |
| vdac23 | 8.848587 | 0.291974572 | 0.597 | 0.324 | 2.2479837 | 21 | vdac2 |
| prdx54 | 9.801620 | 0.253883639 | 0.375 | 0.178 | 2.4901017 | 21 | prdx5 |
| cct72 | 1.092497 | 0.258805796 | 0.455 | 0.227 | 2.7754904 | 21 | cct7 |
| atp5mf3 | 1.106981 | 0.259536465 | 0.551 | 0.287 | 2.8122866 | 21 | atp5mf |
| gapdhs9 | 1.514534 | 0.280356794 | 0.597 | 0.313 | 3.8476743 | 21 | gapdhs |
| cnn25 | 1.657206 | 0.259681612 | 0.335 | 0.156 | 4.2101342 | 21 | cnn2 |
| atp5fa11 | 1.883398 | 0.266593585 | 0.682 | 0.368 | 4.7847726 | 21 | atp5fa1 |
| pnrc211 | 2.720448 | 0.370806852 | 0.938 | 0.595 | 6.9112993 | 21 | pnrc2 |
| anxa2a6 | 2.726107 | 0.363436193 | 0.261 | 0.117 | 6.9256751 | 21 | anxa2a |
| ola11 | 3.078418 | 0.266221942 | 0.466 | 0.244 | 7.8207223 | 21 | ola1 |
| faua8 | 3.262600 | 0.296916722 | 0.943 | 0.716 | 8.2886366 | 21 | faua |
| cox5aa2 | 3.671978 | 0.257280034 | 0.659 | 0.356 | 9.3286614 | 21 | cox5aa |
| mt-atp610 | 5.152172 | 0.343194368 | 0.744 | 0.556 | 0.000131 | 21 | mt-atp6 |
| rplp2l7 | 5.763392 | 0.268582137 | 0.972 | 0.793 | 0.000146 | 21 | rplp2l |
| atp5f1e2 | 7.525719 | 0.315215977 | 0.812 | 0.488 | 0.000191 | 21 | atp5f1e |

| | | | | | | | |
|---|---|---|---|---|---|---|---|
| serbp1a9 | 7.987431 | 0.351679638 | 0.812 | 0.504 | 0.000203 | 21 | serbp1a |
| uqcr101 | 8.195223 | 0.269366424 | 0.631 | 0.346 | 0.000208 | 21 | uqcr10 |
| rpl318 | 9.707088 | 0.297116884 | 0.932 | 0.686 | 0.000247 | 21 | rpl31 |
| snrpb3 | 9.887809 | 0.2616408 | 0.659 | 0.354 | 0.000251 | 21 | snrpb |
| cox4i15 | 1.041592 | 0.279850225 | 0.744 | 0.415 | 0.000265 | 21 | cox4i1 |
| rack15 | 1.686577 | 0.252210638 | 0.955 | 0.762 | 0.000428 | 21 | rack1 |
| naca8 | 2.640422 | 0.307476724 | 0.92 | 0.65 | 0.000671 | 21 | naca |
| rps4x8 | 3.680619 | 0.271555349 | 0.955 | 0.735 | 0.000935 | 21 | rps4x |
| prdx24 | 3.878751 | 0.300480597 | 0.705 | 0.421 | 0.000985 | 21 | prdx2 |
| rps247 | 4.852621 | 0.265298252 | 0.977 | 0.778 | 0.001233 | 21 | rps24 |
| rpl286 | 6.808688 | 0.250835526 | 0.96 | 0.776 | 0.00173 | 21 | rpl28 |
| uba527 | 7.606669 | 0.316814572 | 0.943 | 0.659 | 0.001932 | 21 | uba52 |
| eif3m5 | 9.333799 | 0.279069675 | 0.665 | 0.382 | 0.002371 | 21 | eif3m |
| rps59 | 9.784767 | 0.283384708 | 0.955 | 0.727 | 0.002486 | 21 | rps5 |
| rpl175 | 1.365867 | 0.255652344 | 0.977 | 0.762 | 0.00347 | 21 | rpl17 |
| bsg1 | 1.495165 | 0.271710231 | 0.648 | 0.375 | 0.003798 | 21 | bsg |
| atp5l4 | 4.082494 | 0.253658936 | 0.773 | 0.455 | 0.010372 | 21 | atp5l |
| rps256 | 4.690691 | 0.252032893 | 0.955 | 0.748 | 0.011917 | 21 | rps25 |
| ip6k2b4 | 6.521449 | 0.258815396 | 0.256 | 0.129 | 0.016568 | 21 | ip6k2b |
| hnrnpaba1 | 1.023241 | 0.279522299 | 0.75 | 0.466 | 0.025995 | 21 | hnrnpaba |
| eif3f7 | 1.461965 | 0.275844917 | 0.739 | 0.464 | 0.037141 | 21 | eif3f |
| rps27.110 | 1.661125 | 0.282222358 | 0.96 | 0.708 | 0.042201 | 21 | rps27.1 |
| eef1g8 | 2.434305 | 0.294406421 | 0.926 | 0.651 | 0.061844 | 21 | eef1g |
| hmgn25 | 3.312134 | 0.269128572 | 0.699 | 0.467 | 0.084145 | 21 | hmgn2 |
| khdrbs1a1 | 7.058515 | 0.265105551 | 0.841 | 0.56 | 0.179322 | 21 | khdrbs1a |
| pabpc1a5 | 1.722332 | 0.278074978 | 0.875 | 0.568 | 0.437559 | 21 | pabpc1a |
| cfl16 | 1.802022 | 0.26244338 | 0.875 | 0.598 | 0.457804 | 21 | cfl1 |
| cirbpa1 | 2.816272 | 0.251333814 | 0.881 | 0.622 | 0.715474 | 21 | cirbpa |
| igfbp71 | 0 | 1.311704485 | 0.566 | 0.026 | 0 | 22 | igfbp7 |
| lyve1a | 0 | 1.174004893 | 0.358 | 0.002 | 0 | 22 | lyve1a |
| cldn5b | 0 | 0.990515641 | 0.387 | 0.006 | 0 | 22 | cldn5b |
| she | 0 | 0.953721093 | 0.387 | 0.001 | 0 | 22 | she |
| kdr | 0 | 0.865326359 | 0.376 | 0.001 | 0 | 22 | kdr |
| lyve1b | 0 | 0.844212955 | 0.312 | 0 | 0 | 22 | lyve1b |
| si:dkeyp-97a10.2 | 0 | 0.83833437 | 0.341 | 0.003 | 0 | 22 | si:dkeyp-97a10.2 |

| | | | | | | | |
|---|---|---|---|---|---|---|---|
| mrc1a | 0 | 0.783115358 | 0.295 | 0.005 | 0 | 22 | mrc1a |
| flt4 | 0 | 0.782315825 | 0.312 | 0.003 | 0 | 22 | flt4 |
| tmem88a | 0 | 0.767031487 | 0.329 | 0.006 | 0 | 22 | tmem88a |
| si:ch211-152c2.3 | 0 | 0.72346208 | 0.249 | 0.002 | 0 | 22 | si:ch211-152c2.3 |
| si:dkey-126g1.9 | 0 | 0.613653356 | 0.237 | 0.002 | 0 | 22 | si:dkey-126g1.9 |
| clec14a | 0 | 0.613526079 | 0.243 | 0.002 | 0 | 22 | clec14a |
| pcdh12 | 0 | 0.603702826 | 0.249 | 0.002 | 0 | 22 | pcdh12 |
| rasip1 | 0 | 0.588326942 | 0.243 | 0.001 | 0 | 22 | rasip1 |
| cdh5 | 0 | 0.580272991 | 0.185 | 0.001 | 0 | 22 | cdh5 |
| ldb2a | 0 | 0.567941251 | 0.22 | 0.003 | 0 | 22 | ldb2a |
| sele | 0 | 0.550859843 | 0.202 | 0.002 | 0 | 22 | sele |
| lgi2a | 0 | 0.550782858 | 0.202 | 0.002 | 0 | 22 | lgi2a |
| CT033796.1 | 0 | 0.540859264 | 0.208 | 0.001 | 0 | 22 | CT033796.1 |
| akap12a | 0 | 0.53900535 | 0.208 | 0.001 | 0 | 22 | akap12a |
| rca2.2 | 0 | 0.511760418 | 0.179 | 0 | 0 | 22 | rca2.2 |
| pecam1 | 0 | 0.475795675 | 0.168 | 0.001 | 0 | 22 | pecam1 |
| myct1a | 1.730121 | 0.7474012 | 0.295 | 0.006 | 4.3953738 | 22 | myct1a |
| stab1 | 1.636434 | 0.59566815 | 0.225 | 0.003 | 4.1573620 | 22 | stab1 |
| gpr182 | 8.582969 | 0.376989586 | 0.133 | 0 | 2.1805035 | 22 | gpr182 |
| exoc3l2a | 3.740679 | 0.531867964 | 0.156 | 0.001 | 9.5031969 | 22 | exoc3l2a |
| egfl7 | 5.071433 | 0.609407445 | 0.243 | 0.004 | 1.2883977 | 22 | egfl7 |
| kdrl | 1.033916 | 0.50680977 | 0.162 | 0.001 | 2.6266651 | 22 | kdrl |
| si:ch211-33e4.2 | 3.131350 | 0.450600291 | 0.179 | 0.002 | 7.9551946 | 22 | si:ch211-33e4.2 |
| etv2 | 1.176704 | 0.524718268 | 0.22 | 0.004 | 2.9894165 | 22 | etv2 |
| sox18 | 8.976508 | 0.695859217 | 0.277 | 0.007 | 2.2804818 | 22 | sox18 |
| si:dkey-237i9.8 | 1.052550 | 0.354143588 | 0.133 | 0.001 | 2.6740042 | 22 | si:dkey-237i9.8 |
| parvab | 1.488278 | 0.62231911 | 0.249 | 0.005 | 3.7809715 | 22 | parvab |
| fabp11a4 | 4.000742 | 1.751435774 | 0.786 | 0.08 | 1.0163887 | 22 | fabp11a |
| si:ch211-145b13.6 | 1.165056 | 0.637203691 | 0.243 | 0.005 | 2.9598250 | 22 | si:ch211-145b13.6 |
| si:ch211-248e11.2 | 2.070636 | 0.583846507 | 0.231 | 0.005 | 5.2604515 | 22 | si:ch211-248e11.2 |
| tie1 | 1.346539 | 0.484727182 | 0.185 | 0.003 | 3.4208837 | 22 | tie1 |
| erg | 1.414078 | 0.763157488 | 0.301 | 0.01 | 3.5924670 | 22 | erg |
| LOC100535225 | 1.467554 | 0.379774648 | 0.127 | 0.001 | 3.7283233 | 22 | LOC100535225 |
| plk2b | 7.094431 | 0.696352612 | 0.272 | 0.008 | 1.8023402 | 22 | plk2b |
| tm4sf18 | 5.062335 | 0.554668802 | 0.208 | 0.005 | 1.2860863 | 22 | tm4sf18 |

| | | | | | | | |
|---|---|---|---|---|---|---|---|
| fgd5a1 | 1.101693 | 0.6597596 | 0.289 | 0.011 | 2.7988535 | 22 | fgd5a |
| si:ch211-14k19.8 | 5.449261 | 0.306723368 | 0.121 | 0.001 | 1.3843847 | 22 | si:ch211-14k19.8 |
| notchl | 1.934758 | 0.300561112 | 0.116 | 0.001 | 4.9152535 | 22 | notchl |
| arhgef9b | 9.837302 | 0.33544135 | 0.133 | 0.002 | 2.4991667 | 22 | arhgef9b |
| rab11bb | 4.414520 | 0.803200196 | 0.335 | 0.017 | 1.1215089 | 22 | rab11bb |
| shank3b | 6.055833 | 0.268830749 | 0.11 | 0.001 | 1.5384844 | 22 | shank3b |
| ehd31 | 8.878097 | 0.758000201 | 0.301 | 0.013 | 2.2554805 | 22 | ehd3 |
| ramp22 | 2.221999 | 1.124404642 | 0.486 | 0.039 | 5.6449901 | 22 | ramp2 |
| slc9a3r1b | 1.311242 | 0.466930731 | 0.179 | 0.004 | 3.3312108 | 22 | slc9a3r1b |
| sox7 | 4.493740 | 0.394060999 | 0.156 | 0.003 | 1.1416346 | 22 | sox7 |
| b3gnt7 | 1.726480 | 0.315070037 | 0.121 | 0.002 | 4.3861249 | 22 | b3gnt7 |
| si:ch73-334d15.2 | 1.896560 | 0.47990557 | 0.191 | 0.005 | 4.8182127 | 22 | si:ch73-334d15.2 |
| ppp1r13ba | 2.947815 | 0.407740997 | 0.15 | 0.003 | 7.4889249 | 22 | ppp1r13ba |
| ecscr | 5.048822 | 0.406003948 | 0.162 | 0.004 | 1.2826533 | 22 | ecscr |
| dysf | 3.273367 | 0.250964382 | 0.104 | 0.001 | 8.3159898 | 22 | dysf |
| fam174b | 6.607583 | 0.619353264 | 0.272 | 0.013 | 1.6786565 | 22 | fam174b |
| igf2a | 6.355500 | 0.511710897 | 0.185 | 0.006 | 1.6146150 | 22 | igf2a |
| plk2a | 1.671561 | 0.391143593 | 0.156 | 0.004 | 4.2466019 | 22 | plk2a |
| cdh62 | 9.138797 | 0.895480169 | 0.358 | 0.027 | 2.3217115 | 22 | cdh6 |
| si:dkey-28n18.9 | 1.036565 | 0.308942095 | 0.121 | 0.002 | 2.6333939 | 22 | si:dkey-28n18.9 |
| aqp8a.1 | 4.144028 | 0.521215836 | 0.15 | 0.004 | 1.0527904 | 22 | aqp8a.1 |
| s1pr1 | 8.750003 | 0.441984747 | 0.173 | 0.006 | 2.2229383 | 22 | s1pr1 |
| hapln31 | 2.563640 | 0.77504343 | 0.272 | 0.016 | 6.5129278 | 22 | hapln3 |
| plvapb1 | 1.027732 | 0.803687458 | 0.254 | 0.014 | 2.6109546 | 22 | plvapb |
| rassf9 | 3.843892 | 0.346145056 | 0.133 | 0.003 | 9.7654091 | 22 | rassf9 |
| tmem108 | 6.623591 | 0.426293271 | 0.168 | 0.006 | 1.6827233 | 22 | tmem108 |
| nr5a2 | 1.468032 | 0.311822301 | 0.121 | 0.003 | 3.7295369 | 22 | nr5a2 |
| ednraa1 | 1.012540 | 0.689134069 | 0.295 | 0.02 | 2.5723592 | 22 | ednraa |
| dll4 | 4.543561 | 0.350432595 | 0.145 | 0.004 | 1.1542917 | 22 | dll4 |
| reps2 | 7.333798 | 0.322761973 | 0.127 | 0.003 | 1.8631514 | 22 | reps2 |
| podn | 5.071477 | 0.294656997 | 0.116 | 0.003 | 1.2884089 | 22 | podn |
| ccl38.1 | 1.684733 | 0.62864375 | 0.197 | 0.009 | 4.2800645 | 22 | ccl38.1 |
| nrarpb | 6.191633 | 0.32555245 | 0.133 | 0.004 | 1.5729846 | 22 | nrarpb |
| loxl12 | 2.381392 | 0.843321257 | 0.324 | 0.028 | 6.0499268 | 22 | loxl1 |
| afap1l1b | 2.949002 | 0.500717256 | 0.197 | 0.01 | 7.4919414 | 22 | afap1l1b |

| | | | | | | | |
|---|---|---|---|---|---|---|---|
| cd81b | 5.411311 | 0.45209896 | 0.173 | 0.008 | 1.3747438 | 22 | cd81b |
| cpn15 | 4.46441( | 1.009795943 | 0.497 | 0.073 | 1.1341835 | 22 | cpn1 |
| calcrlb | 3.573314 | 0.492410634 | 0.202 | 0.011 | 9.0780066 | 22 | calcrlb |
| CU929150.12 | 1.037474 | 0.841751285 | 0.289 | 0.025 | 2.6357051 | 22 | CU929150.1 |
| fli1a | 2.560932 | 0.424376451 | 0.168 | 0.008 | 6.5060479 | 22 | fli1a |
| cyth3a | 4.328951 | 0.280165455 | 0.116 | 0.004 | 1.0997701 | 22 | cyth3a |
| marcksl1b8 | 2.214155 | 1.178403041 | 0.63 | 0.137 | 5.6250613 | 22 | marcksl1b |
| LOC101882264 | 7.65845( | 0.453913123 | 0.173 | 0.009 | 1.9456293 | 22 | LOC101882264 |
| gpm6ab1 | 1.529467 | 0.562130721 | 0.225 | 0.017 | 3.8856110 | 22 | gpm6ab |
| piezo2a.2 | 1.435255 | 0.467854636 | 0.191 | 0.012 | 3.6462670 | 22 | piezo2a.2 |
| pgm5 | 1.098893 | 0.348043465 | 0.133 | 0.005 | 2.7917389 | 22 | pgm5 |
| reep1 | 1.077159 | 0.332045183 | 0.139 | 0.007 | 2.7365233 | 22 | reep1 |
| pcdh17 | 5.141268 | 0.28569777 | 0.11 | 0.004 | 1.3061391 | 22 | pcdh17 |
| macf1a1 | 1.574576 | 0.762855277 | 0.382 | 0.057 | 4.0002118 | 22 | macf1a |
| cyp26b11 | 2.111323 | 0.399561725 | 0.133 | 0.007 | 5.3638168 | 22 | cyp26b1 |
| plekhg5a | 2.577189 | 0.363942094 | 0.145 | 0.008 | 6.5473498 | 22 | plekhg5a |
| limch1b | 1.042983 | 0.48180011 | 0.214 | 0.018 | 2.6497004 | 22 | limch1b |
| vat15 | 5.522063 | 0.987038595 | 0.595 | 0.143 | 1.4028801 | 22 | vat1 |
| nova22 | 2.981892 | 0.657297473 | 0.289 | 0.034 | 7.5754967 | 22 | nova2 |
| cxcl12a3 | 4.550931 | 1.078077381 | 0.445 | 0.081 | 1.1561642 | 22 | cxcl12a |
| lfng1 | 2.648098 | 0.369256935 | 0.145 | 0.009 | 6.7274936 | 22 | lfng |
| gas62 | 1.472011 | 0.591845801 | 0.26 | 0.029 | 3.7396448 | 22 | gas6 |
| wasf3b | 1.43326( | 0.516445686 | 0.214 | 0.019 | 3.6411985 | 22 | wasf3b |
| palm2 | 2.538018 | 0.474202852 | 0.197 | 0.017 | 6.4478347 | 22 | palm2 |
| selenop1 | 3.387895 | 0.581598314 | 0.231 | 0.024 | 8.6069481 | 22 | selenop |
| si:ch211-66e2.34 | 2.608356 | 0.753334296 | 0.37 | 0.06 | 6.6265308 | 22 | si:ch211-66e2.3 |
| c1qtnf6a1 | 2.642748 | 0.584236282 | 0.237 | 0.025 | 6.7139018 | 22 | c1qtnf6a |
| hhex | 9.58171( | 0.310496816 | 0.133 | 0.008 | 2.4342335 | 22 | hhex |
| grb10a2 | 1.149701 | 0.690500153 | 0.301 | 0.04 | 2.9208170 | 22 | grb10a |
| anks1b1 | 3.824911 | 0.441495738 | 0.197 | 0.017 | 9.7171874 | 22 | anks1b |
| tspan4b1 | 3.084556 | 0.754850534 | 0.393 | 0.069 | 7.8363151 | 22 | tspan4b |
| scarf1 | 3.790239 | 0.253039545 | 0.104 | 0.005 | 9.6291047 | 22 | scarf1 |
| si:ch211-156j16.12 | 3.745545 | 0.593970653 | 0.237 | 0.026 | 9.5155576 | 22 | si:ch211-156j16.1 |
| krt18a.16 | 1.099657 | 0.933433024 | 0.595 | 0.146 | 2.7936809 | 22 | krt18a.1 |
| fzd41 | 1.249749 | 0.434052075 | 0.185 | 0.016 | 3.1749889 | 22 | fzd4 |

| | | | | | | | |
|---|---|---|---|---|---|---|---|
| fam198b | 2.825346 | 0.271760381 | 0.116 | 0.006 | 7.1777938 | 22 | fam198b |
| cldn11b | 1.040306 | 0.423308977 | 0.162 | 0.013 | 2.6428980 | 22 | cldn11b |
| sema3ga | 2.827437 | 0.288488179 | 0.104 | 0.005 | 7.1831055 | 22 | sema3ga |
| tpm4a7 | 5.185905 | 0.843253961 | 0.486 | 0.11 | 1.3174793 | 22 | tpm4a |
| krt88 | 1.108950 | 0.974970821 | 0.832 | 0.307 | 2.8172880 | 22 | krt8 |
| yrk4 | 2.289328 | 0.70008389 | 0.37 | 0.065 | 5.8160390 | 22 | yrk |
| tcima2 | 1.244144 | 0.882169702 | 0.376 | 0.069 | 3.1607492 | 22 | tcima |
| epas1b | 2.848489 | 0.385994955 | 0.145 | 0.011 | 7.2365867 | 22 | epas1b |
| fkbp1ab1 | 2.081094 | 0.728774171 | 0.341 | 0.058 | 5.2870216 | 22 | fkbp1ab |
| lmo2 | 1.919992 | 0.495309656 | 0.214 | 0.024 | 4.8777403 | 22 | lmo2 |
| akap12b5 | 8.938836 | 0.736497671 | 0.376 | 0.071 | 2.2709114 | 22 | akap12b |
| crip2 | 9.716565 | 0.380155331 | 0.168 | 0.015 | 2.4684934 | 22 | crip2 |
| arl4ab | 1.749326 | 0.374971299 | 0.168 | 0.015 | 4.4441643 | 22 | arl4ab |
| il13ra1 | 8.461778 | 0.489342408 | 0.231 | 0.029 | 2.1497148 | 22 | il13ra1 |
| rbp2a | 1.513630 | 0.672313426 | 0.15 | 0.013 | 3.8453772 | 22 | rbp2a |
| marcksl1a7 | 5.221165 | 0.978914307 | 0.682 | 0.257 | 1.3264370 | 22 | marcksl1a |
| oaz2a | 3.968678 | 0.405061504 | 0.156 | 0.014 | 1.0082428 | 22 | oaz2a |
| si:ch211-1a19.35 | 5.463512 | 0.610029519 | 0.318 | 0.055 | 1.3880053 | 22 | si:ch211-1a19.3 |
| gcnt4a1 | 6.071645 | 0.349592009 | 0.156 | 0.014 | 1.5425016 | 22 | gcnt4a |
| prox1a1 | 1.109485 | 0.410443994 | 0.168 | 0.016 | 2.8186487 | 22 | prox1a |
| adam8a2 | 3.591411 | 0.641036072 | 0.289 | 0.048 | 9.1239812 | 22 | adam8a |
| oaz2b1 | 2.351162 | 0.319015702 | 0.133 | 0.011 | 5.9731289 | 22 | oaz2b |
| CABZ01084131.1 | 7.036389 | 0.366710699 | 0.162 | 0.016 | 1.7875948 | 22 | CABZ01084131.1 |
| npdc1a | 2.704141 | 0.319491658 | 0.145 | 0.013 | 6.8698711 | 22 | npdc1a |
| abcg2a | 5.467880 | 0.321726223 | 0.127 | 0.01 | 1.3891149 | 22 | abcg2a |
| notch1b1 | 6.083829 | 0.270252563 | 0.121 | 0.009 | 1.5455968 | 22 | notch1b |
| si:ch211-207d6.22 | 7.959557 | 0.403918421 | 0.179 | 0.02 | 2.0221256 | 22 | si:ch211-207d6.2 |
| stox2a | 1.650592 | 0.26360141 | 0.116 | 0.009 | 4.1933294 | 22 | stox2a |
| exoc6 | 1.785780 | 0.528073363 | 0.237 | 0.036 | 4.5367752 | 22 | exoc6 |
| ebf3a | 3.276738 | 0.271548814 | 0.116 | 0.009 | 8.3245537 | 22 | ebf3a |
| myo1ca1 | 1.930168 | 0.487744746 | 0.202 | 0.027 | 4.9035922 | 22 | myo1ca |
| spns2 | 1.600417 | 0.264054456 | 0.104 | 0.007 | 4.0658619 | 22 | spns2 |
| ccdc85b6 | 2.412028 | 0.54157689 | 0.272 | 0.049 | 6.1277589 | 22 | ccdc85b |
| kank31 | 2.712685 | 0.338419434 | 0.15 | 0.016 | 6.8915778 | 22 | kank3 |
| tagln28 | 5.763888 | 0.834278301 | 0.561 | 0.207 | 1.4643158 | 22 | tagln2 |

| | | | | | | | |
|---|---|---|---|---|---|---|---|
| atp1b1a5 | 2.1956877 | 0.843250214 | 0.578 | 0.227 | 5.5781428 | 22 | atp1b1a |
| hyal2b | 7.0676664 | 0.303504543 | 0.116 | 0.01 | 1.7955401 | 22 | hyal2b |
| carhsp11 | 7.2642722 | 0.632032942 | 0.324 | 0.073 | 1.8454884 | 22 | carhsp1 |
| lrch1 | 1.3056333 | 0.339512049 | 0.139 | 0.015 | 3.3169607 | 22 | lrch1 |
| fkbp1aa7 | 1.5791899 | 0.714458857 | 0.757 | 0.412 | 4.0119310 | 22 | fkbp1aa |
| rgcc3 | 1.7104822 | 0.777763982 | 0.306 | 0.07 | 4.3454797 | 22 | rgcc |
| lgmn3 | 6.6848300 | 0.639391645 | 0.312 | 0.072 | 1.6982812 | 22 | lgmn |
| ralgapa2 | 2.2012866 | 0.293919372 | 0.121 | 0.012 | 5.5923679 | 22 | ralgapa2 |
| tuba8l35 | 1.9626811 | 0.610543652 | 0.306 | 0.07 | 4.9861929 | 22 | tuba8l3 |
| arhgap311 | 7.4417499 | 0.39342938 | 0.197 | 0.032 | 1.8905763 | 22 | arhgap31 |
| dab2 | 7.7398299 | 0.363644325 | 0.191 | 0.029 | 1.9663037 | 22 | dab2 |
| ushbp1 | 1.3889855 | 0.270290264 | 0.116 | 0.012 | 3.5287183 | 22 | ushbp1 |
| rps6ka3b | 2.3798899 | 0.313835861 | 0.127 | 0.014 | 6.0461099 | 22 | rps6ka3b |
| cavin2b4 | 2.6546588 | 0.606157925 | 0.318 | 0.077 | 6.7441589 | 22 | cavin2b |
| klf2a5 | 9.8191322 | 0.707813772 | 0.358 | 0.098 | 2.4945505 | 22 | klf2a |
| mafba6 | 1.4609733 | 0.599590713 | 0.324 | 0.083 | 3.7116035 | 22 | mafba |
| dab2ipb | 2.2407255 | 0.330029046 | 0.133 | 0.016 | 5.6925642 | 22 | dab2ipb |
| lzts2a1 | 3.3370988 | 0.363576814 | 0.156 | 0.022 | 8.4778977 | 22 | lzts2a |
| CU984579.11 | 9.5226488 | 0.332467479 | 0.139 | 0.017 | 2.4192288 | 22 | CU984579.1 |
| si:ch211-284f22.32 | 1.8334399 | 0.391801717 | 0.197 | 0.033 | 4.6578520 | 22 | si:ch211-284f22.3 |
| rhoca1 | 1.9539577 | 0.532849657 | 0.283 | 0.065 | 4.9640295 | 22 | rhoca |
| arl4aa | 3.0975011 | 0.486301448 | 0.254 | 0.052 | 7.8692014 | 22 | arl4aa |
| cdc42ep5 | 3.1300633 | 0.442087166 | 0.197 | 0.034 | 7.9519258 | 22 | cdc42ep5 |
| grapa2 | 4.7405144 | 0.401143466 | 0.173 | 0.027 | 1.2043276 | 22 | grapa |
| col15a1a2 | 8.0463277 | 0.398965394 | 0.162 | 0.024 | 2.0441694 | 22 | col15a1a |
| fam43a | 1.0065699 | 0.287574627 | 0.11 | 0.011 | 2.5571898 | 22 | fam43a |
| foxp1b | 1.7998422 | 0.452219195 | 0.208 | 0.039 | 4.5724995 | 22 | foxp1b |
| ponzr18 | 4.4459955 | 0.72802375 | 0.618 | 0.287 | 1.1295050 | 22 | ponzr1 |
| acsbg21 | 8.1219344 | 0.402747946 | 0.202 | 0.036 | 2.0633774 | 22 | acsbg2 |
| spry44 | 3.3427900 | 0.478655635 | 0.231 | 0.048 | 8.4923582 | 22 | spry4 |
| pros1 | 6.8780522 | 0.437203948 | 0.231 | 0.047 | 1.7473693 | 22 | pros1 |
| rhocb | 2.8428177 | 0.602474139 | 0.347 | 0.102 | 7.2221769 | 22 | rhocb |
| tmem88b4 | 3.2457155 | 0.452649475 | 0.243 | 0.052 | 8.2457414 | 22 | tmem88b |
| si:ch211-71n6.4 | 5.7851433 | 0.365646978 | 0.168 | 0.028 | 1.4697157 | 22 | si:ch211-71n6.4 |
| hspg23 | 1.4005799 | 0.367015235 | 0.168 | 0.028 | 3.5581733 | 22 | hspg2 |

| | | | | | | | |
|---|---|---|---|---|---|---|---|
| ckba2 | 2.111106 | 0.539678169 | 0.283 | 0.072 | 5.3632651 | 22 | ckba |
| clic23 | 2.214225 | 0.553420092 | 0.277 | 0.07 | 5.6252395 | 22 | clic2 |
| sema4c1 | 4.493914 | 0.279446571 | 0.121 | 0.016 | 1.1416789 | 22 | sema4c |
| gstt1b6 | 7.486593 | 0.513949602 | 0.312 | 0.088 | 1.9019690 | 22 | gstt1b |
| krt946 | 1.591862 | 0.722397643 | 0.451 | 0.164 | 4.0441277 | 22 | krt94 |
| tmsb4x12 | 2.317978 | 0.516620478 | 0.994 | 0.717 | 5.8888231 | 22 | tmsb4x |
| vamp5 | 2.598809 | 0.342639397 | 0.162 | 0.028 | 6.6022754 | 22 | vamp5 |
| tcf125 | 4.121774 | 0.49983813 | 0.26 | 0.067 | 1.0471369 | 22 | tcf12 |
| tpte | 1.169250 | 0.363928066 | 0.162 | 0.029 | 2.9704806 | 22 | tpte |
| sash1a | 1.418720 | 0.367140673 | 0.179 | 0.034 | 3.6042605 | 22 | sash1a |
| pdlim12 | 1.946062 | 0.455117089 | 0.243 | 0.06 | 4.9439717 | 22 | pdlim1 |
| pmp22b2 | 4.544101 | 0.450149193 | 0.208 | 0.046 | 1.1544291 | 22 | pmp22b |
| cyp3c13 | 9.241645 | 0.476365866 | 0.249 | 0.064 | 2.3478401 | 22 | cyp3c1 |
| tbc1d41 | 2.483377 | 0.281814929 | 0.127 | 0.019 | 6.3090197 | 22 | tbc1d4 |
| dlc11 | 3.211808 | 0.340282778 | 0.168 | 0.032 | 8.1596001 | 22 | dlc1 |
| cd276 | 6.202080 | 0.332297688 | 0.15 | 0.027 | 1.5756386 | 22 | cd276 |
| id17 | 6.265449 | 0.635425558 | 0.376 | 0.13 | 1.5917375 | 22 | id1 |
| phlda22 | 6.677760 | 0.475173187 | 0.22 | 0.052 | 1.6964849 | 22 | phlda2 |
| rnd3a2 | 7.803552 | 0.39897303 | 0.173 | 0.035 | 1.9824925 | 22 | rnd3a |
| zgc:1533172 | 9.925758 | 0.394118555 | 0.185 | 0.039 | 2.5216388 | 22 | zgc:153317 |
| cav14 | 1.013017 | 0.48381637 | 0.208 | 0.048 | 2.5735712 | 22 | cav1 |
| gng102 | 1.647117 | 0.444536731 | 0.237 | 0.06 | 4.1845027 | 22 | gng10 |
| mef2cb1 | 1.926781 | 0.337408425 | 0.156 | 0.029 | 4.8949892 | 22 | mef2cb |
| myo1cb2 | 3.762704 | 0.395678262 | 0.173 | 0.036 | 9.5591511 | 22 | myo1cb |
| zgc:1951736 | 3.977453 | 0.573309445 | 0.335 | 0.113 | 1.0104721 | 22 | zgc:195173 |
| smad11 | 6.831531 | 0.286535095 | 0.127 | 0.02 | 1.7355504 | 22 | smad1 |
| slc22a31 | 1.032925 | 0.274914008 | 0.121 | 0.019 | 2.6241477 | 22 | slc22a31 |
| si:ch73-138n13.12 | 3.187395 | 0.374486738 | 0.191 | 0.043 | 8.0975777 | 22 | si:ch73-138n13.1 |
| efna1a | 4.433819 | 0.262891617 | 0.104 | 0.015 | 1.1264119 | 22 | efna1a |
| XLOC-0160945 | 8.811254 | 0.47624142 | 0.277 | 0.082 | 2.2384992 | 22 | XLOC-016094 |
| aplp24 | 1.157710 | 0.469989634 | 0.26 | 0.076 | 2.9411645 | 22 | aplp2 |
| tcf43 | 1.194802 | 0.287240252 | 0.133 | 0.023 | 3.0353951 | 22 | tcf4 |
| srgap1b | 1.510739 | 0.256555638 | 0.104 | 0.015 | 3.8380335 | 22 | srgap1b |
| mmp14b4 | 1.569939 | 0.464277871 | 0.237 | 0.066 | 3.9884315 | 22 | mmp14b |
| adgrl3.12 | 2.759883 | 0.337211411 | 0.15 | 0.03 | 7.0114840 | 22 | adgrl3.1 |

| | | | | | | | |
|---|---|---|---|---|---|---|---|
| klf2b | 2.993032 | 0.312196992 | 0.121 | 0.02 | 7.6037998 | 22 | klf2b |
| asap1b | 3.646800 | 0.352813408 | 0.162 | 0.034 | 9.2646961 | 22 | asap1b |
| ehd41 | 4.629485 | 0.389668157 | 0.162 | 0.034 | 1.1761207 | 22 | ehd4 |
| add3a7 | 5.180852 | 0.584334632 | 0.358 | 0.137 | 1.3161956 | 22 | add3a |
| palm1a | 8.251034 | 0.267483075 | 0.116 | 0.019 | 2.0961752 | 22 | palm1a |
| wwtr12 | 1.667012 | 0.300218957 | 0.145 | 0.028 | 4.2350447 | 22 | wwtr1 |
| si:dkey-9i23.164 | 2.035721 | 0.379252888 | 0.197 | 0.049 | 5.1717494 | 22 | si:dkey-9i23.16 |
| vwa11 | 2.137786 | 0.302460911 | 0.121 | 0.021 | 5.4310470 | 22 | vwa1 |
| arhgef7b | 2.377210 | 0.255639882 | 0.121 | 0.021 | 6.0393038 | 22 | arhgef7b |
| chka2 | 1.348726 | 0.300042482 | 0.145 | 0.029 | 3.4264401 | 22 | chka |
| lima1a | 2.114463 | 0.36971681 | 0.197 | 0.049 | 5.3717938 | 22 | lima1a |
| nos1apa1 | 2.328532 | 0.311568856 | 0.139 | 0.028 | 5.9156361 | 22 | nos1apa |
| ms4a17a.93 | 2.675413 | 0.44525635 | 0.231 | 0.069 | 6.7968890 | 22 | ms4a17a.9 |
| stard8 | 2.990689 | 0.261912415 | 0.127 | 0.023 | 7.5978463 | 22 | stard8 |
| tns2a3 | 5.658908 | 0.32829726 | 0.162 | 0.037 | 1.4376458 | 22 | tns2a |
| ctsla7 | 1.304515 | 0.593384049 | 0.601 | 0.373 | 3.3141219 | 22 | ctsla |
| tjp1a3 | 1.610188 | 0.383971348 | 0.197 | 0.053 | 4.0906839 | 22 | tjp1a |
| gstr1 | 2.102306 | 0.494821369 | 0.329 | 0.127 | 5.3409086 | 22 | gstr |
| snx9b | 2.118105 | 0.402904137 | 0.214 | 0.061 | 5.3810462 | 22 | snx9b |
| hspb15 | 4.090971 | 0.44747273 | 0.254 | 0.081 | 1.0393112 | 22 | hspb1 |
| cnn3a6 | 4.324000 | 0.430159425 | 0.225 | 0.068 | 1.0985122 | 22 | cnn3a |
| anxa134 | 4.394739 | 0.359465468 | 0.197 | 0.053 | 1.1164836 | 22 | anxa13 |
| mrc1b2 | 7.735845 | 0.344130271 | 0.179 | 0.045 | 1.9652914 | 22 | mrc1b |
| rplp17 | 1.431769 | 0.254968129 | 0.988 | 0.882 | 3.6374110 | 22 | rplp1 |
| cald1a6 | 1.508544 | 0.467646708 | 0.318 | 0.119 | 3.8324574 | 22 | cald1a |
| cd81a7 | 1.877664 | 0.546472311 | 0.491 | 0.24 | 4.7702056 | 22 | cd81a |
| pdk2b4 | 4.091983 | 0.504664216 | 0.283 | 0.105 | 1.0395683 | 22 | pdk2b |
| cavin1b4 | 4.180428 | 0.385904308 | 0.173 | 0.045 | 1.0620378 | 22 | cavin1b |
| col4a14 | 4.362051 | 0.531526394 | 0.22 | 0.07 | 1.1081791 | 22 | col4a1 |
| piezo1 | 5.307657 | 0.251705124 | 0.121 | 0.024 | 1.3484103 | 22 | piezo1 |
| rplp2l8 | 6.365937 | 0.315571865 | 0.965 | 0.794 | 1.6172664 | 22 | rplp2l |
| rasgef1bb1 | 6.840650 | 0.273416228 | 0.145 | 0.033 | 1.7378673 | 22 | rasgef1bb |
| sept23 | 7.216561 | 0.49386896 | 0.399 | 0.182 | 1.8333674 | 22 | sept2 |
| mbnl2 | 7.792776 | 0.276688323 | 0.121 | 0.025 | 1.9797548 | 22 | mbnl2 |
| dusp53 | 1.298309 | 0.38097328 | 0.22 | 0.067 | 3.2983557 | 22 | dusp5 |

| | | | | | | | |
|---|---|---|---|---|---|---|---|
| pfn26 | 1.304165 | 0.544445377 | 0.462 | 0.238 | 3.3132312 | 22 | pfn2 |
| myl67 | 1.951301 | 0.439824061 | 0.306 | 0.116 | 4.9572827 | 22 | myl6 |
| appa5 | 2.833323 | 0.401527532 | 0.254 | 0.087 | 7.1980577 | 22 | appa |
| zfp36l2 | 3.647845 | 0.426585868 | 0.283 | 0.101 | 9.2673503 | 22 | zfp36l2 |
| CABZ01076275.11 | 9.012965 | 0.301877893 | 0.104 | 0.02 | 2.2897438 | 22 | CABZ01076275.1 |
| rab8b2 | 1.244707 | 0.395376013 | 0.243 | 0.084 | 3.1621783 | 22 | rab8b |
| tead1b3 | 1.452850 | 0.27993861 | 0.139 | 0.033 | 3.6909668 | 22 | tead1b |
| ms4a17a.113 | 1.550923 | 0.346368697 | 0.179 | 0.052 | 3.9401221 | 22 | ms4a17a.11 |
| sypl2a3 | 2.060394 | 0.35958999 | 0.197 | 0.06 | 5.2344325 | 22 | sypl2a |
| ap1b11 | 2.104171 | 0.346916261 | 0.179 | 0.051 | 5.3456464 | 22 | ap1b1 |
| itm2ba7 | 2.387929 | 0.499633227 | 0.509 | 0.296 | 6.0665349 | 22 | itm2ba |
| npc1 | 3.252643 | 0.272532931 | 0.127 | 0.029 | 8.2633398 | 22 | npc1 |
| pdk4 | 5.430981 | 0.278603131 | 0.127 | 0.03 | 1.3797407 | 22 | pdk4 |
| eva1ba4 | 6.771885 | 0.367062142 | 0.208 | 0.067 | 1.7203975 | 22 | eva1ba |
| lmo4a1 | 7.685073 | 0.271481128 | 0.145 | 0.036 | 1.9523928 | 22 | lmo4a |
| slc25a43 | 9.428621 | 0.284203539 | 0.139 | 0.035 | 2.3953413 | 22 | slc25a43 |
| rpl912 | 1.136562 | 0.372898741 | 0.908 | 0.667 | 2.8874375 | 22 | rpl9 |
| cd6311 | 1.140592 | 0.483645046 | 0.595 | 0.373 | 2.8976739 | 22 | cd63 |
| scarb2a2 | 1.148796 | 0.367415295 | 0.197 | 0.062 | 2.9185177 | 22 | scarb2a |
| fblim12 | 1.175055 | 0.276772637 | 0.145 | 0.037 | 2.9852276 | 22 | fblim1 |
| cdc423 | 1.236652 | 0.49422189 | 0.434 | 0.233 | 3.1417155 | 22 | cdc42 |
| myh103 | 1.656660 | 0.293823885 | 0.156 | 0.042 | 4.2087459 | 22 | myh10 |
| rps248 | 3.354751 | 0.266433109 | 0.925 | 0.778 | 8.5227471 | 22 | rps24 |
| cx433 | 4.520819 | 0.378777546 | 0.185 | 0.057 | 1.1485142 | 22 | cx43 |
| gadd45ba5 | 4.912605 | 0.497869464 | 0.399 | 0.197 | 1.2480473 | 22 | gadd45ba |
| CT030188.12 | 5.956749 | 0.355059986 | 0.179 | 0.055 | 1.5133123 | 22 | CT030188.1 |
| zgc:11418812 | 7.557960 | 0.33691309 | 0.873 | 0.667 | 1.9200997 | 22 | zgc:114188 |
| ftr822 | 8.417741 | 0.271567042 | 0.139 | 0.036 | 2.1385272 | 22 | ftr82 |
| tspan18b2 | 9.743471 | 0.277972001 | 0.15 | 0.041 | 2.4753288 | 22 | tspan18b |
| klf6a6 | 9.979091 | 0.502053158 | 0.364 | 0.18 | 2.5351881 | 22 | klf6a |
| ahnak7 | 1.042506 | 0.437943925 | 0.312 | 0.136 | 2.6484882 | 22 | ahnak |
| krt154 | 1.859253 | 0.483702213 | 0.254 | 0.096 | 4.7234329 | 22 | krt15 |
| spred2b5 | 3.522263 | 0.38560426 | 0.225 | 0.084 | 8.9483112 | 22 | spred2b |
| afdna2 | 5.558217 | 0.266181187 | 0.11 | 0.026 | 1.4120652 | 22 | afdna |
| fstl1b5 | 6.845437 | 0.416870315 | 0.329 | 0.147 | 1.7390835 | 22 | fstl1b |

| | | | | | | | |
|---|---|---|---|---|---|---|---|
| ehd1b2 | 7.878072 | 0.307725407 | 0.168 | 0.052 | 2.0014242 | 22 | ehd1b |
| jund5 | 9.011273 | 0.392198146 | 0.231 | 0.09 | 2.2893140 | 22 | jund |
| qkia6 | 9.599694 | 0.442226737 | 0.382 | 0.199 | 2.4388023 | 22 | qkia |
| rgl11 | 1.075666 | 0.261094927 | 0.139 | 0.039 | 2.7327301 | 22 | rgl1 |
| hsp90ab15 | 1.341271 | 0.251281539 | 0.942 | 0.8 | 3.4075000 | 22 | hsp90ab1 |
| ier2a8 | 1.454495 | 0.485191371 | 0.468 | 0.274 | 3.6951445 | 22 | ier2a |
| mef2aa2 | 1.546540 | 0.369826895 | 0.202 | 0.073 | 3.9289872 | 22 | mef2aa |
| cbx7a3 | 3.417426 | 0.365378474 | 0.162 | 0.052 | 8.6819717 | 22 | cbx7a |
| tshz21 | 5.497246 | 0.251483862 | 0.116 | 0.03 | 1.3965755 | 22 | tshz2 |
| h3f3c9 | 9.715154 | 0.433221038 | 0.486 | 0.304 | 2.4681348 | 22 | h3f3c |
| msna7 | 9.781984 | 0.453947472 | 0.439 | 0.265 | 2.4851130 | 22 | msna |
| si:ch211-204c21.14 | 9.909930 | 0.313598459 | 0.202 | 0.074 | 2.5176177 | 22 | si:ch211-204c21.1 |
| rab5c2 | 1.175961 | 0.367583267 | 0.289 | 0.131 | 2.9875300 | 22 | rab5c |
| tmem176l.16 | 1.374105 | 0.420842866 | 0.382 | 0.201 | 3.4909138 | 22 | tmem176l.1 |
| socs3a8 | 1.435212 | 0.480838209 | 0.52 | 0.314 | 3.6461567 | 22 | socs3a |
| jun7 | 1.973936 | 0.471886137 | 0.445 | 0.25 | 5.0147866 | 22 | jun |
| klf32 | 1.981612 | 0.280280891 | 0.133 | 0.039 | 5.0342876 | 22 | klf3 |
| rpl368 | 2.008995 | 0.264916475 | 0.965 | 0.806 | 5.1038524 | 22 | rpl36 |
| LOC1018844514 | 2.655112 | 0.330811562 | 0.225 | 0.091 | 6.7453140 | 22 | LOC101884451 |
| rplp28 | 2.883784 | 0.268691188 | 0.942 | 0.782 | 7.3262542 | 22 | rplp2 |
| perp3 | 3.848443 | 0.306400817 | 0.173 | 0.061 | 9.7769698 | 22 | perp |
| rap1b3 | 4.341165 | 0.41307917 | 0.393 | 0.22 | 1.1028731 | 22 | rap1b |
| cnn26 | 4.412590 | 0.409021176 | 0.312 | 0.157 | 1.1210185 | 22 | cnn2 |
| lpar13 | 4.668495 | 0.29804792 | 0.191 | 0.07 | 1.1860313 | 22 | lpar1 |
| ctdspla2 | 4.858647 | 0.268395119 | 0.133 | 0.04 | 1.2343393 | 22 | ctdspla |
| rpsa8 | 5.515765 | 0.262781228 | 0.948 | 0.785 | 1.4012803 | 22 | rpsa |
| cox6b16 | 1.336782 | 0.395978158 | 0.26 | 0.122 | 3.3960959 | 22 | cox6b1 |
| myl9a3 | 1.393136 | 0.348375334 | 0.139 | 0.045 | 3.5392639 | 22 | myl9a |
| cdc42se1 | 1.732013 | 0.312034847 | 0.202 | 0.079 | 4.4001790 | 22 | cdc42se1 |
| rps1011 | 2.864034 | 0.272416867 | 0.902 | 0.726 | 7.2760784 | 22 | rps10 |
| junba8 | 3.024665 | 0.429503977 | 0.514 | 0.32 | 7.6841631 | 22 | junba |
| svild5 | 4.524918 | 0.355571673 | 0.202 | 0.084 | 0.000115 | 22 | svild |
| XLOC-0247502 | 5.092957 | 0.327487248 | 0.168 | 0.064 | 0.000129 | 22 | XLOC-024750 |
| CR383676.112 | 5.132720 | 0.309470396 | 0.74 | 0.718 | 0.00013 | 22 | CR383676.1 |
| btg18 | 5.263479 | 0.387568369 | 0.526 | 0.356 | 0.000134 | 22 | btg1 |

| | | | | | | | |
|---|---|---|---|---|---|---|---|
| zfp36l1b6 | 5.462612 | 0.369572525 | 0.289 | 0.14 | 0.000139 | 22 | zfp36l1b |
| sdcbp27 | 9.217495 | 0.391662704 | 0.387 | 0.228 | 0.000234 | 22 | sdcbp2 |
| calm2a1 | 9.319860 | 0.399843718 | 0.37 | 0.209 | 0.000237 | 22 | calm2a |
| foxo1a1 | 1.105526 | 0.260973657 | 0.127 | 0.042 | 0.000281 | 22 | foxo1a |
| pim16 | 1.246645 | 0.369984389 | 0.324 | 0.169 | 0.000317 | 22 | pim1 |
| cox8b3 | 1.401269 | 0.337599792 | 0.225 | 0.101 | 0.000356 | 22 | cox8b |
| dusp6 | 2.342736 | 0.257349841 | 0.15 | 0.055 | 0.000595 | 22 | dusp6 |
| degs1 | 3.247097 | 0.278637116 | 0.162 | 0.063 | 0.000825 | 22 | degs1 |
| CR318588.33 | 3.314427 | 0.28216959 | 0.145 | 0.051 | 0.000842 | 22 | CR318588.3 |
| nr2f23 | 3.423323 | 0.274384784 | 0.145 | 0.052 | 0.00087 | 22 | nr2f2 |
| cyfip11 | 3.558661 | 0.258453866 | 0.162 | 0.062 | 0.000904 | 22 | cyfip1 |
| sept121 | 3.683149 | 0.251339755 | 0.168 | 0.065 | 0.000936 | 22 | sept12 |
| txn3 | 5.070221 | 0.275845293 | 0.145 | 0.052 | 0.001288 | 22 | txn |
| fosb4 | 7.370692 | 0.40365527 | 0.249 | 0.122 | 0.001873 | 22 | fosb |
| rps2112 | 7.488154 | 0.251190592 | 0.942 | 0.775 | 0.001902 | 22 | rps21 |
| kctd5a2 | 1.180068 | 0.336342276 | 0.329 | 0.186 | 0.002998 | 22 | kctd5a |
| aoc25 | 1.408388 | 0.391915429 | 0.179 | 0.077 | 0.003578 | 22 | aoc2 |
| hopx2 | 1.422947 | 0.286856589 | 0.162 | 0.065 | 0.003615 | 22 | hopx |
| atf4b7 | 3.487239 | 0.34788226 | 0.422 | 0.278 | 0.008859 | 22 | atf4b |
| mt-nd111 | 3.735141 | 0.342464684 | 0.538 | 0.397 | 0.009489 | 22 | mt-nd1 |
| rhoaa6 | 4.168420 | 0.327038468 | 0.306 | 0.177 | 0.01059 | 22 | rhoaa |
| btg26 | 5.021920 | 0.372375709 | 0.486 | 0.35 | 0.012758 | 22 | btg2 |
| txnipa5 | 5.090910 | 0.351480648 | 0.387 | 0.244 | 0.012933 | 22 | txnipa |
| hmgn26 | 6.780436 | 0.385867051 | 0.613 | 0.468 | 0.017226 | 22 | hmgn2 |
| pde4ba5 | 8.357236 | 0.311786029 | 0.266 | 0.142 | 0.021232 | 22 | pde4ba |
| si:dkeyp-19e1.32 | 1.357671 | 0.251373652 | 0.15 | 0.063 | 0.034492 | 22 | si:dkeyp-19e1.3 |
| selenot1b1 | 1.405225 | 0.288774035 | 0.179 | 0.083 | 0.0357 | 22 | selenot1b |
| ier2b9 | 1.410014 | 0.363846886 | 0.37 | 0.237 | 0.035821 | 22 | ier2b |
| rpl22l16 | 1.434270 | 0.309226648 | 0.566 | 0.426 | 0.036438 | 22 | rpl22l1 |
| rps27.29 | 1.752210 | 0.254651055 | 0.688 | 0.554 | 0.044515 | 22 | rps27.2 |
| cfd3 | 2.199241 | 0.354628936 | 0.162 | 0.071 | 0.055872 | 22 | cfd |
| vdac34 | 3.155834 | 0.324140816 | 0.289 | 0.169 | 0.080174 | 22 | vdac3 |
| sptan13 | 3.384919 | 0.268618029 | 0.139 | 0.059 | 0.085994 | 22 | sptan1 |
| cltca | 3.503898 | 0.298877359 | 0.185 | 0.091 | 0.089017 | 22 | cltca |
| LOC1039092844 | 3.760101 | 0.264449591 | 0.168 | 0.077 | 0.095525 | 22 | LOC103909284 |

| | | | | | | |
|---|---|---|---|---|---|---|
| rhoab6 | 5.535598 | 0.334010889 | 0.324 | 0.208 | 0.140632 | 22 | rhoab |
| mcl1b7 | 6.284988 | 0.326513621 | 0.382 | 0.251 | 0.15967 | 22 | mcl1b |
| hmgn610 | 6.336262 | 0.314028798 | 0.439 | 0.312 | 0.160973 | 22 | hmgn6 |
| mt-atp611 | 1.008244 | 0.272883433 | 0.665 | 0.557 | 0.256144 | 22 | mt-atp6 |
| crip17 | 1.027077 | 0.316690027 | 0.422 | 0.285 | 0.260929 | 22 | crip1 |
| tspan143 | 1.047078 | 0.298231913 | 0.237 | 0.132 | 0.26601 | 22 | tspan14 |
| csrp1a2 | 1.350191 | 0.257982248 | 0.168 | 0.081 | 0.343016 | 22 | csrp1a |
| gnb1a2 | 1.548720 | 0.298443133 | 0.318 | 0.203 | 0.393452 | 22 | gnb1a |
| myl12.16 | 3.087850 | 0.272797662 | 0.543 | 0.427 | 0.784468 | 22 | myl12.1 |
| sumo2b7 | 3.445329 | 0.290432811 | 0.41 | 0.297 | 0.875286 | 22 | sumo2b |
| si:ch73-234b20.52 | 3.516937 | 0.253566626 | 0.197 | 0.106 | 0.893478 | 22 | si:ch73-234b20.5 |
| arpc1a1 | 3.651892 | 0.25613984 | 0.249 | 0.142 | 0.927763 | 22 | arpc1a |
| ywhabl3 | 4.327940 | 0.320612195 | 0.399 | 0.302 | 1 | 22 | ywhabl |
| mt-nd29 | 4.457248 | 0.295047352 | 0.497 | 0.393 | 1 | 22 | mt-nd2 |
| tpm35 | 4.506447 | 0.299171619 | 0.393 | 0.285 | 1 | 22 | tpm3 |
| aip | 4.882324 | 0.26609993 | 0.191 | 0.103 | 1 | 22 | aip |
| hnrnpa0a2 | 4.901527 | 0.255504536 | 0.162 | 0.082 | 1 | 22 | hnrnpa0a |
| glula4 | 4.943997 | 0.264162425 | 0.225 | 0.125 | 1 | 22 | glula |
| gdi21 | 5.300753 | 0.273044759 | 0.312 | 0.203 | 1 | 22 | gdi2 |
| si:ch211-196l7.410 | 6.025958 | 0.288590468 | 0.434 | 0.331 | 1 | 22 | si:ch211-196l7.4 |
| snx1a1 | 8.155176 | 0.270519735 | 0.214 | 0.123 | 1 | 22 | snx1a |
| fosab8 | 8.221378 | 0.351598734 | 0.462 | 0.362 | 1 | 22 | fosab |
| si:dkey-7j14.66 | 0.0002 | 0.271454782 | 0.295 | 0.197 | 1 | 22 | si:dkey-7j14.6 |
| pfn2l6 | 0.0002 | 0.265776391 | 0.324 | 0.216 | 1 | 22 | pfn2l |
| GCA4 | 0.0002 | 0.271344197 | 0.295 | 0.201 | 1 | 22 | GCA |
| gabarapb7 | 0.0002 | 0.271463495 | 0.399 | 0.306 | 1 | 22 | gabarapb |
| si:ch73-1a9.310 | 0.0004 | 0.250661457 | 0.434 | 0.316 | 1 | 22 | si:ch73-1a9.3 |
| hnrnpa0b6 | 0.0005 | 0.257858044 | 0.434 | 0.342 | 1 | 22 | hnrnpa0b |
| eif4ebp23 | 0.0007 | 0.26666456 | 0.277 | 0.193 | 1 | 22 | eif4ebp2 |
| zgc:1583436 | 0.0008 | 0.252638513 | 0.26 | 0.17 | 1 | 22 | zgc:158343 |
| ywhaqb6 | 0.001 | 0.257519778 | 0.231 | 0.156 | 1 | 22 | ywhaqb |
| tuba8l5 | 0.0088 | 0.253181145 | 0.272 | 0.21 | 1 | 22 | tuba8l |
| hbbe2 | 0 | 3.105083728 | 1 | 0.002 | 0 | 23 | hbbe2 |
| hbae5 | 0 | 3.013773857 | 0.942 | 0.002 | 0 | 23 | hbae5 |
| hbae1.3 | 0 | 2.971472656 | 1 | 0.001 | 0 | 23 | hbae1.3 |

| | | | | | | | |
|---|---|---|---|---|---|---|---|
| hbae1.1 | 0 | 2.797808698 | 0.994 | 0.002 | 0 | 23 | hbae1.1 |
| hbbe1.1 | 0 | 2.665553853 | 0.959 | 0.001 | 0 | 23 | hbbe1.1 |
| hbbe1.2 | 0 | 2.619182441 | 0.971 | 0.001 | 0 | 23 | hbbe1.2 |
| hbbe1.3 | 0 | 2.437364705 | 0.965 | 0 | 0 | 23 | hbbe1.3 |
| si:ch211-250g4.3 | 0 | 2.262208044 | 1 | 0.027 | 0 | 23 | si:ch211-250g4.3 |
| si:ch211-103n10.5 | 0 | 2.233857533 | 0.93 | 0.022 | 0 | 23 | si:ch211-103n10.5 |
| cahz | 0 | 1.937755618 | 0.953 | 0.038 | 0 | 23 | cahz |
| hbae3 | 0 | 1.885458475 | 0.678 | 0 | 0 | 23 | hbae3 |
| creg1 | 0 | 1.746938769 | 0.959 | 0.067 | 0 | 23 | creg1 |
| nt5c2l1 | 0 | 1.744216428 | 0.901 | 0.01 | 0 | 23 | nt5c2l1 |
| zgc:163057 | 0 | 1.570269377 | 0.83 | 0.003 | 0 | 23 | zgc:163057 |
| hemgn | 0 | 1.444829164 | 0.784 | 0.012 | 0 | 23 | hemgn |
| alas2 | 0 | 1.405224766 | 0.754 | 0.008 | 0 | 23 | alas2 |
| blvrb | 0 | 1.22829952 | 0.731 | 0.039 | 0 | 23 | blvrb |
| si:ch211-207c6.2 | 0 | 1.144680937 | 0.567 | 0.005 | 0 | 23 | si:ch211-207c6.2 |
| si:dkey-240n22.3 | 0 | 1.043979643 | 0.48 | 0.003 | 0 | 23 | si:dkey-240n22.3 |
| aqp1a.1 | 0 | 0.958076502 | 0.509 | 0.005 | 0 | 23 | aqp1a.1 |
| epb41b | 0 | 0.859040377 | 0.468 | 0.003 | 0 | 23 | epb41b |
| rfesd | 0 | 0.735876324 | 0.363 | 0.002 | 0 | 23 | rfesd |
| slc4a1a | 0 | 0.674912668 | 0.351 | 0.008 | 0 | 23 | slc4a1a |
| gp1bb | 0 | 0.617444527 | 0.31 | 0.006 | 0 | 23 | gp1bb |
| add2 | 0 | 0.570615785 | 0.298 | 0.006 | 0 | 23 | add2 |
| gp9 | 0 | 0.550986564 | 0.269 | 0.002 | 0 | 23 | gp9 |
| sept3 | 0 | 0.494966229 | 0.257 | 0.001 | 0 | 23 | sept3 |
| klf1 | 0 | 0.483281186 | 0.24 | 0.003 | 0 | 23 | klf1 |
| nmt1b | 0 | 0.475437139 | 0.24 | 0.003 | 0 | 23 | nmt1b |
| XLOC-016167 | 0 | 0.470016082 | 0.246 | 0.004 | 0 | 23 | XLOC-016167 |
| LOC108183423 | 0 | 0.459847177 | 0.193 | 0.002 | 0 | 23 | LOC108183423 |
| slc6a22.2 | 0 | 0.396727933 | 0.193 | 0 | 0 | 23 | slc6a22.2 |
| BX005417.1 | 0 | 0.38627113 | 0.205 | 0.001 | 0 | 23 | BX005417.1 |
| bnip3lb1 | 1.536739 | 1.359641963 | 0.825 | 0.072 | 3.9040877 | 23 | bnip3lb |
| eef2l2 | 5.104119 | 0.410116444 | 0.181 | 0.002 | 1.2967015 | 23 | eef2l2 |
| ak1 | 3.971322 | 1.512171915 | 0.865 | 0.09 | 1.0089145 | 23 | ak1 |
| slc20a1a | 1.912135 | 0.495033704 | 0.263 | 0.006 | 4.8577814 | 23 | slc20a1a |
| TCIM (1 of many)2 | 1.672489 | 1.158940922 | 0.62 | 0.043 | 4.2489606 | 23 | TCIM (1 of many) |

| | | | | | | | |
|---|---|---|---|---|---|---|---|
| tal1 | 2.212565 | 0.387716277 | 0.181 | 0.002 | 5.6210238 | 23 | tal1 |
| prkab1a | 1.754948 | 0.743864349 | 0.386 | 0.016 | 4.4584472 | 23 | prkab1a |
| ptger3 | 3.250166 | 0.347566582 | 0.158 | 0.002 | 8.2570474 | 23 | ptger3 |
| CR293511.1 | 1.018521 | 0.265001 | 0.135 | 0.002 | 2.5875541 | 23 | CR293511.1 |
| tal2 | 2.396744 | 0.279752686 | 0.105 | 0.001 | 6.0889283 | 23 | tal2 |
| dmtn | 7.443129 | 0.382535296 | 0.17 | 0.003 | 1.8909270 | 23 | dmtn |
| tspo | 3.426597 | 1.031393111 | 0.608 | 0.061 | 8.7052698 | 23 | tspo |
| faxdc2 | 8.106714 | 0.540818767 | 0.298 | 0.014 | 2.0595108 | 23 | faxdc2 |
| lmo21 | 9.765148 | 0.707888762 | 0.363 | 0.022 | 2.4808359 | 23 | lmo2 |
| XLOC-020187 | 3.207794 | 0.255980443 | 0.111 | 0.001 | 8.1494016 | 23 | XLOC-020187 |
| si:dkey-93h22.7 | 2.082850 | 0.284764295 | 0.123 | 0.002 | 5.2914823 | 23 | si:dkey-93h22.7 |
| gpx1a4 | 2.570097 | 1.260953026 | 0.901 | 0.229 | 6.5293315 | 23 | gpx1a |
| fth1a | 2.454081 | 1.216649181 | 1 | 0.608 | 6.2345939 | 23 | fth1a |
| zgc:926063 | 6.824730 | 0.875671273 | 0.561 | 0.091 | 1.7338227 | 23 | zgc:92606 |
| si:ch211-161c3.6 | 3.925132 | 0.419357161 | 0.222 | 0.014 | 9.9717989 | 23 | si:ch211-161c3.6 |
| bloc1s61 | 5.071763 | 0.850220739 | 0.556 | 0.091 | 1.2884814 | 23 | bloc1s6 |
| BX649411.2 | 1.289366 | 0.353433971 | 0.193 | 0.012 | 3.2756366 | 23 | BX649411.2 |
| CABZ01086877.1 | 7.746760 | 0.327664321 | 0.175 | 0.01 | 1.9680646 | 23 | CABZ01086877.1 |
| tpd52l2b | 2.190861 | 0.809734396 | 0.561 | 0.105 | 5.5658840 | 23 | tpd52l2b |
| si:ch211-260e23.9 | 2.539397 | 0.64300412 | 0.409 | 0.059 | 6.4513396 | 23 | si:ch211-260e23.9 |
| DIPK2B | 1.459010 | 0.25183464 | 0.135 | 0.006 | 3.7066165 | 23 | DIPK2B |
| purba | 2.072832 | 0.646648506 | 0.444 | 0.071 | 5.2660320 | 23 | purba |
| adrb3a | 2.565351 | 0.342572036 | 0.17 | 0.011 | 6.5172746 | 23 | adrb3a |
| tmem63bb | 6.756078 | 0.26516045 | 0.14 | 0.007 | 1.7163818 | 23 | tmem63bb |
| mibp7 | 2.105560 | 0.929404951 | 0.789 | 0.194 | 5.3491771 | 23 | mibp |
| cox7a12 | 1.098906 | 0.442260495 | 0.304 | 0.037 | 2.7917723 | 23 | cox7a1 |
| abtb1 | 1.212523 | 0.390440907 | 0.228 | 0.022 | 3.0804164 | 23 | abtb1 |
| mibp28 | 4.513372 | 0.820913431 | 0.959 | 0.517 | 1.1466221 | 23 | mibp2 |
| rhag1 | 5.328822 | 0.289565409 | 0.164 | 0.011 | 1.3537873 | 23 | rhag |
| akt1s1 | 2.310664 | 0.596431885 | 0.386 | 0.061 | 5.8702421 | 23 | akt1s1 |
| cdkn1a2 | 9.992718 | 0.588021578 | 0.327 | 0.046 | 2.5386501 | 23 | cdkn1a |
| cxcr4a | 5.390116 | 0.377941447 | 0.199 | 0.017 | 1.3693590 | 23 | cxcr4a |
| txnipa6 | 1.467819 | 0.844976832 | 0.778 | 0.239 | 3.7289949 | 23 | txnipa |
| LOC101885776 | 6.644035 | 0.25870592 | 0.146 | 0.01 | 1.6879171 | 23 | LOC101885776 |
| si:dkey-100n23.3 | 1.391356 | 0.413321889 | 0.211 | 0.021 | 3.5347414 | 23 | si:dkey-100n23.3 |

| | | | | | | | |
|---|---|---|---|---|---|---|---|
| selenoj | 7.899732 | 0.574890636 | 0.357 | 0.06 | 2.0069270 | 23 | selenoj |
| clic24 | 1.404701 | 0.565418083 | 0.392 | 0.068 | 3.5686429 | 23 | clic2 |
| tbcelb | 7.653764 | 0.35704602 | 0.175 | 0.016 | 1.9444387 | 23 | tbcelb |
| si:ch211-227m13.12 | 3.868321 | 0.348058433 | 0.181 | 0.017 | 9.8274715 | 23 | si:ch211-227m13.1 |
| clgn | 5.468896 | 0.326388701 | 0.17 | 0.015 | 1.3893730 | 23 | clgn |
| lrrc8da1 | 4.009010 | 0.36843047 | 0.222 | 0.026 | 1.0184891 | 23 | lrrc8da |
| XLOC-043857 | 3.471607 | 0.605701905 | 0.374 | 0.072 | 8.8196185 | 23 | XLOC-043857 |
| prdx25 | 1.354599 | 0.812337081 | 0.883 | 0.418 | 3.4413593 | 23 | prdx2 |
| jak2a | 7.405583 | 0.363803289 | 0.205 | 0.025 | 1.8813885 | 23 | jak2a |
| rbm24a | 2.097446 | 0.335242597 | 0.199 | 0.024 | 5.3285640 | 23 | rbm24a |
| castor2 | 7.830820 | 0.47206634 | 0.298 | 0.052 | 1.9894200 | 23 | castor2 |
| ctsf2 | 9.353300 | 0.381458302 | 0.234 | 0.035 | 2.3762060 | 23 | ctsf |
| rnf102 | 5.076473 | 0.580658879 | 0.45 | 0.126 | 1.2896780 | 23 | rnf10 |
| gbp1 | 9.020077 | 0.562980104 | 0.444 | 0.12 | 2.2915507 | 23 | gbp |
| gabarapb8 | 1.025729 | 0.652671986 | 0.731 | 0.301 | 2.6058669 | 23 | gabarapb |
| vdac24 | 2.303193 | 0.650223955 | 0.731 | 0.323 | 5.8512631 | 23 | vdac2 |
| FBXO48 | 3.346428 | 0.423895571 | 0.287 | 0.061 | 8.5016004 | 23 | FBXO48 |
| pfkfb4b4 | 3.915035 | 0.525783206 | 0.415 | 0.118 | 9.9461470 | 23 | pfkfb4b |
| gstm.16 | 1.941363 | 0.675027488 | 0.655 | 0.276 | 4.9320335 | 23 | gstm.1 |
| slc16a102 | 5.275794 | 0.307313021 | 0.164 | 0.024 | 1.3403157 | 23 | slc16a10 |
| elocb3 | 1.101652 | 0.556997244 | 0.538 | 0.193 | 2.7987469 | 23 | elocb |
| reep3b1 | 1.116078 | 0.377855258 | 0.287 | 0.065 | 2.8353984 | 23 | reep3b |
| ccni5 | 1.199486 | 0.610110745 | 0.69 | 0.316 | 3.0472958 | 23 | ccni |
| glrx51 | 6.373910 | 0.429566888 | 0.345 | 0.094 | 1.6192919 | 23 | glrx5 |
| histh1l4 | 7.358110 | 0.499764455 | 0.398 | 0.12 | 1.8693278 | 23 | histh1l |
| si:ch211-147a11.3 | 1.971580 | 0.369084612 | 0.316 | 0.081 | 5.0088012 | 23 | si:ch211-147a11.3 |
| ak3 | 3.636910 | 0.308810869 | 0.17 | 0.031 | 9.2395699 | 23 | ak3 |
| fam53b | 4.173071 | 0.371394497 | 0.287 | 0.075 | 1.0601687 | 23 | fam53b |
| gabarapa2 | 9.157209 | 0.472118722 | 0.485 | 0.179 | 2.3263890 | 23 | gabarapa |
| ypel5 | 1.888626 | 0.293849104 | 0.211 | 0.047 | 4.7980560 | 23 | ypel5 |
| pde7a1 | 3.045731 | 0.344461832 | 0.251 | 0.064 | 7.7376818 | 23 | pde7a |
| gabarapl21 | 3.825342 | 0.362656863 | 0.292 | 0.083 | 9.7182830 | 23 | gabarapl2 |
| chmp5b2 | 8.616142 | 0.407679599 | 0.38 | 0.127 | 2.1889311 | 23 | chmp5b |
| bada | 9.439584 | 0.288855551 | 0.164 | 0.033 | 2.3981263 | 23 | bada |
| tcp11l2 | 1.787886 | 0.257236801 | 0.175 | 0.037 | 4.5421244 | 23 | tcp11l2 |

| | | | | | | | |
|---|---|---|---|---|---|---|---|
| zgc:560951 | 1.516146 | 0.393016808 | 0.316 | 0.1 | 3.8517705 | 23 | zgc:56095 |
| isca1 | 4.023693 | 0.256658017 | 0.193 | 0.045 | 1.0222192 | 23 | isca1 |
| ppp1r14bb2 | 1.267574 | 0.277247756 | 0.175 | 0.042 | 3.2202723 | 23 | ppp1r14bb |
| anp32a | 1.313034 | 0.432080419 | 0.462 | 0.195 | 3.3357652 | 23 | anp32a |
| oaz1b6 | 2.703020 | 0.458661884 | 0.696 | 0.358 | 6.8670223 | 23 | oaz1b |
| mthfd21 | 6.176207 | 0.274825293 | 0.146 | 0.032 | 1.5690656 | 23 | mthfd2 |
| fbxo9 | 9.453168 | 0.270151909 | 0.193 | 0.05 | 2.4015775 | 23 | fbxo9 |
| hbp12 | 1.289560 | 0.283238009 | 0.211 | 0.058 | 3.2761276 | 23 | hbp1 |
| nt5dc23 | 3.198643 | 0.377418815 | 0.251 | 0.081 | 8.1261544 | 23 | nt5dc2 |
| map1lc3a6 | 3.891121 | 0.321460951 | 0.251 | 0.08 | 9.8853930 | 23 | map1lc3a |
| zgc:163080 | 1.018166 | 0.288137658 | 0.269 | 0.089 | 2.5866517 | 23 | zgc:163080 |
| pcbp21 | 4.761867 | 0.382766142 | 0.485 | 0.215 | 1.2097523 | 23 | pcbp2 |
| coq8a | 1.204316 | 0.26803846 | 0.181 | 0.05 | 3.0595656 | 23 | coq8a |
| ISCU (1 of many)1 | 6.401003 | 0.354663524 | 0.316 | 0.124 | 1.6261749 | 23 | ISCU (1 of many) |
| tmem591 | 2.230809 | 0.29813683 | 0.31 | 0.126 | 5.6673708 | 23 | tmem59 |
| tspan363 | 1.534273 | 0.275660626 | 0.357 | 0.156 | 3.8978227 | 23 | tspan36 |
| nupr1a | 1.788749 | 0.292898507 | 0.357 | 0.164 | 4.5443182 | 23 | nupr1a |
| irf2bp2b5 | 7.607504 | 0.276222725 | 0.246 | 0.102 | 1.9326865 | 23 | irf2bp2b |
| ndufb83 | 4.533839 | 0.273418949 | 0.491 | 0.265 | 0.000115 | 23 | ndufb8 |
| eif1b6 | 7.119994 | 0.325025369 | 0.673 | 0.405 | 0.000181 | 23 | eif1b |
| atp5if1b3 | 1.509679 | 0.307211319 | 0.386 | 0.207 | 0.000384 | 23 | atp5if1b |
| atp1b1a6 | 3.852149 | 0.285759311 | 0.421 | 0.23 | 0.000979 | 23 | atp1b1a |
| pnrc212 | 4.733510 | 0.318230734 | 0.906 | 0.596 | 0.012025 | 23 | pnrc2 |
| fam32a1 | 8.808439 | 0.258081473 | 0.485 | 0.29 | 0.022378 | 23 | fam32a |
| mbnl15 | 2.023833 | 0.254691041 | 0.333 | 0.187 | 0.051416 | 23 | mbnl1 |
| igl4v8 | 5.641307 | 0.970769793 | 0.413 | 0.025 | 1.4331742 | 24 | igl4v8 |
| LOC101886766 | 1.330790 | 0.589046152 | 0.222 | 0.013 | 3.3808731 | 24 | LOC101886766 |
| zgc:1534996 | 2.338786 | 0.949600576 | 0.97 | 0.418 | 5.9416875 | 24 | zgc:153499 |
| zp36 | 7.989661 | 0.902552175 | 0.988 | 0.477 | 2.0297735 | 24 | zp3 |
| si:dkey-241l7.26 | 5.805337 | 0.875164933 | 0.994 | 0.526 | 1.4748460 | 24 | si:dkey-241l7.2 |
| si:ch211-125e6.126 | 2.358018 | 0.915296866 | 0.952 | 0.408 | 5.9905447 | 24 | si:ch211-125e6.12 |
| zp2.66 | 3.502340 | 0.917504917 | 0.934 | 0.408 | 8.8976970 | 24 | zp2.6 |
| si:zfos-1505d6.36 | 1.185541 | 0.913816205 | 0.982 | 0.418 | 3.0118676 | 24 | si:zfos-1505d6.3 |
| si:dkey-229d11.36 | 1.403891 | 0.92337942 | 0.862 | 0.357 | 3.5665858 | 24 | si:dkey-229d11.3 |
| si:dkeyp-98a7.96 | 9.274157 | 0.89622102 | 0.928 | 0.418 | 2.3560997 | 24 | si:dkeyp-98a7.9 |

| | | | | | | | |
|---|---|---|---|---|---|---|---|
| si:ch73-189n23.16 | 9.804602 | 0.89664532 | 0.958 | 0.428 | 2.4908593 | 24 | si:ch73-189n23.1 |
| h2af1al6 | 2.765654 | 0.87882083 | 1 | 0.486 | 7.0261457 | 24 | h2af1al |
| cldnd6 | 4.372763 | 0.88181778 | 1 | 0.494 | 1.1109005 | 24 | cldnd |
| mt26 | 1.284884 | 0.877084474 | 0.958 | 0.437 | 3.2642498 | 24 | mt2 |
| zp2.36 | 2.679120 | 0.885478117 | 0.934 | 0.423 | 6.8063054 | 24 | zp2.3 |
| zp2.56 | 1.057354 | 0.866031055 | 0.958 | 0.46 | 2.6862081 | 24 | zp2.5 |
| zp2.16 | 1.552167 | 0.873884902 | 0.94 | 0.436 | 3.9432803 | 24 | zp2.1 |
| cldng6 | 2.815797 | 0.877858961 | 0.946 | 0.42 | 7.1535332 | 24 | cldng |
| zp3.26 | 1.118183 | 0.851198712 | 0.946 | 0.456 | 2.8407455 | 24 | zp3.2 |
| tmsb56 | 3.340124 | 0.880445849 | 0.916 | 0.399 | 8.4855851 | 24 | tmsb5 |
| si:ch211-12h2.86 | 3.669454 | 0.840396058 | 0.922 | 0.447 | 9.3222488 | 24 | si:ch211-12h2.8 |
| si:dkey-241l7.46 | 5.056922 | 0.815291605 | 0.964 | 0.498 | 1.2847112 | 24 | si:dkey-241l7.4 |
| si:dkey-240e12.66 | 8.127331 | 0.856463466 | 0.952 | 0.464 | 2.0647485 | 24 | si:dkey-240e12.6 |
| zgc:1751356 | 8.568951 | 0.870034051 | 0.91 | 0.391 | 2.1769421 | 24 | zgc:175135 |
| CR790368.16 | 3.715609 | 0.841547372 | 0.916 | 0.425 | 9.4395069 | 24 | CR790368.1 |
| h1m6 | 1.101603 | 0.836570275 | 0.976 | 0.493 | 2.7986236 | 24 | h1m |
| zp3a.26 | 1.267273 | 0.836313395 | 0.868 | 0.401 | 3.2195084 | 24 | zp3a.2 |
| acp5a6 | 1.547612 | 0.83764891 | 0.952 | 0.442 | 3.9317101 | 24 | acp5a |
| si:dkey-241l7.56 | 4.220474 | 0.816979321 | 0.94 | 0.478 | 1.0722116 | 24 | si:dkey-241l7.5 |
| si:ch211-226h8.146 | 3.842986 | 0.828525602 | 0.952 | 0.464 | 9.7631076 | 24 | si:ch211-226h8.14 |
| si:dkeyp-98a7.86 | 7.008323 | 0.838899844 | 0.85 | 0.373 | 1.7804645 | 24 | si:dkeyp-98a7.8 |
| epcam6 | 1.836607 | 0.824682832 | 0.904 | 0.417 | 4.6659015 | 24 | epcam |
| crp26 | 1.910254 | 0.84042631 | 0.886 | 0.399 | 4.8530012 | 24 | crp2 |
| zgc:1952456 | 3.217095 | 0.835638245 | 0.778 | 0.316 | 8.1730308 | 24 | zgc:195245 |
| zgc:1717796 | 1.773588 | 0.817921313 | 0.88 | 0.418 | 4.5058017 | 24 | zgc:171779 |
| si:dkeyp-20e4.86 | 1.339259 | 0.846447987 | 0.784 | 0.332 | 3.4023880 | 24 | si:dkeyp-20e4.8 |
| ccna16 | 1.760198 | 0.81644622 | 0.868 | 0.388 | 4.4717854 | 24 | ccna1 |
| si:ch211-14a17.76 | 7.651130 | 0.818586875 | 0.784 | 0.333 | 1.9437698 | 24 | si:ch211-14a17.7 |
| mid1ip1l6 | 1.009625 | 0.769286926 | 0.904 | 0.457 | 2.5649528 | 24 | mid1ip1l |
| si:ch211-226h8.46 | 1.544745 | 0.819247419 | 0.808 | 0.36 | 3.9244260 | 24 | si:ch211-226h8.4 |
| zp2.26 | 1.734018 | 0.804789164 | 0.904 | 0.432 | 4.4052740 | 24 | zp2.2 |
| zgc:771186 | 2.260703 | 0.808062465 | 0.844 | 0.374 | 5.7433183 | 24 | zgc:77118 |
| si:dkey-241l7.66 | 7.134561 | 0.787071802 | 0.862 | 0.431 | 1.8125353 | 24 | si:dkey-241l7.6 |
| zar1l6 | 7.412200 | 0.80769181 | 0.802 | 0.324 | 1.8830694 | 24 | zar1l |
| CU571163.16 | 9.966901 | 0.822204737 | 0.778 | 0.327 | 2.5320914 | 24 | CU571163.1 |

| | | | | | | | |
|---|---|---|---|---|---|---|---|
| si:dkeyp-46h3.66 | 1.218448 | 0.811104171 | 0.772 | 0.335 | 3.0954694 | 24 | si:dkeyp-46h3.6 |
| zgc:1734436 | 2.280704 | 0.813257714 | 0.695 | 0.269 | 5.7941296 | 24 | zgc:173443 |
| LOC1005374996 | 7.363234 | 0.803063675 | 0.677 | 0.263 | 1.8706296 | 24 | LOC100537499 |
| lamp26 | 7.974707 | 0.750669216 | 0.874 | 0.481 | 2.0259744 | 24 | lamp2 |
| si:dkeyp-98a7.46 | 1.789202 | 0.79257733 | 0.79 | 0.347 | 4.5454692 | 24 | si:dkeyp-98a7.4 |
| nanog6 | 6.758883 | 0.798704353 | 0.719 | 0.294 | 1.7170942 | 24 | nanog |
| AL929032.16 | 6.816491 | 0.803603737 | 0.677 | 0.258 | 1.7317296 | 24 | AL929032.1 |
| si:dkey-90l23.16 | 9.707317 | 0.791343621 | 0.808 | 0.365 | 2.4661440 | 24 | si:dkey-90l23.1 |
| cth16 | 9.767247 | 0.789092493 | 0.737 | 0.309 | 2.4813692 | 24 | cth1 |
| ccnb26 | 2.328574 | 0.794333078 | 0.749 | 0.314 | 5.9157436 | 24 | ccnb2 |
| zgc:1717766 | 2.933577 | 0.77935196 | 0.856 | 0.404 | 7.4527529 | 24 | zgc:171776 |
| btg46 | 3.616543 | 0.775619944 | 0.731 | 0.301 | 9.1878276 | 24 | btg4 |
| LOC1104378216 | 4.572172 | 0.798974172 | 0.737 | 0.34 | 1.1615603 | 24 | LOC110437821 |
| zp3b6 | 8.131943 | 0.779066313 | 0.749 | 0.317 | 2.0659203 | 24 | zp3b |
| wu:fi42e036 | 1.636484 | 0.772949832 | 0.683 | 0.286 | 4.1574900 | 24 | wu:fi42e03 |
| si:dkeyp-50f7.26 | 6.780991 | 0.760179081 | 0.754 | 0.329 | 1.7227108 | 24 | si:dkeyp-50f7.2 |
| BX649337.16 | 1.736492 | 0.765626333 | 0.796 | 0.357 | 4.4115594 | 24 | BX649337.1 |
| slbp26 | 1.774884 | 0.757141551 | 0.737 | 0.325 | 4.5090930 | 24 | slbp2 |
| zgc:1715176 | 3.364413 | 0.762740723 | 0.707 | 0.317 | 8.5472912 | 24 | zgc:171517 |
| si:dkey-229d11.56 | 3.872391 | 0.762554995 | 0.713 | 0.318 | 9.8378112 | 24 | si:dkey-229d11.5 |
| qdprb26 | 8.382687 | 0.755419147 | 0.766 | 0.354 | 2.1296216 | 24 | qdprb2 |
| kpna76 | 1.836303 | 0.749069623 | 0.647 | 0.269 | 4.6651278 | 24 | kpna7 |
| si:ch1073-263o8.26 | 2.997017 | 0.735909744 | 0.563 | 0.215 | 7.6139238 | 24 | si:ch1073-263o8.2 |
| rrm26 | 3.512113 | 0.714769932 | 0.832 | 0.42 | 8.9225247 | 24 | rrm2 |
| LOC1005351626 | 3.745549 | 0.750579204 | 0.623 | 0.252 | 9.5155696 | 24 | LOC100535162 |
| ccnb16 | 4.268271 | 0.737066371 | 0.707 | 0.324 | 1.0843544 | 24 | ccnb1 |
| npm2b6 | 1.032065 | 0.735306134 | 0.719 | 0.325 | 2.6219623 | 24 | npm2b |
| siva16 | 1.048577 | 0.728811001 | 0.749 | 0.359 | 2.6639104 | 24 | siva1 |
| si:dkeyp-46h3.86 | 1.992957 | 0.740536939 | 0.689 | 0.287 | 5.0631095 | 24 | si:dkeyp-46h3.8 |
| si:ch211-226l4.67 | 4.223967 | 0.737643036 | 0.593 | 0.241 | 1.0730989 | 24 | si:ch211-226l4.6 |
| zar16 | 7.663156 | 0.716027848 | 0.743 | 0.361 | 1.9468250 | 24 | zar1 |
| si:ch211-12h2.66 | 7.965864 | 0.712070983 | 0.79 | 0.387 | 2.0237278 | 24 | si:ch211-12h2.6 |
| si:dkey-152b24.66 | 3.695839 | 0.710148601 | 0.599 | 0.243 | 9.3892798 | 24 | si:dkey-152b24.6 |
| zgc:1630306 | 4.022103 | 0.634237687 | 0.359 | 0.105 | 1.0218153 | 24 | zgc:163030 |
| si:dkey-90l23.26 | 5.108260 | 0.71233796 | 0.641 | 0.282 | 1.2977535 | 24 | si:dkey-90l23.2 |

| | | | | | | | |
|---|---|---|---|---|---|---|---|
| si:dkey-19b23.116 | 7.832653 | 0.713807256 | 0.557 | 0.228 | 1.9898856 | 24 | si:dkey-19b23.11 |
| zgc:1362546 | 9.830590 | 0.70591336 | 0.635 | 0.27 | 2.4974614 | 24 | zgc:136254 |
| s100a16 | 1.547095 | 0.698157141 | 0.665 | 0.298 | 3.9303970 | 24 | s100a1 |
| hnrnpabb7 | 6.850637 | 0.57709652 | 0.826 | 0.557 | 1.7404045 | 24 | hnrnpabb |
| si:ch211-132b12.86 | 7.715662 | 0.697982579 | 0.623 | 0.281 | 1.9601641 | 24 | si:ch211-132b12.8 |
| si:dkeyp-46h3.26 | 1.104489 | 0.68024461 | 0.695 | 0.331 | 2.8059547 | 24 | si:dkeyp-46h3.2 |
| tuba4l6 | 1.811047 | 0.68711904 | 0.707 | 0.332 | 4.6009657 | 24 | tuba4l |
| arhgef10la6 | 1.936173 | 0.685270949 | 0.557 | 0.225 | 4.9188497 | 24 | arhgef10la |
| si:dkey-37o8.16 | 2.224738 | 0.687646722 | 0.587 | 0.251 | 5.6519490 | 24 | si:dkey-37o8.1 |
| LOC1018854776 | 4.449268 | 0.699376319 | 0.545 | 0.228 | 1.1303366 | 24 | LOC101885477 |
| thy16 | 7.163398 | 0.656825637 | 0.743 | 0.376 | 1.8198612 | 24 | thy1 |
| zpcx6 | 1.205825 | 0.676966572 | 0.617 | 0.277 | 3.0634006 | 24 | zpcx |
| zgc:1714746 | 1.817847 | 0.67107865 | 0.563 | 0.249 | 4.6182428 | 24 | zgc:171474 |
| LOC1018838976 | 1.363569 | 0.610682687 | 0.389 | 0.134 | 3.4641484 | 24 | LOC101883897 |
| ctss2.16 | 5.296391 | 0.598702334 | 0.772 | 0.45 | 1.3455483 | 24 | ctss2.1 |
| gtf3ab6 | 5.849761 | 0.640498782 | 0.593 | 0.262 | 1.4861318 | 24 | gtf3ab |
| spink2.17 | 8.097222 | 0.623659185 | 0.713 | 0.372 | 2.0570993 | 24 | spink2.1 |
| ca15b6 | 9.418443 | 0.634160358 | 0.503 | 0.204 | 2.3927555 | 24 | ca15b |
| si:dkey-208k4.26 | 9.810334 | 0.635741783 | 0.557 | 0.242 | 2.4923154 | 24 | si:dkey-208k4.2 |
| cpeb1b6 | 1.277874 | 0.625327837 | 0.539 | 0.221 | 3.2464413 | 24 | cpeb1b |
| zp3a.16 | 1.665999 | 0.63200565 | 0.635 | 0.312 | 4.2324706 | 24 | zp3a.1 |
| si:ch211-146l10.86 | 2.460857 | 0.643526862 | 0.557 | 0.251 | 6.2518097 | 24 | si:ch211-146l10.8 |
| gapdh6 | 4.815576 | 0.626513672 | 0.665 | 0.346 | 1.2233970 | 24 | gapdh |
| slc16a36 | 5.320433 | 0.624131589 | 0.497 | 0.205 | 1.3516561 | 24 | slc16a3 |
| si:ch73-14h10.26 | 5.885713 | 0.626616437 | 0.521 | 0.223 | 1.4952655 | 24 | si:ch73-14h10.2 |
| gdf36 | 1.660288 | 0.619336515 | 0.455 | 0.183 | 4.2179637 | 24 | gdf3 |
| zgc:1526526 | 1.951602 | 0.550955195 | 0.299 | 0.093 | 4.9580460 | 24 | zgc:152652 |
| gbgt1l46 | 2.192484 | 0.562474829 | 0.353 | 0.123 | 5.5700062 | 24 | gbgt1l4 |
| tmem704 | 2.455161 | 0.539555284 | 0.287 | 0.09 | 6.2373389 | 24 | tmem70 |
| AL929192.16 | 2.564117 | 0.563424045 | 0.377 | 0.134 | 6.5141412 | 24 | AL929192.1 |
| uchl16 | 2.668627 | 0.615456776 | 0.593 | 0.289 | 6.7796471 | 24 | uchl1 |
| zgc:554617 | 3.797967 | 0.610421434 | 0.527 | 0.24 | 9.6487368 | 24 | zgc:55461 |
| si:dkeyp-46h3.56 | 5.501621 | 0.610617128 | 0.521 | 0.237 | 1.3976869 | 24 | si:dkeyp-46h3.5 |
| gmnn6 | 5.536608 | 0.610259641 | 0.545 | 0.253 | 1.4065753 | 24 | gmnn |
| si:ch211-5k11.86 | 5.566242 | 0.54194675 | 0.623 | 0.295 | 1.4141039 | 24 | si:ch211-5k11.8 |

| | | | | | | | |
|---|---|---|---|---|---|---|---|
| si:ch211-250e5.166 | 7.47131( | 0.604433051 | 0.575 | 0.284 | 1.8980864 | 24 | si:ch211-250e5.16 |
| si:ch211-251f6.66 | 9.18105[illegible] | 0.607365883 | 0.551 | 0.272 | 2.3324478 | 24 | si:ch211-251f6.6 |
| LOC1005368606 | 1.08849[illegible] | 0.587498937 | 0.581 | 0.287 | 2.7653316 | 24 | LOC100536860 |
| crp36 | 1.08957[illegible] | 0.579786552 | 0.437 | 0.179 | 2.7680629 | 24 | crp3 |
| avd6 | 1.36390[illegible] | 0.596050384 | 0.497 | 0.217 | 3.4650127 | 24 | avd |
| zglp16 | 3.17992[illegible] | 0.571037156 | 0.407 | 0.163 | 8.0786078 | 24 | zglp1 |
| zgc:1034827 | 4.90878[illegible] | 0.568246665 | 0.395 | 0.156 | 1.2470767 | 24 | zgc:103482 |
| hbba16 | 5.64874[illegible] | 0.573849381 | 0.629 | 0.293 | 1.4350639 | 24 | hbba1 |
| nasp7 | 1.20969[illegible] | 0.591635775 | 0.575 | 0.293 | 3.0732388 | 24 | nasp |
| si:dkey-42i9.86 | 1.49122[illegible] | 0.527574601 | 0.305 | 0.105 | 3.7884521 | 24 | si:dkey-42i9.8 |
| sinup6 | 1.868681 | 0.571661162 | 0.407 | 0.167 | 4.7473845 | 24 | sinup |
| pigrl2.36 | 2.12214[illegible] | 0.559692397 | 0.413 | 0.168 | 5.3913132 | 24 | pigrl2.3 |
| pabpn1l7 | 3.11769[illegible] | 0.572620091 | 0.509 | 0.234 | 7.9205150 | 24 | pabpn1l |
| si:ch211-146l10.76 | 1.00148[illegible] | 0.545647936 | 0.383 | 0.155 | 2.5442847 | 24 | si:ch211-146l10.7 |
| birc5b6 | 1.039501 | 0.567979898 | 0.473 | 0.213 | 2.6408544 | 24 | birc5b |
| hnrnpa0l6 | 1.29264[illegible] | 0.312270943 | 0.832 | 0.741 | 3.2839733 | 24 | hnrnpa0l |
| si:dkeyp-98a7.76 | 1.76247[illegible] | 0.535527388 | 0.401 | 0.16 | 4.4775797 | 24 | si:dkeyp-98a7.7 |
| zgc:1737427 | 2.13893[illegible] | 0.547637967 | 0.335 | 0.129 | 5.4339605 | 24 | zgc:173742 |
| larp6a6 | 2.24185[illegible] | 0.567862275 | 0.461 | 0.213 | 5.6954339 | 24 | larp6a |
| sybu6 | 8.05550[illegible] | 0.532857126 | 0.347 | 0.136 | 2.0465008 | 24 | sybu |
| mkrn46 | 8.38061[illegible] | 0.54600882 | 0.431 | 0.193 | 2.1290951 | 24 | mkrn4 |
| mos6 | 1.06384[illegible] | 0.496791156 | 0.311 | 0.114 | 2.7027074 | 24 | mos |
| zgc:1655396 | 1.07239( | 0.500180973 | 0.281 | 0.098 | 2.7244088 | 24 | zgc:165539 |
| bzw1b5 | 1.12502[illegible] | 0.539318502 | 0.593 | 0.377 | 2.8581212 | 24 | bzw1b |
| hbaa16 | 1.19409[illegible] | 0.550103068 | 0.497 | 0.233 | 3.0335993 | 24 | hbaa1 |
| si:ch211-286b5.45 | 1.73255[illegible] | 0.406051461 | 0.204 | 0.057 | 4.4015598 | 24 | si:ch211-286b5.4 |
| plk16 | 1.79851[illegible] | 0.516426081 | 0.365 | 0.148 | 4.5691265 | 24 | plk1 |
| setd41 | 2.45738( | 0.414920277 | 0.204 | 0.059 | 6.2429742 | 24 | setd4 |
| zgc:562316 | 2.920021 | 0.53947405 | 0.425 | 0.187 | 7.4183152 | 24 | zgc:56231 |
| CABZ01059627.15 | 3.50922[illegible] | 0.4788513 | 0.305 | 0.112 | 8.9151785 | 24 | CABZ01059627.1 |
| si:ch1073-190k2.16 | 5.69024( | 0.532363869 | 0.365 | 0.154 | 1.4456056 | 24 | si:ch1073-190k2.1 |
| zgc:1139845 | 6.25261[illegible] | 0.440509729 | 0.251 | 0.083 | 1.5884763 | 24 | zgc:113984 |
| elovl7a6 | 8.25804[illegible] | 0.519944374 | 0.365 | 0.152 | 2.0979567 | 24 | elovl7a |
| zgc:566997 | 8.65226[illegible] | 0.533281024 | 0.449 | 0.209 | 2.1981073 | 24 | zgc:56699 |
| pou5f36 | 9.10647[illegible] | 0.49255944 | 0.317 | 0.121 | 2.3135007 | 24 | pou5f3 |

| | | | | | | | |
|---|---|---|---|---|---|---|---|
| tac3b4 | 1.278126 | 0.45764862 | 0.281 | 0.099 | 3.2470812 | 24 | tac3b |
| si:dkey-188i13.117 | 1.334844 | 0.545209067 | 0.527 | 0.29 | 3.3911723 | 24 | si:dkey-188i13.11 |
| zgc:1738566 | 1.461078 | 0.512343696 | 0.407 | 0.181 | 3.7118700 | 24 | zgc:173856 |
| si:dkey-216e24.36 | 1.650419 | 0.462372337 | 0.281 | 0.101 | 4.1928895 | 24 | si:dkey-216e24.3 |
| tmem54a6 | 2.765254 | 0.484970763 | 0.305 | 0.117 | 7.0251286 | 24 | tmem54a |
| st8sia7.13 | 2.913524 | 0.407212792 | 0.198 | 0.058 | 7.4018086 | 24 | st8sia7.1 |
| ccna26 | 4.006507 | 0.526313768 | 0.479 | 0.238 | 1.0178532 | 24 | ccna2 |
| slbp7 | 7.443384 | 0.52153821 | 0.419 | 0.199 | 1.8909918 | 24 | slbp |
| map1lc3c6 | 1.503301 | 0.479471697 | 0.383 | 0.165 | 3.8191371 | 24 | map1lc3c |
| zgc:1138866 | 1.761618 | 0.465928478 | 0.299 | 0.116 | 4.4753912 | 24 | zgc:113886 |
| cldn7b6 | 2.180391 | 0.485818861 | 0.323 | 0.133 | 5.5392833 | 24 | cldn7b |
| yrdc6 | 2.194227 | 0.448935208 | 0.275 | 0.103 | 5.5744345 | 24 | yrdc |
| spsb13 | 2.520855 | 0.447867857 | 0.228 | 0.078 | 6.4042344 | 24 | spsb1 |
| hsd17b12a6 | 3.640323 | 0.475552176 | 0.329 | 0.136 | 9.2482408 | 24 | hsd17b12a |
| elavl27 | 3.881518 | 0.509107739 | 0.395 | 0.186 | 9.8609979 | 24 | elavl2 |
| si:ch73-160p18.36 | 5.798607 | 0.514026411 | 0.407 | 0.198 | 1.4731361 | 24 | si:ch73-160p18.3 |
| zp2l16 | 7.049705 | 0.485517047 | 0.317 | 0.135 | 1.7909777 | 24 | zp2l1 |
| mibp8 | 7.635240 | 0.49613858 | 0.413 | 0.199 | 1.9397327 | 24 | mibp |
| rcc16 | 7.745850 | 0.500189523 | 0.377 | 0.173 | 1.9678331 | 24 | rcc1 |
| fdx16 | 7.831367 | 0.500339513 | 0.401 | 0.198 | 1.9895589 | 24 | fdx1 |
| ccnh6 | 1.099352 | 0.442692586 | 0.275 | 0.106 | 2.7929048 | 24 | ccnh |
| BX284697.16 | 2.011283 | 0.44375478 | 0.251 | 0.094 | 5.1096649 | 24 | BX284697.1 |
| rdh10b6 | 2.877607 | 0.467252369 | 0.341 | 0.15 | 7.3105614 | 24 | rdh10b |
| sox19b5 | 3.527680 | 0.418566886 | 0.228 | 0.081 | 8.9620721 | 24 | sox19b |
| si:ch211-145c1.16 | 3.669637 | 0.496166337 | 0.401 | 0.192 | 9.3227142 | 24 | si:ch211-145c1.1 |
| zgc:925916 | 3.961733 | 0.474510479 | 0.371 | 0.169 | 1.0064784 | 24 | zgc:92591 |
| rbm4.36 | 4.065771 | 0.457487607 | 0.599 | 0.421 | 1.0329093 | 24 | rbm4.3 |
| cdc206 | 5.182422 | 0.462047268 | 0.305 | 0.13 | 1.3165944 | 24 | cdc20 |
| snx10a6 | 5.368802 | 0.427016951 | 0.269 | 0.105 | 1.3639442 | 24 | snx10a |
| lsm14b6 | 5.839110 | 0.492727595 | 0.467 | 0.247 | 1.4834261 | 24 | lsm14b |
| zgc:1629456 | 9.744485 | 0.44851443 | 0.317 | 0.136 | 2.4755865 | 24 | zgc:162945 |
| mcm3l6 | 1.135527 | 0.387755489 | 0.21 | 0.072 | 2.8848068 | 24 | mcm3l |
| si:dkey-241l7.36 | 1.426439 | 0.490041599 | 0.527 | 0.3 | 3.6238704 | 24 | si:dkey-241l7.3 |
| zgc:660246 | 1.704961 | 0.437944956 | 0.281 | 0.117 | 4.3314552 | 24 | zgc:66024 |
| lin28a3 | 1.970199 | 0.36353005 | 0.18 | 0.057 | 5.0052913 | 24 | lin28a |

| | | | | | | | |
|---|---|---|---|---|---|---|---|
| suv39h1b | 2.673060 | 0.323929441 | 0.144 | 0.041 | 6.7909104 | 24 | suv39h1b |
| si:ch73-90k17.16 | 2.788013 | 0.465556578 | 0.413 | 0.204 | 7.0829476 | 24 | si:ch73-90k17.1 |
| org7 | 3.104570 | 0.482429535 | 0.395 | 0.2 | 7.8871606 | 24 | org |
| CABZ01059627.26 | 3.348493 | 0.429024594 | 0.269 | 0.109 | 8.5068474 | 24 | CABZ01059627.2 |
| si:ch211-241b2.4 | 4.798967 | 0.286150223 | 0.126 | 0.033 | 1.2191776 | 24 | si:ch211-241b2.4 |
| si:dkey-43p13.54 | 5.089050 | 0.383020379 | 0.198 | 0.069 | 1.2928732 | 24 | si:dkey-43p13.5 |
| zp3c6 | 5.177263 | 0.451033827 | 0.335 | 0.156 | 1.3152837 | 24 | zp3c |
| srsf2a5 | 5.255117 | 0.468046788 | 0.521 | 0.349 | 1.3350626 | 24 | srsf2a |
| retsatl6 | 5.255248 | 0.457907714 | 0.353 | 0.167 | 1.3350959 | 24 | retsatl |
| zgc:1134246 | 6.486373 | 0.430109555 | 0.305 | 0.133 | 1.6478632 | 24 | zgc:113424 |
| larp6b6 | 6.805627 | 0.46327931 | 0.341 | 0.162 | 1.7289697 | 24 | larp6b |
| zgc:1628796 | 1.164424 | 0.454189189 | 0.323 | 0.149 | 2.9582195 | 24 | zgc:162879 |
| rpz26 | 1.467800 | 0.458890773 | 0.335 | 0.164 | 3.7289464 | 24 | rpz2 |
| hspa56 | 1.509100 | 0.461079023 | 0.491 | 0.295 | 3.8338702 | 24 | hspa5 |
| si:dkey-68o6.56 | 1.534901 | 0.415105449 | 0.251 | 0.101 | 3.8994174 | 24 | si:dkey-68o6.5 |
| zgc:1714265 | 1.660728 | 0.394273006 | 0.228 | 0.089 | 4.2190805 | 24 | zgc:171426 |
| CR847984.16 | 2.158283 | 0.435253005 | 0.305 | 0.138 | 5.4831198 | 24 | CR847984.1 |
| si:ch211-69g19.26 | 2.396620 | 0.401055988 | 0.251 | 0.104 | 6.0886136 | 24 | si:ch211-69g19.2 |
| gpr1605 | 2.592241 | 0.355263939 | 0.192 | 0.067 | 6.5855898 | 24 | gpr160 |
| BX571780.15 | 3.472329 | 0.334125566 | 0.192 | 0.067 | 8.8214526 | 24 | BX571780.1 |
| cpm5 | 4.283149 | 0.378148722 | 0.228 | 0.09 | 1.0881341 | 24 | cpm |
| h3f3a7 | 4.589075 | 0.404169344 | 0.629 | 0.483 | 1.1658546 | 24 | h3f3a |
| si:ch211-113a14.273 | 4.936328 | 0.337266257 | 0.186 | 0.065 | 1.2540742 | 24 | si:ch211-113a14.27 |
| gstp15 | 5.037643 | 0.447728188 | 0.521 | 0.347 | 1.2798132 | 24 | gstp1 |
| zgc:1719775 | 5.147128 | 0.447004373 | 0.371 | 0.189 | 1.3076278 | 24 | zgc:171977 |
| tmem442 | 5.286184 | 0.317050597 | 0.156 | 0.05 | 1.3429551 | 24 | tmem44 |
| CU467646.26 | 5.820576 | 0.387059903 | 0.251 | 0.104 | 1.4787174 | 24 | CU467646.2 |
| cdca86 | 7.094205 | 0.432848595 | 0.299 | 0.14 | 1.8022828 | 24 | cdca8 |
| eloal | 7.124453 | 0.303258546 | 0.15 | 0.047 | 1.8099674 | 24 | eloal |
| tmem144b6 | 8.083523 | 0.379025918 | 0.204 | 0.078 | 2.0536190 | 24 | tmem144b |
| rbpms2b6 | 1.143549 | 0.439434348 | 0.365 | 0.191 | 2.9051883 | 24 | rbpms2b |
| FP074874.2 | 1.152126 | 0.253043568 | 0.114 | 0.031 | 2.9269768 | 24 | FP074874.2 |
| mad2l17 | 1.284228 | 0.423817019 | 0.323 | 0.159 | 3.2625816 | 24 | mad2l1 |
| dctpp17 | 1.326548 | 0.419290514 | 0.311 | 0.149 | 3.3700968 | 24 | dctpp1 |
| bach2a | 1.498072 | 0.311715291 | 0.138 | 0.043 | 3.8058521 | 24 | bach2a |

| | | | | | | | |
|---|---|---|---|---|---|---|---|
| si:dkey-46g23.26 | 1.667470 | 0.392106572 | 0.269 | 0.119 | 4.2362098 | 24 | si:dkey-46g23.2 |
| CDPF16 | 1.714302 | 0.449351909 | 0.413 | 0.235 | 4.3551867 | 24 | CDPF1 |
| si:ch73-15b2.53 | 1.739702 | 0.362190341 | 0.204 | 0.078 | 4.4197130 | 24 | si:ch73-15b2.5 |
| zgc:868394 | 1.867046 | 0.379996959 | 0.228 | 0.094 | 4.7432308 | 24 | zgc:86839 |
| ankrd226 | 2.218305 | 0.408907924 | 0.269 | 0.122 | 5.6356060 | 24 | ankrd22 |
| si:dkey-189g17.22 | 2.293068 | 0.302155046 | 0.15 | 0.049 | 5.8255402 | 24 | si:dkey-189g17.2 |
| si:ch211-137a8.21 | 2.386432 | 0.318814572 | 0.144 | 0.047 | 6.0627311 | 24 | si:ch211-137a8.2 |
| si:dkeyp-75b4.107 | 2.501129 | 0.373618347 | 0.587 | 0.366 | 6.3541190 | 24 | si:dkeyp-75b4.10 |
| BX548000.11 | 2.983229 | 0.258882429 | 0.12 | 0.034 | 7.5788954 | 24 | BX548000.1 |
| pabpc1l6 | 3.251208 | 0.393414266 | 0.263 | 0.117 | 8.2596947 | 24 | pabpc1l |
| snu13b7 | 3.259088 | 0.428169241 | 0.539 | 0.371 | 8.2797144 | 24 | snu13b |
| BX005461.32 | 3.593969 | 0.323882332 | 0.162 | 0.056 | 9.1304796 | 24 | BX005461.3 |
| kmt5ab4 | 3.985981 | 0.340408357 | 0.198 | 0.076 | 0.000101 | 24 | kmt5ab |
| zgc:1717506 | 3.999351 | 0.405872943 | 0.299 | 0.144 | 0.000102 | 24 | zgc:171750 |
| gyg1a7 | 4.191687 | 0.442717084 | 0.491 | 0.319 | 0.000106 | 24 | gyg1a |
| si:dkey-87o1.22 | 4.831012 | 0.334847591 | 0.186 | 0.069 | 0.000123 | 24 | si:dkey-87o1.2 |
| patl26 | 6.578271 | 0.39018527 | 0.299 | 0.143 | 0.000167 | 24 | patl2 |
| elovl7b6 | 6.836693 | 0.426192246 | 0.383 | 0.205 | 0.000174 | 24 | elovl7b |
| aktip4 | 6.985689 | 0.420196543 | 0.263 | 0.126 | 0.000177 | 24 | aktip |
| asz13 | 7.080991 | 0.331320537 | 0.168 | 0.061 | 0.00018 | 24 | asz1 |
| zgc:922877 | 7.142233 | 0.378562381 | 0.251 | 0.112 | 0.000181 | 24 | zgc:92287 |
| prdx17 | 7.627409 | 0.41967895 | 0.299 | 0.149 | 0.000194 | 24 | prdx1 |
| tph1a6 | 7.701434 | 0.383215089 | 0.257 | 0.117 | 0.000196 | 24 | tph1a |
| cx44.26 | 8.101774 | 0.365618972 | 0.228 | 0.096 | 0.000206 | 24 | cx44.2 |
| nfr6 | 1.005545 | 0.343580544 | 0.222 | 0.092 | 0.000255 | 24 | nfr |
| dynll17 | 1.063604 | 0.380032567 | 0.587 | 0.441 | 0.00027 | 24 | dynll1 |
| si:dkeyp-2e4.2 | 1.072380 | 0.251804123 | 0.108 | 0.03 | 0.000272 | 24 | si:dkeyp-2e4.2 |
| chia.46 | 1.136997 | 0.380042968 | 0.263 | 0.121 | 0.000289 | 24 | chia.4 |
| asf1ba3 | 1.143558 | 0.351135309 | 0.21 | 0.086 | 0.000291 | 24 | asf1ba |
| LOC1003344391 | 1.329073 | 0.282805345 | 0.138 | 0.045 | 0.000338 | 24 | LOC100334439 |
| si:dkey-33c14.7 | 1.485651 | 0.263895106 | 0.12 | 0.037 | 0.000377 | 24 | si:dkey-33c14.7 |
| tmco13 | 1.774888 | 0.396727654 | 0.24 | 0.111 | 0.000451 | 24 | tmco1 |
| h2afva6 | 1.882362 | 0.431847165 | 0.455 | 0.306 | 0.000478 | 24 | h2afva |
| si:ch211-210c8.75 | 1.938559 | 0.397866874 | 0.246 | 0.115 | 0.000492 | 24 | si:ch211-210c8.7 |
| pif15 | 2.137952 | 0.326796212 | 0.216 | 0.089 | 0.000543 | 24 | pif1 |

| | | | | | | | |
|---|---|---|---|---|---|---|---|
| LOC1104377795 | 2.196439 | 0.316416682 | 0.18 | 0.069 | 0.000558 | 24 | LOC110437779 |
| cdca5 | 2.416447 | 0.260226403 | 0.12 | 0.037 | 0.000614 | 24 | cdca5 |
| ubc4 | 2.783794 | 0.256614111 | 0.707 | 0.654 | 0.000707 | 24 | ubc |
| ctsba8 | 2.809851 | 0.396708916 | 0.521 | 0.353 | 0.000714 | 24 | ctsba |
| si:dkey-203a12.23 | 2.978228 | 0.336363855 | 0.383 | 0.209 | 0.000757 | 24 | si:dkey-203a12.2 |
| slmapb | 3.026368 | 0.306168962 | 0.144 | 0.051 | 0.000769 | 24 | slmapb |
| eif4a1a6 | 3.173681 | 0.381946896 | 0.545 | 0.405 | 0.000806 | 24 | eif4a1a |
| dnd16 | 3.372034 | 0.343951344 | 0.216 | 0.092 | 0.000857 | 24 | dnd1 |
| snrpc3 | 4.269762 | 0.417140318 | 0.317 | 0.176 | 0.001085 | 24 | snrpc |
| zgc:1118686 | 4.562297 | 0.374882425 | 0.257 | 0.122 | 0.001159 | 24 | zgc:111868 |
| ypel35 | 4.898560 | 0.365102441 | 0.24 | 0.11 | 0.001244 | 24 | ypel3 |
| cdca95 | 5.180071 | 0.319411421 | 0.18 | 0.071 | 0.001316 | 24 | cdca9 |
| zgc:1017445 | 5.234568 | 0.354488207 | 0.246 | 0.112 | 0.00133 | 24 | zgc:101744 |
| ddx39ab6 | 5.929665 | 0.41057962 | 0.431 | 0.27 | 0.001506 | 24 | ddx39ab |
| stmn1a3 | 6.428783 | 0.38384785 | 0.347 | 0.188 | 0.001633 | 24 | stmn1a |
| zgc:1715661 | 6.611127 | 0.280761272 | 0.162 | 0.06 | 0.00168 | 24 | zgc:171566 |
| kpna27 | 7.302122 | 0.37953809 | 0.275 | 0.137 | 0.001855 | 24 | kpna2 |
| spryd43 | 7.824052 | 0.320079042 | 0.174 | 0.069 | 0.001988 | 24 | spryd4 |
| unga1 | 8.033088 | 0.312958499 | 0.174 | 0.069 | 0.002041 | 24 | unga |
| fam113 | 8.560712 | 0.250083909 | 0.102 | 0.03 | 0.002175 | 24 | fam113 |
| si:ch211-202f5.35 | 1.055935 | 0.319538514 | 0.186 | 0.076 | 0.002683 | 24 | si:ch211-202f5.3 |
| ppp4r2b | 1.152552 | 0.324375683 | 0.168 | 0.067 | 0.002928 | 24 | ppp4r2b |
| si:ch211-157c3.43 | 1.251919 | 0.306375886 | 0.186 | 0.075 | 0.003181 | 24 | si:ch211-157c3.4 |
| rpf13 | 1.311800 | 0.394127733 | 0.257 | 0.131 | 0.003333 | 24 | rpf1 |
| prtfdc1 | 1.394355 | 0.294916442 | 0.144 | 0.053 | 0.003542 | 24 | prtfdc1 |
| COQ10A4 | 1.723568 | 0.30806205 | 0.174 | 0.07 | 0.004379 | 24 | COQ10A |
| CABZ01075618.11 | 1.953559 | 0.260783469 | 0.132 | 0.046 | 0.004963 | 24 | CABZ01075618.1 |
| cfl1l3 | 2.311867 | 0.370582642 | 0.269 | 0.137 | 0.005873 | 24 | cfl1l |
| tatdn23 | 2.741784 | 0.283878671 | 0.156 | 0.06 | 0.006966 | 24 | tatdn2 |
| tbcc | 2.955311 | 0.272381425 | 0.132 | 0.047 | 0.007508 | 24 | tbcc |
| pias4b2 | 3.608784 | 0.268505795 | 0.144 | 0.054 | 0.009168 | 24 | pias4b |
| pttg16 | 3.636595 | 0.359048035 | 0.269 | 0.137 | 0.009239 | 24 | pttg1 |
| zgc:566766 | 3.764113 | 0.409215179 | 0.407 | 0.275 | 0.009563 | 24 | zgc:56676 |
| phex2 | 3.947940 | 0.29820548 | 0.174 | 0.072 | 0.01003 | 24 | phex |
| chtopa5 | 5.961627 | 0.389101856 | 0.341 | 0.207 | 0.015146 | 24 | chtopa |

| | | | | | | | |
|---|---|---|---|---|---|---|---|
| aurka6 | 6.839628 | 0.32200859 | 0.216 | 0.1 | 0.017376 | 24 | aurka |
| cdca7a6 | 6.878015 | 0.377098398 | 0.335 | 0.193 | 0.017474 | 24 | cdca7a |
| BX649350.11 | 7.231980 | 0.257964034 | 0.126 | 0.046 | 0.018373 | 24 | BX649350.1 |
| BX537276.16 | 7.249154 | 0.325263131 | 0.216 | 0.1 | 0.018416 | 24 | BX537276.1 |
| pdzd11 | 7.354372 | 0.339298279 | 0.192 | 0.088 | 0.018684 | 24 | pdzd11 |
| degs25 | 7.563560 | 0.309126174 | 0.198 | 0.089 | 0.019215 | 24 | degs2 |
| ube2c6 | 8.251438 | 0.341329013 | 0.246 | 0.124 | 0.020963 | 24 | ube2c |
| efcab112 | 8.530286 | 0.259703973 | 0.138 | 0.052 | 0.021671 | 24 | efcab11 |
| cdc45 | 8.708314 | 0.254671622 | 0.114 | 0.04 | 0.022123 | 24 | cdc45 |
| itm2cb4 | 9.023485 | 0.347584806 | 0.24 | 0.121 | 0.022924 | 24 | itm2cb |
| orc64 | 9.203722 | 0.314137678 | 0.204 | 0.094 | 0.023382 | 24 | orc6 |
| tspan13a1 | 1.017842 | 0.253638335 | 0.138 | 0.052 | 0.025858 | 24 | tspan13a |
| g3bp15 | 1.052489 | 0.368116577 | 0.329 | 0.194 | 0.026738 | 24 | g3bp1 |
| rnf47 | 1.057228 | 0.344692061 | 0.234 | 0.117 | 0.026859 | 24 | rnf4 |
| gar11 | 1.118284 | 0.356287336 | 0.228 | 0.115 | 0.02841 | 24 | gar1 |
| ddx414 | 1.158641 | 0.276303692 | 0.144 | 0.057 | 0.029435 | 24 | ddx41 |
| buc2l5 | 1.264359 | 0.268965603 | 0.162 | 0.066 | 0.032121 | 24 | buc2l |
| gtf3c6 | 1.274132 | 0.262400195 | 0.132 | 0.05 | 0.032369 | 24 | gtf3c6 |
| zgc:1141236 | 1.355389 | 0.295447041 | 0.192 | 0.086 | 0.034434 | 24 | zgc:114123 |
| cited13 | 1.364720 | 0.264073783 | 0.138 | 0.053 | 0.034671 | 24 | cited1 |
| fbxo325 | 1.570800 | 0.305343697 | 0.192 | 0.088 | 0.039906 | 24 | fbxo32 |
| rhogc | 1.609399 | 0.315154378 | 0.186 | 0.085 | 0.040887 | 24 | rhogc |
| buc6 | 1.695234 | 0.330762914 | 0.281 | 0.148 | 0.043067 | 24 | buc |
| dnajc5ga | 1.900821 | 0.289075642 | 0.156 | 0.066 | 0.04829 | 24 | dnajc5ga |
| fam122b1 | 2.008840 | 0.256616118 | 0.15 | 0.061 | 0.051035 | 24 | fam122b |
| lrp132 | 2.214073 | 0.271152974 | 0.144 | 0.058 | 0.056249 | 24 | lrp13 |
| haus66 | 2.257450 | 0.345853008 | 0.311 | 0.174 | 0.057351 | 24 | haus6 |
| rpain5 | 2.300504 | 0.309730193 | 0.21 | 0.101 | 0.058444 | 24 | rpain |
| faub6 | 2.459887 | 0.365516239 | 0.311 | 0.182 | 0.062493 | 24 | faub |
| bmb6 | 2.488069 | 0.293141538 | 0.168 | 0.073 | 0.063209 | 24 | bmb |
| inka1b6 | 2.690112 | 0.359289236 | 0.281 | 0.157 | 0.068342 | 24 | inka1b |
| hells4 | 2.952949 | 0.340598679 | 0.24 | 0.125 | 0.07502 | 24 | hells |
| vps9d14 | 3.401197 | 0.295336174 | 0.18 | 0.082 | 0.086407 | 24 | vps9d1 |
| chchd107 | 3.407824 | 0.34452335 | 0.251 | 0.134 | 0.086576 | 24 | chchd10 |
| CU655961.46 | 3.615491 | 0.279199695 | 0.168 | 0.073 | 0.091852 | 24 | CU655961.4 |

| | | | | | | | |
|---|---|---|---|---|---|---|---|
| aurkb6 | 3.873119 | 0.312734422 | 0.222 | 0.111 | 0.098397 | 24 | aurkb |
| BX001030.16 | 4.093429 | 0.298078348 | 0.21 | 0.101 | 0.103994 | 24 | BX001030.1 |
| il13ra24 | 4.341972 | 0.318015945 | 0.192 | 0.093 | 0.110308 | 24 | il13ra2 |
| cox177 | 5.699253 | 0.35083525 | 0.461 | 0.342 | 0.14479 | 24 | cox17 |
| gtf2a1l | 6.225932 | 0.252972722 | 0.132 | 0.054 | 0.15817 | 24 | gtf2a1l |
| snrpb25 | 6.389159 | 0.339760256 | 0.251 | 0.139 | 0.162317 | 24 | snrpb2 |
| rnf128a1 | 6.461966 | 0.280577615 | 0.144 | 0.062 | 0.164166 | 24 | rnf128a |
| si:dkey-81e3.23 | 6.904986 | 0.250400894 | 0.132 | 0.053 | 0.175421 | 24 | si:dkey-81e3.2 |
| eloca2 | 7.180076 | 0.310435826 | 0.192 | 0.095 | 0.18241 | 24 | eloca |
| cat | 7.412239 | 0.274420347 | 0.156 | 0.069 | 0.188308 | 24 | cat |
| zgc:1751365 | 8.333308 | 0.295545893 | 0.186 | 0.089 | 0.211708 | 24 | zgc:175136 |
| marcksb6 | 8.341717 | 0.325708511 | 0.281 | 0.161 | 0.211921 | 24 | marcksb |
| prc1a1 | 9.632135 | 0.255496439 | 0.138 | 0.058 | 0.244704 | 24 | prc1a |
| rab35b | 9.955018 | 0.292406642 | 0.15 | 0.067 | 0.252907 | 24 | rab35b |
| npm2a5 | 1.044879 | 0.264696495 | 0.15 | 0.065 | 0.265452 | 24 | npm2a |
| ctsla8 | 1.064470 | 0.328222252 | 0.491 | 0.375 | 0.270429 | 24 | ctsla |
| dynll2a7 | 1.136048 | 0.302222046 | 0.228 | 0.118 | 0.288613 | 24 | dynll2a |
| ccne16 | 1.399909 | 0.283171497 | 0.192 | 0.094 | 0.355647 | 24 | ccne1 |
| rbbp45 | 1.407864 | 0.335538496 | 0.275 | 0.163 | 0.357668 | 24 | rbbp4 |
| prrg24 | 1.429446 | 0.271409532 | 0.168 | 0.078 | 0.363151 | 24 | prrg2 |
| zgc:1529866 | 1.519392 | 0.262498599 | 0.162 | 0.074 | 0.386002 | 24 | zgc:152986 |
| zgc:1010162 | 1.598795 | 0.2887313 | 0.18 | 0.088 | 0.406174 | 24 | zgc:101016 |
| lipg4 | 1.745391 | 0.266466103 | 0.156 | 0.071 | 0.443417 | 24 | lipg |
| abcf2a3 | 1.868657 | 0.338679407 | 0.24 | 0.138 | 0.474733 | 24 | abcf2a |
| khsrp2 | 1.881240 | 0.346168841 | 0.305 | 0.194 | 0.477929 | 24 | khsrp |
| si:ch73-78o10.16 | 1.900437 | 0.303112954 | 0.21 | 0.111 | 0.482806 | 24 | si:ch73-78o10.1 |
| rogdi5 | 1.993508 | 0.291534546 | 0.198 | 0.101 | 0.506451 | 24 | rogdi |
| mcfd21 | 2.132145 | 0.309451067 | 0.204 | 0.108 | 0.541672 | 24 | mcfd2 |
| hmgb3b6 | 2.467386 | 0.33974899 | 0.281 | 0.171 | 0.626839 | 24 | hmgb3b |
| thoc73 | 2.636617 | 0.314561343 | 0.204 | 0.108 | 0.669833 | 24 | thoc7 |
| ivns1abpb3 | 2.636933 | 0.303993636 | 0.198 | 0.104 | 0.669913 | 24 | ivns1abpb |
| rps19bp11 | 2.712291 | 0.300874411 | 0.18 | 0.092 | 0.689058 | 24 | rps19bp1 |
| mrps28 | 2.766211 | 0.258595907 | 0.15 | 0.069 | 0.702756 | 24 | mrps28 |
| pop46 | 3.042906 | 0.302371764 | 0.198 | 0.104 | 0.77305 | 24 | pop4 |
| gpx82 | 3.095570 | 0.259389875 | 0.156 | 0.073 | 0.78643 | 24 | gpx8 |

| | | | | | | | |
|---|---|---|---|---|---|---|---|
| st3gal82 | 3.407907 | 0.258909208 | 0.144 | 0.066 | 0.865779 | 24 | st3gal8 |
| slc48a1b1 | 3.879537 | 0.265880686 | 0.162 | 0.078 | 0.985596 | 24 | slc48a1b |
| zgc:1532843 | 4.160531 | 0.261135656 | 0.174 | 0.085 | 1 | 24 | zgc:153284 |
| mrpl481 | 4.283788 | 0.286050286 | 0.156 | 0.076 | 1 | 24 | mrpl48 |
| hmga1a4 | 4.535562 | 0.280159089 | 0.527 | 0.443 | 1 | 24 | hmga1a |
| kif221 | 4.899246 | 0.261817107 | 0.162 | 0.078 | 1 | 24 | kif22 |
| shmt1 | 5.080290 | 0.262770867 | 0.156 | 0.075 | 1 | 24 | shmt1 |
| si:ch73-340n8.46 | 5.111199 | 0.299758872 | 0.21 | 0.115 | 1 | 24 | si:ch73-340n8.4 |
| cx43.45 | 5.186386 | 0.311627139 | 0.204 | 0.112 | 1 | 24 | cx43.4 |
| tuba8l41 | 5.278458 | 0.292390141 | 0.497 | 0.416 | 1 | 24 | tuba8l4 |
| zgc:1121851 | 5.519144 | 0.264611358 | 0.15 | 0.071 | 1 | 24 | zgc:112185 |
| psme33 | 5.724954 | 0.302371187 | 0.222 | 0.125 | 1 | 24 | psme3 |
| slc25a25a3 | 6.322076 | 0.32857547 | 0.263 | 0.164 | 1 | 24 | slc25a25a |
| fbxw11b4 | 6.890601 | 0.309322787 | 0.228 | 0.132 | 1 | 24 | fbxw11b |
| cycsb4 | 7.310332 | 0.325861573 | 0.407 | 0.315 | 1 | 24 | cycsb |
| dnmt11 | 7.507963 | 0.267765162 | 0.174 | 0.089 | 1 | 24 | dnmt1 |
| hsp90b12 | 8.297294 | 0.320091934 | 0.317 | 0.211 | 1 | 24 | hsp90b1 |
| prpf312 | 8.321729 | 0.251144618 | 0.156 | 0.076 | 1 | 24 | prpf31 |
| sumo3b4 | 8.369297 | 0.285083146 | 0.21 | 0.115 | 1 | 24 | sumo3b |
| rbm4.25 | 8.855305 | 0.307805258 | 0.257 | 0.158 | 1 | 24 | rbm4.2 |
| LOC1039099726 | 9.486443 | 0.318963996 | 0.419 | 0.323 | 1 | 24 | LOC103909972 |
| dap1b4 | 9.818961 | 0.325906725 | 0.347 | 0.25 | 1 | 24 | dap1b |
| ttc323 | 9.901066 | 0.251350569 | 0.174 | 0.089 | 1 | 24 | ttc32 |
| f11r.14 | 0.0001 | 0.311128836 | 0.263 | 0.165 | 1 | 24 | f11r.1 |
| cdc14b2 | 0.0001 | 0.292968651 | 0.216 | 0.125 | 1 | 24 | cdc14b |
| dcun1d51 | 0.0001 | 0.281502969 | 0.162 | 0.084 | 1 | 24 | dcun1d5 |
| c1d3 | 0.0001 | 0.298797137 | 0.21 | 0.12 | 1 | 24 | c1d |
| eif61 | 0.0001 | 0.308373332 | 0.228 | 0.138 | 1 | 24 | eif6 |
| zc3h11a6 | 0.0001 | 0.297266246 | 0.299 | 0.19 | 1 | 24 | zc3h11a |
| pum3 | 0.0001 | 0.258173091 | 0.138 | 0.068 | 1 | 24 | pum3 |
| ndc12 | 0.0001 | 0.263300825 | 0.168 | 0.088 | 1 | 24 | ndc1 |
| BX957362.14 | 0.0001 | 0.280060974 | 0.204 | 0.115 | 1 | 24 | BX957362.1 |
| adipor1a2 | 0.0002 | 0.269454416 | 0.168 | 0.091 | 1 | 24 | adipor1a |
| atp5mc13 | 0.0002 | 0.302873448 | 0.419 | 0.33 | 1 | 24 | atp5mc1 |
| rpp211 | 0.0002 | 0.252315996 | 0.15 | 0.076 | 1 | 24 | rpp21 |

| | | | | | | | |
|---|---|---|---|---|---|---|---|
| cstf12 | 0.0002 | 0.262334408 | 0.168 | 0.09 | 1 | 24 | cstf1 |
| parn6 | 0.0002 | 0.279112983 | 0.222 | 0.131 | 1 | 24 | parn |
| snrpe2 | 0.0003 | 0.319920093 | 0.365 | 0.285 | 1 | 24 | snrpe |
| exosc61 | 0.0003 | 0.251474721 | 0.156 | 0.082 | 1 | 24 | exosc6 |
| eif4g2a | 0.0003 | 0.281803504 | 0.21 | 0.127 | 1 | 24 | eif4g2a |
| si:ch73-281n10.26 | 0.0003 | 0.267200345 | 0.479 | 0.397 | 1 | 24 | si:ch73-281n10.2 |
| dcun1d14 | 0.0004 | 0.257216772 | 0.162 | 0.088 | 1 | 24 | dcun1d1 |
| chrac14 | 0.0004 | 0.281386399 | 0.204 | 0.123 | 1 | 24 | chrac1 |
| mcm51 | 0.0005 | 0.251370981 | 0.168 | 0.093 | 1 | 24 | mcm5 |
| rasl11b6 | 0.0005 | 0.254720609 | 0.198 | 0.114 | 1 | 24 | rasl11b |
| pprc12 | 0.0005 | 0.259355879 | 0.156 | 0.086 | 1 | 24 | pprc1 |
| pcf114 | 0.0006 | 0.309491066 | 0.299 | 0.218 | 1 | 24 | pcf11 |
| eno31 | 0.0007 | 0.26959661 | 0.431 | 0.364 | 1 | 24 | eno3 |
| anp32b6 | 0.0007 | 0.30450957 | 0.335 | 0.254 | 1 | 24 | anp32b |
| prps1a | 0.0008 | 0.286313423 | 0.198 | 0.123 | 1 | 24 | prps1a |
| bud311 | 0.0009 | 0.309693722 | 0.305 | 0.231 | 1 | 24 | bud31 |
| zgc:1102395 | 0.0009 | 0.291574307 | 0.281 | 0.198 | 1 | 24 | zgc:110239 |
| imp36 | 0.0009 | 0.263156743 | 0.204 | 0.125 | 1 | 24 | imp3 |
| sec61a1l3 | 0.0009 | 0.26879282 | 0.192 | 0.116 | 1 | 24 | sec61a1l |
| rpa34 | 0.001 | 0.264098561 | 0.24 | 0.154 | 1 | 24 | rpa3 |
| h2afx6 | 0.001 | 0.253518581 | 0.204 | 0.123 | 1 | 24 | h2afx |
| selenom1 | 0.0011 | 0.259867676 | 0.186 | 0.112 | 1 | 24 | selenom |
| nop161 | 0.0012 | 0.267119775 | 0.18 | 0.109 | 1 | 24 | nop16 |
| mcm73 | 0.0012 | 0.254828853 | 0.228 | 0.148 | 1 | 24 | mcm7 |
| spcs31 | 0.0016 | 0.273552904 | 0.335 | 0.261 | 1 | 24 | spcs3 |
| ccdc583 | 0.0016 | 0.251435698 | 0.168 | 0.1 | 1 | 24 | ccdc58 |
| serp12 | 0.0017 | 0.286312965 | 0.275 | 0.199 | 1 | 24 | serp1 |
| mrps18c2 | 0.0017 | 0.262604163 | 0.186 | 0.117 | 1 | 24 | mrps18c |
| lsm12a4 | 0.0022 | 0.251772167 | 0.186 | 0.117 | 1 | 24 | lsm12a |
| lbh4 | 0.0022 | 0.255640071 | 0.228 | 0.154 | 1 | 24 | lbh |
| rtn4a4 | 0.0024 | 0.289948231 | 0.305 | 0.241 | 1 | 24 | rtn4a |
| polr2l1 | 0.0025 | 0.272105179 | 0.275 | 0.204 | 1 | 24 | polr2l |
| dnajc1 | 0.0032 | 0.260857424 | 0.21 | 0.143 | 1 | 24 | dnajc1 |
| ppifb | 0.0038 | 0.268327052 | 0.228 | 0.162 | 1 | 24 | ppifb |
| STMP14 | 0.0052 | 0.260903085 | 0.353 | 0.295 | 1 | 24 | STMP1 |

| | | | | | | | |
|---|---|---|---|---|---|---|---|
| zfand5b2 | 0.0067 | 0.261691832 | 0.269 | 0.206 | 1 | 24 | zfand5b |
| pcbp22 | 0.0076 | 0.265189845 | 0.275 | 0.218 | 1 | 24 | pcbp2 |
| tomm20a3 | 0.0078 | 0.25782155 | 0.287 | 0.228 | 1 | 24 | tomm20a |
| top1l3 | 0.009 | 0.25407949 | 0.275 | 0.219 | 1 | 24 | top1l |
| lyz | 0 | 2.688858889 | 0.924 | 0.043 | 0 | 25 | lyz |
| lect2l1 | 0 | 2.57303876 | 0.975 | 0.027 | 0 | 25 | lect2l |
| npsn1 | 0 | 2.396211749 | 0.981 | 0.024 | 0 | 25 | npsn |
| BX908782.2 | 0 | 2.276345023 | 0.949 | 0.017 | 0 | 25 | BX908782.2 |
| mpx | 0 | 2.099305347 | 0.956 | 0.005 | 0 | 25 | mpx |
| mmp13a | 0 | 1.795387314 | 0.778 | 0.003 | 0 | 25 | mmp13a |
| ch25hl2 | 0 | 1.768079107 | 0.791 | 0.002 | 0 | 25 | ch25hl2 |
| BX908782.31 | 0 | 1.762939851 | 0.797 | 0.029 | 0 | 25 | BX908782.3 |
| si:ch1073-429i10.1 | 0 | 1.648962995 | 0.639 | 0.004 | 0 | 25 | si:ch1073-429i10.1 |
| scpp8 | 0 | 1.644383507 | 0.759 | 0.004 | 0 | 25 | scpp8 |
| il1b1 | 0 | 1.48482074 | 0.614 | 0.016 | 0 | 25 | il1b |
| cd7al1 | 0 | 1.367006925 | 0.69 | 0.021 | 0 | 25 | cd7al |
| adam8a3 | 0 | 1.355451574 | 0.709 | 0.043 | 0 | 25 | adam8a |
| sult2st1 | 0 | 1.352342072 | 0.652 | 0.023 | 0 | 25 | sult2st1 |
| il6r | 0 | 1.349159027 | 0.627 | 0.007 | 0 | 25 | il6r |
| ly6m5 | 0 | 1.036377335 | 0.5 | 0.008 | 0 | 25 | ly6m5 |
| CR848841.11 | 0 | 1.004856567 | 0.462 | 0.012 | 0 | 25 | CR848841.1 |
| si:ch73-343l4.81 | 0 | 0.997144652 | 0.475 | 0.016 | 0 | 25 | si:ch73-343l4.8 |
| si:ch211-286b5.8 | 0 | 0.99383589 | 0.475 | 0.004 | 0 | 25 | si:ch211-286b5.8 |
| si:ch211-193e13.51 | 0 | 0.920189155 | 0.418 | 0.009 | 0 | 25 | si:ch211-193e13.5 |
| si:ch211-264f5.21 | 0 | 0.889801784 | 0.399 | 0.01 | 0 | 25 | si:ch211-264f5.2 |
| illr41 | 0 | 0.860186562 | 0.405 | 0.006 | 0 | 25 | illr4 |
| si:ch211-9d9.8 | 0 | 0.840303512 | 0.392 | 0.008 | 0 | 25 | si:ch211-9d9.8 |
| si:ch211-284o19.8 | 0 | 0.812923025 | 0.38 | 0.003 | 0 | 25 | si:ch211-284o19.8 |
| chia.6 | 0 | 0.758226585 | 0.348 | 0.003 | 0 | 25 | chia.6 |
| si:ch211-260e23.8 | 0 | 0.735156863 | 0.361 | 0.004 | 0 | 25 | si:ch211-260e23.8 |
| abcb9 | 0 | 0.728509415 | 0.329 | 0.002 | 0 | 25 | abcb9 |
| si:dkey-19b23.14 | 0 | 0.667111337 | 0.31 | 0.002 | 0 | 25 | si:dkey-19b23.14 |
| si:ch211-9d9.1 | 0 | 0.631852972 | 0.272 | 0.001 | 0 | 25 | si:ch211-9d9.1 |
| si:dkey-238m4.4 | 0 | 0.456065277 | 0.171 | 0 | 0 | 25 | si:dkey-238m4.4 |
| cnksr2b | 0 | 0.421732555 | 0.203 | 0.001 | 0 | 25 | cnksr2b |

| | | | | | | | |
|---|---|---|---|---|---|---|---|
| CR354540.1 | 0 | 0.355810582 | 0.158 | 0 | 0 | 25 | CR354540.1 |
| ponzr62 | 5.958556 | 0.89209122 | 0.443 | 0.017 | 1.5137714 | 25 | ponzr6 |
| CT573383.1 | 1.643587 | 0.617695302 | 0.316 | 0.009 | 4.1755328 | 25 | CT573383.1 |
| mmp91 | 1.250771 | 1.094157816 | 0.373 | 0.014 | 3.1775842 | 25 | mmp9 |
| CT030188.13 | 1.582759 | 1.230023428 | 0.665 | 0.049 | 4.0210015 | 25 | CT030188.1 |
| sms1 | 1.966958 | 1.189461532 | 0.563 | 0.034 | 4.9970592 | 25 | sms |
| hbxb | 1.723167 | 0.309551749 | 0.171 | 0.002 | 4.3777082 | 25 | hbxb |
| abcc13 | 3.826651 | 0.302812918 | 0.152 | 0.001 | 9.7216074 | 25 | abcc13 |
| lta4h3 | 3.956904 | 1.448661982 | 0.747 | 0.069 | 1.0052516 | 25 | lta4h |
| ltb4r2 | 5.295720 | 0.9024814 | 0.456 | 0.023 | 1.3453778 | 25 | ltb4r |
| cpa51 | 8.060396 | 1.248037872 | 0.513 | 0.031 | 2.0477438 | 25 | cpa5 |
| ncf11 | 3.490096 | 0.967439045 | 0.494 | 0.029 | 8.8665902 | 25 | ncf1 |
| alox5ap1 | 9.102661 | 0.862093234 | 0.411 | 0.02 | 2.3125311 | 25 | alox5ap |
| hpgd | 2.117955 | 0.534774188 | 0.247 | 0.007 | 5.3806664 | 25 | hpgd |
| si:ch211-8c17.2 | 1.094960 | 0.521581077 | 0.215 | 0.005 | 2.7817474 | 25 | si:ch211-8c17.2 |
| si:ch73-248e21.72 | 1.209567 | 1.086939861 | 0.57 | 0.045 | 3.0729063 | 25 | si:ch73-248e21.7 |
| ryr3 | 4.028471 | 0.35246525 | 0.158 | 0.002 | 1.0234332 | 25 | ryr3 |
| si:ch211-276a23.51 | 4.565761 | 0.709010306 | 0.285 | 0.01 | 1.1599317 | 25 | si:ch211-276a23.5 |
| papss2b | 3.160639 | 0.686120785 | 0.354 | 0.017 | 8.0296041 | 25 | papss2b |
| HACD41 | 1.056425 | 1.03273537 | 0.532 | 0.044 | 2.6838498 | 25 | HACD4 |
| srgn6 | 2.251408 | 1.839362886 | 0.981 | 0.228 | 5.7197042 | 25 | srgn |
| CU499336.21 | 2.787572 | 0.817667031 | 0.43 | 0.029 | 7.0818277 | 25 | CU499336.2 |
| si:ch1073-443f11.21 | 7.225897 | 0.892149612 | 0.468 | 0.037 | 1.8357393 | 25 | si:ch1073-443f11.2 |
| si:ch73-359m17.5 | 2.447843 | 0.424338041 | 0.171 | 0.004 | 6.2187451 | 25 | si:ch73-359m17.5 |
| mmp25b1 | 8.513559 | 0.4726067 | 0.222 | 0.008 | 2.1628697 | 25 | mmp25b |
| vwa111 | 1.498127 | 0.314351997 | 0.127 | 0.002 | 3.8059929 | 25 | vwa11 |
| triobpa | 8.179374 | 0.532677394 | 0.266 | 0.012 | 2.0779700 | 25 | triobpa |
| elovl5 | 1.750719 | 0.540916757 | 0.253 | 0.011 | 4.4477026 | 25 | elovl5 |
| cxcr4b5 | 2.579833 | 1.419598002 | 0.899 | 0.17 | 6.5540666 | 25 | cxcr4b |
| agtrap | 3.690413 | 0.465811478 | 0.222 | 0.008 | 9.3754957 | 25 | agtrap |
| BX548046.2 | 3.317509 | 0.335388241 | 0.158 | 0.004 | 8.4281341 | 25 | BX548046.2 |
| ctss1 | 6.048275 | 0.523005271 | 0.272 | 0.013 | 1.5365644 | 25 | ctss1 |
| lrrc15 | 7.963707 | 0.610944067 | 0.285 | 0.015 | 2.0231797 | 25 | lrrc15 |
| timp2b1 | 1.948069 | 0.874398328 | 0.418 | 0.034 | 4.9490708 | 25 | timp2b |
| jakmip1 | 1.345860 | 0.410490424 | 0.203 | 0.007 | 3.4191595 | 25 | jakmip1 |

| | | | | | | | |
|---|---|---|---|---|---|---|---|
| slc7a111 | 3.392965 | 0.509319421 | 0.209 | 0.008 | 8.6198289 | 25 | slc7a11 |
| tcnbb1 | 5.156493 | 0.852635337 | 0.285 | 0.017 | 1.3100071 | 25 | tcnbb |
| hyal3 | 1.283165 | 0.337657051 | 0.165 | 0.005 | 3.2598828 | 25 | hyal3 |
| glipr1a6 | 2.972692 | 0.972270753 | 0.538 | 0.064 | 7.5521244 | 25 | glipr1a |
| rnaset2 | 9.489736 | 0.795906142 | 0.43 | 0.04 | 2.4108674 | 25 | rnaset2 |
| si:ch211-137j23.6 | 7.933631 | 0.298732022 | 0.101 | 0.002 | 2.0155391 | 25 | si:ch211-137j23.6 |
| alox5a2 | 1.297940 | 0.692996735 | 0.373 | 0.03 | 3.2974187 | 25 | alox5a |
| fcer1gl5 | 1.727158 | 1.341879352 | 0.892 | 0.176 | 4.3878449 | 25 | fcer1gl |
| adora2b | 2.863152 | 0.403243505 | 0.196 | 0.008 | 7.2738391 | 25 | adora2b |
| gpr183b | 8.219814 | 0.37755695 | 0.171 | 0.006 | 2.0882439 | 25 | gpr183b |
| f8 | 4.124371 | 0.301967119 | 0.108 | 0.002 | 1.0477966 | 25 | f8 |
| ppdpfa5 | 2.085281 | 1.073963614 | 0.532 | 0.067 | 5.2976588 | 25 | ppdpfa |
| noctb | 3.030796 | 0.565666849 | 0.241 | 0.013 | 7.6997379 | 25 | noctb |
| s1pr4 | 1.914234 | 0.469935652 | 0.184 | 0.007 | 4.8631137 | 25 | s1pr4 |
| cfbl2 | 4.457536 | 0.963228451 | 0.544 | 0.066 | 1.1324372 | 25 | cfbl |
| si:dkey-27h10.21 | 3.989421 | 0.681561613 | 0.323 | 0.025 | 1.0135124 | 25 | si:dkey-27h10.2 |
| slc7a3a2 | 1.605857 | 0.850729826 | 0.424 | 0.044 | 4.0796809 | 25 | slc7a3a |
| slc2a3b1 | 1.063700 | 0.642187436 | 0.323 | 0.025 | 2.7023315 | 25 | slc2a3b |
| si:ch211-67e16.31 | 1.219888 | 0.693153596 | 0.253 | 0.016 | 3.0991276 | 25 | si:ch211-67e16.3 |
| spi1b3 | 1.637073 | 0.954572803 | 0.519 | 0.068 | 4.1589845 | 25 | spi1b |
| si:dkey-102g19.32 | 1.148390 | 0.798699803 | 0.43 | 0.045 | 2.9174866 | 25 | si:dkey-102g19.3 |
| mir2233 | 5.751716 | 0.674538189 | 0.354 | 0.031 | 1.4612235 | 25 | mir223 |
| scinlb5 | 1.605852 | 0.960177103 | 0.557 | 0.08 | 4.0796692 | 25 | scinlb |
| nccrp12 | 2.185584 | 1.310107211 | 0.778 | 0.191 | 5.5524775 | 25 | nccrp1 |
| LOC1018842674 | 2.946636 | 0.947620196 | 0.43 | 0.048 | 7.4859304 | 25 | LOC101884267 |
| ca61 | 2.213260 | 0.635277783 | 0.266 | 0.018 | 5.6227872 | 25 | ca6 |
| aldh8a1 | 2.650715 | 0.303304422 | 0.158 | 0.006 | 6.7341437 | 25 | aldh8a1 |
| dhrs91 | 2.694165 | 0.686093393 | 0.354 | 0.034 | 6.8445284 | 25 | dhrs9 |
| vamp8 | 2.707052 | 0.887169543 | 0.519 | 0.072 | 6.8772660 | 25 | vamp8 |
| BX908782.1 | 4.582120 | 0.31756413 | 0.101 | 0.002 | 1.1640876 | 25 | BX908782.1 |
| si:dkey-222f2.72 | 5.257518 | 0.67083137 | 0.329 | 0.03 | 1.3356725 | 25 | si:dkey-222f2.7 |
| si:ch211-66e2.35 | 5.379066 | 0.849972142 | 0.462 | 0.059 | 1.3665519 | 25 | si:ch211-66e2.3 |
| CABZ01111959.12 | 6.585153 | 0.602403162 | 0.291 | 0.024 | 1.6729583 | 25 | CABZ01111959.1 |
| LOC1018850651 | 2.140184 | 0.436691576 | 0.222 | 0.014 | 5.4371388 | 25 | LOC101885065 |
| cebpa5 | 3.815524 | 0.856614674 | 0.5 | 0.072 | 9.6933406 | 25 | cebpa |

| | | | | | | | |
|---|---|---|---|---|---|---|---|
| FAR2 | 1.69761[illegible] | 0.520992819 | 0.272 | 0.022 | 4.3127950 | 25 | FAR2 |
| si:ch73-248e21.51 | 1.09502[illegible] | 0.762533988 | 0.373 | 0.043 | 2.7819170 | 25 | si:ch73-248e21.5 |
| itgb24 | 8.82446[illegible] | 0.968049538 | 0.589 | 0.104 | 2.2418560 | 25 | itgb2 |
| odc12 | 2.41148[illegible] | 0.999875898 | 0.614 | 0.117 | 6.1263653 | 25 | odc1 |
| zgc:175264 | 3.11832[illegible] | 0.592602156 | 0.329 | 0.033 | 7.9221132 | 25 | zgc:175264 |
| spns31 | 8.96059[illegible] | 0.415082872 | 0.209 | 0.014 | 2.2764394 | 25 | spns3 |
| capgb4 | 9.51358[illegible] | 0.952402433 | 0.589 | 0.111 | 2.4169258 | 25 | capgb |
| vsir5 | 1.17998[illegible] | 0.919943352 | 0.506 | 0.085 | 2.9977645 | 25 | vsir |
| cebpb9 | 1.32481[illegible] | 1.181969678 | 0.93 | 0.334 | 3.3656905 | 25 | cebpb |
| gadd45aa | 1.47131[illegible] | 0.568845278 | 0.228 | 0.018 | 3.7378738 | 25 | gadd45aa |
| BX323087.1 | 1.22899[illegible] | 0.324714603 | 0.152 | 0.008 | 3.1222709 | 25 | BX323087.1 |
| actb27 | 1.05994[illegible] | 0.645007804 | 1 | 0.866 | 2.6927924 | 25 | actb2 |
| LOC101886073 | 1.67792[illegible] | 0.259671612 | 0.146 | 0.007 | 4.2627654 | 25 | LOC101886073 |
| ank3b2 | 1.15710[illegible] | 0.553544541 | 0.323 | 0.036 | 2.9396204 | 25 | ank3b |
| rab445 | 1.87920[illegible] | 0.781497341 | 0.405 | 0.059 | 4.7741222 | 25 | rab44 |
| bdh1 | 2.99166[illegible] | 0.33607144 | 0.19 | 0.013 | 7.6003272 | 25 | bdh1 |
| cd6312 | 1.23712[illegible] | 1.041829576 | 0.886 | 0.369 | 3.1429276 | 25 | cd63 |
| si:ch211-136m16.82 | 3.96454[illegible] | 0.617598288 | 0.278 | 0.029 | 1.0071924 | 25 | si:ch211-136m16.8 |
| cyp4f31 | 5.87381[illegible] | 0.414587436 | 0.209 | 0.016 | 1.4922418 | 25 | cyp4f3 |
| hlx11 | 8.57975[illegible] | 0.366144458 | 0.215 | 0.017 | 2.1796856 | 25 | hlx1 |
| gsr1 | 1.04388[illegible] | 0.535401152 | 0.31 | 0.036 | 2.6519774 | 25 | gsr |
| mapk111 | 1.79138[illegible] | 0.447384245 | 0.215 | 0.018 | 4.5510092 | 25 | mapk11 |
| mrps153 | 7.10154[illegible] | 0.87429496 | 0.551 | 0.11 | 1.8041463 | 25 | mrps15 |
| zgc:165573 | 1.84748[illegible] | 0.630643226 | 0.38 | 0.052 | 4.6935286 | 25 | zgc:165573 |
| lpcat2 | 2.13753[illegible] | 0.502920082 | 0.234 | 0.021 | 5.4304075 | 25 | lpcat2 |
| cnksr1 | 5.55202[illegible] | 0.313012125 | 0.146 | 0.008 | 1.4104916 | 25 | cnksr1 |
| hopx3 | 6.50775[illegible] | 0.724588063 | 0.405 | 0.062 | 1.6532961 | 25 | hopx |
| si:ch211-243a20.41 | 3.31036[illegible] | 0.440661181 | 0.215 | 0.018 | 8.4099777 | 25 | si:ch211-243a20.4 |
| itgb3a1 | 5.18394[illegible] | 0.39831721 | 0.177 | 0.012 | 1.3169820 | 25 | itgb3a |
| tmsb16 | 1.56442[illegible] | 1.042519916 | 0.633 | 0.156 | 3.9744232 | 25 | tmsb1 |
| alox12 | 1.74671[illegible] | 0.313496869 | 0.177 | 0.013 | 4.4375317 | 25 | alox12 |
| zmp:00000010823 | 1.99226[illegible] | 0.708765253 | 0.361 | 0.053 | 5.0613416 | 25 | zmp:0000001082 |
| CABZ01073834.13 | 4.17081[illegible] | 0.609087331 | 0.323 | 0.042 | 1.0595956 | 25 | CABZ01073834.1 |
| pfn17 | 1.45207[illegible] | 1.002900612 | 0.994 | 0.498 | 3.6890020 | 25 | pfn1 |
| pnp5a3 | 3.74343[illegible] | 0.83937731 | 0.418 | 0.072 | 9.5101893 | 25 | pnp5a |

| | | | | | | | |
|---|---|---|---|---|---|---|---|
| prkcda | 3.20013[illegible] | 0.630171568 | 0.373 | 0.055 | 8.1299387 | 25 | prkcda |
| XLOC-039817 | 6.93347[illegible] | 0.339477767 | 0.152 | 0.01 | 1.7614494 | 25 | XLOC-039817 |
| il343 | 2.02380[illegible] | 0.560994196 | 0.323 | 0.044 | 5.1414848 | 25 | il34 |
| dachd2 | 6.02426[illegible] | 0.297983597 | 0.152 | 0.01 | 1.5304657 | 25 | dachd |
| fam107b3 | 1.29942[illegible] | 0.762507593 | 0.519 | 0.109 | 3.3012000 | 25 | fam107b |
| cfl1l4 | 6.52399[illegible] | 0.908764914 | 0.57 | 0.133 | 1.6574203 | 25 | cfl1l |
| pgp | 2.85598[illegible] | 0.614336939 | 0.38 | 0.061 | 7.2556365 | 25 | pgp |
| CABZ01001434.11 | 2.91861[illegible] | 0.638762564 | 0.171 | 0.014 | 7.4147519 | 25 | CABZ01001434.1 |
| arpc1b7 | 1.74110[illegible] | 0.97570341 | 0.861 | 0.309 | 4.4232807 | 25 | arpc1b |
| slc6a6b4 | 2.05027[illegible] | 0.798978758 | 0.462 | 0.094 | 5.2087144 | 25 | slc6a6b |
| cotl17 | 2.93893[illegible] | 0.963362304 | 0.823 | 0.316 | 7.4663764 | 25 | cotl1 |
| rac27 | 3.02228[illegible] | 0.906455448 | 0.715 | 0.221 | 7.6781219 | 25 | rac2 |
| b3gnt3.41 | 8.40765[illegible] | 0.583757936 | 0.278 | 0.036 | 2.1359640 | 25 | b3gnt3.4 |
| nrros1 | 1.91590[illegible] | 0.595146309 | 0.316 | 0.046 | 4.8673643 | 25 | nrros |
| dub4 | 2.95696[illegible] | 0.624528226 | 0.399 | 0.071 | 7.5121572 | 25 | dub |
| zgc:638313 | 4.72533[illegible] | 0.74360355 | 0.424 | 0.083 | 1.2004715 | 25 | zgc:63831 |
| atf35 | 5.90626[illegible] | 0.736242644 | 0.418 | 0.079 | 1.5004877 | 25 | atf3 |
| il164 | 8.27571[illegible] | 0.558326278 | 0.31 | 0.047 | 2.1024444 | 25 | il16 |
| plek | 1.35123[illegible] | 0.404569652 | 0.234 | 0.027 | 3.4328236 | 25 | plek |
| XLOC-004556 | 9.19489[illegible] | 0.313959821 | 0.171 | 0.015 | 2.3359621 | 25 | XLOC-004556 |
| adam19b2 | 4.82289[illegible] | 0.296025394 | 0.158 | 0.013 | 1.2252563 | 25 | adam19b |
| selenow15 | 1.87240[illegible] | 0.867217631 | 0.81 | 0.413 | 4.7568468 | 25 | selenow1 |
| rgs192 | 4.72114[illegible] | 0.475371822 | 0.253 | 0.034 | 1.1994069 | 25 | rgs19 |
| si:cabz01036022.11 | 6.43300[illegible] | 0.319981043 | 0.139 | 0.011 | 1.6343043 | 25 | si:cabz01036022.1 |
| rhogb4 | 4.79693[illegible] | 0.73926308 | 0.456 | 0.105 | 1.2186623 | 25 | rhogb |
| coro1a7 | 1.13369[illegible] | 0.84900215 | 0.728 | 0.233 | 2.8801605 | 25 | coro1a |
| wasb7 | 1.15789[illegible] | 0.823139446 | 0.595 | 0.17 | 2.9416410 | 25 | wasb |
| lcp15 | 7.54512[illegible] | 0.651147727 | 0.399 | 0.082 | 1.9168398 | 25 | lcp1 |
| apol1 | 7.54565[illegible] | 0.320327756 | 0.152 | 0.013 | 1.9169748 | 25 | apol1 |
| plxnc13 | 2.71906[illegible] | 0.563530083 | 0.285 | 0.045 | 6.9077746 | 25 | plxnc1 |
| myh9a1 | 6.59630[illegible] | 0.545781564 | 0.304 | 0.05 | 1.6757907 | 25 | myh9a |
| si:dkey-163m14.7 | 1.09526[illegible] | 0.29900523 | 0.101 | 0.006 | 2.7825319 | 25 | si:dkey-163m14.7 |
| ccdc88b1 | 1.59843[illegible] | 0.455136103 | 0.234 | 0.031 | 4.0608247 | 25 | ccdc88b |
| spag1a | 1.60490[illegible] | 0.344126152 | 0.171 | 0.017 | 4.0772585 | 25 | spag1a |
| dgkzb | 7.81617[illegible] | 0.277633942 | 0.12 | 0.009 | 1.9856993 | 25 | dgkzb |

| | | | | | | | |
|---|---|---|---|---|---|---|---|
| COX312 | 9.807231 | 0.671382578 | 0.981 | 0.674 | 2.4915272 | 25 | COX3 |
| gpr841 | 1.727839 | 0.332356764 | 0.127 | 0.01 | 4.3895750 | 25 | gpr84 |
| itgae.22 | 2.340526 | 0.441144399 | 0.247 | 0.035 | 5.9461076 | 25 | itgae.2 |
| pgd3 | 2.385274 | 0.606303784 | 0.354 | 0.069 | 6.0597905 | 25 | pgd |
| raraa1 | 3.562807 | 0.566265712 | 0.247 | 0.036 | 9.0513121 | 25 | raraa |
| LOC101882843 | 8.329564 | 0.320355795 | 0.114 | 0.008 | 2.1161258 | 25 | LOC101882843 |
| icn6 | 8.770992 | 0.98931341 | 0.709 | 0.265 | 2.2282707 | 25 | icn |
| antxr2a5 | 1.091213 | 0.779683284 | 0.563 | 0.166 | 2.7722274 | 25 | antxr2a |
| lxn6 | 2.354294 | 0.765466161 | 0.563 | 0.165 | 5.9810846 | 25 | lxn |
| pim17 | 7.955986 | 0.820035344 | 0.557 | 0.166 | 2.0212184 | 25 | pim1 |
| si:ch211-197k17.3 | 2.734394 | 0.468879651 | 0.241 | 0.035 | 6.9467303 | 25 | si:ch211-197k17.3 |
| gapdhs10 | 8.628102 | 0.82084077 | 0.715 | 0.312 | 2.1919693 | 25 | gapdhs |
| si:dkey-262k9.45 | 1.526936 | 0.601922114 | 0.392 | 0.088 | 3.8791822 | 25 | si:dkey-262k9.4 |
| CR381673.14 | 1.555241 | 0.514077136 | 0.304 | 0.055 | 3.9510917 | 25 | CR381673.1 |
| plpp1a2 | 2.655408 | 0.53183013 | 0.367 | 0.076 | 6.7460645 | 25 | plpp1a |
| zdhhc20a | 3.521834 | 0.48992502 | 0.259 | 0.042 | 8.9472200 | 25 | zdhhc20a |
| dusp23 | 5.844711 | 0.774307301 | 0.551 | 0.161 | 1.4848489 | 25 | dusp2 |
| mapre1a5 | 6.786200 | 0.732970358 | 0.608 | 0.193 | 1.7240342 | 25 | mapre1a |
| mt-co211 | 7.806467 | 0.642476367 | 0.943 | 0.669 | 1.9832329 | 25 | mt-co2 |
| si:dkey-53k12.23 | 1.579052 | 0.452122571 | 0.259 | 0.043 | 4.0115819 | 25 | si:dkey-53k12.2 |
| LOC1104382371 | 1.605381 | 0.497842936 | 0.291 | 0.053 | 4.0784729 | 25 | LOC110438237 |
| skap24 | 3.534278 | 0.560973007 | 0.361 | 0.079 | 8.9788342 | 25 | skap2 |
| tmsb4x13 | 7.527962 | 0.598428553 | 1 | 0.717 | 1.9124789 | 25 | tmsb4x |
| gsto2 | 1.881762 | 0.656901338 | 0.392 | 0.094 | 4.7806169 | 25 | gsto2 |
| si:zfos-741a10.34 | 2.440636 | 0.432343567 | 0.209 | 0.03 | 6.2004372 | 25 | si:zfos-741a10.3 |
| ppp1r14aa | 4.372776 | 0.301739485 | 0.228 | 0.034 | 1.1109037 | 25 | ppp1r14aa |
| FP015862.21 | 5.167398 | 0.285190792 | 0.133 | 0.013 | 1.3127776 | 25 | FP015862.2 |
| mrtfab1 | 6.058653 | 0.45931165 | 0.278 | 0.051 | 1.5392008 | 25 | mrtfab |
| cyba4 | 1.012400 | 0.533888847 | 0.361 | 0.08 | 2.5720032 | 25 | cyba |
| laptm57 | 1.029666 | 0.68770803 | 0.57 | 0.173 | 2.6158683 | 25 | laptm5 |
| xbp14 | 1.189253 | 0.779395508 | 0.766 | 0.352 | 3.0212992 | 25 | xbp1 |
| rps6ka13 | 7.400301 | 0.528065072 | 0.323 | 0.068 | 1.8800465 | 25 | rps6ka1 |
| amd1 | 1.107088 | 0.445253581 | 0.272 | 0.049 | 2.8125579 | 25 | amd1 |
| atp1a3a2 | 7.363211 | 0.334244135 | 0.171 | 0.022 | 1.8706237 | 25 | atp1a3a |
| parvg3 | 7.388287 | 0.528661222 | 0.329 | 0.072 | 1.8769945 | 25 | parvg |

| | | | | | | | |
|---|---|---|---|---|---|---|---|
| mt-co111 | 1.292252 | 0.615205919 | 0.981 | 0.638 | 3.2829663 | 25 | mt-co1 |
| trib32 | 1.555432 | 0.739137125 | 0.392 | 0.106 | 3.9515773 | 25 | trib3 |
| lpar5a3 | 2.881299 | 0.458046406 | 0.247 | 0.044 | 7.3199420 | 25 | lpar5a |
| mbnl16 | 4.119958 | 0.68541487 | 0.551 | 0.184 | 1.0466754 | 25 | mbnl1 |
| zgc:640516 | 4.566575 | 0.677448513 | 0.519 | 0.161 | 1.1601385 | 25 | zgc:64051 |
| mt-cyb9 | 5.168733 | 0.659633251 | 0.905 | 0.575 | 1.3131166 | 25 | mt-cyb |
| stk17b4 | 7.050689 | 0.619838802 | 0.418 | 0.112 | 1.7912275 | 25 | stk17b |
| sdr16c5b1 | 1.432678 | 0.413760384 | 0.253 | 0.046 | 3.6397192 | 25 | sdr16c5b |
| smox3 | 2.529267 | 0.57705102 | 0.354 | 0.085 | 6.4256044 | 25 | smox |
| tpd52 | 3.574719 | 0.267383318 | 0.196 | 0.029 | 9.0815759 | 25 | tpd52 |
| arpc5b5 | 6.043892 | 0.618323852 | 0.449 | 0.129 | 1.5354508 | 25 | arpc5b |
| pxk | 1.142050 | 0.384684723 | 0.209 | 0.033 | 2.9013782 | 25 | pxk |
| ptprr | 1.354058 | 0.251006964 | 0.139 | 0.016 | 3.4399850 | 25 | ptprr |
| ncf41 | 2.573796 | 0.264756879 | 0.152 | 0.019 | 6.5387295 | 25 | ncf4 |
| g6pd | 4.168149 | 0.370103323 | 0.19 | 0.029 | 1.0589184 | 25 | g6pd |
| unm-sa16141 | 1.218488 | 0.327653651 | 0.146 | 0.018 | 3.0955708 | 25 | unm-sa1614 |
| sh3bgrl4 | 2.726494 | 0.608640702 | 0.411 | 0.119 | 6.9266585 | 25 | sh3bgrl |
| cox6a22 | 7.047080 | 0.368875241 | 0.215 | 0.037 | 1.7903108 | 25 | cox6a2 |
| arhgdig6 | 7.553716 | 0.685908964 | 0.646 | 0.257 | 1.9190216 | 25 | arhgdig |
| atg16l1 | 5.325989 | 0.360628077 | 0.177 | 0.026 | 1.3530676 | 25 | atg16l1 |
| si:ch211-105j21.92 | 5.416365 | 0.42132516 | 0.203 | 0.034 | 1.3760277 | 25 | si:ch211-105j21.9 |
| syngr2a2 | 6.995316 | 0.460421506 | 0.266 | 0.056 | 1.7771601 | 25 | syngr2a |
| mt-nd411 | 1.371474 | 0.660142353 | 0.848 | 0.445 | 3.4842322 | 25 | mt-nd4 |
| b3gnt2b | 2.917119 | 0.300483406 | 0.152 | 0.02 | 7.4109426 | 25 | b3gnt2b |
| mt-atp612 | 4.244008 | 0.625651633 | 0.899 | 0.554 | 1.0781903 | 25 | mt-atp6 |
| farsa | 4.377236 | 0.350357726 | 0.209 | 0.037 | 1.1120368 | 25 | farsa |
| derl21 | 7.180754 | 0.512532196 | 0.386 | 0.105 | 1.8242706 | 25 | derl2 |
| ets21 | 8.960532 | 0.51117213 | 0.335 | 0.086 | 2.2764233 | 25 | ets2 |
| sema4aa | 1.651919 | 0.268160699 | 0.108 | 0.011 | 4.1967002 | 25 | sema4aa |
| hbegfa1 | 1.983624 | 0.539783685 | 0.373 | 0.1 | 5.0393969 | 25 | hbegfa |
| tnfaip8l2a | 2.105698 | 0.31905288 | 0.165 | 0.024 | 5.3495267 | 25 | tnfaip8l2a |
| cst14b.12 | 3.185635 | 0.44898848 | 0.272 | 0.06 | 8.0931061 | 25 | cst14b.1 |
| fmo5 | 3.708932 | 0.252480304 | 0.152 | 0.021 | 9.4225432 | 25 | fmo5 |
| grk6 | 4.977419 | 0.288838761 | 0.158 | 0.023 | 1.2645134 | 25 | grk6 |
| wbp2nl1 | 5.262604 | 0.517600811 | 0.342 | 0.091 | 1.3369647 | 25 | wbp2nl |

| | | | | | | | |
|---|---|---|---|---|---|---|---|
| arhgef3l2 | 5.643818 | 0.346878605 | 0.184 | 0.03 | 1.4338121 | 25 | arhgef3l |
| itm2bb5 | 6.995606 | 0.646004819 | 0.601 | 0.24 | 1.7772338 | 25 | itm2bb |
| LOC1039091154 | 3.571090 | 0.57836176 | 0.487 | 0.167 | 9.0723543 | 25 | LOC103909115 |
| actb16 | 1.373536 | 0.408895915 | 0.949 | 0.767 | 3.4894684 | 25 | actb1 |
| cited4b1 | 1.618231 | 0.541101955 | 0.405 | 0.119 | 4.1111171 | 25 | cited4b |
| sri1 | 2.606398 | 0.538289021 | 0.468 | 0.159 | 6.6215557 | 25 | sri |
| gpia1 | 2.639096 | 0.549955831 | 0.399 | 0.122 | 6.7046247 | 25 | gpia |
| si:ch73-208g10.14 | 2.878295 | 0.385710505 | 0.209 | 0.04 | 7.3123101 | 25 | si:ch73-208g10.1 |
| selp1 | 3.735716 | 0.258972471 | 0.108 | 0.012 | 9.4905875 | 25 | selp |
| cited4a2 | 4.165128 | 0.393881651 | 0.209 | 0.041 | 1.0581508 | 25 | cited4a |
| itga12 | 4.322633 | 0.343026186 | 0.215 | 0.042 | 1.0981651 | 25 | itga1 |
| si:dkey-24p1.71 | 7.847201 | 0.263006193 | 0.114 | 0.013 | 1.9935815 | 25 | si:dkey-24p1.7 |
| prdx55 | 8.724775 | 0.626448533 | 0.468 | 0.177 | 2.2165293 | 25 | prdx5 |
| cap17 | 9.020559 | 0.674298456 | 0.57 | 0.244 | 2.2916732 | 25 | cap1 |
| crip18 | 1.221276 | 0.669871918 | 0.627 | 0.282 | 3.1026522 | 25 | crip1 |
| mcl1b8 | 1.673816 | 0.645253532 | 0.589 | 0.249 | 4.2523319 | 25 | mcl1b |
| arg2 | 2.146408 | 0.339232741 | 0.133 | 0.018 | 5.4529507 | 25 | arg2 |
| pde4ba6 | 2.837846 | 0.575386776 | 0.424 | 0.141 | 7.2095499 | 25 | pde4ba |
| grasp4 | 3.971406 | 0.506615448 | 0.285 | 0.072 | 1.0089358 | 25 | grasp |
| si:ch211-102c2.45 | 4.256344 | 0.553100869 | 0.449 | 0.144 | 1.0813243 | 25 | si:ch211-102c2.4 |
| zmp:00000009361 | 1.988108 | 0.254261269 | 0.127 | 0.017 | 5.0507905 | 25 | zmp:0000000936 |
| wdr15 | 3.051450 | 0.521396904 | 0.449 | 0.154 | 7.7522088 | 25 | wdr1 |
| rab11fip1b3 | 3.504835 | 0.418518672 | 0.253 | 0.059 | 8.9040354 | 25 | rab11fip1b |
| ptprc6 | 6.666464 | 0.521853496 | 0.392 | 0.123 | 1.6936152 | 25 | ptprc |
| cmtm71 | 1.017501 | 0.349837248 | 0.228 | 0.05 | 2.5849637 | 25 | cmtm7 |
| fermt3b1 | 1.175743 | 0.263626933 | 0.139 | 0.021 | 2.9869756 | 25 | fermt3b |
| CR383676.113 | 1.238245 | 0.62766239 | 0.975 | 0.715 | 3.1457633 | 25 | CR383676.1 |
| thy17 | 1.277166 | 0.63871596 | 0.804 | 0.376 | 3.2446408 | 25 | thy1 |
| fybb4 | 3.427046 | 0.349396254 | 0.209 | 0.043 | 8.7064103 | 25 | fybb |
| sptlc2a1 | 7.103983 | 0.326328093 | 0.215 | 0.046 | 1.8047671 | 25 | sptlc2a |
| nfkbiab2 | 1.901084 | 0.646953635 | 0.481 | 0.199 | 4.8297043 | 25 | nfkbiab |
| zgc:91976 | 3.677301 | 0.263171949 | 0.139 | 0.022 | 9.3421834 | 25 | zgc:91976 |
| nme2b.21 | 3.701913 | 0.288785456 | 0.152 | 0.026 | 9.4047111 | 25 | nme2b.2 |
| vaspa5 | 9.203279 | 0.398469206 | 0.297 | 0.08 | 2.3380931 | 25 | vaspa |
| srd5a2a | 1.014524 | 0.395172871 | 0.304 | 0.084 | 2.5773989 | 25 | srd5a2a |

| | | | | | | | |
|---|---|---|---|---|---|---|---|
| calm1b5 | 3.162227 | 0.446338037 | 0.335 | 0.102 | 8.0336401 | 25 | calm1b |
| tpt13 | 6.393832 | 0.31594127 | 0.962 | 0.811 | 1.6243530 | 25 | tpt1 |
| hsd3b71 | 6.739841 | 0.351341532 | 0.304 | 0.083 | 1.7122568 | 25 | hsd3b7 |
| arpc4l5 | 8.476867 | 0.549342833 | 0.513 | 0.217 | 2.1535480 | 25 | arpc4l |
| myo1f4 | 1.155036 | 0.334123022 | 0.209 | 0.047 | 2.9343692 | 25 | myo1f |
| si:ch73-234b20.53 | 1.425687 | 0.457633437 | 0.329 | 0.104 | 3.6219583 | 25 | si:ch73-234b20.5 |
| elovl1b4 | 1.431556 | 0.483398571 | 0.361 | 0.121 | 3.6368684 | 25 | elovl1b |
| tpd52l2b1 | 1.442118 | 0.441061755 | 0.342 | 0.108 | 3.6637009 | 25 | tpd52l2b |
| zgc:1749043 | 3.115808 | 0.417798629 | 0.215 | 0.051 | 7.9157106 | 25 | zgc:174904 |
| sh3kbp13 | 6.466989 | 0.371978193 | 0.272 | 0.076 | 1.6429385 | 25 | sh3kbp1 |
| rab27a1 | 9.276920 | 0.378982502 | 0.203 | 0.047 | 2.3568016 | 25 | rab27a |
| limd22 | 1.609736 | 0.353520423 | 0.253 | 0.067 | 4.0895349 | 25 | limd2 |
| tmem30ab1 | 1.646174 | 0.355803959 | 0.228 | 0.057 | 4.1821072 | 25 | tmem30ab |
| zgc:77650 | 1.913839 | 0.452236985 | 0.316 | 0.1 | 4.8621090 | 25 | zgc:77650 |
| osgn1 | 2.118906 | 0.319724048 | 0.184 | 0.04 | 5.3830827 | 25 | osgn1 |
| zgc:1528631 | 2.767317 | 0.420021961 | 0.177 | 0.039 | 7.0303690 | 25 | zgc:152863 |
| rab11ba1 | 4.166857 | 0.443728834 | 0.354 | 0.121 | 1.0585901 | 25 | rab11ba |
| asah1a | 6.352564 | 0.268625998 | 0.139 | 0.025 | 1.6138690 | 25 | asah1a |
| kctd12.24 | 9.474933 | 0.402548398 | 0.316 | 0.099 | 2.4071068 | 25 | kctd12.2 |
| sult6b1 | 1.127447 | 0.2542736 | 0.146 | 0.028 | 2.8642801 | 25 | sult6b1 |
| clic15 | 1.186395 | 0.493838707 | 0.354 | 0.128 | 3.0140384 | 25 | clic1 |
| mt-nd38 | 3.154100 | 0.546285813 | 0.684 | 0.37 | 8.0129918 | 25 | mt-nd3 |
| rgs14a | 3.510359 | 0.267840951 | 0.177 | 0.039 | 8.9180682 | 25 | rgs14a |
| arpc26 | 3.533934 | 0.539683115 | 0.582 | 0.298 | 8.9779599 | 25 | arpc2 |
| fmnl1a1 | 4.283492 | 0.332411105 | 0.171 | 0.038 | 1.0882213 | 25 | fmnl1a |
| plekho1b1 | 4.834424 | 0.331439263 | 0.184 | 0.042 | 1.2281855 | 25 | plekho1b |
| hcar1-43 | 5.088229 | 0.28094323 | 0.158 | 0.033 | 1.2926648 | 25 | hcar1-4 |
| cbx7a4 | 6.501339 | 0.296727726 | 0.209 | 0.052 | 1.6516653 | 25 | cbx7a |
| lrrfip1a3 | 8.030656 | 0.312607386 | 0.228 | 0.06 | 2.0401883 | 25 | lrrfip1a |
| rps6kb1b | 9.334749 | 0.357749747 | 0.253 | 0.072 | 2.3714931 | 25 | rps6kb1b |
| pygb1 | 1.072087 | 0.285331649 | 0.19 | 0.045 | 2.7236373 | 25 | pygb |
| elf14 | 1.261042 | 0.416196669 | 0.31 | 0.102 | 3.2036789 | 25 | elf1 |
| capza1b4 | 2.471646 | 0.528573991 | 0.494 | 0.234 | 6.2792188 | 25 | capza1b |
| arpc36 | 3.169115 | 0.53748763 | 0.614 | 0.333 | 8.0511367 | 25 | arpc3 |
| tagln29 | 4.792345 | 0.471690483 | 0.487 | 0.208 | 1.2174953 | 25 | tagln2 |

| | | | | | | | |
|---|---|---|---|---|---|---|---|
| rca2.1 | 5.889043 | 0.397584982 | 0.266 | 0.082 | 1.4961114 | 25 | rca2.1 |
| actr2a6 | 8.321735 | 0.424551464 | 0.361 | 0.132 | 2.1141368 | 25 | actr2a |
| cnp | 9.177836 | 0.481930575 | 0.506 | 0.224 | 2.3316292 | 25 | cnp |
| zgc:771124 | 9.841244 | 0.395342186 | 0.31 | 0.099 | 2.5001680 | 25 | zgc:77112 |
| selplg3 | 1.394202 | 0.319991588 | 0.184 | 0.045 | 3.5419716 | 25 | selplg |
| h1fx5 | 1.407843 | 0.532713159 | 0.443 | 0.19 | 3.5766262 | 25 | h1fx |
| ppt13 | 1.566512 | 0.521360224 | 0.418 | 0.181 | 3.9797260 | 25 | ppt1 |
| iqgap12 | 3.281103 | 0.331447825 | 0.241 | 0.071 | 8.3356430 | 25 | iqgap1 |
| add3a8 | 3.846168 | 0.40679688 | 0.367 | 0.137 | 9.7711912 | 25 | add3a |
| zgc:66427 | 4.542151 | 0.363325969 | 0.253 | 0.077 | 1.1539335 | 25 | zgc:66427 |
| fam49ba4 | 5.141991 | 0.369084223 | 0.31 | 0.104 | 1.3063228 | 25 | fam49ba |
| si:rp71-68n21.93 | 5.427804 | 0.306436135 | 0.184 | 0.046 | 1.3789336 | 25 | si:rp71-68n21.9 |
| si:ch211-226m16.23 | 9.615514 | 0.411388525 | 0.228 | 0.068 | 2.4428214 | 25 | si:ch211-226m16.2 |
| gapdh7 | 9.984728 | 0.555328813 | 0.614 | 0.346 | 2.5366202 | 25 | gapdh |
| tsc22d35 | 1.200965 | 0.467851153 | 0.386 | 0.158 | 3.0510532 | 25 | tsc22d3 |
| lasp15 | 1.895059 | 0.485523389 | 0.513 | 0.252 | 4.8143984 | 25 | lasp1 |
| slc9a3r1a1 | 2.405923 | 0.317332286 | 0.19 | 0.051 | 6.1122494 | 25 | slc9a3r1a |
| dusp15 | 1.003395 | 0.456482683 | 0.367 | 0.148 | 2.5491250 | 25 | dusp1 |
| rab11fip4b1 | 1.050248 | 0.284935538 | 0.146 | 0.033 | 2.6681572 | 25 | rab11fip4b |
| LOC5665872 | 1.704042 | 0.28986103 | 0.171 | 0.044 | 4.3291208 | 25 | LOC566587 |
| acsl4b1 | 2.743218 | 0.326439793 | 0.19 | 0.053 | 6.9691468 | 25 | acsl4b |
| serpine21 | 2.747922 | 0.302646567 | 0.209 | 0.061 | 6.9810967 | 25 | serpine2 |
| itpkcb2 | 3.252465 | 0.34574482 | 0.234 | 0.074 | 8.2628898 | 25 | itpkcb |
| si:ch211-132p1.35 | 4.152808 | 0.395026028 | 0.272 | 0.097 | 1.0550211 | 25 | si:ch211-132p1.3 |
| hmgn27 | 7.045011 | 0.458042032 | 0.734 | 0.467 | 1.7897851 | 25 | hmgn2 |
| CR392341.14 | 1.497687 | 0.269835285 | 0.203 | 0.059 | 3.8048742 | 25 | CR392341.1 |
| arpc45 | 1.527122 | 0.453686948 | 0.481 | 0.237 | 3.8796559 | 25 | arpc4 |
| gpx1a5 | 1.752203 | 0.392091012 | 0.513 | 0.235 | 4.4514737 | 25 | gpx1a |
| samsn1a2 | 2.136464 | 0.292341087 | 0.152 | 0.038 | 5.4276875 | 25 | samsn1a |
| tktb3 | 2.761732 | 0.392291396 | 0.386 | 0.163 | 7.0161810 | 25 | tktb |
| si:ch211-197h24.94 | 2.767540 | 0.306758416 | 0.247 | 0.083 | 7.0309371 | 25 | si:ch211-197h24.9 |
| pkma4 | 3.175809 | 0.394863062 | 0.405 | 0.178 | 8.0681449 | 25 | pkma |
| dbi5 | 3.596655 | 0.48762106 | 0.456 | 0.243 | 9.1373029 | 25 | dbi |
| ptmaa10 | 4.182131 | 0.455631499 | 0.715 | 0.432 | 1.0624705 | 25 | ptmaa |
| si:ch73-1a9.311 | 1.202455 | 0.460442283 | 0.563 | 0.315 | 3.0548393 | 25 | si:ch73-1a9.3 |

| | | | | | | | |
|---|---|---|---|---|---|---|---|
| tpi1b5 | 1.210786 | 0.439779306 | 0.386 | 0.178 | 3.0760039 | 25 | tpi1b |
| pde7a2 | 1.452856 | 0.253640494 | 0.209 | 0.065 | 3.6909828 | 25 | pde7a |
| tpst11 | 1.483479 | 0.264664848 | 0.152 | 0.04 | 3.7687790 | 25 | tpst1 |
| nr4a16 | 2.295456 | 0.438766525 | 0.361 | 0.158 | 5.8316060 | 25 | nr4a1 |
| RASSF54 | 3.057635 | 0.284646977 | 0.196 | 0.061 | 7.7679227 | 25 | RASSF5 |
| lamtor4 | 3.086792 | 0.284145126 | 0.272 | 0.1 | 7.8419952 | 25 | lamtor4 |
| actn42 | 5.025201 | 0.357015244 | 0.259 | 0.097 | 1.2766525 | 25 | actn4 |
| myl12.17 | 5.089841 | 0.424012128 | 0.652 | 0.426 | 1.2930741 | 25 | myl12.1 |
| scarb2a3 | 5.114412 | 0.299336428 | 0.196 | 0.062 | 1.2993164 | 25 | scarb2a |
| capzb6 | 9.095781 | 0.411780628 | 0.506 | 0.271 | 2.3107832 | 25 | capzb |
| aldoaa2 | 9.731300 | 0.41233577 | 0.481 | 0.253 | 2.4722368 | 25 | aldoaa |
| arrb2b5 | 1.030714 | 0.302303107 | 0.259 | 0.096 | 2.6185299 | 25 | arrb2b |
| rgs134 | 1.135386 | 0.363591581 | 0.31 | 0.125 | 2.8844490 | 25 | rgs13 |
| wasa5 | 1.241506 | 0.324061259 | 0.259 | 0.096 | 3.1540467 | 25 | wasa |
| tpm36 | 1.499672 | 0.402915046 | 0.519 | 0.283 | 3.8099189 | 25 | tpm3 |
| tmem592 | 1.836422 | 0.404128268 | 0.297 | 0.126 | 4.6654316 | 25 | tmem59 |
| mt-nd210 | 2.112264 | 0.44695394 | 0.627 | 0.391 | 5.3662089 | 25 | mt-nd2 |
| slc38a22 | 2.594068 | 0.251999843 | 0.158 | 0.046 | 6.5902316 | 25 | slc38a2 |
| crema5 | 2.731428 | 0.428804159 | 0.367 | 0.174 | 6.9391943 | 25 | crema |
| mt-nd112 | 3.140946 | 0.405118097 | 0.652 | 0.396 | 7.9795750 | 25 | mt-nd1 |
| f11r.15 | 4.010583 | 0.407883973 | 0.361 | 0.164 | 1.0188886 | 25 | f11r.1 |
| birc23 | 5.000236 | 0.31055604 | 0.272 | 0.107 | 1.2703101 | 25 | birc2 |
| eno32 | 6.733178 | 0.431556489 | 0.589 | 0.362 | 1.7105640 | 25 | eno3 |
| fam49bb | 8.424373 | 0.26197307 | 0.171 | 0.053 | 2.1402120 | 25 | fam49bb |
| mtpn3 | 8.578948 | 0.357100493 | 0.348 | 0.163 | 2.1794819 | 25 | mtpn |
| ak11 | 1.085609 | 0.305317983 | 0.253 | 0.098 | 2.7579920 | 25 | ak1 |
| LO018102.1 | 1.185217 | 0.285030274 | 0.228 | 0.083 | 3.0110455 | 25 | LO018102.1 |
| scamp2 | 1.548856 | 0.287421898 | 0.209 | 0.074 | 3.9348711 | 25 | scamp2 |
| pnrc213 | 2.764013 | 0.368336346 | 0.848 | 0.597 | 7.0219763 | 25 | pnrc2 |
| cox8a7 | 2.802224 | 0.3629261 | 0.487 | 0.267 | 7.1190510 | 25 | cox8a |
| adgre5b.31 | 2.995278 | 0.255652519 | 0.133 | 0.037 | 7.6095053 | 25 | adgre5b.3 |
| socs3b6 | 7.802030 | 0.348772895 | 0.329 | 0.151 | 1.9821059 | 25 | socs3b |
| ccdc28a | 7.836231 | 0.27919062 | 0.259 | 0.105 | 1.9907945 | 25 | ccdc28a |
| hmha1b3 | 8.582568 | 0.285815215 | 0.203 | 0.074 | 2.1804014 | 25 | hmha1b |
| sh3bgrl33 | 1.996548 | 0.326661007 | 0.329 | 0.156 | 5.0722324 | 25 | sh3bgrl3 |

| | | | | | | | |
|---|---|---|---|---|---|---|---|
| tln14 | 2.225771 | 0.30372613 | 0.31 | 0.142 | 5.6545733 | 25 | tln1 |
| si:dkey-27i16.23 | 2.738130 | 0.281831062 | 0.234 | 0.091 | 6.9562204 | 25 | si:dkey-27i16.2 |
| degs11 | 2.907389 | 0.27133574 | 0.177 | 0.063 | 7.3862229 | 25 | degs1 |
| mcl1a4 | 4.606994 | 0.342432596 | 0.816 | 0.613 | 0.000117 | 25 | mcl1a |
| gpr183a2 | 6.366031 | 0.30064733 | 0.177 | 0.065 | 0.000162 | 25 | gpr183a |
| grap2b6 | 7.960212 | 0.272293603 | 0.285 | 0.124 | 0.000202 | 25 | grap2b |
| zgc:16273010 | 8.236416 | 0.38960039 | 0.639 | 0.402 | 0.000209 | 25 | zgc:162730 |
| abracl4 | 8.966747 | 0.332627909 | 0.424 | 0.221 | 0.000228 | 25 | abracl |
| magt11 | 1.014891 | 0.318403879 | 0.278 | 0.129 | 0.000258 | 25 | magt1 |
| yrk5 | 1.236537 | 0.252349619 | 0.184 | 0.067 | 0.000314 | 25 | yrk |
| pim24 | 3.127311 | 0.357343472 | 0.209 | 0.088 | 0.000794 | 25 | pim2 |
| cdc424 | 4.592414 | 0.333436558 | 0.411 | 0.234 | 0.001167 | 25 | cdc42 |
| msna8 | 5.792889 | 0.339116142 | 0.456 | 0.265 | 0.001472 | 25 | msna |
| aldh9a1a.17 | 6.279945 | 0.277234875 | 0.373 | 0.193 | 0.001595 | 25 | aldh9a1a.1 |
| capza1a4 | 6.442547 | 0.300036922 | 0.405 | 0.221 | 0.001637 | 25 | capza1a |
| svild6 | 6.557984 | 0.290717226 | 0.203 | 0.084 | 0.001666 | 25 | svild |
| si:dkey-112e7.25 | 7.171050 | 0.288107501 | 0.399 | 0.214 | 0.001822 | 25 | si:dkey-112e7.2 |
| btg19 | 1.433671 | 0.337965137 | 0.563 | 0.355 | 0.003642 | 25 | btg1 |
| nfil3-67 | 1.443360 | 0.300125355 | 0.272 | 0.131 | 0.003667 | 25 | nfil3-6 |
| ccni6 | 2.967548 | 0.326269507 | 0.494 | 0.319 | 0.007539 | 25 | ccni |
| si:ch211-212k18.73 | 3.166865 | 0.260726818 | 0.297 | 0.146 | 0.008045 | 25 | si:ch211-212k18.7 |
| s100a10b8 | 4.084227 | 0.336013556 | 0.494 | 0.285 | 0.010376 | 25 | s100a10b |
| chac14 | 7.164702 | 0.324727721 | 0.551 | 0.378 | 0.018202 | 25 | chac1 |
| arhgdia5 | 1.079539 | 0.291904953 | 0.424 | 0.251 | 0.027426 | 25 | arhgdia |
| tmed72 | 1.122106 | 0.280082 | 0.386 | 0.22 | 0.028507 | 25 | tmed7 |
| tnfb3 | 1.260421 | 0.31224107 | 0.133 | 0.05 | 0.032021 | 25 | tnfb |
| rap1b4 | 1.283310 | 0.272234483 | 0.386 | 0.22 | 0.032602 | 25 | rap1b |
| gpx4b2 | 1.627264 | 0.300200854 | 0.373 | 0.22 | 0.041341 | 25 | gpx4b |
| cfl17 | 1.652736 | 0.263410074 | 0.715 | 0.601 | 0.041988 | 25 | cfl1 |
| ucp26 | 1.182184 | 0.273070103 | 0.595 | 0.414 | 0.300334 | 25 | ucp2 |
| serf25 | 3.898922 | 0.255498781 | 0.525 | 0.378 | 0.990521 | 25 | serf2 |
| notch26 | 0.0001 | 0.253797792 | 0.215 | 0.123 | 1 | 25 | notch2 |
| socs3a9 | 0.0006 | 0.260586181 | 0.443 | 0.316 | 1 | 25 | socs3a |
| LOC103908723 | 0 | 2.046966928 | 0.965 | 0.003 | 0 | 26 | LOC103908723 |
| LOC1018867661 | 0 | 1.973738141 | 0.951 | 0.005 | 0 | 26 | LOC101886766 |

| | | | | | | | |
|---|---|---|---|---|---|---|---|
| zgc:194275 | 0 | 1.968368083 | 0.923 | 0.001 | 0 | 26 | zgc:194275 |
| igl4v81 | 0 | 1.893423146 | 0.944 | 0.019 | 0 | 26 | igl4v8 |
| cd37 | 0 | 1.845040656 | 0.894 | 0.002 | 0 | 26 | cd37 |
| CU896602.1 | 0 | 1.664572674 | 0.838 | 0.001 | 0 | 26 | CU896602.1 |
| igic1s1 | 0 | 1.54980338 | 0.803 | 0.001 | 0 | 26 | igic1s1 |
| igl1c3 | 0 | 1.535972513 | 0.754 | 0 | 0 | 26 | igl1c3 |
| igl3v5 | 0 | 1.515732501 | 0.775 | 0.001 | 0 | 26 | igl3v5 |
| XLOC-026705 | 0 | 1.471666132 | 0.718 | 0.001 | 0 | 26 | XLOC-026705 |
| cxcr4a1 | 0 | 1.399343311 | 0.641 | 0.013 | 0 | 26 | cxcr4a |
| ncf12 | 0 | 1.249903206 | 0.648 | 0.027 | 0 | 26 | ncf1 |
| CT573231.12 | 0 | 1.195646934 | 0.585 | 0.021 | 0 | 26 | CT573231.1 |
| LOC1018822641 | 0 | 1.109083506 | 0.5 | 0.006 | 0 | 26 | LOC101882264 |
| BX571825.4 | 0 | 1.091032841 | 0.423 | 0.002 | 0 | 26 | BX571825.4 |
| pax5 | 0 | 1.083384597 | 0.507 | 0.002 | 0 | 26 | pax5 |
| BX571825.5 | 0 | 1.069322261 | 0.5 | 0 | 0 | 26 | BX571825.5 |
| CT573342.2 | 0 | 1.005888532 | 0.486 | 0.001 | 0 | 26 | CT573342.2 |
| epb41a | 0 | 0.978465623 | 0.514 | 0.014 | 0 | 26 | epb41a |
| CU571382.4 | 0 | 0.89204847 | 0.394 | 0.001 | 0 | 26 | CU571382.4 |
| LOC108179343 | 0 | 0.873671678 | 0.387 | 0 | 0 | 26 | LOC108179343 |
| BX571825.3 | 0 | 0.827389563 | 0.345 | 0 | 0 | 26 | BX571825.3 |
| ighv1-4 | 0 | 0.749128438 | 0.303 | 0.003 | 0 | 26 | ighv1-4 |
| ly6m2 | 0 | 0.738127595 | 0.282 | 0.001 | 0 | 26 | ly6m2 |
| LOC101886979 | 0 | 0.711293733 | 0.275 | 0.001 | 0 | 26 | LOC101886979 |
| swap70a | 0 | 0.624744792 | 0.289 | 0.004 | 0 | 26 | swap70a |
| FP016018.1 | 0 | 0.577068501 | 0.246 | 0 | 0 | 26 | FP016018.1 |
| capga | 0 | 0.545281159 | 0.225 | 0.001 | 0 | 26 | capga |
| rtn4rl2b | 0 | 0.496849051 | 0.204 | 0 | 0 | 26 | rtn4rl2b |
| CR944667.2 | 0 | 0.363496252 | 0.148 | 0 | 0 | 26 | CR944667.2 |
| CU571382.1 | 0 | 0.331734003 | 0.127 | 0 | 0 | 26 | CU571382.1 |
| si:ch73-7i4.3 | 2.949062 | 0.604163728 | 0.261 | 0.004 | 7.4920939 | 26 | si:ch73-7i4.3 |
| XLOC-0183561 | 1.588495 | 1.15573137 | 0.535 | 0.025 | 4.0355725 | 26 | XLOC-018356 |
| si:ch211-145b13.61 | 2.859186 | 0.624927373 | 0.282 | 0.006 | 7.2637623 | 26 | si:ch211-145b13.6 |
| pou2f2a1 | 8.431899 | 0.600578694 | 0.296 | 0.006 | 2.1421240 | 26 | pou2f2a |
| sp6 | 7.105110 | 0.335508726 | 0.162 | 0.001 | 1.8050532 | 26 | sp6 |
| LOC110438240 | 2.486667 | 0.377901306 | 0.169 | 0.002 | 6.3173793 | 26 | LOC110438240 |

| | | | | | | | |
|---|---|---|---|---|---|---|---|
| ccr9a5 | 4.593551 | 1.463064901 | 0.789 | 0.082 | 1.1669918 | 26 | ccr9a |
| sftpba2 | 4.177086 | 1.05301264 | 0.507 | 0.031 | 1.0611887 | 26 | sftpba |
| si:ch73-380l3.21 | 3.308153 | 0.937812822 | 0.458 | 0.026 | 8.4043649 | 26 | si:ch73-380l3.2 |
| cxcr4b6 | 7.211531 | 1.725046364 | 0.979 | 0.17 | 1.8320896 | 26 | cxcr4b |
| cobll1a | 4.346824 | 0.392402963 | 0.197 | 0.004 | 1.1043108 | 26 | cobll1a |
| grap2b7 | 3.983262 | 1.362579504 | 0.838 | 0.118 | 1.0119479 | 26 | grap2b |
| illr11 | 4.162618 | 0.709767673 | 0.289 | 0.012 | 1.0575131 | 26 | illr1 |
| si:dkeyp-117h8.2 | 2.471771 | 0.369815765 | 0.148 | 0.003 | 6.2795348 | 26 | si:dkeyp-117h8.2 |
| sh2d4a | 9.434219 | 0.501197007 | 0.232 | 0.008 | 2.3967635 | 26 | sh2d4a |
| CT027980.1 | 5.368963 | 0.306147812 | 0.134 | 0.002 | 1.3639852 | 26 | CT027980.1 |
| rbbp8l1 | 1.256221 | 0.493098239 | 0.225 | 0.009 | 3.1914312 | 26 | rbbp8l |
| irf4b2 | 2.415544 | 0.732669576 | 0.331 | 0.02 | 6.1366917 | 26 | irf4b |
| blnk1 | 9.871758 | 0.656136764 | 0.338 | 0.022 | 2.5079203 | 26 | blnk |
| si:dkey-24p1.1 | 3.197840 | 0.449873456 | 0.19 | 0.006 | 8.1241143 | 26 | si:dkey-24p1.1 |
| tlr19 | 1.903928 | 0.284163979 | 0.127 | 0.003 | 4.8369308 | 26 | tlr19 |
| si:ch211-119e14.13 | 6.806810 | 1.054470232 | 0.563 | 0.066 | 1.7292700 | 26 | si:ch211-119e14.1 |
| si:ch1073-220m6.11 | 7.147135 | 0.605490561 | 0.296 | 0.018 | 1.8157298 | 26 | si:ch1073-220m6.1 |
| btbd11b | 4.411509 | 0.43694713 | 0.197 | 0.008 | 1.1207439 | 26 | btbd11b |
| pacsin2 | 5.173495 | 0.838685314 | 0.444 | 0.044 | 1.3143264 | 26 | pacsin2 |
| XLOC-0425954 | 1.145178 | 1.309791329 | 0.817 | 0.148 | 2.9093260 | 26 | XLOC-042595 |
| tspan2a | 5.184399 | 0.428101447 | 0.211 | 0.01 | 1.3170967 | 26 | tspan2a |
| arhgap15 | 6.083327 | 0.667340823 | 0.352 | 0.028 | 1.5454694 | 26 | arhgap15 |
| csrnp1a | 1.443963 | 0.804613638 | 0.444 | 0.045 | 3.6683897 | 26 | csrnp1a |
| atp11a1 | 2.580193 | 0.459202226 | 0.211 | 0.011 | 6.5549805 | 26 | atp11a |
| si:busm1-266f07.23 | 2.458063 | 1.127424061 | 0.796 | 0.143 | 6.2447096 | 26 | si:busm1-266f07.2 |
| mpeg1.1 | 1.765941 | 0.538284971 | 0.254 | 0.016 | 4.4863742 | 26 | mpeg1.1 |
| cd74a4 | 1.458176 | 1.267654905 | 0.908 | 0.211 | 3.7044973 | 26 | cd74a |
| lmo22 | 2.037206 | 0.571159558 | 0.296 | 0.024 | 5.1755236 | 26 | lmo2 |
| cd74b5 | 1.404665 | 1.209304405 | 0.901 | 0.221 | 3.5685523 | 26 | cd74b |
| syk4 | 9.392768 | 0.876941642 | 0.514 | 0.075 | 2.3862328 | 26 | syk |
| ebf1a | 1.514326 | 0.339682734 | 0.155 | 0.006 | 3.8471467 | 26 | ebf1a |
| XLOC-0004316 | 4.810537 | 1.344212937 | 0.923 | 0.268 | 1.2221170 | 26 | XLOC-000431 |
| dapp11 | 1.440747 | 0.367723147 | 0.162 | 0.008 | 3.6602190 | 26 | dapp1 |
| XLOC-0139644 | 3.061288 | 1.210847082 | 0.732 | 0.165 | 7.7772033 | 26 | XLOC-013964 |
| si:dkey-82o10.4 | 2.007348 | 0.375927986 | 0.155 | 0.007 | 5.0996685 | 26 | si:dkey-82o10.4 |

| | | | | | | | |
|---|---|---|---|---|---|---|---|
| dtx4a | 5.577335 | 0.341064763 | 0.176 | 0.01 | 1.4169219 | 26 | dtx4a |
| bpifcl1 | 2.095765 | 0.430331967 | 0.204 | 0.014 | 5.3242919 | 26 | bpifcl |
| rgs135 | 9.335134 | 0.966432754 | 0.627 | 0.122 | 2.3715908 | 26 | rgs13 |
| derl22 | 1.053845 | 0.861798454 | 0.563 | 0.104 | 2.6772951 | 26 | derl2 |
| cd7al2 | 3.986562 | 0.517107865 | 0.275 | 0.026 | 1.0127861 | 26 | cd7al |
| gpr55a | 9.592149 | 0.325972841 | 0.141 | 0.007 | 2.4368856 | 26 | gpr55a |
| kctd5a3 | 5.244994 | 0.946169909 | 0.711 | 0.182 | 1.3324909 | 26 | kctd5a |
| LOC1039094521 | 1.473229 | 0.489761818 | 0.211 | 0.016 | 3.7427400 | 26 | LOC103909452 |
| cd833 | 2.133960 | 1.035726175 | 0.5 | 0.091 | 5.4213268 | 26 | cd83 |
| bcl11aa | 2.886007 | 0.341239955 | 0.162 | 0.01 | 7.3319025 | 26 | bcl11aa |
| si:dkey-78k11.9 | 5.654351 | 0.311324037 | 0.141 | 0.007 | 1.4364880 | 26 | si:dkey-78k11.9 |
| LOC1039108713 | 4.015261 | 0.972694851 | 0.676 | 0.154 | 1.0200772 | 26 | LOC103910871 |
| gng2 | 1.101975 | 0.621530707 | 0.345 | 0.047 | 2.7995700 | 26 | gng2 |
| s1pr41 | 7.594791 | 0.32816712 | 0.141 | 0.008 | 1.9294566 | 26 | s1pr4 |
| rac28 | 2.645602 | 0.93300285 | 0.775 | 0.221 | 6.7211526 | 26 | rac2 |
| pacsin3 | 4.075679 | 0.451538641 | 0.232 | 0.023 | 1.0354263 | 26 | pacsin3 |
| CABZ01066694.1 | 5.034103 | 0.355526219 | 0.19 | 0.016 | 1.2789139 | 26 | CABZ01066694.1 |
| fam53b1 | 5.172447 | 0.673306621 | 0.423 | 0.074 | 1.3140601 | 26 | fam53b |
| XLOC-0108995 | 3.688413 | 1.072308334 | 0.754 | 0.236 | 9.3704146 | 26 | XLOC-010899 |
| ptmaa11 | 7.015439 | 0.917814439 | 0.915 | 0.43 | 1.7822725 | 26 | ptmaa |
| mhc2dab3 | 7.844646 | 0.666884452 | 0.479 | 0.092 | 1.9929323 | 26 | mhc2dab |
| si:ch73-234b20.54 | 1.293491 | 0.71497251 | 0.479 | 0.103 | 3.2861147 | 26 | si:ch73-234b20.5 |
| ncf22 | 3.535173 | 0.417001442 | 0.211 | 0.022 | 8.9811088 | 26 | ncf2 |
| LOC1005362133 | 3.837338 | 0.809471733 | 0.486 | 0.109 | 9.7487584 | 26 | LOC100536213 |
| CR936442.14 | 6.506038 | 0.869039998 | 0.585 | 0.151 | 1.6528590 | 26 | CR936442.1 |
| spi1b4 | 5.731819 | 0.591168487 | 0.38 | 0.07 | 1.4561688 | 26 | spi1b |
| parvg4 | 3.404329 | 0.616322365 | 0.373 | 0.072 | 8.6486997 | 26 | parvg |
| gadd45ga2 | 3.750993 | 0.670311218 | 0.31 | 0.052 | 9.5293986 | 26 | gadd45ga |
| gpx1a6 | 4.636719 | 0.846601791 | 0.711 | 0.233 | 1.1779585 | 26 | gpx1a |
| rps203 | 1.110966 | 0.338697276 | 1 | 0.91 | 2.8224091 | 26 | rps20 |
| crip19 | 1.265991 | 0.818412466 | 0.803 | 0.281 | 3.2162503 | 26 | crip1 |
| rps27.111 | 2.122313 | 0.551073504 | 1 | 0.709 | 5.3917375 | 26 | rps27.1 |
| rplp18 | 4.683691 | 0.37788322 | 1 | 0.882 | 1.1898919 | 26 | rplp1 |
| cad | 6.054914 | 0.47418415 | 0.218 | 0.027 | 1.5382510 | 26 | cad |
| ccni7 | 7.185593 | 0.815597585 | 0.782 | 0.316 | 1.8255001 | 26 | ccni |

| | | | | | | | |
|---|---|---|---|---|---|---|---|
| rplp29 | 1.759927 | 0.482194822 | 1 | 0.782 | 4.4710955 | 26 | rplp2 |
| ralgps21 | 2.596324 | 0.394160219 | 0.197 | 0.023 | 6.5959631 | 26 | ralgps2 |
| rps2113 | 3.550966 | 0.477084555 | 1 | 0.775 | 9.0212301 | 26 | rps21 |
| h1fx6 | 9.162859 | 0.968649182 | 0.592 | 0.188 | 2.3278243 | 26 | h1fx |
| rpl116 | 4.518515 | 0.415302434 | 1 | 0.829 | 1.1479289 | 26 | rpl11 |
| rsl24d17 | 3.460369 | 0.738098228 | 0.908 | 0.458 | 8.7910685 | 26 | rsl24d1 |
| rps15a9 | 1.722262 | 0.459023881 | 1 | 0.797 | 4.3754068 | 26 | rps15a |
| ikzf23 | 1.901502 | 0.424870871 | 0.232 | 0.033 | 4.8307669 | 26 | ikzf2 |
| hmbsa | 7.399822 | 0.372047958 | 0.218 | 0.029 | 1.8799250 | 26 | hmbsa |
| ascc1 | 7.911370 | 0.524416195 | 0.317 | 0.059 | 2.0098836 | 26 | ascc1 |
| cited4a3 | 8.234233 | 0.474747289 | 0.254 | 0.04 | 2.0919071 | 26 | cited4a |
| znf296 | 1.233585 | 0.267065444 | 0.134 | 0.012 | 3.1339247 | 26 | znf296 |
| mhc2b2 | 1.364135 | 0.468935824 | 0.254 | 0.039 | 3.4655863 | 26 | mhc2b |
| tsc22d36 | 2.892175 | 0.73803657 | 0.521 | 0.157 | 7.3475711 | 26 | tsc22d3 |
| sox121 | 3.519196 | 0.297791015 | 0.134 | 0.012 | 8.9405184 | 26 | sox12 |
| rpl106 | 5.777293 | 0.425710721 | 0.986 | 0.789 | 1.4677213 | 26 | rpl10 |
| pfn18 | 1.653114 | 0.836153708 | 1 | 0.498 | 4.1997380 | 26 | pfn1 |
| rpl369 | 3.800549 | 0.426328086 | 1 | 0.806 | 9.6552969 | 26 | rpl36 |
| rpl328 | 3.819132 | 0.409872427 | 1 | 0.823 | 9.7025053 | 26 | rpl32 |
| rpl194 | 4.612450 | 0.369801155 | 1 | 0.853 | 1.1717930 | 26 | rpl19 |
| si:dkey-183p4.102 | 6.503508 | 0.388911993 | 0.218 | 0.031 | 1.6522163 | 26 | si:dkey-183p4.10 |
| klf132 | 8.027968 | 0.64850365 | 0.408 | 0.102 | 2.0395053 | 26 | klf13 |
| marcksl1a8 | 8.821432 | 0.753408098 | 0.711 | 0.258 | 2.2410848 | 26 | marcksl1a |
| thrap3a | 1.365624 | 0.275891668 | 0.134 | 0.013 | 3.4693699 | 26 | thrap3a |
| rbm38 | 1.374655 | 0.291311786 | 0.162 | 0.018 | 3.4923110 | 26 | rbm38 |
| si:ch211-197h24.95 | 6.449118 | 0.5916831 | 0.359 | 0.082 | 1.6383984 | 26 | si:ch211-197h24.9 |
| rpl86 | 1.388279 | 0.421609796 | 0.993 | 0.794 | 3.5269249 | 26 | rpl8 |
| rpl18a8 | 1.826643 | 0.427110988 | 1 | 0.8 | 4.6405867 | 26 | rpl18a |
| coro1a8 | 2.916431 | 0.700510161 | 0.718 | 0.233 | 7.4091950 | 26 | coro1a |
| wu:fb55g092 | 1.305011 | 0.729499401 | 0.683 | 0.257 | 3.3153809 | 26 | wu:fb55g09 |
| cybb1 | 3.502262 | 0.353579662 | 0.162 | 0.02 | 8.8974978 | 26 | cybb |
| rps510 | 6.823950 | 0.484296498 | 1 | 0.727 | 1.7336247 | 26 | rps5 |
| rpl158 | 7.496551 | 0.449034949 | 0.993 | 0.763 | 1.9044988 | 26 | rpl15 |
| rplp02 | 8.103223 | 0.347156709 | 0.993 | 0.858 | 2.0586238 | 26 | rplp0 |
| cyba5 | 8.939200 | 0.532962652 | 0.352 | 0.081 | 2.2710037 | 26 | cyba |

| | | | | | | | |
|---|---|---|---|---|---|---|---|
| tfdp2 | 1.543366 | 0.398332765 | 0.218 | 0.034 | 3.9209215 | 26 | tfdp2 |
| dap1b5 | 5.162278 | 0.698669262 | 0.669 | 0.247 | 1.3114769 | 26 | dap1b |
| nr4a17 | 1.087656 | 0.669346955 | 0.514 | 0.156 | 2.7631904 | 26 | nr4a1 |
| naca9 | 2.333625 | 0.539305495 | 0.979 | 0.65 | 5.9285766 | 26 | naca |
| ptmab9 | 3.319080 | 0.624580852 | 0.908 | 0.568 | 8.4321232 | 26 | ptmab |
| rpl10a7 | 4.326565 | 0.419835793 | 0.993 | 0.778 | 1.0991639 | 26 | rpl10a |
| arhgdig7 | 7.138716 | 0.708197985 | 0.662 | 0.257 | 1.8135908 | 26 | arhgdig |
| rpl73 | 8.652717 | 0.387110702 | 0.986 | 0.809 | 2.1982229 | 26 | rpl7 |
| rpsa9 | 9.119080 | 0.409345377 | 0.993 | 0.785 | 2.3167023 | 26 | rpsa |
| rplp2l9 | 1.820070 | 0.421413275 | 1 | 0.794 | 4.6238884 | 26 | rplp2l |
| histh1l5 | 2.097524 | 0.655671954 | 0.423 | 0.121 | 5.3287614 | 26 | histh1l |
| ptp4a31 | 4.552214 | 0.467067027 | 0.261 | 0.051 | 1.1564900 | 26 | ptp4a3 |
| rps4x9 | 5.370091 | 0.448773708 | 0.979 | 0.736 | 1.3642717 | 26 | rps4x |
| ppiaa5 | 7.683826 | 0.359107678 | 0.972 | 0.814 | 1.9520760 | 26 | ppiaa |
| pik3r13 | 1.621907 | 0.51693236 | 0.359 | 0.09 | 4.1204569 | 26 | pik3r1 |
| celf29 | 3.622802 | 0.680719517 | 0.634 | 0.255 | 9.2037308 | 26 | celf2 |
| rps26l8 | 4.757275 | 0.483127532 | 0.979 | 0.698 | 1.2085859 | 26 | rps26l |
| rps221 | 5.148392 | 0.38949567 | 1 | 0.841 | 1.3079489 | 26 | rps2 |
| rpl375 | 5.227948 | 0.356532906 | 1 | 0.855 | 1.3281602 | 26 | rpl37 |
| mbnl17 | 6.015021 | 0.594634426 | 0.549 | 0.185 | 1.5281160 | 26 | mbnl1 |
| rps66 | 6.460644 | 0.395750588 | 1 | 0.803 | 1.6413268 | 26 | rps6 |
| cnpy31 | 7.000246 | 0.419388579 | 0.268 | 0.054 | 1.7784125 | 26 | cnpy3 |
| rpl346 | 8.835287 | 0.398068818 | 1 | 0.801 | 2.2446048 | 26 | rpl34 |
| ramp23 | 1.681788 | 0.476030601 | 0.232 | 0.043 | 4.2725845 | 26 | ramp2 |
| tnfaip8l2a1 | 2.363123 | 0.324305478 | 0.169 | 0.024 | 6.0035150 | 26 | tnfaip8l2a |
| hmgn28 | 3.286037 | 0.871834419 | 0.803 | 0.466 | 8.3481789 | 26 | hmgn2 |
| btg110 | 6.812777 | 0.68855012 | 0.732 | 0.354 | 1.7307860 | 26 | btg1 |
| pde4ba7 | 9.455679 | 0.5978635 | 0.451 | 0.141 | 2.4022152 | 26 | pde4ba |
| rps138 | 1.085978 | 0.427035072 | 0.986 | 0.748 | 2.7589286 | 26 | rps13 |
| LOC1039091155 | 1.361054 | 0.600740212 | 0.5 | 0.167 | 3.4577599 | 26 | LOC103909115 |
| rpl356 | 1.688186 | 0.369222798 | 0.993 | 0.813 | 4.2888386 | 26 | rpl35 |
| rpl217 | 2.753399 | 0.390979056 | 0.979 | 0.776 | 6.9950113 | 26 | rpl21 |
| rps298 | 2.914996 | 0.339842681 | 0.993 | 0.844 | 7.4055478 | 26 | rps29 |
| si:ch211-214p16.15 | 3.151852 | 0.770945042 | 0.648 | 0.242 | 8.0072802 | 26 | si:ch211-214p16.1 |
| rpl287 | 3.886748 | 0.395275065 | 0.986 | 0.776 | 9.8742855 | 26 | rpl28 |

| | | | | | | | |
|---|---|---|---|---|---|---|---|
| rpl307 | 1.293921 | 0.44311349 | 0.993 | 0.729 | 3.2872066 | 26 | rpl30 |
| hopx4 | 1.628835 | 0.476993479 | 0.282 | 0.064 | 4.1380575 | 26 | hopx |
| rel2 | 1.642443 | 0.363755868 | 0.197 | 0.034 | 4.1726264 | 26 | rel |
| ccnd2a | 2.819611 | 0.547564058 | 0.345 | 0.094 | 7.1632235 | 26 | ccnd2a |
| rpl247 | 3.715697 | 0.382375874 | 0.993 | 0.793 | 9.4397284 | 26 | rpl24 |
| glo12 | 7.792040 | 0.618445586 | 0.535 | 0.208 | 1.9795679 | 26 | glo1 |
| zgc:165555.3 | 1.054674 | 0.270320996 | 0.141 | 0.019 | 2.6794015 | 26 | zgc:165555.3 |
| tmc83 | 1.957211 | 0.359451575 | 0.19 | 0.033 | 4.9722953 | 26 | tmc8 |
| rpl189 | 2.608236 | 0.393189483 | 1 | 0.792 | 6.6262253 | 26 | rpl18 |
| plk32 | 8.374442 | 0.504580631 | 0.38 | 0.107 | 2.1275271 | 26 | plk3 |
| rps146 | 1.554651 | 0.388733972 | 0.993 | 0.78 | 3.9495926 | 26 | rps14 |
| herpud1 | 3.007643 | 0.445551213 | 0.289 | 0.071 | 7.6409190 | 26 | herpud1 |
| rps174 | 4.975521 | 0.364894963 | 0.965 | 0.778 | 1.2640313 | 26 | rps17 |
| xpc | 5.679380 | 0.324679306 | 0.176 | 0.03 | 1.4428466 | 26 | xpc |
| rps237 | 5.712677 | 0.403122222 | 0.972 | 0.744 | 1.4513056 | 26 | rps23 |
| rack16 | 8.830123 | 0.365921808 | 0.958 | 0.763 | 2.2432929 | 26 | rack1 |
| prkcbb1 | 9.402633 | 0.347850236 | 0.176 | 0.031 | 2.3887390 | 26 | prkcbb |
| rps1610 | 1.018503 | 0.399729753 | 0.986 | 0.765 | 2.5875081 | 26 | rps16 |
| b3gnt3.42 | 1.245519 | 0.367785998 | 0.197 | 0.038 | 3.1642410 | 26 | b3gnt3.4 |
| hsp90ab16 | 1.325352 | 0.35086806 | 0.979 | 0.8 | 3.3670573 | 26 | hsp90ab1 |
| egr33 | 1.632864 | 0.419396226 | 0.239 | 0.053 | 4.1482930 | 26 | egr3 |
| ccl35.13 | 2.150456 | 0.66777631 | 0.31 | 0.082 | 5.4632341 | 26 | ccl35.1 |
| mknk2b7 | 3.389859 | 0.589999802 | 0.514 | 0.199 | 8.6119385 | 26 | mknk2b |
| rpl277 | 3.827703 | 0.383760416 | 0.993 | 0.779 | 9.7242808 | 26 | rpl27 |
| rpl266 | 4.074376 | 0.323422882 | 1 | 0.85 | 1.0350954 | 26 | rpl26 |
| rps189 | 4.695431 | 0.409316633 | 0.993 | 0.753 | 1.1928743 | 26 | rps18 |
| eif3f8 | 4.858794 | 0.570046082 | 0.817 | 0.464 | 1.2343767 | 26 | eif3f |
| rpl7a7 | 5.193408 | 0.340583054 | 0.993 | 0.823 | 1.3193855 | 26 | rpl7a |
| wasa6 | 5.826532 | 0.47193424 | 0.338 | 0.096 | 1.4802304 | 26 | wasa |
| jam2a2 | 7.273678 | 0.36294082 | 0.232 | 0.05 | 1.8478780 | 26 | jam2a |
| rps125 | 8.164282 | 0.332117869 | 1 | 0.841 | 2.0741360 | 26 | rps12 |
| XLOC-0425963 | 1.106006 | 0.35126958 | 0.197 | 0.038 | 2.8098098 | 26 | XLOC-042596 |
| rpl913 | 1.154304 | 0.484537712 | 0.993 | 0.667 | 2.9325109 | 26 | rpl9 |
| dusp24 | 1.162970 | 0.629489748 | 0.458 | 0.162 | 2.9545260 | 26 | dusp2 |
| btk1 | 1.741826 | 0.28052472 | 0.162 | 0.027 | 4.4251104 | 26 | btk |

| | | | | | | | |
|---|---|---|---|---|---|---|---|
| ikzf13 | 1.835950 | 0.277544768 | 0.183 | 0.034 | 4.6642320 | 26 | ikzf1 |
| rpl36a7 | 3.159727 | 0.337645376 | 1 | 0.823 | 8.0272869 | 26 | rpl36a |
| si:ch211-195b13.1 | 5.389643 | 0.344000815 | 0.155 | 0.026 | 1.3692390 | 26 | si:ch211-195b13.1 |
| ptprc7 | 7.253679 | 0.506558817 | 0.387 | 0.123 | 1.8427972 | 26 | ptprc |
| TPGS1 | 8.105080 | 0.370637772 | 0.19 | 0.038 | 2.0590955 | 26 | TPGS1 |
| rpl234 | 1.184650 | 0.275762454 | 0.993 | 0.872 | 3.0096044 | 26 | rpl23 |
| tagapa5 | 1.288599 | 0.516538497 | 0.331 | 0.097 | 3.2736875 | 26 | tagapa |
| faua9 | 1.514382 | 0.426799626 | 0.986 | 0.716 | 3.8472889 | 26 | faua |
| rpl136 | 2.057467 | 0.318531899 | 0.979 | 0.818 | 5.2269974 | 26 | rpl13 |
| rps37 | 2.461272 | 0.394186218 | 0.951 | 0.725 | 6.2528633 | 26 | rps3 |
| ada | 2.664795 | 0.344425663 | 0.204 | 0.044 | 6.7699140 | 26 | ada |
| si:ch1073-429i10.3.1 | 3.030679 | 0.577990234 | 0.528 | 0.225 | 7.6994411 | 26 | si:ch1073-429i10.3.1 |
| rpl23a9 | 3.442529 | 0.458240383 | 0.972 | 0.668 | 8.7457455 | 26 | rpl23a |
| rps249 | 4.061465 | 0.366029415 | 0.986 | 0.778 | 1.0318152 | 26 | rps24 |
| rgcc4 | 5.933593 | 0.389236715 | 0.275 | 0.071 | 1.5074294 | 26 | rgcc |
| unc93b11 | 6.692237 | 0.274711056 | 0.141 | 0.023 | 1.7001628 | 26 | unc93b1 |
| actb28 | 9.974141 | 0.348497 | 1 | 0.866 | 2.5339307 | 26 | actb2 |
| sesn15 | 1.081664 | 0.566316012 | 0.458 | 0.174 | 2.7479678 | 26 | sesn1 |
| rpl35a7 | 1.101757 | 0.36922121 | 0.993 | 0.781 | 2.7990143 | 26 | rpl35a |
| inpp5d2 | 1.103552 | 0.356638947 | 0.176 | 0.035 | 2.8035746 | 26 | inpp5d |
| fli1b2 | 1.663657 | 0.27267617 | 0.176 | 0.034 | 4.2265220 | 26 | fli1b |
| tpt14 | 1.678017 | 0.325710556 | 0.979 | 0.811 | 4.2630024 | 26 | tpt1 |
| rps1012 | 2.354757 | 0.399667417 | 0.972 | 0.726 | 5.9822616 | 26 | rps10 |
| rpl69 | 9.004604 | 0.408854695 | 0.958 | 0.7 | 2.2876197 | 26 | rpl6 |
| ucp27 | 1.075037 | 0.532608568 | 0.803 | 0.412 | 2.7311337 | 26 | ucp2 |
| eif3m6 | 1.391858 | 0.535694931 | 0.711 | 0.382 | 3.5360159 | 26 | eif3m |
| nfkbiab3 | 1.481935 | 0.534473227 | 0.493 | 0.2 | 3.7648567 | 26 | nfkbiab |
| slc25a55a2 | 2.021431 | 0.403227151 | 0.268 | 0.075 | 5.1354463 | 26 | slc25a55a |
| rpl319 | 2.185762 | 0.419972826 | 0.951 | 0.686 | 5.5529307 | 26 | rpl31 |
| rps96 | 2.567831 | 0.347253298 | 0.979 | 0.79 | 6.5235748 | 26 | rps9 |
| nsa27 | 2.781964 | 0.543141825 | 0.732 | 0.387 | 7.0675800 | 26 | nsa2 |
| plcg22 | 3.675270 | 0.334503707 | 0.211 | 0.05 | 9.3370245 | 26 | plcg2 |
| rps196 | 5.021170 | 0.283374643 | 0.993 | 0.868 | 1.2756282 | 26 | rps19 |
| rpl387 | 7.162445 | 0.377885569 | 0.979 | 0.753 | 1.8196193 | 26 | rpl38 |
| zgc:16273011 | 7.928148 | 0.580153544 | 0.796 | 0.401 | 2.0141460 | 26 | zgc:162730 |

| | | | | | | | |
|---|---|---|---|---|---|---|---|
| lyn1 | 1.347147 | 0.315315837 | 0.197 | 0.045 | 3.4224279 | 26 | lyn |
| btf36 | 2.401311 | 0.458654926 | 0.923 | 0.609 | 6.1005328 | 26 | btf3 |
| gm2a3 | 2.892309 | 0.446550728 | 0.359 | 0.12 | 7.3479112 | 26 | gm2a |
| rpl2210 | 2.967241 | 0.402728602 | 0.979 | 0.719 | 7.5382783 | 26 | rpl22 |
| rps1510 | 4.441794 | 0.407508131 | 0.979 | 0.716 | 1.1284378 | 26 | rps15 |
| oaz1b7 | 6.979922 | 0.526780934 | 0.662 | 0.359 | 1.7732493 | 26 | oaz1b |
| si:dkey-27i16.24 | 7.068564 | 0.431874983 | 0.296 | 0.091 | 1.7957687 | 26 | si:dkey-27i16.2 |
| rpl5a10 | 1.681530 | 0.473352502 | 0.845 | 0.535 | 4.2719291 | 26 | rpl5a |
| rpl395 | 2.365246 | 0.278958385 | 1 | 0.877 | 6.0089077 | 26 | rpl39 |
| h2afx7 | 2.757115 | 0.514451295 | 0.352 | 0.122 | 7.0044516 | 26 | h2afx |
| wipf1b2 | 3.201839 | 0.361613013 | 0.232 | 0.064 | 8.1342734 | 26 | wipf1b |
| wasb8 | 3.549204 | 0.493705608 | 0.437 | 0.172 | 9.0167544 | 26 | wasb |
| serf26 | 3.646239 | 0.498933613 | 0.69 | 0.376 | 9.2632718 | 26 | serf2 |
| mbpb | 5.473324 | 0.325748129 | 0.232 | 0.064 | 1.3904981 | 26 | mbpb |
| ran2 | 6.904813 | 0.397732245 | 0.859 | 0.639 | 1.7541679 | 26 | ran |
| tmc6b | 7.245707 | 0.309038088 | 0.162 | 0.035 | 1.8407721 | 26 | tmc6b |
| lmnb2 | 7.917745 | 0.333272904 | 0.232 | 0.064 | 2.0115031 | 26 | lmnb2 |
| rps115 | 8.748311 | 0.320887791 | 0.979 | 0.784 | 2.2225084 | 26 | rps11 |
| rhogb5 | 8.750431 | 0.42594167 | 0.317 | 0.107 | 2.2230470 | 26 | rhogb |
| rps76 | 1.151545 | 0.300518864 | 0.979 | 0.828 | 2.9255007 | 26 | rps7 |
| si:rp71-68n21.94 | 1.359693 | 0.287186386 | 0.19 | 0.046 | 3.4543012 | 26 | si:rp71-68n21.9 |
| gdi22 | 1.439123 | 0.451923467 | 0.472 | 0.202 | 3.6560926 | 26 | gdi2 |
| gpr183a3 | 1.518016 | 0.335795921 | 0.232 | 0.064 | 3.8565202 | 26 | gpr183a |
| arpc1b8 | 1.719989 | 0.506041642 | 0.69 | 0.312 | 4.3696334 | 26 | arpc1b |
| pycard6 | 2.710780 | 0.391710575 | 0.268 | 0.083 | 6.8867384 | 26 | pycard |
| serbp1a10 | 3.461781 | 0.474565146 | 0.824 | 0.505 | 8.7946564 | 26 | serbp1a |
| stk351 | 3.957379 | 0.353673811 | 0.218 | 0.061 | 1.0053723 | 26 | stk35 |
| dub5 | 4.072362 | 0.353627291 | 0.246 | 0.073 | 1.0345836 | 26 | dub |
| mcl1a5 | 6.805832 | 0.436427705 | 0.908 | 0.612 | 1.7290216 | 26 | mcl1a |
| nop533 | 9.150655 | 0.495680043 | 0.535 | 0.266 | 2.3247240 | 26 | nop53 |
| zgc:640517 | 1.041286 | 0.432556015 | 0.415 | 0.162 | 2.6453880 | 26 | zgc:64051 |
| rpl125 | 1.063977 | 0.288218927 | 0.972 | 0.812 | 2.7030343 | 26 | rpl12 |
| rps287 | 1.415286 | 0.302419659 | 1 | 0.84 | 3.5955344 | 26 | rps28 |
| fmnl1a2 | 1.476320 | 0.313452401 | 0.162 | 0.038 | 3.7505913 | 26 | fmnl1a |
| smox4 | 1.960669 | 0.316920378 | 0.275 | 0.087 | 4.9810799 | 26 | smox |

| | | | | | | | |
|---|---|---|---|---|---|---|---|
| skap25 | 2.293556 | 0.394289329 | 0.254 | 0.08 | 5.8267800 | 26 | skap2 |
| rps3a6 | 2.414823 | 0.307868414 | 0.972 | 0.791 | 6.1348590 | 26 | rps3a |
| mki671 | 3.425459 | 0.30269122 | 0.169 | 0.04 | 8.7023805 | 26 | mki67 |
| si:ch211-150o23.2 | 8.707497 | 0.333031767 | 0.225 | 0.066 | 2.2121398 | 26 | si:ch211-150o23.2 |
| CR855337.11 | 9.092033 | 0.258810703 | 0.148 | 0.033 | 2.3098310 | 26 | CR855337.1 |
| cirbpb1 | 1.025620 | 0.3180628 | 0.923 | 0.757 | 2.6055885 | 26 | cirbpb |
| calm1b6 | 1.150851 | 0.319015213 | 0.303 | 0.102 | 2.9237372 | 26 | calm1b |
| nme2b.12 | 1.307724 | 0.252762615 | 0.993 | 0.856 | 3.3222730 | 26 | nme2b.1 |
| CABZ01079427.1 | 1.390139 | 0.255489418 | 0.155 | 0.036 | 3.5316497 | 26 | CABZ01079427.1 |
| foxp1b1 | 1.475785 | 0.265096763 | 0.162 | 0.039 | 3.7492325 | 26 | foxp1b |
| rps27a5 | 1.698677 | 0.319448846 | 0.979 | 0.774 | 4.3154890 | 26 | rps27a |
| arrdc3b1 | 1.929582 | 0.255124104 | 0.155 | 0.037 | 4.9021036 | 26 | arrdc3b |
| eef1g9 | 2.221728 | 0.390011549 | 0.915 | 0.652 | 5.6443022 | 26 | eef1g |
| rps265 | 2.831514 | 0.321950126 | 0.958 | 0.75 | 7.1934624 | 26 | rps26 |
| capzb7 | 3.584383 | 0.450689132 | 0.535 | 0.271 | 9.1061264 | 26 | capzb |
| rps8a5 | 5.067648 | 0.290838753 | 0.965 | 0.801 | 1.2874361 | 26 | rps8a |
| pnrc214 | 6.201118 | 0.442351741 | 0.923 | 0.596 | 1.5753942 | 26 | pnrc2 |
| si:busm1-194e12.112 | 7.863427 | 0.279696097 | 0.127 | 0.027 | 1.9977037 | 26 | si:busm1-194e12.11 |
| slc25a55 | 8.356275 | 0.281625727 | 0.979 | 0.813 | 2.1229118 | 26 | slc25a5 |
| rpl27a8 | 9.237498 | 0.451976847 | 0.866 | 0.521 | 2.3467865 | 26 | rpl27a |
| bcl2l102 | 1.033653 | 0.46206154 | 0.444 | 0.199 | 2.6259962 | 26 | bcl2l10 |
| sypl2b | 1.162431 | 0.271093258 | 0.19 | 0.054 | 2.9531576 | 26 | sypl2b |
| hmha1b4 | 1.778888 | 0.295192274 | 0.232 | 0.074 | 4.5192666 | 26 | hmha1b |
| tefa5 | 1.899153 | 0.389478651 | 0.345 | 0.141 | 4.8247985 | 26 | tefa |
| sept122 | 2.239192 | 0.302112615 | 0.211 | 0.065 | 5.6886675 | 26 | sept12 |
| mych3 | 3.086171 | 0.393496767 | 0.254 | 0.09 | 7.8404174 | 26 | mych |
| tob1a1 | 4.417062 | 0.409097232 | 0.345 | 0.144 | 1.1221546 | 26 | tob1a |
| sftpbb3 | 4.814201 | 0.301685075 | 0.225 | 0.07 | 1.2230478 | 26 | sftpbb |
| CABZ01090890.1 | 5.569757 | 0.300448053 | 0.197 | 0.06 | 1.4149968 | 26 | CABZ01090890.1 |
| romo16 | 7.252697 | 0.430994623 | 0.683 | 0.408 | 1.8425478 | 26 | romo1 |
| uba528 | 7.540366 | 0.374125032 | 0.944 | 0.66 | 1.9156300 | 26 | uba52 |
| eif5a5 | 7.844972 | 0.413551693 | 0.761 | 0.495 | 1.9930152 | 26 | eif5a |
| rpl298 | 8.339145 | 0.289136139 | 0.993 | 0.822 | 2.1185598 | 26 | rpl29 |
| tubb2b1 | 9.329251 | 0.507043554 | 0.5 | 0.259 | 2.3700962 | 26 | tubb2b |
| lbr | 1.000170 | 0.343504394 | 0.232 | 0.077 | 2.5409342 | 26 | lbr |

| | | | | | | | |
|---|---|---|---|---|---|---|---|
| rps257 | 1.267911 | 0.324820815 | 0.972 | 0.748 | 3.2211294 | 26 | rps25 |
| rbm39a4 | 1.494040 | 0.393062973 | 0.352 | 0.153 | 3.7956110 | 26 | rbm39a |
| msna9 | 2.100402 | 0.41517598 | 0.514 | 0.265 | 5.3360722 | 26 | msna |
| tmsb4x14 | 2.773253 | 0.439220779 | 1 | 0.717 | 7.0454514 | 26 | tmsb4x |
| XLOC-0283705 | 2.835182 | 0.395926849 | 0.317 | 0.125 | 7.2027799 | 26 | XLOC-028370 |
| myl12.18 | 2.952420 | 0.424768422 | 0.725 | 0.425 | 7.5006253 | 26 | myl12.1 |
| nrros2 | 3.320434 | 0.274583931 | 0.169 | 0.048 | 8.4355626 | 26 | nrros |
| eif3k3 | 3.491401 | 0.379945084 | 0.486 | 0.244 | 8.8699050 | 26 | eif3k |
| cited4b2 | 5.674576 | 0.345929534 | 0.31 | 0.121 | 1.4416262 | 26 | cited4b |
| eif3ea3 | 6.830311 | 0.412613734 | 0.542 | 0.304 | 1.7352405 | 26 | eif3ea |
| capza1a5 | 7.401865 | 0.395716087 | 0.444 | 0.22 | 1.8804439 | 26 | capza1a |
| arpc5a6 | 9.104089 | 0.321580549 | 0.31 | 0.125 | 2.3128938 | 26 | arpc5a |
| ier2a9 | 9.966183 | 0.435828531 | 0.521 | 0.274 | 2.5319088 | 26 | ier2a |
| fgfr1op2 | 1.028257 | 0.288286743 | 0.197 | 0.064 | 2.6122884 | 26 | fgfr1op2 |
| acin1a5 | 1.484830 | 0.399598068 | 0.451 | 0.228 | 3.7722131 | 26 | acin1a |
| arf6b2 | 1.593663 | 0.342037121 | 0.282 | 0.113 | 4.0487014 | 26 | arf6b |
| btg27 | 1.757832 | 0.398889218 | 0.655 | 0.348 | 4.4657730 | 26 | btg2 |
| ddit36 | 2.079300 | 0.46439606 | 0.366 | 0.178 | 5.2824640 | 26 | ddit3 |
| si:dkey-112e7.26 | 2.234012 | 0.386484406 | 0.43 | 0.214 | 5.6755084 | 26 | si:dkey-112e7.2 |
| sept9b1 | 2.366697 | 0.270261734 | 0.162 | 0.048 | 6.0125945 | 26 | sept9b |
| ero1b1 | 2.396741 | 0.267779447 | 0.141 | 0.038 | 6.0889209 | 26 | ero1b |
| grasp5 | 2.708847 | 0.319932349 | 0.211 | 0.073 | 6.8818280 | 26 | grasp |
| arf6a1 | 3.062119 | 0.349349148 | 0.366 | 0.163 | 7.7793153 | 26 | arf6a |
| XLOC-0140793 | 3.141468 | 0.26389499 | 0.127 | 0.032 | 7.9809013 | 26 | XLOC-014079 |
| zgc:11418813 | 3.782070 | 0.370781842 | 0.937 | 0.667 | 9.6083508 | 26 | zgc:114188 |
| si:ch211-198c19.36 | 3.954904 | 0.544566278 | 0.218 | 0.083 | 1.0047434 | 26 | si:ch211-198c19.3 |
| ahcy1 | 4.241486 | 0.359874022 | 0.352 | 0.157 | 1.0775496 | 26 | ahcy |
| ss181 | 4.659580 | 0.26516886 | 0.211 | 0.073 | 1.1837664 | 26 | ss18 |
| gnb1a3 | 7.612854 | 0.38503034 | 0.401 | 0.202 | 1.9340457 | 26 | gnb1a |
| si:dkeyp-68b7.124 | 1.376155 | 0.260524717 | 0.183 | 0.06 | 3.4961237 | 26 | si:dkeyp-68b7.12 |
| sarnp | 1.547099 | 0.368029295 | 0.366 | 0.176 | 3.9304068 | 26 | sarnp |
| cct24 | 2.106991 | 0.378116014 | 0.465 | 0.254 | 5.3528108 | 26 | cct2 |
| crlf3 | 2.214202 | 0.254812391 | 0.183 | 0.061 | 5.6251810 | 26 | crlf3 |
| eif3c4 | 2.450822 | 0.347150473 | 0.38 | 0.188 | 6.2263141 | 26 | eif3c |
| rpl5b7 | 2.804551 | 0.335210081 | 0.873 | 0.645 | 7.1249626 | 26 | rpl5b |

| | | | | | | | |
|---|---|---|---|---|---|---|---|
| mibp29 | 2.868523 | 0.370618735 | 0.775 | 0.52 | 7.2874848 | 26 | mibp2 |
| COX7A2 (1 of many)2 | 3.073262 | 0.355661859 | 0.493 | 0.266 | 7.8076224 | 26 | COX7A2 (1 of many) |
| ywhaba3 | 4.031249 | 0.376973985 | 0.556 | 0.328 | 0.000102 | 26 | ywhaba |
| imp4 | 4.653915 | 0.252083019 | 0.162 | 0.052 | 0.000118 | 26 | imp4 |
| ppp3ccb | 4.827928 | 0.35773814 | 0.282 | 0.125 | 0.000123 | 26 | ppp3ccb |
| jmjd1cb1 | 5.562812 | 0.272255551 | 0.148 | 0.046 | 0.000141 | 26 | jmjd1cb |
| eef1da5 | 5.937964 | 0.37218448 | 0.796 | 0.505 | 0.000151 | 26 | eef1da |
| irf2bp2b6 | 8.351712 | 0.308343403 | 0.246 | 0.102 | 0.000212 | 26 | irf2bp2b |
| capza1b5 | 8.568050 | 0.342203915 | 0.444 | 0.234 | 0.000218 | 26 | capza1b |
| kdm6bb1 | 1.935030 | 0.266975941 | 0.176 | 0.063 | 0.000492 | 26 | kdm6bb |
| rpl22l17 | 1.953747 | 0.368235132 | 0.69 | 0.425 | 0.000496 | 26 | rpl22l1 |
| CR383676.114 | 2.091392 | 0.518262529 | 0.993 | 0.716 | 0.000531 | 26 | CR383676.1 |
| magt12 | 2.225197 | 0.32977955 | 0.282 | 0.129 | 0.000565 | 26 | magt1 |
| hmgn32 | 2.671377 | 0.251884432 | 0.204 | 0.078 | 0.000679 | 26 | hmgn3 |
| wasf21 | 3.284573 | 0.27710612 | 0.204 | 0.079 | 0.000834 | 26 | wasf2 |
| zfp36l1a6 | 3.292835 | 0.354257077 | 0.465 | 0.255 | 0.000837 | 26 | zfp36l1a |
| CR381673.15 | 3.671396 | 0.295608266 | 0.162 | 0.057 | 0.000933 | 26 | CR381673.1 |
| lasp16 | 3.968696 | 0.343628946 | 0.451 | 0.253 | 0.001008 | 26 | lasp1 |
| rad21a2 | 5.767372 | 0.339693485 | 0.303 | 0.148 | 0.001465 | 26 | rad21a |
| XLOC-0254235 | 6.379575 | 0.349042767 | 0.352 | 0.171 | 0.001621 | 26 | XLOC-025423 |
| sat1a.25 | 6.960840 | 0.340397017 | 0.521 | 0.301 | 0.001768 | 26 | sat1a.2 |
| plekho1b2 | 7.185323 | 0.259148997 | 0.134 | 0.043 | 0.001825 | 26 | plekho1b |
| bhlhe407 | 7.327532 | 0.431403931 | 0.317 | 0.168 | 0.001862 | 26 | bhlhe40 |
| ldha3 | 7.725042 | 0.292050738 | 0.183 | 0.07 | 0.001963 | 26 | ldha |
| hmga1a5 | 1.113781 | 0.384575669 | 0.627 | 0.442 | 0.00283 | 26 | hmga1a |
| si:ch211-260e23.91 | 1.121490 | 0.250142337 | 0.169 | 0.062 | 0.002849 | 26 | si:ch211-260e23.9 |
| zgc:564935 | 1.892540 | 0.316282971 | 0.81 | 0.59 | 0.004808 | 26 | zgc:56493 |
| kmt2bb2 | 1.973612 | 0.267648127 | 0.225 | 0.097 | 0.005014 | 26 | kmt2bb |
| si:ch211-214j24.101 | 2.035190 | 0.27552139 | 0.324 | 0.158 | 0.00517 | 26 | si:ch211-214j24.10 |
| rpl47 | 2.163306 | 0.292360463 | 0.88 | 0.656 | 0.005496 | 26 | rpl4 |
| itm2bb6 | 2.358720 | 0.306920447 | 0.444 | 0.242 | 0.005992 | 26 | itm2bb |
| naa101 | 2.718060 | 0.312643486 | 0.415 | 0.23 | 0.006905 | 26 | naa10 |
| si:dkey-262k9.46 | 2.810776 | 0.302714376 | 0.211 | 0.091 | 0.007141 | 26 | si:dkey-262k9.4 |
| sec61g3 | 3.443695 | 0.276655535 | 0.873 | 0.681 | 0.008749 | 26 | sec61g |
| laptm58 | 4.095652 | 0.274265635 | 0.359 | 0.176 | 0.010405 | 26 | laptm5 |

| | | | | | | | |
|---|---|---|---|---|---|---|---|
| aldoaa3 | 4.126874 | 0.287092034 | 0.458 | 0.253 | 0.010484 | 26 | aldoaa |
| anapc161 | 4.385826 | 0.262883665 | 0.246 | 0.112 | 0.011142 | 26 | anapc16 |
| ifi303 | 5.241482 | 0.28754591 | 0.373 | 0.196 | 0.013316 | 26 | ifi30 |
| ankrd111 | 5.573272 | 0.258560765 | 0.225 | 0.1 | 0.014159 | 26 | ankrd11 |
| dusp16 | 8.175346 | 0.317024816 | 0.296 | 0.15 | 0.020769 | 26 | dusp1 |
| eif3ha1 | 9.795917 | 0.301278891 | 0.401 | 0.231 | 0.024887 | 26 | eif3ha |
| atf4b8 | 1.003641 | 0.309936345 | 0.465 | 0.278 | 0.025498 | 26 | atf4b |
| jpt2 | 1.022395 | 0.269830519 | 0.373 | 0.198 | 0.025974 | 26 | jpt2 |
| eif3g5 | 1.388551 | 0.319628019 | 0.479 | 0.3 | 0.035276 | 26 | eif3g |
| zgc:926064 | 1.477246 | 0.274155806 | 0.211 | 0.096 | 0.037529 | 26 | zgc:92606 |
| arpc27 | 1.536656 | 0.289201796 | 0.5 | 0.299 | 0.039039 | 26 | arpc2 |
| pgam1a4 | 1.615909 | 0.313659131 | 0.296 | 0.155 | 0.041052 | 26 | pgam1a |
| b2m8 | 1.992921 | 0.318858077 | 0.486 | 0.299 | 0.05063 | 26 | b2m |
| arpc4l6 | 2.158883 | 0.297918081 | 0.38 | 0.219 | 0.054846 | 26 | arpc4l |
| prelid3b1 | 2.470588 | 0.296136261 | 0.394 | 0.226 | 0.062765 | 26 | prelid3b |
| sumo2b8 | 2.605653 | 0.30475279 | 0.472 | 0.297 | 0.066197 | 26 | sumo2b |
| tmed73 | 2.901653 | 0.275330931 | 0.387 | 0.221 | 0.073717 | 26 | tmed7 |
| arpc37 | 2.970943 | 0.296516283 | 0.542 | 0.334 | 0.075477 | 26 | arpc3 |
| nfkbiaa4 | 3.478403 | 0.286623023 | 0.296 | 0.153 | 0.088369 | 26 | nfkbiaa |
| tram12 | 3.539865 | 0.280206534 | 0.225 | 0.107 | 0.08993 | 26 | tram1 |
| sept62 | 6.948709 | 0.268936568 | 0.359 | 0.202 | 0.176532 | 26 | sept6 |
| ola12 | 9.423491 | 0.291935928 | 0.394 | 0.245 | 0.239404 | 26 | ola1 |
| eif1b7 | 9.642290 | 0.284124464 | 0.606 | 0.407 | 0.244962 | 26 | eif1b |
| pabpc1a6 | 1.092534 | 0.280885646 | 0.782 | 0.57 | 0.277558 | 26 | pabpc1a |
| dnaja2a | 1.106986 | 0.268115043 | 0.437 | 0.261 | 0.28123 | 26 | dnaja2a |
| irf2bp2a3 | 1.163248 | 0.311417731 | 0.317 | 0.186 | 0.295523 | 26 | irf2bp2a |
| cfl18 | 1.217259 | 0.279062924 | 0.824 | 0.6 | 0.309245 | 26 | cfl1 |
| ier2b10 | 1.263311 | 0.281519989 | 0.394 | 0.237 | 0.320944 | 26 | ier2b |
| cox6b23 | 1.416575 | 0.286946218 | 0.796 | 0.549 | 0.359881 | 26 | cox6b2 |
| arl5c3 | 1.456924 | 0.273672587 | 0.218 | 0.108 | 0.370132 | 26 | arl5c |
| arhgdia6 | 1.689221 | 0.273948993 | 0.408 | 0.252 | 0.429147 | 26 | arhgdia |
| eif4a1b4 | 2.013802 | 0.262117727 | 0.493 | 0.312 | 0.511607 | 26 | eif4a1b |
| litaf3 | 2.357885 | 0.269585851 | 0.268 | 0.147 | 0.599021 | 26 | litaf |
| gng54 | 2.595170 | 0.257276729 | 0.458 | 0.288 | 0.659303 | 26 | gng5 |
| cox8a8 | 2.712157 | 0.254792726 | 0.43 | 0.268 | 0.689024 | 26 | cox8a |

| | | | | | | | |
|---|---|---|---|---|---|---|---|
| ppp1caa | 2.712262 | 0.259820161 | 0.296 | 0.166 | 0.68905 | 26 | ppp1caa |
| seta1 | 3.692403 | 0.29622232 | 0.444 | 0.294 | 0.938055 | 26 | seta |
| dut1 | 5.304316 | 0.26008573 | 0.232 | 0.12 | 1 | 26 | dut |
| rsl1d11 | 5.352589 | 0.265142414 | 0.317 | 0.189 | 1 | 26 | rsl1d1 |
| rps27.210 | 6.465803 | 0.285470321 | 0.796 | 0.553 | 1 | 26 | rps27.2 |
| cox7b4 | 6.496866 | 0.252424172 | 0.627 | 0.427 | 1 | 26 | cox7b |
| hmgn611 | 6.555111 | 0.275956344 | 0.458 | 0.312 | 1 | 26 | hmgn6 |
| si:ch73-1a9.312 | 6.781232 | 0.29368751 | 0.458 | 0.316 | 1 | 26 | si:ch73-1a9.3 |
| XLOC-0430433 | 8.152345 | 0.268977227 | 0.113 | 0.045 | 1 | 26 | XLOC-043043 |
| hmgb2a4 | 0.0004 | 0.321777903 | 0.704 | 0.61 | 1 | 26 | hmgb2a |
| stmn1a4 | 0.003 | 0.31822856 | 0.268 | 0.19 | 1 | 26 | stmn1a |
| hmgb2b5 | 0.0044 | 0.291718699 | 0.401 | 0.338 | 1 | 26 | hmgb2b |
| si:ch211-161c3.61 | 0 | 1.146441859 | 0.515 | 0.011 | 0 | 27 | si:ch211-161c3.6 |
| myb | 0 | 0.780088382 | 0.449 | 0.006 | 0 | 27 | myb |
| maptb | 1.053271 | 0.382292716 | 0.228 | 0.003 | 2.6758362 | 27 | maptb |
| sid1 | 3.934871 | 0.428050888 | 0.132 | 0.001 | 9.9965419 | 27 | sid1 |
| sb:cb811 | 9.091386 | 0.69062769 | 0.426 | 0.026 | 2.3096667 | 27 | sb:cb81 |
| csf1rb | 4.597810 | 0.310014103 | 0.184 | 0.006 | 1.1680738 | 27 | csf1rb |
| XLOC-014336 | 6.088683 | 0.497363977 | 0.36 | 0.026 | 1.5468299 | 27 | XLOC-014336 |
| CU499330.12 | 1.997239 | 0.854391423 | 0.566 | 0.069 | 5.0739870 | 27 | CU499330.1 |
| cd7al3 | 6.445893 | 0.408035625 | 0.331 | 0.026 | 1.6375792 | 27 | cd7al |
| CABZ01066694.11 | 1.691347 | 0.324388676 | 0.243 | 0.015 | 4.2968682 | 27 | CABZ01066694.1 |
| lmo23 | 1.712648 | 0.468961352 | 0.301 | 0.024 | 4.3509822 | 27 | lmo2 |
| cxcr4b7 | 2.807645 | 1.151828484 | 0.816 | 0.173 | 7.1328230 | 27 | cxcr4b |
| si:dkey-217l24.1 | 5.629745 | 0.328769149 | 0.184 | 0.009 | 1.4302369 | 27 | si:dkey-217l24.1 |
| cbfa2t3 | 2.138791 | 0.444395884 | 0.272 | 0.022 | 5.4335990 | 27 | cbfa2t3 |
| ncf23 | 2.252138 | 0.394305468 | 0.265 | 0.022 | 5.7215582 | 27 | ncf2 |
| kdm6bb2 | 1.334526 | 0.611017764 | 0.434 | 0.06 | 3.3903654 | 27 | kdm6bb |
| si:dkey-22o12.21 | 3.506062 | 0.268556637 | 0.162 | 0.009 | 8.9071508 | 27 | si:dkey-22o12.2 |
| pdxka | 5.350557 | 0.351963641 | 0.272 | 0.03 | 1.3593092 | 27 | pdxka |
| si:dkey-261h17.13 | 3.274161 | 0.731050114 | 0.551 | 0.113 | 8.3180084 | 27 | si:dkey-261h17.1 |
| dnmt3bb.11 | 3.488792 | 0.273889336 | 0.176 | 0.014 | 8.8632778 | 27 | dnmt3bb.1 |
| myca1 | 1.614749 | 0.445795302 | 0.301 | 0.039 | 4.1022706 | 27 | myca |
| cables2b1 | 2.554484 | 0.303909907 | 0.176 | 0.014 | 6.4896666 | 27 | cables2b |
| dub6 | 1.304183 | 0.494656933 | 0.419 | 0.071 | 3.3132777 | 27 | dub |

| | | | | | | | |
|---|---|---|---|---|---|---|---|
| mybbp1a1 | 4.357096 | 0.290785049 | 0.25 | 0.028 | 1.1069204 | 27 | mybbp1a |
| msi2a2 | 9.859238 | 0.439061146 | 0.316 | 0.046 | 2.5047394 | 27 | msi2a |
| itgae.23 | 4.119001 | 0.366651106 | 0.272 | 0.035 | 1.0464322 | 27 | itgae.2 |
| si:ch211-214p16.23 | 7.672569 | 0.576803533 | 0.404 | 0.073 | 1.9492163 | 27 | si:ch211-214p16.2 |
| tagapb3 | 2.673889 | 0.443480407 | 0.39 | 0.068 | 6.7930150 | 27 | tagapb |
| MCMDC2 | 2.980720 | 0.417557474 | 0.419 | 0.077 | 7.5725211 | 27 | MCMDC2 |
| si:ch211-197h24.96 | 2.315512 | 0.533071708 | 0.419 | 0.081 | 5.8825583 | 27 | si:ch211-197h24.9 |
| rplp03 | 3.006681 | 0.383656214 | 0.978 | 0.858 | 7.6384743 | 27 | rplp0 |
| fmnl1a3 | 3.881324 | 0.297988644 | 0.265 | 0.037 | 9.8605048 | 27 | fmnl1a |
| si:ch211-222l21.110 | 2.577047 | 0.881446434 | 0.765 | 0.312 | 6.5469901 | 27 | si:ch211-222l21.1 |
| rpl117 | 5.917305 | 0.392804074 | 0.956 | 0.829 | 1.5032913 | 27 | rpl11 |
| slc16a103 | 6.676557 | 0.267475827 | 0.206 | 0.024 | 1.6961793 | 27 | slc16a10 |
| ikzf14 | 1.923599 | 0.323718999 | 0.243 | 0.033 | 4.8869041 | 27 | ikzf1 |
| gtpbp42 | 2.621129 | 0.726745187 | 0.713 | 0.244 | 6.6589796 | 27 | gtpbp4 |
| rplp19 | 2.007774 | 0.333988335 | 0.963 | 0.882 | 5.1007511 | 27 | rplp1 |
| rpl87 | 6.402085 | 0.419659076 | 0.956 | 0.795 | 1.6264497 | 27 | rpl8 |
| rpl610 | 1.018542 | 0.509694083 | 0.963 | 0.7 | 2.5876065 | 27 | rpl6 |
| rps97 | 1.316064 | 0.414915966 | 0.934 | 0.79 | 3.3434606 | 27 | rps9 |
| rps204 | 1.942826 | 0.297688662 | 0.956 | 0.91 | 4.9357500 | 27 | rps20 |
| rpl107 | 2.943269 | 0.413401094 | 0.949 | 0.79 | 7.4773758 | 27 | rpl10 |
| rps15a10 | 9.118934 | 0.420860523 | 0.963 | 0.797 | 2.3166652 | 27 | rps15a |
| eef1a1l12 | 1.166573 | 0.283549924 | 0.978 | 0.915 | 2.9636790 | 27 | eef1a1l1 |
| bahcc1b | 1.181483 | 0.326704635 | 0.228 | 0.032 | 3.0015584 | 27 | bahcc1b |
| rplp2l10 | 1.380499 | 0.430690978 | 0.963 | 0.794 | 3.5071586 | 27 | rplp2l |
| rpl137 | 1.451288 | 0.36592635 | 0.956 | 0.818 | 3.6869977 | 27 | rpl13 |
| rps126 | 1.570323 | 0.38009228 | 0.971 | 0.841 | 3.9894065 | 27 | rps12 |
| rpl248 | 1.715520 | 0.403919677 | 0.956 | 0.794 | 4.3582790 | 27 | rpl24 |
| rpl159 | 2.934019 | 0.448346765 | 0.941 | 0.763 | 7.4538754 | 27 | rpl15 |
| tmsb6 | 2.996848 | 0.348205666 | 0.228 | 0.032 | 7.6134938 | 27 | tmsb |
| rpl1810 | 5.156437 | 0.412277532 | 0.949 | 0.792 | 1.3099929 | 27 | rpl18 |
| gatad2b1 | 1.270106 | 0.380292946 | 0.346 | 0.067 | 3.2267045 | 27 | gatad2b |
| faua10 | 1.797458 | 0.468226065 | 0.926 | 0.717 | 4.5664426 | 27 | faua |
| ahcy2 | 2.983323 | 0.597993228 | 0.559 | 0.155 | 7.5791344 | 27 | ahcy |
| rps27.112 | 5.770564 | 0.484277774 | 0.941 | 0.709 | 1.4660119 | 27 | rps27.1 |
| rbm28 | 8.737863 | 0.413800801 | 0.419 | 0.094 | 2.2198543 | 27 | rbm28 |

| | | | | | | | |
|---|---|---|---|---|---|---|---|
| hsp90ab17 | 1.291756 | 0.390434655 | 0.941 | 0.8 | 3.2817084 | 27 | hsp90ab1 |
| rpl36a8 | 1.294387 | 0.359561634 | 0.949 | 0.823 | 3.2883913 | 27 | rpl36a |
| rpl310 | 1.427035 | 0.316246429 | 0.971 | 0.872 | 3.6253845 | 27 | rpl3 |
| rps3a7 | 1.770154 | 0.40193398 | 0.941 | 0.791 | 4.4970777 | 27 | rps3a |
| rpsa10 | 3.415174 | 0.416120059 | 0.978 | 0.785 | 8.6762505 | 27 | rpsa |
| rpl22l18 | 4.645396 | 0.70331531 | 0.809 | 0.423 | 1.1801629 | 27 | rpl22l1 |
| prr5a1 | 5.802289 | 0.255814267 | 0.184 | 0.023 | 1.4740717 | 27 | prr5a |
| rps2114 | 1.918574 | 0.422501922 | 0.971 | 0.775 | 4.8741392 | 27 | rps21 |
| rpl218 | 2.041349 | 0.399045797 | 0.941 | 0.776 | 5.1860495 | 27 | rpl21 |
| rpl914 | 2.221719 | 0.502831089 | 0.919 | 0.668 | 5.6442786 | 27 | rpl9 |
| rpl7a8 | 2.274169 | 0.354589186 | 0.941 | 0.824 | 5.7775286 | 27 | rpl7a |
| rps511 | 4.517824 | 0.461245393 | 0.934 | 0.728 | 1.1477533 | 27 | rps5 |
| rps67 | 5.255900 | 0.390825206 | 0.956 | 0.803 | 1.3352616 | 27 | rps6 |
| rpl376 | 5.929595 | 0.33013549 | 0.956 | 0.855 | 1.5064137 | 27 | rpl37 |
| ccr9a6 | 8.366736 | 0.522938829 | 0.382 | 0.086 | 2.1255694 | 27 | ccr9a |
| rps4x10 | 1.018834 | 0.440010883 | 0.912 | 0.736 | 2.5883488 | 27 | rps4x |
| rps1013 | 1.021451 | 0.450276176 | 0.956 | 0.726 | 2.5949966 | 27 | rps10 |
| rps38 | 1.071452 | 0.453556857 | 0.949 | 0.725 | 2.7220247 | 27 | rps3 |
| si:ch211-226m16.24 | 1.087812 | 0.348882194 | 0.338 | 0.067 | 2.7635888 | 27 | si:ch211-226m16.2 |
| rpl329 | 1.152563 | 0.378740696 | 0.978 | 0.824 | 2.9280886 | 27 | rpl32 |
| rpl288 | 1.760268 | 0.391924967 | 0.956 | 0.776 | 4.4719621 | 27 | rpl28 |
| mcm61 | 9.068615 | 0.257159971 | 0.257 | 0.043 | 2.3038817 | 27 | mcm6 |
| eef1g10 | 1.145319 | 0.509773333 | 0.897 | 0.652 | 2.9096847 | 27 | eef1g |
| rpl18a9 | 1.823546 | 0.390491061 | 0.963 | 0.801 | 4.6327208 | 27 | rpl18a |
| rpl267 | 3.049639 | 0.330327073 | 0.971 | 0.851 | 7.7476086 | 27 | rpl26 |
| rpl10a8 | 4.126488 | 0.397020149 | 0.934 | 0.779 | 1.0483344 | 27 | rpl10a |
| nr4a18 | 4.739274 | 0.631484584 | 0.529 | 0.156 | 1.2040127 | 27 | nr4a1 |
| rpl278 | 6.791924 | 0.388188789 | 0.956 | 0.78 | 1.7254884 | 27 | rpl27 |
| rps26l9 | 8.939339 | 0.468381648 | 0.934 | 0.699 | 2.2710392 | 27 | rps26l |
| rpl195 | 9.901451 | 0.335688533 | 0.985 | 0.853 | 2.5154637 | 27 | rpl19 |
| rps147 | 1.707220 | 0.386507966 | 0.963 | 0.781 | 4.3371945 | 27 | rps14 |
| rpl235 | 1.979349 | 0.295740906 | 0.963 | 0.873 | 5.0285384 | 27 | rpl23 |
| rps1611 | 2.578130 | 0.401682024 | 0.949 | 0.765 | 6.5497404 | 27 | rps16 |
| rps77 | 4.214782 | 0.344786576 | 0.956 | 0.828 | 1.0707654 | 27 | rps7 |
| rpl308 | 1.666479 | 0.431784318 | 0.949 | 0.73 | 4.2336905 | 27 | rpl30 |

| | | | | | | | |
|---|---|---|---|---|---|---|---|
| rpl396 | 2.451740 | 0.286649117 | 0.949 | 0.877 | 6.2286478 | 27 | rpl39 |
| nop21 | 2.517320 | 0.260691689 | 0.279 | 0.053 | 6.3952534 | 27 | nop2 |
| nop562 | 3.417006 | 0.566950829 | 0.566 | 0.184 | 8.6809054 | 27 | nop56 |
| rack17 | 3.447504 | 0.383557259 | 0.949 | 0.763 | 8.7583860 | 27 | rack1 |
| rps1511 | 4.273346 | 0.453021656 | 0.956 | 0.716 | 1.0856437 | 27 | rps15 |
| rps222 | 6.468050 | 0.366903504 | 0.985 | 0.841 | 1.6432082 | 27 | rps2 |
| si:dkey-27i16.25 | 6.615899 | 0.453702841 | 0.375 | 0.09 | 1.6807693 | 27 | si:dkey-27i16.2 |
| eif3s104 | 8.834171 | 0.482302206 | 0.566 | 0.177 | 2.2443212 | 27 | eif3s10 |
| rps139 | 1.462029 | 0.384518986 | 0.904 | 0.749 | 3.7142860 | 27 | rps13 |
| parvg5 | 2.036317 | 0.340708484 | 0.331 | 0.072 | 5.1732647 | 27 | parvg |
| rpl74 | 2.151057 | 0.354079958 | 0.934 | 0.809 | 5.4647613 | 27 | rpl7 |
| rpl3610 | 2.173144 | 0.341553093 | 0.941 | 0.806 | 5.5208738 | 27 | rpl36 |
| rps238 | 3.365514 | 0.415725842 | 0.956 | 0.744 | 8.5500893 | 27 | rps23 |
| FP074902.21 | 5.537920 | 0.3107594 | 0.338 | 0.076 | 1.4069086 | 27 | FP074902.2 |
| cbfb2 | 5.569598 | 0.277464351 | 0.235 | 0.042 | 1.4149565 | 27 | cbfb |
| rpl357 | 1.071766 | 0.334178427 | 0.941 | 0.814 | 2.7228234 | 27 | rpl35 |
| cops2 | 1.258819 | 0.293438476 | 0.301 | 0.064 | 3.1980315 | 27 | cops2 |
| rps258 | 3.867976 | 0.384305538 | 0.934 | 0.749 | 9.8265931 | 27 | rps25 |
| si:ch211-214p16.16 | 4.496724 | 0.70516181 | 0.691 | 0.242 | 1.1423929 | 27 | si:ch211-214p16.1 |
| mibp210 | 5.218849 | 0.594697567 | 0.86 | 0.52 | 1.3258486 | 27 | mibp2 |
| rps2410 | 5.350798 | 0.356195148 | 0.941 | 0.779 | 1.3593702 | 27 | rps24 |
| rps8a6 | 1.196886 | 0.336423512 | 0.926 | 0.801 | 3.0406909 | 27 | rps8a |
| tnrc182 | 1.885519 | 0.265500481 | 0.235 | 0.044 | 4.7901614 | 27 | tnrc18 |
| rpl347 | 2.095735 | 0.336863597 | 0.926 | 0.802 | 5.3242171 | 27 | rpl34 |
| rps27a6 | 3.097270 | 0.377113098 | 0.978 | 0.774 | 7.8686148 | 27 | rps27a |
| eif3c5 | 3.448759 | 0.46136446 | 0.566 | 0.186 | 8.7615731 | 27 | eif3c |
| gars | 4.108955 | 0.310201801 | 0.316 | 0.072 | 1.0438800 | 27 | gars |
| rps197 | 4.400684 | 0.2898326 | 0.971 | 0.868 | 1.1179940 | 27 | rps19 |
| rpl23a10 | 4.574560 | 0.45084635 | 0.897 | 0.669 | 1.1621671 | 27 | rpl23a |
| rps175 | 6.687983 | 0.364900767 | 0.949 | 0.779 | 1.6990822 | 27 | rps17 |
| rpl126 | 6.987168 | 0.299723675 | 0.904 | 0.813 | 1.7750901 | 27 | rpl12 |
| sept123 | 1.116542 | 0.290683272 | 0.294 | 0.064 | 2.8365763 | 27 | sept12 |
| ddx211 | 1.123256 | 0.53122469 | 0.588 | 0.208 | 2.8536344 | 27 | ddx21 |
| nsa28 | 2.093775 | 0.6166465 | 0.794 | 0.387 | 5.3192365 | 27 | nsa2 |
| rpl299 | 3.675408 | 0.316463734 | 0.949 | 0.823 | 9.3373744 | 27 | rpl29 |

| | | | | | | | |
|---|---|---|---|---|---|---|---|
| rpl35a8 | 4.298586 | 0.351979406 | 0.949 | 0.781 | 1.0920559 | 27 | rpl35a |
| rpl3110 | 5.417315 | 0.428556706 | 0.919 | 0.687 | 1.3762690 | 27 | rpl31 |
| tfr1b | 6.130546 | 0.277690408 | 0.25 | 0.05 | 1.5574653 | 27 | tfr1b |
| rps1810 | 7.638440 | 0.387689181 | 0.949 | 0.754 | 1.9405459 | 27 | rps18 |
| naca10 | 8.029947 | 0.462432216 | 0.897 | 0.651 | 2.0400080 | 27 | naca |
| rps116 | 9.360541 | 0.329765694 | 0.934 | 0.784 | 2.3780455 | 27 | rps11 |
| gnl31 | 1.504685 | 0.257260601 | 0.301 | 0.069 | 3.8226531 | 27 | gnl3 |
| eif3s6ip | 1.665729 | 0.413321265 | 0.507 | 0.161 | 4.2317862 | 27 | eif3s6ip |
| cox6a23 | 1.733823 | 0.264445099 | 0.206 | 0.037 | 4.4047789 | 27 | cox6a2 |
| cxcr3.22 | 1.940870 | 0.270918685 | 0.199 | 0.035 | 4.9307805 | 27 | cxcr3.2 |
| cct42 | 2.268758 | 0.537817292 | 0.632 | 0.243 | 5.7637806 | 27 | cct4 |
| rrp1 | 3.981327 | 0.28893014 | 0.368 | 0.095 | 1.0114561 | 27 | rrp1 |
| metap1 | 4.119905 | 0.292347222 | 0.331 | 0.082 | 1.0466619 | 27 | metap1 |
| si:dkey-239h2.31 | 8.821686 | 0.345945313 | 0.368 | 0.098 | 2.2411494 | 27 | si:dkey-239h2.3 |
| eef1b21 | 8.964025 | 0.366522994 | 0.912 | 0.741 | 2.2773105 | 27 | eef1b2 |
| ppid1 | 9.617482 | 0.262205855 | 0.36 | 0.092 | 2.4433214 | 27 | ppid |
| rpl27a9 | 1.085380 | 0.556575353 | 0.86 | 0.521 | 2.7574081 | 27 | rpl27a |
| rpl13a4 | 1.192440 | 0.316192721 | 0.912 | 0.782 | 3.0293959 | 27 | rpl13a |
| pde7a3 | 3.577570 | 0.314930677 | 0.279 | 0.065 | 9.0888180 | 27 | pde7a |
| serbp1a11 | 4.607986 | 0.536216928 | 0.801 | 0.505 | 1.1706590 | 27 | serbp1a |
| rgs136 | 6.595675 | 0.429195238 | 0.419 | 0.124 | 1.6756313 | 27 | rgs13 |
| rsl24d18 | 6.617482 | 0.575660437 | 0.772 | 0.46 | 1.6811713 | 27 | rsl24d1 |
| eef2b5 | 9.520484 | 0.439509909 | 0.868 | 0.635 | 2.4186792 | 27 | eef2b |
| eif3ha2 | 1.096999 | 0.501014187 | 0.596 | 0.229 | 2.7869262 | 27 | eif3ha |
| tcp13 | 2.371719 | 0.494729752 | 0.625 | 0.244 | 6.0253525 | 27 | tcp1 |
| pdcd4a1 | 2.996812 | 0.264091417 | 0.257 | 0.058 | 7.6134025 | 27 | pdcd4a |
| cct25 | 3.475396 | 0.517728673 | 0.625 | 0.252 | 8.8292437 | 27 | cct2 |
| cct84 | 3.777928 | 0.48872686 | 0.581 | 0.221 | 9.5978280 | 27 | cct8 |
| grap2b8 | 4.235206 | 0.353394903 | 0.419 | 0.123 | 1.0759541 | 27 | grap2b |
| si:ch211-262e15.1 | 4.336111 | 0.263722073 | 0.316 | 0.079 | 1.1015892 | 27 | si:ch211-262e15.1 |
| cct73 | 4.544373 | 0.514745981 | 0.581 | 0.227 | 1.1544981 | 27 | cct7 |
| pinx1 | 4.787817 | 0.384128313 | 0.478 | 0.154 | 1.2163450 | 27 | pinx1 |
| si:ch73-1a9.313 | 5.017318 | 0.590263991 | 0.706 | 0.314 | 1.2746496 | 27 | si:ch73-1a9.3 |
| XLOC-0425955 | 5.158138 | 0.552581453 | 0.463 | 0.152 | 1.3104252 | 27 | XLOC-042595 |
| rpl48 | 7.407992 | 0.421434992 | 0.882 | 0.656 | 1.8820006 | 27 | rpl4 |

| | | | | | | | |
|---|---|---|---|---|---|---|---|
| nop534 | 8.380739 | 0.544993067 | 0.625 | 0.265 | 2.1291267 | 27 | nop53 |
| sox4a5 | 1.034167 | 0.310478142 | 0.265 | 0.062 | 2.6273026 | 27 | sox4a |
| wasb9 | 1.759866 | 0.37923783 | 0.529 | 0.171 | 4.4709404 | 27 | wasb |
| adh56 | 1.792270 | 0.50028887 | 0.61 | 0.24 | 4.5532622 | 27 | adh5 |
| myg11 | 1.809297 | 0.266679455 | 0.287 | 0.071 | 4.5965209 | 27 | myg1 |
| MRPS12 | 2.204717 | 0.26926283 | 0.338 | 0.091 | 5.6010853 | 27 | MRPS12 |
| rsl1d12 | 2.217945 | 0.429228429 | 0.529 | 0.186 | 5.6346910 | 27 | rsl1d1 |
| rpl2211 | 3.189493 | 0.385022215 | 0.934 | 0.72 | 8.1029072 | 27 | rpl22 |
| pho1 | 4.843085 | 0.295733633 | 0.346 | 0.094 | 1.2303857 | 27 | pho |
| rpl388 | 5.999079 | 0.360992982 | 0.941 | 0.753 | 1.5240661 | 27 | rpl38 |
| pfdn41 | 6.010962 | 0.393910672 | 0.529 | 0.186 | 1.5270851 | 27 | pfdn4 |
| rps266 | 7.841660 | 0.339566651 | 0.934 | 0.75 | 1.9921739 | 27 | rps26 |
| rps288 | 8.663562 | 0.305651653 | 0.985 | 0.84 | 2.2009780 | 27 | rps28 |
| usp361 | 1.152847 | 0.263312242 | 0.309 | 0.081 | 2.9288083 | 27 | usp36 |
| eif3ba3 | 1.548488 | 0.498888189 | 0.588 | 0.24 | 3.9339351 | 27 | eif3ba |
| zgc:11418814 | 1.849447 | 0.412742646 | 0.904 | 0.667 | 4.6985214 | 27 | zgc:114188 |
| rpl5a11 | 2.261019 | 0.498652152 | 0.838 | 0.536 | 5.7441199 | 27 | rpl5a |
| eif3ea4 | 2.436214 | 0.546570528 | 0.647 | 0.303 | 6.1892024 | 27 | eif3ea |
| rps299 | 4.014899 | 0.270295116 | 0.956 | 0.844 | 1.0199851 | 27 | rps29 |
| eif2s33 | 6.643528 | 0.510982379 | 0.684 | 0.306 | 1.6877883 | 27 | eif2s3 |
| fabp34 | 6.666821 | 0.497325317 | 0.566 | 0.227 | 1.6937059 | 27 | fabp3 |
| rpl149 | 7.560125 | 0.421166414 | 0.926 | 0.669 | 1.9206498 | 27 | rpl14 |
| ascc11 | 9.645290 | 0.273208049 | 0.25 | 0.06 | 2.4503859 | 27 | ascc1 |
| ccdc251 | 1.365480 | 0.302735914 | 0.346 | 0.101 | 3.4690029 | 27 | ccdc25 |
| eif3k4 | 2.455106 | 0.447019139 | 0.596 | 0.243 | 6.2371969 | 27 | eif3k |
| si:dkey-23i12.5 | 3.230472 | 0.349327517 | 0.463 | 0.158 | 8.2070153 | 27 | si:dkey-23i12.5 |
| nars2 | 3.502574 | 0.251475127 | 0.301 | 0.082 | 8.8982906 | 27 | nars |
| si:dkey-112e7.27 | 3.989420 | 0.431588897 | 0.551 | 0.213 | 1.0135123 | 27 | si:dkey-112e7.2 |
| pa2g4a1 | 4.498592 | 0.31265595 | 0.397 | 0.124 | 1.1428675 | 27 | pa2g4a |
| si:dkey-102m7.3 | 7.171480 | 0.288467249 | 0.426 | 0.138 | 1.8219147 | 27 | si:dkey-102m7.3 |
| si:dkey-250d21.11 | 7.935910 | 0.282772545 | 0.301 | 0.084 | 2.0161181 | 27 | si:dkey-250d21.1 |
| nop582 | 9.544088 | 0.377467622 | 0.5 | 0.179 | 2.4246756 | 27 | nop58 |
| txndc93 | 1.312555 | 0.340335085 | 0.39 | 0.128 | 3.3345460 | 27 | txndc9 |
| eif3m7 | 1.328406 | 0.521683364 | 0.757 | 0.382 | 3.3748159 | 27 | eif3m |
| btf37 | 1.702195 | 0.439559289 | 0.897 | 0.61 | 4.3244276 | 27 | btf3 |

| | | | | | | | |
|---|---|---|---|---|---|---|---|
| pmp22b3 | 1.891233 | 0.29068543 | 0.206 | 0.046 | 4.8046778 | 27 | pmp22b |
| spi1b5 | 2.727569 | 0.355661215 | 0.265 | 0.071 | 6.9293895 | 27 | spi1b |
| ptmaa12 | 3.110177 | 0.558390215 | 0.794 | 0.431 | 7.9014065 | 27 | ptmaa |
| eef1da6 | 3.399241 | 0.49224215 | 0.824 | 0.505 | 8.6357719 | 27 | eef1da |
| uba529 | 4.599083 | 0.385182805 | 0.89 | 0.66 | 1.1683971 | 27 | uba52 |
| pbdc11 | 4.780887 | 0.266184302 | 0.419 | 0.138 | 1.2145845 | 27 | pbdc1 |
| eif3f9 | 6.408908 | 0.522595282 | 0.831 | 0.464 | 1.6281832 | 27 | eif3f |
| rpl176 | 6.764775 | 0.327564407 | 0.949 | 0.763 | 1.7185912 | 27 | rpl17 |
| eif3g6 | 6.964691 | 0.459568713 | 0.669 | 0.298 | 1.7693799 | 27 | eif3g |
| caprin1b1 | 2.051521 | 0.388336038 | 0.603 | 0.244 | 5.2118910 | 27 | caprin1b |
| zbtb205 | 2.764553 | 0.384253047 | 0.331 | 0.107 | 7.0233476 | 27 | zbtb20 |
| twistnb | 2.935160 | 0.258454201 | 0.25 | 0.066 | 7.4567743 | 27 | twistnb |
| wasa7 | 3.557236 | 0.250822904 | 0.324 | 0.096 | 9.0371591 | 27 | wasa |
| rpl5b8 | 3.619961 | 0.387783723 | 0.875 | 0.645 | 9.1965131 | 27 | rpl5b |
| eif2s1b3 | 5.206652 | 0.379186191 | 0.493 | 0.194 | 1.3227501 | 27 | eif2s1b |
| cct31 | 5.672793 | 0.431584864 | 0.537 | 0.224 | 1.4411731 | 27 | cct3 |
| rnf442 | 6.731928 | 0.332109559 | 0.309 | 0.096 | 1.7102464 | 27 | rnf44 |
| cct51 | 7.019800 | 0.445543855 | 0.61 | 0.273 | 1.7833802 | 27 | cct5 |
| eif3i5 | 7.982568 | 0.435035459 | 0.647 | 0.289 | 2.0279715 | 27 | eif3i |
| aldoaa4 | 9.869609 | 0.426058751 | 0.588 | 0.252 | 2.5073744 | 27 | aldoaa |
| slc25a56 | 1.181000 | 0.25797028 | 0.926 | 0.813 | 3.0003318 | 27 | slc25a5 |
| ppiaa6 | 1.776759 | 0.268801223 | 0.949 | 0.814 | 4.5138568 | 27 | ppiaa |
| ipo71 | 2.423443 | 0.281781473 | 0.353 | 0.115 | 6.1567584 | 27 | ipo7 |
| tpt15 | 2.915527 | 0.289961444 | 0.978 | 0.811 | 7.4068973 | 27 | tpt1 |
| elmo13 | 2.928433 | 0.289123889 | 0.25 | 0.07 | 7.4396848 | 27 | elmo1 |
| ak21 | 4.421169 | 0.403636599 | 0.581 | 0.253 | 1.1231980 | 27 | ak2 |
| eif4ebp14 | 5.895399 | 0.388665633 | 0.61 | 0.257 | 1.4977263 | 27 | eif4ebp1 |
| hmgn29 | 7.104911 | 0.525123019 | 0.794 | 0.466 | 1.8050028 | 27 | hmgn2 |
| hnrnpa0b7 | 1.973375 | 0.465283645 | 0.691 | 0.34 | 5.0133594 | 27 | hnrnpa0b |
| hmgn612 | 2.494691 | 0.431241524 | 0.662 | 0.31 | 6.3377641 | 27 | hmgn6 |
| tuba8l6 | 2.609216 | 0.393895967 | 0.5 | 0.208 | 6.6287134 | 27 | tuba8l |
| vdac25 | 3.159668 | 0.430084617 | 0.676 | 0.324 | 8.0271388 | 27 | vdac2 |
| bccip2 | 3.891102 | 0.288778563 | 0.404 | 0.148 | 9.8853457 | 27 | bccip |
| eif5a6 | 3.891865 | 0.449541634 | 0.772 | 0.495 | 9.8872840 | 27 | eif5a |
| metap2b1 | 4.898617 | 0.347834759 | 0.529 | 0.22 | 1.2444937 | 27 | metap2b |

| | | | | | | | |
|---|---|---|---|---|---|---|---|
| si:busm1-266f07.24 | 5.134365 | 0.325932121 | 0.412 | 0.147 | 1.3043854 | 27 | si:busm1-266f07.2 |
| tnrc6c17 | 5.565102 | 0.309336562 | 0.404 | 0.149 | 1.4138143 | 27 | tnrc6c1 |
| atrx6 | 6.198736 | 0.251743128 | 0.368 | 0.127 | 1.5747889 | 27 | atrx |
| laptm59 | 8.013770 | 0.276940885 | 0.478 | 0.175 | 2.0358984 | 27 | laptm5 |
| psme25 | 1.125943 | 0.267593794 | 0.331 | 0.112 | 2.8604585 | 27 | psme2 |
| XLOC-0139645 | 4.531595 | 0.398948627 | 0.426 | 0.169 | 1.1512519 | 27 | XLOC-013964 |
| si:dkey-42i9.43 | 5.058010 | 0.262664767 | 0.412 | 0.155 | 1.2849875 | 27 | si:dkey-42i9.4 |
| arhgdig8 | 7.982610 | 0.398752392 | 0.581 | 0.258 | 2.0279820 | 27 | arhgdig |
| crema6 | 9.625639 | 0.371391221 | 0.426 | 0.173 | 2.4453937 | 27 | crema |
| tefa6 | 1.059087 | 0.250801212 | 0.382 | 0.14 | 2.6906126 | 27 | tefa |
| XLOC-0004317 | 1.063803 | 0.575124877 | 0.544 | 0.272 | 2.7025933 | 27 | XLOC-000431 |
| h1fx7 | 1.280529 | 0.384238044 | 0.456 | 0.19 | 3.2531851 | 27 | h1fx |
| celf210 | 1.321632 | 0.353712518 | 0.581 | 0.256 | 3.3576077 | 27 | celf2 |
| eif4a1b5 | 1.380228 | 0.401364852 | 0.632 | 0.311 | 3.5064704 | 27 | eif4a1b |
| eif3d2 | 1.522632 | 0.39672275 | 0.581 | 0.28 | 3.8682468 | 27 | eif3d |
| atp5pf5 | 2.087719 | 0.354965003 | 0.676 | 0.318 | 5.3038523 | 27 | atp5pf |
| ola13 | 2.112143 | 0.331606064 | 0.551 | 0.243 | 5.3659008 | 27 | ola1 |
| calm2b7 | 2.194915 | 0.373756441 | 0.544 | 0.246 | 5.5761828 | 27 | calm2b |
| si:dkey-11f4.146 | 2.496216 | 0.26232126 | 0.338 | 0.122 | 6.3416392 | 27 | si:dkey-11f4.14 |
| rplp210 | 2.699955 | 0.321575636 | 0.971 | 0.782 | 6.8592363 | 27 | rplp2 |
| cd834 | 2.788383 | 0.40484131 | 0.272 | 0.093 | 7.0838872 | 27 | cd83 |
| hmga1a6 | 3.043295 | 0.453766909 | 0.801 | 0.441 | 7.7314922 | 27 | hmga1a |
| foxp44 | 3.175435 | 0.287054385 | 0.235 | 0.073 | 8.0671927 | 27 | foxp4 |
| tob1a2 | 4.421736 | 0.26987917 | 0.375 | 0.144 | 1.1233421 | 27 | tob1a |
| wu:fb55g093 | 1.183489 | 0.374857838 | 0.551 | 0.258 | 3.0066538 | 27 | wu:fb55g09 |
| rac29 | 1.345502 | 0.316065659 | 0.537 | 0.224 | 3.4182498 | 27 | rac2 |
| stmn1a5 | 1.626063 | 0.326938146 | 0.456 | 0.188 | 4.1310134 | 27 | stmn1a |
| pno12 | 1.991718 | 0.260595819 | 0.419 | 0.171 | 5.0599611 | 27 | pno1 |
| idh22 | 3.007888 | 0.310395356 | 0.5 | 0.226 | 7.6415416 | 27 | idh2 |
| atp5if1b4 | 3.202672 | 0.255507919 | 0.485 | 0.206 | 8.1363882 | 27 | atp5if1b |
| prmt12 | 3.320824 | 0.27200994 | 0.507 | 0.22 | 8.4365546 | 27 | prmt1 |
| ran3 | 3.357477 | 0.346020111 | 0.868 | 0.639 | 8.5296722 | 27 | ran |
| arpc1b9 | 3.915220 | 0.351445074 | 0.699 | 0.312 | 9.9466166 | 27 | arpc1b |
| cnbpb5 | 5.215811 | 0.280950792 | 0.478 | 0.209 | 1.3250768 | 27 | cnbpb |
| hmgn76 | 5.968446 | 0.369766848 | 0.632 | 0.317 | 1.5162838 | 27 | hmgn7 |

| | | | | | | | |
|---|---|---|---|---|---|---|---|
| histh1l6 | 6.888570 | 0.373076677 | 0.309 | 0.122 | 1.7500412 | 27 | histh1l |
| cnbpa1 | 7.085952 | 0.300774044 | 0.485 | 0.216 | 1.8001862 | 27 | cnbpa |
| ldhba2 | 7.841694 | 0.310955592 | 0.559 | 0.26 | 1.9921824 | 27 | ldhba |
| ucp28 | 1.415974 | 0.382802748 | 0.75 | 0.413 | 3.5972835 | 27 | ucp2 |
| sept63 | 2.323559 | 0.252949675 | 0.463 | 0.201 | 5.9030032 | 27 | sept6 |
| prdx56 | 2.780685 | 0.281952238 | 0.412 | 0.178 | 7.0643326 | 27 | prdx5 |
| krt18b7 | 4.681664 | 0.315649229 | 0.404 | 0.176 | 1.1893767 | 27 | krt18b |
| cebpb10 | 5.223622 | 0.429729366 | 0.618 | 0.339 | 1.3270612 | 27 | cebpb |
| sub1a3 | 5.471903 | 0.286188948 | 0.537 | 0.252 | 1.3901369 | 27 | sub1a |
| si:ch211-250g4.31 | 7.811320 | 0.469414323 | 0.14 | 0.039 | 1.9844660 | 27 | si:ch211-250g4.3 |
| gpx1a7 | 8.359924 | 0.35438389 | 0.478 | 0.236 | 2.1238387 | 27 | gpx1a |
| npm1a2 | 1.138699 | 0.370677747 | 0.566 | 0.299 | 2.8928648 | 27 | npm1a |
| ywhaqa2 | 1.221312 | 0.264379516 | 0.463 | 0.209 | 3.1027436 | 27 | ywhaqa |
| sod12 | 1.564553 | 0.325946807 | 0.662 | 0.344 | 3.9747473 | 27 | sod1 |
| ybx16 | 1.814503 | 0.297548918 | 0.897 | 0.695 | 4.6097473 | 27 | ybx1 |
| mdh1aa3 | 1.951307 | 0.263570553 | 0.485 | 0.228 | 4.9572975 | 27 | mdh1aa |
| atp5po1 | 2.218409 | 0.271538499 | 0.566 | 0.276 | 5.6358682 | 27 | atp5po |
| id2a1 | 2.702862 | 0.324300118 | 0.375 | 0.167 | 6.8666229 | 27 | id2a |
| cct6a2 | 3.878237 | 0.256982836 | 0.471 | 0.22 | 9.8526629 | 27 | cct6a |
| crip110 | 6.879855 | 0.292918567 | 0.588 | 0.283 | 0.000175 | 27 | crip1 |
| sumo2b9 | 8.197375 | 0.302848294 | 0.581 | 0.296 | 0.000208 | 27 | sumo2b |
| creg11 | 8.590849 | 0.271183948 | 0.213 | 0.077 | 0.000218 | 27 | creg1 |
| b2m9 | 1.268017 | 0.284422734 | 0.588 | 0.298 | 0.000322 | 27 | b2m |
| uqcrh3 | 1.791434 | 0.264555794 | 0.515 | 0.259 | 0.000455 | 27 | uqcrh |
| romo17 | 2.068012 | 0.346768723 | 0.721 | 0.407 | 0.000525 | 27 | romo1 |
| pabpc1a7 | 2.203655 | 0.342818871 | 0.824 | 0.569 | 0.00056 | 27 | pabpc1a |
| oaz1b8 | 2.995154 | 0.282080598 | 0.676 | 0.359 | 0.000761 | 27 | oaz1b |
| ccnd2a1 | 6.246436 | 0.26729462 | 0.235 | 0.095 | 0.001587 | 27 | ccnd2a |
| slc25a3b5 | 9.237382 | 0.341660663 | 0.772 | 0.457 | 0.002347 | 27 | slc25a3b |
| pfn19 | 1.001844 | 0.394676095 | 0.838 | 0.5 | 0.002545 | 27 | pfn1 |
| arpc28 | 1.695287 | 0.256239496 | 0.574 | 0.298 | 0.004307 | 27 | arpc2 |
| prdx26 | 1.903736 | 0.314286659 | 0.735 | 0.421 | 0.004836 | 27 | prdx2 |
| ddit37 | 2.088325 | 0.268832058 | 0.368 | 0.178 | 0.005305 | 27 | ddit3 |
| atp5mc3b2 | 3.187860 | 0.304188373 | 0.816 | 0.602 | 0.008099 | 27 | atp5mc3b |
| mknk2b8 | 3.924443 | 0.253833662 | 0.397 | 0.2 | 0.00997 | 27 | mknk2b |

| | | | | | | | |
|---|---|---|---|---|---|---|---|
| COX5B4 | 4.945652 | 0.286541648 | 0.654 | 0.376 | 0.012564 | 27 | COX5B |
| hmgb2b6 | 1.815353 | 0.279706131 | 0.574 | 0.336 | 0.046119 | 27 | hmgb2b |
| cox6b24 | 1.951263 | 0.31037224 | 0.846 | 0.549 | 0.049572 | 27 | cox6b2 |
| rps27.211 | 2.287341 | 0.309656856 | 0.794 | 0.553 | 0.05811 | 27 | rps27.2 |
| h2afx8 | 4.455544 | 0.269806171 | 0.257 | 0.123 | 0.113193 | 27 | h2afx |
| zgc:564936 | 6.711745 | 0.289150429 | 0.809 | 0.59 | 0.170512 | 27 | zgc:56493 |
| ptmab10 | 7.706125 | 0.290739539 | 0.824 | 0.569 | 0.195774 | 27 | ptmab |
| CR383676.115 | 2.322611 | 0.438615828 | 0.963 | 0.716 | 0.590059 | 27 | CR383676.1 |
| NC-002333.410 | 0.0001 | 0.272715811 | 0.846 | 0.609 | 1 | 27 | NC-002333.4 |
| cahz1 | 0.0002 | 0.286536966 | 0.118 | 0.05 | 1 | 27 | cahz |
| epdl1 | 0 | 1.866423062 | 0.843 | 0.026 | 0 | 28 | epdl1 |
| spock3 | 0 | 1.12424787 | 0.543 | 0.008 | 0 | 28 | spock3 |
| mpeg1.11 | 0 | 0.947094462 | 0.48 | 0.014 | 0 | 28 | mpeg1.1 |
| syngr3b | 0 | 0.845386679 | 0.417 | 0.002 | 0 | 28 | syngr3b |
| CU928222.3 | 0 | 0.818828024 | 0.362 | 0.001 | 0 | 28 | CU928222.3 |
| cxcl11.6 | 0 | 0.785018655 | 0.291 | 0.004 | 0 | 28 | cxcl11.6 |
| FP085399.2 | 0 | 0.778133121 | 0.37 | 0.001 | 0 | 28 | FP085399.2 |
| cxcl11.5 | 0 | 0.77123378 | 0.299 | 0.003 | 0 | 28 | cxcl11.5 |
| zgc:101810 | 0 | 0.686279236 | 0.331 | 0.004 | 0 | 28 | zgc:101810 |
| p2rx3a | 0 | 0.380479075 | 0.126 | 0 | 0 | 28 | p2rx3a |
| ptprh | 0 | 0.295705857 | 0.15 | 0 | 0 | 28 | ptprh |
| BX323596.21 | 9.220738 | 0.868629324 | 0.354 | 0.008 | 2.3425286 | 28 | BX323596.2 |
| BX323596.1 | 1.437582 | 0.909878088 | 0.331 | 0.007 | 3.6521793 | 28 | BX323596.1 |
| si:ch211-1o7.32 | 1.863199 | 0.903427802 | 0.472 | 0.018 | 4.7334581 | 28 | si:ch211-1o7.3 |
| AL929237.1 | 8.104081 | 0.564911696 | 0.276 | 0.005 | 2.0588418 | 28 | AL929237.1 |
| sftpbb4 | 1.620006 | 1.650410909 | 0.811 | 0.065 | 4.1156258 | 28 | sftpbb |
| mxf | 3.100320 | 0.460322319 | 0.189 | 0.002 | 7.8763630 | 28 | mxf |
| LOC110438030 | 4.829175 | 0.474780809 | 0.205 | 0.003 | 1.2268520 | 28 | LOC110438030 |
| ccl35.14 | 1.023935 | 1.690556334 | 0.811 | 0.078 | 2.6013070 | 28 | ccl35.1 |
| si:dkey-33i11.91 | 7.190628 | 0.9969468 | 0.433 | 0.023 | 1.8267791 | 28 | si:dkey-33i11.9 |
| si:busm1-266f07.25 | 5.863522 | 1.778842412 | 0.969 | 0.142 | 1.4896277 | 28 | si:busm1-266f07.2 |
| grn12 | 7.830552 | 1.739226246 | 0.78 | 0.085 | 1.9893518 | 28 | grn1 |
| si:dkey-148a17.5 | 1.389103 | 0.315627227 | 0.15 | 0.002 | 3.5290179 | 28 | si:dkey-148a17.5 |
| BX649442.1 | 1.437435 | 0.627808442 | 0.165 | 0.003 | 3.6518054 | 28 | BX649442.1 |
| tnfa | 6.125622 | 0.397608877 | 0.189 | 0.004 | 1.5562143 | 28 | tnfa |

| | | | | | | | |
|---|---|---|---|---|---|---|---|
| CR385050.1 | 9.073634 | 0.550344057 | 0.291 | 0.011 | 2.3051568 | 28 | CR385050.1 |
| mhc2dab4 | 8.654768 | 1.459622569 | 0.772 | 0.09 | 2.1987438 | 28 | mhc2dab |
| hepacam2 | 2.332657 | 0.361144627 | 0.189 | 0.004 | 5.9261159 | 28 | hepacam2 |
| si:ch211-79k12.1 | 3.224510 | 0.798240143 | 0.362 | 0.019 | 8.1918686 | 28 | si:ch211-79k12.1 |
| LOC101885445 | 3.367614 | 0.545110069 | 0.268 | 0.01 | 8.5554237 | 28 | LOC101885445 |
| ccl35.2 | 1.524927 | 0.447198758 | 0.173 | 0.004 | 3.8740774 | 28 | ccl35.2 |
| rnaset2l3 | 9.823669 | 1.069744448 | 0.543 | 0.046 | 2.4957033 | 28 | rnaset2l |
| cd74a5 | 1.001077 | 1.871707723 | 1 | 0.211 | 2.5432368 | 28 | cd74a |
| spi1a2 | 3.869162 | 0.933033932 | 0.48 | 0.036 | 9.8296062 | 28 | spi1a |
| si:ch211-214p13.3 | 1.765507 | 0.337551614 | 0.15 | 0.003 | 4.4852725 | 28 | si:ch211-214p13.3 |
| si:ch73-158p21.31 | 2.554140 | 0.728350717 | 0.386 | 0.025 | 6.4887941 | 28 | si:ch73-158p21.3 |
| arap1a | 3.396801 | 0.282350154 | 0.142 | 0.003 | 8.6295733 | 28 | arap1a |
| smcr8b | 5.581669 | 0.27512731 | 0.15 | 0.003 | 1.4180232 | 28 | smcr8b |
| si:dkey-78o7.32 | 3.325060 | 0.777625612 | 0.331 | 0.019 | 8.4473158 | 28 | si:dkey-78o7.3 |
| si:dkey-1h24.6 | 3.745323 | 0.470449118 | 0.22 | 0.008 | 9.5149938 | 28 | si:dkey-1h24.6 |
| cd74b6 | 4.490034 | 1.611905856 | 0.953 | 0.221 | 1.1406933 | 28 | cd74b |
| CU459094.3 | 1.429685 | 0.335715935 | 0.118 | 0.002 | 3.6321160 | 28 | CU459094.3 |
| CR786582.2 | 5.291304 | 0.488174023 | 0.236 | 0.011 | 1.3442559 | 28 | CR786582.2 |
| si:dkey-102c8.3 | 1.821770 | 0.253772297 | 0.134 | 0.003 | 4.6282067 | 28 | si:dkey-102c8.3 |
| arhgef15 | 1.831697 | 0.299026081 | 0.142 | 0.004 | 4.6534276 | 28 | arhgef15 |
| si:ch73-236c18.7 | 4.942321 | 0.340874075 | 0.173 | 0.006 | 1.2555968 | 28 | si:ch73-236c18.7 |
| si:dkey-192g7.3 | 1.373807 | 0.527762621 | 0.283 | 0.017 | 3.4901572 | 28 | si:dkey-192g7.3 |
| zgc:1231071 | 1.154336 | 0.483280344 | 0.236 | 0.012 | 2.9325928 | 28 | zgc:123107 |
| ctss2.22 | 9.167859 | 0.850990056 | 0.465 | 0.048 | 2.3290947 | 28 | ctss2.2 |
| tnfb4 | 1.805293 | 1.030034255 | 0.449 | 0.047 | 4.5863476 | 28 | tnfb |
| ccl39.11 | 7.979506 | 0.738757115 | 0.22 | 0.012 | 2.0271936 | 28 | ccl39.1 |
| si:dkey-27i16.26 | 1.102958 | 1.026104893 | 0.598 | 0.088 | 2.8020662 | 28 | si:dkey-27i16.2 |
| CABZ01078737.1 | 7.108016 | 0.386073739 | 0.205 | 0.01 | 1.8057915 | 28 | CABZ01078737.1 |
| cd835 | 4.124410 | 1.210796145 | 0.591 | 0.09 | 1.0478066 | 28 | cd83 |
| ciita | 4.885436 | 0.362228295 | 0.181 | 0.008 | 1.2411451 | 28 | ciita |
| XLOC-0254236 | 1.507848 | 1.313147568 | 0.772 | 0.167 | 3.8306884 | 28 | XLOC-025423 |
| arpc1b10 | 3.034470 | 1.279363629 | 0.953 | 0.31 | 7.7090721 | 28 | arpc1b |
| grna2 | 4.397123 | 0.822797633 | 0.457 | 0.055 | 1.1170892 | 28 | grna |
| BX548046.21 | 2.723755 | 0.275643702 | 0.134 | 0.005 | 6.9197013 | 28 | BX548046.2 |
| si:ch211-198c19.1 | 6.288418 | 0.397814883 | 0.189 | 0.01 | 1.5975727 | 28 | si:ch211-198c19.1 |

| | | | | | | | |
|---|---|---|---|---|---|---|---|
| abi3a2 | 4.695269 | 0.722900093 | 0.402 | 0.045 | 1.1928331 | 28 | abi3a |
| si:ch211-214p16.24 | 2.489242 | 0.936793885 | 0.512 | 0.072 | 6.3239200 | 28 | si:ch211-214p16.2 |
| kctd12.25 | 6.174418 | 0.933888328 | 0.591 | 0.097 | 1.5686109 | 28 | kctd12.2 |
| BX004816.2 | 1.051350 | 0.313325813 | 0.15 | 0.006 | 2.6709566 | 28 | BX004816.2 |
| zgc:1037003 | 1.901205 | 0.930503154 | 0.417 | 0.052 | 4.8300117 | 28 | zgc:103700 |
| irak3 | 1.113540 | 0.431968605 | 0.22 | 0.015 | 2.8289497 | 28 | irak3 |
| mxc | 5.356281 | 0.280469036 | 0.134 | 0.005 | 1.3607634 | 28 | mxc |
| arpc5b6 | 7.486022 | 0.989672515 | 0.638 | 0.128 | 1.9018240 | 28 | arpc5b |
| si:dkey-53k12.24 | 8.434174 | 0.637031774 | 0.37 | 0.043 | 2.1427021 | 28 | si:dkey-53k12.2 |
| si:busm1-194e12.113 | 6.308516 | 0.658491607 | 0.283 | 0.026 | 1.6026785 | 28 | si:busm1-194e12.11 |
| lygl12 | 1.373787 | 0.933634013 | 0.575 | 0.097 | 3.4901071 | 28 | lygl1 |
| tmsb4x15 | 2.824760 | 0.885206645 | 1 | 0.717 | 7.1763049 | 28 | tmsb4x |
| fgd4b | 4.358024 | 0.54512608 | 0.291 | 0.027 | 1.1071560 | 28 | fgd4b |
| spi1b6 | 5.744055 | 0.740349858 | 0.472 | 0.069 | 1.4592771 | 28 | spi1b |
| cyb561a3a | 1.687085 | 0.527672507 | 0.268 | 0.024 | 4.2860397 | 28 | cyb561a3a |
| gpx1a8 | 1.360653 | 1.157385628 | 0.795 | 0.233 | 3.4567406 | 28 | gpx1a |
| pfn110 | 1.437779 | 1.107010258 | 1 | 0.499 | 3.6526796 | 28 | pfn1 |
| parm1 | 4.951546 | 0.287029273 | 0.15 | 0.008 | 1.2579404 | 28 | parm1 |
| actb29 | 1.761621 | 0.673634336 | 1 | 0.867 | 4.4753987 | 28 | actb2 |
| si:ch211-8c17.32 | 2.114269 | 0.507547285 | 0.276 | 0.026 | 5.3713021 | 28 | si:ch211-8c17.3 |
| mhc2b3 | 8.279678 | 0.839036037 | 0.331 | 0.039 | 2.1034523 | 28 | mhc2b |
| LO018493.12 | 2.692772 | 0.512598825 | 0.268 | 0.025 | 6.8409884 | 28 | LO018493.1 |
| ptmaa13 | 1.309696 | 1.015925566 | 0.945 | 0.43 | 3.3272844 | 28 | ptmaa |
| bpifcl2 | 1.940830 | 0.399709452 | 0.197 | 0.014 | 4.9306806 | 28 | bpifcl |
| plekhf1 | 2.061973 | 0.299086143 | 0.118 | 0.005 | 5.2384435 | 28 | plekhf1 |
| psap4 | 3.636041 | 1.098784913 | 0.835 | 0.284 | 9.2373644 | 28 | psap |
| ccl19a.11 | 1.157623 | 0.645803395 | 0.189 | 0.013 | 2.9409421 | 28 | ccl19a.1 |
| tiam1a | 1.387014 | 0.324082568 | 0.165 | 0.01 | 3.5237094 | 28 | tiam1a |
| ptgs2b | 1.462104 | 0.308483815 | 0.165 | 0.01 | 3.7144766 | 28 | ptgs2b |
| XLOC-0234924 | 1.281415 | 0.954891444 | 0.496 | 0.091 | 3.2554354 | 28 | XLOC-023492 |
| wasb10 | 2.431827 | 0.952014994 | 0.701 | 0.17 | 6.1780581 | 28 | wasb |
| BX005183.35 | 1.678918 | 0.80134708 | 0.449 | 0.074 | 4.2652929 | 28 | BX005183.3 |
| unc93b12 | 2.788369 | 0.452864479 | 0.236 | 0.022 | 7.0838535 | 28 | unc93b1 |
| LOC1018850652 | 9.335544 | 0.356424207 | 0.189 | 0.015 | 2.3716949 | 28 | LOC101885065 |
| lcp16 | 1.188903 | 0.785865807 | 0.457 | 0.082 | 3.0204105 | 28 | lcp1 |

| | | | | | | | |
|---|---|---|---|---|---|---|---|
| CR753876.14 | 3.258068 | 0.916551735 | 0.535 | 0.109 | 8.2771220 | 28 | CR753876.1 |
| nr4a2b | 5.440904 | 0.341155872 | 0.173 | 0.012 | 1.3822618 | 28 | nr4a2b |
| CU499330.13 | 7.719753 | 0.75900128 | 0.433 | 0.07 | 1.9612034 | 28 | CU499330.1 |
| tmsb17 | 2.236767 | 0.933111563 | 0.654 | 0.157 | 5.6825069 | 28 | tmsb1 |
| cotl18 | 2.458418 | 0.987002858 | 0.874 | 0.317 | 6.2456127 | 28 | cotl1 |
| BX901920.14 | 1.009834 | 0.850460233 | 0.409 | 0.072 | 2.5654845 | 28 | BX901920.1 |
| irf83 | 5.868178 | 0.620398914 | 0.331 | 0.047 | 1.4908108 | 28 | irf8 |
| si:ch211-214p16.17 | 2.566425 | 1.009532258 | 0.843 | 0.24 | 6.5200038 | 28 | si:ch211-214p16.1 |
| si:ch211-152p11.41 | 8.748605 | 0.457123676 | 0.244 | 0.027 | 2.2225831 | 28 | si:ch211-152p11.4 |
| p2rx7 | 9.380089 | 0.299671592 | 0.165 | 0.013 | 2.3830118 | 28 | p2rx7 |
| sgk3 | 3.310244 | 0.39710947 | 0.236 | 0.025 | 8.4096769 | 28 | sgk3 |
| si:dkey-7i4.242 | 3.667894 | 0.757493555 | 0.236 | 0.026 | 9.3182848 | 28 | si:dkey-7i4.24 |
| ccl39.31 | 7.225721 | 0.407090686 | 0.134 | 0.008 | 1.8356946 | 28 | ccl39.3 |
| aif1l5 | 7.353214 | 0.903719063 | 0.622 | 0.168 | 1.8680840 | 28 | aif1l |
| XLOC-0283706 | 1.783991 | 0.859883149 | 0.535 | 0.123 | 4.5322307 | 28 | XLOC-028370 |
| il21r.11 | 2.329483 | 0.384191236 | 0.213 | 0.022 | 5.9180520 | 28 | il21r.1 |
| cxcr4b8 | 6.476892 | 1.055869207 | 0.598 | 0.175 | 1.6454545 | 28 | cxcr4b |
| npc22 | 1.253534 | 0.945767754 | 0.795 | 0.34 | 3.1846035 | 28 | npc2 |
| gm2a4 | 3.585999 | 0.760065127 | 0.52 | 0.119 | 9.1102316 | 28 | gm2a |
| hcls14 | 7.278837 | 0.642918125 | 0.402 | 0.075 | 1.8491885 | 28 | hcls1 |
| gabarapb9 | 4.101088 | 0.859091289 | 0.78 | 0.302 | 1.0418815 | 28 | gabarapb |
| XLOC-0054493 | 4.864127 | 0.647473949 | 0.331 | 0.056 | 1.2357315 | 28 | XLOC-005449 |
| LOC1039101362 | 1.466053 | 0.441140045 | 0.22 | 0.026 | 3.7245093 | 28 | LOC103910136 |
| tfe3a2 | 1.931866 | 0.480060008 | 0.276 | 0.04 | 4.9079065 | 28 | tfe3a |
| malt3 | 3.538495 | 0.295892382 | 0.165 | 0.015 | 8.9895488 | 28 | malt3 |
| slc15a4 | 8.040418 | 0.387443089 | 0.205 | 0.023 | 2.0426684 | 28 | slc15a4 |
| actb17 | 2.988923 | 0.601435952 | 0.961 | 0.767 | 7.5933611 | 28 | actb1 |
| arpc29 | 1.180970 | 0.819956154 | 0.772 | 0.297 | 3.0002543 | 28 | arpc2 |
| krt18b8 | 4.850049 | 0.836638379 | 0.598 | 0.174 | 1.2321550 | 28 | krt18b |
| cyba6 | 1.051601 | 0.622073264 | 0.394 | 0.081 | 2.6715938 | 28 | cyba |
| zgc:16273012 | 2.342463 | 0.901348453 | 0.858 | 0.401 | 5.9510285 | 28 | zgc:162730 |
| TPGS11 | 2.686049 | 0.440631719 | 0.26 | 0.038 | 6.8239075 | 28 | TPGS1 |
| gstt1a2 | 3.539451 | 0.50003656 | 0.268 | 0.04 | 8.9919774 | 28 | gstt1a |
| zgc:1368701 | 4.364460 | 0.326655062 | 0.173 | 0.018 | 1.1087912 | 28 | zgc:136870 |
| rac210 | 8.766979 | 0.786751674 | 0.685 | 0.223 | 2.2272511 | 28 | rac2 |

| | | | | | | | |
|---|---|---|---|---|---|---|---|
| tcima3 | 9.350597 | 0.60603854 | 0.362 | 0.07 | 2.3755193 | 28 | tcima |
| rgs182 | 1.154124 | 0.413184739 | 0.236 | 0.033 | 2.9320527 | 28 | rgs18 |
| mbnl32 | 1.395031 | 0.523493311 | 0.331 | 0.061 | 3.5440776 | 28 | mbnl3 |
| ccl39.2 | 9.295459 | 0.28237255 | 0.102 | 0.007 | 2.3615113 | 28 | ccl39.2 |
| zgc:640518 | 1.539425 | 0.733797425 | 0.567 | 0.161 | 3.9109117 | 28 | zgc:64051 |
| id2a2 | 2.451491 | 0.843298793 | 0.543 | 0.166 | 6.2280138 | 28 | id2a |
| ppiab4 | 3.449637 | 0.55631844 | 0.961 | 0.674 | 8.7638043 | 28 | ppiab |
| GNAZ | 6.750149 | 0.351806648 | 0.205 | 0.026 | 1.7148754 | 28 | GNAZ |
| zgc:638314 | 1.259069 | 0.568707265 | 0.386 | 0.084 | 3.1986658 | 28 | zgc:63831 |
| dennd2da | 2.798730 | 0.297409172 | 0.173 | 0.019 | 7.1101739 | 28 | dennd2da |
| mapk112 | 3.263592 | 0.311944939 | 0.165 | 0.019 | 8.2911560 | 28 | mapk11 |
| sept124 | 6.664070 | 0.513412737 | 0.323 | 0.064 | 1.6930070 | 28 | sept12 |
| si:cabz01074946.14 | 7.173626 | 0.488831709 | 0.315 | 0.06 | 1.8224596 | 28 | si:cabz01074946.1 |
| tent5ba1 | 7.200590 | 0.252575468 | 0.134 | 0.012 | 1.8293099 | 28 | tent5ba |
| ifi304 | 7.930612 | 0.796555738 | 0.583 | 0.194 | 2.0147720 | 28 | ifi30 |
| ncf42 | 1.076935 | 0.308096697 | 0.165 | 0.019 | 2.7359544 | 28 | ncf4 |
| cox6b17 | 2.732380 | 0.682380499 | 0.449 | 0.121 | 6.9416131 | 28 | cox6b1 |
| map3k14a1 | 7.801359 | 0.479661996 | 0.228 | 0.036 | 1.9819353 | 28 | map3k14a |
| sh3bgrl5 | 4.251178 | 0.631830702 | 0.441 | 0.12 | 1.0800119 | 28 | sh3bgrl |
| irf22 | 4.458431 | 0.529609717 | 0.339 | 0.074 | 1.1326645 | 28 | irf2 |
| zgc:153733 | 9.386222 | 0.26415184 | 0.157 | 0.018 | 2.3845697 | 28 | zgc:153733 |
| pycard7 | 1.065919 | 0.549215252 | 0.362 | 0.083 | 2.7079693 | 28 | pycard |
| arpc38 | 2.416257 | 0.738998067 | 0.748 | 0.332 | 6.1385009 | 28 | arpc3 |
| plekhj11 | 5.137425 | 0.426112724 | 0.276 | 0.051 | 1.3051628 | 28 | plekhj1 |
| parvg6 | 5.969529 | 0.46113522 | 0.339 | 0.072 | 1.5165590 | 28 | parvg |
| coro1a9 | 9.662558 | 0.712211346 | 0.693 | 0.234 | 2.4547730 | 28 | coro1a |
| si:ch1073-214b20.2 | 1.006611 | 0.273546465 | 0.189 | 0.025 | 2.5572970 | 28 | si:ch1073-214b20.2 |
| vsir6 | 1.842372 | 0.530989064 | 0.37 | 0.087 | 4.6805473 | 28 | vsir |
| lgals2a3 | 2.298881 | 0.866246341 | 0.465 | 0.137 | 5.8403086 | 28 | lgals2a |
| arhgdig9 | 6.450729 | 0.717443226 | 0.654 | 0.258 | 1.6388078 | 28 | arhgdig |
| xaf11 | 7.830456 | 0.460218934 | 0.291 | 0.061 | 1.9893275 | 28 | xaf1 |
| plekho1b3 | 1.871394 | 0.396965234 | 0.236 | 0.042 | 4.7542769 | 28 | plekho1b |
| CR318588.110 | 6.749654 | 0.920248459 | 0.575 | 0.214 | 1.7147497 | 28 | CR318588.1 |
| itgae.24 | 1.532871 | 0.359184035 | 0.213 | 0.036 | 3.8942595 | 28 | itgae.2 |
| sftpba3 | 2.913323 | 0.52867376 | 0.205 | 0.034 | 7.4012978 | 28 | sftpba |

| | | | | | | | |
|---|---|---|---|---|---|---|---|
| mrtfab2 | 3.416102 | 0.39527042 | 0.26 | 0.051 | 8.6786077 | 28 | mrtfab |
| capzb8 | 3.856933 | 0.665961122 | 0.654 | 0.27 | 9.7985398 | 28 | capzb |
| fcer1g2 | 4.468887 | 0.365723138 | 0.236 | 0.043 | 1.1353209 | 28 | fcer1g |
| XLOC-0331372 | 9.718745 | 0.321493781 | 0.205 | 0.034 | 2.4690473 | 28 | XLOC-033137 |
| ppt14 | 2.037180 | 0.618621681 | 0.52 | 0.18 | 5.1754559 | 28 | ppt1 |
| ccr9a7 | 2.305295 | 0.547997059 | 0.346 | 0.087 | 5.8566036 | 28 | ccr9a |
| wipf1a4 | 4.153884 | 0.436044963 | 0.299 | 0.067 | 1.0552943 | 28 | wipf1a |
| XLOC-0268364 | 4.476488 | 0.587549938 | 0.323 | 0.078 | 1.1372520 | 28 | XLOC-026836 |
| rplp211 | 4.731665 | 0.409550195 | 0.984 | 0.782 | 1.2020796 | 28 | rplp2 |
| hdac9b1 | 1.043290 | 0.316366265 | 0.173 | 0.026 | 2.6504801 | 28 | hdac9b |
| XLOC-0203014 | 1.138648 | 0.587153892 | 0.299 | 0.069 | 2.8927358 | 28 | XLOC-020301 |
| il12rb2l1 | 1.956511 | 0.361844049 | 0.22 | 0.041 | 4.9705182 | 28 | il12rb2l |
| pak2b | 2.569063 | 0.442084206 | 0.268 | 0.059 | 6.5267052 | 28 | pak2b |
| itm2bb7 | 3.061185 | 0.619505727 | 0.622 | 0.241 | 7.7769412 | 28 | itm2bb |
| cnn27 | 5.177867 | 0.563560561 | 0.472 | 0.156 | 1.3154372 | 28 | cnn2 |
| XLOC-0004318 | 6.216206 | 0.825671938 | 0.646 | 0.272 | 1.5792273 | 28 | XLOC-000431 |
| XLOC-0089932 | 6.535316 | 0.282240738 | 0.173 | 0.027 | 1.6602971 | 28 | XLOC-008993 |
| arpc46 | 9.555047 | 0.628858887 | 0.598 | 0.236 | 2.4274598 | 28 | arpc4 |
| atp6v1g12 | 1.448389 | 0.548846643 | 0.449 | 0.141 | 3.6796338 | 28 | atp6v1g1 |
| plbd11 | 3.345523 | 0.256989687 | 0.157 | 0.023 | 8.4993030 | 28 | plbd1 |
| dbn14 | 6.889227 | 0.453024244 | 0.276 | 0.065 | 1.7502082 | 28 | dbn1 |
| litaf4 | 1.617213 | 0.535708142 | 0.441 | 0.145 | 4.1085305 | 28 | litaf |
| ncf13 | 1.645665 | 0.322494709 | 0.189 | 0.033 | 4.1808135 | 28 | ncf1 |
| arpc4l7 | 1.853068 | 0.602792864 | 0.551 | 0.218 | 4.7077203 | 28 | arpc4l |
| capza1b6 | 2.088816 | 0.603666037 | 0.575 | 0.233 | 5.3066393 | 28 | capza1b |
| twf1a3 | 3.854690 | 0.503790436 | 0.472 | 0.156 | 9.7928402 | 28 | twf1a |
| wee2 | 4.010732 | 0.416320794 | 0.323 | 0.081 | 1.0189266 | 28 | wee2 |
| anxa3b1 | 1.032420 | 0.344660239 | 0.173 | 0.03 | 2.6228648 | 28 | anxa3b |
| si:dkey-262k9.47 | 1.045549 | 0.446538328 | 0.331 | 0.089 | 2.6562179 | 28 | si:dkey-262k9.4 |
| eif3jb2 | 1.064308 | 0.496381254 | 0.386 | 0.118 | 2.7038760 | 28 | eif3jb |
| myo1f5 | 1.257825 | 0.34242284 | 0.228 | 0.048 | 3.1955050 | 28 | myo1f |
| selplg4 | 1.668423 | 0.338511582 | 0.22 | 0.045 | 4.2386301 | 28 | selplg |
| BX322618.13 | 2.146013 | 0.389449751 | 0.189 | 0.035 | 5.4519466 | 28 | BX322618.1 |
| timm101 | 1.114162 | 0.383754863 | 0.268 | 0.064 | 2.8305297 | 28 | timm10 |
| COX7A2 (1 of many)3 | 1.243819 | 0.581993939 | 0.606 | 0.265 | 3.1599233 | 28 | COX7A2 (1 of many) |

| | | | | | | | |
|---|---|---|---|---|---|---|---|
| arl5c4 | 2.757429 | 0.491744206 | 0.354 | 0.107 | 7.0052501 | 28 | arl5c |
| triqk1 | 3.595595 | 0.392838738 | 0.236 | 0.055 | 9.1346105 | 28 | triqk |
| efhd22 | 4.294689 | 0.369136316 | 0.252 | 0.059 | 1.0910659 | 28 | efhd2 |
| fgl2a4 | 5.505164 | 0.412418751 | 0.291 | 0.075 | 1.3985870 | 28 | fgl2a |
| abracl5 | 6.028451 | 0.582829448 | 0.551 | 0.221 | 1.5315280 | 28 | abracl |
| malt1 | 1.231188 | 0.252375547 | 0.173 | 0.032 | 3.1278348 | 28 | malt1 |
| hcst2 | 1.688918 | 0.318709488 | 0.213 | 0.045 | 4.2906983 | 28 | hcst |
| ncaldb3 | 3.443007 | 0.375386957 | 0.276 | 0.07 | 8.7469601 | 28 | ncaldb |
| rgs137 | 5.297736 | 0.46715984 | 0.394 | 0.125 | 1.3458899 | 28 | rgs13 |
| XLOC-0054483 | 5.840109 | 0.29122696 | 0.157 | 0.028 | 1.4836797 | 28 | XLOC-005448 |
| marcksl1a9 | 6.368070 | 0.595931341 | 0.591 | 0.26 | 1.6178081 | 28 | marcksl1a |
| skap26 | 7.577458 | 0.412590367 | 0.291 | 0.08 | 1.9250532 | 28 | skap2 |
| BX908750.13 | 9.717732 | 0.357345193 | 0.165 | 0.031 | 2.4687900 | 28 | BX908750.1 |
| LOC1039091156 | 1.601924 | 0.491475882 | 0.457 | 0.168 | 4.0696880 | 28 | LOC103909115 |
| myl12.19 | 1.739628 | 0.556361103 | 0.764 | 0.425 | 4.4195254 | 28 | myl12.1 |
| lgals9l12 | 1.894599 | 0.255147461 | 0.157 | 0.028 | 4.8132295 | 28 | lgals9l1 |
| bach1b1 | 4.062905 | 0.276708384 | 0.15 | 0.027 | 1.0321812 | 28 | bach1b |
| gng21 | 6.256687 | 0.307822888 | 0.213 | 0.049 | 1.5895115 | 28 | gng2 |
| LOC1081907174 | 7.291005 | 0.459767678 | 0.291 | 0.082 | 1.8522798 | 28 | LOC108190717 |
| derl23 | 1.660706 | 0.425378655 | 0.339 | 0.106 | 4.2190246 | 28 | derl2 |
| bhlhe408 | 2.592563 | 0.539467843 | 0.433 | 0.167 | 6.5864065 | 28 | bhlhe40 |
| b2m10 | 2.603444 | 0.578676371 | 0.598 | 0.298 | 6.6140511 | 28 | b2m |
| gpr132b1 | 2.737022 | 0.294243763 | 0.173 | 0.035 | 6.9534059 | 28 | gpr132b |
| cfl19 | 2.766444 | 0.483992988 | 0.89 | 0.599 | 7.0281514 | 28 | cfl1 |
| arl8bb | 3.396345 | 0.280859953 | 0.189 | 0.041 | 8.6284144 | 28 | arl8bb |
| si:dkeyp-68b7.125 | 4.179627 | 0.32742381 | 0.236 | 0.06 | 1.0618344 | 28 | si:dkeyp-68b7.12 |
| rab5c3 | 4.248865 | 0.416741934 | 0.378 | 0.131 | 1.0794244 | 28 | rab5c |
| BX248082.14 | 4.426929 | 0.42898572 | 0.173 | 0.036 | 1.1246613 | 28 | BX248082.1 |
| cap18 | 5.067592 | 0.544299158 | 0.551 | 0.245 | 1.2874218 | 28 | cap1 |
| si:dkey-206d17.122 | 5.881889 | 0.288927286 | 0.213 | 0.051 | 1.4942940 | 28 | si:dkey-206d17.12 |
| XLOC-0418702 | 6.214644 | 0.350378017 | 0.157 | 0.031 | 1.5788304 | 28 | XLOC-041870 |
| itgb25 | 7.856309 | 0.415458877 | 0.339 | 0.108 | 1.9958953 | 28 | itgb2 |
| stk17b5 | 1.157541 | 0.418241356 | 0.346 | 0.113 | 2.9407336 | 28 | stk17b |
| sdcbp28 | 3.788500 | 0.505755483 | 0.512 | 0.227 | 9.6246851 | 28 | sdcbp2 |
| cmtm3 | 4.779371 | 0.28694737 | 0.197 | 0.047 | 1.2141993 | 28 | cmtm3 |

| | | | | | | | |
|---|---|---|---|---|---|---|---|
| XLOC-0320023 | 8.058985 | 0.340433903 | 0.189 | 0.044 | 2.0473853 | 28 | XLOC-032002 |
| pak2a1 | 1.368328 | 0.323871654 | 0.244 | 0.068 | 3.4762374 | 28 | pak2a |
| arf6b3 | 4.068546 | 0.364914803 | 0.331 | 0.113 | 1.0336142 | 28 | arf6b |
| tagapa6 | 4.205725 | 0.36639787 | 0.307 | 0.097 | 1.0684645 | 28 | tagapa |
| glrx6 | 4.239051 | 0.510513336 | 0.63 | 0.326 | 1.0769311 | 28 | glrx |
| XLOC-0108996 | 5.136514 | 0.525156962 | 0.535 | 0.239 | 1.3049316 | 28 | XLOC-010899 |
| gdi23 | 6.203851 | 0.470493773 | 0.465 | 0.202 | 1.5760884 | 28 | gdi2 |
| ptmab11 | 6.296232 | 0.474101485 | 0.874 | 0.569 | 1.5995579 | 28 | ptmab |
| asah1b2 | 7.651086 | 0.253971839 | 0.228 | 0.062 | 1.9437585 | 28 | asah1b |
| ostf12 | 7.696740 | 0.309952397 | 0.205 | 0.053 | 1.9553568 | 28 | ostf1 |
| mir21-13 | 9.169228 | 0.390348305 | 0.315 | 0.107 | 2.3294426 | 28 | mir21-1 |
| h1fx8 | 9.725337 | 0.53300961 | 0.449 | 0.19 | 2.4707219 | 28 | h1fx |
| csrp1a3 | 1.574975 | 0.360407924 | 0.26 | 0.081 | 4.0012263 | 28 | csrp1a |
| grap2a3 | 3.897661 | 0.265348015 | 0.189 | 0.048 | 9.9020084 | 28 | grap2a |
| ufc11 | 4.583404 | 0.336409242 | 0.276 | 0.088 | 1.1644139 | 28 | ufc1 |
| capgb5 | 5.102341 | 0.423432351 | 0.323 | 0.115 | 1.2962497 | 28 | capgb |
| crip111 | 9.661911 | 0.502831448 | 0.583 | 0.283 | 2.4546086 | 28 | crip1 |
| histh1l7 | 1.150832 | 0.422252695 | 0.331 | 0.122 | 2.9236911 | 28 | histh1l |
| prdx57 | 1.478409 | 0.427730836 | 0.417 | 0.178 | 3.7559001 | 28 | prdx5 |
| si:dkey-112e7.28 | 1.707432 | 0.453643511 | 0.465 | 0.214 | 4.3377326 | 28 | si:dkey-112e7.2 |
| cdc42l5 | 2.552098 | 0.442930135 | 0.496 | 0.233 | 6.4836058 | 28 | cdc42l |
| BX120005.13 | 2.965243 | 0.263650799 | 0.181 | 0.046 | 7.5332008 | 28 | BX120005.1 |
| rps27.113 | 3.698537 | 0.377781894 | 0.953 | 0.71 | 9.3961350 | 28 | rps27.1 |
| wdr16 | 5.433841 | 0.413035329 | 0.378 | 0.155 | 1.3804673 | 28 | wdr1 |
| atp6ap22 | 7.343731 | 0.329883484 | 0.299 | 0.106 | 1.8656749 | 28 | atp6ap2 |
| ctsk3 | 1.091715 | 0.28463428 | 0.236 | 0.074 | 2.7735043 | 28 | ctsk |
| gabarapa3 | 1.111299 | 0.429532655 | 0.409 | 0.181 | 2.8232566 | 28 | gabarapa |
| si:ch211-212k18.74 | 1.491039 | 0.361077561 | 0.37 | 0.146 | 3.7879870 | 28 | si:ch211-212k18.7 |
| ckba3 | 1.614053 | 0.254363241 | 0.236 | 0.073 | 4.1005022 | 28 | ckba |
| nr4a19 | 1.651244 | 0.450502223 | 0.378 | 0.158 | 4.1949860 | 28 | nr4a1 |
| fam53b2 | 1.676260 | 0.335285136 | 0.236 | 0.077 | 4.2585403 | 28 | fam53b |
| si:ch211-102c2.46 | 2.030368 | 0.40242734 | 0.37 | 0.146 | 5.1581499 | 28 | si:ch211-102c2.4 |
| laptm510 | 3.605784 | 0.404746924 | 0.417 | 0.176 | 9.1604963 | 28 | laptm5 |
| slc6a6b5 | 3.619134 | 0.342946441 | 0.276 | 0.097 | 9.1944122 | 28 | slc6a6b |
| si:ch211-222l21.111 | 4.344743 | 0.459301929 | 0.606 | 0.314 | 1.1037819 | 28 | si:ch211-222l21.1 |

| | | | | | | | |
|---|---|---|---|---|---|---|---|
| tsc22d37 | 4.424021 | 0.407946307 | 0.37 | 0.159 | 1.1239226 | 28 | tsc22d3 |
| pmaip11 | 5.591150 | 0.401040425 | 0.276 | 0.101 | 1.4204318 | 28 | pmaip1 |
| tagapb4 | 5.859853 | 0.30198208 | 0.22 | 0.07 | 1.4886958 | 28 | tagapb |
| RASSF55 | 5.926003 | 0.260246381 | 0.205 | 0.061 | 1.5055011 | 28 | RASSF5 |
| ctsh3 | 7.666653 | 0.304681274 | 0.213 | 0.067 | 1.9477132 | 28 | ctsh |
| GCA5 | 8.553936 | 0.370179589 | 0.441 | 0.2 | 2.1731276 | 28 | GCA |
| ckbb5 | 9.029357 | 0.433013081 | 0.535 | 0.25 | 2.2939081 | 28 | ckbb |
| cnpy32 | 1.039966 | 0.267010567 | 0.189 | 0.055 | 2.6420356 | 28 | cnpy3 |
| mvb12a1 | 1.278799 | 0.261490247 | 0.236 | 0.078 | 3.2487888 | 28 | mvb12a |
| rps15a11 | 1.427995 | 0.318817703 | 0.976 | 0.797 | 3.6278231 | 28 | rps15a |
| rab8b3 | 1.919918 | 0.304853591 | 0.244 | 0.085 | 4.8775531 | 28 | rab8b |
| plekho24 | 3.340591 | 0.28580883 | 0.228 | 0.077 | 8.4867728 | 28 | plekho2 |
| sri2 | 4.334548 | 0.390068117 | 0.362 | 0.161 | 1.1011921 | 28 | sri |
| psme26 | 1.118517 | 0.333465334 | 0.283 | 0.113 | 2.8415926 | 28 | psme2 |
| dusp54 | 1.121607 | 0.338973406 | 0.205 | 0.068 | 2.8494443 | 28 | dusp5 |
| actr2a7 | 1.228918 | 0.330142487 | 0.323 | 0.133 | 3.1220681 | 28 | actr2a |
| naga2 | 1.742530 | 0.252877041 | 0.181 | 0.056 | 4.4268976 | 28 | naga |
| CR392341.15 | 2.059278 | 0.252745121 | 0.189 | 0.06 | 5.2315963 | 28 | CR392341.1 |
| ptprc8 | 2.623639 | 0.322613293 | 0.307 | 0.125 | 6.6653554 | 28 | ptprc |
| ecm1b4 | 2.769777 | 0.420928131 | 0.197 | 0.067 | 7.0366188 | 28 | ecm1b |
| nptna3 | 2.887330 | 0.286395407 | 0.205 | 0.068 | 7.3352624 | 28 | nptna |
| tpi1b6 | 2.987752 | 0.372175346 | 0.378 | 0.179 | 7.5903844 | 28 | tpi1b |
| fcer1gl6 | 3.721729 | 0.283884518 | 0.441 | 0.182 | 9.4550544 | 28 | fcer1gl |
| fam49ba5 | 4.154364 | 0.339911808 | 0.268 | 0.105 | 0.000106 | 28 | fam49ba |
| rps2115 | 5.285180 | 0.313428701 | 0.969 | 0.775 | 0.000134 | 28 | rps21 |
| rpl1811 | 6.678919 | 0.284155805 | 0.961 | 0.792 | 0.00017 | 28 | rpl18 |
| bloc1s62 | 6.826039 | 0.260310521 | 0.252 | 0.096 | 0.000173 | 28 | bloc1s6 |
| rps39 | 7.013080 | 0.339109892 | 0.976 | 0.725 | 0.000178 | 28 | rps3 |
| clic16 | 1.074503 | 0.321646095 | 0.299 | 0.129 | 0.000273 | 28 | clic1 |
| ccng23 | 1.108840 | 0.267228384 | 0.228 | 0.085 | 0.000282 | 28 | ccng2 |
| sod13 | 1.174775 | 0.402902846 | 0.567 | 0.345 | 0.000298 | 28 | sod1 |
| rps512 | 1.267304 | 0.314150328 | 0.945 | 0.728 | 0.000322 | 28 | rps5 |
| si:ch73-1a9.314 | 1.516402 | 0.386626183 | 0.567 | 0.315 | 0.000385 | 28 | si:ch73-1a9.3 |
| commd3 | 1.559542 | 0.255011647 | 0.276 | 0.111 | 0.000396 | 28 | commd3 |
| mtpn4 | 2.165236 | 0.336916732 | 0.346 | 0.164 | 0.00055 | 28 | mtpn |

| | | | | | | | |
|---|---|---|---|---|---|---|---|
| cox8a9 | 2.329784 | 0.387702084 | 0.48 | 0.267 | 0.000592 | 28 | cox8a |
| rps223 | 2.345347 | 0.270641675 | 0.992 | 0.841 | 0.000596 | 28 | rps2 |
| rps176 | 2.987744 | 0.266845744 | 0.953 | 0.779 | 0.000759 | 28 | rps17 |
| sept64 | 3.426442 | 0.328266641 | 0.409 | 0.202 | 0.00087 | 28 | sept6 |
| mcl1a6 | 3.922659 | 0.338195031 | 0.819 | 0.613 | 0.000997 | 28 | mcl1a |
| oaz1b9 | 5.100717 | 0.351807086 | 0.614 | 0.36 | 0.001296 | 28 | oaz1b |
| romo18 | 6.135529 | 0.364214254 | 0.638 | 0.408 | 0.001559 | 28 | romo1 |
| ywhabl4 | 9.660962 | 0.361726534 | 0.512 | 0.301 | 0.002454 | 28 | ywhabl |
| capza1a6 | 1.051345 | 0.334985329 | 0.417 | 0.221 | 0.002671 | 28 | capza1a |
| mapre1a6 | 1.147563 | 0.291732193 | 0.409 | 0.196 | 0.002915 | 28 | mapre1a |
| cox4i16 | 1.193766 | 0.355096448 | 0.661 | 0.417 | 0.003033 | 28 | cox4i1 |
| rps26l10 | 1.283216 | 0.32918138 | 0.945 | 0.699 | 0.00326 | 28 | rps26l |
| rpl18a10 | 1.409482 | 0.254959999 | 0.945 | 0.801 | 0.003581 | 28 | rpl18a |
| eif3f10 | 1.414207 | 0.359432114 | 0.693 | 0.466 | 0.003593 | 28 | eif3f |
| ube2e36 | 1.431244 | 0.285219427 | 0.354 | 0.17 | 0.003636 | 28 | ube2e3 |
| sec61g4 | 1.526125 | 0.287251611 | 0.85 | 0.681 | 0.003877 | 28 | sec61g |
| arhgdia7 | 1.580753 | 0.340780659 | 0.457 | 0.252 | 0.004016 | 28 | arhgdia |
| vma211 | 1.815686 | 0.274426758 | 0.26 | 0.112 | 0.004613 | 28 | vma21 |
| eif3g7 | 2.604223 | 0.334642367 | 0.52 | 0.3 | 0.006616 | 28 | eif3g |
| rpl219 | 2.824801 | 0.264455654 | 0.961 | 0.776 | 0.007176 | 28 | rpl21 |
| mob1a4 | 3.624196 | 0.276556864 | 0.354 | 0.173 | 0.009207 | 28 | mob1a |
| rps98 | 4.020847 | 0.260040882 | 0.961 | 0.79 | 0.010215 | 28 | rps9 |
| glula5 | 4.560864 | 0.281893767 | 0.283 | 0.125 | 0.011587 | 28 | glula |
| ccni8 | 4.703516 | 0.347387795 | 0.52 | 0.319 | 0.011949 | 28 | ccni |
| psma21 | 6.092123 | 0.295621325 | 0.331 | 0.164 | 0.015477 | 28 | psma2 |
| rpl3611 | 8.199683 | 0.276987445 | 0.984 | 0.806 | 0.020831 | 28 | rpl36 |
| tln15 | 8.634024 | 0.26625804 | 0.299 | 0.143 | 0.021935 | 28 | tln1 |
| cox5aa3 | 1.059959 | 0.320853587 | 0.591 | 0.358 | 0.026928 | 28 | cox5aa |
| ak12 | 1.119325 | 0.260137364 | 0.228 | 0.099 | 0.028436 | 28 | ak1 |
| serf27 | 1.849349 | 0.310363472 | 0.614 | 0.378 | 0.046983 | 28 | serf2 |
| atp6v0ca3 | 2.549581 | 0.286177625 | 0.386 | 0.204 | 0.064772 | 28 | atp6v0ca |
| uqcrq4 | 2.953978 | 0.3092331 | 0.48 | 0.287 | 0.075046 | 28 | uqcrq |
| rps1512 | 2.978358 | 0.308361173 | 0.953 | 0.716 | 0.075665 | 28 | rps15 |
| hmgn210 | 2.979965 | 0.360508523 | 0.685 | 0.468 | 0.075706 | 28 | hmgn2 |
| fosl25 | 4.442091 | 0.269615635 | 0.244 | 0.114 | 0.112851 | 28 | fosl2 |

| | | | | | | | |
|---|---|---|---|---|---|---|---|
| fth1a1 | 5.28760[illegible] | 0.297049695 | 0.795 | 0.612 | 0.134332 | 28 | fth1a |
| ponzr19 | 6.84683[illegible] | 0.285856947 | 0.504 | 0.29 | 0.173944 | 28 | ponzr1 |
| XLOC-0425956 | 9.57630[illegible] | 0.348889729 | 0.299 | 0.154 | 0.243286 | 28 | XLOC-042595 |
| naca11 | 9.68900[illegible] | 0.276584122 | 0.843 | 0.652 | 0.246149 | 28 | naca |
| magt13 | 1.00142[illegible] | 0.254520802 | 0.26 | 0.129 | 0.254411 | 28 | magt1 |
| ssr32 | 1.01524[illegible] | 0.288773627 | 0.48 | 0.288 | 0.257922 | 28 | ssr3 |
| rhoab7 | 1.02423[illegible] | 0.28296545 | 0.37 | 0.208 | 0.260206 | 28 | rhoab |
| cdc425 | 1.11770[illegible] | 0.265926422 | 0.409 | 0.234 | 0.283952 | 28 | cdc42 |
| rpl2212 | 1.13725[illegible] | 0.259499178 | 0.913 | 0.72 | 0.28892 | 28 | rpl22 |
| cox7b5 | 1.37432[illegible] | 0.298167754 | 0.638 | 0.427 | 0.349147 | 28 | cox7b |
| rps148 | 1.39994[illegible] | 0.257564664 | 0.976 | 0.781 | 0.355656 | 28 | rps14 |
| b2ml9 | 1.52472[illegible] | 0.275286074 | 0.488 | 0.294 | 0.387357 | 28 | b2ml |
| tagln210 | 1.54438[illegible] | 0.290913169 | 0.37 | 0.21 | 0.392351 | 28 | tagln2 |
| rpl35a9 | 1.619901 | 0.256291586 | 0.976 | 0.781 | 0.411536 | 28 | rpl35a |
| COX5B5 | 1.73277[illegible] | 0.286310331 | 0.591 | 0.377 | 0.44021 | 28 | COX5B |
| rpl389 | 1.77594[illegible] | 0.262029555 | 0.953 | 0.753 | 0.45118 | 28 | rpl38 |
| cst14a.21 | 1.87764[illegible] | 0.275971587 | 0.409 | 0.236 | 0.477016 | 28 | cst14a.2 |
| sec61a12 | 2.11422[illegible] | 0.256314184 | 0.346 | 0.193 | 0.537118 | 28 | sec61a1 |
| zgc:1537132 | 2.41660[illegible] | 0.25720734 | 0.315 | 0.173 | 0.613939 | 28 | zgc:153713 |
| rpl1510 | 2.95240[illegible] | 0.261756764 | 0.969 | 0.763 | 0.750059 | 28 | rpl15 |
| celf211 | 3.20236[illegible] | 0.279011843 | 0.425 | 0.258 | 0.81356 | 28 | celf2 |
| sh3bgrl34 | 3.28424[illegible] | 0.258440231 | 0.291 | 0.156 | 0.834363 | 28 | sh3bgrl3 |
| dusp17 | 4.367041 | 0.252097492 | 0.283 | 0.15 | 1 | 28 | dusp1 |
| ndufb33 | 4.78717[illegible] | 0.268220051 | 0.394 | 0.239 | 1 | 28 | ndufb3 |
| sumo2b10 | 5.10040[illegible] | 0.266502576 | 0.465 | 0.297 | 1 | 28 | sumo2b |
| ybx17 | 5.84513[illegible] | 0.259788279 | 0.913 | 0.695 | 1 | 28 | ybx1 |
| lasp17 | 0.0001 | 0.261886413 | 0.402 | 0.254 | 1 | 28 | lasp1 |
| rps4x11 | 0.0001 | 0.255115893 | 0.945 | 0.736 | 1 | 28 | rps4x |
| rps1612 | 0.0002 | 0.256685136 | 0.969 | 0.765 | 1 | 28 | rps16 |
| zgc:11418815 | 0.0002 | 0.289120597 | 0.913 | 0.668 | 1 | 28 | zgc:114188 |
| pde4ba8 | 0.0002 | 0.251403873 | 0.26 | 0.143 | 1 | 28 | pde4ba |
| rpl27a10 | 0.0003 | 0.275167294 | 0.772 | 0.522 | 1 | 28 | rpl27a |
| rpl915 | 0.0004 | 0.279845784 | 0.906 | 0.668 | 1 | 28 | rpl9 |
| bcl2l103 | 0.0004 | 0.291799452 | 0.323 | 0.201 | 1 | 28 | bcl2l10 |
| mibp211 | 0.0006 | 0.250977274 | 0.709 | 0.521 | 1 | 28 | mibp2 |

| | | | | | | | |
|---|---|---|---|---|---|---|---|
| XLOC-0139646 | 0.0015 | 0.282087194 | 0.276 | 0.171 | 1 | 28 | XLOC-013964 |
| krt5 | 0 | 2.257418537 | 0.827 | 0.007 | 0 | 29 | krt5 |
| si:dkey-248g15.3 | 0 | 1.650148387 | 0.555 | 0.004 | 0 | 29 | si:dkey-248g15.3 |
| si:dkey-193p11.2 | 0 | 1.512085146 | 0.527 | 0.01 | 0 | 29 | si:dkey-193p11.2 |
| icn21 | 0 | 1.490929592 | 0.5 | 0.006 | 0 | 29 | icn2 |
| cldnb | 0 | 1.42611035 | 0.491 | 0.008 | 0 | 29 | cldnb |
| krt97 | 0 | 1.336218008 | 0.436 | 0.004 | 0 | 29 | krt97 |
| spaca4l | 0 | 1.224853456 | 0.582 | 0.007 | 0 | 29 | spaca4l |
| krt17 | 0 | 1.037076738 | 0.409 | 0.006 | 0 | 29 | krt17 |
| CT027832.1 | 0 | 0.940872453 | 0.245 | 0 | 0 | 29 | CT027832.1 |
| apnl | 0 | 0.927866803 | 0.345 | 0.001 | 0 | 29 | apnl |
| pkp3a | 0 | 0.921044036 | 0.491 | 0.007 | 0 | 29 | pkp3a |
| LO017965.1 | 0 | 0.900015351 | 0.345 | 0.001 | 0 | 29 | LO017965.1 |
| cldna | 0 | 0.863856098 | 0.327 | 0.001 | 0 | 29 | cldna |
| zgc:92380 | 0 | 0.807560057 | 0.4 | 0.009 | 0 | 29 | zgc:92380 |
| krtt1c19e | 0 | 0.752038941 | 0.236 | 0.001 | 0 | 29 | krtt1c19e |
| zgc:1018101 | 0 | 0.751961746 | 0.382 | 0.004 | 0 | 29 | zgc:101810 |
| zgc:193505 | 0 | 0.718619731 | 0.327 | 0.005 | 0 | 29 | zgc:193505 |
| si:dkey-33c14.3 | 0 | 0.703438054 | 0.227 | 0.001 | 0 | 29 | si:dkey-33c14.3 |
| ms4a17a.5 | 0 | 0.619899214 | 0.336 | 0.005 | 0 | 29 | ms4a17a.5 |
| cldn1 | 0 | 0.602595614 | 0.227 | 0 | 0 | 29 | cldn1 |
| caspb | 0 | 0.596133233 | 0.309 | 0.001 | 0 | 29 | caspb |
| CU682640.1 | 0 | 0.544602322 | 0.227 | 0 | 0 | 29 | CU682640.1 |
| gpa33a | 0 | 0.533717965 | 0.318 | 0.002 | 0 | 29 | gpa33a |
| cxl34b.11 | 0 | 0.530740448 | 0.264 | 0 | 0 | 29 | cxl34b.11 |
| rhbg | 0 | 0.49990479 | 0.218 | 0.001 | 0 | 29 | rhbg |
| zgc:162941 | 0 | 0.435278931 | 0.182 | 0 | 0 | 29 | zgc:162941 |
| si:ch211-95j8.3 | 0 | 0.417015236 | 0.145 | 0 | 0 | 29 | si:ch211-95j8.3 |
| CU929145.1 | 9.648792 | 0.412582112 | 0.118 | 0 | 2.4512757 | 29 | CU929145.1 |
| RGS13 (1 of many) | 2.109616 | 0.406338286 | 0.182 | 0.001 | 5.3594807 | 29 | RGS13 (1 of many) |
| ms4a17a.4 | 5.292570 | 0.329389357 | 0.227 | 0.002 | 1.3445774 | 29 | ms4a17a.4 |
| LOC110438226 | 6.921370 | 0.320618012 | 0.1 | 0 | 1.7583742 | 29 | LOC110438226 |
| ms4a17a.2 | 1.721183 | 0.273383457 | 0.145 | 0.001 | 4.3726676 | 29 | ms4a17a.2 |
| si:ch211-243g18.2 | 2.166123 | 0.46413098 | 0.145 | 0.001 | 5.5030379 | 29 | si:ch211-243g18.2 |
| LOC103910192 | 1.230792 | 0.809208444 | 0.164 | 0.001 | 3.1268280 | 29 | LOC103910192 |

| | | | | | | | |
|---|---|---|---|---|---|---|---|
| eevs | 2.389447 | 0.256350129 | 0.145 | 0.001 | 6.0703913 | 29 | eevs |
| si:ch73-106l15.4 | 1.412281 | 0.692430962 | 0.145 | 0.001 | 3.5879010 | 29 | si:ch73-106l15.4 |
| cd44a | 8.739400 | 0.332372144 | 0.173 | 0.001 | 2.2202446 | 29 | cd44a |
| si:dkeyp-93a5.2 | 1.567830 | 0.321905172 | 0.109 | 0 | 3.9830735 | 29 | si:dkeyp-93a5.2 |
| ccl25a | 2.648033 | 0.285282271 | 0.1 | 0 | 6.7273294 | 29 | ccl25a |
| fabp1b.2 | 8.070130 | 0.457492558 | 0.127 | 0.001 | 2.0502165 | 29 | fabp1b.2 |
| pou2f3 | 1.734207 | 0.363066656 | 0.155 | 0.001 | 4.4057529 | 29 | pou2f3 |
| pof1b | 9.678532 | 0.645074168 | 0.345 | 0.009 | 2.4588312 | 29 | pof1b |
| col17a1b | 1.953676 | 0.391608533 | 0.145 | 0.001 | 4.9633152 | 29 | col17a1b |
| tp63 | 7.876283 | 0.565071224 | 0.318 | 0.009 | 2.0009698 | 29 | tp63 |
| gsnb | 8.045770 | 0.276503967 | 0.155 | 0.001 | 2.0440279 | 29 | gsnb |
| c6ast1 | 1.158589 | 0.69983327 | 0.2 | 0.003 | 2.9433974 | 29 | c6ast1 |
| si:ch211-241e1.3 | 6.815303 | 0.582202984 | 0.227 | 0.004 | 1.7314278 | 29 | si:ch211-241e1.3 |
| ptgdsb.1 | 1.338359 | 1.205504241 | 0.309 | 0.009 | 3.4001011 | 29 | ptgdsb.1 |
| wu:fb18f06 | 1.943153 | 1.408308921 | 0.655 | 0.046 | 4.9365820 | 29 | wu:fb18f06 |
| si:ch211-264f5.61 | 2.495856 | 0.557856266 | 0.345 | 0.012 | 6.3407237 | 29 | si:ch211-264f5.6 |
| lamc2 | 5.264649 | 0.434847669 | 0.164 | 0.002 | 1.3374841 | 29 | lamc2 |
| aqp3a1 | 4.443633 | 0.997763221 | 0.491 | 0.026 | 1.1289051 | 29 | aqp3a |
| gdpd3a | 9.671006 | 0.333075471 | 0.173 | 0.002 | 2.4569191 | 29 | gdpd3a |
| si:ch1073-406l10.2 | 3.471891 | 0.700761896 | 0.318 | 0.011 | 8.8203394 | 29 | si:ch1073-406l10.2 |
| si:ch211-213d14.2 | 1.355386 | 0.394911213 | 0.236 | 0.005 | 3.4433585 | 29 | si:ch211-213d14.2 |
| krt914 | 1.593402 | 1.943152506 | 0.727 | 0.068 | 4.0480380 | 29 | krt91 |
| si:dkeyp-93a5.3 | 3.164390 | 0.638713824 | 0.118 | 0.001 | 8.0391338 | 29 | si:dkeyp-93a5.3 |
| si:busm1-71b9.3 | 5.123049 | 0.456387674 | 0.218 | 0.005 | 1.3015107 | 29 | si:busm1-71b9.3 |
| scpp71 | 1.400623 | 1.617579604 | 0.418 | 0.02 | 3.5582831 | 29 | scpp7 |
| zgc:77439 | 3.180300 | 0.686850416 | 0.355 | 0.014 | 8.0795545 | 29 | zgc:77439 |
| dsg2.1 | 1.815743 | 0.88229504 | 0.418 | 0.021 | 4.6128959 | 29 | dsg2.1 |
| s100w1 | 2.028033 | 0.675325984 | 0.382 | 0.018 | 5.1522181 | 29 | s100w |
| si:ch211-155m12.1 | 6.357123 | 0.546482818 | 0.109 | 0.001 | 1.6150273 | 29 | si:ch211-155m12.1 |
| aldob1 | 3.125192 | 1.069706623 | 0.5 | 0.033 | 7.9395506 | 29 | aldob |
| bik | 2.944986 | 0.505227365 | 0.264 | 0.009 | 7.4817376 | 29 | bik |
| zgc:153911 | 3.043910 | 0.866187043 | 0.345 | 0.016 | 7.7330534 | 29 | zgc:153911 |
| evpla | 1.155966 | 0.343103306 | 0.2 | 0.005 | 2.9367325 | 29 | evpla |
| cldni | 3.013986 | 0.982298276 | 0.282 | 0.01 | 7.6570335 | 29 | cldni |
| fermt1 | 1.788654 | 0.250329162 | 0.127 | 0.002 | 4.5440776 | 29 | fermt1 |

| | | | | | | | |
|---|---|---|---|---|---|---|---|
| perp4 | 3.031035 | 1.132937396 | 0.618 | 0.058 | 7.7003465 | 29 | perp |
| zgc:158412 | 3.248161 | 0.265007406 | 0.118 | 0.001 | 8.2519540 | 29 | zgc:158412 |
| cldnh | 2.809547 | 0.51201689 | 0.136 | 0.002 | 7.1376550 | 29 | cldnh |
| dsc2l1 | 3.922024 | 0.767551063 | 0.409 | 0.025 | 9.9639036 | 29 | dsc2l |
| FP017215.1 | 1.312589 | 0.522991101 | 0.3 | 0.013 | 3.3346345 | 29 | FP017215.1 |
| anxa1b | 2.678794 | 0.50057678 | 0.218 | 0.006 | 6.8054778 | 29 | anxa1b |
| selenow2b | 4.174385 | 0.345078649 | 0.173 | 0.004 | 1.0605025 | 29 | selenow2b |
| tmem238a1 | 5.132180 | 0.507239472 | 0.327 | 0.016 | 1.3038303 | 29 | tmem238a |
| dspa2 | 9.379846 | 0.73710927 | 0.418 | 0.027 | 2.3829500 | 29 | dspa |
| tmem176l.4 | 9.462602 | 0.490946147 | 0.245 | 0.009 | 2.4039742 | 29 | tmem176l.4 |
| si:ch211-12e13.10 | 1.815790 | 0.350473289 | 0.1 | 0.001 | 4.6130149 | 29 | si:ch211-12e13.10 |
| limk2 | 1.902029 | 0.511370529 | 0.318 | 0.017 | 4.8321047 | 29 | limk2 |
| ppl | 1.204463 | 0.40747953 | 0.236 | 0.009 | 3.0599388 | 29 | ppl |
| slc16a12b | 3.769013 | 0.317090196 | 0.209 | 0.007 | 9.5751782 | 29 | slc16a12b |
| lad1 | 7.430466 | 0.470181867 | 0.136 | 0.003 | 1.8877099 | 29 | lad1 |
| cfd4 | 1.339589 | 1.40232094 | 0.591 | 0.068 | 3.4032268 | 29 | cfd |
| myh11b | 4.744452 | 0.600490615 | 0.155 | 0.004 | 1.2053281 | 29 | myh11b |
| f2rl1.2 | 4.788354 | 0.558611625 | 0.327 | 0.019 | 1.2164813 | 29 | f2rl1.2 |
| cavin2a | 2.107845 | 0.512251673 | 0.291 | 0.016 | 5.3549813 | 29 | cavin2a |
| cfl1l5 | 1.138073 | 1.50730307 | 0.773 | 0.133 | 2.8912766 | 29 | cfl1l |
| icn7 | 9.827225 | 1.801420744 | 0.955 | 0.264 | 2.4966065 | 29 | icn |
| zgc:172339 | 5.210553 | 0.529793182 | 0.136 | 0.003 | 1.3237410 | 29 | zgc:172339 |
| mocs2 | 6.838461 | 0.309250677 | 0.182 | 0.006 | 1.7373112 | 29 | mocs2 |
| bnipl | 8.042859 | 0.271166867 | 0.173 | 0.005 | 2.0432884 | 29 | bnipl |
| gpd1c | 4.240139 | 0.415878398 | 0.291 | 0.017 | 1.0772075 | 29 | gpd1c |
| ak13 | 1.332753 | 1.100300323 | 0.673 | 0.095 | 3.3858611 | 29 | ak1 |
| dnase1l4.1 | 7.133538 | 0.314685144 | 0.191 | 0.007 | 1.8122754 | 29 | dnase1l4.1 |
| gata1a | 1.014402 | 0.316123025 | 0.118 | 0.002 | 2.5770905 | 29 | gata1a |
| elf3 | 2.054193 | 0.416528832 | 0.173 | 0.006 | 5.2186795 | 29 | elf3 |
| mid1ip1a | 1.376088 | 0.476184743 | 0.245 | 0.013 | 3.4959534 | 29 | mid1ip1a |
| LOC100538255 | 2.253849 | 0.880590993 | 0.155 | 0.005 | 5.7259039 | 29 | LOC100538255 |
| si:ch211-69b7.6 | 4.312255 | 0.484160255 | 0.173 | 0.006 | 1.0955285 | 29 | si:ch211-69b7.6 |
| tmsb18 | 7.062335 | 1.621914893 | 0.764 | 0.157 | 1.7941863 | 29 | tmsb1 |
| serpinb1l4 | 1.226797 | 0.253454287 | 0.155 | 0.005 | 3.1166787 | 29 | serpinb1l4 |
| apoeb2 | 3.885382 | 0.784867056 | 0.273 | 0.017 | 9.8708142 | 29 | apoeb |

| | | | | | | | |
|---|---|---|---|---|---|---|---|
| jupa1 | 1.424111 | 0.801715303 | 0.491 | 0.057 | 3.6179556 | 29 | jupa |
| tdh | 1.634311 | 0.945042736 | 0.609 | 0.089 | 4.1519689 | 29 | tdh |
| si:dkey-157l19.2 | 1.855115 | 0.286144819 | 0.191 | 0.009 | 4.7129211 | 29 | si:dkey-157l19.2 |
| cav21 | 3.840993 | 0.469658098 | 0.255 | 0.016 | 9.7580435 | 29 | cav2 |
| krt43 | 8.552983 | 1.044105253 | 0.445 | 0.048 | 2.1728854 | 29 | krt4 |
| fat2 | 2.131331 | 0.266427844 | 0.173 | 0.007 | 5.4146482 | 29 | fat2 |
| p2rx4b | 4.099794 | 0.353854337 | 0.127 | 0.004 | 1.0415528 | 29 | p2rx4b |
| plecb2 | 6.805464 | 0.816529794 | 0.445 | 0.05 | 1.7289281 | 29 | plecb |
| arg21 | 1.461720 | 0.50068094 | 0.264 | 0.017 | 3.7135016 | 29 | arg2 |
| cdh12 | 1.517060 | 0.737148201 | 0.473 | 0.053 | 3.8540910 | 29 | cdh1 |
| spint1a | 3.153205 | 0.678796148 | 0.455 | 0.049 | 8.0107179 | 29 | spint1a |
| ptgs2a | 5.017042 | 0.407018283 | 0.209 | 0.011 | 1.2745796 | 29 | ptgs2a |
| arhgef53 | 2.439203 | 0.54256563 | 0.345 | 0.031 | 6.1967971 | 29 | arhgef5 |
| gcshb1 | 2.639205 | 0.413515365 | 0.236 | 0.014 | 6.7049009 | 29 | gcshb |
| tnks1bp1 | 1.119303 | 0.427972619 | 0.173 | 0.008 | 2.8435900 | 29 | tnks1bp1 |
| casz11 | 3.314975 | 0.474636324 | 0.264 | 0.019 | 8.4216954 | 29 | casz1 |
| zgc:1533173 | 2.198081 | 0.610263271 | 0.373 | 0.038 | 5.5842271 | 29 | zgc:153317 |
| si:ch211-166a6.52 | 2.655855 | 0.577980894 | 0.364 | 0.036 | 6.7472000 | 29 | si:ch211-166a6.5 |
| itgb42 | 9.692660 | 0.669489317 | 0.236 | 0.016 | 2.4624204 | 29 | itgb4 |
| capn2b1 | 2.455311 | 0.503758477 | 0.291 | 0.025 | 6.2377193 | 29 | capn2b |
| wu:fi04e121 | 1.622064 | 0.385187916 | 0.264 | 0.021 | 4.1208554 | 29 | wu:fi04e12 |
| serpinb14 | 4.360427 | 0.378446531 | 0.209 | 0.014 | 1.1077666 | 29 | serpinb14 |
| pon1 | 1.217807 | 0.337876782 | 0.173 | 0.009 | 3.0938409 | 29 | pon1 |
| s100v27 | 1.904454 | 1.026057342 | 0.655 | 0.138 | 4.8382667 | 29 | s100v2 |
| CT573383.11 | 1.625781 | 0.658305336 | 0.182 | 0.011 | 4.1302969 | 29 | CT573383.1 |
| scinla | 5.137968 | 0.319872823 | 0.191 | 0.012 | 1.3053010 | 29 | scinla |
| itga6b | 2.774584 | 0.523098848 | 0.218 | 0.016 | 7.0488320 | 29 | itga6b |
| phlda23 | 2.006796 | 0.857147679 | 0.391 | 0.051 | 5.0982663 | 29 | phlda2 |
| si:dkey-16p21.8 | 2.325986 | 0.752269147 | 0.464 | 0.068 | 5.9091682 | 29 | si:dkey-16p21.8 |
| si:dkey-16l2.201 | 5.052937 | 0.386599062 | 0.255 | 0.022 | 1.2836987 | 29 | si:dkey-16l2.20 |
| dlx3b1 | 5.071296 | 0.252592173 | 0.155 | 0.008 | 1.2883629 | 29 | dlx3b |
| si:ch211-39i22.14 | 7.903456 | 0.828512383 | 0.518 | 0.086 | 2.0078730 | 29 | si:ch211-39i22.1 |
| ptprfb1 | 9.327740 | 0.324407324 | 0.218 | 0.016 | 2.3697125 | 29 | ptprfb |
| tnfrsf11b | 5.222875 | 0.436515044 | 0.127 | 0.006 | 1.3268715 | 29 | tnfrsf11b |
| ano9b | 1.435612 | 0.31385633 | 0.136 | 0.006 | 3.6471745 | 29 | ano9b |

| | | | | | | | |
|---|---|---|---|---|---|---|---|
| tuft1a1 | 4.529451 | 0.450440269 | 0.291 | 0.029 | 1.1507070 | 29 | tuft1a |
| XLOC-041832 | 9.678874 | 0.258388637 | 0.155 | 0.009 | 2.4589180 | 29 | XLOC-041832 |
| CABZ01048956.1 | 4.883739 | 0.25432763 | 0.1 | 0.004 | 1.2407138 | 29 | CABZ01048956.1 |
| znf185 | 1.207393 | 0.258300002 | 0.118 | 0.005 | 3.0673838 | 29 | znf185 |
| ptgs2b1 | 8.537063 | 0.318877414 | 0.164 | 0.01 | 2.1688409 | 29 | ptgs2b |
| cd9b3 | 1.021307 | 0.596702003 | 0.418 | 0.059 | 2.5946323 | 29 | cd9b |
| lgals3b2 | 4.430509 | 0.76569479 | 0.509 | 0.097 | 1.1255709 | 29 | lgals3b |
| cdo11 | 5.020770 | 0.253889925 | 0.136 | 0.007 | 1.2755268 | 29 | cdo1 |
| ptgis1 | 1.089982 | 0.348446205 | 0.191 | 0.015 | 2.7691003 | 29 | ptgis |
| hepacam21 | 2.147023 | 0.292826525 | 0.109 | 0.005 | 5.4545136 | 29 | hepacam2 |
| efna1b2 | 2.009969 | 0.330251981 | 0.209 | 0.019 | 5.1063275 | 29 | efna1b |
| lmna1 | 7.307544 | 0.28798994 | 0.173 | 0.013 | 1.8564815 | 29 | lmna |
| si:dkey-262k9.21 | 2.084371 | 0.48312575 | 0.273 | 0.032 | 5.2953460 | 29 | si:dkey-262k9.2 |
| ahnak8 | 8.802274 | 0.891218081 | 0.564 | 0.135 | 2.2362177 | 29 | ahnak |
| pnp5a4 | 3.871884 | 0.782805467 | 0.409 | 0.074 | 9.8365230 | 29 | pnp5a |
| srd5a2a1 | 6.686719 | 0.639256496 | 0.455 | 0.083 | 1.6987611 | 29 | srd5a2a |
| sult2st11 | 1.513180 | 0.388024106 | 0.255 | 0.029 | 3.8442339 | 29 | sult2st1 |
| cldne | 1.803044 | 0.261375447 | 0.109 | 0.006 | 4.5806336 | 29 | cldne |
| tuba8l23 | 2.428770 | 0.561721542 | 0.4 | 0.067 | 6.1702902 | 29 | tuba8l2 |
| anxa2a7 | 3.711468 | 0.837120686 | 0.527 | 0.116 | 9.4289865 | 29 | anxa2a |
| klf33 | 2.259111 | 0.444351217 | 0.291 | 0.038 | 5.7392717 | 29 | klf3 |
| eppk12 | 1.035596 | 0.488274316 | 0.3 | 0.042 | 2.6309320 | 29 | eppk1 |
| amotl2b | 2.562383 | 0.280728653 | 0.173 | 0.015 | 6.5097355 | 29 | amotl2b |
| gipc2 | 5.680349 | 0.38719219 | 0.255 | 0.032 | 1.4430928 | 29 | gipc2 |
| rab42b | 1.166648 | 0.253877489 | 0.173 | 0.016 | 2.9638695 | 29 | rab42b |
| fthl28 | 1.733002 | 0.296265956 | 0.1 | 0.005 | 4.4026918 | 29 | fthl28 |
| rnd3a3 | 5.670269 | 0.357642313 | 0.264 | 0.035 | 1.4405319 | 29 | rnd3a |
| lsr1 | 7.226379 | 0.30353014 | 0.236 | 0.028 | 1.8358617 | 29 | lsr |
| pycard8 | 1.653775 | 0.591481617 | 0.418 | 0.082 | 4.2014159 | 29 | pycard |
| kremen11 | 2.514837 | 0.261947365 | 0.145 | 0.012 | 6.3889440 | 29 | kremen1 |
| anxa1a6 | 3.908603 | 0.929965681 | 0.573 | 0.161 | 9.9298072 | 29 | anxa1a |
| CYTH2 | 1.309634 | 0.526318809 | 0.209 | 0.024 | 3.3271275 | 29 | CYTH2 |
| prr131 | 4.381010 | 0.334629982 | 0.209 | 0.024 | 1.1129957 | 29 | prr13 |
| zgc:92242 | 1.232711 | 0.307298207 | 0.191 | 0.021 | 3.1317037 | 29 | zgc:92242 |
| pdlim13 | 4.335582 | 0.444584575 | 0.336 | 0.06 | 1.1014546 | 29 | pdlim1 |

| | | | | | | | |
|---|---|---|---|---|---|---|---|
| si:ch211-195b11.3 | 4.388686 | 0.465893743 | 0.245 | 0.034 | 1.1149457 | 29 | si:ch211-195b11.3 |
| jac7 | 2.516246 | 0.608007171 | 0.1 | 0.006 | 6.3925238 | 29 | jac7 |
| mat2ab | 9.587887 | 0.36523332 | 0.227 | 0.031 | 2.4358028 | 29 | mat2ab |
| enam1 | 1.187236 | 0.626116697 | 0.127 | 0.01 | 3.0161731 | 29 | enam |
| fosl1a4 | 1.917956 | 0.665698287 | 0.382 | 0.083 | 4.8725672 | 29 | fosl1a |
| exosc1 | 4.055801 | 0.412828473 | 0.273 | 0.044 | 1.0303762 | 29 | exosc1 |
| FAR21 | 6.742293 | 0.462326666 | 0.191 | 0.023 | 1.7128797 | 29 | FAR2 |
| vaspb1 | 7.553876 | 0.394746791 | 0.3 | 0.052 | 1.9190623 | 29 | vaspb |
| fxyd14 | 8.718655 | 0.496760797 | 0.391 | 0.083 | 2.2149744 | 29 | fxyd1 |
| krt155 | 1.154779 | 0.902332376 | 0.409 | 0.096 | 2.9337172 | 29 | krt15 |
| atp2a31 | 1.771305 | 0.455989398 | 0.236 | 0.035 | 4.5000014 | 29 | atp2a3 |
| cav15 | 2.194318 | 0.47217993 | 0.282 | 0.048 | 5.5746668 | 29 | cav1 |
| fam129ba | 2.875938 | 0.332403238 | 0.191 | 0.024 | 7.3063217 | 29 | fam129ba |
| midn6 | 3.224048 | 0.779452267 | 0.491 | 0.144 | 8.1906948 | 29 | midn |
| hmgn33 | 2.543059 | 0.535416032 | 0.355 | 0.077 | 6.4606423 | 29 | hmgn3 |
| zgc:1532844 | 2.965257 | 0.499300368 | 0.391 | 0.084 | 7.5332357 | 29 | zgc:153284 |
| cast3 | 1.177787 | 0.459210593 | 0.345 | 0.072 | 2.9921680 | 29 | cast |
| tagln211 | 1.213604 | 0.703820655 | 0.618 | 0.208 | 3.0831615 | 29 | tagln2 |
| ywhaz5 | 1.309924 | 0.722466407 | 0.736 | 0.303 | 3.3278636 | 29 | ywhaz |
| nectin1b3 | 4.056312 | 0.324457716 | 0.264 | 0.045 | 1.0305061 | 29 | nectin1b |
| oclna2 | 6.752815 | 0.340183625 | 0.336 | 0.066 | 1.7155527 | 29 | oclna |
| cavin2b5 | 2.789943 | 0.456767493 | 0.355 | 0.078 | 7.0878515 | 29 | cavin2b |
| jund6 | 3.605787 | 0.477096403 | 0.382 | 0.089 | 9.1605026 | 29 | jund |
| cd992 | 4.067099 | 0.324602453 | 0.282 | 0.052 | 1.0332466 | 29 | cd99 |
| lygl13 | 2.248854 | 0.406691306 | 0.418 | 0.099 | 5.7132138 | 29 | lygl1 |
| capns1a4 | 2.476022 | 0.631297517 | 0.509 | 0.158 | 6.2903364 | 29 | capns1a |
| capgb6 | 4.019904 | 0.539530447 | 0.436 | 0.114 | 1.0212566 | 29 | capgb |
| flna6 | 4.428705 | 0.584212025 | 0.436 | 0.121 | 1.1251125 | 29 | flna |
| ptgdsb.21 | 7.196187 | 0.697240381 | 0.218 | 0.037 | 1.8281914 | 29 | ptgdsb.2 |
| s100a10b9 | 7.387247 | 0.853549967 | 0.664 | 0.284 | 1.8767303 | 29 | s100a10b |
| tmc6b1 | 1.402875 | 0.297300202 | 0.218 | 0.035 | 3.5640062 | 29 | tmc6b |
| cst14b.13 | 2.102470 | 0.471813006 | 0.291 | 0.061 | 5.3413260 | 29 | cst14b.1 |
| calm2b8 | 2.260456 | 0.648541713 | 0.664 | 0.245 | 5.7426892 | 29 | calm2b |
| jag1b2 | 2.791449 | 0.267914079 | 0.191 | 0.028 | 7.0916786 | 29 | jag1b |
| slc38a5b6 | 1.318608 | 0.533330991 | 0.445 | 0.128 | 3.3499241 | 29 | slc38a5b |

| | | | | | | | |
|---|---|---|---|---|---|---|---|
| agr21 | 3.212320 | 0.543433198 | 0.191 | 0.03 | 8.1608998 | 29 | agr2 |
| si:ch211-79k12.11 | 3.813304 | 0.35869176 | 0.155 | 0.021 | 9.6876999 | 29 | si:ch211-79k12.1 |
| fbp1a1 | 6.247768 | 0.338813121 | 0.227 | 0.042 | 1.5872456 | 29 | fbp1a |
| myh9a2 | 1.936553 | 0.322405067 | 0.255 | 0.051 | 4.9198143 | 29 | myh9a |
| selenow16 | 2.116071 | 0.657639869 | 0.773 | 0.414 | 5.3758803 | 29 | selenow1 |
| il4r.12 | 3.525966 | 0.341406643 | 0.236 | 0.047 | 8.9577181 | 29 | il4r.1 |
| ctnnd1 | 3.607241 | 0.273160425 | 0.273 | 0.057 | 9.1641967 | 29 | ctnnd1 |
| slc20a1b1 | 5.189608 | 0.478632388 | 0.355 | 0.094 | 1.3184199 | 29 | slc20a1b |
| anxa46 | 7.518350 | 0.506856721 | 0.473 | 0.15 | 1.9100369 | 29 | anxa4 |
| si:ch211-105c13.33 | 1.295069 | 0.424926504 | 0.291 | 0.067 | 3.2901234 | 29 | si:ch211-105c13.3 |
| rarga2 | 2.253481 | 0.320347294 | 0.273 | 0.06 | 5.7249690 | 29 | rarga |
| si:ch73-234b20.55 | 4.318949 | 0.383891538 | 0.382 | 0.105 | 1.0972292 | 29 | si:ch73-234b20.5 |
| entpd2a.11 | 7.177018 | 0.296068468 | 0.191 | 0.033 | 1.8233215 | 29 | entpd2a.1 |
| aldh9a1a.18 | 3.220640 | 0.531576227 | 0.527 | 0.192 | 8.1820363 | 29 | aldh9a1a.1 |
| pmp22b4 | 4.515433 | 0.301715742 | 0.227 | 0.046 | 1.1471458 | 29 | pmp22b |
| prdx58 | 6.952793 | 0.535026583 | 0.491 | 0.178 | 1.7663571 | 29 | prdx5 |
| ccni9 | 1.501489 | 0.58481847 | 0.7 | 0.318 | 3.8145338 | 29 | ccni |
| suco | 1.599219 | 0.289394857 | 0.145 | 0.022 | 4.0628170 | 29 | suco |
| trib33 | 1.606485 | 0.465338032 | 0.364 | 0.108 | 4.0812764 | 29 | trib3 |
| bcl2l13 | 2.084984 | 0.467326131 | 0.236 | 0.053 | 5.2969030 | 29 | bcl2l13 |
| id2b3 | 2.205677 | 0.516967402 | 0.2 | 0.04 | 5.6035227 | 29 | id2b |
| ybx18 | 4.036055 | 0.409614642 | 0.909 | 0.696 | 1.0253599 | 29 | ybx1 |
| ppp3ccb1 | 1.412092 | 0.4598704 | 0.391 | 0.124 | 3.5874221 | 29 | ppp3ccb |
| mpzl2b2 | 5.179893 | 0.322265126 | 0.327 | 0.088 | 1.3159519 | 29 | mpzl2b |
| zfand5a | 7.587184 | 0.348253695 | 0.291 | 0.078 | 1.9275243 | 29 | zfand5a |
| mvp4 | 8.878516 | 0.36736592 | 0.245 | 0.06 | 2.2555872 | 29 | mvp |
| fosb5 | 1.670114 | 0.615847503 | 0.364 | 0.121 | 4.2429254 | 29 | fosb |
| ets22 | 2.032802 | 0.349950544 | 0.309 | 0.087 | 5.1643335 | 29 | ets2 |
| adrb2a | 4.053947 | 0.287962247 | 0.127 | 0.02 | 1.0299052 | 29 | adrb2a |
| atf36 | 7.107550 | 0.405676393 | 0.291 | 0.081 | 1.8056732 | 29 | atf3 |
| itgav3 | 1.367114 | 0.303980454 | 0.245 | 0.062 | 3.4731548 | 29 | itgav |
| tcima4 | 1.940188 | 0.411895968 | 0.264 | 0.071 | 4.9290485 | 29 | tcima |
| ptbp3 | 2.819429 | 0.263422378 | 0.255 | 0.066 | 7.1627594 | 29 | ptbp3 |
| spint21 | 3.906967 | 0.301827479 | 0.264 | 0.069 | 9.9256499 | 29 | spint2 |
| anxa5b5 | 5.713612 | 0.356103156 | 0.336 | 0.105 | 1.4515432 | 29 | anxa5b |

| | | | | | | | |
|---|---|---|---|---|---|---|---|
| rmdn12 | 7.126383 | 0.276504326 | 0.245 | 0.064 | 1.8104576 | 29 | rmdn1 |
| si:dkey-180p18.92 | 8.182656 | 0.374147852 | 0.336 | 0.108 | 2.0788037 | 29 | si:dkey-180p18.9 |
| anxa11a1 | 2.718958 | 0.38925406 | 0.345 | 0.117 | 6.9075148 | 29 | anxa11a |
| fabp35 | 3.476994 | 0.521315943 | 0.518 | 0.228 | 8.8333057 | 29 | fabp3 |
| chmp1b2 | 3.639562 | 0.347414027 | 0.382 | 0.132 | 9.2463075 | 29 | chmp1b |
| ccng24 | 6.154135 | 0.327231572 | 0.282 | 0.085 | 1.5634581 | 29 | ccng2 |
| camk2g21 | 7.846507 | 0.284709101 | 0.173 | 0.039 | 1.9934051 | 29 | camk2g2 |
| rffl | 9.165238 | 0.252554291 | 0.136 | 0.026 | 2.3284287 | 29 | rffl |
| cavin1b5 | 9.338312 | 0.291419804 | 0.191 | 0.045 | 2.3723983 | 29 | cavin1b |
| ier2b11 | 1.716225 | 0.528829869 | 0.518 | 0.237 | 4.3600719 | 29 | ier2b |
| smdt1b3 | 1.783438 | 0.38722396 | 0.4 | 0.152 | 4.5308265 | 29 | smdt1b |
| capns1b5 | 2.073247 | 0.400217934 | 0.391 | 0.145 | 5.2670861 | 29 | capns1b |
| csnk1a11 | 7.251365 | 0.400229003 | 0.545 | 0.237 | 1.8422094 | 29 | csnk1a1 |
| si:ch1073-429i10.3.1 | 8.381085 | 0.398509037 | 0.527 | 0.226 | 2.1292146 | 29 | si:ch1073-429i10.3.1 |
| reep55 | 1.059349 | 0.360622737 | 0.436 | 0.173 | 2.6912769 | 29 | reep5 |
| jdp2b5 | 2.641318 | 0.509833514 | 0.4 | 0.164 | 6.7102693 | 29 | jdp2b |
| atp5if1b5 | 2.698731 | 0.368741177 | 0.491 | 0.207 | 6.8561276 | 29 | atp5if1b |
| ip6k2b5 | 2.741187 | 0.330515346 | 0.355 | 0.129 | 6.9639857 | 29 | ip6k2b |
| myl9a4 | 4.121054 | 0.472219294 | 0.173 | 0.045 | 1.0469537 | 29 | myl9a |
| chmp2bb | 4.303367 | 0.280818573 | 0.227 | 0.066 | 1.0932704 | 29 | chmp2bb |
| smox5 | 6.399513 | 0.271001002 | 0.273 | 0.087 | 1.6257963 | 29 | smox |
| hopx5 | 7.000675 | 0.382877243 | 0.218 | 0.065 | 1.7785216 | 29 | hopx |
| si:ch73-335l21.45 | 1.176013 | 0.517304535 | 0.355 | 0.141 | 2.9876629 | 29 | si:ch73-335l21.4 |
| tob1a3 | 1.944378 | 0.387819586 | 0.364 | 0.145 | 4.9396939 | 29 | tob1a |
| f11r.16 | 2.218615 | 0.330489351 | 0.418 | 0.164 | 5.6363938 | 29 | f11r.1 |
| ppiab5 | 2.238779 | 0.34715897 | 0.9 | 0.674 | 5.6876193 | 29 | ppiab |
| rpl22l19 | 2.390367 | 0.454172503 | 0.736 | 0.425 | 6.0727277 | 29 | rpl22l1 |
| pbxip1b | 2.458679 | 0.330773349 | 0.191 | 0.054 | 6.2462758 | 29 | pbxip1b |
| si:ch73-1a9.315 | 2.500757 | 0.443827818 | 0.618 | 0.315 | 6.3531743 | 29 | si:ch73-1a9.3 |
| wdr17 | 2.704026 | 0.287500879 | 0.4 | 0.156 | 6.8695805 | 29 | wdr1 |
| iqgap13 | 3.292201 | 0.323608393 | 0.227 | 0.071 | 8.3638370 | 29 | iqgap1 |
| cox8a10 | 3.341987 | 0.414026392 | 0.545 | 0.267 | 8.4903187 | 29 | cox8a |
| adh57 | 3.512643 | 0.412532491 | 0.509 | 0.242 | 8.9238719 | 29 | adh5 |
| tuba8l7 | 4.158855 | 0.367480236 | 0.473 | 0.209 | 1.0565573 | 29 | tuba8l |
| eif4a1b6 | 5.185395 | 0.413029824 | 0.609 | 0.311 | 1.3173496 | 29 | eif4a1b |

| | | | | | | | |
|---|---|---|---|---|---|---|---|
| pleca3 | 7.044488 | 0.259123988 | 0.209 | 0.062 | 1.7896522 | 29 | pleca |
| csde12 | 8.389347 | 0.330297155 | 0.473 | 0.207 | 2.1313138 | 29 | csde1 |
| tpm37 | 9.014580 | 0.410636358 | 0.564 | 0.284 | 2.2901542 | 29 | tpm3 |
| sptbn21 | 9.027204 | 0.297736665 | 0.164 | 0.044 | 2.2933613 | 29 | sptbn2 |
| ywhaqb7 | 1.116052 | 0.267818486 | 0.391 | 0.155 | 2.8353308 | 29 | ywhaqb |
| ahcy3 | 1.122889 | 0.310484061 | 0.391 | 0.158 | 2.8526997 | 29 | ahcy |
| pak2a2 | 1.245165 | 0.288043593 | 0.218 | 0.069 | 3.1633425 | 29 | pak2a |
| h3f3c10 | 1.391572 | 0.409120424 | 0.609 | 0.304 | 3.5352894 | 29 | h3f3c |
| higd1a5 | 1.392025 | 0.431322976 | 0.573 | 0.303 | 3.5364410 | 29 | higd1a |
| ccl25b5 | 1.425072 | 0.327361414 | 0.273 | 0.094 | 3.6203961 | 29 | ccl25b |
| ak22 | 1.711441 | 0.380241528 | 0.518 | 0.255 | 4.3479165 | 29 | ak2 |
| pim18 | 1.739741 | 0.36100442 | 0.4 | 0.169 | 4.4198127 | 29 | pim1 |
| si:ch211-270g19.51 | 1.743186 | 0.297535665 | 0.1 | 0.02 | 4.4285655 | 29 | si:ch211-270g19.5 |
| calm1b7 | 1.888273 | 0.510451975 | 0.264 | 0.103 | 4.7971594 | 29 | calm1b |
| atf4b9 | 2.059869 | 0.402512884 | 0.555 | 0.277 | 5.2330988 | 29 | atf4b |
| itm2bb8 | 2.726067 | 0.359984246 | 0.518 | 0.242 | 6.9255755 | 29 | itm2bb |
| znf7061 | 3.270885 | 0.303605589 | 0.427 | 0.183 | 8.3096856 | 29 | znf706 |
| rab1ab2 | 4.262942 | 0.285631691 | 0.345 | 0.138 | 0.000108 | 29 | rab1ab |
| pfn27 | 4.726137 | 0.3551859 | 0.509 | 0.239 | 0.00012 | 29 | pfn2 |
| cebpd7 | 8.021608 | 0.341566532 | 0.364 | 0.152 | 0.000204 | 29 | cebpd |
| rpl5a12 | 8.664262 | 0.402164317 | 0.809 | 0.536 | 0.00022 | 29 | rpl5a |
| myl12.110 | 9.738219 | 0.407727071 | 0.718 | 0.426 | 0.000247 | 29 | myl12.1 |
| junba9 | 1.206011 | 0.52984064 | 0.527 | 0.32 | 0.000306 | 29 | junba |
| ccnd11 | 1.491586 | 0.286025223 | 0.155 | 0.044 | 0.000379 | 29 | ccnd1 |
| tnfaip34 | 1.743630 | 0.260716655 | 0.191 | 0.06 | 0.000443 | 29 | tnfaip3 |
| ier53 | 3.145221 | 0.341952096 | 0.336 | 0.147 | 0.000799 | 29 | ier5 |
| jun8 | 3.364346 | 0.39682459 | 0.5 | 0.25 | 0.000855 | 29 | jun |
| plcg23 | 3.397723 | 0.38654681 | 0.164 | 0.05 | 0.000863 | 29 | plcg2 |
| hdlbpa6 | 3.541038 | 0.307592134 | 0.391 | 0.175 | 0.0009 | 29 | hdlbpa |
| mcl1b9 | 4.206589 | 0.378138537 | 0.5 | 0.251 | 0.001069 | 29 | mcl1b |
| prelid3b2 | 4.411824 | 0.322825049 | 0.473 | 0.226 | 0.001121 | 29 | prelid3b |
| si:dkey-206d17.123 | 9.481087 | 0.267095607 | 0.164 | 0.051 | 0.002409 | 29 | si:dkey-206d17.12 |
| ppib9 | 9.920953 | 0.366703029 | 0.7 | 0.379 | 0.00252 | 29 | ppib |
| ucp29 | 1.241226 | 0.393713577 | 0.7 | 0.414 | 0.003153 | 29 | ucp2 |
| zgc:1538675 | 1.399275 | 0.402106467 | 0.736 | 0.465 | 0.003555 | 29 | zgc:153867 |

| | | | | | | | |
|---|---|---|---|---|---|---|---|
| pfn111 | 1.406488 | 0.439265385 | 0.773 | 0.501 | 0.003573 | 29 | pfn1 |
| epcam7 | 1.536967 | 0.440405893 | 0.636 | 0.422 | 0.003905 | 29 | epcam |
| mibp212 | 2.430276 | 0.375852875 | 0.791 | 0.521 | 0.006174 | 29 | mibp2 |
| vdac26 | 2.492901 | 0.376012112 | 0.564 | 0.326 | 0.006333 | 29 | vdac2 |
| rpsa11 | 2.693839 | 0.261836437 | 0.945 | 0.786 | 0.006844 | 29 | rpsa |
| cct26 | 3.224836 | 0.297947386 | 0.5 | 0.254 | 0.008193 | 29 | cct2 |
| glo13 | 5.095401 | 0.277089992 | 0.427 | 0.209 | 0.012945 | 29 | glo1 |
| klf2a6 | 7.103812 | 0.300722977 | 0.245 | 0.1 | 0.018047 | 29 | klf2a |
| selenok1 | 7.257108 | 0.264993725 | 0.309 | 0.14 | 0.018437 | 29 | selenok |
| eif1b8 | 9.381459 | 0.371546512 | 0.655 | 0.407 | 0.023834 | 29 | eif1b |
| sdf4 | 9.670306 | 0.309734015 | 0.155 | 0.052 | 0.024567 | 29 | sdf4 |
| nsa29 | 9.896397 | 0.347466899 | 0.645 | 0.389 | 0.025142 | 29 | nsa2 |
| ost43 | 1.157060 | 0.31183475 | 0.491 | 0.262 | 0.029395 | 29 | ost4 |
| cnpy1 | 1.283569 | 0.425851896 | 0.245 | 0.108 | 0.032609 | 29 | cnpy1 |
| fosl26 | 1.747584 | 0.265302231 | 0.264 | 0.114 | 0.044397 | 29 | fosl2 |
| rps259 | 1.794644 | 0.251021453 | 0.909 | 0.749 | 0.045593 | 29 | rps25 |
| rplp212 | 1.825261 | 0.259641807 | 0.927 | 0.783 | 0.046371 | 29 | rplp2 |
| atp5pf6 | 1.840494 | 0.298608622 | 0.573 | 0.32 | 0.046758 | 29 | atp5pf |
| eef1da7 | 2.202033 | 0.346213592 | 0.764 | 0.506 | 0.055943 | 29 | eef1da |
| rpl23a11 | 2.533172 | 0.319988031 | 0.909 | 0.669 | 0.064355 | 29 | rpl23a |
| canx2 | 2.712132 | 0.295275076 | 0.236 | 0.102 | 0.068902 | 29 | canx |
| mt-co112 | 2.983451 | 0.311593038 | 0.836 | 0.64 | 0.075795 | 29 | mt-co1 |
| krt89 | 2.992278 | 0.272048395 | 0.591 | 0.312 | 0.076019 | 29 | krt8 |
| tegt4 | 3.062981 | 0.259697227 | 0.464 | 0.238 | 0.077815 | 29 | tegt |
| eef1db7 | 3.189162 | 0.307272649 | 0.509 | 0.285 | 0.081021 | 29 | eef1db |
| zgc:11418816 | 3.275601 | 0.322328953 | 0.909 | 0.668 | 0.083217 | 29 | zgc:114188 |
| btg111 | 3.394932 | 0.327250916 | 0.6 | 0.356 | 0.086248 | 29 | btg1 |
| nfkbiaa5 | 3.473269 | 0.297233694 | 0.318 | 0.153 | 0.088238 | 29 | nfkbiaa |
| itm2ba8 | 4.794331 | 0.30471264 | 0.527 | 0.297 | 0.1218 | 29 | itm2ba |
| irf2bp2a4 | 4.846985 | 0.25655748 | 0.373 | 0.186 | 0.123138 | 29 | irf2bp2a |
| naca12 | 5.005445 | 0.267023892 | 0.818 | 0.653 | 0.127163 | 29 | naca |
| ralbp11 | 5.020128 | 0.301603015 | 0.164 | 0.061 | 0.127536 | 29 | ralbp1 |
| rpl309 | 8.379008 | 0.284842495 | 0.936 | 0.73 | 0.212869 | 29 | rpl30 |
| ddit38 | 1.000655 | 0.304868935 | 0.345 | 0.179 | 0.254217 | 29 | ddit3 |
| capzb9 | 1.042928 | 0.25179258 | 0.5 | 0.272 | 0.264956 | 29 | capzb |

| | | | | | | | |
|---|---|---|---|---|---|---|---|
| rpl27a11 | 1.131297 | 0.329828365 | 0.773 | 0.523 | 0.287406 | 29 | rpl27a |
| fosab9 | 1.243039 | 0.37083701 | 0.573 | 0.362 | 0.315794 | 29 | fosab |
| gapdhs11 | 1.253423 | 0.282083499 | 0.545 | 0.315 | 0.318432 | 29 | gapdhs |
| atp5pd5 | 1.374034 | 0.285470602 | 0.491 | 0.283 | 0.349073 | 29 | atp5pd |
| cotl19 | 1.621689 | 0.293393874 | 0.573 | 0.32 | 0.41199 | 29 | cotl1 |
| fkbp1aa8 | 1.720935 | 0.296550529 | 0.673 | 0.414 | 0.437204 | 29 | fkbp1aa |
| romo19 | 1.816656 | 0.295497174 | 0.655 | 0.409 | 0.461521 | 29 | romo1 |
| lasp18 | 1.959118 | 0.252385745 | 0.464 | 0.254 | 0.497714 | 29 | lasp1 |
| eef1g11 | 2.011666 | 0.286413573 | 0.855 | 0.653 | 0.511064 | 29 | eef1g |
| gadd45ba6 | 2.032935 | 0.25047852 | 0.382 | 0.198 | 0.516467 | 29 | gadd45ba |
| ldhba3 | 2.179894 | 0.267063075 | 0.464 | 0.261 | 0.553802 | 29 | ldhba |
| zgc:16273013 | 2.227408 | 0.334264327 | 0.627 | 0.403 | 0.565873 | 29 | zgc:162730 |
| chac15 | 2.457664 | 0.312858696 | 0.591 | 0.378 | 0.62437 | 29 | chac1 |
| eif2s34 | 2.549745 | 0.259606493 | 0.536 | 0.308 | 0.647763 | 29 | eif2s3 |
| ier2a10 | 2.550833 | 0.295489276 | 0.482 | 0.275 | 0.648039 | 29 | ier2a |
| kctd5a4 | 2.565144 | 0.250774026 | 0.355 | 0.186 | 0.651675 | 29 | kctd5a |
| sdc48 | 3.133230 | 0.343224543 | 0.436 | 0.255 | 0.795997 | 29 | sdc4 |
| csrnp1b5 | 3.372736 | 0.25712946 | 0.318 | 0.165 | 0.856844 | 29 | csrnp1b |
| rpl611 | 7.814472 | 0.265248809 | 0.909 | 0.701 | 1 | 29 | rpl6 |
| arpc210 | 8.521641 | 0.254970914 | 0.5 | 0.3 | 1 | 29 | arpc2 |
| COX5B6 | 0.0001 | 0.268091321 | 0.582 | 0.377 | 1 | 29 | COX5B |
| grna3 | 0.0002 | 0.462378542 | 0.136 | 0.058 | 1 | 29 | grna |
| rps27.114 | 0.0004 | 0.259976603 | 0.909 | 0.71 | 1 | 29 | rps27.1 |
| rpl1410 | 0.0004 | 0.251590002 | 0.882 | 0.669 | 1 | 29 | rpl14 |
| eif3f11 | 0.0004 | 0.273418188 | 0.745 | 0.465 | 1 | 29 | eif3f |
| rsl24d19 | 0.0005 | 0.26319446 | 0.7 | 0.461 | 1 | 29 | rsl24d1 |
| mdka6 | 0.0005 | 0.281843174 | 0.218 | 0.114 | 1 | 29 | mdka |
| COX313 | 0.0006 | 0.250257713 | 0.827 | 0.676 | 1 | 29 | COX3 |
| thy18 | 0.0009 | 0.287088132 | 0.564 | 0.379 | 1 | 29 | thy1 |
| btg28 | 0.0014 | 0.266849278 | 0.482 | 0.351 | 1 | 29 | btg2 |
| myl9b6 | 0.0038 | 0.348518926 | 0.173 | 0.1 | 1 | 29 | myl9b |
| calr3b3 | 0.0068 | 0.273718818 | 0.164 | 0.095 | 1 | 29 | calr3b |
| CR383676.116 | 0.0068 | 0.251809929 | 0.764 | 0.718 | 1 | 29 | CR383676.1 |
| gsdf2 | 0 | 2.455436726 | 0.99 | 0.05 | 0 | 30 | gsdf |
| gstt1a3 | 0 | 1.884150439 | 0.882 | 0.035 | 0 | 30 | gstt1a |

| | | | | | | | |
|---|---|---|---|---|---|---|---|
| dmrt12 | 0 | 1.632765202 | 0.775 | 0.012 | 0 | 30 | dmrt1 |
| igf31 | 0 | 1.609138026 | 0.696 | 0.017 | 0 | 30 | igf3 |
| hspb81 | 0 | 1.452786477 | 0.676 | 0.019 | 0 | 30 | hspb8 |
| gpx3 | 0 | 1.409158422 | 0.569 | 0.002 | 0 | 30 | gpx3 |
| LOC1104380241 | 0 | 1.326119099 | 0.451 | 0.008 | 0 | 30 | LOC110438024 |
| amh | 0 | 1.165348568 | 0.471 | 0.003 | 0 | 30 | amh |
| zgc:92162 | 0 | 1.05965208 | 0.48 | 0.005 | 0 | 30 | zgc:92162 |
| nrgna | 0 | 1.023525855 | 0.402 | 0.005 | 0 | 30 | nrgna |
| scg31 | 0 | 0.974271558 | 0.451 | 0.006 | 0 | 30 | scg3 |
| cyp26a1 | 0 | 0.962234356 | 0.402 | 0.008 | 0 | 30 | cyp26a1 |
| inha1 | 0 | 0.923966411 | 0.412 | 0.006 | 0 | 30 | inha |
| scn1ba | 0 | 0.920279881 | 0.402 | 0.004 | 0 | 30 | scn1ba |
| LOC110437926 | 0 | 0.911664234 | 0.373 | 0.002 | 0 | 30 | LOC110437926 |
| mmel1 | 0 | 0.782713264 | 0.363 | 0.002 | 0 | 30 | mmel1 |
| inhbb | 0 | 0.705071797 | 0.324 | 0.004 | 0 | 30 | inhbb |
| ppp1r1b | 0 | 0.703553312 | 0.324 | 0.004 | 0 | 30 | ppp1r1b |
| kcnj1a.1 | 0 | 0.673132473 | 0.294 | 0 | 0 | 30 | kcnj1a.1 |
| col28a2a | 0 | 0.609985414 | 0.275 | 0.001 | 0 | 30 | col28a2a |
| si:dkeyp-80c12.8 | 0 | 0.608084869 | 0.275 | 0 | 0 | 30 | si:dkeyp-80c12.8 |
| LOC103908925 | 0 | 0.57319953 | 0.255 | 0.002 | 0 | 30 | LOC103908925 |
| cyb5d1 | 0 | 0.55940302 | 0.255 | 0.001 | 0 | 30 | cyb5d1 |
| cnga1a | 0 | 0.530776546 | 0.245 | 0.002 | 0 | 30 | cnga1a |
| CT573799.2 | 0 | 0.361311055 | 0.186 | 0 | 0 | 30 | CT573799.2 |
| XLOC-0422291 | 3.921836 | 0.749923633 | 0.363 | 0.007 | 9.9634250 | 30 | XLOC-042229 |
| LOC110438394 | 1.260297 | 0.470650197 | 0.186 | 0.001 | 3.2017855 | 30 | LOC110438394 |
| nts | 1.888312 | 0.508450369 | 0.186 | 0.001 | 4.7972572 | 30 | nts |
| XLOC-042806 | 5.005829 | 0.360334242 | 0.157 | 0.001 | 1.2717310 | 30 | XLOC-042806 |
| si:ch211-117c9.51 | 2.553908 | 1.113515038 | 0.52 | 0.018 | 6.4882045 | 30 | si:ch211-117c9.5 |
| adm2a | 7.779033 | 0.481409762 | 0.225 | 0.003 | 1.9762634 | 30 | adm2a |
| ctnnd2b | 2.671754 | 0.406482642 | 0.176 | 0.001 | 6.7875911 | 30 | ctnnd2b |
| znf11041 | 3.799345 | 1.089930411 | 0.441 | 0.014 | 9.6522361 | 30 | znf1104 |
| plxnb3 | 1.652161 | 0.610008147 | 0.284 | 0.005 | 4.1973155 | 30 | plxnb3 |
| msx1b | 3.964488 | 0.538379189 | 0.235 | 0.003 | 1.0071783 | 30 | msx1b |
| sv2bb1 | 9.216573 | 0.671323929 | 0.324 | 0.007 | 2.3414706 | 30 | sv2bb |
| ugt5g1 | 7.188653 | 0.785863442 | 0.373 | 0.011 | 1.8262773 | 30 | ugt5g1 |

| | | | | | | | |
|---|---|---|---|---|---|---|---|
| nkx3.3 | 2.77787 | 0.408651471 | 0.167 | 0.002 | 7.0571950 | 30 | nkx3.3 |
| LOC1039109031 | 6.48639 | 1.079413981 | 0.412 | 0.015 | 1.6478698 | 30 | LOC103910903 |
| hpn1 | 3.15499 | 0.654455439 | 0.294 | 0.007 | 8.0152768 | 30 | hpn |
| sox9a1 | 2.28098 | 0.708964436 | 0.343 | 0.01 | 5.7948322 | 30 | sox9a |
| ak41 | 3.94845 | 1.193552397 | 0.549 | 0.031 | 1.0031045 | 30 | ak4 |
| zgc:194990 | 4.37581 | 0.384969051 | 0.167 | 0.002 | 1.1116745 | 30 | zgc:194990 |
| cth2 | 7.74010 | 1.170734424 | 0.569 | 0.035 | 1.9663738 | 30 | cth |
| soul5 | 3.66296 | 0.799780282 | 0.353 | 0.013 | 9.3057636 | 30 | soul5 |
| abtb2a | 2.05676 | 0.290293516 | 0.127 | 0.001 | 5.2252037 | 30 | abtb2a |
| add21 | 1.02723 | 0.608671741 | 0.265 | 0.007 | 2.6096910 | 30 | add2 |
| zgc:1527914 | 1.00880 | 1.512541822 | 0.706 | 0.064 | 2.5628583 | 30 | zgc:152791 |
| slc4a2a | 1.94033 | 0.33040367 | 0.186 | 0.003 | 4.9294294 | 30 | slc4a2a |
| prr15lb | 3.26998 | 0.303399696 | 0.127 | 0.001 | 8.3073982 | 30 | prr15lb |
| fyna | 6.38419 | 0.574900604 | 0.284 | 0.009 | 1.6219051 | 30 | fyna |
| fabp7a1 | 2.45912 | 0.991354398 | 0.412 | 0.02 | 6.2474169 | 30 | fabp7a |
| jam2a3 | 7.66601 | 1.393561429 | 0.608 | 0.048 | 1.9475502 | 30 | jam2a |
| alcamb1 | 1.41099 | 0.805633509 | 0.392 | 0.018 | 3.5846409 | 30 | alcamb |
| kirrel3l | 2.71262 | 0.593411153 | 0.275 | 0.009 | 6.8914122 | 30 | kirrel3l |
| gpm6ab2 | 6.04784 | 0.775615965 | 0.373 | 0.017 | 1.5364547 | 30 | gpm6ab |
| gh1 | 7.13559 | 0.408969603 | 0.147 | 0.002 | 1.8127978 | 30 | gh1 |
| LOC100149700 | 4.96243 | 0.278496866 | 0.118 | 0.001 | 1.2607058 | 30 | LOC100149700 |
| lnx2a | 1.56447 | 0.394921147 | 0.186 | 0.004 | 3.9745386 | 30 | lnx2a |
| cldnc2 | 1.29584 | 1.018517936 | 0.49 | 0.033 | 3.2921016 | 30 | cldnc |
| rhoub | 1.84082 | 0.486434564 | 0.216 | 0.006 | 4.6766084 | 30 | rhoub |
| map1ab | 1.05450 | 0.439069955 | 0.196 | 0.005 | 2.6789820 | 30 | map1ab |
| asb5b | 1.39714 | 0.276405497 | 0.127 | 0.002 | 3.5494434 | 30 | asb5b |
| ftr823 | 1.09943 | 0.904284479 | 0.471 | 0.034 | 2.7931139 | 30 | ftr82 |
| tmem229b | 3.88805 | 0.4056492 | 0.186 | 0.005 | 9.8776085 | 30 | tmem229b |
| large21 | 1.26703 | 0.473652356 | 0.216 | 0.007 | 3.2188977 | 30 | large2 |
| CABZ01021532.11 | 9.00047 | 0.794201595 | 0.324 | 0.016 | 2.2865715 | 30 | CABZ01021532.1 |
| slc2a1b1 | 4.53260 | 0.701403244 | 0.333 | 0.018 | 1.1515070 | 30 | slc2a1b |
| cx34.51 | 1.03926 | 0.543443801 | 0.216 | 0.008 | 2.6402478 | 30 | cx34.5 |
| CDK18 | 2.90796 | 0.258384866 | 0.127 | 0.002 | 7.3876858 | 30 | CDK18 |
| lpar3 | 4.95060 | 0.309655175 | 0.147 | 0.003 | 1.2577005 | 30 | lpar3 |
| rbm47 | 1.88603 | 0.612884135 | 0.284 | 0.014 | 4.7914633 | 30 | rbm47 |

| | | | | | | | |
|---|---|---|---|---|---|---|---|
| XLOC-0405161 | 5.244285 | 0.388410589 | 0.167 | 0.005 | 1.3323106 | 30 | XLOC-040516 |
| fndc5b | 3.062773 | 0.337268248 | 0.147 | 0.004 | 7.7809765 | 30 | fndc5b |
| cx434 | 1.173758 | 1.085973125 | 0.539 | 0.055 | 2.9819339 | 30 | cx43 |
| isg154 | 6.808420 | 1.213175865 | 0.549 | 0.062 | 1.7296792 | 30 | isg15 |
| slc16a7 | 1.326266 | 0.277897612 | 0.147 | 0.004 | 3.3693812 | 30 | slc16a7 |
| dag14 | 1.155622 | 1.019878845 | 0.5 | 0.053 | 2.9358590 | 30 | dag1 |
| si:dkey-188i13.91 | 1.570321 | 0.843231123 | 0.353 | 0.026 | 3.9894005 | 30 | si:dkey-188i13.9 |
| cox6a24 | 1.543620 | 0.809513357 | 0.412 | 0.036 | 3.9215683 | 30 | cox6a2 |
| slc7a3a3 | 4.024487 | 0.897688351 | 0.461 | 0.045 | 1.0224211 | 30 | slc7a3a |
| cahz2 | 4.920435 | 0.871507226 | 0.48 | 0.047 | 1.2500365 | 30 | cahz |
| cldn11a5 | 1.120139 | 1.223616693 | 0.598 | 0.081 | 2.8457139 | 30 | cldn11a |
| ifi466 | 1.310039 | 1.262859843 | 0.725 | 0.129 | 3.3281544 | 30 | ifi46 |
| nfasca | 4.198167 | 0.276599117 | 0.127 | 0.003 | 1.0665444 | 30 | nfasca |
| fam13b1 | 1.845833 | 0.745020826 | 0.382 | 0.035 | 4.6893399 | 30 | fam13b |
| scube11 | 6.165858 | 0.533510686 | 0.294 | 0.02 | 1.5664364 | 30 | scube1 |
| appa6 | 9.343663 | 1.051790316 | 0.588 | 0.086 | 2.3737577 | 30 | appa |
| ndrg3a1 | 1.852788 | 0.61335101 | 0.304 | 0.022 | 4.7070100 | 30 | ndrg3a |
| prss12 | 2.538310 | 0.30661707 | 0.137 | 0.004 | 6.4485782 | 30 | prss12 |
| trim3b | 1.349512 | 0.297182651 | 0.157 | 0.006 | 3.4284375 | 30 | trim3b |
| epdl11 | 1.647822 | 0.730083104 | 0.363 | 0.032 | 4.1862920 | 30 | epdl1 |
| bcam3 | 3.490889 | 0.834152367 | 0.402 | 0.039 | 8.8686060 | 30 | bcam |
| sema7a1 | 1.655698 | 0.429221759 | 0.206 | 0.011 | 4.2063011 | 30 | sema7a |
| elovl51 | 1.933254 | 0.462312152 | 0.216 | 0.012 | 4.9114317 | 30 | elovl5 |
| ifi458 | 1.395968 | 1.354690899 | 0.696 | 0.137 | 3.5464577 | 30 | ifi45 |
| slc31a2 | 1.526511 | 0.37409196 | 0.206 | 0.011 | 3.8781011 | 30 | slc31a2 |
| atp11a2 | 3.964508 | 0.393276122 | 0.206 | 0.011 | 1.0071834 | 30 | atp11a |
| krt810 | 2.696147 | 1.338578678 | 0.941 | 0.309 | 6.8495629 | 30 | krt8 |
| XLOC-040108 | 7.115214 | 0.277494348 | 0.127 | 0.004 | 1.8076201 | 30 | XLOC-040108 |
| blf | 1.070086 | 0.289085071 | 0.127 | 0.004 | 2.7185537 | 30 | blf |
| itga3b1 | 1.605886 | 0.271008244 | 0.137 | 0.005 | 4.0797548 | 30 | itga3b |
| phlda24 | 2.063914 | 0.952256777 | 0.422 | 0.051 | 5.2433735 | 30 | phlda2 |
| si:dkey-6n6.11 | 4.090025 | 0.393694565 | 0.167 | 0.008 | 1.0390710 | 30 | si:dkey-6n6.1 |
| sobpa | 5.211272 | 0.380152689 | 0.186 | 0.01 | 1.3239236 | 30 | sobpa |
| fat21 | 4.900258 | 0.32892174 | 0.157 | 0.007 | 1.2449105 | 30 | fat2 |
| cgnl13 | 1.697002 | 0.586161059 | 0.304 | 0.028 | 4.3112345 | 30 | cgnl1 |

| | | | | | | | |
|---|---|---|---|---|---|---|---|
| rnf24 | 4.612587 | 0.590936818 | 0.304 | 0.029 | 1.1718277 | 30 | rnf24 |
| lgals9l34 | 2.728893 | 0.870258594 | 0.51 | 0.082 | 6.9327532 | 30 | lgals9l3 |
| esr2b3 | 6.873149 | 0.62587467 | 0.343 | 0.038 | 1.7461235 | 30 | esr2b |
| phc2a2 | 8.382713 | 0.90362794 | 0.539 | 0.091 | 2.1296283 | 30 | phc2a |
| lgals173 | 1.835318 | 0.844820488 | 0.402 | 0.054 | 4.6626258 | 30 | lgals17 |
| sparc7 | 2.249691 | 1.119782761 | 0.775 | 0.193 | 5.7153423 | 30 | sparc |
| wfdc2 | 1.318485 | 0.34225275 | 0.167 | 0.009 | 3.3496126 | 30 | wfdc2 |
| ckba4 | 2.455719 | 0.897775656 | 0.461 | 0.072 | 6.2387543 | 30 | ckba |
| marcksl1b9 | 4.992530 | 1.104516102 | 0.637 | 0.14 | 1.2683523 | 30 | marcksl1b |
| krt18a.17 | 7.273204 | 1.096894813 | 0.676 | 0.148 | 1.8477577 | 30 | krt18a.1 |
| prss352 | 8.791390 | 0.616416882 | 0.284 | 0.028 | 2.2334528 | 30 | prss35 |
| cbsb | 9.196739 | 0.625258576 | 0.294 | 0.029 | 2.3364317 | 30 | cbsb |
| aldh1a25 | 4.094959 | 0.993198936 | 0.539 | 0.096 | 1.0403244 | 30 | aldh1a2 |
| csgalnact1a1 | 6.583415 | 0.487606997 | 0.245 | 0.021 | 1.6725166 | 30 | csgalnact1a |
| aplp11 | 8.043974 | 0.468446929 | 0.216 | 0.016 | 2.0435717 | 30 | aplp1 |
| mylka | 2.622165 | 0.424278462 | 0.216 | 0.017 | 6.6616111 | 30 | mylka |
| senp3a | 2.672249 | 0.539531298 | 0.255 | 0.023 | 6.7888491 | 30 | senp3a |
| stard9 | 1.998745 | 0.306711649 | 0.157 | 0.009 | 5.0778122 | 30 | stard9 |
| nhsl1a | 2.060941 | 0.295887497 | 0.147 | 0.008 | 5.2358220 | 30 | nhsl1a |
| ap1m11 | 2.853998 | 0.798225675 | 0.461 | 0.075 | 7.2505822 | 30 | ap1m1 |
| calm1a3 | 4.997289 | 1.063781446 | 0.676 | 0.172 | 1.2695614 | 30 | calm1a |
| si:dkey-188i13.105 | 5.344830 | 0.970982334 | 0.549 | 0.112 | 1.3578542 | 30 | si:dkey-188i13.10 |
| ahcyl2 | 5.781228 | 0.424469572 | 0.216 | 0.017 | 1.4687210 | 30 | ahcyl2 |
| FP245456.1 | 1.177918 | 0.339072287 | 0.147 | 0.008 | 2.9925009 | 30 | FP245456.1 |
| ncam1a2 | 3.601667 | 0.43776216 | 0.235 | 0.02 | 9.1500356 | 30 | ncam1a |
| lpcat21 | 9.645144 | 0.45937666 | 0.245 | 0.022 | 2.4503490 | 30 | lpcat2 |
| tmem45a | 7.897047 | 0.55822062 | 0.265 | 0.027 | 2.0062448 | 30 | tmem45a |
| fhod3b | 2.001638 | 0.323854673 | 0.137 | 0.007 | 5.0851619 | 30 | fhod3b |
| pon11 | 2.271910 | 0.349811666 | 0.157 | 0.01 | 5.7717883 | 30 | pon1 |
| metrn1 | 1.102394 | 0.418470125 | 0.196 | 0.015 | 2.8006328 | 30 | metrn |
| osbpl3a | 3.461687 | 0.437519335 | 0.235 | 0.022 | 8.7944165 | 30 | osbpl3a |
| bag33 | 6.248528 | 0.599293578 | 0.284 | 0.033 | 1.5874387 | 30 | bag3 |
| znfx12 | 8.323328 | 0.472761097 | 0.275 | 0.032 | 2.1145416 | 30 | znfx1 |
| ca7 | 2.004779 | 0.488937193 | 0.275 | 0.031 | 5.0931434 | 30 | ca7 |
| fxyd6l3 | 2.755140 | 0.874391466 | 0.529 | 0.116 | 6.9994336 | 30 | fxyd6l |

| | | | | | | | |
|---|---|---|---|---|---|---|---|
| gadd45aa1 | 4.728976 | 0.45453679 | 0.206 | 0.019 | 1.2013965 | 30 | gadd45aa |
| mfge8b2 | 1.321200 | 0.514147433 | 0.284 | 0.035 | 3.3565101 | 30 | mfge8b |
| zgc:927445 | 1.376914 | 0.985184443 | 0.853 | 0.325 | 3.4980518 | 30 | zgc:92744 |
| si:dkeyp-77h1.4 | 2.543882 | 0.298770974 | 0.127 | 0.007 | 6.4627322 | 30 | si:dkeyp-77h1.4 |
| gdpd4a2 | 3.310426 | 0.292394707 | 0.157 | 0.011 | 8.4101373 | 30 | gdpd4a |
| kank25 | 1.004355 | 0.615901195 | 0.353 | 0.054 | 2.5515639 | 30 | kank2 |
| p2rx71 | 2.027454 | 0.330794274 | 0.167 | 0.013 | 5.1507476 | 30 | p2rx7 |
| nr0b12 | 3.225447 | 0.372823974 | 0.186 | 0.016 | 8.1942482 | 30 | nr0b1 |
| gstt1b7 | 6.043647 | 0.763971185 | 0.451 | 0.088 | 1.5353885 | 30 | gstt1b |
| nos1apa2 | 1.009699 | 0.459540977 | 0.245 | 0.028 | 2.5651403 | 30 | nos1apa |
| tbc1d1 | 2.840202 | 0.361334403 | 0.176 | 0.015 | 7.2155342 | 30 | tbc1d1 |
| ugt1b52 | 1.543976 | 0.56646221 | 0.275 | 0.037 | 3.9224715 | 30 | ugt1b5 |
| angptl44 | 1.809966 | 0.601944082 | 0.343 | 0.055 | 4.5982187 | 30 | angptl4 |
| si:ch211-225b11.11 | 3.473752 | 0.543311684 | 0.275 | 0.037 | 8.8250672 | 30 | si:ch211-225b11.1 |
| selenbp13 | 8.106049 | 0.710566578 | 0.412 | 0.08 | 2.0593418 | 30 | selenbp1 |
| znf326 | 3.354352 | 0.530184981 | 0.294 | 0.041 | 8.5217331 | 30 | znf326 |
| ponzr110 | 5.669836 | 0.925560064 | 0.775 | 0.288 | 1.4404220 | 30 | ponzr1 |
| atf5b2 | 1.188494 | 0.517118429 | 0.275 | 0.038 | 3.0193708 | 30 | atf5b |
| ap2m1a2 | 7.809057 | 0.797731913 | 0.598 | 0.169 | 1.9838909 | 30 | ap2m1a |
| jupa2 | 2.429824 | 0.586607786 | 0.343 | 0.059 | 6.1729684 | 30 | jupa |
| pard3ab | 3.684657 | 0.251703502 | 0.127 | 0.009 | 9.3608713 | 30 | pard3ab |
| CR318588.34 | 5.288096 | 0.677014973 | 0.314 | 0.05 | 1.3434407 | 30 | CR318588.3 |
| myh104 | 1.725076 | 0.458265482 | 0.284 | 0.042 | 4.3825580 | 30 | myh10 |
| ptmaa14 | 2.125530 | 0.867875488 | 0.873 | 0.432 | 5.3999096 | 30 | ptmaa |
| nlgn4xa1 | 4.019218 | 0.271013629 | 0.147 | 0.012 | 1.0210824 | 30 | nlgn4xa |
| CR376783.1 | 5.619012 | 0.34515847 | 0.147 | 0.012 | 1.4275101 | 30 | CR376783.1 |
| slc7a61 | 8.469699 | 0.292270374 | 0.167 | 0.016 | 2.1517272 | 30 | slc7a6 |
| cmbl | 4.406848 | 0.474765014 | 0.245 | 0.034 | 1.1195597 | 30 | cmbl |
| marcksb7 | 5.371790 | 0.760725613 | 0.598 | 0.159 | 1.3647033 | 30 | marcksb |
| b2m11 | 6.880159 | 0.890279796 | 0.716 | 0.297 | 1.7479045 | 30 | b2m |
| mafba7 | 2.266254 | 0.659958163 | 0.392 | 0.084 | 5.7574197 | 30 | mafba |
| podxl4 | 3.162644 | 0.51177322 | 0.333 | 0.057 | 8.0346991 | 30 | podxl |
| clstn12 | 7.519410 | 0.3544204 | 0.196 | 0.023 | 1.9103063 | 30 | clstn1 |
| hmgn613 | 1.579375 | 0.80061118 | 0.765 | 0.31 | 4.0124039 | 30 | hmgn6 |
| adgrl2a1 | 1.608530 | 0.311951158 | 0.176 | 0.019 | 4.0864711 | 30 | adgrl2a |

| | | | | | | | |
|---|---|---|---|---|---|---|---|
| aplp25 | 3.626754 | 0.592399573 | 0.373 | 0.076 | 9.2137698 | 30 | aplp2 |
| krt947 | 8.761365 | 0.833953987 | 0.578 | 0.165 | 2.2258248 | 30 | krt94 |
| plat1 | 9.701033 | 0.280669373 | 0.137 | 0.012 | 2.4645476 | 30 | plat |
| cox5b21 | 1.434243 | 0.341802043 | 0.137 | 0.012 | 3.6436951 | 30 | cox5b2 |
| stat1a5 | 2.103238 | 0.69623108 | 0.471 | 0.122 | 5.3432763 | 30 | stat1a |
| stat23 | 3.665215 | 0.623265667 | 0.324 | 0.063 | 9.3114807 | 30 | stat2 |
| nit2 | 1.007080 | 0.325907142 | 0.167 | 0.018 | 2.5584868 | 30 | nit2 |
| f104 | 1.492180 | 0.568089261 | 0.314 | 0.059 | 3.7908851 | 30 | f10 |
| met1 | 3.424701 | 0.26098627 | 0.118 | 0.009 | 8.7004550 | 30 | met |
| vat16 | 5.663528 | 0.719265493 | 0.52 | 0.146 | 1.4388194 | 30 | vat1 |
| atp1a1a.11 | 2.454162 | 0.739119832 | 0.5 | 0.144 | 6.2347991 | 30 | atp1a1a.1 |
| gata63 | 3.351846 | 0.329166283 | 0.196 | 0.025 | 8.5153647 | 30 | gata6 |
| wu:fc27b112 | 3.723336 | 0.304666027 | 0.157 | 0.017 | 9.4591353 | 30 | wu:fc27b11 |
| si:dkey-204f11.64 | 6.726899 | 0.432953448 | 0.216 | 0.031 | 1.7089689 | 30 | si:dkey-204f11.64 |
| arhgef18a1 | 7.124015 | 0.308511498 | 0.167 | 0.019 | 1.8098562 | 30 | arhgef18a |
| sntb1 | 7.323236 | 0.291219136 | 0.127 | 0.012 | 1.8604682 | 30 | sntb1 |
| gig2e1 | 3.910531 | 0.329994099 | 0.157 | 0.018 | 9.9347065 | 30 | gig2e |
| bmpr2a1 | 4.122522 | 0.330507575 | 0.176 | 0.022 | 1.0473267 | 30 | bmpr2a |
| fthl271 | 8.067223 | 0.569727522 | 0.971 | 0.682 | 2.0494782 | 30 | fthl27 |
| sema3b1 | 9.514881 | 0.299389725 | 0.176 | 0.022 | 2.4172555 | 30 | sema3b |
| si:ch73-1a9.316 | 9.652116 | 0.793944409 | 0.725 | 0.314 | 2.4521200 | 30 | si:ch73-1a9.3 |
| ap1s21 | 1.327525 | 0.701046062 | 0.471 | 0.134 | 3.3725795 | 30 | ap1s2 |
| ndrg22 | 3.052648 | 0.470957863 | 0.275 | 0.051 | 7.7552526 | 30 | ndrg2 |
| FAT42 | 1.473594 | 0.294944697 | 0.167 | 0.021 | 3.7436669 | 30 | FAT4 |
| hspe15 | 1.640248 | 0.77342934 | 0.696 | 0.29 | 4.1670500 | 30 | hspe1 |
| tent5ba2 | 1.731783 | 0.261879481 | 0.127 | 0.013 | 4.3995963 | 30 | tent5ba |
| itm2bb9 | 2.028709 | 0.842546372 | 0.588 | 0.242 | 5.1539377 | 30 | itm2bb |
| prnprs31 | 2.780245 | 0.264195727 | 0.137 | 0.015 | 7.0632148 | 30 | prnprs3 |
| efhd23 | 4.324743 | 0.492885318 | 0.294 | 0.059 | 1.0987011 | 30 | efhd2 |
| atp1b1a7 | 4.391917 | 0.756029598 | 0.588 | 0.229 | 1.1157666 | 30 | atp1b1a |
| fkbp1b | 4.401718 | 0.320234319 | 0.147 | 0.017 | 1.1182565 | 30 | fkbp1b |
| got1 | 5.145457 | 0.356257159 | 0.186 | 0.026 | 1.3072035 | 30 | got1 |
| si:ch211-151g22.1 | 7.245202 | 0.294473259 | 0.157 | 0.019 | 1.8406436 | 30 | si:ch211-151g22.1 |
| enah | 1.141040 | 0.357355471 | 0.176 | 0.024 | 2.8988132 | 30 | enah |
| tjp1a4 | 1.462920 | 0.414391147 | 0.275 | 0.053 | 3.7165485 | 30 | tjp1a |

| | | | | | | | |
|---|---|---|---|---|---|---|---|
| pik3r24 | 2.724550 | 0.546786781 | 0.304 | 0.067 | 6.9217207 | 30 | pik3r2 |
| prr132 | 3.358745 | 0.313492975 | 0.176 | 0.024 | 8.5328936 | 30 | prr13 |
| gnav1 | 3.423326 | 0.279136095 | 0.167 | 0.022 | 8.6969598 | 30 | gnav1 |
| mpped2a | 3.572660 | 0.402675967 | 0.206 | 0.032 | 9.0763434 | 30 | mpped2a |
| fkbp1aa9 | 4.023701 | 0.678735579 | 0.784 | 0.414 | 1.0222213 | 30 | fkbp1aa |
| zbtb10 | 4.824045 | 0.385672896 | 0.206 | 0.033 | 1.2255486 | 30 | zbtb10 |
| LOC110438541 | 6.747104 | 0.255323542 | 0.157 | 0.02 | 1.7141019 | 30 | LOC110438541 |
| cav16 | 9.678268 | 0.522232077 | 0.255 | 0.049 | 2.4587641 | 30 | cav1 |
| CABZ01076275.12 | 1.246719 | 0.291774831 | 0.157 | 0.02 | 3.1672902 | 30 | CABZ01076275.1 |
| ctsf3 | 2.072938 | 0.372753581 | 0.216 | 0.037 | 5.2663010 | 30 | ctsf |
| rhoj2 | 2.544882 | 0.275710829 | 0.137 | 0.016 | 6.4652734 | 30 | rhoj |
| cox6b25 | 2.577634 | 0.600284544 | 0.843 | 0.55 | 6.5484794 | 30 | cox6b2 |
| CR318588.111 | 3.370391 | 0.980069691 | 0.559 | 0.215 | 8.5624793 | 30 | CR318588.1 |
| emilin1a3 | 3.666590 | 0.305422292 | 0.176 | 0.025 | 9.3149719 | 30 | emilin1a |
| zgc:1750881 | 6.830138 | 0.494846001 | 0.294 | 0.065 | 1.7351966 | 30 | zgc:175088 |
| tspan9b4 | 7.909393 | 0.406080063 | 0.255 | 0.05 | 2.0093814 | 30 | tspan9b |
| si:ch211-197h24.97 | 1.131840 | 0.561229538 | 0.333 | 0.083 | 2.8754404 | 30 | si:ch211-197h24.9 |
| actn14 | 1.692451 | 0.435754593 | 0.245 | 0.048 | 4.2996722 | 30 | actn1 |
| fam102ab1 | 2.381827 | 0.37640468 | 0.206 | 0.035 | 6.0510323 | 30 | fam102ab |
| gstm.35 | 2.633027 | 0.640533709 | 0.451 | 0.13 | 6.6892055 | 30 | gstm.3 |
| eif4ebp15 | 4.275633 | 0.71141845 | 0.608 | 0.257 | 1.0862246 | 30 | eif4ebp1 |
| sema4ba5 | 7.435700 | 0.399179422 | 0.225 | 0.042 | 1.8890396 | 30 | sema4ba |
| ptpn13 | 1.183093 | 0.259173045 | 0.137 | 0.017 | 3.0056498 | 30 | ptpn13 |
| grk4 | 2.027234 | 0.360560211 | 0.196 | 0.033 | 5.1501901 | 30 | grk4 |
| ldlrap1a1 | 2.147982 | 0.467387465 | 0.284 | 0.063 | 5.4569505 | 30 | ldlrap1a |
| efhd1 | 3.972811 | 0.26109438 | 0.137 | 0.017 | 1.0092927 | 30 | efhd1 |
| arl4d3 | 4.667433 | 0.420747424 | 0.255 | 0.053 | 1.1857614 | 30 | arl4d |
| fosl1a5 | 8.196602 | 0.543672982 | 0.324 | 0.083 | 2.0823467 | 30 | fosl1a |
| itm2ba9 | 9.993408 | 0.655937242 | 0.637 | 0.296 | 2.5388254 | 30 | itm2ba |
| edf13 | 1.073249 | 0.677799589 | 0.627 | 0.287 | 2.7265898 | 30 | edf1 |
| gmfb2 | 1.190171 | 0.569048493 | 0.382 | 0.112 | 3.0236295 | 30 | gmfb |
| mkln1 | 1.272667 | 0.376132941 | 0.206 | 0.037 | 3.2332113 | 30 | mkln1 |
| lamb1a5 | 1.384551 | 0.398465442 | 0.255 | 0.054 | 3.5174518 | 30 | lamb1a |
| oip5-as11 | 1.562706 | 0.369034302 | 0.186 | 0.032 | 3.9700562 | 30 | oip5-as1 |
| agrn4 | 3.751628 | 0.458209885 | 0.265 | 0.06 | 9.5310130 | 30 | agrn |

| | | | | | | | |
|---|---|---|---|---|---|---|---|
| calm2b9 | 3.842529 | 0.679918169 | 0.578 | 0.246 | 9.7619473 | 30 | calm2b |
| mtpn5 | 4.179978 | 0.585201179 | 0.471 | 0.163 | 1.0619234 | 30 | mtpn |
| jun9 | 5.089934 | 0.786323362 | 0.569 | 0.25 | 1.2930977 | 30 | jun |
| ar5 | 5.449311 | 0.395780078 | 0.284 | 0.067 | 1.3843975 | 30 | ar |
| plekha61 | 5.562837 | 0.277570192 | 0.137 | 0.019 | 1.4132388 | 30 | plekha6 |
| pxnb | 6.184423 | 0.267723721 | 0.127 | 0.016 | 1.5711526 | 30 | pxnb |
| apoeb3 | 8.287342 | 0.271869748 | 0.137 | 0.019 | 2.1053992 | 30 | apoeb |
| pcyt2 | 9.885389 | 0.325821261 | 0.206 | 0.038 | 2.5113832 | 30 | pcyt2 |
| rpl138 | 9.925697 | 0.350510862 | 0.98 | 0.818 | 2.5216234 | 30 | rpl13 |
| cgref1 | 1.314358 | 0.28007807 | 0.147 | 0.022 | 3.3391273 | 30 | cgref1 |
| idh11 | 1.348043 | 0.565018034 | 0.392 | 0.117 | 3.4247041 | 30 | idh1 |
| hsd11b26 | 1.366549 | 0.61362758 | 0.471 | 0.149 | 3.4717200 | 30 | hsd11b2 |
| ppp1r14aa1 | 1.398258 | 0.289612328 | 0.196 | 0.035 | 3.5522768 | 30 | ppp1r14aa |
| dsc2l2 | 1.445806 | 0.298058266 | 0.167 | 0.027 | 3.6730713 | 30 | dsc2l |
| pnrc215 | 1.552374 | 0.522770043 | 0.794 | 0.598 | 3.9438071 | 30 | pnrc2 |
| mov10b.1 | 1.842315 | 0.257017997 | 0.147 | 0.022 | 4.6804036 | 30 | mov10b.1 |
| mbnl33 | 2.997822 | 0.431118209 | 0.265 | 0.062 | 7.6159690 | 30 | mbnl3 |
| ppp3ccb2 | 3.024810 | 0.571562708 | 0.392 | 0.125 | 7.6845298 | 30 | ppp3ccb |
| hmgb3a6 | 4.902313 | 0.461839147 | 0.314 | 0.083 | 1.2454327 | 30 | hmgb3a |
| gstm.21 | 1.094144 | 0.440484717 | 0.255 | 0.061 | 2.7796741 | 30 | gstm.2 |
| igfbp5b3 | 1.862235 | 0.423343574 | 0.333 | 0.088 | 4.7310093 | 30 | igfbp5b |
| si:ch211-286b5.57 | 4.801407 | 0.59563567 | 0.422 | 0.151 | 1.2197974 | 30 | si:ch211-286b5.5 |
| wbp1lb1 | 5.290405 | 0.348737764 | 0.225 | 0.05 | 1.3440273 | 30 | wbp1lb |
| cd1641 | 6.678888 | 0.45648114 | 0.284 | 0.075 | 1.6967715 | 30 | cd164 |
| smad12 | 8.674329 | 0.286106204 | 0.137 | 0.021 | 2.2037133 | 30 | smad1 |
| ctnnb15 | 8.910994 | 0.602534986 | 0.431 | 0.158 | 2.2638381 | 30 | ctnnb1 |
| plxna42 | 1.104298 | 0.265703743 | 0.147 | 0.024 | 2.8054701 | 30 | plxna4 |
| metrnla4 | 1.352264 | 0.426112143 | 0.333 | 0.094 | 3.4354277 | 30 | metrnla |
| erbb23 | 1.629148 | 0.286516359 | 0.176 | 0.033 | 4.1388507 | 30 | erbb2 |
| gadd45ba7 | 2.578128 | 0.674025962 | 0.48 | 0.197 | 6.5497349 | 30 | gadd45ba |
| bambia1 | 2.954355 | 0.298195458 | 0.157 | 0.027 | 7.5055396 | 30 | bambia |
| wu:fj29h11 | 4.221461 | 0.324297489 | 0.147 | 0.025 | 1.0724623 | 30 | wu:fj29h11 |
| tmsb4x16 | 4.849737 | 0.516186273 | 1 | 0.718 | 1.2320758 | 30 | tmsb4x |
| fdxr2 | 7.526368 | 0.345672599 | 0.157 | 0.028 | 1.9120738 | 30 | fdxr |
| CABZ01092156.1 | 7.548223 | 0.252372695 | 0.167 | 0.03 | 1.9176261 | 30 | CABZ01092156.1 |

| | | | | | | | |
|---|---|---|---|---|---|---|---|
| si:ch73-335l21.46 | 9.610678 | 0.710289332 | 0.392 | 0.141 | 2.4415928 | 30 | si:ch73-335l21.4 |
| pi15a4 | 1.275702 | 0.359061069 | 0.235 | 0.055 | 3.2409218 | 30 | pi15a |
| fosb6 | 1.308836 | 0.629500182 | 0.363 | 0.121 | 3.3250988 | 30 | fosb |
| zgc:921402 | 1.466634 | 0.283049185 | 0.167 | 0.031 | 3.7259838 | 30 | zgc:92140 |
| cdv31 | 1.708453 | 0.506636188 | 0.412 | 0.147 | 4.3403271 | 30 | cdv3 |
| psme13 | 2.009162 | 0.504805261 | 0.343 | 0.112 | 5.1042780 | 30 | psme1 |
| myl9a5 | 2.062853 | 0.329811939 | 0.206 | 0.045 | 5.2406797 | 30 | myl9a |
| CABZ01068366.14 | 2.207555 | 0.307555571 | 0.196 | 0.042 | 5.6082956 | 30 | CABZ01068366.1 |
| gstm.17 | 2.551586 | 0.598736244 | 0.598 | 0.279 | 6.4823067 | 30 | gstm.1 |
| CU984600.21 | 2.566129 | 0.267210743 | 0.147 | 0.026 | 6.5192523 | 30 | CU984600.2 |
| pelp1 | 2.898056 | 0.255218183 | 0.157 | 0.028 | 7.3625123 | 30 | pelp1 |
| si:dkey-42i9.44 | 3.309538 | 0.51509101 | 0.422 | 0.156 | 8.4078824 | 30 | si:dkey-42i9.4 |
| cdh13 | 3.634254 | 0.32226755 | 0.235 | 0.056 | 9.2328246 | 30 | cdh1 |
| rab11a1 | 4.630574 | 0.497192354 | 0.373 | 0.127 | 1.1763973 | 30 | rab11a |
| si:ch211-105c13.34 | 5.476028 | 0.461023814 | 0.255 | 0.067 | 1.3911849 | 30 | si:ch211-105c13.3 |
| gamt2 | 1.467152 | 0.290033099 | 0.196 | 0.044 | 3.7273011 | 30 | gamt |
| myl68 | 2.237968 | 0.463582445 | 0.353 | 0.116 | 5.6855582 | 30 | myl6 |
| optn | 7.122943 | 0.252591507 | 0.127 | 0.022 | 1.8095838 | 30 | optn |
| ywhae2 | 7.503270 | 0.274903321 | 0.157 | 0.032 | 1.9062058 | 30 | ywhae2 |
| rpz57 | 7.784117 | 0.479203929 | 0.363 | 0.127 | 1.9775549 | 30 | rpz5 |
| hmgb1b8 | 1.103451 | 0.510706866 | 0.402 | 0.154 | 2.8033197 | 30 | hmgb1b |
| man1b1b1 | 2.280803 | 0.254331459 | 0.157 | 0.032 | 5.7943808 | 30 | man1b1b |
| foxp45 | 3.872598 | 0.371351611 | 0.255 | 0.073 | 9.8383358 | 30 | foxp4 |
| gadd45bb4 | 4.768760 | 0.50019178 | 0.265 | 0.082 | 1.2115035 | 30 | gadd45bb |
| cebpd8 | 5.154376 | 0.46087411 | 0.412 | 0.152 | 1.3094692 | 30 | cebpd |
| ndufa4l8 | 5.581746 | 0.492230821 | 0.706 | 0.422 | 1.4180426 | 30 | ndufa4l |
| cox8b4 | 6.323823 | 0.436186678 | 0.304 | 0.101 | 1.6065672 | 30 | cox8b |
| rcan1a | 6.401751 | 0.276924636 | 0.186 | 0.044 | 1.6263648 | 30 | rcan1a |
| atox12 | 1.484574 | 0.476348418 | 0.333 | 0.123 | 3.7715618 | 30 | atox1 |
| si:ch211-39k3.2 | 1.497662 | 0.286858581 | 0.137 | 0.027 | 3.8048105 | 30 | si:ch211-39k3.2 |
| cited4a4 | 1.584132 | 0.273294496 | 0.176 | 0.042 | 4.0244874 | 30 | cited4a |
| tsg101b | 1.625145 | 0.262069694 | 0.147 | 0.031 | 4.1286816 | 30 | tsg101b |
| slc38a23 | 1.926776 | 0.302600648 | 0.186 | 0.046 | 4.8949753 | 30 | slc38a2 |
| socs3a10 | 1.973330 | 0.688214971 | 0.539 | 0.315 | 5.0132466 | 30 | socs3a |
| cox7c5 | 2.282498 | 0.477223934 | 0.745 | 0.494 | 5.7986884 | 30 | cox7c |

| | | | | | | | |
|---|---|---|---|---|---|---|---|
| tuba8l36 | 2.289433 | 0.377848167 | 0.245 | 0.072 | 5.8163064 | 30 | tuba8l3 |
| ccdc1871 | 2.377519 | 0.275273313 | 0.167 | 0.038 | 6.0400882 | 30 | ccdc187 |
| jdp2b6 | 4.815114 | 0.519965859 | 0.402 | 0.165 | 1.2232798 | 30 | jdp2b |
| cmtm31 | 6.400299 | 0.281320286 | 0.186 | 0.047 | 1.6259960 | 30 | cmtm3 |
| eef1db8 | 6.598856 | 0.520964755 | 0.539 | 0.285 | 1.6764393 | 30 | eef1db |
| pfn2l7 | 6.920792 | 0.463999559 | 0.49 | 0.215 | 1.7582273 | 30 | pfn2l |
| pgrmc15 | 8.319569 | 0.452360619 | 0.382 | 0.152 | 2.1135866 | 30 | pgrmc1 |
| fosab10 | 8.753071 | 0.647558935 | 0.588 | 0.362 | 2.2237178 | 30 | fosab |
| gabarapb10 | 9.328796 | 0.520680619 | 0.559 | 0.305 | 2.3699808 | 30 | gabarapb |
| bin3 | 1.252966 | 0.287658126 | 0.157 | 0.036 | 3.1831605 | 30 | bin3 |
| mdka7 | 1.381670 | 0.404148143 | 0.324 | 0.114 | 3.5101339 | 30 | mdka |
| LOC1039092845 | 1.504062 | 0.374659842 | 0.245 | 0.077 | 3.8210696 | 30 | LOC103909284 |
| yap13 | 1.737490 | 0.312318818 | 0.206 | 0.057 | 4.4140937 | 30 | yap1 |
| zgc:11418817 | 1.826750 | 0.418429874 | 0.961 | 0.668 | 4.6408602 | 30 | zgc:114188 |
| ywhaz6 | 1.991833 | 0.509092785 | 0.559 | 0.305 | 5.0602521 | 30 | ywhaz |
| arpc1a2 | 2.124708 | 0.497864932 | 0.353 | 0.142 | 5.3978221 | 30 | arpc1a |
| h1f06 | 2.246690 | 0.357730102 | 0.235 | 0.072 | 5.7077175 | 30 | h1f0 |
| si:dkey-28b4.77 | 3.009118 | 0.396041531 | 0.304 | 0.11 | 7.6446668 | 30 | si:dkey-28b4.7 |
| ptprga7 | 3.347071 | 0.321160288 | 0.235 | 0.072 | 8.5032346 | 30 | ptprga |
| hist2h2l4 | 3.664359 | 0.384313286 | 0.284 | 0.099 | 9.3093057 | 30 | hist2h2l |
| prkab1b2 | 3.739971 | 0.417868503 | 0.343 | 0.136 | 9.5013965 | 30 | prkab1b |
| s100v28 | 3.797248 | 0.402813025 | 0.363 | 0.14 | 9.6469087 | 30 | s100v2 |
| ppp2cb1 | 3.835893 | 0.445589572 | 0.412 | 0.178 | 9.7450882 | 30 | ppp2cb |
| pfkfb4b5 | 4.093912 | 0.441541617 | 0.314 | 0.121 | 1.0400584 | 30 | pfkfb4b |
| si:dkey-42i9.72 | 4.500270 | 0.25980455 | 0.127 | 0.027 | 1.1432937 | 30 | si:dkey-42i9.7 |
| rpsa12 | 5.126041 | 0.326149734 | 0.99 | 0.785 | 1.3022707 | 30 | rpsa |
| hsp70.31 | 5.222195 | 0.257840954 | 0.108 | 0.02 | 1.3266987 | 30 | hsp70.3 |
| ifi351 | 5.562525 | 0.283416419 | 0.157 | 0.039 | 1.4131596 | 30 | ifi35 |
| tcima5 | 6.160974 | 0.50207521 | 0.225 | 0.072 | 1.5651956 | 30 | tcima |
| tbc1d5 | 7.115292 | 0.253852361 | 0.137 | 0.031 | 1.8076399 | 30 | tbc1d5 |
| ucp210 | 8.531080 | 0.482014763 | 0.647 | 0.415 | 2.1673209 | 30 | ucp2 |
| zgc:1008291 | 8.709153 | 0.296197941 | 0.167 | 0.043 | 2.2125603 | 30 | zgc:100829 |
| id18 | 9.231543 | 0.51441354 | 0.333 | 0.131 | 2.3452737 | 30 | id1 |
| tma73 | 1.075248 | 0.487142546 | 0.588 | 0.331 | 2.7316700 | 30 | tma7 |
| psme27 | 1.079783 | 0.369258201 | 0.304 | 0.113 | 2.7431893 | 30 | psme2 |

| | | | | | | | |
|---|---|---|---|---|---|---|---|
| ccdc85b7 | 1.146792 | 0.262805475 | 0.186 | 0.051 | 2.9134260 | 30 | ccdc85b |
| h2afx19 | 1.226913 | 0.485349822 | 0.461 | 0.227 | 3.1169734 | 30 | h2afx1 |
| oaz1b10 | 1.266283 | 0.456073887 | 0.627 | 0.36 | 3.2169942 | 30 | oaz1b |
| sub1a4 | 1.306276 | 0.470255586 | 0.5 | 0.253 | 3.3185951 | 30 | sub1a |
| sdc49 | 1.388009 | 0.481028599 | 0.52 | 0.254 | 3.5262379 | 30 | sdc4 |
| ppdpfb5 | 1.711326 | 0.434438957 | 0.735 | 0.532 | 4.3476241 | 30 | ppdpfb |
| rps224 | 2.112886 | 0.273547523 | 0.971 | 0.842 | 5.3677893 | 30 | rps2 |
| eif1b9 | 2.171616 | 0.465558711 | 0.647 | 0.407 | 5.5169925 | 30 | eif1b |
| ndufb92 | 2.339312 | 0.456576594 | 0.422 | 0.201 | 5.9430224 | 30 | ndufb9 |
| ap2a1 | 2.436876 | 0.293511407 | 0.206 | 0.062 | 6.1908840 | 30 | ap2a1 |
| si:ch73-42p12.21 | 3.788313 | 0.393555401 | 0.294 | 0.113 | 9.6242095 | 30 | si:ch73-42p12.2 |
| wu:fb16f034 | 3.926800 | 0.362851579 | 0.255 | 0.09 | 9.9760371 | 30 | wu:fb16f03 |
| rps1613 | 4.026171 | 0.320804146 | 0.961 | 0.766 | 0.000102 | 30 | rps16 |
| irf2bp2b7 | 4.076325 | 0.375389363 | 0.275 | 0.102 | 0.000104 | 30 | irf2bp2b |
| ier2b12 | 4.342203 | 0.465978 | 0.471 | 0.237 | 0.00011 | 30 | ier2b |
| ybx19 | 4.937091 | 0.335579611 | 0.892 | 0.696 | 0.000125 | 30 | ybx1 |
| sdcbp29 | 5.688298 | 0.429066525 | 0.461 | 0.228 | 0.000145 | 30 | sdcbp2 |
| ctsk4 | 6.040954 | 0.326952202 | 0.225 | 0.074 | 0.000153 | 30 | ctsk |
| glod51 | 7.123862 | 0.299227815 | 0.147 | 0.038 | 0.000181 | 30 | glod5 |
| gadd45ab2 | 9.328860 | 0.329112869 | 0.216 | 0.07 | 0.000237 | 30 | gadd45ab |
| junba10 | 1.027915 | 0.516789042 | 0.559 | 0.32 | 0.000261 | 30 | junba |
| prkar2aa2 | 1.131337 | 0.353988168 | 0.284 | 0.108 | 0.000287 | 30 | prkar2aa |
| stxbp62 | 1.289561 | 0.264468754 | 0.167 | 0.046 | 0.000328 | 30 | stxbp6 |
| xaf12 | 1.338633 | 0.324975511 | 0.196 | 0.062 | 0.00034 | 30 | xaf1 |
| dap | 1.342785 | 0.322864111 | 0.255 | 0.09 | 0.000341 | 30 | dap |
| lgals3b3 | 1.386871 | 0.345995308 | 0.265 | 0.099 | 0.000352 | 30 | lgals3b |
| rpl916 | 1.446309 | 0.365419971 | 0.912 | 0.668 | 0.000367 | 30 | rpl9 |
| snx121 | 1.463851 | 0.354691142 | 0.235 | 0.083 | 0.000372 | 30 | snx12 |
| eif4a1b7 | 1.704976 | 0.440309821 | 0.559 | 0.312 | 0.000433 | 30 | eif4a1b |
| gpx4b3 | 1.715056 | 0.434643446 | 0.441 | 0.22 | 0.000436 | 30 | gpx4b |
| csde13 | 1.868664 | 0.402029637 | 0.431 | 0.207 | 0.000475 | 30 | csde1 |
| serinc14 | 2.078297 | 0.299042574 | 0.245 | 0.087 | 0.000528 | 30 | serinc1 |
| calm3b2 | 2.281180 | 0.454211026 | 0.51 | 0.278 | 0.00058 | 30 | calm3b |
| gchfr2 | 2.296057 | 0.256931266 | 0.147 | 0.039 | 0.000583 | 30 | gchfr |
| gatad2b2 | 3.071916 | 0.312496465 | 0.206 | 0.069 | 0.00078 | 30 | gatad2b |

| | | | | | | | |
|---|---|---|---|---|---|---|---|
| zfp36l1b7 | 3.249332 | 0.390485918 | 0.333 | 0.14 | 0.000825 | 30 | zfp36l1b |
| sox4a6 | 3.460389 | 0.306058512 | 0.196 | 0.063 | 0.000879 | 30 | sox4a |
| zgc:1951737 | 3.525253 | 0.325425245 | 0.294 | 0.114 | 0.000896 | 30 | zgc:195173 |
| gng76 | 3.609800 | 0.26279401 | 0.196 | 0.062 | 0.000917 | 30 | gng7 |
| txnipa7 | 3.676855 | 0.441799404 | 0.471 | 0.245 | 0.000934 | 30 | txnipa |
| pfn28 | 3.808140 | 0.44959478 | 0.461 | 0.239 | 0.000967 | 30 | pfn2 |
| aif1l6 | 4.212617 | 0.388575301 | 0.392 | 0.17 | 0.00107 | 30 | aif1l |
| cpne13 | 4.238193 | 0.370880616 | 0.314 | 0.134 | 0.001077 | 30 | cpne1 |
| ppib10 | 4.730976 | 0.427120903 | 0.627 | 0.379 | 0.001202 | 30 | ppib |
| rpl3111 | 4.856200 | 0.330437084 | 0.892 | 0.688 | 0.001234 | 30 | rpl31 |
| ecm1b5 | 5.451308 | 0.256867916 | 0.206 | 0.067 | 0.001385 | 30 | ecm1b |
| b2ml10 | 5.763525 | 0.443685644 | 0.5 | 0.294 | 0.001464 | 30 | b2ml |
| map1lc3a7 | 6.086510 | 0.317887005 | 0.225 | 0.081 | 0.001546 | 30 | map1lc3a |
| gng12a | 6.593765 | 0.269204391 | 0.137 | 0.037 | 0.001675 | 30 | gng12a |
| pde4ba9 | 7.856053 | 0.401721011 | 0.324 | 0.143 | 0.001996 | 30 | pde4ba |
| pak2a3 | 8.091107 | 0.262012634 | 0.206 | 0.069 | 0.002056 | 30 | pak2a |
| hmgb1a10 | 8.386198 | 0.449703916 | 0.431 | 0.228 | 0.002131 | 30 | hmgb1a |
| cnn28 | 8.470671 | 0.360898859 | 0.353 | 0.157 | 0.002152 | 30 | cnn2 |
| ywhabl5 | 1.121785 | 0.439515597 | 0.51 | 0.301 | 0.00285 | 30 | ywhabl |
| si:ch73-46j18.58 | 1.176458 | 0.382315536 | 0.441 | 0.209 | 0.002989 | 30 | si:ch73-46j18.5 |
| cox8a11 | 1.418937 | 0.401063106 | 0.49 | 0.268 | 0.003605 | 30 | cox8a |
| myclb1 | 1.475663 | 0.339191542 | 0.235 | 0.087 | 0.003749 | 30 | myclb |
| si:dkey-56m19.53 | 1.814444 | 0.257880959 | 0.176 | 0.056 | 0.00461 | 30 | si:dkey-56m19.5 |
| tegt5 | 1.818929 | 0.394387227 | 0.441 | 0.238 | 0.004621 | 30 | tegt |
| uqcrh4 | 1.890332 | 0.415182466 | 0.471 | 0.26 | 0.004802 | 30 | uqcrh |
| aqp3a2 | 1.921196 | 0.252103648 | 0.118 | 0.029 | 0.004881 | 30 | aqp3a |
| crema7 | 2.185693 | 0.364850731 | 0.373 | 0.175 | 0.005553 | 30 | crema |
| rpl348 | 2.379111 | 0.278637123 | 0.971 | 0.802 | 0.006044 | 30 | rpl34 |
| atp6v1e1a1 | 2.392152 | 0.30523901 | 0.176 | 0.058 | 0.006077 | 30 | atp6v1e1a |
| CU694197.11 | 2.625521 | 0.34893023 | 0.324 | 0.145 | 0.00667 | 30 | CU694197.1 |
| ier2a11 | 2.778060 | 0.479004503 | 0.461 | 0.275 | 0.007058 | 30 | ier2a |
| psmf1 | 3.035917 | 0.288745595 | 0.147 | 0.044 | 0.007713 | 30 | psmf1 |
| rdh10a4 | 3.351482 | 0.29249978 | 0.157 | 0.048 | 0.008514 | 30 | rdh10a |
| copb2 | 3.667133 | 0.257043921 | 0.186 | 0.063 | 0.009316 | 30 | copb2 |
| arl5c5 | 3.989150 | 0.322866367 | 0.265 | 0.108 | 0.010134 | 30 | arl5c |

| | | | | | | | |
|---|---|---|---|---|---|---|---|
| rps27.212 | 4.094671 | 0.344254807 | 0.735 | 0.554 | 0.010403 | 30 | rps27.2 |
| tpm4a8 | 4.354458 | 0.328797771 | 0.275 | 0.114 | 0.011063 | 30 | tpm4a |
| rps2411 | 5.310139 | 0.270876831 | 0.941 | 0.779 | 0.01349 | 30 | rps24 |
| tnfrsf9a2 | 5.318121 | 0.291032575 | 0.157 | 0.049 | 0.013511 | 30 | tnfrsf9a |
| btg29 | 5.969083 | 0.470517537 | 0.5 | 0.351 | 0.015164 | 30 | btg2 |
| rpl3210 | 6.063658 | 0.251276408 | 0.98 | 0.824 | 0.015405 | 30 | rpl32 |
| rgs5b3 | 6.508700 | 0.269658339 | 0.206 | 0.073 | 0.016535 | 30 | rgs5b |
| rps27l7 | 6.686964 | 0.406217587 | 0.402 | 0.218 | 0.016988 | 30 | rps27l |
| cald1a7 | 6.944971 | 0.319346713 | 0.284 | 0.12 | 0.017644 | 30 | cald1a |
| si:dkey-222f8.3 | 6.948747 | 0.281910677 | 0.186 | 0.066 | 0.017653 | 30 | si:dkey-222f8.3 |
| ppp1cb2 | 7.190343 | 0.263071236 | 0.196 | 0.07 | 0.018267 | 30 | ppp1cb |
| ccdc32 | 7.578149 | 0.286089599 | 0.157 | 0.05 | 0.019252 | 30 | ccdc32 |
| jak14 | 7.751959 | 0.318578178 | 0.245 | 0.1 | 0.019694 | 30 | jak1 |
| rps289 | 8.872867 | 0.252402478 | 0.99 | 0.84 | 0.022542 | 30 | rps28 |
| h3f3c11 | 9.743447 | 0.390217473 | 0.52 | 0.305 | 0.024753 | 30 | h3f3c |
| hmgn77 | 9.990090 | 0.405484189 | 0.51 | 0.318 | 0.02538 | 30 | hmgn7 |
| rpl3810 | 1.030716 | 0.28979434 | 0.951 | 0.754 | 0.026185 | 30 | rpl38 |
| tgif14 | 1.086165 | 0.302999854 | 0.245 | 0.1 | 0.027594 | 30 | tgif1 |
| rps2116 | 1.138270 | 0.283418322 | 0.971 | 0.776 | 0.028918 | 30 | rps21 |
| vamp31 | 1.190455 | 0.38588618 | 0.373 | 0.196 | 0.030244 | 30 | vamp3 |
| rpl249 | 1.217076 | 0.259502192 | 0.961 | 0.794 | 0.03092 | 30 | rpl24 |
| ier54 | 1.230778 | 0.394427079 | 0.304 | 0.147 | 0.031268 | 30 | ier5 |
| eif4ebp24 | 1.300385 | 0.382923144 | 0.373 | 0.193 | 0.033036 | 30 | eif4ebp2 |
| egr18 | 1.358824 | 0.390312218 | 0.363 | 0.184 | 0.034521 | 30 | egr1 |
| ywhaqb8 | 1.445002 | 0.338480745 | 0.324 | 0.155 | 0.03671 | 30 | ywhaqb |
| tob1a4 | 1.465608 | 0.322186591 | 0.314 | 0.145 | 0.037234 | 30 | tob1a |
| rpl10a9 | 1.605095 | 0.267484097 | 0.961 | 0.779 | 0.040777 | 30 | rpl10a |
| hp1bp36 | 2.080977 | 0.354652945 | 0.265 | 0.121 | 0.052867 | 30 | hp1bp3 |
| txndc171 | 2.170473 | 0.276373209 | 0.167 | 0.058 | 0.055141 | 30 | txndc17 |
| midn7 | 2.701440 | 0.307946763 | 0.314 | 0.146 | 0.06863 | 30 | midn |
| wbp2nl2 | 2.961117 | 0.293269931 | 0.225 | 0.093 | 0.075227 | 30 | wbp2nl |
| rpl612 | 3.154870 | 0.309581948 | 0.931 | 0.701 | 0.080149 | 30 | rpl6 |
| lamc13 | 3.203513 | 0.25090392 | 0.167 | 0.058 | 0.081385 | 30 | lamc1 |
| rps149 | 3.379682 | 0.258616793 | 0.941 | 0.781 | 0.085861 | 30 | rps14 |
| litaf5 | 3.625122 | 0.338516298 | 0.304 | 0.147 | 0.092096 | 30 | litaf |

| | | | | | | | |
|---|---|---|---|---|---|---|---|
| akt1s11 | 3.730514 | 0.27769481 | 0.176 | 0.065 | 0.094774 | 30 | akt1s1 |
| iscub1 | 4.180460 | 0.325938836 | 0.196 | 0.078 | 0.106205 | 30 | iscub |
| naca13 | 5.045089 | 0.312099564 | 0.863 | 0.652 | 0.12817 | 30 | naca |
| selenof3 | 5.048355 | 0.355290991 | 0.461 | 0.268 | 0.128253 | 30 | selenof |
| s100a10b10 | 5.711214 | 0.340403666 | 0.529 | 0.286 | 0.145093 | 30 | s100a10b |
| glrx52 | 5.799502 | 0.347924391 | 0.225 | 0.097 | 0.147336 | 30 | glrx5 |
| kctd12.26 | 6.077817 | 0.302203908 | 0.235 | 0.101 | 0.154407 | 30 | kctd12.2 |
| dynlt32 | 6.573753 | 0.293591862 | 0.324 | 0.158 | 0.167006 | 30 | dynlt3 |
| fundc23 | 7.669156 | 0.283190103 | 0.225 | 0.097 | 0.194835 | 30 | fundc2 |
| anxa47 | 8.470683 | 0.322359208 | 0.304 | 0.151 | 0.215198 | 30 | anxa4 |
| rpl279 | 9.524686 | 0.254840359 | 0.951 | 0.78 | 0.241975 | 30 | rpl27 |
| fth1a2 | 9.676876 | 0.3032283 | 0.833 | 0.612 | 0.245841 | 30 | fth1a |
| mcl1b10 | 1.081793 | 0.333249874 | 0.451 | 0.252 | 0.27483 | 30 | mcl1b |
| adssl | 1.133603 | 0.259993728 | 0.255 | 0.113 | 0.287992 | 30 | adssl |
| rps1513 | 1.147294 | 0.283812274 | 0.922 | 0.717 | 0.29147 | 30 | rps15 |
| arf2b1 | 1.164536 | 0.31342086 | 0.333 | 0.169 | 0.295851 | 30 | arf2b |
| gabarapa4 | 1.280438 | 0.323038935 | 0.343 | 0.182 | 0.325295 | 30 | gabarapa |
| psmc52 | 1.332308 | 0.265245229 | 0.235 | 0.103 | 0.338473 | 30 | psmc5 |
| chchd22 | 1.416410 | 0.251227555 | 0.196 | 0.08 | 0.359839 | 30 | chchd2 |
| map1lc3b3 | 1.520402 | 0.270381622 | 0.265 | 0.125 | 0.386258 | 30 | map1lc3b |
| ndufa14 | 1.832216 | 0.324502342 | 0.422 | 0.243 | 0.465475 | 30 | ndufa1 |
| capns1b6 | 1.895192 | 0.355927635 | 0.284 | 0.146 | 0.481474 | 30 | capns1b |
| sat1a.26 | 1.914643 | 0.365581177 | 0.471 | 0.302 | 0.486415 | 30 | sat1a.2 |
| rnaseka1 | 2.143030 | 0.276405284 | 0.275 | 0.13 | 0.544437 | 30 | rnaseka |
| rpl3010 | 2.186428 | 0.273754061 | 0.931 | 0.73 | 0.555462 | 30 | rpl30 |
| rplp213 | 2.221957 | 0.258381799 | 0.951 | 0.783 | 0.564488 | 30 | rplp2 |
| elmo14 | 2.419011 | 0.271678908 | 0.176 | 0.071 | 0.61455 | 30 | elmo1 |
| ppp2r1ba1 | 2.759835 | 0.270277643 | 0.216 | 0.096 | 0.701136 | 30 | ppp2r1ba |
| csrnp1b6 | 3.073331 | 0.319085162 | 0.314 | 0.165 | 0.78078 | 30 | csrnp1b |
| rpl2213 | 3.220908 | 0.257542498 | 0.892 | 0.721 | 0.818272 | 30 | rpl22 |
| vps28 | 3.378291 | 0.263394921 | 0.225 | 0.102 | 0.858255 | 30 | vps28 |
| vdac35 | 3.889171 | 0.285446981 | 0.324 | 0.169 | 0.988044 | 30 | vdac3 |
| chmp5b3 | 4.470505 | 0.260320427 | 0.265 | 0.129 | 1 | 30 | chmp5b |
| atp5meb3 | 5.619759 | 0.331777765 | 0.51 | 0.34 | 1 | 30 | atp5meb |
| atp5mea2 | 6.538431 | 0.294509064 | 0.294 | 0.154 | 1 | 30 | atp5mea |

| | | | | | | | |
|---|---|---|---|---|---|---|---|
| rplp2l11 | 7.416019 | 0.261851638 | 0.98 | 0.794 | 1 | 30 | rplp2l |
| mif2 | 0.0001 | 0.324298257 | 0.353 | 0.208 | 1 | 30 | mif |
| rpl27a12 | 0.0001 | 0.307923376 | 0.716 | 0.523 | 1 | 30 | rpl27a |
| atf4b10 | 0.0001 | 0.30654263 | 0.441 | 0.278 | 1 | 30 | atf4b |
| elocb4 | 0.0001 | 0.296145532 | 0.343 | 0.197 | 1 | 30 | elocb |
| rps1014 | 0.0001 | 0.28371807 | 0.951 | 0.727 | 1 | 30 | rps10 |
| rhoaa7 | 0.0001 | 0.264838411 | 0.324 | 0.177 | 1 | 30 | rhoaa |
| si:dkey-51e6.14 | 0.0001 | 0.309465513 | 0.235 | 0.12 | 1 | 30 | si:dkey-51e6.1 |
| myl12.111 | 0.0001 | 0.301044718 | 0.608 | 0.427 | 1 | 30 | myl12.1 |
| atp5f1e3 | 0.0001 | 0.291961753 | 0.637 | 0.492 | 1 | 30 | atp5f1e |
| tpm38 | 0.0001 | 0.291405845 | 0.461 | 0.285 | 1 | 30 | tpm3 |
| hif1ab2 | 0.0001 | 0.256473764 | 0.245 | 0.123 | 1 | 30 | hif1ab |
| rps4x12 | 0.0001 | 0.261462367 | 0.941 | 0.737 | 1 | 30 | rps4x |
| prdx33 | 0.0002 | 0.272697667 | 0.275 | 0.143 | 1 | 30 | prdx3 |
| csnk1a12 | 0.0002 | 0.272535706 | 0.402 | 0.238 | 1 | 30 | csnk1a1 |
| serinc24 | 0.0002 | 0.267408713 | 0.245 | 0.125 | 1 | 30 | serinc2 |
| rab5c4 | 0.0002 | 0.271316988 | 0.255 | 0.132 | 1 | 30 | rab5c |
| tmem50a2 | 0.0002 | 0.256347299 | 0.284 | 0.152 | 1 | 30 | tmem50a |
| ppiab6 | 0.0002 | 0.253469579 | 0.843 | 0.675 | 1 | 30 | ppiab |
| cirbpa2 | 0.0002 | 0.250362178 | 0.775 | 0.624 | 1 | 30 | cirbpa |
| ost44 | 0.0002 | 0.280215275 | 0.431 | 0.262 | 1 | 30 | ost4 |
| rab106 | 0.0002 | 0.275746302 | 0.363 | 0.215 | 1 | 30 | rab10 |
| pim19 | 0.0003 | 0.284638814 | 0.304 | 0.17 | 1 | 30 | pim1 |
| eif5a7 | 0.0003 | 0.275696306 | 0.667 | 0.496 | 1 | 30 | eif5a |
| pomp3 | 0.0003 | 0.288746901 | 0.412 | 0.261 | 1 | 30 | pomp |
| sod14 | 0.0003 | 0.292829896 | 0.5 | 0.346 | 1 | 30 | sod1 |
| prdx59 | 0.0004 | 0.285770865 | 0.304 | 0.18 | 1 | 30 | prdx5 |
| gnb1b3 | 0.0004 | 0.275968635 | 0.422 | 0.267 | 1 | 30 | gnb1b |
| nucks1b | 0.0004 | 0.258503884 | 0.225 | 0.115 | 1 | 30 | nucks1b |
| cdc42l6 | 0.0004 | 0.265886903 | 0.382 | 0.235 | 1 | 30 | cdc42l |
| btf38 | 0.0005 | 0.268575269 | 0.814 | 0.611 | 1 | 30 | btf3 |
| etfa2 | 0.0005 | 0.260895711 | 0.196 | 0.099 | 1 | 30 | etfa |
| sap30l1 | 0.0007 | 0.272844298 | 0.265 | 0.152 | 1 | 30 | sap30l |
| taf131 | 0.0007 | 0.262053264 | 0.382 | 0.239 | 1 | 30 | taf13 |
| ywhae13 | 0.0007 | 0.274441771 | 0.441 | 0.296 | 1 | 30 | ywhae1 |

| | | | | | | | |
|---|---|---|---|---|---|---|---|
| prdx27 | 0.0013 | 0.267424085 | 0.578 | 0.423 | 1 | 30 | prdx2 |
| mgst3b4 | 0.0013 | 0.261042962 | 0.314 | 0.194 | 1 | 30 | mgst3b |
| cox5ab2 | 0.0016 | 0.28835242 | 0.333 | 0.219 | 1 | 30 | cox5ab |
| ywhaba4 | 0.0018 | 0.25994591 | 0.471 | 0.33 | 1 | 30 | ywhaba |
| atp5f1d5 | 0.0024 | 0.266249123 | 0.392 | 0.275 | 1 | 30 | atp5f1d |
| ndfip11 | 0.0027 | 0.250732353 | 0.275 | 0.172 | 1 | 30 | ndfip1 |
| uba5210 | 0.003 | 0.250875444 | 0.863 | 0.661 | 1 | 30 | uba52 |
| ctsla9 | 0.0032 | 0.252254737 | 0.52 | 0.375 | 1 | 30 | ctsla |
| mt-nd412 | 0.0055 | 0.264718657 | 0.51 | 0.45 | 1 | 30 | mt-nd4 |
| si:dkey-21e2.13 | 0 | 2.531088288 | 0.946 | 0.003 | 0 | 31 | si:dkey-21e2.13 |
| si:dkey-21e2.15 | 0 | 2.290181862 | 0.839 | 0.005 | 0 | 31 | si:dkey-21e2.15 |
| si:dkey-21e2.16 | 0 | 2.228518734 | 0.731 | 0.003 | 0 | 31 | si:dkey-21e2.16 |
| si:ch211-212d10.11 | 0 | 1.997010096 | 0.903 | 0.019 | 0 | 31 | si:ch211-212d10.1 |
| si:dkey-21e2.14 | 0 | 1.699772343 | 0.71 | 0.002 | 0 | 31 | si:dkey-21e2.14 |
| si:dkey-21e2.10 | 0 | 1.558637031 | 0.398 | 0.001 | 0 | 31 | si:dkey-21e2.10 |
| si:dkey-78l4.2 | 0 | 1.52940346 | 0.484 | 0.005 | 0 | 31 | si:dkey-78l4.2 |
| il13 | 0 | 1.521958098 | 0.581 | 0.001 | 0 | 31 | il13 |
| il4 | 0 | 1.486955 | 0.581 | 0.004 | 0 | 31 | il4 |
| il11b | 0 | 1.403464986 | 0.581 | 0.006 | 0 | 31 | il11b |
| csf3a | 0 | 1.38847585 | 0.581 | 0.003 | 0 | 31 | csf3a |
| si:dkey-21e2.8 | 0 | 1.181232428 | 0.29 | 0.001 | 0 | 31 | si:dkey-21e2.8 |
| CR847844.1 | 0 | 0.990911665 | 0.462 | 0 | 0 | 31 | CR847844.1 |
| si:dkey-78l4.1 | 0 | 0.891251514 | 0.237 | 0.001 | 0 | 31 | si:dkey-78l4.1 |
| LOC103911745 | 0 | 0.834604012 | 0.419 | 0 | 0 | 31 | LOC103911745 |
| si:dkey-12f6.5 | 0 | 0.830242622 | 0.398 | 0.002 | 0 | 31 | si:dkey-12f6.5 |
| si:dkey-8e10.2 | 0 | 0.627203044 | 0.323 | 0.001 | 0 | 31 | si:dkey-8e10.2 |
| BX571675.1 | 0 | 0.462289624 | 0.226 | 0.001 | 0 | 31 | BX571675.1 |
| kcnc1b | 0 | 0.286832764 | 0.151 | 0 | 0 | 31 | kcnc1b |
| cldni1 | 6.211966 | 0.709037462 | 0.355 | 0.01 | 1.5781500 | 31 | cldni |
| sox13 | 3.095039 | 0.664363554 | 0.333 | 0.011 | 7.8629476 | 31 | sox13 |
| si:dkeyp-28d2.4 | 4.341200 | 0.262191111 | 0.151 | 0.002 | 1.1028821 | 31 | si:dkeyp-28d2.4 |
| faslg1 | 2.266420 | 0.715363284 | 0.366 | 0.015 | 5.7578405 | 31 | faslg |
| si:dkey-33i11.92 | 1.466537 | 0.810073476 | 0.43 | 0.024 | 3.7257386 | 31 | si:dkey-33i11.9 |
| LOC1001482542 | 1.884351 | 0.765978269 | 0.419 | 0.023 | 4.7871958 | 31 | LOC100148254 |
| si:ch211-112c15.8 | 1.742745 | 0.307756973 | 0.14 | 0.002 | 4.4274458 | 31 | si:ch211-112c15.8 |

| | | | | | | | |
|---|---|---|---|---|---|---|---|
| ccl34b.41 | 1.190604 | 1.067644433 | 0.473 | 0.032 | 3.0247310 | 31 | ccl34b.4 |
| gata3 | 3.102808 | 0.456240491 | 0.237 | 0.008 | 7.8826850 | 31 | gata3 |
| slc35f4 | 6.720660 | 0.318477169 | 0.161 | 0.003 | 1.7073838 | 31 | slc35f4 |
| si:ch211-67e16.32 | 8.545280 | 0.646894367 | 0.333 | 0.016 | 2.1709285 | 31 | si:ch211-67e16.3 |
| LOC103911004 | 5.393422 | 0.294599144 | 0.108 | 0.001 | 1.3701989 | 31 | LOC103911004 |
| ca22 | 5.128806 | 1.047507542 | 0.548 | 0.051 | 1.3029732 | 31 | ca2 |
| XLOC-024867 | 1.156733 | 0.449664973 | 0.237 | 0.01 | 2.9386822 | 31 | XLOC-024867 |
| si:cabz01036022.12 | 1.521008 | 0.456563502 | 0.226 | 0.011 | 3.8641213 | 31 | si:cabz01036022.1 |
| laptm511 | 6.349284 | 1.43605795 | 0.817 | 0.173 | 1.6130356 | 31 | laptm5 |
| CU075735.1 | 1.588251 | 0.304745332 | 0.14 | 0.004 | 4.0349517 | 31 | CU075735.1 |
| tnfrsf9a3 | 8.810785 | 1.015450651 | 0.441 | 0.047 | 2.2383799 | 31 | tnfrsf9a |
| CABZ01073834.14 | 1.090549 | 0.728127024 | 0.419 | 0.043 | 2.7705404 | 31 | CABZ01073834.1 |
| blk | 7.240346 | 0.383511967 | 0.183 | 0.008 | 1.8394100 | 31 | blk |
| si:ch211-147k10.5 | 9.814286 | 0.270795518 | 0.129 | 0.004 | 2.4933194 | 31 | si:ch211-147k10.5 |
| samsn1a3 | 4.754303 | 0.683262396 | 0.376 | 0.037 | 1.2078307 | 31 | samsn1a |
| scgn2 | 9.340863 | 0.542985327 | 0.29 | 0.022 | 2.3730463 | 31 | scgn |
| samsn1b1 | 8.569098 | 0.460028667 | 0.269 | 0.022 | 2.1769795 | 31 | samsn1b |
| CABZ01041494.1 | 2.132862 | 0.255739637 | 0.108 | 0.004 | 5.4185371 | 31 | CABZ01041494.1 |
| si:dkey-276j7.2 | 6.812817 | 0.299293312 | 0.14 | 0.006 | 1.7307962 | 31 | si:dkey-276j7.2 |
| si:dkey-27h10.22 | 5.382205 | 0.475780143 | 0.269 | 0.027 | 1.3673493 | 31 | si:dkey-27h10.2 |
| tfa | 3.312683 | 0.567775647 | 0.172 | 0.012 | 8.4158719 | 31 | tfa |
| MFAP4 (1 of many)1 | 6.621181 | 0.662662019 | 0.409 | 0.06 | 1.6821110 | 31 | MFAP4 (1 of many) |
| CR392341.16 | 3.210749 | 0.581957411 | 0.387 | 0.059 | 8.1569085 | 31 | CR392341.1 |
| pfn112 | 4.028597 | 1.002630905 | 1 | 0.5 | 1.0234653 | 31 | pfn1 |
| si:ch211-207n23.21 | 1.607955 | 0.49233984 | 0.247 | 0.026 | 4.0850099 | 31 | si:ch211-207n23.2 |
| rgs138 | 1.927013 | 0.861670888 | 0.559 | 0.124 | 4.8955787 | 31 | rgs13 |
| sult5a11 | 5.705139 | 0.362265579 | 0.151 | 0.01 | 1.4493907 | 31 | sult5a1 |
| cd247l2 | 1.383056 | 0.372989263 | 0.204 | 0.019 | 3.5136545 | 31 | cd247l |
| rplp214 | 1.701529 | 0.512865635 | 0.978 | 0.783 | 4.3227365 | 31 | rplp2 |
| wasb11 | 2.563018 | 0.882691677 | 0.645 | 0.172 | 6.5113484 | 31 | wasb |
| nbeal21 | 2.543187 | 0.351386363 | 0.215 | 0.022 | 6.4609666 | 31 | nbeal2 |
| si:dkey-151g10.61 | 6.074965 | 0.250582741 | 0.989 | 0.976 | 1.5433450 | 31 | si:dkey-151g10.6 |
| CU855821.11 | 3.798720 | 0.320371535 | 0.161 | 0.013 | 9.6506483 | 31 | CU855821.1 |
| zbtb16a1 | 3.933332 | 0.488289471 | 0.301 | 0.043 | 9.9926299 | 31 | zbtb16a |
| si:dkey-262k9.22 | 8.145710 | 0.475725362 | 0.258 | 0.033 | 2.0694177 | 31 | si:dkey-262k9.2 |

| | | | | | | |
|---|---|---|---|---|---|---|
| tnfb5 | 9.076344 | 0.606004883 | 0.323 | 0.049 | 2.3058452 | 31 | tnfb |
| si:ch1073-110a20.31 | 2.086132 | 0.551232848 | 0.323 | 0.048 | 5.2998208 | 31 | si:ch1073-110a20.3 |
| srgn7 | 3.430538 | 0.928602544 | 0.71 | 0.234 | 8.7152832 | 31 | srgn |
| gpr183a4 | 5.985696 | 0.606016344 | 0.366 | 0.064 | 1.5206660 | 31 | gpr183a |
| fosl27 | 1.485862 | 0.789284505 | 0.484 | 0.112 | 3.7748334 | 31 | fosl2 |
| rplp110 | 3.309727 | 0.382716987 | 0.978 | 0.882 | 8.4083614 | 31 | rplp1 |
| rplp2l12 | 3.580111 | 0.476693805 | 0.989 | 0.795 | 9.0952720 | 31 | rplp2l |
| zmp:00000013232 | 6.935379 | 0.943283275 | 0.516 | 0.126 | 1.7619332 | 31 | zmp:0000001323 |
| rps205 | 6.030061 | 0.333429283 | 0.968 | 0.91 | 1.5319369 | 31 | rps20 |
| ndufa4 | 6.805293 | 0.339112811 | 0.172 | 0.017 | 1.7288847 | 31 | ndufa4 |
| si:ch211-119e14.14 | 6.811983 | 0.609175683 | 0.376 | 0.07 | 1.7305845 | 31 | si:ch211-119e14.1 |
| XLOC-0430434 | 1.418405 | 0.574375951 | 0.29 | 0.044 | 3.6034594 | 31 | XLOC-043043 |
| tnip13 | 2.609749 | 0.60068145 | 0.398 | 0.079 | 6.6300698 | 31 | tnip1 |
| si:dkey-25e12.32 | 7.563639 | 0.464923831 | 0.301 | 0.048 | 1.9215427 | 31 | si:dkey-25e12.3 |
| h1fx9 | 1.822478 | 0.816984466 | 0.645 | 0.189 | 4.6300066 | 31 | h1fx |
| rpl349 | 6.236007 | 0.44004268 | 0.968 | 0.802 | 1.5842576 | 31 | rpl34 |
| abca2 | 1.229241 | 0.378664902 | 0.215 | 0.027 | 3.1228878 | 31 | abca2 |
| sla23 | 1.589343 | 0.624687558 | 0.333 | 0.06 | 4.0377282 | 31 | sla2 |
| rpl3811 | 4.296448 | 0.490640484 | 0.989 | 0.754 | 1.0915127 | 31 | rpl38 |
| si:ch211-239m17.1 | 9.922312 | 0.269511817 | 0.151 | 0.014 | 2.5207634 | 31 | si:ch211-239m17.1 |
| rpl397 | 3.513358 | 0.332692332 | 0.946 | 0.878 | 8.9256877 | 31 | rpl39 |
| arhgdig10 | 7.262594 | 0.784856396 | 0.72 | 0.259 | 1.8450621 | 31 | arhgdig |
| rps127 | 1.115325 | 0.378366136 | 0.957 | 0.842 | 2.8334856 | 31 | rps12 |
| FP236356.12 | 1.204582 | 0.474914406 | 0.28 | 0.046 | 3.0602412 | 31 | FP236356.1 |
| raraa2 | 2.299722 | 0.38219068 | 0.247 | 0.037 | 5.8424451 | 31 | raraa |
| rps177 | 5.325710 | 0.450723596 | 0.968 | 0.779 | 1.3529966 | 31 | rps17 |
| grap2b9 | 5.750619 | 0.596506799 | 0.495 | 0.123 | 1.4609449 | 31 | grap2b |
| ms4a17a.63 | 7.382636 | 0.513366219 | 0.29 | 0.051 | 1.8755587 | 31 | ms4a17a.6 |
| nr4a110 | 1.070820 | 0.723170376 | 0.548 | 0.157 | 2.7204199 | 31 | nr4a1 |
| arpc1b11 | 1.723610 | 0.846564658 | 0.785 | 0.312 | 4.3788331 | 31 | arpc1b |
| rps2910 | 4.281800 | 0.370591718 | 0.978 | 0.844 | 1.0877913 | 31 | rps29 |
| prkcq | 6.592234 | 0.274950363 | 0.129 | 0.012 | 1.6747570 | 31 | prkcq |
| rpl35a10 | 8.019064 | 0.418961905 | 0.957 | 0.782 | 2.0372432 | 31 | rpl35a |
| rpl377 | 1.652843 | 0.359673135 | 0.968 | 0.855 | 4.1990498 | 31 | rpl37 |
| CABZ01084575.13 | 3.055781 | 0.415454721 | 0.247 | 0.04 | 7.7632140 | 31 | CABZ01084575.1 |

| | | | | | | | |
|---|---|---|---|---|---|---|---|
| ada1 | 5.544879 | 0.41064518 | 0.258 | 0.044 | 1.4086766 | 31 | ada |
| rps1811 | 3.709828 | 0.449545421 | 0.968 | 0.754 | 9.4248184 | 31 | rps18 |
| rps2810 | 5.498792 | 0.359862561 | 0.968 | 0.841 | 1.3969681 | 31 | rps28 |
| rpl268 | 1.418330 | 0.343649056 | 0.978 | 0.851 | 3.6032695 | 31 | rpl26 |
| rps1614 | 1.732184 | 0.432859289 | 0.989 | 0.766 | 4.4006147 | 31 | rps16 |
| rps225 | 2.512384 | 0.3789169 | 0.968 | 0.842 | 6.3827127 | 31 | rps2 |
| dennd2da1 | 3.503782 | 0.261376247 | 0.161 | 0.02 | 8.9013583 | 31 | dennd2da |
| rpl3211 | 5.433729 | 0.386514353 | 0.978 | 0.824 | 1.3804389 | 31 | rpl32 |
| unc119b1 | 7.913448 | 0.360315431 | 0.226 | 0.037 | 2.0104115 | 31 | unc119b |
| cebpb11 | 1.150065 | 0.802874428 | 0.753 | 0.339 | 2.9217423 | 31 | cebpb |
| tnfaip35 | 2.161383 | 0.463433134 | 0.29 | 0.06 | 5.4909938 | 31 | tnfaip3 |
| si:zfos-2330d3.11 | 4.193859 | 0.306095486 | 0.194 | 0.028 | 1.0654499 | 31 | si:zfos-2330d3.1 |
| rps26l11 | 4.738143 | 0.473864096 | 0.935 | 0.7 | 1.2037254 | 31 | rps26l |
| rps27.115 | 2.147689 | 0.460164853 | 0.935 | 0.71 | 5.4562043 | 31 | rps27.1 |
| grap2a4 | 4.425636 | 0.423512538 | 0.247 | 0.048 | 1.1243329 | 31 | grap2a |
| rpl118 | 4.896750 | 0.335776917 | 0.946 | 0.83 | 1.2440193 | 31 | rpl11 |
| rpl3011 | 6.135846 | 0.427992418 | 0.946 | 0.73 | 1.5588117 | 31 | rpl30 |
| capgb7 | 7.722376 | 0.540371605 | 0.419 | 0.115 | 1.9618696 | 31 | capgb |
| rps15a12 | 8.968989 | 0.395180763 | 0.968 | 0.798 | 2.2785717 | 31 | rps15a |
| bhlhe409 | 9.895885 | 0.668873216 | 0.495 | 0.167 | 2.5140496 | 31 | bhlhe40 |
| coro1a10 | 1.078229 | 0.667065748 | 0.656 | 0.236 | 2.7392418 | 31 | coro1a |
| rpl1411 | 1.356356 | 0.486233566 | 0.957 | 0.669 | 3.4458227 | 31 | rpl14 |
| calm1b8 | 1.534914 | 0.501027027 | 0.387 | 0.102 | 3.8994497 | 31 | calm1b |
| rpl1812 | 1.741561 | 0.384850226 | 0.978 | 0.793 | 4.4244375 | 31 | rpl18 |
| zgc:1366051 | 2.520096 | 0.307664678 | 0.151 | 0.02 | 6.4023062 | 31 | zgc:136605 |
| rps1410 | 3.079211 | 0.381068218 | 0.957 | 0.781 | 7.8227370 | 31 | rps14 |
| rps1310 | 3.790085 | 0.409368443 | 0.946 | 0.749 | 9.6287114 | 31 | rps13 |
| rpl2710 | 4.335128 | 0.377588011 | 0.946 | 0.78 | 1.1013395 | 31 | rpl27 |
| glipr1a7 | 6.402265 | 0.390384493 | 0.301 | 0.068 | 1.6264956 | 31 | glipr1a |
| rpl7a9 | 6.436620 | 0.346948287 | 0.978 | 0.824 | 1.6352234 | 31 | rpl7a |
| tnfrsf9b1 | 6.556551 | 0.275313186 | 0.129 | 0.016 | 1.6656919 | 31 | tnfrsf9b |
| si:ch211-226m16.25 | 8.744057 | 0.403298605 | 0.301 | 0.069 | 2.2214278 | 31 | si:ch211-226m16.2 |
| rpl3612 | 1.026547 | 0.376384706 | 0.968 | 0.807 | 2.6079450 | 31 | rpl36 |
| arl5c6 | 1.146204 | 0.519500095 | 0.387 | 0.108 | 2.9119334 | 31 | arl5c |
| nfkbiaa6 | 1.278157 | 0.67262624 | 0.462 | 0.152 | 3.2471602 | 31 | nfkbiaa |

| | | | | | | | |
|---|---|---|---|---|---|---|---|
| rpl236 | 1.332405 | 0.276155965 | 0.968 | 0.873 | 3.3849771 | 31 | rpl23 |
| rps2412 | 1.339665 | 0.378209803 | 0.968 | 0.779 | 3.4034189 | 31 | rps24 |
| rpsa13 | 1.385033 | 0.378538926 | 0.978 | 0.786 | 3.5186777 | 31 | rpsa |
| rpl18a11 | 1.699478 | 0.352133119 | 0.935 | 0.801 | 4.3175240 | 31 | rpl18a |
| rpl1511 | 1.721976 | 0.393346284 | 0.957 | 0.764 | 4.3746813 | 31 | rpl15 |
| rps239 | 2.335125 | 0.399775029 | 0.946 | 0.745 | 5.9323853 | 31 | rps23 |
| rps2117 | 2.350940 | 0.394607151 | 0.957 | 0.776 | 5.9725638 | 31 | rps21 |
| XLOC-0397572 | 2.995950 | 0.270218404 | 0.172 | 0.026 | 7.6112121 | 31 | XLOC-039757 |
| rps78 | 5.441851 | 0.330020109 | 0.957 | 0.829 | 1.3825024 | 31 | rps7 |
| rps68 | 5.874390 | 0.340086496 | 0.935 | 0.804 | 1.4923889 | 31 | rps6 |
| ppp1r184 | 7.652604 | 0.384166592 | 0.258 | 0.056 | 1.9441442 | 31 | ppp1r18 |
| adcy71 | 9.790062 | 0.339284557 | 0.172 | 0.028 | 2.4871653 | 31 | adcy7 |
| wipf1a5 | 1.014487 | 0.412818525 | 0.29 | 0.068 | 2.5773046 | 31 | wipf1a |
| hmha1b5 | 3.498028 | 0.406002626 | 0.301 | 0.074 | 8.8867405 | 31 | hmha1b |
| rps198 | 4.193191 | 0.295394316 | 0.978 | 0.869 | 1.0652804 | 31 | rps19 |
| rpl36a9 | 7.026159 | 0.319708858 | 0.957 | 0.824 | 1.7849957 | 31 | rpl36a |
| nme2b.13 | 8.349322 | 0.288355648 | 0.978 | 0.857 | 2.1211452 | 31 | nme2b.1 |
| tagln212 | 8.833291 | 0.580387962 | 0.548 | 0.209 | 2.2440976 | 31 | tagln2 |
| irf4b3 | 9.474091 | 0.284832284 | 0.151 | 0.023 | 2.4068928 | 31 | irf4b |
| ccl34b.12 | 9.661360 | 0.475856238 | 0.172 | 0.029 | 2.4544686 | 31 | ccl34b.1 |
| rpl88 | 1.102896 | 0.351114281 | 0.957 | 0.795 | 2.8019074 | 31 | rpl8 |
| crip112 | 1.217291 | 0.612640804 | 0.677 | 0.284 | 3.0925297 | 31 | crip1 |
| rpl358 | 1.322584 | 0.331227541 | 0.968 | 0.814 | 3.3600250 | 31 | rpl35 |
| zgc:11418818 | 1.499195 | 0.470535728 | 0.957 | 0.668 | 3.8087071 | 31 | zgc:114188 |
| rps513 | 1.503126 | 0.403236721 | 0.935 | 0.729 | 3.8186931 | 31 | rps5 |
| rps4x13 | 1.540263 | 0.4101457 | 0.968 | 0.737 | 3.9130388 | 31 | rps4x |
| ptpn64 | 2.431259 | 0.440187737 | 0.355 | 0.099 | 6.1766156 | 31 | ptpn6 |
| rpl23a12 | 5.014569 | 0.440539805 | 0.925 | 0.67 | 1.2739514 | 31 | rpl23a |
| rpl2910 | 5.684920 | 0.330695076 | 0.968 | 0.823 | 1.4442540 | 31 | rpl29 |
| rasal32 | 7.704246 | 0.298001355 | 0.161 | 0.027 | 1.9572638 | 31 | rasal3 |
| rpl75 | 8.151402 | 0.335191447 | 0.968 | 0.809 | 2.0708638 | 31 | rpl7 |
| lpar5a4 | 9.755028 | 0.335732046 | 0.215 | 0.045 | 2.4782648 | 31 | lpar5a |
| n4bp1 | 1.481994 | 0.347377662 | 0.204 | 0.042 | 3.7650078 | 31 | n4bp1 |
| rpl127 | 1.543715 | 0.309289399 | 0.946 | 0.813 | 3.9218086 | 31 | rpl12 |
| si:ch211-191c10.13 | 1.602320 | 0.352723172 | 0.226 | 0.049 | 4.0706944 | 31 | si:ch211-191c10.1 |

| | | | | | | | |
|---|---|---|---|---|---|---|---|
| si:ch73-226l13.21 | 1.607022 | 0.254531934 | 0.118 | 0.016 | 4.0826406 | 31 | si:ch73-226l13.2 |
| selplg5 | 1.727181 | 0.325838742 | 0.215 | 0.046 | 4.3879037 | 31 | selplg |
| cox8b5 | 1.903778 | 0.43558272 | 0.344 | 0.101 | 4.8365503 | 31 | cox8b |
| si:ch211-241f5.33 | 2.044331 | 0.379958832 | 0.258 | 0.062 | 5.1936232 | 31 | si:ch211-241f5.3 |
| rpl139 | 3.882211 | 0.310618975 | 0.968 | 0.818 | 9.8627580 | 31 | rpl13 |
| si:ch211-66e2.36 | 4.051931 | 0.38312807 | 0.258 | 0.063 | 1.0293931 | 31 | si:ch211-66e2.3 |
| rplp04 | 4.642906 | 0.284278143 | 0.968 | 0.859 | 1.1795303 | 31 | rplp0 |
| rps1514 | 5.478449 | 0.42063203 | 0.957 | 0.717 | 1.3918000 | 31 | rps15 |
| rps267 | 9.503034 | 0.363030363 | 0.957 | 0.751 | 2.4142459 | 31 | rps26 |
| cd6313 | 1.132648 | 0.581187992 | 0.742 | 0.373 | 2.8774943 | 31 | cd63 |
| rpl177 | 1.265349 | 0.347857374 | 0.957 | 0.764 | 3.2146203 | 31 | rpl17 |
| rpl2110 | 1.434117 | 0.334703749 | 0.946 | 0.777 | 3.6433748 | 31 | rpl21 |
| wipf1b3 | 2.431279 | 0.30164284 | 0.258 | 0.065 | 6.1766654 | 31 | wipf1b |
| rab446 | 2.594092 | 0.358121233 | 0.247 | 0.061 | 6.5902920 | 31 | rab44 |
| rpl2410 | 3.936857 | 0.326162576 | 0.957 | 0.794 | 1.0001587 | 31 | rpl24 |
| faua11 | 5.967667 | 0.395838019 | 0.946 | 0.718 | 1.5160858 | 31 | faua |
| rpl2214 | 7.641996 | 0.372382876 | 0.914 | 0.721 | 1.9414491 | 31 | rpl22 |
| si:ch211-222l21.112 | 1.230246 | 0.557227038 | 0.677 | 0.314 | 3.1254407 | 31 | si:ch211-222l21.1 |
| il344 | 1.372577 | 0.31822311 | 0.204 | 0.046 | 3.4870319 | 31 | il34 |
| rpl3112 | 1.734563 | 0.409085901 | 0.935 | 0.687 | 4.4066577 | 31 | rpl31 |
| si:dkeyp-68b7.126 | 2.263727 | 0.35265223 | 0.237 | 0.06 | 5.7509985 | 31 | si:dkeyp-68b7.12 |
| sept125 | 2.274321 | 0.366807029 | 0.247 | 0.065 | 5.7779126 | 31 | sept12 |
| rac211 | 2.305922 | 0.515538694 | 0.559 | 0.225 | 5.8581967 | 31 | rac2 |
| mrps154 | 2.361157 | 0.408517242 | 0.355 | 0.113 | 5.9985204 | 31 | mrps15 |
| rps27a7 | 2.564296 | 0.306445581 | 0.925 | 0.775 | 6.5145948 | 31 | rps27a |
| rpl196 | 2.708425 | 0.253678284 | 0.935 | 0.854 | 6.8807539 | 31 | rpl19 |
| itga44 | 2.873723 | 0.341539728 | 0.247 | 0.065 | 7.3006938 | 31 | itga4 |
| rpl10a10 | 3.719574 | 0.332500017 | 0.957 | 0.779 | 9.4495778 | 31 | rpl10a |
| rpl917 | 4.423257 | 0.422581369 | 0.935 | 0.668 | 1.1237285 | 31 | rpl9 |
| rps1015 | 5.967147 | 0.364791123 | 0.946 | 0.727 | 1.5159539 | 31 | rps10 |
| rps3a8 | 6.037626 | 0.302330351 | 0.935 | 0.792 | 1.5338590 | 31 | rps3a |
| dusp18 | 6.427267 | 0.476761533 | 0.409 | 0.149 | 1.6328473 | 31 | dusp1 |
| tagapa7 | 8.157327 | 0.459648838 | 0.312 | 0.098 | 2.0723690 | 31 | tagapa |
| CU499336.22 | 1.962839 | 0.256333912 | 0.161 | 0.033 | 4.9865945 | 31 | CU499336.2 |
| eno1a5 | 2.331791 | 0.426370672 | 0.387 | 0.142 | 5.9239175 | 31 | eno1a |

| | | | | | | | |
|---|---|---|---|---|---|---|---|
| id33 | 4.141497 | 0.537704949 | 0.473 | 0.198 | 1.0521475 | 31 | id3 |
| ldha4 | 4.747427 | 0.336814427 | 0.247 | 0.07 | 1.2060840 | 31 | ldha |
| batf3 | 8.341899 | 0.26294181 | 0.172 | 0.039 | 2.1192595 | 31 | batf3 |
| cotl110 | 9.500296 | 0.52690884 | 0.667 | 0.32 | 2.4135502 | 31 | cotl1 |
| si:ch73-234b20.56 | 1.029021 | 0.395982447 | 0.312 | 0.106 | 2.6142281 | 31 | si:ch73-234b20.5 |
| cox6b18 | 1.408277 | 0.398934621 | 0.344 | 0.122 | 3.5777301 | 31 | cox6b1 |
| sh2b32 | 1.413714 | 0.308690848 | 0.226 | 0.062 | 3.5915417 | 31 | sh2b3 |
| rpl108 | 1.434466 | 0.292742136 | 0.935 | 0.79 | 3.6442610 | 31 | rpl10 |
| rbp54 | 1.443226 | 0.415319761 | 0.323 | 0.111 | 3.6665175 | 31 | rbp5 |
| rps2510 | 1.497525 | 0.338386697 | 0.957 | 0.749 | 3.8044633 | 31 | rps25 |
| zfp36l1a7 | 1.788041 | 0.473075061 | 0.559 | 0.255 | 4.5425197 | 31 | zfp36l1a |
| kdm6ba | 1.874394 | 0.310997834 | 0.194 | 0.049 | 4.7618983 | 31 | kdm6ba |
| CR381673.16 | 2.053345 | 0.275921145 | 0.215 | 0.057 | 5.2165254 | 31 | CR381673.1 |
| rsl24d110 | 2.364851 | 0.481049519 | 0.753 | 0.461 | 6.0079054 | 31 | rsl24d1 |
| cfbl3 | 2.392271 | 0.353939084 | 0.247 | 0.071 | 6.0775657 | 31 | cfbl |
| XLOC-0140794 | 2.882139 | 0.270051744 | 0.151 | 0.032 | 7.3220760 | 31 | XLOC-014079 |
| si:ch73-359m17.22 | 3.044546 | 0.397397125 | 0.43 | 0.164 | 7.7346709 | 31 | si:ch73-359m17.2 |
| si:ch1073-443f11.22 | 3.749027 | 0.277796677 | 0.172 | 0.041 | 9.5244046 | 31 | si:ch1073-443f11.2 |
| si:ch211-102c2.47 | 4.367516 | 0.377867519 | 0.398 | 0.146 | 1.1095675 | 31 | si:ch211-102c2.4 |
| scinlb6 | 4.902591 | 0.35495778 | 0.269 | 0.084 | 1.2455034 | 31 | scinlb |
| ndfip21 | 5.135170 | 0.318519677 | 0.215 | 0.06 | 1.3045901 | 31 | ndfip2 |
| zgc:640519 | 5.642944 | 0.370926665 | 0.43 | 0.163 | 1.4335899 | 31 | zgc:64051 |
| ptmaa15 | 6.039340 | 0.49704488 | 0.774 | 0.433 | 1.5342944 | 31 | ptmaa |
| gapdhs12 | 7.438659 | 0.475732684 | 0.613 | 0.315 | 1.8897913 | 31 | gapdhs |
| chac16 | 8.626884 | 0.459050791 | 0.699 | 0.378 | 2.1916600 | 31 | chac1 |
| derl24 | 1.004440 | 0.342218045 | 0.312 | 0.107 | 2.5517815 | 31 | derl2 |
| uba5211 | 1.107918 | 0.382684082 | 0.925 | 0.661 | 2.8146669 | 31 | uba52 |
| btg112 | 1.845167 | 0.470625759 | 0.656 | 0.356 | 4.6876481 | 31 | btg1 |
| rpl27a13 | 2.058188 | 0.432599925 | 0.785 | 0.523 | 5.2288287 | 31 | rpl27a |
| tmsb4x17 | 2.097830 | 0.452132728 | 1 | 0.718 | 5.3295388 | 31 | tmsb4x |
| rps99 | 2.781928 | 0.290333913 | 0.946 | 0.791 | 7.0674886 | 31 | rps9 |
| cd28l2 | 3.692275 | 0.267937045 | 0.161 | 0.04 | 9.3802271 | 31 | cd28l |
| ets23 | 3.719999 | 0.382165067 | 0.258 | 0.087 | 9.4506598 | 31 | ets2 |
| sik14 | 3.830263 | 0.334876751 | 0.312 | 0.111 | 9.7307835 | 31 | sik1 |
| LOC5665873 | 4.343406 | 0.263611219 | 0.172 | 0.044 | 0.00011 | 31 | LOC566587 |

| | | | | | | | |
|---|---|---|---|---|---|---|---|
| eif3f12 | 5.022758 | 0.448496515 | 0.796 | 0.465 | 0.000128 | 31 | eif3f |
| mbpb1 | 5.758078 | 0.281347745 | 0.215 | 0.065 | 0.000146 | 31 | mbpb |
| rpl5a13 | 5.789342 | 0.411874437 | 0.806 | 0.537 | 0.000147 | 31 | rpl5a |
| gnai11 | 6.357605 | 0.286724336 | 0.226 | 0.07 | 0.000162 | 31 | gnai1 |
| rps8a7 | 7.148939 | 0.281979401 | 0.968 | 0.801 | 0.000182 | 31 | rps8a |
| rps117 | 1.364174 | 0.267632931 | 0.935 | 0.785 | 0.000347 | 31 | rps11 |
| si:dkey-206d17.124 | 1.572249 | 0.272057233 | 0.183 | 0.051 | 0.000399 | 31 | si:dkey-206d17.12 |
| mbnl18 | 1.933244 | 0.393380038 | 0.419 | 0.187 | 0.000491 | 31 | mbnl1 |
| sec61g5 | 3.431422 | 0.326656884 | 0.871 | 0.682 | 0.000872 | 31 | sec61g |
| rnf1301 | 3.594967 | 0.266370716 | 0.183 | 0.054 | 0.000913 | 31 | rnf130 |
| egr34 | 4.259848 | 0.315685593 | 0.183 | 0.054 | 0.001082 | 31 | egr3 |
| rack18 | 6.512274 | 0.268306668 | 0.925 | 0.764 | 0.001654 | 31 | rack1 |
| tagapb5 | 7.381841 | 0.289776339 | 0.215 | 0.07 | 0.001875 | 31 | tagapb |
| rpl613 | 8.475686 | 0.302928285 | 0.892 | 0.702 | 0.002153 | 31 | rpl6 |
| pim25 | 8.594873 | 0.3247506 | 0.247 | 0.089 | 0.002184 | 31 | pim2 |
| msna10 | 1.112478 | 0.413816576 | 0.505 | 0.266 | 0.002826 | 31 | msna |
| cap19 | 1.112828 | 0.382281037 | 0.505 | 0.246 | 0.002827 | 31 | cap1 |
| crema8 | 1.373557 | 0.419902208 | 0.376 | 0.175 | 0.00349 | 31 | crema |
| aldocb3 | 1.381504 | 0.265272298 | 0.204 | 0.066 | 0.00351 | 31 | aldocb |
| tefa7 | 1.420774 | 0.341291137 | 0.333 | 0.141 | 0.003609 | 31 | tefa |
| rpl289 | 1.776214 | 0.260729126 | 0.925 | 0.777 | 0.004512 | 31 | rpl28 |
| naca14 | 2.299152 | 0.317722068 | 0.849 | 0.653 | 0.005841 | 31 | naca |
| eef1g12 | 2.430185 | 0.342700869 | 0.903 | 0.653 | 0.006174 | 31 | eef1g |
| actb210 | 2.522947 | 0.252692705 | 0.989 | 0.867 | 0.00641 | 31 | actb2 |
| csrnp1b7 | 3.888710 | 0.34292798 | 0.366 | 0.165 | 0.009879 | 31 | csrnp1b |
| elf15 | 4.240004 | 0.278516856 | 0.269 | 0.104 | 0.010772 | 31 | elf1 |
| pgam1a5 | 4.723326 | 0.316392987 | 0.355 | 0.156 | 0.012 | 31 | pgam1a |
| pnrc216 | 8.144538 | 0.362379084 | 0.828 | 0.598 | 0.020691 | 31 | pnrc2 |
| rpl22l110 | 8.408468 | 0.386415026 | 0.677 | 0.426 | 0.021362 | 31 | rpl22l1 |
| myl12.112 | 9.817725 | 0.391073116 | 0.667 | 0.427 | 0.024942 | 31 | myl12.1 |
| nsa210 | 1.448714 | 0.382169434 | 0.634 | 0.389 | 0.036805 | 31 | nsa2 |
| dusp25 | 1.641866 | 0.376153173 | 0.355 | 0.164 | 0.041712 | 31 | dusp2 |
| arrb2b6 | 1.976449 | 0.255816989 | 0.247 | 0.097 | 0.050212 | 31 | arrb2b |
| ssr33 | 2.316431 | 0.361350692 | 0.527 | 0.288 | 0.058849 | 31 | ssr3 |
| irf2bp2a5 | 2.637755 | 0.277010699 | 0.398 | 0.186 | 0.067012 | 31 | irf2bp2a |

| | | | | | | | |
|---|---|---|---|---|---|---|---|
| CR855311.15 | 2.835461 | 0.41801586 | 0.226 | 0.089 | 0.072035 | 31 | CR855311.1 |
| LOC1039091157 | 5.210440 | 0.26276714 | 0.366 | 0.17 | 0.132371 | 31 | LOC103909115 |
| rpl5b9 | 5.545158 | 0.329720003 | 0.882 | 0.646 | 0.140875 | 31 | rpl5b |
| si:dkey-11f4.147 | 7.066036 | 0.284648139 | 0.28 | 0.123 | 0.179513 | 31 | si:dkey-11f4.14 |
| notch27 | 7.218577 | 0.285343306 | 0.28 | 0.123 | 0.183388 | 31 | notch2 |
| ppp3ccb3 | 1.248942 | 0.271842884 | 0.28 | 0.126 | 0.317294 | 31 | ppp3ccb |
| arpc5a7 | 1.314899 | 0.284998727 | 0.28 | 0.126 | 0.33405 | 31 | arpc5a |
| tnrc6c18 | 1.981202 | 0.287349373 | 0.312 | 0.151 | 0.503325 | 31 | tnrc6c1 |
| arpc39 | 2.348715 | 0.340209062 | 0.548 | 0.335 | 0.596691 | 31 | arpc3 |
| xbp15 | 3.323761 | 0.355277876 | 0.559 | 0.356 | 0.844402 | 31 | xbp1 |
| reep56 | 3.637194 | 0.307129147 | 0.333 | 0.174 | 0.924029 | 31 | reep5 |
| zgc:564937 | 3.857621 | 0.310721415 | 0.828 | 0.591 | 0.980029 | 31 | zgc:56493 |
| txnipa8 | 4.416466 | 0.302566664 | 0.441 | 0.245 | 1 | 31 | txnipa |
| hmgn211 | 4.429118 | 0.363526526 | 0.688 | 0.468 | 1 | 31 | hmgn2 |
| ddx212 | 4.965847 | 0.294298061 | 0.387 | 0.21 | 1 | 31 | ddx21 |
| sat1a.27 | 5.284044 | 0.295730963 | 0.527 | 0.302 | 1 | 31 | sat1a.2 |
| rps27.213 | 5.519343 | 0.324151304 | 0.839 | 0.553 | 1 | 31 | rps27.2 |
| serf28 | 5.745411 | 0.312316068 | 0.591 | 0.378 | 1 | 31 | serf2 |
| sesn16 | 6.353896 | 0.274645134 | 0.344 | 0.176 | 1 | 31 | sesn1 |
| btf39 | 7.250148 | 0.28549078 | 0.817 | 0.611 | 1 | 31 | btf3 |
| midn8 | 0.0001 | 0.262249032 | 0.29 | 0.146 | 1 | 31 | midn |
| fabp36 | 0.0001 | 0.314748178 | 0.398 | 0.229 | 1 | 31 | fabp3 |
| tsc22d38 | 0.0001 | 0.288488638 | 0.301 | 0.16 | 1 | 31 | tsc22d3 |
| arpc211 | 0.0002 | 0.283250963 | 0.495 | 0.3 | 1 | 31 | arpc2 |
| pabpc1a8 | 0.0002 | 0.277474342 | 0.774 | 0.57 | 1 | 31 | pabpc1a |
| GCA6 | 0.0003 | 0.268903429 | 0.355 | 0.201 | 1 | 31 | GCA |
| cox8a12 | 0.0003 | 0.25579462 | 0.452 | 0.268 | 1 | 31 | cox8a |
| capza1a7 | 0.0004 | 0.270618084 | 0.376 | 0.222 | 1 | 31 | capza1a |
| caprin1b2 | 0.0005 | 0.262008038 | 0.409 | 0.247 | 1 | 31 | caprin1b |
| ssr43 | 0.0006 | 0.262690083 | 0.527 | 0.332 | 1 | 31 | ssr4 |
| mcl1a7 | 0.0007 | 0.274738584 | 0.817 | 0.614 | 1 | 31 | mcl1a |
| dbi6 | 0.0008 | 0.278865909 | 0.387 | 0.244 | 1 | 31 | dbi |
| col10a1a | 0 | 2.812779031 | 0.91 | 0.007 | 0 | 32 | col10a1a |
| scpp72 | 0 | 2.652330529 | 0.899 | 0.018 | 0 | 32 | scpp7 |
| ifitm5 | 0 | 2.324811563 | 0.978 | 0.002 | 0 | 32 | ifitm5 |

| | | | | | | | |
|---|---|---|---|---|---|---|---|
| cd81b1 | 0 | 2.215311883 | 0.899 | 0.004 | 0 | 32 | cd81b |
| enam2 | 0 | 2.097433061 | 0.787 | 0.006 | 0 | 32 | enam |
| tmem119a | 0 | 1.830010309 | 0.888 | 0.002 | 0 | 32 | tmem119a |
| panx3 | 0 | 1.752652921 | 0.865 | 0.001 | 0 | 32 | panx3 |
| si:ch211-106n13.3 | 0 | 1.752529213 | 0.876 | 0.001 | 0 | 32 | si:ch211-106n13.3 |
| sgms2a | 0 | 1.729713159 | 0.865 | 0.004 | 0 | 32 | sgms2a |
| si:dkey-12l12.11 | 0 | 1.598489858 | 0.854 | 0.017 | 0 | 32 | si:dkey-12l12.1 |
| phospho11 | 0 | 1.557867799 | 0.843 | 0.008 | 0 | 32 | phospho1 |
| sod3b1 | 0 | 1.556881308 | 0.921 | 0.004 | 0 | 32 | sod3b |
| cthrc1a | 0 | 1.553790517 | 0.876 | 0.004 | 0 | 32 | cthrc1a |
| bgna | 0 | 1.529397063 | 0.831 | 0.012 | 0 | 32 | bgna |
| col10a1b | 0 | 1.441352423 | 0.528 | 0.006 | 0 | 32 | col10a1b |
| col11a1a1 | 0 | 1.434990603 | 0.809 | 0.004 | 0 | 32 | col11a1a |
| loxl4 | 0 | 1.430518991 | 0.809 | 0.002 | 0 | 32 | loxl4 |
| col11a2 | 0 | 1.377934716 | 0.708 | 0.001 | 0 | 32 | col11a2 |
| fgfbp2a | 0 | 1.365456559 | 0.416 | 0.001 | 0 | 32 | fgfbp2a |
| cdh112 | 0 | 1.247181352 | 0.831 | 0.025 | 0 | 32 | cdh11 |
| slc8a4b1 | 0 | 1.215371948 | 0.742 | 0.011 | 0 | 32 | slc8a4b |
| spp1 | 0 | 1.150409356 | 0.618 | 0.002 | 0 | 32 | spp1 |
| zgc:921621 | 0 | 1.148533999 | 0.607 | 0.005 | 0 | 32 | zgc:92162 |
| smpd3 | 0 | 1.126288611 | 0.685 | 0.007 | 0 | 32 | smpd3 |
| chst11 | 0 | 1.113106776 | 0.674 | 0.005 | 0 | 32 | chst11 |
| itga10 | 0 | 1.068053762 | 0.708 | 0.002 | 0 | 32 | itga10 |
| LO018513.1 | 0 | 1.028818821 | 0.539 | 0.001 | 0 | 32 | LO018513.1 |
| bambib | 0 | 0.997896443 | 0.685 | 0.011 | 0 | 32 | bambib |
| ano5a | 0 | 0.977561741 | 0.652 | 0.006 | 0 | 32 | ano5a |
| bmp8a | 0 | 0.971131179 | 0.618 | 0.007 | 0 | 32 | bmp8a |
| runx3 | 0 | 0.969099984 | 0.663 | 0.017 | 0 | 32 | runx3 |
| col5a2b | 0 | 0.965875507 | 0.652 | 0.001 | 0 | 32 | col5a2b |
| qpct | 0 | 0.963678222 | 0.584 | 0.005 | 0 | 32 | qpct |
| klf2b1 | 0 | 0.955671029 | 0.629 | 0.017 | 0 | 32 | klf2b |
| entpd5a1 | 0 | 0.936058625 | 0.382 | 0.002 | 0 | 32 | entpd5a |
| pcolce2b | 0 | 0.91891123 | 0.629 | 0.018 | 0 | 32 | pcolce2b |
| dlx3b2 | 0 | 0.855220764 | 0.539 | 0.006 | 0 | 32 | dlx3b |
| kazald3 | 0 | 0.855197813 | 0.584 | 0.003 | 0 | 32 | kazald3 |

| | | | | | | | |
|---|---|---|---|---|---|---|---|
| sp7 | 0 | 0.808969788 | 0.562 | 0.001 | 0 | 32 | sp7 |
| si:ch211-170d8.81 | 0 | 0.802967251 | 0.584 | 0.009 | 0 | 32 | si:ch211-170d8.8 |
| bmp32 | 0 | 0.779074213 | 0.517 | 0.01 | 0 | 32 | bmp3 |
| COLGALT1 (1 of many) | 0 | 0.774190752 | 0.449 | 0.007 | 0 | 32 | COLGALT1 (1 of many) |
| tnfrsf11a | 0 | 0.738395362 | 0.483 | 0.01 | 0 | 32 | tnfrsf11a |
| myadml2 | 0 | 0.706209434 | 0.461 | 0 | 0 | 32 | myadml2 |
| thrap3a1 | 0 | 0.704214956 | 0.562 | 0.01 | 0 | 32 | thrap3a |
| prp | 0 | 0.666693831 | 0.438 | 0.001 | 0 | 32 | prp |
| CABZ01059119.1 | 0 | 0.656950236 | 0.517 | 0.011 | 0 | 32 | CABZ01059119.1 |
| satb2 | 0 | 0.635981104 | 0.449 | 0.002 | 0 | 32 | satb2 |
| bmp2a | 0 | 0.633692419 | 0.494 | 0.002 | 0 | 32 | bmp2a |
| ism11 | 0 | 0.61302805 | 0.449 | 0.008 | 0 | 32 | ism1 |
| dlx5a1 | 0 | 0.56683042 | 0.371 | 0.002 | 0 | 32 | dlx5a |
| pltp | 0 | 0.554825654 | 0.348 | 0.002 | 0 | 32 | pltp |
| plcd3b | 0 | 0.52492194 | 0.382 | 0.001 | 0 | 32 | plcd3b |
| si:ch211-106h11.3 | 0 | 0.509885374 | 0.371 | 0.002 | 0 | 32 | si:ch211-106h11.3 |
| ihha | 0 | 0.492907221 | 0.213 | 0 | 0 | 32 | ihha |
| dlx2b | 0 | 0.46705608 | 0.337 | 0.001 | 0 | 32 | dlx2b |
| wnt10a | 0 | 0.46017473 | 0.247 | 0.001 | 0 | 32 | wnt10a |
| cspg41 | 0 | 0.451883651 | 0.36 | 0.005 | 0 | 32 | cspg4 |
| zgc:162171 | 0 | 0.430997314 | 0.36 | 0.002 | 0 | 32 | zgc:162171 |
| scpp1 | 0 | 0.412595197 | 0.326 | 0 | 0 | 32 | scpp1 |
| megf10 | 0 | 0.343518631 | 0.281 | 0.001 | 0 | 32 | megf10 |
| ppfibp2a | 0 | 0.300927968 | 0.191 | 0 | 0 | 32 | ppfibp2a |
| chodl | 0 | 0.282481095 | 0.191 | 0 | 0 | 32 | chodl |
| mxra5a | 4.844807 | 0.406665827 | 0.281 | 0.003 | 1.2308234 | 32 | mxra5a |
| nid2b | 1.089845 | 0.356912216 | 0.258 | 0.002 | 2.7687518 | 32 | nid2b |
| phlda3 | 1.950852 | 0.543651086 | 0.393 | 0.007 | 4.9561417 | 32 | phlda3 |
| nrp2b | 2.270469 | 0.689159104 | 0.449 | 0.01 | 5.7681289 | 32 | nrp2b |
| myh11b1 | 3.633622 | 0.398183487 | 0.281 | 0.003 | 9.2312184 | 32 | myh11b |
| si:ch211-110e21.3 | 2.343091 | 0.565984875 | 0.348 | 0.006 | 5.9526244 | 32 | si:ch211-110e21.3 |
| zgc:1626121 | 1.229974 | 0.552042583 | 0.326 | 0.005 | 3.1247501 | 32 | zgc:162612 |
| ndnf2 | 7.130145 | 1.232578411 | 0.764 | 0.033 | 1.8114133 | 32 | ndnf |
| mfsd2b | 5.033178 | 0.278542728 | 0.191 | 0.001 | 1.2786789 | 32 | mfsd2b |
| serpinh1a | 9.362199 | 0.663900386 | 0.494 | 0.013 | 2.3784667 | 32 | serpinh1a |

| | | | | | | | |
|---|---|---|---|---|---|---|---|
| nav2a | 1.112653 | 0.307887906 | 0.191 | 0.001 | 2.8266957 | 32 | nav2a |
| rcn1 | 1.702843 | 0.847208803 | 0.573 | 0.019 | 4.3260751 | 32 | rcn1 |
| heyl1 | 6.954487 | 0.405180081 | 0.202 | 0.002 | 1.7667875 | 32 | heyl |
| crip21 | 1.197210 | 0.887040172 | 0.483 | 0.014 | 3.0415129 | 32 | crip2 |
| si:dkey-234i14.3 | 1.319997 | 0.304410199 | 0.135 | 0 | 3.3534547 | 32 | si:dkey-234i14.3 |
| rnd2 | 1.422551 | 0.348883632 | 0.135 | 0 | 3.6139911 | 32 | rnd2 |
| zbtb7c | 1.397529 | 0.385286976 | 0.247 | 0.003 | 3.5504234 | 32 | zbtb7c |
| si:cabz01100188.1 | 1.693227 | 0.335014588 | 0.213 | 0.002 | 4.3016438 | 32 | si:cabz01100188.1 |
| sema3e1 | 4.768832 | 0.48562767 | 0.371 | 0.008 | 1.2115219 | 32 | sema3e |
| xylt1 | 3.419468 | 0.461070864 | 0.326 | 0.006 | 8.6871601 | 32 | xylt1 |
| igfbp1a | 1.910553 | 1.110153849 | 0.483 | 0.015 | 4.8537599 | 32 | igfbp1a |
| lmo7a | 2.094651 | 0.475271967 | 0.348 | 0.007 | 5.3214614 | 32 | lmo7a |
| golim4b | 7.228226 | 0.320059667 | 0.247 | 0.003 | 1.8363309 | 32 | golim4b |
| pcdh171 | 5.358684 | 0.397246612 | 0.258 | 0.004 | 1.3613738 | 32 | pcdh17 |
| tfap2c | 2.911909 | 0.456496717 | 0.236 | 0.003 | 7.3977069 | 32 | tfap2c |
| serpinh1b1 | 4.310114 | 0.958971297 | 0.618 | 0.027 | 1.0949846 | 32 | serpinh1b |
| s1pr11 | 7.904186 | 0.475755386 | 0.303 | 0.006 | 2.0080584 | 32 | s1pr1 |
| plod1a | 7.988199 | 0.76163805 | 0.551 | 0.023 | 2.0294021 | 32 | plod1a |
| fam20a | 7.236876 | 0.32831247 | 0.27 | 0.005 | 1.8385283 | 32 | fam20a |
| XLOC-005794 | 6.751915 | 0.346554635 | 0.191 | 0.002 | 1.7153242 | 32 | XLOC-005794 |
| fkbp92 | 1.554083 | 0.903315704 | 0.652 | 0.033 | 3.9481494 | 32 | fkbp9 |
| si:dkey-193p11.21 | 1.810967 | 1.286715452 | 0.393 | 0.012 | 4.6007619 | 32 | si:dkey-193p11.2 |
| nupr1b3 | 2.865526 | 1.064642527 | 0.719 | 0.042 | 7.2798695 | 32 | nupr1b |
| pcolceb | 2.299931 | 0.298481879 | 0.202 | 0.002 | 5.8429750 | 32 | pcolceb |
| fpr12 | 1.742530 | 0.754446377 | 0.494 | 0.019 | 4.4268994 | 32 | fpr1 |
| fgfr32 | 4.244847 | 0.723067142 | 0.483 | 0.018 | 1.0784035 | 32 | fgfr3 |
| col14a1b | 9.182962 | 0.500675377 | 0.27 | 0.005 | 2.3329316 | 32 | col14a1b |
| olfml3b5 | 6.534546 | 1.135303514 | 0.787 | 0.052 | 1.6601015 | 32 | olfml3b |
| p4ha2 | 2.194041 | 0.535669362 | 0.416 | 0.013 | 5.5739620 | 32 | p4ha2 |
| msx2b | 3.994982 | 0.29024776 | 0.18 | 0.002 | 1.0149251 | 32 | msx2b |
| thbs1b1 | 2.503895 | 0.506217671 | 0.348 | 0.01 | 6.3611470 | 32 | thbs1b |
| twist21 | 1.019265 | 0.625184964 | 0.348 | 0.01 | 2.5894444 | 32 | twist2 |
| cdc42ep3 | 1.214421 | 0.493058961 | 0.371 | 0.011 | 3.0852371 | 32 | cdc42ep3 |
| LOC1018861141 | 2.557695 | 0.601457838 | 0.416 | 0.015 | 6.4978246 | 32 | LOC101886114 |
| btbd6a | 2.500481 | 0.491666251 | 0.292 | 0.007 | 6.3524739 | 32 | btbd6a |

| | | | | | | | |
|---|---|---|---|---|---|---|---|
| ccdc801 | 7.235349 | 0.759021083 | 0.584 | 0.03 | 1.8381406 | 32 | ccdc80 |
| tcf72 | 7.800354 | 0.816011211 | 0.483 | 0.021 | 1.9816800 | 32 | tcf7 |
| her62 | 2.340766 | 0.814698138 | 0.506 | 0.023 | 5.9467182 | 32 | her6 |
| serpine22 | 6.182692 | 0.987502907 | 0.798 | 0.058 | 1.5707131 | 32 | serpine2 |
| pdgfbb | 9.270502 | 0.448610918 | 0.236 | 0.004 | 2.3551712 | 32 | pdgfbb |
| cd82b1 | 1.168987 | 1.264541465 | 0.865 | 0.072 | 2.9698131 | 32 | cd82b |
| mef2cb2 | 1.278003 | 0.802204983 | 0.539 | 0.027 | 3.2467675 | 32 | mef2cb |
| p3h1 | 1.314122 | 0.436017119 | 0.382 | 0.013 | 3.3385269 | 32 | p3h1 |
| clec3bb | 1.924095 | 0.625705671 | 0.225 | 0.004 | 4.8881646 | 32 | clec3bb |
| nradd1 | 1.010266 | 0.433647664 | 0.292 | 0.008 | 2.5665822 | 32 | nradd |
| sox11a1 | 5.068137 | 0.78377274 | 0.517 | 0.027 | 1.2875602 | 32 | sox11a |
| acvrl11 | 1.992352 | 0.526388186 | 0.404 | 0.016 | 5.0615703 | 32 | acvrl1 |
| col11a1b1 | 3.948765 | 0.612825764 | 0.393 | 0.016 | 1.0031839 | 32 | col11a1b |
| plekhh1 | 2.257901 | 0.352144978 | 0.247 | 0.006 | 5.7361977 | 32 | plekhh1 |
| tox1 | 2.278512 | 0.569721503 | 0.438 | 0.02 | 5.7885615 | 32 | tox |
| rcn3 | 2.653345 | 0.356667984 | 0.27 | 0.007 | 6.7408238 | 32 | rcn3 |
| zgc:110796 | 4.565794 | 0.399768921 | 0.27 | 0.007 | 1.1599401 | 32 | zgc:110796 |
| fkbp141 | 1.690674 | 0.757553189 | 0.607 | 0.04 | 4.2951584 | 32 | fkbp14 |
| plxna3 | 8.010746 | 0.536714591 | 0.393 | 0.017 | 2.0351301 | 32 | plxna3 |
| LOC100535130 | 1.511792 | 0.38338274 | 0.101 | 0.001 | 3.8407086 | 32 | LOC100535130 |
| farp2 | 1.599341 | 0.518472189 | 0.416 | 0.019 | 4.0631269 | 32 | farp2 |
| FAM163A | 2.701051 | 0.29116235 | 0.157 | 0.002 | 6.8620218 | 32 | FAM163A |
| stc2a | 1.455868 | 0.286147997 | 0.18 | 0.003 | 3.6986346 | 32 | stc2a |
| p4ha1b | 4.065237 | 0.805732811 | 0.719 | 0.056 | 1.0327736 | 32 | p4ha1b |
| ccdc1872 | 6.612583 | 0.719534136 | 0.562 | 0.036 | 1.6799268 | 32 | ccdc187 |
| cplx2 | 4.366713 | 0.35759579 | 0.247 | 0.007 | 1.1093635 | 32 | cplx2 |
| rarga3 | 1.353176 | 0.848180142 | 0.697 | 0.058 | 3.4377437 | 32 | rarga |
| LOC108179036 | 1.548856 | 0.665881256 | 0.506 | 0.031 | 3.9348695 | 32 | LOC108179036 |
| cpe | 1.670350 | 0.385951544 | 0.247 | 0.007 | 4.2435247 | 32 | cpe |
| vkorc1 | 2.804136 | 0.488393298 | 0.36 | 0.015 | 7.1239079 | 32 | vkorc1 |
| fam20cb | 1.227451 | 0.342973462 | 0.27 | 0.009 | 3.1183405 | 32 | fam20cb |
| htra3b1 | 2.767406 | 0.458869111 | 0.326 | 0.013 | 7.0305969 | 32 | htra3b |
| mef2aa3 | 2.970329 | 1.084061676 | 0.73 | 0.07 | 7.5461219 | 32 | mef2aa |
| BX322661.2 | 2.943106 | 0.255486013 | 0.135 | 0.002 | 7.4769622 | 32 | BX322661.2 |
| alcama2 | 2.475105 | 0.723571432 | 0.483 | 0.03 | 6.2880064 | 32 | alcama |

| | | | | | | | |
|---|---|---|---|---|---|---|---|
| zgc:1533174 | 6.316315 | 0.728527405 | 0.539 | 0.038 | 1.6046598 | 32 | zgc:153317 |
| creb3l1 | 7.725337 | 0.267588364 | 0.225 | 0.006 | 1.9626220 | 32 | creb3l1 |
| serpinf15 | 9.065671 | 0.801055743 | 0.573 | 0.043 | 2.3031339 | 32 | serpinf1 |
| mef2ca1 | 8.731506 | 0.380276111 | 0.258 | 0.008 | 2.2182391 | 32 | mef2ca |
| ak5 | 1.116162 | 0.291987489 | 0.225 | 0.006 | 2.8356107 | 32 | ak5 |
| tmem198b | 9.682768 | 0.857667347 | 0.64 | 0.054 | 2.4599073 | 32 | tmem198b |
| crtap3 | 6.431292 | 0.662087599 | 0.528 | 0.038 | 1.6338697 | 32 | crtap |
| cdh7a | 1.137786 | 0.376531234 | 0.213 | 0.006 | 2.8905458 | 32 | cdh7a |
| rbfox11 | 5.596620 | 0.577817839 | 0.449 | 0.027 | 1.4218213 | 32 | rbfox1 |
| foxd1 | 6.409496 | 0.295450285 | 0.191 | 0.005 | 1.6283326 | 32 | foxd1 |
| fkbp11 | 1.080180 | 0.549703245 | 0.404 | 0.023 | 2.7441994 | 32 | fkbp11 |
| frmd61 | 8.266338 | 0.397032664 | 0.348 | 0.017 | 2.1000631 | 32 | frmd6 |
| tnfaip8l3 | 1.243771 | 0.310150492 | 0.191 | 0.005 | 3.1598004 | 32 | tnfaip8l3 |
| rtn1a | 1.675255 | 0.412627798 | 0.36 | 0.018 | 4.2559873 | 32 | rtn1a |
| tmem45a1 | 1.305121 | 0.469895129 | 0.427 | 0.026 | 3.3156604 | 32 | tmem45a |
| fabp7a2 | 4.561811 | 0.57939478 | 0.382 | 0.021 | 1.1589282 | 32 | fabp7a |
| cgref11 | 5.585916 | 0.615329539 | 0.371 | 0.02 | 1.4191021 | 32 | cgref1 |
| col1a1a5 | 9.150415 | 1.641171354 | 0.989 | 0.163 | 2.3246631 | 32 | col1a1a |
| cx435 | 5.689833 | 0.898207755 | 0.618 | 0.055 | 1.4455020 | 32 | cx43 |
| nrgna1 | 1.106397 | 0.674751612 | 0.213 | 0.007 | 2.8108037 | 32 | nrgna |
| zeb2a2 | 1.063228 | 0.676327898 | 0.551 | 0.044 | 2.7011319 | 32 | zeb2a |
| marveld1 | 1.689182 | 0.35886016 | 0.27 | 0.011 | 4.2913690 | 32 | marveld1 |
| MDFI1 | 4.389872 | 0.506267074 | 0.382 | 0.022 | 1.1152472 | 32 | MDFI |
| twist1a1 | 3.722681 | 0.502835156 | 0.337 | 0.018 | 9.4574724 | 32 | twist1a |
| sparc8 | 3.282158 | 1.570402136 | 1 | 0.192 | 8.3383225 | 32 | sparc |
| tbx2b1 | 3.047539 | 0.502466162 | 0.337 | 0.018 | 7.7422751 | 32 | tbx2b |
| bambia2 | 5.631825 | 0.458028131 | 0.404 | 0.025 | 1.4307651 | 32 | bambia |
| col1a25 | 8.824087 | 1.636516006 | 0.989 | 0.179 | 2.2417593 | 32 | col1a2 |
| VIT1 | 1.429847 | 0.348043621 | 0.213 | 0.007 | 3.6325279 | 32 | VIT |
| bcl11aa1 | 1.827668 | 0.337926805 | 0.247 | 0.01 | 4.6431916 | 32 | bcl11aa |
| jdp2b7 | 2.129388 | 1.414683587 | 0.944 | 0.161 | 5.4097109 | 32 | jdp2b |
| pdlim4 | 5.647529 | 0.259626232 | 0.18 | 0.005 | 1.4347548 | 32 | pdlim4 |
| asip11 | 1.071094 | 0.429360492 | 0.337 | 0.019 | 2.7211166 | 32 | asip1 |
| pleca4 | 2.551364 | 0.744705538 | 0.607 | 0.06 | 6.4817408 | 32 | pleca |
| kremen12 | 4.172993 | 0.414301647 | 0.258 | 0.011 | 1.0601490 | 32 | kremen1 |

| | | | | | | | |
|---|---|---|---|---|---|---|---|
| znf385a4 | 8.399613 | 0.706029342 | 0.562 | 0.052 | 2.1339217 | 32 | znf385a |
| col1a1b5 | 1.010352 | 1.467395946 | 0.966 | 0.166 | 2.5667999 | 32 | col1a1b |
| nrp2a3 | 1.994780 | 0.476212703 | 0.337 | 0.019 | 5.0677410 | 32 | nrp2a |
| fscn1a2 | 3.308864 | 0.610960932 | 0.562 | 0.051 | 8.4061694 | 32 | fscn1a |
| hsp70l | 1.092938 | 0.266055065 | 0.202 | 0.007 | 2.7766095 | 32 | hsp70l |
| phex3 | 6.444945 | 0.807366022 | 0.652 | 0.069 | 1.6373383 | 32 | phex |
| timp2a5 | 6.628415 | 0.725641065 | 0.607 | 0.063 | 1.6839489 | 32 | timp2a |
| tspan4a3 | 9.866112 | 0.627030141 | 0.449 | 0.036 | 2.5064859 | 32 | tspan4a |
| pard6gb | 1.891054 | 0.312108038 | 0.303 | 0.016 | 4.8042250 | 32 | pard6gb |
| reck1 | 1.547654 | 0.470490962 | 0.393 | 0.028 | 3.9318156 | 32 | reck |
| paplnb | 2.664951 | 0.502790015 | 0.135 | 0.003 | 6.7703085 | 32 | paplnb |
| fosb7 | 2.905119 | 1.084999341 | 0.82 | 0.118 | 7.3804563 | 32 | fosb |
| XLOC-035413 | 3.942756 | 0.491734641 | 0.438 | 0.035 | 1.0016572 | 32 | XLOC-035413 |
| kdelr3 | 1.012484 | 0.380770366 | 0.303 | 0.017 | 2.5722156 | 32 | kdelr3 |
| bmp62 | 5.427587 | 0.397339162 | 0.27 | 0.014 | 1.3788787 | 32 | bmp6 |
| fxyd15 | 1.758124 | 0.765838712 | 0.674 | 0.081 | 4.4665144 | 32 | fxyd1 |
| hoxb7a1 | 3.473054 | 0.355162042 | 0.281 | 0.015 | 8.8232942 | 32 | hoxb7a |
| plod31 | 3.623810 | 0.481746689 | 0.438 | 0.037 | 9.2062908 | 32 | plod3 |
| sox63 | 3.949714 | 0.605509529 | 0.393 | 0.031 | 1.0034249 | 32 | sox6 |
| cbfb3 | 5.523726 | 0.546970499 | 0.461 | 0.041 | 1.4033026 | 32 | cbfb |
| fkbp75 | 5.838482 | 0.706386486 | 0.494 | 0.048 | 1.4832664 | 32 | fkbp7 |
| FP102018.11 | 8.941638 | 0.447360207 | 0.315 | 0.02 | 2.2716231 | 32 | FP102018.1 |
| cpamd8 | 2.224067 | 0.288609179 | 0.157 | 0.005 | 5.6502429 | 32 | cpamd8 |
| tgfb2l1 | 7.556204 | 0.341596592 | 0.202 | 0.008 | 1.9196538 | 32 | tgfb2l |
| aqp3a3 | 1.145681 | 0.606770539 | 0.371 | 0.028 | 2.9106044 | 32 | aqp3a |
| ncam1a3 | 1.568628 | 0.394332824 | 0.315 | 0.02 | 3.9850994 | 32 | ncam1a |
| cav17 | 1.095061 | 0.770062317 | 0.483 | 0.047 | 2.7820039 | 32 | cav1 |
| fhl1b1 | 1.209532 | 0.397659775 | 0.303 | 0.019 | 3.0728172 | 32 | fhl1b |
| mmp16a5 | 4.923494 | 0.772779208 | 0.629 | 0.076 | 1.2508136 | 32 | mmp16a |
| zgc:162939 | 9.250843 | 0.400422141 | 0.281 | 0.017 | 2.3501767 | 32 | zgc:162939 |
| col5a15 | 9.776031 | 1.097840433 | 0.809 | 0.129 | 2.4836007 | 32 | col5a1 |
| notch3 | 1.177866 | 0.576013311 | 0.337 | 0.024 | 2.9923710 | 32 | notch3 |
| krt18a.18 | 1.268138 | 1.095781532 | 0.876 | 0.147 | 3.2217051 | 32 | krt18a.1 |
| adgra2 | 2.682963 | 0.255920828 | 0.18 | 0.007 | 6.8160696 | 32 | adgra2 |
| lhfpl62 | 4.462194 | 0.478817061 | 0.326 | 0.023 | 1.1336204 | 32 | lhfpl6 |

| | | | | | | | |
|---|---|---|---|---|---|---|---|
| copz2 | 5.202651 | 0.272007132 | 0.225 | 0.011 | 1.3217336 | 32 | copz2 |
| surf4l | 1.146285 | 0.60043576 | 0.494 | 0.052 | 2.9121393 | 32 | surf4l |
| hsp70.32 | 1.644758 | 0.498830945 | 0.292 | 0.019 | 4.1785090 | 32 | hsp70.3 |
| sgcd | 5.297219 | 0.251794547 | 0.191 | 0.008 | 1.3457585 | 32 | sgcd |
| itgb1a4 | 3.683486 | 0.583682241 | 0.483 | 0.051 | 9.3578976 | 32 | itgb1a |
| srgap3 | 1.037543 | 0.438867152 | 0.337 | 0.026 | 2.6358781 | 32 | srgap3 |
| eva1ba5 | 2.320077 | 0.675192038 | 0.551 | 0.065 | 5.8941572 | 32 | eva1ba |
| atp1b3a | 8.442419 | 0.287464727 | 0.225 | 0.012 | 2.1447965 | 32 | atp1b3a |
| sox181 | 9.196977 | 0.295784912 | 0.202 | 0.01 | 2.3364922 | 32 | sox18 |
| efna1a1 | 1.067273 | 0.354333561 | 0.247 | 0.014 | 2.7114086 | 32 | efna1a |
| id2b4 | 1.093789 | 0.642685877 | 0.416 | 0.039 | 2.7787718 | 32 | id2b |
| prkd21 | 6.980084 | 0.446009258 | 0.382 | 0.033 | 1.7732905 | 32 | prkd2 |
| CR376783.11 | 4.484421 | 0.284603941 | 0.225 | 0.012 | 1.1392673 | 32 | CR376783.1 |
| bckdhbl1 | 7.031083 | 0.367541427 | 0.36 | 0.03 | 1.7862467 | 32 | bckdhbl |
| anxa135 | 1.609021 | 0.484741004 | 0.483 | 0.052 | 4.0877183 | 32 | anxa13 |
| serinc54 | 2.167037 | 0.730500602 | 0.618 | 0.086 | 5.5053586 | 32 | serinc5 |
| tmeff1b3 | 5.578318 | 0.496841041 | 0.416 | 0.041 | 1.4171717 | 32 | tmeff1b |
| pmp22a5 | 1.064282 | 0.925366687 | 0.91 | 0.164 | 2.7038084 | 32 | pmp22a |
| bhmt2 | 1.805971 | 0.54568637 | 0.416 | 0.041 | 4.5880718 | 32 | bhmt |
| ctnnb16 | 2.801883 | 0.930352502 | 0.831 | 0.156 | 7.1181847 | 32 | ctnnb1 |
| robo13 | 3.075033 | 0.420359264 | 0.326 | 0.026 | 7.8121217 | 32 | robo1 |
| fgfr1a1 | 3.208050 | 0.387592146 | 0.292 | 0.021 | 8.1500530 | 32 | fgfr1a |
| icn8 | 6.005786 | 1.402989504 | 0.955 | 0.265 | 1.5257701 | 32 | icn |
| fstl35 | 2.561035 | 0.550854062 | 0.303 | 0.024 | 6.5063118 | 32 | fstl3 |
| hoxb8a1 | 2.950211 | 0.295498984 | 0.191 | 0.01 | 7.4950114 | 32 | hoxb8a |
| mmp14a4 | 9.221532 | 0.497806074 | 0.483 | 0.058 | 2.3427303 | 32 | mmp14a |
| ckap45 | 1.043355 | 0.517688274 | 0.483 | 0.059 | 2.6506446 | 32 | ckap4 |
| sh3pxd2aa3 | 1.784447 | 0.457683956 | 0.348 | 0.033 | 4.5333882 | 32 | sh3pxd2aa |
| adgrl3.13 | 5.434887 | 0.495152148 | 0.326 | 0.029 | 1.3807331 | 32 | adgrl3.1 |
| si:ch211-133n4.43 | 5.880353 | 0.538811182 | 0.472 | 0.057 | 1.4939037 | 32 | si:ch211-133n4.4 |
| ier5l1 | 1.627929 | 0.403388401 | 0.303 | 0.025 | 4.1357550 | 32 | ier5l |
| egr19 | 6.278435 | 0.94423379 | 0.854 | 0.18 | 1.5950364 | 32 | egr1 |
| zgc:153675 | 6.522871 | 0.567736144 | 0.506 | 0.066 | 1.6571355 | 32 | zgc:153675 |
| fosab11 | 1.122584 | 1.26609157 | 1 | 0.359 | 2.8519250 | 32 | fosab |
| fkbp10b | 7.051924 | 0.338882561 | 0.303 | 0.026 | 1.7915414 | 32 | fkbp10b |

| | | | | | | | |
|---|---|---|---|---|---|---|---|
| SCARF22 | 2.355223 | 0.394202915 | 0.303 | 0.027 | 5.9834446 | 32 | SCARF2 |
| nuak1b | 9.747192 | 0.295978957 | 0.202 | 0.012 | 2.4762742 | 32 | nuak1b |
| atf37 | 1.328901 | 0.745590863 | 0.539 | 0.08 | 3.3760732 | 32 | atf3 |
| rpz58 | 1.463834 | 0.700742667 | 0.708 | 0.124 | 3.7188726 | 32 | rpz5 |
| igfbp31 | 1.770664 | 0.288458545 | 0.202 | 0.012 | 4.4983729 | 32 | igfbp3 |
| mrc22 | 4.271579 | 0.370473531 | 0.315 | 0.029 | 1.0851947 | 32 | mrc2 |
| schip12 | 1.091129 | 0.320458364 | 0.27 | 0.022 | 2.7720137 | 32 | schip1 |
| prkab1b3 | 2.087458 | 0.706932426 | 0.708 | 0.134 | 5.3031879 | 32 | prkab1b |
| id19 | 2.493760 | 0.883817672 | 0.697 | 0.129 | 6.3353991 | 32 | id1 |
| b3glcta | 3.528652 | 0.287057802 | 0.236 | 0.017 | 8.9645416 | 32 | b3glcta |
| angptl45 | 3.574456 | 0.487238928 | 0.438 | 0.054 | 9.0809076 | 32 | angptl4 |
| inhbab1 | 1.834605 | 0.277827322 | 0.169 | 0.009 | 4.6608140 | 32 | inhbab |
| cd993 | 5.614216 | 0.51785751 | 0.416 | 0.051 | 1.4262916 | 32 | cd99 |
| jun10 | 6.710632 | 1.13491289 | 0.899 | 0.248 | 1.7048363 | 32 | jun |
| cntfr2 | 9.210562 | 0.442904299 | 0.247 | 0.02 | 2.3399434 | 32 | cntfr |
| dpt1 | 1.135754 | 0.544216578 | 0.225 | 0.016 | 2.8853846 | 32 | dpt |
| cavin1b6 | 8.328314 | 0.483324447 | 0.382 | 0.044 | 2.1158082 | 32 | cavin1b |
| slc39a71 | 1.135411 | 0.403512887 | 0.371 | 0.042 | 2.8845135 | 32 | slc39a7 |
| perp5 | 1.136134 | 0.496576765 | 0.449 | 0.06 | 2.8863501 | 32 | perp |
| ppfibp1b | 1.187154 | 0.28372361 | 0.247 | 0.02 | 3.0159668 | 32 | ppfibp1b |
| fat1a2 | 7.768330 | 0.395067568 | 0.303 | 0.03 | 1.9735443 | 32 | fat1a |
| myo1b | 1.090781 | 0.252052425 | 0.18 | 0.011 | 2.7711296 | 32 | myo1b |
| yap14 | 2.844511 | 0.488758313 | 0.427 | 0.056 | 7.2264807 | 32 | yap1 |
| sox4b | 3.096069 | 0.352875141 | 0.191 | 0.013 | 7.8655652 | 32 | sox4b |
| elovl1b5 | 3.324961 | 0.74802707 | 0.629 | 0.121 | 8.4470649 | 32 | elovl1b |
| cnn3a7 | 4.057383 | 0.49672042 | 0.472 | 0.068 | 1.0307782 | 32 | cnn3a |
| ier2b13 | 7.732713 | 0.936846307 | 0.876 | 0.235 | 1.9644958 | 32 | ier2b |
| irf2bpl2 | 1.762292 | 0.443056143 | 0.371 | 0.045 | 4.4771043 | 32 | irf2bpl |
| endouc3 | 9.067737 | 0.614239416 | 0.472 | 0.07 | 2.3036587 | 32 | endouc |
| si:ch211-156j16.13 | 1.645447 | 0.404872811 | 0.281 | 0.027 | 4.1802582 | 32 | si:ch211-156j16.1 |
| ecrg4a3 | 1.691385 | 0.525617965 | 0.326 | 0.036 | 4.2969640 | 32 | ecrg4a |
| hmgb1b9 | 4.864670 | 0.6988741 | 0.73 | 0.152 | 1.2358694 | 32 | hmgb1b |
| klf2a7 | 5.333400 | 0.694932556 | 0.562 | 0.098 | 1.3549504 | 32 | klf2a |
| junba11 | 8.195015 | 1.078995164 | 0.944 | 0.318 | 2.0819437 | 32 | junba |
| calua1 | 8.464930 | 0.334859282 | 0.303 | 0.032 | 2.1505155 | 32 | calua |

| | | | | | | | |
|---|---|---|---|---|---|---|---|
| s100v29 | 1.019834 | 0.6797249 | 0.697 | 0.138 | 2.5908901 | 32 | s100v2 |
| CU929237.12 | 1.147294 | 0.287386107 | 0.247 | 0.022 | 2.9147008 | 32 | CU929237.1 |
| dap1 | 1.245143 | 0.524389012 | 0.539 | 0.088 | 3.1632877 | 32 | dap |
| creb3l2 | 1.799484 | 0.408598211 | 0.337 | 0.039 | 4.5715896 | 32 | creb3l2 |
| fn1a2 | 2.103701 | 0.639662695 | 0.506 | 0.081 | 5.3444527 | 32 | fn1a |
| midn9 | 1.285140 | 0.724818314 | 0.685 | 0.143 | 3.2649004 | 32 | midn |
| mob2b | 1.330893 | 0.349529036 | 0.281 | 0.028 | 3.3811344 | 32 | mob2b |
| pcolcea2 | 2.768412 | 0.598926961 | 0.449 | 0.066 | 7.0331522 | 32 | pcolcea |
| si:ch211-286o17.13 | 5.816116 | 0.488750672 | 0.371 | 0.048 | 1.4775843 | 32 | si:ch211-286o17.1 |
| metrnlb | 7.722441 | 0.323523662 | 0.258 | 0.024 | 1.9618863 | 32 | metrnlb |
| slc31a1 | 1.308233 | 0.268342342 | 0.247 | 0.023 | 3.3235673 | 32 | slc31a1 |
| chsy1 | 9.342063 | 0.292456954 | 0.236 | 0.022 | 2.3733513 | 32 | chsy1 |
| s100a10b11 | 9.698697 | 0.947201909 | 0.989 | 0.283 | 2.4639540 | 32 | s100a10b |
| anxa2a8 | 2.300193 | 0.600052406 | 0.607 | 0.116 | 5.8436409 | 32 | anxa2a |
| arg22 | 2.369549 | 0.305285206 | 0.213 | 0.018 | 6.0198416 | 32 | arg2 |
| ptprfb2 | 2.564452 | 0.288361165 | 0.202 | 0.017 | 6.5149907 | 32 | ptprfb |
| nucb2a1 | 3.495819 | 0.438725283 | 0.404 | 0.059 | 8.8811300 | 32 | nucb2a |
| emid11 | 5.236058 | 0.405312093 | 0.27 | 0.029 | 1.3302207 | 32 | emid1 |
| si:ch73-335l21.47 | 5.365512 | 0.929029247 | 0.629 | 0.14 | 1.3631083 | 32 | si:ch73-335l21.4 |
| LOC1018835071 | 8.749687 | 0.319617277 | 0.213 | 0.019 | 2.2228581 | 32 | LOC101883507 |
| mych4 | 1.120189 | 0.557477587 | 0.506 | 0.089 | 2.8458419 | 32 | mych |
| gadd45ga3 | 1.417604 | 0.532810325 | 0.371 | 0.052 | 3.6014245 | 32 | gadd45ga |
| tuba8l37 | 1.695774 | 0.474253102 | 0.449 | 0.071 | 4.3081144 | 32 | tuba8l3 |
| ptbp31 | 5.417932 | 0.50482846 | 0.416 | 0.065 | 1.3764257 | 32 | ptbp3 |
| tsc22d39 | 1.585065 | 0.716795708 | 0.663 | 0.157 | 4.0268589 | 32 | tsc22d3 |
| lhfpl2b | 2.075253 | 0.303597241 | 0.225 | 0.022 | 5.2721826 | 32 | lhfpl2b |
| ccnd12 | 4.645896 | 0.373017054 | 0.326 | 0.043 | 1.1802899 | 32 | ccnd1 |
| meis1b6 | 6.524707 | 0.440224019 | 0.438 | 0.072 | 1.6576018 | 32 | meis1b |
| kdelr2a | 7.000177 | 0.465506006 | 0.449 | 0.076 | 1.7783951 | 32 | kdelr2a |
| atp2b1a2 | 8.749858 | 0.423318638 | 0.438 | 0.071 | 2.2229015 | 32 | atp2b1a |
| txndc5 | 9.761484 | 0.398839916 | 0.438 | 0.071 | 2.4799050 | 32 | txndc5 |
| kank1a | 1.651673 | 0.250605849 | 0.191 | 0.016 | 4.1960772 | 32 | kank1a |
| mgst3a1 | 1.951769 | 0.416619876 | 0.438 | 0.073 | 4.9584695 | 32 | mgst3a |
| prrx1a1 | 2.327355 | 0.251900513 | 0.169 | 0.013 | 5.9126475 | 32 | prrx1a |
| h1f07 | 2.364272 | 0.487499261 | 0.427 | 0.07 | 6.0064333 | 32 | h1f0 |

| | | | | | | | |
|---|---|---|---|---|---|---|---|
| gadd45ba8 | 6.160441 | 0.801832134 | 0.742 | 0.196 | 1.5650601 | 32 | gadd45ba |
| prdx510 | 7.948202 | 0.677264508 | 0.708 | 0.177 | 2.0192409 | 32 | prdx5 |
| anxa5b6 | 9.689633 | 0.550965548 | 0.528 | 0.104 | 2.4616513 | 32 | anxa5b |
| si:busm1-57f23.11 | 1.028421 | 0.387740944 | 0.258 | 0.029 | 2.6127043 | 32 | si:busm1-57f23.1 |
| plod23 | 1.191526 | 0.3374209 | 0.258 | 0.029 | 3.0270742 | 32 | plod2 |
| meis1a2 | 1.196363 | 0.254714926 | 0.213 | 0.02 | 3.0393612 | 32 | meis1a |
| fstl1b6 | 1.377055 | 0.576888992 | 0.674 | 0.146 | 3.4984090 | 32 | fstl1b |
| asph2 | 4.690806 | 0.366843364 | 0.315 | 0.042 | 1.1916992 | 32 | asph |
| lgals3b4 | 6.124455 | 0.473218656 | 0.506 | 0.097 | 1.5559178 | 32 | lgals3b |
| socs3a11 | 6.850765 | 0.975291089 | 0.899 | 0.313 | 1.7404370 | 32 | socs3a |
| tsc22d1 | 7.565110 | 0.363168525 | 0.315 | 0.042 | 1.9219163 | 32 | tsc22d1 |
| snai1a2 | 8.189446 | 0.335543947 | 0.371 | 0.055 | 2.0805288 | 32 | snai1a |
| fam168a7 | 9.008693 | 0.423043618 | 0.427 | 0.073 | 2.2886586 | 32 | fam168a |
| efna1b3 | 1.149302 | 0.306267572 | 0.202 | 0.019 | 2.9198035 | 32 | efna1b |
| vgll4b | 4.416630 | 0.319035207 | 0.236 | 0.026 | 1.1220449 | 32 | vgll4b |
| tns3.2 | 7.845611 | 0.302860256 | 0.225 | 0.024 | 1.9931777 | 32 | tns3.2 |
| ier2a12 | 1.358199 | 0.823189126 | 0.876 | 0.272 | 3.4505065 | 32 | ier2a |
| bcar1 | 1.640821 | 0.25159885 | 0.202 | 0.02 | 4.1685062 | 32 | bcar1 |
| tent5ba3 | 1.061715 | 0.270943698 | 0.157 | 0.013 | 2.6972887 | 32 | tent5ba |
| slc38a5b7 | 1.089891 | 0.560827138 | 0.573 | 0.127 | 2.7688696 | 32 | slc38a5b |
| fzd23 | 1.113226 | 0.323306854 | 0.303 | 0.042 | 2.8281522 | 32 | fzd2 |
| sdc410 | 5.991598 | 0.766808126 | 0.843 | 0.252 | 1.5221655 | 32 | sdc4 |
| klf133 | 9.616392 | 0.46107832 | 0.506 | 0.103 | 2.4430443 | 32 | klf13 |
| h3f3c12 | 1.027651 | 0.792429096 | 0.876 | 0.302 | 2.6107488 | 32 | h3f3c |
| sept15 | 1.295188 | 0.29586965 | 0.292 | 0.04 | 3.2904273 | 32 | sept15 |
| ptprga8 | 3.080278 | 0.39539583 | 0.404 | 0.071 | 7.8254464 | 32 | ptprga |
| frzb2 | 4.633257 | 0.415761998 | 0.191 | 0.019 | 1.1770790 | 32 | frzb |
| si:ch73-46j18.59 | 5.023108 | 0.600703795 | 0.809 | 0.207 | 1.2761205 | 32 | si:ch73-46j18.5 |
| selenok2 | 8.076812 | 0.489713878 | 0.596 | 0.138 | 2.0519141 | 32 | selenok |
| calm2a2 | 1.649890 | 0.618538907 | 0.764 | 0.208 | 4.1915462 | 32 | calm2a |
| ssr34 | 2.264935 | 0.719806137 | 0.888 | 0.286 | 5.7540674 | 32 | ssr3 |
| klf11a | 2.567499 | 0.369407312 | 0.303 | 0.045 | 6.5227331 | 32 | klf11a |
| tpm4a9 | 2.593667 | 0.474977817 | 0.528 | 0.112 | 6.5892133 | 32 | tpm4a |
| klf34 | 3.343333 | 0.286344067 | 0.281 | 0.039 | 8.4937387 | 32 | klf3 |
| sgce2 | 6.634465 | 0.304334959 | 0.247 | 0.032 | 1.6854860 | 32 | sgce |

| | | | | | | | |
|---|---|---|---|---|---|---|---|
| fosl1a6 | 6.939186 | 0.489521445 | 0.427 | 0.083 | 1.7629003 | 32 | fosl1a |
| map1lc3b4 | 7.294686 | 0.448937042 | 0.551 | 0.123 | 1.8532151 | 32 | map1lc3b |
| neo1a3 | 8.850334 | 0.327858172 | 0.247 | 0.032 | 2.2484275 | 32 | neo1a |
| sdc22 | 1.049748 | 0.413918427 | 0.427 | 0.082 | 2.6668859 | 32 | sdc2 |
| zdhhc20a1 | 1.346953 | 0.358495629 | 0.292 | 0.043 | 3.4219344 | 32 | zdhhc20a |
| tspan72 | 1.352374 | 0.402898208 | 0.427 | 0.081 | 3.4357076 | 32 | tspan7 |
| triqk2 | 1.374589 | 0.325799149 | 0.337 | 0.055 | 3.4921434 | 32 | triqk |
| ier55 | 1.569758 | 0.540148931 | 0.596 | 0.145 | 3.9879725 | 32 | ier5 |
| f13a1b3 | 2.086685 | 0.285776558 | 0.247 | 0.032 | 5.3012241 | 32 | f13a1b |
| capns1b7 | 4.965561 | 0.495908645 | 0.596 | 0.144 | 1.2615009 | 32 | capns1b |
| mdka8 | 6.988515 | 0.665544442 | 0.494 | 0.113 | 1.7754322 | 32 | mdka |
| supt3h1 | 7.206398 | 0.30955685 | 0.27 | 0.038 | 1.8307856 | 32 | supt3h |
| twist31 | 1.440523 | 0.27901218 | 0.236 | 0.03 | 3.6596500 | 32 | twist3 |
| sumo11 | 1.443018 | 0.424112854 | 0.551 | 0.126 | 3.6659895 | 32 | sumo1 |
| col5a2a5 | 1.499988 | 0.517196774 | 0.472 | 0.1 | 3.8107217 | 32 | col5a2a |
| irf2bp2b8 | 1.998814 | 0.454601741 | 0.472 | 0.101 | 5.0779875 | 32 | irf2bp2b |
| zgc:16273014 | 2.744508 | 0.865462515 | 0.921 | 0.402 | 6.9724245 | 32 | zgc:162730 |
| pkig1 | 3.213038 | 0.389554907 | 0.472 | 0.098 | 8.1627236 | 32 | pkig |
| tspan9a1 | 3.819852 | 0.277176498 | 0.202 | 0.023 | 9.7043363 | 32 | tspan9a |
| btg210 | 5.592074 | 0.770260901 | 0.944 | 0.348 | 1.4206664 | 32 | btg2 |
| anxa1a7 | 7.892140 | 0.558056696 | 0.64 | 0.161 | 2.0049982 | 32 | anxa1a |
| sar1b2 | 9.650790 | 0.371837994 | 0.438 | 0.089 | 2.4517833 | 32 | sar1b |
| cdkn1a3 | 1.216409 | 0.335683334 | 0.303 | 0.048 | 3.0902887 | 32 | cdkn1a |
| rrbp1a1 | 1.975892 | 0.336554377 | 0.337 | 0.058 | 5.0197542 | 32 | rrbp1a |
| loxa1 | 3.057298 | 0.350674313 | 0.157 | 0.015 | 7.7670677 | 32 | loxa |
| mmp26 | 3.283212 | 0.388746836 | 0.506 | 0.11 | 8.3410025 | 32 | mmp2 |
| antxr2a6 | 3.456673 | 0.551130981 | 0.64 | 0.168 | 8.7816791 | 32 | antxr2a |
| enpp12 | 5.167461 | 0.469495013 | 0.258 | 0.039 | 1.3127935 | 32 | enpp1 |
| sec61a13 | 5.748558 | 0.576893742 | 0.674 | 0.191 | 1.4604213 | 32 | sec61a1 |
| net12 | 5.947754 | 0.28360611 | 0.236 | 0.032 | 1.5110270 | 32 | net1 |
| arl4d4 | 1.193916 | 0.346703123 | 0.315 | 0.053 | 3.0331437 | 32 | arl4d |
| ppib11 | 1.350552 | 0.768517305 | 0.921 | 0.378 | 3.4310790 | 32 | ppib |
| pmp22b5 | 1.548968 | 0.300724458 | 0.292 | 0.046 | 3.9351557 | 32 | pmp22b |
| vdac36 | 2.527495 | 0.511241634 | 0.629 | 0.167 | 6.4211011 | 32 | vdac3 |
| hdlbpa7 | 2.862833 | 0.532326976 | 0.64 | 0.174 | 7.2730296 | 32 | hdlbpa |

| | | | | | | | |
|---|---|---|---|---|---|---|---|
| tdh1 | 1.390102 | 0.38362114 | 0.427 | 0.091 | 3.5315564 | 32 | tdh |
| sdf41 | 1.440934 | 0.316156538 | 0.303 | 0.051 | 3.6606950 | 32 | sdf4 |
| znf704 | 2.281856 | 0.288802732 | 0.236 | 0.033 | 5.7970558 | 32 | znf704 |
| metrnla5 | 2.328114 | 0.478279477 | 0.427 | 0.094 | 5.9145756 | 32 | metrnla |
| lrrc174 | 2.720850 | 0.337178391 | 0.326 | 0.058 | 6.9123198 | 32 | lrrc17 |
| calr2 | 5.511653 | 0.347302225 | 0.348 | 0.066 | 1.4002356 | 32 | calr |
| tacc1 | 7.188359 | 0.302319392 | 0.247 | 0.037 | 1.8262027 | 32 | tacc1 |
| serf29 | 7.640496 | 0.705729926 | 0.91 | 0.376 | 1.9410680 | 32 | serf2 |
| ip6k2b6 | 7.799803 | 0.523321864 | 0.506 | 0.128 | 1.9815400 | 32 | ip6k2b |
| h2afy25 | 8.627900 | 0.413309201 | 0.506 | 0.122 | 2.1919180 | 32 | h2afy2 |
| gng22 | 9.436014 | 0.265324151 | 0.292 | 0.049 | 2.3972194 | 32 | gng2 |
| hmgn78 | 1.122950 | 0.667717676 | 0.865 | 0.316 | 2.8528557 | 32 | hmgn7 |
| atf4b11 | 2.046208 | 0.623573076 | 0.82 | 0.276 | 5.1983925 | 32 | atf4b |
| kdelr2b | 3.450202 | 0.446325056 | 0.494 | 0.121 | 8.7652389 | 32 | kdelr2b |
| CABZ01080702.17 | 5.319252 | 0.537384267 | 0.416 | 0.097 | 1.3513562 | 32 | CABZ01080702.1 |
| hmgn614 | 9.707955 | 0.647751602 | 0.876 | 0.309 | 2.4663060 | 32 | hmgn6 |
| klf11b5 | 4.348722 | 0.37277246 | 0.382 | 0.082 | 1.1047929 | 32 | klf11b |
| prss353 | 4.890372 | 0.432726688 | 0.202 | 0.029 | 1.2423991 | 32 | prss35 |
| si:dkey-28b4.78 | 5.893698 | 0.350421482 | 0.461 | 0.109 | 1.4972942 | 32 | si:dkey-28b4.7 |
| ldlrap1a2 | 7.170058 | 0.34407242 | 0.326 | 0.062 | 1.8215533 | 32 | ldlrap1a |
| spry23 | 1.212312 | 0.361093673 | 0.348 | 0.072 | 3.0798806 | 32 | spry2 |
| arf51 | 1.733646 | 0.498598685 | 0.685 | 0.21 | 4.4043283 | 32 | arf5 |
| nme35 | 3.813508 | 0.358426449 | 0.393 | 0.088 | 9.6882179 | 32 | nme3 |
| gadd45ab3 | 5.167373 | 0.34905439 | 0.337 | 0.069 | 1.3127713 | 32 | gadd45ab |
| evla3 | 5.282575 | 0.332601893 | 0.404 | 0.09 | 1.3420382 | 32 | evla |
| id34 | 7.279151 | 0.555971213 | 0.652 | 0.196 | 1.8492683 | 32 | id3 |
| nr1d2a3 | 8.295153 | 0.34043084 | 0.393 | 0.089 | 2.1073837 | 32 | nr1d2a |
| gng77 | 1.253782 | 0.268451139 | 0.315 | 0.062 | 3.1852354 | 32 | gng7 |
| pfkfb4b6 | 2.850122 | 0.36695408 | 0.472 | 0.12 | 7.2407371 | 32 | pfkfb4b |
| si:ch211-194k22.8 | 3.345166 | 0.275857015 | 0.281 | 0.052 | 8.4983943 | 32 | si:ch211-194k22.8 |
| vamp32 | 3.967436 | 0.416741867 | 0.652 | 0.194 | 1.0079271 | 32 | vamp3 |
| spred2b6 | 7.620896 | 0.356309758 | 0.371 | 0.084 | 1.9360886 | 32 | spred2b |
| lrpap12 | 9.759294 | 0.254624134 | 0.315 | 0.063 | 2.4793487 | 32 | lrpap1 |
| emp23 | 1.053315 | 0.275213028 | 0.292 | 0.056 | 2.6759487 | 32 | emp2 |
| cish2 | 1.136738 | 0.368648683 | 0.281 | 0.055 | 2.8878840 | 32 | cish |

| | | | | | | | |
|---|---|---|---|---|---|---|---|
| tpm13 | 1.217627 | 0.499594007 | 0.652 | 0.202 | 3.0933826 | 32 | tpm1 |
| myca2 | 1.860549 | 0.283231594 | 0.236 | 0.04 | 4.7267248 | 32 | myca |
| arl1 | 2.226219 | 0.280511168 | 0.348 | 0.076 | 5.6557100 | 32 | arl1 |
| map1lc3a8 | 3.353613 | 0.280364734 | 0.36 | 0.08 | 8.5198559 | 32 | map1lc3a |
| selenof4 | 4.450082 | 0.542539695 | 0.753 | 0.266 | 1.1305434 | 32 | selenof |
| chd61 | 5.835115 | 0.267492097 | 0.258 | 0.048 | 1.4824109 | 32 | chd6 |
| cited4a5 | 8.721402 | 0.268108574 | 0.236 | 0.041 | 2.2156723 | 32 | cited4a |
| pgrmc16 | 1.095461 | 0.316184916 | 0.551 | 0.151 | 2.7830189 | 32 | pgrmc1 |
| klf6a7 | 1.478910 | 0.487191006 | 0.584 | 0.179 | 3.7571720 | 32 | klf6a |
| rab2a2 | 1.635124 | 0.413672514 | 0.584 | 0.176 | 4.1540333 | 32 | rab2a |
| plecb3 | 1.730665 | 0.27921674 | 0.27 | 0.052 | 4.3967557 | 32 | plecb |
| rhoab8 | 2.301157 | 0.440748064 | 0.652 | 0.206 | 5.8460917 | 32 | rhoab |
| tomm20b4 | 3.561012 | 0.46021463 | 0.685 | 0.227 | 9.0467532 | 32 | tomm20b |
| atp2b25 | 4.043887 | 0.270894306 | 0.36 | 0.083 | 1.0273495 | 32 | atp2b2 |
| gpc1b | 4.305291 | 0.257335284 | 0.213 | 0.036 | 1.0937591 | 32 | gpc1b |
| mgst3b5 | 5.547745 | 0.438717723 | 0.607 | 0.192 | 1.4094048 | 32 | mgst3b |
| col6a15 | 7.422928 | 0.350503286 | 0.393 | 0.097 | 1.8857950 | 32 | col6a1 |
| tfg | 7.724146 | 0.309308759 | 0.393 | 0.098 | 1.9623195 | 32 | tfg |
| svild7 | 9.310136 | 0.292903388 | 0.36 | 0.084 | 2.3652401 | 32 | svild |
| junbb4 | 1.154228 | 0.303217511 | 0.315 | 0.07 | 2.9323171 | 32 | junbb |
| her91 | 1.206023 | 0.264198198 | 0.169 | 0.024 | 3.0639021 | 32 | her9 |
| itga52 | 1.249875 | 0.323453698 | 0.191 | 0.031 | 3.1753084 | 32 | itga5 |
| cald1a8 | 1.424558 | 0.351091559 | 0.449 | 0.119 | 3.6190916 | 32 | cald1a |
| grpel23 | 1.975950 | 0.27410556 | 0.416 | 0.106 | 5.0199016 | 32 | grpel2 |
| hnrnpa0a3 | 3.380993 | 0.250113177 | 0.348 | 0.081 | 8.5894130 | 32 | hnrnpa0a |
| mmp14b5 | 3.778868 | 0.274236854 | 0.303 | 0.066 | 9.6002149 | 32 | mmp14b |
| chd4b1 | 4.276044 | 0.291491719 | 0.416 | 0.108 | 1.0863292 | 32 | chd4b |
| tpt16 | 4.862947 | 0.355293061 | 0.989 | 0.812 | 1.2354317 | 32 | tpt1 |
| tgif15 | 6.568934 | 0.274870202 | 0.393 | 0.1 | 1.6688379 | 32 | tgif1 |
| ciarta4 | 6.981378 | 0.366588828 | 0.427 | 0.117 | 1.7736192 | 32 | ciarta |
| cd1642 | 7.800075 | 0.262982693 | 0.326 | 0.075 | 1.9816092 | 32 | cd164 |
| ephb35 | 8.713524 | 0.259066633 | 0.258 | 0.052 | 2.2136710 | 32 | ephb3 |
| osr24 | 8.714713 | 0.537878482 | 0.247 | 0.052 | 2.2139729 | 32 | osr2 |
| histh1l8 | 9.117517 | 0.47585693 | 0.427 | 0.122 | 2.3163053 | 32 | histh1l |
| mafba8 | 9.332464 | 0.304549597 | 0.348 | 0.084 | 2.3709127 | 32 | mafba |

| | | | | | | | |
|---|---|---|---|---|---|---|---|
| cyr615 | 9.561178 | 0.316563961 | 0.303 | 0.067 | 2.4290172 | 32 | cyr61 |
| snd11 | 1.036775 | 0.315461412 | 0.427 | 0.117 | 2.6339285 | 32 | snd1 |
| ssr44 | 1.607433 | 0.596717135 | 0.775 | 0.331 | 4.0836842 | 32 | ssr4 |
| spcs12 | 2.206934 | 0.440329473 | 0.663 | 0.23 | 5.6067181 | 32 | spcs1 |
| jund7 | 2.581862 | 0.28849438 | 0.36 | 0.09 | 6.5592211 | 32 | jund |
| sept7a | 2.817469 | 0.286116482 | 0.326 | 0.077 | 7.1577803 | 32 | sept7a |
| wls6 | 3.019172 | 0.339828841 | 0.461 | 0.131 | 7.6702074 | 32 | wls |
| rgcc5 | 3.097400 | 0.400108101 | 0.303 | 0.071 | 7.8689452 | 32 | rgcc |
| appb4 | 3.994726 | 0.259505736 | 0.236 | 0.047 | 1.0148602 | 32 | appb |
| ostc2 | 4.094218 | 0.38921531 | 0.584 | 0.189 | 1.0401362 | 32 | ostc |
| flot2a6 | 4.793662 | 0.266048222 | 0.315 | 0.074 | 1.2178299 | 32 | flot2a |
| tmem167a | 5.095227 | 0.272931166 | 0.404 | 0.108 | 1.2944424 | 32 | tmem167a |
| ssr11 | 5.902002 | 0.445492546 | 0.584 | 0.2 | 1.4994036 | 32 | ssr1 |
| myl9b7 | 6.360760 | 0.292057 | 0.382 | 0.098 | 1.6159511 | 32 | myl9b |
| krt811 | 7.361600 | 0.526934102 | 0.831 | 0.31 | 1.8702145 | 32 | krt8 |
| notch28 | 1.626944 | 0.348083944 | 0.427 | 0.122 | 4.1332528 | 32 | notch2 |
| tmem2582 | 4.189400 | 0.446018104 | 0.798 | 0.304 | 1.0643172 | 32 | tmem258 |
| LOC1018844515 | 4.411173 | 0.30096355 | 0.348 | 0.091 | 1.1206586 | 32 | LOC101884451 |
| mt-nd211 | 4.887582 | 0.550392698 | 0.91 | 0.391 | 1.2416903 | 32 | mt-nd2 |
| slc3a2b4 | 5.266483 | 0.327642812 | 0.438 | 0.128 | 1.3379501 | 32 | slc3a2b |
| ier3ip11 | 6.330160 | 0.322784164 | 0.517 | 0.163 | 1.6081773 | 32 | ier3ip1 |
| cops91 | 7.329433 | 0.269823267 | 0.404 | 0.114 | 1.8620425 | 32 | cops9 |
| ywhaqb9 | 8.699483 | 0.335222489 | 0.494 | 0.154 | 2.2101038 | 32 | ywhaqb |
| si:ch1073-325m22.23 | 9.867598 | 0.416437544 | 0.674 | 0.249 | 2.5068635 | 32 | si:ch1073-325m22.2 |
| ssr21 | 9.993924 | 0.500898205 | 0.82 | 0.332 | 2.5389565 | 32 | ssr2 |
| glud1a2 | 1.030085 | 0.26856678 | 0.371 | 0.099 | 2.6169325 | 32 | glud1a |
| cd81a8 | 1.370959 | 0.354637779 | 0.708 | 0.24 | 3.4829229 | 32 | cd81a |
| ost45 | 1.378759 | 0.441139367 | 0.697 | 0.261 | 3.5027380 | 32 | ost4 |
| ptmaa16 | 1.527520 | 0.558767743 | 0.955 | 0.432 | 3.8806669 | 32 | ptmaa |
| si:ch211-286b5.58 | 1.539505 | 0.281117392 | 0.494 | 0.151 | 3.9111141 | 32 | si:ch211-286b5.5 |
| pim26 | 1.933454 | 0.266729096 | 0.337 | 0.088 | 4.9119412 | 32 | pim2 |
| krtcap22 | 2.451808 | 0.354151812 | 0.573 | 0.195 | 6.2288191 | 32 | krtcap2 |
| p4hb1 | 2.486448 | 0.494838505 | 0.573 | 0.212 | 6.3168211 | 32 | p4hb |
| atf4a3 | 2.695734 | 0.353278681 | 0.551 | 0.185 | 6.8485145 | 32 | atf4a |
| dusp26 | 3.222716 | 0.399738695 | 0.506 | 0.163 | 8.1873103 | 32 | dusp2 |

| | | | | | | | |
|---|---|---|---|---|---|---|---|
| ahnak9 | 4.103973 | 0.320467544 | 0.449 | 0.136 | 1.0426145 | 32 | ahnak |
| nfia6 | 5.664729 | 0.283711332 | 0.36 | 0.099 | 1.4391244 | 32 | nfia |
| eif4a1b8 | 9.100180 | 0.474297702 | 0.764 | 0.311 | 2.3119007 | 32 | eif4a1b |
| ywhabb2 | 9.460376 | 0.396568135 | 0.674 | 0.252 | 2.4034087 | 32 | ywhabb |
| eif1b10 | 1.158894 | 0.528690321 | 0.876 | 0.406 | 2.9441723 | 32 | eif1b |
| CR383676.117 | 1.438514 | 0.643619011 | 1 | 0.717 | 3.6545457 | 32 | CR383676.1 |
| serp13 | 1.520182 | 0.376861877 | 0.562 | 0.197 | 3.8620239 | 32 | serp1 |
| zgc:77849 | 1.540239 | 0.308308516 | 0.539 | 0.18 | 3.9129778 | 32 | zgc:77849 |
| mtdha1 | 2.009310 | 0.253221952 | 0.472 | 0.147 | 5.1046545 | 32 | mtdha |
| pfn2l8 | 2.894343 | 0.341010241 | 0.618 | 0.215 | 7.3530802 | 32 | pfn2l |
| qkia7 | 3.050776 | 0.347483314 | 0.573 | 0.199 | 7.7504988 | 32 | qkia |
| mt-nd39 | 4.434781 | 0.468345007 | 0.865 | 0.37 | 1.1266562 | 32 | mt-nd3 |
| ccnd2a2 | 5.011816 | 0.297466449 | 0.337 | 0.095 | 1.2732519 | 32 | ccnd2a |
| atrx7 | 6.204859 | 0.269023194 | 0.416 | 0.128 | 1.5763444 | 32 | atrx |
| si:dkey-79d12.53 | 7.165875 | 0.266964112 | 0.27 | 0.068 | 1.8204906 | 32 | si:dkey-79d12.5 |
| socs3b7 | 7.299712 | 0.305598802 | 0.472 | 0.151 | 1.8544920 | 32 | socs3b |
| mycb5 | 7.993601 | 0.419267582 | 0.551 | 0.2 | 2.0307745 | 32 | mycb |
| si:zfos-169g10.32 | 8.273518 | 0.255512376 | 0.382 | 0.115 | 2.1018872 | 32 | si:zfos-169g10.3 |
| elocb5 | 1.190235 | 0.332585897 | 0.551 | 0.195 | 3.0237940 | 32 | elocb |
| si:dkey-56m19.54 | 1.297685 | 0.266058916 | 0.236 | 0.056 | 3.2967709 | 32 | si:dkey-56m19.5 |
| nfkbiab4 | 1.481397 | 0.330307015 | 0.562 | 0.2 | 3.7634906 | 32 | nfkbiab |
| mcl1b11 | 1.578937 | 0.398047691 | 0.663 | 0.25 | 4.0112895 | 32 | mcl1b |
| idh23 | 2.469166 | 0.313274523 | 0.618 | 0.226 | 6.2729171 | 32 | idh2 |
| ptp4a14 | 2.748255 | 0.261397429 | 0.416 | 0.132 | 6.9819432 | 32 | ptp4a1 |
| calm3b3 | 2.839914 | 0.399401906 | 0.697 | 0.277 | 7.2148024 | 32 | calm3b |
| reep57 | 2.949040 | 0.293817093 | 0.506 | 0.173 | 7.4920371 | 32 | reep5 |
| smim151 | 2.949846 | 0.267828804 | 0.461 | 0.153 | 7.4940854 | 32 | smim15 |
| zfp36l21 | 4.083257 | 0.263507937 | 0.348 | 0.101 | 1.0373516 | 32 | zfp36l2 |
| fkbp1aa10 | 4.321042 | 0.503687588 | 0.888 | 0.413 | 1.0977608 | 32 | fkbp1aa |
| jpt1b6 | 5.161918 | 0.379123862 | 0.719 | 0.276 | 1.3113854 | 32 | jpt1b |
| mfap25 | 6.028641 | 0.25095618 | 0.393 | 0.12 | 1.5315763 | 32 | mfap2 |
| rpl22l111 | 6.107977 | 0.532132152 | 0.865 | 0.424 | 1.5517316 | 32 | rpl22l1 |
| nsa211 | 6.958816 | 0.500566685 | 0.854 | 0.388 | 1.7678872 | 32 | nsa2 |
| ddit39 | 8.818555 | 0.309745403 | 0.506 | 0.178 | 2.2403539 | 32 | ddit3 |
| marcksl1b10 | 9.186200 | 0.321307475 | 0.427 | 0.142 | 2.3337543 | 32 | marcksl1b |

| | | | | | | | |
|---|---|---|---|---|---|---|---|
| mt-nd113 | 1.659765 | 0.462346474 | 0.865 | 0.396 | 4.2166344 | 32 | mt-nd1 |
| sem12 | 1.968603 | 0.341371514 | 0.685 | 0.272 | 5.0012371 | 32 | sem1 |
| cebpb12 | 2.512386 | 0.518609574 | 0.719 | 0.339 | 6.3827179 | 32 | cebpb |
| pno13 | 5.398338 | 0.29834417 | 0.472 | 0.171 | 1.3714480 | 32 | pno1 |
| tnrc6c19 | 6.185994 | 0.260570306 | 0.438 | 0.15 | 1.5715518 | 32 | tnrc6c1 |
| rpl398 | 8.827715 | 0.277776623 | 1 | 0.877 | 2.2426811 | 32 | rpl39 |
| mibp213 | 9.598915 | 0.538026343 | 0.899 | 0.521 | 2.4386045 | 32 | mibp2 |
| cnn29 | 1.077639 | 0.263884541 | 0.449 | 0.157 | 2.7377429 | 32 | cnn2 |
| si:dkey-42i9.45 | 1.183510 | 0.288748901 | 0.438 | 0.156 | 3.0067087 | 32 | si:dkey-42i9.4 |
| kctd12.27 | 1.255444 | 0.251275376 | 0.326 | 0.1 | 3.1894568 | 32 | kctd12.2 |
| ndufa15 | 1.281805 | 0.305721434 | 0.607 | 0.242 | 3.2564279 | 32 | ndufa1 |
| mt-cyb10 | 1.290663 | 0.496992172 | 0.989 | 0.576 | 3.2789316 | 32 | mt-cyb |
| chd4a3 | 1.292053 | 0.266579507 | 0.337 | 0.107 | 3.2824622 | 32 | chd4a |
| higd1a6 | 1.315078 | 0.388122781 | 0.708 | 0.303 | 3.3409581 | 32 | higd1a |
| tcf126 | 1.373354 | 0.267291158 | 0.247 | 0.068 | 3.4890064 | 32 | tcf12 |
| cnbpb6 | 1.433263 | 0.321296969 | 0.539 | 0.209 | 3.6412065 | 32 | cnbpb |
| tmed101 | 1.832154 | 0.297128152 | 0.483 | 0.181 | 4.6545885 | 32 | tmed10 |
| si:ch73-1a9.317 | 2.404077 | 0.389036651 | 0.742 | 0.315 | 6.1075580 | 32 | si:ch73-1a9.3 |
| mt-nd413 | 3.146353 | 0.460371612 | 0.921 | 0.447 | 7.9933098 | 32 | mt-nd4 |
| cox6a14 | 3.370889 | 0.429776278 | 0.831 | 0.389 | 8.5637437 | 32 | cox6a1 |
| XLOC-0234925 | 5.043564 | 0.261874922 | 0.303 | 0.094 | 1.2813176 | 32 | XLOC-023492 |
| gtpbp43 | 5.345622 | 0.303396305 | 0.618 | 0.246 | 1.3580553 | 32 | gtpbp4 |
| gnb1a4 | 5.958407 | 0.31917271 | 0.517 | 0.202 | 1.5137334 | 32 | gnb1a |
| csrnp1b8 | 6.730609 | 0.252618315 | 0.449 | 0.165 | 1.7099112 | 32 | csrnp1b |
| eif3ea5 | 7.235505 | 0.348879582 | 0.719 | 0.304 | 1.8381800 | 32 | eif3ea |
| tob1b3 | 9.716070 | 0.285787591 | 0.449 | 0.168 | 2.4683676 | 32 | tob1b |
| ptmab12 | 2.067121 | 0.447096803 | 0.978 | 0.569 | 5.2515213 | 32 | ptmab |
| cox7a2a2 | 2.088260 | 0.281096729 | 0.607 | 0.251 | 5.3052254 | 32 | cox7a2a |
| ybx110 | 2.434715 | 0.393700045 | 0.989 | 0.695 | 6.1853938 | 32 | ybx1 |
| rhoaa8 | 2.600839 | 0.269947055 | 0.461 | 0.177 | 6.6074322 | 32 | rhoaa |
| COX314 | 2.943949 | 0.463706857 | 1 | 0.676 | 7.4791046 | 32 | COX3 |
| ndufa4l9 | 3.243224 | 0.42209147 | 0.888 | 0.421 | 8.2394119 | 32 | ndufa4l |
| cdc426 | 5.555170 | 0.294643257 | 0.562 | 0.233 | 0.000141 | 32 | cdc42 |
| sec61b2 | 6.581448 | 0.440828455 | 0.876 | 0.469 | 0.000167 | 32 | sec61b |
| mt-atp613 | 7.310854 | 0.470283941 | 0.966 | 0.555 | 0.000186 | 32 | mt-atp6 |

| | | | | | | | |
|---|---|---|---|---|---|---|---|
| rab107 | 8.07519( | 0.258232234 | 0.528 | 0.214 | 0.000205 | 32 | rab10 |
| tpm39 | 8.78276 | 0.274071774 | 0.674 | 0.284 | 0.000223 | 32 | tpm3 |
| mt-co113 | 9.36118 | 0.453829306 | 1 | 0.639 | 0.000238 | 32 | mt-co1 |
| itm2ba10 | 1.21957( | 0.292014277 | 0.697 | 0.296 | 0.00031 | 32 | itm2ba |
| gng55 | 1.44279 | 0.261161385 | 0.685 | 0.287 | 0.000367 | 32 | gng5 |
| ddx52 | 2.63259 | 0.348027812 | 0.764 | 0.354 | 0.000669 | 32 | ddx5 |
| ubb3 | 2.80149 | 0.343805106 | 0.697 | 0.325 | 0.000712 | 32 | ubb |
| rpl2911 | 3.49256 | 0.30102659 | 1 | 0.823 | 0.000887 | 32 | rpl29 |
| rps226 | 3.57977 | 0.311588444 | 1 | 0.842 | 0.000909 | 32 | rps2 |
| gabarapb11 | 6.04769 | 0.298852913 | 0.674 | 0.305 | 0.001536 | 32 | gabarapb |
| tma74 | 7.32624 | 0.310928388 | 0.719 | 0.331 | 0.001861 | 32 | tma7 |
| dad12 | 1.32855 | 0.316438002 | 0.764 | 0.358 | 0.003375 | 32 | dad1 |
| eef1g13 | 1.56386 | 0.416712825 | 1 | 0.652 | 0.003973 | 32 | eef1g |
| mt-co212 | 1.59734 | 0.44913787 | 0.989 | 0.671 | 0.004058 | 32 | mt-co2 |
| sec61g6 | 2.16854 | 0.374139353 | 0.989 | 0.681 | 0.005509 | 32 | sec61g |
| rpl76 | 2.29319 | 0.284522813 | 0.989 | 0.809 | 0.005826 | 32 | rpl7 |
| rpl5a14 | 2.37835 | 0.458591075 | 0.955 | 0.536 | 0.006042 | 32 | rpl5a |
| si:ch211-196l7.411 | 2.43182( | 0.289465328 | 0.697 | 0.33 | 0.006178 | 32 | si:ch211-196l7.4 |
| rps27.116 | 2.89768 | 0.381731878 | 0.989 | 0.71 | 0.007362 | 32 | rps27.1 |
| lum1 | 3.72052 | 0.278936012 | 0.101 | 0.022 | 0.009452 | 32 | lum |
| xbp16 | 4.76583 | 0.309769813 | 0.742 | 0.355 | 0.012108 | 32 | xbp1 |
| rpl27a14 | 6.01944 | 0.4545243 | 0.966 | 0.522 | 0.015292 | 32 | rpl27a |
| rps27.214 | 7.23752 | 0.441415835 | 0.978 | 0.552 | 0.018387 | 32 | rps27.2 |
| sumo2b11 | 1.31107 | 0.253622147 | 0.629 | 0.296 | 0.033308 | 32 | sumo2b |
| rpl1813 | 1.47437 | 0.303386222 | 0.989 | 0.793 | 0.037457 | 32 | rpl18 |
| oaz1b11 | 2.131001 | 0.254832791 | 0.764 | 0.36 | 0.054138 | 32 | oaz1b |
| dynll18 | 2.75668 | 0.317381139 | 0.854 | 0.44 | 0.070034 | 32 | dynll1 |
| hnrnpa0b8 | 2.83657 | 0.276561086 | 0.708 | 0.341 | 0.072063 | 32 | hnrnpa0b |
| rpl3212 | 3.10122( | 0.289854521 | 1 | 0.824 | 0.078787 | 32 | rpl32 |
| bsg2 | 3.20704 | 0.256824748 | 0.798 | 0.375 | 0.081475 | 32 | bsg |
| zgc:1935413 | 4.35201 | 0.263399199 | 0.775 | 0.381 | 0.110563 | 32 | zgc:193541 |
| rpl18a12 | 8.76605 | 0.292020659 | 0.989 | 0.801 | 0.222702 | 32 | rpl18a |
| eef2b6 | 8.96536 | 0.374949964 | 0.978 | 0.636 | 0.227765 | 32 | eef2b |
| rpl109 | 1.25795( | 0.275131146 | 0.978 | 0.79 | 0.319582 | 32 | rpl10 |
| rpl10a11 | 1.769051 | 0.288518025 | 0.989 | 0.779 | 0.449427 | 32 | rpl10a |

| | | | | | | | |
|---|---|---|---|---|---|---|---|
| rps128 | 1.975346 | 0.250268066 | 1 | 0.841 | 0.501837 | 32 | rps12 |
| rpl918 | 2.147957 | 0.384617459 | 1 | 0.668 | 0.545689 | 32 | rpl9 |
| ucp211 | 2.361047 | 0.260189361 | 0.831 | 0.414 | 0.599824 | 32 | ucp2 |
| rpl3613 | 2.738481 | 0.288250672 | 0.989 | 0.806 | 0.695711 | 32 | rpl36 |
| rps79 | 2.776230 | 0.260513294 | 1 | 0.828 | 0.705301 | 32 | rps7 |
| rpl119 | 3.242533 | 0.267967551 | 0.989 | 0.83 | 0.823766 | 32 | rpl11 |
| rpl378 | 3.842087 | 0.252472466 | 1 | 0.855 | 0.976082 | 32 | rpl37 |
| h3f3d1 | 4.156444 | 0.305688877 | 0.955 | 0.676 | 1 | 32 | h3f3d |
| eef1da8 | 4.766154 | 0.357175334 | 0.876 | 0.506 | 1 | 32 | eef1da |
| rps69 | 7.807675 | 0.262829293 | 0.989 | 0.804 | 1 | 32 | rps6 |
| rps1812 | 0.0001 | 0.298490687 | 0.978 | 0.754 | 1 | 32 | rps18 |
| rps8a8 | 0.0001 | 0.26157307 | 0.989 | 0.801 | 1 | 32 | rps8a |
| faua12 | 0.0002 | 0.317398274 | 0.989 | 0.717 | 1 | 32 | faua |
| rps15a13 | 0.0002 | 0.298199503 | 1 | 0.798 | 1 | 32 | rps15a |
| rps4x14 | 0.0002 | 0.308422301 | 1 | 0.736 | 1 | 32 | rps4x |
| rpl89 | 0.0002 | 0.274894551 | 1 | 0.795 | 1 | 32 | rpl8 |
| rps2118 | 0.0002 | 0.297067878 | 0.989 | 0.776 | 1 | 32 | rps21 |
| rpl1512 | 0.0002 | 0.286376194 | 0.989 | 0.764 | 1 | 32 | rpl15 |
| eif5a8 | 0.0003 | 0.310047603 | 0.888 | 0.495 | 1 | 32 | eif5a |
| rpl3410 | 0.0003 | 0.265801789 | 0.989 | 0.802 | 1 | 32 | rpl34 |
| NC-002333.411 | 0.0003 | 0.316817171 | 0.978 | 0.608 | 1 | 32 | NC-002333.4 |
| rps268 | 0.0004 | 0.284743631 | 0.989 | 0.751 | 1 | 32 | rps26 |
| rps1615 | 0.0004 | 0.301821583 | 1 | 0.766 | 1 | 32 | rps16 |
| rpl35a11 | 0.0004 | 0.283478201 | 1 | 0.782 | 1 | 32 | rpl35a |
| atp5f1e4 | 0.0005 | 0.262564645 | 0.91 | 0.49 | 1 | 32 | atp5f1e |
| rps1515 | 0.0005 | 0.326881422 | 0.989 | 0.717 | 1 | 32 | rps15 |
| slc25a3b6 | 0.0006 | 0.2757702 | 0.831 | 0.458 | 1 | 32 | slc25a3b |
| rps1016 | 0.0006 | 0.315834479 | 0.989 | 0.727 | 1 | 32 | rps10 |
| rps310 | 0.0007 | 0.302609902 | 1 | 0.726 | 1 | 32 | rps3 |
| zgc:11418819 | 0.0008 | 0.336699916 | 0.966 | 0.668 | 1 | 32 | zgc:114188 |
| rps1311 | 0.0009 | 0.288417873 | 0.989 | 0.749 | 1 | 32 | rps13 |
| rpl2411 | 0.0009 | 0.262492897 | 1 | 0.794 | 1 | 32 | rpl24 |
| eef1b22 | 0.0009 | 0.260829107 | 0.966 | 0.742 | 1 | 32 | eef1b2 |
| rpsa14 | 0.0024 | 0.257958368 | 0.989 | 0.786 | 1 | 32 | rpsa |
| rplp2l13 | 0.0043 | 0.253912217 | 1 | 0.795 | 1 | 32 | rplp2l |

| | | | | | | | |
|---|---|---|---|---|---|---|---|
| rpl614 | 0.0044 | 0.308713264 | 1 | 0.701 | 1 | 32 | rpl6 |
| rpl3012 | 0.0047 | 0.288337609 | 0.989 | 0.73 | 1 | 32 | rpl30 |
| rps514 | 0.0048 | 0.280428452 | 0.978 | 0.729 | 1 | 32 | rps5 |
| rps27a8 | 0.0069 | 0.251371819 | 0.989 | 0.775 | 1 | 32 | rps27a |
| rpl2711 | 0.0069 | 0.256337745 | 1 | 0.78 | 1 | 32 | rpl27 |
| si:dkey-78l4.6 | 0 | 2.592495886 | 0.885 | 0.006 | 0 | 33 | si:dkey-78l4.6 |
| si:dkey-9i23.4 | 0 | 2.469931763 | 0.885 | 0.013 | 0 | 33 | si:dkey-9i23.4 |
| si:dkey-78l4.42 | 0 | 2.011301148 | 0.862 | 0.026 | 0 | 33 | si:dkey-78l4.4 |
| LOC100150882 | 0 | 1.991489029 | 0.678 | 0.008 | 0 | 33 | LOC100150882 |
| si:dkey-78l4.7 | 0 | 1.653986171 | 0.529 | 0.001 | 0 | 33 | si:dkey-78l4.7 |
| spink4 | 0 | 1.102346968 | 0.448 | 0.004 | 0 | 33 | spink4 |
| si:dkey-21e2.31 | 0 | 0.960631899 | 0.425 | 0.004 | 0 | 33 | si:dkey-21e2.3 |
| il11b1 | 0 | 0.938769867 | 0.437 | 0.007 | 0 | 33 | il11b |
| gata31 | 0 | 0.898494992 | 0.448 | 0.006 | 0 | 33 | gata3 |
| CR847936.10 | 0 | 0.656388176 | 0.31 | 0.003 | 0 | 33 | CR847936.10 |
| si:dkey-78l4.8 | 0 | 0.472504233 | 0.184 | 0 | 0 | 33 | si:dkey-78l4.8 |
| cfbl4 | 1.714599 | 1.876013564 | 0.943 | 0.066 | 4.3559390 | 33 | cfbl |
| ctss11 | 1.011195 | 0.81918108 | 0.414 | 0.014 | 2.5689416 | 33 | ctss1 |
| cpa52 | 2.403319 | 1.387799077 | 0.621 | 0.033 | 6.1056328 | 33 | cpa5 |
| AL954142.11 | 1.606069 | 0.975471421 | 0.517 | 0.022 | 4.0802207 | 33 | AL954142.1 |
| tnfrsf9a4 | 2.450088 | 1.471797666 | 0.713 | 0.045 | 6.2244508 | 33 | tnfrsf9a |
| eomesa2 | 1.280277 | 1.162308459 | 0.667 | 0.038 | 3.2525460 | 33 | eomesa |
| adgre131 | 2.187857 | 0.592205927 | 0.299 | 0.007 | 5.5582526 | 33 | adgre13 |
| CR396595.1 | 3.571206 | 0.347446211 | 0.161 | 0.002 | 9.0726492 | 33 | CR396595.1 |
| ogfrl1 | 1.372839 | 0.573201201 | 0.276 | 0.007 | 3.4876992 | 33 | ogfrl1 |
| ptprjb.1 | 3.484271 | 0.384418007 | 0.184 | 0.003 | 8.8517906 | 33 | ptprjb.1 |
| acp5b | 9.341940 | 0.671286457 | 0.368 | 0.013 | 2.3733198 | 33 | acp5b |
| cox6a25 | 4.880056 | 1.024553738 | 0.586 | 0.035 | 1.2397783 | 33 | cox6a2 |
| rab36 | 4.414883 | 0.288929883 | 0.149 | 0.002 | 1.1216012 | 33 | rab36 |
| si:dkey-78l4.21 | 1.022114 | 0.654012525 | 0.253 | 0.007 | 2.5966828 | 33 | si:dkey-78l4.2 |
| dusp55 | 8.005052 | 1.266575864 | 0.713 | 0.065 | 2.0336835 | 33 | dusp5 |
| nansb | 4.605181 | 0.276379847 | 0.115 | 0.001 | 1.1699464 | 33 | nansb |
| CABZ01073834.15 | 1.765789 | 0.967209461 | 0.575 | 0.042 | 4.4859871 | 33 | CABZ01073834.1 |
| capgb8 | 2.643942 | 1.467819486 | 0.874 | 0.112 | 6.7169356 | 33 | capgb |
| kctd7 | 3.108631 | 0.348202811 | 0.184 | 0.004 | 7.8974782 | 33 | kctd7 |

| | | | | | | | |
|---|---|---|---|---|---|---|---|
| LOC101886102 | 9.880279 | 0.360487361 | 0.172 | 0.004 | 2.5100849 | 33 | LOC101886102 |
| prodha | 2.032816 | 0.596474393 | 0.299 | 0.013 | 5.1643700 | 33 | prodha |
| laptm512 | 4.565754 | 1.536723271 | 0.931 | 0.173 | 1.1599298 | 33 | laptm5 |
| tnfrsf9b2 | 2.037722 | 0.555117963 | 0.299 | 0.014 | 5.1768348 | 33 | tnfrsf9b |
| hhex1 | 2.048127 | 0.443897637 | 0.218 | 0.008 | 5.2032677 | 33 | hhex |
| si:ch211-241f5.34 | 6.414651 | 1.061448418 | 0.575 | 0.06 | 1.6296422 | 33 | si:ch211-241f5.3 |
| rnaset2l4 | 3.673283 | 0.85252358 | 0.517 | 0.048 | 9.3319772 | 33 | rnaset2l |
| fcer1gl7 | 4.636309 | 1.487444409 | 0.943 | 0.179 | 1.1778543 | 33 | fcer1gl |
| XLOC-0183562 | 2.204061 | 0.757460755 | 0.391 | 0.028 | 5.5994185 | 33 | XLOC-018356 |
| grap2b10 | 2.846104 | 1.22217109 | 0.793 | 0.122 | 7.2305275 | 33 | grap2b |
| parvb | 1.913746 | 0.398933353 | 0.207 | 0.008 | 4.8618737 | 33 | parvb |
| yrk6 | 5.468952 | 0.973480165 | 0.586 | 0.065 | 1.3893873 | 33 | yrk |
| adgre14 | 2.724327 | 0.323317049 | 0.126 | 0.003 | 6.9211540 | 33 | adgre14 |
| si:dkey-33i11.93 | 8.138141 | 0.628810911 | 0.356 | 0.024 | 2.0674947 | 33 | si:dkey-33i11.9 |
| ptprc9 | 1.255149 | 1.201751015 | 0.77 | 0.122 | 3.1887068 | 33 | ptprc |
| dnajb9b | 3.778988 | 0.60460006 | 0.333 | 0.023 | 9.6005192 | 33 | dnajb9b |
| itgb26 | 6.446863 | 1.114633816 | 0.701 | 0.106 | 1.6378256 | 33 | itgb2 |
| vaspa6 | 8.581542 | 0.967313906 | 0.621 | 0.079 | 2.1801409 | 33 | vaspa |
| CABZ01039863.1 | 1.951931 | 0.315534463 | 0.126 | 0.003 | 4.9588824 | 33 | CABZ01039863.1 |
| scgn3 | 1.340020 | 0.562372989 | 0.322 | 0.022 | 3.4043218 | 33 | scgn |
| BX511132.1 | 2.854747 | 0.26581807 | 0.126 | 0.003 | 7.2524853 | 33 | BX511132.1 |
| hck3 | 1.151788 | 0.690218525 | 0.379 | 0.032 | 2.9261190 | 33 | hck |
| ndufa41 | 8.866561 | 0.498711213 | 0.264 | 0.016 | 2.2525499 | 33 | ndufa4 |
| mylk5 | 2.863901 | 0.503086933 | 0.276 | 0.018 | 7.2757422 | 33 | mylk5 |
| si:ch211-198c19.11 | 5.074296 | 0.380246826 | 0.207 | 0.01 | 1.2891250 | 33 | si:ch211-198c19.1 |
| nptna4 | 1.010642 | 0.880213881 | 0.54 | 0.066 | 2.5675375 | 33 | nptna |
| cremb | 1.159780 | 0.524238094 | 0.276 | 0.018 | 2.9464214 | 33 | cremb |
| ccr9a8 | 3.316184 | 0.970259747 | 0.609 | 0.086 | 8.4247661 | 33 | ccr9a |
| wasa8 | 1.778723 | 1.020651472 | 0.621 | 0.095 | 4.5188483 | 33 | wasa |
| fermt3b2 | 6.959783 | 0.546644293 | 0.287 | 0.02 | 1.7681329 | 33 | fermt3b |
| SLA (1 of many) | 2.952108 | 0.303903188 | 0.126 | 0.004 | 7.4998315 | 33 | SLA (1 of many) |
| osmr1 | 1.931859 | 0.356554205 | 0.184 | 0.008 | 4.9078903 | 33 | osmr |
| si:ch211-191c10.14 | 2.701507 | 0.792038026 | 0.437 | 0.048 | 6.8631803 | 33 | si:ch211-191c10.1 |
| fybb5 | 1.040109 | 0.69030425 | 0.414 | 0.043 | 2.6423972 | 33 | fybb |
| si:ch211-102c2.48 | 4.894480 | 1.117794767 | 0.759 | 0.144 | 1.2434428 | 33 | si:ch211-102c2.4 |

| | | | | | | | |
|---|---|---|---|---|---|---|---|
| alox5ap2 | 1.456929 | 0.562097069 | 0.299 | 0.023 | 3.7013294 | 33 | alox5ap |
| CU207241.1 | 2.270673 | 0.476946634 | 0.253 | 0.017 | 5.7686456 | 33 | CU207241.1 |
| cyba7 | 4.131820 | 0.845485063 | 0.563 | 0.08 | 1.0496891 | 33 | cyba |
| mir2234 | 1.302285 | 0.602247264 | 0.356 | 0.033 | 3.3084566 | 33 | mir223 |
| sik15 | 1.436504 | 0.992415041 | 0.644 | 0.109 | 3.6494403 | 33 | sik1 |
| CU855821.12 | 3.524689 | 0.409004804 | 0.218 | 0.013 | 8.9544738 | 33 | CU855821.1 |
| ca23 | 3.695747 | 0.757064176 | 0.448 | 0.052 | 9.3890464 | 33 | ca2 |
| bhlhe4010 | 7.992534 | 1.149968499 | 0.759 | 0.166 | 2.0305034 | 33 | bhlhe40 |
| il41 | 1.035938 | 0.488586801 | 0.161 | 0.007 | 2.6318025 | 33 | il4 |
| lcp2b1 | 2.411906 | 0.466170516 | 0.23 | 0.014 | 6.1274483 | 33 | lcp2b |
| ccr24 | 4.937494 | 0.756432579 | 0.402 | 0.044 | 1.2543704 | 33 | ccr2 |
| cyb5r2 | 4.990482 | 0.392292305 | 0.195 | 0.011 | 1.2678321 | 33 | cyb5r2 |
| zgc:6405110 | 1.493926 | 1.097441337 | 0.759 | 0.161 | 3.7953215 | 33 | zgc:64051 |
| skap27 | 6.500155 | 0.78485323 | 0.54 | 0.079 | 1.6513645 | 33 | skap2 |
| si:zfos-741a10.35 | 3.172447 | 0.552149354 | 0.322 | 0.03 | 8.0596038 | 33 | si:zfos-741a10.3 |
| dgkzb1 | 9.417349 | 0.359952243 | 0.172 | 0.009 | 2.3924776 | 33 | dgkzb |
| si:dkey-1c7.11 | 1.264225 | 0.441409993 | 0.253 | 0.019 | 3.2117653 | 33 | si:dkey-1c7.1 |
| arpc1b12 | 1.187898 | 1.183109393 | 0.954 | 0.312 | 3.0178570 | 33 | arpc1b |
| coro1a11 | 1.426009 | 1.118584221 | 0.862 | 0.235 | 3.6227764 | 33 | coro1a |
| lpcat22 | 2.762919 | 0.514755486 | 0.264 | 0.022 | 7.0191973 | 33 | lpcat2 |
| atr | 2.966264 | 0.281484258 | 0.126 | 0.005 | 7.5357953 | 33 | atr |
| srgn8 | 5.660190 | 1.055854724 | 0.874 | 0.233 | 1.4379715 | 33 | srgn |
| pfn113 | 2.660903 | 1.104593216 | 1 | 0.5 | 6.7600247 | 33 | pfn1 |
| csf3a1 | 2.774802 | 0.299155589 | 0.138 | 0.006 | 7.0493855 | 33 | csf3a |
| si:dkey-237j10.2 | 3.782943 | 0.35963119 | 0.184 | 0.011 | 9.6105689 | 33 | si:dkey-237j10.2 |
| arap3 | 8.210136 | 0.28960419 | 0.161 | 0.009 | 2.0857851 | 33 | arap3 |
| nitr5 | 1.512252 | 0.332819987 | 0.149 | 0.008 | 3.8418786 | 33 | nitr5 |
| pik3cd | 7.668602 | 0.432885369 | 0.23 | 0.018 | 1.9482085 | 33 | pik3cd |
| il165 | 2.262587 | 0.590517152 | 0.379 | 0.048 | 5.7481039 | 33 | il16 |
| ppp1r185 | 4.842938 | 0.711271736 | 0.402 | 0.055 | 1.2303485 | 33 | ppp1r18 |
| viml1 | 5.800390 | 0.347358233 | 0.161 | 0.009 | 1.4735891 | 33 | viml |
| sort1b2 | 1.954355 | 0.566056619 | 0.287 | 0.029 | 4.9650406 | 33 | sort1b |
| sox51 | 2.730493 | 0.318983471 | 0.184 | 0.012 | 6.9368199 | 33 | sox5 |
| ccl33.31 | 1.106391 | 0.960736287 | 0.425 | 0.06 | 2.8107875 | 33 | ccl33.3 |
| si:dkey-21e2.151 | 2.495975 | 0.623683987 | 0.161 | 0.01 | 6.3410264 | 33 | si:dkey-21e2.15 |

| | | | | | | | |
|---|---|---|---|---|---|---|---|
| def6c1 | 5.477134 | 0.356342445 | 0.184 | 0.012 | 1.3914660 | 33 | def6c |
| parvg7 | 1.163440 | 0.699733545 | 0.46 | 0.072 | 2.9557198 | 33 | parvg |
| fam49ba6 | 1.377525 | 0.781188567 | 0.563 | 0.104 | 3.4996042 | 33 | fam49ba |
| il15l | 7.071114 | 0.528030821 | 0.264 | 0.026 | 1.7964166 | 33 | il15l |
| spi1b7 | 4.805111 | 0.696125904 | 0.448 | 0.07 | 1.2207385 | 33 | spi1b |
| XLOC-0430435 | 8.754452 | 0.710978013 | 0.345 | 0.044 | 2.2240685 | 33 | XLOC-043043 |
| spns32 | 1.088178 | 0.383861596 | 0.195 | 0.015 | 2.7645185 | 33 | spns3 |
| tln16 | 1.962105 | 0.85378411 | 0.621 | 0.141 | 4.9847301 | 33 | tln1 |
| ccl35.15 | 4.387363 | 0.926858503 | 0.483 | 0.082 | 1.1146096 | 33 | ccl35.1 |
| ubash3ba1 | 1.233914 | 0.472015036 | 0.241 | 0.023 | 3.1347590 | 33 | ubash3ba |
| rgs139 | 1.343418 | 0.90801216 | 0.586 | 0.124 | 3.4129558 | 33 | rgs13 |
| btk2 | 1.557269 | 0.480294238 | 0.264 | 0.027 | 3.9562430 | 33 | btk |
| rhogb6 | 2.821915 | 0.807405594 | 0.529 | 0.107 | 7.1690759 | 33 | rhogb |
| ldha5 | 3.418882 | 0.664057845 | 0.425 | 0.069 | 8.6856700 | 33 | ldha |
| hexa | 4.267473 | 0.365981502 | 0.207 | 0.018 | 1.0841515 | 33 | hexa |
| tagapb6 | 4.753736 | 0.65318308 | 0.425 | 0.069 | 1.2076867 | 33 | tagapb |
| ptpn65 | 2.101008 | 0.806980129 | 0.506 | 0.098 | 5.3376123 | 33 | ptpn6 |
| slc41a12 | 8.246155 | 0.320558152 | 0.172 | 0.013 | 2.0949358 | 33 | slc41a1 |
| limd23 | 9.532398 | 0.66374365 | 0.414 | 0.067 | 2.4217058 | 33 | limd2 |
| kctd12.28 | 3.640648 | 0.753043913 | 0.506 | 0.099 | 9.2490665 | 33 | kctd12.2 |
| atp2a32 | 1.165874 | 0.457282478 | 0.287 | 0.035 | 2.9619030 | 33 | atp2a3 |
| zgc:565851 | 1.166786 | 0.288201265 | 0.161 | 0.012 | 2.9642220 | 33 | zgc:56585 |
| glipr1a8 | 4.026386 | 0.668650303 | 0.402 | 0.067 | 1.0229035 | 33 | glipr1a |
| il10ra3 | 7.892997 | 0.529355267 | 0.31 | 0.041 | 2.0052160 | 33 | il10ra |
| pla2g4aa1 | 1.796816 | 0.323120977 | 0.195 | 0.017 | 4.5648135 | 33 | pla2g4aa |
| sla24 | 2.415871 | 0.67213826 | 0.379 | 0.06 | 6.1375215 | 33 | sla2 |
| cd6314 | 3.296658 | 0.910566678 | 0.92 | 0.372 | 8.3751609 | 33 | cd63 |
| sla13 | 3.996281 | 0.382246265 | 0.23 | 0.024 | 1.0152552 | 33 | sla1 |
| nkl.31 | 8.252592 | 0.521535636 | 0.126 | 0.008 | 2.0965710 | 33 | nkl.3 |
| tnikb | 8.549339 | 0.42497064 | 0.218 | 0.022 | 2.1719598 | 33 | tnikb |
| CABZ01084575.14 | 1.510415 | 0.544175586 | 0.299 | 0.039 | 3.8372103 | 33 | CABZ01084575.1 |
| myl12.113 | 6.678835 | 0.856822274 | 0.92 | 0.425 | 1.6967580 | 33 | myl12.1 |
| rps6ka14 | 8.913796 | 0.576273777 | 0.402 | 0.069 | 2.2645499 | 33 | rps6ka1 |
| gpr132b2 | 1.171009 | 0.503117875 | 0.276 | 0.035 | 2.9749506 | 33 | gpr132b |
| si:ch211-136m16.83 | 3.023571 | 0.445716861 | 0.253 | 0.03 | 7.6813827 | 33 | si:ch211-136m16.8 |

| | | | | | | | |
|---|---|---|---|---|---|---|---|
| itm2bb10 | 3.157743 | 0.87897012 | 0.759 | 0.241 | 8.0222471 | 33 | itm2bb |
| CU499336.23 | 6.034387 | 0.439145002 | 0.264 | 0.033 | 1.5330360 | 33 | CU499336.2 |
| arhgap20b1 | 8.046516 | 0.39443473 | 0.207 | 0.021 | 2.0442176 | 33 | arhgap20b |
| cd74b7 | 8.846132 | 0.920644589 | 0.77 | 0.225 | 2.2473599 | 33 | cd74b |
| si:ch211-66e2.37 | 1.012560 | 0.664390085 | 0.368 | 0.062 | 2.5724087 | 33 | si:ch211-66e2.3 |
| aldocb4 | 1.516761 | 0.587281431 | 0.379 | 0.065 | 3.8533337 | 33 | aldocb |
| calm1b9 | 2.479670 | 0.717294215 | 0.483 | 0.102 | 6.2996040 | 33 | calm1b |
| si:dkey-172h23.2 | 3.718261 | 0.500015762 | 0.299 | 0.043 | 9.4462429 | 33 | si:dkey-172h23.2 |
| sema7a2 | 6.659187 | 0.271661911 | 0.149 | 0.011 | 1.6917664 | 33 | sema7a |
| crip113 | 7.579550 | 0.904122643 | 0.816 | 0.283 | 1.9255848 | 33 | crip1 |
| si:dkey-21o19.21 | 1.351632 | 0.324732857 | 0.172 | 0.015 | 3.4338215 | 33 | si:dkey-21o19.2 |
| b3gnt3.43 | 2.387867 | 0.510721398 | 0.276 | 0.038 | 6.0663768 | 33 | b3gnt3.4 |
| actb211 | 9.760492 | 0.515623042 | 1 | 0.867 | 2.4796532 | 33 | actb2 |
| actr2a8 | 1.182445 | 0.736958529 | 0.54 | 0.132 | 3.0040019 | 33 | actr2a |
| rac212 | 1.342511 | 0.846309378 | 0.724 | 0.224 | 3.4106500 | 33 | rac2 |
| cd74a6 | 1.677373 | 0.835038674 | 0.747 | 0.215 | 4.2613680 | 33 | cd74a |
| RASSF56 | 1.707464 | 0.534497007 | 0.356 | 0.061 | 4.3378145 | 33 | RASSF5 |
| csf2rb3 | 1.932263 | 0.488813587 | 0.276 | 0.038 | 4.9089145 | 33 | csf2rb |
| rorab | 4.858983 | 0.303445288 | 0.138 | 0.011 | 1.2344246 | 33 | rorab |
| cass4 | 5.691282 | 0.262674455 | 0.126 | 0.009 | 1.4458704 | 33 | cass4 |
| rnf1302 | 7.066948 | 0.544369954 | 0.322 | 0.053 | 1.7953581 | 33 | rnf130 |
| sult5a12 | 7.859882 | 0.291131446 | 0.138 | 0.011 | 1.9968031 | 33 | sult5a1 |
| si:ch73-264p11.14 | 1.029872 | 0.388712205 | 0.23 | 0.028 | 2.6163915 | 33 | si:ch73-264p11.1 |
| elf16 | 1.915950 | 0.686246344 | 0.46 | 0.103 | 4.8674716 | 33 | elf1 |
| mbnl19 | 3.622979 | 0.785776522 | 0.632 | 0.186 | 9.2041782 | 33 | mbnl1 |
| FP236356.13 | 6.773403 | 0.496581085 | 0.299 | 0.045 | 1.7207832 | 33 | FP236356.1 |
| edaradd2 | 8.686861 | 0.352877661 | 0.195 | 0.021 | 2.2068971 | 33 | edaradd |
| si:dkey-262k9.48 | 9.417068 | 0.627957531 | 0.425 | 0.09 | 2.3924062 | 33 | si:dkey-262k9.4 |
| si:dkey-21e2.141 | 2.112206 | 0.304021562 | 0.103 | 0.006 | 5.3660596 | 33 | si:dkey-21e2.14 |
| jak2a1 | 2.361602 | 0.335076631 | 0.218 | 0.026 | 5.9996515 | 33 | jak2a |
| crema9 | 2.694087 | 0.839008819 | 0.586 | 0.173 | 6.8443285 | 33 | crema |
| si:dkey-53k12.25 | 2.866731 | 0.470944714 | 0.287 | 0.044 | 7.2829304 | 33 | si:dkey-53k12.2 |
| si:ch211-243a20.42 | 8.151551 | 0.332210924 | 0.184 | 0.02 | 2.0709016 | 33 | si:ch211-243a20.4 |
| BX120005.14 | 1.486954 | 0.614649649 | 0.287 | 0.046 | 3.7776082 | 33 | BX120005.1 |
| syk5 | 1.776149 | 0.572760958 | 0.391 | 0.077 | 4.5123071 | 33 | syk |

| | | | | | | | |
|---|---|---|---|---|---|---|---|
| arhgdia8 | 3.312726 | 0.784442638 | 0.713 | 0.25 | 8.4159804 | 33 | arhgdia |
| csk4 | 1.738637 | 0.627463703 | 0.448 | 0.103 | 4.4170081 | 33 | csk |
| arpc310 | 2.496436 | 0.807111782 | 0.782 | 0.333 | 6.3421963 | 33 | arpc3 |
| csrnp1a1 | 8.013746 | 0.512151851 | 0.287 | 0.048 | 2.0358921 | 33 | csrnp1a |
| LOC1018842675 | 1.746220 | 0.490251874 | 0.299 | 0.051 | 4.4362724 | 33 | LOC101884267 |
| selplg6 | 1.997631 | 0.45961024 | 0.276 | 0.045 | 5.0749819 | 33 | selplg |
| cfl110 | 2.855185 | 0.665158554 | 0.931 | 0.6 | 7.2535990 | 33 | cfl1 |
| wasb12 | 3.355514 | 0.715454897 | 0.598 | 0.172 | 8.5246837 | 33 | wasb |
| gnsb2 | 3.777896 | 0.452254086 | 0.253 | 0.039 | 9.5977461 | 33 | gnsb |
| mrps155 | 3.965917 | 0.657119142 | 0.46 | 0.113 | 1.0075412 | 33 | mrps15 |
| si:dkey-222f2.73 | 4.206405 | 0.366424223 | 0.23 | 0.032 | 1.0686373 | 33 | si:dkey-222f2.7 |
| arpc5a8 | 5.693193 | 0.633616551 | 0.483 | 0.125 | 1.4463558 | 33 | arpc5a |
| il345 | 6.429732 | 0.494121902 | 0.276 | 0.045 | 1.6334735 | 33 | il34 |
| si:ch211-215e19.32 | 6.884799 | 0.316575575 | 0.172 | 0.019 | 1.7490832 | 33 | si:ch211-215e19.3 |
| rasal33 | 9.532434 | 0.357863916 | 0.207 | 0.027 | 2.4217148 | 33 | rasal3 |
| far12 | 1.672739 | 0.416447005 | 0.241 | 0.036 | 4.2495936 | 33 | far1 |
| grasp6 | 2.699035 | 0.51027582 | 0.356 | 0.072 | 6.8568995 | 33 | grasp |
| zgc:1538676 | 2.815685 | 0.707968629 | 0.908 | 0.464 | 7.1532493 | 33 | zgc:153867 |
| rasgrp41 | 3.524682 | 0.371669951 | 0.207 | 0.028 | 8.9544559 | 33 | rasgrp4 |
| zmp:00000010824 | 7.270232 | 0.51275139 | 0.299 | 0.055 | 1.8470024 | 33 | zmp:0000001082 |
| si:ch73-208g10.15 | 1.289110 | 0.407807828 | 0.253 | 0.041 | 3.2749861 | 33 | si:ch73-208g10.1 |
| cotl111 | 1.813724 | 0.801843571 | 0.793 | 0.319 | 4.6077670 | 33 | cotl1 |
| gapdhs13 | 3.538454 | 0.735099218 | 0.77 | 0.314 | 8.9894443 | 33 | gapdhs |
| atp5if1b6 | 4.272781 | 0.715090325 | 0.609 | 0.206 | 1.0855000 | 33 | atp5if1b |
| hif1ab3 | 4.495475 | 0.606071639 | 0.46 | 0.122 | 1.1420755 | 33 | hif1ab |
| wipf1a6 | 4.932458 | 0.508542812 | 0.333 | 0.067 | 1.2530911 | 33 | wipf1a |
| hcls15 | 6.093020 | 0.508185708 | 0.356 | 0.076 | 1.5479318 | 33 | hcls1 |
| capzb10 | 6.341213 | 0.711024814 | 0.713 | 0.271 | 1.6109852 | 33 | capzb |
| LOC1039091158 | 7.676469 | 0.671619589 | 0.552 | 0.168 | 1.9502069 | 33 | LOC103909115 |
| si:ch211-222l21.113 | 9.665783 | 0.771680822 | 0.77 | 0.313 | 2.4555924 | 33 | si:ch211-222l21.1 |
| faslg2 | 1.244220 | 0.302443801 | 0.149 | 0.016 | 3.1609419 | 33 | faslg |
| dicp3.1 | 1.377713 | 0.251615442 | 0.138 | 0.014 | 3.5000820 | 33 | dicp3.1 |
| arrb2b7 | 1.492657 | 0.527106124 | 0.402 | 0.096 | 3.7920970 | 33 | arrb2b |
| si:ch211-132b12.71 | 1.541509 | 0.423671946 | 0.287 | 0.053 | 3.9162039 | 33 | si:ch211-132b12.7 |
| arhgap4a | 1.687087 | 0.263120709 | 0.149 | 0.016 | 4.2860456 | 33 | arhgap4a |

| | | | | | | | |
|---|---|---|---|---|---|---|---|
| btg113 | 2.195208 | 0.74486618 | 0.793 | 0.355 | 5.5769268 | 33 | btg1 |
| actb18 | 4.836885 | 0.481725588 | 0.954 | 0.768 | 1.2288106 | 33 | actb1 |
| myd88 | 5.087661 | 0.473773634 | 0.299 | 0.059 | 1.2925203 | 33 | myd88 |
| sh3bgrl6 | 6.701599 | 0.57504214 | 0.448 | 0.121 | 1.7025412 | 33 | sh3bgrl |
| sh2d3ca1 | 1.310499 | 0.271291869 | 0.172 | 0.022 | 3.3293246 | 33 | sh2d3ca |
| rab27a2 | 1.448218 | 0.415221592 | 0.264 | 0.048 | 3.6791996 | 33 | rab27a |
| rgl2 | 5.690210 | 0.269778082 | 0.172 | 0.023 | 1.4455979 | 33 | rgl2 |
| rgs3a | 7.063097 | 0.323457383 | 0.184 | 0.026 | 1.7943799 | 33 | rgs3a |
| vsir7 | 1.078809 | 0.531965916 | 0.368 | 0.088 | 2.7407146 | 33 | vsir |
| XLOC-0140795 | 1.279619 | 0.405013602 | 0.207 | 0.032 | 3.2508728 | 33 | XLOC-014079 |
| rplp215 | 2.349389 | 0.434329192 | 1 | 0.783 | 5.9686235 | 33 | rplp2 |
| cox6b19 | 3.515539 | 0.574890994 | 0.437 | 0.122 | 8.9312284 | 33 | cox6b1 |
| pak1 | 3.854931 | 0.258209663 | 0.149 | 0.018 | 9.7934533 | 33 | pak1 |
| arpc212 | 3.888639 | 0.684279494 | 0.713 | 0.299 | 9.8790886 | 33 | arpc2 |
| ptpn222 | 4.132995 | 0.330619797 | 0.172 | 0.024 | 1.0499875 | 33 | ptpn22 |
| tmsb4x18 | 4.143079 | 0.574980964 | 1 | 0.718 | 1.0525493 | 33 | tmsb4x |
| pik3ap1 | 5.837998 | 0.278139362 | 0.149 | 0.018 | 1.4831435 | 33 | pik3ap1 |
| sept7b1 | 6.097877 | 0.507445572 | 0.414 | 0.11 | 1.5491657 | 33 | sept7b |
| cers11 | 1.163977 | 0.313381221 | 0.161 | 0.021 | 2.9570843 | 33 | cers1 |
| fam210b | 1.227001 | 0.466613068 | 0.276 | 0.056 | 3.1171972 | 33 | fam210b |
| lcp17 | 1.990595 | 0.542628695 | 0.345 | 0.084 | 5.0571066 | 33 | lcp1 |
| pdxdc1 | 3.062512 | 0.275494842 | 0.149 | 0.019 | 7.7803122 | 33 | pdxdc1 |
| raraa3 | 3.340320 | 0.357128577 | 0.218 | 0.038 | 8.4860854 | 33 | raraa |
| anks1b2 | 3.412291 | 0.267733337 | 0.149 | 0.019 | 8.6689263 | 33 | anks1b |
| lrrfip1a4 | 3.746374 | 0.414793974 | 0.287 | 0.061 | 9.5176634 | 33 | lrrfip1a |
| rab447 | 4.510304 | 0.460763518 | 0.287 | 0.061 | 1.1458428 | 33 | rab44 |
| dtnbp1a3 | 6.786465 | 0.479545383 | 0.356 | 0.088 | 1.7241015 | 33 | dtnbp1a |
| fscn1a3 | 9.545356 | 0.419147583 | 0.264 | 0.053 | 2.4249977 | 33 | fscn1a |
| lasp19 | 1.146340 | 0.637638698 | 0.644 | 0.253 | 2.9122792 | 33 | lasp1 |
| ostf13 | 1.272620 | 0.382907132 | 0.264 | 0.054 | 3.2330935 | 33 | ostf1 |
| prkcbb2 | 1.320166 | 0.296505952 | 0.195 | 0.031 | 3.3538825 | 33 | prkcbb |
| arl113 | 1.380027 | 0.30377672 | 0.172 | 0.025 | 3.5059604 | 33 | arl11 |
| plxnc14 | 1.723953 | 0.401237477 | 0.241 | 0.046 | 4.3797031 | 33 | plxnc1 |
| mtpn6 | 2.438982 | 0.615172817 | 0.483 | 0.163 | 6.1962356 | 33 | mtpn |
| ucp212 | 3.750969 | 0.658114563 | 0.782 | 0.414 | 9.5293390 | 33 | ucp2 |

| | | | | | | | |
|---|---|---|---|---|---|---|---|
| tpi1b7 | 6.773412 | 0.584450561 | 0.517 | 0.178 | 1.7207855 | 33 | tpi1b |
| dbnlb | 7.056995 | 0.352350988 | 0.23 | 0.044 | 1.7928298 | 33 | dbnlb |
| gsk3b | 1.001439 | 0.287792805 | 0.184 | 0.03 | 2.5441566 | 33 | gsk3b |
| XLOC-0397573 | 1.302163 | 0.333167082 | 0.172 | 0.026 | 3.3081472 | 33 | XLOC-039757 |
| arpc4l8 | 2.856974 | 0.602713665 | 0.575 | 0.219 | 7.2581425 | 33 | arpc4l |
| pygb2 | 3.454807 | 0.384284266 | 0.23 | 0.045 | 8.7769388 | 33 | pygb |
| tmc84 | 4.757198 | 0.313901768 | 0.195 | 0.034 | 1.2085662 | 33 | tmc8 |
| mef2d2 | 5.941223 | 0.409459055 | 0.31 | 0.076 | 1.5093678 | 33 | mef2d |
| si:rp71-36a1.3 | 6.154956 | 0.262625998 | 0.138 | 0.019 | 1.5636665 | 33 | si:rp71-36a1.3 |
| fli1b3 | 9.603523 | 0.312632157 | 0.195 | 0.035 | 2.4397750 | 33 | fli1b |
| sat1a.28 | 1.035012 | 0.63192488 | 0.678 | 0.301 | 2.6294491 | 33 | sat1a.2 |
| nfkbie3 | 1.399391 | 0.449252594 | 0.356 | 0.097 | 3.5551531 | 33 | nfkbie |
| arpc5b7 | 2.426838 | 0.511074016 | 0.425 | 0.131 | 6.1653831 | 33 | arpc5b |
| LOC1104382372 | 3.207914 | 0.345218728 | 0.253 | 0.055 | 8.1497058 | 33 | LOC110438237 |
| rpl3411 | 3.979885 | 0.35780323 | 0.977 | 0.802 | 1.0110899 | 33 | rpl34 |
| hic13 | 4.378774 | 0.437029913 | 0.287 | 0.07 | 1.1124277 | 33 | hic1 |
| elmo15 | 5.671494 | 0.41388588 | 0.287 | 0.07 | 1.4408430 | 33 | elmo1 |
| sqstm1 | 7.147078 | 0.40772392 | 0.31 | 0.079 | 1.8157152 | 33 | sqstm1 |
| arhgdig11 | 8.943720 | 0.603338895 | 0.632 | 0.259 | 2.2721522 | 33 | arhgdig |
| oaz1b12 | 9.422272 | 0.576481393 | 0.759 | 0.36 | 2.3937284 | 33 | oaz1b |
| BX649485.11 | 1.356159 | 0.285308626 | 0.161 | 0.026 | 3.4453228 | 33 | BX649485.1 |
| CR855389.11 | 1.613084 | 0.319598842 | 0.172 | 0.03 | 4.0980414 | 33 | CR855389.1 |
| si:ch1073-443f11.23 | 2.225589 | 0.308129105 | 0.207 | 0.041 | 5.6541113 | 33 | si:ch1073-443f11.2 |
| clic17 | 3.468937 | 0.486600325 | 0.402 | 0.129 | 8.8128359 | 33 | clic1 |
| myh11a5 | 3.995889 | 0.383732773 | 0.287 | 0.071 | 1.0151557 | 33 | myh11a |
| unm-hu7910 | 4.645674 | 0.29035051 | 0.184 | 0.034 | 1.1802335 | 33 | unm-hu7910 |
| mir21-14 | 4.691947 | 0.501467682 | 0.356 | 0.108 | 1.1919892 | 33 | mir21-1 |
| srgap31 | 5.012632 | 0.2500077 | 0.161 | 0.027 | 1.2734593 | 33 | srgap3 |
| nme2b.14 | 5.018882 | 0.28498518 | 0.989 | 0.857 | 1.2750471 | 33 | nme2b.1 |
| FP101882.11 | 5.610749 | 0.292291784 | 0.195 | 0.038 | 1.4254109 | 33 | FP101882.1 |
| cebpb13 | 5.695579 | 0.64580868 | 0.736 | 0.339 | 1.4469620 | 33 | cebpb |
| tes2 | 6.786960 | 0.397069352 | 0.264 | 0.065 | 1.7242272 | 33 | tes |
| mcl1a8 | 9.758901 | 0.503882885 | 0.908 | 0.613 | 2.4792488 | 33 | mcl1a |
| zfp36l1a8 | 1.208324 | 0.594594429 | 0.598 | 0.255 | 3.0697486 | 33 | zfp36l1a |
| batf31 | 1.334027 | 0.317805586 | 0.195 | 0.038 | 3.3890958 | 33 | batf3 |

| | | | | | | | |
|---|---|---|---|---|---|---|---|
| si:rp71-36a1.1 | 1.781812 | 0.310794124 | 0.184 | 0.036 | 4.5266956 | 33 | si:rp71-36a1.1 |
| cap110 | 1.806723 | 0.53666886 | 0.609 | 0.246 | 4.5899805 | 33 | cap1 |
| ches1 | 1.859568 | 0.258003253 | 0.161 | 0.028 | 4.7242331 | 33 | ches1 |
| tagapa8 | 2.045269 | 0.521291147 | 0.333 | 0.098 | 5.1960083 | 33 | tagapa |
| ets24 | 2.871125 | 0.423494836 | 0.31 | 0.087 | 7.2940931 | 33 | ets2 |
| txn4 | 3.441482 | 0.379855625 | 0.23 | 0.052 | 8.7430868 | 33 | txn |
| si:zfos-2330d3.12 | 3.559750 | 0.40439705 | 0.161 | 0.029 | 9.0435457 | 33 | si:zfos-2330d3.1 |
| LOC1005362134 | 4.006888 | 0.459238165 | 0.368 | 0.112 | 1.0179500 | 33 | LOC100536213 |
| gstk1 | 4.210978 | 0.358659816 | 0.241 | 0.058 | 1.0697991 | 33 | gstk1 |
| sec61g7 | 4.730297 | 0.44586968 | 0.931 | 0.681 | 1.2017321 | 33 | sec61g |
| rpl269 | 4.969345 | 0.299915964 | 0.977 | 0.851 | 1.2624622 | 33 | rpl26 |
| plek1 | 5.276050 | 0.293874252 | 0.161 | 0.029 | 1.3403805 | 33 | plek |
| gng56 | 5.549745 | 0.564377792 | 0.621 | 0.287 | 1.4099129 | 33 | gng5 |
| LOC1001511962 | 5.684015 | 0.305009264 | 0.195 | 0.04 | 1.4440242 | 33 | LOC100151196 |
| ccdc88b2 | 5.796656 | 0.28112391 | 0.172 | 0.033 | 1.4726406 | 33 | ccdc88b |
| mapre1a7 | 6.979209 | 0.501886 | 0.529 | 0.196 | 1.7730682 | 33 | mapre1a |
| si:busm1-266f07.26 | 7.042071 | 0.423081433 | 0.46 | 0.148 | 1.7890382 | 33 | si:busm1-266f07.2 |
| rps2911 | 7.954189 | 0.308806669 | 1 | 0.844 | 2.0207618 | 33 | rps29 |
| rarga4 | 9.608057 | 0.42162023 | 0.241 | 0.061 | 2.4409271 | 33 | rarga |
| CR753876.15 | 1.022355 | 0.449700052 | 0.368 | 0.111 | 2.5972934 | 33 | CR753876.1 |
| CR855337.12 | 1.128524 | 0.328358418 | 0.172 | 0.034 | 2.8670158 | 33 | CR855337.1 |
| rhoab9 | 1.525125 | 0.564174606 | 0.494 | 0.207 | 3.8745822 | 33 | rhoab |
| si:ch211-226m16.26 | 1.943463 | 0.369719717 | 0.264 | 0.069 | 4.9373696 | 33 | si:ch211-226m16.2 |
| capza1a8 | 2.164890 | 0.526958813 | 0.529 | 0.221 | 5.4999044 | 33 | capza1a |
| pnrc217 | 2.290488 | 0.488185347 | 0.897 | 0.598 | 5.8189862 | 33 | pnrc2 |
| cmtm72 | 2.876323 | 0.316684132 | 0.218 | 0.051 | 7.3072988 | 33 | cmtm7 |
| BX901920.15 | 4.712315 | 0.359936118 | 0.276 | 0.074 | 1.1971638 | 33 | BX901920.1 |
| anxa48 | 5.533462 | 0.458221309 | 0.414 | 0.151 | 1.4057762 | 33 | anxa4 |
| XLOC-0268365 | 6.021689 | 0.397324319 | 0.287 | 0.079 | 1.5298103 | 33 | XLOC-026836 |
| ccni10 | 6.802424 | 0.520310459 | 0.667 | 0.319 | 1.7281559 | 33 | ccni |
| map4k5 | 8.310084 | 0.263857367 | 0.195 | 0.043 | 2.1111769 | 33 | map4k5 |
| larp4ab1 | 8.792207 | 0.338547769 | 0.276 | 0.077 | 2.2336603 | 33 | larp4ab |
| zgc:1629444 | 9.374029 | 0.524949612 | 0.598 | 0.275 | 2.3814722 | 33 | zgc:162944 |
| pgam1a6 | 9.624689 | 0.469547934 | 0.425 | 0.155 | 2.4451524 | 33 | pgam1a |
| twf1a4 | 9.733196 | 0.43114195 | 0.437 | 0.158 | 2.4727185 | 33 | twf1a |

| | | | | | | | |
|---|---|---|---|---|---|---|---|
| mob1a5 | 1.346181 | 0.439797839 | 0.46 | 0.173 | 3.4199734 | 33 | mob1a |
| tescb | 1.394408 | 0.253183656 | 0.149 | 0.028 | 3.5424952 | 33 | tescb |
| mhc2dab5 | 1.627131 | 0.35923222 | 0.322 | 0.095 | 4.1337283 | 33 | mhc2dab |
| gng23 | 2.108253 | 0.309814456 | 0.207 | 0.05 | 5.3560177 | 33 | gng2 |
| BX322787.12 | 2.129953 | 0.465888493 | 0.368 | 0.122 | 5.4111477 | 33 | BX322787.1 |
| mt-co114 | 2.131259 | 0.471723516 | 0.908 | 0.64 | 5.4144642 | 33 | mt-co1 |
| pde4ba10 | 2.287407 | 0.486324364 | 0.391 | 0.142 | 5.8111582 | 33 | pde4ba |
| RASA21 | 3.295441 | 0.295854062 | 0.195 | 0.045 | 8.3720696 | 33 | RASA2 |
| CR392341.17 | 4.117298 | 0.350259301 | 0.23 | 0.06 | 1.0459996 | 33 | CR392341.1 |
| rps15a14 | 4.883001 | 0.358419631 | 0.989 | 0.798 | 1.2405264 | 33 | rps15a |
| si:dkeyp-68b7.127 | 5.204087 | 0.350931727 | 0.23 | 0.061 | 1.3220984 | 33 | si:dkeyp-68b7.12 |
| rpl1513 | 7.566349 | 0.362886901 | 0.977 | 0.764 | 1.9222311 | 33 | rpl15 |
| iqgap14 | 1.059140 | 0.328983286 | 0.253 | 0.071 | 2.6907463 | 33 | iqgap1 |
| cxcr4b9 | 1.093268 | 0.508262928 | 0.46 | 0.177 | 2.7774486 | 33 | cxcr4b |
| ccdc28a1 | 1.255233 | 0.385216686 | 0.322 | 0.105 | 3.1889207 | 33 | ccdc28a |
| CU467110.11 | 1.473652 | 0.297989008 | 0.276 | 0.079 | 3.7438138 | 33 | CU467110.1 |
| BX005340.1 | 1.734198 | 0.272359895 | 0.184 | 0.043 | 4.4057310 | 33 | BX005340.1 |
| snap23.1 | 1.772844 | 0.355902827 | 0.253 | 0.073 | 4.5039112 | 33 | snap23.1 |
| wdr18 | 1.962403 | 0.464339274 | 0.402 | 0.156 | 4.9854861 | 33 | wdr1 |
| myo1f6 | 2.097967 | 0.316749232 | 0.195 | 0.048 | 5.3298856 | 33 | myo1f |
| arhgef3l3 | 2.605089 | 0.284121185 | 0.149 | 0.031 | 6.6182296 | 33 | arhgef3l |
| sh2b33 | 2.782204 | 0.297994814 | 0.23 | 0.062 | 7.0681917 | 33 | sh2b3 |
| cdkn1bb3 | 3.376153 | 0.460136699 | 0.391 | 0.154 | 8.5771176 | 33 | cdkn1bb |
| stxbp21 | 4.554868 | 0.321911589 | 0.218 | 0.059 | 1.1571644 | 33 | stxbp2 |
| sec11a2 | 4.883287 | 0.42774584 | 0.46 | 0.19 | 1.2405991 | 33 | sec11a |
| si:ch211-105j21.93 | 4.919011 | 0.253834957 | 0.161 | 0.035 | 1.2496748 | 33 | si:ch211-105j21.9 |
| arl4aa1 | 5.952297 | 0.326230175 | 0.207 | 0.054 | 1.5121812 | 33 | arl4aa |
| ptmab13 | 6.367575 | 0.456859998 | 0.874 | 0.57 | 1.6176825 | 33 | ptmab |
| rpl3213 | 6.412840 | 0.317246684 | 0.989 | 0.824 | 1.6291820 | 33 | rpl32 |
| cox8b6 | 9.644529 | 0.389540537 | 0.299 | 0.102 | 2.4501927 | 33 | cox8b |
| dusp19 | 9.761238 | 0.429569905 | 0.391 | 0.15 | 2.4798425 | 33 | dusp1 |
| lamp1a2 | 1.186412 | 0.369836697 | 0.287 | 0.094 | 3.0140822 | 33 | lamp1a |
| rps26l12 | 1.242645 | 0.389717752 | 0.954 | 0.7 | 3.1569421 | 33 | rps26l |
| tsc22d310 | 1.298752 | 0.45677993 | 0.391 | 0.159 | 3.2994798 | 33 | tsc22d3 |
| slc25a33 | 1.435081 | 0.328254988 | 0.287 | 0.092 | 3.6458244 | 33 | slc25a33 |

| | | | | | | | |
|---|---|---|---|---|---|---|---|
| bcl2l104 | 1.597178 | 0.486237734 | 0.46 | 0.2 | 4.0576332 | 33 | bcl2l10 |
| plekho25 | 1.703049 | 0.336815648 | 0.253 | 0.077 | 4.3265960 | 33 | plekho2 |
| rpl18a13 | 1.861991 | 0.321096344 | 0.977 | 0.801 | 4.7303904 | 33 | rpl18a |
| si:dkey-118j18.21 | 1.951417 | 0.306182202 | 0.207 | 0.057 | 4.9575771 | 33 | si:dkey-118j18.2 |
| sh3kbp14 | 2.109340 | 0.296567198 | 0.253 | 0.077 | 5.3587791 | 33 | sh3kbp1 |
| FO203432.11 | 2.192526 | 0.355890894 | 0.172 | 0.041 | 5.5701129 | 33 | FO203432.1 |
| rps2811 | 2.218873 | 0.281953408 | 0.977 | 0.841 | 5.6370486 | 33 | rps28 |
| h1fx10 | 2.554323 | 0.448831277 | 0.46 | 0.191 | 6.4892583 | 33 | h1fx |
| nfe2l2a2 | 2.809667 | 0.328342598 | 0.23 | 0.069 | 7.1379595 | 33 | nfe2l2a |
| ncaldb4 | 2.976287 | 0.298885394 | 0.241 | 0.071 | 7.5612590 | 33 | ncaldb |
| stk17b6 | 3.506351 | 0.371815007 | 0.322 | 0.114 | 8.9078856 | 33 | stk17b |
| si:dkeyp-19e1.33 | 4.051577 | 0.314625244 | 0.218 | 0.063 | 0.000103 | 33 | si:dkeyp-19e1.3 |
| tec2 | 4.951049 | 0.269044912 | 0.149 | 0.035 | 0.000126 | 33 | tec |
| rpl399 | 5.164961 | 0.253032401 | 0.989 | 0.877 | 0.000131 | 33 | rpl39 |
| rpl2215 | 5.474265 | 0.368306348 | 0.954 | 0.721 | 0.000139 | 33 | rpl22 |
| gabarapb12 | 6.211796 | 0.464163756 | 0.586 | 0.305 | 0.000158 | 33 | gabarapb |
| rps2119 | 6.627960 | 0.346853163 | 0.977 | 0.776 | 0.000168 | 33 | rps21 |
| dub7 | 7.171996 | 0.286454341 | 0.241 | 0.074 | 0.000182 | 33 | dub |
| ncalda | 7.259101 | 0.273977151 | 0.195 | 0.054 | 0.000184 | 33 | ncalda |
| rps27a9 | 8.036399 | 0.312811074 | 0.966 | 0.775 | 0.000204 | 33 | rps27a |
| rpl1412 | 8.903573 | 0.374196972 | 0.931 | 0.67 | 0.000226 | 33 | rpl14 |
| rps1516 | 1.012976 | 0.373342584 | 0.977 | 0.717 | 0.000257 | 33 | rps15 |
| rel3 | 1.076027 | 0.260044055 | 0.149 | 0.035 | 0.000273 | 33 | rel |
| msna11 | 1.236446 | 0.461541572 | 0.529 | 0.266 | 0.000314 | 33 | msna |
| MYADM1 | 1.320774 | 0.306729015 | 0.161 | 0.041 | 0.000336 | 33 | MYADM |
| dok24 | 1.398743 | 0.274942373 | 0.161 | 0.04 | 0.000355 | 33 | dok2 |
| efhd24 | 1.436079 | 0.303063517 | 0.207 | 0.06 | 0.000365 | 33 | efhd2 |
| COX315 | 1.491110 | 0.423066589 | 0.931 | 0.676 | 0.000379 | 33 | COX3 |
| rpl5a15 | 1.973433 | 0.427930025 | 0.851 | 0.537 | 0.000501 | 33 | rpl5a |
| serf210 | 2.053031 | 0.436658894 | 0.678 | 0.378 | 0.000522 | 33 | serf2 |
| rps227 | 2.216483 | 0.28657761 | 0.977 | 0.842 | 0.000563 | 33 | rps2 |
| zgc:1537133 | 2.704175 | 0.370498233 | 0.402 | 0.173 | 0.000687 | 33 | zgc:153713 |
| cox8a13 | 3.015796 | 0.427978237 | 0.529 | 0.268 | 0.000766 | 33 | cox8a |
| cyb5r3 | 3.270277 | 0.264272789 | 0.172 | 0.047 | 0.000831 | 33 | cyb5r3 |
| cdc42l7 | 4.163124 | 0.414143908 | 0.483 | 0.234 | 0.001058 | 33 | cdc42l |

| | | | | | | | |
|---|---|---|---|---|---|---|---|
| eno1a6 | 4.369707 | 0.337758608 | 0.356 | 0.142 | 0.00111 | 33 | eno1a |
| prelid3b3 | 5.080369 | 0.397088122 | 0.483 | 0.227 | 0.001291 | 33 | prelid3b |
| arpc47 | 5.684419 | 0.394017774 | 0.506 | 0.238 | 0.001444 | 33 | arpc4 |
| rps515 | 6.433567 | 0.322479295 | 0.931 | 0.729 | 0.001634 | 33 | rps5 |
| ywhabl6 | 6.573604 | 0.453975642 | 0.563 | 0.301 | 0.00167 | 33 | ywhabl |
| rpl2810 | 6.982780 | 0.314730823 | 0.977 | 0.777 | 0.001774 | 33 | rpl28 |
| fam107b4 | 8.209949 | 0.353132923 | 0.299 | 0.113 | 0.002086 | 33 | fam107b |
| sept65 | 8.304901 | 0.408924883 | 0.437 | 0.202 | 0.00211 | 33 | sept6 |
| stk17al1 | 9.601263 | 0.31421236 | 0.299 | 0.109 | 0.002439 | 33 | stk17al |
| slc6a6b6 | 1.008432 | 0.285226982 | 0.276 | 0.097 | 0.002562 | 33 | slc6a6b |
| hmha1b6 | 1.019139 | 0.256189796 | 0.23 | 0.074 | 0.002589 | 33 | hmha1b |
| plp22 | 1.105118 | 0.266076238 | 0.184 | 0.054 | 0.002808 | 33 | plp2 |
| eif3f13 | 1.248462 | 0.405180975 | 0.724 | 0.466 | 0.003172 | 33 | eif3f |
| higd1a7 | 1.270884 | 0.412644059 | 0.586 | 0.304 | 0.003229 | 33 | higd1a |
| dnajb6a | 1.397007 | 0.256625159 | 0.184 | 0.054 | 0.003549 | 33 | dnajb6a |
| rpl379 | 1.426611 | 0.269032719 | 0.989 | 0.855 | 0.003624 | 33 | rpl37 |
| rab11fip1b4 | 1.434458 | 0.306594089 | 0.195 | 0.061 | 0.003644 | 33 | rab11fip1b |
| pnp5a5 | 1.444246 | 0.253776726 | 0.23 | 0.075 | 0.003669 | 33 | pnp5a |
| rpl3812 | 1.726333 | 0.343286565 | 1 | 0.754 | 0.004386 | 33 | rpl38 |
| prkar2aa3 | 1.924251 | 0.31188178 | 0.287 | 0.109 | 0.004889 | 33 | prkar2aa |
| rpl2912 | 2.043385 | 0.287970047 | 1 | 0.823 | 0.005191 | 33 | rpl29 |
| pef1 | 2.109379 | 0.290103198 | 0.218 | 0.073 | 0.005359 | 33 | pef1 |
| rps1813 | 2.501846 | 0.308558218 | 0.931 | 0.754 | 0.006356 | 33 | rps18 |
| prkar1ab | 3.092179 | 0.2879394 | 0.218 | 0.074 | 0.007856 | 33 | prkar1ab |
| si:dkey-112e7.29 | 3.686586 | 0.358563367 | 0.448 | 0.215 | 0.009366 | 33 | si:dkey-112e7.2 |
| si:ch211-196l7.412 | 4.804252 | 0.389566581 | 0.609 | 0.33 | 0.012205 | 33 | si:ch211-196l7.4 |
| sptssa | 5.481968 | 0.264456152 | 0.241 | 0.087 | 0.013927 | 33 | sptssa |
| nr4a33 | 5.643892 | 0.282365565 | 0.161 | 0.047 | 0.014338 | 33 | nr4a3 |
| si:ch73-234b20.57 | 5.920918 | 0.301466648 | 0.276 | 0.106 | 0.015042 | 33 | si:ch73-234b20.5 |
| eif3s105 | 6.105462 | 0.325742882 | 0.391 | 0.179 | 0.015511 | 33 | eif3s10 |
| rpl35a12 | 6.336418 | 0.300571282 | 0.977 | 0.782 | 0.016098 | 33 | rpl35a |
| xbp17 | 7.971044 | 0.417908185 | 0.609 | 0.356 | 0.02025 | 33 | xbp1 |
| anxa11a2 | 8.763137 | 0.309530942 | 0.287 | 0.118 | 0.022263 | 33 | anxa11a |
| pdk2b5 | 8.773815 | 0.351058731 | 0.264 | 0.106 | 0.02229 | 33 | pdk2b |
| rnf443 | 8.983597 | 0.309879886 | 0.253 | 0.097 | 0.022823 | 33 | rnf44 |

| | | | | | | | |
|---|---|---|---|---|---|---|---|
| rpl919 | 9.262111 | 0.362585795 | 0.943 | 0.669 | 0.02353 | 33 | rpl9 |
| uqcrq5 | 9.569134 | 0.386018989 | 0.529 | 0.287 | 0.02431 | 33 | uqcrq |
| rps27.117 | 9.603271 | 0.340878834 | 0.943 | 0.71 | 0.024397 | 33 | rps27.1 |
| bcl2l11 | 9.751603 | 0.27961105 | 0.218 | 0.077 | 0.024774 | 33 | bcl2l1 |
| lypla2 | 1.036050 | 0.252095506 | 0.184 | 0.059 | 0.026321 | 33 | lypla2 |
| rpl3013 | 1.190242 | 0.305790747 | 0.931 | 0.731 | 0.030238 | 33 | rpl30 |
| vamp81 | 1.197640 | 0.266215751 | 0.218 | 0.076 | 0.030426 | 33 | vamp8 |
| metrnla6 | 1.279544 | 0.285713224 | 0.253 | 0.095 | 0.032507 | 33 | metrnla |
| tnfaip36 | 1.356351 | 0.268109288 | 0.184 | 0.06 | 0.034458 | 33 | tnfaip3 |
| nrros3 | 1.361611 | 0.26238948 | 0.161 | 0.049 | 0.034592 | 33 | nrros |
| eef1da9 | 1.434913 | 0.377764445 | 0.747 | 0.507 | 0.036454 | 33 | eef1da |
| ndufb93 | 1.716832 | 0.290974915 | 0.425 | 0.201 | 0.043616 | 33 | ndufb9 |
| gnai2a3 | 1.977162 | 0.324519397 | 0.333 | 0.149 | 0.05023 | 33 | gnai2a |
| rpl3614 | 2.151824 | 0.294466215 | 1 | 0.806 | 0.054667 | 33 | rpl36 |
| zgc:11418820 | 2.168011 | 0.354583607 | 0.92 | 0.668 | 0.055078 | 33 | zgc:114188 |
| tmx1 | 3.053390 | 0.255120804 | 0.241 | 0.092 | 0.077571 | 33 | tmx1 |
| ap2m1a3 | 3.066665 | 0.305605035 | 0.368 | 0.171 | 0.077909 | 33 | ap2m1a |
| ptp4a15 | 3.289850 | 0.274801799 | 0.31 | 0.132 | 0.083579 | 33 | ptp4a1 |
| rpl1110 | 3.361995 | 0.265807453 | 0.977 | 0.83 | 0.085411 | 33 | rpl11 |
| si:dkey-206d17.125 | 3.424334 | 0.257817495 | 0.161 | 0.051 | 0.086995 | 33 | si:dkey-206d17.12 |
| ndufa61 | 3.609552 | 0.338051991 | 0.494 | 0.26 | 0.091701 | 33 | ndufa6 |
| rps178 | 3.786732 | 0.260873755 | 0.954 | 0.779 | 0.096202 | 33 | rps17 |
| tnip14 | 3.842807 | 0.260254511 | 0.218 | 0.08 | 0.097627 | 33 | tnip1 |
| cct52 | 3.994556 | 0.35162687 | 0.506 | 0.275 | 0.101482 | 33 | cct5 |
| reep58 | 4.048629 | 0.312562669 | 0.368 | 0.174 | 0.102855 | 33 | reep5 |
| uba5212 | 4.418565 | 0.319491121 | 0.885 | 0.661 | 0.112254 | 33 | uba52 |
| zgc:564938 | 4.678202 | 0.333040993 | 0.793 | 0.591 | 0.11885 | 33 | zgc:56493 |
| magt14 | 5.050637 | 0.262966199 | 0.299 | 0.13 | 0.128311 | 33 | magt1 |
| rpl23a13 | 5.171943 | 0.350513935 | 0.943 | 0.67 | 0.131393 | 33 | rpl23a |
| slc3a2b5 | 5.547483 | 0.33886695 | 0.287 | 0.129 | 0.140934 | 33 | slc3a2b |
| ufc12 | 6.469963 | 0.262305088 | 0.23 | 0.089 | 0.164369 | 33 | ufc1 |
| rpl359 | 6.548387 | 0.256667881 | 0.966 | 0.814 | 0.166362 | 33 | rpl35 |
| abracl6 | 6.583575 | 0.320302954 | 0.448 | 0.222 | 0.167256 | 33 | abracl |
| b2ml11 | 6.777544 | 0.348054779 | 0.529 | 0.294 | 0.172184 | 33 | b2ml |
| rps1411 | 7.275366 | 0.284308106 | 0.966 | 0.781 | 0.184831 | 33 | rps14 |

| | | | | | | | |
|---|---|---|---|---|---|---|---|
| XLOC-0234926 | 7.353679 | 0.262158407 | 0.241 | 0.094 | 0.18682 | 33 | XLOC-023492 |
| fkbp55 | 7.997981 | 0.304966355 | 0.287 | 0.128 | 0.203189 | 33 | fkbp5 |
| rpl2712 | 8.700458 | 0.263816685 | 0.943 | 0.78 | 0.221035 | 33 | rpl27 |
| rps1616 | 8.707767 | 0.315407078 | 0.989 | 0.766 | 0.221221 | 33 | rps16 |
| gpr183a5 | 8.734893 | 0.260759602 | 0.184 | 0.065 | 0.22191 | 33 | gpr183a |
| capza1b7 | 8.875129 | 0.34895748 | 0.437 | 0.235 | 0.225473 | 33 | capza1b |
| nop563 | 8.945527 | 0.313166783 | 0.379 | 0.187 | 0.227261 | 33 | nop56 |
| BX005183.36 | 1.116108 | 0.252590869 | 0.207 | 0.077 | 0.283547 | 33 | BX005183.3 |
| gm2a5 | 1.147603 | 0.255715173 | 0.287 | 0.122 | 0.291549 | 33 | gm2a |
| spcs13 | 1.164645 | 0.296615286 | 0.448 | 0.232 | 0.295878 | 33 | spcs1 |
| rps1017 | 1.269710 | 0.305663457 | 0.954 | 0.727 | 0.32257 | 33 | rps10 |
| litaf6 | 1.316857 | 0.316931411 | 0.31 | 0.147 | 0.334548 | 33 | litaf |
| dbi7 | 1.339254 | 0.314312174 | 0.46 | 0.244 | 0.340238 | 33 | dbi |
| rpl7a10 | 1.390712 | 0.254911642 | 0.989 | 0.824 | 0.353311 | 33 | rpl7a |
| si:ch1073-325m22.24 | 1.573290 | 0.310367331 | 0.46 | 0.25 | 0.399694 | 33 | si:ch1073-325m22.2 |
| arf6a2 | 1.595668 | 0.291668964 | 0.345 | 0.164 | 0.40538 | 33 | arf6a |
| btf310 | 1.697306 | 0.32252779 | 0.839 | 0.611 | 0.431201 | 33 | btf3 |
| rplp2l14 | 1.847350 | 0.26710265 | 0.954 | 0.795 | 0.469319 | 33 | rplp2l |
| ndufa4l10 | 1.944791 | 0.336440651 | 0.701 | 0.423 | 0.494074 | 33 | ndufa4l |
| nr4a111 | 2.135648 | 0.32885137 | 0.333 | 0.159 | 0.542562 | 33 | nr4a1 |
| hsp90ab18 | 2.315249 | 0.250600099 | 0.977 | 0.801 | 0.588189 | 33 | hsp90ab1 |
| GCA7 | 2.517323 | 0.327558895 | 0.379 | 0.201 | 0.639526 | 33 | GCA |
| selenok3 | 2.528356 | 0.26486002 | 0.299 | 0.14 | 0.642329 | 33 | selenok |
| ppt15 | 2.600761 | 0.318234597 | 0.356 | 0.182 | 0.660723 | 33 | ppt1 |
| commd31 | 3.283341 | 0.275790048 | 0.253 | 0.111 | 0.834133 | 33 | commd3 |
| eif4ebp16 | 4.290875 | 0.280137252 | 0.483 | 0.259 | 1 | 33 | eif4ebp1 |
| eif4ebp25 | 4.373128 | 0.304128152 | 0.368 | 0.193 | 1 | 33 | eif4ebp2 |
| tomm7 | 4.763185 | 0.283930975 | 0.414 | 0.218 | 1 | 33 | tomm7 |
| rpl810 | 4.886150 | 0.274027619 | 0.989 | 0.795 | 1 | 33 | rpl8 |
| rps27.215 | 5.102105 | 0.350120971 | 0.874 | 0.553 | 1 | 33 | rps27.2 |
| rpl615 | 5.746814 | 0.289374227 | 0.92 | 0.702 | 1 | 33 | rpl6 |
| pkma5 | 6.024289 | 0.265891533 | 0.356 | 0.179 | 1 | 33 | pkma |
| tmed74 | 7.313788 | 0.265996007 | 0.414 | 0.221 | 1 | 33 | tmed7 |
| ube2d1b1 | 7.391589 | 0.252164444 | 0.299 | 0.144 | 1 | 33 | ube2d1b |
| rps1312 | 7.776551 | 0.28257504 | 0.966 | 0.749 | 1 | 33 | rps13 |

| | | | | | | | |
|---|---|---|---|---|---|---|---|
| id35 | 8.355163 | 0.280107214 | 0.391 | 0.198 | 1 | 33 | id3 |
| naca15 | 9.208801 | 0.295439954 | 0.874 | 0.652 | 1 | 33 | naca |
| atp5pd6 | 9.596571 | 0.27818743 | 0.494 | 0.284 | 1 | 33 | atp5pd |
| rhoaa9 | 0.0001 | 0.251058787 | 0.345 | 0.177 | 1 | 33 | rhoaa |
| rpl2111 | 0.0001 | 0.256124341 | 0.966 | 0.777 | 1 | 33 | rpl21 |
| mibp214 | 0.0001 | 0.285825682 | 0.713 | 0.522 | 1 | 33 | mibp2 |
| ak14 | 0.0001 | 0.280034104 | 0.218 | 0.099 | 1 | 33 | ak1 |
| ak23 | 0.0001 | 0.288156516 | 0.437 | 0.256 | 1 | 33 | ak2 |
| rps4x15 | 0.0002 | 0.277694061 | 0.954 | 0.737 | 1 | 33 | rps4x |
| faua13 | 0.0002 | 0.286513926 | 0.931 | 0.718 | 1 | 33 | faua |
| idh24 | 0.0002 | 0.265784986 | 0.402 | 0.228 | 1 | 33 | idh2 |
| mt-nd310 | 0.0002 | 0.297671386 | 0.586 | 0.372 | 1 | 33 | mt-nd3 |
| mt-cyb11 | 0.0002 | 0.296272475 | 0.77 | 0.577 | 1 | 33 | mt-cyb |
| mt-atp614 | 0.0002 | 0.30722852 | 0.793 | 0.557 | 1 | 33 | mt-atp6 |
| rap1b5 | 0.0003 | 0.272110216 | 0.391 | 0.221 | 1 | 33 | rap1b |
| CR383676.118 | 0.0003 | 0.369058195 | 0.851 | 0.718 | 1 | 33 | CR383676.1 |
| cox7a2a3 | 0.0004 | 0.252203751 | 0.437 | 0.253 | 1 | 33 | cox7a2a |
| mt-co213 | 0.0006 | 0.265425712 | 0.793 | 0.672 | 1 | 33 | mt-co2 |
| hmgn615 | 0.0006 | 0.251357291 | 0.529 | 0.312 | 1 | 33 | hmgn6 |
| ddx213 | 0.0006 | 0.252519417 | 0.368 | 0.211 | 1 | 33 | ddx21 |
| romo110 | 0.0007 | 0.273338494 | 0.598 | 0.409 | 1 | 33 | romo1 |
| rpsa15 | 0.0007 | 0.253271741 | 0.966 | 0.786 | 1 | 33 | rpsa |
| cdc427 | 0.0007 | 0.257515194 | 0.391 | 0.235 | 1 | 33 | cdc42 |
| eef1db9 | 0.0008 | 0.278528545 | 0.448 | 0.286 | 1 | 33 | eef1db |
| nsa212 | 0.0008 | 0.267632797 | 0.621 | 0.389 | 1 | 33 | nsa2 |
| MFAP4 (1 of many)2 | 0.0009 | 0.270547247 | 0.149 | 0.062 | 1 | 33 | MFAP4 (1 of many) |
| ckbb6 | 0.001 | 0.254282687 | 0.437 | 0.252 | 1 | 33 | ckbb |
| rpl49 | 0.001 | 0.258652997 | 0.851 | 0.658 | 1 | 33 | rpl4 |
| rpl3113 | 0.0013 | 0.262329198 | 0.908 | 0.688 | 1 | 33 | rpl31 |
| ssr45 | 0.0013 | 0.254814751 | 0.517 | 0.332 | 1 | 33 | ssr4 |
| rpl27a15 | 0.0013 | 0.287331178 | 0.77 | 0.523 | 1 | 33 | rpl27a |
| hmgn212 | 0.0015 | 0.270186523 | 0.69 | 0.468 | 1 | 33 | hmgn2 |
| slc25a3b7 | 0.0016 | 0.266895939 | 0.667 | 0.459 | 1 | 33 | slc25a3b |
| atp5f1e5 | 0.0017 | 0.252872179 | 0.678 | 0.492 | 1 | 33 | atp5f1e |
| rsl24d111 | 0.0019 | 0.275857846 | 0.667 | 0.461 | 1 | 33 | rsl24d1 |

| | | | | | | | |
|---|---|---|---|---|---|---|---|
| rps311 | 0.0021 | 0.260062858 | 0.943 | 0.726 | 1 | 33 | rps3 |
| pabpc1a9 | 0.0022 | 0.263761785 | 0.782 | 0.571 | 1 | 33 | pabpc1a |
| ctrb11 | 0 | 1.872668132 | 0.737 | 0.016 | 0 | 34 | ctrb1 |
| prss59.1 | 0 | 1.690187872 | 0.579 | 0.011 | 0 | 34 | prss59.1 |
| CELA1 (1 of many) | 0 | 1.688118745 | 0.592 | 0.012 | 0 | 34 | CELA1 (1 of many) |
| zgc:136461 | 0 | 1.313105608 | 0.474 | 0.008 | 0 | 34 | zgc:136461 |
| ctrl | 0 | 1.265874274 | 0.447 | 0.007 | 0 | 34 | ctrl |
| c6ast4 | 5.258296 | 1.043399144 | 0.316 | 0.004 | 1.3358702 | 34 | c6ast4 |
| prss11 | 6.407808 | 1.74337091 | 0.645 | 0.02 | 1.6279036 | 34 | prss1 |
| ela3l | 8.192733 | 1.343646509 | 0.5 | 0.014 | 2.0813639 | 34 | ela3l |
| cpb1 | 3.669891 | 0.997550224 | 0.382 | 0.008 | 9.3233595 | 34 | cpb1 |
| si:dkey-14d8.6 | 4.016882 | 0.94904912 | 0.382 | 0.008 | 1.0204889 | 34 | si:dkey-14d8.6 |
| si:ch211-240l19.8 | 8.586144 | 1.33047953 | 0.526 | 0.018 | 2.1813099 | 34 | si:ch211-240l19.8 |
| ela2 | 2.738661 | 1.247258911 | 0.434 | 0.012 | 6.9575687 | 34 | ela2 |
| cpa1 | 7.438796 | 0.743090885 | 0.25 | 0.005 | 1.8898262 | 34 | cpa1 |
| zgc:112160 | 1.867006 | 1.025671729 | 0.342 | 0.011 | 4.7431303 | 34 | zgc:112160 |
| ela2l | 5.168426 | 0.979152719 | 0.316 | 0.01 | 1.3130388 | 34 | ela2l |
| zgc:92590 | 4.493527 | 0.570711351 | 0.184 | 0.003 | 1.1415805 | 34 | zgc:92590 |
| cel.1 | 1.508766 | 0.735750883 | 0.25 | 0.006 | 3.8330224 | 34 | cel.1 |
| si:dkey-14d8.7 | 1.092092 | 0.526230425 | 0.211 | 0.005 | 2.7744607 | 34 | si:dkey-14d8.7 |
| c6ast3 | 1.522293 | 0.658355123 | 0.237 | 0.008 | 3.8673872 | 34 | c6ast3 |
| erp27 | 2.108419 | 0.395153988 | 0.158 | 0.003 | 5.3564397 | 34 | erp27 |
| cpa53 | 6.310126 | 1.151741781 | 0.447 | 0.034 | 1.6030876 | 34 | cpa5 |
| pla2g1b | 7.184379 | 0.502883658 | 0.184 | 0.006 | 1.8251915 | 34 | pla2g1b |
| spink41 | 9.101413 | 0.550772798 | 0.184 | 0.006 | 2.3122141 | 34 | spink4 |
| si:ch211-240l19.6 | 3.387340 | 0.528758917 | 0.184 | 0.006 | 8.6055395 | 34 | si:ch211-240l19.6 |
| cel.2 | 5.669693 | 0.44146781 | 0.184 | 0.006 | 1.4403855 | 34 | cel.2 |
| sycn.2 | 7.687715 | 0.565992259 | 0.184 | 0.007 | 1.9530641 | 34 | sycn.2 |
| si:ch211-240l19.5 | 1.998169 | 0.564167451 | 0.171 | 0.006 | 5.0763485 | 34 | si:ch211-240l19.5 |
| prss59.21 | 5.165290 | 0.970779432 | 0.263 | 0.016 | 1.3122419 | 34 | prss59.2 |
| cpa4 | 5.070130 | 0.363418723 | 0.145 | 0.005 | 1.2880667 | 34 | cpa4 |
| amy2a | 4.199317 | 0.539619328 | 0.171 | 0.011 | 1.0668367 | 34 | amy2a |
| apoda.2 | 3.487890 | 0.637925135 | 0.237 | 0.034 | 8.8609850 | 34 | apoda.2 |
| tmsb4x19 | 4.381261 | 0.546283816 | 0.947 | 0.719 | 1.1130594 | 34 | tmsb4x |
| gamt3 | 6.997377 | 0.516188134 | 0.224 | 0.044 | 1.7776836 | 34 | gamt |

| | | | | | | | |
|---|---|---|---|---|---|---|---|
| pdia21 | 1.369958 | 0.25810931 | 0.118 | 0.017 | 3.4803786 | 34 | pdia2 |
| pfn114 | 2.465747 | 0.65634231 | 0.776 | 0.502 | 6.2642306 | 34 | pfn1 |
| actb212 | 1.052700 | 0.354599285 | 0.947 | 0.867 | 2.6743866 | 34 | actb2 |
| rpl178 | 1.566760 | 0.282766041 | 0.829 | 0.765 | 3.9803558 | 34 | rpl17 |
| rpl2112 | 2.184467 | 0.253545297 | 0.842 | 0.778 | 0.000555 | 34 | rpl21 |
| rps312 | 1.581168 | 0.267539361 | 0.803 | 0.727 | 0.004017 | 34 | rps3 |
| rpl1514 | 3.939892 | 0.296739696 | 0.908 | 0.764 | 0.010009 | 34 | rpl15 |
| grn13 | 5.236936 | 0.636041591 | 0.25 | 0.091 | 0.013304 | 34 | grn1 |
| fcer1gl8 | 5.454151 | 0.60470671 | 0.382 | 0.183 | 0.013856 | 34 | fcer1gl |
| bhmt3 | 6.244401 | 0.333643616 | 0.158 | 0.043 | 0.015864 | 34 | bhmt |
| rpl3412 | 7.733651 | 0.255508213 | 0.908 | 0.803 | 0.019647 | 34 | rpl34 |
| rps27.118 | 1.157847 | 0.295263244 | 0.855 | 0.711 | 0.029415 | 34 | rps27.1 |
| rps26l13 | 1.269370 | 0.284681765 | 0.803 | 0.701 | 0.032248 | 34 | rps26l |
| rps1617 | 1.298102 | 0.26148291 | 0.868 | 0.767 | 0.032978 | 34 | rps16 |
| rpl23a14 | 1.380171 | 0.304338631 | 0.776 | 0.671 | 0.035063 | 34 | rpl23a |
| faua14 | 1.576591 | 0.278514671 | 0.829 | 0.719 | 0.040053 | 34 | faua |
| si:dkey-5n18.12 | 1.797895 | 0.496261728 | 0.197 | 0.065 | 0.045676 | 34 | si:dkey-5n18.1 |
| rpl616 | 2.187813 | 0.258708929 | 0.763 | 0.703 | 0.055581 | 34 | rpl6 |
| c1qb2 | 4.131682 | 0.490210554 | 0.211 | 0.074 | 0.104965 | 34 | c1qb |
| lgals3bpb2 | 1.905261 | 0.398112094 | 0.145 | 0.044 | 0.484032 | 34 | lgals3bpb |
| CU929150.13 | 4.131979 | 0.279871155 | 0.105 | 0.028 | 1 | 34 | CU929150.1 |
| lgals2a4 | 6.577179 | 0.547010444 | 0.276 | 0.139 | 1 | 34 | lgals2a |
| arpc1b13 | 9.932005 | 0.461393367 | 0.474 | 0.315 | 1 | 34 | arpc1b |
| zgc:11418821 | 0.0001 | 0.256320311 | 0.776 | 0.669 | 1 | 34 | zgc:114188 |
| cfl111 | 0.0002 | 0.277930157 | 0.632 | 0.602 | 1 | 34 | cfl1 |
| cd74b8 | 0.0003 | 0.467540387 | 0.382 | 0.227 | 1 | 34 | cd74b |
| ccl33.32 | 0.0004 | 0.574689247 | 0.158 | 0.062 | 1 | 34 | ccl33.3 |
| rpl920 | 0.0004 | 0.253624307 | 0.803 | 0.67 | 1 | 34 | rpl9 |
| c1qc2 | 0.0005 | 0.380686154 | 0.171 | 0.071 | 1 | 34 | c1qc |
| CR318588.112 | 0.0005 | 0.542041316 | 0.355 | 0.217 | 1 | 34 | CR318588.1 |
| marco2 | 0.0006 | 0.356268389 | 0.118 | 0.043 | 1 | 34 | marco |
| slc40a12 | 0.0006 | 0.291811571 | 0.118 | 0.042 | 1 | 34 | slc40a1 |
| NPC2 (1 of many)2 | 0.0007 | 0.316134679 | 0.118 | 0.042 | 1 | 34 | NPC2 (1 of many) |
| aldob2 | 0.001 | 0.294751904 | 0.105 | 0.037 | 1 | 34 | aldob |
| cotl112 | 0.0016 | 0.405741259 | 0.421 | 0.322 | 1 | 34 | cotl1 |

| | | | | | | | |
|---|---|---|---|---|---|---|---|
| si:busm1-266f07.27 | 0.0033 | 0.353616675 | 0.263 | 0.149 | 1 | 34 | si:busm1-266f07.2 |
| icn22 | 0 | 2.502608321 | 0.984 | 0.005 | 0 | 35 | icn2 |
| cldnb1 | 0 | 2.45954331 | 0.967 | 0.008 | 0 | 35 | cldnb |
| zgc:1935051 | 0 | 2.42617627 | 0.984 | 0.003 | 0 | 35 | zgc:193505 |
| si:dkey-7c18.24 | 0 | 2.400450448 | 0.967 | 0.002 | 0 | 35 | si:dkey-7c18.24 |
| zgc:111983 | 0 | 2.345686003 | 1 | 0.001 | 0 | 35 | zgc:111983 |
| si:dkey-193p11.22 | 0 | 2.314913878 | 0.934 | 0.01 | 0 | 35 | si:dkey-193p11.2 |
| cldne1 | 0 | 2.293193347 | 0.984 | 0.002 | 0 | 35 | cldne |
| krt171 | 0 | 2.213901295 | 0.934 | 0.005 | 0 | 35 | krt17 |
| krt51 | 0 | 2.194873862 | 0.918 | 0.01 | 0 | 35 | krt5 |
| mid1ip1a1 | 0 | 2.191567549 | 0.967 | 0.01 | 0 | 35 | mid1ip1a |
| si:dkey-247k7.2 | 0 | 2.184705734 | 0.967 | 0.001 | 0 | 35 | si:dkey-247k7.2 |
| si:ch211-69b7.61 | 0 | 2.08569387 | 0.902 | 0.003 | 0 | 35 | si:ch211-69b7.6 |
| si:ch211-207n23.22 | 0 | 2.030079521 | 0.951 | 0.024 | 0 | 35 | si:ch211-207n23.2 |
| si:ch211-217k17.9 | 0 | 2.024190699 | 0.934 | 0.001 | 0 | 35 | si:ch211-217k17.9 |
| lye | 0 | 1.934637767 | 0.902 | 0.001 | 0 | 35 | lye |
| ppl1 | 0 | 1.855661506 | 0.967 | 0.006 | 0 | 35 | ppl |
| krt1-19d | 0 | 1.80884878 | 0.787 | 0.001 | 0 | 35 | krt1-19d |
| anxa1c | 0 | 1.731959163 | 0.82 | 0.001 | 0 | 35 | anxa1c |
| dsg2.11 | 0 | 1.727993889 | 0.869 | 0.021 | 0 | 35 | dsg2.1 |
| abcb5 | 0 | 1.716848988 | 0.852 | 0.022 | 0 | 35 | abcb5 |
| pkp3a1 | 0 | 1.706078759 | 0.918 | 0.007 | 0 | 35 | pkp3a |
| dsc2l3 | 0 | 1.697101032 | 0.885 | 0.024 | 0 | 35 | dsc2l |
| sec23b | 0 | 1.690880006 | 0.967 | 0.031 | 0 | 35 | sec23b |
| zgc:1539111 | 0 | 1.680900633 | 0.689 | 0.015 | 0 | 35 | zgc:153911 |
| scel | 0 | 1.666940367 | 0.869 | 0.004 | 0 | 35 | scel |
| evpla1 | 0 | 1.660708803 | 0.902 | 0.002 | 0 | 35 | evpla |
| soul2 | 0 | 1.652867109 | 0.852 | 0.001 | 0 | 35 | soul2 |
| si:dkey-248g15.31 | 0 | 1.630694807 | 0.721 | 0.006 | 0 | 35 | si:dkey-248g15.3 |
| cldn23a | 0 | 1.584348811 | 0.82 | 0.002 | 0 | 35 | cldn23a |
| tnks1bp11 | 0 | 1.574048785 | 0.82 | 0.005 | 0 | 35 | tnks1bp1 |
| si:ch211-125o16.4 | 0 | 1.557164193 | 0.836 | 0 | 0 | 35 | si:ch211-125o16.4 |
| CABZ01048956.11 | 0 | 1.52630313 | 0.738 | 0.001 | 0 | 35 | CABZ01048956.1 |
| zgc:153665 | 0 | 1.524003935 | 0.77 | 0.016 | 0 | 35 | zgc:153665 |
| anxa1b1 | 0 | 1.51894866 | 0.77 | 0.005 | 0 | 35 | anxa1b |

| | | | | | | | |
|---|---|---|---|---|---|---|---|
| rhcgl1 | 0 | 1.505304258 | 0.705 | 0.001 | 0 | 35 | rhcgl1 |
| krt971 | 0 | 1.498436773 | 0.623 | 0.005 | 0 | 35 | krt97 |
| LOC108190590 | 0 | 1.474096258 | 0.77 | 0 | 0 | 35 | LOC108190590 |
| znf1851 | 0 | 1.439490796 | 0.803 | 0.002 | 0 | 35 | znf185 |
| zmp:0000000906 | 0 | 1.364741936 | 0.705 | 0.002 | 0 | 35 | zmp:0000000906 |
| tgm5l | 0 | 1.325696309 | 0.623 | 0.003 | 0 | 35 | tgm5l |
| si:ch211-98n17.5 | 0 | 1.321144483 | 0.705 | 0.001 | 0 | 35 | si:ch211-98n17.5 |
| si:ch211-79k12.12 | 0 | 1.315342514 | 0.803 | 0.019 | 0 | 35 | si:ch211-79k12.1 |
| AL929237.11 | 0 | 1.305643758 | 0.623 | 0.005 | 0 | 35 | AL929237.1 |
| hrc | 0 | 1.301180514 | 0.689 | 0.006 | 0 | 35 | hrc |
| agr1 | 0 | 1.298261945 | 0.639 | 0.001 | 0 | 35 | agr1 |
| lmo7a1 | 0 | 1.277189671 | 0.705 | 0.006 | 0 | 35 | lmo7a |
| zgc:110333 | 0 | 1.232783544 | 0.721 | 0.001 | 0 | 35 | zgc:110333 |
| LOC101882729 | 0 | 1.181011155 | 0.656 | 0.001 | 0 | 35 | LOC101882729 |
| tmem176l.41 | 0 | 1.166790699 | 0.607 | 0.008 | 0 | 35 | tmem176l.4 |
| CR854839.1 | 0 | 1.160522818 | 0.656 | 0.002 | 0 | 35 | CR854839.1 |
| chst6 | 0 | 1.158489223 | 0.639 | 0.005 | 0 | 35 | chst6 |
| stx11b.1 | 0 | 1.145867725 | 0.623 | 0.001 | 0 | 35 | stx11b.1 |
| LOC100330250 | 0 | 1.143150349 | 0.541 | 0.002 | 0 | 35 | LOC100330250 |
| si:dkeyp-86h10.3 | 0 | 1.113042201 | 0.541 | 0.002 | 0 | 35 | si:dkeyp-86h10.3 |
| ftr83 | 0 | 1.108559681 | 0.656 | 0.005 | 0 | 35 | ftr83 |
| sts | 0 | 1.096747936 | 0.607 | 0.009 | 0 | 35 | sts |
| bicdl2l | 0 | 1.094107249 | 0.574 | 0.001 | 0 | 35 | bicdl2l |
| ptgs2b2 | 0 | 1.087626842 | 0.525 | 0.009 | 0 | 35 | ptgs2b |
| evplb | 0 | 1.068953144 | 0.623 | 0 | 0 | 35 | evplb |
| cyldb | 0 | 1.054509795 | 0.607 | 0.001 | 0 | 35 | cyldb |
| mir203a | 0 | 1.049910215 | 0.557 | 0 | 0 | 35 | mir203a |
| slc16a12b1 | 0 | 1.036381086 | 0.574 | 0.006 | 0 | 35 | slc16a12b |
| si:ch73-52f15.5 | 0 | 0.99941176 | 0.541 | 0.001 | 0 | 35 | si:ch73-52f15.5 |
| myzap | 0 | 0.982294326 | 0.525 | 0.006 | 0 | 35 | myzap |
| sox3 | 0 | 0.967804753 | 0.508 | 0.004 | 0 | 35 | sox3 |
| fam129aa | 0 | 0.960911956 | 0.525 | 0 | 0 | 35 | fam129aa |
| apodb | 0 | 0.940857726 | 0.492 | 0.002 | 0 | 35 | apodb |
| capn9 | 0 | 0.929128823 | 0.525 | 0.002 | 0 | 35 | capn9 |
| ANXA1 (1 of many) | 0 | 0.919172464 | 0.443 | 0.001 | 0 | 35 | ANXA1 (1 of many) |

| | | | | | | | |
|---|---|---|---|---|---|---|---|
| zgc:100868 | 0 | 0.906480812 | 0.525 | 0 | 0 | 35 | zgc:100868 |
| FO681323.1 | 0 | 0.901244922 | 0.508 | 0 | 0 | 35 | FO681323.1 |
| bik1 | 0 | 0.883576377 | 0.508 | 0.008 | 0 | 35 | bik |
| si:dkeyp-1h4.6 | 0 | 0.881279084 | 0.459 | 0.003 | 0 | 35 | si:dkeyp-1h4.6 |
| papss2a | 0 | 0.862291297 | 0.459 | 0.005 | 0 | 35 | papss2a |
| pkp2 | 0 | 0.859086691 | 0.443 | 0.004 | 0 | 35 | pkp2 |
| CABZ01084230.1 | 0 | 0.841514973 | 0.459 | 0.001 | 0 | 35 | CABZ01084230.1 |
| LOC100333235 | 0 | 0.839018968 | 0.443 | 0.001 | 0 | 35 | LOC100333235 |
| cldnh1 | 0 | 0.82755951 | 0.295 | 0.002 | 0 | 35 | cldnh |
| ccl20b | 0 | 0.826742121 | 0.475 | 0.003 | 0 | 35 | ccl20b |
| CABZ01067657.1 | 0 | 0.825644182 | 0.426 | 0.001 | 0 | 35 | CABZ01067657.1 |
| cx35.4 | 0 | 0.824080375 | 0.459 | 0.003 | 0 | 35 | cx35.4 |
| CU914622.2 | 0 | 0.819270777 | 0.377 | 0 | 0 | 35 | CU914622.2 |
| XLOC-041945 | 0 | 0.803678627 | 0.443 | 0.001 | 0 | 35 | XLOC-041945 |
| si:ch211-182p11.1 | 0 | 0.801767381 | 0.41 | 0.002 | 0 | 35 | si:ch211-182p11.1 |
| eps8l1a | 0 | 0.796991975 | 0.443 | 0.002 | 0 | 35 | eps8l1a |
| btr12 | 0 | 0.794833185 | 0.459 | 0.003 | 0 | 35 | btr12 |
| cldnf | 0 | 0.791632122 | 0.459 | 0.001 | 0 | 35 | cldnf |
| LOC100149234 | 0 | 0.784673761 | 0.459 | 0 | 0 | 35 | LOC100149234 |
| tmem176l.3b | 0 | 0.774851084 | 0.41 | 0.001 | 0 | 35 | tmem176l.3b |
| ponzr2 | 0 | 0.774812685 | 0.426 | 0 | 0 | 35 | ponzr2 |
| si:dkeyp-51b9.3 | 0 | 0.744138776 | 0.377 | 0 | 0 | 35 | si:dkeyp-51b9.3 |
| st14b | 0 | 0.738899737 | 0.426 | 0.002 | 0 | 35 | st14b |
| hmgcra | 0 | 0.731371127 | 0.41 | 0.003 | 0 | 35 | hmgcra |
| ponzr10 | 0 | 0.723712612 | 0.328 | 0 | 0 | 35 | ponzr10 |
| si:dkey-221j11.3 | 0 | 0.721038963 | 0.393 | 0 | 0 | 35 | si:dkey-221j11.3 |
| CABZ01015815.1 | 0 | 0.710999139 | 0.393 | 0.003 | 0 | 35 | CABZ01015815.1 |
| si:rp71-1h20.9 | 0 | 0.708969762 | 0.41 | 0.002 | 0 | 35 | si:rp71-1h20.9 |
| si:dkey-175d9.2 | 0 | 0.702745671 | 0.41 | 0 | 0 | 35 | si:dkey-175d9.2 |
| wu:fc34e06 | 0 | 0.689772068 | 0.377 | 0 | 0 | 35 | wu:fc34e06 |
| CABZ01068248.1 | 0 | 0.686965121 | 0.393 | 0.002 | 0 | 35 | CABZ01068248.1 |
| CABZ01068469.1 | 0 | 0.686466943 | 0.377 | 0.001 | 0 | 35 | CABZ01068469.1 |
| CR388148.2 | 0 | 0.681241296 | 0.344 | 0 | 0 | 35 | CR388148.2 |
| rassf7b | 0 | 0.67891868 | 0.344 | 0.003 | 0 | 35 | rassf7b |
| selenow2b1 | 0 | 0.678174782 | 0.41 | 0.003 | 0 | 35 | selenow2b |

| | | | | | | | |
|---|---|---|---|---|---|---|---|
| si:ch211-197e7.1 | 0 | 0.669823284 | 0.377 | 0 | 0 | 35 | si:ch211-197e7.1 |
| BX005027.2 | 0 | 0.668097909 | 0.328 | 0 | 0 | 35 | BX005027.2 |
| dbnla | 0 | 0.667229167 | 0.426 | 0.003 | 0 | 35 | dbnla |
| gpa33a1 | 0 | 0.662640785 | 0.393 | 0.003 | 0 | 35 | gpa33a |
| anxa3a | 0 | 0.662145136 | 0.393 | 0.001 | 0 | 35 | anxa3a |
| erbb3a | 0 | 0.661969281 | 0.377 | 0.002 | 0 | 35 | erbb3a |
| si:ch73-347e22.8 | 0 | 0.657935014 | 0.361 | 0 | 0 | 35 | si:ch73-347e22.8 |
| rnf222 | 0 | 0.652243038 | 0.344 | 0 | 0 | 35 | rnf222 |
| si:dkey-73p2.1 | 0 | 0.63683049 | 0.328 | 0 | 0 | 35 | si:dkey-73p2.1 |
| adgrf6 | 0 | 0.628670414 | 0.361 | 0.001 | 0 | 35 | adgrf6 |
| CR589947.2 | 0 | 0.628394145 | 0.344 | 0 | 0 | 35 | CR589947.2 |
| CR926130.1 | 0 | 0.614895485 | 0.295 | 0 | 0 | 35 | CR926130.1 |
| zgc:136892 | 0 | 0.605659994 | 0.344 | 0.001 | 0 | 35 | zgc:136892 |
| paqr5b | 0 | 0.601126959 | 0.328 | 0 | 0 | 35 | paqr5b |
| foxq1a | 0 | 0.594209662 | 0.311 | 0.002 | 0 | 35 | foxq1a |
| CR677617.1 | 0 | 0.592479291 | 0.311 | 0.002 | 0 | 35 | CR677617.1 |
| LOC110438610 | 0 | 0.584064234 | 0.344 | 0.001 | 0 | 35 | LOC110438610 |
| si:ch73-184c24.1 | 0 | 0.576895734 | 0.328 | 0.001 | 0 | 35 | si:ch73-184c24.1 |
| vsig8b | 0 | 0.564762332 | 0.328 | 0 | 0 | 35 | vsig8b |
| LOC407643 | 0 | 0.563054016 | 0.311 | 0.001 | 0 | 35 | LOC407643 |
| esrp1 | 0 | 0.556829946 | 0.295 | 0.002 | 0 | 35 | esrp1 |
| XLOC-027129 | 0 | 0.554229905 | 0.328 | 0 | 0 | 35 | XLOC-027129 |
| LOC100331080 | 0 | 0.548575912 | 0.23 | 0 | 0 | 35 | LOC100331080 |
| tmem45b | 0 | 0.547631806 | 0.311 | 0.001 | 0 | 35 | tmem45b |
| LOC110438215 | 0 | 0.546068925 | 0.279 | 0.001 | 0 | 35 | LOC110438215 |
| LOC103909616 | 0 | 0.543944604 | 0.295 | 0 | 0 | 35 | LOC103909616 |
| si:dkey-223p19.1 | 0 | 0.539769041 | 0.23 | 0.001 | 0 | 35 | si:dkey-223p19.1 |
| CABZ01039863.11 | 0 | 0.536148952 | 0.328 | 0.002 | 0 | 35 | CABZ01039863.1 |
| si:dkey-27n6.4 | 0 | 0.529619323 | 0.311 | 0 | 0 | 35 | si:dkey-27n6.4 |
| noxa1 | 0 | 0.525400751 | 0.246 | 0.001 | 0 | 35 | noxa1 |
| LOC100001846 | 0 | 0.523279978 | 0.295 | 0.001 | 0 | 35 | LOC100001846 |
| CABZ01027551.1 | 0 | 0.50134775 | 0.197 | 0.001 | 0 | 35 | CABZ01027551.1 |
| nectin4a | 0 | 0.495397211 | 0.279 | 0.001 | 0 | 35 | nectin4a |
| si:ch73-196i15.3 | 0 | 0.493933311 | 0.262 | 0.001 | 0 | 35 | si:ch73-196i15.3 |
| CU062645.1 | 0 | 0.493141488 | 0.262 | 0 | 0 | 35 | CU062645.1 |

| | | | | | | | |
|---|---|---|---|---|---|---|---|
| zgc:194839 | 0 | 0.489312918 | 0.23 | 0.001 | 0 | 35 | zgc:194839 |
| LOC100149931 | 0 | 0.487807957 | 0.246 | 0 | 0 | 35 | LOC100149931 |
| gna14 | 0 | 0.483446742 | 0.246 | 0 | 0 | 35 | gna14 |
| si:dkey-165e24.1 | 0 | 0.482862716 | 0.246 | 0 | 0 | 35 | si:dkey-165e24.1 |
| tgm1 | 0 | 0.478233085 | 0.246 | 0 | 0 | 35 | tgm1 |
| pkp1b | 0 | 0.466566568 | 0.279 | 0.001 | 0 | 35 | pkp1b |
| CU693379.1 | 0 | 0.465123674 | 0.246 | 0.001 | 0 | 35 | CU693379.1 |
| si:ch211-226h7.5 | 0 | 0.463465991 | 0.197 | 0 | 0 | 35 | si:ch211-226h7.5 |
| si:busm1-194e12.8 | 0 | 0.459689148 | 0.246 | 0 | 0 | 35 | si:busm1-194e12.8 |
| zgc:113229 | 0 | 0.445927511 | 0.23 | 0 | 0 | 35 | zgc:113229 |
| styk1b | 0 | 0.439010408 | 0.246 | 0.001 | 0 | 35 | styk1b |
| tlcd1 | 0 | 0.437980483 | 0.279 | 0.002 | 0 | 35 | tlcd1 |
| si:ch211-149a19.3 | 0 | 0.428907498 | 0.246 | 0 | 0 | 35 | si:ch211-149a19.3 |
| adgrg2a | 0 | 0.42676866 | 0.23 | 0 | 0 | 35 | adgrg2a |
| BX004785.3 | 0 | 0.425564925 | 0.246 | 0.001 | 0 | 35 | BX004785.3 |
| ttc22 | 0 | 0.424056003 | 0.213 | 0 | 0 | 35 | ttc22 |
| il20ra | 0 | 0.423677725 | 0.262 | 0.001 | 0 | 35 | il20ra |
| sp8a | 0 | 0.421022531 | 0.23 | 0 | 0 | 35 | sp8a |
| si:dkey-11o15.7 | 0 | 0.412970673 | 0.246 | 0 | 0 | 35 | si:dkey-11o15.7 |
| zgc:165423 | 0 | 0.412332036 | 0.164 | 0 | 0 | 35 | zgc:165423 |
| il17rc | 0 | 0.410081408 | 0.246 | 0.001 | 0 | 35 | il17rc |
| abca12 | 0 | 0.401042722 | 0.23 | 0.001 | 0 | 35 | abca12 |
| LOC101882034 | 0 | 0.39928673 | 0.213 | 0 | 0 | 35 | LOC101882034 |
| si:dkey-19b23.12 | 0 | 0.398624155 | 0.213 | 0.001 | 0 | 35 | si:dkey-19b23.12 |
| klf5l | 0 | 0.395621736 | 0.23 | 0.001 | 0 | 35 | klf5l |
| zgc:123295 | 0 | 0.393802065 | 0.197 | 0 | 0 | 35 | zgc:123295 |
| card14 | 0 | 0.385120543 | 0.23 | 0.001 | 0 | 35 | card14 |
| XLOC-011066 | 0 | 0.38373564 | 0.197 | 0 | 0 | 35 | XLOC-011066 |
| si:ch1073-340i21.3 | 0 | 0.361444421 | 0.18 | 0 | 0 | 35 | si:ch1073-340i21.3 |
| CABZ01046954.1 | 0 | 0.355331583 | 0.197 | 0 | 0 | 35 | CABZ01046954.1 |
| BX901957.1 | 0 | 0.340062638 | 0.197 | 0 | 0 | 35 | BX901957.1 |
| adgrf8 | 0 | 0.333046385 | 0.164 | 0 | 0 | 35 | adgrf8 |
| muc13a | 0 | 0.329151942 | 0.148 | 0 | 0 | 35 | muc13a |
| CABZ01049361.1 | 0 | 0.321464558 | 0.148 | 0 | 0 | 35 | CABZ01049361.1 |
| LOC100333218 | 0 | 0.318576605 | 0.164 | 0 | 0 | 35 | LOC100333218 |

| | | | | | | | |
|---|---|---|---|---|---|---|---|
| si:dkey-61p9.6 | 0 | 0.315061229 | 0.148 | 0 | 0 | 35 | si:dkey-61p9.6 |
| sparta | 0 | 0.311133437 | 0.164 | 0 | 0 | 35 | sparta |
| LOC100537138 | 0 | 0.306917431 | 0.18 | 0 | 0 | 35 | LOC100537138 |
| mala | 0 | 0.306306402 | 0.148 | 0 | 0 | 35 | mala |
| si:ch73-338o16.4 | 0 | 0.305236858 | 0.164 | 0 | 0 | 35 | si:ch73-338o16.4 |
| si:ch211-229n2.6 | 0 | 0.293074042 | 0.164 | 0 | 0 | 35 | si:ch211-229n2.6 |
| plekhs1 | 0 | 0.29107162 | 0.148 | 0 | 0 | 35 | plekhs1 |
| zgc:172075 | 0 | 0.280205973 | 0.115 | 0 | 0 | 35 | zgc:172075 |
| fam177b | 0 | 0.265992561 | 0.164 | 0 | 0 | 35 | fam177b |
| si:ch211-250e5.2 | 0 | 0.265923226 | 0.131 | 0 | 0 | 35 | si:ch211-250e5.2 |
| si:ch211-232i5.1 | 0 | 0.26234812 | 0.131 | 0 | 0 | 35 | si:ch211-232i5.1 |
| pof1b1 | 4.628813 | 0.967052768 | 0.525 | 0.009 | 1.1759501 | 35 | pof1b |
| ttc39a | 1.383765 | 0.45770188 | 0.246 | 0.001 | 3.5154572 | 35 | ttc39a |
| nkx3-1 | 3.937479 | 0.894528737 | 0.393 | 0.005 | 1.0003165 | 35 | nkx3-1 |
| adgrg11 | 7.583894 | 0.513854662 | 0.23 | 0.001 | 1.9266883 | 35 | adgrg11 |
| zgc:922421 | 1.058141 | 1.296410385 | 0.705 | 0.019 | 2.6882081 | 35 | zgc:92242 |
| si:dkey-228d14.5 | 2.137500 | 0.575974745 | 0.328 | 0.003 | 5.4303193 | 35 | si:dkey-228d14.5 |
| ano1 | 8.011458 | 0.676683696 | 0.426 | 0.006 | 2.0353110 | 35 | ano1 |
| si:ch73-208h1.1 | 9.463678 | 0.320659716 | 0.148 | 0 | 2.4042474 | 35 | si:ch73-208h1.1 |
| LOC110440078 | 1.134693 | 0.252555054 | 0.148 | 0 | 2.8826877 | 35 | LOC110440078 |
| LO017816.1 | 3.209596 | 0.344999974 | 0.213 | 0.001 | 8.1539799 | 35 | LO017816.1 |
| dhrs13a.21 | 3.175266 | 1.998516424 | 0.967 | 0.04 | 8.0667645 | 35 | dhrs13a.2 |
| crfb16 | 1.076618 | 0.290747147 | 0.115 | 0 | 2.7351504 | 35 | crfb16 |
| f2rl1.21 | 1.943277 | 1.304840734 | 0.689 | 0.019 | 4.9368952 | 35 | f2rl1.2 |
| zgc:153932 | 2.765699 | 0.4685033 | 0.246 | 0.002 | 7.0262589 | 35 | zgc:153932 |
| elf31 | 3.245798 | 0.738050251 | 0.393 | 0.005 | 8.2459505 | 35 | elf3 |
| frmd4ba | 2.359379 | 1.009415618 | 0.574 | 0.013 | 5.9940047 | 35 | frmd4ba |
| si:dkeyp-11e3.1 | 3.440675 | 0.267839756 | 0.131 | 0 | 8.7410362 | 35 | si:dkeyp-11e3.1 |
| rarres3 | 4.863299 | 0.535864313 | 0.295 | 0.003 | 1.2355212 | 35 | rarres3 |
| BX548046.22 | 2.664421 | 0.609886764 | 0.344 | 0.004 | 6.7689622 | 35 | BX548046.2 |
| CU914622.1 | 3.465389 | 0.332131077 | 0.148 | 0 | 8.8038223 | 35 | CU914622.1 |
| dspa3 | 1.957009 | 1.369702968 | 0.77 | 0.026 | 4.9717837 | 35 | dspa |
| s100w2 | 3.352734 | 1.15213901 | 0.639 | 0.018 | 8.5176214 | 35 | s100w |
| ltc4s | 5.273342 | 0.346846419 | 0.164 | 0.001 | 1.3396926 | 35 | ltc4s |
| LOC108180802 | 1.362024 | 0.350830739 | 0.197 | 0.001 | 3.4602223 | 35 | LOC108180802 |

| | | | | | | | |
|---|---|---|---|---|---|---|---|
| rnf183 | 3.335347 | 0.339107125 | 0.18 | 0.001 | 8.4734493 | 35 | rnf183 |
| si:dkey-242g16.2 | 9.688704 | 0.438346292 | 0.262 | 0.002 | 2.4614153 | 35 | si:dkey-242g16.2 |
| wu:fb18f061 | 1.225478 | 2.12836725 | 0.951 | 0.047 | 3.1133287 | 35 | wu:fb18f06 |
| si:ch211-195b11.31 | 3.393517 | 1.701728849 | 0.803 | 0.032 | 8.6212315 | 35 | si:ch211-195b11.3 |
| CABZ01070852.1 | 3.148303 | 0.522712122 | 0.262 | 0.003 | 7.9982642 | 35 | CABZ01070852.1 |
| ccl20a.3 | 6.328289 | 0.392870152 | 0.246 | 0.002 | 1.6077019 | 35 | ccl20a.3 |
| stard141 | 4.424912 | 1.706468258 | 0.918 | 0.043 | 1.1241490 | 35 | stard14 |
| krt44 | 1.196873 | 2.274502354 | 0.934 | 0.048 | 3.0406560 | 35 | krt4 |
| lgals3bpa | 1.337654 | 0.271037432 | 0.148 | 0.001 | 3.3983117 | 35 | lgals3bpa |
| xkr91 | 3.090615 | 1.022216645 | 0.557 | 0.015 | 7.8517083 | 35 | xkr9 |
| si:dkey-61f9.2 | 1.389893 | 0.272853673 | 0.131 | 0 | 3.5310241 | 35 | si:dkey-61f9.2 |
| zgc:91849 | 1.677324 | 0.379584584 | 0.23 | 0.002 | 4.2612430 | 35 | zgc:91849 |
| nox1 | 4.013598 | 0.445715567 | 0.262 | 0.003 | 1.0196547 | 35 | nox1 |
| tmem30b | 2.331193 | 0.854117738 | 0.508 | 0.012 | 5.9223967 | 35 | tmem30b |
| wsb2 | 9.607189 | 0.898396353 | 0.541 | 0.014 | 2.4407065 | 35 | wsb2 |
| b3gnt7l | 1.746453 | 0.447789487 | 0.213 | 0.002 | 4.4368650 | 35 | b3gnt7l |
| aldob3 | 6.687069 | 1.597701659 | 0.77 | 0.034 | 1.6988500 | 35 | aldob |
| erbb3b | 1.703677 | 0.332708096 | 0.164 | 0.001 | 4.3281923 | 35 | erbb3b |
| nqo1 | 1.008219 | 0.881560007 | 0.508 | 0.014 | 2.5613819 | 35 | nqo1 |
| LOC108181504 | 1.075730 | 0.280252591 | 0.148 | 0.001 | 2.7328930 | 35 | LOC108181504 |
| golph3l | 1.491849 | 0.281157171 | 0.148 | 0.001 | 3.7900428 | 35 | golph3l |
| macc1 | 8.142211 | 0.689862966 | 0.393 | 0.008 | 2.0685288 | 35 | macc1 |
| zgc:162184 | 1.035485 | 0.40574697 | 0.23 | 0.002 | 2.6306515 | 35 | zgc:162184 |
| phlda25 | 4.367041 | 1.722642207 | 0.902 | 0.05 | 1.1094467 | 35 | phlda2 |
| rgs21 | 7.452574 | 1.347431656 | 0.689 | 0.028 | 1.8933265 | 35 | rgs2 |
| zgc:194398 | 2.635625 | 0.338414528 | 0.197 | 0.002 | 6.6958072 | 35 | zgc:194398 |
| si:dkey-61p9.7 | 5.145978 | 0.300845871 | 0.164 | 0.001 | 1.3073358 | 35 | si:dkey-61p9.7 |
| cdh26.1 | 7.825736 | 0.290461335 | 0.164 | 0.001 | 1.9881282 | 35 | cdh26.1 |
| oclnb | 2.346287 | 1.244471683 | 0.738 | 0.032 | 5.9607443 | 35 | oclnb |
| si:dkey-160o24.3 | 3.873485 | 0.251010657 | 0.115 | 0 | 9.8405900 | 35 | si:dkey-160o24.3 |
| tfap2c1 | 2.813845 | 0.484364068 | 0.262 | 0.004 | 7.1485753 | 35 | tfap2c |
| oat | 3.848530 | 0.539240646 | 0.295 | 0.005 | 9.7771913 | 35 | oat |
| perp6 | 1.002986 | 1.714231219 | 0.918 | 0.058 | 2.5480870 | 35 | perp |
| cyp8b2 | 2.754753 | 0.428036363 | 0.197 | 0.002 | 6.9984520 | 35 | cyp8b2 |
| fam83b | 1.225791 | 0.365550279 | 0.18 | 0.002 | 3.1141222 | 35 | fam83b |

| | | | | | | | |
|---|---|---|---|---|---|---|---|
| BX005421.3 | 3.597120 | 0.306785929 | 0.18 | 0.002 | 9.1384856 | 35 | BX005421.3 |
| si:ch73-22a13.3 | 4.229447 | 0.363422396 | 0.18 | 0.002 | 1.0744911 | 35 | si:ch73-22a13.3 |
| elovl1a | 1.925480 | 1.38919497 | 0.787 | 0.041 | 4.8916842 | 35 | elovl1a |
| krt96 | 4.188906 | 0.301122788 | 0.131 | 0.001 | 1.0641916 | 35 | krt96 |
| barx2 | 5.038282 | 0.253173251 | 0.131 | 0.001 | 1.2799757 | 35 | barx2 |
| elovl6l | 1.588541 | 0.789104557 | 0.475 | 0.014 | 4.0356906 | 35 | elovl6l |
| fam169aa | 5.835000 | 0.378333564 | 0.213 | 0.002 | 1.4823819 | 35 | fam169aa |
| si:busm1-104n07.3 | 1.942127 | 0.304776123 | 0.164 | 0.001 | 4.9339753 | 35 | si:busm1-104n07.3 |
| gcnt4a2 | 5.348881 | 0.834415013 | 0.459 | 0.014 | 1.3588833 | 35 | gcnt4a |
| capn2l | 1.019654 | 0.27279295 | 0.148 | 0.001 | 2.5904315 | 35 | capn2l |
| tuft1a2 | 4.562881 | 1.126811637 | 0.639 | 0.028 | 1.1591999 | 35 | tuft1a |
| tmprss13b | 2.660212 | 0.269440593 | 0.131 | 0.001 | 6.7582692 | 35 | tmprss13b |
| FO704779.1 | 2.200794 | 0.32289284 | 0.148 | 0.001 | 5.5911184 | 35 | FO704779.1 |
| LOC110438236 | 5.895152 | 0.399570546 | 0.197 | 0.002 | 1.4976634 | 35 | LOC110438236 |
| bbox1 | 1.342540 | 0.670548574 | 0.426 | 0.013 | 3.4107248 | 35 | bbox1 |
| pik3c2g | 2.452357 | 0.261997041 | 0.164 | 0.001 | 6.2302141 | 35 | pik3c2g |
| krt915 | 2.533884 | 2.098218832 | 0.918 | 0.069 | 6.4373335 | 35 | krt91 |
| CABZ01056052.1 | 2.225669 | 0.268911971 | 0.148 | 0.001 | 5.6543139 | 35 | CABZ01056052.1 |
| si:dkey-30c15.2 | 2.632852 | 0.405041342 | 0.23 | 0.003 | 6.6887628 | 35 | si:dkey-30c15.2 |
| ildr1a | 1.802136 | 1.025163841 | 0.607 | 0.027 | 4.5783288 | 35 | ildr1a |
| si:ch211-153b23.4 | 2.406528 | 0.292537015 | 0.115 | 0.001 | 6.1137845 | 35 | si:ch211-153b23.4 |
| limk21 | 4.303929 | 0.801971402 | 0.475 | 0.017 | 1.0934134 | 35 | limk2 |
| rhpn22 | 2.426907 | 1.094551842 | 0.656 | 0.033 | 6.1655587 | 35 | rhpn2 |
| nansb1 | 3.100217 | 0.279350776 | 0.148 | 0.001 | 7.8761015 | 35 | nansb |
| si:ch211-157c3.44 | 6.059263 | 1.778538624 | 0.902 | 0.073 | 1.5393558 | 35 | si:ch211-157c3.4 |
| rassf10a | 8.930429 | 0.315309103 | 0.18 | 0.002 | 2.2687756 | 35 | rassf10a |
| rnh1 | 5.039641 | 0.37455804 | 0.213 | 0.003 | 1.2803209 | 35 | rnh1 |
| spint22 | 1.178145 | 1.574435962 | 0.885 | 0.067 | 2.9930781 | 35 | spint2 |
| arrdc1a | 3.281892 | 0.813239989 | 0.541 | 0.023 | 8.3376471 | 35 | arrdc1a |
| si:dkey-262k9.23 | 1.030956 | 1.044505663 | 0.623 | 0.031 | 2.6191442 | 35 | si:dkey-262k9.2 |
| trpm4b.2 | 1.174681 | 0.28274938 | 0.148 | 0.001 | 2.9842773 | 35 | trpm4b.2 |
| xkr8.2 | 2.085685 | 0.427928199 | 0.197 | 0.003 | 5.2986835 | 35 | xkr8.2 |
| homer2 | 3.175323 | 0.673925377 | 0.377 | 0.011 | 8.0669087 | 35 | homer2 |
| si:ch1073-385f13.3 | 4.840945 | 0.533649189 | 0.262 | 0.005 | 1.2298422 | 35 | si:ch1073-385f13.3 |
| cd9b4 | 5.225601 | 1.58848024 | 0.82 | 0.058 | 1.3275640 | 35 | cd9b |

| | | | | | | | |
|---|---|---|---|---|---|---|---|
| zgc:112433 | 3.219275 | 0.322509099 | 0.164 | 0.002 | 8.1785689 | 35 | zgc:112433 |
| SEMA4F | 1.333359 | 0.402383807 | 0.246 | 0.005 | 3.3873996 | 35 | SEMA4F |
| CT027980.11 | 4.534284 | 0.384253156 | 0.197 | 0.003 | 1.1519350 | 35 | CT027980.1 |
| trib1 | 4.590639 | 0.454662785 | 0.246 | 0.005 | 1.1662518 | 35 | trib1 |
| jupa3 | 5.193026 | 1.311648999 | 0.803 | 0.058 | 1.3192884 | 35 | jupa |
| wu:fi04e122 | 4.718710 | 0.903158183 | 0.492 | 0.021 | 1.1987883 | 35 | wu:fi04e12 |
| LO018605.2 | 1.861584 | 0.275681009 | 0.131 | 0.001 | 4.7293546 | 35 | LO018605.2 |
| ch25hl1.1 | 7.193556 | 0.291583676 | 0.197 | 0.003 | 1.8275231 | 35 | ch25hl1.1 |
| plekhf11 | 8.262082 | 0.538107118 | 0.246 | 0.005 | 2.0989820 | 35 | plekhf1 |
| tlcd2 | 3.944919 | 0.547461503 | 0.311 | 0.008 | 1.0022068 | 35 | tlcd2 |
| arhgef54 | 1.112661 | 0.936836034 | 0.59 | 0.031 | 2.8267178 | 35 | arhgef5 |
| iqsec2b | 1.670076 | 0.357613989 | 0.197 | 0.003 | 4.2428287 | 35 | iqsec2b |
| cx34.4 | 6.604975 | 0.355336389 | 0.23 | 0.004 | 1.6779939 | 35 | cx34.4 |
| gdpd3a1 | 7.476869 | 0.328096678 | 0.197 | 0.003 | 1.8994985 | 35 | gdpd3a |
| myh14 | 1.480770 | 0.259047178 | 0.131 | 0.001 | 3.7618983 | 35 | myh14 |
| casz12 | 4.391532 | 0.777257863 | 0.459 | 0.019 | 1.1156688 | 35 | casz1 |
| si:ch211-161h7.5 | 3.552643 | 0.277048878 | 0.115 | 0.001 | 9.0254907 | 35 | si:ch211-161h7.5 |
| CABZ01072077.1 | 1.446745 | 0.308380523 | 0.148 | 0.002 | 3.6754558 | 35 | CABZ01072077.1 |
| ms4a17a.51 | 4.662465 | 0.42863199 | 0.262 | 0.006 | 1.1844992 | 35 | ms4a17a.5 |
| si:dkey-204f11.641 | 5.180791 | 0.863458867 | 0.557 | 0.03 | 1.3161800 | 35 | si:dkey-204f11.64 |
| epdl12 | 1.688327 | 0.979346512 | 0.574 | 0.032 | 4.2891959 | 35 | epdl1 |
| plekhn1 | 9.989016 | 0.302382754 | 0.164 | 0.002 | 2.5377095 | 35 | plekhn1 |
| zgc:158263 | 2.239776 | 0.260465699 | 0.164 | 0.002 | 5.6901533 | 35 | zgc:158263 |
| ehf | 1.900399 | 0.293918843 | 0.148 | 0.002 | 4.8279643 | 35 | ehf |
| si:ch73-204p21.2 | 4.218475 | 0.991578111 | 0.639 | 0.039 | 1.0717037 | 35 | si:ch73-204p21.2 |
| slc2a1b2 | 3.640025 | 0.742188034 | 0.426 | 0.018 | 9.2474844 | 35 | slc2a1b |
| tcnbb2 | 2.564796 | 0.967362333 | 0.426 | 0.018 | 6.5158655 | 35 | tcnbb |
| cast4 | 4.304458 | 1.220994413 | 0.82 | 0.071 | 1.0935477 | 35 | cast |
| zgc:162193 | 7.026522 | 0.36954583 | 0.213 | 0.004 | 1.7850880 | 35 | zgc:162193 |
| atp8b1 | 8.150942 | 0.368637952 | 0.23 | 0.005 | 2.0707468 | 35 | atp8b1 |
| net13 | 1.960000 | 0.91088677 | 0.541 | 0.031 | 4.9793805 | 35 | net1 |
| si:dkey-157l19.21 | 3.398691 | 0.507785913 | 0.295 | 0.009 | 8.6343747 | 35 | si:dkey-157l19.2 |
| zgc:154054 | 7.432540 | 0.429683437 | 0.262 | 0.007 | 1.8882368 | 35 | zgc:154054 |
| si:dkey-38p12.3 | 8.644884 | 0.404580916 | 0.246 | 0.006 | 2.1962328 | 35 | si:dkey-38p12.3 |
| klf35 | 1.010465 | 0.987480862 | 0.59 | 0.038 | 2.5670876 | 35 | klf3 |

| | | | | | | | |
|---|---|---|---|---|---|---|---|
| kremen13 | 5.050961 | 0.543974585 | 0.328 | 0.011 | 1.2831967 | 35 | kremen1 |
| gna15.1 | 1.646084 | 0.377647268 | 0.18 | 0.003 | 4.1818787 | 35 | gna15.1 |
| krt222 | 1.874068 | 0.272067888 | 0.148 | 0.002 | 4.7610704 | 35 | krt222 |
| CABZ01039859.1 | 2.387482 | 0.265654812 | 0.148 | 0.002 | 6.0654005 | 35 | CABZ01039859.1 |
| cdh14 | 7.822938 | 1.123172559 | 0.705 | 0.054 | 1.9874175 | 35 | cdh1 |
| si:dkey-33i11.94 | 3.112116 | 0.803103349 | 0.475 | 0.024 | 7.9063330 | 35 | si:dkey-33i11.9 |
| plcd1a | 4.378357 | 0.388226186 | 0.213 | 0.005 | 1.1123216 | 35 | plcd1a |
| si:ch211-264f5.62 | 1.111887 | 0.564617057 | 0.344 | 0.013 | 2.8247514 | 35 | si:ch211-264f5.6 |
| tnfrsf11b1 | 1.602382 | 0.628763425 | 0.23 | 0.006 | 4.0708520 | 35 | tnfrsf11b |
| gsnb1 | 1.651244 | 0.281098454 | 0.148 | 0.002 | 4.1949860 | 35 | gsnb |
| osbpl7 | 3.867110 | 0.841468784 | 0.525 | 0.03 | 9.8243937 | 35 | osbpl7 |
| xiap | 7.123718 | 0.994831146 | 0.607 | 0.042 | 1.8097805 | 35 | xiap |
| dnmbp | 7.570251 | 0.643169859 | 0.377 | 0.016 | 1.9232223 | 35 | dnmbp |
| myo1ea | 1.266734 | 0.408375287 | 0.197 | 0.004 | 3.2181387 | 35 | myo1ea |
| kras | 2.432700 | 0.980696296 | 0.607 | 0.043 | 6.1802748 | 35 | kras |
| si:ch211-158d24.4 | 1.096343 | 0.433231505 | 0.18 | 0.003 | 2.7852595 | 35 | si:ch211-158d24.4 |
| si:ch211-217k17.10 | 1.325658 | 0.455592205 | 0.197 | 0.004 | 3.3678364 | 35 | si:ch211-217k17.10 |
| chmp4c | 6.976072 | 0.720556422 | 0.475 | 0.025 | 1.7722713 | 35 | chmp4c |
| ptpn131 | 1.072855 | 0.580172112 | 0.377 | 0.016 | 2.7255896 | 35 | ptpn13 |
| FP017215.11 | 1.555613 | 0.586662132 | 0.344 | 0.014 | 3.9520358 | 35 | FP017215.1 |
| dlx3b3 | 1.521802 | 0.463384299 | 0.262 | 0.008 | 3.8661396 | 35 | dlx3b |
| tead3b | 2.051575 | 0.744560892 | 0.41 | 0.02 | 5.2120272 | 35 | tead3b |
| fam160a1a | 1.531402 | 0.316889457 | 0.197 | 0.004 | 3.8905268 | 35 | fam160a1a |
| zbtb16b2 | 1.226667 | 0.745459926 | 0.41 | 0.021 | 3.1163499 | 35 | zbtb16b |
| mpp6a | 1.146622 | 0.565945493 | 0.344 | 0.015 | 2.9129948 | 35 | mpp6a |
| zgc:113279 | 2.434356 | 0.32978894 | 0.131 | 0.002 | 6.1844837 | 35 | zgc:113279 |
| spint1a1 | 4.079386 | 0.937938808 | 0.639 | 0.05 | 1.0363682 | 35 | spint1a |
| si:dkey-87o1.23 | 7.854046 | 1.190404442 | 0.738 | 0.068 | 1.9953204 | 35 | si:dkey-87o1.2 |
| samd10a | 1.296062 | 0.405010743 | 0.246 | 0.007 | 3.2926464 | 35 | samd10a |
| hopx6 | 2.838969 | 1.163785952 | 0.689 | 0.064 | 7.2124012 | 35 | hopx |
| bicdl2 | 7.968371 | 0.540721093 | 0.328 | 0.014 | 2.0243646 | 35 | bicdl2 |
| ssh2a | 8.300170 | 0.782580461 | 0.475 | 0.029 | 2.1086583 | 35 | ssh2a |
| paqr6 | 7.864664 | 0.629218234 | 0.361 | 0.017 | 1.9980179 | 35 | paqr6 |
| myo5b | 7.999198 | 0.28938118 | 0.164 | 0.003 | 2.0321962 | 35 | myo5b |
| btr25 | 3.153056 | 0.423911752 | 0.23 | 0.007 | 8.0103402 | 35 | btr25 |

| | | | | | | | |
|---|---|---|---|---|---|---|---|
| ugt8 | 6.807829 | 0.267809907 | 0.131 | 0.002 | 1.7295290 | 35 | ugt8 |
| tiam1a1 | 8.452489 | 0.478532207 | 0.279 | 0.01 | 2.1473550 | 35 | tiam1a |
| zgc:92481 | 2.338414 | 0.625872215 | 0.41 | 0.023 | 5.9407427 | 35 | zgc:92481 |
| lmbr1l1 | 1.307485 | 0.566834865 | 0.344 | 0.016 | 3.3216679 | 35 | lmbr1l |
| si:ch211-152p11.42 | 6.197645 | 0.775030671 | 0.443 | 0.027 | 1.5745117 | 35 | si:ch211-152p11.4 |
| usp2b | 1.277785 | 0.350884944 | 0.18 | 0.004 | 3.2462138 | 35 | usp2b |
| si:ch73-359m17.51 | 7.219273 | 0.570997146 | 0.197 | 0.005 | 1.8340564 | 35 | si:ch73-359m17.5 |
| CT027702.1 | 1.063240 | 0.299656363 | 0.148 | 0.003 | 2.7011616 | 35 | CT027702.1 |
| pard6b | 1.746449 | 0.417345351 | 0.246 | 0.008 | 4.4368554 | 35 | pard6b |
| zgc:1532845 | 1.932180 | 1.269113663 | 0.77 | 0.083 | 4.9087045 | 35 | zgc:153284 |
| galnt7 | 2.797683 | 0.831313456 | 0.525 | 0.039 | 7.1075145 | 35 | galnt7 |
| nfkbiz | 9.782826 | 0.585910789 | 0.41 | 0.023 | 2.4853269 | 35 | nfkbiz |
| cfl1l6 | 4.623516 | 1.543255942 | 0.902 | 0.135 | 1.1746044 | 35 | cfl1l |
| eppk13 | 5.195940 | 0.904856335 | 0.541 | 0.042 | 1.3200286 | 35 | eppk1 |
| rab11al | 5.101064 | 0.401594019 | 0.246 | 0.009 | 1.2959254 | 35 | rab11al |
| ptgs2a1 | 2.685911 | 0.559829872 | 0.279 | 0.011 | 6.8235571 | 35 | ptgs2a |
| mgst1.24 | 7.534784 | 1.319593872 | 0.836 | 0.115 | 1.9142120 | 35 | mgst1.2 |
| si:ch211-285f17.1 | 4.859260 | 0.283553283 | 0.164 | 0.004 | 1.2344952 | 35 | si:ch211-285f17.1 |
| si:dkey-125i10.3 | 7.735209 | 0.28428866 | 0.148 | 0.003 | 1.9651299 | 35 | si:dkey-125i10.3 |
| b4galnt1a | 8.875803 | 0.386231768 | 0.213 | 0.007 | 2.2548977 | 35 | b4galnt1a |
| si:ch211-166a6.53 | 1.119126 | 0.770893958 | 0.492 | 0.037 | 2.8431411 | 35 | si:ch211-166a6.5 |
| alox121 | 4.320084 | 0.46483006 | 0.295 | 0.013 | 1.0975173 | 35 | alox12 |
| ptpn212 | 7.854518 | 0.839091438 | 0.475 | 0.036 | 1.9954403 | 35 | ptpn21 |
| dusp56 | 2.964520 | 1.091193136 | 0.639 | 0.067 | 7.5313642 | 35 | dusp5 |
| fam169ab | 3.177429 | 0.287144858 | 0.164 | 0.004 | 8.0722599 | 35 | fam169ab |
| zmp:0000001114 | 8.137701 | 0.396448199 | 0.246 | 0.01 | 2.0673829 | 35 | zmp:0000001114 |
| kdm6ba1 | 1.499436 | 0.899514099 | 0.541 | 0.048 | 3.8093179 | 35 | kdm6ba |
| ccng25 | 2.010554 | 1.096718694 | 0.705 | 0.083 | 5.1078148 | 35 | ccng2 |
| prkcdb | 1.177513 | 0.539772017 | 0.295 | 0.014 | 2.9914736 | 35 | prkcdb |
| ms4a17a.41 | 6.728105 | 0.265195358 | 0.148 | 0.003 | 1.7092752 | 35 | ms4a17a.4 |
| rhov | 1.411052 | 0.411242355 | 0.262 | 0.012 | 3.5847781 | 35 | rhov |
| si:dkey-16l2.202 | 1.420829 | 0.623556713 | 0.361 | 0.022 | 3.6096163 | 35 | si:dkey-16l2.20 |
| ngef | 1.513916 | 0.275261895 | 0.148 | 0.003 | 3.8461059 | 35 | ngef |
| apnl1 | 1.555065 | 0.28034236 | 0.148 | 0.003 | 3.9506431 | 35 | apnl |
| cavin2a1 | 4.123845 | 0.509701918 | 0.311 | 0.016 | 1.0476630 | 35 | cavin2a |

| | | | | | | | |
|---|---|---|---|---|---|---|---|
| ptgis2 | 5.601548 | 0.583489527 | 0.295 | 0.015 | 1.4230733 | 35 | ptgis |
| pld2 | 1.530445 | 0.287734398 | 0.18 | 0.005 | 3.8880979 | 35 | pld2 |
| pdlim2 | 1.714471 | 0.378387153 | 0.23 | 0.009 | 4.3556150 | 35 | pdlim2 |
| ripk4 | 3.332012 | 0.296680501 | 0.197 | 0.007 | 8.4649780 | 35 | ripk4 |
| si:ch211-221f10.2 | 9.842533 | 0.298475052 | 0.18 | 0.006 | 2.5004956 | 35 | si:ch211-221f10.2 |
| ahnak10 | 1.518913 | 1.264474712 | 0.852 | 0.135 | 3.8588002 | 35 | ahnak |
| igf2bp2b | 9.372409 | 0.520867561 | 0.377 | 0.025 | 2.3810607 | 35 | igf2bp2b |
| ftr36 | 1.333746 | 0.397134447 | 0.18 | 0.006 | 3.3883835 | 35 | ftr36 |
| si:dkey-30g5.1 | 9.655977 | 0.376325174 | 0.213 | 0.008 | 2.4531011 | 35 | si:dkey-30g5.1 |
| tob1b4 | 3.400309 | 1.359626309 | 0.902 | 0.166 | 8.6384860 | 35 | tob1b |
| cst14b.14 | 5.777818 | 0.904323434 | 0.574 | 0.061 | 1.4678547 | 35 | cst14b.1 |
| cdaa3 | 1.568824 | 0.680395936 | 0.426 | 0.033 | 3.9855998 | 35 | cdaa |
| si:ch1073-345a8.1 | 2.064938 | 0.36894887 | 0.246 | 0.011 | 5.2459763 | 35 | si:ch1073-345a8.1 |
| fuca1.1 | 7.517319 | 0.785715914 | 0.475 | 0.042 | 1.9097749 | 35 | fuca1.1 |
| osbpl3a1 | 1.049663 | 0.562480447 | 0.344 | 0.022 | 2.6666689 | 35 | osbpl3a |
| nipa2 | 5.789726 | 0.800126389 | 0.508 | 0.048 | 1.4708799 | 35 | nipa2 |
| anxa1a8 | 1.453346 | 1.421100236 | 0.869 | 0.161 | 3.6922273 | 35 | anxa1a |
| sqstm11 | 2.467469 | 0.980813555 | 0.639 | 0.078 | 6.2686067 | 35 | sqstm1 |
| fam43a1 | 2.500303 | 0.46403488 | 0.246 | 0.012 | 6.3520212 | 35 | fam43a |
| c1galt1b | 3.089396 | 0.712244198 | 0.459 | 0.04 | 7.8486108 | 35 | c1galt1b |
| grhl13 | 3.196987 | 0.669987929 | 0.393 | 0.03 | 8.1219478 | 35 | grhl1 |
| cpeb4b | 4.417895 | 0.710036208 | 0.492 | 0.046 | 1.1223663 | 35 | cpeb4b |
| lmna2 | 6.954406 | 0.444409348 | 0.262 | 0.013 | 1.7667669 | 35 | lmna |
| s100v210 | 2.463627 | 1.296499825 | 0.803 | 0.139 | 6.2588448 | 35 | s100v2 |
| si:ch73-141c7.1 | 2.659581 | 0.257384175 | 0.18 | 0.006 | 6.7566676 | 35 | si:ch73-141c7.1 |
| klc1b | 4.682150 | 0.411554016 | 0.279 | 0.015 | 1.1895003 | 35 | klc1b |
| s100v1 | 7.862184 | 0.489301419 | 0.246 | 0.012 | 1.9973878 | 35 | s100v1 |
| midn10 | 8.692473 | 1.362510398 | 0.803 | 0.144 | 2.2083229 | 35 | midn |
| fam83hb | 3.104203 | 0.322203711 | 0.197 | 0.008 | 7.8862292 | 35 | fam83hb |
| hsd11b1la | 4.056229 | 0.456997419 | 0.262 | 0.014 | 1.0304849 | 35 | hsd11b1la |
| cpeb2 | 8.682870 | 0.402223332 | 0.18 | 0.007 | 2.2058832 | 35 | cpeb2 |
| prodha1 | 1.446058 | 0.396127978 | 0.262 | 0.014 | 3.6737103 | 35 | prodha |
| pnp5a6 | 2.774007 | 0.994228704 | 0.607 | 0.074 | 7.0473652 | 35 | pnp5a |
| cdc42ep51 | 1.202053 | 0.694587772 | 0.41 | 0.034 | 3.0538168 | 35 | cdc42ep5 |
| ldlrb | 8.118950 | 0.287229607 | 0.164 | 0.006 | 2.0626192 | 35 | ldlrb |

| | | | | | | | |
|---|---|---|---|---|---|---|---|
| cx30.9 | 1.040489 | 0.276040665 | 0.164 | 0.006 | 2.6433625 | 35 | cx30.9 |
| ifngr1 | 2.220149 | 0.277888018 | 0.164 | 0.006 | 5.6402902 | 35 | ifngr1 |
| tjp3 | 2.235059 | 0.521802608 | 0.311 | 0.02 | 5.6781681 | 35 | tjp3 |
| XLOC-044191 | 2.276185 | 0.438994194 | 0.262 | 0.015 | 5.7826484 | 35 | XLOC-044191 |
| zgc:174164 | 7.182033 | 0.294566276 | 0.164 | 0.006 | 1.8245955 | 35 | zgc:174164 |
| slc7a3b1 | 8.980163 | 0.626104087 | 0.361 | 0.028 | 2.2814104 | 35 | slc7a3b |
| stk10 | 4.418584 | 0.659902333 | 0.459 | 0.044 | 1.1225414 | 35 | stk10 |
| cxadr1 | 4.900903 | 0.670511905 | 0.393 | 0.033 | 1.2450746 | 35 | cxadr |
| oclna3 | 7.799664 | 0.84529695 | 0.574 | 0.066 | 1.9815046 | 35 | oclna |
| hunk | 9.538854 | 0.259530001 | 0.148 | 0.005 | 2.4233460 | 35 | hunk |
| chmp1b3 | 1.235411 | 1.034671328 | 0.77 | 0.131 | 3.1385619 | 35 | chmp1b |
| LO017965.11 | 1.865080 | 0.255489581 | 0.131 | 0.004 | 4.7382359 | 35 | LO017965.1 |
| si:ch211-202f5.36 | 2.178901 | 0.935120937 | 0.607 | 0.075 | 5.5354995 | 35 | si:ch211-202f5.3 |
| BX664618.1 | 2.999579 | 0.305815823 | 0.18 | 0.007 | 7.6204315 | 35 | BX664618.1 |
| cbx3b | 1.148013 | 0.623925871 | 0.377 | 0.031 | 2.9165281 | 35 | cbx3b |
| bnipl1 | 3.656340 | 0.307686951 | 0.164 | 0.006 | 9.2889331 | 35 | bnipl |
| XLOC-0418321 | 3.924343 | 0.414353975 | 0.197 | 0.009 | 9.9697951 | 35 | XLOC-041832 |
| rnf128a2 | 8.516719 | 0.689288476 | 0.541 | 0.061 | 2.1636725 | 35 | rnf128a |
| si:busm1-57f23.12 | 1.272922 | 0.602114916 | 0.361 | 0.029 | 3.2338596 | 35 | si:busm1-57f23.1 |
| icn9 | 1.979087 | 1.512722887 | 0.967 | 0.267 | 5.0278724 | 35 | icn |
| vsig10 | 2.425084 | 0.462299934 | 0.295 | 0.02 | 6.1609268 | 35 | vsig10 |
| zgc:113625 | 3.549987 | 0.330268297 | 0.164 | 0.006 | 9.0187430 | 35 | zgc:113625 |
| slc38a5a | 5.995626 | 0.481830806 | 0.279 | 0.018 | 1.5231889 | 35 | slc38a5a |
| serpinb141 | 6.378465 | 0.434775133 | 0.246 | 0.014 | 1.6204491 | 35 | serpinb14 |
| tnfaip37 | 9.673015 | 0.817856023 | 0.508 | 0.059 | 2.4574296 | 35 | tnfaip3 |
| zgc:774861 | 1.099822 | 0.830826745 | 0.541 | 0.066 | 2.7940982 | 35 | zgc:77486 |
| mboat1 | 3.511806 | 0.294969758 | 0.148 | 0.005 | 8.9217432 | 35 | mboat1 |
| ckap46 | 4.962179 | 0.769668209 | 0.508 | 0.06 | 1.2606417 | 35 | ckap4 |
| sh3d21 | 5.956516 | 0.630408776 | 0.393 | 0.035 | 1.5132530 | 35 | sh3d21 |
| zfand5a1 | 1.045253 | 0.881087945 | 0.574 | 0.078 | 2.6554658 | 35 | zfand5a |
| ptprfb3 | 2.059607 | 0.445021257 | 0.262 | 0.017 | 5.2324326 | 35 | ptprfb |
| chac17 | 2.386311 | 1.412058537 | 0.984 | 0.377 | 6.0624247 | 35 | chac1 |
| ovol1a | 7.529331 | 0.332888743 | 0.148 | 0.005 | 1.9128266 | 35 | ovol1a |
| pleca5 | 1.372906 | 0.829206963 | 0.508 | 0.061 | 3.4878700 | 35 | pleca |
| itprid2 | 1.915384 | 0.368186854 | 0.246 | 0.015 | 4.8660336 | 35 | itprid2 |

| | | | | | | | |
|---|---|---|---|---|---|---|---|
| FAM83G | 2.625405 | 0.389967429 | 0.213 | 0.012 | 6.6698434 | 35 | FAM83G |
| cd994 | 5.655098 | 0.729637778 | 0.459 | 0.052 | 1.4366778 | 35 | cd99 |
| gfpt2 | 1.067965 | 0.316241244 | 0.18 | 0.008 | 2.7131660 | 35 | gfpt2 |
| lipia | 1.108879 | 0.428537732 | 0.279 | 0.019 | 2.8171080 | 35 | lipia |
| slc20a1b2 | 1.464339 | 0.875745067 | 0.623 | 0.094 | 3.7201551 | 35 | slc20a1b |
| tdh2 | 3.231098 | 0.895995163 | 0.607 | 0.091 | 8.2086052 | 35 | tdh |
| nt5c2b3 | 3.536989 | 0.703693131 | 0.426 | 0.046 | 8.9857223 | 35 | nt5c2b |
| anxa2a9 | 3.803797 | 0.984113288 | 0.705 | 0.117 | 9.6635473 | 35 | anxa2a |
| hlfa | 1.236580 | 0.314780368 | 0.213 | 0.012 | 3.1415328 | 35 | hlfa |
| baiap2l1a1 | 1.338636 | 0.446350121 | 0.377 | 0.035 | 3.4008051 | 35 | baiap2l1a |
| st14a | 1.746202 | 0.48379923 | 0.361 | 0.032 | 4.4362264 | 35 | st14a |
| zgc:1533175 | 2.064723 | 0.583291038 | 0.393 | 0.039 | 5.2454291 | 35 | zgc:153317 |
| ier2b14 | 6.596009 | 1.306637581 | 0.852 | 0.236 | 1.6757161 | 35 | ier2b |
| snphb | 2.887459 | 0.561396105 | 0.328 | 0.029 | 7.3355913 | 35 | snphb |
| esrp2 | 3.197484 | 0.308368692 | 0.164 | 0.007 | 8.1232095 | 35 | esrp2 |
| si:dkey-105i14.1 | 3.248268 | 0.258822167 | 0.131 | 0.005 | 8.2522260 | 35 | si:dkey-105i14.1 |
| si:ch73-335l21.1 | 5.381799 | 0.436331987 | 0.23 | 0.015 | 1.3672461 | 35 | si:ch73-335l21.1 |
| ezrb | 8.525329 | 0.798398691 | 0.525 | 0.071 | 2.1658599 | 35 | ezrb |
| zgc:153759 | 1.196790 | 0.294386221 | 0.164 | 0.007 | 3.0404471 | 35 | zgc:153759 |
| sh3d19 | 1.433468 | 0.367437926 | 0.213 | 0.012 | 3.6417264 | 35 | sh3d19 |
| btbd10b | 9.170715 | 0.752896245 | 0.541 | 0.076 | 2.3298203 | 35 | btbd10b |
| akap9 | 1.136348 | 0.340589646 | 0.18 | 0.009 | 2.8868930 | 35 | akap9 |
| efna1b4 | 1.137498 | 0.449617006 | 0.262 | 0.019 | 2.8898145 | 35 | efna1b |
| selenow17 | 1.651867 | 1.209249541 | 0.951 | 0.415 | 4.1965693 | 35 | selenow1 |
| bcl6ab | 4.541284 | 0.440111927 | 0.23 | 0.015 | 1.1537133 | 35 | bcl6ab |
| epcam8 | 5.925386 | 1.210928875 | 0.984 | 0.421 | 1.5053443 | 35 | epcam |
| sorl1 | 8.358600 | 0.456422892 | 0.279 | 0.022 | 2.1235023 | 35 | sorl1 |
| spns21 | 2.651511 | 0.33949005 | 0.164 | 0.008 | 6.7361649 | 35 | spns2 |
| etnk11 | 3.947916 | 0.618271239 | 0.361 | 0.037 | 1.0029682 | 35 | etnk1 |
| chmp2bb1 | 5.374927 | 0.650929396 | 0.492 | 0.066 | 1.3655002 | 35 | chmp2bb |
| tmem593 | 6.732122 | 0.880782746 | 0.672 | 0.126 | 1.7102956 | 35 | tmem59 |
| b2ml12 | 7.347410 | 1.149855011 | 0.918 | 0.293 | 1.8666096 | 35 | b2ml |
| scpp73 | 1.161335 | 0.455331751 | 0.279 | 0.022 | 2.9503738 | 35 | scpp7 |
| fosb8 | 2.264674 | 1.161990946 | 0.639 | 0.121 | 5.7534051 | 35 | fosb |
| kalrnb1 | 3.049656 | 0.3088042 | 0.18 | 0.01 | 7.7476534 | 35 | kalrnb |

| | | | | | | | |
|---|---|---|---|---|---|---|---|
| mhc2dab6 | 5.088707 | 0.908547224 | 0.59 | 0.094 | 1.2927860 | 35 | mhc2dab |
| acaca | 5.67124( | 0.308158553 | 0.18 | 0.01 | 1.4407785 | 35 | acaca |
| nfe2l2a3 | 7.88535( | 0.651274958 | 0.492 | 0.068 | 2.0032733 | 35 | nfe2l2a |
| tiparp | 1.817094 | 0.611118835 | 0.377 | 0.042 | 4.6163282 | 35 | tiparp |
| b2m12 | 2.940682 | 1.224415844 | 0.902 | 0.298 | 7.4708039 | 35 | b2m |
| sh3yl1 | 3.319195 | 0.528605804 | 0.311 | 0.029 | 8.4324173 | 35 | sh3yl1 |
| tmem79b | 3.664349 | 0.362099256 | 0.197 | 0.012 | 9.3092808 | 35 | tmem79b |
| epha2a | 3.697351 | 0.35738483 | 0.164 | 0.008 | 9.3931207 | 35 | epha2a |
| dhrs13a.11 | 4.170453 | 0.440295096 | 0.279 | 0.024 | 1.0595037 | 35 | dhrs13a.1 |
| gnb1a5 | 5.110793 | 1.029261591 | 0.803 | 0.202 | 1.2983971 | 35 | gnb1a |
| uba7 | 5.689439 | 0.257690955 | 0.148 | 0.007 | 1.4454021 | 35 | uba7 |
| arid3a | 1.747924 | 0.279676739 | 0.18 | 0.01 | 4.4406019 | 35 | arid3a |
| si:ch1073-429i10.3.1 | 1.993222 | 1.020927877 | 0.852 | 0.226 | 5.0637820 | 35 | si:ch1073-429i10.3.1 |
| reep3b2 | 2.083519 | 0.673940216 | 0.475 | 0.066 | 5.2931802 | 35 | reep3b |
| csrp2 | 2.160003 | 0.327691668 | 0.213 | 0.014 | 5.4874894 | 35 | csrp2 |
| creb3l11 | 3.830711 | 0.251863004 | 0.148 | 0.007 | 9.7319236 | 35 | creb3l1 |
| zbtb7b | 5.481648 | 0.27171688 | 0.164 | 0.009 | 1.3926128 | 35 | zbtb7b |
| cldn7b7 | 6.758458 | 0.944053498 | 0.689 | 0.133 | 1.7169864 | 35 | cldn7b |
| jdp2b8 | 1.803259 | 1.258924611 | 0.689 | 0.164 | 4.5811801 | 35 | jdp2b |
| nfkbiaa7 | 2.08764( | 1.037232345 | 0.705 | 0.151 | 5.3036510 | 35 | nfkbiaa |
| adrb2a1 | 2.38851( | 0.376366706 | 0.246 | 0.02 | 6.0680100 | 35 | adrb2a |
| zmp:0000000991 | 2.882214 | 0.369344836 | 0.197 | 0.013 | 7.3222655 | 35 | zmp:0000000991 |
| ap1m3 | 2.90149( | 0.252010069 | 0.148 | 0.007 | 7.3712356 | 35 | ap1m3 |
| tmsb19 | 2.938853 | 1.206405447 | 0.689 | 0.16 | 7.4661580 | 35 | tmsb1 |
| crema10 | 3.13296( | 0.954401234 | 0.754 | 0.173 | 7.9592850 | 35 | crema |
| daam1a1 | 3.367652 | 0.455794432 | 0.311 | 0.031 | 8.5555210 | 35 | daam1a |
| errfi1a1 | 9.098414 | 0.402226371 | 0.246 | 0.02 | 2.3114522 | 35 | errfi1a |
| si:ch211-251p5.5 | 9.490164 | 0.511635182 | 0.295 | 0.029 | 2.4109764 | 35 | si:ch211-251p5.5 |
| evi5b | 1.076613 | 0.502624576 | 0.295 | 0.029 | 2.7351363 | 35 | evi5b |
| adipor21 | 1.296368 | 0.734103063 | 0.525 | 0.084 | 3.2934231 | 35 | adipor2 |
| fosl1a7 | 1.495169 | 0.841491235 | 0.508 | 0.083 | 3.7984778 | 35 | fosl1a |
| rnd3a4 | 1.610556 | 0.5147739 | 0.328 | 0.035 | 4.0916185 | 35 | rnd3a |
| erbb24 | 2.479598 | 0.537511155 | 0.311 | 0.033 | 6.2994198 | 35 | erbb2 |
| trib34 | 6.642124 | 0.857746571 | 0.574 | 0.108 | 1.6874317 | 35 | trib3 |
| nrg1 | 6.914218 | 0.365107149 | 0.23 | 0.018 | 1.7565572 | 35 | nrg1 |

| | | | | | | | |
|---|---|---|---|---|---|---|---|
| sipa1l3 | 9.546038 | 0.376011134 | 0.23 | 0.018 | 2.4251711 | 35 | sipa1l3 |
| mgst1.13 | 1.236111 | 0.604397279 | 0.41 | 0.055 | 3.1403403 | 35 | mgst1.1 |
| CR847566.13 | 2.646590 | 0.559879685 | 0.393 | 0.05 | 6.7236622 | 35 | CR847566.1 |
| fxyd16 | 3.577497 | 0.702982617 | 0.508 | 0.083 | 9.0886335 | 35 | fxyd1 |
| tuba8l24 | 4.239514 | 0.634678566 | 0.459 | 0.068 | 1.0770487 | 35 | tuba8l2 |
| arg23 | 4.600607 | 0.380114723 | 0.23 | 0.019 | 1.1687842 | 35 | arg2 |
| si:ch73-138n13.13 | 7.362700 | 0.506113691 | 0.361 | 0.044 | 1.8704940 | 35 | si:ch73-138n13.1 |
| ankrd227 | 9.919425 | 0.75765684 | 0.656 | 0.121 | 2.5200301 | 35 | ankrd22 |
| slc37a1 | 1.779815 | 0.335971536 | 0.197 | 0.014 | 4.5216216 | 35 | slc37a1 |
| si:ch211-197g15.6 | 2.243397 | 0.310989822 | 0.164 | 0.01 | 5.6993507 | 35 | si:ch211-197g15.6 |
| crybg1a1 | 2.340186 | 0.358403554 | 0.23 | 0.019 | 5.9452427 | 35 | crybg1a |
| rasgef1ba2 | 3.124572 | 0.607022516 | 0.426 | 0.06 | 7.9379753 | 35 | rasgef1ba |
| mcl1b12 | 3.553634 | 1.024595979 | 0.836 | 0.25 | 9.0280077 | 35 | mcl1b |
| zgc:1527915 | 3.723108 | 0.589043398 | 0.459 | 0.067 | 9.4585574 | 35 | zgc:152791 |
| tjp1b | 4.091420 | 0.447038738 | 0.279 | 0.028 | 1.0394254 | 35 | tjp1b |
| si:ch73-22o12.12 | 5.148807 | 0.417722263 | 0.295 | 0.031 | 1.3080544 | 35 | si:ch73-22o12.1 |
| gpatch81 | 5.794087 | 0.640390561 | 0.426 | 0.063 | 1.4719879 | 35 | gpatch8 |
| rhoca2 | 6.501429 | 0.630180313 | 0.443 | 0.066 | 1.6516882 | 35 | rhoca |
| vamp82 | 8.838166 | 0.677505708 | 0.475 | 0.075 | 2.2453362 | 35 | vamp8 |
| ier56 | 1.069747 | 0.911204134 | 0.639 | 0.146 | 2.7176934 | 35 | ier5 |
| jag1b3 | 1.283483 | 0.452600845 | 0.279 | 0.028 | 3.2606889 | 35 | jag1b |
| f11r.17 | 1.806620 | 0.924476715 | 0.721 | 0.163 | 4.5897194 | 35 | f11r.1 |
| si:dkey-79d12.54 | 2.176486 | 0.669733316 | 0.443 | 0.068 | 5.5293633 | 35 | si:dkey-79d12.5 |
| porb1 | 2.466915 | 0.573274385 | 0.344 | 0.043 | 6.2671983 | 35 | porb |
| ccdc50 | 2.678529 | 0.445060588 | 0.279 | 0.028 | 6.8048042 | 35 | ccdc50 |
| chmp4bb1 | 3.551494 | 0.622825327 | 0.508 | 0.085 | 9.0225713 | 35 | chmp4bb |
| AL954191.12 | 4.371631 | 0.441701522 | 0.213 | 0.017 | 1.1106129 | 35 | AL954191.1 |
| ddit310 | 5.571133 | 1.034439109 | 0.689 | 0.178 | 1.4153463 | 35 | ddit3 |
| zbtb16a2 | 9.249441 | 0.546024369 | 0.344 | 0.043 | 2.3498205 | 35 | zbtb16a |
| chmp5b4 | 9.263397 | 0.726337194 | 0.623 | 0.128 | 2.3533660 | 35 | chmp5b |
| cishb | 1.403760 | 0.276305096 | 0.148 | 0.009 | 3.5662533 | 35 | cishb |
| si:dkey-85k7.71 | 1.450837 | 0.311584265 | 0.148 | 0.009 | 3.6858532 | 35 | si:dkey-85k7.7 |
| elovl7a7 | 1.728093 | 0.808587605 | 0.738 | 0.152 | 4.3902204 | 35 | elovl7a |
| si:ch73-335l21.48 | 2.638311 | 1.089719522 | 0.623 | 0.141 | 6.7026310 | 35 | si:ch73-335l21.4 |
| tent5ba4 | 3.556411 | 0.261798583 | 0.18 | 0.013 | 9.0350636 | 35 | tent5ba |

| | | | | | | | |
|---|---|---|---|---|---|---|---|
| CABZ01029938.1 | 3.902554 | 0.44943378 | 0.295 | 0.033 | 9.9144399 | 35 | CABZ01029938.1 |
| s100a10b12 | 5.936451 | 1.074346145 | 0.902 | 0.285 | 1.5081555 | 35 | s100a10b |
| gipc21 | 7.185094 | 0.438373154 | 0.295 | 0.032 | 1.8253732 | 35 | gipc2 |
| capns1a5 | 7.392955 | 0.905983951 | 0.656 | 0.158 | 1.8781803 | 35 | capns1a |
| abca3b | 9.827389 | 0.341080618 | 0.213 | 0.018 | 2.4966484 | 35 | abca3b |
| efnb2a1 | 1.194417 | 0.347880719 | 0.197 | 0.015 | 3.0344182 | 35 | efnb2a |
| cavin2b6 | 1.822765 | 0.60279362 | 0.475 | 0.078 | 4.6307348 | 35 | cavin2b |
| strbp2 | 2.242966 | 0.347409189 | 0.213 | 0.018 | 5.6982560 | 35 | strbp |
| zgc:1748634 | 3.443156 | 0.789163036 | 0.541 | 0.111 | 8.7473386 | 35 | zgc:174863 |
| zgc:113333 | 4.748432 | 0.276137794 | 0.148 | 0.009 | 1.2063392 | 35 | zgc:113333 |
| itpkb | 5.334164 | 0.329662617 | 0.197 | 0.016 | 1.3551444 | 35 | itpkb |
| si:ch1073-443f11.24 | 7.083204 | 0.404901374 | 0.328 | 0.041 | 1.7994881 | 35 | si:ch1073-443f11.2 |
| zgc:923801 | 9.492416 | 0.291983771 | 0.164 | 0.011 | 2.4115483 | 35 | zgc:92380 |
| zgc:92107 | 1.357475 | 0.311830957 | 0.18 | 0.014 | 3.4486676 | 35 | zgc:92107 |
| fuca22 | 1.623039 | 0.635296823 | 0.426 | 0.068 | 4.1233308 | 35 | fuca2 |
| mgst2 | 1.661490 | 0.370298256 | 0.213 | 0.019 | 4.2210158 | 35 | mgst2 |
| fam129ba1 | 1.665761 | 0.38214675 | 0.246 | 0.024 | 4.2318678 | 35 | fam129ba |
| stap2b | 2.686315 | 0.306308725 | 0.164 | 0.011 | 6.8245844 | 35 | stap2b |
| si:ch211-207d6.23 | 2.875447 | 0.34658882 | 0.23 | 0.021 | 7.3050750 | 35 | si:ch211-207d6.2 |
| jun11 | 3.357627 | 1.07644657 | 0.803 | 0.25 | 8.5300527 | 35 | jun |
| XLOC-018182 | 3.739950 | 0.367722943 | 0.18 | 0.014 | 9.5013443 | 35 | XLOC-018182 |
| b3gnt5a1 | 4.026074 | 0.447814338 | 0.295 | 0.035 | 1.0228241 | 35 | b3gnt5a |
| itsn2b | 4.536527 | 0.422917612 | 0.295 | 0.035 | 1.1525048 | 35 | itsn2b |
| stk352 | 4.674461 | 0.611820888 | 0.393 | 0.061 | 1.1875470 | 35 | stk35 |
| fkbp56 | 7.502763 | 0.726002321 | 0.59 | 0.127 | 1.9060770 | 35 | fkbp5 |
| ifi459 | 8.710910 | 0.78023416 | 0.639 | 0.139 | 2.2130069 | 35 | ifi45 |
| plpp2b | 1.015412 | 0.403961566 | 0.295 | 0.034 | 2.5796549 | 35 | plpp2b |
| pvrl2l | 1.114931 | 0.286867264 | 0.18 | 0.014 | 2.8324840 | 35 | pvrl2l |
| ucp213 | 1.186197 | 0.924348311 | 0.951 | 0.414 | 3.0135359 | 35 | ucp2 |
| zgc:174935 | 1.255288 | 0.385545768 | 0.246 | 0.025 | 3.1890602 | 35 | zgc:174935 |
| gadd45aa2 | 1.519623 | 0.349999899 | 0.213 | 0.019 | 3.8606025 | 35 | gadd45aa |
| si:ch211-286b5.59 | 2.016334 | 0.777712746 | 0.639 | 0.151 | 5.1224980 | 35 | si:ch211-286b5.5 |
| si:ch211-39i22.15 | 2.410713 | 0.734610565 | 0.475 | 0.088 | 6.1244175 | 35 | si:ch211-39i22.1 |
| cds1 | 2.678816 | 0.268323811 | 0.131 | 0.008 | 6.8055332 | 35 | cds1 |
| rabac1 | 3.206118 | 0.476710565 | 0.344 | 0.048 | 8.1451445 | 35 | rabac1 |

| | | | | | | | |
|---|---|---|---|---|---|---|---|
| myo6a | 3.802572 | 0.254432222 | 0.148 | 0.01 | 9.6604357 | 35 | myo6a |
| esyt2a | 6.449061 | 0.285403787 | 0.18 | 0.014 | 1.6383841 | 35 | esyt2a |
| csnk1da | 6.652040 | 0.68796831 | 0.557 | 0.114 | 1.6899509 | 35 | csnk1da |
| arfgap3 | 7.403407 | 0.377742804 | 0.23 | 0.023 | 1.8808357 | 35 | arfgap3 |
| mboat2a1 | 9.037690 | 0.386138452 | 0.246 | 0.026 | 2.2960252 | 35 | mboat2a |
| s100u1 | 1.897994 | 0.411433432 | 0.311 | 0.04 | 4.8218540 | 35 | s100u |
| CABZ01021599.1 | 2.730935 | 0.292959834 | 0.164 | 0.012 | 6.9379421 | 35 | CABZ01021599.1 |
| gabarapl22 | 3.694512 | 0.575510957 | 0.459 | 0.084 | 9.3859078 | 35 | gabarapl2 |
| dcun1d2b | 4.866940 | 0.290813092 | 0.18 | 0.015 | 1.2364463 | 35 | dcun1d2b |
| arl4d5 | 5.334544 | 0.491663958 | 0.361 | 0.053 | 1.3552409 | 35 | arl4d |
| gmds | 7.198319 | 0.48282175 | 0.344 | 0.05 | 1.8287329 | 35 | gmds |
| prdx511 | 1.313277 | 0.791926229 | 0.672 | 0.178 | 3.3363804 | 35 | prdx5 |
| fosab12 | 1.507263 | 1.04909877 | 0.918 | 0.361 | 3.8292024 | 35 | fosab |
| si:ch73-103b11.2 | 1.734377 | 0.330033406 | 0.213 | 0.021 | 4.4061860 | 35 | si:ch73-103b11.2 |
| ubb4 | 1.841427 | 1.047021658 | 0.803 | 0.325 | 4.6781471 | 35 | ubb |
| zyg11 | 2.337393 | 0.414731988 | 0.262 | 0.031 | 5.9381487 | 35 | zyg11 |
| plcb3 | 2.954926 | 0.326579913 | 0.213 | 0.021 | 7.5069914 | 35 | plcb3 |
| fut9b | 3.684584 | 0.360865412 | 0.213 | 0.021 | 9.3606875 | 35 | fut9b |
| rpz4 | 8.590255 | 0.326450529 | 0.23 | 0.024 | 2.1823543 | 35 | rpz4 |
| atxn1a | 1.079044 | 0.331114931 | 0.23 | 0.025 | 2.7413136 | 35 | atxn1a |
| atp1a1b | 1.143364 | 0.292324485 | 0.164 | 0.013 | 2.9047187 | 35 | atp1a1b |
| nmi | 1.800187 | 0.580029068 | 0.459 | 0.089 | 4.5733765 | 35 | nmi |
| waslb | 1.803134 | 0.390698028 | 0.262 | 0.031 | 4.5808643 | 35 | waslb |
| mpc11 | 2.197085 | 0.58841204 | 0.41 | 0.074 | 5.5816960 | 35 | mpc1 |
| jund8 | 2.601843 | 0.640433776 | 0.459 | 0.09 | 6.6099827 | 35 | jund |
| zdhhc16a | 2.909094 | 0.346107867 | 0.213 | 0.022 | 7.3905548 | 35 | zdhhc16a |
| krt156 | 3.287434 | 1.068641514 | 0.459 | 0.097 | 8.3517264 | 35 | krt15 |
| plekha62 | 3.771008 | 0.271743844 | 0.197 | 0.019 | 9.5802464 | 35 | plekha6 |
| cmah2 | 3.892260 | 0.279331637 | 0.18 | 0.016 | 9.8882875 | 35 | cmah |
| tmod21 | 3.982627 | 0.503746615 | 0.328 | 0.049 | 1.0117864 | 35 | tmod2 |
| slc38a5b8 | 6.734826 | 0.674031695 | 0.557 | 0.128 | 1.7109826 | 35 | slc38a5b |
| mcl1a9 | 1.018868 | 0.759181742 | 0.934 | 0.614 | 2.5884360 | 35 | mcl1a |
| atf4b12 | 1.091978 | 0.835663214 | 0.803 | 0.277 | 2.7741706 | 35 | atf4b |
| pim110 | 1.126554 | 0.795336876 | 0.639 | 0.169 | 2.8620127 | 35 | pim1 |
| CR318588.35 | 1.611200 | 0.733053449 | 0.328 | 0.051 | 4.0932547 | 35 | CR318588.3 |

| | | | | | | | |
|---|---|---|---|---|---|---|---|
| prelid3b4 | 1.687902 | 0.785266144 | 0.754 | 0.226 | 4.2881174 | 35 | prelid3b |
| cx30.3 | 2.014306 | 0.348374421 | 0.197 | 0.019 | 5.1173464 | 35 | cx30.3 |
| GGT7 | 2.390024 | 0.28417491 | 0.18 | 0.016 | 6.0718577 | 35 | GGT7 |
| jmjd1cb2 | 2.513824 | 0.472914469 | 0.311 | 0.046 | 6.3863701 | 35 | jmjd1cb |
| selenot21 | 3.374771 | 0.549647072 | 0.377 | 0.066 | 8.5736077 | 35 | selenot2 |
| crtc3 | 3.558080 | 0.257274026 | 0.18 | 0.017 | 9.0393047 | 35 | crtc3 |
| mark2b | 4.005909 | 0.345821257 | 0.213 | 0.023 | 1.0177014 | 35 | mark2b |
| btg211 | 4.416566 | 0.892933354 | 0.869 | 0.349 | 1.1220288 | 35 | btg2 |
| pdcd6ip | 5.493467 | 0.397995205 | 0.344 | 0.054 | 1.3956155 | 35 | pdcd6ip |
| zgc:123258 | 7.436817 | 0.515486561 | 0.328 | 0.052 | 1.8893235 | 35 | zgc:123258 |
| aste1a | 7.517484 | 0.264451344 | 0.197 | 0.02 | 1.9098169 | 35 | aste1a |
| ippk | 7.541112 | 0.262010516 | 0.18 | 0.017 | 1.9158197 | 35 | ippk |
| higd1a8 | 8.454351 | 0.849458087 | 0.803 | 0.303 | 2.1478279 | 35 | higd1a |
| adam28 | 1.088455 | 0.275646634 | 0.131 | 0.009 | 2.7652203 | 35 | adam28 |
| itpkcb3 | 1.188888 | 0.495689625 | 0.41 | 0.074 | 3.0203716 | 35 | itpkcb |
| vps4b1 | 1.317546 | 0.571725688 | 0.492 | 0.105 | 3.3472270 | 35 | vps4b |
| pycard9 | 1.385202 | 0.618859533 | 0.426 | 0.084 | 3.5191059 | 35 | pycard |
| per33 | 1.419183 | 0.573874272 | 0.41 | 0.078 | 3.6054355 | 35 | per3 |
| plecb4 | 2.171874 | 0.484917843 | 0.328 | 0.052 | 5.5176465 | 35 | plecb |
| tax1bp33 | 2.346605 | 0.603094716 | 0.475 | 0.101 | 5.9615519 | 35 | tax1bp3 |
| mt-co115 | 2.742968 | 0.716818919 | 0.984 | 0.64 | 6.9685104 | 35 | mt-co1 |
| cd82b2 | 3.027475 | 0.515043839 | 0.41 | 0.076 | 7.6913002 | 35 | cd82b |
| lgals3b5 | 3.247513 | 0.594392558 | 0.459 | 0.098 | 8.2503072 | 35 | lgals3b |
| smox6 | 5.487928 | 0.511978614 | 0.443 | 0.087 | 1.3942082 | 35 | smox |
| fxyd6l4 | 6.747315 | 0.599716971 | 0.525 | 0.117 | 1.7141556 | 35 | fxyd6l |
| ccdc1711 | 7.106079 | 0.326083229 | 0.197 | 0.021 | 1.8052993 | 35 | ccdc171 |
| reep59 | 7.108389 | 0.76526254 | 0.607 | 0.173 | 1.8058862 | 35 | reep5 |
| zgc:112148 | 9.180508 | 0.307722725 | 0.18 | 0.018 | 2.3323082 | 35 | zgc:112148 |
| degs26 | 9.365390 | 0.541809873 | 0.459 | 0.088 | 2.3792773 | 35 | degs2 |
| fgd1 | 1.029439 | 0.257264827 | 0.164 | 0.015 | 2.6152912 | 35 | fgd1 |
| prrg25 | 1.031456 | 0.412564938 | 0.426 | 0.077 | 2.6204156 | 35 | prrg2 |
| baiap2a | 1.178529 | 0.348342025 | 0.262 | 0.035 | 2.9940535 | 35 | baiap2a |
| exosc11 | 1.532023 | 0.410418698 | 0.295 | 0.045 | 3.8921052 | 35 | exosc1 |
| rab5ab | 1.544234 | 0.568980254 | 0.443 | 0.092 | 3.9231281 | 35 | rab5ab |
| ehbp1l1a | 1.963225 | 0.3004383 | 0.18 | 0.018 | 4.9875741 | 35 | ehbp1l1a |

| | | | | | | | |
|---|---|---|---|---|---|---|---|
| si:ch211-212k18.75 | 1.973276 | 0.702892067 | 0.574 | 0.146 | 5.0131101 | 35 | si:ch211-212k18.7 |
| kitlga1 | 2.143652 | 0.297820673 | 0.18 | 0.018 | 5.4459480 | 35 | kitlga |
| tjp1a5 | 2.409749 | 0.427450466 | 0.328 | 0.053 | 6.1219680 | 35 | tjp1a |
| dusp61 | 2.422283 | 0.510192967 | 0.328 | 0.055 | 6.1538123 | 35 | dusp6 |
| plce1 | 2.530143 | 0.251280803 | 0.131 | 0.01 | 6.4278284 | 35 | plce1 |
| sox131 | 2.613989 | 0.292242065 | 0.148 | 0.012 | 6.6408400 | 35 | sox13 |
| ptdss2 | 2.705143 | 0.278881474 | 0.148 | 0.012 | 6.8724177 | 35 | ptdss2 |
| si:dkey-71l4.4 | 3.115845 | 0.25211484 | 0.148 | 0.013 | 7.9158060 | 35 | si:dkey-71l4.4 |
| gadd45ba9 | 3.236608 | 0.748839328 | 0.689 | 0.197 | 8.2226032 | 35 | gadd45ba |
| hsd17b12a7 | 3.417797 | 0.670886993 | 0.59 | 0.137 | 8.6829148 | 35 | hsd17b12a |
| cebpd9 | 5.523503 | 0.714817076 | 0.59 | 0.152 | 1.4032461 | 35 | cebpd |
| znfx13 | 6.985216 | 0.362702023 | 0.246 | 0.033 | 1.7745943 | 35 | znfx1 |
| zgc:1017151 | 7.508085 | 0.326002753 | 0.213 | 0.025 | 1.9074289 | 35 | zgc:101715 |
| mef2d3 | 7.850485 | 0.512426331 | 0.393 | 0.076 | 1.9944159 | 35 | mef2d |
| ms4a17a.102 | 8.390948 | 0.508129784 | 0.328 | 0.054 | 2.1317203 | 35 | ms4a17a.10 |
| slc25a55a3 | 8.604858 | 0.446640993 | 0.393 | 0.075 | 2.1860644 | 35 | slc25a55a |
| irf2bp2a6 | 1.084350 | 0.753518304 | 0.623 | 0.186 | 2.7547934 | 35 | irf2bp2a |
| bada1 | 2.210894 | 0.387961198 | 0.246 | 0.034 | 5.6167778 | 35 | bada |
| pls3 | 3.376097 | 0.267213484 | 0.148 | 0.013 | 8.5769761 | 35 | pls3 |
| calm2b10 | 6.309024 | 0.752170817 | 0.738 | 0.246 | 1.6028077 | 35 | calm2b |
| taok1b | 8.906557 | 0.321977046 | 0.213 | 0.027 | 2.2627109 | 35 | taok1b |
| tmem176l.3a1 | 9.952704 | 0.398651183 | 0.295 | 0.048 | 2.5284845 | 35 | tmem176l.3a |
| irs2b1 | 1.130461 | 0.268478917 | 0.164 | 0.016 | 2.8719371 | 35 | irs2b |
| si:ch211-213a13.2 | 1.153897 | 0.313042449 | 0.197 | 0.023 | 2.9314755 | 35 | si:ch211-213a13.2 |
| calcoco2 | 1.230193 | 0.264998272 | 0.164 | 0.017 | 3.1253076 | 35 | calcoco2 |
| rnf213a6 | 1.304256 | 0.616201526 | 0.426 | 0.096 | 3.3134644 | 35 | rnf213a |
| actn43 | 1.515519 | 0.530617721 | 0.443 | 0.098 | 3.8501774 | 35 | actn4 |
| si:busm1-266f07.28 | 1.548585 | 0.736745388 | 0.574 | 0.148 | 3.9341818 | 35 | si:busm1-266f07.2 |
| si:ch211-132b12.72 | 1.849035 | 0.445035124 | 0.311 | 0.054 | 4.6974740 | 35 | si:ch211-132b12.7 |
| dhrs13l13 | 2.620749 | 0.388057751 | 0.262 | 0.04 | 6.6580138 | 35 | dhrs13l1 |
| cd74a7 | 2.678082 | 0.800507698 | 0.705 | 0.216 | 6.8036678 | 35 | cd74a |
| cd74b9 | 3.075288 | 0.893591406 | 0.689 | 0.226 | 7.8127692 | 35 | cd74b |
| micu2 | 3.092240 | 0.401510157 | 0.246 | 0.036 | 7.8558364 | 35 | micu2 |
| bach1b2 | 3.411438 | 0.285472725 | 0.213 | 0.027 | 8.6667600 | 35 | bach1b |
| si:ch211-191a24.4 | 3.465109 | 0.4090677 | 0.279 | 0.044 | 8.8031117 | 35 | si:ch211-191a24.4 |

| | | | | | | | |
|---|---|---|---|---|---|---|---|
| CR383676.119 | 3.974732 | 0.742797745 | 1 | 0.717 | 1.0097807 | 35 | CR383676.1 |
| klf2a8 | 4.677790 | 0.776048908 | 0.426 | 0.1 | 1.1883927 | 35 | klf2a |
| slc25a24 | 5.143657 | 0.344601821 | 0.246 | 0.036 | 1.3067462 | 35 | slc25a24 |
| si:dkey-177p2.62 | 6.946909 | 0.530091242 | 0.393 | 0.083 | 1.7648622 | 35 | si:dkey-177p2.6 |
| wbp1lb2 | 8.237935 | 0.387282825 | 0.295 | 0.05 | 2.0928476 | 35 | wbp1lb |
| COX316 | 8.981443 | 0.651110266 | 0.984 | 0.676 | 2.2817357 | 35 | COX3 |
| tnfrsfa2 | 1.097230 | 0.311920295 | 0.23 | 0.032 | 2.7875131 | 35 | tnfrsfa |
| papss2b1 | 1.395131 | 0.286607054 | 0.18 | 0.021 | 3.5443320 | 35 | papss2b |
| ca24 | 1.455729 | 0.339125321 | 0.311 | 0.053 | 3.6982796 | 35 | ca2 |
| mvp5 | 1.473480 | 0.42926308 | 0.328 | 0.061 | 3.7433764 | 35 | mvp |
| rsrp12 | 1.626851 | 0.584391925 | 0.492 | 0.122 | 4.1330162 | 35 | rsrp1 |
| fgfr33 | 1.881170 | 0.259589976 | 0.18 | 0.021 | 4.7791130 | 35 | fgfr3 |
| naga3 | 2.182889 | 0.436357867 | 0.311 | 0.056 | 5.5456317 | 35 | naga |
| tax1bp1b1 | 2.202933 | 0.47591057 | 0.361 | 0.073 | 5.5965530 | 35 | tax1bp1b |
| stk17al2 | 2.233186 | 0.50746291 | 0.475 | 0.108 | 5.6734113 | 35 | stk17al |
| CU633745.1 | 2.327807 | 0.259982761 | 0.18 | 0.021 | 5.9137953 | 35 | CU633745.1 |
| taok1a | 2.578288 | 0.303324714 | 0.18 | 0.021 | 6.5501416 | 35 | taok1a |
| ist11 | 2.811799 | 0.523246393 | 0.426 | 0.096 | 7.1433771 | 35 | ist1 |
| rin21 | 3.402864 | 0.251261659 | 0.197 | 0.025 | 8.6449765 | 35 | rin2 |
| golgb11 | 3.439902 | 0.379257852 | 0.295 | 0.051 | 8.7390728 | 35 | golgb1 |
| psph | 4.074838 | 0.302343402 | 0.213 | 0.028 | 1.0352128 | 35 | psph |
| prr133 | 4.370898 | 0.271515117 | 0.197 | 0.025 | 1.1104266 | 35 | prr13 |
| abhd2a | 5.240898 | 0.307834189 | 0.213 | 0.029 | 1.3314502 | 35 | abhd2a |
| tprg1 | 8.083656 | 0.312739089 | 0.262 | 0.041 | 2.0536528 | 35 | tprg1 |
| map1lc3a9 | 1.116038 | 0.459696343 | 0.377 | 0.081 | 2.8352970 | 35 | map1lc3a |
| wipf2a | 1.509659 | 0.326544339 | 0.23 | 0.034 | 3.8352889 | 35 | wipf2a |
| rhoab10 | 1.798767 | 0.656460423 | 0.639 | 0.207 | 4.5697693 | 35 | rhoab |
| zswim51 | 1.823686 | 0.372298026 | 0.262 | 0.043 | 4.6330752 | 35 | zswim5 |
| mat2aa2 | 1.896496 | 0.61268289 | 0.59 | 0.168 | 4.8180502 | 35 | mat2aa |
| dusp110 | 2.120895 | 0.550768194 | 0.557 | 0.149 | 5.3881348 | 35 | dusp1 |
| CABZ01055347.1 | 2.944532 | 0.333043033 | 0.23 | 0.034 | 7.4805860 | 35 | CABZ01055347.1 |
| kmt2ca | 3.150189 | 0.262241948 | 0.213 | 0.03 | 8.0030555 | 35 | kmt2ca |
| fam199x | 3.565188 | 0.260317103 | 0.164 | 0.019 | 9.0573623 | 35 | fam199x |
| slc10a3 | 4.516363 | 0.25806414 | 0.164 | 0.019 | 1.1473821 | 35 | slc10a3 |
| atl3 | 4.875716 | 0.468353049 | 0.328 | 0.065 | 1.2386757 | 35 | atl3 |

| | | | | | | | |
|---|---|---|---|---|---|---|---|
| afdna3 | 5.360027 | 0.276180375 | 0.197 | 0.026 | 1.3617149 | 35 | afdna |
| emd4 | 7.425347 | 0.400037092 | 0.344 | 0.071 | 1.8864095 | 35 | emd |
| zgc:923136 | 7.726841 | 0.519092808 | 0.459 | 0.108 | 1.9630041 | 35 | zgc:92313 |
| gnpnat1 | 9.298343 | 0.304762804 | 0.213 | 0.031 | 2.3622441 | 35 | gnpnat1 |
| si:ch211-107o10.3 | 1.162947 | 0.380830674 | 0.246 | 0.041 | 2.9544677 | 35 | si:ch211-107o10.3 |
| pcdh111 | 1.333885 | 0.267236345 | 0.148 | 0.016 | 3.3887369 | 35 | pcdh11 |
| ciartb | 1.749135 | 0.362532467 | 0.262 | 0.045 | 4.4436780 | 35 | ciartb |
| btg114 | 1.918238 | 0.696718445 | 0.852 | 0.355 | 4.8732840 | 35 | btg1 |
| pnp5b1 | 2.036687 | 0.255064089 | 0.18 | 0.023 | 5.1742057 | 35 | pnp5b |
| arhgef43 | 2.186382 | 0.352926198 | 0.23 | 0.036 | 5.5545040 | 35 | arhgef4 |
| sdcbp210 | 2.395855 | 0.652034151 | 0.656 | 0.228 | 6.0866715 | 35 | sdcbp2 |
| far13 | 3.434233 | 0.405408229 | 0.23 | 0.037 | 8.7246703 | 35 | far1 |
| eif1b11 | 4.469311 | 0.677788423 | 0.869 | 0.407 | 1.1354285 | 35 | eif1b |
| CABZ01079427.11 | 5.880736 | 0.303470013 | 0.23 | 0.036 | 1.4940010 | 35 | CABZ01079427.1 |
| wbp2nl3 | 6.599151 | 0.430187002 | 0.393 | 0.093 | 1.6765145 | 35 | wbp2nl |
| abcc31 | 6.803455 | 0.349697097 | 0.18 | 0.024 | 1.7284179 | 35 | abcc3 |
| ube2d1b2 | 8.193805 | 0.522143916 | 0.508 | 0.144 | 2.0816363 | 35 | ube2d1b |
| cflara3 | 8.854671 | 0.417075344 | 0.311 | 0.064 | 2.2495293 | 35 | cflara |
| per2 | 9.796260 | 0.358538173 | 0.213 | 0.033 | 2.4887401 | 35 | per2 |
| mycbp21 | 1.002604 | 0.414000825 | 0.246 | 0.043 | 2.5471167 | 35 | mycbp2 |
| nr4a34 | 1.123075 | 0.346336759 | 0.262 | 0.047 | 2.8531740 | 35 | nr4a3 |
| egr110 | 1.264780 | 0.673698508 | 0.574 | 0.183 | 3.2131754 | 35 | egr1 |
| rap2b | 1.303051 | 0.376711104 | 0.279 | 0.053 | 3.3104026 | 35 | rap2b |
| alox5a3 | 1.468149 | 0.37050841 | 0.213 | 0.033 | 3.7298336 | 35 | alox5a |
| furina1 | 1.616223 | 0.453765279 | 0.311 | 0.065 | 4.1060169 | 35 | furina |
| klf8 | 1.652210 | 0.314744757 | 0.213 | 0.033 | 4.1974401 | 35 | klf8 |
| clint1a | 1.746538 | 0.393432312 | 0.279 | 0.054 | 4.4370800 | 35 | clint1a |
| hexb1 | 2.018425 | 0.343367534 | 0.246 | 0.043 | 5.1278106 | 35 | hexb |
| klf2b2 | 2.366691 | 0.258108194 | 0.164 | 0.021 | 6.0125787 | 35 | klf2b |
| itm2bb11 | 2.758663 | 0.597441594 | 0.705 | 0.243 | 7.0083835 | 35 | itm2bb |
| uckl1b | 3.484165 | 0.253100264 | 0.164 | 0.021 | 8.8515228 | 35 | uckl1b |
| LOC1039092846 | 4.272371 | 0.429001452 | 0.344 | 0.077 | 1.0853960 | 35 | LOC103909284 |
| epn2 | 4.316509 | 0.257787464 | 0.197 | 0.029 | 1.0966091 | 35 | epn2 |
| CU929259.1 | 4.988366 | 0.269995932 | 0.18 | 0.025 | 1.2672945 | 35 | CU929259.1 |
| lsr2 | 5.488899 | 0.264137923 | 0.197 | 0.029 | 1.3944548 | 35 | lsr |

| | | | | | | | |
|---|---|---|---|---|---|---|---|
| cebpa6 | 6.867681 | 0.380777797 | 0.344 | 0.076 | 1.7447345 | 35 | cebpa |
| cnn210 | 7.272864 | 0.588447112 | 0.508 | 0.157 | 1.8476711 | 35 | cnn2 |
| arl114 | 8.681215 | 0.253558009 | 0.18 | 0.025 | 2.2054628 | 35 | arl11 |
| rab18a | 8.713592 | 0.327191771 | 0.213 | 0.035 | 2.2136880 | 35 | rab18a |
| plpp1a3 | 1.091127 | 0.426054718 | 0.344 | 0.078 | 2.7720100 | 35 | plpp1a |
| utrn | 1.236693 | 0.272670539 | 0.213 | 0.035 | 3.1418199 | 35 | utrn |
| capns1b8 | 1.306446 | 0.499047326 | 0.492 | 0.146 | 3.3190270 | 35 | capns1b |
| rtn31 | 1.432705 | 0.415047169 | 0.344 | 0.079 | 3.6397875 | 35 | rtn3 |
| nktr1 | 1.627274 | 0.324309359 | 0.23 | 0.04 | 4.1340914 | 35 | nktr |
| arpc1a3 | 1.731004 | 0.508904591 | 0.492 | 0.142 | 4.3976169 | 35 | arpc1a |
| ier2a13 | 3.769493 | 0.666306518 | 0.705 | 0.275 | 9.5763982 | 35 | ier2a |
| chmp1a | 3.776599 | 0.374810239 | 0.279 | 0.057 | 9.5944500 | 35 | chmp1a |
| arl5c7 | 4.291805 | 0.457171734 | 0.41 | 0.108 | 1.0903332 | 35 | arl5c |
| skia | 6.966928 | 0.328732333 | 0.246 | 0.047 | 1.7699480 | 35 | skia |
| LOC103910015 | 8.876467 | 0.282412492 | 0.164 | 0.023 | 2.2550665 | 35 | LOC103910015 |
| susd61 | 9.806935 | 0.31531587 | 0.279 | 0.058 | 2.4914518 | 35 | susd6 |
| ftr824 | 1.036997 | 0.277184333 | 0.213 | 0.037 | 2.6344918 | 35 | ftr82 |
| csnk1a13 | 1.058812 | 0.550253296 | 0.656 | 0.238 | 2.6899128 | 35 | csnk1a1 |
| ctss2.23 | 1.137347 | 0.286909666 | 0.262 | 0.052 | 2.8894316 | 35 | ctss2.2 |
| hmgn34 | 1.180705 | 0.418307908 | 0.328 | 0.078 | 2.9995812 | 35 | hmgn3 |
| tp53inp13 | 2.233701 | 0.35897225 | 0.279 | 0.06 | 5.6747181 | 35 | tp53inp1 |
| ptbp32 | 2.277706 | 0.380417581 | 0.295 | 0.066 | 5.7865130 | 35 | ptbp3 |
| fosl28 | 2.336772 | 0.468114482 | 0.41 | 0.114 | 5.9365715 | 35 | fosl2 |
| tpd52l2a | 2.795896 | 0.312585071 | 0.262 | 0.054 | 7.1029759 | 35 | tpd52l2a |
| il154 | 2.800051 | 0.379737101 | 0.279 | 0.06 | 7.1135297 | 35 | il15 |
| si:ch211-196l7.413 | 3.038962 | 0.626086667 | 0.738 | 0.33 | 7.7204848 | 35 | si:ch211-196l7.4 |
| gbgt1l47 | 3.802460 | 0.481760391 | 0.459 | 0.125 | 9.6601502 | 35 | gbgt1l4 |
| mt-co214 | 4.828875 | 0.5734004 | 0.967 | 0.671 | 1.2267758 | 35 | mt-co2 |
| ak15 | 5.356727 | 0.359470725 | 0.377 | 0.099 | 1.3608767 | 35 | ak1 |
| CABZ01090890.11 | 5.977890 | 0.300708675 | 0.279 | 0.06 | 1.5186831 | 35 | CABZ01090890.1 |
| atf38 | 6.008297 | 0.503535603 | 0.328 | 0.082 | 1.5264079 | 35 | atf3 |
| cdc42bpb1 | 6.925644 | 0.288125194 | 0.18 | 0.029 | 1.7594599 | 35 | cdc42bpb |
| vps281 | 7.756067 | 0.411016341 | 0.377 | 0.101 | 1.9704290 | 35 | vps28 |
| junba12 | 8.076890 | 0.717362871 | 0.705 | 0.32 | 2.0519339 | 35 | junba |
| zgc:1583437 | 1.026649 | 0.586962923 | 0.508 | 0.17 | 2.6082042 | 35 | zgc:158343 |

| | | | | | | | |
|---|---|---|---|---|---|---|---|
| tmbim1a | 1.179458 | 0.253762872 | 0.164 | 0.025 | 2.9964149 | 35 | tmbim1a |
| SBNO21 | 1.206925 | 0.263543602 | 0.197 | 0.035 | 3.0661941 | 35 | SBNO2 |
| fam102ab2 | 1.876285 | 0.31663326 | 0.197 | 0.035 | 4.7667034 | 35 | fam102ab |
| galnt12 | 2.187822 | 0.27585648 | 0.164 | 0.026 | 5.5581626 | 35 | galnt12 |
| ftr86 | 2.323083 | 0.321166274 | 0.213 | 0.039 | 5.9017936 | 35 | ftr86 |
| ponzr63 | 2.777533 | 0.256106585 | 0.148 | 0.021 | 7.0563243 | 35 | ponzr6 |
| alas12 | 2.821160 | 0.414290316 | 0.459 | 0.138 | 7.1671589 | 35 | alas1 |
| pde4ba11 | 3.906753 | 0.464657423 | 0.459 | 0.142 | 9.9251078 | 35 | pde4ba |
| ppp2r2d1 | 5.782256 | 0.305297957 | 0.262 | 0.059 | 1.4689823 | 35 | ppp2r2d |
| smdt1b4 | 6.026757 | 0.46966768 | 0.459 | 0.153 | 1.5310978 | 35 | smdt1b |
| mt-cyb12 | 6.807696 | 0.586952908 | 0.934 | 0.577 | 1.7294953 | 35 | mt-cyb |
| fbp1a2 | 8.201019 | 0.333240697 | 0.213 | 0.043 | 2.0834689 | 35 | fbp1a |
| psme28 | 1.162701 | 0.443984521 | 0.377 | 0.113 | 2.9538419 | 35 | psme2 |
| gstt1b8 | 1.275202 | 0.460436979 | 0.328 | 0.09 | 3.2396520 | 35 | gstt1b |
| mvb12a2 | 1.299640 | 0.306238569 | 0.311 | 0.078 | 3.3017369 | 35 | mvb12a |
| adipor1b | 1.564427 | 0.326659205 | 0.23 | 0.049 | 3.9744290 | 35 | adipor1b |
| tpm310 | 1.576311 | 0.55769414 | 0.672 | 0.284 | 4.0046192 | 35 | tpm3 |
| si:ch1073-190k2.17 | 1.783386 | 0.514420231 | 0.492 | 0.155 | 4.5306944 | 35 | si:ch1073-190k2.1 |
| zgc:16273015 | 1.827435 | 0.633769002 | 0.852 | 0.403 | 4.6426000 | 35 | zgc:162730 |
| nfil3-68 | 2.042097 | 0.522692073 | 0.41 | 0.132 | 5.1879486 | 35 | nfil3-6 |
| ddx3xa | 2.415739 | 0.350305091 | 0.344 | 0.094 | 6.1371850 | 35 | ddx3xa |
| ppm1aa | 2.638107 | 0.315400632 | 0.279 | 0.068 | 6.7021117 | 35 | ppm1aa |
| klf6a8 | 2.785047 | 0.499260168 | 0.508 | 0.181 | 7.0754132 | 35 | klf6a |
| elf17 | 2.801445 | 0.406075611 | 0.361 | 0.104 | 7.1170727 | 35 | elf1 |
| ccdc1873 | 2.993940 | 0.279475326 | 0.197 | 0.038 | 7.6061057 | 35 | ccdc187 |
| ctsk5 | 3.137252 | 0.311932992 | 0.295 | 0.074 | 7.9701889 | 35 | ctsk |
| pmaip12 | 3.221520 | 0.355668504 | 0.361 | 0.101 | 8.1842724 | 35 | pmaip1 |
| zbtb101 | 3.582253 | 0.28054102 | 0.18 | 0.034 | 9.1007159 | 35 | zbtb10 |
| tegt6 | 4.161120 | 0.482358998 | 0.607 | 0.238 | 1.0571327 | 35 | tegt |
| h3f3c13 | 4.388446 | 0.534527372 | 0.721 | 0.304 | 1.1148847 | 35 | h3f3c |
| egr2a1 | 6.094785 | 0.265523608 | 0.148 | 0.024 | 1.5483802 | 35 | egr2a |
| nr1d2b | 7.379742 | 0.261838379 | 0.197 | 0.039 | 1.8748235 | 35 | nr1d2b |
| tnrc6c110 | 7.885267 | 0.437008703 | 0.443 | 0.15 | 2.0032522 | 35 | tnrc6c1 |
| sdf42 | 8.209940 | 0.341859521 | 0.23 | 0.052 | 2.0857353 | 35 | sdf4 |
| csrnp1b9 | 8.741928 | 0.519449571 | 0.459 | 0.165 | 2.2208869 | 35 | csrnp1b |

| | | | | | | | |
|---|---|---|---|---|---|---|---|
| plekhf2 | 1.197987 | 0.267828381 | 0.18 | 0.035 | 3.0434874 | 35 | plekhf2 |
| tmem54a7 | 1.263124 | 0.310815984 | 0.41 | 0.118 | 3.2089685 | 35 | tmem54a |
| herpud2 | 1.318931 | 0.280118554 | 0.18 | 0.035 | 3.3507447 | 35 | herpud2 |
| vaspb2 | 1.677308 | 0.284352967 | 0.23 | 0.053 | 4.2612017 | 35 | vaspb |
| ar6 | 2.392879 | 0.370660714 | 0.262 | 0.067 | 6.0791112 | 35 | ar |
| stag2b | 2.468980 | 0.304991172 | 0.246 | 0.06 | 6.2724443 | 35 | stag2b |
| mid1ip1b1 | 2.770005 | 0.339058176 | 0.23 | 0.054 | 7.0371983 | 35 | mid1ip1b |
| sar1b3 | 2.922615 | 0.373304056 | 0.311 | 0.09 | 7.4249045 | 35 | sar1b |
| meis1b7 | 3.309051 | 0.319242917 | 0.279 | 0.073 | 8.4066443 | 35 | meis1b |
| foxp46 | 3.378855 | 0.342486014 | 0.279 | 0.074 | 8.5839820 | 35 | foxp4 |
| zgc:162698 | 3.519093 | 0.281066381 | 0.23 | 0.054 | 8.9402569 | 35 | zgc:162698 |
| u2surp2 | 3.886354 | 0.356185099 | 0.344 | 0.104 | 9.8732829 | 35 | u2surp |
| ptp4a2b3 | 4.695795 | 0.386850024 | 0.443 | 0.15 | 0.000119 | 35 | ptp4a2b |
| cish3 | 8.310110 | 0.278803773 | 0.23 | 0.055 | 0.000211 | 35 | cish |
| qkia8 | 8.470608 | 0.426235001 | 0.525 | 0.2 | 0.000215 | 35 | qkia |
| pitpnb | 1.038058 | 0.285615169 | 0.213 | 0.05 | 0.000264 | 35 | pitpnb |
| si:ch211-260e23.92 | 1.256851 | 0.253228516 | 0.246 | 0.062 | 0.000319 | 35 | si:ch211-260e23.9 |
| yap15 | 1.452399 | 0.336855532 | 0.23 | 0.058 | 0.000369 | 35 | yap1 |
| cnn3a8 | 1.552576 | 0.269942741 | 0.262 | 0.07 | 0.000394 | 35 | cnn3a |
| CR318588.113 | 1.808515 | 0.561504533 | 0.541 | 0.216 | 0.000459 | 35 | CR318588.1 |
| txnrd3 | 2.375313 | 0.290549748 | 0.328 | 0.099 | 0.000603 | 35 | txnrd3 |
| nupr1a1 | 2.411993 | 0.423744043 | 0.443 | 0.165 | 0.000613 | 35 | nupr1a |
| dedd15 | 2.504954 | 0.358634949 | 0.443 | 0.157 | 0.000636 | 35 | dedd1 |
| itm2ba11 | 2.677978 | 0.450449915 | 0.672 | 0.297 | 0.00068 | 35 | itm2ba |
| mt-nd414 | 2.878459 | 0.493303812 | 0.869 | 0.448 | 0.000731 | 35 | mt-nd4 |
| tpp13 | 3.034641 | 0.266165639 | 0.279 | 0.077 | 0.000771 | 35 | tpp1 |
| zgc:1584634 | 3.862859 | 0.314216837 | 0.279 | 0.081 | 0.000981 | 35 | zgc:158463 |
| ppp4cb | 3.902153 | 0.29759881 | 0.361 | 0.116 | 0.000991 | 35 | ppp4cb |
| elovl7b7 | 4.091270 | 0.535312951 | 0.508 | 0.206 | 0.001039 | 35 | elovl7b |
| ccnl1b | 4.196205 | 0.337807641 | 0.311 | 0.095 | 0.001066 | 35 | ccnl1b |
| pmp22b6 | 4.833530 | 0.328267499 | 0.197 | 0.047 | 0.001228 | 35 | pmp22b |
| si:ch211-191j22.32 | 5.217709 | 0.364405699 | 0.361 | 0.124 | 0.001326 | 35 | si:ch211-191j22.3 |
| cyp3c14 | 5.403673 | 0.253565125 | 0.246 | 0.065 | 0.001373 | 35 | cyp3c1 |
| NC-002333.412 | 1.265643 | 0.482694445 | 0.984 | 0.609 | 0.003215 | 35 | NC-002333.4 |
| gnal | 1.714995 | 0.284445749 | 0.164 | 0.037 | 0.004357 | 35 | gnal |

| | | | | | | | |
|---|---|---|---|---|---|---|---|
| si:ch211-39i2.21 | 1.717088 | 0.288041346 | 0.23 | 0.062 | 0.004362 | 35 | si:ch211-39i2.2 |
| junbb5 | 1.776234 | 0.299387572 | 0.246 | 0.071 | 0.004513 | 35 | junbb |
| cdc42l8 | 1.786917 | 0.421420164 | 0.541 | 0.235 | 0.00454 | 35 | cdc42l |
| rab5aa | 1.998496 | 0.253378447 | 0.279 | 0.084 | 0.005077 | 35 | rab5aa |
| nfixb4 | 2.200753 | 0.260260969 | 0.279 | 0.083 | 0.005591 | 35 | nfixb |
| mapk3 | 2.227567 | 0.309551135 | 0.295 | 0.093 | 0.005659 | 35 | mapk3 |
| tmem106a | 2.259791 | 0.279837684 | 0.279 | 0.084 | 0.005741 | 35 | tmem106a |
| rab5c5 | 2.743741 | 0.277922198 | 0.377 | 0.132 | 0.00697 | 35 | rab5c |
| slc3a2b6 | 2.911882 | 0.340030233 | 0.361 | 0.129 | 0.007398 | 35 | slc3a2b |
| sik16 | 3.348342 | 0.338411018 | 0.328 | 0.111 | 0.008506 | 35 | sik1 |
| ddx53 | 4.246387 | 0.463492043 | 0.705 | 0.355 | 0.010788 | 35 | ddx5 |
| sdc411 | 4.562476 | 0.463376029 | 0.557 | 0.255 | 0.011591 | 35 | sdc4 |
| tcima6 | 4.681878 | 0.313074281 | 0.246 | 0.072 | 0.011894 | 35 | tcima |
| zgc:1630801 | 5.411780 | 0.295740185 | 0.279 | 0.09 | 0.013749 | 35 | zgc:163080 |
| notch29 | 5.846446 | 0.329398071 | 0.344 | 0.123 | 0.014853 | 35 | notch2 |
| litaf7 | 6.468594 | 0.305207088 | 0.393 | 0.147 | 0.016433 | 35 | litaf |
| egfra5 | 7.669090 | 0.250950717 | 0.148 | 0.033 | 0.019483 | 35 | egfra |
| scaf1 | 9.692596 | 0.260393829 | 0.164 | 0.04 | 0.024624 | 35 | scaf1 |
| sptlc2b | 1.054857 | 0.25447228 | 0.18 | 0.047 | 0.026799 | 35 | sptlc2b |
| lrrfip1a5 | 1.063114 | 0.297636398 | 0.213 | 0.061 | 0.027008 | 35 | lrrfip1a |
| dnajb1b | 1.252597 | 0.275712165 | 0.197 | 0.053 | 0.031822 | 35 | dnajb1b |
| anxa11a3 | 1.260779 | 0.296646944 | 0.328 | 0.118 | 0.03203 | 35 | anxa11a |
| irf23 | 1.278313 | 0.253333058 | 0.246 | 0.076 | 0.032476 | 35 | irf2 |
| creb3l3l1 | 1.302049 | 0.267242209 | 0.213 | 0.062 | 0.033079 | 35 | creb3l3l |
| arhgdig12 | 1.355686 | 0.408413433 | 0.574 | 0.26 | 0.034441 | 35 | arhgdig |
| psme14 | 1.355972 | 0.339191371 | 0.311 | 0.113 | 0.034448 | 35 | psme1 |
| syap1 | 1.415159 | 0.268678248 | 0.246 | 0.076 | 0.035952 | 35 | syap1 |
| si:dkey-42i9.46 | 1.417786 | 0.335887752 | 0.393 | 0.157 | 0.036019 | 35 | si:dkey-42i9.4 |
| pfn29 | 1.540678 | 0.375627954 | 0.541 | 0.24 | 0.039141 | 35 | pfn2 |
| cnot11 | 2.393689 | 0.284059496 | 0.213 | 0.064 | 0.060812 | 35 | cnot1 |
| sf3b1 | 2.399116 | 0.254920063 | 0.295 | 0.101 | 0.06095 | 35 | sf3b1 |
| znf7062 | 2.538791 | 0.327052356 | 0.443 | 0.184 | 0.064498 | 35 | znf706 |
| brd4 | 2.744170 | 0.258761293 | 0.262 | 0.086 | 0.069716 | 35 | brd4 |
| dazap2 | 3.088031 | 0.303194956 | 0.279 | 0.097 | 0.078451 | 35 | dazap2 |
| rab108 | 3.505672 | 0.337605837 | 0.492 | 0.215 | 0.089062 | 35 | rab10 |

| | | | | | | | |
|---|---|---|---|---|---|---|---|
| smim152 | 3.736005 | 0.331169383 | 0.377 | 0.154 | 0.094913 | 35 | smim15 |
| ndufv1 | 4.073801 | 0.258220957 | 0.23 | 0.072 | 0.103495 | 35 | ndufv1 |
| ets25 | 4.251599 | 0.266310931 | 0.262 | 0.088 | 0.108012 | 35 | ets2 |
| zgc:110339 | 4.839803 | 0.28191981 | 0.295 | 0.106 | 0.122955 | 35 | zgc:110339 |
| calm1b10 | 5.974843 | 0.254006942 | 0.295 | 0.104 | 0.151791 | 35 | calm1b |
| atp5if1b7 | 6.124351 | 0.306470249 | 0.475 | 0.208 | 0.155589 | 35 | atp5if1b |
| atf4a4 | 6.574527 | 0.346870587 | 0.426 | 0.186 | 0.167026 | 35 | atf4a |
| ankrd112 | 7.545739 | 0.289164309 | 0.279 | 0.101 | 0.1917 | 35 | ankrd11 |
| sptssa1 | 9.669534 | 0.333816343 | 0.246 | 0.087 | 0.245655 | 35 | sptssa |
| ywhabl7 | 1.101954 | 0.377126566 | 0.59 | 0.302 | 0.279951 | 35 | ywhabl |
| chmp2a4 | 1.129708 | 0.333698938 | 0.344 | 0.145 | 0.287003 | 35 | chmp2a |
| si:ch211-105c13.35 | 1.152468 | 0.296163321 | 0.213 | 0.068 | 0.292785 | 35 | si:ch211-105c13.3 |
| glo14 | 1.574320 | 0.321298478 | 0.459 | 0.21 | 0.399956 | 35 | glo1 |
| ctsz2 | 1.645303 | 0.251338069 | 0.262 | 0.094 | 0.417989 | 35 | ctsz |
| zgc:778491 | 1.721280 | 0.257686889 | 0.426 | 0.181 | 0.437291 | 35 | zgc:77849 |
| mt-nd311 | 1.797337 | 0.39620698 | 0.672 | 0.372 | 0.456614 | 35 | mt-nd3 |
| cd836 | 2.002754 | 0.312166375 | 0.262 | 0.094 | 0.5088 | 35 | cd83 |
| gng57 | 2.536807 | 0.315910188 | 0.59 | 0.288 | 0.644476 | 35 | gng5 |
| cd6315 | 2.888927 | 0.360087634 | 0.721 | 0.374 | 0.733932 | 35 | cd63 |
| cers5 | 2.933047 | 0.256416153 | 0.18 | 0.055 | 0.745141 | 35 | cers5 |
| gabarapa5 | 3.378691 | 0.271750911 | 0.41 | 0.182 | 0.858357 | 35 | gabarapa |
| h2afx110 | 3.725962 | 0.26289406 | 0.508 | 0.228 | 0.946581 | 35 | h2afx1 |
| yipf4 | 4.985644 | 0.267738844 | 0.246 | 0.093 | 1 | 35 | yipf4 |
| cst14a.22 | 6.641887 | 0.339530302 | 0.475 | 0.237 | 1 | 35 | cst14a.2 |
| eif3s106 | 7.292529 | 0.260695319 | 0.393 | 0.18 | 1 | 35 | eif3s10 |
| hdlbpa8 | 7.786049 | 0.254141431 | 0.393 | 0.176 | 1 | 35 | hdlbpa |
| si:dkey-11f4.148 | 0.0001 | 0.269110761 | 0.295 | 0.123 | 1 | 35 | si:dkey-11f4.14 |
| ifi305 | 0.0002 | 0.287286403 | 0.41 | 0.197 | 1 | 35 | ifi30 |
| birc24 | 0.0002 | 0.267581775 | 0.262 | 0.108 | 1 | 35 | birc2 |
| nfkbiab5 | 0.0002 | 0.380257375 | 0.393 | 0.202 | 1 | 35 | nfkbiab |
| gnb1b4 | 0.0002 | 0.289455897 | 0.508 | 0.267 | 1 | 35 | gnb1b |
| si:ch73-1a9.318 | 0.0002 | 0.334646863 | 0.574 | 0.317 | 1 | 35 | si:ch73-1a9.3 |
| ddx614 | 0.0003 | 0.251946928 | 0.475 | 0.236 | 1 | 35 | ddx61 |
| mt-nd114 | 0.0003 | 0.310717225 | 0.689 | 0.398 | 1 | 35 | mt-nd1 |
| sumo2b12 | 0.0003 | 0.291809688 | 0.557 | 0.297 | 1 | 35 | sumo2b |

| | | | | | | | |
|---|---|---|---|---|---|---|---|
| rplp216 | 0.0003 | 0.268121503 | 0.951 | 0.784 | 1 | 35 | rplp2 |
| bhlhe4011 | 0.0003 | 0.259342332 | 0.361 | 0.169 | 1 | 35 | bhlhe40 |
| ppp3ccb4 | 0.0004 | 0.285136389 | 0.279 | 0.126 | 1 | 35 | ppp3ccb |
| ccni11 | 0.0005 | 0.267353159 | 0.59 | 0.32 | 1 | 35 | ccni |
| cox8a14 | 0.0006 | 0.254807377 | 0.508 | 0.268 | 1 | 35 | cox8a |
| ybx111 | 0.0006 | 0.268143987 | 0.869 | 0.697 | 1 | 35 | ybx1 |
| mt-atp615 | 0.0015 | 0.323901288 | 0.852 | 0.557 | 1 | 35 | mt-atp6 |
| myl12.114 | 0.0016 | 0.307100534 | 0.656 | 0.427 | 1 | 35 | myl12.1 |
| mt-nd212 | 0.0017 | 0.254175477 | 0.689 | 0.393 | 1 | 35 | mt-nd2 |
| bzw1b6 | 0.0018 | 0.273087677 | 0.639 | 0.379 | 1 | 35 | bzw1b |
| si:ch211-243a20.31 | 0 | 1.717956594 | 0.855 | 0.011 | 0 | 36 | si:ch211-243a20.3 |
| lum2 | 0 | 1.551231855 | 0.8 | 0.019 | 0 | 36 | lum |
| wfdc21 | 0 | 1.540862236 | 0.745 | 0.007 | 0 | 36 | wfdc2 |
| sfrp2 | 0 | 1.164206246 | 0.527 | 0.005 | 0 | 36 | sfrp2 |
| tnmd | 0 | 1.145109821 | 0.473 | 0.002 | 0 | 36 | tnmd |
| CR855375.3 | 0 | 0.394396611 | 0.145 | 0 | 0 | 36 | CR855375.3 |
| trabd2b | 1.099088 | 0.485304159 | 0.273 | 0.002 | 2.7922333 | 36 | trabd2b |
| cyp26b12 | 1.112647 | 1.040576839 | 0.418 | 0.007 | 2.8266819 | 36 | cyp26b1 |
| thbs2b | 2.095708 | 0.797127181 | 0.382 | 0.005 | 5.3241482 | 36 | thbs2b |
| CABZ01085139.1 | 2.092868 | 0.545348925 | 0.255 | 0.002 | 5.3169316 | 36 | CABZ01085139.1 |
| pbx3b1 | 5.059792 | 1.291635186 | 0.673 | 0.019 | 1.2854403 | 36 | pbx3b |
| twist1a2 | 1.207594 | 1.181045485 | 0.6 | 0.017 | 3.0678926 | 36 | twist1a |
| pcolce2b1 | 1.183915 | 1.141970881 | 0.6 | 0.019 | 3.0077365 | 36 | pcolce2b |
| CU929237.13 | 8.524299 | 1.142921427 | 0.618 | 0.021 | 2.1655981 | 36 | CU929237.1 |
| sfrp1b2 | 3.120478 | 1.082320565 | 0.527 | 0.015 | 7.9275751 | 36 | sfrp1b |
| col2a1b1 | 2.846634 | 1.045122698 | 0.527 | 0.015 | 7.2318752 | 36 | col2a1b |
| apcdd1l1 | 4.754989 | 0.77757842 | 0.4 | 0.009 | 1.2080050 | 36 | apcdd1l |
| tmem26b | 7.332365 | 0.333363894 | 0.127 | 0.001 | 1.8627875 | 36 | tmem26b |
| col11a1b2 | 1.579515 | 1.036970813 | 0.527 | 0.016 | 4.0127581 | 36 | col11a1b |
| thbs4b | 1.438266 | 0.552305995 | 0.255 | 0.003 | 3.6539166 | 36 | thbs4b |
| abi3bpb | 8.959197 | 0.997857544 | 0.509 | 0.015 | 2.2760841 | 36 | abi3bpb |
| cd248b | 3.438950 | 0.60192029 | 0.273 | 0.004 | 8.7366530 | 36 | cd248b |
| pdgfrl3 | 4.505182 | 1.273487977 | 0.709 | 0.031 | 1.1445415 | 36 | pdgfrl |
| cav22 | 1.397319 | 1.032963754 | 0.509 | 0.015 | 3.5498907 | 36 | cav2 |
| si:dkey-12l12.12 | 1.730394 | 1.105016636 | 0.582 | 0.021 | 4.3960664 | 36 | si:dkey-12l12.1 |

| | | | | | | | |
|---|---|---|---|---|---|---|---|
| chrna5 | 3.430227 | 0.435480354 | 0.145 | 0.001 | 8.7144920 | 36 | chrna5 |
| wnt11r2 | 7.456317 | 0.879676827 | 0.382 | 0.009 | 1.8942773 | 36 | wnt11r |
| crabp2b2 | 2.668240 | 0.954641603 | 0.545 | 0.019 | 6.7786641 | 36 | crabp2b |
| ism12 | 1.612934 | 0.850249356 | 0.382 | 0.009 | 4.0976610 | 36 | ism1 |
| enpp21 | 1.768712 | 0.538243775 | 0.309 | 0.006 | 4.4934130 | 36 | enpp2 |
| LO018188.13 | 3.007847 | 1.210532212 | 0.6 | 0.027 | 7.6414369 | 36 | LO018188.1 |
| chrdl2 | 6.919710 | 0.49047667 | 0.218 | 0.003 | 1.7579524 | 36 | chrdl2 |
| rasl12 | 3.910384 | 0.626282994 | 0.273 | 0.005 | 9.9343312 | 36 | rasl12 |
| efnb2a2 | 3.783797 | 0.970889655 | 0.436 | 0.015 | 9.6127378 | 36 | efnb2a |
| mkxa | 1.198636 | 0.364867184 | 0.145 | 0.001 | 3.0451367 | 36 | mkxa |
| hhipl21 | 5.357001 | 0.471628183 | 0.218 | 0.003 | 1.3609463 | 36 | hhipl2 |
| igsf10 | 1.615905 | 0.30087437 | 0.127 | 0.001 | 4.1052071 | 36 | igsf10 |
| itga11a | 3.319751 | 0.462456607 | 0.2 | 0.003 | 8.4338285 | 36 | itga11a |
| coch | 8.717685 | 0.711750563 | 0.309 | 0.007 | 2.2147279 | 36 | coch |
| sfrp1a3 | 1.644789 | 1.297408624 | 0.636 | 0.035 | 4.1785864 | 36 | sfrp1a |
| alx4a1 | 7.278941 | 0.537262632 | 0.255 | 0.005 | 1.8492150 | 36 | alx4a |
| adh8a4 | 1.623838 | 1.658265122 | 0.727 | 0.046 | 4.1253605 | 36 | adh8a |
| si:dkey-282h22.51 | 3.453553 | 0.673377507 | 0.255 | 0.005 | 8.7737539 | 36 | si:dkey-282h22.5 |
| zfhx4 | 2.061603 | 0.69534709 | 0.345 | 0.01 | 5.2375025 | 36 | zfhx4 |
| timp2a6 | 8.306070 | 1.318382374 | 0.836 | 0.063 | 2.1101571 | 36 | timp2a |
| col16a13 | 1.321732 | 1.036192113 | 0.636 | 0.039 | 3.3578624 | 36 | col16a1 |
| cyr616 | 1.876051 | 1.441821581 | 0.8 | 0.066 | 4.7661093 | 36 | cyr61 |
| fndc1 | 1.232622 | 0.414425738 | 0.182 | 0.003 | 3.1314766 | 36 | fndc1 |
| aqp3a4 | 2.399964 | 1.435527935 | 0.509 | 0.028 | 6.0971098 | 36 | aqp3a |
| tpm4b | 4.419092 | 0.858999961 | 0.418 | 0.018 | 1.1226705 | 36 | tpm4b |
| vwa12 | 5.432178 | 0.871901428 | 0.436 | 0.021 | 1.3800448 | 36 | vwa1 |
| fibinb1 | 1.812799 | 0.844252869 | 0.455 | 0.022 | 4.6054168 | 36 | fibinb |
| cntfr3 | 1.314043 | 0.883575737 | 0.418 | 0.019 | 3.3383282 | 36 | cntfr |
| crabp1a1 | 2.253209 | 0.645316067 | 0.291 | 0.009 | 5.7242788 | 36 | crabp1a |
| asip12 | 1.506505 | 1.101614141 | 0.4 | 0.019 | 3.8272776 | 36 | asip1 |
| calhm2 | 2.249068 | 0.550604877 | 0.309 | 0.011 | 5.7137586 | 36 | calhm2 |
| cpe1 | 1.412088 | 0.522064518 | 0.255 | 0.008 | 3.5874102 | 36 | cpe |
| ecrg4a4 | 3.717658 | 1.18648555 | 0.545 | 0.036 | 9.4447110 | 36 | ecrg4a |
| ucmaa1 | 2.626809 | 0.949041537 | 0.345 | 0.015 | 6.6734095 | 36 | ucmaa |
| tnc | 3.083554 | 0.36065531 | 0.145 | 0.002 | 7.8337692 | 36 | tnc |

| | | | | | | | |
|---|---|---|---|---|---|---|---|
| FP102018.12 | 7.307901 | 0.828169415 | 0.4 | 0.02 | 1.8565722 | 36 | FP102018.1 |
| XLOC-0429171 | 1.144612 | 0.678803333 | 0.327 | 0.013 | 2.9078890 | 36 | XLOC-042917 |
| mmp27 | 1.289741 | 1.480070206 | 0.891 | 0.11 | 3.2765889 | 36 | mmp2 |
| loxa2 | 4.465912 | 0.708093947 | 0.345 | 0.015 | 1.1345649 | 36 | loxa |
| ctgfa2 | 1.342560 | 1.096655086 | 0.509 | 0.033 | 3.4107757 | 36 | ctgfa |
| col12a1b3 | 1.402318 | 1.200249698 | 0.527 | 0.036 | 3.5625896 | 36 | col12a1b |
| efcc1 | 1.729957 | 0.444658454 | 0.218 | 0.006 | 4.3949562 | 36 | efcc1 |
| chad | 7.533579 | 0.624335308 | 0.236 | 0.007 | 1.9139058 | 36 | chad |
| nuak1a | 7.784408 | 0.437285931 | 0.236 | 0.007 | 1.9776290 | 36 | nuak1a |
| cacna1g | 9.292106 | 0.418590969 | 0.2 | 0.005 | 2.3606596 | 36 | cacna1g |
| palm1a1 | 2.944885 | 0.685365332 | 0.382 | 0.018 | 7.4814822 | 36 | palm1a |
| endouc4 | 3.210048 | 1.420297151 | 0.727 | 0.07 | 8.1551274 | 36 | endouc |
| BX640522.23 | 4.773742 | 1.116081401 | 0.564 | 0.041 | 1.2127692 | 36 | BX640522.2 |
| sh3pxd2ab | 2.092691 | 0.388708581 | 0.218 | 0.006 | 5.3164832 | 36 | sh3pxd2ab |
| lhfpl63 | 3.889424 | 0.784999033 | 0.418 | 0.023 | 9.8810827 | 36 | lhfpl6 |
| glsl | 5.473552 | 0.266447833 | 0.109 | 0.001 | 1.3905560 | 36 | glsl |
| mfap53 | 3.128234 | 1.267279113 | 0.673 | 0.062 | 7.9472802 | 36 | mfap5 |
| loxl13 | 1.933144 | 0.840202574 | 0.473 | 0.03 | 4.9111539 | 36 | loxl1 |
| boc5 | 2.266347 | 1.14273908 | 0.691 | 0.066 | 5.7576557 | 36 | boc |
| emx22 | 4.183883 | 0.754912762 | 0.418 | 0.024 | 1.0629155 | 36 | emx2 |
| zgc:113307 | 1.243623 | 0.319024293 | 0.109 | 0.001 | 3.1594247 | 36 | zgc:113307 |
| notum1a | 1.674790 | 0.265364599 | 0.109 | 0.001 | 4.2548048 | 36 | notum1a |
| mfap26 | 3.856965 | 1.458113293 | 0.891 | 0.118 | 9.7986203 | 36 | mfap2 |
| myl9a6 | 5.605651 | 1.027600142 | 0.564 | 0.044 | 1.4241158 | 36 | myl9a |
| si:ch1073-459j12.15 | 7.590766 | 1.280902446 | 0.764 | 0.083 | 1.9284341 | 36 | si:ch1073-459j12.1 |
| col1a1a6 | 1.172092 | 1.798944077 | 1 | 0.165 | 2.9776999 | 36 | col1a1a |
| cdon4 | 4.510571 | 1.191255501 | 0.691 | 0.07 | 1.1459107 | 36 | cdon |
| lrp1aa4 | 3.426728 | 1.085083407 | 0.673 | 0.065 | 8.7056034 | 36 | lrp1aa |
| pcsk61 | 2.927496 | 0.510226124 | 0.273 | 0.01 | 7.4373043 | 36 | pcsk6 |
| efna2a | 6.290860 | 0.445144948 | 0.255 | 0.009 | 1.5981930 | 36 | efna2a |
| col1a26 | 7.624307 | 1.839091102 | 1 | 0.181 | 1.9369552 | 36 | col1a2 |
| vipr1a | 1.171489 | 0.399645267 | 0.2 | 0.006 | 2.9761688 | 36 | vipr1a |
| meis2b3 | 2.128208 | 0.787310002 | 0.436 | 0.028 | 5.4067125 | 36 | meis2b |
| cemip | 2.980406 | 0.656081405 | 0.291 | 0.013 | 7.5717229 | 36 | cemip |
| pitx2 | 3.296265 | 0.511941033 | 0.255 | 0.01 | 8.3741635 | 36 | pitx2 |

| | | | | | | | |
|---|---|---|---|---|---|---|---|
| adgra21 | 1.693083 | 0.439095688 | 0.218 | 0.007 | 4.3012798 | 36 | adgra2 |
| LOC1005373421 | 7.540351 | 0.556597946 | 0.291 | 0.013 | 1.9156264 | 36 | LOC100537342 |
| col5a16 | 2.509229 | 1.322208433 | 0.909 | 0.131 | 6.3746987 | 36 | col5a1 |
| col5a2a6 | 1.842385 | 1.221587947 | 0.782 | 0.099 | 4.6805791 | 36 | col5a2a |
| gulp1a4 | 2.095497 | 0.832656688 | 0.473 | 0.036 | 5.3236114 | 36 | gulp1a |
| si:ch211-37e10.2 | 1.111259 | 0.258995078 | 0.127 | 0.002 | 2.8231555 | 36 | si:ch211-37e10.2 |
| fstl1b7 | 1.391957 | 1.37919617 | 0.909 | 0.146 | 3.5362684 | 36 | fstl1b |
| dcn5 | 1.585422 | 1.412280406 | 0.909 | 0.142 | 4.0277651 | 36 | dcn |
| anos1b | 2.360000 | 0.419363017 | 0.218 | 0.008 | 5.9955823 | 36 | anos1b |
| pigr1 | 3.816951 | 0.695481881 | 0.364 | 0.023 | 9.6969646 | 36 | pigr |
| igf1 | 4.970806 | 0.317647465 | 0.127 | 0.003 | 1.2628333 | 36 | igf1 |
| serpinf16 | 3.731892 | 0.854594272 | 0.509 | 0.044 | 9.4808734 | 36 | serpinf1 |
| thbs2a | 4.181243 | 0.55796374 | 0.236 | 0.01 | 1.0622449 | 36 | thbs2a |
| aoc26 | 1.284261 | 1.226213109 | 0.655 | 0.076 | 3.2626652 | 36 | aoc2 |
| sparc9 | 1.100204 | 1.542650284 | 0.982 | 0.194 | 2.7950684 | 36 | sparc |
| col17a1b1 | 4.832322 | 0.263478071 | 0.109 | 0.002 | 1.2276514 | 36 | col17a1b |
| rbfox12 | 2.114281 | 0.732292581 | 0.4 | 0.029 | 5.3713333 | 36 | rbfox1 |
| tpm4a10 | 4.645897 | 1.23091323 | 0.782 | 0.112 | 1.1802901 | 36 | tpm4a |
| efna5b1 | 5.292709 | 0.560913956 | 0.236 | 0.01 | 1.3446129 | 36 | efna5b |
| lamc3 | 2.312919 | 0.287951253 | 0.145 | 0.004 | 5.8759720 | 36 | lamc3 |
| col6a35 | 2.569394 | 1.212235301 | 0.782 | 0.113 | 6.5275464 | 36 | col6a3 |
| omd | 1.412270 | 0.341897959 | 0.109 | 0.002 | 3.5878736 | 36 | omd |
| srpx21 | 1.597580 | 0.424203475 | 0.218 | 0.009 | 4.0586526 | 36 | srpx2 |
| zgc:153704 | 1.712954 | 0.490991181 | 0.2 | 0.007 | 4.3517607 | 36 | zgc:153704 |
| pcsk9 | 4.435226 | 0.267348795 | 0.127 | 0.003 | 1.1267692 | 36 | pcsk9 |
| ecm1b6 | 1.603927 | 0.893444015 | 0.6 | 0.066 | 4.0747788 | 36 | ecm1b |
| tmeff2a | 1.700529 | 0.337602124 | 0.145 | 0.004 | 4.3201945 | 36 | tmeff2a |
| col1a1b6 | 1.848564 | 1.572069404 | 0.891 | 0.169 | 4.6962789 | 36 | col1a1b |
| cdh113 | 4.762169 | 0.872148605 | 0.382 | 0.029 | 1.2098290 | 36 | cdh11 |
| col6a25 | 9.938385 | 1.004728028 | 0.618 | 0.074 | 2.5248468 | 36 | col6a2 |
| fam69c | 1.457834 | 0.605931372 | 0.291 | 0.017 | 3.7036282 | 36 | fam69c |
| sema3d2 | 1.506305 | 0.53883596 | 0.309 | 0.018 | 3.8267680 | 36 | sema3d |
| hmbox1a1 | 5.312332 | 0.690497453 | 0.436 | 0.039 | 1.3495981 | 36 | hmbox1a |
| fbn2b5 | 1.845272 | 0.889381376 | 0.491 | 0.05 | 4.6879138 | 36 | fbn2b |
| ehd2b2 | 2.170405 | 0.740161119 | 0.4 | 0.033 | 5.5139149 | 36 | ehd2b |

| | | | | | | | |
|---|---|---|---|---|---|---|---|
| plpp32 | 2.467467 | 1.05803033 | 0.564 | 0.065 | 6.2686022 | 36 | plpp3 |
| efemp2a | 3.837918 | 0.5014955 | 0.273 | 0.016 | 9.7502325 | 36 | efemp2a |
| col12a1a2 | 1.940847 | 0.743881495 | 0.382 | 0.031 | 4.9307224 | 36 | col12a1a |
| wnt16 | 2.379868 | 0.321604064 | 0.145 | 0.004 | 6.0460556 | 36 | wnt16 |
| olfml3b6 | 3.886967 | 0.884862675 | 0.509 | 0.055 | 9.8748411 | 36 | olfml3b |
| LOC101885478 | 3.408477 | 0.263515182 | 0.109 | 0.002 | 8.6592371 | 36 | LOC101885478 |
| ccl441 | 9.380791 | 0.367460196 | 0.145 | 0.005 | 2.3831900 | 36 | ccl44 |
| pmp22a6 | 1.269223 | 1.292856332 | 0.873 | 0.166 | 3.2244617 | 36 | pmp22a |
| col6a16 | 1.816525 | 1.061479114 | 0.673 | 0.097 | 4.6148819 | 36 | col6a1 |
| csf1b | 5.998066 | 0.295573937 | 0.109 | 0.003 | 1.5238086 | 36 | csf1b |
| sostdc1a2 | 7.210313 | 0.771113662 | 0.327 | 0.024 | 1.8317801 | 36 | sostdc1a |
| meox1 | 1.277201 | 0.286391567 | 0.127 | 0.004 | 3.2447305 | 36 | meox1 |
| cpn16 | 4.857891 | 1.01828422 | 0.582 | 0.077 | 1.2341473 | 36 | cpn1 |
| si:ch211-195b15.8 | 1.238218 | 0.355608842 | 0.164 | 0.006 | 3.1456938 | 36 | si:ch211-195b15.8 |
| mmp14a5 | 4.747466 | 0.818047493 | 0.509 | 0.059 | 1.2060938 | 36 | mmp14a |
| ebf3a1 | 8.083665 | 0.39279222 | 0.2 | 0.009 | 2.0536552 | 36 | ebf3a |
| tmem88a1 | 1.086293 | 0.421696536 | 0.2 | 0.009 | 2.7597291 | 36 | tmem88a |
| prss354 | 1.151147 | 0.821501474 | 0.345 | 0.028 | 2.9244899 | 36 | prss35 |
| zgc:1626122 | 3.744984 | 0.351503297 | 0.164 | 0.006 | 9.5141318 | 36 | zgc:162612 |
| s100v211 | 1.027819 | 1.124493468 | 0.764 | 0.14 | 2.6111749 | 36 | s100v2 |
| fbln51 | 1.231528 | 0.807063754 | 0.455 | 0.048 | 3.1286970 | 36 | fbln5 |
| ckba5 | 2.712124 | 0.854979415 | 0.564 | 0.073 | 6.8901530 | 36 | ckba |
| marcksl1b11 | 8.148656 | 1.250376198 | 0.745 | 0.141 | 2.0701662 | 36 | marcksl1b |
| filip1l1 | 2.784529 | 0.479169216 | 0.273 | 0.018 | 7.0740975 | 36 | filip1l |
| svep1 | 4.003585 | 0.41248906 | 0.2 | 0.01 | 1.0171109 | 36 | svep1 |
| zgc:1135311 | 5.842014 | 0.599999241 | 0.309 | 0.024 | 1.4841636 | 36 | zgc:113531 |
| cavin2b7 | 1.298648 | 0.965991013 | 0.564 | 0.078 | 3.2992173 | 36 | cavin2b |
| efna5a | 1.584947 | 0.279068876 | 0.109 | 0.003 | 4.0265601 | 36 | efna5a |
| pcdh7b | 1.431872 | 0.420576813 | 0.182 | 0.009 | 3.6376727 | 36 | pcdh7b |
| krt916 | 2.752917 | 0.963993364 | 0.545 | 0.071 | 6.9937871 | 36 | krt91 |
| col8a2 | 3.015468 | 0.271653673 | 0.127 | 0.004 | 7.6607969 | 36 | col8a2 |
| bcl11aa2 | 4.245574 | 0.397502818 | 0.2 | 0.011 | 1.0785880 | 36 | bcl11aa |
| rnd3b4 | 1.474860 | 0.762515259 | 0.436 | 0.049 | 3.7468819 | 36 | rnd3b |
| twist1b | 1.496694 | 0.331761477 | 0.164 | 0.007 | 3.8023516 | 36 | twist1b |
| fkbp76 | 1.771740 | 0.735408874 | 0.436 | 0.05 | 4.5011075 | 36 | fkbp7 |

| | | | | | | | |
|---|---|---|---|---|---|---|---|
| si:ch73-46j18.510 | 2.937113 | 1.187431344 | 0.873 | 0.208 | 7.4617365 | 36 | si:ch73-46j18.5 |
| hmcn1 | 1.163016 | 0.325754032 | 0.164 | 0.007 | 2.9546439 | 36 | hmcn1 |
| ogna1 | 1.234646 | 0.431687254 | 0.182 | 0.009 | 3.1366194 | 36 | ogna |
| emilin1b | 1.368883 | 0.482323682 | 0.236 | 0.015 | 3.4776495 | 36 | emilin1b |
| raph1a | 2.756675 | 0.31060808 | 0.164 | 0.007 | 7.0033351 | 36 | raph1a |
| twsg1a | 3.430052 | 0.412654987 | 0.236 | 0.015 | 8.7140478 | 36 | twsg1a |
| ntn1b | 5.132093 | 0.275961746 | 0.109 | 0.003 | 1.3038084 | 36 | ntn1b |
| cald1a9 | 6.194551 | 1.007898052 | 0.673 | 0.119 | 1.5737257 | 36 | cald1a |
| zfhx3 | 1.407478 | 0.601048627 | 0.345 | 0.033 | 3.5757000 | 36 | zfhx3 |
| si:ch211-39i22.16 | 2.253060 | 1.03300519 | 0.564 | 0.088 | 5.7238998 | 36 | si:ch211-39i22.1 |
| si:dkey-238c7.161 | 2.638005 | 0.881847963 | 0.509 | 0.07 | 6.7018532 | 36 | si:dkey-238c7.16 |
| tnfsf122 | 2.960271 | 0.586528961 | 0.309 | 0.027 | 7.5205692 | 36 | tnfsf12 |
| hsp70.1 | 3.687234 | 0.251005055 | 0.109 | 0.003 | 9.3674195 | 36 | hsp70.1 |
| mmp14b6 | 9.166820 | 0.802859902 | 0.491 | 0.066 | 2.3288307 | 36 | mmp14b |
| cav18 | 1.579169 | 0.813703808 | 0.418 | 0.049 | 4.0118798 | 36 | cav1 |
| gas1a5 | 1.844963 | 0.759826376 | 0.455 | 0.058 | 4.6871300 | 36 | gas1a |
| paplna2 | 1.856080 | 0.652354985 | 0.309 | 0.027 | 4.7153719 | 36 | paplna |
| serpinh1b2 | 2.914926 | 0.602960819 | 0.327 | 0.03 | 7.4053697 | 36 | serpinh1b |
| cldn11a6 | 3.155025 | 1.038960676 | 0.545 | 0.083 | 8.0153427 | 36 | cldn11a |
| pabpc1b5 | 6.387133 | 0.803781912 | 0.527 | 0.077 | 1.6226512 | 36 | pabpc1b |
| cd81a9 | 7.146597 | 1.157757745 | 0.909 | 0.24 | 1.8155931 | 36 | cd81a |
| si:ch211-105c13.36 | 1.960912 | 0.844043667 | 0.491 | 0.067 | 4.9816991 | 36 | si:ch211-105c13.3 |
| fsta1 | 2.152901 | 0.369911719 | 0.145 | 0.006 | 5.4694468 | 36 | fsta |
| cercam | 2.223817 | 0.451582856 | 0.255 | 0.019 | 5.6496091 | 36 | cercam |
| si:dkey-261h17.14 | 2.365266 | 1.172312029 | 0.636 | 0.115 | 6.0089589 | 36 | si:dkey-261h17.1 |
| fzd7a | 6.625012 | 0.451002144 | 0.236 | 0.017 | 1.6830844 | 36 | fzd7a |
| cavin2a2 | 1.412786 | 0.437141317 | 0.236 | 0.017 | 3.5891837 | 36 | cavin2a |
| net14 | 2.038566 | 0.545476 | 0.327 | 0.032 | 5.1789789 | 36 | net1 |
| si:dkey-16l2.203 | 2.394081 | 0.50025622 | 0.273 | 0.023 | 6.0821633 | 36 | si:dkey-16l2.20 |
| cfd5 | 3.175272 | 1.005414884 | 0.491 | 0.07 | 8.0667798 | 36 | cfd |
| grm4 | 3.936508 | 0.392758078 | 0.182 | 0.01 | 1.0000701 | 36 | grm4 |
| auts2a | 4.005832 | 0.4295842 | 0.218 | 0.015 | 1.0176816 | 36 | auts2a |
| rbp55 | 9.796922 | 0.834433441 | 0.636 | 0.11 | 2.4889082 | 36 | rbp5 |
| plxdc21 | 1.178253 | 0.390235259 | 0.2 | 0.013 | 2.9933531 | 36 | plxdc2 |
| col21a12 | 1.394288 | 0.547546354 | 0.255 | 0.02 | 3.5421901 | 36 | col21a1 |

| | | | | | | | |
|---|---|---|---|---|---|---|---|
| cspg42 | 4.252811 | 0.350436824 | 0.145 | 0.007 | 1.0804266 | 36 | cspg4 |
| metrnla7 | 4.354240 | 0.891055707 | 0.564 | 0.094 | 1.1061948 | 36 | metrnla |
| qsox11 | 7.125399 | 0.656928656 | 0.327 | 0.034 | 1.8102077 | 36 | qsox1 |
| si:ch73-217n20.1 | 8.448731 | 0.38287264 | 0.182 | 0.011 | 2.1464003 | 36 | si:ch73-217n20.1 |
| fgf18a | 1.308883 | 0.278579175 | 0.127 | 0.005 | 3.3252178 | 36 | fgf18a |
| CABZ01074363.11 | 1.663079 | 0.652214341 | 0.291 | 0.027 | 4.2250544 | 36 | CABZ01074363.1 |
| nfixb5 | 2.263102 | 0.780858564 | 0.527 | 0.082 | 5.7494118 | 36 | nfixb |
| ahnak11 | 4.203933 | 0.959166646 | 0.673 | 0.136 | 1.0680092 | 36 | ahnak |
| cd1512 | 4.502918 | 0.560395677 | 0.255 | 0.021 | 1.1439664 | 36 | cd151 |
| tfpia1 | 8.535158 | 0.494291029 | 0.255 | 0.021 | 2.1683570 | 36 | tfpia |
| glis2a2 | 9.957520 | 0.498525664 | 0.291 | 0.027 | 2.5297081 | 36 | glis2a |
| cnpy4 | 1.351754 | 0.473557071 | 0.327 | 0.034 | 3.4341326 | 36 | cnpy4 |
| chd37 | 5.530593 | 0.847485899 | 0.636 | 0.125 | 1.4050473 | 36 | chd3 |
| RAMP1 | 7.515912 | 0.288283673 | 0.127 | 0.006 | 1.9094174 | 36 | RAMP1 |
| meis1b8 | 1.093548 | 0.77042277 | 0.473 | 0.073 | 2.7781596 | 36 | meis1b |
| lrp5 | 1.686312 | 0.375296765 | 0.2 | 0.014 | 4.2840770 | 36 | lrp5 |
| tmem119b2 | 1.766645 | 0.442475151 | 0.255 | 0.022 | 4.4881639 | 36 | tmem119b |
| actn15 | 2.860478 | 0.616775008 | 0.382 | 0.048 | 7.2670450 | 36 | actn1 |
| mex3b2 | 2.959414 | 0.441324086 | 0.236 | 0.019 | 7.5183917 | 36 | mex3b |
| si:ch211-251b21.12 | 3.581600 | 0.347479625 | 0.182 | 0.011 | 9.0990564 | 36 | si:ch211-251b21.1 |
| fkbp93 | 3.694360 | 0.57747005 | 0.327 | 0.036 | 9.3855226 | 36 | fkbp9 |
| si:ch211-195m9.3 | 4.458710 | 0.45829912 | 0.2 | 0.014 | 1.1327353 | 36 | si:ch211-195m9.3 |
| si:ch211-133n4.44 | 5.428593 | 0.705375012 | 0.418 | 0.059 | 1.3791340 | 36 | si:ch211-133n4.4 |
| crispld21 | 1.153371 | 0.547123305 | 0.236 | 0.02 | 2.9301397 | 36 | crispld2 |
| myl69 | 1.669654 | 0.889269894 | 0.6 | 0.116 | 4.2417568 | 36 | myl6 |
| stom1 | 1.988717 | 0.467293095 | 0.255 | 0.023 | 5.0523380 | 36 | stom |
| igf2a1 | 2.255811 | 0.326264018 | 0.145 | 0.008 | 5.7308883 | 36 | igf2a |
| sema3c | 2.273588 | 0.292894099 | 0.145 | 0.008 | 5.7760507 | 36 | sema3c |
| adamts15a5 | 2.475939 | 0.675779781 | 0.418 | 0.058 | 6.2901247 | 36 | adamts15a |
| tgfb33 | 2.861739 | 0.562744884 | 0.309 | 0.033 | 7.2702480 | 36 | tgfb3 |
| nrp2b1 | 6.146818 | 0.352258562 | 0.182 | 0.012 | 1.5615992 | 36 | nrp2b |
| fhod3b1 | 7.820148 | 0.309329783 | 0.145 | 0.008 | 1.9867086 | 36 | fhod3b |
| tril1 | 8.064160 | 0.492277681 | 0.273 | 0.026 | 2.0486999 | 36 | tril |
| agmo | 8.134681 | 0.304294775 | 0.164 | 0.01 | 2.0666157 | 36 | agmo |
| mir214a | 1.116760 | 0.281121322 | 0.127 | 0.006 | 2.8371308 | 36 | mir214a |

| | | | | | | | |
|---|---|---|---|---|---|---|---|
| hoxa9b | 1.213596 | 0.356588546 | 0.182 | 0.012 | 3.0831413 | 36 | hoxa9b |
| c1qtnf6a2 | 4.055571 | 0.465187863 | 0.273 | 0.027 | 1.0303179 | 36 | c1qtnf6a |
| hoxc8a | 7.600087 | 0.434091666 | 0.236 | 0.021 | 1.9308021 | 36 | hoxc8a |
| LOC1039092847 | 8.595300 | 0.696713845 | 0.473 | 0.077 | 2.1836360 | 36 | LOC103909284 |
| parvaa2 | 8.724615 | 0.509897793 | 0.309 | 0.035 | 2.2164886 | 36 | parvaa |
| SCARF23 | 1.413338 | 0.462449026 | 0.273 | 0.028 | 3.5905862 | 36 | SCARF2 |
| p4ha21 | 1.420240 | 0.386231039 | 0.2 | 0.015 | 3.6081202 | 36 | p4ha2 |
| vwde | 1.618294 | 0.369570033 | 0.109 | 0.005 | 4.1112778 | 36 | vwde |
| edil3a3 | 1.801634 | 0.54704802 | 0.255 | 0.025 | 4.5770526 | 36 | edil3a |
| sema6e | 2.335023 | 0.287104978 | 0.164 | 0.01 | 5.9321266 | 36 | sema6e |
| calua2 | 4.776196 | 0.557587217 | 0.291 | 0.033 | 1.2133926 | 36 | calua |
| ckap47 | 9.437589 | 0.681557006 | 0.4 | 0.06 | 2.3976197 | 36 | ckap4 |
| bmp33 | 1.075299 | 0.41453373 | 0.182 | 0.013 | 2.7317984 | 36 | bmp3 |
| dbn15 | 1.313165 | 0.705478327 | 0.418 | 0.066 | 3.3360957 | 36 | dbn1 |
| pawr | 1.379569 | 0.391903642 | 0.236 | 0.022 | 3.5047957 | 36 | pawr |
| aebp13 | 2.016722 | 0.513327948 | 0.364 | 0.048 | 5.1234836 | 36 | aebp1 |
| myl9b8 | 6.489421 | 0.752038208 | 0.527 | 0.099 | 1.6486376 | 36 | myl9b |
| itgb3b1 | 7.335322 | 0.372837062 | 0.182 | 0.014 | 1.8635386 | 36 | itgb3b |
| hsd11b27 | 7.472768 | 1.059969214 | 0.655 | 0.149 | 1.8984569 | 36 | hsd11b2 |
| reck2 | 1.085705 | 0.486783234 | 0.273 | 0.03 | 2.7582338 | 36 | reck |
| si:ch211-51c14.1 | 1.369373 | 0.25795095 | 0.127 | 0.007 | 3.4788932 | 36 | si:ch211-51c14.1 |
| sgcb | 1.806817 | 0.344568242 | 0.182 | 0.014 | 4.5902188 | 36 | sgcb |
| sdc23 | 2.128282 | 0.668153334 | 0.473 | 0.082 | 5.4069013 | 36 | sdc2 |
| egfl71 | 2.956220 | 0.297908627 | 0.127 | 0.007 | 7.5102769 | 36 | egfl7 |
| cald1b | 3.684031 | 0.343098706 | 0.145 | 0.009 | 9.3592826 | 36 | cald1b |
| tgfbr33 | 4.248776 | 0.571866435 | 0.345 | 0.046 | 1.0794017 | 36 | tgfbr3 |
| cd99l22 | 6.900402 | 0.50182112 | 0.291 | 0.035 | 1.7530471 | 36 | cd99l2 |
| akap12b6 | 7.484339 | 0.790787917 | 0.436 | 0.074 | 1.9013963 | 36 | akap12b |
| itgb1a5 | 7.619467 | 0.594305718 | 0.364 | 0.053 | 1.9357256 | 36 | itgb1a |
| si:ch211-106j24.11 | 7.904202 | 0.381321829 | 0.218 | 0.02 | 2.0080627 | 36 | si:ch211-106j24.1 |
| gstr2 | 8.954254 | 0.689093284 | 0.6 | 0.127 | 2.2748283 | 36 | gstr |
| cldn11b1 | 9.696019 | 0.301524201 | 0.182 | 0.014 | 2.4632736 | 36 | cldn11b |
| pleca6 | 1.135009 | 0.612921075 | 0.4 | 0.062 | 2.8834914 | 36 | pleca |
| hmgb3a7 | 1.259552 | 0.788557904 | 0.455 | 0.083 | 3.1998926 | 36 | hmgb3a |
| cnn3a9 | 1.771037 | 0.64280643 | 0.418 | 0.069 | 4.4993203 | 36 | cnn3a |

| | | | | | | | |
|---|---|---|---|---|---|---|---|
| fhl1b2 | 1.915614 | 0.328206064 | 0.218 | 0.02 | 4.8666198 | 36 | fhl1b |
| jun12 | 2.150919 | 1.075399329 | 0.8 | 0.25 | 5.4644118 | 36 | jun |
| nfia7 | 2.372434 | 0.733024039 | 0.509 | 0.099 | 6.0271703 | 36 | nfia |
| itgb43 | 2.845919 | 0.367155175 | 0.2 | 0.017 | 7.2300575 | 36 | itgb4 |
| marcksl1a10 | 3.786757 | 1.04046174 | 0.8 | 0.261 | 9.6202566 | 36 | marcksl1a |
| loxl3b | 4.572099 | 0.379769036 | 0.182 | 0.015 | 1.1615419 | 36 | loxl3b |
| htra1a2 | 4.787988 | 0.557468202 | 0.236 | 0.024 | 1.2163883 | 36 | htra1a |
| jcada2 | 6.816114 | 0.428290628 | 0.218 | 0.021 | 1.7316337 | 36 | jcada |
| tgfb21 | 1.190817 | 0.330327486 | 0.182 | 0.015 | 3.0252717 | 36 | tgfb2 |
| fzd1 | 1.229509 | 0.30194672 | 0.145 | 0.01 | 3.1235692 | 36 | fzd1 |
| bnc22 | 1.233158 | 0.54845902 | 0.309 | 0.04 | 3.1328386 | 36 | bnc2 |
| jag1b4 | 1.346845 | 0.551063275 | 0.255 | 0.029 | 3.4216606 | 36 | jag1b |
| hic14 | 1.682281 | 0.633144562 | 0.418 | 0.07 | 4.2738366 | 36 | hic1 |
| FQ323119.11 | 2.898192 | 0.286634625 | 0.127 | 0.008 | 7.3628575 | 36 | FQ323119.1 |
| pag1 | 3.079306 | 0.336796416 | 0.2 | 0.018 | 7.8229784 | 36 | pag1 |
| slit31 | 4.261543 | 0.531153512 | 0.291 | 0.036 | 1.0826451 | 36 | slit3 |
| kank4 | 4.318150 | 0.329342279 | 0.145 | 0.01 | 1.0970261 | 36 | kank4 |
| CR847566.14 | 6.337481 | 0.590092959 | 0.345 | 0.051 | 1.6100370 | 36 | CR847566.1 |
| sox11a2 | 6.375770 | 0.547755039 | 0.255 | 0.029 | 1.6197644 | 36 | sox11a |
| vmo1a1 | 6.991106 | 0.313954802 | 0.109 | 0.006 | 1.7760907 | 36 | vmo1a |
| LO018340.11 | 7.584965 | 0.479795452 | 0.255 | 0.029 | 1.9269606 | 36 | LO018340.1 |
| krt18b9 | 8.268477 | 0.795582092 | 0.709 | 0.176 | 2.1006067 | 36 | krt18b |
| lgals2a5 | 1.043261 | 1.015443061 | 0.582 | 0.138 | 2.6504067 | 36 | lgals2a |
| si:ch211-250c4.42 | 1.070128 | 0.398568956 | 0.2 | 0.018 | 2.7186607 | 36 | si:ch211-250c4.4 |
| gng78 | 1.404244 | 0.572058839 | 0.382 | 0.062 | 3.5674828 | 36 | gng7 |
| slc43a3b3 | 1.626710 | 0.477292973 | 0.255 | 0.029 | 4.1326569 | 36 | slc43a3b |
| amotl2b1 | 1.908889 | 0.37356697 | 0.182 | 0.016 | 4.8495332 | 36 | amotl2b |
| adamts15b1 | 2.097578 | 0.487755734 | 0.236 | 0.025 | 5.3288969 | 36 | adamts15b |
| fbln1 | 2.733306 | 0.296905083 | 0.109 | 0.006 | 6.9439661 | 36 | fbln1 |
| mrc23 | 3.251210 | 0.487811527 | 0.255 | 0.03 | 8.2597001 | 36 | mrc2 |
| tmem2041 | 3.263400 | 0.310338085 | 0.145 | 0.01 | 8.2906698 | 36 | tmem204 |
| wnt5b2 | 3.914187 | 0.367704238 | 0.182 | 0.016 | 9.9439929 | 36 | wnt5b |
| lepr | 5.033979 | 0.307547049 | 0.127 | 0.008 | 1.2788825 | 36 | lepr |
| lrrc175 | 5.725832 | 0.648287709 | 0.364 | 0.058 | 1.4546476 | 36 | lrrc17 |
| ppib12 | 6.272549 | 0.929721013 | 0.855 | 0.379 | 1.5935411 | 36 | ppib |

| | | | | | | | |
|---|---|---|---|---|---|---|---|
| tuft1a3 | 6.934064 | 0.491499433 | 0.255 | 0.03 | 1.7615989 | 36 | tuft1a |
| myh105 | 7.864621 | 0.493367455 | 0.309 | 0.043 | 1.9980071 | 36 | myh10 |
| utrn1 | 1.157542 | 0.474441465 | 0.273 | 0.034 | 2.9407364 | 36 | utrn |
| angptl1a3 | 1.172203 | 0.608355371 | 0.273 | 0.035 | 2.9779828 | 36 | angptl1a |
| s100a10b13 | 1.478516 | 0.995110934 | 0.836 | 0.285 | 3.7561721 | 36 | s100a10b |
| znf385a5 | 2.024828 | 0.620709403 | 0.345 | 0.054 | 5.1440768 | 36 | znf385a |
| ddr2a | 2.438434 | 0.284925526 | 0.127 | 0.008 | 6.1948420 | 36 | ddr2a |
| BX088707.31 | 4.321989 | 0.306536441 | 0.127 | 0.008 | 1.0980014 | 36 | BX088707.3 |
| si:ch211-67e16.11 | 4.495193 | 0.263493836 | 0.127 | 0.008 | 1.1420039 | 36 | si:ch211-67e16.11 |
| ccdc3b3 | 4.746368 | 0.449463839 | 0.255 | 0.031 | 1.2058148 | 36 | ccdc3b |
| fam20a1 | 5.259664 | 0.263005355 | 0.109 | 0.006 | 1.3362176 | 36 | fam20a |
| si:ch211-212k18.94 | 5.379823 | 0.515500171 | 0.327 | 0.05 | 1.3667440 | 36 | si:ch211-212k18.9 |
| prr12b | 9.073529 | 0.316483837 | 0.164 | 0.014 | 2.3051301 | 36 | prr12b |
| dedd16 | 9.716791 | 0.782660895 | 0.6 | 0.157 | 2.4685507 | 36 | dedd1 |
| CABZ01092746.14 | 1.056036 | 0.726372226 | 0.455 | 0.089 | 2.6828597 | 36 | CABZ01092746.1 |
| sgce3 | 1.076297 | 0.470141602 | 0.255 | 0.032 | 2.7343337 | 36 | sgce |
| CR383676.120 | 1.150803 | 0.794932065 | 1 | 0.717 | 2.9236150 | 36 | CR383676.1 |
| mgst1.25 | 1.650780 | 0.672712307 | 0.527 | 0.116 | 4.1938085 | 36 | mgst1.2 |
| cygb15 | 2.017011 | 0.656022055 | 0.364 | 0.062 | 5.1242165 | 36 | cygb1 |
| vat17 | 2.253801 | 0.747703757 | 0.6 | 0.147 | 5.7257835 | 36 | vat1 |
| ackr3b3 | 5.924036 | 0.47986212 | 0.255 | 0.032 | 1.5050015 | 36 | ackr3b |
| fermt21 | 8.062838 | 0.450552729 | 0.255 | 0.033 | 2.0483640 | 36 | fermt2 |
| pcolcea3 | 8.915534 | 0.701248672 | 0.382 | 0.068 | 2.2649914 | 36 | pcolcea |
| arl6ip5b3 | 8.942076 | 0.678134786 | 0.473 | 0.103 | 2.2717344 | 36 | arl6ip5b |
| fzd24 | 9.114033 | 0.485418221 | 0.291 | 0.043 | 2.3154202 | 36 | fzd2 |
| ccdc802 | 1.069704 | 0.477545245 | 0.255 | 0.033 | 2.7175835 | 36 | ccdc80 |
| ccl19a.12 | 1.087946 | 0.420851035 | 0.164 | 0.014 | 2.7639275 | 36 | ccl19a.1 |
| myocd1 | 1.094426 | 0.294781612 | 0.127 | 0.009 | 2.7803915 | 36 | myocd |
| krt45 | 1.337799 | 0.526929268 | 0.327 | 0.051 | 3.3986798 | 36 | krt4 |
| meis1a3 | 1.823793 | 0.296240488 | 0.2 | 0.021 | 4.6333461 | 36 | meis1a |
| tmem176l.17 | 2.013135 | 0.791331678 | 0.691 | 0.202 | 5.1143715 | 36 | tmem176l.1 |
| cd248a3 | 2.266587 | 0.526731254 | 0.309 | 0.048 | 5.7582655 | 36 | cd248a |
| anxa1a9 | 2.363602 | 0.829991245 | 0.618 | 0.163 | 6.0047324 | 36 | anxa1a |
| hmgb1b10 | 3.482299 | 0.623817972 | 0.618 | 0.154 | 8.8467815 | 36 | hmgb1b |
| quo | 3.587522 | 0.258179219 | 0.127 | 0.009 | 9.1141000 | 36 | quo |

| | | | | | | | |
|---|---|---|---|---|---|---|---|
| slc38a5b9 | 5.29927[illegible] | 0.685905093 | 0.527 | 0.129 | 1.3462820 | 36 | slc38a5b |
| epha31 | 6.767132 | 0.36264495 | 0.2 | 0.022 | 1.7191901 | 36 | epha3 |
| lpl1 | 7.837492 | 0.290207218 | 0.2 | 0.021 | 1.9911149 | 36 | lpl |
| crtap4 | 9.457460 | 0.530587726 | 0.273 | 0.04 | 2.4026678 | 36 | crtap |
| nucb2a2 | 1.151941 | 0.505746698 | 0.345 | 0.061 | 2.9265081 | 36 | nucb2a |
| roraa | 1.542829 | 0.271136773 | 0.182 | 0.018 | 3.9195594 | 36 | roraa |
| postnb5 | 1.591104 | 0.950806591 | 0.436 | 0.096 | 4.0422019 | 36 | postnb |
| fat1a3 | 1.711099 | 0.445291588 | 0.236 | 0.031 | 4.3470470 | 36 | fat1a |
| calm2b11 | 2.528684 | 0.764127275 | 0.764 | 0.247 | 6.4241219 | 36 | calm2b |
| mgst1.14 | 2.610445 | 0.508748672 | 0.327 | 0.056 | 6.6318367 | 36 | mgst1.1 |
| cd28l3 | 4.633967 | 0.438733503 | 0.273 | 0.04 | 1.1772593 | 36 | cd28l |
| ptbp1b2 | 6.639013 | 0.420849938 | 0.273 | 0.04 | 1.6866413 | 36 | ptbp1b |
| ptprk2 | 7.166531 | 0.479476309 | 0.255 | 0.036 | 1.8206573 | 36 | ptprk |
| fbln22 | 1.259412 | 0.483801835 | 0.218 | 0.028 | 3.1995374 | 36 | fbln2 |
| efna1b5 | 1.356652 | 0.379341216 | 0.182 | 0.02 | 3.4465762 | 36 | efna1b |
| lsp11 | 1.516192 | 0.355171911 | 0.182 | 0.02 | 3.8518875 | 36 | lsp1 |
| pdcd61 | 2.216804 | 0.555019113 | 0.4 | 0.083 | 5.6317911 | 36 | pdcd6 |
| klf7b | 2.887990 | 0.41526207 | 0.255 | 0.037 | 7.3369411 | 36 | klf7b |
| tmem45a2 | 3.521317 | 0.339326008 | 0.218 | 0.028 | 8.9459059 | 36 | tmem45a |
| XLOC-0045561 | 3.841657 | 0.313678248 | 0.164 | 0.016 | 9.7597307 | 36 | XLOC-004556 |
| fam107b5 | 3.912479 | 0.782800956 | 0.455 | 0.113 | 9.9396534 | 36 | fam107b |
| hoxc9a | 6.543150 | 0.286497751 | 0.164 | 0.017 | 1.6622873 | 36 | hoxc9a |
| thrap3a2 | 7.037037 | 0.29949772 | 0.145 | 0.013 | 1.7877593 | 36 | thrap3a |
| col6a4a4 | 9.910736 | 0.64273104 | 0.345 | 0.066 | 2.5178226 | 36 | col6a4a |
| flot2a7 | 1.192981 | 0.537640955 | 0.364 | 0.074 | 3.0307688 | 36 | flot2a |
| atrn | 1.557322 | 0.294325909 | 0.164 | 0.017 | 3.9563782 | 36 | atrn |
| fam198a1 | 1.786011 | 0.372996624 | 0.182 | 0.021 | 4.5373622 | 36 | fam198a |
| calr3 | 1.950214 | 0.516418206 | 0.345 | 0.067 | 4.9545199 | 36 | calr |
| kirrel1a2 | 2.411323 | 0.453399779 | 0.236 | 0.034 | 6.1259682 | 36 | kirrel1a |
| rasgef1bb2 | 2.413123 | 0.374982269 | 0.236 | 0.033 | 6.1305410 | 36 | rasgef1bb |
| ghra3 | 2.766369 | 0.451211796 | 0.236 | 0.034 | 7.0279611 | 36 | ghra |
| slc22a18 | 2.832780 | 0.255231491 | 0.145 | 0.014 | 7.1966792 | 36 | slc22a18 |
| tenm34 | 3.012453 | 0.550104365 | 0.327 | 0.062 | 7.6531387 | 36 | tenm3 |
| arhgap10 | 3.016757 | 0.315188546 | 0.2 | 0.025 | 7.6640726 | 36 | arhgap10 |
| tspan9b5 | 3.228126 | 0.495847482 | 0.291 | 0.051 | 8.2010560 | 36 | tspan9b |

| | | | | | | | |
|---|---|---|---|---|---|---|---|
| flna7 | 3.249384 | 0.724289673 | 0.473 | 0.122 | 8.2550622 | 36 | flna |
| nhsl2 | 3.728434 | 0.288298538 | 0.127 | 0.011 | 9.4720888 | 36 | nhsl2 |
| ccnd13 | 4.008969 | 0.399174015 | 0.273 | 0.044 | 1.0184785 | 36 | ccnd1 |
| fhl1a | 4.200664 | 0.250677308 | 0.109 | 0.008 | 1.0671786 | 36 | fhl1a |
| tpte1 | 4.456150 | 0.39830232 | 0.218 | 0.03 | 1.1320849 | 36 | tpte |
| lrp1ab4 | 4.843457 | 0.501553015 | 0.382 | 0.078 | 1.2304804 | 36 | lrp1ab |
| klf6a9 | 6.081997 | 0.720517937 | 0.6 | 0.18 | 1.5451314 | 36 | klf6a |
| si:busm1-57f23.13 | 7.458589 | 0.45066901 | 0.218 | 0.03 | 1.8948546 | 36 | si:busm1-57f23.1 |
| gpc6a1 | 7.555338 | 0.27844772 | 0.145 | 0.014 | 1.9194337 | 36 | gpc6a |
| si:dkey-7j14.67 | 7.686252 | 0.682500259 | 0.636 | 0.196 | 1.9526924 | 36 | si:dkey-7j14.6 |
| pdia51 | 8.578332 | 0.445075563 | 0.291 | 0.05 | 2.1793254 | 36 | pdia5 |
| h3f3c14 | 8.708312 | 0.763669752 | 0.764 | 0.304 | 2.2123468 | 36 | h3f3c |
| csad | 9.801883 | 0.256313638 | 0.164 | 0.018 | 2.4901685 | 36 | csad |
| gadd45ba10 | 1.007294 | 0.796236464 | 0.618 | 0.198 | 2.5590321 | 36 | gadd45ba |
| BX663503.32 | 1.140657 | 0.382643514 | 0.236 | 0.034 | 2.8978415 | 36 | BX663503.3 |
| MMP23B | 1.195369 | 0.318036803 | 0.182 | 0.022 | 3.0368358 | 36 | MMP23B |
| plod1a1 | 1.219627 | 0.333219629 | 0.2 | 0.026 | 3.0984647 | 36 | plod1a |
| scara32 | 1.269003 | 0.454218342 | 0.236 | 0.035 | 3.2239023 | 36 | scara3 |
| p3h4 | 1.296930 | 0.286475452 | 0.145 | 0.014 | 3.2948530 | 36 | p3h4 |
| cfap45 | 1.812189 | 0.301936678 | 0.127 | 0.011 | 4.6038678 | 36 | cfap45 |
| igfbp5b4 | 2.081971 | 0.66495705 | 0.4 | 0.089 | 5.2892486 | 36 | igfbp5b |
| appb5 | 2.092157 | 0.479300699 | 0.273 | 0.047 | 5.3151261 | 36 | appb |
| ephb36 | 2.344249 | 0.478769193 | 0.291 | 0.052 | 5.9555650 | 36 | ephb3 |
| marcksa | 2.352714 | 0.311894852 | 0.182 | 0.022 | 5.9770699 | 36 | marcksa |
| esyt2a1 | 2.438972 | 0.267344691 | 0.145 | 0.015 | 6.1962103 | 36 | esyt2a |
| lrpap13 | 2.729283 | 0.476560793 | 0.327 | 0.064 | 6.9337448 | 36 | lrpap1 |
| pdgfrb4 | 3.491883 | 0.486801876 | 0.291 | 0.052 | 8.8711311 | 36 | pdgfrb |
| hsp70.2 | 3.523383 | 0.265412648 | 0.127 | 0.011 | 8.9511554 | 36 | hsp70.2 |
| lpar14 | 4.279546 | 0.525536726 | 0.345 | 0.07 | 1.0872187 | 36 | lpar1 |
| tmeff1b4 | 6.372795 | 0.398002507 | 0.255 | 0.042 | 1.6190086 | 36 | tmeff1b |
| aff21 | 7.330430 | 0.270471051 | 0.127 | 0.012 | 1.8622959 | 36 | aff2 |
| mgp2 | 8.396926 | 0.312536291 | 0.182 | 0.022 | 2.1332391 | 36 | mgp |
| ywhah2 | 9.829252 | 0.485832146 | 0.309 | 0.06 | 2.4971214 | 36 | ywhah |
| adamts22 | 1.069661 | 0.358731102 | 0.2 | 0.028 | 2.7174742 | 36 | adamts2 |
| tspan4b2 | 1.432862 | 0.512771707 | 0.345 | 0.072 | 3.6401875 | 36 | tspan4b |

| | | | | | | | |
|---|---|---|---|---|---|---|---|
| adh8b1 | 1.703303 | 0.375603299 | 0.273 | 0.048 | 4.3272413 | 36 | adh8b |
| fkbp10b1 | 1.787860 | 0.306642004 | 0.2 | 0.027 | 4.5420601 | 36 | fkbp10b |
| plod32 | 2.362617 | 0.458866189 | 0.236 | 0.039 | 6.0022299 | 36 | plod3 |
| tcf127 | 2.513485 | 0.524903065 | 0.327 | 0.068 | 6.3855111 | 36 | tcf12 |
| tuba8l38 | 2.638501 | 0.469668875 | 0.345 | 0.072 | 6.7031128 | 36 | tuba8l3 |
| h2afx111 | 3.070154 | 0.691864628 | 0.655 | 0.227 | 7.7997269 | 36 | h2afx1 |
| cbfa2t31 | 3.574844 | 0.296306956 | 0.182 | 0.024 | 9.0818927 | 36 | cbfa2t3 |
| magixa | 4.773210 | 0.269135528 | 0.127 | 0.012 | 1.2126341 | 36 | magixa |
| myo1ca2 | 4.933143 | 0.357396402 | 0.2 | 0.029 | 1.2532651 | 36 | myo1ca |
| LOC1018844516 | 6.013110 | 0.567648151 | 0.382 | 0.091 | 1.5276307 | 36 | LOC101884451 |
| si:ch211-71n6.41 | 6.020117 | 0.341555631 | 0.2 | 0.029 | 1.5294107 | 36 | si:ch211-71n6.4 |
| palld | 6.767602 | 0.351689687 | 0.218 | 0.033 | 1.7193093 | 36 | palld |
| gsna1 | 1.055317 | 0.355650731 | 0.218 | 0.034 | 2.6810330 | 36 | gsna |
| tagln5 | 1.395158 | 0.785132591 | 0.327 | 0.075 | 3.5444001 | 36 | tagln |
| zgc:770861 | 1.478688 | 0.428524002 | 0.309 | 0.062 | 3.7566092 | 36 | zgc:77086 |
| ier2b15 | 1.522603 | 0.763108185 | 0.618 | 0.238 | 3.8681749 | 36 | ier2b |
| LOC1018823341 | 1.778839 | 0.331571424 | 0.145 | 0.017 | 4.5191424 | 36 | LOC101882334 |
| ppfibp1a1 | 2.111879 | 0.380566432 | 0.255 | 0.044 | 5.3652288 | 36 | ppfibp1a |
| ppfibp1b1 | 2.376890 | 0.278710594 | 0.164 | 0.021 | 6.0384913 | 36 | ppfibp1b |
| spred2b7 | 3.060341 | 0.47766217 | 0.364 | 0.085 | 7.7747973 | 36 | spred2b |
| fosb9 | 3.147553 | 0.762768505 | 0.436 | 0.122 | 7.9963588 | 36 | fosb |
| hmgb1a11 | 4.019124 | 0.645279468 | 0.636 | 0.228 | 1.0210586 | 36 | hmgb1a |
| rhoab11 | 4.102852 | 0.671444221 | 0.582 | 0.208 | 1.0423296 | 36 | rhoab |
| dhrs13l14 | 4.269042 | 0.302431114 | 0.236 | 0.04 | 1.0845502 | 36 | dhrs13l1 |
| vkorc11 | 4.673267 | 0.279360798 | 0.145 | 0.017 | 1.1872436 | 36 | vkorc1 |
| BX323797.4 | 6.441944 | 0.264730943 | 0.145 | 0.017 | 1.6365758 | 36 | BX323797.4 |
| sertad41 | 6.543019 | 0.276622569 | 0.145 | 0.017 | 1.6622541 | 36 | sertad4 |
| hmgn616 | 6.723785 | 0.667758729 | 0.764 | 0.311 | 1.7081775 | 36 | hmgn6 |
| FAT43 | 7.375029 | 0.263451373 | 0.164 | 0.021 | 1.8736263 | 36 | FAT4 |
| krt812 | 8.546642 | 0.712103633 | 0.8 | 0.312 | 2.1712744 | 36 | krt8 |
| ctbp11 | 9.654118 | 0.523843013 | 0.491 | 0.14 | 2.4526287 | 36 | ctbp1 |
| pitpnc1b | 1.228706 | 0.260250255 | 0.164 | 0.022 | 3.1215289 | 36 | pitpnc1b |
| tmem1765 | 1.235270 | 0.488725638 | 0.345 | 0.081 | 3.1382056 | 36 | tmem176 |
| zyx1 | 1.261473 | 0.500776162 | 0.327 | 0.076 | 3.2047727 | 36 | zyx |
| ptpn132 | 1.379354 | 0.262274623 | 0.145 | 0.018 | 3.5042505 | 36 | ptpn13 |

| | | | | | | | |
|---|---|---|---|---|---|---|---|
| selenof5 | 1.429257 | 0.619310775 | 0.709 | 0.267 | 3.6310276 | 36 | selenof |
| thbs1a1 | 1.494216 | 0.293927615 | 0.127 | 0.014 | 3.7960568 | 36 | thbs1a |
| ctsk6 | 1.743341 | 0.456187731 | 0.327 | 0.074 | 4.4289599 | 36 | ctsk |
| si:ch211-198c19.37 | 1.848888 | 0.629798656 | 0.345 | 0.083 | 4.6971021 | 36 | si:ch211-198c19.3 |
| XLOC-0354131 | 1.973101 | 0.296308078 | 0.218 | 0.037 | 5.0126648 | 36 | XLOC-035413 |
| nova23 | 2.128100 | 0.3498744 | 0.218 | 0.037 | 5.4064400 | 36 | nova2 |
| puraa1 | 2.588318 | 0.349933131 | 0.236 | 0.043 | 6.5756221 | 36 | puraa |
| hmga21 | 2.860907 | 0.277147935 | 0.127 | 0.014 | 7.2681356 | 36 | hmga2 |
| ubtd21 | 3.059618 | 0.320765533 | 0.2 | 0.032 | 7.7729597 | 36 | ubtd2 |
| rbp45 | 3.067630 | 0.667684041 | 0.491 | 0.142 | 7.7933140 | 36 | rbp4 |
| emp24 | 3.145396 | 0.495053968 | 0.273 | 0.057 | 7.9908790 | 36 | emp2 |
| kank26 | 3.245749 | 0.408493301 | 0.273 | 0.055 | 8.2458275 | 36 | kank2 |
| amotl2a | 3.340238 | 0.280444576 | 0.145 | 0.018 | 8.4858761 | 36 | amotl2a |
| tsku1 | 3.424495 | 0.411456586 | 0.218 | 0.038 | 8.6999299 | 36 | tsku |
| abl11 | 3.959483 | 0.348485036 | 0.218 | 0.038 | 1.0059068 | 36 | abl1 |
| kitlga2 | 4.631145 | 0.261659692 | 0.145 | 0.018 | 1.1765426 | 36 | kitlga |
| wu:fi46h04 | 5.036196 | 0.275307599 | 0.145 | 0.018 | 1.2794458 | 36 | wu:fi46h04 |
| dnmt3ab5 | 5.123654 | 0.47924389 | 0.364 | 0.093 | 1.3016644 | 36 | dnmt3ab |
| si:ch73-1a9.319 | 5.308443 | 0.669980767 | 0.745 | 0.316 | 1.3486100 | 36 | si:ch73-1a9.3 |
| cdkn1a4 | 5.356623 | 0.366438976 | 0.255 | 0.049 | 1.3608501 | 36 | cdkn1a |
| kdelr31 | 5.780059 | 0.254351592 | 0.145 | 0.018 | 1.4684241 | 36 | kdelr3 |
| dip2ca | 5.880300 | 0.278269661 | 0.145 | 0.018 | 1.4938903 | 36 | dip2ca |
| col4a15 | 6.932540 | 0.529815394 | 0.309 | 0.071 | 1.7612120 | 36 | col4a1 |
| mir199-32 | 7.026667 | 0.293969661 | 0.164 | 0.023 | 1.7851248 | 36 | mir199-3 |
| kdelr2a1 | 7.292749 | 0.460954596 | 0.327 | 0.077 | 1.8527229 | 36 | kdelr2a |
| igflr11 | 1.039413 | 0.275391168 | 0.145 | 0.019 | 2.6406295 | 36 | igflr1 |
| ms4a17a.81 | 1.044850 | 0.289021385 | 0.145 | 0.019 | 2.6544438 | 36 | ms4a17a.8 |
| colec124 | 1.314531 | 0.401190255 | 0.291 | 0.063 | 3.3395674 | 36 | colec12 |
| tnrc6c111 | 1.510643 | 0.56235726 | 0.473 | 0.15 | 3.8377887 | 36 | tnrc6c1 |
| hdlbpa9 | 1.580413 | 0.554991567 | 0.527 | 0.175 | 4.0150397 | 36 | hdlbpa |
| bmp1a | 1.598494 | 0.293125913 | 0.145 | 0.019 | 4.0609750 | 36 | bmp1a |
| hist2h2l5 | 1.640615 | 0.436722361 | 0.382 | 0.099 | 4.1679841 | 36 | hist2h2l |
| ptgdsb.22 | 1.672610 | 0.279281566 | 0.218 | 0.038 | 4.2492679 | 36 | ptgdsb.2 |
| p3h11 | 1.749960 | 0.257683304 | 0.127 | 0.015 | 4.4457734 | 36 | p3h1 |
| hspg24 | 2.337073 | 0.33343032 | 0.182 | 0.029 | 5.9373352 | 36 | hspg2 |

| | | | | | | | |
|---|---|---|---|---|---|---|---|
| hoxc3a3 | 2.572788 | 0.331125342 | 0.182 | 0.029 | 6.5361688 | 36 | hoxc3a |
| tmem165 | 3.111501 | 0.341584758 | 0.255 | 0.052 | 7.9047700 | 36 | tmem165 |
| antxr2a7 | 3.391428 | 0.565927398 | 0.509 | 0.17 | 8.6159239 | 36 | antxr2a |
| phyh | 3.423322 | 0.286492101 | 0.164 | 0.024 | 8.6969501 | 36 | phyh |
| skib1 | 3.482386 | 0.255604859 | 0.164 | 0.024 | 8.8470021 | 36 | skib |
| tbx2b2 | 3.622069 | 0.265754095 | 0.145 | 0.02 | 9.2018686 | 36 | tbx2b |
| grb10b3 | 3.765974 | 0.338516087 | 0.218 | 0.04 | 9.5674578 | 36 | grb10b |
| tpst12 | 4.068329 | 0.336850356 | 0.218 | 0.041 | 1.0335590 | 36 | tpst1 |
| dhrs3b1 | 4.647988 | 0.337956229 | 0.164 | 0.025 | 1.1808214 | 36 | dhrs3b |
| pef11 | 4.666786 | 0.380690057 | 0.309 | 0.073 | 1.1855970 | 36 | pef1 |
| tjp1a6 | 5.376056 | 0.400930652 | 0.255 | 0.054 | 1.3657870 | 36 | tjp1a |
| tspan73 | 6.392723 | 0.462024902 | 0.327 | 0.082 | 1.6240713 | 36 | tspan7 |
| sar1b4 | 6.571026 | 0.438024164 | 0.345 | 0.09 | 1.6693692 | 36 | sar1b |
| gli33 | 7.137100 | 0.352829834 | 0.2 | 0.036 | 1.8131802 | 36 | gli3 |
| gstt1b9 | 7.954248 | 0.444112059 | 0.345 | 0.09 | 2.0207767 | 36 | gstt1b |
| creg21 | 8.236692 | 0.31901621 | 0.2 | 0.036 | 2.0925317 | 36 | creg2 |
| lims1 | 8.700441 | 0.368268496 | 0.236 | 0.048 | 2.2103472 | 36 | lims1 |
| pcdh112 | 9.635850 | 0.294520202 | 0.127 | 0.016 | 2.4479877 | 36 | pcdh11 |
| si:ch211-286o17.14 | 1.021409 | 0.479605526 | 0.236 | 0.05 | 2.5948910 | 36 | si:ch211-286o17.1 |
| atp2b26 | 1.118560 | 0.428786647 | 0.327 | 0.084 | 2.8417027 | 36 | atp2b2 |
| cebpd10 | 1.306897 | 0.590824886 | 0.473 | 0.152 | 3.3201720 | 36 | cebpd |
| ip6k2b7 | 1.384528 | 0.557791384 | 0.418 | 0.129 | 3.5173943 | 36 | ip6k2b |
| rap1b6 | 2.705095 | 0.589399907 | 0.564 | 0.221 | 6.8722951 | 36 | rap1b |
| ehd32 | 2.920594 | 0.279430551 | 0.127 | 0.017 | 7.4197692 | 36 | ehd3 |
| sept7a1 | 2.968052 | 0.398431962 | 0.309 | 0.078 | 7.5403376 | 36 | sept7a |
| h2afy26 | 3.088703 | 0.477804659 | 0.4 | 0.123 | 7.8468510 | 36 | h2afy2 |
| dhrs3a4 | 3.131780 | 0.420032808 | 0.2 | 0.038 | 7.9562883 | 36 | dhrs3a |
| gpc43 | 3.138996 | 0.309269487 | 0.182 | 0.032 | 7.9746199 | 36 | gpc4 |
| slc7a14 | 3.583522 | 0.335942035 | 0.2 | 0.037 | 9.1039397 | 36 | slc7a1 |
| grb10a3 | 3.812725 | 0.332318971 | 0.218 | 0.043 | 9.6862286 | 36 | grb10a |
| c64 | 3.840575 | 0.596314633 | 0.382 | 0.114 | 9.7569820 | 36 | c6 |
| ier57 | 4.073802 | 0.554698947 | 0.436 | 0.147 | 1.0349494 | 36 | ier5 |
| map1lc3a10 | 4.247624 | 0.457218612 | 0.309 | 0.081 | 1.0791090 | 36 | map1lc3a |
| gsta.13 | 5.516690 | 0.39238821 | 0.236 | 0.051 | 1.4015151 | 36 | gsta.1 |
| ppm1la | 5.700537 | 0.291242407 | 0.2 | 0.038 | 1.4482214 | 36 | ppm1la |

| | | | | | | | |
|---|---|---|---|---|---|---|---|
| slc20a22 | 6.350381 | 0.274197543 | 0.164 | 0.027 | 1.6133142 | 36 | slc20a2 |
| fpr13 | 6.410720 | 0.274835067 | 0.145 | 0.022 | 1.6286435 | 36 | fpr1 |
| homer3b | 6.532570 | 0.289468605 | 0.182 | 0.033 | 1.6595996 | 36 | homer3b |
| kras1 | 6.753687 | 0.336211607 | 0.218 | 0.045 | 1.7157744 | 36 | kras |
| ptprsa2 | 6.865346 | 0.332069979 | 0.182 | 0.033 | 1.7441412 | 36 | ptprsa |
| serinc15 | 7.437937 | 0.38496234 | 0.327 | 0.087 | 1.8896079 | 36 | serinc1 |
| qkia9 | 8.139771 | 0.514429969 | 0.545 | 0.2 | 2.0679090 | 36 | qkia |
| mylka1 | 9.232203 | 0.265484793 | 0.127 | 0.018 | 2.3454413 | 36 | mylka |
| prss233 | 9.740485 | 0.353098815 | 0.182 | 0.033 | 2.4745702 | 36 | prss23 |
| zgc:1530182 | 9.923582 | 0.264879512 | 0.164 | 0.027 | 2.5210860 | 36 | zgc:153018 |
| foxp47 | 1.073909 | 0.415474054 | 0.291 | 0.074 | 2.7282676 | 36 | foxp4 |
| vasna3 | 1.083786 | 0.406553293 | 0.218 | 0.046 | 2.7533588 | 36 | vasna |
| tead1b4 | 1.554171 | 0.327731173 | 0.182 | 0.034 | 3.9483733 | 36 | tead1b |
| crip114 | 1.690234 | 0.751261459 | 0.6 | 0.285 | 4.2940411 | 36 | crip1 |
| id110 | 1.764372 | 0.513674689 | 0.418 | 0.132 | 4.4823875 | 36 | id1 |
| rsu12 | 2.100370 | 0.335256638 | 0.236 | 0.053 | 5.3359900 | 36 | rsu1 |
| mt-nd213 | 2.119443 | 0.594959926 | 0.782 | 0.392 | 5.3844451 | 36 | mt-nd2 |
| pgrmc17 | 2.329523 | 0.402760365 | 0.473 | 0.153 | 5.9181556 | 36 | pgrmc1 |
| cnn211 | 2.461040 | 0.490553552 | 0.455 | 0.158 | 6.2522726 | 36 | cnn2 |
| rarga5 | 2.471733 | 0.377705863 | 0.255 | 0.061 | 6.2794378 | 36 | rarga |
| ches11 | 2.498592 | 0.257929441 | 0.164 | 0.028 | 6.3476734 | 36 | ches1 |
| cyb5r31 | 2.700976 | 0.294725173 | 0.218 | 0.047 | 6.8618320 | 36 | cyb5r3 |
| rhoca3 | 2.703823 | 0.342175865 | 0.273 | 0.067 | 6.8690630 | 36 | rhoca |
| tmem106ba | 2.797360 | 0.316559166 | 0.218 | 0.047 | 7.1066948 | 36 | tmem106ba |
| fosab13 | 2.798078 | 0.699197813 | 0.727 | 0.362 | 7.1085193 | 36 | fosab |
| BX663516.2 | 2.874649 | 0.255047704 | 0.109 | 0.014 | 7.3030461 | 36 | BX663516.2 |
| txnipa9 | 3.308285 | 0.510154502 | 0.618 | 0.245 | 8.4046986 | 36 | txnipa |
| c1qtnf53 | 3.394713 | 0.385452123 | 0.182 | 0.035 | 8.6242687 | 36 | c1qtnf5 |
| rer11 | 3.424483 | 0.401172932 | 0.4 | 0.125 | 8.6998994 | 36 | rer1 |
| pbx1a3 | 3.587378 | 0.361567475 | 0.236 | 0.055 | 9.1137342 | 36 | pbx1a |
| sc:d02021 | 3.665342 | 0.351815324 | 0.218 | 0.046 | 9.3118021 | 36 | sc:d0202 |
| ywhabb3 | 3.833122 | 0.510761724 | 0.618 | 0.254 | 9.7380487 | 36 | ywhabb |
| surf4l1 | 3.924073 | 0.330508317 | 0.236 | 0.054 | 9.9691077 | 36 | surf4l |
| laptm4a3 | 5.041721 | 0.449006635 | 0.382 | 0.123 | 0.000128 | 36 | laptm4a |
| loxl2a4 | 5.103202 | 0.330391937 | 0.218 | 0.047 | 0.00013 | 36 | loxl2a |

| | | | | | | | |
|---|---|---|---|---|---|---|---|
| gpm6ab3 | 5.195945 | 0.273698824 | 0.127 | 0.019 | 0.000132 | 36 | gpm6ab |
| degs12 | 5.581731 | 0.363113664 | 0.255 | 0.063 | 0.000142 | 36 | degs1 |
| ddit311 | 6.055996 | 0.573531133 | 0.473 | 0.179 | 0.000154 | 36 | ddit3 |
| fzd7b3 | 6.228226 | 0.394539765 | 0.218 | 0.049 | 0.000158 | 36 | fzd7b |
| bcl2l12 | 6.677411 | 0.347982085 | 0.291 | 0.077 | 0.00017 | 36 | bcl2l1 |
| CABZ01068366.15 | 7.493813 | 0.326118652 | 0.2 | 0.042 | 0.00019 | 36 | CABZ01068366.1 |
| socs3a12 | 7.630219 | 0.61415484 | 0.709 | 0.315 | 0.000194 | 36 | socs3a |
| wls7 | 7.655133 | 0.465233748 | 0.4 | 0.132 | 0.000194 | 36 | wls |
| ccl39.73 | 7.973190 | 0.323320756 | 0.255 | 0.062 | 0.000203 | 36 | ccl39.7 |
| si:ch211-212k18.84 | 8.160567 | 0.314500951 | 0.218 | 0.049 | 0.000207 | 36 | si:ch211-212k18.8 |
| ptk2aa | 8.237978 | 0.250793978 | 0.127 | 0.019 | 0.000209 | 36 | ptk2aa |
| crhbp1 | 8.704659 | 0.340431749 | 0.127 | 0.019 | 0.000221 | 36 | crhbp |
| jpt1b7 | 8.752233 | 0.556959165 | 0.636 | 0.278 | 0.000222 | 36 | jpt1b |
| angptl46 | 9.101603 | 0.36958855 | 0.236 | 0.056 | 0.000231 | 36 | angptl4 |
| krt948 | 9.486107 | 0.564365631 | 0.473 | 0.167 | 0.000241 | 36 | krt94 |
| si:dkey-6n21.124 | 1.031720 | 0.306330856 | 0.2 | 0.042 | 0.000262 | 36 | si:dkey-6n21.12 |
| rtn4b | 1.095814 | 0.268274138 | 0.164 | 0.03 | 0.000278 | 36 | rtn4b |
| ppp3r1b | 1.133469 | 0.322117432 | 0.255 | 0.064 | 0.000288 | 36 | ppp3r1b |
| mgst3b6 | 1.342695 | 0.467572526 | 0.509 | 0.194 | 0.000341 | 36 | mgst3b |
| eif2s23 | 1.345341 | 0.511880233 | 0.709 | 0.323 | 0.000342 | 36 | eif2s2 |
| tegt7 | 1.572213 | 0.507640147 | 0.564 | 0.239 | 0.000399 | 36 | tegt |
| ctnnb2 | 1.744618 | 0.403036701 | 0.345 | 0.105 | 0.000443 | 36 | ctnnb2 |
| rab109 | 1.872656 | 0.506694727 | 0.527 | 0.215 | 0.000476 | 36 | rab10 |
| asph3 | 1.912480 | 0.309386361 | 0.2 | 0.044 | 0.000486 | 36 | asph |
| serf211 | 1.941031 | 0.532312417 | 0.745 | 0.378 | 0.000493 | 36 | serf2 |
| socs3b8 | 2.046278 | 0.556052882 | 0.418 | 0.152 | 0.00052 | 36 | socs3b |
| fn1a3 | 2.080708 | 0.499086005 | 0.291 | 0.083 | 0.000529 | 36 | fn1a |
| caprin1a1 | 2.123007 | 0.27950287 | 0.236 | 0.057 | 0.000539 | 36 | caprin1a |
| sept9a | 2.137453 | 0.282178389 | 0.218 | 0.049 | 0.000543 | 36 | sept9a |
| tns1b2 | 2.174353 | 0.253195545 | 0.182 | 0.037 | 0.000552 | 36 | tns1b |
| jund9 | 2.506819 | 0.414497239 | 0.309 | 0.091 | 0.000637 | 36 | jund |
| rsrp13 | 2.800792 | 0.398578466 | 0.382 | 0.123 | 0.000712 | 36 | rsrp1 |
| si:ch211-222l21.114 | 2.835555 | 0.523122685 | 0.709 | 0.315 | 0.00072 | 36 | si:ch211-222l21.1 |
| ablim1b3 | 3.037282 | 0.261378262 | 0.2 | 0.043 | 0.000772 | 36 | ablim1b |
| clic43 | 3.271549 | 0.33573264 | 0.236 | 0.059 | 0.000831 | 36 | clic4 |

| | | | | | | | |
|---|---|---|---|---|---|---|---|
| hopx7 | 3.702891 | 0.336071827 | 0.255 | 0.066 | 0.000941 | 36 | hopx |
| eva1ba6 | 3.740629 | 0.410326134 | 0.255 | 0.068 | 0.00095 | 36 | eva1ba |
| rcan1a1 | 3.748827 | 0.285418826 | 0.2 | 0.044 | 0.000952 | 36 | rcan1a |
| si:ch211-106h4.125 | 4.054044 | 0.433468016 | 0.236 | 0.059 | 0.00103 | 36 | si:ch211-106h4.12 |
| tuba8l8 | 4.230980 | 0.411462448 | 0.545 | 0.21 | 0.001075 | 36 | tuba8l |
| rrbp1a2 | 4.783837 | 0.30438267 | 0.236 | 0.059 | 0.001215 | 36 | rrbp1a |
| junba13 | 4.991089 | 0.600301721 | 0.673 | 0.321 | 0.001268 | 36 | junba |
| mt-nd415 | 5.048174 | 0.497445601 | 0.836 | 0.448 | 0.001282 | 36 | mt-nd4 |
| tmem2583 | 5.578636 | 0.48242147 | 0.673 | 0.306 | 0.001417 | 36 | tmem258 |
| si:dkey-177p2.63 | 6.148940 | 0.349697622 | 0.291 | 0.084 | 0.001562 | 36 | si:dkey-177p2.6 |
| bcl2a1 | 6.946308 | 0.274892354 | 0.164 | 0.033 | 0.001765 | 36 | bcl2a |
| sept102 | 6.963164 | 0.304571436 | 0.218 | 0.053 | 0.001769 | 36 | sept10 |
| ssr12 | 6.978854 | 0.42857657 | 0.509 | 0.201 | 0.001773 | 36 | ssr1 |
| selenow18 | 7.124508 | 0.522813911 | 0.818 | 0.416 | 0.00181 | 36 | selenow1 |
| tpm311 | 8.303197 | 0.486859712 | 0.636 | 0.285 | 0.002109 | 36 | tpm3 |
| reep510 | 8.470468 | 0.430056586 | 0.455 | 0.174 | 0.002152 | 36 | reep5 |
| pard3bb5 | 8.828389 | 0.288834201 | 0.164 | 0.034 | 0.002243 | 36 | pard3bb |
| leng8 | 8.958648 | 0.324212008 | 0.2 | 0.046 | 0.002276 | 36 | leng8 |
| brd3b4 | 8.963634 | 0.322138517 | 0.255 | 0.069 | 0.002277 | 36 | brd3b |
| rgs5b4 | 9.102297 | 0.621526837 | 0.255 | 0.073 | 0.002312 | 36 | rgs5b |
| ybx112 | 9.266577 | 0.410652306 | 0.964 | 0.696 | 0.002354 | 36 | ybx1 |
| creb3l3l2 | 9.428298 | 0.320631747 | 0.236 | 0.062 | 0.002395 | 36 | creb3l3l |
| ndufc23 | 9.619821 | 0.429065409 | 0.418 | 0.154 | 0.002444 | 36 | ndufc2 |
| si:ch211-132b12.73 | 1.089730 | 0.30276226 | 0.218 | 0.054 | 0.002768 | 36 | si:ch211-132b12.7 |
| itm2ba12 | 1.288797 | 0.515293078 | 0.618 | 0.298 | 0.003274 | 36 | itm2ba |
| tob1a5 | 1.355904 | 0.42684408 | 0.4 | 0.145 | 0.003445 | 36 | tob1a |
| stau27 | 1.371219 | 0.372546998 | 0.309 | 0.097 | 0.003484 | 36 | stau2 |
| pho2 | 1.439316 | 0.383063478 | 0.309 | 0.096 | 0.003657 | 36 | pho |
| chd62 | 1.449528 | 0.315370684 | 0.2 | 0.049 | 0.003683 | 36 | chd6 |
| fcho2 | 1.509463 | 0.28776749 | 0.164 | 0.034 | 0.003835 | 36 | fcho2 |
| zfyve211 | 1.524599 | 0.264170924 | 0.236 | 0.062 | 0.003873 | 36 | zfyve21 |
| vim4 | 1.625204 | 0.251026869 | 0.2 | 0.046 | 0.004129 | 36 | vim |
| capns1b9 | 1.635729 | 0.4060242 | 0.4 | 0.146 | 0.004156 | 36 | capns1b |
| cmtm32 | 1.652604 | 0.278993674 | 0.2 | 0.048 | 0.004198 | 36 | cmtm3 |
| creb3l21 | 1.682601 | 0.254464994 | 0.182 | 0.04 | 0.004275 | 36 | creb3l2 |

| | | | | | | | |
|---|---|---|---|---|---|---|---|
| anxa5b7 | 1.974163 | 0.375394234 | 0.327 | 0.106 | 0.005015 | 36 | anxa5b |
| s100u2 | 2.107033 | 0.251788903 | 0.182 | 0.041 | 0.005353 | 36 | s100u |
| ier2a14 | 2.170998 | 0.605608599 | 0.564 | 0.275 | 0.005515 | 36 | ier2a |
| cyp2n134 | 2.354077 | 0.31757631 | 0.182 | 0.042 | 0.005981 | 36 | cyp2n13 |
| crim15 | 2.568218 | 0.286688679 | 0.182 | 0.042 | 0.006525 | 36 | crim1 |
| lmnb21 | 3.213616 | 0.330243414 | 0.236 | 0.065 | 0.008164 | 36 | lmnb2 |
| si:dkey-180p18.93 | 3.316233 | 0.358752856 | 0.327 | 0.109 | 0.008425 | 36 | si:dkey-180p18.9 |
| si:dkey-239h2.32 | 3.429693 | 0.37815863 | 0.309 | 0.1 | 0.008713 | 36 | si:dkey-239h2.3 |
| bnip3lb2 | 3.720040 | 0.318245063 | 0.273 | 0.081 | 0.009451 | 36 | bnip3lb |
| fkbp1aa11 | 3.984729 | 0.471995859 | 0.818 | 0.415 | 0.010123 | 36 | fkbp1aa |
| spcs14 | 4.454683 | 0.455373575 | 0.527 | 0.232 | 0.011317 | 36 | spcs1 |
| napab | 4.776647 | 0.276827999 | 0.291 | 0.089 | 0.012135 | 36 | napab |
| vkorc1l1 | 5.039381 | 0.284826222 | 0.218 | 0.057 | 0.012803 | 36 | vkorc1l1 |
| fkbp142 | 5.079086 | 0.256970225 | 0.182 | 0.043 | 0.012903 | 36 | fkbp14 |
| ndfip1l4 | 5.378888 | 0.355896331 | 0.255 | 0.076 | 0.013665 | 36 | ndfip1l |
| tgfbr2b1 | 5.642144 | 0.257241391 | 0.127 | 0.024 | 0.014334 | 36 | tgfbr2b |
| COX317 | 5.674145 | 0.444785971 | 0.964 | 0.677 | 0.014415 | 36 | COX3 |
| pdia39 | 6.277672 | 0.445427455 | 0.691 | 0.314 | 0.015948 | 36 | pdia3 |
| ppap2d3 | 6.471121 | 0.292019384 | 0.164 | 0.037 | 0.01644 | 36 | ppap2d |
| gna11b1 | 6.631249 | 0.291152037 | 0.236 | 0.066 | 0.016847 | 36 | gna11b |
| vasnb4 | 6.951046 | 0.273371316 | 0.164 | 0.037 | 0.017659 | 36 | vasnb |
| ctsla10 | 6.997542 | 0.461991073 | 0.782 | 0.374 | 0.017777 | 36 | ctsla |
| ptp4a16 | 7.155348 | 0.371945654 | 0.364 | 0.133 | 0.018178 | 36 | ptp4a1 |
| ndrg23 | 7.295175 | 0.328306543 | 0.2 | 0.052 | 0.018533 | 36 | ndrg2 |
| XLOC-0268366 | 8.334961 | 0.539225376 | 0.255 | 0.079 | 0.021175 | 36 | XLOC-026836 |
| ssr46 | 8.469668 | 0.45525557 | 0.673 | 0.332 | 0.021517 | 36 | ssr4 |
| ndnf3 | 8.783847 | 0.315117142 | 0.164 | 0.037 | 0.022315 | 36 | ndnf |
| capns1a6 | 9.412178 | 0.361333196 | 0.418 | 0.16 | 0.023912 | 36 | capns1a |
| ccl25b6 | 9.423737 | 0.498543811 | 0.291 | 0.094 | 0.023941 | 36 | ccl25b |
| BX284696.12 | 9.593087 | 0.294704543 | 0.218 | 0.06 | 0.024371 | 36 | BX284696.1 |
| tubb51 | 9.622459 | 0.317365811 | 0.218 | 0.06 | 0.024446 | 36 | tubb5 |
| ms4a17a.114 | 1.011301 | 0.298664659 | 0.2 | 0.053 | 0.025692 | 36 | ms4a17a.11 |
| rab6a | 1.023010 | 0.25291065 | 0.218 | 0.059 | 0.02599 | 36 | rab6a |
| sdc412 | 1.050822 | 0.459105173 | 0.582 | 0.255 | 0.026696 | 36 | sdc4 |
| trappc12 | 1.062499 | 0.388352207 | 0.327 | 0.118 | 0.026993 | 36 | trappc1 |

| | | | | | | | |
|---|---|---|---|---|---|---|---|
| cd82b3 | 1.072516 | 0.342821956 | 0.255 | 0.077 | 0.027247 | 36 | cd82b |
| ilk | 1.112816 | 0.315649375 | 0.273 | 0.085 | 0.028271 | 36 | ilk |
| wu:fi37e093 | 1.117223 | 0.339529447 | 0.364 | 0.13 | 0.028383 | 36 | wu:fi37e09 |
| cd6316 | 1.128982 | 0.465386638 | 0.764 | 0.374 | 0.028682 | 36 | cd63 |
| hmgn79 | 1.144740 | 0.477785316 | 0.636 | 0.319 | 0.029082 | 36 | hmgn7 |
| serinc25 | 1.190715 | 0.370278789 | 0.345 | 0.125 | 0.03025 | 36 | serinc2 |
| entpd14 | 1.194571 | 0.279247639 | 0.273 | 0.083 | 0.030348 | 36 | entpd1 |
| mcl1b13 | 1.237563 | 0.442171876 | 0.564 | 0.252 | 0.03144 | 36 | mcl1b |
| atp10a6 | 1.277607 | 0.262267318 | 0.218 | 0.06 | 0.032458 | 36 | atp10a |
| XLOC-0320024 | 1.313975 | 0.326009671 | 0.182 | 0.045 | 0.033382 | 36 | XLOC-032002 |
| zbtb206 | 1.380208 | 0.390022906 | 0.309 | 0.109 | 0.035064 | 36 | zbtb20 |
| tax1bp34 | 1.471198 | 0.285888595 | 0.309 | 0.102 | 0.037376 | 36 | tax1bp3 |
| per34 | 1.632164 | 0.31688522 | 0.255 | 0.078 | 0.041465 | 36 | per3 |
| itga94 | 1.670683 | 0.272416407 | 0.164 | 0.038 | 0.042444 | 36 | itga9 |
| dkk3b4 | 1.702695 | 0.321091141 | 0.218 | 0.062 | 0.043257 | 36 | dkk3b |
| NC-002333.413 | 1.803597 | 0.442983884 | 0.927 | 0.61 | 0.04582 | 36 | NC-002333.4 |
| zfp36l22 | 1.887139 | 0.352413247 | 0.309 | 0.102 | 0.047943 | 36 | zfp36l2 |
| fgfr21 | 1.933904 | 0.253513501 | 0.145 | 0.032 | 0.049131 | 36 | fgfr2 |
| rhocb1 | 1.951740 | 0.287769837 | 0.309 | 0.104 | 0.049584 | 36 | rhocb |
| pdlim14 | 1.953197 | 0.27787075 | 0.218 | 0.061 | 0.049621 | 36 | pdlim1 |
| irf2bpl3 | 2.227686 | 0.274400287 | 0.182 | 0.046 | 0.056594 | 36 | irf2bpl |
| ube2v11 | 2.247759 | 0.381674773 | 0.509 | 0.219 | 0.057104 | 36 | ube2v1 |
| lamb1a6 | 2.340878 | 0.319924256 | 0.2 | 0.055 | 0.05947 | 36 | lamb1a |
| mdka9 | 2.367643 | 0.379191573 | 0.327 | 0.114 | 0.06015 | 36 | mdka |
| her63 | 2.389297 | 0.250859921 | 0.127 | 0.026 | 0.0607 | 36 | her6 |
| copg2 | 2.436904 | 0.265096288 | 0.2 | 0.054 | 0.06191 | 36 | copg2 |
| vdac37 | 2.482708 | 0.314164617 | 0.436 | 0.169 | 0.063073 | 36 | vdac3 |
| pdgfra3 | 2.658635 | 0.314831222 | 0.218 | 0.062 | 0.067543 | 36 | pdgfra |
| krt18a.19 | 2.791782 | 0.378181813 | 0.4 | 0.151 | 0.070925 | 36 | krt18a.1 |
| sox4a7 | 2.931633 | 0.324708939 | 0.218 | 0.063 | 0.074478 | 36 | sox4a |
| hnrnpa0b9 | 2.950518 | 0.433498493 | 0.691 | 0.342 | 0.074958 | 36 | hnrnpa0b |
| aldh9a1a.19 | 2.999019 | 0.345484176 | 0.473 | 0.194 | 0.07619 | 36 | aldh9a1a.1 |
| CABZ01080702.18 | 3.153569 | 0.329872788 | 0.291 | 0.099 | 0.080116 | 36 | CABZ01080702.1 |
| sost2 | 3.154315 | 0.25787753 | 0.109 | 0.02 | 0.080135 | 36 | sost |
| clu3 | 3.408726 | 0.255918991 | 0.145 | 0.032 | 0.086599 | 36 | clu |

| | | | | | | | |
|---|---|---|---|---|---|---|---|
| h1f08 | 3.571635 | 0.333133511 | 0.236 | 0.072 | 0.090737 | 36 | h1f0 |
| higd1a9 | 3.605002 | 0.448187624 | 0.618 | 0.304 | 0.091585 | 36 | higd1a |
| gpatch82 | 3.654824 | 0.279868804 | 0.218 | 0.064 | 0.092851 | 36 | gpatch8 |
| emd5 | 3.679154 | 0.258261692 | 0.236 | 0.072 | 0.093469 | 36 | emd |
| b2ml13 | 4.341468 | 0.440053923 | 0.582 | 0.295 | 0.110295 | 36 | b2ml |
| flot1b3 | 4.375732 | 0.325820911 | 0.291 | 0.101 | 0.111165 | 36 | flot1b |
| mgst3a2 | 4.874808 | 0.331263051 | 0.236 | 0.075 | 0.123845 | 36 | mgst3a |
| map1lc3b5 | 4.917598 | 0.363115729 | 0.327 | 0.125 | 0.124932 | 36 | map1lc3b |
| cux1a4 | 5.002371 | 0.261896657 | 0.2 | 0.056 | 0.127085 | 36 | cux1a |
| cdca4 | 5.063142 | 0.286778485 | 0.218 | 0.065 | 0.128629 | 36 | cdca4 |
| tpd52l2b2 | 5.604161 | 0.317055975 | 0.309 | 0.11 | 0.142374 | 36 | tpd52l2b |
| dynlt33 | 6.014276 | 0.303339648 | 0.4 | 0.158 | 0.152793 | 36 | dynlt3 |
| emc2 | 6.088547 | 0.3136345 | 0.236 | 0.076 | 0.15468 | 36 | emc2 |
| ptena1 | 6.338011 | 0.264917161 | 0.218 | 0.066 | 0.161017 | 36 | ptena |
| pfn2l9 | 6.484670 | 0.351373565 | 0.509 | 0.216 | 0.164743 | 36 | pfn2l |
| eif2s35 | 6.768539 | 0.413229117 | 0.618 | 0.309 | 0.171955 | 36 | eif2s3 |
| ssr22 | 7.739798 | 0.375252335 | 0.691 | 0.334 | 0.19663 | 36 | ssr2 |
| bsg3 | 8.014316 | 0.451677572 | 0.691 | 0.377 | 0.203604 | 36 | bsg |
| lgals9l35 | 8.160351 | 0.299400391 | 0.255 | 0.084 | 0.207314 | 36 | lgals9l3 |
| rbfox23 | 8.642115 | 0.26705925 | 0.164 | 0.043 | 0.219553 | 36 | rbfox2 |
| klf2a9 | 8.776789 | 0.30832229 | 0.291 | 0.101 | 0.222974 | 36 | klf2a |
| celf212 | 8.939934 | 0.404132073 | 0.545 | 0.258 | 0.227119 | 36 | celf2 |
| mt-nd115 | 9.318529 | 0.442047957 | 0.691 | 0.398 | 0.236737 | 36 | mt-nd1 |
| u2surp3 | 9.796606 | 0.308467434 | 0.291 | 0.105 | 0.248883 | 36 | u2surp |
| rab2a3 | 9.884002 | 0.356791494 | 0.418 | 0.178 | 0.251103 | 36 | rab2a |
| park71 | 1.011224 | 0.263269324 | 0.255 | 0.084 | 0.256902 | 36 | park7 |
| plpp2b1 | 1.204959 | 0.283029408 | 0.145 | 0.035 | 0.30612 | 36 | plpp2b |
| agrn5 | 1.229933 | 0.335395009 | 0.2 | 0.061 | 0.312465 | 36 | agrn |
| gbp2 | 1.261192 | 0.293165218 | 0.327 | 0.123 | 0.320406 | 36 | gbp |
| zgc:1538677 | 1.389238 | 0.424769184 | 0.782 | 0.466 | 0.352936 | 36 | zgc:153867 |
| ppm1aa1 | 1.401606 | 0.270017728 | 0.218 | 0.069 | 0.356078 | 36 | ppm1aa |
| irf2bp2a7 | 1.435470 | 0.33564246 | 0.436 | 0.187 | 0.364681 | 36 | irf2bp2a |
| ccdc85b8 | 1.544615 | 0.259724958 | 0.182 | 0.052 | 0.39241 | 36 | ccdc85b |
| nucks1b1 | 1.608126 | 0.308255558 | 0.309 | 0.115 | 0.408545 | 36 | nucks1b |
| gatad2b3 | 1.621956 | 0.260751854 | 0.218 | 0.069 | 0.412058 | 36 | gatad2b |

| | | | | | | | |
|---|---|---|---|---|---|---|---|
| pfn210 | 1.628296 | 0.350294961 | 0.527 | 0.24 | 0.413669 | 36 | pfn2 |
| cxcl12a4 | 1.701676 | 0.32021927 | 0.255 | 0.085 | 0.432311 | 36 | cxcl12a |
| ccnd2a3 | 1.779625 | 0.281386343 | 0.273 | 0.096 | 0.452114 | 36 | ccnd2a |
| tln17 | 1.841148 | 0.36050935 | 0.345 | 0.143 | 0.467744 | 36 | tln1 |
| ciarta5 | 1.858456 | 0.332622336 | 0.309 | 0.119 | 0.472141 | 36 | ciarta |
| hp1bp37 | 2.068185 | 0.336519571 | 0.309 | 0.122 | 0.525423 | 36 | hp1bp3 |
| plecb5 | 2.208002 | 0.26745955 | 0.182 | 0.053 | 0.560943 | 36 | plecb |
| elocb6 | 2.273448 | 0.339374384 | 0.436 | 0.197 | 0.57757 | 36 | elocb |
| atf4b13 | 2.315631 | 0.401391232 | 0.545 | 0.279 | 0.588286 | 36 | atf4b |
| s1pr22 | 2.348393 | 0.256742475 | 0.145 | 0.038 | 0.596609 | 36 | s1pr2 |
| zgc:919101 | 2.503560 | 0.362410965 | 0.564 | 0.271 | 0.63603 | 36 | zgc:91910 |
| si:ch73-335l21.49 | 2.511180 | 0.445141442 | 0.345 | 0.142 | 0.637965 | 36 | si:ch73-335l21.4 |
| taf12 | 2.541739 | 0.288190206 | 0.327 | 0.13 | 0.645729 | 36 | taf12 |
| ndufa4l11 | 2.580116 | 0.401333636 | 0.8 | 0.423 | 0.655479 | 36 | ndufa4l |
| add3a9 | 2.593286 | 0.323860191 | 0.345 | 0.139 | 0.658824 | 36 | add3a |
| cnbpb7 | 2.683119 | 0.357062371 | 0.455 | 0.211 | 0.681646 | 36 | cnbpb |
| mtpn7 | 2.691131 | 0.333388374 | 0.382 | 0.164 | 0.683682 | 36 | mtpn |
| rps27.216 | 2.853375 | 0.406243271 | 0.873 | 0.554 | 0.7249 | 36 | rps27.2 |
| add11 | 2.858065 | 0.293590411 | 0.2 | 0.063 | 0.726092 | 36 | add1 |
| ylpm1 | 2.927776 | 0.25830459 | 0.182 | 0.053 | 0.743802 | 36 | ylpm1 |
| anxa136 | 3.022288 | 0.288115784 | 0.182 | 0.055 | 0.767812 | 36 | anxa13 |
| tmem167a1 | 3.429587 | 0.2614995 | 0.291 | 0.109 | 0.871287 | 36 | tmem167a |
| capn2a2 | 3.534878 | 0.268510885 | 0.182 | 0.055 | 0.898036 | 36 | capn2a |
| fam168a8 | 3.632153 | 0.300859103 | 0.218 | 0.075 | 0.922749 | 36 | fam168a |
| psmb72 | 3.683959 | 0.32691439 | 0.4 | 0.174 | 0.93591 | 36 | psmb7 |
| cavin1b7 | 3.964726 | 0.262847123 | 0.164 | 0.046 | 1 | 36 | cavin1b |
| ginm1 | 4.339654 | 0.276771798 | 0.273 | 0.101 | 1 | 36 | ginm1 |
| hspb16 | 4.475014 | 0.327859826 | 0.236 | 0.083 | 1 | 36 | hspb1 |
| zgc:11418822 | 4.572843 | 0.373587947 | 0.945 | 0.669 | 1 | 36 | zgc:114188 |
| srsf9 | 4.793324 | 0.28562653 | 0.345 | 0.141 | 1 | 36 | srsf9 |
| psmc53 | 4.896160 | 0.287471561 | 0.273 | 0.104 | 1 | 36 | psmc5 |
| mt-nd55 | 4.953854 | 0.270539861 | 0.327 | 0.13 | 1 | 36 | mt-nd5 |
| FO082781.17 | 4.963577 | 0.26288816 | 0.182 | 0.056 | 1 | 36 | FO082781.1 |
| slc44a21 | 5.005567 | 0.255618239 | 0.182 | 0.056 | 1 | 36 | slc44a2 |
| cdc428 | 5.061107 | 0.366127506 | 0.473 | 0.235 | 1 | 36 | cdc42 |

| | | | | | | | |
|---|---|---|---|---|---|---|---|
| enpp13 | 5.100619 | 0.291607312 | 0.145 | 0.04 | 1 | 36 | enpp1 |
| rbm39a5 | 5.656006 | 0.293713069 | 0.364 | 0.154 | 1 | 36 | rbm39a |
| atrx8 | 5.760797 | 0.330032623 | 0.309 | 0.129 | 1 | 36 | atrx |
| furina2 | 6.309004 | 0.27865703 | 0.2 | 0.066 | 1 | 36 | furina |
| lman21 | 6.332832 | 0.308932011 | 0.345 | 0.146 | 1 | 36 | lman2 |
| calr3a1 | 6.393676 | 0.365571864 | 0.364 | 0.159 | 1 | 36 | calr3a |
| kctd12.29 | 6.637916 | 0.262781487 | 0.273 | 0.101 | 1 | 36 | kctd12.2 |
| ndfip12 | 6.660079 | 0.330566989 | 0.382 | 0.172 | 1 | 36 | ndfip1 |
| rps4x16 | 6.663449 | 0.329188124 | 1 | 0.737 | 1 | 36 | rps4x |
| si:ch211-196l7.414 | 6.838293 | 0.381514817 | 0.618 | 0.331 | 1 | 36 | si:ch211-196l7.4 |
| rhoaa10 | 7.009305 | 0.348630197 | 0.382 | 0.178 | 1 | 36 | rhoaa |
| pik3r25 | 7.127171 | 0.314695175 | 0.2 | 0.068 | 1 | 36 | pik3r2 |
| eif3d3 | 7.576757 | 0.334226615 | 0.564 | 0.282 | 1 | 36 | eif3d |
| kmt2bb3 | 7.769331 | 0.301228779 | 0.255 | 0.098 | 1 | 36 | kmt2bb |
| atox13 | 8.031565 | 0.256856109 | 0.309 | 0.124 | 1 | 36 | atox1 |
| CR753876.16 | 8.710024 | 0.292760822 | 0.291 | 0.112 | 1 | 36 | CR753876.1 |
| acin1a6 | 8.720966 | 0.340071992 | 0.473 | 0.23 | 1 | 36 | acin1a |
| notch31 | 8.777357 | 0.256824375 | 0.109 | 0.026 | 1 | 36 | notch3 |
| rbm281 | 9.598562 | 0.269852136 | 0.255 | 0.097 | 1 | 36 | rbm28 |
| cxcl18b6 | 0.0001 | 0.261218572 | 0.291 | 0.111 | 1 | 36 | cxcl18b |
| pomp4 | 0.0001 | 0.347827617 | 0.509 | 0.261 | 1 | 36 | pomp |
| rpl2811 | 0.0001 | 0.28113948 | 0.982 | 0.777 | 1 | 36 | rpl28 |
| rpl921 | 0.0001 | 0.383078479 | 1 | 0.669 | 1 | 36 | rpl9 |
| ssr35 | 0.0001 | 0.295142331 | 0.582 | 0.289 | 1 | 36 | ssr3 |
| ywhaqb10 | 0.0001 | 0.322216732 | 0.345 | 0.156 | 1 | 36 | ywhaqb |
| grinab6 | 0.0001 | 0.333620701 | 0.291 | 0.124 | 1 | 36 | grinab |
| rab1ab3 | 0.0001 | 0.25176843 | 0.327 | 0.139 | 1 | 36 | rab1ab |
| ctnnb17 | 0.0001 | 0.279469923 | 0.364 | 0.16 | 1 | 36 | ctnnb1 |
| capza1a9 | 0.0001 | 0.316583957 | 0.455 | 0.222 | 1 | 36 | capza1a |
| rps1618 | 0.0001 | 0.28932001 | 0.945 | 0.766 | 1 | 36 | rps16 |
| mt-atp616 | 0.0002 | 0.384957938 | 0.764 | 0.557 | 1 | 36 | mt-atp6 |
| mif3 | 0.0002 | 0.292003301 | 0.436 | 0.208 | 1 | 36 | mif |
| eif4a1b9 | 0.0002 | 0.362408917 | 0.564 | 0.313 | 1 | 36 | eif4a1b |
| zfp36l1b8 | 0.0002 | 0.299698155 | 0.327 | 0.141 | 1 | 36 | zfp36l1b |
| zgc:1583438 | 0.0002 | 0.379098447 | 0.364 | 0.17 | 1 | 36 | zgc:158343 |

| | | | | | | | |
|---|---|---|---|---|---|---|---|
| arf52 | 0.0002 | 0.314178786 | 0.436 | 0.213 | 1 | 36 | arf5 |
| nupr1a2 | 0.0002 | 0.294201785 | 0.364 | 0.166 | 1 | 36 | nupr1a |
| nenf | 0.0002 | 0.260787015 | 0.2 | 0.071 | 1 | 36 | nenf |
| wu:fb16f035 | 0.0002 | 0.26003582 | 0.236 | 0.091 | 1 | 36 | wu:fb16f03 |
| notch210 | 0.0002 | 0.272220981 | 0.291 | 0.124 | 1 | 36 | notch2 |
| tagln213 | 0.0003 | 0.310344351 | 0.436 | 0.211 | 1 | 36 | tagln2 |
| ppdpfb6 | 0.0003 | 0.356196335 | 0.8 | 0.533 | 1 | 36 | ppdpfb |
| zgc:16273016 | 0.0003 | 0.38869424 | 0.709 | 0.404 | 1 | 36 | zgc:162730 |
| sec11a3 | 0.0003 | 0.284669631 | 0.4 | 0.191 | 1 | 36 | sec11a |
| fkbp31 | 0.0003 | 0.26996984 | 0.382 | 0.179 | 1 | 36 | fkbp3 |
| rnf103 | 0.0003 | 0.288152516 | 0.291 | 0.129 | 1 | 36 | rnf10 |
| xbp18 | 0.0004 | 0.341519454 | 0.636 | 0.356 | 1 | 36 | xbp1 |
| calm1a4 | 0.0004 | 0.30887811 | 0.364 | 0.175 | 1 | 36 | calm1a |
| mtdha2 | 0.0004 | 0.261186611 | 0.327 | 0.148 | 1 | 36 | mtdha |
| anxa11b4 | 0.0005 | 0.273858913 | 0.364 | 0.175 | 1 | 36 | anxa11b |
| glo15 | 0.0005 | 0.292522617 | 0.418 | 0.21 | 1 | 36 | glo1 |
| mt-co116 | 0.0005 | 0.34977633 | 0.927 | 0.641 | 1 | 36 | mt-co1 |
| grpel24 | 0.0006 | 0.259375492 | 0.255 | 0.108 | 1 | 36 | grpel2 |
| mycb6 | 0.0007 | 0.330513373 | 0.4 | 0.202 | 1 | 36 | mycb |
| cst14a.23 | 0.0008 | 0.289122635 | 0.455 | 0.237 | 1 | 36 | cst14a.2 |
| tmed75 | 0.0008 | 0.255009558 | 0.436 | 0.222 | 1 | 36 | tmed7 |
| nsa213 | 0.0008 | 0.327723975 | 0.655 | 0.39 | 1 | 36 | nsa2 |
| nucks1a2 | 0.0008 | 0.274313187 | 0.382 | 0.189 | 1 | 36 | nucks1a |
| b4galt61 | 0.0008 | 0.255027421 | 0.182 | 0.069 | 1 | 36 | b4galt6 |
| plscr3b | 0.0009 | 0.261184899 | 0.255 | 0.11 | 1 | 36 | plscr3b |
| si:ch73-376l24.31 | 0.001 | 0.271781069 | 0.273 | 0.124 | 1 | 36 | si:ch73-376l24.3 |
| pno14 | 0.0012 | 0.26075996 | 0.345 | 0.173 | 1 | 36 | pno1 |
| anxa49 | 0.0014 | 0.259978673 | 0.309 | 0.152 | 1 | 36 | anxa4 |
| calm2a3 | 0.0014 | 0.29119721 | 0.4 | 0.211 | 1 | 36 | calm2a |
| mt-nd312 | 0.0015 | 0.285504855 | 0.636 | 0.373 | 1 | 36 | mt-nd3 |
| sem13 | 0.0016 | 0.274683259 | 0.491 | 0.274 | 1 | 36 | sem1 |
| zgc:1119865 | 0.0017 | 0.261358749 | 0.364 | 0.187 | 1 | 36 | zgc:111986 |
| mt-cyb13 | 0.0018 | 0.318915666 | 0.745 | 0.578 | 1 | 36 | mt-cyb |
| pdk2b6 | 0.0019 | 0.258423754 | 0.236 | 0.107 | 1 | 36 | pdk2b |
| rpl5a16 | 0.0024 | 0.323137553 | 0.836 | 0.537 | 1 | 36 | rpl5a |

| | | | | | | | |
|---|---|---|---|---|---|---|---|
| cox8a15 | 0.0024 | 0.252099821 | 0.491 | 0.269 | 1 | 36 | cox8a |
| crebzf4 | 0.0026 | 0.251989277 | 0.236 | 0.11 | 1 | 36 | crebzf |
| cdh24 | 0.0034 | 0.259637605 | 0.164 | 0.068 | 1 | 36 | cdh2 |
| krtcap23 | 0.0037 | 0.253681783 | 0.364 | 0.197 | 1 | 36 | krtcap2 |
| syncrip3 | 0.0039 | 0.264964998 | 0.527 | 0.314 | 1 | 36 | syncrip |
| sec61b3 | 0.0045 | 0.277911638 | 0.745 | 0.47 | 1 | 36 | sec61b |
| uba5213 | 0.0047 | 0.284087069 | 0.909 | 0.662 | 1 | 36 | uba52 |
| tubb4b2 | 0.005 | 0.274001427 | 0.8 | 0.593 | 1 | 36 | tubb4b |
| mt-co215 | 0.005 | 0.271283011 | 0.8 | 0.672 | 1 | 36 | mt-co2 |
| eef1g14 | 0.0053 | 0.275853517 | 0.909 | 0.653 | 1 | 36 | eef1g |
| rps1517 | 0.0077 | 0.264122524 | 0.945 | 0.718 | 1 | 36 | rps15 |
| rpl27a16 | 0.0098 | 0.302409456 | 0.836 | 0.524 | 1 | 36 | rpl27a |
| gp1bb1 | 0 | 1.954633235 | 0.961 | 0.006 | 0 | 37 | gp1bb |
| thbs1b2 | 0 | 1.624096683 | 0.706 | 0.009 | 0 | 37 | thbs1b |
| hbegfb | 0 | 1.601440051 | 0.686 | 0.009 | 0 | 37 | hbegfb |
| itgb3b2 | 0 | 1.437674379 | 0.667 | 0.012 | 0 | 37 | itgb3b |
| apln1 | 0 | 1.263833024 | 0.471 | 0.005 | 0 | 37 | apln |
| rac3a | 0 | 1.076646805 | 0.451 | 0.005 | 0 | 37 | rac3a |
| itga2b | 0 | 1.052286577 | 0.431 | 0 | 0 | 37 | itga2b |
| f5 | 0 | 0.985332452 | 0.353 | 0.003 | 0 | 37 | f5 |
| si:ch211-33e4.21 | 0 | 0.952855699 | 0.392 | 0.003 | 0 | 37 | si:ch211-33e4.2 |
| tnr | 0 | 0.805662649 | 0.333 | 0.002 | 0 | 37 | tnr |
| mpl | 0 | 0.473724184 | 0.176 | 0 | 0 | 37 | mpl |
| tubb1 | 2.391814 | 0.700224326 | 0.275 | 0.002 | 6.0764050 | 37 | tubb1 |
| sele1 | 1.102741 | 0.878342085 | 0.353 | 0.003 | 2.8015160 | 37 | sele |
| parvb1 | 1.803988 | 1.038018479 | 0.471 | 0.007 | 4.5830326 | 37 | parvb |
| vegfc | 1.517035 | 0.72562661 | 0.275 | 0.002 | 3.8540274 | 37 | vegfc |
| rpgra | 5.915373 | 0.62996477 | 0.196 | 0.001 | 1.5028005 | 37 | rpgra |
| blf1 | 3.518825 | 0.816871188 | 0.314 | 0.004 | 8.9395753 | 37 | blf |
| zgc:171686 | 1.198839 | 0.892688062 | 0.353 | 0.006 | 3.0456508 | 37 | zgc:171686 |
| bmp161 | 1.074694 | 0.984598515 | 0.373 | 0.007 | 2.7302609 | 37 | bmp16 |
| zgc:162989 | 8.280876 | 0.36129828 | 0.137 | 0.001 | 2.1037566 | 37 | zgc:162989 |
| p2ry13 | 8.647906 | 0.49266601 | 0.196 | 0.002 | 2.1970005 | 37 | p2ry13 |
| zmat4b | 2.319590 | 0.376962671 | 0.157 | 0.001 | 5.8929192 | 37 | zmat4b |
| adra2a | 8.845325 | 0.461970831 | 0.157 | 0.001 | 2.2471548 | 37 | adra2a |

| | | | | | | | |
|---|---|---|---|---|---|---|---|
| hacd31 | 3.624253 | 1.376709698 | 0.686 | 0.036 | 9.2074159 | 37 | hacd3 |
| tal11 | 3.512647 | 0.488026046 | 0.235 | 0.004 | 8.9238813 | 37 | tal1 |
| plac8l1 | 1.314140 | 0.703395604 | 0.235 | 0.004 | 3.3385743 | 37 | plac8l1 |
| reps21 | 3.913765 | 0.544767365 | 0.235 | 0.004 | 9.9429203 | 37 | reps2 |
| arhgef9b1 | 5.195659 | 0.482374509 | 0.196 | 0.003 | 1.3199573 | 37 | arhgef9b |
| si:ch211-195b11.32 | 1.419037 | 1.584554819 | 0.627 | 0.033 | 3.6050652 | 37 | si:ch211-195b11.3 |
| gata1a1 | 4.901177 | 0.409719971 | 0.196 | 0.003 | 1.2451442 | 37 | gata1a |
| fgl2a5 | 1.140181 | 1.796160896 | 0.902 | 0.074 | 2.8966299 | 37 | fgl2a |
| wnt2 | 7.512704 | 0.54373115 | 0.176 | 0.002 | 1.9086025 | 37 | wnt2 |
| plek2 | 3.564998 | 1.196261265 | 0.549 | 0.028 | 9.0568778 | 37 | plek |
| spns22 | 1.098620 | 0.760053914 | 0.294 | 0.008 | 2.7910465 | 37 | spns2 |
| GARNL3 | 8.799412 | 0.501155384 | 0.196 | 0.003 | 2.2354908 | 37 | GARNL3 |
| myocd2 | 3.567309 | 0.66712342 | 0.294 | 0.008 | 9.0627491 | 37 | myocd |
| celf41 | 4.714743 | 0.647602544 | 0.235 | 0.006 | 1.1977806 | 37 | celf4 |
| tpm4a11 | 2.136630 | 1.638584207 | 0.902 | 0.112 | 5.4281098 | 37 | tpm4a |
| fli1a1 | 1.216398 | 0.6644238 | 0.275 | 0.009 | 3.0902608 | 37 | fli1a |
| mir2235 | 7.829298 | 1.18682263 | 0.51 | 0.034 | 1.9890332 | 37 | mir223 |
| admb | 1.500078 | 0.674992234 | 0.196 | 0.005 | 3.8109501 | 37 | admb |
| lpp | 4.327796 | 1.225343209 | 0.51 | 0.035 | 1.0994766 | 37 | lpp |
| cald1a10 | 6.847955 | 1.662502703 | 0.882 | 0.119 | 1.7397231 | 37 | cald1a |
| myh11a6 | 1.042085 | 1.355472867 | 0.706 | 0.07 | 2.6474182 | 37 | myh11a |
| susd1 | 4.456357 | 0.682694117 | 0.275 | 0.01 | 1.1321376 | 37 | susd1 |
| lmo24 | 4.525069 | 0.949208208 | 0.431 | 0.025 | 1.1495938 | 37 | lmo2 |
| XLOC-019062 | 3.127663 | 0.548696344 | 0.196 | 0.005 | 7.9458300 | 37 | XLOC-019062 |
| rap1gap2a | 9.261234 | 0.680312045 | 0.275 | 0.01 | 2.3528166 | 37 | rap1gap2a |
| boka | 3.036466 | 0.7237252 | 0.294 | 0.012 | 7.7141438 | 37 | boka |
| si:ch211-214p13.91 | 6.784803 | 0.91037962 | 0.333 | 0.016 | 1.7236792 | 37 | si:ch211-214p13.9 |
| pdgfd | 1.578429 | 0.571919907 | 0.235 | 0.008 | 4.0100010 | 37 | pdgfd |
| fhl2a1 | 6.223993 | 0.975730066 | 0.333 | 0.017 | 1.5812055 | 37 | fhl2a |
| si:ch211-153b23.5 | 7.439569 | 0.336507861 | 0.137 | 0.003 | 1.8900226 | 37 | si:ch211-153b23.5 |
| bmp63 | 2.074761 | 0.821944866 | 0.294 | 0.014 | 5.2709312 | 37 | bmp6 |
| rhag2 | 7.190668 | 0.644393724 | 0.275 | 0.012 | 1.8267892 | 37 | rhag |
| arhgap20b2 | 1.047940 | 0.889289327 | 0.353 | 0.021 | 2.6622936 | 37 | arhgap20b |
| sept126 | 2.262256 | 1.185535102 | 0.608 | 0.064 | 5.7472622 | 37 | sept12 |
| dennd2c | 3.712795 | 0.611145537 | 0.275 | 0.013 | 9.4323567 | 37 | dennd2c |

| | | | | | | | |
|---|---|---|---|---|---|---|---|
| gcnt4a3 | 3.468883 | 0.78348377 | 0.294 | 0.015 | 8.8126996 | 37 | gcnt4a |
| derl25 | 3.707094 | 1.355767426 | 0.765 | 0.106 | 9.4178735 | 37 | derl2 |
| XLOC-0268367 | 3.175790 | 1.26689596 | 0.667 | 0.078 | 8.0680967 | 37 | XLOC-026836 |
| blvrb1 | 4.020632 | 1.095178977 | 0.51 | 0.047 | 1.0214417 | 37 | blvrb |
| tln18 | 3.526904 | 1.436286751 | 0.824 | 0.141 | 8.9601013 | 37 | tln1 |
| si:dkey-217l24.11 | 8.473028 | 0.585599693 | 0.235 | 0.01 | 2.1525728 | 37 | si:dkey-217l24.1 |
| nfe21 | 2.002565 | 0.687003067 | 0.294 | 0.016 | 5.0875188 | 37 | nfe2 |
| ctgfa3 | 1.934883 | 1.1807665 | 0.412 | 0.033 | 4.9155708 | 37 | ctgfa |
| AL954142.12 | 6.004287 | 0.829495666 | 0.353 | 0.024 | 1.5253891 | 37 | AL954142.1 |
| zgc:171506 | 3.006962 | 0.545784308 | 0.216 | 0.009 | 7.6391872 | 37 | zgc:171506 |
| mef2aa4 | 3.395758 | 1.179446275 | 0.588 | 0.073 | 8.6269246 | 37 | mef2aa |
| dera | 9.916925 | 0.762105659 | 0.333 | 0.023 | 2.5193948 | 37 | dera |
| fhl1a1 | 1.457011 | 0.502266198 | 0.196 | 0.008 | 3.7015381 | 37 | fhl1a |
| pmp22b7 | 1.640518 | 1.093629908 | 0.471 | 0.046 | 4.1677377 | 37 | pmp22b |
| LOC1081907175 | 1.904127 | 1.225736923 | 0.627 | 0.082 | 4.8374368 | 37 | LOC108190717 |
| cdkn2c | 1.926222 | 0.394799143 | 0.157 | 0.005 | 4.8935682 | 37 | cdkn2c |
| hpca | 4.579706 | 0.646287736 | 0.275 | 0.016 | 1.1634744 | 37 | hpca |
| fermt3b3 | 1.604395 | 0.771717649 | 0.314 | 0.021 | 4.0759677 | 37 | fermt3b |
| lrrc8da2 | 8.411270 | 0.78758915 | 0.353 | 0.027 | 2.1368832 | 37 | lrrc8da |
| b3gnt3.44 | 8.083795 | 0.931876642 | 0.412 | 0.038 | 2.0536882 | 37 | b3gnt3.4 |
| mocs21 | 1.586527 | 0.517707078 | 0.176 | 0.007 | 4.0305734 | 37 | mocs2 |
| ptger2b | 1.872112 | 0.412693219 | 0.157 | 0.005 | 4.7561028 | 37 | ptger2b |
| rap1b7 | 4.132790 | 1.475356469 | 0.882 | 0.219 | 1.0499354 | 37 | rap1b |
| CR753876.17 | 7.764197 | 1.223592742 | 0.706 | 0.111 | 1.9724943 | 37 | CR753876.1 |
| XLOC-0203015 | 9.081095 | 1.286194542 | 0.549 | 0.07 | 2.3070523 | 37 | XLOC-020301 |
| nuak1a1 | 2.664382 | 0.558519534 | 0.176 | 0.007 | 6.7688638 | 37 | nuak1a |
| tpte2 | 7.293753 | 0.751398083 | 0.353 | 0.029 | 1.8529782 | 37 | tpte |
| si:ch211-222l21.115 | 1.815411 | 1.365410107 | 0.98 | 0.314 | 4.6120524 | 37 | si:ch211-222l21.1 |
| myl9b9 | 6.389437 | 1.114215758 | 0.627 | 0.098 | 1.6232366 | 37 | myl9b |
| slc5a8l | 1.318148 | 0.365178662 | 0.157 | 0.006 | 3.3487561 | 37 | slc5a8l |
| ophn1 | 9.625788 | 0.787875731 | 0.373 | 0.037 | 2.4454316 | 37 | ophn1 |
| itprid21 | 2.519058 | 0.518540086 | 0.235 | 0.015 | 6.3996677 | 37 | itprid2 |
| pim27 | 7.289940 | 1.24623609 | 0.549 | 0.088 | 1.8520093 | 37 | pim2 |
| p2rx72 | 4.784981 | 0.546634433 | 0.216 | 0.013 | 1.2156245 | 37 | p2rx7 |
| aatka1 | 4.192791 | 0.50955341 | 0.196 | 0.011 | 1.0651787 | 37 | aatka |

| | | | | | | | |
|---|---|---|---|---|---|---|---|
| srgn9 | 5.153001 | 1.270887539 | 0.882 | 0.235 | 1.3091200 | 37 | srgn |
| hif1aa | 8.81088( | 0.291236825 | 0.118 | 0.004 | 2.2384041 | 37 | hif1aa |
| rdh8a | 1.838011 | 0.339602086 | 0.137 | 0.006 | 4.6694680 | 37 | rdh8a |
| si:ch211-214p16.25 | 1.058105 | 0.976137229 | 0.51 | 0.075 | 2.6881168 | 37 | si:ch211-214p16.2 |
| si:ch211-284f22.33 | 5.137689 | 0.732104988 | 0.333 | 0.034 | 1.3052300 | 37 | si:ch211-284f22.3 |
| ptmaa17 | 7.39358( | 1.164005967 | 0.961 | 0.433 | 1.8783392 | 37 | ptmaa |
| si:ch211-226m16.27 | 1.951149 | 0.917146019 | 0.471 | 0.069 | 4.9568952 | 37 | si:ch211-226m16.2 |
| rbpms | 1.118923 | 0.581920953 | 0.235 | 0.018 | 2.8426253 | 37 | rbpms |
| si:ch73-234b20.58 | 1.978852 | 0.996797868 | 0.569 | 0.105 | 5.0272738 | 37 | si:ch73-234b20.5 |
| btk3 | 2.361507 | 0.61584812 | 0.294 | 0.028 | 5.9994093 | 37 | btk |
| btg115 | 4.561949 | 1.179366119 | 0.902 | 0.356 | 1.1589633 | 37 | btg1 |
| cldn5b1 | 7.629402 | 0.424339279 | 0.176 | 0.01 | 1.9382496 | 37 | cldn5b |
| XLOC-0283707 | 1.290188 | 1.272467061 | 0.608 | 0.126 | 3.2777239 | 37 | XLOC-028370 |
| limd24 | 1.411559 | 0.972213308 | 0.451 | 0.068 | 3.5860659 | 37 | limd2 |
| cavin2b8 | 1.45376( | 0.857065151 | 0.49 | 0.079 | 3.6932775 | 37 | cavin2b |
| nup88 | 1.54499( | 0.440898638 | 0.157 | 0.009 | 3.9250480 | 37 | nup88 |
| actn16 | 1.657144 | 0.837414594 | 0.373 | 0.048 | 4.2099758 | 37 | actn1 |
| mgst1.26 | 1.744088 | 1.001631434 | 0.588 | 0.116 | 4.4308561 | 37 | mgst1.2 |
| ano5b | 2.899867 | 0.629775639 | 0.255 | 0.023 | 7.3671127 | 37 | ano5b |
| svila | 7.640615 | 0.664276036 | 0.294 | 0.03 | 1.9410984 | 37 | svila |
| si:dkey-238c7.162 | 2.861196 | 0.879166858 | 0.451 | 0.071 | 7.2688693 | 37 | si:dkey-238c7.16 |
| iqgap2 | 5.33787( | 0.402979278 | 0.137 | 0.007 | 1.3560858 | 37 | iqgap2 |
| tmsb4x20 | 6.143764 | 0.790332826 | 1 | 0.719 | 1.5608232 | 37 | tmsb4x |
| blvra | 1.32652( | 0.388465529 | 0.157 | 0.009 | 3.3700246 | 37 | blvra |
| adgrg1 | 1.013131 | 0.446049621 | 0.176 | 0.012 | 2.5738611 | 37 | adgrg1 |
| fosab14 | 6.285006 | 1.184565286 | 0.902 | 0.362 | 1.5967057 | 37 | fosab |
| fn1b4 | 8.985174 | 0.97302233 | 0.529 | 0.102 | 2.2826835 | 37 | fn1b |
| mtss1 | 1.530396 | 0.55364864 | 0.255 | 0.026 | 3.8879717 | 37 | mtss1 |
| ptpn11b1 | 1.691138 | 0.55991719 | 0.235 | 0.022 | 4.2963364 | 37 | ptpn11b |
| tagln214 | 1.838494 | 1.078472472 | 0.706 | 0.21 | 4.6706951 | 37 | tagln2 |
| pbx1b2 | 1.902563 | 0.591904093 | 0.294 | 0.034 | 4.8334614 | 37 | pbx1b |
| asap1b1 | 1.954721 | 0.699194458 | 0.294 | 0.034 | 4.9659704 | 37 | asap1b |
| lzts2a2 | 3.993253 | 0.494836416 | 0.235 | 0.023 | 1.0144859 | 37 | lzts2a |
| pfn115 | 5.434088 | 1.029874652 | 1 | 0.502 | 1.3805300 | 37 | pfn1 |
| rpz59 | 1.654566 | 0.960489233 | 0.569 | 0.127 | 4.2034252 | 37 | rpz5 |

| | | | | | | | |
|---|---|---|---|---|---|---|---|
| zgc:921403 | 2.019462 | 0.618435381 | 0.275 | 0.031 | 5.1304443 | 37 | zgc:92140 |
| lpar15 | 2.862378 | 0.865714641 | 0.412 | 0.07 | 7.2718734 | 37 | lpar1 |
| CU639469.1 | 8.644985 | 0.367012595 | 0.118 | 0.006 | 2.1962584 | 37 | CU639469.1 |
| tmem35 | 1.427980 | 0.352491512 | 0.137 | 0.008 | 3.6277849 | 37 | tmem35 |
| f13a1b4 | 2.794151 | 0.556333237 | 0.275 | 0.032 | 7.0985430 | 37 | f13a1b |
| mylka2 | 3.798868 | 0.498894691 | 0.196 | 0.017 | 9.6510251 | 37 | mylka |
| stk26 | 2.200893 | 0.473001656 | 0.216 | 0.021 | 5.5913691 | 37 | stk26 |
| bcl2l105 | 3.337014 | 0.960021598 | 0.686 | 0.2 | 8.4776845 | 37 | bcl2l10 |
| meis1b9 | 3.834393 | 0.717773926 | 0.412 | 0.073 | 9.7412775 | 37 | meis1b |
| mcl1a10 | 3.838673 | 0.768481763 | 0.941 | 0.614 | 9.7521495 | 37 | mcl1a |
| rgs22 | 6.440135 | 0.580930783 | 0.255 | 0.03 | 1.6361164 | 37 | rgs2 |
| atp2a33 | 2.446670 | 0.604407483 | 0.275 | 0.035 | 6.2157657 | 37 | atp2a3 |
| ppiab7 | 2.707195 | 0.635695838 | 0.98 | 0.675 | 6.8776302 | 37 | ppiab |
| ap1s11 | 3.020004 | 0.822208343 | 0.471 | 0.099 | 7.6723221 | 37 | ap1s1 |
| soga1 | 6.760810 | 0.522040229 | 0.216 | 0.023 | 1.7175840 | 37 | soga1 |
| zgc:1625091 | 1.312816 | 0.545399961 | 0.255 | 0.031 | 3.3352110 | 37 | zgc:162509 |
| zgc:1538678 | 1.641236 | 0.875118876 | 0.882 | 0.465 | 4.1695619 | 37 | zgc:153867 |
| CR383676.121 | 2.000208 | 0.793223083 | 1 | 0.718 | 5.0815284 | 37 | CR383676.1 |
| zgc:153409 | 2.870839 | 0.391851654 | 0.176 | 0.016 | 7.2933679 | 37 | zgc:153409 |
| actb213 | 3.352571 | 0.552870179 | 1 | 0.867 | 8.5172075 | 37 | actb2 |
| myl12.115 | 3.679166 | 0.869862108 | 0.863 | 0.427 | 9.3469234 | 37 | myl12.1 |
| gng103 | 5.145963 | 0.747668327 | 0.353 | 0.061 | 1.3073319 | 37 | gng10 |
| gapdhs14 | 1.850226 | 0.954294268 | 0.745 | 0.316 | 4.7005009 | 37 | gapdhs |
| ccnd2a4 | 2.369468 | 0.892868414 | 0.431 | 0.095 | 6.0196357 | 37 | ccnd2a |
| dusp57 | 3.047271 | 0.678173567 | 0.373 | 0.068 | 7.7415919 | 37 | dusp5 |
| XLOC-0320025 | 4.569150 | 0.650318799 | 0.294 | 0.044 | 1.1607927 | 37 | XLOC-032002 |
| flna8 | 4.582361 | 0.904800025 | 0.49 | 0.122 | 1.1641489 | 37 | flna |
| vash2 | 5.485976 | 0.343605612 | 0.118 | 0.008 | 1.3937123 | 37 | vash2 |
| creg12 | 6.626429 | 0.797177974 | 0.392 | 0.078 | 1.6834445 | 37 | creg1 |
| arpc1b14 | 1.148433 | 0.955482844 | 0.804 | 0.314 | 2.9175946 | 37 | arpc1b |
| zeb2a3 | 1.155225 | 0.679710708 | 0.294 | 0.047 | 2.9348510 | 37 | zeb2a |
| si:ch211-105j21.94 | 1.318128 | 0.563919769 | 0.255 | 0.035 | 3.3487065 | 37 | si:ch211-105j21.9 |
| gna11b2 | 1.521639 | 0.747207309 | 0.353 | 0.065 | 3.8657243 | 37 | gna11b |
| tfpia2 | 2.687977 | 0.391048901 | 0.196 | 0.022 | 6.8288061 | 37 | tfpia |
| vaspb3 | 2.837710 | 0.610231618 | 0.314 | 0.053 | 7.2092047 | 37 | vaspb |

| | | | | | | | |
|---|---|---|---|---|---|---|---|
| si:dkey-51e6.15 | 6.154128 | 0.833580867 | 0.471 | 0.119 | 1.5634564 | 37 | si:dkey-51e6.1 |
| tspan74 | 6.192932 | 0.775032336 | 0.392 | 0.082 | 1.5733144 | 37 | tspan7 |
| LOC101886015 | 1.411329 | 0.491902042 | 0.196 | 0.023 | 3.5854837 | 37 | LOC101886015 |
| si:cabz01074946.15 | 2.804629 | 0.638104233 | 0.333 | 0.061 | 7.1251605 | 37 | si:cabz01074946.1 |
| ehd1a1 | 6.182687 | 0.55426538 | 0.235 | 0.033 | 1.5707116 | 37 | ehd1a |
| cyr617 | 6.524333 | 0.663751297 | 0.353 | 0.068 | 1.6575068 | 37 | cyr61 |
| sb:cb812 | 2.055232 | 0.511501455 | 0.216 | 0.029 | 5.2213192 | 37 | sb:cb81 |
| itpr3 | 2.139376 | 0.326479657 | 0.118 | 0.009 | 5.4350853 | 37 | itpr3 |
| zgc:6405111 | 2.975072 | 0.80217774 | 0.549 | 0.163 | 7.5581717 | 37 | zgc:64051 |
| si:ch211-102c2.49 | 3.100385 | 0.867542428 | 0.51 | 0.147 | 7.8765283 | 37 | si:ch211-102c2.4 |
| zgc:91860 | 3.303225 | 0.298848327 | 0.118 | 0.009 | 8.3918433 | 37 | zgc:91860 |
| zgc:664332 | 4.313629 | 0.564393354 | 0.235 | 0.035 | 1.0958775 | 37 | zgc:66433 |
| cotl113 | 4.409849 | 0.868488787 | 0.765 | 0.32 | 1.1203223 | 37 | cotl1 |
| plxna2 | 5.277951 | 0.298059315 | 0.118 | 0.009 | 1.3408634 | 37 | plxna2 |
| ilk1 | 6.844914 | 0.756542987 | 0.373 | 0.085 | 1.7389504 | 37 | ilk |
| RASSF57 | 8.608499 | 0.691377511 | 0.314 | 0.062 | 2.1869892 | 37 | RASSF5 |
| suv39h1b1 | 1.723301 | 0.623736099 | 0.255 | 0.041 | 4.3780480 | 37 | suv39h1b |
| XLOC-0254237 | 1.966002 | 0.908077417 | 0.549 | 0.171 | 4.9946294 | 37 | XLOC-025423 |
| pnrc218 | 4.510571 | 0.635553247 | 0.922 | 0.598 | 1.1459107 | 37 | pnrc2 |
| actb19 | 5.866039 | 0.488724604 | 0.941 | 0.768 | 1.4902673 | 37 | actb1 |
| pacs1a | 1.037382 | 0.344376818 | 0.118 | 0.01 | 2.6354697 | 37 | pacs1a |
| glcci1a | 1.087918 | 0.351134767 | 0.137 | 0.014 | 2.7638564 | 37 | glcci1a |
| alox5a4 | 2.272395 | 0.553842705 | 0.216 | 0.033 | 5.7730203 | 37 | alox5a |
| rabgap1l | 2.995376 | 0.610816964 | 0.235 | 0.04 | 7.6097547 | 37 | rabgap1l |
| rfx5 | 3.329616 | 0.514326613 | 0.216 | 0.034 | 8.4588895 | 37 | rfx5 |
| dub8 | 3.388984 | 0.626387587 | 0.333 | 0.074 | 8.6097146 | 37 | dub |
| XLOC-0437603 | 5.609433 | 0.616911524 | 0.255 | 0.046 | 1.4250765 | 37 | XLOC-043760 |
| yrk7 | 7.068628 | 0.633557781 | 0.314 | 0.068 | 1.7957850 | 37 | yrk |
| macf1a2 | 7.603520 | 0.561597842 | 0.294 | 0.06 | 1.9316744 | 37 | macf1a |
| zfpm11 | 8.657371 | 0.269683947 | 0.137 | 0.015 | 2.1994052 | 37 | zfpm1 |
| bmp2k | 9.478394 | 0.364299349 | 0.157 | 0.019 | 2.4079861 | 37 | bmp2k |
| LOC1039091159 | 1.382485 | 0.749318352 | 0.51 | 0.17 | 3.5122035 | 37 | LOC103909115 |
| rhof | 1.502522 | 0.464681287 | 0.176 | 0.024 | 3.8171580 | 37 | rhof |
| kdm6bb3 | 1.612358 | 0.60983075 | 0.294 | 0.063 | 4.0961962 | 37 | kdm6bb |
| si:ch211-196l7.415 | 1.614586 | 0.773989511 | 0.706 | 0.331 | 4.1018558 | 37 | si:ch211-196l7.4 |

| | | | | | | | |
|---|---|---|---|---|---|---|---|
| tspan5a | 1.683726 | 0.383457813 | 0.137 | 0.015 | 4.2775082 | 37 | tspan5a |
| hdr1 | 1.742183 | 0.569335664 | 0.255 | 0.046 | 4.4260182 | 37 | hdr |
| fhl1b3 | 7.363137 | 0.337521158 | 0.157 | 0.02 | 1.8706050 | 37 | fhl1b |
| atp2b1a3 | 8.952524 | 0.622634946 | 0.314 | 0.073 | 2.2743888 | 37 | atp2b1a |
| znfl2a | 1.050423 | 0.261087102 | 0.118 | 0.012 | 2.6686002 | 37 | znfl2a |
| prkacba1 | 2.373331 | 0.387316381 | 0.157 | 0.021 | 6.0294485 | 37 | prkacba |
| tpm312 | 3.815836 | 0.760265584 | 0.608 | 0.285 | 9.6941315 | 37 | tpm3 |
| si:dkey-52l18.4 | 3.960067 | 0.598444082 | 0.255 | 0.053 | 1.0060550 | 37 | si:dkey-52l18.4 |
| jdp2b9 | 9.051068 | 0.82881629 | 0.471 | 0.165 | 2.2994240 | 37 | jdp2b |
| fkbp1b1 | 1.133976 | 0.275620939 | 0.137 | 0.018 | 2.8808683 | 37 | fkbp1b |
| LOC5533161 | 1.543559 | 0.409031051 | 0.157 | 0.023 | 3.9214125 | 37 | LOC553316 |
| dusp62 | 2.188256 | 0.492855196 | 0.255 | 0.055 | 5.5592655 | 37 | dusp6 |
| ier2a15 | 3.794970 | 0.752232993 | 0.608 | 0.275 | 9.6411216 | 37 | ier2a |
| BX005183.37 | 7.187108 | 0.626531631 | 0.294 | 0.077 | 1.8258850 | 37 | BX005183.3 |
| si:ch211-203d1.3 | 8.400287 | 0.327760648 | 0.137 | 0.019 | 2.1340930 | 37 | si:ch211-203d1.3 |
| selenow19 | 1.167341 | 0.652272428 | 0.725 | 0.416 | 2.9656314 | 37 | selenow1 |
| zgc:560952 | 1.211345 | 0.590435054 | 0.353 | 0.102 | 3.0774220 | 37 | zgc:56095 |
| XLOC-0437612 | 1.255725 | 0.374468234 | 0.137 | 0.02 | 3.1901717 | 37 | XLOC-043761 |
| shq1 | 1.742588 | 0.440217838 | 0.176 | 0.031 | 4.4270472 | 37 | shq1 |
| mtus1b1 | 1.810055 | 0.353811066 | 0.157 | 0.025 | 4.5984453 | 37 | mtus1b |
| abhd31 | 2.810106 | 0.330238822 | 0.137 | 0.02 | 7.1390746 | 37 | abhd3 |
| b3galt2 | 3.143992 | 0.289626948 | 0.176 | 0.03 | 7.9873119 | 37 | b3galt2 |
| bambib1 | 3.708702 | 0.25607088 | 0.118 | 0.015 | 9.4219597 | 37 | bambib |
| lrrfip1a6 | 3.815223 | 0.490772205 | 0.255 | 0.061 | 9.6925765 | 37 | lrrfip1a |
| actn44 | 5.289375 | 0.535393376 | 0.333 | 0.098 | 0.000134 | 37 | actn4 |
| ubl3a | 8.538021 | 0.465370588 | 0.216 | 0.048 | 0.000217 | 37 | ubl3a |
| bloc1s5 | 1.013631 | 0.372776088 | 0.157 | 0.027 | 0.000258 | 37 | bloc1s5 |
| gsna2 | 1.171090 | 0.42603449 | 0.176 | 0.034 | 0.000298 | 37 | gsna |
| cap111 | 1.180825 | 0.71849524 | 0.529 | 0.247 | 0.0003 | 37 | cap1 |
| si:zfos-464b6.22 | 1.415484 | 0.490552871 | 0.216 | 0.049 | 0.00036 | 37 | si:zfos-464b6.2 |
| palld1 | 1.910051 | 0.306326344 | 0.176 | 0.033 | 0.000485 | 37 | palld |
| cmtm61 | 2.988328 | 0.478049343 | 0.235 | 0.058 | 0.000759 | 37 | cmtm6 |
| snx5 | 3.048565 | 0.558768003 | 0.255 | 0.069 | 0.000774 | 37 | snx5 |
| slc25a55a4 | 3.170530 | 0.514957126 | 0.275 | 0.076 | 0.000805 | 37 | slc25a55a |
| trib35 | 3.252943 | 0.58737073 | 0.333 | 0.109 | 0.000826 | 37 | trib3 |

| | | | | | | | |
|---|---|---|---|---|---|---|---|
| msna12 | 4.559224 | 0.614904707 | 0.549 | 0.267 | 0.001158 | 37 | msna |
| ing5a | 4.805022 | 0.322574481 | 0.137 | 0.023 | 0.001221 | 37 | ing5a |
| trip10a | 5.394181 | 0.316120044 | 0.118 | 0.017 | 0.00137 | 37 | trip10a |
| vamp51 | 5.971332 | 0.355741464 | 0.157 | 0.029 | 0.001517 | 37 | vamp5 |
| fosb10 | 6.202141 | 0.690128939 | 0.353 | 0.122 | 0.001576 | 37 | fosb |
| fam53b3 | 6.535458 | 0.472868553 | 0.275 | 0.077 | 0.00166 | 37 | fam53b |
| rab11fip1b5 | 7.033841 | 0.515482082 | 0.235 | 0.061 | 0.001787 | 37 | rab11fip1b |
| dnase1l12 | 8.685382 | 0.507611134 | 0.216 | 0.054 | 0.002207 | 37 | dnase1l1 |
| cfl112 | 1.036129 | 0.510052613 | 0.804 | 0.601 | 0.002632 | 37 | cfl1 |
| myh9a3 | 1.166637 | 0.431729364 | 0.216 | 0.052 | 0.002964 | 37 | myh9a |
| phlda26 | 1.330867 | 0.531256352 | 0.216 | 0.053 | 0.003381 | 37 | phlda2 |
| ehbp1l1a1 | 1.387461 | 0.319691633 | 0.118 | 0.018 | 0.003525 | 37 | ehbp1l1a |
| arpc5a9 | 1.521995 | 0.558634028 | 0.353 | 0.126 | 0.003867 | 37 | arpc5a |
| prkcaa1 | 1.574467 | 0.316465009 | 0.118 | 0.019 | 0.004 | 37 | prkcaa |
| arpc213 | 1.739588 | 0.601375298 | 0.569 | 0.3 | 0.004419 | 37 | arpc2 |
| crema11 | 1.862860 | 0.653844335 | 0.412 | 0.175 | 0.004733 | 37 | crema |
| capza1b8 | 2.309813 | 0.606152782 | 0.49 | 0.236 | 0.005868 | 37 | capza1b |
| dyrk1b | 2.752382 | 0.31727291 | 0.137 | 0.025 | 0.006992 | 37 | dyrk1b |
| rassf2b | 2.881824 | 0.272650187 | 0.118 | 0.019 | 0.007321 | 37 | rassf2b |
| cyth1b | 3.257916 | 0.346894008 | 0.157 | 0.032 | 0.008277 | 37 | cyth1b |
| sh3kbp15 | 3.658745 | 0.532796302 | 0.255 | 0.077 | 0.009295 | 37 | sh3kbp1 |
| ppp1cbl1 | 3.687662 | 0.523910042 | 0.235 | 0.068 | 0.009369 | 37 | ppp1cbl |
| mxd4 | 4.331514 | 0.320434808 | 0.137 | 0.026 | 0.011004 | 37 | mxd4 |
| cpn17 | 4.693488 | 0.564754374 | 0.255 | 0.078 | 0.011924 | 37 | cpn1 |
| sept9b2 | 4.956028 | 0.408567911 | 0.196 | 0.049 | 0.012591 | 37 | sept9b |
| hmgn617 | 5.120364 | 0.568548331 | 0.569 | 0.312 | 0.013008 | 37 | hmgn6 |
| mob1a6 | 5.827273 | 0.564852728 | 0.412 | 0.174 | 0.014804 | 37 | mob1a |
| kctd5a5 | 5.869453 | 0.551498184 | 0.431 | 0.187 | 0.014911 | 37 | kctd5a |
| badb | 6.959448 | 0.270477365 | 0.157 | 0.032 | 0.01768 | 37 | badb |
| inka1b7 | 7.805293 | 0.536488977 | 0.412 | 0.157 | 0.019829 | 37 | inka1b |
| selenok4 | 7.994777 | 0.572183803 | 0.353 | 0.14 | 0.020311 | 37 | selenok |
| plk33 | 8.189767 | 0.549842596 | 0.314 | 0.11 | 0.020806 | 37 | plk3 |
| vclb | 8.788506 | 0.275456775 | 0.118 | 0.02 | 0.022327 | 37 | vclb |
| arhgef12a | 1.005640 | 0.275755581 | 0.118 | 0.02 | 0.025548 | 37 | arhgef12a |
| eef2k | 1.251680 | 0.316920191 | 0.118 | 0.021 | 0.031799 | 37 | eef2k |

| | | | | | | | |
|---|---|---|---|---|---|---|---|
| ier2b16 | 1.521625 | 0.625443117 | 0.471 | 0.238 | 0.038657 | 37 | ier2b |
| pdlim15 | 1.966442 | 0.435019522 | 0.216 | 0.061 | 0.049957 | 37 | pdlim1 |
| rhoab12 | 2.142456 | 0.522294705 | 0.451 | 0.208 | 0.054429 | 37 | rhoab |
| nr4a112 | 2.145530 | 0.620175668 | 0.373 | 0.159 | 0.054507 | 37 | nr4a1 |
| hmgb1a12 | 2.321854 | 0.550863231 | 0.471 | 0.229 | 0.058987 | 37 | hmgb1a |
| dynll2b2 | 2.699536 | 0.450757741 | 0.353 | 0.131 | 0.068582 | 37 | dynll2b |
| sdc413 | 3.454617 | 0.527610383 | 0.529 | 0.255 | 0.087765 | 37 | sdc4 |
| ddx54 | 3.549101 | 0.534405032 | 0.588 | 0.356 | 0.090165 | 37 | ddx5 |
| ralgapa1 | 3.625455 | 0.373081551 | 0.137 | 0.029 | 0.092105 | 37 | ralgapa1 |
| pbx1a4 | 4.327471 | 0.434457197 | 0.196 | 0.055 | 0.109939 | 37 | pbx1a |
| mob2b1 | 4.532717 | 0.358207671 | 0.137 | 0.03 | 0.115154 | 37 | mob2b |
| cd6317 | 4.918824 | 0.557418375 | 0.608 | 0.375 | 0.124963 | 37 | cd63 |
| add12 | 5.573361 | 0.368533778 | 0.216 | 0.063 | 0.141591 | 37 | add1 |
| rgl21 | 6.511478 | 0.292365833 | 0.118 | 0.023 | 0.165424 | 37 | rgl2 |
| tnikb1 | 6.648287 | 0.260484011 | 0.118 | 0.023 | 0.1689 | 37 | tnikb |
| sept24 | 8.219359 | 0.540537561 | 0.392 | 0.184 | 0.208813 | 37 | sept2 |
| tes3 | 8.363170 | 0.417799664 | 0.216 | 0.065 | 0.212466 | 37 | tes |
| nup931 | 9.652417 | 0.315519143 | 0.157 | 0.038 | 0.24522 | 37 | nup93 |
| h3f3c15 | 9.709058 | 0.530180085 | 0.529 | 0.306 | 0.246659 | 37 | h3f3c |
| si:dkey-185m8.22 | 1.067069 | 0.357477966 | 0.176 | 0.048 | 0.271089 | 37 | si:dkey-185m8.2 |
| orai1a2 | 1.179198 | 0.274391127 | 0.118 | 0.024 | 0.299575 | 37 | orai1a |
| dynlrb12 | 1.259109 | 0.474132014 | 0.451 | 0.214 | 0.319877 | 37 | dynlrb1 |
| si:ch211-166a6.54 | 1.288626 | 0.267615261 | 0.157 | 0.039 | 0.327376 | 37 | si:ch211-166a6.5 |
| ip6k2b8 | 1.290162 | 0.396739745 | 0.333 | 0.13 | 0.327766 | 37 | ip6k2b |
| erbin1 | 1.398039 | 0.370259885 | 0.196 | 0.058 | 0.355172 | 37 | erbin |
| serf212 | 1.719120 | 0.467769069 | 0.588 | 0.379 | 0.436742 | 37 | serf2 |
| CR848749.21 | 1.945561 | 0.305325811 | 0.118 | 0.025 | 0.49427 | 37 | CR848749.2 |
| btg212 | 2.054984 | 0.510426661 | 0.588 | 0.351 | 0.522069 | 37 | btg2 |
| wu:fb55g094 | 2.432121 | 0.555157194 | 0.471 | 0.261 | 0.617881 | 37 | wu:fb55g09 |
| si:ch211-260e23.93 | 2.710842 | 0.483876762 | 0.196 | 0.063 | 0.68869 | 37 | si:ch211-260e23.9 |
| gbp3 | 2.831943 | 0.515517388 | 0.294 | 0.123 | 0.719455 | 37 | gbp |
| CR318588.36 | 3.202781 | 0.47450015 | 0.176 | 0.052 | 0.813667 | 37 | CR318588.3 |
| gmds1 | 3.321358 | 0.326400458 | 0.176 | 0.05 | 0.843791 | 37 | gmds |
| emilin1a4 | 3.422844 | 0.29539961 | 0.118 | 0.026 | 0.869574 | 37 | emilin1a |
| rhocb2 | 3.758895 | 0.40315412 | 0.275 | 0.104 | 0.954947 | 37 | rhocb |

| | | | | | | | |
|---|---|---|---|---|---|---|---|
| gadd45ba11 | 4.087237 | 0.504560021 | 0.412 | 0.199 | 1 | 37 | gadd45ba |
| tubb4b3 | 4.447257 | 0.378036235 | 0.765 | 0.593 | 1 | 37 | tubb4b |
| si:dkey-85k15.4 | 5.480221 | 0.315741661 | 0.137 | 0.034 | 1 | 37 | si:dkey-85k15.4 |
| si:ch211-214p16.18 | 6.923749 | 0.439060294 | 0.529 | 0.245 | 1 | 37 | si:ch211-214p16.1 |
| jak2a2 | 6.996572 | 0.256734528 | 0.118 | 0.027 | 1 | 37 | jak2a |
| GNAZ1 | 7.254283 | 0.267146769 | 0.118 | 0.028 | 1 | 37 | GNAZ |
| tfdp21 | 8.640301 | 0.288975879 | 0.137 | 0.036 | 1 | 37 | tfdp2 |
| anp32e | 8.848539 | 0.405988855 | 0.314 | 0.132 | 1 | 37 | anp32e |
| mfng3 | 0.0001 | 0.318159456 | 0.157 | 0.046 | 1 | 37 | mfng |
| hmgn213 | 0.0001 | 0.546612683 | 0.627 | 0.469 | 1 | 37 | hmgn2 |
| lyn2 | 0.0001 | 0.303534053 | 0.157 | 0.046 | 1 | 37 | lyn |
| CABZ01079663.13 | 0.0001 | 0.333997291 | 0.137 | 0.038 | 1 | 37 | CABZ01079663.1 |
| ankrd10b | 0.0001 | 0.263225637 | 0.118 | 0.029 | 1 | 37 | ankrd10b |
| zgc:1537134 | 0.0002 | 0.440845221 | 0.353 | 0.174 | 1 | 37 | zgc:153713 |
| crebrf | 0.0002 | 0.34024925 | 0.137 | 0.039 | 1 | 37 | crebrf |
| twf1a5 | 0.0002 | 0.458272068 | 0.333 | 0.159 | 1 | 37 | twf1a |
| rbms1b | 0.0002 | 0.316147541 | 0.137 | 0.039 | 1 | 37 | rbms1b |
| vps36 | 0.0002 | 0.25054667 | 0.137 | 0.038 | 1 | 37 | vps36 |
| calm2a4 | 0.0002 | 0.473613543 | 0.392 | 0.211 | 1 | 37 | calm2a |
| mt-co117 | 0.0002 | 0.366397397 | 0.902 | 0.641 | 1 | 37 | mt-co1 |
| rap2c | 0.0002 | 0.421260587 | 0.275 | 0.118 | 1 | 37 | rap2c |
| sh3bgrl7 | 0.0003 | 0.435403701 | 0.275 | 0.122 | 1 | 37 | sh3bgrl |
| pim111 | 0.0003 | 0.492087623 | 0.333 | 0.17 | 1 | 37 | pim1 |
| spred2b8 | 0.0003 | 0.435054609 | 0.216 | 0.086 | 1 | 37 | spred2b |
| ppp3r1a | 0.0003 | 0.279312662 | 0.157 | 0.049 | 1 | 37 | ppp3r1a |
| capza1a10 | 0.0003 | 0.413038164 | 0.412 | 0.222 | 1 | 37 | capza1a |
| sept9a1 | 0.0003 | 0.32712648 | 0.157 | 0.05 | 1 | 37 | sept9a |
| lbr1 | 0.0004 | 0.266421466 | 0.216 | 0.079 | 1 | 37 | lbr |
| histh1l9 | 0.0004 | 0.440249715 | 0.275 | 0.123 | 1 | 37 | histh1l |
| ndfip22 | 0.0004 | 0.316523054 | 0.176 | 0.061 | 1 | 37 | ndfip2 |
| TCIM (1 of many)3 | 0.0005 | 0.317021157 | 0.157 | 0.05 | 1 | 37 | TCIM (1 of many) |
| elmo16 | 0.0005 | 0.288704164 | 0.196 | 0.071 | 1 | 37 | elmo1 |
| sh2b34 | 0.0005 | 0.370420883 | 0.176 | 0.063 | 1 | 37 | sh2b3 |
| kmt5aa | 0.0005 | 0.42007544 | 0.314 | 0.152 | 1 | 37 | kmt5aa |
| lrrfip2 | 0.0005 | 0.272853572 | 0.137 | 0.041 | 1 | 37 | lrrfip2 |

| | | | | | | | |
|---|---|---|---|---|---|---|---|
| sh3bgrl35 | 0.0006 | 0.400318896 | 0.314 | 0.157 | 1 | 37 | sh3bgrl3 |
| gabarapa6 | 0.0008 | 0.454108206 | 0.333 | 0.182 | 1 | 37 | gabarapa |
| CR855337.13 | 0.0008 | 0.273709663 | 0.118 | 0.034 | 1 | 37 | CR855337.1 |
| ppt16 | 0.0009 | 0.461649594 | 0.333 | 0.183 | 1 | 37 | ppt1 |
| arpc4l9 | 0.0009 | 0.390119615 | 0.392 | 0.22 | 1 | 37 | arpc4l |
| kif5ba1 | 0.001 | 0.275540016 | 0.157 | 0.055 | 1 | 37 | kif5ba |
| ywhaba5 | 0.0011 | 0.390608874 | 0.49 | 0.33 | 1 | 37 | ywhaba |
| hopx8 | 0.0011 | 0.305611401 | 0.176 | 0.066 | 1 | 37 | hopx |
| mt-cyb14 | 0.0011 | 0.332844587 | 0.745 | 0.578 | 1 | 37 | mt-cyb |
| siah1 | 0.0012 | 0.311644501 | 0.137 | 0.045 | 1 | 37 | siah1 |
| mibp215 | 0.0012 | 0.336469482 | 0.647 | 0.523 | 1 | 37 | mibp2 |
| skap28 | 0.0013 | 0.376677895 | 0.196 | 0.082 | 1 | 37 | skap2 |
| jun13 | 0.0013 | 0.462468585 | 0.431 | 0.251 | 1 | 37 | jun |
| csrp1a4 | 0.0013 | 0.375966359 | 0.196 | 0.082 | 1 | 37 | csrp1a |
| ucp214 | 0.0017 | 0.356117418 | 0.588 | 0.416 | 1 | 37 | ucp2 |
| klf7b1 | 0.0018 | 0.28627766 | 0.118 | 0.037 | 1 | 37 | klf7b |
| rhogb7 | 0.0021 | 0.344520121 | 0.235 | 0.109 | 1 | 37 | rhogb |
| irf2bp2a8 | 0.0021 | 0.377371193 | 0.333 | 0.187 | 1 | 37 | irf2bp2a |
| wdr110 | 0.0022 | 0.390179818 | 0.294 | 0.157 | 1 | 37 | wdr1 |
| mir21-15 | 0.0022 | 0.324243599 | 0.235 | 0.109 | 1 | 37 | mir21-1 |
| thrap3b | 0.0022 | 0.341251943 | 0.216 | 0.097 | 1 | 37 | thrap3b |
| lims11 | 0.0023 | 0.278490929 | 0.137 | 0.049 | 1 | 37 | lims1 |
| snx122 | 0.0023 | 0.327883061 | 0.196 | 0.084 | 1 | 37 | snx12 |
| txnipa10 | 0.0023 | 0.401626969 | 0.392 | 0.246 | 1 | 37 | txnipa |
| rnf444 | 0.0023 | 0.342595751 | 0.216 | 0.098 | 1 | 37 | rnf44 |
| si:ch211-225b11.12 | 0.0024 | 0.255427378 | 0.118 | 0.038 | 1 | 37 | si:ch211-225b11.1 |
| si:dkey-261h17.15 | 0.0025 | 0.313982742 | 0.255 | 0.117 | 1 | 37 | si:dkey-261h17.1 |
| taf71 | 0.0026 | 0.307627512 | 0.157 | 0.061 | 1 | 37 | taf7 |
| cnn212 | 0.0027 | 0.375076952 | 0.294 | 0.158 | 1 | 37 | cnn2 |
| klf6a10 | 0.0028 | 0.446156934 | 0.314 | 0.182 | 1 | 37 | klf6a |
| qkia10 | 0.0028 | 0.36376727 | 0.353 | 0.201 | 1 | 37 | qkia |
| rab8b4 | 0.0029 | 0.319351886 | 0.196 | 0.086 | 1 | 37 | rab8b |
| fkbp57 | 0.0029 | 0.382828066 | 0.255 | 0.129 | 1 | 37 | fkbp5 |
| fthl272 | 0.0029 | 0.255423744 | 0.784 | 0.684 | 1 | 37 | fthl27 |
| si:ch211-119e14.15 | 0.003 | 0.283559572 | 0.176 | 0.072 | 1 | 37 | si:ch211-119e14.1 |

| | | | | | | | |
|---|---|---|---|---|---|---|---|
| mt-co216 | 0.0031 | 0.303140033 | 0.843 | 0.672 | 1 | 37 | mt-co2 |
| jupa4 | 0.0033 | 0.256658576 | 0.157 | 0.061 | 1 | 37 | jupa |
| oaz1b13 | 0.004 | 0.33345515 | 0.51 | 0.362 | 1 | 37 | oaz1b |
| cdc261 | 0.0043 | 0.342830198 | 0.196 | 0.091 | 1 | 37 | cdc26 |
| sat1a.29 | 0.0047 | 0.332714288 | 0.471 | 0.303 | 1 | 37 | sat1a.2 |
| nr1d2a4 | 0.0049 | 0.301707221 | 0.196 | 0.091 | 1 | 37 | nr1d2a |
| irf24 | 0.0051 | 0.265552939 | 0.176 | 0.077 | 1 | 37 | irf2 |
| COX318 | 0.0057 | 0.281282281 | 0.843 | 0.677 | 1 | 37 | COX3 |
| reep511 | 0.0061 | 0.392761847 | 0.294 | 0.175 | 1 | 37 | reep5 |
| arf6a3 | 0.0061 | 0.337961692 | 0.294 | 0.164 | 1 | 37 | arf6a |
| il10ra4 | 0.0063 | 0.260794906 | 0.118 | 0.043 | 1 | 37 | il10ra |
| nfyba | 0.0065 | 0.335056296 | 0.314 | 0.182 | 1 | 37 | nfyba |
| chmp1b4 | 0.0072 | 0.304009998 | 0.255 | 0.134 | 1 | 37 | chmp1b |
| mtdha3 | 0.0081 | 0.289884105 | 0.275 | 0.149 | 1 | 37 | mtdha |
| eif1b12 | 0.0083 | 0.304657207 | 0.529 | 0.408 | 1 | 37 | eif1b |
| ube2l3b | 0.0084 | 0.295072105 | 0.294 | 0.166 | 1 | 37 | ube2l3b |
| sik17 | 0.0085 | 0.358319334 | 0.216 | 0.112 | 1 | 37 | sik1 |
| itm2ba13 | 0.0086 | 0.332459851 | 0.431 | 0.299 | 1 | 37 | itm2ba |
| arpc311 | 0.0099 | 0.31986633 | 0.471 | 0.336 | 1 | 37 | arpc3 |
| kmt2bb4 | 0.01 | 0.313047444 | 0.196 | 0.098 | 1 | 37 | kmt2bb |
| lum3 | 0 | 2.100585105 | 1 | 0.019 | 0 | 38 | lum |
| ogna2 | 0 | 1.813196308 | 0.976 | 0.007 | 0 | 38 | ogna |
| cgref12 | 0 | 1.587633042 | 0.952 | 0.019 | 0 | 38 | cgref1 |
| fgl2b2 | 0 | 1.449580529 | 0.929 | 0.013 | 0 | 38 | fgl2b |
| mettl24 | 0 | 1.408330137 | 0.905 | 0.01 | 0 | 38 | mettl24 |
| erg1 | 0 | 1.400253692 | 0.905 | 0.011 | 0 | 38 | erg |
| tmem119b3 | 0 | 1.387056064 | 0.905 | 0.02 | 0 | 38 | tmem119b |
| htra3b2 | 0 | 1.347357879 | 0.929 | 0.012 | 0 | 38 | htra3b |
| igfbp32 | 0 | 1.31919495 | 0.786 | 0.011 | 0 | 38 | igfbp3 |
| mmp23bb | 0 | 1.300409539 | 0.833 | 0.006 | 0 | 38 | mmp23bb |
| si:ch211-243a20.32 | 0 | 1.276761106 | 0.881 | 0.012 | 0 | 38 | si:ch211-243a20.3 |
| ak51 | 0 | 1.133811601 | 0.81 | 0.005 | 0 | 38 | ak5 |
| olfml11 | 0 | 1.099840046 | 0.738 | 0.005 | 0 | 38 | olfml1 |
| col5a3a1 | 0 | 1.003248079 | 0.714 | 0.008 | 0 | 38 | col5a3a |
| creb3l12 | 0 | 0.868327851 | 0.619 | 0.006 | 0 | 38 | creb3l1 |

| | | | | | | | |
|---|---|---|---|---|---|---|---|
| omd1 | 0 | 0.866442453 | 0.405 | 0.001 | 0 | 38 | omd |
| apcdd1l2 | 0 | 0.860254151 | 0.619 | 0.008 | 0 | 38 | apcdd1l |
| npnt | 0 | 0.807871408 | 0.595 | 0.006 | 0 | 38 | npnt |
| wisp1b | 0 | 0.779054231 | 0.524 | 0.001 | 0 | 38 | wisp1b |
| fgl1 | 0 | 0.725741526 | 0.429 | 0.001 | 0 | 38 | fgl1 |
| si:ch211-37e10.21 | 0 | 0.561439412 | 0.429 | 0.002 | 0 | 38 | si:ch211-37e10.2 |
| c1qtnf6b | 0 | 0.525460297 | 0.381 | 0.001 | 0 | 38 | c1qtnf6b |
| wnt21 | 0 | 0.435941681 | 0.333 | 0.002 | 0 | 38 | wnt2 |
| XLOC-008935 | 0 | 0.317863915 | 0.238 | 0.001 | 0 | 38 | XLOC-008935 |
| olfml2bb | 2.812472 | 0.298271199 | 0.19 | 0.001 | 7.1450863 | 38 | olfml2bb |
| scinla1 | 2.266881 | 1.261933878 | 0.643 | 0.011 | 5.7590128 | 38 | scinla |
| twist32 | 7.751258 | 1.550935842 | 0.976 | 0.028 | 1.9692072 | 38 | twist3 |
| col7a1 | 1.321041 | 0.352556661 | 0.238 | 0.001 | 3.3561057 | 38 | col7a1 |
| cdo12 | 1.846426 | 0.834823246 | 0.5 | 0.007 | 4.6908457 | 38 | cdo1 |
| twist22 | 1.147928 | 0.94484069 | 0.595 | 0.01 | 2.9163132 | 38 | twist2 |
| CABZ01074363.12 | 6.434119 | 1.448508708 | 0.905 | 0.026 | 1.6345881 | 38 | CABZ01074363.1 |
| fam19a1a | 2.034238 | 0.380139882 | 0.262 | 0.002 | 5.1679831 | 38 | fam19a1a |
| wnt10b | 2.844615 | 0.439081683 | 0.333 | 0.003 | 7.2267451 | 38 | wnt10b |
| irx1b | 1.501656 | 0.434450773 | 0.286 | 0.002 | 3.8149588 | 38 | irx1b |
| col16a14 | 1.764125 | 1.959516907 | 1 | 0.039 | 4.4817614 | 38 | col16a1 |
| si:dkeyp-59c12.1 | 9.965030 | 0.364969877 | 0.238 | 0.002 | 2.5316160 | 38 | si:dkeyp-59c12.1 |
| srpx22 | 6.657053 | 0.74485919 | 0.476 | 0.008 | 1.6912244 | 38 | srpx2 |
| fkbp111 | 1.396963 | 1.11124148 | 0.762 | 0.023 | 3.5489850 | 38 | fkbp11 |
| vwa13 | 7.627093 | 0.977106528 | 0.714 | 0.02 | 1.9376632 | 38 | vwa1 |
| serpinf17 | 6.100380 | 1.546954235 | 0.976 | 0.043 | 1.5498015 | 38 | serpinf1 |
| cspg5a1 | 4.413402 | 0.755582398 | 0.5 | 0.01 | 1.1212249 | 38 | cspg5a |
| dnm3a | 2.037933 | 0.502341711 | 0.381 | 0.006 | 5.1773700 | 38 | dnm3a |
| scara33 | 2.070569 | 1.201756979 | 0.881 | 0.033 | 5.2602821 | 38 | scara3 |
| megf6a | 3.309045 | 0.54800434 | 0.405 | 0.007 | 8.4066293 | 38 | megf6a |
| rcn31 | 4.561958 | 0.554309587 | 0.429 | 0.008 | 1.1589656 | 38 | rcn3 |
| si:dkey-12l12.13 | 1.583438 | 0.879844746 | 0.667 | 0.021 | 4.0227254 | 38 | si:dkey-12l12.1 |
| kdelr32 | 1.540316 | 0.816735049 | 0.595 | 0.017 | 3.9131742 | 38 | kdelr3 |
| copz21 | 1.249392 | 0.660235292 | 0.476 | 0.011 | 3.1740812 | 38 | copz2 |
| col15a1a3 | 1.526197 | 1.047167866 | 0.69 | 0.024 | 3.8773039 | 38 | col15a1a |
| pigr2 | 6.090836 | 0.838911067 | 0.667 | 0.022 | 1.5473771 | 38 | pigr |

| | | | | | | | |
|---|---|---|---|---|---|---|---|
| bhmt4 | 4.076375 | 1.438711785 | 0.881 | 0.041 | 1.0356031 | 38 | bhmt |
| gtdc1 | 3.653394 | 0.529350749 | 0.429 | 0.009 | 9.2814488 | 38 | gtdc1 |
| nradd2 | 5.039815 | 0.507075757 | 0.405 | 0.008 | 1.2803650 | 38 | nradd |
| casz13 | 1.644048 | 0.804432185 | 0.595 | 0.019 | 4.1767051 | 38 | casz1 |
| abi3bpa1 | 5.555421 | 0.598034226 | 0.476 | 0.012 | 1.4113548 | 38 | abi3bpa |
| cd248b1 | 1.226485 | 0.459541629 | 0.286 | 0.004 | 3.1158860 | 38 | cd248b |
| CMKLR1 (1 of many) | 5.397672 | 0.49843659 | 0.357 | 0.007 | 1.3712786 | 38 | CMKLR1 (1 of many) |
| pdgfrl4 | 7.350894 | 1.122391198 | 0.738 | 0.032 | 1.8674948 | 38 | pdgfrl |
| hpdb1 | 2.557939 | 0.672979024 | 0.476 | 0.013 | 6.4984461 | 38 | hpdb |
| zgc:1133071 | 4.873470 | 0.272064215 | 0.167 | 0.001 | 1.2381051 | 38 | zgc:113307 |
| asip13 | 5.989676 | 0.781980209 | 0.571 | 0.019 | 1.5216773 | 38 | asip1 |
| srpx | 9.074887 | 1.024619536 | 0.69 | 0.028 | 2.3054752 | 38 | srpx |
| clu4 | 2.805112 | 1.130777237 | 0.714 | 0.031 | 7.1263876 | 38 | clu |
| c1qtnf22 | 1.100597 | 0.826994315 | 0.643 | 0.025 | 2.7960691 | 38 | c1qtnf2 |
| timp2a7 | 4.240446 | 1.431304878 | 0.952 | 0.064 | 1.0772854 | 38 | timp2a |
| FQ378018.1 | 1.199108 | 0.429125418 | 0.262 | 0.004 | 3.0463349 | 38 | FQ378018.1 |
| hhipl22 | 1.098405 | 0.332655883 | 0.238 | 0.003 | 2.7904979 | 38 | hhipl2 |
| rgma1 | 1.235036 | 0.508744742 | 0.405 | 0.011 | 3.1376089 | 38 | rgma |
| fkbp77 | 1.167803 | 1.138460889 | 0.833 | 0.049 | 2.9668035 | 38 | fkbp7 |
| TENT5A | 1.756038 | 0.462448708 | 0.31 | 0.006 | 4.4612154 | 38 | TENT5A |
| si:ch211-106h4.126 | 1.477911 | 1.605818569 | 0.881 | 0.057 | 3.7546334 | 38 | si:ch211-106h4.12 |
| nupr1b4 | 3.155521 | 1.131777685 | 0.786 | 0.044 | 8.0166017 | 38 | nupr1b |
| krt917 | 5.395089 | 1.719220473 | 0.976 | 0.071 | 1.3706225 | 38 | krt91 |
| cxcl8a1 | 1.938733 | 0.732712366 | 0.429 | 0.013 | 4.9253527 | 38 | cxcl8a |
| prelp1 | 2.929985 | 0.64102624 | 0.357 | 0.009 | 7.4436269 | 38 | prelp |
| irx1a1 | 4.826920 | 0.455032508 | 0.262 | 0.005 | 1.2262790 | 38 | irx1a |
| ccdc803 | 2.637257 | 0.878320473 | 0.667 | 0.032 | 6.6999520 | 38 | ccdc80 |
| twist1a3 | 7.477660 | 0.738311406 | 0.5 | 0.018 | 1.8996997 | 38 | twist1a |
| fkbp143 | 4.153829 | 1.045733773 | 0.738 | 0.041 | 1.0552803 | 38 | fkbp14 |
| fkbp94 | 1.817540 | 0.88556164 | 0.69 | 0.035 | 4.6174606 | 38 | fkbp9 |
| rgs5a3 | 2.012720 | 0.943482416 | 0.714 | 0.037 | 5.1133168 | 38 | rgs5a |
| ecrg4a5 | 1.170358 | 1.137660409 | 0.69 | 0.036 | 2.9732957 | 38 | ecrg4a |
| nkd2a | 1.697606 | 0.463233896 | 0.381 | 0.011 | 4.3127700 | 38 | nkd2a |
| atoh82 | 1.558964 | 0.704145375 | 0.571 | 0.025 | 3.9605500 | 38 | atoh8 |
| cygb16 | 6.195456 | 1.381197621 | 0.857 | 0.06 | 1.5739558 | 38 | cygb1 |

| | | | | | | | |
|---|---|---|---|---|---|---|---|
| XLOC-041451 | 7.425207 | 0.263824331 | 0.167 | 0.002 | 1.8863738 | 38 | XLOC-041451 |
| ccdc3b4 | 1.570025 | 0.818451753 | 0.619 | 0.03 | 3.9886491 | 38 | ccdc3b |
| vkorc12 | 4.470107 | 0.576096117 | 0.452 | 0.016 | 1.1356309 | 38 | vkorc1 |
| p4ha22 | 2.250371 | 0.601758665 | 0.429 | 0.015 | 5.7170679 | 38 | p4ha2 |
| col5a2a7 | 4.275357 | 1.882712041 | 1 | 0.099 | 1.0861545 | 38 | col5a2a |
| serpinh1b3 | 7.498121 | 1.147556006 | 0.595 | 0.03 | 1.9048978 | 38 | serpinh1b |
| reck3 | 1.067176 | 0.784000836 | 0.595 | 0.029 | 2.7111611 | 38 | reck |
| si:ch211-51c14.11 | 2.317298 | 0.34605493 | 0.286 | 0.006 | 5.8870955 | 38 | si:ch211-51c14.1 |
| ckap48 | 6.689402 | 1.006482113 | 0.833 | 0.059 | 1.6994426 | 38 | ckap4 |
| col21a13 | 5.610637 | 0.723891764 | 0.476 | 0.02 | 1.4253823 | 38 | col21a1 |
| si:ch211-133n4.45 | 3.200605 | 1.187600857 | 0.786 | 0.058 | 8.1311395 | 38 | si:ch211-133n4.4 |
| cd82a3 | 1.688875 | 1.194606802 | 0.857 | 0.068 | 4.2905893 | 38 | cd82a |
| dpt2 | 2.951298 | 0.70556233 | 0.429 | 0.016 | 7.4977728 | 38 | dpt |
| ctgfb1 | 1.414687 | 0.518399486 | 0.333 | 0.01 | 3.5940132 | 38 | ctgfb |
| postnb6 | 3.397761 | 1.726365913 | 0.976 | 0.094 | 8.6320134 | 38 | postnb |
| hkdc11 | 3.748842 | 0.285762821 | 0.214 | 0.004 | 9.5239355 | 38 | hkdc1 |
| pabpc1b6 | 6.806449 | 1.259592663 | 0.881 | 0.076 | 1.7291785 | 38 | pabpc1b |
| klf12a | 3.406024 | 0.55454095 | 0.452 | 0.019 | 8.6530051 | 38 | klf12a |
| ehd2b3 | 6.851350 | 0.767617199 | 0.595 | 0.033 | 1.7405855 | 38 | ehd2b |
| pvalb6 | 7.462351 | 0.289219553 | 0.143 | 0.002 | 1.8958103 | 38 | pvalb6 |
| glis2a3 | 4.528429 | 0.652057732 | 0.524 | 0.027 | 1.1504474 | 38 | glis2a |
| olfml3b7 | 1.649661 | 0.936157008 | 0.738 | 0.055 | 4.1909651 | 38 | olfml3b |
| cercam1 | 4.710936 | 0.576621695 | 0.429 | 0.019 | 1.1968134 | 38 | cercam |
| ecm21 | 1.284091 | 0.430580021 | 0.31 | 0.01 | 3.2622357 | 38 | ecm2 |
| nucb2a3 | 3.264201 | 0.883428237 | 0.762 | 0.06 | 8.2927047 | 38 | nucb2a |
| col5a17 | 7.810226 | 1.808716227 | 1 | 0.131 | 1.9841881 | 38 | col5a1 |
| tns2b | 8.405264 | 0.440738856 | 0.333 | 0.011 | 2.1353575 | 38 | tns2b |
| colec125 | 8.694528 | 0.923650028 | 0.786 | 0.062 | 2.2088450 | 38 | colec12 |
| aebp14 | 1.880704 | 0.977654881 | 0.667 | 0.048 | 4.7779293 | 38 | aebp1 |
| ms4a17a.115 | 1.562486 | 0.872383166 | 0.69 | 0.051 | 3.9694976 | 38 | ms4a17a.11 |
| nfixb6 | 6.546577 | 1.068579768 | 0.881 | 0.081 | 1.6631581 | 38 | nfixb |
| fkbp10b2 | 6.850810 | 0.568906955 | 0.5 | 0.027 | 1.7404483 | 38 | fkbp10b |
| traf4a | 3.455319 | 0.35952233 | 0.286 | 0.009 | 8.7782396 | 38 | traf4a |
| fam198a2 | 2.083356 | 0.52477343 | 0.429 | 0.02 | 5.2927661 | 38 | fam198a |
| CABZ01039304.11 | 5.358525 | 0.479561396 | 0.381 | 0.017 | 1.3613335 | 38 | CABZ01039304.1 |

| | | | | | | | |
|---|---|---|---|---|---|---|---|
| calua3 | 6.152246 | 0.641188592 | 0.524 | 0.032 | 1.5629781 | 38 | calua |
| wnt11r3 | 2.465134 | 0.419296614 | 0.286 | 0.01 | 6.2626744 | 38 | wnt11r |
| col6a36 | 7.400943 | 1.29381174 | 0.976 | 0.113 | 1.8802097 | 38 | col6a3 |
| dcn6 | 8.978498 | 1.666751254 | 1 | 0.143 | 2.2809875 | 38 | dcn |
| zgc:565852 | 2.013580 | 0.427620939 | 0.31 | 0.012 | 5.1155020 | 38 | zgc:56585 |
| epha4b1 | 3.920522 | 0.496386385 | 0.31 | 0.012 | 9.9600881 | 38 | epha4b |
| lamb1b2 | 4.576755 | 0.616756675 | 0.452 | 0.025 | 1.1627247 | 38 | lamb1b |
| col1a1a7 | 5.942991 | 2.262891599 | 1 | 0.166 | 1.5098170 | 38 | col1a1a |
| si:ch1073-459j12.16 | 6.514843 | 1.162477038 | 0.833 | 0.084 | 1.6550960 | 38 | si:ch1073-459j12.1 |
| col1a1b7 | 4.439687 | 2.289162756 | 1 | 0.169 | 1.1279027 | 38 | col1a1b |
| plpp1a4 | 5.920055 | 0.981750598 | 0.81 | 0.077 | 1.5039900 | 38 | plpp1a |
| sdc24 | 7.243278 | 0.930913886 | 0.833 | 0.081 | 1.8401548 | 38 | sdc2 |
| pdgfra4 | 3.510489 | 0.877292604 | 0.714 | 0.061 | 8.9183980 | 38 | pdgfra |
| sgcd1 | 5.863361 | 0.36880783 | 0.262 | 0.009 | 1.4895869 | 38 | sgcd |
| p3h41 | 9.549230 | 0.418157387 | 0.333 | 0.014 | 2.4259819 | 38 | p3h4 |
| tgfbi2 | 1.134823 | 0.797341092 | 0.5 | 0.032 | 2.8830200 | 38 | tgfbi |
| apbb2b | 6.696663 | 0.518560742 | 0.381 | 0.019 | 1.7012873 | 38 | apbb2b |
| crhbp2 | 2.485266 | 0.693121623 | 0.381 | 0.019 | 6.3138207 | 38 | crhbp |
| mmp28 | 8.204981 | 1.175172299 | 0.929 | 0.111 | 2.0844756 | 38 | mmp2 |
| col1a27 | 1.645026 | 2.312814842 | 1 | 0.182 | 4.1791909 | 38 | col1a2 |
| rcn11 | 1.852242 | 0.547925218 | 0.405 | 0.021 | 4.7056221 | 38 | rcn1 |
| XLOC-0418322 | 2.061025 | 0.37286399 | 0.262 | 0.009 | 5.2360359 | 38 | XLOC-041832 |
| htra1a3 | 2.563594 | 0.630559981 | 0.429 | 0.024 | 6.5128122 | 38 | htra1a |
| fam69c1 | 1.017686 | 0.463507666 | 0.357 | 0.017 | 2.5854334 | 38 | fam69c |
| CABZ01092746.15 | 1.323638 | 1.154349195 | 0.833 | 0.088 | 3.3627040 | 38 | CABZ01092746.1 |
| atf5b3 | 1.467947 | 0.610644723 | 0.548 | 0.038 | 3.7293195 | 38 | atf5b |
| olfml2ba | 6.785214 | 0.314581327 | 0.214 | 0.006 | 1.7237838 | 38 | olfml2ba |
| sparc10 | 2.173873 | 2.181259624 | 1 | 0.195 | 5.5227252 | 38 | sparc |
| irx3a | 5.645618 | 0.330696072 | 0.238 | 0.008 | 1.4342694 | 38 | irx3a |
| glis2b | 2.421575 | 0.344070302 | 0.31 | 0.014 | 6.1520127 | 38 | glis2b |
| BX957346.1 | 2.876650 | 0.276700572 | 0.238 | 0.008 | 7.3081301 | 38 | BX957346.1 |
| tcf44 | 3.352019 | 0.483765732 | 0.405 | 0.024 | 8.5158065 | 38 | tcf4 |
| tmeff1b5 | 3.574850 | 0.578518009 | 0.548 | 0.042 | 9.0819088 | 38 | tmeff1b |
| crtap5 | 9.168168 | 0.676997499 | 0.524 | 0.04 | 2.3291732 | 38 | crtap |
| antxr2a8 | 3.147632 | 1.43890477 | 0.976 | 0.169 | 7.9965594 | 38 | antxr2a |

| | | | | | | | |
|---|---|---|---|---|---|---|---|
| fxyd17 | 3.288418 | 0.942563666 | 0.762 | 0.083 | 8.3542260 | 38 | fxyd1 |
| smoc22 | 7.159678 | 0.438916732 | 0.286 | 0.012 | 1.8189163 | 38 | smoc2 |
| si:ch73-138n13.14 | 1.262873 | 0.610873038 | 0.548 | 0.044 | 3.2083312 | 38 | si:ch73-138n13.1 |
| selenop2 | 1.050295 | 0.511184956 | 0.405 | 0.025 | 2.6682752 | 38 | selenop |
| angptl47 | 9.807717 | 0.872378242 | 0.595 | 0.055 | 2.4916506 | 38 | angptl4 |
| bcl11ba2 | 2.552820 | 0.384256249 | 0.31 | 0.015 | 6.4854414 | 38 | bcl11ba |
| ier5l2 | 3.996427 | 0.574711942 | 0.405 | 0.026 | 1.0152924 | 38 | ier5l |
| klf7a | 4.528009 | 0.321551671 | 0.238 | 0.009 | 1.1503409 | 38 | klf7a |
| slc7a10a | 6.007116 | 0.390408102 | 0.31 | 0.015 | 1.5261080 | 38 | slc7a10a |
| plpp33 | 7.913936 | 0.728810826 | 0.667 | 0.065 | 2.0105355 | 38 | plpp3 |
| zgc:194981 | 9.519878 | 0.786882033 | 0.714 | 0.076 | 2.4185251 | 38 | zgc:194981 |
| p3h3 | 1.837130 | 0.35152581 | 0.262 | 0.011 | 4.6672309 | 38 | p3h3 |
| zgc:1533176 | 2.079106 | 0.64832708 | 0.5 | 0.04 | 5.2819704 | 38 | zgc:153317 |
| FAT44 | 2.538385 | 0.517890747 | 0.357 | 0.021 | 6.4487691 | 38 | FAT4 |
| dag15 | 2.911614 | 0.709727335 | 0.595 | 0.055 | 7.3969554 | 38 | dag1 |
| pmp22a7 | 1.269171 | 1.355187047 | 1 | 0.166 | 3.2243306 | 38 | pmp22a |
| f3b5 | 5.309262 | 1.093610815 | 0.786 | 0.098 | 1.3488182 | 38 | f3b |
| cyr618 | 6.108251 | 1.026459728 | 0.643 | 0.067 | 1.5518013 | 38 | cyr61 |
| ephb37 | 7.279136 | 0.659031177 | 0.571 | 0.052 | 1.8492646 | 38 | ephb3 |
| rrbp1a3 | 6.859827 | 0.742678408 | 0.595 | 0.058 | 1.7427392 | 38 | rrbp1a |
| mfap27 | 7.647911 | 1.029112905 | 0.881 | 0.119 | 1.9429520 | 38 | mfap2 |
| nt5e | 1.327384 | 0.418491313 | 0.286 | 0.014 | 3.3722200 | 38 | nt5e |
| nfia8 | 6.676788 | 0.876165476 | 0.786 | 0.098 | 1.6962381 | 38 | nfia |
| fndc3ba1 | 2.834355 | 0.41339081 | 0.31 | 0.017 | 7.2006800 | 38 | fndc3ba |
| appb6 | 3.637533 | 0.586385273 | 0.524 | 0.046 | 9.2411545 | 38 | appb |
| col11a1b3 | 8.822343 | 0.455283105 | 0.31 | 0.017 | 2.2413163 | 38 | col11a1b |
| p3h12 | 1.279798 | 0.367968051 | 0.286 | 0.015 | 3.2513289 | 38 | p3h1 |
| wipi1 | 4.096514 | 0.338700444 | 0.262 | 0.013 | 1.0407194 | 38 | wipi1 |
| smoc1 | 1.365881 | 0.272159193 | 0.214 | 0.009 | 3.4700217 | 38 | smoc1 |
| c1qtnf6a3 | 1.643071 | 0.483400833 | 0.381 | 0.027 | 4.1742222 | 38 | c1qtnf6a |
| fosl1a8 | 2.145500 | 0.827011503 | 0.69 | 0.083 | 5.4506427 | 38 | fosl1a |
| si:ch73-46j18.511 | 3.432822 | 1.333655111 | 1 | 0.208 | 8.7210857 | 38 | si:ch73-46j18.5 |
| klf6a11 | 5.263573 | 1.253722961 | 0.952 | 0.18 | 1.3372109 | 38 | klf6a |
| arhgap36 | 2.538174 | 0.253248618 | 0.214 | 0.009 | 6.4482335 | 38 | arhgap36 |
| p4ha1b1 | 2.685295 | 0.700279237 | 0.571 | 0.059 | 6.8219932 | 38 | p4ha1b |

| | | | | | | | |
|---|---|---|---|---|---|---|---|
| zgc:1536751 | 3.956733 | 0.654149875 | 0.619 | 0.067 | 1.0052080 | 38 | zgc:153675 |
| yaf2 | 8.948234 | 0.589639943 | 0.5 | 0.047 | 2.2732990 | 38 | yaf2 |
| tuba1b2 | 1.031254 | 0.536469598 | 0.452 | 0.039 | 2.6199022 | 38 | tuba1b |
| ca8 | 1.649350 | 0.320076537 | 0.262 | 0.014 | 4.1901753 | 38 | ca8 |
| zgc:1101581 | 2.003885 | 0.506358776 | 0.429 | 0.035 | 5.0908704 | 38 | zgc:110158 |
| fam102ab3 | 2.060836 | 0.473506656 | 0.429 | 0.035 | 5.2355549 | 38 | fam102ab |
| LOC1018823342 | 3.229479 | 0.420540003 | 0.286 | 0.016 | 8.2044922 | 38 | LOC101882334 |
| CU929237.14 | 3.458857 | 0.524685071 | 0.333 | 0.022 | 8.7872281 | 38 | CU929237.1 |
| fosb11 | 1.707898 | 1.0407544 | 0.81 | 0.121 | 4.3389165 | 38 | fosb |
| ehd1b3 | 3.937800 | 0.581142805 | 0.524 | 0.052 | 1.0003981 | 38 | ehd1b |
| dnase1l13 | 3.989201 | 0.559174793 | 0.524 | 0.053 | 1.0134565 | 38 | dnase1l1 |
| sertad42 | 1.579336 | 0.35020031 | 0.286 | 0.017 | 4.0123051 | 38 | sertad4 |
| col6a17 | 1.680618 | 0.928864797 | 0.714 | 0.097 | 4.2696109 | 38 | col6a1 |
| XLOC-026514 | 3.702274 | 0.256357818 | 0.167 | 0.006 | 9.4056290 | 38 | XLOC-026514 |
| mrc24 | 3.977725 | 0.486331818 | 0.381 | 0.03 | 1.0105412 | 38 | mrc2 |
| sh3pxd2aa4 | 4.939717 | 0.506256023 | 0.405 | 0.034 | 1.2549353 | 38 | sh3pxd2aa |
| znf7041 | 5.815459 | 0.517832301 | 0.405 | 0.034 | 1.4774175 | 38 | znf704 |
| flrt2 | 6.746897 | 0.29208217 | 0.238 | 0.012 | 1.7140492 | 38 | flrt2 |
| mmp14b7 | 4.784253 | 0.722429642 | 0.571 | 0.066 | 1.2154396 | 38 | mmp14b |
| CR847566.15 | 5.925571 | 0.592407304 | 0.5 | 0.051 | 1.5053914 | 38 | CR847566.1 |
| si:ch211-284f22.34 | 6.029715 | 0.505882018 | 0.405 | 0.034 | 1.5318491 | 38 | si:ch211-284f22.3 |
| soul51 | 1.447612 | 0.306117269 | 0.262 | 0.015 | 3.6776598 | 38 | soul5 |
| sept103 | 7.857059 | 0.565724608 | 0.5 | 0.052 | 1.9960860 | 38 | sept10 |
| lrrc58b1 | 1.445192 | 0.378032709 | 0.31 | 0.021 | 3.6715105 | 38 | lrrc58b |
| cd28l4 | 2.987884 | 0.514238154 | 0.429 | 0.04 | 7.5907195 | 38 | cd28l |
| marcksl1a11 | 3.123395 | 1.297375527 | 1 | 0.261 | 7.9349871 | 38 | marcksl1a |
| prrx1a2 | 8.500738 | 0.315126305 | 0.238 | 0.013 | 2.1596127 | 38 | prrx1a |
| wfdc1 | 9.588795 | 0.292043058 | 0.214 | 0.011 | 2.4360335 | 38 | wfdc1 |
| egr111 | 1.088115 | 1.071458691 | 0.929 | 0.183 | 2.7643563 | 38 | egr1 |
| tcf128 | 2.224965 | 0.586268356 | 0.571 | 0.068 | 5.6525259 | 38 | tcf12 |
| LOC1018835072 | 2.380674 | 0.374033311 | 0.286 | 0.019 | 6.0481044 | 38 | LOC101883507 |
| dnmt3ab6 | 3.294249 | 0.690881383 | 0.667 | 0.092 | 8.3690418 | 38 | dnmt3ab |
| crip22 | 3.431352 | 0.404203938 | 0.262 | 0.016 | 8.7173506 | 38 | crip2 |
| jdp2b10 | 3.473295 | 1.369402637 | 0.81 | 0.164 | 8.8239059 | 38 | jdp2b |
| foxo1a2 | 4.059084 | 0.536572514 | 0.429 | 0.041 | 1.0312104 | 38 | foxo1a |

| | | | | | | | |
|---|---|---|---|---|---|---|---|
| gstcd | 4.971581 | 0.315767243 | 0.19 | 0.009 | 1.2630302 | 38 | gstcd |
| arhgef1b4 | 8.074892 | 0.776103946 | 0.595 | 0.079 | 2.0514264 | 38 | arhgef1b |
| ppm1la1 | 9.524179 | 0.475314496 | 0.405 | 0.037 | 2.4196177 | 38 | ppm1la |
| si:dkey-101k6.52 | 1.110794 | 0.331540032 | 0.286 | 0.019 | 2.8219739 | 38 | si:dkey-101k6.5 |
| ak42 | 1.410840 | 0.443385095 | 0.381 | 0.034 | 3.5842408 | 38 | ak4 |
| vat18 | 1.736515 | 0.894555191 | 0.857 | 0.147 | 4.4116163 | 38 | vat1 |
| rasgef1bb3 | 1.803581 | 0.420600287 | 0.381 | 0.033 | 4.5819992 | 38 | rasgef1bb |
| mych5 | 2.364003 | 0.763123468 | 0.643 | 0.09 | 6.0057508 | 38 | mych |
| rgcc6 | 2.991568 | 0.704665954 | 0.571 | 0.071 | 7.6000794 | 38 | rgcc |
| col2a1b2 | 3.391589 | 0.447921869 | 0.262 | 0.017 | 8.6163334 | 38 | col2a1b |
| cebpd11 | 5.195029 | 0.962040282 | 0.857 | 0.152 | 1.3197971 | 38 | cebpd |
| fam114a1 | 1.138914 | 0.267467465 | 0.214 | 0.011 | 2.8934114 | 38 | fam114a1 |
| atf39 | 4.118352 | 0.848997001 | 0.595 | 0.082 | 1.0462674 | 38 | atf3 |
| marveld11 | 2.120525 | 0.264589 | 0.214 | 0.012 | 5.3871942 | 38 | marveld1 |
| si:ch211-39i22.17 | 5.538393 | 0.745770037 | 0.619 | 0.088 | 1.4070288 | 38 | si:ch211-39i22.1 |
| serf213 | 8.484755 | 1.15205247 | 0.976 | 0.378 | 2.1555521 | 38 | serf2 |
| znf385a6 | 1.192503 | 0.542646974 | 0.476 | 0.054 | 3.0295555 | 38 | znf385a |
| fpr14 | 3.796466 | 0.41160781 | 0.286 | 0.022 | 9.6449227 | 38 | fpr1 |
| mycb7 | 4.523941 | 1.093702226 | 0.881 | 0.2 | 1.1493072 | 38 | mycb |
| calm3a | 7.282588 | 0.569866189 | 0.548 | 0.071 | 1.8501416 | 38 | calm3a |
| LOC1018860301 | 7.900815 | 0.302189081 | 0.238 | 0.015 | 2.0072022 | 38 | LOC101886030 |
| sdc414 | 8.419784 | 1.187867851 | 0.929 | 0.254 | 2.1390462 | 38 | sdc4 |
| fam110b | 9.152255 | 0.346074744 | 0.262 | 0.018 | 2.3251304 | 38 | fam110b |
| s100v212 | 1.036851 | 0.756301284 | 0.81 | 0.14 | 2.6341218 | 38 | s100v2 |
| loxl2b2 | 1.388474 | 0.465713339 | 0.429 | 0.046 | 3.5274183 | 38 | loxl2b |
| lrp1aa5 | 1.646739 | 0.563675125 | 0.524 | 0.066 | 4.1835406 | 38 | lrp1aa |
| sulf2b1 | 2.154811 | 0.43514799 | 0.405 | 0.042 | 5.4742987 | 38 | sulf2b |
| eef1da10 | 2.510180 | 1.022373624 | 1 | 0.507 | 6.3771136 | 38 | eef1da |
| ddit312 | 3.035762 | 0.965816654 | 0.857 | 0.178 | 7.7123552 | 38 | ddit3 |
| sec61a14 | 4.641059 | 0.934257919 | 0.881 | 0.192 | 1.1790610 | 38 | sec61a1 |
| tgfbr34 | 4.709164 | 0.46365863 | 0.429 | 0.046 | 1.1963631 | 38 | tgfbr3 |
| slc38a5b10 | 8.185491 | 0.756569755 | 0.738 | 0.129 | 2.0795241 | 38 | slc38a5b |
| lzts2a3 | 1.261247 | 0.318349759 | 0.286 | 0.023 | 3.2041992 | 38 | lzts2a |
| sema6e1 | 1.429360 | 0.299926851 | 0.19 | 0.01 | 3.6312901 | 38 | sema6e |
| sept7a2 | 1.891309 | 0.677767813 | 0.548 | 0.077 | 4.8048719 | 38 | sept7a |

| | | | | | | | |
|---|---|---|---|---|---|---|---|
| gas1a6 | 2.025765 | 0.540803824 | 0.476 | 0.058 | 5.1464580 | 38 | gas1a |
| ift20 | 3.748321 | 0.342820486 | 0.286 | 0.023 | 9.5226098 | 38 | ift20 |
| adh8a5 | 4.699090 | 0.553310826 | 0.429 | 0.048 | 1.1938038 | 38 | adh8a |
| vdac38 | 1.280824 | 0.831454712 | 0.833 | 0.168 | 3.2539349 | 38 | vdac3 |
| lysmd3 | 1.297313 | 0.401140601 | 0.381 | 0.04 | 3.2958240 | 38 | lysmd3 |
| st7 | 1.712194 | 0.410964377 | 0.333 | 0.032 | 4.3498306 | 38 | st7 |
| loxl14 | 1.903219 | 0.376964762 | 0.333 | 0.031 | 4.8351285 | 38 | loxl1 |
| cd2761 | 2.547434 | 0.390499545 | 0.31 | 0.028 | 6.4717574 | 38 | cd276 |
| fosab15 | 2.859419 | 1.290827386 | 0.976 | 0.362 | 7.2643549 | 38 | fosab |
| zgc:1583439 | 3.649304 | 0.916669033 | 0.833 | 0.169 | 9.2710585 | 38 | zgc:158343 |
| adrb2a2 | 3.744072 | 0.343487926 | 0.262 | 0.02 | 9.5118156 | 38 | adrb2a |
| mafba9 | 4.192670 | 0.620913174 | 0.571 | 0.084 | 1.0651480 | 38 | mafba |
| SCARF24 | 4.753472 | 0.391249263 | 0.31 | 0.028 | 1.2076197 | 38 | SCARF2 |
| tmem45a3 | 5.966219 | 0.356711569 | 0.31 | 0.028 | 1.5157180 | 38 | tmem45a |
| ip6k2b9 | 6.111473 | 0.759535372 | 0.714 | 0.129 | 1.5526198 | 38 | ip6k2b |
| si:dkey-56m19.55 | 1.103158 | 0.511551378 | 0.452 | 0.056 | 2.8025732 | 38 | si:dkey-56m19.5 |
| cyp2n135 | 1.488745 | 0.478455504 | 0.381 | 0.041 | 3.7821572 | 38 | cyp2n13 |
| zgc:1716791 | 1.792645 | 0.371289578 | 0.31 | 0.028 | 4.5542158 | 38 | zgc:171679 |
| emp25 | 2.757287 | 0.511578861 | 0.452 | 0.057 | 7.0048891 | 38 | emp2 |
| cxcl18b7 | 3.471076 | 1.103346411 | 0.619 | 0.11 | 8.8182696 | 38 | cxcl18b |
| ms4a17a.94 | 3.495986 | 0.593453715 | 0.5 | 0.069 | 8.8815548 | 38 | ms4a17a.9 |
| ier2a16 | 3.905155 | 1.143389089 | 0.929 | 0.274 | 9.9210480 | 38 | ier2a |
| fstl1b8 | 4.194288 | 0.793053976 | 0.786 | 0.147 | 1.0655590 | 38 | fstl1b |
| meis1a4 | 5.245898 | 0.316707446 | 0.262 | 0.021 | 1.3327203 | 38 | meis1a |
| hcar1-44 | 9.115846 | 0.421890172 | 0.333 | 0.033 | 2.3158807 | 38 | hcar1-4 |
| hdlbpa10 | 1.340151 | 0.795254396 | 0.833 | 0.175 | 3.4046541 | 38 | hdlbpa |
| reep512 | 2.286001 | 0.749971242 | 0.833 | 0.173 | 5.8075871 | 38 | reep5 |
| tspan75 | 3.125497 | 0.571536381 | 0.548 | 0.082 | 7.9403263 | 38 | tspan7 |
| rbfox13 | 3.685438 | 0.36814458 | 0.31 | 0.029 | 9.3628577 | 38 | rbfox1 |
| calhm21 | 4.258648 | 0.258825604 | 0.19 | 0.012 | 1.0819095 | 38 | calhm2 |
| ugp2b | 4.533607 | 0.513296258 | 0.5 | 0.07 | 1.1517628 | 38 | ugp2b |
| jun14 | 5.368622 | 1.075376375 | 0.952 | 0.25 | 1.3638985 | 38 | jun |
| ppib13 | 7.481733 | 1.019039934 | 1 | 0.379 | 1.9007343 | 38 | ppib |
| dusp58 | 8.268959 | 0.707470959 | 0.476 | 0.068 | 2.1007292 | 38 | dusp5 |
| ghra4 | 9.440284 | 0.387295238 | 0.333 | 0.034 | 2.3983041 | 38 | ghra |

| | | | | | | | |
|---|---|---|---|---|---|---|---|
| plod1a2 | 1.013914 | 0.346422529 | 0.286 | 0.026 | 2.5758500 | 38 | plod1a |
| dip2ca1 | 1.099699 | 0.273460466 | 0.238 | 0.018 | 2.7937853 | 38 | dip2ca |
| marcksl1b12 | 1.285780 | 0.657925649 | 0.762 | 0.141 | 3.2665259 | 38 | marcksl1b |
| si:dkey-100n23.5 | 1.816161 | 0.377302768 | 0.381 | 0.044 | 4.6139570 | 38 | si:dkey-100n23.5 |
| wbp1lb3 | 1.987020 | 0.475584014 | 0.405 | 0.05 | 5.0480254 | 38 | wbp1lb |
| kdelr2b1 | 2.041091 | 0.698590012 | 0.667 | 0.122 | 5.1853917 | 38 | kdelr2b |
| efnb2a3 | 9.474473 | 0.294241499 | 0.214 | 0.016 | 2.4069899 | 38 | efnb2a |
| tuba8l39 | 1.004673 | 0.506612892 | 0.5 | 0.072 | 2.5523717 | 38 | tuba8l3 |
| ecm1b7 | 1.240749 | 0.49794473 | 0.476 | 0.067 | 3.1521238 | 38 | ecm1b |
| ptk2aa1 | 1.270429 | 0.262695568 | 0.238 | 0.019 | 3.2275267 | 38 | ptk2aa |
| anxa410 | 1.502004 | 0.7155712 | 0.738 | 0.151 | 3.8158431 | 38 | anxa4 |
| pdk2b7 | 2.509911 | 0.687858363 | 0.595 | 0.105 | 6.3764306 | 38 | pdk2b |
| junba14 | 2.942816 | 1.132063279 | 0.929 | 0.32 | 7.4762243 | 38 | junba |
| map1lc3a11 | 3.218591 | 0.502574836 | 0.524 | 0.081 | 8.1768306 | 38 | map1lc3a |
| tnfsf123 | 3.373515 | 0.333588366 | 0.286 | 0.027 | 8.5704160 | 38 | tnfsf12 |
| krt813 | 4.507012 | 1.081295959 | 1 | 0.312 | 1.1450065 | 38 | krt8 |
| gpc44 | 5.088766 | 0.309978631 | 0.31 | 0.032 | 1.2928010 | 38 | gpc4 |
| ANKRD50 | 6.415938 | 0.323576776 | 0.262 | 0.024 | 1.6299690 | 38 | ANKRD50 |
| cbfa2t32 | 7.038355 | 0.3035627 | 0.262 | 0.024 | 1.7880942 | 38 | cbfa2t3 |
| tmem39a | 7.603385 | 0.365025994 | 0.357 | 0.042 | 1.9316399 | 38 | tmem39a |
| susd62 | 1.061136 | 0.434523444 | 0.429 | 0.058 | 2.6958163 | 38 | susd6 |
| pdgfrb5 | 1.138156 | 0.473588596 | 0.405 | 0.052 | 2.8914853 | 38 | pdgfrb |
| id111 | 1.420038 | 0.685737767 | 0.69 | 0.131 | 3.6076077 | 38 | id1 |
| tob1a6 | 1.438835 | 0.692253038 | 0.714 | 0.145 | 3.6553624 | 38 | tob1a |
| stau28 | 1.748126 | 0.552966227 | 0.571 | 0.096 | 4.4411151 | 38 | stau2 |
| zgc:16273017 | 2.046007 | 1.116741435 | 0.952 | 0.403 | 5.1978809 | 38 | zgc:162730 |
| adgre5b.32 | 3.325816 | 0.3934627 | 0.333 | 0.037 | 8.4492372 | 38 | adgre5b.3 |
| map7d1b | 3.507057 | 0.317332948 | 0.262 | 0.025 | 8.9096789 | 38 | map7d1b |
| zgc:1533722 | 6.229557 | 0.377292948 | 0.31 | 0.033 | 1.5826191 | 38 | zgc:153372 |
| foxo3b5 | 6.782648 | 0.460339121 | 0.476 | 0.071 | 1.7231318 | 38 | foxo3b |
| dusp63 | 1.064200 | 0.474687095 | 0.405 | 0.055 | 2.7036020 | 38 | dusp6 |
| rab2a4 | 1.095194 | 0.727407707 | 0.786 | 0.177 | 2.7823427 | 38 | rab2a |
| her64 | 1.699958 | 0.457138264 | 0.262 | 0.026 | 4.3187438 | 38 | her6 |
| si:ch211-286o17.15 | 2.431692 | 0.429764875 | 0.381 | 0.049 | 6.1777138 | 38 | si:ch211-286o17.1 |
| epdl13 | 2.786992 | 0.278319657 | 0.31 | 0.034 | 7.0803542 | 38 | epdl1 |

| | | | | | | | |
|---|---|---|---|---|---|---|---|
| triqk3 | 2.901201 | 0.41527894 | 0.405 | 0.056 | 7.3705018 | 38 | triqk |
| snd12 | 3.365127 | 0.56974276 | 0.619 | 0.118 | 8.5491066 | 38 | snd1 |
| zgc:922422 | 4.782280 | 0.276271809 | 0.238 | 0.021 | 1.2149382 | 38 | zgc:92242 |
| selenom2 | 5.092259 | 0.502504239 | 0.619 | 0.112 | 1.2936886 | 38 | selenom |
| laptm4a4 | 1.621830 | 0.60692917 | 0.619 | 0.122 | 4.1202609 | 38 | laptm4a |
| dhrs13l15 | 1.918130 | 0.342072095 | 0.333 | 0.04 | 4.8730092 | 38 | dhrs13l1 |
| socs3b9 | 3.171262 | 0.680171753 | 0.714 | 0.152 | 8.0565932 | 38 | socs3b |
| tmem594 | 3.638364 | 0.547871718 | 0.643 | 0.127 | 9.2432653 | 38 | tmem59 |
| XLOC-0354132 | 3.803956 | 0.401726218 | 0.31 | 0.037 | 9.6639520 | 38 | XLOC-035413 |
| arf53 | 4.166809 | 0.712173065 | 0.857 | 0.211 | 1.0585780 | 38 | arf5 |
| fxyd6l5 | 6.087109 | 0.545016364 | 0.619 | 0.118 | 1.5464301 | 38 | fxyd6l |
| zfp36l1b9 | 6.631413 | 0.708335079 | 0.667 | 0.14 | 1.6847105 | 38 | zfp36l1b |
| nfic6 | 6.907092 | 0.501020143 | 0.5 | 0.084 | 1.7547467 | 38 | nfic |
| ssr36 | 7.362738 | 0.844447066 | 0.976 | 0.288 | 1.8705036 | 38 | ssr3 |
| smco4 | 7.823281 | 0.304417774 | 0.262 | 0.027 | 1.9875046 | 38 | smco4 |
| prdx41 | 9.142438 | 0.661384555 | 0.738 | 0.168 | 2.3226365 | 38 | prdx4 |
| nova24 | 2.131614 | 0.30558023 | 0.31 | 0.036 | 5.4153672 | 38 | nova2 |
| nanos13 | 2.383330 | 0.365089846 | 0.286 | 0.032 | 6.0548511 | 38 | nanos1 |
| XLOC-0441911 | 2.716537 | 0.255067739 | 0.19 | 0.015 | 6.9013642 | 38 | XLOC-044191 |
| rassf1 | 2.949449 | 0.291917108 | 0.262 | 0.028 | 7.4930757 | 38 | rassf1 |
| fbp1a3 | 3.072762 | 0.374584683 | 0.333 | 0.043 | 7.8063525 | 38 | fbp1a |
| pgrmc18 | 4.221273 | 0.575250356 | 0.714 | 0.152 | 1.0724144 | 38 | pgrmc1 |
| eif4a1b10 | 4.626553 | 0.865994952 | 0.929 | 0.312 | 1.1753757 | 38 | eif4a1b |
| clstn13 | 5.820968 | 0.265855771 | 0.238 | 0.023 | 1.4788169 | 38 | clstn1 |
| spry45 | 6.110313 | 0.464921968 | 0.357 | 0.049 | 1.5523251 | 38 | spry4 |
| capns1b10 | 6.116013 | 0.537315781 | 0.69 | 0.146 | 1.5537731 | 38 | capns1b |
| col27a1b3 | 9.738551 | 0.345814926 | 0.262 | 0.028 | 2.4740790 | 38 | col27a1b |
| ssr47 | 1.120797 | 0.881431033 | 0.929 | 0.332 | 2.8473864 | 38 | ssr4 |
| eef1a1l13 | 1.205330 | 0.349883487 | 1 | 0.916 | 3.0621410 | 38 | eef1a1l1 |
| ablim1b4 | 1.416115 | 0.324279594 | 0.333 | 0.043 | 3.5976422 | 38 | ablim1b |
| cers2a | 1.712775 | 0.335667121 | 0.286 | 0.034 | 4.3513054 | 38 | cers2a |
| grb10b4 | 1.716157 | 0.503069951 | 0.31 | 0.04 | 4.3598982 | 38 | grb10b |
| atf4b14 | 1.753029 | 0.827231899 | 0.905 | 0.278 | 4.4535717 | 38 | atf4b |
| nme36 | 1.775912 | 0.486920519 | 0.5 | 0.089 | 4.5117051 | 38 | nme3 |
| gpx83 | 1.794889 | 0.456083708 | 0.452 | 0.073 | 4.5599168 | 38 | gpx8 |

| | | | | | | | |
|---|---|---|---|---|---|---|---|
| fstl36 | 4.020507 | 0.35816578 | 0.238 | 0.025 | 1.0214098 | 38 | fstl3 |
| yif1a | 8.752145 | 0.340574825 | 0.405 | 0.062 | 2.2234825 | 38 | yif1a |
| errfi1a2 | 1.188378 | 0.255525208 | 0.214 | 0.021 | 3.0190767 | 38 | errfi1a |
| selenow110 | 1.544132 | 0.846042196 | 0.976 | 0.416 | 3.9228686 | 38 | selenow1 |
| dap2 | 1.774872 | 0.401071805 | 0.5 | 0.09 | 4.5090639 | 38 | dap |
| cybrd11 | 1.984000 | 0.367261993 | 0.357 | 0.052 | 5.0403536 | 38 | cybrd1 |
| sh3gl1b2 | 2.388451 | 0.381452276 | 0.31 | 0.041 | 6.0678610 | 38 | sh3gl1b |
| myl9b10 | 3.178466 | 0.506069697 | 0.524 | 0.099 | 8.0748945 | 38 | myl9b |
| wls8 | 4.622491 | 0.514265424 | 0.619 | 0.131 | 1.1743438 | 38 | wls |
| rarga6 | 4.833346 | 0.454456459 | 0.381 | 0.061 | 1.2279116 | 38 | rarga |
| rimkla | 5.801174 | 0.293906579 | 0.31 | 0.041 | 1.4737884 | 38 | rimkla |
| serp14 | 8.490636 | 0.61174256 | 0.786 | 0.198 | 2.1570461 | 38 | serp1 |
| ier2b17 | 9.180425 | 0.799200969 | 0.81 | 0.237 | 2.3322870 | 38 | ier2b |
| gars1 | 1.121581 | 0.382436528 | 0.429 | 0.074 | 2.8493781 | 38 | gars |
| midn11 | 1.218038 | 0.650823425 | 0.619 | 0.145 | 3.0944272 | 38 | midn |
| sdr16c5b2 | 1.261629 | 0.313142425 | 0.333 | 0.048 | 3.2051703 | 38 | sdr16c5b |
| rbfox24 | 1.725522 | 0.318524137 | 0.31 | 0.042 | 4.3836899 | 38 | rbfox2 |
| rabac11 | 1.733802 | 0.321634963 | 0.333 | 0.048 | 4.4047254 | 38 | rabac1 |
| LO018493.13 | 1.744730 | 0.323109588 | 0.238 | 0.027 | 4.4324867 | 38 | LO018493.1 |
| mylipb | 1.809228 | 0.331977825 | 0.238 | 0.027 | 4.5963444 | 38 | mylipb |
| anxa137 | 1.814710 | 0.354147713 | 0.357 | 0.054 | 4.6102730 | 38 | anxa13 |
| zgc:1132084 | 2.274090 | 0.399026846 | 0.333 | 0.049 | 5.7773257 | 38 | zgc:113208 |
| tpd52l2b3 | 2.456151 | 0.40919076 | 0.548 | 0.11 | 6.2398531 | 38 | tpd52l2b |
| BX640522.24 | 2.633979 | 0.255597057 | 0.31 | 0.042 | 6.6916254 | 38 | BX640522.2 |
| pleca7 | 2.705384 | 0.436838024 | 0.381 | 0.063 | 6.8730304 | 38 | pleca |
| anxa2a10 | 2.735123 | 0.487393472 | 0.571 | 0.118 | 6.9485804 | 38 | anxa2a |
| gulp1a5 | 2.935337 | 0.306594499 | 0.286 | 0.037 | 7.4572256 | 38 | gulp1a |
| ctsk7 | 3.141298 | 0.381914619 | 0.429 | 0.074 | 7.9804689 | 38 | ctsk |
| CABZ01058650.1 | 3.638757 | 0.318487671 | 0.31 | 0.043 | 9.2442634 | 38 | CABZ01058650.1 |
| slc39a72 | 3.807790 | 0.323229176 | 0.31 | 0.044 | 9.6736924 | 38 | slc39a7 |
| tnfrsfa3 | 4.555944 | 0.257763349 | 0.262 | 0.032 | 1.1574377 | 38 | tnfrsfa |
| sgce4 | 5.059545 | 0.284513685 | 0.262 | 0.032 | 1.2853775 | 38 | sgce |
| slc3a2b7 | 7.188496 | 0.571319948 | 0.571 | 0.129 | 1.8262375 | 38 | slc3a2b |
| rrbp1b | 7.802455 | 0.29455948 | 0.31 | 0.044 | 1.9822139 | 38 | rrbp1b |
| si:dkey-238c7.163 | 8.062653 | 0.477473809 | 0.405 | 0.071 | 2.0483172 | 38 | si:dkey-238c7.16 |

| | | | | | | | |
|---|---|---|---|---|---|---|---|
| sept9a2 | 8.619983 | 0.309571194 | 0.333 | 0.049 | 2.1899068 | 38 | sept9a |
| klf11b6 | 9.406826 | 0.400608661 | 0.452 | 0.083 | 2.3898041 | 38 | klf11b |
| ccnd14 | 1.113371 | 0.293039821 | 0.31 | 0.044 | 2.8285203 | 38 | ccnd1 |
| cd82b4 | 1.466330 | 0.375798881 | 0.429 | 0.076 | 3.7252132 | 38 | cd82b |
| mgst3b7 | 2.317925 | 0.588885952 | 0.738 | 0.194 | 5.8886899 | 38 | mgst3b |
| spred2b9 | 2.485794 | 0.403598485 | 0.452 | 0.085 | 6.3151609 | 38 | spred2b |
| emd6 | 2.502369 | 0.368383405 | 0.405 | 0.071 | 6.3572692 | 38 | emd |
| jag1b5 | 2.890312 | 0.315832774 | 0.238 | 0.029 | 7.3428392 | 38 | jag1b |
| ATP11B | 3.010431 | 0.298859787 | 0.238 | 0.029 | 7.6480009 | 38 | ATP11B |
| ier58 | 3.078223 | 0.545673496 | 0.619 | 0.147 | 7.8202271 | 38 | ier5 |
| cdon5 | 3.121097 | 0.398382589 | 0.405 | 0.071 | 7.9291483 | 38 | cdon |
| alg5 | 3.288906 | 0.338524667 | 0.405 | 0.071 | 8.3554665 | 38 | alg5 |
| fgl2a6 | 4.617641 | 0.597076449 | 0.405 | 0.076 | 1.1731118 | 38 | fgl2a |
| klf11a1 | 5.174508 | 0.338022189 | 0.31 | 0.046 | 1.3145840 | 38 | klf11a |
| spry24 | 5.199969 | 0.404722619 | 0.405 | 0.073 | 1.3210521 | 38 | spry2 |
| si:ch211-140m22.72 | 6.803366 | 0.4224263 | 0.381 | 0.067 | 1.7283952 | 38 | si:ch211-140m22.7 |
| aldh9a1a.110 | 6.905055 | 0.627228268 | 0.714 | 0.193 | 1.7542294 | 38 | aldh9a1a.1 |
| socs3a13 | 7.079684 | 0.879766936 | 0.905 | 0.315 | 1.7985939 | 38 | socs3a |
| CABZ01075242.1 | 7.298387 | 0.262352787 | 0.19 | 0.02 | 1.8541552 | 38 | CABZ01075242.1 |
| hspg25 | 7.395901 | 0.273773577 | 0.238 | 0.029 | 1.8789287 | 38 | hspg2 |
| vasna4 | 8.951280 | 0.305342372 | 0.31 | 0.046 | 2.2740727 | 38 | vasna |
| skia1 | 9.582302 | 0.334623265 | 0.31 | 0.047 | 2.4343840 | 38 | skia |
| trib36 | 1.013425 | 0.546754196 | 0.5 | 0.109 | 2.5746065 | 38 | trib3 |
| sar1b5 | 1.084116 | 0.434054469 | 0.452 | 0.09 | 2.7541984 | 38 | sar1b |
| gadd45ga4 | 1.231608 | 0.379385716 | 0.333 | 0.054 | 3.1289022 | 38 | gadd45ga |
| mibp216 | 1.263720 | 0.728683387 | 0.976 | 0.522 | 3.2104820 | 38 | mibp2 |
| atox14 | 1.430114 | 0.470203097 | 0.548 | 0.124 | 3.6332051 | 38 | atox1 |
| f105 | 1.627319 | 0.391482048 | 0.357 | 0.06 | 4.1342039 | 38 | f10 |
| ctdspla3 | 1.784336 | 0.274905899 | 0.286 | 0.041 | 4.5331079 | 38 | ctdspla |
| calm1a5 | 1.841726 | 0.515175277 | 0.69 | 0.175 | 4.6789053 | 38 | calm1a |
| bnip3lb3 | 1.892980 | 0.354953573 | 0.429 | 0.081 | 4.8091157 | 38 | bnip3lb |
| fhl1b4 | 2.296283 | 0.250339534 | 0.19 | 0.02 | 5.8337092 | 38 | fhl1b |
| ostc3 | 3.280184 | 0.549760114 | 0.714 | 0.19 | 8.3333098 | 38 | ostc |
| arl6ip5b4 | 3.466259 | 0.513288317 | 0.476 | 0.104 | 8.8060310 | 38 | arl6ip5b |
| s100a10b14 | 3.598040 | 0.707029914 | 0.952 | 0.285 | 9.1408231 | 38 | s100a10b |

| | | | | | | | |
|---|---|---|---|---|---|---|---|
| ubtfl | 3.889141 | 0.250013428 | 0.238 | 0.03 | 9.8803639 | 38 | ubtfl |
| kdelr2a2 | 4.623378 | 0.416254884 | 0.405 | 0.077 | 1.1745693 | 38 | kdelr2a |
| boc6 | 4.637826 | 0.361611647 | 0.381 | 0.068 | 1.1782399 | 38 | boc |
| pno15 | 4.696813 | 0.513941252 | 0.667 | 0.172 | 1.1932254 | 38 | pno1 |
| sec61g8 | 7.325051 | 0.561250311 | 1 | 0.682 | 1.8609292 | 38 | sec61g |
| lhfpl2a1 | 7.608456 | 0.254413919 | 0.262 | 0.036 | 1.9329284 | 38 | lhfpl2a |
| zgc:1008292 | 7.729543 | 0.329842875 | 0.286 | 0.043 | 1.9636904 | 38 | zgc:100829 |
| gng24 | 8.486120 | 0.345020679 | 0.31 | 0.05 | 2.1558990 | 38 | gng2 |
| si:dkey-9i23.165 | 8.578969 | 0.376468531 | 0.31 | 0.05 | 2.1794873 | 38 | si:dkey-9i23.16 |
| rgs3a1 | 8.853416 | 0.281451163 | 0.214 | 0.026 | 2.2492104 | 38 | rgs3a |
| yipf5 | 1.155684 | 0.311435129 | 0.405 | 0.076 | 2.9360162 | 38 | yipf5 |
| rpl3910 | 1.236013 | 0.367528517 | 1 | 0.878 | 3.1400934 | 38 | rpl39 |
| h3f3c16 | 1.383410 | 0.705062826 | 0.905 | 0.304 | 3.5145540 | 38 | h3f3c |
| cox4i23 | 1.525280 | 0.271539269 | 0.262 | 0.037 | 3.8749743 | 38 | cox4i2 |
| si:dkey-7j14.68 | 1.598640 | 0.578995651 | 0.714 | 0.197 | 4.0613467 | 38 | si:dkey-7j14.6 |
| fam114a2 | 1.757800 | 0.254929163 | 0.262 | 0.037 | 4.4656912 | 38 | fam114a2 |
| nsa214 | 3.139778 | 0.741097834 | 0.976 | 0.389 | 7.9766060 | 38 | nsa2 |
| txndc51 | 3.696879 | 0.374304188 | 0.381 | 0.073 | 9.3919213 | 38 | txndc5 |
| ramp24 | 4.727325 | 0.289439243 | 0.286 | 0.044 | 1.2009770 | 38 | ramp2 |
| slc38a101 | 5.073527 | 0.275556031 | 0.238 | 0.033 | 1.2889296 | 38 | slc38a10 |
| btg213 | 5.219978 | 0.767526577 | 0.905 | 0.35 | 1.3261356 | 38 | btg2 |
| csrnp1b10 | 6.470260 | 0.552775852 | 0.619 | 0.165 | 1.6437696 | 38 | csrnp1b |
| arhgap312 | 7.058943 | 0.261412359 | 0.238 | 0.033 | 1.7933246 | 38 | arhgap31 |
| rps228 | 7.366830 | 0.425920221 | 1 | 0.842 | 1.8715433 | 38 | rps2 |
| nfkbib | 7.950751 | 0.271560561 | 0.286 | 0.045 | 2.0198883 | 38 | nfkbib |
| eef2b7 | 8.475611 | 0.564661673 | 0.976 | 0.637 | 2.1532291 | 38 | eef2b |
| eif4ebp3l6 | 8.844306 | 0.675577287 | 0.69 | 0.206 | 2.2468959 | 38 | eif4ebp3l |
| sec23a | 9.201174 | 0.269299309 | 0.19 | 0.023 | 2.3375583 | 38 | sec23a |
| hmbox1a2 | 9.912599 | 0.290024528 | 0.262 | 0.04 | 2.5182957 | 38 | hmbox1a |
| nupr1a3 | 1.083755 | 0.504723985 | 0.619 | 0.165 | 2.7532797 | 38 | nupr1a |
| unc50 | 1.231158 | 0.256303156 | 0.286 | 0.046 | 3.1277584 | 38 | unc50 |
| tead1b5 | 1.339369 | 0.294263441 | 0.238 | 0.034 | 3.4026693 | 38 | tead1b |
| ube2e37 | 1.372847 | 0.472762954 | 0.643 | 0.17 | 3.4877179 | 38 | ube2e3 |
| gstz11 | 1.442596 | 0.280579801 | 0.238 | 0.034 | 3.6649152 | 38 | gstz1 |
| tmem167a2 | 1.605764 | 0.398496309 | 0.476 | 0.109 | 4.0794454 | 38 | tmem167a |

| | | | | | | | |
|---|---|---|---|---|---|---|---|
| plecb6 | 2.625070 | 0.258415481 | 0.31 | 0.053 | 6.6689914 | 38 | plecb |
| bhlhe4012 | 2.670929 | 0.616439215 | 0.595 | 0.168 | 6.7854954 | 38 | bhlhe40 |
| ssr23 | 2.758160 | 0.649595219 | 0.952 | 0.333 | 7.0071073 | 38 | ssr2 |
| prelid3b5 | 3.646835 | 0.614467612 | 0.714 | 0.227 | 9.2647857 | 38 | prelid3b |
| rpl1010 | 4.873004 | 0.429604421 | 1 | 0.791 | 1.2379867 | 38 | rpl10 |
| serpine23 | 5.727669 | 0.304340452 | 0.333 | 0.062 | 1.4551145 | 38 | serpine2 |
| FO082781.18 | 7.571730 | 0.325704077 | 0.31 | 0.056 | 1.9235982 | 38 | FO082781.1 |
| sec621 | 7.832768 | 0.297307015 | 0.381 | 0.077 | 1.9899147 | 38 | sec62 |
| inka1a2 | 8.386211 | 0.256811539 | 0.214 | 0.03 | 2.1305171 | 38 | inka1a |
| gpatch83 | 8.609225 | 0.356179236 | 0.333 | 0.064 | 2.1871736 | 38 | gpatch8 |
| nid1b4 | 8.668537 | 0.337042115 | 0.357 | 0.069 | 2.2022420 | 38 | nid1b |
| ddx214 | 1.051941 | 0.549178267 | 0.69 | 0.21 | 2.6724571 | 38 | ddx21 |
| asph4 | 1.081662 | 0.358037132 | 0.262 | 0.044 | 2.7479624 | 38 | asph |
| tnrc6c112 | 1.164513 | 0.457456628 | 0.571 | 0.15 | 2.9584456 | 38 | tnrc6c1 |
| ifi306 | 1.257574 | 0.461105553 | 0.69 | 0.196 | 3.1948689 | 38 | ifi30 |
| hbegfa2 | 1.337644 | 0.396849955 | 0.452 | 0.103 | 3.3982847 | 38 | hbegfa |
| si:ch73-335l21.410 | 1.344334 | 0.665631978 | 0.524 | 0.142 | 3.4152828 | 38 | si:ch73-335l21.4 |
| tsc22d311 | 1.391397 | 0.452891593 | 0.595 | 0.16 | 3.5348460 | 38 | tsc22d3 |
| fzd25 | 1.784163 | 0.267384685 | 0.262 | 0.043 | 4.5326670 | 38 | fzd2 |
| anxa5b8 | 1.986756 | 0.389550554 | 0.452 | 0.105 | 5.0473544 | 38 | anxa5b |
| cx436 | 1.999736 | 0.408180887 | 0.31 | 0.058 | 5.0803313 | 38 | cx43 |
| ybx113 | 2.117445 | 0.50731688 | 1 | 0.696 | 5.3793696 | 38 | ybx1 |
| cald1a11 | 2.475506 | 0.370104351 | 0.5 | 0.12 | 6.2890233 | 38 | cald1a |
| cdca41 | 2.607884 | 0.305322553 | 0.333 | 0.065 | 6.6253313 | 38 | cdca4 |
| chd38 | 3.494079 | 0.414775511 | 0.5 | 0.126 | 8.8767092 | 38 | chd3 |
| mmp16a6 | 3.606439 | 0.271144695 | 0.381 | 0.079 | 9.1621591 | 38 | mmp16a |
| sec61b4 | 3.908623 | 0.67718362 | 0.952 | 0.47 | 9.9298584 | 38 | sec61b |
| ghrb4 | 4.081511 | 0.301119312 | 0.262 | 0.044 | 1.0369080 | 38 | ghrb |
| yap16 | 4.654180 | 0.264718586 | 0.31 | 0.058 | 1.1823945 | 38 | yap1 |
| mfap54 | 4.669805 | 0.26310215 | 0.333 | 0.064 | 1.1863640 | 38 | mfap5 |
| si:dkey-28b4.79 | 6.609333 | 0.392331419 | 0.452 | 0.11 | 1.6791011 | 38 | si:dkey-28b4.7 |
| rpl128 | 6.755369 | 0.364372445 | 0.976 | 0.813 | 1.7162015 | 38 | rpl12 |
| tpt17 | 1.056104 | 0.385654519 | 1 | 0.812 | 2.6830327 | 38 | tpt1 |
| zfyve212 | 1.086842 | 0.349890047 | 0.31 | 0.061 | 2.7611234 | 38 | zfyve21 |
| zfand31 | 1.133283 | 0.288519803 | 0.333 | 0.068 | 2.8791061 | 38 | zfand3 |

| | | | | | | | |
|---|---|---|---|---|---|---|---|
| LOC1081802691 | 1.138454 | 0.274878439 | 0.31 | 0.059 | 2.8922425 | 38 | LOC108180269 |
| map1lc3b6 | 1.263049 | 0.429818657 | 0.476 | 0.125 | 3.2087769 | 38 | map1lc3b |
| nr4a113 | 1.371593 | 0.465566627 | 0.571 | 0.159 | 3.4845337 | 38 | nr4a1 |
| selenof6 | 1.411998 | 0.554731816 | 0.786 | 0.267 | 3.5871826 | 38 | selenof |
| sec13 | 1.505297 | 0.281368209 | 0.31 | 0.061 | 3.8242076 | 38 | sec13 |
| tdh3 | 1.524134 | 0.288096722 | 0.405 | 0.092 | 3.8720646 | 38 | tdh |
| glo16 | 1.615604 | 0.442047422 | 0.69 | 0.21 | 4.1044424 | 38 | glo1 |
| serpine12 | 2.048635 | 0.376591384 | 0.238 | 0.04 | 5.2045590 | 38 | serpine1 |
| spcs15 | 2.057301 | 0.549564401 | 0.69 | 0.232 | 5.2265739 | 38 | spcs1 |
| vapb1 | 2.083879 | 0.329433858 | 0.381 | 0.086 | 5.2940971 | 38 | vapb |
| pik3r26 | 2.104687 | 0.254386778 | 0.333 | 0.068 | 5.3469578 | 38 | pik3r2 |
| LOC1039092848 | 2.227622 | 0.334873914 | 0.357 | 0.078 | 5.6592746 | 38 | LOC103909284 |
| rhoaa11 | 3.865557 | 0.456352471 | 0.595 | 0.177 | 9.8204488 | 38 | rhoaa |
| ciarta6 | 4.546182 | 0.288538947 | 0.476 | 0.118 | 1.1549577 | 38 | ciarta |
| pbx1a5 | 4.569056 | 0.251702328 | 0.286 | 0.055 | 1.1607688 | 38 | pbx1a |
| si:dkey-239h2.33 | 4.857761 | 0.396761474 | 0.405 | 0.1 | 1.2341144 | 38 | si:dkey-239h2.3 |
| ssr13 | 5.316341 | 0.451006262 | 0.643 | 0.201 | 1.3506166 | 38 | ssr1 |
| zgc:926065 | 5.377234 | 0.310004902 | 0.405 | 0.096 | 1.3660864 | 38 | zgc:92606 |
| ldlrad4a2 | 5.582724 | 0.26664668 | 0.214 | 0.035 | 1.4182910 | 38 | ldlrad4a |
| dcaf7 | 6.347551 | 0.293845735 | 0.452 | 0.113 | 1.6125955 | 38 | dcaf7 |
| sox4a8 | 7.727070 | 0.2682407 | 0.31 | 0.063 | 1.9630621 | 38 | sox4a |
| anxa11b5 | 7.899910 | 0.399712012 | 0.595 | 0.174 | 2.0069723 | 38 | anxa11b |
| tram13 | 8.362168 | 0.339029054 | 0.429 | 0.107 | 2.1244088 | 38 | tram1 |
| rps27l8 | 8.705777 | 0.428107933 | 0.69 | 0.218 | 2.2117027 | 38 | rps27l |
| crip115 | 1.136246 | 0.576859675 | 0.81 | 0.285 | 2.8866331 | 38 | crip1 |
| flot2a8 | 1.302452 | 0.321900102 | 0.333 | 0.074 | 3.3088804 | 38 | flot2a |
| eif2s36 | 1.377020 | 0.537674813 | 0.857 | 0.308 | 3.4983209 | 38 | eif2s3 |
| arpc1a4 | 1.704512 | 0.303436008 | 0.524 | 0.143 | 4.3303134 | 38 | arpc1a |
| pim34 | 2.043504 | 0.347573825 | 0.452 | 0.117 | 5.1915240 | 38 | pim3 |
| rnd3b5 | 2.303200 | 0.260375209 | 0.262 | 0.05 | 5.8512802 | 38 | rnd3b |
| pim112 | 2.581559 | 0.406894296 | 0.571 | 0.17 | 6.5584517 | 38 | pim1 |
| fbxo91 | 2.884027 | 0.263560365 | 0.262 | 0.051 | 7.3268723 | 38 | fbxo9 |
| slc25a331 | 2.894045 | 0.307572546 | 0.381 | 0.092 | 7.3523230 | 38 | slc25a33 |
| zgc:927446 | 3.017849 | 0.574317175 | 0.881 | 0.327 | 7.6668471 | 38 | zgc:92744 |
| srprb | 3.575642 | 0.358099826 | 0.548 | 0.162 | 9.0839195 | 38 | srprb |

| | | | | | | | |
|---|---|---|---|---|---|---|---|
| rps1518 | 3.624806 | 0.471708517 | 1 | 0.718 | 9.2088209 | 38 | rps15 |
| si:ch211-204c21.15 | 4.353718 | 0.274124502 | 0.333 | 0.075 | 0.000111 | 38 | si:ch211-204c21.1 |
| lgals2a6 | 5.720090 | 0.378465058 | 0.5 | 0.139 | 0.000145 | 38 | lgals2a |
| ndfip13 | 6.192908 | 0.352512322 | 0.571 | 0.172 | 0.000157 | 38 | ndfip1 |
| ipo72 | 7.097750 | 0.354420123 | 0.429 | 0.117 | 0.00018 | 38 | ipo7 |
| tob1b5 | 7.421512 | 0.427785229 | 0.548 | 0.169 | 0.000189 | 38 | tob1b |
| surf4l2 | 7.764826 | 0.299779431 | 0.262 | 0.054 | 0.000197 | 38 | surf4l |
| gstr3 | 8.125530 | 0.361100748 | 0.452 | 0.128 | 0.000206 | 38 | gstr |
| junbb6 | 8.611383 | 0.307493122 | 0.31 | 0.071 | 0.000219 | 38 | junbb |
| rps129 | 8.625413 | 0.347640871 | 1 | 0.842 | 0.000219 | 38 | rps12 |
| rap1b8 | 9.020495 | 0.378274524 | 0.69 | 0.221 | 0.000229 | 38 | rap1b |
| tmem2584 | 1.136580 | 0.495561282 | 0.81 | 0.306 | 0.000289 | 38 | tmem258 |
| cyb5r32 | 1.218420 | 0.26801934 | 0.238 | 0.047 | 0.00031 | 38 | cyb5r3 |
| mtdha4 | 1.394098 | 0.34488102 | 0.5 | 0.148 | 0.000354 | 38 | mtdha |
| rab1ab4 | 1.595388 | 0.340491229 | 0.476 | 0.139 | 0.000405 | 38 | rab1ab |
| b2ml14 | 2.108739 | 0.455659544 | 0.833 | 0.294 | 0.000536 | 38 | b2ml |
| maff4 | 2.302724 | 0.281956573 | 0.333 | 0.081 | 0.000585 | 38 | maff |
| mxra8a4 | 2.396719 | 0.256018157 | 0.286 | 0.063 | 0.000609 | 38 | mxra8a |
| rpl2216 | 2.429038 | 0.449793587 | 1 | 0.721 | 0.000617 | 38 | rpl22 |
| cnbpb8 | 2.803524 | 0.3696183 | 0.643 | 0.21 | 0.000712 | 38 | cnbpb |
| s100u3 | 2.833251 | 0.28179796 | 0.214 | 0.041 | 0.00072 | 38 | s100u |
| rps610 | 2.883196 | 0.376083868 | 1 | 0.804 | 0.000732 | 38 | rps6 |
| rpl1111 | 3.609251 | 0.363770671 | 1 | 0.83 | 0.000917 | 38 | rpl11 |
| add3a10 | 3.615799 | 0.323568839 | 0.476 | 0.139 | 0.000919 | 38 | add3a |
| rps3a9 | 3.908874 | 0.37449593 | 1 | 0.792 | 0.000993 | 38 | rps3a |
| ier3ip12 | 3.936842 | 0.343301923 | 0.524 | 0.164 | 0.001 | 38 | ier3ip1 |
| col6a26 | 4.795437 | 0.340134453 | 0.31 | 0.075 | 0.001218 | 38 | col6a2 |
| romo111 | 4.896260 | 0.560841875 | 0.929 | 0.409 | 0.001244 | 38 | romo1 |
| prdx512 | 5.254150 | 0.323411099 | 0.571 | 0.18 | 0.001335 | 38 | prdx5 |
| yipf41 | 5.343719 | 0.266210743 | 0.357 | 0.093 | 0.001358 | 38 | yipf4 |
| ndufa4l12 | 5.575462 | 0.558962566 | 0.952 | 0.423 | 0.001416 | 38 | ndufa4l |
| rps206 | 5.780576 | 0.269228906 | 1 | 0.911 | 0.001469 | 38 | rps20 |
| rpl18a14 | 6.171098 | 0.384043416 | 1 | 0.802 | 0.001568 | 38 | rpl18a |
| ap2m1a4 | 7.094560 | 0.297733561 | 0.548 | 0.172 | 0.001802 | 38 | ap2m1a |
| cd1643 | 7.131435 | 0.277528996 | 0.31 | 0.076 | 0.001812 | 38 | cd164 |

| | | | | | | | |
|---|---|---|---|---|---|---|---|
| rpl617 | 7.350603 | 0.454376902 | 1 | 0.702 | 0.001867 | 38 | rpl6 |
| nck1b | 8.298695 | 0.280192675 | 0.262 | 0.059 | 0.002108 | 38 | nck1b |
| eef1g15 | 8.834511 | 0.477421257 | 0.976 | 0.653 | 0.002244 | 38 | eef1g |
| rps1313 | 1.099603 | 0.414830991 | 1 | 0.75 | 0.002794 | 38 | rps13 |
| selenok5 | 1.274387 | 0.346684189 | 0.452 | 0.14 | 0.003238 | 38 | selenok |
| XLOC-0004319 | 1.580465 | 0.357750857 | 0.762 | 0.274 | 0.004015 | 38 | XLOC-000431 |
| syf2 | 1.768403 | 0.292355664 | 0.5 | 0.157 | 0.004493 | 38 | syf2 |
| rpl35a13 | 1.843241 | 0.383430946 | 1 | 0.783 | 0.004683 | 38 | rpl35a |
| rpl27a17 | 2.226440 | 0.566566521 | 1 | 0.523 | 0.005656 | 38 | rpl27a |
| p4hb2 | 2.308569 | 0.399334947 | 0.595 | 0.213 | 0.005865 | 38 | p4hb |
| si:ch1073-325m22.2! | 2.567888 | 0.365819607 | 0.69 | 0.25 | 0.006524 | 38 | si:ch1073-325m22.2 |
| chmp5b5 | 2.686797 | 0.282549937 | 0.429 | 0.129 | 0.006826 | 38 | chmp5b |
| ahnak12 | 2.995928 | 0.287296318 | 0.452 | 0.137 | 0.007611 | 38 | ahnak |
| zgc:11418823 | 3.096838 | 0.486976257 | 1 | 0.669 | 0.007868 | 38 | zgc:114188 |
| rhoab13 | 3.103991 | 0.371762851 | 0.595 | 0.208 | 0.007886 | 38 | rhoab |
| xbp19 | 3.304410 | 0.508243343 | 0.833 | 0.356 | 0.008395 | 38 | xbp1 |
| h2afx112 | 3.455945 | 0.376140209 | 0.643 | 0.228 | 0.00878 | 38 | h2afx1 |
| CR383676.122 | 4.078493 | 0.641599914 | 1 | 0.718 | 0.010361 | 38 | CR383676.1 |
| rpl2913 | 4.278686 | 0.340940889 | 1 | 0.823 | 0.01087 | 38 | rpl29 |
| ginm11 | 4.540862 | 0.275272167 | 0.357 | 0.101 | 0.011536 | 38 | ginm1 |
| h2afy27 | 4.625154 | 0.307391363 | 0.405 | 0.123 | 0.01175 | 38 | h2afy2 |
| rpl179 | 5.227776 | 0.380993311 | 1 | 0.764 | 0.013281 | 38 | rpl17 |
| cops92 | 5.300708 | 0.315857661 | 0.381 | 0.115 | 0.013466 | 38 | cops9 |
| ptprga9 | 5.330805 | 0.265065263 | 0.286 | 0.072 | 0.013543 | 38 | ptprga |
| rpl77 | 5.578170 | 0.325429798 | 0.976 | 0.81 | 0.014171 | 38 | rpl7 |
| irf2bp2a9 | 5.853731 | 0.337570108 | 0.548 | 0.186 | 0.014871 | 38 | irf2bp2a |
| faua15 | 5.915933 | 0.415326174 | 0.976 | 0.718 | 0.015029 | 38 | faua |
| rps1018 | 6.367388 | 0.421000487 | 1 | 0.728 | 0.016176 | 38 | rps10 |
| celf213 | 6.695839 | 0.360481204 | 0.714 | 0.258 | 0.017011 | 38 | celf2 |
| si:ch211-66k16.271 | 6.896918 | 0.251922504 | 0.333 | 0.093 | 0.017522 | 38 | si:ch211-66k16.27 |
| rpl3014 | 7.010663 | 0.416838693 | 1 | 0.731 | 0.017811 | 38 | rpl30 |
| cd81a10 | 7.386595 | 0.328069844 | 0.69 | 0.242 | 0.018766 | 38 | cd81a |
| gadd45ba12 | 8.581235 | 0.402435296 | 0.548 | 0.198 | 0.021801 | 38 | gadd45ba |
| aimp2 | 8.672688 | 0.292213029 | 0.429 | 0.137 | 0.022033 | 38 | aimp2 |
| rab5aa1 | 8.706407 | 0.268384286 | 0.31 | 0.084 | 0.022119 | 38 | rab5aa |

| | | | | | | | |
|---|---|---|---|---|---|---|---|
| ifrd1 | 8.747221 | 0.265651479 | 0.357 | 0.105 | 0.022222 | 38 | ifrd1 |
| rps27a10 | 8.944539 | 0.368048613 | 1 | 0.776 | 0.022724 | 38 | rps27a |
| zgc:1102396 | 9.244718 | 0.317269861 | 0.571 | 0.197 | 0.023486 | 38 | zgc:110239 |
| rpl3413 | 1.082804 | 0.357793272 | 1 | 0.803 | 0.027509 | 38 | rpl34 |
| gtpbp44 | 1.191603 | 0.358128172 | 0.667 | 0.247 | 0.030273 | 38 | gtpbp4 |
| gabarapb13 | 1.305548 | 0.412912016 | 0.762 | 0.306 | 0.033167 | 38 | gabarapb |
| rpl320 | 1.402787 | 0.268583408 | 1 | 0.873 | 0.035638 | 38 | rpl3 |
| gnb1a6 | 1.485375 | 0.325900378 | 0.571 | 0.203 | 0.037736 | 38 | gnb1a |
| rpl36a10 | 1.581347 | 0.326565394 | 1 | 0.824 | 0.040174 | 38 | rpl36a |
| rps2120 | 1.667113 | 0.394476608 | 1 | 0.777 | 0.042353 | 38 | rps21 |
| rps1619 | 1.853509 | 0.392747979 | 1 | 0.766 | 0.047088 | 38 | rps16 |
| eif3ba4 | 1.882460 | 0.348122379 | 0.643 | 0.242 | 0.047824 | 38 | eif3ba |
| histh1l10 | 1.932069 | 0.385308579 | 0.381 | 0.123 | 0.049084 | 38 | histh1l |
| rps27.217 | 2.087024 | 0.538752603 | 1 | 0.554 | 0.053021 | 38 | rps27.2 |
| rps1814 | 2.088128 | 0.39409543 | 1 | 0.755 | 0.053049 | 38 | rps18 |
| rpl922 | 2.162855 | 0.472636603 | 1 | 0.669 | 0.054947 | 38 | rpl9 |
| mcl1b14 | 2.165107 | 0.360962829 | 0.667 | 0.252 | 0.055005 | 38 | mcl1b |
| eef1b23 | 2.348211 | 0.369878649 | 0.976 | 0.742 | 0.059656 | 38 | eef1b2 |
| eif2s1b4 | 2.375319 | 0.288774821 | 0.548 | 0.196 | 0.060345 | 38 | eif2s1b |
| rpl5a17 | 2.384581 | 0.546723294 | 1 | 0.537 | 0.06058 | 38 | rpl5a |
| pfn211 | 2.479585 | 0.312046355 | 0.643 | 0.24 | 0.062994 | 38 | pfn2 |
| eif1b13 | 2.550754 | 0.468682374 | 0.929 | 0.407 | 0.064802 | 38 | eif1b |
| tmem176l.18 | 2.584787 | 0.314674835 | 0.571 | 0.203 | 0.065667 | 38 | tmem176l.1 |
| dbi8 | 2.741292 | 0.378684957 | 0.619 | 0.244 | 0.069643 | 38 | dbi |
| zgc:778492 | 3.196636 | 0.330197312 | 0.5 | 0.181 | 0.081211 | 38 | zgc:77849 |
| rpl3710 | 3.352042 | 0.315500371 | 1 | 0.856 | 0.085159 | 38 | rpl37 |
| ptp4a17 | 3.561917 | 0.289162631 | 0.405 | 0.133 | 0.090491 | 38 | ptp4a1 |
| itm2ba14 | 3.799057 | 0.361801068 | 0.762 | 0.298 | 0.096515 | 38 | itm2ba |
| rpl7a11 | 4.766824 | 0.323597442 | 1 | 0.824 | 0.121101 | 38 | rpl7a |
| adh58 | 5.084888 | 0.301439911 | 0.643 | 0.243 | 0.129182 | 38 | adh5 |
| vdac27 | 5.961884 | 0.358740311 | 0.81 | 0.327 | 0.151462 | 38 | vdac2 |
| rack19 | 6.682536 | 0.353281105 | 1 | 0.764 | 0.16977 | 38 | rack1 |
| rplp111 | 7.516348 | 0.287286548 | 1 | 0.883 | 0.190953 | 38 | rplp1 |
| rpsa16 | 8.842490 | 0.352757544 | 1 | 0.786 | 0.224643 | 38 | rpsa |
| rps2511 | 8.868462 | 0.368170155 | 1 | 0.75 | 0.225303 | 38 | rps25 |

| | | | | | | | |
|---|---|---|---|---|---|---|---|
| rpl2812 | 9.772654 | 0.356706095 | 1 | 0.778 | 0.248274 | 38 | rpl28 |
| sumo2b13 | 9.777708 | 0.366612183 | 0.714 | 0.297 | 0.248403 | 38 | sumo2b |
| cnn213 | 1.085869 | 0.256395866 | 0.452 | 0.158 | 0.275865 | 38 | cnn2 |
| nme2b.15 | 1.180059 | 0.258155568 | 1 | 0.857 | 0.299794 | 38 | nme2b.1 |
| rpl811 | 1.187956 | 0.348607351 | 1 | 0.796 | 0.3018 | 38 | rpl8 |
| ccl25b7 | 1.207013 | 0.2774657 | 0.31 | 0.095 | 0.306642 | 38 | ccl25b |
| rpl1814 | 1.237682 | 0.351574174 | 1 | 0.793 | 0.314433 | 38 | rpl18 |
| rps199 | 1.353947 | 0.280114207 | 1 | 0.869 | 0.34397 | 38 | rps19 |
| edf14 | 1.413878 | 0.304877982 | 0.714 | 0.288 | 0.359196 | 38 | edf1 |
| btg116 | 1.421354 | 0.386906422 | 0.833 | 0.356 | 0.361095 | 38 | btg1 |
| rpl1515 | 1.426585 | 0.372594878 | 1 | 0.765 | 0.362424 | 38 | rpl15 |
| rps710 | 1.427570 | 0.315213359 | 1 | 0.829 | 0.362674 | 38 | rps7 |
| rpl3214 | 1.480380 | 0.332566139 | 1 | 0.825 | 0.376091 | 38 | rpl32 |
| rpl3114 | 1.481102 | 0.418004084 | 1 | 0.688 | 0.376274 | 38 | rpl31 |
| mdh1aa4 | 1.483462 | 0.271926681 | 0.595 | 0.23 | 0.376874 | 38 | mdh1aa |
| rps4x17 | 1.508870 | 0.385807613 | 1 | 0.737 | 0.383329 | 38 | rps4x |
| rpl3510 | 1.684327 | 0.320616981 | 1 | 0.815 | 0.427903 | 38 | rpl35 |
| eif3c6 | 1.724874 | 0.280684383 | 0.5 | 0.189 | 0.438204 | 38 | eif3c |
| nop564 | 1.807208 | 0.281726352 | 0.5 | 0.187 | 0.459121 | 38 | nop56 |
| ctsla11 | 1.843071 | 0.39049795 | 0.881 | 0.374 | 0.468232 | 38 | ctsla |
| rpl23a15 | 1.932845 | 0.424551279 | 0.976 | 0.67 | 0.491039 | 38 | rpl23a |
| rplp05 | 1.984318 | 0.278573666 | 1 | 0.859 | 0.504116 | 38 | rplp0 |
| rplp2l15 | 2.089492 | 0.356071035 | 1 | 0.795 | 0.530836 | 38 | rplp2l |
| mknk2b9 | 2.481222 | 0.348816174 | 0.5 | 0.201 | 0.630354 | 38 | mknk2b |
| tomm54 | 2.540598 | 0.252046862 | 0.595 | 0.231 | 0.645439 | 38 | tomm5 |
| eif5a9 | 2.546648 | 0.493804016 | 0.976 | 0.496 | 0.646976 | 38 | eif5a |
| rpl2610 | 3.357047 | 0.288353853 | 1 | 0.852 | 0.852858 | 38 | rpl26 |
| rps118 | 3.434475 | 0.325943831 | 1 | 0.785 | 0.872529 | 38 | rps11 |
| rpl1310 | 3.471266 | 0.302951706 | 1 | 0.819 | 0.881875 | 38 | rpl13 |
| hmgn618 | 4.177469 | 0.354609784 | 0.738 | 0.312 | 1 | 38 | hmgn6 |
| rpl197 | 4.190407 | 0.282651836 | 1 | 0.854 | 1 | 38 | rpl19 |
| jpt1b8 | 4.774166 | 0.286600281 | 0.69 | 0.278 | 1 | 38 | jpt1b |
| ak24 | 5.455601 | 0.306944897 | 0.595 | 0.256 | 1 | 38 | ak2 |
| rpl10a12 | 5.608719 | 0.340814121 | 1 | 0.78 | 1 | 38 | rpl10a |
| rpl2713 | 6.091236 | 0.345297111 | 1 | 0.781 | 1 | 38 | rpl27 |

| | | | | | | | |
|---|---|---|---|---|---|---|---|
| txndc94 | 6.423901 | 0.25253543 | 0.357 | 0.13 | 1 | 38 | txndc9 |
| fabp37 | 6.614155 | 0.25402761 | 0.571 | 0.23 | 1 | 38 | fabp3 |
| srgn10 | 7.024090 | 0.255133231 | 0.595 | 0.236 | 1 | 38 | srgn |
| cdc429 | 8.240778 | 0.266426922 | 0.571 | 0.235 | 1 | 38 | cdc42 |
| rps2413 | 8.722812 | 0.338489791 | 1 | 0.78 | 1 | 38 | rps24 |
| si:ch211-196l7.416 | 0.0001 | 0.339526339 | 0.714 | 0.331 | 1 | 38 | si:ch211-196l7.4 |
| pdia310 | 0.0001 | 0.306528251 | 0.738 | 0.314 | 1 | 38 | pdia3 |
| ccng12 | 0.0001 | 0.363787291 | 0.976 | 0.444 | 1 | 38 | ccng1 |
| rps2310 | 0.0001 | 0.358359267 | 1 | 0.745 | 1 | 38 | rps23 |
| rps516 | 0.0001 | 0.378676192 | 1 | 0.729 | 1 | 38 | rps5 |
| cd6318 | 0.0001 | 0.337185637 | 0.857 | 0.374 | 1 | 38 | cd63 |
| rps15a15 | 0.0001 | 0.351791479 | 1 | 0.799 | 1 | 38 | rps15a |
| rpl1413 | 0.0001 | 0.400514347 | 1 | 0.67 | 1 | 38 | rpl14 |
| dap1b6 | 0.0002 | 0.313346131 | 0.571 | 0.25 | 1 | 38 | dap1b |
| si:dkey-11f4.149 | 0.0002 | 0.27214804 | 0.333 | 0.124 | 1 | 38 | si:dkey-11f4.14 |
| eif3i6 | 0.0002 | 0.288202568 | 0.667 | 0.292 | 1 | 38 | eif3i |
| rps27.119 | 0.0002 | 0.3963443 | 1 | 0.711 | 1 | 38 | rps27.1 |
| wu:fb55g095 | 0.0003 | 0.269490364 | 0.595 | 0.26 | 1 | 38 | wu:fb55g09 |
| rps8a9 | 0.0003 | 0.287199428 | 0.976 | 0.802 | 1 | 38 | rps8a |
| si:ch73-1a9.320 | 0.0004 | 0.282400692 | 0.69 | 0.317 | 1 | 38 | si:ch73-1a9.3 |
| serbp1a12 | 0.0004 | 0.454248271 | 0.976 | 0.507 | 1 | 38 | serbp1a |
| rps1412 | 0.0004 | 0.323308239 | 1 | 0.782 | 1 | 38 | rps14 |
| nfil3-69 | 0.0004 | 0.267893501 | 0.333 | 0.132 | 1 | 38 | nfil3-6 |
| higd1a10 | 0.0005 | 0.31618725 | 0.643 | 0.304 | 1 | 38 | higd1a |
| eif3ea6 | 0.0005 | 0.255779072 | 0.69 | 0.306 | 1 | 38 | eif3ea |
| rpl3615 | 0.0005 | 0.319359589 | 1 | 0.807 | 1 | 38 | rpl36 |
| rpl2113 | 0.0005 | 0.303134582 | 0.976 | 0.777 | 1 | 38 | rpl21 |
| eif3m8 | 0.0007 | 0.337172535 | 0.786 | 0.385 | 1 | 38 | eif3m |
| rps26l14 | 0.0007 | 0.387896829 | 1 | 0.701 | 1 | 38 | rps26l |
| rpl2412 | 0.0009 | 0.309369128 | 1 | 0.795 | 1 | 38 | rpl24 |
| rps313 | 0.001 | 0.353849628 | 1 | 0.727 | 1 | 38 | rps3 |
| uba5214 | 0.0012 | 0.394131988 | 1 | 0.662 | 1 | 38 | uba52 |
| eif3f14 | 0.0012 | 0.398971108 | 0.905 | 0.466 | 1 | 38 | eif3f |
| rps2912 | 0.0014 | 0.266257223 | 1 | 0.845 | 1 | 38 | rps29 |
| ddx55 | 0.0017 | 0.262319364 | 0.738 | 0.355 | 1 | 38 | ddx5 |

| | | | | | | | |
|---|---|---|---|---|---|---|---|
| ppiaa7 | 0.0018 | 0.267904951 | 1 | 0.815 | 1 | 38 | ppiaa |
| rsl24d112 | 0.002 | 0.384276031 | 0.952 | 0.461 | 1 | 38 | rsl24d1 |
| rplp217 | 0.0025 | 0.313715475 | 1 | 0.784 | 1 | 38 | rplp2 |
| rps910 | 0.0027 | 0.292735006 | 1 | 0.791 | 1 | 38 | rps9 |
| rps269 | 0.0033 | 0.314079469 | 1 | 0.752 | 1 | 38 | rps26 |
| ppdpfb7 | 0.0033 | 0.36522831 | 0.929 | 0.533 | 1 | 38 | ppdpfb |
| rps2812 | 0.0039 | 0.265129557 | 1 | 0.841 | 1 | 38 | rps28 |
| rpl3813 | 0.0048 | 0.321504988 | 1 | 0.755 | 1 | 38 | rpl38 |
| zgc:193726 | 0 | 2.666533795 | 0.9 | 0 | 0 | 39 | zgc:193726 |
| lgals2b | 0 | 2.398127003 | 0.95 | 0.001 | 0 | 39 | lgals2b |
| si:dkey-96g2.1 | 0 | 2.392942323 | 1 | 0 | 0 | 39 | si:dkey-96g2.1 |
| BX855618.1 | 0 | 2.323715902 | 0.9 | 0 | 0 | 39 | BX855618.1 |
| si:ch211-137i24.101 | 0 | 2.081031186 | 1 | 0.009 | 0 | 39 | si:ch211-137i24.10 |
| aqp8a.11 | 0 | 2.043392374 | 0.9 | 0.004 | 0 | 39 | aqp8a.1 |
| tm4sf4 | 0 | 1.874956729 | 0.9 | 0.001 | 0 | 39 | tm4sf4 |
| cldn15a | 0 | 1.829041709 | 0.85 | 0.001 | 0 | 39 | cldn15a |
| cftr | 0 | 1.817454739 | 0.85 | 0.001 | 0 | 39 | cftr |
| zgc:113142 | 0 | 1.808415538 | 0.9 | 0 | 0 | 39 | zgc:113142 |
| onecut1 | 0 | 1.44261968 | 0.8 | 0.007 | 0 | 39 | onecut1 |
| entpd8 | 0 | 1.420541817 | 0.8 | 0.006 | 0 | 39 | entpd8 |
| nr5a21 | 0 | 1.299197027 | 0.7 | 0.003 | 0 | 39 | nr5a2 |
| mgll | 0 | 1.278846625 | 0.75 | 0.006 | 0 | 39 | mgll |
| gprc5c | 0 | 1.182076141 | 0.65 | 0.003 | 0 | 39 | gprc5c |
| noxo1a | 0 | 1.175480428 | 0.55 | 0.001 | 0 | 39 | noxo1a |
| serpinf2b | 0 | 1.174666218 | 0.6 | 0 | 0 | 39 | serpinf2b |
| ptprjb.11 | 0 | 1.129101969 | 0.65 | 0.003 | 0 | 39 | ptprjb.1 |
| rassf7b1 | 0 | 1.12068799 | 0.65 | 0.004 | 0 | 39 | rassf7b |
| c5 | 0 | 1.045969112 | 0.65 | 0.001 | 0 | 39 | c5 |
| si:ch211-25g7.5 | 0 | 1.038789251 | 0.6 | 0.001 | 0 | 39 | si:ch211-25g7.5 |
| zgc:198329 | 0 | 1.014030085 | 0.6 | 0 | 0 | 39 | zgc:198329 |
| cdh17 | 0 | 0.991774886 | 0.5 | 0 | 0 | 39 | cdh17 |
| erbb3a1 | 0 | 0.964852457 | 0.55 | 0.002 | 0 | 39 | erbb3a |
| si:dkeyp-72e1.6 | 0 | 0.898116222 | 0.55 | 0.001 | 0 | 39 | si:dkeyp-72e1.6 |
| cldn15lb | 0 | 0.897288278 | 0.3 | 0 | 0 | 39 | cldn15lb |
| gpa33b | 0 | 0.808182482 | 0.45 | 0.002 | 0 | 39 | gpa33b |

| | | | | | | | |
|---|---|---|---|---|---|---|---|
| ppp1r9bb | 0 | 0.798753923 | 0.5 | 0.002 | 0 | 39 | ppp1r9bb |
| LOC1081808021 | 0 | 0.792334571 | 0.45 | 0.001 | 0 | 39 | LOC108180802 |
| foxa2 | 0 | 0.707588717 | 0.35 | 0 | 0 | 39 | foxa2 |
| st3gal7 | 0 | 0.700418088 | 0.35 | 0.001 | 0 | 39 | st3gal7 |
| tmem54b | 0 | 0.682585699 | 0.4 | 0.001 | 0 | 39 | tmem54b |
| gpr39 | 0 | 0.666013337 | 0.3 | 0 | 0 | 39 | gpr39 |
| mnx2a | 0 | 0.624522648 | 0.35 | 0 | 0 | 39 | mnx2a |
| cbln17 | 0 | 0.599501995 | 0.35 | 0 | 0 | 39 | cbln17 |
| tmem30c | 0 | 0.587387461 | 0.35 | 0.001 | 0 | 39 | tmem30c |
| pla1a | 0 | 0.561227817 | 0.35 | 0 | 0 | 39 | pla1a |
| si:dkeyp-82a1.4 | 0 | 0.548824213 | 0.25 | 0 | 0 | 39 | si:dkeyp-82a1.4 |
| si:dkey-208c12.2 | 0 | 0.538362613 | 0.3 | 0.001 | 0 | 39 | si:dkey-208c12.2 |
| slc5a9 | 0 | 0.535073646 | 0.3 | 0 | 0 | 39 | slc5a9 |
| CR855996.2 | 0 | 0.498322147 | 0.3 | 0.001 | 0 | 39 | CR855996.2 |
| clic5b | 3.72666[illegible] | 0.713113851 | 0.4 | 0.002 | 9.4676015 | 39 | clic5b |
| nkx2.2a | 2.79702[illegible] | 0.515643765 | 0.25 | 0.001 | 7.1058389 | 39 | nkx2.2a |
| itga6l | 3.30392[illegible] | 0.487100442 | 0.25 | 0.001 | 8.3936216 | 39 | itga6l |
| si:ch73-206d17.1 | 6.66727[illegible] | 0.355692615 | 0.2 | 0 | 1.6938218 | 39 | si:ch73-206d17.1 |
| sox9b | 1.98534[illegible] | 0.714541553 | 0.4 | 0.002 | 5.0437813 | 39 | sox9b |
| sb:cb1058 | 1.76877[illegible] | 0.80590095 | 0.4 | 0.002 | 4.4935791 | 39 | sb:cb1058 |
| nkx6.1 | 1.95849[illegible] | 0.451346592 | 0.25 | 0.001 | 4.9755577 | 39 | nkx6.1 |
| si:ch211-248a14.8 | 1.26063[illegible] | 0.281924159 | 0.2 | 0 | 3.2026489 | 39 | si:ch211-248a14.8 |
| pdzk1ip1 | 1.73693[illegible] | 0.857416017 | 0.4 | 0.002 | 4.4126881 | 39 | pdzk1ip1 |
| ccl19a.13 | 2.17940[illegible] | 1.781488435 | 0.9 | 0.014 | 5.5367746 | 39 | ccl19a.1 |
| ccl20b1 | 1.10384[illegible] | 1.047759604 | 0.5 | 0.004 | 2.8043279 | 39 | ccl20b |
| GUCY1A2 | 2.91708[illegible] | 0.536078175 | 0.25 | 0.001 | 7.4108498 | 39 | GUCY1A2 |
| tmem45b1 | 1.58997[illegible] | 0.639366129 | 0.35 | 0.002 | 4.0393347 | 39 | tmem45b |
| eps8l1a1 | 8.99518[illegible] | 0.69524116 | 0.4 | 0.003 | 2.2852261 | 39 | eps8l1a |
| slc3a1 | 3.66291[illegible] | 0.470281832 | 0.2 | 0.001 | 9.3056325 | 39 | slc3a1 |
| LOC101886880 | 7.53703[illegible] | 0.27197946 | 0.15 | 0 | 1.9147844 | 39 | LOC101886880 |
| si:dkey-234i14.6 | 9.74618[illegible] | 0.311093489 | 0.15 | 0 | 2.4760194 | 39 | si:dkey-234i14.6 |
| rassf10a1 | 4.57738[illegible] | 0.689780383 | 0.35 | 0.002 | 1.1628845 | 39 | rassf10a |
| foxa3 | 6.36362[illegible] | 0.396226989 | 0.25 | 0.001 | 1.6166786 | 39 | foxa3 |
| slc5a1 | 1.58221[illegible] | 0.490891595 | 0.25 | 0.001 | 4.0196276 | 39 | slc5a1 |
| cobll1a1 | 4.30170[illegible] | 0.929934895 | 0.5 | 0.006 | 1.0928476 | 39 | cobll1a |

| | | | | | | | |
|---|---|---|---|---|---|---|---|
| gdpd2 | 4.867108 | 0.714816813 | 0.4 | 0.004 | 1.2364890 | 39 | gdpd2 |
| si:dkey-45k15.1 | 2.804300 | 0.405429557 | 0.2 | 0.001 | 7.1243257 | 39 | si:dkey-45k15.1 |
| ehf1 | 5.045833 | 0.573605501 | 0.3 | 0.002 | 1.2818939 | 39 | ehf |
| CABZ01068469.11 | 1.134663 | 0.473072768 | 0.3 | 0.002 | 2.8826138 | 39 | CABZ01068469.1 |
| abcc2 | 1.397934 | 0.446341551 | 0.25 | 0.001 | 3.5514531 | 39 | abcc2 |
| shank3a | 1.684475 | 0.536311083 | 0.3 | 0.002 | 4.2794098 | 39 | shank3a |
| chn2 | 1.202006 | 0.540741856 | 0.3 | 0.002 | 3.0536972 | 39 | chn2 |
| CR396595.11 | 2.659768 | 0.656962942 | 0.3 | 0.002 | 6.7571408 | 39 | CR396595.1 |
| igf2a2 | 1.238834 | 0.772181521 | 0.5 | 0.008 | 3.1472597 | 39 | igf2a |
| stox2a1 | 1.712003 | 0.867884746 | 0.55 | 0.009 | 4.3493450 | 39 | stox2a |
| cers3b | 1.702783 | 0.283459257 | 0.15 | 0.001 | 4.3259219 | 39 | cers3b |
| vegfab | 1.462118 | 1.038074896 | 0.6 | 0.011 | 3.7145112 | 39 | vegfab |
| bbox11 | 9.981377 | 1.219511075 | 0.65 | 0.014 | 2.5357689 | 39 | bbox1 |
| zgc:194392 | 1.878896 | 0.309159849 | 0.2 | 0.001 | 4.7733368 | 39 | zgc:194392 |
| si:dkey-14d8.61 | 2.407607 | 1.433691577 | 0.55 | 0.01 | 6.1165260 | 39 | si:dkey-14d8.6 |
| dab2ipb1 | 2.464933 | 1.188994497 | 0.7 | 0.016 | 6.2621631 | 39 | dab2ipb |
| kalrnb2 | 2.492765 | 0.848098511 | 0.55 | 0.01 | 6.3328706 | 39 | kalrnb |
| arhgap29a | 9.378568 | 0.944672176 | 0.5 | 0.008 | 2.3826253 | 39 | arhgap29a |
| lsp12 | 9.348506 | 1.161617894 | 0.75 | 0.019 | 2.3749880 | 39 | lsp1 |
| cx30.31 | 1.566819 | 1.272164676 | 0.75 | 0.019 | 3.9805054 | 39 | cx30.3 |
| tnks1bp12 | 3.130893 | 0.85099372 | 0.5 | 0.008 | 7.9540341 | 39 | tnks1bp1 |
| f11r.2 | 5.795611 | 0.550742741 | 0.3 | 0.003 | 1.4723752 | 39 | f11r.2 |
| cldn2 | 7.728692 | 0.435510587 | 0.1 | 0 | 1.9634743 | 39 | cldn2 |
| b3gnt2l | 5.771681 | 1.290066814 | 0.75 | 0.02 | 1.4662955 | 39 | b3gnt2l |
| pbx3b2 | 5.559893 | 1.263454339 | 0.75 | 0.021 | 1.4124908 | 39 | pbx3b |
| vangl1 | 3.109437 | 0.688918377 | 0.35 | 0.005 | 7.8995268 | 39 | vangl1 |
| eps8l2 | 1.759017 | 0.447121728 | 0.25 | 0.002 | 4.4687837 | 39 | eps8l2 |
| EML5 | 1.073278 | 0.516512379 | 0.25 | 0.002 | 2.7266650 | 39 | EML5 |
| frmd4ba1 | 1.956207 | 0.967241204 | 0.6 | 0.015 | 4.9697439 | 39 | frmd4ba |
| si:dkey-163m14.21 | 2.868526 | 1.375113786 | 0.7 | 0.021 | 7.2874920 | 39 | si:dkey-163m14.2 |
| elf32 | 1.003707 | 0.787704888 | 0.4 | 0.007 | 2.5499189 | 39 | elf3 |
| il17rc1 | 2.785577 | 0.343537593 | 0.2 | 0.002 | 7.0767588 | 39 | il17rc |
| klc4 | 8.473797 | 0.6728474 | 0.4 | 0.007 | 2.1527683 | 39 | klc4 |
| ank3b3 | 1.032993 | 1.483943157 | 0.9 | 0.038 | 2.6243197 | 39 | ank3b |
| pvrl2l1 | 1.835238 | 0.830570477 | 0.55 | 0.014 | 4.6624246 | 39 | pvrl2l |

| | | | | | | | |
|---|---|---|---|---|---|---|---|
| sall1a | 3.315378 | 0.513231269 | 0.3 | 0.004 | 8.4227182 | 39 | sall1a |
| enpp6 | 4.435248 | 0.528864961 | 0.25 | 0.003 | 1.1267749 | 39 | enpp6 |
| ezra | 9.637788 | 0.421288989 | 0.25 | 0.003 | 2.4484801 | 39 | ezra |
| prox1a2 | 8.200427 | 0.996380945 | 0.6 | 0.018 | 2.0833186 | 39 | prox1a |
| cldnc3 | 1.655397 | 1.453424787 | 0.85 | 0.035 | 4.2055383 | 39 | cldnc |
| palm21 | 2.279390 | 1.089709173 | 0.6 | 0.018 | 5.7907923 | 39 | palm2 |
| cldnk | 7.063118 | 0.252354706 | 0.15 | 0.001 | 1.7943853 | 39 | cldnk |
| tmem196b | 7.754811 | 0.278984119 | 0.15 | 0.001 | 1.9701098 | 39 | tmem196b |
| cxcl20 | 8.188998 | 0.429860917 | 0.3 | 0.004 | 2.0804150 | 39 | cxcl20 |
| cdh15 | 5.159551 | 1.78709175 | 1 | 0.055 | 1.3107841 | 39 | cdh1 |
| mmp15b | 2.086692 | 0.850263939 | 0.5 | 0.013 | 5.3012415 | 39 | mmp15b |
| hnf1ba | 4.918092 | 0.623976241 | 0.35 | 0.006 | 1.2494412 | 39 | hnf1ba |
| xirp2a3 | 4.994888 | 1.222606603 | 0.8 | 0.035 | 1.2689513 | 39 | xirp2a |
| rab11fip1a | 1.446556 | 0.553207249 | 0.3 | 0.005 | 3.6749756 | 39 | rab11fip1a |
| si:ch73-308m11.1 | 2.723995 | 0.304347952 | 0.15 | 0.001 | 6.9203098 | 39 | si:ch73-308m11.1 |
| sigirr | 3.027198 | 0.252445076 | 0.15 | 0.001 | 7.6905988 | 39 | sigirr |
| itpk1a | 3.305412 | 0.264899436 | 0.15 | 0.001 | 8.3973993 | 39 | itpk1a |
| si:dkey-205h13.1 | 5.946440 | 0.509192511 | 0.25 | 0.003 | 1.5106932 | 39 | si:dkey-205h13.1 |
| frzb3 | 7.892043 | 1.089808097 | 0.6 | 0.02 | 2.0049736 | 39 | frzb |
| plk2b1 | 1.354305 | 1.04628971 | 0.45 | 0.011 | 3.4406121 | 39 | plk2b |
| sorbs2a | 7.093217 | 0.872335959 | 0.45 | 0.011 | 1.8020319 | 39 | sorbs2a |
| CR762483.1 | 4.613864 | 0.513836814 | 0.2 | 0.002 | 1.1721523 | 39 | CR762483.1 |
| mir125a-2 | 5.178808 | 0.335491481 | 0.2 | 0.002 | 1.3156763 | 39 | mir125a-2 |
| slc27a2a | 5.521940 | 0.34070994 | 0.2 | 0.002 | 1.4028489 | 39 | slc27a2a |
| si:ch211-266g18.6 | 5.812691 | 0.415945927 | 0.2 | 0.002 | 1.4767143 | 39 | si:ch211-266g18.6 |
| patj | 2.039064 | 0.778500146 | 0.5 | 0.014 | 5.1802440 | 39 | patj |
| cers3a | 4.613412 | 0.284407326 | 0.15 | 0.001 | 1.1720375 | 39 | cers3a |
| ckmt1 | 3.771824 | 0.379525433 | 0.25 | 0.003 | 9.5823206 | 39 | ckmt1 |
| f2rl1.22 | 4.739602 | 1.139399413 | 0.6 | 0.021 | 1.2040960 | 39 | f2rl1.2 |
| cadm2a2 | 5.871267 | 1.013517393 | 0.6 | 0.021 | 1.4915955 | 39 | cadm2a |
| traf4a1 | 7.441966 | 0.705260488 | 0.4 | 0.009 | 1.8906316 | 39 | traf4a |
| pdlim21 | 8.293147 | 0.703569153 | 0.4 | 0.009 | 2.1068740 | 39 | pdlim2 |
| cgnb | 1.017276 | 0.580040618 | 0.35 | 0.007 | 2.5843918 | 39 | cgnb |
| cxadr2 | 1.071647 | 1.303111522 | 0.75 | 0.034 | 2.7225214 | 39 | cxadr |
| pdx1 | 2.037369 | 0.341570283 | 0.2 | 0.002 | 5.1759365 | 39 | pdx1 |

| | | | | | | | |
|---|---|---|---|---|---|---|---|
| myo5b1 | 1.314107 | 0.441657637 | 0.25 | 0.004 | 3.3384901 | 39 | myo5b |
| c9 | 2.082141 | 0.254826393 | 0.15 | 0.001 | 5.2896799 | 39 | c9 |
| crb3a | 1.042857 | 0.417715092 | 0.2 | 0.002 | 2.6493784 | 39 | crb3a |
| ugt81 | 1.364785 | 0.332963831 | 0.2 | 0.002 | 3.4672383 | 39 | ugt8 |
| fn3krp | 1.849345 | 0.353941967 | 0.2 | 0.002 | 4.6982633 | 39 | fn3krp |
| her15.1 | 2.659081 | 0.666082771 | 0.25 | 0.004 | 6.7553959 | 39 | her15.1 |
| krt92 | 1.040321 | 0.300739198 | 0.1 | 0.001 | 2.6429375 | 39 | krt92 |
| UNC13B | 7.117353 | 0.414576299 | 0.2 | 0.002 | 1.8081637 | 39 | UNC13B |
| si:ch1073-209e23.1 | 4.248228 | 0.270959325 | 0.15 | 0.001 | 1.0792623 | 39 | si:ch1073-209e23.1 |
| igsf9a | 5.566257 | 0.258400773 | 0.15 | 0.001 | 1.4141076 | 39 | igsf9a |
| pam1 | 1.500772 | 1.134623556 | 0.6 | 0.025 | 3.8127130 | 39 | pam |
| slc4a4b | 1.416738 | 0.646461133 | 0.35 | 0.009 | 3.5992251 | 39 | slc4a4b |
| chia.61 | 4.598260 | 0.721725143 | 0.3 | 0.006 | 1.1681880 | 39 | chia.6 |
| cd9b5 | 1.433568 | 1.628504588 | 0.9 | 0.061 | 3.6419805 | 39 | cd9b |
| myo9bl | 1.746187 | 0.701072608 | 0.45 | 0.015 | 4.4361898 | 39 | myo9bl |
| prxl2b | 6.163027 | 1.006544849 | 0.55 | 0.022 | 1.5657171 | 39 | prxl2b |
| rapgef2 | 1.193951 | 0.667286268 | 0.4 | 0.012 | 3.0332333 | 39 | rapgef2 |
| si:ch73-281f12.4 | 1.732317 | 0.54365519 | 0.25 | 0.005 | 4.4009519 | 39 | si:ch73-281f12.4 |
| alcama3 | 3.269931 | 0.960589549 | 0.65 | 0.032 | 8.3072615 | 39 | alcama |
| tspan11 | 1.060197 | 0.684941223 | 0.35 | 0.01 | 2.6934313 | 39 | tspan11 |
| s100v11 | 4.648914 | 0.64281994 | 0.4 | 0.012 | 1.1810568 | 39 | s100v1 |
| gpx1b1 | 1.644692 | 1.054300148 | 0.6 | 0.028 | 4.1783404 | 39 | gpx1b |
| LO016987.2 | 5.635740 | 0.788922506 | 0.45 | 0.016 | 1.4317597 | 39 | LO016987.2 |
| nectin1b4 | 7.286890 | 1.206101263 | 0.75 | 0.046 | 1.8512344 | 39 | nectin1b |
| jupa5 | 7.948281 | 1.323529966 | 0.85 | 0.06 | 2.0192608 | 39 | jupa |
| si:ch211-39i22.18 | 8.358044 | 1.641575134 | 1 | 0.088 | 2.1233613 | 39 | si:ch211-39i22.1 |
| FO704777.1 | 2.737674 | 0.314732081 | 0.2 | 0.003 | 6.9550630 | 39 | FO704777.1 |
| notch32 | 8.122878 | 1.041522957 | 0.55 | 0.025 | 2.0636173 | 39 | notch3 |
| noxa11 | 1.924655 | 0.306477628 | 0.15 | 0.002 | 4.8895873 | 39 | noxa1 |
| si:rp71-1h20.91 | 2.362960 | 0.491755258 | 0.2 | 0.003 | 6.0031008 | 39 | si:rp71-1h20.9 |
| spint23 | 9.212951 | 1.432534485 | 0.9 | 0.069 | 2.3405503 | 39 | spint2 |
| si:ch73-61d6.31 | 2.813210 | 0.578590246 | 0.4 | 0.014 | 7.1469609 | 39 | si:ch73-61d6.3 |
| traf4b | 3.870810 | 0.396876627 | 0.2 | 0.003 | 9.8337938 | 39 | traf4b |
| serpinb1 | 1.542112 | 1.020383863 | 0.7 | 0.044 | 3.9177359 | 39 | serpinb1 |
| sema3b2 | 1.791175 | 0.861759559 | 0.5 | 0.023 | 4.5504822 | 39 | sema3b |

| | | | | | | | |
|---|---|---|---|---|---|---|---|
| msmo1 | 3.748851 | 0.503060823 | 0.35 | 0.011 | 9.5239576 | 39 | msmo1 |
| tanc1b | 1.588426 | 0.422027857 | 0.25 | 0.006 | 4.0353971 | 39 | tanc1b |
| si:ch73-184c24.11 | 2.154975 | 0.270249826 | 0.15 | 0.002 | 5.4747164 | 39 | si:ch73-184c24.1 |
| sh3pxd2aa5 | 5.943098 | 0.962839244 | 0.6 | 0.034 | 1.5098441 | 39 | sh3pxd2aa |
| bpifcl3 | 2.227763 | 0.645342796 | 0.4 | 0.015 | 5.6596333 | 39 | bpifcl |
| LOC100537265 | 7.270672 | 0.269388653 | 0.15 | 0.002 | 1.8471143 | 39 | LOC100537265 |
| itsn2b1 | 7.516675 | 1.063749908 | 0.6 | 0.035 | 1.9096113 | 39 | itsn2b |
| pkp21 | 1.863653 | 0.504327873 | 0.25 | 0.006 | 4.7346119 | 39 | pkp2 |
| rhpn23 | 5.693813 | 1.088089689 | 0.6 | 0.035 | 1.4465134 | 39 | rhpn2 |
| mpp7a | 1.591905 | 0.534806672 | 0.35 | 0.012 | 4.0442357 | 39 | mpp7a |
| p2rx1 | 3.497754 | 0.550740942 | 0.3 | 0.009 | 8.8860460 | 39 | p2rx1 |
| hsd20b2 | 9.639005 | 0.4196093 | 0.25 | 0.006 | 2.4487893 | 39 | hsd20b2 |
| ftr825 | 1.147489 | 0.989809685 | 0.6 | 0.036 | 2.9151969 | 39 | ftr82 |
| CU462878.1 | 1.157542 | 0.383925381 | 0.25 | 0.006 | 2.9407367 | 39 | CU462878.1 |
| fat1a4 | 1.658115 | 0.858055053 | 0.55 | 0.031 | 4.2124425 | 39 | fat1a |
| metrn2 | 1.993931 | 0.675801823 | 0.4 | 0.016 | 5.0655817 | 39 | metrn |
| cmah3 | 2.150949 | 0.633993944 | 0.4 | 0.016 | 5.4644861 | 39 | cmah |
| basp1 | 2.339498 | 0.290085607 | 0.15 | 0.002 | 5.9434970 | 39 | basp1 |
| slc16a104 | 5.503044 | 0.698466803 | 0.5 | 0.026 | 1.3980484 | 39 | slc16a10 |
| c1qtnf121 | 1.611348 | 0.42544972 | 0.25 | 0.007 | 4.0936307 | 39 | c1qtnf12 |
| st51 | 8.876690 | 0.688451531 | 0.4 | 0.017 | 2.2551231 | 39 | st5 |
| pard3ab1 | 1.069651 | 0.436070556 | 0.3 | 0.01 | 2.7174503 | 39 | pard3ab |
| zgc:158482 | 1.723764 | 0.411464766 | 0.25 | 0.007 | 4.3792244 | 39 | zgc:158482 |
| mrtfba | 4.087918 | 0.605130231 | 0.35 | 0.013 | 1.0385356 | 39 | mrtfba |
| bves | 5.926079 | 0.289102039 | 0.15 | 0.002 | 1.5055205 | 39 | bves |
| pcdh1b1 | 6.573636 | 0.664568575 | 0.35 | 0.013 | 1.6700322 | 39 | pcdh1b |
| hkdc12 | 3.596068 | 0.332785444 | 0.2 | 0.004 | 9.1358123 | 39 | hkdc1 |
| bik2 | 8.976710 | 0.479766397 | 0.3 | 0.01 | 2.2805332 | 39 | bik |
| mid1ip1a2 | 9.203082 | 0.711133645 | 0.35 | 0.014 | 2.3380431 | 39 | mid1ip1a |
| nos1apa3 | 9.457288 | 0.802192314 | 0.5 | 0.029 | 2.4026242 | 39 | nos1apa |
| si:dkey-266m15.5 | 1.725341 | 0.343260691 | 0.2 | 0.005 | 4.3832296 | 39 | si:dkey-266m15.5 |
| si:dkey-74k8.3 | 5.341274 | 0.411834353 | 0.2 | 0.005 | 1.3569507 | 39 | si:dkey-74k8.3 |
| nectin4b | 6.941534 | 0.287434497 | 0.15 | 0.003 | 1.7634969 | 39 | nectin4b |
| muc5e | 8.553557 | 0.27081682 | 0.15 | 0.003 | 2.1730311 | 39 | muc5e |
| btr251 | 1.075661 | 0.4434363 | 0.25 | 0.007 | 2.7327183 | 39 | btr25 |

| | | | | | | | |
|---|---|---|---|---|---|---|---|
| anxa411 | 2.257545 | 1.802301402 | 1 | 0.151 | 5.7352943 | 39 | anxa4 |
| irg1l | 3.734653 | 0.297311254 | 0.1 | 0.001 | 9.4878881 | 39 | irg1l |
| blnk2 | 5.785673 | 0.766425156 | 0.45 | 0.024 | 1.4698503 | 39 | blnk |
| plekhg5b | 1.533073 | 0.608243096 | 0.35 | 0.015 | 3.8947741 | 39 | plekhg5b |
| sts1 | 3.122926 | 0.526511047 | 0.3 | 0.011 | 7.9337950 | 39 | sts |
| gdpd5a | 3.344729 | 0.335210971 | 0.2 | 0.005 | 8.4972840 | 39 | gdpd5a |
| hsp70.33 | 3.673282 | 1.002554185 | 0.4 | 0.02 | 9.3319753 | 39 | hsp70.3 |
| arl4ca | 4.024423 | 0.312714949 | 0.15 | 0.003 | 1.0224049 | 39 | arl4ca |
| zgc:1582631 | 4.310066 | 0.289168029 | 0.15 | 0.003 | 1.0949725 | 39 | zgc:158263 |
| kng1 | 7.349833 | 0.380213927 | 0.25 | 0.008 | 1.8672251 | 39 | kng1 |
| pof1b2 | 1.016661 | 0.523852857 | 0.3 | 0.011 | 2.5828289 | 39 | pof1b |
| fxyd18 | 2.247670 | 1.230384065 | 0.8 | 0.084 | 5.7102057 | 39 | fxyd1 |
| si:ch211-117c9.52 | 1.941705 | 0.79647342 | 0.4 | 0.021 | 4.9329037 | 39 | si:ch211-117c9.5 |
| crb3b | 2.772018 | 0.526812021 | 0.3 | 0.012 | 7.0423123 | 39 | crb3b |
| dvl1a | 7.893337 | 0.318231346 | 0.15 | 0.003 | 2.0053025 | 39 | dvl1a |
| TBC1D8B | 9.417219 | 0.437961201 | 0.2 | 0.005 | 2.3924444 | 39 | TBC1D8B |
| myzap1 | 1.115255 | 0.437962851 | 0.25 | 0.008 | 2.8333071 | 39 | myzap |
| cryl1 | 1.248168 | 0.723929305 | 0.45 | 0.027 | 3.1709732 | 39 | cryl1 |
| wdr37 | 1.309601 | 0.498874803 | 0.3 | 0.012 | 3.3270415 | 39 | wdr37 |
| itgb1b.1 | 2.005737 | 0.405906124 | 0.2 | 0.006 | 5.0955769 | 39 | itgb1b.1 |
| sytl4 | 3.972095 | 0.475378695 | 0.3 | 0.012 | 1.0091108 | 39 | sytl4 |
| GAN | 4.281571 | 0.296529449 | 0.15 | 0.003 | 1.0877333 | 39 | GAN |
| ccl20a.31 | 4.371189 | 0.389879239 | 0.15 | 0.003 | 1.1105007 | 39 | ccl20a.3 |
| capn1a | 5.790355 | 0.369267002 | 0.25 | 0.009 | 1.4710397 | 39 | capn1a |
| cav23 | 8.373117 | 0.634743398 | 0.35 | 0.017 | 2.1271904 | 39 | cav2 |
| sgk1 | 9.811962 | 1.084090013 | 0.7 | 0.067 | 2.4927291 | 39 | sgk1 |
| tnk1 | 1.061162 | 0.567740306 | 0.35 | 0.017 | 2.6958824 | 39 | tnk1 |
| si:ch211-153b23.51 | 1.756790 | 0.44565567 | 0.15 | 0.003 | 4.4631252 | 39 | si:ch211-153b23.5 |
| znf1083 | 2.038044 | 0.258743169 | 0.15 | 0.003 | 5.1776518 | 39 | znf1083 |
| slc8a2a | 3.206585 | 0.290510273 | 0.1 | 0.001 | 8.1463300 | 39 | slc8a2a |
| si:dkey-88e18.2 | 7.626096 | 0.316034975 | 0.15 | 0.003 | 1.9374098 | 39 | si:dkey-88e18.2 |
| celsr1a | 7.934581 | 0.343132693 | 0.2 | 0.006 | 2.0157804 | 39 | celsr1a |
| ptprfa | 1.096893 | 0.338462301 | 0.2 | 0.006 | 2.7866588 | 39 | ptprfa |
| foxp48 | 1.259101 | 1.237741873 | 0.7 | 0.074 | 3.1987478 | 39 | foxp4 |
| clic25 | 3.805723 | 1.120769828 | 0.7 | 0.072 | 9.6684417 | 39 | clic2 |

| | | | | | | | |
|---|---|---|---|---|---|---|---|
| ahcyl21 | 4.01422[illegible] | 0.592618054 | 0.35 | 0.018 | 1.0198148 | 39 | ahcyl2 |
| myo1cb3 | 6.03674[illegible] | 0.841159239 | 0.5 | 0.037 | 1.5336359 | 39 | myo1cb |
| macc11 | 1.41858[illegible] | 0.404425277 | 0.25 | 0.009 | 3.6039252 | 39 | macc1 |
| alkal2b | 1.67643[illegible] | 0.288317472 | 0.15 | 0.003 | 4.2589850 | 39 | alkal2b |
| aldh3a2b | 3.98273[illegible] | 0.401932081 | 0.25 | 0.01 | 1.0118150 | 39 | aldh3a2b |
| spint1a2 | 6.02324[illegible] | 0.89985293 | 0.6 | 0.052 | 1.5302064 | 39 | spint1a |
| myl9b11 | 8.83997[illegible] | 1.198415912 | 0.8 | 0.099 | 2.2457946 | 39 | myl9b |
| gsap | 1.06944[illegible] | 0.333912861 | 0.2 | 0.006 | 2.7169329 | 39 | gsap |
| si:ch73-22o12.13 | 1.17233[illegible] | 0.785662583 | 0.45 | 0.031 | 2.9783264 | 39 | si:ch73-22o12.1 |
| b3gnt3.45 | 2.38385[illegible] | 0.853902931 | 0.5 | 0.039 | 6.0561923 | 39 | b3gnt3.4 |
| daam1a2 | 3.31241[illegible] | 0.803478702 | 0.45 | 0.032 | 8.4151964 | 39 | daam1a |
| slc9a3r1b1 | 3.81716[illegible] | 0.384687302 | 0.2 | 0.006 | 9.6975033 | 39 | slc9a3r1b |
| XLOC-026010 | 7.42534[illegible] | 0.388672864 | 0.25 | 0.01 | 1.8864099 | 39 | XLOC-026010 |
| her65 | 1.28424[illegible] | 0.73670946 | 0.4 | 0.026 | 3.2626333 | 39 | her6 |
| prodha2 | 1.46497[illegible] | 0.462263578 | 0.3 | 0.015 | 3.7217798 | 39 | prodha |
| gdpd3a2 | 2.08170[illegible] | 0.256673635 | 0.15 | 0.004 | 5.2885638 | 39 | gdpd3a |
| fam84a | 2.57594[illegible] | 0.403555838 | 0.25 | 0.01 | 6.5441957 | 39 | fam84a |
| fa2h1 | 3.20278[illegible] | 0.460197294 | 0.25 | 0.01 | 8.1366628 | 39 | fa2h |
| stard142 | 3.90098[illegible] | 0.813011156 | 0.55 | 0.046 | 9.9104610 | 39 | stard14 |
| erbb25 | 7.76238[illegible] | 0.700162916 | 0.45 | 0.033 | 1.9720327 | 39 | erbb2 |
| s100u4 | 1.29727[illegible] | 0.717165185 | 0.5 | 0.041 | 3.2957252 | 39 | s100u |
| XLOC-038073 | 3.38270[illegible] | 0.379758376 | 0.2 | 0.007 | 8.5937727 | 39 | XLOC-038073 |
| ccdc85cb | 5.37999[illegible] | 0.275087789 | 0.15 | 0.004 | 1.3667876 | 39 | ccdc85cb |
| pkn2 | 5.38568[illegible] | 0.407080978 | 0.25 | 0.011 | 1.3682343 | 39 | pkn2 |
| NAV1 (1 of many)1 | 6.94123[illegible] | 0.715930321 | 0.4 | 0.028 | 1.7634203 | 39 | NAV1 (1 of many) |
| grb10a4 | 7.29070[illegible] | 0.922077689 | 0.5 | 0.043 | 1.8522044 | 39 | grb10a |
| nbeal1 | 7.47088[illegible] | 0.419569292 | 0.25 | 0.011 | 1.8979794 | 39 | nbeal1 |
| sypl2a4 | 9.00844[illegible] | 0.914109279 | 0.6 | 0.061 | 2.2885958 | 39 | sypl2a |
| XLOC-023879 | 1.23800[illegible] | 0.278669216 | 0.15 | 0.004 | 3.1451519 | 39 | XLOC-023879 |
| AL590151.1 | 1.93071[illegible] | 0.332105094 | 0.2 | 0.007 | 4.9049844 | 39 | AL590151.1 |
| cmbl1 | 2.89869[illegible] | 0.603421849 | 0.45 | 0.035 | 7.3641247 | 39 | cmbl |
| ryr31 | 3.09018[illegible] | 0.341156483 | 0.15 | 0.004 | 7.8506034 | 39 | ryr3 |
| PLEKHH3 | 3.11290[illegible] | 0.452469062 | 0.3 | 0.016 | 7.9083290 | 39 | PLEKHH3 |
| cdc42ep1a | 3.47109[illegible] | 0.288286355 | 0.15 | 0.004 | 8.8183225 | 39 | cdc42ep1a |
| CABZ01074363.13 | 4.23154[illegible] | 0.601795696 | 0.4 | 0.028 | 1.0750241 | 39 | CABZ01074363.1 |

| | | | | | | | |
|---|---|---|---|---|---|---|---|
| XLOC-0352021 | 4.990212 | 0.446258316 | 0.25 | 0.011 | 1.2677634 | 39 | XLOC-035202 |
| tjp1b1 | 6.130311 | 0.612995508 | 0.4 | 0.028 | 1.5574055 | 39 | tjp1b |
| mink1 | 6.180102 | 0.596444768 | 0.35 | 0.022 | 1.5700549 | 39 | mink1 |
| tex2 | 6.225359 | 0.470578401 | 0.35 | 0.022 | 1.5815525 | 39 | tex2 |
| kalrna | 1.000150 | 0.421154278 | 0.25 | 0.011 | 2.5408830 | 39 | kalrna |
| hivep3b | 1.028632 | 0.354051986 | 0.2 | 0.007 | 2.6132400 | 39 | hivep3b |
| stm | 1.332880 | 0.268185225 | 0.1 | 0.002 | 3.3861825 | 39 | stm |
| wwtr13 | 2.233116 | 0.676558283 | 0.4 | 0.029 | 5.6732331 | 39 | wwtr1 |
| ugt2a7 | 2.345537 | 0.453655204 | 0.3 | 0.017 | 5.9588382 | 39 | ugt2a7 |
| si:ch1073-345a8.11 | 2.672128 | 0.577249242 | 0.25 | 0.012 | 6.7885423 | 39 | si:ch1073-345a8.1 |
| lsr3 | 2.796798 | 0.747297054 | 0.4 | 0.029 | 7.1052666 | 39 | lsr |
| frmd4bb | 3.575722 | 0.414355633 | 0.15 | 0.004 | 9.0841222 | 39 | frmd4bb |
| dnmbp1 | 7.934214 | 0.482005779 | 0.3 | 0.017 | 2.0156871 | 39 | dnmbp |
| krt18a.110 | 2.197245 | 1.318502086 | 0.9 | 0.151 | 5.5821009 | 39 | krt18a.1 |
| rgs12b | 2.710278 | 0.431562432 | 0.25 | 0.012 | 6.8854613 | 39 | rgs12b |
| mylka3 | 3.700475 | 0.479601339 | 0.3 | 0.018 | 9.4010580 | 39 | mylka |
| slc34a2b | 4.015749 | 0.42122241 | 0.25 | 0.012 | 1.0202011 | 39 | slc34a2b |
| elmsan1a | 5.120265 | 0.423744001 | 0.25 | 0.012 | 1.3008035 | 39 | elmsan1a |
| krt949 | 6.066747 | 1.597755546 | 0.85 | 0.167 | 1.5412572 | 39 | krt94 |
| fgl2a7 | 6.735506 | 0.883921473 | 0.65 | 0.076 | 1.7111553 | 39 | fgl2a |
| si:ch73-234b20.59 | 1.372160 | 0.944111302 | 0.75 | 0.106 | 3.4859727 | 39 | si:ch73-234b20.5 |
| phactr4a4 | 1.859681 | 0.835947623 | 0.45 | 0.041 | 4.7245220 | 39 | phactr4a |
| btr121 | 2.156029 | 0.382621477 | 0.15 | 0.005 | 5.4773941 | 39 | btr12 |
| myo1ea1 | 2.299941 | 0.285904588 | 0.15 | 0.005 | 5.8430005 | 39 | myo1ea |
| pacsin31 | 2.838006 | 0.504556068 | 0.35 | 0.024 | 7.2099543 | 39 | pacsin3 |
| meis1b10 | 4.221157 | 0.951437394 | 0.6 | 0.074 | 1.0723851 | 39 | meis1b |
| ephb38 | 4.533680 | 0.971719522 | 0.5 | 0.053 | 1.1517815 | 39 | ephb3 |
| prr134 | 4.586582 | 0.566038209 | 0.35 | 0.025 | 1.1652213 | 39 | prr13 |
| sdk1a | 5.286666 | 0.34795463 | 0.2 | 0.009 | 1.3430776 | 39 | sdk1a |
| arhgef55 | 5.316495 | 0.682733012 | 0.4 | 0.033 | 1.3506557 | 39 | arhgef5 |
| pdlim16 | 5.529679 | 0.861163354 | 0.55 | 0.061 | 1.4048151 | 39 | pdlim1 |
| btbd6a1 | 8.129070 | 0.352563129 | 0.2 | 0.009 | 2.0651902 | 39 | btbd6a |
| mylk51 | 8.357075 | 0.589465764 | 0.3 | 0.019 | 2.1231149 | 39 | mylk5 |
| bmp162 | 9.503505 | 0.396351977 | 0.2 | 0.009 | 2.4143656 | 39 | bmp16 |
| efhd25 | 2.022141 | 0.789473346 | 0.55 | 0.061 | 5.1372512 | 39 | efhd2 |

| | | | | | | | |
|---|---|---|---|---|---|---|---|
| clmn | 3.673518 | 0.274449953 | 0.15 | 0.005 | 9.3325738 | 39 | clmn |
| llgl2 | 3.752412 | 0.6165307 | 0.35 | 0.026 | 9.5330032 | 39 | llgl2 |
| plecb7 | 4.410779 | 0.78615441 | 0.5 | 0.053 | 1.1205584 | 39 | plecb |
| glsb | 5.129777 | 0.564597933 | 0.3 | 0.02 | 1.3032200 | 39 | glsb |
| si:ch211-87m7.2 | 5.281141 | 0.559925456 | 0.4 | 0.034 | 1.3416739 | 39 | si:ch211-87m7.2 |
| st14a1 | 5.580973 | 0.608790097 | 0.4 | 0.033 | 1.4178463 | 39 | st14a |
| hpn2 | 9.807723 | 0.356926309 | 0.2 | 0.009 | 2.4916521 | 39 | hpn |
| sema6bb | 1.122533 | 0.305407156 | 0.2 | 0.009 | 2.8517970 | 39 | sema6bb |
| ap1m2 | 1.678022 | 0.487728559 | 0.35 | 0.026 | 4.2630152 | 39 | ap1m2 |
| cdr2a | 3.157991 | 0.376020887 | 0.2 | 0.009 | 8.0228785 | 39 | cdr2a |
| cttnbp2nlb | 3.361221 | 0.359040452 | 0.2 | 0.009 | 8.5391822 | 39 | cttnbp2nlb |
| apol11 | 3.682316 | 0.425330254 | 0.25 | 0.014 | 9.3549238 | 39 | apol1 |
| lmo7a2 | 4.685250 | 0.362168955 | 0.2 | 0.009 | 1.1902878 | 39 | lmo7a |
| tjap1 | 6.072029 | 0.292074662 | 0.2 | 0.009 | 1.5425990 | 39 | tjap1 |
| SEMA4F1 | 1.074952 | 0.338018298 | 0.15 | 0.006 | 2.7309179 | 39 | SEMA4F |
| fgfr34 | 1.421196 | 0.457182843 | 0.3 | 0.021 | 3.6105497 | 39 | fgfr3 |
| XLOC-0021291 | 1.585440 | 0.393639435 | 0.15 | 0.006 | 4.0278113 | 39 | XLOC-002129 |
| icn10 | 1.807441 | 1.554423695 | 0.95 | 0.269 | 4.5918050 | 39 | icn |
| b3gnt2b1 | 1.930298 | 0.527886238 | 0.3 | 0.022 | 4.9039229 | 39 | b3gnt2b |
| si:dkey-157l19.22 | 2.199964 | 0.310744214 | 0.2 | 0.01 | 5.5890103 | 39 | si:dkey-157l19.2 |
| si:ch73-103b11.21 | 2.216805 | 0.436291912 | 0.3 | 0.021 | 5.6317942 | 39 | si:ch73-103b11.2 |
| CABZ01086029.11 | 4.458260 | 0.496582575 | 0.3 | 0.022 | 1.1326209 | 39 | CABZ01086029.1 |
| znf1852 | 1.841397 | 0.254998348 | 0.15 | 0.006 | 4.6780696 | 39 | znf185 |
| adamts12 | 1.887365 | 0.373068513 | 0.15 | 0.006 | 4.7948529 | 39 | adamts12 |
| cltb | 2.201805 | 0.526947774 | 0.3 | 0.023 | 5.5936872 | 39 | cltb |
| gcshb2 | 2.203551 | 0.383403653 | 0.25 | 0.016 | 5.5981234 | 39 | gcshb |
| arhgap17a | 2.260782 | 0.375843223 | 0.2 | 0.01 | 5.7435172 | 39 | arhgap17a |
| rhoua3 | 2.613515 | 0.588027502 | 0.35 | 0.031 | 6.6396372 | 39 | rhoua |
| sowahd1 | 2.650521 | 0.356923325 | 0.25 | 0.016 | 6.7336492 | 39 | sowahd |
| ubl3a1 | 3.581940 | 0.545965265 | 0.45 | 0.048 | 9.0999193 | 39 | ubl3a |
| mpp5a1 | 3.957462 | 0.427934621 | 0.25 | 0.016 | 1.0053934 | 39 | mpp5a |
| ywhah3 | 5.443131 | 0.671363134 | 0.5 | 0.061 | 1.3828276 | 39 | ywhah |
| si:ch211-221f10.21 | 5.559714 | 0.311637521 | 0.15 | 0.006 | 1.4124453 | 39 | si:ch211-221f10.2 |
| pgd4 | 6.441486 | 0.68310122 | 0.55 | 0.072 | 1.6364596 | 39 | pgd |
| ripor1.1 | 6.600607 | 0.330438167 | 0.25 | 0.016 | 1.6768843 | 39 | ripor1.1 |

| | | | | | | | |
|---|---|---|---|---|---|---|---|
| rgl22 | 6.635045 | 0.406539451 | 0.3 | 0.023 | 1.6856332 | 39 | rgl2 |
| magi3a | 8.312625 | 0.394067121 | 0.25 | 0.017 | 2.1118223 | 39 | magi3a |
| XLOC-0045562 | 9.459215 | 0.474174567 | 0.25 | 0.017 | 2.4031138 | 39 | XLOC-004556 |
| lnx2b | 1.057053 | 0.70593329 | 0.45 | 0.05 | 2.6854451 | 39 | lnx2b |
| zgc:193711 | 1.098609 | 0.427805929 | 0.3 | 0.023 | 2.7910185 | 39 | zgc:193711 |
| dapk1 | 1.118781 | 0.281351292 | 0.2 | 0.011 | 2.8422651 | 39 | dapk1 |
| f3b6 | 1.261522 | 0.88902738 | 0.65 | 0.1 | 3.2048967 | 39 | f3b |
| bmpr1bb1 | 1.748970 | 0.362786925 | 0.2 | 0.011 | 4.4432582 | 39 | bmpr1bb |
| kifc3 | 2.157162 | 0.487901555 | 0.3 | 0.024 | 5.4802719 | 39 | kifc3 |
| cobll1b | 2.430857 | 0.302928375 | 0.2 | 0.011 | 6.1755936 | 39 | cobll1b |
| CU459056.1 | 3.221896 | 0.609314912 | 0.35 | 0.033 | 8.1852280 | 39 | CU459056.1 |
| dhdhl | 4.260186 | 0.426980797 | 0.35 | 0.032 | 1.0823003 | 39 | dhdhl |
| mfsd12a | 5.664196 | 0.362714139 | 0.25 | 0.017 | 1.4389892 | 39 | mfsd12a |
| zgc:923137 | 6.289890 | 0.789288236 | 0.7 | 0.109 | 1.5979467 | 39 | zgc:92313 |
| cebpd12 | 7.096491 | 0.983947091 | 0.8 | 0.153 | 1.8028637 | 39 | cebpd |
| hsp70.21 | 7.148072 | 0.589273803 | 0.2 | 0.012 | 1.8159678 | 39 | hsp70.2 |
| ptpn133 | 9.067990 | 0.387968684 | 0.25 | 0.018 | 2.3037230 | 39 | ptpn13 |
| zgc:923802 | 1.064889 | 0.371887846 | 0.2 | 0.012 | 2.7053513 | 39 | zgc:92380 |
| pxylp1 | 1.080260 | 0.251532363 | 0.15 | 0.007 | 2.7444026 | 39 | pxylp1 |
| srebf2 | 1.160281 | 0.457542937 | 0.3 | 0.025 | 2.9476944 | 39 | srebf2 |
| CABZ01055347.11 | 1.204345 | 0.644910015 | 0.35 | 0.035 | 3.0596402 | 39 | CABZ01055347.1 |
| inhbb1 | 1.403488 | 0.326165698 | 0.15 | 0.007 | 3.5655636 | 39 | inhbb |
| epha4a | 1.532401 | 0.25301903 | 0.15 | 0.007 | 3.8930658 | 39 | epha4a |
| tmem238a2 | 1.745040 | 0.345061781 | 0.25 | 0.018 | 4.4332754 | 39 | tmem238a |
| sptbn22 | 2.269241 | 0.585837622 | 0.4 | 0.044 | 5.7650076 | 39 | sptbn2 |
| krt814 | 2.349616 | 1.232211844 | 1 | 0.313 | 5.9692011 | 39 | krt8 |
| cldnb2 | 2.490053 | 0.519094633 | 0.2 | 0.012 | 6.3259820 | 39 | cldnb |
| kctd9a | 2.540436 | 0.37138155 | 0.2 | 0.012 | 6.4539798 | 39 | kctd9a |
| lygl14 | 2.668611 | 0.714270326 | 0.65 | 0.101 | 6.7796075 | 39 | lygl1 |
| gsna3 | 2.683101 | 0.48022339 | 0.35 | 0.034 | 6.8164193 | 39 | gsna |
| cldnh2 | 2.967450 | 0.268461227 | 0.1 | 0.003 | 7.5388091 | 39 | cldnh |
| emilin1a5 | 3.735096 | 0.47014007 | 0.3 | 0.026 | 9.4890135 | 39 | emilin1a |
| tnksb | 5.869954 | 0.358075775 | 0.25 | 0.019 | 1.4912618 | 39 | tnksb |
| ctnnd11 | 5.996485 | 0.816487024 | 0.45 | 0.058 | 1.5234070 | 39 | ctnnd1 |
| cnn214 | 7.363834 | 0.946530109 | 0.75 | 0.158 | 1.8707822 | 39 | cnn2 |

| | | | | | | | |
|---|---|---|---|---|---|---|---|
| atp2a34 | 7.946635 | 0.56496759 | 0.35 | 0.036 | 2.0188427 | 39 | atp2a3 |
| stard13b1 | 8.106827 | 0.266241555 | 0.2 | 0.012 | 2.0595395 | 39 | stard13b |
| si:ch211-213d14.21 | 9.670496 | 0.381105535 | 0.15 | 0.007 | 2.4567895 | 39 | si:ch211-213d14.2 |
| LOC103911313 | 9.900590 | 0.253211111 | 0.1 | 0.003 | 2.5152451 | 39 | LOC103911313 |
| slc10a31 | 9.929882 | 0.418807716 | 0.25 | 0.019 | 2.5226867 | 39 | slc10a3 |
| jun15 | 9.930657 | 1.144850325 | 0.95 | 0.251 | 2.5228834 | 39 | jun |
| dbpb1 | 1.164606 | 0.514448441 | 0.4 | 0.045 | 2.9586831 | 39 | dbpb |
| XLOC-0152912 | 1.175837 | 0.319688646 | 0.2 | 0.013 | 2.9872139 | 39 | XLOC-015291 |
| taok1b1 | 1.202072 | 0.419163599 | 0.3 | 0.027 | 3.0538656 | 39 | taok1b |
| s100v213 | 1.444699 | 1.006993652 | 0.7 | 0.141 | 3.6702583 | 39 | s100v2 |
| stxbp63 | 1.751710 | 0.621895832 | 0.4 | 0.047 | 4.4502201 | 39 | stxbp6 |
| cald1a12 | 1.846452 | 0.932073275 | 0.65 | 0.121 | 4.6909119 | 39 | cald1a |
| smurf2 | 2.008105 | 0.465274271 | 0.3 | 0.027 | 5.1015918 | 39 | smurf2 |
| zgc:1752641 | 2.012224 | 0.528929912 | 0.35 | 0.036 | 5.1120564 | 39 | zgc:175264 |
| h1f09 | 2.643175 | 0.756318447 | 0.5 | 0.072 | 6.7149876 | 39 | h1f0 |
| perp7 | 4.161053 | 0.712203288 | 0.45 | 0.062 | 1.0571155 | 39 | perp |
| tent5ba5 | 5.623589 | 0.376275194 | 0.2 | 0.013 | 1.4286728 | 39 | tent5ba |
| agpat32 | 6.753619 | 0.542751492 | 0.35 | 0.039 | 1.7157569 | 39 | agpat3 |
| PDE3B1 | 7.282899 | 0.439912937 | 0.25 | 0.021 | 1.8502205 | 39 | PDE3B |
| dspa4 | 9.206913 | 0.558907839 | 0.3 | 0.029 | 2.3390163 | 39 | dspa |
| nipa21 | 9.906258 | 0.646736278 | 0.4 | 0.049 | 2.5166848 | 39 | nipa2 |
| camk2g22 | 1.098212 | 0.599044979 | 0.35 | 0.039 | 2.7900089 | 39 | camk2g2 |
| RNF14 (1 of many) | 1.226418 | 0.266508218 | 0.15 | 0.008 | 3.1157163 | 39 | RNF14 (1 of many) |
| nr4a2b1 | 1.230768 | 0.448988367 | 0.2 | 0.014 | 3.1267669 | 39 | nr4a2b |
| esrp21 | 1.454901 | 0.255822173 | 0.15 | 0.008 | 3.6961781 | 39 | esrp2 |
| capzb11 | 1.579726 | 1.017486144 | 0.9 | 0.273 | 4.0132956 | 39 | capzb |
| tmod22 | 1.742733 | 0.534622618 | 0.4 | 0.049 | 4.4274155 | 39 | tmod2 |
| kif13ba | 1.785491 | 0.417272656 | 0.25 | 0.021 | 4.5360403 | 39 | kif13ba |
| syne2b | 1.946992 | 0.281123635 | 0.2 | 0.014 | 4.9463341 | 39 | syne2b |
| leng9 | 1.978797 | 0.337335659 | 0.15 | 0.008 | 5.0271339 | 39 | leng9 |
| furina3 | 2.223833 | 0.793468754 | 0.45 | 0.066 | 5.6496500 | 39 | furina |
| dbn16 | 2.332022 | 0.732311927 | 0.45 | 0.067 | 5.9245020 | 39 | dbn1 |
| fosl1a9 | 2.509919 | 1.001454139 | 0.5 | 0.084 | 6.3764497 | 39 | fosl1a |
| XLOC-024446 | 3.996424 | 0.250625724 | 0.1 | 0.004 | 1.0152917 | 39 | XLOC-024446 |
| abcd3a | 4.194876 | 0.460619393 | 0.3 | 0.031 | 1.0657084 | 39 | abcd3a |

| | | | | | | | |
|---|---|---|---|---|---|---|---|
| tjp31 | 4.272341 | 0.439867293 | 0.25 | 0.021 | 1.0853884 | 39 | tjp3 |
| taf8 | 4.322539 | 0.28973157 | 0.15 | 0.008 | 1.0981410 | 39 | taf8 |
| golgb12 | 5.078258 | 0.506863186 | 0.4 | 0.052 | 1.2901315 | 39 | golgb1 |
| nfil3-610 | 5.667343 | 0.875073513 | 0.65 | 0.132 | 1.4397886 | 39 | nfil3-6 |
| hipk22 | 6.910953 | 0.477767286 | 0.25 | 0.022 | 1.7557278 | 39 | hipk2 |
| mcu | 7.663862 | 0.398283214 | 0.3 | 0.031 | 1.9470043 | 39 | mcu |
| gna11b3 | 7.695843 | 0.656591255 | 0.45 | 0.066 | 1.9551289 | 39 | gna11b |
| ddt1 | 8.038323 | 0.60186025 | 0.6 | 0.106 | 2.0421361 | 39 | ddt |
| spns23 | 8.077496 | 0.358909894 | 0.15 | 0.008 | 2.0520880 | 39 | spns2 |
| sdcbp211 | 9.506582 | 0.946849149 | 0.85 | 0.229 | 2.4151471 | 39 | sdcbp2 |
| bdh11 | 9.724134 | 0.295392252 | 0.2 | 0.015 | 2.4704164 | 39 | bdh1 |
| ch25hl31 | 1.135405 | 0.648144541 | 0.4 | 0.054 | 2.8844964 | 39 | ch25hl3 |
| myh9a4 | 1.309566 | 0.511205302 | 0.4 | 0.052 | 3.3269529 | 39 | myh9a |
| tmsb4x21 | 1.344345 | 0.779523535 | 1 | 0.72 | 3.4153109 | 39 | tmsb4x |
| pstpip21 | 1.371052 | 0.255008713 | 0.25 | 0.022 | 3.4831598 | 39 | pstpip2 |
| nectin3b | 1.393929 | 0.409504602 | 0.25 | 0.022 | 3.5412781 | 39 | nectin3b |
| eml2 | 1.413102 | 0.510601715 | 0.35 | 0.042 | 3.5899878 | 39 | eml2 |
| rpz510 | 1.598063 | 0.77372933 | 0.65 | 0.128 | 4.0598798 | 39 | rpz5 |
| fgfr22 | 2.135802 | 0.442978942 | 0.3 | 0.032 | 5.4260067 | 39 | fgfr2 |
| myl610 | 2.155082 | 0.841149751 | 0.6 | 0.118 | 5.4749868 | 39 | myl6 |
| si:ch211-202f5.37 | 2.231875 | 0.635592854 | 0.5 | 0.077 | 5.6700805 | 39 | si:ch211-202f5.3 |
| ptbp2b | 2.300175 | 0.336298365 | 0.2 | 0.015 | 5.8435964 | 39 | ptbp2b |
| capn2a3 | 2.332958 | 0.58810733 | 0.4 | 0.055 | 5.9268813 | 39 | capn2a |
| gdpd5b | 2.696613 | 0.423515528 | 0.3 | 0.032 | 6.8507470 | 39 | gdpd5b |
| si:dkey-204f11.642 | 2.888948 | 0.400976112 | 0.3 | 0.032 | 7.3393738 | 39 | si:dkey-204f11.64 |
| hipk3b1 | 3.301813 | 0.471032826 | 0.35 | 0.043 | 8.3882560 | 39 | hipk3b |
| ecm1b8 | 3.321598 | 0.616786863 | 0.45 | 0.068 | 8.4385217 | 39 | ecm1b |
| ak16 | 3.833949 | 0.685947413 | 0.55 | 0.1 | 9.7401481 | 39 | ak1 |
| zgc:924811 | 4.897372 | 0.487229771 | 0.25 | 0.024 | 1.2441774 | 39 | zgc:92481 |
| als2a | 5.473734 | 0.310805877 | 0.15 | 0.009 | 1.3906023 | 39 | als2a |
| cyp11c11 | 5.877023 | 0.432884123 | 0.25 | 0.023 | 1.4930577 | 39 | cyp11c1 |
| canx3 | 6.110052 | 0.71683773 | 0.55 | 0.103 | 1.5522588 | 39 | canx |
| azin1b | 7.070953 | 0.531113319 | 0.3 | 0.034 | 1.7963758 | 39 | azin1b |
| gipc22 | 7.084304 | 0.471811402 | 0.3 | 0.033 | 1.7997675 | 39 | gipc2 |
| tspan9a2 | 7.290637 | 0.388150075 | 0.25 | 0.024 | 1.8521864 | 39 | tspan9a |

| | | | | | | | |
|---|---|---|---|---|---|---|---|
| id2a3 | 7.424734 | 0.969895669 | 0.7 | 0.169 | 1.8862538 | 39 | id2a |
| ptmaa18 | 7.983774 | 0.99299317 | 0.95 | 0.434 | 2.0282778 | 39 | ptmaa |
| tubb6 | 9.054385 | 0.299915722 | 0.15 | 0.009 | 2.3002666 | 39 | tubb6 |
| dsg2.12 | 9.965115 | 0.415935717 | 0.25 | 0.024 | 2.5316375 | 39 | dsg2.1 |
| mafk | 1.050962 | 0.253731768 | 0.1 | 0.004 | 2.6699699 | 39 | mafk |
| pgpep1 | 1.111890 | 0.551274816 | 0.3 | 0.035 | 2.8247586 | 39 | pgpep1 |
| msrb3 | 1.167891 | 0.388123945 | 0.2 | 0.016 | 2.9670289 | 39 | msrb3 |
| picalma | 1.185976 | 0.435388462 | 0.3 | 0.034 | 3.0129729 | 39 | picalma |
| rbms2a | 1.273789 | 0.375720762 | 0.3 | 0.034 | 3.2360618 | 39 | rbms2a |
| tns3.21 | 1.485607 | 0.410475405 | 0.25 | 0.025 | 3.7741855 | 39 | tns3.2 |
| socs3a14 | 1.491235 | 1.114430402 | 0.9 | 0.316 | 3.7884827 | 39 | socs3a |
| rbms2b | 1.547123 | 0.391272957 | 0.25 | 0.025 | 3.9304681 | 39 | rbms2b |
| ezrb1 | 1.733247 | 0.628883834 | 0.45 | 0.073 | 4.4033142 | 39 | ezrb |
| pdgfc1 | 1.792898 | 0.25024789 | 0.15 | 0.01 | 4.5548596 | 39 | pdgfc |
| ddr1 | 1.834610 | 0.263447386 | 0.15 | 0.01 | 4.6608291 | 39 | ddr1 |
| adh8b2 | 1.879689 | 0.645548099 | 0.35 | 0.049 | 4.7753522 | 39 | adh8b |
| flii | 1.950598 | 0.398912441 | 0.3 | 0.035 | 4.9554956 | 39 | flii |
| cyb5r33 | 2.047939 | 0.505227246 | 0.35 | 0.047 | 5.2027909 | 39 | cyb5r3 |
| lrrc176 | 2.054771 | 0.591928953 | 0.4 | 0.059 | 5.2201469 | 39 | lrrc17 |
| tuba8l25 | 2.112383 | 0.395842436 | 0.45 | 0.07 | 5.3665093 | 39 | tuba8l2 |
| padi2 | 2.944398 | 0.256209682 | 0.2 | 0.017 | 7.4802435 | 39 | padi2 |
| fkbp58 | 3.016228 | 0.734749211 | 0.6 | 0.128 | 7.6627280 | 39 | fkbp5 |
| ctsk8 | 3.050000 | 0.613986346 | 0.45 | 0.075 | 7.7485251 | 39 | ctsk |
| per1b1 | 3.188953 | 0.528383384 | 0.45 | 0.073 | 8.1015360 | 39 | per1b |
| rpz | 3.275274 | 0.298362041 | 0.2 | 0.017 | 8.3208353 | 39 | rpz |
| CU929259.11 | 3.868141 | 0.3782856 | 0.25 | 0.026 | 9.8270130 | 39 | CU929259.1 |
| XLOC-040069 | 4.117946 | 0.342568499 | 0.2 | 0.017 | 1.0461644 | 39 | XLOC-040069 |
| apcdd1l3 | 4.381212 | 0.360307738 | 0.15 | 0.01 | 1.1130471 | 39 | apcdd1l |
| gas1b | 4.385451 | 0.295070922 | 0.15 | 0.01 | 1.1141238 | 39 | gas1b |
| afdna4 | 5.334449 | 0.521731264 | 0.25 | 0.027 | 1.3552168 | 39 | afdna |
| sh3bgrl8 | 5.801928 | 0.622455283 | 0.6 | 0.122 | 1.4739798 | 39 | sh3bgrl |
| selenol | 5.826028 | 0.450725011 | 0.25 | 0.027 | 1.4801026 | 39 | selenol |
| alpl | 5.863026 | 0.308518884 | 0.15 | 0.01 | 1.4895018 | 39 | alpl |
| zgc:77158 | 5.983769 | 0.305082949 | 0.2 | 0.017 | 1.5201765 | 39 | zgc:77158 |
| lrrfip1a7 | 7.161167 | 0.536749296 | 0.4 | 0.062 | 1.8192945 | 39 | lrrfip1a |

| | | | | | | | |
|---|---|---|---|---|---|---|---|
| wbp1lb4 | 8.45558( | 0.534240187 | 0.35 | 0.051 | 2.1481401 | 39 | wbp1lb |
| rab11fip1b6 | 8.470489 | 0.490874911 | 0.4 | 0.061 | 2.1519279 | 39 | rab11fip1b |
| prkacaa | 1.117021 | 0.46241681 | 0.3 | 0.038 | 2.8377920 | 39 | prkacaa |
| rc3h1a | 1.176903 | 0.378390767 | 0.2 | 0.018 | 2.9899223 | 39 | rc3h1a |
| lpin1 | 1.234576 | 0.288613646 | 0.2 | 0.018 | 3.1364423 | 39 | lpin1 |
| CABZ01079427.12 | 1.302472 | 0.352651127 | 0.3 | 0.037 | 3.3089315 | 39 | CABZ01079427.1 |
| rassf91 | 1.311026 | 0.271332054 | 0.1 | 0.005 | 3.3306639 | 39 | rassf9 |
| cnot6b | 1.539421 | 0.285500996 | 0.15 | 0.011 | 3.9109007 | 39 | cnot6b |
| CABZ01072614.18 | 1.577684 | 0.618706735 | 0.5 | 0.096 | 4.0081085 | 39 | CABZ01072614.1 |
| lrrc11 | 1.593222 | 0.333529189 | 0.25 | 0.027 | 4.0475811 | 39 | lrrc1 |
| dsc2l4 | 1.633072 | 0.444202989 | 0.25 | 0.028 | 4.1488207 | 39 | dsc2l |
| LOC110439449 | 1.677251 | 0.27678367 | 0.15 | 0.011 | 4.2610585 | 39 | LOC110439449 |
| RALGDS | 1.728527 | 0.296884583 | 0.2 | 0.018 | 4.3913230 | 39 | RALGDS |
| si:ch73-193i22.1 | 2.035002 | 0.295896318 | 0.15 | 0.011 | 5.1699244 | 39 | si:ch73-193i22.1 |
| cyr619 | 2.158940 | 0.882344894 | 0.4 | 0.068 | 5.4847876 | 39 | cyr61 |
| inpp5b | 2.262852 | 0.456888923 | 0.25 | 0.028 | 5.7487766 | 39 | inpp5b |
| usp43b2 | 2.274952 | 0.42000196 | 0.2 | 0.019 | 5.7795175 | 39 | usp43b |
| bambia3 | 2.291354 | 0.439802572 | 0.25 | 0.028 | 5.8211854 | 39 | bambia |
| man2b2 | 2.368200 | 0.298253706 | 0.2 | 0.018 | 6.0164134 | 39 | man2b2 |
| CT737123.1 | 2.502612 | 0.252692436 | 0.15 | 0.011 | 6.3578879 | 39 | CT737123.1 |
| gadd45ba13 | 2.748445 | 0.972294602 | 0.7 | 0.199 | 6.9824269 | 39 | gadd45ba |
| cyb5a1 | 2.892482 | 0.488770574 | 0.4 | 0.065 | 7.3483522 | 39 | cyb5a |
| gpd1c1 | 2.896138 | 0.3047186 | 0.2 | 0.019 | 7.3576393 | 39 | gpd1c |
| nrarpa | 2.991244 | 0.332398052 | 0.15 | 0.011 | 7.5992577 | 39 | nrarpa |
| mgst3b8 | 3.105408 | 0.810017909 | 0.7 | 0.195 | 7.8892913 | 39 | mgst3b |
| gabarapl23 | 3.329677 | 0.640221681 | 0.45 | 0.085 | 8.4590468 | 39 | gabarapl2 |
| limk22 | 3.435541 | 0.379312655 | 0.2 | 0.019 | 8.7279937 | 39 | limk2 |
| cd151l2 | 3.470725 | 0.456398101 | 0.4 | 0.064 | 8.8173793 | 39 | cd151l |
| cyp3c15 | 3.501071 | 0.527475121 | 0.4 | 0.066 | 8.8944727 | 39 | cyp3c1 |
| zfand32 | 3.835175 | 0.550234703 | 0.4 | 0.068 | 9.7432646 | 39 | zfand3 |
| cdc42se11 | 4.130409 | 0.567142786 | 0.45 | 0.08 | 0.000105 | 39 | cdc42se1 |
| serinc26 | 4.505463 | 0.742974873 | 0.55 | 0.125 | 0.000114 | 39 | serinc2 |
| irf6 | 4.601380 | 0.578335001 | 0.35 | 0.051 | 0.000117 | 39 | irf6 |
| zgc:1716792 | 4.729423 | 0.372561828 | 0.25 | 0.029 | 0.00012 | 39 | zgc:171679 |
| actn45 | 4.741964 | 0.604490954 | 0.5 | 0.099 | 0.00012 | 39 | actn4 |

| | | | | | | | |
|---|---|---|---|---|---|---|---|
| ap1b12 | 5.006434 | 0.468265454 | 0.35 | 0.052 | 0.000127 | 39 | ap1b1 |
| sult6b11 | 5.274129 | 0.321908001 | 0.25 | 0.029 | 0.000134 | 39 | sult6b1 |
| crtap6 | 6.166702 | 0.449580908 | 0.3 | 0.041 | 0.000157 | 39 | crtap |
| setd1a | 6.299610 | 0.299312417 | 0.2 | 0.019 | 0.00016 | 39 | setd1a |
| arg24 | 6.500815 | 0.362141753 | 0.2 | 0.019 | 0.000165 | 39 | arg2 |
| mical3a | 6.660860 | 0.30810475 | 0.15 | 0.011 | 0.000169 | 39 | mical3a |
| si:ch211-107o10.31 | 6.912759 | 0.46782646 | 0.3 | 0.042 | 0.000176 | 39 | si:ch211-107o10.3 |
| ankrd10b1 | 8.542422 | 0.309476726 | 0.25 | 0.029 | 0.000217 | 39 | ankrd10b |
| chmp4bb2 | 1.050440 | 0.60240605 | 0.45 | 0.087 | 0.000267 | 39 | chmp4bb |
| LOC108190056 | 1.055498 | 0.312027639 | 0.15 | 0.012 | 0.000268 | 39 | LOC108190056 |
| ier59 | 1.141478 | 0.721987057 | 0.6 | 0.148 | 0.00029 | 39 | ier5 |
| nudt4a1 | 1.320974 | 0.362756302 | 0.2 | 0.02 | 0.000336 | 39 | nudt4a |
| efna1b6 | 1.386769 | 0.357711947 | 0.2 | 0.02 | 0.000352 | 39 | efna1b |
| tollip | 1.851082 | 0.379709022 | 0.3 | 0.042 | 0.00047 | 39 | tollip |
| vclb1 | 1.997628 | 0.32011818 | 0.2 | 0.021 | 0.000507 | 39 | vclb |
| si:ch211-222k6.3 | 2.051533 | 0.314736951 | 0.2 | 0.021 | 0.000521 | 39 | si:ch211-222k6.3 |
| appa7 | 2.156095 | 0.605479683 | 0.45 | 0.089 | 0.000548 | 39 | appa |
| lama54 | 2.294646 | 0.394694616 | 0.25 | 0.031 | 0.000583 | 39 | lama5 |
| gnb2 | 2.418401 | 0.252116259 | 0.15 | 0.012 | 0.000614 | 39 | gnb2 |
| qkia11 | 2.455434 | 0.770011688 | 0.7 | 0.201 | 0.000624 | 39 | qkia |
| tuft1a4 | 2.598250 | 0.348319353 | 0.25 | 0.031 | 0.00066 | 39 | tuft1a |
| bhmt5 | 2.689400 | 0.46806514 | 0.3 | 0.043 | 0.000683 | 39 | bhmt |
| porb2 | 2.876860 | 0.470045837 | 0.3 | 0.044 | 0.000731 | 39 | porb |
| zgc:1630802 | 2.971906 | 0.565489764 | 0.45 | 0.09 | 0.000755 | 39 | zgc:163080 |
| CABZ01092156.11 | 3.055220 | 0.375610662 | 0.25 | 0.031 | 0.000776 | 39 | CABZ01092156.1 |
| sntb11 | 3.132581 | 0.267183204 | 0.15 | 0.012 | 0.000796 | 39 | sntb1 |
| atp1b1a8 | 3.137096 | 0.770720449 | 0.75 | 0.231 | 0.000797 | 39 | atp1b1a |
| fosb12 | 3.267502 | 1.043324824 | 0.5 | 0.123 | 0.00083 | 39 | fosb |
| errfi1a3 | 3.521380 | 0.334349354 | 0.2 | 0.021 | 0.000895 | 39 | errfi1a |
| crema12 | 3.852754 | 0.737890592 | 0.65 | 0.175 | 0.000979 | 39 | crema |
| si:ch1073-392o20.21 | 4.043478 | 0.41884177 | 0.35 | 0.057 | 0.001027 | 39 | si:ch1073-392o20.2 |
| chd1l | 4.149798 | 0.304264292 | 0.15 | 0.012 | 0.001054 | 39 | chd1l |
| epcam9 | 4.494151 | 0.908687019 | 1 | 0.423 | 0.001142 | 39 | epcam |
| hiat1a | 5.164620 | 0.386770035 | 0.35 | 0.056 | 0.001312 | 39 | hiat1a |
| rbm25a | 5.174943 | 0.304428777 | 0.2 | 0.021 | 0.001315 | 39 | rbm25a |

| | | | | | | | |
|---|---|---|---|---|---|---|---|
| smad13 | 5.318832 | 0.338728838 | 0.2 | 0.022 | 0.001351 | 39 | smad1 |
| nae1 | 6.453134 | 0.265156603 | 0.2 | 0.022 | 0.001639 | 39 | nae1 |
| stk261 | 6.465414 | 0.326210916 | 0.2 | 0.022 | 0.001643 | 39 | stk26 |
| slc38a24 | 7.179769 | 0.503622531 | 0.3 | 0.047 | 0.001824 | 39 | slc38a2 |
| arhgap51 | 7.421772 | 0.419657466 | 0.25 | 0.033 | 0.001886 | 39 | arhgap5 |
| myo10 | 7.698450 | 0.311835338 | 0.15 | 0.013 | 0.001956 | 39 | myo10 |
| pcyox12 | 8.038867 | 0.498115938 | 0.35 | 0.061 | 0.002042 | 39 | pcyox1 |
| MMP23B1 | 8.251311 | 0.330192421 | 0.2 | 0.022 | 0.002096 | 39 | MMP23B |
| LOC1018844517 | 9.550945 | 0.559220813 | 0.45 | 0.092 | 0.002426 | 39 | LOC101884451 |
| cyb5r1 | 9.606923 | 0.47717653 | 0.35 | 0.061 | 0.002441 | 39 | cyb5r1 |
| tmed76 | 9.964567 | 0.669164607 | 0.75 | 0.222 | 0.002531 | 39 | tmed7 |
| prkar1aa1 | 1.096262 | 0.424415991 | 0.35 | 0.06 | 0.002785 | 39 | prkar1aa |
| pak4 | 1.106528 | 0.33218315 | 0.25 | 0.033 | 0.002811 | 39 | pak4 |
| pawr1 | 1.110570 | 0.295473456 | 0.2 | 0.022 | 0.002821 | 39 | pawr |
| hnrnpul11 | 1.120965 | 0.367415218 | 0.25 | 0.034 | 0.002848 | 39 | hnrnpul1 |
| mcl1b15 | 1.123296 | 0.836026018 | 0.75 | 0.252 | 0.002854 | 39 | mcl1b |
| lipf3 | 1.170248 | 0.552575817 | 0.3 | 0.048 | 0.002973 | 39 | lipf |
| aplp26 | 1.174009 | 0.544272976 | 0.4 | 0.078 | 0.002983 | 39 | aplp2 |
| trioa | 1.265942 | 0.343593548 | 0.2 | 0.023 | 0.003216 | 39 | trioa |
| tpm4a12 | 1.266085 | 0.627098228 | 0.5 | 0.115 | 0.003216 | 39 | tpm4a |
| atf310 | 1.301538 | 0.754086405 | 0.4 | 0.083 | 0.003307 | 39 | atf3 |
| vdrb2 | 1.331857 | 0.41034965 | 0.3 | 0.047 | 0.003384 | 39 | vdrb |
| fosab16 | 1.455585 | 1.02489875 | 0.8 | 0.363 | 0.003698 | 39 | fosab |
| cd6319 | 1.483976 | 0.798748285 | 0.95 | 0.375 | 0.00377 | 39 | cd63 |
| cers2a1 | 1.491455 | 0.380390957 | 0.25 | 0.034 | 0.003789 | 39 | cers2a |
| ndufa53 | 1.520154 | 0.64963183 | 0.6 | 0.161 | 0.003862 | 39 | ndufa5 |
| dop1b | 1.546568 | 0.395353097 | 0.2 | 0.023 | 0.003929 | 39 | dop1b |
| mtmr111 | 1.567998 | 0.262020562 | 0.15 | 0.013 | 0.003984 | 39 | mtmr11 |
| acsl1a1 | 1.569851 | 0.334986539 | 0.2 | 0.023 | 0.003988 | 39 | acsl1a |
| igfbp33 | 1.578163 | 0.296332368 | 0.15 | 0.013 | 0.004009 | 39 | igfbp3 |
| pcbd11 | 1.582045 | 0.337536904 | 0.35 | 0.059 | 0.004019 | 39 | pcbd1 |
| XLOC-0242481 | 1.812747 | 0.329891006 | 0.2 | 0.023 | 0.004605 | 39 | XLOC-024248 |
| oaz1b14 | 1.814496 | 0.768050025 | 0.9 | 0.361 | 0.00461 | 39 | oaz1b |
| ndufa32 | 1.912682 | 0.563987188 | 0.7 | 0.193 | 0.004859 | 39 | ndufa3 |
| cygb17 | 1.922461 | 0.540183176 | 0.35 | 0.062 | 0.004884 | 39 | cygb1 |

| | | | | | | | |
|---|---|---|---|---|---|---|---|
| pdlim5b | 1.944637 | 0.367950689 | 0.2 | 0.023 | 0.00494 | 39 | pdlim5b |
| enc3 | 1.995967 | 0.270800088 | 0.15 | 0.014 | 0.005071 | 39 | enc3 |
| selenow111 | 2.062508 | 0.777425235 | 0.9 | 0.417 | 0.00524 | 39 | selenow1 |
| ccdc186 | 2.066208 | 0.255222403 | 0.15 | 0.014 | 0.005249 | 39 | ccdc186 |
| schip13 | 2.151246 | 0.312887548 | 0.2 | 0.023 | 0.005465 | 39 | schip1 |
| hmgn619 | 2.225849 | 0.767294567 | 0.85 | 0.313 | 0.005655 | 39 | hmgn6 |
| ildr1b | 2.366072 | 0.359874823 | 0.25 | 0.034 | 0.006011 | 39 | ildr1b |
| utrn2 | 2.743927 | 0.339975567 | 0.25 | 0.035 | 0.006971 | 39 | utrn |
| jdp2b11 | 2.963799 | 0.966883349 | 0.55 | 0.166 | 0.00753 | 39 | jdp2b |
| itgb3b3 | 3.016489 | 0.289328375 | 0.15 | 0.014 | 0.007663 | 39 | itgb3b |
| hist2h2l6 | 3.025596 | 0.558999184 | 0.45 | 0.1 | 0.007687 | 39 | hist2h2l |
| ewsr1a2 | 3.428621 | 0.496526363 | 0.3 | 0.051 | 0.00871 | 39 | ewsr1a |
| zgc:101583 | 3.757444 | 0.302493121 | 0.2 | 0.024 | 0.009546 | 39 | zgc:101583 |
| im:6904045 | 3.888075 | 0.260657433 | 0.15 | 0.014 | 0.009878 | 39 | im:6904045 |
| cnn3b | 4.081495 | 0.403827211 | 0.3 | 0.049 | 0.010369 | 39 | cnn3b |
| gadd45bb5 | 4.142172 | 0.566023167 | 0.4 | 0.083 | 0.010523 | 39 | gadd45bb |
| slc38a5b11 | 4.421016 | 0.685818901 | 0.5 | 0.13 | 0.011232 | 39 | slc38a5b |
| mbpb2 | 4.466120 | 0.42523832 | 0.35 | 0.065 | 0.011346 | 39 | mbpb |
| notch211 | 4.673381 | 0.59571084 | 0.5 | 0.124 | 0.011873 | 39 | notch2 |
| calm2a5 | 4.793792 | 0.675223688 | 0.7 | 0.211 | 0.012179 | 39 | calm2a |
| micu21 | 4.826990 | 0.356351556 | 0.25 | 0.036 | 0.012263 | 39 | micu2 |
| per1a | 4.908442 | 0.322752311 | 0.2 | 0.025 | 0.01247 | 39 | per1a |
| si:ch211-153b23.71 | 4.917616 | 0.288040085 | 0.2 | 0.024 | 0.012493 | 39 | si:ch211-153b23.7 |
| hmga22 | 4.920115 | 0.264446773 | 0.15 | 0.014 | 0.0125 | 39 | hmga2 |
| rell1 | 5.124804 | 0.494441095 | 0.3 | 0.05 | 0.01302 | 39 | rell1 |
| mkrn1 | 5.190473 | 0.3568325 | 0.3 | 0.05 | 0.013186 | 39 | mkrn1 |
| chpt1 | 5.983727 | 0.255328951 | 0.15 | 0.015 | 0.015202 | 39 | chpt1 |
| tmem56a | 6.466893 | 0.281988876 | 0.2 | 0.025 | 0.016429 | 39 | tmem56a |
| dhrs92 | 6.480379 | 0.38666695 | 0.25 | 0.037 | 0.016463 | 39 | dhrs9 |
| osbpl9 | 6.630750 | 0.311509185 | 0.2 | 0.025 | 0.016845 | 39 | osbpl9 |
| ctsf4 | 6.685617 | 0.430419406 | 0.25 | 0.038 | 0.016985 | 39 | ctsf |
| nfkbiz1 | 7.487679 | 0.312651756 | 0.2 | 0.025 | 0.019022 | 39 | nfkbiz |
| kctd12.210 | 7.507404 | 0.522619885 | 0.45 | 0.102 | 0.019073 | 39 | kctd12.2 |
| zfpm12 | 7.712888 | 0.262598881 | 0.15 | 0.015 | 0.019595 | 39 | zfpm1 |
| nudt3b | 8.009000 | 0.293761742 | 0.2 | 0.025 | 0.020347 | 39 | nudt3b |

| | | | | | | | |
|---|---|---|---|---|---|---|---|
| sept104 | 8.735699 | 0.491697863 | 0.3 | 0.053 | 0.022193 | 39 | sept10 |
| prkar2aa4 | 9.661913 | 0.595911185 | 0.45 | 0.109 | 0.024546 | 39 | prkar2aa |
| sptssa2 | 9.708805 | 0.510575721 | 0.4 | 0.087 | 0.024665 | 39 | sptssa |
| mprip | 9.776110 | 0.390705545 | 0.2 | 0.026 | 0.024836 | 39 | mprip |
| tmbim1a1 | 9.974181 | 0.31627951 | 0.2 | 0.025 | 0.025339 | 39 | tmbim1a |
| smdt1b5 | 1.012460 | 0.612046973 | 0.55 | 0.154 | 0.025722 | 39 | smdt1b |
| ywhaqb11 | 1.077905 | 0.627227907 | 0.55 | 0.156 | 0.027384 | 39 | ywhaqb |
| mthfs | 1.126419 | 0.262480835 | 0.2 | 0.025 | 0.028617 | 39 | mthfs |
| elovl1b6 | 1.157903 | 0.534931842 | 0.5 | 0.124 | 0.029417 | 39 | elovl1b |
| ier2b18 | 1.173319 | 0.828161654 | 0.65 | 0.238 | 0.029808 | 39 | ier2b |
| pigf | 1.189648 | 0.352293814 | 0.2 | 0.026 | 0.030223 | 39 | pigf |
| si:ch211-74m13.32 | 1.301981 | 0.463559024 | 0.35 | 0.069 | 0.033077 | 39 | si:ch211-74m13.3 |
| mtmr8 | 1.422233 | 0.351974671 | 0.2 | 0.026 | 0.036132 | 39 | mtmr8 |
| tjp1a7 | 1.477304 | 0.447927167 | 0.3 | 0.054 | 0.037531 | 39 | tjp1a |
| lrig2 | 1.520182 | 0.261355012 | 0.15 | 0.016 | 0.03862 | 39 | lrig2 |
| phlda27 | 1.560852 | 0.483460911 | 0.3 | 0.054 | 0.039653 | 39 | phlda2 |
| enc14 | 1.597996 | 0.406237444 | 0.25 | 0.04 | 0.040597 | 39 | enc1 |
| vaspb4 | 1.630155 | 0.373191973 | 0.3 | 0.053 | 0.041414 | 39 | vaspb |
| cnn3a10 | 1.667492 | 0.425894876 | 0.35 | 0.07 | 0.042363 | 39 | cnn3a |
| magi1b | 1.761422 | 0.269813474 | 0.15 | 0.016 | 0.044749 | 39 | magi1b |
| chchd6a | 1.794200 | 0.285469642 | 0.2 | 0.026 | 0.045582 | 39 | chchd6a |
| si:dkey-56m19.56 | 1.874967 | 0.521299416 | 0.3 | 0.057 | 0.047634 | 39 | si:dkey-56m19.5 |
| ccdc28a2 | 1.936031 | 0.473241161 | 0.45 | 0.106 | 0.049185 | 39 | ccdc28a |
| mrpl20 | 2.103652 | 0.456520866 | 0.4 | 0.088 | 0.053443 | 39 | mrpl20 |
| hectd3 | 2.123142 | 0.381394854 | 0.25 | 0.04 | 0.053938 | 39 | hectd3 |
| cox141 | 2.128336 | 0.539971163 | 0.45 | 0.111 | 0.05407 | 39 | cox14 |
| mfsd13a | 2.128414 | 0.261664212 | 0.15 | 0.016 | 0.054072 | 39 | mfsd13a |
| sycn.21 | 2.138533 | 0.334120586 | 0.1 | 0.008 | 0.054329 | 39 | sycn.2 |
| aga | 2.242915 | 0.309197337 | 0.25 | 0.039 | 0.056981 | 39 | aga |
| XLOC-006268 | 2.463515 | 0.283973795 | 0.15 | 0.016 | 0.062586 | 39 | XLOC-006268 |
| serpinh1a1 | 2.540442 | 0.259860714 | 0.15 | 0.016 | 0.06454 | 39 | serpinh1a |
| si:ch211-132b12.74 | 2.855247 | 0.340265638 | 0.3 | 0.054 | 0.072538 | 39 | si:ch211-132b12.7 |
| top2b1 | 2.863147 | 0.328498323 | 0.25 | 0.041 | 0.072738 | 39 | top2b |
| zgc:927447 | 2.883074 | 0.810616592 | 0.75 | 0.328 | 0.073245 | 39 | zgc:92744 |
| nup214 | 2.927268 | 0.257430943 | 0.15 | 0.016 | 0.074367 | 39 | nup214 |

| | | | | | | | |
|---|---|---|---|---|---|---|---|
| cltca1 | 3.026122 | 0.468281135 | 0.4 | 0.091 | 0.076879 | 39 | cltca |
| bhlhe4013 | 3.072147 | 0.543441518 | 0.6 | 0.169 | 0.078048 | 39 | bhlhe40 |
| gps1 | 3.226544 | 0.312470915 | 0.2 | 0.028 | 0.08197 | 39 | gps1 |
| echs12 | 3.422306 | 0.397560519 | 0.4 | 0.089 | 0.086944 | 39 | echs1 |
| cd2762 | 3.496126 | 0.337404479 | 0.2 | 0.028 | 0.088819 | 39 | cd276 |
| psme29 | 3.651304 | 0.509984735 | 0.45 | 0.114 | 0.092761 | 39 | psme2 |
| dusp8a | 3.684735 | 0.366541195 | 0.2 | 0.028 | 0.093611 | 39 | dusp8a |
| foxo4 | 3.727899 | 0.504118085 | 0.25 | 0.043 | 0.094707 | 39 | foxo4 |
| robo24 | 4.061099 | 0.347446419 | 0.3 | 0.055 | 0.103172 | 39 | robo2 |
| rgcc7 | 4.286630 | 0.430532479 | 0.35 | 0.072 | 0.108902 | 39 | rgcc |
| junba15 | 4.597290 | 0.790746578 | 0.8 | 0.321 | 0.116794 | 39 | junba |
| cst14a.24 | 4.811282 | 0.742699777 | 0.65 | 0.237 | 0.122231 | 39 | cst14a.2 |
| tpm313 | 5.024052 | 0.670981013 | 0.75 | 0.286 | 0.127636 | 39 | tpm3 |
| h2afx113 | 5.226324 | 0.699612196 | 0.65 | 0.229 | 0.132775 | 39 | h2afx1 |
| zgc:774391 | 5.289365 | 0.26778757 | 0.15 | 0.017 | 0.134376 | 39 | zgc:77439 |
| ubb5 | 5.850035 | 0.728609972 | 0.75 | 0.327 | 0.14862 | 39 | ubb |
| oxsr1a | 6.494235 | 0.438430787 | 0.25 | 0.044 | 0.164986 | 39 | oxsr1a |
| tegt8 | 6.766811 | 0.662665834 | 0.65 | 0.239 | 0.171911 | 39 | tegt |
| susd63 | 6.921861 | 0.367336748 | 0.3 | 0.058 | 0.17585 | 39 | susd6 |
| timm17a1 | 6.997199 | 0.391612442 | 0.35 | 0.076 | 0.177764 | 39 | timm17a |
| dbi9 | 7.361895 | 0.697822796 | 0.65 | 0.245 | 0.187029 | 39 | dbi |
| fkbp1b2 | 7.512834 | 0.307597183 | 0.15 | 0.018 | 0.190864 | 39 | fkbp1b |
| sept7a3 | 7.815150 | 0.468549638 | 0.35 | 0.079 | 0.198544 | 39 | sept7a |
| LOC5665874 | 8.692607 | 0.490692698 | 0.25 | 0.045 | 0.220836 | 39 | LOC566587 |
| bpnt1 | 8.707478 | 0.321353781 | 0.2 | 0.03 | 0.221213 | 39 | bpnt1 |
| ist12 | 8.821136 | 0.483478388 | 0.4 | 0.097 | 0.224101 | 39 | ist1 |
| trip11 | 8.867197 | 0.256058049 | 0.15 | 0.018 | 0.225271 | 39 | trip11 |
| cox6b110 | 9.228398 | 0.56045952 | 0.45 | 0.123 | 0.234447 | 39 | cox6b1 |
| ralbp12 | 1.014621 | 0.430831179 | 0.3 | 0.062 | 0.257765 | 39 | ralbp1 |
| ralgapa11 | 1.030010 | 0.26325511 | 0.2 | 0.03 | 0.261674 | 39 | ralgapa1 |
| bri31 | 1.132016 | 0.366560075 | 0.3 | 0.061 | 0.287589 | 39 | bri3 |
| g6pd1 | 1.133047 | 0.301628616 | 0.2 | 0.03 | 0.287851 | 39 | g6pd |
| supt4h11 | 1.139150 | 0.40986941 | 0.4 | 0.097 | 0.289401 | 39 | supt4h1 |
| snrkb1 | 1.152030 | 0.267560826 | 0.15 | 0.018 | 0.292673 | 39 | snrkb |
| nat9 | 1.161975 | 0.348545633 | 0.3 | 0.06 | 0.2952 | 39 | nat9 |

| | | | | | | | |
|---|---|---|---|---|---|---|---|
| cyfip12 | 1.170315 | 0.466374574 | 0.3 | 0.063 | 0.297319 | 39 | cyfip1 |
| ywhabl8 | 1.229169 | 0.60553125 | 0.8 | 0.302 | 0.312271 | 39 | ywhabl |
| hexb2 | 1.258383 | 0.252964485 | 0.25 | 0.044 | 0.319692 | 39 | hexb |
| brd7 | 1.260490 | 0.346174583 | 0.3 | 0.061 | 0.320228 | 39 | brd7 |
| plod24 | 1.261663 | 0.328462844 | 0.2 | 0.031 | 0.320526 | 39 | plod2 |
| ppp1caa1 | 1.268379 | 0.548389991 | 0.55 | 0.166 | 0.322232 | 39 | ppp1caa |
| klf6a12 | 1.350206 | 0.664648236 | 0.55 | 0.182 | 0.34302 | 39 | klf6a |
| dynll2b3 | 1.392969 | 0.383881483 | 0.5 | 0.131 | 0.353884 | 39 | dynll2b |
| mpc2 | 1.471721 | 0.392914159 | 0.4 | 0.098 | 0.373891 | 39 | mpc2 |
| man2b1 | 1.488614 | 0.287608479 | 0.2 | 0.031 | 0.378182 | 39 | man2b1 |
| myl9a7 | 1.498542 | 0.422461575 | 0.25 | 0.046 | 0.380705 | 39 | myl9a |
| zdhhc13 | 1.529760 | 0.382323182 | 0.25 | 0.046 | 0.388636 | 39 | zdhhc13 |
| rsrp14 | 1.557701 | 0.530008121 | 0.45 | 0.124 | 0.395734 | 39 | rsrp1 |
| mrpl15 | 1.650954 | 0.332068618 | 0.25 | 0.045 | 0.419425 | 39 | mrpl15 |
| arpc1a5 | 1.654095 | 0.512052008 | 0.5 | 0.143 | 0.420223 | 39 | arpc1a |
| ctso | 1.739972 | 0.260532453 | 0.15 | 0.019 | 0.44204 | 39 | ctso |
| dazap21 | 1.772314 | 0.377737332 | 0.4 | 0.097 | 0.450256 | 39 | dazap2 |
| tax1bp35 | 1.819727 | 0.477599845 | 0.4 | 0.102 | 0.462302 | 39 | tax1bp3 |
| kansl1b | 1.840267 | 0.299650831 | 0.25 | 0.045 | 0.46752 | 39 | kansl1b |
| cox8b7 | 1.842790 | 0.445341166 | 0.4 | 0.103 | 0.468161 | 39 | cox8b |
| rhocb3 | 1.899162 | 0.499327242 | 0.4 | 0.104 | 0.482482 | 39 | rhocb |
| ctr9 | 1.901644 | 0.296204035 | 0.2 | 0.032 | 0.483113 | 39 | ctr9 |
| pik3cb | 1.925930 | 0.272355573 | 0.15 | 0.019 | 0.489283 | 39 | pik3cb |
| usp8 | 1.959657 | 0.404102631 | 0.2 | 0.032 | 0.497851 | 39 | usp8 |
| cbr1 | 2.041673 | 0.296467975 | 0.15 | 0.019 | 0.518687 | 39 | cbr1 |
| arpin | 2.042764 | 0.275294887 | 0.25 | 0.046 | 0.518964 | 39 | arpin |
| relch | 2.058779 | 0.267639069 | 0.15 | 0.019 | 0.523033 | 39 | relch |
| cldn7b8 | 2.135598 | 0.516126752 | 0.5 | 0.135 | 0.542549 | 39 | cldn7b |
| cry21 | 2.189220 | 0.380010756 | 0.35 | 0.081 | 0.556171 | 39 | cry2 |
| xpc1 | 2.223130 | 0.295823339 | 0.2 | 0.032 | 0.564786 | 39 | xpc |
| ppiab8 | 2.264939 | 0.479861235 | 0.95 | 0.676 | 0.575408 | 39 | ppiab |
| rca2.11 | 2.447398 | 0.450835235 | 0.35 | 0.084 | 0.621762 | 39 | rca2.1 |
| cyp2ad6 | 2.458513 | 0.283621165 | 0.15 | 0.019 | 0.624585 | 39 | cyp2ad6 |
| NDUFB14 | 2.462722 | 0.577017771 | 0.6 | 0.213 | 0.625655 | 39 | NDUFB1 |
| nt5c2b4 | 2.483646 | 0.368202021 | 0.25 | 0.048 | 0.63097 | 39 | nt5c2b |

| | | | | | | | |
|---|---|---|---|---|---|---|---|
| rnf128a3 | 2.585191 | 0.402556854 | 0.3 | 0.062 | 0.656768 | 39 | rnf128a |
| cavin2b9 | 2.600098 | 0.354872402 | 0.35 | 0.08 | 0.660555 | 39 | cavin2b |
| u2surp4 | 2.689293 | 0.449246622 | 0.4 | 0.105 | 0.683215 | 39 | u2surp |
| rnf167 | 2.775224 | 0.268778211 | 0.15 | 0.02 | 0.705046 | 39 | rnf167 |
| rhoab14 | 2.836112 | 0.581848939 | 0.6 | 0.209 | 0.720514 | 39 | rhoab |
| sdhc2 | 2.901136 | 0.484616139 | 0.5 | 0.149 | 0.737034 | 39 | sdhc |
| oip5-as12 | 2.985542 | 0.304944087 | 0.2 | 0.033 | 0.758477 | 39 | oip5-as1 |
| hspa4b | 2.985719 | 0.324563501 | 0.3 | 0.064 | 0.758522 | 39 | hspa4b |
| srpra1 | 2.998612 | 0.311024068 | 0.3 | 0.063 | 0.761798 | 39 | srpra |
| calua4 | 3.025518 | 0.338488845 | 0.2 | 0.033 | 0.768633 | 39 | calua |
| sdc415 | 3.028311 | 0.638624686 | 0.7 | 0.256 | 0.769342 | 39 | sdc4 |
| ypel51 | 3.045043 | 0.398859185 | 0.25 | 0.049 | 0.773593 | 39 | ypel5 |
| col4a3bpb | 3.193328 | 0.25619468 | 0.2 | 0.032 | 0.811265 | 39 | col4a3bpb |
| dctn5 | 3.242605 | 0.37016802 | 0.25 | 0.049 | 0.823784 | 39 | dctn5 |
| rab1ba | 3.467327 | 0.365691362 | 0.25 | 0.049 | 0.880875 | 39 | rab1ba |
| hmgb3a8 | 3.631179 | 0.403282998 | 0.35 | 0.084 | 0.922501 | 39 | hmgb3a |
| gpx4b4 | 3.643123 | 0.492489379 | 0.65 | 0.222 | 0.925535 | 39 | gpx4b |
| ctdsp26 | 3.834536 | 0.412376503 | 0.35 | 0.087 | 0.974164 | 39 | ctdsp2 |
| ppt17 | 3.836333 | 0.565305276 | 0.55 | 0.183 | 0.97462 | 39 | ppt1 |
| pmp22b8 | 3.854574 | 0.297680363 | 0.25 | 0.048 | 0.979255 | 39 | pmp22b |
| pacsin21 | 3.861732 | 0.32946237 | 0.25 | 0.048 | 0.981073 | 39 | pacsin2 |
| foxk1 | 3.960477 | 0.322150907 | 0.2 | 0.033 | 1 | 39 | foxk1 |
| lxn7 | 4.019256 | 0.4812744 | 0.55 | 0.169 | 1 | 39 | lxn |
| zgc:16273018 | 4.092799 | 0.673843589 | 0.85 | 0.404 | 1 | 39 | zgc:162730 |
| fkbp32 | 4.140444 | 0.508632811 | 0.55 | 0.179 | 1 | 39 | fkbp3 |
| ccdc804 | 4.176260 | 0.328243994 | 0.2 | 0.034 | 1 | 39 | ccdc80 |
| tmem176l.3a2 | 4.400629 | 0.282716937 | 0.25 | 0.049 | 1 | 39 | tmem176l.3a |
| si:ch73-269m14.2 | 4.445290 | 0.304985014 | 0.25 | 0.049 | 1 | 39 | si:ch73-269m14.2 |
| nnt | 4.479503 | 0.25649335 | 0.25 | 0.048 | 1 | 39 | nnt |
| si:ch211-191j22.33 | 4.526039 | 0.386449994 | 0.45 | 0.124 | 1 | 39 | si:ch211-191j22.3 |
| zgc:1748635 | 4.606468 | 0.491219064 | 0.4 | 0.112 | 1 | 39 | zgc:174863 |
| ifrd11 | 4.841550 | 0.363735836 | 0.4 | 0.105 | 1 | 39 | ifrd1 |
| actb214 | 5.141431 | 0.400368417 | 1 | 0.868 | 1 | 39 | actb2 |
| si:dkey-253i9.4 | 5.258574 | 0.299224725 | 0.15 | 0.021 | 1 | 39 | si:dkey-253i9.4 |
| slmapa | 5.376672 | 0.258899017 | 0.2 | 0.034 | 1 | 39 | slmapa |

| | | | | | | | |
|---|---|---|---|---|---|---|---|
| tjp2b4 | 5.401626 | 0.285765471 | 0.3 | 0.065 | 1 | 39 | tjp2b |
| fam172a | 5.559768 | 0.420940687 | 0.3 | 0.068 | 1 | 39 | fam172a |
| uqcrh5 | 5.628635 | 0.521798375 | 0.7 | 0.261 | 1 | 39 | uqcrh |
| ndufa22 | 5.692027 | 0.472731264 | 0.5 | 0.157 | 1 | 39 | ndufa2 |
| zfp36l23 | 6.142726 | 0.379837866 | 0.4 | 0.103 | 1 | 39 | zfp36l2 |
| cav19 | 6.338871 | 0.367051362 | 0.25 | 0.05 | 1 | 39 | cav1 |
| selenos | 6.365402 | 0.270026238 | 0.25 | 0.049 | 1 | 39 | selenos |
| klf2b3 | 6.376442 | 0.380853395 | 0.15 | 0.021 | 1 | 39 | klf2b |
| tspo1 | 6.494699 | 0.312518199 | 0.3 | 0.067 | 1 | 39 | tspo |
| ccdc85b9 | 7.047201 | 0.366058498 | 0.25 | 0.052 | 1 | 39 | ccdc85b |
| ucp215 | 7.452848 | 0.630743475 | 0.85 | 0.416 | 1 | 39 | ucp2 |
| cpb11 | 7.838451 | 0.360594895 | 0.1 | 0.01 | 1 | 39 | cpb1 |
| tmc6b2 | 8.023579 | 0.397781801 | 0.2 | 0.037 | 1 | 39 | tmc6b |
| tspan144 | 8.396824 | 0.445519409 | 0.45 | 0.133 | 1 | 39 | tspan14 |
| itm2bb12 | 8.482676 | 0.480397498 | 0.7 | 0.244 | 1 | 39 | itm2bb |
| bsdc1 | 8.542650 | 0.305218757 | 0.3 | 0.069 | 1 | 39 | bsdc1 |
| hbs1l | 8.951791 | 0.328734387 | 0.2 | 0.036 | 1 | 39 | hbs1l |
| pmp22a8 | 9.276132 | 0.437801024 | 0.55 | 0.168 | 1 | 39 | pmp22a |
| lgals3a | 9.279800 | 0.401717141 | 0.4 | 0.111 | 1 | 39 | lgals3a |
| flot2a9 | 9.282285 | 0.45500682 | 0.3 | 0.075 | 1 | 39 | flot2a |
| rsu13 | 9.320522 | 0.386931246 | 0.25 | 0.054 | 1 | 39 | rsu1 |
| cdc42ep52 | 9.720918 | 0.283126213 | 0.2 | 0.036 | 1 | 39 | cdc42ep5 |
| psap5 | 9.743177 | 0.54601302 | 0.75 | 0.289 | 1 | 39 | psap |
| cd1513 | 0.0001 | 0.282587808 | 0.15 | 0.022 | 1 | 39 | cd151 |
| zfp36l1b10 | 0.0001 | 0.570322246 | 0.45 | 0.141 | 1 | 39 | zfp36l1b |
| wdr111 | 0.0001 | 0.434181225 | 0.5 | 0.157 | 1 | 39 | wdr1 |
| sptlc1 | 0.0001 | 0.260526302 | 0.2 | 0.036 | 1 | 39 | sptlc1 |
| irf25 | 0.0001 | 0.496802911 | 0.3 | 0.077 | 1 | 39 | irf2 |
| pafah1b1b | 0.0001 | 0.443598083 | 0.3 | 0.072 | 1 | 39 | pafah1b1b |
| clk4a | 0.0001 | 0.451296103 | 0.35 | 0.093 | 1 | 39 | clk4a |
| cnpy11 | 0.0001 | 0.337695458 | 0.4 | 0.109 | 1 | 39 | cnpy1 |
| ndufs52 | 0.0001 | 0.471323053 | 0.5 | 0.166 | 1 | 39 | ndufs5 |
| COX5B7 | 0.0001 | 0.593519238 | 0.8 | 0.378 | 1 | 39 | COX5B |
| si:dkey-188i13.106 | 0.0001 | 0.441090337 | 0.4 | 0.115 | 1 | 39 | si:dkey-188i13.10 |
| sdf43 | 0.0001 | 0.297353483 | 0.25 | 0.053 | 1 | 39 | sdf4 |

| | | | | | | | |
|---|---|---|---|---|---|---|---|
| ndufa71 | 0.0001 | 0.345976433 | 0.4 | 0.111 | 1 | 39 | ndufa7 |
| lpp1 | 0.0001 | 0.252272345 | 0.2 | 0.036 | 1 | 39 | lpp |
| pef12 | 0.0001 | 0.380700185 | 0.3 | 0.074 | 1 | 39 | pef1 |
| dedd17 | 0.0001 | 0.445830589 | 0.5 | 0.158 | 1 | 39 | dedd1 |
| ubqln4 | 0.0001 | 0.414490743 | 0.25 | 0.055 | 1 | 39 | ubqln4 |
| CR383676.123 | 0.0001 | 0.663208818 | 1 | 0.718 | 1 | 39 | CR383676.1 |
| si:dkey-187e18.12 | 0.0001 | 0.344568466 | 0.3 | 0.072 | 1 | 39 | si:dkey-187e18.1 |
| ckba6 | 0.0001 | 0.423871867 | 0.3 | 0.075 | 1 | 39 | ckba |
| npc23 | 0.0001 | 0.588758755 | 0.8 | 0.344 | 1 | 39 | npc2 |
| map3k14a2 | 0.0001 | 0.27621346 | 0.2 | 0.037 | 1 | 39 | map3k14a |
| ndufb84 | 0.0002 | 0.553383141 | 0.65 | 0.267 | 1 | 39 | ndufb8 |
| pigk | 0.0002 | 0.25926163 | 0.1 | 0.011 | 1 | 39 | pigk |
| s100a10b15 | 0.0002 | 0.550015272 | 0.75 | 0.287 | 1 | 39 | s100a10b |
| coq9 | 0.0002 | 0.351898056 | 0.2 | 0.039 | 1 | 39 | coq9 |
| aldh6a11 | 0.0002 | 0.323815108 | 0.35 | 0.094 | 1 | 39 | aldh6a1 |
| kpna1 | 0.0002 | 0.299445513 | 0.15 | 0.023 | 1 | 39 | kpna1 |
| marcksl1b13 | 0.0002 | 0.510590881 | 0.45 | 0.143 | 1 | 39 | marcksl1b |
| pim113 | 0.0002 | 0.520761634 | 0.5 | 0.17 | 1 | 39 | pim1 |
| dynlrb13 | 0.0002 | 0.450019192 | 0.6 | 0.214 | 1 | 39 | dynlrb1 |
| btg214 | 0.0002 | 0.711846741 | 0.65 | 0.351 | 1 | 39 | btg2 |
| s1pr23 | 0.0002 | 0.258456401 | 0.2 | 0.038 | 1 | 39 | s1pr2 |
| mef2aa5 | 0.0002 | 0.367229661 | 0.3 | 0.075 | 1 | 39 | mef2aa |
| pdcd6ip1 | 0.0002 | 0.289667789 | 0.25 | 0.055 | 1 | 39 | pdcd6ip |
| hdlbpa11 | 0.0002 | 0.515333976 | 0.5 | 0.176 | 1 | 39 | hdlbpa |
| txndc52 | 0.0002 | 0.329047071 | 0.3 | 0.073 | 1 | 39 | txndc5 |
| si:dkey-16l2.204 | 0.0002 | 0.260278388 | 0.15 | 0.023 | 1 | 39 | si:dkey-16l2.20 |
| si:ch73-46j18.512 | 0.0002 | 0.510614669 | 0.6 | 0.21 | 1 | 39 | si:ch73-46j18.5 |
| ela2l1 | 0.0002 | 0.43186635 | 0.1 | 0.011 | 1 | 39 | ela2l |
| XLOC-0438571 | 0.0002 | 0.31756644 | 0.3 | 0.075 | 1 | 39 | XLOC-043857 |
| ralbb | 0.0002 | 0.297049076 | 0.2 | 0.039 | 1 | 39 | ralbb |
| maff5 | 0.0002 | 0.544006108 | 0.3 | 0.082 | 1 | 39 | maff |
| si:ch211-166a6.55 | 0.0002 | 0.29108528 | 0.2 | 0.039 | 1 | 39 | si:ch211-166a6.5 |
| becn1 | 0.0002 | 0.373123747 | 0.15 | 0.024 | 1 | 39 | becn1 |
| pora | 0.0002 | 0.275740687 | 0.2 | 0.039 | 1 | 39 | pora |
| slc44a22 | 0.0002 | 0.301324358 | 0.25 | 0.056 | 1 | 39 | slc44a2 |

| | | | | | | | |
|---|---|---|---|---|---|---|---|
| psmb42 | 0.0002 | 0.439007068 | 0.55 | 0.194 | 1 | 39 | psmb4 |
| cd1644 | 0.0002 | 0.345809227 | 0.3 | 0.077 | 1 | 39 | cd164 |
| ndfip1l5 | 0.0002 | 0.36730431 | 0.3 | 0.076 | 1 | 39 | ndfip1l |
| mgst3a3 | 0.0003 | 0.292348308 | 0.3 | 0.075 | 1 | 39 | mgst3a |
| glg1a | 0.0003 | 0.320745164 | 0.2 | 0.039 | 1 | 39 | glg1a |
| nr1d2b1 | 0.0003 | 0.306534748 | 0.2 | 0.04 | 1 | 39 | nr1d2b |
| chmp1a1 | 0.0003 | 0.350966496 | 0.25 | 0.058 | 1 | 39 | chmp1a |
| eif3ja1 | 0.0003 | 0.466008707 | 0.55 | 0.198 | 1 | 39 | eif3ja |
| lamc14 | 0.0003 | 0.415122273 | 0.25 | 0.059 | 1 | 39 | lamc1 |
| arrb2b8 | 0.0003 | 0.3569525 | 0.35 | 0.098 | 1 | 39 | arrb2b |
| hnrnpul1l | 0.0003 | 0.300200684 | 0.25 | 0.057 | 1 | 39 | hnrnpul1l |
| calm3b4 | 0.0003 | 0.491581666 | 0.7 | 0.28 | 1 | 39 | calm3b |
| si:ch211-286b5.510 | 0.0003 | 0.474881276 | 0.45 | 0.153 | 1 | 39 | si:ch211-286b5.5 |
| GCA8 | 0.0003 | 0.473129017 | 0.55 | 0.202 | 1 | 39 | GCA |
| chmp2a5 | 0.0003 | 0.402243621 | 0.45 | 0.145 | 1 | 39 | chmp2a |
| naga4 | 0.0003 | 0.28757225 | 0.25 | 0.057 | 1 | 39 | naga |
| rrbp1a4 | 0.0003 | 0.412070425 | 0.25 | 0.06 | 1 | 39 | rrbp1a |
| reep513 | 0.0003 | 0.447877672 | 0.5 | 0.175 | 1 | 39 | reep5 |
| gsto21 | 0.0003 | 0.361695244 | 0.35 | 0.098 | 1 | 39 | gsto2 |
| dpm1 | 0.0003 | 0.266486431 | 0.2 | 0.04 | 1 | 39 | dpm1 |
| atp6v1e1a2 | 0.0003 | 0.321517526 | 0.25 | 0.059 | 1 | 39 | atp6v1e1a |
| tmem2585 | 0.0003 | 0.562375208 | 0.7 | 0.307 | 1 | 39 | tmem258 |
| gchfr3 | 0.0003 | 0.266602022 | 0.2 | 0.04 | 1 | 39 | gchfr |
| lysmd31 | 0.0003 | 0.319036816 | 0.2 | 0.041 | 1 | 39 | lysmd3 |
| ctsa3 | 0.0003 | 0.327522984 | 0.35 | 0.098 | 1 | 39 | ctsa |
| rtn32 | 0.0003 | 0.398406488 | 0.3 | 0.08 | 1 | 39 | rtn3 |
| per35 | 0.0003 | 0.362834893 | 0.3 | 0.079 | 1 | 39 | per3 |
| arpc5a10 | 0.0003 | 0.449878478 | 0.4 | 0.127 | 1 | 39 | arpc5a |
| ppp1r10 | 0.0003 | 0.351773321 | 0.25 | 0.06 | 1 | 39 | ppp1r10 |
| rpz41 | 0.0003 | 0.310613605 | 0.15 | 0.025 | 1 | 39 | rpz4 |
| hp1bp38 | 0.0003 | 0.364935868 | 0.4 | 0.122 | 1 | 39 | hp1bp3 |
| tp53inp14 | 0.0003 | 0.401764566 | 0.25 | 0.061 | 1 | 39 | tp53inp1 |
| atf4b15 | 0.0003 | 0.543882713 | 0.65 | 0.279 | 1 | 39 | atf4b |
| scaf11 | 0.0003 | 0.273209712 | 0.2 | 0.04 | 1 | 39 | scaf1 |
| elf18 | 0.0004 | 0.460221279 | 0.35 | 0.105 | 1 | 39 | elf1 |

| | | | | | | | |
|---|---|---|---|---|---|---|---|
| cap112 | 0.0004 | 0.465440504 | 0.65 | 0.247 | 1 | 39 | cap1 |
| dhrs13l16 | 0.0004 | 0.266452406 | 0.2 | 0.041 | 1 | 39 | dhrs13l1 |
| stk35l2 | 0.0004 | 0.328721679 | 0.2 | 0.041 | 1 | 39 | stk35l |
| aip1 | 0.0004 | 0.411064161 | 0.35 | 0.104 | 1 | 39 | aip |
| psmc54 | 0.0004 | 0.400702692 | 0.35 | 0.104 | 1 | 39 | psmc5 |
| srp541 | 0.0004 | 0.296370848 | 0.25 | 0.059 | 1 | 39 | srp54 |
| wdr26b | 0.0004 | 0.358674843 | 0.2 | 0.042 | 1 | 39 | wdr26b |
| usp362 | 0.0004 | 0.370668899 | 0.3 | 0.083 | 1 | 39 | usp36 |
| galnt71 | 0.0004 | 0.265940669 | 0.2 | 0.041 | 1 | 39 | galnt7 |
| ywhaz7 | 0.0004 | 0.523321785 | 0.7 | 0.306 | 1 | 39 | ywhaz |
| ptbp1b3 | 0.0004 | 0.264854581 | 0.2 | 0.041 | 1 | 39 | ptbp1b |
| vps282 | 0.0004 | 0.352743099 | 0.35 | 0.102 | 1 | 39 | vps28 |
| tnrc6a2 | 0.0004 | 0.36527867 | 0.3 | 0.081 | 1 | 39 | tnrc6a |
| cry1aa1 | 0.0004 | 0.324477269 | 0.25 | 0.06 | 1 | 39 | cry1aa |
| laptm4a5 | 0.0004 | 0.345411579 | 0.4 | 0.123 | 1 | 39 | laptm4a |
| fosl29 | 0.0005 | 0.605408493 | 0.35 | 0.115 | 1 | 39 | fosl2 |
| ap2a11 | 0.0005 | 0.427850468 | 0.25 | 0.063 | 1 | 39 | ap2a1 |
| mt-nd56 | 0.0005 | 0.443839796 | 0.4 | 0.131 | 1 | 39 | mt-nd5 |
| mir21-16 | 0.0005 | 0.474241237 | 0.35 | 0.109 | 1 | 39 | mir21-1 |
| ppp3ccb5 | 0.0005 | 0.373298017 | 0.4 | 0.126 | 1 | 39 | ppp3ccb |
| srsf5b | 0.0005 | 0.362207296 | 0.2 | 0.043 | 1 | 39 | srsf5b |
| sptan14 | 0.0005 | 0.258095417 | 0.25 | 0.059 | 1 | 39 | sptan1 |
| timm8a | 0.0005 | 0.27155446 | 0.25 | 0.06 | 1 | 39 | timm8a |
| snx123 | 0.0005 | 0.409649879 | 0.3 | 0.084 | 1 | 39 | snx12 |
| si:ch73-376l24.32 | 0.0005 | 0.351446553 | 0.4 | 0.124 | 1 | 39 | si:ch73-376l24.3 |
| slc6a6b7 | 0.0005 | 0.288549526 | 0.35 | 0.098 | 1 | 39 | slc6a6b |
| zgc:162396 | 0.0005 | 0.26319041 | 0.15 | 0.026 | 1 | 39 | zgc:162396 |
| lsm14ab1 | 0.0005 | 0.316756913 | 0.3 | 0.081 | 1 | 39 | lsm14ab |
| zgc:1121601 | 0.0005 | 0.419386233 | 0.1 | 0.013 | 1 | 39 | zgc:112160 |
| p4hb3 | 0.0006 | 0.48927635 | 0.55 | 0.214 | 1 | 39 | p4hb |
| snrnp2001 | 0.0006 | 0.319034149 | 0.2 | 0.043 | 1 | 39 | snrnp200 |
| foxo1a3 | 0.0006 | 0.252325134 | 0.2 | 0.042 | 1 | 39 | foxo1a |
| itgav4 | 0.0006 | 0.371177825 | 0.25 | 0.064 | 1 | 39 | itgav |
| si:cabz01101003.1 | 0.0006 | 0.252054648 | 0.15 | 0.026 | 1 | 39 | si:cabz01101003.1 |
| si:dkey-177p2.64 | 0.0006 | 0.365227716 | 0.3 | 0.085 | 1 | 39 | si:dkey-177p2.6 |

| | | | | | | | |
|---|---|---|---|---|---|---|---|
| gata64 | 0.0006 | 0.286048374 | 0.15 | 0.027 | 1 | 39 | gata6 |
| tmem50a3 | 0.0007 | 0.37015259 | 0.45 | 0.152 | 1 | 39 | tmem50a |
| ppp1r14bb3 | 0.0007 | 0.260085002 | 0.2 | 0.043 | 1 | 39 | ppp1r14bb |
| syncripl | 0.0007 | 0.354735067 | 0.3 | 0.085 | 1 | 39 | syncripl |
| asah1b3 | 0.0007 | 0.313241854 | 0.25 | 0.063 | 1 | 39 | asah1b |
| mt-co118 | 0.0007 | 0.465719514 | 0.95 | 0.641 | 1 | 39 | mt-co1 |
| cib11 | 0.0007 | 0.34280897 | 0.3 | 0.084 | 1 | 39 | cib1 |
| cyp2ad34 | 0.0007 | 0.337072102 | 0.25 | 0.063 | 1 | 39 | cyp2ad3 |
| slc39a73 | 0.0007 | 0.274232939 | 0.2 | 0.044 | 1 | 39 | slc39a7 |
| ppp3r1b1 | 0.0008 | 0.317850306 | 0.25 | 0.065 | 1 | 39 | ppp3r1b |
| jund10 | 0.0008 | 0.481533975 | 0.3 | 0.091 | 1 | 39 | jund |
| ndufv21 | 0.0008 | 0.256284566 | 0.3 | 0.082 | 1 | 39 | ndufv2 |
| lgals9l36 | 0.0008 | 0.330937937 | 0.3 | 0.085 | 1 | 39 | lgals9l3 |
| ankrd228 | 0.0008 | 0.297482583 | 0.4 | 0.123 | 1 | 39 | ankrd22 |
| hnrnpm | 0.0008 | 0.338427921 | 0.3 | 0.085 | 1 | 39 | hnrnpm |
| tsc22d11 | 0.0008 | 0.255280858 | 0.2 | 0.044 | 1 | 39 | tsc22d1 |
| calm2b12 | 0.0008 | 0.486724422 | 0.6 | 0.248 | 1 | 39 | calm2b |
| rnaset21 | 0.0008 | 0.304359749 | 0.2 | 0.045 | 1 | 39 | rnaset2 |
| slirp1 | 0.0009 | 0.329971494 | 0.4 | 0.13 | 1 | 39 | slirp |
| rab5aa2 | 0.0009 | 0.308886917 | 0.3 | 0.085 | 1 | 39 | rab5aa |
| ccnd15 | 0.0009 | 0.275867588 | 0.2 | 0.045 | 1 | 39 | ccnd1 |
| lrpap14 | 0.0009 | 0.306343733 | 0.25 | 0.065 | 1 | 39 | lrpap1 |
| ilk2 | 0.0009 | 0.320774326 | 0.3 | 0.086 | 1 | 39 | ilk |
| abcf12 | 0.0009 | 0.375816605 | 0.35 | 0.111 | 1 | 39 | abcf1 |
| cnot12 | 0.001 | 0.272448666 | 0.25 | 0.064 | 1 | 39 | cnot1 |
| wu:fb16f036 | 0.001 | 0.439412843 | 0.3 | 0.091 | 1 | 39 | wu:fb16f03 |
| si:dkey-28b4.710 | 0.001 | 0.365478321 | 0.35 | 0.111 | 1 | 39 | si:dkey-28b4.7 |
| lmnb22 | 0.001 | 0.320164752 | 0.25 | 0.066 | 1 | 39 | lmnb2 |
| midn12 | 0.001 | 0.508979818 | 0.4 | 0.147 | 1 | 39 | midn |
| st13 | 0.001 | 0.287076167 | 0.4 | 0.129 | 1 | 39 | st13 |
| si:ch73-335l21.411 | 0.001 | 0.568595042 | 0.4 | 0.143 | 1 | 39 | si:ch73-335l21.4 |
| ube2h | 0.001 | 0.289093106 | 0.25 | 0.065 | 1 | 39 | ube2h |
| sik2b3 | 0.001 | 0.278605928 | 0.2 | 0.046 | 1 | 39 | sik2b |
| ubap1 | 0.0011 | 0.264285607 | 0.15 | 0.028 | 1 | 39 | ubap1 |
| atp5pf7 | 0.0011 | 0.524875083 | 0.65 | 0.322 | 1 | 39 | atp5pf |

| | | | | | | | |
|---|---|---|---|---|---|---|---|
| sik18 | 0.0011 | 0.411360371 | 0.35 | 0.112 | 1 | 39 | sik1 |
| mt-nd416 | 0.0011 | 0.504636674 | 0.85 | 0.449 | 1 | 39 | mt-nd4 |
| ctnnb18 | 0.0011 | 0.386942171 | 0.45 | 0.16 | 1 | 39 | ctnnb1 |
| si:dkey-79d12.55 | 0.0012 | 0.429000813 | 0.25 | 0.069 | 1 | 39 | si:dkey-79d12.5 |
| mgat1a | 0.0012 | 0.277769386 | 0.2 | 0.045 | 1 | 39 | mgat1a |
| sptlc2b1 | 0.0012 | 0.315232348 | 0.2 | 0.047 | 1 | 39 | sptlc2b |
| csnk1db | 0.0013 | 0.412797885 | 0.4 | 0.139 | 1 | 39 | csnk1db |
| mif4 | 0.0013 | 0.345448014 | 0.55 | 0.209 | 1 | 39 | mif |
| hadh | 0.0013 | 0.314659545 | 0.25 | 0.067 | 1 | 39 | hadh |
| prss59.11 | 0.0013 | 0.346911216 | 0.1 | 0.014 | 1 | 39 | prss59.1 |
| ndfip14 | 0.0014 | 0.421859882 | 0.45 | 0.173 | 1 | 39 | ndfip1 |
| slc39a3 | 0.0014 | 0.254556209 | 0.1 | 0.015 | 1 | 39 | slc39a3 |
| ptenb1 | 0.0014 | 0.29570602 | 0.3 | 0.088 | 1 | 39 | ptenb |
| dusp59 | 0.0014 | 0.420217407 | 0.25 | 0.069 | 1 | 39 | dusp5 |
| ctsh4 | 0.0015 | 0.310232255 | 0.25 | 0.068 | 1 | 39 | ctsh |
| ndufs42 | 0.0015 | 0.291696861 | 0.35 | 0.112 | 1 | 39 | ndufs4 |
| cry1ab | 0.0016 | 0.298062383 | 0.2 | 0.048 | 1 | 39 | cry1ab |
| arhgap42a | 0.0016 | 0.285859681 | 0.2 | 0.047 | 1 | 39 | arhgap42a |
| stx12 | 0.0016 | 0.35816014 | 0.25 | 0.069 | 1 | 39 | stx12 |
| tmem30aa3 | 0.0016 | 0.348365149 | 0.25 | 0.07 | 1 | 39 | tmem30aa |
| ipo73 | 0.0016 | 0.372167616 | 0.35 | 0.117 | 1 | 39 | ipo7 |
| pik3r27 | 0.0016 | 0.296386556 | 0.25 | 0.068 | 1 | 39 | pik3r2 |
| ppm1aa2 | 0.0016 | 0.298442395 | 0.25 | 0.069 | 1 | 39 | ppm1aa |
| nmt1a | 0.0017 | 0.271619567 | 0.25 | 0.067 | 1 | 39 | nmt1a |
| si:ch1073-325m22.2( | 0.0017 | 0.476137126 | 0.55 | 0.251 | 1 | 39 | si:ch1073-325m22.2 |
| nfe2l2a4 | 0.0017 | 0.309923868 | 0.25 | 0.07 | 1 | 39 | nfe2l2a |
| b2m13 | 0.0018 | 0.485758149 | 0.65 | 0.3 | 1 | 39 | b2m |
| si:ch211-105c13.37 | 0.0018 | 0.327994629 | 0.25 | 0.068 | 1 | 39 | si:ch211-105c13.3 |
| pomp5 | 0.0018 | 0.423147154 | 0.6 | 0.261 | 1 | 39 | pomp |
| trim25 | 0.0018 | 0.340394429 | 0.2 | 0.049 | 1 | 39 | trim25 |
| zgc:1538679 | 0.0018 | 0.502456642 | 0.8 | 0.467 | 1 | 39 | zgc:153867 |
| arpc4l10 | 0.0019 | 0.38712511 | 0.55 | 0.221 | 1 | 39 | arpc4l |
| stat1a6 | 0.0019 | 0.447184557 | 0.35 | 0.125 | 1 | 39 | stat1a |
| erp441 | 0.0019 | 0.286081256 | 0.3 | 0.092 | 1 | 39 | erp44 |
| ip6k2a1 | 0.002 | 0.286628788 | 0.25 | 0.07 | 1 | 39 | ip6k2a |

| | | | | | | | |
|---|---|---|---|---|---|---|---|
| si:ch211-212k18.76 | 0.002 | 0.427473262 | 0.4 | 0.148 | 1 | 39 | si:ch211-212k18.7 |
| zgc:1132085 | 0.0021 | 0.304991048 | 0.2 | 0.05 | 1 | 39 | zgc:113208 |
| fam32a2 | 0.0021 | 0.42530384 | 0.65 | 0.292 | 1 | 39 | fam32a |
| NC-002333.414 | 0.0021 | 0.473891488 | 0.95 | 0.61 | 1 | 39 | NC-002333.4 |
| fkbp1aa12 | 0.0021 | 0.49340044 | 0.8 | 0.416 | 1 | 39 | fkbp1aa |
| rdh10a5 | 0.0021 | 0.332335639 | 0.2 | 0.049 | 1 | 39 | rdh10a |
| zgc:1231052 | 0.0022 | 0.259762662 | 0.2 | 0.049 | 1 | 39 | zgc:123105 |
| f11r.18 | 0.0022 | 0.40359748 | 0.45 | 0.166 | 1 | 39 | f11r.1 |
| cdipt | 0.0023 | 0.299048575 | 0.25 | 0.071 | 1 | 39 | cdipt |
| gmnn7 | 0.0024 | 0.421355697 | 0.65 | 0.257 | 1 | 39 | gmnn |
| smg7 | 0.0024 | 0.321987953 | 0.2 | 0.051 | 1 | 39 | smg7 |
| mt-nd116 | 0.0025 | 0.506096402 | 0.75 | 0.399 | 1 | 39 | mt-nd1 |
| mapk12a | 0.0025 | 0.284200296 | 0.1 | 0.016 | 1 | 39 | mapk12a |
| ap1s3b2 | 0.0025 | 0.268528607 | 0.25 | 0.068 | 1 | 39 | ap1s3b |
| crip116 | 0.0025 | 0.348354079 | 0.7 | 0.286 | 1 | 39 | crip1 |
| pdia311 | 0.0025 | 0.483733791 | 0.65 | 0.315 | 1 | 39 | pdia3 |
| yipf3 | 0.0027 | 0.259552721 | 0.2 | 0.05 | 1 | 39 | yipf3 |
| tob1a7 | 0.0027 | 0.340721766 | 0.4 | 0.146 | 1 | 39 | tob1a |
| ndufv11 | 0.0027 | 0.288104466 | 0.25 | 0.073 | 1 | 39 | ndufv1 |
| mrps35 | 0.0028 | 0.27514217 | 0.2 | 0.051 | 1 | 39 | mrps35 |
| mt-co217 | 0.0028 | 0.426451975 | 0.85 | 0.672 | 1 | 39 | mt-co2 |
| rnf104 | 0.0028 | 0.413351944 | 0.35 | 0.129 | 1 | 39 | rnf10 |
| itpa | 0.0028 | 0.25318225 | 0.15 | 0.032 | 1 | 39 | itpa |
| tspan9b6 | 0.0029 | 0.261964635 | 0.2 | 0.051 | 1 | 39 | tspan9b |
| CR847566.16 | 0.0029 | 0.315651344 | 0.2 | 0.052 | 1 | 39 | CR847566.1 |
| msna13 | 0.003 | 0.359060159 | 0.65 | 0.267 | 1 | 39 | msna |
| snap23.11 | 0.0031 | 0.305690346 | 0.25 | 0.074 | 1 | 39 | snap23.1 |
| si:ch211-196l7.417 | 0.0032 | 0.434627566 | 0.7 | 0.332 | 1 | 39 | si:ch211-196l7.4 |
| ogt.12 | 0.0032 | 0.332465376 | 0.25 | 0.075 | 1 | 39 | ogt.1 |
| eif3s107 | 0.0033 | 0.354905912 | 0.45 | 0.18 | 1 | 39 | eif3s10 |
| atp5if1b8 | 0.0033 | 0.372438241 | 0.5 | 0.209 | 1 | 39 | atp5if1b |
| spry25 | 0.0033 | 0.282868959 | 0.25 | 0.073 | 1 | 39 | spry2 |
| cpsf6 | 0.0034 | 0.279293925 | 0.3 | 0.097 | 1 | 39 | cpsf6 |
| pnp5a7 | 0.0035 | 0.380705367 | 0.25 | 0.076 | 1 | 39 | pnp5a |
| slc25a55a5 | 0.0035 | 0.325326538 | 0.25 | 0.076 | 1 | 39 | slc25a55a |

| | | | | | | | |
|---|---|---|---|---|---|---|---|
| ywhaqa3 | 0.0036 | 0.403829578 | 0.5 | 0.211 | 1 | 39 | ywhaqa |
| stau29 | 0.0037 | 0.27668409 | 0.3 | 0.097 | 1 | 39 | stau2 |
| metrnla8 | 0.0037 | 0.269603298 | 0.3 | 0.096 | 1 | 39 | metrnla |
| mt-nd214 | 0.0041 | 0.491731693 | 0.75 | 0.394 | 1 | 39 | mt-nd2 |
| si:dkey-262k9.24 | 0.0041 | 0.300343025 | 0.15 | 0.034 | 1 | 39 | si:dkey-262k9.2 |
| scamp21 | 0.0042 | 0.265128523 | 0.25 | 0.076 | 1 | 39 | scamp2 |
| sh3kbp16 | 0.0043 | 0.31463678 | 0.25 | 0.078 | 1 | 39 | sh3kbp1 |
| helz | 0.0043 | 0.345707559 | 0.2 | 0.056 | 1 | 39 | helz |
| pgrmc19 | 0.0044 | 0.353272859 | 0.4 | 0.154 | 1 | 39 | pgrmc1 |
| zgc:1119866 | 0.0045 | 0.377489217 | 0.45 | 0.188 | 1 | 39 | zgc:111986 |
| tent4b | 0.0045 | 0.254510375 | 0.2 | 0.054 | 1 | 39 | tent4b |
| ppib14 | 0.0045 | 0.469747738 | 0.75 | 0.381 | 1 | 39 | ppib |
| khdrbs1b1 | 0.0045 | 0.392959835 | 0.3 | 0.107 | 1 | 39 | khdrbs1b |
| skp13 | 0.0045 | 0.379134319 | 0.6 | 0.27 | 1 | 39 | skp1 |
| pfn212 | 0.0045 | 0.384326971 | 0.55 | 0.241 | 1 | 39 | pfn2 |
| atrn1 | 0.0046 | 0.262021927 | 0.1 | 0.018 | 1 | 39 | atrn |
| irf2bp2b9 | 0.0046 | 0.323625543 | 0.3 | 0.103 | 1 | 39 | irf2bp2b |
| supt16h1 | 0.0047 | 0.298763668 | 0.25 | 0.077 | 1 | 39 | supt16h |
| arl12 | 0.0047 | 0.270606079 | 0.25 | 0.078 | 1 | 39 | arl1 |
| vps4b2 | 0.0049 | 0.380893018 | 0.3 | 0.107 | 1 | 39 | vps4b |
| fam53b4 | 0.0049 | 0.293734466 | 0.25 | 0.078 | 1 | 39 | fam53b |
| flot1b4 | 0.0049 | 0.282604132 | 0.3 | 0.102 | 1 | 39 | flot1b |
| fscn1a4 | 0.005 | 0.254825961 | 0.2 | 0.054 | 1 | 39 | fscn1a |
| abi2b1 | 0.0051 | 0.283292367 | 0.2 | 0.055 | 1 | 39 | abi2b |
| dusp111 | 0.0051 | 0.320078394 | 0.4 | 0.151 | 1 | 39 | dusp1 |
| atf4a5 | 0.0052 | 0.358003068 | 0.45 | 0.187 | 1 | 39 | atf4a |
| ier2a17 | 0.0054 | 0.499099539 | 0.55 | 0.276 | 1 | 39 | ier2a |
| cyc11 | 0.0054 | 0.282932919 | 0.4 | 0.154 | 1 | 39 | cyc1 |
| chmp1b5 | 0.0055 | 0.341247157 | 0.35 | 0.134 | 1 | 39 | chmp1b |
| birc25 | 0.0055 | 0.370016803 | 0.3 | 0.109 | 1 | 39 | birc2 |
| zgc:1584635 | 0.0057 | 0.337353764 | 0.25 | 0.082 | 1 | 39 | zgc:158463 |
| uqcrq6 | 0.0057 | 0.379285072 | 0.6 | 0.288 | 1 | 39 | uqcrq |
| si:ch211-235e9.62 | 0.006 | 0.345610762 | 0.4 | 0.162 | 1 | 39 | si:ch211-235e9.6 |
| vps29 | 0.006 | 0.33941575 | 0.35 | 0.135 | 1 | 39 | vps29 |
| pfdn5 | 0.0061 | 0.302925272 | 0.4 | 0.159 | 1 | 39 | pfdn5 |

| | | | | | | | |
|---|---|---|---|---|---|---|---|
| si:rp71-36a1.11 | 0.0062 | 0.260925882 | 0.15 | 0.037 | 1 | 39 | si:rp71-36a1.1 |
| wasf22 | 0.0062 | 0.290224902 | 0.25 | 0.081 | 1 | 39 | wasf2 |
| CABZ01113812.1 | 0.0062 | 0.257918936 | 0.15 | 0.036 | 1 | 39 | CABZ01113812.1 |
| arf2a1 | 0.0062 | 0.327352067 | 0.35 | 0.133 | 1 | 39 | arf2a |
| hmgn710 | 0.0062 | 0.405121073 | 0.65 | 0.319 | 1 | 39 | hmgn7 |
| ahnak13 | 0.0064 | 0.403750474 | 0.35 | 0.138 | 1 | 39 | ahnak |
| larp1b | 0.0065 | 0.302005801 | 0.15 | 0.037 | 1 | 39 | larp1b |
| ndufs61 | 0.0066 | 0.357784064 | 0.4 | 0.163 | 1 | 39 | ndufs6 |
| sqstm12 | 0.0067 | 0.278584433 | 0.25 | 0.08 | 1 | 39 | sqstm1 |
| pdcd4a2 | 0.0068 | 0.351564188 | 0.2 | 0.06 | 1 | 39 | pdcd4a |
| selenof7 | 0.0068 | 0.331745098 | 0.6 | 0.269 | 1 | 39 | selenof |
| tsc22d312 | 0.0071 | 0.318594835 | 0.4 | 0.161 | 1 | 39 | tsc22d3 |
| mtpn8 | 0.0071 | 0.335499748 | 0.4 | 0.165 | 1 | 39 | mtpn |
| CABZ01045617.11 | 0.0071 | 0.290188216 | 0.2 | 0.059 | 1 | 39 | CABZ01045617.1 |
| atp5fa12 | 0.0073 | 0.376132974 | 0.75 | 0.372 | 1 | 39 | atp5fa1 |
| vdac28 | 0.0075 | 0.390798028 | 0.65 | 0.328 | 1 | 39 | vdac2 |
| b2ml15 | 0.0075 | 0.390451246 | 0.6 | 0.296 | 1 | 39 | b2ml |
| cox7c6 | 0.0077 | 0.434802522 | 0.85 | 0.495 | 1 | 39 | cox7c |
| cfl113 | 0.0077 | 0.399733231 | 0.9 | 0.602 | 1 | 39 | cfl1 |
| rtf1 | 0.0078 | 0.303809919 | 0.25 | 0.084 | 1 | 39 | rtf1 |
| gnal1 | 0.0078 | 0.252587315 | 0.15 | 0.038 | 1 | 39 | gnal |
| capza1b9 | 0.0078 | 0.296525849 | 0.55 | 0.236 | 1 | 39 | capza1b |
| usp10 | 0.0082 | 0.268349765 | 0.25 | 0.082 | 1 | 39 | usp10 |
| sfpq1 | 0.0082 | 0.352912992 | 0.3 | 0.112 | 1 | 39 | sfpq |
| si:dkey-180p18.94 | 0.0082 | 0.288515546 | 0.3 | 0.11 | 1 | 39 | si:dkey-180p18.9 |
| fundc11 | 0.0082 | 0.260666006 | 0.25 | 0.082 | 1 | 39 | fundc1 |
| krtcap24 | 0.0083 | 0.352114589 | 0.45 | 0.197 | 1 | 39 | krtcap2 |
| tagln215 | 0.0085 | 0.29599697 | 0.5 | 0.211 | 1 | 39 | tagln2 |
| abl12 | 0.0086 | 0.259650544 | 0.15 | 0.038 | 1 | 39 | abl1 |
| h1fx11 | 0.0086 | 0.391729991 | 0.45 | 0.192 | 1 | 39 | h1fx |
| skap29 | 0.0087 | 0.253838831 | 0.25 | 0.082 | 1 | 39 | skap2 |
| plscr3b1 | 0.0087 | 0.33593784 | 0.3 | 0.111 | 1 | 39 | plscr3b |
| sept7b2 | 0.0089 | 0.301216194 | 0.3 | 0.111 | 1 | 39 | sept7b |
| ndufs3 | 0.0091 | 0.324577163 | 0.3 | 0.114 | 1 | 39 | ndufs3 |
| ythdf1 | 0.0091 | 0.251144398 | 0.2 | 0.06 | 1 | 39 | ythdf1 |

| | | | | | | | |
|---|---|---|---|---|---|---|---|
| arl5c8 | 0.0092 | 0.288201453 | 0.3 | 0.109 | 1 | 39 | arl5c |
| add3a11 | 0.0096 | 0.339518444 | 0.35 | 0.139 | 1 | 39 | add3a |
| tnfrsf1a3 | 0.0097 | 0.255057958 | 0.2 | 0.061 | 1 | 39 | tnfrsf1a |
| COX319 | 0.0099 | 0.365470654 | 0.9 | 0.677 | 1 | 39 | COX3 |
| arf2b2 | 0.0099 | 0.344819561 | 0.4 | 0.17 | 1 | 39 | arf2b |
