## Supplementary material for "Direct Male Development in Chromosomally ZZ Zebrafish": SuppMethods Scripts Wilson

### Supplementary Methods

**SuppMethods 19dpf Seurat**

```
#!/usr/bin/env Rscript


library(Seurat)
library(ggplot2)
library(patchwork)
library(dplyr)
library(sctransform)
library(cowplot)

ZZ.data <- Read10X(data.dir =
"/pathwayTo/19zz/strainedCounts_clustersFALSE")
ZZ <- CreateSeuratObject(counts = ZZ.data, project = "ZZ", min.cells = 3,
min.features = 200)
ZW.data <- Read10X(data.dir =
"/pahthwayTo/19zw/strainedCounts_clustersFALSE")
ZW <- CreateSeuratObject(counts = ZW.data, project = "ZW", min.cells = 3,
min.features = 200)
ZZ[["percent.mt"]] <- PercentageFeatureSet(ZZ, pattern = "^mt-")
ZW[["percent.mt"]] <- PercentageFeatureSet(ZW, pattern = "^mt-")
mitoplot <- VlnPlot(ZZ, features = c("nFeature_RNA", "nCount_RNA",
"percent.mt"), ncol = 3)
ggsave(paste("NA19zz_mitoplot.pdf", sep=""), width=20, height=10,
mitoplot, device = "pdf", limitsize = FALSE)
mitoplot <- VlnPlot(ZW, features = c("nFeature_RNA", "nCount_RNA",
"percent.mt"), ncol = 3)
ggsave(paste("NA19zw_mitoplot.pdf", sep=""), width=20, height=10,
mitoplot, device = "pdf", limitsize = FALSE)
getwd()
ZZ <- subset(ZZ, subset = nFeature_RNA > 200 & nFeature_RNA < 2000 &
percent.mt < 8)
ZW <- subset(ZW, subset = nFeature_RNA > 200 & nFeature_RNA < 2000 &
percent.mt < 8)
ZZ <- NormalizeData(ZZ)
ZW <- NormalizeData(ZW)
gonadcombined <- merge(x=ZZ, y=ZW, merge.data = TRUE)
gonadcombined <- FindVariableFeatures(gonadcombined, selection.method =
"vst", nfeatures = 2000)
top10 <- head(VariableFeatures(gonadcombined), 10)
top10
all.genes <- rownames(gonadcombined)
gonadcombined <- ScaleData(gonadcombined, features = all.genes)
gonadcombined <- RunPCA(gonadcombined, features = VariableFeatures(object
= gonadcombined))
elbow <- ElbowPlot(gonadcombined, ndims = 50)
ggsave(paste("elbow.pdf", sep=""), width=20, height=10, elbow, device =
"pdf", limitsize = FALSE)

gonadcombined <- FindNeighbors(gonadcombined, dims = 1:40)
```

```
gonadcombined <- FindClusters(gonadcombined, resolution = 1)
gonadcombined <- RunUMAP(gonadcombined, dims = 1:40)

testplot2 <- DimPlot(gonadcombined, reduction = "umap", label = TRUE)
ggsave(paste("ALLcells_Nadia19dpf_ndims40res1.pdf", sep=""), width=10,
height=10, testplot2, device = "pdf", limitsize = FALSE)
p3 <- DimPlot(gonadcombined, reduction = "umap", group.by = "orig.ident",
raster=FALSE)
ggsave(paste("ALL_cells_gonadumapreplicat19dpf.pdf", sep=""), width=10,
height=10, p3, device = "pdf", limitsize = FALSE)

custom_theme <- theme(legend.position = c(0.1,0.1), axis.title =
element_blank(),plot.title = element_text(size=20, face="bold.italic"),
axis.text = element_blank(), axis.ticks = element_blank(), axis.line =
element_blank(), panel.border = element_rect(colour = "black", fill=NA,
size=1))

p1 <- FeaturePlot(gonadcombined, features = c("cpa2", "alas2",
"lypc"),label.size = 8, order=TRUE,pt.size=1.5, label=FALSE, ncol=3, cols
= c("gray90","red2"))
p3 <- p1 & custom_theme
ggsave(paste("ALL_cells_MarkerGenes_GonadCombined.jpeg", sep=""),
width=30, height=10, p3, device = "jpeg", limitsize = FALSE)


saveRDS(gonadcombined, file =
"./Nadia19dpf_allcells_SoupXclusterFalse_40dims_res1.rds")

gonadsubset <- subset(gonadcombined, idents = c(2,8, 10,21,6), invert =
TRUE)
gonadsubset <- FindVariableFeatures(gonadsubset, selection.method = "vst",
nfeatures = 2000)
top10 <- head(VariableFeatures(gonadsubset), 10)
top10
all.genes <- rownames(gonadsubset)
gonadsubset <- ScaleData(gonadsubset, features = all.genes)
gonadsubset <- RunPCA(gonadsubset, features = VariableFeatures(object =
gonadsubset))
elbow <- ElbowPlot(gonadsubset, ndims = 50)
ggsave(paste("elbowsubset.pdf", sep=""), width=20, height=10, elbow,
device = "pdf", limitsize = FALSE)
gonadsubset <- FindNeighbors(gonadsubset, dims = 1:50)
gonadsubset <- FindClusters(gonadsubset, resolution = 1.5)
gonadsubset <- RunUMAP(gonadsubset, dims = 1:50)

testplot2 <- DimPlot(gonadsubset, reduction = "umap", label = TRUE)
ggsave(paste("Subset1_Nadia19dpfsubset1_ndims15res075.pdf", sep=""),
width=10, height=10, testplot2, device = "pdf", limitsize = FALSE)
p3 <- DimPlot(gonadsubset, reduction = "umap", group.by = "orig.ident",
raster=FALSE)
ggsave(paste("Subset1_gonadumapreplicat19dpfsubset1_nopanciridrbcs.pdf",
sep=""), width=10, height=10, p3, device = "pdf", limitsize = FALSE)
custom_theme <- theme(legend.position = c(0.1,0.1), axis.title =
element_blank(),plot.title = element_text(size=20, face="bold.italic"),
```

```
axis.text = element_blank(), axis.ticks = element_blank(), axis.line =
element_blank(), panel.border = element_rect(colour = "black", fill=NA,
size=1))
p1 <- FeaturePlot(gonadsubset, features = c("neu3.5", "nid2a","egfl6",
"lhx9", "cpa2", "rspo1", "islr2", "s100a10a", "ddx4", "slc29a1b",
"cyp11a1", "eya1", "lypc","gsdf", "amh", "alas2", "hbba1", "cpa5",
"amy2a", "prss59.2"),label.size = 8, order=TRUE,pt.size=1, label=TRUE,
ncol=4, cols = c("gray90","red2"))
p3 <- p1 & custom_theme
ggsave(paste("Subset1_MarkerGenes.jpeg", sep=""), width=40, height=50, p3,
device = "jpeg", limitsize = FALSE)



gonadsubset2 <- subset(gonadsubset, idents = c(5), invert = TRUE)
gonadsubset2 <- FindVariableFeatures(gonadsubset2, selection.method =
"vst", nfeatures = 2000)
top10 <- head(VariableFeatures(gonadsubset2), 10)
top10
all.genes <- rownames(gonadsubset2)
gonadsubset <- ScaleData(gonadsubset2, features = all.genes)
gonadsubset <- RunPCA(gonadsubset2, features = VariableFeatures(object =
gonadsubset2))
elbow <- ElbowPlot(gonadsubset2, ndims = 50)
ggsave(paste("elbowsubset2.pdf", sep=""), width=20, height=10, elbow,
device = "pdf", limitsize = FALSE)



gonadsubset2 <- FindNeighbors(gonadsubset2, dims = 1:25)
gonadsubset2 <- FindClusters(gonadsubset2, resolution = 2.25)
gonadsubset2 <- RunUMAP(gonadsubset2, dims = 1:25)

testplot2 <- DimPlot(gonadsubset2, reduction = "umap", label = TRUE)
ggsave(paste("Subset2_Nadia19dpfsubset2_ndims25res2-25.pdf", sep=""),
width=10, height=10, testplot2, device = "pdf", limitsize = FALSE)


saveRDS(gonadsubset2, file =
"./Nadia19dpf_allcells_gonadSubset2_SoupXclusterFalse_25dims_res2-25.rds")


all.markers2 <- FindAllMarkers(gonadsubset2, only.pos = TRUE, min.pct =
0.25, logfc.threshold = 0.25)
all.markers2 %>% group_by(cluster) %>% top_n(n = 2, wt = avg_log2FC)
write.csv(all.markers2,
file="19dpf_SoupxNoCluster_Subset2_allMarkers_subset2ndims25res2-
25_minpct25.csv")


all.markers2 <- FindAllMarkers(gonadsubset2, only.pos = TRUE, min.pct =
0.1, logfc.threshold = 0.25)
all.markers2 %>% group_by(cluster) %>% top_n(n = 2, wt = avg_log2FC)
```

```
write.csv(all.markers2,
file="19dpf_SoupxNoCluster_Subset2_allMarkers_subset2ndims25res2-
25_minpct10.csv")
```

≈≈≈≈≈≈≈≈≈≈≈≈≈≈≈≈≈≈≈

**2. SuppMethods 30dpf Seurat**

```
#!/usr/bin/env Rscript

library(Seurat)
library(ggplot2)
library(patchwork)
library(dplyr)
library(sctransform)
library(cowplot)


ZZ.data <- Read10X(data.dir = "/pathTo/30zw/strainedCounts_clustersFalse")
ZZ <- CreateSeuratObject(counts = ZZ.data, project = "ZZ", min.cells = 3,
min.features = 200)

ZW.data <- Read10X(data.dir = "/pathTo/30ZZ/strainedCounts_clustersFalse")
ZW <- CreateSeuratObject(counts = ZW.data, project = "ZW", min.cells = 3,
min.features = 200)

ZZ[["percent.mt"]] <- PercentageFeatureSet(ZZ, pattern = "^mt-")
ZW[["percent.mt"]] <- PercentageFeatureSet(ZW, pattern = "^mt-")
mitoplot <- VlnPlot(ZZ, features = c("nFeature_RNA", "nCount_RNA",
"percent.mt"), ncol = 3)
ggsave(paste("NA30-ZZ_mitoplot.pdf", sep=""), width=20, height=10,
mitoplot, device = "pdf", limitsize = FALSE)
mitoplot <- VlnPlot(ZW, features = c("nFeature_RNA", "nCount_RNA",
"percent.mt"), ncol = 3)
ggsave(paste("NA30-ZW_mitoplot.pdf", sep=""), width=20, height=10,
mitoplot, device = "pdf", limitsize = FALSE)
ZZ <- subset(ZZ, subset = nFeature_RNA > 200 & nFeature_RNA < 2000 &
percent.mt < 8)
ZW <- subset(ZW, subset = nFeature_RNA > 200 & nFeature_RNA < 2000 &
percent.mt < 8)
ZZ <- NormalizeData(ZZ)
ZW <- NormalizeData(ZW)

gonadcombined <- merge(x=ZW, y=ZZ, merge.data = TRUE)
gonadcombined <- FindVariableFeatures(gonadcombined, selection.method =
"vst", nfeatures = 2000)
top10 <- head(VariableFeatures(gonadcombined), 10)
top10
all.genes <- rownames(gonadcombined)
gonadcombined <- ScaleData(gonadcombined, features = all.genes)
gonadcombined <- RunPCA(gonadcombined, features = VariableFeatures(object
= gonadcombined))
elbow <- ElbowPlot(gonadcombined, ndims = 50)
```

```
ggsave(paste("elbow.pdf", sep=""), width=20, height=10, elbow, device =
"pdf", limitsize = FALSE)
gonadcombined <- FindNeighbors(gonadcombined, dims = 1:30)
gonadcombined <- FindClusters(gonadcombined, resolution = 0.75)
gonadcombined <- RunUMAP(gonadcombined, dims = 1:30)
testplot2 <- DimPlot(gonadcombined, reduction = "umap", label = TRUE)
ggsave(paste("Nadia30dpfumapRes075ndims30.pdf", sep=""), width=10,
height=10, testplot2, device = "pdf", limitsize = FALSE)
saveRDS(gonadcombined, file = "./Nadia30dpf_30dims_res075_20230203.rds")


p1 <- FeaturePlot(gonadcombined, features = c("ambp", "hpda","neu3.5",
"nid2a","egfl6", "lhx9", "cpa2","prss59.2","prss1","amy2a","hbba1","hbaa1"
, "rspo1", "islr2", "s100a10a", "ddx4", "slc29a1b", "cyp11a1", "eya1",
"lypc","gsdf", "amh", "alas2", "coro1a", "mpx", "star",
"cyp19a1a"),label.size = 8, order=TRUE,pt.size=1, label=FALSE, ncol=4,
cols = c("gray90","red2"))
ggsave(paste("gonadcombined30dpf_Markers.jpeg", sep=""),width=40,
height=70, p1, device = "jpeg", limitsize = FALSE)


gonadsubset <- subset(gonadcombined, idents = c(6, 4, 7, 29, 31, 19),
invert = TRUE)
table(Idents(gonadsubset))

gonadsubset <- FindVariableFeatures(gonadsubset, selection.method = "vst",
nfeatures = 2000)
top10 <- head(VariableFeatures(gonadsubset), 10)
top10
all.genes <- rownames(gonadsubset)
gonadsubset <- ScaleData(gonadsubset, features = all.genes)
gonadsubset <- RunPCA(gonadsubset, features = VariableFeatures(object =
gonadsubset))
elbow <- ElbowPlot(gonadsubset, ndims = 50)
ggsave(paste("subsetelbow.pdf", sep=""), width=20, height=10, elbow,
device = "pdf", limitsize = FALSE)



gonadsubset <- FindNeighbors(gonadsubset, dims = 1:35)
gonadsubset <- FindClusters(gonadsubset, resolution = 2.25)
gonadsubset <- RunUMAP(gonadsubset, dims = 1:35)




gonadsubset <- FindClusters(gonadsubset, resolution = 2.25)
testplot2 <- DimPlot(gonadsubset, reduction = "umap", label = TRUE)
ggsave(paste("Nadia30dpfLiverPancreasErythExcludeumap_ndims35_res2-
25.pdf", sep=""), width=10, height=10, testplot2, device = "pdf",
limitsize = FALSE)
```

```
saveRDS(gonadsubset, file =
"./Nadia30dpf_gonadSUBSET_SoupXclusterFalse_35dims_res2-25.rds")


all.markers2 <- FindAllMarkers(gonadsubset, only.pos = TRUE, min.pct =
0.25, logfc.threshold = 0.25)
all.markers2 %>% group_by(cluster) %>% top_n(n = 2, wt = avg_log2FC)
write.csv(all.markers2,
file="30dpf_SoupxNoCluster_Subset_allMarkers_subset2ndims35res2-
25_minpct25.csv")


all.markers2 <- FindAllMarkers(gonadsubset, only.pos = TRUE, min.pct =
0.1, logfc.threshold = 0.25)
all.markers2 %>% group_by(cluster) %>% top_n(n = 2, wt = avg_log2FC)
```

≈≈≈≈≈≈≈≈≈≈≈≈≈≈≈≈≈≈≈

**3. SuppMethods 19dpf 30dpf Merged Seurat**

```
#!/usr/bin/env Rscript

library(Seurat)
library(ggplot2)
library(patchwork)
library(dplyr)
library(sctransform)
library(cowplot)

ZW_30.data <- Read10X(data.dir =
"/pathTo/30ZZ/strainedCounts_clustersFalse")
ZW_30 <- CreateSeuratObject(counts = ZW_30.data, project = "ZW_30",
min.cells = 3, min.features = 200)
ZZ_30.data <- Read10X(data.dir =
"/pathTo/30zw/strainedCounts_clustersFalse")
ZZ_30 <- CreateSeuratObject(counts = ZZ_30.data, project = "ZZ_30",
min.cells = 3, min.features = 200)
ZZ_30[["percent.mt"]] <- PercentageFeatureSet(ZZ_30, pattern = "^mt-")
ZW_30[["percent.mt"]] <- PercentageFeatureSet(ZW_30, pattern = "^mt-")
ZZ_30 <- subset(ZZ_30, subset = nFeature_RNA > 200 & nFeature_RNA < 2000 &
percent.mt < 8)
ZW_30 <- subset(ZW_30, subset = nFeature_RNA > 200 & nFeature_RNA < 2000 &
percent.mt < 8)
ZZ_30 <- NormalizeData(ZZ_30)
ZW_30 <- NormalizeData(ZW_30)

ZZ_19.data <- Read10X(data.dir =
"/pathTo/19zz/strainedCounts_clustersFALSE")
ZZ_19 <- CreateSeuratObject(counts = ZZ_19.data, project = "ZZ_19",
min.cells = 3, min.features = 200)
```

```
ZW_19.data <- Read10X(data.dir =
"/pathTo/19zw/strainedCounts_clustersFALSE")
ZW_19 <- CreateSeuratObject(counts = ZW_19.data, project = "ZW_19",
min.cells = 3, min.features = 200)
ZZ_19[["percent.mt"]] <- PercentageFeatureSet(ZZ_19, pattern = "^mt-")
ZW_19[["percent.mt"]] <- PercentageFeatureSet(ZW_19, pattern = "^mt-")
ZZ_19 <- subset(ZZ_19, subset = nFeature_RNA > 200 & nFeature_RNA < 2000 &
percent.mt < 8)
ZW_19 <- subset(ZW_19, subset = nFeature_RNA > 200 & nFeature_RNA < 2000 &
percent.mt < 8)
ZZ_19 <- NormalizeData(ZZ_19)
ZW_19 <- NormalizeData(ZW_19)



ZZ_30 <- AddMetaData(ZZ_30, metadata = "male", col.name = "sex")
ZW_30 <- AddMetaData(ZW_30, metadata = "female", col.name = "sex")
ZZ_19  <- AddMetaData(ZZ_19, metadata = "male", col.name = "sex")
ZW_19 <- AddMetaData(ZW_19, metadata = "female", col.name = "sex")

ZZ_30 <- AddMetaData(ZZ_30, metadata = "30dpf", col.name = "age")
ZW_30 <- AddMetaData(ZW_30, metadata = "30dpf", col.name = "age")
ZZ_19  <- AddMetaData(ZZ_19, metadata = "19dpf", col.name = "age")
ZW_19 <- AddMetaData(ZW_19, metadata = "19dpf", col.name = "age")

gonadcombined <- merge(ZW_30, y = c(ZZ_30, ZZ_19, ZW_19), add.cell.ids =
c("ZZ_30", "ZW_30", "ZZ_19", "ZW_19"), project = "Nadia", merge.data =
TRUE)
gonadcombined <- FindVariableFeatures(gonadcombined, selection.method =
"vst", nfeatures = 2000)
top10 <- head(VariableFeatures(gonadcombined), 10)
top10
all.genes <- rownames(gonadcombined)
gonadcombined <- ScaleData(gonadcombined, features = all.genes)
gonadcombined <- RunPCA(gonadcombined, features = VariableFeatures(object
= gonadcombined))
elbow <- ElbowPlot(gonadcombined, ndims = 100)
ggsave(paste("elbow.pdf", sep=""), width=20, height=10, elbow, device =
"pdf", limitsize = FALSE)



gonadcombined <- FindNeighbors(gonadcombined, dims = 1:40)
gonadcombined <- FindClusters(gonadcombined, resolution = 0.75)
gonadcombined <- RunUMAP(gonadcombined, dims = 1:40)
testplot2 <- DimPlot(gonadcombined, reduction = "umap", label = TRUE)
ggsave(paste("Nadia_19dpf_30dpfCOMBINED_ndims40res075.pdf", sep=""),
width=10, height=10, testplot2, device = "pdf", limitsize = FALSE)

p3 <- DimPlot(gonadcombined, reduction = "umap", group.by = "orig.ident",
raster=FALSE)
ggsave(paste("Nadia_19dpf_30dpfCOMBINED_ndims40res075_origIdent.pdf",
sep=""), width=10, height=10, p3, device = "pdf", limitsize = FALSE)
```

```
p3 <- DimPlot(gonadcombined, reduction = "umap", split.by = "orig.ident",
raster=FALSE)
ggsave(paste("Nadia_19dpf_30dpfCOMBINED_ndims40res075_origIdentsplit.pdf",
sep=""), width=40, height=10, p3, device = "pdf", limitsize = FALSE)

custom_theme <- theme(legend.position = c(0.1,0.1), axis.title =
element_blank(),plot.title = element_text(size=20, face="bold.italic"),
axis.text = element_blank(), axis.ticks = element_blank(), axis.line =
element_blank(), panel.border = element_rect(colour = "black", fill=NA,
size=1))

p1 <- FeaturePlot(gonadcombined, features = c("ambp", "hpda","neu3.5",
"nid2a","egfl6", "lhx9", "cpa2","prss59.2","prss1","amy2a","hbba1","hbaa1"
, "rspo1", "islr2", "s100a10a", "ddx4", "slc29a1b", "cyp11a1", "eya1",
"lypc","gsdf", "amh", "alas2", "coro1a", "mpx", "star",
"cyp19a1a"),label.size = 8, order=TRUE,pt.size=1, label=FALSE, ncol=4,
cols = c("gray90","red2"))
p3 <- p1 & custom_theme
ggsave(paste("gonadcombined_Markers.jpeg", sep=""),width=40, height=70,
p3, device = "jpeg", limitsize = FALSE)



#all.markers <- FindAllMarkers(gonadcombined, only.pos = TRUE, min.pct =
0.25, logfc.threshold = 0.25)
#all.markers %>% group_by(cluster) %>% top_n(n = 2, wt = avg_log2FC)
#write.csv(all.markers, file="allMarkers_gonadcombinedTest.csv")


gonadsubset <- subset(gonadcombined, idents = c(10, 2, 8, 15, 30, 29),
invert = TRUE)
table(Idents(gonadsubset))

gonadsubset <- FindVariableFeatures(gonadsubset, selection.method = "vst",
nfeatures = 2000)
top10 <- head(VariableFeatures(gonadsubset), 10)
top10
all.genes <- rownames(gonadsubset)
gonadsubset <- ScaleData(gonadsubset, features = all.genes)
gonadsubset <- RunPCA(gonadsubset, features = VariableFeatures(object =
gonadsubset), npcs = 200)
elbow <- ElbowPlot(gonadsubset, ndims = 100)
ggsave(paste("subsetelbow.pdf", sep=""), width=20, height=10, elbow,
device = "pdf", limitsize = FALSE)



gonadsubset <- FindNeighbors(gonadsubset, dims = 1:45)
gonadsubset <- FindClusters(gonadsubset, resolution = 1.5)
gonadsubset <- RunUMAP(gonadsubset, dims = 1:45)
testplot2 <- DimPlot(gonadsubset, reduction = "umap", label = TRUE)
```

```
ggsave(paste("Nadia_19dpf_30dpfCOMBINED_SUBSET___ndims45res1-
5_20230203.pdf", sep=""), width=10, height=10, testplot2, device = "pdf",
limitsize = FALSE)




###############################
#
#

saveRDS(gonadsubset, file = "./Nadia30dpf_19dpf_SUBSET2_45dims_res1-
5_20230203.rds")



p3 <- DimPlot(gonadsubset, reduction = "umap", group.by = "orig.ident",
raster=FALSE)
ggsave(paste("Nadia_19dpf_30dpfCOMBINED_SUBSET_45dims_res1-
5_origIdent_20230203.pdf", sep=""), width=10, height=10, p3, device =
"pdf", limitsize = FALSE)

p3 <- DimPlot(gonadsubset, reduction = "umap", split.by = "orig.ident",
raster=FALSE)
ggsave(paste("Nadia_19dpf_30dpfCOMBINED_SUBSET_45dims_res1-
55_origIdentsplit_20230203.pdf", sep=""), width=40, height=10, p3, device
= "pdf", limitsize = FALSE)

p3 <- DimPlot(gonadsubset, reduction = "umap", group.by = "sex",
raster=FALSE)
ggsave(paste("Nadia_19dpf_30dpfCOMBINED_SUBSET_n45dims_res1-
5_sex_20230203.pdf", sep=""), width=10, height=10, p3, device = "pdf",
limitsize = FALSE)

p3 <- DimPlot(gonadsubset, reduction = "umap", group.by = "age",
raster=FALSE)
ggsave(paste("Nadia_19dpf_30dpfCOMBINED_SUBSET_45dims_res1-
5_age_20230203.pdf", sep=""), width=10, height=10, p3, device = "pdf",
limitsize = FALSE)



all.markers <- FindAllMarkers(gonadsubset, only.pos = TRUE, min.pct =
0.25, logfc.threshold = 0.25)
all.markers %>% group_by(cluster) %>% top_n(n = 2, wt = avg_log2FC)
write.csv(all.markers, file="allMarkers_gonadCOMBINED_SUBSET_ndims45_res1-
5_20230203_minpct25.csv")


all.markers <- FindAllMarkers(gonadsubset, only.pos = TRUE, min.pct = 0.1,
logfc.threshold = 0.25)
all.markers %>% group_by(cluster) %>% top_n(n = 2, wt = avg_log2FC)
```

```
write.csv(all.markers, file="allMarkers_gonadCOMBINED_SUBSET_ndims45_res1-5_20230203_minpct10.csv")
```

≈≈≈≈≈≈≈≈≈≈≈≈≈≈≈≈≈≈≈≈

**4. SuppMethods soupx commands**

```
##30dpf ZW

library(DropletUtils)
library(SoupX)
sc = load10X("/pathTo/30dpf_zw/1/outs/")

gcGenes = c("hbaa1", "hbaa2", "hbba2", "hbba1", "hbae5", "ddx4", "dnd1", "sycp1", "sycp3", "dmc1")

useToEst = estimateNonExpressingCells(sc, nonExpressedGeneList = list(gcGenes))

sc = calculateContaminationFraction(sc, list(gcGenes), useToEst = useToEst)

out = adjustCounts(sc)

DropletUtils:::write10xCounts("./strainedCounts_clusters", out)


##30dpf ZZ
library(DropletUtils)
library(SoupX)
sc = load10X("/pathTo/30dpf_zz/2/outs/")

gcGenes = c("hbaa1", "hbaa2", "hbba2", "hbba1", "hbae5", "ddx4", "dnd1", "sycp1", "sycp3", "dmc1")

useToEst = estimateNonExpressingCells(sc, nonExpressedGeneList = list(gcGenes), clusters=FALSE)

sc = calculateContaminationFraction(sc, list(gcGenes), useToEst = useToEst)

out = adjustCounts(sc)

DropletUtils:::write10xCounts("./strainedCounts_clustersFalse", out)

##19dpf ZW
library(DropletUtils)
library(SoupX)
sc = load10X("/pathTo/19dpf_zw/zw_19NAfem/outs/")

gcGenes = c("hbaa1", "cpa5", "cpa1", "cel.2", "cel.1", "amy2a", "hbba1", "ela2l", "ela2", "prss1", "hbae5", "prss59.2")
```

```
useToEst = estimateNonExpressingCells(sc, nonExpressedGeneList =
list(gcGenes),clusters=FALSE)

sc = calculateContaminationFraction(sc, list(gcGenes), useToEst =
useToEst)

out = adjustCounts(sc)

DropletUtils:::write10xCounts("./strainedCounts_clustersFALSE", out)


##19dpf ZZ
library(DropletUtils)
library(SoupX)
sc = load10X("/pathTo/19dpf_zz/zz_19NAmale/outs/")

gcGenes = c("hbaa1", "cpa5", "cpa1", "cel.2", "cel.1", "amy2a", "hbba1",
"ela2l", "ela2", "prss1", "hbae5", "prss59.2")

useToEst = estimateNonExpressingCells(sc, nonExpressedGeneList =
list(gcGenes),clusters=FALSE)

sc = calculateContaminationFraction(sc, list(gcGenes), useToEst =
useToEst)

out = adjustCounts(sc)

DropletUtils:::write10xCounts("./strainedCounts_clustersFALSE", out)
```
